# Burkholderia PglL enzymes are serine preferring oligosaccharidetransferases which target conserved proteins across the Burkholderia genus

**DOI:** 10.1101/2021.04.20.440587

**Authors:** Andrew J. Hayes, Jessica M. Lewis, Mark R. Davies, Nichollas E. Scott

## Abstract

Glycosylation is increasingly recognised as a common protein modification within bacterial proteomes. While great strides have been made in identifying species that contain glycosylation systems, our understanding of the proteins and sites targeted by these enzymes is far more limited. Within this work we explore the conservation of glycoproteins and *O*-linked glycosylation sites across the pan-*Burkholderia* glycoproteome. Using a multi-protease glycoproteomic approach we generate high-confidence glycoproteomes and associated glycosylation sites in two widely utilized *B. cenocepacia* strains, K56-2 and H111. This resource reveals glycosylation occurs exclusively at serine residues and that glycoproteins/glycosylation sites are highly conserved across 294 publicly available *B. cenocepacia* genomes. Consistent with this we demonstrate that the substitution of Serine for Threonine residues in a model protein results in a dramatic decrease in glycosylation efficiency by the oligosaccharidetransferase pglL_BC_ even when pglL_BC_ is overexpressed. This preference for glycosylation at Serine residues is observed across at least 9 *Burkholderia* glycoproteomes supporting that Serine is the dominant residue targeted by pglL-mediated glycosylation across the *Burkholderia* genus. Using population genomics we observe that pglL targeted glycosylated proteins are common across *Burkholderia* species. Combined, this work demonstrates that PglL enzymes of the Burkholderia genus are Serine-preferring oligosaccharidetransferases that target conserved and shared protein substrates across the *Burkholderia* genus.

## Introduction

Glycosylation is a common class of protein modifications increasingly recognised within bacterial proteomes ^1,2^. Within bacterial proteomes, the installation of glycans has been shown to influence protein functions by enhancing protein stability ^3–5^, stabilizing protein complexes ^6^, masking antigenic sites ^7^ and modulating protein enzymatic activities ^8,9^. As these events fine-tune the proteome, identifying novel glycosylation systems has been a key goal of microbial glycoproteomics. Over the last decade, with the aid of mass spectrometry-based proteomics, a range of bacterial glycosylation systems have been identified ^1,2,10,11^. Although these studies have demonstrated the commonality of bacterial glycosylation, the majority of follow up studies have focused on understanding the biology of protein glycosylation glycans, including elucidating their biosynthetic pathways ^12–14^ and defining their diversity ^15–17^. This focus has resulted in a limited understanding of the specific glycoproteins and glycosylation sites within the majority of known bacterial glycoproteomes.

To date arguably the best characterised bacterial glycoproteome is that of the *N*-linked glycosylation system of *Campylobacter jejuni*, where 134 glycosylation sites have been experimentally identified ^18^. The success in mapping glycosylation across the *C. jejuni* glycoproteome has largely been due to two features of this system; i) glycosylation events within the Campylobacter genus occurring within a unique glycosylation sequon, D/E-X-N-X-S/T ^19^, which restricts the possible glycosylation sites within a protein and ii) the chemical nature of *N*-linked glycosylation which can allow localisation information to be obtained using collision-based Mass Spectrometry (MS) fragmentation approaches ^20^. For bacterial mediated *O*-linked glycosylation systems, far less is known about the sites of glycosylation ^1,2^. This limited understanding of bacterial *O*-linked glycosylation sites is largely driven by technical limitations associated with the analysis of *O*-linked glycopeptides, where the chemical nature of *O*-linked glycosylation requires approaches such as Electron-transfer dissociation (ETD) or Electron-Transfer/Higher-Energy Collision Dissociation (EThcD) to localise glycosylation sites ^20^. Defining the glycosylation sites are critical for enabling functional characterisation of glycoproteins as well as for understanding the conservation of sites across species. From previous studies it has become clear that bacterial glycosylation sites can vary even within closely related strains/species ^21,22^ and that understanding glycosylation site heterogeneity can provide critical insights into functionally important nuances within glycoproteins. An example of this can be seen in the pilin of *Neisseria* species where high levels of glycosylation are correlated with masking conserved antigenic regions while forms of pilin prone to amino acid sequence variation possess limited pilin glycosylation ^7,14,23,24^.

One of the most widespread families of *O*-linked glycosylating enzymes are the PglL oligosaccharidetransferases ^25^ which have been experimentally shown to *O*-glycosylate proteins within *Acinetobacter* ^26,27^, *Neisseria* ^14,17,28^, *Burkholderia* ^12,16,29^, *Francisella* ^30^, *Pseudomonas* ^31^, and *Ralstonia* ^32^ species. Within these species, PglL mediates *O*-linked glycosylation of substrates within the periplasmic space, and can be responsible for the modification of a single protein or multiple proteins depending on the enzyme ^10,11^. To date, no rigid glycosylation sequon has been observed within PglL substrates, with glycosylation predominately occurring in disordered regions rich in Alanine and Prolines ^15,29,33,34^. For PglL enzymes responsible for the glycosylation of multiple proteins, also known as general glycosylation systems, the abolishment of PglL has been shown to result in profound effects such as the attenuation of virulence ^27,29,32^. Within these systems the lack of a detailed understanding of the glycoproteomes limits our ability to understand how glycosylation influences virulence and the glycosylation sites which are critical for virulence associated processes.

Recently we began exploring the pan-glycoproteome of *Burkholderia* species focusing on the conservation of the glycans used for glycosylation ^12,16^ and the properties of glycopeptides which can influence their enrichment using zwitterionic hydrophilic interaction liquid chromatography (ZIC-HILIC) ^35^. These studies have highlighted that the glycoproteomes of *Burkholderia* species appear larger than initially thought, yet, due to the glycosylation sites associated with these proteins not being defined, this has limited our ability to understand the general trends within site utilisation across the *Burkholderia* pan-glycoproteome. To improve our understanding of *Burkholderia* glycosylation we have undertaken a site focused glycoproteomic study of *Burkholderia cenocepacia* within two widely utilised strains, K56-2 ^36^ and H111 ^37^. Leveraging this curated resource, we gain and experimentally confirm a previously unrecognised preference in the specificity of *O*-linked glycosylation as well as demonstrate that *Burkholderia* glycosylation targets conserved protein substrates across this genus. Combined this work further demonstrates a previously underappreciated aspect of *O*-linked glycosylation conservation within *Burkholderia* species, the conservation of proteins substrates.

## Methods

### Bacterial strains and growth conditions

Strains and plasmids used in this study are listed in Tables 1 and 2, respectively. Strains of *Escherichia coli* and *B. cenocepacia* were grown at 37°C on Lysogeny Broth (LB) medium. When required, antibiotics were added to a final concentration of 50 μg/ml trimethoprim for *E. coli* and 100 μg/ml for *B. cenocepacia*, 20 μg/ml tetracycline for *E. coli* and 150 μg/ml for *B. cenocepacia* and 40 μg/ml kanamycin for *E. coli*. Ampicillin was used at 100 μg/ml and polymyxin B at 25 μg/ml for triparental mating to select against donor and helper *E. coli* strains as previously described ^38^. Induction of tagged glycoproteins within *Burkholderia* strains was undertaken by the addition of either Rhamnose (final concentration 0.1%, for pSCrhaB2-based plasmids) or Arabinose (final concentration 0.5%, for pKM4-based plasmids) to overnight cultures. Antibiotics were purchased from Thermo Fisher Scientific while all other chemicals, unless otherwise stated, were provided by Sigma-Aldrich.

**Table 1:** Strains list

| Strain name | Description | Source/<br>Reference |
| --- | --- | --- |
| *E. coli* strain |  |  |
| *E. coli* pir2 | F<sup>-</sup> Δlac169 rpoS(Am) robA1 creC510 hsdR514 endA recA1 uidA(ΔMlul)::pir-116 | Thermo Scientific |
| *B. cenocepacia* strains |  |  |
| *B. cenocepacia* K56-2 | Clinical isolate of the ET12 lineage (Darling P 1998, Mahenthiralingam E *et al* 2005) | Canadian *B. cepacia* research and referral repository <sup>36</sup> |
| *B. cenocepacia* K56-2 ΔpglL | ΔpglL (BCAL0960) derivative of K56-2 created using pYM8 | 5 |
| *B. cenocepacia* K56-2 ΔpglL amrAB::S7-pglL-his<sub>10</sub> | amrAB::S7-pglL-his<sub>10</sub> chromosomal complement derivative of ΔpglL (BCAL0960) overexpressing pglL from the S7 promoter | 5 |
| *B. cenocepacia* H111 | Clinically non-ET12 lineage isolate | 37 |
| *B. cenocepacia* H111 ΔpglL candidate 1 | ΔpglL (I35_RS13570) derivative of H111 created using pYM8 candidate | This study |
| *B. cenocepacia* H111 ΔpglL candidate 2 | ΔpglL (I35_RS13570) derivative of H111 created using pYM8 candidate | This study |

**Table 2:**
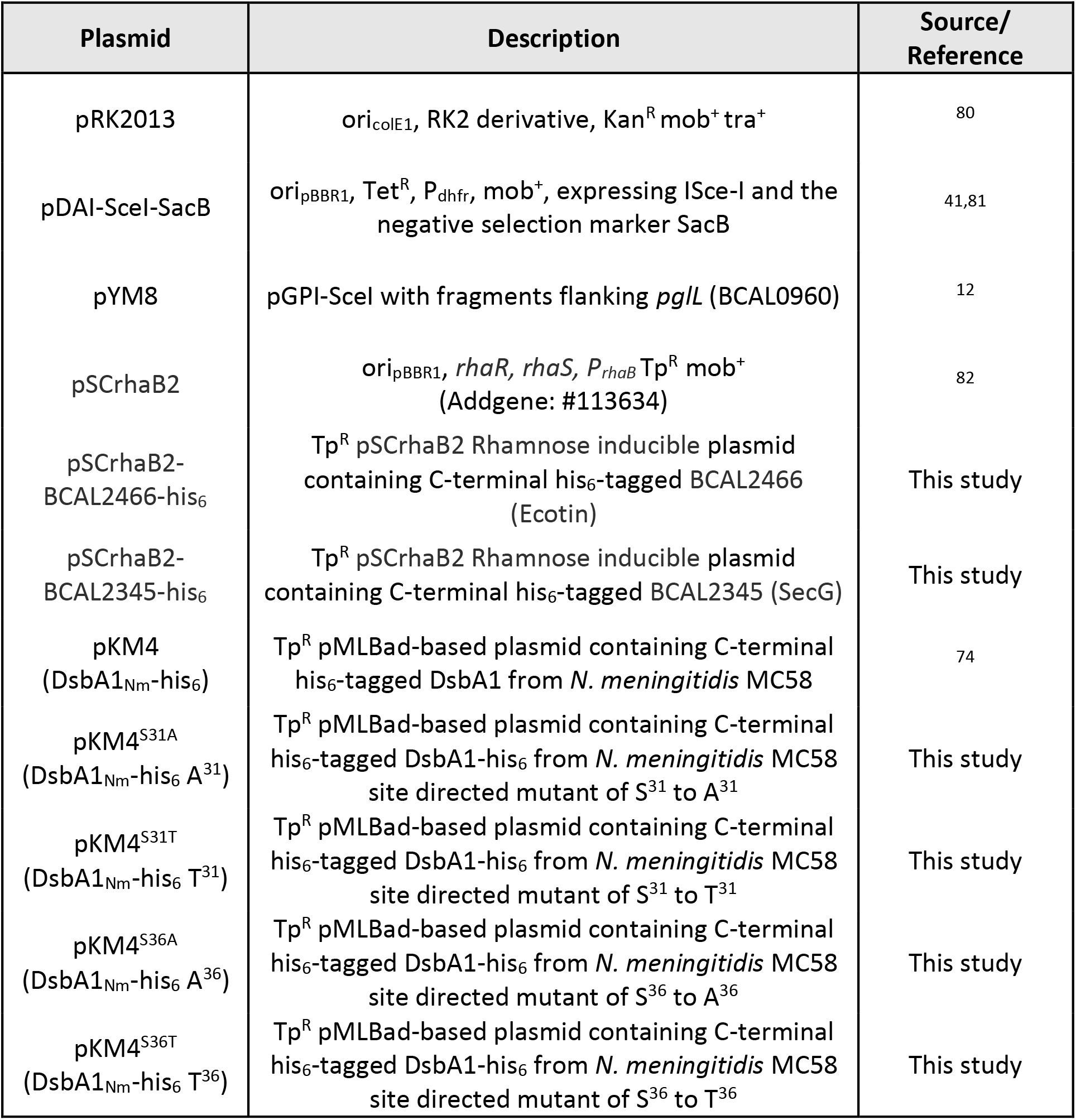
Plasmid list

### Recombinant DNA methods

Oligonucleotides used in this study are provided in Table 3. The inducible pSCrhaB2-BCAL2345-his_6_ and pSCrhaB2-BCAL2466-his_6_ constructs were generated using Gibson assembly ^39^ by inserting PCR amplified fragments into *Nde*I and *Xba*I linearized pSCrhaB2 using NEBuilder^®^ HiFi DNA master mix according to the manufacturer’s instructions (New England Biolabs). pKM4 Site directed mutagenesis was undertaken using PCR-based site replacement and *Dpn*I digestion ^40^. All restriction endonuclease digestions, and agarose gel electrophoresis were performed using standard molecular biology techniques ^40^. All restriction enzymes were used according to the manufacturer’s instructions (New England Biolabs). Chemically competent *E. coli* pir2 cells were transformed using heat shock-based transformation ^40^. PCR amplifications were carried out using Phusion DNA polymerase (Thermo Fisher Scientific) according to the manufacturer’s recommendations with the addition of 2.5% DMSO for the amplification of *B. cenocepacia* DNA, due to its high GC content. Genomic DNA isolations were performed using genomic DNA clean-up Kits (Zmyo research), while PCR recoveries and restriction digest purifications were performed using DNA Clean & Concentrator Kits (Zmyo research). Colony and screening PCRs were performed using GoTaq DNA polymerase (Promega; supplemented with 10% DMSO when screening *B. cenocepacia* gDNA). DNA sequencing was undertaken at the Australian Genome Research Facility (Melbourne, Australia).

**Table 3:**
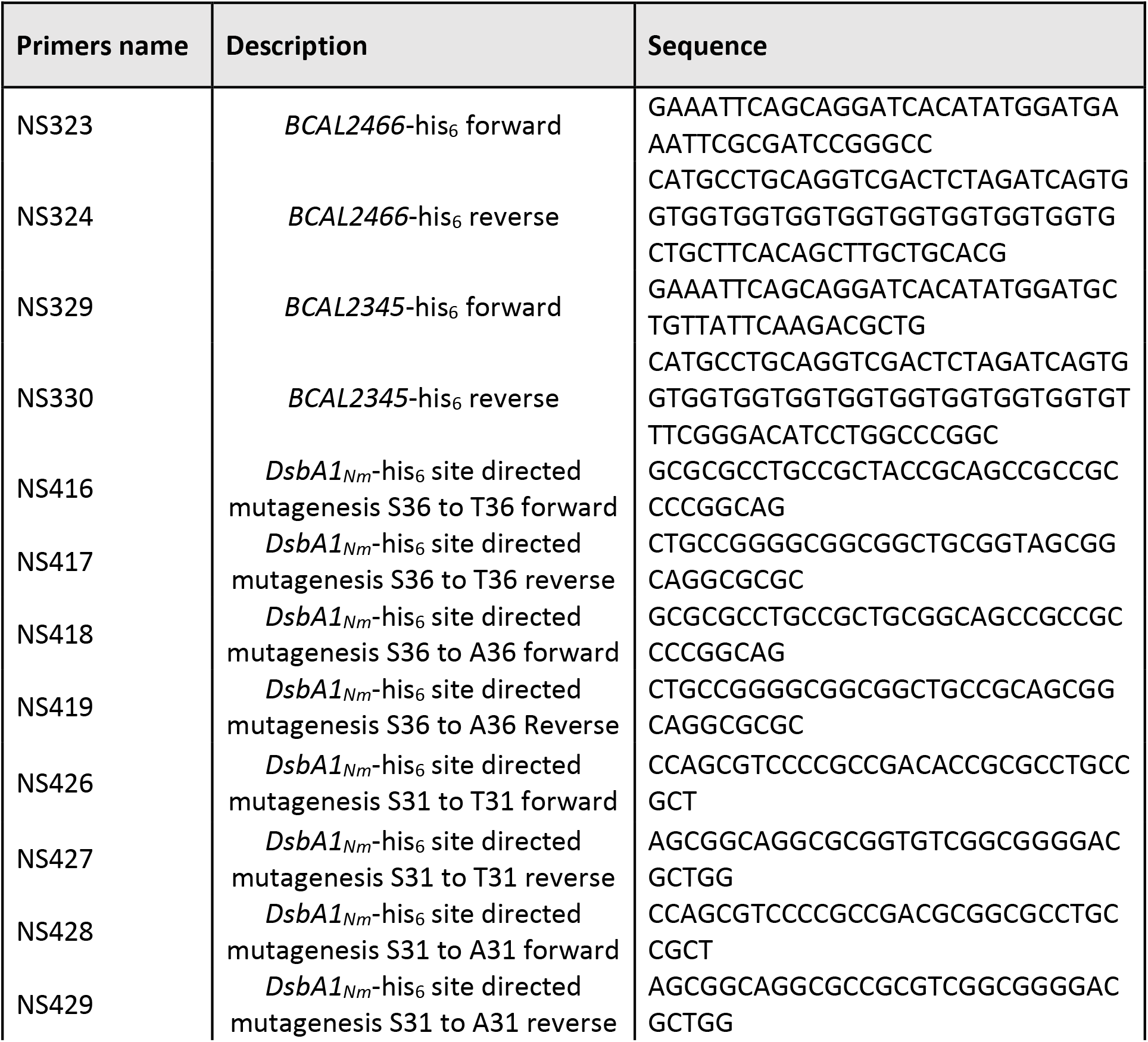
Primer list

### Construction of unmarked H111 Δ*pglL* deletion mutants

Deletions of H111 *pglL* (Gene accession: I35_RS13570) were undertaken using the approach of Flannagan *et al*. for the construction of unmarked, non-polar deletions in *B. cenocepacia* ^41^ using the plasmid pYM8 ^12^.

### Protein Immunoblotting

Bacterial whole-cell lysates were prepared from overnight LB cultures of *Burkholderia* strains. 1ml of overnight cultures at an OD_600_ of 1.0 were pelleted, resuspended in 1X Laemmli loading buffer [24.8 mM Tris, 10 mM glycerol, 0.5% (w/v) SDS, 3.6 mM β-mercaptoethanol and 0.001% (w/v) of bromophenol blue (pH 6.8)] and heated for 10 minutes at 95°C. Lysates were then subjected to SDS-PAGE using pre-cast 4-12% gels (Invitrogen) and transferred to nitrocellulose membranes. Membranes were blocked for 1 hour in 5% skim milk in TBS-T (20 mM Tris, 150 mM NaCl and 0.1% Tween 20) and then incubated for at least 16 hours at 4°C with either mouse monoclonal anti-His (1:2,000; AD1.1.10, Biorad) or mouse anti-RNA pol (1:5,000; 4RA2, Neoclone). Proteins were detected using anti-mouse IgG horseradish peroxidase (HRP)-conjugated secondary antibodies (1:3,000; catalog number NEF822001EA, Perkin-Elmer) and developed with Clarity Western ECL Substrates (BioRad). All antibodies were diluted in TBS-T with 1% bovine serum albumin (BSA; Sigma-Aldrich). Images were obtained using an Amersham imager 600 (GE life sciences) or a Biorad ChemiDoc imaging station (Biorad).

### Preparation of cell lysates for proteomic analysis

*B. cenocepacia* strains were grown overnight on LB plates as previously described ^5^. Plates were flooded with 5 ml of pre-chilled sterile phosphate-buffered saline (PBS) and cells collected with a cell scraper. Cells were washed 3 times in PBS and collected by centrifugation at 10,000 x *g* at 4°C then snap frozen. Frozen whole cell samples were resuspended in 4% SDS, 100 mM Tris pH 8.0, 20 mM DTT and boiled at 95°C with shaking for 10 minutes. Samples were then clarified by centrifugation at 17,000 x *g* for 10 minutes, the supernatant collected, and protein concentration determined by bicinchoninic acid assays (Thermo Fisher Scientific). For glycoproteomic analysis, 1mg of protein for each sample (three biological replicates per strain/per enzyme) was acetone precipitated by mixing 4 volumes of ice-cold acetone with one volume of sample. For quantitative proteomic comparisons of H111 strains, 200μg of protein for each biological replicate (four biological replicates per strain) were acetone precipitated by mixing 4 volumes of ice-cold acetone with one volume of sample. Samples were precipitated overnight at −20°C and then centrifuged at 10,000 x *g* for 10 minutes at 0°C. The precipitated protein pellets were resuspended in 80% ice-cold acetone and precipitated again for an additional 4 hours at −20°C. Following incubation, samples were spun down at 17,000 x *g* for 10 minutes at 0°C to pellet precipitated protein, the supernatant discarded, and excess acetone evaporated at 65°C for 5 minutes.

### Digestion of Proteome samples for glycoproteomic analysis

Dried protein pellets were resuspended in 6 M urea, 2 M thiourea, 50 mM NH_4_HCO_3_ and reduced / alkylated as previously described ^42^. Samples were then digested with one of three different protease combinations; 1) Trypsin (Promega) and Lys-c (Wako); 2) Thermolysin (Promega) or 3) Pepsin (Promega). 1) For the Trypsin / Lys-C digests; Lys-C (1/200 w/w) was added to reduced / alkylated samples for 4 hours at room temperature before the sample was diluted with 100mM NH_4_HCO_3_ four-fold to reduce the urea/thiourea concentration below 2M and trypsin (1/50 w/w) added. 2) For Thermolysin digestions reduced / alkylated samples were diluted with 100mM NH_4_HCO_3_ four-fold to reduce the urea/thiourea concentration below 2M and Thermolysin (1/25 w/w) added. 3) For Pepsin digests reduced / alkylated samples were diluted with 0.1% TFA four-fold to reduce the urea/thiourea concentration below 2M, the pH was confirmed to be pH ^~^2 and Pepsin (1/25 w/w) added. Digests were allowed to proceed overnight at room temperature with shaking Digested samples were acidified to a final concentration of 0.5% formic acid and desalted on 50 mg tC18 Sep-Pak columns (Waters corporation) according to the manufacturer’s instructions. tC18 Sep-Pak columns were conditioned with 10 bed volumes of Buffer B (0.1% formic acid, 80% acetonitrile), then equilibrated with 10 bed volumes of Buffer A* (0.1% TFA, 2% acetonitrile) before use. Samples were loaded on to equilibrated columns then columns washed with at least 10 bed volumes of Buffer A* before bound peptides were eluted with Buffer B. Eluted peptides were dried by vacuum centrifugation at room temperature and stored at −20 °C.

### ZIC-HILIC enrichment of glycopeptides

ZIC-HILIC enrichment was performed as previously described with minor modifications ^43^. ZIC-HILIC Stage-tips ^44^ were created by packing 0.5cm of 10 μm ZIC-HILIC resin (Millipore/Sigma) into p200 tips containing a frit of C8 Empore™ (Sigma) material. Prior to use, the columns were washed with Milli-Q water, followed by 95% acetonitrile and then equilibrated with 80% acetonitrile and 1% TFA. Digested proteome samples were resuspended in 80% acetonitrile and 1% TFA. Samples were adjusted to a concentration of 5 μg/μL (a total of 500 μg of peptide used for each enrichment) then loaded onto equilibrated ZIC-HILIC columns. ZIC-HILIC columns were washed with 20 bed volumes of 80% acetonitrile, 1% TFA to remove non-glycosylated peptides and bound peptides eluted with 10 bed volumes of Milli-Q water. Eluted peptides were dried by vacuum centrifugation at room temperature and stored at −20°C.

### LC-MS analysis of glycopeptide enrichments

ZIC-HILIC enriched samples were re-suspended in Buffer A* and separated using a two-column chromatography set up composed of a PepMap100 C18 20 mm x 75 μm trap and a PepMap C18 500 mm x 75 μm analytical column (Thermo Fisher Scientific). Samples were concentrated onto the trap column at 5 μL/min for 5 minutes with Buffer A (0.1% formic acid, 2% DMSO) and then infused into an Orbitrap Fusion™ Lumos™ Tribrid™ Mass Spectrometer (Thermo Fisher Scientific) at 300 nl/minute via the analytical column using a Dionex Ultimate 3000 UPLC (Thermo Fisher Scientific). 185-minute analytical runs were undertaken by altering the buffer composition from 2% Buffer B (0.1% formic acid, 77.9% acetonitrile, 2% DMSO) to 28% B over 150 minutes, then from 28% B to 40% B over 10 minutes, then from 40% B to 100% B over 2 minutes. The composition was held at 100% B for 3 minutes, and then dropped to 2% B over 5 minutes before being held at 2% B for another 15 minutes. The Lumos™ Mass Spectrometer was operated in a data-dependent mode automatically switching between the acquisition of a single Orbitrap MS scan (350-1800 m/z, maximal injection time of 50 ms, an Automated Gain Control (AGC) set to a maximum of 1*10^6^ ions and a resolution of 120k) every 3 seconds and Orbitrap MS/MS HCD scans of precursors (NCE 28% with 5% Stepping, maximal injection time of 60 ms, an AGC set to a maximum of 1*10^5^ ions and a resolution of 15k). Scans containing the oxonium ions (204.0867; 138.0545 or 366.1396 m/z) triggered three additional product-dependent MS/MS scans ^45^ of potential glycopeptides; a Orbitrap EThcD scan (NCE 15%, maximal injection time of 250 ms, AGC set to a maximum of 2*10^5^ ions with a resolution of 30k and using the extended mass range setting to improve the detection of high mass glycopeptide fragment ions ^46^); a ion trap CID scan (NCE 35%, maximal injection time of 40 ms, an AGC set to a maximum of 5*10^4^ ions) and a stepped collision energy HCD scan (using NCE 35% with 8% Stepping, maximal injection time of 150 ms, an AGC set to a maximum of 2*10^5^ ions and a resolution of 30k).

### Glycopeptide identifications using Byonic

Raw data files were processed using Byonic v3.5.3 (Protein Metrics Inc. ^47^). Tryptic samples were searched with a n-ragged semi-tryptic specificity allowing a maximum of two missed cleavage events while Pepsin and Thermolysin samples were searched with non-specific specificity. Carbamidomethyl was set as a fixed modification of cysteine while oxidation of methionine was included as a variable modification. The Burkholderia glycans HexHexNAc_2_ (elemental composition: C_22_O_15_H_36_N_2_, mass: 568.2115) and Suc-HexHexNAc_2_ (elemental composition: C_26_O_18_H_40_N_2_, mass: 668.2276) were included as variable modifications at serine and threonine residues. K56-2 samples were searched against the K56-2 proteome ^48^ (Uniprot accession: UP000011196, 7,467 proteins) while H111 samples were searched against the H111 proteome ^49^ (Uniprot accession: UP000245426, 8111 proteins). A maximum mass precursor tolerance of 5 ppm was allowed while a mass tolerance of up to 10 ppm was set for HCD fragments and 20 ppm for EThcD fragments. Separate datasets from the same strain were combined using R and only glycopeptides with a Byonic score >300 used for further analysis. This score cut-off is in line with previous reports highlighting that score thresholds greater than at least 150 are required for robust tryptic glycopeptide assignments within Byonic ^50,51^. Manual inspection of glycopeptides to assess the correctness of assignments was undertaken using the guidelines of Chen *et al*. ^52^ with the additional requirement that HCD glycopeptide spectra should contain evidence for glycan fragments such as oxonium ions or the presence of Y_0_ to Y_2_ ions. Glycosylation sites were defined as localised if EThcD scans enabled the unambiguous assignment to a specific serine / threonine residue based on c and z ions or a HCD scan contained only a single serine / threonine residue. Partial localisation is defined as a EThcD spectra containing fragmentation information which allows the ruling out of potential serine / threonine residues yet does not provide evidence enabling the exact site of glycosylation to be assigned. All glycopeptide spectra deemed correct, as well as spectra which provided either partial or complete localisation of glycosylation sites, are provided within Supplementary Data 1 to 4.

### Analysis of Glycosylation sites using O-Pair

Automated glycosylation site analysis was undertaken using O-Pair within MetaMorpheus (non-public release version MM0523 ^53^). Glycopeptide enriched samples were first processed using the Freestyle Viewer (1.7 SP1, Thermo Fisher Scientific) to remove ion-trap CID scans and then searched allowing for a maximum of 4 glycans. The *Burkholderia* glycans were defined as HexHexNAc_2_ (elemental composition: C_22_O_15_H_36_N_2_, mass: 568.2115) or Suc-HexHexNAc_2_ (elemental composition: C_26_O_18_H_40_N_2_, mass: 668.2276). K56-2 samples were searched against the K56-2 proteome ^48^ (Uniprot accession: UP000011196) while H111 samples were searched against the H111 proteome ^49^ (Uniprot accession: UP000245426). Tryptic samples were searched using the default settings while Pepsin and Thermolysin samples were searched with non-specific specificity allowing 5 to 55 amino acids and the search partitioned into 100 partitions. Analysis of the tryptic glycoproteome samples of *B. pseudomallei* K96243; *B. multivorans* MSMB2008; *B. dolosa* AU0158; *B. humptydooensis* MSMB43; *B. ubonensis* MSMB22; *B. anthina* MSMB649; *B. diffusa* MSMB375 and *B. pseudomultivorans* MSMB2199 (PRIDE ProteomeXchange Consortium accession: PXD018587) were undertaken as above using the proteome databases as outlined in ^16^. Only class “level 1” glycopeptides with a Q-value less than 0.01 and a site-specific probability of >0.75 were considered as localised and used for further analysis.

### Digestion of whole cell samples for proteomic comparisons

Dried protein pellets of H111 strains were resuspended in 6 M urea, 2 M thiourea, 40 mM NH_4_HCO_3_ and reduced / alkylated prior to digestion with Lys-C then trypsin overnight as described above. Digested samples were acidified to a final concentration of 0.5% formic acid and desalted with home-made high-capacity StageTips composed of 1 mg Empore™ C18 material (3M) and 5 mg of OLIGO R3 reverse phase resin (Thermo Fisher Scientific) as described ^44,54^. Columns were wet with Buffer B (0.1% formic acid, 80% acetonitrile) and conditioned with Buffer A* prior to use. Acidified samples were loaded onto conditioned columns, washed with 10 bed volumes of Buffer A*, and bound peptides were eluted with Buffer B, before being dried via vacuum centrifugation at room temperature and stored at −20°C.

### LC-MS analysis of H111 proteome samples

Stagetip cleaned-up H111 proteome samples were resuspended in Buffer A* and separated using a two-column chromatography set up comprised of a PepMap100 C18 20 mm x 75 μm trap and a PepMap C18 500mm x 75μm analytical column (Thermo Scientific). Samples were concentrated onto the trap column at 5 μl/minute with Buffer A for 5 minutes and infused into an Orbitrap Elite™ Mass Spectrometer (Thermo Scientific) at 300 nl/minute via the analytical column using an Dionex Ultimate 3000 UPLC (Thermo Scientific). The peptides were separated using a 125-minute gradient altering the buffer composition from 1% buffer B to 23% B over 95 minutes, then from 23% B to 40% B over 10 minutes, then from 40% B to 100% B over 5 minutes, the composition was held at 100% B for 5 minutes, and then dropped to 3% B over 2 minutes and held at 3% B for another 8 minutes. The Orbitrap Elite™ was operated in a data-dependent mode automatically switching between the acquisition of a single Orbitrap MS scan (300-1650 m/z, maximal injection time of 50 ms, an AGC set to a maximum of 1*10^6^ ions and a resolution of 60k) followed by 20 data-dependent CID MS-MS events (NCE 30%, maximal injection time of 80 ms, an AGC set to a maximum of 2*10^4^ ions).

### In-gel digestion of DsbA1_Nm_-his_6_ within Burkholderia lysates

Whole cell lysates of induced *Burkholderia* strains prepared as above for immunoblotting were separated on pre-cast 4-12% gels then fixed in fixative buffer (10% methanol, 7% acetic acid) for 1hr before being stained with Coomassie G-250 overnight. The region corresponding to ^~^25-35kDa was then excised and processed as previously described (38). Briefly, the excised gel regions were sectioned into ^~^2 mm^2^ pieces then destained in a solution of 100 mM NH_4_HCO_3_ / 50% ethanol for 15 minutes at room temperature with shaking. Destaining was repeated twice to ensure the removal of excess Coomassie. Destained bands were dehydrated with 100% ethanol for 5 minutes and then rehydrated in 50 mM NH_4_HCO_3_ containing 10 mM DTT. Samples were reduced for 60 minutes at 56 °C with shaking and washed twice in 100% ethanol for 10 minutes to remove DTT. Reduced ethanol washed samples were sequentially alkylated with 55 mM of Iodoacetamide in 50 mM NH_4_HCO_3_ in the dark for 45 minutes at room temperature. Alkylated samples were then washed with Milli-Q water followed by 100% ethanol twice for 5 minutes to remove residual Iodoacetamide then vacuum-dried for 10 minutes. Alkylated samples were then rehydrated with 20 ng/μl trypsin (Promega) in 50 mM NH_4_HCO_3_ at 4 °C for 1 hour. Excess trypsin was removed, the gel pieces covered in 40 mM NH_4_HCO_3_, and incubated overnight at 37 °C. Peptides were concentrated and desalted using C18 stage tips (64, 65) before analysis by LC-MS.

### LC-MS analysis of DsbA1_Nm_-his_6_ glycopeptides

Stagetip cleaned-up samples were resuspended in Buffer A* and separated using a two-column set up as described above, coupled to a Orbitrap Lumos™ Mass Spectrometer equipped with a FAIMS Pro interface (Thermo Fisher Scientific). 125-minute gradients were run for each sample altering the buffer composition from 3% Buffer B to 28% B over 95 minutes, then from 28% B to 40% B over 10 minutes, then from 40% B to 80% B over 7 minutes, the composition was held at 80% B for 3 minutes, and then dropped to 3% B over 1 minutes and held at 3% B for another 9 minutes. The Lumos™ Mass Spectrometer was operated in a stepped FAIMS data-dependent mode at three different FAIMS CVs −25, −45 and −65 as previously described ^35^. For each FAIMS CV a single Orbitrap MS scan (350-2000 m/z, maximal injection time of 50 ms, an AGC of maximum of 1*10^6^ ions and a resolution of 120k) was acquired every 2 seconds followed by Orbitrap MS/MS HCD scans of precursors (NCE 30%, maximal injection time of 80 ms, an AGC set to a maximum of 1*10^5^ ions and a resolution of 15k). HCD scans containing the oxonium ions (204.0867; 138.0545 or 366.1396 m/z) triggered an additional Orbitrap EThcD scan, an ion-trap CID scan and a Orbitrap HCD scan of potential glycopeptides with scan parameters described above. For PRM experiments, 65-minute gradients were run for each sample altering the buffer composition from 3% buffer B to 28% B over 35 minutes, then from 28% B to 40% B over 10 minutes, then from 40% B to 80% B over 7 minutes, the composition was held at 80% B for 3 minutes, and then dropped to 3% B over 1 minutes and held at 3% B for another 9 minutes. The Lumos™ Mass Spectrometer was operated at a FAIMS CV of −25 in a data-dependent mode automatically switching between the acquisition of a single Orbitrap MS scan (350-2000 m/z, maximal injection time of 50 ms, an AGC set to a maximum of 1*10^6^ ions and a resolution of 60k) every 3 seconds followed by ion-trap EThcD MS2 events (NCE 25%, maximal injection time of 200 ms, an AGC set to a maximum of 6*10^4^ ions) of precursors and then a Orbitrap EThcD PRM scan (NCE 25%, maximal injection time of 450 ms, an AGC set to a maximum of 2*10^5^ ions) of the m/z 1547.77, 1581.12 and 1543.10 which correspond to the +3 charge state of the HexHexNAc_2_ glycosylated ^23^VQTSVPADSAPAATAAAAPAGLVEGQNYTVLANPIPQQQAGK^64^; the Suc-HexHexNAc_2_ glycosylated ^23^VQTSVPADSAPAATAAAAPAGLVEGQNYTVLANPIPQQQAGK^64^ and the HexHexNAc_2_ glycosylated ^23^VQTSVPADSAPAASAAAAPAGLVEGQNYTVLANPIPQQQAGK^64^ peptides.

### Proteomic analysis

H111 Proteome and in-gel datasets were processed using MaxQuant (v1.5.5.1 or 1.6.3.4. ^55^). The H111 proteome dataset was searched against the H111 proteome ^49^ (Uniprot accession: UP000245426) and *B. cenocepacia* strain J2315 (Uniprot accession: UP000001035, 6,993 proteins) to enable the matching of J2315 gene accessions. In-gel digests were searched against either *B. cenocepacia* strain J2315 (Uniprot accession: UP000001035), *B. ubonensis* MSMB22 (Burkholderia Genome Database ^56^, Strain number: 3404) or *B. humptydooensis* MSMB43 (Burkholderia Genome Database ^56^, Strain number: 4072) depending on the sample type and a custom database of DsbA1_Nm_-his_6_ containing the desired point mutations at position 31 and 36 within Uniprot entry Q9K189. All searches were undertaken using “Trypsin” enzyme specificity with carbamidomethylation of cysteine as a fixed modification. Oxidation of methionine and acetylation of protein N-termini were included as variable modifications and a maximum of 2 missed cleavages allowed. For in-gel samples HexHexNAc_2_ (elemental composition: C_22_O_15_H_36_N_2_, mass: 568.2115) and Suc-HexHexNAc_2_ (elemental composition: C_26_O_18_H_40_N_2_, mass: 668.2276) were also included as variable modifications. To enhance the identification of peptides between samples, the Match between Runs option was enabled with a precursor match window set to 2 minutes and an alignment window of 10 minutes. For label free quantitation (LFQ) the MaxLFQ option in Maxquant ^57^ was enabled. The resulting outputs were processed within the Perseus (v1.5.0.9) ^58^ analysis environment to remove reverse matches and common protein contaminates prior to further analysis. For LFQ comparisons missing values were imputed based on the observed total peptide intensities with a range of 0.3σ and a downshift of 2.5σ using Perseus. Enrichment analysis was undertaken using Fisher exact tests in Perseus with PglL altered proteins defined as those proteins which were previously reported by Oppy *et al*. as differentially altered in K56-2 Δ*pglL* compared to K56-2 Wt ^5^. To compare the relative abundance of glycosylated and non-glycosylated peptides from DsbA1_Nm_-his_6_ point mutants the area under the curve of these peptides were extracted using the FreeStyle viewer and the resulting data provided within the supplementary document. Statistically analysis of the area under the curve was undertaken in Prism (version 7.0e) using a two-side t-test.

### Visualization of glycoproteome and proteome datasets

Data visualization was undertaken using ggplot2 ^59^ within R with all scripts included in the PRIDE uploaded datasets. To aid in the analysis of the MS/MS data the Interactive Peptide Spectral Annotator ^60^ (http://www.interactivepeptidespectralannotator.com/PeptideAnnotator.html) was used. Mass spectrometry data (Raw data files, Byonic/Maxquant/O-pair search outputs, R Scripts and output tables) have been deposited into the PRIDE ProteomeXchange Consortium repository ^61,62^. The glycoproteomic datasets are available with the identifier: PXD024090 and can be accessed using the Username: and Password: HDbdkjb6; the H111 proteome analysis is available with the identifier: PXD023755 and can be accessed using the username, Password: QMdQyTe2; the DsbA1_Nm_-his_6_ (*B. cenocepacia* K56-2) associated analysis is available with the identifier: PXD023955 and can be accessed using the Username: Password: vAQRDw5X; the DsbA1_Nm_-his_6_ (*B. humptydooensis* MSMB43 and *B. ubonensis* MSMB22) associated analysis is available with the identifier: PXD024056 and can be accessed using the Username: Password: Bj3dwihm.

### Bioinformatic analysis of glycoproteins and glycosylation sites

Draft and complete genome sequences of *B. cenocepacia* (n=294) and complete genome sequences of other *Burkholderia* species, subsetted to contain a maximum of 20 of any individual species (n=174), were obtained from the *Burkholderia* database ^56^. Isolates were screened for the presence of the glycosylated motif containing genes from *B. cenocepacia* J2315 using an 80% identity and length BlastN thresholds using screen_assembly3.py v1.2.7 ^63^. Gene hits were translated to protein sequences and the translated hits screened for the presence of the motif using seqkit locate ^64^, allowing for up to 4 mismatched amino acids (> 80% conservation of the 21 amino acid motif). The identity and coverage of the genes at 80% identity as well as the motif coverage at 80% identity were visualised using ggplot2. Sequence logos were generated using ggseqlogo ^65^. To compare glycoprotein homologues and pglL sequences between *Burkholderia* species the *Burkholderia* Orthologous Groups information provided by the *Burkholderia* database was used ^56^.

## Results

### *O*-linked glycosylation targets similar proteins and glycosylation disruption impacts the proteome in a similar manner across *B. cenocepacia* strains

Recently we noted that the physiochemical properties of peptides heavily influences the ability of ZIC-HILIC enrichment to isolate bacterial glycopeptides ^35^.. This observation suggests numerous glycoproteins have likely been overlooked using trypsin alone to assess the glycoproteome of *B. cenocepacia* in previous studies ^16,29^. To increase the coverage of the *B. cenocepacia* glycoproteome we undertook glycoproteomic analysis using multiple proteases ^66,67^ and two widely used *B. cenocepacia* strains, K56-2 and H111. Glycopeptide enrichments of Trypsin, Thermolysin and Pepsin digested samples enabled the identification of 584 and 666 unique glycopeptides from K56-2 and H111 strains (Figure 1 A, Supplementary Tables 1 and 2). Although these glycopeptide datasets identified more glycoproteins than identified within previously published studies (Supplementary Figure 1A ^16,29^) the majority of identified glycoproteins were unique to a single strain (Supplementary Figure 1B). This high degree of heterogeneity suggested the presence of erroneous assignments within the datasets, a known issue associated with the assignment of a relatively limited population of modified peptides within large database searches ^68^. Consistent with this, examination of the glycopeptide scores reveal a bias toward lower scores (Supplementary Figure 2A and B) supporting a higher than ideal false discovery rate despite stringent filtering ^69^. To improve the quality of these glycopeptide datasets we manually curated the Byonic assigned glycopeptides, identifying 256 and 328 high-quality unique glycopeptides in K56-2 and H111 (Figure 1A; Supplementary Data 1 and 2). Of these unique glycopeptides, 167 and 175 glycopeptides within K56-2 and H111 respectively provided partial or complete site localisation (Figure 1A; Supplementary Data 3 and 4). Consistent with improving the data quality, the curated glycopeptides revealed a gaussian distribution within the scores, removing predominately low scoring identifications (Supplementary Figure 2C-F). Of these manually assessed glycopeptides, we noted >65% of glycoproteins were identified within both strains (Figure 1B) supporting a conserved glycoproteome across *B. cenocepacia* strains.

**Figure 1.**
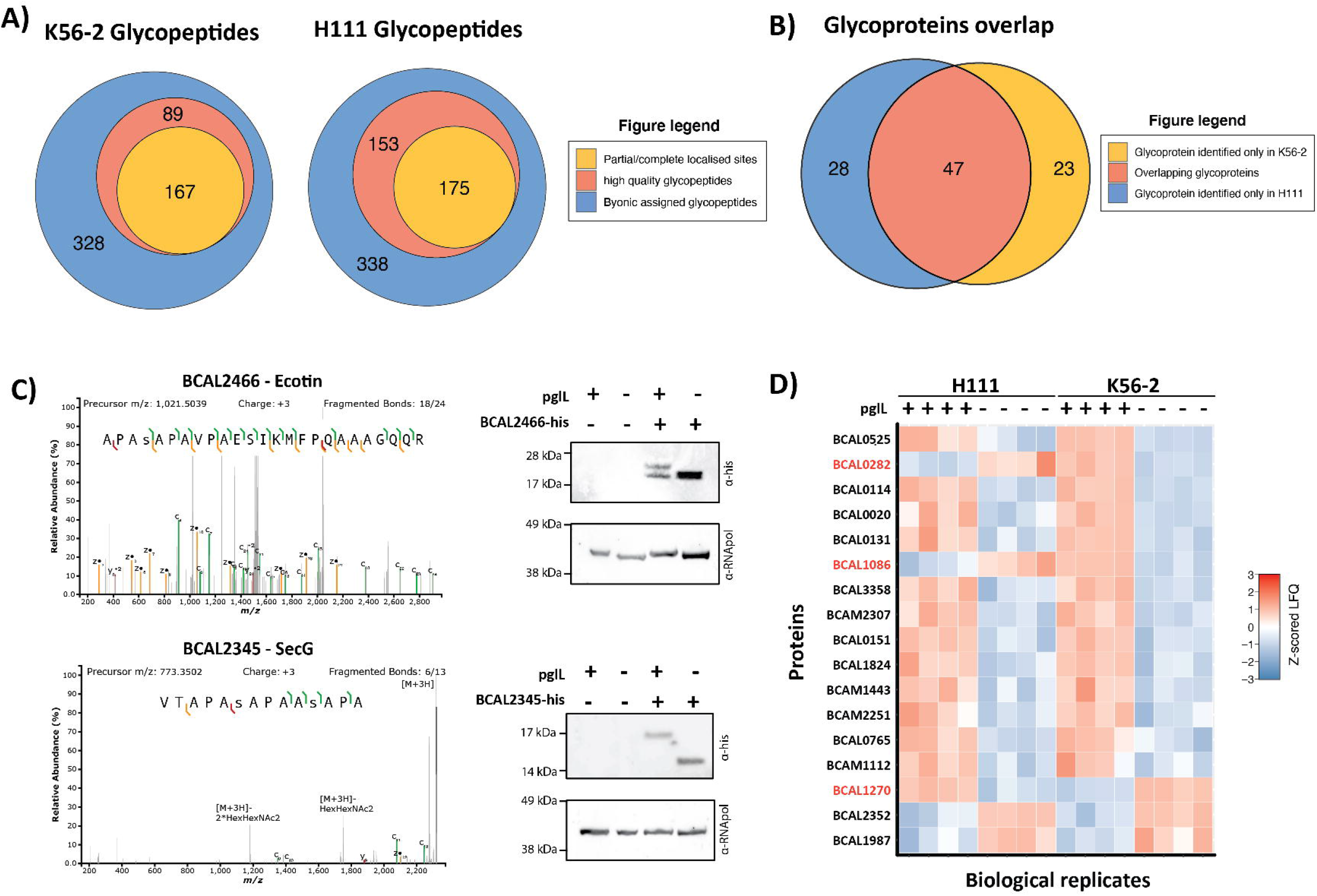
*B. cenocepacia* glycoproteomic analysis reveals similar substrates and functional impacts on *B. cenocepacia* strains. **A)** Within *B. cenocepacia* strains K56-2 and H111 584 and 666 unique glycopeptides were identified respectively (Byonic scores >300). Within these glycopeptides manual curation identified 256 and 328 glycopeptides as high-quality unique glycopeptides of which 167 and 175 provided partial or complete glycosylation site information. **B)** Across the manually curated glycopeptides a total of 98 unique glycoproteins were identified of which 47 were identified within both strains. **C)** Western blotting of two potential glycoproteins, His_6_ tagged BCAL2466 and BCAL2345, reveals alterations in protein banding when expressed within the glycosylation null strain K56-2 Δ*pglL* compared to the K56-2 WT. **D)** LFQ analysis of the most altered proteins observed between H111 Δ*pglL* compared to H111 WT reveals similar proteome alterations as seen previously within K56-2 ^5^.

To validate the accuracy of these manually filtered assignments two small novel glycoproteins, BCAL2466 and BCAL2345, were His_6_ epitope tagged and expressed within K56-2 WT and K56-2 Δ*pglL* (Figure 1C). Consistent with a single glycosylation event on BCAL2466 and multiple glycosylation events on BCAL2345, the expression of these proteins within K56-2 Δ*pglL* resulted in reduced gel mobility supporting their glycosylation status. To functionally support the observed similarities within the glycoproteomes of *B. cenocepacia* strains, we generated two independent H111 Δ*pglL* mutants and assessed the proteome changes observed in the absence of glycosylation within H111, compared to our previously published K56-2 findings (Figure 1D ^5^). Although the proteomic alterations show congruent behaviour, it should be noted strain differences are observed within some proteins such as BCAL1086, BCAL0282 and BCAL1270 (Figure 1D; Supplementary Table 3). Regardless of these differences between K56-2 and H111 the proteins observed to undergo statistically significant alterations within H111 Δ*pglL* were enriched for proteins observed to be altered within K56-2 Δ*pglL* (Fisher exact test p-value = 0.0006 Supplementary table 4). Independent H111 Δ*pglL* mutants also showed similar alterations with the proteome and were highly similar (Fisher exact test p-value = 5.67 x 10^−15^, Supplementary table 3 and 4). Taken together these findings support that the *O*-linked glycoproteome of *B. cenocepacia* strains are similar in substrates and functional consequences.

### Glycosylation site analysis reveals that *B. cenocepacia* glycosylation occurs solely on serine residues

Our manually curated glycoproteomic data revealed ^~^50% of unique glycopeptides provided at least partial site localisation information (Figure 1A, Supplementary Data 3 and 4) of which 88 sites could be precisely localised across 70 glycoproteins (Supplementary Table 5). Examination of the sites of glycosylation revealed 69 out of 70 sites (>98%) were observed on Serine residues and that glycosylation favoured Alanine at the −1 position, yet this was not a strict requirement (Figure 2A). As only a single Threonine modification was identified on a single glycopeptide derived from the protein BCAM0996, we sought to confirm the accuracy of this assignment. Examination of glycopeptides from BCAM0996 revealed multiple serine residues are modified within the same peptide observed to be modified at T^159^ (Supplementary Data 3 and 4). The close proximity of this sole threonine modification to multiple serine modification events further raised concerns of miss-localisation. Manual annotation of the assigned glycopeptide supported the incorrect localisation of the glycosylation site due to the incorrect assignment of the glycan composition and peptide caused by a secondary modification within the peptide sequence revealing S^167^ to be the correct localisation site (Supplementary Figure 3). This finding suggested all localised glycosylation sites within *B. cenocepacia* were observed on Serine residues. To independently validate this preference for Serine, we re-analysed these glycoproteomic datasets with O-Pair, a glycosylation site localisation focused glycoproteomic tool ^53^, supporting that the majority of high quality localisable glycosylation events occurred on Serine residues (Supplementary Figure 4; Supplementary Table 6). Taken together, these findings support that glycosylation within the *B. cenocepacia* glycoproteome localises exclusively to Serine residues.

**Figure 2.**
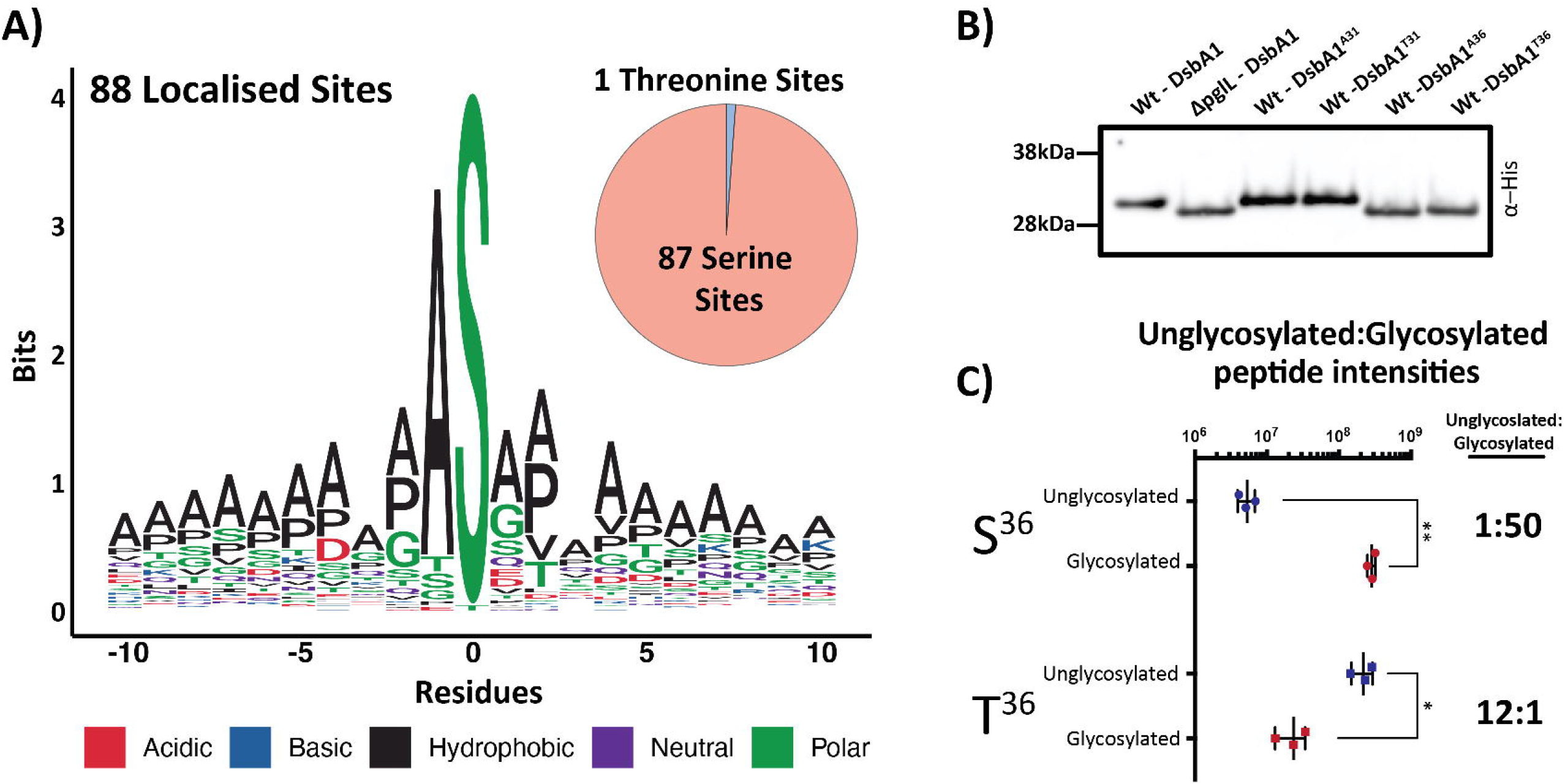
*O*-linked glycosylation predominately occurs on Serine residues across the *B. cenocepacia* glycoproteome. **A)** Sequence analysis of localised glycosylation sites across *B. cenocepacia* strains reveal the majority of assigned sites are Serine. **B)** Western analysis of *DsbA1_Nm_*-his_6_ variants reveals substitution of only S^36^ to Alanine or Threonine results in reduced glycosylation. **C)** Relative amounts of the *DsbA1_Nm_*-his_6_ glycosylated and non-glycosylated peptides ^23^VQTSVPADSAPAASAAAAPAGLVEGQNYTVLANPIPQQQAGK^64^ and ^23^VQTSVPADSAPAATAAAAPAGLVEGQNYTVLANPIPQQQAGK^64^ observed within K56-2 WT reveals the alteration of S^36^ to T^36^ dramatically reduces glycosylation.

### Threonine residues undergo poor glycosylation in *B. cenocepacia*

To experimentally explore this preference for Serine, we utilised the glycoprotein DsbA1_Nm_-his_6_ that we have previously observed is predominately glycosylated by a single glycosylation event when expressed in *B. cenocepacia* ^5,29^. Past work showed that glycosylation occurs at an unknown site within the peptide ^23^VQTSVPADSAPAASAAAAPAGLVEGQNYTVLANPIPQQQAGK^64^, suggesting that one of the three Serine residues (S^26^, S^31^ and S^36^) was the site of glycosylation within DsbA1_Nm_-his_6_. Of these sites, S^31^ and S^36^ are flanked by Alanine residues, consistent with the preferred glycosylation sites within native substrates (Figure 2A), yet only substitution of S^36^ with either Alanine or Threonine resulted in gel mobility shifts consistent with the loss of glycosylation (Figure 2B). Glycopeptide analysis further confirmed S^36^ was the sole residue modified within this peptide (Supplementary Figure 5 and Supplementary Table 7). As MS analysis provides greater dynamic range then western blotting, we manually investigated if glycosylation still occurred at T^36^ with analysis supporting the presence of a glycosylated form of the ^23^VQTSVPADSAPAATAAAAPAGLVEGQNYTVLANPIPQQQAGK^64^ at <10% of the abundance of the unmodified form (Figure 2C, Supplementary Figure 6A). Targeted MS analysis supported the identity of this glycopeptide yet could not provide definitive site localisation to T^36^ (Supplementary Figure 6B). To further assess if Threonine could be modified, we introduced the DsbA1_Nm_-his_6_ point mutants T^36^ and A^36^ into K56-2 ΔpglL AmrAB::S7-pglL-his, an overexpressing PglL_BC_ strain ^5^, to assess if increasing PglL_BC_ levels enhanced Threonine glycosylation. Western analysis demonstrates the majority of DsbA1_Nm_-his_6_ T^36^ remained unglycosylated within K56-2 ΔpglL AmrAB::S7-pglL-his. Yet, in contrast to K56-2 WT, a faint band potentially corresponding to glycosylated DsbA1_Nm_-his_6_ T^36^ was observed (Figure 3A). Targeted MS analysis confirms the glycosylation of T^36^ in DsbA1_Nm_-his_6_ T^36^ when expressed in K56-2 ΔpglL AmrAB::S7-pglL-his (Figure 3B and C). Combined this data supports that it is possible to glycosylate Threonine in *B. cenocepacia*, but the preferred residue for glycosylation is Serine, even when PglL_BC_ is overexpressed.

**Figure 3.**
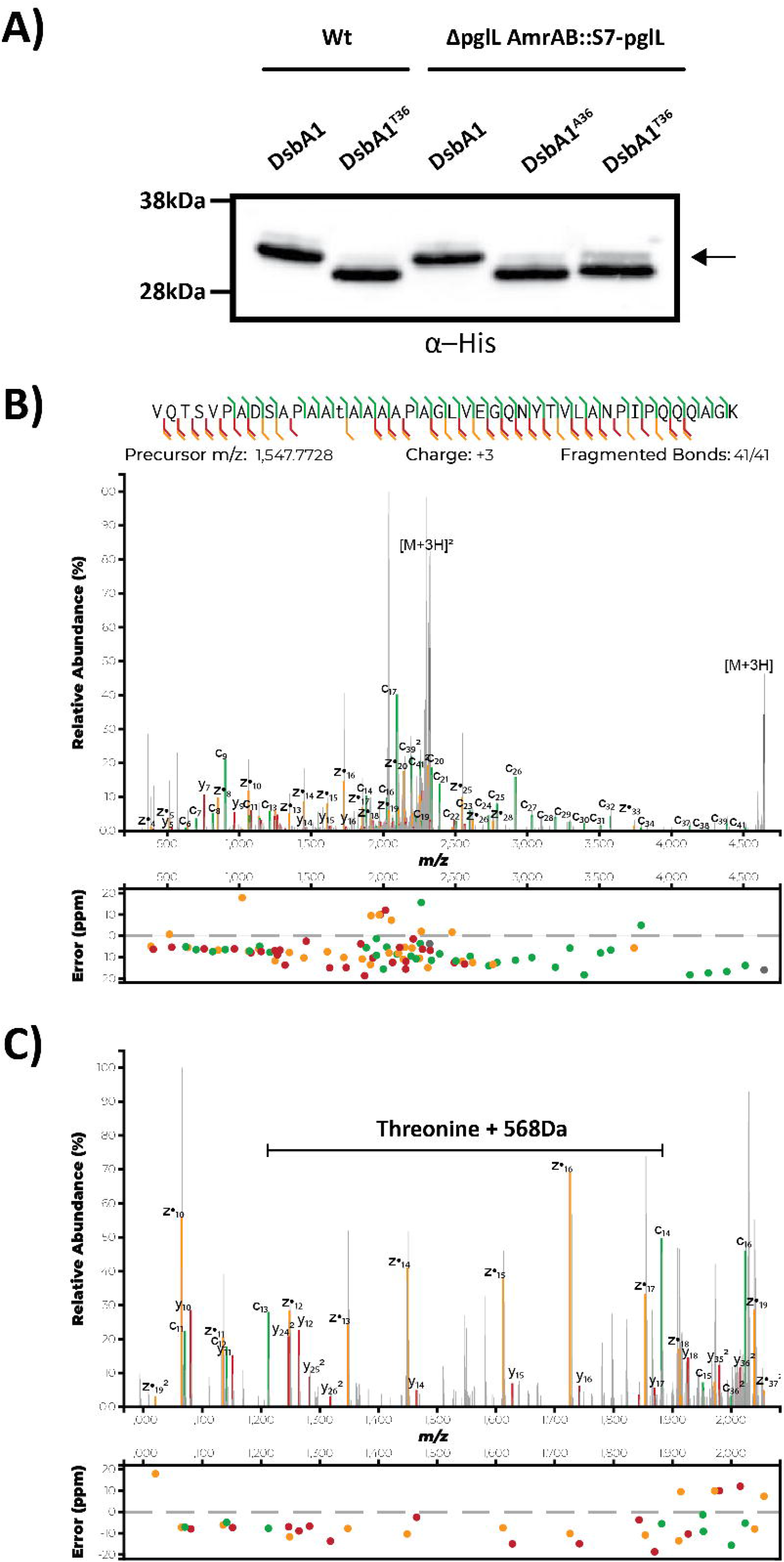
Threonine residues undergo poor glycosylation in *B. cenocepacia* even with overexpression of *pglL*_BC_. **A)** Western analysis of *DsbA1_Nm_*-his_6_ variants reveals a minor product, indicated with an arrow, is observed within DsbA1_Nm_-his_6_ T^36^ when expressed within K56-2 Δ*pglL* AmrAB::S7-*pglL*-his. **B and C)** Targeted MS analysis using EThcD fragmentation enabled the confirmation of glycosylation at residue T^36^ within ^23^VQTSVPADSAPAATAAAAPAGLVEGQNYTV LANPIPQQQAGK^64^.

### Glycoproteins / glycosylation sites are highly conserved across *B. cenocepacia* strains

With our proteome and site-directed analysis supporting Serine as the favoured residue for glycosylation, we next addressed whether these glycosylation proteins/sites are conserved across a diverse set of *B. cenocepacia* strains. Leveraging our curated glycoprotein dataset (Supplementary Table 5) we assessed the conservation of these 70 glycoproteins across 294 publicly available *B. cenocepacia* genomes ^56^. Across strains, the frequency of observation, defined as the presence of proteins with a minimum blast identity of >80%, revealed 65 glycoproteins are present in >80% of available genomes and that these glycoproteins are highly similar, with an average amino acid sequence identity >95% (Figure 4A, Supplementary Table 8). This high level of conservation is also reflected at the glycosylation site (±10 amino acids) where the majority of glycosylation sites are highly conserved at the sequence level (Figure 4B, Supplementary Table 8). It should be noted that although the majority of glycosylation sites are conserved (Supplementary Figure 7 to 12), variations are observed; such as in S^140^ of BCAL1746, which is present in only a fraction of strains (Figure 4C,); S^404^ of BCAL1674, where multiple alterations within the flanking sequences were noted (Figure 4E); and S^262^ of BCAL0163, where the serine required for glycosylation is lost across multiple strains (Figure 4F). This data supports that both *B. cenocepacia* glycoproteins and glycosylation sites are highly conserved across *B. cenocepacia* strains.

**Figure 4.**
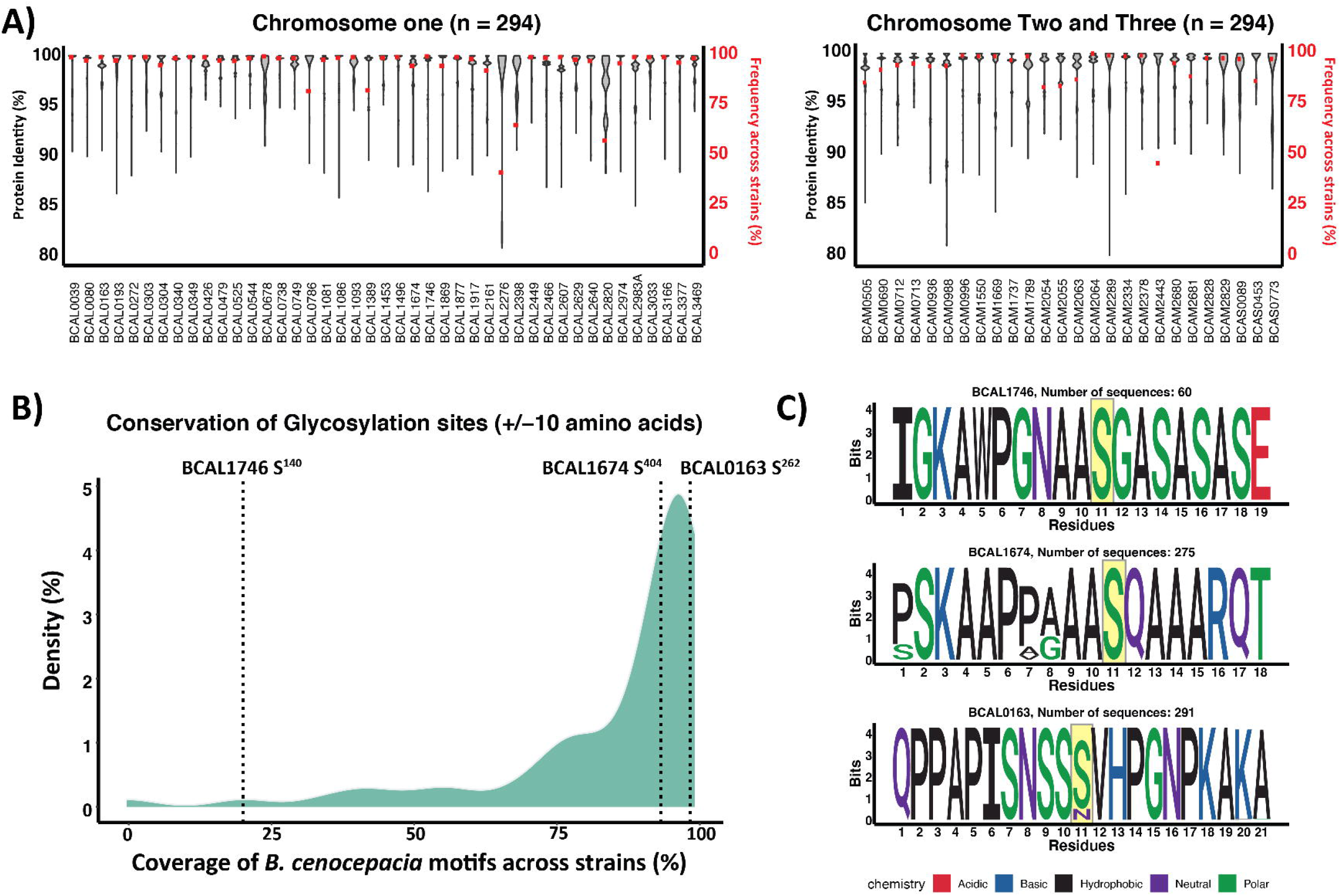
Glycoproteins and glycosylation sites are conserved across *B. cenocepacia*. **A)** Violin plot showing the amino acid sequence variation and frequency of 69 glycoproteins across 294 *B. cenocepacia* genomes. Relative frequency is shown in red (right axis) and protein sequence identity (>80% minimum) is shown in black (violin plot, left axis). **B)** Density plot representing the conservation of the glycosylation site sequence across 69 glycoproteins within 294 *B. cenocepacia* genomes **C)** Sequence logos of glycoproteins denoted within the density plot; BCAL1746 S^140^; BCAL1674 S^404^ and BCAL0163 S^262^.

### Across *Burkholderia* species similar glycoproteins are targeted for glycosylation at Serine residues

In light of the high sequence identity of PglL across *Burkholderia* species (Supplementary Figure 13) we sought to assess if the preference for serine glycosylation was a common feature across *Burkholderia* glycoproteomes. To investigate this, we utilized our recently published glycoproteomic datasets of eight Burkholderia species (*B. pseudomallei* K96243; *B. multivorans* MSMB2008; *B. dolosa* AU0158; *B. humptydooensis* MSMB43; *B. ubonensis* MSMB22; *B. anthina* MSMB649; *B. diffusa* MSMB375 and *B. pseudomultivorans* MSMB2199 ^16^) and re-analysed these datasets using O-Pair. As with *B. cenocepacia*, Serine was the dominant residue targeted for glycosylation with only 11 out of 440 high confidence sites localised to Threonine (Figure 5A; Supplementary table 9). To experimentally support this preference, we introduced DsbA1_Nm_-his_6_ variants into two strains, *B. humptydooensis* MSMB43 and *B. ubonensis* MSMB22, to compare the preference for glycosylation at Serine over Threonine residues. Similar to *B. cenocepacia* we find DsbA1_Nm_-his_6_ T^36^ is predominantly unable to be glycosylated within both *B. humptydooensis* MSMB43 and *B. ubonensis* MSMB22 (Figure 5B and C). As with *B. cenocepacia*, the glycosylated form of the peptide ^23^VQTSVPADSAPAATAAAAPAGLVEGQNYTVLANPIPQQQAGK^64^ was observed at low levels within both *B. humptydooensis* MSMB43 and *B. ubonensis* MSMB22 (Supplementary Figure 14; Supplementary Table 10). Finally, to survey the similarities between Burkholderia glycoproteomes we assessed the sequence identity of glycoproteins revealing the majority of glycoproteins are homologues of confirmed glycoproteins within *B. cenocepacia* (Figure 5D, Supplementary Table 11). Enrichment analysis supports that for *B. anthina* MSMB649; *B. ubonensis* MSMB22; *B. multivorans* MSMB2008; *B. pseudomultivorans* MSMB2199, and *B. dolosa* AU0158, this overlap represents a statistically significant enrichment (Figure 5E Supplementary Table 11). Combined these results support that Serine is the preferred target of glycosylation across *Burkholderia* glycoproteomes and that similar glycoproteins were targeted across the *Burkholderia* genus.

**Figure 5.**
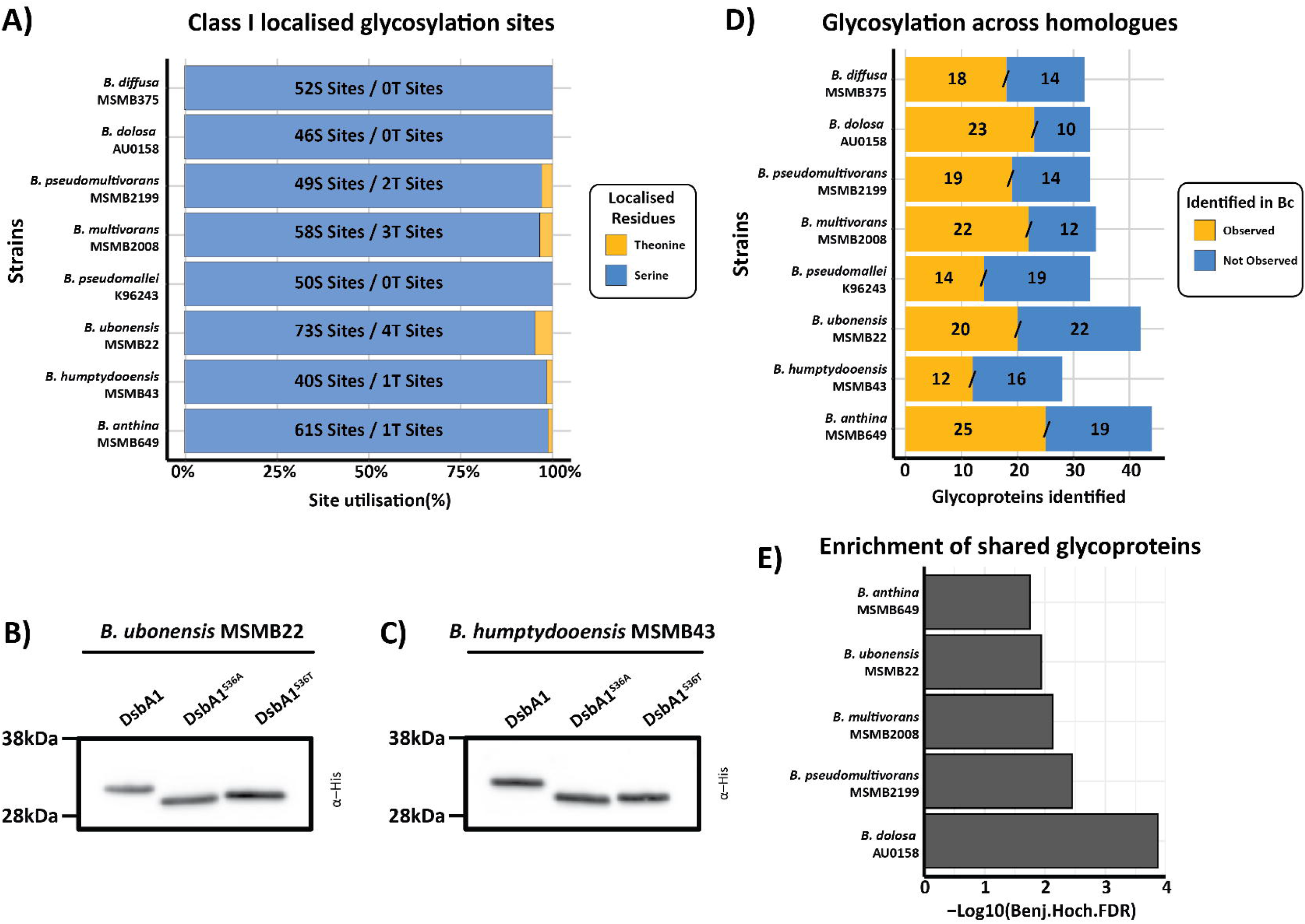
Serine glycosylation and the conservation of glycoproteins across *Burkholderia* species. **A)** The re-analysis of *Burkholderia* glycopeptide enrichment datasets reveals the majority of assigned glycosylation sites are localised to Serine residues. **B)** and **C)** Western analysis of DsbA1_Nm_-his_6_ variants expressed within *B. ubonensis* MSMB22 (B) and *B. humptydooensis* MSMB43 (C) supports that Threonine is disfavoured for glycosylation similar to *B. cenocepacia*. **D)** Analysis of glycoproteins observed across *Burkholderia* species supports multiple *B. cenocepacia* glycoprotein homologues are observed glycosylated within other species. **E)** Enrichment analysis of glycoproteins shows an enrichment of shared glycoproteins in *B. anthina* MSMB649; *B. ubonensis* MSMB22; *B. multivorans* MSMB2008; *B. pseudomultivorans* MSMB2199 and *B. dolosa* AU0158 compared to *B. cenocepacia*

## Discussion

The conservation of glycosylation systems across bacterial species is increasingly being recognised as a common phenomenon in bacterial genera ^17,70,71^. Although previous studies have sought to confirm the presence of protein glycosylation as well as highlight differences within the glycans utilised across species, little attention has been given to the protein’s substrates and glycosylation sites themselves. Within this work we undertook a glycosylation-site-focused analysis of *Burkholderia* species, revealing both experimentally (Figure 1B and 5D) and bioinformatically (Figure 4A) that proteins subjected to glycosylation are conserved across *Burkholderia* strains / species and that *O*-linked glycosylation within this genus is overwhelmingly restricted to Serine residues (Figure 2A and 5A). Our findings here highlight that protein glycosylation targets nearly 100 protein substrates, the majority of which appear conserved and targeted for glycosylation across multiple *Burkholderia* species, supporting that these glycosylation events are responsible for shared, yet still largely unknown, functions across the *Burkholderia* genus. Consistent with this concept, phenotypic studies have previously shown the loss of *O*-linked glycosylation leads to similar impacts on *Burkholderia* species, such as in the case of glycosylation null *B. cenocepacia, B. pseudomallei* and *B. thailandensis* strains, which all possess biofilm defects ^5,72^. Thus, taken together this highlights that *Burkholderia* glycoproteomes are far more similar and conserved then previously noted.

Similar to the optimal motif proposed for the *N. meningitidis* PglL, WPAAASAG ^73^, our proteomic analysis supports that Serine residues flanked by Alanine are predominately targeted for glycosylation within *B. cenocepacia*, yet glycosylation can still occur within divergent sequences lacking consensus residues (Figure 2A). An unexpected finding within this study is that Serine is strongly targeted for glycosylation across *Burkholderia* glycoproteomes, and that a dramatic difference in the extent by which threonine/serine residues can be glycosylated exists. In fact, although multiple studies have examined the glycan promiscuity and general ability of PglL enzymes to glycosylate proteins ^74–76^, no previous reports have suggested that threonine/serine residues are not equivalent targets for glycosylation. This difference in preference between these residues is striking with semi-quantitative comparisons of the unglycosylated and glycosylated peptides suggesting Serine is glycosylated ^~^500 times more efficiently than Threonine within *B. cenocepacia* (Figure 2B and C), with a similar trend also noted within *B. humptydooensis* MSMB43 and *B. ubonensis* MSMB22 (Supplementary Figure 14). Although we confirmed that Threonine can be glycosylated at a low occupancy, this dramatic difference in preference supports that Threonine glycosylation events are likely rare in *Burkholderia* glycoproteomes, or if observed, are likely found at modest levels of occupancy. Interestingly, within other PglL glycosylation systems where only a limited number of glycosylation sites have been confirmed to date, such as in *N. gonorrhoeae* ^33,34^, *A. baumannii* ^15^ and *F. tularensis* ^30^, all confirmed glycosylation sites appear to occur on serine residues. These trends suggest that Serine specificity may be a more general feature of the PglL oligosaccharidetransferases, yet further work is required to confirm this preference across known PglL enzymes.

At the genus level our analysis supports that across *Burkholderia* species glycoproteins are conserved (Figure 5D and Supplementary Figure 15). Although the limited number of known glycosylation sites within other bacterial glycosylation systems has hampered the analysis of glycosylation site conservation, the identification of varying numbers of glycosylation sites in proteins such as pilin of *Neisseria* species ^7,14,23,24^ has suggested that glycosylation sites maybe highly variable across strains/species. In contrast, our analysis highlights that the majority of glycosylation sites, within *B. cenocepacia* at least, appear conserved and that loss of glycosylation sites appear rare. This said, we did note examples of glycosylation sites which were only observed within a subset of genomes, such as S^140^ of BCAL1746, yet this level of variability was the exception, not the rule, across *B. cenocepacia*. Although our glycoproteome of *B. cenocepacia* significantly expands the known number of glycosylation sites from six sites in 23 glycoproteins ^29^ to 88 sites in 70 glycoproteins, it is important to note these glycosylation events likely still only represent a subset of the complete *B. cenocepacia* glycoproteome. It is well known that growth under different conditions, such as nutrient limitation ^77^ and or in the presence of chemical queues ^78,79^ can dramatically alter the observable proteome of *B. cenocepacia*. As the growth of *B. cenocepacia* strains were undertaken under laboratory conditions with growth on rich media, it is possible multiple glycoproteins not expressed under these conditions have been missed. Thus, to better understand glycoproteins associated with virulence or survival under specific environmental conditions further studies may be required.

In summary, this work furthers our understanding of both *B. cenocepacia* and the *Burkholderia* pan-glycoproteome as a whole and represents the first step in understanding the roles and specific functions of glycoproteins across the *Burkholderia* genus. The insights into the preference for glycosylation at Serine residues improves our ability to predict the specific residues likely to be modified within a glycopeptide, while the identification that glycoproteins identified within one species are likely glycosylated within another provides a new opportunity to dissect the conserved roles of glycoproteins across this genus. These insights will aid in future studies to understand why glycosylation events have been maintained within different *Burkholderia* species.

## ACKNOWLEDGEMENTS

N.E.S is supported by an Australian Research Council Future Fellowship (FT200100270) and an ARC Discovery Project Grant (DP210100362). M.R.D is supported by a University of Melbourne CR Roper fellowship. We thank the Melbourne Mass Spectrometry and Proteomics Facility of The Bio21 Molecular Science and Biotechnology Institute for access to MS instrumentation and Byonic. We thank Lei Lu, Nick Riley and Lloyd M. Smith for providing the custom version of O-Pair for the analysis of *Burkholderia* glycopeptides. pSCrhaB2 was a gift from Miguel Valvano (Addgene plasmid # 113634; http://n2t.net/addgene:113634; RRID:Addgene_113634)

## Supplementary Data and Tables

**Supplementary Data 1: Manually curated *B. cenocepacia* H111 glycopeptides (Best scoring unique glycopeptides)**. For each of the best scoring unique glycopeptide, the Byonic assigned spectra is provided in addition to the assigned J2315 gene number, protein name, peptide assignment, glycan assignment, if the spectra enables localisation of the glycan within the sequence, assignment associated metrics (m/z; mass error, score, delta score, delta mod score, scan number within the LC-MS run and scan time), enzymatic digest for which the best scoring glycopeptide was observed in, replicate for which the best scoring glycopeptide was observed in and page within the pdf the spectra can be found.

**Supplementary Data 2: Manually curated *B. cenocepacia* K56-2 glycopeptides (Best scoring unique glycopeptides)**. For each of the best scoring unique glycopeptide, the Byonic assigned spectra is provided in addition to the assigned J2315 gene number, protein name, peptide assignment, glycan assignment, if the spectra enables localisation of the glycan within the sequence, assignment associated metrics (m/z; mass error, score, delta score, delta mod score, scan number within the LC-MS run and scan time), enzymatic digest for which the best scoring glycopeptide was observed in, replicate for which the best scoring glycopeptide was observed in and page within the pdf the spectra can be found.

**Supplementary Data 3: Manually curated *B. cenocepacia* H111 glycopeptides (Best localised unique glycopeptides)**. For each of the best localising unique glycopeptide, the Byonic assigned spectra is provided in addition to the assigned J2315 gene number, protein name, peptide assignment, glycan assignment, if the spectra enables localisation of the glycan within the sequence, assignment associated metrics (m/z; mass error, score, delta score, delta mod score, scan number within the LC-MS run and scan time), enzymatic digest for which the best scoring glycopeptide was observed in, replicate for which the best scoring glycopeptide was observed in and page within the pdf the spectra can be found.

**Supplementary Data 4: Manually curated *B. cenocepacia* K56-2 glycopeptides (Best localised unique glycopeptides)**. For each of the best localising unique glycopeptide, the Byonic assigned spectra is provided in addition to the assigned J2315 gene number, protein name, peptide assignment, glycan assignment, if the spectra enables localisation of the glycan within the sequence, assignment associated metrics (m/z; mass error, score, delta score, delta mod score, scan number within the LC-MS run and scan time), enzymatic digest for which the best scoring glycopeptide was observed in, replicate for which the best scoring glycopeptide was observed in and page within the pdf the spectra can be found.

**Supplementary Table 1. Combined Byonic search results of *B. cenocepacia* H111 glycopeptide enrichments**. The combined Byonic search results from all proteases across each of the three biological replicates of *B. cenocepacia* H111 are provided. The complete list of all glycopeptides identified, the best unique glycopeptides filtered for glycopeptides with a Byonic score over 300 and the curated list of glycopeptides associated with Supplementary data 1 are provided.

**Supplementary Table 2. Combined Byonic search results of *B. cenocepacia* K56-2 glycopeptide enrichments**. The combined Byonic search results from all proteases across each of the three biological replicates of *B. cenocepacia* K56-2 are provided. The complete list of all glycopeptides identified, the best unique glycopeptides filtered for glycopeptides with a Byonic score over 300 and the curated list of glycopeptides associated with supplementary data 2 are provided.

**Supplementary Table 3. LFQ proteome analysis of *B. cenocepacia* strains H111 WT vs H111 Δ*pglL* candidate 1 vs H111 Δ*pglL* candidate 8**. The Perseus processed MaxQuant search results for the protein analysis of four biological replicates of strains H111 WT, H111 Δ*pglL* candidate 1 and H111 Δ*pglL* candidate 8 are provided. For each identified protein, the log2 LFQ protein values, the T-test information including the −log_10_(*p*-value), difference in the mean between the groups and if the resulting *p*-values is below the multiple hypothesis corrected *p*-value are provided. Categorical information used for enrichment analysis including GO terms and if proteins were considered to be altered within K56-2 Δ*pglL* within Oppy *et al*. 2019 ^5^ are provided. For each protein the protein score, number of MS/MS for the corresponding protein, expected molecular weight, number of peptides identified, peptide sequence coverage and corresponding J2315 gene number for the protein are provided.

**Supplementary Table 4: Enrichment analysis of global proteome changes in response to loss of glycosylation in *B. cenocepacia* H111**. Using Perseus, proteins determined to undergo statistically significant changes were assessed for co-occurrence of statistically significant changes across strains as well as enrichment for proteins considered to be altered within K56-2 Δ*pglL* within Oppy *et al*. 2019 ^5^.

**Supplementary Table 5. Curated *B. cenocepacia* glycosylation sites**. The identified glycosylation sites from *B. cenocepacia* K56-2 and H111 are provided. For each site the J315 Gene name, position of the site within the J2315 version of the protein, residue the glycan was localised to, amino acid sequence +/- 10 amino acid either side of the localised site and which strain these sites were observed in are provided.

**Supplementary Table 6. O-Pair analysis of *B. cenocepacia* glycosylation sites**. O-pair site localisation information is provided with only class “level 1” glycopeptides with a Q-value less then 0.01 and a site-specific probability of >0.75 considered as localised and used for data visualisation.

**Supplementary Table 7. Proteome analysis of the 25-35kDa of *B. cenocepacia* lysates expressing DsbA1_Nm_-his_6_ variants**. MaxQuant peptide search results of induced K56-2 expressing DsbA1_Nm_-his_6_ WT or T^36^. For each peptide the mass, Protein Group, Modification status, identification type (by MS/MS or matching), Retention time, Charge, PEP, MS/MS scan number for the best scoring peptide spectral match, Raw file of the best scoring peptide spectral match, Score, Delta Score, Intensity and number of MS/MS are provided.

**Supplementary Table 8. Glycoprotein and glycosylation site conservation across Burkholderia genomes**. Summary of sequence identity percentages of glycoproteins and glycosylation motifs identified across *B. cenocepacia* strains and representative Burkholderia species.

**Supplementary Table 9. O-Pair analysis of multiple *Burkholderia* species**. O-pair site localisation information is provided with only class “level 1” glycopeptides with a Q-value less than 0.01 and a site-specific probability of >0.75 considered as localised and used for data visualisation.

**Supplementary Table 10. Proteome analysis of the 25-35kDa of *B. ubonensis* MSMB22 and *B. humptydooensis* MSMB43 lysates expressing DsbA1_Nm_-his_6_ variants**. MaxQuant peptide search results of induced K56-2 expressing DsbA1_Nm_-his_6_ WT or T^36^. For each peptide the mass, Protein Group, Modification status, identification type (by MS/MS or matching), Retention time, Charge, PEP, MS/MS scan number for the best scoring peptide spectral match, Raw file of the best scoring peptide spectral match, Score, Delta Score, Intensity and number of MS/MS are provided.

**Supplementary Table 11. Glycoprotein homologues identified across Burkholderia species**. Summary of glycoprotein homologues identified, and enrichment analysis of glycoproteins identified across *Burkholderia* species.

