## Supplementary Data 1 for "Burkholderia PglL enzymes are serine preferring oligosaccharidetransferases which target conserved proteins across the Burkholderia genus"

### *Burkholderia cenocepacia* H111 Best Scoring Unique glycopeptides

| J2315 numbers | Protein Name | Peptide < ProteinMetrics Confidential > | Glycans NHFAGNa | Localised site (Yes/No) | Observed m/z | z | Observed (M+H) | Calc. mass (M+H) | Mass error (ppm) | Cleavage | Score | Delta | Delta Mod | Comment | Scan Time | Enzyme | Replicate | Page |
|---|---|---|---|---|---|---|---|---|---|---|---|---|---|---|---|---|---|---|
| BCAL0080 | >tr\|A0A1V2WZJ2\|A0A1V2WZJ2_9BURK Class I cytochrome c OS=Burkholderia cenocepacia OX=95486 GN=A8E72_32710 PE=4 SV=1 | Y.MANNDGANFPEPAAPAANAAQPAS[+568.212]GAPASGADASNAQA.A | HexNAc(2)Hex(1) | Yes | 1345.2598 | 3 | 4033.7649 | 4033.7476 | 4.3 | Non | 661.09 | 592.27 | 106.51 | Nsco_20191108_BC_ZIC_HILIC_H111_thermolysin_B3.42890.42890.3 | 84.7311 | Thermolysin | B3 | 28 |
| BCAL0080 | >tr\|A0A1V2WZJ2\|A0A1V2WZJ2_9BURK Class I cytochrome c OS=Burkholderia cenocepacia OX=95486 GN=A8E72_32710 PE=4 SV=1 | Y.MANNDGANFPEPAAPAANAAQPAS[+568.212]GAPASGADASNAQAA.A | HexNAc(2)Hex(1) | Yes | 1369.2733 | 3 | 4105.8054 | 4104.7847 | 4.2 | Non | 466.66 | 430.12 | 35.3 | Nsco_20191108_BC_ZIC_HILIC_H111_thermolysin_B3.43275.43275.3 | 85.4568 | Thermolysin | B3 | 29 |
| BCAL0080 | >tr\|A0A1V2WZJ2\|A0A1V2WZJ2_9BURK Class I cytochrome c OS=Burkholderia cenocepacia OX=95486 GN=A8E72_32710 PE=4 SV=1 | Y.MANNDGANFPEPAAPAANAAQPAS[+568.212][+100.064]GAPAS[+568.212]GADASNAQAAAA.M | HexNAc(2)Hex(1) 100.064,HexNAc(2)Hex(1) | No | 1639.0439 | 3 | 4915.1171 | 4915.1345 | -3.5 | Non | 268.13 | 247.1 | 2.69 | Nsco_20191108_BC_ZIC_HILIC_H111_thermolysin_B3.44017.44017.3 | 86.817 | Thermolysin | B3 | 30 |
| BCAL0080 | >tr\|A0A1V2WZJ2\|A0A1V2WZJ2_9BURK Class I cytochrome c OS=Burkholderia cenocepacia OX=95486 GN=A8E72_32710 PE=4 SV=1 | R.AIVYMANNDGANFPEPAAPAANAAQPAS[+568.212]GAPAS[+568.212][+100.064]GADASNAQAAAAMAAIAAIPK.A | HexNAc(2)Hex(1), HexNAc(2)Hex(1) 100.064 | No | 1557.7234 | 4 | 6227.8717 | 6227.8922 | -3.3 | Specific | 806.09 | 806.09 | 0 | Nsco_20191108_BC_ZIC_HILIC_H111_typsin_B2.71129.71129.4 | 140.2105 | Trypsin | B2 | 31 |
| BCAL0080 | >tr\|A0A1V2WZJ2\|A0A1V2WZJ2_9BURK Class I cytochrome c OS=Burkholderia cenocepacia OX=95486 GN=A8E72_32710 PE=4 SV=1 | A.IVYMANNDGANFPEPAAPAANAAQPAS[+568.212][+100.064]GAPAS[+568.212]GADASNAQAAAAMAAIAAIPK.A | HexNAc(2)Hex(1) 100.064,HexNAc(2)Hex(1) | No | 1540.4741 | 4 | 6158.8747 | 6156.8551 | 2.1 | NRagged | 464.47 | 464.47 | 0 | Nsco_20191108_BC_ZIC_HILIC_H111_typsin_B3.66894.66894.4 | 139.2643 | Trypsin | B3 | 32 |
| BCAL0163 | >tr\|A0A144S4M8\|A0A144S4M8_9BURK BON domain protein OS=Burkholderia cenocepacia OX=95486 GN=A8E72_32405 PE=4 SV=1 | A.LQDASPASGAS[+568.212]GAQAAAAPADN.A | HexNAc(2)Hex(1) | No | 1255.0607 | 2 | 2509.1141 | 2509.106 | 3.2 | Non | 554.16 | 266.54 | 0 | Nsco_20191108_BC_ZIC_HILIC_H111_pepsin_B1.22640.22640.2 | 48.7976 | Pepsin | B1 | 33 |
| BCAL0163 | >tr\|A0A144S4M8\|A0A144S4M8_9BURK BON domain protein OS=Burkholderia cenocepacia OX=95486 GN=A8E72_32405 PE=4 SV=1 | S.VQSTPLQPPAPISNSS[+568.212]SVHPGNPKAKAQ.- | HexNAc(2)Hex(1) | Yes | 852.1834 | 4 | 3405.7117 | 3405.702 | 2.8 | Non | 783.02 | 702.53 | 22.83 | Nsco_20191108_BC_ZIC_HILIC_H111_pepsin_B2.21640.21640.4 | 45.2027 | Pepsin | B2 | 34 |
| BCAL0163 | >tr\|A0A144S4M8\|A0A144S4M8_9BURK BON domain protein OS=Burkholderia cenocepacia OX=95486 GN=A8E72_32405 PE=4 SV=1 | A.SVQSTPLQPPAPISNS[+568.212]SSVHPGNPKAKAQ.- | HexNAc(2)Hex(1) | No | 873.9426 | 4 | 3492.7486 | 3492.734 | 4.2 | Non | 404.08 | 404.08 | 0 | Nsco_20191108_BC_ZIC_HILIC_H111_pepsin_B3.22640.22640.4 | 46.758 | Pepsin | B3 | 35 |
| BCAL0163 | >tr\|A0A144S4M8\|A0A144S4M8_9BURK BON domain protein OS=Burkholderia cenocepacia OX=95486 GN=A8E72_32405 PE=4 SV=1 | A.LQDASPASGASGAQAAAAPADNATVGAVPDASVQSTPLQPPAPIS[+568.212]NSSSVHPGNPKAKAQ.- | HexNAc(2)Hex(1) | No | 1549.7634 | 4 | 6196.0319 | 6196.0076 | 3.9 | Non | 830.97 | 830.97 | 0 | Nsco_20191108_BC_ZIC_HILIC_H111_thermolysin_B1.44267.44267.4 | 83.5595 | Thermolysin | B1 | 36 |
| BCAL0163 | >tr\|A0A144S4M8\|A0A144S4M8_9BURK BON domain protein OS=Burkholderia cenocepacia OX=95486 GN=A8E72_32405 PE=4 SV=1 | Y.VKPQDAQALQDASPASGASGAQAAAAPADNAT[+568.212]VGAVPDASVQSTPLQPPAP.I | HexNAc(2)Hex(1) | No | 1778.5364 | 3 | 5333.5946 | 5330.567 | 3.3 | Non | 148.88 | 148.88 | 0 | Nsco_20191108_BC_ZIC_HILIC_H111_thermolysin_B1.57277.57277.3 | 107.1961 | Thermolysin | B1 | 37 |

| | | | | | | | | | | | | | | | | | | |
|---|---|---|---|---|---|---|---|---|---|---|---|---|---|---|---|---|---|---|
| BCAL0163 | >tr\|A0A144S4M8\|A0A144S4M8_9BURK BON domain protein OS=Burkholderia cenocepacia OX=95486 GN=A8E72_32405 PE=4 SV=1 | Y.VKPQDAQALQDASPAS[+568.212]GASGAQA.A | HexNAc(2)Hex(1) | No | 1368.6446 | 2 | 2736.2819 | 2736.2694 | 4.6 | Non | 539.84 | 480.2 | 4.11 | Nsco_20191108_BC_ZIC_HILIC_H111_thermolysin_B2.25007.25007.2 | 52.3677 | Thermolysin | B2 | 38 |
| BCAL0163 | >tr\|A0A144S4M8\|A0A144S4M8_9BURK BON domain protein OS=Burkholderia cenocepacia OX=95486 GN=A8E72_32405 PE=4 SV=1 | A.LQDASPASGAS[+568.212]GAQAAAAPADNAT.V | HexNAc(2)Hex(1) | No | 1341.606 | 2 | 2682.2047 | 2681.1908 | 3.9 | Non | 545.39 | 207.92 | 2.4 | Nsco_20191108_BC_ZIC_HILIC_H111_thermolysin_B2.26277.26277.2 | 54.5842 | Thermolysin | B2 | 39 |
| BCAL0163 | >tr\|A0A144S4M8\|A0A144S4M8_9BURK BON domain protein OS=Burkholderia cenocepacia OX=95486 GN=A8E72_32405 PE=4 SV=1 | A.LQDASPASGAS[+568.212]GAQAAAAPADNATVGAVPDASVQSTP.L | HexNAc(2)Hex(1) | Yes | 1297.2759 | 3 | 3889.8133 | 3889.7945 | 4.8 | Non | 260.83 | 253.64 | 45.47 | Nsco_20191108_BC_ZIC_HILIC_H111_thermolysin_B2.46669.46669.3 | 92.8997 | Thermolysin | B2 | 40 |
| BCAL0163 | >tr\|A0A144S4M8\|A0A144S4M8_9BURK BON domain protein OS=Burkholderia cenocepacia OX=95486 GN=A8E72_32405 PE=4 SV=1 | Y.VKPQDAQALQDAS[+568.212][+100.064]PASGASGAQAAAAPADNA.T | HexNAc(2)Hex(1) 100.064 | No | 1759.8258 | 2 | 3518.6443 | 3517.6416 | -0.2 | Non | 276.03 | 115.52 | 2.12 | Nsco_20191108_BC_ZIC_HILIC_H111_thermolysin_B3.32267.32267.2 | 64.9265 | Thermolysin | B3 | 41 |
| BCAL0163 | >tr\|A0A144S4M8\|A0A144S4M8_9BURK BON domain protein OS=Burkholderia cenocepacia OX=95486 GN=A8E72_32405 PE=4 SV=1 | Y.VKPQDAQALQDASPAS[+568.212]GASGAQ.A | HexNAc(2)Hex(1) | No | 1333.1246 | 2 | 2665.242 | 2665.2323 | 3.6 | Non | 494.43 | 461.19 | 4.68 | Nsco_20191108_BC_ZIC_HILIC_H111_thermolysin_B3.23059.23059.2 | 49.2567 | Thermolysin | B3 | 42 |
| BCAL0163 | >tr\|A0A144S4M8\|A0A144S4M8_9BURK BON domain protein OS=Burkholderia cenocepacia OX=95486 GN=A8E72_32405 PE=4 SV=1 | Y.VKPQDAQALQDASPAS[+568.212]GASGAQAAAAPADNAT.V | HexNAc(2)Hex(1) | No | 1759.825 | 2 | 3518.6428 | 3518.6253 | 5 | Non | 579.7 | 300.08 | 3.43 | Nsco_20191108_BC_ZIC_HILIC_H111_thermolysin_B3.32243.32243.2 | 64.8849 | Thermolysin | B3 | 43 |
| BCAL0163 | >tr\|A0A144S4M8\|A0A144S4M8_9BURK BON domain protein OS=Burkholderia cenocepacia OX=95486 GN=A8E72_32405 PE=4 SV=1 | S.GASGAQAAAAPADNATVGAVPDASVQST[+568.212][+100.064]PLQPPAPIS[+568.212][+100.064]NSSSVHPGNPKAKAQ.- | HexNAc(2)Hex(1) 100.064,HexNAc(2)Hex(1) 100.064 | No | 1549.5074 | 4 | 6195.008 | 6194.9866 | 3.5 | Non | 457.38 | 457.38 | 0 | Nsco_20191108_BC_ZIC_HILIC_H111_thermolysin_B3.42433.42433.4 | 83.785 | Thermolysin | B3 | 44 |
| BCAL0163 | >tr\|A0A144S4M8\|A0A144S4M8_9BURK BON domain protein OS=Burkholderia cenocepacia OX=95486 GN=A8E72_32405 PE=4 SV=1 | Y.VKPQDAQALQDASPASGAS[+568.212]GAQAAAAPADNATVGAVPDASVQSTPLQPPAPISNSSSVHPGNPK.A | HexNAc(2)Hex(1) | Yes | 1659.8156 | 4 | 6636.2404 | 6635.2143 | 3.4 | NRagged | 901.28 | 550.13 | 85.05 | Nsco_20191108_BC_ZIC_HILIC_H111_typsin_B2.47144.47144.4 | 91.2822 | Trypsin | B2 | 45 |
| BCAL0163 | >tr\|A0A144S4M8\|A0A144S4M8_9BURK BON domain protein OS=Burkholderia cenocepacia OX=95486 GN=A8E72_32405 PE=4 SV=1 | K.PQDAQALQDASPASGASGAQAAAAPADNATVGAVPDASVQST[+568.212]PLQPPAPISNSSSVHPGNPK.A | HexNAc(2)Hex(1) | No | 1603.0234 | 4 | 6409.0719 | 6408.051 | 2.7 | Specific | 1091.36 | 1068.39 | 0 | Nsco_20191108_BC_ZIC_HILIC_H111_typsin_B2.50885.50885.4 | 97.7338 | Trypsin | B2 | 46 |
| BCAL0163 | >tr\|A0A144S4M8\|A0A144S4M8_9BURK BON domain protein OS=Burkholderia cenocepacia OX=95486 GN=A8E72_32405 PE=4 SV=1 | V.KPQDAQALQDASPASGAS[+568.212]GAQAAAAPADNAT[+568.212][+100.064]VGAVPDASVQST[+568.212]PLQPPAPISNSSSVHPGNPK.A | HexNAc(2)Hex(1), HexNAc(2)Hex(1) 100.064,HexNAc(2)Hex(1) | No | 1297.1181 | 6 | 7777.6722 | 7772.6331 | 2.9 | NRagged | 287.53 | 287.53 | 0 | Nsco_20191108_BC_ZIC_HILIC_H111_typsin_B2.51562.51562.6 | 98.9045 | Trypsin | B2 | 47 |
| BCAL0163 | >tr\|A0A144S4M8\|A0A144S4M8_9BURK BON domain protein OS=Burkholderia cenocepacia OX=95486 GN=A8E72_32405 PE=4 SV=1 | V.FQYVKPQDAQALQDAS[+568.212]PASGASGAQAAAAPADNATVGAVPDASVQSTPLQPPAPISNSSSVHPGNPK.A | HexNAc(2)Hex(1) | No | 1769.8647 | 4 | 7076.4372 | 7073.4047 | 3.2 | NRagged | 590.81 | 590.81 | 1.2 | Nsco_20191108_BC_ZIC_HILIC_H111_typsin_B2.51800.51800.4 | 99.3283 | Trypsin | B2 | 48 |
| BCAL0163 | >tr\|A0A144S4M8\|A0A144S4M8_9BURK BON domain protein OS=Burkholderia cenocepacia OX=95486 GN=A8E72_32405 PE=4 SV=1 | K.VFQYVKPQDAQALQDASPASGASGAQAAAAPADNATVGAVPDASVQST[+568.212]PLQPPAPISNSSSVHPGNPK.A | HexNAc(2)Hex(1) | No | 1793.8806 | 4 | 7172.5005 | 7172.4731 | 3.8 | Specific | 1198.94 | 1198.94 | 0 | Nsco_20191108_BC_ZIC_HILIC_H111_typsin_B2.54057.54057.4 | 103.675 | Trypsin | B2 | 49 |

| | | | | | | | | | | | | | | | | | | |
|---|---|---|---|---|---|---|---|---|---|---|---|---|---|---|---|---|---|---|
| BCAL0163 | >tr\|A0A144S4M8\|A0A144S4M8_9BURK BON domain protein OS=Burkholderia cenocepacia OX=95486 GN=A8E72_32405 PE=4 SV=1 | V.KVFQYVKPQDAQALQDASPASGAS[+568.212]GAQAAAAPADNATVGAVPDASVQSTPLQPPAPISNSSSVHPGNPK.A | HexNAc(2)Hex(1) | Yes | 1460.9267 | 5 | 7300.6043 | 7300.568 | 5 | NRagged | 1341.58 | 1341.58 | 22.45 | Nsco_20191108_BC_ZIC_HILIC_H111_typsin_B3.45311.45311.5 | 93.2228 | Trypsin | B3 | 50 |
| BCAL0193 | >tr\|A0A071MDE2\|A0A071MDE2_9BURK Uncharacterized protein OS=Burkholderia cenocepacia OX=95486 GN=A8E72_00390 PE=4 SV=1 | Q.LKHHGSKKGQAKAAAAS[+568.212]AAGTNDAGTQN.- | HexNAc(2)Hex(1) | Yes | 652.5254 | 5 | 3258.5977 | 3258.5833 | 4.4 | Non | 882.97 | 882.97 | 102.81 | Nsco_20191108_BC_ZIC_HILIC_H111_thermolysin_B1.2860.2860.5 | 16.0864 | Thermolysin | B1 | 51 |
| BCAL0193 | >tr\|A0A071MDE2\|A0A071MDE2_9BURK Uncharacterized protein OS=Burkholderia cenocepacia OX=95486 GN=A8E72_00390 PE=4 SV=1 | P.AADTSAAAPAPAKKDHS[+568.212][+100.064]KPKHQ.L | HexNAc(2)Hex(1) 100.064 | No | 966.1519 | 3 | 2896.4413 | 2895.4334 | 1.6 | Non | 360.56 | 293.32 | 9.6 | Nsco_20191108_BC_ZIC_HILIC_H111_thermolysin_B3.5940.5940.3 | 21.4368 | Thermolysin | B3 | 52 |
| BCAL0193 | >tr\|A0A071MDE2\|A0A071MDE2_9BURK Uncharacterized protein OS=Burkholderia cenocepacia OX=95486 GN=A8E72_00390 PE=4 SV=1 | Q.ASAPAADT[+568.212]SAAAPAPAKK.D | HexNAc(2)Hex(1) | No | 1082.5309 | 2 | 2164.0546 | 2164.0503 | 2 | NRagged | 769.06 | 615.58 | 0 | Nsco_20191108_BC_ZIC_HILIC_H111_typsin_B1.6903.6903.2 | 22.6365 | Trypsin | B1 | 53 |
| BCAL0272 | >tr\|A0A142PJB0\|A0A142PJB0_9BURK Lipoprotein OS=Burkholderia cenocepacia OX=95486 GN=A8E72_20920 PE=4 SV=1 | K.PIQPQNTPPSDVKPTDENASSDESPDTSGSPLTLSPELSSTSTMPPAS[+568.212]GPAATK.- | HexNAc(2)Hex(1) | Yes | 1515.9615 | 4 | 6060.8244 | 6058.798 | 3.2 | Specific | 944.57 | 888.07 | 96.92 | Nsco_20191108_BC_ZIC_HILIC_H111_typsin_B2.51007.51007.4 | 97.9364 | Trypsin | B2 | 54 |
| BCAL0303 | >tr\|A0A144S0T9\|A0A144S0T9_9BURK Outer membrane lipid asymmetry maintenance protein MlaD OS=Burkholderia cenocepacia OX=95486 GN=yrbD PE=4 SV=1 | F.LYS[+568.212]KAADAGGAKPAAGASAAPAAPAPVAV.P | HexNAc(2)Hex(1) | No | 1030.5266 | 3 | 3089.5654 | 3089.5525 | 4.2 | CRagged | 222.88 | 189.14 | 2.91 | Nsco_20191108_BC_ZIC_HILIC_H111_pepsin_B1.28060.28060.3 | 59.3181 | Pepsin | B1 | 55 |
| BCAL0303 | >tr\|A0A144S0T9\|A0A144S0T9_9BURK Outer membrane lipid asymmetry maintenance protein MlaD OS=Burkholderia cenocepacia OX=95486 GN=yrbD PE=4 SV=1 | L.YSKAADAGGAKPAAGAS[+568.212]AAPAAPAPVAVPASAVSGSAGQ.- | HexNAc(2)Hex(1) | No | 1296.9756 | 3 | 3888.9124 | 3888.8985 | 3.6 | CRagged | 785.85 | 782.53 | 7.13 | Nsco_20191108_BC_ZIC_HILIC_H111_pepsin_B1.31112.31112.3 | 65.6963 | Pepsin | B1 | 56 |
| BCAL0303 | >tr\|A0A144S0T9\|A0A144S0T9_9BURK Outer membrane lipid asymmetry maintenance protein MlaD OS=Burkholderia cenocepacia OX=95486 GN=yrbD PE=4 SV=1 | S.KAADAGGAKPAAGAS[+568.212]AAPAAPAPVAV.P | HexNAc(2)Hex(1) | Yes | 909.4655 | 3 | 2726.3819 | 2726.3731 | 3.2 | Non | 741.46 | 731.06 | 731.06 | Nsco_20191108_BC_ZIC_HILIC_H111_pepsin_B2.23564.23564.3 | 48.9597 | Pepsin | B2 | 57 |
| BCAL0303 | >tr\|A0A144S0T9\|A0A144S0T9_9BURK Outer membrane lipid asymmetry maintenance protein MlaD OS=Burkholderia cenocepacia OX=95486 GN=yrbD PE=4 SV=1 | Y.SKAADAGGAKPAAGAS[+568.212]AAPAAPAPVAV.P | HexNAc(2)Hex(1) | No | 938.4753 | 3 | 2813.4114 | 2813.4051 | 2.2 | Non | 621.17 | 600 | 47.88 | Nsco_20191108_BC_ZIC_HILIC_H111_pepsin_B2.23611.23611.3 | 49.0427 | Pepsin | B2 | 58 |
| BCAL0303 | >tr\|A0A144S0T9\|A0A144S0T9_9BURK Outer membrane lipid asymmetry maintenance protein MlaD OS=Burkholderia cenocepacia OX=95486 GN=yrbD PE=4 SV=1 | L.YSKAADAGGAKPAAGAS[+568.212]AAPAAPAPVAV.P | HexNAc(2)Hex(1) | Yes | 744.8754 | 4 | 2976.48 | 2976.4684 | 3.9 | CRagged | 843.54 | 806.59 | 509.24 | Nsco_20191108_BC_ZIC_HILIC_H111_pepsin_B3.25168.25168.4 | 51.9446 | Pepsin | B3 | 59 |
| BCAL0303 | >tr\|A0A144S0T9\|A0A144S0T9_9BURK Outer membrane lipid asymmetry maintenance protein MlaD OS=Burkholderia cenocepacia OX=95486 GN=yrbD PE=4 SV=1 | F.LYSKAADAGGAKPAAGAS[+568.212]AAPAAPAPVAVPASAVSGSAGQ.- | HexNAc(2)Hex(1) | No | 1334.671 | 3 | 4001.9983 | 4001.9826 | 3.9 | CRagged | 752 | 752 | 18 | Nsco_20191108_BC_ZIC_HILIC_H111_thermolysin_B1.36046.36046.3 | 69.1968 | Thermolysin | B1 | 60 |

| | | | | | | | | | | | | | | | | | | |
|---|---|---|---|---|---|---|---|---|---|---|---|---|---|---|---|---|---|---|
| BCAL0303 | >tr\|A0A144S0T9\|A0A144S0T9_9BURK Outer membrane lipid asymmetry maintenance protein MlaD OS=Burkholderia cenocepacia OX=95486 GN=yrbD PE=4 SV=1 | K.AADAGGAKPAAGASAAPAAPAPVAVPAS[+568.212]AVSGSAGQ.- | HexNAc(2)Hex(1) | No | 1755.8662 | 2 | 3510.7251 | 3510.7082 | 4.8 | CRagged | 800.05 | 686.18 | 1.17 | Nsco_20191108_BC_ZIC_HILIC_H111_typsin_B1.36885.36885.2 | 74.4191 | Trypsin | B1 | 61 |
| BCAL0303 | >tr\|A0A144S0T9\|A0A144S0T9_9BURK Outer membrane lipid asymmetry maintenance protein MlaD OS=Burkholderia cenocepacia OX=95486 GN=yrbD PE=4 SV=1 | A.ADAGGAKPAAGAS[+568.212][+100.064]AAPAAPAPVAVPASAVSGSAGQ.- | HexNAc(2)Hex(1) 100.064 | No | 1181.246 | 3 | 3541.7234 | 3539.7351 | -5.2 | Non | 279.51 | 279.51 | 13.16 | Nsco_20191108_BC_ZIC_HILIC_H111_typsin_B2.39186.39186.3 | 77.2324 | Trypsin | B2 | 62 |
| BCAL0303 | >tr\|A0A144S0T9\|A0A144S0T9_9BURK Outer membrane lipid asymmetry maintenance protein MlaD OS=Burkholderia cenocepacia OX=95486 GN=yrbD PE=4 SV=1 | K.PAAGASAAPAAPAPVAVPAS[+568.212][+100.064]AVS[+568.212]GSAGQ.- | HexNAc(2)Hex(1) 100.064,HexNAc(2)Hex(1) | Yes | 1180.2254 | 3 | 3538.6616 | 3537.6705 | -3.4 | CRagged | 247.45 | 206.29 | 1.76 | Nsco_20191108_BC_ZIC_HILIC_H111_typsin_B2.42457.42457.3 | 82.9779 | Trypsin | B2 | 63 |
| BCAL0304 | >tr\|A0A2N9CYS8\|A0A2N9CYS8_9BURK Surface lipoprotein OS=Burkholderia cenocepacia OX=95486 GN=F01_50090 PE=4 SV=1 | A.PASGTAESPNPAS[+568.212]ET[+568.212]NV.P | HexNAc(2)Hex(1), HexNAc(2)Hex(1) | No | 1383.091 | 2 | 2765.1747 | 2765.163 | 4.2 | Non | 683.52 | 496.16 | 9.94 | Nsco_20191108_BC_ZIC_HILIC_H111_pepsin_B1.21155.21155.2 | 45.8986 | Pepsin | B1 | 64 |
| BCAL0304 | >tr\|A0A2N9CYS8\|A0A2N9CYS8_9BURK Surface lipoprotein OS=Burkholderia cenocepacia OX=95486 GN=F01_50090 PE=4 SV=1 | A.VAAPASGTAESPNPAS[+568.212]ET[+568.212]NV.P | HexNAc(2)Hex(1), HexNAc(2)Hex(1) | No | 1002.7762 | 3 | 3006.3141 | 3006.3056 | 2.8 | Non | 549.86 | 421.72 | 8.83 | Nsco_20191108_BC_ZIC_HILIC_H111_pepsin_B1.26846.26846.3 | 56.911 | Pepsin | B1 | 65 |
| BCAL0304 | >tr\|A0A2N9CYS8\|A0A2N9CYS8_9BURK Surface lipoprotein OS=Burkholderia cenocepacia OX=95486 GN=F01_50090 PE=4 SV=1 | A.PASGTAESPNPAS[+568.212][+100.064]ET[+568.212]NVPA.M | HexNAc(2)Hex(1) 100.064,HexNAc(2)Hex(1) | No | 1518.1598 | 2 | 3035.3124 | 3033.3169 | -3.7 | Non | 314.5 | 153.64 | 8.22 | Nsco_20191108_BC_ZIC_HILIC_H111_pepsin_B2.29530.29530.2 | 60.8298 | Pepsin | B2 | 66 |
| BCAL0304 | >tr\|A0A2N9CYS8\|A0A2N9CYS8_9BURK Surface lipoprotein OS=Burkholderia cenocepacia OX=95486 GN=F01_50090 PE=4 SV=1 | A.VAAPASGTAESPNPAS[+568.212]ET[+568.212]NVPAM.Q | HexNAc(2)Hex(1), HexNAc(2)Hex(1) | No | 1102.4886 | 3 | 3305.4511 | 3305.436 | 4.6 | Non | 480.43 | 256.57 | 2.4 | Nsco_20191108_BC_ZIC_HILIC_H111_pepsin_B3.35544.35544.3 | 76.6557 | Pepsin | B3 | 67 |
| BCAL0304 | >tr\|A0A2N9CYS8\|A0A2N9CYS8_9BURK Surface lipoprotein OS=Burkholderia cenocepacia OX=95486 GN=F01_50090 PE=4 SV=1 | A.PASGTAESPNPAS[+568.212]ET[+568.212]NVPAMQ.V | HexNAc(2)Hex(1), HexNAc(2)Hex(1) | No | 1064.792 | 3 | 3192.3615 | 3192.3519 | 3 | Non | 334.39 | 334.39 | 1.2 | Nsco_20191108_BC_ZIC_HILIC_H111_pepsin_B3.31204.31204.3 | 65.5756 | Pepsin | B3 | 68 |
| BCAL0304 | >tr\|A0A2N9CYS8\|A0A2N9CYS8_9BURK Surface lipoprotein OS=Burkholderia cenocepacia OX=95486 GN=F01_50090 PE=4 SV=1 | A.VAAPASGTAESPNPAS[+568.212]ET[+568.212]NVPA.M | HexNAc(2)Hex(1), HexNAc(2)Hex(1) | No | 1587.7082 | 2 | 3174.4091 | 3174.3955 | 4.3 | Non | 472.31 | 305.68 | 6.36 | Nsco_20191108_BC_ZIC_HILIC_H111_thermolysin_B2.30194.30194.2 | 61.2273 | Thermolysin | B2 | 69 |
| BCAL0304 | >tr\|A0A2N9CYS8\|A0A2N9CYS8_9BURK Surface lipoprotein OS=Burkholderia cenocepacia OX=95486 GN=F01_50090 PE=4 SV=1 | A.VAAPASGTAES[+568.212][+100.064]PNPAS[+568.212]ETNVP.A | HexNAc(2)Hex(1) 100.064,HexNAc(2)Hex(1) | No | 1602.2084 | 2 | 3203.4095 | 3203.4224 | -4 | Non | 349.49 | 208.59 | 3.63 | Nsco_20191108_BC_ZIC_HILIC_H111_thermolysin_B2.32625.32625.2 | 65.2288 | Thermolysin | B2 | 70 |
| BCAL0304 | >tr\|A0A2N9CYS8\|A0A2N9CYS8_9BURK Surface lipoprotein OS=Burkholderia cenocepacia OX=95486 GN=F01_50090 PE=4 SV=1 | A.APASGTAESPNPAS[+568.212][+100.064]ET[+568.212][+100.064]NVPAMQ.V | HexNAc(2)Hex(1) 100.064,HexNAc(2)Hex(1) 100.064 | No | 1732.26 | 2 | 3463.5127 | 3463.517 | -1.2 | Non | 338.9 | 87.16 | 2.4 | Nsco_20191108_BC_ZIC_HILIC_H111_thermolysin_B2.38282.38282.2 | 75.6943 | Thermolysin | B2 | 71 |
| BCAL0304 | >tr\|A0A2N9CYS8\|A0A2N9CYS8_9BURK Surface lipoprotein OS=Burkholderia cenocepacia OX=95486 GN=F01_50090 PE=4 SV=1 | A.VAAPASGT[+568.212]AES[+568.212]PNPASETNVPAMQ.V | HexNAc(2)Hex(1), HexNAc(2)Hex(1) | No | 1717.2554 | 2 | 3433.5036 | 3433.4946 | 2.6 | Non | 482.57 | 394.87 | 2.47 | Nsco_20191108_BC_ZIC_HILIC_H111_thermolysin_B2.36740.36740.2 | 72.7165 | Thermolysin | B2 | 72 |

| | | | | | | | | | | | | | | | | | |
|---|---|---|---|---|---|---|---|---|---|---|---|---|---|---|---|---|---|
| BCAL0304 | >tr\|A0A2N9CYS8\|A0A2N9CYS8_9BURK Surface lipoprotein OS=Burkholderia cenocepacia OX=95486 GN=F01_50090 PE=4 SV=1 | L.PEDGAAPAAAGAAGT[+568.212][+100.064]AGAAAVGGAAPAGAAVAAPASGT[+568.212]AES[+568.212]PNPASETNVPAMQVAPPSPGGFRFPSIR.L | HexNAc(2)Hex(1)100.064,HexNAc(2)Hex(1),HexNAc(2)Hex(1) | No | 1589.3426 | 5 | 7942.6838 | 7938.7048 | -4.3 | NRagged | 316.83 | 316.83 | 0 | Nsco_20191108_BC_ZIC_HILIC_H111_typsin_B2.65020.65020.5 | 126.0041 | Trypsin | B2 | 73 |
| BCAL0332 | >tr\|A0A2N9CYY5\|A0A2N9CYY5_9BURK ClpXP protease specificity-enhancing factor OS=Burkholderia cenocepacia OX=95486 GN=sspB PE=4 SV=1 | E.DSGAFDDEQADDAQRDES[+568.212]VSPLAPVADSGANEEPSEGADEPPKTDGDGSK.G | HexNAc(2)Hex(1) | No | 1412.1012 | 4 | 5645.383 | 5643.3656 | 1.9 | NRagged | 572.55 | 572.55 | 0 | Nsco_20191108_BC_ZIC_HILIC_H111_typsin_B2.43714.43714.4 | 85.2165 | Trypsin | B2 | 74 |
| BCAL0332 | >tr\|A0A2N9CYY5\|A0A2N9CYY5_9BURK ClpXP protease specificity-enhancing factor OS=Burkholderia cenocepacia OX=95486 GN=sspB PE=4 SV=1 | A.DDAQRDESVSPLAPVADS[+568.212]GANEEPSEGADEPPKTDGDGSK.G | HexNAc(2)Hex(1) | No | 1153.5083 | 4 | 4611.0115 | 4607.9875 | 3 | NRagged | 634.53 | 634.53 | 1.2 | Nsco_20191108_BC_ZIC_HILIC_H111_typsin_B3.35283.35283.4 | 73.8531 | Trypsin | B3 | 75 |
| BCAL0340 | >tr\|A0A1V2W5P4\|A0A1V2W5P4_9BURK Uncharacterized protein OS=Burkholderia cenocepacia OX=95486 GN=A8E72_11900 PE=4 SV=1 | K.AAAAPAAEAAASAAPT[+568.212]PPAAQK.G | HexNAc(2)Hex(1) | No | 1236.6064 | 2 | 2472.2055 | 2472.1988 | 2.7 | Specific | 817.86 | 747.77 | 19.34 | Nsco_20191108_BC_ZIC_HILIC_H111_typsin_B1.21382.21382.2 | 47.1614 | Trypsin | B1 | 76 |
| BCAL0349 (typ | >tr\|A0A2N9D2R1\|A0A2N9D2R1_9BURK Outer membrane protein OS=Burkholderia cenocepacia OX=95486 GN=F01_70026 PE=3 SV=1 | T.TALPQANPNAGGAS[+568.212]GTVVHGTAGTLTPPPANAAPGQ.V | HexNAc(2)Hex(1) | Yes | 1277.9608 | 3 | 3831.8677 | 3831.8519 | 4.1 | Non | 548.84 | 334.26 | 39.38 | Nsco_20191108_BC_ZIC_HILIC_H111_pepsin_B1.36350.36350.3 | 77.6999 | Pepsin | B1 | 77 |
| BCAL0349 (typ | >tr\|A0A2N9D2R1\|A0A2N9D2R1_9BURK Outer membrane protein OS=Burkholderia cenocepacia OX=95486 GN=F01_70026 PE=3 SV=1 | L.PQANPNAGGASGTVVHGT[+568.212]AGTLTPPPANAAPGQ.V | HexNAc(2)Hex(1) | No | 1182.9037 | 3 | 3546.6967 | 3546.6831 | 3.8 | CRagged | 556.85 | 505.61 | 3.6 | Nsco_20191108_BC_ZIC_HILIC_H111_pepsin_B2.30910.30910.3 | 63.7214 | Pepsin | B2 | 78 |
| BCAL0349 (typ | >tr\|A0A2N9D2R1\|A0A2N9D2R1_9BURK Outer membrane protein OS=Burkholderia cenocepacia OX=95486 GN=F01_70026 PE=3 SV=1 | A.TTALPQANPNAGGASGT[+568.212]VVHGTAGTLTPPPANAAPGQ.V | HexNAc(2)Hex(1) | No | 1311.6434 | 3 | 3932.9155 | 3932.8996 | 4.1 | Non | 327.12 | 308.22 | 1.2 | Nsco_20191108_BC_ZIC_HILIC_H111_pepsin_B2.37363.37363.3 | 78.598 | Pepsin | B2 | 79 |
| BCAL0349 (typ | >tr\|A0A2N9D2R1\|A0A2N9D2R1_9BURK Outer membrane protein OS=Burkholderia cenocepacia OX=95486 GN=F01_70026 PE=3 SV=1 | T.ALPQANPNAGGAS[+568.212]GTVVHGTAGTLTPPPANAAPGQVVVGGKVPDEATKA.A | HexNAc(2)Hex(1) | Yes | 1271.3992 | 4 | 5082.5749 | 5081.5549 | 3.3 | Non | 1019.7 | 1019.7 | 77.15 | Nsco_20191108_BC_ZIC_HILIC_H111_thermolysin_B1.41139.41139.4 | 78.3066 | Thermolysin | B1 | 80 |
| BCAL0349 (typ | >tr\|A0A2N9D2R1\|A0A2N9D2R1_9BURK Outer membrane protein OS=Burkholderia cenocepacia OX=95486 GN=F01_70026 PE=3 SV=1 | T.ALPQANPNAGGAS[+568.212][+100.064]GTVVHGTAGTLTPPPANAAPGQVVVGGKVPDEATKAAVLQK.L | HexNAc(2)Hex(1) 100.064 | Yes | 1145.1952 | 5 | 5721.9468 | 5720.9621 | -3.2 | Non | 803.05 | 467.38 | 82.36 | Nsco_20191108_BC_ZIC_HILIC_H111_thermolysin_B1.44302.44302.5 | 83.627 | Thermolysin | B1 | 81 |
| BCAL0349 (typ | >tr\|A0A2N9D2R1\|A0A2N9D2R1_9BURK Outer membrane protein OS=Burkholderia cenocepacia OX=95486 GN=F01_70026 PE=3 SV=1 | T.VATTALPQANPNAGGAS[+568.212]GTVVHGTAGTLTPPPANAAPGQVVVGGKVPDEATKAAVLQK.L | HexNAc(2)Hex(1) | No | 1199.4313 | 5 | 5993.1273 | 5993.099 | 4.7 | Non | 896.04 | 896.04 | 29.65 | Nsco_20191108_BC_ZIC_HILIC_H111_thermolysin_B1.45401.45401.5 | 85.4774 | Thermolysin | B1 | 82 |
| BCAL0349 (typ | >tr\|A0A2N9D2R1\|A0A2N9D2R1_9BURK Outer membrane protein OS=Burkholderia cenocepacia OX=95486 GN=F01_70026 PE=3 SV=1 | T.VATTALPQANPNAGGAS[+568.212][+100.064]GTVVHGTAGTLTPPPANAAPGQVVVGGKVPDEATKAA.V | HexNAc(2)Hex(1) 100.064 | No | 1406.964 | 4 | 5624.8341 | 5624.8569 | -4.1 | Non | 832.16 | 832.16 | 76.86 | Nsco_20191108_BC_ZIC_HILIC_H111_thermolysin_B1.45794.45794.4 | 86.1307 | Thermolysin | B1 | 83 |
| BCAL0349 (typ | >tr\|A0A2N9D2R1\|A0A2N9D2R1_9BURK Outer membrane protein OS=Burkholderia cenocepacia OX=95486 GN=F01_70026 PE=3 SV=1 | A.TVTPVGNGTVATTALPQANPNAGGASGTVVHGT[+568.212]AGTLTPPPANAAPGQVVVGGKVPDEATKA.A | HexNAc(2)Hex(1) | No | 1571.046 | 4 | 6281.1623 | 6280.1743 | -2.5 | Non | 717.57 | 717.57 | 0 | Nsco_20191108_BC_ZIC_HILIC_H111_thermolysin_B1.52394.52394.4 | 97.6529 | Thermolysin | B1 | 84 |

| | | | | | | | | | | | | | | | | | | |
|---|---|---|---|---|---|---|---|---|---|---|---|---|---|---|---|---|---|---|
| BCAL0349 (typ | >tr\|A0A2N9D2R1\|A0A2N9D2R1_9BUR K Outer membrane protein OS=Burkholderia cenocepacia OX=95486 GN=F01_70026 PE=3 SV=1 | T.VTPVGNGTVATTALPQANPNAGGASGTVVHGTAGTLT[+568.212][+100.064]PPPANAAPGQVVVGGKVPDEATKAAVLQK.L | HexNAc(2)Hex(1) 100.064 | No | 1365.1086 | 5 | 6821.5141 | 6818.5338 | -4.4 | Non | 905.81 | 176.65 | 0 | Nsco_20191108_BC_ZIC_HILIC_H111_thermolysin_B1.50091.50091.5 | 93.6016 | Thermolysin | B1 | 85 |
| BCAL0349 (typ | >tr\|A0A2N9D2R1\|A0A2N9D2R1_9BUR K Outer membrane protein OS=Burkholderia cenocepacia OX=95486 GN=F01_70026 PE=3 SV=1 | T.VTPVGNGTVATTALPQANPNAGGAS[+568.212]GTVVHGTAGTLTPPPANAAPGQVVVGGKVPDEATKA.A | HexNAc(2)Hex(1) | No | 1545.5436 | 4 | 6179.1525 | 6179.1266 | 4.2 | Non | 947.6 | 947.6 | 0 | Nsco_20191108_BC_ZIC_HILIC_H111_thermolysin_B1.48566.48566.4 | 91.0348 | Thermolysin | B1 | 86 |
| BCAL0349 (typ | >tr\|A0A2N9D2R1\|A0A2N9D2R1_9BUR K Outer membrane protein OS=Burkholderia cenocepacia OX=95486 GN=F01_70026 PE=3 SV=1 | T.TALPQANPNAGGAS[+568.212]GTVVHGTAGTLTPPPANAAPGQVVVGGKVPDEATKA.A | HexNAc(2)Hex(1) | Yes | 1297.1617 | 4 | 5185.6251 | 5182.6026 | 2.4 | Non | 851.32 | 245.93 | 84.48 | Nsco_20191108_BC_ZIC_HILIC_H111_thermolysin_B1.42189.42189.4 | 80.0561 | Thermolysin | B1 | 87 |
| BCAL0349 (typ | >tr\|A0A2N9D2R1\|A0A2N9D2R1_9BUR K Outer membrane protein OS=Burkholderia cenocepacia OX=95486 GN=F01_70026 PE=3 SV=1 | T.VATTALPQANPNAGGAS[+568.212]GTVVHGTAGTLTPPPANAAPGQVVVGGKVPDEATKA.A | HexNAc(2)Hex(1) | Yes | 1364.1989 | 4 | 5453.7736 | 5453.7558 | 3.3 | Non | 1069.56 | 1032.4 | 116.78 | Nsco_20191108_BC_ZIC_HILIC_H111_thermolysin_B1.43972.43972.4 | 83.1087 | Thermolysin | B1 | 88 |
| BCAL0349 (typ | >tr\|A0A2N9D2R1\|A0A2N9D2R1_9BUR K Outer membrane protein OS=Burkholderia cenocepacia OX=95486 GN=F01_70026 PE=3 SV=1 | T.VATTALPQANPNAGGAS[+568.212]GTVVHGTAGTLTPPPAN.A | HexNAc(2)Hex(1) | Yes | 1227.2776 | 3 | 3679.8184 | 3678.7981 | 4.6 | Non | 367.42 | 345.16 | 47.16 | Nsco_20191108_BC_ZIC_HILIC_H111_thermolysin_B2.39792.39792.3 | 78.7213 | Thermolysin | B2 | 89 |
| BCAL0349 (typ | >tr\|A0A2N9D2R1\|A0A2N9D2R1_9BUR K Outer membrane protein OS=Burkholderia cenocepacia OX=95486 GN=F01_70026 PE=3 SV=1 | A.LPQANPNAGGASGTVVHGT[+568.212][+100.064]AGTLTPPPANAAPGQVVVGGKVPDEATKA.A | HexNAc(2)Hex(1) 100.064 | No | 1278.6484 | 4 | 5111.5719 | 5110.5818 | -2.6 | Non | 335.82 | 323.31 | 0 | Nsco_20191108_BC_ZIC_HILIC_H111_thermolysin_B2.40376.40376.4 | 79.8677 | Thermolysin | B2 | 90 |
| BCAL0349 (typ | >tr\|A0A2N9D2R1\|A0A2N9D2R1_9BUR K Outer membrane protein OS=Burkholderia cenocepacia OX=95486 GN=F01_70026 PE=3 SV=1 | T.VTPVGNGTVATTALPQANPNAGGAS[+568.212]GTVVHGTAGTLTPPPAN.A | HexNAc(2)Hex(1) | No | 1469.0619 | 3 | 4405.1711 | 4404.1689 | -0.3 | Non | 299.6 | 138.55 | 2.4 | Nsco_20191108_BC_ZIC_HILIC_H111_thermolysin_B2.46162.46162.3 | 91.7994 | Thermolysin | B2 | 91 |
| BCAL0349 (typ | >tr\|A0A2N9D2R1\|A0A2N9D2R1_9BUR K Outer membrane protein OS=Burkholderia cenocepacia OX=95486 GN=F01_70026 PE=3 SV=1 | T.ALPQANPNAGGASGTVVHGTAGT[+568.212]LTPPPANAAPGQVVVGGK.V | HexNAc(2)Hex(1) | No | 1424.0583 | 3 | 4270.1603 | 4270.1474 | 3 | NRagged | 598.25 | 598.25 | 0 | Nsco_20191108_BC_ZIC_HILIC_H111_typsin_B1.38428.38428.3 | 77.2536 | Trypsin | B1 | 92 |
| BCAL0349 (typ | >tr\|A0A2N9D2R1\|A0A2N9D2R1_9BUR K Outer membrane protein OS=Burkholderia cenocepacia OX=95486 GN=F01_70026 PE=3 SV=1 | V.GNGTVATTALPQANPNAGGAS[+568.212]GTVVHGTAGTLTPPPANAAPGQVVVGGKVPDEATK.A | HexNAc(2)Hex(1) | No | 1428.7255 | 4 | 5711.88 | 5711.8523 | 4.9 | NRagged | 1116.29 | 1116.29 | 0 | Nsco_20191108_BC_ZIC_HILIC_H111_typsin_B1.42452.42452.4 | 83.9668 | Trypsin | B1 | 93 |
| BCAL0349 (typ | >tr\|A0A2N9D2R1\|A0A2N9D2R1_9BUR K Outer membrane protein OS=Burkholderia cenocepacia OX=95486 GN=F01_70026 PE=3 SV=1 | T.ALPQANPNAGGAS[+568.212]GTVVHGTAGTLTPPPANAAPGQVVVGGKVPDEATK.A | HexNAc(2)Hex(1) | Yes | 1253.3899 | 4 | 5010.5378 | 5010.5178 | 4 | NRagged | 974.71 | 974.71 | 93.18 | Nsco_20191108_BC_ZIC_HILIC_H111_typsin_B2.39145.39145.4 | 77.1581 | Trypsin | B2 | 94 |
| BCAL0349 (typ | >tr\|A0A2N9D2R1\|A0A2N9D2R1_9BUR K Outer membrane protein OS=Burkholderia cenocepacia OX=95486 GN=F01_70026 PE=3 SV=1 | A.QNTGATVTPVGNGTVATTAL[illegible]QANPNAGGASGTVVHGTAGTL[illegible][+568.212]PPPANAAPGQVVVGGKVPDEATK.A | HexNAc(2)Hex(1) | No | 1670.8411 | 4 | 6680.3428 | 6680.345 | -0.3 | NRagged | 615.65 | 198.73 | 0 | Nsco_20191108_BC_ZIC_HILIC_H111_typsin_B2.55798.55798.4 | 106.9982 | Trypsin | B2 | 95 |
| BCAL0349 (typ | >tr\|A0A2N9D2R1\|A0A2N9D2R1_9BUR K Outer membrane protein OS=Burkholderia cenocepacia OX=95486 GN=F01_70026 PE=3 SV=1 | V.TPVGNGTVATTALPQANPNAGGAS[+568.212][+100.064]GTVVHGTAGTLT[+568.212][+100.064]PPPANAAPGQVVVGGKVPDEATK.A | HexNAc(2)Hex(1) 100.064,HexNAc(2)Hex(1) 100.064 | No | 1695.8493 | 4 | 6780.3755 | 6777.3607 | 0.7 | NRagged | 458.14 | 63.6 | 0 | Nsco_20191108_BC_ZIC_HILIC_H111_typsin_B2.56710.56710.4 | 108.7073 | Trypsin | B2 | 96 |

| | | | | | | | | | | | | | | | | | | |
|---|---|---|---|---|---|---|---|---|---|---|---|---|---|---|---|---|---|---|
| BCAL0358 | >tr\|A0A142PJ25\|A0A142PJ25_9BURK Aminopeptidase N OS=Burkholderia cenocepacia OX=95486 GN=pepN_1 PE=4 SV=1 | L.SSAGAPHTGSSPT[+568.212]VAAPPSAS.N | HexNAc(2)Hex(1) | No | 1203.049 | 2 | 2405.0906 | 2405.0838 | 2.8 | CRagged | 388.69 | 354.52 | 0 | Nsco_20191108_BC_ZIC_HILIC_H111_pepsin_B2.16836.16836.2 | 37.0932 | Pepsin | B2 | 97 |
| BCAL0358 | >tr\|A0A142PJ25\|A0A142PJ25_9BURK Aminopeptidase N OS=Burkholderia cenocepacia OX=95486 GN=pepN_1 PE=4 SV=1 | L.SSAGAPHTGS[+568.212]SPTVAAPPSASN.V | HexNAc(2)Hex(1) | No | 1260.0722 | 2 | 2519.1371 | 2519.1267 | 4.1 | CRagged | 635.65 | 631.94 | 0 | Nsco_20191108_BC_ZIC_HILIC_H111_pepsin_B2.16231.16231.2 | 36.2053 | Pepsin | B2 | 98 |
| BCAL0426 | >tr\|A0A1V6KWZ1\|A0A1V6KWZ1_9BURK Membrane protein insertase YidC OS=Burkholderia cenocepacia OX=95486 GN=yidC PE=3 SV=1 | T.HTAPAAAGGASGTGATTT[+568.212]AGDVPAAAAGAAPSTTAPAAQ.A | HexNAc(2)Hex(1) | No | 1272.2641 | 3 | 3814.7777 | 3814.7737 | 1.1 | Non | 172.9 | 172.9 | 1.2 | Nsco_20191108_BC_ZIC_HILIC_H111_pepsin_B2.31114.31114.3 | 64.1861 | Pepsin | B2 | 99 |
| BCAL0426 | >tr\|A0A1V6KWZ1\|A0A1V6KWZ1_9BURK Membrane protein insertase YidC OS=Burkholderia cenocepacia OX=95486 GN=yidC PE=3 SV=1 | S.ATHTAPAAAGGASGT[+568.212]GATTTAGDVPAA.A | HexNAc(2)Hex(1) | No | 1411.1507 | 2 | 2821.2941 | 2821.2858 | 2.9 | Non | 732.51 | 500.51 | 0 | Nsco_20191108_BC_ZIC_HILIC_H111_thermolysin_B2.19464.19464.2 | 42.59 | Thermolysin | B2 | 100 |
| BCAL0426 | >tr\|A0A1V6KWZ1\|A0A1V6KWZ1_9BURK Membrane protein insertase YidC OS=Burkholderia cenocepacia OX=95486 GN=yidC PE=3 SV=1 | S.ATHTAPAAAGGASGTGATTT[568.212]AGDVPA.A | HexNAc(2)Hex(1) | No | 1375.6329 | 2 | 2750.2584 | 2750.2486 | 3.6 | Non | 685.64 | 508.9 | 1.67 | Nsco_20191108_BC_ZIC_HILIC_H111_thermolysin_B3.17670.17670.2 | 39.8571 | Thermolysin | B3 | 101 |
| BCAL0426 | >tr\|A0A1V6KWZ1\|A0A1V6KWZ1_9BURK Membrane protein insertase YidC OS=Burkholderia cenocepacia OX=95486 GN=yidC PE=3 SV=1 | S.ATHTAPAAAGGASGTGATTT[568.212]AGDVPAAAAGAAPSTTAPAAQAQ.L | HexNAc(2)Hex(1) | No | 1396.6617 | 3 | 4187.9707 | 4185.9542 | 2.3 | Non | 631.07 | 574.7 | 1.2 | Nsco_20191108_BC_ZIC_HILIC_H111_thermolysin_B3.32236.32236.3 | 64.8714 | Thermolysin | B3 | 102 |
| BCAL0426 | >tr\|A0A1V6KWZ1\|A0A1V6KWZ1_9BURK Membrane protein insertase YidC OS=Burkholderia cenocepacia OX=95486 GN=yidC PE=3 SV=1 | R.DHGRPSMFFPSATHTAPAAAGGAS[+568.212]GTGATTTAGDVPAAAAGAAPSTTAPAAQAQLVK.F | HexNAc(2)Hex(1) | Yes | 1157.7617 | 5 | 5784.7795 | 5784.757 | 3.9 | Specific | 1146.79 | 1145.41 | 48.09 | Nsco_20191108_BC_ZIC_HILIC_H111_typsin_B2.43258.43258.5 | 84.3869 | Trypsin | B2 | 103 |
| BCAL0479 | >tr\|A0A2N9CSF4\|A0A2N9CSF4_9BURK Peptidoglycan D,D-transpeptidase MrdA OS=Burkholderia cenocepacia OX=95486 GN=mrdA PE=3 SV=1 | A.VAAAASATEPVSAPVVGDT[+568.212]S[+568.212]KPAT.I | HexNAc(2)Hex(1), HexNAc(2)Hex(1) | No | 1667.287 | 2 | 3333.5667 | 3333.5578 | 2.7 | Non | 660.44 | 357.82 | 9.42 | Nsco_20191108_BC_ZIC_HILIC_H111_thermolysin_B1.30870.30870.2 | 60.3488 | Thermolysin | B1 | 104 |
| BCAL0479 | >tr\|A0A2N9CSF4\|A0A2N9CSF4_9BURK Peptidoglycan D,D-transpeptidase MrdA OS=Burkholderia cenocepacia OX=95486 GN=mrdA PE=3 SV=1 | R.RPAS[+568.212]DAQPVVATPR.D | HexNAc(2)Hex(1) | Yes | 1017.0083 | 2 | 2033.0094 | 2033.0033 | 3 | Specific | 632.91 | 349.99 | 39.47 | Nsco_20191108_BC_ZIC_HILIC_H111_typsin_B1.9404.9404.2 | 27.4075 | Trypsin | B1 | 105 |
| BCAL0481 | >tr\|A0A2N9CS00\|A0A2N9CS00_9BURK Rod shape-determining protein MreC OS=Burkholderia cenocepacia OX=95486 GN=DFS07_10564 PE=4 SV=1 | K.PAAAAPPAGANPAPAPAAPAKPAAAPAKPAAGQS[+568.212][+100.064]GAQR.- | HexNAc(2)Hex(1) 100.064 | No | 1326.3508 | 3 | 3977.0379 | 3977.0366 | 0.3 | Specific | 474.1 | 474.1 | 474.1 | Nsco_20191108_BC_ZIC_HILIC_H111_typsin_B2.16250.16250.3 | 36.756 | Trypsin | B2 | 106 |
| BCAL0525 | >tr\|A0A119PFA4\|A0A119PFA4_9BURK Flagellar M-ring protein OS=Burkholderia cenocepacia OX=95486 GN=fliF PE=3 SV=1 | L.SNTPPQPAS[+568.212]APIVAGNGQNAPQTTPVSD.R | HexNAc(2)Hex(1) | Yes | 1095.5194 | 3 | 3284.5438 | 3284.5288 | 4.5 | CRagged | 539.46 | 429.01 | 14.55 | Nsco_20191108_BC_ZIC_HILIC_H111_pepsin_B1.36617.36617.3 | 78.3735 | Pepsin | B1 | 107 |
| BCAL0525 | >tr\|A0A119PFA4\|A0A119PFA4_9BURK Flagellar M-ring protein OS=Burkholderia cenocepacia OX=95486 GN=fliF PE=3 SV=1 | A.LSNTPPQPASAPIVAGNGQNAPQTT[+568.212]PVSDRKDQTTNYE.L | HexNAc(2)Hex(1) | No | 1512.0564 | 3 | 4534.1546 | 4533.1387 | 2.8 | Non | 753.41 | 704.3 | 0 | Nsco_20191108_BC_ZIC_HILIC_H111_thermolysin_B2.33278.33278.3 | 66.396 | Thermolysin | B2 | 108 |
| BCAL0525 | >tr\|A0A119PFA4\|A0A119PFA4_9BURK Flagellar M-ring protein OS=Burkholderia cenocepacia OX=95486 GN=fliF PE=3 SV=1 | R.SQQTSSATELAQGGASGVPGALS[+568.212]NTPPQPAS[+568.212][+100.064]APIVAGNGQNAPQTTPVSDRK.D | HexNAc(2)Hex(1), HexNAc(2)Hex(1) 100.064 | No | 1559.7397 | 4 | 6235.9372 | 6234.9659 | -5.1 | Specific | 695.91 | 695.91 | 0 | Nsco_20191108_BC_ZIC_HILIC_H111_typsin_B3.41487.41487.4 | 86.0175 | Trypsin | B3 | 109 |

| | | | | | | | | | | | | | | | | | | |
|---|---|---|---|---|---|---|---|---|---|---|---|---|---|---|---|---|---|---|
| BCAL0544 | >tr\|A0A2N9CS86\|A0A2N9CS86_9BURK Dipeptide transport system substrate-binding protein OS=Burkholderia cenocepacia OX=95486 GN=dppA PE=4 SV=1 | S.VYQGAGQAASAPMPPT[+568.212]QWSYDKN.L | HexNAc(2)Hex(1) | No | 1518.1818 | 2 | 3035.3563 | 3035.3463 | 3.3 | Non | 479.1 | 418.9 | 6 | Nsco_20191108_BC_ZIC_HILIC_H111_thermolysin_B2.39343.39343.2 | 77.7918 | Thermolysin | B2 | 110 |
| BCAL0544 | >tr\|A0A2N9CS86\|A0A2N9CS86_9BURK Dipeptide transport system substrate-binding protein OS=Burkholderia cenocepacia OX=95486 GN=dppA PE=4 SV=1 | K.AILESVYQGAGQAAS[+568.212]APMPPTQWSYDK.N | HexNAc(2)Hex(1) | No | 1718.3034 | 2 | 3435.5995 | 3434.5832 | 3.8 | Specific | 782.03 | 749.56 | 8.4 | Nsco_20191108_BC_ZIC_HILIC_H111_typsin_B1.57958.57958.2 | 109.4793 | Trypsin | B1 | 111 |
| BCAL0544 | >tr\|A0A2N9CS86\|A0A2N9CS86_9BURK Dipeptide transport system substrate-binding protein OS=Burkholderia cenocepacia OX=95486 GN=dppA PE=4 SV=1 | K.KAILESVYQGAGQAAS[+568.212]APMPPTQWSYDK.N | HexNAc(2)Hex(1) | No | 1188.2358 | 3 | 3562.6928 | 3562.6782 | 4.1 | Specific | 754.77 | 754.77 | 9.6 | Nsco_20191108_BC_ZIC_HILIC_H111_typsin_B2.50150.50150.3 | 96.4702 | Trypsin | B2 | 112 |
| BCAL0678 | >tr\|A0A142PL18\|A0A142PL18_9BURK Cell division protein OS=Burkholderia cenocepacia OX=95486 GN=A8E72_03155 PE=4 SV=1 | S.KVAPPPADNGAS[+568.212]QPQQFDPNRALQ.G | HexNAc(2)Hex(1) | Yes | 1038.8367 | 3 | 3114.4955 | 3114.4862 | 3 | Non | 609.94 | 549.65 | 549.65 | Nsco_20191108_BC_ZIC_HILIC_H111_pepsin_B1.27083.27083.3 | 57.3842 | Pepsin | B1 | 113 |
| BCAL0678 | >tr\|A0A142PL18\|A0A142PL18_9BURK Cell division protein OS=Burkholderia cenocepacia OX=95486 GN=A8E72_03155 PE=4 SV=1 | S.KVAPPPADNGAS[+568.212]QPQQFDPNRAL.Q | HexNAc(2)Hex(1) | Yes | 996.151 | 3 | 2986.4384 | 2986.4276 | 3.6 | NRagged | 652.56 | 631.5 | 631.5 | Nsco_20191108_BC_ZIC_HILIC_H111_pepsin_B1.27669.27669.3 | 58.5354 | Pepsin | B1 | 114 |
| BCAL0678 | >tr\|A0A142PL18\|A0A142PL18_9BURK Cell division protein OS=Burkholderia cenocepacia OX=95486 GN=A8E72_03155 PE=4 SV=1 | F.VSKVAPPPADNGASQPQQFDPNRALQGKT[+568.212]PGQPVPQAAQPAPPNTAPGQA.A | HexNAc(2)Hex(1) | No | 1391.4462 | 4 | 5562.7629 | 5561.7419 | 3.2 | CRagged | 263.81 | 247.51 | 3.6 | Nsco_20191108_BC_ZIC_HILIC_H111_pepsin_B1.35328.35328.4 | 75.1743 | Pepsin | B1 | 115 |
| BCAL0678 | >tr\|A0A142PL18\|A0A142PL18_9BURK Cell division protein OS=Burkholderia cenocepacia OX=95486 GN=A8E72_03155 PE=4 SV=1 | S.KVAPPPADNGAS[+568.212]QPQQFDPNRALQGKTPGQPVPQAAQPAPPNTAPGQA.A | HexNAc(2)Hex(1) | Yes | 1344.6699 | 4 | 5375.6579 | 5375.6415 | 3.1 | Non | 753.23 | 753.23 | 297.94 | Nsco_20191108_BC_ZIC_HILIC_H111_pepsin_B1.34700.34700.4 | 73.7034 | Pepsin | B1 | 116 |
| BCAL0678 | >tr\|A0A142PL18\|A0A142PL18_9BURK Cell division protein OS=Burkholderia cenocepacia OX=95486 GN=A8E72_03155 PE=4 SV=1 | S.KVAPPPADNGASQPQQFDPNRALQGKT[+568.212][+100.064]PGQPVPQAAQPAPPNTAPGQAANQ.T | HexNAc(2)Hex(1) 100.064 | No | 1447.9617 | 4 | 5788.8249 | 5788.8441 | -3.3 | Non | 616.03 | 481.95 | 9.3 | Nsco_20191108_BC_ZIC_HILIC_H111_pepsin_B1.35407.35407.4 | 75.3781 | Pepsin | B1 | 117 |
| BCAL0678 | >tr\|A0A142PL18\|A0A142PL18_9BURK Cell division protein OS=Burkholderia cenocepacia OX=95486 GN=A8E72_03155 PE=4 SV=1 | S.KVAPPPADNGASQPQQFDPNRALQGKT[+568.212]PGQPVPQAAQPAPPNTAP.G | HexNAc(2)Hex(1) | No | 1281.3821 | 4 | 5122.5065 | 5119.5243 | -5.4 | Non | 253.73 | 253.73 | 10.07 | Nsco_20191108_BC_ZIC_HILIC_H111_pepsin_B2.35513.35513.4 | 74.1341 | Pepsin | B2 | 118 |
| BCAL0678 | >tr\|A0A142PL18\|A0A142PL18_9BURK Cell division protein OS=Burkholderia cenocepacia OX=95486 GN=A8E72_03155 PE=4 SV=1 | S.KVAPPPADNGAS[+568.212]QPQQFDPNRA.L | HexNAc(2)Hex(1) | Yes | 958.4571 | 3 | 2873.3566 | 2873.3436 | 4.6 | Non | 589.42 | 589.42 | 589.42 | Nsco_20191108_BC_ZIC_HILIC_H111_pepsin_B2.18898.18898.3 | 40.4584 | Pepsin | B2 | 119 |
| BCAL0678 | >tr\|A0A142PL18\|A0A142PL18_9BURK Cell division protein OS=Burkholderia cenocepacia OX=95486 GN=A8E72_03155 PE=4 SV=1 | S.KVAPPPADNGAS[+568.212]QPQQFDPNRALQG.K | HexNAc(2)Hex(1) | Yes | 1057.8449 | 3 | 3171.5201 | 3171.5077 | 3.9 | Non | 509.23 | 509.23 | 509.23 | Nsco_20191108_BC_ZIC_HILIC_H111_pepsin_B2.28821.28821.3 | 59.3228 | Pepsin | B2 | 120 |
| BCAL0678 | >tr\|A0A142PL18\|A0A142PL18_9BURK Cell division protein OS=Burkholderia cenocepacia OX=95486 GN=A8E72_03155 PE=4 SV=1 | S.KVAPPPADNGAS[+568.212]QPQQFDPNRALQGKTPGQPVPQA.A | HexNAc(2)Hex(1) | Yes | 1044.523 | 4 | 4175.0701 | 4175.0527 | 4.2 | Non | 776.89 | 776.89 | 543.75 | Nsco_20191108_BC_ZIC_HILIC_H111_pepsin_B2.31728.31728.4 | 65.5889 | Pepsin | B2 | 121 |

| | | | | | | | | | | | | | | | | | |
|---|---|---|---|---|---|---|---|---|---|---|---|---|---|---|---|---|---|
| BCAL0678 | >tr\|A0A142PL18\|A0A142PL18_9BURK Cell division protein OS=Burkholderia cenocepacia OX=95486 GN=A8E72_03155 PE=4 SV=1 | S.KVAPPPADNGAS[+568.212][+100.064]QPQQFDPNRALQGKTPGQPVPQAAQ.P | HexNAc(2)Hex(1) 100.064 | No | 1119.5551 | 4 | 4475.1986 | 4474.2124 | -3.9 | Non | 366.59 | 282.54 | 5.72 | Nsco_20191108_BC_ZIC_HILIC_H111_pepsin_B2.32825.32825.2 | 67.8781 | Pepsin | B2 | 122 |
| BCAL0678 | >tr\|A0A142PL18\|A0A142PL18_9BURK Cell division protein OS=Burkholderia cenocepacia OX=95486 GN=A8E72_03155 PE=4 SV=1 | K.VAPPPADNGAS[+568.212][+100.064]QPQQFDPNRALQGKTPGQPVPQAAQPAPPNTAPGQA.A | HexNAc(2)Hex(1) 100.064 | Yes | 1337.6516 | 4 | 5347.5846 | 5347.6105 | -4.8 | Non | 560.62 | 560.62 | 232.52 | Nsco_20191108_BC_ZIC_HILIC_H111_pepsin_B2.39476.39476.4 | 83.9714 | Pepsin | B2 | 123 |
| BCAL0678 | >tr\|A0A142PL18\|A0A142PL18_9BURK Cell division protein OS=Burkholderia cenocepacia OX=95486 GN=A8E72_03155 PE=4 SV=1 | S.KVAPPPADNGAS[+568.212]QPQQFDPNRALQGKTPGQPVPQAAQPAPPNTAPGQAAN.Q | HexNAc(2)Hex(1) | Yes | 1390.9406 | 4 | 5560.7404 | 5560.7215 | 3.4 | Non | 753.49 | 753.49 | 363.84 | Nsco_20191108_BC_ZIC_HILIC_H111_pepsin_B2.34965.34965.4 | 72.7835 | Pepsin | B2 | 124 |
| BCAL0678 | >tr\|A0A142PL18\|A0A142PL18_9BURK Cell division protein OS=Burkholderia cenocepacia OX=95486 GN=A8E72_03155 PE=4 SV=1 | S.KVAPPPADNGAS[+568.212]QPQQFDPNRALQGKTPGQPVPQAAQPAPPNTAPGQ.A | HexNAc(2)Hex(1) | Yes | 1326.9128 | 4 | 5304.6296 | 5304.6044 | 4.7 | Non | 726.93 | 726.93 | 266.35 | Nsco_20191108_BC_ZIC_HILIC_H111_pepsin_B3.34094.34094.4 | 72.7935 | Pepsin | B3 | 125 |
| BCAL0678 | >tr\|A0A142PL18\|A0A142PL18_9BURK Cell division protein OS=Burkholderia cenocepacia OX=95486 GN=A8E72_03155 PE=4 SV=1 | S.KVAPPPADNGAS[+568.212]QPQQFDPNRALQGKTPGQPVPQAAQPAPPNTA.P | HexNAc(2)Hex(1) | Yes | 1256.3777 | 4 | 5022.4889 | 5022.4716 | 3.5 | Non | 602.85 | 602.85 | 265.96 | Nsco_20191108_BC_ZIC_HILIC_H111_pepsin_B3.33636.33636.4 | 71.5919 | Pepsin | B3 | 126 |
| BCAL0678 | >tr\|A0A142PL18\|A0A142PL18_9BURK Cell division protein OS=Burkholderia cenocepacia OX=95486 GN=A8E72_03155 PE=4 SV=1 | S.KVAPPPADNGAS[+568.212]QPQQFDPNRALQGKTPGQPVPQAAQPAPPNTAPGQAA.N | HexNAc(2)Hex(1) | Yes | 1362.4308 | 4 | 5446.7013 | 5446.6786 | 4.2 | Non | 780.89 | 780.89 | 370.32 | Nsco_20191108_BC_ZIC_HILIC_H111_pepsin_B3.34360.34360.4 | 73.5177 | Pepsin | B3 | 127 |
| BCAL0678 | >tr\|A0A142PL18\|A0A142PL18_9BURK Cell division protein OS=Burkholderia cenocepacia OX=95486 GN=A8E72_03155 PE=4 SV=1 | F.VSKVAPPPADNGAS[+568.212][+100.064]QPQQFDPNRALQGKTPGQPVPQAAQPAPPNTAPGQAA.N | HexNAc(2)Hex(1) 100.064 | No | 1434.4614 | 4 | 5734.8239 | 5732.843 | -4.5 | CRagged | 477.51 | 477.51 | 229.78 | Nsco_20191108_BC_ZIC_HILIC_H111_pepsin_B3.35710.35710.4 | 77.0909 | Pepsin | B3 | 128 |
| BCAL0678 | >tr\|A0A142PL18\|A0A142PL18_9BURK Cell division protein OS=Burkholderia cenocepacia OX=95486 GN=A8E72_03155 PE=4 SV=1 | A.QKQQQQQQAANTPKPTSSAT[+568.212]AAA.A | HexNAc(2)Hex(1) | No | 1476.7124 | 2 | 2952.4176 | 2951.4076 | 2.3 | Non | 498.87 | 408.91 | 2.47 | Nsco_20191108_BC_ZIC_HILIC_H111_pepsin_B3.7831.7831.2 | 22.0979 | Pepsin | B3 | 129 |
| BCAL0678 | >tr\|A0A142PL18\|A0A142PL18_9BURK Cell division protein OS=Burkholderia cenocepacia OX=95486 GN=A8E72_03155 PE=4 SV=1 | A.AAQKQQQQQQAANT[+568.212]PKPTSSATAAA.A | HexNAc(2)Hex(1) | No | 1032.1702 | 3 | 3094.496 | 3093.4818 | 3.5 | Non | 378.39 | 378.39 | 14.7 | Nsco_20191108_BC_ZIC_HILIC_H111_pepsin_B3.8210.8210.3 | 22.633 | Pepsin | B3 | 130 |
| BCAL0678 | >tr\|A0A142PL18\|A0A142PL18_9BURK Cell division protein OS=Burkholderia cenocepacia OX=95486 GN=A8E72_03155 PE=4 SV=1 | R.FAAQKQAQQAAAQKQQQQQQAANTPKPT[+568.212][+100.064]SSA.T | HexNAc(2)Hex(1) 100.064 | No | 1318.3182 | 3 | 3952.9402 | 3950.933 | 0.1 | Non | 170.22 | 145.54 | 0.88 | Nsco_20191108_BC_ZIC_HILIC_H111_thermolysin_B1.12441.12441.3 | 30.8095 | Thermolysin | B1 | 131 |
| BCAL0678 | >tr\|A0A142PL18\|A0A142PL18_9BURK Cell division protein OS=Burkholderia cenocepacia OX=95486 GN=A8E72_03155 PE=4 SV=1 | K.VAPPPADNGAS[+568.212]QPQQFDPNRA.L | HexNAc(2)Hex(1) | No | 1373.1332 | 2 | 2745.2591 | 2745.2486 | 3.8 | Non | 317.04 | 254.66 | 254.66 | Nsco_20191108_BC_ZIC_HILIC_H111_thermolysin_B1.25881.25881.2 | 51.6772 | Thermolysin | B1 | 132 |
| BCAL0678 | >tr\|A0A142PL18\|A0A142PL18_9BURK Cell division protein OS=Burkholderia cenocepacia OX=95486 GN=A8E72_03155 PE=4 SV=1 | Q.AAAQKQQQQQQAANT[+568.212]PKPTSSATAA.A | HexNAc(2)Hex(1) | No | 1031.8346 | 3 | 3093.4893 | 3093.4818 | 2.4 | Non | 385.59 | 283.64 | 3.62 | Nsco_20191108_BC_ZIC_HILIC_H111_thermolysin_B1.6531.6531.3 | 21.9265 | Thermolysin | B1 | 133 |
| BCAL0678 | >tr\|A0A142PL18\|A0A142PL18_9BURK Cell division protein OS=Burkholderia cenocepacia OX=95486 GN=A8E72_03155 PE=4 SV=1 | Q.AAAQKQQQQQQAANT[+568.212]PKPTSSATAAA.A | HexNAc(2)Hex(1) | No | 1055.5145 | 3 | 3164.529 | 3164.519 | 3.2 | Non | 458.21 | 366.96 | 1.56 | Nsco_20191108_BC_ZIC_HILIC_H111_thermolysin_B2.7398.7398.3 | 23.3115 | Thermolysin | B2 | 134 |

| | | | | | | | | | | | | | | | | | | |
|---|---|---|---|---|---|---|---|---|---|---|---|---|---|---|---|---|---|---|
| BCAL0678 | >tr\|A0A142PL18\|A0A142PL18_9BURK Cell division protein OS=Burkholderia cenocepacia OX=95486 GN=A8E72_03155 PE=4 SV=1 | K.QQQQQQAANTPKPTS[+568.212]SATAAAAAKPPTANDANTGYFLQVGAYK.T | HexNAc(2)Hex(1) | No | 1244.1078 | 4 | 4973.4093 | 4973.3923 | 3.4 | Specific | 1270 | 1207.06 | 17.78 | Nsco_20191108_BC_ZIC_HILIC_H111_typsin_B1.35724.35724.4 | 72.5132 | Trypsin | B1 | 135 |
| BCAL0678 | >tr\|A0A142PL18\|A0A142PL18_9BURK Cell division protein OS=Burkholderia cenocepacia OX=95486 GN=A8E72_03155 PE=4 SV=1 | Q.QAANTPKPTS[+568.212]SATAAAAAKPPT[+568.212][+100.064]ANDANTGYFLQVGAYKTEGDAEQQR.A | HexNAc(2)Hex(1), HexNAc(2)Hex(1) 100.064 | No | 1204.7749 | 5 | 6019.8454 | 6015.8116 | 3.4 | NRagged | 738.44 | 738.44 | 2.58 | Nsco_20191108_BC_ZIC_HILIC_H111_typsin_B1.34157.34157.5 | 69.8041 | Trypsin | B1 | 136 |
| BCAL0678 | >tr\|A0A142PL18\|A0A142PL18_9BURK Cell division protein OS=Burkholderia cenocepacia OX=95486 GN=A8E72_03155 PE=4 SV=1 | K.QQQQQQAANTPKPTSS[+568.212][+100.064]ATAAAAAKPPTANDANTGYFLQVGAYKTEGDAEQQR.A | HexNAc(2)Hex(1) 100.064 | No | 1218.3798 | 5 | 6087.8697 | 6087.893 | -3.8 | Specific | 1260.89 | 1260.89 | 0 | Nsco_20191108_BC_ZIC_HILIC_H111_typsin_B2.35579.35579.5 | 70.8298 | Trypsin | B2 | 137 |
| BCAL0678 | >tr\|A0A142PL18\|A0A142PL18_9BURK Cell division protein OS=Burkholderia cenocepacia OX=95486 GN=A8E72_03155 PE=4 SV=1 | S.KVAPPPADNGAS[+568.212]QPQQFDPNR.A | HexNAc(2)Hex(1) | Yes | 934.7765 | 3 | 2802.3148 | 2802.3064 | 3 | NRagged | 428.78 | 428.78 | 428.78 | Nsco_20191108_BC_ZIC_HILIC_H111_typsin_B2.16578.16578.3 | 37.2989 | Trypsin | B2 | 138 |
| BCAL0678 | >tr\|A0A142PL18\|A0A142PL18_9BURK Cell division protein OS=Burkholderia cenocepacia OX=95486 GN=A8E72_03155 PE=4 SV=1 | K.VAPPPADNGAS[+568.212]QPQQFDPNRALQGK.T | HexNAc(2)Hex(1) | Yes | 1057.8446 | 3 | 3171.5193 | 3171.5077 | 3.7 | Specific | 671.21 | 671.21 | 671.21 | Nsco_20191108_BC_ZIC_HILIC_H111_typsin_B2.29596.29596.3 | 60.2503 | Trypsin | B2 | 139 |
| BCAL0678 | >tr\|A0A142PL18\|A0A142PL18_9BURK Cell division protein OS=Burkholderia cenocepacia OX=95486 GN=A8E72_03155 PE=4 SV=1 | Q.QQAANTPKPTSSATAAAAAKPPT[+568.212]ANDANT[+568.212][+100.064]GYFLQVGAYKTEGDAEQQR.A | HexNAc(2)Hex(1), HexNAc(2)Hex(1) 100.064 | No | 1229.9755 | 5 | 6145.8482 | 6143.8702 | -4.7 | NRagged | 523.48 | 523.48 | 0 | Nsco_20191108_BC_ZIC_HILIC_H111_typsin_B2.37796.37796.5 | 74.6971 | Trypsin | B2 | 140 |
| BCAL0678 | >tr\|A0A142PL18\|A0A142PL18_9BURK Cell division protein OS=Burkholderia cenocepacia OX=95486 GN=A8E72_03155 PE=4 SV=1 | Q.KQQQQQQAANTPKPTS[+568.212]SATAAAAAKPPTANDANTGYFLQVGAYKTEGDAEQQR.[illegible] | HexNAc(2)Hex(1) | Yes | 1020.6626 | 6 | 6118.9391 | 6115.9239 | 0.8 | NRagged | 1389.56 | 1103.5 | 15.81 | Nsco_20191108_BC_ZIC_HILIC_H111_typsin_B3.29736.29736.6 | 63.1584 | Trypsin | B3 | 141 |
| BCAL0678 | >tr\|A0A142PL18\|A0A142PL18_9BURK Cell division protein OS=Burkholderia cenocepacia OX=95486 GN=A8E72_03155 PE=4 SV=1 | K.VAPPPADNGAS[+568.212]QPQQFDPNR.A | HexNAc(2)Hex(1) | Yes | 1337.6127 | 2 | 2674.2182 | 2674.2115 | 2.5 | Specific | 557.94 | 510.14 | 510.14 | Nsco_20191108_BC_ZIC_HILIC_H111_typsin_B3.22139.22139.2 | 48.3419 | Trypsin | B3 | 142 |
| BCAL0678 | >tr\|A0A142PL18\|A0A142PL18_9BURK Cell division protein OS=Burkholderia cenocepacia OX=95486 GN=A8E72_03155 PE=4 SV=1 | Q.KQQQQQQAANTPKPTS[+568.212]SATAAAAAKPPTANDANTGYFLQVGAYK.T | HexNAc(2)Hex(1) | Yes | 1021.5052 | 5 | 5103.4971 | 5101.4873 | 0.6 | NRagged | 1343.42 | 1328.26 | 42.28 | Nsco_20191108_BC_ZIC_HILIC_H111_typsin_B3.30947.30947.5 | 65.4326 | Trypsin | B3 | 143 |
| BCAL0738 | >tr\|A0A142PKY4\|A0A142PKY4_9BURK C-terminal processing protease-3 OS=Burkholderia cenocepacia OX=95486 GN=A8E72_10915 PE=3 SV=1 | A.T[+568.212]ASKAS[+568.212][+100.064]GASAAKPAS[+568.212]APKPASAPK.- | HexNAc(2)Hex(1), HexNAc(2)Hex(1) 100.064,HexNAc(2)Hex(1) | No | 1319.6238 | 3 | 3956.8568 | 3956.8708 | -3.5 | Non | 156.06 | 156.06 | 1.2 | Nsco_20191108_BC_ZIC_HILIC_H111_thermolysin_B1.4774.4774.3 | 19.118 | Thermolysin | B1 | 144 |
| BCAL0749 | >tr\|A0A2N9D2V0\|A0A2N9D2V0_9BURK Cytochrome c oxidase subunit 2 OS=Burkholderia cenocepacia OX=95486 GN=ctaC_1 PE=3 SV=1 | Q.VADARNGKLPEDTAGAAT[+568.212][+100.064]AAAPAEAAS[+568.212][+100.064]A.P | HexNAc(2)Hex(1) 100.064,HexNAc(2)Hex(1) 100.064 | No | 1301.9514 | 3 | 3903.8395 | 3903.8207 | 4.8 | Non | 163.27 | 152.83 | 5.6 | Nsco_20191108_BC_ZIC_HILIC_H111_thermolysin_B2.33883.33883.3 | 67.38 | Thermolysin | B2 | 145 |
| BCAL0749 | >tr\|A0A2N9D2V0\|A0A2N9D2V0_9BURK Cytochrome c oxidase subunit 2 OS=Burkholderia cenocepacia OX=95486 GN=ctaC_1 PE=3 SV=1 | G.AATAAAPAEAASAPAQAASGAEQPAAAASAALS[+568.212][+100.064]T[+568.212][+100.064]IYFETGK.S | HexNAc(2)Hex(1) 100.064,HexNAc(2)Hex(1) 100.064 | No | 1691.1403 | 3 | 5071.4065 | 5069.3863 | 2.7 | NRagged | 567.51 | 545.2 | 1.2 | Nsco_20191108_BC_ZIC_HILIC_H111_typsin_B1.66797.66797.3 | 123.8368 | Trypsin | B1 | 146 |
| BCAL0749 | >tr\|A0A2N9D2V0\|A0A2N9D2V0_9BURK Cytochrome c oxidase subunit 2 OS=Burkholderia cenocepacia OX=95486 GN=ctaC_1 PE=3 SV=1 | A.GAATAAAPAEAASAPAQAAS[+568.212]GAEQPAAAAS[+568.212][+100.064]AALSTIYFETGK.S | HexNAc(2)Hex(1), HexNAc(2)Hex(1) 100.064 | No | 1676.4614 | 3 | 5027.3696 | 5026.3437 | 4.5 | NRagged | 433.97 | 433.97 | 0 | Nsco_20191108_BC_ZIC_HILIC_H111_typsin_B1.67138.67138.3 | 124.4222 | Trypsin | B1 | 147 |

| | | | | | | | | | | | | | | | | | | |
|---|---|---|---|---|---|---|---|---|---|---|---|---|---|---|---|---|---|---|
| BCAL0749 | >tr\|A0A2N9D2V0\|A0A2N9D2V0_9BURK Cytochrome c oxidase subunit 2 OS=Burkholderia cenocepacia OX=95486 GN=ctaC_1 PE=3 SV=1 | E.DTAGAATAAAPAEAASAPAQAAS[+568.212][+100.064]GAEQPAAAAS[+568.212]AALSTIYFETGK.S | HexNAc(2)Hex(1) 100.064,HexNAc(2)Hex(1) | No | 1771.8249 | 3 | 5313.4602 | 5313.4555 | 0.9 | NRagged | 396.18 | 396.18 | 0 | Nsco_20191108_BC_ZIC_HILIC_H111_typsin_B1.72265.72265.3 | 133.0596 | Trypsin | B1 | 148 |
| BCAL0749 | >tr\|A0A2N9D2V0\|A0A2N9D2V0_9BURK Cytochrome c oxidase subunit 2 OS=Burkholderia cenocepacia OX=95486 GN=ctaC_1 PE=3 SV=1 | G.KLPEDTAGAAT[+568.212][+100.064]AAAPAEAASAPAQAAS[+568.212]GAEQPAAAASAALSTIYFETGK.S | HexNAc(2)Hex(1) 100.064,HexNAc(2)Hex(1) | No | 1447.1931 | 4 | 5785.7506 | 5780.7299 | 0.7 | NRagged | 243.78 | 243.78 | 0 | Nsco_20191108_BC_ZIC_HILIC_H111_typsin_B2.63829.63829.4 | 123.3676 | Trypsin | B2 | 149 |
| BCAL0749 | >tr\|A0A2N9D2V0\|A0A2N9D2V0_9BURK Cytochrome c oxidase subunit 2 OS=Burkholderia cenocepacia OX=95486 GN=ctaC_1 PE=3 SV=1 | L.PEDTAGAATAAAPAEAASAPAQAASGAEQPAAAASAALS[+568.212][+100.064]T[+568.212][+100.064]IYFETGK.S | HexNAc(2)Hex(1) 100.064,HexNAc(2)Hex(1) 100.064 | No | 1410.6657 | 4 | 5639.6408 | 5639.6148 | 4.6 | NRagged | 493.11 | 493.11 | 1.2 | Nsco_20191108_BC_ZIC_HILIC_H111_typsin_B2.62442.62442.4 | 120.3245 | Trypsin | B2 | 150 |
| BCAL0749 | >tr\|A0A2N9D2V0\|A0A2N9D2V0_9BURK Cytochrome c oxidase subunit 2 OS=Burkholderia cenocepacia OX=95486 GN=ctaC_1 PE=3 SV=1 | Q.PAAAASAALS[+568.212]T[+568.212]IYFETGK.S | HexNAc(2)Hex(1), HexNAc(2)Hex(1) | No | 1454.1756 | 2 | 2907.3439 | 2905.3347 | 0.8 | NRagged | 388.67 | 378.93 | 15.6 | Nsco_20191108_BC_ZIC_HILIC_H111_typsin_B3.59776.59776.4 | 122.6454 | Trypsin | B3 | 151 |
| BCAL0786 | >tr\|A0A1V2XQK6\|A0A1V2XQK6_9BURK Membrane protein OS=Burkholderia cenocepacia OX=95486 GN=A8E72_26595 PE=4 SV=1 | K.LLSVPAPAST[+568.212]EGDHHDK.- | HexNAc(2)Hex(1) | No | 781.3706 | 3 | 2342.0973 | 2342.0882 | 3.9 | Specific | 542.13 | 542.13 | 0 | Nsco_20191108_BC_ZIC_HILIC_H111_typsin_B2.15440.15440.3 | 35.4194 | Trypsin | B2 | 152 |
| BCAL1081 | >tr\|A0A1V2XQC3\|A0A1V2XQC3_9BURK Multidrug efflux system membrane fusion protein OS=Burkholderia cenocepacia OX=95486 GN=mdtA PE=3 SV=1 | K.RYQTLLSQDSIAS[+568.212]QTVDTQASLVK.Q | HexNAc(2)Hex(1) | No | 1074.5406 | 3 | 3221.6074 | 3220.5955 | 2.7 | Specific | 440.52 | 357.48 | 1.2 | Nsco_20191108_BC_ZIC_HILIC_H111_typsin_B3.39618.39618.3 | 82.4359 | Trypsin | B3 | 153 |
| BCAL1086 | >tr\|A0A107L3U1\|A0A107L3U1_9BURK Putative lipoprotein OS=Burkholderia cenocepacia OX=95486 GN=A8E72_07105 PE=4 SV=1 | S.AAQAQIDAAAS[+568.212]AVVAHAAS[+568.212][+100.064]EAGAK.I | HexNAc(2)Hex(1), HexNAc(2)Hex(1) 100.064 | Yes | 1139.5326 | 3 | 3416.5832 | 3415.5973 | -5.1 | Non | 482.86 | 468.7 | 0 | Nsco_20191108_BC_ZIC_HILIC_H111_thermolysin_B1.44403.44403.3 | 83.8155 | Thermolysin | B1 | 154 |
| BCAL1086 | >tr\|A0A107L3U1\|A0A107L3U1_9BURK Putative lipoprotein OS=Burkholderia cenocepacia OX=95486 GN=A8E72_07105 PE=4 SV=1 | K.LQQWSQQSAAGAKPAS[+568.212]GE.- | HexNAc(2)Hex(1) | No | 1206.5597 | 2 | 2412.1122 | 2412.1049 | 3 | CRagged | 771.99 | 632.32 | 2.87 | Nsco_20191108_BC_ZIC_HILIC_H111_typsin_B1.15216.15216.2 | 36.5522 | Trypsin | B1 | 155 |
| BCAL1086 | >tr\|A0A107L3U1\|A0A107L3U1_9BURK Putative lipoprotein OS=Burkholderia cenocepacia OX=95486 GN=A8E72_07105 PE=4 SV=1 | K.KLQQWSQQS[+568.212]AAGAKPASGE.- | HexNAc(2)Hex(1) | No | 1270.6077 | 2 | 2540.2081 | 2540.1998 | 3.3 | CRagged | 820.17 | 701.91 | 2.83 | Nsco_20191108_BC_ZIC_HILIC_H111_typsin_B1.11185.11185.2 | 30.1493 | Trypsin | B1 | 156 |
| BCAL1086 | >tr\|A0A107L3U1\|A0A107L3U1_9BURK Putative lipoprotein OS=Burkholderia cenocepacia OX=95486 GN=A8E72_07105 PE=4 SV=1 | K.ALDQVAST[+568.212]VNQQINAAK.A | HexNAc(2)Hex(1) | No | 1170.0812 | 2 | 2339.1552 | 2339.146 | 3.9 | Specific | 757.76 | 635.78 | 3.22 | Nsco_20191108_BC_ZIC_HILIC_H111_typsin_B3.35315.35315.2 | 73.9304 | Trypsin | B3 | 157 |
| BCAL1093 | >tr\|A0A2N9CGE7\|A0A2N9CGE7_9BURK Signal peptide transmembrane protein OS=Burkholderia cenocepacia OX=95486 GN=DFS07_119133 PE=4 SV=1 | K.HAYDEAHPAEAAS[+568.212]AASH.- | HexNAc(2)Hex(1) | No | 768.3269 | 3 | 2302.9661 | 2302.9582 | 3.4 | CRagged | 733.24 | 729.7 | 1.54 | Nsco_20191108_BC_ZIC_HILIC_H111_typsin_B1.7502.7502.3 | 24.0483 | Trypsin | B1 | 158 |
| BCAL1389 | >tr\|A0A2N9CG52\|A0A2N9CG52_9BURK Cyclic di-GMP-binding protein OS=Burkholderia cenocepacia OX=95486 GN=bcsB PE=3 SV=1 | P.AVPAAAS[+568.212]T[+568.212][+100.064]AAAGHAAAADVAPAAGPVAAPAAASAPAAGLPATTVHVPFASLGAFDPLR.L | HexNAc(2)Hex(1), HexNAc(2)Hex(1) 100.064 | No | 1598.0516 | 4 | 6389.1847 | 6389.1838 | 0.1 | NRagged | 954.87 | 371.84 | 0 | Nsco_20191108_BC_ZIC_HILIC_H111_typsin_B2.65506.65506.4 | 127.0975 | Trypsin | B2 | 159 |

| | | | | | | | | | | | | | | | | | | |
|---|---|---|---|---|---|---|---|---|---|---|---|---|---|---|---|---|---|---|
| BCAL1389 | >tr\|A0A2N9CG52\|A0A2N9CG52_9BURK Cyclic di-GMP-binding protein OS=Burkholderia cenocepacia OX=95486 GN=bcsB PE=3 SV=1 | A.APMPAVPAAAST[+568.212]AAAGHAAAADVAPAAGPVAAPAAAS[+568.212]APAAGLPATTVHVPFASLGAFDPLR.L | HexNAc(2)Hex(1), HexNAc(2)Hex(1) | Yes (1 of 2) | 1672.0869 | 4 | 6685.3257 | 6685.3029 | 3.4 | NRagged | 1064.73 | 1061.81 | 159.46 | Nsco_20191108_BC_ZIC_HILIC_H111_typsin_B2.67298.67298.4 | 131.0906 | Trypsin | B2 | 160 |
| BCAL1496 | >tr\|A0A0M1IFG9\|A0A0M1IFG9_9BURK Signal peptide protein OS=Burkholderia cenocepacia OX=95486 GN=A8F55_16750 PE=4 SV=1 | K.KAGKKAKAADAAS[+568.212]Q.- | HexNAc(2)Hex(1) | Yes | 638.3309 | 3 | 1912.9781 | 1912.971 | 3.8 | Non | 431.43 | 251.88 | 251.88 | Nsco_20191108_BC_ZIC_HILIC_H111_pepsin_B1.2102.2102.3 | 13.2946 | Pepsin | B1 | 161 |
| BCAL1496 | >tr\|A0A0M1IFG9\|A0A0M1IFG9_9BURK Signal peptide protein OS=Burkholderia cenocepacia OX=95486 GN=A8F55_16750 PE=4 SV=1 | H.KAAAKKAGKKAKAADAAS[+568.212]Q.- | HexNAc(2)Hex(1) | Yes | 596.3257 | 4 | 2382.281 | 2382.2722 | 3.7 | Non | 638.78 | 549.42 | 549.42 | Nsco_20191108_BC_ZIC_HILIC_H111_pepsin_B2.2867.2867.4 | 14.0627 | Pepsin | B2 | 162 |
| BCAL1496 | >tr\|A0A0M1IFG9\|A0A0M1IFG9_9BURK Signal peptide protein OS=Burkholderia cenocepacia OX=95486 GN=A8F55_16750 PE=4 SV=1 | G.KKAKAADAAS[+568.212]Q.- | HexNAc(2)Hex(1) | Yes | 552.9459 | 3 | 1656.823 | 1656.8174 | 3.4 | Non | 326.6 | 179.24 | 179.24 | Nsco_20191108_BC_ZIC_HILIC_H111_pepsin_B2.2605.2605.3 | 13.2624 | Pepsin | B2 | 163 |
| BCAL1496 | >tr\|A0A0M1IFG9\|A0A0M1IFG9_9BURK Signal peptide protein OS=Burkholderia cenocepacia OX=95486 GN=A8F55_16750 PE=4 SV=1 | A.KKAGKKAKAADAAS[+568.212]Q.- | HexNAc(2)Hex(1) | Yes | 511.0242 | 4 | 2041.0748 | 2041.0659 | 4.4 | Non | 432.15 | 280.15 | 280.15 | Nsco_20191108_BC_ZIC_HILIC_H111_pepsin_B2.2745.2745.4 | 13.726 | Pepsin | B2 | 164 |
| BCAL1496 | >tr\|A0A0M1IFG9\|A0A0M1IFG9_9BURK Signal peptide protein OS=Burkholderia cenocepacia OX=95486 GN=A8F55_16750 PE=4 SV=1 | A.AAKKAGKKAKAADAAS[+568.212]Q.- | HexNAc(2)Hex(1) | Yes | 546.5425 | 4 | 2183.1482 | 2183.1401 | 3.7 | Non | 645.31 | 568.86 | 568.86 | Nsco_20191108_BC_ZIC_HILIC_H111_thermolysin_B1.2030.2030.4 | 13.2481 | Thermolysin | B1 | 165 |
| BCAL1496 | >tr\|A0A0M1IFG9\|A0A0M1IFG9_9BURK Signal peptide protein OS=Burkholderia cenocepacia OX=95486 GN=A8F55_16750 PE=4 SV=1 | K.AAAKKAGKKAKAADAAS[+568.212]Q.- | HexNAc(2)Hex(1) | Yes | 564.302 | 4 | 2254.186 | 2254.1773 | 3.9 | Non | 657.05 | 583.97 | 583.97 | Nsco_20191108_BC_ZIC_HILIC_H111_thermolysin_B3.2327.2327.4 | 14.0219 | Thermolysin | B3 | 166 |
| BCAL1496 | >tr\|A0A0M1IFG9\|A0A0M1IFG9_9BURK Signal peptide protein OS=Burkholderia cenocepacia OX=95486 GN=A8F55_16750 PE=4 SV=1 | K.AKAADAAS[+568.212]Q.- | HexNAc(2)Hex(1) | Yes | 700.819 | 2 | 1400.6308 | 1400.6275 | 2.4 | CRagged | 314.51 | 138.87 | 138.87 | Nsco_20191108_BC_ZIC_HILIC_H111_typsin_B1.2165.2165.2 | 14.2832 | Trypsin | B1 | 167 |
| BCAL1537 | >tr\|A0A1V6KNH3\|A0A1V6KNH3_9BURK Lipoprotein OS=Burkholderia cenocepacia OX=95486 GN=A8E72_13475 PE=4 SV=1 | S.AVQQASAPVAAIPVIDSQAQT[+568.212]SVQPQAGETTGPSTVDDLQRQ.I | HexNAc(2)Hex(1) | No | 1610.1277 | 3 | 4828.3685 | 4828.3494 | 3.9 | Non | 364.19 | 364.19 | 0 | Nsco_20191108_BC_ZIC_HILIC_H111_thermolysin_B1.57459.57459.3 | 107.5788 | Thermolysin | B1 | 168 |
| BCAL1649 | >tr\|A0A1V6KP99\|A0A1V6KP99_9BURK Membrane protein OS=Burkholderia cenocepacia OX=95486 GN=A8E72_01255 PE=4 SV=1 | V.ASAASSPAPAPAAPASPS[+568.212][+100.064]EAT[+568.212][+100.064]QAPHQQQTSAGYRN.I | HexNAc(2)Hex(1) 100.064,HexNAc(2)Hex(1) 100.064 | No | 1571.7136 | 3 | 4713.1263 | 4712.146 | -4.9 | Non | 230.26 | 29.55 | 2.4 | Nsco_20191108_BC_ZIC_HILIC_H111_thermolysin_B1.23209.23209.3 | 47.0602 | Thermolysin | B1 | 169 |
| BCAL1674 | >tr\|A0A2N9CH15\|A0A2N9CH15_9BURK Multidrug efflux system OS=Burkholderia cenocepacia OX=95486 GN=acrA PE=3 SV=1 | Q.FAPDTAVKPVEKAPPS[+568.212]KAAPPAAASQA.A | HexNAc(2)Hex(1) | No | 1062.5435 | 3 | 3185.6159 | 3185.61 | 1.9 | Non | 748.99 | 695.75 | 16.78 | Nsco_20191108_BC_ZIC_HILIC_H111_thermolysin_B3.20755.20755.3 | 45.2678 | Thermolysin | B3 | 170 |
| BCAL1746 | >tr\|A0A1V2Y428\|A0A1V2Y428_9BURK Uncharacterized protein OS=Burkholderia cenocepacia OX=95486 GN=A8F55_03700 PE=4 SV=1 | L.IDHIGKAWPGNAAS[+568.212]GASASE.- | HexNAc(2)Hex(1) | No | 836.3886 | 3 | 2507.1514 | 2507.142 | 3.7 | CRagged | 734.05 | 680.07 | 6 | Nsco_20191108_BC_ZIC_HILIC_H111_pepsin_B2.22428.22428.3 | 46.795 | Pepsin | B2 | 171 |

| | | | | | | | | | | | | | | | | | | |
|---|---|---|---|---|---|---|---|---|---|---|---|---|---|---|---|---|---|---|
| BCAL1877 | >tr\|A0A2N9CHN6\|A0A2N9CHN6_9BURK Protein HflK OS=Burkholderia cenocepacia OX=95486 GN=F01_210017 PE=3 SV=1 | A.AAST[+568.212]GAS[+568.212]AADAASAPAATVPSAA.A | HexNAc(2)Hex(1), HexNAc(2)Hex(1) | No | 1008.4515 | 3 | 3023.34 | 3023.3322 | 2.6 | Non | 342.23 | 273.47 | 0 | Nsco_20191108_BC_ZIC_HILIC_H111_pepsin_B2.29475.29475.3 | 60.7221 | Pepsin | B2 | 172 |
| BCAL1877 | >tr\|A0A2N9CHN6\|A0A2N9CHN6_9BURK Protein HflK OS=Burkholderia cenocepacia OX=95486 GN=F01_210017 PE=3 SV=1 | P.LDKLVEQGRQNAAAS[+568.212]T[+568.212]GASAADAASAPAATVPSA.A | HexNAc(2)Hex(1), HexNAc(2)Hex(1) | No | 1435.6845 | 3 | 4305.039 | 4304.0159 | 4.6 | Non | 247.74 | 203.21 | 5.6 | Nsco_20191108_BC_ZIC_HILIC_H111_thermolysin_B1.33074.33074.3 | 64.0234 | Thermolysin | B1 | 173 |
| BCAL1877 | >tr\|A0A2N9CHN6\|A0A2N9CHN6_9BURK Protein HflK OS=Burkholderia cenocepacia OX=95486 GN=F01_210017 PE=3 SV=1 | A.AASTGASAADAASAPAAT[+568.212]VPS[+568.212].A | HexNAc(2)Hex(1), HexNAc(2)Hex(1) | No | 1441.1368 | 2 | 2881.2662 | 2881.2579 | 2.9 | Non | 621.81 | 258.83 | 0.76 | Nsco_20191108_BC_ZIC_HILIC_H111_thermolysin_B2.26699.26699.2 | 55.2555 | Thermolysin | B2 | 174 |
| BCAL1877 | >tr\|A0A2N9CHN6\|A0A2N9CHN6_9BURK Protein HflK OS=Burkholderia cenocepacia OX=95486 GN=F01_210017 PE=3 SV=1 | P.LDKLVEQGRQNAAAST[+568.212]GAS[+568.212]AADAASAPAATVPS.A | HexNAc(2)Hex(1), HexNAc(2)Hex(1) | No | 1411.6694 | 3 | 4232.9936 | 4232.9787 | 3.5 | Non | 306.31 | 283.72 | 6.13 | Nsco_20191108_BC_ZIC_HILIC_H111_thermolysin_B2.30688.30688.3 | 62.0296 | Thermolysin | B2 | 175 |
| BCAL1885 | >tr\|A0A1V2XJM4\|A0A1V2XJM4_9BURK Helix-turn-helix domain-containing protein OS=Burkholderia cenocepacia OX=95486 GN=A8F55_38750 PE=4 SV=1 | A.AAST[+568.212]AAPAQPAS[+568.212]VVVA.A | HexNAc(2)Hex(1), HexNAc(2)Hex(1) | No | 1274.099 | 2 | 2547.1908 | 2547.1819 | 3.5 | Non | 500.63 | 283.69 | 94.58 | Nsco_20191108_BC_ZIC_HILIC_H111_pepsin_B1.29311.29311.2 | 61.8555 | Pepsin | B1 | 176 |
| BCAL1885 | >tr\|A0A1V2XJM4\|A0A1V2XJM4_9BURK Helix-turn-helix domain-containing protein OS=Burkholderia cenocepacia OX=95486 GN=A8F55_38750 PE=4 SV=1 | T.AAAST[+568.212]AAPAQPAS[+568.212]VVVAAGQS.M | HexNAc(2)Hex(1), HexNAc(2)Hex(1) | No | 1482.1934 | 2 | 2963.3794 | 2961.3682 | 1.5 | Non | 320.1 | 240.96 | 20.27 | Nsco_20191108_BC_ZIC_HILIC_H111_thermolysin_B3.33206.33206.2 | 66.5721 | Thermolysin | B3 | 177 |
| BCAL1893 | >tr\|A0A1V2VRW6\|A0A1V2VRW6_9BURK Family M23 peptidase OS=Burkholderia cenocepacia OX=95486 GN=nlpD_1 PE=4 SV=1 | R.VAPPGGAAVAGAPAAAPIVGGAVAT[+568.212]APLSSGPAAPAAGTSSALAATPPAAATGSSDTAAAPSGPVTFAWPAR.G | HexNAc(2)Hex(1) | No | 1692.3637 | 4 | 6766.4331 | 6766.401 | 4.7 | Specific | 960.85 | 960.85 | 7.56 | Nsco_20191108_BC_ZIC_HILIC_H111_typsin_B1.77573.77573.4 | 142.8025 | Trypsin | B1 | 178 |
| BCAL1917 | >tr\|A0A2N9CHU5\|A0A2N9CHU5_9BURK Uncharacterized protein OS=Burkholderia cenocepacia OX=95486 GN=F01_210059 PE=4 SV=1 | T.ATAGTT[+568.212]TAAPAPTASAPEA.A | HexNAc(2)Hex(1) | No | 1113.0168 | 2 | 2225.0263 | 2225.0191 | 3.2 | Non | 479.56 | 271.22 | 0 | Nsco_20191108_BC_ZIC_HILIC_H111_pepsin_B2.24784.24784.2 | 51.3388 | Pepsin | B2 | 179 |
| BCAL1917 | >tr\|A0A2N9CHU5\|A0A2N9CHU5_9BURK Uncharacterized protein OS=Burkholderia cenocepacia OX=95486 GN=F01_210059 PE=4 SV=1 | T.TATAGTTTAAPAPTAS[+568.212]APEAAAKPAKTKR.A | HexNAc(2)Hex(1) | No | 1093.223 | 3 | 3277.6545 | 3277.6646 | -3.1 | Non | 380.66 | 380.66 | 1.2 | Nsco_20191108_BC_ZIC_HILIC_H111_thermolysin_B1.12081.12081.3 | 30.2274 | Thermolysin | B1 | 180 |
| BCAL1917 | >tr\|A0A2N9CHU5\|A0A2N9CHU5_9BURK Uncharacterized protein OS=Burkholderia cenocepacia OX=95486 GN=F01_210059 PE=4 SV=1 | K.LSKPAATTSATTSTTTSAGTASTTTATAGTTTAAPAPTAS[+568.212]APEAAAKPAKTKR.A | HexNAc(2)Hex(1) | No | 1112.9653 | 5 | 5560.7976 | 5560.7723 | 4.5 | Non | 1523.41 | 1442.53 | 37.43 | Nsco_20191108_BC_ZIC_HILIC_H111_thermolysin_B1.18180.18180.5 | 39.1099 | Thermolysin | B1 | 181 |
| BCAL1917 | >tr\|A0A2N9CHU5\|A0A2N9CHU5_9BURK Uncharacterized protein OS=Burkholderia cenocepacia OX=95486 GN=F01_210059 PE=4 SV=1 | K.LSKPAATTSATTSTTTSAGTASTTTATAGTTTAAPAPTAS[+568.212][+100.064]APEAAAKPA.K | HexNAc(2)Hex(1) 100.064 | No | 1717.4998 | 3 | 5150.4847 | 5147.4976 | -4.5 | Non | 390.69 | 378.24 | 5.94 | Nsco_20191108_BC_ZIC_HILIC_H111_thermolysin_B1.30680.30680.3 | 60.0593 | Thermolysin | B1 | 182 |
| BCAL1917 | >tr\|A0A2N9CHU5\|A0A2N9CHU5_9BURK Uncharacterized protein OS=Burkholderia cenocepacia OX=95486 GN=F01_210059 PE=4 SV=1 | T.AAPAPTAS[+568.212]APEAAAKPAKTKRASKKEK.A | HexNAc(2)Hex(1) | Yes | 649.7541 | 5 | 3244.7412 | 3244.7271 | 4.4 | Non | 796.81 | 784.32 | 121.44 | Nsco_20191108_BC_ZIC_HILIC_H111_thermolysin_B1.3500.3500.5 | 17.3463 | Thermolysin | B1 | 183 |

| | | | | | | | | | | | | | | | | | | |
|---|---|---|---|---|---|---|---|---|---|---|---|---|---|---|---|---|---|---|
| BCAL1917 | >tr\|A0A2N9CHU5\|A0A2N9CHU5_9BURK Uncharacterized protein OS=Burkholderia cenocepacia OX=95486 GN=F01_210059 PE=4 SV=1 | G.TTTAAPAPT[+568.212]AS[+568.212][+100.064]APEAAAKPAKTKRASKKEKAA.A | HexNAc(2)Hex(1), HexNAc(2)Hex(1) 100.064 | No | 872.8513 | 5 | 4360.2276 | 4358.2199 | 0.2 | Non | 386.81 | 386.81 | 0 | Nsco_20191108_BC_ZIC_HILIC_H111_thermolysin_B1.4588.4588.4 | 18.8915 | Thermolysin | B1 | 184 |
| BCAL1917 | >tr\|A0A2N9CHU5\|A0A2N9CHU5_9BURK Uncharacterized protein OS=Burkholderia cenocepacia OX=95486 GN=F01_210059 PE=4 SV=1 | T.TAAPAPTASAPEAAAKPAKT[+568.212]KRASKKEKA.A | HexNAc(2)Hex(1) | No | 1139.9388 | 3 | 3417.802 | 3416.8119 | -3.9 | Non | 161.58 | 161.58 | 9.6 | Nsco_20191108_BC_ZIC_HILIC_H111_thermolysin_B1.5091.5091.3 | 19.6085 | Thermolysin | B1 | 185 |
| BCAL1917 | >tr\|A0A2N9CHU5\|A0A2N9CHU5_9BURK Uncharacterized protein OS=Burkholderia cenocepacia OX=95486 GN=F01_210059 PE=4 SV=1 | T.AAPAPTAS[+568.212]APEAAAKPAKTKRASKKEKA.A | HexNAc(2)Hex(1) | Yes | 663.961 | 5 | 3315.7758 | 3315.7642 | 3.5 | Non | 1153.47 | 1053.48 | 140.09 | Nsco_20191108_BC_ZIC_HILIC_H111_thermolysin_B1.3693.3693.5 | 17.6616 | Thermolysin | B1 | 186 |
| BCAL1917 | >tr\|A0A2N9CHU5\|A0A2N9CHU5_9BURK Uncharacterized protein OS=Burkholderia cenocepacia OX=95486 GN=F01_210059 PE=4 SV=1 | T.AAPAPTAS[+568.212]APEAAAKPAKTKRASKKEKAAAA.A | HexNAc(2)Hex(1) | Yes | 882.9785 | 4 | 3528.8922 | 3528.8755 | 4.7 | Non | 761.25 | 761.25 | 127.74 | Nsco_20191108_BC_ZIC_HILIC_H111_thermolysin_B1.4372.4372.4 | 18.6162 | Thermolysin | B1 | 187 |
| BCAL1917 | >tr\|A0A2N9CHU5\|A0A2N9CHU5_9BURK Uncharacterized protein OS=Burkholderia cenocepacia OX=95486 GN=F01_210059 PE=4 SV=1 | T.AGTTTAAPAPTAS[+568.212]APEAAAKPAKTKR.A | HexNAc(2)Hex(1) | No | 1002.5198 | 3 | 3005.545 | 3004.5321 | 3.2 | Non | 486.62 | 400.84 | 2.4 | Nsco_20191108_BC_ZIC_HILIC_H111_thermolysin_B1.8726.8726.3 | 24.9449 | Thermolysin | B1 | 188 |
| BCAL1917 | >tr\|A0A2N9CHU5\|A0A2N9CHU5_9BURK Uncharacterized protein OS=Burkholderia cenocepacia OX=95486 GN=F01_210059 PE=4 SV=1 | A.ATTSATTSTTTSAGTASTSTTATGTTT[+568.212][+100.064]AAPAPTASAPEAAAKPAKTKR.A | HexNAc(2)Hex(1) 100.064 | No | 1033.7095 | 5 | 5164.5183 | 5164.5354 | -3.3 | Non | 298.51 | 298.51 | 0 | Nsco_20191108_BC_ZIC_HILIC_H111_thermolysin_B2.17991.17991.5 | 40.013 | Thermolysin | B2 | 189 |
| BCAL1917 | >tr\|A0A2N9CHU5\|A0A2N9CHU5_9BURK Uncharacterized protein OS=Burkholderia cenocepacia OX=95486 GN=F01_210059 PE=4 SV=1 | T.AAPAPTAS[+568.212]APEAAAKPAKTKRAS.K | HexNAc(2)Hex(1) | Yes | 683.6077 | 4 | 2731.4092 | 2731.3996 | 3.5 | Non | 695.41 | 663.67 | 96.64 | Nsco_20191108_BC_ZIC_HILIC_H111_thermolysin_B2.5786.5786.4 | 20.9413 | Thermolysin | B2 | 190 |
| BCAL1917 | >tr\|A0A2N9CHU5\|A0A2N9CHU5_9BURK Uncharacterized protein OS=Burkholderia cenocepacia OX=95486 GN=F01_210059 PE=4 SV=1 | T.ATAGTTTAAPAPTAS[+568.212]APEAAAKPAKTKR.A | HexNAc(2)Hex(1) | No | 1059.8811 | 3 | 3177.6288 | 3176.6169 | 2.7 | Non | 724.14 | 537.25 | 2.4 | Nsco_20191108_BC_ZIC_HILIC_H111_thermolysin_B2.9225.9225.3 | 25.9324 | Thermolysin | B2 | 191 |
| BCAL1917 | >tr\|A0A2N9CHU5\|A0A2N9CHU5_9BURK Uncharacterized protein OS=Burkholderia cenocepacia OX=95486 GN=F01_210059 PE=4 SV=1 | T.AAPAPTAS[+568.212]APEAAAKPAKTKRASKKEKAAA.A | HexNAc(2)Hex(1) | Yes | 692.3754 | 5 | 3457.8481 | 3457.8384 | 2.8 | Non | 1074.62 | 1074.62 | 140.46 | Nsco_20191108_BC_ZIC_HILIC_H111_thermolysin_B3.4136.4136.5 | 18.4944 | Thermolysin | B3 | 192 |
| BCAL1917 | >tr\|A0A2N9CHU5\|A0A2N9CHU5_9BURK Uncharacterized protein OS=Burkholderia cenocepacia OX=95486 GN=F01_210059 PE=4 SV=1 | T.AAPAPTAS[+568.212]APEAAAKPAKTKRASKKEKAA.A | HexNAc(2)Hex(1) | Yes | 678.1687 | 5 | 3386.8143 | 3386.8013 | 3.8 | Non | 1151.4 | 1103.63 | 130.25 | Nsco_20191108_BC_ZIC_HILIC_H111_thermolysin_B3.3893.3893.5 | 18.0785 | Thermolysin | B3 | 193 |
| BCAL1917 | >tr\|A0A2N9CHU5\|A0A2N9CHU5_9BURK Uncharacterized protein OS=Burkholderia cenocepacia OX=95486 GN=F01_210059 PE=4 SV=1 | T.AAPAPTAS[+568.212]APEAAAKPAKTKR.A | HexNAc(2)Hex(1) | Yes | 644.0909 | 4 | 2573.3418 | 2573.3305 | 4.4 | Non | 917.07 | 748.87 | 124.03 | Nsco_20191108_BC_ZIC_HILIC_H111_thermolysin_B3.5571.5571.4 | 20.9475 | Thermolysin | B3 | 194 |
| BCAL1917 | >tr\|A0A2N9CHU5\|A0A2N9CHU5_9BURK Uncharacterized protein OS=Burkholderia cenocepacia OX=95486 GN=F01_210059 PE=4 SV=1 | P.AATTSATTSTTTSAGTASTS[+568.212]TTATAGTTTAAPAPTASAPEAAAKPAKTKR.A | HexNAc(2)Hex(1) | No | 1285.1395 | 4 | 5137.5363 | 5135.5085 | 4.1 | Non | 698.47 | 689.16 | 0 | Nsco_20191108_BC_ZIC_HILIC_H111_thermolysin_B3.17015.17015.4 | 38.7465 | Thermolysin | B3 | 195 |
| BCAL1917 | >tr\|A0A2N9CHU5\|A0A2N9CHU5_9BURK Uncharacterized protein OS=Burkholderia cenocepacia OX=95486 GN=F01_210059 PE=4 SV=1 | T.TAAPAPTASAPEAAAKPAKT[+568.212]KRASKKEKAAA.A | HexNAc(2)Hex(1) | No | 712.5804 | 5 | 3558.8729 | 3558.8861 | -3.7 | Non | 715.88 | 715.88 | 9.6 | Nsco_20191108_BC_ZIC_HILIC_H111_thermolysin_B3.5230.5230.5 | 20.3847 | Thermolysin | B3 | 196 |

<table>
<tr><td>BCAL1917</td><td>>tr|A0A2N9CHU5|A0A2N9CHU5_9BURK Uncharacterized protein OS=Burkholderia cenocepacia OX=95486 GN=F01_210059 PE=4 SV=1</td><td>T.TAAPAPTAS[+568.212]APEAAAKPAKTKR.A</td><td>HexNAc(2)Hex(1)</td><td>No</td><td>1337.6868</td><td>2</td><td>2674.3663</td><td>2674.3781</td><td>-4.4</td><td>Non</td><td>490.81</td><td>380.09</td><td>2.4</td><td>Nsco_20191108_BC_ZIC_HILIC_H111_thermolysin_B3.7211.7211.2</td><td>23.2425</td><td>Thermolysin</td><td>B3</td><td>197</td></tr>
<tr><td>BCAL1917</td><td>>tr|A0A2N9CHU5|A0A2N9CHU5_9BURK Uncharacterized protein OS=Burkholderia cenocepacia OX=95486 GN=F01_210059 PE=4 SV=1</td><td>A.AAAAAAASADAGASAPAAASS[+568.212]T[+568.212]K.A</td><td>HexNAc(2)Hex(1), HexNAc(2)Hex(1)</td><td>No</td><td>1498.6727</td><td>2</td><td>2996.338</td><td>2996.3325</td><td>1.8</td><td>NRagged</td><td>855.07</td><td>790.62</td><td>1.2</td><td>Nsco_20191108_BC_ZIC_HILIC_H111_typsin_B1.11090.11090.2</td><td>30.0018</td><td>Trypsin</td><td>B1</td><td>198</td></tr>
<tr><td>BCAL1917</td><td>>tr|A0A2N9CHU5|A0A2N9CHU5_9BURK Uncharacterized protein OS=Burkholderia cenocepacia OX=95486 GN=F01_210059 PE=4 SV=1</td><td>T.STTATAGTTTAAPAPT[+568.212]ASAPEAAAKPAK.T</td><td>HexNAc(2)Hex(1)</td><td>No</td><td>1027.5093</td><td>3</td><td>3080.5134</td><td>3080.5005</td><td>4.2</td><td>NRagged</td><td>679.67</td><td>588.94</td><td>2.4</td><td>Nsco_20191108_BC_ZIC_HILIC_H111_typsin_B1.15161.15161.3</td><td>36.4723</td><td>Trypsin</td><td>B1</td><td>199</td></tr>
<tr><td>BCAL1917</td><td>>tr|A0A2N9CHU5|A0A2N9CHU5_9BURK Uncharacterized protein OS=Burkholderia cenocepacia OX=95486 GN=F01_210059 PE=4 SV=1</td><td>A.TAGTTTAAPAPTAS[+568.212]APEAAAKPAK.T</td><td>HexNAc(2)Hex(1)</td><td>No</td><td>907.453</td><td>3</td><td>2720.3444</td><td>2720.336</td><td>3.1</td><td>NRagged</td><td>708.75</td><td>145.82</td><td>7.02</td><td>Nsco_20191108_BC_ZIC_HILIC_H111_typsin_B1.12720.12720.3</td><td>32.3796</td><td>Trypsin</td><td>B1</td><td>200</td></tr>
<tr><td>BCAL1917</td><td>>tr|A0A2N9CHU5|A0A2N9CHU5_9BURK Uncharacterized protein OS=Burkholderia cenocepacia OX=95486 GN=F01_210059 PE=4 SV=1</td><td>T.SAGTASTSTTATAGTTTAAPAPAS[+568.212]APEAAAKPAK.T</td><td>HexNAc(2)Hex(1)</td><td>No</td><td>1219.262</td><td>3</td><td>3655.7715</td><td>3655.7556</td><td>4.3</td><td>NRagged</td><td>934.29</td><td>918.95</td><td>4.53</td><td>Nsco_20191108_BC_ZIC_HILIC_H111_typsin_B2.19051.19051.3</td><td>41.4334</td><td>Trypsin</td><td>B2</td><td>201</td></tr>
<tr><td>BCAL1917</td><td>>tr|A0A2N9CHU5|A0A2N9CHU5_9BURK Uncharacterized protein OS=Burkholderia cenocepacia OX=95486 GN=F01_210059 PE=4 SV=1</td><td>S.KPAATTSATTSTTTSAGTASTSTATAGTTTAAPAPTAS[+568.212]APEAAAKPAK.T</td><td>HexNAc(2)Hex(1)</td><td>No</td><td>1244.8641</td><td>4</td><td>4976.4347</td><td>4975.4125</td><td>3.8</td><td>NRagged</td><td>454.51</td><td>454.51</td><td>25.96</td><td>Nsco_20191108_BC_ZIC_HILIC_H111_typsin_B2.20241.20241.4</td><td>43.5398</td><td>Trypsin</td><td>B2</td><td>202</td></tr>
<tr><td>BCAL1917</td><td>>tr|A0A2N9CHU5|A0A2N9CHU5_9BURK Uncharacterized protein OS=Burkholderia cenocepacia OX=95486 GN=F01_210059 PE=4 SV=1</td><td>T.TSATTSTTTSAGTASTS[+568.212][+100.064]TTATAGTTTAAPAPT[+568.212][+100.064]ASAPEAAAKPAK.T</td><td>HexNAc(2)Hex(1) 100.064,HexNAc(2)Hex(1) 100.064</td><td>No</td><td>1319.631</td><td>4</td><td>5275.5021</td><td>5275.4824</td><td>3.7</td><td>NRagged</td><td>366.32</td><td>366.32</td><td>2.4</td><td>Nsco_20191108_BC_ZIC_HILIC_H111_typsin_B2.43076.43076.4</td><td>84.0457</td><td>Trypsin</td><td>B2</td><td>203</td></tr>
<tr><td>BCAL1917</td><td>>tr|A0A2N9CHU5|A0A2N9CHU5_9BURK Uncharacterized protein OS=Burkholderia cenocepacia OX=95486 GN=F01_210059 PE=4 SV=1</td><td>A.GTASTSTTATAGTTTAAPAPT[568.212]ASAPEAAAKPAK.T</td><td>HexNAc(2)Hex(1)</td><td>No</td><td>1166.5699</td><td>3</td><td>3497.6951</td><td>3497.6865</td><td>2.5</td><td>NRagged</td><td>863.61</td><td>803.48</td><td>3.18</td><td>Nsco_20191108_BC_ZIC_HILIC_H111_typsin_B2.18137.18137.3</td><td>39.9377</td><td>Trypsin</td><td>B2</td><td>204</td></tr>
<tr><td>BCAL1917</td><td>>tr|A0A2N9CHU5|A0A2N9CHU5_9BURK Uncharacterized protein OS=Burkholderia cenocepacia OX=95486 GN=F01_210059 PE=4 SV=1</td><td>K.LSKPAATTSATTSTTTSAGTASTTTATAGTTTAAPAPTAS[+568.212]APEAAAK.P</td><td>HexNAc(2)Hex(1)</td><td>No</td><td>1220.8451</td><td>4</td><td>4880.3587</td><td>4879.3437</td><td>2.4</td><td>Specific</td><td>915.76</td><td>906.55</td><td>60.93</td><td>Nsco_20191108_BC_ZIC_HILIC_H111_typsin_B3.24861.24861.4</td><td>53.4965</td><td>Trypsin</td><td>B3</td><td>205</td></tr>
<tr><td>BCAL1917</td><td>>tr|A0A2N9CHU5|A0A2N9CHU5_9BURK Uncharacterized protein OS=Burkholderia cenocepacia OX=95486 GN=F01_210059 PE=4 SV=1</td><td>K.PAATTSATTSTTTSAGTASTSTTTAGTTTAAPAPT[+568.212]ASAPEAAAKPAK.T</td><td>HexNAc(2)Hex(1)</td><td>No</td><td>1213.089</td><td>4</td><td>4849.3341</td><td>4847.3175</td><td>2</td><td>Specific</td><td>1213.9</td><td>1213.9</td><td>4.8</td><td>Nsco_20191108_BC_ZIC_HILIC_H111_typsin_B3.24719.24719.4</td><td>53.2254</td><td>Trypsin</td><td>B3</td><td>206</td></tr>
<tr><td>BCAL1917</td><td>>tr|A0A2N9CHU5|A0A2N9CHU5_9BURK Uncharacterized protein OS=Burkholderia cenocepacia OX=95486 GN=F01_210059 PE=4 SV=1</td><td>A.AAAAAASADAGASAPAAAS[+568.212]S[+568.212][+100.064]T[+568.212]K.A</td><td>HexNAc(2)Hex(1), HexNAc(2)Hex(1) 100.064,HexNAc(2)Hex(1)</td><td>No</td><td>1797.2834</td><td>2</td><td>3593.5596</td><td>3593.571</td><td>-3.2</td><td>NRagged</td><td>582.16</td><td>537.26</td><td>0</td><td>Nsco_20191108_BC_ZIC_HILIC_H111_typsin_B3.12030.12030.2</td><td>30.1645</td><td>Trypsin</td><td>B3</td><td>207</td></tr>
<tr><td>BCAL1917</td><td>>tr|A0A2N9CHU5|A0A2N9CHU5_9BURK Uncharacterized protein OS=Burkholderia cenocepacia OX=95486 GN=F01_210059 PE=4 SV=1</td><td>A.STSTTATAGTTTAAPAPT[+568.212]ASAPEAAAKPAK.T</td><td>HexNAc(2)Hex(1)</td><td>No</td><td>1090.2005</td><td>3</td><td>3268.5869</td><td>3268.5802</td><td>2</td><td>NRagged</td><td>567.87</td><td>567.87</td><td>2.77</td><td>Nsco_20191108_BC_ZIC_HILIC_H111_typsin_B3.16581.16581.3</td><td>37.8916</td><td>Trypsin</td><td>B3</td><td>208</td></tr>
<tr><td>BCAL1917</td><td>>tr|A0A2N9CHU5|A0A2N9CHU5_9BURK Uncharacterized protein OS=Burkholderia cenocepacia OX=95486 GN=F01_210059 PE=4 SV=1</td><td>T.STTTSAGTASTSTTATAGTTTAAPT[+568.212]ASAPEAAAKPAK.T</td><td>HexNAc(2)Hex(1)</td><td>No</td><td>1349.3188</td><td>3</td><td>4045.9418</td><td>4045.9307</td><td>2.7</td><td>NRagged</td><td>720.97</td><td>718.16</td><td>4.8</td><td>Nsco_20191108_BC_ZIC_HILIC_H111_typsin_B3.20204.20204.3</td><td>44.5606</td><td>Trypsin</td><td>B3</td><td>209</td></tr>
</table>

| | | | | | | | | | | | | | | | | | | |
|---|---|---|---|---|---|---|---|---|---|---|---|---|---|---|---|---|---|---|
| BCAL1917 | >tr\|A0A2N9CHU5\|A0A2N9CHU5_9BURK Uncharacterized protein OS=Burkholderia cenocepacia OX=95486 GN=F01_210059 PE=4 SV=1 | A.TTSTTTSAGTASTSTTATAGTTAPAPT[+568.212]ASAPEAAAKPAK.T | HexNAc(2)Hex(1) | No | 1417.6876 | 3 | 4251.0483 | 4248.026 | 2.9 | NRagged | 842.67 | 186.29 | 0.29 | Nsco_20191108_BC_ZIC_HILIC_H111_typsin_B3.20809.20809.3 | 45.7375 | Trypsin | B3 | 210 |
| BCAL1917 | >tr\|A0A2N9CHU5\|A0A2N9CHU5_9BURK Uncharacterized protein OS=Burkholderia cenocepacia OX=95486 GN=F01_210059 PE=4 SV=1 | S.ATTSTTTSAGTASTSTTATAGTAAPAPT[+568.212]ASAPEAAAKPAK.T | HexNAc(2)Hex(1) | No | 1440.3658 | 3 | 4319.0828 | 4319.0631 | 4.5 | NRagged | 892.62 | 884.51 | 4.8 | Nsco_20191108_BC_ZIC_HILIC_H111_typsin_B3.21061.21061.3 | 46.2839 | Trypsin | B3 | 211 |
| BCAL1917 | >tr\|A0A2N9CHU5\|A0A2N9CHU5_9BURK Uncharacterized protein OS=Burkholderia cenocepacia OX=95486 GN=F01_210059 PE=4 SV=1 | T.TSAGTASTSTTATAGTTTAAPATAS[+568.212]APEAAAKPAK.T | HexNAc(2)Hex(1) | No | 1252.9331 | 3 | 3756.7848 | 3756.8033 | -4.9 | NRagged | 608.21 | 601.73 | 3.99 | Nsco_20191108_BC_ZIC_HILIC_H111_typsin_B3.21338.21338.3 | 46.7666 | Trypsin | B3 | 212 |
| BCAL1917 | >tr\|A0A2N9CHU5\|A0A2N9CHU5_9BURK Uncharacterized protein OS=Burkholderia cenocepacia OX=95486 GN=F01_210059 PE=4 SV=1 | K.SVGHLEENGLT[+568.212]IGGASTPPK.G | HexNAc(2)Hex(1) | No | 844.7475 | 3 | 2532.2279 | 2532.2199 | 3.2 | Specific | 575.83 | 517.48 | 4.8 | Nsco_20191108_BC_ZIC_HILIC_H111_typsin_B3.25309.25309.3 | 54.4126 | Trypsin | B3 | 213 |
| BCAL1917 | >tr\|A0A2N9CHU5\|A0A2N9CHU5_9BURK Uncharacterized protein OS=Burkholderia cenocepacia OX=95486 GN=F01_210059 PE=4 SV=1 | P.AATTSATTSTTTSAGTASTSTTAGTTTAAPAPT[+568.212][+100.064]ASAPEAAAKPAK.T | HexNAc(2)Hex(1) 100.064 | No | 1617.4548 | 3 | 4850.3499 | 4850.3288 | 4.4 | NRagged | 594.2 | 582.41 | 4.8 | Nsco_20191108_BC_ZIC_HILIC_H111_typsin_B3.25793.25793.3 | 55.4594 | Trypsin | B3 | 214 |
| BCAL1917 | >tr\|A0A2N9CHU5\|A0A2N9CHU5_9BURK Uncharacterized protein OS=Burkholderia cenocepacia OX=95486 GN=F01_210059 PE=4 SV=1 | K.PAATTSATTSTTTSAGTASTSTTAGTTTAAPAPT[+568.212]ASAPEAAAK.P | HexNAc(2)Hex(1) | No | 1518.0553 | 3 | 4552.1513 | 4551.1327 | 3.4 | Specific | 679.17 | 679.17 | 4.8 | Nsco_20191108_BC_ZIC_HILIC_H111_typsin_B3.27783.27783.3 | 59.2997 | Trypsin | B3 | 215 |
| BCAL1917 | >tr\|A0A2N9CHU5\|A0A2N9CHU5_9BURK Uncharacterized protein OS=Burkholderia cenocepacia OX=95486 GN=F01_210059 PE=4 SV=1 | K.LSKPAATTSATTSTTTSAGTASTTTATAGTTTAAPAPTAS[+568.212]APEAAAKPAK.T | HexNAc(2)Hex(1) | yes | 1294.6423 | 4 | 5175.5475 | 5175.5286 | 3.7 | Specific | 1605.91 | 1543.27 | 67.29 | Nsco_20191108_BC_ZIC_HILIC_H111_typsin_B3.22646.22646.4 | 49.3785 | Trypsin | B3 | 216 |
| BCAL2097 | >tr\|A0A2N9CIF7\|A0A2N9CIF7_9BURK Cell division protein ZipA OS=Burkholderia cenocepacia OX=95486 GN=F01_210252 PE=3 SV=1 | T.PADLQAEATGVDTS[+568.212]AESSEPAAGEEAVPAATHEAAAEPAEPT[+568.212][+100.064]EPVMPAATTVSSAPPAIVDR.R | HexNAc(2)Hex(1), HexNAc(2)Hex(1) 100.064 | No | 1455.4792 | 5 | 7273.3667 | 7272.3334 | 4.1 | NRagged | 234.05 | 234.05 | 0 | Nsco_20191108_BC_ZIC_HILIC_H111_typsin_B2.65630.65630.3 | 127.3905 | Trypsin | B2 | 217 |
| BCAL2097 | >tr\|A0A2N9CIF7\|A0A2N9CIF7_9BURK Cell division protein ZipA OS=Burkholderia cenocepacia OX=95486 GN=F01_210252 PE=3 SV=1 | G.GAAPADTPADLQAEATGVDTSAESS[+568.212][+100.064]EPAAGEEAVPAATHEAAAEPAEPTEPVMPAATTVSSAPPAIVDR.R | HexNAc(2)Hex(1) 100.064 | No | 1458.4841 | 5 | 7288.3912 | 7287.382 | 0.8 | NRagged | 243.47 | 243.47 | 0 | Nsco_20191108_BC_ZIC_HILIC_H111_typsin_B2.65723.65723.5 | 127.6274 | Trypsin | B2 | 218 |
| BCAL2107 | >tr\|A0A2N9CIG6\|A0A2N9CIG6_9BURK Fused putative transporter subunits of ABC superfamily: ATP-binding components OS=Burkholderia cenocepacia OX=95486 GN=yheS PE=4 SV=1 | A.DS[+568.212]AAAK.A | HexNAc(2)Hex(1) | Yes | 565.7523 | 2 | 1130.4974 | 1130.4947 | 2.4 | NRagged | 241.82 | 23.12 | 23.12 | Nsco_20191108_BC_ZIC_HILIC_H111_typsin_B1.1961.1961.2 | 13.168 | Trypsin | B1 | 219 |
| BCAL2161 | >tr\|A0A2N9CIP1\|A0A2N9CIP1_9BURK Putative exported protein OS=Burkholderia cenocepacia OX=95486 GN=F01_230051 PE=4 SV=1 | S.VAPPLQGDGAAPGGAS[+568.212]WPAPPPASGPAPGLPASSVQGT[+568.212]P.- | HexNAc(2)Hex(1), HexNAc(2)Hex(1) | Yes (1 of 2) | 1541.0685 | 3 | 4621.1911 | 4621.1727 | 4 | Non | 465.76 | 361.02 | 34.77 | Nsco_20191108_BC_ZIC_HILIC_H111_thermolysin_B1.62166.62166.3 | 117.7316 | Thermolysin | B1 | 220 |
| BCAL2191 | >tr\|A0A119PBH5\|A0A119PBH5_9BURK Glycine zipper 2TM protein OS=Burkholderia cenocepacia OX=95486 GN=A8E72_00900 PE=4 SV=1 | K.ATSSNEQNGQTQAALIAS[+568.212]QPAVDT[+568.212][+100.064]AAAASAALAAQAQR.Q | HexNAc(2)Hex(1), HexNAc(2)Hex(1) 100.064 | No | 1631.4314 | 3 | 4892.2796 | 4890.2985 | -5.2 | Specific | 294.5 | 294.5 | 0 | Nsco_20191108_BC_ZIC_HILIC_H111_typsin_B3.50609.50609.3 | 103.5296 | Trypsin | B3 | 221 |

| | | | | | | | | | | | | | | | | | | |
|---|---|---|---|---|---|---|---|---|---|---|---|---|---|---|---|---|---|---|
| BCAL2276 | >tr\|A0A2N9CJ70\|A0A2N9CJ70_9BURK Putative membrane protein OS=Burkholderia cenocepacia OX=95486 GN=DFS07_11429 PE=4 SV=1 | K.STIDTAASNAGVPVS[+568.212]SVNYIVHDAGK.G | HexNAc(2)Hex(1) | No | 1047.841 | 3 | 3141.5084 | 3141.4958 | 4 | Specific | 598.71 | 598.71 | 1.2 | Nsco_20191108_BC_ZIC_HILIC_H111_typsin_B2.46258.46258.3 | 89.8439 | Trypsin | B2 | 222 |
| BCAL2398 | >tr\|A0A2N9CKD9\|A0A2N9CKD9_9BURK Periplasmic protein TonB, links inner and outer membranes OS=Burkholderia cenocepacia OX=95486 GN=F01_260105 PE=4 SV=1 | K.ASPPYAADKPIVAVFPPVVPAAPAS[+568.212]SASATR.- | HexNAc(2)Hex(1) | No | 1191.9495 | 3 | 3573.834 | 3573.8211 | 3.6 | Specific | 861.49 | 861.49 | 2.4 | Nsco_20191108_BC_ZIC_HILIC_H111_typsin_B2.54229.54229.3 | 104.0211 | Trypsin | B2 | 223 |
| BCAL2449 | >tr\|A0A144SYR9\|A0A144SYR9_9BURK Diguanylate cyclase (GGDEF) domain protein OS=Burkholderia cenocepacia OX=95486 GN=gmr_4 PE=4 SV=1 | R.RTGSVNNAPGAFS[+568.212]ASGVYPIAER.V | HexNAc(2)Hex(1) | No | 963.7993 | 3 | 2889.3835 | 2889.3749 | 3 | Specific | 465.63 | 396.16 | 2.4 | Nsco_20191108_BC_ZIC_HILIC_H111_typsin_B1.31667.31667.3 | 65.5806 | Trypsin | B1 | 224 |
| BCAL2463 | >tr\|A0A1V2XLQ9\|A0A1V2XLQ9_9BURK Putative exported protein OS=Burkholderia cenocepacia OX=95486 GN=A8F55_33885 PE=4 SV=1 | V.SPGIANVT[+568.212]AS[+568.212]AAPRPEST.A | HexNAc(2)Hex(1), HexNAc(2)Hex(1) | No | 1431.6581 | 2 | 2862.309 | 2862.2997 | 3.2 | Non | 312.22 | 144.26 | 0 | Nsco_20191108_BC_ZIC_HILIC_H111_thermolysin_B3.16208.16208.2 | 37.5219 | Thermolysin | B3 | 225 |
| BCAL2466 | >tr\|A0A144SXW2\|A0A144SXW2_9BURK Ecotin OS=Burkholderia cenocepacia OX=95486 GN=eco PE=4 SV=1 | A.APASAPAVPAES[+568.212]IKMFPQ.A | HexNAc(2)Hex(1) | No | 1190.5817 | 2 | 2380.1561 | 2380.1476 | 3.6 | Non | 620.21 | 426.33 | 65.03 | Nsco_20191108_BC_ZIC_HILIC_H111_thermolysin_B1.47927.47927.2 | 89.9114 | Thermolysin | B1 | 226 |
| BCAL2466 | >tr\|A0A144SXW2\|A0A144SXW2_9BURK Ecotin OS=Burkholderia cenocepacia OX=95486 GN=eco PE=4 SV=1 | A.APASAPAVPAES[+568.212]IKMFPQA.A | HexNAc(2)Hex(1) | No | 1226.5984 | 2 | 2452.1896 | 2451.1847 | 0.6 | Non | 552.79 | 324.07 | 39.88 | Nsco_20191108_BC_ZIC_HILIC_H111_thermolysin_B1.49468.49468.2 | 92.5878 | Thermolysin | B1 | 227 |
| BCAL2466 | >tr\|A0A144SXW2\|A0A144SXW2_9BURK Ecotin OS=Burkholderia cenocepacia OX=95486 GN=eco PE=4 SV=1 | A.APASAPAVPAES[+568.212]IKMFPQAAAGQQR.V | HexNAc(2)Hex(1) | No | 1021.8416 | 3 | 3063.5102 | 3062.4987 | 2.7 | NRagged | 702.19 | 648.84 | 88.78 | Nsco_20191108_BC_ZIC_HILIC_H111_typsin_B2.38634.38634.3 | 76.2715 | Trypsin | B2 | 228 |
| BCAL2607 | >tr\|A0A1V2WZX0\|A0A1V2WZX0_9BURK DNA repair ATPase OS=Burkholderia cenocepacia OX=95486 GN=F01_260195 PE=4 SV=1 | M.KERWEQHRAAAKGAS[+568.212]APAQ.- | HexNAc(2)Hex(1) | Yes | 665.8308 | 4 | 2660.3013 | 2660.2911 | 3.8 | Non | 659.69 | 477.63 | 477.63 | Nsco_20191108_BC_ZIC_HILIC_H111_pepsin_B2.4597.4597.4 | 17.4269 | Pepsin | B2 | 229 |
| BCAL2640 | >tr\|A0A144SWF6\|A0A144SWF6_9BURK Putative exported protein | A.LRGAADRYAPPPAAVPVAAT[+568.212]SGAQGGA.A | HexNAc(2)Hex(1) | No | 1021.1807 | 3 | 3061.5276 | 3060.512 | 4 | Non | 155.97 | 107.63 | 2.4 | Nsco_20191108_BC_ZIC_HILIC_H111_thermolysin_B2.29749.29749.3 | 60.5034 | Thermolysin | B2 | 230 |
| BCAL2640 | >tr\|A0A144SWF6\|A0A144SWF6_9BURK Putative exported protein OS=Burkholderia cenocepacia OX=95486 GN=A8E72_18590 PE=4 SV=1 | A.LRGAADRYAPPPAAVPVAAT[+568.212]SGAQGGAAA.A | HexNAc(2)Hex(1) | No | 1068.2055 | 3 | 3202.6019 | 3202.5862 | 4.9 | Non | 472.56 | 280.84 | 2.4 | Nsco_20191108_BC_ZIC_HILIC_H111_thermolysin_B2.30511.30511.3 | 61.7489 | Thermolysin | B2 | 231 |
| BCAL2640 | >tr\|A0A144SWF6\|A0A144SWF6_9BURK Putative exported protein OS=Burkholderia cenocepacia OX=95486 GN=A8E72_18590 PE=4 SV=1 | A.LRGAADRYAPPPAAVPVAAT[+568.212]SGAQGGAA.A | HexNAc(2)Hex(1) | No | 1044.5257 | 3 | 3131.5624 | 3131.5491 | 4.2 | Non | 406.84 | 276.6 | 2.4 | Nsco_20191108_BC_ZIC_HILIC_H111_thermolysin_B3.29780.29780.3 | 60.9241 | Thermolysin | B3 | 232 |
| BCAL2640 | >tr\|A0A144SWF6\|A0A144SWF6_9BURK Putative exported protein OS=Burkholderia cenocepacia OX=95486 GN=A8E72_18590 PE=4 SV=1 | R.GAADRYAPPPAAVPVAAT[+568.212]SGAQGGAAAAAAPAGTKPANAPR.E | HexNAc(2)Hex(1) | No | 1052.53 | 4 | 4207.0981 | 4207.0902 | 1.9 | Specific | 1111.07 | 1111.07 | 0.8 | Nsco_20191108_BC_ZIC_HILIC_H111_typsin_B1.29076.29076.3 | 60.9225 | Trypsin | B1 | 233 |
| BCAL2640 | >tr\|A0A144SWF6\|A0A144SWF6_9BURK Putative exported protein OS=Burkholderia cenocepacia OX=95486 GN=A8E72_18590 PE=4 SV=1 | R.YAPPPAAVPVAATSGAQGGAAAAAAPAGT[+568.212]KPANAPREPAVRR.V | HexNAc(2)Hex(1) | No | 1112.5768 | 4 | 4447.2853 | 4445.2695 | 2 | Specific | 661.91 | 661.91 | 3.6 | Nsco_20191108_BC_ZIC_HILIC_H111_typsin_B1.25357.25357.4 | 54.6382 | Trypsin | B1 | 234 |
| BCAL2640 | >tr\|A0A144SWF6\|A0A144SWF6_9BURK Putative exported protein OS=Burkholderia cenocepacia OX=95486 GN=A8E72_18590 PE=4 SV=1 | R.GAADRYAPPPAAVPVAAT[+568.212]SGAQGGAAAAAAPAGTKPANAPREPAVR.R | HexNAc(2)Hex(1) | No | 1191.105 | 4 | 4761.3981 | 4759.3922 | -0.2 | Specific | 1118.94 | 1118.94 | 0.8 | Nsco_20191108_BC_ZIC_HILIC_H111_typsin_B1.28719.28719.4 | 60.3221 | Trypsin | B1 | 235 |

| | | | | | | | | | | | | | | | | | | |
|---|---|---|---|---|---|---|---|---|---|---|---|---|---|---|---|---|---|---|
| BCAL2640 | >tr\|A0A144SWF6\|A0A144SWF6_9BURK Putative exported protein OS=Burkholderia cenocepacia OX=95486 GN=A8E72_18590 PE=4 SV=1 | R.YAPPPAAVPVAATSGAQGGAAAAAAPAGT[+568.212]KPANAPREPAVR.R | HexNAc(2)Hex(1) | No | 1073.0513 | 4 | 4289.1832 | 4289.1684 | 3.5 | Specific | 1032.58 | 1032.58 | 1.2 | Nsco_20191108_BC_ZIC_HILIC_H111_typsin_B2.30521.30521.4 | 62.0121 | Trypsin | B2 | 236 |
| BCAL2640 | >tr\|A0A144SWF6\|A0A144SWF6_9BURK Putative exported protein OS=Burkholderia cenocepacia OX=95486 GN=A8E72_18590 PE=4 SV=1 | R.GAADRYAPPPAAVPVAAT[+568.212]SGAQGGAAAAAAPAGTK.P | HexNAc(2)Hex(1) | No | 1200.9278 | 3 | 3600.7688 | 3600.7664 | 0.7 | Specific | 491.08 | 469.86 | 2 | Nsco_20191108_BC_ZIC_HILIC_H111_typsin_B2.32989.32989.5 | 66.322 | Trypsin | B2 | 237 |
| BCAL2640 | >tr\|A0A144SWF6\|A0A144SWF6_9BURK Putative exported protein OS=Burkholderia cenocepacia OX=95486 GN=A8E72_18590 PE=4 SV=1 | R.YAPPPAAVPVAATSGAQGGAAAAAAPAGT[+568.212]KPANAPR.E | HexNAc(2)Hex(1) | No | 1246.2967 | 3 | 3736.8755 | 3736.8664 | 2.4 | Specific | 879.44 | 857.22 | 2.21 | Nsco_20191108_BC_ZIC_HILIC_H111_typsin_B2.31307.31307.3 | 63.3668 | Trypsin | B2 | 238 |
| BCAL2820 | >tr\|A0A1V2WC66\|A0A1V2WC66_9BURK Multidrug transporter OS=Burkholderia cenocepacia OX=95486 GN=oprM PE=3 SV=1 | D.YDKAAAPAPASAT[+568.212]ATNG.- | HexNAc(2)Hex(1) | No | 1072.9942 | 2 | 2144.9811 | 2144.9717 | 4.4 | Non | 551.21 | 411.92 | 1.2 | Nsco_20191108_BC_ZIC_HILIC_H111_pepsin_B1.12913.12913.2 | 31.746 | Pepsin | B1 | 239 |
| BCAL2820 | >tr\|A0A1V2WC66\|A0A1V2WC66_9BURK Multidrug transporter OS=Burkholderia cenocepacia OX=95486 GN=oprM PE=3 SV=1 | A.LGGGWIQRAGETPRAPDAPVDYDKAAAPAPAS[+568.212]ATATNG.- | HexNAc(2)Hex(1) | Yes | 1073.021 | 4 | 4289.0623 | 4289.048 | 3.3 | Non | 572.87 | 533.13 | 60.25 | Nsco_20191108_BC_ZIC_HILIC_H111_thermolysin_B1.40111.40111.4 | 76.5565 | Thermolysin | B1 | 240 |
| BCAL2820 | >tr\|A0A1V2WC66\|A0A1V2WC66_9BURK Multidrug transporter OS=Burkholderia cenocepacia OX=95486 GN=oprM PE=3 SV=1 | R.AGETPRAPDAPVDYDKAAAPAPAS[+568.212]ATATNG.- | HexNAc(2)Hex(1) | No | 1141.2008 | 3 | 3421.5879 | 3421.5765 | 3.3 | Non | 729.11 | 628.95 | 4.8 | Nsco_20191108_BC_ZIC_HILIC_H111_thermolysin_B2.29377.29377.3 | 59.8556 | Thermolysin | B2 | 241 |
| BCAL2820 | >tr\|A0A1V2WC66\|A0A1V2WC66_9BURK Multidrug transporter OS=Burkholderia cenocepacia OX=95486 GN=oprM PE=3 SV=1 | P.VDYDKAAAPAPAS[+568.212]ATATNG.- | HexNAc(2)Hex(1) | No | 1180.5431 | 2 | 2360.0789 | 2359.0671 | 3.6 | Non | 597.71 | 502.43 | 4 | Nsco_20191108_BC_ZIC_HILIC_H111_thermolysin_B2.20925.20925.2 | 45.2202 | Thermolysin | B2 | 242 |
| BCAL2820 | >tr\|A0A1V2WC66\|A0A1V2WC66_9BURK Multidrug transporter OS=Burkholderia cenocepacia OX=95486 GN=oprM PE=3 SV=1 | W.IQRAGETPRAPDAPVDYDKAAAPAPAS[+568.212]ATATNG.- | HexNAc(2)Hex(1) | No | 1273.6166 | 3 | 3818.8353 | 3818.8203 | 3.9 | Non | 676.21 | 638.85 | 4.8 | Nsco_20191108_BC_ZIC_HILIC_H111_thermolysin_B2.25629.25629.3 | 53.4603 | Thermolysin | B2 | 243 |
| BCAL2820 | >tr\|A0A1V2WC66\|A0A1V2WC66_9BURK Multidrug transporter OS=Burkholderia cenocepacia OX=95486 GN=oprM PE=3 SV=1 | K.AAAPAPASAT[+568.212]ATNG.- | HexNAc(2)Hex(1) | No | 869.9 | 2 | 1738.7927 | 1738.7865 | 3.5 | CRagged | 423.92 | 286.45 | 0 | Nsco_20191108_BC_ZIC_HILIC_H111_typsin_B1.11032.11032.2 | 29.9075 | Trypsin | B1 | 244 |
| BCAL2820 | >tr\|A0A1V2WC66\|A0A1V2WC66_9BURK Multidrug transporter OS=Burkholderia cenocepacia OX=95486 GN=oprM PE=3 SV=1 | K.RPDAPVAQAYPAS[+568.212]GVYATQPGAAGAR.S | HexNAc(2)Hex(1) | No | 1037.5055 | 3 | 3110.502 | 3110.4913 | 3.5 | Specific | 699.97 | 637.21 | 12.09 | Nsco_20191108_BC_ZIC_HILIC_H111_typsin_B1.26451.26451.3 | 56.5269 | Trypsin | B1 | 245 |
| BCAL2974 | >tr\|A0A144U3Q1\|A0A144U3Q1_9BURK Uncharacterized protein OS=Burkholderia cenocepacia OX=95486 GN=A8E72_12460 PE=4 SV=1 | M.PFAASAPS[+568.212]QKYQGSKKSAL.R | HexNAc(2)Hex(1) | No | 845.4247 | 3 | 2534.2597 | 2534.2508 | 3.5 | NRagged | 641.1 | 567.25 | 1.2 | Nsco_20191108_BC_ZIC_HILIC_H111_pepsin_B1.12679.12679.3 | 31.416 | Pepsin | B1 | 246 |
| BCAL2974 | >tr\|A0A144U3Q1\|A0A144U3Q1_9BURK Uncharacterized protein OS=Burkholderia cenocepacia OX=95486 GN=A8E72_12460 PE=4 SV=1 | F.RLNEHPQMPFAAS[+568.212]APSQ.K | HexNAc(2)Hex(1) | No | 817.0482 | 3 | 2449.1302 | 2449.1188 | 4.7 | CRagged | 435.18 | 421.83 | 3.39 | Nsco_20191108_BC_ZIC_HILIC_H111_pepsin_B1.26120.26120.3 | 55.4949 | Pepsin | B1 | 247 |
| BCAL2974 | >tr\|A0A144U3Q1\|A0A144U3Q1_9BURK Uncharacterized protein OS=Burkholderia cenocepacia OX=95486 GN=A8E72_12460 PE=4 SV=1 | L.NEHPQMPFAASAPS[+568.212]QKYQG.S | HexNAc(2)Hex(1) | No | 886.0625 | 3 | 2656.1728 | 2656.1719 | 0.4 | CRagged | 463.46 | 463.46 | 9.76 | Nsco_20191108_BC_ZIC_HILIC_H111_pepsin_B2.24698.24698.3 | 51.1852 | Pepsin | B2 | 248 |

| | | | | | | | | | | | | | | | | | | |
|---|---|---|---|---|---|---|---|---|---|---|---|---|---|---|---|---|---|---|
| BCAL2974 | >tr\|A0A144U3Q1\|A0A144U3Q1_9BURK Uncharacterized protein OS=Burkholderia cenocepacia OX=95486 GN=A8E72_12460 PE=4 SV=1 | F.RLNEHPQMPFAAS[+568.212]APSQKYQGSKKSAL.R | HexNAc(2)Hex(1) | Yes | 708.7547 | 5 | 3539.7445 | 3539.7323 | 3.5 | Specific | 1130.77 | 1103.56 | 104.29 | Nsco_20191108_BC_ZIC_HILIC_H111_pepsin_B3.19692.19692.5 | 40.9669 | Pepsin | B3 | 249 |
| BCAL2974 | >tr\|A0A144U3Q1\|A0A144U3Q1_9BURK Uncharacterized protein OS=Burkholderia cenocepacia OX=95486 GN=A8E72_12460 PE=4 SV=1 | L.NEHPQMPFAAS[+568.212]APSQKYQGSKKSAL.R | HexNAc(2)Hex(1) | Yes | 818.3957 | 4 | 3270.5608 | 3270.5471 | 4.2 | Specific | 1164.9 | 1137.25 | 171.47 | Nsco_20191108_BC_ZIC_HILIC_H111_pepsin_B3.19176.19176.4 | 40.0011 | Pepsin | B3 | 250 |
| BCAL2974 | >tr\|A0A144U3Q1\|A0A144U3Q1_9BURK Uncharacterized protein OS=Burkholderia cenocepacia OX=95486 GN=A8E72_12460 PE=4 SV=1 | P.FAASAPSQKYQGS[+568.212]KKSA.L | HexNAc(2)Hex(1) | No | 1162.5658 | 2 | 2324.1244 | 2324.114 | 4.5 | Non | 611.86 | 452.14 | 0.09 | Nsco_20191108_BC_ZIC_HILIC_H111_thermolysin_B2.5849.5849.2 | 21.0158 | Thermolysin | B2 | 251 |
| BCAL2974 | >tr\|A0A144U3Q1\|A0A144U3Q1_9BURK Uncharacterized protein OS=Burkholderia cenocepacia OX=95486 GN=A8E72_12460 PE=4 SV=1 | R.LNEHPQMPFAASAPS[+568.212]QK.Y | HexNAc(2)Hex(1) | No | 1211.0638 | 2 | 2421.1203 | 2421.1126 | 3.2 | Specific | 870.93 | 774.64 | 4.79 | Nsco_20191108_BC_ZIC_HILIC_H111_typsin_B1.22975.22975.2 | 50.0902 | Trypsin | B1 | 252 |
| BCAL2983A | >tr\|A0A2N9CL86\|A0A2N9CL86_9BURK Putative lipoprotein OS=Burkholderia cenocepacia OX=95486 GN=F01_310136 PE=4 SV=1 | S.ADASAPVAGTRPAVTSLSGGASSAASGAVAT[+568.212]DAAAQGNVAELTQMLHDGR.I | HexNAc(2)Hex(1) | No | 1301.8806 | 4 | 5204.5006 | 5204.4772 | 4.5 | NRagged | 604.47 | 604.47 | 0 | Nsco_20191108_BC_ZIC_HILIC_H111_typsin_B3.53186.53186.5 | 108.5861 | Trypsin | B3 | 253 |
| BCAL3033 | >tr\|A0A2N9CM16\|A0A2N9CM16_9BURK Outer-membrane lipoprotein carrier protein OS=Burkholderia cenocepacia OX=95486 GN=lolA PE=3 SV=1 | Q.IVKAPAKGAS[+568.212]AAQAAPKPTDN.S | HexNAc(2)Hex(1) | Yes | 644.5867 | 4 | 2575.3249 | 2574.3145 | 2.8 | Non | 729.64 | 648.17 | 408.82 | Nsco_20191108_BC_ZIC_HILIC_H111_pepsin_B3.10005.10005.4 | 25.4729 | Pepsin | B3 | 254 |
| BCAL3033 | >tr\|A0A2N9CM16\|A0A2N9CM16_9BURK Outer-membrane lipoprotein carrier protein OS=Burkholderia cenocepacia OX=95486 GN=lolA PE=3 SV=1 | Q.IVKAPAKGAS[+568.212]AAQAAPKPTDNSSGTF.V | HexNAc(2)Hex(1) | Yes | 764.1379 | 4 | 3053.5298 | 3053.5161 | 4.5 | NRagged | 1082.94 | 1032.62 | 506.95 | Nsco_20191108_BC_ZIC_HILIC_H111_pepsin_B3.16451.16451.4 | 35.4703 | Pepsin | B3 | 255 |
| BCAL3033 | >tr\|A0A2N9CM16\|A0A2N9CM16_9BURK Outer-membrane lipoprotein carrier protein OS=Burkholderia cenocepacia OX=95486 GN=lolA PE=3 SV=1 | Q.IVKAPAKGAS[+568.212]AAQAAPKPTDNSSGTFV.F | HexNAc(2)Hex(1) | Yes | 788.9051 | 4 | 3152.5986 | 3152.5845 | 4.5 | Non | 805.9 | 805.9 | 378.72 | Nsco_20191108_BC_ZIC_HILIC_H111_pepsin_B3.19759.19759.4 | 41.0925 | Pepsin | B3 | 256 |
| BCAL3033 | >tr\|A0A2N9CM16\|A0A2N9CM16_9BURK Outer-membrane lipoprotein carrier protein OS=Burkholderia cenocepacia OX=95486 GN=lolA PE=3 SV=1 | Q.IVKAPAKGAS[+568.212]AAQAAPKPTDNSSGT.F | HexNAc(2)Hex(1) | Yes | 727.371 | 4 | 2906.4621 | 2906.4477 | 5 | Non | 727.77 | 543.13 | 403.44 | Nsco_20191108_BC_ZIC_HILIC_H111_thermolysin_B1.9738.9738.4 | 26.6693 | Thermolysin | B1 | 257 |
| BCAL3033 | >tr\|A0A2N9CM16\|A0A2N9CM16_9BURK Outer-membrane lipoprotein carrier protein OS=Burkholderia cenocepacia OX=95486 GN=lolA PE=3 SV=1 | Q.IVKAPAKGAS[+568.212]AAQAAPKPTDNSSGTFVF.A | HexNAc(2)Hex(1) | Yes | 825.6715 | 4 | 3299.6643 | 3299.6529 | 3.4 | NRagged | 451.66 | 424.44 | 313.42 | Nsco_20191108_BC_ZIC_HILIC_H111_thermolysin_B3.26971.26971.4 | 55.9849 | Thermolysin | B3 | 258 |
| BCAL3033 | >tr\|A0A2N9CM16\|A0A2N9CM16_9BURK Outer-membrane lipoprotein carrier protein OS=Burkholderia cenocepacia OX=95486 GN=lolA PE=3 SV=1 | K.GAS[+568.212]AAQAAPKPTDNSSGTFVFARPGK.F | HexNAc(2)Hex(1) | Yes | 776.1301 | 4 | 3101.4988 | 3101.4909 | 2.5 | Specific | 1141.01 | 1022.65 | 329.49 | Nsco_20191108_BC_ZIC_HILIC_H111_typsin_B1.21425.21425.4 | 47.2372 | Trypsin | B1 | 259 |
| BCAL3033 | >tr\|A0A2N9CM16\|A0A2N9CM16_9BURK Outer-membrane lipoprotein carrier protein OS=Burkholderia cenocepacia OX=95486 GN=lolA PE=3 SV=1 | K.GASAAQAAPKPT[+568.212]DNSSGTFVFAR.P | HexNAc(2)Hex(1) | No | 940.448 | 3 | 2819.3293 | 2819.3218 | 2.7 | Specific | 945.35 | 771.26 | 9.1 | Nsco_20191108_BC_ZIC_HILIC_H111_typsin_B3.26104.26104.3 | 56.044 | Trypsin | B3 | 260 |
| BCAL3166 | >tr\|A0A1V2WZ65\|A0A1V2WZ65_9BURK BON domain-containing protein OS=Burkholderia cenocepacia OX=95486 GN=A8E72_33585 PE=4 SV=1 | A.TEAPAAATSAPKAAAKT[+568.212]AKKANRKLGY.A | HexNAc(2)Hex(1) | No | 1085.9139 | 3 | 3255.7273 | 3254.7114 | 3.8 | Non | 772.23 | 364.05 | 9.6 | Nsco_20191108_BC_ZIC_HILIC_H111_pepsin_B3.12083.12083.3 | 28.6806 | Pepsin | B3 | 261 |

| | | | | | | | | | | | | | | | | | | |
|---|---|---|---|---|---|---|---|---|---|---|---|---|---|---|---|---|---|---|
| BCAL3377 | >tr\|A0A142PIH6\|A0A142PIH6_9BURK Outer membrane protein assembly factor BamE OS=Burkholderia cenocepacia OX=95486 GN=bamE PE=3 SV=1 | D.IDGDRGGKKAKAAAAAAKKAS[+568.212]EAAA.A | HexNAc(2)Hex(1) | Yes | 565.6979 | 5 | 2824.4605 | 2824.4534 | 2.5 | Non | 649.03 | 597.74 | 597.74 | Nsco_20191108_BC_ZIC_HILIC_H111_pepsin_B2.4588.4588.5 | 17.4183 | Pepsin | B2 | 262 |
| BCAL3377 | >tr\|A0A142PIH6\|A0A142PIH6_9BURK Outer membrane protein assembly factor BamE OS=Burkholderia cenocepacia OX=95486 GN=bamE PE=3 SV=1 | A.AAAASPATVPAS[+568.212]GAAVDQDANAQAARAANRATNQ.V | HexNAc(2)Hex(1) | No | 1249.5988 | 3 | 3746.7819 | 3746.77 | 3.2 | Non | 766.21 | 698.32 | 9.6 | Nsco_20191108_BC_ZIC_HILIC_H111_thermolysin_B1.28242.28242.3 | 55.8564 | Thermolysin | B1 | 263 |
| BCAL3377 | >tr\|A0A142PIH6\|A0A142PIH6_9BURK Outer membrane protein assembly factor BamE OS=Burkholderia cenocepacia OX=95486 GN=bamE PE=3 SV=1 | A.ASPATVPAS[+568.212]GAAVDQDANAQAARAANRATNQ.V | HexNAc(2)Hex(1) | No | 1178.8954 | 3 | 3534.6716 | 3533.6586 | 2.7 | Non | 624.68 | 580.63 | 3.86 | Nsco_20191108_BC_ZIC_HILIC_H111_thermolysin_B1.26493.26493.3 | 52.7385 | Thermolysin | B1 | 264 |
| BCAL3377 | >tr\|A0A142PIH6\|A0A142PIH6_9BURK Outer membrane protein assembly factor BamE OS=Burkholderia cenocepacia OX=95486 GN=bamE PE=3 SV=1 | A.AAAAKKASEAAAAASAAQAAAAAS[+568.212]PAT[+568.212]VPASGAAVDQDANAQ.A | HexNAc(2)Hex(1), HexNAc(2)Hex(1) | No | 1601.7587 | 3 | 4803.2614 | 4801.2393 | 3.2 | Non | 375.64 | 169.53 | 4.8 | Nsco_20191108_BC_ZIC_HILIC_H111_thermolysin_B1.38320.38320.3 | 73.5381 | Thermolysin | B1 | 265 |
| BCAL3377 | >tr\|A0A142PIH6\|A0A142PIH6_9BURK Outer membrane protein assembly factor BamE OS=Burkholderia cenocepacia OX=95486 GN=bamE PE=3 SV=1 | A.AASPATVPAS[+568.212]GAAVDQDANAQAARAANRATNQ.V | HexNAc(2)Hex(1) | No | 1202.2415 | 3 | 3604.71 | 3604.6957 | 3.9 | Non | 754.14 | 628.56 | 9.6 | Nsco_20191108_BC_ZIC_HILIC_H111_thermolysin_B1.27168.27168.3 | 53.8751 | Thermolysin | B1 | 266 |
| BCAL3377 | >tr\|A0A142PIH6\|A0A142PIH6_9BURK Outer membrane protein assembly factor BamE OS=Burkholderia cenocepacia OX=95486 GN=bamE PE=3 SV=1 | A.AAASPATVPAS[+568.212]GAAVDQDANAQAARAANR.A | HexNAc(2)Hex(1) | No | 1088.1919 | 3 | 3262.5612 | 3261.5466 | 3.5 | Non | 670.85 | 352.84 | 9.6 | Nsco_20191108_BC_ZIC_HILIC_H111_thermolysin_B2.22544.22544.3 | 48.0203 | Thermolysin | B2 | 267 |
| BCAL3377 | >tr\|A0A142PIH6\|A0A142PIH6_9BURK Outer membrane protein assembly factor BamE OS=Burkholderia cenocepacia OX=95486 GN=bamE PE=3 SV=1 | A.AAASPATVPAS[+568.212]GAAVDQDANAQ.A | HexNAc(2)Hex(1) | No | 1276.0848 | 2 | 2551.1622 | 2551.153 | 3.6 | Non | 310.38 | 216.69 | 2.4 | Nsco_20191108_BC_ZIC_HILIC_H111_thermolysin_B2.27766.27766.2 | 57.1181 | Thermolysin | B2 | 268 |
| BCAL3377 | >tr\|A0A142PIH6\|A0A142PIH6_9BURK Outer membrane protein assembly factor BamE OS=Burkholderia cenocepacia OX=95486 GN=bamE PE=3 SV=1 | Q.AAAAASPATVPAS[+568.212]GAAVDQDANAQAARAANRATNQ.V | HexNAc(2)Hex(1) | No | 1273.9446 | 3 | 3819.8192 | 3817.8071 | 1.4 | Non | 725.88 | 651.03 | 2.45 | Nsco_20191108_BC_ZIC_HILIC_H111_thermolysin_B3.27596.27596.3 | 57.259 | Thermolysin | B3 | 269 |
| BCAL3377 | >tr\|A0A142PIH6\|A0A142PIH6_9BURK Outer membrane protein assembly factor BamE OS=Burkholderia cenocepacia OX=95486 GN=bamE PE=3 SV=1 | A.AAAASPATVPAS[+568.212]GAAVDQDANAQ.A | HexNAc(2)Hex(1) | No | 1311.6047 | 2 | 2622.2022 | 2622.1901 | 4.6 | Non | 398.76 | 278.3 | 12.86 | Nsco_20191108_BC_ZIC_HILIC_H111_thermolysin_B3.28288.28288.2 | 58.4224 | Thermolysin | B3 | 270 |
| BCAL3377 | >tr\|A0A142PIH6\|A0A142PIH6_9BURK Outer membrane protein assembly factor BamE OS=Burkholderia cenocepacia OX=95486 GN=bamE PE=3 SV=1 | A.AAASPATVPAS[+568.212]GAAVDQDANAQAARAANRATNQ.V | HexNAc(2)Hex(1) | No | 1225.9203 | 3 | 3675.7465 | 3675.7329 | 3.7 | Non | 710.7 | 638.62 | 5.75 | Nsco_20191108_BC_ZIC_HILIC_H111_thermolysin_B3.26468.26468.3 | 55.226 | Thermolysin | B3 | 271 |

| | | | | | | | | | | | | | | | | | | |
|---|---|---|---|---|---|---|---|---|---|---|---|---|---|---|---|---|---|---|
| BCAL3377 | >tr\|A0A142PIH6\|A0A142PIH6_9BURK Outer membrane protein assembly factor BamE OS=Burkholderia cenocepacia OX=95486 GN=bamE PE=3 SV=1 | K.KASEAAAAAS[+568.212]AAQAAAAASPATVPASGAAVDQDANAQAAR.A | HexNAc(2)Hex(1) | Yes | 1030.498 | 4 | 4118.97 | 4118.9596 | 2.5 | Specific | 1133.19 | 1029.66 | 156.35 | Nsco_20191108_BC_ZIC_HILIC_H111_typsin_B2.39556.39556.4 | 77.8852 | Trypsin | B2 | 272 |
| BCAL3377 | >tr\|A0A142PIH6\|A0A142PIH6_9BURK Outer membrane protein assembly factor BamE OS=Burkholderia cenocepacia OX=95486 GN=bamE PE=3 SV=1 | A.AAAAKKASEAAAAASAAQAAAAAS[+568.212][+100.064]PAT[+568.212][+100.064]VPASGAAVDQDANAQAAR.A | HexNAc(2)Hex(1) 100.064,HexNAc(2)Hex(1) 100.064 | No | 1326.395 | 4 | 5302.5583 | 5299.5426 | 1.1 | NRagged | 149.57 | 149.57 | 2.4 | Nsco_20191108_BC_ZIC_HILIC_H111_typsin_B2.34416.34416.4 | 68.7503 | Trypsin | B2 | 273 |
| BCAL3377 | >tr\|A0A142PIH6\|A0A142PIH6_9BURK Outer membrane protein assembly factor BamE OS=Burkholderia cenocepacia OX=95486 GN=bamE PE=3 SV=1 | K.ASEAAAAASAAQAAAAASPAT[+568.212]VPAS[+568.212]GAAVDQDANAQAAR.A | HexNAc(2)Hex(1), HexNAc(2)Hex(1) | No | 1520.3664 | 3 | 4559.0846 | 4559.0762 | 1.8 | Specific | 1089.29 | 1044.6 | 0 | Nsco_20191108_BC_ZIC_HILIC_H111_typsin_B2.43338.43338.3 | 84.5161 | Trypsin | B2 | 274 |
| BCAL3377 | >tr\|A0A142PIH6\|A0A142PIH6_9BURK Outer membrane protein assembly factor BamE OS=Burkholderia cenocepacia OX=95486 GN=bamE PE=3 SV=1 | A.SEAAAAASAAQAAAAASPAT[+568.212][+100.064]VPAS[+568.212]GAAVDQDANAQAAR.A | HexNAc(2)Hex(1) 100.064,HexNAc(2)Hex(1) | No | 1530.0365 | 3 | 4588.0949 | 4588.1031 | -1.8 | NRagged | 801.32 | 801.32 | 0 | Nsco_20191108_BC_ZIC_HILIC_H111_typsin_B2.44897.44897.3 | 87.3539 | Trypsin | B2 | 275 |
| BCAL3377 | >tr\|A0A142PIH6\|A0A142PIH6_9BURK Outer membrane protein assembly factor BamE OS=Burkholderia cenocepacia OX=95486 GN=bamE PE=3 SV=1 | A.KKASEAAAAAS[+568.212]AAQAAAAASPAT[+568.212]VPASGAAVDQDANAQAAR.A | HexNAc(2)Hex(1), HexNAc(2)Hex(1) | No | 1204.5761 | 4 | 4815.2825 | 4815.2662 | 3.4 | NRagged | 879.73 | 879.73 | 6.08 | Nsco_20191108_BC_ZIC_HILIC_H111_typsin_B3.30338.30338.4 | 64.2638 | Trypsin | B3 | 276 |
| BCAL3377 | >tr\|A0A142PIH6\|A0A142PIH6_9BURK Outer membrane protein assembly factor BamE OS=Burkholderia cenocepacia OX=95486 GN=bamE PE=3 SV=1 | A.SAAQAAAAAS[+568.212]PAT[+568.212]VPAS[+568.212][+100.064]GAAVDQDANAQAAR.A | HexNAc(2)Hex(1), HexNAc(2)Hex(1), HexNAc(2)Hex(1) 100.064 | No | 1529.6973 | 3 | 4587.0774 | 4585.0545 | 3.5 | NRagged | 374.52 | 374.52 | 0 | Nsco_20191108_BC_ZIC_HILIC_H111_typsin_B3.42023.42023.3 | 87.0473 | Trypsin | B3 | 277 |
| BCAL3469 | >tr\|A0A142PJ18\|A0A142PJ18_9BURK Cell division protein FtsL OS=Burkholderia cenocepacia OX=95486 GN=ftsL PE=3 SV=1 | A.IDAPIPAS[+568.212]ADTAGKGKGGAR.- | HexNAc(2)Hex(1) | No | 606.057 | 4 | 2421.206 | 2421.1991 | 2.9 | Non | 579.19 | 487.06 | 2.4 | Nsco_20191108_BC_ZIC_HILIC_H111_thermolysin_B2.12797.12797.4 | 31.8597 | Thermolysin | B2 | 278 |
| BCAM0157 | >tr\|A0A2N9CQK4\|A0A2N9CQK4_9BURK Putative exported protein OS=Burkholderia cenocepacia OX=95486 GN=F01_420142 PE=4 SV=1 | R.AVDC[+57.021]GQLDAATSGPDDNFRPPAS[+568.212]GTVIGTGR.A | HexNAc(2)Hex(1) | No | 1224.2327 | 3 | 3670.6834 | 3670.6661 | 4.7 | Specific | 746.57 | 746.57 | 4.8 | Nsco_20191108_BC_ZIC_HILIC_H111_typsin_B1.39845.39845.3 | 79.8131 | Trypsin | B1 | 279 |
| BCAM0264 | >tr\|A0A2N9CNS4\|A0A2N9CNS4_9BURK Receptor family ligand-binding protein OS=Burkholderia cenocepacia OX=95486 GN=F01_390005 PE=4 SV=1 | I.ATAGAS[+568.212]T[+568.212][+100.064]GAIGVSSYF.M | HexNAc(2)Hex(1), HexNAc(2)Hex(1) 100.064 | No | 1348.6044 | 2 | 2696.2015 | 2696.1935 | 3 | NRagged | 244.27 | 96.45 | 0 | Nsco_20191108_BC_ZIC_HILIC_H111_pepsin_B2.17819.17819.2 | 38.7448 | Pepsin | B2 | 280 |
| BCAM0505 | >tr\|A0A1V2WB29\|A0A1V2WB29_9BURK BON domain protein OS=Burkholderia cenocepacia OX=95486 GN=osmY_2 PE=4 SV=1 | D.SGMASES[+568.212]NQPVT[+568.212]DTWITTKVKGE.L | HexNAc(2)Hex(1), HexNAc(2)Hex(1) | No | 1202.5412 | 3 | 3605.609 | 3602.6048 | -1.6 | Non | 253.84 | 253.84 | 4.8 | Nsco_20191108_BC_ZIC_HILIC_H111_thermolysin_B1.49324.49324.3 | 92.3335 | Thermolysin | B1 | 281 |
| BCAM0505 | >tr\|A0A1V2WB29\|A0A1V2WB29_9BURK BON domain protein OS=Burkholderia cenocepacia OX=95486 GN=osmY_2 PE=4 SV=1 | A.QAS[+568.212]STDSGMASESNQPVTDTWITTKVKGELATTDCVKSTDISVKT.V | HexNAc(2)Hex(1) | No | 1311.1262 | 4 | 5241.4831 | 5240.4897 | -1.9 | Non | 319.81 | 319.81 | 1.4 | Nsco_20191108_BC_ZIC_HILIC_H111_thermolysin_B1.55291.55291.4 | 103.1577 | Thermolysin | B1 | 282 |

| | | | | | | | | | | | | | | | | | | |
|---|---|---|---|---|---|---|---|---|---|---|---|---|---|---|---|---|---|---|
| BCAM0505 | >tr\|A0A1V2WB29\|A0A1V2WB29_9BURK BON domain protein OS=Burkholderia cenocepacia OX=95486 GN=osmY_2 PE=4 SV=1 | D.SGMASESNQPVTDTWIT[+568.212]TKVKGELATTDGVKS[+568.212]TDISVKT.V | HexNAc(2)Hex(1), HexNAc(2)Hex(1) | No | 1306.3752 | 4 | 5222.4792 | 5219.4669 | 0.4 | Non | 555.79 | 555.79 | 0 | Nsco_20191108_BC_ZIC_HILIC_H111_thermolysin_B3.52234.52234.4 | 103.1762 | Thermolysin | B3 | 283 |
| BCAM0505 | >tr\|A0A1V2WB29\|A0A1V2WB29_9BURK BON domain protein OS=Burkholderia cenocepacia OX=95486 GN=osmY_2 PE=4 SV=1 | A.QASSTDSGMAS[+568.212]ESNQPVTDTWITTK.V | HexNAc(2)Hex(1) | No | 1607.2037 | 2 | 3213.4002 | 3210.4002 | -3.1 | NRagged | 537.66 | 537.66 | 2.59 | Nsco_20191108_BC_ZIC_HILIC_H111_typsin_B1.40204.40204.2 | 80.4063 | Trypsin | B1 | 284 |
| BCAM0505 | >tr\|A0A1V2WB29\|A0A1V2WB29_9BURK BON domain protein OS=Burkholderia cenocepacia OX=95486 GN=osmY_2 PE=4 SV=1 | Q.ASSTDSGMASES[+568.212]NQPVTDTWITTK.V | HexNAc(2)Hex(1) | No | 1541.6788 | 2 | 3082.3502 | 3082.3416 | 2.8 | NRagged | 715.39 | 715.39 | 1.66 | Nsco_20191108_BC_ZIC_HILIC_H111_typsin_B2.39570.39570.2 | 77.9153 | Trypsin | B2 | 285 |
| BCAM0505 | >tr\|A0A1V2WB29\|A0A1V2WB29_9BURK BON domain protein OS=Burkholderia cenocepacia OX=95486 GN=osmY_2 PE=4 SV=1 | A.HAQASSTDSGMASES[+568.212]NQPVTDTWITTK.V | HexNAc(2)Hex(1) | No | 1140.1741 | 3 | 3418.5077 | 3418.4962 | 3.4 | NRagged | 759.04 | 759.04 | 2.4 | Nsco_20191108_BC_ZIC_HILIC_H111_typsin_B3.29909.29909.3 | 63.4467 | Trypsin | B3 | 286 |
| BCAM0690 | >tr\|A0A1V2Y2A4\|A0A1V2Y2A4_9BURK Flagellar motor protein OS=Burkholderia cenocepacia OX=95486 GN=A8F55_06330 PE=4 SV=1 | R.ALIDAGVPASSVFAAAFGS[+568.212]EQPVSSNADDEGRAK.N | HexNAc(2)Hex(1) | No | 1301.6159 | 3 | 3902.8332 | 3902.8302 | 0.8 | Specific | 635.27 | 600.91 | 3.7 | Nsco_20191108_BC_ZIC_HILIC_H111_typsin_B1.61806.61806.3 | 115.5056 | Trypsin | B1 | 287 |
| BCAM0712 | >tr\|A0A2N9CPU2\|A0A2N9CPU2_9BURK Cobalt-zinc-cadmium resistance protein CzcB OS=Burkholderia cenocepacia OX=95486 GN=czcB PE=3 SV=1 | G.GGGASAPT[+568.212]AAEVAQPAAGAGAR.G | HexNAc(2)Hex(1) | No | 1204.0726 | 2 | 2407.138 | 2406.1267 | 3.3 | NRagged | 657.6 | 562.44 | 48.38 | Nsco_20191108_BC_ZIC_HILIC_H111_typsin_B2.19920.19920.2 | 42.9078 | Trypsin | B2 | 288 |
| BCAM0936 | >tr\|A0A1V2XJK6\|A0A1V2XJK6_9BURK Mechanosensitive ion channel protein OS=Burkholderia cenocepacia OX=95486 GN=A8F55_37840 PE=4 SV=1 | A.AAPAPAAASAAS[+568.212]DAAPALT[+568.212]PQQAR.Q | HexNAc(2)Hex(1), HexNAc(2)Hex(1) | No | 1104.5209 | 3 | 3311.5482 | 3311.5384 | 2.9 | NRagged | 767.39 | 758.8 | 31.72 | Nsco_20191108_BC_ZIC_HILIC_H111_typsin_B3.27082.27082.3 | 57.9011 | Trypsin | B3 | 289 |
| BCAM0988 | >tr\|A0A2N9CRH5\|A0A2N9CRH5_9BURK Putative exported protein OS=Burkholderia cenocepacia OX=95486 GN=F01_420505 PE=4 SV=1 | Q.PAAPTAGAS[+568.212]GPHVWSGA.I | HexNAc(2)Hex(1) | No | 1051.4849 | 2 | 2101.9625 | 2101.956 | 3.1 | Non | 557.2 | 394.13 | 5.76 | Nsco_20191108_BC_ZIC_HILIC_H111_pepsin_B2.25604.25604.2 | 52.8615 | Pepsin | B2 | 290 |
| BCAM0996 | >tr\|A0A2N9CRL1\|A0A2N9CRL1_9BURK Sporulation related OS=Burkholderia cenocepacia OX=95486 GN=F01_420513 PE=4 SV=1 | Q.AVAPRDDDVS[+568.212]DVQAGVAHDEPPASDTT.V | HexNAc(2)Hex(1) | No | 1101.8256 | 3 | 3303.4622 | 3303.4507 | 3.5 | Non | 511.96 | 384.22 | 7.61 | Nsco_20191108_BC_ZIC_HILIC_H111_thermolysin_B2.28438.28438.3 | 58.3006 | Thermolysin | B2 | 291 |
| BCAM0996 | >tr\|A0A2N9CRL1\|A0A2N9CRL1_9BURK Sporulation related OS=Burkholderia cenocepacia OX=95486 GN=F01_420513 PE=4 SV=1 | A.VAPRDDDVSDVQAGVAHDEPPAS[+568.212]DTT.V | HexNAc(2)Hex(1) | No | 1078.1468 | 3 | 3232.4257 | 3232.4135 | 3.8 | Non | 609.61 | 485.16 | 1.1 | Nsco_20191108_BC_ZIC_HILIC_H111_thermolysin_B2.27181.27181.3 | 56.0176 | Thermolysin | B2 | 292 |
| BCAM0996 | >tr\|A0A2N9CRL1\|A0A2N9CRL1_9BURK Sporulation related OS=Burkholderia cenocepacia OX=95486 GN=F01_420513 PE=4 SV=1 | K.DAAKPAAKPDTTTTASVTPPKPAPKPAAPAAKPAAPKPAPATVANAGPAS[+568.212]PDSGDASSPASPAGAR.F | HexNAc(2)Hex(1) | No | 1106.2348 | 6 | 6632.3723 | 6628.3541 | 0.7 | Specific | 431.46 | 429.91 | 31.05 | Nsco_20191108_BC_ZIC_HILIC_H111_typsin_B1.20631.20631.6 | 45.7781 | Trypsin | B1 | 293 |
| BCAM0996 | >tr\|A0A2N9CRL1\|A0A2N9CRL1_9BURK Sporulation related OS=Burkholderia cenocepacia OX=95486 GN=F01_420513 PE=4 SV=1 | P.APATVANAGPASPDS[+568.212][+100.064]GDASSPASPAGAR.F | HexNAc(2)Hex(1) 100.064 | No | 1560.214 | 2 | 3119.4208 | 3118.4298 | -4 | NRagged | 426.55 | 417.46 | 2.4 | Nsco_20191108_BC_ZIC_HILIC_H111_typsin_B1.22179.22179.2 | 48.6827 | Trypsin | B1 | 294 |

| | | | | | | | | | | | | | | | | | | |
|---|---|---|---|---|---|---|---|---|---|---|---|---|---|---|---|---|---|---|
| BCAM0996 | >tr\|A0A2N9CRL1\|A0A2N9CRL1_9BURK Sporulation related OS=Burkholderia cenocepacia OX=95486 GN=F01_420513 PE=4 SV=1 | K.PAAPAAKPAAPKPAPAT[+568.212]VANAGPASPDSGDASSPASPAGAR.F | HexNAc(2)Hex(1) | No | 1048.2685 | 4 | 4190.0521 | 4186.0422 | -0.9 | Specific | 550.99 | 288.16 | 8 | Nsco_20191108_BC_ZIC_HILIC_H111_typsin_B2.20495.20495.4 | 44.0185 | Trypsin | B2 | 295 |
| BCAM0996 | >tr\|A0A2N9CRL1\|A0A2N9CRL1_9BURK Sporulation related OS=Burkholderia cenocepacia OX=95486 GN=F01_420513 PE=4 SV=1 | R.DDDVSDVQAGVAHDEPPAS[+568.212]DTTVAAAPAPAPK.D | HexNAc(2)Hex(1) | No | 1228.2275 | 3 | 3682.6679 | 3682.6614 | 1.8 | Specific | 1148.33 | 1148.33 | 4.8 | Nsco_20191108_BC_ZIC_HILIC_H111_typsin_B2.37298.37298.3 | 73.8101 | Trypsin | B2 | 296 |
| BCAM0996 | >tr\|A0A2N9CRL1\|A0A2N9CRL1_9BURK Sporulation related OS=Burkholderia cenocepacia OX=95486 GN=F01_420513 PE=4 SV=1 | P.AAPKPAPAT[+568.212][+100.064]VANAGPASPDSGDASSPASPAGAR.F | HexNAc(2)Hex(1) 100.064 | No | 1195.2413 | 3 | 3583.7093 | 3582.7045 | 0.4 | NRagged | 293.98 | 293.98 | 2.1 | Nsco_20191108_BC_ZIC_HILIC_H111_typsin_B2.21349.21349.3 | 45.5733 | Trypsin | B2 | 297 |
| BCAM0996 | >tr\|A0A2N9CRL1\|A0A2N9CRL1_9BURK Sporulation related OS=Burkholderia cenocepacia OX=95486 GN=F01_420513 PE=4 SV=1 | P.AAPAAKPAAPKPAPAT[+568.212][+100.064]VANAGPASPDSGDASSPASPAGAR.F | HexNAc(2)Hex(1) 100.064 | No | 1048.7671 | 4 | 4192.0467 | 4189.0535 | -4 | NRagged | 299.3 | 299.3 | 7.6 | Nsco_20191108_BC_ZIC_HILIC_H111_typsin_B3.19856.19856.2 | 43.877 | Trypsin | B3 | 298 |
| BCAM1550 | >tr\|A0A142PC85\|A0A142PC85_9BURK Peptidoglycan-associated protein OS=Burkholderia cenocepacia OX=95486 GN=pal PE=3 SV=1 | A.AKTPENAGAAPEPS[+568.212]SETVA.T | HexNAc(2)Hex(1) | No | 1198.0492 | 2 | 2395.0911 | 2395.0882 | 1.2 | Non | 443.15 | 275.96 | 1.2 | Nsco_20191108_BC_ZIC_HILIC_H111_pepsin_B1.16299.16299.2 | 37.1808 | Pepsin | B1 | 299 |
| BCAM1550 | >tr\|A0A142PC85\|A0A142PC85_9BURK Peptidoglycan-associated protein OS=Burkholderia cenocepacia OX=95486 GN=pal PE=3 SV=1 | H.NAAKTPENAGAAPEPS[+568.212]SETVA.T | HexNAc(2)Hex(1) | No | 1290.5912 | 2 | 2580.1752 | 2580.1683 | 2.7 | Non | 468 | 427.26 | 1.2 | Nsco_20191108_BC_ZIC_HILIC_H111_pepsin_B1.17269.17269.2 | 38.6954 | Pepsin | B1 | 300 |
| BCAM1550 | >tr\|A0A142PC85\|A0A142PC85_9BURK Peptidoglycan-associated protein OS=Burkholderia cenocepacia OX=95486 GN=pal PE=3 SV=1 | T.PENAGAAPEPSSETVAT[+568.212][+100.064]VTADDLNNPNSPLAK.R | HexNAc(2)Hex(1) 100.064 | No | 1282.9384 | 3 | 3846.8007 | 3845.7938 | 0.9 | NRagged | 581.29 | 581.29 | 2.4 | Nsco_20191108_BC_ZIC_HILIC_H111_typsin_B1.47137.47137.3 | 92.0418 | Trypsin | B1 | 301 |
| BCAM1550 | >tr\|A0A142PC85\|A0A142PC85_9BURK Peptidoglycan-associated protein OS=Burkholderia cenocepacia OX=95486 GN=pal PE=3 SV=1 | K.TPENAGAAPEPSSETVATVT[+568.212]ADDLNNPNSPLAKR.S | HexNAc(2)Hex(1) | No | 1335.6364 | 3 | 4004.8945 | 4002.8786 | 2.3 | Specific | 858 | 198.34 | 0 | Nsco_20191108_BC_ZIC_HILIC_H111_typsin_B2.42072.42072.3 | 82.3448 | Trypsin | B2 | 302 |
| BCAM1550 | >tr\|A0A142PC85\|A0A142PC85_9BURK Peptidoglycan-associated protein OS=Burkholderia cenocepacia OX=95486 GN=pal PE=3 SV=1 | K.TPENAGAAPEPSSETVAT[+568.212]VTADDLNNPNSPLAK.R | HexNAc(2)Hex(1) | No | 1282.9346 | 3 | 3846.7893 | 3846.7775 | 3.1 | Specific | 950.71 | 313.89 | 2.4 | Nsco_20191108_BC_ZIC_HILIC_H111_typsin_B2.47780.47780.3 | 92.3752 | Trypsin | B2 | 303 |
| BCAM1550 | >tr\|A0A142PC85\|A0A142PC85_9BURK Peptidoglycan-associated protein OS=Burkholderia cenocepacia OX=95486 GN=pal PE=3 SV=1 | T.PENAGAAPEPSSETVAT[+568.212][+100.064]VTADDLNNPNSPLAKR.S | HexNAc(2)Hex(1) 100.064 | No | 1334.9743 | 3 | 4002.9083 | 4001.8949 | 2.5 | NRagged | 412.57 | 412.57 | 0 | Nsco_20191108_BC_ZIC_HILIC_H111_typsin_B3.39599.39599.3 | 82.3936 | Trypsin | B3 | 304 |
| BCAM1669 | >tr\|A0A1V2X7H1\|A0A1V2X7H1_9BURK Purine nucleoside phosphorylase OS=Burkholderia cenocepacia OX=95486 GN=A8E72_24650 PE=4 SV=1 | A.AQARVHGIDNSGAGSQPAAT[+568.212]VEGGAPVVRAQNPRDSVY.F | HexNAc(2)Hex(1) | No | 875.0314 | 5 | 4371.1279 | 4371.1084 | 4.5 | Non | 554.26 | 554.26 | 2.4 | Nsco_20191108_BC_ZIC_HILIC_H111_pepsin_B1.26985.26985.5 | 57.1927 | Pepsin | B1 | 305 |
| BCAM1669 | >tr\|A0A1V2X7H1\|A0A1V2X7H1_9BURK Purine nucleoside phosphorylase OS=Burkholderia cenocepacia OX=95486 GN=A8E72_24650 PE=4 SV=1 | A.AQARVHGIDNS[+568.212]GAGSQPAA.T | HexNAc(2)Hex(1) | No | 1188.5564 | 2 | 2376.1055 | 2375.0957 | 2.7 | Non | 375.97 | 339.11 | 5.6 | Nsco_20191108_BC_ZIC_HILIC_H111_pepsin_B1.9852.9852.2 | 26.9269 | Pepsin | B1 | 306 |
| BCAM1669 | >tr\|A0A1V2X7H1\|A0A1V2X7H1_9BURK Purine nucleoside phosphorylase OS=Burkholderia cenocepacia OX=95486 GN=A8E72_24650 PE=4 SV=1 | Q.ARVHGIDNS[+568.212]GAGSQPAAT.V | HexNAc(2)Hex(1) | No | 1139.0313 | 2 | 2277.0553 | 2277.0477 | 3.3 | Non | 473.51 | 147.67 | 0.63 | Nsco_20191108_BC_ZIC_HILIC_H111_pepsin_B2.9736.9736.2 | 25.8538 | Pepsin | B2 | 307 |

| | | | | | | | | | | | | | | | | | | |
|---|---|---|---|---|---|---|---|---|---|---|---|---|---|---|---|---|---|---|
| BCAM1669 | >tr\|A0A1V2X7H1\|A0A1V2X7H1_9BURK Purine nucleoside phosphorylase OS=Burkholderia cenocepacia OX=95486 GN=A8E72_24650 PE=4 SV=1 | A.AQARVHGIDNSGAGS[+568.212]QPAAT.V | HexNAc(2)Hex(1) | No | 1238.5804 | 2 | 2476.1535 | 2476.1434 | 4.1 | Non | 500.02 | 157.54 | 1.63 | Nsco_20191108_BC_ZIC_HILIC_H111_pepsin_B2.10761.10761.2 | 27.4903 | Pepsin | B2 | 308 |
| BCAM1669 | >tr\|A0A1V2X7H1\|A0A1V2X7H1_9BURK Purine nucleoside phosphorylase OS=Burkholderia cenocepacia OX=95486 GN=A8E72_24650 PE=4 SV=1 | A.RVHGIDNSGAGS[+568.212]QPAAT.V | HexNAc(2)Hex(1) | No | 1103.5121 | 2 | 2206.0169 | 2206.0106 | 2.9 | Non | 465.26 | 365.35 | 3.12 | Nsco_20191108_BC_ZIC_HILIC_H111_pepsin_B2.8843.8843.2 | 24.4525 | Pepsin | B2 | 309 |
| BCAM1669 | >tr\|A0A1V2X7H1\|A0A1V2X7H1_9BURK Purine nucleoside phosphorylase OS=Burkholderia cenocepacia OX=95486 GN=A8E72_24650 PE=4 SV=1 | A.RVHGIDNSGAGS[+568.212]QPAA.T | HexNAc(2)Hex(1) | No | 1052.9897 | 2 | 2104.9721 | 2104.9629 | 4.3 | Non | 416.48 | 285.03 | 5.5 | Nsco_20191108_BC_ZIC_HILIC_H111_pepsin_B3.8746.8746.2 | 23.5508 | Pepsin | B3 | 310 |
| BCAM1669 | >tr\|A0A1V2X7H1\|A0A1V2X7H1_9BURK Purine nucleoside phosphorylase OS=Burkholderia cenocepacia OX=95486 GN=A8E72_24650 PE=4 SV=1 | R.VHGIDNSGAGSQPAAT[+568.212]VEGGAPVVRAQNPRDSVYF.G | HexNAc(2)Hex(1) | No | 1024.2471 | 4 | 4093.9665 | 4091.9429 | 4.1 | NRagged | 785.56 | 696.72 | 9.6 | Nsco_20191108_BC_ZIC_HILIC_H111_thermolysin_B1.39135.39135.4 | 74.9032 | Thermolysin | B1 | 311 |
| BCAM1669 | >tr\|A0A1V2X7H1\|A0A1V2X7H1_9BURK Purine nucleoside phosphorylase OS=Burkholderia cenocepacia OX=95486 GN=A8E72_24650 PE=4 SV=1 | L.VQLENAGYKPSQSSPYYPADICAAQARVHGIDNSGAGS[+568.212]QPAAT[+568.212]VEGGAPVVRAQNPRDSVYF.G | HexNAc(2)Hex(1), HexNAc(2)Hex(1) | No | 1266.6069 | 6 | 7594.6048 | 7593.5852 | 2.1 | Specific | 347.94 | 142.68 | 0 | Nsco_20191108_BC_ZIC_HILIC_H111_thermolysin_B1.43288.43288.3 | 81.9145 | Thermolysin | B1 | 312 |
| BCAM1669 | >tr\|A0A1V2X7H1\|A0A1V2X7H1_9BURK Purine nucleoside phosphorylase OS=Burkholderia cenocepacia OX=95486 GN=A8E72_24650 PE=4 SV=1 | R.VHGIDNSGAGSQPAAT[+568.212]VEGGAPVVRAQNPRDSVY.F | HexNAc(2)Hex(1) | No | 1316.637 | 3 | 3947.8963 | 3944.8744 | 3 | Non | 922.88 | 906.89 | 9.6 | Nsco_20191108_BC_ZIC_HILIC_H111_thermolysin_B2.30783.30783.3 | 62.1659 | Thermolysin | B2 | 313 |
| BCAM1669 | >tr\|A0A1V2X7H1\|A0A1V2X7H1_9BURK Purine nucleoside phosphorylase OS=Burkholderia cenocepacia OX=95486 GN=A8E72_24650 PE=4 SV=1 | G.IDNSGAGSQPAAT[+568.212]VEGGAPVVRAQNPRDSVYF.G | HexNAc(2)Hex(1) | No | 1266.9399 | 3 | 3798.8051 | 3798.7941 | 2.9 | NRagged | 755.68 | 683.26 | 12 | Nsco_20191108_BC_ZIC_HILIC_H111_thermolysin_B2.41526.41526.3 | 82.134 | Thermolysin | B2 | 314 |
| BCAM1669 | >tr\|A0A1V2X7H1\|A0A1V2X7H1_9BURK Purine nucleoside phosphorylase OS=Burkholderia cenocepacia OX=95486 GN=A8E72_24650 PE=4 SV=1 | R.VHGIDNSGAGS[+568.212]QPAATVEGGAP.V | HexNAc(2)Hex(1) | No | 1280.5865 | 2 | 2560.1657 | 2560.1533 | 4.8 | Non | 589.46 | 497.26 | 1.2 | Nsco_20191108_BC_ZIC_HILIC_H111_thermolysin_B3.24499.24499.2 | 51.8666 | Thermolysin | B3 | 315 |
| BCAM1669 | >tr\|A0A1V2X7H1\|A0A1V2X7H1_9BURK Purine nucleoside phosphorylase OS=Burkholderia cenocepacia OX=95486 GN=A8E72_24650 PE=4 SV=1 | G.IDNSGAGSQPAAT[+568.212]VEGGAPVVRAQNPRD.S | HexNAc(2)Hex(1) | No | 1101.8645 | 3 | 3303.579 | 3302.5619 | 4.2 | Non | 510.46 | 430.49 | 12 | Nsco_20191108_BC_ZIC_HILIC_H111_thermolysin_B3.24567.24567.3 | 51.9771 | Thermolysin | B3 | 316 |
| BCAM1669 | >tr\|A0A1V2X7H1\|A0A1V2X7H1_9BURK Purine nucleoside phosphorylase OS=Burkholderia cenocepacia OX=95486 GN=A8E72_24650 PE=4 SV=1 | G.IDNSGAGSQPAAT[+568.212]VEGGAPVVRAQNPRDS.V | HexNAc(2)Hex(1) | No | 1130.5406 | 3 | 3389.6071 | 3389.5939 | 3.9 | Non | 698.9 | 666.63 | 12 | Nsco_20191108_BC_ZIC_HILIC_H111_thermolysin_B3.24570.24570.3 | 51.9841 | Thermolysin | B3 | 317 |
| BCAM1669 | >tr\|A0A1V2X7H1\|A0A1V2X7H1_9BURK Purine nucleoside phosphorylase OS=Burkholderia cenocepacia OX=95486 GN=A8E72_24650 PE=4 SV=1 | H.GIDNSGAGSQPAAT[+568.212]VEGGAPVVRAQNPRDS.V | HexNAc(2)Hex(1) | No | 1149.5388 | 3 | 3446.6019 | 3446.6154 | -3.9 | Non | 284.76 | 129.46 | 1.44 | Nsco_20191108_BC_ZIC_HILIC_H111_thermolysin_B3.27776.27776.3 | 57.5507 | Thermolysin | B3 | 318 |
| BCAM1669 | >tr\|A0A1V2X7H1\|A0A1V2X7H1_9BURK Purine nucleoside phosphorylase OS=Burkholderia cenocepacia OX=95486 GN=A8E72_24650 PE=4 SV=1 | G.IDNSGAGSQPAAT[+568.212]VEGGAPVVRAQNPRDSVY.F | HexNAc(2)Hex(1) | No | 1217.9178 | 3 | 3651.7388 | 3651.7256 | 3.6 | Non | 809.25 | 796.29 | 12 | Nsco_20191108_BC_ZIC_HILIC_H111_thermolysin_B3.33078.33078.3 | 66.3496 | Thermolysin | B3 | 319 |
| BCAM1669 | >tr\|A0A1V2X7H1\|A0A1V2X7H1_9BURK Purine nucleoside phosphorylase OS=Burkholderia cenocepacia OX=95486 GN=A8E72_24650 PE=4 SV=1 | R.VHGIDNSGAGSQPAAT[+568.212]VEGGAPVVR.A | HexNAc(2)Hex(1) | No | 1457.7007 | 2 | 2914.3942 | 2914.3912 | 1 | Specific | 974.46 | 925.6 | 0.05 | Nsco_20191108_BC_ZIC_HILIC_H111_typsin_B1.25669.25669.2 | 55.2181 | Trypsin | B1 | 320 |

| | | | | | | | | | | | | | | | | | | |
|---|---|---|---|---|---|---|---|---|---|---|---|---|---|---|---|---|---|---|
| BCAM1737 | >tr\|A0A2N9CVE8\|A0A2N9CVE8_9BURK<br>Putative exported glycoprotein<br>OS=Burkholderia cenocepacia OX=95486<br>GN=F01_460192 PE=4 SV=1 | W.SQAGSAAPADTPASAAPSAS<br>[+568.212]AT[+568.212]P<br>ATRA.A | HexNAc(2)Hex(1),<br>HexNAc(2)Hex(1) | No | 1174.2 | 3 | 3520.5853 | 3519.5716 | 3 | Non | 468.72 | 468.72 | 3.01 | Nsco_20191108_BC_ZIC_HILIC_H111_pepsin_B1.19<br>245.19245.3 | 42.0253 | Pepsin | B1 | 321 |
| BCAM1737 | >tr\|A0A2N9CVE8\|A0A2N9CVE8_9BURK<br>Putative exported glycoprotein<br>OS=Burkholderia cenocepacia OX=95486<br>GN=F01_460192 PE=4 SV=1 | E.RAISWSQAGSAAPADT[+568.<br>212]PASAAPSASATPATRA.A | HexNAc(2)Hex(1) | No | 1189.2407 | 3 | 3565.7074 | 3564.6936 | 2.9 | Non | 331.43 | 311.67 | 2.4 | Nsco_20191108_BC_ZIC_HILIC_H111_pepsin_B1.28<br>994.28994.3 | 61.2309 | Pepsin | B1 | 322 |
| BCAM1737 | >tr\|A0A2N9CVE8\|A0A2N9CVE8_9BURK<br>Putative exported glycoprotein<br>OS=Burkholderia cenocepacia OX=95486<br>GN=F01_460192 PE=4 SV=1 | W.SQAGSAAPADTPASAAPSAS<br>[+568.212]AT[+568.212]P<br>ATRAAA.A | HexNAc(2)Hex(1),<br>HexNAc(2)Hex(1) | No | 1221.5586 | 3 | 3662.6612 | 3661.6458 | 3.3 | Non | 478.39 | 444.26 | 3.6 | Nsco_20191108_BC_ZIC_HILIC_H111_pepsin_B2.21<br>264.21264.3 | 44.5412 | Pepsin | B2 | 323 |
| BCAM1737 | >tr\|A0A2N9CVE8\|A0A2N9CVE8_9BURK<br>Putative exported glycoprotein<br>OS=Burkholderia cenocepacia OX=95486<br>GN=F01_460192 PE=4 SV=1 | P.ADTPASAAPSAS[+568.212<br>][+100.064]AT[+568.212]P<br>AT[+568.212][+100.064]R<br>A.A | HexNAc(2)Hex(1)<br>100.064,HexNAc(<br>2)Hex(1),HexNAc(<br>2)Hex(1) 100.064 | No | 1207.2055 | 3 | 3619.602 | 3618.6029 | -1.2 | Non | 196.8 | 196.8 | 3.44 | Nsco_20191108_BC_ZIC_HILIC_H111_pepsin_B3.23<br>406.23406.3 | 48.2958 | Pepsin | B3 | 324 |
| BCAM1737 | >tr\|A0A2N9CVE8\|A0A2N9CVE8_9BURK<br>Putative exported glycoprotein<br>OS=Burkholderia cenocepacia OX=95486<br>GN=F01_460192 PE=4 SV=1 | S.WSQAGSAAPADTPASAAPSA<br>S[+568.212]AT[+568.212]P<br>ATRA.A | HexNAc(2)Hex(1),<br>HexNAc(2)Hex(1) | No | 1236.2268 | 3 | 3706.6659 | 3705.6509 | 3.1 | Non | 180.07 | 180.07 | 2.4 | Nsco_20191108_BC_ZIC_HILIC_H111_pepsin_B3.25<br>874.25874.3 | 53.4267 | Pepsin | B3 | 325 |
| BCAM1737 | >tr\|A0A2N9CVE8\|A0A2N9CVE8_9BURK<br>Putative exported glycoprotein<br>OS=Burkholderia cenocepacia OX=95486<br>GN=F01_460192 PE=4 SV=1 | A.ISWSQAGSAAPADT[+568.2<br>12]PASAAPS[+568.212]ASA<br>TPATR.A | HexNAc(2)Hex(1),<br>HexNAc(2)Hex(1) | No | 1279.2545 | 3 | 3835.749 | 3834.7299 | 4.1 | Non | 514.82 | 230.7 | 2.27 | Nsco_20191108_BC_ZIC_HILIC_H111_thermolysin_B<br>2.34280.34280.3 | 68.0681 | Thermolysin | B2 | 326 |
| BCAM1737 | >tr\|A0A2N9CVE8\|A0A2N9CVE8_9BURK<br>Putative exported glycoprotein<br>OS=Burkholderia cenocepacia OX=95486<br>GN=F01_460192 PE=4 SV=1 | S.WSQAGSAAPADTPASAAPSA<br>S[+568.212][+100.064]AT[<br>+568.212][+100.064]PATR<br>AA.A | HexNAc(2)Hex(1)<br>100.064,HexNAc(<br>2)Hex(1) 100.064 | No | 1326.2803 | 3 | 3976.8264 | 3976.816 | 2.6 | Non | 399.36 | 399.36 | 2.4 | Nsco_20191108_BC_ZIC_HILIC_H111_thermolysin_B<br>2.35161.35161.3 | 69.6204 | Thermolysin | B2 | 327 |
| BCAM1737 | >tr\|A0A2N9CVE8\|A0A2N9CVE8_9BURK<br>Putative exported glycoprotein<br>OS=Burkholderia cenocepacia OX=95486<br>GN=F01_460192 PE=4 SV=1 | A.GSAAPADTPASAAPS[+568.<br>212][+100.064]AS[+568.2<br>12]AT[+568.212][+100.06<br>4]PATRAA.A | HexNAc(2)Hex(1)<br>100.064,HexNAc(<br>2)Hex(1),HexNAc(<br>2)Hex(1) 100.064 | No | 1359.616 | 3 | 4076.8335 | 4072.8205 | -0.1 | Non | 324.49 | 324.49 | 0 | Nsco_20191108_BC_ZIC_HILIC_H111_thermolysin_B<br>3.37010.37010.3 | 73.8046 | Thermolysin | B3 | 328 |
| BCAM1737 | >tr\|A0A2N9CVE8\|A0A2N9CVE8_9BURK<br>Putative exported glycoprotein<br>OS=Burkholderia cenocepacia OX=95486<br>GN=F01_460192 PE=4 SV=1 | S.WSQAGSAAPADTPASAAPSA<br>S[+568.212][+100.064]AT[<br>+568.212][+100.064]PATR.<br>A | HexNAc(2)Hex(1)<br>100.064,HexNAc(<br>2)Hex(1) 100.064 | No | 1279.2556 | 3 | 3835.7523 | 3834.7418 | 1.9 | Non | 319.11 | 319.11 | 2.4 | Nsco_20191108_BC_ZIC_HILIC_H111_thermolysin_B<br>3.34011.34011.3 | 67.9246 | Thermolysin | B3 | 329 |
| BCAM1737 | >tr\|A0A2N9CVE8\|A0A2N9CVE8_9BURK<br>Putative exported glycoprotein<br>OS=Burkholderia cenocepacia OX=95486<br>GN=F01_460192 PE=4 SV=1 | A.ISWSQAGSAAPADTPAS[+56<br>8.212]AAPSASAT[+568.212<br>]PATRAA.A | HexNAc(2)Hex(1),<br>HexNAc(2)Hex(1) | No | 1326.6125 | 3 | 3977.8231 | 3976.8041 | 3.9 | Non | 684.89 | 302.89 | 5.88 | Nsco_20191108_BC_ZIC_HILIC_H111_thermolysin_B<br>3.34894.34894.3 | 69.4909 | Thermolysin | B3 | 330 |
| BCAM1737 | >tr\|A0A2N9CVE8\|A0A2N9CVE8_9BURK<br>Putative exported glycoprotein<br>OS=Burkholderia cenocepacia OX=95486<br>GN=F01_460192 PE=4 SV=1 | A.GSAAPADTPASAAPSAS[+56<br>8.212][+100.064]AT[+568.<br>212]PAT[+568.212][+100.<br>064]R.A | HexNAc(2)Hex(1)<br>100.064,HexNAc(<br>2)Hex(1),HexNAc(<br>2)Hex(1) 100.064 | No | 1311.9205 | 3 | 3933.7468 | 3930.7463 | -2.4 | NRagged | 321.68 | 321.68 | 1.76 | Nsco_20191108_BC_ZIC_HILIC_H111_typsin_B1.345<br>74.34574.3 | 70.5281 | Trypsin | B1 | 331 |
| BCAM1737 | >tr\|A0A2N9CVE8\|A0A2N9CVE8_9BURK<br>Putative exported glycoprotein<br>OS=Burkholderia cenocepacia OX=95486<br>GN=F01_460192 PE=4 SV=1 | Q.AGSAAPADTPASAAPS[+56<br>8.212][+100.064]AS[+568.<br>212]AT[+568.212][+100.0<br>64]PATR.A | HexNAc(2)Hex(1)<br>100.064,HexNAc(<br>2)Hex(1),HexNAc(<br>2)Hex(1) 100.064 | No | 1335.935 | 3 | 4005.7904 | 4001.7834 | -1.6 | NRagged | 475.79 | 475.79 | 0 | Nsco_20191108_BC_ZIC_HILIC_H111_typsin_B1.362<br>12.36212.3 | 73.2799 | Trypsin | B1 | 332 |

| | | | | | | | | | | | | | | | | | | |
|---|---|---|---|---|---|---|---|---|---|---|---|---|---|---|---|---|---|---|
| BCAM1737 | >tr\|A0A2N9CVE8\|A0A2N9CVE8_9BURK Putative exported glycoprotein OS=Burkholderia cenocepacia OX=95486 GN=F01_460192 PE=4 SV=1 | R.AISWSQAGSAAPADTPASAAPSAS[+568.212]ATPATR.A | HexNAc(2)Hex(1) | No | 1669.2892 | 2 | 3337.5712 | 3337.5554 | 4.7 | Specific | 851.85 | 798.79 | 4.8 | Nsco_20191108_BC_ZIC_HILIC_H111_typsin_B2.37171.37171.2 | 73.5891 | Trypsin | B2 | 333 |
| BCAM1737 | >tr\|A0A2N9CVE8\|A0A2N9CVE8_9BURK Putative exported glycoprotein OS=Burkholderia cenocepacia OX=95486 GN=F01_460192 PE=4 SV=1 | S.QAGSAAPADTPASAAPSAS[+568.212]AT[+568.212]PAT[+568.212][+100.064]R.A | HexNAc(2)Hex(1), HexNAc(2)Hex(1), HexNAc(2)Hex(1) 100.064 | No | 1345.2692 | 3 | 4033.7929 | 4029.778 | 0.4 | NRagged | 254.86 | 254.86 | 1.55 | Nsco_20191108_BC_ZIC_HILIC_H111_typsin_B3.35430.35430.3 | 74.1491 | Trypsin | B3 | 334 |
| BCAM2054 (ty | >tr\|A0A1V2XAF9\|A0A1V2XAF9_9BURK Peptide-binding protein OS=Burkholderia cenocepacia OX=95486 GN=bcscD PE=4 SV=1 | R.APEQAPAPPVAPLAS[+568.212]GAAAGVAQPVPGPTLLPREPAAGVSTK.E | HexNAc(2)Hex(1) | No | 1493.1235 | 3 | 4477.3559 | 4477.3348 | 4.7 | Specific | 1112.68 | 1112.68 | 3.6 | Nsco_20191108_BC_ZIC_HILIC_H111_typsin_B1.55514.55514.3 | 105.5981 | Trypsin | B1 | 335 |
| BCAM2055 | >tr\|A0A1V2VXC7\|A0A1V2VXC7_9BURK EscC/YscC/HrcC family type III secretion system outer membrane ring protein OS=Burkholderia cenocepacia OX=95486 GN=A8E72_28425 PE=3 SV=1 | D.VTVPLTPKPLPGTKFGPPALPPPKDTAAS[+568.212]QPAA.T | HexNAc(2)Hex(1) | No | 958.2687 | 4 | 3830.053 | 3830.0361 | 4.4 | Non | 296.76 | 296.76 | 44.15 | Nsco_20191108_BC_ZIC_HILIC_H111_pepsin_B1.40237.40237.4 | 88.1894 | Pepsin | B1 | 336 |
| BCAM2055 | >tr\|A0A1V2VXC7\|A0A1V2VXC7_9BURK EscC/YscC/HrcC family type III secretion system outer membrane ring protein OS=Burkholderia cenocepacia OX=95486 GN=A8E72_28425 PE=3 SV=1 | D.VTVPLTPKPLPGTKFGPPALPPPKDT[+568.212]AASQPAATTAG.V | HexNAc(2)Hex(1) | No | 1040.8079 | 4 | 4160.2098 | 4160.1901 | 4.7 | Non | 439.15 | 398.15 | 6.47 | Nsco_20191108_BC_ZIC_HILIC_H111_pepsin_B2.40975.40975.4 | 87.9263 | Pepsin | B2 | 337 |
| BCAM2055 | >tr\|A0A1V2VXC7\|A0A1V2VXC7_9BURK EscC/YscC/HrcC family type III secretion system outer membrane ring protein OS=Burkholderia cenocepacia OX=95486 GN=A8E72_28425 PE=3 SV=1 | K.DTAASQPAATT[+568.212]AGVVTHVDEHH.- | HexNAc(2)Hex(1) | No | 928.7593 | 3 | 2784.2635 | 2783.249 | 4 | CRagged | 324.81 | 324.81 | 1.2 | Nsco_20191108_BC_ZIC_HILIC_H111_typsin_B2.20983.20983.3 | 44.8364 | Trypsin | B2 | 338 |
| BCAM2063 | >tr\|A0A2N9CW94\|A0A2N9CW94_9BURK Porin OS=Burkholderia cenocepacia OX=95486 GN=F01_460501 PE=3 SV=1 | A.SSPAAAEPAAGASDAAAPAQQAADAAAPAPT[+568.212]GFWERSN.L | HexNAc(2)Hex(1) | No | 1374.963 | 3 | 4122.8745 | 4119.8538 | 2.6 | Non | 635.81 | 612.9 | 4.83 | Nsco_20191108_BC_ZIC_HILIC_H111_thermolysin_B3.49038.49038.3 | 96.7602 | Thermolysin | B3 | 339 |
| BCAM2064 | >tr\|A0A2N9CWA7\|A0A2N9CWA7_9BURK Periplasmic trehalase OS=Burkholderia cenocepacia OX=95486 GN=treA PE=3 SV=1 | A.DNANQAAAQAAGQSAIPATTAAAAAPASGT[+568.212]LPPPSQLYGDLFVAVQTAQLYPDQK.T | HexNAc(2)Hex(1) | No | 1494.9893 | 4 | 5976.9352 | 5976.9149 | 3.4 | NRagged | 729.06 | 729.06 | 0 | Nsco_20191108_BC_ZIC_HILIC_H111_typsin_B2.79285.79285.4 | 160.566 | Trypsin | B2 | 340 |
| BCAM2289 | >tr\|A0A142PAR8\|A0A142PAR8_9BURK Purine nucleoside phosphorylase OS=Burkholderia cenocepacia OX=95486 GN=A8E72_27535 PE=4 SV=1 | D.TSGYGAQPAPLVHSGAPAAAS[+568.212]SNARD.S | HexNAc(2)Hex(1) | No | 1007.8045 | 3 | 3021.3989 | 3021.392 | 2.3 | Non | 921.88 | 333.53 | 2.4 | Nsco_20191108_BC_ZIC_HILIC_H111_pepsin_B1.19535.19535.3 | 42.5368 | Pepsin | B1 | 341 |
| BCAM2289 | >tr\|A0A142PAR8\|A0A142PAR8_9BURK Purine nucleoside phosphorylase OS=Burkholderia cenocepacia OX=95486 GN=A8E72_27535 PE=4 SV=1 | R.VHGADTSGYGAQPAPLVHSGAPAAAS[+568.212]SNAR.D | HexNAc(2)Hex(1) | Yes | 847.1534 | 4 | 3385.5918 | 3385.5779 | 4.1 | Specific | 1294.2 | 1230.17 | 71.23 | Nsco_20191108_BC_ZIC_HILIC_H111_typsin_B3.17844.17844.4 | 40.1436 | Trypsin | B3 | 342 |
| BCAM2301 | >tr\|A0A1V2VWJ7\|A0A1V2VWJ7_9BURK Amine dehydrogenase OS=Burkholderia cenocepacia OX=95486 GN=aauA PE=4 SV=1 | A.DAASGPSGASGAAAAGAS[+568.212][+100.064]DDPMSC[+57.021]DYWK.Y | HexNAc(2)Hex(1) 100.064 | No | 1671.6795 | 2 | 3342.3517 | 3341.3584 | -3 | NRagged | 447.12 | 447.12 | 6 | Nsco_20191108_BC_ZIC_HILIC_H111_typsin_B2.40371.40371.2 | 79.3924 | Trypsin | B2 | 343 |

| | | | | | | | | | | | | | | | | | | |
|---|---|---|---|---|---|---|---|---|---|---|---|---|---|---|---|---|---|---|
| BCAM2307 | >tr\|A0A2N9CXU5\|A0A2N9CXU5_9BURK Bacillolysin OS=Burkholderia cenocepacia OX=95486 GN=F01_480197 PE=4 SV=1 | F.EAVNVYYHIDTFLRYVNQT[+568.212][+100.064]LGIKALPYQYTGGVQYDPHGES[+568.212][+100.064]GDDNSSYSSSSGR.L | HexNAc(2)Hex(1) 100.064,HexNAc(2)Hex(1) 100.064 | No | 1472.6734 | 5 | 7359.3378 | 7355.3629 | -5.2 | NRagged | 516.72 | 516.72 | 0 | Nsco_20191108_BC_ZIC_HILIC_H111_typsin_B3.33778.33778.5 | 70.9602 | Trypsin | B3 | 344 |
| BCAM2334 | >tr\|A0A2N9CXX4\|A0A2N9CXX4_9BURK Multidrug efflux system OS=Burkholderia cenocepacia OX=95486 GN=emrA PE=4 SV=1 | R.VHDGVAS[+568.212]DAEAAAAAIIRENQGG.- | HexNAc(2)Hex(1) | Yes | 930.7722 | 3 | 2790.302 | 2790.2912 | 3.9 | CRagged | 667.4 | 634.86 | 634.86 | Nsco_20191108_BC_ZIC_HILIC_H111_typsin_B2.51636.51636.3 | 99.0316 | Trypsin | B2 | 345 |
| BCAM2443 | >tr\|A0A2N9CXV2\|A0A2N9CXV2_9BURK Cytochrome c family protein OS=Burkholderia cenocepacia OX=95486 GN=F01_480355 PE=4 SV=1 | G.ATPQDAPAAAS[+568.212]APPPAPAAAAAPAAKPFTPPPES.A | HexNAc(2)Hex(1) | No | 1224.9368 | 3 | 3672.7959 | 3672.7803 | 4.3 | Non | 446.78 | 188.36 | 2.38 | Nsco_20191108_BC_ZIC_HILIC_H111_pepsin_B2.38609.38609.3 | 81.695 | Pepsin | B2 | 346 |
| BCAM2443 | >tr\|A0A2N9CXV2\|A0A2N9CXV2_9BURK Cytochrome c family protein OS=Burkholderia cenocepacia OX=95486 GN=F01_480355 PE=4 SV=1 | A.TPQDAPAAASAPPPAPAAAAAPAAKPFT[+568.212]PPPESAIPADDFGK.T | HexNAc(2)Hex(1) | No | 1506.4057 | 3 | 4517.2025 | 4516.193 | 1.4 | NRagged | 425.1 | 46.73 | 8.4 | Nsco_20191108_BC_ZIC_HILIC_H111_typsin_B2.49021.49021.3 | 94.5171 | Trypsin | B2 | 347 |
| BCAM2443 | >tr\|A0A2N9CXV2\|A0A2N9CXV2_9BURK Cytochrome c family protein OS=Burkholderia cenocepacia OX=95486 GN=F01_480355 PE=4 SV=1 | T.PQDAPAAASAPPPAPAAAAAPAAKPFT[+568.212][+100.064]PPPESAIPADDFGK.T | HexNAc(2)Hex(1) 100.064 | No | 1506.0768 | 3 | 4516.216 | 4515.2093 | 0.7 | NRagged | 521.44 | 501.79 | 7.2 | Nsco_20191108_BC_ZIC_HILIC_H111_typsin_B2.49077.49077.3 | 94.6144 | Trypsin | B2 | 348 |
| BCAM2681 | >tr\|A0A1V6L0G1\|A0A1V6L0G1_9BURK Putative lipoprotein OS=Burkholderia cenocepacia OX=95486 GN=F01_490209 PE=4 SV=1 | R.GASVAVHAGSAPSEAVGGGT[+568.212]PAEQVAALDPK.A | HexNAc(2)Hex(1) | No | 1124.2145 | 3 | 3370.6289 | 3369.618 | 2.2 | Specific | 900.38 | 819.7 | 4.27 | Nsco_20191108_BC_ZIC_HILIC_H111_typsin_B1.35986.35986.3 | 72.9199 | Trypsin | B1 | 349 |
| BCAM2828 | >tr\|A0A2N9CZJ3\|A0A2N9CZJ3_9BURK Hopanoid biosynthesis associated RND transporter like protein HpnN OS=Burkholderia cenocepacia OX=95486 GN=hpnN PE=4 SV=1 | K.RAAIAAAAS[+568.212]ELLPALTQPAAPPATDAQR.V | HexNAc(2)Hex(1) | No | 1104.2425 | 3 | 3310.7128 | 3310.7013 | 3.5 | Specific | 662.67 | 648.76 | 36.7 | Nsco_20191108_BC_ZIC_HILIC_H111_typsin_B2.51893.51893.3 | 99.4892 | Trypsin | B2 | 350 |
| BCAS0453 | >tr\|A0A142PSQ3\|A0A142PSQ3_9BURK Uncharacterized protein OS=Burkholderia cenocepacia OX=95486 GN=A8E72_07995 PE=4 SV=1 | R.DSLGNGVALDWPAS[+568.212]GVGGVADERQK.L | HexNAc(2)Hex(1) | No | 1023.1548 | 3 | 3067.4498 | 3066.4386 | 2.6 | Specific | 547.29 | 535.04 | 61.99 | Nsco_20191108_BC_ZIC_HILIC_H111_typsin_B1.47980.47980.3 | 93.4226 | Trypsin | B1 | 351 |
| BCAS0453 | >tr\|A0A142PSQ3\|A0A142PSQ3_9BURK Uncharacterized protein OS=Burkholderia cenocepacia OX=95486 GN=A8E72_07995 PE=4 SV=1 | R.DSLGNGVALDWPAS[+568.212]GVGGVADER.Q | HexNAc(2)Hex(1) | No | 1405.6471 | 2 | 2810.287 | 2810.285 | 0.7 | Specific | 399.7 | 376.4 | 53.27 | Nsco_20191108_BC_ZIC_HILIC_H111_typsin_B1.59228.59228.2 | 111.4013 | Trypsin | B1 | 352 |
| BCAS0773 | >tr\|A0A2N9CZW2\|A0A2N9CZW2_9BURK Uncharacterized protein OS=Burkholderia cenocepacia OX=95486 GN=DFS07_107251 PE=4 SV=1 | A.QTDAAS[+568.212]APAAAAAQDAK.A | HexNAc(2)Hex(1) | No | 1063.4866 | 2 | 2125.9659 | 2125.9619 | 1.9 | NRagged | 491.58 | 432.38 | 29.17 | Nsco_20191108_BC_ZIC_HILIC_H111_typsin_B1.11041.11041.2 | 29.9258 | Trypsin | B1 | 353 |
| I35_6143 | >tr\|A0A2N9CX34\|A0A2N9CX34_9BURK Uncharacterized protein OS=Burkholderia cenocepacia OX=95486 GN=F01_480128 PE=4 SV=1 | A.PAERSPFDTPDTRPLRVTS[+568.212]DAL.V | HexNAc(2)Hex(1) | No | 753.1206 | 4 | 3009.4607 | 3009.4535 | 2.4 | NRagged | 684.21 | 615.43 | 14.88 | Nsco_20191108_BC_ZIC_HILIC_H111_pepsin_B1.34820.34820.4 | 73.9991 | Pepsin | B1 | 354 |
| I35_6143 | >tr\|A0A2N9CX34\|A0A2N9CX34_9BURK Uncharacterized protein OS=Burkholderia cenocepacia OX=95486 GN=F01_480128 PE=4 SV=1 | F.AAPAERSPFDTPDTRPLRVT[+568.212]SDAL.V | HexNAc(2)Hex(1) | No | 788.6396 | 4 | 3151.5366 | 3151.5277 | 2.8 | Specific | 550.31 | 537.55 | 3.39 | Nsco_20191108_BC_ZIC_HILIC_H111_pepsin_B1.35214.35214.4 | 74.8912 | Pepsin | B1 | 355 |

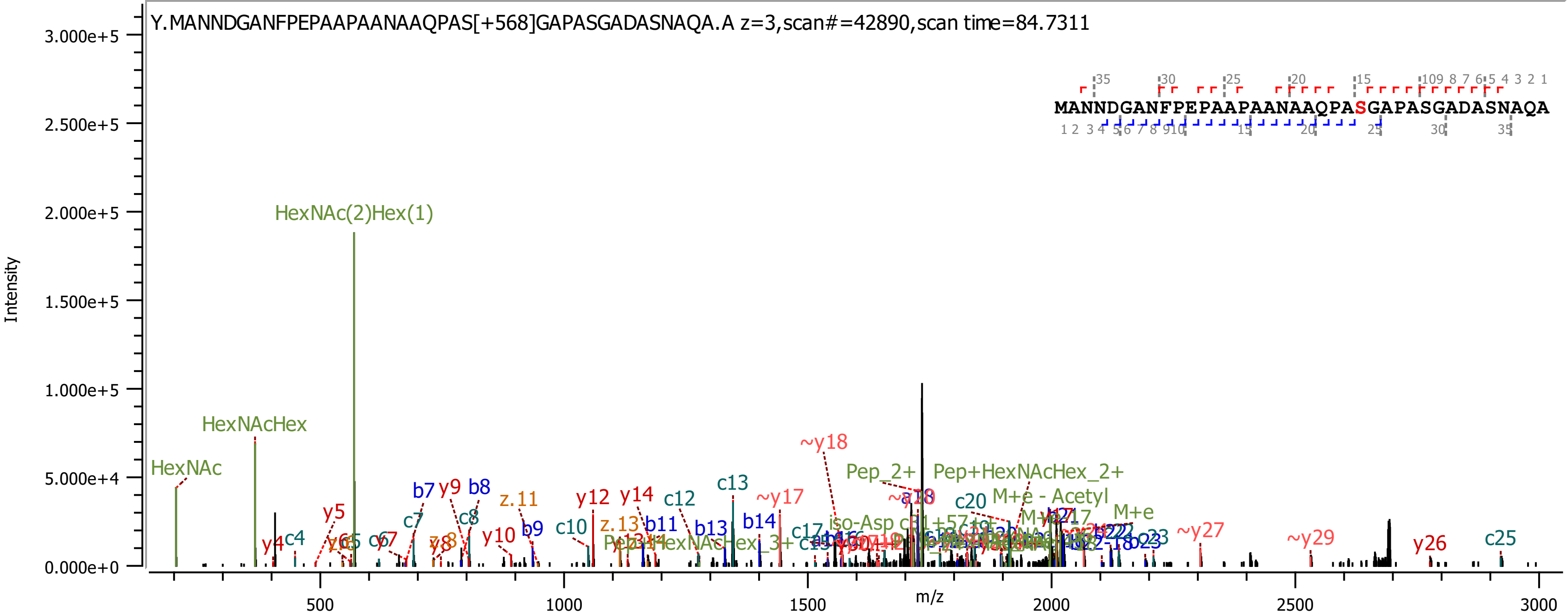
Y.MANNDGANFPEPAAPAANAAQPAS[+568]GAPASGADASNAQA.A z=3,scan#=42890,scan time=84.7311
MANNDGANFPEPAAPAANAAQPASGAPASGADASNAQA
Intensity
m/z
3.000e+5
2.500e+5
2.000e+5
1.500e+5
1.000e+5
5.000e+4
0.000e+0
500
1000
1500
2000
2500
3000
HexNAc(2)Hex(1)
HexNAcHex
HexNAc
~y18
Pep_2+
Pep+HexNAcHex_2+
M+e - Acetyl
M+e
c20
c13
~y17
c12
y12
y14
c10
b11
b13
b14
z.11
z.13
b7
y9
b8
c8
c7
y10
b9
y5
y4
c4
c25
y26
~y29
~y27
c23

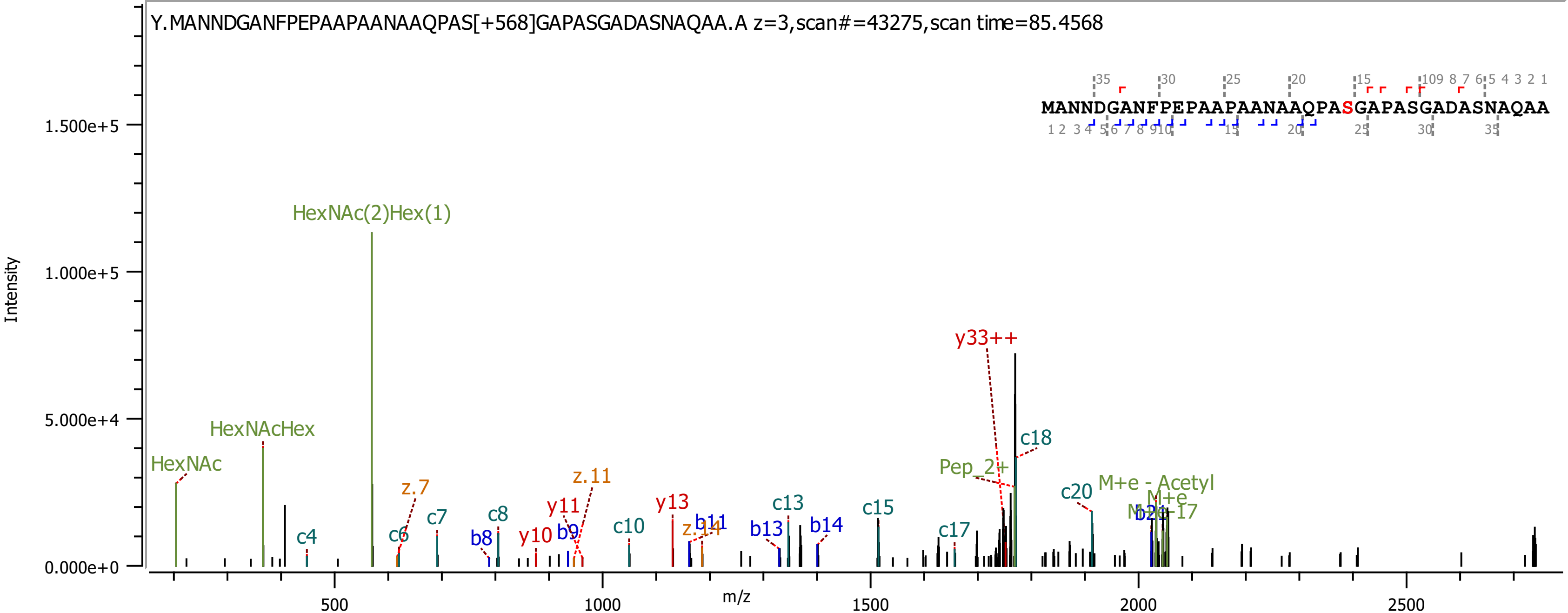

Y.MANNDGANFPEPAAPAANAAQPAS[+568]GAPASGADASNAQAA.A z=3,scan#=43275,scan time=85.4568
MANNDGANFPEPAAPAANAAQPASGAPASGADASNAQAA
Intensity
m/z
HexNAc
HexNAcHex
HexNAc(2)Hex(1)
c4
c6
z.7
c7
b8
c8
y10
b9
y11
z.11
c10
y13
z.14
b11
b13
c13
b14
c15
c17
Pep_2+
y33++
c18
c20
M+e - Acetyl
M+e
M+e-17
b21

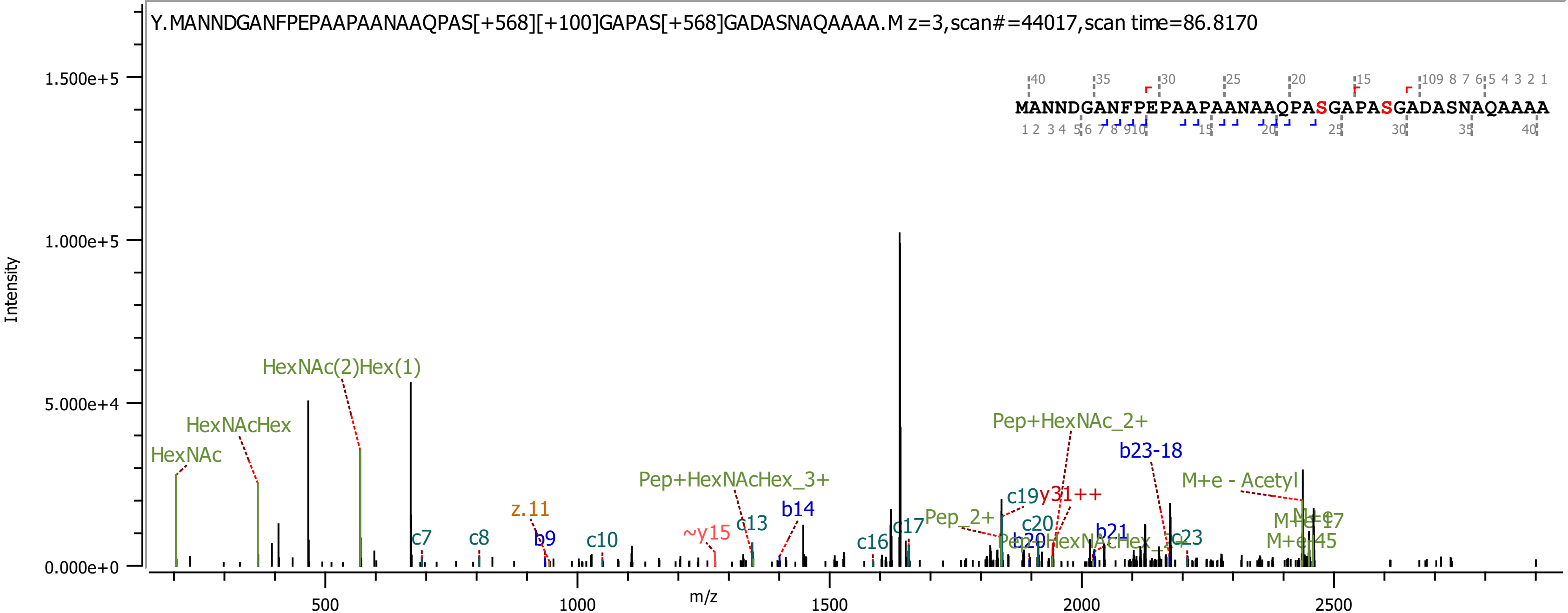

Y.MANNDGANFPEPAAPAANAAQPAS[+568][+100]GAPAS[+568]GADASNAQAAAA.M z=3,scan#=44017,scan time=86.8170
MANNDGANFPEPAAPAANAAQPASGAPASGADASNAQAAAA
Intensity
1.500e+5
1.000e+5
5.000e+4
0.000e+0
m/z
500
1000
1500
2000
2500
HexNAc
HexNAcHex
HexNAc(2)Hex(1)
c7
c8
z.11
b9
c10
~y15
Pep+HexNAcHex_3+
c13
b14
c16
c17
Pep_2+
c19
y31++
Pep+HexNAc_2+
c20
b20
Pep+HexNAcHex_2+
b21
b23-18
c23
M+e - Acetyl
M+e
M+e-17
M+e-45

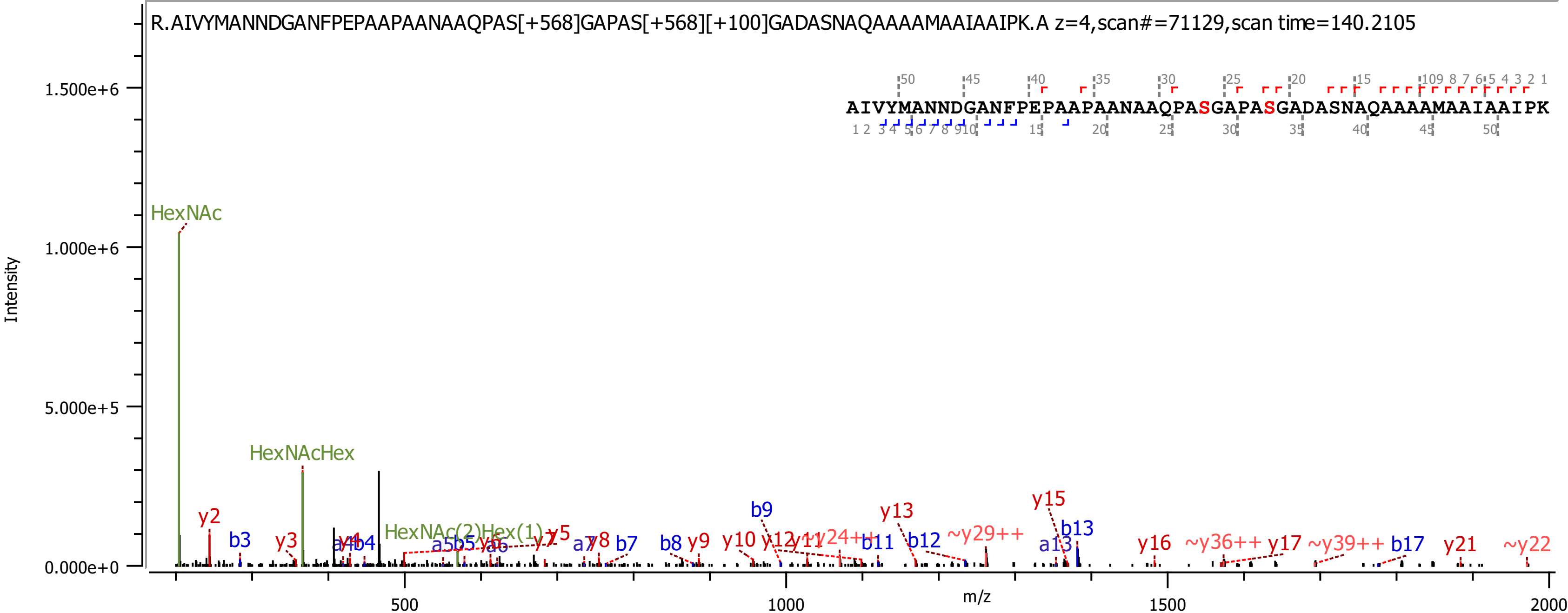
R.AIVYMANNDGANFPEPAAPAANAAQPAS[+568]GAPAS[+568][+100]GADASNAQAAAAMAAIAAIPK.A z=4,scan#=71129,scan time=140.2105
AIVYMANNDGANFPEPAAPAANAAQPASGAPASGADASNAQAAAAMAAIAAIPK
Intensity
m/z
1.500e+6
1.000e+6
5.000e+5
0.000e+0
500
1000
1500
2000
HexNAc
HexNAcHex
HexNAc(2)Hex(1)
y2
b3
y3
a4
y4
b4
a5
b5
y5
y6
a6
y7
a7
y8
b7
b8
y9
b9
y10
y12
y11
~y24++
b11
y13
b12
~y29++
y15
b13
a13
y16
~y36++
y17
~y39++
b17
y21
~y22

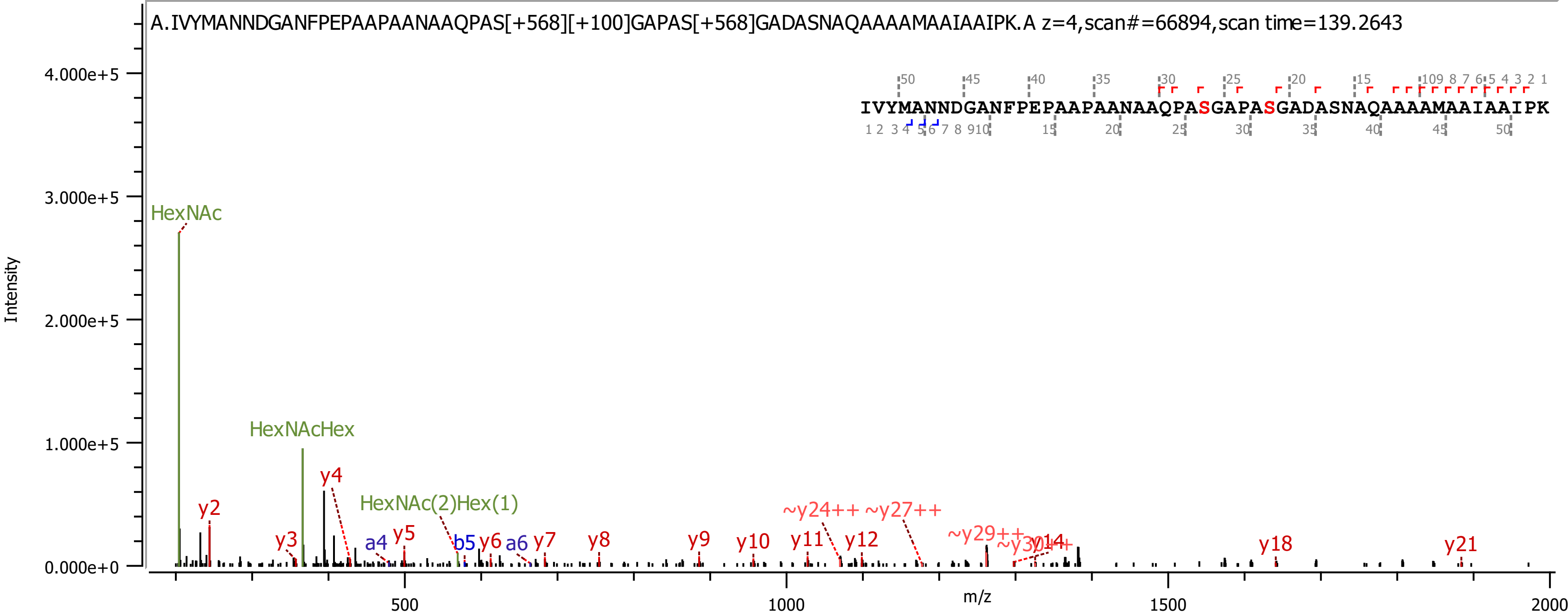

A.IVYMANNDGANFPEPAAPAANAAQPAS[+568][+100]GAPAS[+568]GADASNAQAAAAMAAIAAIPK.A z=4,scan#=66894,scan time=139.2643
IVYMANNDGANFPEPAAPAANAAQPASGAPASGADASNAQAAAAMAAIAAIPK
Intensity
m/z
HexNAc
HexNAcHex
HexNAc(2)Hex(1)
y2
y3
y4
y5
y6
y7
y8
y9
y10
y11
y12
y18
y21
a4
a6
b5
~y24++
~y27++
~y29++

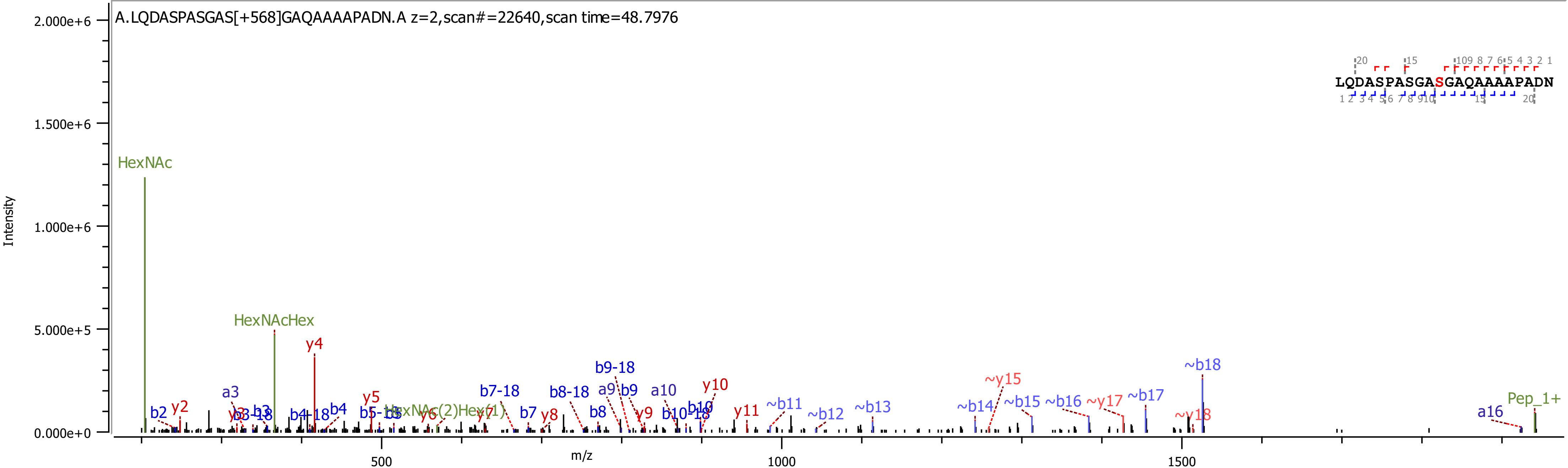
A.LQDASPASGAS[+568]GAQAAAAPADN.A z=2,scan#=22640,scan time=48.7976
LQDASPASGASGAQAAAAPADN
Intensity
m/z
HexNAc
HexNAcHex
y4
~b18
Pep_1+
a16
~y18
~b17
~y17
~b16
~b15
~b14
~y15
~b13
~b12
~b11
y11
y10
b10
b10-18
a10
b9
a9
b9-18
y9
b8
b8-18
y8
b7
b7-18
y7
y6
HexNAc(2)Hex(1)
b5
b5-18
y5
b4
b4-18
b3
b3-18
y3
a3
y2
b2

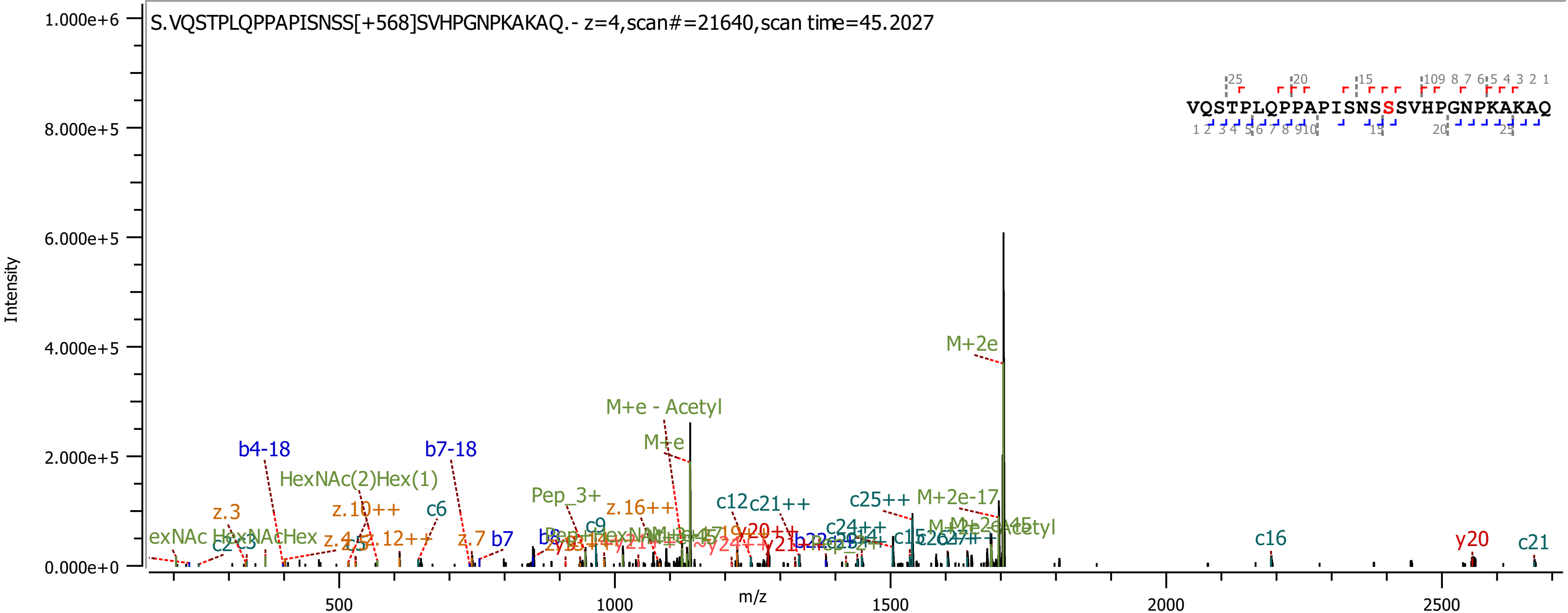

S.VQSTPLQPPAPISNSS[+568]SVHPGNPKAKAQ.- z=4,scan#=21640,scan time=45.2027
Intensity
m/z
1.000e+6
8.000e+5
6.000e+5
4.000e+5
2.000e+5
0.000e+0
500
1000
1500
2000
2500
VQSTPLQPPAPISNSSSVHPGNPKAKAQ
M+2e
M+e - Acetyl
M+e
b4-18
b7-18
HexNAc(2)Hex(1)
Pep_3+
z.3
z.10++
c6
z.16++
c12
c21++
c25++
M+2e-17
z.12++
z.7
b7
c16
y20
c21

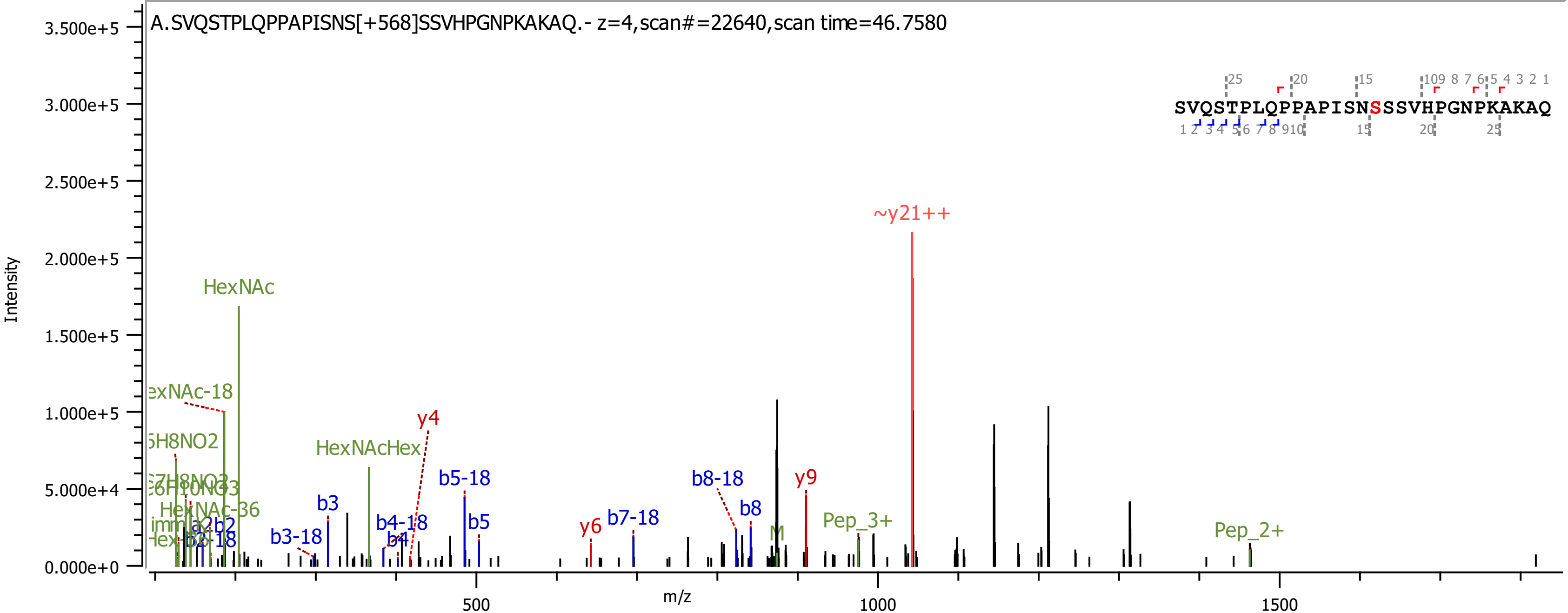

A.SVQSTPLQPPAPISNS[+568]SSVHPGNPKAKAQ.- z=4,scan#=22640,scan time=46.7580
SVQSTPLQPPAPISNSSSVHPGNPKAKAQ
Intensity
m/z
0.000e+0
5.000e+4
1.000e+5
1.500e+5
2.000e+5
2.500e+5
3.000e+5
3.500e+5
500
1000
1500
~y21++
HexNAc
HexNAcHex
y4
y6
y9
b3
b3-18
b4-18
b4
b5-18
b5
b7-18
b8-18
b8
HexNAc-36
Pep_3+
Pep_2+
M

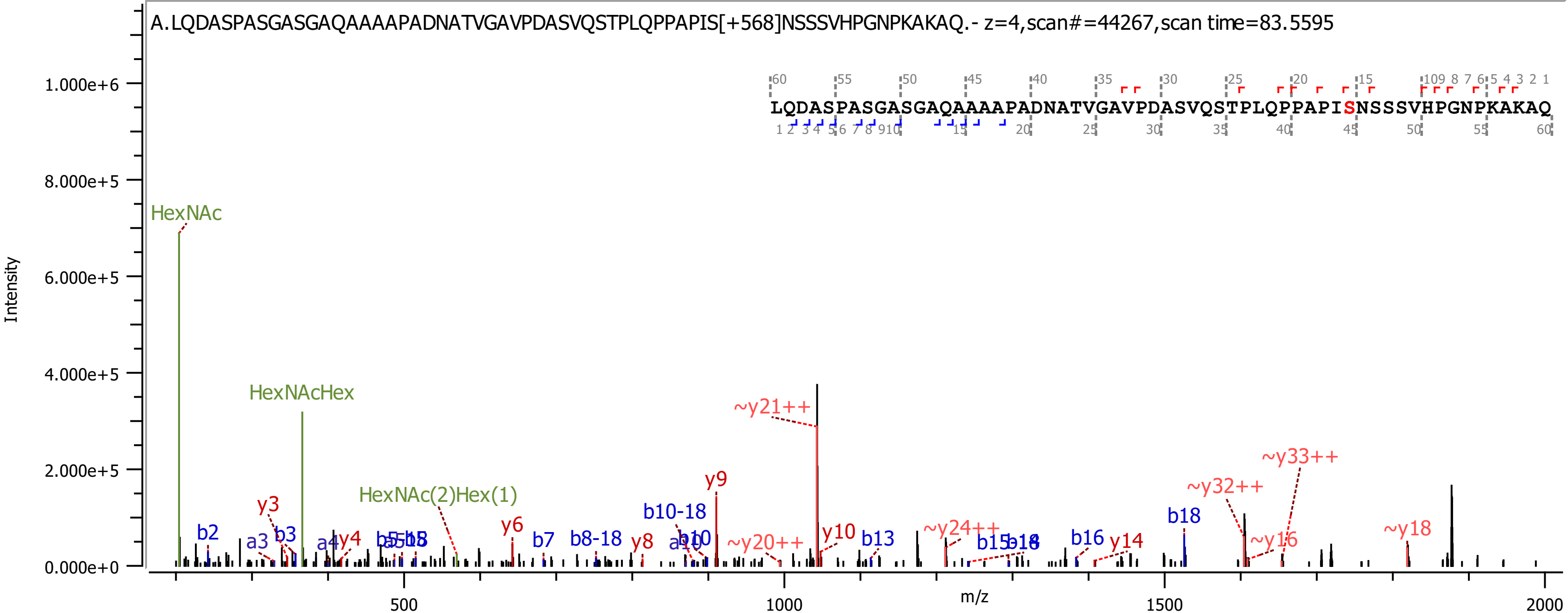

A.LQDASPASGASGAQAAAAPADNATVGAVPDASVQSTPLQPPAPIS[+568]NSSSVHPGNPKAKAQ.- z=4,scan#=44267,scan time=83.5595
LQDASPASGASGAQAAAAPADNATVGAVPDASVQSTPLQPPAPISNSSSVHPGNPKAKAQ
Intensity
m/z
HexNAc
HexNAcHex
HexNAc(2)Hex(1)
~y21++
~y33++
~y32++
y9
b18
~y18
~y24++
~y20++
~y16
y14
b16
b13
y10
b10-18
b8-18
b7
y6
y8
y3
y4
b2
a3
b3
a4
b10
a10

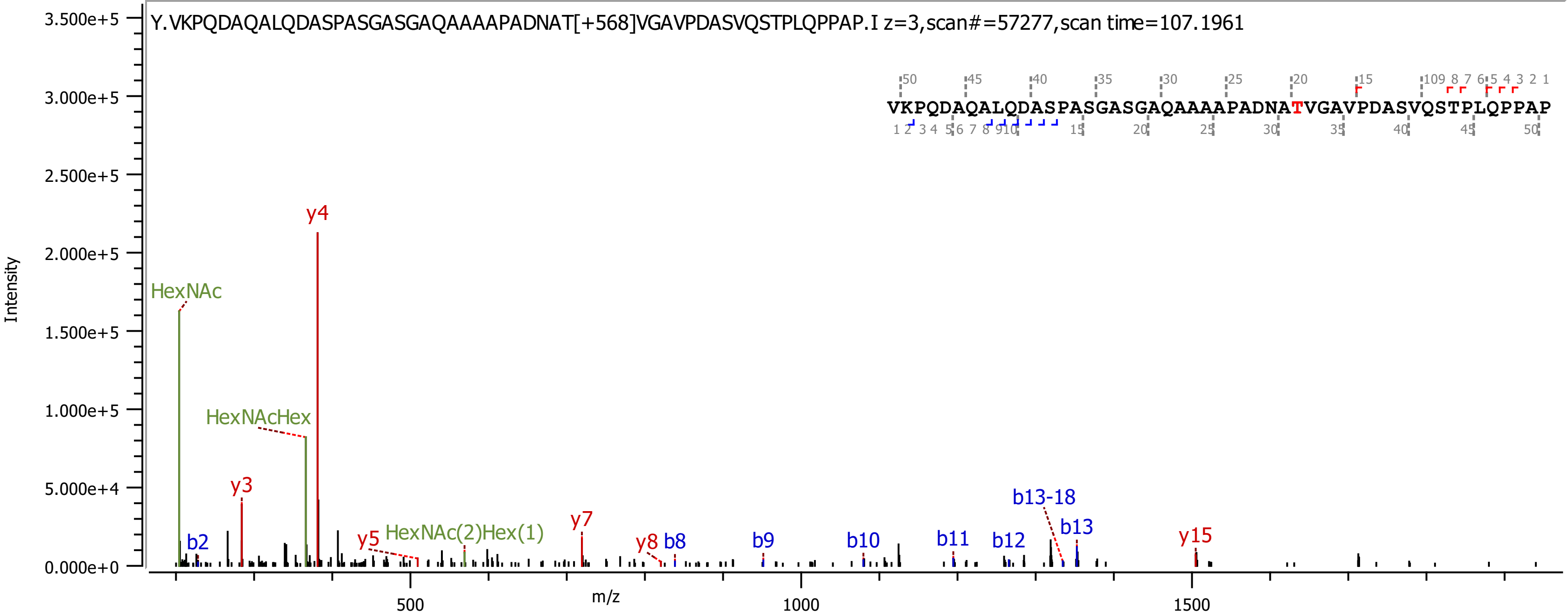
Y.VKPQDAQALQDASPASGASGAQAAAAPADNAT[+568]VGAVPDASVQSTPLQPPAP.I z=3,scan#=57277,scan time=107.1961
VKPQDAQALQDASPASGASGAQAAAAPADNATVGAVPDASVQSTPLQPPAP
Intensity
m/z
HexNAc
HexNAcHex
HexNAc(2)Hex(1)
y3
y4
y5
y7
y8
y15
b2
b8
b9
b10
b11
b12
b13-18
b13
3.500e+5
3.000e+5
2.500e+5
2.000e+5
1.500e+5
1.000e+5
5.000e+4
0.000e+0
500
1000
1500

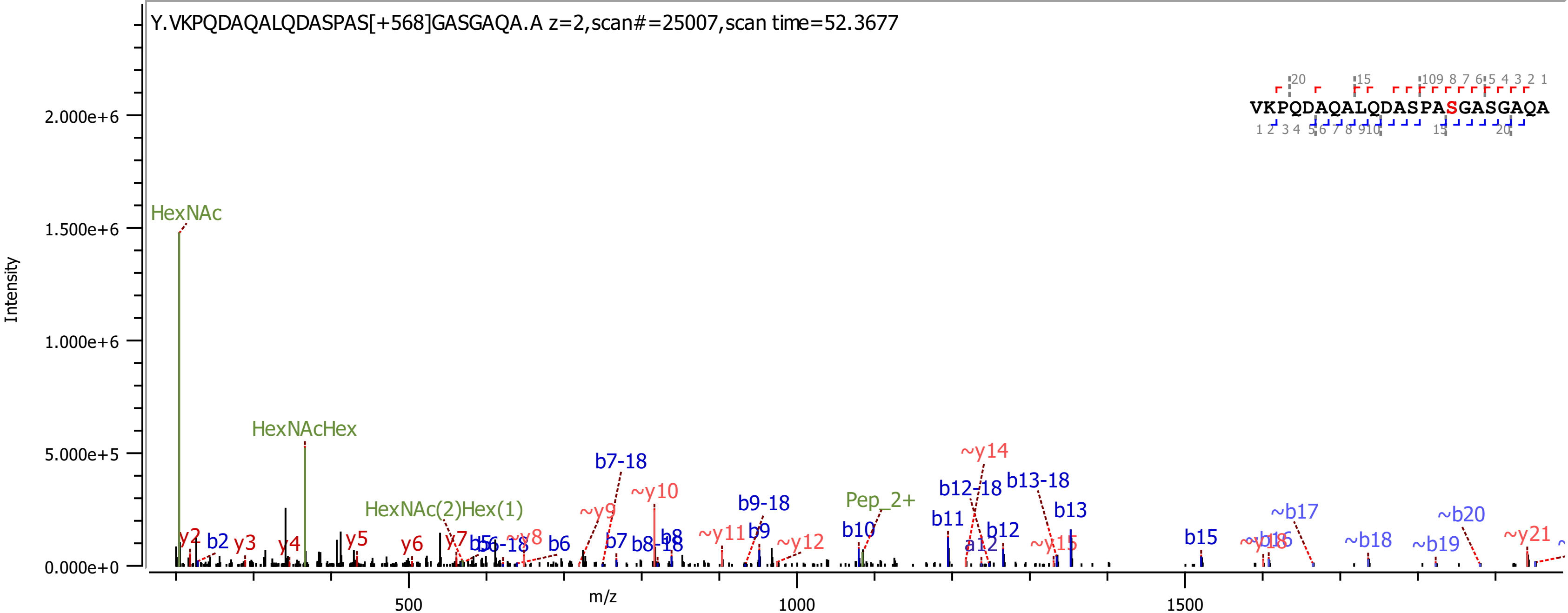

Y.VKPQDAQALQDASPAS[+568]GASGAQA.A z=2,scan#=25007,scan time=52.3677
VKPQDAQALQDASPASGASGAQA
Intensity
2.000e+6
1.500e+6
1.000e+6
5.000e+5
0.000e+0
m/z
500
1000
1500
HexNAc
HexNAcHex
HexNAc(2)Hex(1)
Pep_2+
y2
b2
y3
y4
y5
y6
y7
b5
b6-18
~y8
b6
~y9
b7-18
b7
b8-18
b8
~y10
~y11
b9
b9-18
~y12
b10
b11
b12-18
~y14
a12
b12
b13-18
b13
~y15
b15
~y16
~b16
~b17
~b18
~b19
~b20
~y21

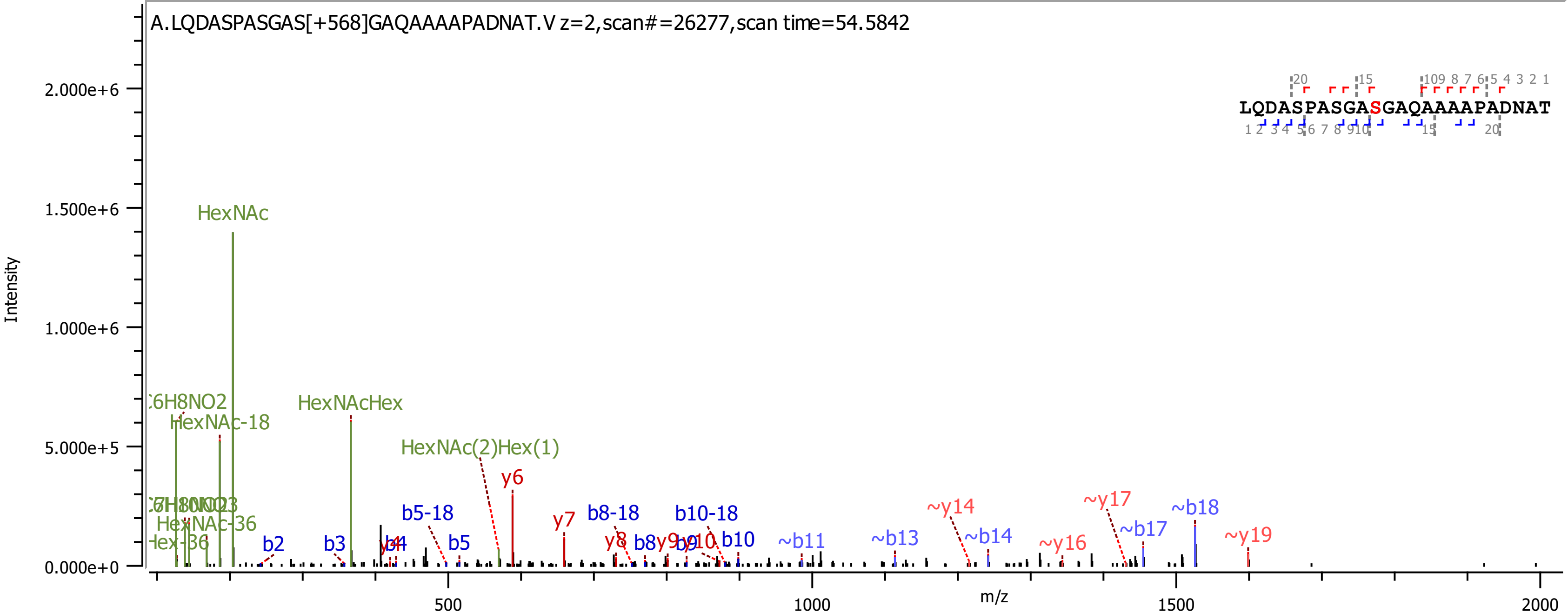
A.LQDASPASGAS[+568]GAQAAAAPADNAT.V z=2,scan#=26277,scan time=54.5842
LQDASPASGASGAQAAAAPADNAT
Intensity
m/z
2.000e+6
1.500e+6
1.000e+6
5.000e+5
0.000e+0
500
1000
1500
2000
HexNAc
HexNAc-18
HexNAc-36
HexNAcHex
HexNAc(2)Hex(1)
y6
y7
y8
b2
b3
b5
b5-18
b8-18
b8
b10-18
b10
~b11
~b13
~b14
~y14
~y16
~y17
~b17
~b18
~y19

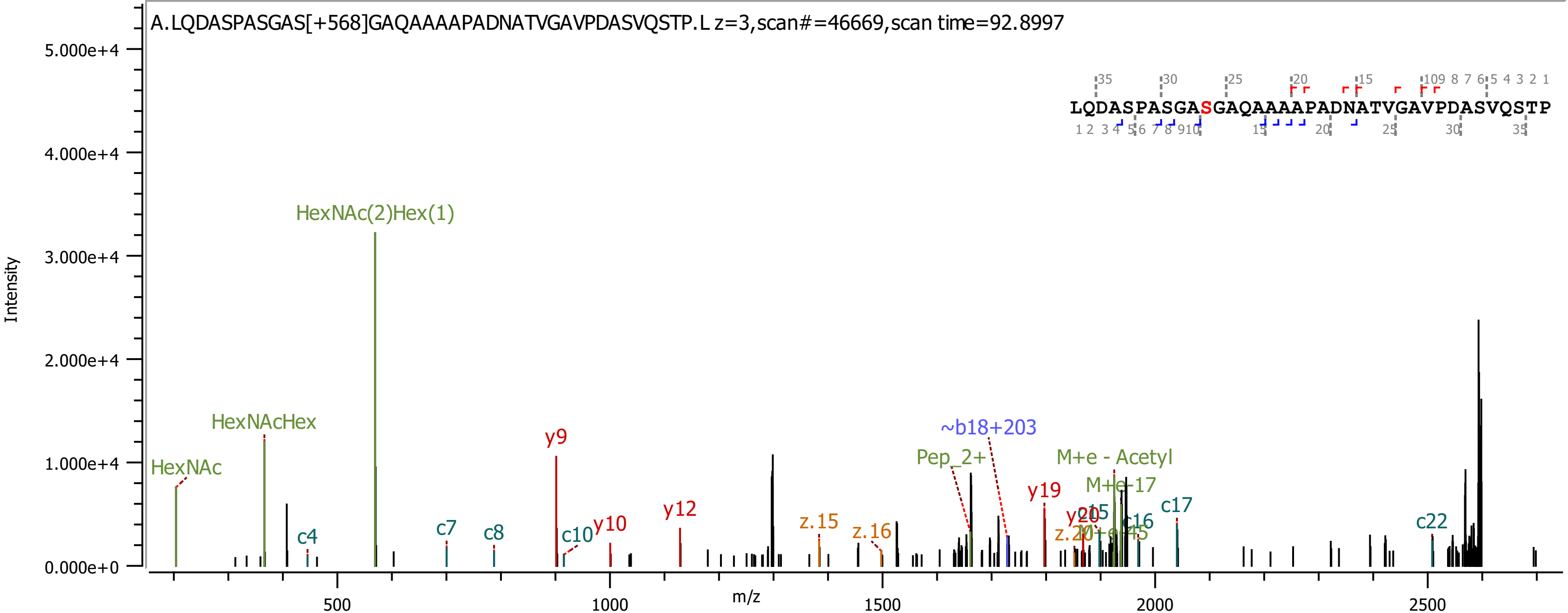
A.LQDASPASGAS[+568]GAQAAAAPADNATVGAVPDASVQSTP.L z=3,scan#=46669,scan time=92.8997
LQDASPASGASGAQAAAAPADNATVGAVPDASVQSTP
Intensity
m/z
HexNAc
HexNAcHex
HexNAc(2)Hex(1)
c4
c7
c8
y9
c10
y10
y12
z.15
z.16
Pep_2+
~b18+203
y19
y20
c15
M+e - Acetyl
M+e-17
M+e-45
c16
c17
c22

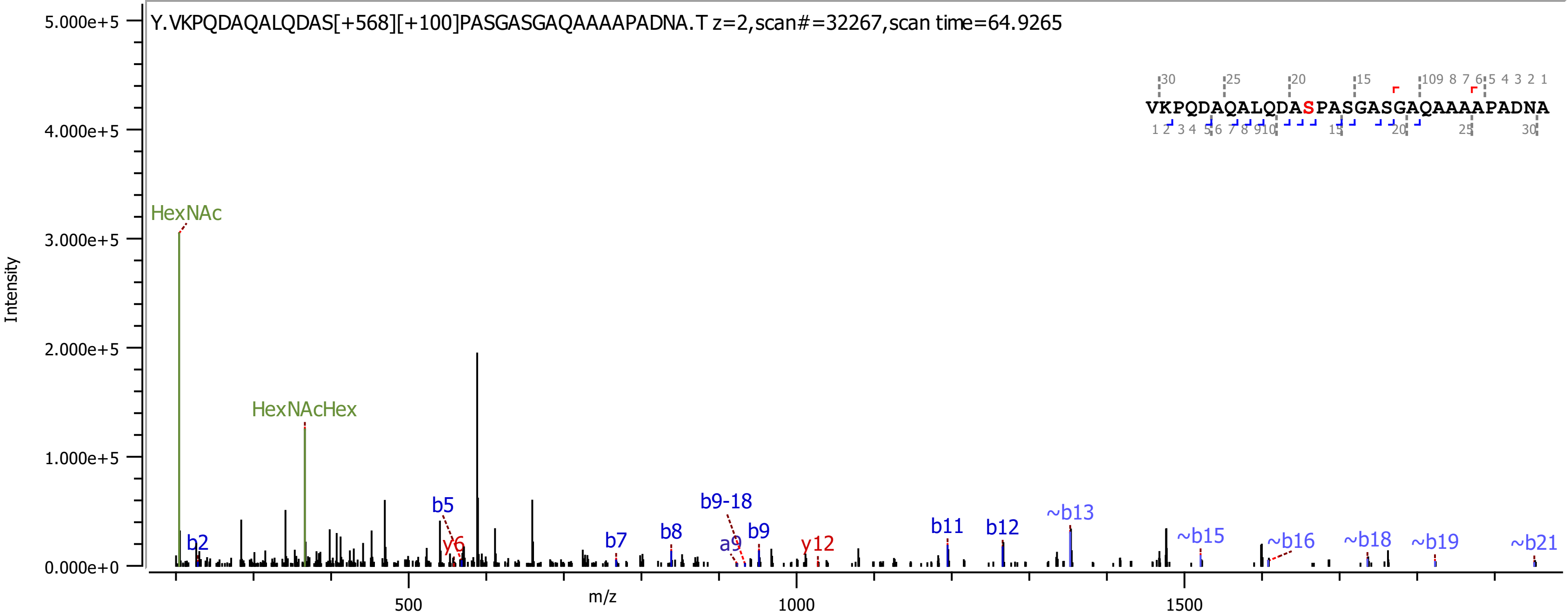
Y.VKPQDAQALQDAS[+568][+100]PASGASGAQAAAAPADNA.T z=2,scan#=32267,scan time=64.9265
VKPQDAQALQDASPASGASGAQAAAAPADNA
Intensity
m/z
5.000e+5
4.000e+5
3.000e+5
2.000e+5
1.000e+5
0.000e+0
500
1000
1500
HexNAc
HexNAcHex
b2
b5
y6
b7
b8
b9-18
a9
b9
y12
b11
b12
~b13
~b15
~b16
~b18
~b19
~b21

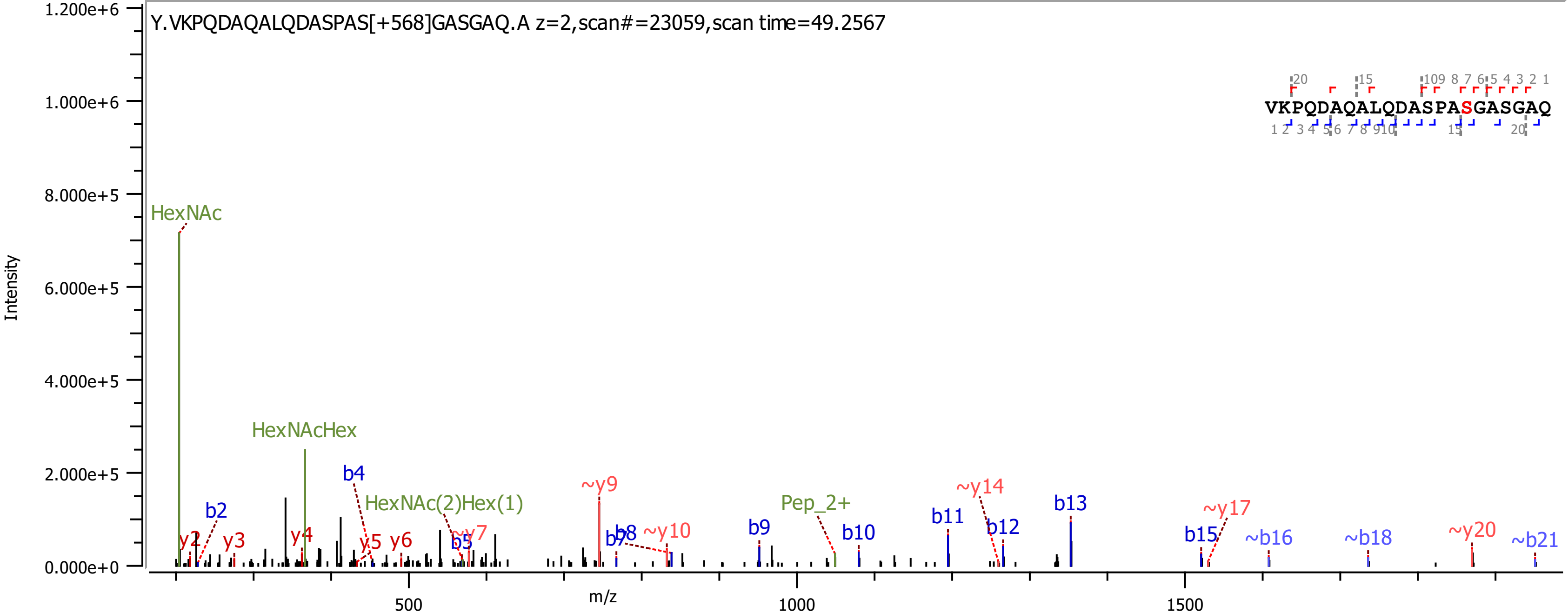
Y.VKPQDAQALQDASPAS[+568]GASGAQ.A z=2,scan#=23059,scan time=49.2567
VKPQDAQALQDASPASGASGAQ
Intensity
m/z
HexNAc
HexNAcHex
HexNAc(2)Hex(1)
Pep_2+
b2
b4
b5
b7
b8
b9
b10
b11
b12
b13
b15
~b16
~b18
~b21
y2
y3
y4
y5
y6
~y7
~y9
~y10
~y14
~y17
~y20
0.000e+0
2.000e+5
4.000e+5
6.000e+5
8.000e+5
1.000e+6
1.200e+6
500
1000
1500

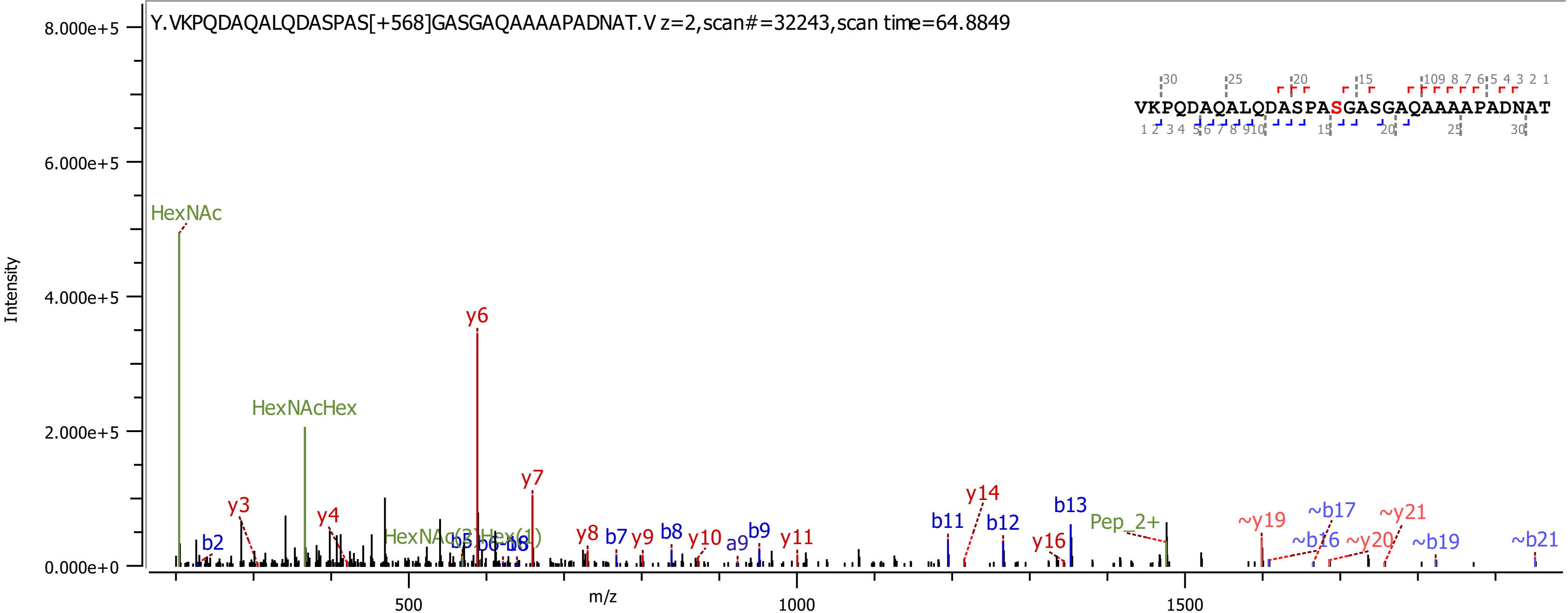

Y.VKPQDAQALQDASPAS[+568]GASGAQAAAAPADNAT.V z=2,scan#=32243,scan time=64.8849
VKPQDAQALQDASPASGASGAQAAAAPADNAT
Intensity
m/z
HexNAc
HexNAcHex
y6
y7
y3
y4
b2
y8
b7
y9
b8
y10
a9
b9
y11
b11
y14
b12
b13
y16
Pep_2+
~y19
~b17
~y21
~b16
~y20
~b19
~b21

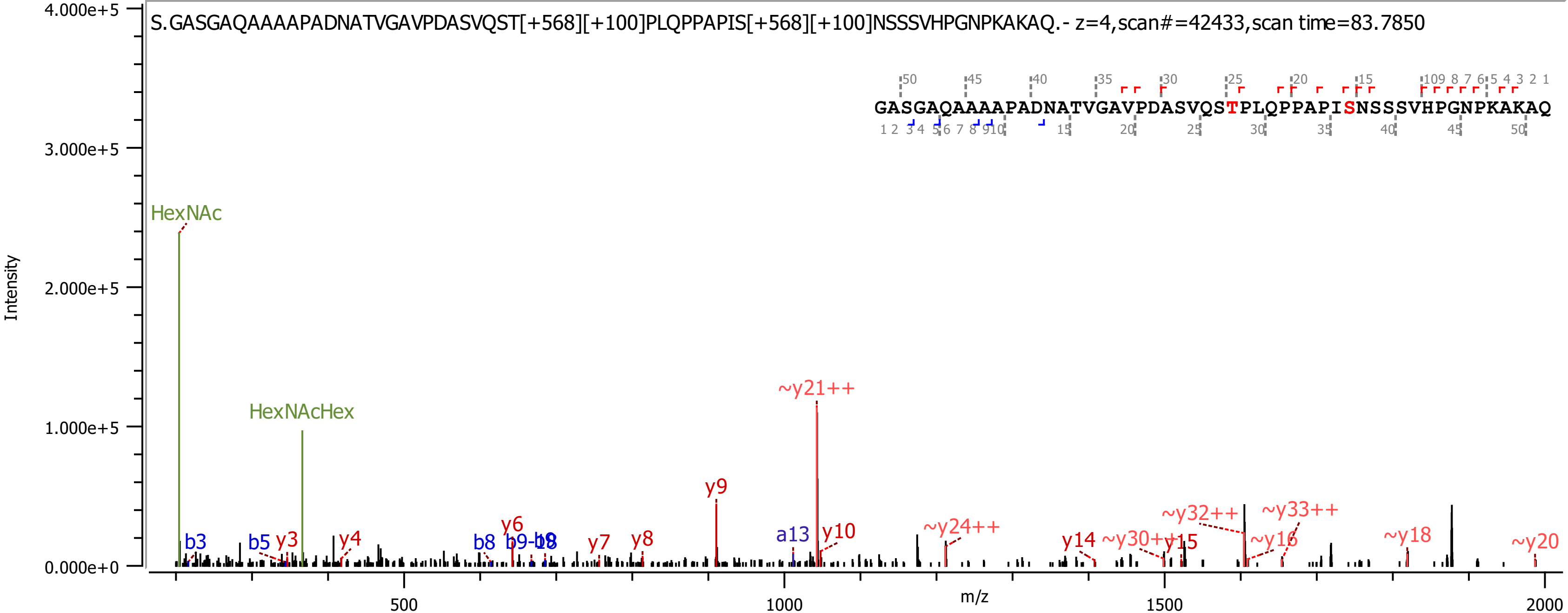

S.GASGAQAAAAPADNATVGAVPDASVQST[+568][+100]PLQPPAPIS[+568][+100]NSSSVHPGNPKAKAQ.- z=4,scan#=42433,scan time=83.7850
GASGAQAAAAPADNATVGAVPDASVQSTPLQPPAPISNSSSVHPGNPKAKAQ
HexNAc
HexNAcHex
~y21++
y9
~y24++
~y32++
~y33++
~y18
~y20
b3
b5
y3
y4
b8
y6
y7
y8
a13
y10
y14
~y30++
y15
~y16
Intensity
m/z
4.000e+5
3.000e+5
2.000e+5
1.000e+5
0.000e+0
500
1000
1500
2000

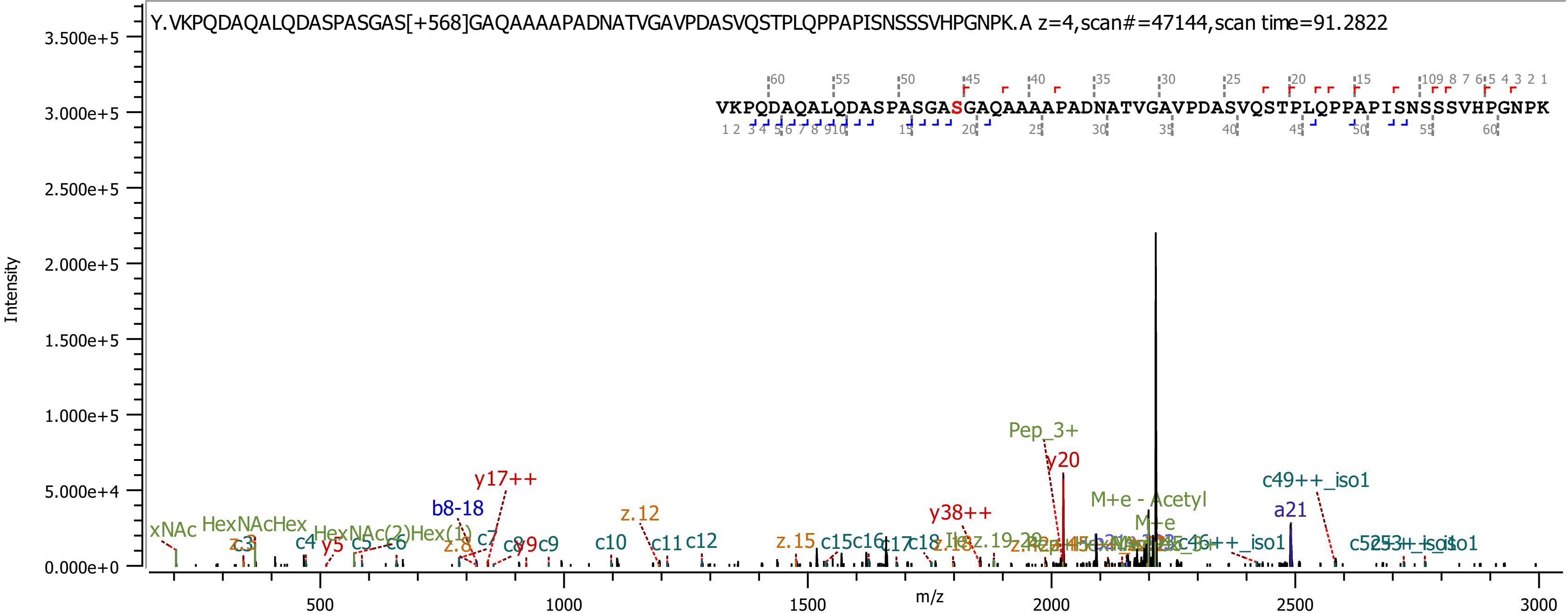
Y.VKPQDAQALQDASPASGAS[+568]GAQAAAAPADNATVGAVPDASVQSTPLQPPAPISNSSSVHPGNPK.A z=4,scan#=47144,scan time=91.2822
VKPQDAQALQDASPASGASGAQAAAAPADNATVGAVPDASVQSTPLQPPAPISNSSSVHPGNPK
Intensity
m/z
Pep_3+
y20
y17++
b8-18
z.12
y38++
M+e - Acetyl
M+e
a21
c49++_iso1
HexNAcHex
HexNAc(2)Hex(1)
c4
c5
c6
c7
c9
c10
c11
c12
z.15
c15
c16
c17
c18
z.18
y5
y9
z.8
xNAc

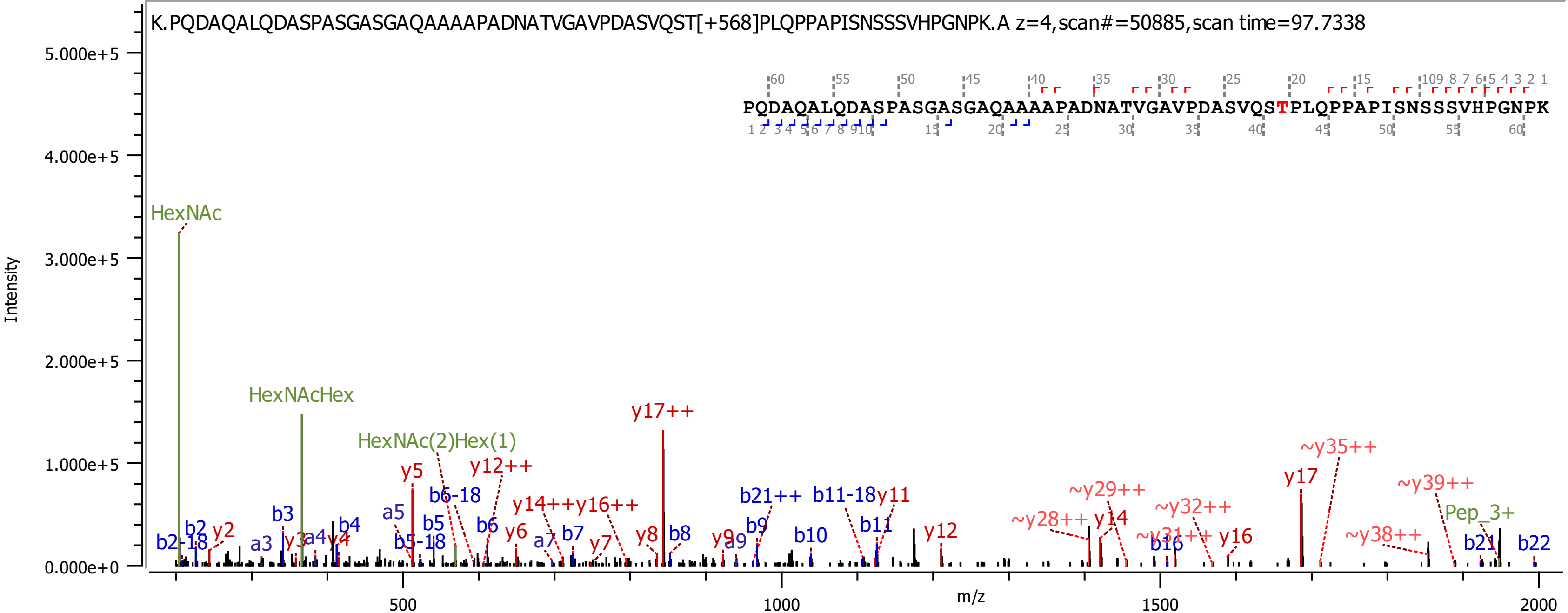
K.PQDAQALQDASPASGASGAQAAAAPADNATVGAVPDASVQST[+568]PLQPPAPISNSSSVHPGNPK.A z=4,scan#=50885,scan time=97.7338
PQDAQALQDASPASGASGAQAAAAPADNATVGAVPDASVQSTPLQPPAPISNSSSVHPGNPK
Intensity
m/z
5.000e+5
4.000e+5
3.000e+5
2.000e+5
1.000e+5
0.000e+0
500
1000
1500
2000
HexNAc
HexNAcHex
HexNAc(2)Hex(1)
y17++
y5
y12++
b6-18
b5
b5-18
b6
y6
a5
b3
b2
b2-18
y2
a3
y3
a4
y4
b4
y14++
y16++
a7
b7
y7
y8
b8
y9
a9
b9
b21++
b11-18
y11
b10
b11
y12
~y28++
~y29++
y14
~y31++
b16
~y32++
y16
y17
~y35++
~y38++
~y39++
Pep_3+
b21
b22

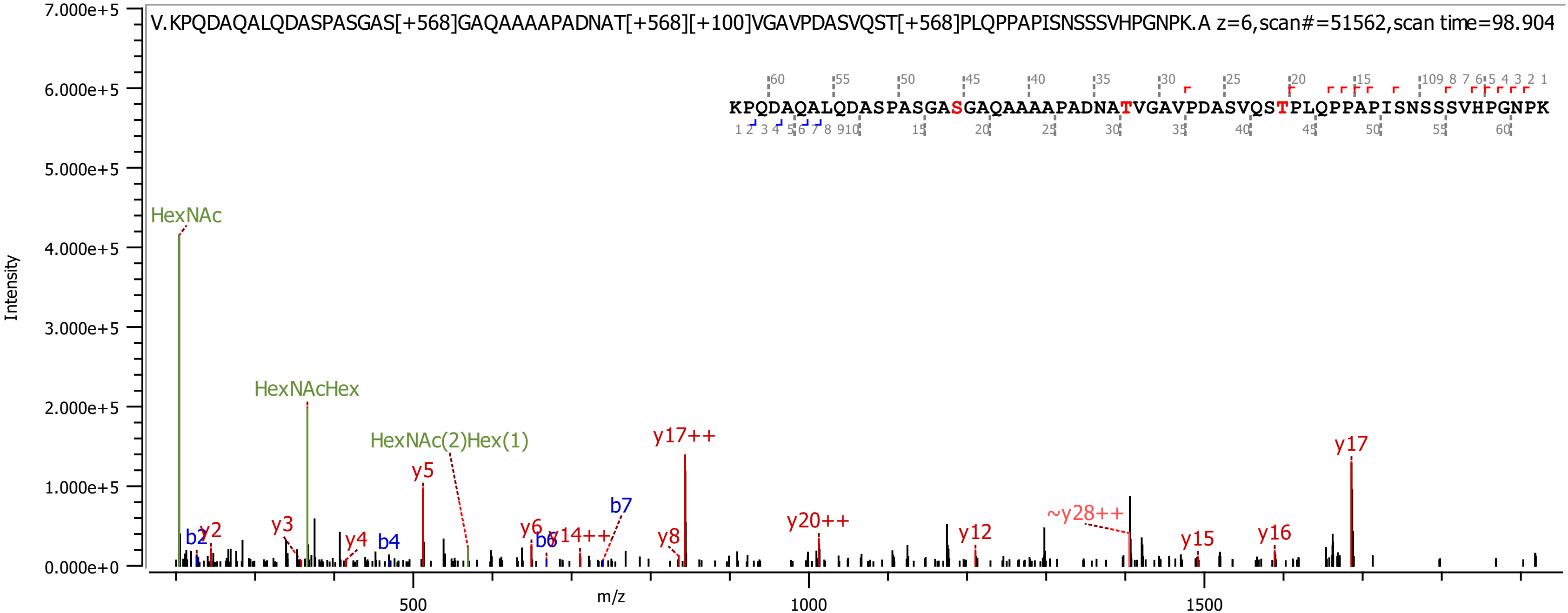
V.KPQDAQALQDASPASGAS[+568]GAQAAAAPADNAT[+568][+100]VGAVPDASVQST[+568]PLQPPAPISNSSSVHPGNPK.A z=6,scan#=51562,scan time=98.904
KPQDAQALQDASPASGASGAQAAAAPADNATVGAVPDASVQSTPLQPPAPISNSSSVHPGNPK
Intensity
m/z
HexNAc
HexNAcHex
HexNAc(2)Hex(1)
b2
y2
y3
y4
b4
y5
y6
b6
y14++
b7
y8
y17++
y20++
y12
~y28++
y15
y16
y17
7.000e+5
6.000e+5
5.000e+5
4.000e+5
3.000e+5
2.000e+5
1.000e+5
0.000e+0
500
1000
1500

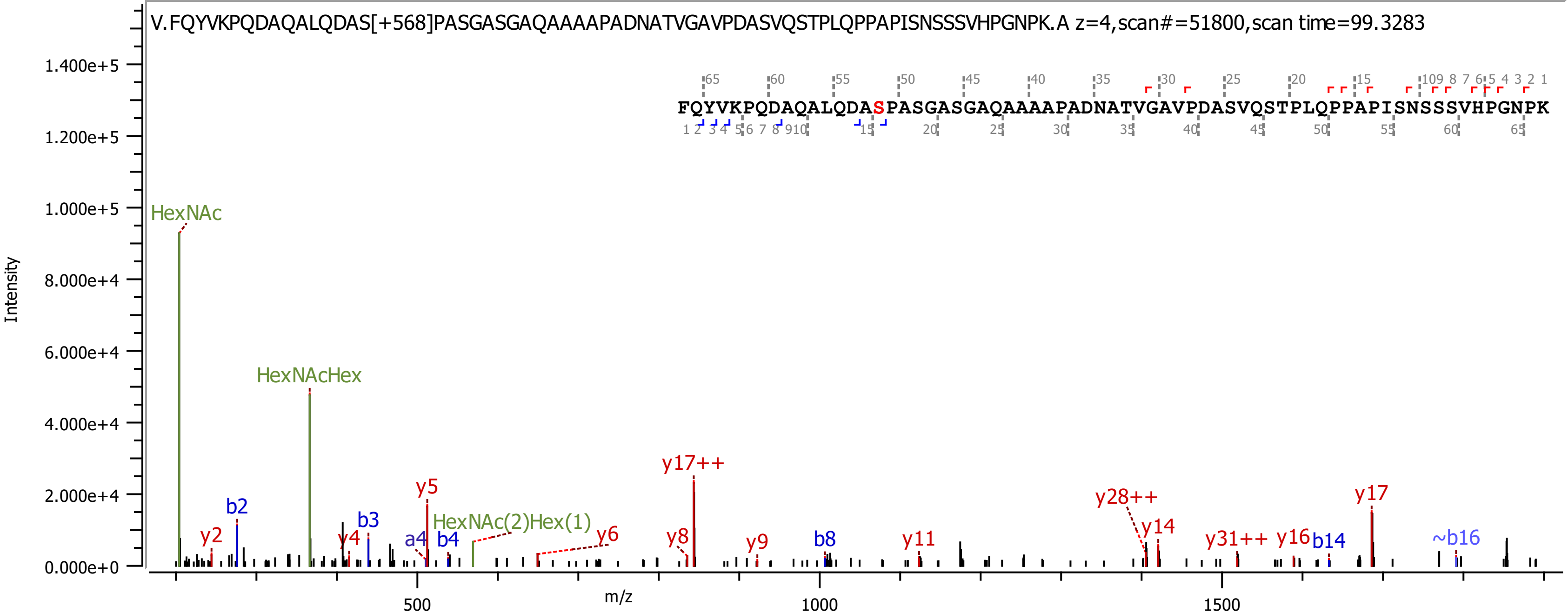
V.FQYVKPQDAQALQDAS[+568]PASGASGAQAAAAPADNATVGAVPDASVQSTPLQPPAPISNSSSVHPGNPK.A z=4,scan#=51800,scan time=99.3283
FQYVKPQDAQALQDASPASGASGAQAAAAPADNATVGAVPDASVQSTPLQPPAPISNSSSVHPGNPK
Intensity
m/z
HexNAc
HexNAcHex
HexNAc(2)Hex(1)
b2
b3
b4
b8
b14
~b16
a4
y2
y4
y5
y6
y8
y9
y11
y14
y16
y17
y17++
y28++
y31++

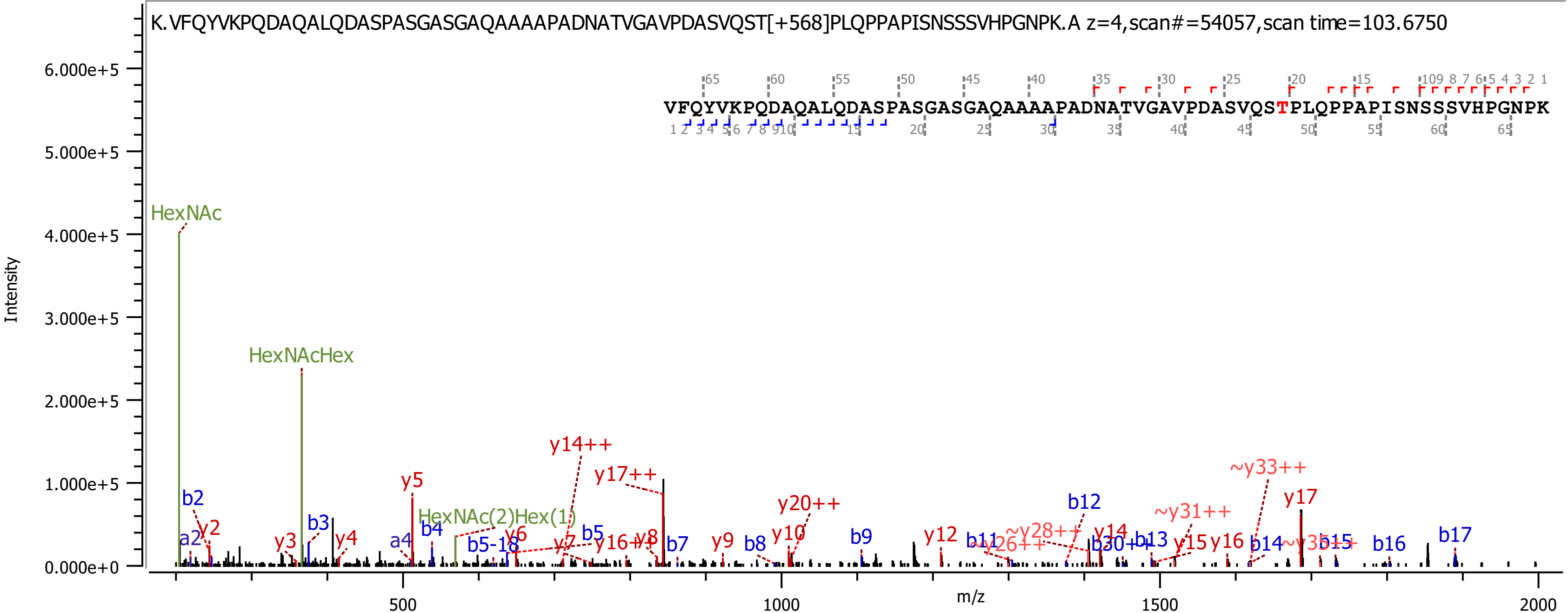
K.VFQYVKPQDAQALQDASPASGASGAQAAAAPADNATVGAVPDASVQST[+568]PLQPPAPISNSSSVHPGNPK.A z=4,scan#=54057,scan time=103.6750
VFQYVKPQDAQALQDASPASGASGAQAAAAPADNATVGAVPDASVQSTPLQPPAPISNSSSVHPGNPK
Intensity
m/z
HexNAc
HexNAcHex
HexNAc(2)Hex(1)
y14++
y17++
y5
y20++
~y33++
y17
b12
~y31++
b2
y2
a2
y3
b3
y4
a4
b4
b5-18
y6
y7
b5
y16++
y8
b7
y9
b8
y10
b9
y12
b11
~y26++
~y28++
y14
b30++
b13
y15
y16
b14
~y35++
b15
b16
b17

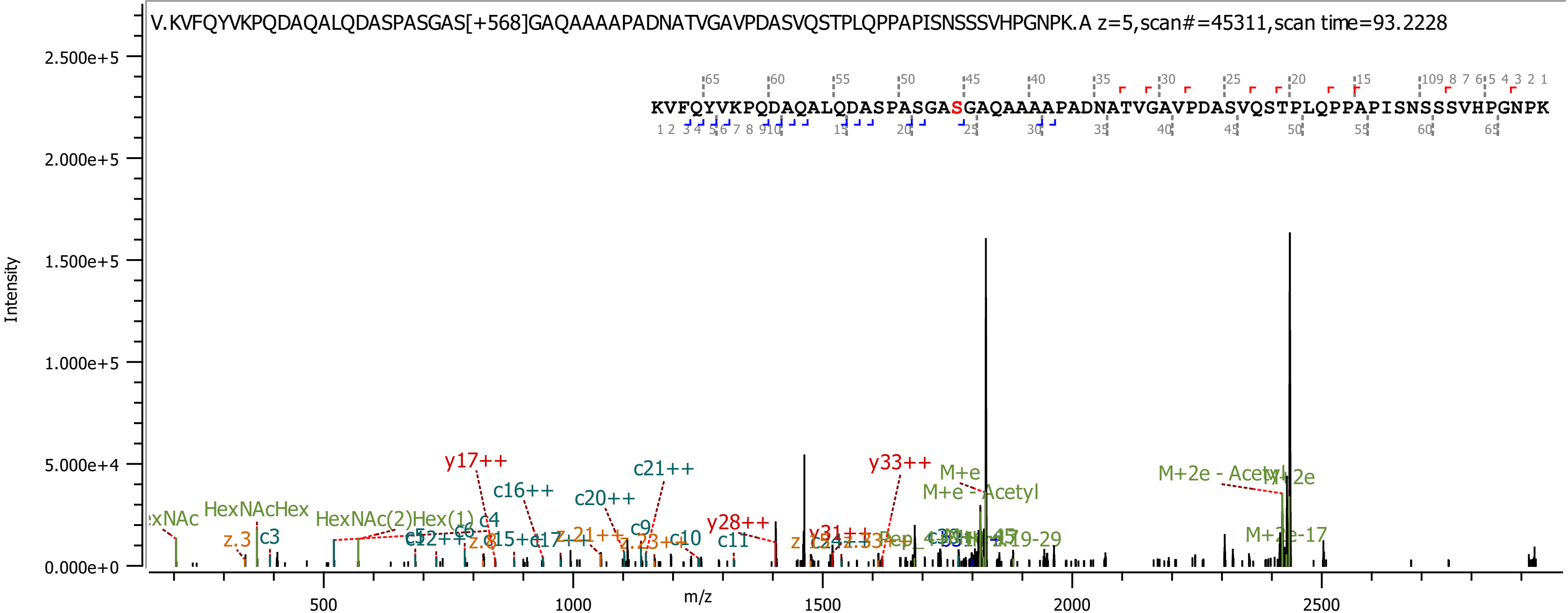
V.KVFQYVKPQDAQALQDASPASGAS[+568]GAQAAAAPADNATVGAVPDASVQSTPLQPPAPISNSSSVHPGNPK.A z=5,scan#=45311,scan time=93.2228
KVFQYVKPQDAQALQDASPASGASGAQAAAAPADNATVGAVPDASVQSTPLQPPAPISNSSSVHPGNPK
Intensity
m/z
2.500e+5
2.000e+5
1.500e+5
1.000e+5
5.000e+4
0.000e+0
500
1000
1500
2000
2500
y17++
c16++
c20++
c21++
y28++
y33++
M+e
M+e - Acetyl
M+2e - Acetyl
M+2e
M+2e-17
HexNAcHex
HexNAc(2)Hex(1)
c3
c4
c6
c9
c10
c11
z.3
z.8
z.21++
z.23++

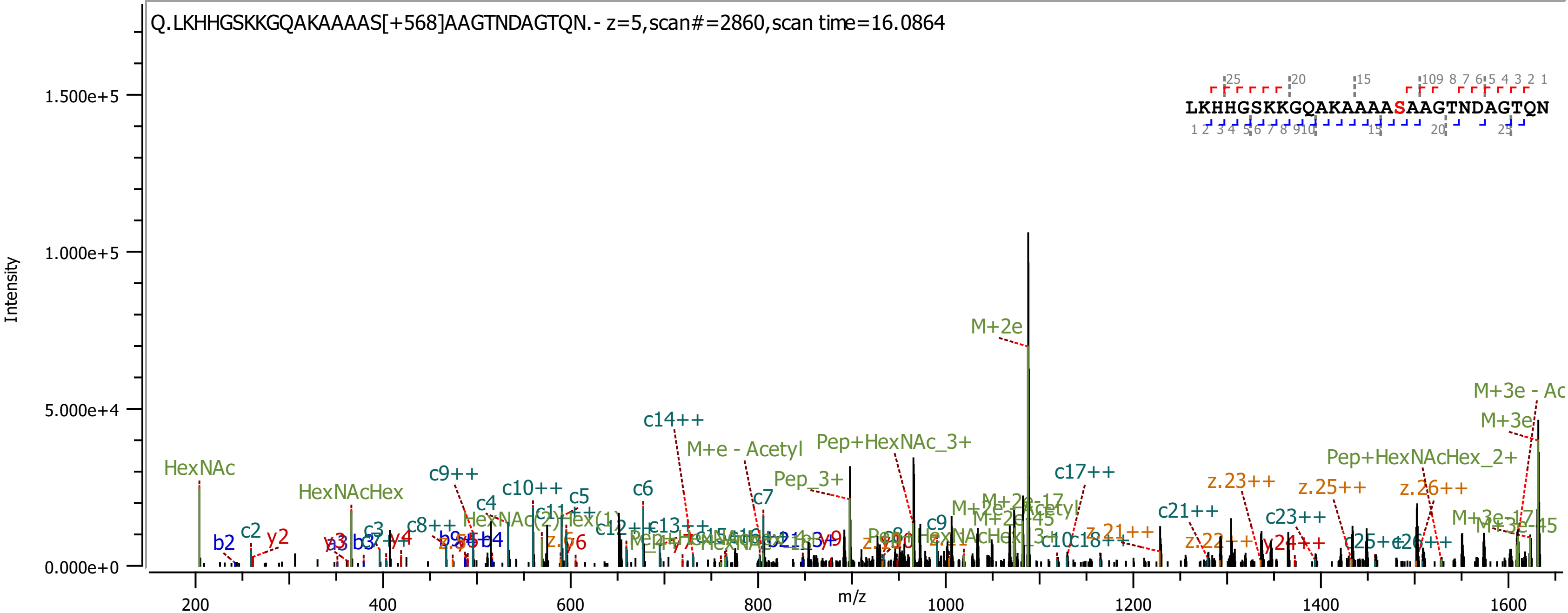

Q.LKHHGSKKGQAKAAAAS[+568]AAGTNDAGTQN.- z=5,scan#=2860,scan time=16.0864
LKHHGSKKGQAKAAAASAAGTNDAGTQN
Intensity
m/z
1.500e+5
1.000e+5
5.000e+4
0.000e+0
200
400
600
800
1000
1200
1400
1600
M+2e
HexNAc
HexNAcHex
c14++
M+e - Acetyl
Pep+HexNAc_3+
Pep_3+
c17++
Pep+HexNAcHex_2+
M+3e - Ac
M+3e
z.23++
z.25++
z.26++
c21++
c23++
c9++
c10++
c5
c6
c4
c7
c9
c2
y2
b2
c3
y4
c8++
z.y6
c13++
z.21++
z.22++
y24++
c25++
c26++

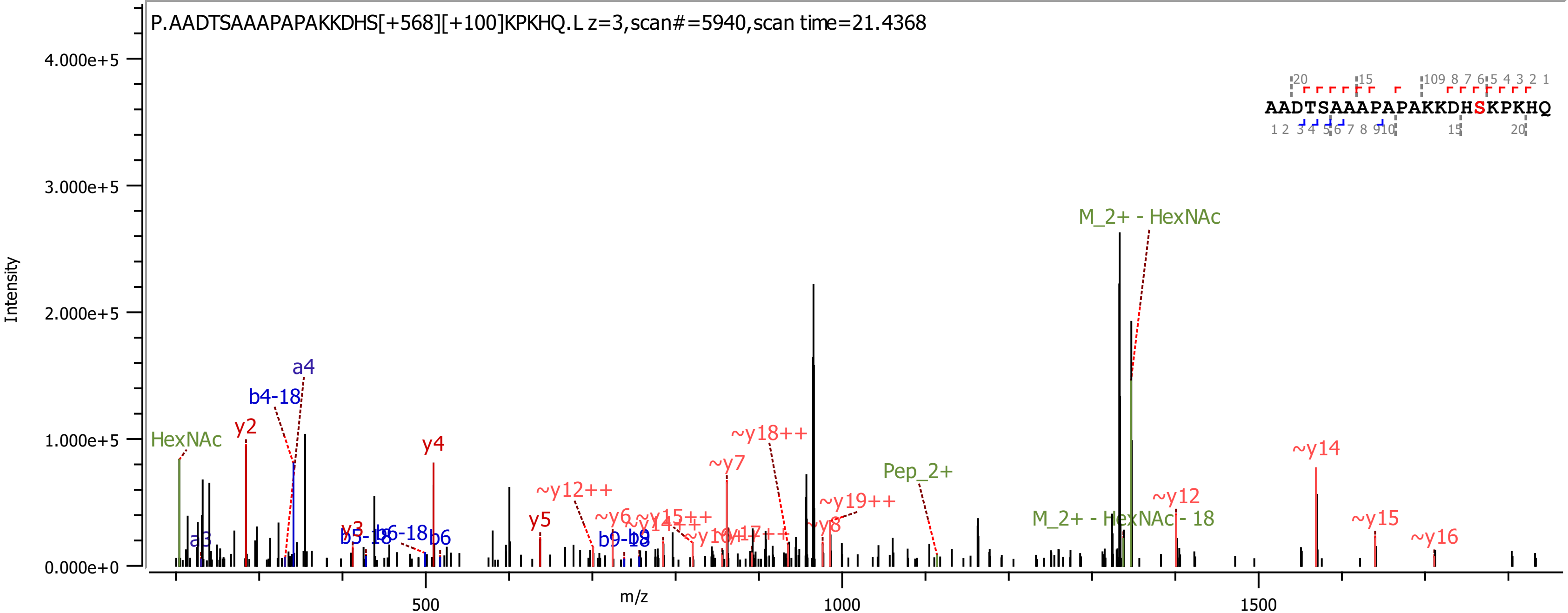
P.AADTSAAAPAPAKKDHS[+568][+100]KPKHQ.L z=3,scan#=5940,scan time=21.4368
AADTSAAAPAPAKKDHSKPKHQ
Intensity
m/z
4.000e+5
3.000e+5
2.000e+5
1.000e+5
0.000e+0
500
1000
1500
HexNAc
y2
b4-18
a4
y4
y5
~y12++
~y7
~y18++
~y19++
Pep_2+
M_2+ - HexNAc
M_2+ - HexNAc - 18
~y12
~y14
~y15
~y16
b6
b6-18

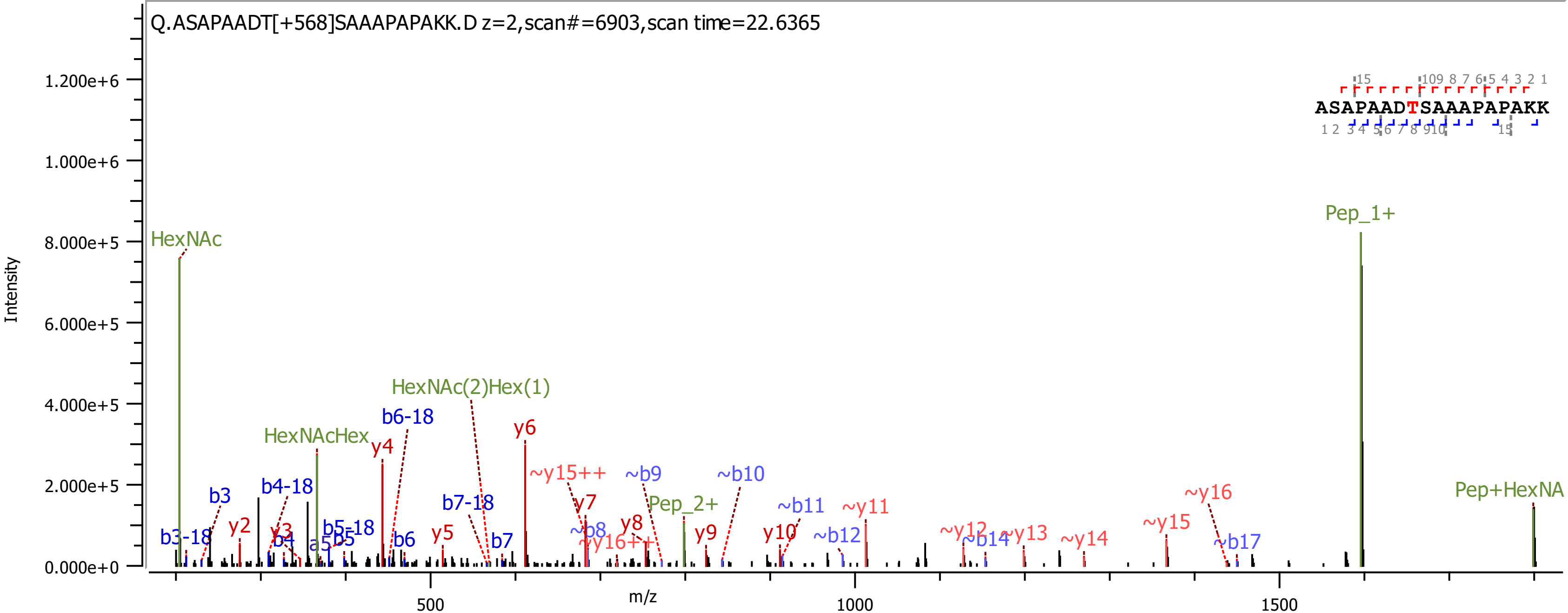

Q.ASAPAADT[+568]SAAAPAPAKK.D z=2,scan#=6903,scan time=22.6365
ASAPAADTSAAAPAPAKK
Intensity
m/z
0.000e+0
2.000e+5
4.000e+5
6.000e+5
8.000e+5
1.000e+6
1.200e+6
500
1000
1500
HexNAc
HexNAcHex
HexNAc(2)Hex(1)
Pep_1+
Pep_2+
Pep+HexNA
b3-18
b3
y2
b4-18
y3
b4
a5
b5
b5-18
y4
b6-18
b6
y5
b7-18
b7
y6
~y15++
y7
~b8
~y16++
y8
~b9
y9
~b10
y10
~b11
~b12
~y11
~y12
~b14
~y13
~y14
~y15
~y16
~b17

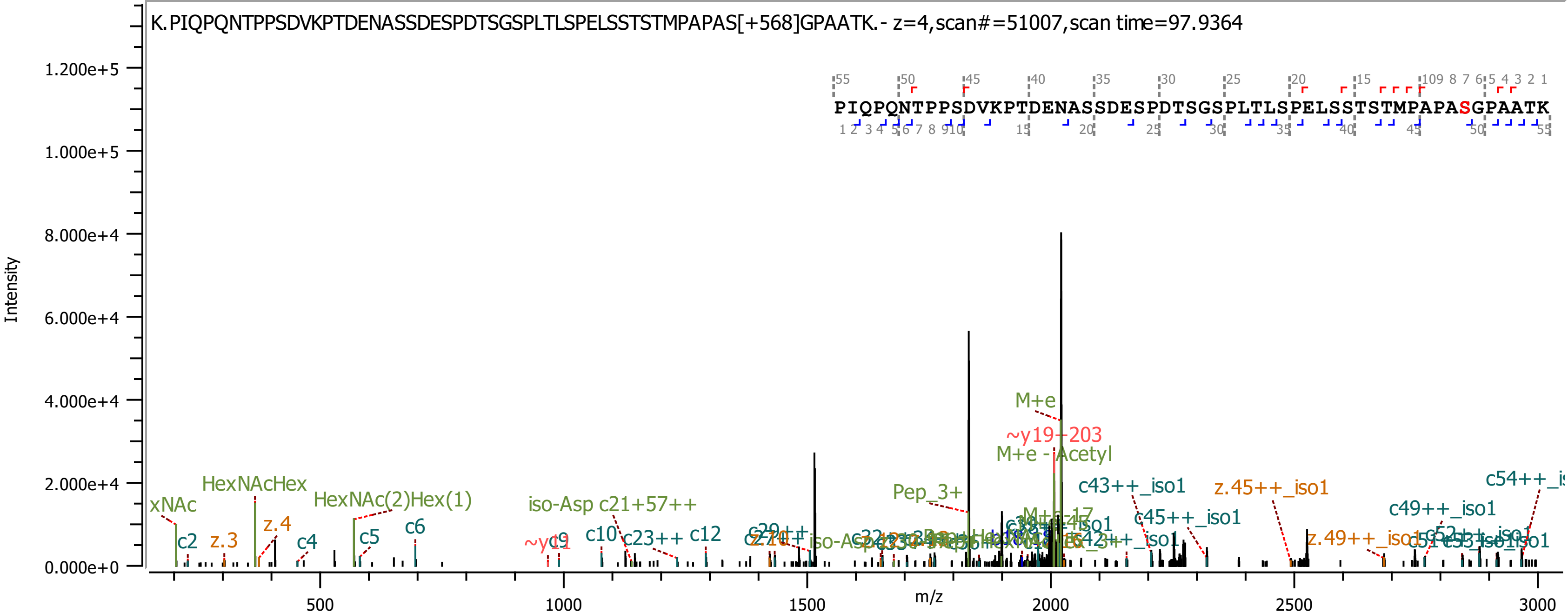
K.PIQPQNTPPSDVKPTDENASSDESPDTSGSPLTLSPELSSTSTMPAPAS[+568]GPAATK.- z=4,scan#=51007,scan time=97.9364
PIQPQNTPPSDVKPTDENASSDESPDTSGSPLTLSPELSSTSTMPAPASGPAATK
Intensity
m/z
1.200e+5
1.000e+5
8.000e+4
6.000e+4
4.000e+4
2.000e+4
0.000e+0
500
1000
1500
2000
2500
3000
M+e
~y19-203
M+e - Acetyl
Pep_3+
c43++_iso1
z.45++_iso1
c54++_is
c49++_iso1
c45++_iso1
z.49++_iso1
xNAc
HexNAcHex
HexNAc(2)Hex(1)
iso-Asp c21+57++
c2
z.3
z.4
c4
c5
c6
~y11
c9
c10
c23++
c12

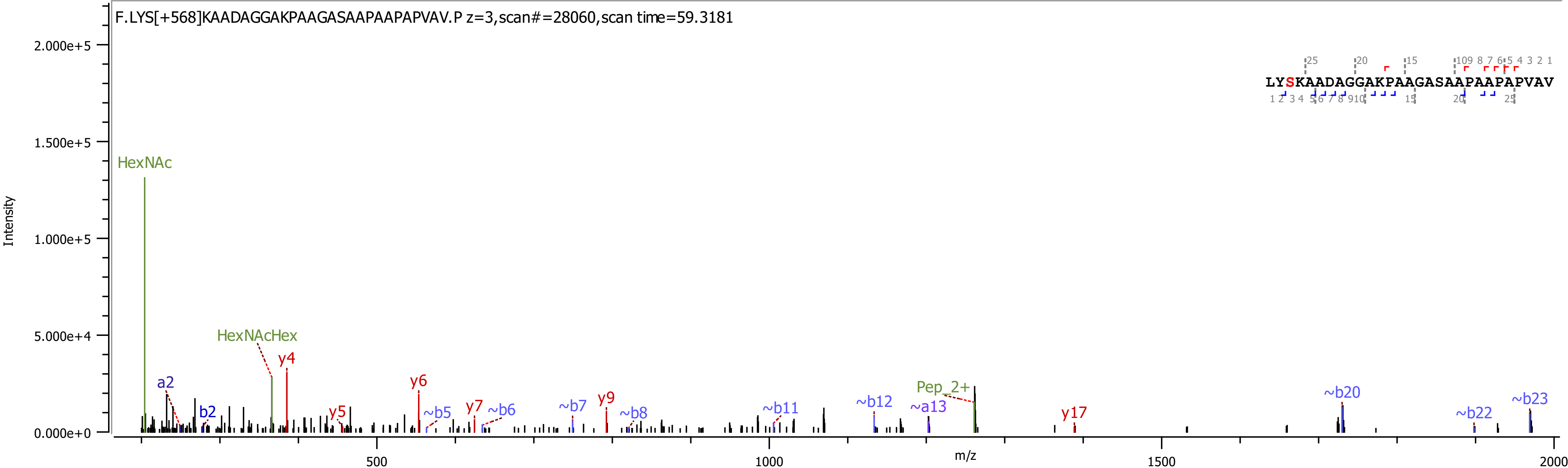
F.LYS[+568]KAADAGGAKPAAGASAAPAAPAPVAV.P z=3,scan#=28060,scan time=59.3181
Intensity
m/z
HexNAc
HexNAcHex
a2
b2
y4
y5
y6
~b5
y7
~b6
~b7
y9
~b8
~b11
~b12
~a13
Pep_2+
y17
~b20
~b22
~b23
LYSKAADAGGAKPAAGASAAPAAPAPVAV

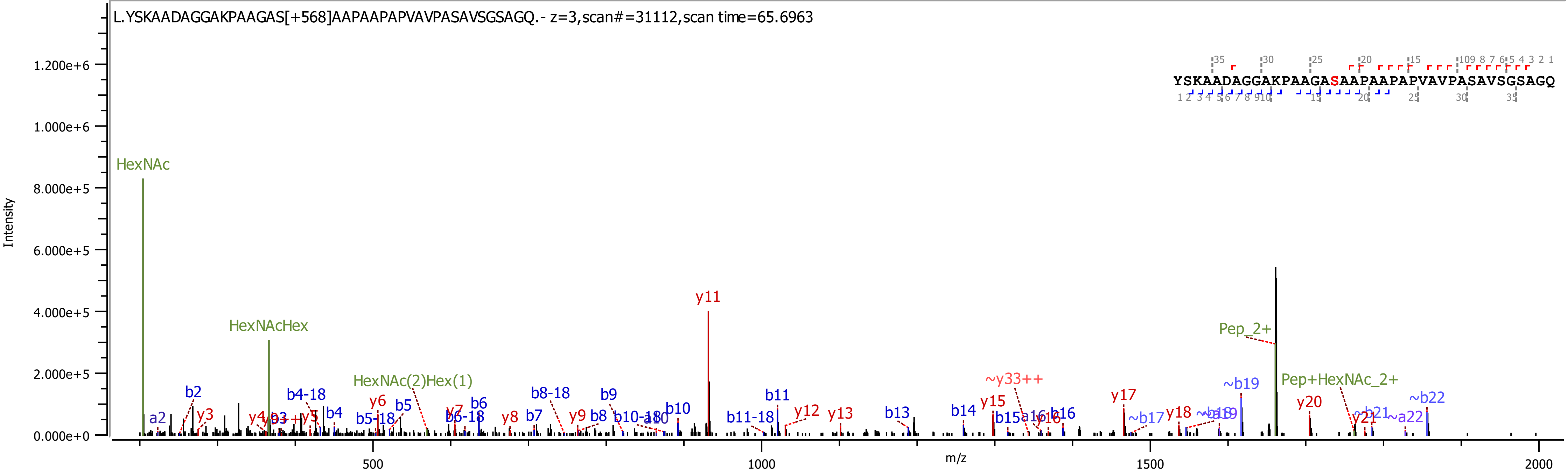
L.YSKAADAGGAKPAAGAS[+568]AAPAAPAPVAVPASAVSGSAGQ.- z=3,scan#=31112,scan time=65.6963
YSKAADAGGAKPAAGASAAPAAPAPVAVPASAVSGSAGQ
Intensity
m/z
HexNAc
HexNAcHex
HexNAc(2)Hex(1)
Pep_2+
Pep+HexNAc_2+
y11
~y33++
~b19
~b22

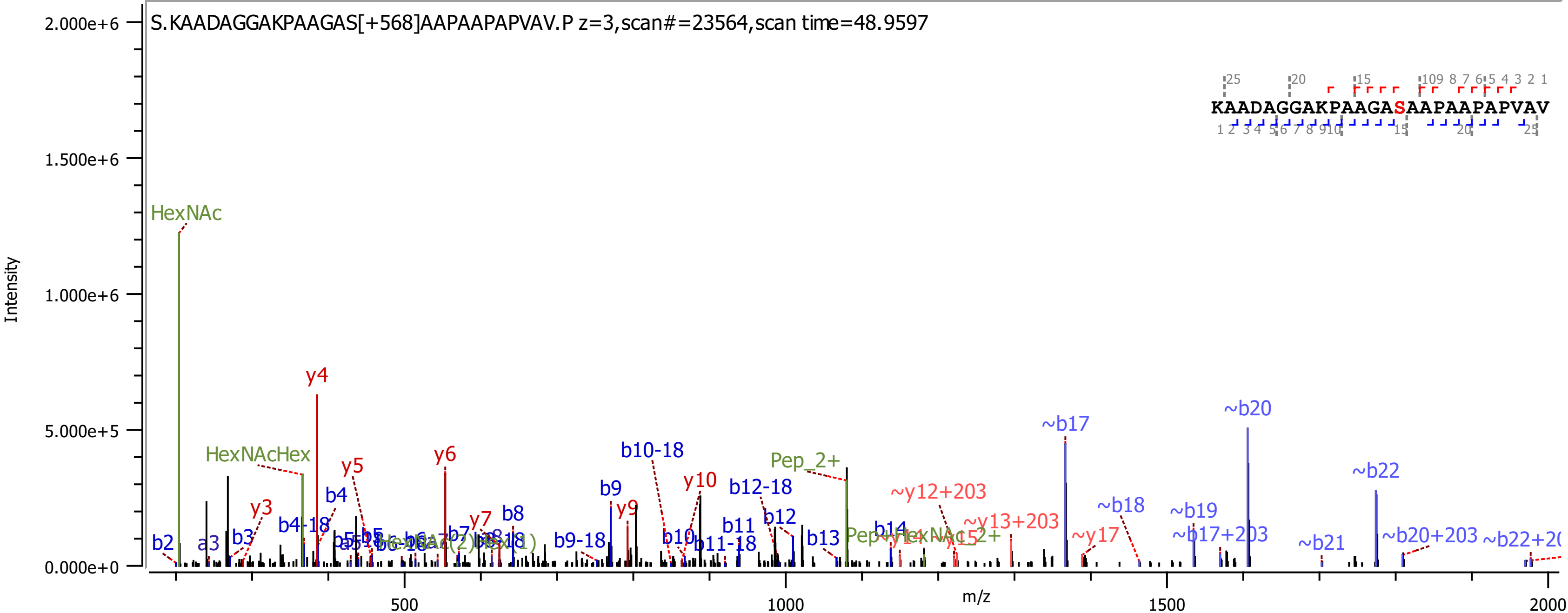
S.KAADAGGAKPAAGAS[+568]AAPAAPAPVAV.P z=3,scan#=23564,scan time=48.9597
KAADAGGAKPAAGASAAPAAPAPVAV
Intensity
m/z
2.000e+6
1.500e+6
1.000e+6
5.000e+5
0.000e+0
500
1000
1500
2000
HexNAc
HexNAcHex
Pep_2+
y3
y4
y5
y6
y7
y9
y10
b2
a3
b3
b4
b4-18
b8
b9
b9-18
b10
b10-18
b11
b11-18
b12
b12-18
b13
b14
~y12+203
~y13+203
~y17
~b17
~b18
~b19
~b17+203
~b20
~b21
~b22
~b20+203

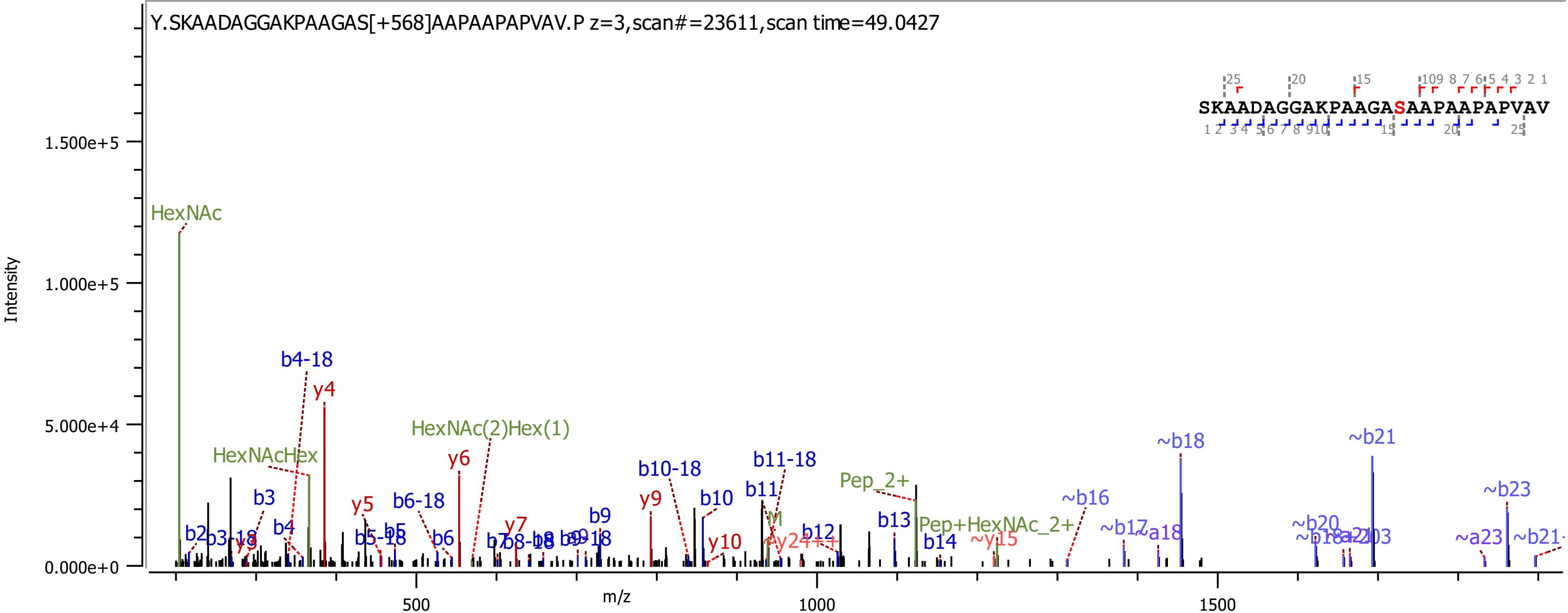

Y.SKAADAGGAKPAAGAS[+568]AAPAAPAPVAV.P z=3,scan#=23611,scan time=49.0427
SKAADAGGAKPAAGASAAPAAPAPVAV
Intensity
1.500e+5
1.000e+5
5.000e+4
0.000e+0
m/z
500
1000
1500
HexNAc
HexNAcHex
HexNAc(2)Hex(1)
b4-18
y4
y6
b3
b4
y5
b6-18
b5
b6
y7
b9
y9
b10-18
b10
b11-18
b11
y10
Pep_2+
Pep+HexNAc_2+
b12
b13
b14
~y15
~b16
~b17
~a18
~b18
~b20
~b21
~b23
~a23

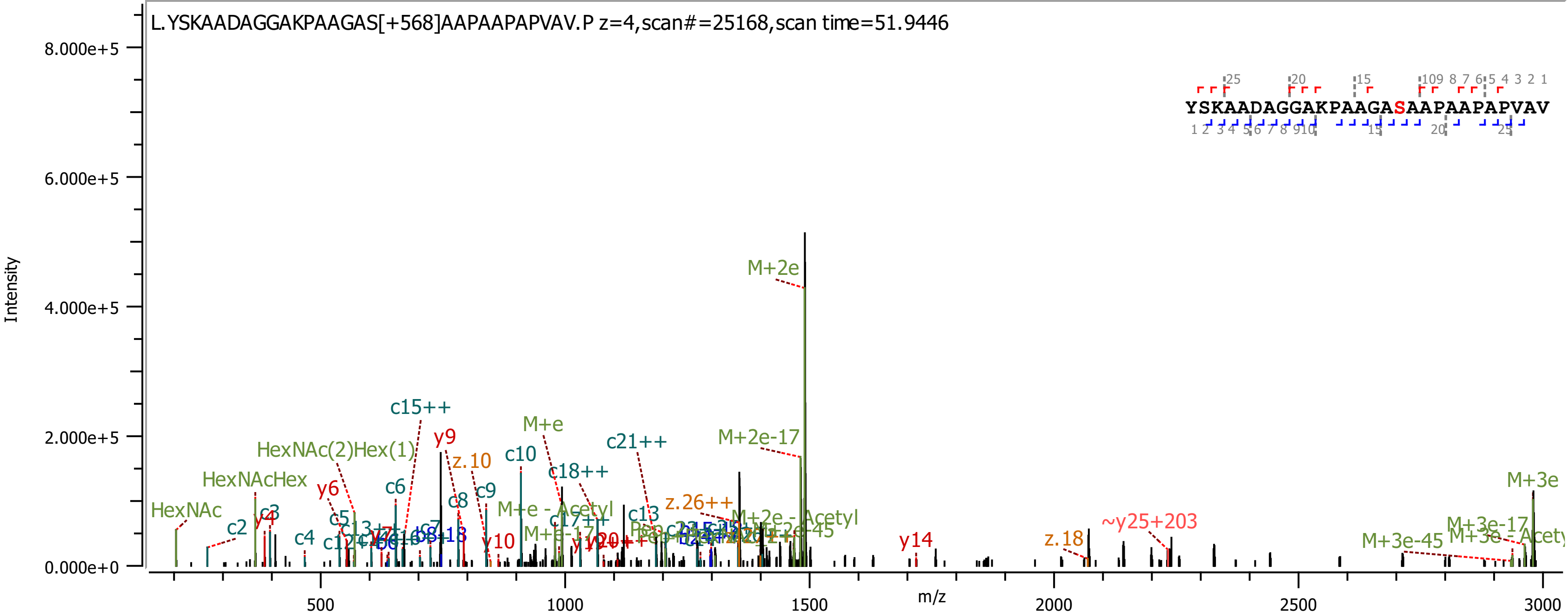
L.YSKAADAGGAKPAAGAS[+568]AAPAAPAPVAV.P z=4,scan#=25168,scan time=51.9446
YSKAADAGGAKPAAGASAAPAAPAPVAV
Intensity
m/z
8.000e+5
6.000e+5
4.000e+5
2.000e+5
0.000e+0
500
1000
1500
2000
2500
3000
M+2e
M+2e-17
c15++
y9
z.10
c10
M+e
c21++
HexNAc(2)Hex(1)
HexNAcHex
HexNAc
c18++
c9
c8
c6
y6
c5
c2
c3
y4
c4
c13
z.26++
M+e - Acetyl
M+2e - Acetyl
c17++
y10
y14
z.18
~y25+203
M+3e
M+3e-17
M+3e-45
M+3e - Acety

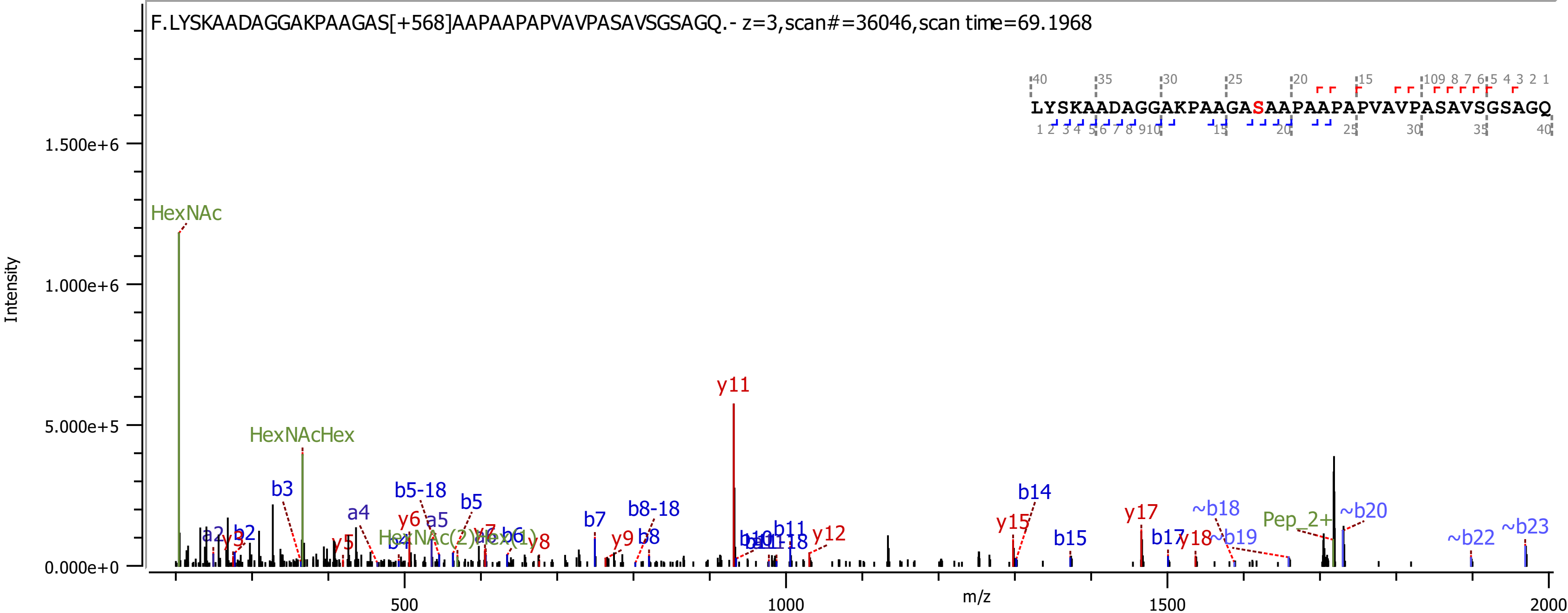
F.LYSKAADAGGAKPAAGAS[+568]AAPAAPAPVAVPASAVSGSAGQ.- z=3,scan#=36046,scan time=69.1968
LYSKAADAGGAKPAAGASAAPAAPAPVAVPASAVSGSAGQ
Intensity
m/z
0.000e+0
5.000e+5
1.000e+6
1.500e+6
500
1000
1500
2000
HexNAc
HexNAcHex
HexNAc(2)Hex(1)
Pep_2+
a2
a4
a5
b2
b3
b4
b5
b5-18
b6
b7
b8
b8-18
b10
b11
b14
b15
b17
~b18
~b19
~b20
~b22
~b23
y3
y5
y6
y7
y8
y9
y11
y12
y15
y17
y18

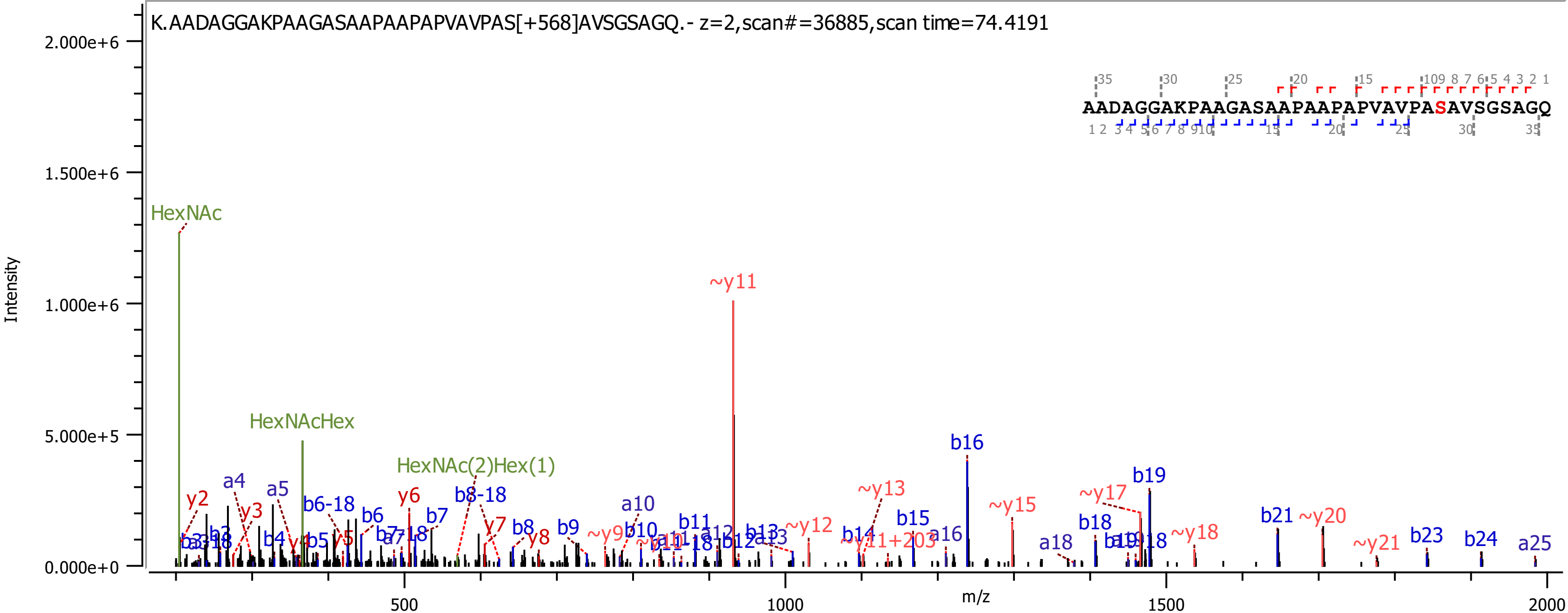
K.AADAGGAKPAAGASAAPAAPAPVAVPAS[+568]AVSGSAGQ.- z=2,scan#=36885,scan time=74.4191
AADAGGAKPAAGASAAPAAPAPVAVPASAVSGSAGQ
Intensity
m/z
2.000e+6
1.500e+6
1.000e+6
5.000e+5
0.000e+0
500
1000
1500
2000
HexNAc
HexNAcHex
HexNAc(2)Hex(1)
~y11
b16
b19
~y17
~y15
~y13
~y12
b21
~y20
~y18
~y21
b23
b24
a25
b18
a18
a16
b15
b14
~y11+203
b13
b12
a12
b11
a10
b10
~y9
b9
y8
b8
y7
b8-18
b7
y6
b6
b6-18
a5
a4
y3
y2
b4
y5
b5

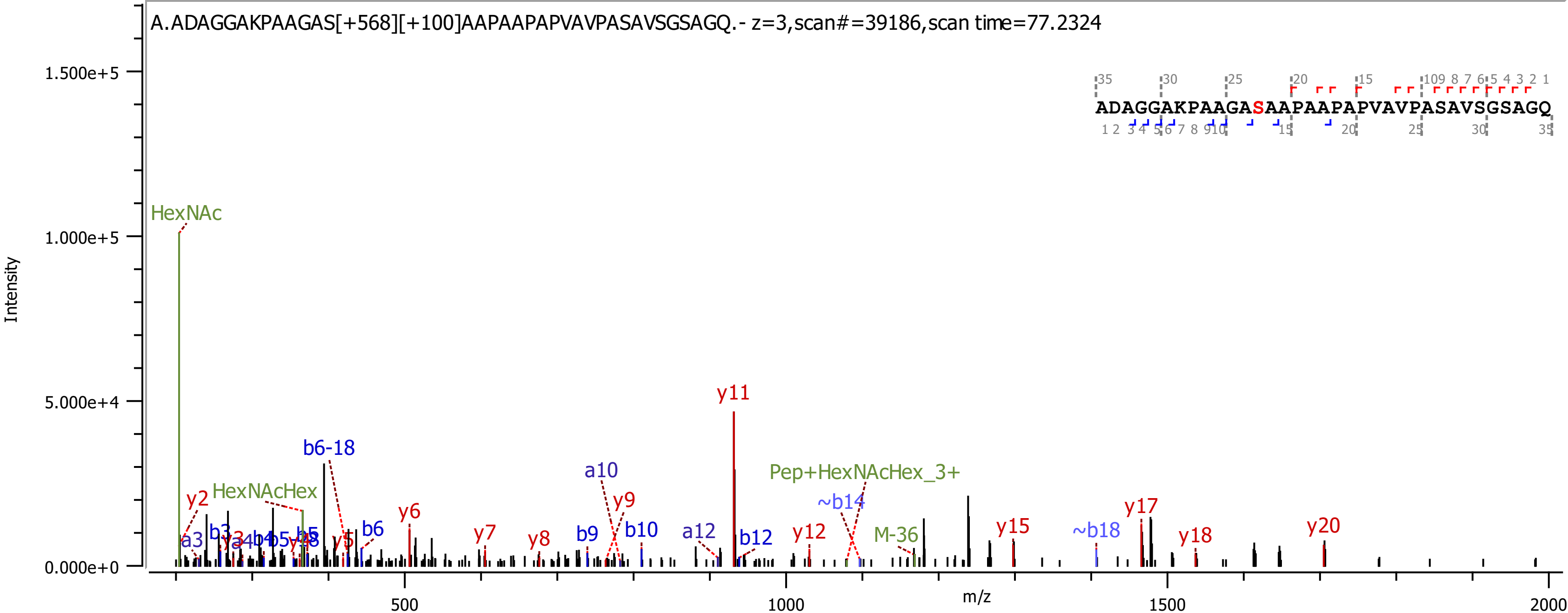

A.ADAGGAKPAAGAS[+568][+100]AAPAAPAPVAVPASAVSGSAGQ.- z=3,scan#=39186,scan time=77.2324
ADAGGAKPAAGASAAPAAPAPVAVPASAVSGSAGQ
Intensity
m/z
1.500e+5
1.000e+5
5.000e+4
0.000e+0
500
1000
1500
2000
HexNAc
HexNAcHex
b6-18
a10
y9
y11
Pep+HexNAcHex_3+
~b14
M-36
y2
a3
b3
y3
b4
b5
y4
b5
y5
b6
y6
y7
y8
b9
b10
a12
b12
y12
y15
~b18
y17
y18
y20

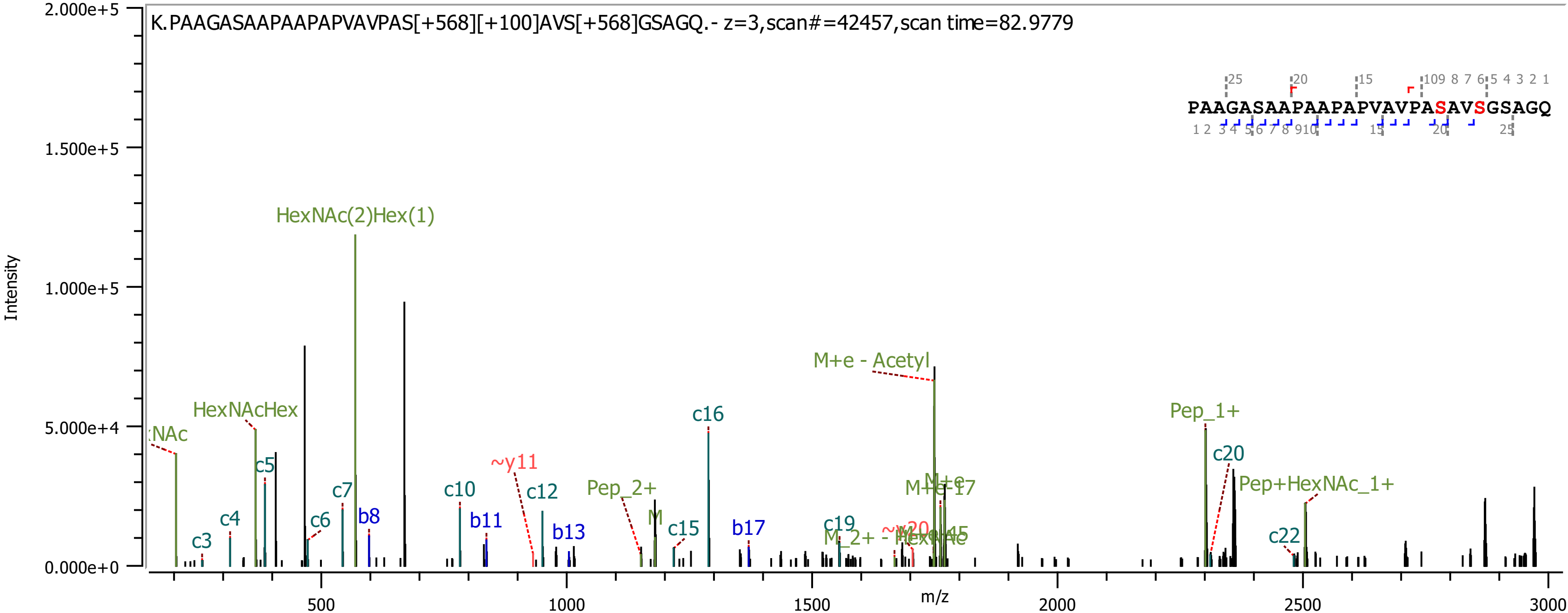

K.PAAGASAAPAAPAPVAVPAS[+568][+100]AVS[+568]GSAGQ.- z=3,scan#=42457,scan time=82.9779
PAAGASAAPAAPAPVAVPASAVSGSAGQ
Intensity
m/z
2.000e+5
1.500e+5
1.000e+5
5.000e+4
0.000e+0
500
1000
1500
2000
2500
3000
HexNAc(2)Hex(1)
HexNAcHex
M+e - Acetyl
Pep_1+
c20
Pep+HexNAc_1+
c16
c5
c4
c3
c6
c7
b8
c10
b11
~y11
c12
b13
Pep_2+
M
c15
b17
c19
c22

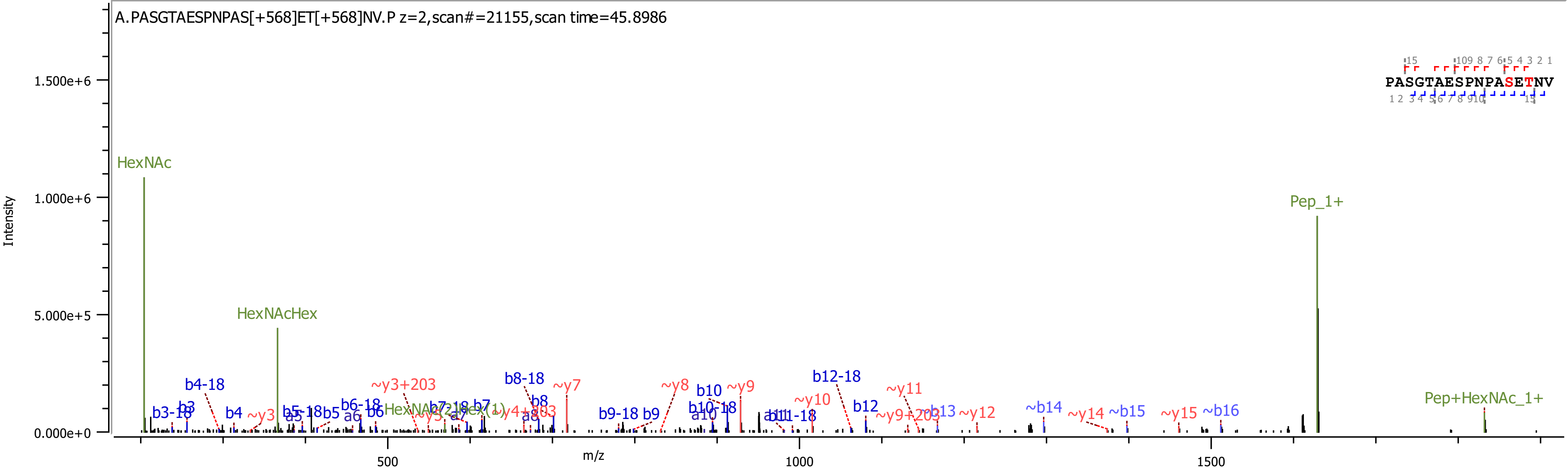

A.PASGTAESPNPAS[+568]ET[+568]NV.P z=2,scan#=21155,scan time=45.8986
Intensity
m/z
HexNAc
HexNAcHex
Pep_1+
Pep+HexNAc_1+
PASGTAESPNPASETNV

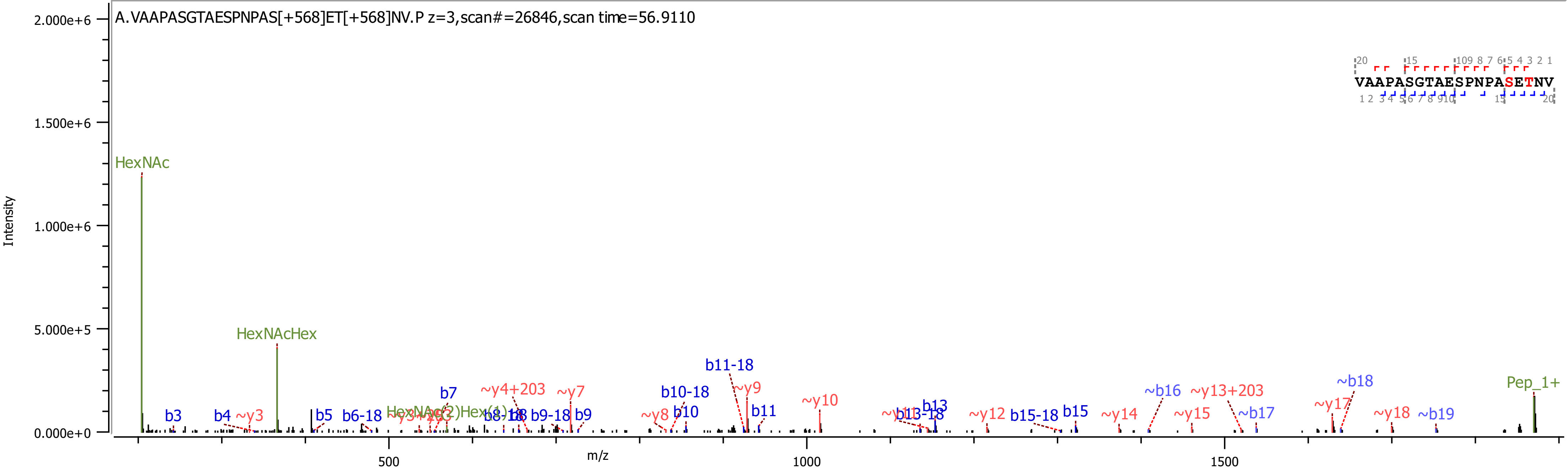
A.VAAPASGTAESPNPAS[+568]ET[+568]NV.P z=3,scan#=26846,scan time=56.9110
VAAPASGTAESPNPASETNV
Intensity
m/z
HexNAc
HexNAcHex
Pep_1+
b3
b4
~y3
b5
b6-18
b7
~y4+203
~y7
b9-18
b9
~y8
b10
b10-18
b11-18
~y9
b11
~y10
b13
b13-18
~y12
b15-18
b15
~y14
~b16
~y13+203
~y15
~b17
~y17
~b18
~y18
~b19
2.000e+6
1.500e+6
1.000e+6
5.000e+5
0.000e+0
500
1000
1500

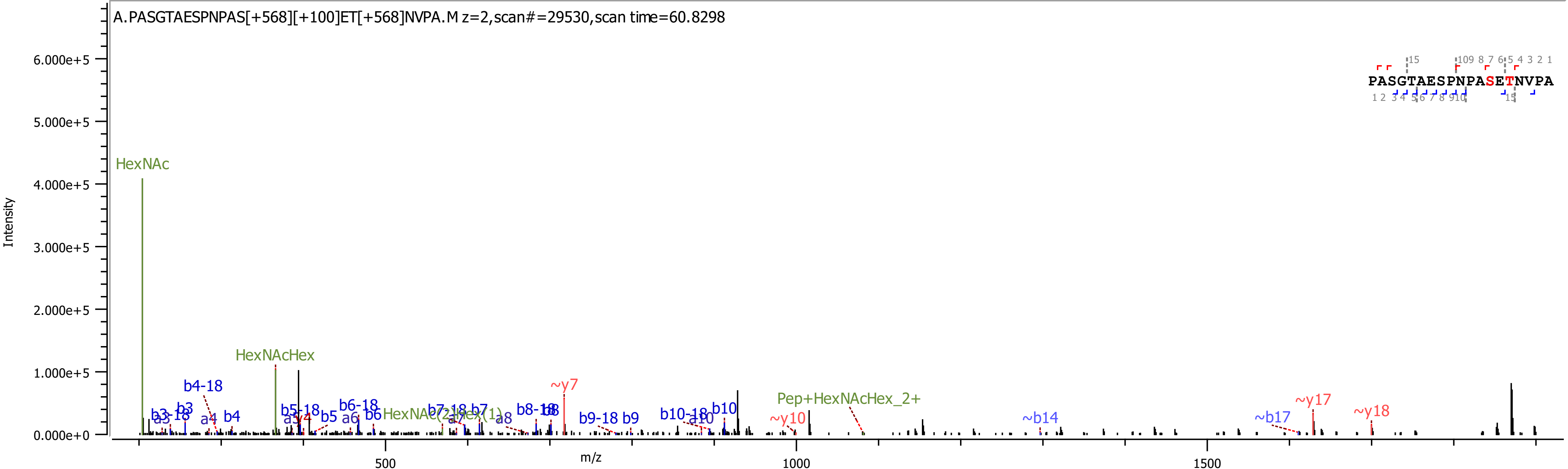
A.PASGTAESPNPAS[+568][+100]ET[+568]NVPA.M z=2,scan#=29530,scan time=60.8298
PASGTAESPNPASETNVPA
Intensity
m/z
0.000e+0
1.000e+5
2.000e+5
3.000e+5
4.000e+5
5.000e+5
6.000e+5
500
1000
1500
HexNAc
HexNAcHex
b4-18
b3
b4
a4
b5-18
b5
b6-18
b6
a6
b7-18
b7
a8
b8-18
~y7
b9-18 b9
b10-18
a10
b10
~y10
Pep+HexNAcHex_2+
~b14
~b17
~y17
~y18

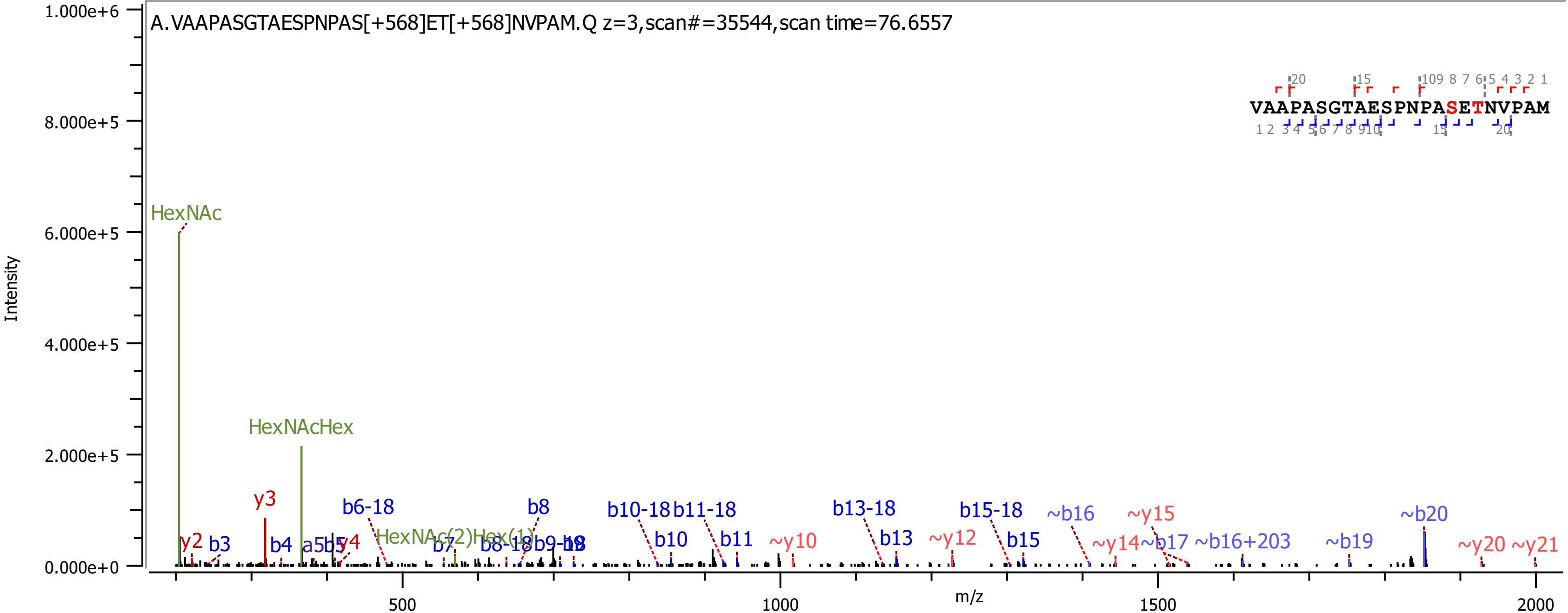

A.VAAPASGTAESPNPAS[+568]ET[+568]NVPAM.Q z=3,scan#=35544,scan time=76.6557
VAAPASGTAESPNPASETNVPAM
Intensity
m/z
HexNAc
HexNAcHex
HexNAc(2)Hex(1)
y2
b3
y3
b4
a5
b5
y4
b6-18
b7
b8
b8-18
b9-18
b10-18
b10
b11-18
b11
~y10
b13-18
b13
~y12
b15-18
b15
~b16
~y14
~b17
~y15
~b16+203
~b19
~b20
~y20
~y21

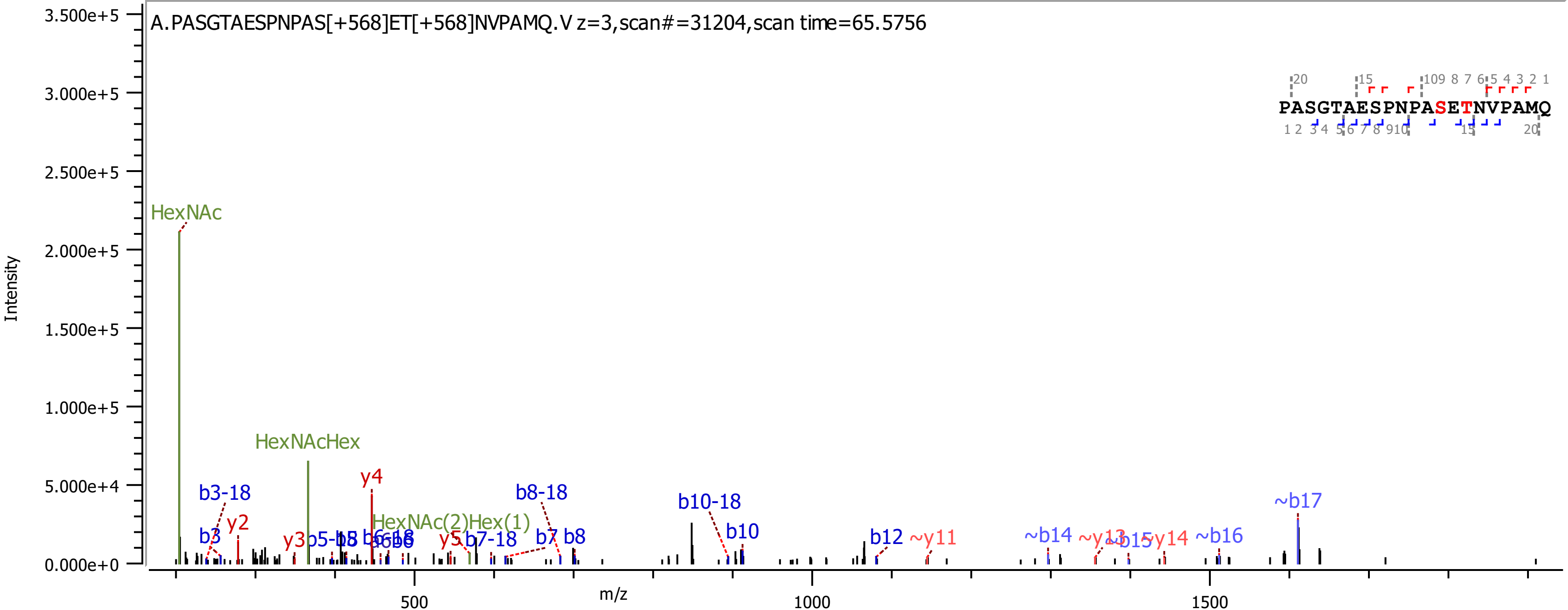
A.PASGTAESPNPAS[+568]ET[+568]NVPAMQ.V z=3,scan#=31204,scan time=65.5756
PASGTAESPNPASETNVPAMQ
Intensity
m/z
HexNAc
HexNAcHex
HexNAc(2)Hex(1)
b3-18
b3
y2
y3
y4
y5
b7-18
b8-18
b7
b8
b10-18
b10
b12
~y11
~b14
~y14
~b16
~b17
3.500e+5
3.000e+5
2.500e+5
2.000e+5
1.500e+5
1.000e+5
5.000e+4
0.000e+0
500
1000
1500

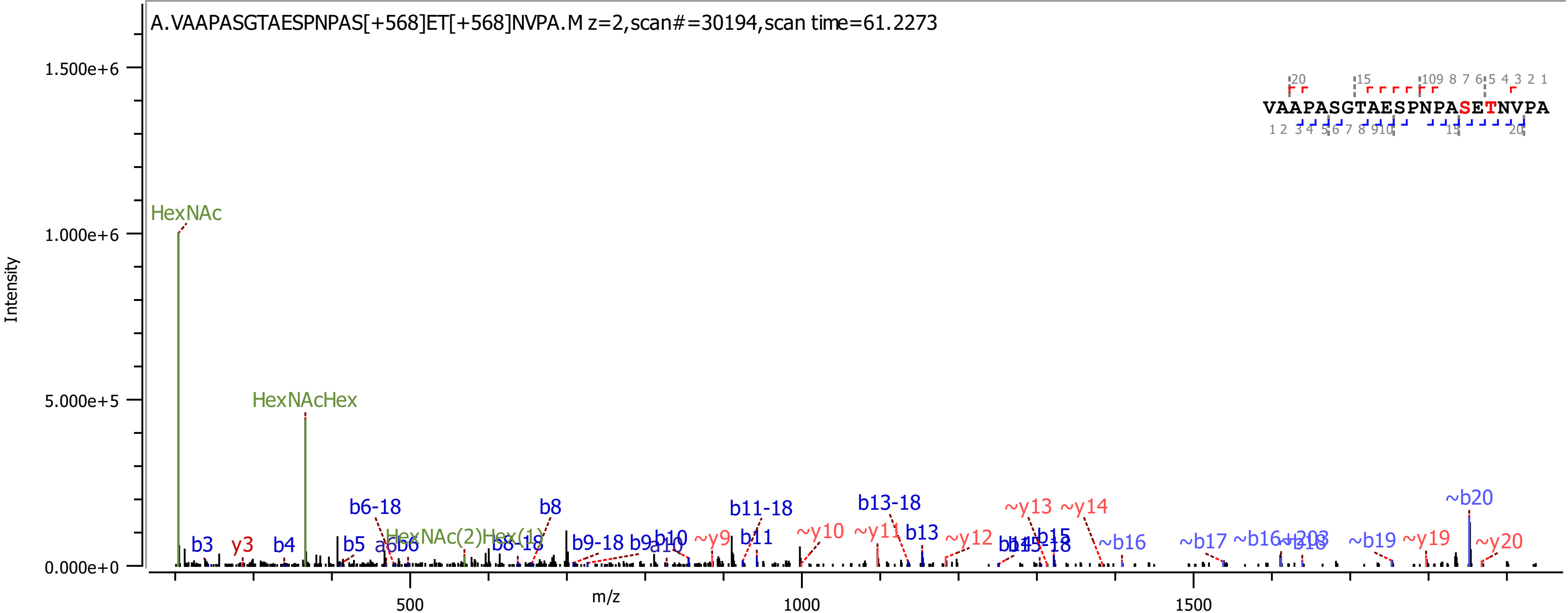

A.VAAPASGTAESPNPAS[+568]ET[+568]NVPA.M z=2,scan#=30194,scan time=61.2273
VAAPASGTAESPNPASETNVPA
Intensity
m/z
HexNAc
HexNAcHex
HexNAc(2)Hex(1)
b3
y3
b4
b5
b6-18
b8
b8-18
b9-18
b10
~y9
b11
b11-18
~y10
~y11
b13
b13-18
~y12
~y13
~y14
b15
~b16
~b17
~b19
~y19
~b20
~y20

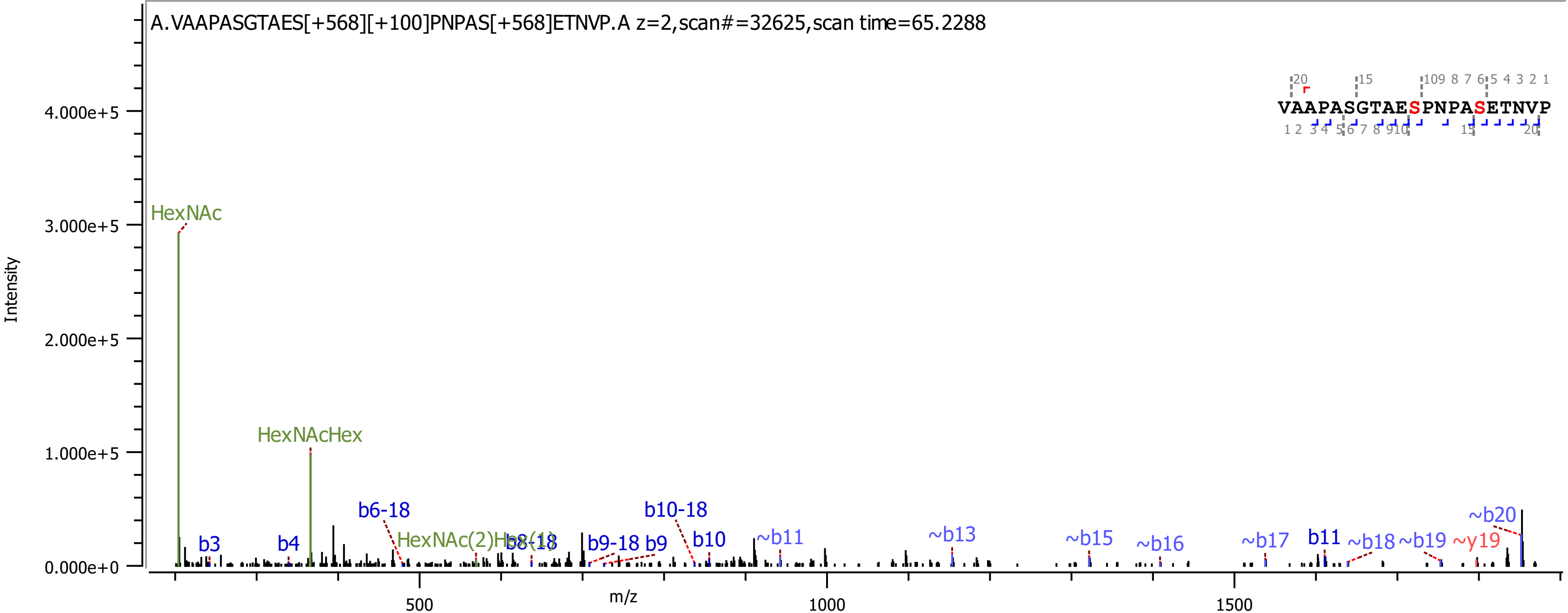

A.VAAPASGTAES[+568][+100]PNPAS[+568]ETNVP.A z=2,scan#=32625,scan time=65.2288
VAAPASGTAESPNPASETNVP
Intensity
m/z
HexNAc
HexNAcHex
HexNAc(2)Hex(1)
b3
b4
b6-18
b8-18
b9-18
b9
b10-18
b10
~b11
~b13
~b15
~b16
~b17
b11
~b18
~b19
~y19
~b20

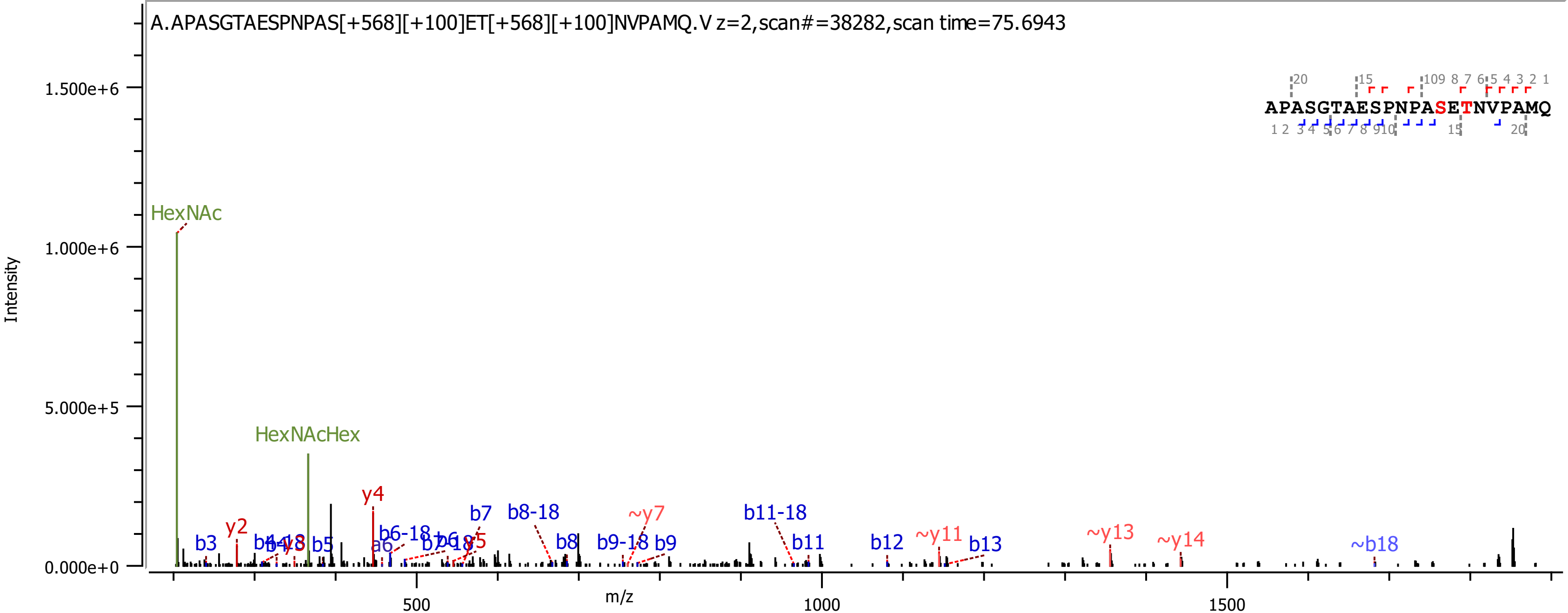
A.APASGTAESPNPAS[+568][+100]ET[+568][+100]NVPAMQ.V z=2,scan#=38282,scan time=75.6943
APASGTAESPNPASETNVPAMQ
Intensity
1.500e+6
1.000e+6
5.000e+5
0.000e+0
m/z
500
1000
1500
HexNAc
HexNAcHex
y2
b3
b4-18
b4
y3
b5
y4
a6
b6-18
b6
b7-18
y5
b7
b8-18
b8
~y7
b9-18
b9
b11-18
b11
b12
~y11
b13
~y13
~y14
~b18

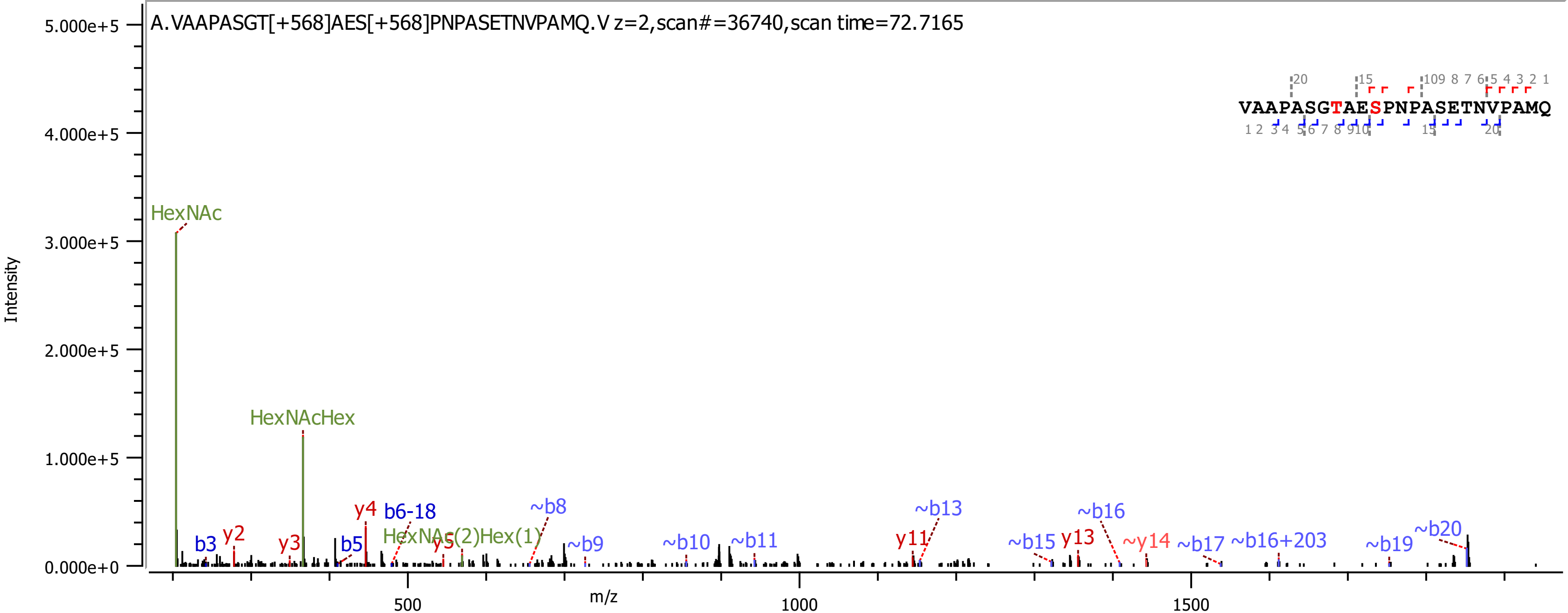

A.VAAPASGT[+568]AES[+568]PNPASETNVPAMQ.V z=2,scan#=36740,scan time=72.7165
VAAPASGTAESPNPASETNVPAMQ
Intensity
5.000e+5
4.000e+5
3.000e+5
2.000e+5
1.000e+5
0.000e+0
m/z
500
1000
1500
HexNAc
HexNAcHex
HexNAc(2)Hex(1)
b3
y2
y3
y4
b5
b6-18
y5
~b8
~b9
~b10
~b11
y11
~b13
~b15
y13
~b16
~y14
~b17
~b16+203
~b19
~b20

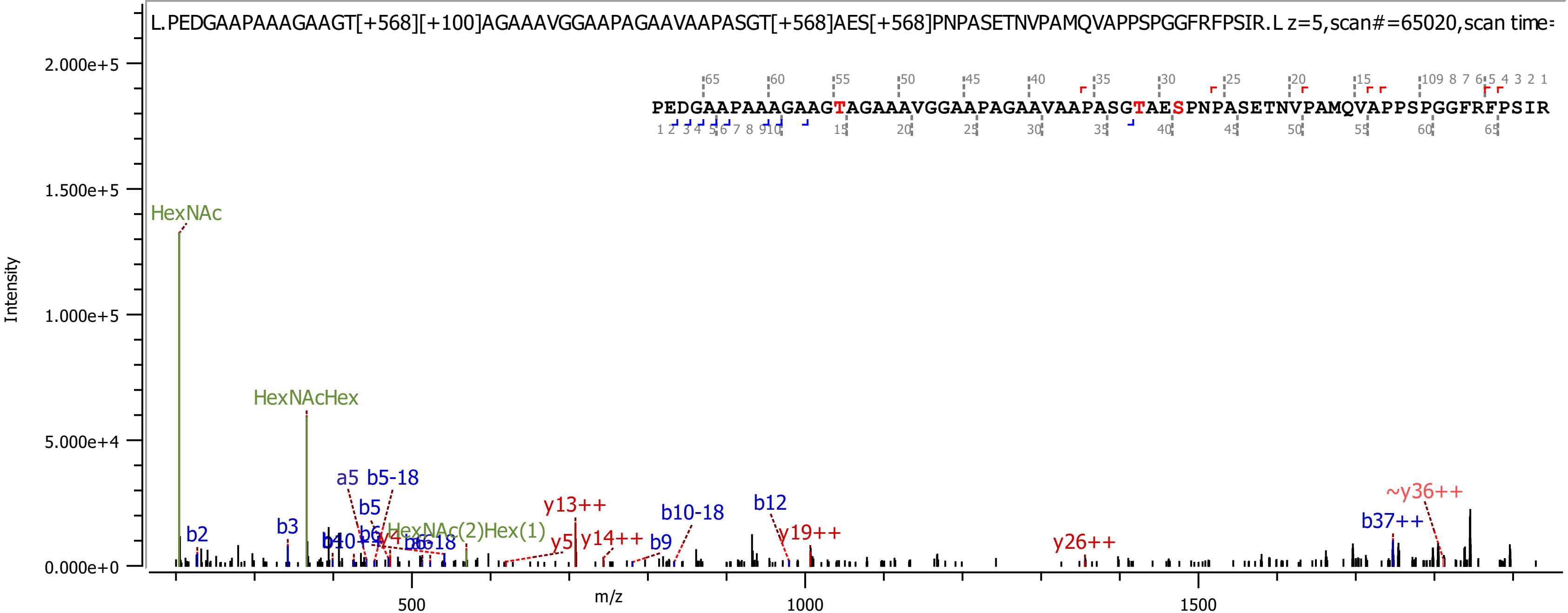
L.PEDGAAPAAAGAAGT[+568][+100]AGAAAVGGAAPAGAAVAAPASGT[+568]AES[+568]PNPASETNVPAMQVAPPSPGGFRFPSIR.L z=5,scan#=65020,scan time=
PEDGAAPAAAGAAGTAGAAAVGGAAPAGAAVAAPASGTAESPNPASETNVPAMQVAPPSPGGFRFPSIR
Intensity
m/z
2.000e+5
1.500e+5
1.000e+5
5.000e+4
0.000e+0
500
1000
1500
HexNAc
HexNAcHex
HexNAc(2)Hex(1)
a5
b5-18
b5
b2
b3
b12
b10-18
b9
b37++
y13++
y5
y14++
y19++
y26++
~y36++

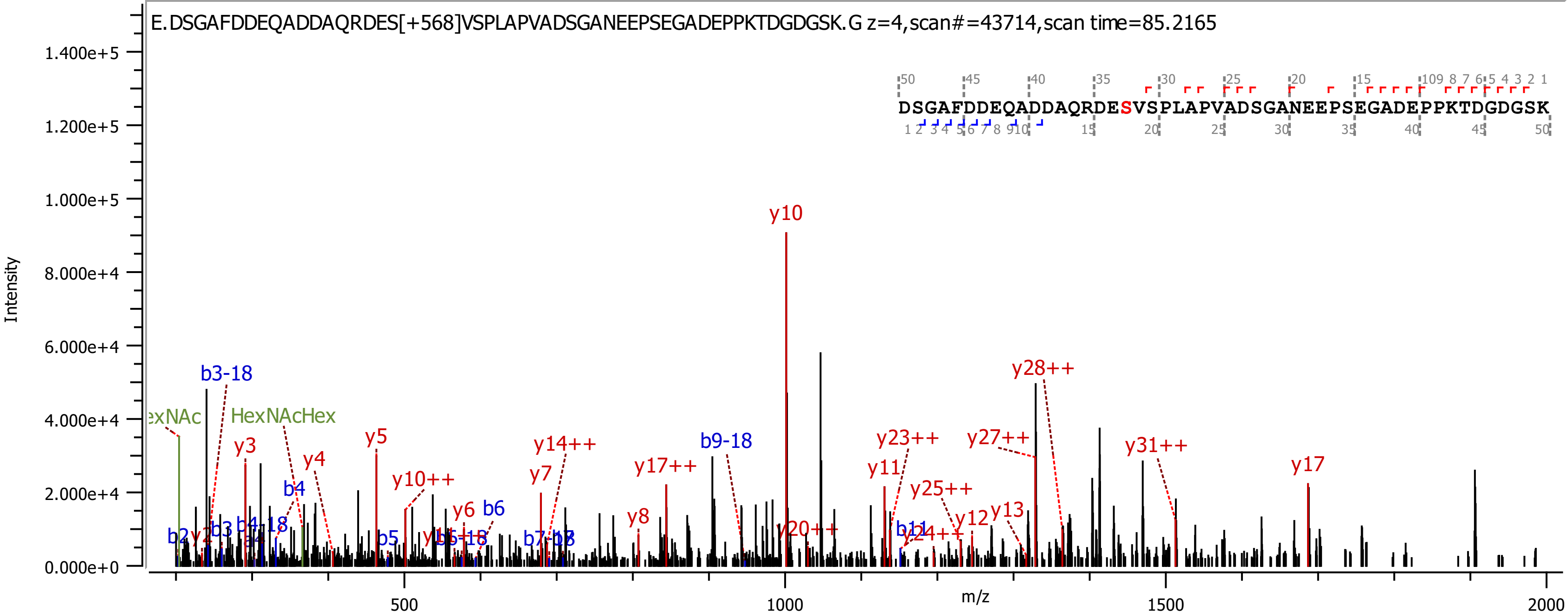

E.DSGAFDDEQADDAQRDES[+568]VSPLAPVADSGANEEPSEGADEPPKTDGDGSK.G z=4,scan#=43714,scan time=85.2165
DSGAFDDEQADDAQRDESVSPLAPVADSGANEEPSEGADEPPKTDGDGSK
Intensity
m/z
y10
b3-18
y28++
HexNAcHex
y5
y3
y4
y14++
b9-18
y23++
y27++
y31++
y10++
y17++
y11
y17
y7
b4
y25++
y6
b6
y13
y12
y8
y20++
b2
b3
b5

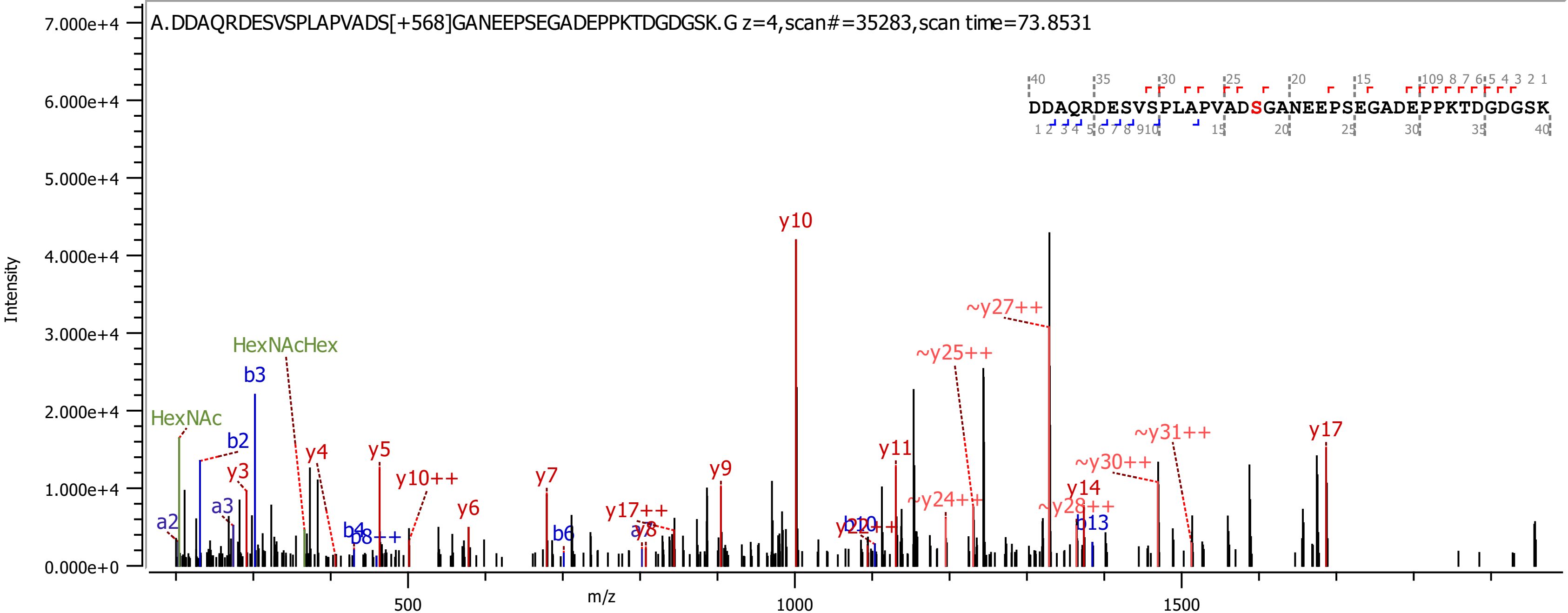
A.DDAQRDESVSPLAPVADS[+568]GANEEPSEGADEPPKTDGDGSK.G z=4,scan#=35283,scan time=73.8531
DDAQRDESVSPLAPVADSGANEEPSEGADEPPKTDGDGSK
Intensity
m/z
y10
~y27++
~y25++
HexNAcHex
b3
HexNAc
b2
y3
y4
y5
y10++
a3
a2
y7
y6
b4
b8++
b6
y17++
a8
y9
y11
~y24++
b10
y22++
~y31++
y17
~y30++
y14
~y28++
b13

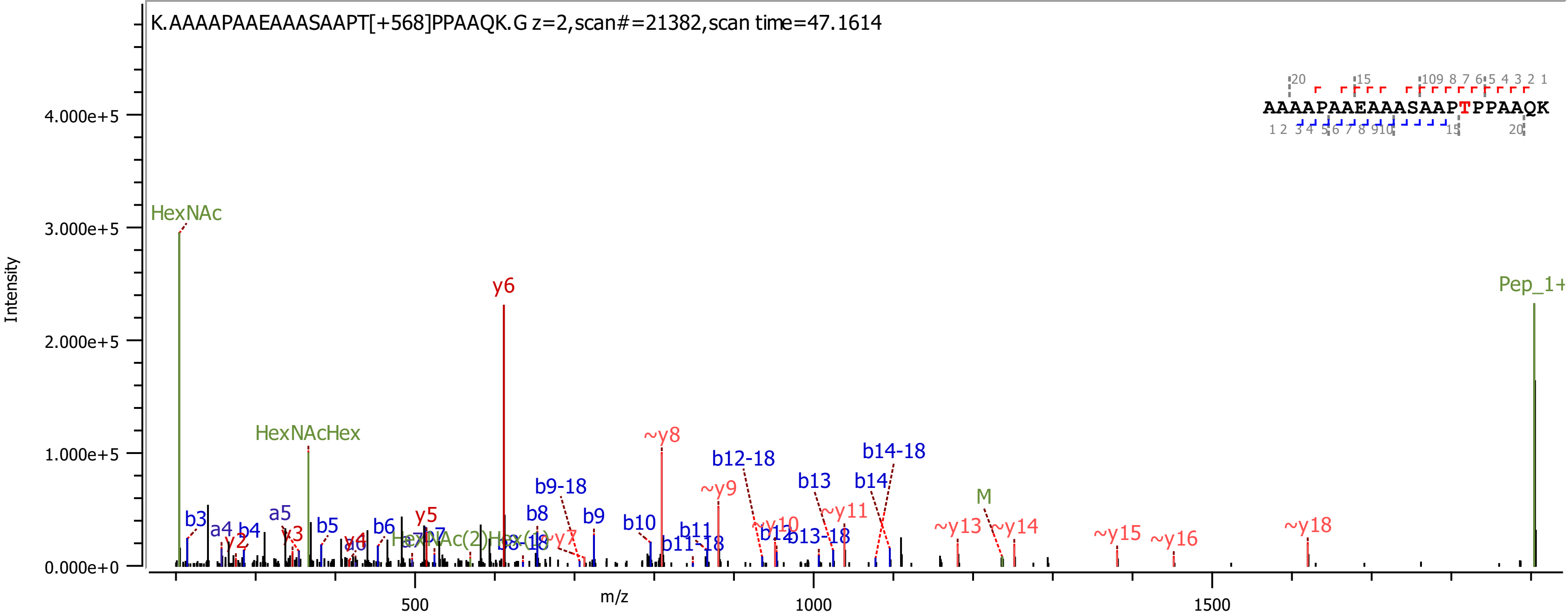
K.AAAAPAAEAAASAAPT[+568]PPAAQK.G z=2,scan#=21382,scan time=47.1614
AAAAPAAEAAASAAPTPPAAQK
Intensity
4.000e+5
3.000e+5
2.000e+5
1.000e+5
0.000e+0
m/z
500
1000
1500
HexNAc
HexNAcHex
y6
Pep_1+
~y8
~y9
b12-18
b14-18
b13
b14
b9-18
b8
b9
b10
b11
b11-18
~y10
b12
b13-18
~y11
M
~y13
~y14
~y15
~y16
~y18
b3
a4
b4
a5
b5
b6
y5
y2
y3
y4

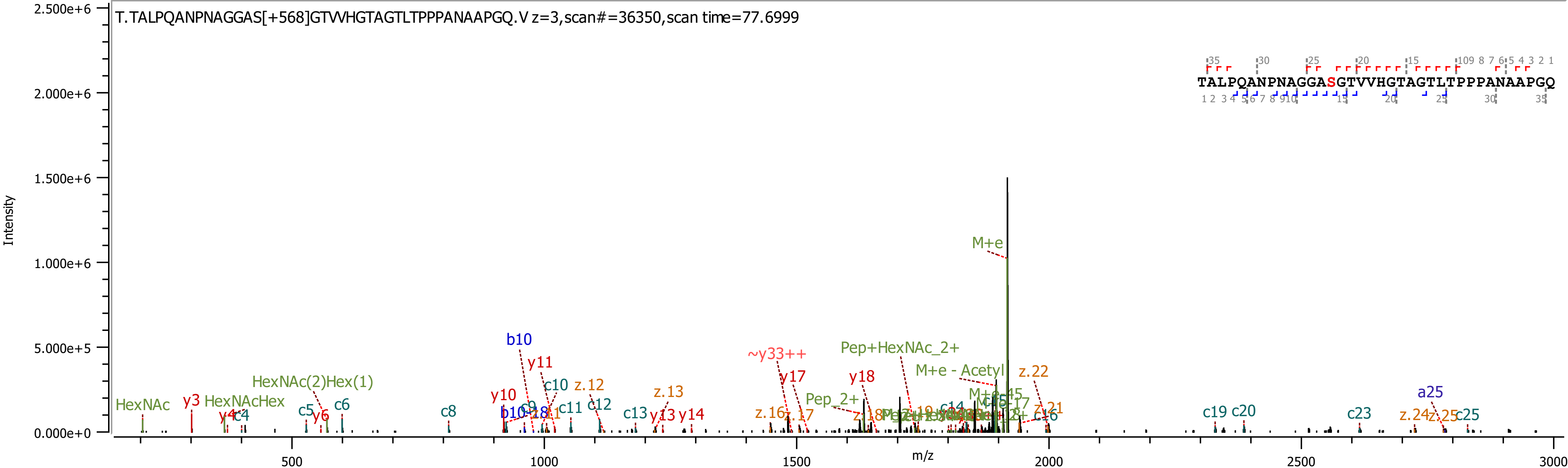

T.TALPQANPNAGGAS[+568]GTVVHGTAGTLTPPPANAAPGQ.V z=3,scan#=36350,scan time=77.6999
TALPQANPNAGGASGTVVHGTAGTLTPPPANAAPGQ
Intensity
m/z
2.500e+6
2.000e+6
1.500e+6
1.000e+6
5.000e+5
0.000e+0
500
1000
1500
2000
2500
3000
M+e
Pep+HexNAc_2+
M+e - Acetyl
~y33++
b10
y11
c10
z.12
z.13
y17
y18
z.22
HexNAc(2)Hex(1)
HexNAc
y3
HexNAcHex
y10
c6
c5
y6
y4
c4
c8
c11
c12
c13
y13
y14
z.16
z.17
Pep_2+
c19
c20
c23
a25
z.24
z.25
c25

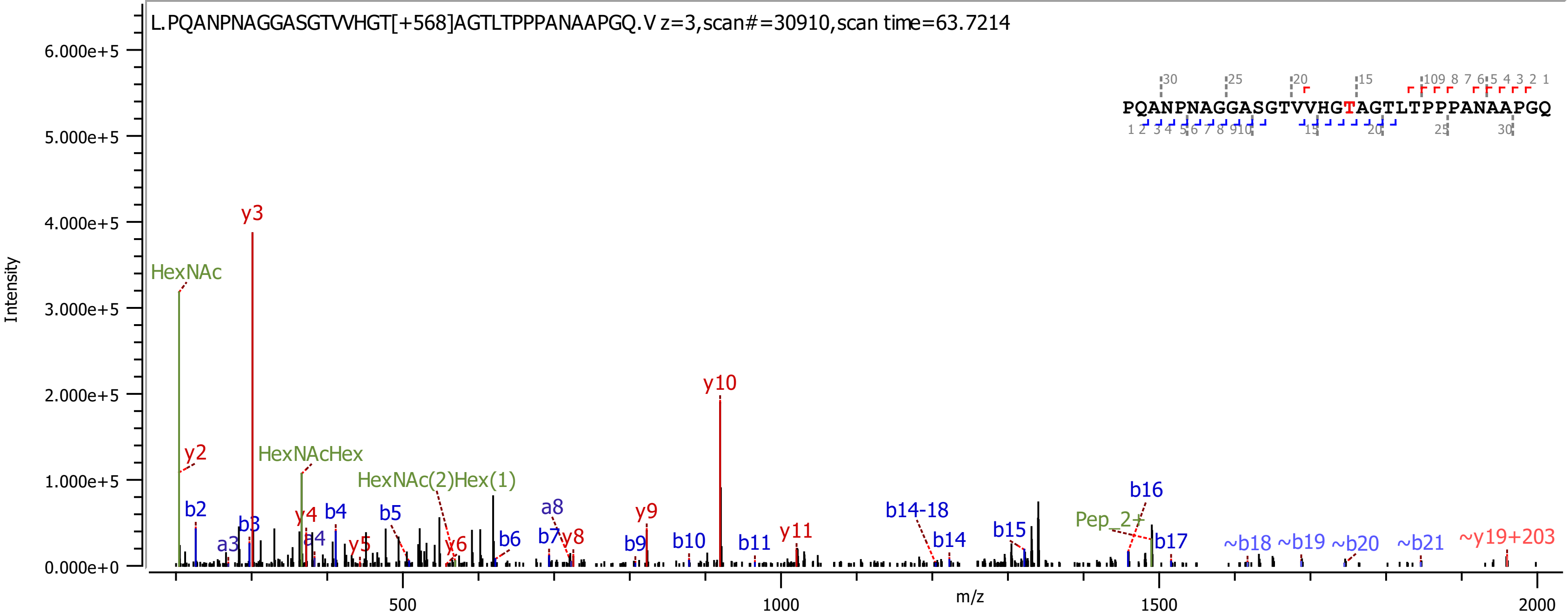

L.PQANPNAGGASGTVVHGT[+568]AGTLTPPPANAAPGQ.V z=3,scan#=30910,scan time=63.7214
PQANPNAGGASGTVVHGTAGTLTPPPANAAPGQ
Intensity
m/z
HexNAc
y2
y3
b2
b3
a3
HexNAcHex
y4
a4
b4
y5
HexNAc(2)Hex(1)
b5
y6
b6
a8
b7
y8
y9
b9
b10
y10
b11
y11
b14-18
b14
b15
Pep_2+
b16
b17
~b18
~b19
~b20
~b21
~y19+203

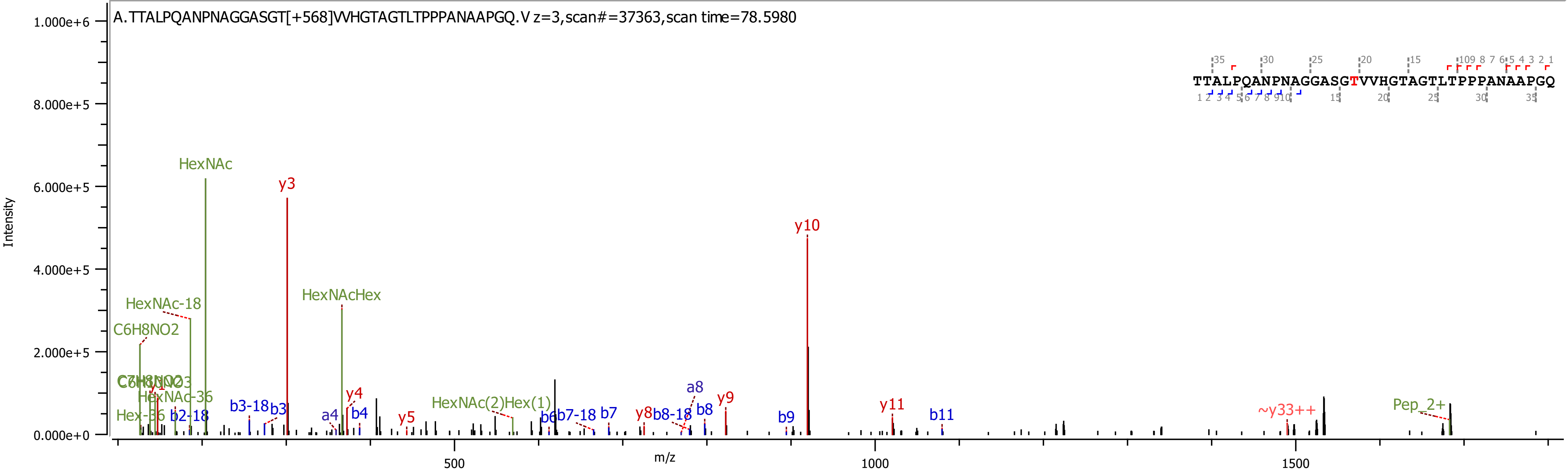
A.TTALPQANPNAGGASGT[+568]VVHGTAGTLTPPPANAAPGQ.V z=3,scan#=37363,scan time=78.5980
TTALPQANPNAGGASGTVVHGTAGTLTPPPANAAPGQ
Intensity
m/z
HexNAc
y3
y10
HexNAc-18
HexNAcHex
C6H8NO2
HexNAc-36
Hex-36
b2-18
b3-18
b3
a4
y4
b4
y5
HexNAc(2)Hex(1)
b6
b7-18
b7
y8
b8-18
a8
b8
y9
b9
y11
b11
~y33++
Pep_2+

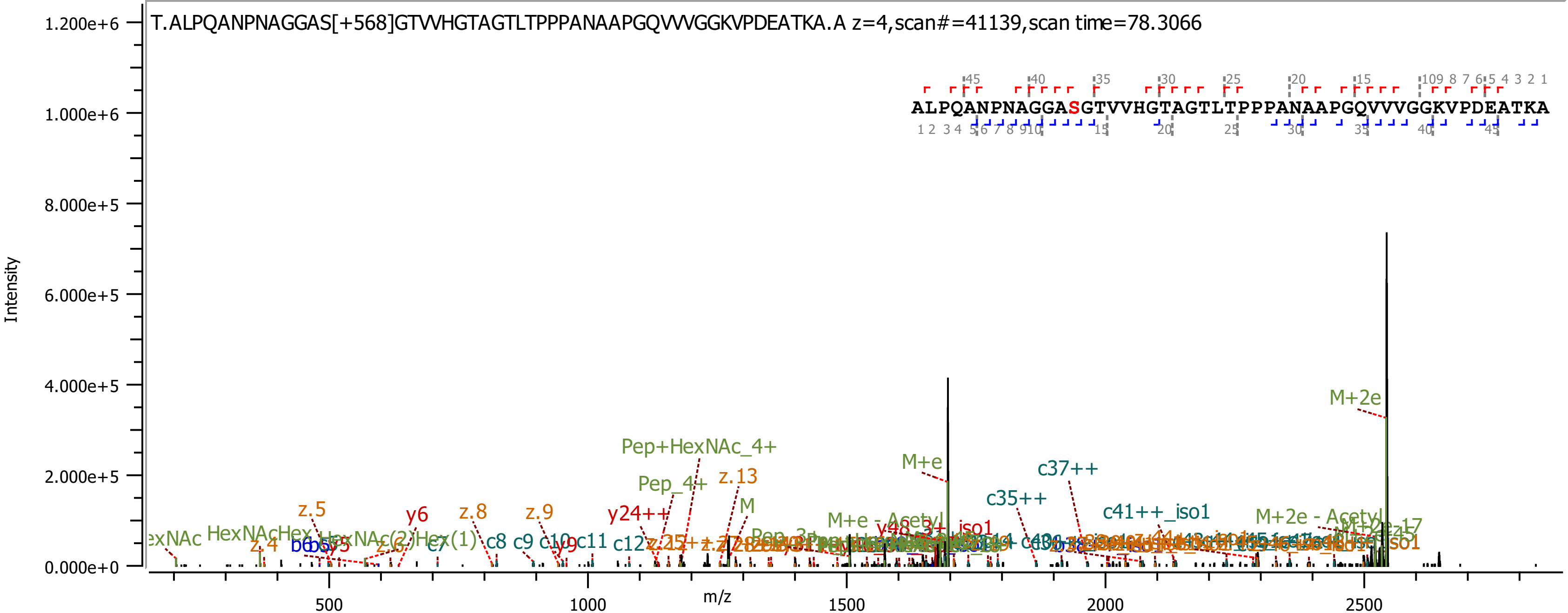
T.ALPQANPNAGGAS[+568]GTVVHGTAGTLTPPPANAAPGQVVVGGKVPDEATKA.A z=4,scan#=41139,scan time=78.3066
ALPQANPNAGGASGTVVHGTAGTLTPPPANAAPGQVVVGGKVPDEATKA
Intensity
m/z
1.200e+6
1.000e+6
8.000e+5
6.000e+5
4.000e+5
2.000e+5
0.000e+0
500
1000
1500
2000
2500
M+2e
Pep+HexNAc_4+
Pep_4+
z.13
M+e
c37++
c35++
c41++_iso1
M+2e - Acetyl
M+e - Acetyl
z.5
y6
z.8
z.9
y24++
M
HexNAcHex
HexNAc(2)Hex(1)
z.4
c7
c8
c9
c11
c12
y9
z.6

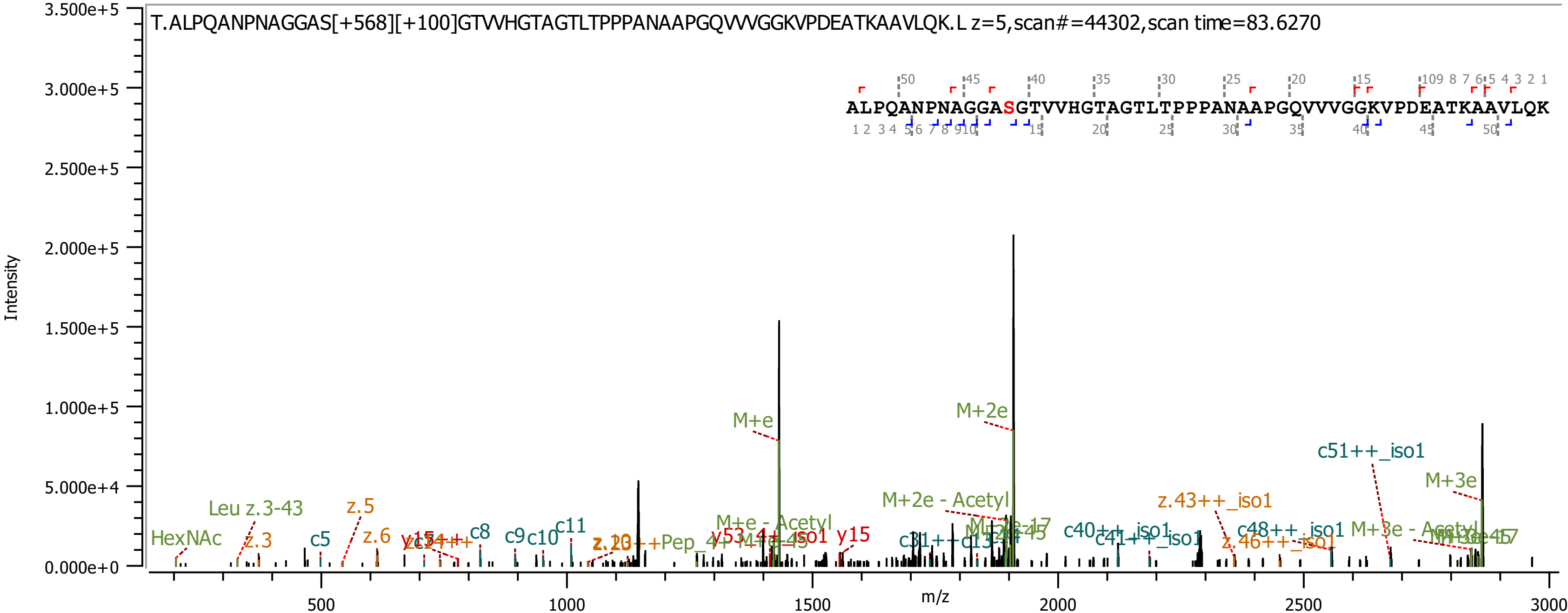
T.ALPQANPNAGGAS[+568][+100]GTVVHGTAGTLTPPPANAAPGQVVVGGKVPDEATKAAVLQK.L z=5,scan#=44302,scan time=83.6270
ALPQANPNAGGASGTVVHGTAGTLTPPPANAAPGQVVVGGKVPDEATKAAVLQK
Intensity
m/z
M+e
M+2e
M+3e
c51++_iso1
M+2e - Acetyl
M+e - Acetyl
z.43++_iso1
c48++_iso1
M+3e - Acetyl
c40++_iso1
c41++_iso1
z.46++_iso1
HexNAc
Leu z.3-43
z.3
z.5
z.6
c5
c8
c9
c10
c11
y15
c31++
c13

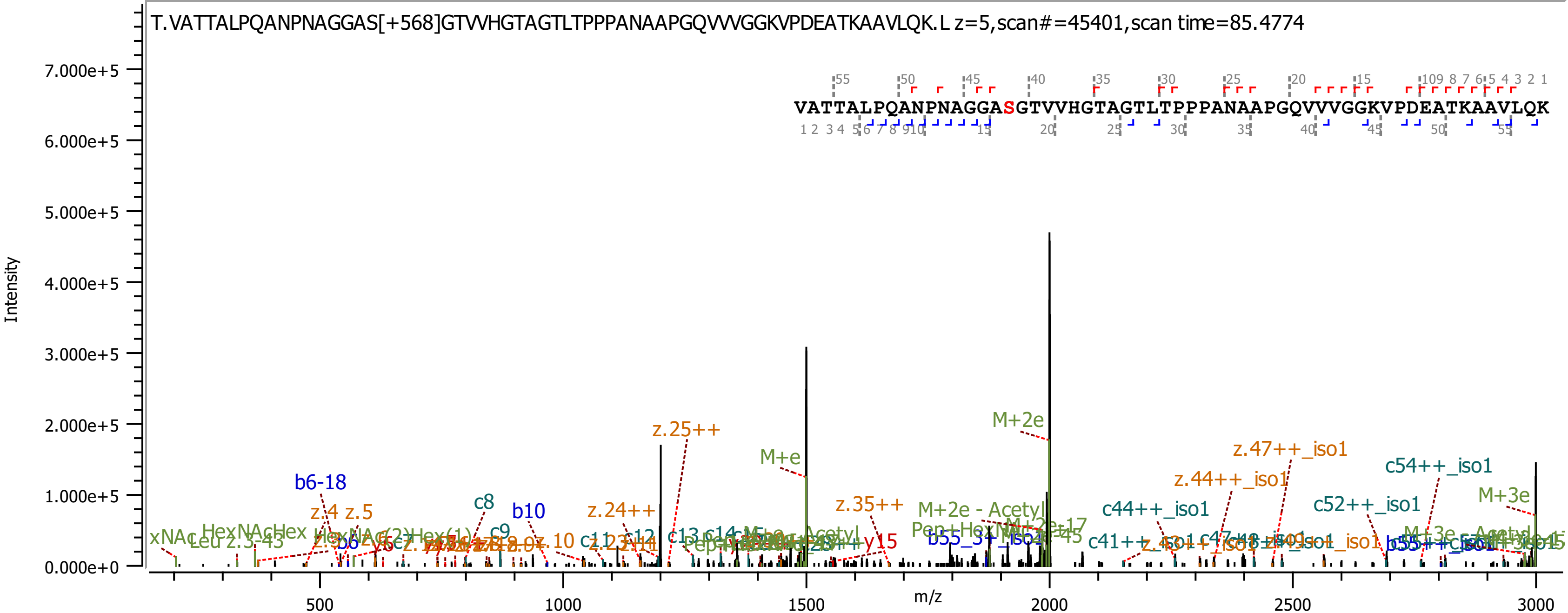

T.VATTALPQANPNAGGAS[+568]GTVVHGTAGTLTPPPANAAPGQVVVGGKVPDEATKAAVLQK.L z=5,scan#=45401,scan time=85.4774
VATTALPQANPNAGGASGTVVHGTAGTLTPPPANAAPGQVVVGGKVPDEATKAAVLQK
Intensity
m/z
7.000e+5
6.000e+5
5.000e+5
4.000e+5
3.000e+5
2.000e+5
1.000e+5
0.000e+0
500
1000
1500
2000
2500
3000
b6-18
z.4
z.5
c8
c9
b10
z.24++
z.25++
c11
c12
c13
M+e
z.35++
y15
M+2e
M+2e - Acetyl
c41++
c44++_iso1
z.44++_iso1
z.47++_iso1
c52++_iso1
c54++_iso1
M+3e

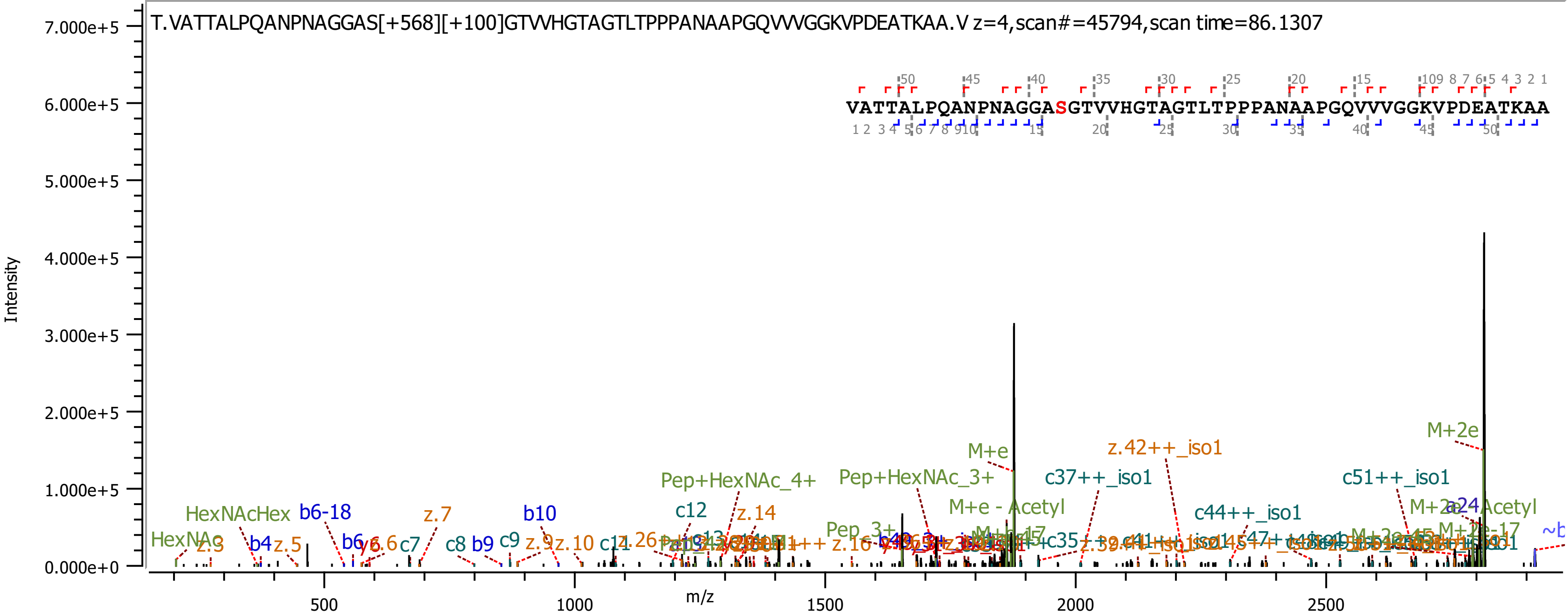
T.VATTALPQANPNAGGAS[+568][+100]GTVVHGTAGTLTPPPANAAPGQVVVGGKVPDEATKAA.V z=4,scan#=45794,scan time=86.1307
VATTALPQANPNAGGASGTVVHGTAGTLTPPPANAAPGQVVVGGKVPDEATKAA
Intensity
m/z
M+2e
M+e
z.42++_iso1
c37++_iso1
c51++_iso1
Pep+HexNAc_4+
Pep+HexNAc_3+
M+e - Acetyl
c44++_iso1
HexNAcHex
b6-18
z.7
b10
c12
z.14
HexNAc
z.3
b4
z.5
b6
c7
c8
b9
c9
z.9
z.10
Pep_3+
c35

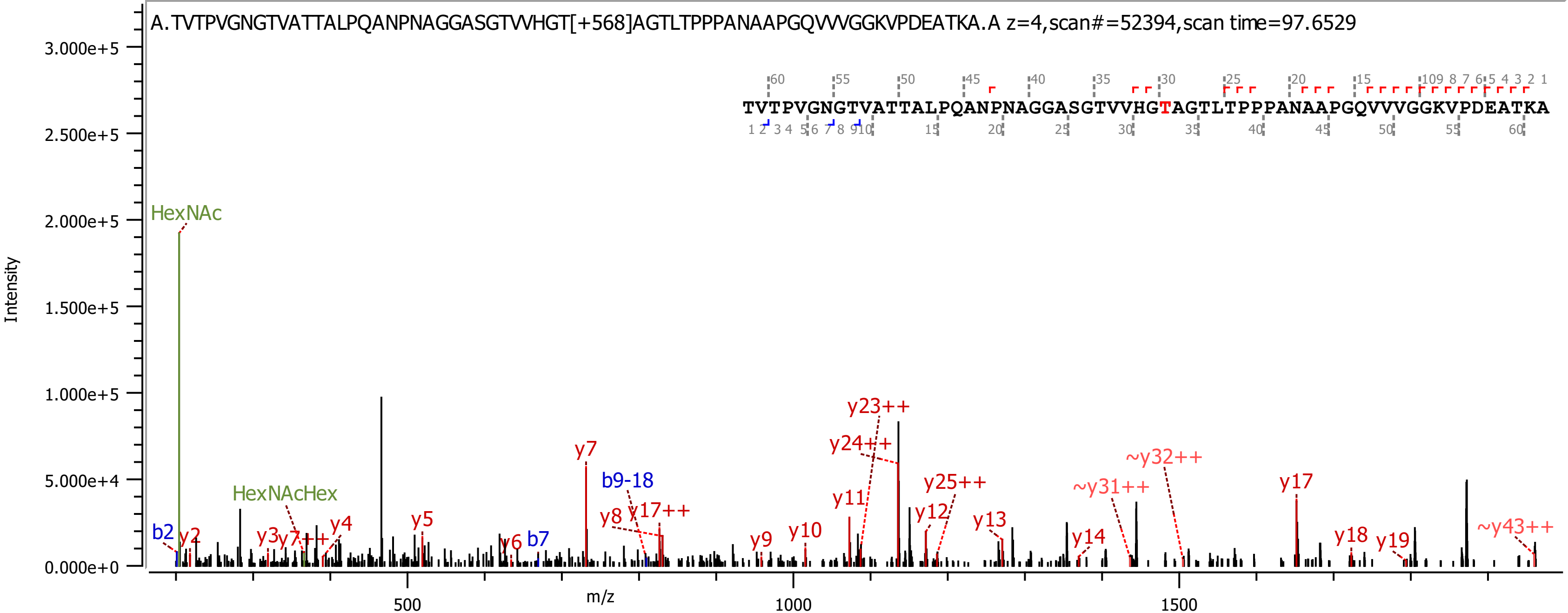
A.TVTPVGNGTVATTALPQANPNAGGASGTVVHGT[+568]AGTLTPPPANAAPGQVVVGGKVPDEATKA.A z=4,scan#=52394,scan time=97.6529
TVTPVGNGTVATTALPQANPNAGGASGTVVHGTAGTLTPPPANAAPGQVVVGGKVPDEATKA
Intensity
m/z
HexNAc
HexNAcHex
b2
y2
y3
y7++
y4
y5
y6
b7
y7
b9-18
y8
y17++
y9
y10
y11
y23++
y24++
y12
y25++
y13
~y31++
~y32++
y14
y17
y18
y19
~y43++

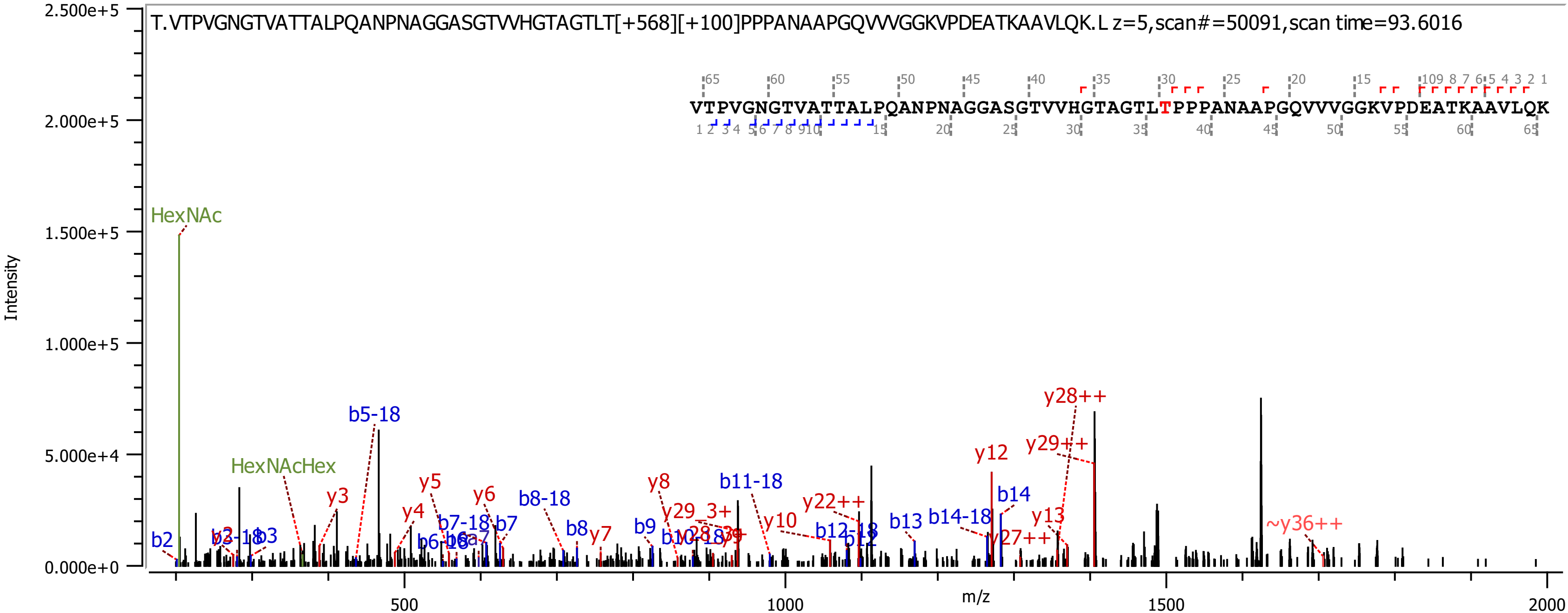

T.VTPVGNGTVATTALPQANPNAGGASGTVVHGTAGTLT[+568][+100]PPPANAAPGQVVVGGKVPDEATKAAVLQK.L z=5,scan#=50091,scan time=93.6016
VTPVGNGTVATTALPQANPNAGGASGTVVHGTAGTLTPPPANAAPGQVVVGGKVPDEATKAAVLQK
HexNAc
HexNAcHex
b5-18
y28++
y29++
y12
b14
b14-18
y13
~y36++
b13
y22++
b11-18
b12-18
b12
y10
y8
y29_3+
b9
b10-18
y7
b8
b8-18
y6
b7
b7-18
y5
y4
b6
y3
b3
b2
Intensity
m/z

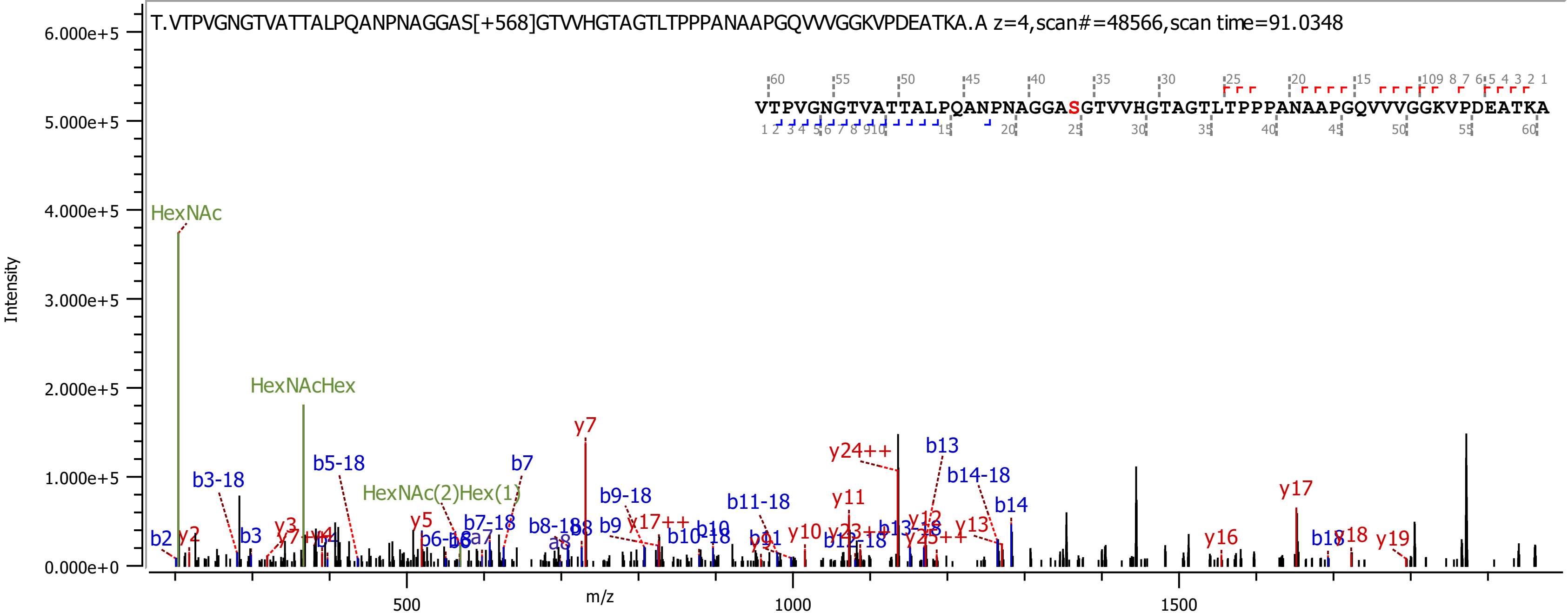
T.VTPVGNGTVATTALPQANPNAGGAS[+568]GTVVHGTAGTLTPPPANAAPGQVVVGGKVPDEATKA.A z=4,scan#=48566,scan time=91.0348
VTPVGNGTVATTALPQANPNAGGASGTVVHGTAGTLTPPPANAAPGQVVVGGKVPDEATKA
Intensity
m/z
HexNAc
HexNAcHex
HexNAc(2)Hex(1)
y7
y24++
b13
b14-18
b14
y17
b3-18
b5-18
b7
b9-18
b11-18
y11
y16
y19
y18
b18

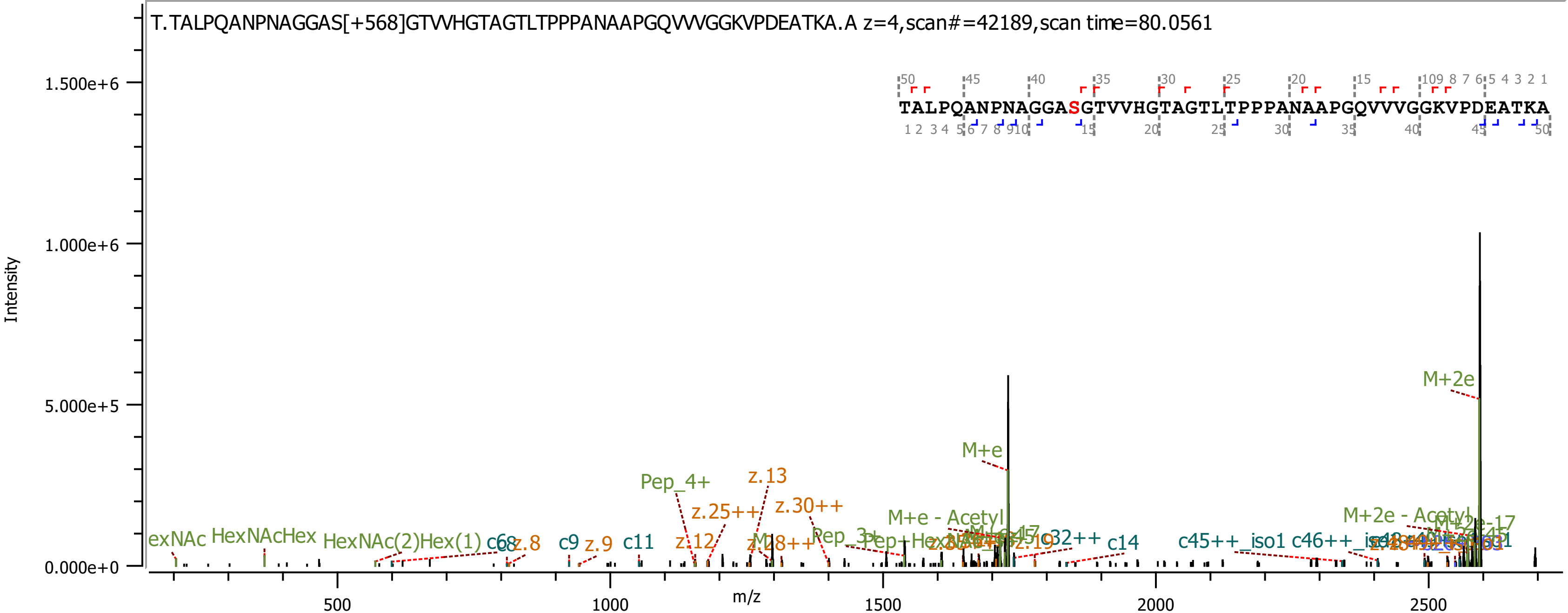

T.TALPQANPNAGGAS[+568]GTVVHGTAGTLTPPPANAAPGQVVVGGKVPDEATKA.A z=4,scan#=42189,scan time=80.0561
TALPQANPNAGGASGTVVHGTAGTLTPPPANAAPGQVVVGGKVPDEATKA
Intensity
m/z
1.500e+6
1.000e+6
5.000e+5
0.000e+0
500
1000
1500
2000
2500
M+2e
M+e
Pep_4+
z.13
z.25++
z.30++
z.12
M+e - Acetyl
M+2e - Acetyl
HexNAcHex
HexNAc(2)Hex(1)
c9
z.9
c11
z.8
c14
c45++_iso1
z.19

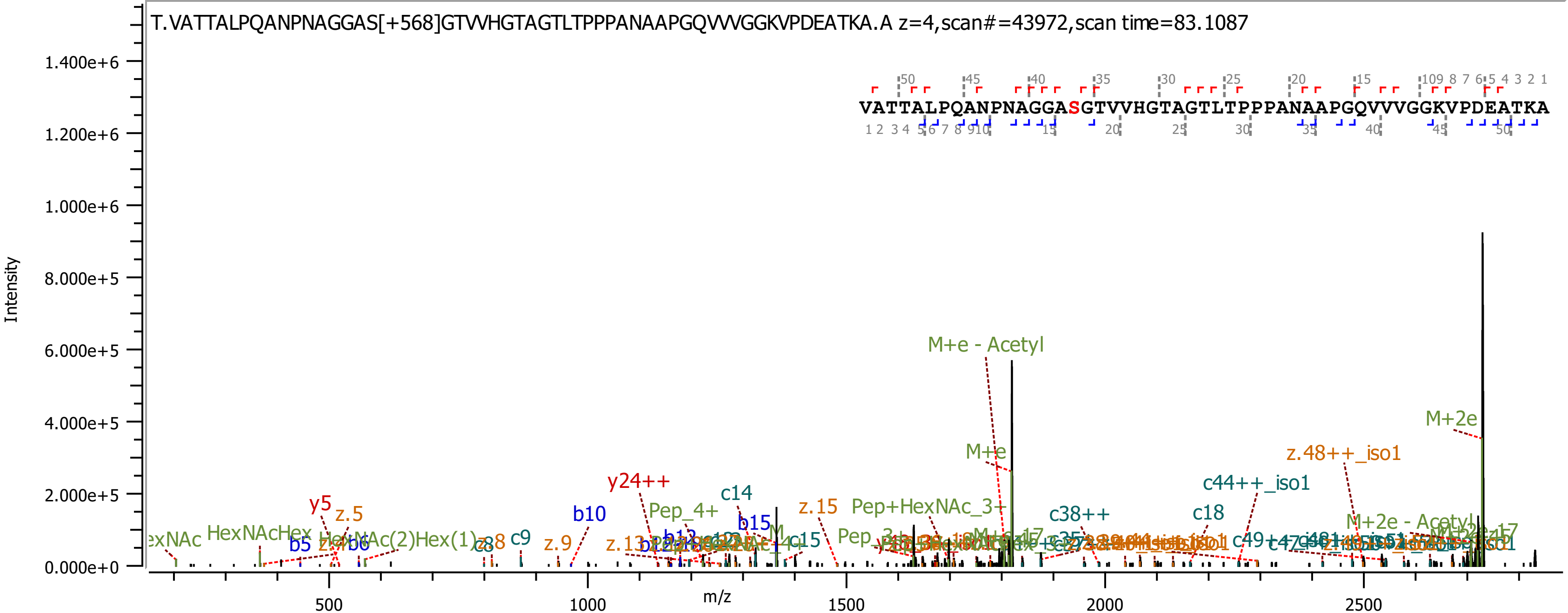
T.VATTALPQANPNAGGAS[+568]GTVVHGTAGTLTPPPANAAPGQVVVGGKVPDEATKA.A z=4,scan#=43972,scan time=83.1087
VATTALPQANPNAGGASGTVVHGTAGTLTPPPANAAPGQVVVGGKVPDEATKA
Intensity
m/z
1.400e+6
1.200e+6
1.000e+6
8.000e+5
6.000e+5
4.000e+5
2.000e+5
0.000e+0
500
1000
1500
2000
2500
M+e - Acetyl
M+2e
M+e
z.48++_iso1
c44++_iso1
y24++
c14
c38++
c18
y5
z.5
b10
Pep_4+
z.15
Pep+HexNAc_3+
b15
M+2e - Acetyl
HexNAcHex
b5
b6
c9
z.9
c15
Pep_3+

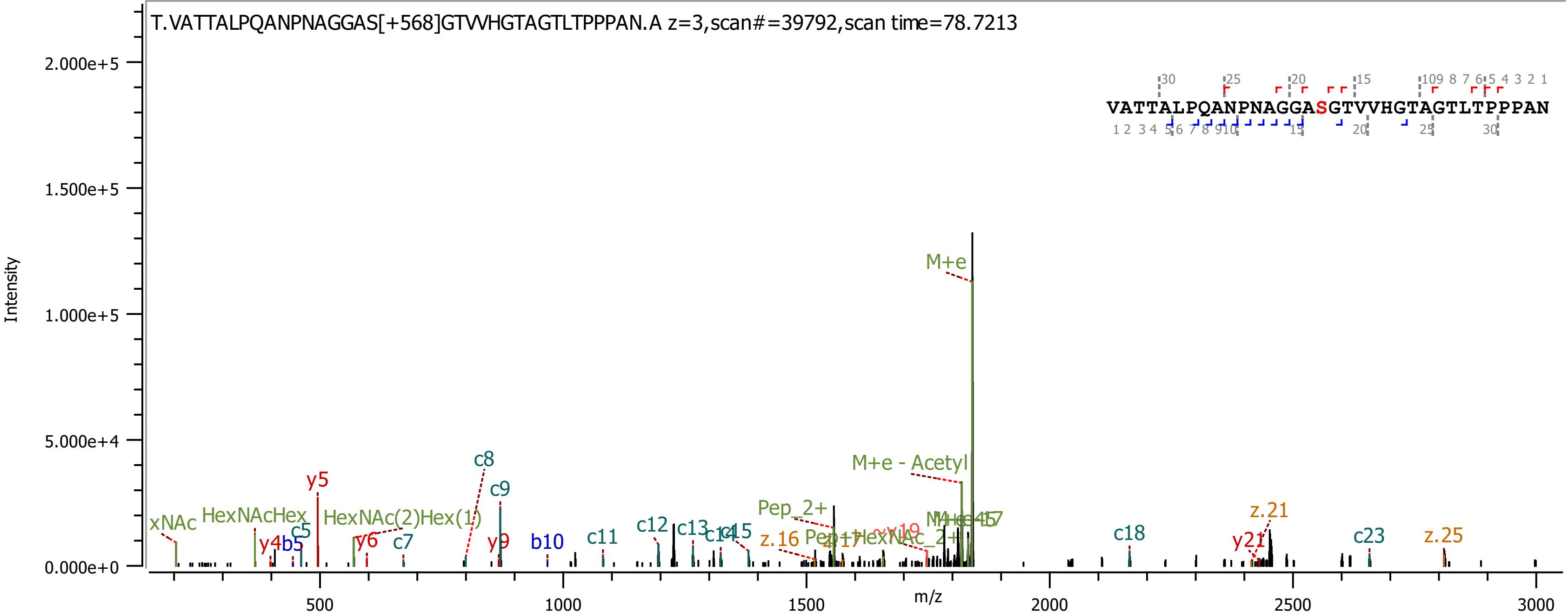
T.VATTALPQANPNAGGAS[+568]GTVVHGTAGTLTPPPAN.A z=3,scan#=39792,scan time=78.7213
VATTALPQANPNAGGASGTVVHGTAGTLTPPPAN
Intensity
2.000e+5
1.500e+5
1.000e+5
5.000e+4
0.000e+0
m/z
500
1000
1500
2000
2500
3000
M+e
M+e - Acetyl
Pep_2+
z.16
z.21
z.25
c5
c7
c8
c9
c11
c12
c13
c18
c23
y4
y5
y6
y9
y21
b5
b10
xNAc
HexNAcHex
HexNAc(2)Hex(1)

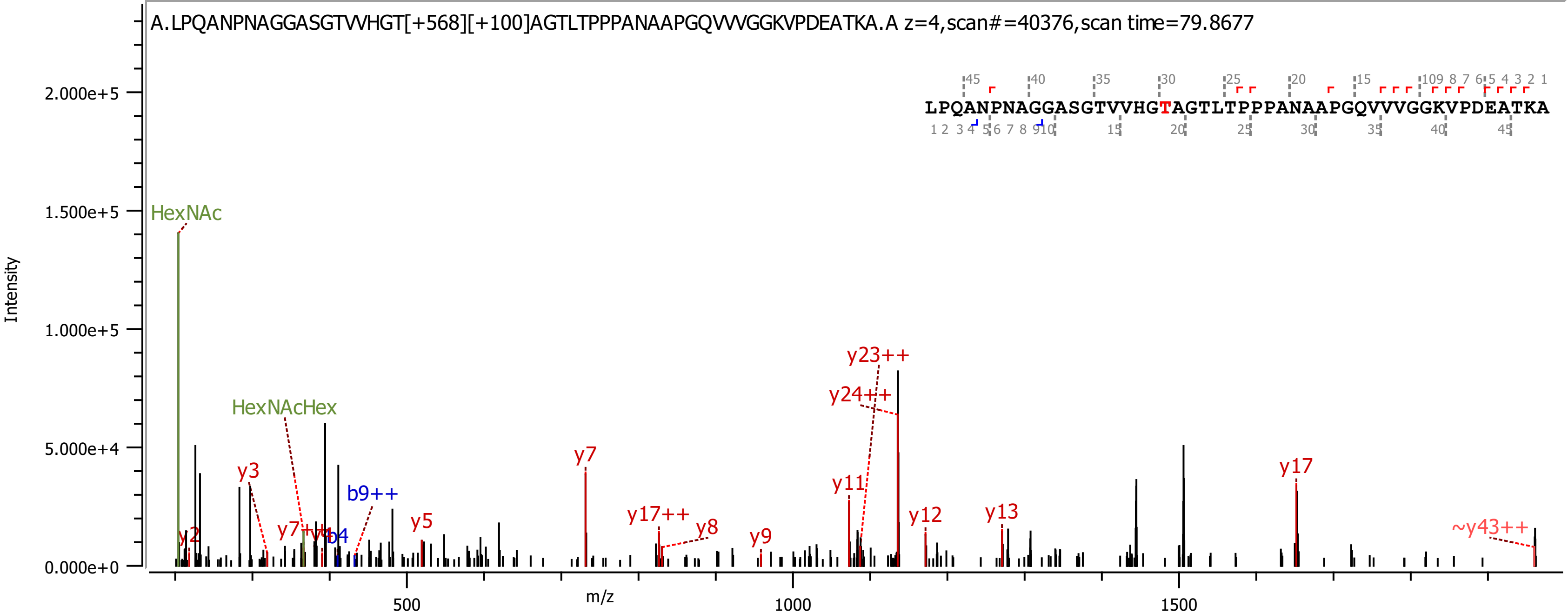
A.LPQANPNAGGASGTVVHGT[+568][+100]AGTLTPPPANAAPGQVVVGGKVPDEATKA.A z=4,scan#=40376,scan time=79.8677
LPQANPNAGGASGTVVHGTAGTLTPPPANAAPGQVVVGGKVPDEATKA
Intensity
m/z
HexNAc
HexNAcHex
y2
y3
y7++
y4
b4
b9++
y5
y7
y17++
y8
y9
y11
y23++
y24++
y12
y13
y17
~y43++
2.000e+5
1.500e+5
1.000e+5
5.000e+4
0.000e+0
500
1000
1500

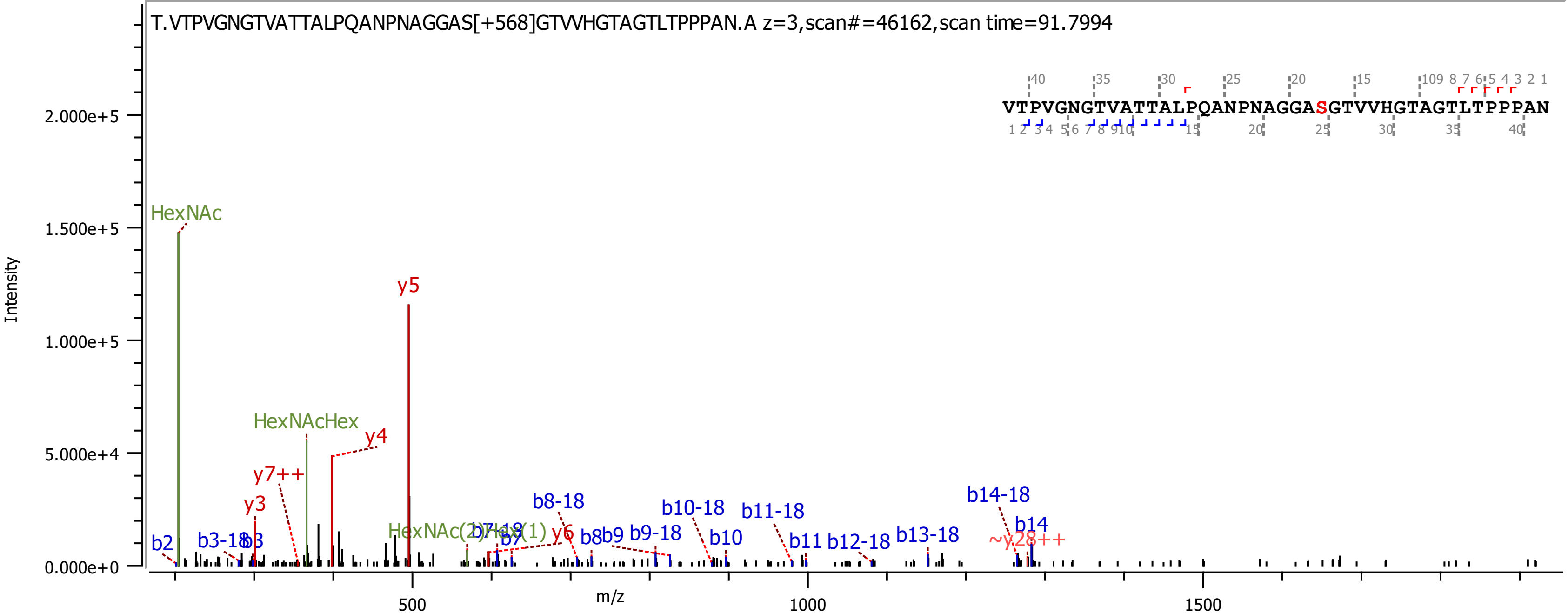
T.VTPVGNGTVATTALPQANPNAGGAS[+568]GTVVHGTAGTLTPPPAN.A z=3,scan#=46162,scan time=91.7994
VTPVGNGTVATTALPQANPNAGGASGTVVHGTAGTLTPPPAN
Intensity
m/z
HexNAc
HexNAcHex
y5
y4
y7++
y3
y6
b2
b3-18
b3
b7
b8-18
b8
b9
b9-18
b10-18
b10
b11-18
b11
b12-18
b13-18
b14-18
b14
~y28++

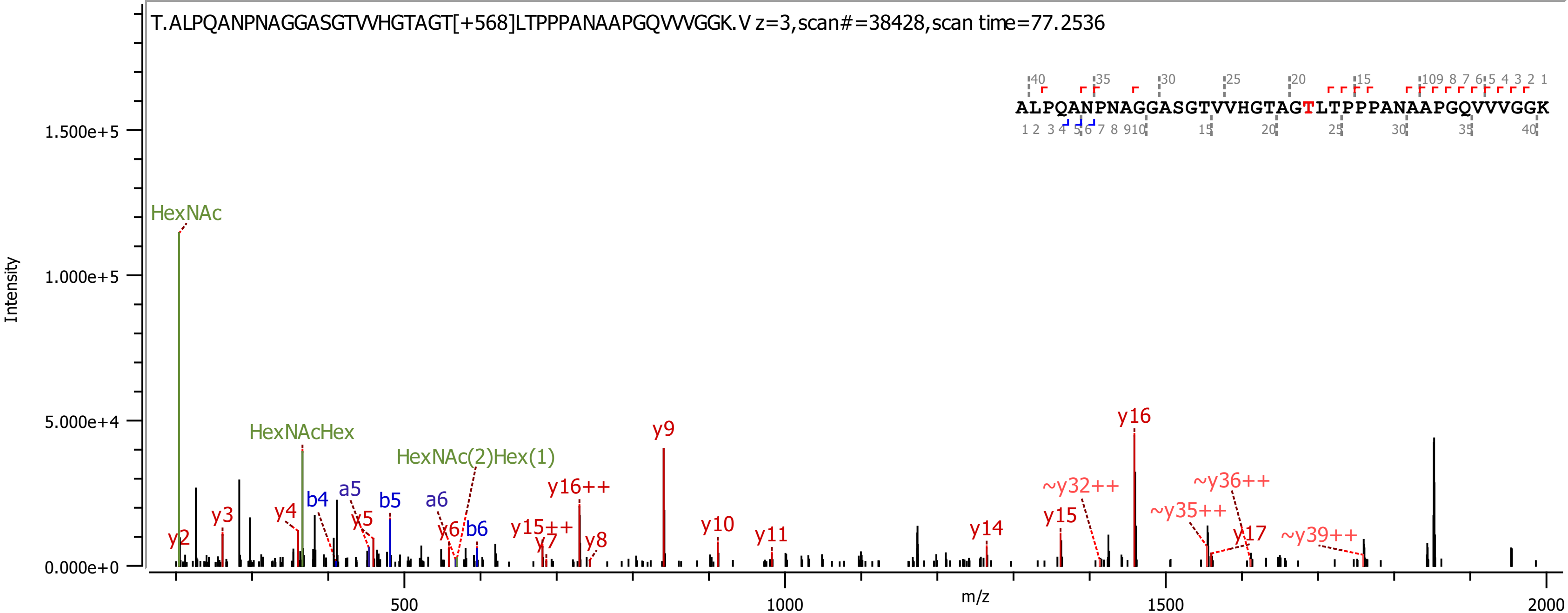

T.ALPQANPNAGGASGTVVHGTAGT[+568]LTPPPANAAPGQVVVGGK.V z=3,scan#=38428,scan time=77.2536
ALPQANPNAGGASGTVVHGTAGTLTPPPANAAPGQVVVGGK
Intensity
m/z
1.500e+5
1.000e+5
5.000e+4
0.000e+0
500
1000
1500
2000
HexNAc
HexNAcHex
HexNAc(2)Hex(1)
y2
y3
y4
b4
a5
y5
b5
a6
y6
b6
y15++
y7
y16++
y8
y9
y10
y11
y14
y15
~y32++
y16
~y35++
~y36++
y17
~y39++

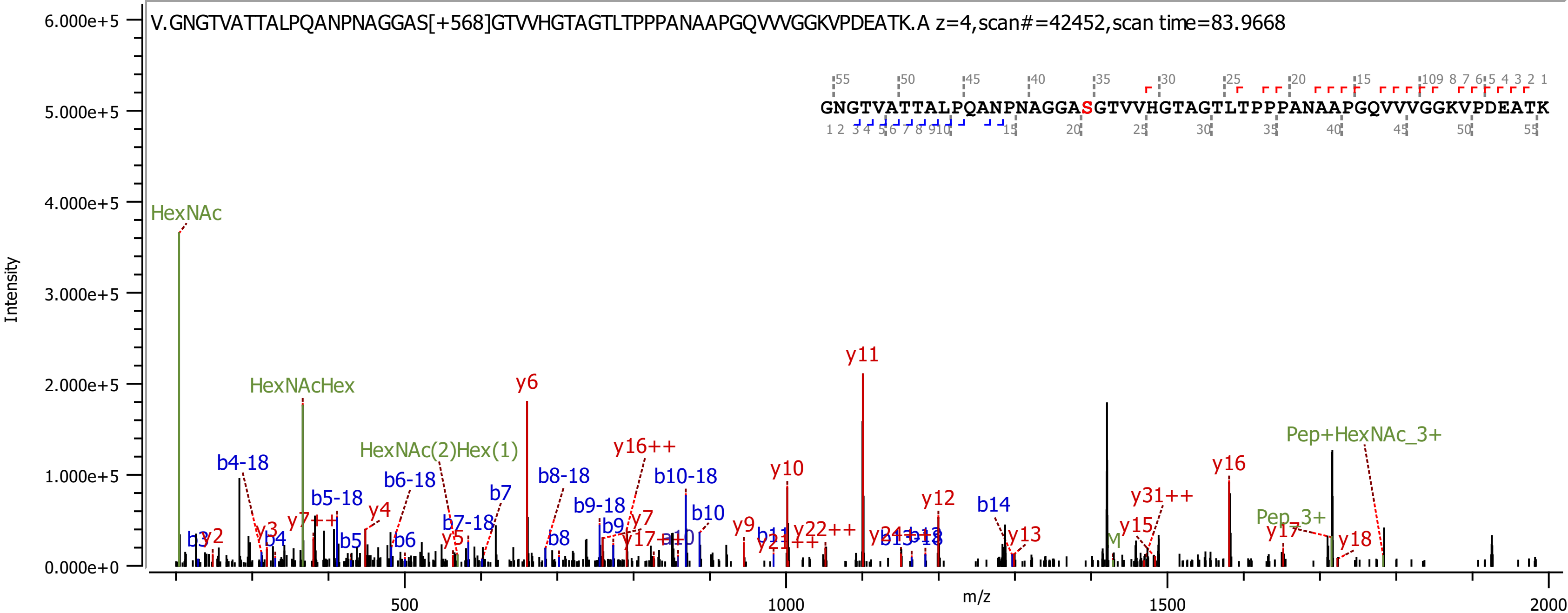
V.GNGTVATTALPQANPNAGGAS[+568]GTVVHGTAGTLTPPPANAAPGQVVVGGKVPDEATK.A z=4,scan#=42452,scan time=83.9668
GNGTVATTALPQANPNAGGASGTVVHGTAGTLTPPPANAAPGQVVVGGKVPDEATK
Intensity
m/z
6.000e+5
5.000e+5
4.000e+5
3.000e+5
2.000e+5
1.000e+5
0.000e+0
500
1000
1500
2000
HexNAc
HexNAcHex
HexNAc(2)Hex(1)
Pep+HexNAc_3+
Pep_3+
y11
y6
y16++
y10
y12
y16
y31++
y15
y13
y17
y18
y9
y7
y4
y2
y3
y5
b4-18
b5-18
b6-18
b7-18
b8-18
b9-18
b10-18
b7
b10
b14
b3
b4
b5
b6
b8
b9

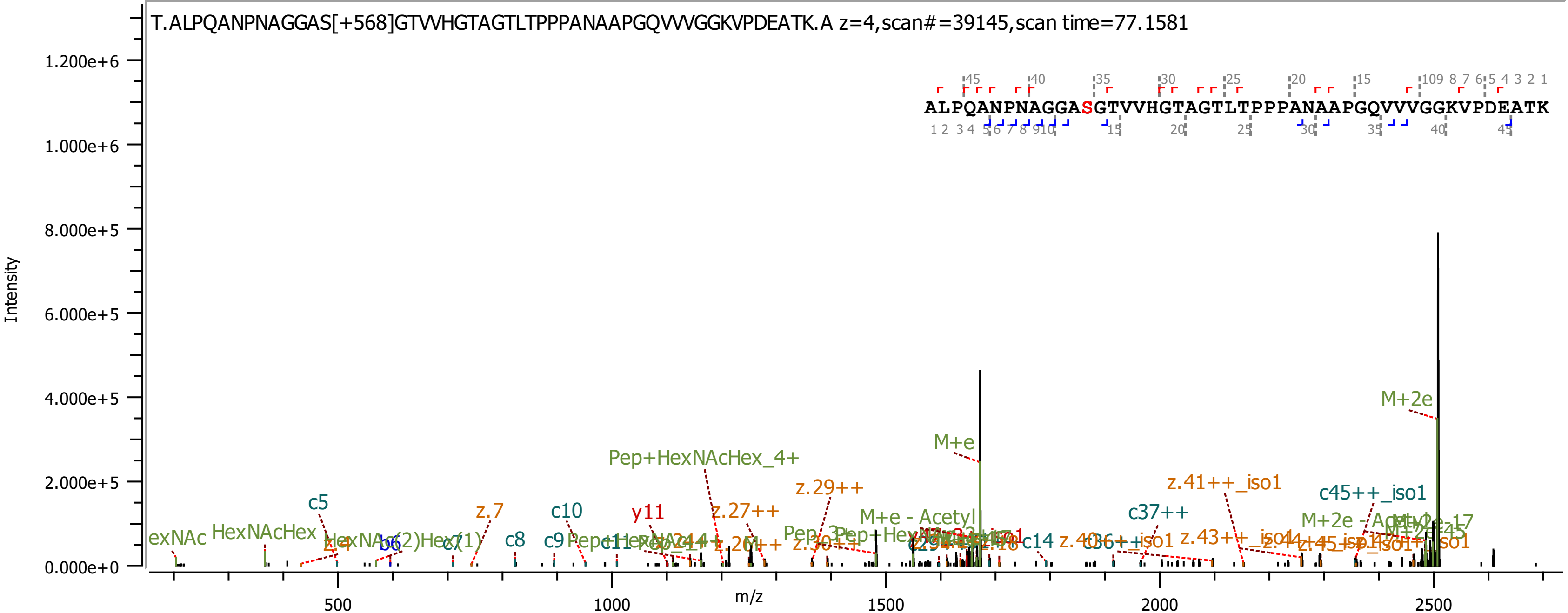
T.ALPQANPNAGGAS[+568]GTVVHGTAGTLTPPPANAAPGQVVVGGKVPDEATK.A z=4,scan#=39145,scan time=77.1581
ALPQANPNAGGASGTVVHGTAGTLTPPPANAAPGQVVVGGKVPDEATK
Intensity
m/z
1.200e+6
1.000e+6
8.000e+5
6.000e+5
4.000e+5
2.000e+5
0.000e+0
500
1000
1500
2000
2500
M+2e
M+e
Pep+HexNAcHex_4+
z.29++
z.41++_iso1
c45++_iso1
c37++
M+e - Acetyl
M+2e - Acetyl
c5
z.7
c10
y11
z.27++
HexNAcHex
c8
c9
c14
b6
c7

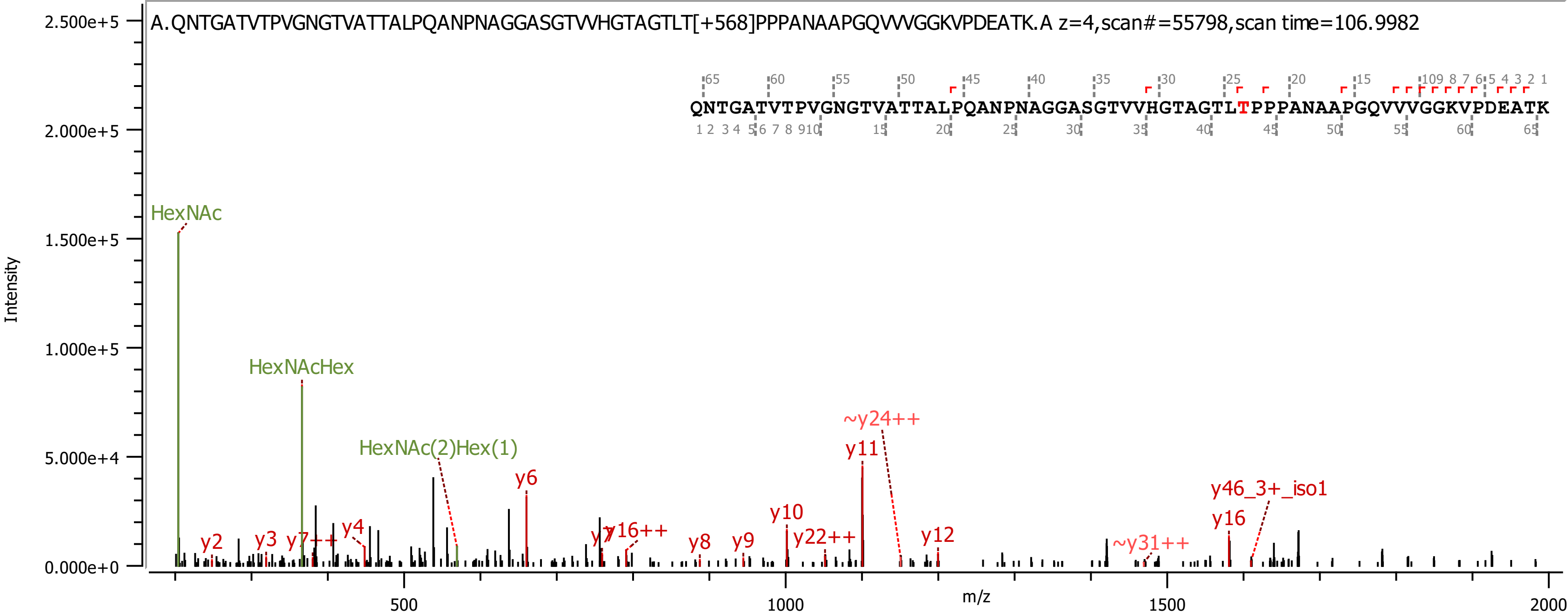

A.QNTGATVTPVGNGTVATTALPQANPNAGGASGTVVHGTAGTLT[+568]PPPANAAPGQVVVGGKVPDEATK.A z=4,scan#=55798,scan time=106.9982
QNTGATVTPVGNGTVATTALPQANPNAGGASGTVVHGTAGTLTPPPANAAPGQVVVGGKVPDEATK
Intensity
m/z
HexNAc
HexNAcHex
HexNAc(2)Hex(1)
y2
y3
y7++
y4
y6
y7
y16++
y8
y9
y10
y22++
~y24++
y11
y12
~y31++
y16
y46_3+_iso1
2.500e+5
2.000e+5
1.500e+5
1.000e+5
5.000e+4
0.000e+0
500
1000
1500
2000

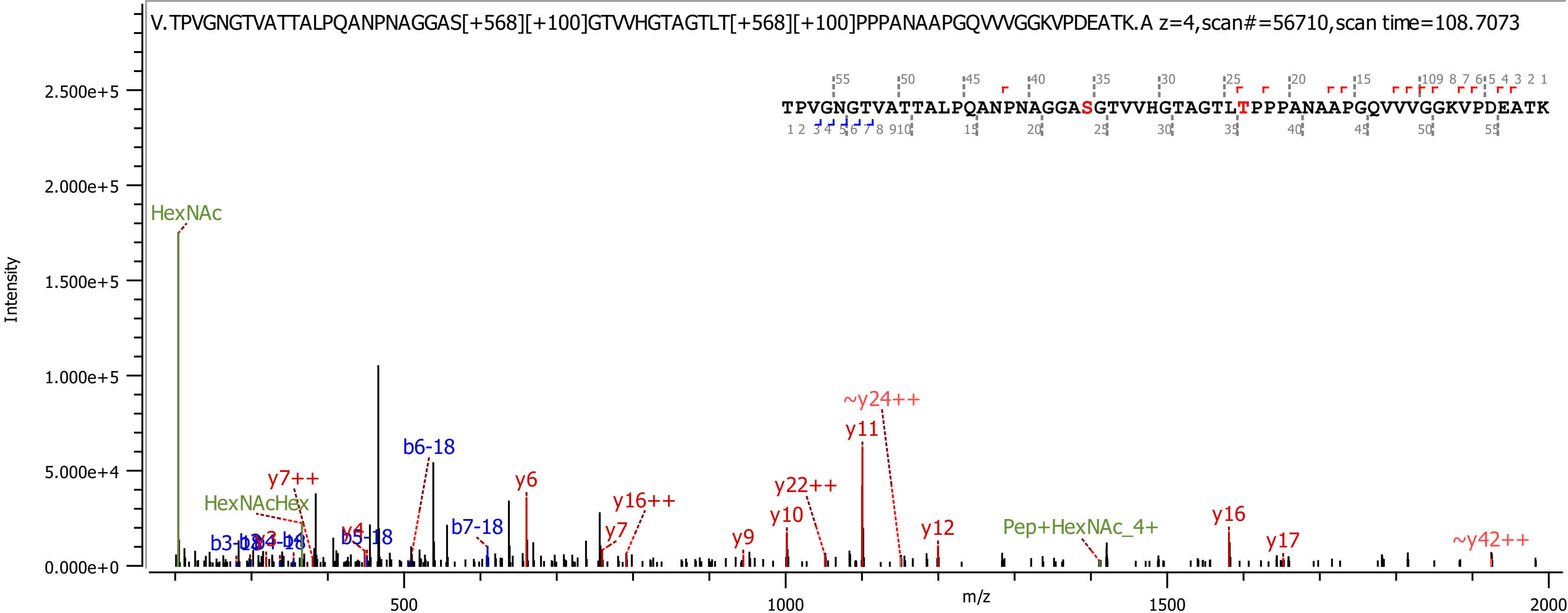
V.TPVGNGTVATTALPQANPNAGGAS[+568][+100]GTVVHGTAGTLT[+568][+100]PPPANAAPGQVVVGGKVPDEATK.A z=4,scan#=56710,scan time=108.7073
TPVGNGTVATTALPQANPNAGGASGTVVHGTAGTLTPPPANAAPGQVVVGGKVPDEATK
Intensity
m/z
HexNAc
HexNAcHex
y7++
b6-18
y6
b7-18
y16++
y7
y4
b5-18
b3-18
y9
y10
y22++
y11
~y24++
y12
Pep+HexNAc_4+
y16
y17
~y42++
2.500e+5
2.000e+5
1.500e+5
1.000e+5
5.000e+4
0.000e+0
500
1000
1500
2000

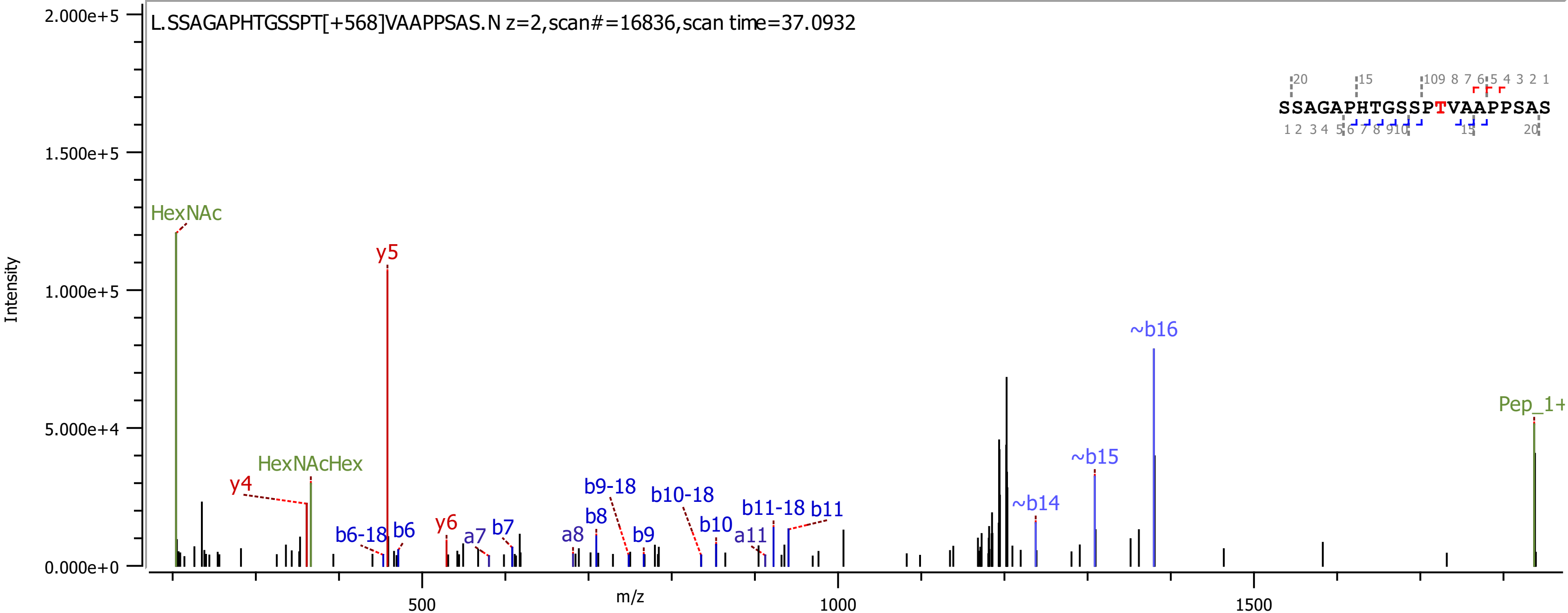
L.SSAGAPHTGSSPT[+568]VAAPPSAS.N z=2,scan#=16836,scan time=37.0932
SSAGAPHTGSSPTVAAPPSAS
Intensity
m/z
2.000e+5
1.500e+5
1.000e+5
5.000e+4
0.000e+0
500
1000
1500
HexNAc
y5
~b16
Pep_1+
~b15
~b14
HexNAcHex
y4
b9-18
b10-18
b11-18
b11
b8
b10
a11
b9
a8
b7
a7
y6
b6
b6-18

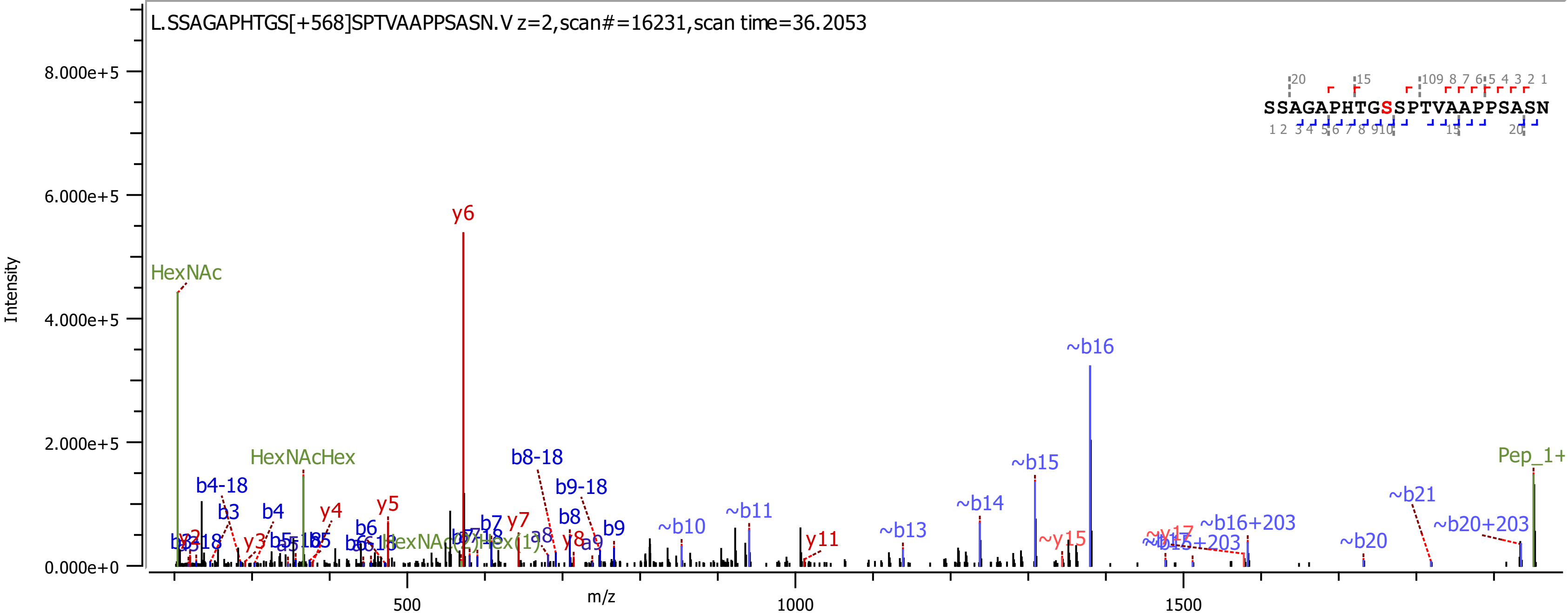

L.SSAGAPHTGS[+568]SPTVAAPPSASN.V z=2,scan#=16231,scan time=36.2053
SSAGAPHTGSSPTVAAPPSASN
Intensity
m/z
8.000e+5
6.000e+5
4.000e+5
2.000e+5
0.000e+0
500
1000
1500
HexNAc
HexNAcHex
y6
~b16
~b15
~b14
~b13
~b11
~b10
~b21
~b20
~b20+203
~b16+203
~y15
b8-18
b9-18
b4-18
b3
b4
y4
y5
b6
b7
y7
b8
b9
y11
Pep_1+

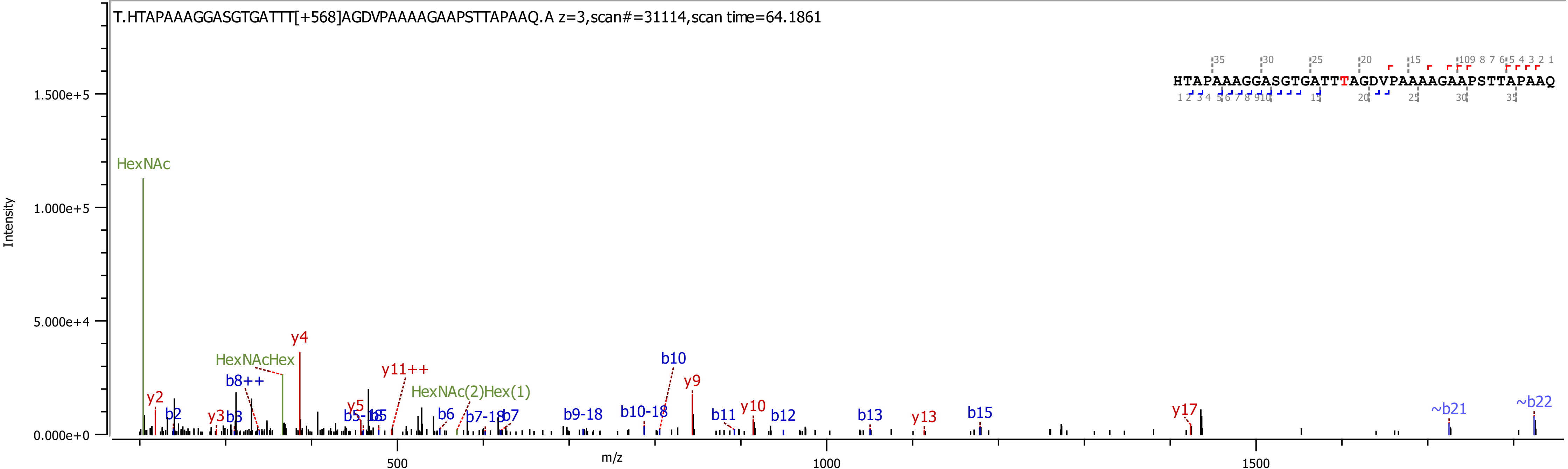
T.HTAPAAAGGASGTGATTT[+568]AGDVPAAAAGAAPSTTAPAAQ.A z=3,scan#=31114,scan time=64.1861
HTAPAAAGGASGTGATTTAGDVPAAAAGAAPSTTAPAAQ
Intensity
m/z
HexNAc
HexNAcHex
HexNAc(2)Hex(1)
y2
y3
y4
y5
y9
y10
y11++
y13
y17
b2
b3
b5-18
b5
b6
b7-18
b7
b8++
b9-18
b10-18
b10
b11
b12
b13
b15
~b21
~b22
1.500e+5
1.000e+5
5.000e+4
0.000e+0
500
1000
1500

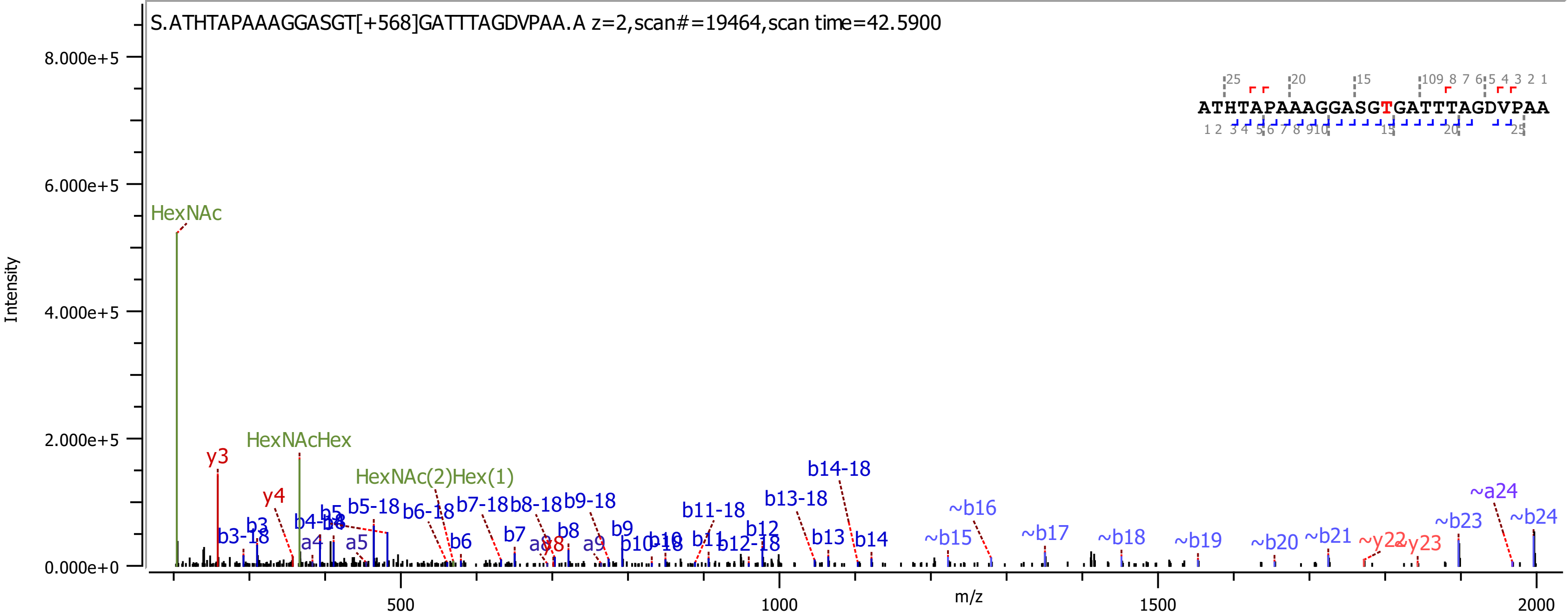
S.ATHTAPAAAGGASGT[+568]GATTTAGDVPAA.A z=2,scan#=19464,scan time=42.5900
ATHTAPAAAGGASGTGATTTAGDVPAA
Intensity
m/z
HexNAc
HexNAcHex
HexNAc(2)Hex(1)
y3
y4
y8
~y22
~y23
b3-18
b3
b4-18
b5
b5-18
b6-18
b6
b7-18
b7
b8-18
b8
b9-18
b9
b10-18
b10
b11-18
b11
b12-18
b12
b13-18
b13
b14-18
b14
~b15
~b16
~b17
~b18
~b19
~b20
~b21
~b23
~b24
~a24
a4
a5
a8
a9

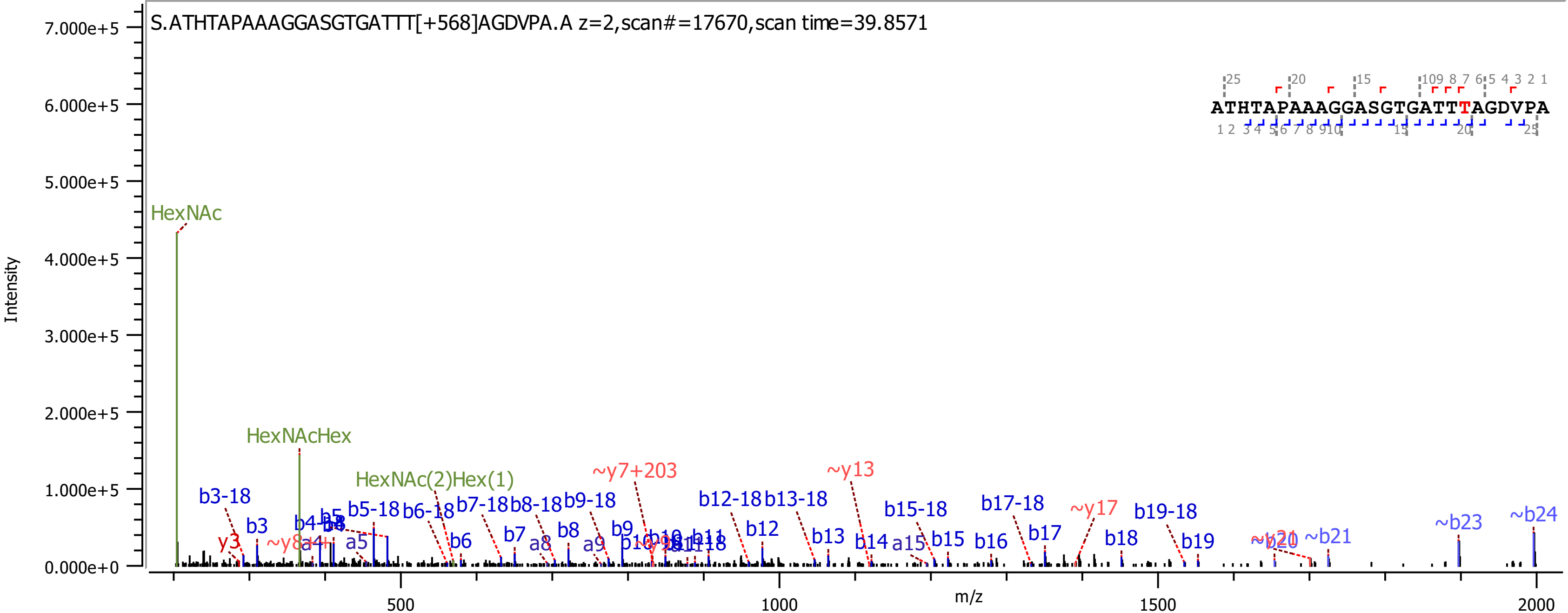
S.ATHTAPAAAGGASGTGATTT[+568]AGDVPA.A z=2,scan#=17670,scan time=39.8571
ATHTAPAAAGGASGTGATTTAGDVPA
Intensity
m/z
0.000e+0
1.000e+5
2.000e+5
3.000e+5
4.000e+5
5.000e+5
6.000e+5
7.000e+5
500
1000
1500
2000
HexNAc
HexNAcHex
HexNAc(2)Hex(1)
~y7+203
~y13
~y17
~b23
~b24
~b21
b3-18
b3
y3
b5-18
b6-18
b7-18
b8-18
b9-18
b12-18
b13-18
b15-18
b17-18
b19-18
b6
b7
a8
b8
a9
b9
b12
b13
b14
a15
b15
b16
b17
b18
b19
a5

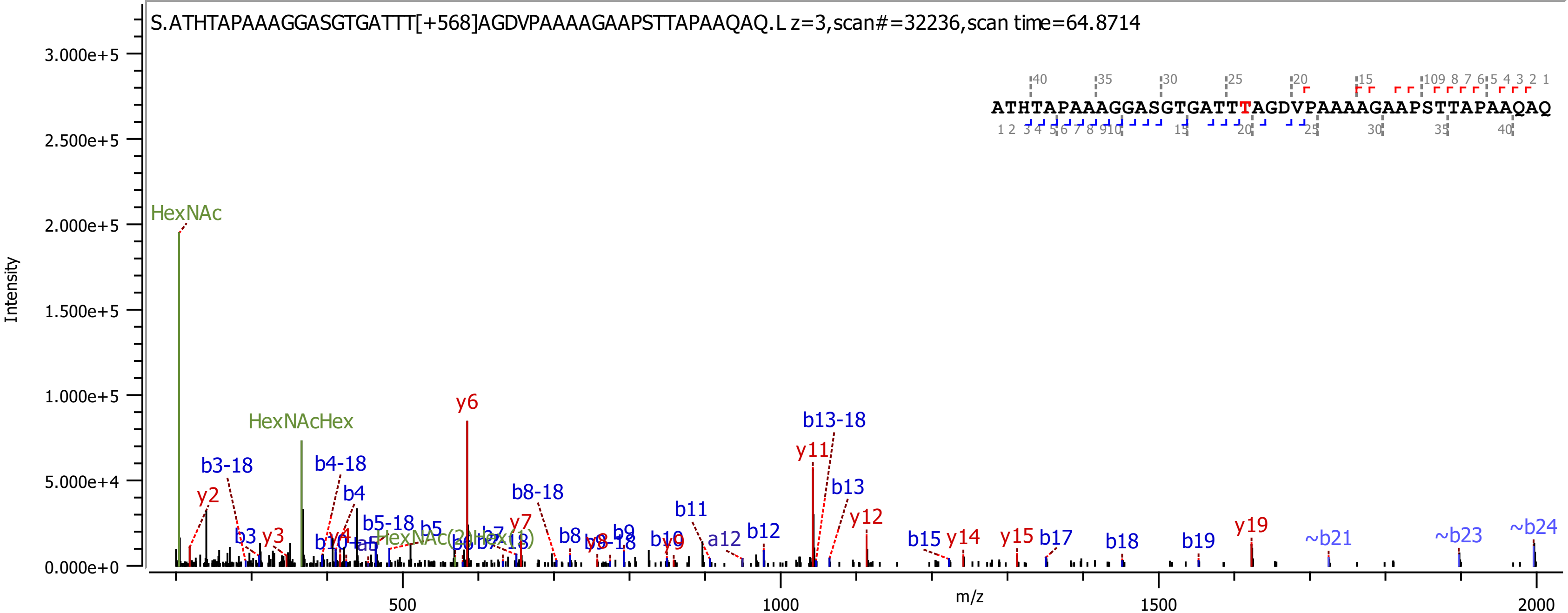
S.ATHTAPAAAGGASGTGATTT[+568]AGDVPAAAAGAAPSTTAPAAQAQ.L z=3,scan#=32236,scan time=64.8714
ATHTAPAAAGGASGTGATTTAGDVPAAAAGAAPSTTAPAAQAQ
Intensity
m/z
HexNAc
HexNAcHex
y6
b13-18
y11
b3-18
b4-18
y2
b4
b8-18
b11
b13
y12
b5-18
b5
y7
y3
b3
b8
b9
b10
y9
a12
b12
b15
y14
y15
b17
b18
b19
y19
~b21
~b23
~b24

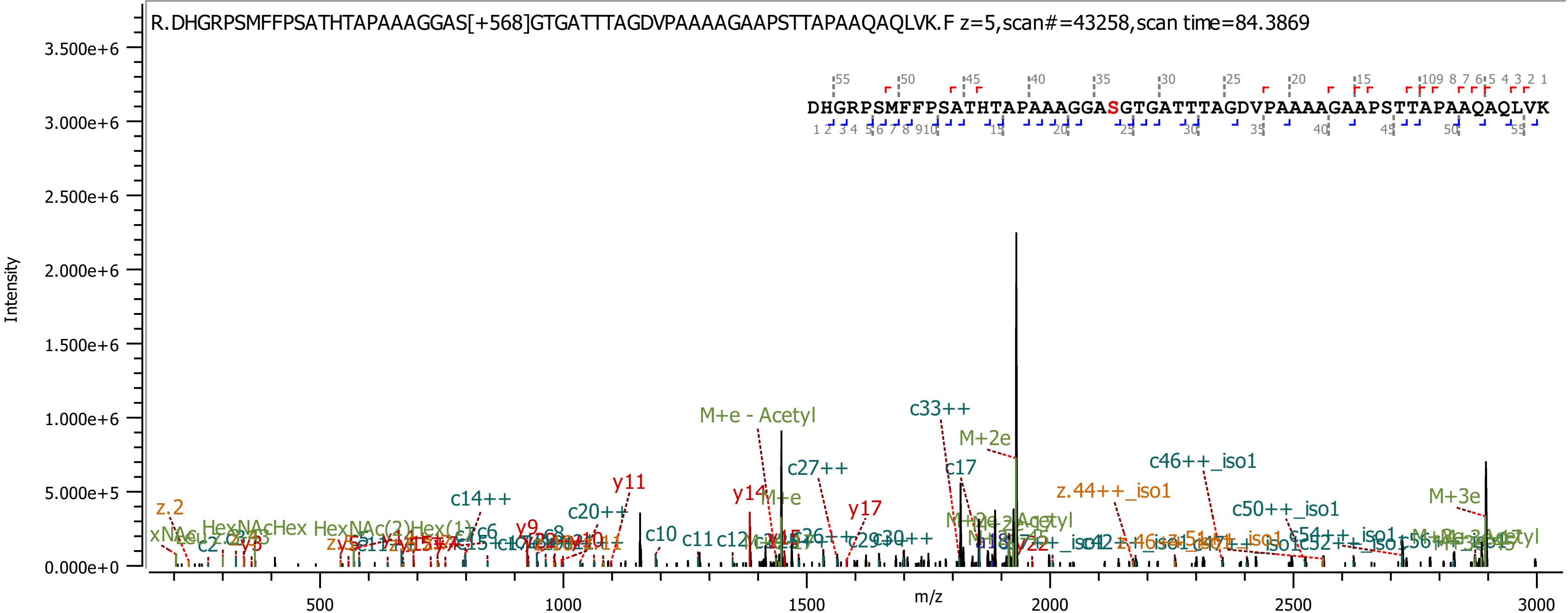

R.DHGRPSMFFPSATHTAPAAAGGAS[+568]GTGATTTAGDVPAAAAGAAPSTTAPAAQAQLVK.F z=5,scan#=43258,scan time=84.3869
DHGRPSMFFPSATHTAPAAAGGASGTGATTTAGDVPAAAAGAAPSTTAPAAQAQLVK
Intensity
m/z
3.500e+6
3.000e+6
2.500e+6
2.000e+6
1.500e+6
1.000e+6
5.000e+5
0.000e+0
500
1000
1500
2000
2500
3000
M+e - Acetyl
c33++
M+2e
c27++
c17
c46++_iso1
y11
y14
M+e
y17
z.44++_iso1
c50++_iso1
M+3e
c14++
c20++
z.2
c10
c11
c12
y9
c8
y3
c2

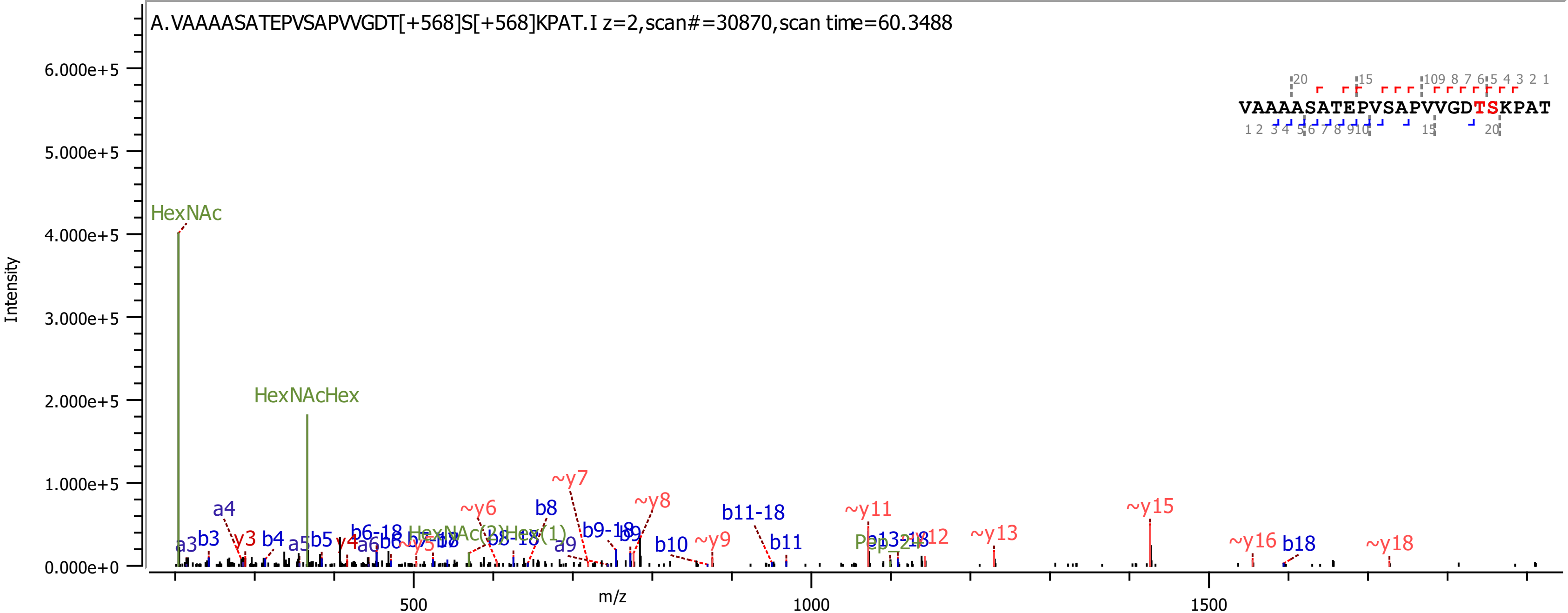
A.VAAAASATEPVSAPVVGDT[+568]S[+568]KPAT.I z=2,scan#=30870,scan time=60.3488
Intensity
m/z
VAAAASATEPVSAPVVGDTSKPAT
HexNAc
HexNAcHex
a3
b3
a4
y3
b4
a5
b5
y4
a6
b6
b8
b9
b10
b11
b18
b6-18
b8-18
b9-18
b11-18
a9
~y6
~y7
~y8
~y9
~y11
~y13
~y15
~y16
~y18
0.000e+0
1.000e+5
2.000e+5
3.000e+5
4.000e+5
5.000e+5
6.000e+5
500
1000
1500

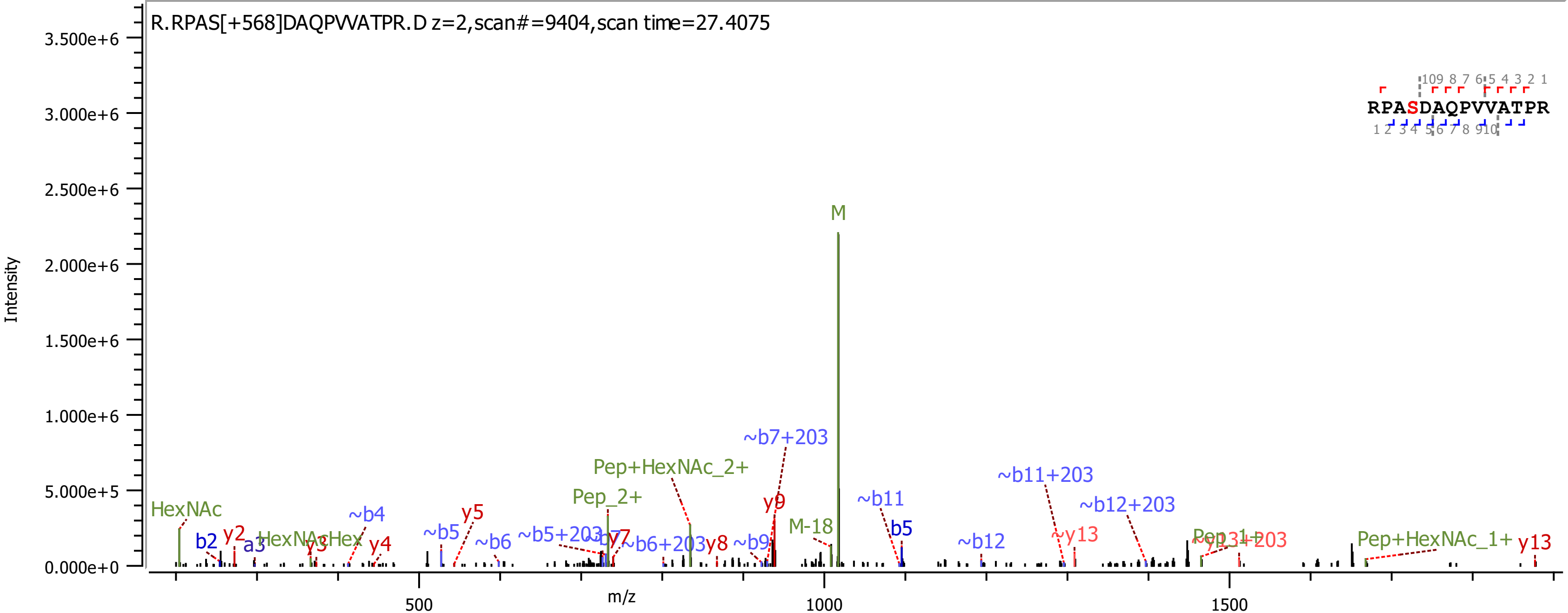
R.RPAS[+568]DAQPVVATPR.D z=2,scan#=9404,scan time=27.4075
Intensity
m/z
M
~b7+203
Pep+HexNAc_2+
~b11+203
~b12+203
Pep_2+
~b11
y9
M-18
b5
HexNAc
~b4
y5
~b5
~b6
~b5+203
~b7
y7
~b6+203
y8
~b9
~b12
~y13
Pep_1+
~y13+203
Pep+HexNAc_1+
y13
b2
y2
a3
y3
HexNAcHex
y4
RPASDAQPVVATPR

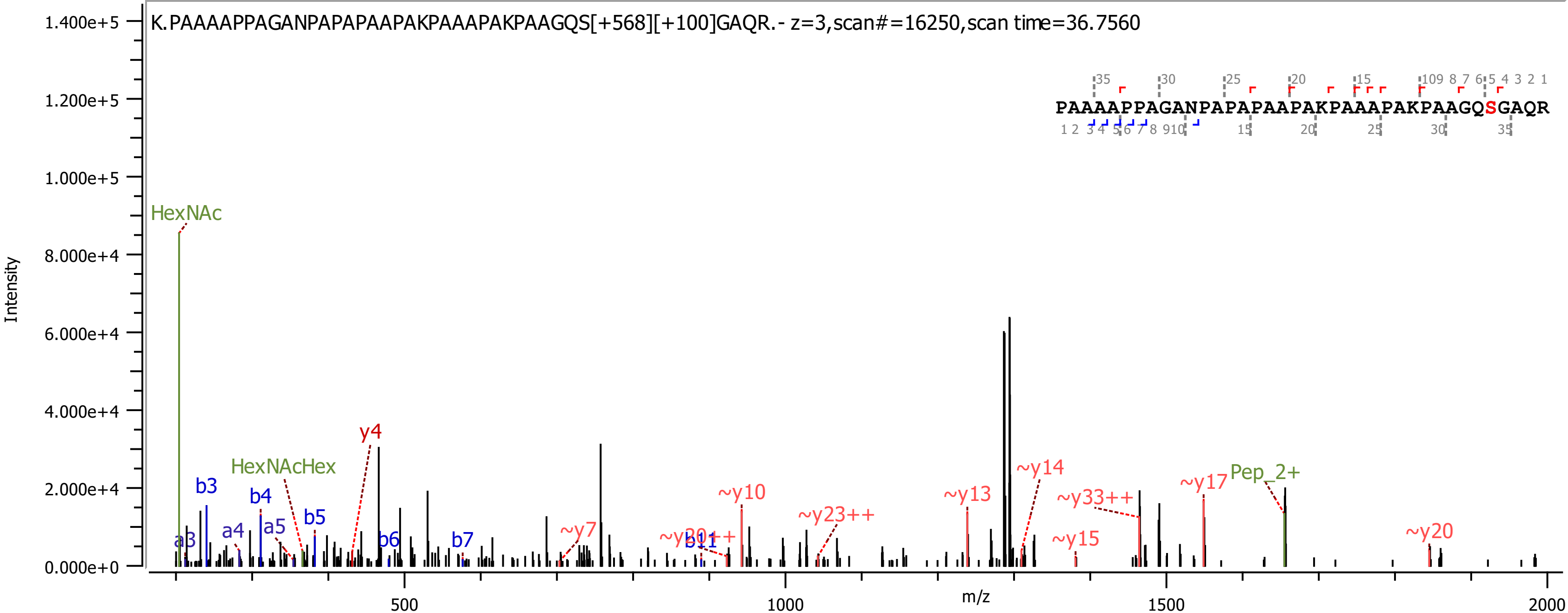
K.PAAAAPPAGANPAPAPAAPAKPAAAPAKPAAGQS[+568][+100]GAQR.- z=3,scan#=16250,scan time=36.7560
PAAAAPPAGANPAPAPAAPAKPAAAPAKPAAGQSGAQR
Intensity
m/z
HexNAc
HexNAcHex
Pep_2+
y4
~y7
~y10
~y13
~y14
~y15
~y17
~y20
~y23++
~y33++
b3
b4
b5
b6
b7
b11
a3
a4
a5

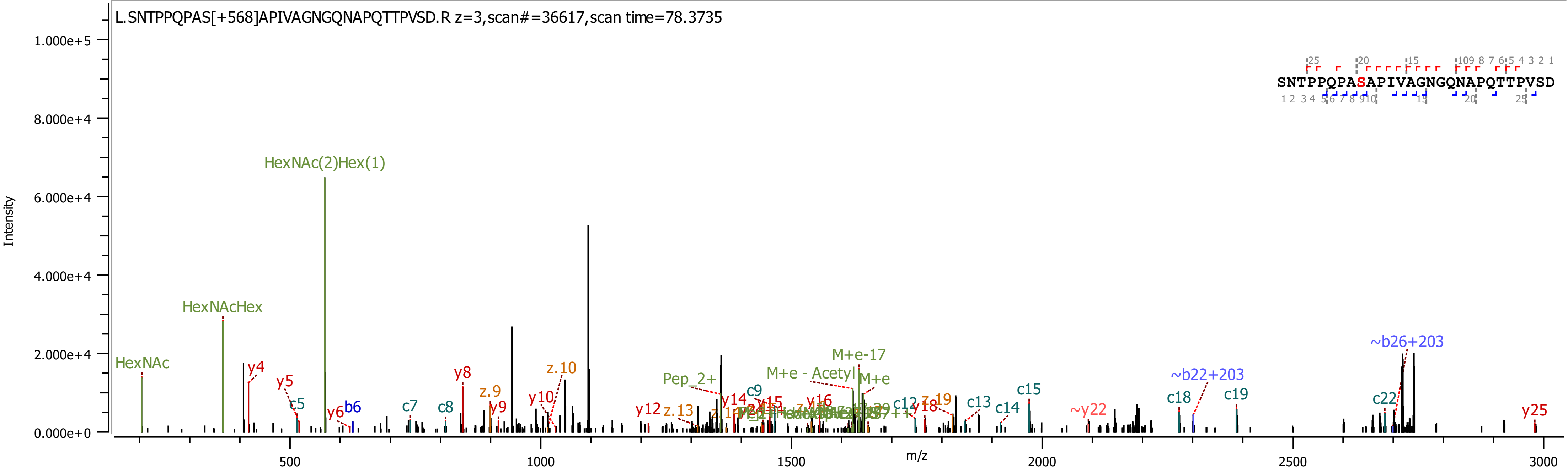

L.SNTPPQPAS[+568]APIVAGNGQNAPQTTPVSD.R z=3,scan#=36617,scan time=78.3735
SNTPPQPASAPIVAGNGQNAPQTTPVSD
Intensity
m/z
HexNAc
HexNAcHex
HexNAc(2)Hex(1)
y4
y5
c5
y6
b6
c7
c8
y8
z.9
y9
y10
z.10
y12
z.13
Pep_2+
y14
c9
y15
y16
M+e - Acetyl
M+e-17
M+e
c12
y18
z.19
c13
c14
c15
~y22
~b22+203
c18
c19
c22
~b26+203
y25

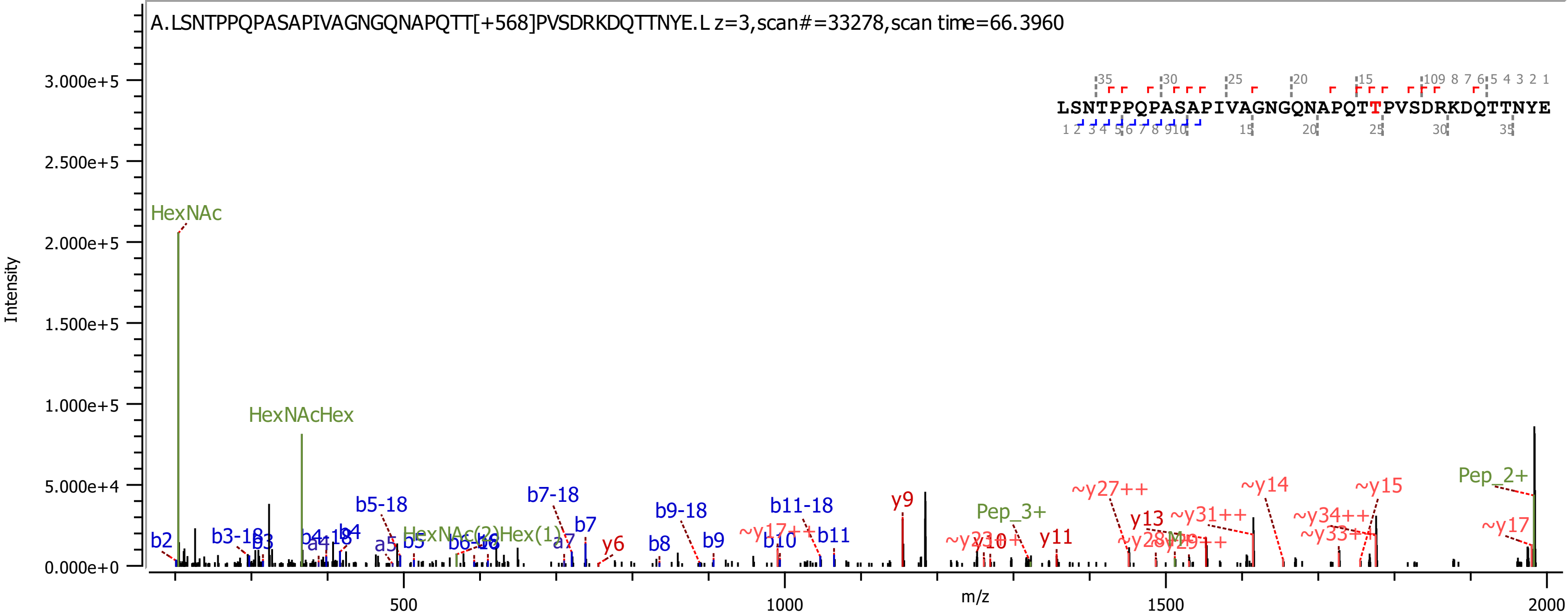
A.LSNTPPQPASAPIVAGNGQNAPQTT[+568]PVSDRKDQTTNYE.L z=3,scan#=33278,scan time=66.3960
LSNTPPQPASAPIVAGNGQNAPQTTPVSDRKDQTTNYE
Intensity
3.000e+5
2.500e+5
2.000e+5
1.500e+5
1.000e+5
5.000e+4
0.000e+0
m/z
500
1000
1500
2000
HexNAc
HexNAcHex
b2
b3-18
b3
b4-18
b4
a4
a5
b5
b5-18
b6-18
HexNAc(2)Hex(1)
a7
b7
b7-18
y6
b8
b9-18
b9
~y17++
b10
b11-18
b11
y9
Pep_3+
y10
y11
~y27++
y13
~y31++
~y14
~y34++
~y33++
~y15
Pep_2+
~y17

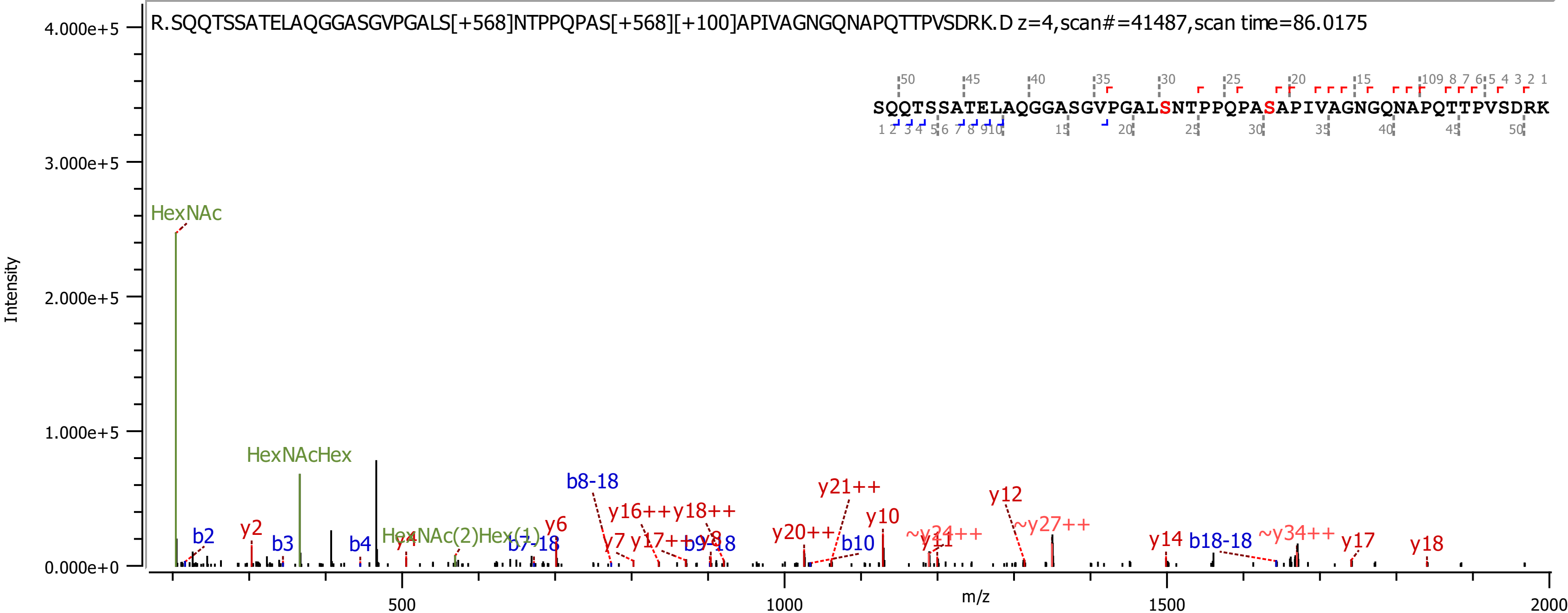
R.SQQTSSATELAQGGASGVPGALS[+568]NTPPQPAS[+568][+100]APIVAGNGQNAPQTTPVSDRK.D z=4,scan#=41487,scan time=86.0175
SQQTSSATELAQGGASGVPGALSNTPPQPASAPIVAGNGQNAPQTTPVSDRK
Intensity
m/z
HexNAc
HexNAcHex
HexNAc(2)Hex(1)
b2
y2
b3
b4
y4
b7-18
y6
b8-18
y7
y16++
y17++
y18++
y8
b9-18
y20++
y21++
b10
y10
~y24++
y11
y12
~y27++
y14
b18-18
~y34++
y17
y18
4.000e+5
3.000e+5
2.000e+5
1.000e+5
0.000e+0
500
1000
1500
2000

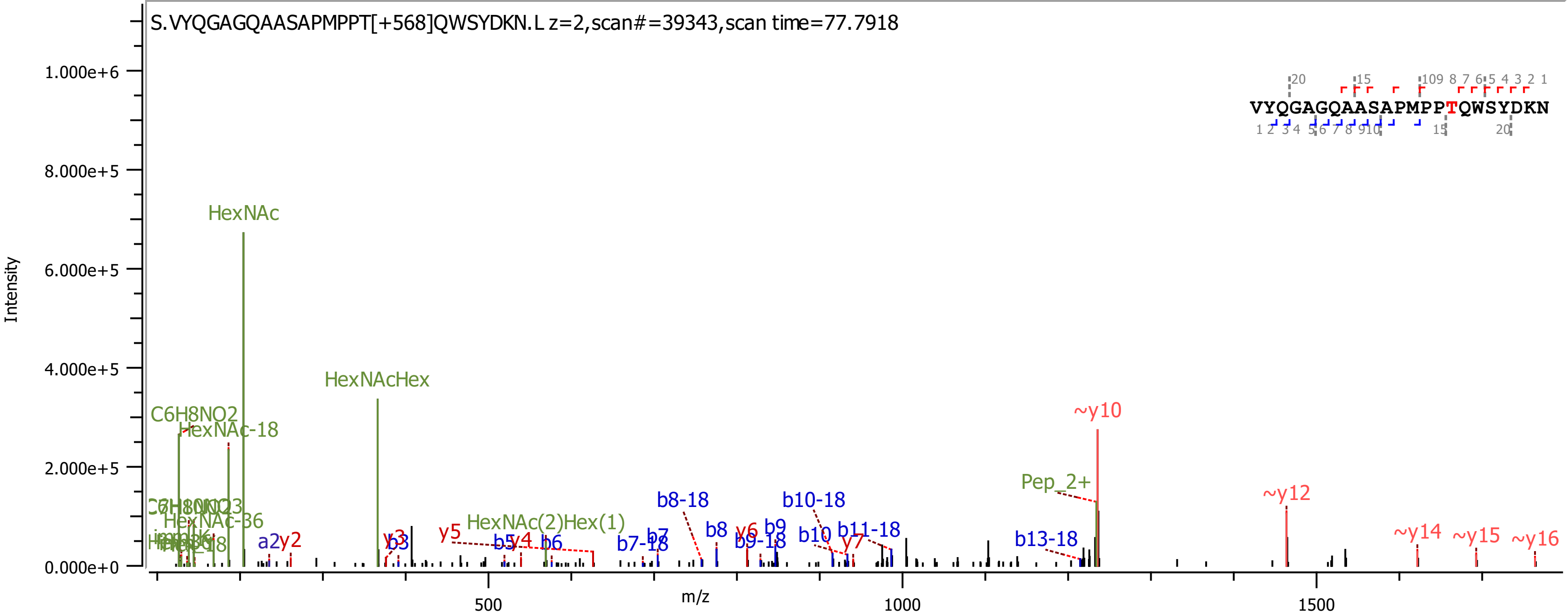
S.VYQGAGQAASAPMPPT[+568]QWSYDKN.L z=2,scan#=39343,scan time=77.7918
VYQGAGQAASAPMPPTQWSYDKN
Intensity
m/z
1.000e+6
8.000e+5
6.000e+5
4.000e+5
2.000e+5
0.000e+0
500
1000
1500
HexNAc
HexNAcHex
C6H8NO2
HexNAc-18
HexNAc-36
HexNAc(2)Hex(1)
Pep_2+
~y10
~y12
~y14
~y15
~y16
a2
y2
y3
b3
y5
b5
y4
b6
b7
b7-18
b8-18
b8
y6
b9
b9-18
b10-18
b10
y7
b11-18
b13-18

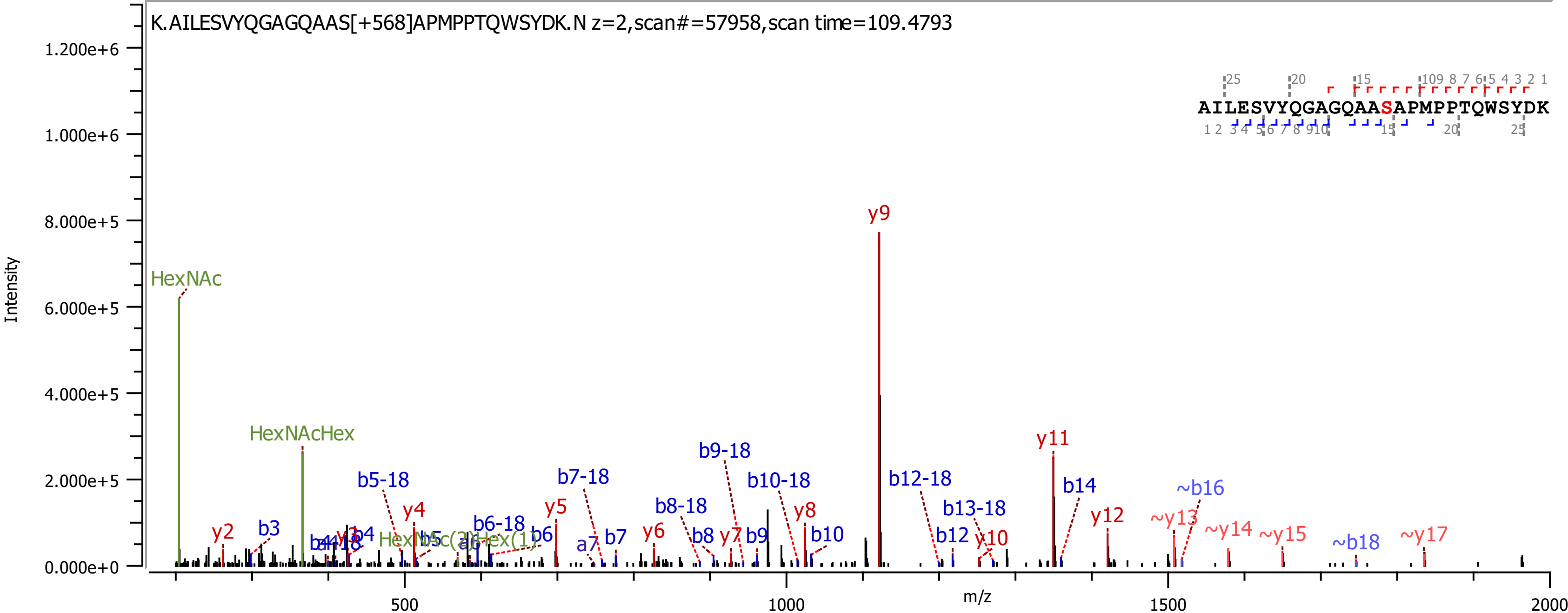

K.AILESVYQGAGQAAS[+568]APMPPTQWSYDK.N z=2,scan#=57958,scan time=109.4793
AILESVYQGAGQAASAPMPPTQWSYDK
Intensity
m/z
HexNAc
HexNAcHex
y9
y11
b14
~b16
y12
~y13
~y14
~y15
~b18
~y17
b9-18
b10-18
b12-18
b13-18
b5-18
b7-18
b8-18
b6-18
y2
b3
y4
y5
y6
y7
y8
b7
b8
b9
b10
b12
y10
a7
b6
b4

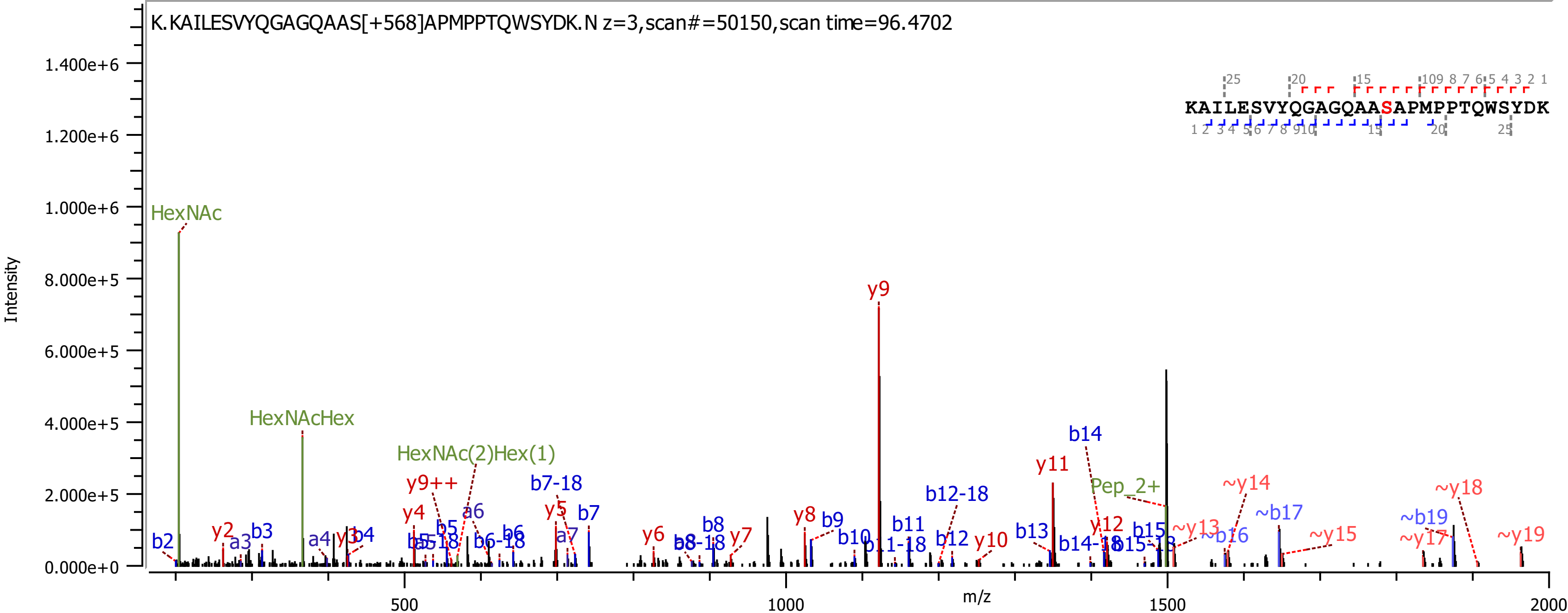

K.KAILESVYQGAGQAAS[+568]APMPPTQWSYDK.N z=3,scan#=50150,scan time=96.4702
KAILESVYQGAGQAASAPMPPTQWSYDK
Intensity
m/z
HexNAc
HexNAcHex
HexNAc(2)Hex(1)
y9
y9++
b7-18
b14
y11
Pep_2+
~y14
~y18
b12-18
~b17
~b19

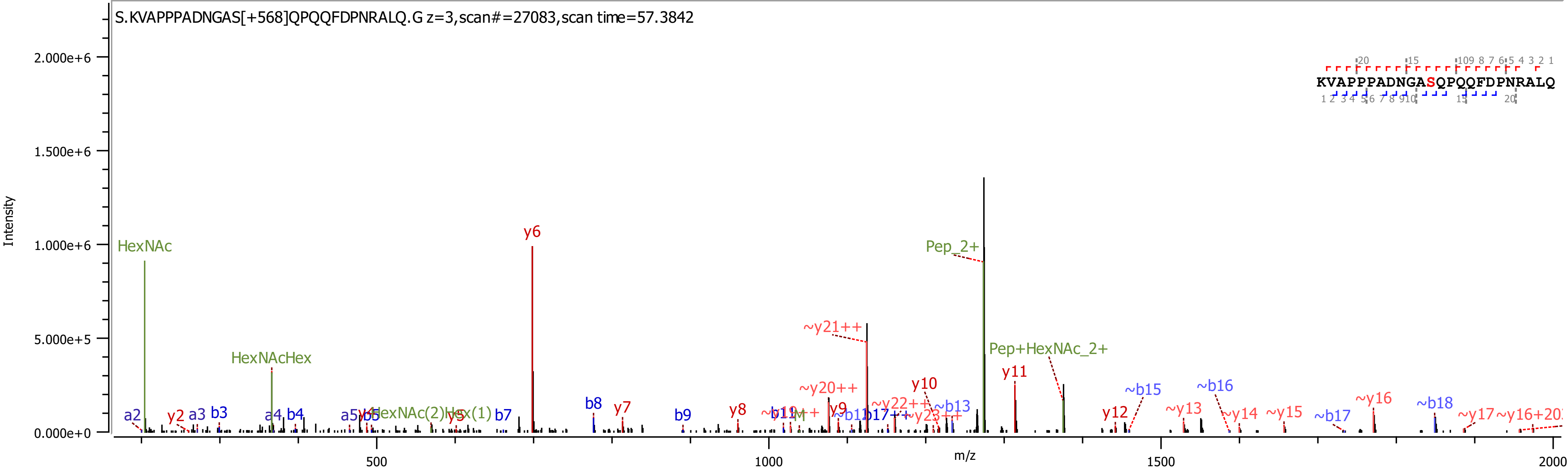

S.KVAPPPADNGAS[+568]QPQQFDPNRALQ.G z=3,scan#=27083,scan time=57.3842
KVAPPPADNGASQPQQFDPNRALQ
Intensity
m/z
HexNAc
HexNAcHex
y6
Pep_2+
~y21++
Pep+HexNAc_2+
y11
~y20++
y10
~b15
~b16
~y16
~b18
b8
y7
y8
y12
~y13
~y14
~y15
~b17
~y17
a2
y2
a3
b3
a4
b4
a5
b5
y5
b7
b9
y9
~b13

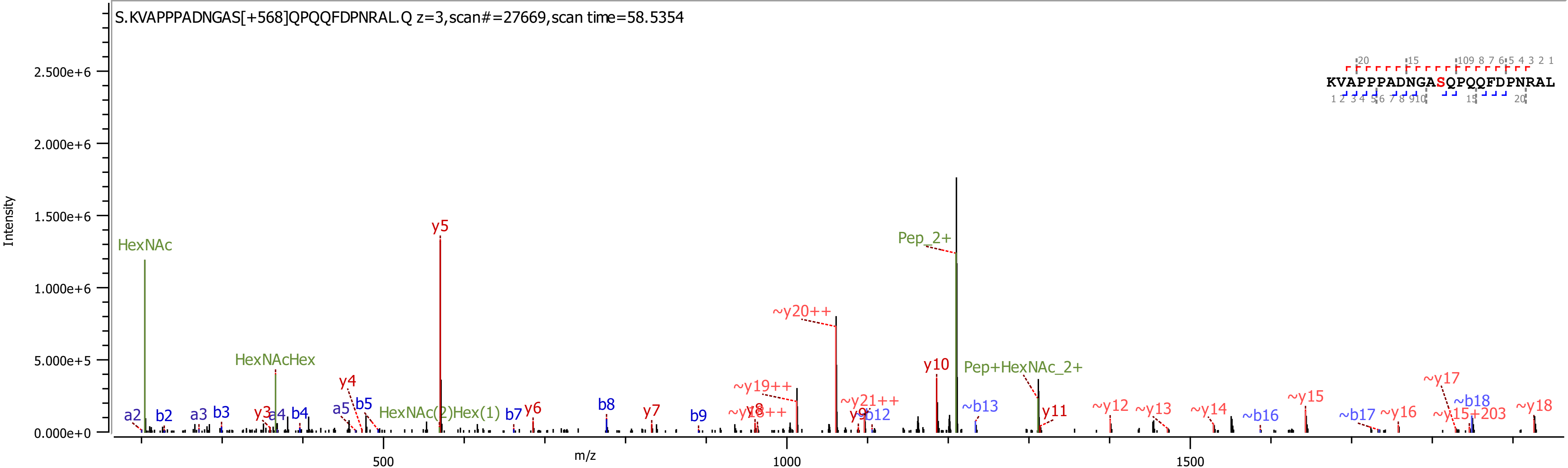

S.KVAPPPADNGAS[+568]QPQQFDPNRAL.Q z=3,scan#=27669,scan time=58.5354
KVAPPPADNGASQPQQFDPNRAL
Intensity
m/z
HexNAc
HexNAcHex
HexNAc(2)Hex(1)
Pep_2+
Pep+HexNAc_2+
y5
~y20++
y10
~y19++
~y21++
y4
y3
y6
y7
y8
y9
y11
a2
b2
a3
b3
a4
b4
a5
b5
b7
b8
b9
~y18++
b12
~b13
~y12
~y13
~y14
~b16
~y15
~b17
~y16
~y17
~b18
~y15+203
~y18

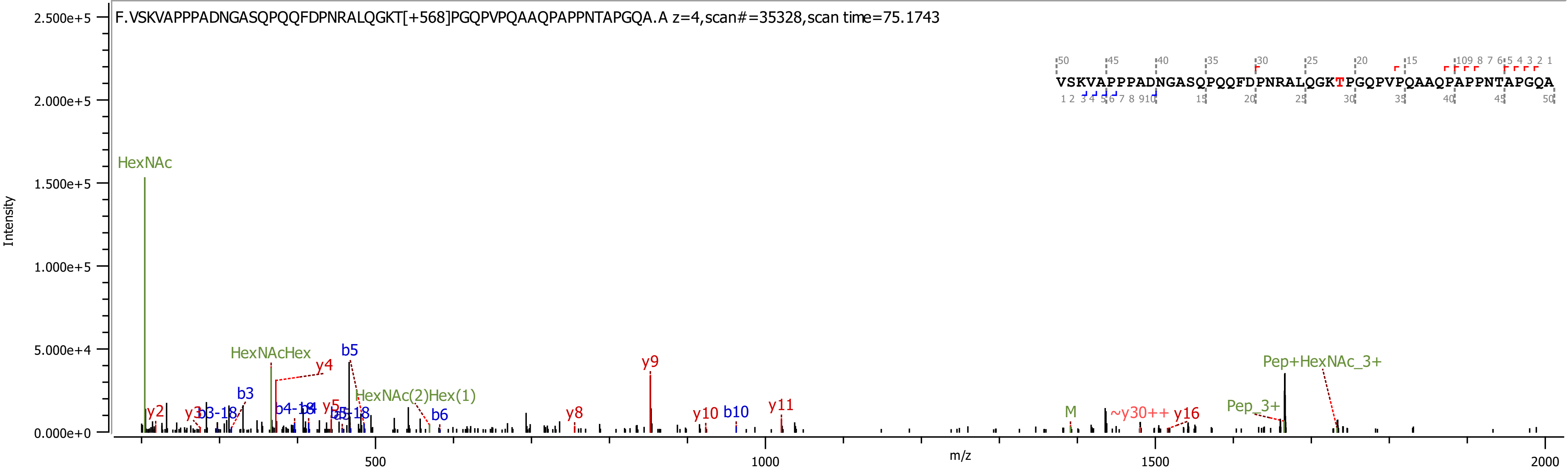
F.VSKVAPPPADNGASQPQQFDPNRALQGKT[+568]PGQPVPQAAQPAPPNTAPGQA.A z=4,scan#=35328,scan time=75.1743
VSKVAPPPADNGASQPQQFDPNRALQGKTPGQPVPQAAQPAPPNTAPGQA
Intensity
m/z
HexNAc
HexNAcHex
HexNAc(2)Hex(1)
Pep+HexNAc_3+
Pep_3+
M
~y30++
y2
y3
y4
y5
y8
y9
y10
y11
y16
b3
b3-18
b4-18
b5
b5-18
b6
b10
2.500e+5
2.000e+5
1.500e+5
1.000e+5
5.000e+4
0.000e+0
500
1000
1500
2000

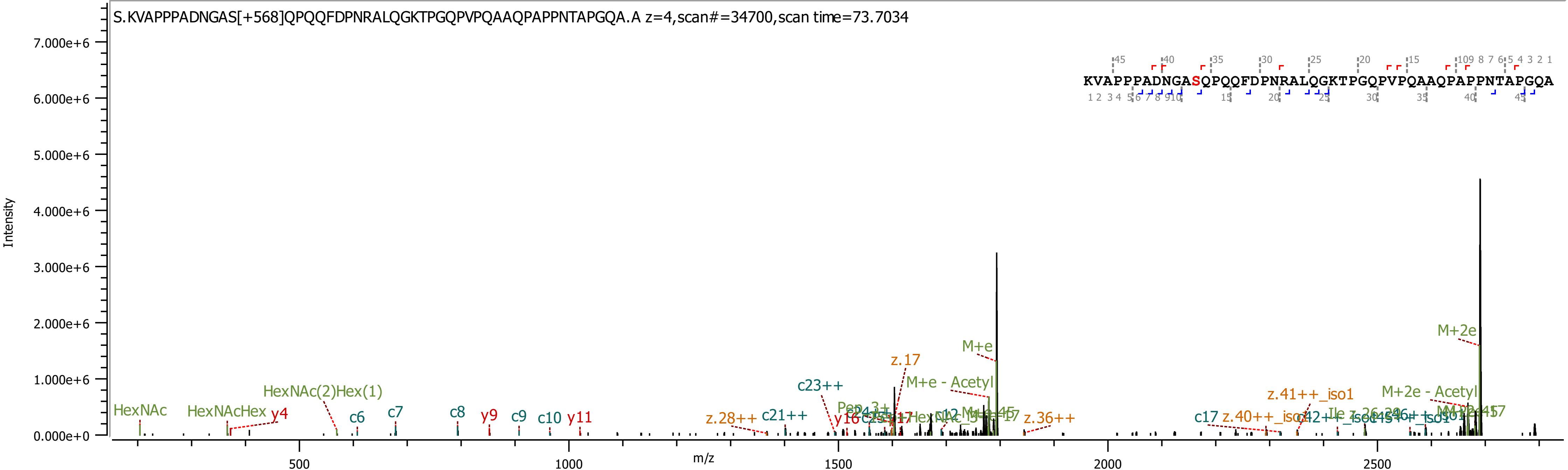

S.KVAPPPADNGAS[+568]QPQQFDPNRALQGKTPGQPVPQAAQPAPPNTAPGQA.A z=4,scan#=34700,scan time=73.7034
KVAPPPADNGASQPQQFDPNRALQGKTPGQPVPQAAQPAPPNTAPGQA
Intensity
m/z
HexNAc
HexNAcHex
y4
HexNAc(2)Hex(1)
c6
c7
c8
y9
c9
c10
y11
z.28++
c21++
c23++
z.17
M+e - Acetyl
M+e
z.36++
c17
z.41++_iso1
M+2e - Acetyl
M+2e

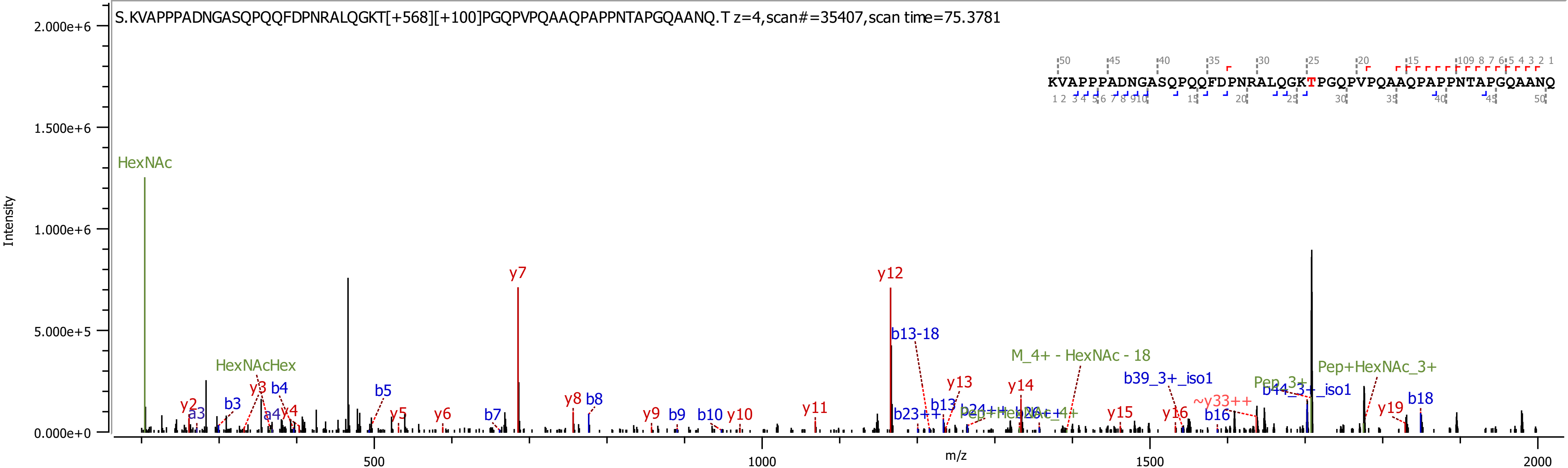

S.KVAPPPADNGASQPQQFDPNRALQGKT[+568][+100]PGQPVPQAAQPAPPNTAPGQAANQ.T z=4,scan#=35407,scan time=75.3781
KVAPPPADNGASQPQQFDPNRALQGKTPGQPVPQAAQPAPPNTAPGQAANQ
Intensity
m/z
HexNAc
HexNAcHex
y7
y12
b13-18
M_4+ - HexNAc - 18
b39_3+_iso1
Pep+HexNAc_3+
b44_3+_iso1
y13
y14
b18
y19
~y33++
y2
a3
b3
y3
b4
a4
y4
b5
y5
y6
b7
y8
b8
y9
b9
b10
y10
y11
b23++
b13
b24++
y15
y16
b16

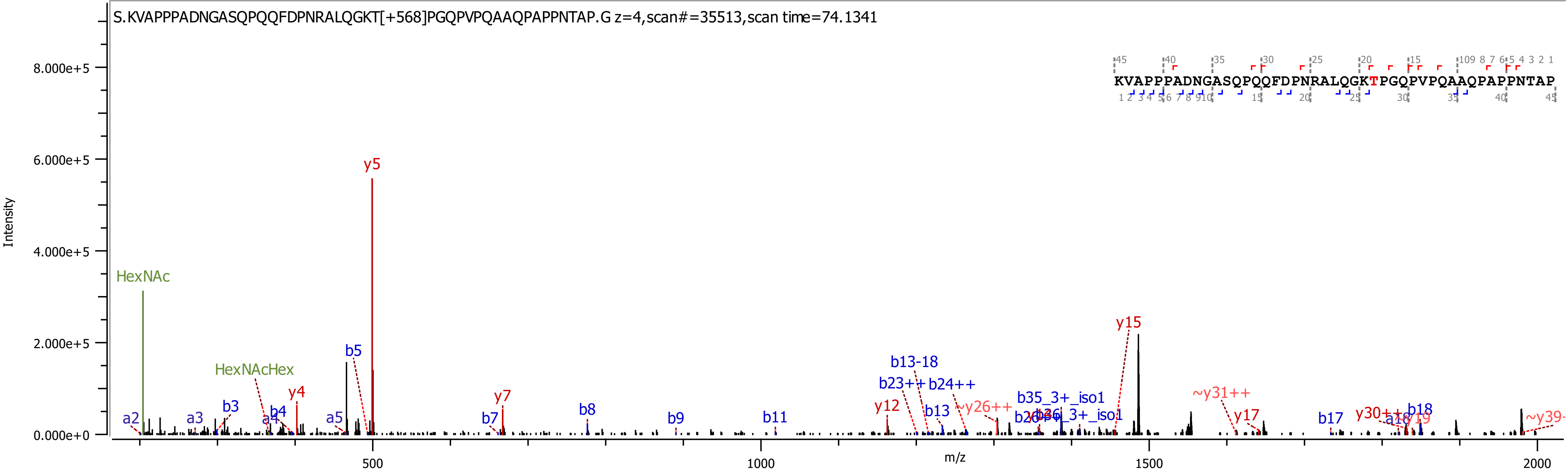
S.KVAPPPADNGASQPQQFDPNRALQGKT[+568]PGQPVPQAAQPAPPNTAP.G z=4,scan#=35513,scan time=74.1341
KVAPPPADNGASQPQQFDPNRALQGKTPGQPVPQAAQPAPPNTAP
Intensity
8.000e+5
6.000e+5
4.000e+5
2.000e+5
0.000e+0
m/z
500
1000
1500
2000
HexNAc
HexNAcHex
y5
y4
y7
y12
y15
y17
b5
b3
b4
b7
b8
b9
b11
b13
b13-18
b23++
b24++
b17
b18
a2
a3
a4
a5
~y26++
~y31++
b35_3+_iso1
y30++
y19

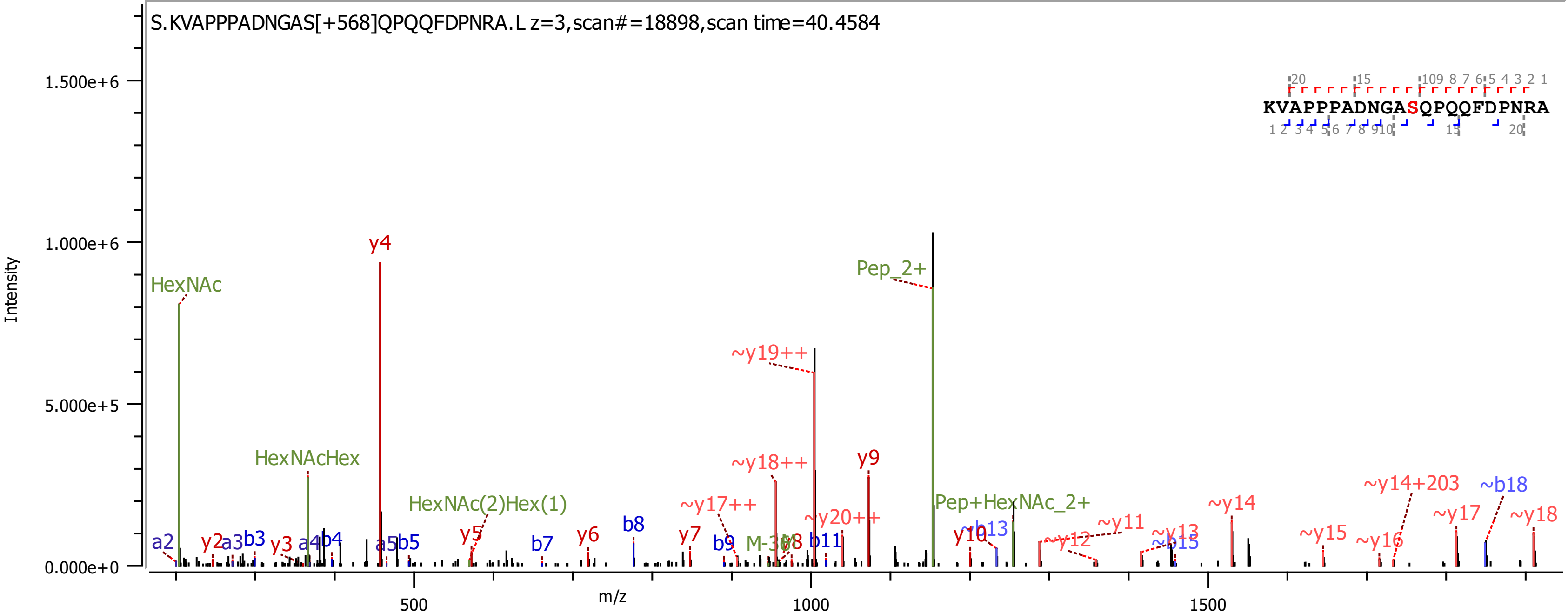
S.KVAPPPADNGAS[+568]QPQQFDPNRA.L z=3,scan#=18898,scan time=40.4584
KVAPPPADNGASQPQQFDPNRA
Intensity
m/z
1.500e+6
1.000e+6
5.000e+5
0.000e+0
500
1000
1500
HexNAc
HexNAcHex
HexNAc(2)Hex(1)
y4
Pep_2+
Pep+HexNAc_2+
~y19++
~y18++
~y17++
~y20++
y9
y10
~y14+203
~b18
~y14
~y17
~y18
~y11
~y12
~b13
~y13
~b15
~y15
~y16
a2
y2
a3
b3
y3
a4
b4
a5
b5
y5
b7
y6
b8
y7
b9
b11

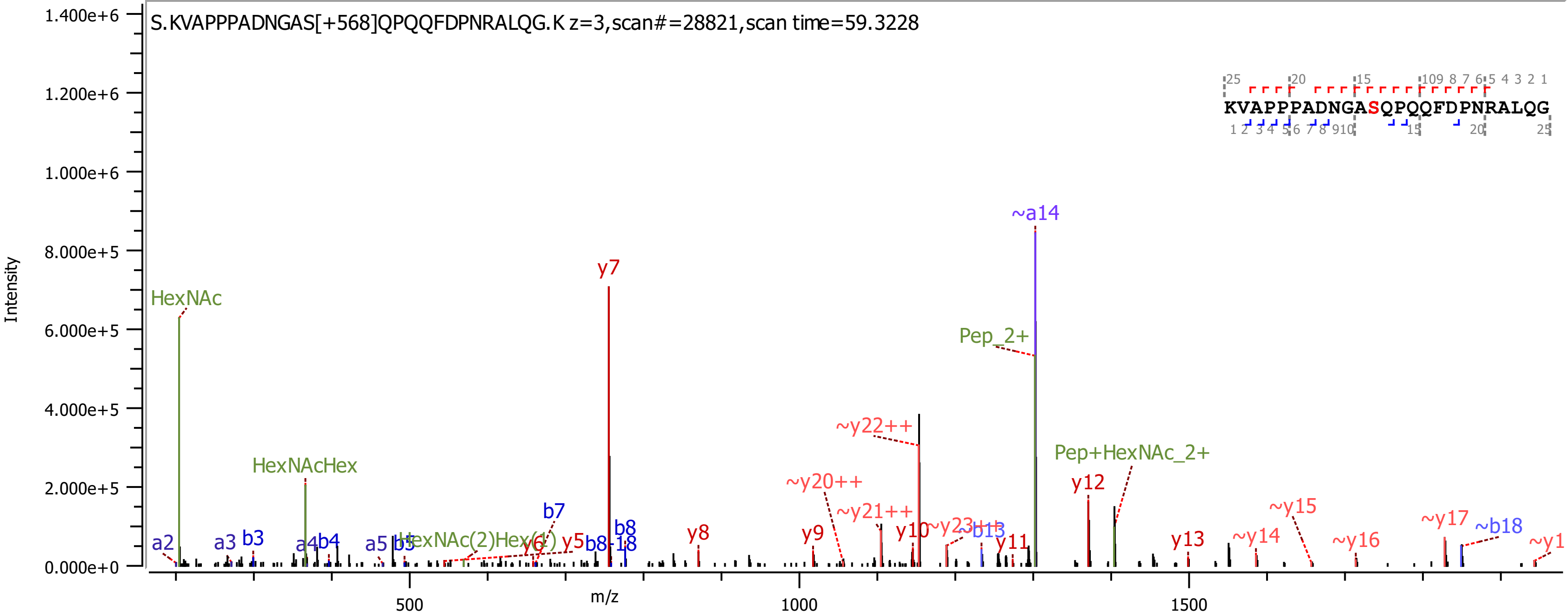
S.KVAPPPADNGAS[+568]QPQQFDPNRALQG.K z=3,scan#=28821,scan time=59.3228
KVAPPPADNGASQPQQFDPNRALQG
Intensity
m/z
HexNAc
HexNAcHex
HexNAc(2)Hex(1)
Pep_2+
Pep+HexNAc_2+
~a14
y7
y8
y9
y10
y11
y12
y13
~y14
~y15
~y16
~y17
~y20++
~y21++
~y22++
~y23++
a2
a3
b3
a4
b4
a5
b5
y6
y5
b7
b8
b8-18
~b13
~b18
0.000e+0
2.000e+5
4.000e+5
6.000e+5
8.000e+5
1.000e+6
1.200e+6
1.400e+6
500
1000
1500

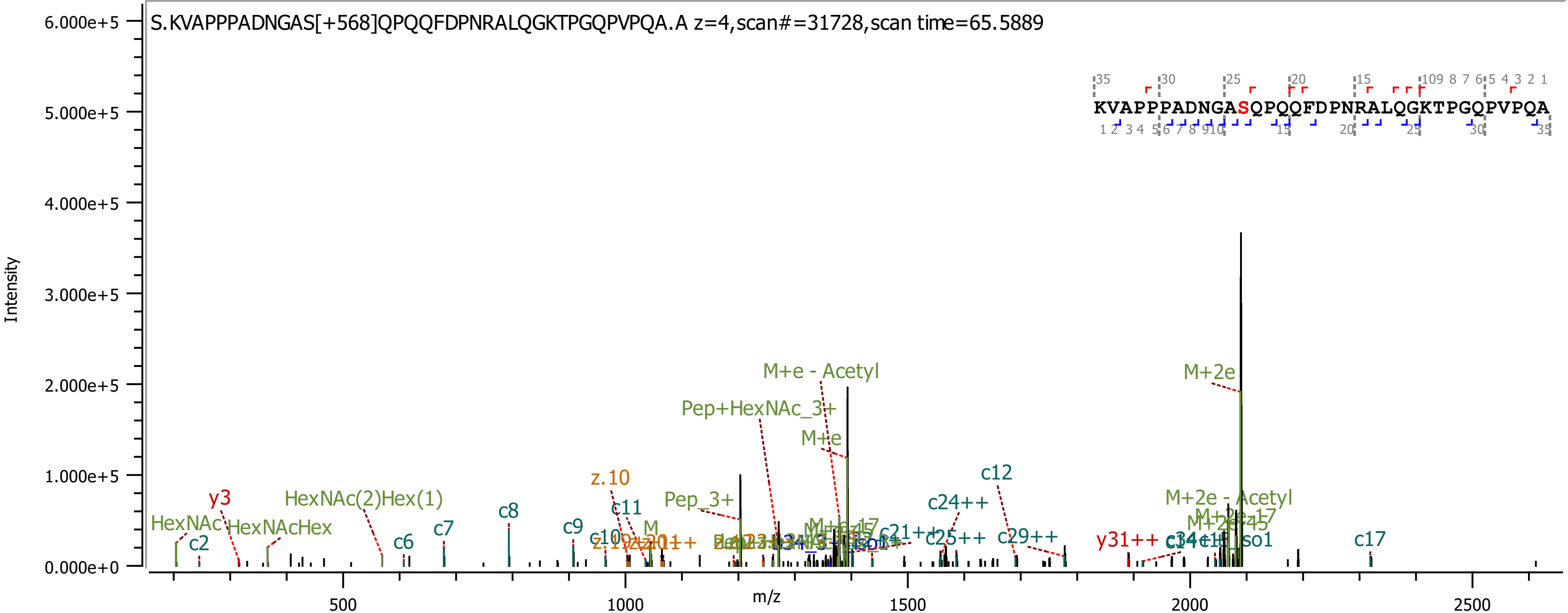
S.KVAPPPADNGAS[+568]QPQQFDPNRALQGKTPGQPVPQA.A z=4,scan#=31728,scan time=65.5889
KVAPPPADNGASQPQQFDPNRALQGKTPGQPVPQA
Intensity
m/z
6.000e+5
5.000e+5
4.000e+5
3.000e+5
2.000e+5
1.000e+5
0.000e+0
500
1000
1500
2000
2500
M+e - Acetyl
Pep+HexNAc_3+
M+e
M+2e
c12
z.10
c24++
M+2e - Acetyl
y3
HexNAc(2)Hex(1)
c8
c11
Pep_3+
HexNAc
HexNAcHex
c2
c6
c7
c9
c10
c25++
c29++
y31++
c17

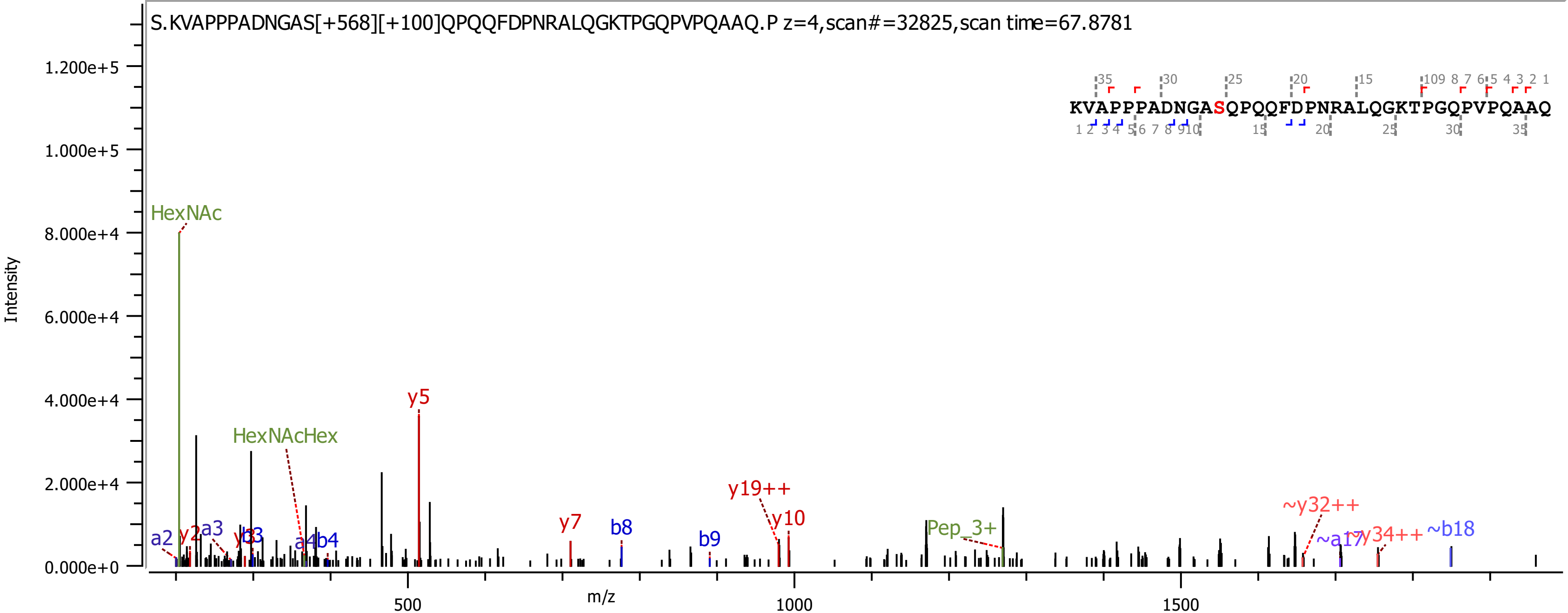
S.KVAPPPADNGAS[+568][+100]QPQQFDPNRALQGKTPGQPVPQAAQ.P z=4,scan#=32825,scan time=67.8781
KVAPPPADNGASQPQQFDPNRALQGKTPGQPVPQAAQ
Intensity
m/z
HexNAc
HexNAcHex
y5
y7
b8
b9
y19++
y10
Pep_3+
~y32++
~a17
~y34++
~b18
a2
y2
a3
y3
b3
a4
b4

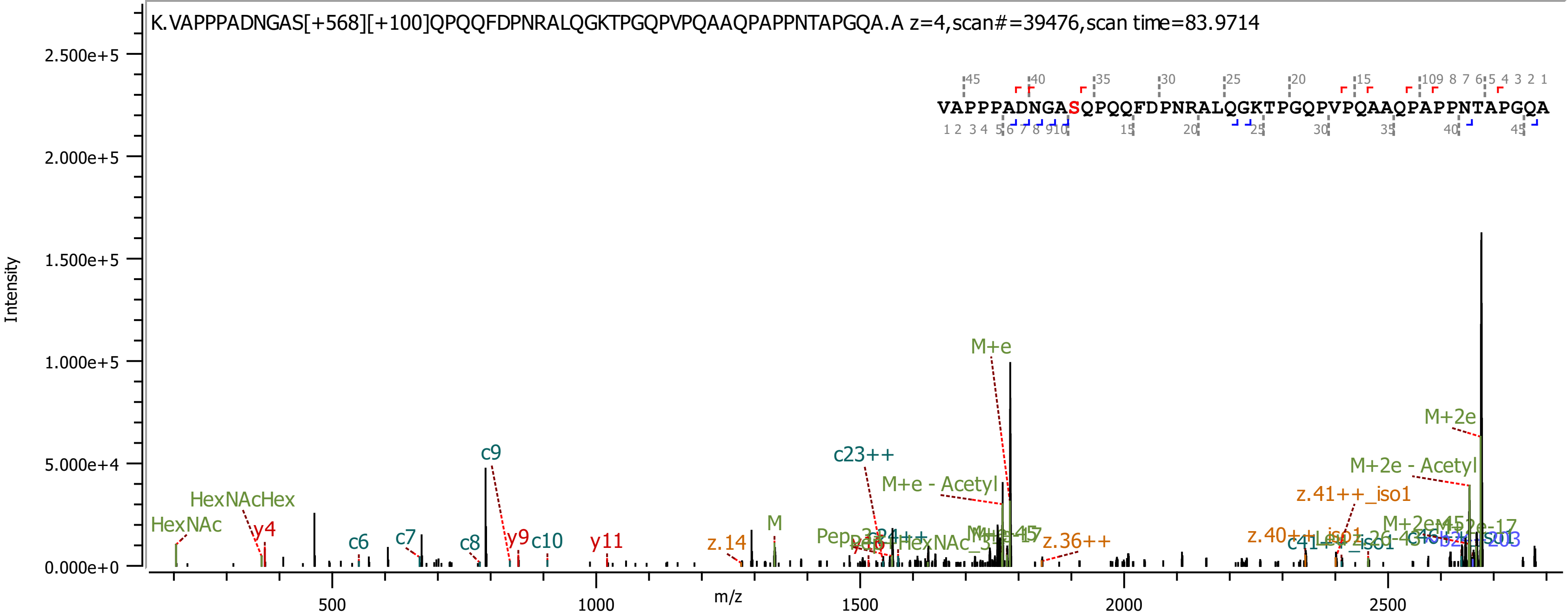
K.VAPPPADNGAS[+568][+100]QPQQFDPNRALQGKTPGQPVPQAAQPAPPNTAPGQA.A z=4,scan#=39476,scan time=83.9714
VAPPPADNGASQPQQFDPNRALQGKTPGQPVPQAAQPAPPNTAPGQA
Intensity
m/z
2.500e+5
2.000e+5
1.500e+5
1.000e+5
5.000e+4
0.000e+0
500
1000
1500
2000
2500
HexNAc
HexNAcHex
y4
c6
c7
c8
c9
y9
c10
y11
z.14
M
c23++
M+e
M+e - Acetyl
z.36++
z.41++_iso1
M+2e
M+2e - Acetyl

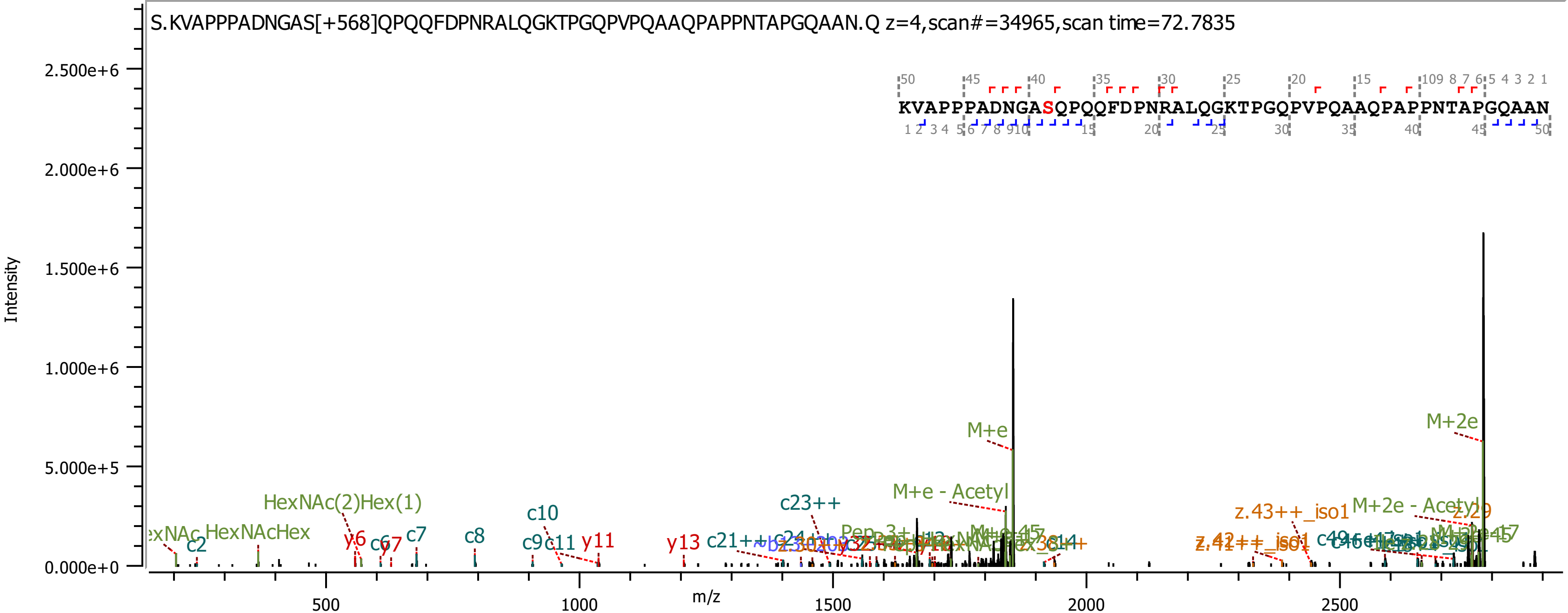

S.KVAPPPADNGAS[+568]QPQQFDPNRALQGKTPGQPVPQAAQPAPPNTAPGQAAN.Q z=4,scan#=34965,scan time=72.7835
KVAPPPADNGASQPQQFDPNRALQGKTPGQPVPQAAQPAPPNTAPGQAAN
Intensity
m/z
2.500e+6
2.000e+6
1.500e+6
1.000e+6
5.000e+5
0.000e+0
500
1000
1500
2000
2500
M+e
M+2e
M+e - Acetyl
M+2e - Acetyl
HexNAc(2)Hex(1)
HexNAcHex
c2
c7
c8
c10
c9
c11
y6
y11
y13
c23++
c21++
z.43++_iso1

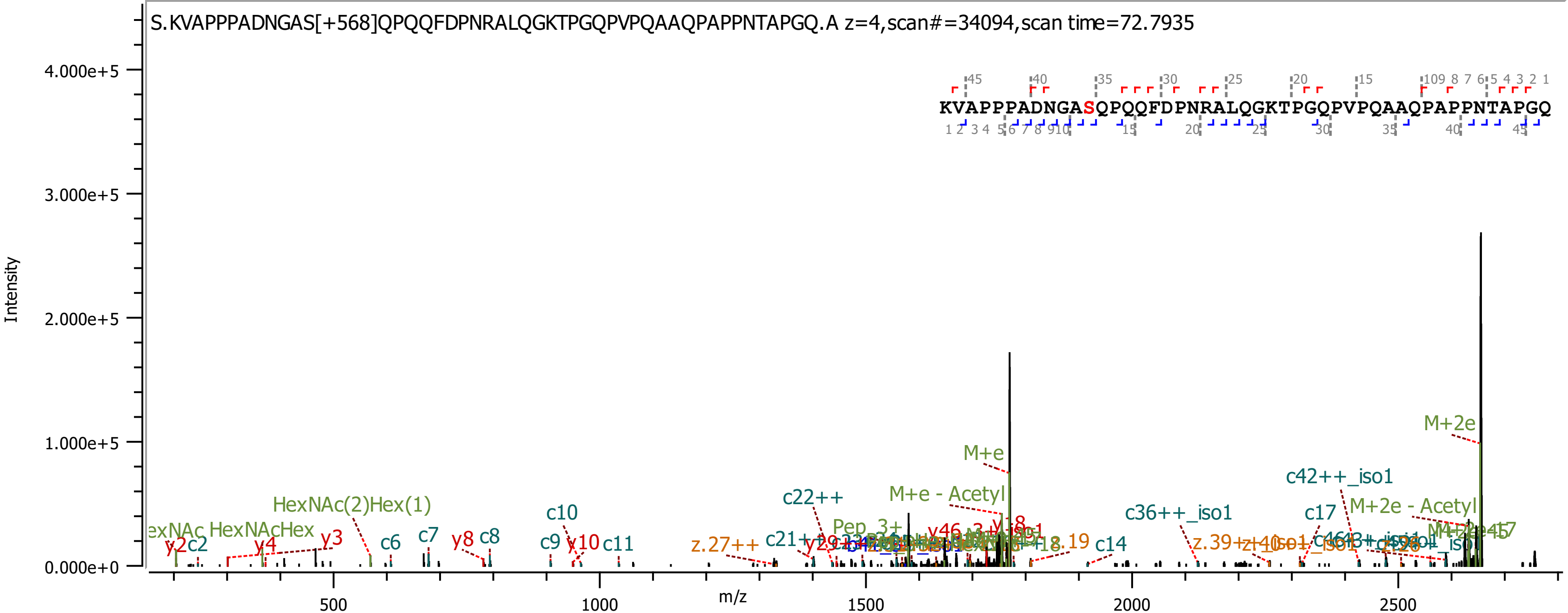
S.KVAPPPADNGAS[+568]QPQQFDPNRALQGKTPGQPVPQAAQPAPPNTAPGQ.A z=4,scan#=34094,scan time=72.7935
Intensity
4.000e+5
3.000e+5
2.000e+5
1.000e+5
0.000e+0
m/z
500
1000
1500
2000
2500
KVAPPPADNGASQPQQFDPNRALQGKTPGQPVPQAAQPAPPNTAPGQ
M+2e
M+e
c42++_iso1
M+e - Acetyl
M+2e - Acetyl
c22++
c36++_iso1
c17
c10
HexNAc(2)Hex(1)
HexNAcHex
y3
y4
y2
c2
c6
c7
y8
c8
c9
y10
c11
z.27++
c14
Pep_3+

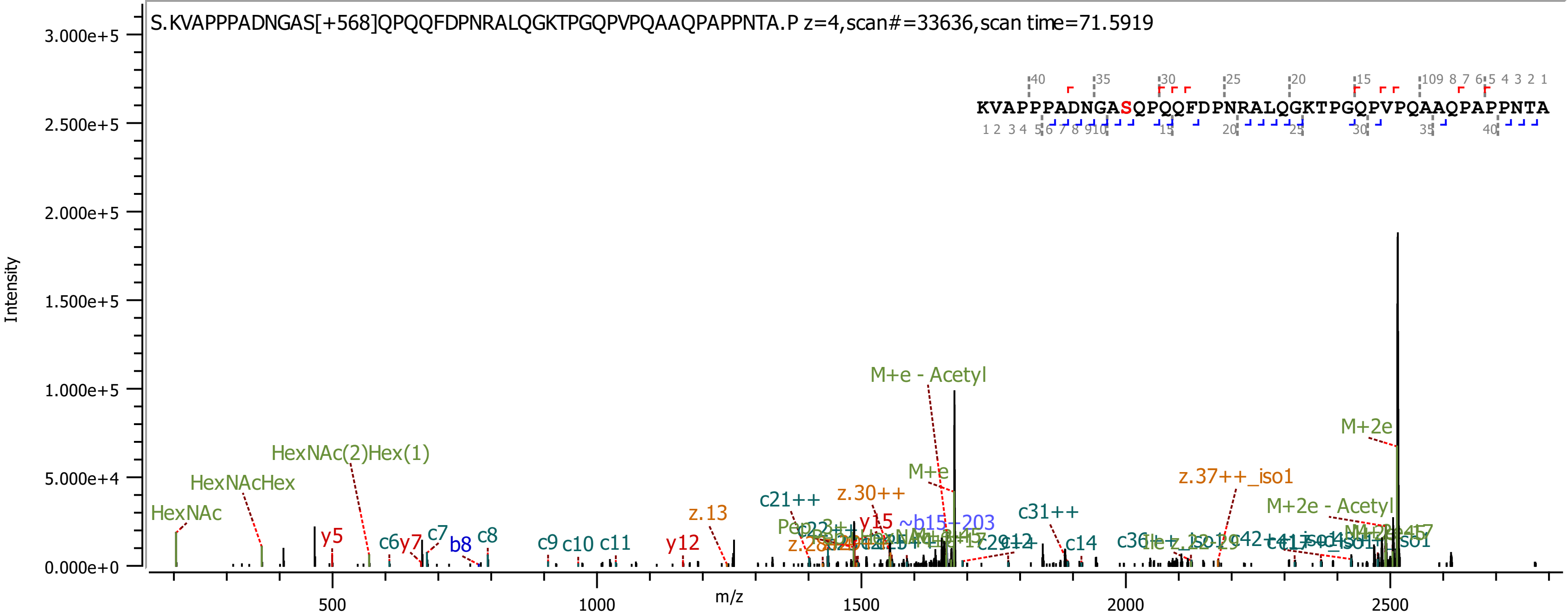
S.KVAPPPADNGAS[+568]QPQQFDPNRALQGKTPGQPVPQAAQPAPPNTA.P z=4,scan#=33636,scan time=71.5919
KVAPPPADNGASQPQQFDPNRALQGKTPGQPVPQAAQPAPPNTA
Intensity
m/z
M+e - Acetyl
M+2e
HexNAc(2)Hex(1)
HexNAcHex
HexNAc
M+e
z.37++_iso1
z.30++
c21++
M+2e - Acetyl
z.13
c31++
y15
~b15-203
y5
c6
y7
c7
b8
c8
c9
c10
c11
y12
c14

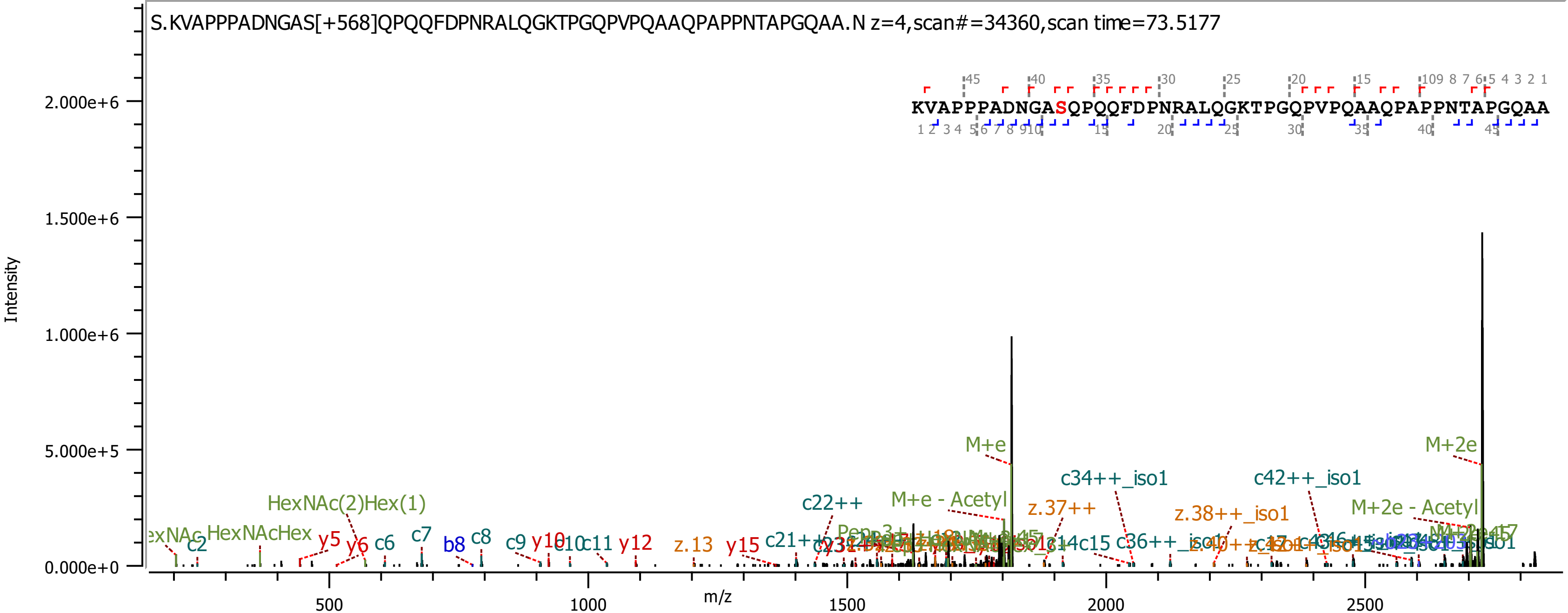

S.KVAPPPADNGAS[+568]QPQQFDPNRALQGKTPGQPVPQAAQPAPPNTAPGQAA.N z=4,scan#=34360,scan time=73.5177
KVAPPPADNGASQPQQFDPNRALQGKTPGQPVPQAAQPAPPNTAPGQAA
Intensity
m/z
M+e
M+2e
M+e - Acetyl
M+2e - Acetyl
c34++_iso1
c42++_iso1
z.37++
z.38++_iso1
c22++
HexNAc(2)Hex(1)
HexNAcHex
y5
y6
c6
c7
b8
c8
c9
y10
c11
y12
z.13
y15
c15

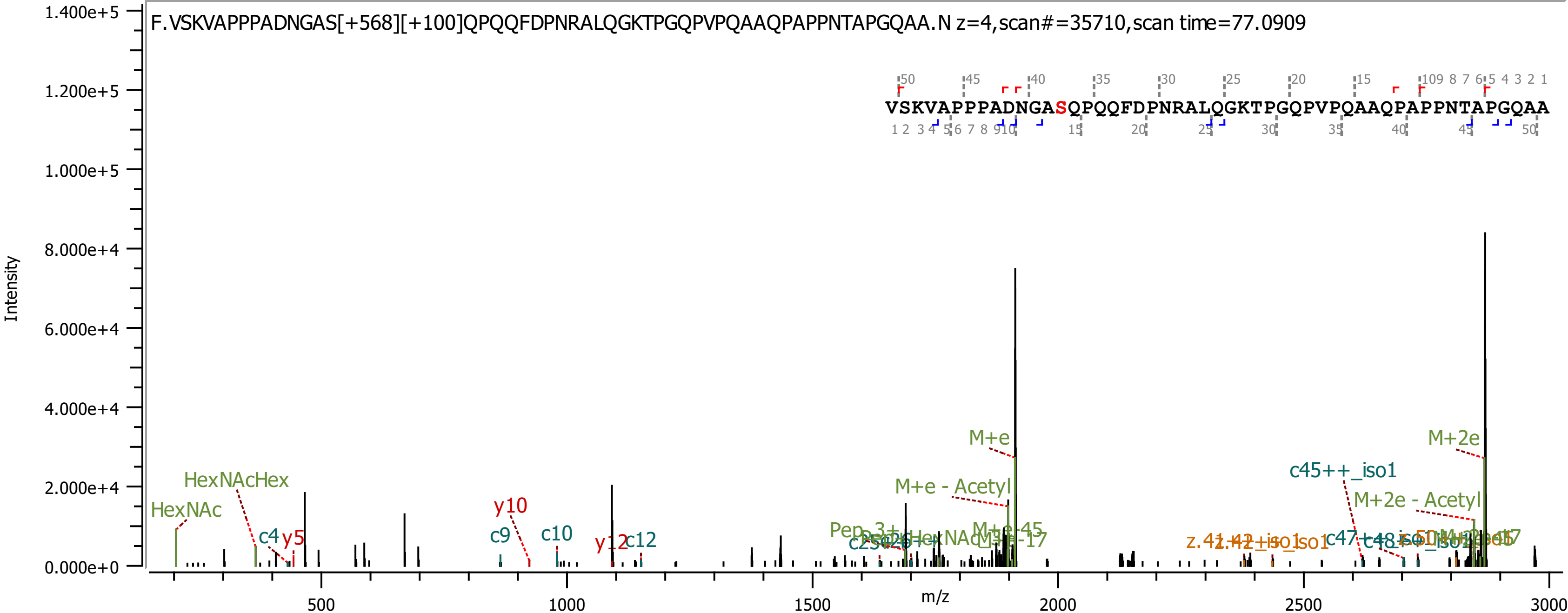

F.VSKVAPPPADNGAS[+568][+100]QPQQFDPNRALQGKTPGQPVPQAAQPAPPNTAPGQAA.N z=4,scan#=35710,scan time=77.0909
VSKVAPPPADNGASQPQQFDPNRALQGKTPGQPVPQAAQPAPPNTAPGQAA
Intensity
m/z
HexNAc
HexNAcHex
c4
y5
y10
c9
c10
y12
c12
M+e
M+e - Acetyl
M+2e
M+2e - Acetyl
c45++_iso1

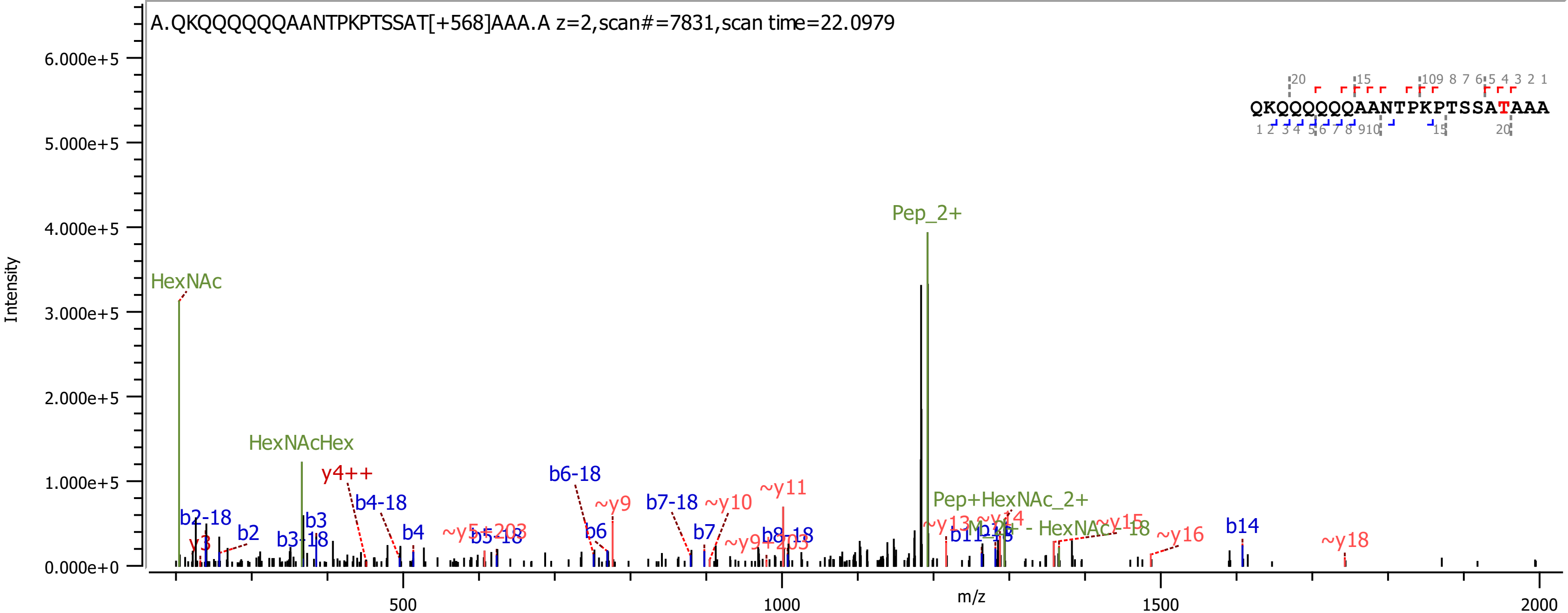

A.QKQQQQQQAANTPKPTSSAT[+568]AAA.A z=2,scan#=7831,scan time=22.0979
QKQQQQQQAANTPKPTSSATAAA
Intensity
m/z
6.000e+5
5.000e+5
4.000e+5
3.000e+5
2.000e+5
1.000e+5
0.000e+0
500
1000
1500
2000
HexNAc
HexNAcHex
Pep_2+
Pep+HexNAc_2+
y4++
b2-18
b2
y3
b3
b3-18
b4-18
b4
~y5+203
b5-18
b6-18
b6
~y9
b7-18
b7
~y10
~y11
~y9+203
b8-18
~y13
~y14
b11-18
b12
~y15
~y16
b14
~y18

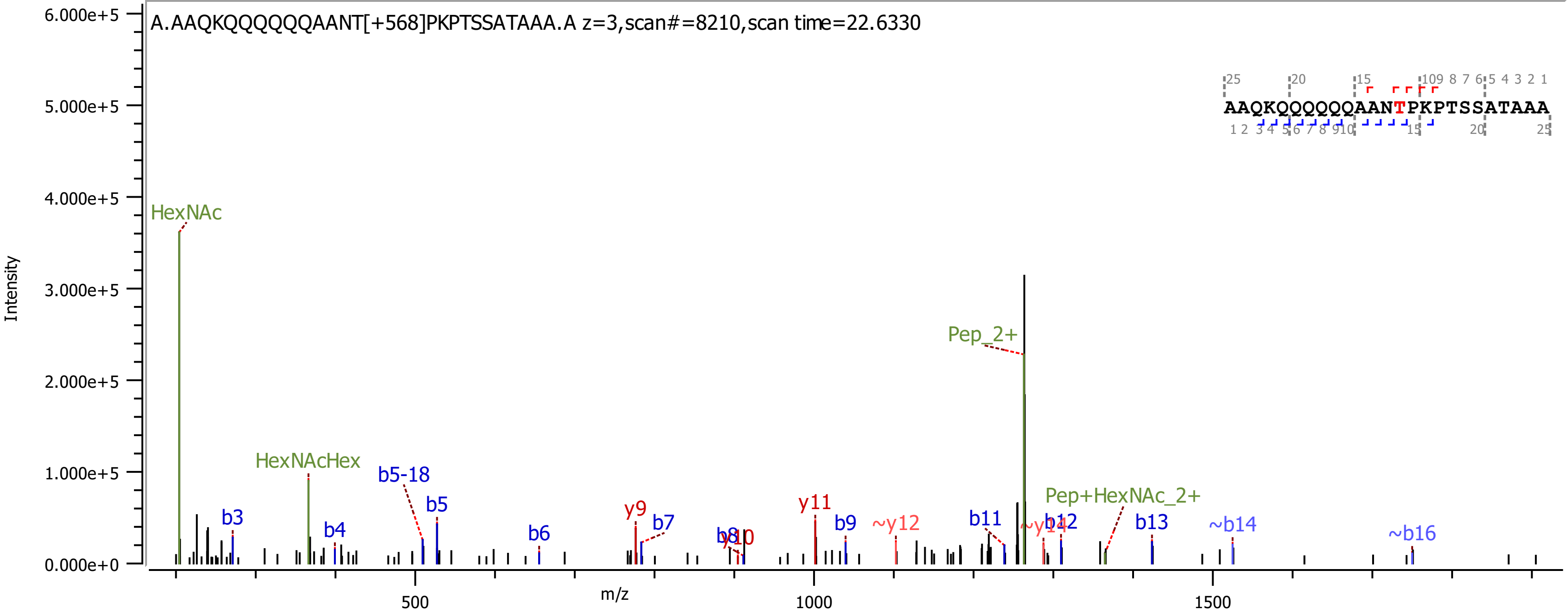

A.AAQKQQQQQQAANT[+568]PKPTSSATAAA.A z=3,scan#=8210,scan time=22.6330
AAQKQQQQQQAANTPKPTSSATAAA
Intensity
m/z
HexNAc
HexNAcHex
Pep_2+
Pep+HexNAc_2+
b3
b4
b5-18
b5
b6
y9
b7
b8
y10
y11
b9
~y12
b11
~y14
b12
b13
~b14
~b16

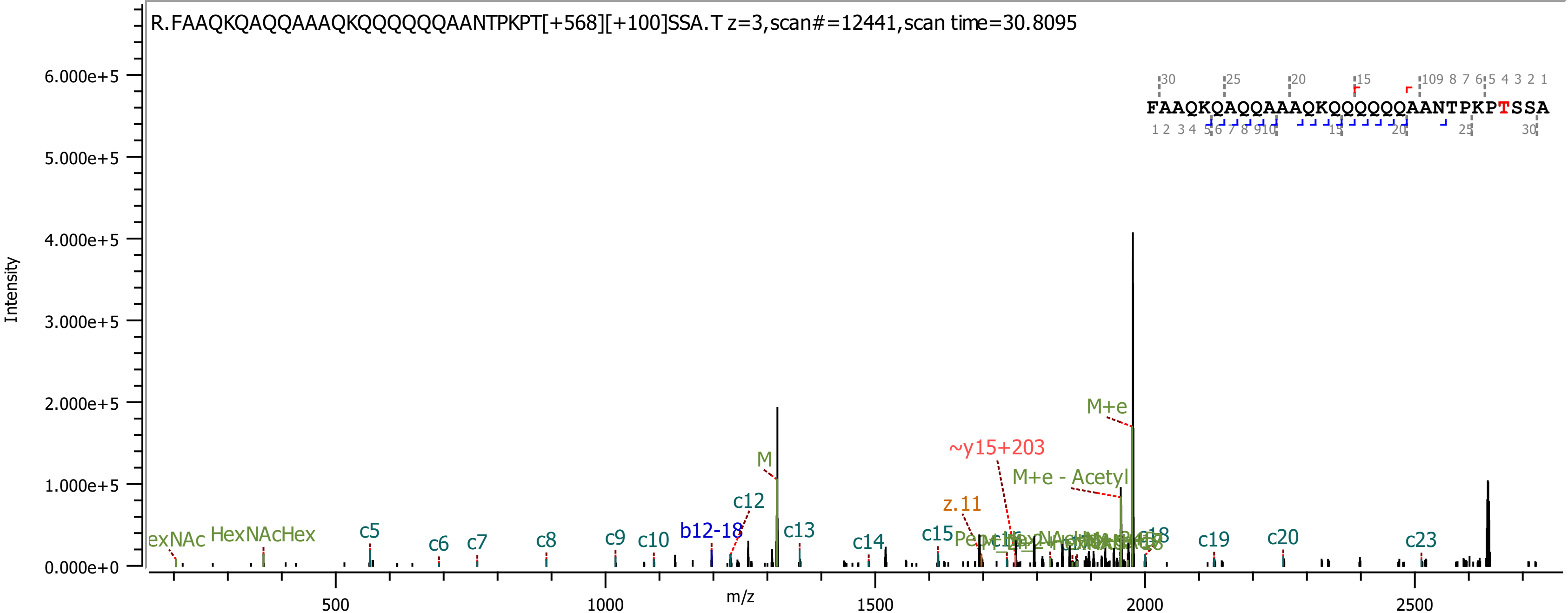
R.FAAQKQAQQAAAQKQQQQQQAANTPKPT[+568][+100]SSA.T z=3,scan#=12441,scan time=30.8095
FAAQKQAQQAAAQKQQQQQQAANTPKPTSSA
Intensity
6.000e+5
5.000e+5
4.000e+5
3.000e+5
2.000e+5
1.000e+5
0.000e+0
m/z
500
1000
1500
2000
2500
exNAc
HexNAcHex
c5
c6
c7
c8
c9
c10
b12-18
c12
M
c13
c14
c15
z.11
~y15+203
M+e - Acetyl
M+e
c19
c20
c23

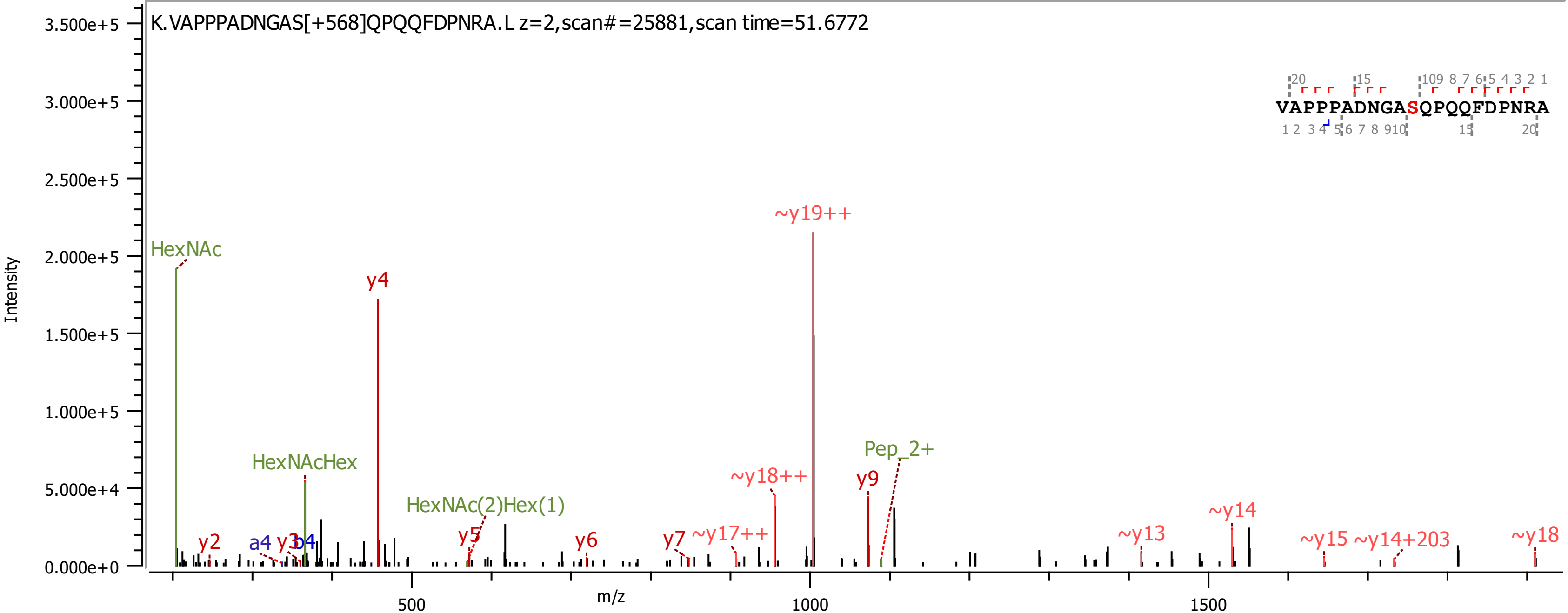
K.VAPPPADNGAS[+568]QPQQFDPNRA.L z=2,scan#=25881,scan time=51.6772
VAPPPADNGASQPQQFDPNRA
Intensity
m/z
HexNAc
HexNAcHex
HexNAc(2)Hex(1)
Pep_2+
y2
a4
y3
b4
y4
y5
y6
y7
~y17++
~y18++
~y19++
y9
~y13
~y14
~y15
~y14+203
~y18
0.000e+0
5.000e+4
1.000e+5
1.500e+5
2.000e+5
2.500e+5
3.000e+5
3.500e+5
500
1000
1500

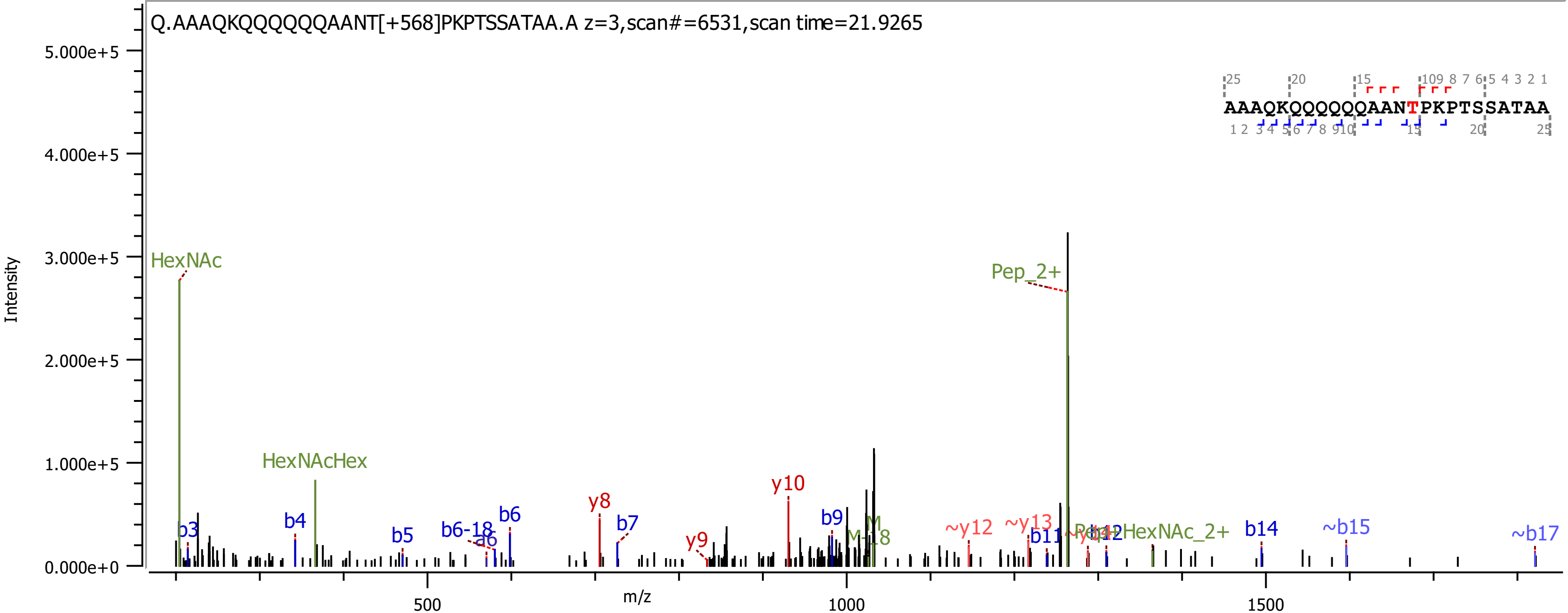

Q.AAAQKQQQQQQAANT[+568]PKPTSSATAA.A z=3,scan#=6531,scan time=21.9265
AAAQKQQQQQQAANTPKPTSSATAA
Intensity
m/z
HexNAc
HexNAcHex
Pep_2+
b3
b4
b5
b6-18
a6
b6
y8
b7
y9
y10
b9
M-18
~y12
~y13
b11
b14
~b15
~b17

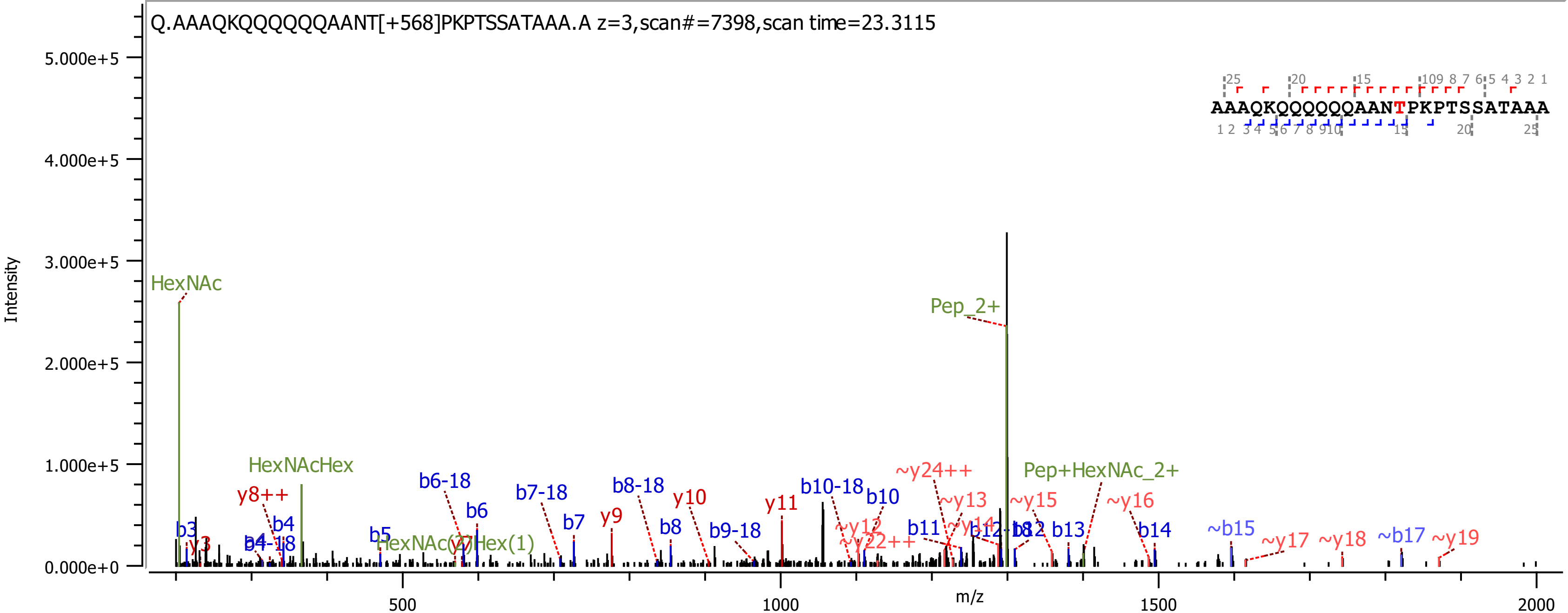

Q.AAAQKQQQQQQAANT[+568]PKPTSSATAAA.A z=3,scan#=7398,scan time=23.3115
AAAQKQQQQQQAANTPKPTSSATAAA
Intensity
m/z
HexNAc
HexNAcHex
Pep_2+
Pep+HexNAc_2+
HexNAc(2)Hex(1)
y8++
y9
y10
y11
b3
b4
b4-18
b5
b6
b6-18
b7
b7-18
b8
b8-18
b9-18
b10
b10-18
b11
b12
b12-18
b13
b14
~b15
~b17
~y12
~y13
~y14
~y15
~y16
~y17
~y18
~y19
~y22++
~y24++

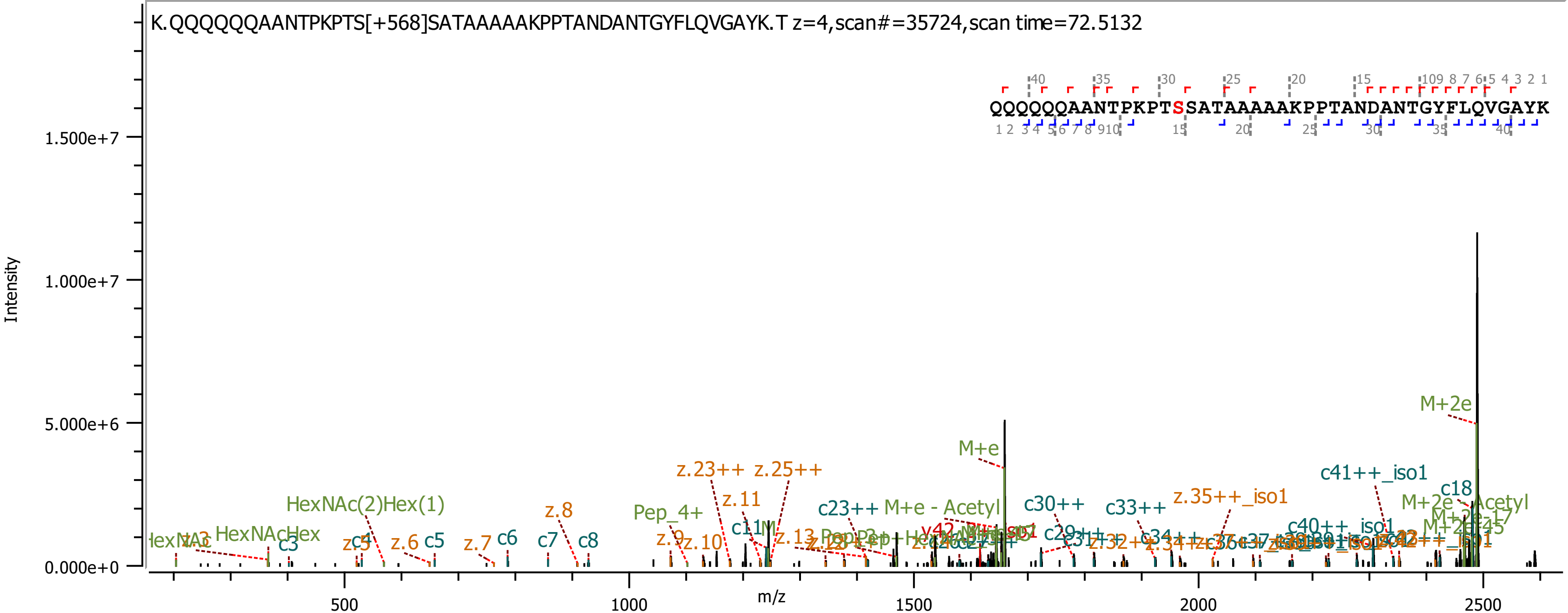

K.QQQQQQAANTPKPTS[+568]SATAAAAAKPPTANDANTGYFLQVGAYK.T z=4,scan#=35724,scan time=72.5132
QQQQQQAANTPKPTSSATAAAAAKPPTANDANTGYFLQVGAYK
Intensity
m/z
0.000e+0
5.000e+6
1.000e+7
1.500e+7
500
1000
1500
2000
2500
M+2e
M+e
c41++_iso1
c18
z.35++_iso1
c33++
c30++
c23++
M+e - Acetyl
z.23++
z.25++
z.11
HexNAc(2)Hex(1)
z.8
Pep_4+
HexNAcHex
c3
c4
z.5
z.6
c5
z.7
c6
c7
c8
z.9
z.10
c40++_iso1

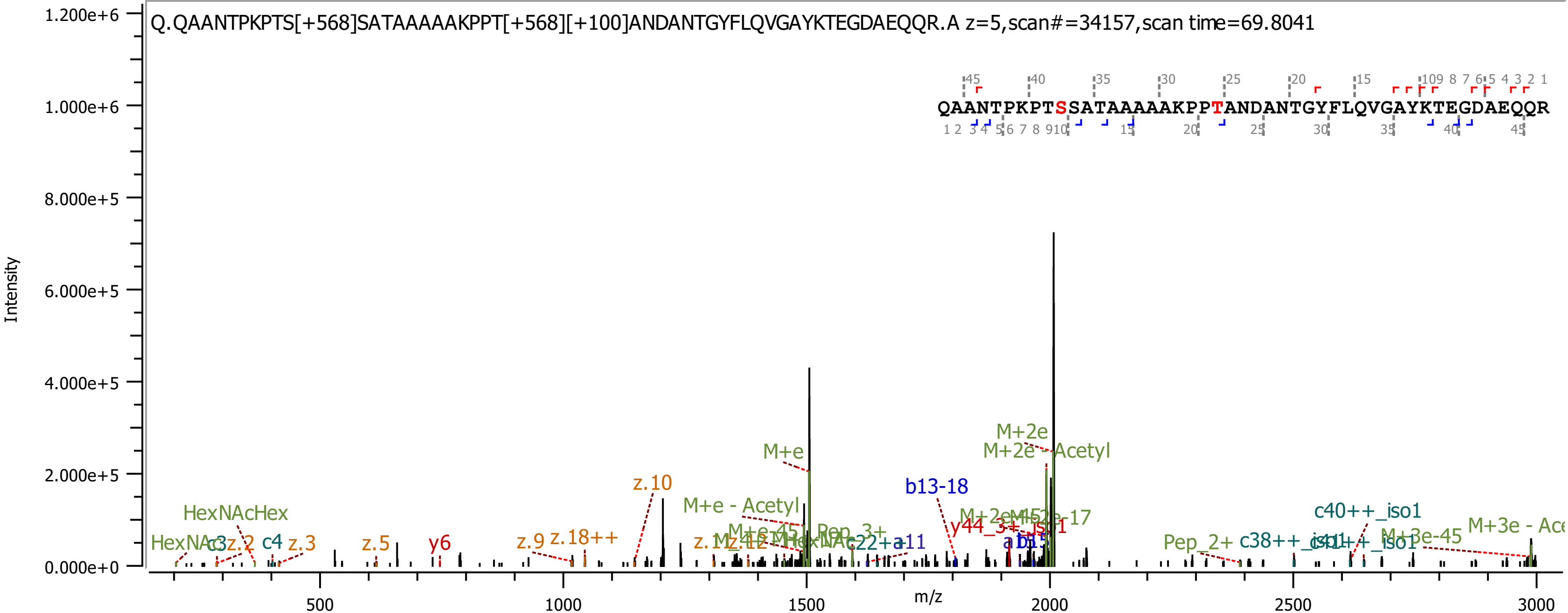

Q.QAANTPKPTS[+568]SATAAAAAKPPT[+568][+100]ANDANTGYFLQVGAYKTEGDAEQQR.A z=5,scan#=34157,scan time=69.8041
QAANTPKPTSSATAAAAAKPPTANDANTGYFLQVGAYKTEGDAEQQR
Intensity
m/z
1.200e+6
1.000e+6
8.000e+5
6.000e+5
4.000e+5
2.000e+5
0.000e+0
500
1000
1500
2000
2500
3000
HexNAcHex
HexNAc
c3
z.2
c4
z.3
z.5
y6
z.9
z.18++
z.10
M+e
M+e - Acetyl
M+2e
M+2e - Acetyl
b13-18
c22+a11
Pep_3+
Pep_2+
c40++_iso1
M+3e-45
M+3e - Ace

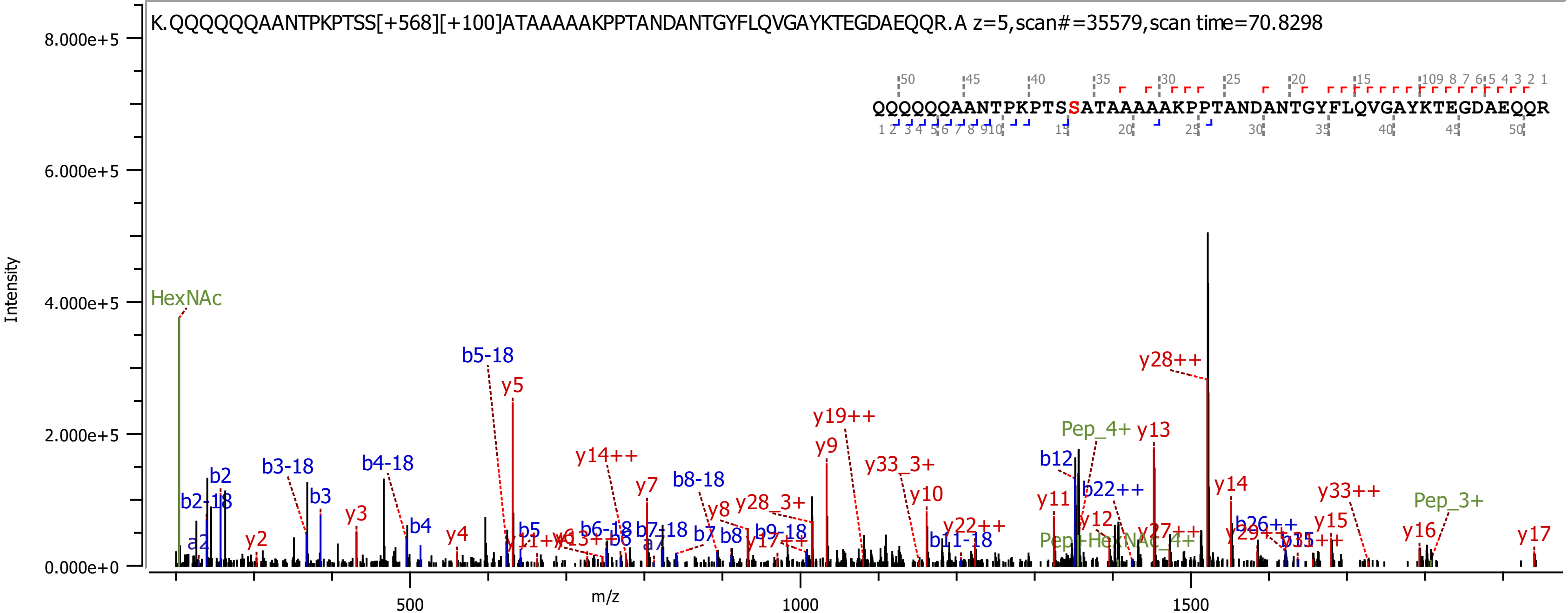
K.QQQQQQAANTPKPTSS[+568][+100]ATAAAAAKPPTANDANTGYFLQVGAYKTEGDAEQQR.A z=5,scan#=35579,scan time=70.8298
QQQQQQAANTPKPTSSATAAAAAKPPTANDANTGYFLQVGAYKTEGDAEQQR
Intensity
m/z
8.000e+5
6.000e+5
4.000e+5
2.000e+5
0.000e+0
500
1000
1500
HexNAc
b5-18
y5
y19++
y9
y28++
Pep_4+
y13
b12
y14
y33++
Pep_3+
y17

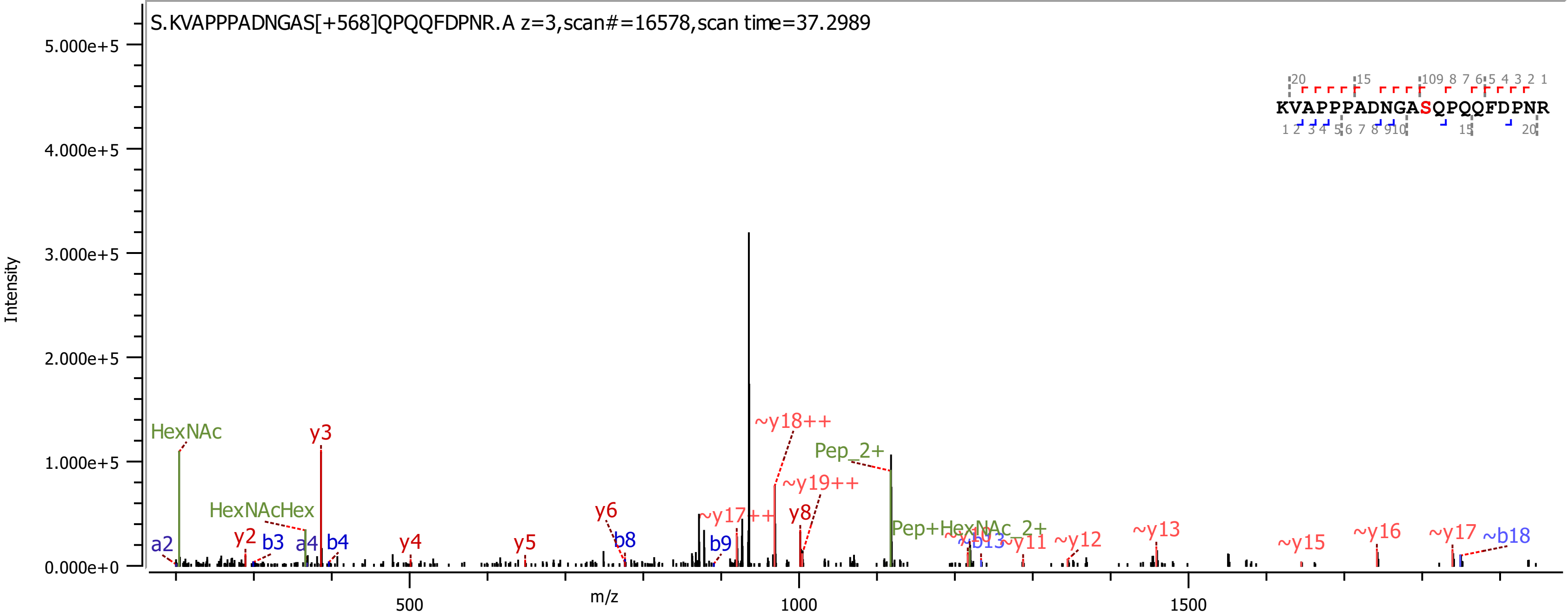
S.KVAPPPADNGAS[+568]QPQQFDPNR.A z=3,scan#=16578,scan time=37.2989
KVAPPPADNGASQPQQFDPNR
Intensity
5.000e+5
4.000e+5
3.000e+5
2.000e+5
1.000e+5
0.000e+0
m/z
500
1000
1500
HexNAc
y3
~y18++
Pep_2+
~y19++
HexNAcHex
y6
~y17++
y8
a2
y2
b3
a4
b4
y4
y5
b8
b9
Pep+HexNAc_2+
~y11
~y12
~y13
~y15
~y16
~y17
~b18

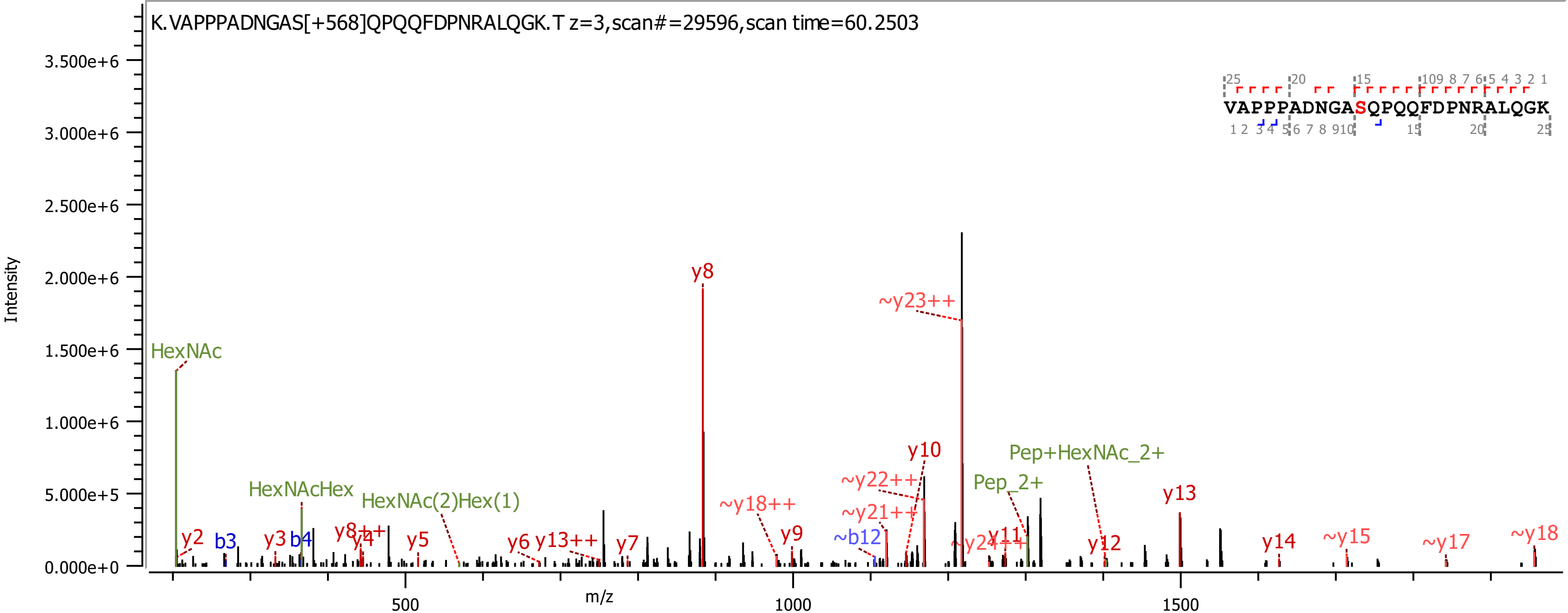
K.VAPPPADNGAS[+568]QPQQFDPNRALQGK.T z=3,scan#=29596,scan time=60.2503
Intensity
m/z
VAPPPADNGASQPQQFDPNRALQGK
HexNAc
HexNAcHex
HexNAc(2)Hex(1)
y8
~y23++
y10
~y22++
~y21++
~y18++
Pep+HexNAc_2+
Pep_2+
y13
y2
b3
y3
b4
y8++
y4
y5
y6
y13++
y7
y9
~b12
~y24++
y11
y12
y14
~y15
~y17
~y18

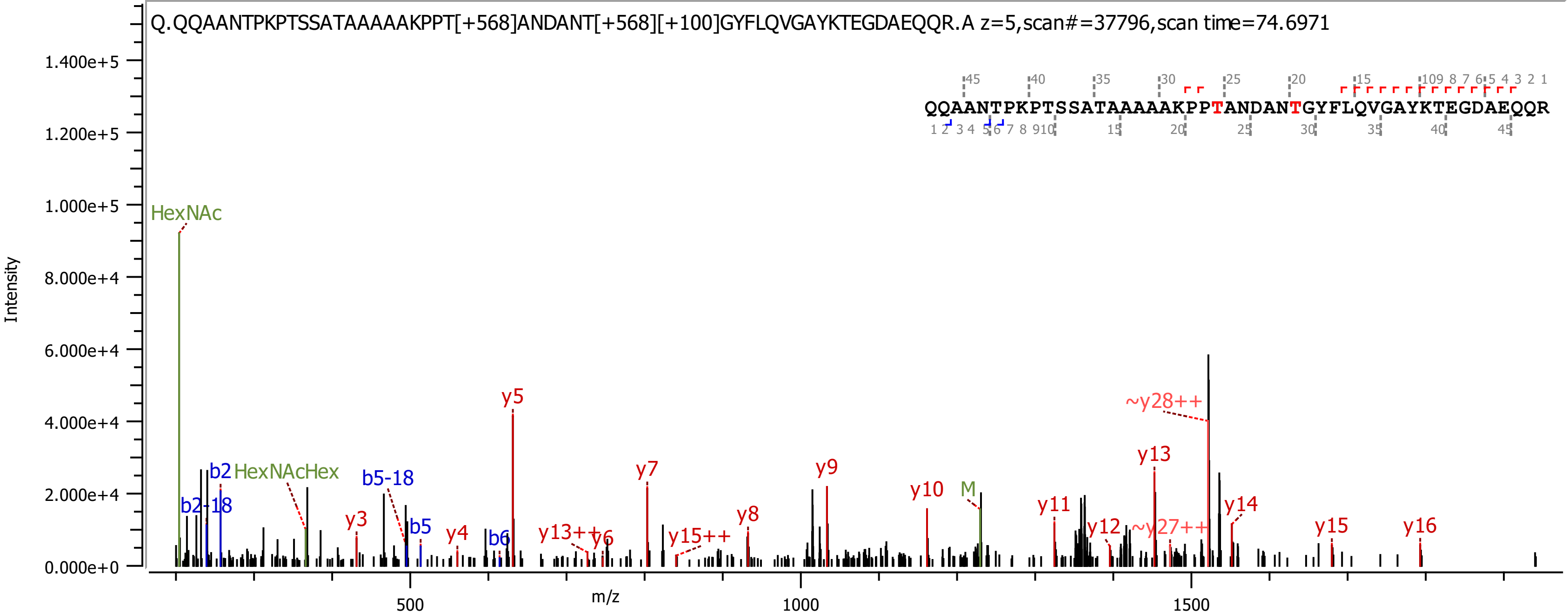

Q.QQAANTPKPTSSATAAAAAKPPT[+568]ANDANT[+568][+100]GYFLQVGAYKTEGDAEQQR.A z=5,scan#=37796,scan time=74.6971
QQAANTPKPTSSATAAAAAKPPTANDANTGYFLQVGAYKTEGDAEQQR
Intensity
m/z
HexNAc
HexNAcHex
b2
b2-18
b5-18
b5
b6
y3
y4
y5
y13++
y6
y7
y15++
y8
y9
y10
M
y11
y12
~y27++
y13
~y28++
y14
y15
y16

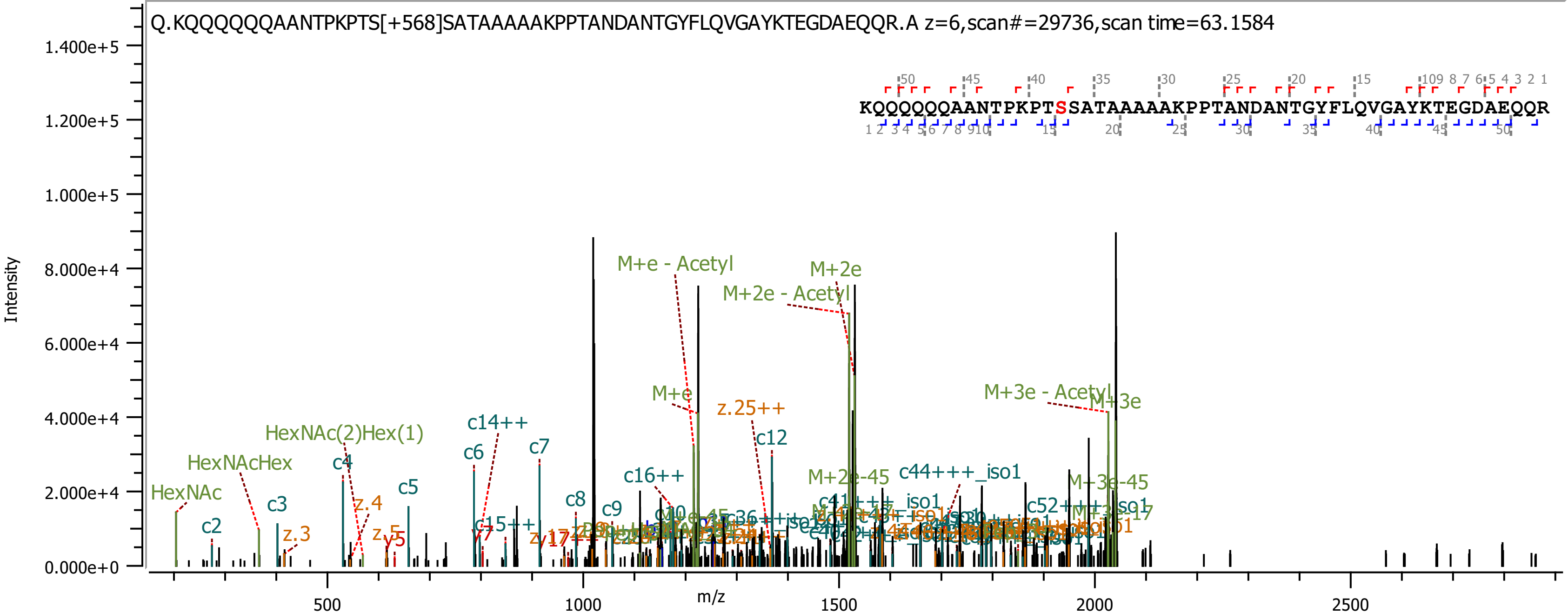

Q.KQQQQQQAANTPKPTS[+568]SATAAAAAKPPTANDANTGYFLQVGAYKTEGDAEQQR.A z=6,scan#=29736,scan time=63.1584
KQQQQQQAANTPKPTSSATAAAAAKPPTANDANTGYFLQVGAYKTEGDAEQQR
Intensity
m/z
1.400e+5
1.200e+5
1.000e+5
8.000e+4
6.000e+4
4.000e+4
2.000e+4
0.000e+0
500
1000
1500
2000
2500
M+e - Acetyl
M+2e
M+2e - Acetyl
M+e
M+3e - Acetyl
M+3e
HexNAc(2)Hex(1)
HexNAcHex
HexNAc
c14++
c6
c7
c4
c5
c3
c2
z.4
z.3
z.5
y5
y7
c8
c9
c12
c16++
c15++
z.25++
M+2e-45
c44+++_iso1
M+3e-45
c52+++_iso1

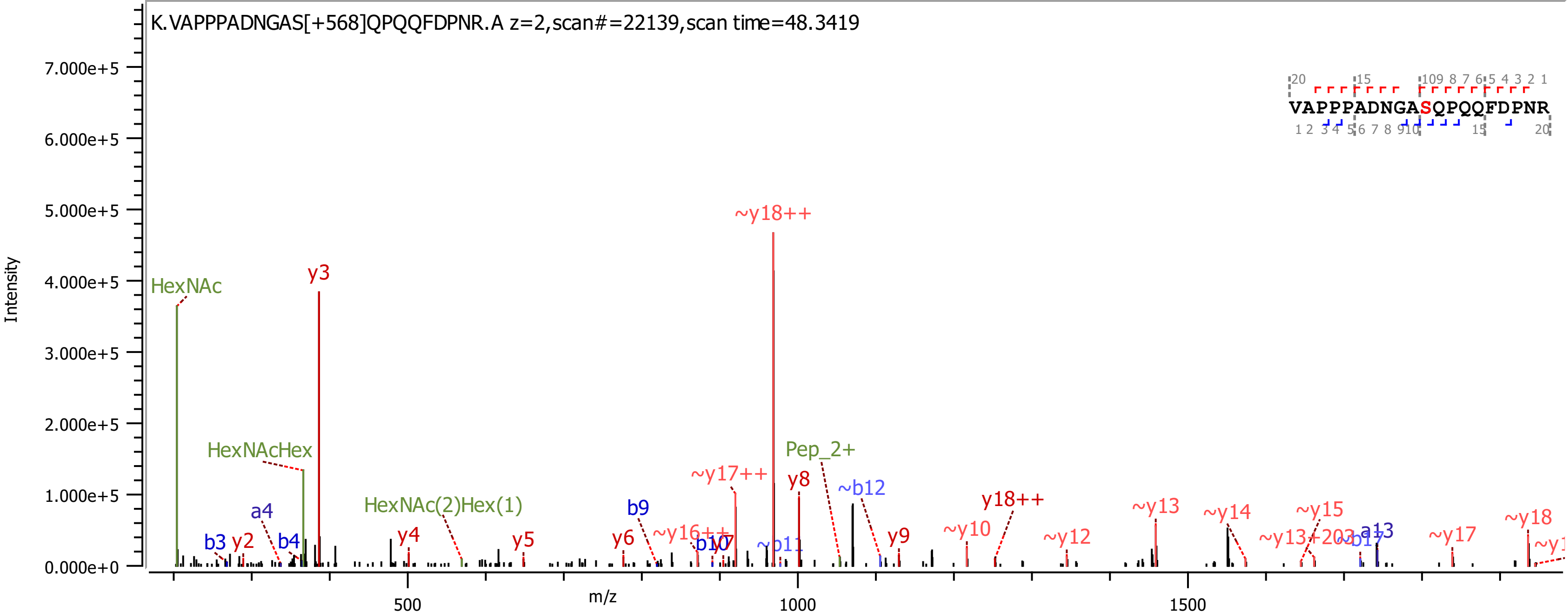

K.VAPPPADNGAS[+568]QPQQFDPNR.A z=2,scan#=22139,scan time=48.3419
VAPPPADNGASQPQQFDPNR
Intensity
m/z
7.000e+5
6.000e+5
5.000e+5
4.000e+5
3.000e+5
2.000e+5
1.000e+5
0.000e+0
500
1000
1500
HexNAc
y3
~y18++
HexNAcHex
Pep_2+
~y17++
y8
~b12
HexNAc(2)Hex(1)
a4
b9
y18++
~y13
~y14
~y15
~y18
~y10
b3
y2
b4
y4
y5
y6
~y16++
b10
~b11
y9
~y12
~y13+203
a13
~b17
~y17

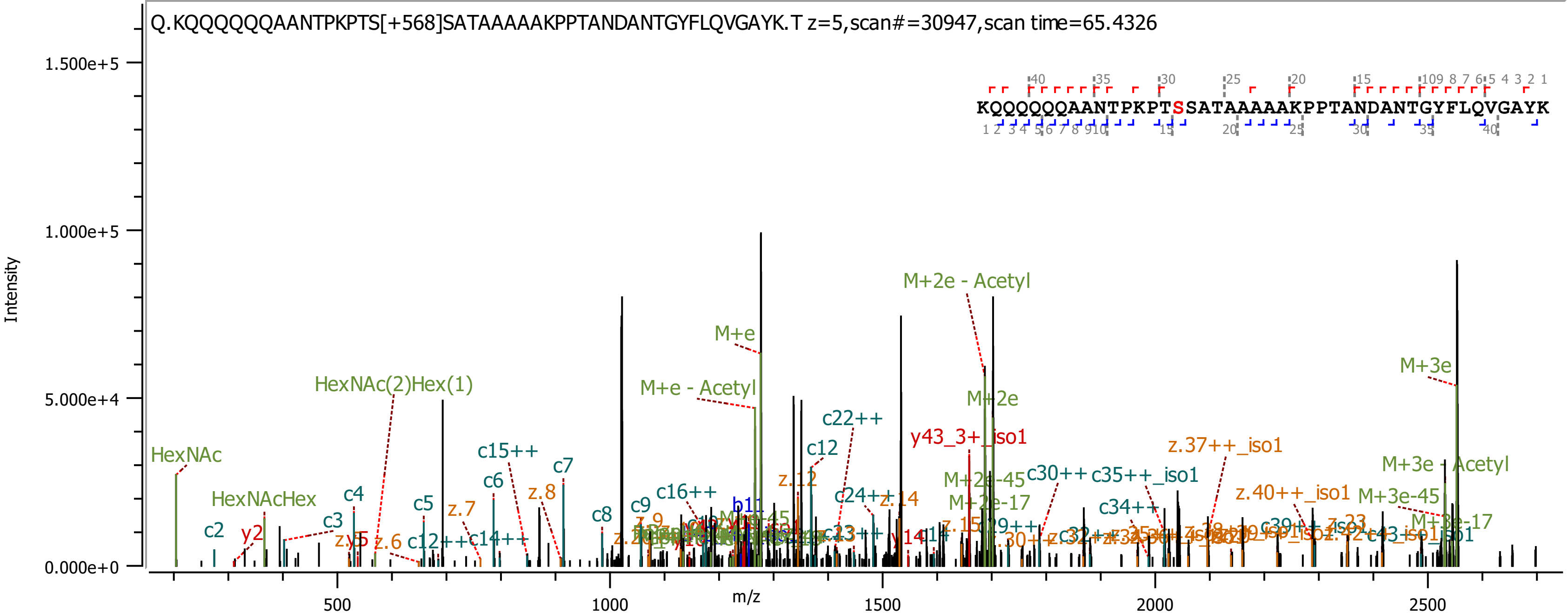

Q.KQQQQQQAANTPKPTS[+568]SATAAAAAKPPTANDANTGYFLQVGAYK.T z=5,scan#=30947,scan time=65.4326
KQQQQQQAANTPKPTSSATAAAAAKPPTANDANTGYFLQVGAYK
Intensity
m/z
1.500e+5
1.000e+5
5.000e+4
0.000e+0
500
1000
1500
2000
2500
HexNAc
HexNAcHex
HexNAc(2)Hex(1)
M+e
M+e - Acetyl
M+2e - Acetyl
M+2e
M+3e
M+3e - Acetyl
M+3e-45
M+3e-17
M+2e-45
M+2e-17
y43_3+_iso1
z.37++_iso1
z.40++_iso1
c22++
c12
c15++
c16++
c24++
c30++
c35++_iso1
c34++
c2
c3
c4
c5
c6
c7
c8
c9
y2
z.6
z.7
z.8
z.12
z.14
c12++
c14++
c13++
b11
y14

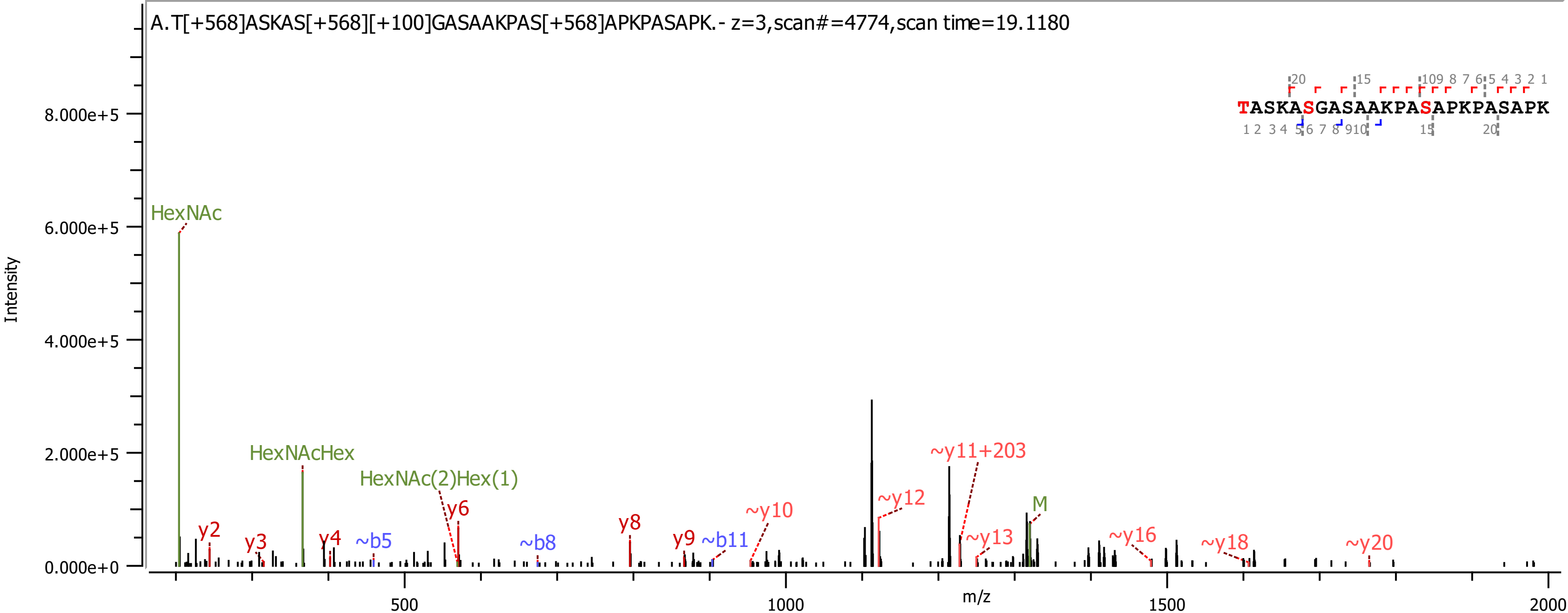

A.T[+568]ASKAS[+568][+100]GASAAKPAS[+568]APKPASAPK.- z=3,scan#=4774,scan time=19.1180
TASKASGASAAKPASAPKPASAPK
Intensity
m/z
HexNAc
HexNAcHex
HexNAc(2)Hex(1)
y2
y3
y4
~b5
y6
~b8
y8
y9
~b11
~y10
~y12
~y11+203
~y13
M
~y16
~y18
~y20

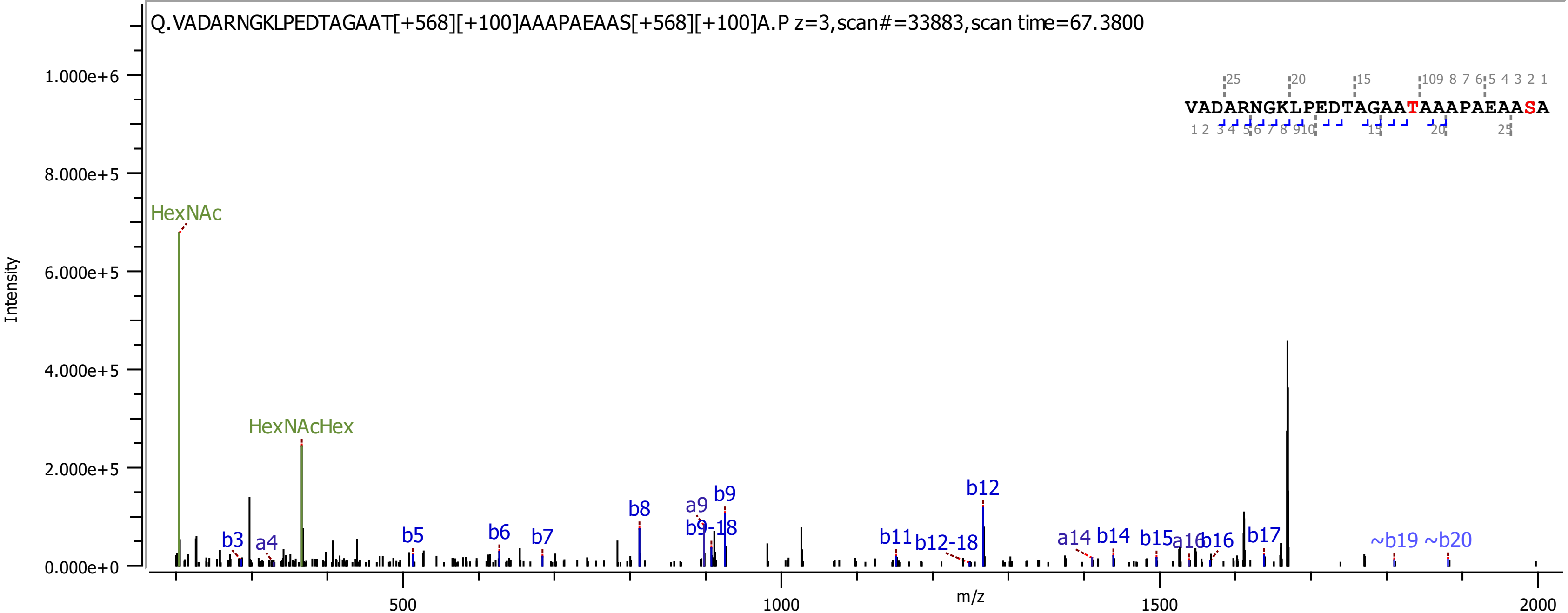
Q.VADARNGKLPEDTAGAAT[+568][+100]AAAPAEAAS[+568][+100]A.P z=3,scan#=33883,scan time=67.3800
Intensity
m/z
VADARNGKLPEDTAGAATAAAPAEAASA
HexNAc
HexNAcHex
b3
a4
b5
b6
b7
b8
a9
b9-18
b9
b11
b12-18
b12
a14
b14
b15
a16
b16
b17
~b19
~b20

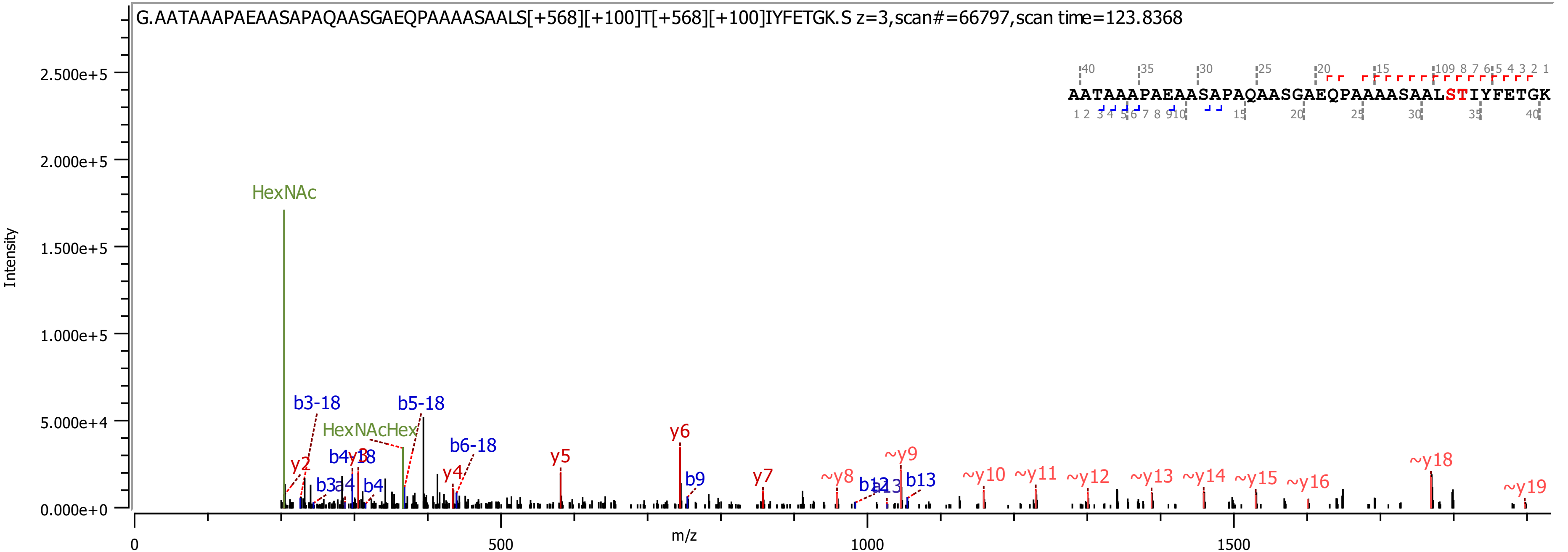

G.AATAAAPAEAASAPAQAASGAEQPAAAASAALS[+568][+100]T[+568][+100]IYFETGK.S z=3,scan#=66797,scan time=123.8368
AATAAAPAEAASAPAQAASGAEQPAAAASAALSTIYFETGK
Intensity
m/z
HexNAc
HexNAcHex
b3-18
b5-18
b6-18
b4-18
y2
y3
y4
y5
y6
y7
b9
b12
b13
~y8
~y9
~y10
~y11
~y12
~y13
~y14
~y15
~y16
~y18
~y19

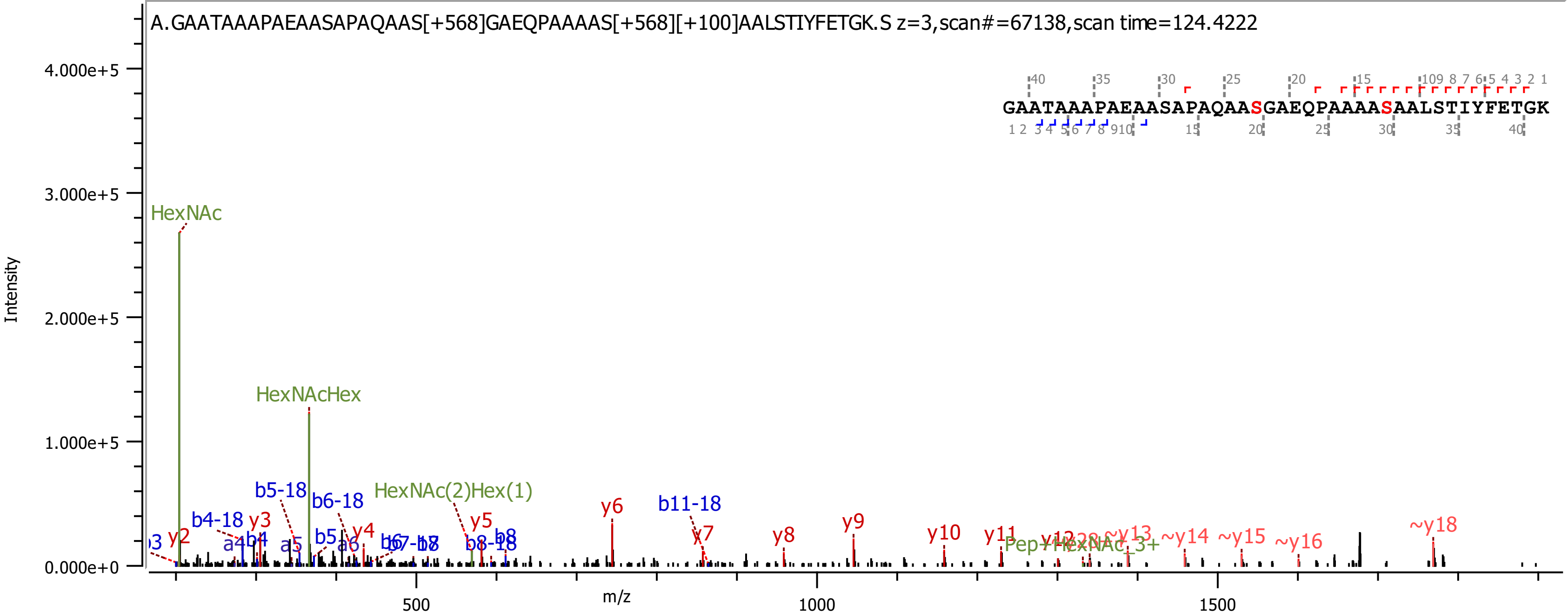

A.GAATAAAPAEAASAPAQAAS[+568]GAEQPAAAAS[+568][+100]AALSTIYFETGK.S z=3,scan#=67138,scan time=124.4222
GAATAAAPAEAASAPAQAASGAEQPAAAASAALSTIYFETGK
Intensity
4.000e+5
3.000e+5
2.000e+5
1.000e+5
0.000e+0
m/z
500
1000
1500
HexNAc
HexNAcHex
HexNAc(2)Hex(1)
b4-18
b5-18
b6-18
b11-18
y2
y3
y4
y5
y6
y7
y8
y9
y10
y11
~y13
~y14
~y15
~y16
~y18

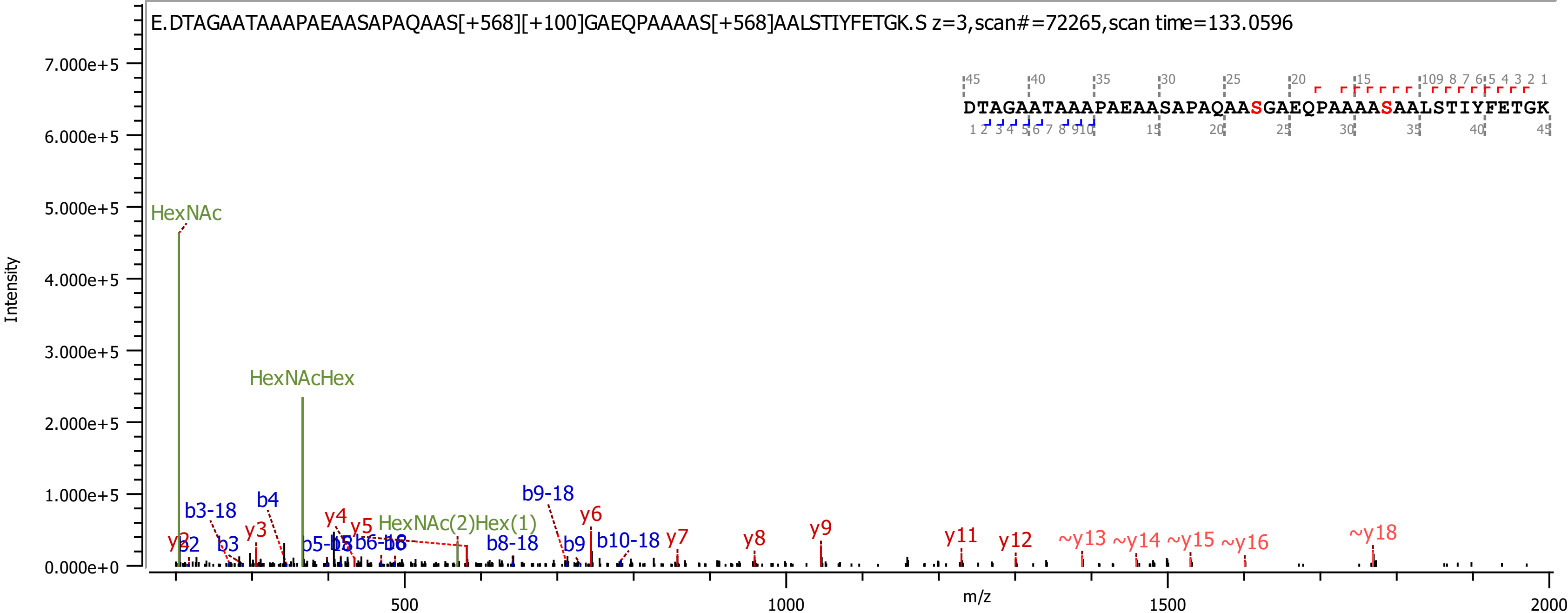
E.DTAGAATAAAPAEAASAPAQAAS[+568][+100]GAEQPAAAAS[+568]AALSTIYFETGK.S z=3,scan#=72265,scan time=133.0596
DTAGAATAAAPAEAASAPAQAASGAEQPAAAASAALSTIYFETGK
Intensity
m/z
HexNAc
HexNAcHex
HexNAc(2)Hex(1)
b3-18
b4
b3
b9-18
b8-18
b9
b10-18
y2
y3
y4
y5
y6
y7
y8
y9
y11
y12
~y13
~y14
~y15
~y16
~y18
7.000e+5
6.000e+5
5.000e+5
4.000e+5
3.000e+5
2.000e+5
1.000e+5
0.000e+0
500
1000
1500
2000

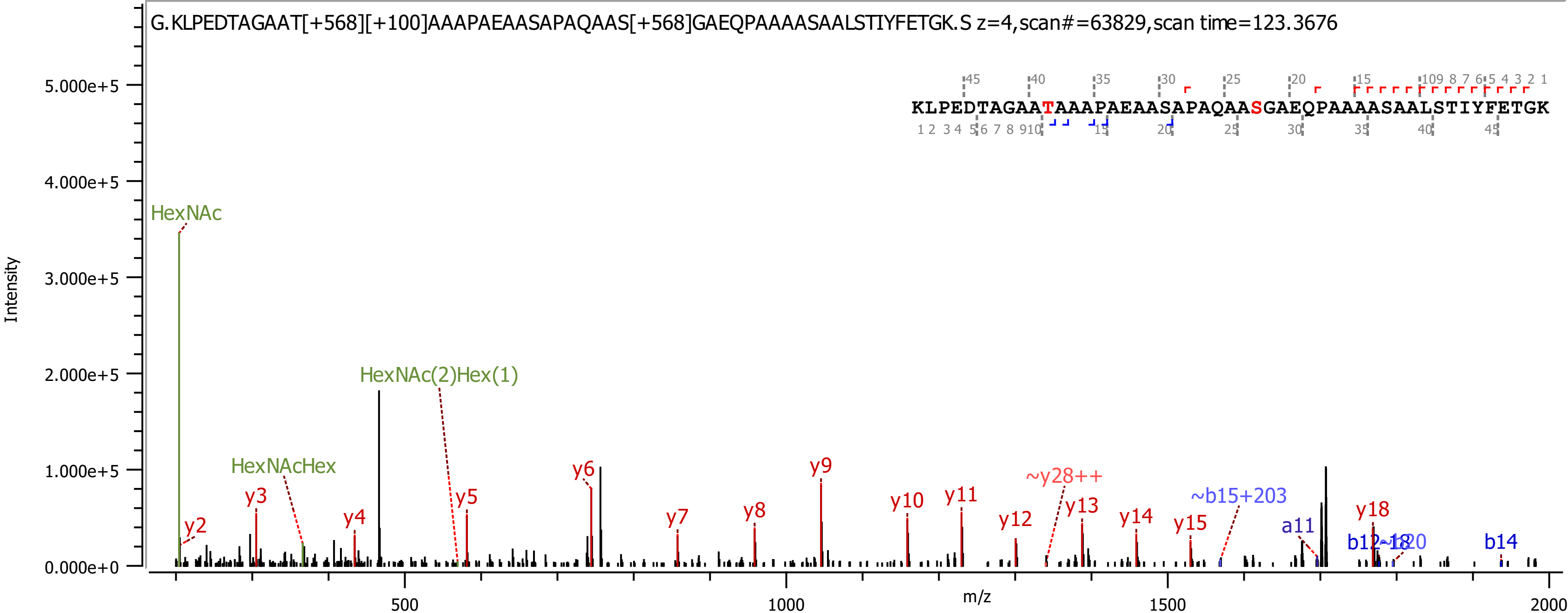

G.KLPEDTAGAAT[+568][+100]AAAPAEAASAPAQAAS[+568]GAEQPAAAASAALSTIYFETGK.S z=4,scan#=63829,scan time=123.3676
KLPEDTAGAATAAAPAEAASAPAQAASGAEQPAAAASAALSTIYFETGK
Intensity
m/z
HexNAc
HexNAcHex
HexNAc(2)Hex(1)
y2
y3
y4
y5
y6
y7
y8
y9
y10
y11
y12
~y28++
y13
y14
y15
~b15+203
a11
b12
y18
b14

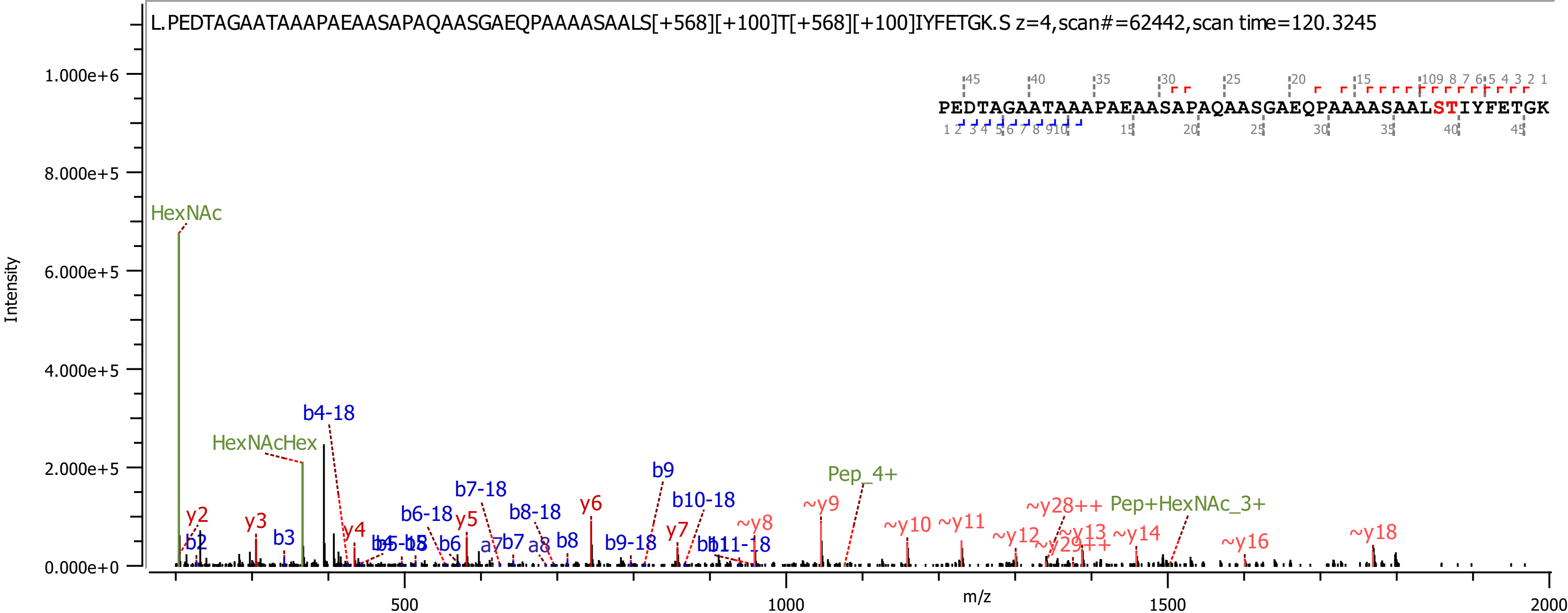
L.PEDTAGAATAAAPAEAASAPAQAASGAEQPAAAASAALS[+568][+100]T[+568][+100]IYFETGK.S z=4,scan#=62442,scan time=120.3245
PEDTAGAATAAAPAEAASAPAQAASGAEQPAAAASAALSTIYFETGK
Intensity
m/z
1.000e+6
8.000e+5
6.000e+5
4.000e+5
2.000e+5
0.000e+0
500
1000
1500
2000
HexNAc
HexNAcHex
b4-18
y2
b2
y3
b3
y4
b5-18
b6-18
y5
b6
b7-18
a7
b7
b8-18
a8
b8
y6
b9-18
b9
y7
b10-18
b11-18
~y8
~y9
Pep_4+
~y10
~y11
~y12
~y28++
~y29++
~y13
Pep+HexNAc_3+
~y14
~y16
~y18

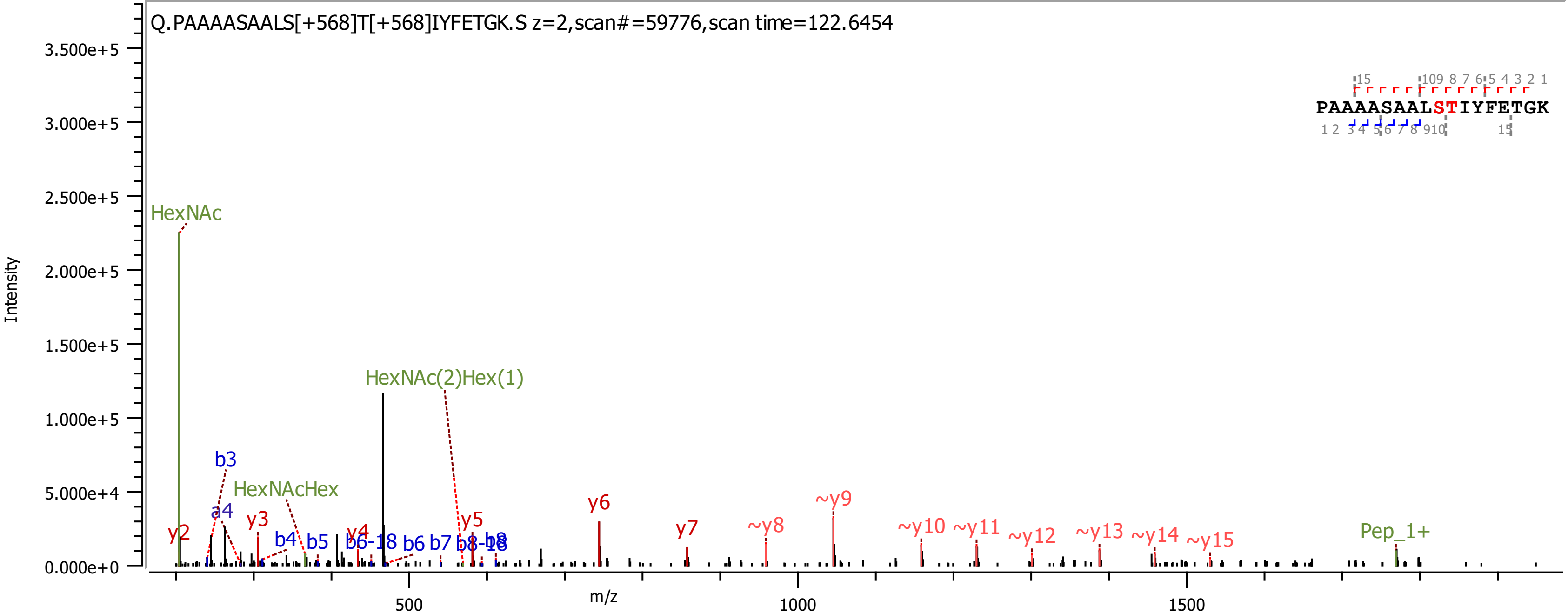

Q.PAAAASAALS[+568]T[+568]IYFETGK.S z=2,scan#=59776,scan time=122.6454
PAAAASAALSTIYFETGK
Intensity
m/z
HexNAc
HexNAc(2)Hex(1)
HexNAcHex
Pep_1+
b3
a4
y2
y3
b4
b5
y4
b6-18
b6
b7
b8-18
b8
y5
y6
y7
~y8
~y9
~y10
~y11
~y12
~y13
~y14
~y15

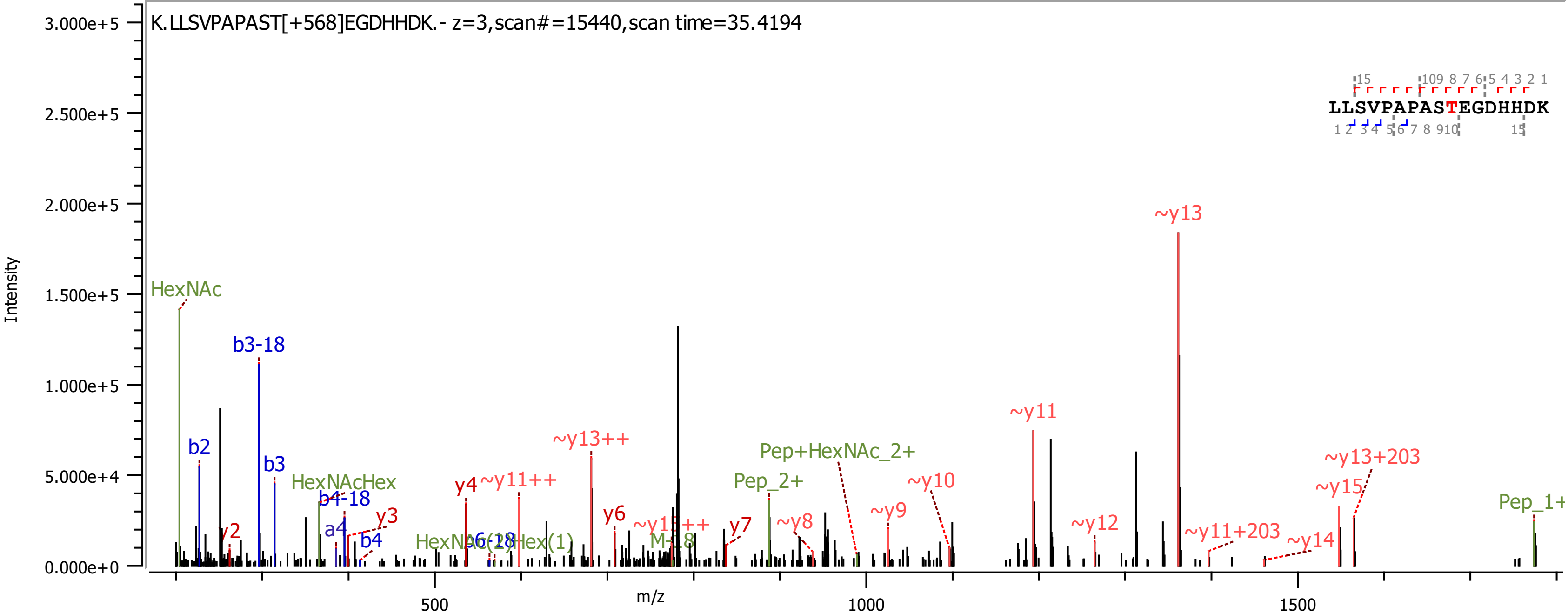

K.LLSVPAPAST[+568]EGDHHDK.- z=3,scan#=15440,scan time=35.4194
LLSVPAPASTEGDHHDK
Intensity
m/z
3.000e+5
2.500e+5
2.000e+5
1.500e+5
1.000e+5
5.000e+4
0.000e+0
500
1000
1500
HexNAc
b3-18
b2
b3
HexNAcHex
b4-18
a4
b4
y2
y3
y4
~y11++
~y13++
y6
~y15++
y7
M-18
Pep_2+
Pep+HexNAc_2+
~y8
~y9
~y10
~y11
~y12
~y13
~y11+203
~y14
~y15
~y13+203
Pep_1+

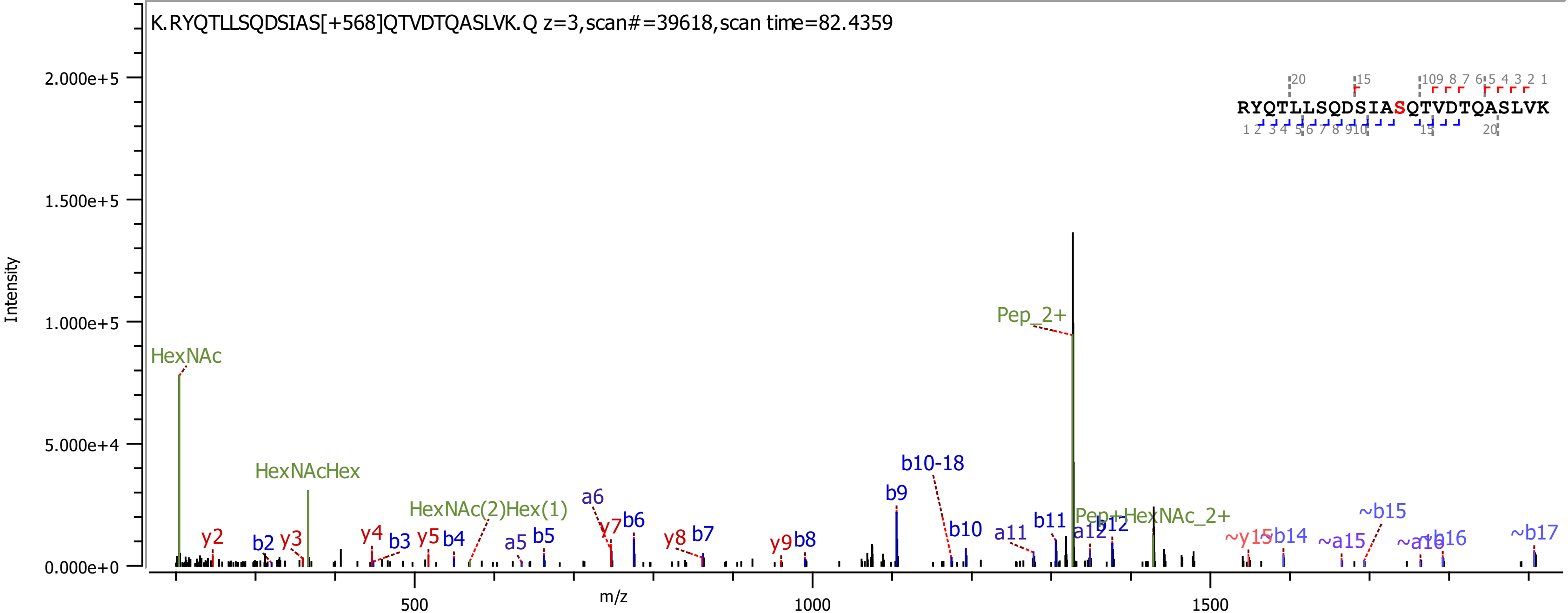
K.RYQTLLSQDSIAS[+568]QTVDTQASLVK.Q z=3,scan#=39618,scan time=82.4359
RYQTLLSQDSIASQTVDTQASLVK
Intensity
m/z
2.000e+5
1.500e+5
1.000e+5
5.000e+4
0.000e+0
500
1000
1500
HexNAc
HexNAcHex
HexNAc(2)Hex(1)
Pep_2+
Pep+HexNAc_2+
y2
y3
y4
y5
y7
y8
y9
b2
b3
b4
b5
b6
b7
b8
b9
b10
b10-18
b11
b12
a5
a6
a11
a12
~y15
~b14
~a15
~b15
~a16
~b16
~b17

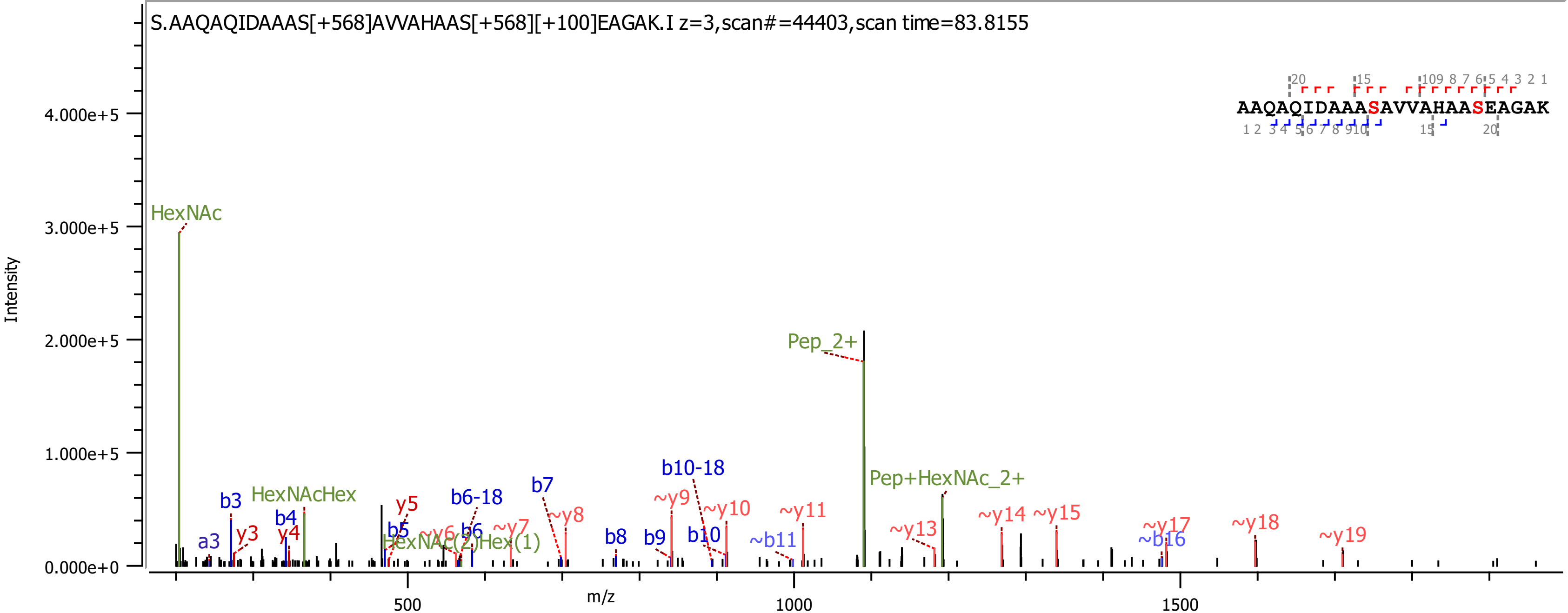
S.AAQAQIDAAAS[+568]AVVAHAAS[+568][+100]EAGAK.I z=3,scan#=44403,scan time=83.8155
AAQAQIDAAASAVVAHAASEAGAK
Intensity
m/z
HexNAc
HexNAcHex
HexNAc(2)Hex(1)
Pep_2+
Pep+HexNAc_2+
a3
b3
b4
b5
b6
b6-18
b7
b8
b9
b10
b10-18
~b11
~b16
y3
y4
y5
~y6
~y7
~y8
~y9
~y10
~y11
~y13
~y14
~y15
~y17
~y18
~y19
0.000e+0
1.000e+5
2.000e+5
3.000e+5
4.000e+5
500
1000
1500

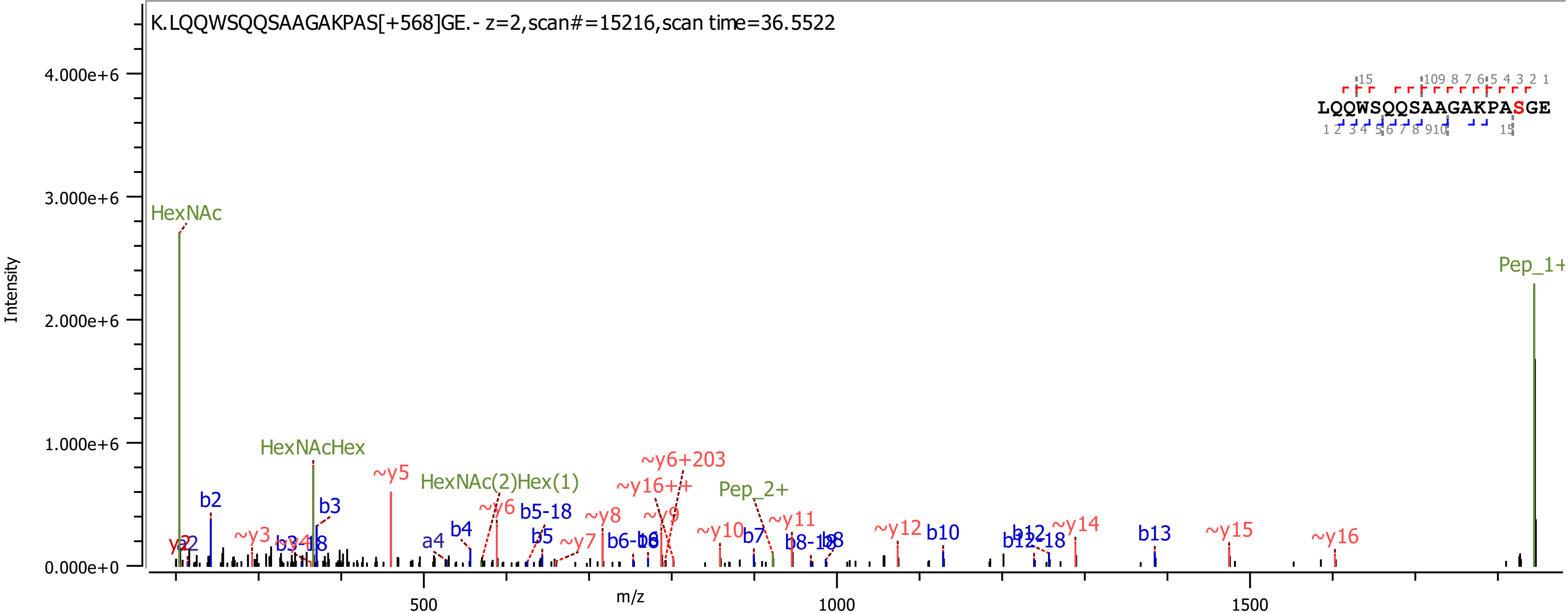
K.LQQWSQQSAAGAKPAS[+568]GE.- z=2,scan#=15216,scan time=36.5522
LQQWSQQSAAGAKPASGE
Intensity
m/z
HexNAc
HexNAcHex
HexNAc(2)Hex(1)
Pep_2+
Pep_1+
b2
b3
b3-18
a4
b4
b5-18
b5
b6-18
b7
b8-18
b10
b12-18
b13
~y3
~y5
~y6
~y7
~y8
~y9
~y10
~y11
~y12
~y14
~y15
~y16
~y16++
~y6+203

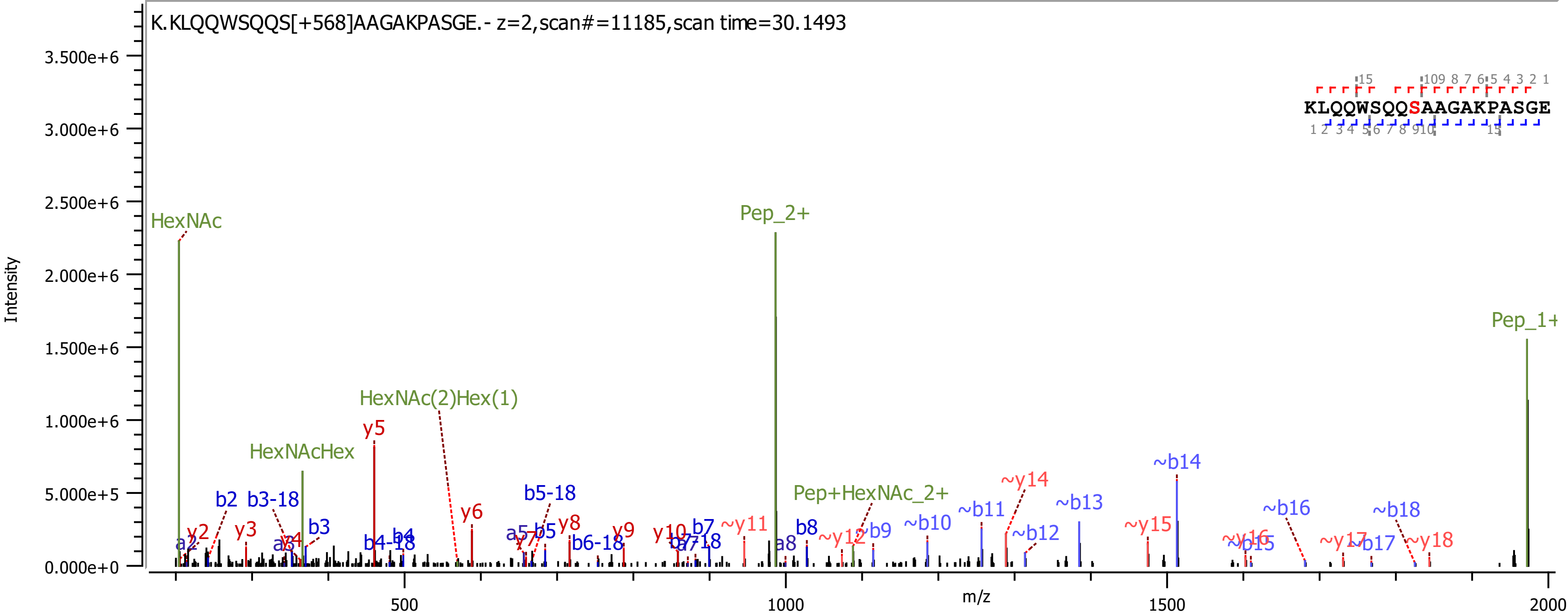

K.KLQQWSQQS[+568]AAGAKPASGE.- z=2,scan#=11185,scan time=30.1493
KLQQWSQQSAAGAKPASGE
Intensity
3.500e+6
3.000e+6
2.500e+6
2.000e+6
1.500e+6
1.000e+6
5.000e+5
0.000e+0
m/z
500
1000
1500
2000
HexNAc
Pep_2+
Pep_1+
HexNAc(2)Hex(1)
HexNAcHex
Pep+HexNAc_2+
y2
y3
y5
y6
y8
y9
y10
~y11
~y12
~y14
~y15
~y16
~y17
~y18
a2
a3
a5
a8
b2
b3-18
b3
b4-18
b4
b5-18
b5
b6-18
b7
b7-18
b8
~b9
~b10
~b11
~b12
~b13
~b14
~b15
~b16
~b17
~b18

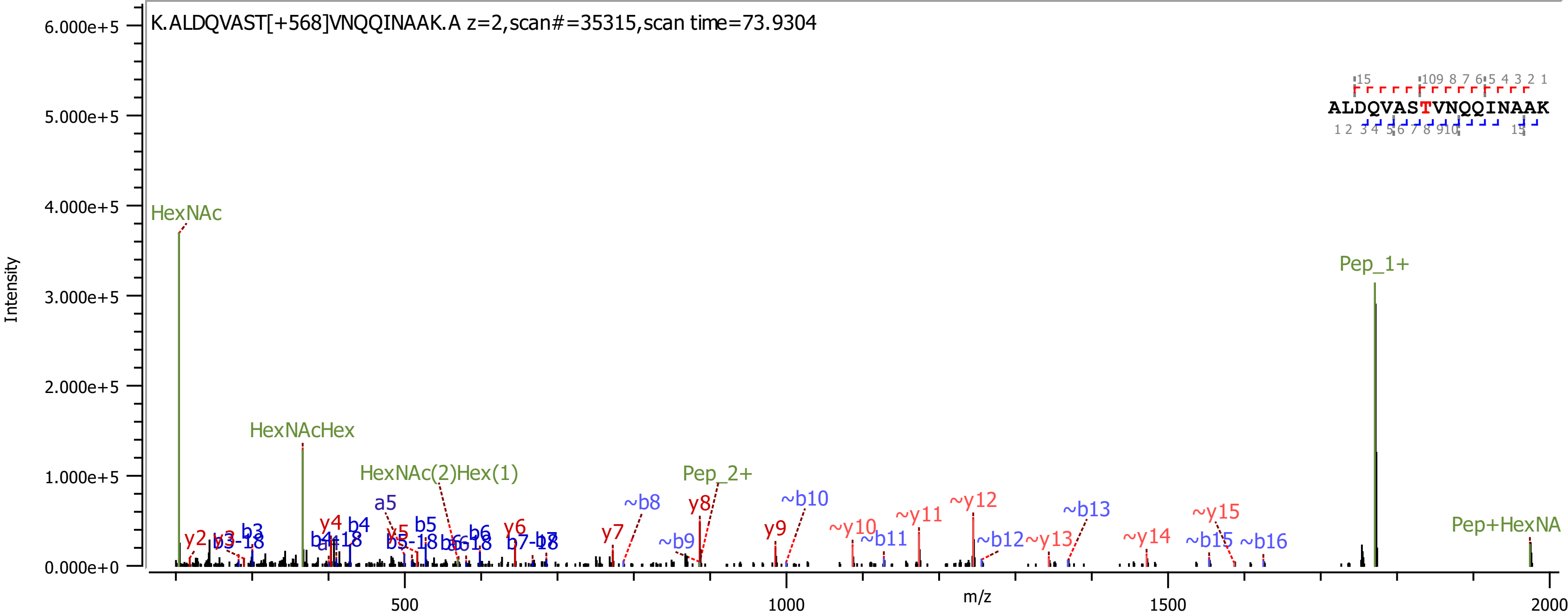
K.ALDQVAST[+568]VNQQINAAK.A z=2,scan#=35315,scan time=73.9304
ALDQVASTVNQQINAAK
Intensity
6.000e+5
5.000e+5
4.000e+5
3.000e+5
2.000e+5
1.000e+5
0.000e+0
m/z
500
1000
1500
2000
HexNAc
HexNAcHex
HexNAc(2)Hex(1)
Pep_2+
Pep_1+
Pep+HexNA
a5
y2
y3
b3
b3-18
y4
b4
b4-18
y5
b5
b5-18
b6
b6-18
y6
b7
b7-18
y7
~b8
y8
~b9
y9
~b10
~y10
~b11
~y11
~y12
~b12
~y13
~b13
~y14
~y15
~b15
~b16

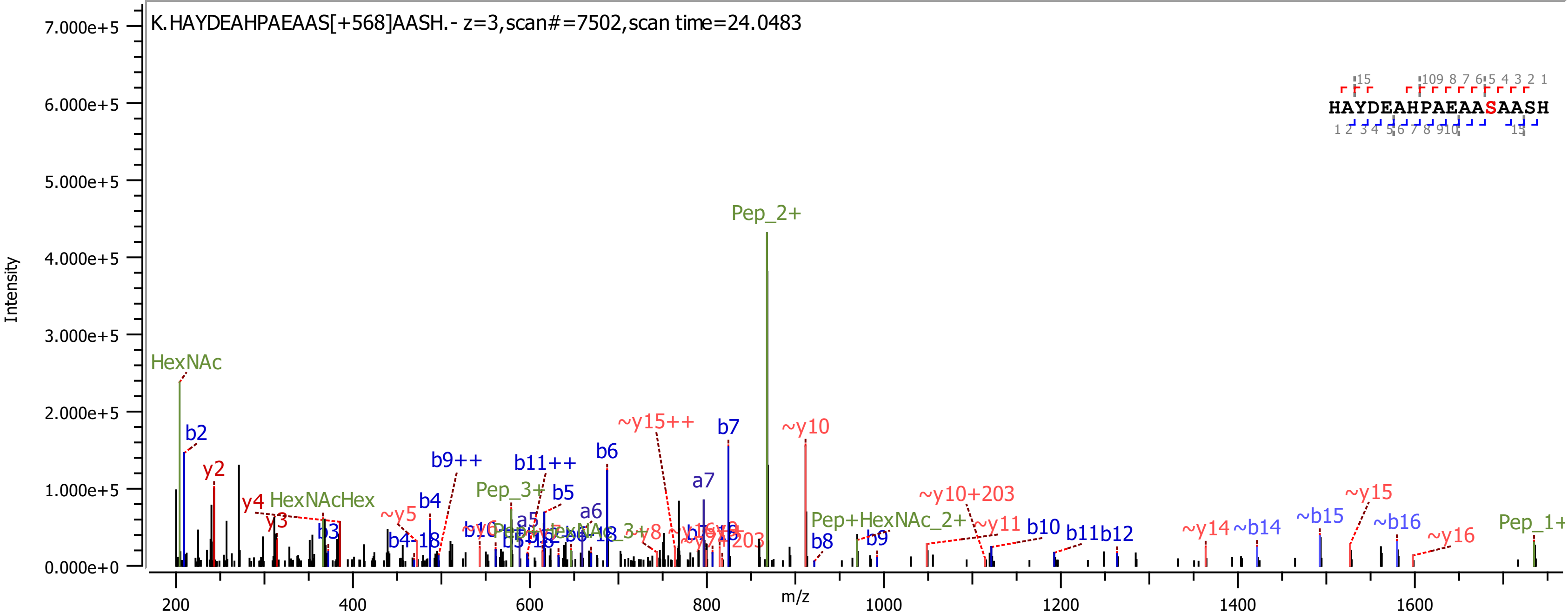

K.HAYDEAHPAEAAS[+568]AASH.- z=3,scan#=7502,scan time=24.0483
HAYDEAHPAEAASAASH
Intensity
m/z
7.000e+5
6.000e+5
5.000e+5
4.000e+5
3.000e+5
2.000e+5
1.000e+5
0.000e+0
200
400
600
800
1000
1200
1400
1600
Pep_2+
HexNAc
b2
y2
y4
HexNAcHex
y3
b3
~y5
b4
b4-18
b9++
b11++
Pep_3+
b5
a5
a6
b6
~y15++
a7
b7
~y10
Pep+HexNAc_2+
b8
b9
~y10+203
~y11
b10
b11
b12
~y14
~b14
~b15
~y15
~b16
~y16
Pep_1+

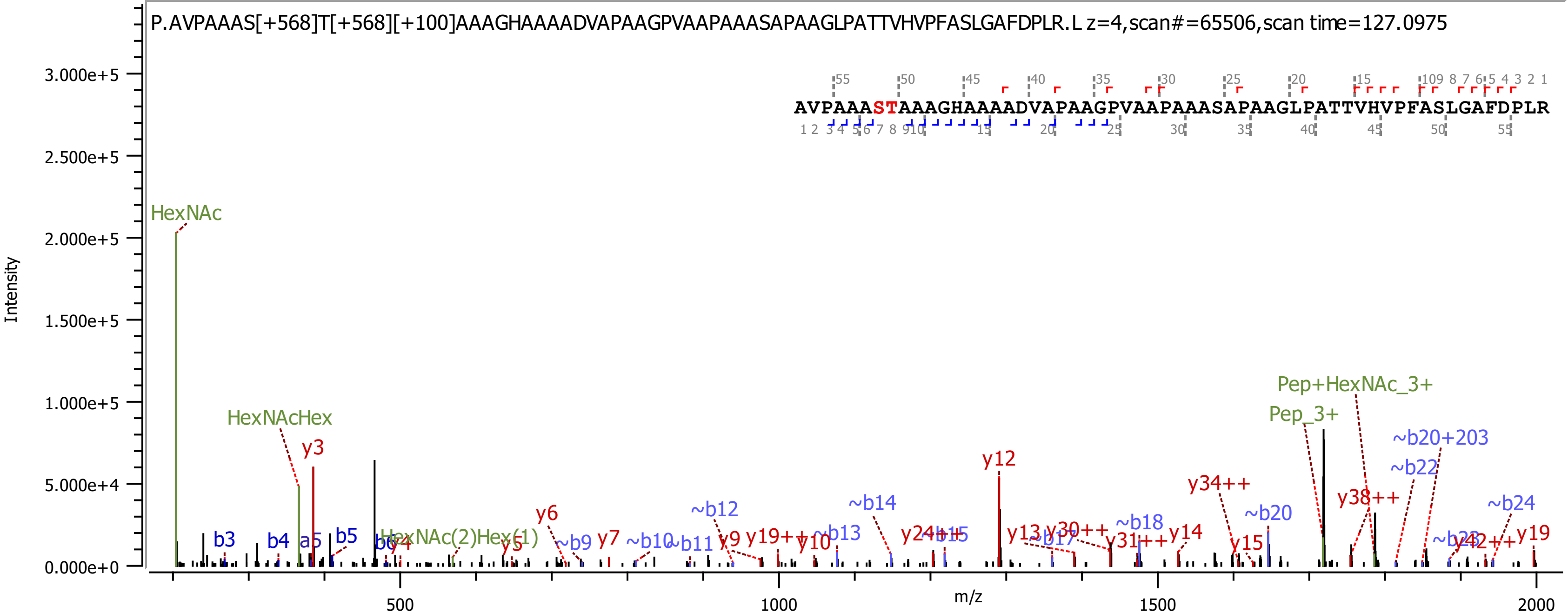

P.AVPAAAS[+568]T[+568][+100]AAAGHAAAADVAPAAGPVAAPAAASAPAAGLPATTVHVPFASLGAFDPLR.L z=4,scan#=65506,scan time=127.0975
AVPAAASTAAAGHAAAADVAPAAGPVAAPAAASAPAAGLPATTVHVPFASLGAFDPLR
Intensity
m/z
HexNAc
HexNAcHex
y3
y12
Pep+HexNAc_3+
Pep_3+
~b20+203
~b22
y34++
y38++
~b20
~b24
~b12
~b14
y6
y7
~b18
y14
y19
~b13
~b9
~b10
~b11
y9
y10
y13
y15
b3
b4
a5
b5

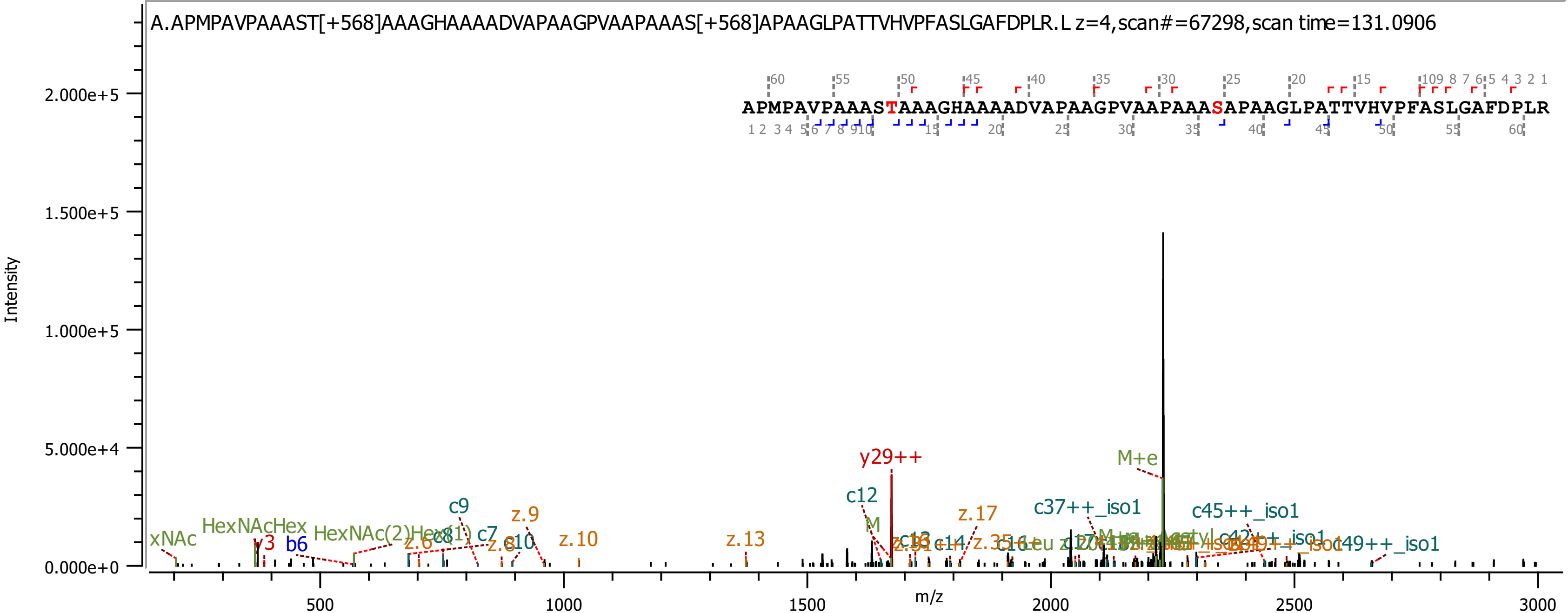
A.APMPAVPAAAST[+568]AAAGHAAAADVAPAAGPVAAPAAAS[+568]APAAGLPATTVHVPFASLGAFDPLR.L z=4,scan#=67298,scan time=131.0906
APMPAVPAAASTAAAGHAAAADVAPAAGPVAAPAAASAPAAGLPATTVHVPFASLGAFDPLR
Intensity
m/z
2.000e+5
1.500e+5
1.000e+5
5.000e+4
0.000e+0
500
1000
1500
2000
2500
3000
y29++
M+e
c12
c9
z.9
z.17
c37++_iso1
c45++_iso1
c49++_iso1
z.10
z.13
xNAc
HexNAcHex
y3
b6
c7
z.8
c10
M

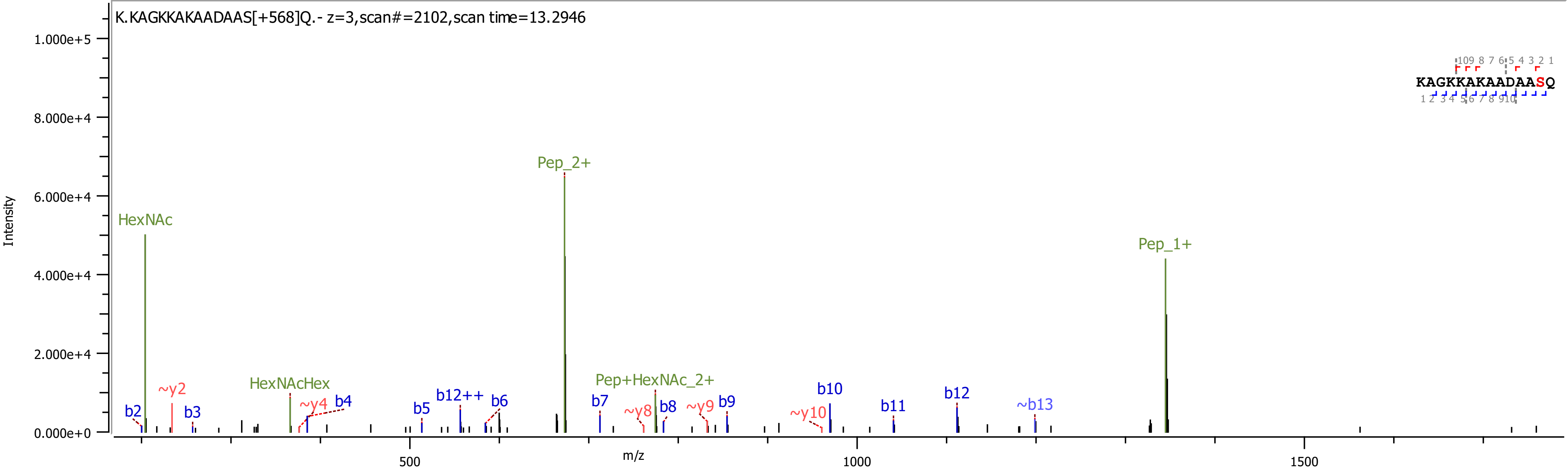

K.KAGKKAKAADAAS[+568]Q.- z=3,scan#=2102,scan time=13.2946
Intensity
m/z
HexNAc
Pep_2+
Pep_1+
HexNAcHex
Pep+HexNAc_2+
b2
~y2
b3
~y4
b4
b5
b12++
b6
b7
~y8
b8
~y9
b9
~y10
b10
b11
b12
~b13
KAGKKAKAADAASQ

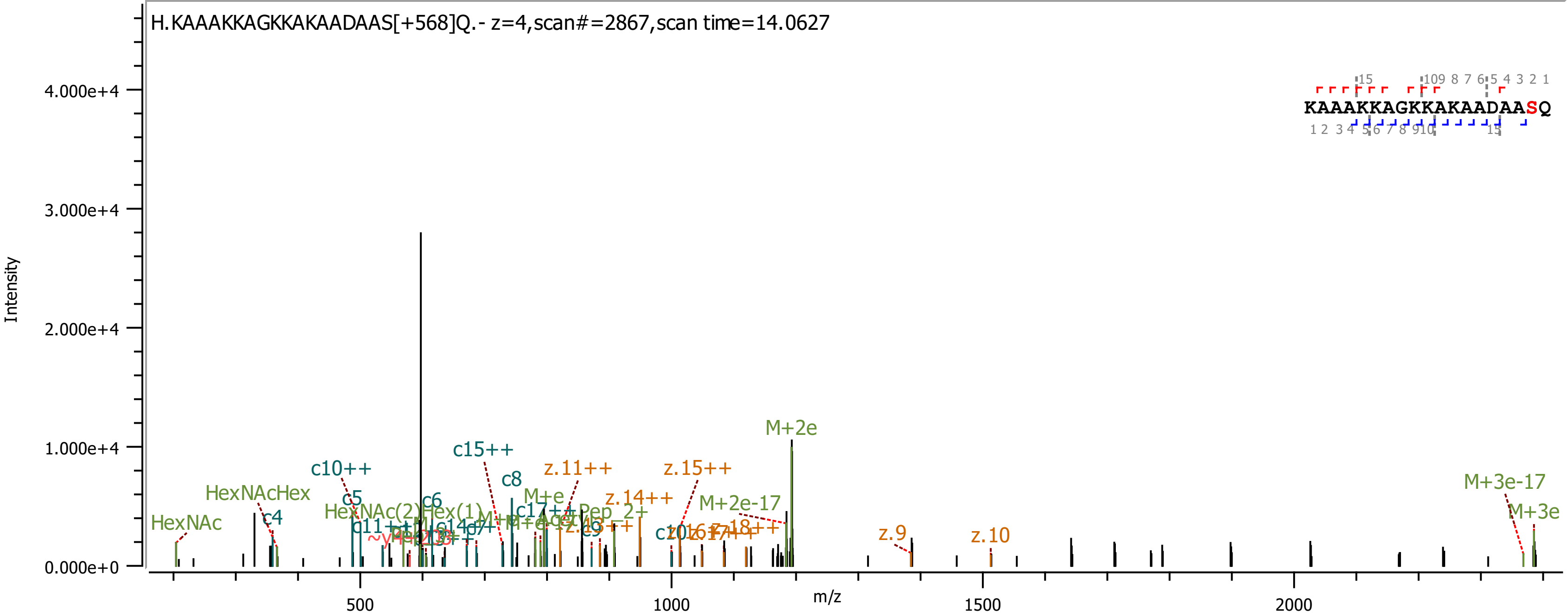
H.KAAAKKAGKKAKAADAAS[+568]Q.- z=4,scan#=2867,scan time=14.0627
KAAAKKAGKKAKAADAASQ
Intensity
4.000e+4
3.000e+4
2.000e+4
1.000e+4
0.000e+0
m/z
500
1000
1500
2000
HexNAc
HexNAcHex
c4
c10++
c5
c6
c15++
c8
M+e
z.11++
z.15++
z.14++
M+2e-17
M+2e
z.9
z.10
M+3e-17
M+3e

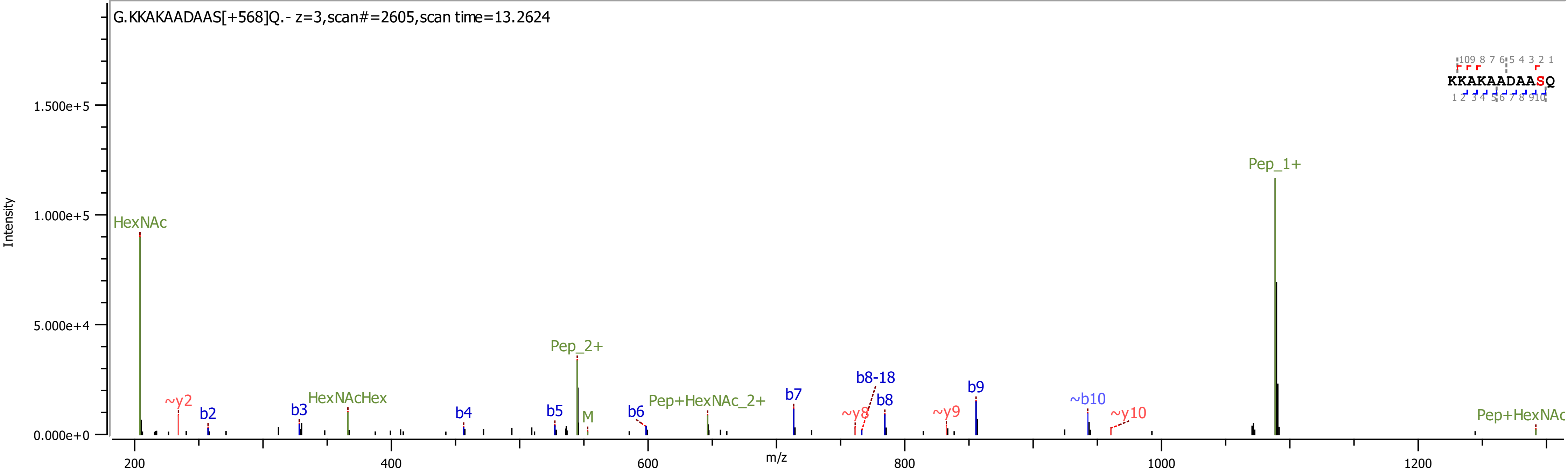

G.KKAKAADAAS[+568]Q.- z=3,scan#=2605,scan time=13.2624
Intensity
m/z
KKAKAADAASQ
HexNAc
~y2
b2
b3
HexNAcHex
b4
b5
Pep_2+
M
b6
Pep+HexNAc_2+
b7
~y8
b8-18
b8
~y9
b9
~b10
~y10
Pep_1+
Pep+HexNAc

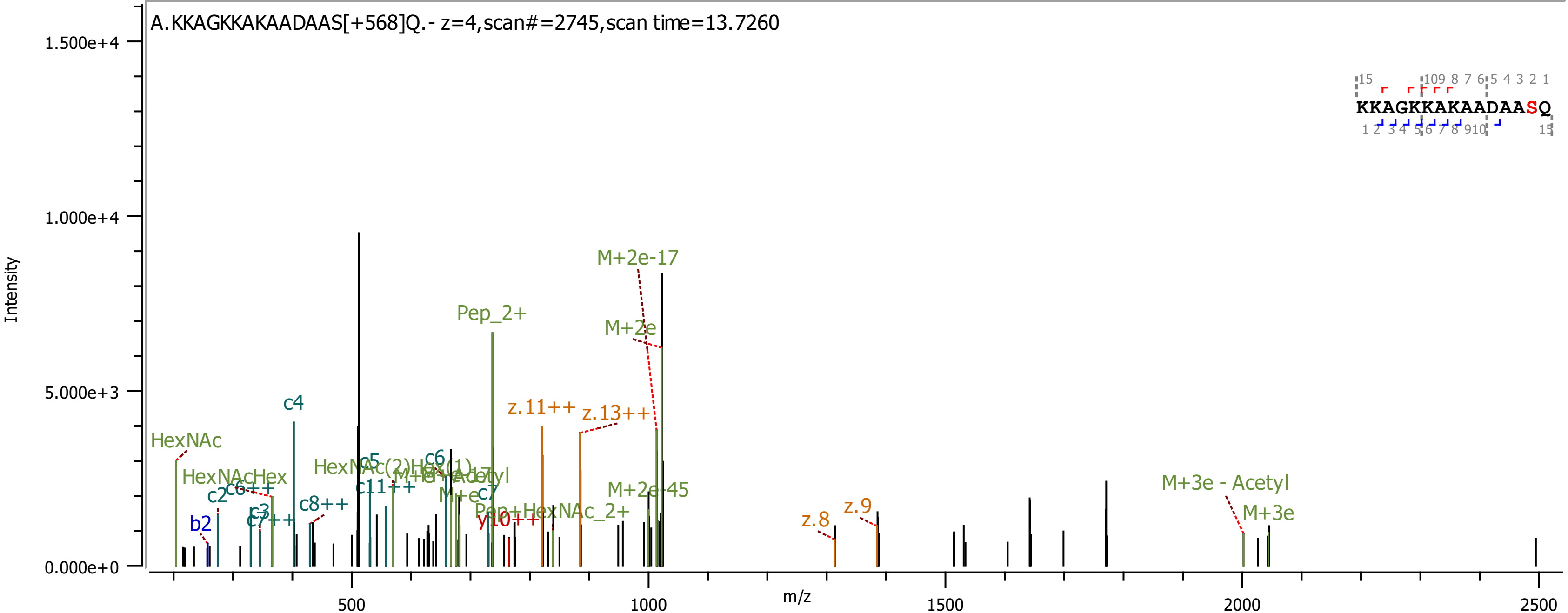

A.KKAGKKAKAADAAS[+568]Q.- z=4,scan#=2745,scan time=13.7260
Intensity
m/z
1.500e+4
1.000e+4
5.000e+3
0.000e+0
500
1000
1500
2000
2500
KKAGKKAKAADAASQ
M+2e-17
Pep_2+
M+2e
z.11++
z.13++
c4
HexNAc
HexNAcHex
c2
c6++
c8++
c3
c7++
b2
c5
c6
c11++
c7
y10++
Pep+HexNAc_2+
M+2e-45
z.8
z.9
M+3e - Acetyl
M+3e

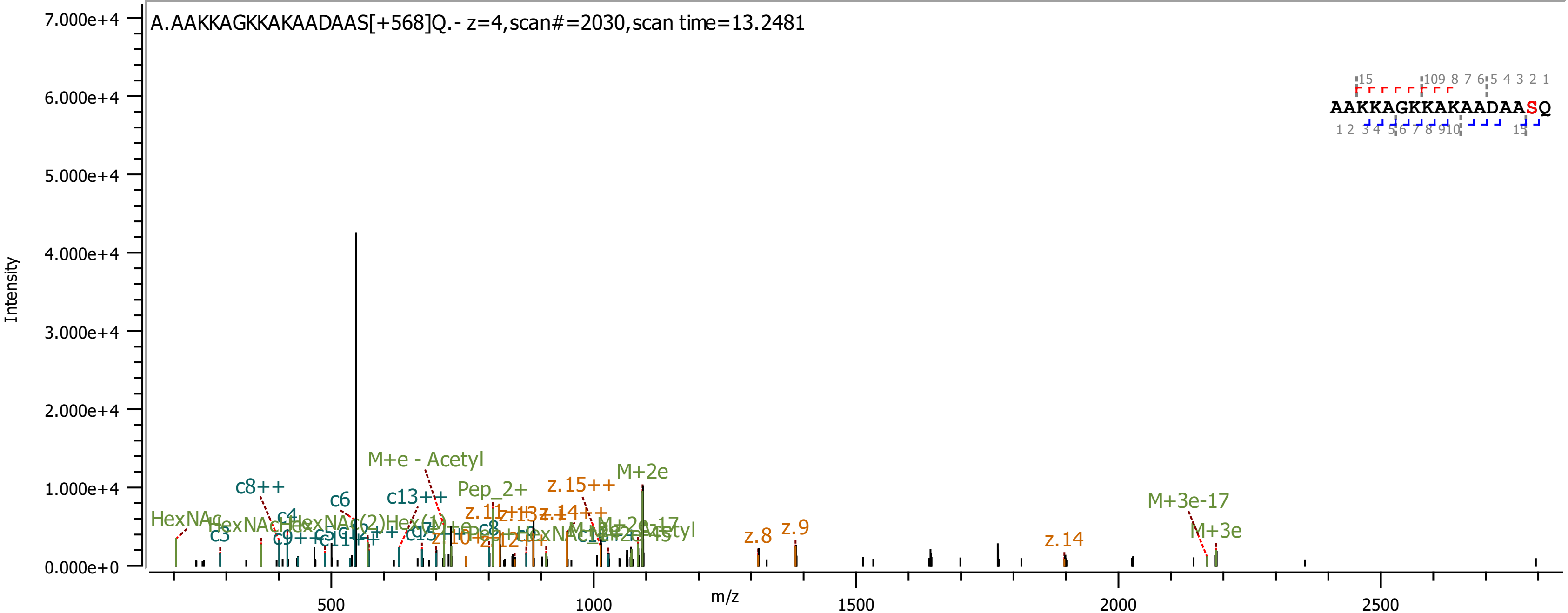

A.AAKKAGKKAKAADAAS[+568]Q.- z=4,scan#=2030,scan time=13.2481
Intensity
m/z
7.000e+4
6.000e+4
5.000e+4
4.000e+4
3.000e+4
2.000e+4
1.000e+4
0.000e+0
500
1000
1500
2000
2500
AAKKAGKKAKAADAASQ
HexNAc
c3
c8++
c6
M+e - Acetyl
c13++
Pep_2+
z.15++
M+2e
z.8
z.9
z.14
M+3e-17
M+3e

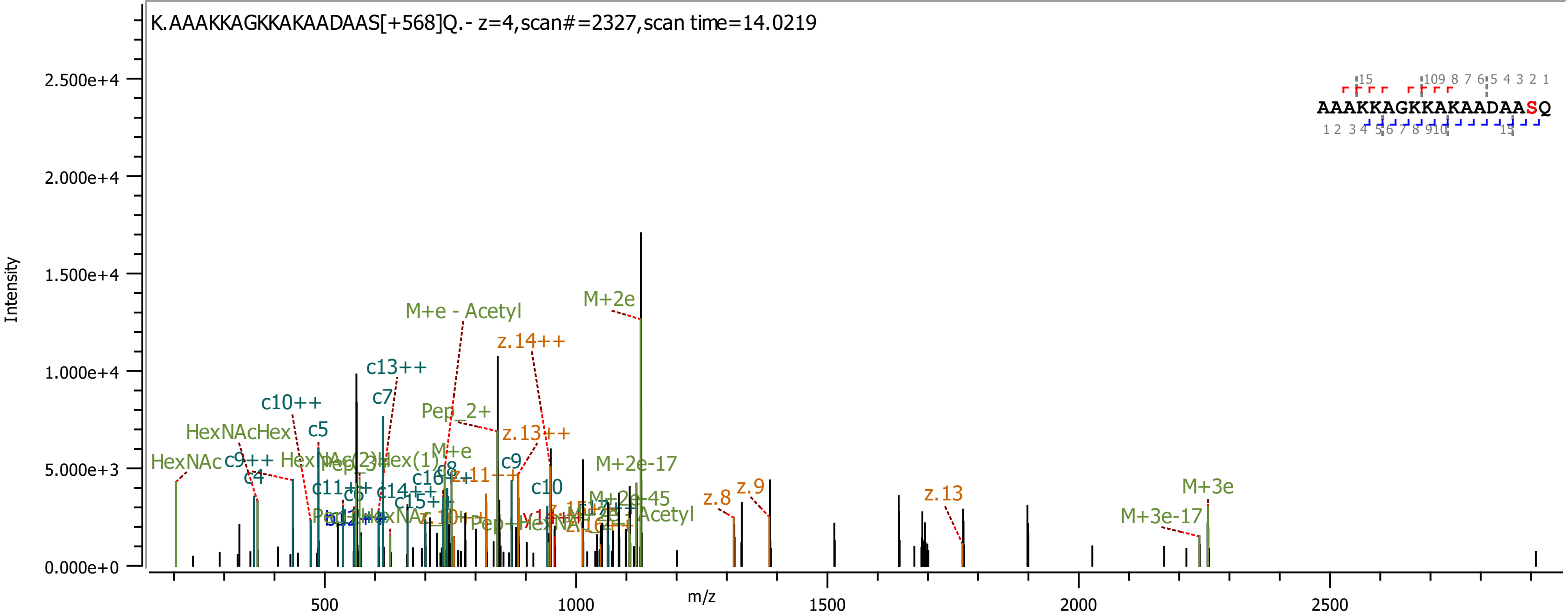
K.AAAKKAGKKAKAADAAS[+568]Q.- z=4,scan#=2327,scan time=14.0219
AAAKKAGKKAKAADAASQ
Intensity
2.500e+4
2.000e+4
1.500e+4
1.000e+4
5.000e+3
0.000e+0
m/z
500
1000
1500
2000
2500
M+2e
M+e - Acetyl
z.14++
c13++
c10++
c7
Pep_2+
z.13++
HexNAcHex
c5
M+e
HexNAc
c9++
c9
c4
z.11++
c10
c11++
c14++
c15++
M+2e-17
M+2e-45
Acetyl
z.8
z.9
z.13
M+3e
M+3e-17

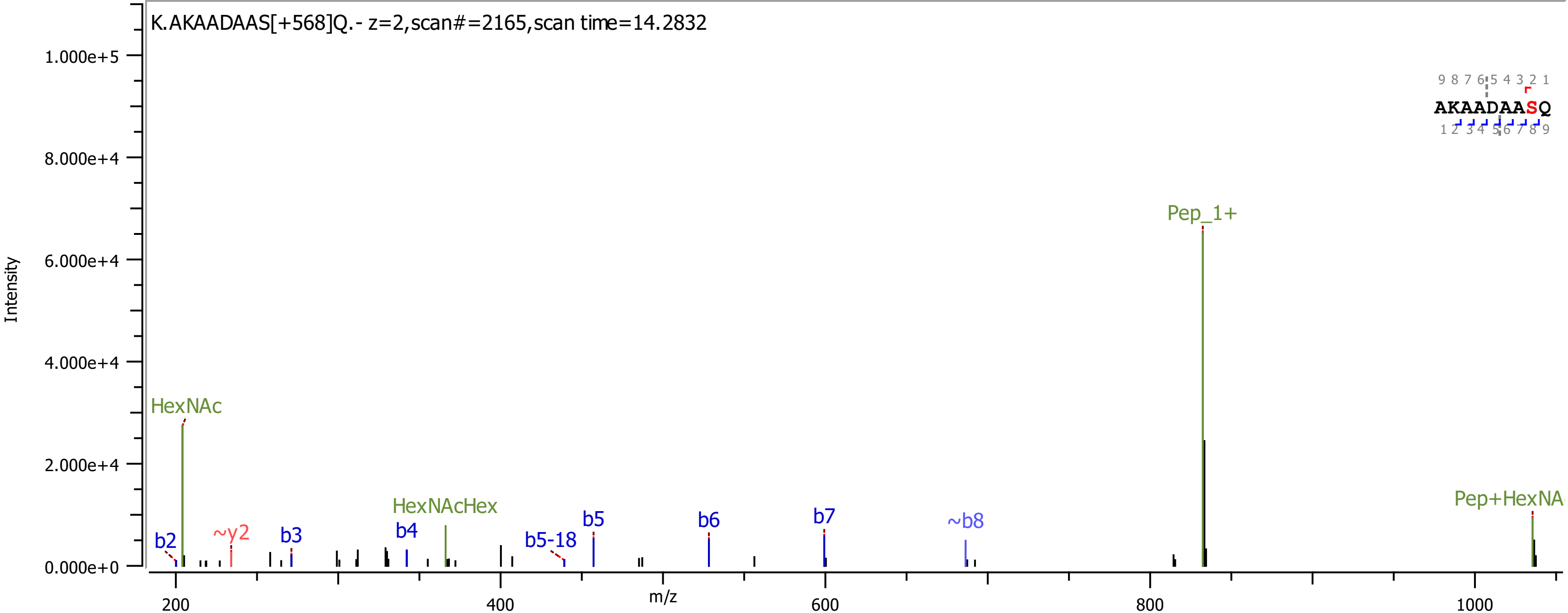

K.AKAADAAS[+568]Q.- z=2,scan#=2165,scan time=14.2832
Intensity
1.000e+5
8.000e+4
6.000e+4
4.000e+4
2.000e+4
0.000e+0
m/z
200
400
600
800
1000
9 8 7 6 5 4 3 2 1
AKAADAASQ
1 2 3 4 5 6 7 8 9
HexNAc
b2
~y2
b3
b4
HexNAcHex
b5-18
b5
b6
b7
~b8
Pep_1+

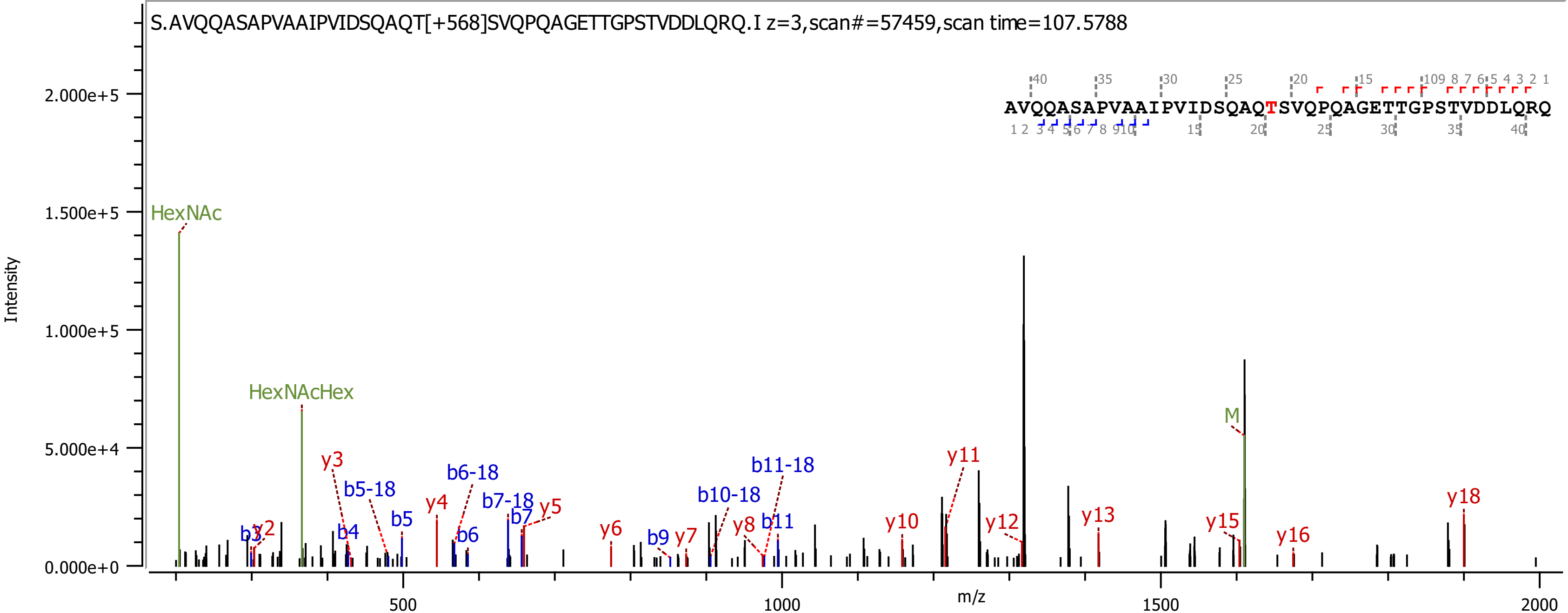
S.AVQQASAPVAAIPVIDSQAQT[+568]SVQPQAGETTGPSTVDDLQRQ.I z=3,scan#=57459,scan time=107.5788
AVQQASAPVAAIPVIDSQAQTSVQPQAGETTGPSTVDDLQRQ
Intensity
m/z
0.000e+0
5.000e+4
1.000e+5
1.500e+5
2.000e+5
500
1000
1500
2000
HexNAc
HexNAcHex
M
b3
b4
b5
b5-18
b6
b6-18
b7
b7-18
b9
b10-18
b11
b11-18
y2
y3
y4
y5
y6
y7
y8
y10
y11
y12
y13
y15
y16
y18

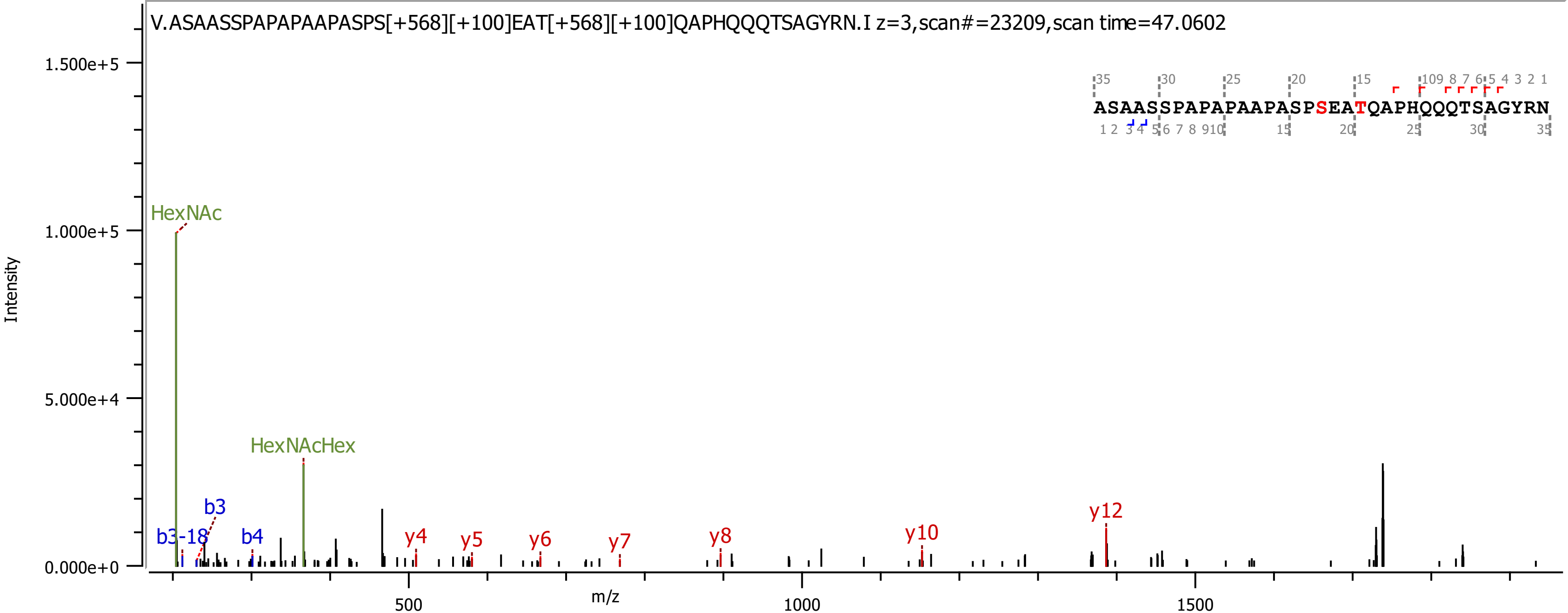
V.ASAASSPAPAPAAPASPS[+568][+100]EAT[+568][+100]QAPHQQQTSAGYRN.I z=3,scan#=23209,scan time=47.0602
ASAASSPAPAPAAPASPSEATQAPHQQQTSAGYRN
Intensity
m/z
1.500e+5
1.000e+5
5.000e+4
0.000e+0
500
1000
1500
HexNAc
HexNAcHex
b3
b3-18
b4
y4
y5
y6
y7
y8
y10
y12

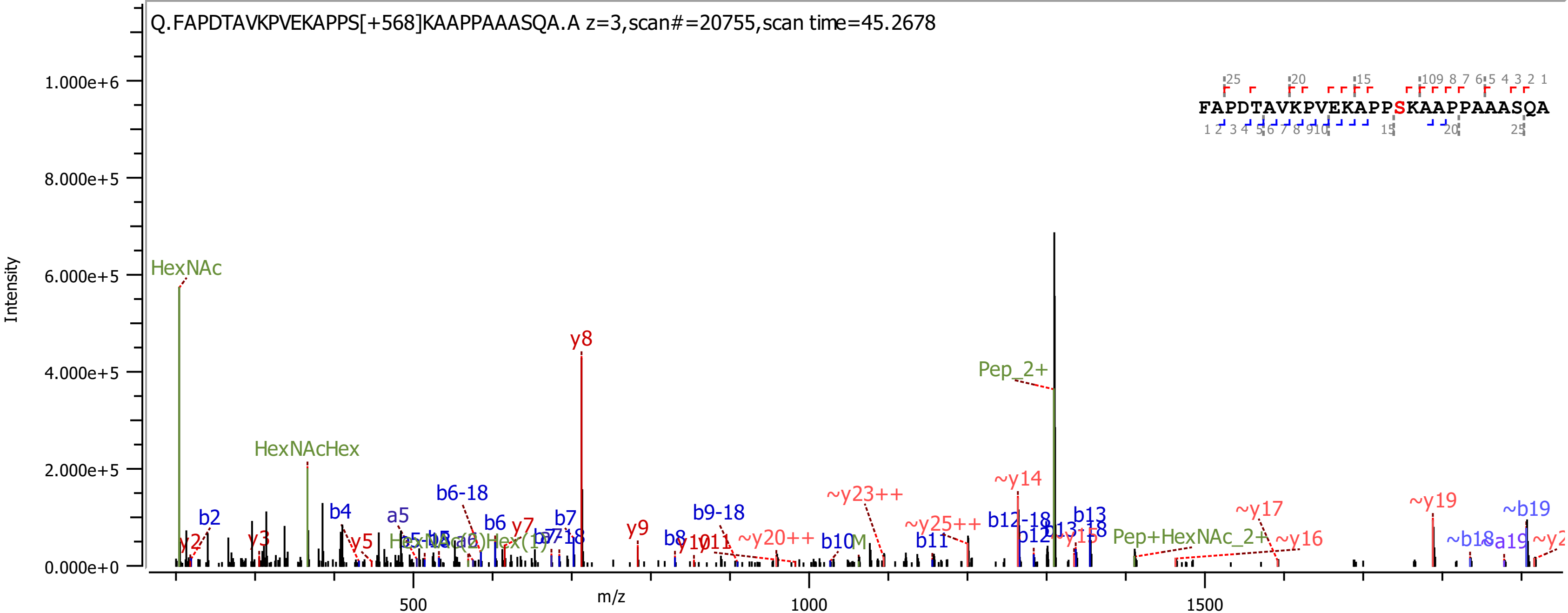
Q.FAPDTAVKPVEKAPPS[+568]KAAPPAAASQA.A z=3,scan#=20755,scan time=45.2678
FAPDTAVKPVEKAPPSKAAPPAAASQA
Intensity
m/z
HexNAc
HexNAcHex
y8
Pep_2+
~y14
~y23++
~y25++
~y19
~b19
~y17
~y16
Pep+HexNAc_2+
b2
b4
a5
b6-18
b6
y7
b7
y9
b8
b9-18
~y20++
b10
b11
b12-18
b12
b13
y2
y3
y5
~b18
~a19
1.000e+6
8.000e+5
6.000e+5
4.000e+5
2.000e+5
0.000e+0
500
1000
1500

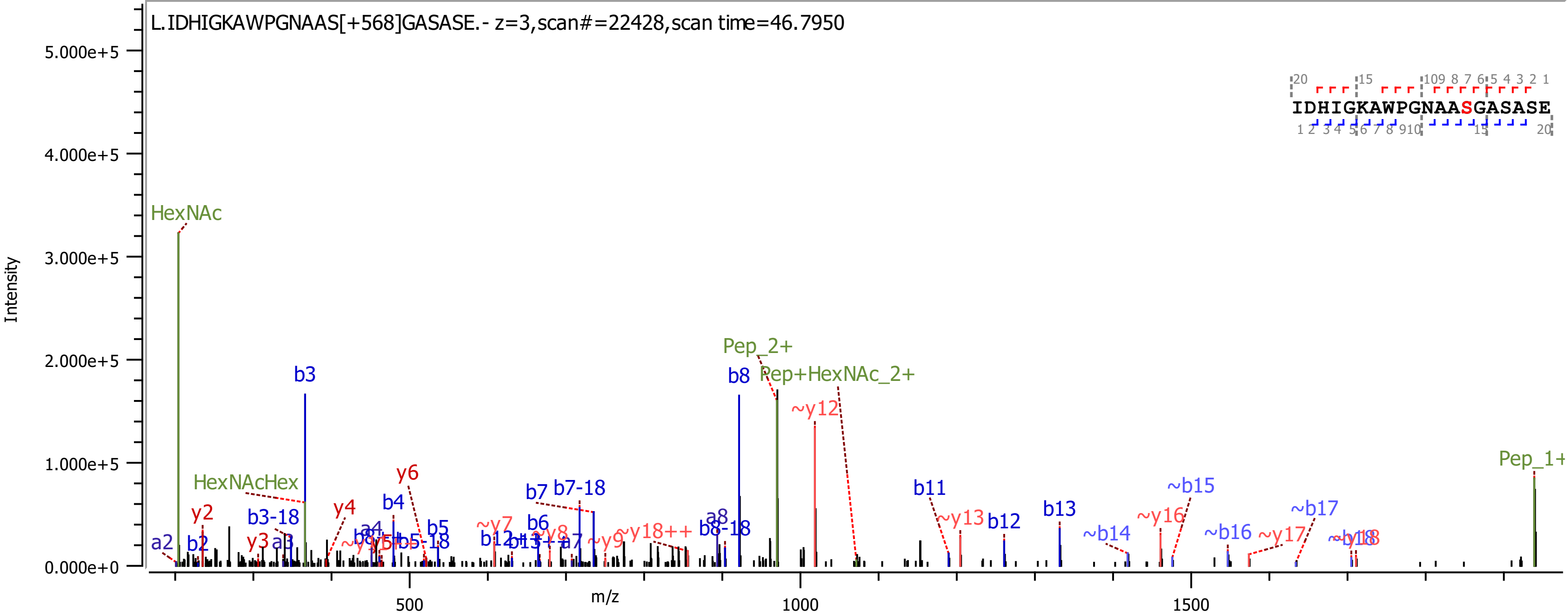

L.IDHIGKAWPGNAAS[+568]GASASE.- z=3,scan#=22428,scan time=46.7950
IDHIGKAWPGNAAS GASASE
Intensity
m/z
5.000e+5
4.000e+5
3.000e+5
2.000e+5
1.000e+5
0.000e+0
500
1000
1500
HexNAc
HexNAcHex
Pep_2+
Pep+HexNAc_2+
Pep_1+
b3
b8
~y12
b11
b13
~b15
~b17
y6
b7
b7-18
y4
b4
y2
b3-18
a2
b2
y3
a3
b5
b5-18
~y7
b6
b12
~y13
~y16
~b14
~b16
~y17
a8
b8-18
~y18++
~y9
~y8
a7

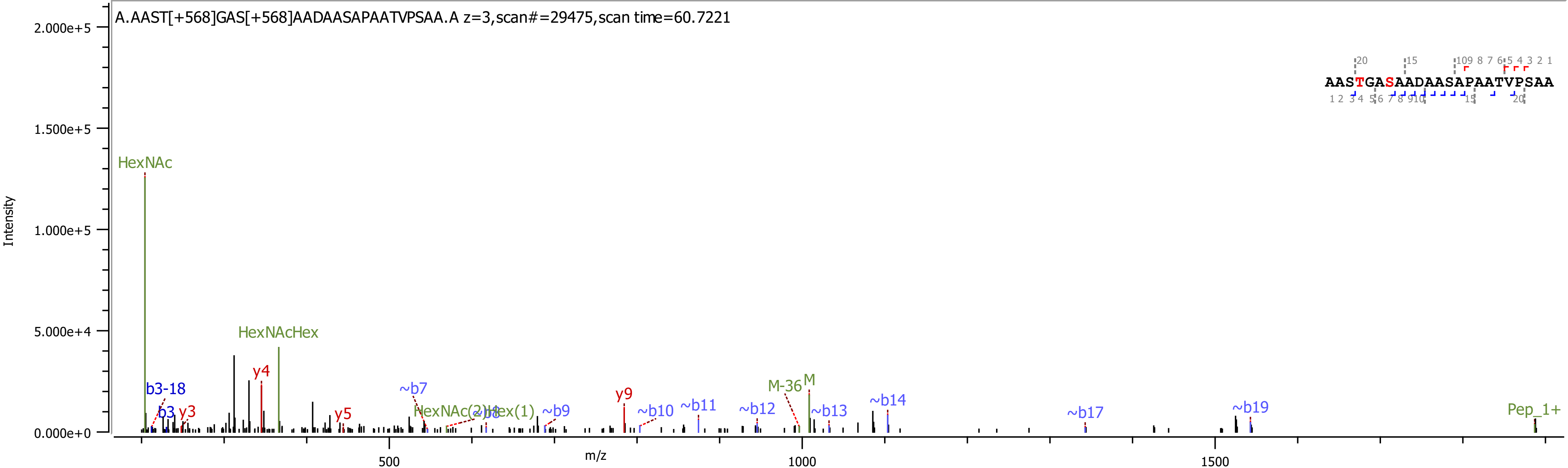

A.AAST[+568]GAS[+568]AADAASAPAATVPSAA.A z=3,scan#=29475,scan time=60.7221
AASTGASAADAASAPAATVPSAA
Intensity
2.000e+5
1.500e+5
1.000e+5
5.000e+4
0.000e+0
m/z
500
1000
1500
HexNAc
HexNAcHex
b3-18
b3
y3
y4
y5
~b7
HexNAc(2)Hex(1)
~b8
~b9
y9
~b10
~b11
~b12
M-36
M
~b13
~b14
~b17
~b19
Pep_1+

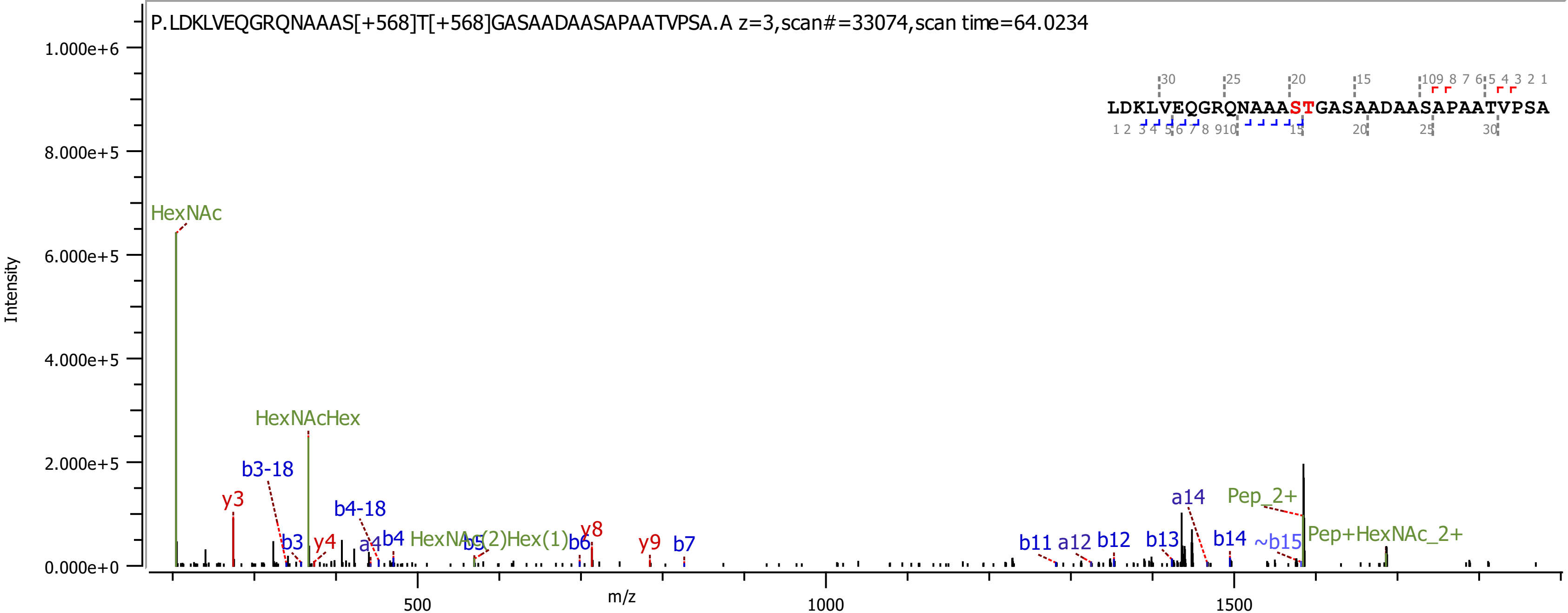

P.LDKLVEQGRQNAAAS[+568]T[+568]GASAADAASAPAATVPSA.A z=3,scan#=33074,scan time=64.0234
LDKLVEQGRQNAAASTGASAADAASAPAATVPSA
Intensity
m/z
HexNAc
HexNAcHex
b3-18
y3
b4-18
b3
y4
a4
b4
HexNAc(2)Hex(1)
b5
b6
y8
y9
b7
b11
a12
b12
b13
a14
b14
~b15
Pep_2+
Pep+HexNAc_2+

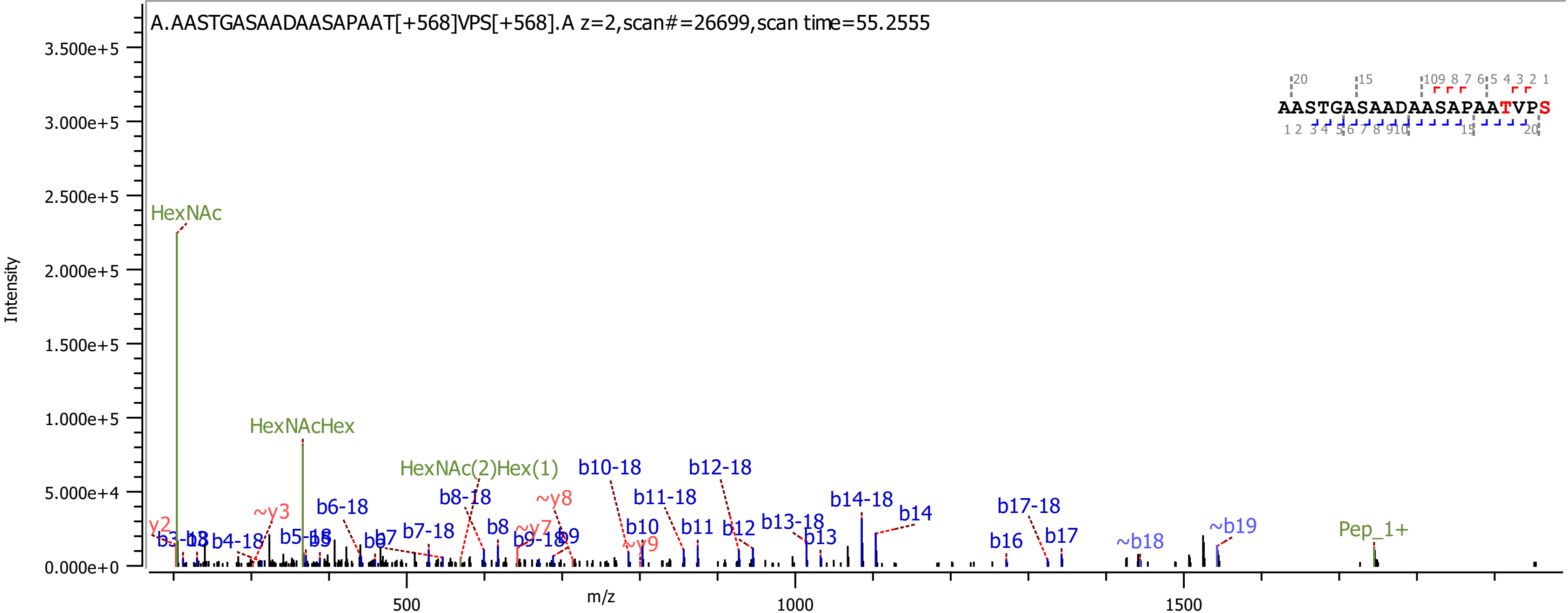
A.AASTGASAADAASAPAAT[+568]VPS[+568].A z=2,scan#=26699,scan time=55.2555
AASTGASAADAASAPAATVPS
Intensity
m/z
HexNAc
HexNAcHex
HexNAc(2)Hex(1)
Pep_1+
b10-18
b12-18
b11-18
b14-18
b14
b17-18
b8-18
b6-18
~y8
~y3
b13-18
b13
b8
b10
b11
b12
b16
b17
~b18
~b19
b7-18
b7
b9
y2
b4-18
b5-18
~y9
~y7

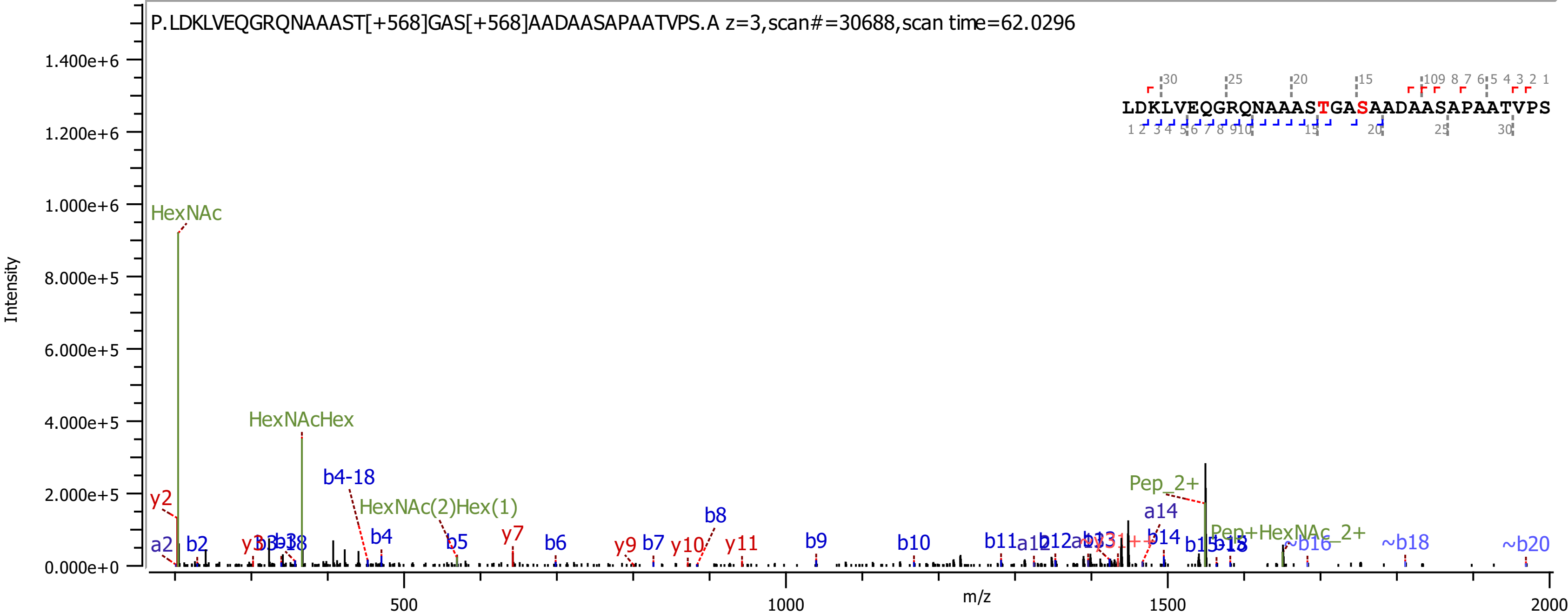

P.LDKLVEQGRQNAAAST[+568]GAS[+568]AADAASAPAATVPS.A z=3,scan#=30688,scan time=62.0296
LDKLVEQGRQNAAASTGASAADAASAPAATVPS
Intensity
m/z
HexNAc
HexNAcHex
HexNAc(2)Hex(1)
Pep_2+
Pep+HexNAc_2+
y2
a2
b2
b4-18
b4
b5
y7
b6
y9
b7
y10
b8
y11
b9
b10
b11
a14
b14
~b16
~b18
~b20

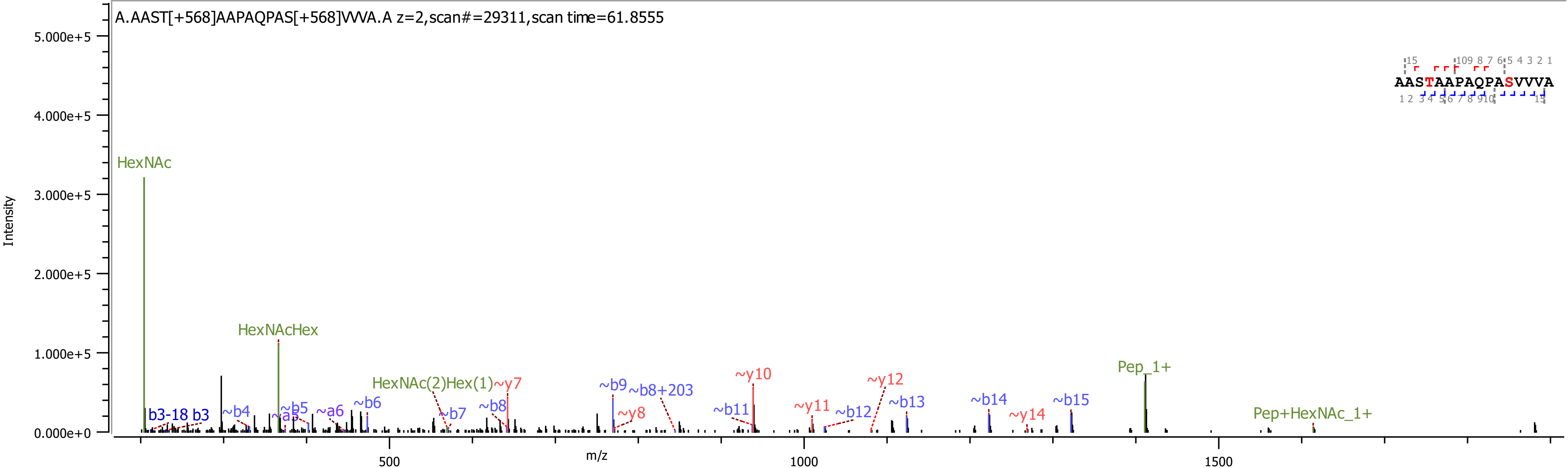
A.AAST[+568]AAPAQPAS[+568]VVVA.A z=2,scan#=29311,scan time=61.8555
AASTAAPAQPASVVVA
Intensity
5.000e+5
4.000e+5
3.000e+5
2.000e+5
1.000e+5
0.000e+0
m/z
500
1000
1500
HexNAc
HexNAcHex
HexNAc(2)Hex(1)
b3-18 b3
~b4
~a5
~b5
~a6
~b6
~b7
~b8
~y7
~b9
~b8+203
~y8
~b11
~y10
~y11
~b12
~y12
~b13
~b14
~y14
~b15
Pep_1+
Pep+HexNAc_1+

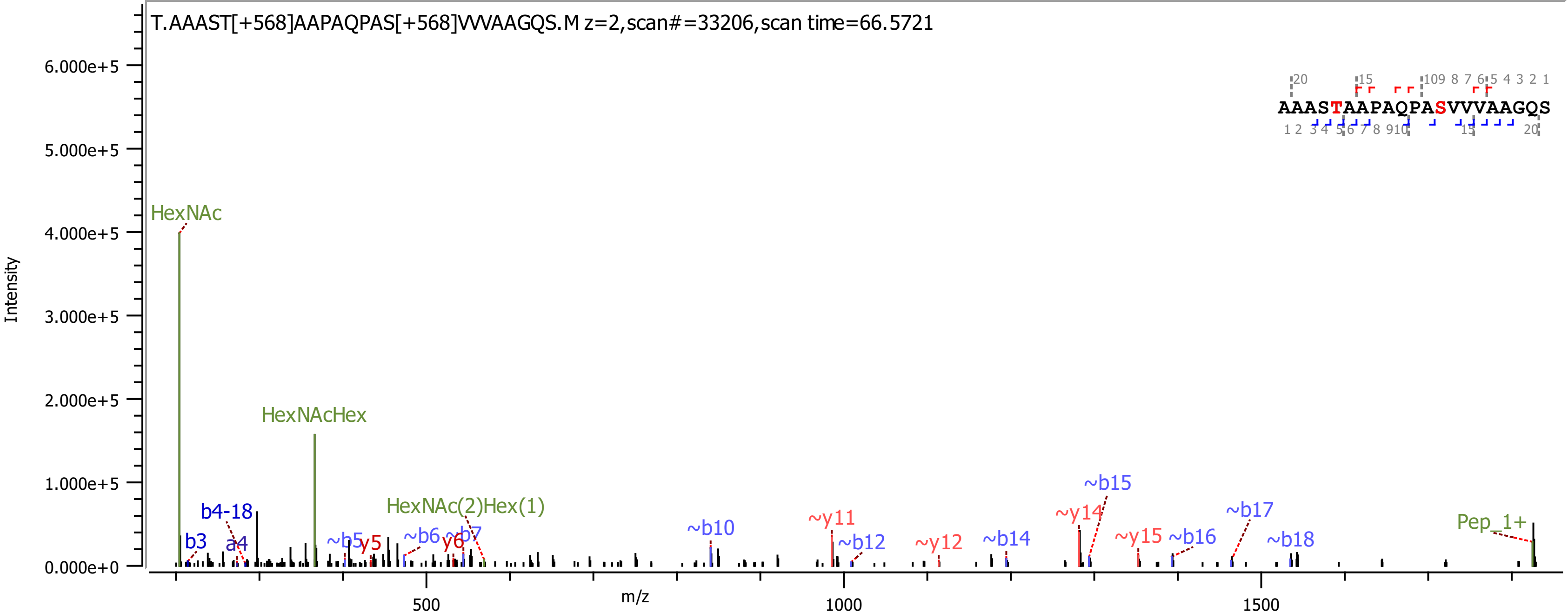

T.AAAST[+568]AAPAQPAS[+568]VVVAAGQS.M z=2,scan#=33206,scan time=66.5721
Intensity
m/z
6.000e+5
5.000e+5
4.000e+5
3.000e+5
2.000e+5
1.000e+5
0.000e+0
500
1000
1500
HexNAc
HexNAcHex
HexNAc(2)Hex(1)
b4-18
b3
a4
~b5
y5
~b6
y6
~b7
~b10
~y11
~b12
~y12
~b14
~y14
~b15
~y15
~b16
~b17
~b18
Pep_1+
AAASTAAPAQPASVVVAAGQS

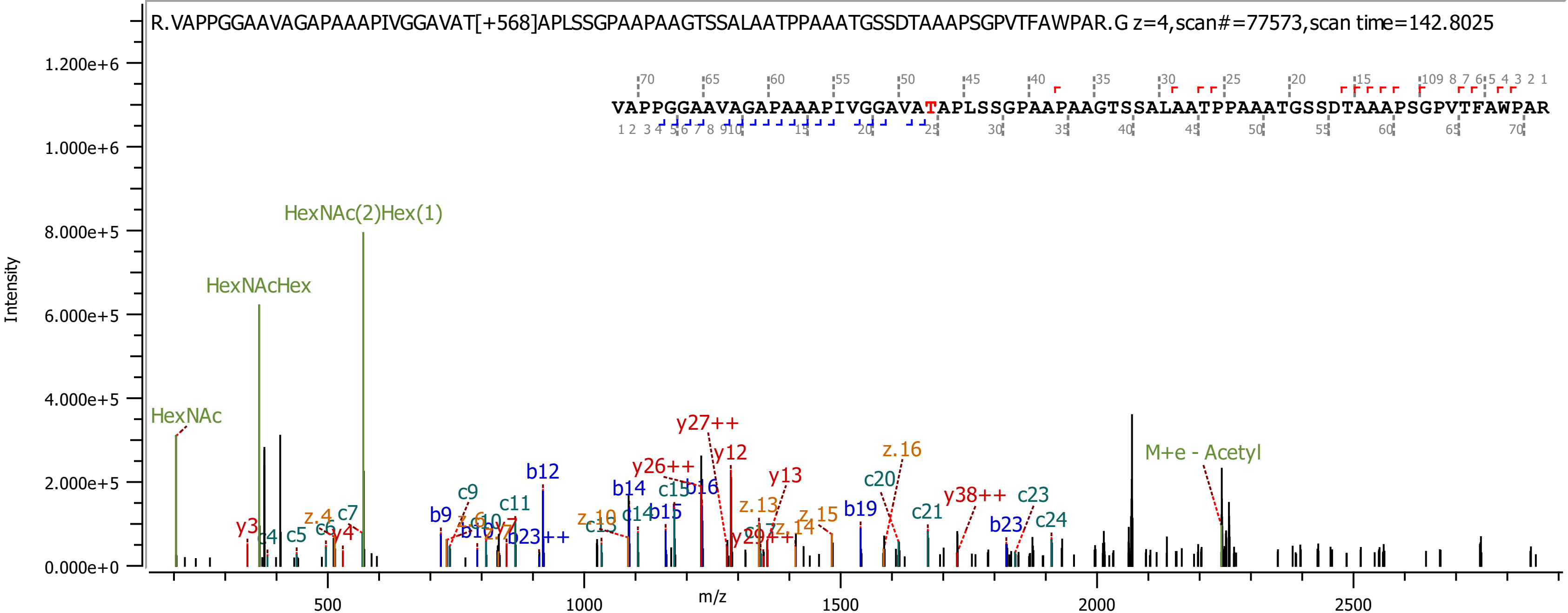

R.VAPPGGAAVAGAPAAAPIVGGAVAT[+568]APLSSGPAAPAAGTSSALAATPPAAATGSSDTAAAPSGPVTFAWPAR.G z=4,scan#=77573,scan time=142.8025
VAPPGGAAVAGAPAAAPIVGGAVATAPLSSGPAAPAAGTSSALAATPPAAATGSSDTAAAPSGPVTFAWPAR
Intensity
m/z
HexNAc
HexNAcHex
HexNAc(2)Hex(1)
M+e - Acetyl
y27++
y26++
y12
y13
y38++
b12
b14
b16
b19
b23
c20
c21
c23
c24
z.16
z.15
z.13

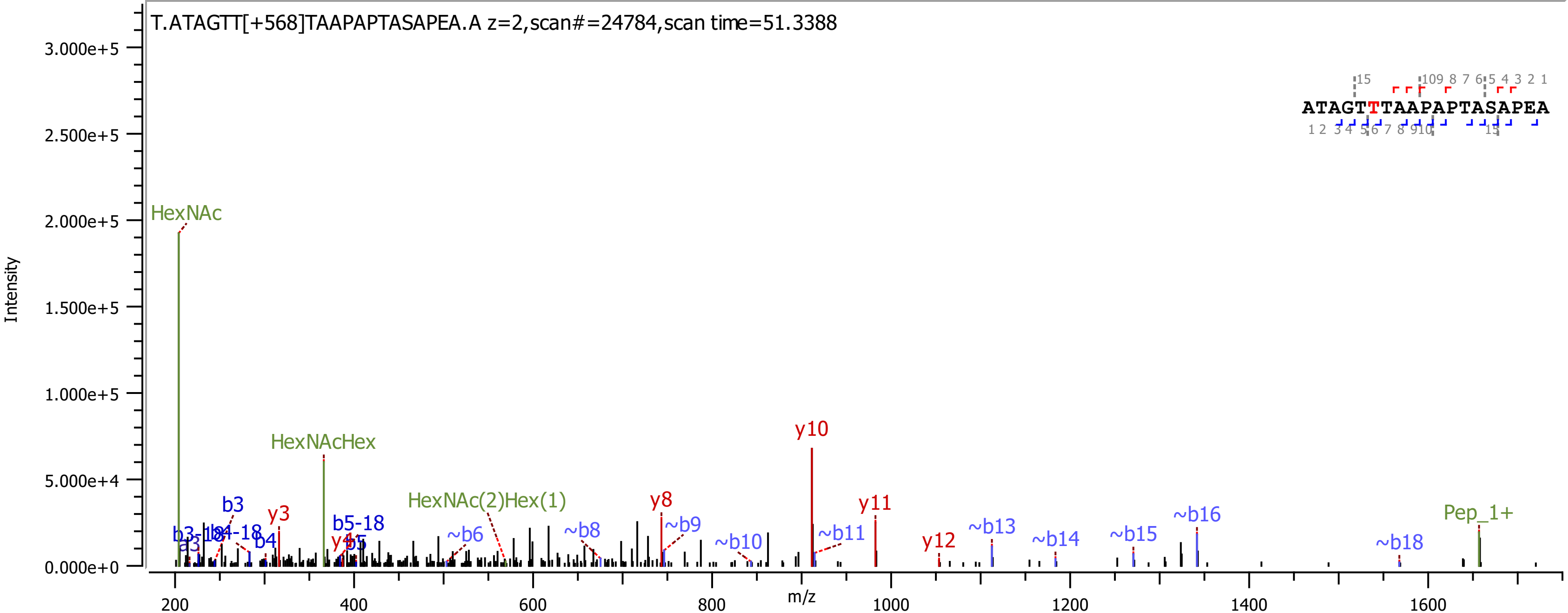

T.ATAGTT[+568]TAAPAPTASAPEA.A z=2,scan#=24784,scan time=51.3388
ATAGTTTAAPAPTASAPEA
Intensity
m/z
HexNAc
HexNAcHex
HexNAc(2)Hex(1)
Pep_1+
y3
y8
y10
y11
y12
b3
b4
b5-18
~b6
~b8
~b9
~b10
~b11
~b13
~b14
~b15
~b16
~b18

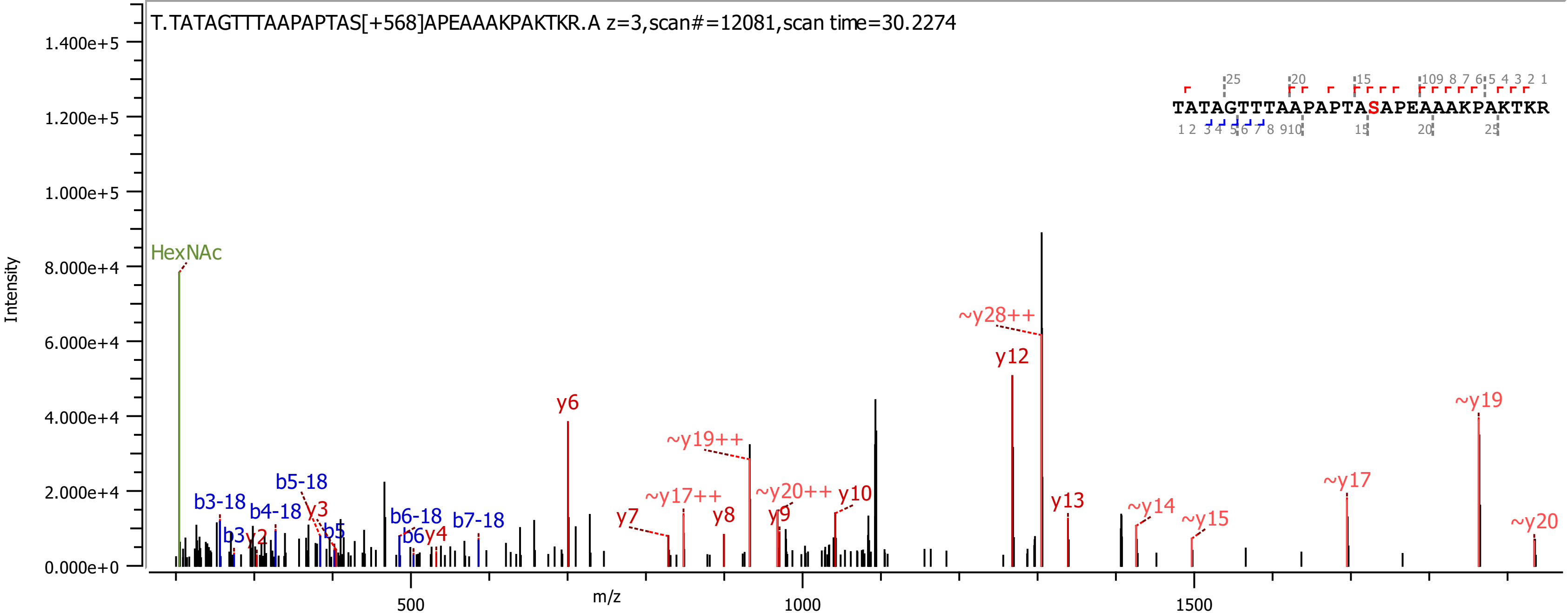

T.TATAGTTTAAPAPTAS[+568]APEAAAKPAKTKR.A z=3,scan#=12081,scan time=30.2274
TATAGTTTAAPAPTASAPEAAAKPAKTKR
Intensity
m/z
1.400e+5
1.200e+5
1.000e+5
8.000e+4
6.000e+4
4.000e+4
2.000e+4
0.000e+0
500
1000
1500
HexNAc
b3-18
b3
y2
b4-18
y3
b5-18
b5
b6-18
b6
y4
b7-18
y6
y7
~y17++
~y19++
y8
y9
~y20++
y10
~y28++
y12
y13
~y14
~y15
~y17
~y19
~y20

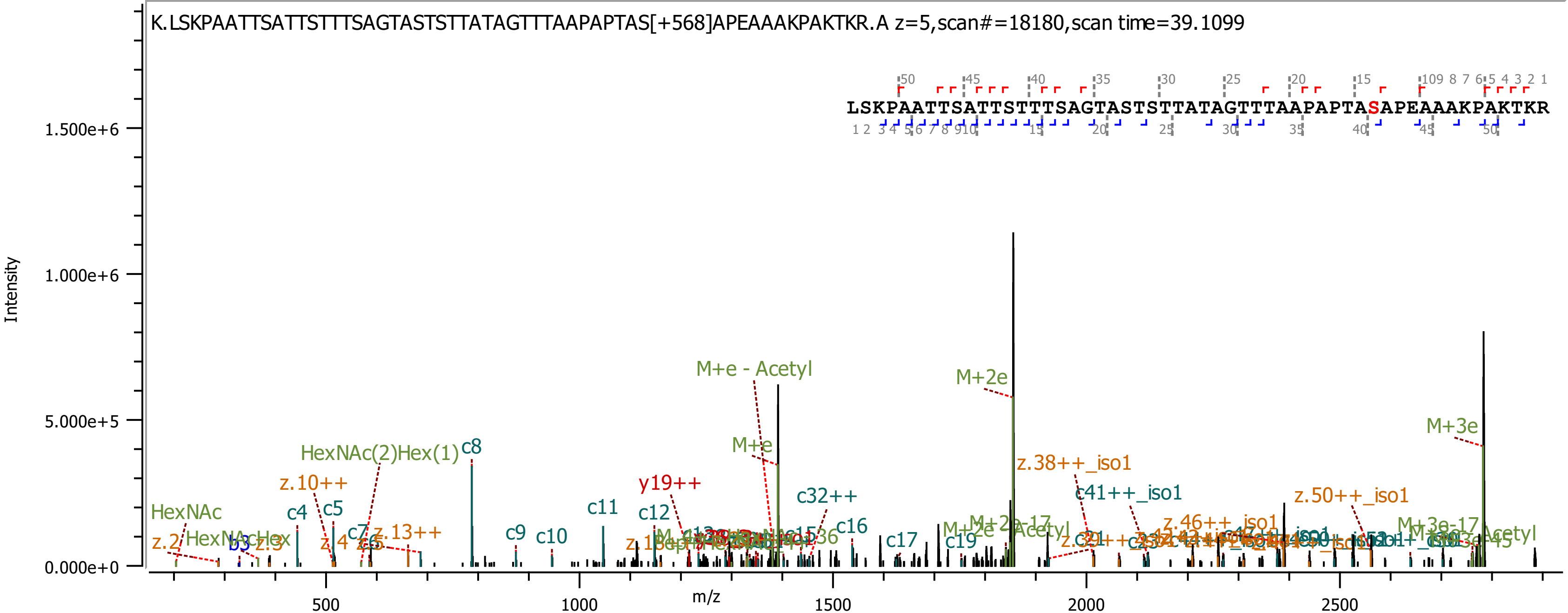
K.LSKPAATTSATTSTTTSAGTASTSTTATAGTTTAAPAPTAS[+568]APEAAAKPAKTKR.A z=5,scan#=18180,scan time=39.1099
LSKPAATTSATTSTTTSAGTASTSTTATAGTTTAAPAPTASAPEAAAKPAKTKR
Intensity
m/z
1.500e+6
1.000e+6
5.000e+5
0.000e+0
500
1000
1500
2000
2500
M+e - Acetyl
M+2e
M+3e
M+e
HexNAc(2)Hex(1)
c8
z.10++
c4
c5
HexNAc
z.2
z.13++
c9
c10
c11
c12
y19++
c32++
c16
c17
c19
z.38++_iso1
c41++_iso1
z.50++_iso1
z.46++_iso1
M+3e-17
M+2e-17
Acetyl

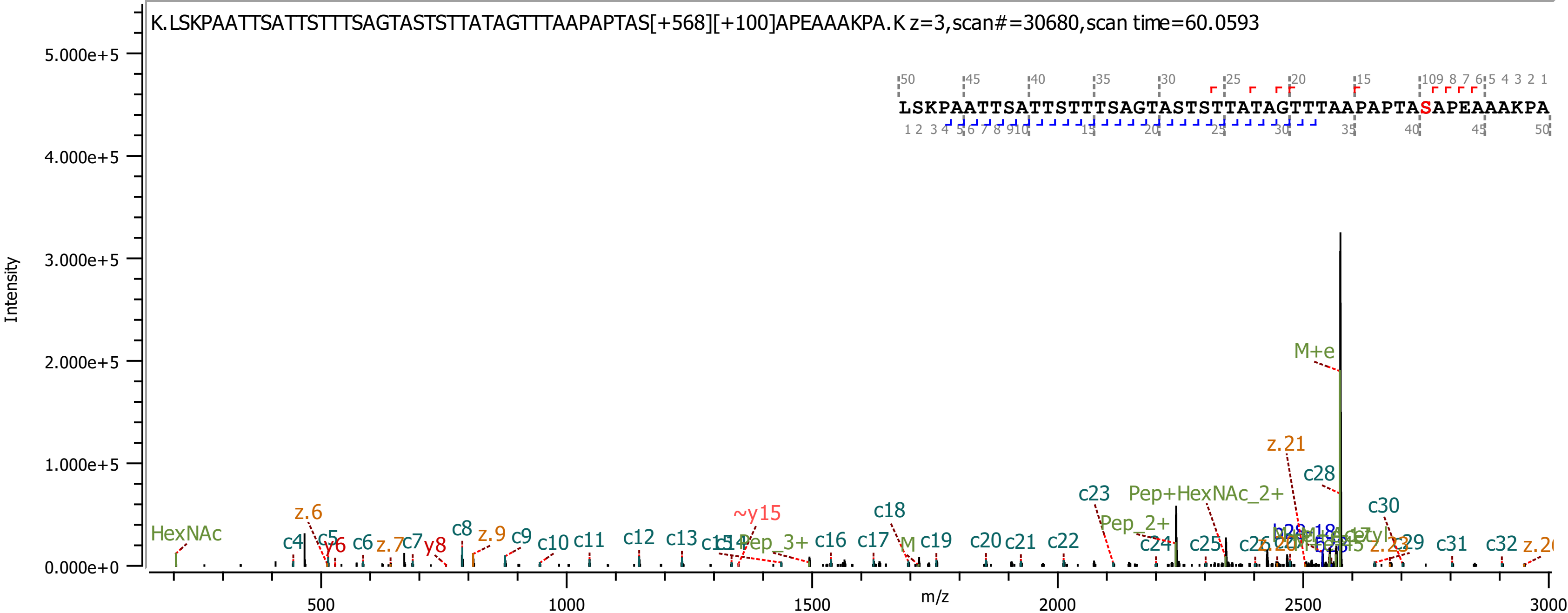
K.LSKPAATTSATTSTTTSAGTASTSTTATAGTTTAAPAPTAS[+568][+100]APEAAAKPA.K z=3,scan#=30680,scan time=60.0593
LSKPAATTSATTSTTTSAGTASTSTTATAGTTTAAPAPTASAPEAAAKPA
Intensity
m/z
5.000e+5
4.000e+5
3.000e+5
2.000e+5
1.000e+5
0.000e+0
500
1000
1500
2000
2500
3000
HexNAc
z.6
c4
c5
y6
c6
z.7
c7
y8
c8
z.9
c9
c10
c11
c12
c13
~y15
Pep_3+
c16
c17
c18
M
c19
c20
c21
c22
c23
Pep_2+
c24
Pep+HexNAc_2+
c25
M+e
z.21
c28
c30
c29
c31
c32

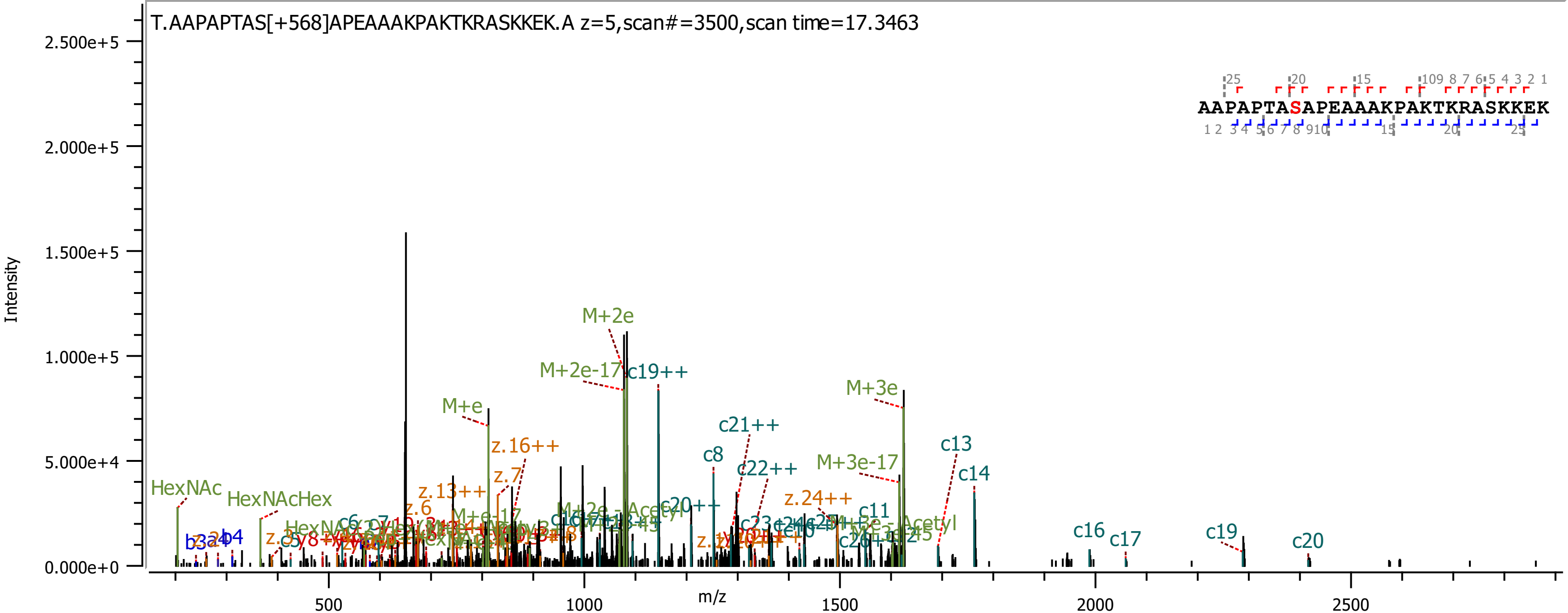
T.AAPAPTAS[+568]APEAAAKPAKTKRASKKEK.A z=5,scan#=3500,scan time=17.3463
AAPAPTASAPEAAAKPAKTKRASKKEK
Intensity
m/z
2.500e+5
2.000e+5
1.500e+5
1.000e+5
5.000e+4
0.000e+0
500
1000
1500
2000
2500
M+2e
M+2e-17
c19++
M+3e
M+e
c21++
z.16++
c8
c22++
M+3e-17
c13
c14
z.7
HexNAc
HexNAcHex
z.13++
z.24++
c11
c16
c17
c19
c20
z.6
c20++
b4
b3
c6

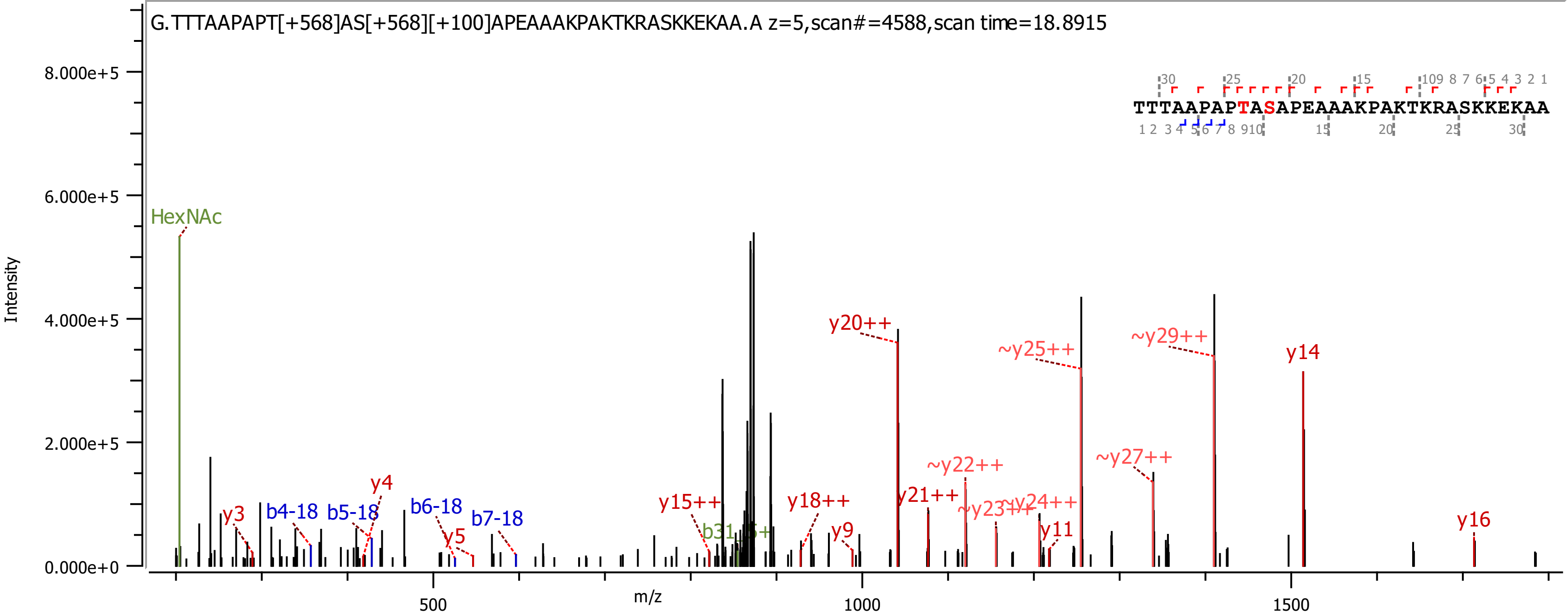

G.TTTAAPAPT[+568]AS[+568][+100]APEAAAKPAKTKRASKKEKAA.A z=5,scan#=4588,scan time=18.8915
TTTAAPAPTASAPEAAAKPAKTKRASKKEKAA
Intensity
m/z
HexNAc
y3
b4-18
b5-18
y4
b6-18
y5
b7-18
y15++
y18++
y9
y20++
y21++
~y22++
~y23++
~y24++
y11
~y25++
~y27++
~y29++
y14
y16

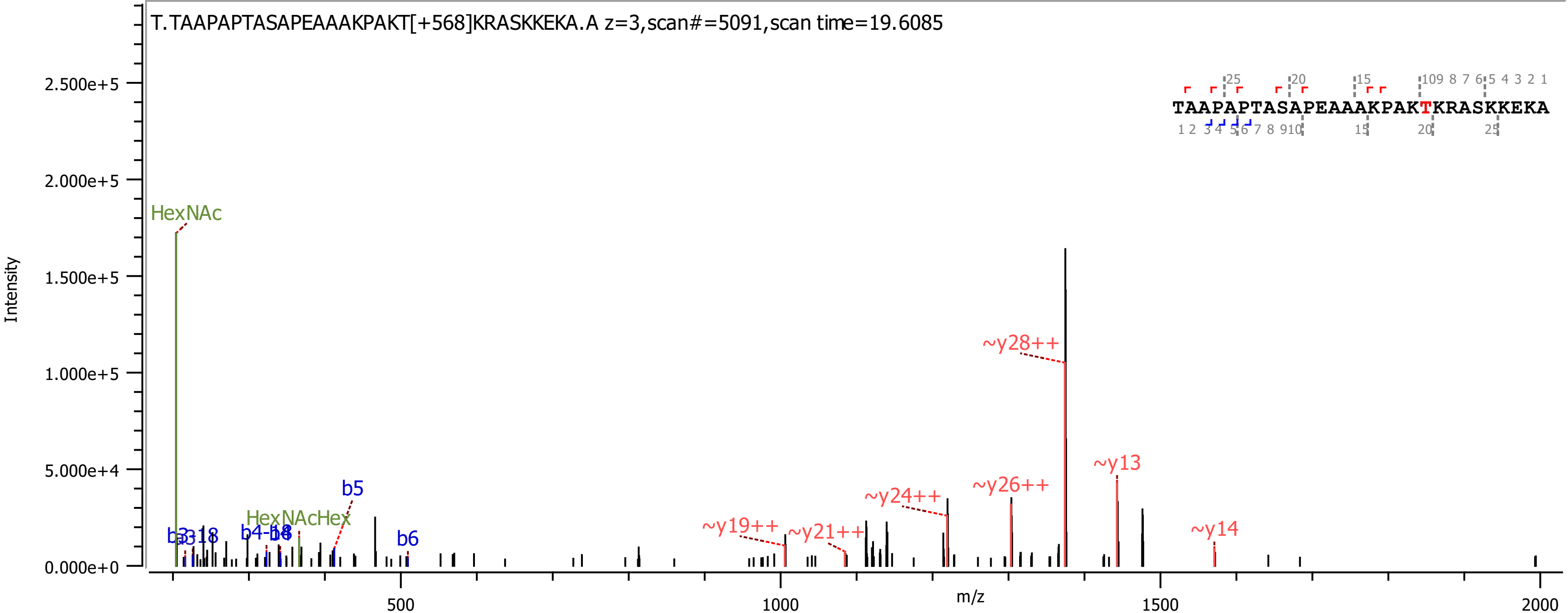

T.TAAPAPTASAPEAAAKPAKT[+568]KRASKKEKA.A z=3,scan#=5091,scan time=19.6085
TAAPAPTASAPEAAAKPAKTKRASKKEKA
Intensity
m/z
HexNAc
HexNAcHex
b5
b6
~y19++
~y21++
~y24++
~y26++
~y28++
~y13
~y14

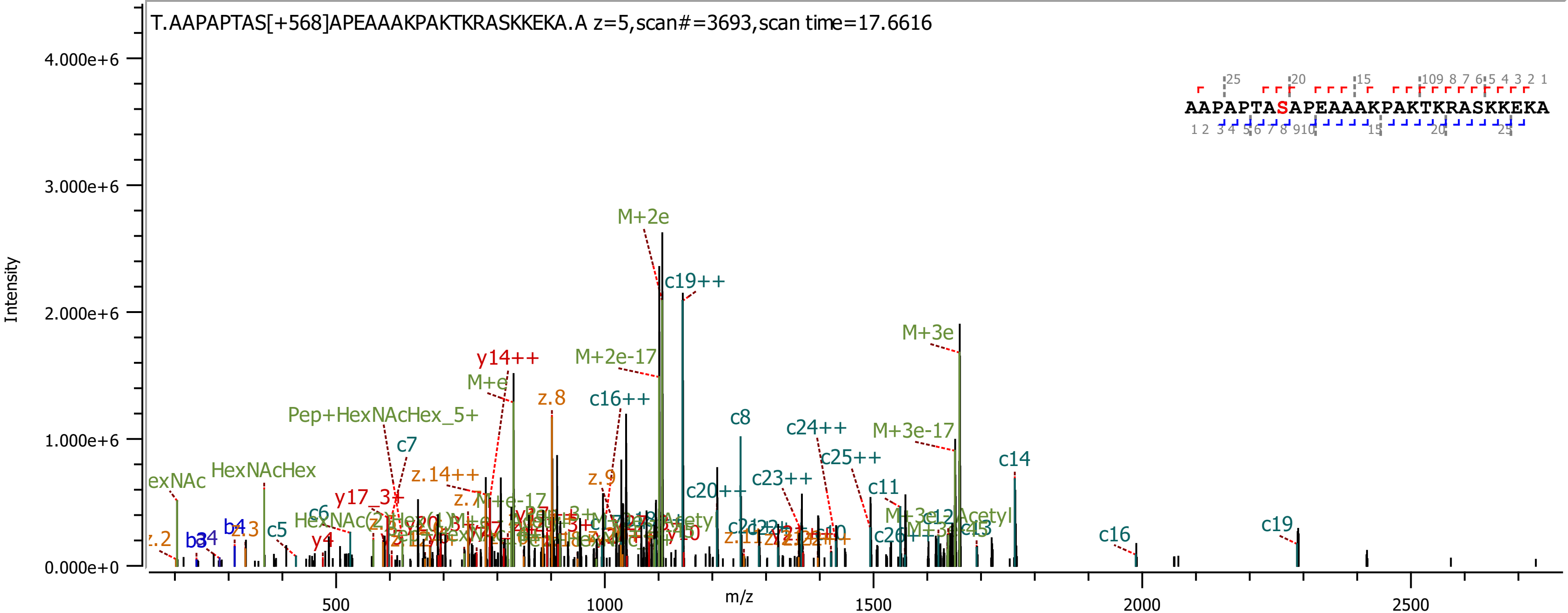
T.AAPAPTAS[+568]APEAAAKPAKTKRASKKEKA.A z=5,scan#=3693,scan time=17.6616
AAPAPTASAPEAAAKPAKTKRASKKEKA
Intensity
4.000e+6
3.000e+6
2.000e+6
1.000e+6
0.000e+0
m/z
500
1000
1500
2000
2500
M+2e
c19++
M+3e
M+2e-17
y14++
M+e
z.8
c16++
Pep+HexNAcHex_5+
c7
c8
c24++
M+3e-17
c25++
c14
HexNAcHex
z.14++
z.9
c20++
c23++
c11
y17_3+
c6
c5
b4
z.3
y4
c16
c19

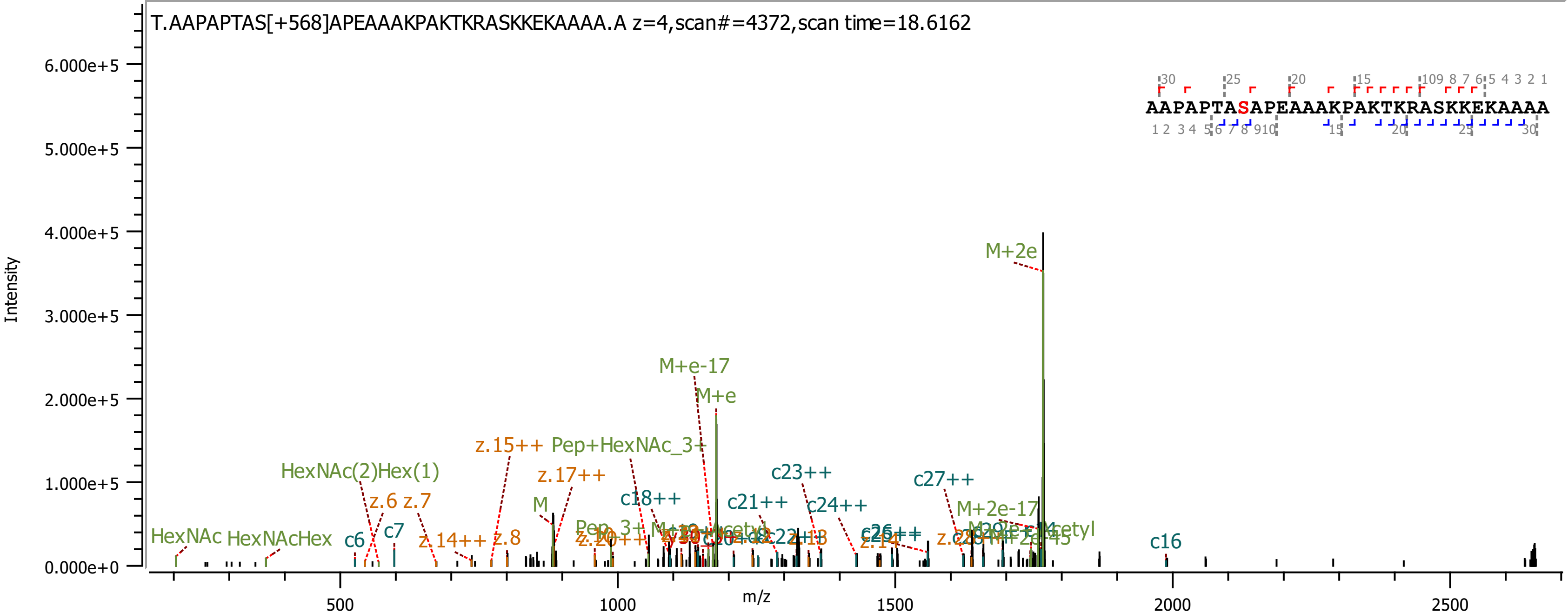
T.AAPAPTAS[+568]APEAAAKPAKTKRASKKEKAAAA.A z=4,scan#=4372,scan time=18.6162
AAPAPTASAPEAAAKPAKTKRASKKEKAAAA
Intensity
6.000e+5
5.000e+5
4.000e+5
3.000e+5
2.000e+5
1.000e+5
0.000e+0
m/z
500
1000
1500
2000
2500
M+2e
M+e-17
M+e
z.15++
Pep+HexNAc_3+
z.17++
HexNAc(2)Hex(1)
z.6
z.7
M
c18++
c21++
c23++
c24++
c27++
M+2e-17
HexNAc
HexNAcHex
c6
c7
z.14++
z.8
c16

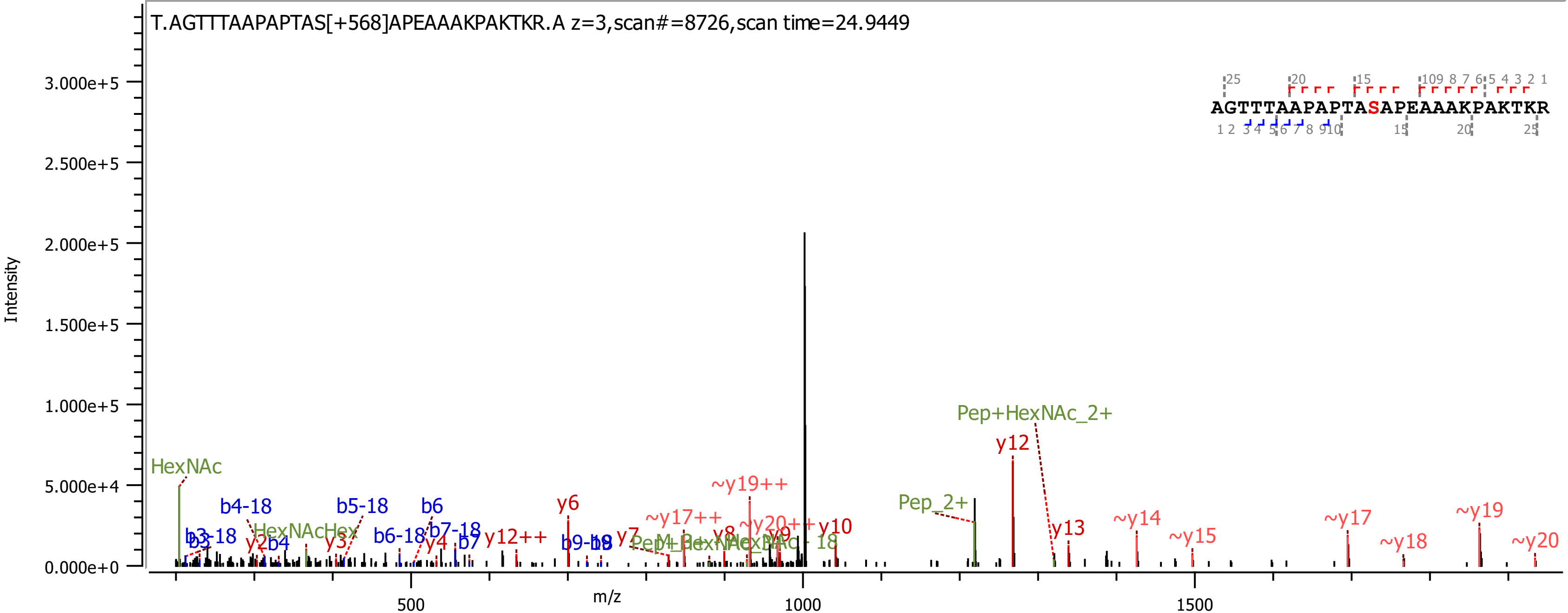

T.AGTTTAAPAPTAS[+568]APEAAAKPAKTKR.A z=3,scan#=8726,scan time=24.9449
AGTTTAAPAPTASAPEAAAKPAKTKR
Intensity
m/z
HexNAc
b4-18
b5-18
b6
b3-18
b3
y2
b4
HexNAcHex
y3
b6-18
b7-18
y4
b7
y12++
y6
b9-18
y7
~y17++
~y19++
~y20++
y8
y9
y10
Pep_2+
Pep+HexNAc_2+
y12
y13
~y14
~y15
~y17
~y18
~y19
~y20

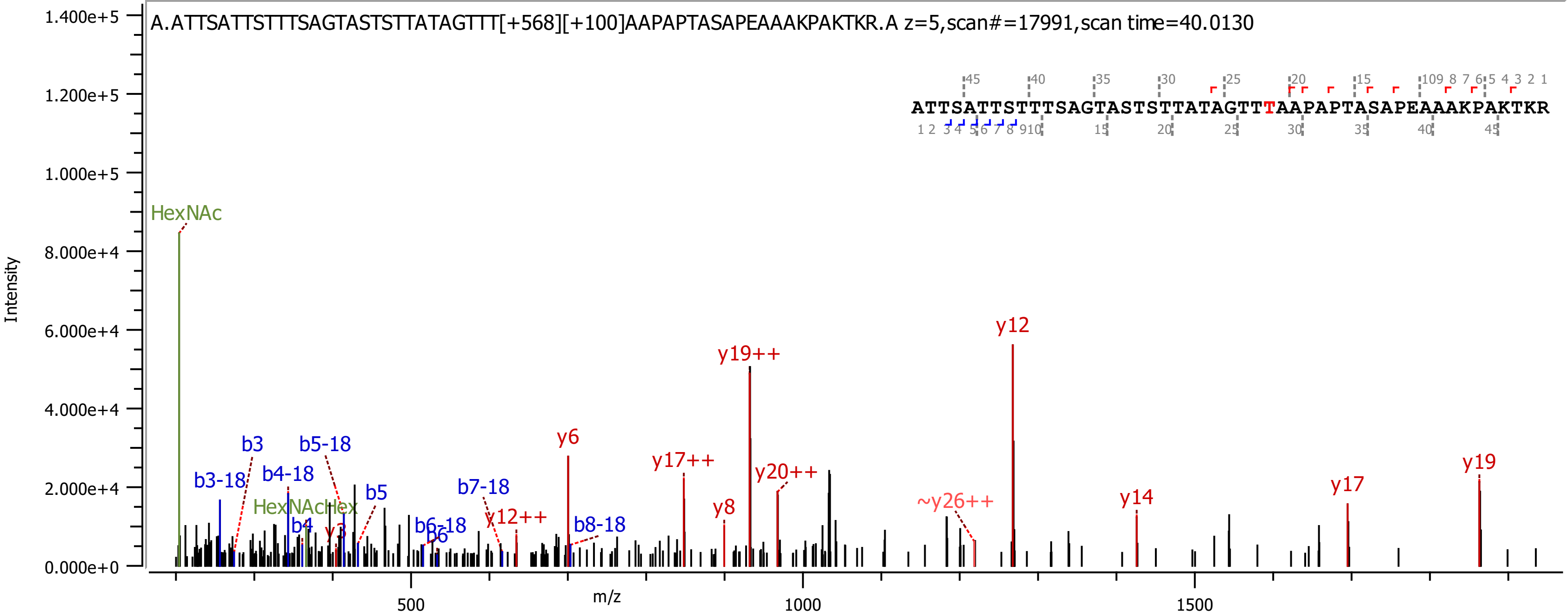
A.ATTSATTSTTTSAGTASTSTTATAGTTT[+568][+100]AAPAPTASAPEAAAKPAKTKR.A z=5,scan#=17991,scan time=40.0130
ATTSATTSTTTSAGTASTSTTATAGTTTAAPAPTASAPEAAAKPAKTKR
Intensity
m/z
HexNAc
HexNAcHex
b3-18
b3
b4-18
b4
b5-18
b5
b6-18
b6
b7-18
b8-18
y3
y6
y8
y12
y12++
y14
y17
y17++
y19
y19++
y20++
~y26++

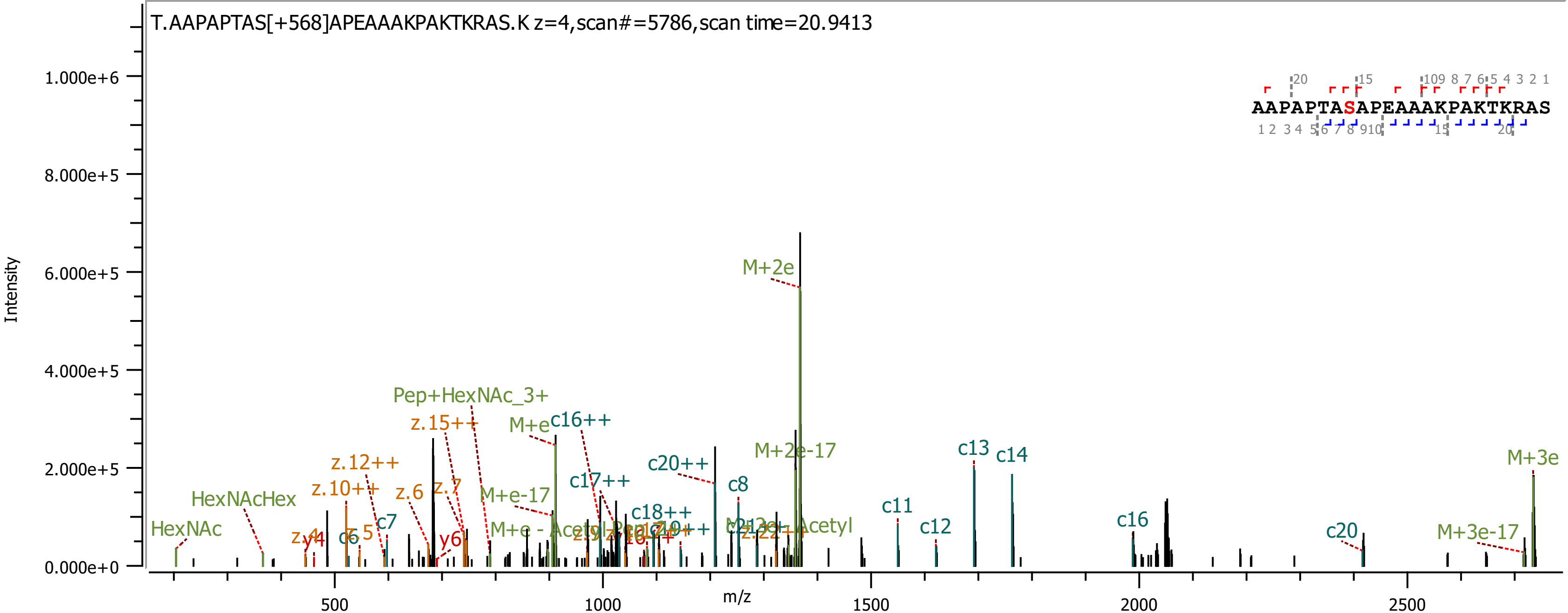

T.AAPAPTAS[+568]APEAAAKPAKTKRAS.K z=4,scan#=5786,scan time=20.9413
AAPAPTASAPEAAAKPAKTKRAS
Intensity
0.000e+0
2.000e+5
4.000e+5
6.000e+5
8.000e+5
1.000e+6
m/z
500
1000
1500
2000
2500
HexNAc
HexNAcHex
z.12++
z.10++
z.6
z.7
z.15++
Pep+HexNAc_3+
M+e
c16++
M+e-17
c17++
c20++
c18++
c8
M+2e
M+2e-17
c7
c6
z.5
y6
c11
c12
c13
c14
c16
c20
M+3e-17
M+3e

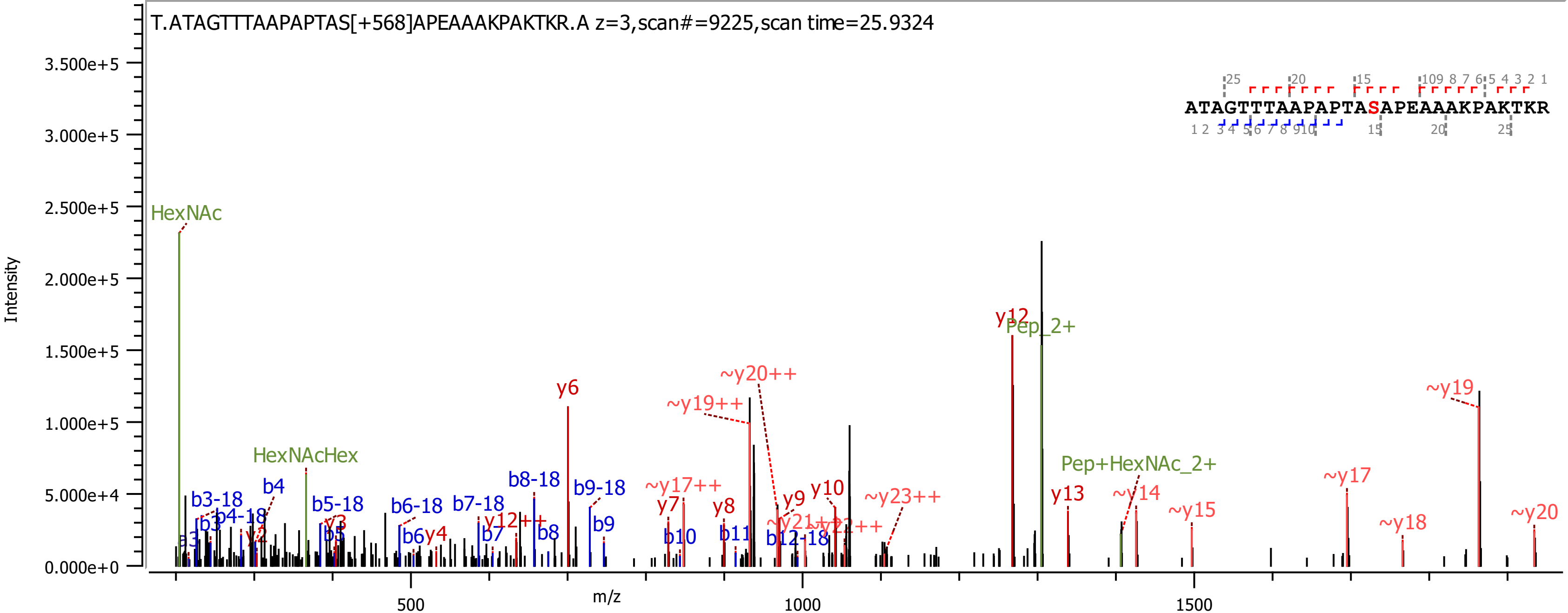
T.ATAGTTTAAPAPTAS[+568]APEAAAKPAKTKR.A z=3,scan#=9225,scan time=25.9324
ATAGTTTAAPAPTASAPEAAAKPAKTKR
Intensity
m/z
0.000e+0
5.000e+4
1.000e+5
1.500e+5
2.000e+5
2.500e+5
3.000e+5
3.500e+5
500
1000
1500
HexNAc
HexNAcHex
Pep_2+
Pep+HexNAc_2+
b3-18
b3
b4-18
b4
b5-18
b5
b6-18
b6
b7-18
b7
b8-18
b8
b9-18
b9
b10
b11
b12-18
y2
y3
y4
y6
y7
y8
y9
y10
y12
y12++
y13
~y14
~y15
~y17
~y17++
~y18
~y19
~y19++
~y20
~y20++
~y21++
~y23++

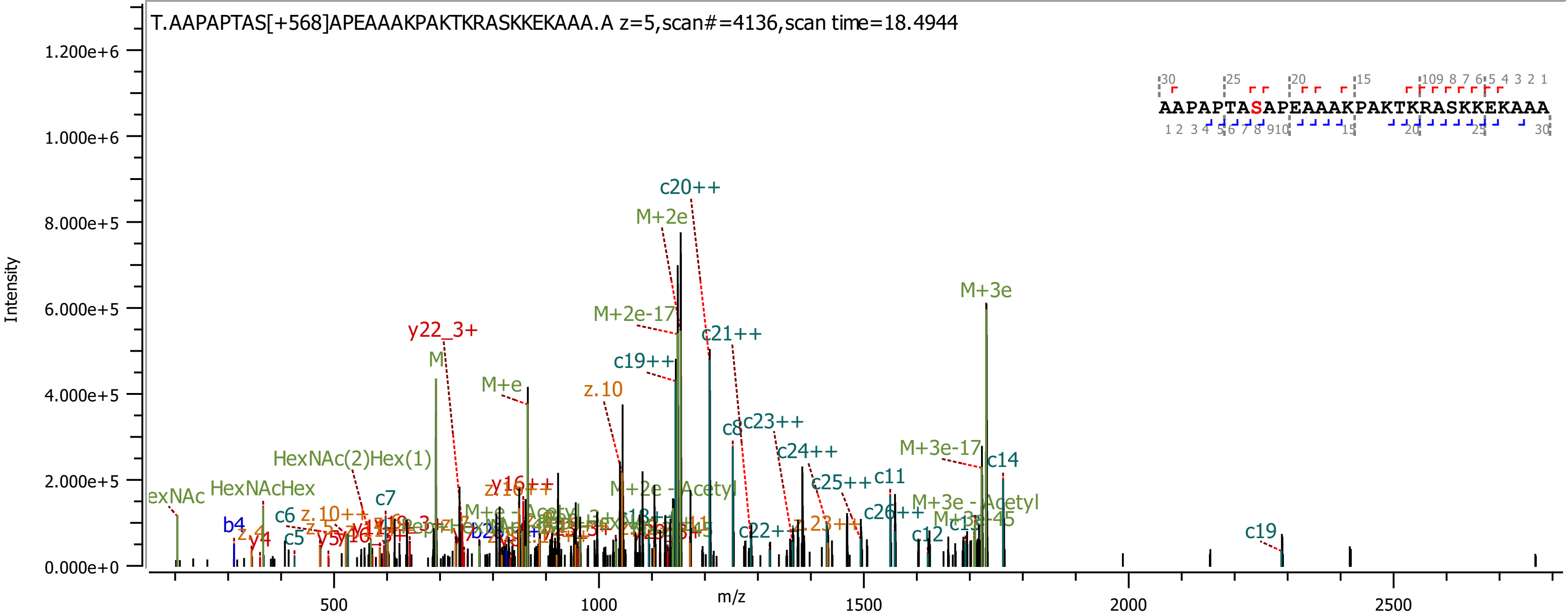

T.AAPAPTAS[+568]APEAAAKPAKTKRASKKEKAAA.A z=5,scan#=4136,scan time=18.4944
AAPAPTASAPEAAAKPAKTKRASKKEKAAA
Intensity
m/z
1.200e+6
1.000e+6
8.000e+5
6.000e+5
4.000e+5
2.000e+5
0.000e+0
500
1000
1500
2000
2500
c20++
M+2e
M+3e
M+2e-17
c21++
y22_3+
M
c19++
M+e
z.10
c8
c23++
c24++
M+3e-17
c14
HexNAc(2)Hex(1)
c11
c25++
HexNAcHex
M+2e - Acetyl
c7
M+3e - Acetyl
c26++
c6
b4
c5
c22++
c12
c19

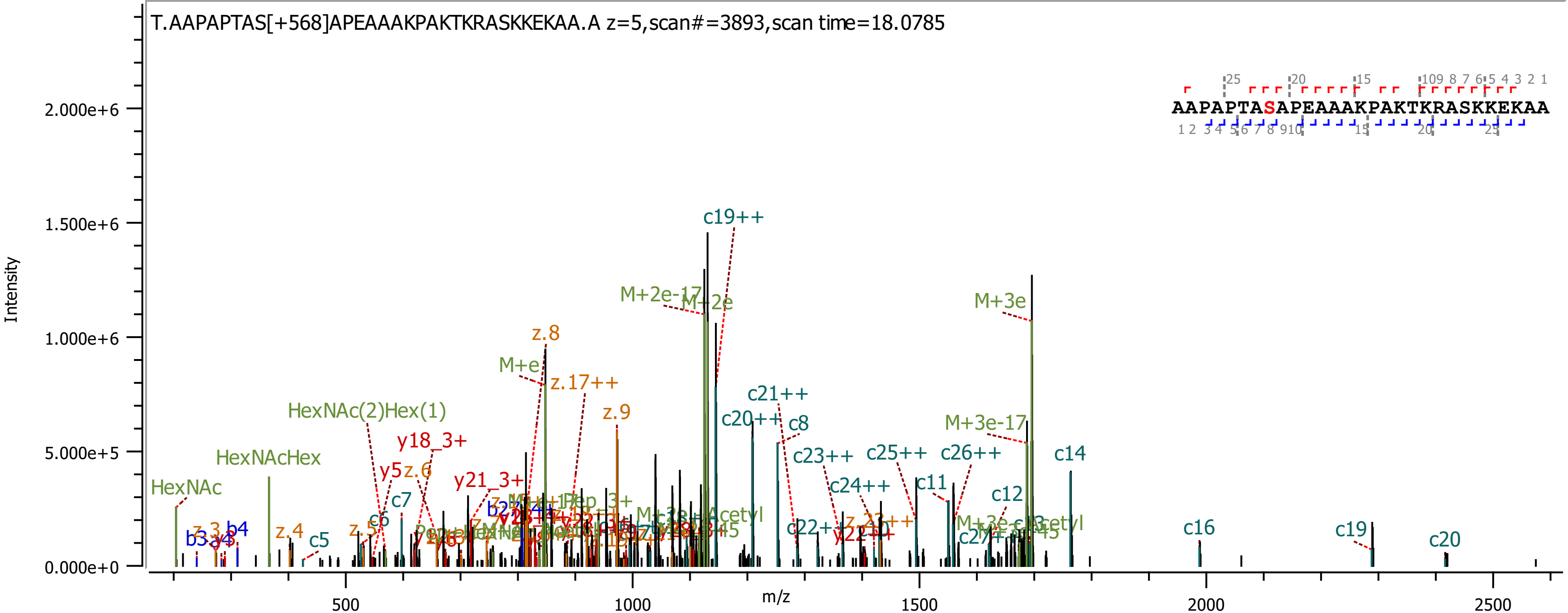
T.AAPAPTAS[+568]APEAAAKPAKTKRASKKEKAA.A z=5,scan#=3893,scan time=18.0785
AAPAPTASAPEAAAKPAKTKRASKKEKAA
Intensity
m/z
2.000e+6
1.500e+6
1.000e+6
5.000e+5
0.000e+0
500
1000
1500
2000
2500
c19++
M+2e-17
M+2e
M+3e
z.8
M+e
z.17++
z.9
c21++
c20++
c8
HexNAc(2)Hex(1)
y18_3+
M+3e-17
c14
HexNAcHex
c23++
c25++
c26++
y5
z.6
y21_3+
c24++
c11
HexNAc
c7
c12
c6
c16
c19
c20
z.3
b4
b3
z.4
c5
z.5

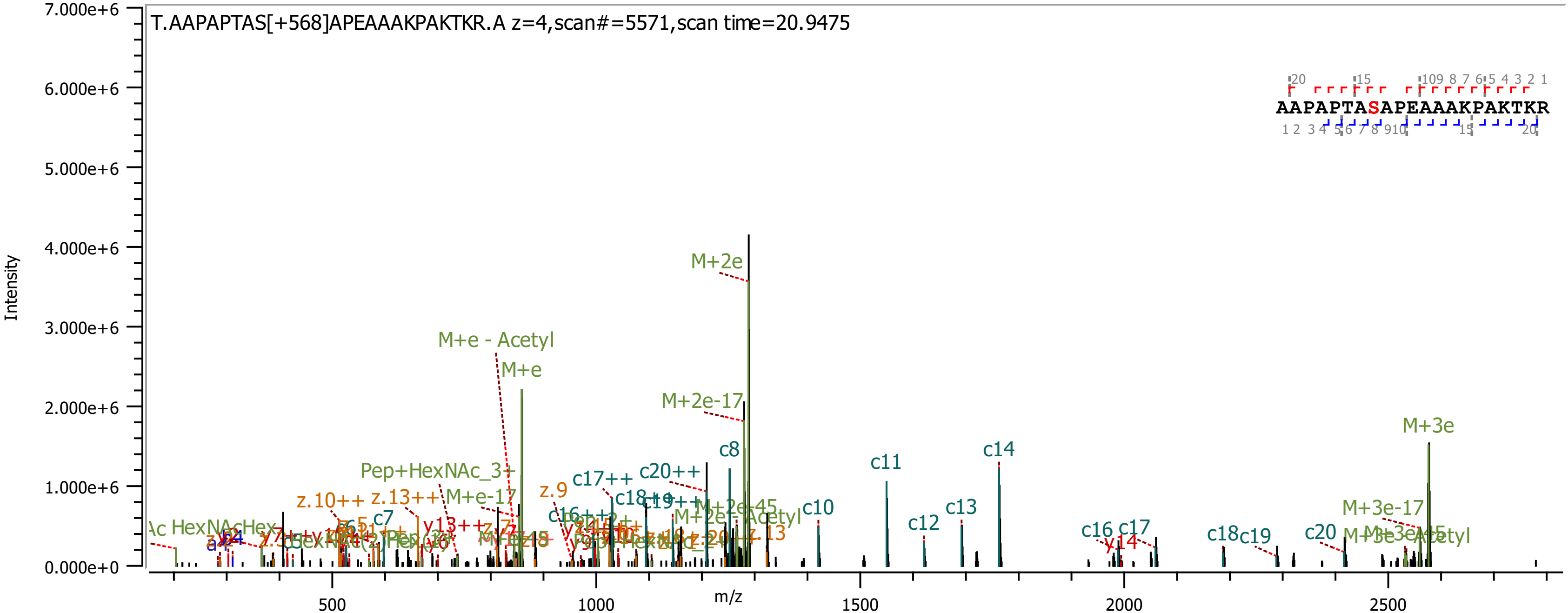
T.AAPAPTAS[+568]APEAAAKPAKTKR.A z=4,scan#=5571,scan time=20.9475
AAPAPTASAPEAAAKPAKTKR
Intensity
m/z
7.000e+6
6.000e+6
5.000e+6
4.000e+6
3.000e+6
2.000e+6
1.000e+6
0.000e+0
500
1000
1500
2000
2500
M+2e
M+e - Acetyl
M+e
M+2e-17
M+3e
c8
c11
c14
c10
c12
c13
c16
c17
c18
c19
c20
c7
c17++
c20++
z.9
z.10++
z.13++
M+e-17
Pep+HexNAc_3+
M+3e-17
y14
z.13

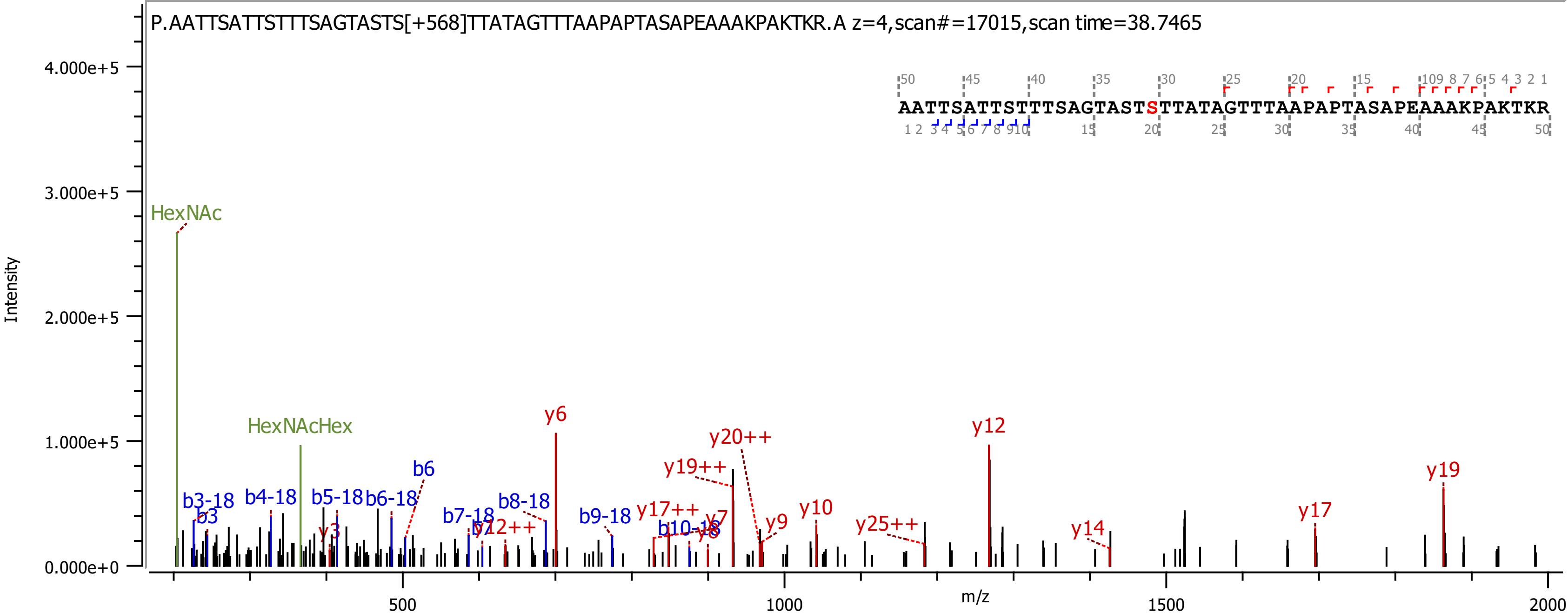

P.AATTSATTSTTTSAGTASTS[+568]TTATAGTTTAAPAPTASAPEAAAKPAKTKR.A z=4,scan#=17015,scan time=38.7465
AATTSATTSTTTSAGTASTSTTATAGTTTAAPAPTASAPEAAAKPAKTKR
HexNAc
HexNAcHex
b3-18
b3
b4-18
b5-18
b6-18
b6
y3
b7-18
b8-18
y12++
y6
b9-18
y17++
b10-18
y7
y8
y19++
y20++
y9
y10
y25++
y12
y14
y17
y19
Intensity
m/z
4.000e+5
3.000e+5
2.000e+5
1.000e+5
0.000e+0
500
1000
1500
2000

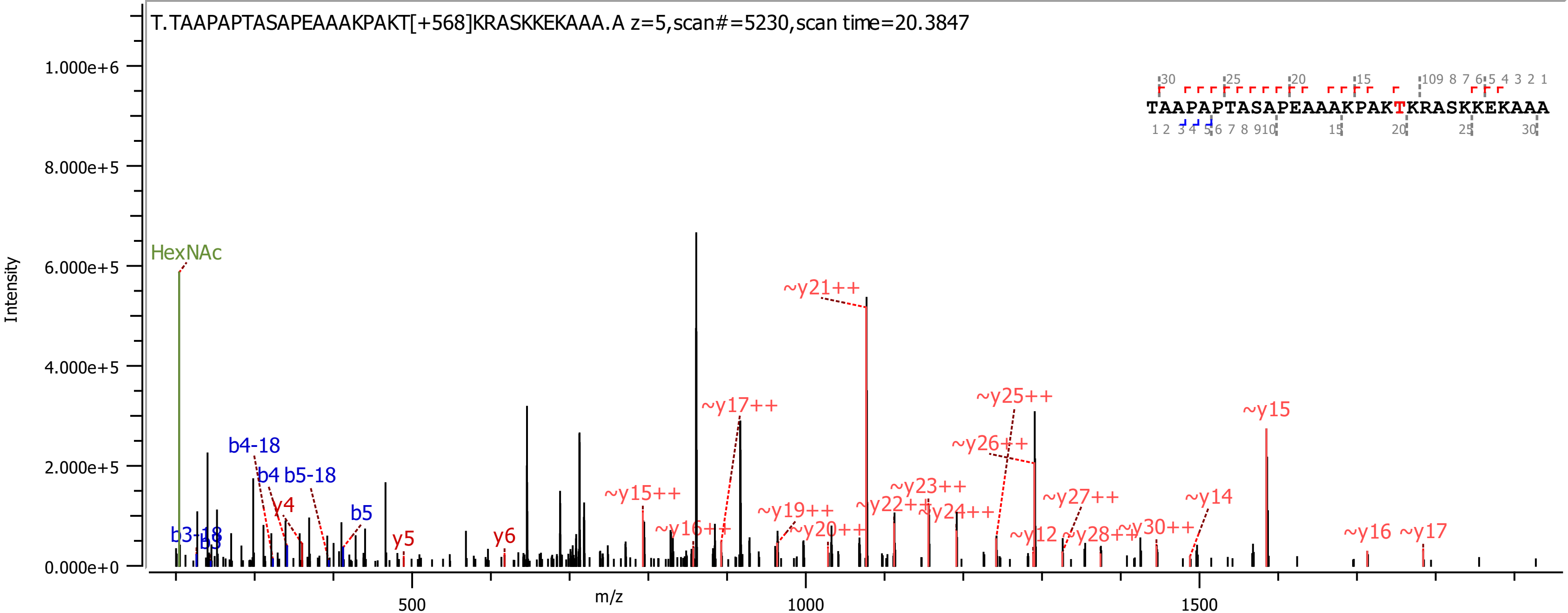
T.TAAPAPTASAPEAAAKPAKT[+568]KRASKKEKAAA.A z=5,scan#=5230,scan time=20.3847
TAAPAPTASAPEAAAKPAKTKRASKKEKAAA
Intensity
m/z
HexNAc
b4-18
b4 b5-18
b5
b3-18
b3
y4
y5
y6
~y15++
~y16++
~y17++
~y19++
~y20++
~y21++
~y22++
~y23++
~y24++
~y25++
~y26++
~y27++
~y28++
~y30++
~y12
~y14
~y15
~y16
~y17
1.000e+6
8.000e+5
6.000e+5
4.000e+5
2.000e+5
0.000e+0
500
1000
1500

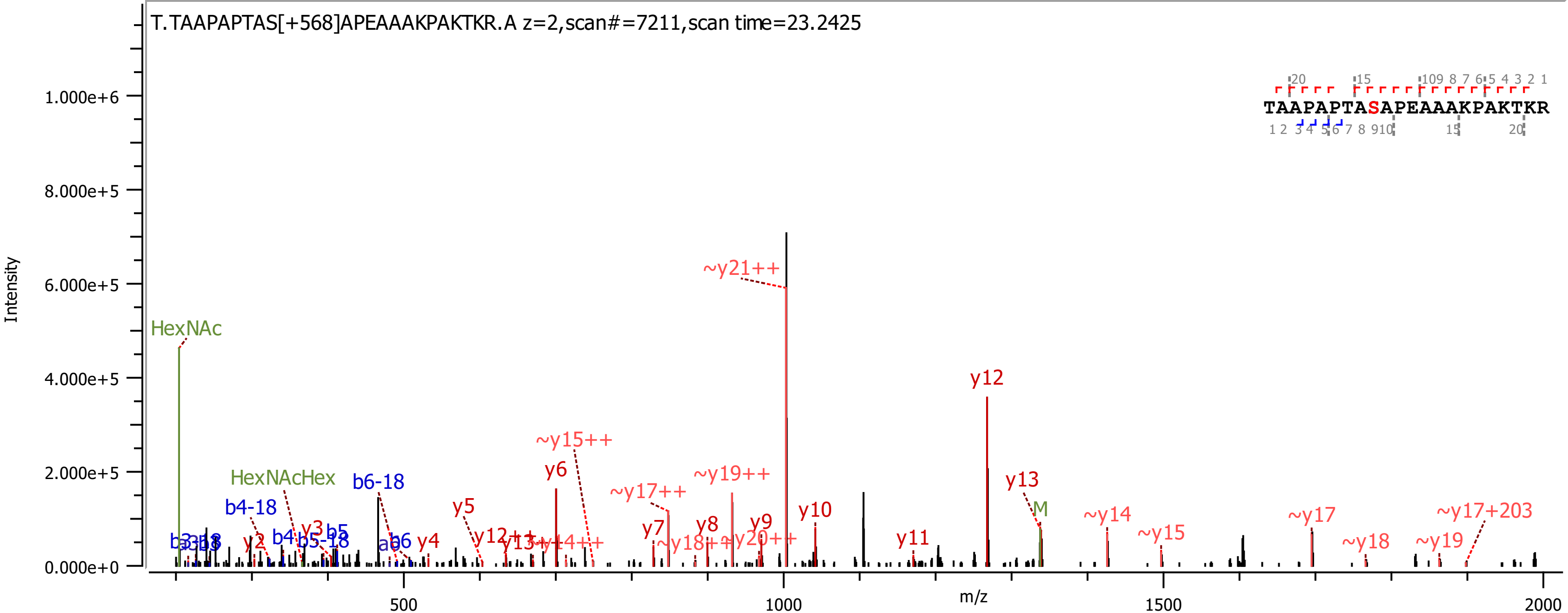
T.TAAPAPTAS[+568]APEAAAKPAKTKR.A z=2,scan#=7211,scan time=23.2425
TAAPAPTASAPEAAAKPAKTKR
Intensity
m/z
0.000e+0
2.000e+5
4.000e+5
6.000e+5
8.000e+5
1.000e+6
500
1000
1500
2000
HexNAc
HexNAcHex
b4-18
b6-18
b4
b5-18
b5
b6
y2
y3
y4
y5
y6
y7
y8
y9
y10
y11
y12
y13
M
~y21++
~y15++
~y17++
~y19++
y12++
~y18++
~y20++
~y14
~y15
~y17
~y18
~y19
~y17+203

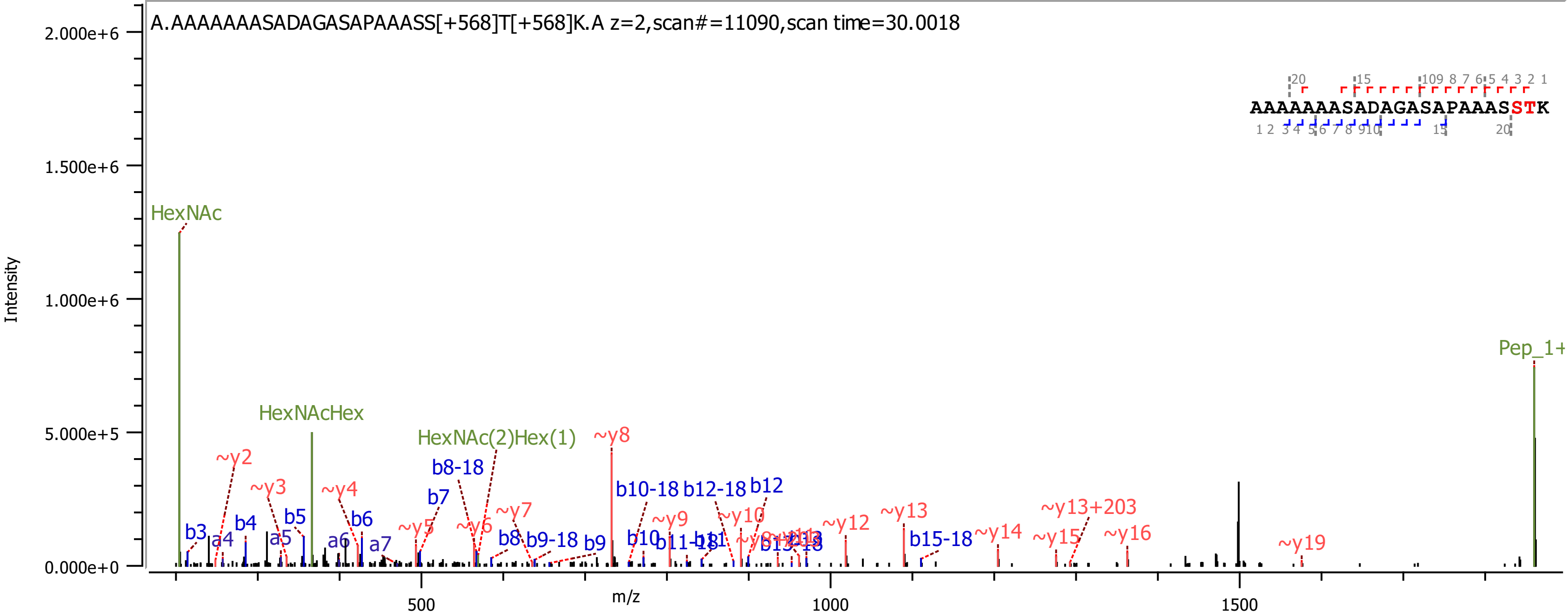

A.AAAAAAASADAGASAPAAASS[+568]T[+568]K.A z=2,scan#=11090,scan time=30.0018
AAAAAAASADAGASAPAAASSTK
Intensity
m/z
2.000e+6
1.500e+6
1.000e+6
5.000e+5
0.000e+0
500
1000
1500
HexNAc
HexNAcHex
HexNAc(2)Hex(1)
Pep_1+
~y2
~y3
~y4
~y5
~y6
~y7
~y8
~y9
~y10
~y12
~y13
~y13+203
~y14
~y15
~y16
~y19
b3
a4
b4
a5
b5
a6
b6
a7
b7
b8-18
b8
b9-18
b9
b10-18
b12-18
b12
b10
b15-18

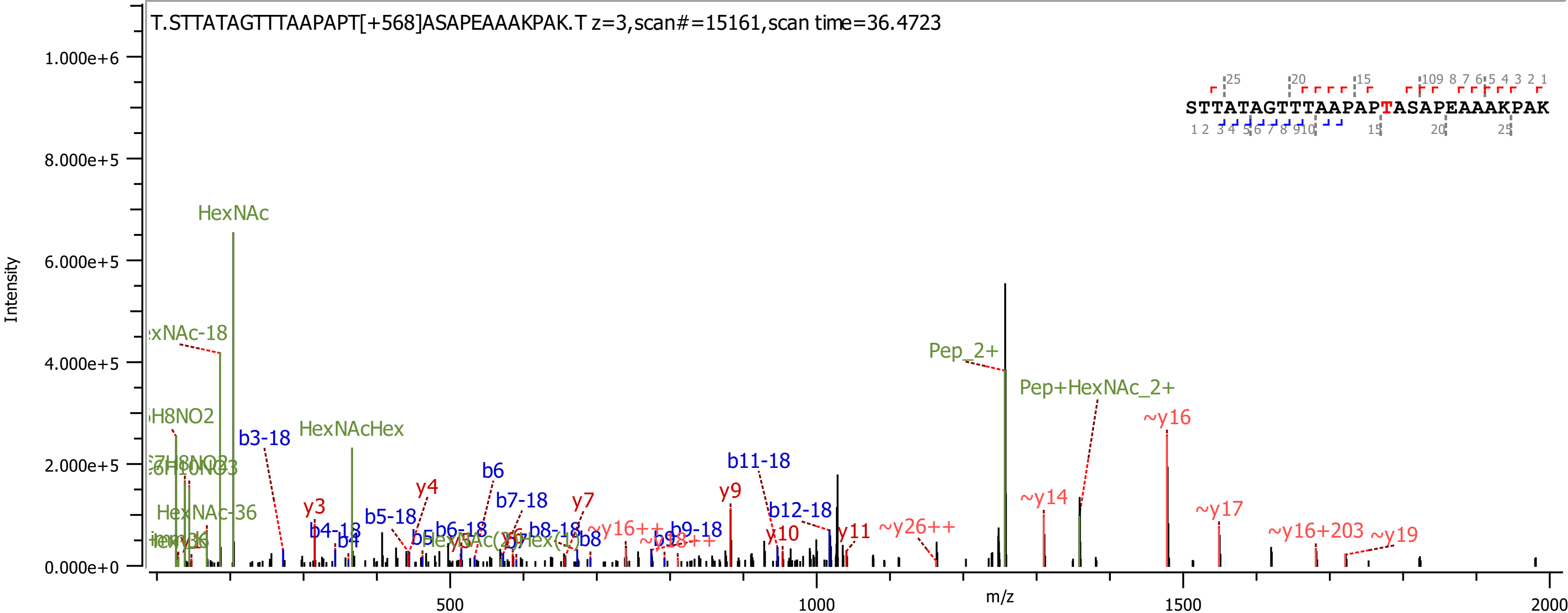
T.STTATAGTTTAAPAPT[+568]ASAPEAAAKPAK.T z=3,scan#=15161,scan time=36.4723
STTATAGTTTAAPAPTASAPEAAAKPAK
Intensity
m/z
1.000e+6
8.000e+5
6.000e+5
4.000e+5
2.000e+5
0.000e+0
500
1000
1500
2000
HexNAc
HexNAcHex
HexNAc-36
Pep_2+
Pep+HexNAc_2+
b3-18
b4-18
b4
b5-18
b6
b6-18
b7-18
b8-18
b8
b9-18
b11-18
b12-18
y3
y4
y7
y9
y10
y11
~y14
~y16
~y17
~y16+203
~y19
~y26++
~y16++

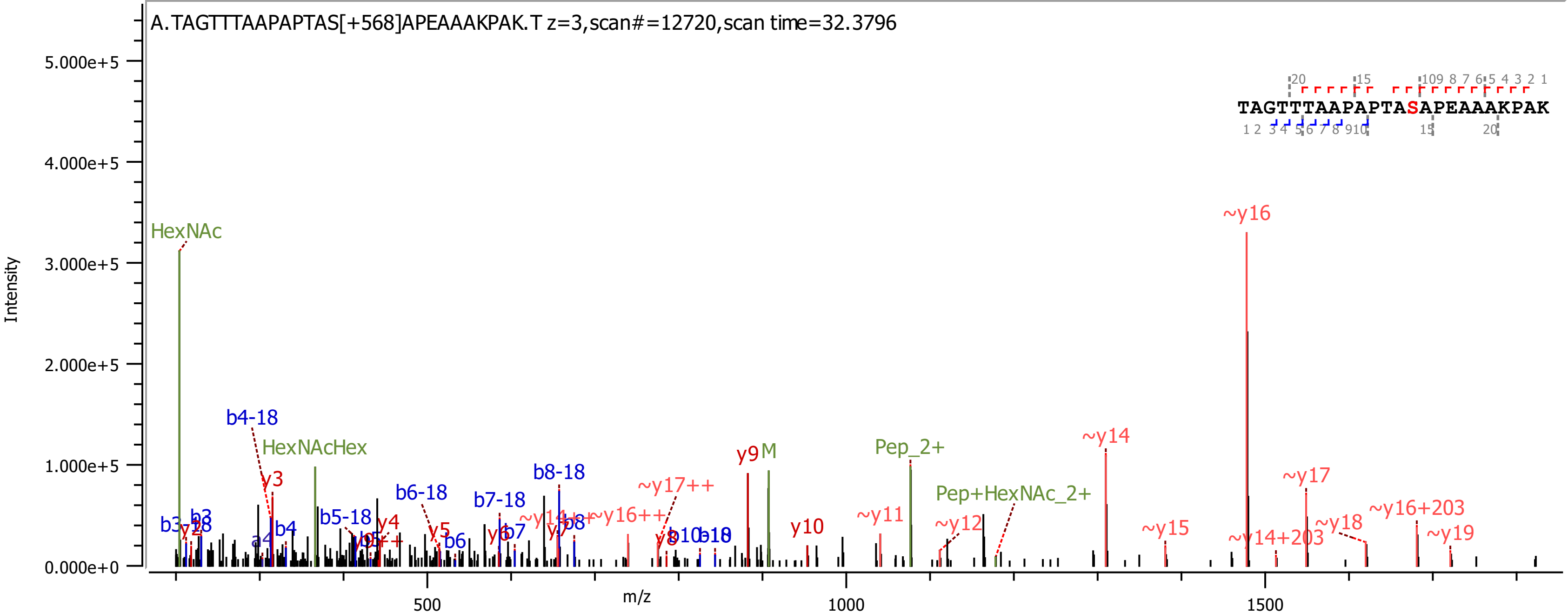
A.TAGTTTAAPAPTAS[+568]APEAAAKPAK.T z=3,scan#=12720,scan time=32.3796
TAGTTTAAPAPTASAPEAAAKPAK
Intensity
m/z
HexNAc
HexNAcHex
b4-18
y3
b3
y2
b4
a4
b5-18
b6-18
y4
y5
b6
b7-18
y6
b7
b8-18
y7
b8
~y14++
~y16++
~y17++
y8
b10
y9
M
y10
~y11
Pep_2+
~y12
Pep+HexNAc_2+
~y14
~y15
~y16
~y14+203
~y17
~y18
~y16+203
~y19

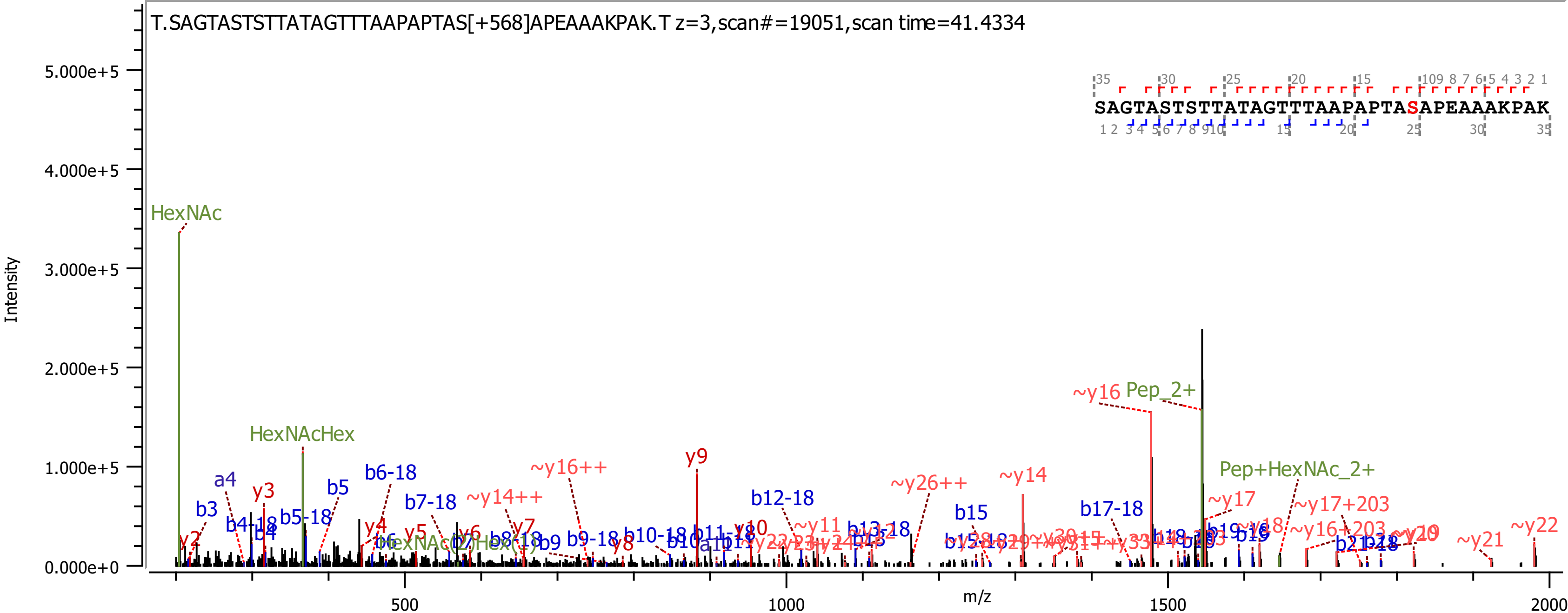

T.SAGTASTSTTATAGTTTAAPAPTAS[+568]APEAAAKPAK.T z=3,scan#=19051,scan time=41.4334
SAGTASTSTTATAGTTTAAPAPTASAPEAAAKPAK
Intensity
m/z
5.000e+5
4.000e+5
3.000e+5
2.000e+5
1.000e+5
0.000e+0
500
1000
1500
2000
HexNAc
HexNAcHex
Pep_2+
Pep+HexNAc_2+
~y16
~y14
~y16++
~y14++
~y26++
~y17
~y17+203
~y16+203
~y21
~y22
y2
y3
y4
y5
y6
y7
y8
y9
y10
a4
b3
b4
b4-18
b5
b5-18
b6-18
b7-18
b9
b9-18
b10-18
b12-18
b13-18
b15
b17-18
b19

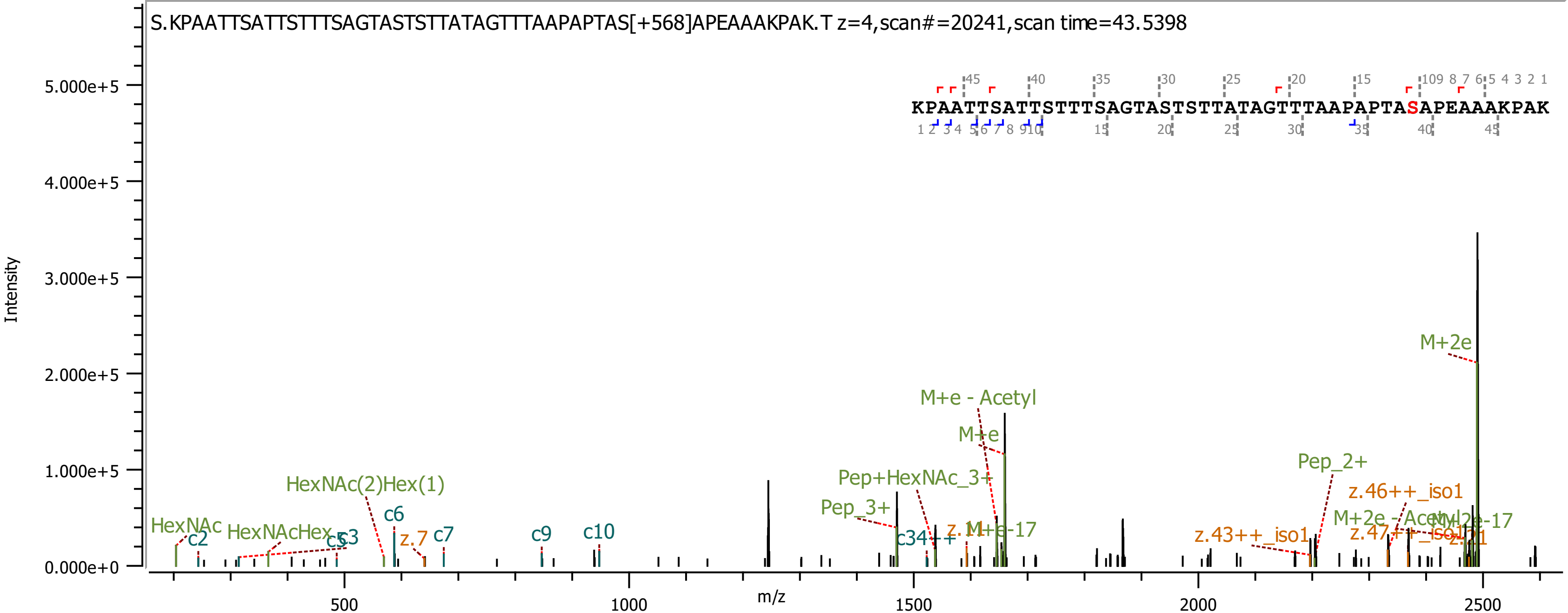

S.KPAATTSATTSTTTSAGTASTSTTATAGTTTAAPAPTAS[+568]APEAAAKPAK.T z=4,scan#=20241,scan time=43.5398
KPAATTSATTSTTTSAGTASTSTTATAGTTTAAPAPTASAPEAAAKPAK
Intensity
m/z
5.000e+5
4.000e+5
3.000e+5
2.000e+5
1.000e+5
0.000e+0
500
1000
1500
2000
2500
HexNAc
c2
HexNAcHex
c3
c5
HexNAc(2)Hex(1)
c6
z.7
c7
c9
c10
Pep_3+
Pep+HexNAc_3+
c34++
M+e - Acetyl
M+e
z.1
M+e-17
Pep_2+
z.43++_iso1
z.46++_iso1
M+2e - Acetyl
M+2e-17
z.47++_iso1
z.11
M+2e

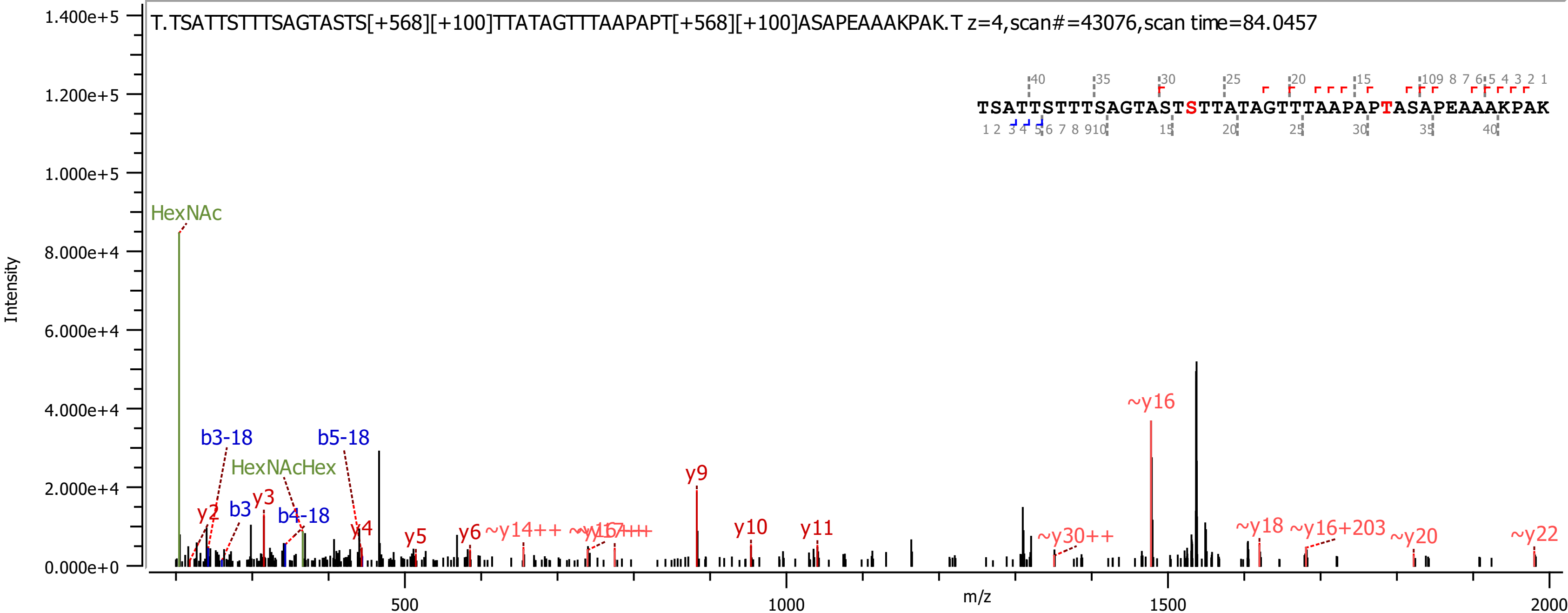
T.TSATTSTTTSAGTASTS[+568][+100]TTATAGTTTAAPAPT[+568][+100]ASAPEAAAKPAK.T z=4,scan#=43076,scan time=84.0457
TSATTSTTTSAGTASTSTTATAGTTTAAPAPTASAPEAAAKPAK
Intensity
m/z
1.400e+5
1.200e+5
1.000e+5
8.000e+4
6.000e+4
4.000e+4
2.000e+4
0.000e+0
500
1000
1500
2000
HexNAc
HexNAcHex
b3-18
b5-18
b3
b4-18
y2
y3
y4
y5
y6
~y14++
y9
y10
y11
~y30++
~y16
~y18
~y16+203
~y20
~y22

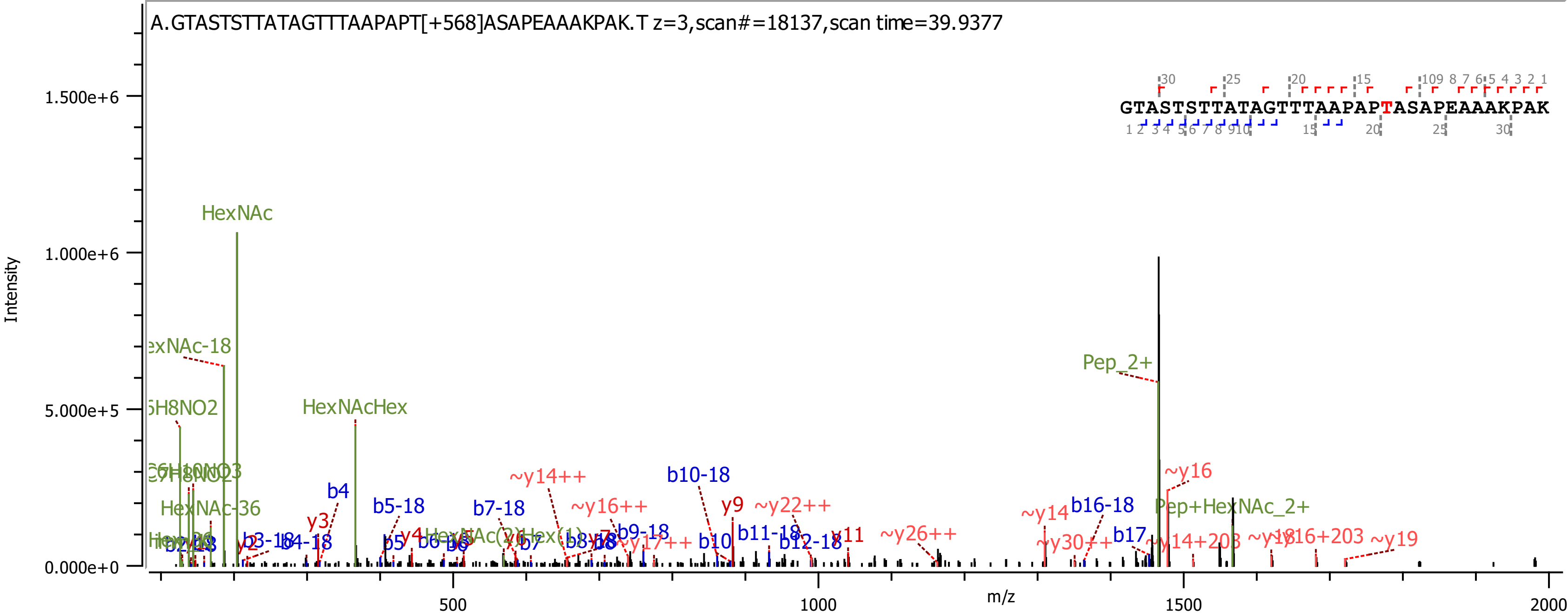
A.GTASTSTTATAGTTTAAPAPT[+568]ASAPEAAAKPAK.T z=3,scan#=18137,scan time=39.9377
GTASTSTTATAGTTTAAPAPTASAPEAAAKPAK
Intensity
m/z
1.500e+6
1.000e+6
5.000e+5
0.000e+0
500
1000
1500
2000
HexNAc
HexNAcHex
Pep_2+
Pep+HexNAc_2+
~y16
~y14++
~y16++
~y22++
~y26++
~y14
~y30++
~y19
b4
b5-18
b7-18
b10-18
b16-18
b17
y3
y9
y11
b10
b11-18
b12-18
b9-18

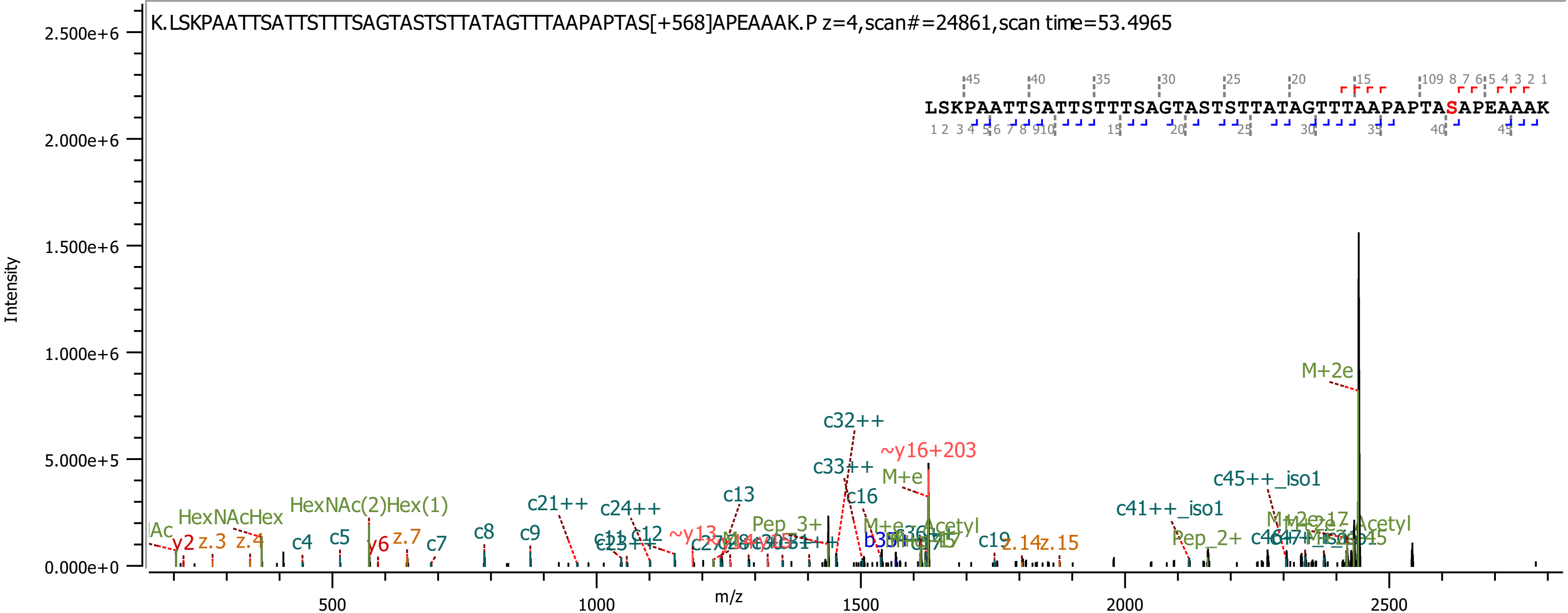

K.LSKPAATTSATTSTTTSAGTASTSTTATAGTTTAAPAPTAS[+568]APEAAAK.P z=4,scan#=24861,scan time=53.4965
LSKPAATTSATTSTTTSAGTASTSTTATAGTTTAAPAPTASAPEAAAK
Intensity
m/z
M+2e
c32++
~y16+203
c33++
M+e
c45++_iso1
c13
c16
c41++_iso1
HexNAc(2)Hex(1)
c21++
c24++
HexNAcHex
Pep_3+
c8
c9
c12
c19
z.14
z.15
Pep_2+
y2
z.3
z.4
c4
c5
y6
z.7
c7

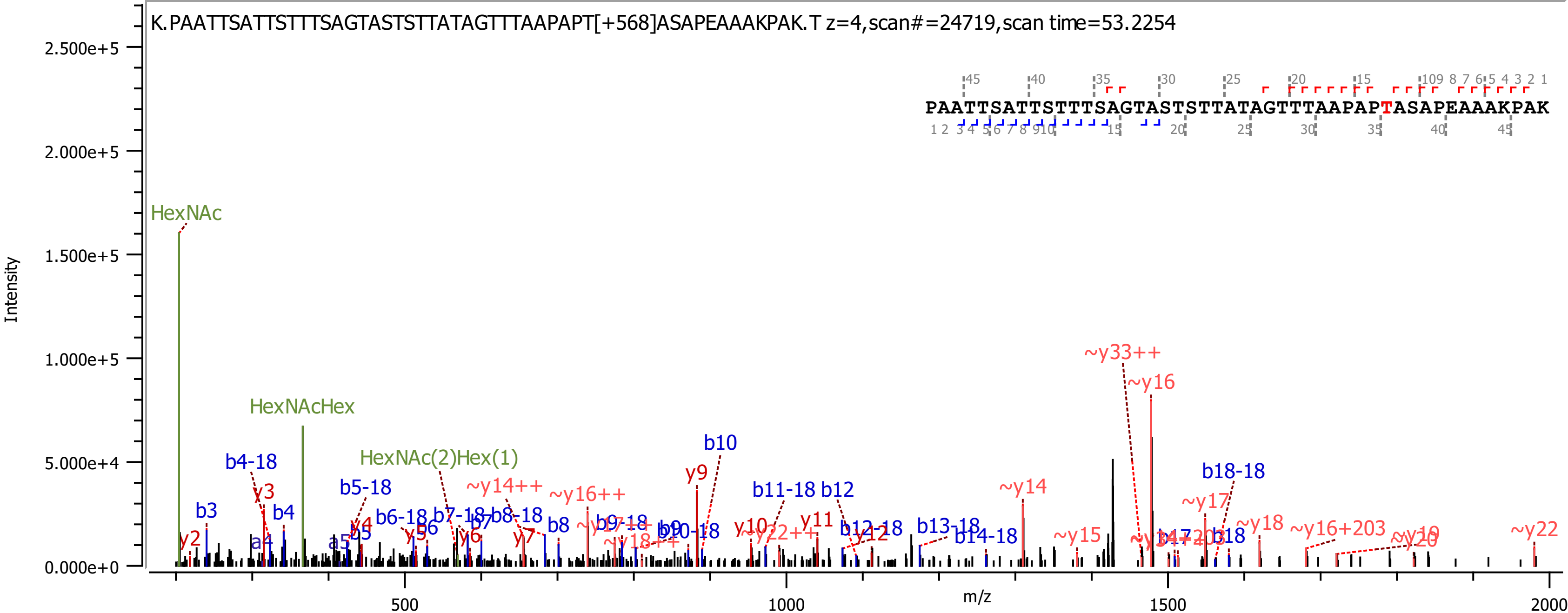

K.PAATTSATTSTTTSAGTASTSTTATAGTTTAAPAPT[+568]ASAPEAAAKPAK.T z=4,scan#=24719,scan time=53.2254
PAATTSATTSTTTSAGTASTSTTATAGTTTAAPAPTASAPEAAAKPAK
Intensity
m/z
HexNAc
HexNAcHex
HexNAc(2)Hex(1)
~y33++
~y16
b10
y9
b18-18
~y14
~y17
b4-18
y3
b3
b4
b5-18
~y14++
~y16++
b11-18 b12
~y18
~y16+203
~y22
~y15
b13-18
b14-18
y2
a4
a5
b5
y4
b6-18
b6
y5
b7-18
b7
y6
b8-18
y7
b8
y10
y11
~y12
b12-18
b18
0.000e+0
5.000e+4
1.000e+5
1.500e+5
2.000e+5
2.500e+5
500
1000
1500
2000

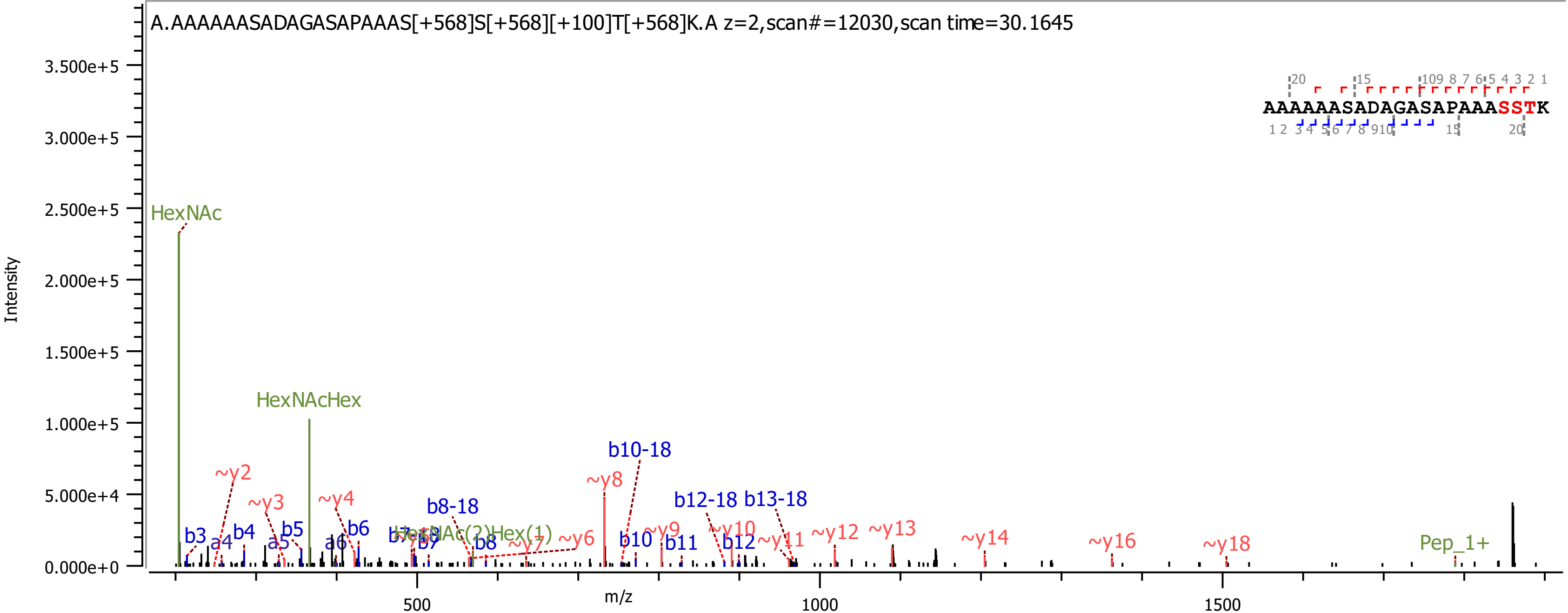

A.AAAAAASADAGASAPAAAS[+568]S[+568][+100]T[+568]K.A z=2,scan#=12030,scan time=30.1645
AAAAAASADAGASAPAAASSTK
Intensity
m/z
HexNAc
HexNAcHex
HexNAc(2)Hex(1)
Pep_1+
b10-18
b8-18
b12-18 b13-18
~y2
~y3
~y4
~y6
~y7
~y8
~y9
~y10
~y11
~y12
~y13
~y14
~y16
~y18
b3
a4
b4
a5
b5
a6
b6
b7
b8
b10
b11
b12

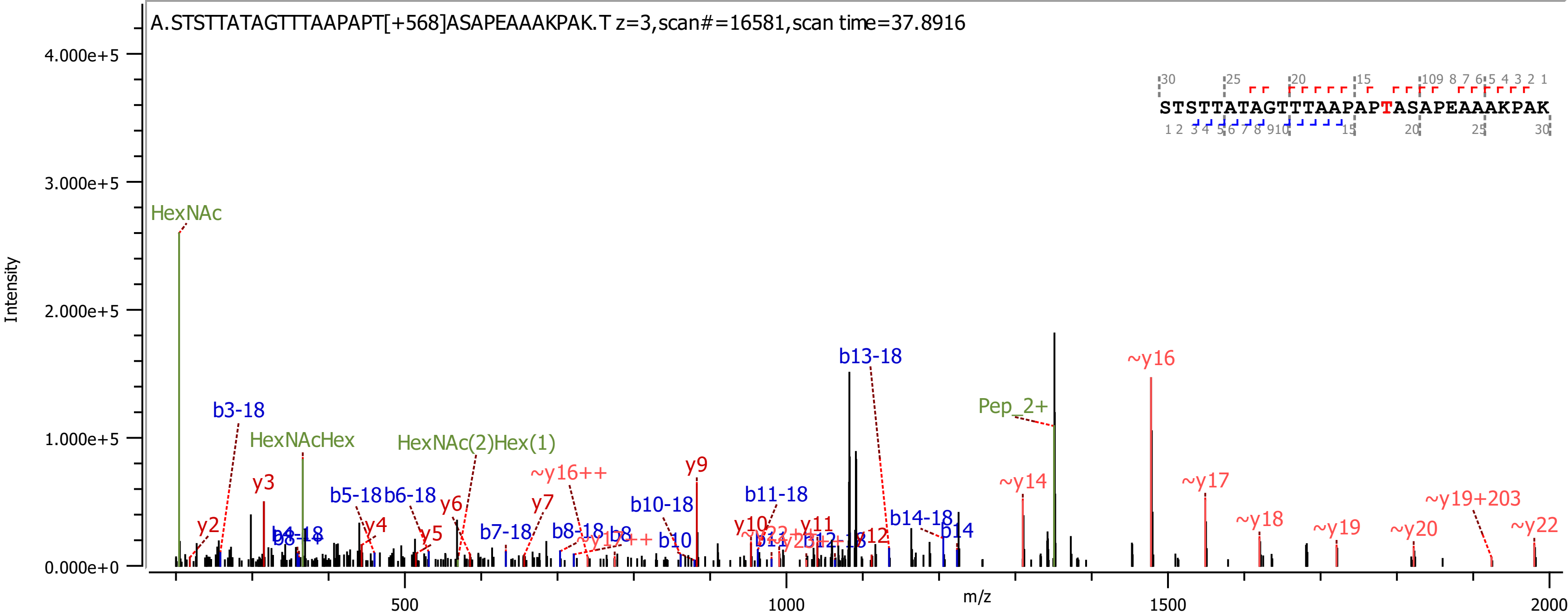

A.STSTTATAGTTTAAPAPT[+568]ASAPEAAAKPAK.T z=3,scan#=16581,scan time=37.8916
STSTTATAGTTTAAPAPTASAPEAAAKPAK
Intensity
m/z
4.000e+5
3.000e+5
2.000e+5
1.000e+5
0.000e+0
500
1000
1500
2000
HexNAc
HexNAcHex
HexNAc(2)Hex(1)
Pep_2+
b3-18
b5-18
b6-18
b7-18
b8-18
b8
b10-18
b10
b11-18
b13-18
b14-18
b14
y2
y3
y4
y5
y6
y7
y9
y10
y11
y12
~y16++
~y14
~y16
~y17
~y18
~y19
~y19+203
~y20
~y22

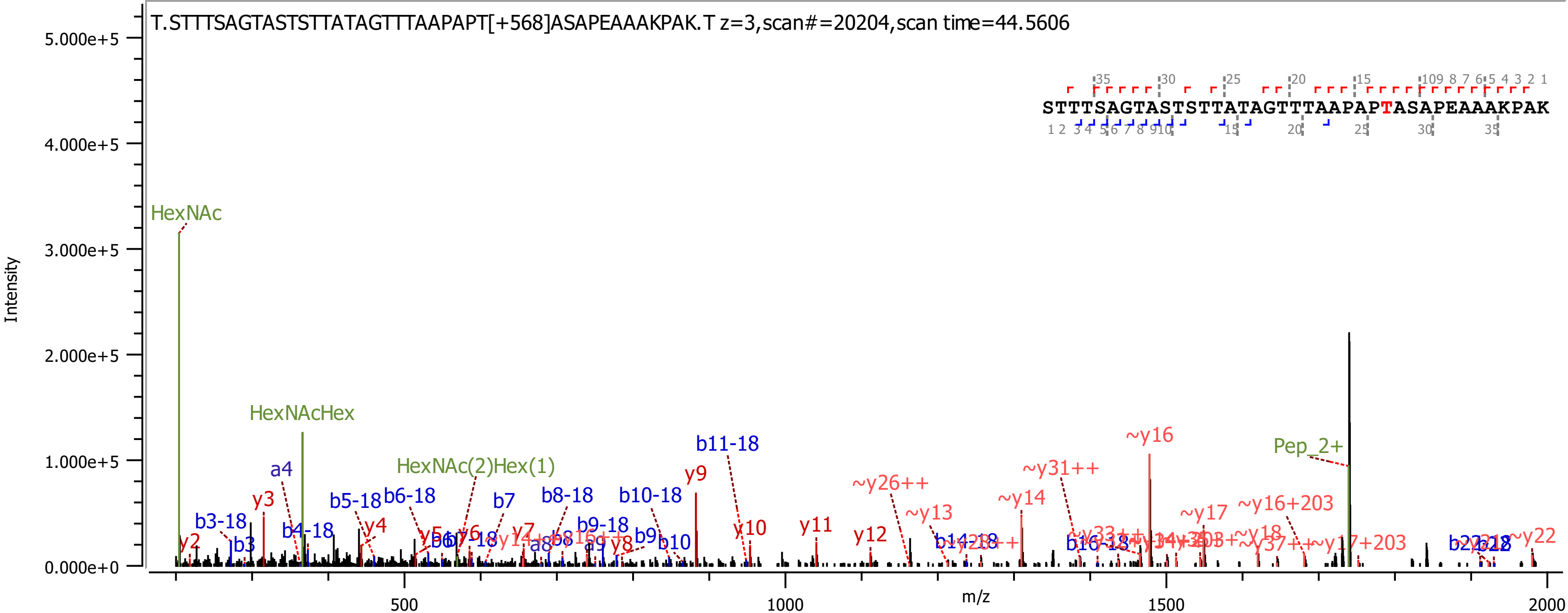
T.STTTSAGTASTSTTATAGTTTAAPAPT[+568]ASAPEAAAKPAK.T z=3,scan#=20204,scan time=44.5606
STTTSAGTASTSTTATAGTTTAAPAPTASAPEAAAKPAK
Intensity
m/z
5.000e+5
4.000e+5
3.000e+5
2.000e+5
1.000e+5
0.000e+0
500
1000
1500
2000
HexNAc
HexNAcHex
HexNAc(2)Hex(1)
Pep_2+
a4
b3-18
b3
b4-18
b5-18
b6-18
b7
b8-18
b9-18
b9
b10-18
b10
b11-18
y2
y3
y4
y5
y6
y7
y8
y9
y10
y11
y12
~y26++
~y13
~y14
~y31++
~y16
~y17
~y16+203
~y18
~y22
~y37++

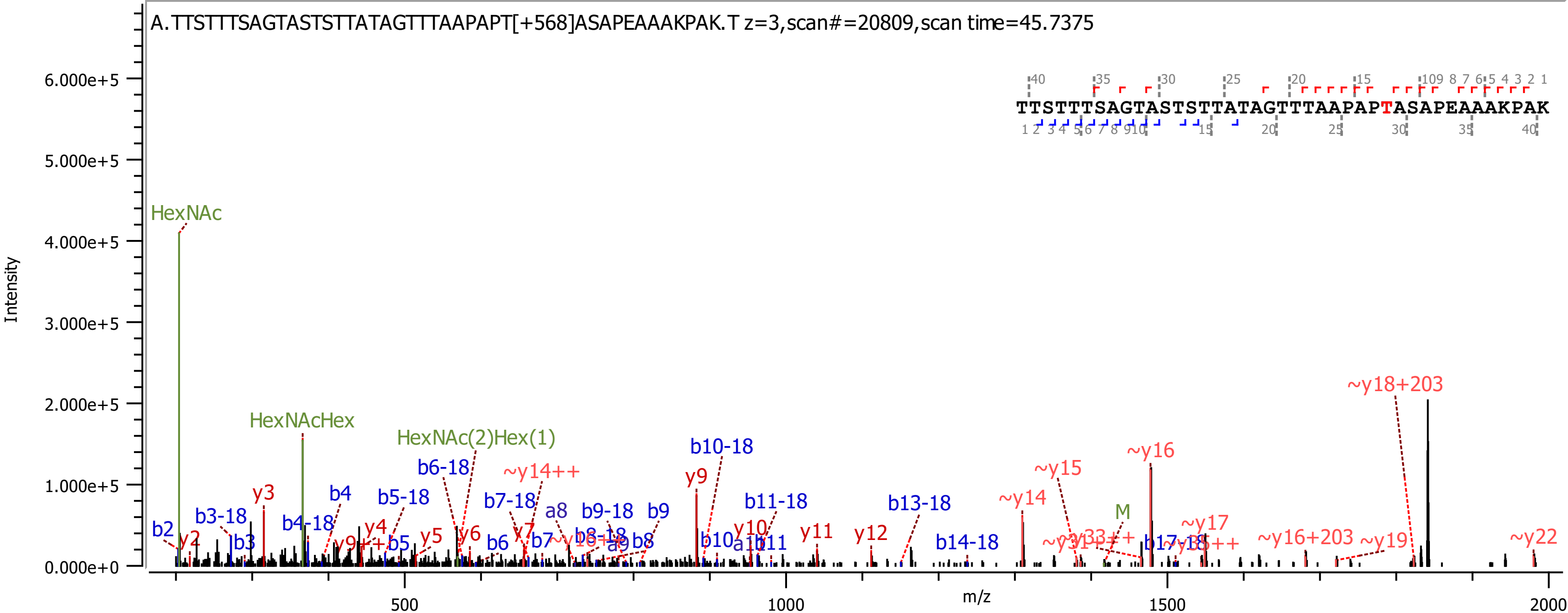

A.TTSTTTSAGTASTSTTATAGTTTAAPAPT[+568]ASAPEAAAKPAK.T z=3,scan#=20809,scan time=45.7375
TTSTTTSAGTASTSTTATAGTTTAAPAPTASAPEAAAKPAK
Intensity
6.000e+5
5.000e+5
4.000e+5
3.000e+5
2.000e+5
1.000e+5
0.000e+0
m/z
500
1000
1500
2000
HexNAc
HexNAcHex
HexNAc(2)Hex(1)
b2
y2
b3-18
b3
y3
b4-18
b4
y4
b5-18
b5
y5
b6-18
y6
b6
~y14++
b7-18
y7
b7
a8
b8-18
b9-18
a9
b9
b8
y9
b10-18
b10
y10
a11
b11
b11-18
y11
y12
b13-18
b14-18
~y14
~y15
M
~y16
~y17
~y18+203
~y16+203
~y19
~y22

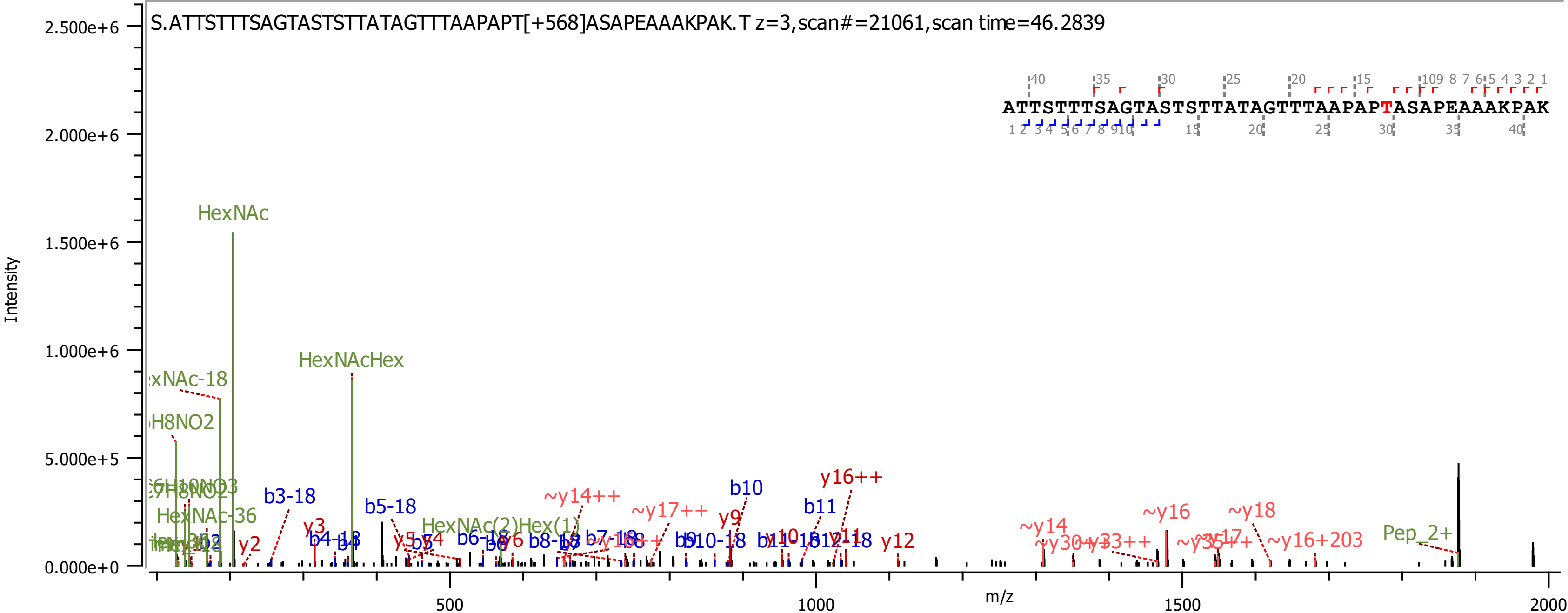

S.ATTSTTTSAGTASTSTTATAGTTTAAPAPT[+568]ASAPEAAAKPAK.T z=3,scan#=21061,scan time=46.2839
ATTSTTTSAGTASTSTTATAGTTTAAPAPTASAPEAAAKPAK
Intensity
m/z
2.500e+6
2.000e+6
1.500e+6
1.000e+6
5.000e+5
0.000e+0
500
1000
1500
2000
HexNAc
HexNAcHex
HexNAc-36
HexNAc(2)Hex(1)
Pep_2+
b3-18
b5-18
b6-18
b7-18
b10
b11
y2
y3
y6
y9
y12
y16++
~y14++
~y17++
~y14
~y16
~y18
~y16+203

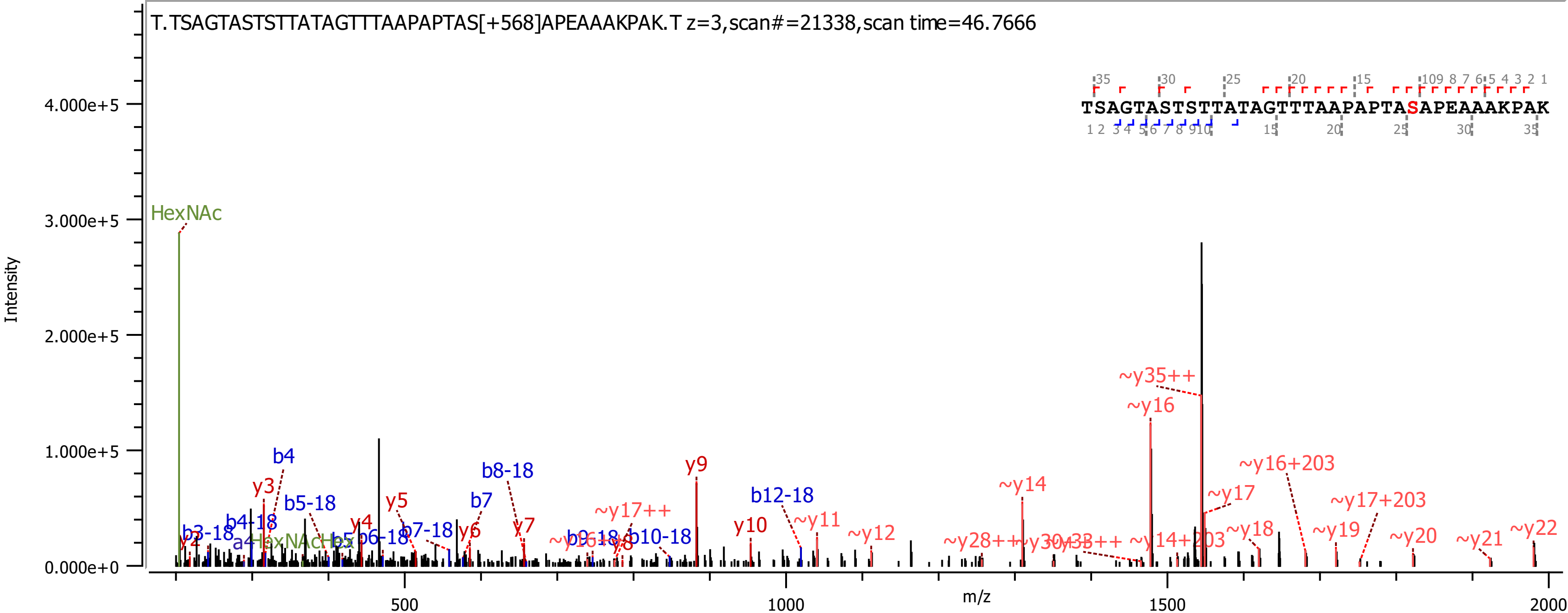

T.TSAGTASTSTTATAGTTTAAPAPTAS[+568]APEAAAKPAK.T z=3,scan#=21338,scan time=46.7666
TSAGTASTSTTATAGTTTAAPAPTASAPEAAAKPAK
Intensity
m/z
HexNAc
b4
y3
b5-18
b8-18
y9
b12-18
~y14
~y35++
~y16
~y16+203
~y17
~y17+203
~y18
~y19
~y20
~y21
~y22
y5
b7
y4
y6
y7
~y17++
y10
~y11
~y12
b10-18
b3-18
b4-18
b6-18
b7-18
~y28++
~y14+203

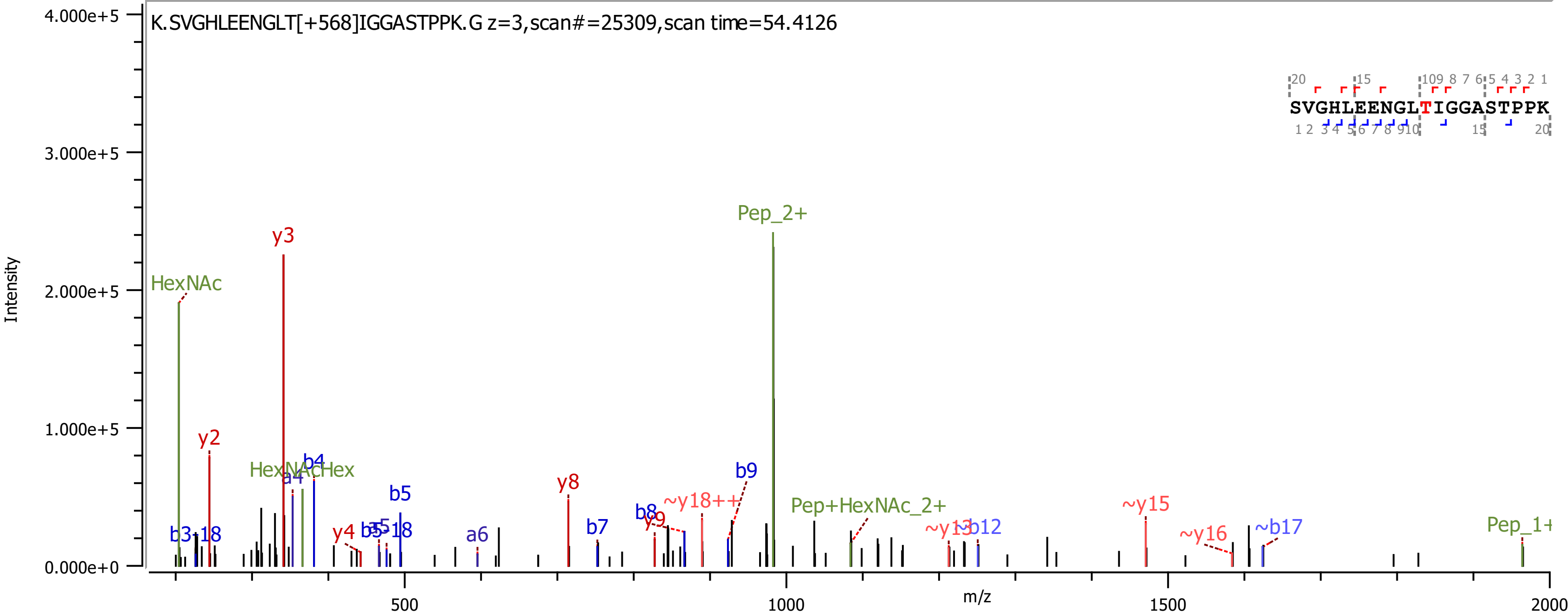

K.SVGHLEENGLT[+568]IGGASTPPK.G z=3,scan#=25309,scan time=54.4126
SVGHLEENGLTIGGASTPPK
Intensity
m/z
4.000e+5
3.000e+5
2.000e+5
1.000e+5
0.000e+0
500
1000
1500
2000
HexNAc
y3
y2
Pep_2+
HexNAcHex
a4
b4
b5
b3-18
y4
b5-18
a6
y8
b7
b8
y9
~y18++
b9
Pep+HexNAc_2+
~y13
~b12
~y15
~y16
~b17

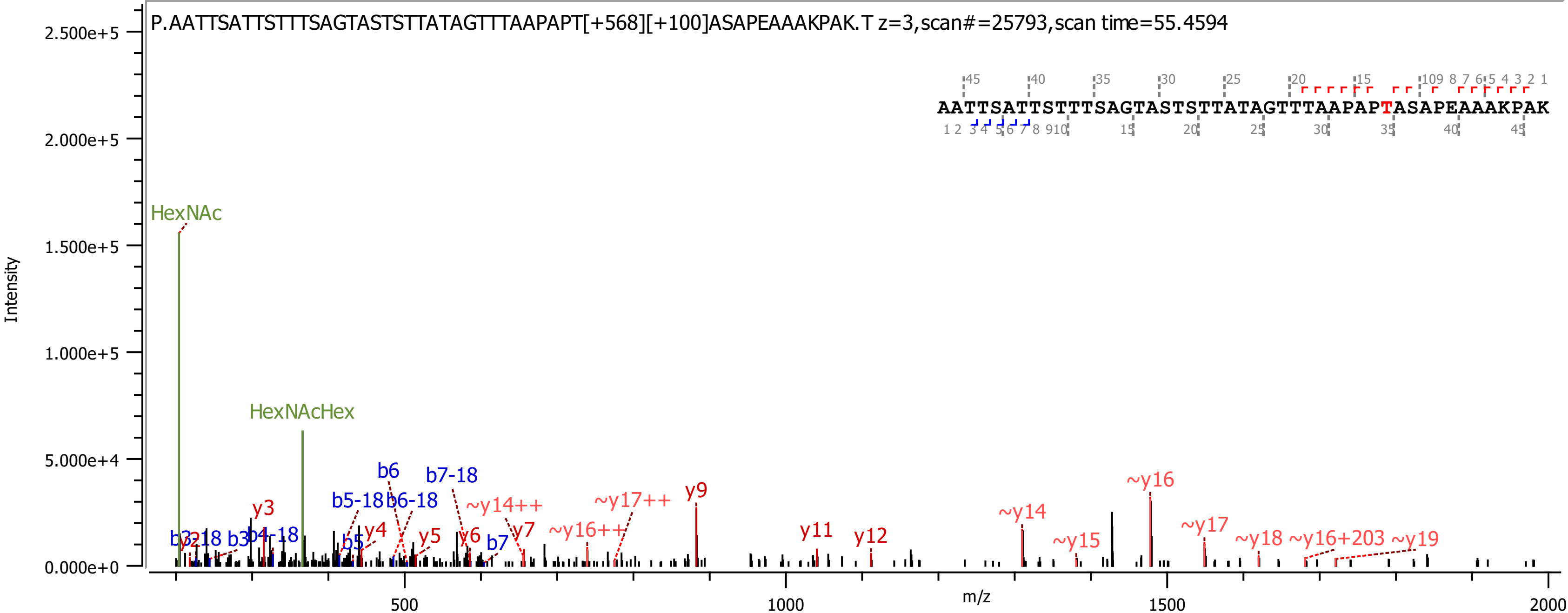
P.AATTSATTSTTTSAGTASTSTTATAGTTTAAPAPT[+568][+100]ASAPEAAAKPAK.T z=3,scan#=25793,scan time=55.4594
AATTSATTSTTTSAGTASTSTTATAGTTTAAPAPTASAPEAAAKPAK
Intensity
m/z
2.500e+5
2.000e+5
1.500e+5
1.000e+5
5.000e+4
0.000e+0
500
1000
1500
2000
HexNAc
HexNAcHex
b6
b7-18
b5-18
b6-18
y3
y4
y5
y6
y7
b7
b5
~y14++
~y16++
~y17++
y9
y11
y12
~y14
~y15
~y16
~y17
~y18
~y16+203
~y19

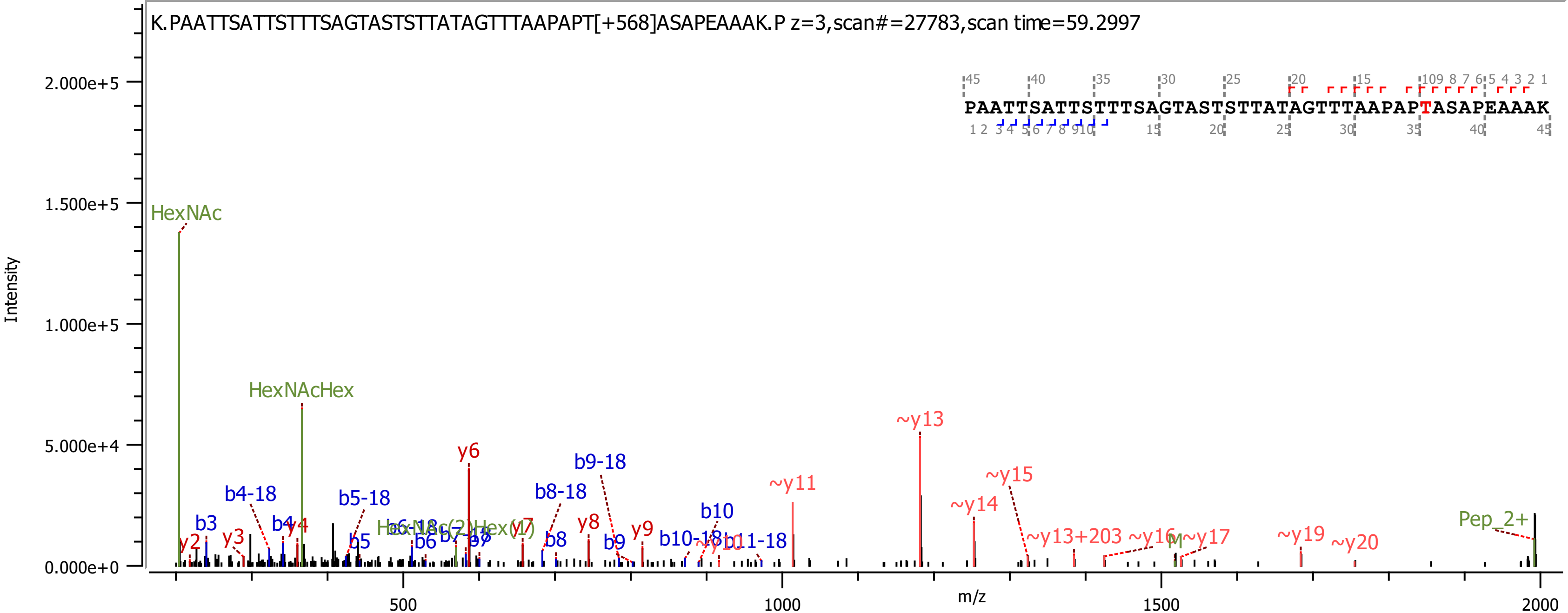

K.PAATTSATTSTTTSAGTASTSTTATAGTTTAAPAPT[+568]ASAPEAAAK.P z=3,scan#=27783,scan time=59.2997
PAATTSATTSTTTSAGTASTSTTATAGTTTAAPAPTASAPEAAAK
Intensity
m/z
HexNAc
HexNAcHex
Pep_2+
y6
~y13
~y11
~y15
~y14
~y19
~y20
b9-18
b8-18
b4-18
b5-18
b10
b10-18
b11-18
~y13+203

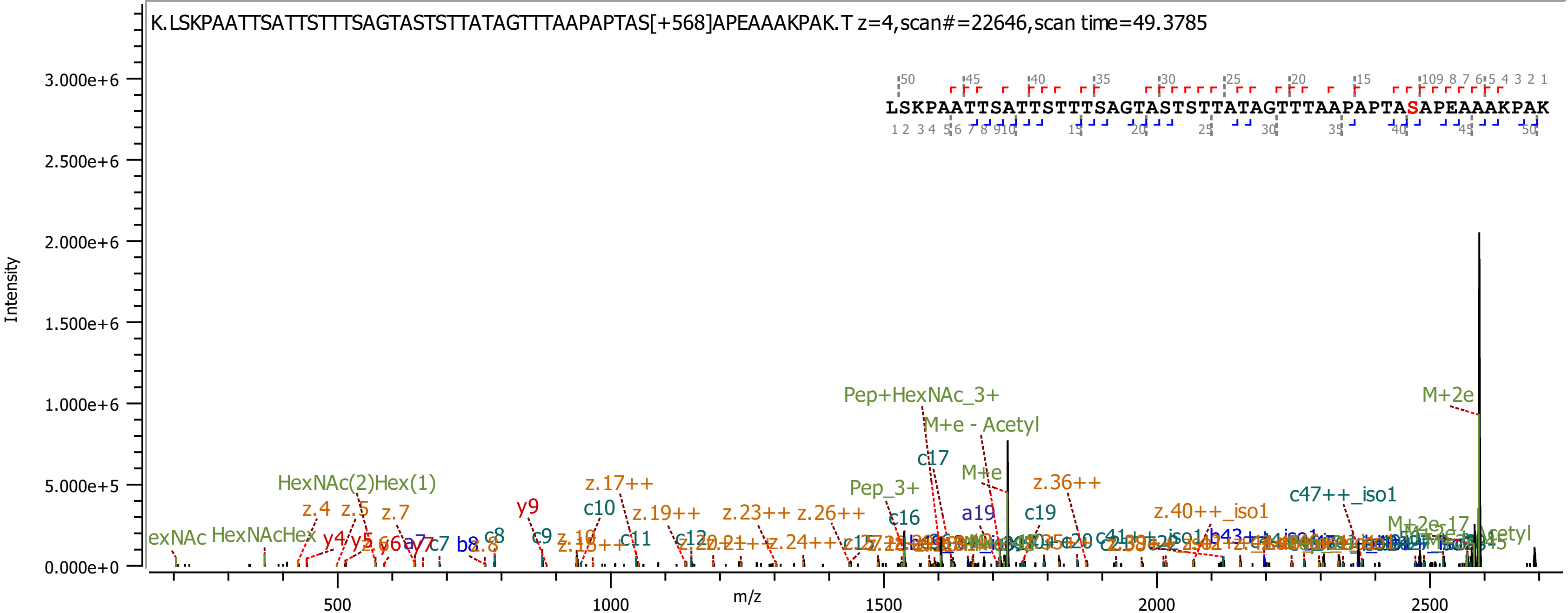
K.LSKPAATTSATTSTTTSAGTASTSTTATAGTTTAAPAPTAS[+568]APEAAAKPAK.T z=4,scan#=22646,scan time=49.3785
LSKPAATTSATTSTTTSAGTASTSTTATAGTTTAAPAPTASAPEAAAKPAK
Intensity
3.000e+6
2.500e+6
2.000e+6
1.500e+6
1.000e+6
5.000e+5
0.000e+0
m/z
500
1000
1500
2000
2500
Pep+HexNAc_3+
M+e - Acetyl
M+2e
c17
M+e
z.36++
Pep_3+
HexNAc(2)Hex(1)
z.17++
c47++_iso1
z.4
z.5
z.7
y9
c10
z.19++
z.23++
z.26++
c16
a19
c19
z.40++_iso1
HexNAcHex
y4
c8
c9
c11
c12
M+2e-17

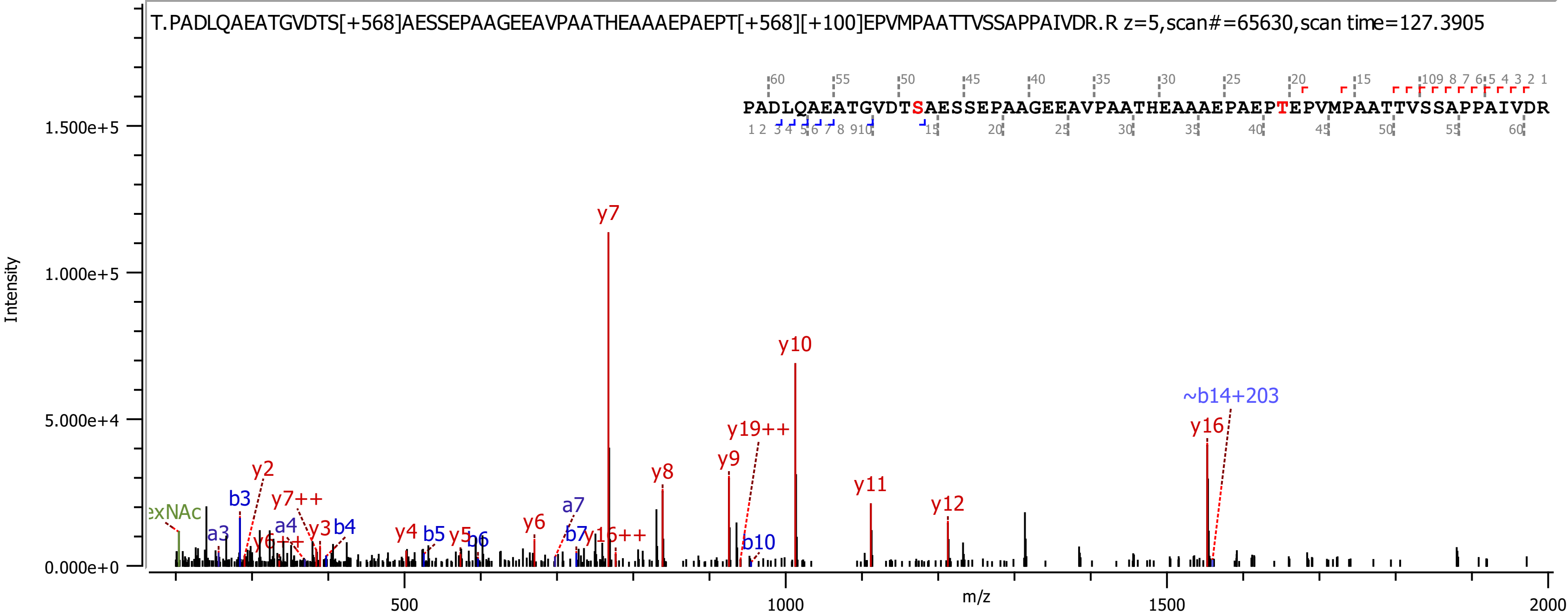
T.PADLQAEATGVDTS[+568]AESSEPAAGEEAVPAATHEAAAEPAEPT[+568][+100]EPVMPAATTVSSAPPAIVDR.R z=5,scan#=65630,scan time=127.3905
PADLQAEATGVDTSAESSEPAAGEEAVPAATHEAAAEPAEPTEPVMPAATTVSSAPPAIVDR
Intensity
m/z
y7
y10
~b14+203
y16
y19++
y9
y8
y11
y12
y2
b3
y7++
exNAc
a4
y3
b4
a3
y6++
y4
b5
y5
b6
y6
a7
b7
y16++
b10
1.500e+5
1.000e+5
5.000e+4
0.000e+0
500
1000
1500
2000

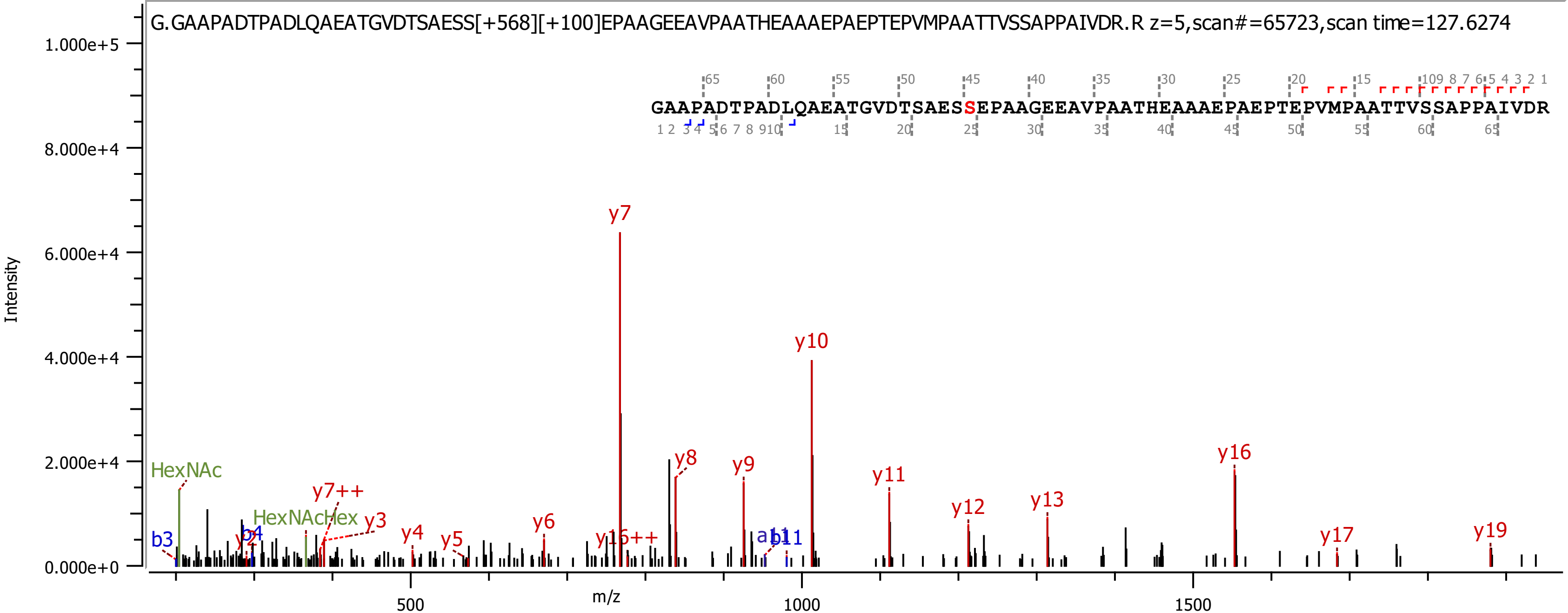

G.GAAPADTPADLQAEATGVDTSAESS[+568][+100]EPAAGEEAVPAATHEAAAEPAEPTEPVMPAATTVSSAPPAIVDR.R z=5,scan#=65723,scan time=127.6274
GAAPADTPADLQAEATGVDTSAESSEPAAGEEAVPAATHEAAAEPAEPTEPVMPAATTVSSAPPAIVDR
Intensity
m/z
y7
y10
y8
y9
y11
y12
y13
y16
y17
y19
HexNAc
HexNAcHex
y7++
y3
y4
y5
y6
y16++
b3
b4
y2
a11
b11

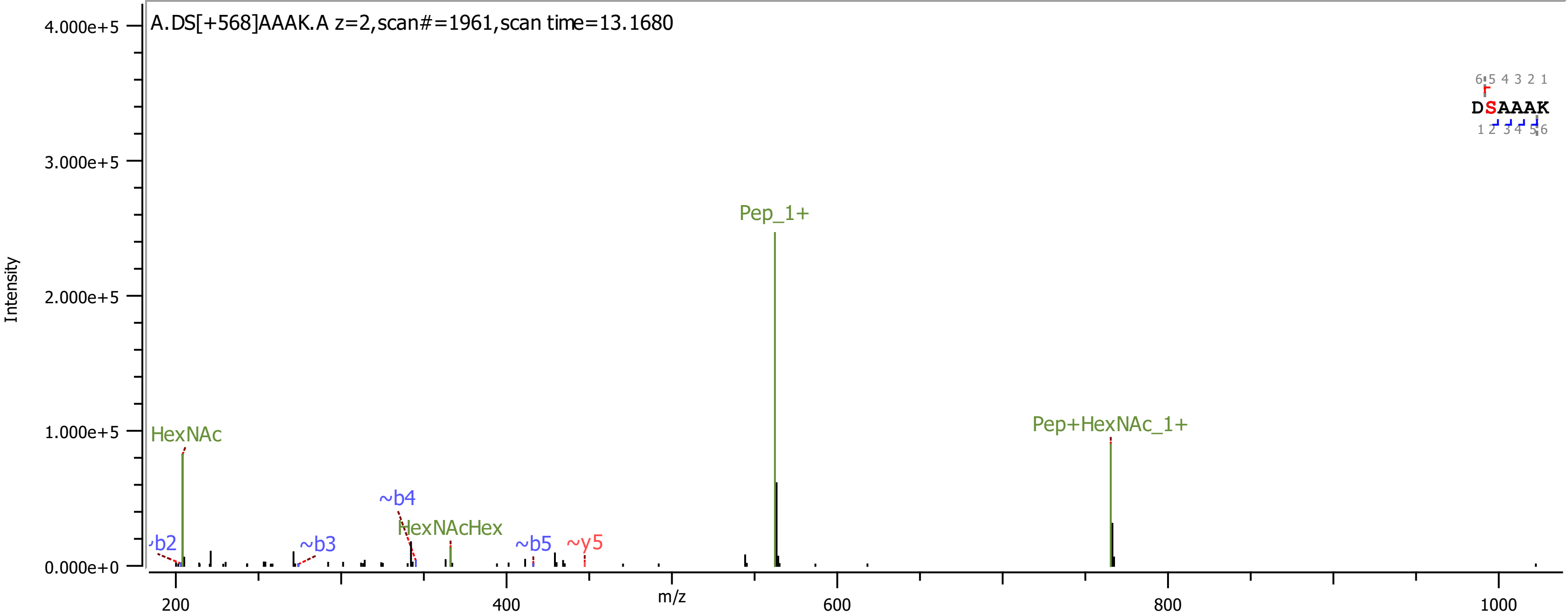
A.DS[+568]AAAK.A z=2,scan#=1961,scan time=13.1680
Intensity
m/z
HexNAc
Pep_1+
Pep+HexNAc_1+
HexNAcHex
~b2
~b3
~b4
~b5
~y5
DSAAAK

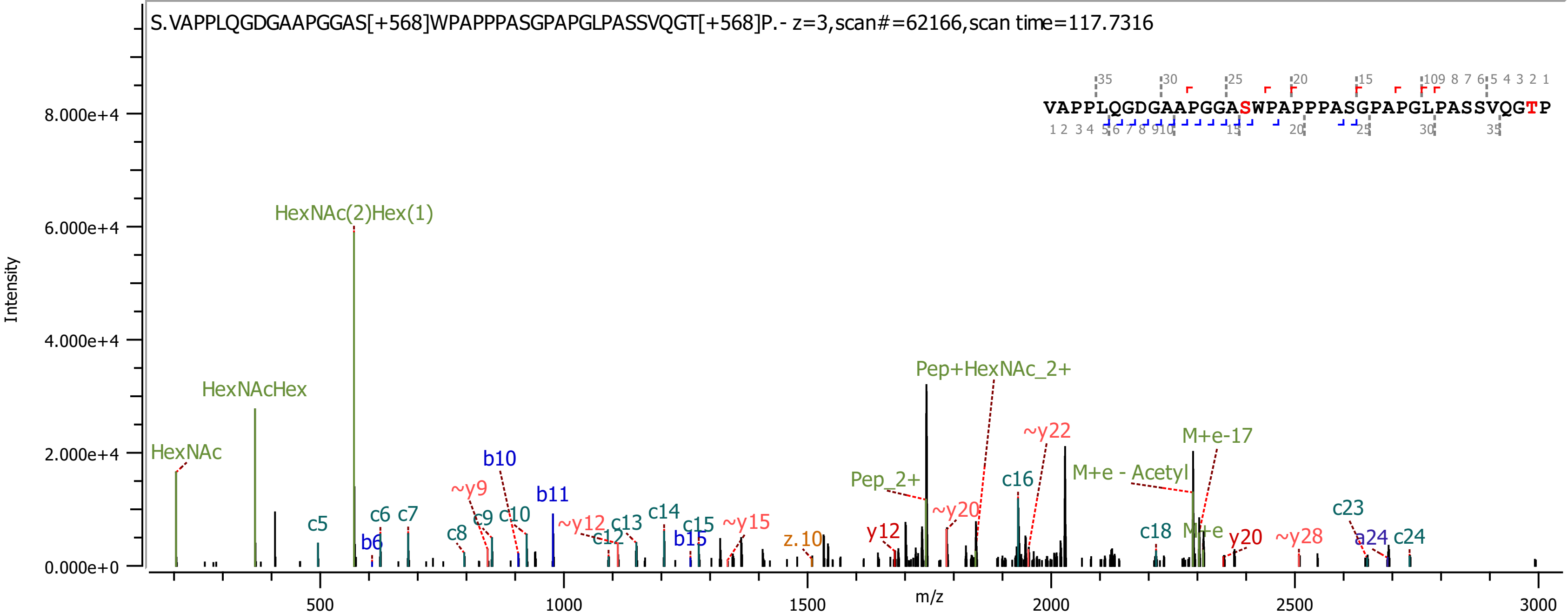
S.VAPPLQGDGAAPGGAS[+568]WPAPPPASGPAPGLPASSVQGT[+568]P.- z=3,scan#=62166,scan time=117.7316
VAPPLQGDGAAPGGASWPAPPPASGPAPGLPASSVQGTP
Intensity
m/z
HexNAc
HexNAcHex
HexNAc(2)Hex(1)
Pep+HexNAc_2+
Pep_2+
M+e-17
M+e - Acetyl
M+e
c5
c6
c7
c8
c9
c10
c12
c13
c14
c15
c16
c18
c23
c24
b6
b10
b11
b15
a24
~y9
~y12
~y15
~y20
~y22
~y28
y12
y20
z.10
0.000e+0
2.000e+4
4.000e+4
6.000e+4
8.000e+4
500
1000
1500
2000
2500
3000

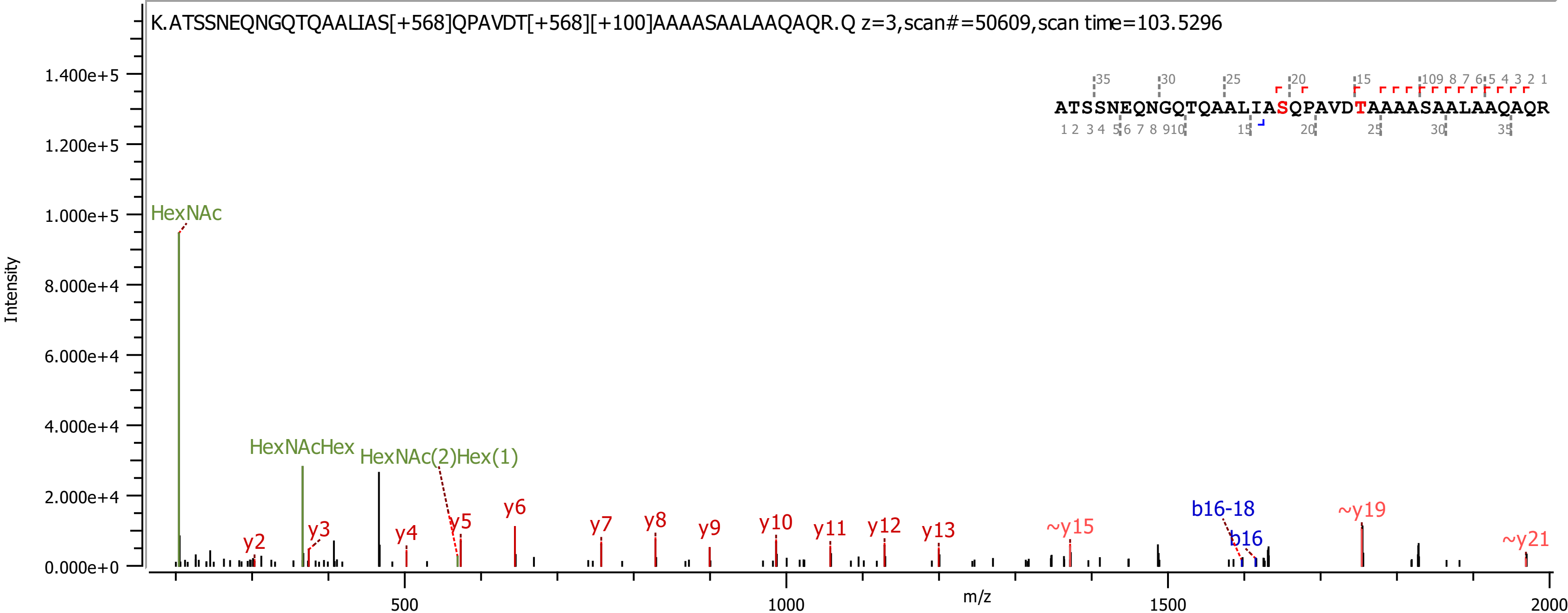

K.ATSSNEQNGQTQAALIAS[+568]QPAVDT[+568][+100]AAAASAALAAQAQR.Q z=3,scan#=50609,scan time=103.5296
ATSSNEQNGQTQAALIASQPAVDTAAAASAALAAQAQR
Intensity
m/z
HexNAc
HexNAcHex
HexNAc(2)Hex(1)
y2
y3
y4
y5
y6
y7
y8
y9
y10
y11
y12
y13
~y15
b16-18
b16
~y19
~y21

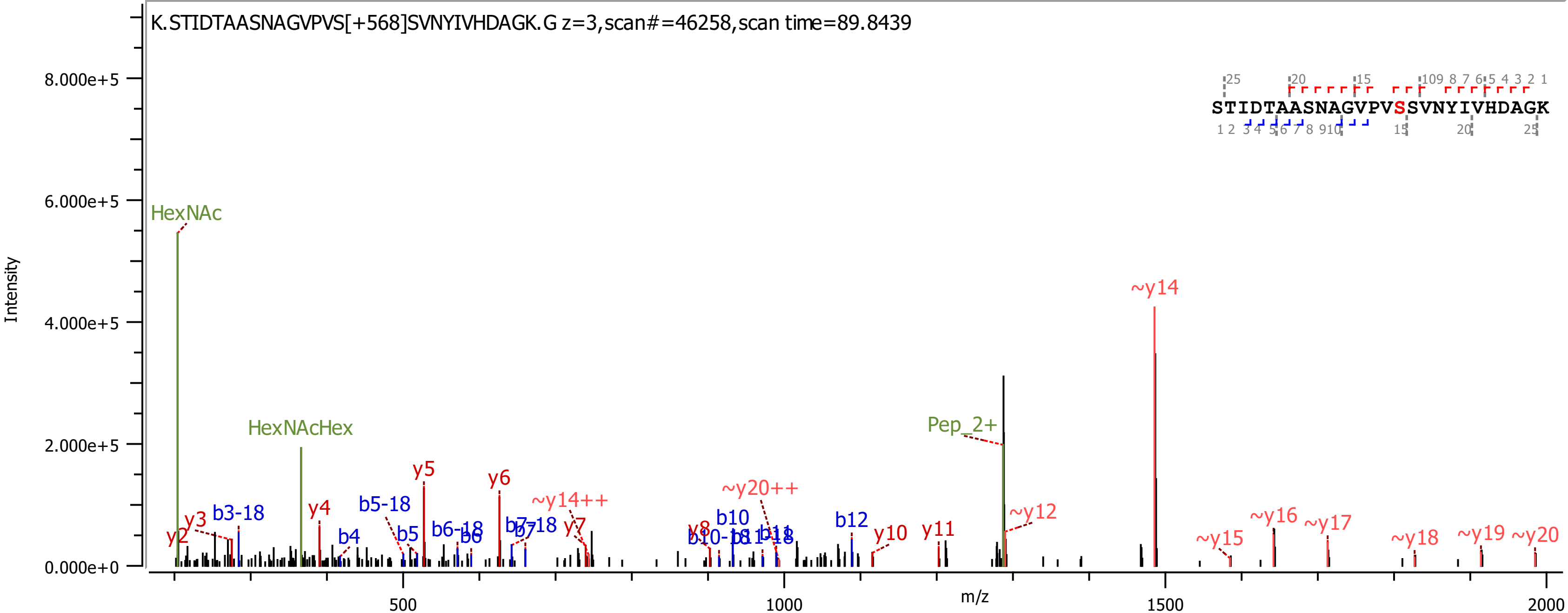
K.STIDTAASNAGVPVS[+568]SVNYIVHDAGK.G z=3,scan#=46258,scan time=89.8439
STIDTAASNAGVPVSSVNYIVHDAGK
Intensity
m/z
HexNAc
HexNAcHex
Pep_2+
y2
y3
y4
y5
y6
y7
y8
y10
y11
b3-18
b4
b5-18
b5
b6-18
b6
b7-18
b7
b10
b10-18
b11
b11-18
b12
~y14
~y14++
~y20++
~y12
~y15
~y16
~y17
~y18
~y19
~y20

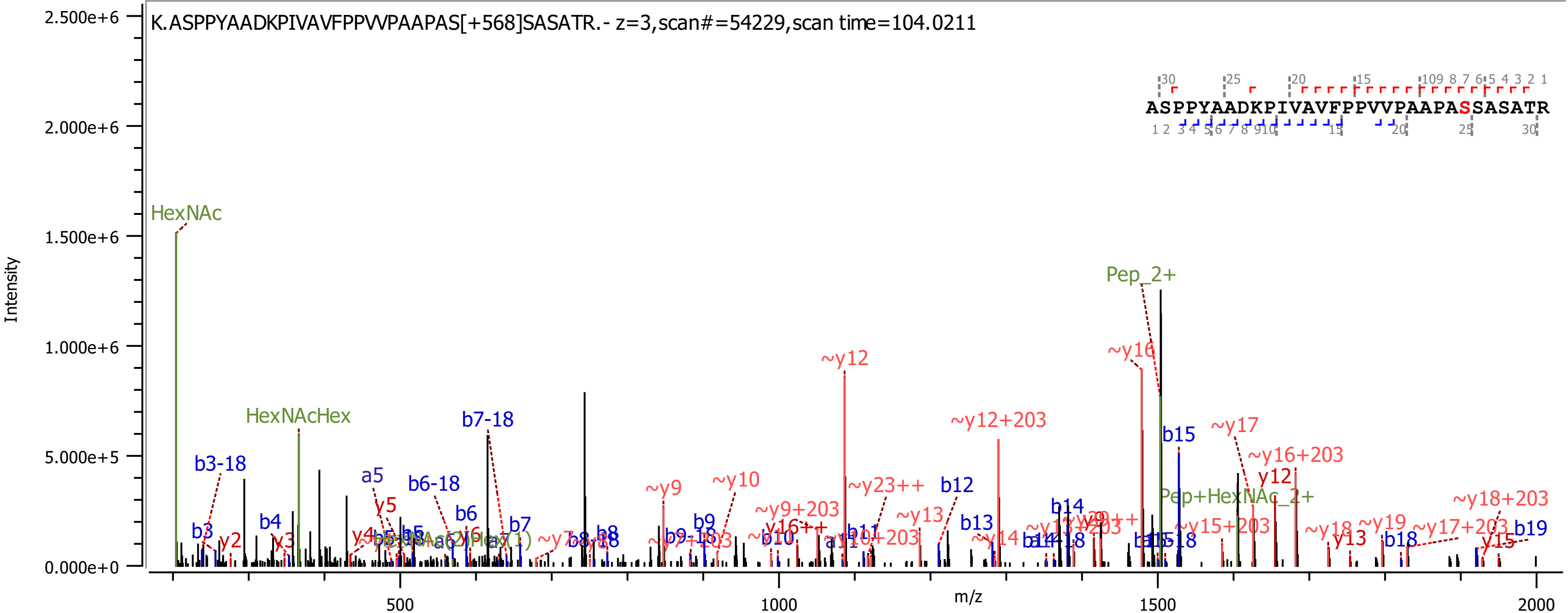
K.ASPPYAADKPIVAVFPPVVPAAPAS[+568]SASATR.- z=3,scan#=54229,scan time=104.0211
ASPPYAADKPIVAVFPPVVPAAPASSASATR
Intensity
m/z
2.500e+6
2.000e+6
1.500e+6
1.000e+6
5.000e+5
0.000e+0
500
1000
1500
2000
HexNAc
HexNAcHex
Pep_2+
Pep+HexNAc_2+
b3-18
b3
y2
b4
y3
a5
y5
y4
b6-18
b6
b7-18
b7
b5
b8
b9
~y9
~y10
~y9+203
~y12
~y23++
~y13
b10
b11
~y10+203
b12
b13
~y14
~y12+203
b14
~y16
b15
~y17
~y16+203
y12
~y15+203
~y18
y13
~y19
b18
~y17+203
~y18+203
y15
b19

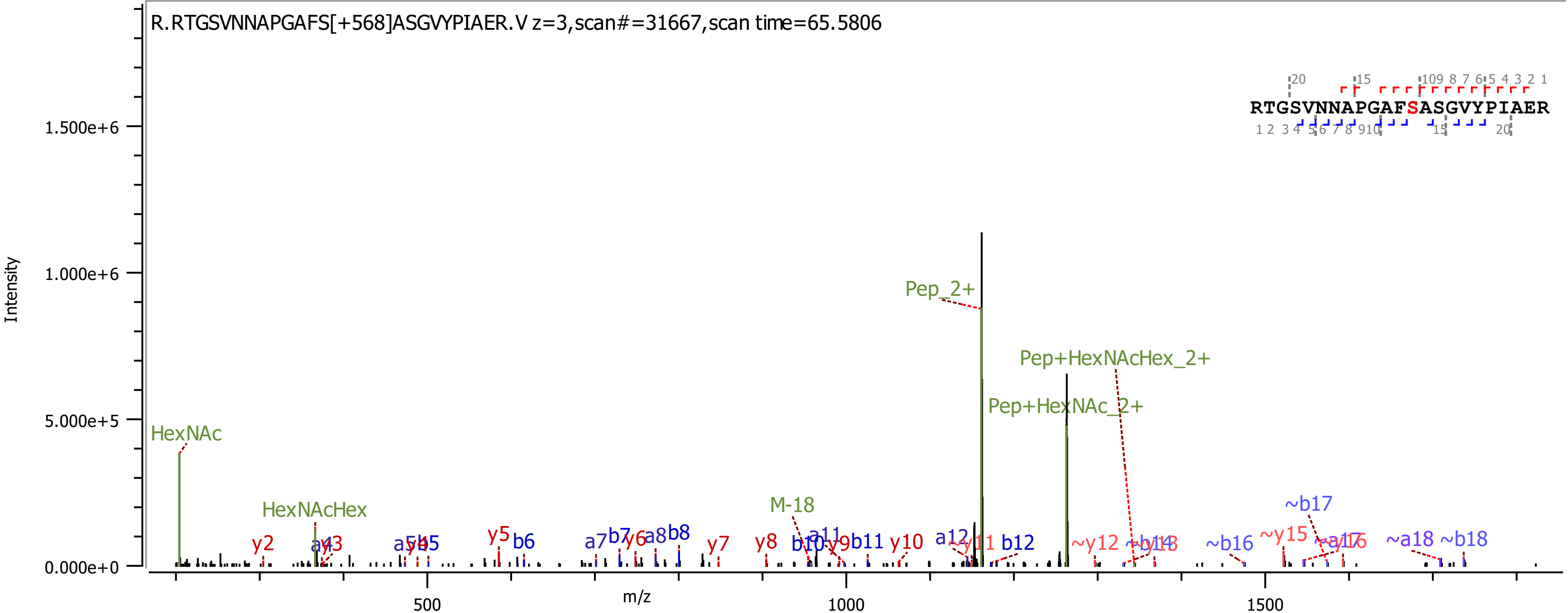
R.RTGSVNNAPGAFS[+568]ASGVYPIAER.V z=3,scan#=31667,scan time=65.5806
RTGSVNNAPGAFSASGVYPIAER
Intensity
m/z
0.000e+0
5.000e+5
1.000e+6
1.500e+6
500
1000
1500
HexNAc
HexNAcHex
M-18
Pep_2+
Pep+HexNAcHex_2+
Pep+HexNAc_2+
y2
y5
b6
a7
b7
y6
a8
b8
y7
y8
b10
a11
y9
b11
y10
a12
b12
~y12
~b16
~y15
~b17
~a18
~b18

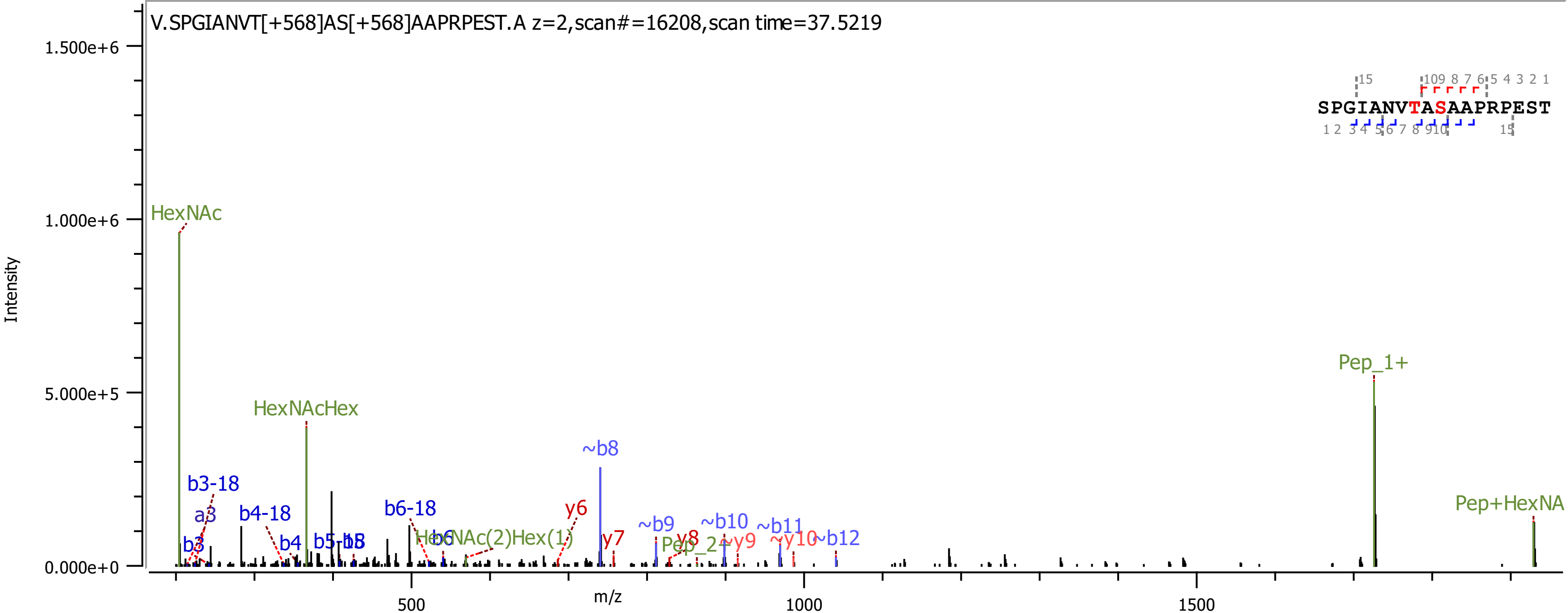

V.SPGIANVT[+568]AS[+568]AAPRPEST.A z=2,scan#=16208,scan time=37.5219
Intensity
1.500e+6
1.000e+6
5.000e+5
0.000e+0
m/z
500
1000
1500
HexNAc
HexNAcHex
b3-18
a3
b3
b4-18
b4
b5-18
b5
b6-18
b6
HexNAc(2)Hex(1)
~b8
y6
y7
~b9
y8
Pep_2+
~b10
~y9
~b11
~y10
~b12
Pep_1+
Pep+HexNA
SPGIANVTASAAPRPEST

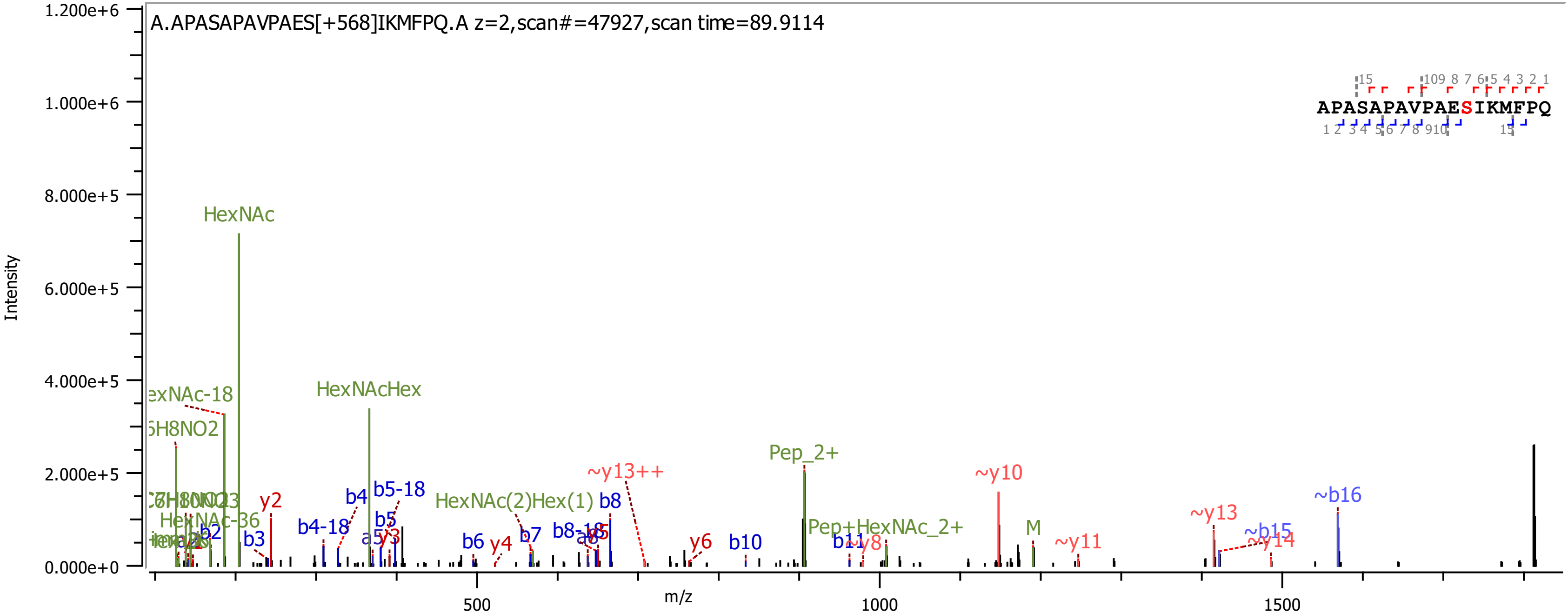

A.APASAPAVPAES[+568]IKMFPQ.A z=2,scan#=47927,scan time=89.9114
APASAPAVPAESIKMFPQ
Intensity
m/z
1.200e+6
1.000e+6
8.000e+5
6.000e+5
4.000e+5
2.000e+5
0.000e+0
500
1000
1500
HexNAc
HexNAcHex
Pep_2+
~y10
~y13++
~b16
~y13
~b15
~y14
HexNAc(2)Hex(1)
Pep+HexNAc_2+
M
~y11
b8
b5-18
b4
b4-18
b5
y2
b2
b3
y3
b6
y4
b7
b8-18
y5
y6
b10
b11
~y8
HexNAc-36

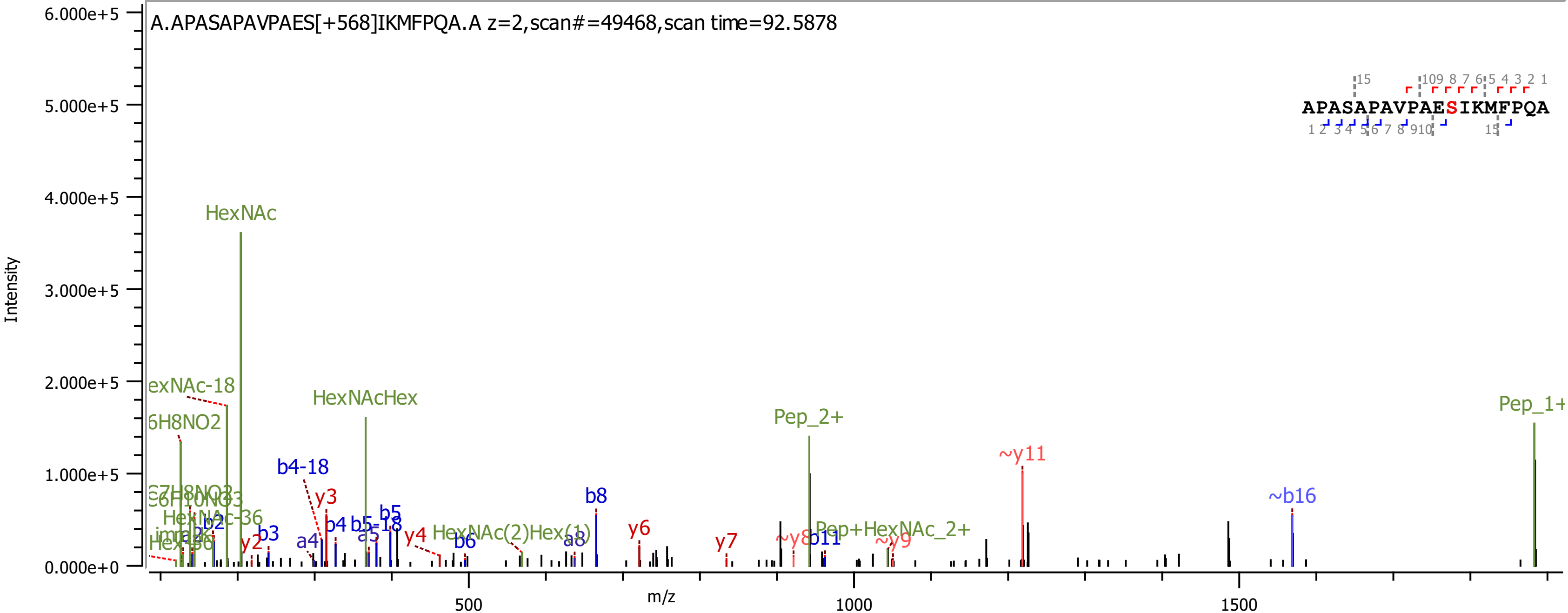
A.APASAPAVPAES[+568]IKMFPQA.A z=2,scan#=49468,scan time=92.5878
APASAPAVPAESIKMFPQA
Intensity
6.000e+5
5.000e+5
4.000e+5
3.000e+5
2.000e+5
1.000e+5
0.000e+0
m/z
500
1000
1500
HexNAc
HexNAcHex
Pep_2+
Pep_1+
~y11
~b16
b4-18
b8
y3
y6
y7
b5
b4
b5-18
a5
a4
b3
y2
y4
b6
a8
HexNAc(2)Hex(1)
Pep+HexNAc_2+
b11
~y8
~y9
HexNAc-36
b2

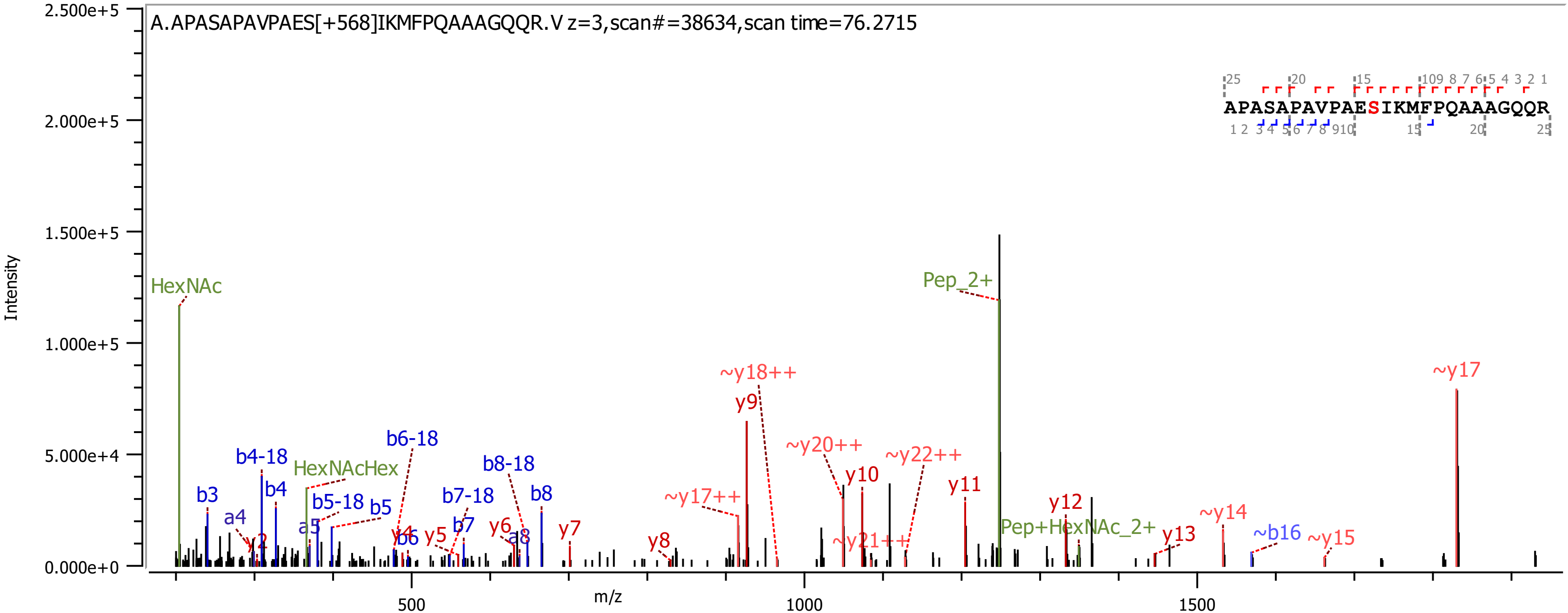
A.APASAPAVPAES[+568]IKMFPQAAAGQQR.V z=3,scan#=38634,scan time=76.2715
APASAPAVPAESIKMFPQAAAGQQR
Intensity
m/z
HexNAc
HexNAcHex
Pep_2+
Pep+HexNAc_2+
b3
a4
b4-18
b4
y2
a5
b5-18
b5
b6-18
y4
b6
y5
b7-18
b7
y6
b8-18
a8
b8
y7
y8
~y17++
y9
~y18++
~y20++
y10
~y21++
~y22++
y11
y12
y13
~y14
~b16
~y15
~y17

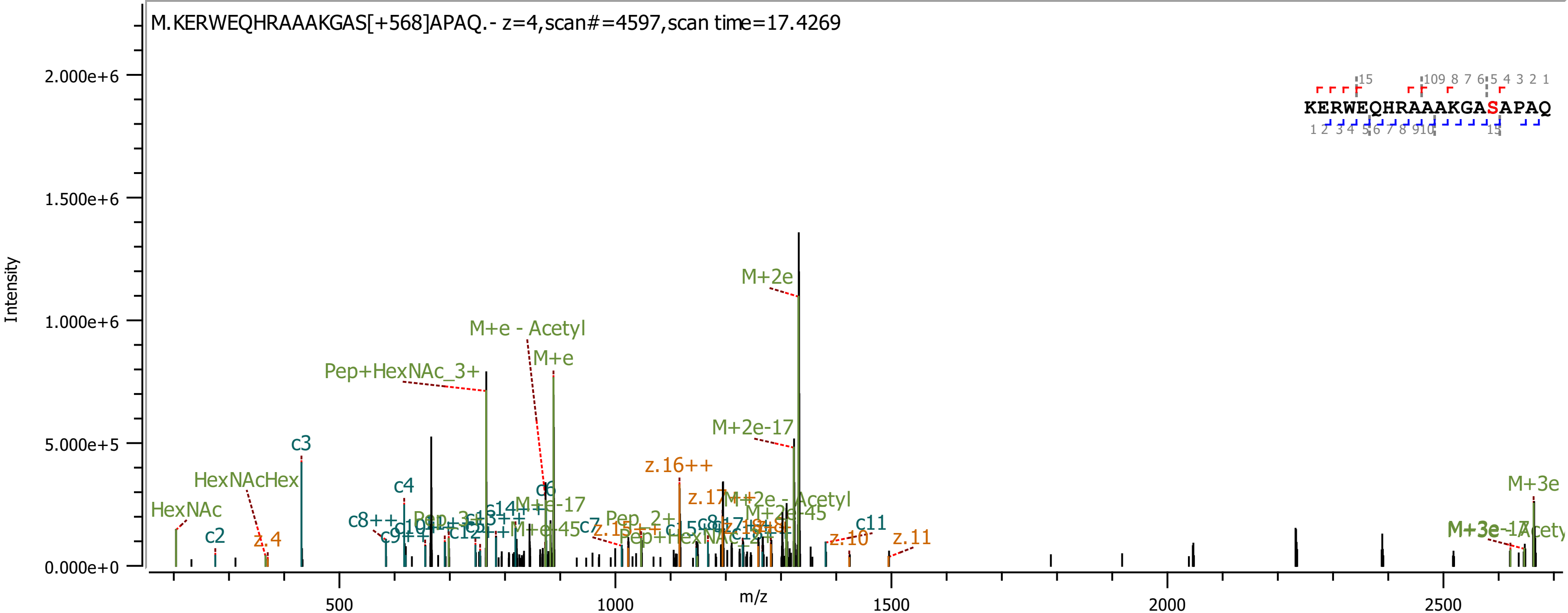

M.KERWEQHRAAAKGAS[+568]APAQ.- z=4,scan#=4597,scan time=17.4269
KERWEQHRAAAKGASAPAQ
Intensity
m/z
2.000e+6
1.500e+6
1.000e+6
5.000e+5
0.000e+0
500
1000
1500
2000
2500
M+2e
M+e - Acetyl
M+e
Pep+HexNAc_3+
M+2e-17
c3
HexNAcHex
HexNAc
c2
z.4
c4
c6
z.16++
M+e-17
M+2e-45
c11
z.10
z.11
M+3e
c7
c8++

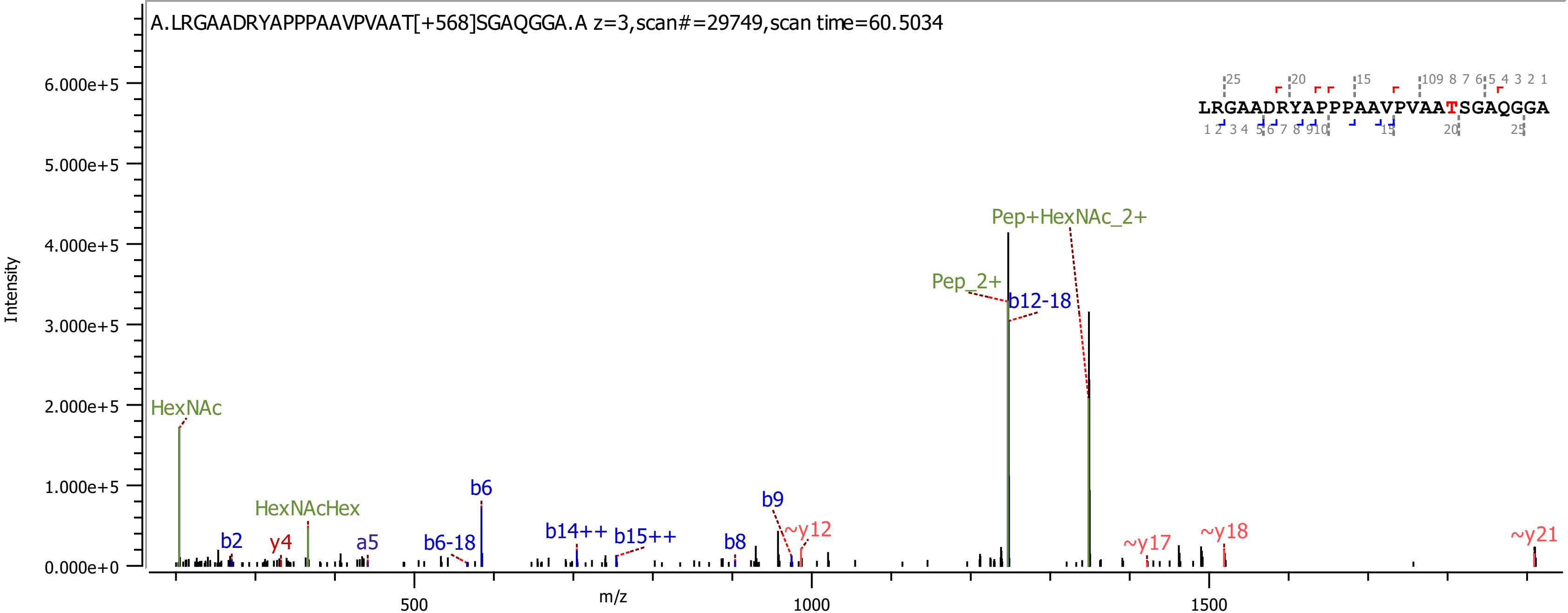
A.LRGAADRYAPPPAAVPVAAT[+568]SGAQGGA.A z=3,scan#=29749,scan time=60.5034
LRGAADRYAPPPAAVPVAATSGAQGGA
Intensity
m/z
HexNAc
HexNAcHex
b2
y4
a5
b6-18
b6
b14++
b15++
b8
b9
~y12
Pep_2+
b12-18
Pep+HexNAc_2+
~y17
~y18
~y21

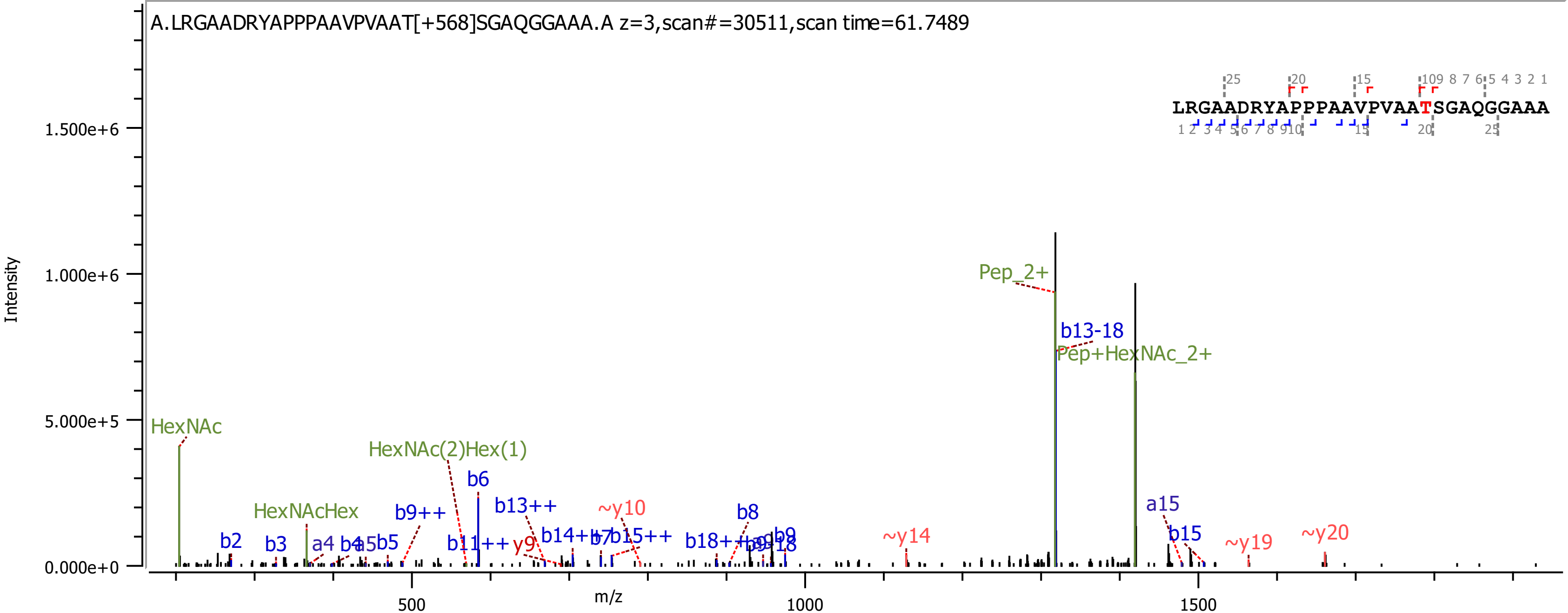
A.LRGAADRYAPPPAAVPVAAT[+568]SGAQGGAAA.A z=3,scan#=30511,scan time=61.7489
LRGAADRYAPPPAAVPVAATSGAQGGAAA
Intensity
m/z
HexNAc
HexNAcHex
HexNAc(2)Hex(1)
Pep_2+
Pep+HexNAc_2+
b13-18
b6
b9++
b13++
b11++
y9
b14++
b15++
~y10
b8
b18++
b9
~y14
a15
b15
~y19
~y20
b2
b3
a4
b4
a5
b5
0.000e+0
5.000e+5
1.000e+6
1.500e+6
500
1000
1500

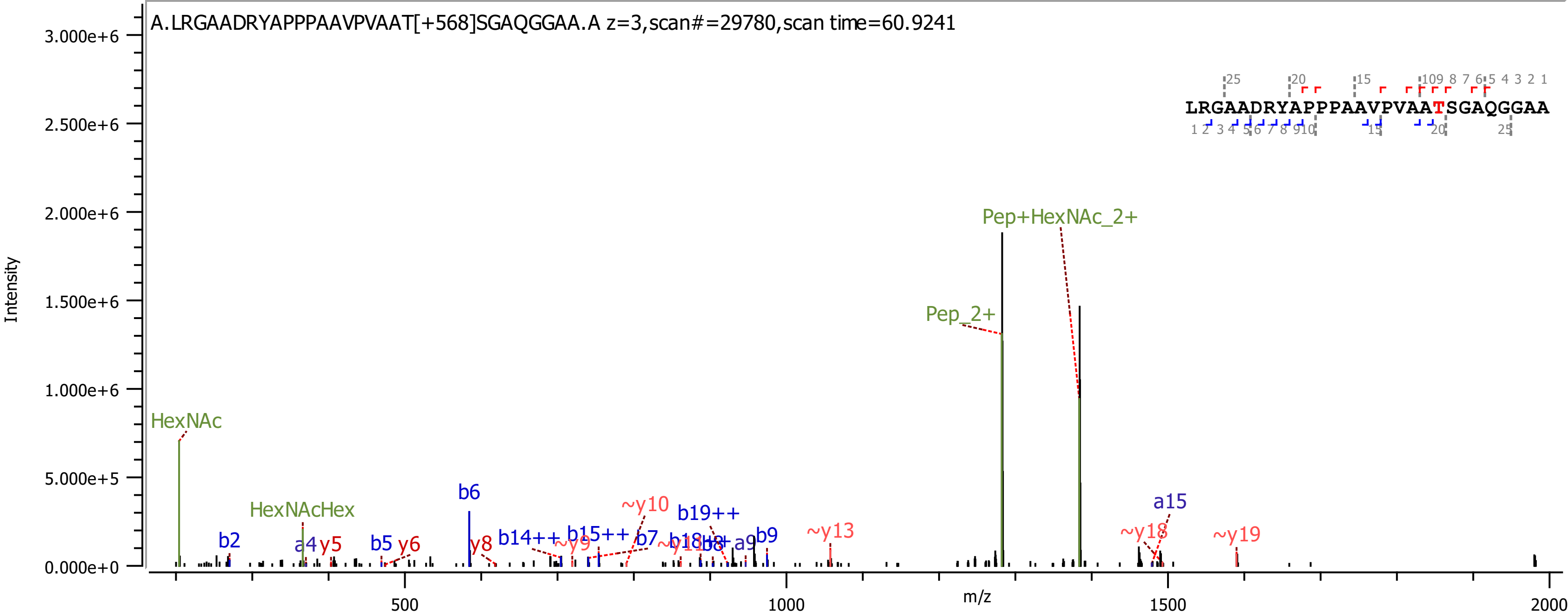
A.LRGAADRYAPPPAAVPVAAT[+568]SGAQGGAA.A z=3,scan#=29780,scan time=60.9241
LRGAADRYAPPPAAVPVAATSGAQGGAA
Pep+HexNAc_2+
Pep_2+
HexNAc
HexNAcHex
b2
a4
y5
b5
y6
b6
y8
b14++
~y9
b15++
~y10
b7
b18++
b8
a9
b19++
b9
~y13
a15
~y18
~y19
Intensity
m/z

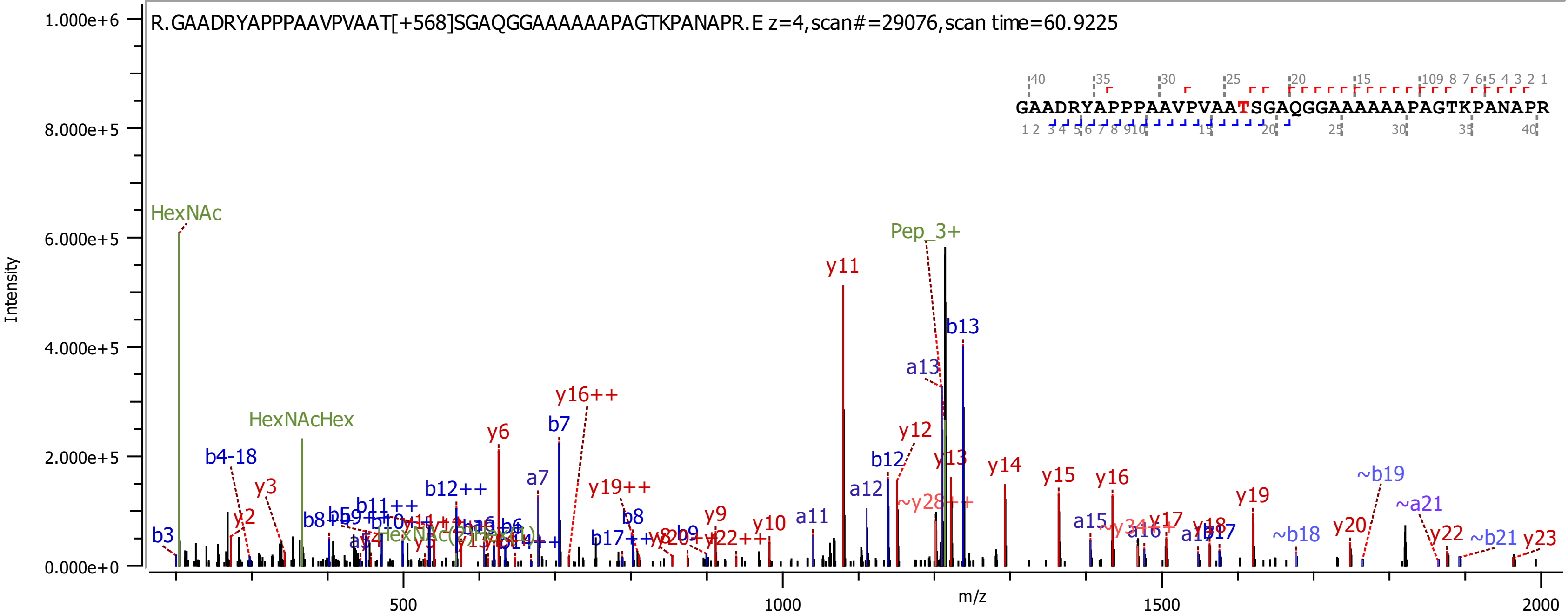
R.GAADRYAPPPAAVPVAAT[+568]SGAQGGAAAAAAPAGTKPANAPR.E z=4,scan#=29076,scan time=60.9225
GAADRYAPPPAAVPVAATSGAQGGAAAAAAPAGTKPANAPR
Intensity
m/z
1.000e+6
8.000e+5
6.000e+5
4.000e+5
2.000e+5
0.000e+0
500
1000
1500
2000
HexNAc
HexNAcHex
Pep_3+
y11
b13
a13
y16++
b7
y6
y12
b12
y13
y14
y15
y16
~b19
a7
y19++
b12++
b4-18
y3
y2
b3
a12
~y28++
y19
~a21
y20
y17
y18
b17
a17
a16
a15
~y34++
~b18
y22
~b21
y23
a11
y10
y9
b9
b8
y22++
b11++
b17++

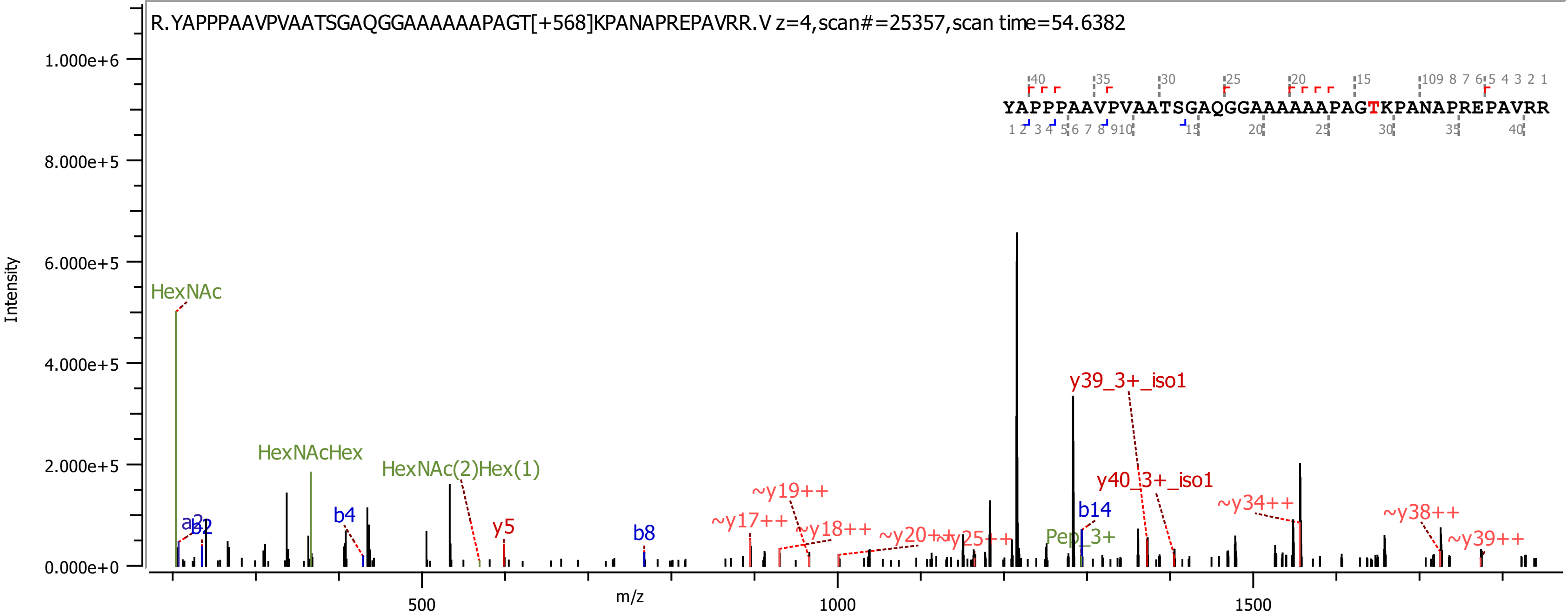

R.YAPPPAAVPVAATSGAQGGAAAAAAPAGT[+568]KPANAPREPAVRR.V z=4,scan#=25357,scan time=54.6382
YAPPPAAVPVAATSGAQGGAAAAAAPAGTKPANAPREPAVRR
Intensity
m/z
HexNAc
HexNAcHex
HexNAc(2)Hex(1)
a2
b2
b4
y5
b8
~y17++
~y19++
~y18++
~y20++
~y25++
Pep_3+
b14
y39_3+_iso1
y40_3+_iso1
~y34++
~y38++
~y39++

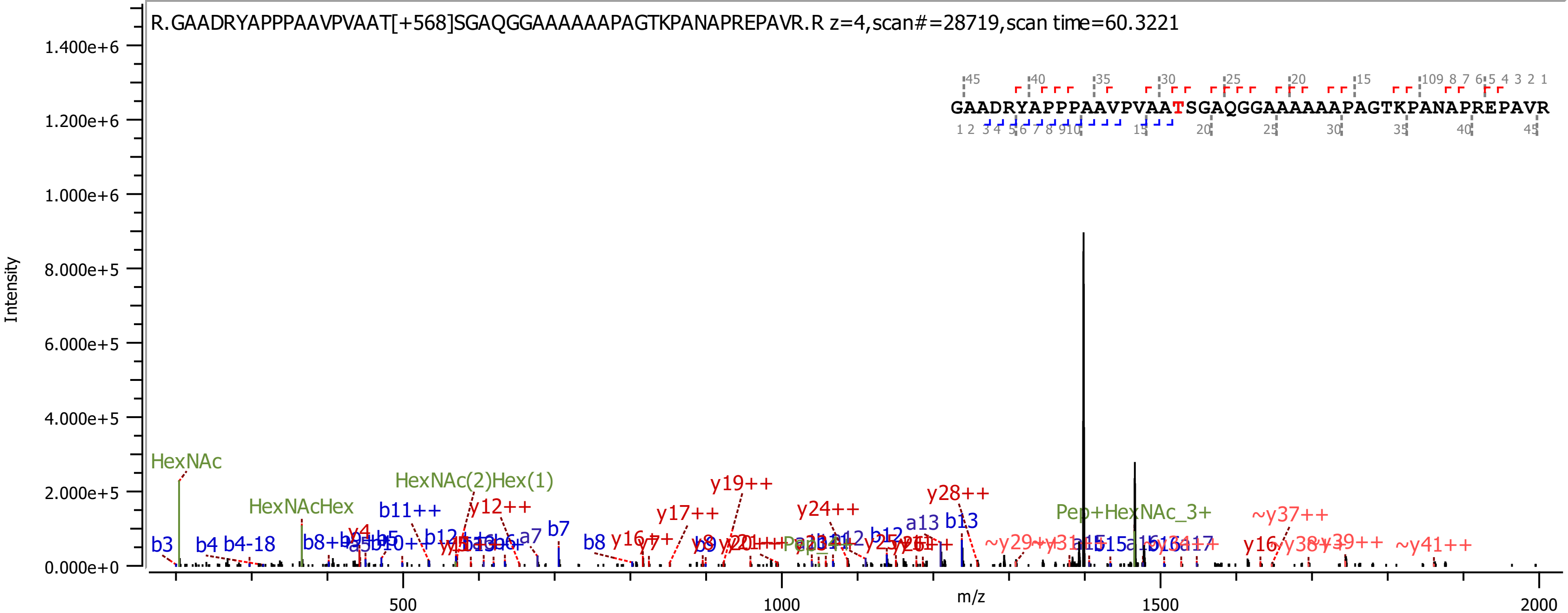
R.GAADRYAPPPAAVPVAAT[+568]SGAQGGAAAAAAPAGTKPANAPREPAVR.R z=4,scan#=28719,scan time=60.3221
GAADRYAPPPAAVPVAATSGAQGGAAAAAAPAGTKPANAPREPAVR
Intensity
m/z
1.400e+6
1.200e+6
1.000e+6
8.000e+5
6.000e+5
4.000e+5
2.000e+5
0.000e+0
500
1000
1500
2000
HexNAc
HexNAcHex
HexNAc(2)Hex(1)
Pep+HexNAc_3+
b3
b4 b4-18
b11++
y12++
a7
b7
b8
y16++
y17++
y19++
y24++
y28++
a13 b13
~y37++
~y29++
~y41++
y16

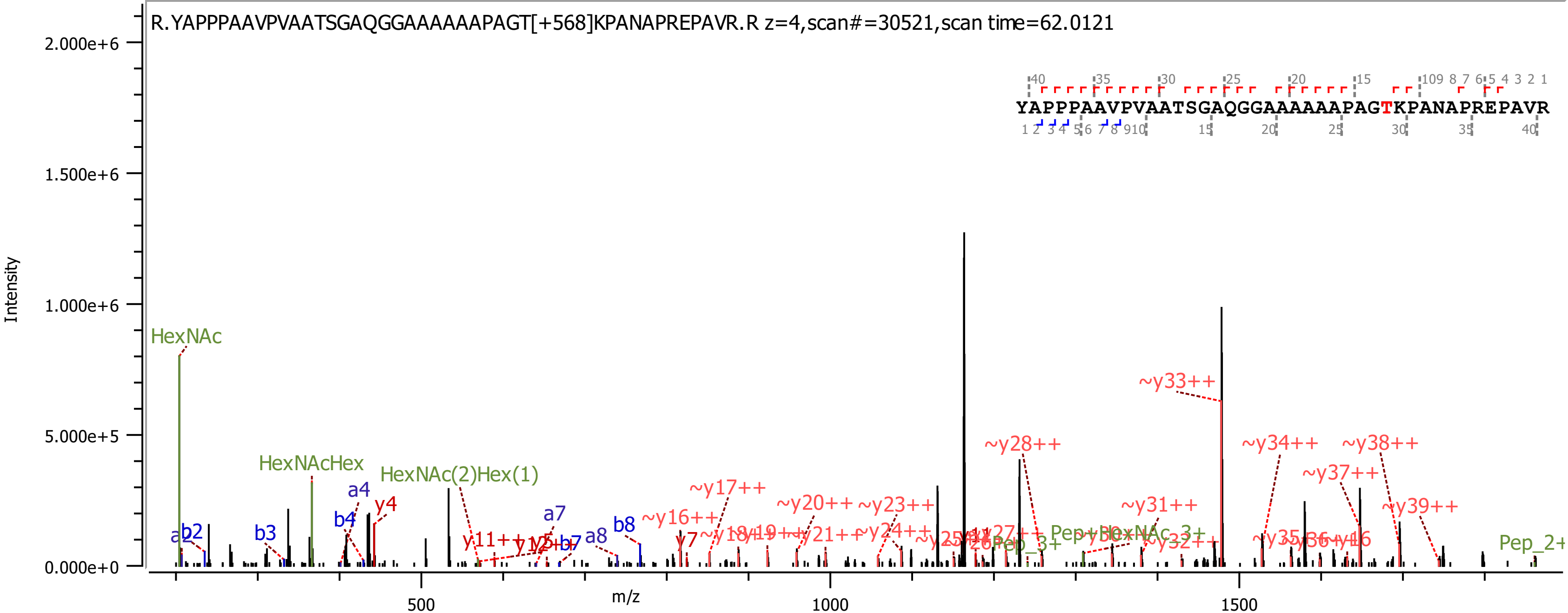
R.YAPPPAAVPVAATSGAQGGAAAAAAPAGT[+568]KPANAPREPAVR.R z=4,scan#=30521,scan time=62.0121
YAPPPAAVPVAATSGAQGGAAAAAAPAGTKPANAPREPAVR
Intensity
m/z
2.000e+6
1.500e+6
1.000e+6
5.000e+5
0.000e+0
500
1000
1500
HexNAc
HexNAcHex
HexNAc(2)Hex(1)
a2
b2
b3
a4
b4
y4
a7
a8
b8
y7
~y16++
~y17++
~y20++
~y21++
~y23++
~y24++
~y28++
~y31++
~y33++
~y34++
~y37++
~y38++
~y39++
Pep_3+
Pep_2+

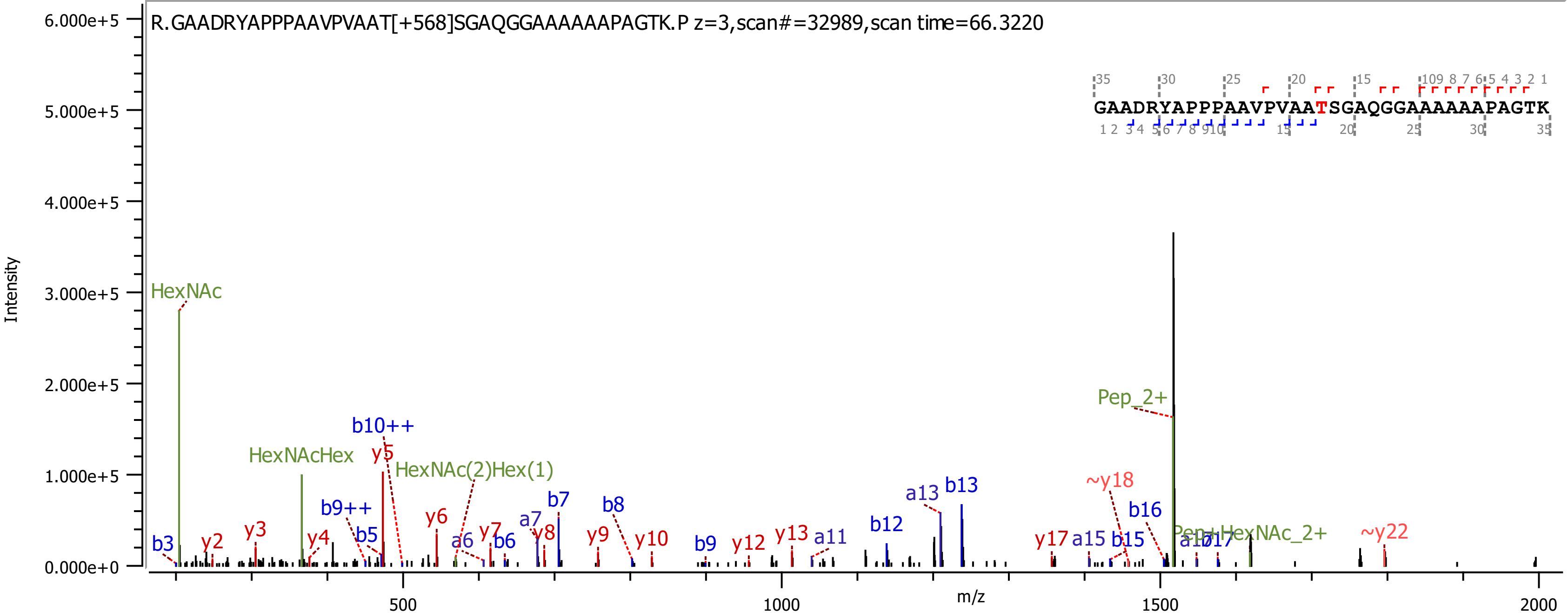

R.GAADRYAPPPAAVPVAAT[+568]SGAQGGAAAAAAPAGTK.P z=3,scan#=32989,scan time=66.3220
GAADRYAPPPAAVPVAATSGAQGGAAAAAAPAGTK
Intensity
m/z
HexNAc
HexNAcHex
HexNAc(2)Hex(1)
Pep_2+
Pep+HexNAc_2+
b10++
b9++
y5
~y18
~y22

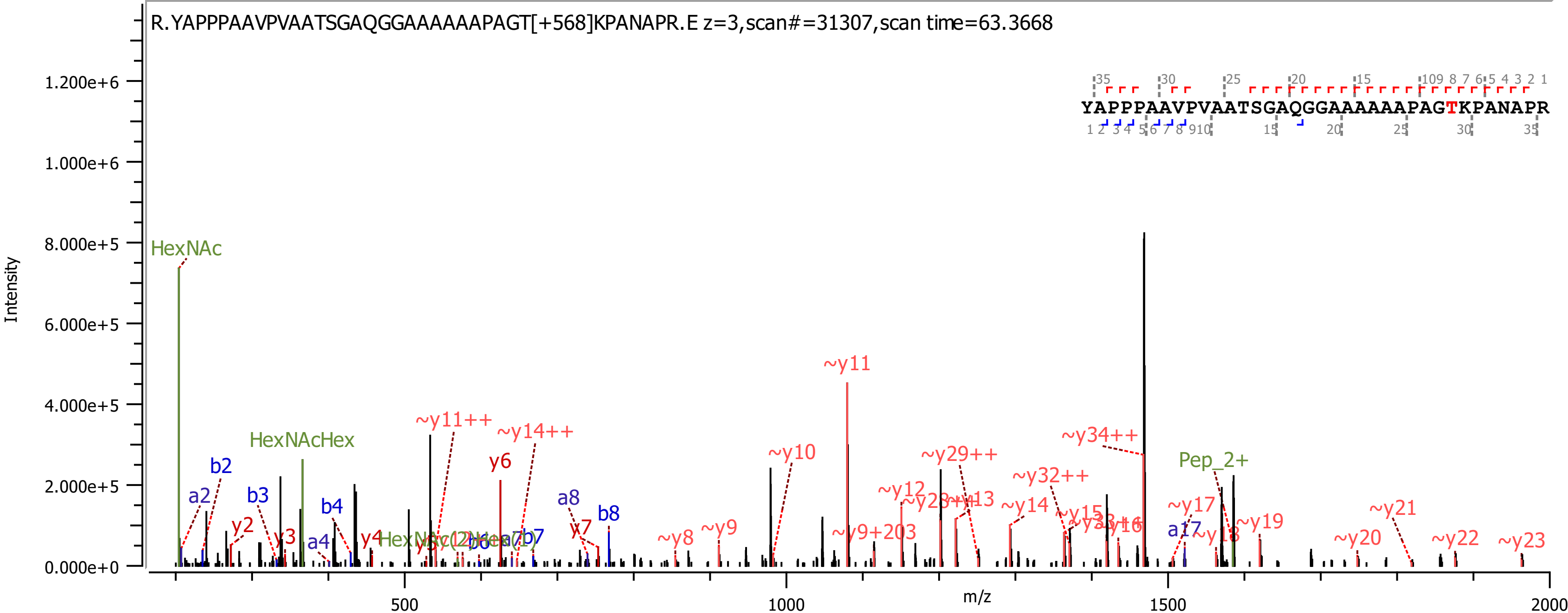
R.YAPPPAAVPVAATSGAQGGAAAAAAPAGT[+568]KPANAPR.E z=3,scan#=31307,scan time=63.3668
YAPPPAAVPVAATSGAQGGAAAAAAPAGTKPANAPR
Intensity
m/z
0.000e+0
2.000e+5
4.000e+5
6.000e+5
8.000e+5
1.000e+6
1.200e+6
500
1000
1500
2000
HexNAc
HexNAcHex
Pep_2+
a2
b2
b3
y2
y3
a4
b4
y4
y6
a8
b8
y7
b7
~y8
~y9
~y10
~y11
~y11++
~y14++
~y9+203
~y12
~y29++
~y34++
~y32++
~y14
~y15
~y17
a17
~y19
~y20
~y21
~y22
~y23

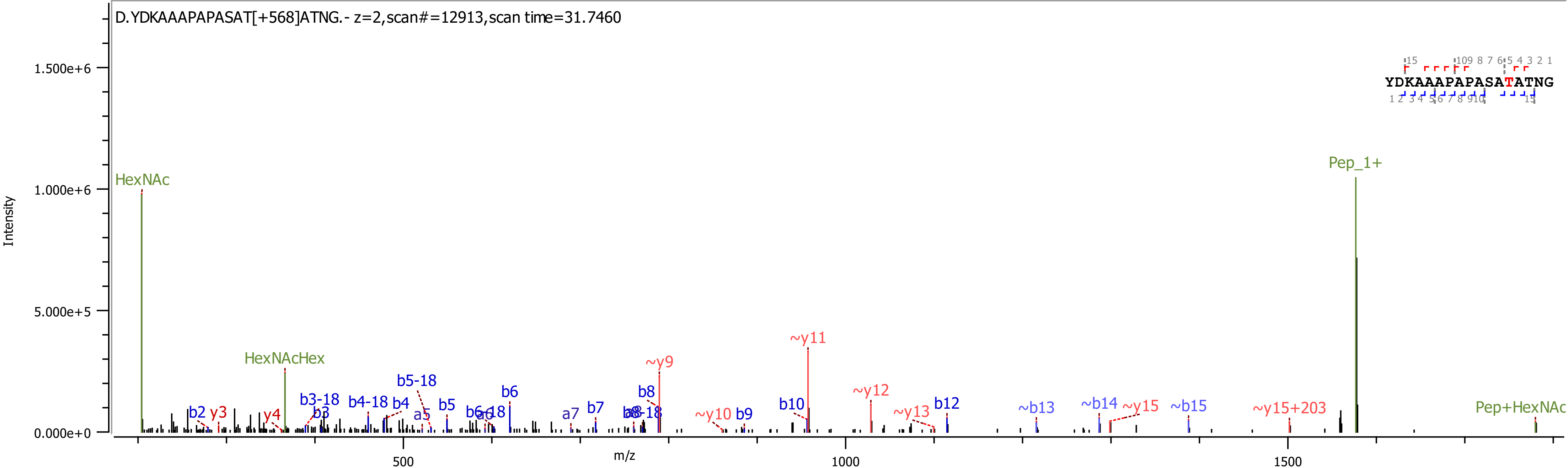

D.YDKAAAPAPASAT[+568]ATNG.- z=2,scan#=12913,scan time=31.7460
YDKAAAPAPASATATNG
Intensity
m/z
HexNAc
HexNAcHex
Pep_1+
Pep+HexNAc
b2
y3
y4
b3-18
b3
b4-18
b4
b5-18
a5
b5
b6-18
a6
b6
a7
b7
b8
b8-18
~y9
~y10
b9
b10
~y11
~y12
~y13
b12
~b13
~b14
~y15
~b15
~y15+203
1.500e+6
1.000e+6
5.000e+5
0.000e+0
500
1000
1500

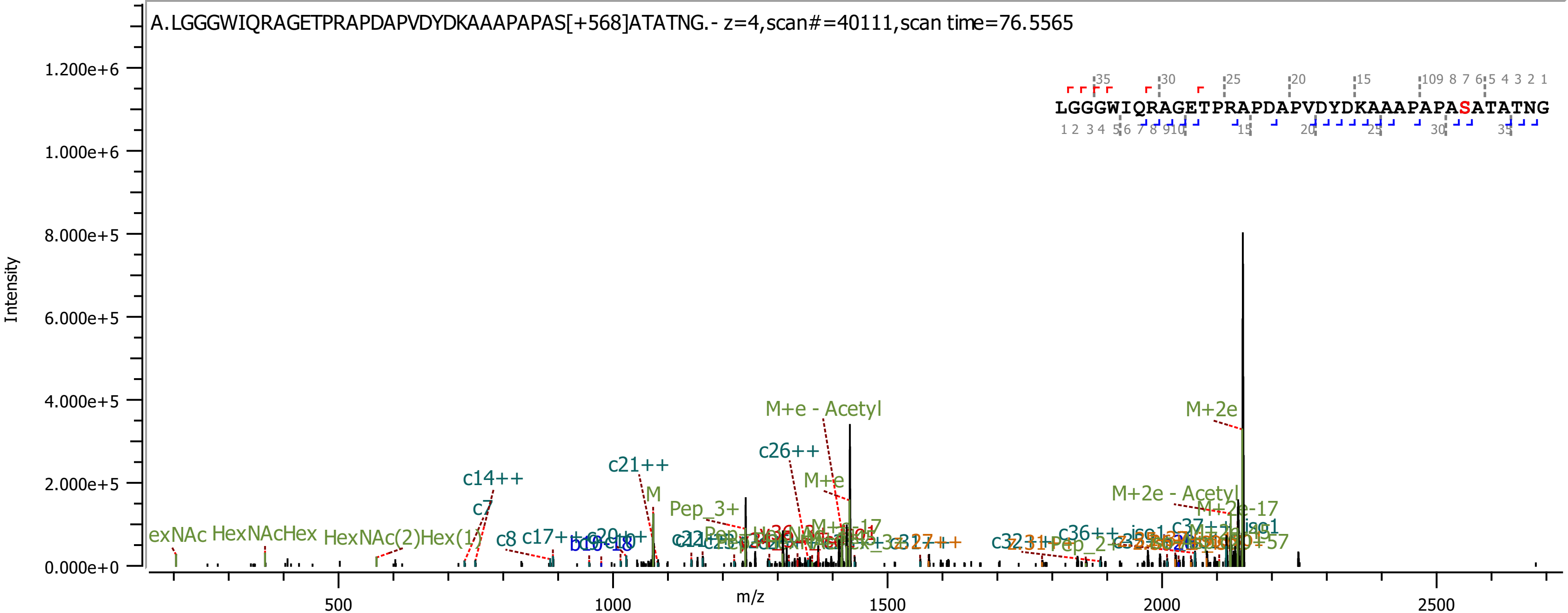
A.LGGGWIQRAGETPRAPDAPVDYDKAAAPAPAS[+568]ATATNG.- z=4,scan#=40111,scan time=76.5565
LGGGWIQRAGETPRAPDAPVDYDKAAAPAPASATATNG
Intensity
m/z
1.200e+6
1.000e+6
8.000e+5
6.000e+5
4.000e+5
2.000e+5
0.000e+0
500
1000
1500
2000
2500
M+e - Acetyl
M+2e
c26++
c21++
c14++
M+e
M
c7
Pep_3+
M+2e - Acetyl
M+2e-17
HexNAcHex
HexNAc(2)Hex(1)
c8
c17++
c36++
c37++

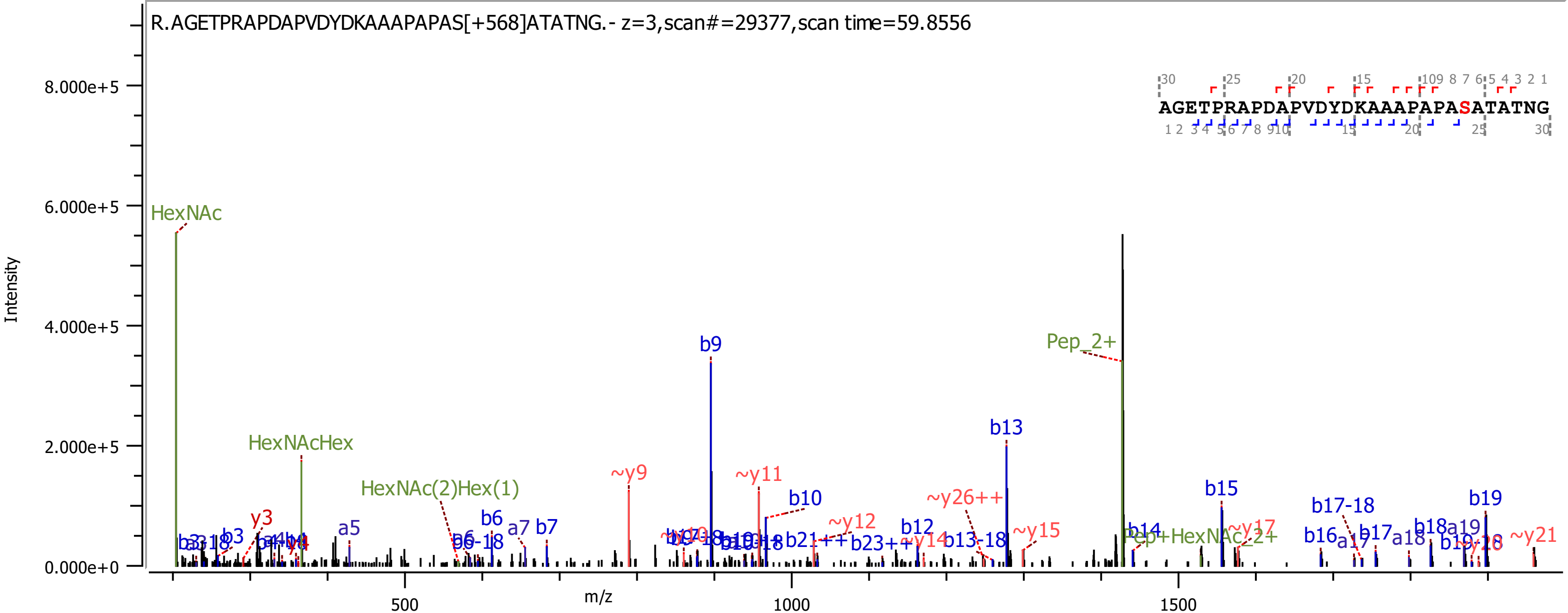
R.AGETPRAPDAPVDYDKAAAPAPAS[+568]ATATNG.- z=3,scan#=29377,scan time=59.8556
AGETPRAPDAPVDYDKAAAPAPASATATNG
Intensity
m/z
8.000e+5
6.000e+5
4.000e+5
2.000e+5
0.000e+0
500
1000
1500
HexNAc
HexNAcHex
HexNAc(2)Hex(1)
Pep_2+
Pep+HexNAc_2+
b9
b13
b10
b15
b19
~y9
~y11
~y12
~y26++
~y15
~y17
~y21
b17-18
b6
b7
a7
a5
y3
b12
b14
b16
a17
b17
a18
b18
a19
b13-18
b21++
b23++
~y14

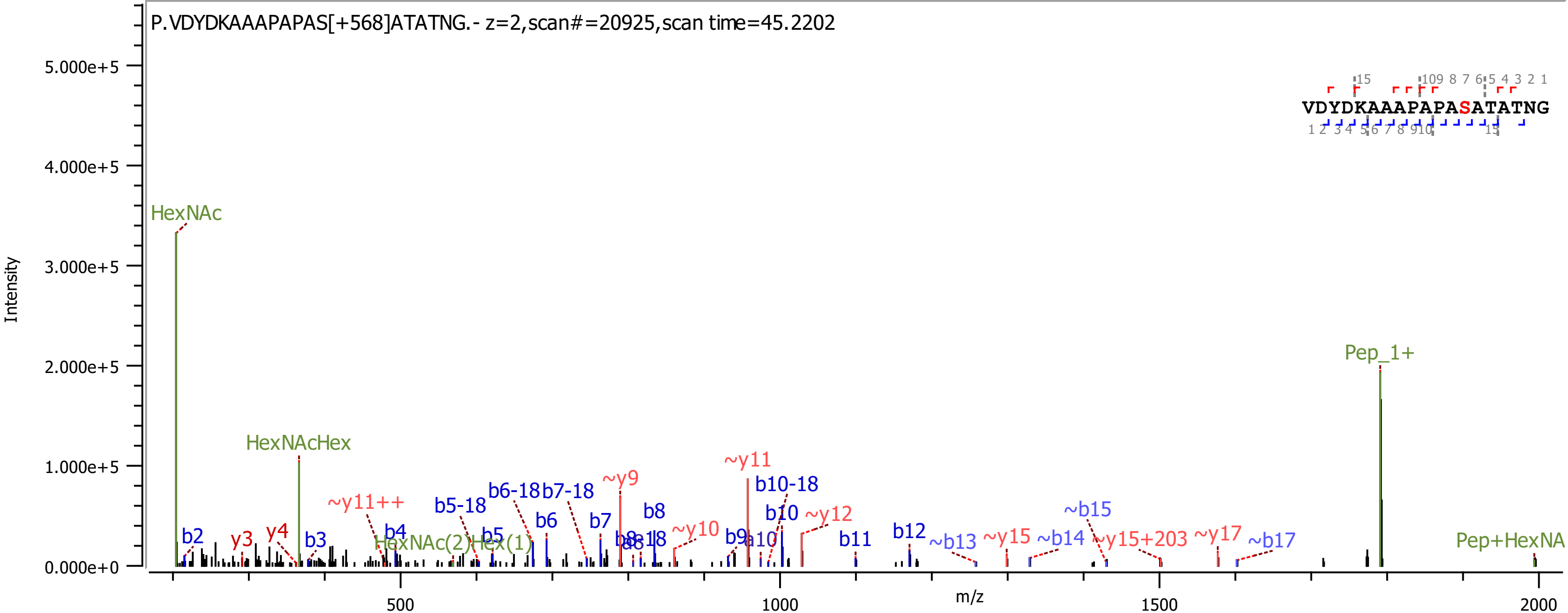
P.VDYDKAAAPAPAS[+568]ATATNG.- z=2,scan#=20925,scan time=45.2202
VDYDKAAAPAPASATATNG
Intensity
5.000e+5
4.000e+5
3.000e+5
2.000e+5
1.000e+5
0.000e+0
m/z
500
1000
1500
2000
HexNAc
HexNAcHex
HexNAc(2)Hex(1)
Pep_1+
Pep+HexNA
b2
y3
y4
b3
~y11++
b4
b5-18
b5
b6-18
b7-18
b6
b7
~y9
b8
~y10
b9
a10
~y11
b10-18
b10
~y12
b11
b12
~b13
~y15
~b14
~b15
~y15+203
~y17
~b17

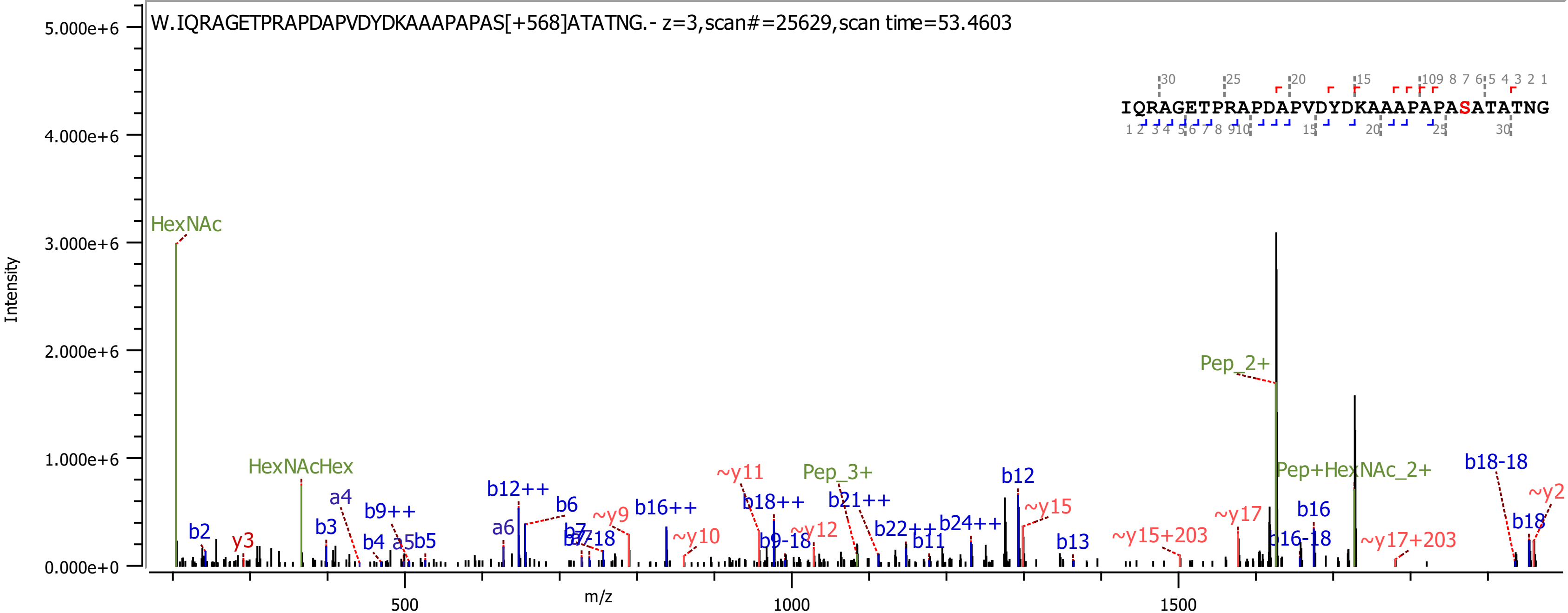
W.IQRAGETPRAPDAPVDYDKAAAPAPAS[+568]ATATNG.- z=3,scan#=25629,scan time=53.4603
IQRAGETPRAPDAPVDYDKAAAPAPASATATNG
Intensity
m/z
5.000e+6
4.000e+6
3.000e+6
2.000e+6
1.000e+6
0.000e+0
500
1000
1500
HexNAc
HexNAcHex
Pep_2+
Pep+HexNAc_2+
Pep_3+
b2
y3
b3
a4
b9++
b4
a5
b5
a6
b12++
b6
b7
~y9
b16++
~y10
~y11
b18++
b9-18
~y12
b21++
b22++
b11
b24++
b12
~y15
b13
~y15+203
~y17
b16
b16-18
~y17+203
b18-18
b18
~y2

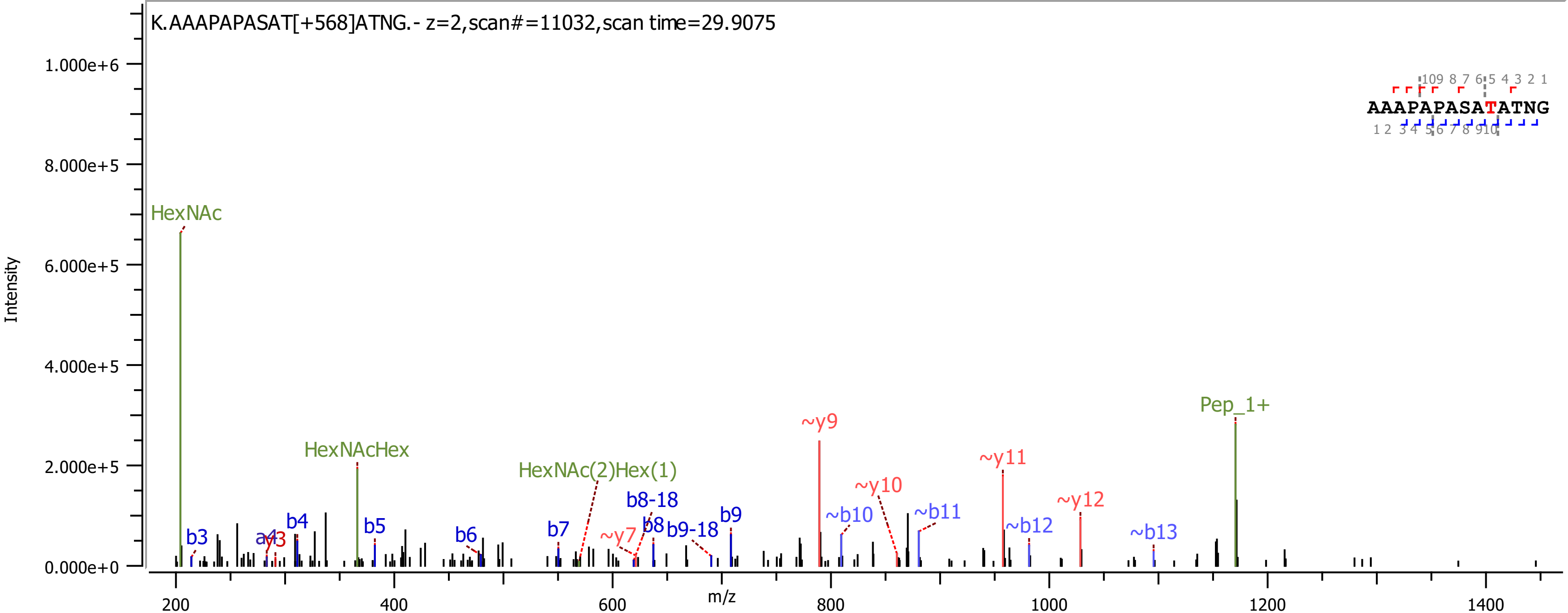
K.AAAPAPASAT[+568]ATNG.- z=2,scan#=11032,scan time=29.9075
AAAPAPASATATNG
HexNAc
HexNAcHex
HexNAc(2)Hex(1)
Pep_1+
b3
a4
y3
b4
b5
b6
b7
~y7
b8-18
b8
b9-18
b9
~y9
~b10
~y10
~b11
~y11
~b12
~y12
~b13
Intensity
m/z

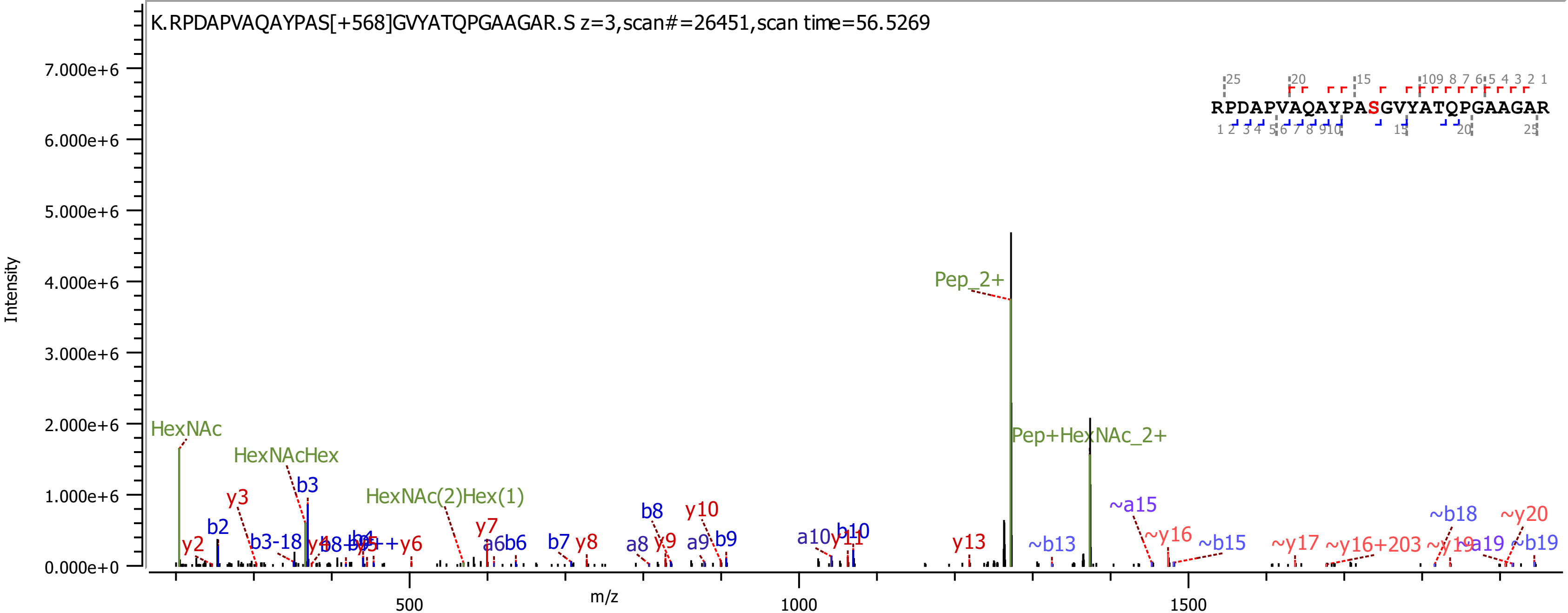
K.RPDAPVAQAYPAS[+568]GVYATQPGAAGAR.S z=3,scan#=26451,scan time=56.5269
RPDAPVAQAYPASGVYATQPGAAGAR
Intensity
m/z
Pep_2+
Pep+HexNAc_2+
HexNAc
HexNAcHex
HexNAc(2)Hex(1)
b2
b3
b3-18
b7
b8
b9
b10
y2
y3
y6
y7
y8
y9
y10
y11
y13
a6b6
a8
a9
a10
~b13
~a15
~y16
~b15
~y17
~y16+203
~b18
~y20
~a19
~b19

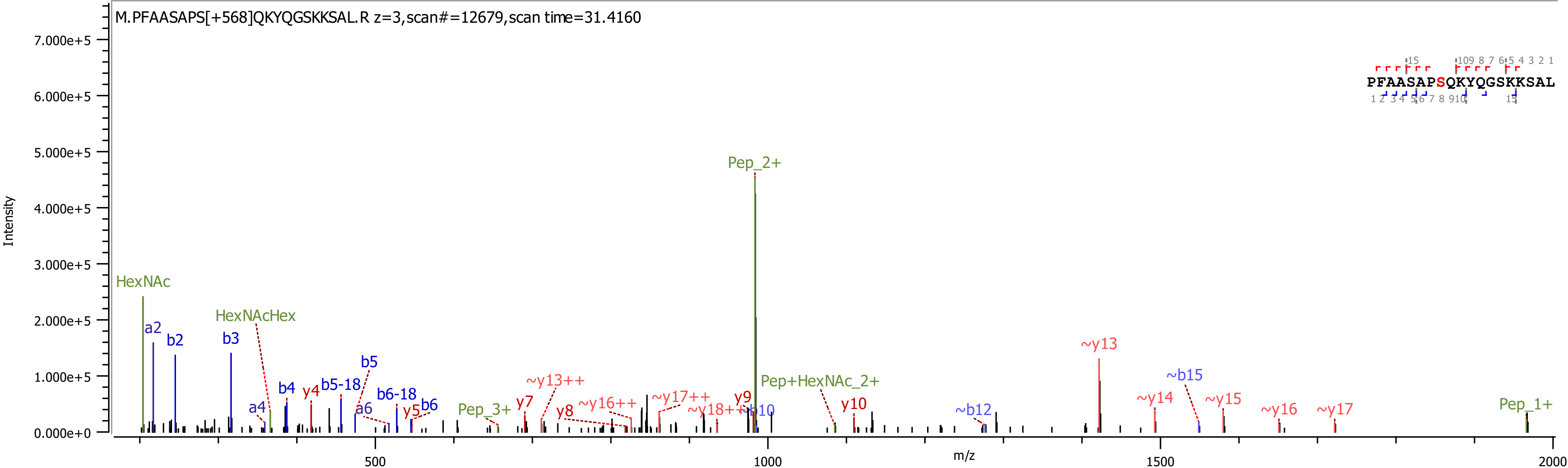
M.PFAASAPS[+568]QKYQGSKKSAL.R z=3,scan#=12679,scan time=31.4160
PFAASAPSQKYQGSKKSAL
Intensity
m/z
HexNAc
a2
b2
b3
HexNAcHex
a4
b4
y4
b5-18
b5
a6
b6-18
y5
b6
Pep_3+
y7
~y13++
y8
~y16++
~y17++
~y18++
y9
b10
Pep_2+
Pep+HexNAc_2+
y10
~b12
~y13
~y14
~b15
~y15
~y16
~y17
Pep_1+

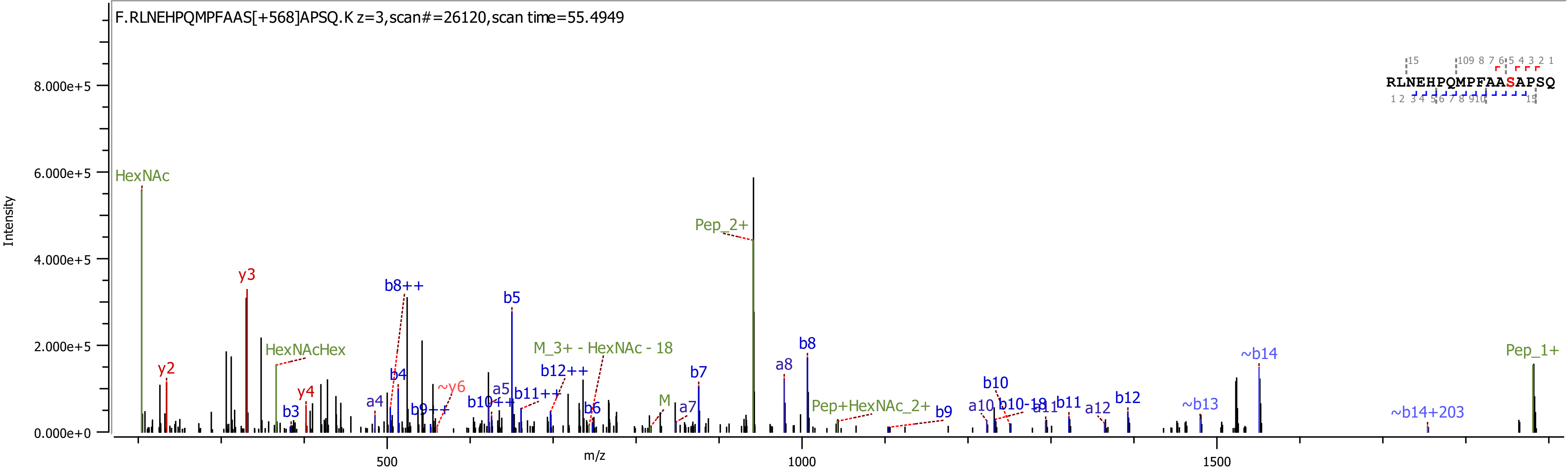

F.RLNEHPQMPFAAS[+568]APSQ.K z=3,scan#=26120,scan time=55.4949
Intensity
m/z
HexNAc
y2
y3
HexNAcHex
b3
y4
a4
b8++
b4
b9++
~y6
b10++
a5
b5
b11++
b12++
M_3+ - HexNAc - 18
b6
M
a7
b7
Pep_2+
a8
b8
Pep+HexNAc_2+
b9
a10
b10
b10-18
a11
b11
a12
b12
~b13
~b14
~b14+203
Pep_1+
RLNEHPQMPFAASAPSQ

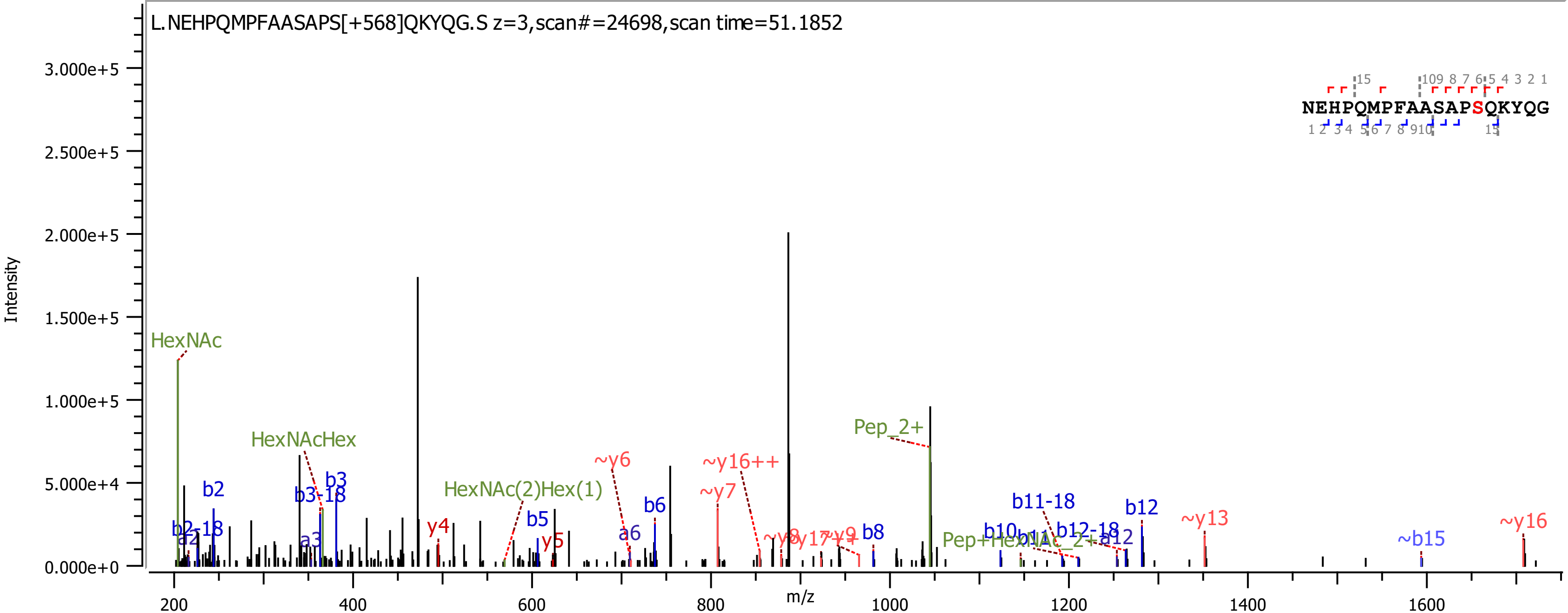
L.NEHPQMPFAASAPS[+568]QKYQG.S z=3,scan#=24698,scan time=51.1852
NEHPQMPFAASAPSQKYQG
Intensity
m/z
HexNAc
HexNAcHex
HexNAc(2)Hex(1)
Pep_2+
Pep+HexNAc_2+
b2
b2-18
a2
b3
b3-18
a3
b5
b6
a6
b8
b10
b11
b11-18
b12
b12-18
a12
~b15
y4
y5
~y6
~y7
~y8
~y9
~y13
~y16
~y16++
~y17++

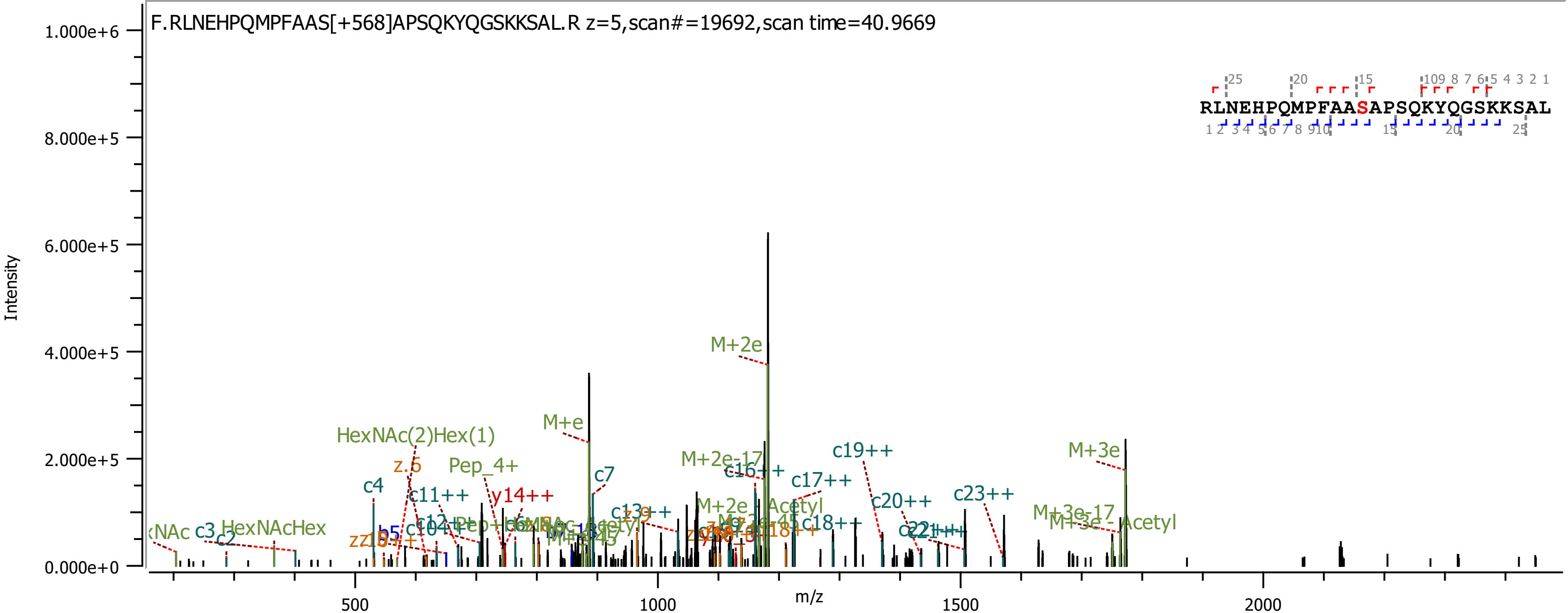

F.RLNEHPQMPFAAS[+568]APSQKYQGSKKSAL.R z=5,scan#=19692,scan time=40.9669
RLNEHPQMPFAASAPSQKYQGSKKSAL
Intensity
m/z
1.000e+6
8.000e+5
6.000e+5
4.000e+5
2.000e+5
0.000e+0
500
1000
1500
2000
M+2e
M+e
HexNAc(2)Hex(1)
Pep_4+
c4
c7
c11++
y14++
M+2e-17
c16++
c17++
c19++
M+2e - Acetyl
c18++
c20++
c23++
M+3e
M+3e-17
M+3e - Acetyl
HexNAcHex
c3
c2
z.6
b5

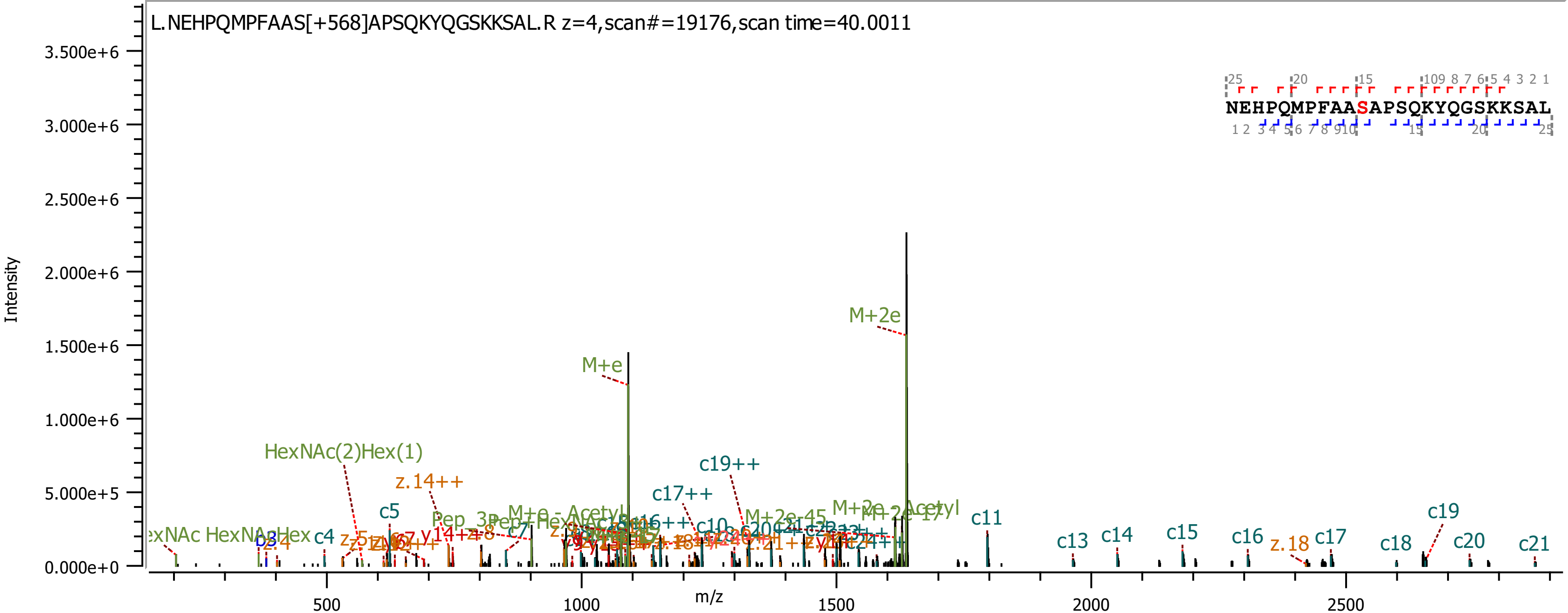

L.NEHPQMPFAAS[+568]APSQKYQGSKKSAL.R z=4,scan#=19176,scan time=40.0011
NEHPQMPFAASAPSQKYQGSKKSAL
Intensity
m/z
M+2e
M+e
HexNAc(2)Hex(1)
z.14++
c19++
c17++
c5
c4
c7
c11
c13
c14
c15
c16
z.18
c17
c18
c19
c20
c21
M+2e-45
M+2e - Acetyl

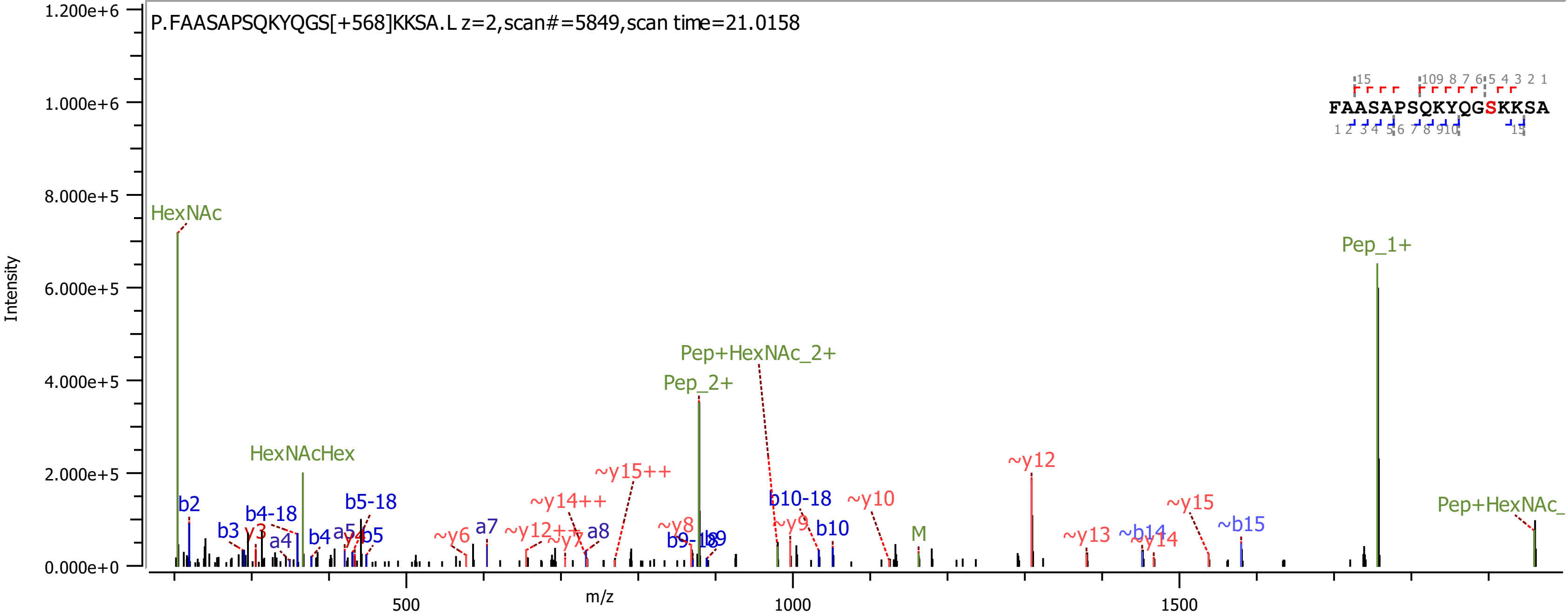
P.FAASAPSQKYQGS[+568]KKSA.L z=2,scan#=5849,scan time=21.0158
FAASAPSQKYQGSKKSA
HexNAc
HexNAcHex
Pep_2+
Pep+HexNAc_2+
Pep_1+
Pep+HexNAc_
b2
b3
y3
b4-18
a4
b4
a5
y4
b5
b5-18
~y6
a7
~y12++
~y7
a8
~y14++
~y15++
~y8
b9-18
b9
b10-18
~y9
b10
~y10
M
~y12
~y13
~b14
~y14
~y15
~b15
Intensity
m/z
0.000e+0
2.000e+5
4.000e+5
6.000e+5
8.000e+5
1.000e+6
1.200e+6
500
1000
1500

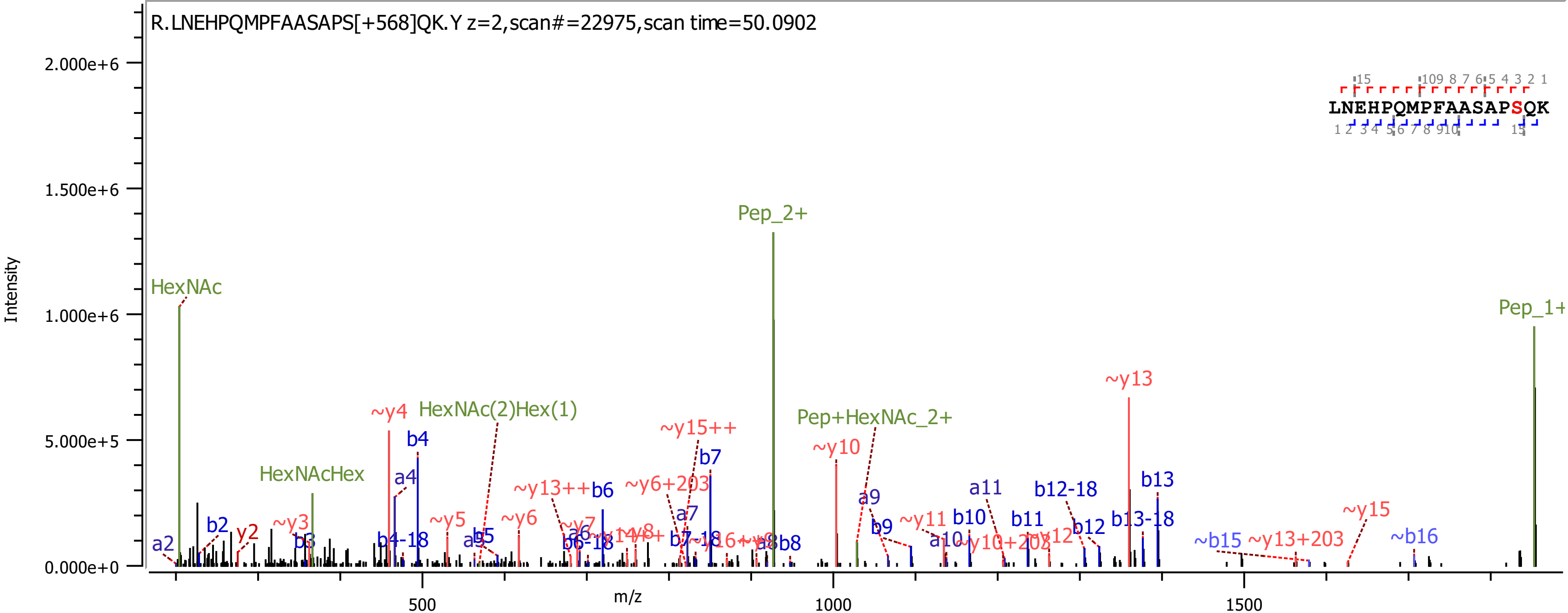
R.LNEHPQMPFAASAPS[+568]QK.Y z=2,scan#=22975,scan time=50.0902
LNEHPQMPFAASAPSQK
Intensity
2.000e+6
1.500e+6
1.000e+6
5.000e+5
0.000e+0
m/z
500
1000
1500
HexNAc
HexNAcHex
HexNAc(2)Hex(1)
Pep_2+
Pep+HexNAc_2+
Pep_1+
a2
b2
y2
~y3
b3
~y4
b4
a4
b4-18
~y5
a5
b5
~y6
~y13++
b6
b6-18
a6
~y7
~y6+203
~y15++
b7
a7
b7-18
b8
~y10
a9
b9
~y11
a10
b10
~y10+203
a11
b11
~y12
b12-18
b12
~y13
b13
b13-18
~b15
~y13+203
~y15
~b16

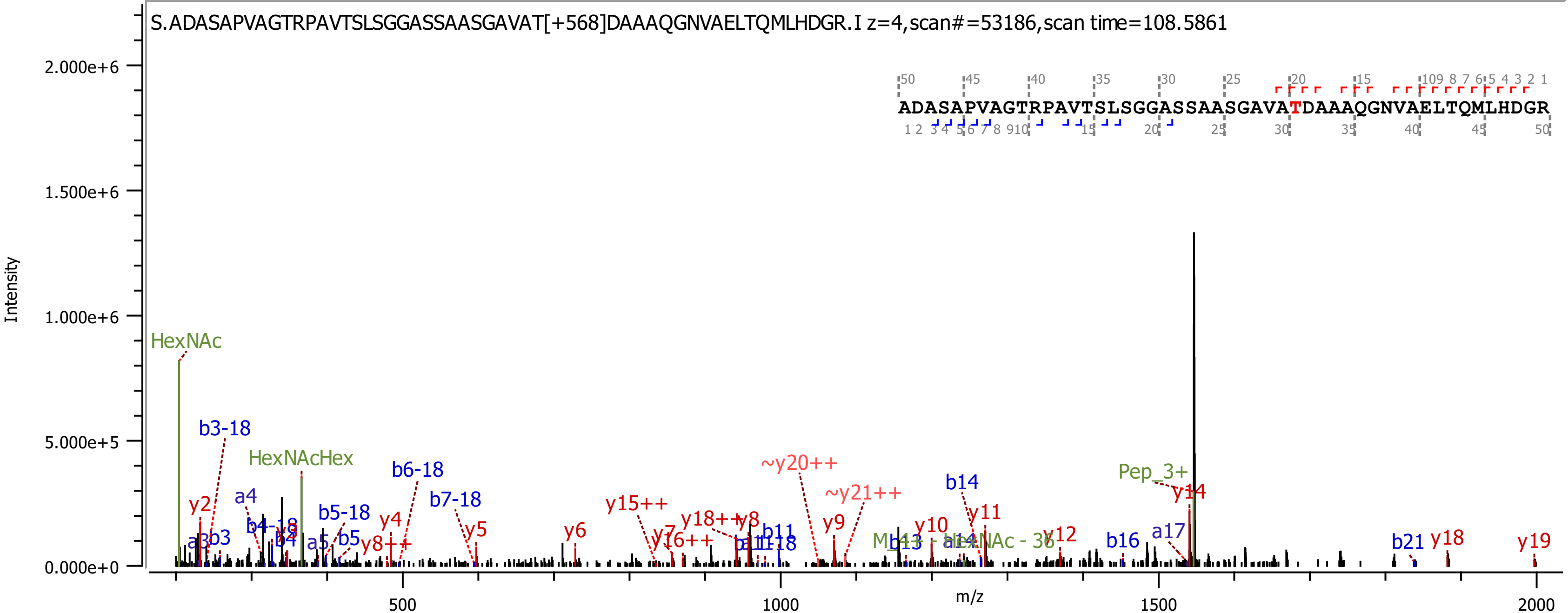
S.ADASAPVAGTRPAVTSLSGGASSAASGAVAT[+568]DAAAQGNVAELTQMLHDGR.I z=4,scan#=53186,scan time=108.5861
ADASAPVAGTRPAVTSLSGGASSAASGAVATDAAAQGNVAELTQMLHDGR
Intensity
m/z
HexNAc
HexNAcHex
Pep_3+
b3-18
b6-18
b7-18
b5-18
b14
b16
b21
a4
a17
y2
y4
y5
y6
y9
y10
y11
y12
y14
y18
y19
y15++
~y20++
~y21++
y18++
y8++
y16++

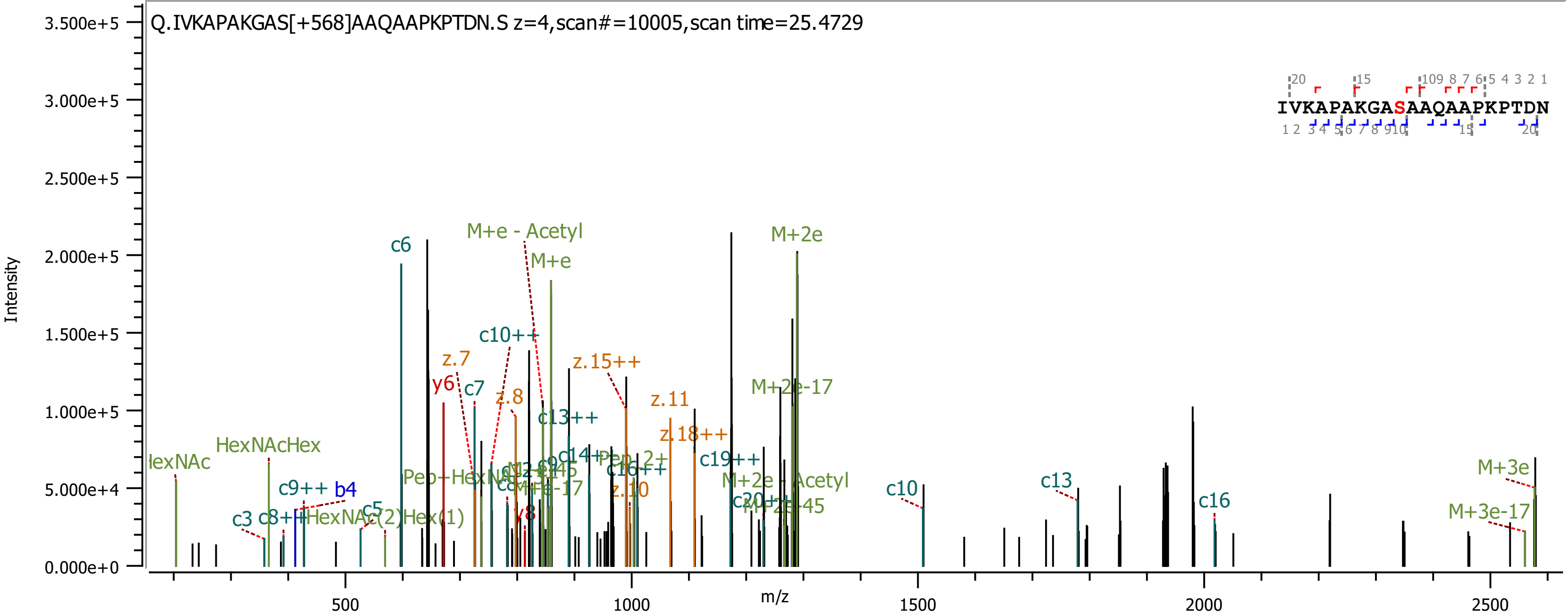

Q.IVKAPAKGAS[+568]AAQAAPKPTDN.S z=4,scan#=10005,scan time=25.4729
IVKAPAKGASAAQAAPKPTDN
Intensity
m/z
c6
M+e - Acetyl
M+e
M+2e
c10++
z.7
y6
c7
z.8
z.15++
z.11
z.18++
c13++
M+2e-17
c14++
c16++
c19++
M+2e - Acetyl
HexNAcHex
c9++
b4
c3
c5
c10
c13
c16
M+3e
M+3e-17
z.10
y8

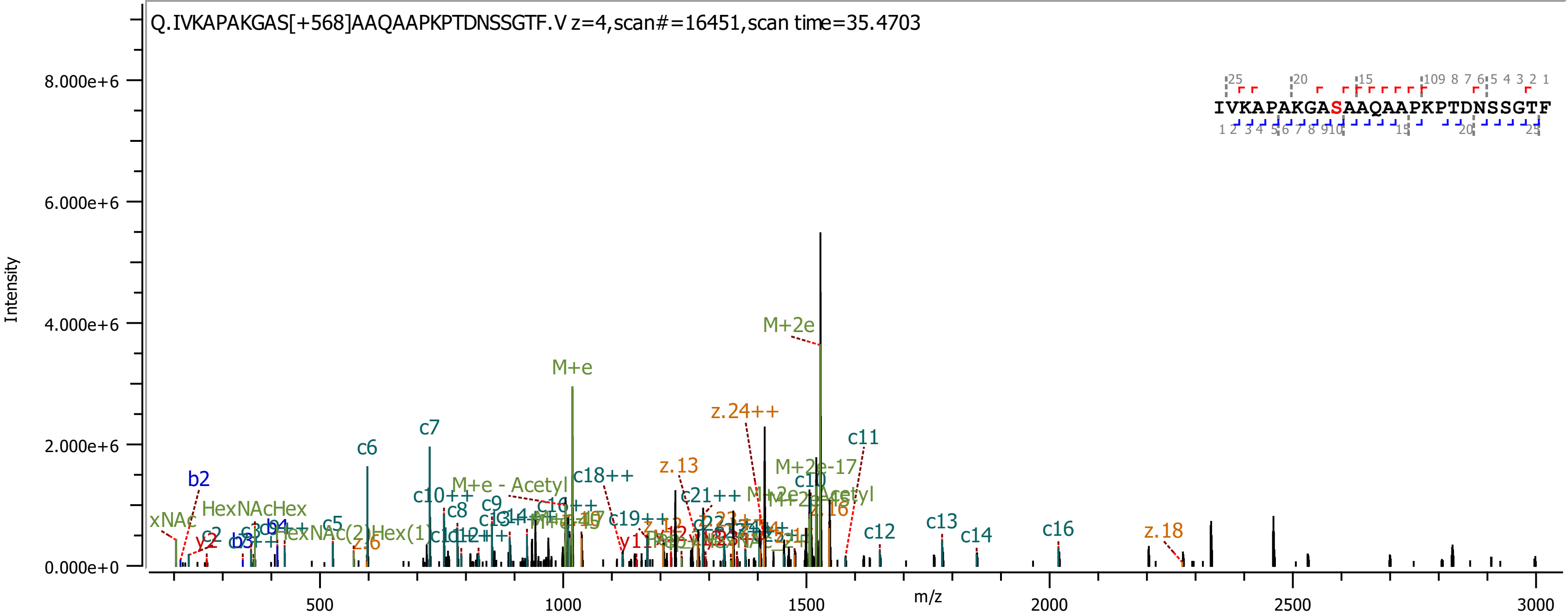

Q.IVKAPAKGAS[+568]AAQAAPKPTDNSSGTF.V z=4,scan#=16451,scan time=35.4703
IVKAPAKGASAAQAAPKPTDNSSGTF
Intensity
m/z
8.000e+6
6.000e+6
4.000e+6
2.000e+6
0.000e+0
500
1000
1500
2000
2500
3000
M+2e
M+e
z.24++
c11
c7
c6
z.13
M+2e-17
c18++
M+e - Acetyl
b2
c10++
c10
c8
c9
HexNAcHex
c21++
z.16
xNAc
c5
c12
c13
c14
c16
z.18
c2
y2
b3
z.6

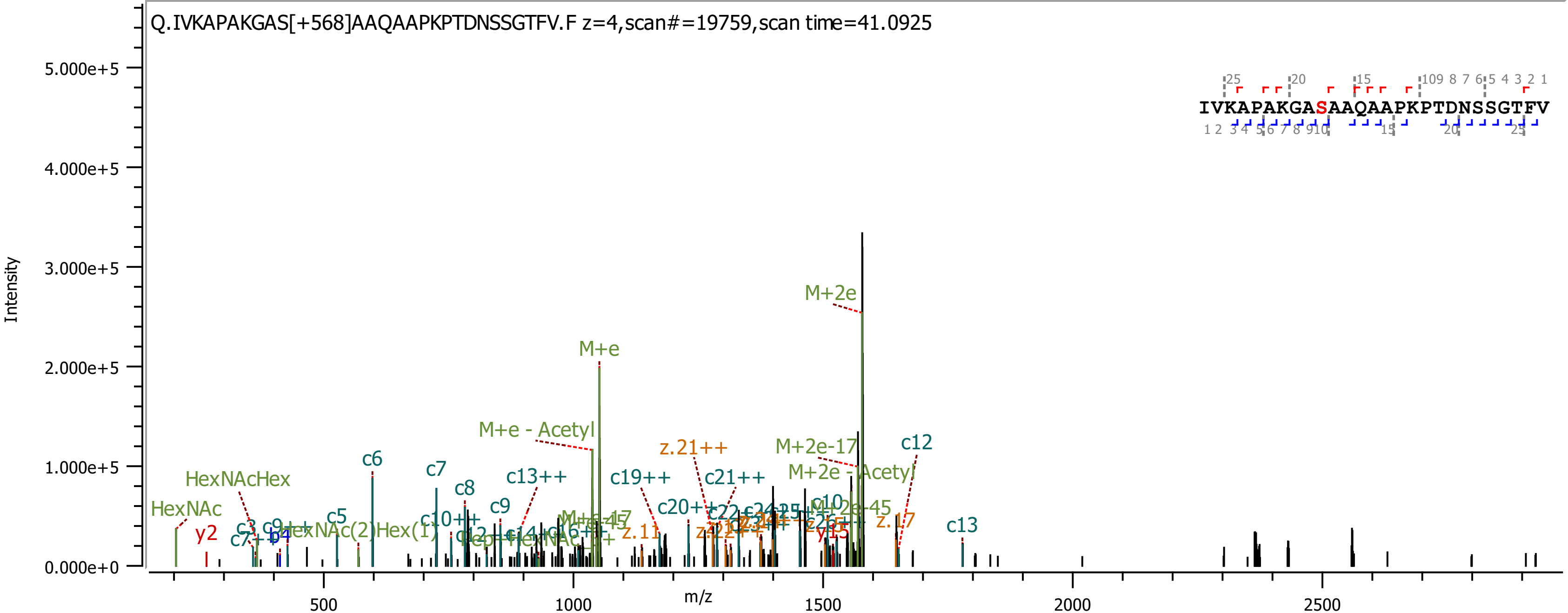
Q.IVKAPAKGAS[+568]AAQAAPKPTDNSSGTFV.F z=4,scan#=19759,scan time=41.0925
IVKAPAKGASAAQAAPKPTDNSSGTFV
Intensity
m/z
5.000e+5
4.000e+5
3.000e+5
2.000e+5
1.000e+5
0.000e+0
500
1000
1500
2000
2500
M+2e
M+e
M+e - Acetyl
z.21++
M+2e-17
c12
M+2e - Acetyl
c6
c7
HexNAcHex
c8
c13++
c19++
c21++
c9
HexNAc
c20++
c5
c10++
M+2e-45
z.17
c13
y2
z.11
y15
b4

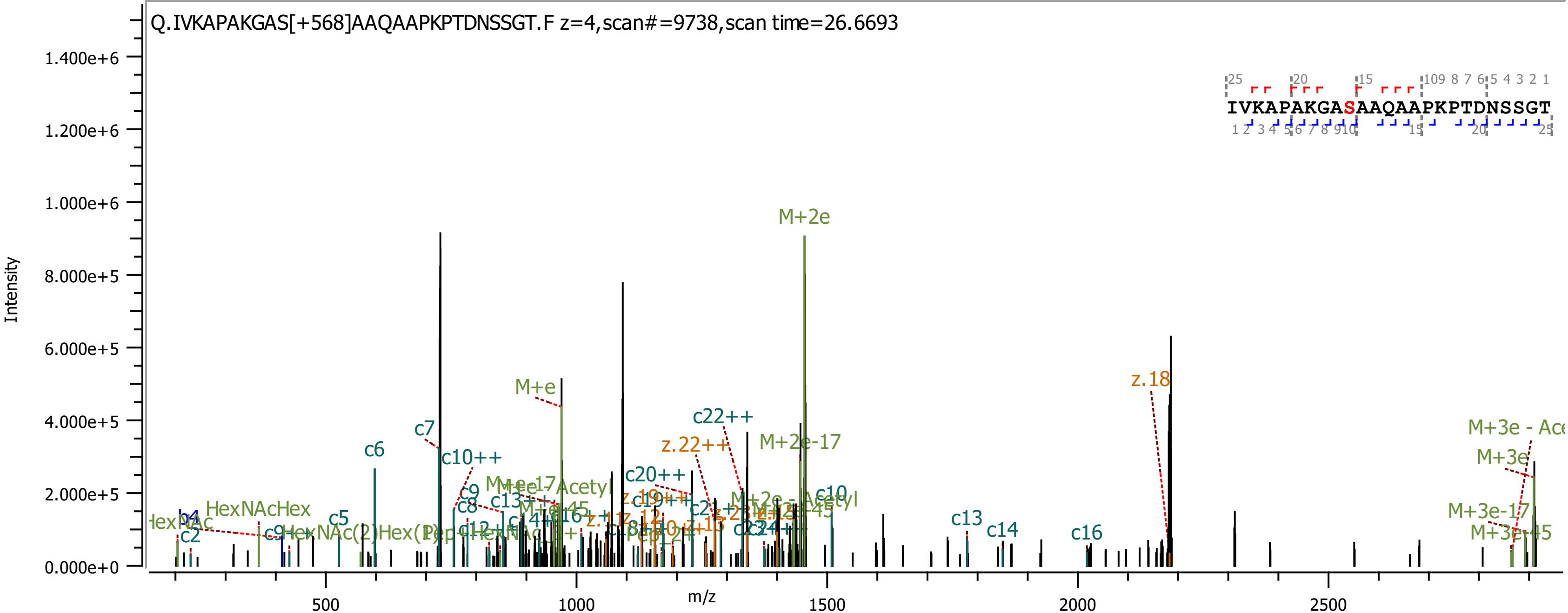

Q.IVKAPAKGAS[+568]AAQAAPKPTDNSSGT.F z=4,scan#=9738,scan time=26.6693
IVKAPAKGASAAQAAPKPTDNSSGT
Intensity
m/z
1.400e+6
1.200e+6
1.000e+6
8.000e+5
6.000e+5
4.000e+5
2.000e+5
0.000e+0
500
1000
1500
2000
2500
M+2e
M+e
z.18
c22++
z.22++
M+2e-17
c6
c7
c10++
c20++
M+3e
M+3e-17
M+3e-45
HexNAcHex
c5
c2
c8
c9
c13
c14
c16

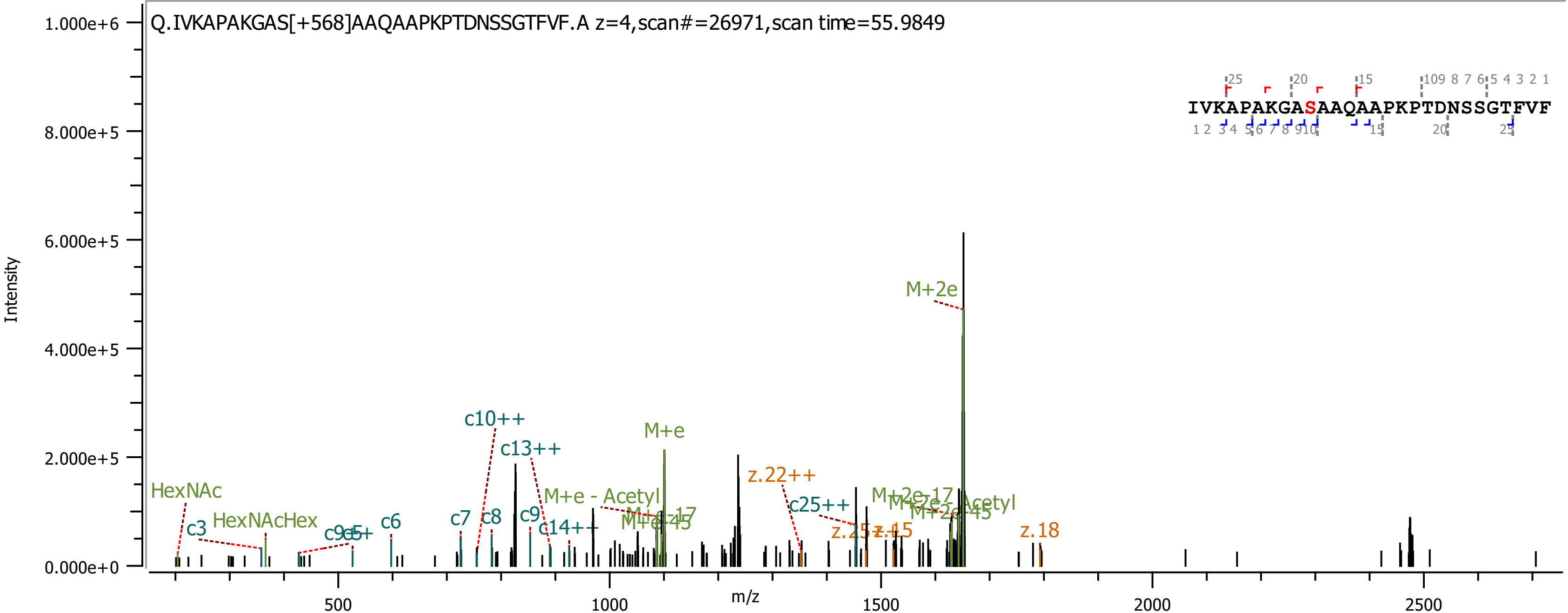
Q.IVKAPAKGAS[+568]AAQAAPKPTDNSSGTFVF.A z=4,scan#=26971,scan time=55.9849
IVKAPAKGASAAQAAPKPTDNSSGTFVF
Intensity
m/z
1.000e+6
8.000e+5
6.000e+5
4.000e+5
2.000e+5
0.000e+0
500
1000
1500
2000
2500
HexNAc
HexNAcHex
c3
c6
c7
c8
c9
c10++
c13++
c14++
c25++
M+e
M+e - Acetyl
M+2e
z.22++
z.15
z.18

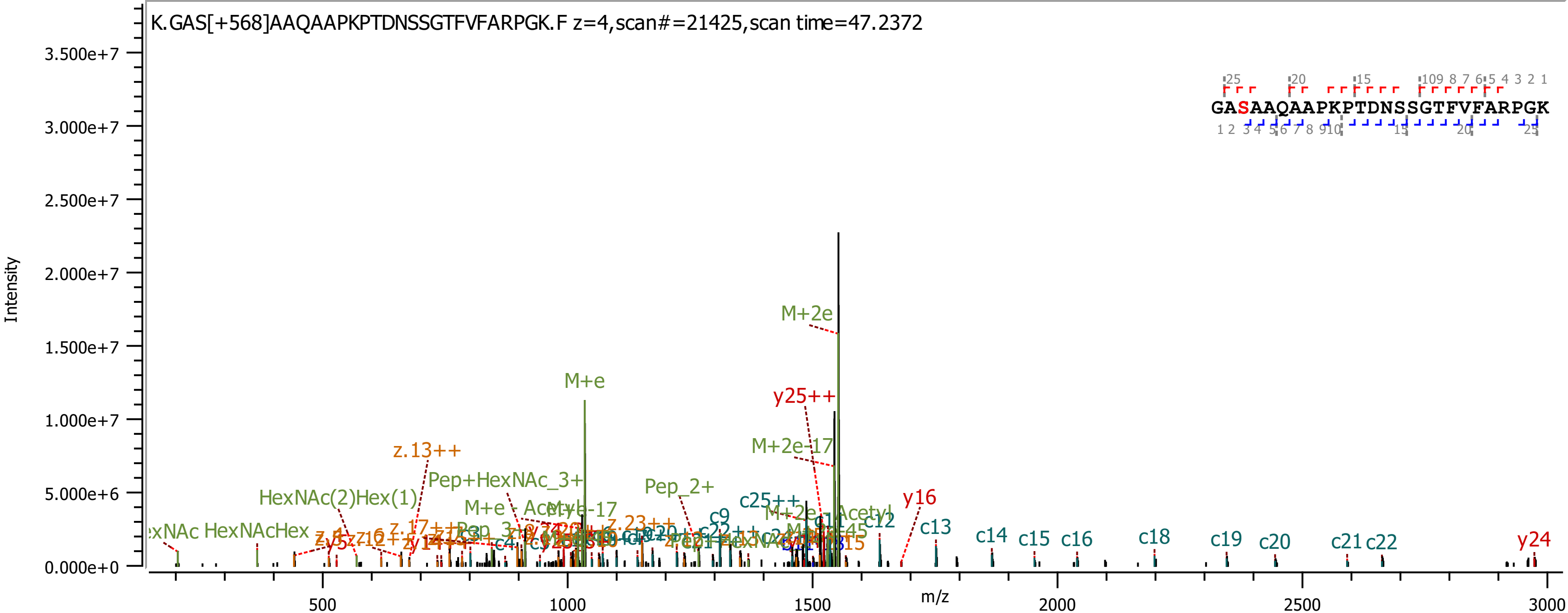

K.GAS[+568]AAQAAPKPTDNSSGTFVFARPGK.F z=4,scan#=21425,scan time=47.2372
GASAAQAAPKPTDNSSGTFVFARPGK
Intensity
m/z
3.500e+7
3.000e+7
2.500e+7
2.000e+7
1.500e+7
1.000e+7
5.000e+6
0.000e+0
500
1000
1500
2000
2500
3000
M+2e
M+e
y25++
M+2e-17
z.13++
Pep+HexNAc_3+
Pep_2+
HexNAc(2)Hex(1)
HexNAcHex
c25++
y16
c13
c12
c9
c14
c15
c16
c18
c19
c20
c21
c22
y24

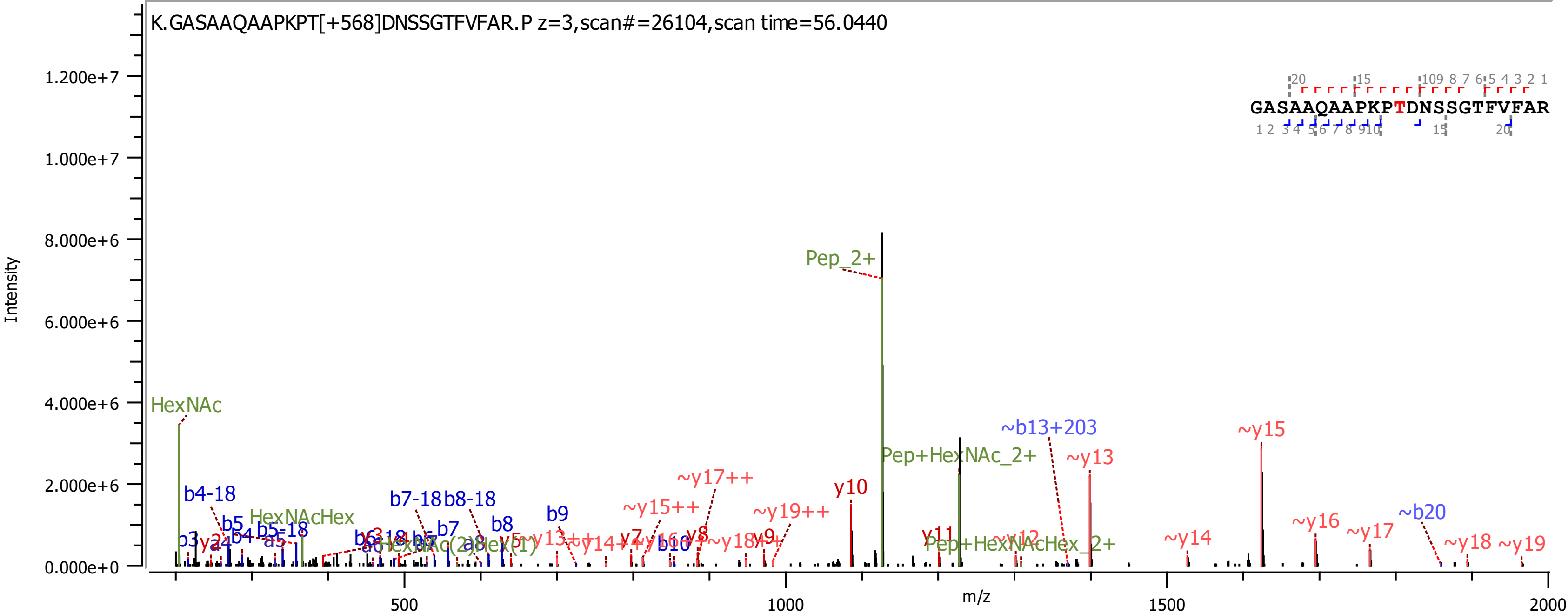

K.GASAAQAAPKPT[+568]DNSSGTFVFAR.P z=3,scan#=26104,scan time=56.0440
GASAAQAAPKPTDNSSGTFVFAR
Intensity
m/z
0.000e+0
2.000e+6
4.000e+6
6.000e+6
8.000e+6
1.000e+7
1.200e+7
500
1000
1500
2000
HexNAc
HexNAcHex
Pep_2+
Pep+HexNAc_2+
Pep+HexNAcHex_2+
b3
b4-18
b5
b4
b5-18
a5
b7-18
b8-18
b7
b8
b9
b10
~b13+203
~b20
y10
y11
y7
y8
y9
~y13
~y14
~y15
~y16
~y17
~y18
~y19
~y17++
~y15++
~y19++
~y12

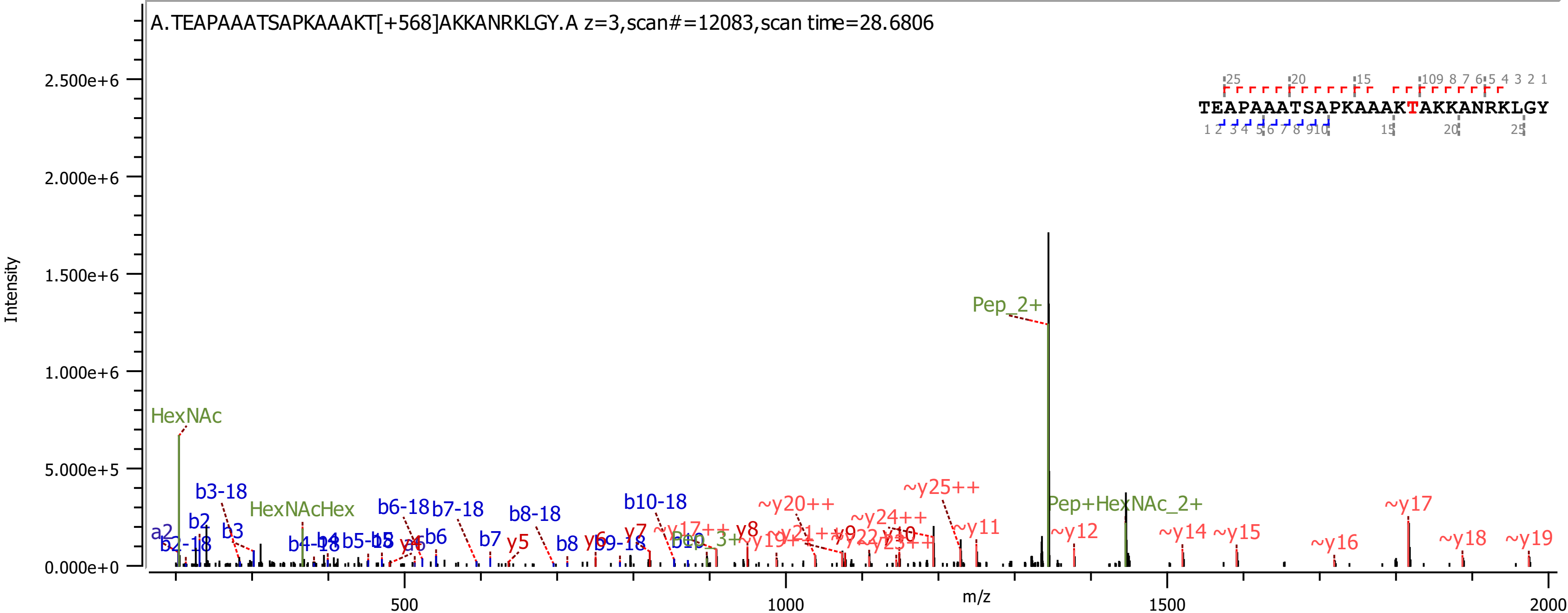
A.TEAPAAATSAPKAAAKT[+568]AKKANRKLGY.A z=3,scan#=12083,scan time=28.6806
TEAPAAATSAPKAAAKTAKKANRKLGY
Intensity
m/z
0.000e+0
5.000e+5
1.000e+6
1.500e+6
2.000e+6
2.500e+6
500
1000
1500
2000
HexNAc
HexNAcHex
Pep_2+
Pep+HexNAc_2+
a2
b2
b2-18
b3
b3-18
b4-18
b6
b6-18
b7
b7-18
b8
b8-18
b9-18
b10-18
y5
y7
y8
~y11
~y12
~y14
~y15
~y16
~y17
~y18
~y19
~y20++
~y24++
~y25++

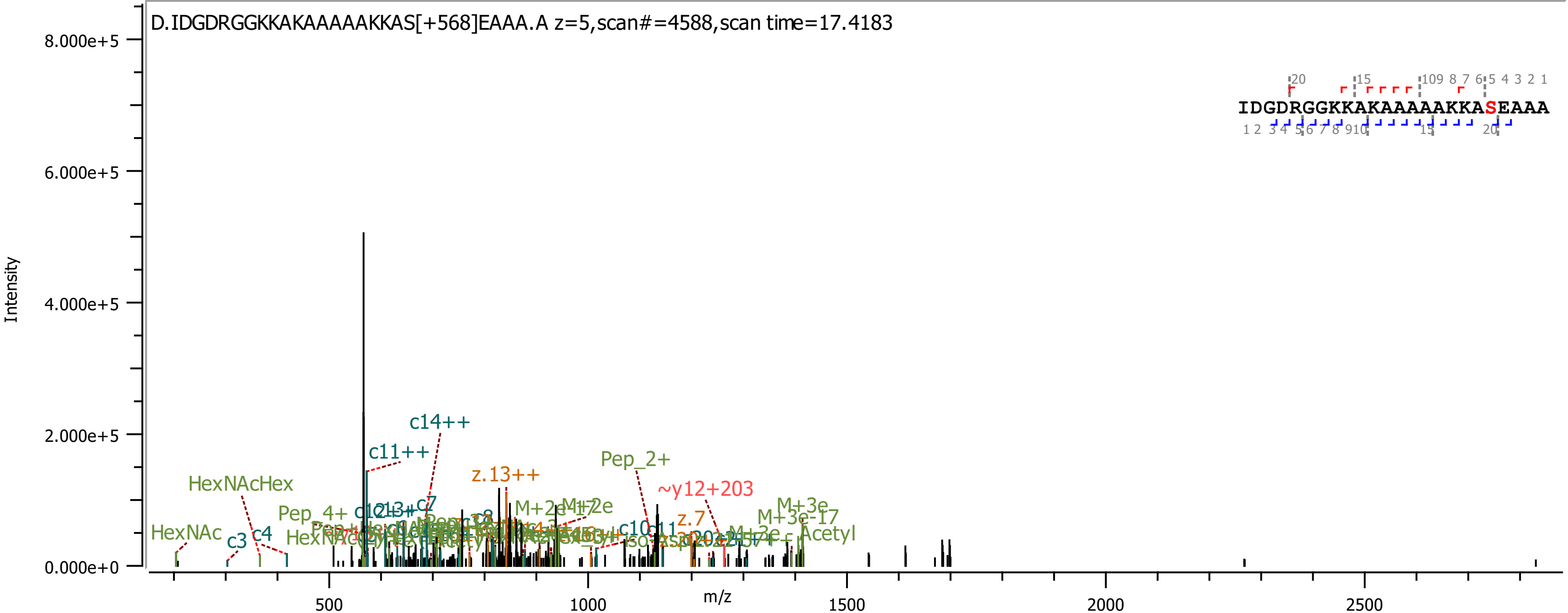

D.IDGDRGGKKAKAAAAAKKAS[+568]EAAA.A z=5,scan#=4588,scan time=17.4183
IDGDRGGKKAKAAAAAKKASEAAA
Intensity
m/z
8.000e+5
6.000e+5
4.000e+5
2.000e+5
0.000e+0
500
1000
1500
2000
2500
c14++
c11++
z.13++
Pep_2+
~y12+203
HexNAcHex
HexNAc
c3
c4
Pep_4+
c7
z.7
M+3e
M+3e-17
Acetyl

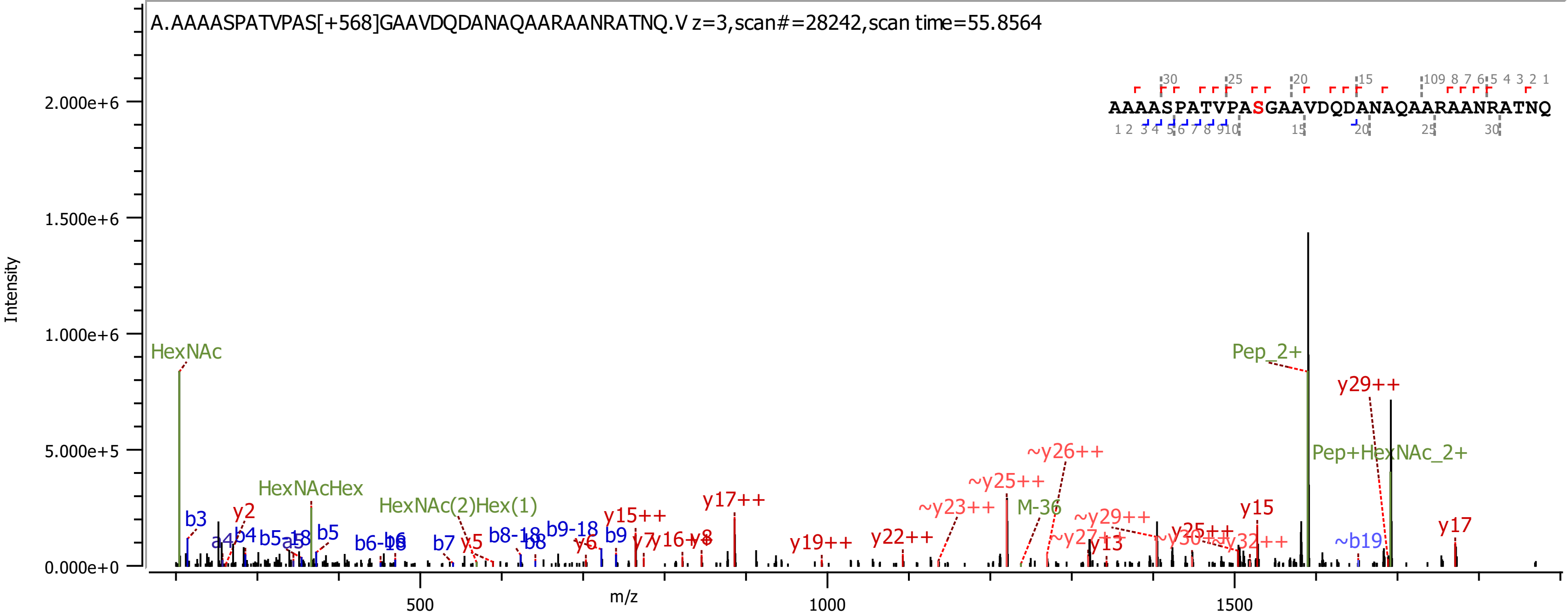

A.AAAASPATVPAS[+568]GAAVDQDANAQAARAANRATNQ.V z=3,scan#=28242,scan time=55.8564
AAAASPATVPASGAAVDQDANAQAARAANRATNQ
Intensity
m/z
HexNAc
HexNAcHex
HexNAc(2)Hex(1)
Pep_2+
Pep+HexNAc_2+
M-36
y29++
~y26++
~y25++
~y23++
y17++
y15++
y22++
y19++
~y29++
y15
y17
~b19

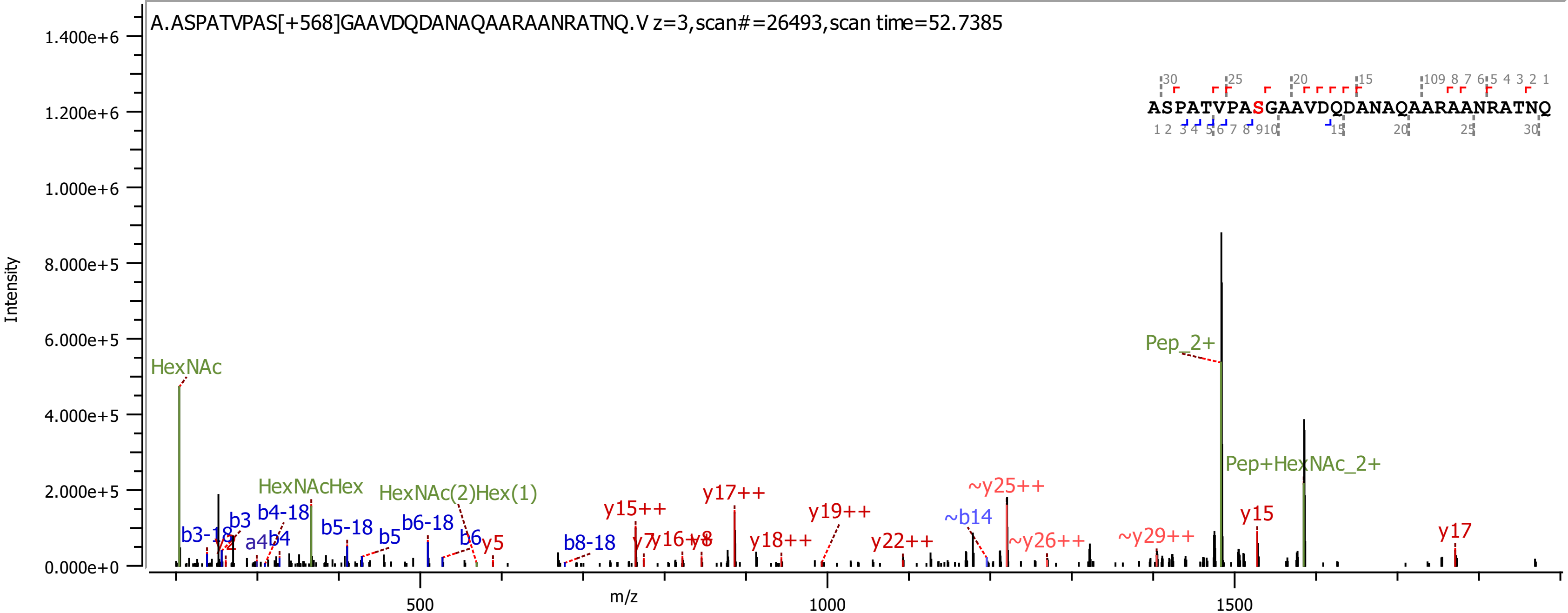
A.ASPATVPAS[+568]GAAVDQDANAQAARAANRATNQ.V z=3,scan#=26493,scan time=52.7385
ASPATVPASGAAVDQDANAQAARAANRATNQ
Intensity
m/z
1.400e+6
1.200e+6
1.000e+6
8.000e+5
6.000e+5
4.000e+5
2.000e+5
0.000e+0
500
1000
1500
HexNAc
HexNAcHex
HexNAc(2)Hex(1)
Pep_2+
Pep+HexNAc_2+
b3-18
b3
b4-18
a4
b4
b5-18
b5
b6-18
b6
y5
b8-18
y15++
y7
y16++
y8
y17++
y18++
y19++
y22++
~b14
~y25++
~y26++
~y29++
y15
y17

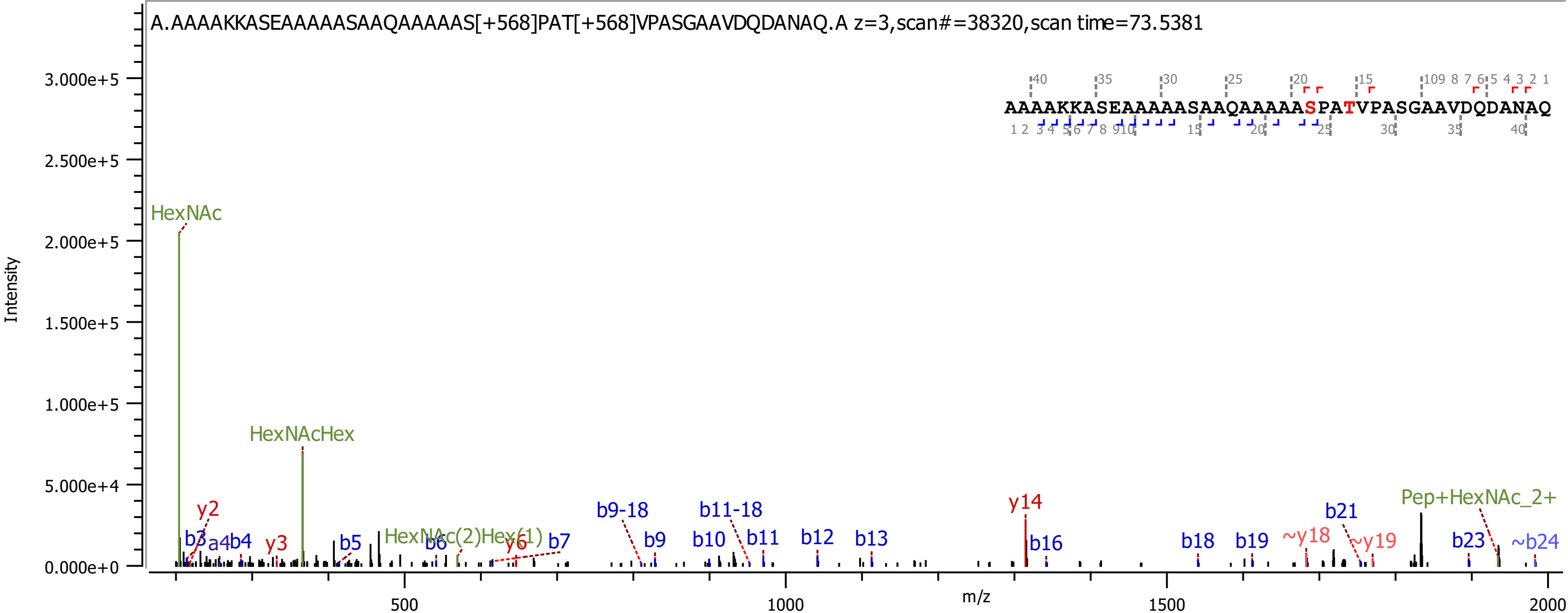

A.AAAAKKASEAAAAASAAQAAAAAS[+568]PAT[+568]VPASGAAVDQDANAQ.A z=3,scan#=38320,scan time=73.5381
AAAAKKASEAAAAASAAQAAAAASPATVPASGAAVDQDANAQ
Intensity
m/z
HexNAc
HexNAcHex
HexNAc(2)Hex(1)
Pep+HexNAc_2+
y2
y3
y6
y14
~y18
~y19
b3
a4
b4
b5
b6
b7
b9-18
b9
b10
b11-18
b11
b12
b13
b16
b18
b19
b21
b23
~b24

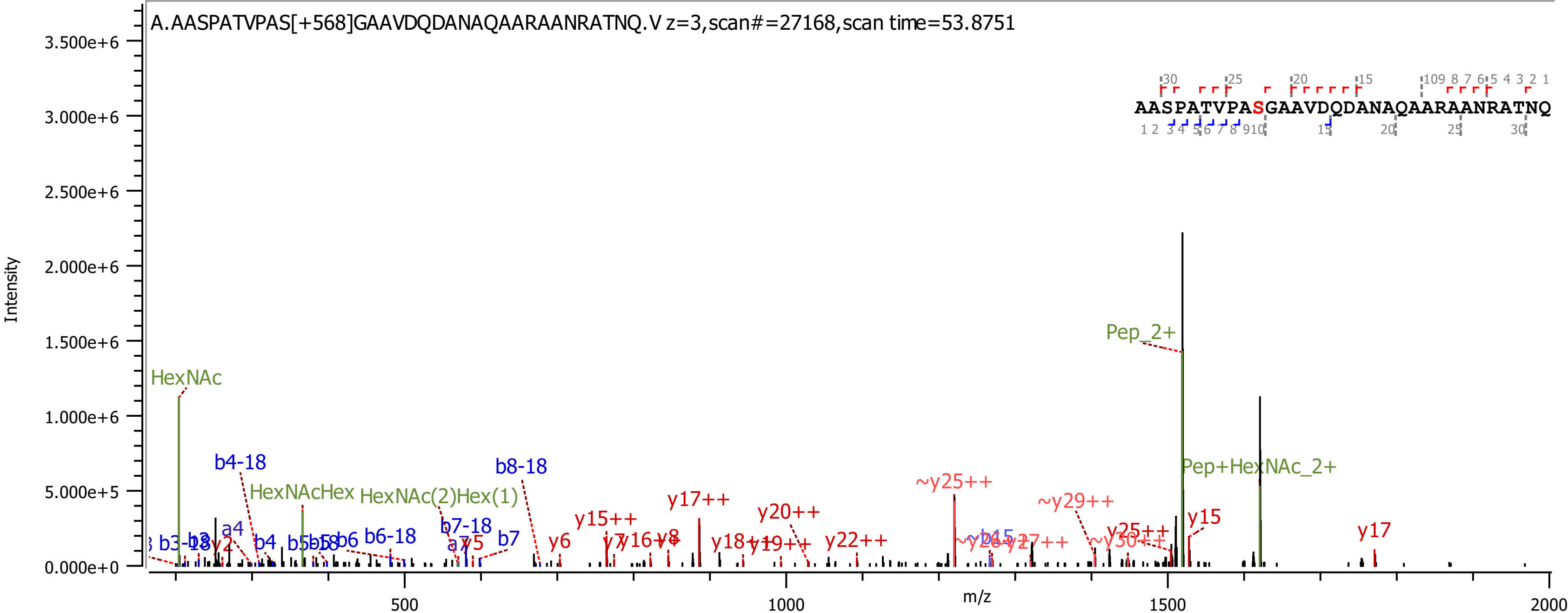

A.AASPATVPAS[+568]GAAVDQDANAQAARAANRATNQ.V z=3,scan#=27168,scan time=53.8751
AASPATVPASGAAVDQDANAQAARAANRATNQ
Intensity
m/z
HexNAc
HexNAcHex
HexNAc(2)Hex(1)
Pep_2+
Pep+HexNAc_2+
b4-18
b8-18
b6-18
b7-18
b7
b6
b4
a4
y2
y5
y6
y8
y15++
y17++
y20++
y22++
~y25++
~y29++
y25++
y15
y17

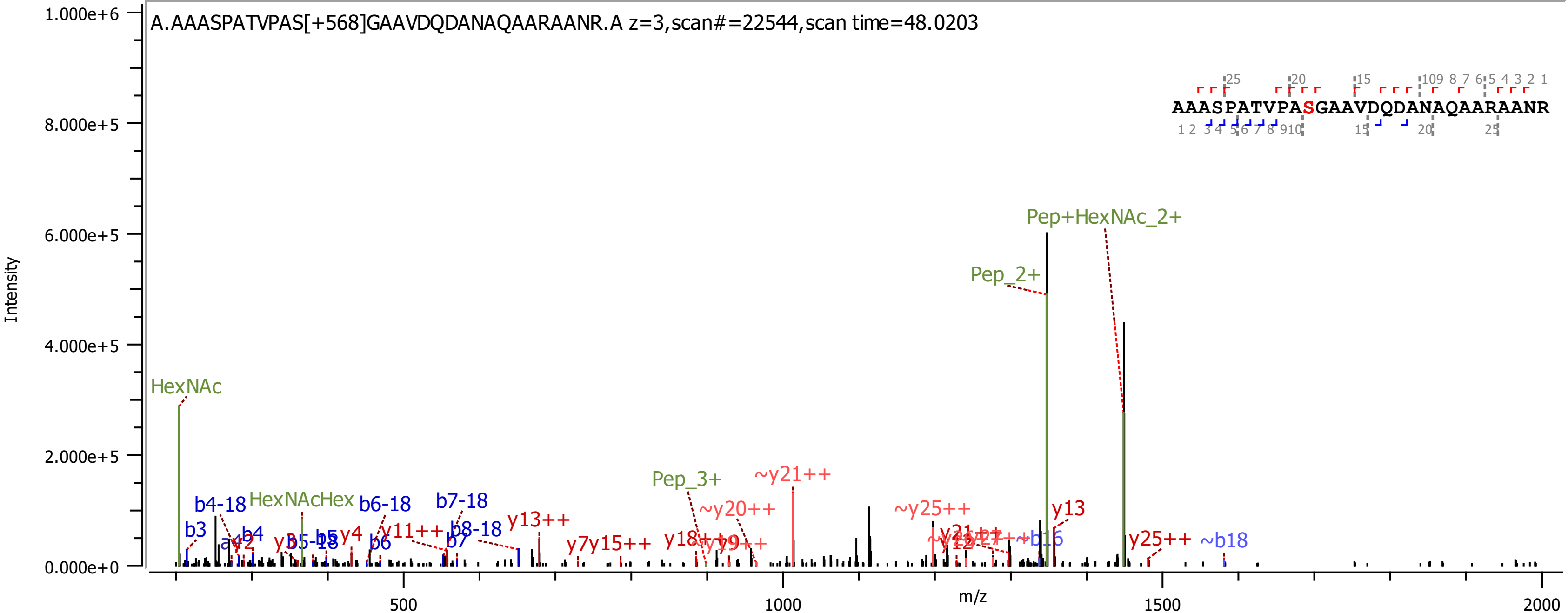

A.AAASPATVPAS[+568]GAAVDQDANAQAARAANR.A z=3,scan#=22544,scan time=48.0203
AAASPATVPASGAAVDQDANAQAARAANR
Intensity
m/z
1.000e+6
8.000e+5
6.000e+5
4.000e+5
2.000e+5
0.000e+0
500
1000
1500
2000
Pep+HexNAc_2+
Pep_2+
HexNAc
HexNAcHex
Pep_3+
~y21++
~y20++
~y25++
y13
y25++
~b18
b3
b4-18
b4
b5-18
b5
y4
b6-18
b6
b7-18
b7
b8-18
y11++
y13++
y7
y15++
y18++
y9++

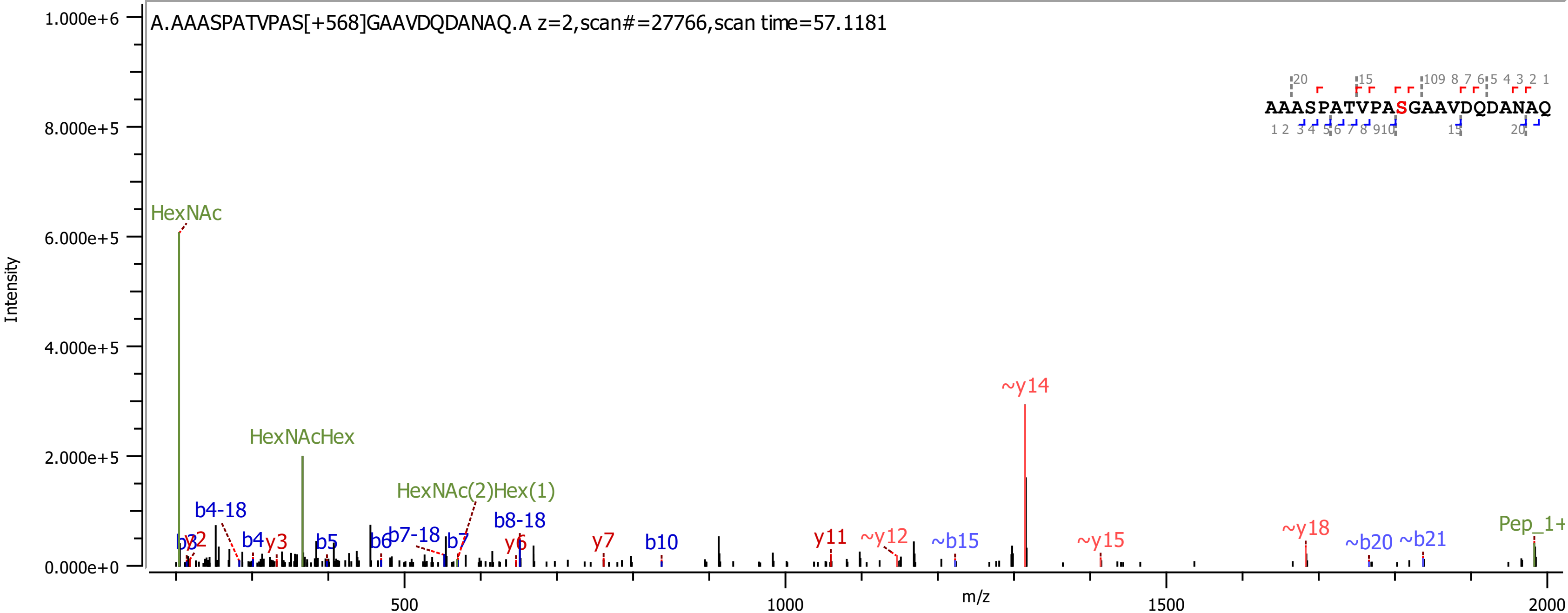
A.AAASPATVPAS[+568]GAAVDQDANAQ.A z=2,scan#=27766,scan time=57.1181
AAASPATVPASGAAVDQDANAQ
Intensity
m/z
HexNAc
HexNAcHex
HexNAc(2)Hex(1)
b4-18
b7-18
b8-18
~y14
~y12
~y15
~y18
~b15
~b20
~b21
Pep_1+

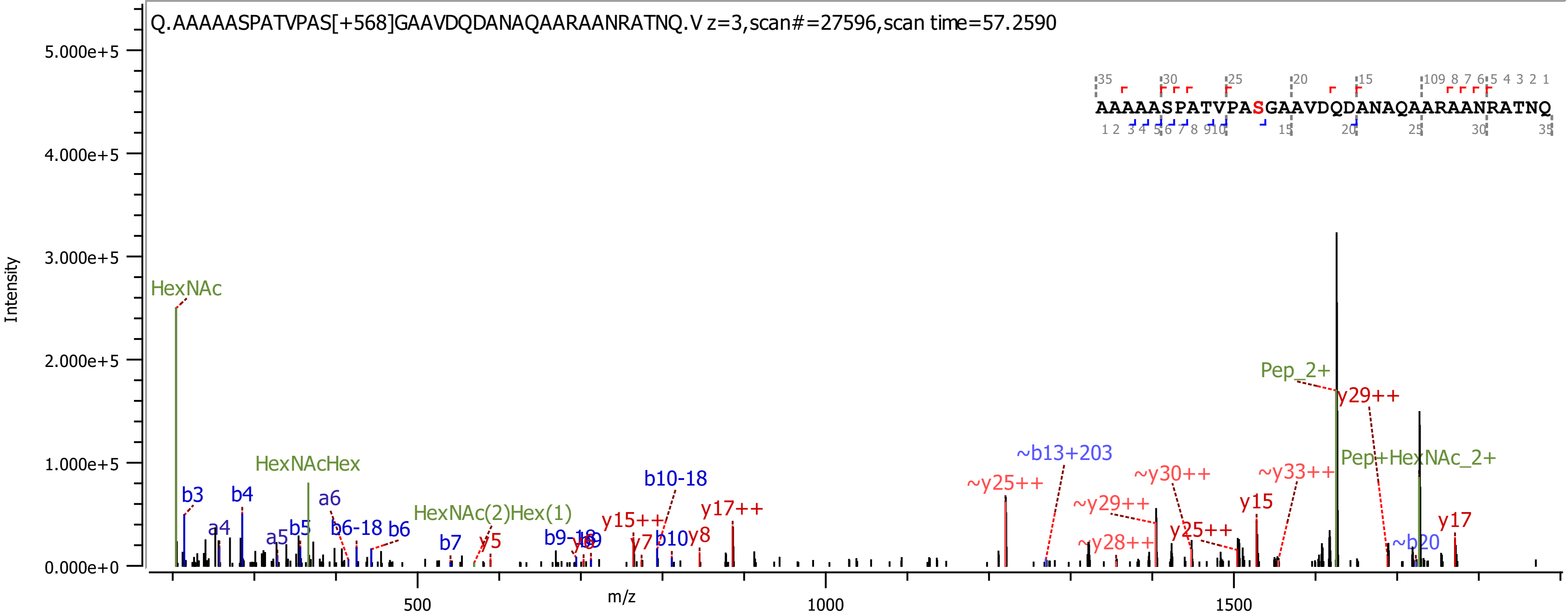
Q.AAAAASPATVPAS[+568]GAAVDQDANAQAARAANRATNQ.V z=3,scan#=27596,scan time=57.2590
AAAAASPATVPASGAAVDQDANAQAARAANRATNQ
Intensity
m/z
HexNAc
HexNAcHex
HexNAc(2)Hex(1)
Pep_2+
Pep+HexNAc_2+
b3
b4
a4
a5
b5
a6
b6-18
b6
b7
y5
b9-18
b10-18
y15++
y7
b10
y8
y17++
~y25++
~b13+203
~y29++
~y28++
~y30++
y25++
y15
~y33++
y29++
~b20
y17
0.000e+0
1.000e+5
2.000e+5
3.000e+5
4.000e+5
5.000e+5
500
1000
1500

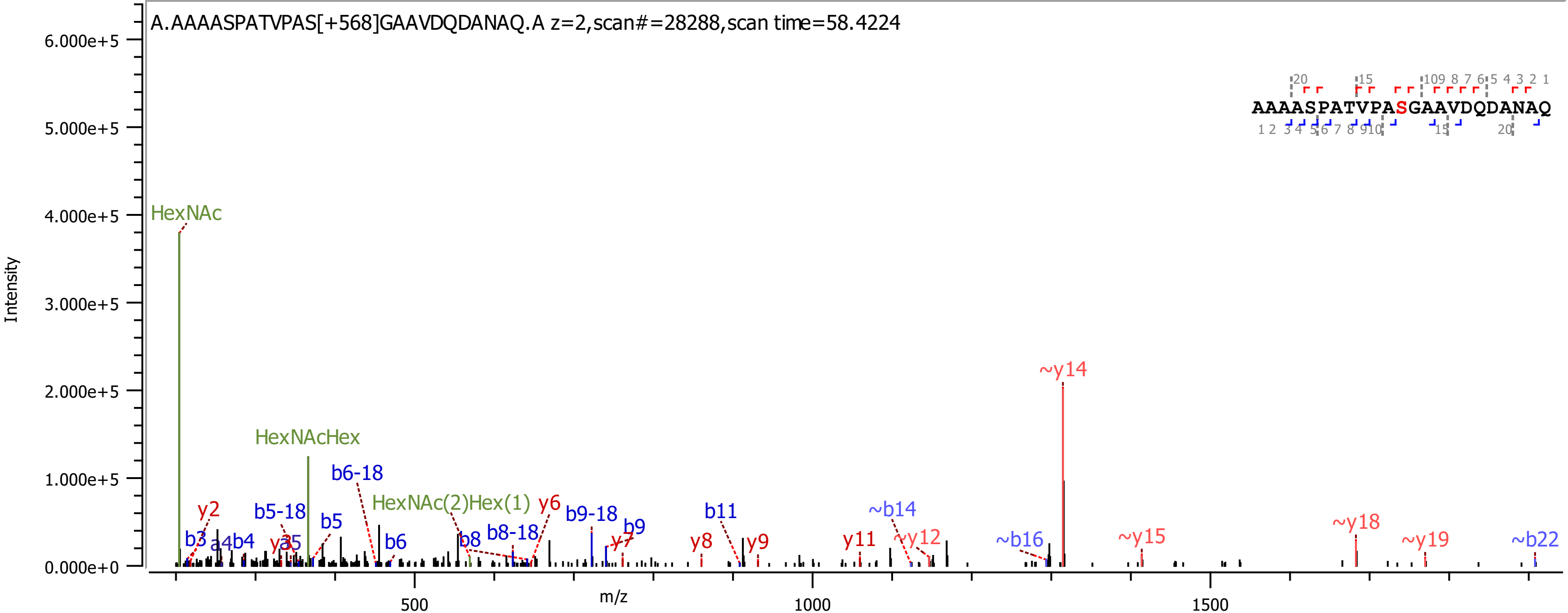
A.AAAASPATVPAS[+568]GAAVDQDANAQ.A z=2,scan#=28288,scan time=58.4224
AAAASPATVPASGAAVDQDANAQ
Intensity
m/z
HexNAc
HexNAcHex
HexNAc(2)Hex(1)
b6-18
b5-18
b9-18
b8-18
y2
y6
b3
a4
b4
y3
y5
b5
b6
b8
b9
y7
b11
y8
y9
y11
~b14
~y12
~y14
~b16
~y15
~y18
~y19
~b22

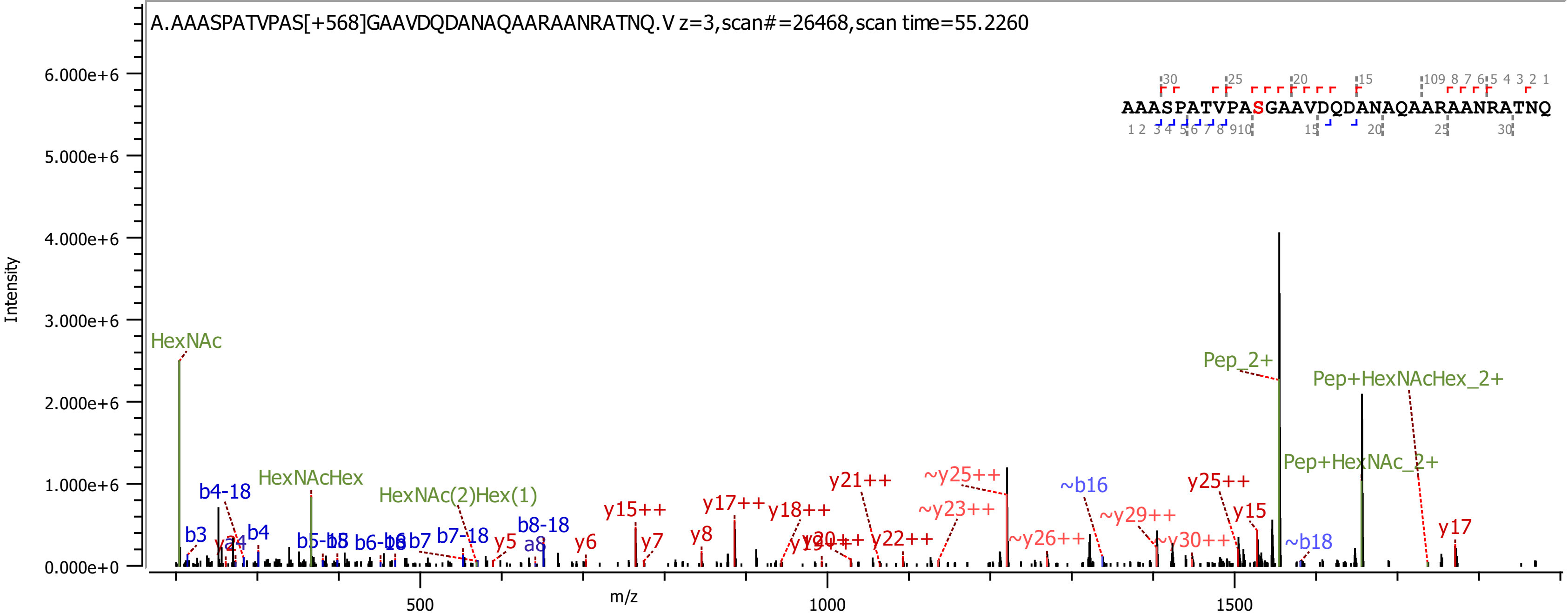
A.AAASPATVPAS[+568]GAAVDQDANAQAARAANRATNQ.V z=3,scan#=26468,scan time=55.2260
AAASPATVPASGAAVDQDANAQAARAANRATNQ
Intensity
m/z
HexNAc
HexNAcHex
HexNAc(2)Hex(1)
Pep_2+
Pep+HexNAcHex_2+
Pep+HexNAc_2+
b3
b4
b4-18
b7
b7-18
b8-18
a8
y5
y6
y7
y8
y15++
y17++
y18++
y21++
y22++
y25++
y15
y17
~y23++
~y25++
~y26++
~y29++
~y30++
~b16
~b18

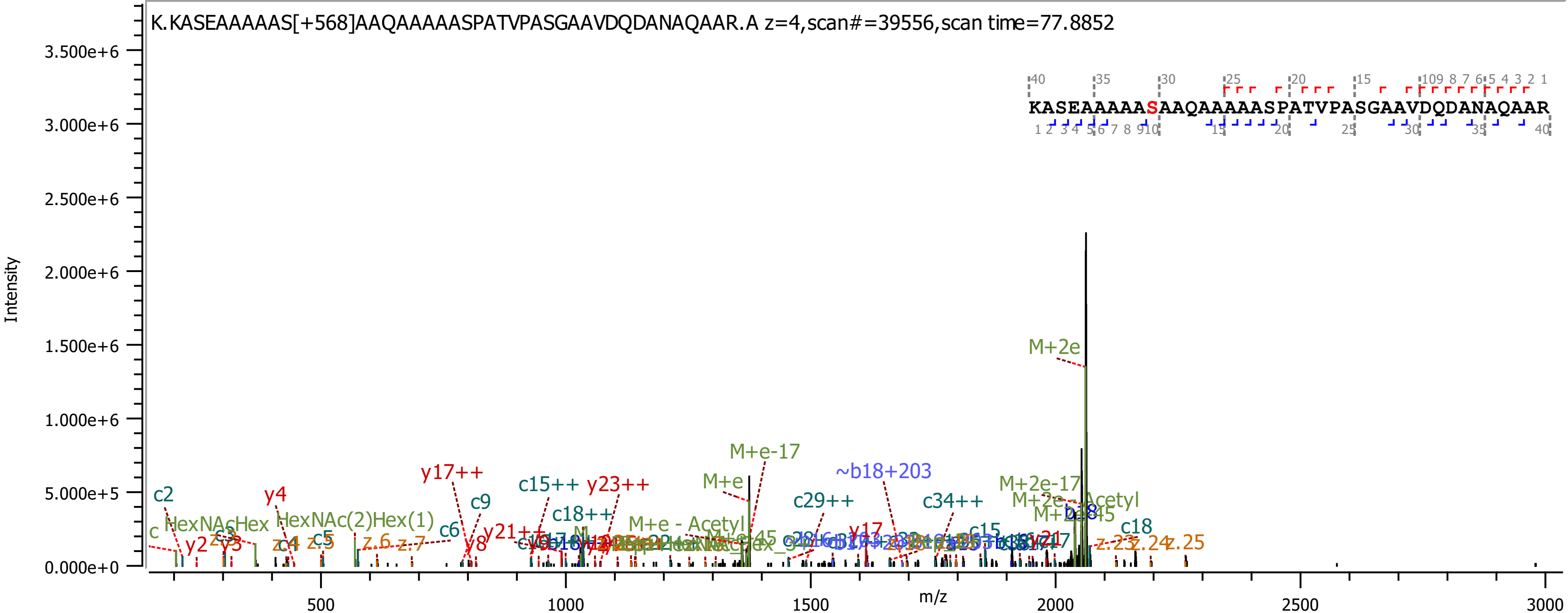

K.KASEAAAAAS[+568]AAQAAAAASPATVPASGAAVDQDANAQAAR.A z=4,scan#=39556,scan time=77.8852
KASEAAAAASAAQAAAAASPATVPASGAAVDQDANAQAAR
Intensity
m/z
3.500e+6
3.000e+6
2.500e+6
2.000e+6
1.500e+6
1.000e+6
5.000e+5
0.000e+0
500
1000
1500
2000
2500
3000
M+2e
M+e-17
~b18+203
M+e
y17++
c15++
y23++
c2
y4
c9
c29++
c34++
M+2e-17
M+2e - Acetyl
c18++
HexNAcHex
HexNAc(2)Hex(1)
c6
y21++
M+e - Acetyl
y2
y3
z.4
c5
z.6
z.7
y8
y17
c15
c18
z.23
z.24
z.25

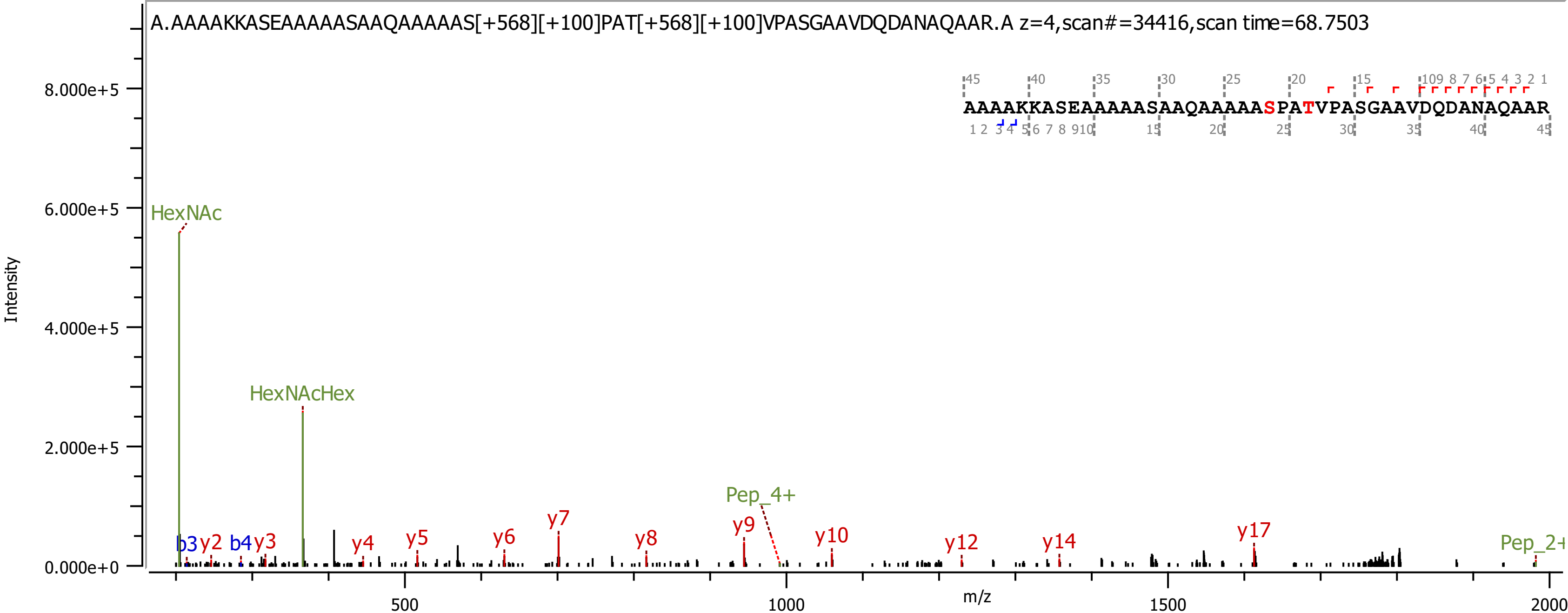

A.AAAAKKASEAAAAASAAQAAAAAS[+568][+100]PAT[+568][+100]VPASGAAVDQDANAQAAR.A z=4,scan#=34416,scan time=68.7503
AAAAKKASEAAAAASAAQAAAAASPATVPASGAAVDQDANAQAAR
HexNAc
HexNAcHex
Pep_4+
Pep_2+
b3
b4
y2
y3
y4
y5
y6
y7
y8
y9
y10
y12
y14
y17
Intensity
m/z

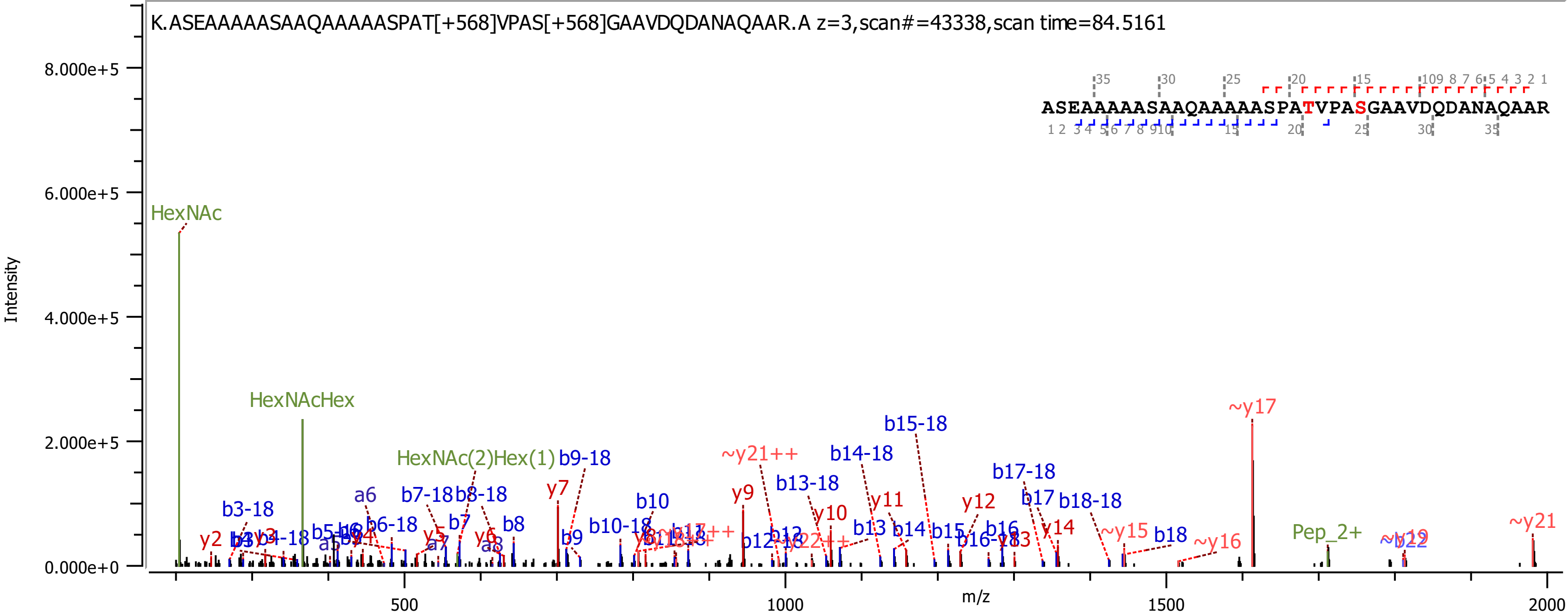

K.ASEAAAAASAAQAAAAASPAT[+568]VPAS[+568]GAAVDQDANAQAAR.A z=3,scan#=43338,scan time=84.5161
ASEAAAAASAAQAAAAASPATVPASGAAVDQDANAQAAR
Intensity
m/z
8.000e+5
6.000e+5
4.000e+5
2.000e+5
0.000e+0
500
1000
1500
2000
HexNAc
HexNAcHex
HexNAc(2)Hex(1)
b9-18
~y17
b15-18
~y21++
b14-18
b17-18
b13-18
b3-18
a6
b7-18
b8-18
y7
b10
y9
y11
y12
b17
b18-18
y10
b13
b14
b15
b16
y14
~y15
b18
~y16
Pep_2+
~y21
b10-18
b7
b8
b9
b6-18
y2
y5
y6
a7
a8

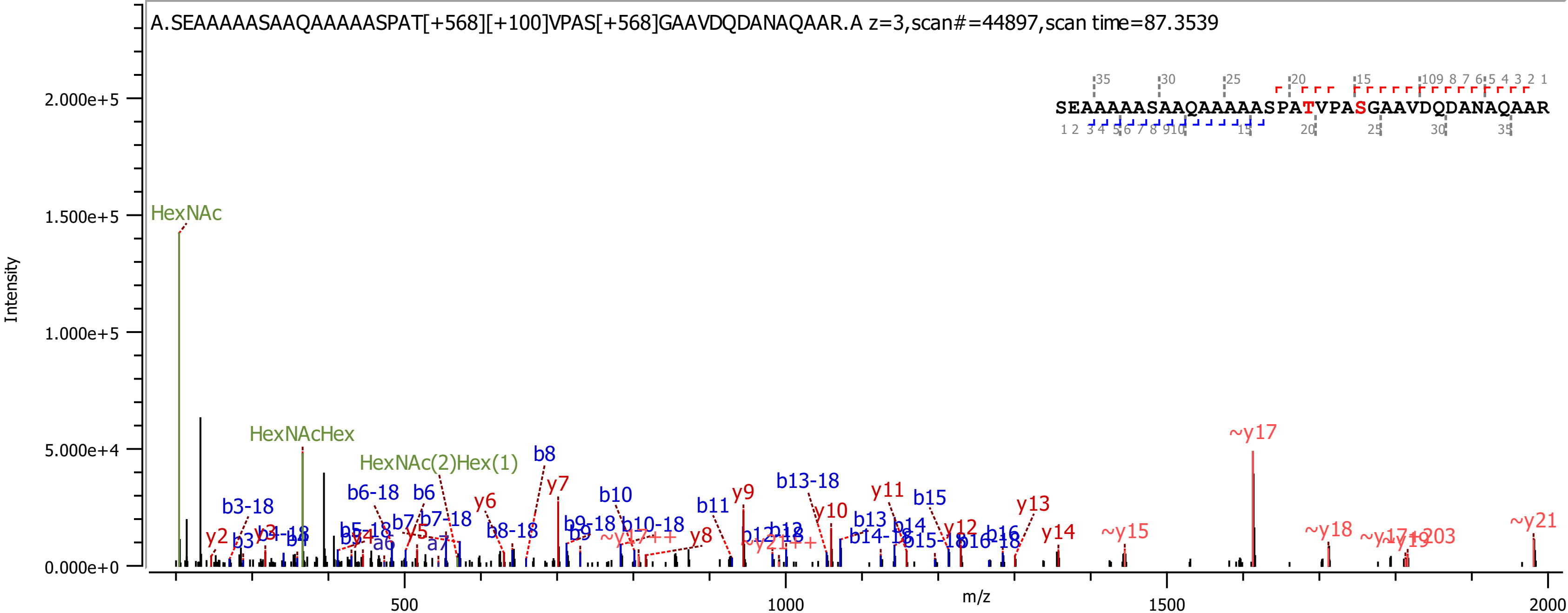
A.SEAAAAASAAQAAAAASPAT[+568][+100]VPAS[+568]GAAVDQDANAQAAR.A z=3,scan#=44897,scan time=87.3539
SEAAAAASAAQAAAAASPATVPASGAAVDQDANAQAAR
Intensity
m/z
HexNAc
HexNAcHex
HexNAc(2)Hex(1)
~y17
~y21
~y18
~y15
y14
y13
y12
y11
y10
y9
y8
y7
y6
y5
y2
y3
b3-18
b6-18
b6
b7-18
b8
b8-18
b9-18
b10
b10-18
b11
b13-18
b13
b14-18
b15
b15-18
b16
b16-18
b4
b3
a6
a7
0.000e+0
5.000e+4
1.000e+5
1.500e+5
2.000e+5
500
1000
1500
2000

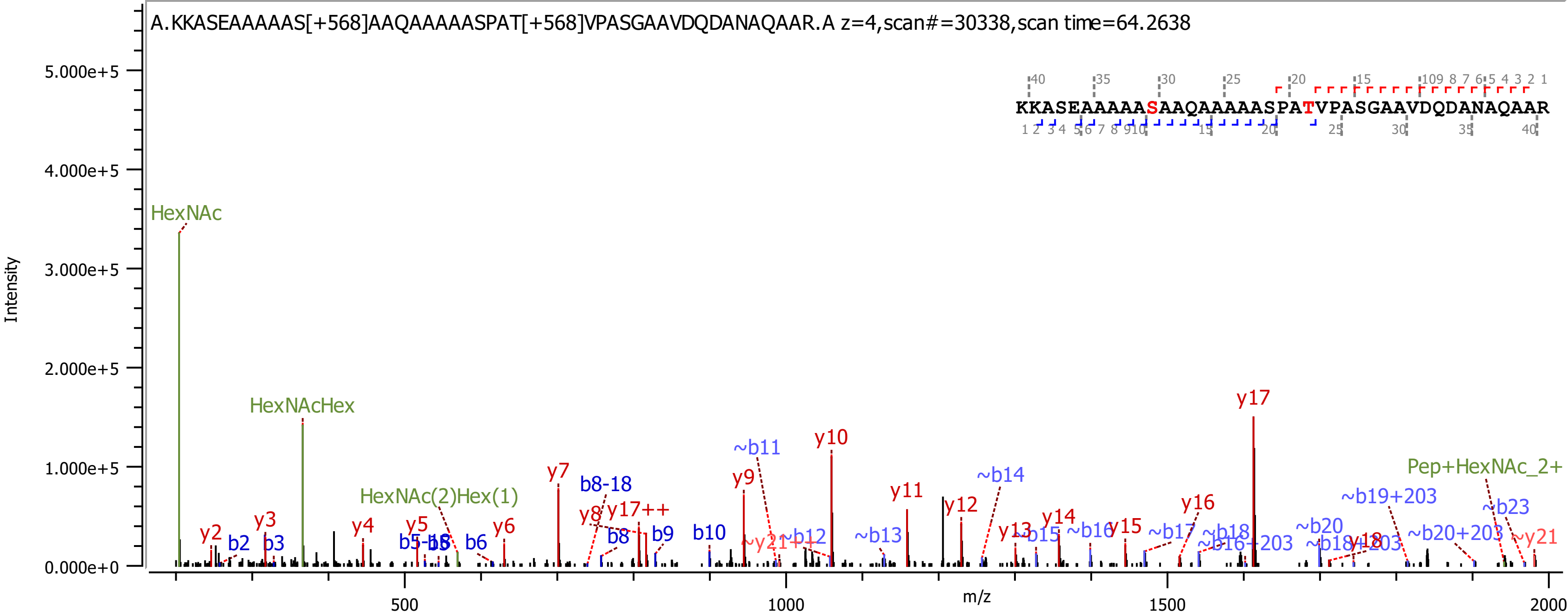

A.KKASEAAAAAS[+568]AAQAAAAASPAT[+568]VPASGAAVDQDANAQAAR.A z=4,scan#=30338,scan time=64.2638
KKASEAAAAASAAQAAAAASPATVPASGAAVDQDANAQAAR
Intensity
m/z
HexNAc
HexNAcHex
HexNAc(2)Hex(1)
Pep+HexNAc_2+
y2
y3
y4
y5
y6
y7
y8
y9
y10
y11
y12
y13
y14
y15
y16
y17
y17++
y18
~y21
~y21++
b2
b3
b6
b8
b8-18
b9
b10
~b11
~b12
~b13
~b14
~b15
~b16
~b17
~b18
~b20
~b23
~b16+203
~b18+203
~b19+203
~b20+203

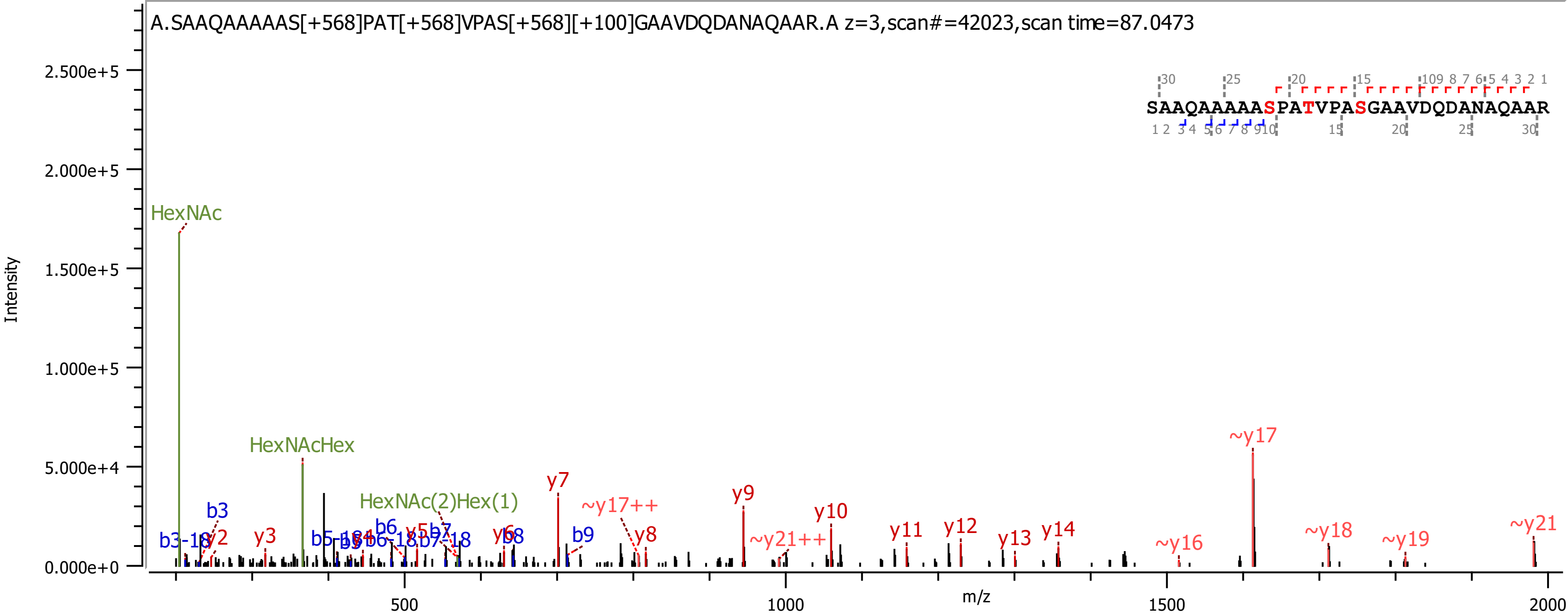
A.SAAQAAAAAS[+568]PAT[+568]VPAS[+568][+100]GAAVDQDANAQAAR.A z=3,scan#=42023,scan time=87.0473
SAAQAAAAASPATVPASGAAVDQDANAQAAR
Intensity
m/z
2.500e+5
2.000e+5
1.500e+5
1.000e+5
5.000e+4
0.000e+0
500
1000
1500
2000
HexNAc
HexNAcHex
HexNAc(2)Hex(1)
b3
b3-18
y2
y3
b6
y7
~y17++
y8
y9
~y21++
y10
y11
y12
y13
y14
~y16
~y17
~y18
~y19
~y21
b8
b9

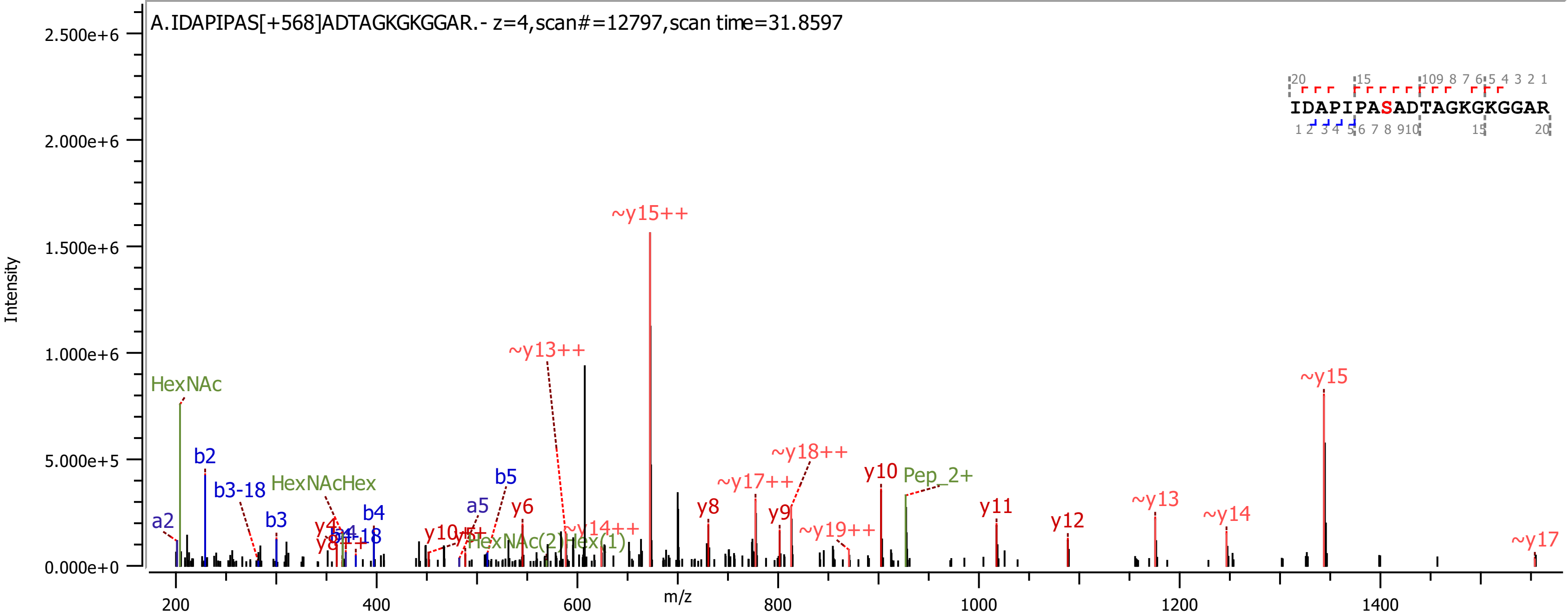

A.IDAPIPAS[+568]ADTAGKGKGGAR.- z=4,scan#=12797,scan time=31.8597
IDAPIPASADTAGKGKGGAR
Intensity
m/z
HexNAc
b2
b3-18
HexNAcHex
a2
b3
b4
a5
b5
y6
y8
y9
y10
y11
y12
~y13++
~y15++
~y14++
~y17++
~y18++
~y19++
~y13
~y14
~y15
~y17
Pep_2+
HexNAc(2)Hex(1)

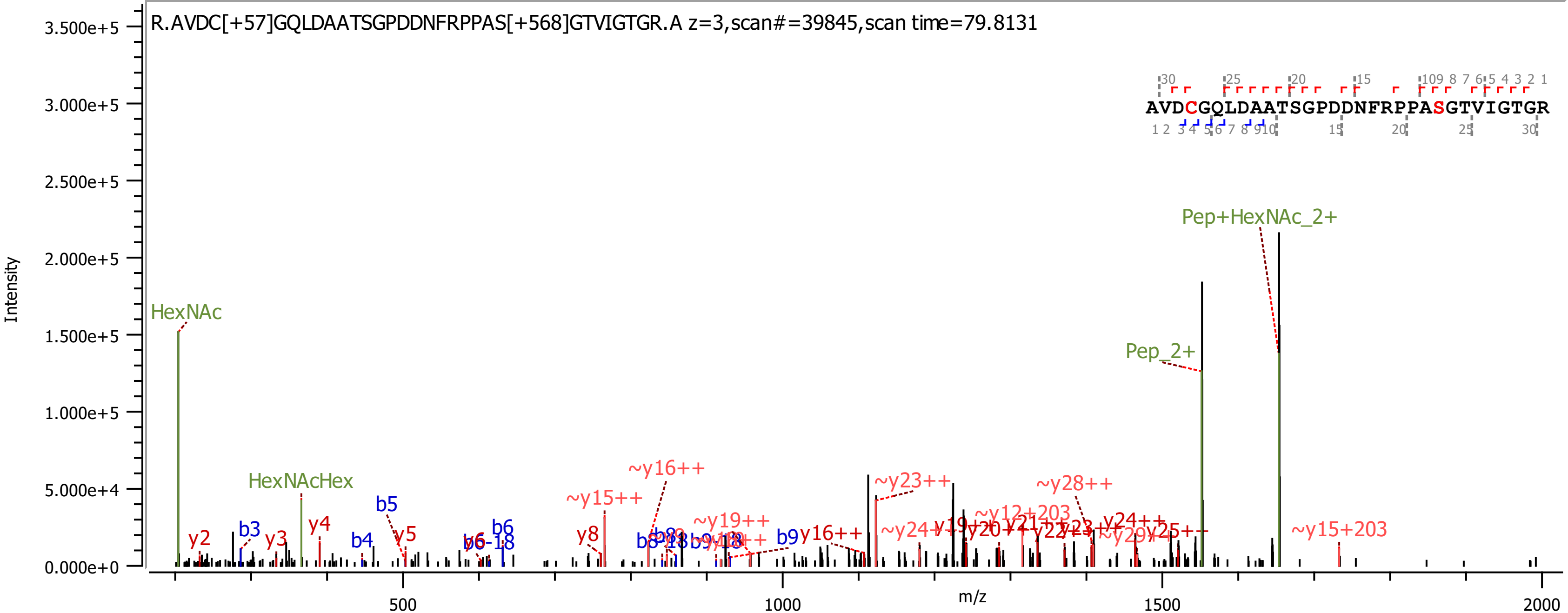
R.AVDC[+57]GQLDAATSGPDDNFRPPAS[+568]GTVIGTGR.A z=3,scan#=39845,scan time=79.8131
AVDCGQLDAATSGPDDNFRPPASGTVIGTGR
Intensity
m/z
HexNAc
HexNAcHex
Pep+HexNAc_2+
Pep_2+
~y15+203
~y12+203
~y23++
~y28++
~y16++
~y15++
~y19++
y2
y3
y4
y5
y8
b3
b4
b5
b6
b6-18
b9
y16++
y24++
~y24++
y25++
3.500e+5
3.000e+5
2.500e+5
2.000e+5
1.500e+5
1.000e+5
5.000e+4
0.000e+0
500
1000
1500
2000

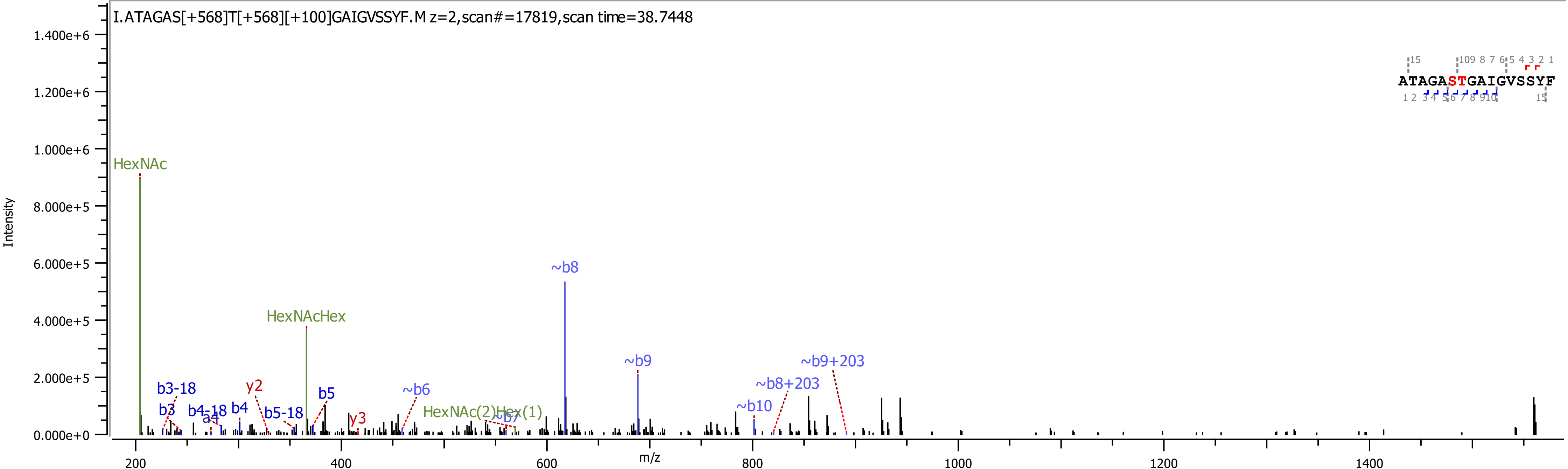
I.ATAGAS[+568]T[+568][+100]GAIGVSSYF.M z=2,scan#=17819,scan time=38.7448
Intensity
m/z
HexNAc
HexNAcHex
HexNAc(2)Hex(1)
b3-18
b3
b4-18
a4
b4
y2
b5-18
b5
y3
~b6
~b7
~b8
~b9
~b10
~b8+203
~b9+203
ATAGASTGAIGVSSYF

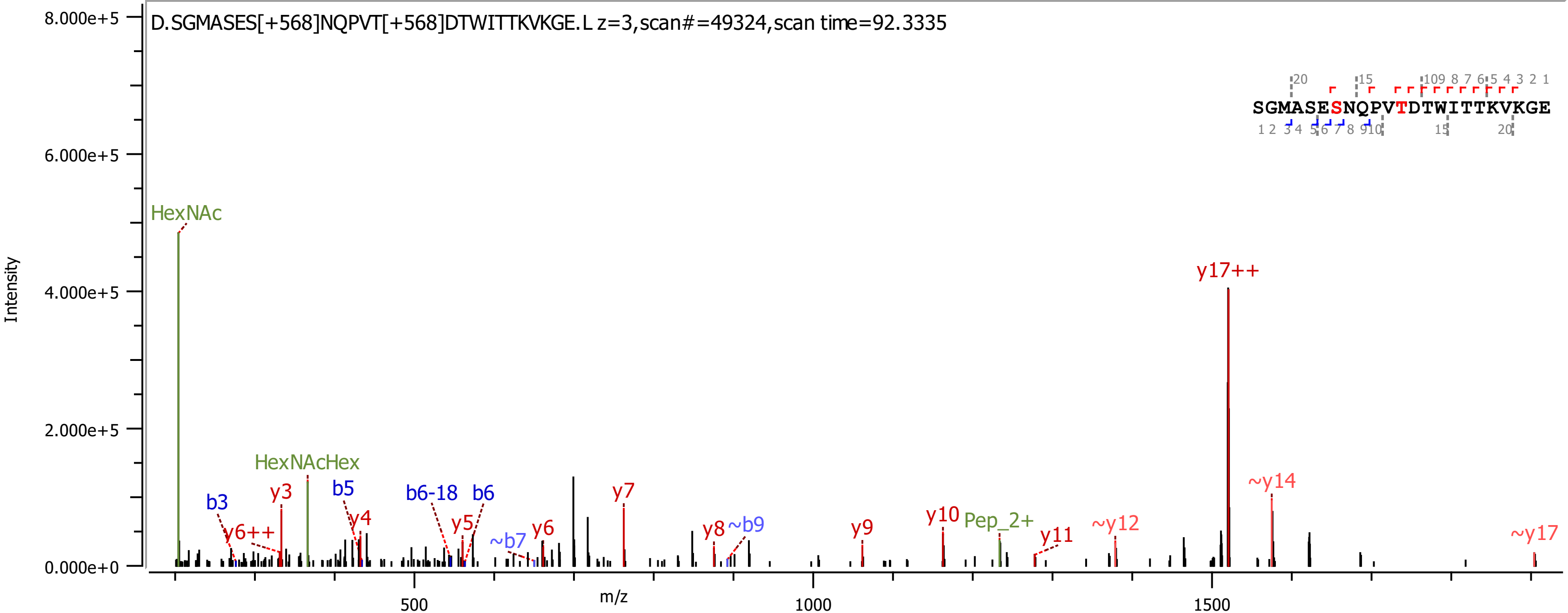

D.SGMASES[+568]NQPVT[+568]DTWITTKVKGE.L z=3,scan#=49324,scan time=92.3335
SGMASESNQPVTDTWITTKVKGE
Intensity
m/z
8.000e+5
6.000e+5
4.000e+5
2.000e+5
0.000e+0
500
1000
1500
HexNAc
HexNAcHex
b3
y6++
y3
b5
y4
b6-18
b6
y5
~b7
y6
y7
y8
~b9
y9
y10
Pep_2+
y11
~y12
y17++
~y14
~y17

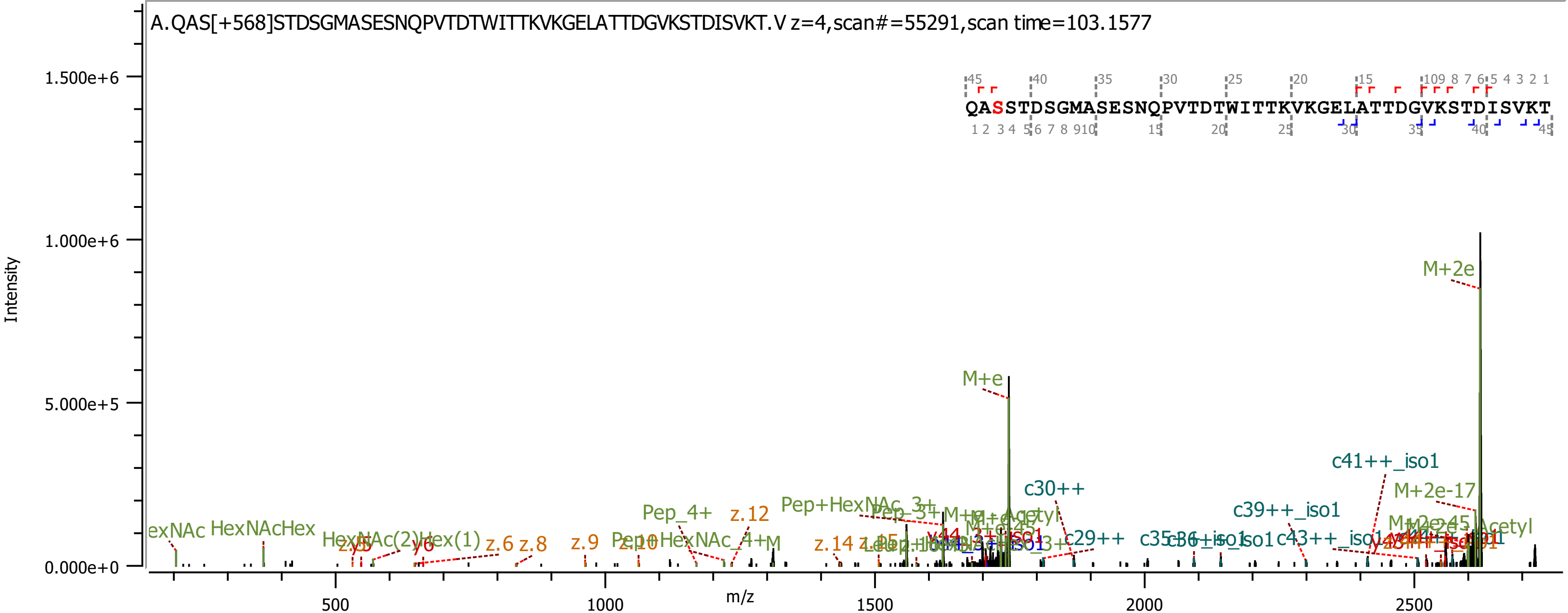
A.QAS[+568]STDSGMASESNQPVTDTWITTKVKGELATTDGVKSTDISVKT.V z=4,scan#=55291,scan time=103.1577
QASSTDSGMASESNQPVTDTWITTKVKGELATTDGVKSTDISVKT
Intensity
m/z
1.500e+6
1.000e+6
5.000e+5
0.000e+0
500
1000
1500
2000
2500
M+2e
M+e
c41++_iso1
M+2e-17
c30++
c39++_iso1
c29++
c43++_iso1
Pep_4+
z.12
Pep+HexNAc_3+
HexNAcHex
HexNAc(2)Hex(1)
z.6
z.8
z.9
z.10
z.14
y6
y5
Pep+HexNAc_4+

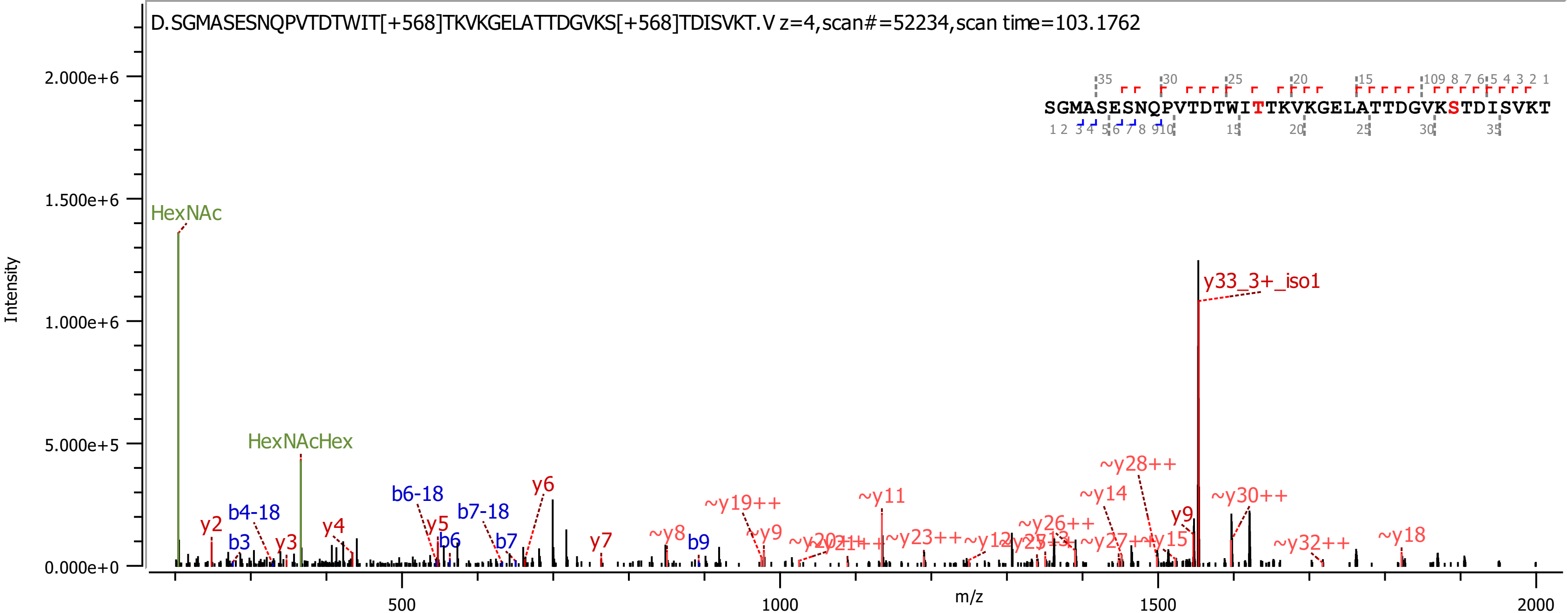

D.SGMASESNQPVTDTWIT[+568]TKVKGELATTDGVKS[+568]TDISVKT.V z=4,scan#=52234,scan time=103.1762
SGMASESNQPVTDTWITTKVKGELATTDGVKSTDISVKT
Intensity
m/z
HexNAc
HexNAcHex
y33_3+_iso1
~y28++
~y30++
~y14
~y11
~y19++
y6
b6-18
b7-18
b4-18
y2
b3
y3
y4
y5
b6
b7
y7
~y8
b9
~y9
y9
~y23++
~y26++
~y15
~y32++
~y18

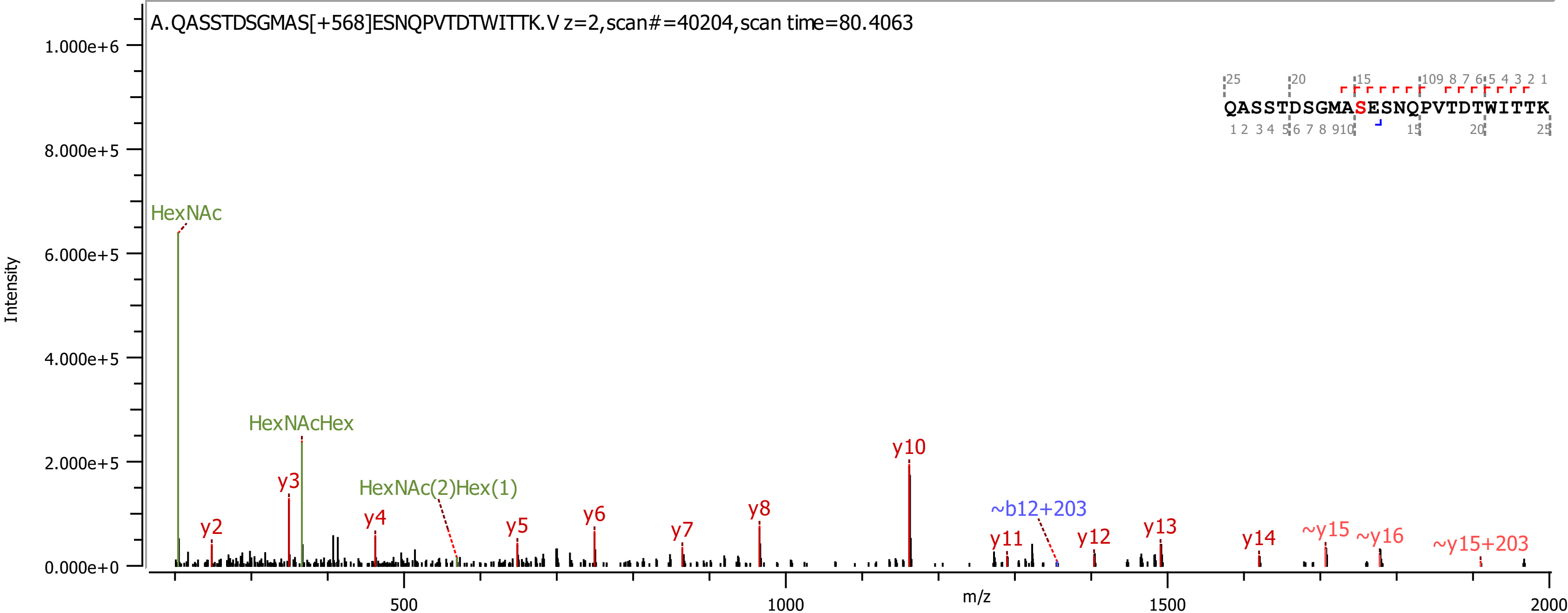

A.QASSTDSGMAS[+568]ESNQPVTDTWITTK.V z=2,scan#=40204,scan time=80.4063
QASSTDSGMASESNQPVTDTWITTK
Intensity
m/z
HexNAc
HexNAcHex
HexNAc(2)Hex(1)
y2
y3
y4
y5
y6
y7
y8
y10
~b12+203
y11
y12
y13
y14
~y15
~y16
~y15+203

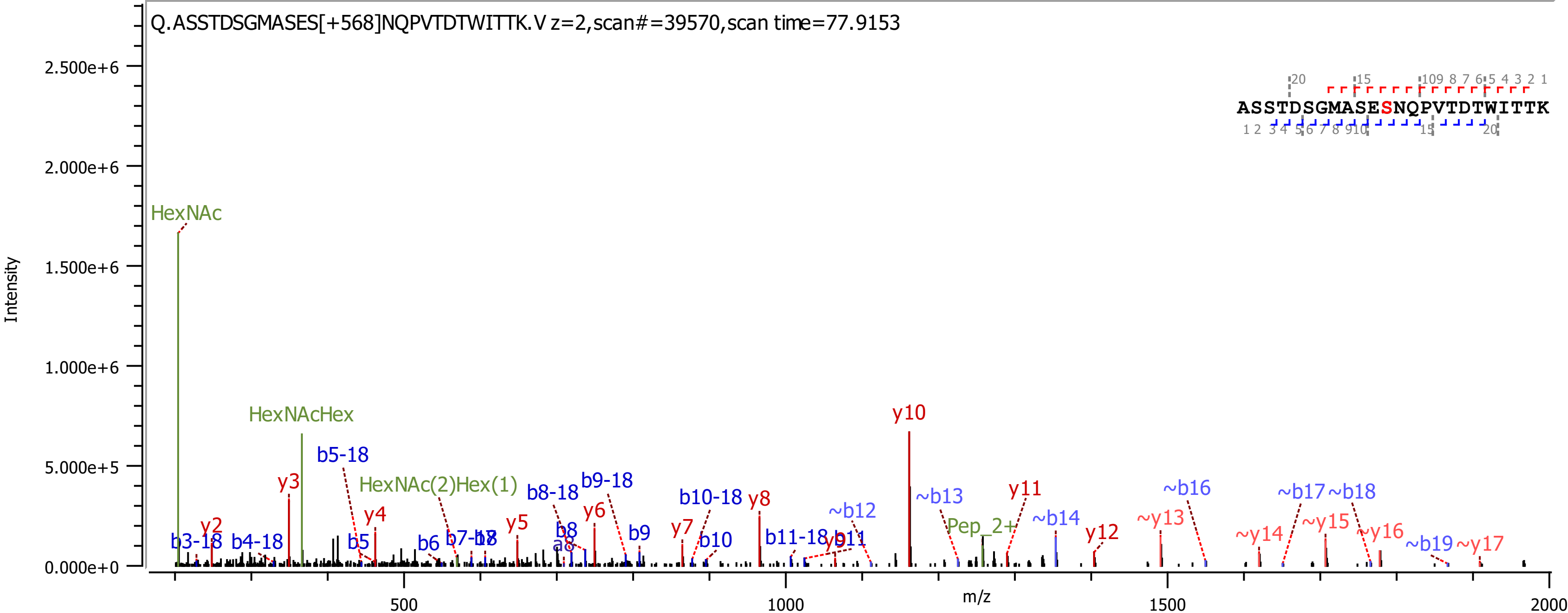
Q.ASSTDSGMASES[+568]NQPVTDTWITTK.V z=2,scan#=39570,scan time=77.9153
ASSTDSGMASESNQPVTDTWITTK
Intensity
m/z
HexNAc
HexNAcHex
HexNAc(2)Hex(1)
Pep_2+
y10
y3
y2
y4
y5
y6
y7
y8
y11
y12
~y13
~y14
~y15
~y16
~y17
b3-18
b4-18
b5-18
b5
b6
b8-18
b9-18
b8
a8
b9
b10-18
b10
b11-18
~b12
~b13
~b14
~b16
~b17
~b18
~b19
0.000e+0
5.000e+5
1.000e+6
1.500e+6
2.000e+6
2.500e+6
500
1000
1500
2000

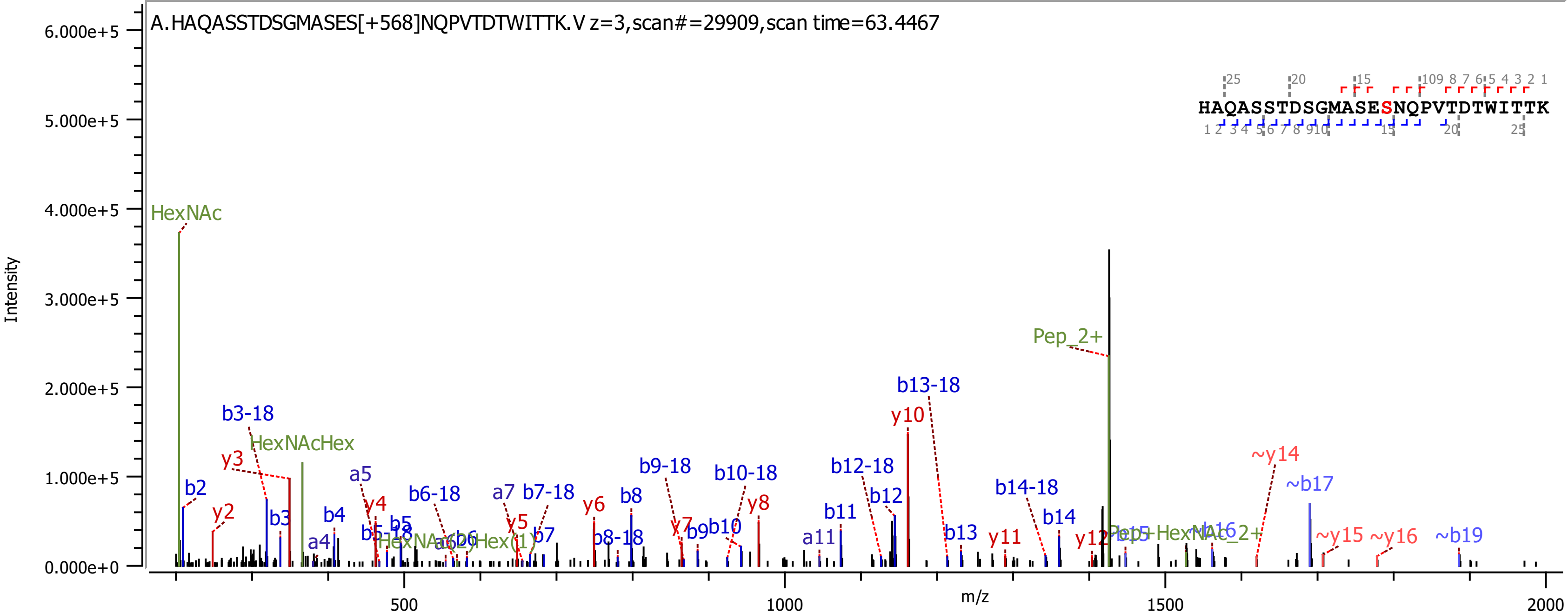

A.HAQASSTDSGMASES[+568]NQPVTDTWITTK.V z=3,scan#=29909,scan time=63.4467
HAQASSTDSGMASESNQPVTDTWITTK
Intensity
m/z
6.000e+5
5.000e+5
4.000e+5
3.000e+5
2.000e+5
1.000e+5
0.000e+0
500
1000
1500
2000
HexNAc
HexNAcHex
Pep_2+
b2
b3-18
y3
y2
b3
a4
b4
a5
y4
b5-18
b5
b6-18
a7
b7-18
y5
b7
y6
b8
b8-18
b9-18
y7
b9
b10
b10-18
y8
a11
b11
b12-18
b12
y10
b13-18
b13
y11
b14-18
b14
y12
~y14
~b17
~y15
~y16
~b19

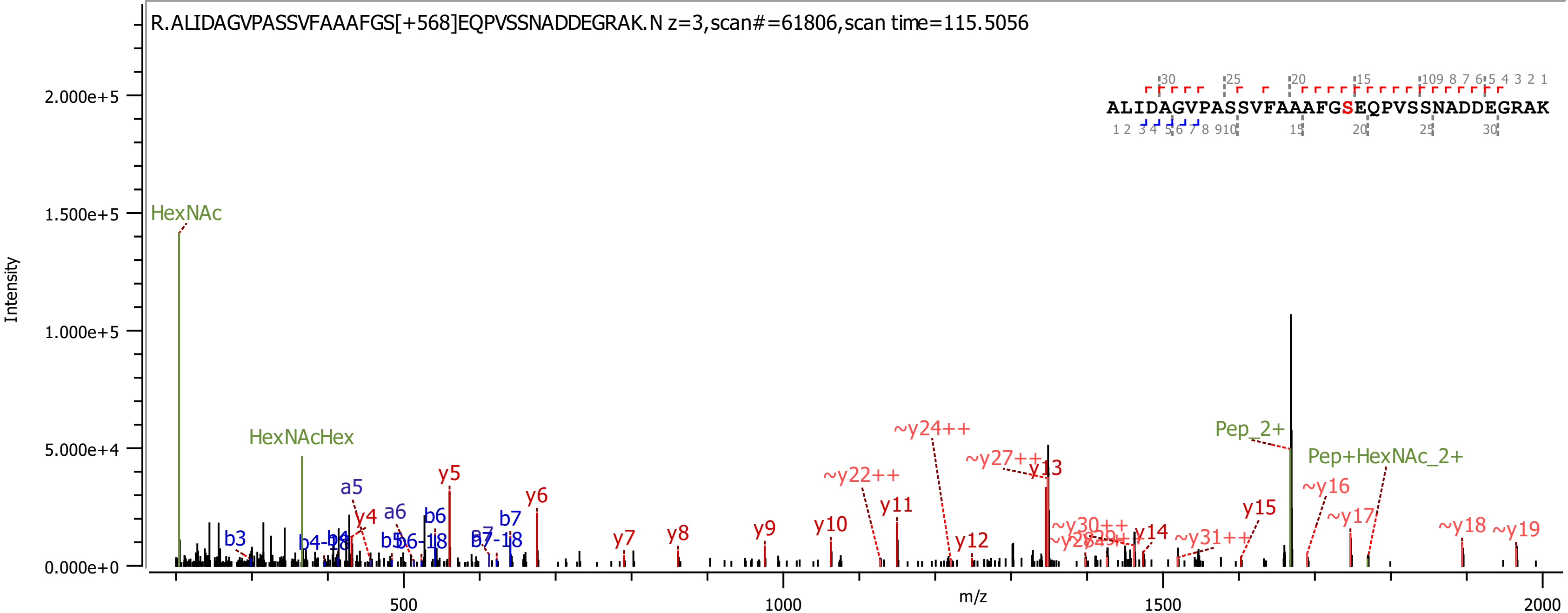
R.ALIDAGVPASSVFAAAFGS[+568]EQPVSSNADDEGRAK.N z=3,scan#=61806,scan time=115.5056
ALIDAGVPASSVFAAAFGSEQPVSSNADDEGRAK
Intensity
m/z
2.000e+5
1.500e+5
1.000e+5
5.000e+4
0.000e+0
500
1000
1500
2000
HexNAc
HexNAcHex
Pep_2+
Pep+HexNAc_2+
b3
b4-18
b4
a5
a6
b5
b6-18
b6
b7
b7-18
y4
y5
y6
y7
y8
y9
y10
y11
y12
y13
y14
y15
~y16
~y17
~y18
~y19
~y22++
~y24++
~y27++
~y30++
~y31++

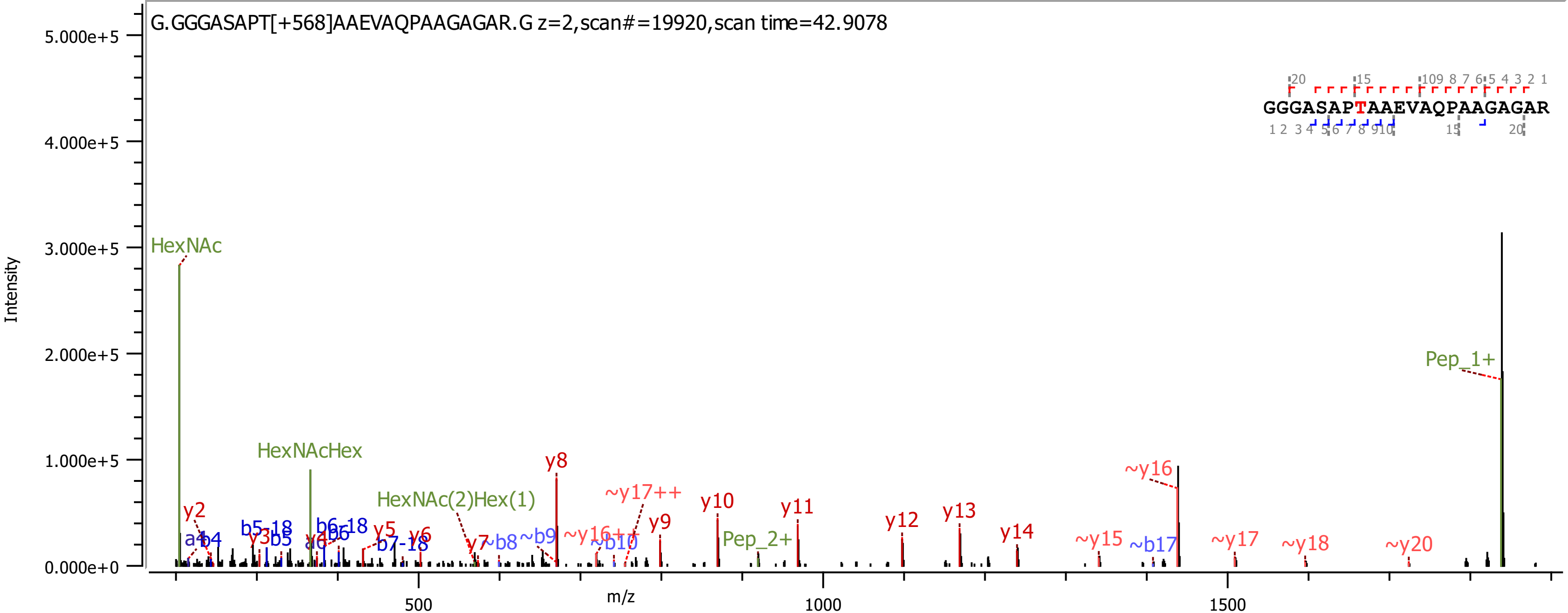
G.GGGASAPT[+568]AAEVAQPAAGAGAR.G z=2,scan#=19920,scan time=42.9078
GGGASAPTAAEVAQPAAGAGAR
Intensity
m/z
HexNAc
HexNAcHex
HexNAc(2)Hex(1)
Pep_2+
Pep_1+
y2
y3
y5
y6
y7
y8
y9
y10
y11
y12
y13
y14
~y15
~y16
~y17
~y18
~y20
~y16++
~y17++
a4
b4
b5
b5-18
a6
b6
b6-18
b7-18
~b8
~b9
~b10
~b17
0.000e+0
1.000e+5
2.000e+5
3.000e+5
4.000e+5
5.000e+5
500
1000
1500

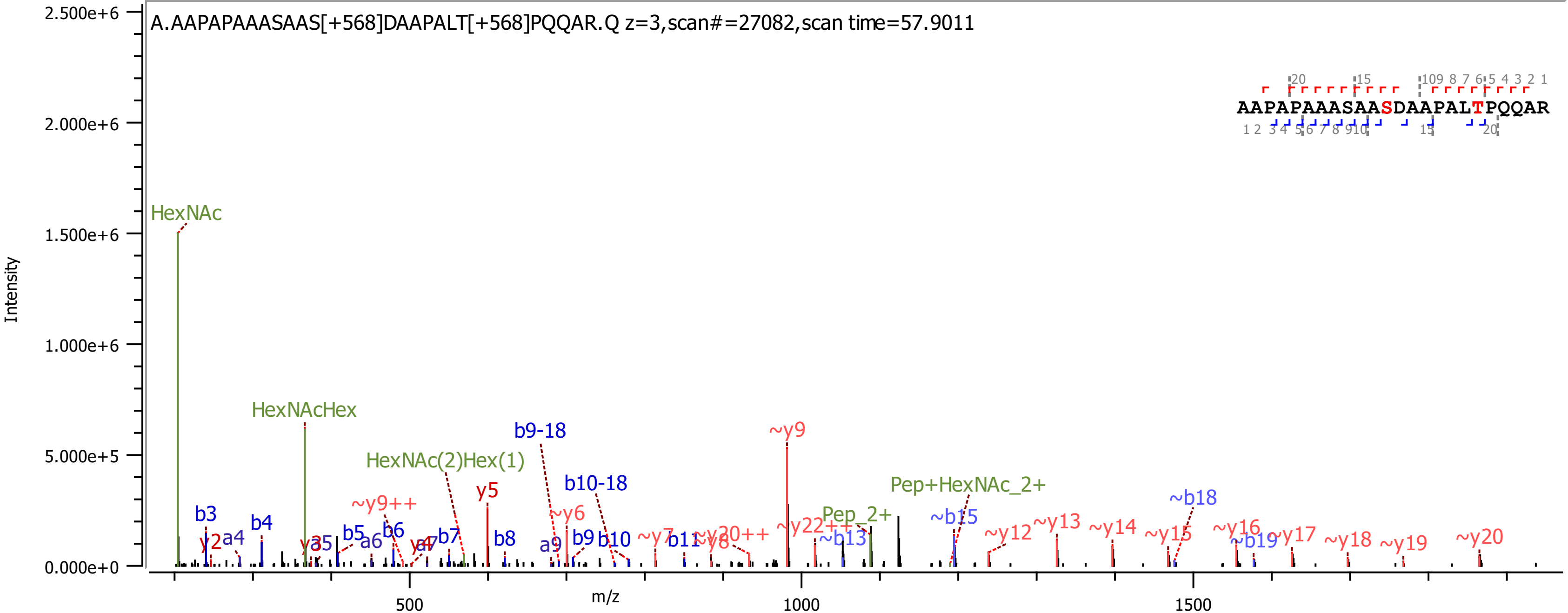
A.AAPAPAAASAAS[+568]DAAPALT[+568]PQQAR.Q z=3,scan#=27082,scan time=57.9011
AAPAPAAASAASDAAPALTPQQAR
Intensity
m/z
2.500e+6
2.000e+6
1.500e+6
1.000e+6
5.000e+5
0.000e+0
500
1000
1500
HexNAc
HexNAcHex
HexNAc(2)Hex(1)
Pep_2+
Pep+HexNAc_2+
b3
b4
b5
b6
b7
b8
b9
b10
b11
b9-18
b10-18
a4
a6
a9
y2
y5
~y6
~y7
~y9
~y9++
~y20++
~y22++
~y12
~y13
~y14
~y15
~y16
~y17
~y18
~y19
~y20
~b13
~b15
~b18
~b19

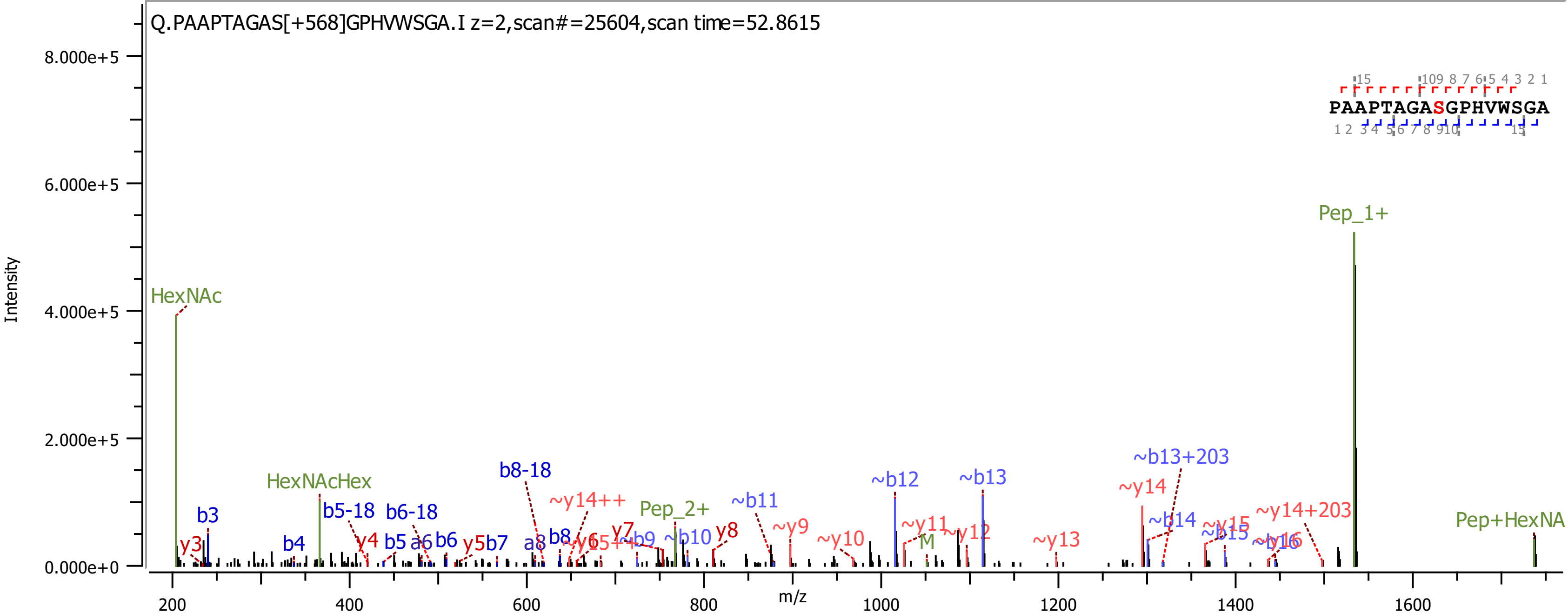
Q.PAAPTAGAS[+568]GPHVWSGA.I z=2,scan#=25604,scan time=52.8615
PAAPTAGASGPHVWSGA
Intensity
m/z
8.000e+5
6.000e+5
4.000e+5
2.000e+5
0.000e+0
200
400
600
800
1000
1200
1400
1600
HexNAc
HexNAcHex
Pep_1+
Pep_2+
Pep+HexNA
b3
b4
b5
b6
b8
a6
a8
b5-18
b6-18
b8-18
y3
y4
y5
y7
y8
b7
~y14++
~b9
~b10
~b11
~b12
~b13
~b13+203
~b14
~b15
~y9
~y10
~y11
~y12
~y13
~y14
~y15
~y16
~y14+203
M

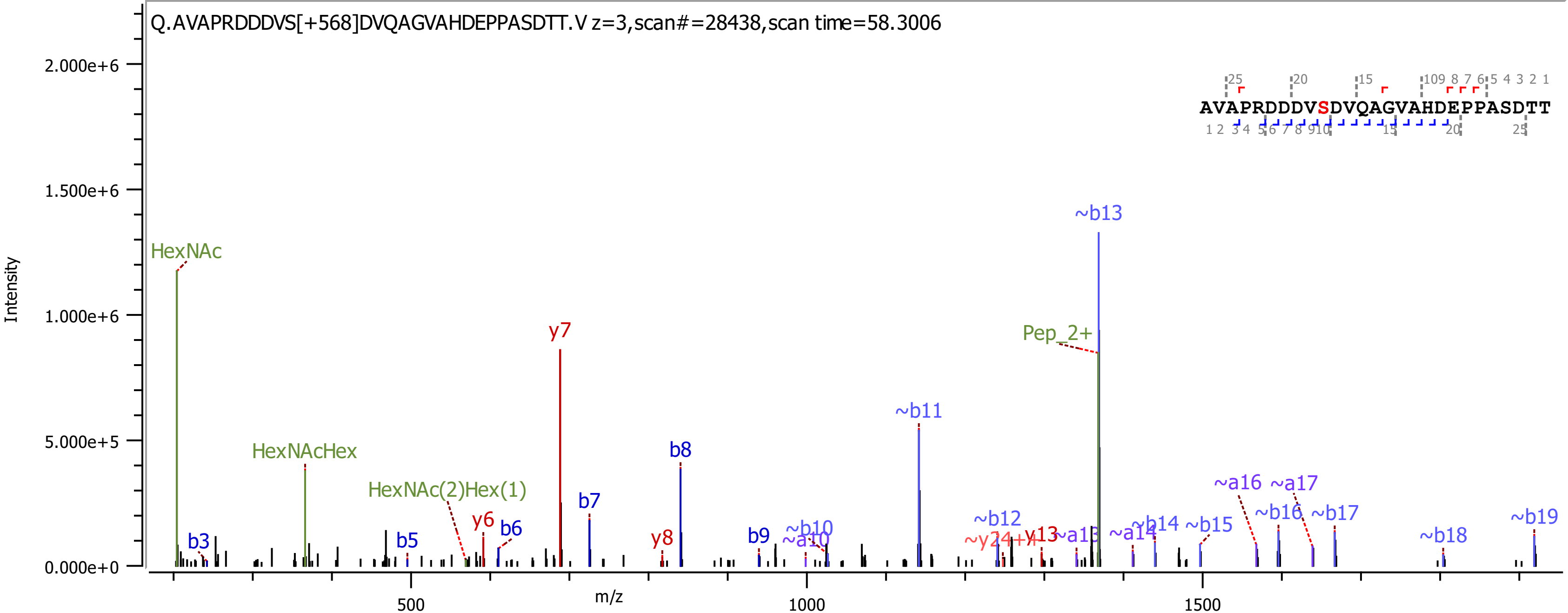

Q.AVAPRDDDVS[+568]DVQAGVAHDEPPASDTT.V z=3,scan#=28438,scan time=58.3006
AVAPRDDDVSDVQAGVAHDEPPASDTT
Intensity
m/z
2.000e+6
1.500e+6
1.000e+6
5.000e+5
0.000e+0
500
1000
1500
HexNAc
HexNAcHex
HexNAc(2)Hex(1)
y6
y7
y8
y13
b3
b5
b6
b7
b8
b9
~b11
~b12
~b13
~b14
~b15
~b16
~b17
~b18
~b19
~a16
~a17
~a14
Pep_2+

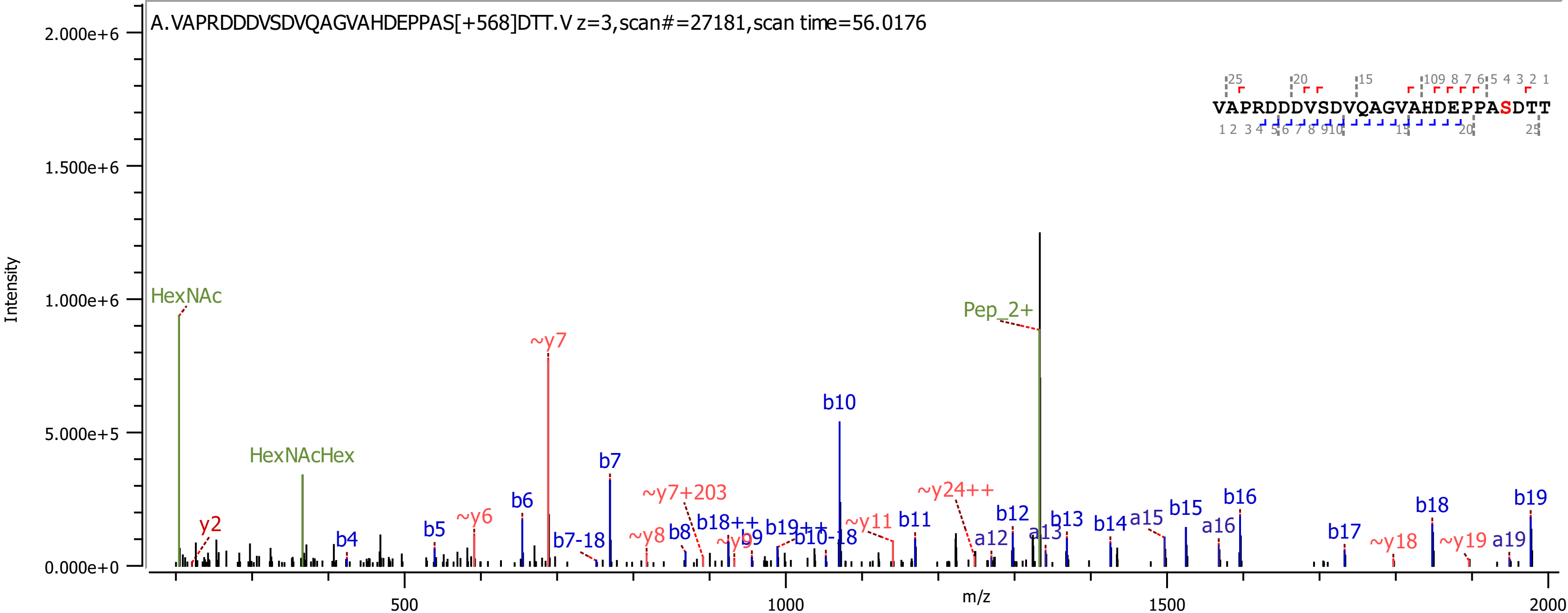

A.VAPRDDDVSDVQAGVAHDEPPAS[+568]DTT.V z=3,scan#=27181,scan time=56.0176
VAPRDDDVSDVQAGVAHDEPPASDTT
Intensity
m/z
HexNAc
HexNAcHex
Pep_2+
~y7
b10
b7
b6
b5
b4
y2
~y6
b7-18
~y7+203
~y8
b8
b18++
~y9
b9
b19++
b10-18
~y11
b11
~y24++
a12
b12
a13
b13
b14
a15
b15
a16
b16
b17
~y18
b18
~y19
a19
b19

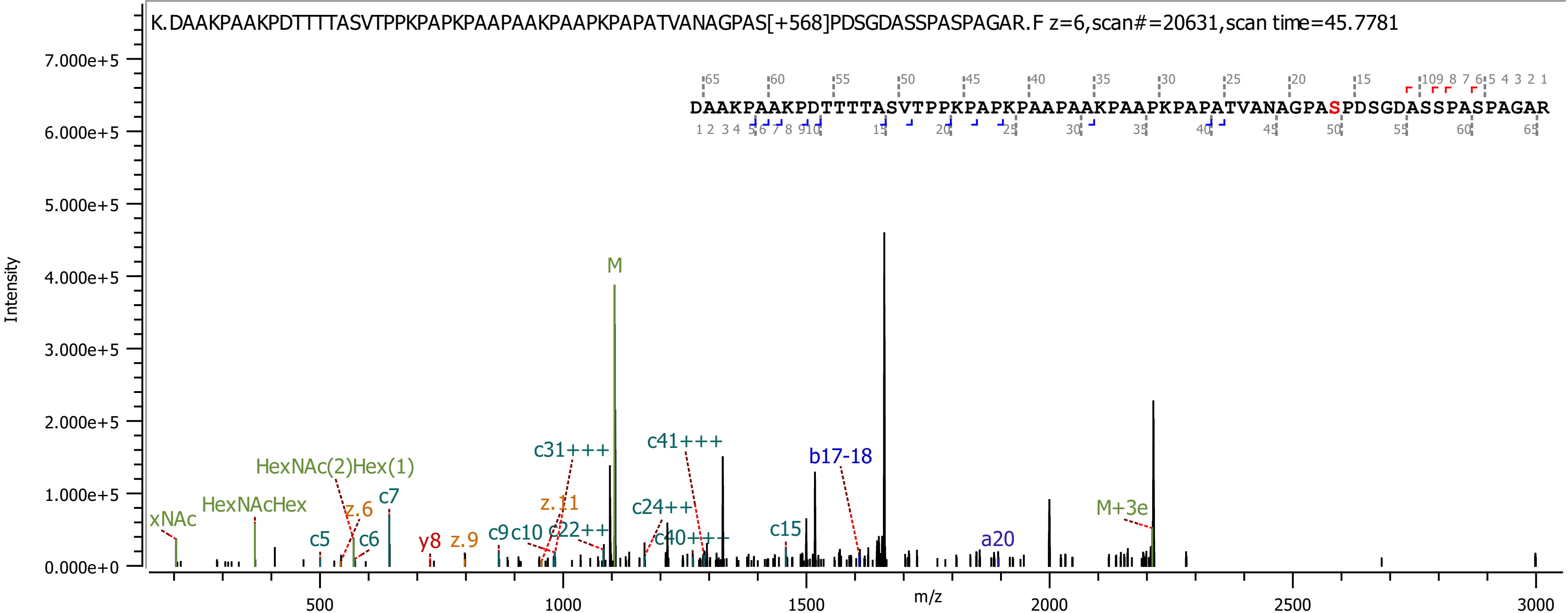

K.DAAKPAAKPDTTTTASVTPPKPAPKPAAPAAKPAAPKPAPATVANAGPAS[+568]PDSGDASSPASPAGAR.F z=6,scan#=20631,scan time=45.7781
DAAKPAAKPDTTTTASVTPPKPAPKPAAPAAKPAAPKPAPATVANAGPASPDSGDASSPASPAGAR
Intensity
m/z
M
M+3e
xNAc
HexNAcHex
HexNAc(2)Hex(1)
c5
c6
c7
z.6
y8
z.9
c9
c10
z.11
c22++
c31+++
c24++
c40+++
c41+++
c15
b17-18
a20

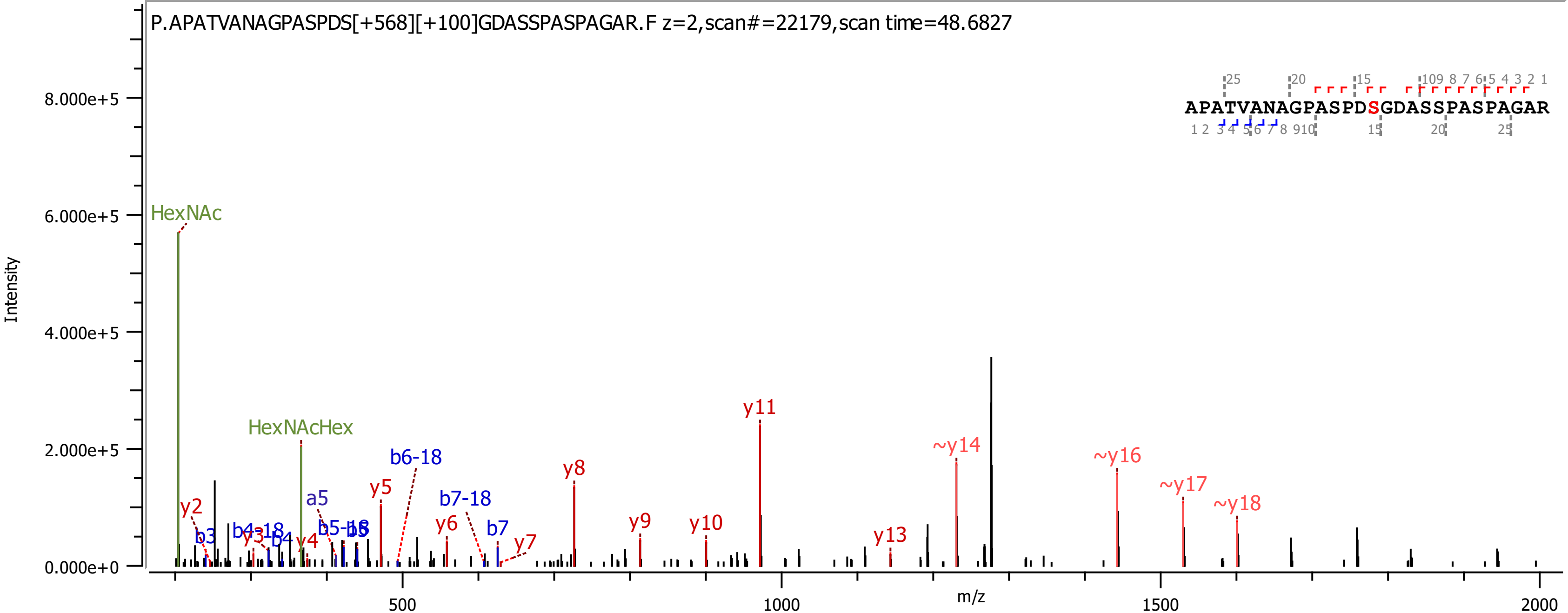

P.APATVANAGPASPDS[+568][+100]GDASSPASPAGAR.F z=2,scan#=22179,scan time=48.6827
APATVANAGPASPDSGDASSPASPAGAR
Intensity
8.000e+5
6.000e+5
4.000e+5
2.000e+5
0.000e+0
m/z
500
1000
1500
2000
HexNAc
HexNAcHex
y2
b3
b4-18
y3
b4
y4
a5
b5-18
b5
y5
b6-18
y6
b7-18
b7
y7
y8
y9
y10
y11
y13
~y14
~y16
~y17
~y18

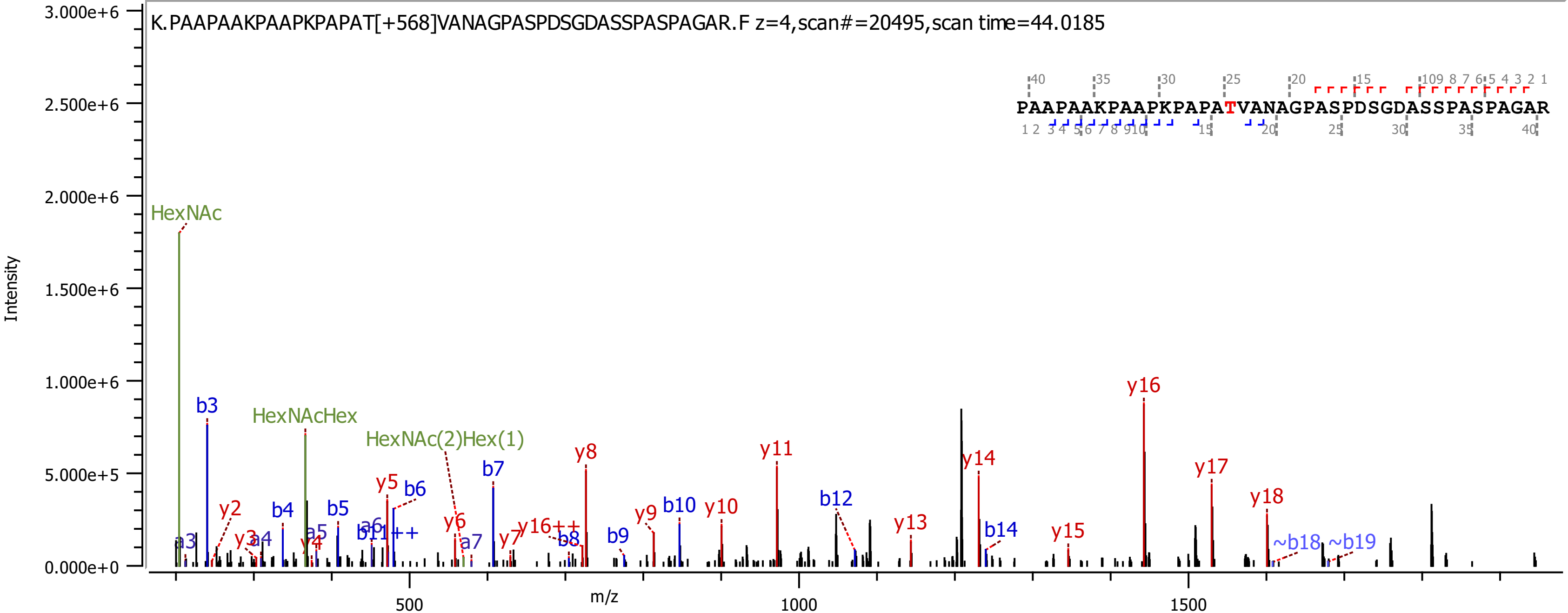
K.PAAPAAKPAAPKPAPAT[+568]VANAGPASPDSGDASSPASPAGAR.F z=4,scan#=20495,scan time=44.0185
PAAPAAKPAAPKPAPATVANAGPASPDSGDASSPASPAGAR
Intensity
m/z
HexNAc
HexNAcHex
HexNAc(2)Hex(1)
b3
b4
b5
b6
b7
b8
b9
b10
b12
b14
~b18
~b19
y2
y3
y4
y5
y6
y7
y8
y9
y10
y11
y13
y14
y15
y16
y17
y18
y16++
b11++
a3
a4
a5
a6
a7

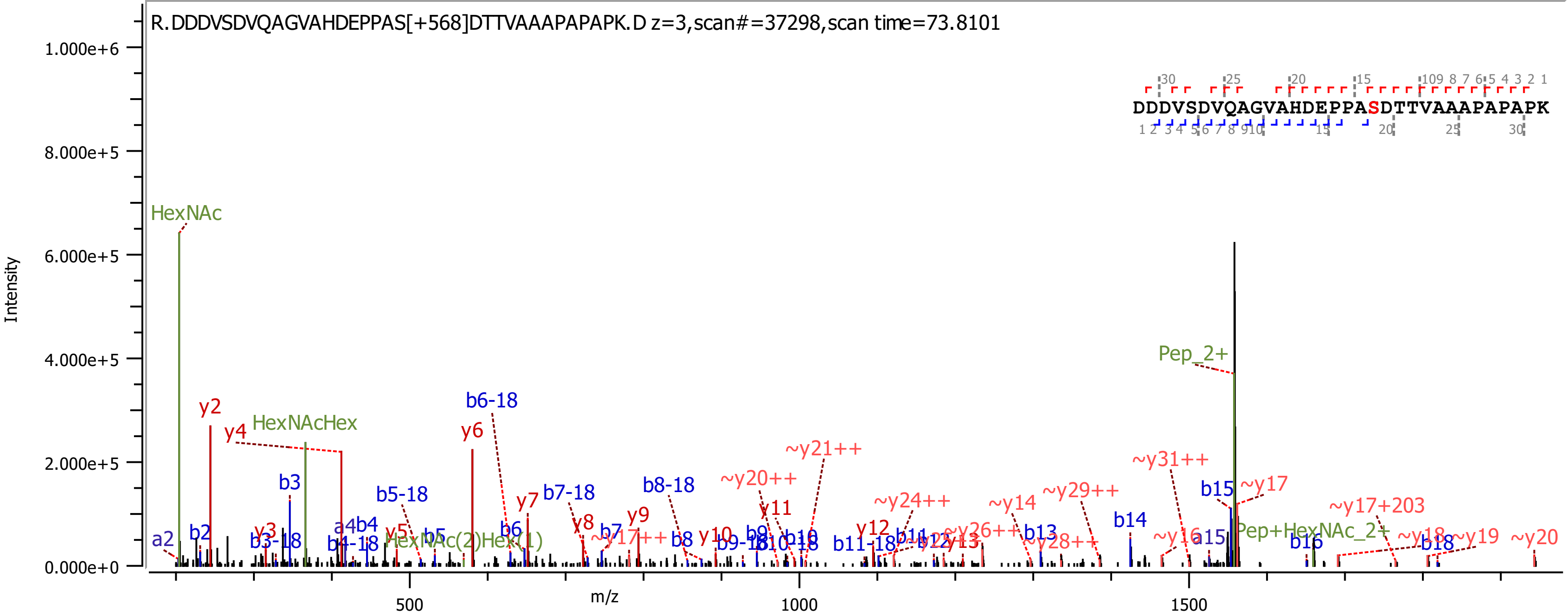
R.DDDVSDVQAGVAHDEPPAS[+568]DTTVAAAPAPAPK.D z=3,scan#=37298,scan time=73.8101
DDDVSDVQAGVAHDEPPASDTTVAAAPAPAPK
Intensity
m/z
1.000e+6
8.000e+5
6.000e+5
4.000e+5
2.000e+5
0.000e+0
500
1000
1500
HexNAc
HexNAcHex
HexNAc(2)Hex(1)
Pep_2+
Pep+HexNAc_2+
y2
y4
y6
y7
b6-18
b3
b5-18
b7-18
b8-18
~y20++
~y21++
~y31++
~y17
b15
~y17+203
~y24++
~y14
~y29++
b14

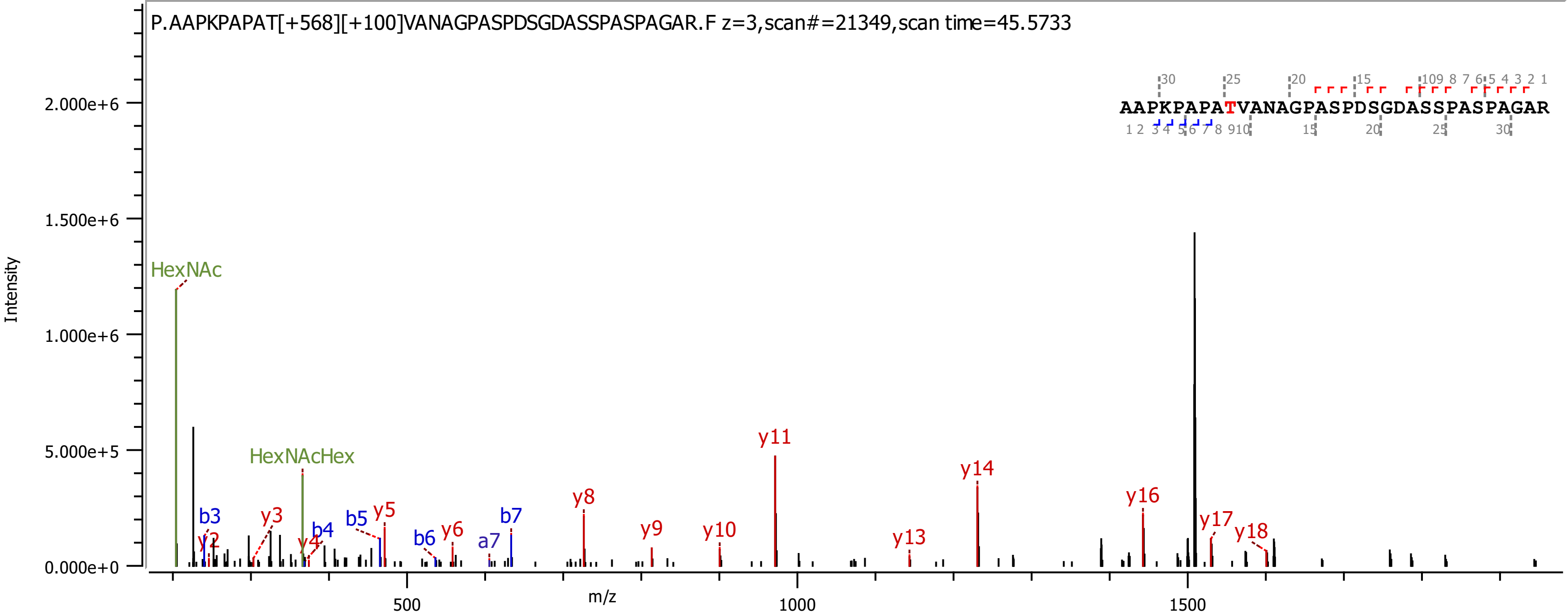

P.AAPKPAPAT[+568][+100]VANAGPASPDSGDASSPASPAGAR.F z=3,scan#=21349,scan time=45.5733
AAPKPAPATVANAGPASPDSGDASSPASPAGAR
Intensity
m/z
2.000e+6
1.500e+6
1.000e+6
5.000e+5
0.000e+0
500
1000
1500
HexNAc
HexNAcHex
b3
b4
b5
b6
b7
a7
y2
y3
y4
y5
y6
y8
y9
y10
y11
y13
y14
y16
y17
y18

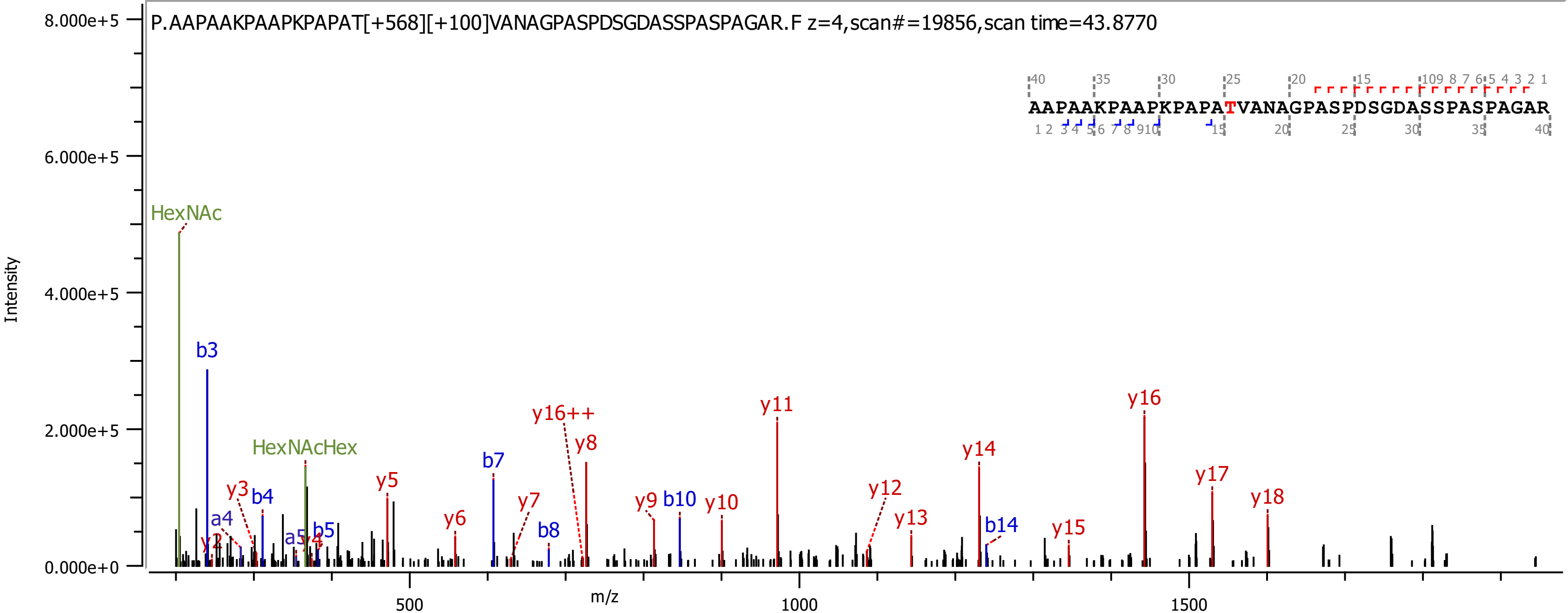

P.AAPAAKPAAPKPAPAT[+568][+100]VANAGPASPDSGDASSPASPAGAR.F z=4,scan#=19856,scan time=43.8770
AAPAAKPAAPKPAPATVANAGPASPDSGDASSPASPAGAR
Intensity
m/z
8.000e+5
6.000e+5
4.000e+5
2.000e+5
0.000e+0
500
1000
1500
HexNAc
HexNAcHex
b3
b4
b5
b7
b8
b10
b14
a4
a5
y2
y3
y4
y5
y6
y7
y8
y9
y10
y11
y12
y13
y14
y15
y16
y16++
y17
y18

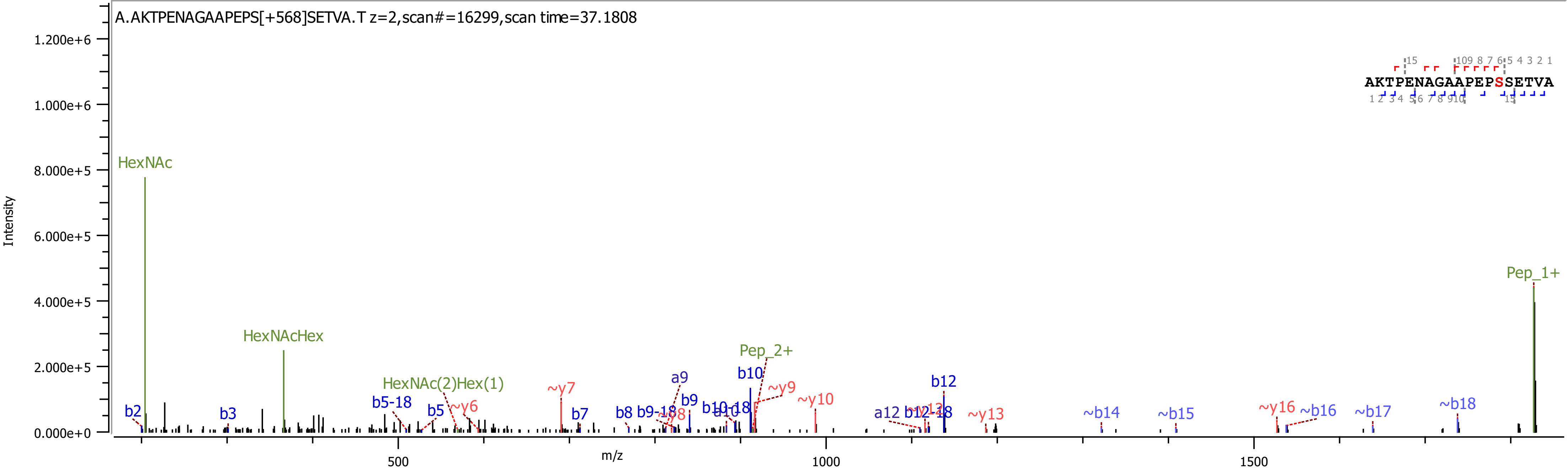
A.AKTPENAGAAPEPS[+568]SETVA.T z=2,scan#=16299,scan time=37.1808
AKTPENAGAAPEPSSETVA
Intensity
m/z
HexNAc
HexNAcHex
HexNAc(2)Hex(1)
Pep_2+
Pep_1+
b2
b3
b5-18
b5
~y6
~y7
b7
b8
b9-18
a9
b9
b10
b10-18
a10
~y9
~y10
a12
b12-18
b12
~y13
~b14
~b15
~y16
~b16
~b17
~b18

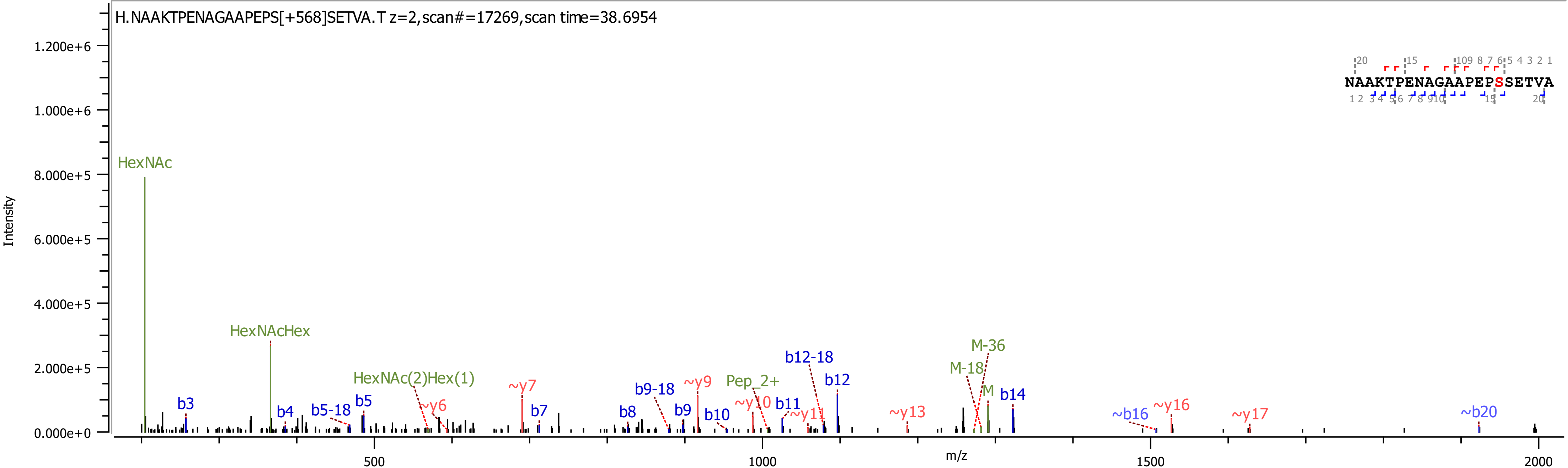
H.NAAKTPENAGAAPEPS[+568]SETVA.T z=2,scan#=17269,scan time=38.6954
NAAKTPENAGAAPEPSSETVA
Intensity
m/z
HexNAc
HexNAcHex
HexNAc(2)Hex(1)
Pep_2+
M-36
M-18
M
b3
b4
b5-18
b5
b7
b8
b9-18
b9
b10
b11
b12-18
b12
b14
~b16
~b20
~y6
~y7
~y9
~y10
~y11
~y13
~y16
~y17

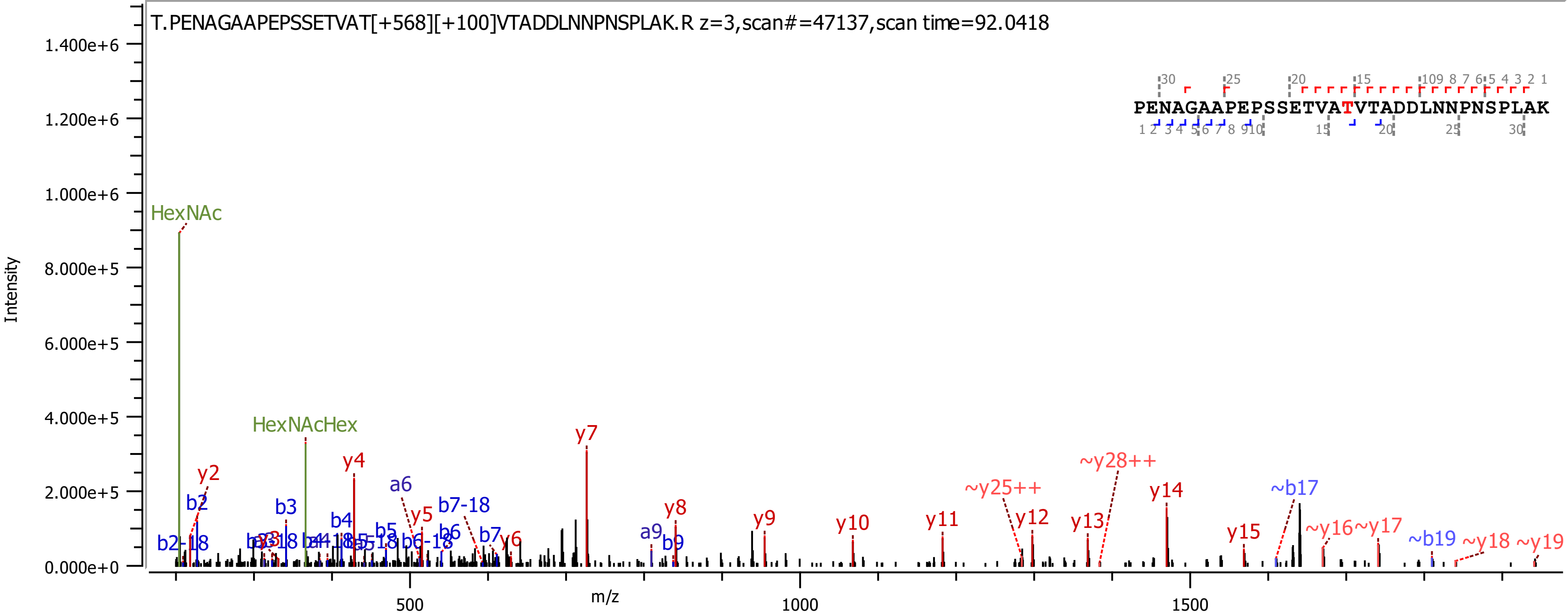

T.PENAGAAPEPSSETVAT[+568][+100]VTADDLNNPNSPLAK.R z=3,scan#=47137,scan time=92.0418
PENAGAAPEPSSETVATVTADDLNNPNSPLAK
Intensity
m/z
HexNAc
HexNAcHex
y2
y4
y5
y6
y7
y8
y9
y10
y11
y12
y13
y14
y15
~y16
~y17
~y18
~y19
~y25++
~y28++
b2
b2-18
b3
b4
b5
b6
b6-18
b7
b7-18
b9
a6
a9
~b17
~b19

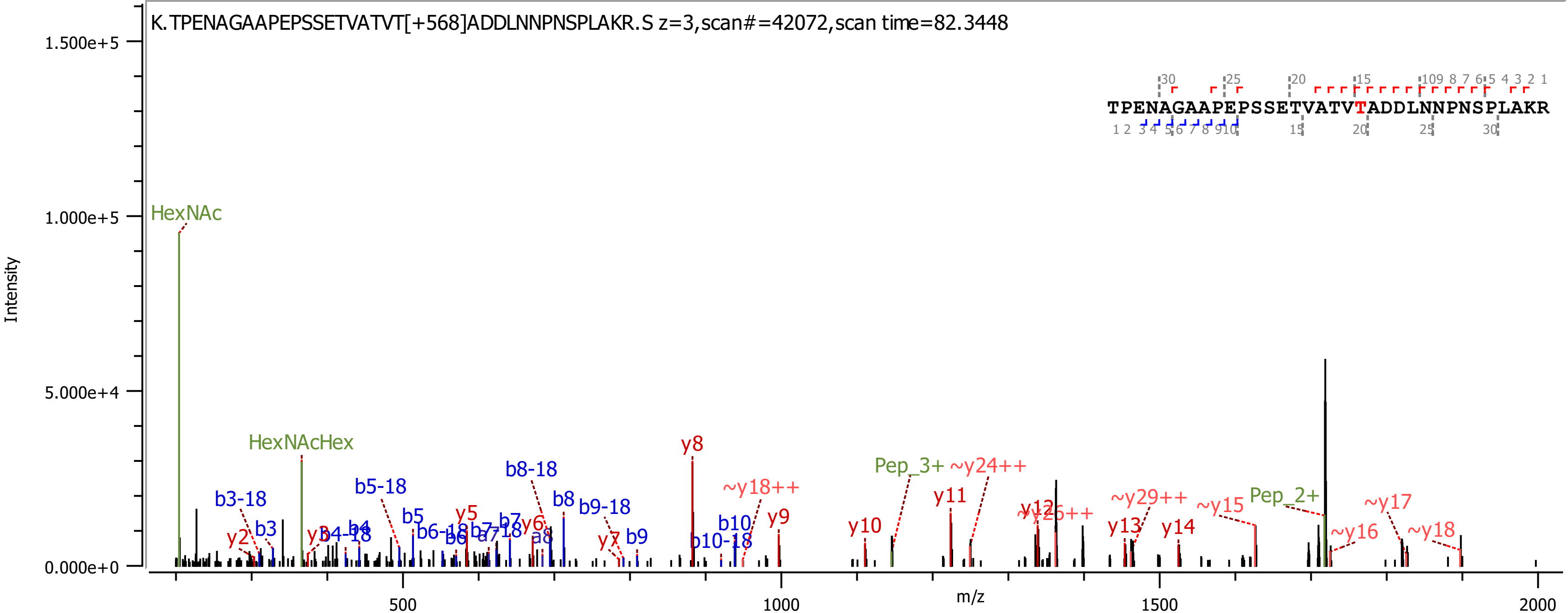

K.TPENAGAAPEPSSETVATVT[+568]ADDLNNPNSPLAKR.S z=3,scan#=42072,scan time=82.3448
TPENAGAAPEPSSETVATVTADDLNNPNSPLAKR
Intensity
m/z
1.500e+5
1.000e+5
5.000e+4
0.000e+0
500
1000
1500
2000
HexNAc
HexNAcHex
y8
Pep_3+
~y24++
~y18++
y11
y9
y10
b10
b10-18
Pep_2+
~y15
~y29++
y13
y14
~y17
~y16
~y18
y12
b8-18
b8
b9-18
b9
b5-18
b5
b3-18
b3
y2
b4
b4-18
y5
y6
b7
a8
y7

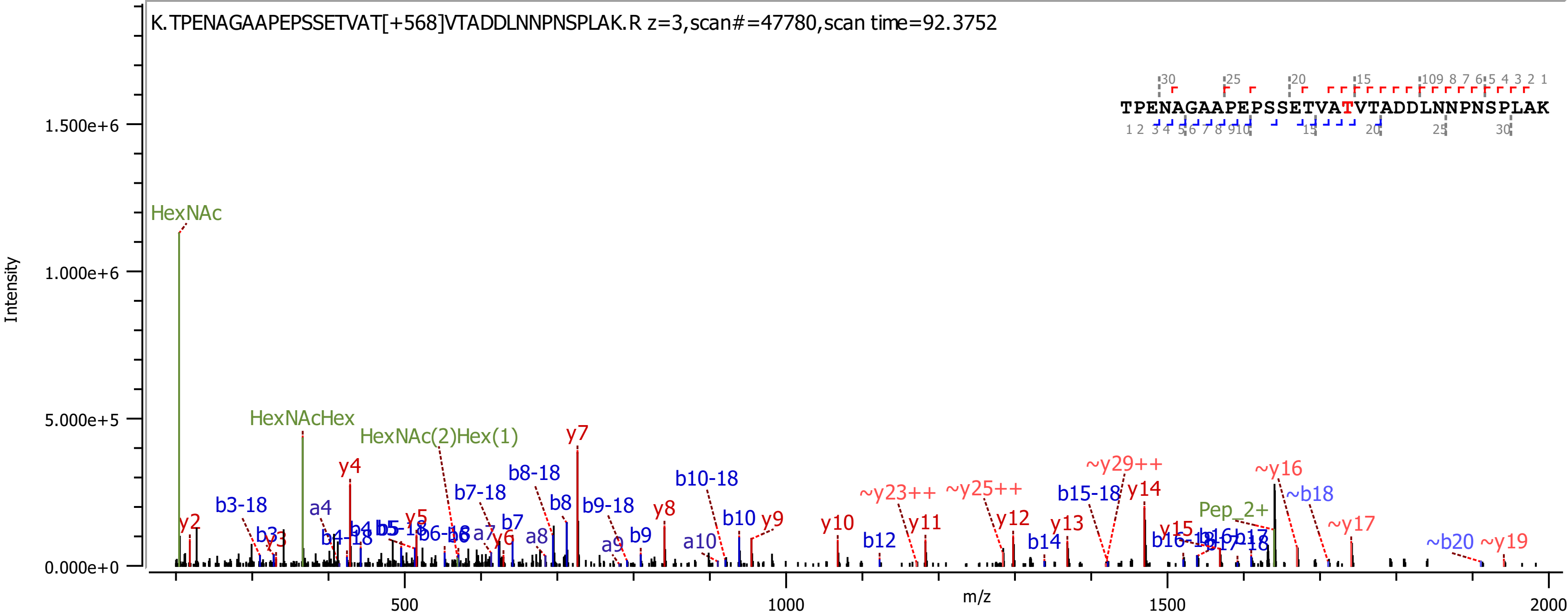

K.TPENAGAAPEPSSETVAT[+568]VTADDLNNPNSPLAK.R z=3,scan#=47780,scan time=92.3752
TPENAGAAPEPSSETVATVTADDLNNPNSPLAK
Intensity
m/z
HexNAc
HexNAcHex
HexNAc(2)Hex(1)
y2
y3
y4
y5
y6
y7
y8
y9
y10
y11
y12
y13
y14
y15
~y16
~y17
~y19
~y23++
~y25++
~y29++
b3-18
b3
a4
b4-18
b4
b5
b5-18
b6-18
a7
b7-18
b7
a8
b8-18
b8
b9-18
a9
b9
b10-18
a10
b10
b12
b14
b15-18
b16-18
b16
b17-18
b17
~b18
~b20
Pep_2+
0.000e+0
5.000e+5
1.000e+6
1.500e+6
500
1000
1500
2000

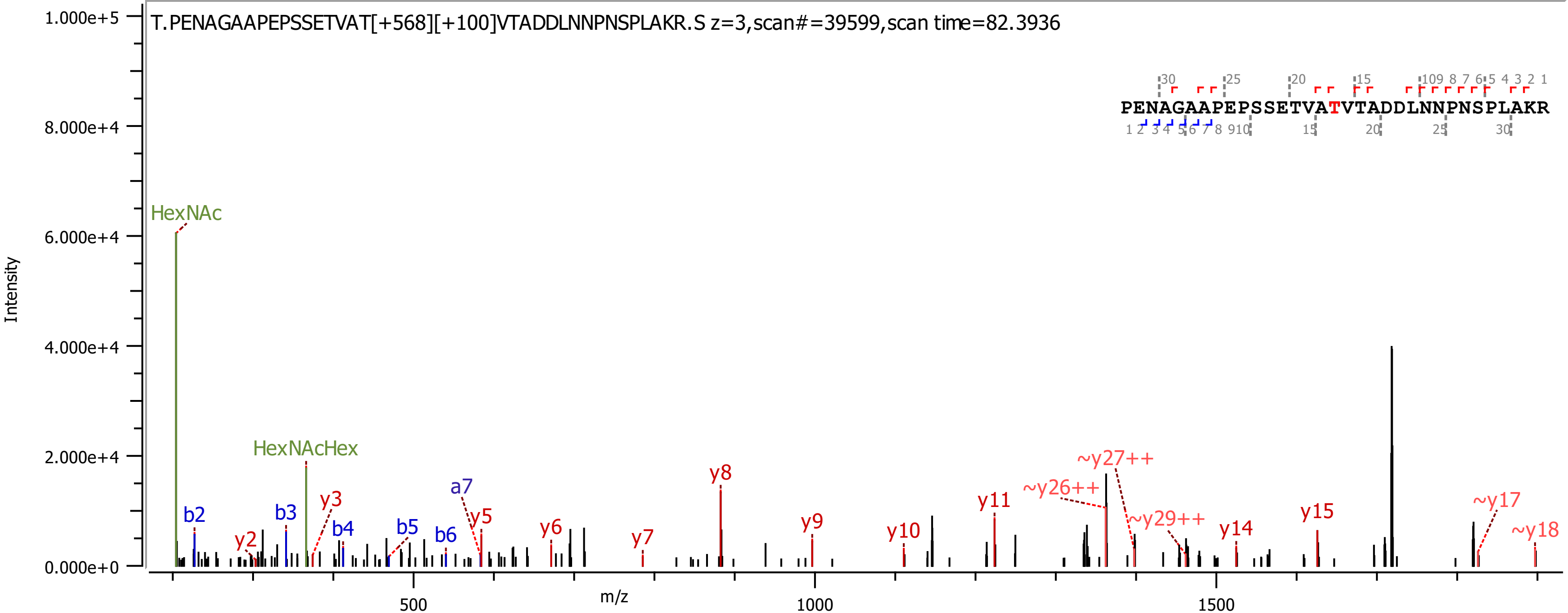
T.PENAGAAPEPSSETVAT[+568][+100]VTADDLNNPNSPLAKR.S z=3,scan#=39599,scan time=82.3936
PENAGAAPEPSSETVATVTADDLNNPNSPLAKR
Intensity
m/z
1.000e+5
8.000e+4
6.000e+4
4.000e+4
2.000e+4
0.000e+0
500
1000
1500
HexNAc
HexNAcHex
b2
b3
b4
b5
b6
a7
y2
y3
y5
y6
y7
y8
y9
y10
y11
~y26++
~y27++
~y29++
y14
y15
~y17
~y18

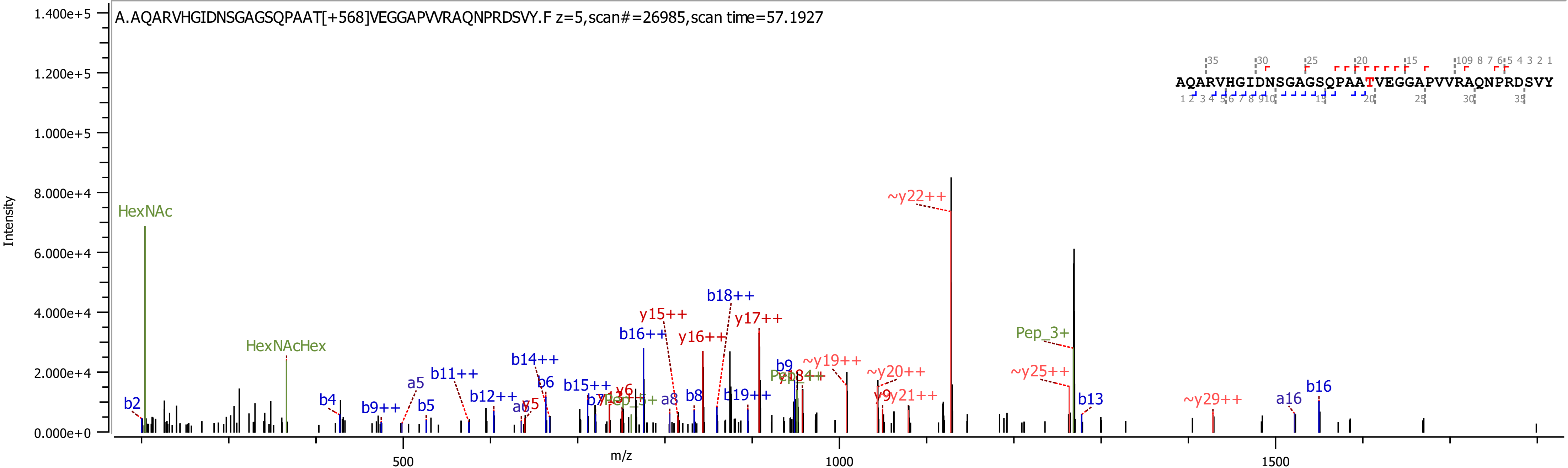
A.AQARVHGIDNSGAGSQPAAT[+568]VEGGAPVVRAQNPRDSVY.F z=5,scan#=26985,scan time=57.1927
AQARVHGIDNSGAGSQPAATVEGGAPVVRAQNPRDSVY
Intensity
m/z
1.400e+5
1.200e+5
1.000e+5
8.000e+4
6.000e+4
4.000e+4
2.000e+4
0.000e+0
500
1000
1500
HexNAc
HexNAcHex
Pep_3+
~y22++
y15++
y16++
y17++
b16++
b18++
b14++
b11++
b12++
b15++
b19++
~y19++
~y20++
~y25++
~y29++
y9
y21++
b2
b4
b9++
a5
b5
b6
b8
b9
b13
a16
b16
a8

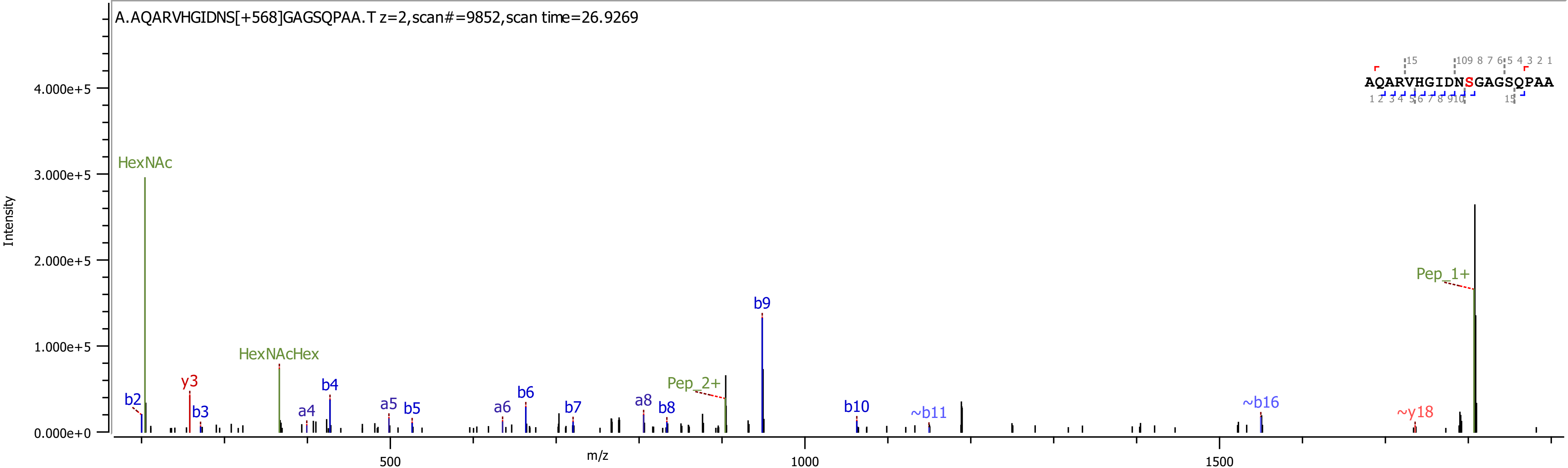

A.AQARVHGIDNS[+568]GAGSQPAA.T z=2,scan#=9852,scan time=26.9269
AQARVHGIDNSGAGSQPAA
Intensity
4.000e+5
3.000e+5
2.000e+5
1.000e+5
0.000e+0
HexNAc
HexNAcHex
b2
y3
b3
a4
b4
a5
b5
a6
b6
b7
a8
b8
Pep_2+
b9
b10
~b11
~b16
~y18
Pep_1+
500
1000
1500
m/z

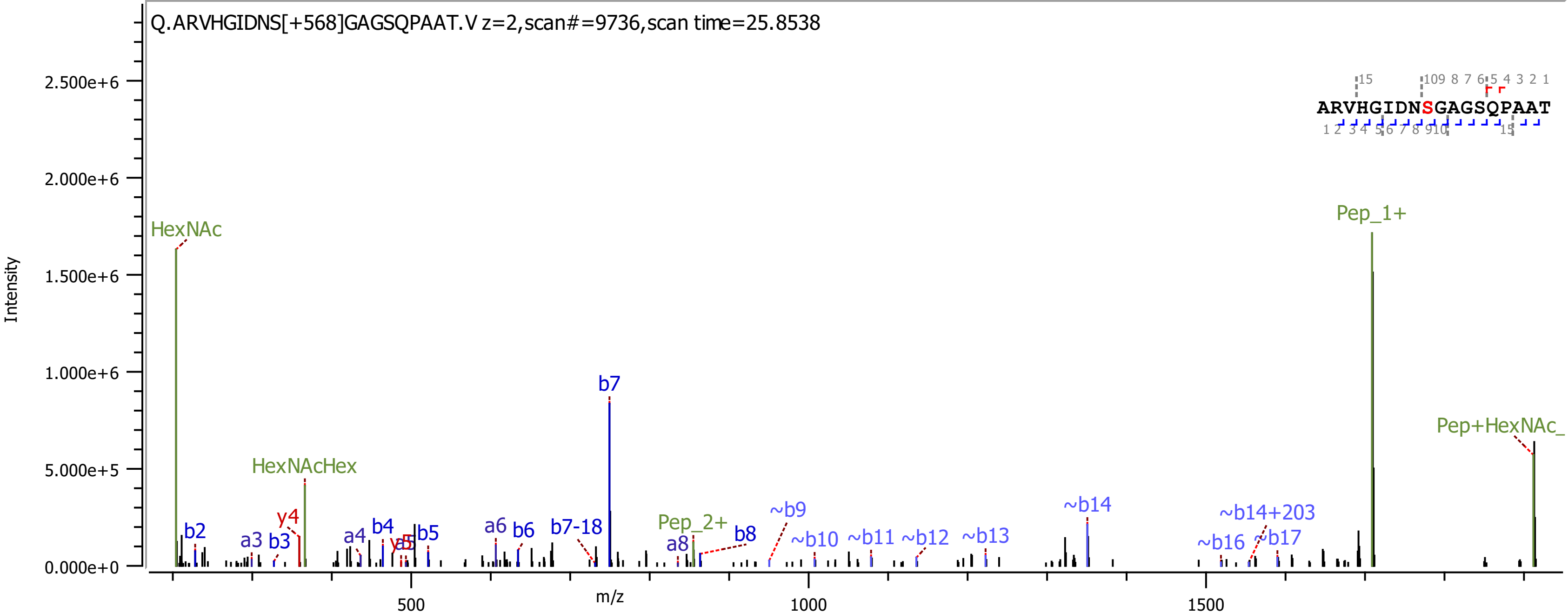

Q.ARVHGIDNS[+568]GAGSQPAAT.V z=2,scan#=9736,scan time=25.8538
ARVHGIDNSGAGSQPAAT
Intensity
m/z
2.500e+6
2.000e+6
1.500e+6
1.000e+6
5.000e+5
0.000e+0
500
1000
1500
HexNAc
HexNAcHex
b2
a3
b3
y4
a4
b4
y5
b5
a6
b6
b7-18
b7
Pep_2+
a8
b8
~b9
~b10
~b11
~b12
~b13
~b14
~b16
~b14+203
~b17
Pep_1+
Pep+HexNAc_

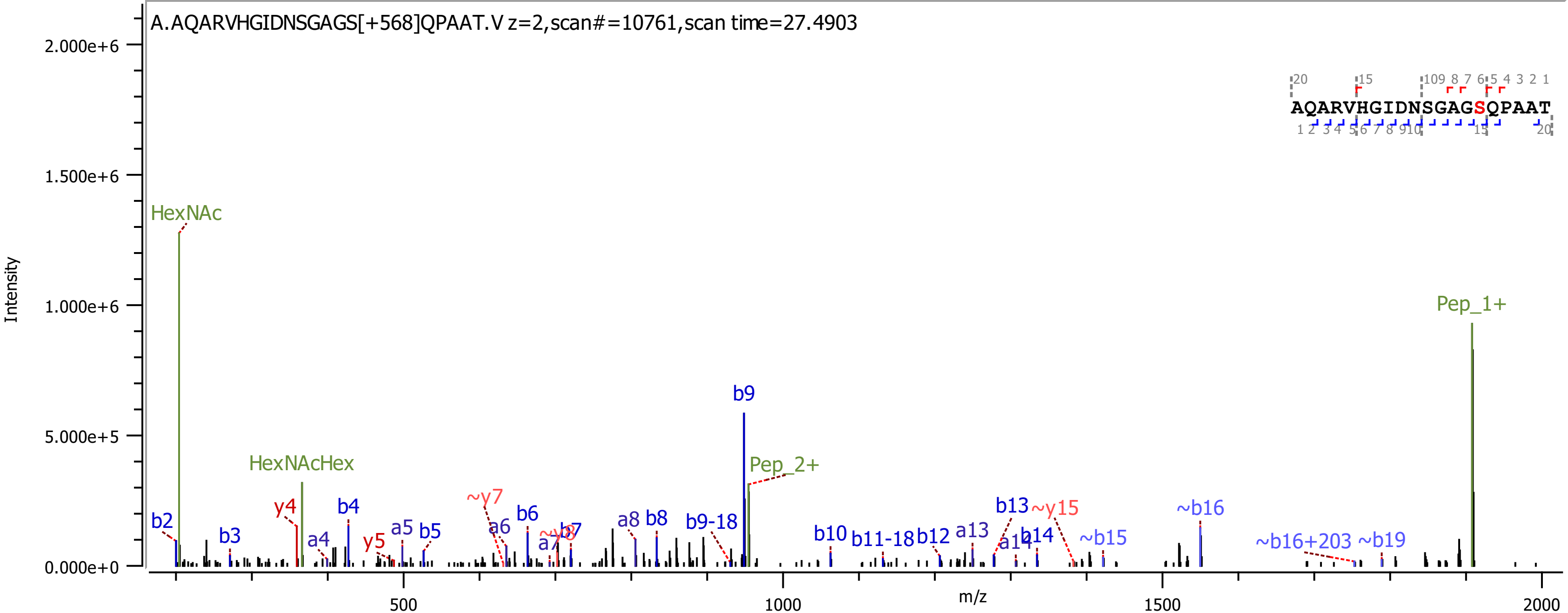

A.AQARVHGIDNSGAGS[+568]QPAAT.V z=2,scan#=10761,scan time=27.4903
AQARVHGIDNSGAGSQPAAT
Intensity
2.000e+6
1.500e+6
1.000e+6
5.000e+5
0.000e+0
m/z
500
1000
1500
2000
HexNAc
HexNAcHex
Pep_2+
Pep_1+
b2
b3
y4
a4
b4
y5
a5
b5
~y7
a6
b6
b7
a8
b8
b9
b9-18
b10
b11-18
b12
a13
b13
~y15
a14
b14
~b15
~b16
~b16+203
~b19

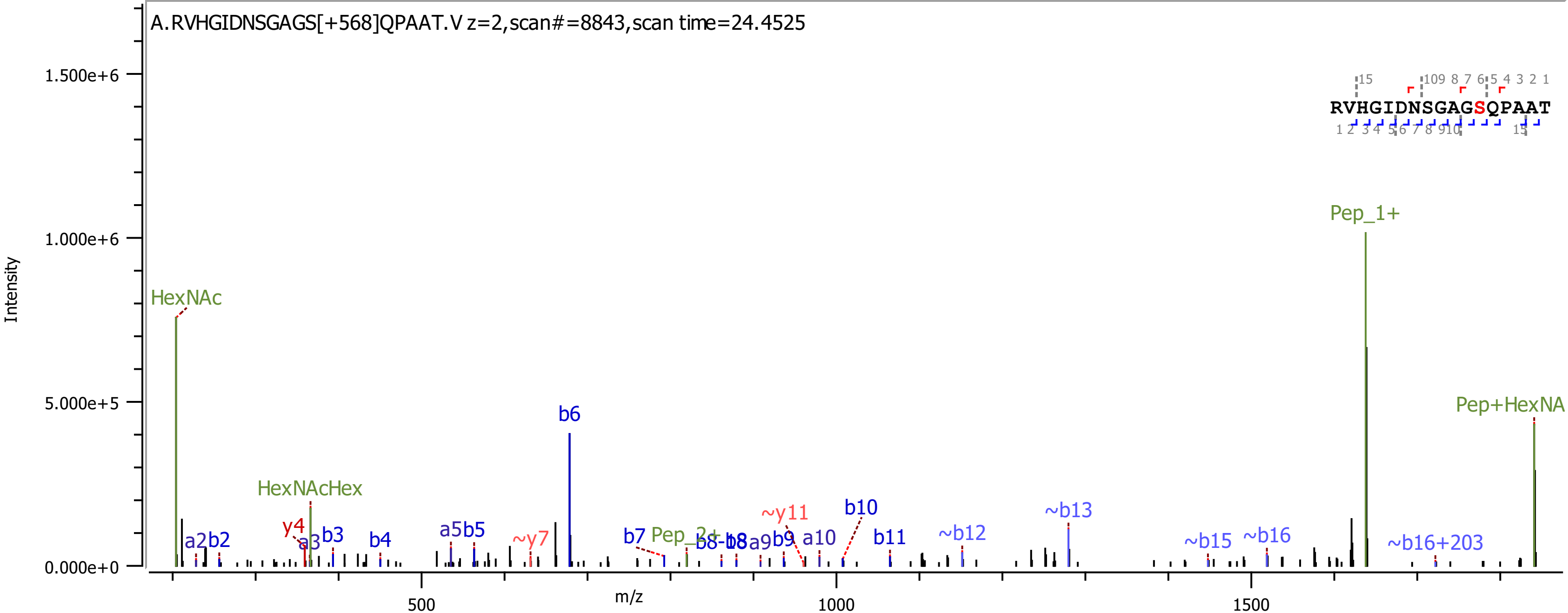
A.RVHGIDNSGAGS[+568]QPAAT.V z=2,scan#=8843,scan time=24.4525
RVHGIDNSGAGSQPAAT
Intensity
m/z
HexNAc
HexNAcHex
Pep_1+
Pep+HexNA
b6
b10
~y11
~y7
~b13
~b12
~b15
~b16
~b16+203
b11
a10
b7
b4
b3
a5b5
a2b2
y4
a3
1.500e+6
1.000e+6
5.000e+5
0.000e+0
500
1000
1500

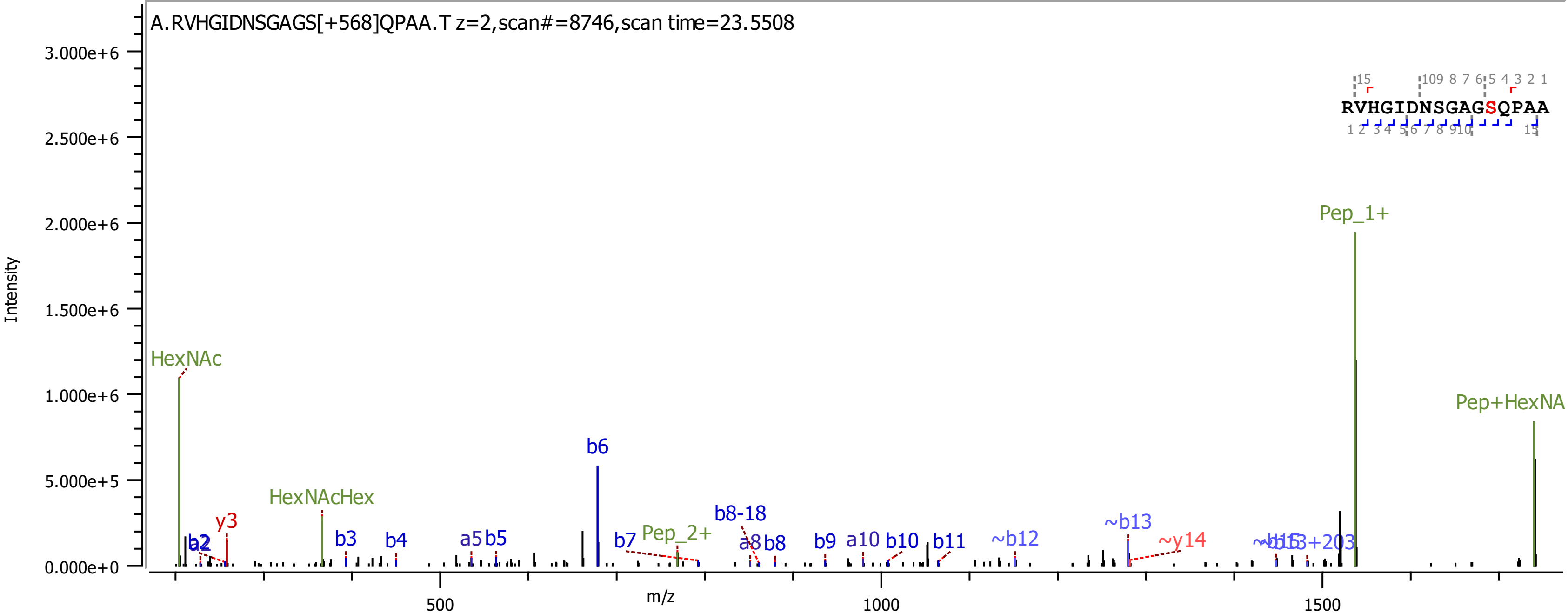

A.RVHGIDNSGAGS[+568]QPAA.T z=2,scan#=8746,scan time=23.5508
RVHGIDNSGAGSQPAA
Intensity
m/z
HexNAc
HexNAcHex
Pep_2+
Pep_1+
Pep+HexNA
b2
a2
y3
b3
b4
a5
b5
b6
b7
b8-18
a8
b8
b9
a10
b10
b11
~b12
~b13
~y14
~b15
~b13+203

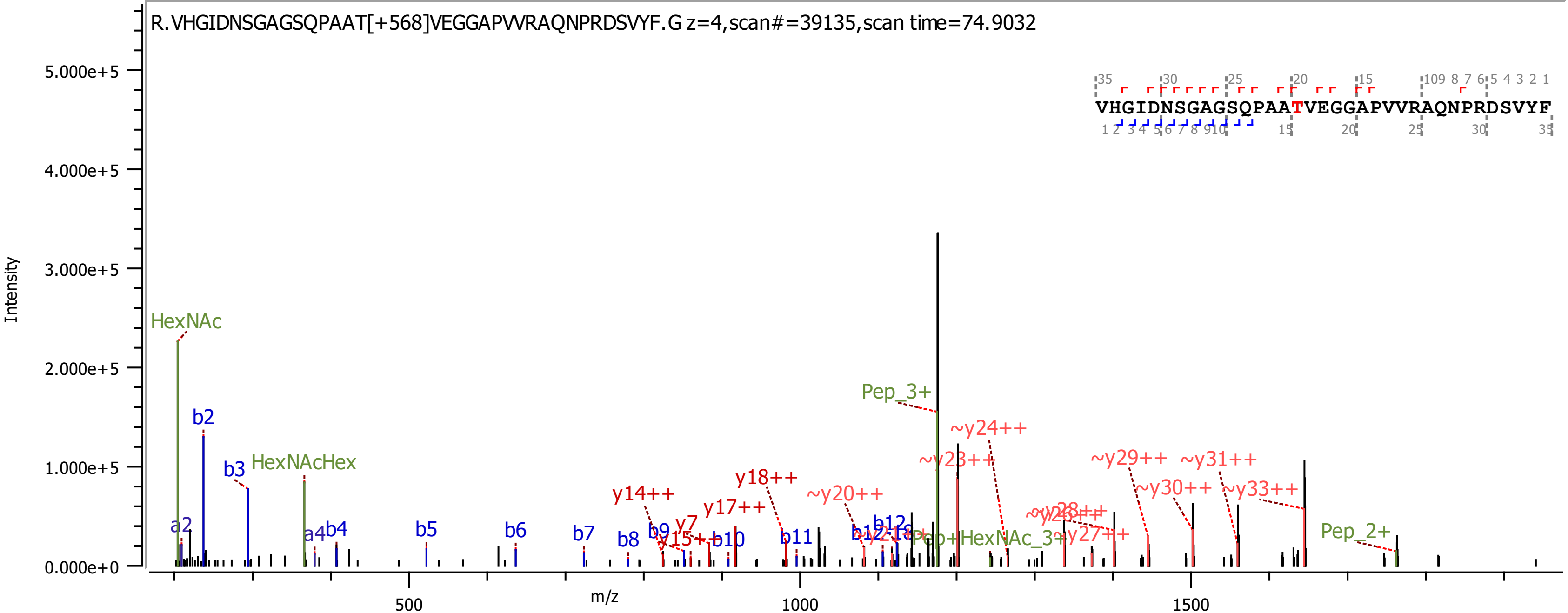
R.VHGIDNSGAGSQPAAT[+568]VEGGAPVVRAQNPRDSVYF.G z=4,scan#=39135,scan time=74.9032
VHGIDNSGAGSQPAATVEGGAPVVRAQNPRDSVYF
Intensity
m/z
HexNAc
HexNAcHex
Pep_3+
Pep_2+
~y24++
~y23++
~y29++
~y31++
~y30++
~y33++
~y20++
y14++
y17++
y18++
b2
b3
a2
a4
b4
b5
b6
b7
b8
b9
y7
b10
b11
b12
~y27++

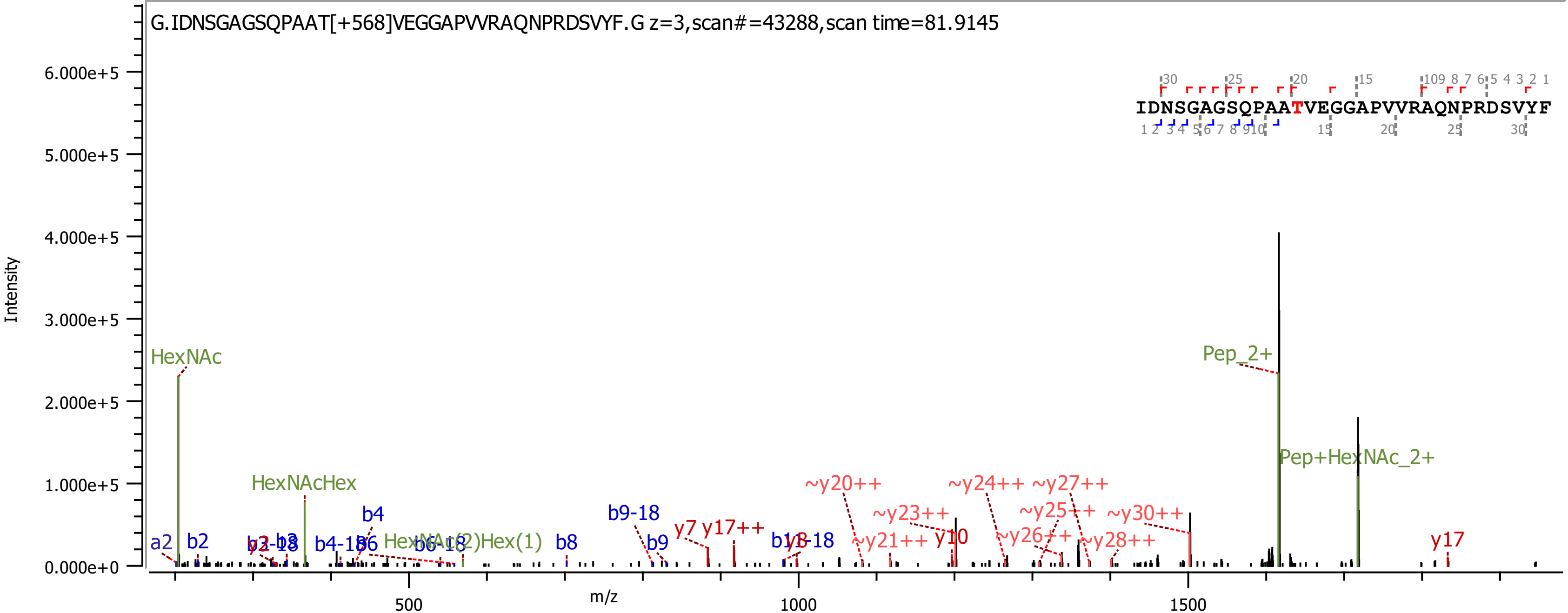
G.IDNSGAGSQPAAT[+568]VEGGAPVVRAQNPRDSVYF.G z=3,scan#=43288,scan time=81.9145
Intensity
6.000e+5
5.000e+5
4.000e+5
3.000e+5
2.000e+5
1.000e+5
0.000e+0
m/z
500
1000
1500
IDNSGAGSQPAATVEGGAPVVRAQNPRDSVYF
HexNAc
HexNAcHex
Pep_2+
Pep+HexNAc_2+
a2
b2
b4
b8
b9
b9-18
y7
y17++
~y20++
~y23++
~y21++
y10
~y24++
~y27++
~y25++
~y26++
~y28++
~y30++
y17

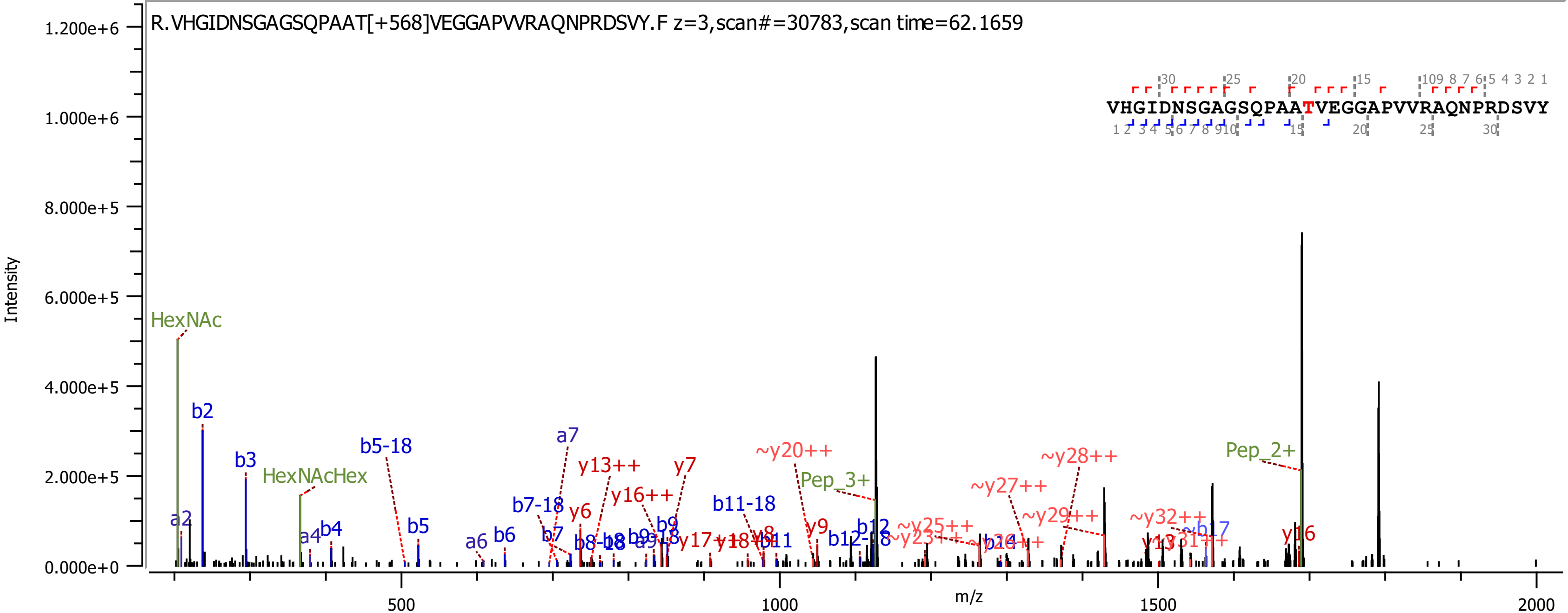

R.VHGIDNSGAGSQPAAT[+568]VEGGAPVVRAQNPRDSVY.F z=3,scan#=30783,scan time=62.1659
VHGIDNSGAGSQPAATVEGGAPVVRAQNPRDSVY
Intensity
m/z
1.200e+6
1.000e+6
8.000e+5
6.000e+5
4.000e+5
2.000e+5
0.000e+0
500
1000
1500
2000
HexNAc
HexNAcHex
Pep_3+
Pep_2+
a2
b2
b3
a4
b4
b5-18
b5
a6
b6
a7
b7-18
b7
y6
y13++
y16++
y7
b8-18
b9
b9-18
b11-18
b11
y8
y9
~y20++
b12
b12-18
~y25++
~y23++
~y27++
~y28++
~y29++
~y32++
~b17
y16

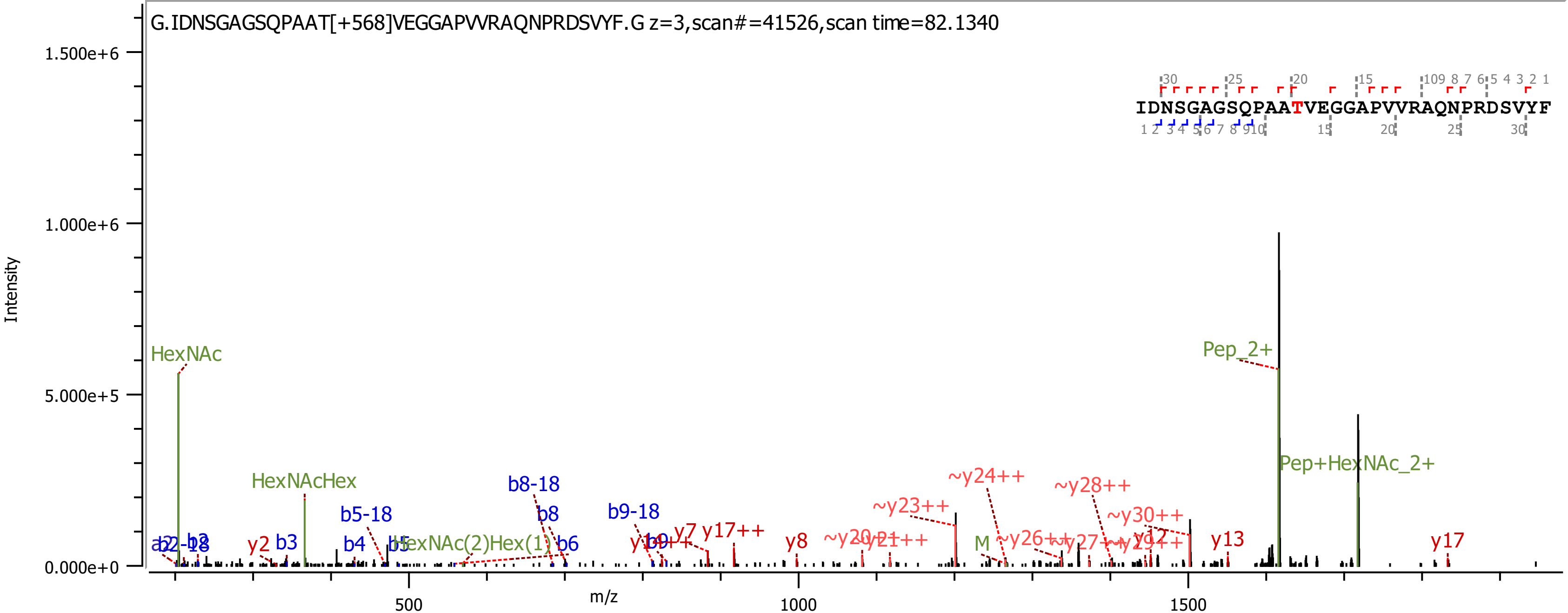

G.IDNSGAGSQPAAT[+568]VEGGAPVVRAQNPRDSVYF.G z=3,scan#=41526,scan time=82.1340
IDNSGAGSQPAATVEGGAPVVRAQNPRDSVYF
Intensity
m/z
1.500e+6
1.000e+6
5.000e+5
0.000e+0
500
1000
1500
HexNAc
HexNAcHex
HexNAc(2)Hex(1)
Pep_2+
Pep+HexNAc_2+
M
b3
b4
b5-18
b6
b8
b8-18
b9-18
y2
y7
y8
y13
y17
y17++
~y23++
~y24++
~y28++
~y30++

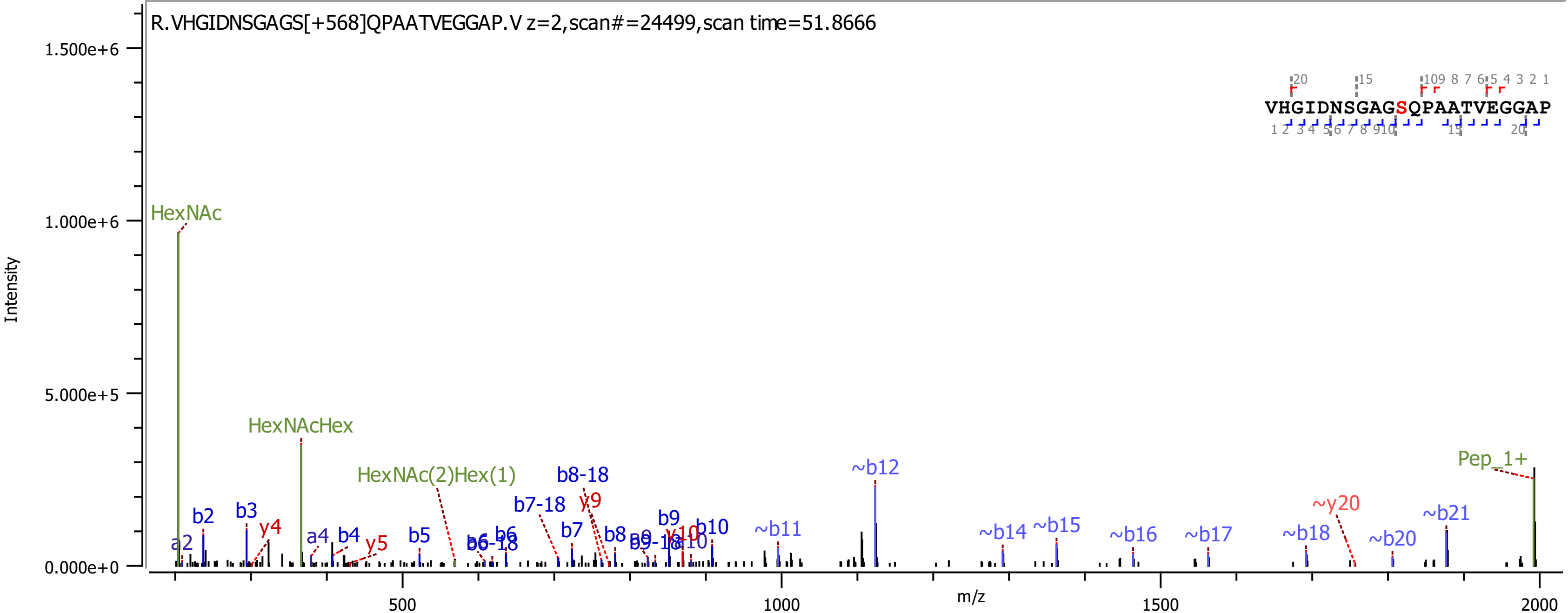
R.VHGIDNSGAGS[+568]QPAATVEGGAP.V z=2,scan#=24499,scan time=51.8666
VHGIDNSGAGSQPAATVEGGAP
Intensity
m/z
HexNAc
HexNAcHex
HexNAc(2)Hex(1)
Pep_1+
~b12
~y20
~b21
b8-18
b7-18
y9
b2
b3
y4
a2
a4
b4
y5
b5
b6
b7
b8
b9
b10
~b11
~b14
~b15
~b16
~b17
~b18
~b20

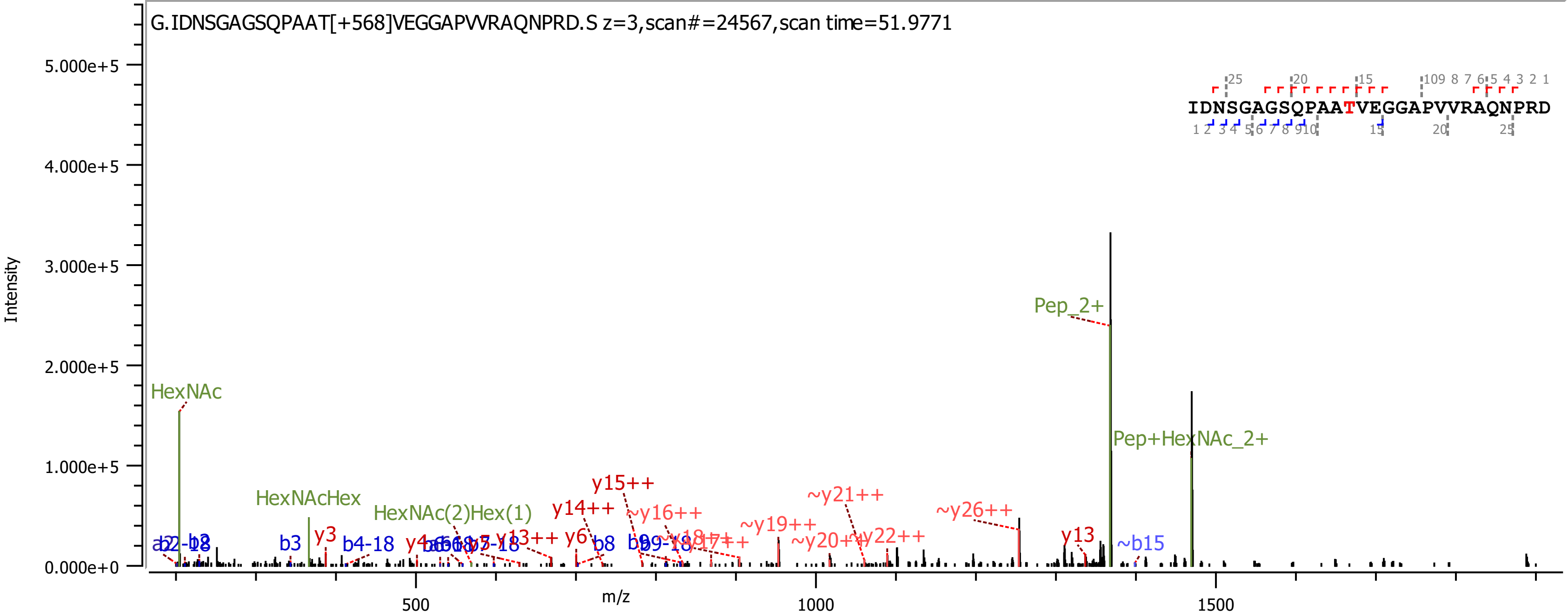
G.IDNSGAGSQPAAT[+568]VEGGAPVVRAQNPRD.S z=3,scan#=24567,scan time=51.9771
IDNSGAGSQPAATVEGGAPVVRAQNPRD
Intensity
5.000e+5
4.000e+5
3.000e+5
2.000e+5
1.000e+5
0.000e+0
m/z
500
1000
1500
HexNAc
HexNAcHex
HexNAc(2)Hex(1)
Pep_2+
Pep+HexNAc_2+
b3
y3
b4-18
y15++
y14++
y13++
y6
b8
~y16++
~y19++
~y21++
~y20++
~y22++
~y26++
y13
~b15

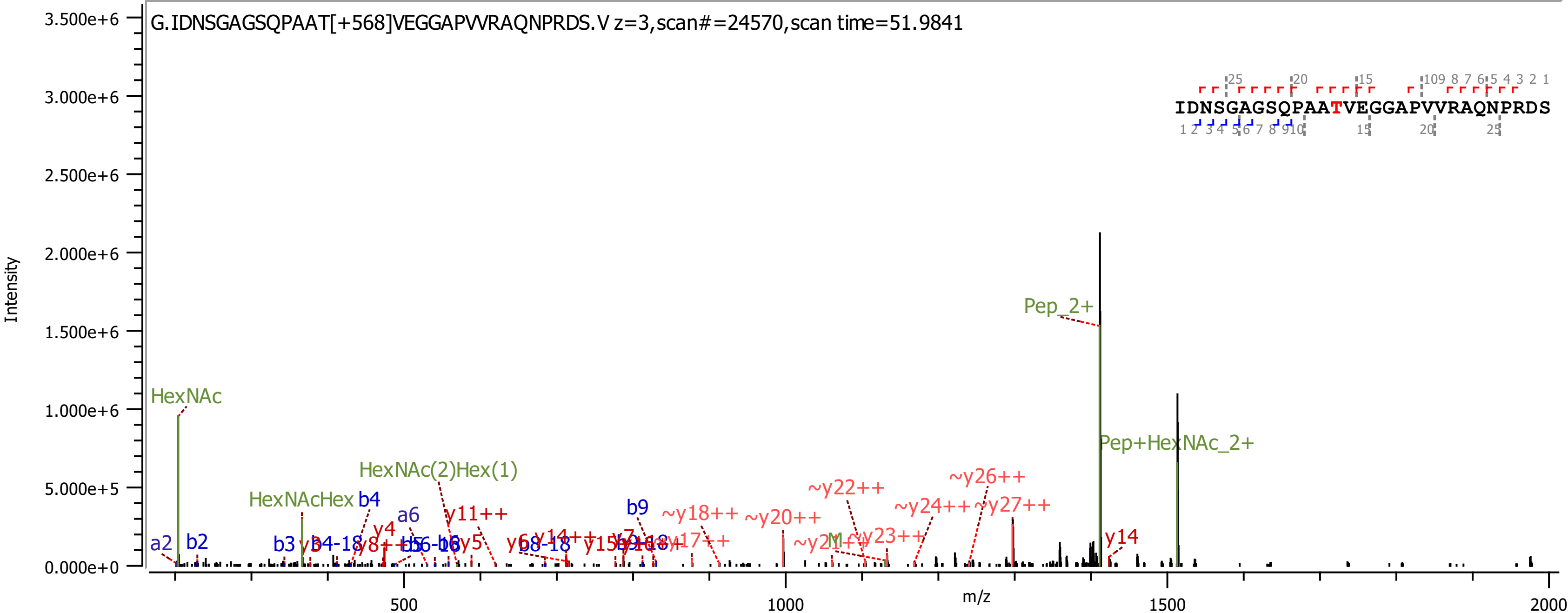

G.IDNSGAGSQPAAT[+568]VEGGAPVVRAQNPRDS.V z=3,scan#=24570,scan time=51.9841
IDNSGAGSQPAATVEGGAPVVRAQNPRDS
Intensity
m/z
HexNAc
HexNAcHex
HexNAc(2)Hex(1)
Pep_2+
Pep+HexNAc_2+
b2
b3
b4
b9
a2
a6
y4
y5
y11++
y14
~y18++
~y20++
~y22++
~y23++
~y24++
~y26++
~y27++

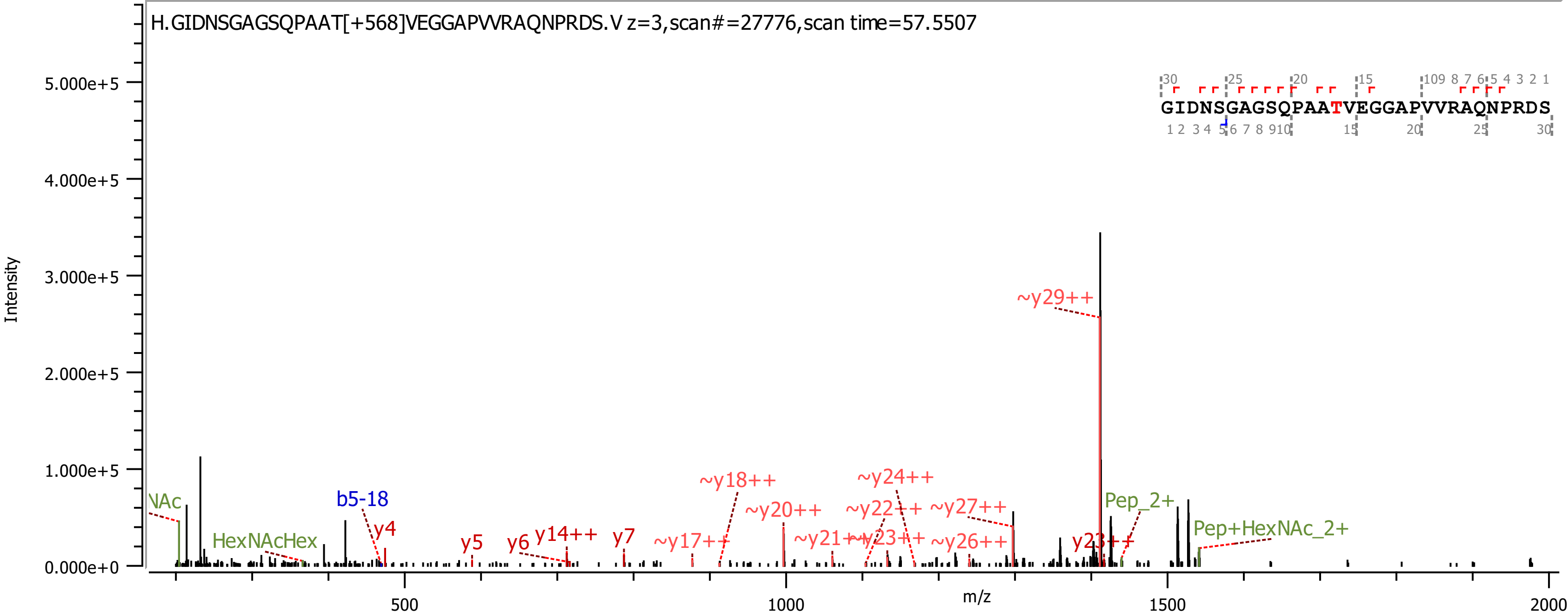

H.GIDNSGAGSQPAAT[+568]VEGGAPVVRAQNPRDS.V z=3,scan#=27776,scan time=57.5507
GIDNSGAGSQPAAT VEGGAPVVRAQNPRDS
Intensity
m/z
5.000e+5
4.000e+5
3.000e+5
2.000e+5
1.000e+5
0.000e+0
500
1000
1500
2000
~y29++
NAc
HexNAcHex
b5-18
y4
y5
y6
y14++
y7
~y17++
~y18++
~y20++
~y21++
~y22++
~y23++
~y24++
~y26++
~y27++
y23++
Pep_2+
Pep+HexNAc_2+

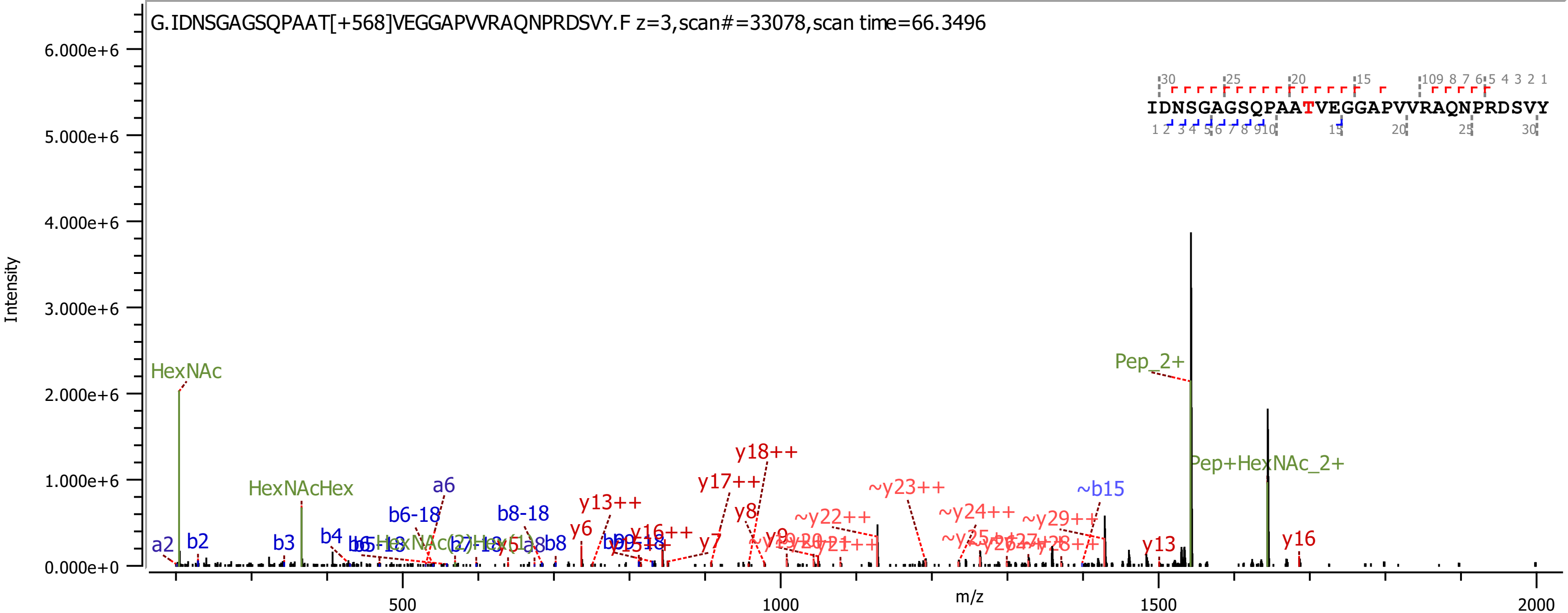

G.IDNSGAGSQPAAT[+568]VEGGAPVVRAQNPRDSVY.F z=3,scan#=33078,scan time=66.3496
IDNSGAGSQPAATVEGGAPVVRAQNPRDSVY
Intensity
m/z
6.000e+6
5.000e+6
4.000e+6
3.000e+6
2.000e+6
1.000e+6
0.000e+0
500
1000
1500
2000
HexNAc
HexNAcHex
Pep_2+
Pep+HexNAc_2+
a2
b2
b3
b4
a6
b6-18
b8-18
b8
y6
y13++
y16++
y7
y17++
y18++
y8
~y22++
~y23++
~y24++
~y29++
~b15
y13
y16

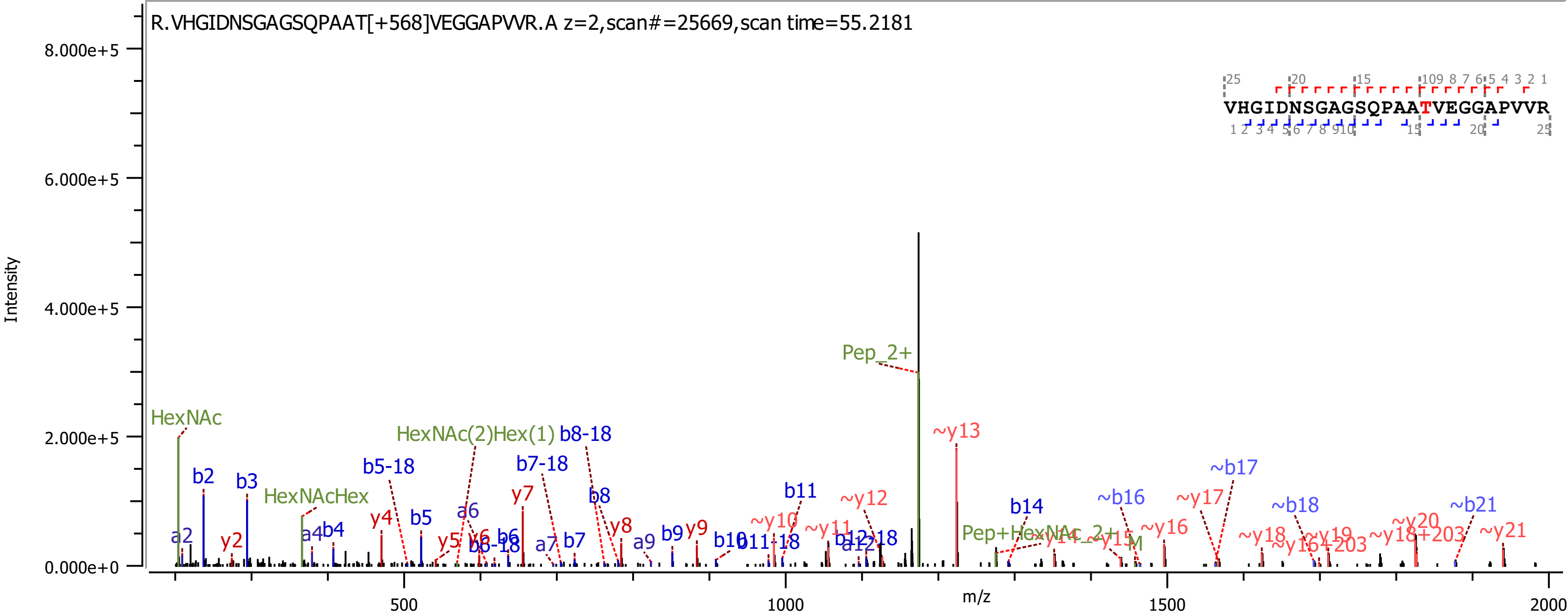

R.VHGIDNSGAGSQPAAT[+568]VEGGAPVVR.A z=2,scan#=25669,scan time=55.2181
VHGIDNSGAGSQPAATVEGGAPVVR
Intensity
8.000e+5
6.000e+5
4.000e+5
2.000e+5
0.000e+0
m/z
500
1000
1500
2000
Pep_2+
~y13
HexNAc
HexNAc(2)Hex(1)
b8-18
b5-18
b7-18
~b17
b2
b3
HexNAcHex
b8
b11
~y12
~b16
~y17
~b18
~b21
y7
a6
b14
y4
b5
y8
~y10
~y16
a2
y2
a4
b4
y5
y6
b6-18
b6
a7
b7
a9
b9
y9
b10
b11-18
~y11
b12-18
a12
Pep+HexNAc_2+
~y14
~y15
M
~y18
~y19
~y16+203
~y20
~y18+203
~y21

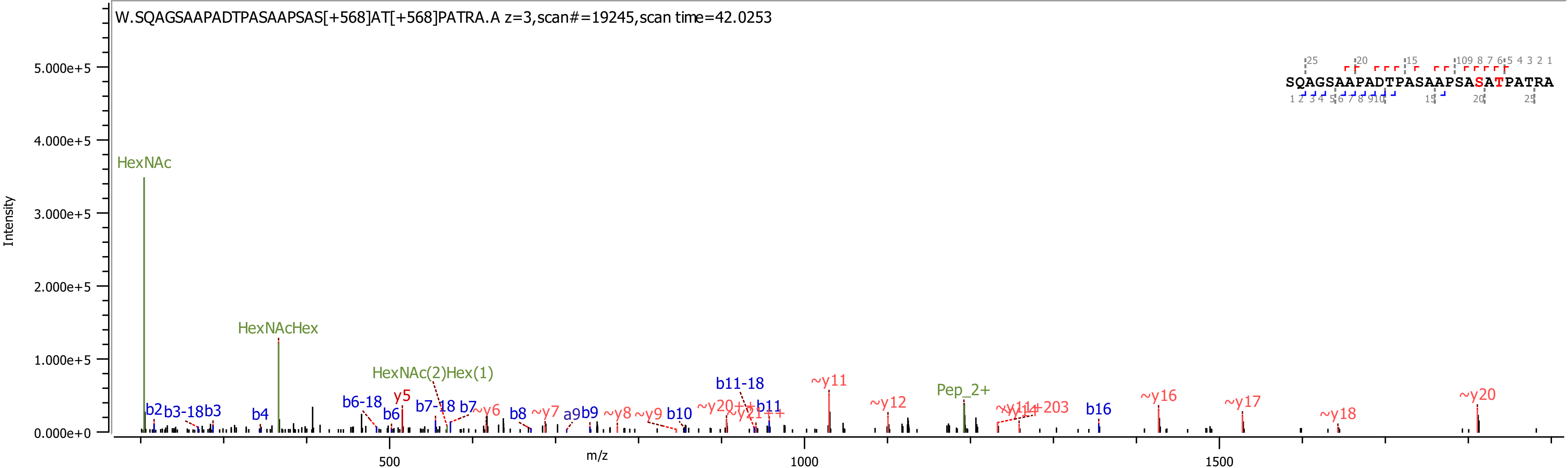
W.SQAGSAAPADTPASAAPSAS[+568]AT[+568]PATRA.A z=3,scan#=19245,scan time=42.0253
SQAGSAAPADTPASAAPSASATPATRA
Intensity
m/z
HexNAc
HexNAcHex
HexNAc(2)Hex(1)
Pep_2+
b2
b3-18
b3
b4
b6-18
b6
y5
b7-18
b7
~y6
b8
~y7
a9
b9
~y8
~y9
b10
b11-18
b11
~y11
~y12
~y16
~y17
~y18
~y20
b16
5.000e+5
4.000e+5
3.000e+5
2.000e+5
1.000e+5
0.000e+0
500
1000
1500

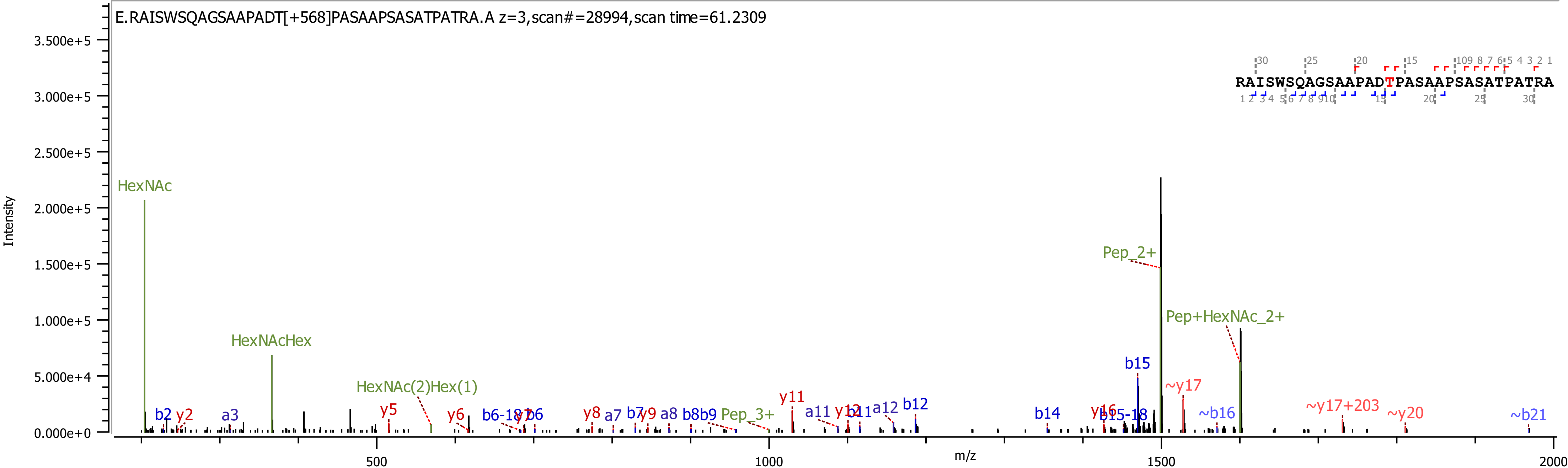

E.RAISWSQAGSAAPADT[+568]PASAAPSASATPATRA.A z=3,scan#=28994,scan time=61.2309
RAISWSQAGSAAPADTPASAAPSASATPATRA
Intensity
m/z
HexNAc
HexNAcHex
HexNAc(2)Hex(1)
Pep_2+
Pep+HexNAc_2+
Pep_3+
b2
y2
a3
y5
y6
b6-18
b6
y8
a7
b7
y9
a8
b8
b9
y11
a11
y12
b11
a12
b12
b14
y16
b15-18
b15
~y17
~b16
~y17+203
~y20
~b21

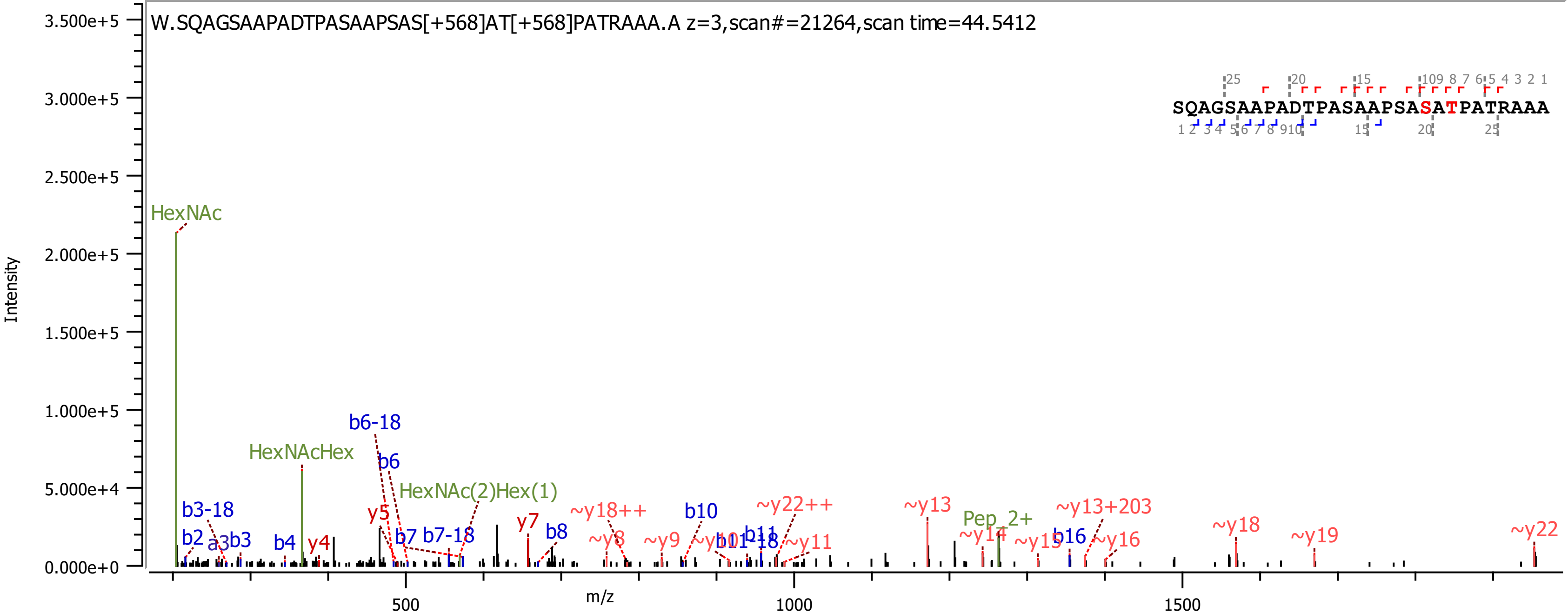
W.SQAGSAAPADTPASAAPSAS[+568]AT[+568]PATRAAA.A z=3,scan#=21264,scan time=44.5412
SQAGSAAPADTPASAAPSASATPATRAAA
Intensity
m/z
HexNAc
HexNAcHex
HexNAc(2)Hex(1)
b6-18
b6
b3-18
b2
a3
b3
b4
y4
y5
b7
b7-18
y7
b8
~y18++
~y8
~y9
b10
~y10
b11-18
b11
~y22++
~y11
~y13
Pep_2+
~y14
~y15
b16
~y13+203
~y16
~y18
~y19
~y22
3.500e+5
3.000e+5
2.500e+5
2.000e+5
1.500e+5
1.000e+5
5.000e+4
0.000e+0
500
1000
1500

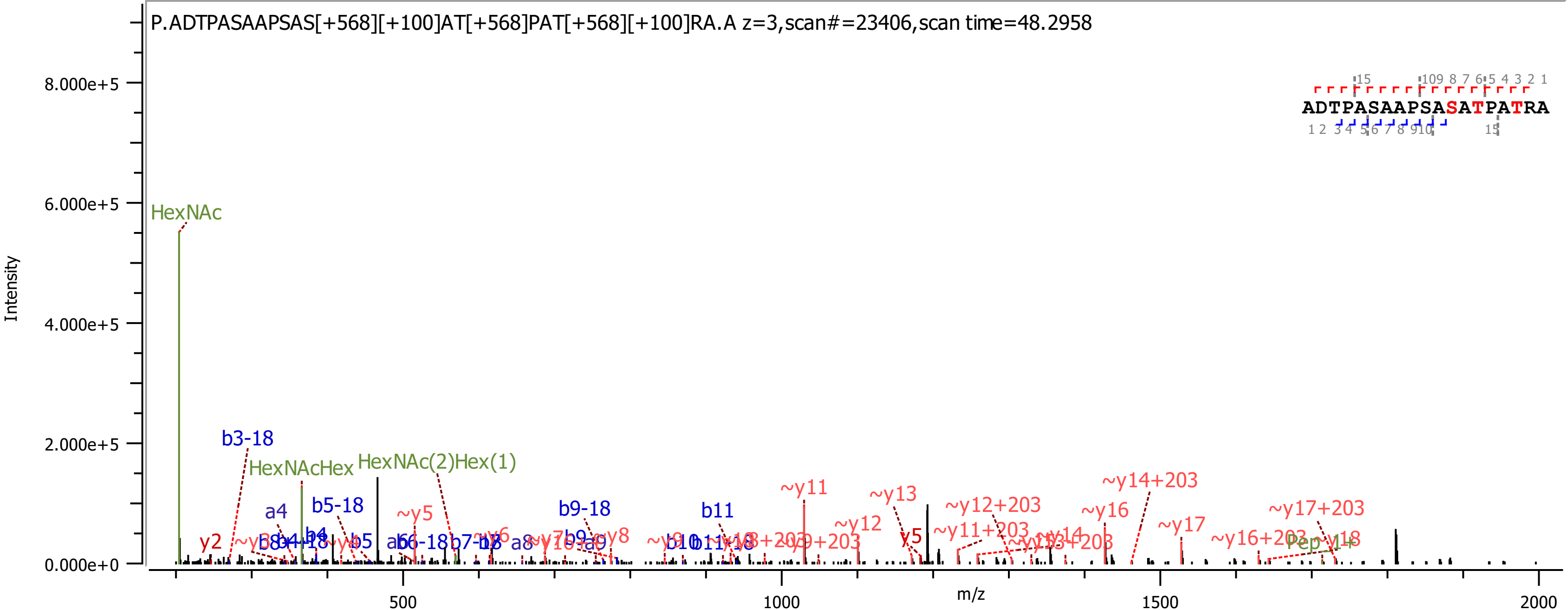

P.ADTPASAAPSAS[+568][+100]AT[+568]PAT[+568][+100]RA.A z=3,scan#=23406,scan time=48.2958
ADTPASAAPSASATPATRA
Intensity
m/z
HexNAc
b3-18
HexNAcHex
HexNAc(2)Hex(1)
a4
b5-18
~y5
b9-18
b11
~y11
~y13
~y12
~y12+203
~y14+203
~y16
~y17
~y11+203
~y17+203
~y16+203
y5
y2

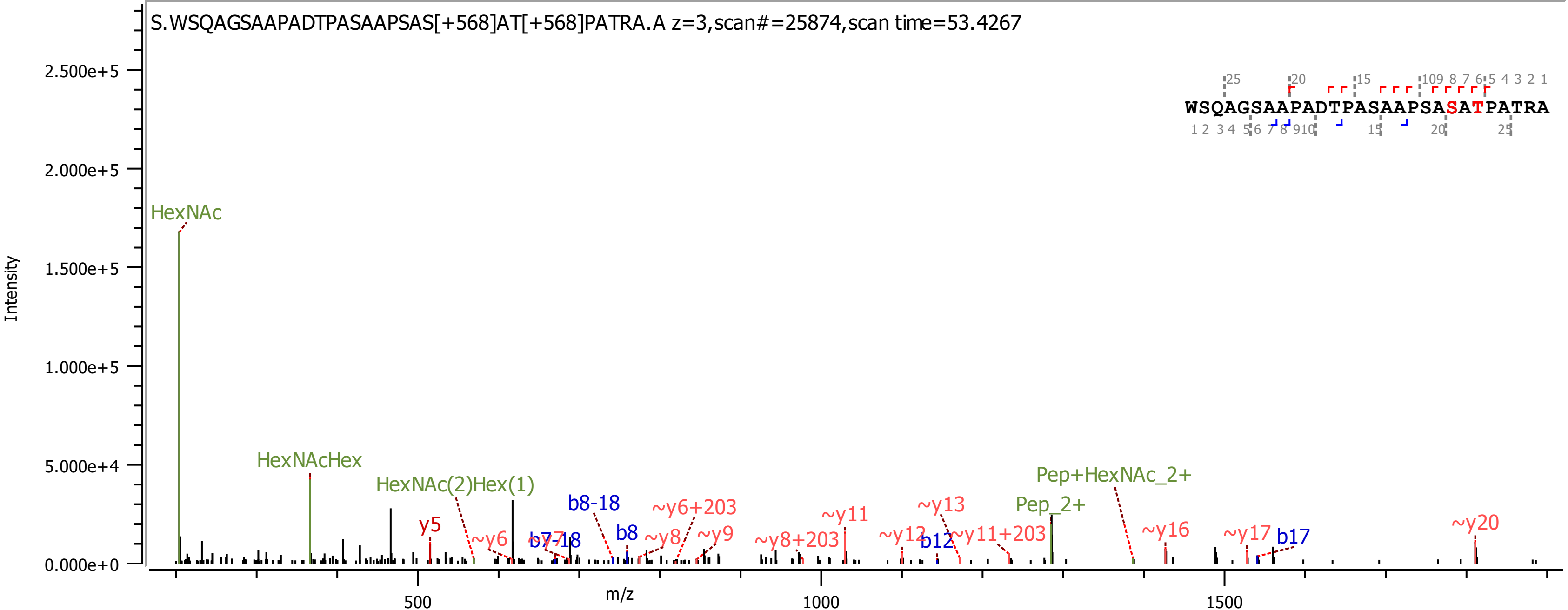
S.WSQAGSAAPADTPASAAPSAS[+568]AT[+568]PATRA.A z=3,scan#=25874,scan time=53.4267
WSQAGSAAPADTPASAAPSASATPATRA
Intensity
m/z
HexNAc
HexNAcHex
HexNAc(2)Hex(1)
y5
~y6
b7-18
b8-18
b8
~y6+203
~y8
~y9
~y8+203
~y11
~y12
b12
~y13
~y11+203
Pep_2+
Pep+HexNAc_2+
~y16
~y17
b17
~y20

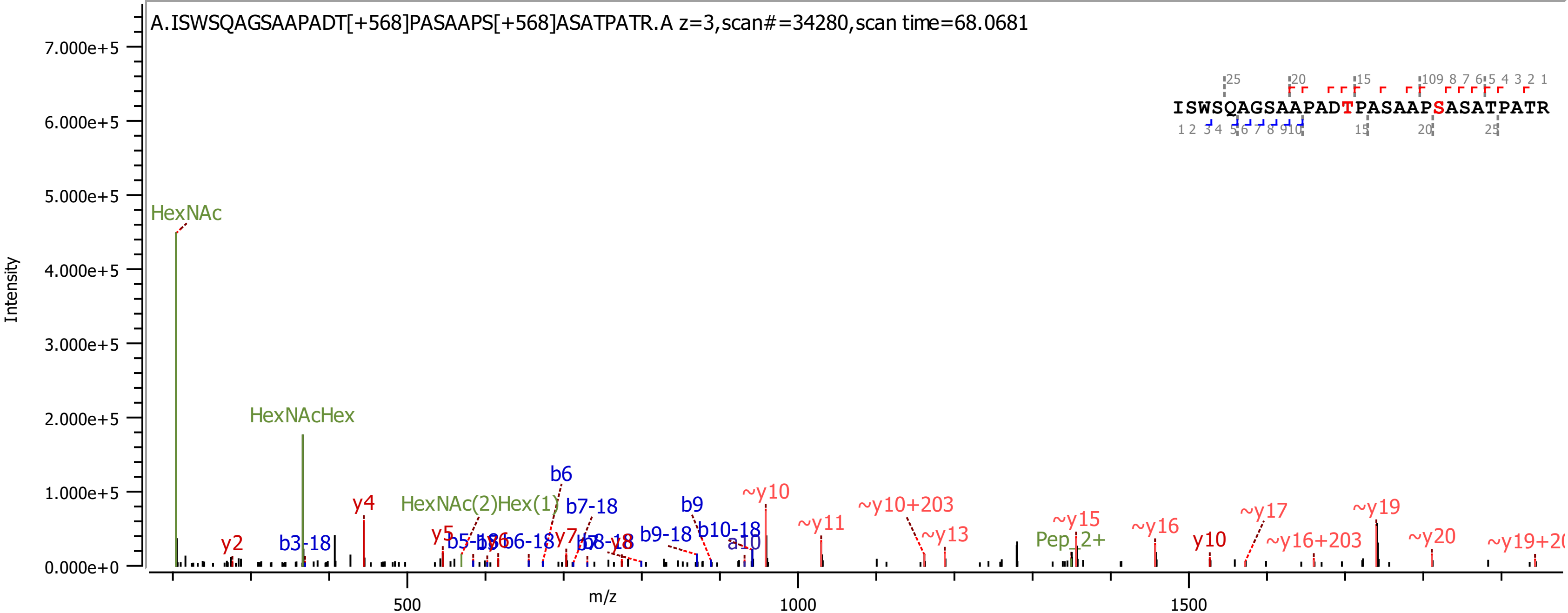

A.ISWSQAGSAAPADT[+568]PASAAPS[+568]ASATPATR.A z=3,scan#=34280,scan time=68.0681
ISWSQAGSAAPADTPASAAPSASATPATR
Intensity
m/z
HexNAc
HexNAcHex
HexNAc(2)Hex(1)
Pep_2+
y2
y4
y5
y7
y10
b3-18
b6
b7-18
b9
b9-18
b10-18
a10
~y10
~y11
~y10+203
~y13
~y15
~y16
~y17
~y16+203
~y19
~y20

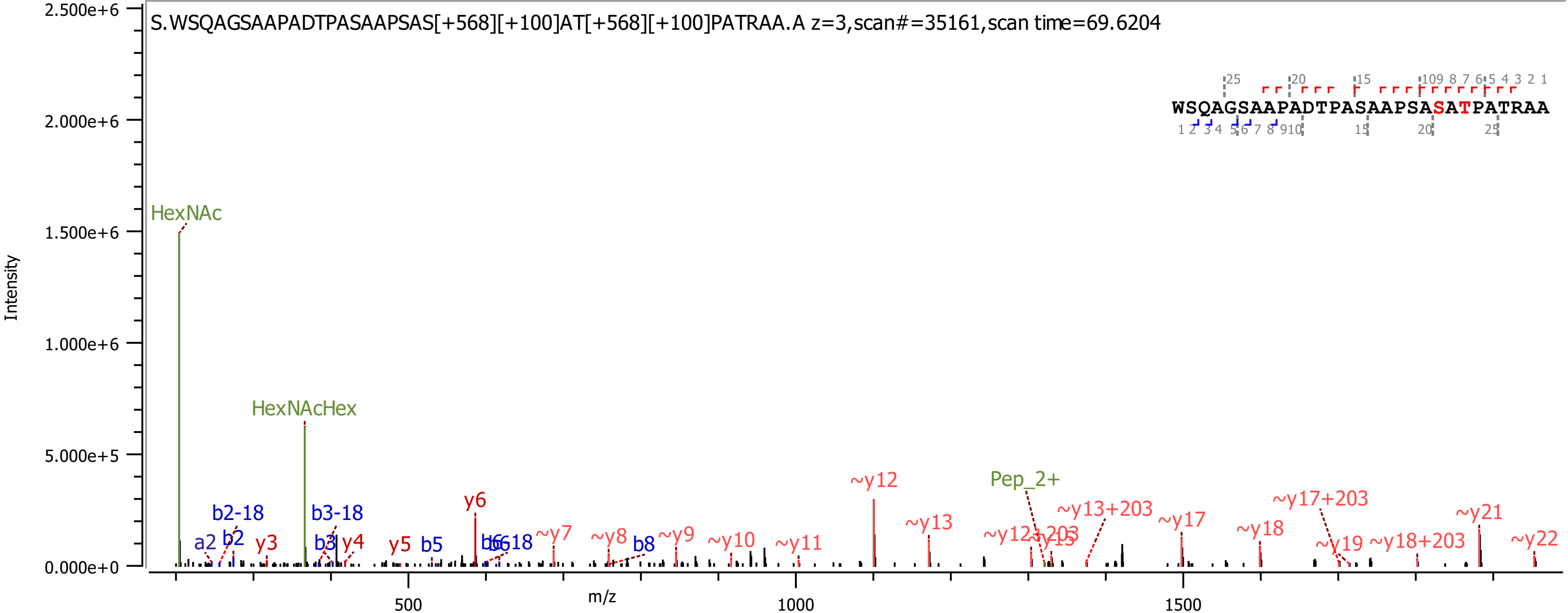

S.WSQAGSAAPADTPASAAPSAS[+568][+100]AT[+568][+100]PATRAA.A z=3,scan#=35161,scan time=69.6204
WSQAGSAAPADTPASAAPSASATPATRAA
Intensity
m/z
2.500e+6
2.000e+6
1.500e+6
1.000e+6
5.000e+5
0.000e+0
500
1000
1500
HexNAc
HexNAcHex
Pep_2+
a2
b2
b2-18
y3
b3
b3-18
y4
y5
b5
y6
b6
b6-18
~y7
~y8
b8
~y9
~y10
~y11
~y12
~y13
~y12+203
~y15
~y13+203
~y17
~y18
~y17+203
~y19
~y18+203
~y21
~y22

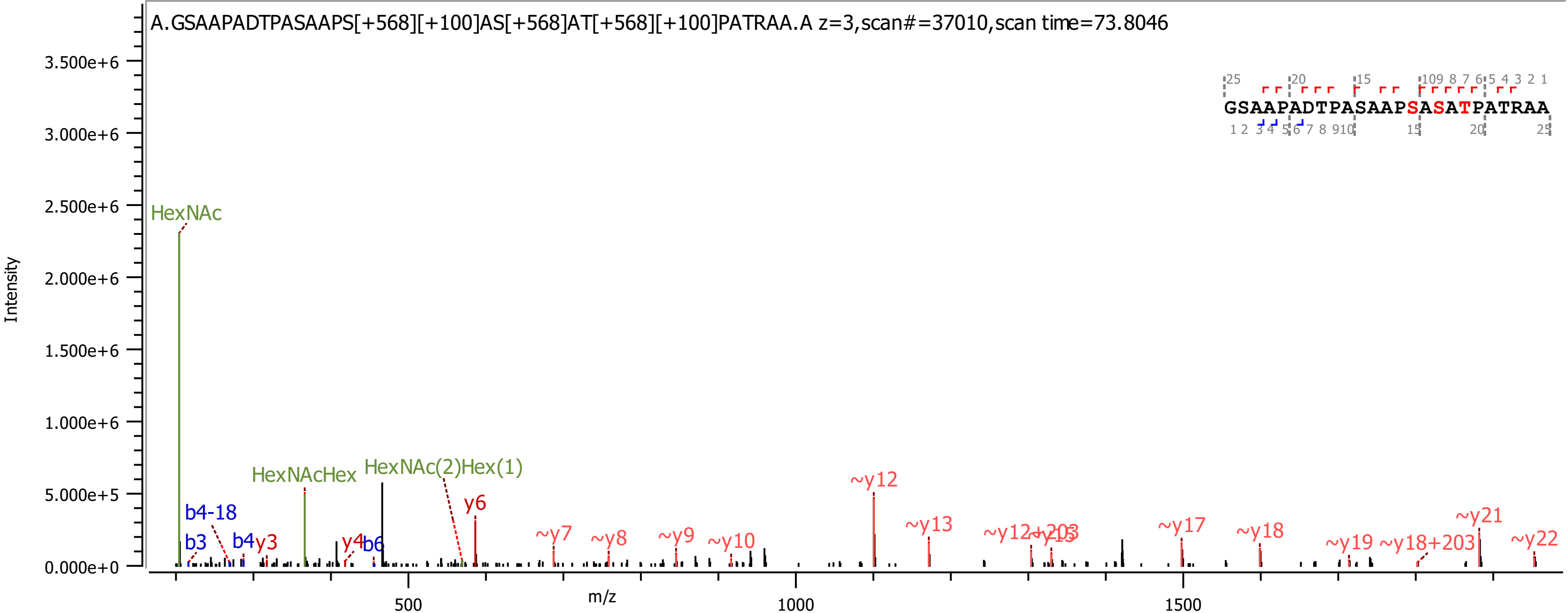
A.GSAAPADTPASAAPS[+568][+100]AS[+568]AT[+568][+100]PATRAA.A z=3,scan#=37010,scan time=73.8046
Intensity
3.500e+6
3.000e+6
2.500e+6
2.000e+6
1.500e+6
1.000e+6
5.000e+5
0.000e+0
m/z
500
1000
1500
HexNAc
HexNAcHex
HexNAc(2)Hex(1)
b4-18
b3
b4
y3
y4
b6
y6
~y7
~y8
~y9
~y10
~y12
~y13
~y12+203
~y15
~y17
~y18
~y19
~y18+203
~y21
~y22
GSAAPADTPASAAPSASATPATRAA

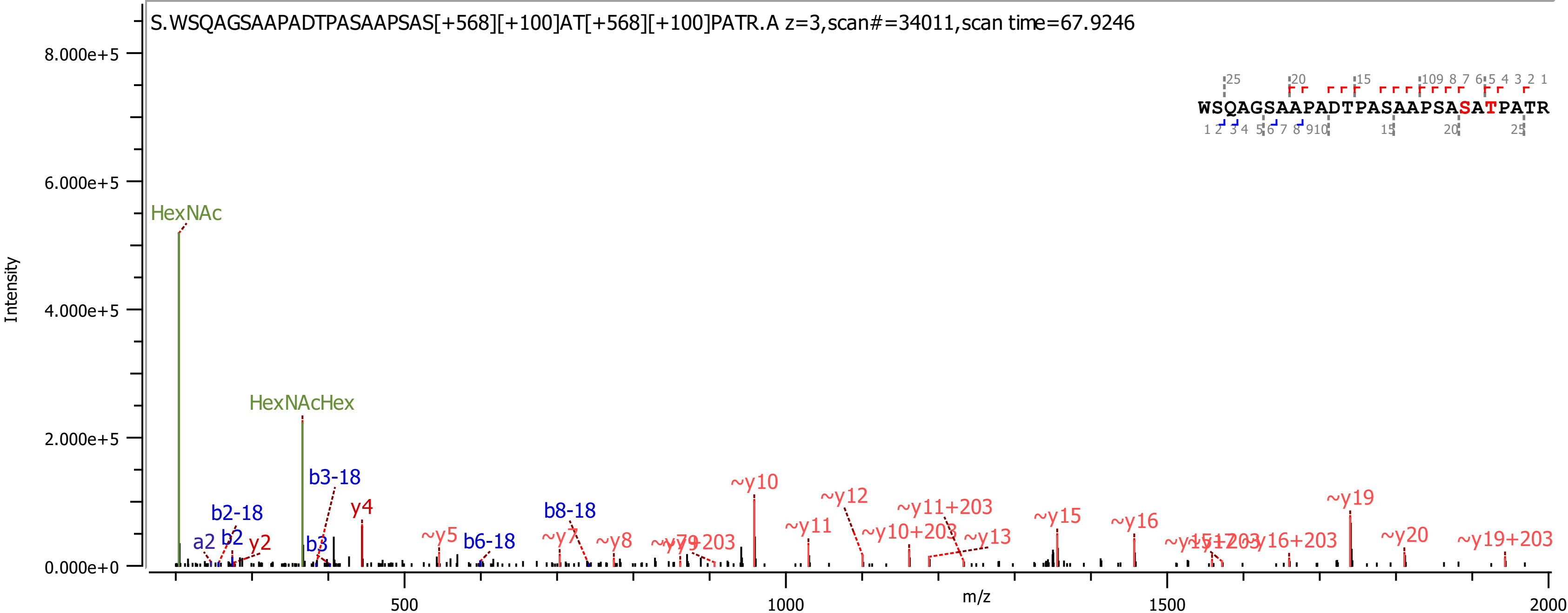

S.WSQAGSAAPADTPASAAPSAS[+568][+100]AT[+568][+100]PATR.A z=3,scan#=34011,scan time=67.9246
WSQAGSAAPADTPASAAPSASATPATR
Intensity
m/z
HexNAc
HexNAcHex
b3-18
b2-18
a2
b2
y2
b3
y4
~y5
b6-18
b8-18
~y7
~y8
~y10
~y11
~y12
~y11+203
~y10+203
~y13
~y15
~y16
~y16+203
~y19
~y20
~y19+203

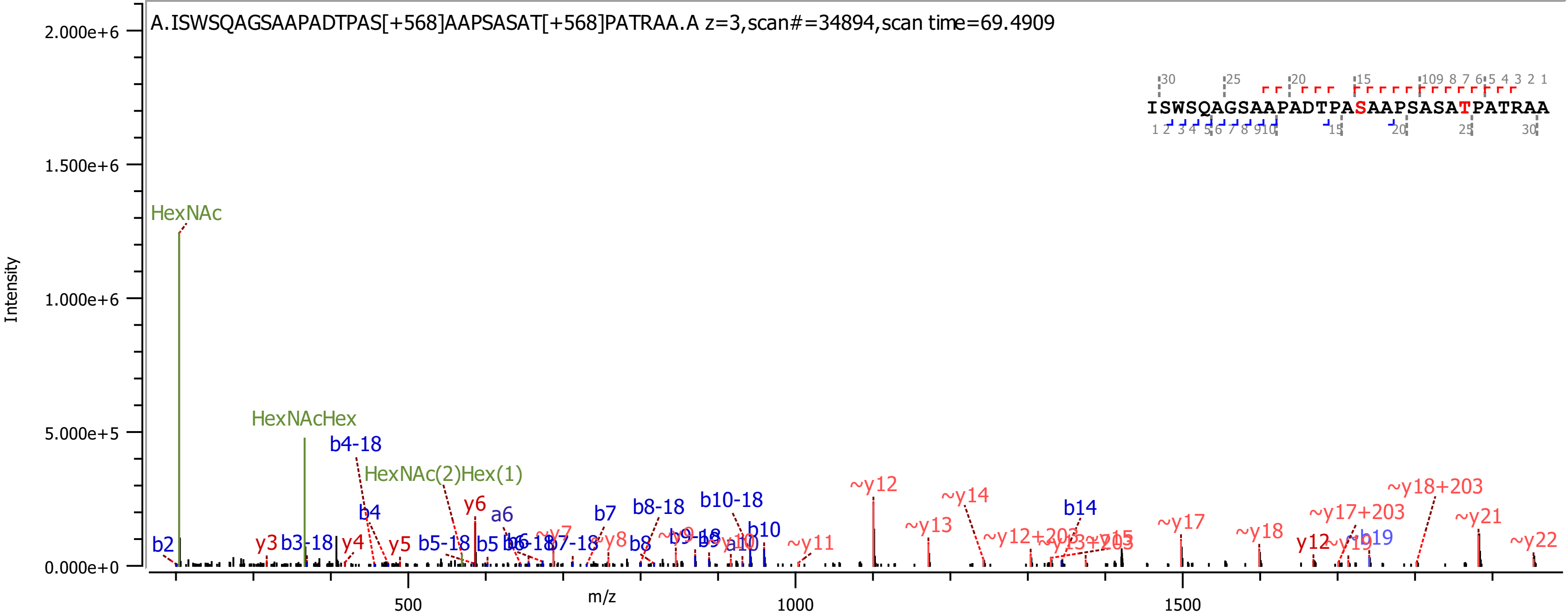
A.ISWSQAGSAAPADTPAS[+568]AAPSASAT[+568]PATRAA.A z=3,scan#=34894,scan time=69.4909
ISWSQAGSAAPADTPASAAPSASATPATRAA
Intensity
m/z
HexNAc
HexNAcHex
HexNAc(2)Hex(1)
b4-18
b4
y6
a6
b7
b8-18
b10-18
b10
~y12
~y14
~y13
b14
~y17
~y18
~y17+203
~y18+203
~y21
~y22
~y11
y12
b19
b2
y3
b3-18
y4
y5
b5-18
b5
b8

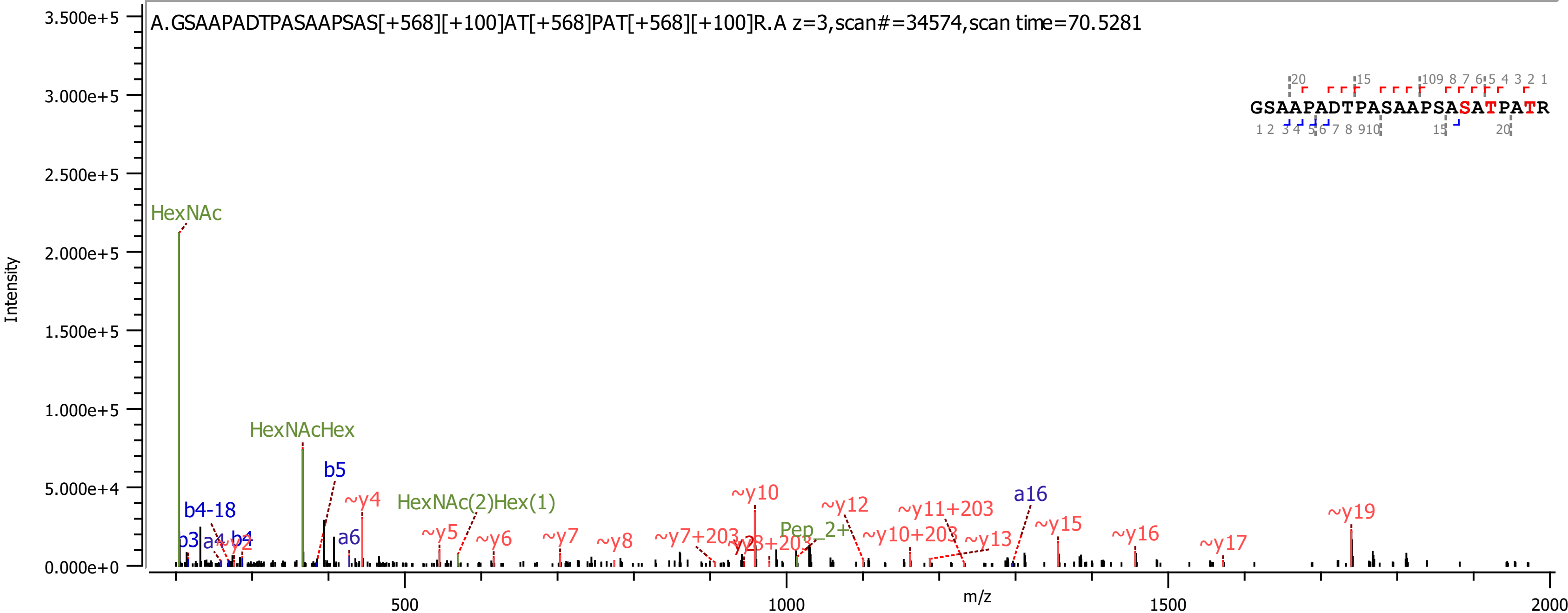

A.GSAAPADTPASAAPSAS[+568][+100]AT[+568]PAT[+568][+100]R.A z=3,scan#=34574,scan time=70.5281
GSAAPADTPASAAPSASATPATR
Intensity
m/z
HexNAc
HexNAcHex
HexNAc(2)Hex(1)
Pep_2+
b5
b4-18
a6
a16
~y4
~y5
~y6
~y7
~y8
~y7+203
~y10
~y12
~y11+203
~y10+203
~y13
~y15
~y16
~y17
~y19

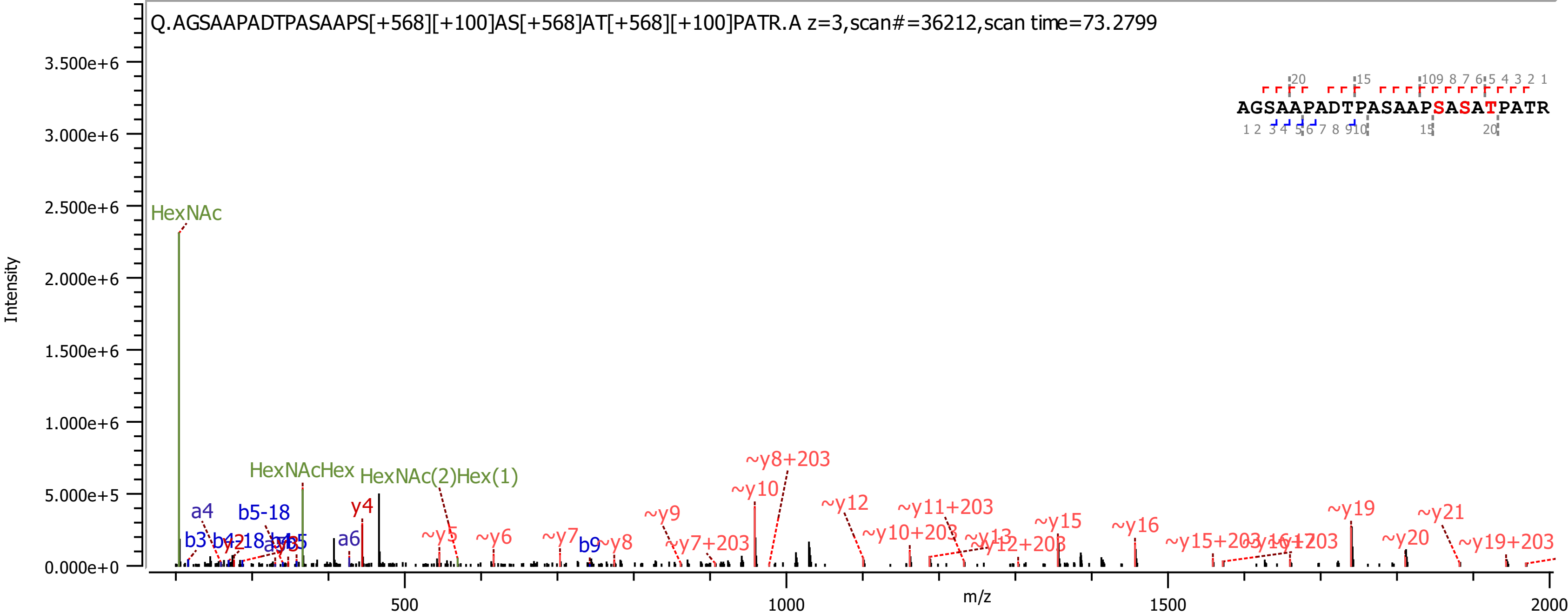
Q.AGSAAPADTPASAAPS[+568][+100]AS[+568]AT[+568][+100]PATR.A z=3,scan#=36212,scan time=73.2799
AGSAAPADTPASAAPSASATPATR
Intensity
m/z
HexNAc
HexNAcHex
HexNAc(2)Hex(1)
~y8+203
~y10
~y12
~y11+203
~y19
~y21
~y15
~y16
~y10+203
~y9
~y5
~y6
~y7
~y8
~y7+203
~y13
~y12+203
~y15+203
~y20
~y19+203
y4
a4
b5-18
a6
b3
b9

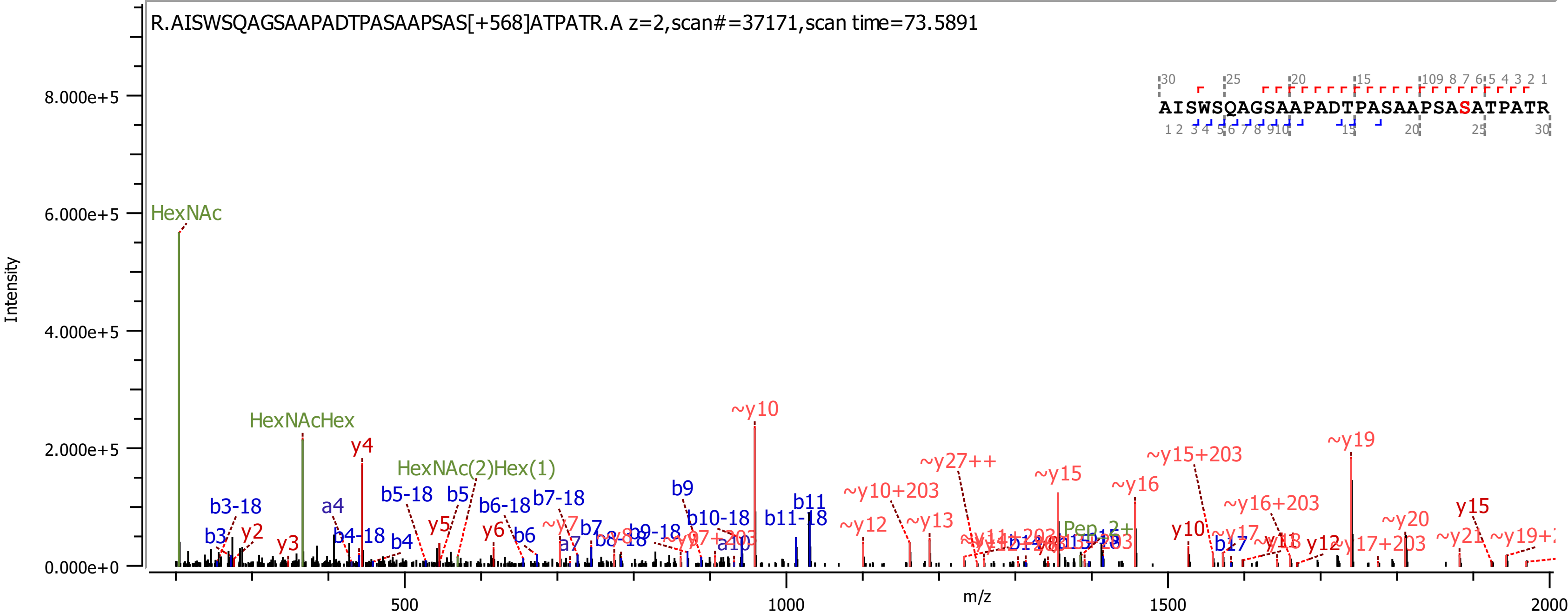
R.AISWSQAGSAAPADTPASAAPSAS[+568]ATPATR.A z=2,scan#=37171,scan time=73.5891
AISWSQAGSAAPADTPASAAPSASATPATR
Intensity
8.000e+5
6.000e+5
4.000e+5
2.000e+5
0.000e+0
m/z
500
1000
1500
2000
HexNAc
HexNAcHex
HexNAc(2)Hex(1)
y4
~y10
~y19
~y27++
~y15+203
~y15
~y16
~y10+203
~y16+203
y15
b3-18
b3
y2
y3
a4
b4-18
b4
b5-18
b5
y5
b6-18
b6
y6
b7-18
b7
~y7
a7
b8-18
b9
b9-18
b10-18
a10
b11
b11-18
~y12
~y13
Pep-2+
y10
~y17
b17
y12
~y20
~y21

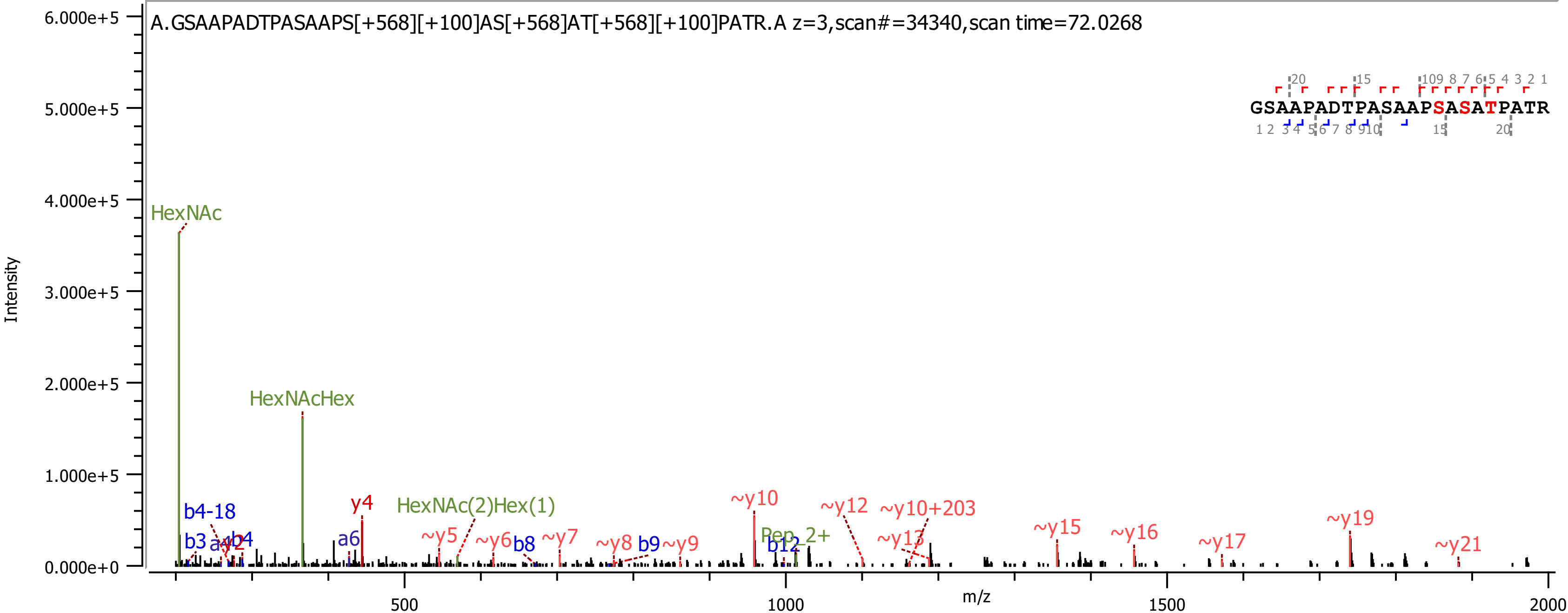
A.GSAAPADTPASAAPS[+568][+100]AS[+568]AT[+568][+100]PATR.A z=3,scan#=34340,scan time=72.0268
GSAAPADTPASAAPSASATPATR
Intensity
m/z
6.000e+5
5.000e+5
4.000e+5
3.000e+5
2.000e+5
1.000e+5
0.000e+0
500
1000
1500
2000
HexNAc
HexNAcHex
HexNAc(2)Hex(1)
Pep_2+
b4-18
b3
b4
y2
a6
y4
~y5
~y6
b8
~y7
~y8
b9
~y9
~y10
b12
~y12
~y10+203
~y13
~y15
~y16
~y17
~y19
~y21

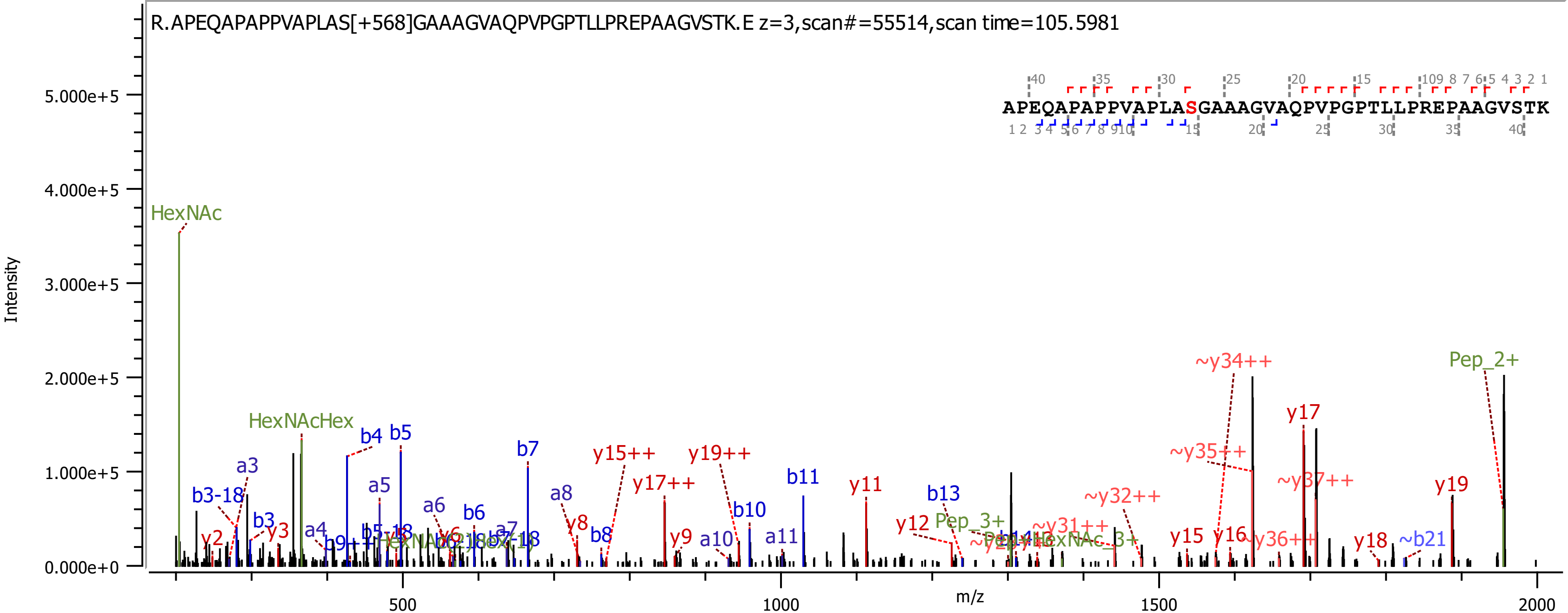
R.APEQAPAPPVAPLAS[+568]GAAAGVAQPVPGPTLLPREPAAGVSTK.E z=3,scan#=55514,scan time=105.5981
APEQAPAPPVAPLASGAAAGVAQPVPGPTLLPREPAAGVSTK
Intensity
m/z
5.000e+5
4.000e+5
3.000e+5
2.000e+5
1.000e+5
0.000e+0
500
1000
1500
2000
HexNAc
HexNAcHex
b3-18
a3
b3
y2
y3
a4
b4
b5
a5
a6
b6
a7
b7
a8
y8
b8
y15++
y17++
y19++
y9
a10
b10
a11
b11
y11
y12
b13
Pep_3+
~y31++
~y32++
~y34++
~y35++
y15
y16
~y36++
y17
~y37++
y18
~b21
y19
Pep_2+

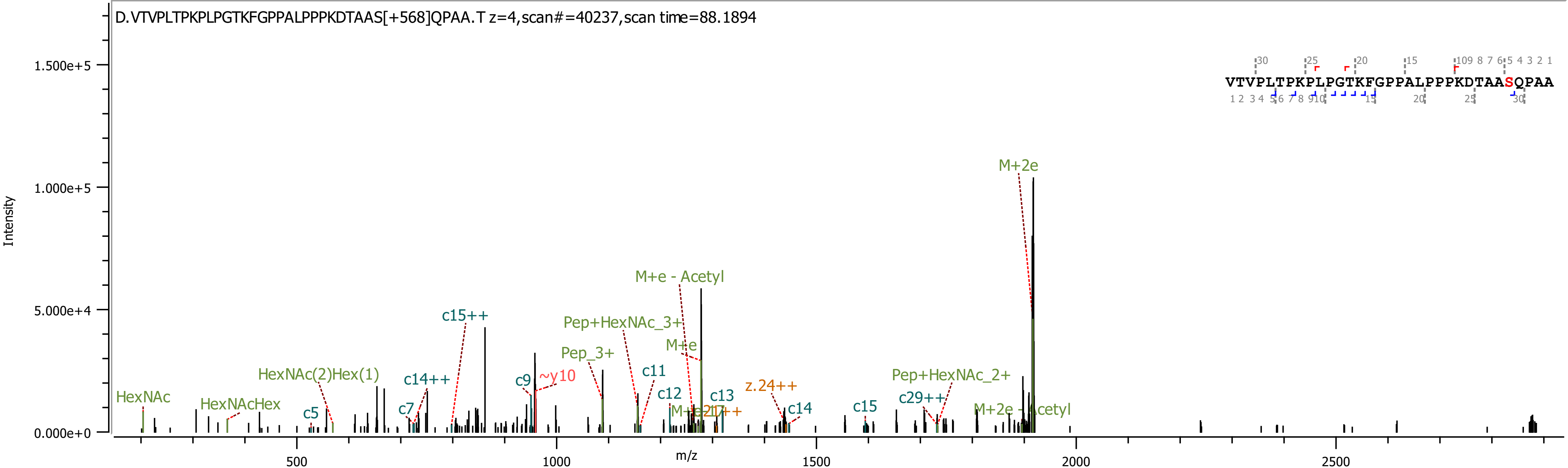

D.VTVPLTPKPLPGTKFGPPALPPPKDTAAS[+568]QPAA.T z=4,scan#=40237,scan time=88.1894
VTVPLTPKPLPGTKFGPPALPPPKDTAASQPAA
Intensity
m/z
1.500e+5
1.000e+5
5.000e+4
0.000e+0
500
1000
1500
2000
2500
M+2e
M+e - Acetyl
Pep+HexNAc_3+
M+e
Pep_3+
c15++
c11
c12
c13
z.24++
c14
c15
c29++
Pep+HexNAc_2+
M+2e - Acetyl
HexNAc(2)Hex(1)
c14++
c9
~y10
c7
c5
HexNAc
HexNAcHex

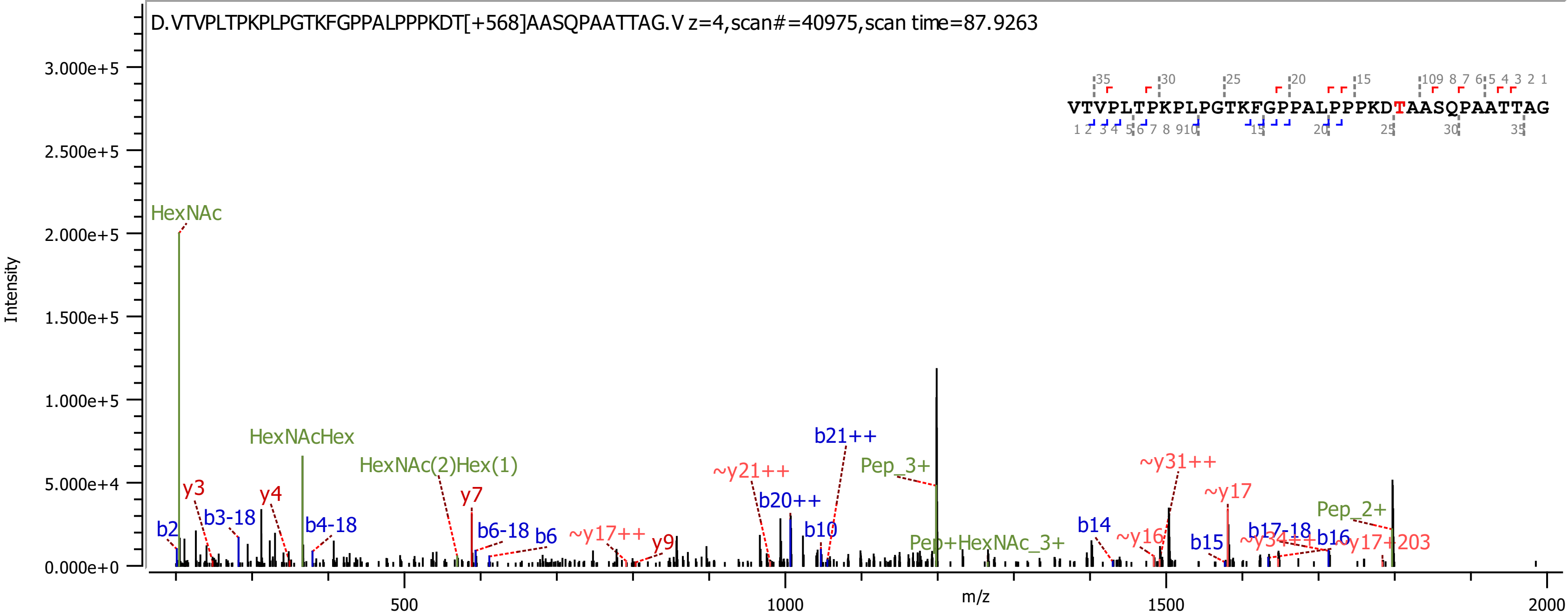

D.VTVPLTPKPLPGTKFGPPALPPPKDT[+568]AASQPAATTAG.V z=4,scan#=40975,scan time=87.9263
VTVPLTPKPLPGTKFGPPALPPPKDTAASQPAATTAG
Intensity
m/z
HexNAc
HexNAcHex
HexNAc(2)Hex(1)
Pep_3+
Pep+HexNAc_3+
Pep_2+
b2
b3-18
b4-18
b6-18
b6
b10
b14
b15
b16
b17-18
b20++
b21++
y3
y4
y7
y9
~y16
~y17
~y17++
~y17+203
~y21++
~y31++
~y34++

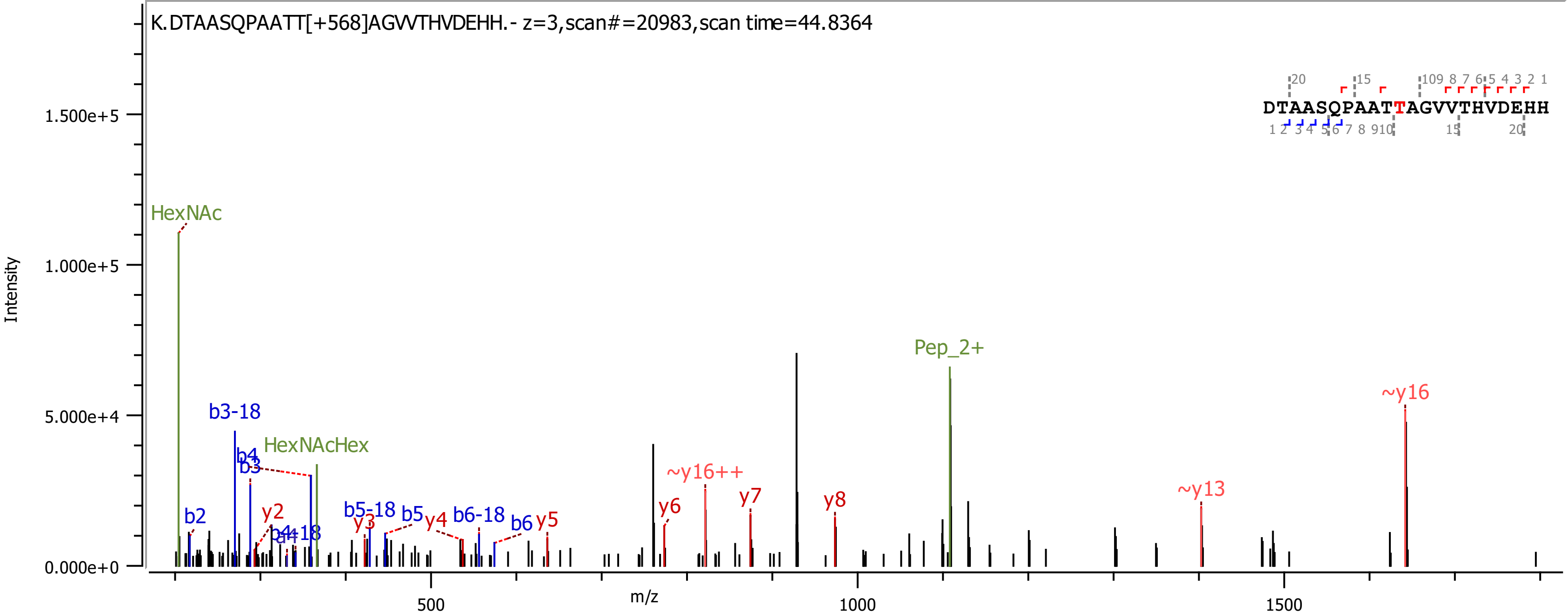

K.DTAASQPAATT[+568]AGVVTHVDEHH.- z=3,scan#=20983,scan time=44.8364
DTAASQPAATTAGVVTHVDEHH
Intensity
m/z
HexNAc
HexNAcHex
Pep_2+
b2
b3-18
b4
b3
y2
b4-18
b5-18
y3
b5
y4
b6-18
b6
y5
~y16++
y6
y7
y8
~y13
~y16

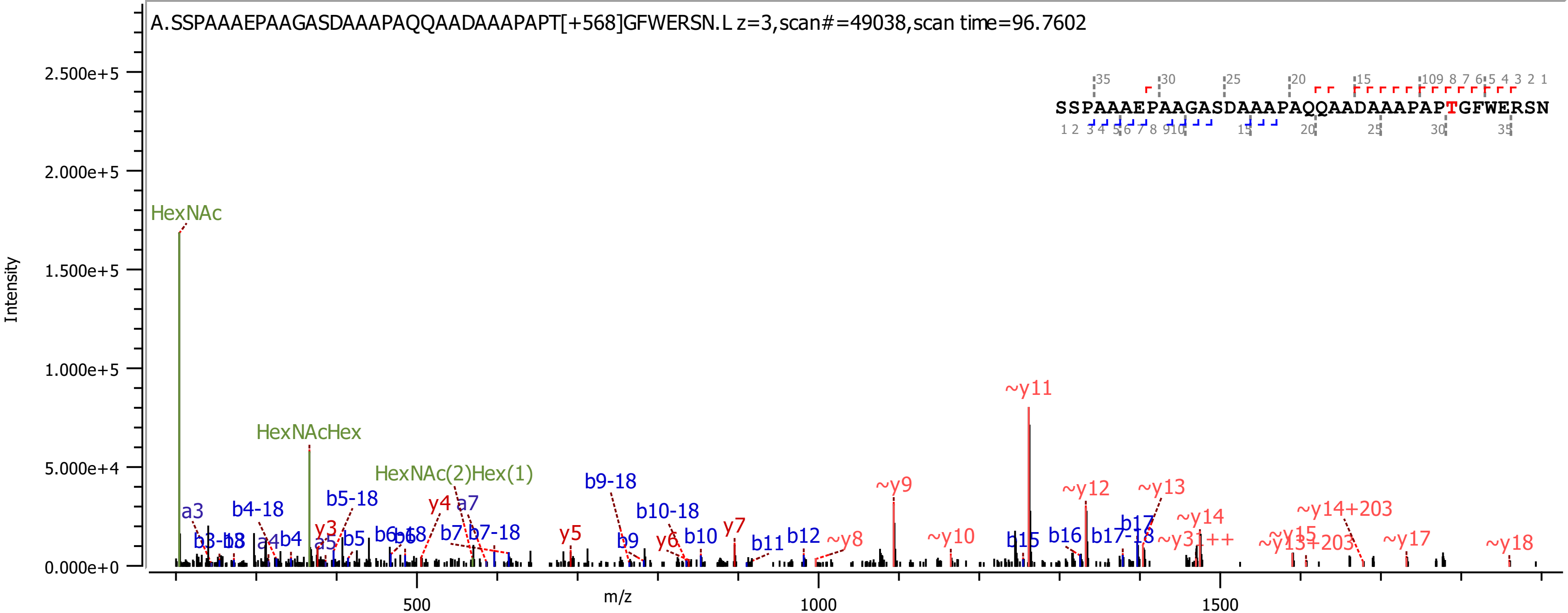

A.SSPAAAEPAAGASDAAAPAQQAADAAAPAPT[+568]GFWERSN.L z=3,scan#=49038,scan time=96.7602
SSPAAAEPAAGASDAAAPAQQAADAAAPAPTGFWERSN
Intensity
m/z
HexNAc
HexNAcHex
HexNAc(2)Hex(1)
~y11
~y9
~y12
~y13
~y14
~y14+203
b9-18
b10-18
b4-18
b5-18
y4
a7
a3
y3
y5
y6
y7
b10
b11
b12
~y8
~y10
b15
b16
b17-18
b17
~y31++
~y15
~y17
~y18

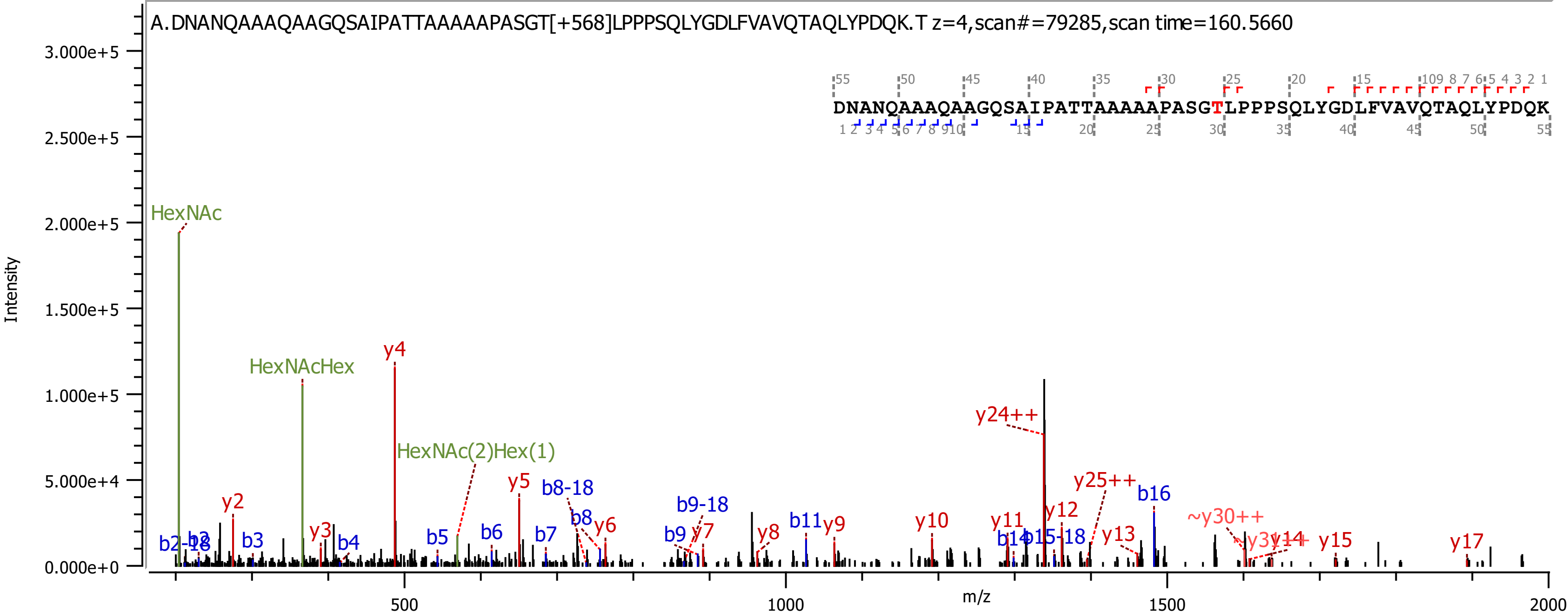

A.DNANQAAAQAAGQSAIPATTAAAAAPASGT[+568]LPPPSQLYGDLFVAVQTAQLYPDQK.T z=4,scan#=79285,scan time=160.5660
DNANQAAAQAAGQSAIPATTAAAAAPASGTLPPPSQLYGDLFVAVQTAQLYPDQK
Intensity
m/z
HexNAc
HexNAcHex
HexNAc(2)Hex(1)
y2
y3
y4
y5
y6
y7
y8
y9
y10
y11
y12
y13
y14
y15
y17
y24++
y25++
~y30++
b2-18
b3
b4
b5
b6
b7
b8
b8-18
b9
b9-18
b11
b14
b15-18
b16

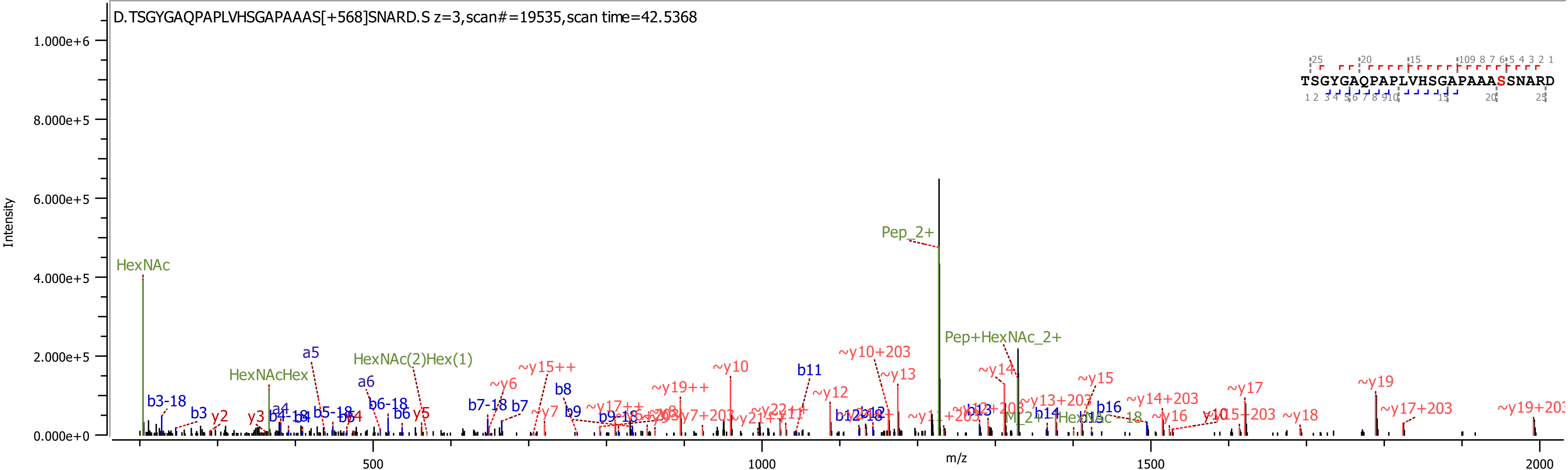
D.TSGYGAQPAPLVHSGAPAAAS[+568]SNARD.S z=3,scan#=19535,scan time=42.5368
TSGYGAQPAPLVHSGAPAAASSNARD
Intensity
1.000e+6
8.000e+5
6.000e+5
4.000e+5
2.000e+5
0.000e+0
m/z
500
1000
1500
2000
HexNAc
HexNAcHex
HexNAc(2)Hex(1)
Pep_2+
Pep+HexNAc_2+
b3-18
b3
y2
y3
a4
a5
a6
b6-18
b6
y5
b7-18
b7
b8
b9
~y6
~y7
~y15++
~y17++
~y19++
~y10
b11
~y10+203
~y12
~y13
~y14
~y13+203
~y15
b14
b16
~y14+203
~y16
~y17
~y15+203
~y18
~y19
~y17+203

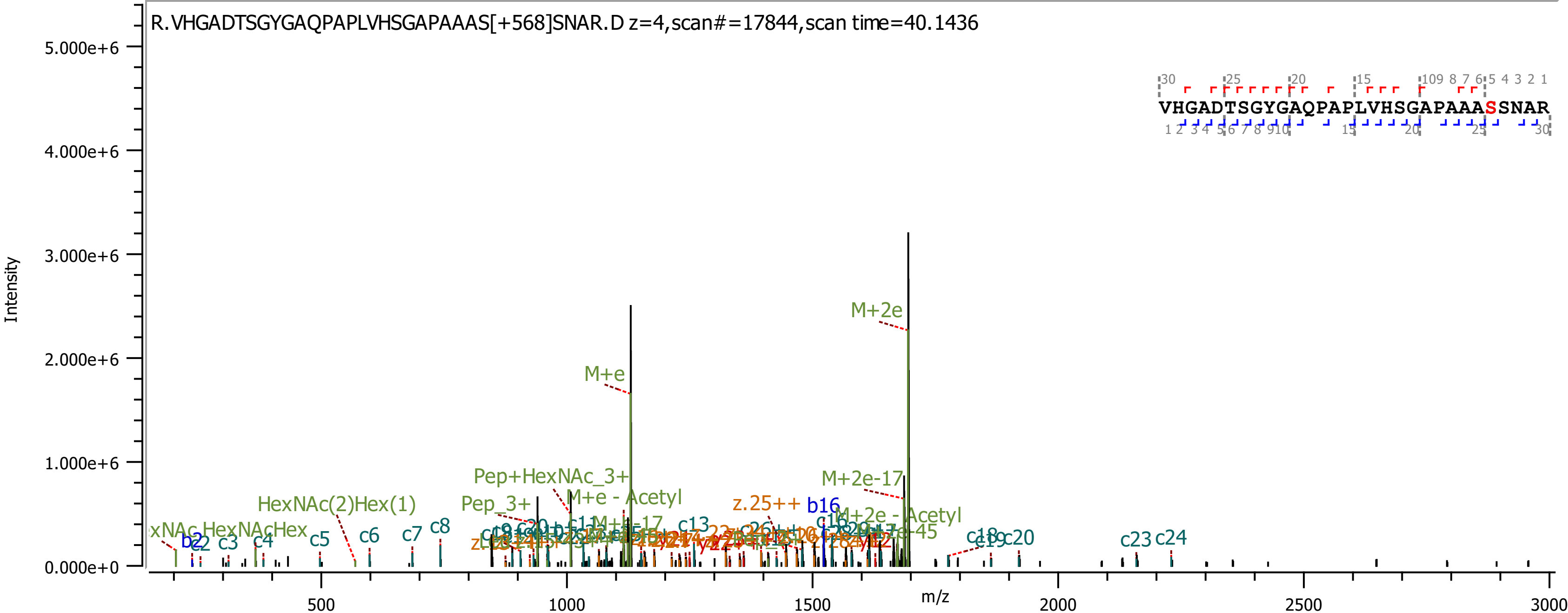

R.VHGADTSGYGAQPAPLVHSGAPAAAS[+568]SNAR.D z=4,scan#=17844,scan time=40.1436
VHGADTSGYGAQPAPLVHSGAPAAASSNAR
Intensity
m/z
M+2e
M+e
M+2e-17
Pep+HexNAc_3+
M+e - Acetyl
Pep_3+
HexNAc(2)Hex(1)
z.25++
b16
M+2e - Acetyl
b2
c2
c3
c4
c5
c6
c7
c8
c13
c18
c19
c20
c23
c24

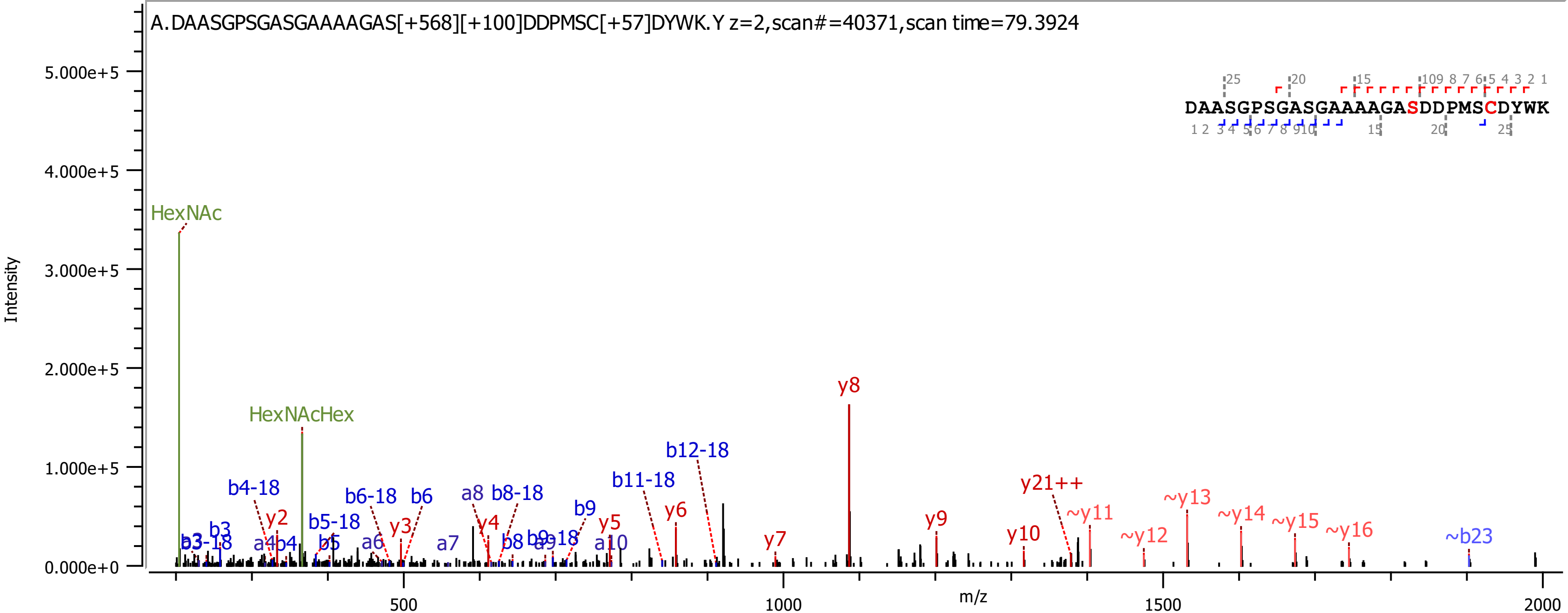
A.DAASGPSGASGAAAAGAS[+568][+100]DDPMSC[+57]DYWK.Y z=2,scan#=40371,scan time=79.3924
DAASGPSGASGAAAAGASDDPMSCDYWK
Intensity
m/z
HexNAc
HexNAcHex
y8
b12-18
b11-18
y21++
~y11
~y13
~y14
~y15
~y16
~y12
~b23
y10
y9
y7
y6
y5
a10
b9
a9-18
b9
b8
y4
a8
b8-18
a7
a6
b6
b6-18
y3
b5-18
b5
y2
b4-18
b4
a4
b3
b3-18
5.000e+5
4.000e+5
3.000e+5
2.000e+5
1.000e+5
0.000e+0
500
1000
1500
2000

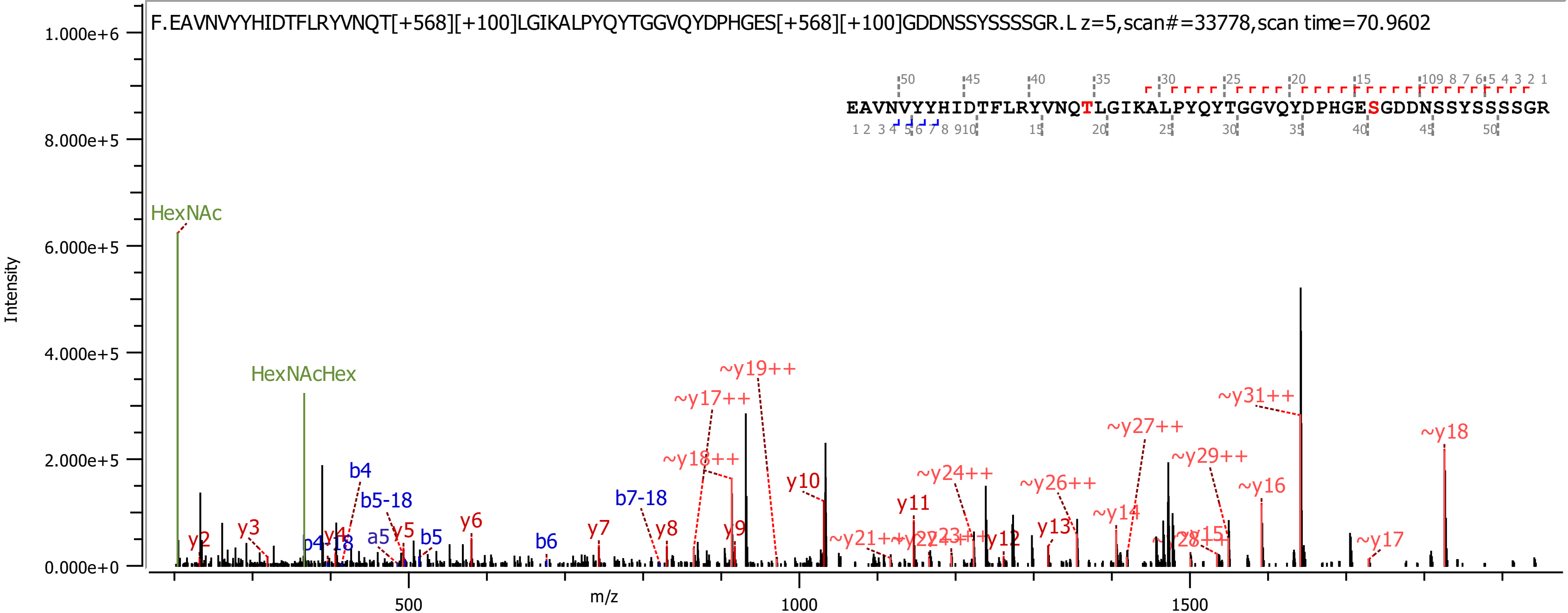

F.EAVNVYYHIDTFLRYVNQT[+568][+100]LGIKALPYQYTGGVQYDPHGES[+568][+100]GDDNSSYSSSSGR.L z=5,scan#=33778,scan time=70.9602
EAVNVYYHIDTFLRYVNQTLGIKALPYQYTGGVQYDPHGESGDDNSSYSSSSGR
Intensity
m/z
HexNAc
HexNAcHex
y2
y3
b4-18
y4
b4
b5-18
a5
y5
b5
y6
b6
y7
b7-18
y8
~y18++
~y17++
y9
~y19++
y10
~y21++
y11
~y24++
y12
y13
~y26++
~y14
~y27++
~y29++
~y15
~y16
~y31++
~y17
~y18
1.000e+6
8.000e+5
6.000e+5
4.000e+5
2.000e+5
0.000e+0
500
1000
1500

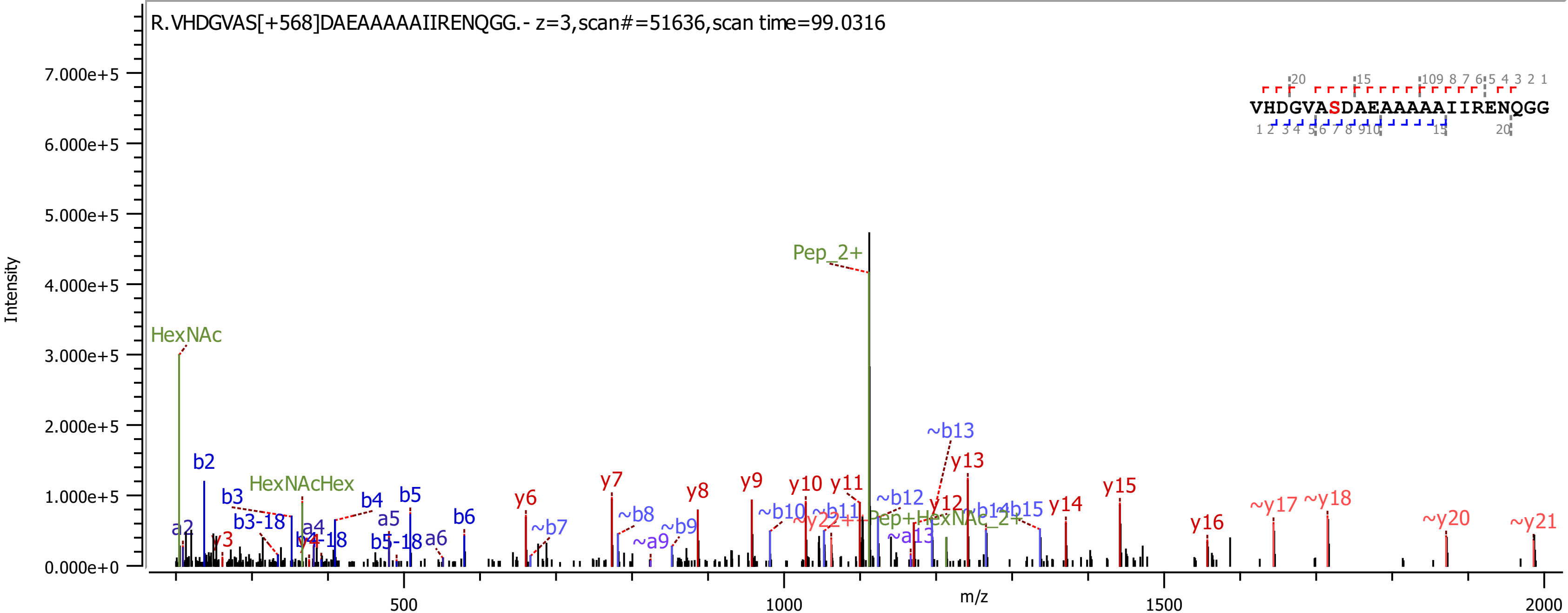
R.VHDGVAS[+568]DAEAAAAAIIRENQGG.- z=3,scan#=51636,scan time=99.0316
VHDGVASDAEAAAAAIIRENQGG
Intensity
m/z
7.000e+5
6.000e+5
5.000e+5
4.000e+5
3.000e+5
2.000e+5
1.000e+5
0.000e+0
500
1000
1500
2000
Pep_2+
HexNAc
HexNAcHex
Pep+HexNAc_2+
a2
b2
y3
b3
b3-18
a4
b4-18
b4
a5
b5
b5-18
a6
b6
y6
~b7
y7
~b8
~a9
~b9
y8
y9
~b10
y10
~b11
y11
~y22++
~b12
~a13
y12
~b13
y13
~b14
~b15
y14
y15
y16
~y17
~y18
~y20
~y21

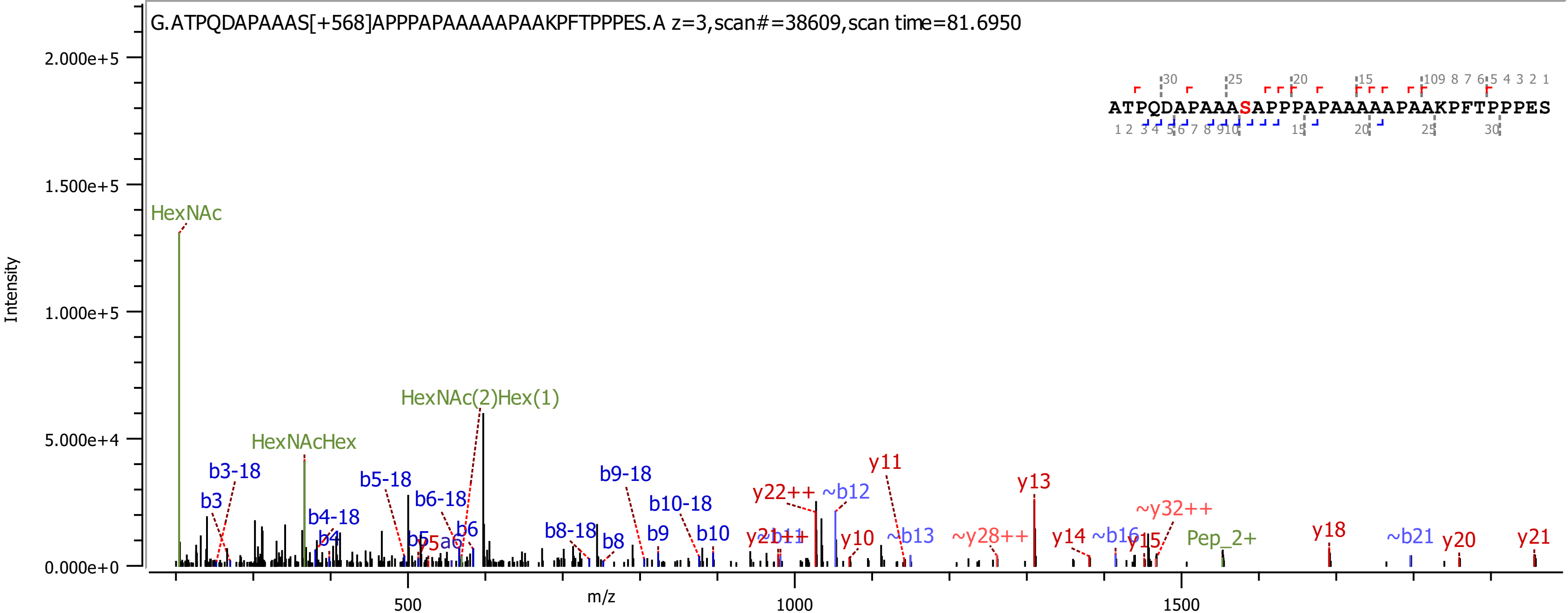

G.ATPQDAPAAAS[+568]APPPAPAAAAAPAAKPFTPPPES.A z=3,scan#=38609,scan time=81.6950
ATPQDAPAAASAPPPAPAAAAAPAAKPFTPPPES
Intensity
m/z
2.000e+5
1.500e+5
1.000e+5
5.000e+4
0.000e+0
500
1000
1500
HexNAc
HexNAc(2)Hex(1)
HexNAcHex
b3-18
b3
b4-18
b4
b5-18
b5
b6-18
b6
y5a6
b8-18
b8
b9-18
b9
b10-18
b10
y21++
~b11
y22++
~b12
y11
y10
~b13
y13
~y28++
y14
~b16
y15
~y32++
Pep_2+
y18
~b21
y20
y21

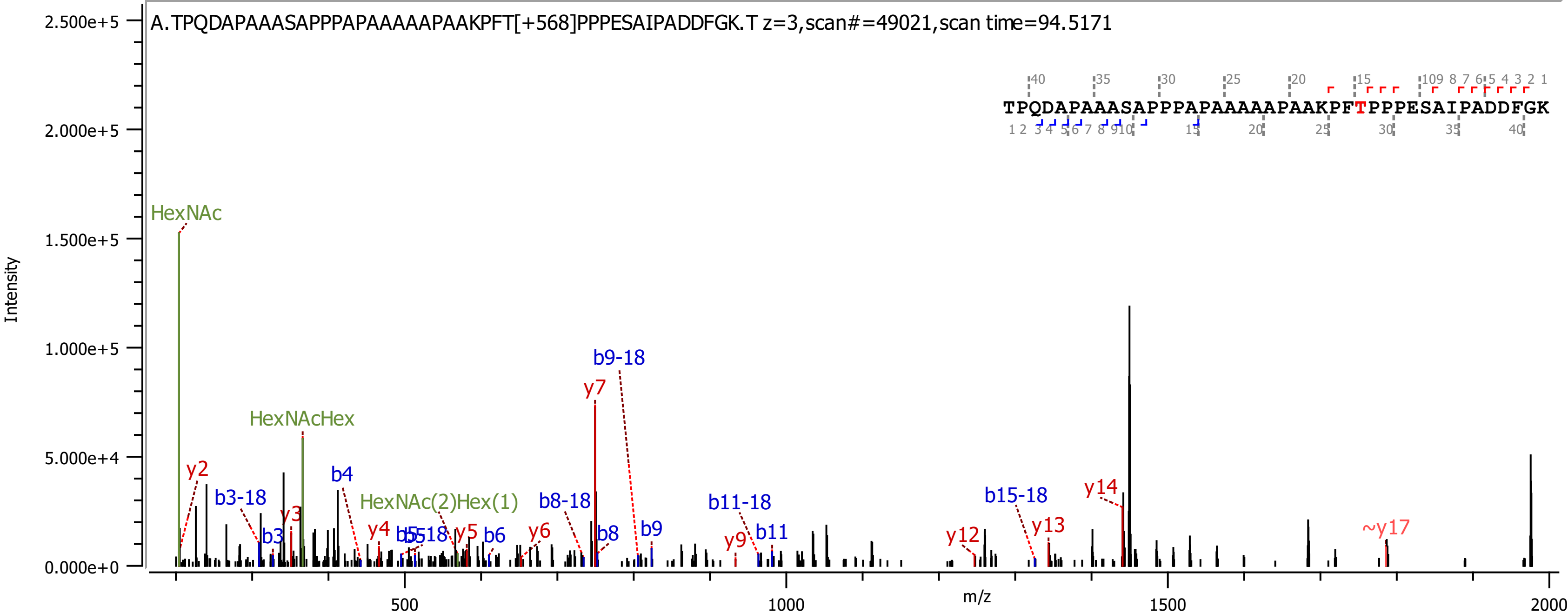
A.TPQDAPAAASAPPPAPAAAAAPAAKPFT[+568]PPPESAIPADDFGK.T z=3,scan#=49021,scan time=94.5171
TPQDAPAAASAPPPAPAAAAAPAAKPFTPPPESAIPADDFGK
Intensity
m/z
HexNAc
HexNAcHex
HexNAc(2)Hex(1)
y2
y3
y4
y5
y6
y7
y9
y12
y13
y14
~y17
b3
b3-18
b4
b5
b5-18
b6
b8
b8-18
b9
b9-18
b11
b11-18
b15-18
2.500e+5
2.000e+5
1.500e+5
1.000e+5
5.000e+4
0.000e+0
500
1000
1500
2000

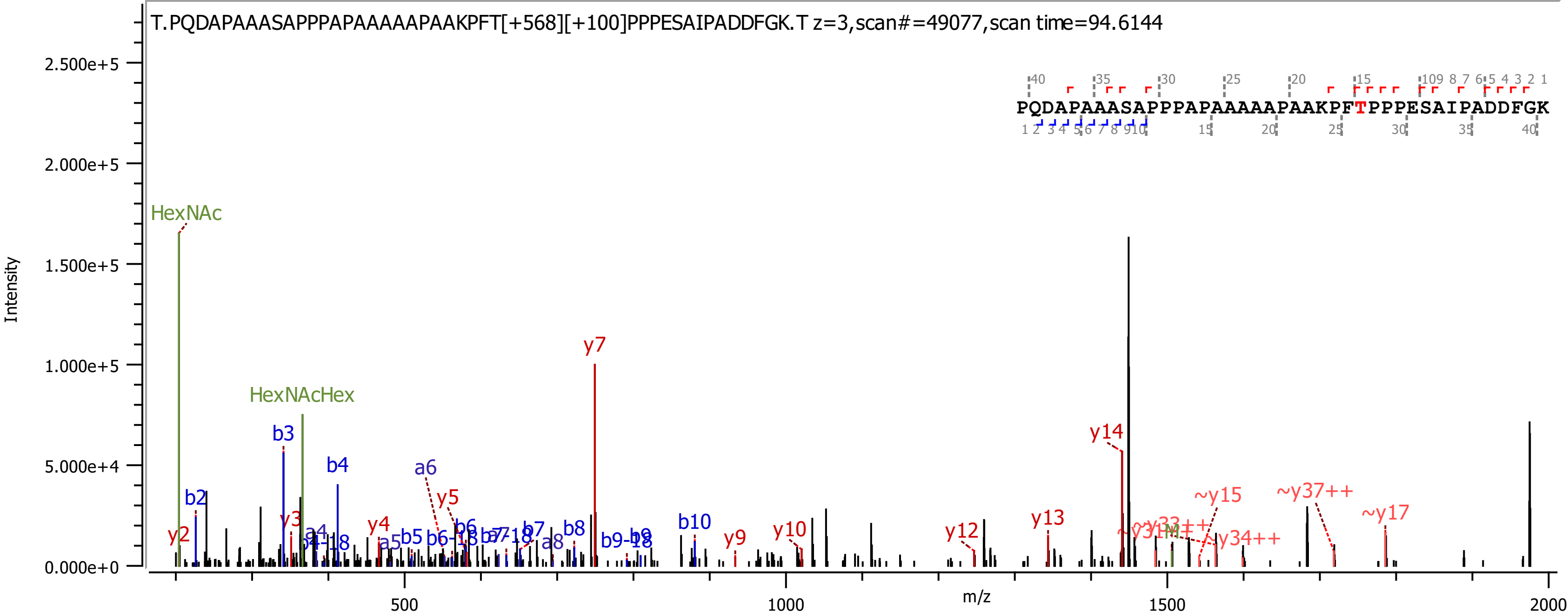

T.PQDAPAAASAPPPAPAAAAAPAAKPFT[+568][+100]PPPESAIPADDFGK.T z=3,scan#=49077,scan time=94.6144
PQDAPAAASAPPPAPAAAAAPAAKPFTPPPESAIPADDFGK
Intensity
m/z
HexNAc
HexNAcHex
y7
y14
b3
b4
a6
y5
b2
y3
y2
y4
a5
b5
b6
b7
a8
b8
b10
y9
y10
y12
y13
~y15
~y37++
~y17
~y34++

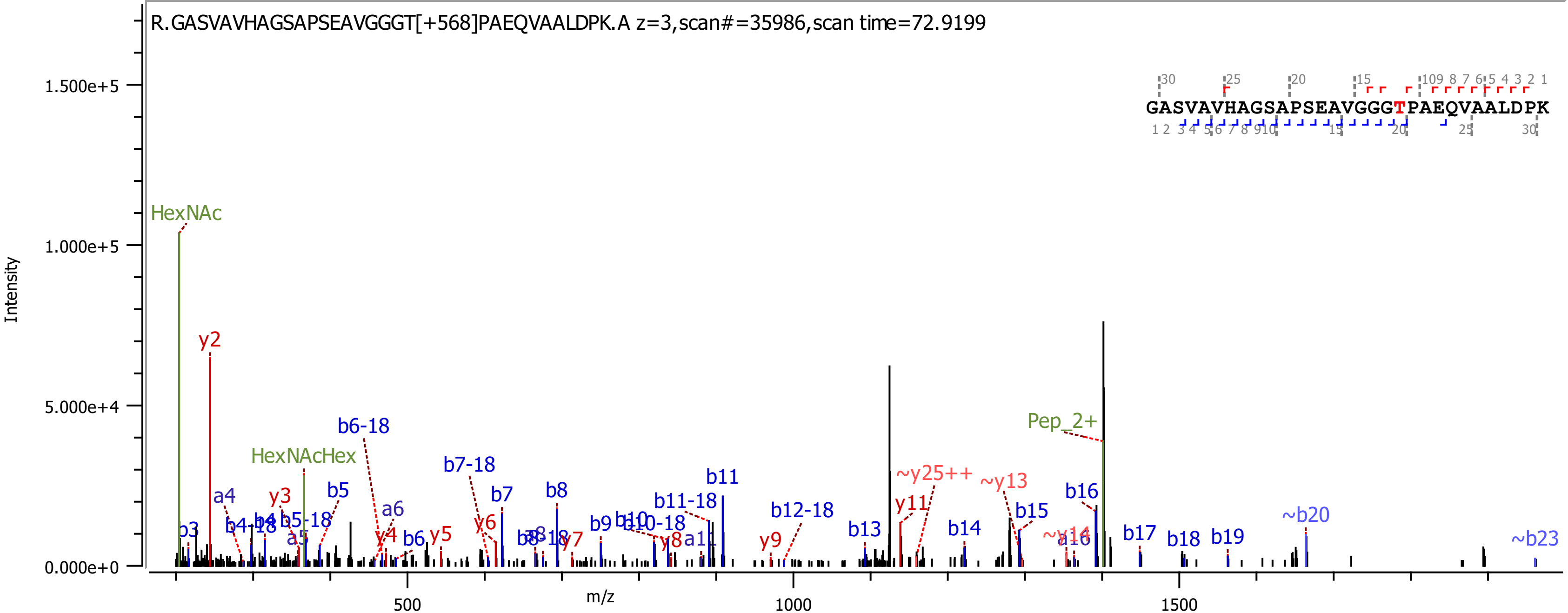
R.GASVAVHAGSAPSEAVGGGT[+568]PAEQVAALDPK.A z=3,scan#=35986,scan time=72.9199
GASVAVHAGSAPSEAVGGGTPAEQVAALDPK
Intensity
m/z
1.500e+5
1.000e+5
5.000e+4
0.000e+0
500
1000
1500
HexNAc
y2
HexNAcHex
b6-18
b7-18
b11
~y25++
~y13
Pep_2+
b16
~b20
~b23
b3
a4
y3
b5
a6
b7
b8
b9
b11-18
b12-18
y11
b15
b13
b14
b17
b18
b19
y9
y5
y6
y7
y8
b6
y4
a5
b4-18
b5-18
b10-18
~y14
a16

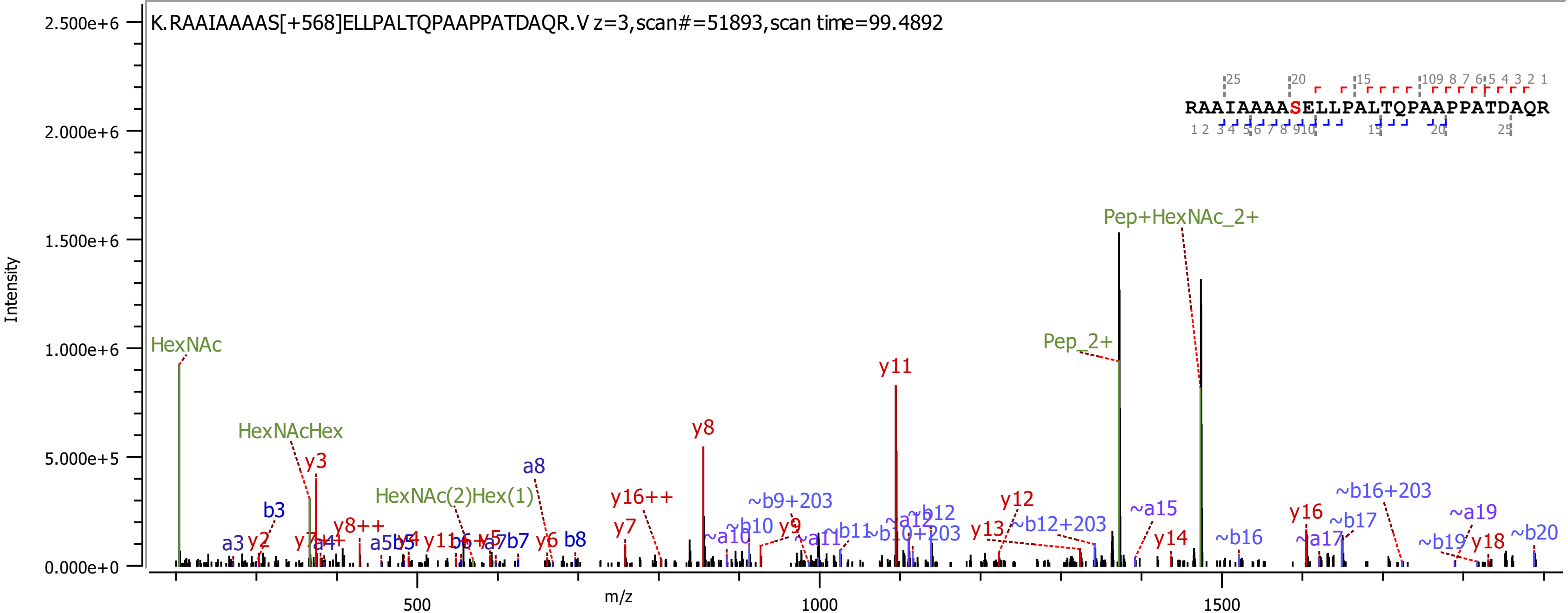
K.RAAIAAAAS[+568]ELLPALTQPAAPPATDAQR.V z=3,scan#=51893,scan time=99.4892
RAAIAAAASELLPALTQPAAPPATDAQR
Intensity
2.500e+6
2.000e+6
1.500e+6
1.000e+6
5.000e+5
0.000e+0
m/z
500
1000
1500
HexNAc
HexNAcHex
HexNAc(2)Hex(1)
Pep_2+
Pep+HexNAc_2+
y3
y8
y11
y16++
y7
y12
y13
y14
y16
y18
y6
b3
b8
a3
a8
~b9+203
~b10
~a10
~b12
~a12
~b12+203
~a15
~b16
~b16+203
~b17
~a17
~a19
~b19
~b20

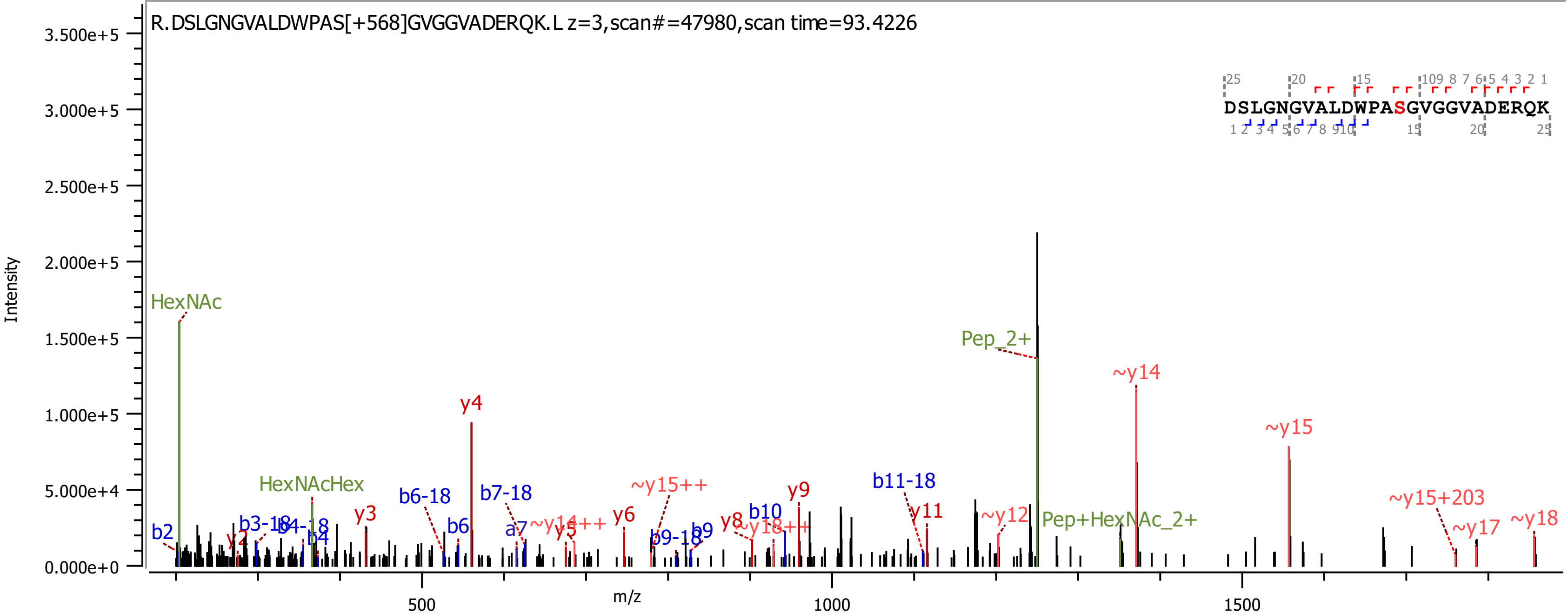
R.DSLGNGVALDWPAS[+568]GVGGVADERQK.L z=3,scan#=47980,scan time=93.4226
DSLGNGVALDWPASGVGGVADERQK
Intensity
m/z
HexNAc
HexNAcHex
Pep_2+
Pep+HexNAc_2+
y4
~y14
~y15
~y15+203
~y17
~y18
b2
y2
b3-18
b4-18
b4
y3
b6-18
b6
b7-18
a7
~y14++
y5
y6
~y15++
b9-18
b9
y8
~y18++
b10
y9
b11-18
y11
~y12

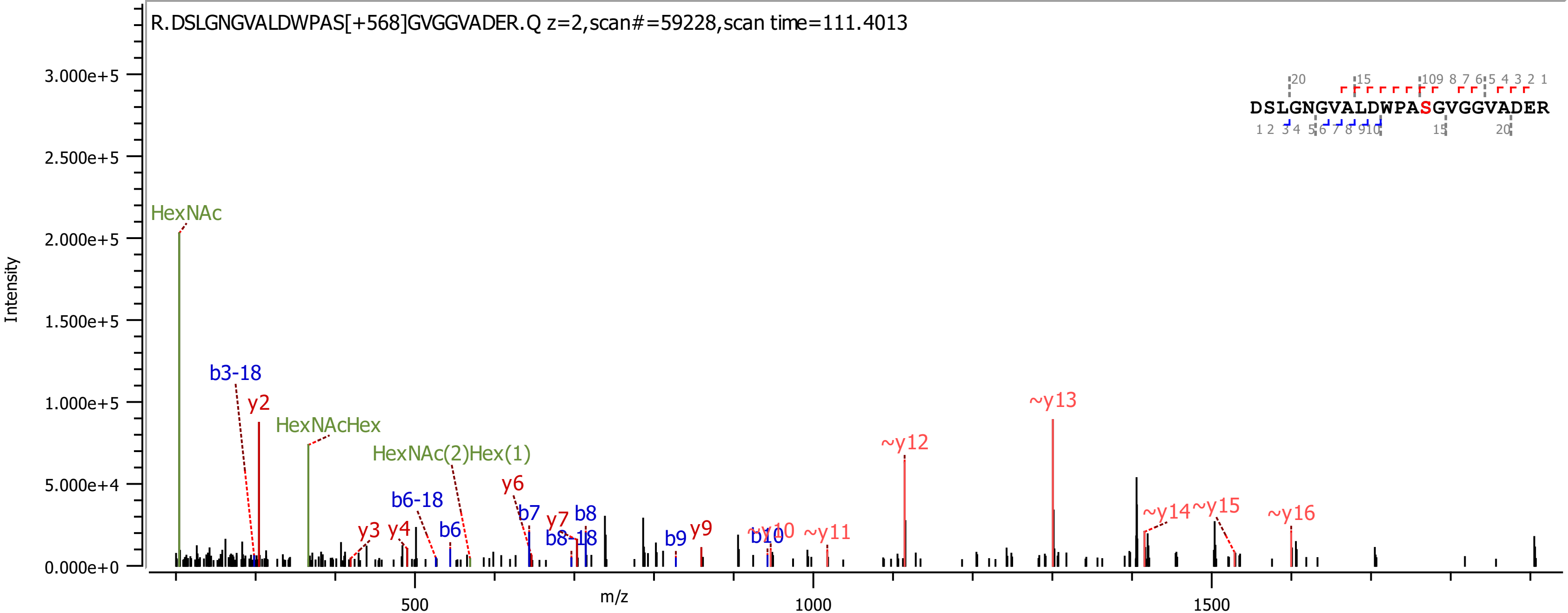

R.DSLGNGVALDWPAS[+568]GVGGVADER.Q z=2,scan#=59228,scan time=111.4013
DSLGNGVALDWPASGVGGVADER
Intensity
m/z
3.000e+5
2.500e+5
2.000e+5
1.500e+5
1.000e+5
5.000e+4
0.000e+0
500
1000
1500
HexNAc
b3-18
y2
HexNAcHex
HexNAc(2)Hex(1)
y3
y4
b6-18
b6
y6
b7
y7
b8
b8-18
b9
y9
~y10
b10
~y11
~y12
~y13
~y14
~y15
~y16

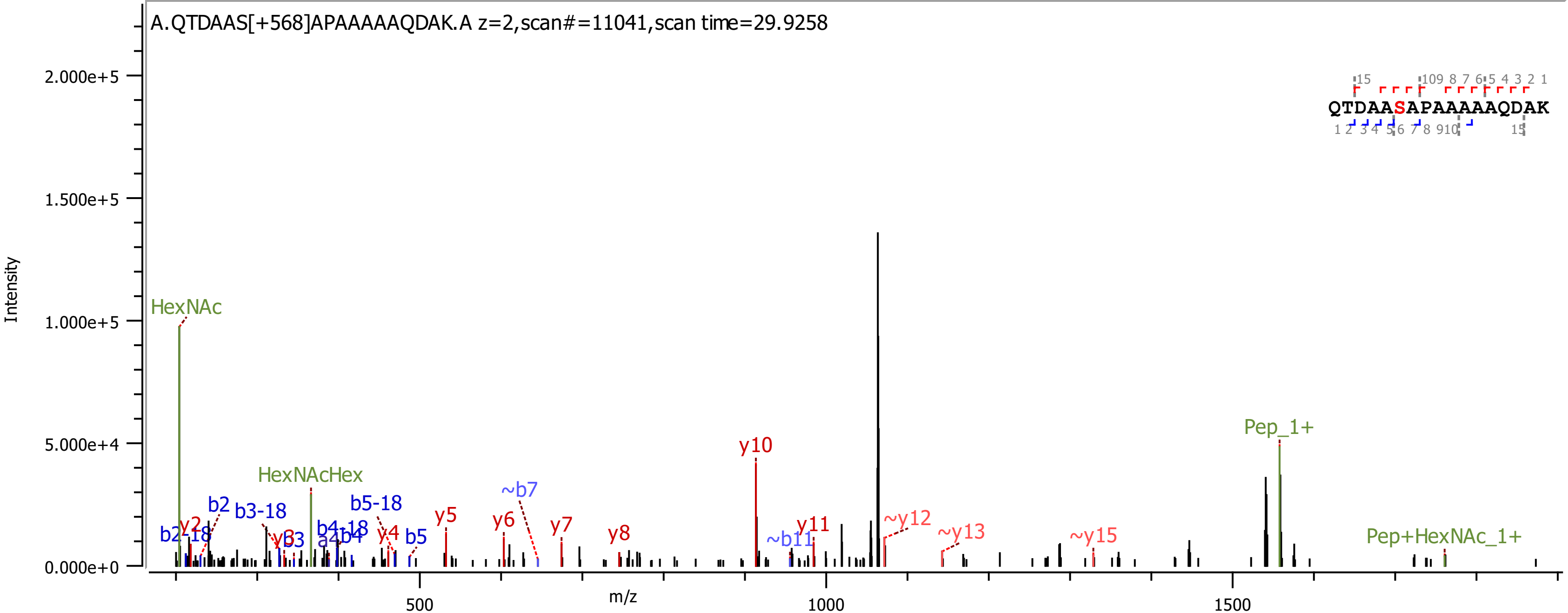
A.QTDAAS[+568]APAAAAAQDAK.A z=2,scan#=11041,scan time=29.9258
QTDAASAPAAAAAQDAK
Intensity
m/z
HexNAc
HexNAcHex
b2-18
y2
b2
b3-18
y3
b3
a4
b4-18
b4
b5-18
y4
b5
y5
y6
~b7
y7
y8
y10
~b11
y11
~y12
~y13
~y15
Pep_1+
Pep+HexNAc_1+
2.000e+5
1.500e+5
1.000e+5
5.000e+4
0.000e+0
500
1000
1500

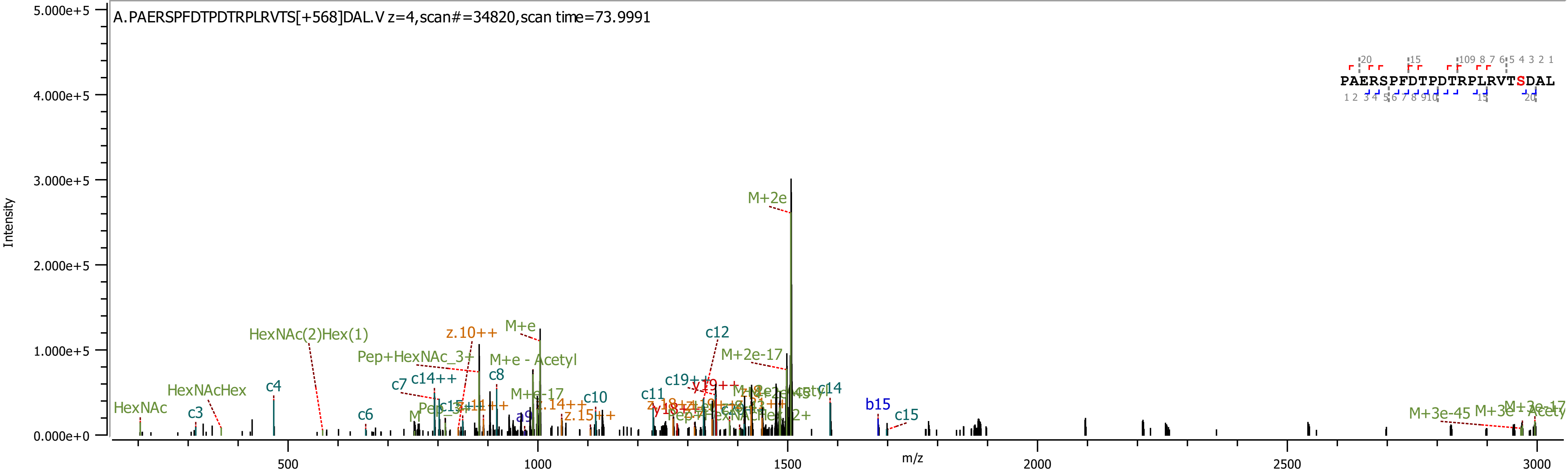
A.PAERSPFDTPDTRPLRVTS[+568]DAL.V z=4,scan#=34820,scan time=73.9991
PAERSPFDTPDTRPLRVTSDAL
Intensity
m/z
5.000e+5
4.000e+5
3.000e+5
2.000e+5
1.000e+5
0.000e+0
500
1000
1500
2000
2500
3000
M+2e
M+e
HexNAc(2)Hex(1)
z.10++
Pep+HexNAc_3+
M+e - Acetyl
c12
M+2e-17
HexNAcHex
c4
c7
c14++
c8
c3
HexNAc
c6
c10
c11
c14
b15
c15
M+3e-45

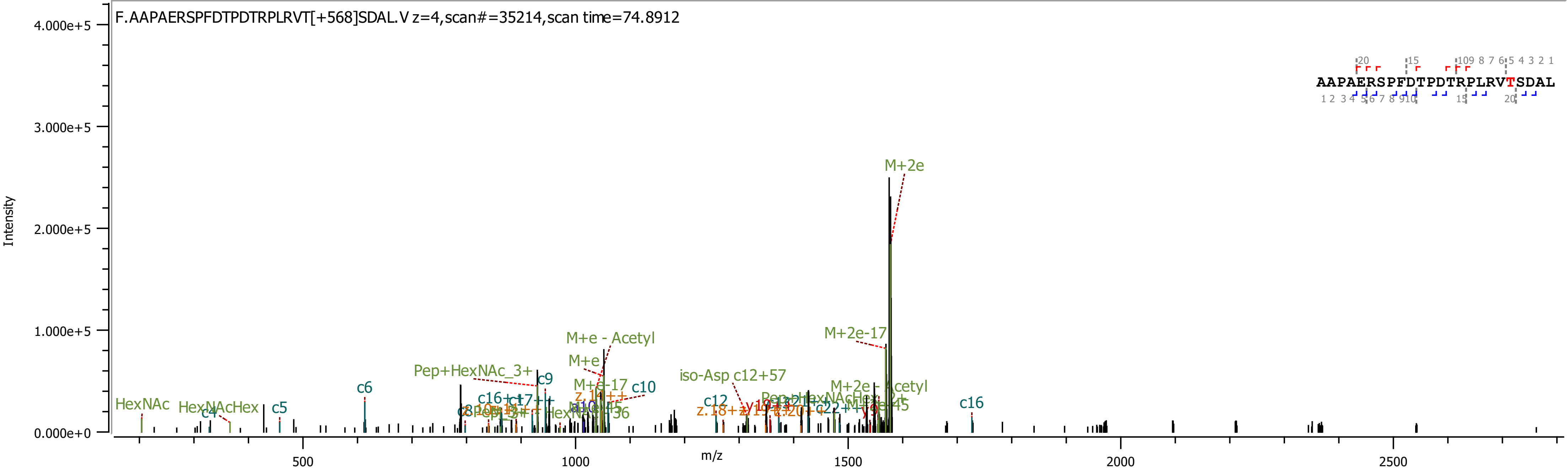
F.AAPAERSPFDTPDTRPLRVT[+568]SDAL.V z=4,scan#=35214,scan time=74.8912
AAPAERSPFDTPDTRPLRVTSDAL
Intensity
m/z
4.000e+5
3.000e+5
2.000e+5
1.000e+5
0.000e+0
500
1000
1500
2000
2500
M+2e
M+2e-17
M+e - Acetyl
M+e
M+e-17
Pep+HexNAc_3+
iso-Asp c12+57
M+2e - Acetyl
HexNAc
HexNAcHex
c4
c5
c6
c9
c10
c12
c16
