## Supplementary Data 2 for "Burkholderia PglL enzymes are serine preferring oligosaccharidetransferases which target conserved proteins across the Burkholderia genus"

### *Burkholderia cenocepacia* K56-2 Best Scoring Unique glycopeptides

<table>
<tr><th>J2315 Gene name</th><th>Protein Name</th><th>Peptide < ProteinMetrics Confidential ></th><th>Glycans NHFAGNa</th><th>Localised site (Yes/No)</th><th>Observed m/z</th><th>z</th><th>Observed (M+H)</th><th>Calc. mass (M+H)</th><th>Mass error (ppm)</th><th>Cleavage</th><th>Score</th><th>Delta</th><th>Delta Mod</th><th>Comment</th><th>Scan Time</th><th>Enzyme</th><th>Replicate</th><th>Page</th></tr>
<tr><td>BCAL0039</td><td>>gi|443605781|gb|ELT73605.1| ABC transporter, substrate-binding protein, family 3 [Burkholderia cenocepacia K56-2Valvano]</td><td>T.AAPASAATAGS[+568.212]RLDDVLARGA.L</td><td>HexNAc(2)Hex(1)</td><td>No</td><td>837.0821</td><td>3</td><td>2509.2316</td><td>2509.2264</td><td>2.1</td><td>Non</td><td>412.99</td><td>336.37</td><td>1.2</td><td>Nsco_20191108_BC_ZIC_HILIC_K56_2_thermolysin_B1.32915.32915.3</td><td>66.6456</td><td>Thermolysin</td><td>B1</td><td>20</td></tr>
<tr><td>BCAL0080</td><td>>gi|443593207|gb|ELT61961.1| cytochrome C domain protein, partial [Burkholderia cenocepacia K56-2Valvano]</td><td>N.AAQPASGAPASGADAS[+568.212]NAQA.-</td><td>HexNAc(2)Hex(1)</td><td>No</td><td>1141.0048</td><td>2</td><td>2281.0023</td><td>2280.995</td><td>3.2</td><td>Non</td><td>421.08</td><td>283.51</td><td>0.87</td><td>Nsco_20191108_BC_ZIC_HILIC_K56_2_thermolysin_B1.12891.12891.2</td><td>31.0195</td><td>Thermolysin</td><td>B1</td><td>21</td></tr>
<tr><td>BCAL0080</td><td>>gi|443593207|gb|ELT61961.1| cytochrome C domain protein, partial [Burkholderia cenocepacia K56-2Valvano]</td><td>Y.MANNDGANFPEPAAPAANAAQPAS[+568.212]GAPASGADASNAQ.A</td><td>HexNAc(2)Hex(1)</td><td>No</td><td>1321.5789</td><td>3</td><td>3962.7222</td><td>3962.7105</td><td>3</td><td>Non</td><td>547.71</td><td>520.06</td><td>44.52</td><td>Nsco_20191108_BC_ZIC_HILIC_K56_2_thermolysin_B1.40978.40978.3</td><td>83.9361</td><td>Thermolysin</td><td>B1</td><td>22</td></tr>
<tr><td>BCAL0080</td><td>>gi|443593207|gb|ELT61961.1| cytochrome C domain protein, partial [Burkholderia cenocepacia K56-2Valvano]</td><td>Y.MANNDGANFPEPAAPAANAAQPAS[+568.212]GAPASGADASNAQA.-</td><td>HexNAc(2)Hex(1)</td><td>Yes (1 of 2)</td><td>1345.2585</td><td>3</td><td>4033.7609</td><td>4033.7476</td><td>3.3</td><td>Non</td><td>273.98</td><td>227.31</td><td>41.79</td><td>Nsco_20191108_BC_ZIC_HILIC_K56_2_thermolysin_B2_.46558.46558.3</td><td>85.0541</td><td>Thermolysin</td><td>B2</td><td>23</td></tr>
<tr><td>BCAL0163</td><td>>gi|443595598|gb|ELT64173.1| BON domain protein [Burkholderia cenocepacia K56-2Valvano]</td><td>V.KPQDAQALQDASPASGASGAQAAAAPADNATVGAVPDASVQSTPLQPPAPISNS[+568.212]S[+568.212][+100.064]SVHPGNPK.A</td><td>HexNAc(2)Hex(1),<br>HexNAc(2)Hex(1)<br>100.064</td><td>No</td><td>1202.579</td><td>6</td><td>7210.4378</td><td>7204.4215</td><td>-0.5</td><td>Non</td><td>176.52</td><td>176.52</td><td>0</td><td>Nsco_20191108_BC_ZIC_HILIC_K56_2_thermolysin_B3.50702.50702.6</td><td>101.3486</td><td>Thermolysin</td><td>B3</td><td>24</td></tr>
<tr><td>BCAL0163</td><td>>gi|443595598|gb|ELT64173.1| BON domain protein [Burkholderia cenocepacia K56-2Valvano]</td><td>V.KVFQYVKPQDAQALQDASPASGAS[+568.212]GAQAAAAPADNATVGAVPDASVQSTPLQPPAPISNSSSVHPGNPK.A</td><td>HexNAc(2)Hex(1)</td><td>No</td><td>1460.9239</td><td>5</td><td>7300.5903</td><td>7300.568</td><td>3</td><td>NRagged</td><td>1310.09</td><td>1268.2</td><td>36.68</td><td>Nsco_20191108_BC_ZIC_HILIC_K56_2_typsin_B2.46071.46071.5</td><td>93.6647</td><td>Trypsin</td><td>B2</td><td>25</td></tr>
<tr><td>BCAL0163</td><td>>gi|443595598|gb|ELT64173.1| BON domain protein [Burkholderia cenocepacia K56-2Valvano]</td><td>A.LQDASPAS[+568.212]GASGAQAAAAPADN.A</td><td>HexNAc(2)Hex(1)</td><td>No</td><td>1255.0619</td><td>2</td><td>2509.1166</td><td>2509.106</td><td>4.2</td><td>Non</td><td>395.8</td><td>267.22</td><td>1.16</td><td>Nsco_20191108_BC_ZIC_HILIC_K56_2_pepsin_B2.22491.22491.2</td><td>49.0361</td><td>Pepsin</td><td>B2</td><td>26</td></tr>
<tr><td>BCAL0163</td><td>>gi|443595598|gb|ELT64173.1| BON domain protein [Burkholderia cenocepacia K56-2Valvano]</td><td>K.PQDAQALQDASPASGASGAQAAAAPADNATVGAVPDASVQST[+568.212]PLQPPAPISNSSSVHPGNPK.A</td><td>HexNAc(2)Hex(1)</td><td>No</td><td>1602.7728</td><td>4</td><td>6408.0695</td><td>6408.051</td><td>2.9</td><td>Specific</td><td>1162.92</td><td>1162.9</td><td>0</td><td>Nsco_20191108_BC_ZIC_HILIC_K56_2_typsin_B2.48573.48573.4</td><td>97.8402</td><td>Trypsin</td><td>B2</td><td>27</td></tr>
<tr><td>BCAL0163</td><td>>gi|443595598|gb|ELT64173.1| BON domain protein [Burkholderia cenocepacia K56-2Valvano]</td><td>K.VFQYVKPQDAQALQDASPASGAS[+568.212]GAQAAAAPADNATVGAVPDASVQSTPLQPPAPISNSSSVHPGNPK.A</td><td>HexNAc(2)Hex(1)</td><td>Yes</td><td>1435.303</td><td>5</td><td>7172.486</td><td>7172.4731</td><td>1.8</td><td>Specific</td><td>1194.18</td><td>1194.2</td><td>111.85</td><td>Nsco_20191108_BC_ZIC_HILIC_K56_2_typsin_B1.51502.51502.5</td><td>101.1388</td><td>Trypsin</td><td>B1</td><td>28</td></tr>
<tr><td>BCAL0163</td><td>>gi|443595598|gb|ELT64173.1| BON domain protein [Burkholderia cenocepacia K56-2Valvano]</td><td>Y.VKPQDAQALQDASPASGAS[+568.212]GAQAAAAPADNATVGAVPDASVQSTPLQPPAPISNSSSVHPGNPK.A</td><td>HexNAc(2)Hex(1)</td><td>Yes</td><td>1659.8165</td><td>4</td><td>6636.2443</td><td>6635.2143</td><td>4</td><td>NRagged</td><td>963.66</td><td>963.66</td><td>93.72</td><td>Nsco_20191108_BC_ZIC_HILIC_K56_2_typsin_B3.45475.45475.4</td><td>91.0302</td><td>Trypsin</td><td>B3</td><td>29</td></tr>
<tr><td>BCAL0163</td><td>>gi|443595598|gb|ELT64173.1| BON domain protein [Burkholderia cenocepacia K56-2Valvano]</td><td>S.VQSTPLQPPAPISNSS[+568.212]SVHPGNPKAKAQ.-</td><td>HexNAc(2)Hex(1)</td><td>Yes</td><td>852.1833</td><td>4</td><td>3405.7114</td><td>3405.702</td><td>2.8</td><td>Non</td><td>717.1</td><td>717.1</td><td>0</td><td>Nsco_20191108_BC_ZIC_HILIC_K56_2_pepsin_B1.20579.20579.4</td><td>45.1081</td><td>Pepsin</td><td>B1</td><td>30</td></tr>
<tr><td>BCAL0193</td><td>>gi|443590944|gb|ELT59881.1| hypothetical protein BURCENK562V_2433 [Burkholderia cenocepacia K56-2Valvano]</td><td>S.AAAPAPAKKDHSKPKHQLKHHGSKKGQAKAAAAS[+568.212][+100.064]AAGTNDAGTQN.-</td><td>HexNAc(2)Hex(1)<br>100.064</td><td>No</td><td>1025.3224</td><td>5</td><td>5122.5828</td><td>5121.6064</td><td>-5.3</td><td>Non</td><td>459.47</td><td>459.47</td><td>51.91</td><td>Nsco_20191108_BC_ZIC_HILIC_K56_2_pepsin_B1.3536.3536.5</td><td>16.5137</td><td>Pepsin</td><td>B1</td><td>31</td></tr>
</table>

| | | | | | | | | | | | | | | | | | | |
|---|---|---|---|---|---|---|---|---|---|---|---|---|---|---|---|---|---|---|
| BCAL0193 | >gi\|443590944\|gb\|ELT59881.1\| hypothetical protein BURCENK562V_2433 [Burkholderia cenocepacia K56-2Valvano] | P.AADTSAAAPAPAKKDHS[+568.212][+100.064]KPKHQ.L | HexNAc(2)Hex(1) 100.064 | No | 965.8129 | 3 | 2895.4241 | 2895.4334 | -3.2 | Non | 491.4 | 491.4 | 7.58 | Nsco_20191108_BC_ZIC_HILIC_K56_2_thermolysin_B2_.5992.5992.3 | 21.3403 | Thermolysin | B2 | 32 |
| BCAL0193 | >gi\|443590944\|gb\|ELT59881.1\| hypothetical protein BURCENK562V_2433 [Burkholderia cenocepacia K56-2Valvano] | A.AAPAPAKKDHSKPKHQLKHHGSKKGQAKAAAAS[+568.212]AAGT[+568.212]ND.A | HexNAc(2)Hex(1), HexNAc(2)Hex(1) | No | 842.5956 | 6 | 5050.537 | 5047.5091 | 3.5 | Non | 299.37 | 299.37 | 137.37 | Nsco_20191108_BC_ZIC_HILIC_K56_2_pepsin_B3_reinject.3333.3333.6 | 16.2103 | Pepsin | B3 | 33 |
| BCAL0193 | >gi\|443590944\|gb\|ELT59881.1\| hypothetical protein BURCENK562V_2433 [Burkholderia cenocepacia K56-2Valvano] | Q.ASAPAADT[+568.212]SAAAPAPAKK.D | HexNAc(2)Hex(1) | No | 1082.5332 | 2 | 2164.0591 | 2164.0503 | 4.1 | NRagged | 781.84 | 637.11 | 0 | Nsco_20191108_BC_ZIC_HILIC_K56_2_typsin_B1.6825.6825.2 | 22.7563 | Trypsin | B1 | 34 |
| BCAL0193 | >gi\|443590944\|gb\|ELT59881.1\| hypothetical protein BURCENK562V_2433 [Burkholderia cenocepacia K56-2Valvano] | A.DTSAAAPAPAKKDHSKPKHQLKHHGSKKGQAKAAAASAAGTNDAGT[+568.212]QN.- | HexNAc(2)Hex(1) | No | 1066.1395 | 5 | 5326.6685 | 5324.649 | 2.4 | Non | 785.04 | 785.04 | 785.04 | Nsco_20191108_BC_ZIC_HILIC_K56_2_pepsin_B3_reinject.3443.3443.5 | 16.382 | Pepsin | B3 | 35 |
| BCAL0193 | >gi\|443590944\|gb\|ELT59881.1\| hypothetical protein BURCENK562V_2433 [Burkholderia cenocepacia K56-2Valvano] | A.PAADTSAAAPAPAKKDHS[+568.212]KPKHQ.L | HexNAc(2)Hex(1) | Yes (1 of 2) | 724.3645 | 4 | 2894.4362 | 2892.4221 | 2.5 | Non | 283.84 | 283.84 | 8.4 | Nsco_20191108_BC_ZIC_HILIC_K56_2_thermolysin_B2_.6117.6117.4 | 21.5289 | Thermolysin | B2 | 36 |
| BCAL0193 | >gi\|443590944\|gb\|ELT59881.1\| hypothetical protein BURCENK562V_2433 [Burkholderia cenocepacia K56-2Valvano] | T.SAAAPAPAKKDHSKPKHQLKHHGSKKGQAKAAAAS[+568.212]AAGT[+568.212]ND.A | HexNAc(2)Hex(1), HexNAc(2)Hex(1) | No | 868.9396 | 6 | 5208.601 | 5205.5782 | 2.4 | Non | 303.13 | 303.13 | 145.33 | Nsco_20191108_BC_ZIC_HILIC_K56_2_pepsin_B3_reinject.3757.3757.6 | 16.838 | Pepsin | B3 | 37 |
| BCAL0193 | >gi\|443590944\|gb\|ELT59881.1\| hypothetical protein BURCENK562V_2433 [Burkholderia cenocepacia K56-2Valvano] | T.SAAAPAPAKKDHSKPKHQLKHHGSKKGQAKAAAAS[+568.212]AAGTNDAGTQN.- | HexNAc(2)Hex(1) | No | 639.4553 | 8 | 5108.5912 | 5108.5744 | 3.3 | Non | 467.89 | 467.89 | 16.28 | Nsco_20191108_BC_ZIC_HILIC_K56_2_pepsin_B3_reinject.3233.3233.8 | 16.0795 | Pepsin | B3 | 38 |
| BCAL0303 | >gi\|443596149\|gb\|ELT64670.1\| hypothetical protein BURCENK562V_3632 [Burkholderia cenocepacia K56-2Valvano] | K.AADAGGAKPAAGAS[+568.212]AAPAAPAPVAVPASAVSGSAGQ.- | HexNAc(2)Hex(1) | No | 1755.8598 | 2 | 3510.7124 | 3510.7082 | 1.2 | CRagged | 830.5 | 764.54 | 9.2 | Nsco_20191108_BC_ZIC_HILIC_K56_2_typsin_B1.35749.35749.2 | 74.5524 | Trypsin | B1 | 39 |
| BCAL0303 | >gi\|443596149\|gb\|ELT64670.1\| hypothetical protein BURCENK562V_3632 [Burkholderia cenocepacia K56-2Valvano] | A.ADAGGAKPAAGASAAPAAPAPVAVPAS[+568.212]AVS[+568.212][+100.064]GSAGQ.- | HexNAc(2)Hex(1), HexNAc(2)Hex(1) 100.064 | No | 1369.9816 | 3 | 4107.9303 | 4107.9467 | -4 | Non | 410.05 | 410.05 | 0 | Nsco_20191108_BC_ZIC_HILIC_K56_2_typsin_B2.33198.33198.3 | 70.2605 | Trypsin | B2 | 40 |
| BCAL0303 | >gi\|443596149\|gb\|ELT64670.1\| hypothetical protein BURCENK562V_3632 [Burkholderia cenocepacia K56-2Valvano] | S.KAADAGGAKPAAGAS[+568.212]AAPAAPAPVAV.P | HexNAc(2)Hex(1) | Yes | 909.4652 | 3 | 2726.381 | 2726.3731 | 2.9 | Non | 745.72 | 742.58 | 742.58 | Nsco_20191108_BC_ZIC_HILIC_K56_2_pepsin_B1.22517.22517.3 | 48.7792 | Pepsin | B1 | 41 |
| BCAL0303 | >gi\|443596149\|gb\|ELT64670.1\| hypothetical protein BURCENK562V_3632 [Burkholderia cenocepacia K56-2Valvano] | S.KAADAGGAKPAAGAS[+568.212]AAPAAPAPVAVPASAVSGSAGQ.- | HexNAc(2)Hex(1) | No | 1213.9411 | 3 | 3639.8087 | 3638.8032 | 0.6 | Non | 673.04 | 647.56 | 0.38 | Nsco_20191108_BC_ZIC_HILIC_K56_2_pepsin_B1.29892.29892.3 | 63.7666 | Pepsin | B1 | 42 |
| BCAL0303 | >gi\|443596149\|gb\|ELT64670.1\| hypothetical protein BURCENK562V_3632 [Burkholderia cenocepacia K56-2Valvano] | F.LYSKAADAGGAKPAAGASAAPAAPAPVAVPAS[+568.212]AVSGSAGQ.- | HexNAc(2)Hex(1) | No | 1334.6696 | 3 | 4001.9941 | 4001.9826 | 2.9 | CRagged | 289.83 | 289.83 | 7.2 | Nsco_20191108_BC_ZIC_HILIC_K56_2_pepsin_B2.33169.33169.3 | 70.6148 | Pepsin | B2 | 43 |
| BCAL0303 | >gi\|443596149\|gb\|ELT64670.1\| hypothetical protein BURCENK562V_3632 [Burkholderia cenocepacia K56-2Valvano] | L.YSKAADAGGAKPAAGAS[+568.212]AAPAAPAPVAV.P | HexNAc(2)Hex(1) | No | 992.8305 | 3 | 2976.4769 | 2976.4684 | 2.8 | CRagged | 794.53 | 737.06 | 49.7 | Nsco_20191108_BC_ZIC_HILIC_K56_2_pepsin_B3_reinject.24724.24724.3 | 51.6664 | Pepsin | B3 | 44 |

<table>
<tr><td>BCAL0303</td><td>>gi|443596149|gb|ELT64670.1| hypothetical protein BURCENK562V_3632 [Burkholderia cenocepacia K56-2Valvano]</td><td>L.YSKAADAGGAKPAAGAS[+568.212]AAPAAPAPVAVPASAVSG.S</td><td>HexNAc(2)Hex(1)</td><td>No</td><td>1182.9243</td><td>3</td><td>3546.7582</td><td>3545.7493</td><td>1.6</td><td>CRagged</td><td>783.83</td><td>744.61</td><td>16.57</td><td>Nsco_20191108_BC_ZIC_HILIC_K56_2_pepsin_B3_reinject.31280.31280.3</td><td>64.1422</td><td>Pepsin</td><td>B3</td><td>45</td></tr>
<tr><td>BCAL0303</td><td>>gi|443596149|gb|ELT64670.1| hypothetical protein BURCENK562V_3632 [Burkholderia cenocepacia K56-2Valvano]</td><td>L.YSKAADAGGAKPAAGASAAPAAPAPVAVPAS[+568.212]AVSGSAGQ.-</td><td>HexNAc(2)Hex(1)</td><td>No</td><td>1296.9753</td><td>3</td><td>3888.9113</td><td>3888.8985</td><td>3.3</td><td>CRagged</td><td>1045.52</td><td>937.06</td><td>1.2</td><td>Nsco_20191108_BC_ZIC_HILIC_K56_2_pepsin_B3_reinject.31719.31719.3</td><td>64.9573</td><td>Pepsin</td><td>B3</td><td>46</td></tr>
<tr><td>BCAL0332</td><td>>gi|443603144|gb|ELT71169.1| stringent starvation protein B [Burkholderia cenocepacia K56-2Valvano]</td><td>A.FQVDAVAGEGEDSGAFDDDAAQADDAQRDESVSPLAPVADS[+568.212][+100.064]GANEEPSEGADEPPKTDGDGSK.G</td><td>HexNAc(2)Hex(1) 100.064</td><td>No</td><td>1744.5016</td><td>4</td><td>6974.9844</td><td>6973.9813</td><td>0</td><td>NRagged</td><td>822.93</td><td>822.93</td><td>0</td><td>Nsco_20191108_BC_ZIC_HILIC_K56_2_typsin_B2.56297.56297.4</td><td>111.8126</td><td>Trypsin</td><td>B2</td><td>47</td></tr>
<tr><td>BCAL0332</td><td>>gi|443603144|gb|ELT71169.1| stringent starvation protein B [Burkholderia cenocepacia K56-2Valvano]</td><td>D.SGAFDDDAAQADDAQRDES[+568.212]VSPLAPVADSGANEEPS[+568.212][+100.064]EGADEPPKT[+568.212][+100.064]DGDGSK.G</td><td>HexNAc(2)Hex(1), HexNAc(2)Hex(1) 100.064,HexNAc(2)Hex(1) 100.064</td><td>No</td><td>1748.9992</td><td>4</td><td>6992.9751</td><td>6992.9484</td><td>3.8</td><td>Non</td><td>593.37</td><td>593.37</td><td>0</td><td>Nsco_20191108_BC_ZIC_HILIC_K56_2_thermolysin_B3.58745.58745.4</td><td>116.544</td><td>Thermolysin</td><td>B3</td><td>48</td></tr>
<tr><td>BCAL0348</td><td>>gi|443597060|gb|ELT65512.1| type VI secretion-associated protein, ImpA family [Burkholderia cenocepacia K56-2Valvano]</td><td>Q.AVQQAQPERIEPVFGQPIQT[+568.212]EET[+568.212][+100.064]HVQQQTASRPPVTQTIAGIQNR.A</td><td>HexNAc(2)Hex(1), HexNAc(2)Hex(1) 100.064</td><td>No</td><td>1250.6227</td><td>5</td><td>6249.0843</td><td>6243.0702</td><td>-1</td><td>Non</td><td>729.84</td><td>729.84</td><td>0</td><td>Nsco_20191108_BC_ZIC_HILIC_K56_2_thermolysin_B3.39945.39945.5</td><td>82.0146</td><td>Thermolysin</td><td>B3</td><td>49</td></tr>
<tr><td>BCAL0348</td><td>>gi|443597060|gb|ELT65512.1| type VI secretion-associated protein, ImpA family [Burkholderia cenocepacia K56-2Valvano]</td><td>T.GSAPHTQAVQQAQPERIEPVFGQPIQTEET[+568.212]HVQQQTASRPPVTQTIAGIQNR.A</td><td>HexNAc(2)Hex(1)</td><td>No</td><td>1251.6292</td><td>5</td><td>6254.1166</td><td>6253.1032</td><td>1.6</td><td>Non</td><td>469.57</td><td>469.57</td><td>0</td><td>Nsco_20191108_BC_ZIC_HILIC_K56_2_thermolysin_B3.44121.44121.5</td><td>89.5488</td><td>Thermolysin</td><td>B3</td><td>50</td></tr>
<tr><td>BCAL0358</td><td>>gi|443598967|gb|ELT67284.1| peptidase family M1 [Burkholderia cenocepacia K56-2Valvano]</td><td>S.SAGAPHTGSSPT[+568.212]VAAPPSAS[+568.212]NVDK.S</td><td>HexNAc(2)Hex(1), HexNAc(2)Hex(1)</td><td>No</td><td>1115.1758</td><td>3</td><td>3343.5128</td><td>3342.4966</td><td>3.8</td><td>Non</td><td>708.56</td><td>554.71</td><td>3.78</td><td>Nsco_20191108_BC_ZIC_HILIC_K56_2_thermolysin_B3.13859.13859.3</td><td>34.0672</td><td>Thermolysin</td><td>B3</td><td>51</td></tr>
<tr><td>BCAL0426</td><td>>gi|443602087|gb|ELT70191.1| membrane protein insertase, YidC/Oxa1 family [Burkholderia cenocepacia K56-2Valvano]</td><td>S.ATHTAPAAAGGASGT[+568.212]GATTAGDVPAA.A</td><td>HexNAc(2)Hex(1)</td><td>No</td><td>1360.6226</td><td>2</td><td>2720.2378</td><td>2720.2381</td><td>-0.1</td><td>Non</td><td>616.62</td><td>434.57</td><td>0</td><td>Nsco_20191108_BC_ZIC_HILIC_K56_2_thermolysin_B1.19339.19339.2</td><td>41.3403</td><td>Thermolysin</td><td>B1</td><td>52</td></tr>
<tr><td>BCAL0426</td><td>>gi|443602087|gb|ELT70191.1| membrane protein insertase, YidC/Oxa1 family [Burkholderia cenocepacia K56-2Valvano]</td><td>R.DHGRPSMFFPSATHTAPAAAGGAS[+568.212]GTGATTAGDVPAAAAGAAPSTTAPAAQAQLVK.F</td><td>HexNAc(2)Hex(1)</td><td>Yes</td><td>1137.5507</td><td>5</td><td>5683.7242</td><td>5683.7093</td><td>2.6</td><td>Specific</td><td>1336.17</td><td>1307.6</td><td>53.87</td><td>Nsco_20191108_BC_ZIC_HILIC_K56_2_typsin_B1.41182.41182.5</td><td>84.0593</td><td>Trypsin</td><td>B1</td><td>53</td></tr>
<tr><td>BCAL0426</td><td>>gi|443602087|gb|ELT70191.1| membrane protein insertase, YidC/Oxa1 family [Burkholderia cenocepacia K56-2Valvano]</td><td>F.FPSATHTAPAAAGGASGTGATT[+568.212][+100.064]AGDVPAAAAGAAPSTTAPAAQ.A</td><td>HexNAc(2)Hex(1) 100.064</td><td>No</td><td>1080.2585</td><td>4</td><td>4318.0124</td><td>4317.028</td><td>-4.4</td><td>CRagged</td><td>596.73</td><td>516.78</td><td>0</td><td>Nsco_20191108_BC_ZIC_HILIC_K56_2_pepsin_B2.36657.36657.4</td><td>78.7405</td><td>Pepsin</td><td>B2</td><td>54</td></tr>
<tr><td>BCAL0426</td><td>>gi|443602087|gb|ELT70191.1| membrane protein insertase, YidC/Oxa1 family [Burkholderia cenocepacia K56-2Valvano]</td><td>T.HTAPAAAGGASGTGATT[+568.212]AGDVPAAAAGAAPSTTAPAAQ.A</td><td>HexNAc(2)Hex(1)</td><td>No</td><td>1238.5833</td><td>3</td><td>3713.7354</td><td>3713.726</td><td>2.5</td><td>Non</td><td>804.69</td><td>742.1</td><td>1.2</td><td>Nsco_20191108_BC_ZIC_HILIC_K56_2_pepsin_B3_reinject.30629.30629.3</td><td>62.8704</td><td>Pepsin</td><td>B3</td><td>55</td></tr>
<tr><td>BCAL0479</td><td>>gi|443597239|gb|ELT65680.1| penicillin-binding protein 2, partial [Burkholderia cenocepacia K56-2Valvano]</td><td>A.AASAT[+568.212]EPVS[+568.212]AP.V</td><td>HexNAc(2)Hex(1), HexNAc(2)Hex(1)</td><td>No</td><td>1068.9668</td><td>2</td><td>2136.9264</td><td>2136.9177</td><td>4.1</td><td>Non</td><td>375.8</td><td>192.35</td><td>62.46</td><td>Nsco_20191108_BC_ZIC_HILIC_K56_2_thermolysin_B1.18700.18700.2</td><td>40.3138</td><td>Thermolysin</td><td>B1</td><td>56</td></tr>
<tr><td>BCAL0520</td><td>>gi|443592535|gb|ELT61330.1| hypothetical protein BURCENK562V_3076, partial [Burkholderia cenocepacia K56-2Valvano]</td><td>A.ARGSGAS[+568.212]AAGADAAAAT[+568.212]SAVP.F</td><td>HexNAc(2)Hex(1), HexNAc(2)Hex(1)</td><td>No</td><td>1433.6428</td><td>2</td><td>2866.2784</td><td>2866.2695</td><td>3.1</td><td>Non</td><td>357.63</td><td>283.69</td><td>2.4</td><td>Nsco_20191108_BC_ZIC_HILIC_K56_2_thermolysin_B1.20407.20407.2</td><td>43.3508</td><td>Thermolysin</td><td>B1</td><td>57</td></tr>
</table>

| | | | | | | | | | | | | | | | | | | |
|---|---|---|---|---|---|---|---|---|---|---|---|---|---|---|---|---|---|---|
| BCAL0520 | >gi\|443592535\|gb\|ELT61330.1\| hypothetical protein BURCENK562V_3076, partial [Burkholderia cenocepacia K56-2Valvano] | R.GSGASAAGADAAAAT[+568.212]S[+568.212]AVPFAQTLK.Q | HexNAc(2)Hex(1), HexNAc(2)Hex(1) | No | 1664.2688 | 2 | 3327.5304 | 3327.5221 | 2.5 | Specific | 968.77 | 869.46 | 14.4 | Nsco_20191108_BC_ZIC_HILIC_K56_2_typsin_B2.39446.39446.2 | 81.8591 | Trypsin | B2 | 58 |
| BCAL0525 | >gi\|443597648\|gb\|ELT66063.1\| flagellar M-ring protein FliF, partial [Burkholderia cenocepacia K56-2Valvano] | L.SNTPPQPAS[+568.212]APIVAGNGQNAPQTTPVSD.R | HexNAc(2)Hex(1) | Yes | 1095.5191 | 3 | 3284.5427 | 3284.5288 | 4.2 | CRagged | 535.25 | 449.78 | 35.51 | Nsco_20191108_BC_ZIC_HILIC_K56_2_pepsin_B3_reinject.37980.37980.3 | 78.3801 | Pepsin | B3 | 59 |
| BCAL0525 | >gi\|443597648\|gb\|ELT66063.1\| flagellar M-ring protein FliF, partial [Burkholderia cenocepacia K56-2Valvano] | R.SQQTSSATELAQGGASGVPGALSNT[+68.212]PPQPAS[+568.212]APIVAGNGQNAPQTTPVSDRK.D | HexNAc(2)Hex(1), HexNAc(2)Hex(1) | No | 1534.984 | 4 | 6136.9142 | 6134.9019 | 0.9 | Specific | 711.45 | 711.45 | 0 | Nsco_20191108_BC_ZIC_HILIC_K56_2_typsin_B2.40293.40293.4 | 83.3257 | Trypsin | B2 | 60 |
| BCAL0544 | >gi\|443601095\|gb\|ELT69255.1\| ABC transporter, substrate-binding protein, family 5 [Burkholderia cenocepacia K56-2Valvano] | K.KAILESVYQGAGQAAS[+568.212]APMPPTQWSYDK.N | HexNAc(2)Hex(1) | No | 1188.2355 | 3 | 3562.692 | 3562.6782 | 3.9 | Specific | 898.13 | 898.13 | 6.78 | Nsco_20191108_BC_ZIC_HILIC_K56_2_typsin_B2.47878.47878.3 | 96.7111 | Trypsin | B2 | 61 |
| BCAL0678 | >gi\|443605648\|gb\|ELT73487.1\| sporulation and cell division repeat protein [Burkholderia cenocepacia K56-2Valvano] | Q.AAAQKQQQQQQAANT[+568.212]PKPTSSAT.A | HexNAc(2)Hex(1) | No | 984.477 | 3 | 2951.4163 | 2951.4076 | 3 | Non | 463.13 | 421.62 | 5.62 | Nsco_20191108_BC_ZIC_HILIC_K56_2_thermolysin_B1.4818.4818.3 | 18.5162 | Thermolysin | B1 | 62 |
| BCAL0678 | >gi\|443605648\|gb\|ELT73487.1\| sporulation and cell division repeat protein [Burkholderia cenocepacia K56-2Valvano] | Q.AAAQKQQQQQQAANT[+568.212]PKPTSSATA.A | HexNAc(2)Hex(1) | No | 1008.1566 | 3 | 3022.4553 | 3022.4447 | 3.5 | Non | 603.81 | 562.68 | 1.48 | Nsco_20191108_BC_ZIC_HILIC_K56_2_thermolysin_B2_.5231.5231.3 | 20.1389 | Thermolysin | B2 | 63 |
| BCAL0678 | >gi\|443605648\|gb\|ELT73487.1\| sporulation and cell division repeat protein [Burkholderia cenocepacia K56-2Valvano] | F.AAQKQAQQAAAQKQQQQQQAANT[568.212]PKPTSSATAAA.A | HexNAc(2)Hex(1) | No | 1339.9922 | 3 | 4017.962 | 4017.9596 | 0.6 | CRagged | 527.89 | 389.69 | 9.6 | Nsco_20191108_BC_ZIC_HILIC_K56_2_pepsin_B3_reinject.9448.9448.3 | 26.3294 | Pepsin | B3 | 64 |
| BCAL0678 | >gi\|443605648\|gb\|ELT73487.1\| sporulation and cell division repeat protein [Burkholderia cenocepacia K56-2Valvano] | A.AAQKQQQQQQAANT[+568.212]PKPTSSATAAA.A | HexNAc(2)Hex(1) | No | 1032.1697 | 3 | 3094.4945 | 3093.4818 | 3 | Non | 516.81 | 460.86 | 14.7 | Nsco_20191108_BC_ZIC_HILIC_K56_2_pepsin_B2.6966.6966.3 | 22.9774 | Pepsin | B2 | 65 |
| BCAL0678 | >gi\|443605648\|gb\|ELT73487.1\| sporulation and cell division repeat protein [Burkholderia cenocepacia K56-2Valvano] | Q.KQQQQQQAANTPKPTS[+568.212]SATAAAAAKPPTANDANTGYFLQVGAY.K | HexNAc(2)Hex(1) | No | 1244.6036 | 4 | 4975.3927 | 4973.3923 | -1.3 | Non | 586.51 | 547.75 | 15.42 | Nsco_20191108_BC_ZIC_HILIC_K56_2_thermolysin_B3.37078.37078.4 | 76.7191 | Thermolysin | B3 | 66 |
| BCAL0678 | >gi\|443605648\|gb\|ELT73487.1\| sporulation and cell division repeat protein [Burkholderia cenocepacia K56-2Valvano] | Q.KQQQQQQAANTPKPTS[+568.212]SATAAAAAKPPTANDANTGYFLQVGAYK.T | HexNAc(2)Hex(1) | No | 1021.106 | 5 | 5101.501 | 5101.4873 | 2.7 | NRagged | 625.67 | 625.67 | 4.2 | Nsco_20191108_BC_ZIC_HILIC_K56_2_typsin_B1.30503.30503.5 | 65.5598 | Trypsin | B1 | 67 |
| BCAL0678 | >gi\|443605648\|gb\|ELT73487.1\| sporulation and cell division repeat protein [Burkholderia cenocepacia K56-2Valvano] | Q.KQQQQQQAANTPKPTS[+568.212]SATAAAAAKPPTANDANTGYFLQVGAYKTEGDAEQQR.A | HexNAc(2)Hex(1) | No | 1020.3292 | 6 | 6116.939 | 6115.9239 | 1.9 | NRagged | 740.63 | 729.18 | 5 | Nsco_20191108_BC_ZIC_HILIC_K56_2_typsin_B2.29453.29453.6 | 63.378 | Trypsin | B2 | 68 |
| BCAL0678 | >gi\|443605648\|gb\|ELT73487.1\| sporulation and cell division repeat protein [Burkholderia cenocepacia K56-2Valvano] | S.KVAPPPADNGAS[+568.212]QPQQFDPNRA.L | HexNAc(2)Hex(1) | Yes | 958.4554 | 3 | 2873.3517 | 2873.3436 | 2.8 | Non | 630.58 | 630.58 | 630.58 | Nsco_20191108_BC_ZIC_HILIC_K56_2_pepsin_B3_reinject.18300.18300.3 | 40.8167 | Pepsin | B3 | 69 |
| BCAL0678 | >gi\|443605648\|gb\|ELT73487.1\| sporulation and cell division repeat protein [Burkholderia cenocepacia K56-2Valvano] | S.KVAPPPADNGAS[+568.212]QPQQFDPNRAL.Q | HexNAc(2)Hex(1) | Yes | 996.151 | 3 | 2986.4384 | 2986.4276 | 3.6 | NRagged | 681.76 | 616.02 | 616.02 | Nsco_20191108_BC_ZIC_HILIC_K56_2_pepsin_B1.27173.27173.3 | 57.9729 | Pepsin | B1 | 70 |
| BCAL0678 | >gi\|443605648\|gb\|ELT73487.1\| sporulation and cell division repeat protein [Burkholderia cenocepacia K56-2Valvano] | S.KVAPPPADNGAS[+568.212]QPQQFDPNRALQ.G | HexNAc(2)Hex(1) | No | 1038.8365 | 3 | 3114.4948 | 3114.4862 | 2.8 | Non | 684.78 | 621.45 | 621.45 | Nsco_20191108_BC_ZIC_HILIC_K56_2_pepsin_B1.26704.26704.3 | 56.8912 | Pepsin | B1 | 71 |
| BCAL0678 | >gi\|443605648\|gb\|ELT73487.1\| sporulation and cell division repeat protein [Burkholderia cenocepacia K56-2Valvano] | S.KVAPPPADNGAS[+568.212]QPQQFDPNRALQG.K | HexNAc(2)Hex(1) | Yes | 1057.8449 | 3 | 3171.5201 | 3171.5077 | 3.9 | Non | 621.38 | 611.92 | 611.92 | Nsco_20191108_BC_ZIC_HILIC_K56_2_pepsin_B1.27694.27694.3 | 59.0443 | Pepsin | B1 | 72 |
| BCAL0678 | >gi\|443605648\|gb\|ELT73487.1\| sporulation and cell division repeat protein [Burkholderia cenocepacia K56-2Valvano] | S.KVAPPPADNGAS[+568.212]QPQQFDPNRALQGKTPGQPVPQA.A | HexNAc(2)Hex(1) | Yes | 1044.5223 | 4 | 4175.0672 | 4175.0527 | 3.5 | Non | 743.18 | 743.18 | 488.41 | Nsco_20191108_BC_ZIC_HILIC_K56_2_pepsin_B3_reinject.31852.31852.4 | 65.2064 | Pepsin | B3 | 73 |

| | | | | | | | | | | | | | | | | | | |
|---|---|---|---|---|---|---|---|---|---|---|---|---|---|---|---|---|---|---|
| BCAL0678 | >gi\|443605648\|gb\|ELT73487.1\| sporulation and cell division repeat protein [Burkholderia cenocepacia K56-2Valvano] | S.KVAPPPADNGAS[+568.212]QPQQFDPNRALQGKTPGQPVPQAAQPAPPNTA.P | HexNAc(2)Hex(1) | Yes | 1256.3777 | 4 | 5022.4889 | 5022.4716 | 3.5 | Non | 665.41 | 665.41 | 320.9 | Nsco_20191108_BC_ZIC_HILIC_K56_2_pepsin_B1.33145.33145.4 | 71.3551 | Pepsin | B1 | 74 |
| BCAL0678 | >gi\|443605648\|gb\|ELT73487.1\| sporulation and cell division repeat protein [Burkholderia cenocepacia K56-2Valvano] | S.KVAPPPADNGAS[+568.212]QPQQFDPNRALQGKTPGQPVPQAAQPAPPNTAPGQ.A | HexNAc(2)Hex(1) | Yes | 1326.9113 | 4 | 5304.6232 | 5304.6044 | 3.6 | Non | 836.81 | 836.81 | 393.47 | Nsco_20191108_BC_ZIC_HILIC_K56_2_pepsin_B3_reinject.35229.35229.4 | 72.3622 | Pepsin | B3 | 75 |
| BCAL0678 | >gi\|443605648\|gb\|ELT73487.1\| sporulation and cell division repeat protein [Burkholderia cenocepacia K56-2Valvano] | S.KVAPPPADNGAS[+568.212]QPQQFDPNRALQGKTPGQPVPQAAQPAPPNTAPGQA.A | HexNAc(2)Hex(1) | Yes | 1075.9376 | 5 | 5375.659 | 5375.6415 | 3.3 | Non | 957.7 | 957.7 | 499.25 | Nsco_20191108_BC_ZIC_HILIC_K56_2_pepsin_B2.34545.34545.5 | 73.7842 | Pepsin | B2 | 76 |
| BCAL0678 | >gi\|443605648\|gb\|ELT73487.1\| sporulation and cell division repeat protein [Burkholderia cenocepacia K56-2Valvano] | S.KVAPPPADNGAS[+568.212]QPQQFDPNRALQGKTPGQPVPQAAQPAPPNTAPGQAA.N | HexNAc(2)Hex(1) | Yes | 1362.4287 | 4 | 5446.693 | 5446.6786 | 2.7 | Non | 807.9 | 807.9 | 435.58 | Nsco_20191108_BC_ZIC_HILIC_K56_2_pepsin_B1.33981.33981.4 | 73.5497 | Pepsin | B1 | 77 |
| BCAL0678 | >gi\|443605648\|gb\|ELT73487.1\| sporulation and cell division repeat protein [Burkholderia cenocepacia K56-2Valvano] | S.KVAPPPADNGAS[+568.212]QPQQFDPNRALQGKTPGQPVPQAAQPAPPNTAPGQAAN.Q | HexNAc(2)Hex(1) | Yes | 1113.3529 | 5 | 5562.7354 | 5560.7215 | 1.3 | Non | 1014.59 | 1000.8 | 483.05 | Nsco_20191108_BC_ZIC_HILIC_K56_2_pepsin_B1.33588.33588.5 | 72.4426 | Pepsin | B1 | 78 |
| BCAL0678 | >gi\|443605648\|gb\|ELT73487.1\| sporulation and cell division repeat protein [Burkholderia cenocepacia K56-2Valvano] | S.KVAPPPADNGAS[+568.212]QPQQFDPNRALQGKTPGQPVPQAAQPAPPNTAPGQAANQ.T | HexNAc(2)Hex(1) | Yes | 1422.9518 | 4 | 5688.7853 | 5688.7801 | 0.9 | Non | 680.89 | 672.05 | 302.36 | Nsco_20191108_BC_ZIC_HILIC_K56_2_pepsin_B3_reinject.35184.35184.4 | 72.264 | Pepsin | B3 | 79 |
| BCAL0678 | >gi\|443605648\|gb\|ELT73487.1\| sporulation and cell division repeat protein [Burkholderia cenocepacia K56-2Valvano] | S.KVAPPPADNGASQPQQFDPNRALQGKT[+568.212][+100.064]PGQPVPQAAQPAPPNTAPGQAANQTQGGL.L | HexNAc(2)Hex(1) 100.064 | No | 1562.0194 | 4 | 6245.0558 | 6245.0773 | -3.4 | NRagged | 730.82 | 730.82 | 3.6 | Nsco_20191108_BC_ZIC_HILIC_K56_2_pepsin_B3_reinject.39257.39257.4 | 81.4775 | Pepsin | B3 | 80 |
| BCAL0678 | >gi\|443605648\|gb\|ELT73487.1\| sporulation and cell division repeat protein [Burkholderia cenocepacia K56-2Valvano] | T.PKPT[+568.212]S[+568.212]S[+568.212][+100.064]ATAAAAAKPPTANDANTGYFLQVGAYKTEGDAEQQR.A | HexNAc(2)Hex(1), HexNAc(2)Hex(1), HexNAc(2)Hex(1) 100.064 | No | 1525.4624 | 4 | 6098.8278 | 6098.7998 | 4.6 | Non | 496.51 | 496.51 | 0 | Nsco_20191108_BC_ZIC_HILIC_K56_2_thermolysin_B3.40382.40382.4 | 82.8135 | Thermolysin | B3 | 81 |
| BCAL0678 | >gi\|443605648\|gb\|ELT73487.1\| sporulation and cell division repeat protein [Burkholderia cenocepacia K56-2Valvano] | K.PTSSATAAAAAKPPT[+568.212]ANDANTGYFLQVGAYK.T | HexNAc(2)Hex(1) | No | 1208.9186 | 3 | 3624.7412 | 3622.7283 | 1.7 | Specific | 915.91 | 852.98 | 1.2 | Nsco_20191108_BC_ZIC_HILIC_K56_2_typsin_B1.39532.39532.3 | 81.3052 | Trypsin | B1 | 82 |
| BCAL0678 | >gi\|443605648\|gb\|ELT73487.1\| sporulation and cell division repeat protein [Burkholderia cenocepacia K56-2Valvano] | Q.QAANTPKPTSSATAAAAAKPPT[+568.212][+100.064]ANDANT[+568.212]GYFLQVGAYK.T | HexNAc(2)Hex(1) 100.064,HexNAc(2)Hex(1) | No | 1251.8569 | 4 | 5004.4059 | 5001.375 | 4.2 | NRagged | 594.1 | 594.1 | 0 | Nsco_20191108_BC_ZIC_HILIC_K56_2_typsin_B2.34826.34826.4 | 73.3709 | Trypsin | B2 | 83 |
| BCAL0678 | >gi\|443605648\|gb\|ELT73487.1\| sporulation and cell division repeat protein [Burkholderia cenocepacia K56-2Valvano] | Q.QAANTPKPTSS[+568.212][+100.064]AT[+568.212][+100.064]AAAAAKPPTANDANTGYFLQVGAYKTEGDAEQQR.A | HexNAc(2)Hex(1) 100.064,HexNAc(2)Hex(1) 100.064 | No | 1223.9764 | 5 | 6115.8531 | 6115.8756 | -3.7 | NRagged | 820.45 | 820.45 | 0 | Nsco_20191108_BC_ZIC_HILIC_K56_2_typsin_B2.34844.34844.5 | 73.4024 | Trypsin | B2 | 84 |
| BCAL0678 | >gi\|443605648\|gb\|ELT73487.1\| sporulation and cell division repeat protein [Burkholderia cenocepacia K56-2Valvano] | A.QKQQQQQQAANTPKPT[+568.212]SSATAAA.A | HexNAc(2)Hex(1) | No | 1476.2091 | 2 | 2951.4109 | 2951.4076 | 1.1 | Non | 588.4 | 506.86 | 0.59 | Nsco_20191108_BC_ZIC_HILIC_K56_2_pepsin_B1.7372.7372.2 | 22.8943 | Pepsin | B1 | 85 |
| BCAL0678 | >gi\|443605648\|gb\|ELT73487.1\| sporulation and cell division repeat protein [Burkholderia cenocepacia K56-2Valvano] | Q.QQAANT[+568.212]PKPTS[+568.212]SATAAAAAKPPTANDANTGYFLQVGAYK.T | HexNAc(2)Hex(1), HexNAc(2)Hex(1) | No | 1258.1035 | 4 | 5029.3922 | 5029.3696 | 4.5 | NRagged | 490.84 | 465.88 | 4.32 | Nsco_20191108_BC_ZIC_HILIC_K56_2_typsin_B2.36809.36809.4 | 77.0609 | Trypsin | B2 | 86 |
| BCAL0678 | >gi\|443605648\|gb\|ELT73487.1\| sporulation and cell division repeat protein [Burkholderia cenocepacia K56-2Valvano] | Q.QQAANTPKPTSS[+568.212]ATAAAAAKPPT[+568.212]ANDANTGYFLQVGAYKTEGDAEQQR.A | HexNAc(2)Hex(1), HexNAc(2)Hex(1) | No | 1209.9703 | 5 | 6045.8226 | 6043.8062 | 1.6 | NRagged | 854.96 | 854.96 | 0 | Nsco_20191108_BC_ZIC_HILIC_K56_2_typsin_B2.33550.33550.5 | 70.9351 | Trypsin | B2 | 87 |
| BCAL0678 | >gi\|443605648\|gb\|ELT73487.1\| sporulation and cell division repeat protein [Burkholderia cenocepacia K56-2Valvano] | K.QQQQQQAANTPKPTS[+568.212][+100.064]SAT[+568.212][+100.064]AAAAAKPPTANDANTGYFL.Q | HexNAc(2)Hex(1) 100.064,HexNAc(2)Hex(1) 100.064 | No | 1275.8588 | 4 | 5100.4132 | 5095.388 | 1.7 | NRagged | 189.07 | 189.07 | 0 | Nsco_20191108_BC_ZIC_HILIC_K56_2_thermolysin_B3.37083.37083.4 | 76.7329 | Thermolysin | B3 | 88 |
| BCAL0678 | >gi\|443605648\|gb\|ELT73487.1\| sporulation and cell division repeat protein [Burkholderia cenocepacia K56-2Valvano] | K.QQQQQQAANTPKPTSS[+568.212]ATAAAAAKPPTANDANTGYFLQV.G | HexNAc(2)Hex(1) | No | 1519.0698 | 3 | 4555.1947 | 4554.1754 | 3.5 | Non | 650.23 | 592.01 | 0 | Nsco_20191108_BC_ZIC_HILIC_K56_2_thermolysin_B3.38672.38672.3 | 79.7172 | Thermolysin | B3 | 89 |

| | | | | | | | | | | | | | | | | | | |
|---|---|---|---|---|---|---|---|---|---|---|---|---|---|---|---|---|---|---|
| BCAL0678 | >gi\|443605648\|gb\|ELT73487.1\| sporulation and cell division repeat protein [Burkholderia cenocepacia K56-2Valvano] | K.QQQQQQAANTPKPTS[+568.212]SATAAAAAKPPTANDANTGYFLQVGAYK.T | HexNAc(2)Hex(1) | Yes | 1244.1068 | 4 | 4973.4054 | 4973.3923 | 2.6 | Specific | 1261.39 | 1149.3 | 13.15 | Nsco_20191108_BC_ZIC_HILIC_K56_2_typsin_B3.34982.34982.4 | 72.4325 | Trypsin | B3 | 90 |
| BCAL0678 | >gi\|443605648\|gb\|ELT73487.1\| sporulation and cell division repeat protein [Burkholderia cenocepacia K56-2Valvano] | K.QQQQQQAANTPKPTS[+568.212]SATAAAAAKPPTANDANTGYFLQVGAYKT[+568.212][+100.064]EGDA.E | HexNAc(2)Hex(1), HexNAc(2)Hex(1) 100.064 | No | 1223.7767 | 5 | 6114.8546 | 6114.8437 | 1.8 | Non | 321.1 | 321.1 | 4.94 | Nsco_20191108_BC_ZIC_HILIC_K56_2_thermolysin_B3.36187.36187.5 | 74.9504 | Thermolysin | B3 | 91 |
| BCAL0678 | >gi\|443605648\|gb\|ELT73487.1\| sporulation and cell division repeat protein [Burkholderia cenocepacia K56-2Valvano] | K.QQQQQQAANTPKPTSSATAAAAAKPPTANDANT[+568.212]GYFLQVGAYKTEGDAEQQR.A | HexNAc(2)Hex(1) | No | 1199.1738 | 5 | 5991.84 | 5987.829 | -0.4 | Specific | 1367.58 | 458.19 | 0 | Nsco_20191108_BC_ZIC_HILIC_K56_2_typsin_B2.33883.33883.5 | 71.5446 | Trypsin | B2 | 92 |
| BCAL0678 | >gi\|443605648\|gb\|ELT73487.1\| sporulation and cell division repeat protein [Burkholderia cenocepacia K56-2Valvano] | K.VAPPPADNGAS[+568.212]QPQQFDPNR.A | HexNAc(2)Hex(1) | No | 1337.6141 | 2 | 2674.2208 | 2674.2115 | 3.5 | Specific | 607.46 | 540.04 | 540.04 | Nsco_20191108_BC_ZIC_HILIC_K56_2_typsin_B2.21592.21592.2 | 48.5971 | Trypsin | B2 | 93 |
| BCAL0678 | >gi\|443605648\|gb\|ELT73487.1\| sporulation and cell division repeat protein [Burkholderia cenocepacia K56-2Valvano] | K.VAPPPADNGAS[+568.212]QPQQFDPNRA.L | HexNAc(2)Hex(1) | Yes | 1373.1335 | 2 | 2745.2597 | 2745.2486 | 4 | Non | 430.75 | 319.55 | 319.55 | Nsco_20191108_BC_ZIC_HILIC_K56_2_thermolysin_B1.25115.25115.2 | 52.1185 | Thermolysin | B1 | 94 |
| BCAL0678 | >gi\|443605648\|gb\|ELT73487.1\| sporulation and cell division repeat protein [Burkholderia cenocepacia K56-2Valvano] | K.VAPPPADNGAS[+568.212]QPQQFDPNRALQGK.T | HexNAc(2)Hex(1) | No | 1057.8439 | 3 | 3171.5171 | 3171.5077 | 3 | Specific | 699.57 | 699.57 | 699.57 | Nsco_20191108_BC_ZIC_HILIC_K56_2_typsin_B3.27804.27804.3 | 60.0678 | Trypsin | B3 | 95 |
| BCAL0678 | >gi\|443605648\|gb\|ELT73487.1\| sporulation and cell division repeat protein [Burkholderia cenocepacia K56-2Valvano] | K.VAPPPADNGASQPQQFDPNRALQGKT[+568.212][+100.064]PGQPVPQAAQPAPPNTAPGQA.A | HexNAc(2)Hex(1) 100.064 | No | 1337.9092 | 4 | 5348.6149 | 5347.6105 | 0.2 | Non | 207.24 | 207.24 | 2.56 | Nsco_20191108_BC_ZIC_HILIC_K56_2_pepsin_B3_reinject.40181.40181.4 | 83.8184 | Pepsin | B3 | 96 |
| BCAL0678 | >gi\|443605648\|gb\|ELT73487.1\| sporulation and cell division repeat protein [Burkholderia cenocepacia K56-2Valvano] | F.VSKVAPPPADNGAS[+568.212]QPQQFDPNRA.L | HexNAc(2)Hex(1) | No | 1020.8231 | 3 | 3060.4548 | 3059.444 | 2.4 | CRagged | 557.21 | 180.84 | 80.42 | Nsco_20191108_BC_ZIC_HILIC_K56_2_pepsin_B2.20692.20692.3 | 45.6549 | Pepsin | B2 | 97 |
| BCAL0678 | >gi\|443605648\|gb\|ELT73487.1\| sporulation and cell division repeat protein [Burkholderia cenocepacia K56-2Valvano] | F.VSKVAPPPADNGAS[+568.212]QPQQFDPNRALQGKTPGQPVPQAAQPAPPNTAPGQA.A | HexNAc(2)Hex(1) | No | 1391.1888 | 4 | 5561.7336 | 5561.7419 | -1.5 | CRagged | 554.99 | 554.99 | 202.81 | Nsco_20191108_BC_ZIC_HILIC_K56_2_pepsin_B2.35137.35137.4 | 75.2139 | Pepsin | B2 | 98 |
| BCAL0678 | >gi\|443605648\|gb\|ELT73487.1\| sporulation and cell division repeat protein [Burkholderia cenocepacia K56-2Valvano] | F.VSKVAPPPADNGASQPQQFDPNRALQGKT[+568.212]PGQPVPQAAQPAPPNTAPGQAAN.Q | HexNAc(2)Hex(1) | No | 1437.7139 | 4 | 5747.8337 | 5746.822 | 1.5 | CRagged | 357.31 | 287.71 | 3.6 | Nsco_20191108_BC_ZIC_HILIC_K56_2_pepsin_B3_reinject.35940.35940.4 | 73.9241 | Pepsin | B3 | 99 |
| BCAL0738 | >gi\|443604531\|gb\|ELT72458.1\| peptidase, S41 family [Burkholderia cenocepacia K56-2Valvano] | K.ASGASAAKPASAPKPAS[+568.212]APK.- | HexNAc(2)Hex(1) | No | 778.3971 | 3 | 2333.1768 | 2333.1718 | 2.1 | Specific | 812.76 | 587.88 | 4.9 | Nsco_20191108_BC_ZIC_HILIC_K56_2_typsin_B3.4955.4955.3 | 19.5407 | Trypsin | B3 | 100 |
| BCAL0749 | >gi\|443604502\|gb\|ELT72429.1\| cytochrome c oxidase, subunit II [Burkholderia cenocepacia K56-2Valvano] | G.AATAAAPAEAASAPAQAASGAEQPAAAASAALS[+568.212][+100.064]T[+568.212][+100.064]IYFETGK.S | HexNAc(2)Hex(1) 100.064,HexNAc(2)Hex(1) 100.064 | No | 1691.4734 | 3 | 5072.4055 | 5069.3863 | 1.8 | Non | 287.24 | 225.55 | 1.2 | Nsco_20191108_BC_ZIC_HILIC_K56_2_thermolysin_B3.63162.63162.3 | 125.8637 | Thermolysin | B3 | 101 |
| BCAL0749 | >gi\|443604502\|gb\|ELT72429.1\| cytochrome c oxidase, subunit II [Burkholderia cenocepacia K56-2Valvano] | T.AGAATAAAPAEAASAPAQAAS[+568.212][+100.064]GAEQPAAAAS[+568.212]AALST[+568.212]IYFETGK.S | HexNAc(2)Hex(1) 100.064,HexNAc(2)Hex(1),HexNAc(2)Hex(1) | No | 1418.1528 | 4 | 5669.5895 | 5665.5924 | -2.9 | Non | 168.77 | 168.77 | 0 | Nsco_20191108_BC_ZIC_HILIC_K56_2_thermolysin_B3.61428.61428.4 | 122.1804 | Thermolysin | B3 | 102 |
| BCAL0749 | >gi\|443604502\|gb\|ELT72429.1\| cytochrome c oxidase, subunit II [Burkholderia cenocepacia K56-2Valvano] | R.NGKLPEDTAGAAT[+568.212]AAAPAEAASAPAQAAS[+568.212]GAEQPAAAASAALSTIYFETGK.S | HexNAc(2)Hex(1), HexNAc(2)Hex(1) | Yes (1 of 2) | 1463.693 | 4 | 5851.7502 | 5851.7303 | 3.4 | Specific | 1107.43 | 1017.6 | 111.99 | Nsco_20191108_BC_ZIC_HILIC_K56_2_typsin_B3.61271.61271.4 | 117.8329 | Trypsin | B3 | 103 |
| BCAL0749 | >gi\|443604502\|gb\|ELT72429.1\| cytochrome c oxidase, subunit II [Burkholderia cenocepacia K56-2Valvano] | L.PEDTAGAATAAAPAEAASAPAQAASGAEQPAAAASAALS[+568.212][+100.064]T[+568.212][+100.064]IYFETGK.S | HexNAc(2)Hex(1) 100.064,HexNAc(2)Hex(1) 100.064 | No | 1410.6604 | 4 | 5639.6198 | 5639.6148 | 0.9 | CRagged | 510.79 | 70.27 | 1.2 | Nsco_20191108_BC_ZIC_HILIC_K56_2_thermolysin_B3.60731.60731.4 | 120.695 | Thermolysin | B3 | 104 |
| BCAL0749 | >gi\|443604502\|gb\|ELT72429.1\| cytochrome c oxidase, subunit II [Burkholderia cenocepacia K56-2Valvano] | K.QVADARNGKLPEDT[+568.212]AGAATAAAPAEAASAPAQAASGAEQPAAAASAALSTIYFETGK.S | HexNAc(2)Hex(1) | No | 1481.9689 | 4 | 5924.8537 | 5923.848 | 0.4 | Specific | 880.62 | 880.62 | 1.6 | Nsco_20191108_BC_ZIC_HILIC_K56_2_typsin_B3.62336.62336.4 | 119.6719 | Trypsin | B3 | 105 |

| | | | | | | | | | | | | | | | | | | |
|---|---|---|---|---|---|---|---|---|---|---|---|---|---|---|---|---|---|---|
| BCAL0749 | >gi\|443604502\|gb\|ELT72429.1\| cytochrome c oxidase, subunit II [Burkholderia cenocepacia K56-2Valvano] | Q.VADARNGKLPEDTAGAAT[+568.212]AAAPAEAAS[+568.212]APAQAASGAEQPA.A | HexNAc(2)Hex(1), HexNAc(2)Hex(1) | Yes (1 of 2) | 1595.0779 | 3 | 4783.2191 | 4782.1971 | 3.9 | Non | 226.43 | 204.84 | 0.8 | Nsco_20191108_BC_ZIC_HILIC_K56_2_thermolysin_B2_.33508.33508.3 | 63.5736 | Thermolysin | B2 | 106 |
| BCAL0786 | >gi\|443598725\|gb\|ELT67055.1\| hypothetical protein BURCENK562V_1882 [Burkholderia cenocepacia K56-2Valvano] | K.LLSVPAPAS[+568.212]TEGDHHDK.- | HexNAc(2)Hex(1) | No | 1171.5508 | 2 | 2342.0944 | 2342.0882 | 2.6 | Specific | 726.94 | 522.22 | 0 | Nsco_20191108_BC_ZIC_HILIC_K56_2_typsin_B3.14451.14451.4 | 36.0021 | Trypsin | B3 | 107 |
| BCAL1086 | >gi\|443597491\|gb\|ELT65913.1\| putative lipoprotein [Burkholderia cenocepacia K56-2Valvano] | K.ALDQVAS[+568.212]TVNQQINAAK.A | HexNAc(2)Hex(1) | No | 1170.0798 | 2 | 2339.1522 | 2339.146 | 2.7 | Specific | 684.32 | 642.21 | 0.8 | Nsco_20191108_BC_ZIC_HILIC_K56_2_typsin_B1.35456.35456.2 | 74.0097 | Trypsin | B1 | 108 |
| BCAL1086 | >gi\|443597491\|gb\|ELT65913.1\| putative lipoprotein [Burkholderia cenocepacia K56-2Valvano] | Q.IDAAAS[+568.212]AVVAH.A | HexNAc(2)Hex(1) | Yes | 796.8835 | 2 | 1592.7597 | 1592.7538 | 3.7 | Non | 394.42 | 147.31 | 147.31 | Nsco_20191108_BC_ZIC_HILIC_K56_2_thermolysin_B2_.18308.18308.2 | 39.2035 | Thermolysin | B2 | 109 |
| BCAL1086 | >gi\|443597491\|gb\|ELT65913.1\| putative lipoprotein [Burkholderia cenocepacia K56-2Valvano] | K.KLQQWSQQSAAGAKPAS[+568.212]GE.- | HexNAc(2)Hex(1) | No | 1270.6071 | 2 | 2540.2069 | 2540.1998 | 2.8 | CRagged | 814.72 | 675.6 | 2.34 | Nsco_20191108_BC_ZIC_HILIC_K56_2_typsin_B2.11123.11123.2 | 29.9123 | Trypsin | B2 | 110 |
| BCAL1086 | >gi\|443597491\|gb\|ELT65913.1\| putative lipoprotein [Burkholderia cenocepacia K56-2Valvano] | E.KSGTPVAQPDTAAS[+568.212]AAAD.A | HexNAc(2)Hex(1) | No | 1113.5132 | 2 | 2226.0191 | 2226.0143 | 2.1 | Non | 524.82 | 394.52 | 0 | Nsco_20191108_BC_ZIC_HILIC_K56_2_pepsin_B1.13985.13985.2 | 33.9556 | Pepsin | B1 | 111 |
| BCAL1086 | >gi\|443597491\|gb\|ELT65913.1\| putative lipoprotein [Burkholderia cenocepacia K56-2Valvano] | E.KSGT[+568.212]PVAQPDTAASAAADAANN.A | HexNAc(2)Hex(1) | No | 1298.5951 | 2 | 2596.183 | 2596.1744 | 3.3 | Non | 583.11 | 522.96 | 1.89 | Nsco_20191108_BC_ZIC_HILIC_K56_2_pepsin_B2.20376.20376.2 | 44.9604 | Pepsin | B2 | 112 |
| BCAL1086 | >gi\|443597491\|gb\|ELT65913.1\| putative lipoprotein [Burkholderia cenocepacia K56-2Valvano] | E.KSGTPVAQPDTAAS[+568.212]AAADAANNAAKAL.D | HexNAc(2)Hex(1) | No | 1017.4953 | 3 | 3050.4713 | 3050.4648 | 2.1 | NRagged | 456.51 | 268.96 | 3.6 | Nsco_20191108_BC_ZIC_HILIC_K56_2_pepsin_B1.37651.37651.4 | 82.7525 | Pepsin | B1 | 113 |
| BCAL1086 | >gi\|443597491\|gb\|ELT65913.1\| putative lipoprotein [Burkholderia cenocepacia K56-2Valvano] | K.LQQWSQQSAAGAKPAS[+568.212]GE.- | HexNAc(2)Hex(1) | No | 1206.5588 | 2 | 2412.1102 | 2412.1049 | 2.2 | CRagged | 750.26 | 616.49 | 1.06 | Nsco_20191108_BC_ZIC_HILIC_K56_2_typsin_B2.15236.15236.2 | 36.5826 | Trypsin | B2 | 114 |
| BCAL1086 | >gi\|443597491\|gb\|ELT65913.1\| putative lipoprotein [Burkholderia cenocepacia K56-2Valvano] | K.SGTPVAQPDTAAS[+568.212]AAADAANNAAK.A | HexNAc(2)Hex(1) | No | 1369.6319 | 2 | 2738.2565 | 2738.2486 | 2.9 | Specific | 650.93 | 618.24 | 9.86 | Nsco_20191108_BC_ZIC_HILIC_K56_2_typsin_B3.28764.28764.2 | 61.6522 | Trypsin | B3 | 115 |
| BCAL1093 | >gi\|443599421\|gb\|ELT67702.1\| hypothetical protein BURCENK562V_2687 [Burkholderia cenocepacia K56-2Valvano] | K.HAYDEAHPAEAAS[+568.212]AASH.- | HexNAc(2)Hex(1) | No | 768.3271 | 3 | 2302.9666 | 2302.9582 | 3.6 | CRagged | 642.3 | 642.3 | 6 | Nsco_20191108_BC_ZIC_HILIC_K56_2_typsin_B3.7342.7342.3 | 24.2457 | Trypsin | B3 | 116 |
| BCAL1357 | >gi\|443602613\|gb\|ELT70680.1\| PF11745 family protein [Burkholderia cenocepacia K56-2Valvano] | A.TDMSPAAS[+568.212]T[+568.212]AAGASSGAGAAAASGTMGLK.L | HexNAc(2)Hex(1), HexNAc(2)Hex(1) | No | 1202.196 | 3 | 3604.5736 | 3604.5623 | 3.1 | NRagged | 493.66 | 231.83 | 7.8 | Nsco_20191108_BC_ZIC_HILIC_K56_2_typsin_B3.30818.30818.3 | 65.4053 | Trypsin | B3 | 117 |
| BCAL1453 | >gi\|443604751\|gb\|ELT72660.1\| HlyD family secretion protein [Burkholderia cenocepacia K56-2Valvano] | R.ENAAHSASS[+568.212]ADAQYQQAIAALDAAK.L | HexNAc(2)Hex(1) | No | 1024.4731 | 3 | 3071.4046 | 3070.3971 | 1.3 | Specific | 915.41 | 915.41 | 2.4 | Nsco_20191108_BC_ZIC_HILIC_K56_2_typsin_B1.39870.39870.3 | 81.8189 | Trypsin | B1 | 118 |
| BCAL1453 | >gi\|443604751\|gb\|ELT72660.1\| HlyD family secretion protein [Burkholderia cenocepacia K56-2Valvano] | K.ENRENAAHSAS[+568.212]SADAQYQQAIAALDAAK.L | HexNAc(2)Hex(1) | Yes | 868.1537 | 4 | 3469.5931 | 3469.5837 | 2.7 | Specific | 1111.53 | 987.06 | 56.26 | Nsco_20191108_BC_ZIC_HILIC_K56_2_typsin_B3.34661.34661.4 | 71.9269 | Trypsin | B3 | 119 |
| BCAL1496 | >gi\|443597624\|gb\|ELT66041.1\| hypothetical protein BURCENK562V_3510 [Burkholderia cenocepacia K56-2Valvano] | K.AAAKKAGKKAKAADAAS[+568.212]Q.- | HexNAc(2)Hex(1) | Yes | 564.3015 | 4 | 2254.1841 | 2254.1773 | 3 | Non | 480.3 | 448.6 | 448.6 | Nsco_20191108_BC_ZIC_HILIC_K56_2_thermolysin_B2_.2250.2250.4 | 13.9237 | Thermolysin | B2 | 120 |
| BCAL1496 | >gi\|443597624\|gb\|ELT66041.1\| hypothetical protein BURCENK562V_3510 [Burkholderia cenocepacia K56-2Valvano] | A.AAKKAGKKAKAADAAS[+568.212]Q.- | HexNAc(2)Hex(1) | Yes | 546.5424 | 4 | 2183.1477 | 2183.1401 | 3.5 | Non | 455.21 | 455.21 | 455.21 | Nsco_20191108_BC_ZIC_HILIC_K56_2_thermolysin_B2_.2204.2204.4 | 13.7129 | Thermolysin | B2 | 121 |

| | | | | | | | | | | | | | | | | | | |
|---|---|---|---|---|---|---|---|---|---|---|---|---|---|---|---|---|---|---|
| BCAL1496 | >gi\|443597624\|gb\|ELT66041.1\| hypothetical protein BURCENK562V_3510 [Burkholderia cenocepacia K56-2Valvano] | K.AGKKAKAADAAS[+568.212]Q.- | HexNAc(2)Hex(1) | Yes | 595.632 | 3 | 1784.8815 | 1784.876 | 3.1 | Non | 195.28 | 64.78 | 64.78 | Nsco_20191108_BC_ZIC_HILIC_K56_2_thermolysin_B1.2214.2214.3 | 13.3883 | Thermolysin | B1 | 122 |
| BCAL1496 | >gi\|443597624\|gb\|ELT66041.1\| hypothetical protein BURCENK562V_3510 [Burkholderia cenocepacia K56-2Valvano] | K.AKAADAAS[+568.212]Q.- | HexNAc(2)Hex(1) | Yes | 700.8192 | 2 | 1400.631 | 1400.6275 | 2.5 | CRagged | 403.43 | 208.72 | 208.72 | Nsco_20191108_BC_ZIC_HILIC_K56_2_typsin_B3.2167.2167.2 | 14.4123 | Trypsin | B3 | 123 |
| BCAL1496 | >gi\|443597624\|gb\|ELT66041.1\| hypothetical protein BURCENK562V_3510 [Burkholderia cenocepacia K56-2Valvano] | H.KAAAKKAGKKAKAADAAS[+568.212]Q.- | HexNAc(2)Hex(1) | Yes | 596.3253 | 4 | 2382.2795 | 2382.2722 | 3.1 | Non | 709.68 | 648.15 | 648.15 | Nsco_20191108_BC_ZIC_HILIC_K56_2_pepsin_B3_reinject.2004.2004.4 | 13.5708 | Pepsin | B3 | 124 |
| BCAL1496 | >gi\|443597624\|gb\|ELT66041.1\| hypothetical protein BURCENK562V_3510 [Burkholderia cenocepacia K56-2Valvano] | K.KAGKKAKAADAAS[+568.212]Q.- | HexNAc(2)Hex(1) | Yes | 638.3307 | 3 | 1912.9774 | 1912.971 | 3.4 | Non | 349.29 | 249.11 | 249.11 | Nsco_20191108_BC_ZIC_HILIC_K56_2_pepsin_B2.1928.1928.3 | 13.3268 | Pepsin | B2 | 125 |
| BCAL1496 | >gi\|443597624\|gb\|ELT66041.1\| hypothetical protein BURCENK562V_3510 [Burkholderia cenocepacia K56-2Valvano] | A.KKAGKKAKAADAAS[+568.212]Q.- | HexNAc(2)Hex(1) | Yes | 511.0234 | 4 | 2041.0719 | 2041.0659 | 2.9 | Non | 349.79 | 268.41 | 268.41 | Nsco_20191108_BC_ZIC_HILIC_K56_2_pepsin_B2.1914.1914.4 | 13.2702 | Pepsin | B2 | 126 |
| BCAL1496 | >gi\|443597624\|gb\|ELT66041.1\| hypothetical protein BURCENK562V_3510 [Burkholderia cenocepacia K56-2Valvano] | G.KKAKAADAAS[+568.212]Q.- | HexNAc(2)Hex(1) | Yes | 552.9457 | 3 | 1656.8225 | 1656.8174 | 3.1 | Non | 367.41 | 216.13 | 216.13 | Nsco_20191108_BC_ZIC_HILIC_K56_2_pepsin_B1.1926.1926.3 | 13.1059 | Pepsin | B1 | 127 |
| BCAL1503 | >gi\|443597627\|gb\|ELT66044.1\| segregation and condensation protein B [Burkholderia cenocepacia K56-2Valvano] | P.AGESAEASGDAAADVASTQEMPSRDT [+568.212][+100.064]LDADRGQTE QTEQVGAEAVPADVQPEPQPADWSEED RKPAAE.A | HexNAc(2)Hex(1) 100.064 | No | 1548.6949 | 5 | 7739.4452 | 7733.4351 | -1.3 | Non | 437.87 | 437.87 | 0 | Nsco_20191108_BC_ZIC_HILIC_K56_2_thermolysin_B1.39793.39793.5 | 81.2183 | Thermolysin | B1 | 128 |
| BCAL1503 | >gi\|443597627\|gb\|ELT66044.1\| segregation and condensation protein B [Burkholderia cenocepacia K56-2Valvano] | P.AGESAEASGDAAADVASTQEMPSRDT [+568.212][+100.064]LDADRGQTE QTEQVGAEAVPADVQPEPQPADWSEED RKPAAEA.A | HexNAc(2)Hex(1) 100.064 | No | 1562.7001 | 5 | 7809.4715 | 7804.4722 | -2.2 | Non | 274.45 | 274.45 | 0 | Nsco_20191108_BC_ZIC_HILIC_K56_2_thermolysin_B1.40060.40060.5 | 81.8258 | Thermolysin | B1 | 129 |
| BCAL1503 | >gi\|443597627\|gb\|ELT66044.1\| segregation and condensation protein B [Burkholderia cenocepacia K56-2Valvano] | T.EQVGAEAVPADVQPEPQPADWS[+568.212][+100.064]EEDRKPAAEAA AGDGAEAAGDMPGEADQADASR.S | HexNAc(2)Hex(1) 100.064 | No | 1551.9427 | 4 | 6204.7492 | 6198.724 | 0.8 | NRagged | 510.38 | 510.38 | 22.15 | Nsco_20191108_BC_ZIC_HILIC_K56_2_typsin_B2.45231.45231.4 | 92.148 | Trypsin | B2 | 130 |
| BCAL1503 | >gi\|443597627\|gb\|ELT66044.1\| segregation and condensation protein B [Burkholderia cenocepacia K56-2Valvano] | T.LDADRGQTEQTEQVGAEAVPADVQPE PQPADWSEEDRKPAAEAAAGDGAEAA GDM[+15.995]PGEADQADAS[+568.212][+100.064]RSRPADDET[+568.212][+100.064]L.D | HexNAc(2)Hex(1) 100.064,HexNAc(2)Hex(1) 100.064 | No | 1515.848 | 6 | 9090.0514 | 9081.9984 | 2.9 | NRagged | 383.98 | 337.77 | 7.84 | Nsco_20191108_BC_ZIC_HILIC_K56_2_thermolysin_B2_.53771.53771.6 | 99.391 | Thermolysin | B2 | 131 |
| BCAL1503 | >gi\|443597627\|gb\|ELT66044.1\| segregation and condensation protein B [Burkholderia cenocepacia K56-2Valvano] | D.VASTQEMPSRDTLDADRGQTEQTEQV GAEAVPADVQPEPQPADWSEEDRKPA AEAAAGDGAEAAGDMPGEADQADASR S[+568.212][+100.064]RP.A | HexNAc(2)Hex(1) 100.064 | No | 1295.1584 | 7 | 9060.0649 | 9056.05 | 0.2 | Non | 374.51 | 226.21 | 17.98 | Nsco_20191108_BC_ZIC_HILIC_K56_2_thermolysin_B2_.52190.52190.7 | 96.1447 | Thermolysin | B2 | 132 |
| BCAL1649 | >gi\|443605315\|gb\|ELT73173.1\| PF11159 family protein [Burkholderia cenocepacia K56-2Valvano] | A.LMSGMPPRGNPGERPPEWS[+568.212]NQPPAGNAPGTTPAAPPANPAP ASAG.V | HexNAc(2)Hex(1) | No | 1647.7775 | 3 | 4941.3178 | 4938.2945 | 2.7 | Non | 386.6 | 386.6 | 3.6 | Nsco_20191108_BC_ZIC_HILIC_K56_2_thermolysin_B2_.42757.42757.3 | 78.5914 | Thermolysin | B2 | 133 |
| BCAL1674 | >gi\|443605593\|gb\|ELT73435.1\| periplasmic multidrug efflux lipoprotein [Burkholderia cenocepacia K56-2Valvano] | T.AVKPVEKAPSSKAAPPAAAS[+568.212]QAAARQT.- | HexNAc(2)Hex(1) | No | 1057.8885 | 3 | 3171.6508 | 3171.6379 | 4.1 | Non | 172.85 | 172.85 | 16.72 | Nsco_20191108_BC_ZIC_HILIC_K56_2_pepsin_B3_reinject.10109.10109.3 | 27.5306 | Pepsin | B3 | 134 |

| | | | | | | | | | | | | | | | | | | |
|---|---|---|---|---|---|---|---|---|---|---|---|---|---|---|---|---|---|---|
| BCAL1674 | >gi\|443605593\|gb\|ELT73435.1\| periplasmic multidrug efflux lipoprotein [Burkholderia cenocepacia K56-2Valvano] | E.KAPSSKAAPPAAAS[+568.212]QAAARQT.- | HexNAc(2)Hex(1) | No | 850.0991 | 3 | 2548.2826 | 2548.2737 | 3.5 | Non | 370.47 | 370.47 | 10.8 | Nsco_20191108_BC_ZIC_HILIC_K56_2_pepsin_B1.6180.6180.3 | 20.969 | Pepsin | B1 | 135 |
| BCAL1674 | >gi\|443605593\|gb\|ELT73435.1\| periplasmic multidrug efflux lipoprotein [Burkholderia cenocepacia K56-2Valvano] | D.TAVKPVEKAPSSKAAPPAAAS[+568.212]QAAARQT.- | HexNAc(2)Hex(1) | No | 1091.5619 | 3 | 3272.6711 | 3272.6856 | -4.4 | Non | 433.93 | 417.23 | 16.8 | Nsco_20191108_BC_ZIC_HILIC_K56_2_pepsin_B2.11787.11787.3 | 30.799 | Pepsin | B2 | 136 |
| BCAL1674 | >gi\|443605593\|gb\|ELT73435.1\| periplasmic multidrug efflux lipoprotein [Burkholderia cenocepacia K56-2Valvano] | P.VEKAPSSKAAPPAAAS[+568.212]QA.A | HexNAc(2)Hex(1) | No | 1125.0602 | 2 | 2249.1132 | 2249.1031 | 4.5 | Non | 632.4 | 364.09 | 7.47 | Nsco_20191108_BC_ZIC_HILIC_K56_2_thermolysin_B2_.7412.7412.2 | 23.3896 | Thermolysin | B2 | 137 |
| BCAL1746 | >gi\|443592790\|gb\|ELT61568.1\| putative lipoprotein [Burkholderia cenocepacia K56-2Valvano] | L.IDHIGKAWPGNAASGAS[+568.212]ASASE.- | HexNAc(2)Hex(1) | No | 889.0777 | 3 | 2665.2186 | 2665.2111 | 2.8 | CRagged | 722.83 | 722.83 | 0.53 | Nsco_20191108_BC_ZIC_HILIC_K56_2_pepsin_B2.22484.22484.3 | 49.0176 | Pepsin | B2 | 138 |
| BCAL1798 | >gi\|443598613\|gb\|ELT66949.1\| hypothetical protein BURCENK562V_2315 [Burkholderia cenocepacia K56-2Valvano] | T.AATPAAASAPS[+568.212]AEQRAARHEARIEQ.R | HexNAc(2)Hex(1) | No | 1043.5105 | 3 | 3128.517 | 3128.5091 | 2.5 | Non | 610.85 | 521.74 | 6.51 | Nsco_20191108_BC_ZIC_HILIC_K56_2_pepsin_B2.12045.12045.3 | 31.1659 | Pepsin | B2 | 139 |
| BCAL1798 | >gi\|443598613\|gb\|ELT66949.1\| hypothetical protein BURCENK562V_2315 [Burkholderia cenocepacia K56-2Valvano] | S.AQAPATTAATPAAAS[+568.212]APSAEQR.A | HexNAc(2)Hex(1) | No | 1304.1209 | 2 | 2607.2345 | 2607.2268 | 3 | Non | 684.62 | 339.9 | 3.6 | Nsco_20191108_BC_ZIC_HILIC_K56_2_thermolysin_B1.17795.17795.2 | 38.7969 | Thermolysin | B1 | 140 |
| BCAL1798 | >gi\|443598613\|gb\|ELT66949.1\| hypothetical protein BURCENK562V_2315 [Burkholderia cenocepacia K56-2Valvano] | S.AQAPATTAATPAAAS[+568.212]APSAEQRA.A | HexNAc(2)Hex(1) | No | 1339.6401 | 2 | 2678.2728 | 2678.2639 | 3.3 | Non | 562.38 | 328.57 | 3.6 | Nsco_20191108_BC_ZIC_HILIC_K56_2_thermolysin_B1.19018.19018.2 | 40.7989 | Thermolysin | B1 | 141 |
| BCAL1798 | >gi\|443598613\|gb\|ELT66949.1\| hypothetical protein BURCENK562V_2315 [Burkholderia cenocepacia K56-2Valvano] | A.SAQAPATTAATPAAAS[+568.212]APSAEQR.A | HexNAc(2)Hex(1) | No | 1348.1365 | 2 | 2695.2657 | 2694.2588 | 1.3 | NRagged | 892.33 | 839.31 | 3.6 | Nsco_20191108_BC_ZIC_HILIC_K56_2_typsin_B3.16665.16665.2 | 39.6691 | Trypsin | B3 | 142 |
| BCAL1869 | >gi\|443596292\|gb\|ELT64810.1\| hypothetical protein BURCENK562V_2581 [Burkholderia cenocepacia K56-2Valvano] | A.DASAPDAVAS[+568.212][+100.064]APVAAS[+568.212]AVPDFDARQKV.L | HexNAc(2)Hex(1) 100.064,HexNAc(2)Hex(1) | Yes (1 of 2) | 1288.9348 | 3 | 3864.7899 | 3862.7979 | -3.8 | Non | 293.29 | 293.29 | 2.4 | Nsco_20191108_BC_ZIC_HILIC_K56_2_thermolysin_B2_.45532.45532.3 | 83.2144 | Thermolysin | B2 | 143 |
| BCAL1869 | >gi\|443596292\|gb\|ELT64810.1\| hypothetical protein BURCENK562V_2581 [Burkholderia cenocepacia K56-2Valvano] | P.VAAS[+568.212]AVPDFDARQK.V | HexNAc(2)Hex(1) | Yes | 681.6658 | 3 | 2042.9828 | 2042.9764 | 3.1 | Non | 394.43 | 175.06 | 175.06 | Nsco_20191108_BC_ZIC_HILIC_K56_2_thermolysin_B1.18482.18482.3 | 39.9444 | Thermolysin | B1 | 144 |
| BCAL1877 | >gi\|443594658\|gb\|ELT63293.1\| HflK protein [Burkholderia cenocepacia K56-2Valvano] | A.AAST[+568.212]GAS[+568.212]AADAASAPAATVPS.A | HexNAc(2)Hex(1), HexNAc(2)Hex(1) | No | 1441.1372 | 2 | 2881.2672 | 2881.2579 | 3.2 | Non | 655.09 | 344.29 | 12.39 | Nsco_20191108_BC_ZIC_HILIC_K56_2_thermolysin_B1.26761.26761.2 | 55.2382 | Thermolysin | B1 | 145 |
| BCAL1877 | >gi\|443594658\|gb\|ELT63293.1\| HflK protein [Burkholderia cenocepacia K56-2Valvano] | A.AASTGASAADAASAPAAT[+568.212]VPS[+568.212][+100.064]A.A | HexNAc(2)Hex(1), HexNAc(2)Hex(1) 100.064 | No | 1526.6785 | 2 | 3052.3497 | 3052.359 | -3.1 | Non | 372.17 | 372.17 | 0 | Nsco_20191108_BC_ZIC_HILIC_K56_2_pepsin_B2.29721.29721.2 | 63.226 | Pepsin | B2 | 146 |
| BCAL1877 | >gi\|443594658\|gb\|ELT63293.1\| HflK protein [Burkholderia cenocepacia K56-2Valvano] | A.AAST[+568.212]GAS[+568.212]AADAASAPAATVPSAA.A | HexNAc(2)Hex(1), HexNAc(2)Hex(1) | No | 1512.1733 | 2 | 3023.3392 | 3023.3322 | 2.3 | Non | 829.49 | 314.77 | 16.52 | Nsco_20191108_BC_ZIC_HILIC_K56_2_pepsin_B1.28514.28514.2 | 60.7583 | Pepsin | B1 | 147 |
| BCAL1877 | >gi\|443594658\|gb\|ELT63293.1\| HflK protein [Burkholderia cenocepacia K56-2Valvano] | A.ATAS[+568.212]GVDVLRSRE.A | HexNAc(2)Hex(1) | No | 964.9725 | 2 | 1928.9376 | 1928.9295 | 4.2 | Non | 483.57 | 209.95 | 0 | Nsco_20191108_BC_ZIC_HILIC_K56_2_thermolysin_B1.14431.14431.2 | 33.5436 | Thermolysin | B1 | 148 |
| BCAL1877 | >gi\|443594658\|gb\|ELT63293.1\| HflK protein [Burkholderia cenocepacia K56-2Valvano] | L.VEQGRQNAAAST[+568.212]GAS[+568.212]AADAASAPAATVPSAA.A | HexNAc(2)Hex(1), HexNAc(2)Hex(1) | No | 1302.5969 | 3 | 3905.776 | 3905.7629 | 3.3 | CRagged | 458.8 | 458.8 | 1.53 | Nsco_20191108_BC_ZIC_HILIC_K56_2_pepsin_B2.26028.26028.3 | 55.7646 | Pepsin | B2 | 149 |

| | | | | | | | | | | | | | | | | | | |
|---|---|---|---|---|---|---|---|---|---|---|---|---|---|---|---|---|---|---|
| BCAL1917 | >gi\|443605668\|gb\|ELT73505.1\| hypothetical protein BURCENK562V_0636 [Burkholderia cenocepacia K56-2Valvano] | K.AAAAAASADAGASAPAAASS[+568.212]T[+568.212]K.A | HexNAc(2)Hex(1), HexNAc(2)Hex(1) | No | 1463.1529 | 2 | 2925.2985 | 2925.2954 | 1.1 | Specific | 871.39 | 776.45 | 1.2 | Nsco_20191108_BC_ZIC_HILIC_K56_2_typsin_B1.9657.9657.2 | 28.075 | Trypsin | B1 | 150 |
| BCAL1917 | >gi\|443605668\|gb\|ELT73505.1\| hypothetical protein BURCENK562V_0636 [Burkholderia cenocepacia K56-2Valvano] | T.AAPAPTAS[+568.212]APEAAAKPAKTKR.A | HexNAc(2)Hex(1) | Yes | 644.0907 | 4 | 2573.341 | 2573.3305 | 4.1 | Non | 928.61 | 779.55 | 128.8 | Nsco_20191108_BC_ZIC_HILIC_K56_2_thermolysin_B1.5828.5828.4 | 20.4309 | Thermolysin | B1 | 151 |
| BCAL1917 | >gi\|443605668\|gb\|ELT73505.1\| hypothetical protein BURCENK562V_0636 [Burkholderia cenocepacia K56-2Valvano] | T.AAPAPTAS[+568.212]APEAAAKPAKTKRA.S | HexNAc(2)Hex(1) | No | 1323.1912 | 2 | 2645.3751 | 2644.3676 | 1.6 | Non | 766.18 | 588.28 | 2.4 | Nsco_20191108_BC_ZIC_HILIC_K56_2_thermolysin_B1.6283.6283.2 | 21.1178 | Thermolysin | B1 | 152 |
| BCAL1917 | >gi\|443605668\|gb\|ELT73505.1\| hypothetical protein BURCENK562V_0636 [Burkholderia cenocepacia K56-2Valvano] | T.AAPAPTASAPEAAAKPAKT[+568.212]KRASKKEK.A | HexNAc(2)Hex(1) | No | 649.7533 | 5 | 3244.7373 | 3244.7271 | 3.1 | Non | 526.01 | 499.99 | 9.6 | Nsco_20191108_BC_ZIC_HILIC_K56_2_thermolysin_B2_.3798.3798.4 | 17.7541 | Thermolysin | B2 | 153 |
| BCAL1917 | >gi\|443605668\|gb\|ELT73505.1\| hypothetical protein BURCENK562V_0636 [Burkholderia cenocepacia K56-2Valvano] | T.AAPAPTAS[+568.212]APEAAAKPAKTKRASKKEKA.A | HexNAc(2)Hex(1) | Yes | 663.9601 | 5 | 3315.7712 | 3315.7642 | 2.1 | Non | 1029.46 | 993.3 | 140.99 | Nsco_20191108_BC_ZIC_HILIC_K56_2_thermolysin_B2_.3999.3999.5 | 18.067 | Thermolysin | B2 | 154 |
| BCAL1917 | >gi\|443605668\|gb\|ELT73505.1\| hypothetical protein BURCENK562V_0636 [Burkholderia cenocepacia K56-2Valvano] | T.AAPAPTAS[+568.212]APEAAAKPAKTKRASKKEKAA.A | HexNAc(2)Hex(1) | Yes | 678.167 | 5 | 3386.8058 | 3386.8013 | 1.3 | Non | 940.1 | 902.62 | 127.47 | Nsco_20191108_BC_ZIC_HILIC_K56_2_thermolysin_B2_.4052.4052.4 | 18.147 | Thermolysin | B2 | 155 |
| BCAL1917 | >gi\|443605668\|gb\|ELT73505.1\| hypothetical protein BURCENK562V_0636 [Burkholderia cenocepacia K56-2Valvano] | A.AASADAGASAPAAASS[+568.212]T[+568.212]KA.T | HexNAc(2)Hex(1), HexNAc(2)Hex(1) | No | 1356.5999 | 2 | 2712.1924 | 2712.184 | 3.1 | Non | 765.09 | 360.69 | 1.39 | Nsco_20191108_BC_ZIC_HILIC_K56_2_thermolysin_B1.8292.8292.2 | 24.0281 | Thermolysin | B1 | 156 |
| BCAL1917 | >gi\|443605668\|gb\|ELT73505.1\| hypothetical protein BURCENK562V_0636 [Burkholderia cenocepacia K56-2Valvano] | P.AATTSATTSTTTSAGTASTSTTATAGT[+68.212]TTAAPAPTASAPEAAAKPAKTKR.A | HexNAc(2)Hex(1) | No | 1027.9116 | 5 | 5135.529 | 5135.5085 | 4 | Non | 904.63 | 904.63 | 0 | Nsco_20191108_BC_ZIC_HILIC_K56_2_thermolysin_B2_.17868.17868.5 | 38.5085 | Thermolysin | B2 | 157 |
| BCAL1917 | >gi\|443605668\|gb\|ELT73505.1\| hypothetical protein BURCENK562V_0636 [Burkholderia cenocepacia K56-2Valvano] | T.AGTTTAAPAPTAS[+568.212]APEAAAKPAKTKR.A | HexNAc(2)Hex(1) | Yes | 751.8917 | 4 | 3004.5449 | 3004.5321 | 4.3 | Non | 907.53 | 830.57 | 99.82 | Nsco_20191108_BC_ZIC_HILIC_K56_2_thermolysin_B2_.8205.8205.4 | 24.6392 | Thermolysin | B2 | 158 |
| BCAL1917 | >gi\|443605668\|gb\|ELT73505.1\| hypothetical protein BURCENK562V_0636 [Burkholderia cenocepacia K56-2Valvano] | A.ASADAGASAPAAASS[+568.212]T[+568.212]KA.T | HexNAc(2)Hex(1), HexNAc(2)Hex(1) | No | 1321.0827 | 2 | 2641.1581 | 2641.1469 | 4.2 | Non | 735.79 | 376.97 | 2.4 | Nsco_20191108_BC_ZIC_HILIC_K56_2_thermolysin_B1.7704.7704.2 | 23.1603 | Thermolysin | B1 | 159 |
| BCAL1917 | >gi\|443605668\|gb\|ELT73505.1\| hypothetical protein BURCENK562V_0636 [Burkholderia cenocepacia K56-2Valvano] | A.ASADAGASAPAAAS[+568.212]S[+568.212][+100.064]T[+568.212]KATKGSKKKSKKDKAASAA.A | HexNAc(2)Hex(1), HexNAc(2)Hex(1) 100.064,HexNAc(2)Hex(1) | No | 718.6356 | 7 | 5024.4053 | 5024.4279 | -4.5 | Non | 374.67 | 374.67 | 0 | Nsco_20191108_BC_ZIC_HILIC_K56_2_thermolysin_B1.2759.2759.7 | 14.9823 | Thermolysin | B1 | 160 |
| BCAL1917 | >gi\|443605668\|gb\|ELT73505.1\| hypothetical protein BURCENK562V_0636 [Burkholderia cenocepacia K56-2Valvano] | T.ATAGTTTAAPAPTAS[+568.212]APEAAAKPAKTKR.A | HexNAc(2)Hex(1) | No | 1059.5459 | 3 | 3176.6232 | 3176.6169 | 2 | Non | 967.99 | 614.32 | 2.4 | Nsco_20191108_BC_ZIC_HILIC_K56_2_thermolysin_B2_.8998.8998.3 | 25.7506 | Thermolysin | B2 | 161 |
| BCAL1917 | >gi\|443605668\|gb\|ELT73505.1\| hypothetical protein BURCENK562V_0636 [Burkholderia cenocepacia K56-2Valvano] | A.ATTSATTSTTTSAGTASTSTTAT[+568.212]AGTTTAAPAPTASAPEAAAKPAKTKR.A | HexNAc(2)Hex(1) | No | 1013.9047 | 5 | 5065.4942 | 5064.4714 | 3.8 | Non | 666.61 | 666.61 | 0 | Nsco_20191108_BC_ZIC_HILIC_K56_2_thermolysin_B2_.17348.17348.5 | 37.743 | Thermolysin | B2 | 162 |

| | | | | | | | | | | | | | | | | | | |
|---|---|---|---|---|---|---|---|---|---|---|---|---|---|---|---|---|---|---|
| BCAL1917 | >gi\|443605668\|gb\|ELT73505.1\| hypothetical protein BURCENK562V_0636 [Burkholderia cenocepacia K56-2Valvano] | S.ATTSTTTSAGTASTSTTATAGTTTAAPA[+568.212]ASAPEAAAK.P | HexNAc(2)Hex(1) | No | 1342.3036 | 3 | 4024.8962 | 4022.8783 | 2.8 | Non | 702.39 | 604.38 | 4.8 | Nsco_20191108_BC_ZIC_HILIC_K56_2_thermolysin_B3.24355.24355.3 | 53.0549 | Thermolysin | B3 | 163 |
| BCAL1917 | >gi\|443605668\|gb\|ELT73505.1\| hypothetical protein BURCENK562V_0636 [Burkholderia cenocepacia K56-2Valvano] | S.ATTSTTTSAGTASTSTTATAGTTTAAPA[+568.212]ASAPEAAAKPAK.T | HexNAc(2)Hex(1) | No | 1440.3653 | 3 | 4319.0813 | 4319.0631 | 4.2 | NRagged | 885.76 | 816.33 | 4.8 | Nsco_20191108_BC_ZIC_HILIC_K56_2_typsin_B1.20175.20175.3 | 46.4565 | Trypsin | B1 | 164 |
| BCAL1917 | >gi\|443605668\|gb\|ELT73505.1\| hypothetical protein BURCENK562V_0636 [Burkholderia cenocepacia K56-2Valvano] | A.GTASTSTTATAGTTTAAPAPT[+568.212]ASAPEAAAKPAK.T | HexNAc(2)Hex(1) | No | 1166.5695 | 3 | 3497.694 | 3497.6865 | 2.2 | NRagged | 873.9 | 854.34 | 4.8 | Nsco_20191108_BC_ZIC_HILIC_K56_2_typsin_B2.17188.17188.3 | 40.2245 | Trypsin | B2 | 165 |
| BCAL1917 | >gi\|443605668\|gb\|ELT73505.1\| hypothetical protein BURCENK562V_0636 [Burkholderia cenocepacia K56-2Valvano] | K.LSKPAATTSATTSTTTSAGTASTSTTATATTTAAPAPTAS[+568.212]APEAAAK.P | HexNAc(2)Hex(1) | No | 1220.5935 | 4 | 4879.3524 | 4879.3437 | 1.8 | Non | 1001.69 | 1001.7 | 63.47 | Nsco_20191108_BC_ZIC_HILIC_K56_2_thermolysin_B3.24738.24738.4 | 53.7309 | Thermolysin | B3 | 166 |
| BCAL1917 | >gi\|443605668\|gb\|ELT73505.1\| hypothetical protein BURCENK562V_0636 [Burkholderia cenocepacia K56-2Valvano] | K.LSKPAATTSATTSTTTSAGTASTSTTATATTTAAPAPTAS[+568.212]APEAAAKP.A | HexNAc(2)Hex(1) | No | 1245.3497 | 4 | 4978.3771 | 4976.3965 | -5.2 | Non | 269.42 | 269.42 | 30.56 | Nsco_20191108_BC_ZIC_HILIC_K56_2_thermolysin_B3.26131.26131.4 | 56.4092 | Thermolysin | B3 | 167 |
| BCAL1917 | >gi\|443605668\|gb\|ELT73505.1\| hypothetical protein BURCENK562V_0636 [Burkholderia cenocepacia K56-2Valvano] | K.LSKPAATTSATTSTTTSAGTASTSTTATATTTAAPAPTAS[+568.212]APEAAAKPA.K | HexNAc(2)Hex(1) | Yes | 1262.6184 | 4 | 5047.4518 | 5047.4336 | 3.6 | Non | 1201.96 | 1202 | 99.55 | Nsco_20191108_BC_ZIC_HILIC_K56_2_thermolysin_B3.26332.26332.4 | 56.7591 | Thermolysin | B3 | 168 |
| BCAL1917 | >gi\|443605668\|gb\|ELT73505.1\| hypothetical protein BURCENK562V_0636 [Burkholderia cenocepacia K56-2Valvano] | K.LSKPAATTSATTSTTTSAGTASTSTTATATTTAAPAPTAS[+568.212]APEAAAKPAK.T | HexNAc(2)Hex(1) | Yes | 1036.1147 | 5 | 5176.5446 | 5175.5286 | 2.5 | Specific | 1639.98 | 1640 | 78.23 | Nsco_20191108_BC_ZIC_HILIC_K56_2_typsin_B2.21791.21791.5 | 48.9606 | Trypsin | B2 | 169 |
| BCAL1917 | >gi\|443605668\|gb\|ELT73505.1\| hypothetical protein BURCENK562V_0636 [Burkholderia cenocepacia K56-2Valvano] | K.LSKPAATTSATTSTTTSAGTASTSTTATATTTAAPAPTASAPEAAAKPAKT[+568.212].K | HexNAc(2)Hex(1) | No | 1056.3196 | 5 | 5277.5688 | 5276.5763 | -2.1 | Non | 1202.73 | 1202.7 | 118.24 | Nsco_20191108_BC_ZIC_HILIC_K56_2_thermolysin_B3.23031.23031.5 | 50.7231 | Thermolysin | B3 | 170 |
| BCAL1917 | >gi\|443605668\|gb\|ELT73505.1\| hypothetical protein BURCENK562V_0636 [Burkholderia cenocepacia K56-2Valvano] | K.PAATTSATTSTTTSAGTASTSTTATAGTAAPAPT[+568.212]ASAPEAAAKPAK.T | HexNAc(2)Hex(1) | No | 1616.4483 | 3 | 4847.3303 | 4847.3175 | 2.6 | Specific | 1160.31 | 1160.3 | 4.8 | Nsco_20191108_BC_ZIC_HILIC_K56_2_typsin_B2.25468.25468.3 | 55.8308 | Trypsin | B2 | 171 |
| BCAL1917 | >gi\|443605668\|gb\|ELT73505.1\| hypothetical protein BURCENK562V_0636 [Burkholderia cenocepacia K56-2Valvano] | K.PAATTSATTSTTTSAGTASTSTTATAGTAAPAPTASAPEAAAKPAKT[+568.212].K | HexNAc(2)Hex(1) | No | 1237.8422 | 4 | 4948.3468 | 4948.3652 | -3.7 | Non | 667.31 | 667.31 | 75.16 | Nsco_20191108_BC_ZIC_HILIC_K56_2_thermolysin_B3.26231.26231.4 | 56.5788 | Thermolysin | B3 | 172 |
| BCAL1917 | >gi\|443605668\|gb\|ELT73505.1\| hypothetical protein BURCENK562V_0636 [Burkholderia cenocepacia K56-2Valvano] | A.SADAGASAPAAASST[+568.212][+100.064]KAT[+568.212]KGS[+568.212][+100.064]KKKSKKDKAASAAAASGA.- | HexNAc(2)Hex(1) 100.064,HexNAc(2)Hex(1),HexNAc(2)Hex(1) 100.064 | No | 903.1068 | 6 | 5413.6044 | 5410.6196 | -4.7 | Non | 193.86 | 193.86 | 0 | Nsco_20191108_BC_ZIC_HILIC_K56_2_pepsin_B1.2469.2469.6 | 14.4722 | Pepsin | B1 | 173 |
| BCAL1917 | >gi\|443605668\|gb\|ELT73505.1\| hypothetical protein BURCENK562V_0636 [Burkholderia cenocepacia K56-2Valvano] | T.SAGTASTSTTATAGTTTAAPAPTAS[+568.212]APEAAAKPAK.T | HexNAc(2)Hex(1) | No | 1219.9271 | 3 | 3657.7668 | 3655.7556 | 1.2 | NRagged | 758.67 | 715.82 | 4.22 | Nsco_20191108_BC_ZIC_HILIC_K56_2_typsin_B1.17651.17651.3 | 41.5742 | Trypsin | B1 | 174 |

| | | | | | | | | | | | | | | | | | | |
|---|---|---|---|---|---|---|---|---|---|---|---|---|---|---|---|---|---|---|
| BCAL1917 | >gi\|443605668\|gb\|ELT73505.1\|<br>hypothetical protein BURCENK562V_0636<br>[Burkholderia cenocepacia K56-2Valvano] | L.SKPAATTSATTSTTTSAGTASTSTTATA[illegible]<br>TTAAPAPTAS[+568.212]APEAAAKP<br>AK.T | HexNAc(2)Hex(1) | Yes | 1266.371 | 4 | 5062.4621 | 5062.4445 | 3.5 | NRagged | 1229.78 | 1229.8 | 53.78 | Nsco_20191108_BC_ZIC_HILIC_K56_2_typsi<br>n_B2.19359.19359.4 | 44.3527 | Trypsin | B2 | 175 |
| BCAL1917 | >gi\|443605668\|gb\|ELT73505.1\|<br>hypothetical protein BURCENK562V_0636<br>[Burkholderia cenocepacia K56-2Valvano] | T.STTATAGTTTAAPAPT[+568.212]AS<br>APEAAAK.P | HexNAc(2)Hex(1) | No | 1392.665 | 2 | 2784.3228 | 2784.3157 | 2.6 | NRagged | 603.53 | 510.41 | 2.4 | Nsco_20191108_BC_ZIC_HILIC_K56_2_typsi<br>n_B3.18227.18227.2 | 42.3733 | Trypsin | B3 | 176 |
| BCAL1917 | >gi\|443605668\|gb\|ELT73505.1\|<br>hypothetical protein BURCENK562V_0636<br>[Burkholderia cenocepacia K56-2Valvano] | T.STTATAGTTTAAPAPT[+568.212]AS<br>APEAAAKPAK.T | HexNAc(2)Hex(1) | No | 1027.5076 | 3 | 3080.5082 | 3080.5005 | 2.5 | NRagged | 740.42 | 737.18 | 2.4 | Nsco_20191108_BC_ZIC_HILIC_K56_2_typsi<br>n_B1.14937.14937.3 | 36.647 | Trypsin | B1 | 177 |
| BCAL1917 | >gi\|443605668\|gb\|ELT73505.1\|<br>hypothetical protein BURCENK562V_0636<br>[Burkholderia cenocepacia K56-2Valvano] | T.STTTSAGTASTSTTATAGTTTAAPAPTA[illegible]<br>+568.212]APEAAAKPAK.T | HexNAc(2)Hex(1) | No | 1349.652 | 3 | 4046.9414 | 4045.9307 | 1.8 | NRagged | 830.7 | 823.79 | 2.99 | Nsco_20191108_BC_ZIC_HILIC_K56_2_typsi<br>n_B2.19554.19554.3 | 44.715 | Trypsin | B2 | 178 |
| BCAL1917 | >gi\|443605668\|gb\|ELT73505.1\|<br>hypothetical protein BURCENK562V_0636<br>[Burkholderia cenocepacia K56-2Valvano] | K.SVGHLEENGLT[+568.212]IGGASTP<br>PK.G | HexNAc(2)Hex(1) | No | 844.7462 | 3 | 2532.224 | 2532.2199 | 1.6 | Specific | 550.22 | 550.22 | 4.8 | Nsco_20191108_BC_ZIC_HILIC_K56_2_typsi<br>n_B2.24679.24679.3 | 54.2878 | Trypsin | B2 | 179 |
| BCAL1917 | >gi\|443605668\|gb\|ELT73505.1\|<br>hypothetical protein BURCENK562V_0636<br>[Burkholderia cenocepacia K56-2Valvano] | A.TAGTTTAAPAPTAS[+568.212]APE<br>AAAKPAK.T | HexNAc(2)Hex(1) | No | 907.4528 | 3 | 2720.3437 | 2720.336 | 2.8 | NRagged | 413.69 | 118.08 | 0 | Nsco_20191108_BC_ZIC_HILIC_K56_2_typsi<br>n_B2.12755.12755.3 | 32.393 | Trypsin | B2 | 180 |
| BCAL1917 | >gi\|443605668\|gb\|ELT73505.1\|<br>hypothetical protein BURCENK562V_0636<br>[Burkholderia cenocepacia K56-2Valvano] | A.TAGTTTAAPAPTAS[+568.212]APE<br>AAAKPAKTKR.A | HexNAc(2)Hex(1) | No | 1036.1936 | 3 | 3106.5663 | 3105.5798 | -5.4 | Non | 436.37 | 436.37 | 2.4 | Nsco_20191108_BC_ZIC_HILIC_K56_2_ther<br>molysin_B2_.10450.10450.3 | 28.0854 | Thermolysin | B2 | 181 |
| BCAL1917 | >gi\|443605668\|gb\|ELT73505.1\|<br>hypothetical protein BURCENK562V_0636<br>[Burkholderia cenocepacia K56-2Valvano] | T.TSTTTSAGTASTSTTATAGTTTAAPAPT<br>[+568.212][+100.064]APEAAAKP<br>AK.T | HexNAc(2)Hex(1)<br>100.064 | No | 1416.3502 | 3 | 4247.0359 | 4247.0424 | -1.5 | NRagged | 628.08 | 628.08 | 4.43 | Nsco_20191108_BC_ZIC_HILIC_K56_2_typsi<br>n_B2.20176.20176.3 | 45.9778 | Trypsin | B2 | 182 |
| BCAL1917 | >gi\|443605668\|gb\|ELT73505.1\|<br>hypothetical protein BURCENK562V_0636<br>[Burkholderia cenocepacia K56-2Valvano] | A.TTSTTTSAGTASTSTTATAGTTTAAPAP[illegible]<br>AS[+568.212]APEAAAKPAK.T | HexNAc(2)Hex(1) | Yes | 1417.3538 | 3 | 4250.0467 | 4248.026 | 3.3 | NRagged | 828.7 | 167.61 | 2.23 | Nsco_20191108_BC_ZIC_HILIC_K56_2_typsi<br>n_B1.19806.19806.3 | 45.8123 | Trypsin | B1 | 183 |
| BCAL2009 | >gi\|443598395\|gb\|ELT66759.1\| PF08811<br>family protein [Burkholderia cenocepacia<br>K56-2Valvano] | R.AHAMPVTDAAS[+568.212]AAGLR<br>DTPAKPAR.G | HexNAc(2)Hex(1) | No | 736.6168 | 4 | 2943.4452 | 2943.4364 | 3 | Specific | 321.81 | 274.8 | 1.6 | Nsco_20191108_BC_ZIC_HILIC_K56_2_typsi<br>n_B1.16553.16553.4 | 39.5274 | Trypsin | B1 | 184 |
| BCAL2161 | >gi\|443595154\|gb\|ELT63756.1\|<br>hypothetical protein BURCENK562V_1535<br>[Burkholderia cenocepacia K56-2Valvano] | S.SVAPPLQGDGAAPGGAS[+568.212<br>]WPAPPPASGPAPGLPASSVQGT[+56<br>8.212]P.- | HexNAc(2)Hex(1),<br>HexNAc(2)Hex(1) | Yes | 1570.0799 | 3 | 4708.2251 | 4708.2047 | 4.3 | Non | 281.68 | 234.86 | 31.41 | Nsco_20191108_BC_ZIC_HILIC_K56_2_pepsi<br>n_B3_reinject.52747.52747.3 | 119.4034 | Pepsin | B3 | 185 |
| BCAL2161 | >gi\|443595154\|gb\|ELT63756.1\|<br>hypothetical protein BURCENK562V_1535<br>[Burkholderia cenocepacia K56-2Valvano] | S.VAPPLQGDGAAPGGASWPAPPPAS[<br>+568.212]GPAPGLPASSVQGTP.- | HexNAc(2)Hex(1) | No | 1351.6608 | 3 | 4052.9679 | 4052.9611 | 1.7 | Non | 405.9 | 362.09 | 14.46 | Nsco_20191108_BC_ZIC_HILIC_K56_2_ther<br>molysin_B1.55174.55174.3 | 123.2653 | Thermolysin | B1 | 186 |
| BCAL2276 | >gi\|443595650\|gb\|ELT64220.1\|<br>hypothetical protein BURCENK562V_2512<br>[Burkholderia cenocepacia K56-2Valvano] | K.STIDTAASNAGVPVS[+568.212]SVN<br>YIVHDAGK.G | HexNAc(2)Hex(1) | No | 1047.8413 | 3 | 3141.5095 | 3141.4958 | 4.4 | Non | 723.57 | 685.92 | 1.2 | Nsco_20191108_BC_ZIC_HILIC_K56_2_ther<br>molysin_B3.44475.44475.3 | 90.1581 | Thermolysin | B3 | 187 |

| | | | | | | | | | | | | | | | | | | |
|---|---|---|---|---|---|---|---|---|---|---|---|---|---|---|---|---|---|---|
| BCAL2345 | >gi\|443599338\|gb\|ELT67627.1\| preprotein translocase, SecG subunit [Burkholderia cenocepacia K56-2Valvano] | A.VT[+568.212]APASAPAAS[+568.212]APA.V | HexNAc(2)Hex(1), HexNAc(2)Hex(1) | No | 1159.5265 | 2 | 2318.0458 | 2318.0392 | 2.8 | Non | 493.43 | 149.42 | 59.06 | Nsco_20191108_BC_ZIC_HILIC_K56_2_thermolysin_B1.20443.20443.2 | 43.4074 | Thermolysin | B1 | 188 |
| BCAL2398 | >gi\|443599855\|gb\|ELT68098.1\| PRC-barrel domain protein [Burkholderia cenocepacia K56-2Valvano] | K.ASPPYAADKPIVAVFPPVVPAAPAS[+568.212]SASATR.- | HexNAc(2)Hex(1) | No | 1191.9501 | 3 | 3573.8358 | 3573.8211 | 4.1 | Non | 708.62 | 708.62 | 2.4 | Nsco_20191108_BC_ZIC_HILIC_K56_2_thermolysin_B3.52294.52294.3 | 104.3649 | Thermolysin | B3 | 189 |
| BCAL2449 | >gi\|443591735\|gb\|ELT60603.1\| diguanylate cyclase (GGDEF) domain protein [Burkholderia cenocepacia K56-2Valvano] | R.RTGSVNNAPGAFSAS[+568.212]GVYPIAER.V | HexNAc(2)Hex(1) | Yes | 963.7997 | 3 | 2889.3846 | 2889.3749 | 3.4 | Specific | 447.58 | 341.35 | 22.33 | Nsco_20191108_BC_ZIC_HILIC_K56_2_typsin_B3.31131.31131.3 | 65.9643 | Trypsin | B3 | 190 |
| BCAL2466 | >gi\|443591728\|gb\|ELT60596.1\| ecotin [Burkholderia cenocepacia K56-2Valvano] | A.APASAPAVPAES[+568.212]IKMFPQAAAGQQR.V | HexNAc(2)Hex(1) | No | 1021.8398 | 3 | 3063.5048 | 3062.4987 | 0.9 | NRagged | 751.55 | 672.06 | 81.86 | Nsco_20191108_BC_ZIC_HILIC_K56_2_typsin_B2.36444.36444.3 | 76.3838 | Trypsin | B2 | 191 |
| BCAL2629 | >gi\|443603168\|gb\|ELT71192.1\| bifunctional uroporphyrinogen-III synthetase/uroporphyrin-III C-methyltransferase, partial [Burkholderia cenocepacia K56-2Valvano] | K.AAPADAASS[+568.212]VAAGEPR.W | HexNAc(2)Hex(1) | Yes | 1004.9679 | 2 | 2008.9285 | 2008.9193 | 4.6 | Non | 702.03 | 575.5 | 5.99 | Nsco_20191108_BC_ZIC_HILIC_K56_2_thermolysin_B3.16957.16957.2 | 39.3801 | Thermolysin | B3 | 192 |
| BCAL2629 | >gi\|443603168\|gb\|ELT71192.1\| bifunctional uroporphyrinogen-III synthetase/uroporphyrin-III C-methyltransferase, partial [Burkholderia cenocepacia K56-2Valvano] | K.AAPADAAS[+568.212]SVAAGEPRWK.V | HexNAc(2)Hex(1) | Yes | 775.0395 | 3 | 2323.1038 | 2323.0936 | 4.4 | Non | 570.64 | 464.89 | 0.8 | Nsco_20191108_BC_ZIC_HILIC_K56_2_thermolysin_B1.24528.24528.3 | 51.0012 | Thermolysin | B1 | 193 |
| BCAL2629 | >gi\|443603168\|gb\|ELT71192.1\| bifunctional uroporphyrinogen-III synthetase/uroporphyrin-III C-methyltransferase, partial [Burkholderia cenocepacia K56-2Valvano] | H.AAPKAAPADAAS[+568.212]SVAAGEPRWK.V | HexNAc(2)Hex(1) | No | 897.4465 | 3 | 2690.325 | 2690.3155 | 3.5 | Non | 540.27 | 387.31 | 0.8 | Nsco_20191108_BC_ZIC_HILIC_K56_2_thermolysin_B1.20915.20915.3 | 44.2319 | Thermolysin | B1 | 194 |
| BCAL2629 | >gi\|443603168\|gb\|ELT71192.1\| bifunctional uroporphyrinogen-III synthetase/uroporphyrin-III C-methyltransferase, partial [Burkholderia cenocepacia K56-2Valvano] | E.AIVPHAAPKAAPADAAS[+568.212]SVAAGEPRW.K | HexNAc(2)Hex(1) | No | 770.6379 | 4 | 3079.5298 | 3079.5219 | 2.6 | Non | 291.36 | 291.36 | 2 | Nsco_20191108_BC_ZIC_HILIC_K56_2_pepsin_B3_reinject.31321.31321.4 | 64.2187 | Pepsin | B3 | 195 |
| BCAL2820 | >gi\|443594108\|gb\|ELT62784.1\| outer membrane multidrug efflux protein OprB [Burkholderia cenocepacia K56-2Valvano] | K.RPDAPVAQAYPAS[+568.212]GVYATQPGAAGAR.S | HexNAc(2)Hex(1) | No | 1037.838 | 3 | 3111.4995 | 3110.4913 | 1.6 | Specific | 649.41 | 645.11 | 1.41 | Nsco_20191108_BC_ZIC_HILIC_K56_2_typsin_B1.25608.25608.3 | 56.8 | Trypsin | B1 | 196 |
| BCAL2820 | >gi\|443594108\|gb\|ELT62784.1\| outer membrane multidrug efflux protein OprB [Burkholderia cenocepacia K56-2Valvano] | D.YDKAAAPAPAS[+568.212]ATATNG.- | HexNAc(2)Hex(1) | No | 1072.9926 | 2 | 2144.9779 | 2144.9717 | 2.9 | Non | 455.71 | 364.43 | 1.6 | Nsco_20191108_BC_ZIC_HILIC_K56_2_pepsin_B2.12674.12674.2 | 32.2169 | Pepsin | B2 | 197 |
| BCAL2974 | >gi\|443594432\|gb\|ELT63086.1\| hypothetical protein BURCENK562V_0304 [Burkholderia cenocepacia K56-2Valvano] | P.FAASAPS[+568.212]QKYQGSKK.S | HexNAc(2)Hex(1) | No | 1083.5312 | 2 | 2166.0551 | 2166.0449 | 4.7 | Non | 594.76 | 434.49 | 0.25 | Nsco_20191108_BC_ZIC_HILIC_K56_2_thermolysin_B1.5609.5609.2 | 20.0631 | Thermolysin | B1 | 198 |
| BCAL2974 | >gi\|443594432\|gb\|ELT63086.1\| hypothetical protein BURCENK562V_0304 [Burkholderia cenocepacia K56-2Valvano] | P.FAASAPSQKYQGS[+568.212]KKSA.L | HexNAc(2)Hex(1) | No | 1162.5644 | 2 | 2324.1215 | 2324.114 | 3.2 | Non | 743.28 | 499.05 | 1.89 | Nsco_20191108_BC_ZIC_HILIC_K56_2_thermolysin_B1.6072.6072.2 | 20.8198 | Thermolysin | B1 | 199 |
| BCAL2974 | >gi\|443594432\|gb\|ELT63086.1\| hypothetical protein BURCENK562V_0304 [Burkholderia cenocepacia K56-2Valvano] | R.LNEHPQMPFAASAPS[+568.212]QK.Y | HexNAc(2)Hex(1) | No | 807.711 | 3 | 2421.1185 | 2421.1126 | 2.4 | Specific | 861.79 | 837.7 | 11.02 | Nsco_20191108_BC_ZIC_HILIC_K56_2_typsin_B2.22692.22692.3 | 50.5873 | Trypsin | B2 | 200 |

| | | | | | | | | | | | | | | | | | | |
|---|---|---|---|---|---|---|---|---|---|---|---|---|---|---|---|---|---|---|
| BCAL2974 | >gi\|443594432\|gb\|ELT63086.1\| hypothetical protein BURCENK562V_0304 [Burkholderia cenocepacia K56-2Valvano] | L.NEHPQMPFAASAPS[+568.212]QKYQG.S | HexNAc(2)Hex(1) | No | 886.0642 | 3 | 2656.178 | 2656.1719 | 2.3 | CRagged | 660.53 | 660.53 | 8.32 | Nsco_20191108_BC_ZIC_HILIC_K56_2_pepsin_B2.23936.23936.3 | 51.6854 | Pepsin | B2 | 201 |
| BCAL2974 | >gi\|443594432\|gb\|ELT63086.1\| hypothetical protein BURCENK562V_0304 [Burkholderia cenocepacia K56-2Valvano] | L.NEHPQMPFAAS[+568.212]APSQKYQGSKKSAL.R | HexNAc(2)Hex(1) | Yes | 818.3941 | 4 | 3270.5544 | 3270.5471 | 2.2 | Specific | 1159.51 | 1149.4 | 137.04 | Nsco_20191108_BC_ZIC_HILIC_K56_2_pepsin_B1.17741.17741.4 | 39.9659 | Pepsin | B1 | 202 |
| BCAL2974 | >gi\|443594432\|gb\|ELT63086.1\| hypothetical protein BURCENK562V_0304 [Burkholderia cenocepacia K56-2Valvano] | M.PFAASAPS[+568.212]QKYQGSKKSAL.R | HexNAc(2)Hex(1) | No | 845.4256 | 3 | 2534.2623 | 2534.2508 | 4.5 | NRagged | 705.99 | 676.5 | 1.2 | Nsco_20191108_BC_ZIC_HILIC_K56_2_pepsin_B2.12496.12496.3 | 31.9139 | Pepsin | B2 | 203 |
| BCAL2974 | >gi\|443594432\|gb\|ELT63086.1\| hypothetical protein BURCENK562V_0304 [Burkholderia cenocepacia K56-2Valvano] | F.RLNEHPQMPFAAS[+568.212]APSQ.K | HexNAc(2)Hex(1) | Yes | 817.0473 | 3 | 2449.1274 | 2449.1188 | 3.5 | CRagged | 547.35 | 498.47 | 9.62 | Nsco_20191108_BC_ZIC_HILIC_K56_2_pepsin_B1.25396.25396.3 | 54.2559 | Pepsin | B1 | 204 |
| BCAL2974 | >gi\|443594432\|gb\|ELT63086.1\| hypothetical protein BURCENK562V_0304 [Burkholderia cenocepacia K56-2Valvano] | F.RLNEHPQMPFAAS[+568.212]APSQKYQGSKKSAL.R | HexNAc(2)Hex(1) | Yes | 708.7539 | 5 | 3539.7405 | 3539.7323 | 2.3 | Specific | 1167.14 | 1140.8 | 122.97 | Nsco_20191108_BC_ZIC_HILIC_K56_2_pepsin_B2.18532.18532.5 | 41.5782 | Pepsin | B2 | 205 |
| BCAL2983A | >gi\|443599451\|gb\|ELT67730.1\| PF11180 family protein [Burkholderia cenocepacia K56-2Valvano] | S.ADASAPVAGTRPAVTSLSGGASSAASGAVAT[+568.212]DAAAQGNVAELTQMLHDGR.I | HexNAc(2)Hex(1) | No | 1301.8783 | 4 | 5204.4914 | 5204.4772 | 2.7 | NRagged | 898.91 | 855.7 | 0 | Nsco_20191108_BC_ZIC_HILIC_K56_2_typsin_B3.55819.55819.4 | 108.6091 | Trypsin | B3 | 206 |
| BCAL3017 | >gi\|443591015\|gb\|ELT59947.1\| hypothetical protein BURCENK562V_1182 [Burkholderia cenocepacia K56-2Valvano] | A.DTASGSDAQAS[+568.212]C[+57.021]AIAYVTGVGGSPR.G | HexNAc(2)Hex(1) | No | 1483.6628 | 2 | 2966.3184 | 2966.3055 | 4.3 | NRagged | 792.59 | 792.59 | 3.6 | Nsco_20191108_BC_ZIC_HILIC_K56_2_typsin_B1.50784.50784.2 | 100.0398 | Trypsin | B1 | 207 |
| BCAL3033 | >gi\|443591005\|gb\|ELT59937.1\| outer-membrane lipoprotein carrier protein [Burkholderia cenocepacia K56-2Valvano] | K.GASAAQAAPKPT[+568.212]DNSSGTFVFAR.P | HexNAc(2)Hex(1) | No | 940.4492 | 3 | 2819.3332 | 2819.3218 | 4.1 | Specific | 934.51 | 784.38 | 8.88 | Nsco_20191108_BC_ZIC_HILIC_K56_2_typsin_B2.25616.25616.3 | 56.1071 | Trypsin | B2 | 208 |
| BCAL3033 | >gi\|443591005\|gb\|ELT59937.1\| outer-membrane lipoprotein carrier protein [Burkholderia cenocepacia K56-2Valvano] | K.GAS[+568.212]AAQAAPKPTDNSSGTFVFARPGK.F | HexNAc(2)Hex(1) | Yes | 776.1301 | 4 | 3101.4988 | 3101.4909 | 2.5 | Non | 1048.85 | 995.34 | 369.44 | Nsco_20191108_BC_ZIC_HILIC_K56_2_thermolysin_B3.21596.21596.4 | 48.0474 | Thermolysin | B3 | 209 |
| BCAL3033 | >gi\|443591005\|gb\|ELT59937.1\| outer-membrane lipoprotein carrier protein [Burkholderia cenocepacia K56-2Valvano] | Q.IVKAPAKGAS[+568.212]AAQAAPKPTDN.S | HexNAc(2)Hex(1) | No | 859.1134 | 3 | 2575.3257 | 2574.3145 | 3.1 | Non | 407.82 | 407.82 | 23.8 | Nsco_20191108_BC_ZIC_HILIC_K56_2_pepsin_B2.8774.8774.3 | 25.7362 | Pepsin | B2 | 210 |
| BCAL3033 | >gi\|443591005\|gb\|ELT59937.1\| outer-membrane lipoprotein carrier protein [Burkholderia cenocepacia K56-2Valvano] | Q.IVKAPAKGAS[+568.212]AAQAAPKPTDNSSGT.F | HexNAc(2)Hex(1) | Yes | 727.3692 | 4 | 2906.4548 | 2906.4477 | 2.5 | Non | 907.13 | 606.48 | 449.61 | Nsco_20191108_BC_ZIC_HILIC_K56_2_thermolysin_B2_.9118.9118.4 | 25.9654 | Thermolysin | B2 | 211 |
| BCAL3033 | >gi\|443591005\|gb\|ELT59937.1\| outer-membrane lipoprotein carrier protein [Burkholderia cenocepacia K56-2Valvano] | Q.IVKAPAKGAS[+568.212]AAQAAPKPTDNSSGTF.V | HexNAc(2)Hex(1) | Yes | 764.1371 | 4 | 3053.5266 | 3053.5161 | 3.4 | NRagged | 1208.22 | 1174.3 | 519.79 | Nsco_20191108_BC_ZIC_HILIC_K56_2_thermolysin_B1.15310.15310.4 | 34.8469 | Thermolysin | B1 | 212 |
| BCAL3033 | >gi\|443591005\|gb\|ELT59937.1\| outer-membrane lipoprotein carrier protein [Burkholderia cenocepacia K56-2Valvano] | Q.IVKAPAKGAS[+568.212]AAQAAPKPTDNSSGTFV.F | HexNAc(2)Hex(1) | Yes | 788.9039 | 4 | 3152.5937 | 3152.5845 | 2.9 | Non | 855.15 | 855.15 | 397.28 | Nsco_20191108_BC_ZIC_HILIC_K56_2_thermolysin_B2_.19314.19314.4 | 40.6759 | Thermolysin | B2 | 213 |
| BCAL3166 | >gi\|443602480\|gb\|ELT70558.1\| BON domain protein [Burkholderia cenocepacia K56-2Valvano] | A.ATSAPKAAAKT[+568.212]AKKANRKLGY.A | HexNAc(2)Hex(1) | No | 905.826 | 3 | 2715.4635 | 2714.4571 | 1.1 | Non | 536.47 | 356.4 | 13.2 | Nsco_20191108_BC_ZIC_HILIC_K56_2_pepsin_B2.4421.4421.3 | 18.4718 | Pepsin | B2 | 214 |
| BCAL3166 | >gi\|443602480\|gb\|ELT70558.1\| BON domain protein [Burkholderia cenocepacia K56-2Valvano] | A.PAAATSAPKAAAKT[+568.212]AKKANRKLGY.A | HexNAc(2)Hex(1) | No | 739.1546 | 4 | 2953.5967 | 2953.5841 | 4.3 | Non | 424.76 | 424.76 | 13.2 | Nsco_20191108_BC_ZIC_HILIC_K56_2_pepsin_B2.6997.6997.4 | 23.0272 | Pepsin | B2 | 215 |

| | | | | | | | | | | | | | | | | | | |
|---|---|---|---|---|---|---|---|---|---|---|---|---|---|---|---|---|---|---|
| BCAL3166 | >gi\|443602480\|gb\|ELT70558.1\| BON domain protein [Burkholderia cenocepacia K56-2Valvano] | A.TEAPAAATSAPKAAAKT[+568.212]AKKANRKLGY.A | HexNAc(2)Hex(1) | No | 1085.9101 | 3 | 3255.7156 | 3254.7114 | 0.3 | Non | 939.76 | 370.14 | 9.06 | Nsco_20191108_BC_ZIC_HILIC_K56_2_pepsin_B3_reinject.11245.11245.3 | 29.3064 | Pepsin | B3 | 216 |
| BCAL3377 | >gi\|443595379\|gb\|ELT63971.1\| SmpA/OmlA family protein, partial [Burkholderia cenocepacia K56-2Valvano] | A.AAKKAS[+568.212]EAAAAASAAQA.A | HexNAc(2)Hex(1) | No | 1028.5033 | 2 | 2055.9993 | 2055.9928 | 3.2 | Non | 364.7 | 73.61 | 0.09 | Nsco_20191108_BC_ZIC_HILIC_K56_2_thermolysin_B2_.6539.6539.2 | 22.1025 | Thermolysin | B2 | 217 |
| BCAL3377 | >gi\|443595379\|gb\|ELT63971.1\| SmpA/OmlA family protein, partial [Burkholderia cenocepacia K56-2Valvano] | D.IDGDRGGKKAKAAAAAKKAS[+568.212]EAAA.A | HexNAc(2)Hex(1) | Yes | 706.8718 | 4 | 2824.4656 | 2824.4534 | 4.3 | Non | 240.3 | 240.3 | 240.3 | Nsco_20191108_BC_ZIC_HILIC_K56_2_pepsin_B2.3463.3463.3 | 17.0527 | Pepsin | B2 | 218 |
| BCAL3469 | >gi\|443597454\|gb\|ELT65879.1\| cell division protein FtsL [Burkholderia cenocepacia K56-2Valvano] | A.IDAPIPAS[+568.212]ADTAGKGKGGAR.- | HexNAc(2)Hex(1) | No | 807.7407 | 3 | 2421.2075 | 2421.1991 | 3.5 | Non | 796.3 | 567.41 | 2.4 | Nsco_20191108_BC_ZIC_HILIC_K56_2_thermolysin_B1.12694.12694.3 | 30.7374 | Thermolysin | B1 | 219 |
| BCAM0157 | >gi\|443592579\|gb\|ELT61370.1\| hypothetical protein BURCENK562V_A2231 [Burkholderia cenocepacia K56-2Valvano] | R.AVDC[+57.021]GQLDAATSGPDDNFRPPAS[+568.212]GTVIGTGR.A | HexNAc(2)Hex(1) | No | 1224.2299 | 3 | 3670.6752 | 3670.6661 | 2.5 | Specific | 715.98 | 715.98 | 4.8 | Nsco_20191108_BC_ZIC_HILIC_K56_2_trypsin_B1.38785.38785.3 | 80.0577 | Trypsin | B1 | 220 |
| BCAM0175a | >gi\|443598439\|gb\|ELT66798.1\| putative lipoprotein [Burkholderia cenocepacia K56-2Valvano] | A.TPSSTPANASAPADT[+568.212]AAQAAPPR.V | HexNAc(2)Hex(1) | No | 1359.6371 | 2 | 2718.2669 | 2718.2588 | 3 | NRagged | 795.66 | 290.44 | 0 | Nsco_20191108_BC_ZIC_HILIC_K56_2_trypsin_B3.18096.18096.2 | 42.1515 | Trypsin | B3 | 221 |
| BCAM0345 | >gi\|443597793\|gb\|ELT66203.1\| hypothetical protein BURCENK562V_A1193 [Burkholderia cenocepacia K56-2Valvano] | R.SAPAGMQDGASS[+568.212]APQPALDR.L | HexNAc(2)Hex(1) | No | 1248.0628 | 2 | 2495.1183 | 2495.109 | 3.7 | Specific | 804.86 | 763.61 | 6.85 | Nsco_20191108_BC_ZIC_HILIC_K56_2_trypsin_B3.21707.21707.2 | 48.7322 | Trypsin | B3 | 222 |
| BCAM0505 | >gi\|443591075\|gb\|ELT60005.1\| BON domain protein [Burkholderia cenocepacia K56-2Valvano] | Q.ASSTDSGMASES[+568.212]NQPVTDTWITTK.V | HexNAc(2)Hex(1) | No | 1541.681 | 2 | 3082.3546 | 3082.3416 | 4.2 | NRagged | 653.93 | 653.93 | 1.2 | Nsco_20191108_BC_ZIC_HILIC_K56_2_trypsin_B1.37726.37726.2 | 78.0391 | Trypsin | B1 | 223 |
| BCAM0505 | >gi\|443591075\|gb\|ELT60005.1\| BON domain protein [Burkholderia cenocepacia K56-2Valvano] | A.HAQASSTDSGMASES[+568.212]NQPVTDTWITTK.V | HexNAc(2)Hex(1) | No | 1140.1741 | 3 | 3418.5077 | 3418.4962 | 3.4 | Non | 850.7 | 850.7 | 2.4 | Nsco_20191108_BC_ZIC_HILIC_K56_2_thermolysin_B3.30162.30162.3 | 63.8622 | Thermolysin | B3 | 224 |
| BCAM0505 | >gi\|443591075\|gb\|ELT60005.1\| BON domain protein [Burkholderia cenocepacia K56-2Valvano] | A.QAS[+568.212]STDSGMASESNQPVTDTW.I | HexNAc(2)Hex(1) | Yes | 1384.0723 | 2 | 2767.1373 | 2767.1258 | 4.2 | Non | 269.64 | 185.63 | 21.49 | Nsco_20191108_BC_ZIC_HILIC_K56_2_thermolysin_B1.53831.53831.2 | 119.557 | Thermolysin | B1 | 225 |
| BCAM0505 | >gi\|443591075\|gb\|ELT60005.1\| BON domain protein [Burkholderia cenocepacia K56-2Valvano] | A.QASSTDSGMASES[+568.212]NQPVTDTWITTK.V | HexNAc(2)Hex(1) | No | 1606.7017 | 2 | 3212.396 | 3210.4002 | -3.4 | NRagged | 518.98 | 518.98 | 3.6 | Nsco_20191108_BC_ZIC_HILIC_K56_2_trypsin_B3.39328.39328.2 | 80.5455 | Trypsin | B3 | 226 |
| BCAM0505 | >gi\|443591075\|gb\|ELT60005.1\| BON domain protein [Burkholderia cenocepacia K56-2Valvano] | D.SGMASES[+568.212]NQPVT[+568.212]DTWITTK.V | HexNAc(2)Hex(1), HexNAc(2)Hex(1) | No | 1064.7974 | 3 | 3192.3776 | 3189.3774 | -3.1 | NRagged | 347.67 | 347.67 | 4.8 | Nsco_20191108_BC_ZIC_HILIC_K56_2_trypsin_B3.50916.50916.3 | 100.1249 | Trypsin | B3 | 227 |
| BCAM0505 | >gi\|443591075\|gb\|ELT60005.1\| BON domain protein [Burkholderia cenocepacia K56-2Valvano] | A.SSTDSGMASES[+568.212]NQPVTDTWITTK.V | HexNAc(2)Hex(1) | No | 1506.6624 | 2 | 3012.3174 | 3011.3045 | 3.2 | Non | 542.49 | 525.03 | 1.2 | Nsco_20191108_BC_ZIC_HILIC_K56_2_thermolysin_B3.38287.38287.2 | 78.9073 | Thermolysin | B3 | 228 |
| BCAM0690 | >gi\|443592239\|gb\|ELT61058.1\| OmpA family protein [Burkholderia cenocepacia K56-2Valvano] | R.ALIDAGVPASSVFAAAFGS[+568.212]EQPVSSNADDEGRAK.N | HexNAc(2)Hex(1) | No | 1301.9484 | 3 | 3903.8307 | 3902.8302 | -0.7 | Specific | 506.19 | 469.64 | 6 | Nsco_20191108_BC_ZIC_HILIC_K56_2_trypsin_B1.60279.60279.3 | 115.7062 | Trypsin | B1 | 229 |
| BCAM0713 | >gi\|443605703\|gb\|ELT73537.1\| cation efflux system protein CzcA [Burkholderia cenocepacia K56-2Valvano] | K.RLPIALPAPAAGAS[+568.212]APLAAAPYVPLAELATIDVAPGPNQISREDGK.R | HexNAc(2)Hex(1) | Yes | 1275.4307 | 4 | 5098.7008 | 5098.6834 | 3.4 | Specific | 1131.96 | 1132 | 379.2 | Nsco_20191108_BC_ZIC_HILIC_K56_2_trypsin_B2.70574.70574.4 | 143.2396 | Trypsin | B2 | 230 |
| BCAM0996 | >gi\|443592743\|gb\|ELT61524.1\| sporulation and cell division repeat protein [Burkholderia cenocepacia K56-2Valvano] | R.DDDVSDVQAGVAHDEPPAS[+568.212]DTTVAAAPAPAPK.D | HexNAc(2)Hex(1) | No | 1228.2274 | 3 | 3682.6675 | 3682.6614 | 1.7 | Specific | 1134.46 | 1134.5 | 4.8 | Nsco_20191108_BC_ZIC_HILIC_K56_2_trypsin_B1.35376.35376.3 | 73.8871 | Trypsin | B1 | 231 |

<table>
<tr><td>BCAM0996</td><td>>gi|443592743|gb|ELT61524.1| sporulation and cell division repeat protein [Burkholderia cenocepacia K56-2Valvano]</td><td>K.PAAPAAKPAAPKPAPATVANAGAASPDS[+568.212]GDASSPASPAGAR.F</td><td>HexNAc(2)Hex(1)</td><td>No</td><td>1040.7653</td><td>4</td><td>4160.0394</td><td>4160.0266</td><td>3.1</td><td>Specific</td><td>1138.91</td><td>1138.9</td><td>0.4</td><td>Nsco_20191108_BC_ZIC_HILIC_K56_2_typsin_B2.19311.19311.4</td><td>44.2627</td><td>Trypsin</td><td>B2</td><td>232</td></tr>
<tr><td>BCAM0996</td><td>>gi|443592743|gb|ELT61524.1| sporulation and cell division repeat protein [Burkholderia cenocepacia K56-2Valvano]</td><td>K.PAAPKPAPATVANAGAASPDSGDAS[+568.212]SPASPAGAR.F</td><td>HexNAc(2)Hex(1)</td><td>No</td><td>1185.2342</td><td>3</td><td>3553.688</td><td>3553.6776</td><td>2.9</td><td>Specific</td><td>906.46</td><td>906.46</td><td>2.4</td><td>Nsco_20191108_BC_ZIC_HILIC_K56_2_typsin_B1.19901.19901.3</td><td>45.9826</td><td>Trypsin</td><td>B1</td><td>233</td></tr>
<tr><td>BCAM0996</td><td>>gi|443592743|gb|ELT61524.1| sporulation and cell division repeat protein [Burkholderia cenocepacia K56-2Valvano]</td><td>P.VTDDIAIDIPNRPAHQAVAPRDDDVSDVQAGVAHDEPPASDT[+568.212]TVAAAPAPAPK.D</td><td>HexNAc(2)Hex(1)</td><td>No</td><td>1187.7791</td><td>5</td><td>5934.8662</td><td>5932.8483</td><td>1.9</td><td>NRagged</td><td>903.05</td><td>347.07</td><td>2.4</td><td>Nsco_20191108_BC_ZIC_HILIC_K56_2_typsin_B3.38825.38825.5</td><td>79.6066</td><td>Trypsin</td><td>B3</td><td>234</td></tr>
<tr><td>BCAM1398</td><td>>gi|443598009|gb|ELT66406.1| gram-negative porin [Burkholderia cenocepacia K56-2Valvano]</td><td>L.GGGAWGDNSTVTSGGQPAVS[+568.212][+100.064]AINNGYQTAQSQQR.I</td><td>HexNAc(2)Hex(1) 100.064</td><td>No</td><td>1346.2901</td><td>3</td><td>4036.8557</td><td>4032.8293</td><td>3.2</td><td>CRagged</td><td>373.27</td><td>373.27</td><td>6</td><td>Nsco_20191108_BC_ZIC_HILIC_K56_2_thermolysin_B3.42045.42045.3</td><td>85.7221</td><td>Thermolysin</td><td>B3</td><td>235</td></tr>
<tr><td>BCAM1550</td><td>>gi|443602195|gb|ELT70283.1| peptidoglycan-associated lipoprotein [Burkholderia cenocepacia K56-2Valvano]</td><td>T.PENAGAAPEPSSETVAT[+568.212][+100.064]VTADDLNNPNSPLAK.R</td><td>HexNAc(2)Hex(1) 100.064</td><td>No</td><td>1282.6044</td><td>3</td><td>3845.7987</td><td>3845.7938</td><td>1.3</td><td>NRagged</td><td>631.68</td><td>631.68</td><td>2.4</td><td>Nsco_20191108_BC_ZIC_HILIC_K56_2_typsin_B1.45909.45909.3</td><td>92.3259</td><td>Trypsin</td><td>B1</td><td>236</td></tr>
<tr><td>BCAM1550</td><td>>gi|443602195|gb|ELT70283.1| peptidoglycan-associated lipoprotein [Burkholderia cenocepacia K56-2Valvano]</td><td>T.PENAGAAPEPSSET[+568.212][+100.064]VATVTADDLNNPNSPLAKR.S</td><td>HexNAc(2)Hex(1) 100.064</td><td>No</td><td>1334.9717</td><td>3</td><td>4002.9007</td><td>4001.8949</td><td>0.6</td><td>NRagged</td><td>737.92</td><td>705.13</td><td>0.82</td><td>Nsco_20191108_BC_ZIC_HILIC_K56_2_typsin_B1.40248.40248.3</td><td>82.4319</td><td>Trypsin</td><td>B1</td><td>237</td></tr>
<tr><td>BCAM1550</td><td>>gi|443602195|gb|ELT70283.1| peptidoglycan-associated lipoprotein [Burkholderia cenocepacia K56-2Valvano]</td><td>K.TPENAGAAPEPSSETVAT[+568.212]VTADDLNNPNSPLAK.R</td><td>HexNAc(2)Hex(1)</td><td>No</td><td>1282.9327</td><td>3</td><td>3846.7835</td><td>3846.7775</td><td>1.6</td><td>Specific</td><td>1037.88</td><td>260.78</td><td>2.4</td><td>Nsco_20191108_BC_ZIC_HILIC_K56_2_typsin_B3.46141.46141.3</td><td>92.2169</td><td>Trypsin</td><td>B3</td><td>238</td></tr>
<tr><td>BCAM1550</td><td>>gi|443602195|gb|ELT70283.1| peptidoglycan-associated lipoprotein [Burkholderia cenocepacia K56-2Valvano]</td><td>K.TPENAGAAPEPSSETVATVT[+568.212]ADDLNNPNSPLAKR.S</td><td>HexNAc(2)Hex(1)</td><td>No</td><td>1334.9677</td><td>3</td><td>4002.8884</td><td>4002.8786</td><td>2.5</td><td>Specific</td><td>1109.66</td><td>253.74</td><td>0</td><td>Nsco_20191108_BC_ZIC_HILIC_K56_2_typsin_B1.40339.40339.3</td><td>82.5901</td><td>Trypsin</td><td>B1</td><td>239</td></tr>
<tr><td>BCAM1669</td><td>>gi|443598083|gb|ELT66474.1| hypothetical protein BURCENK562V_A2079 [Burkholderia cenocepacia K56-2Valvano]</td><td>R.VHGIDNSGAGS[+568.212]QPAATVEGGAPVV.R</td><td>HexNAc(2)Hex(1)</td><td>No</td><td>1380.1554</td><td>2</td><td>2759.3035</td><td>2758.2901</td><td>3.6</td><td>Non</td><td>698.88</td><td>650.18</td><td>2.4</td><td>Nsco_20191108_BC_ZIC_HILIC_K56_2_thermolysin_B3.34070.34070.2</td><td>70.9495</td><td>Thermolysin</td><td>B3</td><td>240</td></tr>
<tr><td>BCAM1669</td><td>>gi|443598083|gb|ELT66474.1| hypothetical protein BURCENK562V_A2079 [Burkholderia cenocepacia K56-2Valvano]</td><td>R.VHGIDNSGAGS[+568.212]QPAATVEGGAPVVR.A</td><td>HexNAc(2)Hex(1)</td><td>No</td><td>972.1373</td><td>3</td><td>2914.3973</td><td>2914.3912</td><td>2.1</td><td>Specific</td><td>894.84</td><td>894.84</td><td>1.2</td><td>Nsco_20191108_BC_ZIC_HILIC_K56_2_typsin_B3.23965.23965.3</td><td>52.8858</td><td>Trypsin</td><td>B3</td><td>241</td></tr>
<tr><td>BCAM1789</td><td>>gi|443592202|gb|ELT61024.1| putative esterase [Burkholderia cenocepacia K56-2Valvano]</td><td>A.TPPAPAPAASPAQPPAVQTATTPS[+568.212]TAQEPSVNPGSSVVLR.T</td><td>HexNAc(2)Hex(1)</td><td>No</td><td>1473.0701</td><td>3</td><td>4417.1957</td><td>4416.194</td><td>-0.4</td><td>Non</td><td>975.65</td><td>248.91</td><td>0</td><td>Nsco_20191108_BC_ZIC_HILIC_K56_2_thermolysin_B3.44674.44674.3</td><td>90.4706</td><td>Thermolysin</td><td>B3</td><td>242</td></tr>
<tr><td>BCAM2055</td><td>>gi|443591388|gb|ELT60286.1| type III secretion outer membrane pore, YscC/HrcC-like family protein [Burkholderia cenocepacia K56-2Valvano]</td><td>K.DTAASQPAATT[+568.212]AGVVTHVDEHH.-</td><td>HexNAc(2)Hex(1)</td><td>No</td><td>696.5697</td><td>4</td><td>2783.2568</td><td>2783.249</td><td>2.8</td><td>Non</td><td>629.35</td><td>629.35</td><td>1.2</td><td>Nsco_20191108_BC_ZIC_HILIC_K56_2_thermolysin_B3.20174.20174.4</td><td>45.2282</td><td>Thermolysin</td><td>B3</td><td>243</td></tr>
<tr><td>BCAM2063</td><td>>gi|443595086|gb|ELT63692.1| carbohydrate-selective porin, OprB family [Burkholderia cenocepacia K56-2Valvano]</td><td>A.SSPAAAEPAAGAS[+568.212]DAAAPAQQAADAAAPAPTGFWER.S</td><td>HexNAc(2)Hex(1)</td><td>No</td><td>1306.9322</td><td>3</td><td>3918.782</td><td>3918.7788</td><td>0.8</td><td>NRagged</td><td>755.5</td><td>755.5</td><td>8.4</td><td>Nsco_20191108_BC_ZIC_HILIC_K56_2_typsin_B2.47912.47912.3</td><td>96.7611</td><td>Trypsin</td><td>B2</td><td>244</td></tr>
<tr><td>BCAM2064</td><td>>gi|443595102|gb|ELT63708.1| alpha,alpha-trehalase [Burkholderia cenocepacia K56-2Valvano]</td><td>Q.AAAQAVGQSAIPATTAAAAAPAS[+568.212]GT[+568.212]LPPPPSQLYGDLFVAVQTAQLYPDQK.T</td><td>HexNAc(2)Hex(1), HexNAc(2)Hex(1)</td><td>No</td><td>1533.5116</td><td>4</td><td>6131.0246</td><td>6128.0021</td><td>2</td><td>NRagged</td><td>367.41</td><td>367.41</td><td>0</td><td>Nsco_20191108_BC_ZIC_HILIC_K56_2_typsin_B2.77678.77678.4</td><td>160.9177</td><td>Trypsin</td><td>B2</td><td>245</td></tr>
<tr><td>BCAM2064</td><td>>gi|443595102|gb|ELT63708.1| alpha,alpha-trehalase [Burkholderia cenocepacia K56-2Valvano]</td><td>A.DNANQAAAQAVGQSAIPATTAAAAAPAS[+568.212]GTLPPPPSQLYGDLFVAVQTAQLYPDQK.T</td><td>HexNAc(2)Hex(1)</td><td>No</td><td>1526.2625</td><td>4</td><td>6102.028</td><td>6101.999</td><td>4.8</td><td>NRagged</td><td>932.43</td><td>887.63</td><td>5.77</td><td>Nsco_20191108_BC_ZIC_HILIC_K56_2_typsin_B3.85867.85867.4</td><td>160.5315</td><td>Trypsin</td><td>B3</td><td>246</td></tr>
<tr><td>BCAM2289</td><td>>gi|443599704|gb|ELT67963.1| hypothetical protein BURCENK562V_A2967 [Burkholderia cenocepacia K56-2Valvano]</td><td>R.VHGADTSGYGAQPAPLVHSGAPAAAS[+568.212]SNAR.D</td><td>HexNAc(2)Hex(1)</td><td>Yes</td><td>1129.1995</td><td>3</td><td>3385.5839</td><td>3385.5779</td><td>1.8</td><td>Specific</td><td>1258.46</td><td>1100.4</td><td>2.4</td><td>Nsco_20191108_BC_ZIC_HILIC_K56_2_typsin_B3.17140.17140.3</td><td>40.4898</td><td>Trypsin</td><td>B3</td><td>247</td></tr>
</table>

| | | | | | | | | | | | | | | | | | | |
|---|---|---|---|---|---|---|---|---|---|---|---|---|---|---|---|---|---|---|
| BCAM2334 | >gi\|443606050\|gb\|ELT73855.1\| auxiliary transport protein, membrane fusion protein (MFP) domain protein, partial [Burkholderia cenocepacia K56-2Valvano] | R.VHDGVAS[+568.212]DAEAAAAAII RENQGG.- | HexNAc(2)Hex(1) | Yes | 930.7708 | 3 | 2790.298 | 2790.2912 | 2.4 | Non | 592.51 | 592.51 | 592.51 | Nsco_20191108_BC_ZIC_HILIC_K56_2_ther molysin_B3.49647.49647.3 | 99.379 | Thermolysin | B3 | 248 |
| BCAM2378 | >gi\|443594879\|gb\|ELT63498.1\| X-Pro dipeptidyl-peptidase (S15 family) [Burkholderia cenocepacia K56-2Valvano] | L.AQGQQEGASAAAGT[+568.212]ANT PSL.A | HexNAc(2)Hex(1) | No | 1199.5435 | 2 | 2398.0797 | 2398.074 | 2.4 | Specific | 333.01 | 260.81 | 2.4 | Nsco_20191108_BC_ZIC_HILIC_K56_2_pepsi n_B2.27030.27030.2 | 57.6522 | Pepsin | B2 | 249 |
| BCAM2378 | >gi\|443594879\|gb\|ELT63498.1\| X-Pro dipeptidyl-peptidase (S15 family) [Burkholderia cenocepacia K56-2Valvano] | A.LAQGQQEGASAAAGT[+568.212]A NTPS.L | HexNAc(2)Hex(1) | No | 1199.5439 | 2 | 2398.0806 | 2398.074 | 2.8 | Non | 452.69 | 317.95 | 3.6 | Nsco_20191108_BC_ZIC_HILIC_K56_2_ther molysin_B1.16414.16414.2 | 36.6046 | Thermolysin | B1 | 250 |
| BCAM2378 | >gi\|443594879\|gb\|ELT63498.1\| X-Pro dipeptidyl-peptidase (S15 family) [Burkholderia cenocepacia K56-2Valvano] | T.PVIVLASPYYAGLADSPNHDVDVELDG TPHPAATAGAAAS[+568.212][+100. 064]ASAR.I | HexNAc(2)Hex(1) 100.064 | No | 1246.6063 | 4 | 4983.4035 | 4983.4021 | 0.3 | NRagged | 511.21 | 511.21 | 4.8 | Nsco_20191108_BC_ZIC_HILIC_K56_2_typsi n_B1.56834.56834.4 | 110.1277 | Trypsin | B1 | 251 |
| BCAM2378 | >gi\|443594879\|gb\|ELT63498.1\| X-Pro dipeptidyl-peptidase (S15 family) [Burkholderia cenocepacia K56-2Valvano] | R.TPVIVLASPYYAGLADSPNHDVDVELD GTPHPAATAGAAAS[+568.212]ASAR .I | HexNAc(2)Hex(1) | No | 1246.8569 | 4 | 4984.4059 | 4984.3858 | 4 | Specific | 1206.74 | 471.98 | 2.36 | Nsco_20191108_BC_ZIC_HILIC_K56_2_typsi n_B3.56510.56510.4 | 109.7632 | Trypsin | B3 | 252 |
| BCAM2443 | >gi\|443594187\|gb\|ELT62858.1\| cytochrome C [Burkholderia cenocepacia K56-2Valvano] | G.ATPQDAPAAASAPPPAPAAAPAAK PFT[+568.212]PPPESAIPADDFGK.T | HexNAc(2)Hex(1) | No | 1482.3965 | 3 | 4445.1751 | 4445.1559 | 4.3 | NRagged | 918.54 | 918.54 | 8.4 | Nsco_20191108_BC_ZIC_HILIC_K56_2_typsi n_B1.46671.46671.3 | 93.5713 | Trypsin | B1 | 253 |
| BCAM2680 | >gi\|443603690\|gb\|ELT71680.1\| hypothetical protein BURCENK562V_A2523, partial [Burkholderia cenocepacia K56-2Valvano] | A.AAQQASVPAAAS[+568.212]AT[+5 68.212]VVVK.A | HexNAc(2)Hex(1), HexNAc(2)Hex(1) | No | 1403.1841 | 2 | 2805.3609 | 2805.351 | 3.5 | Non | 722.03 | 483.86 | 93.02 | Nsco_20191108_BC_ZIC_HILIC_K56_2_ther molysin_B2_.25792.25792.2 | 51.1424 | Thermolysin | B2 | 254 |
| BCAM2680 | >gi\|443603690\|gb\|ELT71680.1\| hypothetical protein BURCENK562V_A2523, partial [Burkholderia cenocepacia K56-2Valvano] | A.AAQQASVPAAAS[+568.212]AT[+5 68.212]VVVKAAPQPQNP.V | HexNAc(2)Hex(1), HexNAc(2)Hex(1) | No | 1203.9231 | 3 | 3609.7547 | 3608.7436 | 2.1 | Non | 385.63 | 385.63 | 72.79 | Nsco_20191108_BC_ZIC_HILIC_K56_2_ther molysin_B2_.36532.36532.3 | 68.4457 | Thermolysin | B2 | 255 |
| BCAM2680 | >gi\|443603690\|gb\|ELT71680.1\| hypothetical protein BURCENK562V_A2523, partial [Burkholderia cenocepacia K56-2Valvano] | Q.AASAAAAQQAS[+568.212]VPAAA S[+568.212][+100.064]AT[+568.2 12]V.V | HexNAc(2)Hex(1), HexNAc(2)Hex(1) 100.064,HexNAc( 2)Hex(1) | No | 1173.8671 | 3 | 3519.5868 | 3518.5753 | 2.3 | Non | 242.35 | 44.1 | 0 | Nsco_20191108_BC_ZIC_HILIC_K56_2_pepsi n_B3_reinject.32149.32149.3 | 65.8129 | Pepsin | B3 | 256 |
| BCAM2829 | >gi\|443591146\|gb\|ELT60069.1\| hypothetical protein BURCENK562V_A2847 [Burkholderia cenocepacia K56-2Valvano] | D.SAMPASAPAAAS[+568.212][+10 0.064]APASSGATTA.P | HexNAc(2)Hex(1) 100.064 | No | 1258.0699 | 2 | 2515.1325 | 2514.1403 | -4.4 | Non | 211.02 | 170.46 | 2.4 | Nsco_20191108_BC_ZIC_HILIC_K56_2_pepsi n_B3_reinject.38438.38438.3 | 79.4583 | Pepsin | B3 | 257 |
| BCAM2829 | >gi\|443591146\|gb\|ELT60069.1\| hypothetical protein BURCENK562V_A2847 [Burkholderia cenocepacia K56-2Valvano] | D.SAMPASAPAAASAPASS[+568.212 ]GAT[+568.212]TAPAANGAPIA.P | HexNAc(2)Hex(1), HexNAc(2)Hex(1) | No | 1248.9059 | 3 | 3744.7033 | 3744.6903 | 3.5 | Non | 482.2 | 259.04 | 1.2 | Nsco_20191108_BC_ZIC_HILIC_K56_2_pepsi n_B2.36147.36147.3 | 77.5477 | Pepsin | B2 | 258 |
| BCAM2829 | >gi\|443591146\|gb\|ELT60069.1\| hypothetical protein BURCENK562V_A2847 [Burkholderia cenocepacia K56-2Valvano] | D.S[+568.212]AMPASAPAAASAPAS SGAT[+568.212]TAPAANGAPIAPAM .P | HexNAc(2)Hex(1), HexNAc(2)Hex(1) | No | 1348.6181 | 3 | 4043.8397 | 4043.8207 | 4.7 | Non | 283.55 | 225.63 | 1.18 | Nsco_20191108_BC_ZIC_HILIC_K56_2_pepsi n_B3_reinject.43487.43487.3 | 92.7742 | Pepsin | B3 | 259 |
| BCAS0089 | >gi\|443598997\|gb\|ELT67309.1\| hypothetical protein BURCENK562V_B0207 [Burkholderia cenocepacia K56-2Valvano] | N.AVQQAPATKAAPAVPAS[+568.21 2]GQ.- | HexNAc(2)Hex(1) | No | 1166.0856 | 2 | 2331.164 | 2331.1562 | 3.3 | Non | 576.42 | 463 | 48.07 | Nsco_20191108_BC_ZIC_HILIC_K56_2_ther molysin_B1.15804.15804.2 | 35.6316 | Thermolysin | B1 | 260 |
| BCAS0089 | >gi\|443598997\|gb\|ELT67309.1\| hypothetical protein BURCENK562V_B0207 [Burkholderia cenocepacia K56-2Valvano] | A.VQQAPATKAAPAVPAS[+568.212] GQ.- | HexNAc(2)Hex(1) | No | 1130.5662 | 2 | 2260.1251 | 2260.1191 | 2.7 | Non | 626.38 | 388.77 | 65.96 | Nsco_20191108_BC_ZIC_HILIC_K56_2_ther molysin_B1.14541.14541.2 | 33.7178 | Thermolysin | B1 | 261 |

| | | | | | | | | | | | | | | | | | | |
|---|---|---|---|---|---|---|---|---|---|---|---|---|---|---|---|---|---|---|
| BCAS0321 | >gi\|443594043\|gb\|ELT62726.1\| autotransporter-associated beta strand repeat (3 repeats) [Burkholderia cenocepacia K56-2Valvano] | W.NIT[+568.212]GASS[+568.212]AADASLA.G | HexNAc(2)Hex(1), HexNAc(2)Hex(1) | No | 1193.0223 | 2 | 2385.0372 | 2385.0298 | 3.1 | Non | 210.12 | 1.04 | 1.04 | Nsco_20191108_BC_ZIC_HILIC_K56_2_pepsin_B1.25683.25683.2 | 54.848 | Pepsin | B1 | 262 |
| BURCENK562V_3104 | >gi\|443598112\|gb\|ELT66499.1\| hypothetical protein BURCENK562V_3104 [Burkholderia cenocepacia K56-2Valvano] | G.AAADGAKDTTSSAVHST[+568.212]KKHTKHAAKSAKSHAGSA.K | HexNAc(2)Hex(1) | No | 797.6005 | 5 | 3983.9733 | 3983.9541 | 4.8 | Non | 652.05 | 652.05 | 1.2 | Nsco_20191108_BC_ZIC_HILIC_K56_2_pepsin_B3_reinject.3845.3845.5 | 16.9543 | Pepsin | B3 | 263 |
| BURCENK562V_3104 | >gi\|443598112\|gb\|ELT66499.1\| hypothetical protein BURCENK562V_3104 [Burkholderia cenocepacia K56-2Valvano] | S.AAFAQTGTT[+568.212]AQGAAGAGVQAQTPAAGVGAGAQGGAGANAS[+568.212]GNATGAATDAVGAAADGAK.D | HexNAc(2)Hex(1), HexNAc(2)Hex(1) | No | 1513.7054 | 4 | 6051.8 | 6048.7672 | 3.8 | NRagged | 392.33 | 276.15 | 5.06 | Nsco_20191108_BC_ZIC_HILIC_K56_2_typsin_B2.61923.61923.3 | 123.5184 | Trypsin | B2 | 264 |
| BURCENK562V_3104 | >gi\|443598112\|gb\|ELT66499.1\| hypothetical protein BURCENK562V_3104 [Burkholderia cenocepacia K56-2Valvano] | K.AKADEATEGGAS[+568.212]VGAQGGASAQGATQ.- | HexNAc(2)Hex(1) | No | 1429.6388 | 2 | 2858.2704 | 2858.2658 | 1.6 | CRagged | 834.52 | 757.98 | 2.47 | Nsco_20191108_BC_ZIC_HILIC_K56_2_typsin_B1.11369.11369.2 | 30.6907 | Trypsin | B1 | 265 |
| BURCENK562V_3104 | >gi\|443598112\|gb\|ELT66499.1\| hypothetical protein BURCENK562V_3104 [Burkholderia cenocepacia K56-2Valvano] | N.ASGNATGAATDAVGAAADGAKDT[+568.212]TSSAVHS.T | HexNAc(2)Hex(1) | No | 1067.4822 | 3 | 3200.4321 | 3200.4197 | 3.9 | Non | 959.33 | 875.29 | 2.4 | Nsco_20191108_BC_ZIC_HILIC_K56_2_thermolysin_B1.29679.29679.3 | 60.5536 | Thermolysin | B1 | 266 |
| BURCENK562V_3104 | >gi\|443598112\|gb\|ELT66499.1\| hypothetical protein BURCENK562V_3104 [Burkholderia cenocepacia K56-2Valvano] | N.ASGNATGAATDAVGAAADGAKDTTS[+568.212]AVHSTKKH.T | HexNAc(2)Hex(1) | Yes | 739.751 | 5 | 3694.7259 | 3694.7162 | 2.6 | Non | 493.63 | 493.63 | 20.41 | Nsco_20191108_BC_ZIC_HILIC_K56_2_thermolysin_B1.19879.19879.5 | 42.3129 | Thermolysin | B1 | 267 |
| BURCENK562V_3104 | >gi\|443598112\|gb\|ELT66499.1\| hypothetical protein BURCENK562V_3104 [Burkholderia cenocepacia K56-2Valvano] | K.DT[+568.212]TSSAVHSTKK.H | HexNAc(2)Hex(1) | No | 610.6228 | 3 | 1829.8539 | 1829.8498 | 2.2 | Specific | 250.3 | 191.7 | 0.8 | Nsco_20191108_BC_ZIC_HILIC_K56_2_typsin_B1.2097.2097.3 | 14.3265 | Trypsin | B1 | 268 |
| BURCENK562V_3104 | >gi\|443598112\|gb\|ELT66499.1\| hypothetical protein BURCENK562V_3104 [Burkholderia cenocepacia K56-2Valvano] | Q.GAAGAGVQAQTPAAGVGAGAQGGAGANAS[+568.212][+100.064]GNATGAATDAVGAAADGAK.D | HexNAc(2)Hex(1) 100.064 | No | 1512.0524 | 3 | 4534.1427 | 4533.1211 | 4 | NRagged | 867 | 854.17 | 4.8 | Nsco_20191108_BC_ZIC_HILIC_K56_2_typsin_B2.63789.63789.3 | 127.5147 | Trypsin | B2 | 269 |
| BURCENK562V_3104 | >gi\|443598112\|gb\|ELT66499.1\| hypothetical protein BURCENK562V_3104 [Burkholderia cenocepacia K56-2Valvano] | G.VGAGAQGGAGANASGNATGAAT[+568.212]DAVGAAADGAKDTTSSAVHS.T | HexNAc(2)Hex(1) | No | 1371.2926 | 3 | 4111.8633 | 4110.8454 | 3.5 | Non | 959.2 | 917.8 | 3.6 | Nsco_20191108_BC_ZIC_HILIC_K56_2_thermolysin_B1.33767.33767.3 | 68.324 | Thermolysin | B1 | 270 |
| BURCENK562V_3104 | >gi\|443598112\|gb\|ELT66499.1\| hypothetical protein BURCENK562V_3104 [Burkholderia cenocepacia K56-2Valvano] | G.VGAGAQGGAGANASGNAT[+568.212]GAATDAVGAAADGAKDTTSSAVHST.K | HexNAc(2)Hex(1) | No | 1405.3107 | 3 | 4213.9175 | 4211.8931 | 4.2 | Non | 539.37 | 424.76 | 0.81 | Nsco_20191108_BC_ZIC_HILIC_K56_2_thermolysin_B2_.37230.37230.3 | 69.5539 | Thermolysin | B2 | 271 |
| BURCENK562V_3104 | >gi\|443598112\|gb\|ELT66499.1\| hypothetical protein BURCENK562V_3104 [Burkholderia cenocepacia K56-2Valvano] | G.VGAGAQGGAGANASGNATGAATDAVGAAADGAKDTTSS[+568.212]AVHSTKK.H | HexNAc(2)Hex(1) | Yes | 894.4257 | 5 | 4468.0996 | 4468.083 | 3.7 | Non | 1192.63 | 979.42 | 27.83 | Nsco_20191108_BC_ZIC_HILIC_K56_2_thermolysin_B1.26977.26977.5 | 55.6311 | Thermolysin | B1 | 272 |
| BURCENK562V_3104 | >gi\|443598112\|gb\|ELT66499.1\| hypothetical protein BURCENK562V_3104 [Burkholderia cenocepacia K56-2Valvano] | G.VGAGAQGGAGANASGNATGAATDAVGAAADGAKDTTSS[+568.212]AVHSTKKH.T | HexNAc(2)Hex(1) | No | 922.0375 | 5 | 4606.1584 | 4605.1419 | 2.9 | Non | 992.62 | 905.74 | 10.81 | Nsco_20191108_BC_ZIC_HILIC_K56_2_thermolysin_B2_.25510.25510.5 | 50.6999 | Thermolysin | B2 | 273 |
| BURCENK562V_3104 | >gi\|443598112\|gb\|ELT66499.1\| hypothetical protein BURCENK562V_3104 [Burkholderia cenocepacia K56-2Valvano] | G.VQAQTPAAGVGAGAQGGAGANASGNATGAATDAVGAAADGAKDTTS[+568.212]SAVHSTKKH.T | HexNAc(2)Hex(1) | No | 1086.9199 | 5 | 5430.5705 | 5428.5607 | 0.6 | Non | 1011.88 | 195.27 | 0.88 | Nsco_20191108_BC_ZIC_HILIC_K56_2_thermolysin_B2_.31008.31008.5 | 59.674 | Thermolysin | B2 | 274 |

| | | | | | | | | | | | | | | | | | | |
|---|---|---|---|---|---|---|---|---|---|---|---|---|---|---|---|---|---|---|
| BCAS0773 | >gi\|443593562\|gb\|ELT62293.1\| hypothetical protein BURCENK562V_B0326 [Burkholderia cenocepacia K56-2Valvano] | A.QTDAAS[+568.212]APAAAAAQDAK.A | HexNAc(2)Hex(1) | No | 1063.4875 | 2 | 2125.9678 | 2125.9619 | 2.8 | Non | 808.63 | 642.97 | 33.43 | Nsco_20191108_BC_ZIC_HILIC_K56_2_thermolysin_B3.10973.10973.2 | 29.3941 | Thermolysin | B3 | 275 |

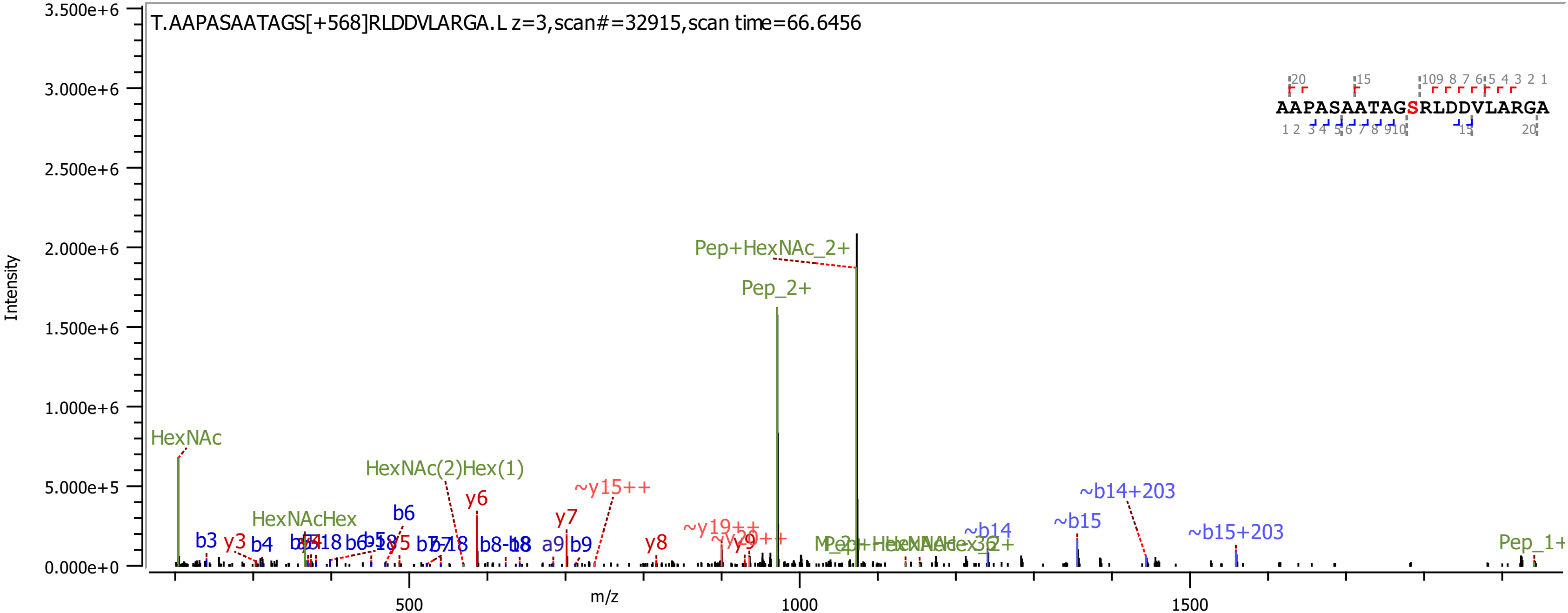
T.AAPASAATAGS[+568]RLDDVLARGA.L z=3,scan#=32915,scan time=66.6456
AAPASAATAGSRLDDVLARGA
Intensity
m/z
3.500e+6
3.000e+6
2.500e+6
2.000e+6
1.500e+6
1.000e+6
5.000e+5
0.000e+0
500
1000
1500
Pep+HexNAc_2+
Pep_2+
HexNAc
HexNAc(2)Hex(1)
HexNAcHex
~y15++
~y19++
~b14+203
~b15+203
~b14
~b15
Pep_1+
b3
y3
b4
b6
y6
y7
y8
y9
b8
a9
b9

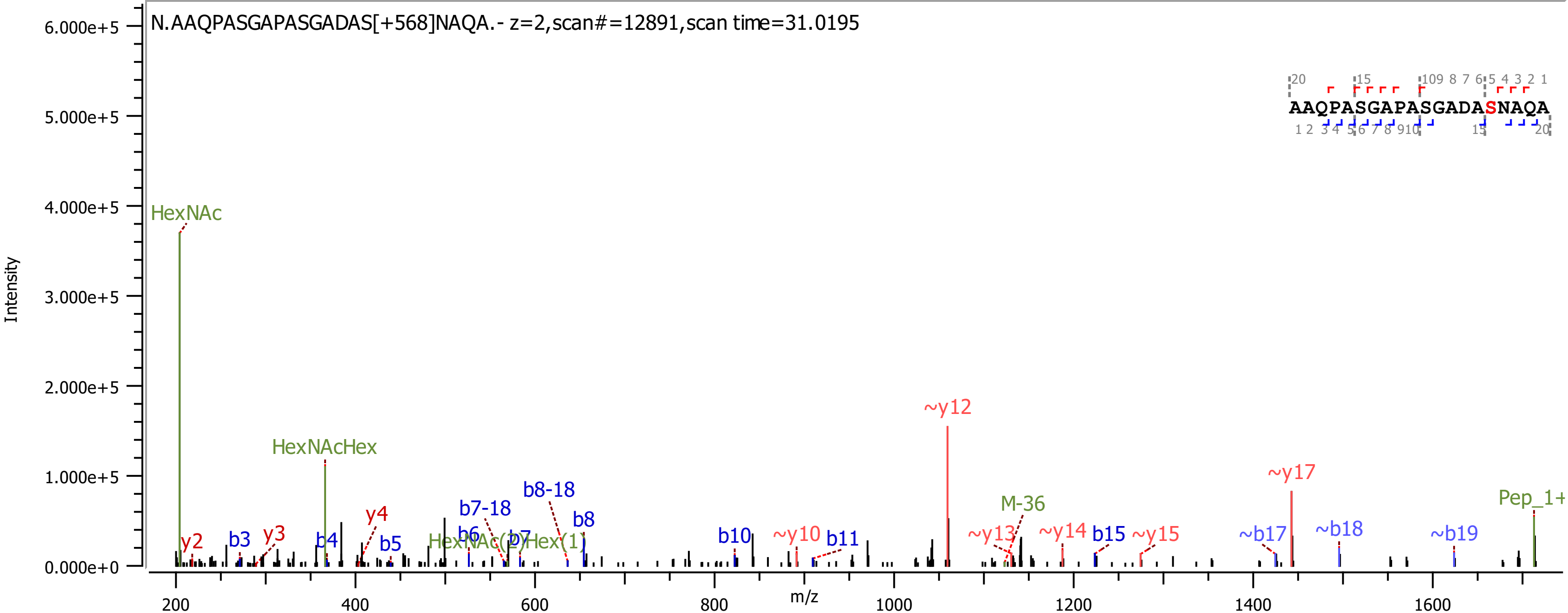

N.AAQPASGAPASGADAS[+568]NAQA.- z=2,scan#=12891,scan time=31.0195
AAQPASGAPASGADASNAQA
Intensity
m/z
HexNAc
HexNAcHex
HexNAc(2)Hex(1)
M-36
Pep_1+
y2
y3
y4
b3
b4
b5
b6
b7
b7-18
b8-18
b8
b10
b11
b15
~y10
~y12
~y13
~y14
~y15
~y17
~b17
~b18
~b19

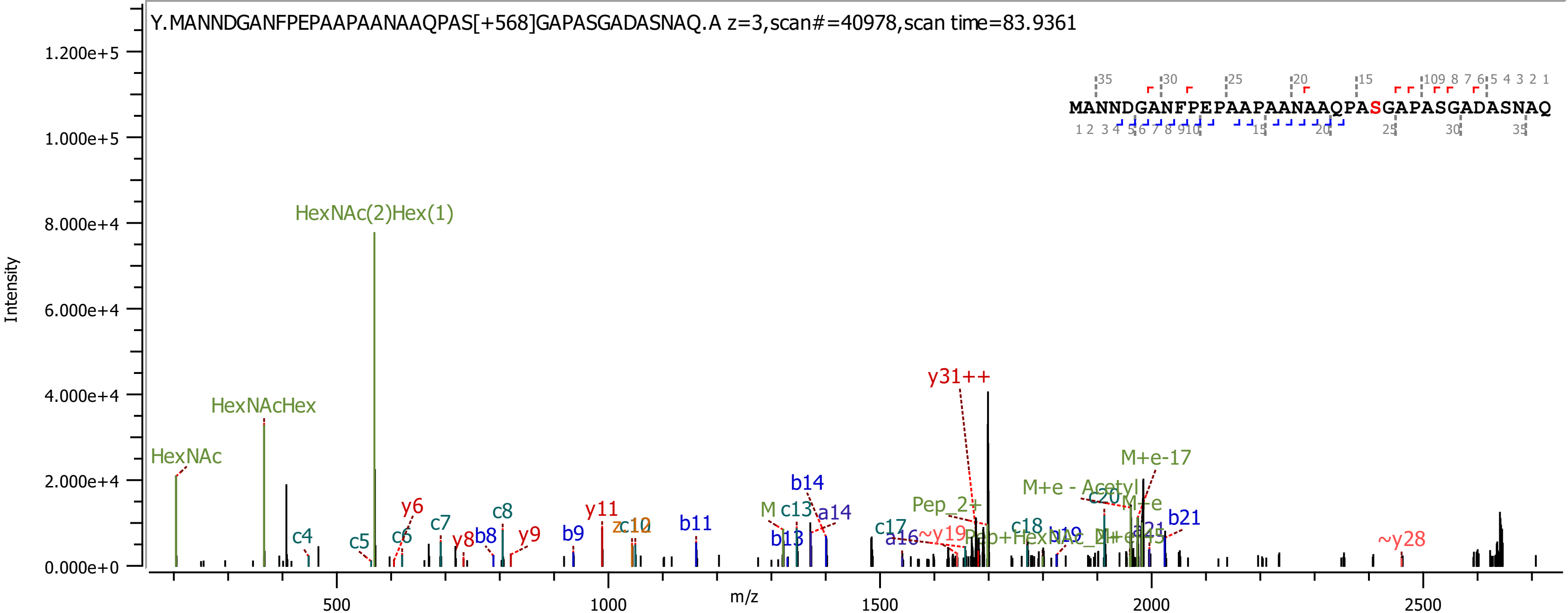

Y.MANNDGANFPEPAAPAANAAQPAS[+568]GAPASGADASNAQ.A z=3,scan#=40978,scan time=83.9361
MANNDGANFPEPAAPAANAAQPASGAPASGADASNAQ
Intensity
m/z
HexNAc(2)Hex(1)
HexNAcHex
HexNAc
y31++
M+e-17
M+e - Acetyl
b14
a14
c13
Pep_2+
b21
~y28
c17
~y19
a16
c18
b13
b11
c8
y11
c10
y6
c7
b8
y8
y9
b9
c4
c5
c6

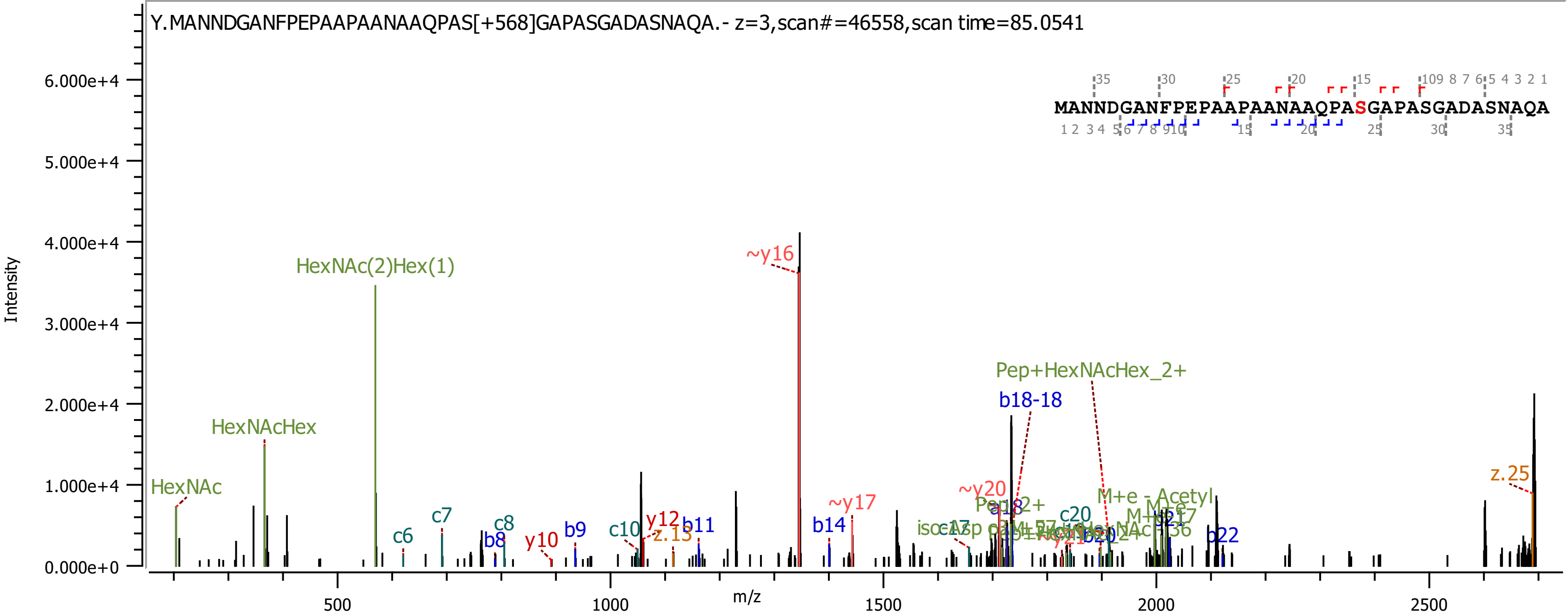
Y.MANNDGANFPEPAAPAANAAQPAS[+568]GAPASGADASNAQA.- z=3,scan#=46558,scan time=85.0541
MANNDGANFPEPAAPAANAAQPASGAPASGADASNAQA
Intensity
m/z
HexNAc
HexNAcHex
HexNAc(2)Hex(1)
~y16
~y17
~y20
Pep+HexNAcHex_2+
b18-18
z.25
c6
c7
c8
b8
y10
b9
c10
y12
z.13
b11
b14
c20
b22
M+e - Acetyl

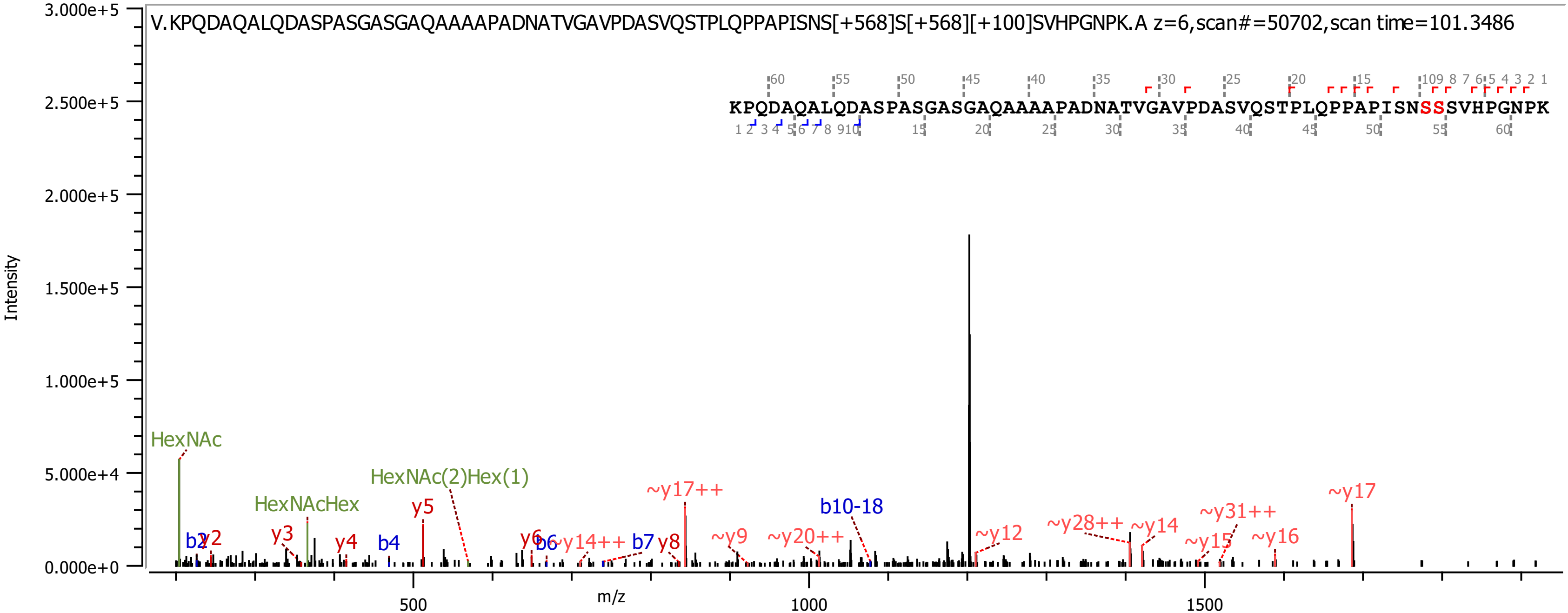
V.KPQDAQALQDASPASGASGAQAAAAPADNATVGAVPDASVQSTPLQPPAPISNS[+568]S[+568][+100]SVHPGNPK.A z=6,scan#=50702,scan time=101.3486
KPQDAQALQDASPASGASGAQAAAAPADNATVGAVPDASVQSTPLQPPAPISNSSSVHPGNPK
Intensity
m/z
3.000e+5
2.500e+5
2.000e+5
1.500e+5
1.000e+5
5.000e+4
0.000e+0
500
1000
1500
HexNAc
HexNAcHex
HexNAc(2)Hex(1)
b2
y2
y3
y4
b4
y5
y6
b6
~y14++
b7
y8
~y17++
~y9
~y20++
b10-18
~y12
~y28++
~y14
~y15
~y31++
~y16
~y17

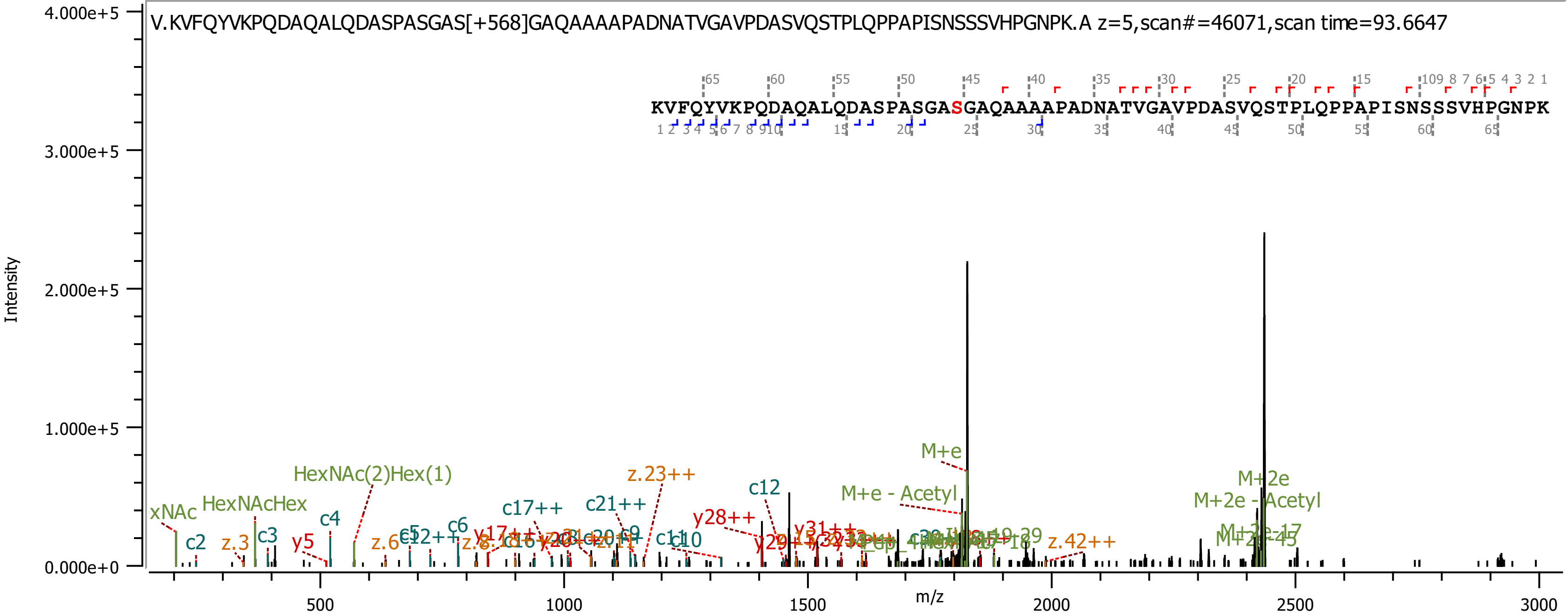
V.KVFQYVKPQDAQALQDASPASGAS[+568]GAQAAAAPADNATVGAVPDASVQSTPLQPPAPISNSSSVHPGNPK.A z=5,scan#=46071,scan time=93.6647
KVFQYVKPQDAQALQDASPASGASGAQAAAAPADNATVGAVPDASVQSTPLQPPAPISNSSSVHPGNPK
Intensity
4.000e+5
3.000e+5
2.000e+5
1.000e+5
0.000e+0
m/z
500
1000
1500
2000
2500
3000
xNAc
HexNAcHex
HexNAc(2)Hex(1)
c2
z.3
c3
y5
c4
z.6
c6
c17++
c21++
z.23++
c12
y28++
y31++
M+e
M+e - Acetyl
z.42++
M+2e
M+2e - Acetyl
M+2e-17
M+2e-45

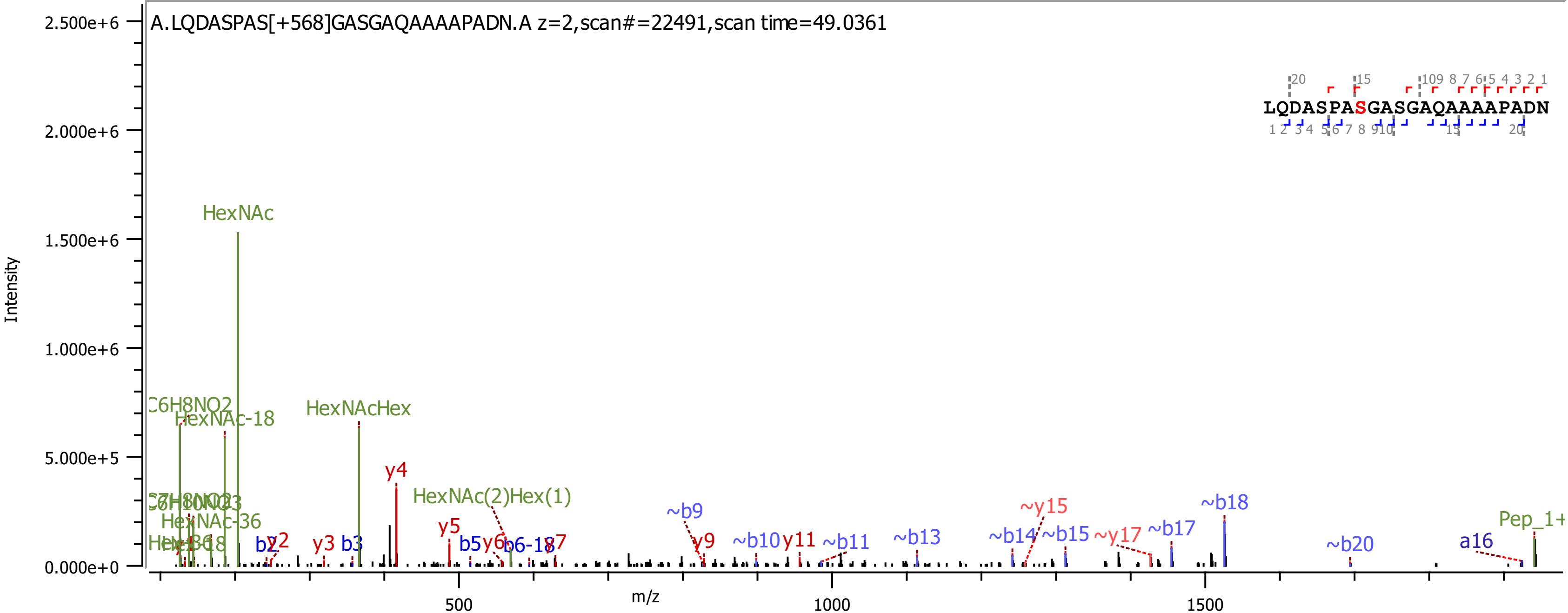
A.LQDASPAS[+568]GASGAQAAAAPADN.A z=2,scan#=22491,scan time=49.0361
LQDASPASGASGAQAAAAPADN
Intensity
m/z
HexNAc
HexNAcHex
HexNAc-18
C6H8NO2
HexNAc(2)Hex(1)
HexNAc-36
y4
y5
~b9
~y15
~b18
Pep_1+
~b14
~b15
~y17
~b17
~b20
a16
~b13
~b11
y11
~b10
y9
b5
b3
y3
b2
y2

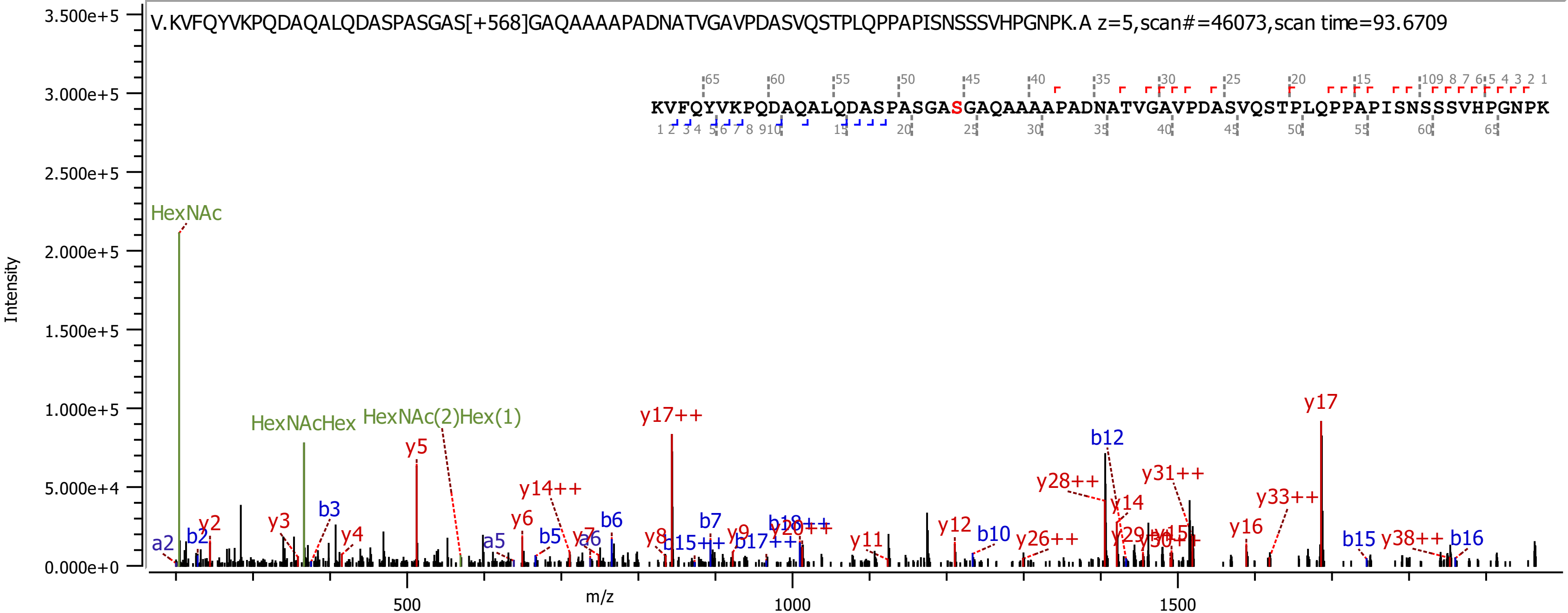
V.KVFQYVKPQDAQALQDASPASGAS[+568]GAQAAAAPADNATVGAVPDASVQSTPLQPPAPISNSSSVHPGNPK.A z=5,scan#=46073,scan time=93.6709
KVFQYVKPQDAQALQDASPASGASGAQAAAAPADNATVGAVPDASVQSTPLQPPAPISNSSSVHPGNPK
Intensity
m/z
HexNAc
HexNAcHex
HexNAc(2)Hex(1)
y17++
y17
y5
b12
y31++
y28++
y14++
y33++
y14
y6
b6
b3
y2
y3
y4
a2
b2
a5
b5
y7
a6
b7
y8
b15++
y9
b18++
y20++
b17++
y11
y12
b10
y26++
y29++
y30++
y15
y16
b15
y38++
b16

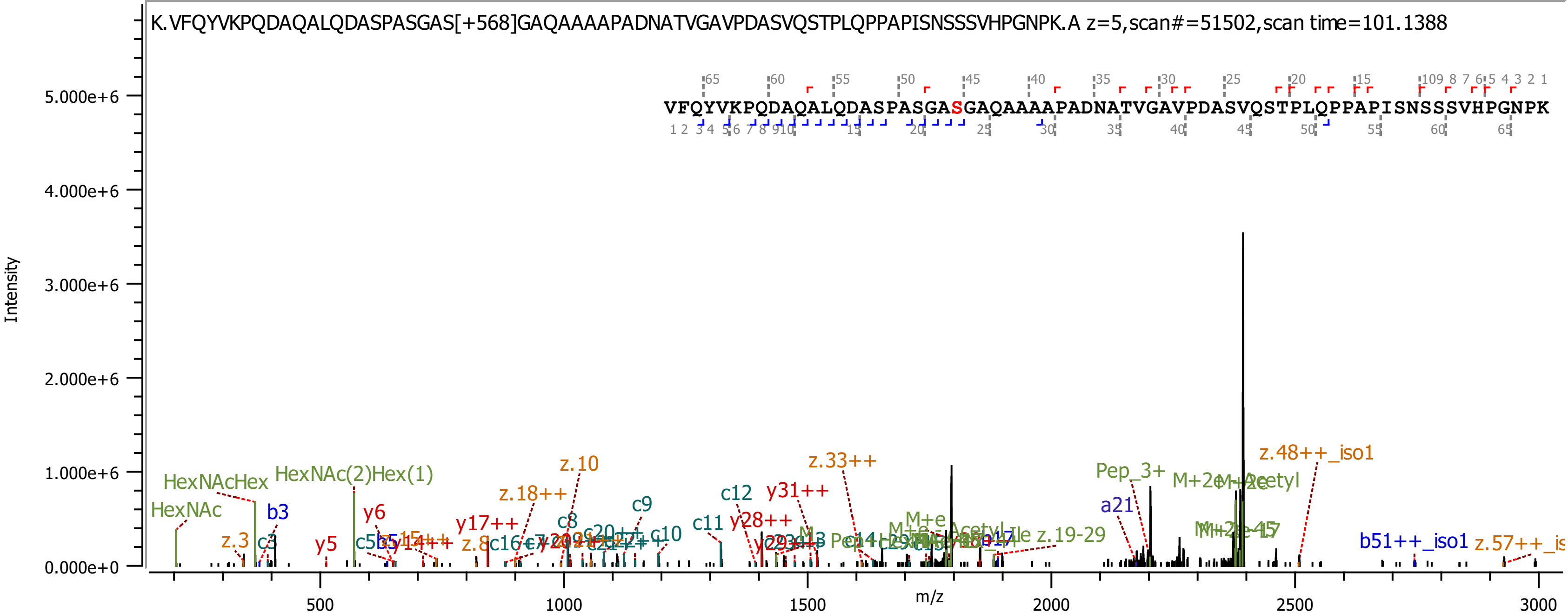
K.VFQYVKPQDAQALQDASPASGAS[+568]GAQAAAAPADNATVGAVPDASVQSTPLQPPAPISNSSSVHPGNPK.A z=5,scan#=51502,scan time=101.1388
VFQYVKPQDAQALQDASPASGASGAQAAAAPADNATVGAVPDASVQSTPLQPPAPISNSSSVHPGNPK
Intensity
m/z
5.000e+6
4.000e+6
3.000e+6
2.000e+6
1.000e+6
0.000e+0
500
1000
1500
2000
2500
3000
HexNAc
HexNAcHex
HexNAc(2)Hex(1)
b3
c3
z.3
y5
y6
y17++
z.18++
z.10
z.8
c8
c9
c10
c11
c12
y28++
y31++
z.33++
M+e
Pep_3+
a21
z.19-29
M+2e-Acetyl
z.48++_iso1
b51++_iso1
z.57++_is

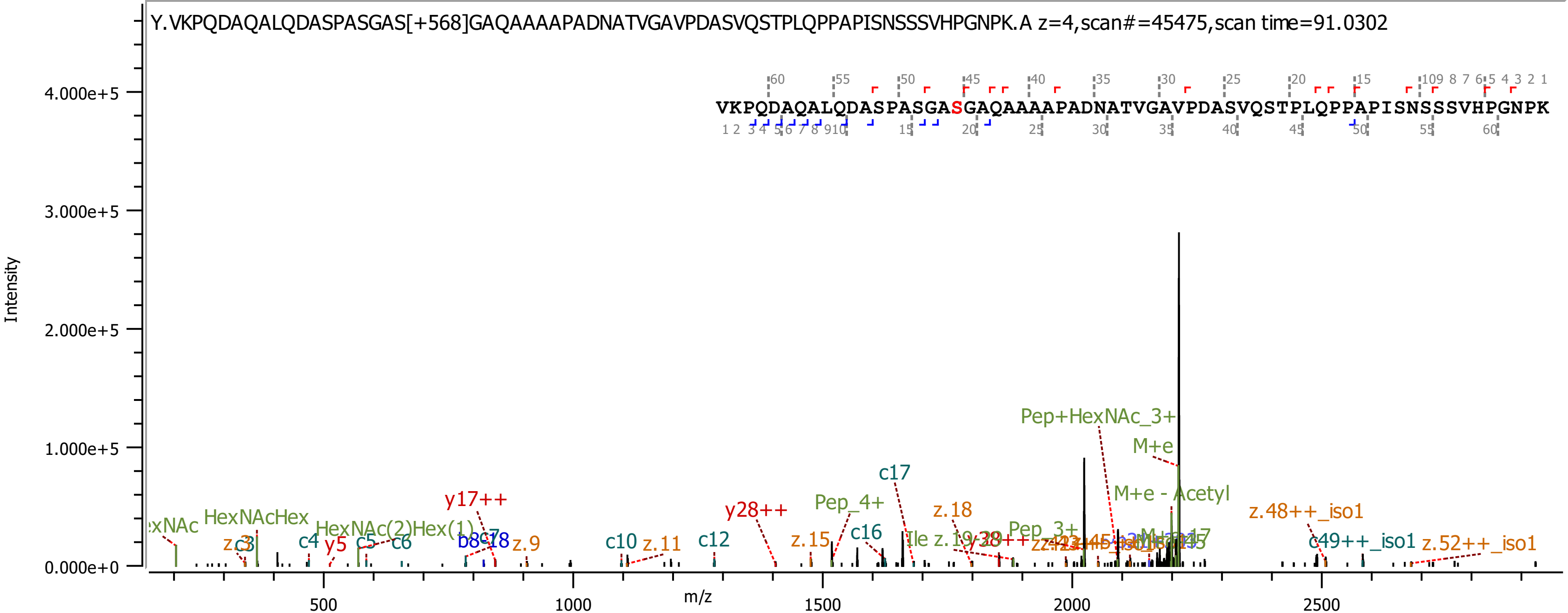
Y.VKPQDAQALQDASPASGAS[+568]GAQAAAAPADNATVGAVPDASVQSTPLQPPAPISNSSSVHPGNPK.A z=4,scan#=45475,scan time=91.0302
VKPQDAQALQDASPASGASGAQAAAAPADNATVGAVPDASVQSTPLQPPAPISNSSSVHPGNPK
Intensity
4.000e+5
3.000e+5
2.000e+5
1.000e+5
0.000e+0
m/z
500
1000
1500
2000
2500
Pep+HexNAc_3+
M+e
M+e - Acetyl
c17
Pep_4+
z.18
z.48++_iso1
y17++
y28++
c16
HexNAcHex
HexNAc(2)Hex(1)
c4
y5
c5
c6
b8
z.9
c10
z.11
c12
z.15
Pep_3+
c49++_iso1
z.52++_iso1

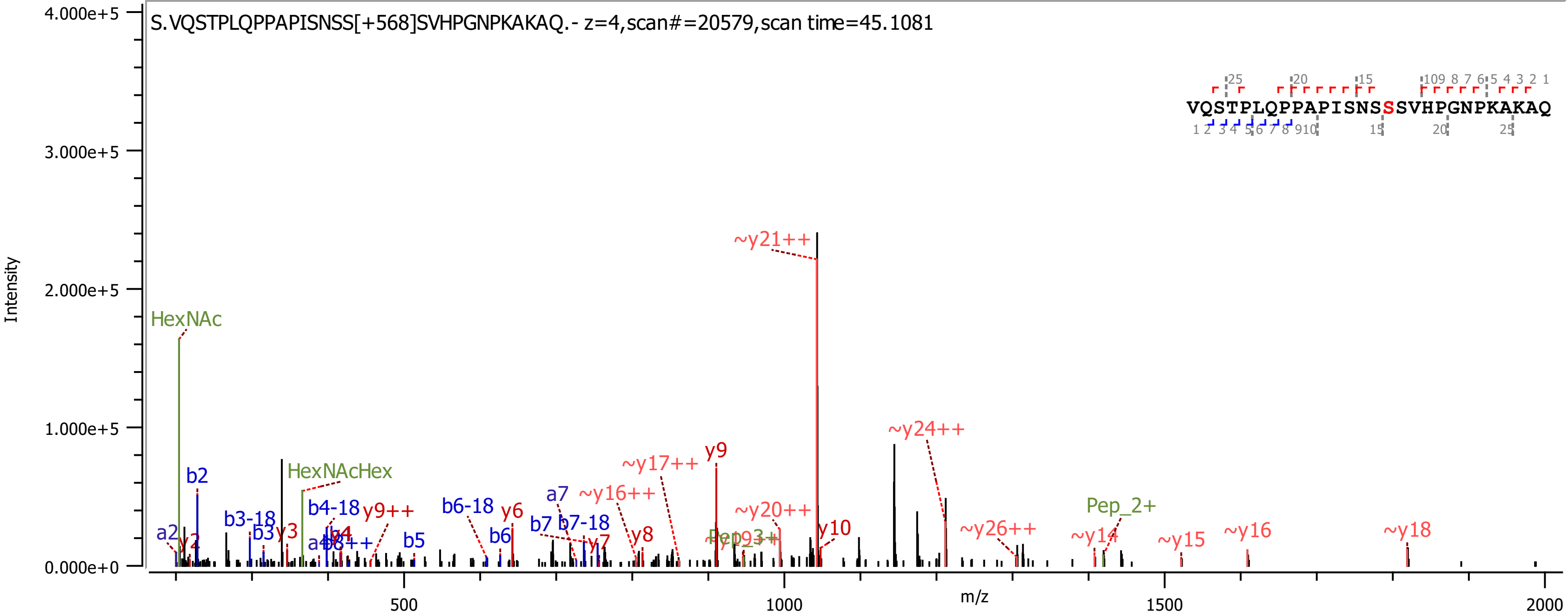
S.VQSTPLQPPAPISNSS[+568]SVHPGNPKAKAQ.- z=4,scan#=20579,scan time=45.1081
VQSTPLQPPAPISNSSSVHPGNPKAKAQ
Intensity
m/z
HexNAc
HexNAcHex
~y21++
~y24++
~y17++
y9
~y16++
~y20++
y10
~y26++
Pep_2+
~y14
~y15
~y16
~y18
b2
b3-18
b4-18
y9++
b6-18
y6
a7
b7
b7-18
y7
y8
b6
b5
a2
y2
b3
y3
y4
Pep_3+

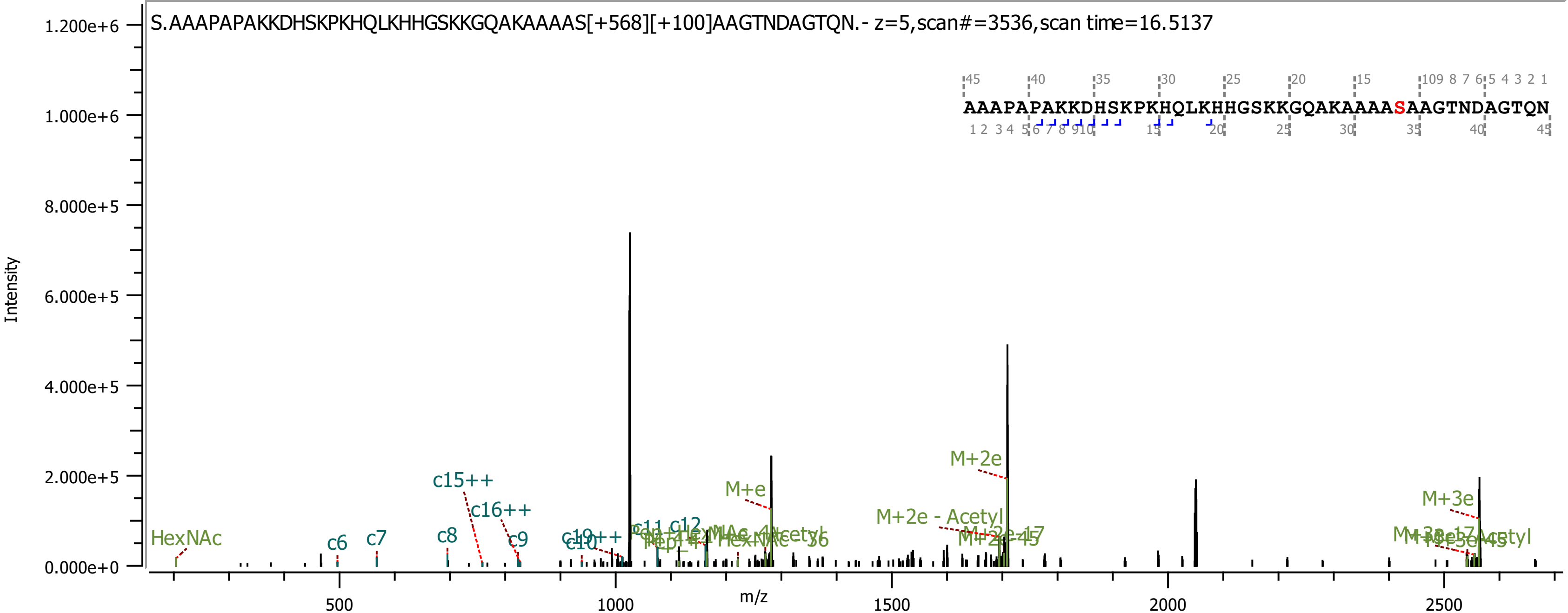
S.AAAPAPAKKDHSKPKHQLKHHGSKKGQAKAAAAS[+568][+100]AAGTNDAGTQN.- z=5,scan#=3536,scan time=16.5137
AAAPAPAKKDHSKPKHQLKHHGSKKGQAKAAAASAAGTNDAGTQN
Intensity
m/z
HexNAc
c6
c7
c8
c9
c10
c11
c12
c15++
c16++
M+e
M+2e
M+2e - Acetyl
M+3e

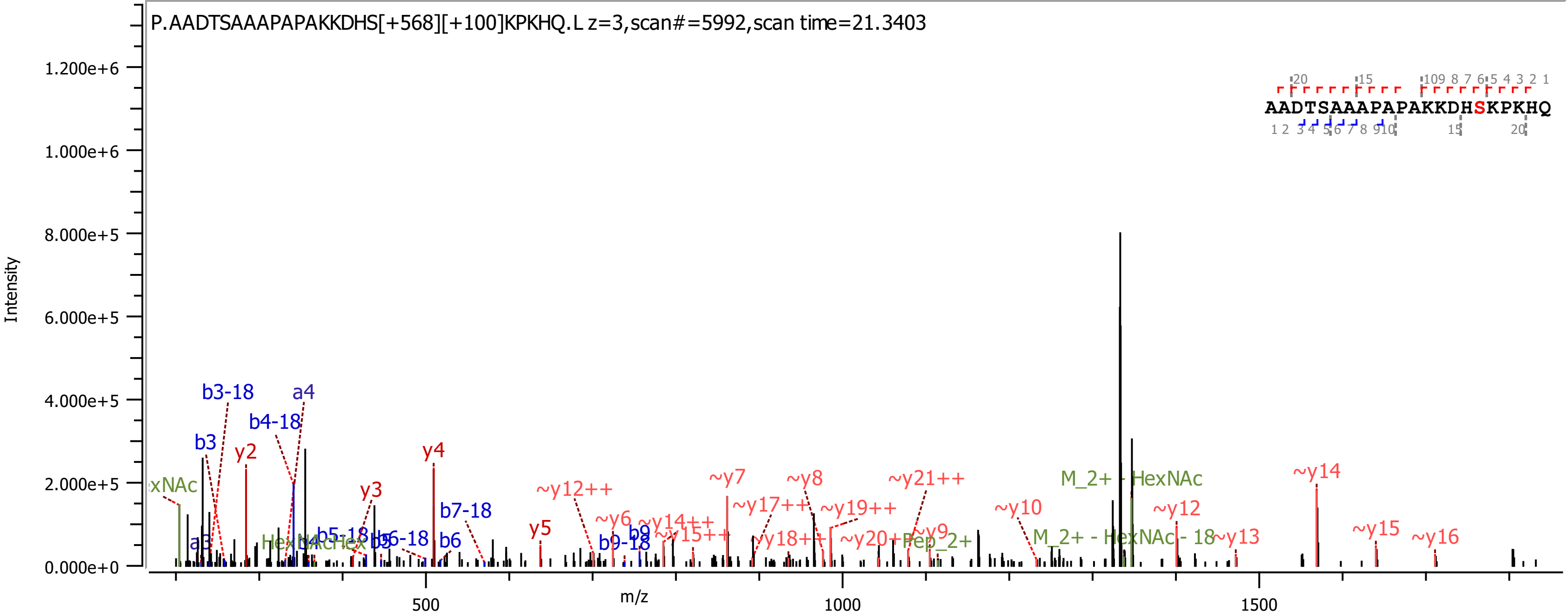
P.AADTSAAAPAPAKKDHS[+568][+100]KPKHQ.L z=3,scan#=5992,scan time=21.3403
Intensity
m/z
AADTSAAAPAPAKKDHSKPKHQ
b3-18
a4
b4-18
b3
y2
y4
y3
b7-18
~y12++
~y7
~y8
~y21++
M_2+ - HexNAc
~y14
~y17++
~y19++
~y10
~y12
~y6
~y14++
~y15++
~y18++
~y20++
Pep_2+
~y9
M_2+ - HexNAc - 18
~y13
~y15
~y16
y5
b6
b9
b9-18
a3
b5-18
b6-18
b5

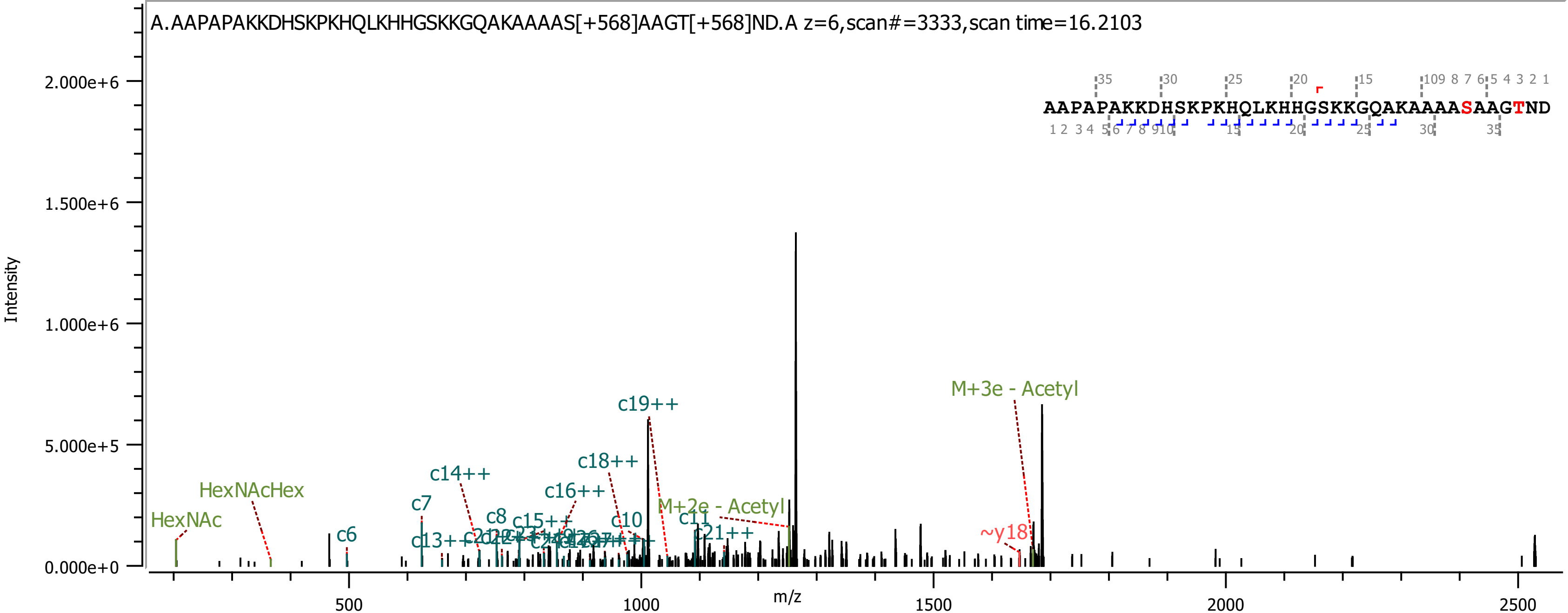
A.AAPAPAKKDHSKPKHQLKHHGSKKGQAKAAAAS[+568]AAGT[+568]ND.A z=6,scan#=3333,scan time=16.2103
AAPAPAKKDHSKPKHQLKHHGSKKGQAKAAAASAAGTND
Intensity
2.000e+6
1.500e+6
1.000e+6
5.000e+5
0.000e+0
m/z
500
1000
1500
2000
2500
HexNAc
HexNAcHex
c6
c7
c8
c10
c11
c13++
c14++
c15++
c16++
c18++
c19++
c21++
M+2e - Acetyl
M+3e - Acetyl
~y18

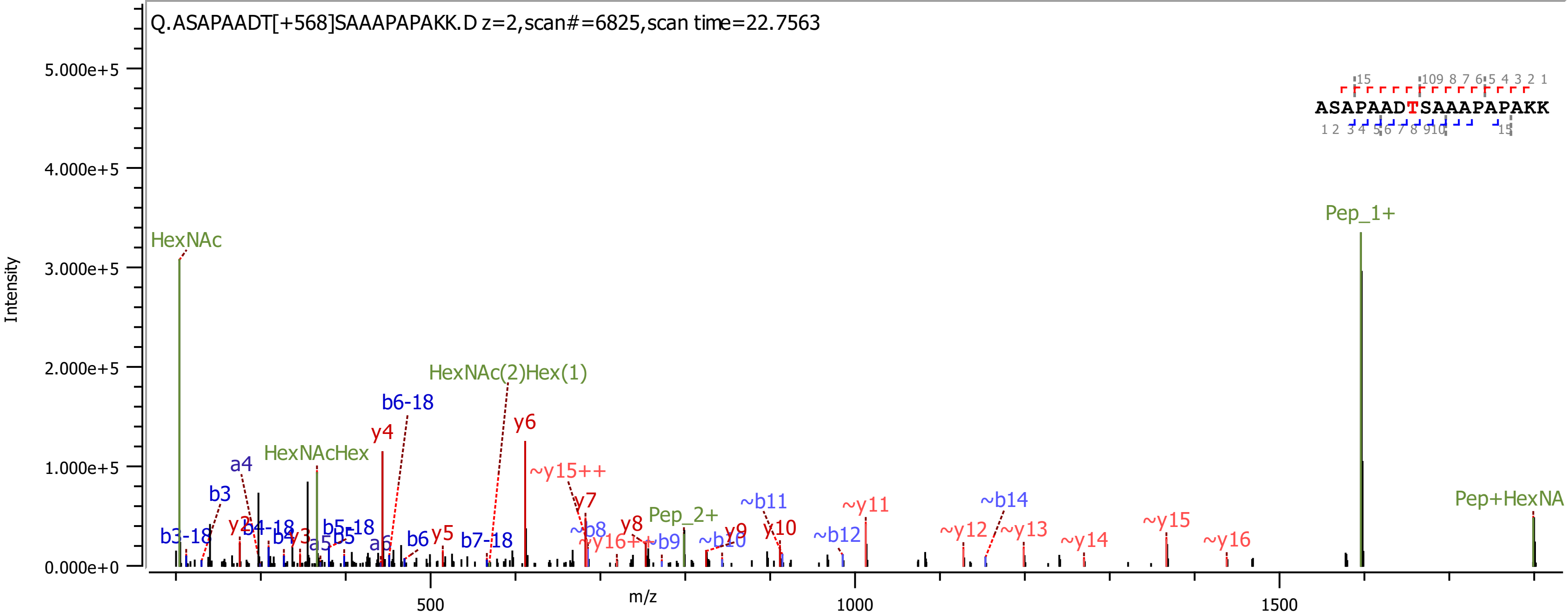
Q.ASAPAADT[+568]SAAAPAPAKK.D z=2,scan#=6825,scan time=22.7563
ASAPAADTSAAAPAPAKK
Intensity
m/z
0.000e+0
1.000e+5
2.000e+5
3.000e+5
4.000e+5
5.000e+5
500
1000
1500
HexNAc
HexNAcHex
HexNAc(2)Hex(1)
Pep_1+
Pep_2+
Pep+HexNA
b3-18
b3
a4
y2
b4-18
b4
y3
a5
b5
b5-18
y4
a6
b6-18
b6
y5
b7-18
y6
~y15++
y7
~b8
~y16++
y8
~b9
~b10
y9
y10
~b11
~b12
~y11
~y12
~b14
~y13
~y14
~y15
~y16

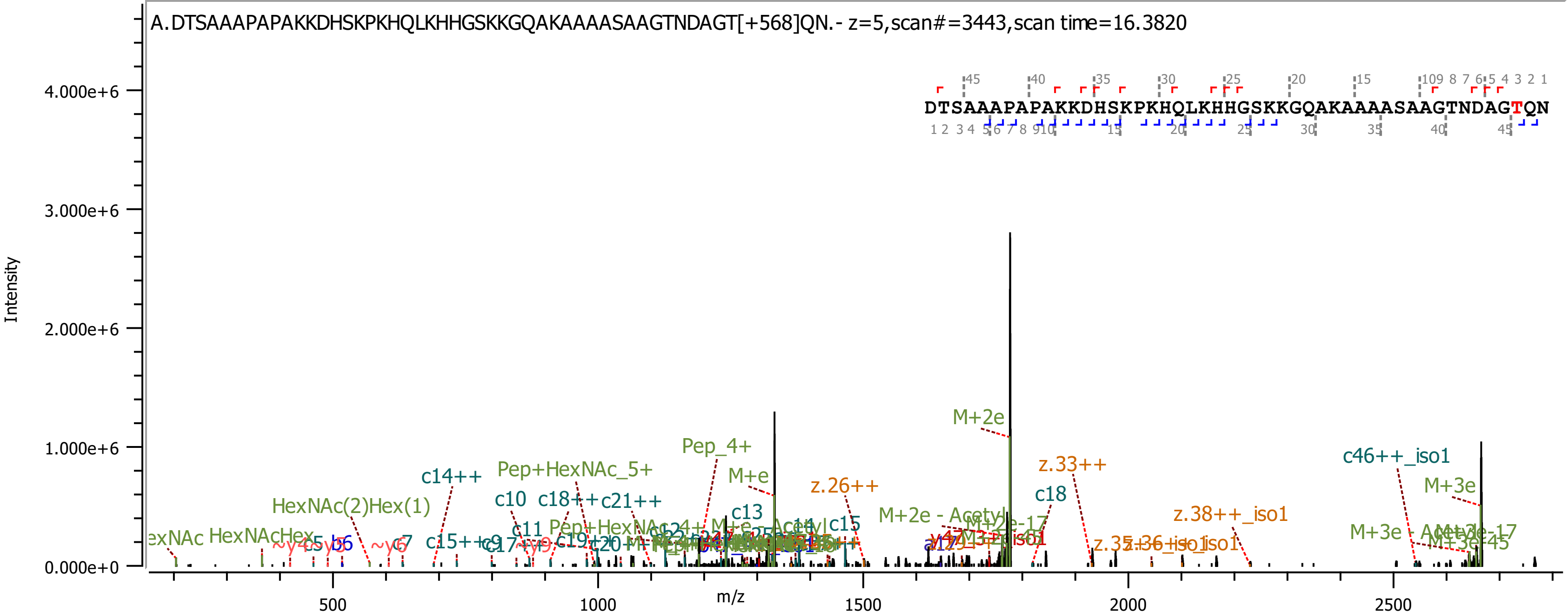

A.DTSAAAPAPAKKDHSKPKHQLKHHGSKKGQAKAAAASAAGTNDAGT[+568]QN.- z=5,scan#=3443,scan time=16.3820
DTSAAAPAPAKKDHSKPKHQLKHHGSKKGQAKAAAASAAGTNDAGTQN
Intensity
4.000e+6
3.000e+6
2.000e+6
1.000e+6
0.000e+0
m/z
500
1000
1500
2000
2500
M+2e
Pep_4+
M+e
c14++
Pep+HexNAc_5+
HexNAc(2)Hex(1)
c10
c18++
c21++
c13
z.26++
z.33++
c18
c46++_iso1
M+3e
z.38++_iso1
M+2e - Acetyl
M+3e - Acetyl
c15
c11
c7
HexNAcHex

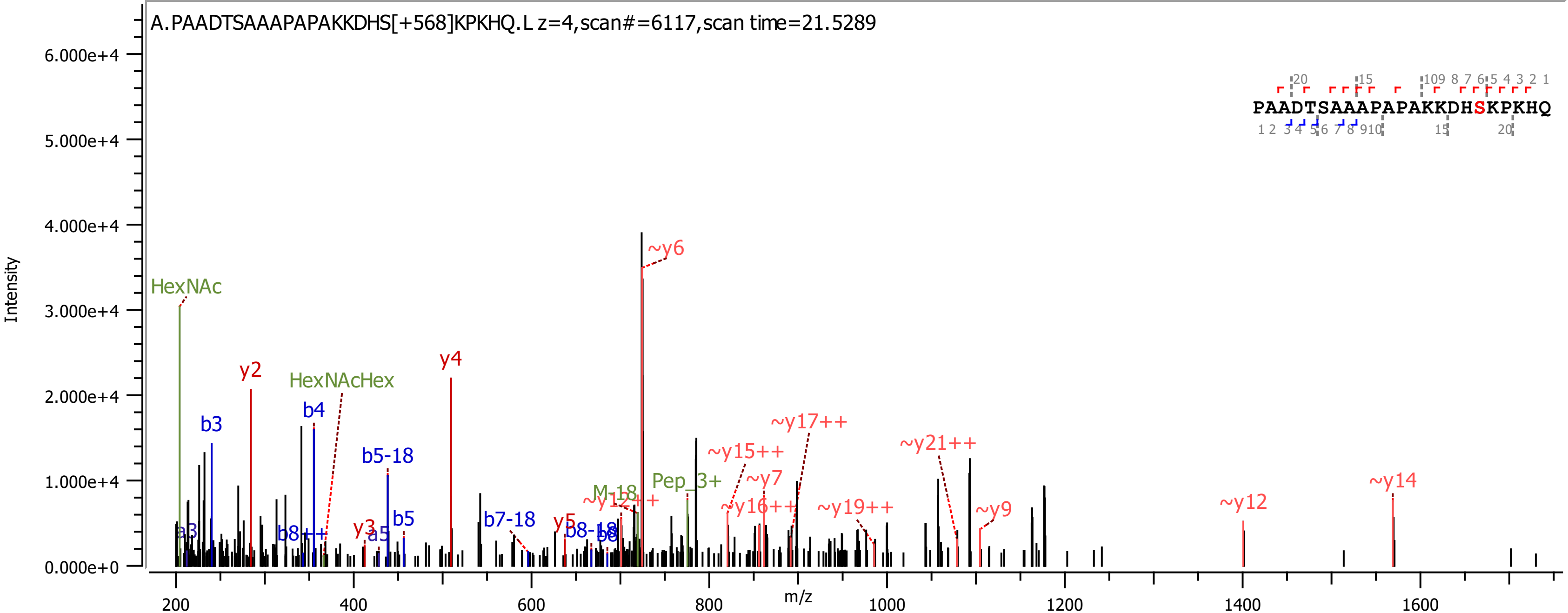
A.PAADTSAAAPAPAKKDHS[+568]KPKHQ.L z=4,scan#=6117,scan time=21.5289
PAADTSAAAPAPAKKDHSKPKHQ
Intensity
m/z
HexNAc
HexNAcHex
y2
y3
y4
y5
b3
b4
b5
b5-18
b7-18
b8++
a3
a5
~y6
~y7
~y9
~y12
~y14
~y15++
~y16++
~y17++
~y19++
~y21++
Pep_3+
M-18

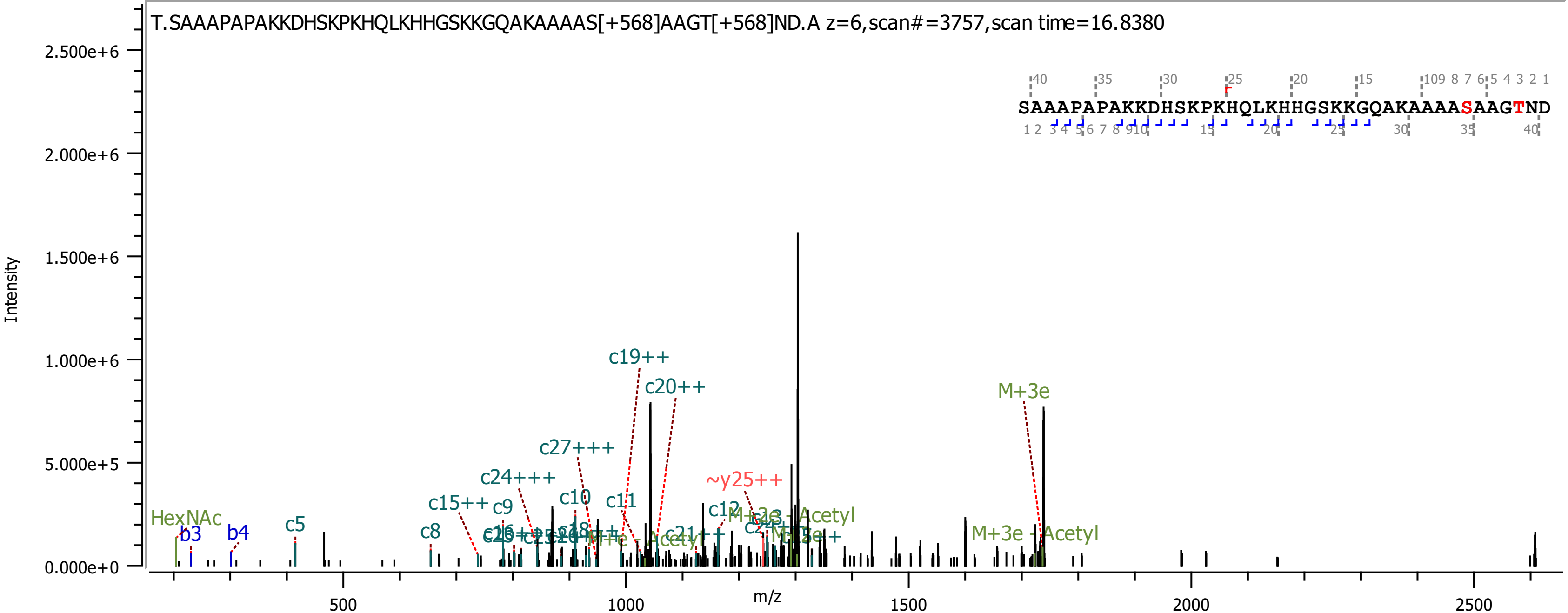
T.SAAAPAPAKKDHSKPKHQLKHHGSKKGQAKAAAAS[+568]AAGT[+568]ND.A z=6,scan#=3757,scan time=16.8380
SAAAPAPAKKDHSKPKHQLKHHGSKKGQAKAAAASAAGTND
Intensity
m/z
2.500e+6
2.000e+6
1.500e+6
1.000e+6
5.000e+5
0.000e+0
500
1000
1500
2000
2500
HexNAc
b3
b4
c5
c8
c15++
c9
c24+++
c27+++
c10
c11
c19++
c20++
~y25++
c12
M+3e - Acetyl
M+3e
M+3e - Acetyl
M+e - Acetyl

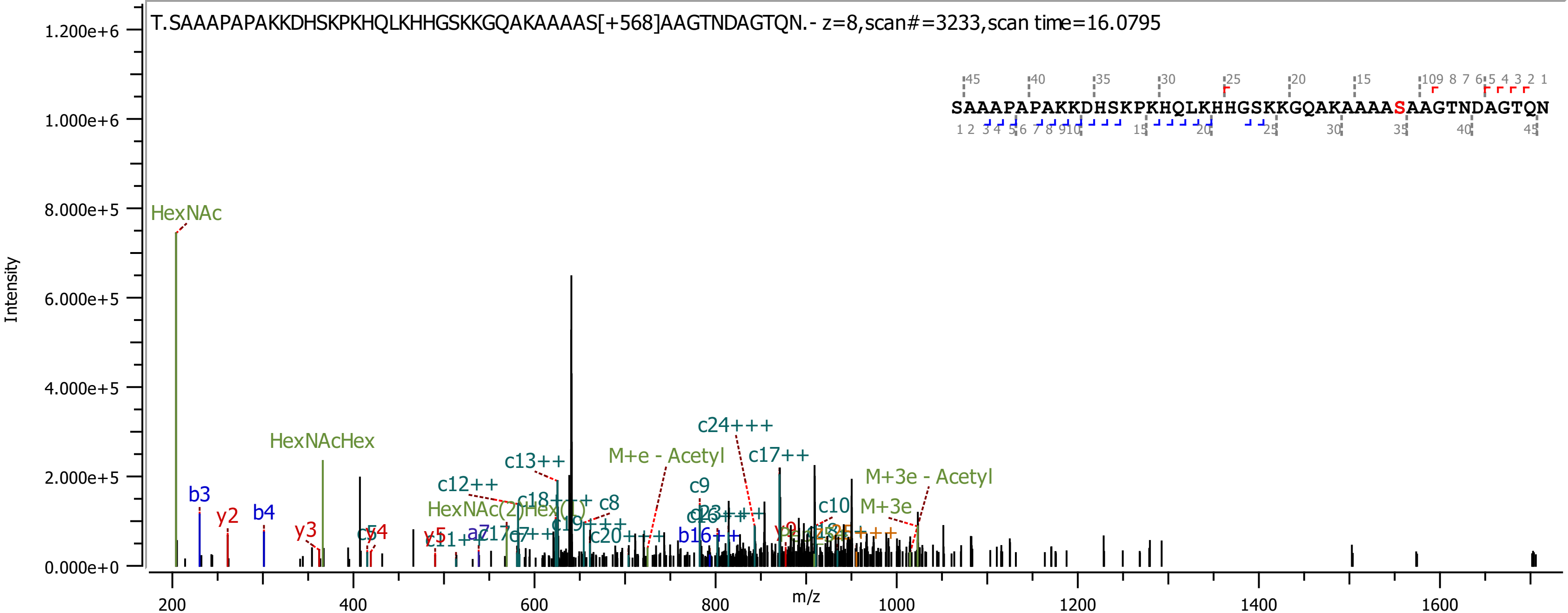
T.SAAAPAPAKKDHSKPKHQLKHHGSKKGQAKAAAAS[+568]AAGTNDAGTQN.- z=8,scan#=3233,scan time=16.0795
SAAAPAPAKKDHSKPKHQLKHHGSKKGQAKAAAASAAGTNDAGTQN
Intensity
m/z
HexNAc
HexNAcHex
HexNAc(2)Hex(1)
M+e - Acetyl
M+3e - Acetyl
M+3e
c24+++
c17++
c13++
c12++
c9
c10
c8
b3
b4
y2
y3
y5
b16++
c20+++

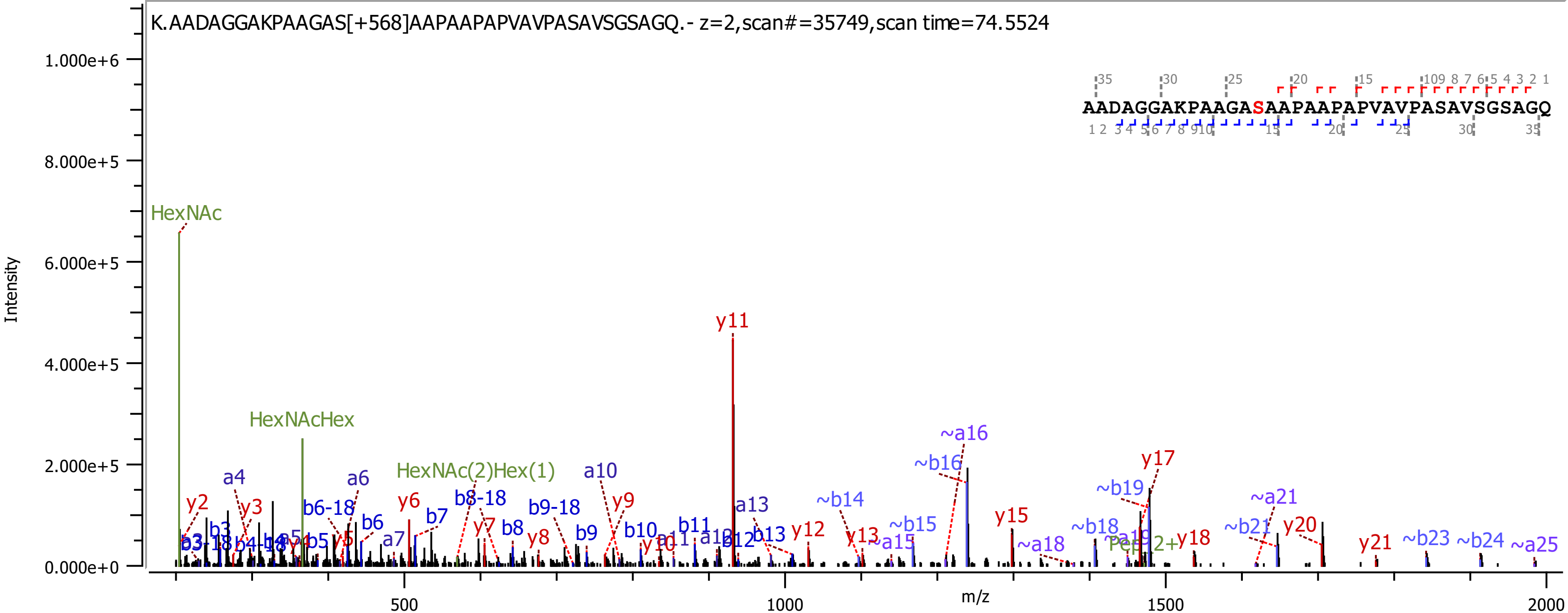
K.AADAGGAKPAAGAS[+568]AAPAAPAPVAVPASAVSGSAGQ.- z=2,scan#=35749,scan time=74.5524
AADAGGAKPAAGASAAPAAPAPVAVPASAVSGSAGQ
Intensity
m/z
HexNAc
HexNAcHex
HexNAc(2)Hex(1)
y11
~a16
~b16
y17
~b19
~a21
y20
~b14
~b15
~a15
y15
~a18
~b18
y18
~b21
y21
~b23
~b24
~a25
y12
y13
a13
b13
b12
a10
y9
b10
y10
b11
a11
a12
b9-18
b9
b8-18
y7
b8
y8
a6
b6
b6-18
y6
b7
a7
a4
y2
y3
b3
b3-18
b4-18
b5
a5
y5

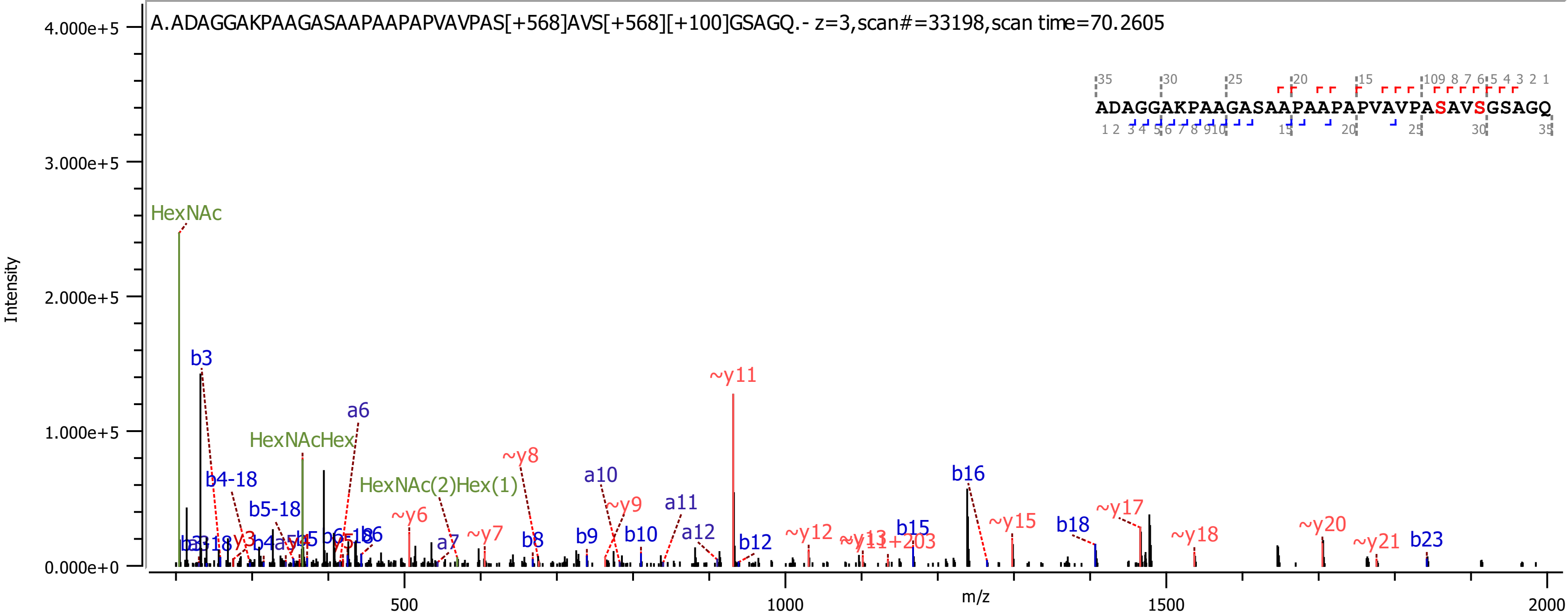
A.ADAGGAKPAAGASAAPAAPAPVAVPAS[+568]AVS[+568][+100]GSAGQ.- z=3,scan#=33198,scan time=70.2605
ADAGGAKPAAGASAAPAAPAPVAVPASAVSGSAGQ
Intensity
m/z
4.000e+5
3.000e+5
2.000e+5
1.000e+5
0.000e+0
500
1000
1500
2000
HexNAc
HexNAcHex
HexNAc(2)Hex(1)
b3
b4-18
b5-18
a6
~y6
a7
~y7
~y8
b8
b9
a10
~y9
b10
a11
a12
~y11
b12
~y12
~y13
b15
b16
~y15
b18
~y17
~y18
~y20
~y21
b23

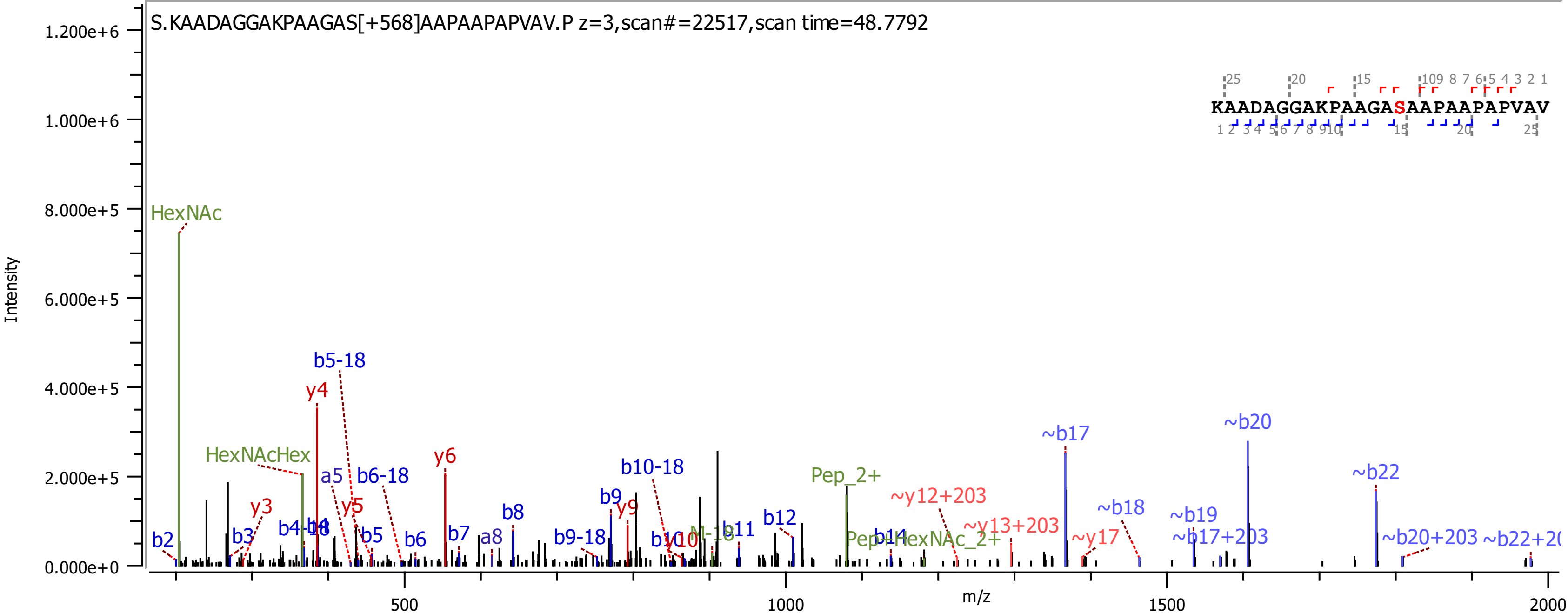
S.KAADAGGAKPAAGAS[+568]AAPAAPAPVAV.P z=3,scan#=22517,scan time=48.7792
KAADAGGAKPAAGASAAPAAPAPVAV
Intensity
m/z
HexNAc
HexNAcHex
b5-18
y4
y6
b10-18
Pep_2+
~y12+203
~y13+203
~b17
~b20
~b22
~b18
~b19
~b17+203
~b20+203
~y17
b2
b3
y3
y5
a5
b6-18
b5
b6
b7
a8
b8
b9
y9
b9-18
b11
b12
b14
Pep+HexNAc_2+
M-18
1.200e+6
1.000e+6
8.000e+5
6.000e+5
4.000e+5
2.000e+5
0.000e+0
500
1000
1500
2000

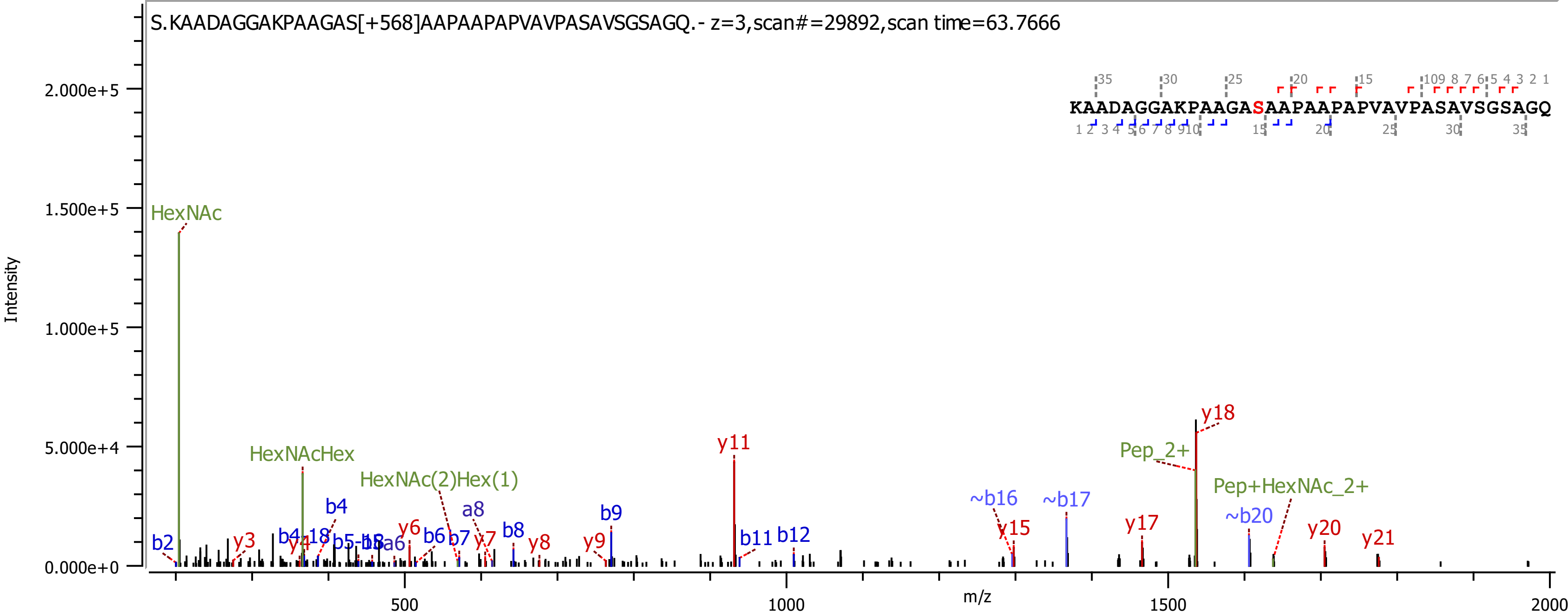

S.KAADAGGAKPAAGAS[+568]AAPAAPAPVAVPASAVSGSAGQ.- z=3,scan#=29892,scan time=63.7666
KAADAGGAKPAAGASAAPAAPAPVAVPASAVSGSAGQ
Intensity
m/z
HexNAc
HexNAcHex
HexNAc(2)Hex(1)
Pep_2+
Pep+HexNAc_2+
y11
y18
~b16
~b17
~b20
y15
y17
y20
y21
b2
y3
b4
a8
b8
b9
b11
b12
y6
b6
b7
y7
y8
y9

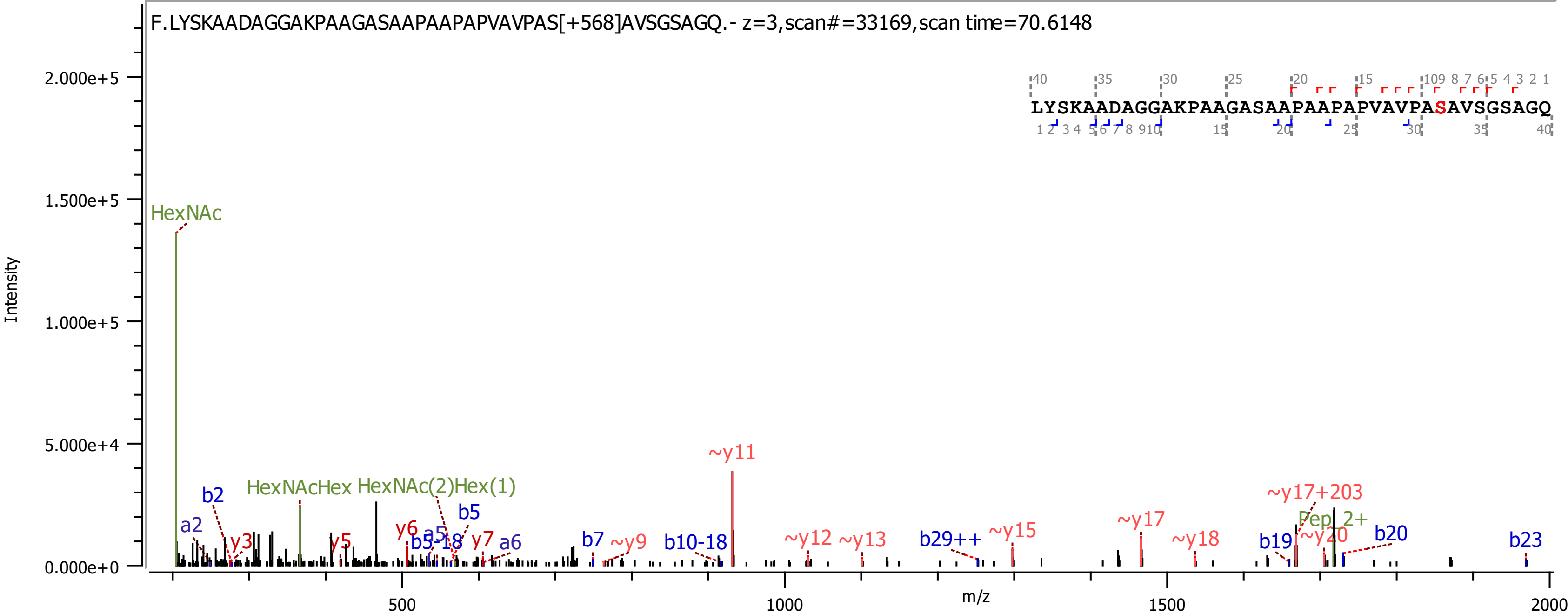
F.LYSKAADAGGAKPAAGASAAPAAPAPVAVPAS[+568]AVSGSAGQ.- z=3,scan#=33169,scan time=70.6148
LYSKAADAGGAKPAAGASAAPAAPAPVAVPASAVSGSAGQ
Intensity
m/z
2.000e+5
1.500e+5
1.000e+5
5.000e+4
0.000e+0
500
1000
1500
2000
HexNAc
HexNAcHex
HexNAc(2)Hex(1)
a2
b2
y3
y5
y6
a5
b5
b5-18
y7
a6
b7
~y9
b10-18
~y11
~y12
~y13
b29++
~y15
~y17
~y17+203
~y18
b19
Pep 2+
~y20
b20
b23

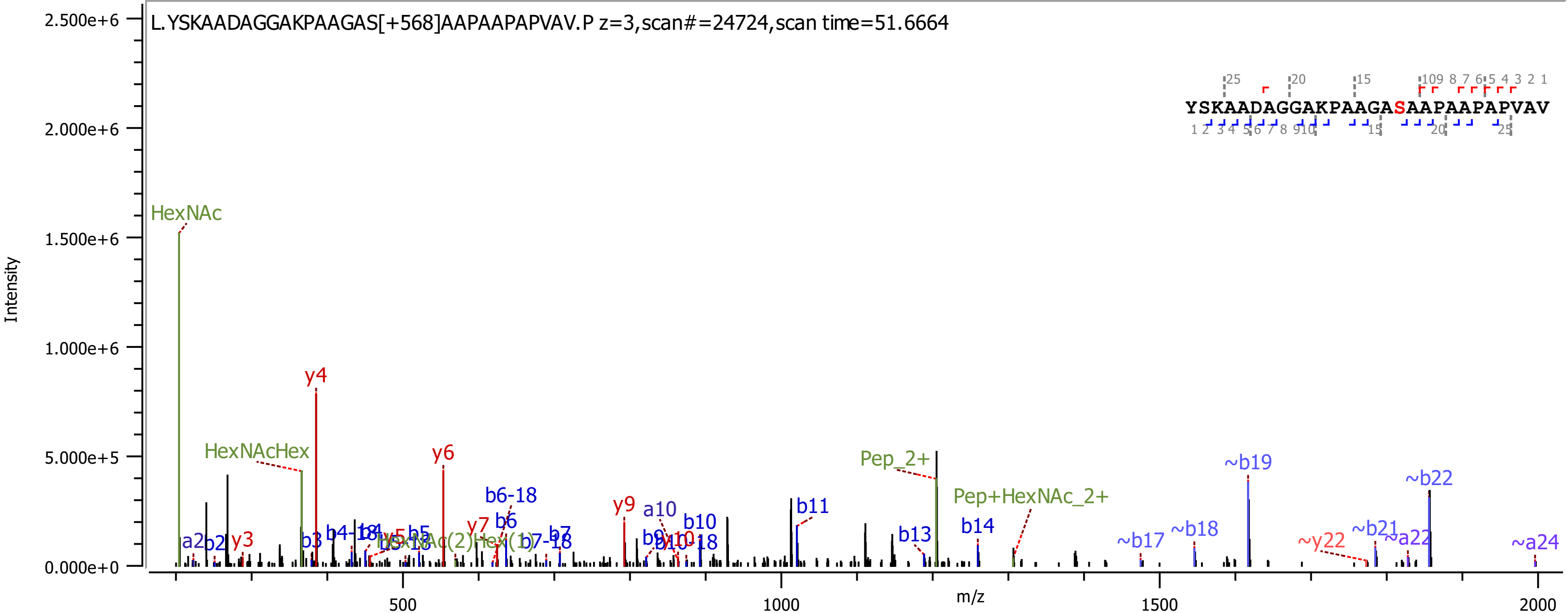

L.YSKAADAGGAKPAAGAS[+568]AAPAAPAPVAV.P z=3,scan#=24724,scan time=51.6664
YSKAADAGGAKPAAGASAAPAAPAPVAV
Intensity
m/z
HexNAc
HexNAcHex
Pep_2+
Pep+HexNAc_2+
y4
y6
y9
b11
~b19
~b22

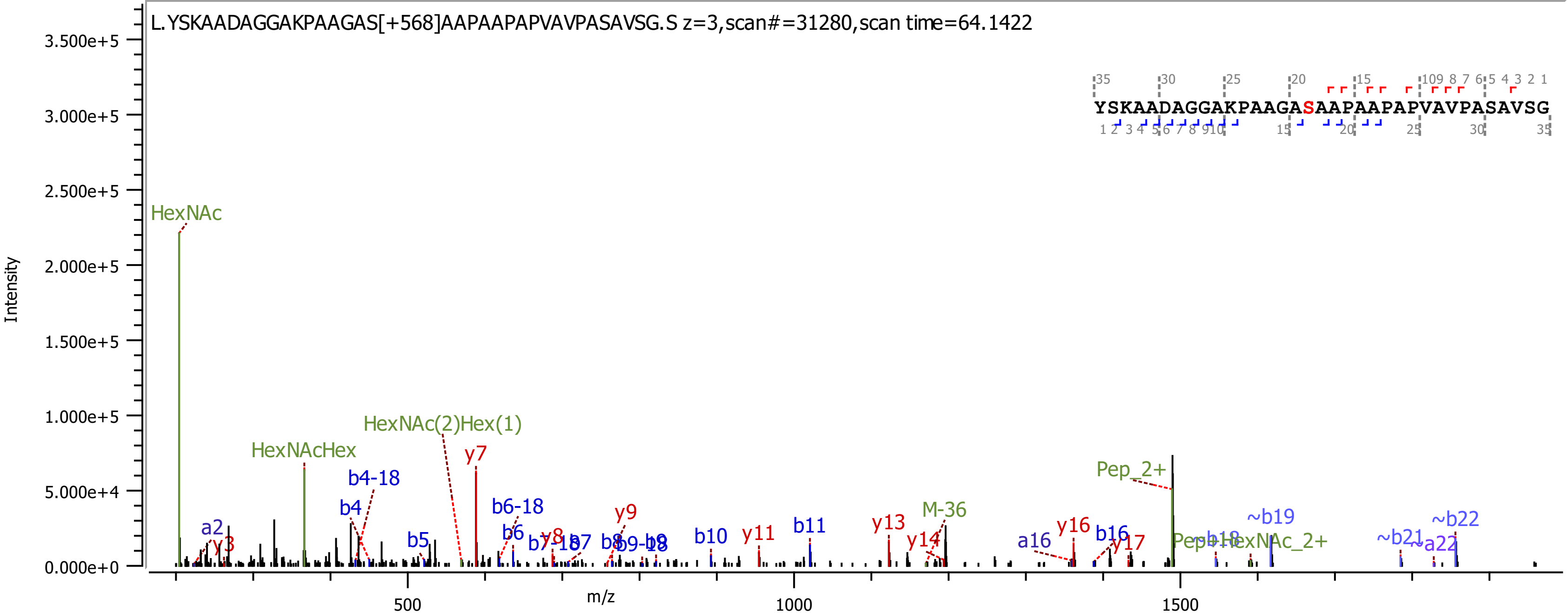
L.YSKAADAGGAKPAAGAS[+568]AAPAAPAPVAVPASAVSG.S z=3,scan#=31280,scan time=64.1422
YSKAADAGGAKPAAGASAAPAAPAPVAVPASAVSG
Intensity
m/z
3.500e+5
3.000e+5
2.500e+5
2.000e+5
1.500e+5
1.000e+5
5.000e+4
0.000e+0
500
1000
1500
HexNAc
HexNAcHex
HexNAc(2)Hex(1)
y7
b4-18
b4
a2
y3
b5
b6-18
b6
y8
b7
y9
b10
y11
b11
y13
M-36
y14
Pep_2+
a16
y16
b16
y17
~b19
~b21
~a22
~b22

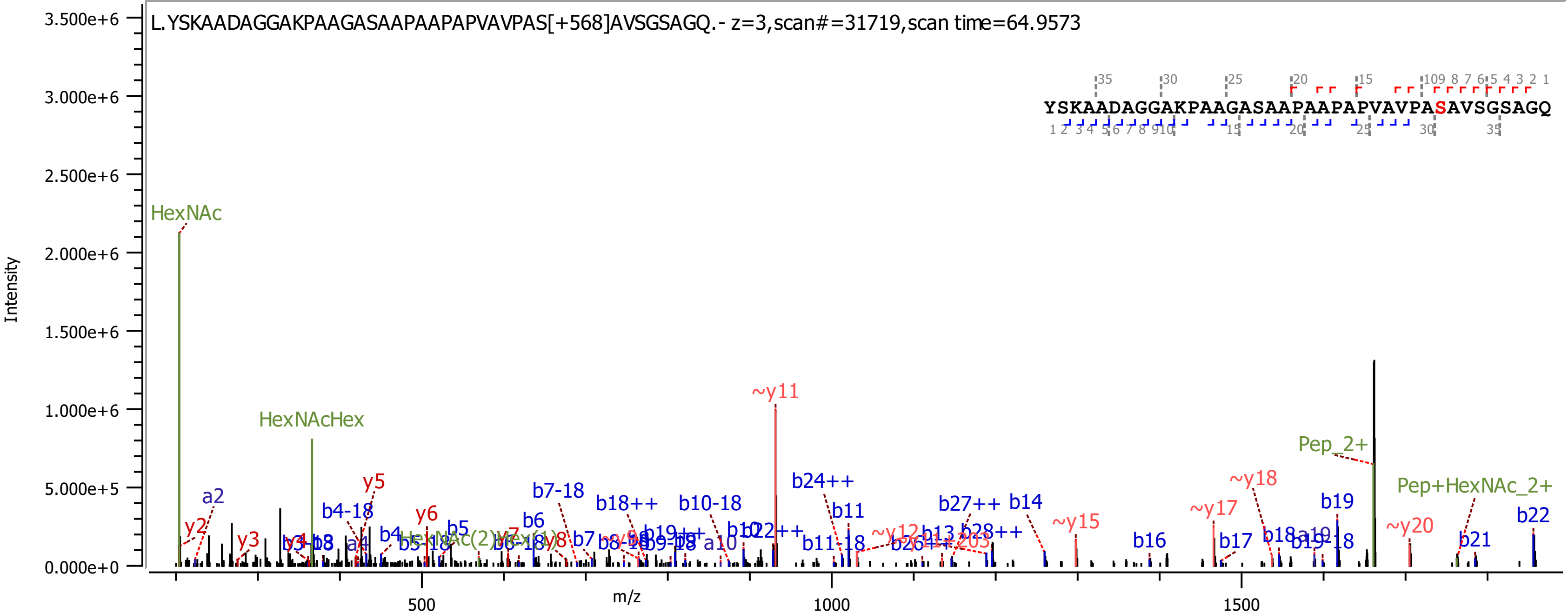

L.YSKAADAGGAKPAAGASAAPAAPAPVAVPAS[+568]AVSGSAGQ.- z=3,scan#=31719,scan time=64.9573
YSKAADAGGAKPAAGASAAPAAPAPVAVPASAVSGSAGQ
Intensity
m/z
HexNAc
HexNAcHex
~y11
Pep_2+
Pep+HexNAc_2+
~y18
~y17
~y15
~y20
b24++
b11
b27++
b14
b19
b22
b21
b16
b17
b7-18
b18++
b10-18
b4-18
y5
y6
a2
y2
y3
b5
b6
b11-18

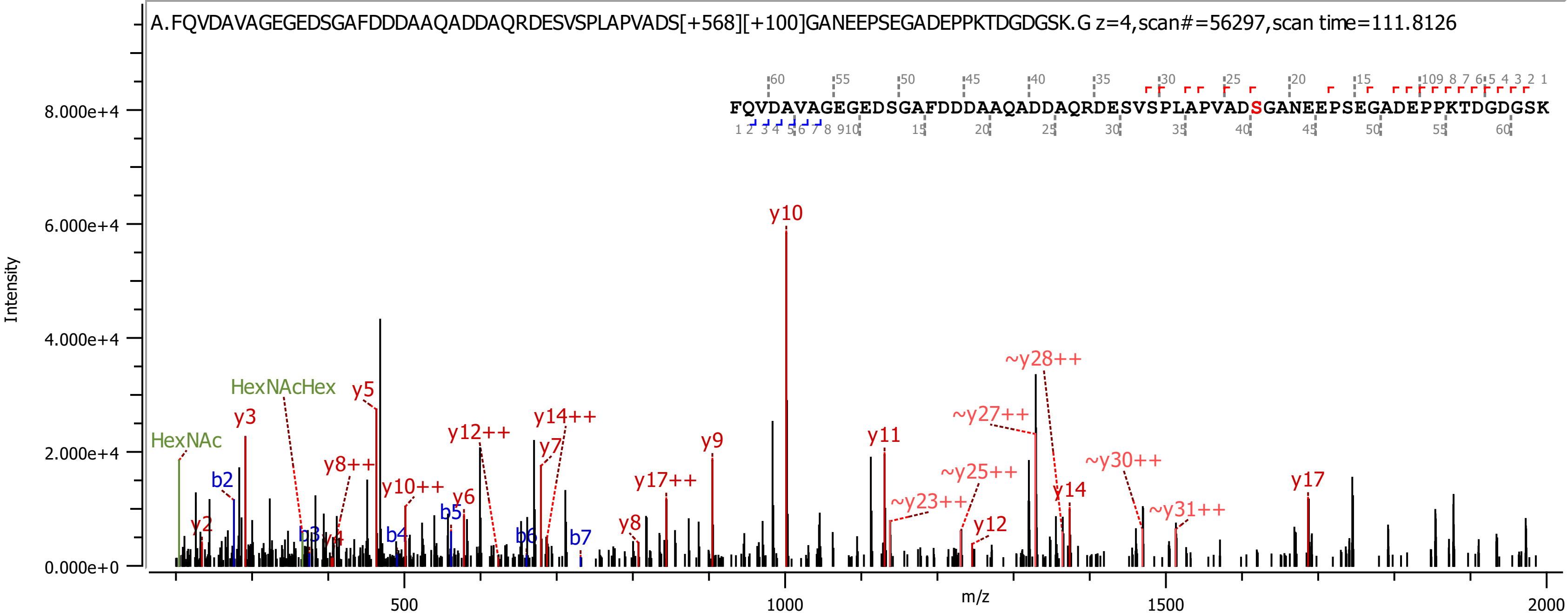
A.FQVDAVAGEGEDSGAFDDDAAQADDAQRDESVSPLAPVADS[+568][+100]GANEEPSEGADEPPKTDGDGSK.G z=4,scan#=56297,scan time=111.8126
FQVDAVAGEGEDSGAFDDDAAQADDAQRDESVSPLAPVADSGANEEPSEGADEPPKTDGDGSK
Intensity
m/z
HexNAc
HexNAcHex
y10
y5
y3
y12++
y14++
y7
y8++
y10++
y6
y9
y17++
y8
y2
b2
b3
y4
b4
b5
b6
b7
y11
~y28++
~y27++
~y25++
~y23++
~y30++
y14
y12
~y31++
y17
8.000e+4
6.000e+4
4.000e+4
2.000e+4
0.000e+0
500
1000
1500
2000

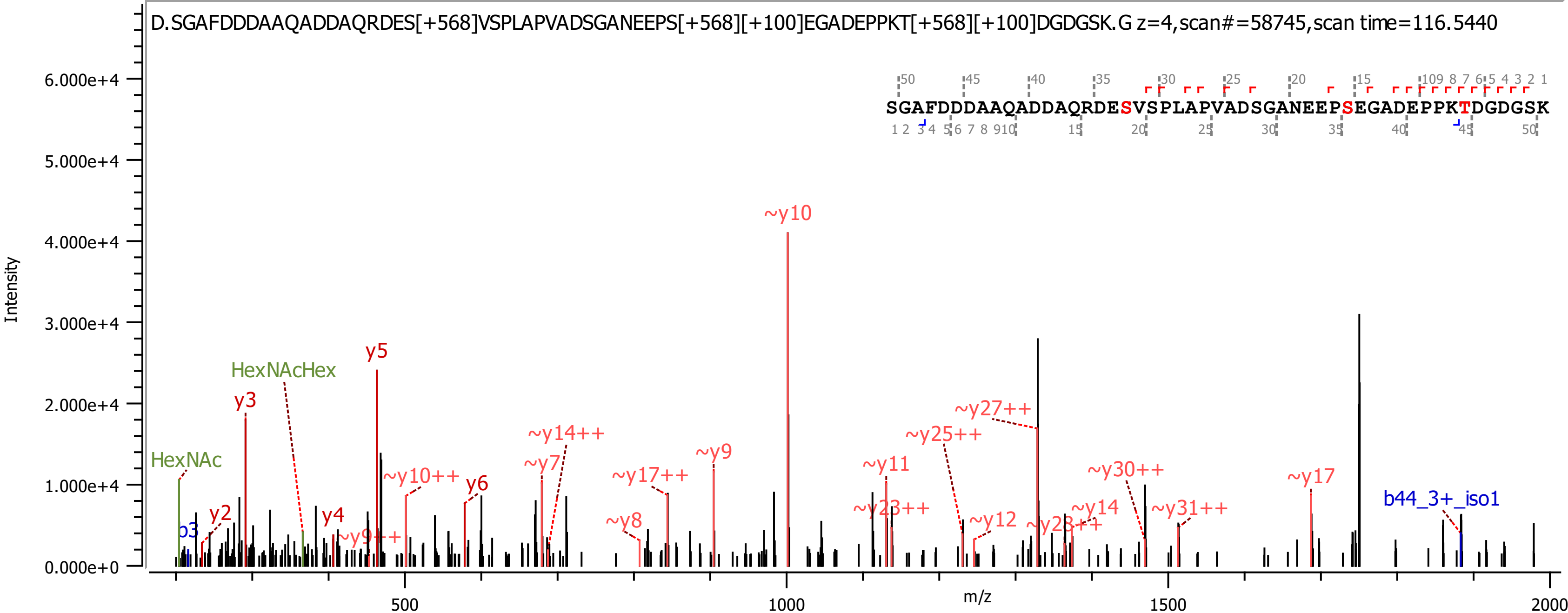
D.SGAFDDDAAQADDAQRDES[+568]VSPLAPVADSGANEEPS[+568][+100]EGADEPPKT[+568][+100]DGDGSK.G z=4,scan#=58745,scan time=116.5440
SGAFDDDAAQADDAQRDESVSPLAPVADSGANEEPSEGADEPPKTDGDGSK
Intensity
6.000e+4
5.000e+4
4.000e+4
3.000e+4
2.000e+4
1.000e+4
0.000e+0
m/z
500
1000
1500
2000
HexNAc
HexNAcHex
b3
y2
y3
y4
y5
~y9++
~y10++
y6
~y7
~y14++
~y8
~y17++
~y9
~y10
~y11
~y23++
~y25++
~y27++
~y12
~y28++
~y14
~y30++
~y31++
~y17
b44_3+_iso1

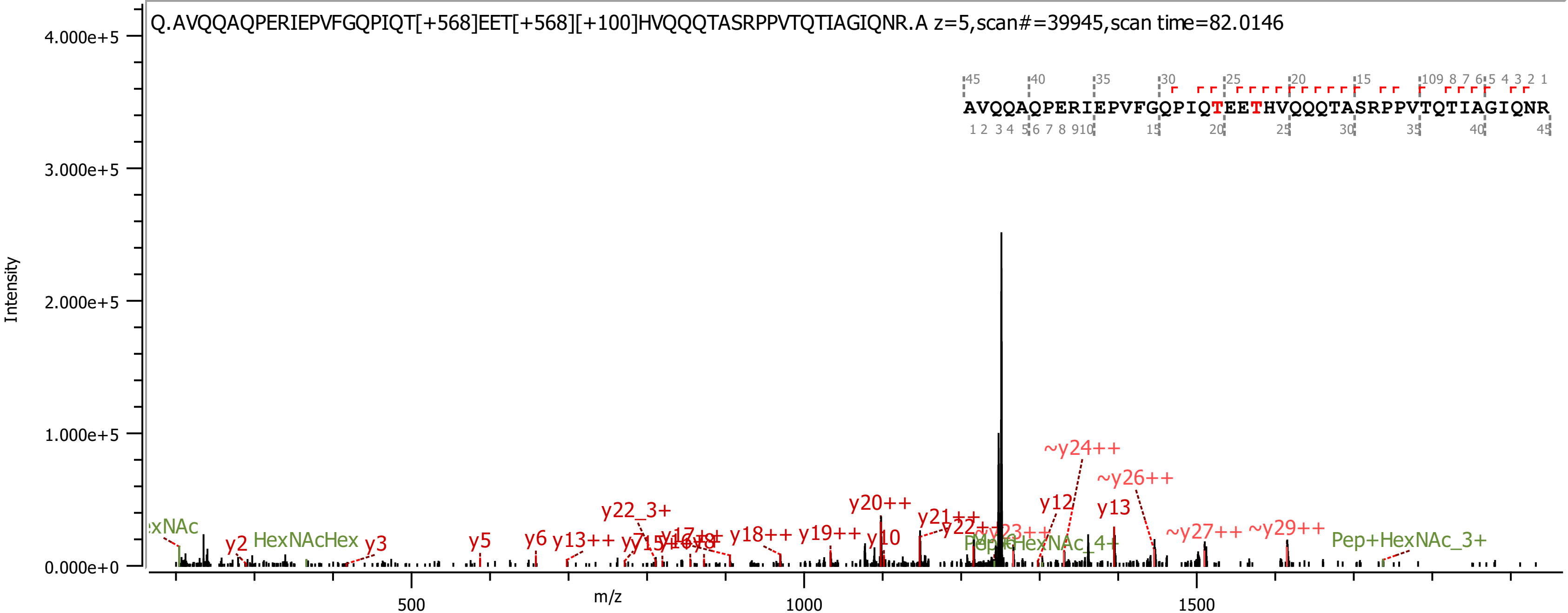

Q.AVQQAQPERIEPVFGQPIQT[+568]EET[+568][+100]HVQQQTASRPPVTQTIAGIQNR.A z=5,scan#=39945,scan time=82.0146
AVQQAQPERIEPVFGQPIQTEETHVQQQTASRPPVTQTIAGIQNR
Intensity
4.000e+5
3.000e+5
2.000e+5
1.000e+5
0.000e+0
m/z
500
1000
1500
y2
HexNAcHex
y3
y5
y6
y13++
y22_3+
y18++
y19++
y20++
y10
y21++
~y24++
y12
y13
~y26++
~y27++
~y29++
Pep+HexNAc_3+

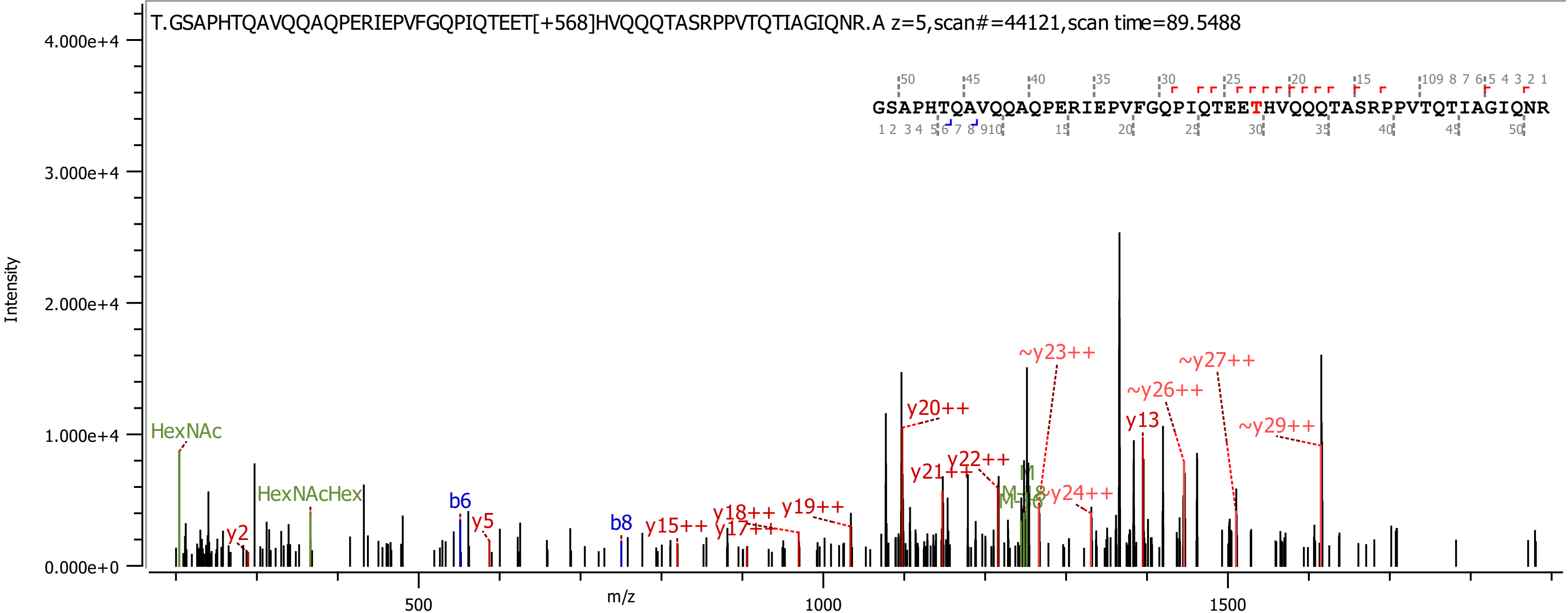
T.GSAPHTQAVQQAQPERIEPVFGQPIQTEET[+568]HVQQQTASRPPVTQTIAGIQNR.A z=5,scan#=44121,scan time=89.5488
GSAPHTQAVQQAQPERIEPVFGQPIQTEETHVQQQTASRPPVTQTIAGIQNR
Intensity
m/z
HexNAc
HexNAcHex
y2
b6
y5
b8
y15++
y17++
y18++
y19++
y20++
y21++
y22++
~y23++
~y24++
y13
~y26++
~y27++
~y29++

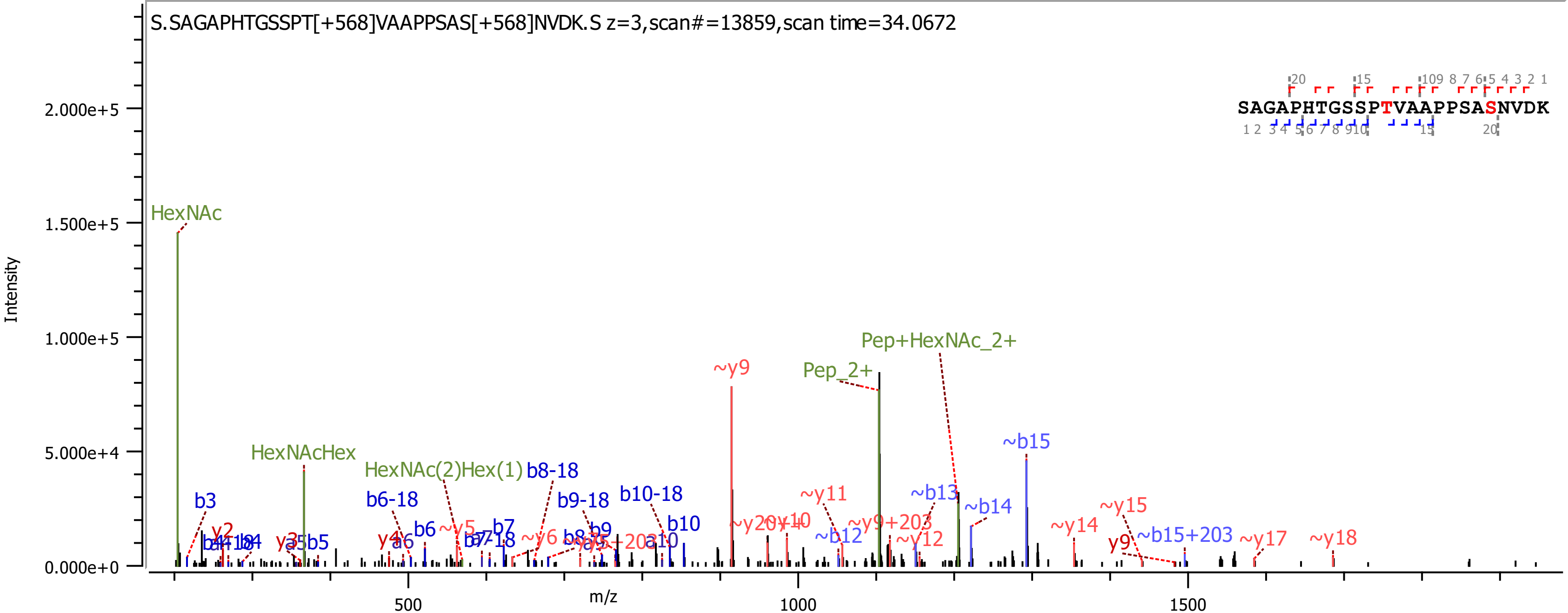

S.SAGAPHTGSSPT[+568]VAAPPSAS[+568]NVDK.S z=3,scan#=13859,scan time=34.0672
SAGAPHTGSSPTVAAPPSASNVDK
Intensity
m/z
0.000e+0
5.000e+4
1.000e+5
1.500e+5
2.000e+5
500
1000
1500
HexNAc
HexNAcHex
HexNAc(2)Hex(1)
Pep_2+
Pep+HexNAc_2+
b3
b5
b6
b6-18
b7
b7-18
b8-18
b9
b9-18
b10
b10-18
~y9
~y11
~b12
~b13
~b14
~b15
~y12
~y14
~y15
~y9+203
~b15+203
~y17
~y18
y9
y2
y3
y4

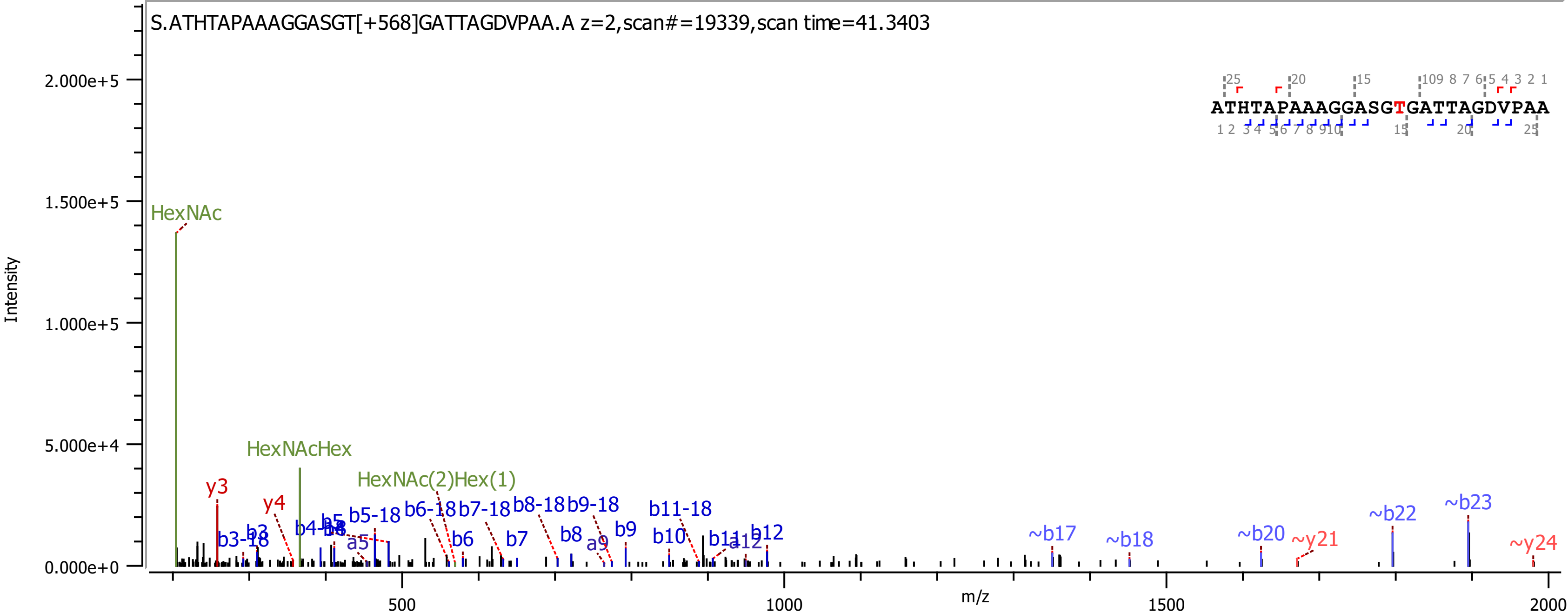

S.ATHTAPAAAGGASGT[+568]GATTAGDVPAA.A z=2,scan#=19339,scan time=41.3403
ATHTAPAAAGGASGTGATTAGDVPAA
Intensity
m/z
HexNAc
HexNAcHex
HexNAc(2)Hex(1)
y3
y4
b3-18
b4-18
b5-18
b6-18
b7-18
b8-18
b9-18
b11-18
b6
b7
b8
a9
b9
b10
b12
a5
~b17
~b18
~b20
~y21
~b22
~b23
~y24

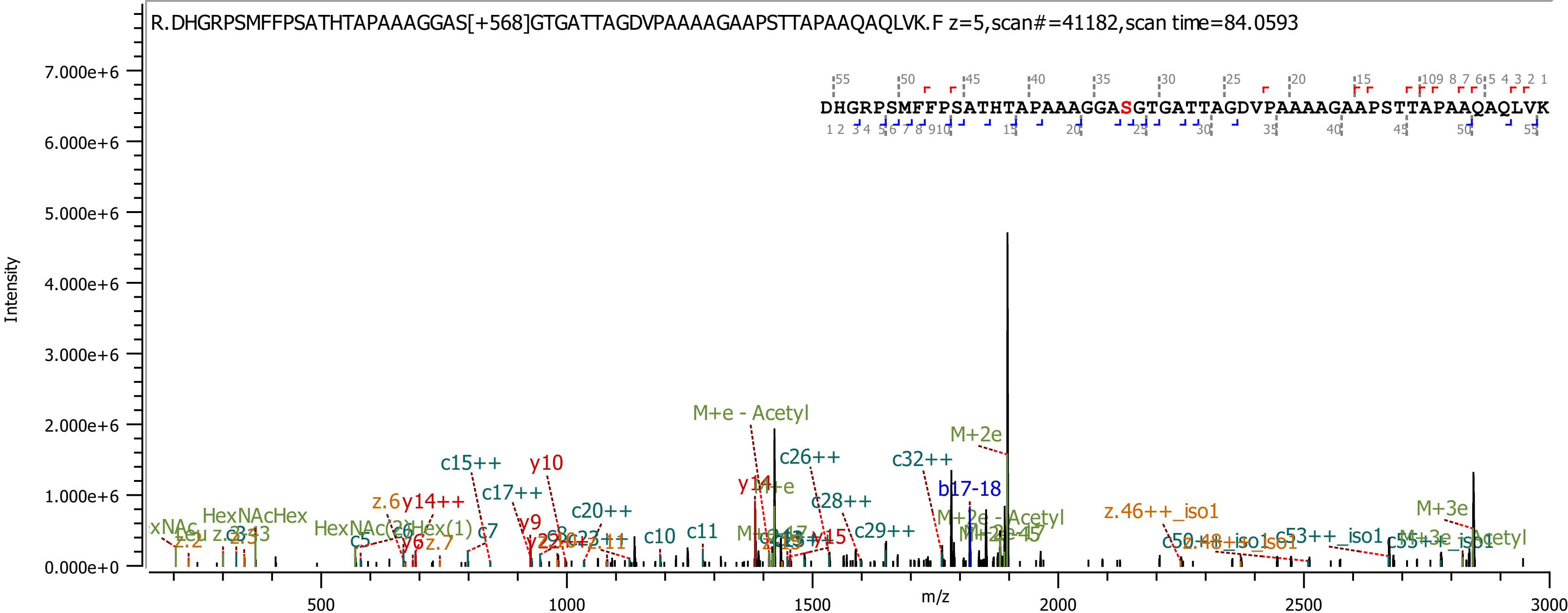

R.DHGRPSMFFPSATHTAPAAAGGAS[+568]GTGATTAGDVPAAAAGAAPSTTAPAAQAQLVK.F z=5,scan#=41182,scan time=84.0593
DHGRPSMFFPSATHTAPAAAGGASGTGATTAGDVPAAAAGAAPSTTAPAAQAQLVK
Intensity
m/z
M+e - Acetyl
M+2e
c15++
y10
c17++
c20++
c26++
c32++
y14
b17-18
c28++
z.46++_iso1
M+3e
c29++
c10
c11
y9
c7
c5
y6
z.7
z.6
y14++
HexNAcHex
HexNAc(2)Hex(1)
z.2
z.11
y15
M+3e - Acetyl
M+2e - Acetyl
c53++_iso1

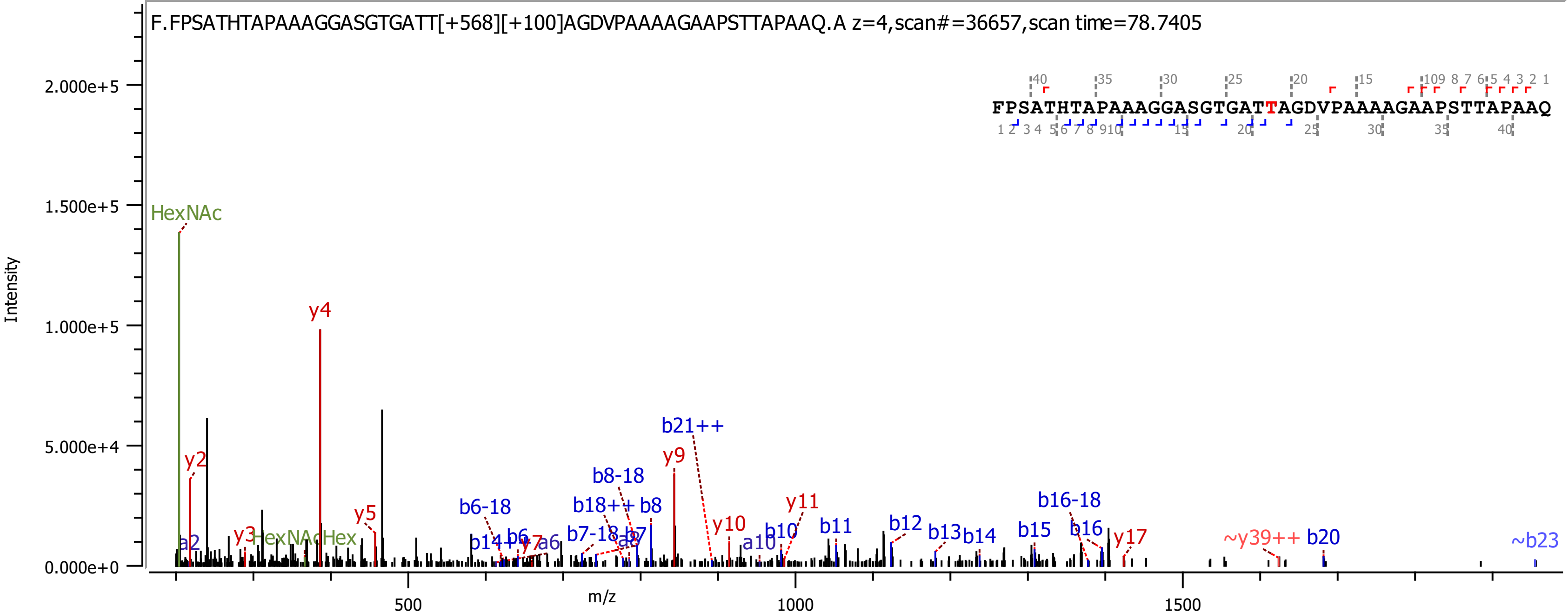

F.FPSATHTAPAAAGGASGTGATT[+568][+100]AGDVPAAAAGAAPSTTAPAAQ.A z=4,scan#=36657,scan time=78.7405
FPSATHTAPAAAGGASGTGATTAGDVPAAAAGAAPSTTAPAAQ
Intensity
m/z
HexNAc
y4
y2
y3
HexNAcHex
y5
a2
b21++
y9
b8-18
b6-18
b18++
b8
b7-18
a8
b7
b14++
b6
y7
a6
y10
y11
a10
b10
b11
b12
b13
b14
b15
b16-18
b16
y17
~y39++
b20
~b23

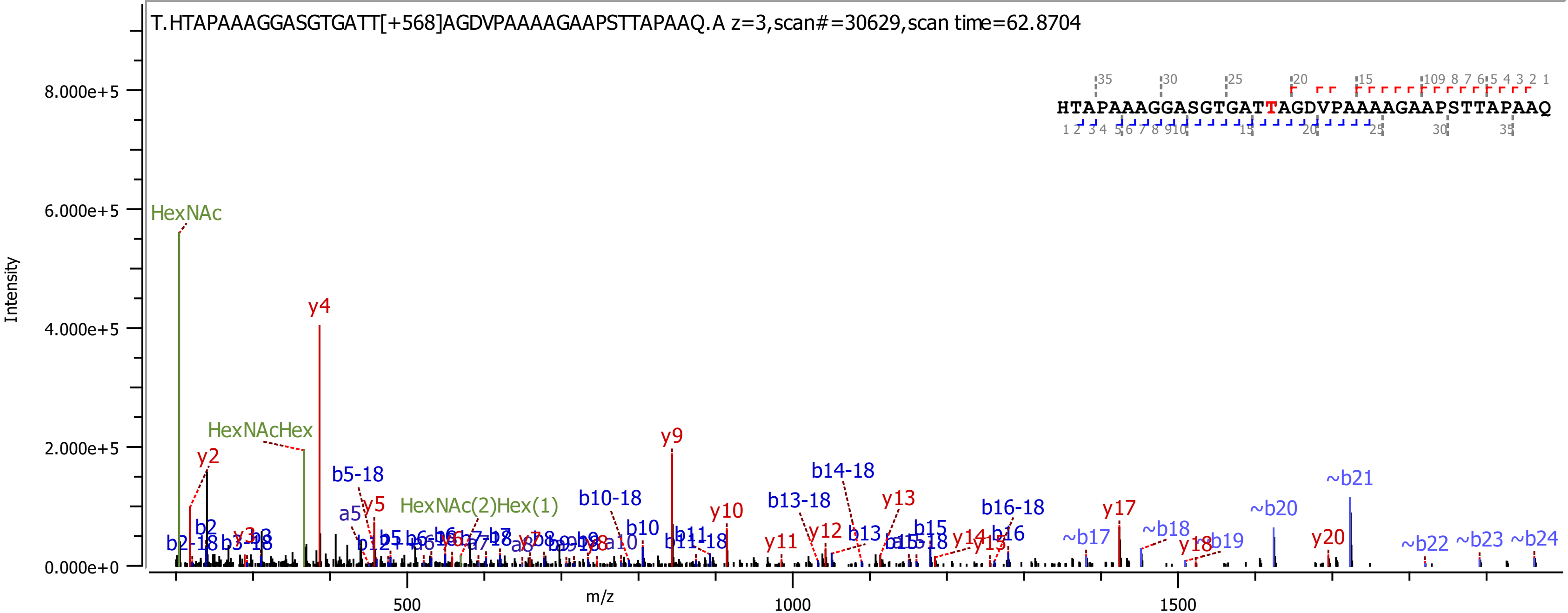
T.HTAPAAAGGASGTGATT[+568]AGDVPAAAAGAAPSTTAPAAQ.A z=3,scan#=30629,scan time=62.8704
HTAPAAAGGASGTGATTAGDVPAAAAGAAPSTTAPAAQ
Intensity
m/z
HexNAc
HexNAcHex
HexNAc(2)Hex(1)
y2
y4
y5
y9
y10
y11
y12
y13
y17
y20
b5-18
b10-18
b13-18
b14-18
b16-18
~b17
~b18
~b20
~b21
~b22
~b23
~b24

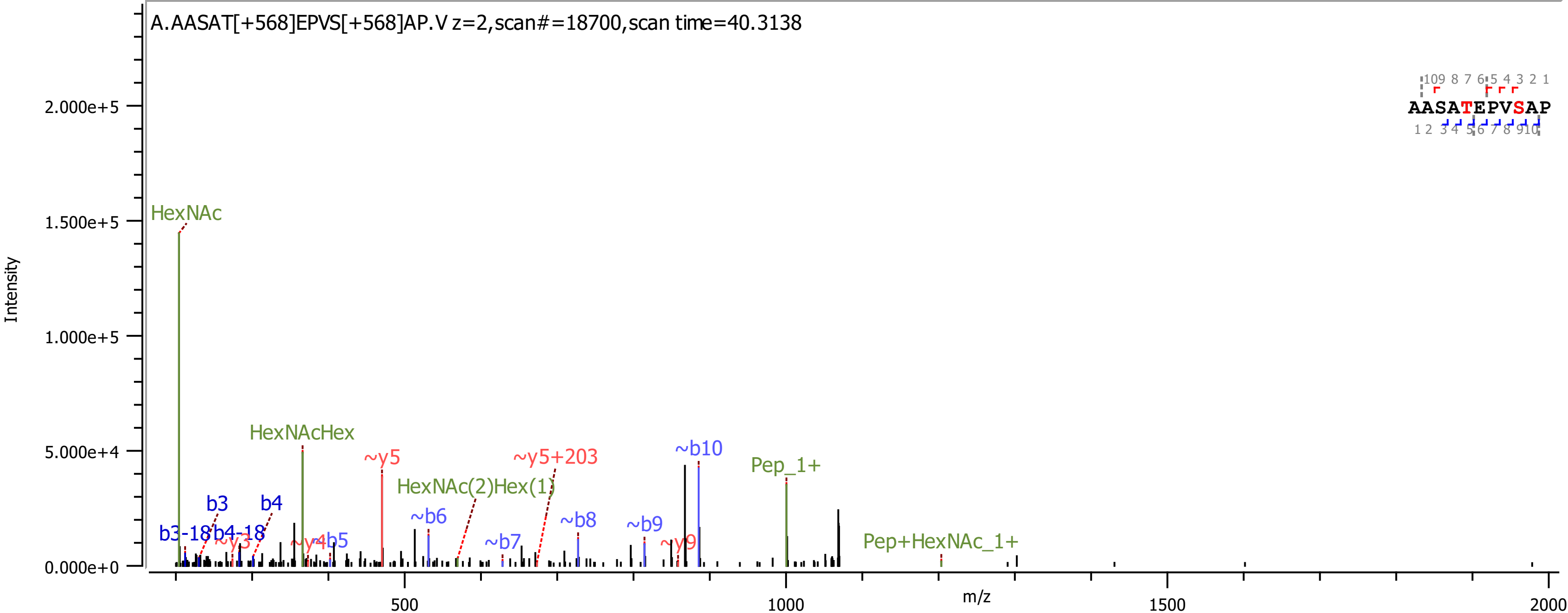

A.AASAT[+568]EPVS[+568]AP.V z=2,scan#=18700,scan time=40.3138
AASATEPVSAP
Intensity
m/z
HexNAc
HexNAcHex
~y5
~y5+203
HexNAc(2)Hex(1)
~b10
Pep_1+
Pep+HexNAc_1+
b3
b4
b3-18
b4-18
~y3
~y4
~b5
~b6
~b7
~b8
~b9
~y9

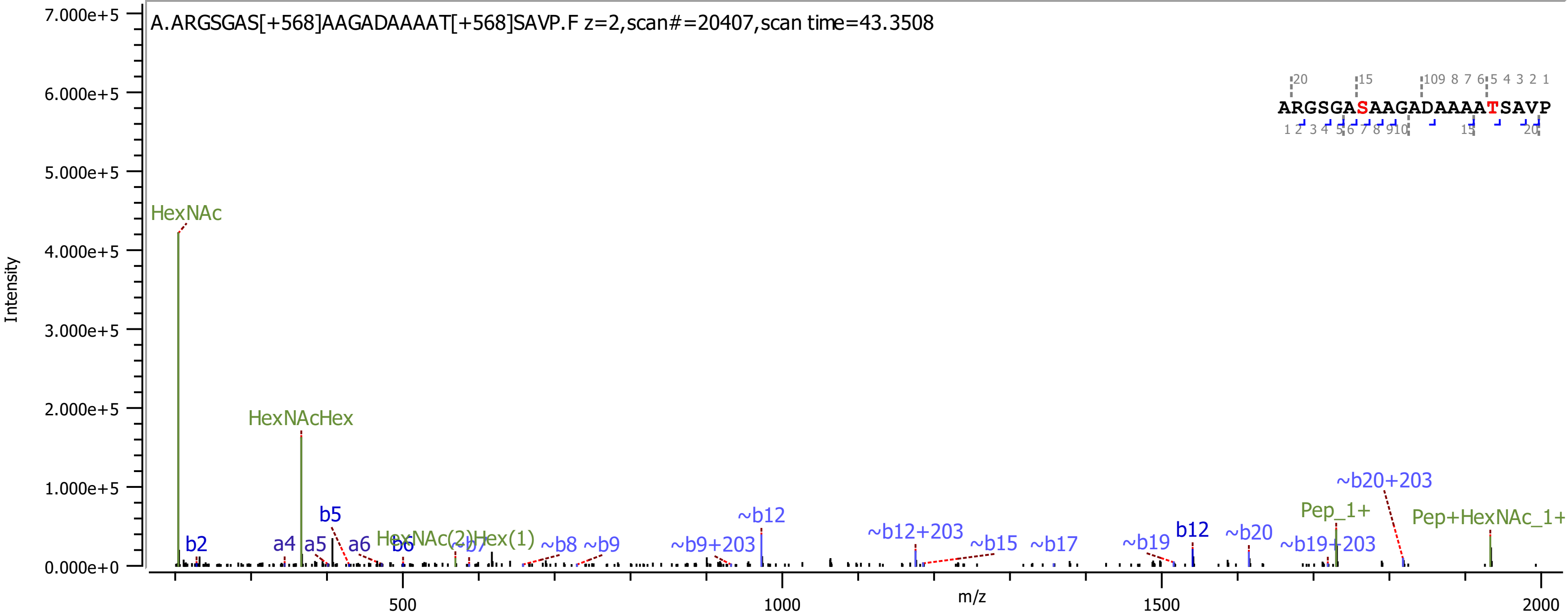

A.ARGSGAS[+568]AAGADAAAAT[+568]SAVP.F z=2,scan#=20407,scan time=43.3508
ARGSGASAAGADAAAATSAVP
Intensity
m/z
HexNAc
HexNAcHex
HexNAc(2)Hex(1)
b2
a4
a5
b5
a6
b6
~b7
~b8
~b9
~b9+203
~b12
~b12+203
~b15
~b17
~b19
b12
~b20
~b19+203
Pep_1+
~b20+203
Pep+HexNAc_1+
0.000e+0
1.000e+5
2.000e+5
3.000e+5
4.000e+5
5.000e+5
6.000e+5
7.000e+5
500
1000
1500
2000

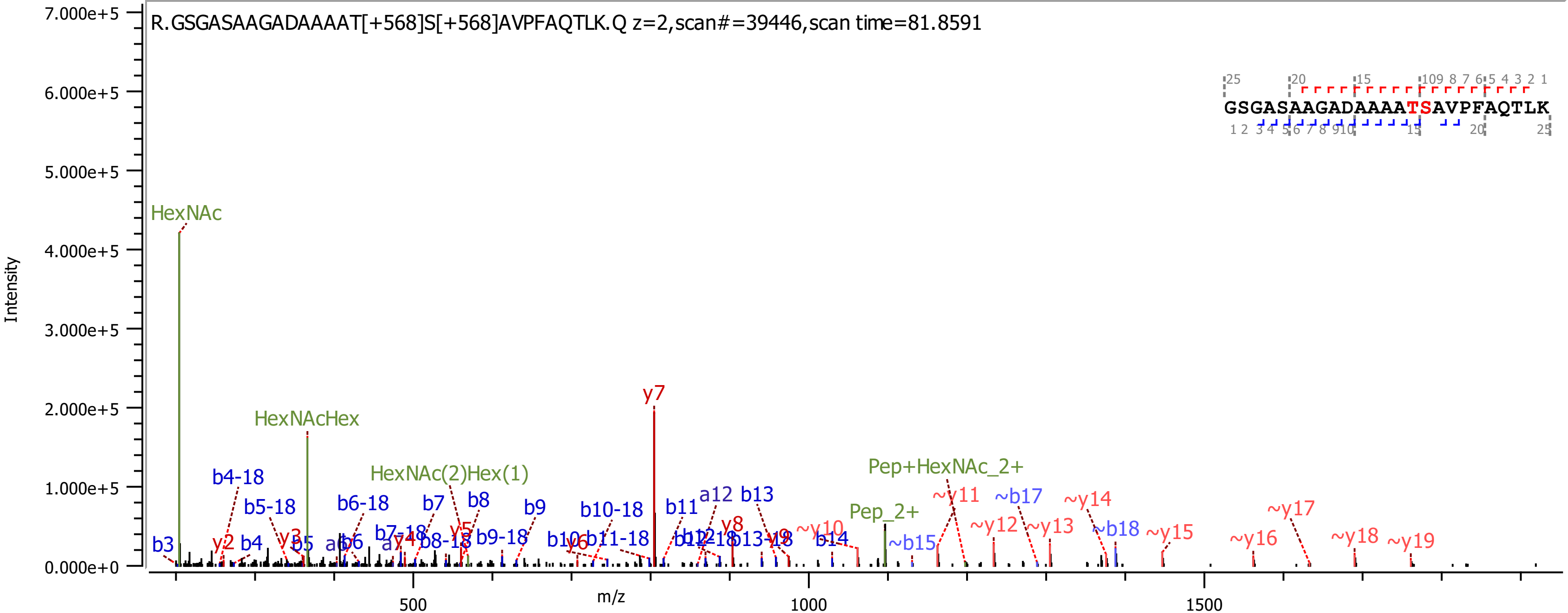
R.GSGASAAGADAAAAT[+568]S[+568]AVPFAQTLK.Q z=2,scan#=39446,scan time=81.8591
Intensity
m/z
7.000e+5
6.000e+5
5.000e+5
4.000e+5
3.000e+5
2.000e+5
1.000e+5
0.000e+0
500
1000
1500
HexNAc
HexNAcHex
HexNAc(2)Hex(1)
Pep+HexNAc_2+
Pep_2+
y7
GSGASAAGADAAAATSAVPFAQTLK

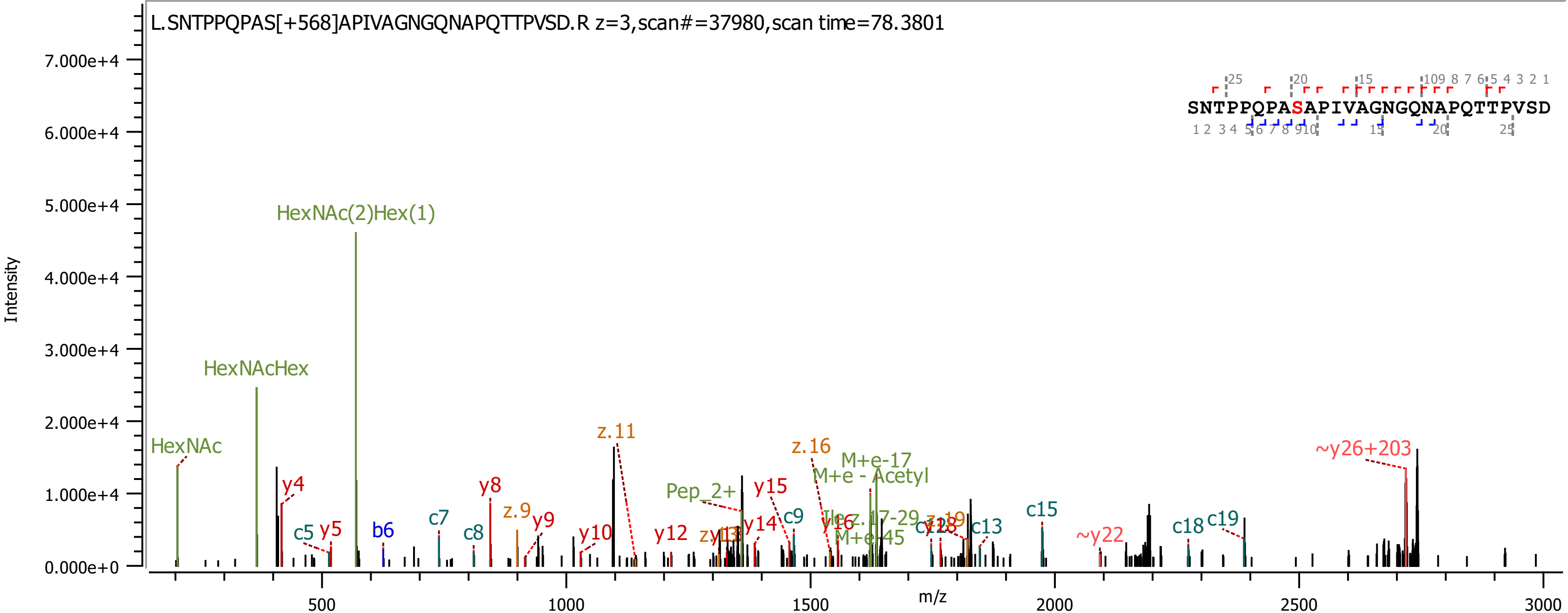

L.SNTPPQPAS[+568]APIVAGNGQNAPQTTPVSD.R z=3,scan#=37980,scan time=78.3801
SNTPPQPASAPIVAGNGQNAPQTTPVSD
Intensity
m/z
HexNAc
HexNAcHex
HexNAc(2)Hex(1)
y4
c5
y5
b6
c7
c8
y8
z.9
y9
y10
z.11
y12
Pep_2+
y14
y15
c9
z.16
M+e-17
M+e - Acetyl
M+e-45
c12
c13
c15
~y22
c18
c19
~y26+203

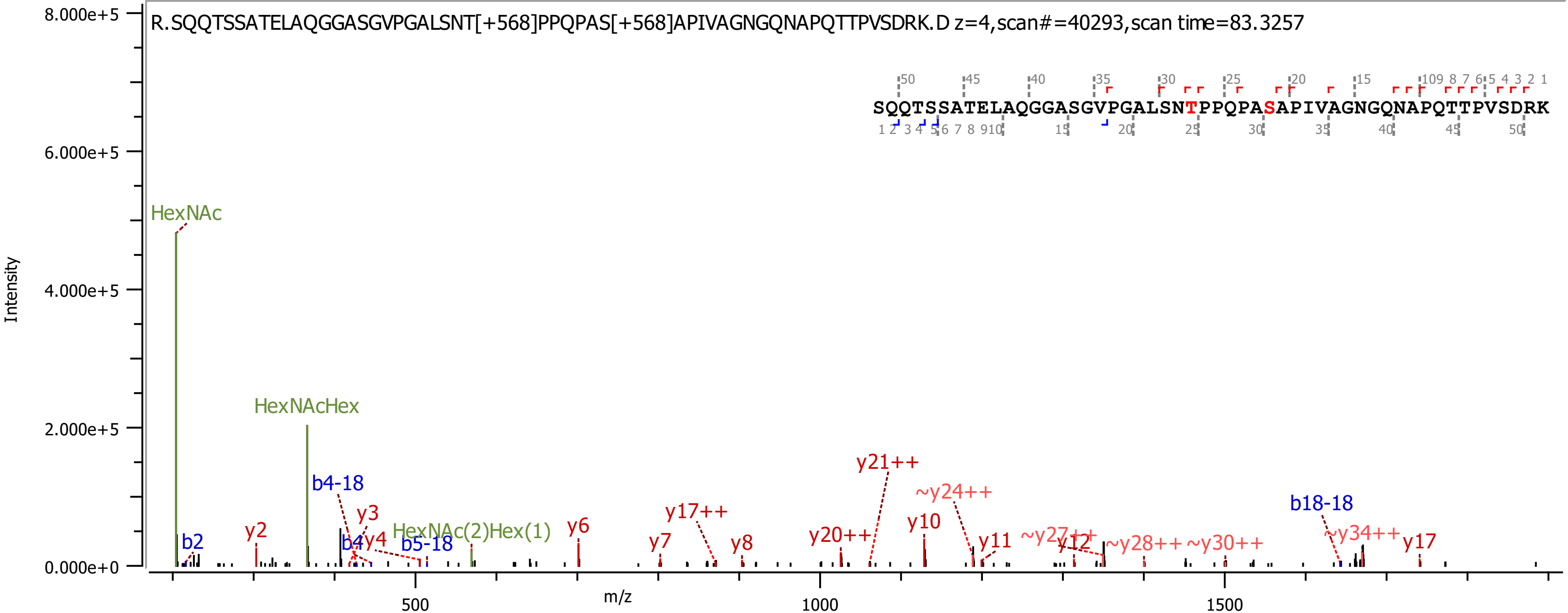
R.SQQTSSATELAQGGASGVPGALSNT[+568]PPQPAS[+568]APIVAGNGQNAPQTTPVSDRK.D z=4,scan#=40293,scan time=83.3257
SQQTSSATELAQGGASGVPGALSNTPPQPASAPIVAGNGQNAPQTTPVSDRK
Intensity
m/z
8.000e+5
6.000e+5
4.000e+5
2.000e+5
0.000e+0
500
1000
1500
HexNAc
HexNAcHex
HexNAc(2)Hex(1)
b2
b4-18
b4
b5-18
b18-18
y2
y3
y4
y6
y7
y8
y10
y11
y12
y17
y17++
y20++
y21++
~y24++
~y27++
~y28++
~y30++
~y34++

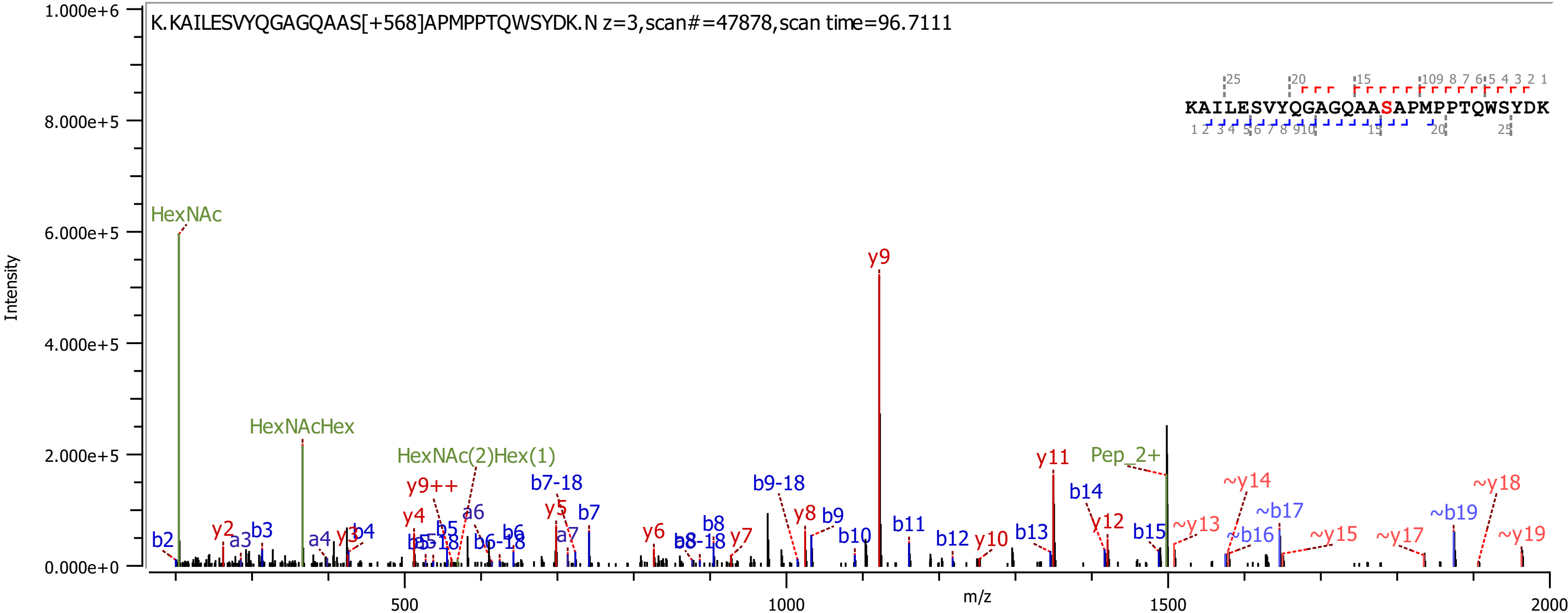
K.KAILESVYQGAGQAAS[+568]APMPPTQWSYDK.N z=3,scan#=47878,scan time=96.7111
KAILESVYQGAGQAASAPMPPTQWSYDK
Intensity
m/z
1.000e+6
8.000e+5
6.000e+5
4.000e+5
2.000e+5
0.000e+0
500
1000
1500
2000
HexNAc
HexNAcHex
HexNAc(2)Hex(1)
Pep_2+
y9
y9++
y2
y3
y4
y5
y6
y7
y8
y10
y11
y12
~y13
~y14
~y15
~y17
~y18
~y19
b2
b3
b4
b5
b6
b7
b8
b9
b10
b11
b12
b13
b14
b15
~b16
~b17
~b19
a3
a4
a6
a7
b5-18
b6-18
b7-18
b8-18
b9-18

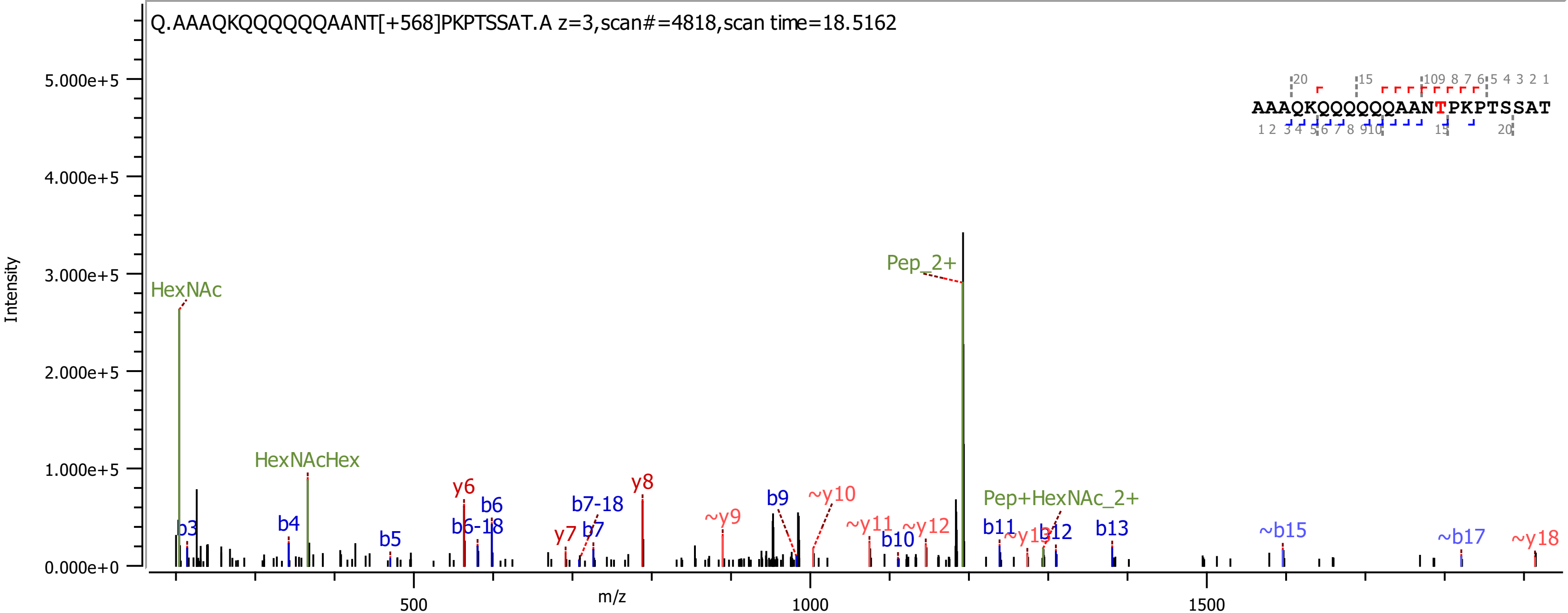
Q.AAAQKQQQQQQAANT[+568]PKPTSSAT.A z=3,scan#=4818,scan time=18.5162
AAAQKQQQQQQAANTPKPTSSAT
Intensity
m/z
HexNAc
HexNAcHex
Pep_2+
Pep+HexNAc_2+
b3
b4
b5
b6
b6-18
b7-18
b7
b9
b10
b11
b12
b13
~b15
~b17
y6
y7
y8
~y9
~y10
~y11
~y12
~y18

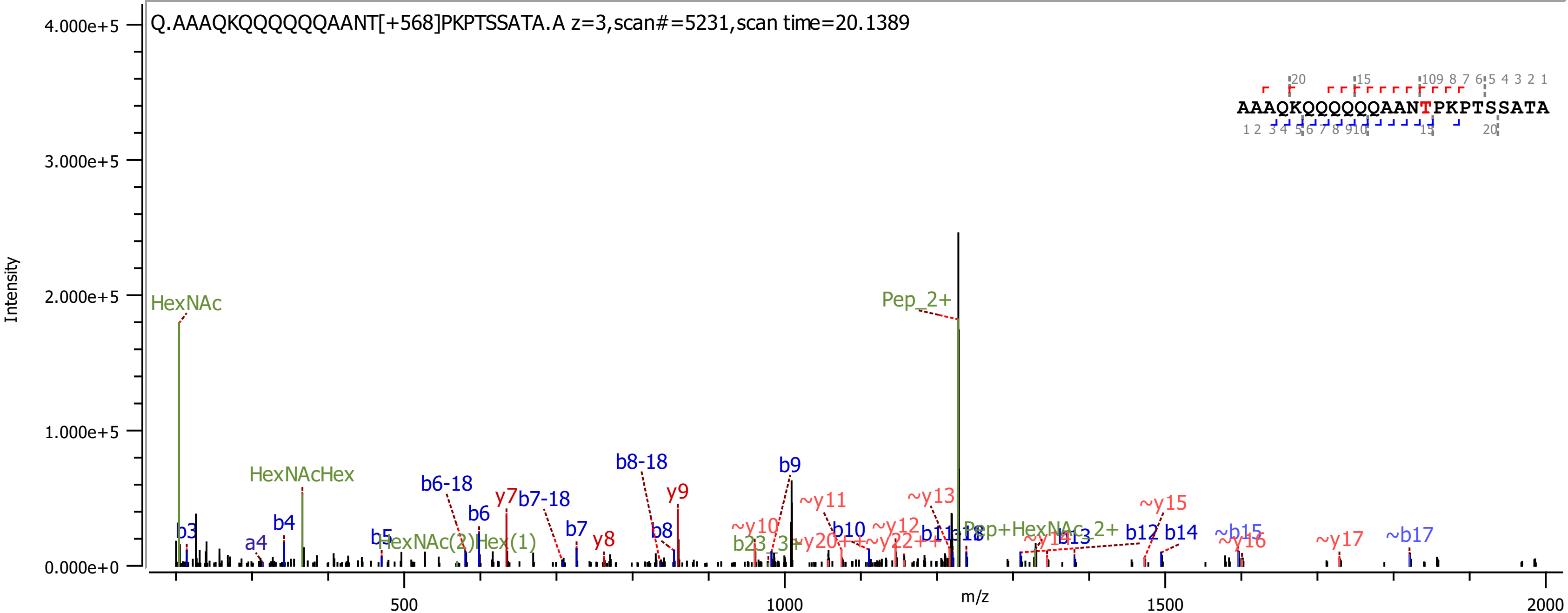

Q.AAAQKQQQQQQAANT[+568]PKPTSSATA.A z=3,scan#=5231,scan time=20.1389
AAAQKQQQQQQAANTPKPTSSATA
Intensity
4.000e+5
3.000e+5
2.000e+5
1.000e+5
0.000e+0
m/z
500
1000
1500
2000
HexNAc
HexNAcHex
Pep_2+
b3
a4
b4
b5
HexNAc(2)Hex(1)
b6-18
b6
y7
b7-18
b7
y8
b8-18
b8
y9
b9
~y10
~y11
b10
~y12
~y13
~y15
b12
b14
~b15
~y16
~y17
~b17

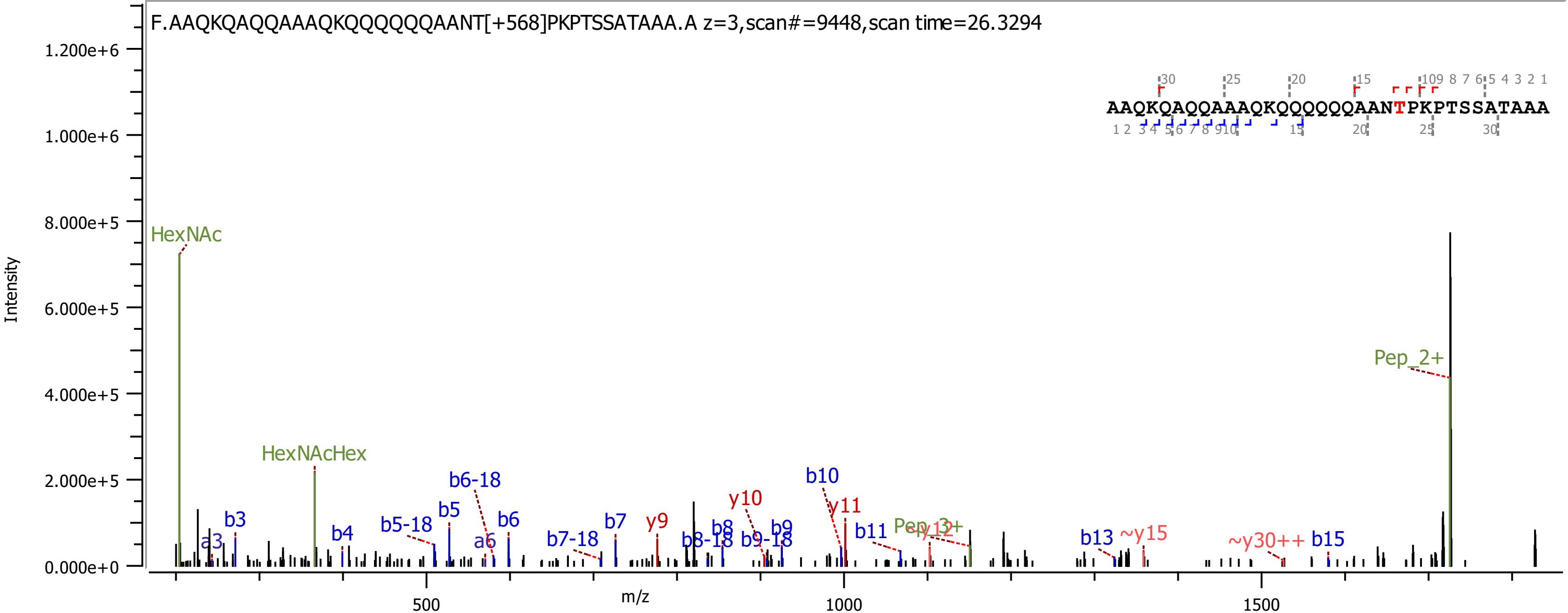

F.AAQKQAQQAAAQKQQQQQQAANT[+568]PKPTSSATAAA.A z=3,scan#=9448,scan time=26.3294
AAQKQAQQAAAQKQQQQQQAANTPKPTSSATAAA
Intensity
m/z
1.200e+6
1.000e+6
8.000e+5
6.000e+5
4.000e+5
2.000e+5
0.000e+0
500
1000
1500
HexNAc
HexNAcHex
b3
a3
b4
b5-18
b5
b6-18
a6
b6
b7-18
b7
y9
b8-18
b8
b9-18
b9
y10
b10
y11
b11
Pep_3+
~y12
b13
~y15
~y30++
b15
Pep_2+

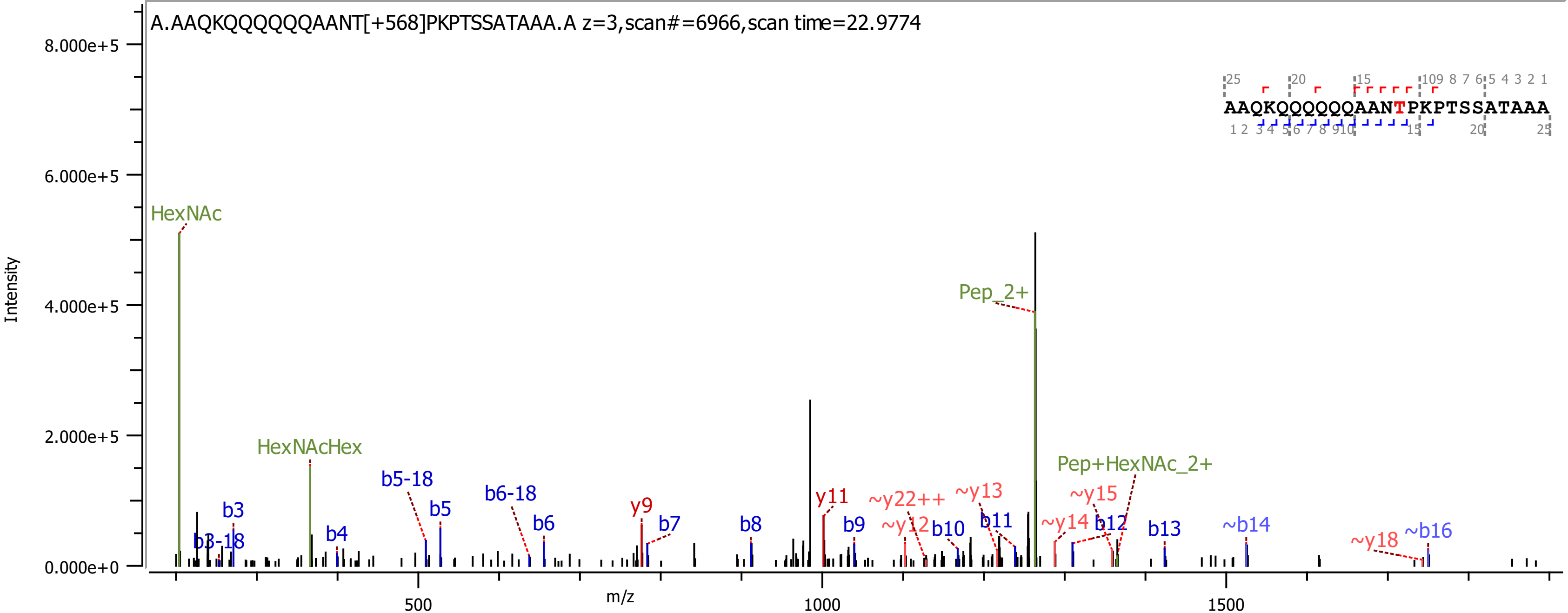

A.AAQKQQQQQQAANT[+568]PKPTSSATAAA.A z=3,scan#=6966,scan time=22.9774
AAQKQQQQQQAANTPKPTSSATAAA
Intensity
m/z
8.000e+5
6.000e+5
4.000e+5
2.000e+5
0.000e+0
500
1000
1500
HexNAc
HexNAcHex
Pep_2+
Pep+HexNAc_2+
b3
b3-18
b4
b5-18
b5
b6-18
b6
y9
b7
b8
y11
b9
~y22++
~y12
b10
~y13
b11
~y14
~y15
b12
b13
~b14
~y18
~b16

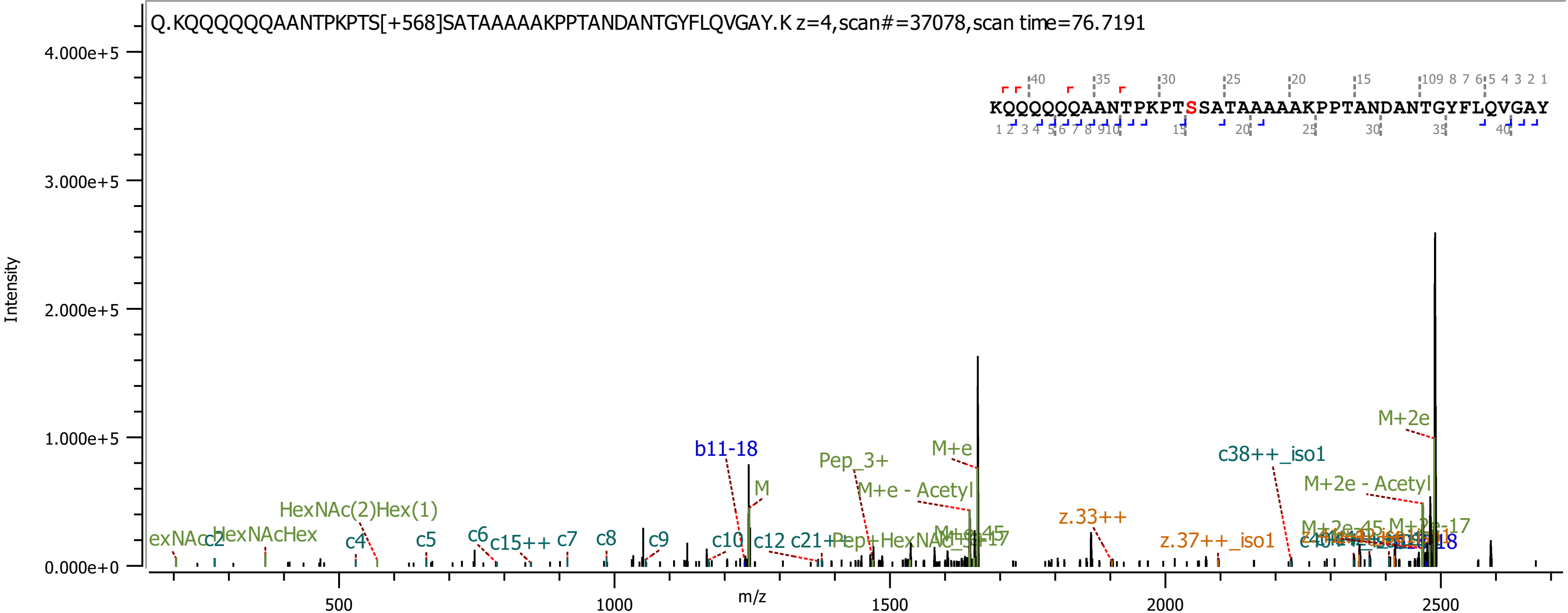

Q.KQQQQQQAANTPKPTS[+568]SATAAAAAKPPTANDANTGYFLQVGAY.K z=4,scan#=37078,scan time=76.7191
KQQQQQQAANTPKPTSSATAAAAAKPPTANDANTGYFLQVGAY
Intensity
m/z
4.000e+5
3.000e+5
2.000e+5
1.000e+5
0.000e+0
500
1000
1500
2000
2500
M+2e
b11-18
Pep_3+
M+e
c38++_iso1
M
M+e - Acetyl
M+2e - Acetyl
HexNAc(2)Hex(1)
z.33++
HexNAcHex
c2
c4
c5
c6
c15++
c7
c8
c9
c10
c12
z.37++_iso1

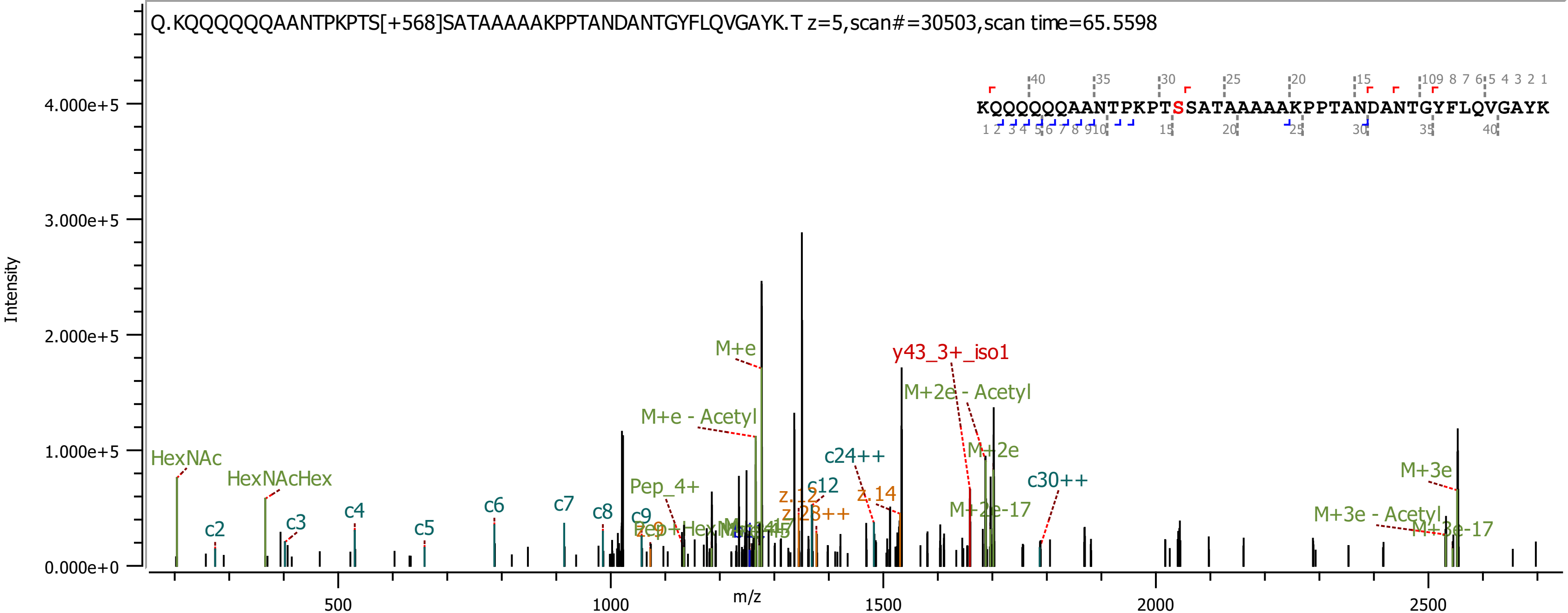
Q.KQQQQQQAANTPKPTS[+568]SATAAAAAKPPTANDANTGYFLQVGAYK.T z=5,scan#=30503,scan time=65.5598
KQQQQQQAANTPKPTSSATAAAAAKPPTANDANTGYFLQVGAYK
Intensity
m/z
4.000e+5
3.000e+5
2.000e+5
1.000e+5
0.000e+0
500
1000
1500
2000
2500
HexNAc
HexNAcHex
c2
c3
c4
c5
c6
c7
c8
c9
M+e
M+e - Acetyl
Pep_4+
c12
c24++
z.14
y43_3+_iso1
M+2e - Acetyl
M+2e
M+2e-17
c30++
M+3e
M+3e - Acetyl
M+3e-17

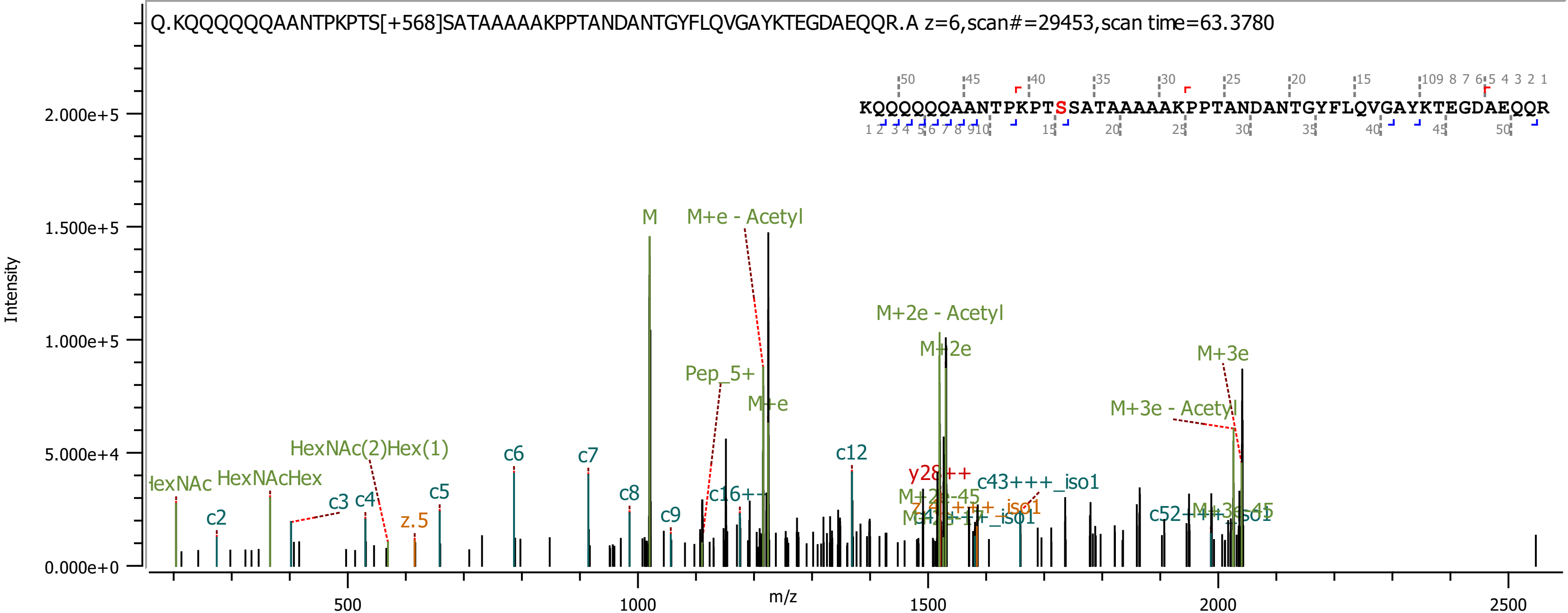
Q.KQQQQQQAANTPKPTS[+568]SATAAAAAKPPTANDANTGYFLQVGAYKTEGDAEQQR.A z=6,scan#=29453,scan time=63.3780
KQQQQQQAANTPKPTSSATAAAAAKPPTANDANTGYFLQVGAYKTEGDAEQQR
Intensity
m/z
0.000e+0
5.000e+4
1.000e+5
1.500e+5
2.000e+5
500
1000
1500
2000
2500
M
M+e - Acetyl
M+2e - Acetyl
M+2e
M+3e
M+3e - Acetyl
Pep_5+
M+e
HexNAc(2)Hex(1)
HexNAcHex
c2
c3
c4
c5
c6
c7
c8
c9
c12
z.5
y28++
c43+++_iso1
c16++

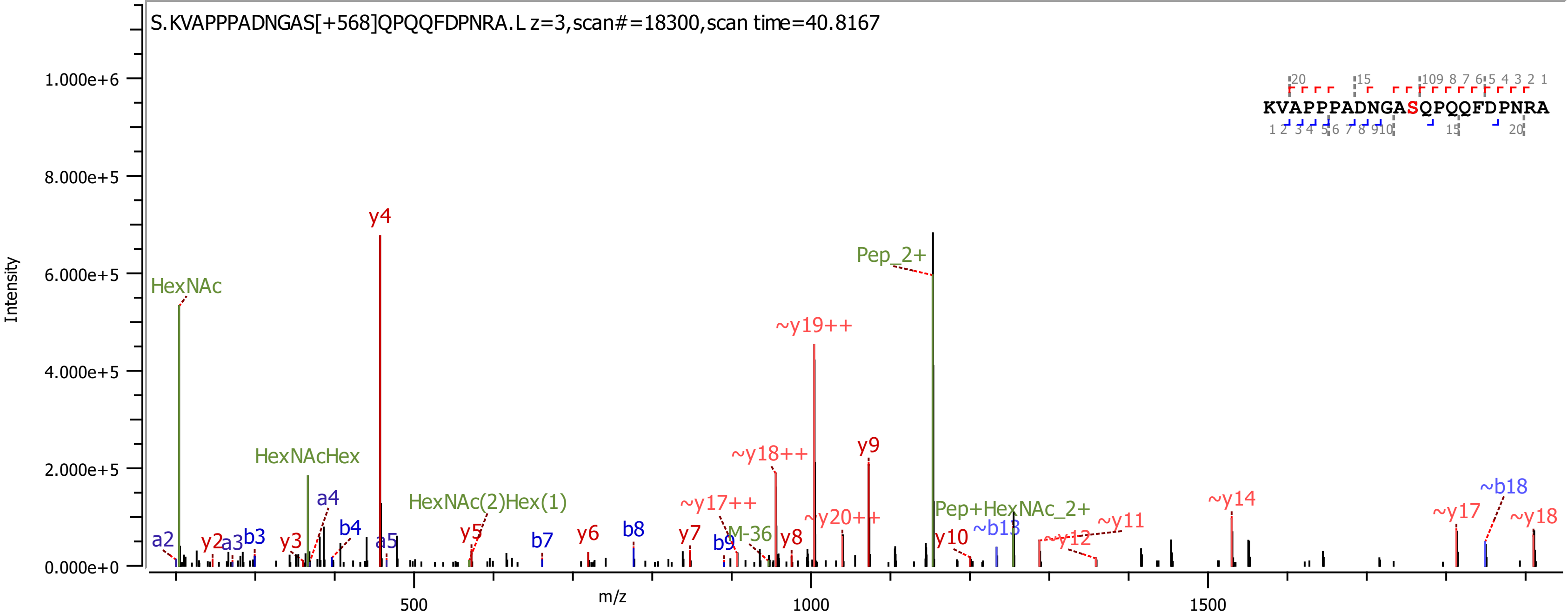

S.KVAPPPADNGAS[+568]QPQQFDPNRA.L z=3,scan#=18300,scan time=40.8167
Intensity
m/z
1.000e+6
8.000e+5
6.000e+5
4.000e+5
2.000e+5
0.000e+0
500
1000
1500
KVAPPPADNGASQPQQFDPNRA
HexNAc
HexNAcHex
HexNAc(2)Hex(1)
Pep_2+
Pep+HexNAc_2+
M-36
y4
~y19++
~y18++
y9
~y17++
~y20++
~y14
~y17
~b18
~y18
~y11
~y12
~b13
y10
y8
y7
y6
y5
y3
y2
b9
b8
b7
b4
b3
a5
a4
a3
a2

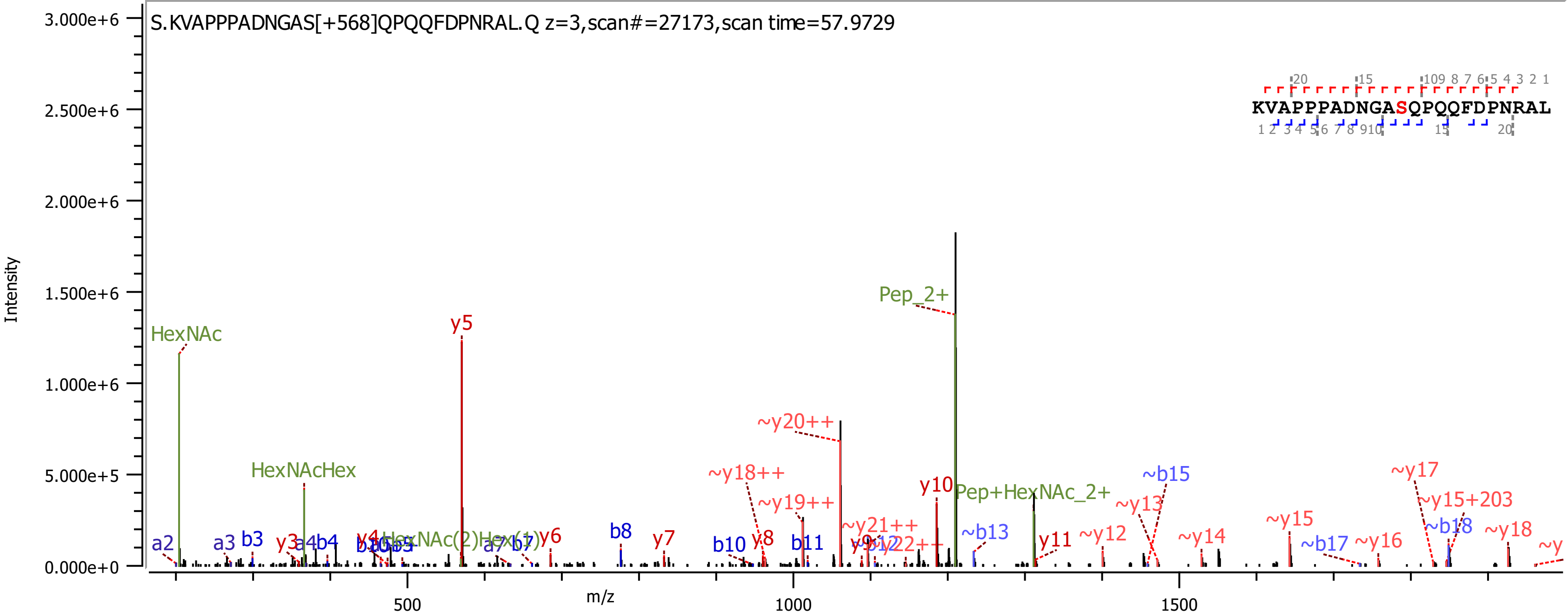

S.KVAPPPADNGAS[+568]QPQQFDPNRAL.Q z=3,scan#=27173,scan time=57.9729
KVAPPPADNGASQPQQFDPNRAL
Intensity
m/z
3.000e+6
2.500e+6
2.000e+6
1.500e+6
1.000e+6
5.000e+5
0.000e+0
500
1000
1500
HexNAc
HexNAcHex
y5
Pep_2+
~y20++
~y18++
~y19++
y10
Pep+HexNAc_2+
~b15
~y13
~y17
~y15+203
~y21++
~b13
~y12
~y14
~y15
~b18
~y18
~b17
~y16
a2
a3
b3
y3
a4
b4
y4
HexNAc(2)Hex(1)
a7
b7
y6
b8
y7
b10
y8
b11
y11

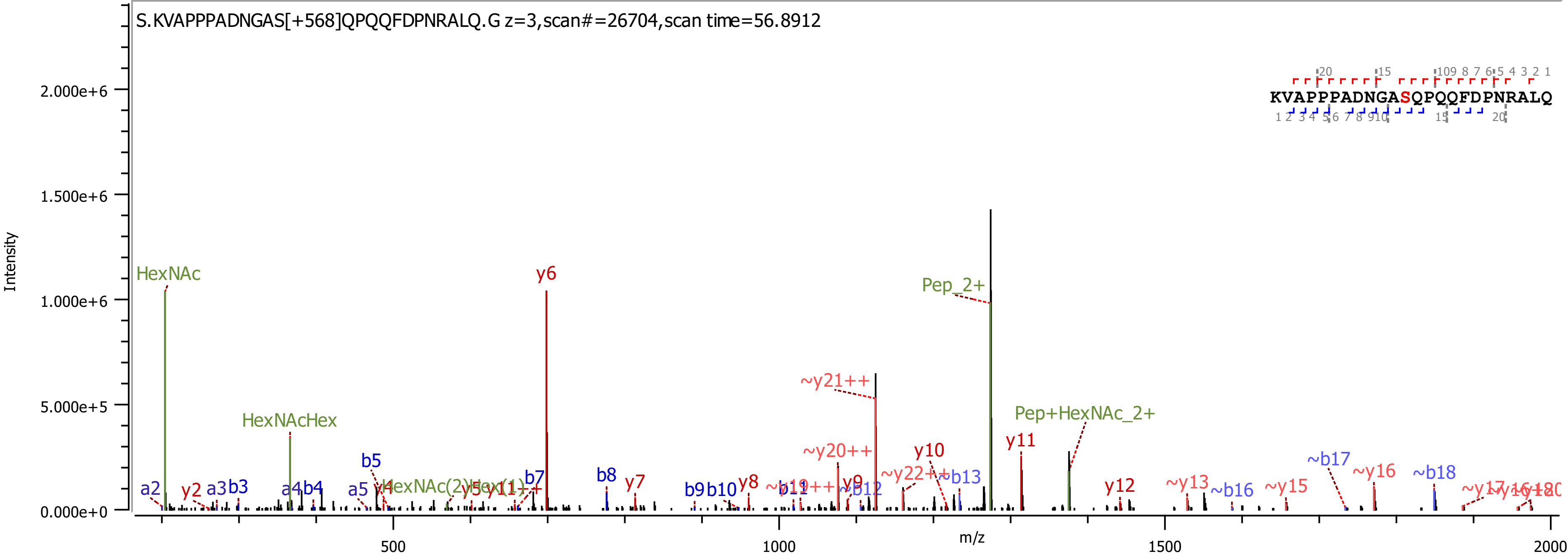

S.KVAPPPADNGAS[+568]QPQQFDPNRALQ.G z=3,scan#=26704,scan time=56.8912
KVAPPPADNGASQPQQFDPNRALQ
Intensity
m/z
HexNAc
HexNAcHex
y6
Pep_2+
~y21++
Pep+HexNAc_2+
~y20++
y10
y11
~y22++
~b13
b5
b7
b8
y7
y8
b9
b10
~b17
~y16
~b18
y12
~y13
~b16
~y15
a2
y2
a3
b3
a4
b4
a5

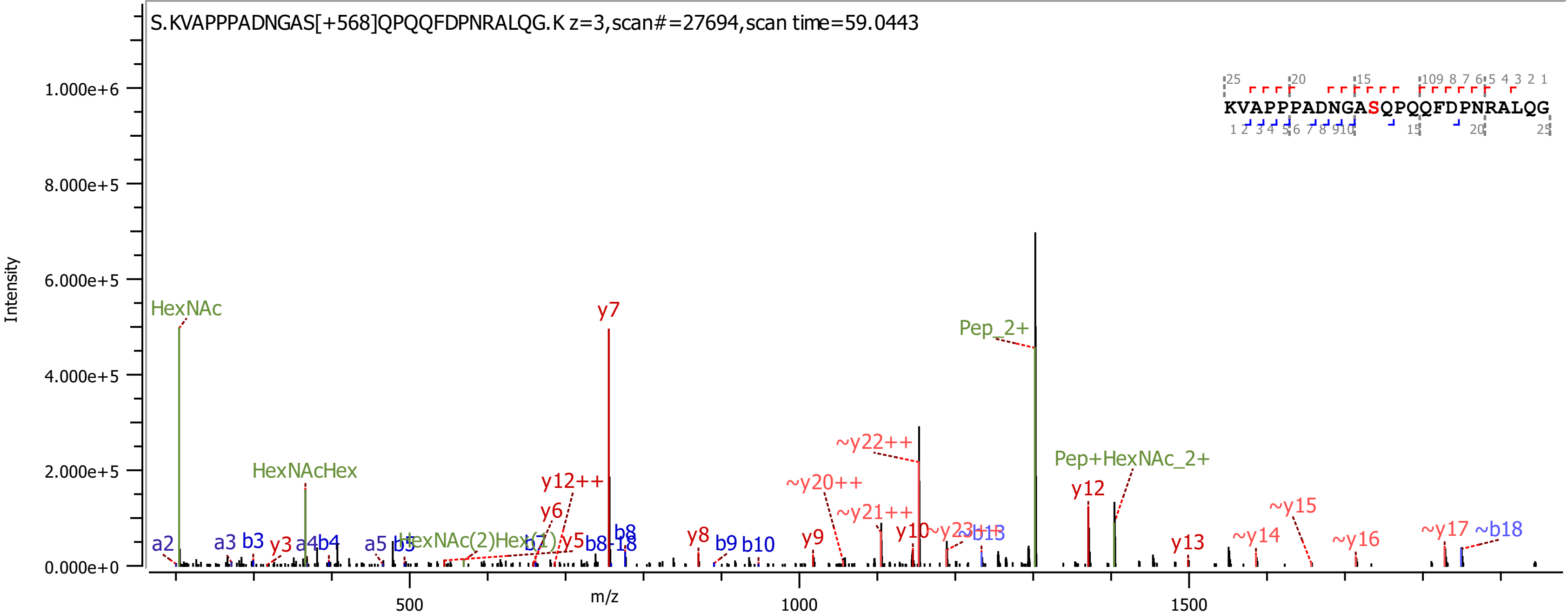
S.KVAPPPADNGAS[+568]QPQQFDPNRALQG.K z=3,scan#=27694,scan time=59.0443
KVAPPPADNGASQPQQFDPNRALQG
Intensity
m/z
HexNAc
HexNAcHex
y7
Pep_2+
~y22++
Pep+HexNAc_2+
y12++
~y20++
y12
~y15
y6
~y21++
y10
~y23
~b13
~y14
~y17
~b18
HexNAc(2)Hex(1)
y13
~y16
y8
y9
a2
a3
b3
y3
a4
b4
a5
b5
y5
b8-18
b8
b9
b10
0.000e+0
2.000e+5
4.000e+5
6.000e+5
8.000e+5
1.000e+6
500
1000
1500

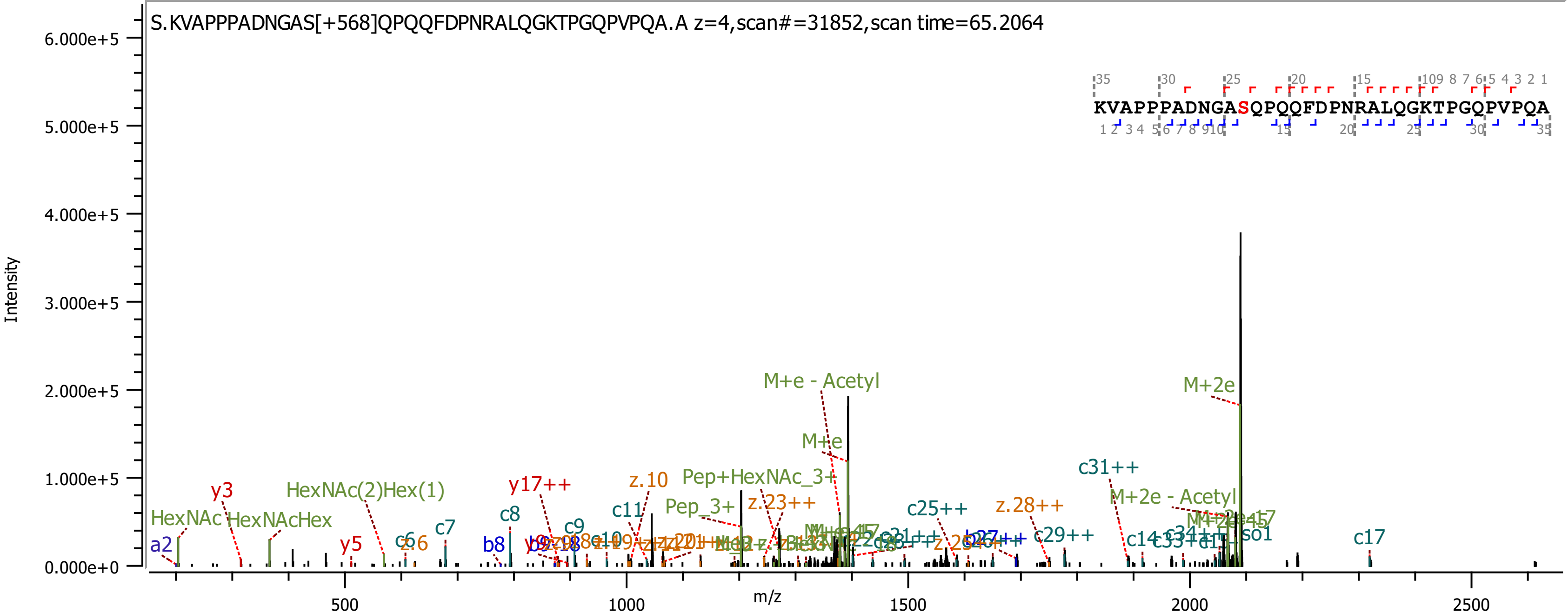
S.KVAPPPADNGAS[+568]QPQQFDPNRALQGKTPGQPVPQA.A z=4,scan#=31852,scan time=65.2064
KVAPPPADNGASQPQQFDPNRALQGKTPGQPVPQA
Intensity
m/z
M+e - Acetyl
M+2e
M+e
c31++
Pep+HexNAc_3+
z.10
y17++
HexNAc(2)Hex(1)
y3
c11
Pep_3+
z.23++
c25++
z.28++
M+2e - Acetyl
HexNAc
HexNAcHex
c8
c7
c9
c29++
c17
a2
y5
z.6
b8
c14

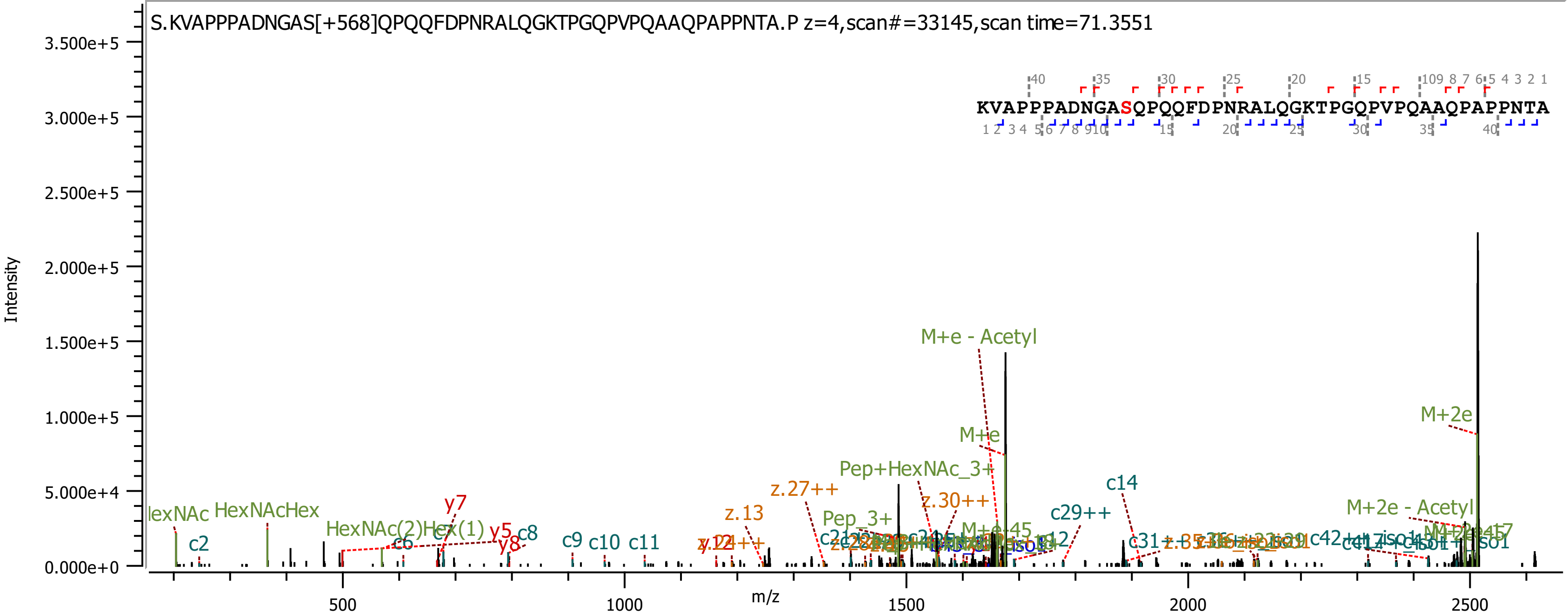
S.KVAPPPADNGAS[+568]QPQQFDPNRALQGKTPGQPVPQAAQPAPPNTA.P z=4,scan#=33145,scan time=71.3551
KVAPPPADNGASQPQQFDPNRALQGKTPGQPVPQAAQPAPPNTA
Intensity
m/z
M+e - Acetyl
M+2e
M+e
Pep+HexNAc_3+
M+2e - Acetyl
c14
c29++
z.30++
z.27++
z.13
Pep_3+
HexNAcHex
HexNAc(2)Hex(1)
y7
y5
y8
c8
c9
c10
c11
c2
c6
c12
c31++
c42++

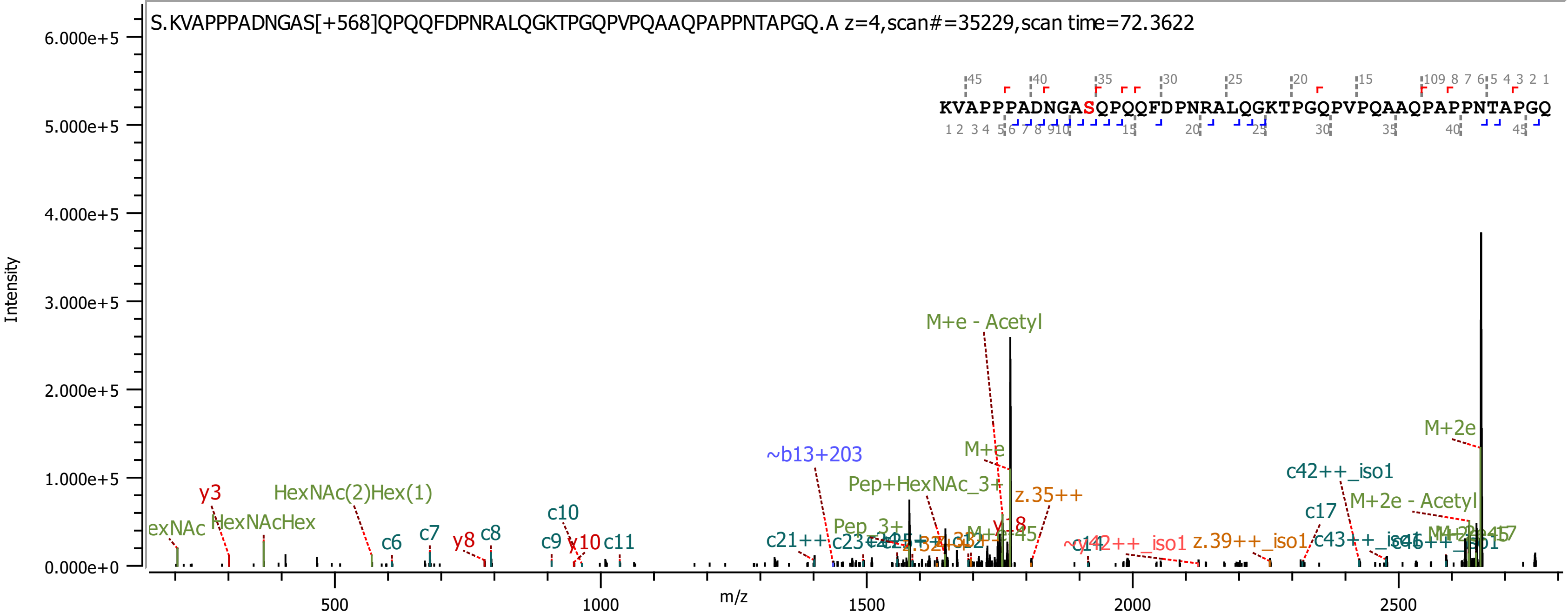
S.KVAPPPADNGAS[+568]QPQQFDPNRALQGKTPGQPVPQAAQPAPPNTAPGQ.A z=4,scan#=35229,scan time=72.3622
KVAPPPADNGASQPQQFDPNRALQGKTPGQPVPQAAQPAPPNTAPGQ
Intensity
m/z
M+e - Acetyl
M+2e
M+e
~b13+203
c42++_iso1
Pep+HexNAc_3+
z.35++
M+2e - Acetyl
y3
HexNAc(2)Hex(1)
c10
c17
HexNAcHex
Pep_3+
y18
c7
c8
y8
c6
c9
y10
c11
c21++
z.39++_iso1
c43++_iso1
c46++_iso1

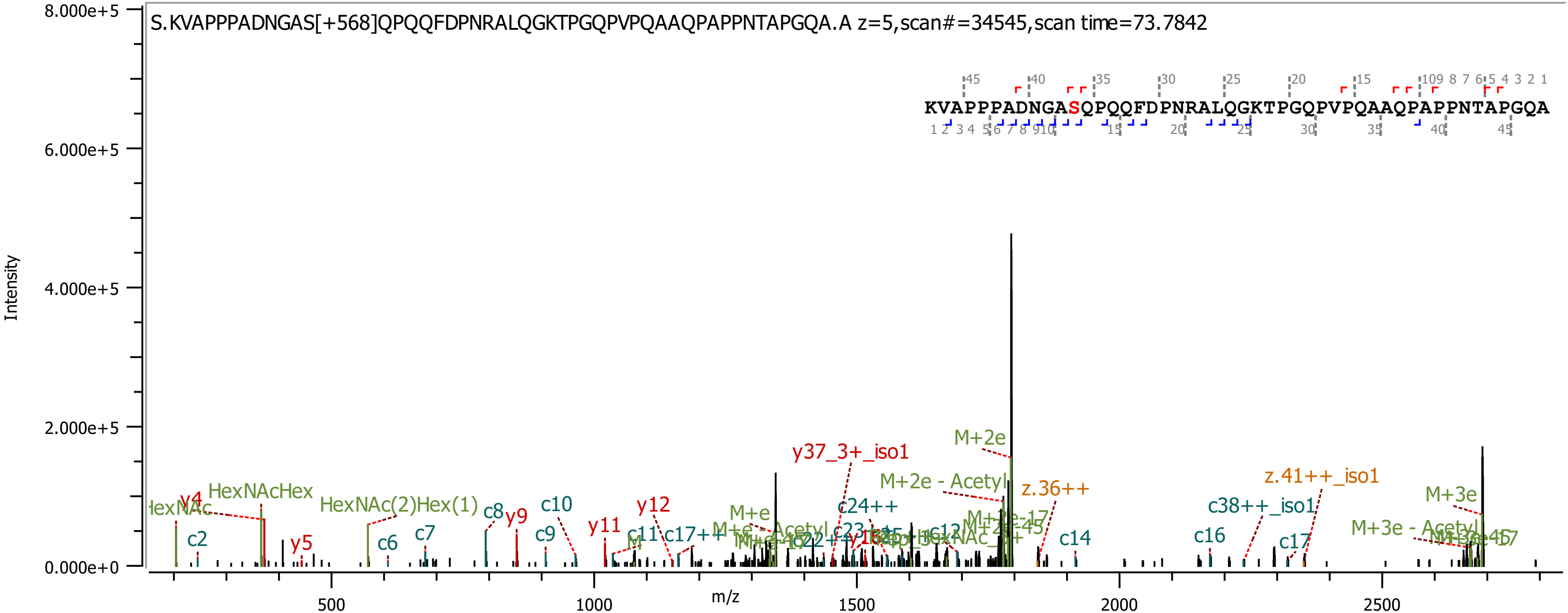
S.KVAPPPADNGAS[+568]QPQQFDPNRALQGKTPGQPVPQAAQPAPPNTAPGQA.A z=5,scan#=34545,scan time=73.7842
KVAPPPADNGASQPQQFDPNRALQGKTPGQPVPQAAQPAPPNTAPGQA
Intensity
m/z
8.000e+5
6.000e+5
4.000e+5
2.000e+5
0.000e+0
500
1000
1500
2000
2500
HexNAcHex
HexNAc(2)Hex(1)
y4
y5
y9
y11
y12
c2
c6
c7
c8
c9
c10
c11
c14
c16
c17
c24++
y37_3+_iso1
M+2e
M+2e - Acetyl
z.36++
z.41++_iso1
c38++_iso1
M+3e
M+3e - Acetyl
M+e

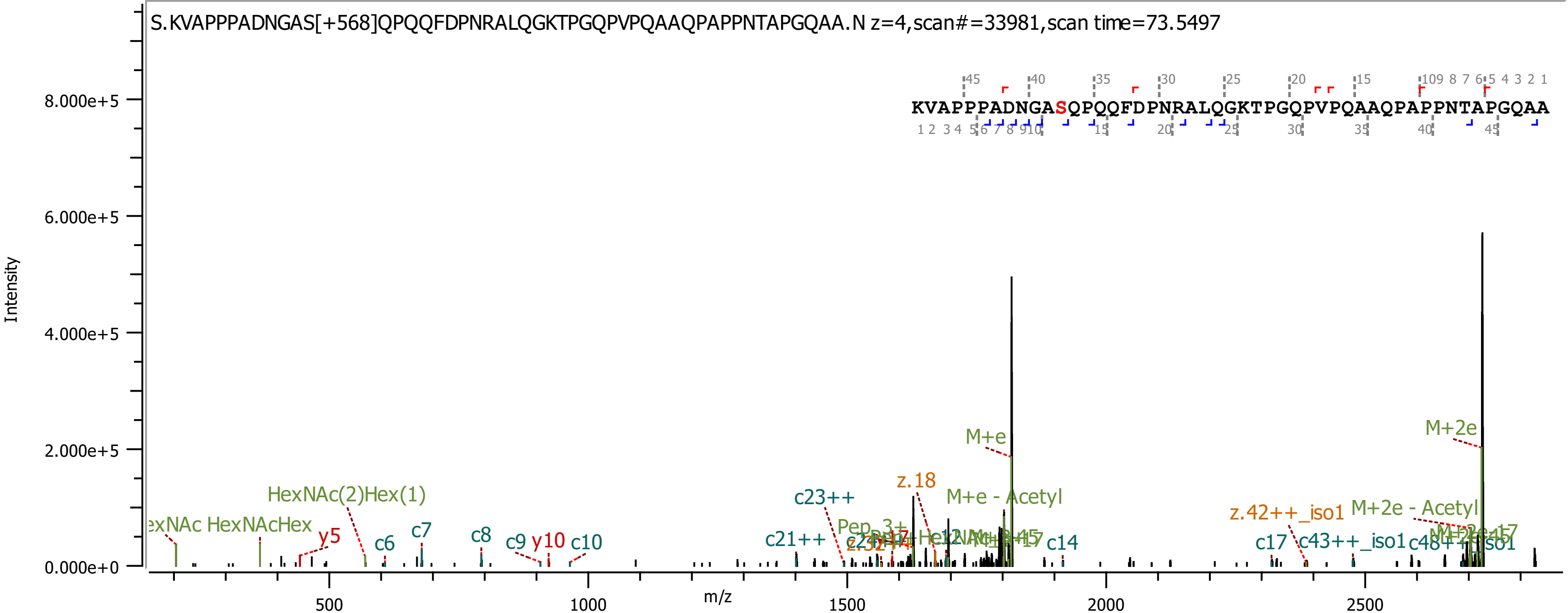
S.KVAPPPADNGAS[+568]QPQQFDPNRALQGKTPGQPVPQAAQPAPPNTAPGQAA.N z=4,scan#=33981,scan time=73.5497
KVAPPPADNGASQPQQFDPNRALQGKTPGQPVPQAAQPAPPNTAPGQAA
Intensity
0.000e+0
2.000e+5
4.000e+5
6.000e+5
8.000e+5
m/z
500
1000
1500
2000
2500
HexNAc(2)Hex(1)
HexNAcHex
y5
c6
c7
c8
c9
y10
c10
c21++
c23++
z.18
M+e - Acetyl
M+e
c14
z.42++_iso1
M+2e - Acetyl
c17
c43++_iso1
M+2e

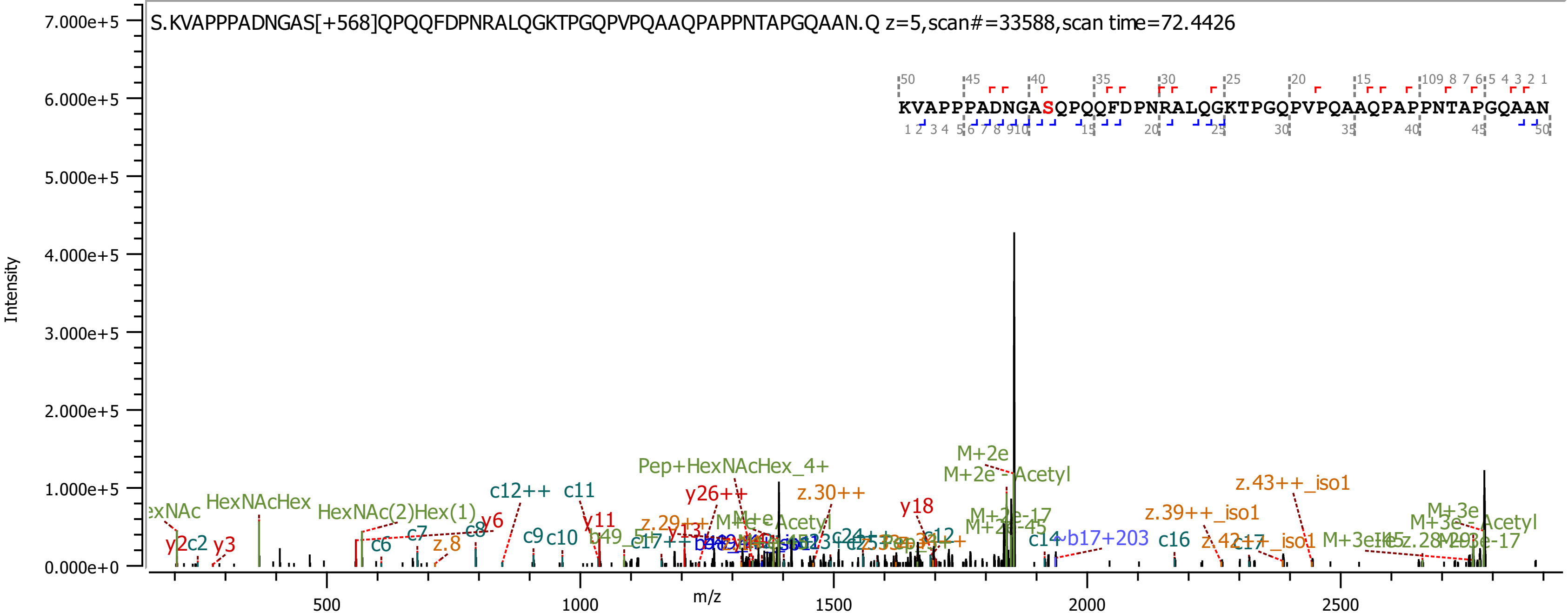
S.KVAPPPADNGAS[+568]QPQQFDPNRALQGKTPGQPVPQAAQPAPPNTAPGQAAN.Q z=5,scan#=33588,scan time=72.4426
KVAPPPADNGASQPQQFDPNRALQGKTPGQPVPQAAQPAPPNTAPGQAAN
Intensity
m/z
7.000e+5
6.000e+5
5.000e+5
4.000e+5
3.000e+5
2.000e+5
1.000e+5
0.000e+0
500
1000
1500
2000
2500
HexNAcHex
HexNAc(2)Hex(1)
Pep+HexNAcHex_4+
M+2e
M+2e - Acetyl
z.43++_iso1
z.39++_iso1
z.30++
y26++
y18
y11
y6
y3
c12++
c11
c10
c9
c8
c7
c6
z.8
c14
b17+203
c16
M+3e
M+3e - Acetyl

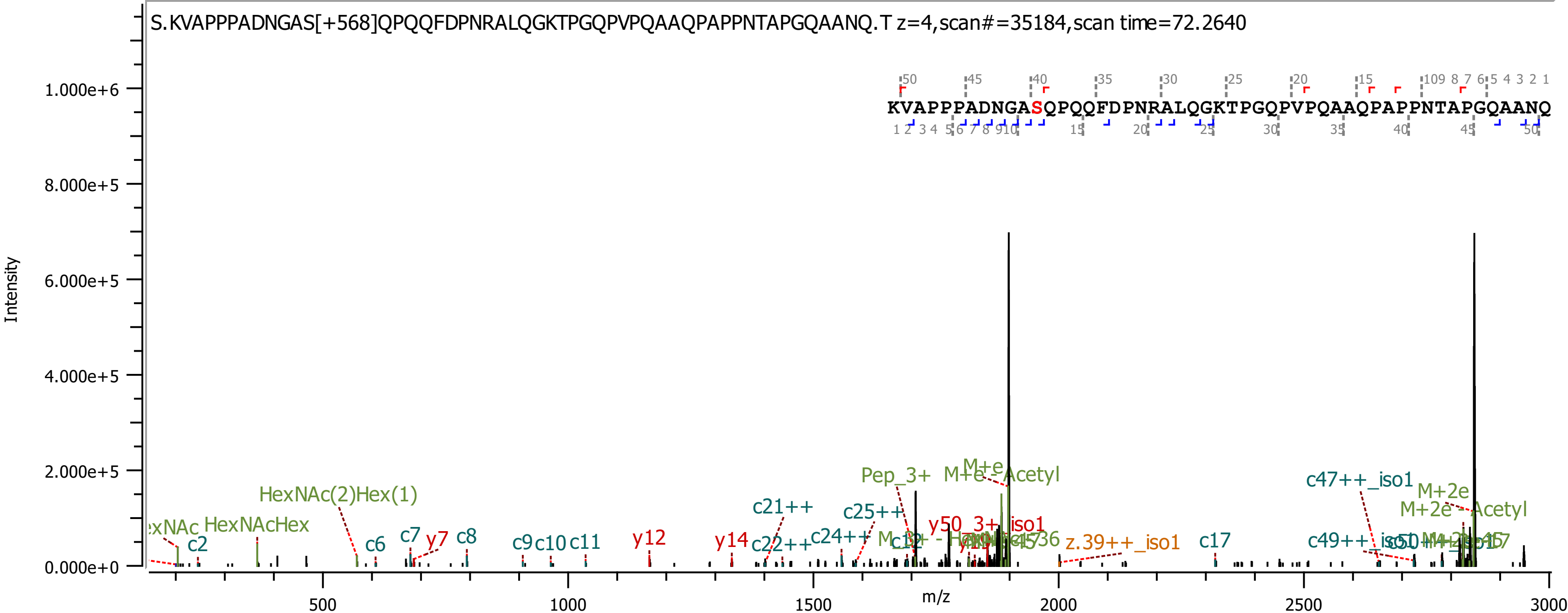

S.KVAPPPADNGAS[+568]QPQQFDPNRALQGKTPGQPVPQAAQPAPPNTAPGQAANQ.T z=4,scan#=35184,scan time=72.2640
KVAPPPADNGASQPQQFDPNRALQGKTPGQPVPQAAQPAPPNTAPGQAANQ
Intensity
m/z
HexNAc(2)Hex(1)
HexNAcHex
c2
c6
c7
y7
c8
c9
c10
c11
y12
y14
c21++
c22++
c24++
c25++
Pep_3+
M+e
M+e - Acetyl
y50_3+_iso1
z.39++_iso1
c17
c47++_iso1
c49++_iso1
M+2e
M+2e - Acetyl

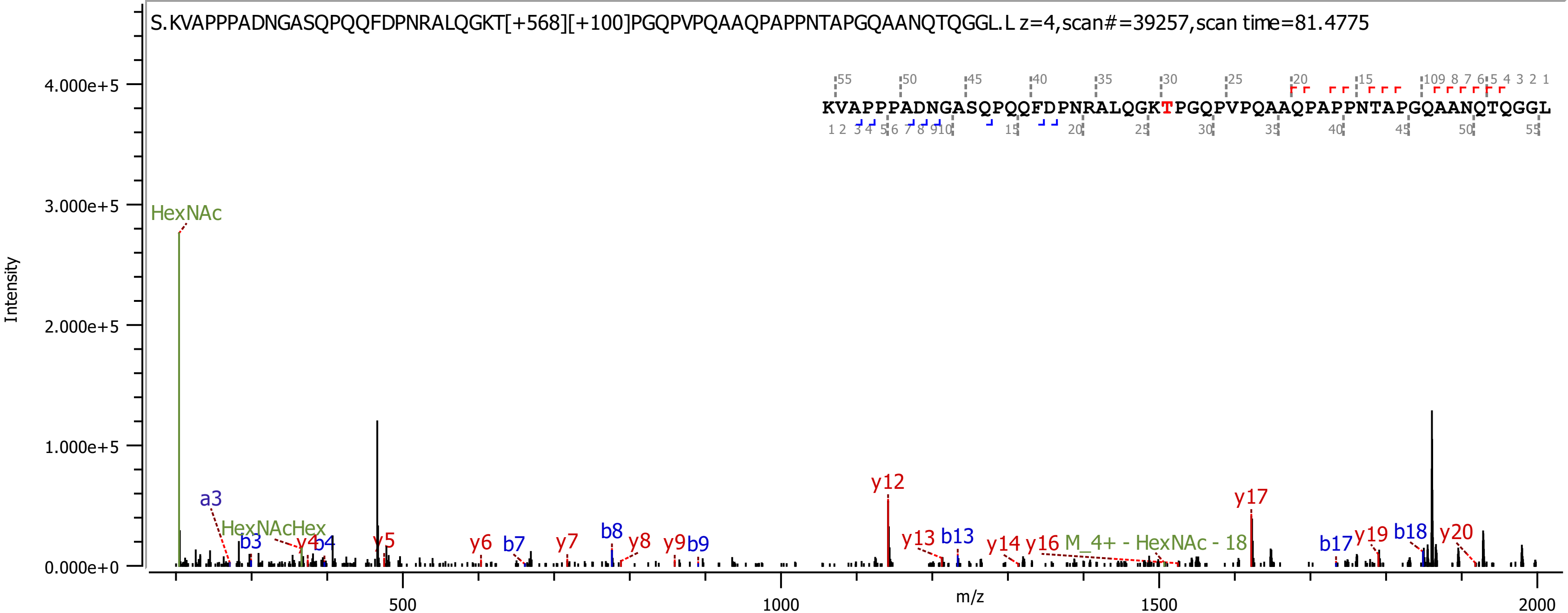

S.KVAPPPADNGASQPQQFDPNRALQGKT[+568][+100]PGQPVPQAAQPAPPNTAPGQAANQTQGGL.L z=4,scan#=39257,scan time=81.4775
KVAPPPADNGASQPQQFDPNRALQGKTPGQPVPQAAQPAPPNTAPGQAANQTQGGL
Intensity
m/z
4.000e+5
3.000e+5
2.000e+5
1.000e+5
0.000e+0
500
1000
1500
2000
HexNAc
a3
HexNAcHex
b3
y4
b4
y5
y6
b7
y7
b8
y8
y9
b9
y12
y13
b13
y14
y16
M_4+ - HexNAc - 18
y17
b17
y19
b18
y20

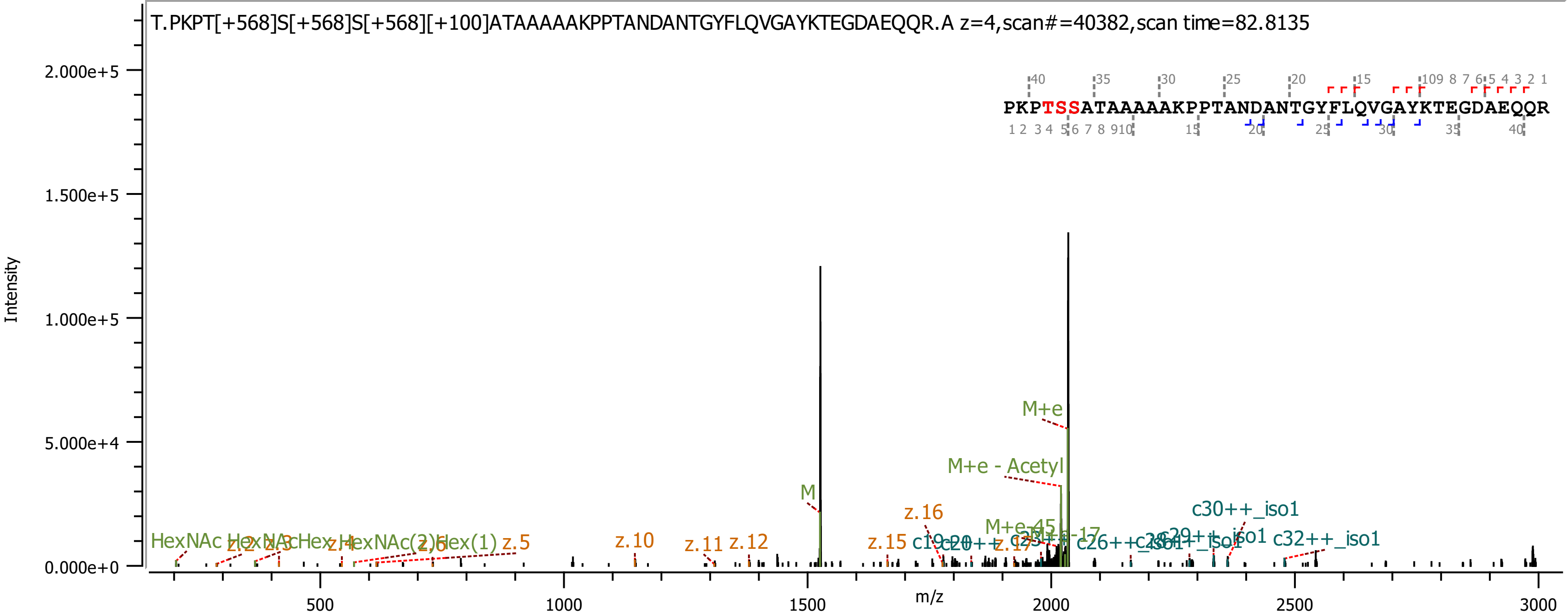
T.PKPT[+568]S[+568]S[+568][+100]ATAAAAAKPPTANDANTGYFLQVGAYKTEGDAEQQR.A z=4,scan#=40382,scan time=82.8135
PKPTSSATAAAAAKPPTANDANTGYFLQVGAYKTEGDAEQQR
Intensity
m/z
M+e
M+e - Acetyl
M
z.16
c30++_iso1
c32++_iso1
z.10
z.11
z.12
z.15
z.5
HexNAc
z.2
z.3
z.4
z.6
2.000e+5
1.500e+5
1.000e+5
5.000e+4
0.000e+0
500
1000
1500
2000
2500
3000

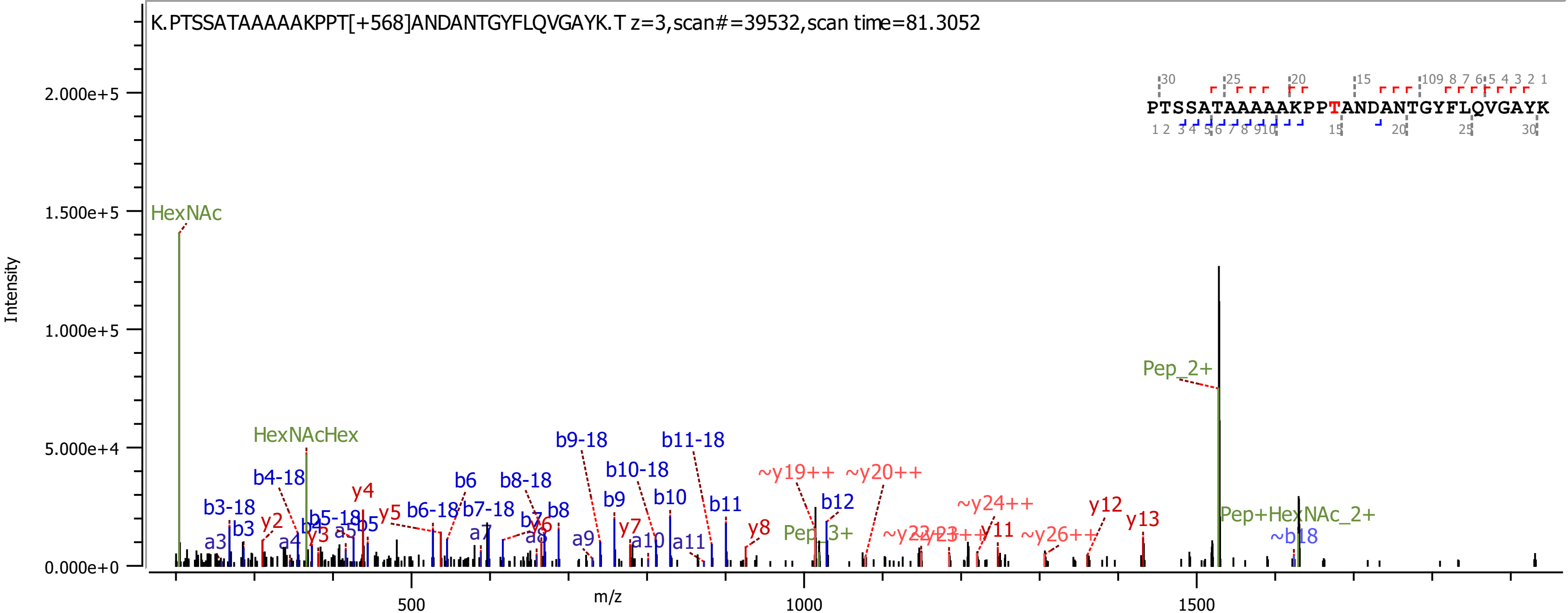
K.PTSSATAAAAAKPPT[+568]ANDANTGYFLQVGAYK.T z=3,scan#=39532,scan time=81.3052
PTSSATAAAAAKPPTANDANTGYFLQVGAYK
Intensity
m/z
HexNAc
HexNAcHex
Pep_2+
Pep_3+
Pep+HexNAc_2+
~b18
b3-18
b4-18
b5-18
b6-18
b7-18
b8-18
b9-18
b10-18
b11-18
b3
b4
b5
b6
b7
b8
b9
b10
b11
b12
a3
a4
a5
a7
a8
a9
a10
a11
y2
y3
y4
y5
y6
y7
y8
y11
y12
y13
~y19++
~y20++
~y24++
~y26++
2.000e+5
1.500e+5
1.000e+5
5.000e+4
0.000e+0
500
1000
1500

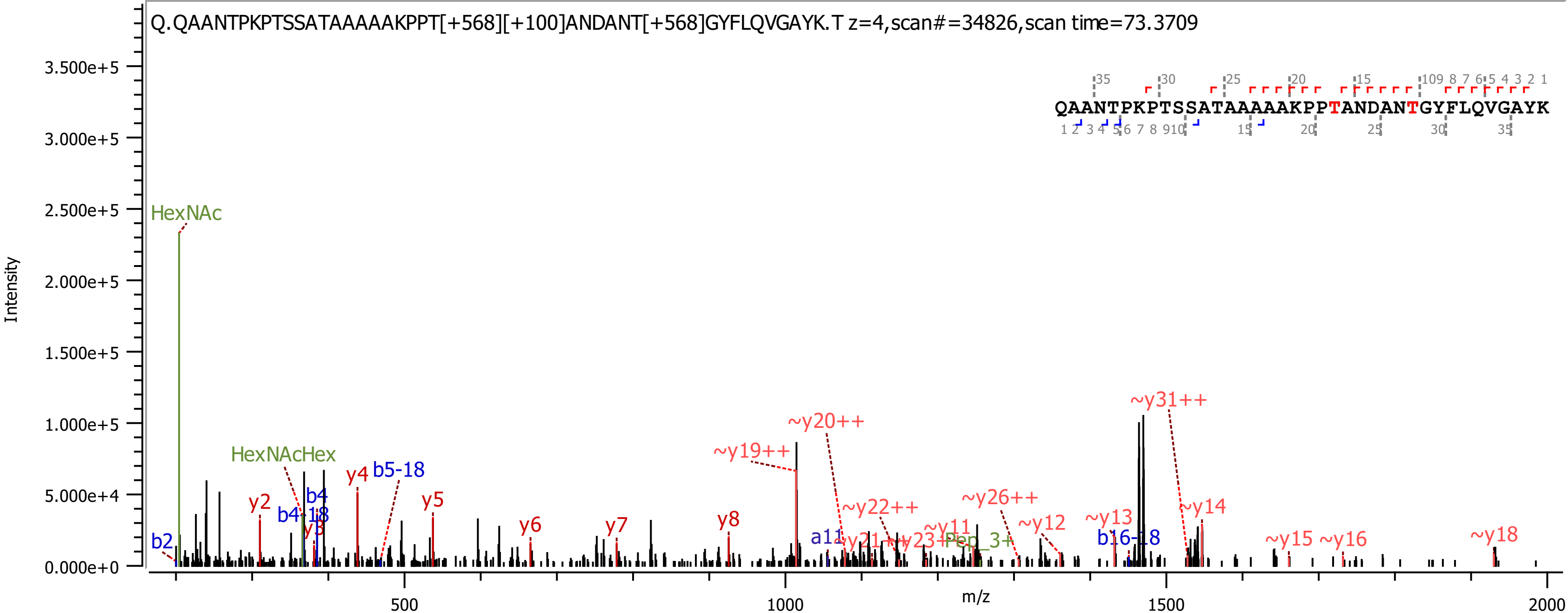

Q.QAANTPKPTSSATAAAAAKPPT[+568][+100]ANDANT[+568]GYFLQVGAYK.T z=4,scan#=34826,scan time=73.3709
QAANTPKPTSSATAAAAAKPPTANDANTGYFLQVGAYK
Intensity
m/z
HexNAc
HexNAcHex
b2
y2
b4-18
b4
y3
y4
b5-18
y5
y6
y7
y8
~y19++
~y20++
a11
~y21++
~y22++
~y23++
~y11
Pep_3+
~y26++
~y12
~y13
b16-18
~y31++
~y14
~y15
~y16
~y18

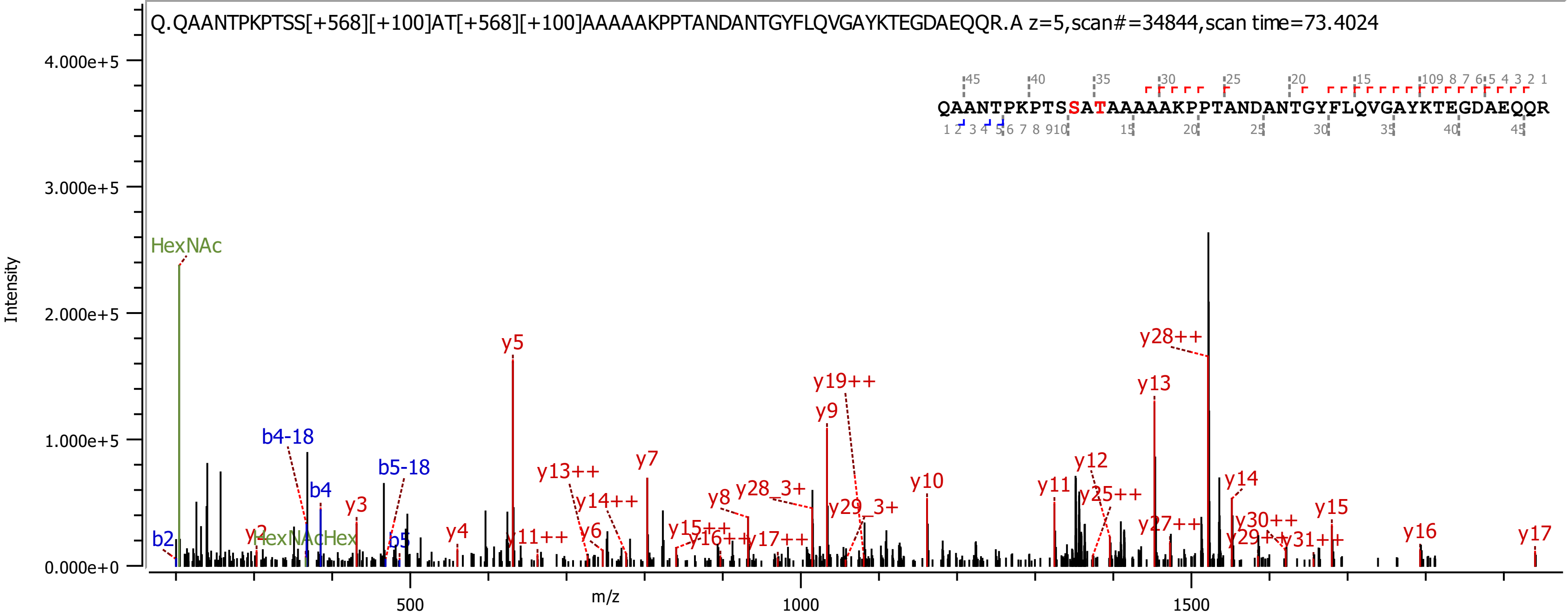

Q.QAANTPKPTSS[+568][+100]AT[+568][+100]AAAAAKPPTANDANTGYFLQVGAYKTEGDAEQQR.A z=5,scan#=34844,scan time=73.4024
QAANTPKPTSSATAAAAAKPPTANDANTGYFLQVGAYKTEGDAEQQR
Intensity
4.000e+5
3.000e+5
2.000e+5
1.000e+5
0.000e+0
m/z
500
1000
1500
HexNAc
HexNAcHex
b2
b4-18
b4
b5-18
b5
y2
y3
y4
y5
y6
y7
y8
y9
y10
y11
y12
y13
y14
y15
y16
y17
y11++
y13++
y14++
y15++
y16++
y17++
y19++
y25++
y27++
y28++
y29++
y30++
y31++
y28_3+
y29_3+

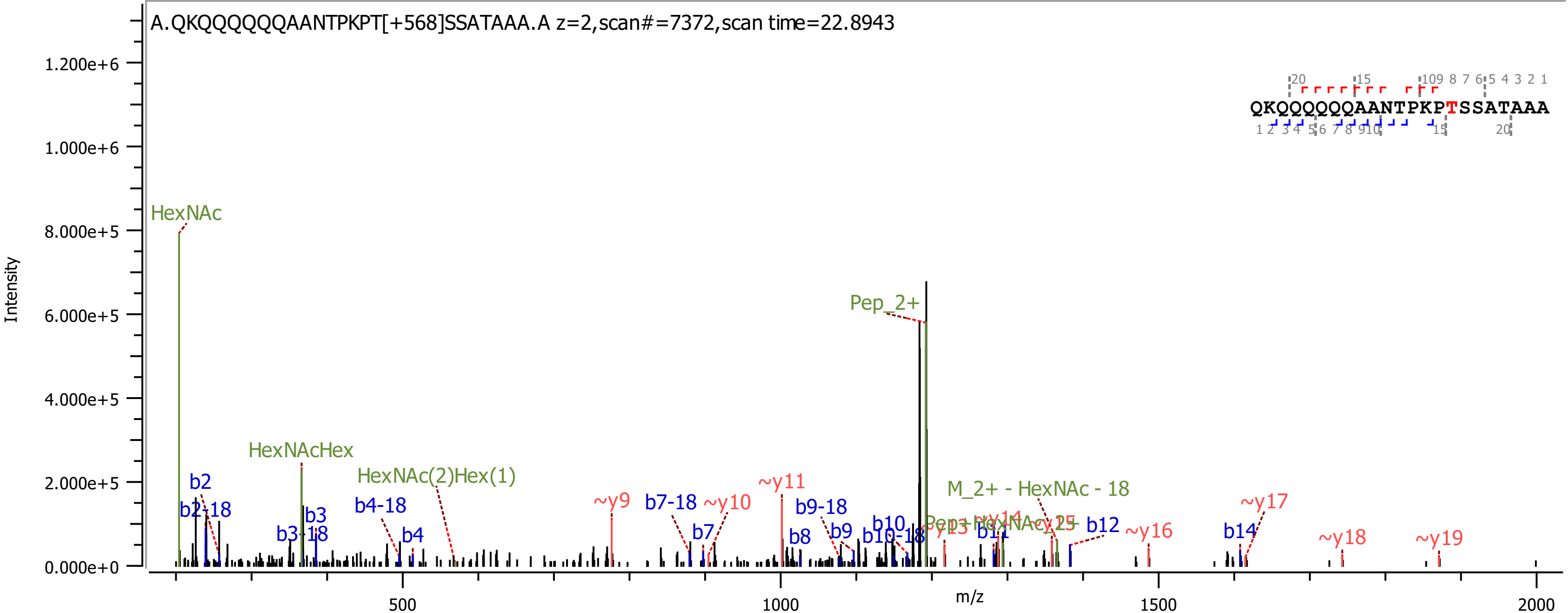

A.QKQQQQQQAANTPKPT[+568]SSATAAA.A z=2,scan#=7372,scan time=22.8943
QKQQQQQQAANTPKPTSSATAAA
Intensity
m/z
1.200e+6
1.000e+6
8.000e+5
6.000e+5
4.000e+5
2.000e+5
0.000e+0
500
1000
1500
2000
HexNAc
HexNAcHex
HexNAc(2)Hex(1)
Pep_2+
M_2+ - HexNAc - 18
b2
b2-18
b3
b3-18
b4-18
b4
b7-18
b7
b8
b9-18
b9
b10
b10-18
b11
b12
b14
~y9
~y10
~y11
~y13
~y14
~y15
~y16
~y17
~y18
~y19

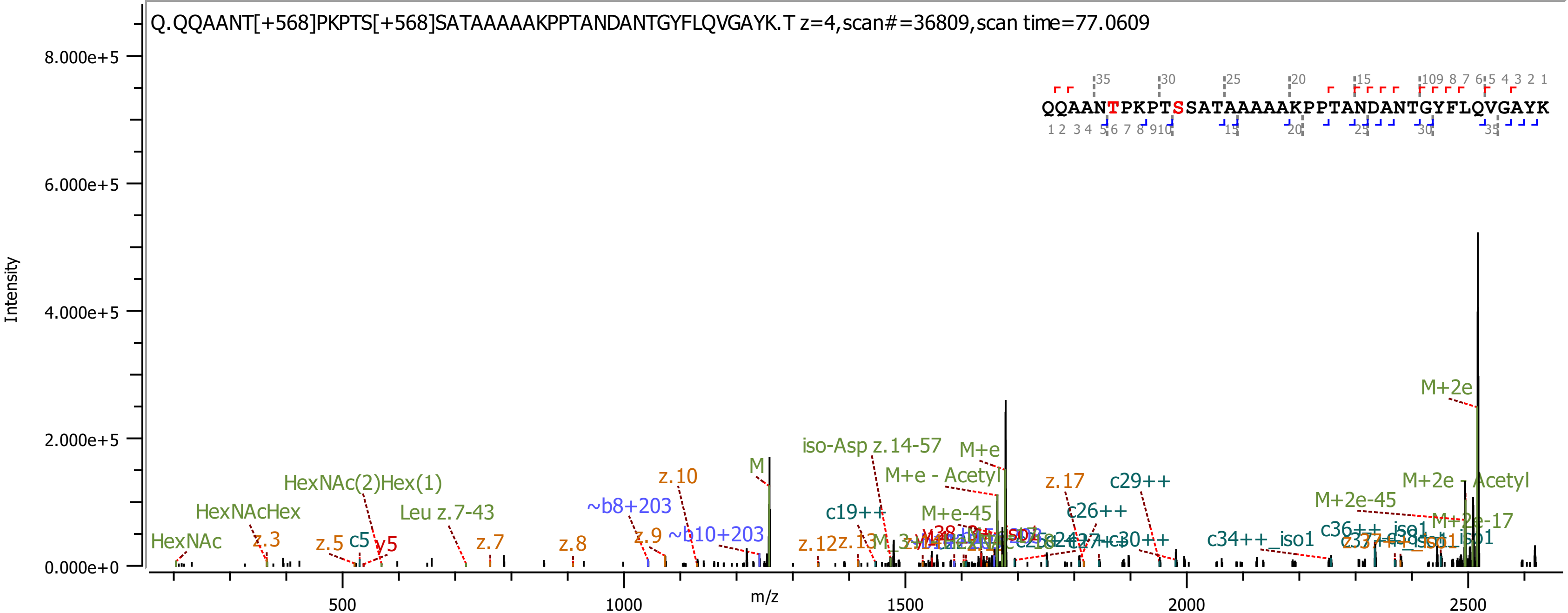

Q.QQAANT[+568]PKPTS[+568]SATAAAAAKPPTANDANTGYFLQVGAYK.T z=4,scan#=36809,scan time=77.0609
QQAANTPKPTSSATAAAAAKPPTANDANTGYFLQVGAYK
Intensity
m/z
8.000e+5
6.000e+5
4.000e+5
2.000e+5
0.000e+0
500
1000
1500
2000
2500
M+2e
M+2e - Acetyl
M+2e-45
M+2e-17
M
M+e
M+e - Acetyl
M+e-45
iso-Asp z.14-57
HexNAc
HexNAcHex
HexNAc(2)Hex(1)
Leu z.7-43
z.3
z.5
c5
y5
z.7
z.8
z.9
z.10
~b8+203
~b10+203
c19++
z.12
z.13
z.17
c26++
c29++
c30++
c34++_iso1
c36++_iso1

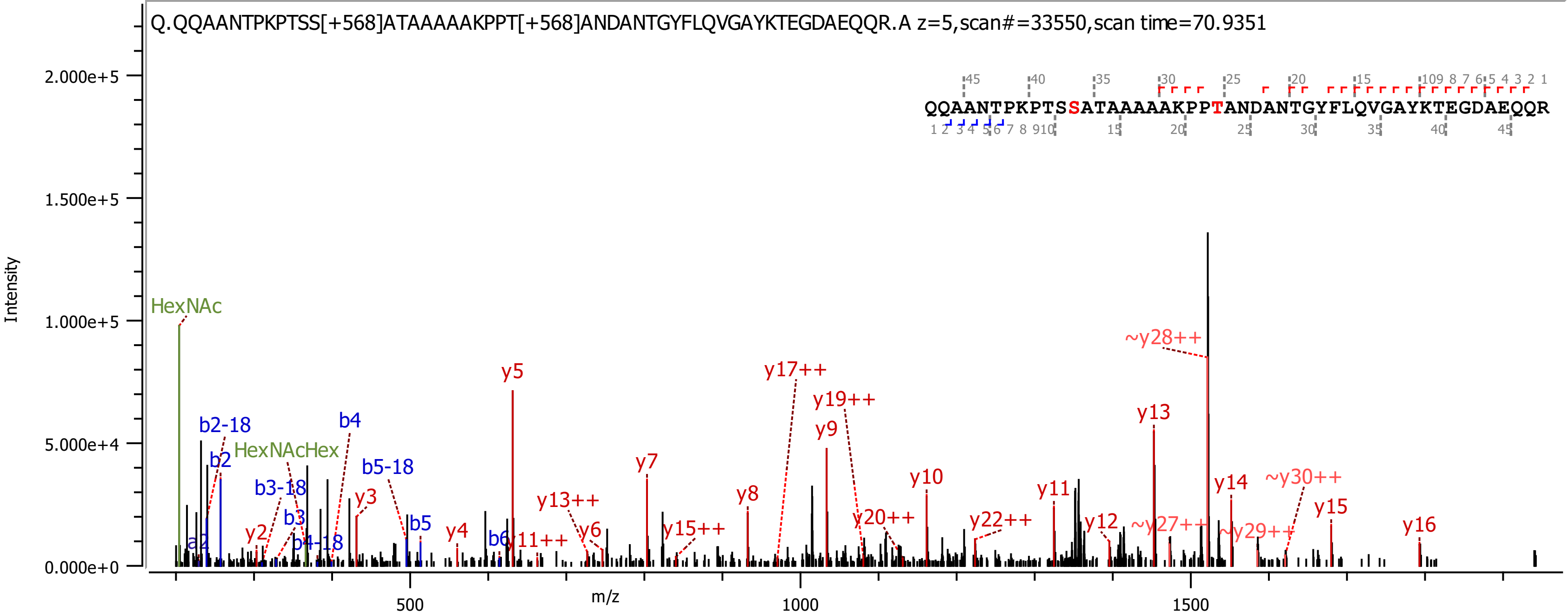

Q.QQAANTPKPTSS[+568]ATAAAAAKPPT[+568]ANDANTGYFLQVGAYKTEGDAEQQR.A z=5,scan#=33550,scan time=70.9351
QQAANTPKPTSSATAAAAAKPPTANDANTGYFLQVGAYKTEGDAEQQR
Intensity
2.000e+5
1.500e+5
1.000e+5
5.000e+4
0.000e+0
m/z
500
1000
1500
HexNAc
HexNAcHex
a2
b2-18
b2
b3-18
b3
b4-18
b4
b5-18
b5
b6
y2
y3
y4
y5
y6
y7
y8
y9
y10
y11
y12
y13
y14
y15
y16
y11++
y13++
y15++
y17++
y19++
y20++
y22++
~y27++
~y28++
~y29++
~y30++

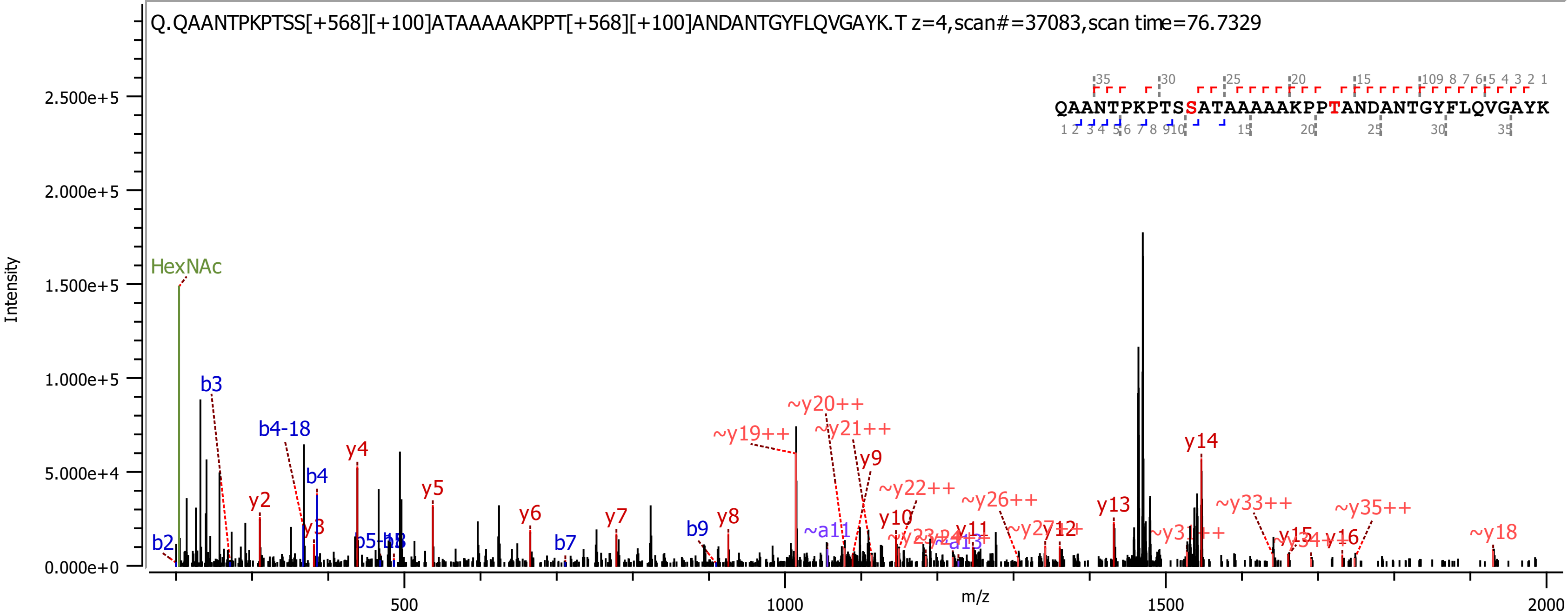

Q.QAANTPKPTSS[+568][+100]ATAAAAAKPPT[+568][+100]ANDANTGYFLQVGAYK.T z=4,scan#=37083,scan time=76.7329
QAANTPKPTSSATAAAAAKPPTANDANTGYFLQVGAYK
Intensity
m/z
HexNAc
b2
b3
b4-18
b4
b7
b9
y2
y3
y4
y5
y6
y7
y8
y9
y10
y11
y12
y13
y14
y15
y16
~y18
~y19++
~y20++
~y21++
~y22++
~y26++
~y33++
~y35++
~a11
0.000e+0
5.000e+4
1.000e+5
1.500e+5
2.000e+5
2.500e+5
500
1000
1500
2000

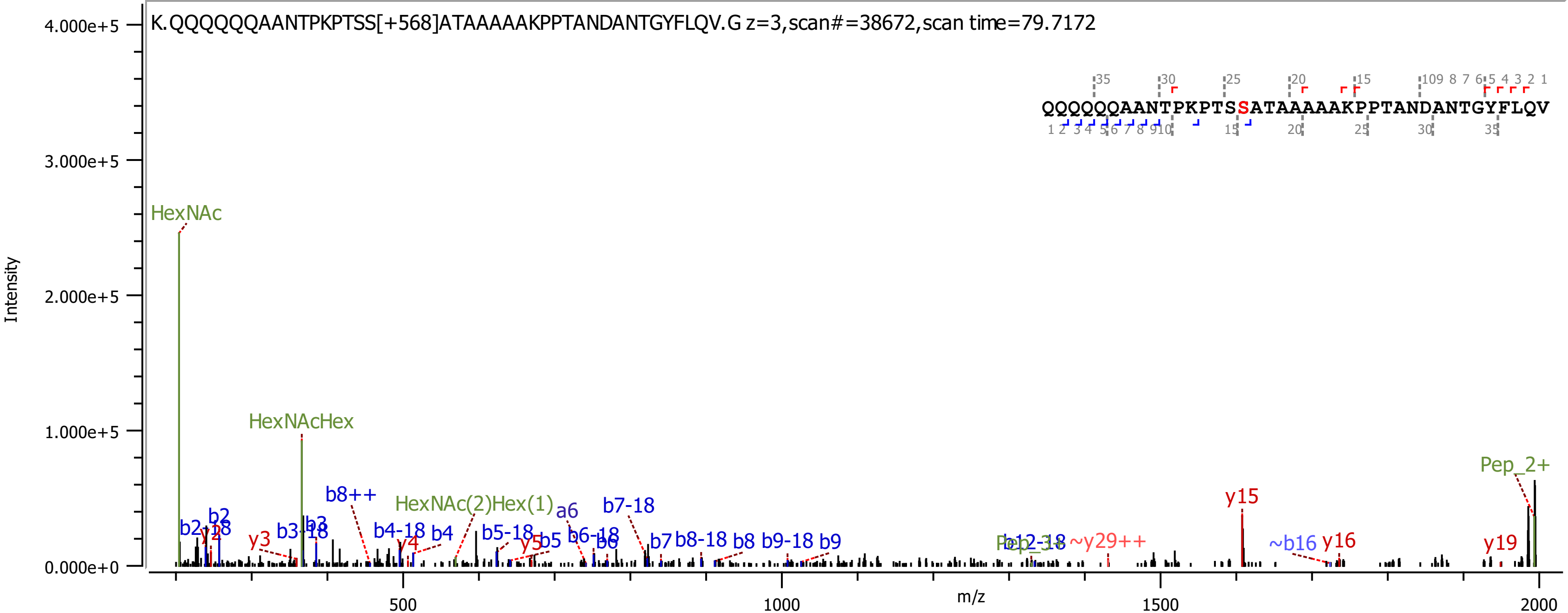
K.QQQQQQAANTPKPTSS[+568]ATAAAAAKPPTANDANTGYFLQV.G z=3,scan#=38672,scan time=79.7172
QQQQQQAANTPKPTSSATAAAAAKPPTANDANTGYFLQV
Intensity
m/z
HexNAc
HexNAcHex
HexNAc(2)Hex(1)
Pep_2+
y15
y16
y19
~y29++
~b16
b8++
a6
b7-18
b7
b8-18
b8
b9-18
b9
b4-18
b4
b5-18
b5
b6-18
b6
y3
y4
y5
b3-18
b3
b2-18
b2
y2

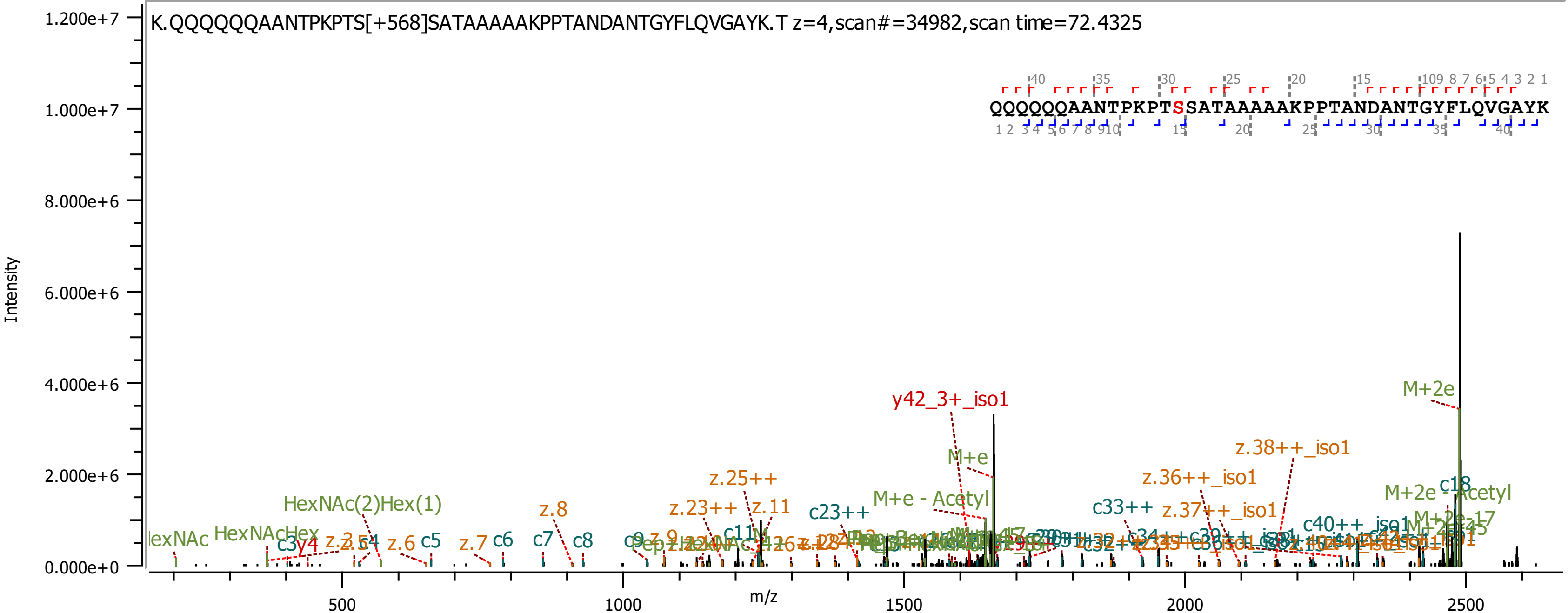

K.QQQQQQAANTPKPTS[+568]SATAAAAAKPPTANDANTGYFLQVGAYK.T z=4,scan#=34982,scan time=72.4325
QQQQQQAANTPKPTSSATAAAAAKPPTANDANTGYFLQVGAYK
Intensity
m/z
y42_3+_iso1
M+2e
M+e
z.38++_iso1
z.36++_iso1
z.37++_iso1
z.25++
z.23++
z.11
c23++
M+e - Acetyl
c33++
HexNAc(2)Hex(1)
z.8
M+2e - Acetyl
c18
c40++_iso1
HexNAcHex
c5
c6
c7
c8
z.6
z.7

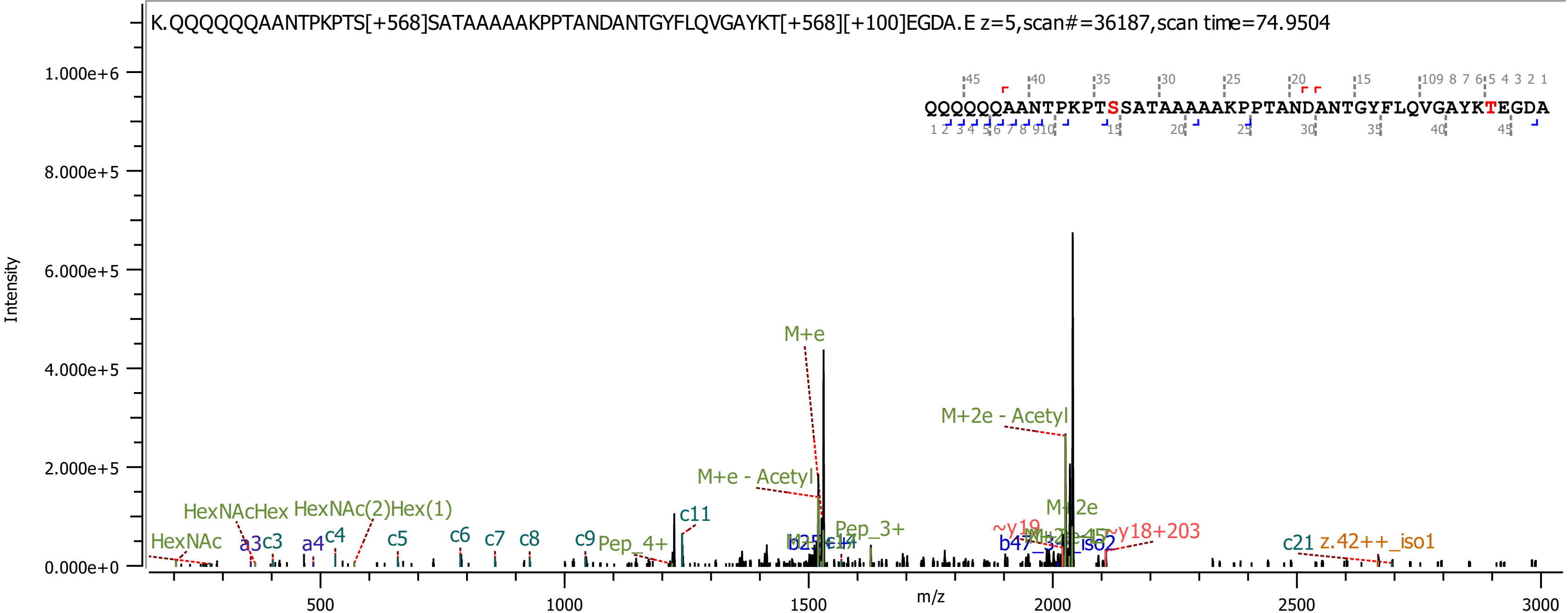
K.QQQQQQAANTPKPTS[+568]SATAAAAAKPPTANDANTGYFLQVGAYKT[+568][+100]EGDA.E z=5,scan#=36187,scan time=74.9504
QQQQQQAANTPKPTSSATAAAAAKPPTANDANTGYFLQVGAYKTEGDA
Intensity
m/z
M+e
M+2e - Acetyl
M+e - Acetyl
M+2e
HexNAcHex
HexNAc(2)Hex(1)
HexNAc
a3
c3
a4
c4
c5
c6
c7
c8
c9
Pep_4+
c11
Pep_3+
~y19
~y18+203
c21
z.42++_iso1

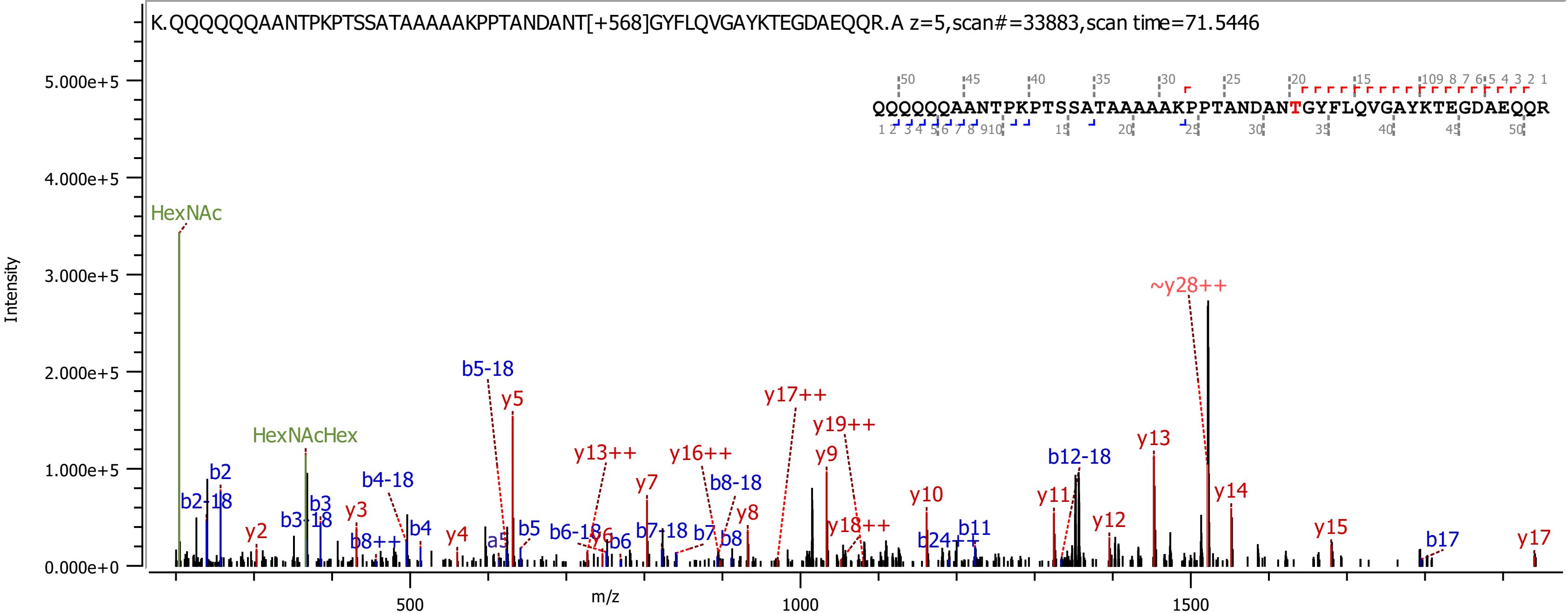
K.QQQQQQAANTPKPTSSATAAAAAKPPTANDANT[+568]GYFLQVGAYKTEGDAEQQR.A z=5,scan#=33883,scan time=71.5446
QQQQQQAANTPKPTSSATAAAAAKPPTANDANTGYFLQVGAYKTEGDAEQQR
Intensity
m/z
5.000e+5
4.000e+5
3.000e+5
2.000e+5
1.000e+5
0.000e+0
500
1000
1500
HexNAc
HexNAcHex
~y28++
b5-18
y5
y17++
y19++
y13++
y16++
y9
y7
b8-18
b12-18
y13
y14
y10
y11
y8
y12
b2
b2-18
y2
b3
b3-18
y3
b4-18
b4
b8++
y4
a5
b5
b6-18
y6
b6
b7-18
b7
b8
y18++
b24++
b11
y15
b17
y17

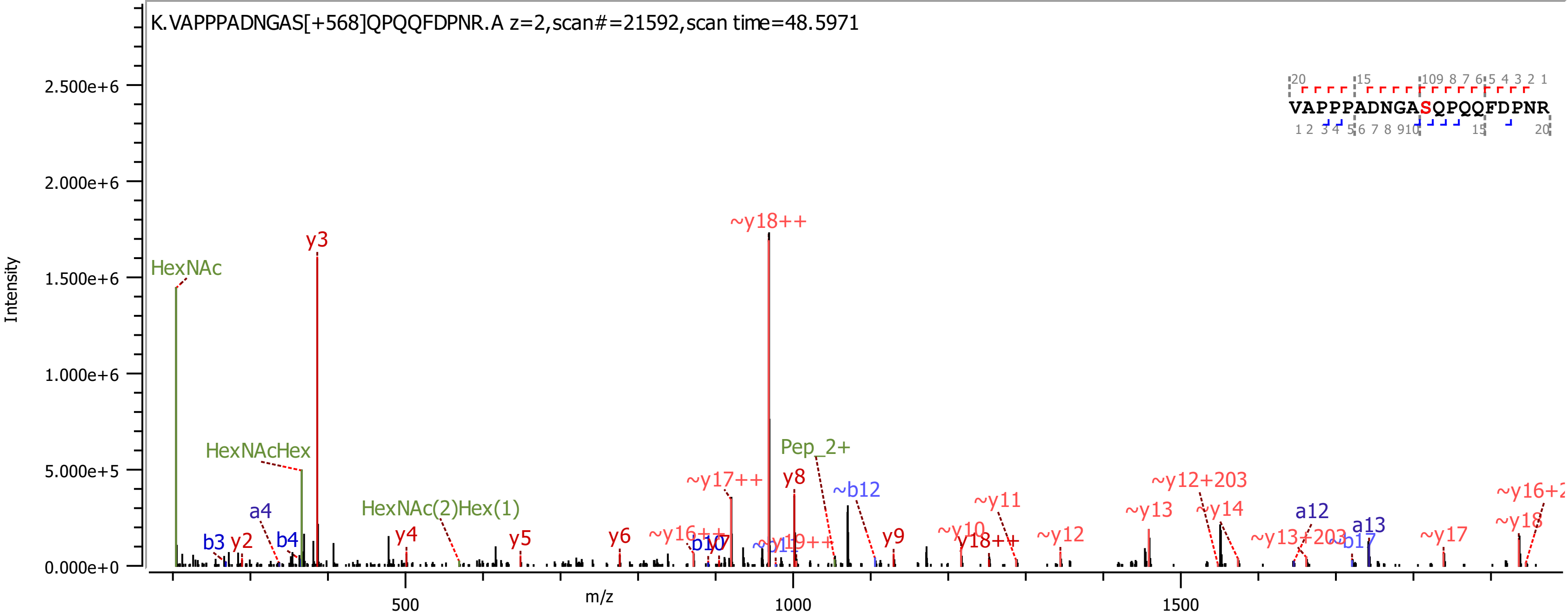
K.VAPPPADNGAS[+568]QPQQFDPNR.A z=2,scan#=21592,scan time=48.5971
VAPPPADNGASQPQQFDPNR
Intensity
m/z
HexNAc
HexNAcHex
HexNAc(2)Hex(1)
Pep_2+
y3
~y18++
y8
~b12
y9
~y11
~y12
~y13
~y14
~y12+203
a12
a13
~y17
~y18
~y17++
b3
y2
a4
b4
y4
y5
y6
b10
y18++
~b17
~y13+203

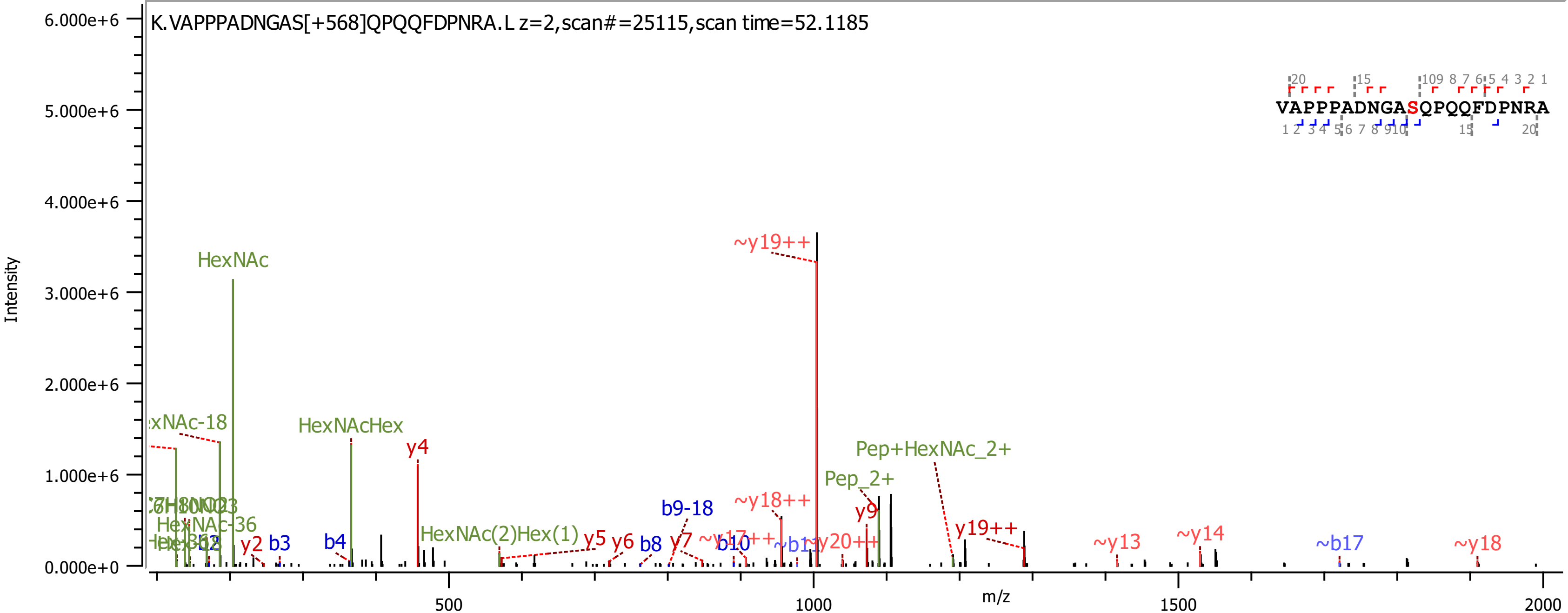
K.VAPPPADNGAS[+568]QPQQFDPNRA.L z=2,scan#=25115,scan time=52.1185
VAPPPADNGASQPQQFDPNRA
Intensity
m/z
6.000e+6
5.000e+6
4.000e+6
3.000e+6
2.000e+6
1.000e+6
0.000e+0
500
1000
1500
2000
HexNAc
~y19++
HexNAcHex
y4
Pep+HexNAc_2+
Pep_2+
~y18++
y9
b9-18
HexNAc-36
HexNAc(2)Hex(1)
y5
y6
b8
y7
b10
y19++
~y13
~y14
~b17
~y18
y2
b3
b4

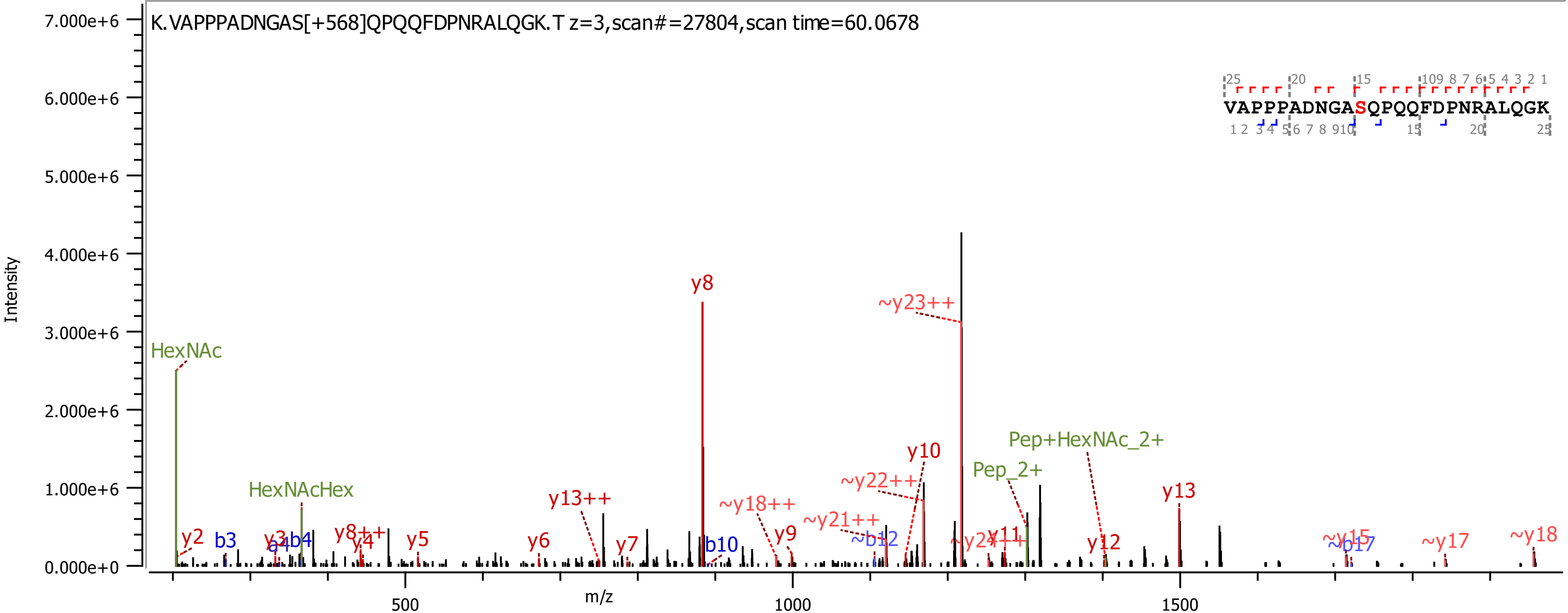
K.VAPPPADNGAS[+568]QPQQFDPNRALQGK.T z=3,scan#=27804,scan time=60.0678
VAPPPADNGASQPQQFDPNRALQGK
Intensity
m/z
7.000e+6
6.000e+6
5.000e+6
4.000e+6
3.000e+6
2.000e+6
1.000e+6
0.000e+0
500
1000
1500
HexNAc
HexNAcHex
y2
b3
y3
a4
b4
y8++
y4
y5
y6
y13++
y7
y8
b10
~y18++
y9
~y21++
~b12
~y22++
y10
~y23++
~y24++
y11
Pep_2+
Pep+HexNAc_2+
y12
y13
~y15
~b17
~y17
~y18

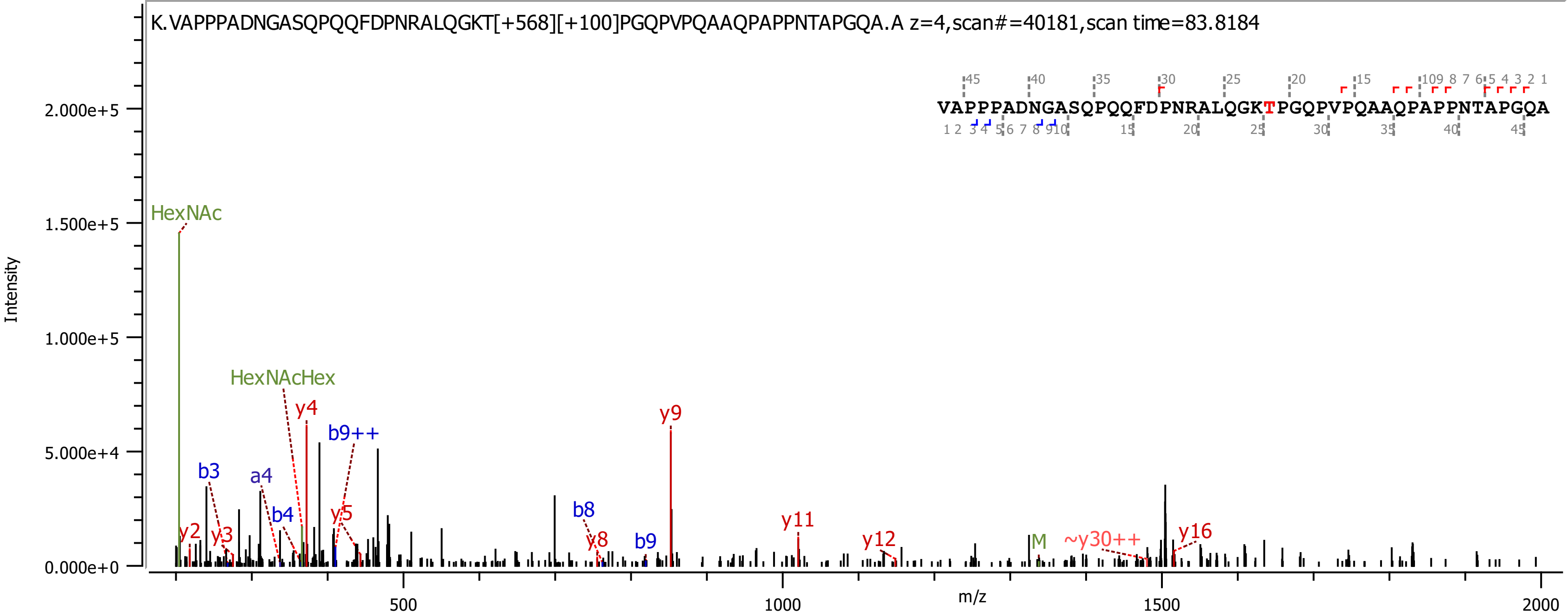

K.VAPPPADNGASQPQQFDPNRALQGKT[+568][+100]PGQPVPQAAQPAPPNTAPGQA.A z=4,scan#=40181,scan time=83.8184
VAPPPADNGASQPQQFDPNRALQGKTPGQPVPQAAQPAPPNTAPGQA
HexNAc
HexNAcHex
y2
y3
b3
a4
b4
y4
b9++
y5
b8
y8
b9
y9
y11
y12
M
~y30++
y16
Intensity
m/z
2.000e+5
1.500e+5
1.000e+5
5.000e+4
0.000e+0
500
1000
1500
2000

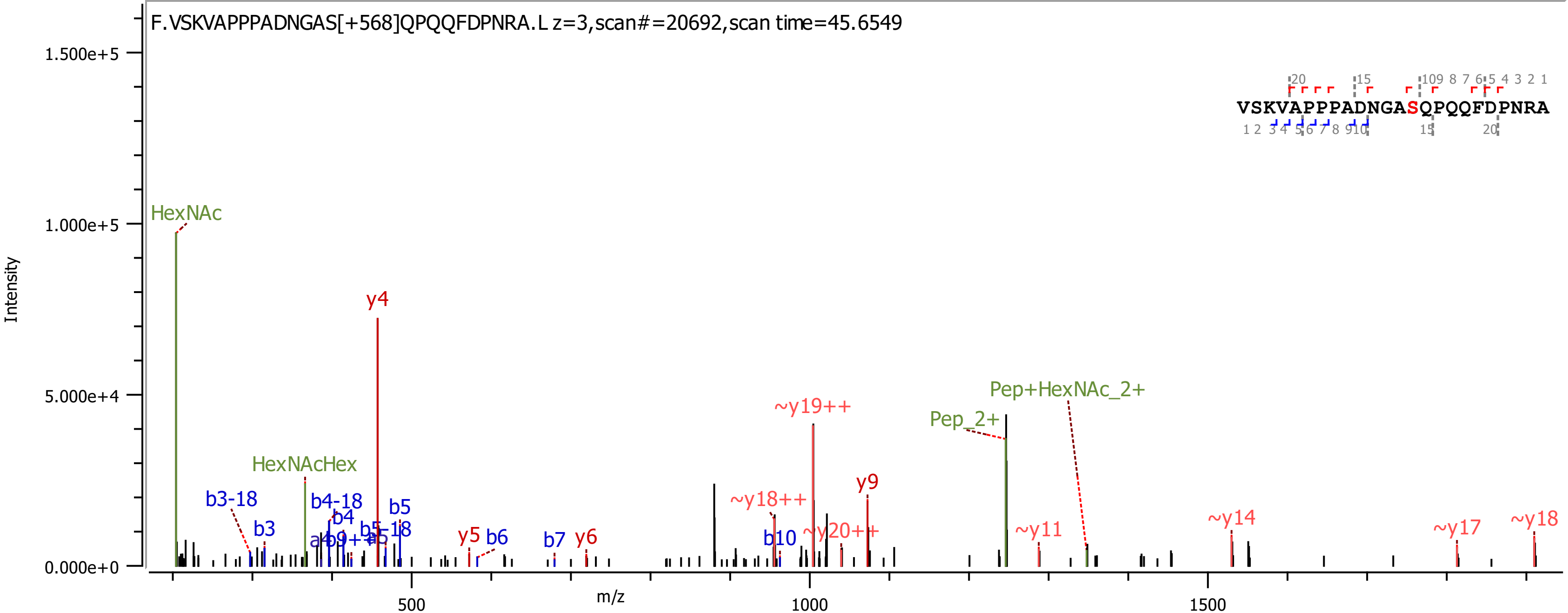

F.VSKVAPPPADNGAS[+568]QPQQFDPNRA.L z=3,scan#=20692,scan time=45.6549
VSKVAPPPADNGASQPQQFDPNRA
Intensity
m/z
1.500e+5
1.000e+5
5.000e+4
0.000e+0
500
1000
1500
HexNAc
HexNAcHex
y4
b3-18
b3
b4-18
b4
b5
b5-18
y5
b6
b7
y6
~y18++
~y19++
b10
~y20++
y9
Pep_2+
Pep+HexNAc_2+
~y11
~y14
~y17
~y18

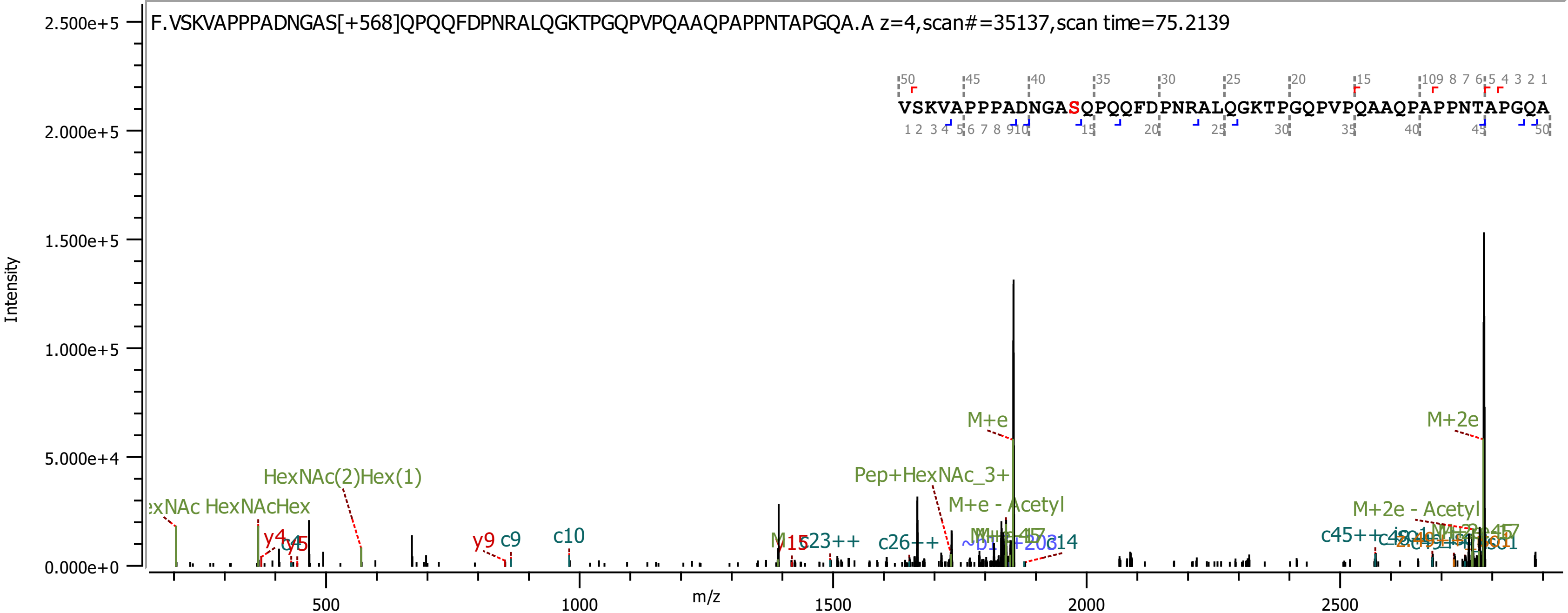
F.VSKVAPPPADNGAS[+568]QPQQFDPNRALQGKTPGQPVPQAAQPAPPNTAPGQA.A z=4,scan#=35137,scan time=75.2139
VSKVAPPPADNGASQPQQFDPNRALQGKTPGQPVPQAAQPAPPNTAPGQA
Intensity
m/z
2.500e+5
2.000e+5
1.500e+5
1.000e+5
5.000e+4
0.000e+0
500
1000
1500
2000
2500
M+e
M+2e
Pep+HexNAc_3+
M+e - Acetyl
M+2e - Acetyl
HexNAc(2)Hex(1)
HexNAcHex
y4
y5
y9
c9
c10
c23++
c26++
c45++

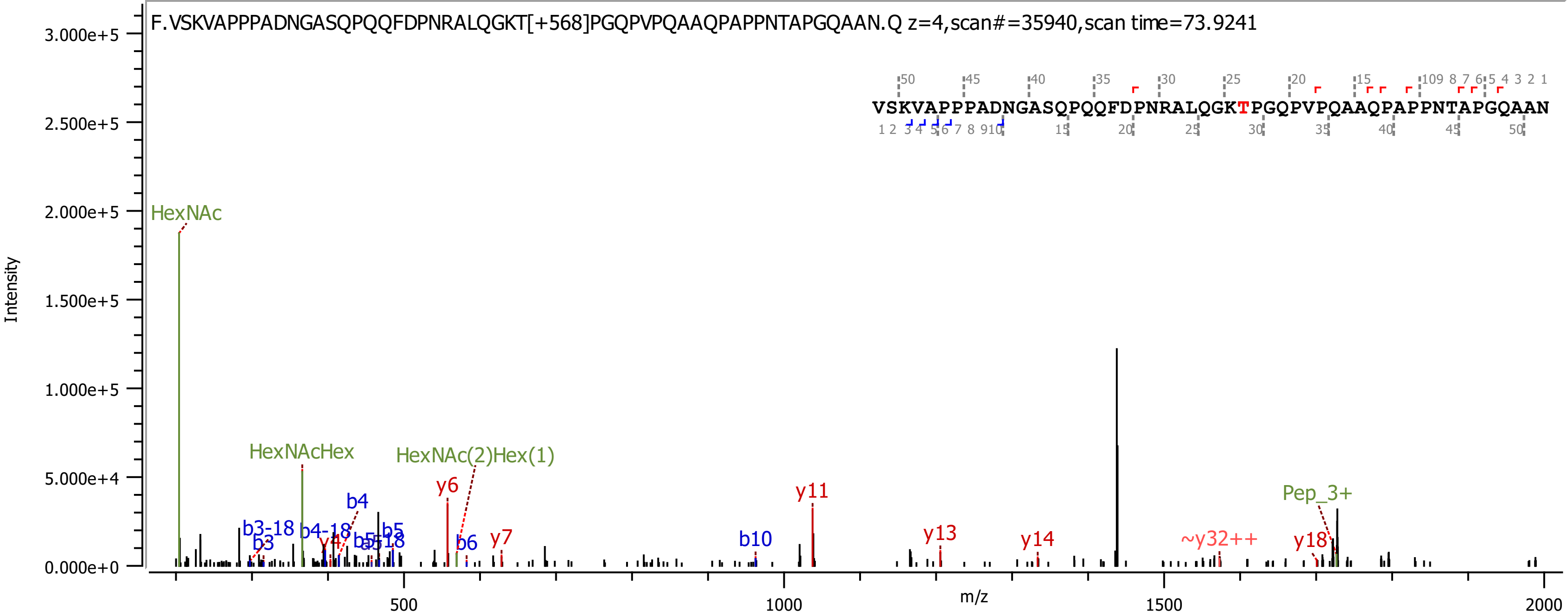

F.VSKVAPPPADNGASQPQQFDPNRALQGKT[+568]PGQPVPQAAQPAPPNTAPGQAAN.Q z=4,scan#=35940,scan time=73.9241
VSKVAPPPADNGASQPQQFDPNRALQGKTPGQPVPQAAQPAPPNTAPGQAAN
HexNAc
HexNAcHex
HexNAc(2)Hex(1)
b3-18
b3
b4-18
b4
y4
b5
b5-18
y6
b6
y7
b10
y11
y13
y14
~y32++
y18
Pep_3+
Intensity
m/z
3.000e+5
2.500e+5
2.000e+5
1.500e+5
1.000e+5
5.000e+4
0.000e+0
500
1000
1500
2000

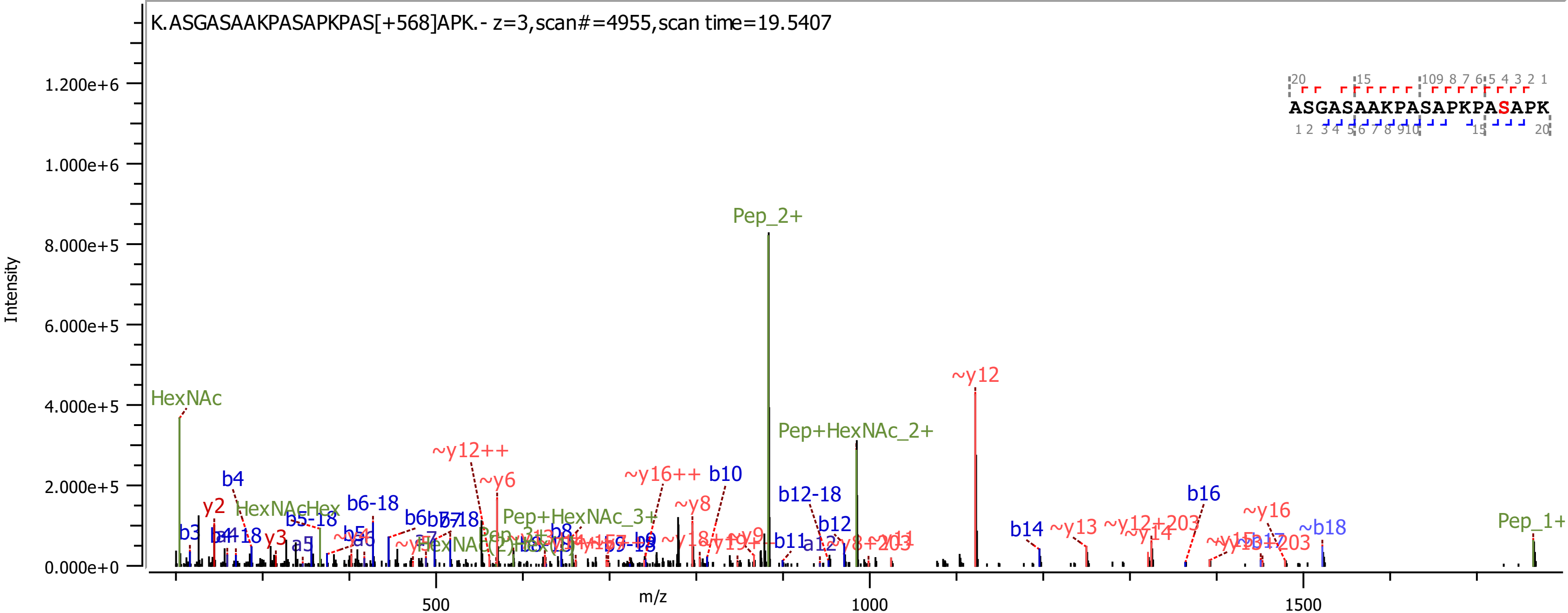
K.ASGASAAKPASAPKPAS[+568]APK.- z=3,scan#=4955,scan time=19.5407
Intensity
m/z
ASGASAAKPASAPKPASAPK
Pep_2+
~y12
Pep+HexNAc_2+
HexNAc
~y12++
~y6
~y16++
b10
b12-18
b16
~y16
~b18
Pep_1+
b4
y2
HexNAcHex
b5-18
b6-18
Pep+HexNAc_3+
~y8
b12
b14
~y13
~y12+203
~y14
b3
y3
a5
b8
b9
b11
a12

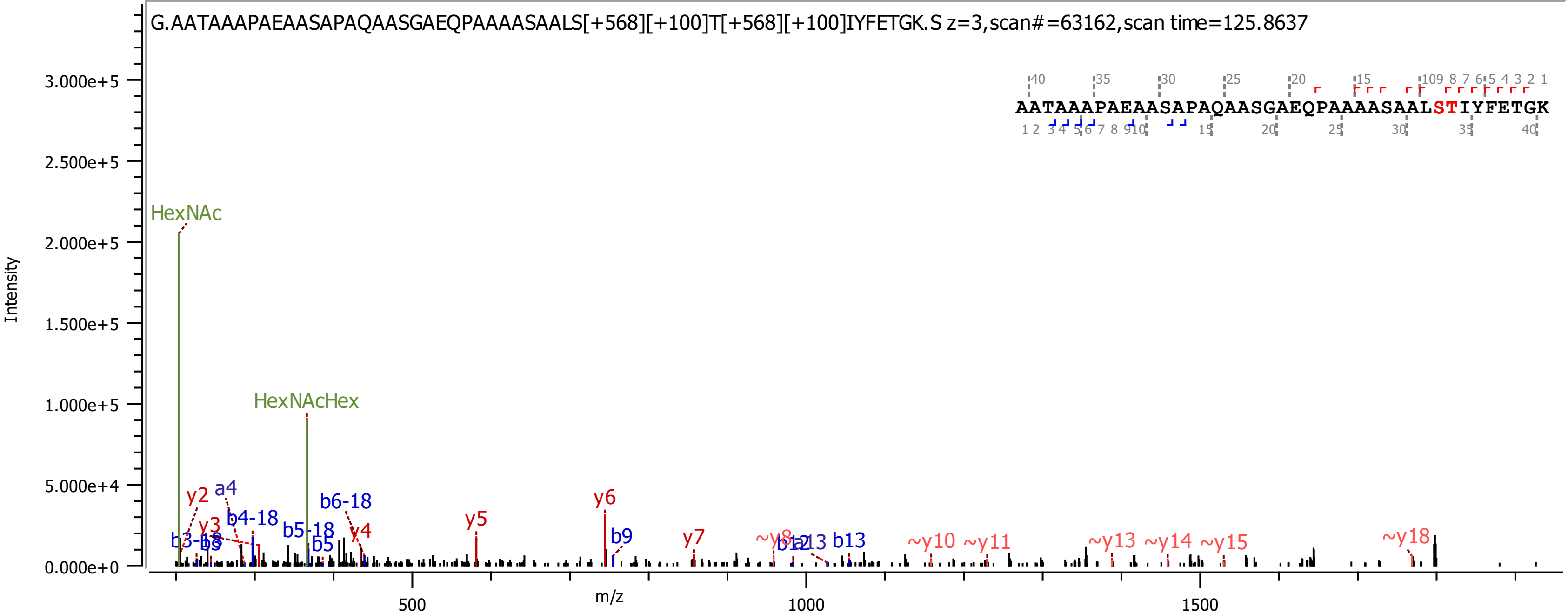

G.AATAAAPAEAASAPAQAASGAEQPAAAASAALS[+568][+100]T[+568][+100]IYFETGK.S z=3,scan#=63162,scan time=125.8637
AATAAAPAEAASAPAQAASGAEQPAAAASAALSTIYFETGK
Intensity
m/z
HexNAc
HexNAcHex
y2
a4
y3
b4-18
b3-18
b3
b5-18
b5
b6-18
y4
y5
y6
b9
y7
~y8
b12
b13
~y10
~y11
~y13
~y14
~y15
~y18

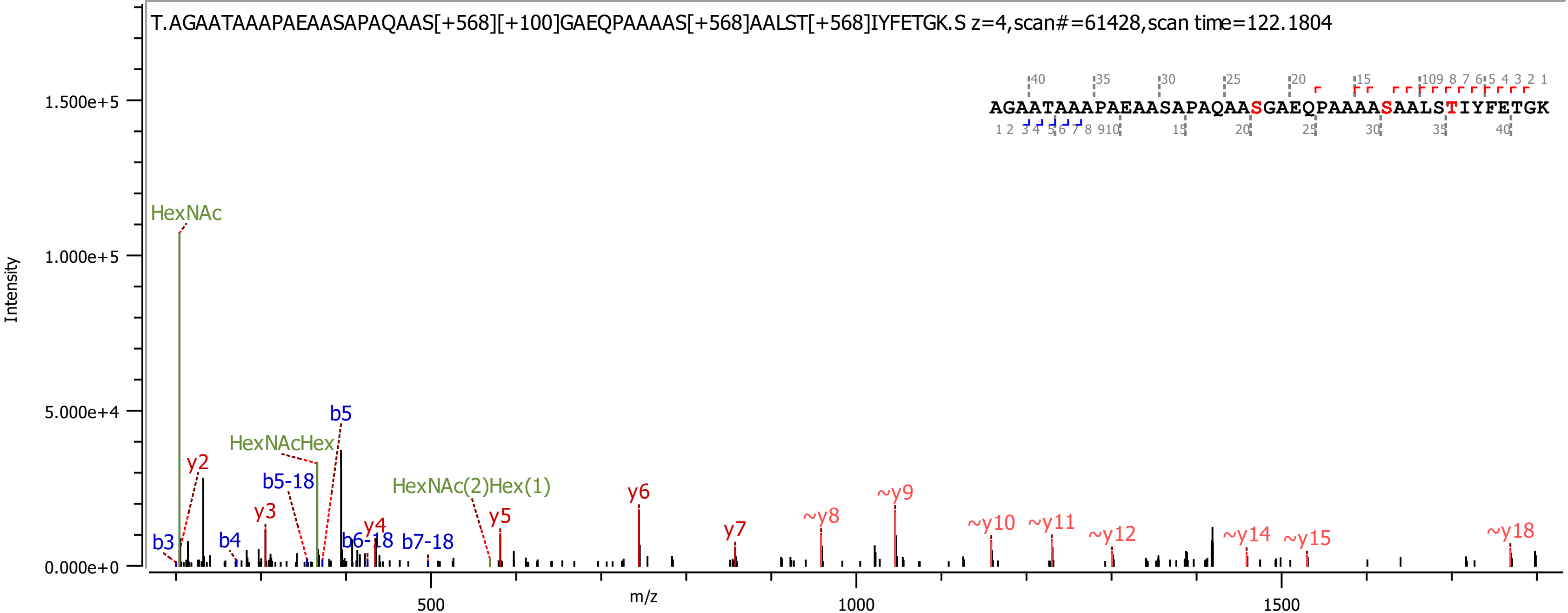

T.AGAATAAAPAEAASAPAQAAS[+568][+100]GAEQPAAAAS[+568]AALST[+568]IYFETGK.S z=4,scan#=61428,scan time=122.1804
AGAATAAAPAEAASAPAQAASGAEQPAAAASAALSTIYFETGK
Intensity
m/z
HexNAc
HexNAcHex
HexNAc(2)Hex(1)
b3
b4
b5
b5-18
b6-18
b7-18
y2
y3
y4
y5
y6
y7
~y8
~y9
~y10
~y11
~y12
~y14
~y15
~y18

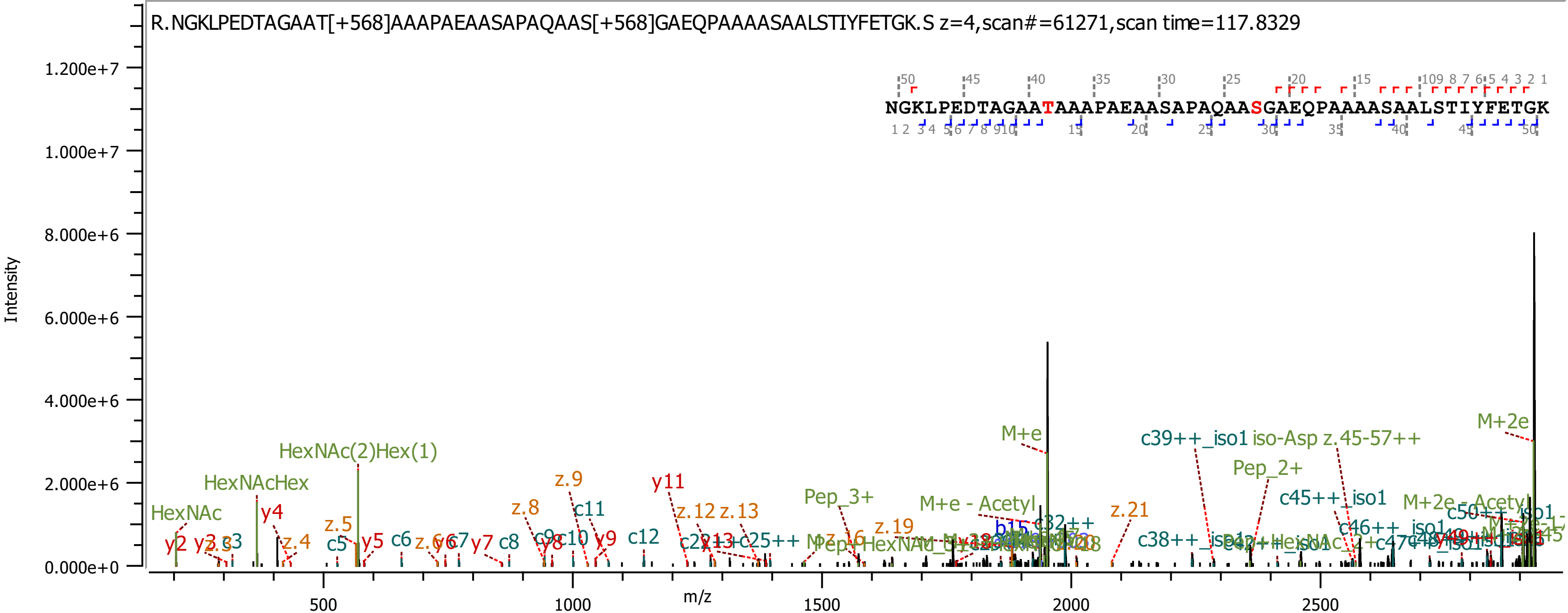
R.NGKLPEDTAGAAT[+568]AAAPAEAASAPAQAAS[+568]GAEQPAAAASAALSTIYFETGK.S z=4,scan#=61271,scan time=117.8329
NGKLPEDTAGAATAAAPAEAASAPAQAASGAEQPAAAASAALSTIYFETGK
Intensity
m/z
1.200e+7
1.000e+7
8.000e+6
6.000e+6
4.000e+6
2.000e+6
0.000e+0
500
1000
1500
2000
2500
HexNAc
HexNAcHex
HexNAc(2)Hex(1)
M+e
M+2e
M+e - Acetyl
M+2e - Acetyl
Pep_2+
Pep_3+
c39++_iso1 iso-Asp z.45-57++
c45++_iso1
c46++_iso1
c38++_iso1
y11
z.9
z.8
c11
z.12 z.13
z.21
z.19
y4
z.5
z.4
y5
c6
c5
y9
c12
c8
y7
y2
c3

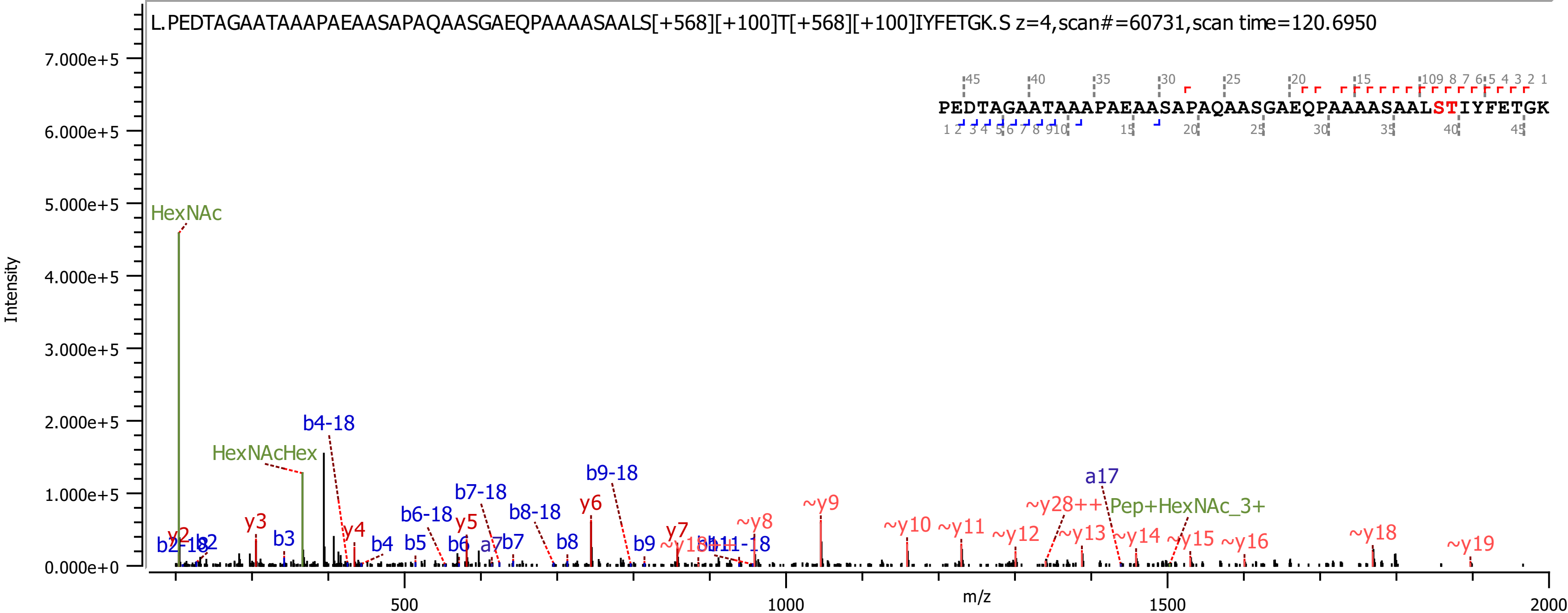

L.PEDTAGAATAAAPAEAASAPAQAASGAEQPAAAASAALS[+568][+100]T[+568][+100]IYFETGK.S z=4,scan#=60731,scan time=120.6950
PEDTAGAATAAAPAEAASAPAQAASGAEQPAAAASAALSTIYFETGK
Intensity
m/z
HexNAc
HexNAcHex
Pep+HexNAc_3+
b4-18
b6-18
b7-18
b8-18
b9-18
y2
y3
y4
y5
y6
y7
~y8
~y9
~y10
~y11
~y12
~y13
~y14
~y15
~y16
~y18
~y19
~y28++
a17
b3
b4
b5
b6
a7
b7
b8
b9

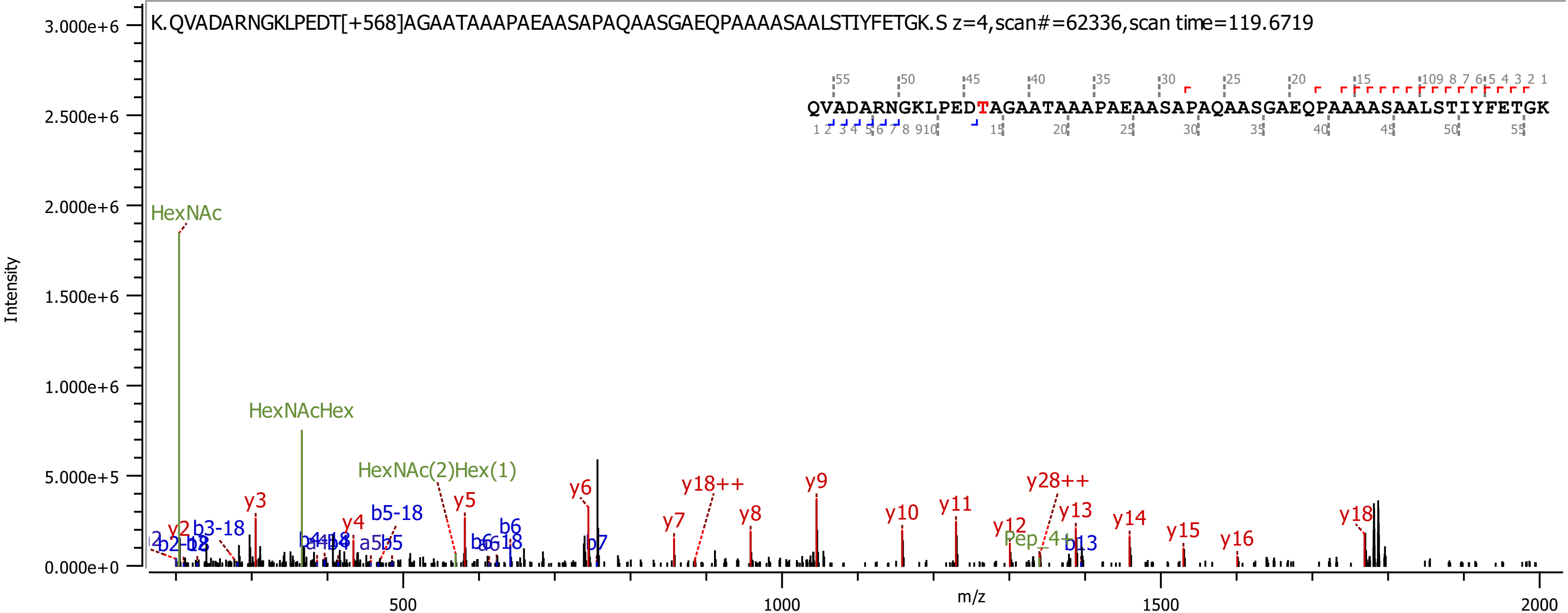
K.QVADARNGKLPEDT[+568]AGAATAAAPAEAASAPAQAASGAEQPAAAASAALSTIYFETGK.S z=4,scan#=62336,scan time=119.6719
QVADARNGKLPEDTAGAATAAAPAEAASAPAQAASGAEQPAAAASAALSTIYFETGK
Intensity
m/z
3.000e+6
2.500e+6
2.000e+6
1.500e+6
1.000e+6
5.000e+5
0.000e+0
500
1000
1500
2000
HexNAc
HexNAcHex
HexNAc(2)Hex(1)
Pep_4+
y2
y3
y4
y5
y6
y7
y8
y9
y10
y11
y12
y13
y14
y15
y16
y18
y18++
y28++
b3-18
b5-18
a5
b5
b6
b6-18
b7
b13

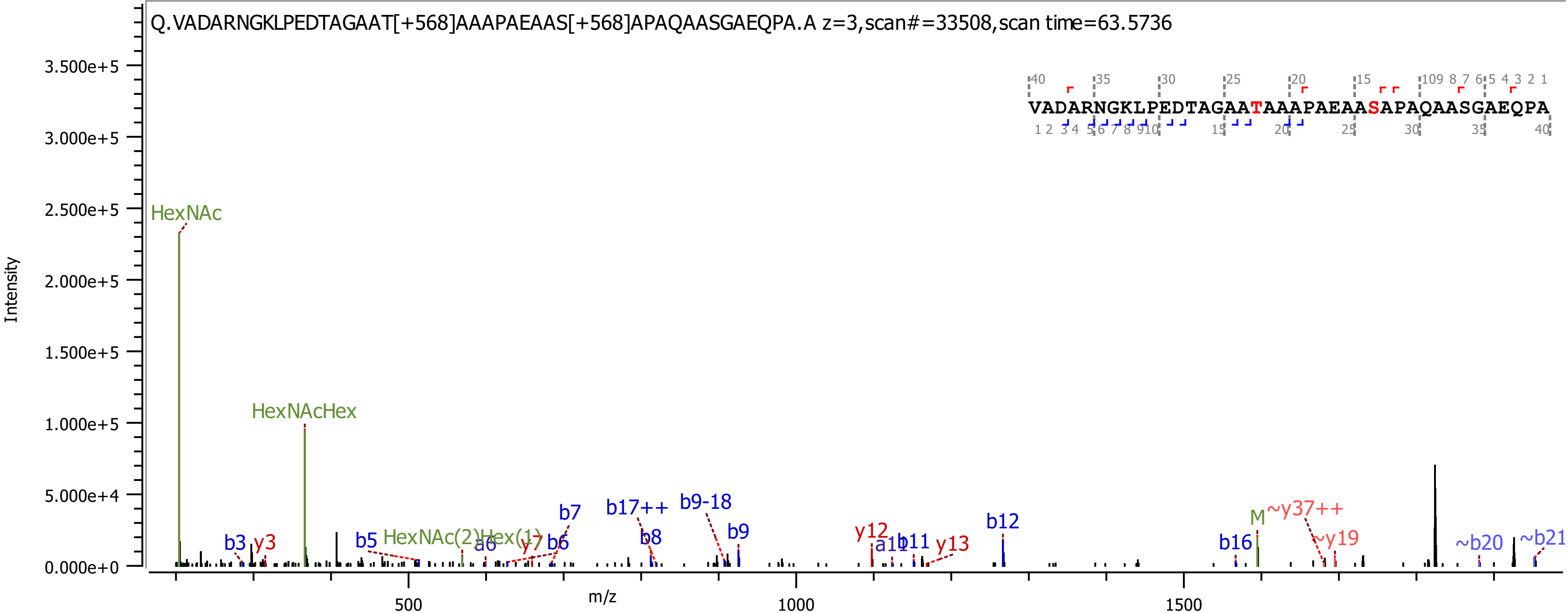
Q.VADARNGKLPEDTAGAAT[+568]AAAPAEAAS[+568]APAQAASGAEQPA.A z=3,scan#=33508,scan time=63.5736
VADARNGKLPEDTAGAATAAAPAEAASAPAQAASGAEQPA
Intensity
m/z
3.500e+5
3.000e+5
2.500e+5
2.000e+5
1.500e+5
1.000e+5
5.000e+4
0.000e+0
500
1000
1500
HexNAc
HexNAcHex
HexNAc(2)Hex(1)
b3
y3
b5
a6
y7
b6
b7
b17++
b8
b9-18
b9
y12
a11
b11
y13
b12
b16
M
~y37++
~y19
~b20
~b21

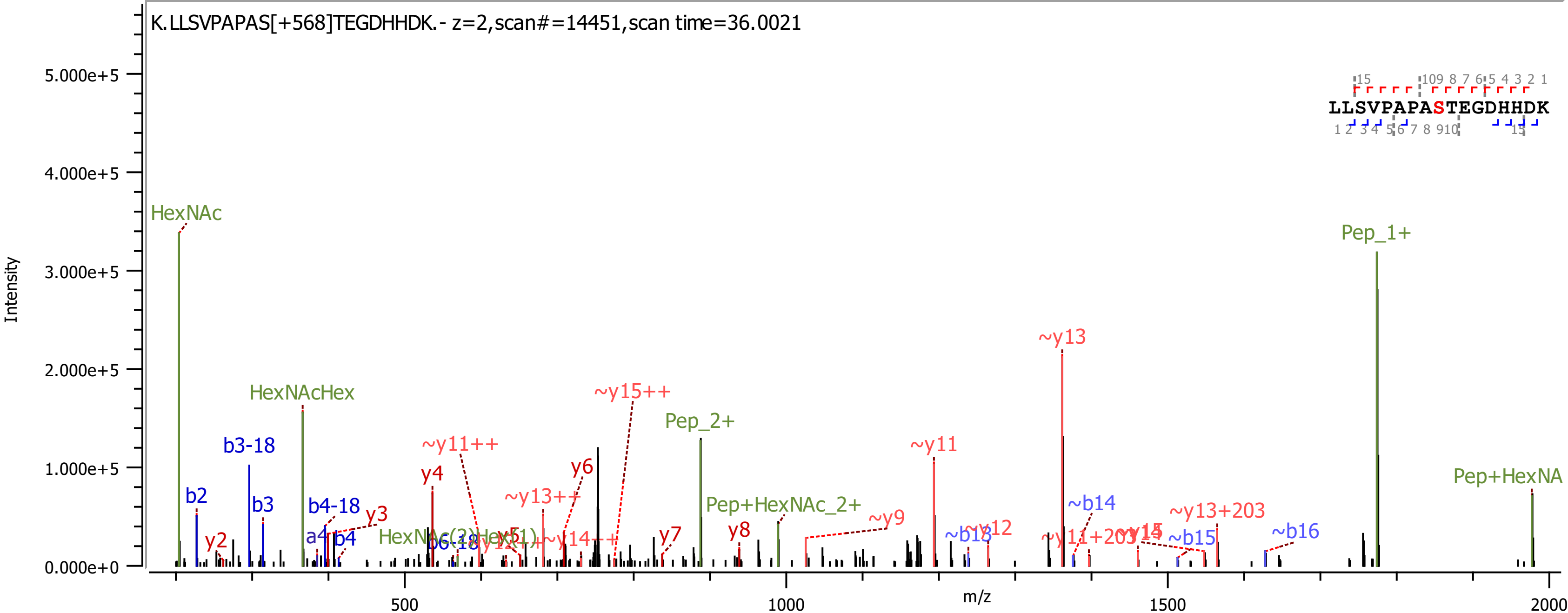
K.LLSVPAPAS[+568]TEGDHHDK.- z=2,scan#=14451,scan time=36.0021
Intensity
m/z
LLSVPAPASTEGDHHDK
HexNAc
HexNAcHex
Pep_1+
Pep_2+
Pep+HexNAc_2+
Pep+HexNA
HexNAc(2)Hex(1)
b2
b3-18
b3
b4-18
a4
b4
y2
y3
y4
y6
y7
y8
~y11++
~y13++
~y14++
~y15++
~y9
~y11
~y12
~y13
~y11+203
~y13+203
~b13
~b14
~b15
~b16
5.000e+5
4.000e+5
3.000e+5
2.000e+5
1.000e+5
0.000e+0
500
1000
1500
2000

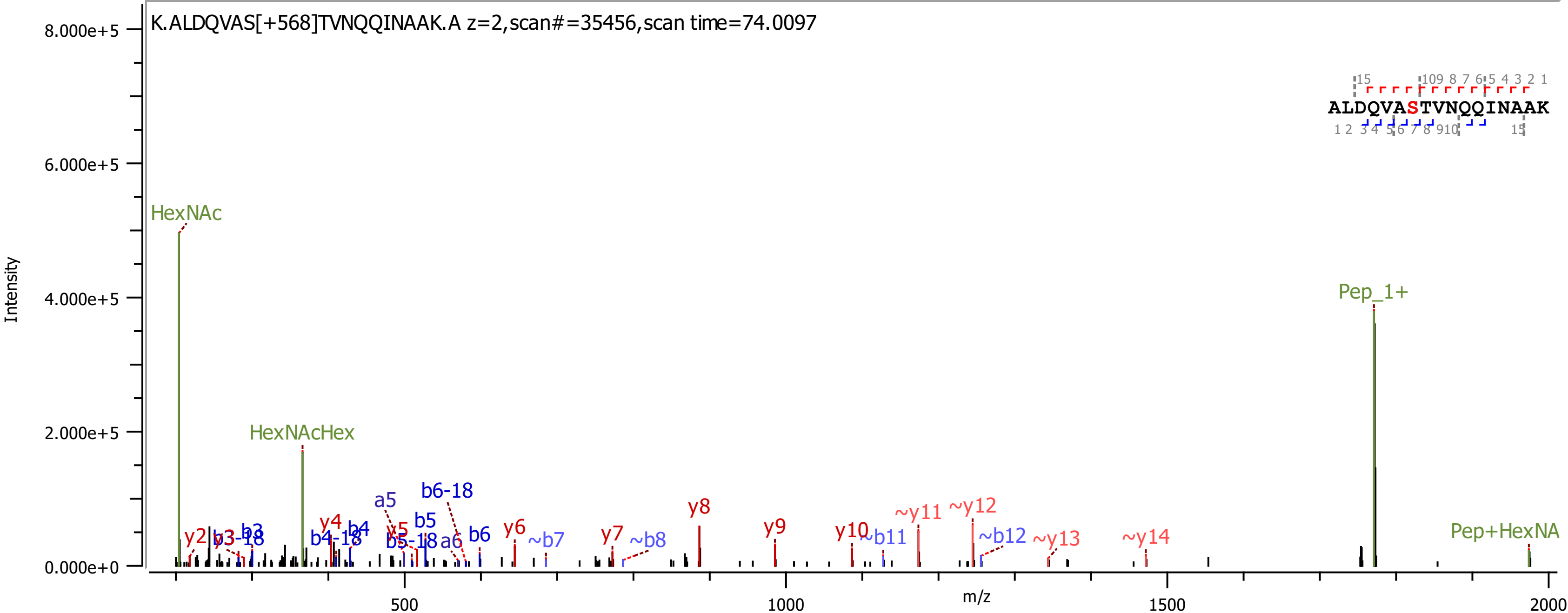
K.ALDQVAS[+568]TVNQQINAAK.A z=2,scan#=35456,scan time=74.0097
ALDQVASTVNQQINAAK
Intensity
m/z
8.000e+5
6.000e+5
4.000e+5
2.000e+5
0.000e+0
500
1000
1500
2000
HexNAc
HexNAcHex
Pep_1+
Pep+HexNA
y2
y3
b3
b3-18
y4
b4
b4-18
a5
y5
b5
b5-18
b6-18
a6
b6
y6
~b7
y7
~b8
y8
y9
y10
~b11
~y11
~y12
~b12
~y13
~y14

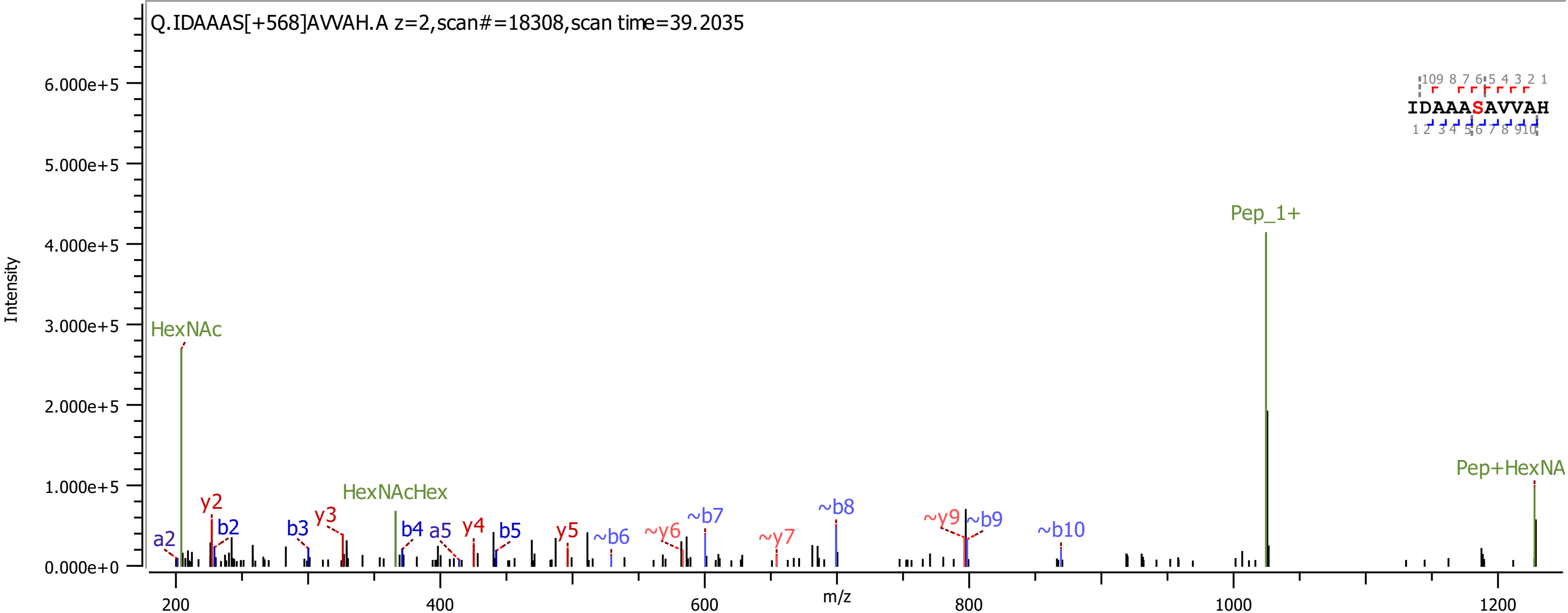

Q.IDAAAS[+568]AVVAH.A z=2,scan#=18308,scan time=39.2035
Intensity
6.000e+5
5.000e+5
4.000e+5
3.000e+5
2.000e+5
1.000e+5
0.000e+0
m/z
200
400
600
800
1000
1200
HexNAc
HexNAcHex
Pep_1+
Pep+HexNA
a2
y2
b2
b3
y3
b4
a5
y4
b5
y5
~b6
~y6
~b7
~y7
~b8
~y9
~b9
~b10
IDAAASAVVAH

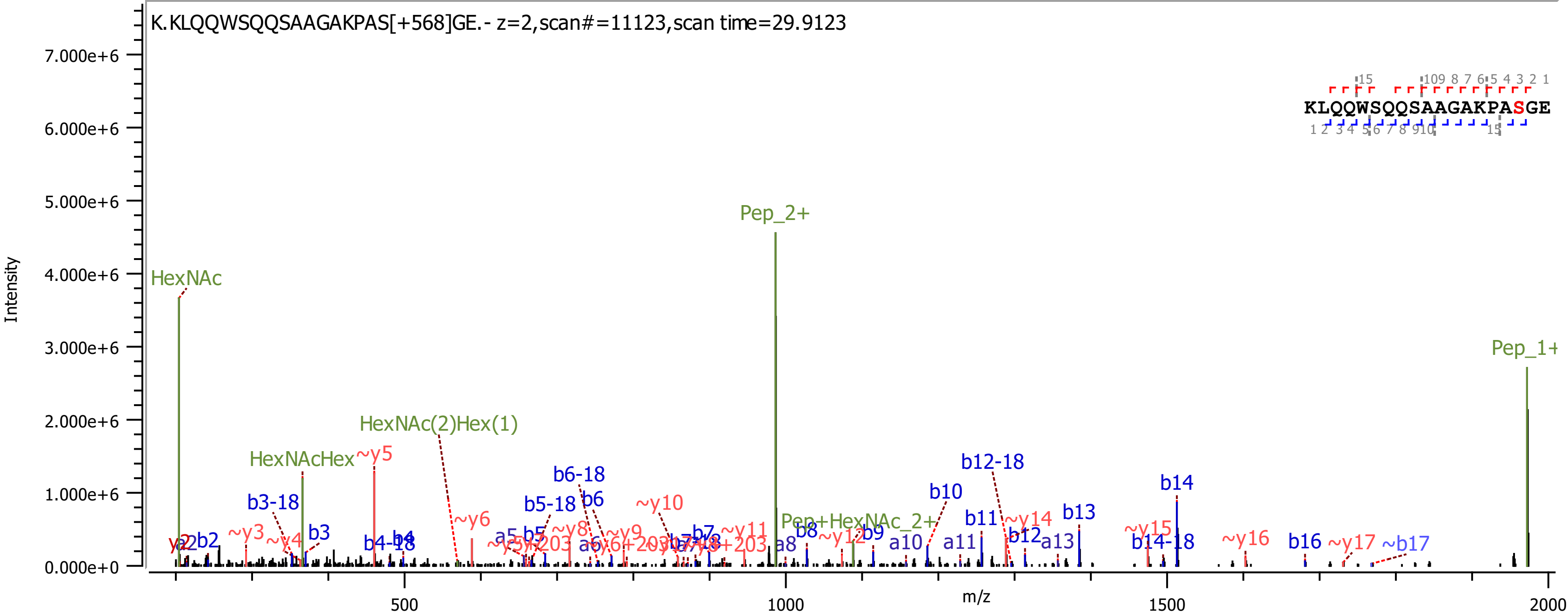

K.KLQQWSQQSAAGAKPAS[+568]GE.- z=2,scan#=11123,scan time=29.9123
KLQQWSQQSAAGAKPASGE
Intensity
m/z
HexNAc
Pep_2+
Pep_1+
HexNAcHex
HexNAc(2)Hex(1)
Pep+HexNAc_2+
b3-18
b6-18
b5-18
b12-18
b14-18
b4-18
b2
b3
b4
b5
b6
b7
b8
b9
b10
b11
b12
b13
b14
b16
~b17
~y3
~y4
~y5
~y6
~y8
~y9
~y10
~y11
~y12
~y14
~y15
~y16
~y17
a5
a6
a8
a10
a11
a13

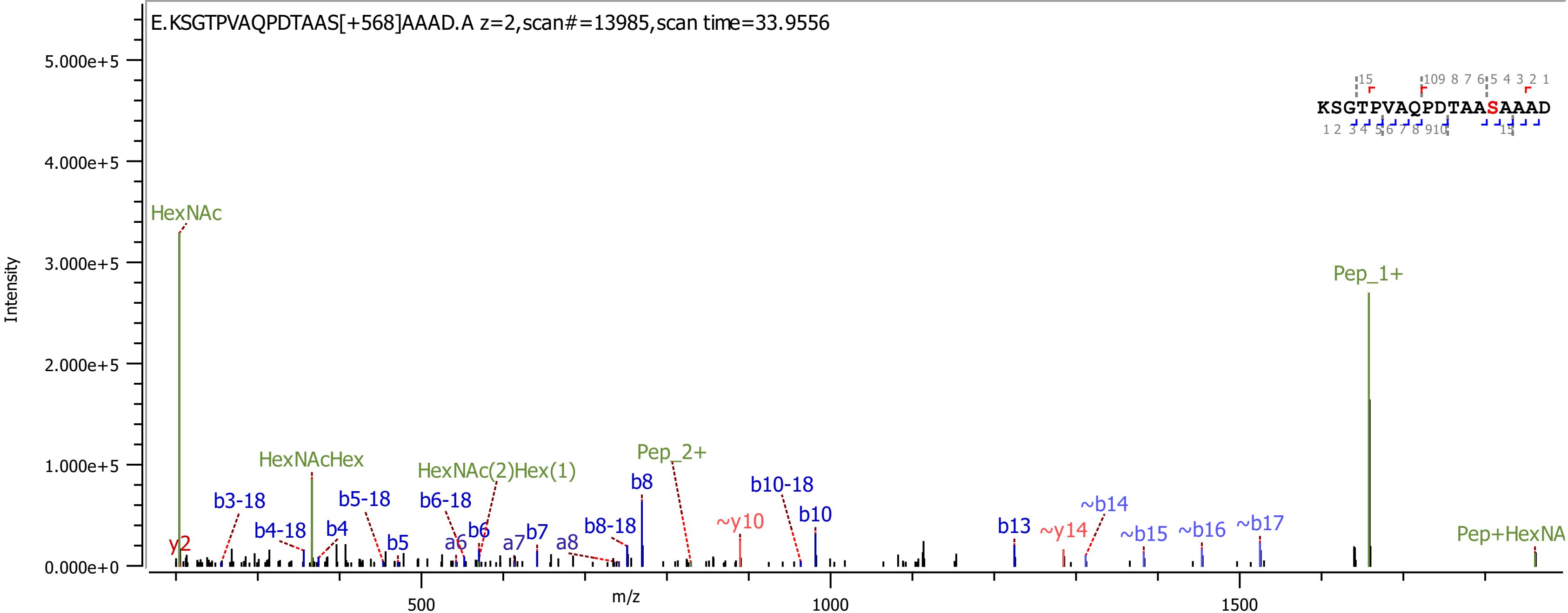
E.KSGTPVAQPDTAAS[+568]AAAD.A z=2,scan#=13985,scan time=33.9556
KSGTPVAQPDTAASAAAD
Intensity
m/z
HexNAc
Pep_1+
HexNAcHex
HexNAc(2)Hex(1)
Pep_2+
Pep+HexNA
y2
b3-18
b4-18
b4
b5-18
b5
b6-18
a6
b6
a7
b7
a8
b8-18
b8
~y10
b10-18
b10
b13
~y14
~b14
~b15
~b16
~b17

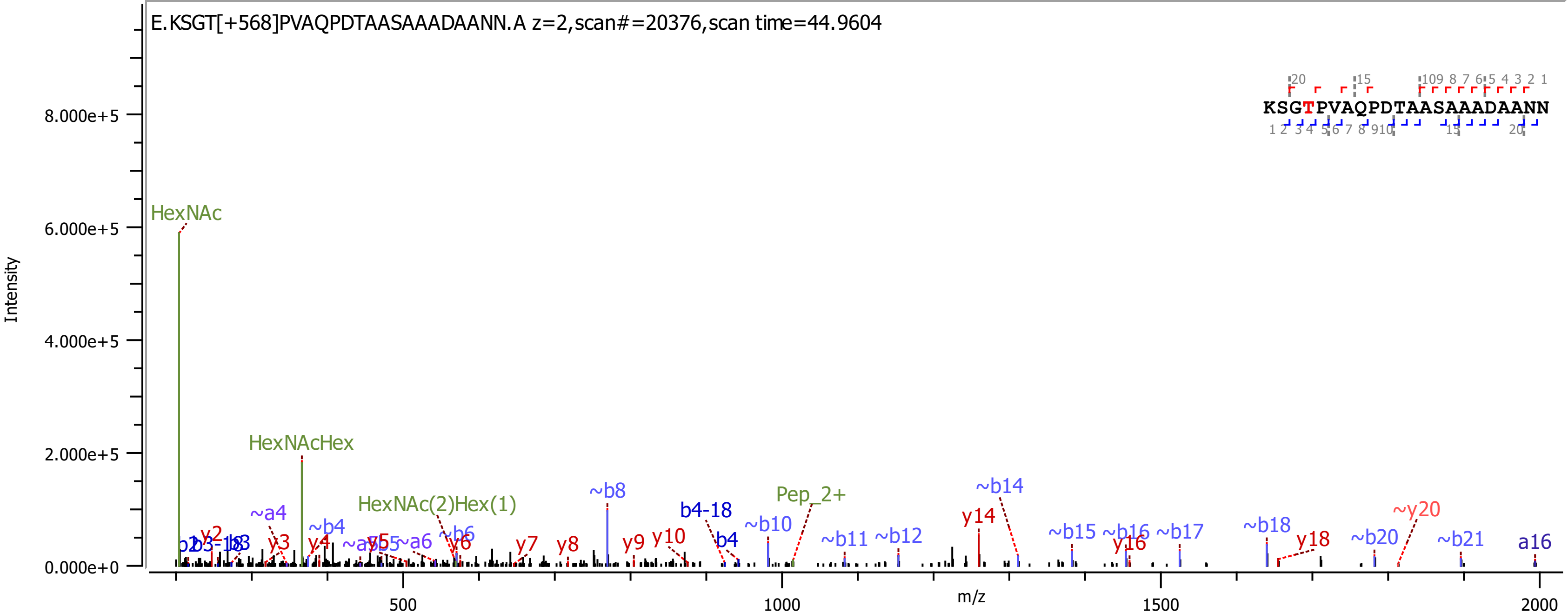
E.KSGT[+568]PVAQPDTAASAAADAANN.A z=2,scan#=20376,scan time=44.9604
KSGTPVAQPDTAASAAADAANN
Intensity
m/z
HexNAc
HexNAcHex
HexNAc(2)Hex(1)
Pep_2+
~b8
b4-18
~b10
~b14
y14
~b15
~b16
~b17
~b18
y18
~b20
~y20
~b21
a16
~b11
~b12
y16
y7
y8
y9
y10
b4
~a4
~b4
y3
y4
y5
~a6
~b6
y6
y2

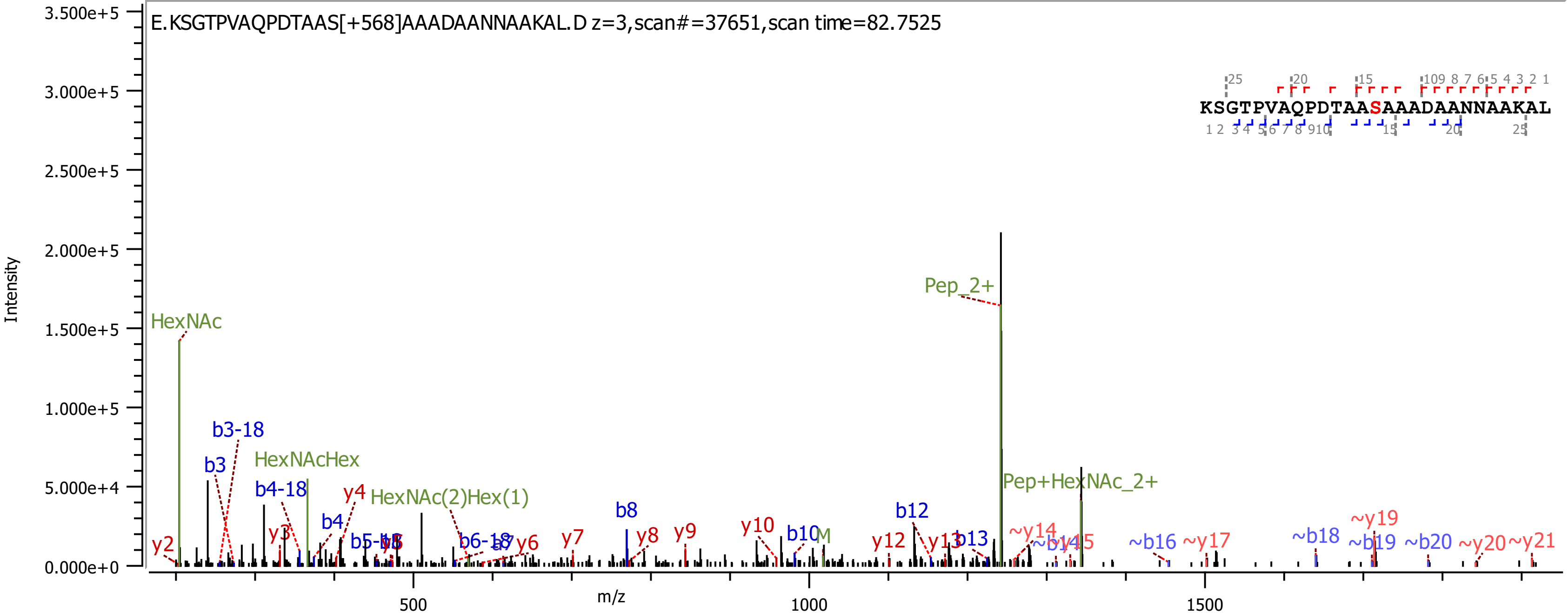
E.KSGTPVAQPDTAAS[+568]AAADAANNAAKAL.D z=3,scan#=37651,scan time=82.7525
KSGTPVAQPDTAASAAADAANNAAKAL
Intensity
m/z
HexNAc
b3-18
b3
HexNAcHex
b4-18
y4
HexNAc(2)Hex(1)
b4
y3
y2
b5-18
b5
b6-18
a7
y6
y7
b8
y8
y9
y10
b10
M
b12
y12
y13
b13
Pep_2+
Pep+HexNAc_2+
~y14
~b14
~y15
~b16
~y17
~b18
~y19
~b19
~b20
~y20
~y21

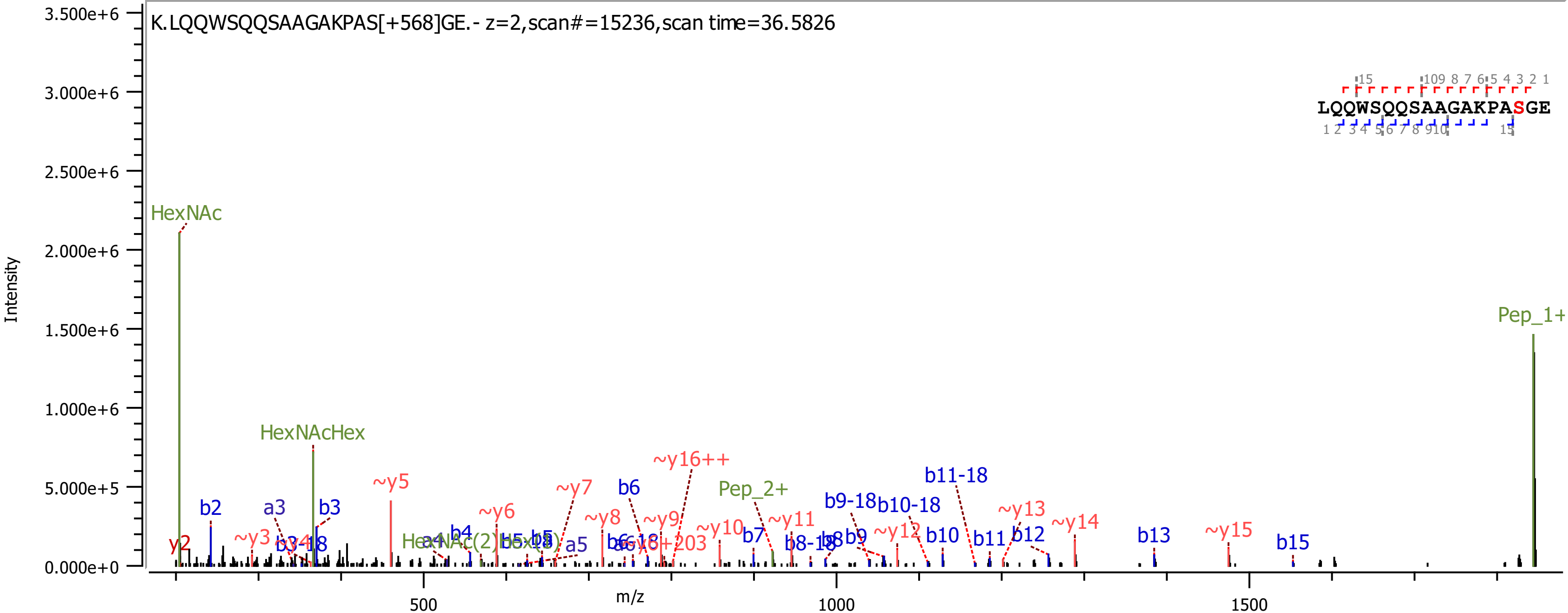
K.LQQWSQQSAAGAKPAS[+568]GE.- z=2,scan#=15236,scan time=36.5826
LQQWSQQSAAGAKPASGE
Intensity
m/z
HexNAc
HexNAcHex
Pep_1+
Pep_2+
~y16++
b11-18
b9-18
b10-18
~y5
~y6
~y7
~y8
~y9
~y10
~y11
~y12
~y13
~y14
~y15
b2
b3
b6
b7
b10
b11
b12
b13
b15
a3
a5
y2
~y3
3.500e+6
3.000e+6
2.500e+6
2.000e+6
1.500e+6
1.000e+6
5.000e+5
0.000e+0
500
1000
1500

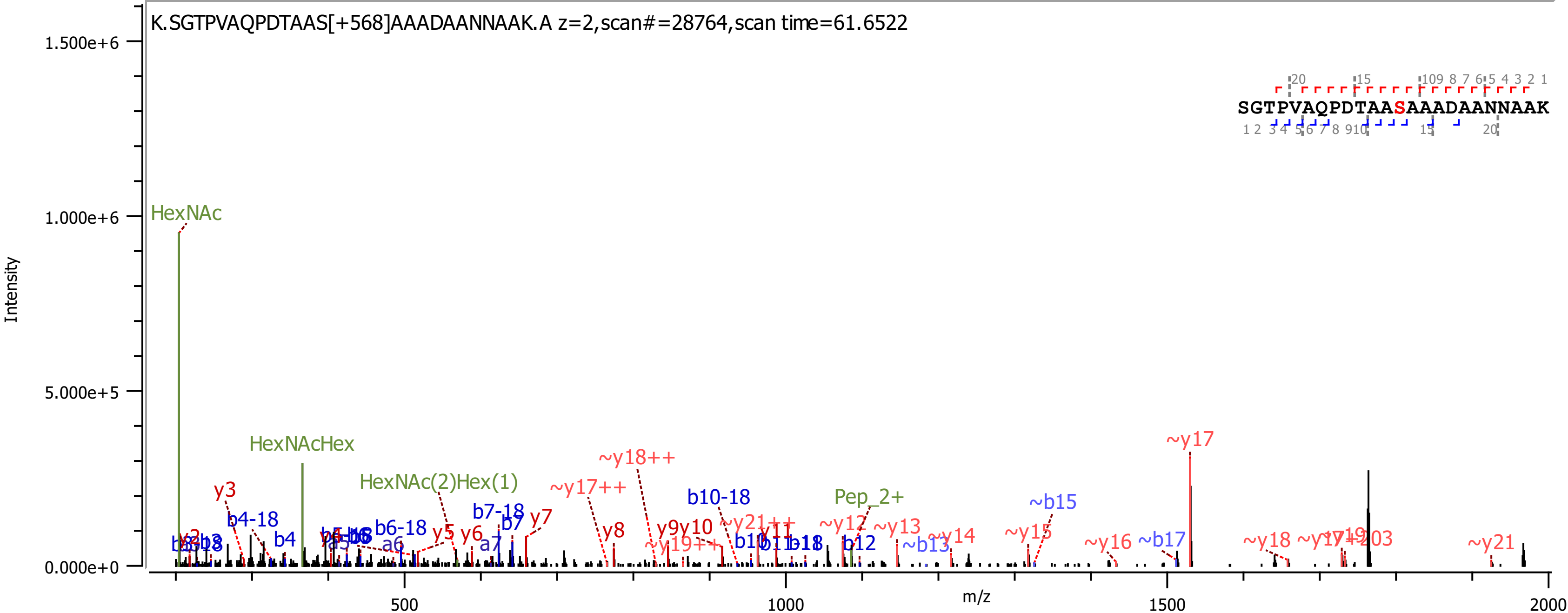

K.SGTPVAQPDTAAS[+568]AAADAANNAAK.A z=2,scan#=28764,scan time=61.6522
SGTPVAQPDTAASAAADAANNAAK
Intensity
m/z
1.500e+6
1.000e+6
5.000e+5
0.000e+0
500
1000
1500
2000
HexNAc
HexNAcHex
HexNAc(2)Hex(1)
Pep_2+
y3
b4-18
b4
b6-18
y5
y6
a6
b7-18
b7
a7
y7
~y17++
y8
~y18++
y9y10
~y19++
b10-18
~y21++
b10
y11
b12
~y12
~y13
~b13
~y14
~y15
~b15
~y16
~b17
~y17
~y18
~y19
~y21

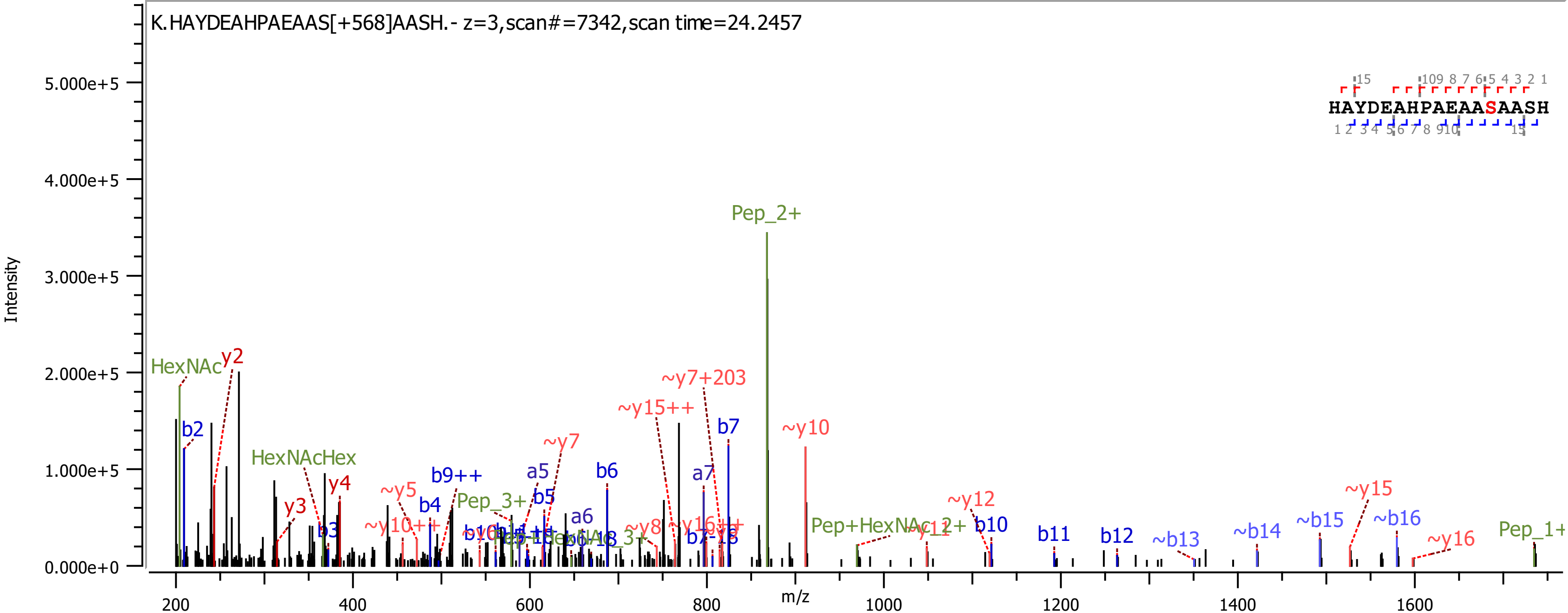
K.HAYDEAHPAEAAS[+568]AASH.- z=3,scan#=7342,scan time=24.2457
HAYDEAHPAEAASAASH
Intensity
m/z
5.000e+5
4.000e+5
3.000e+5
2.000e+5
1.000e+5
0.000e+0
200
400
600
800
1000
1200
1400
1600
Pep_2+
HexNAc
y2
b2
HexNAcHex
y3
y4
b3
~y5
~y10++
b9++
b4
Pep_3+
a5
b5
~y7
b6
a6
~y15++
~y7+203
b7
a7
~y8
~y16++
~y10
Pep+HexNAc_2+
~y11
~y12
b10
b11
b12
~b13
~b14
~b15
~y15
~b16
~y16
Pep_1+

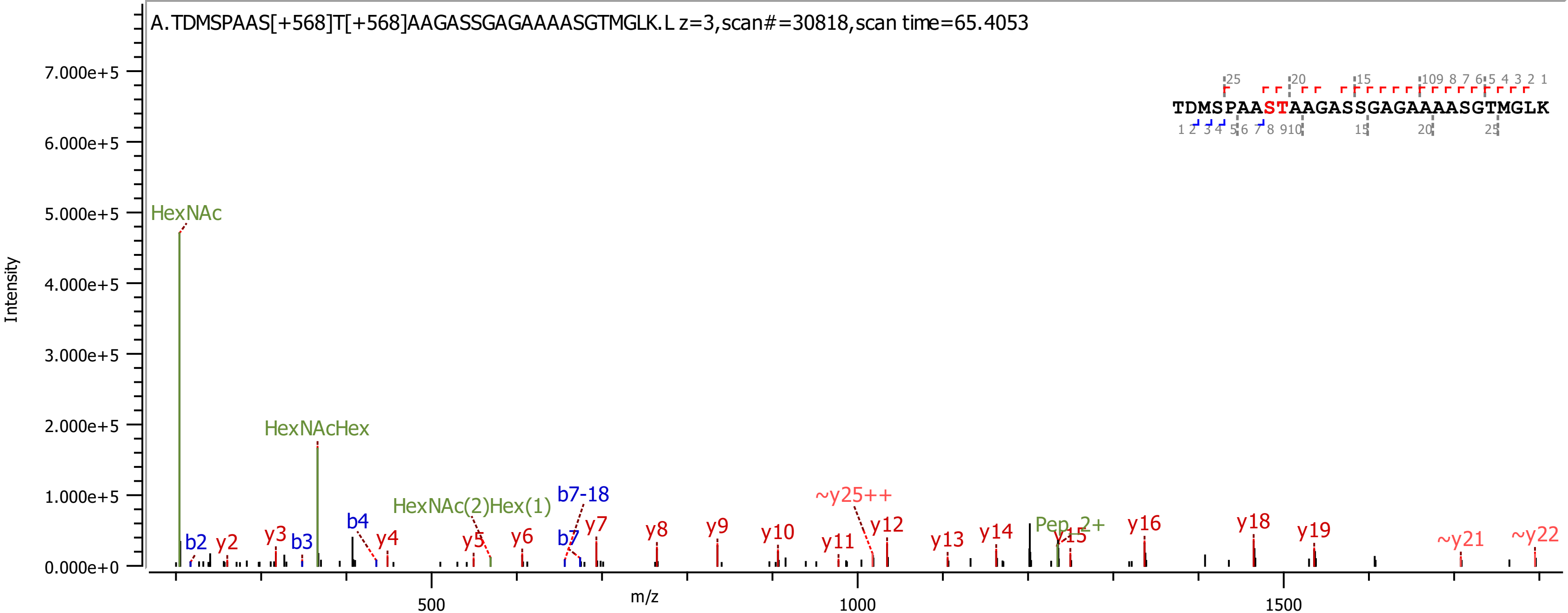
A.TDMSPAAS[+568]T[+568]AAGASSGAGAAAASGTMGLK.L z=3,scan#=30818,scan time=65.4053
TDMSPAASTAAGASSGAGAAAASGTMGLK
Intensity
m/z
HexNAc
HexNAcHex
HexNAc(2)Hex(1)
b2
b3
b4
b7
b7-18
y2
y3
y4
y5
y6
y7
y8
y9
y10
y11
y12
y13
y14
y15
y16
y18
y19
~y21
~y22
~y25++
Pep_2+

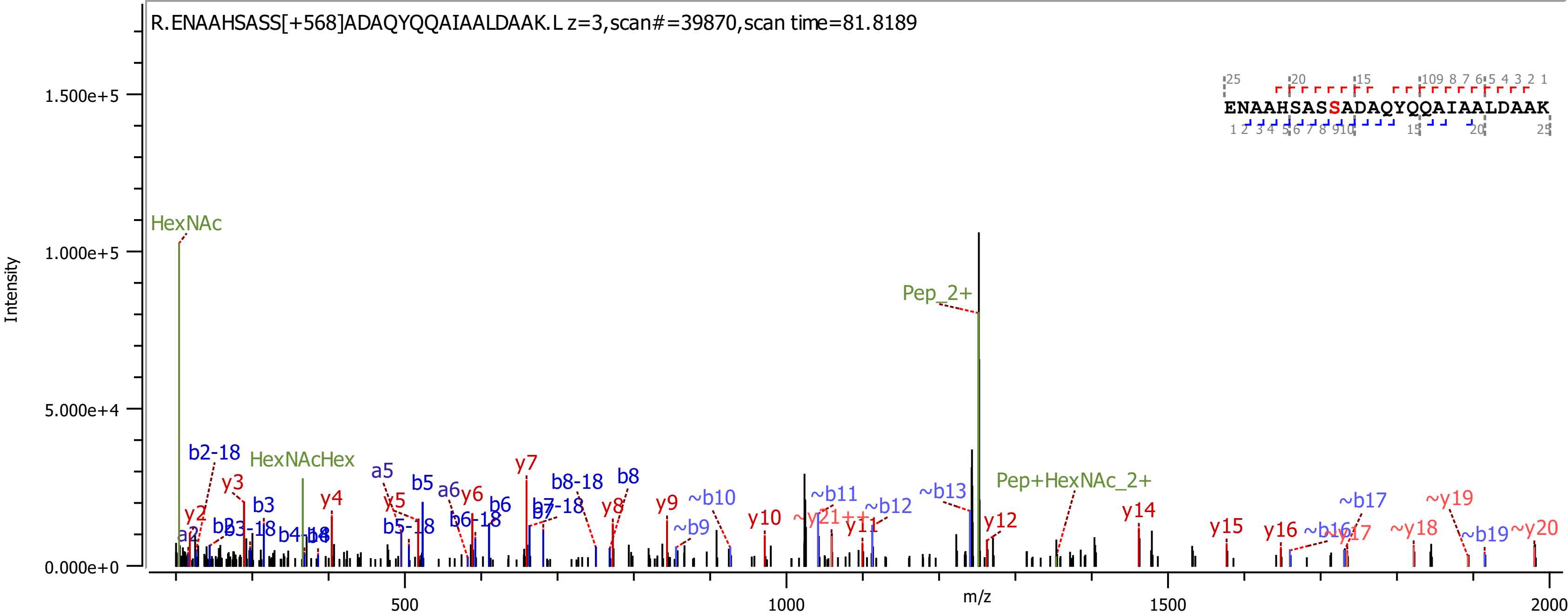
R.ENAAHSASS[+568]ADAQYQQAIAALDAAK.L z=3,scan#=39870,scan time=81.8189
ENAAHSASSADAQYQQAIAALDAAK
Intensity
m/z
HexNAc
HexNAcHex
Pep_2+
Pep+HexNAc_2+
0.000e+0
5.000e+4
1.000e+5
1.500e+5
500
1000
1500
2000

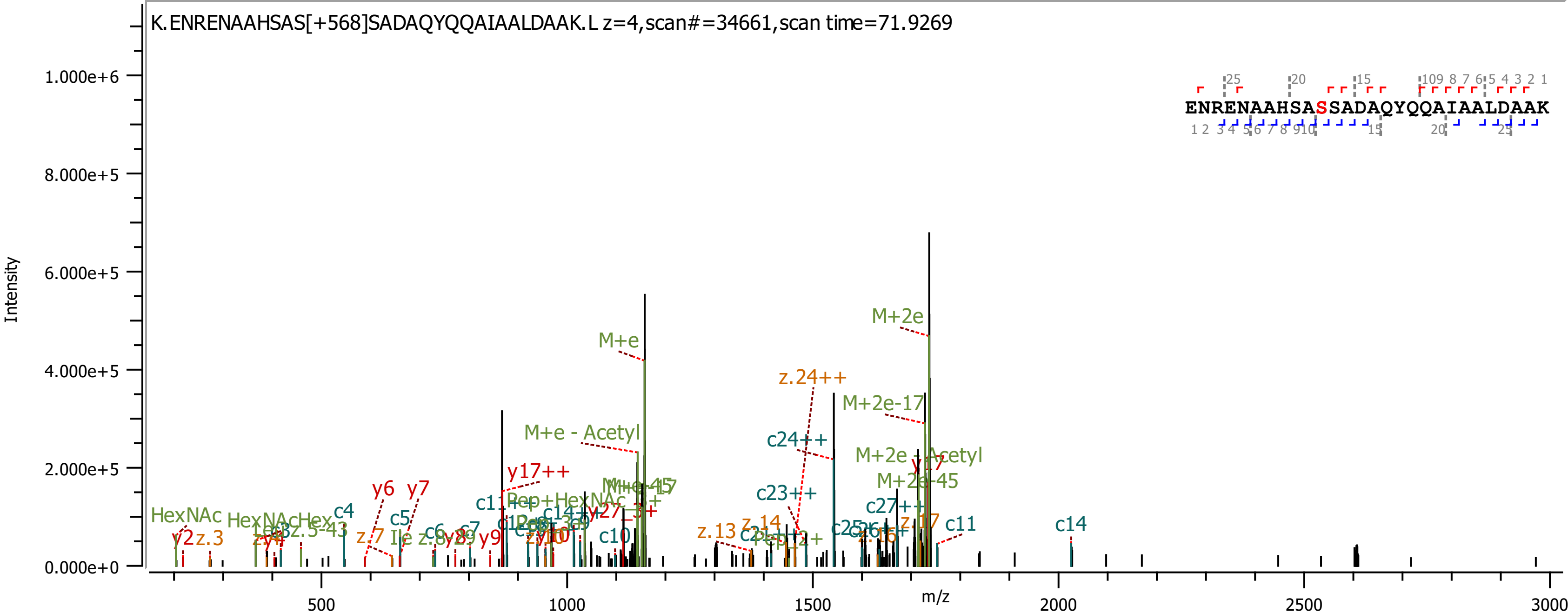
K.ENRENAAHSAS[+568]SADAQYQQAIAALDAAK.L z=4,scan#=34661,scan time=71.9269
ENRENAAHSASSADAQYQQAIAALDAAK
Intensity
m/z
M+e
M+2e
M+2e-17
M+e - Acetyl
z.24++
c24++
c23++
y17++
y6
y7
c4
c5
c10
c11
c14
HexNAc
c27++
M+2e-45

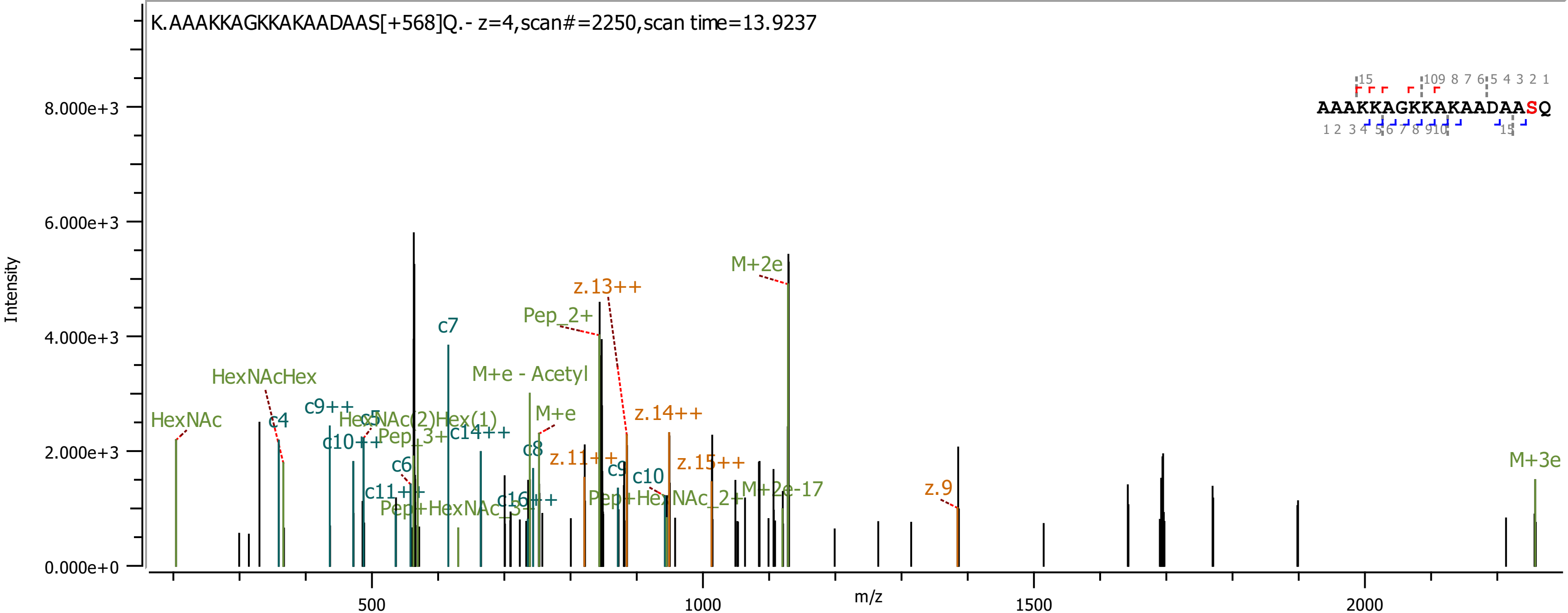
K.AAAKKAGKKAKAADAAS[+568]Q.- z=4,scan#=2250,scan time=13.9237
AAAKKAGKKAKAADAASQ
Intensity
m/z
HexNAc
HexNAcHex
c4
c9++
c10++
HexNAc(2)Hex(1)
Pep_3+
c14++
c7
c6
c11++
Pep+HexNAc_3+
c16++
M+e - Acetyl
M+e
c8
Pep_2+
z.13++
z.11++
c9
c10
z.14++
z.15++
Pep+HexNAc_2+
M+2e
M+2e-17
z.9
M+3e

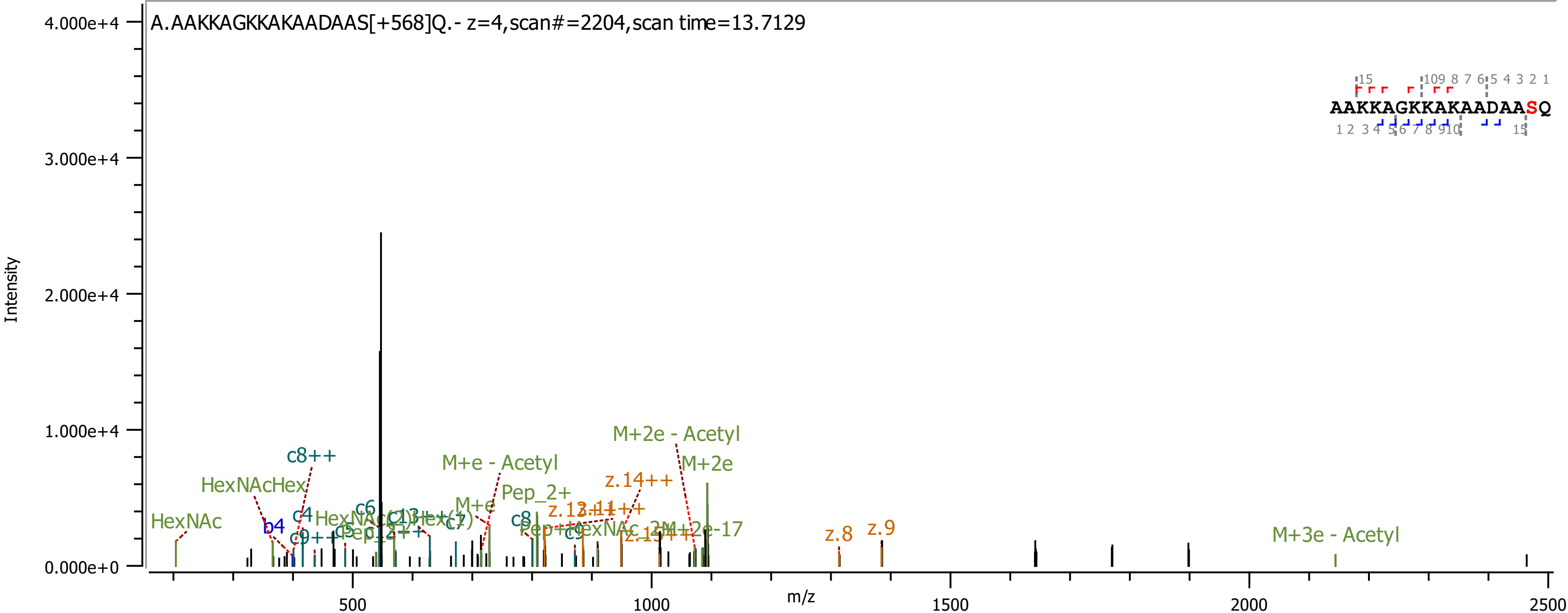
A.AAKKAGKKAKAADAAS[+568]Q.- z=4,scan#=2204,scan time=13.7129
Intensity
m/z
4.000e+4
3.000e+4
2.000e+4
1.000e+4
0.000e+0
500
1000
1500
2000
2500
AAKKAGKKAKAADAASQ
HexNAc
HexNAcHex
b4
c4
c8++
c6
M+e - Acetyl
M+e
Pep_2+
c8
M+2e - Acetyl
M+2e
z.14++
z.8
z.9
M+3e - Acetyl

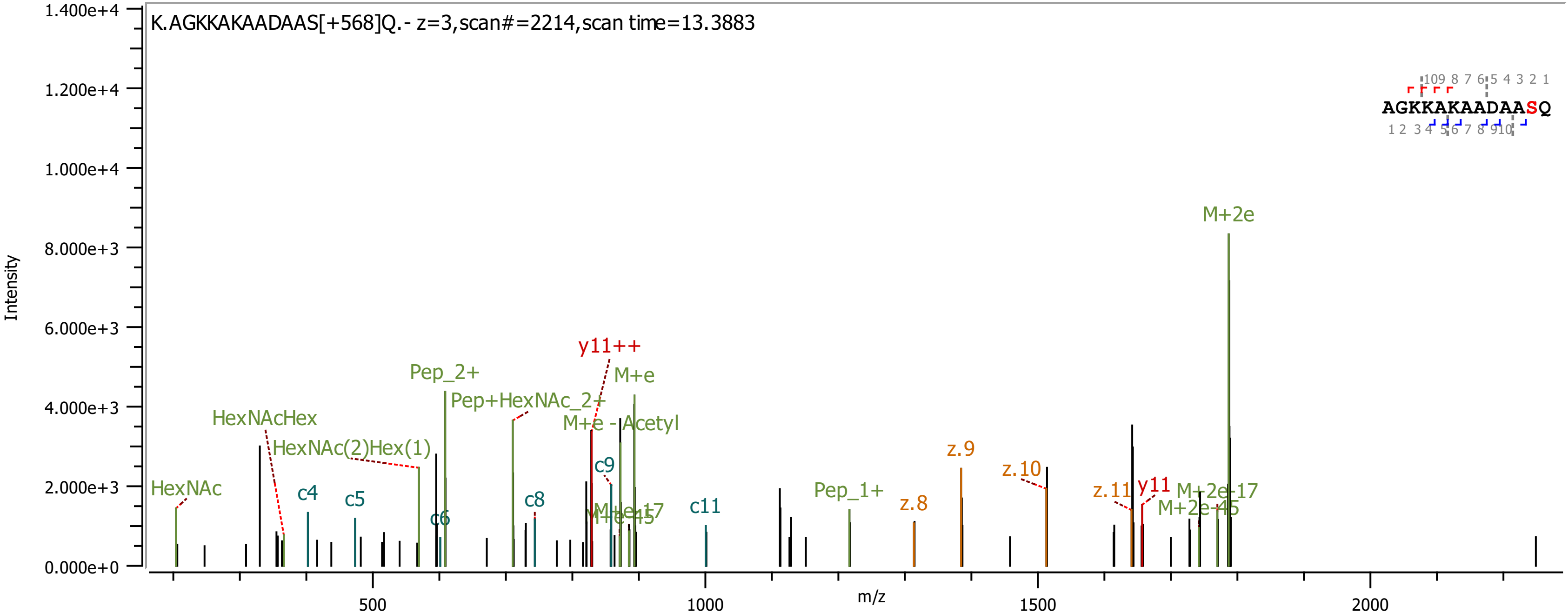
K.AGKKAKAADAAS[+568]Q.- z=3,scan#=2214,scan time=13.3883
AGKKAKAADAASQ
Intensity
m/z
HexNAc
HexNAcHex
HexNAc(2)Hex(1)
c4
c5
c6
Pep_2+
Pep+HexNAc_2+
c8
y11++
M+e
M+e -Acetyl
c9
M+e-17
M+e-45
c11
Pep_1+
z.8
z.9
z.10
z.11
y11
M+2e
M+2e-17
M+2e-45
0.000e+0
2.000e+3
4.000e+3
6.000e+3
8.000e+3
1.000e+4
1.200e+4
1.400e+4
500
1000
1500
2000

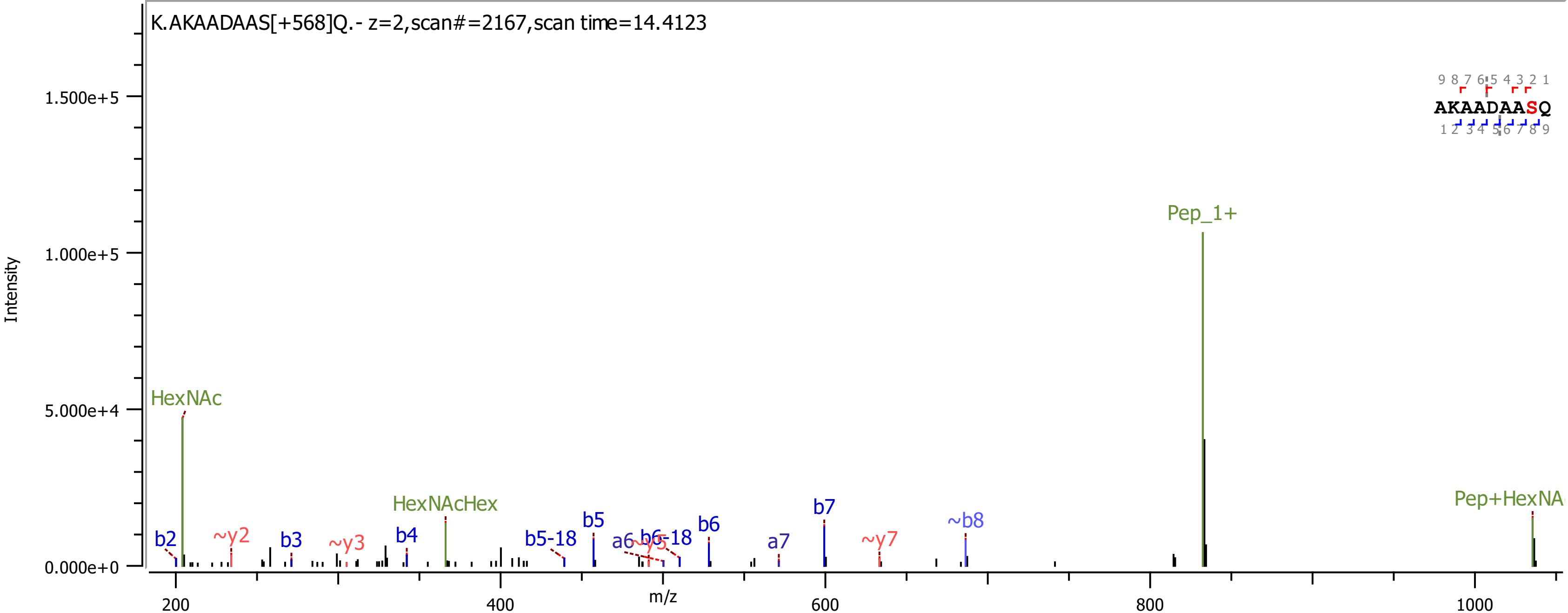

K.AKAADAAS[+568]Q.- z=2,scan#=2167,scan time=14.4123
Intensity
1.500e+5
1.000e+5
5.000e+4
0.000e+0
m/z
200
400
600
800
1000
HexNAc
HexNAcHex
Pep_1+
Pep+HexNA
b2
~y2
b3
~y3
b4
b5-18
b5
a6
~y5
b6-18
b6
a7
b7
~y7
~b8
AKAADAASQ
9 8 7 6 5 4 3 2 1
1 2 3 4 5 6 7 8 9

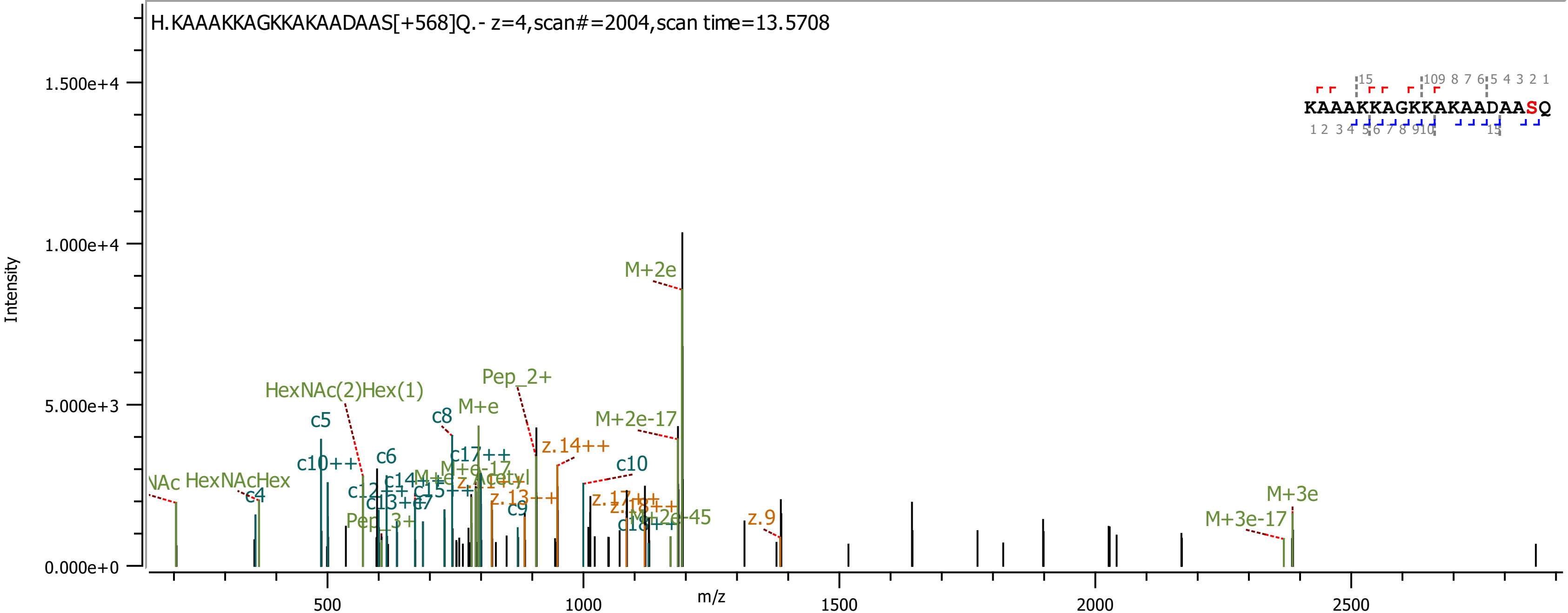
H.KAAAKKAGKKAKAADAAS[+568]Q.- z=4,scan#=2004,scan time=13.5708
Intensity
m/z
1.500e+4
1.000e+4
5.000e+3
0.000e+0
500
1000
1500
2000
2500
KAAAKKAGKKAKAADAASQ
M+2e
Pep_2+
HexNAc(2)Hex(1)
M+e
c5
c8
M+2e-17
z.14++
c6
c10++
c17++
c10
HexNAcHex
c4
z.9
M+3e
M+3e-17
M+2e-45
Pep_3+
c9

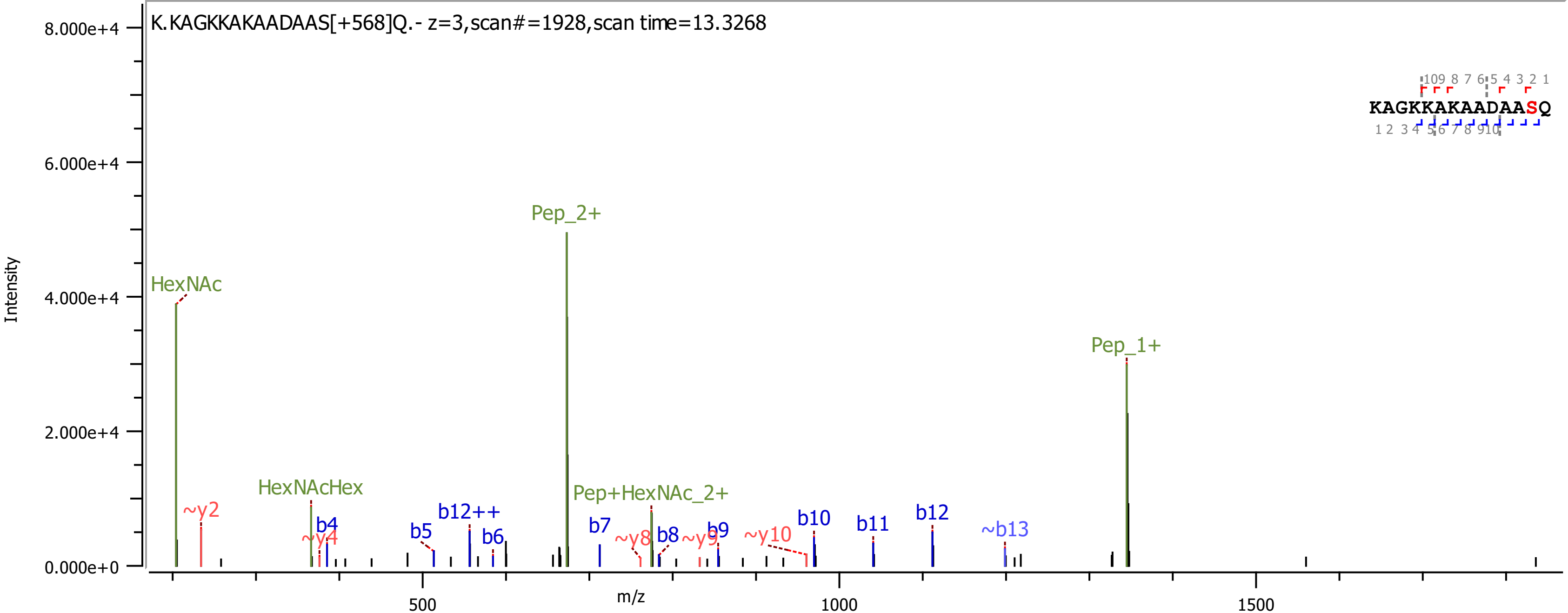
K.KAGKKAKAADAAS[+568]Q.- z=3,scan#=1928,scan time=13.3268
KAGKKAKAADAASQ
Intensity
m/z
HexNAc
HexNAcHex
Pep_2+
Pep+HexNAc_2+
Pep_1+
~y2
~y4
~y8
~y9
~y10
b4
b5
b6
b7
b8
b9
b10
b11
b12
b12++
~b13

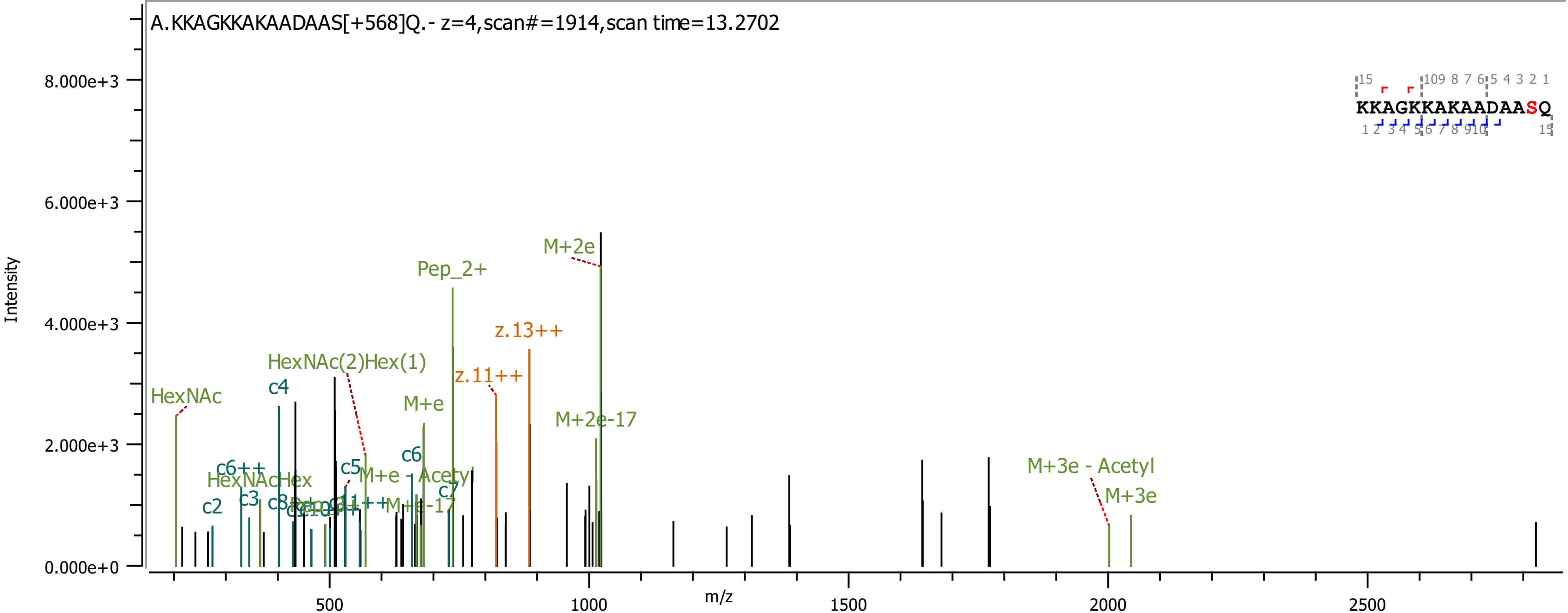

A.KKAGKKAKAADAAS[+568]Q.- z=4,scan#=1914,scan time=13.2702
Intensity
m/z
KKAGKKAKAADAASQ
HexNAc
c2
c6++
HexNAcHex
c3
c4
HexNAc(2)Hex(1)
c5
M+e - Acetyl
Pep_2+
M+e
c6
c7
M+e-17
z.11++
z.13++
M+2e
M+2e-17
M+3e - Acetyl
M+3e

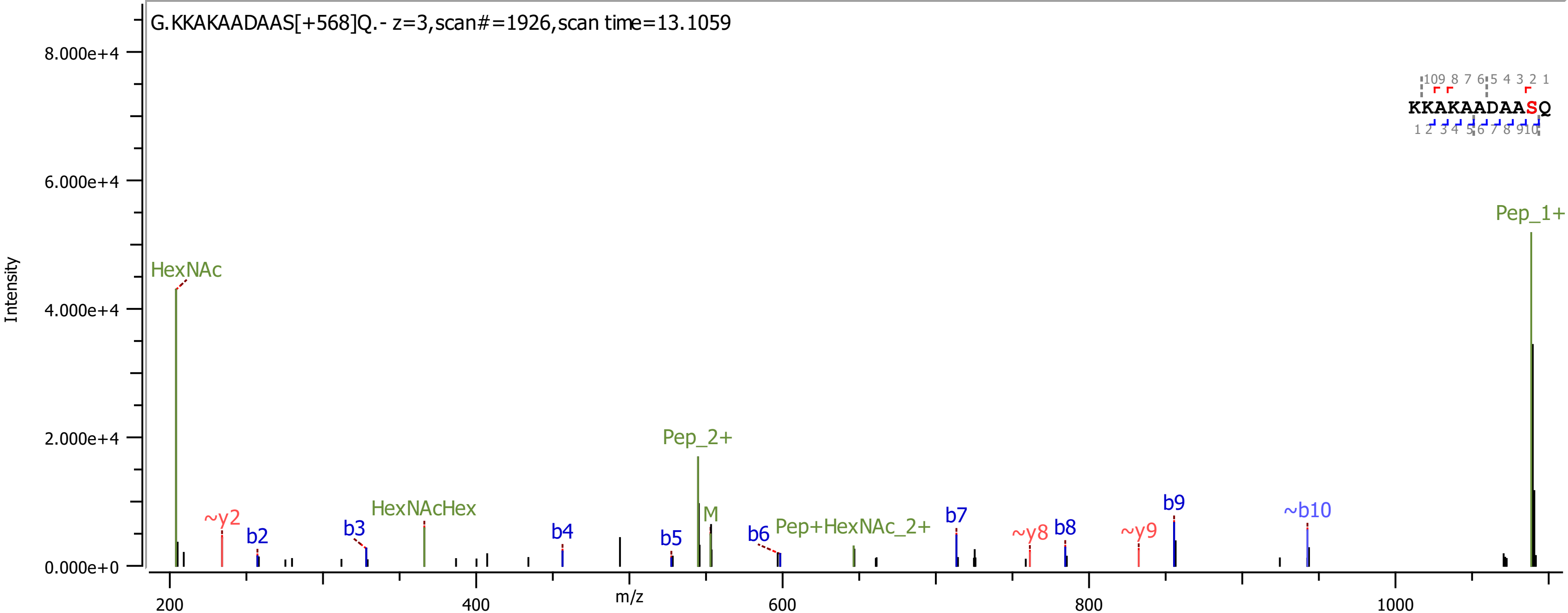

G.KKAKAADAAS[+568]Q.- z=3,scan#=1926,scan time=13.1059
KKAKAADAASQ
HexNAc
~y2
b2
b3
HexNAcHex
b4
b5
Pep_2+
M
b6
Pep+HexNAc_2+
b7
~y8
b8
~y9
b9
~b10
Pep_1+
Intensity
m/z
8.000e+4
6.000e+4
4.000e+4
2.000e+4
0.000e+0
200
400
600
800
1000

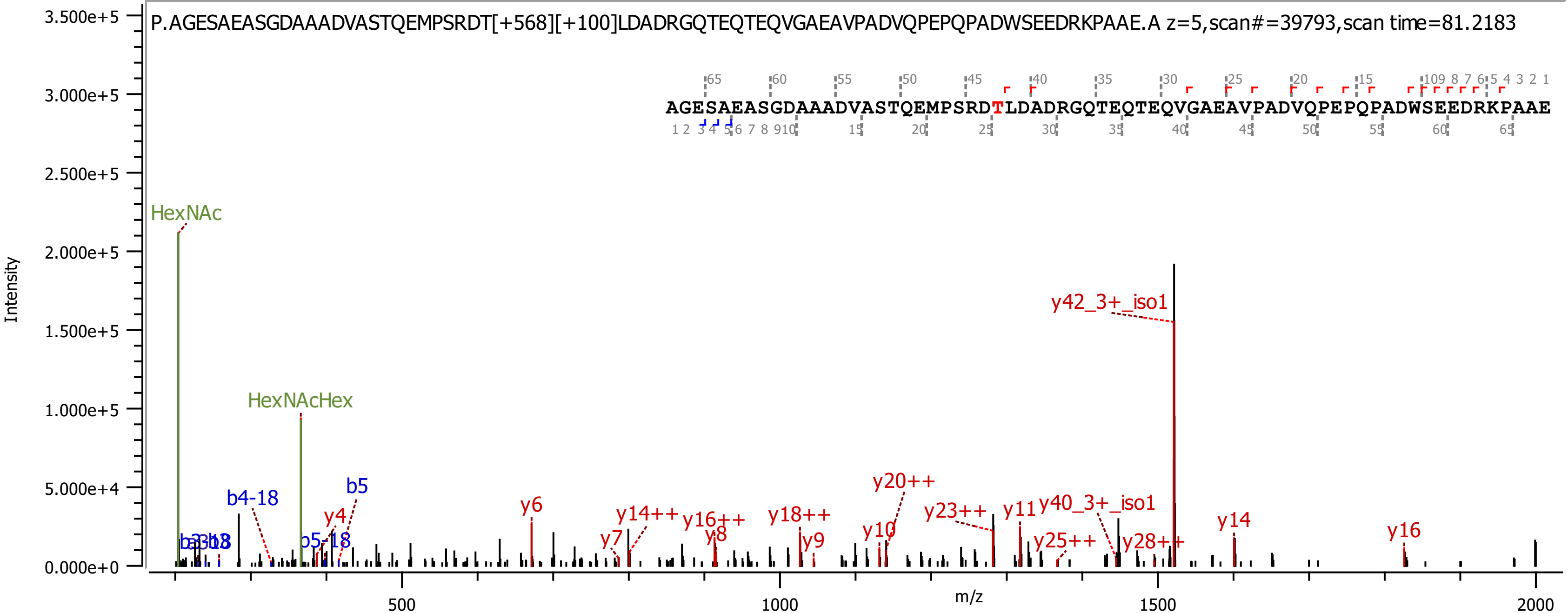

P.AGESAEASGDAAADVASTQEMPSRDT[+568][+100]LDADRGQTEQTEQVGAEAVPADVQPEPQPADWSEEDRKPAAE.A z=5,scan#=39793,scan time=81.2183
AGESAEASGDAAADVASTQEMPSRDTLDADRGQTEQTEQVGAEAVPADVQPEPQPADWSEEDRKPAAE
Intensity
m/z
HexNAc
HexNAcHex
b4-18
b5
b5-18
y4
y6
y7
y14++
y16++
y8
y18++
y9
y10
y20++
y23++
y11
y40_3+_iso1
y42_3+_iso1
y25++
y28++
y14
y16

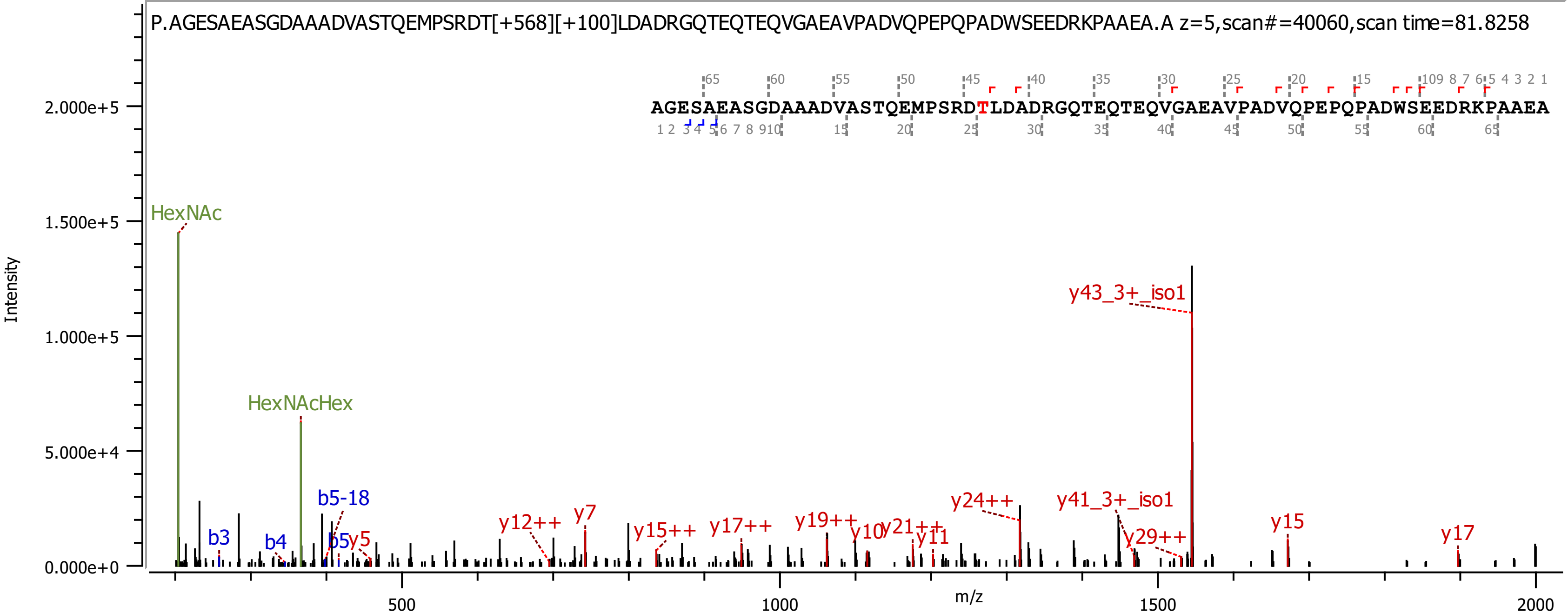
P.AGESAEASGDAAADVASTQEMPSRDT[+568][+100]LDADRGQTEQTEQVGAEAVPADVQPEPQPADWSEEDRKPAAEA.A z=5,scan#=40060,scan time=81.8258
AGESAEASGDAAADVASTQEMPSRDTLDADRGQTEQTEQVGAEAVPADVQPEPQPADWSEEDRKPAAEA
Intensity
m/z
HexNAc
HexNAcHex
b3
b4
b5-18
b5
y5
y12++
y7
y15++
y17++
y19++
y10
y21++
y11
y24++
y43_3+_iso1
y41_3+_iso1
y29++
y15
y17

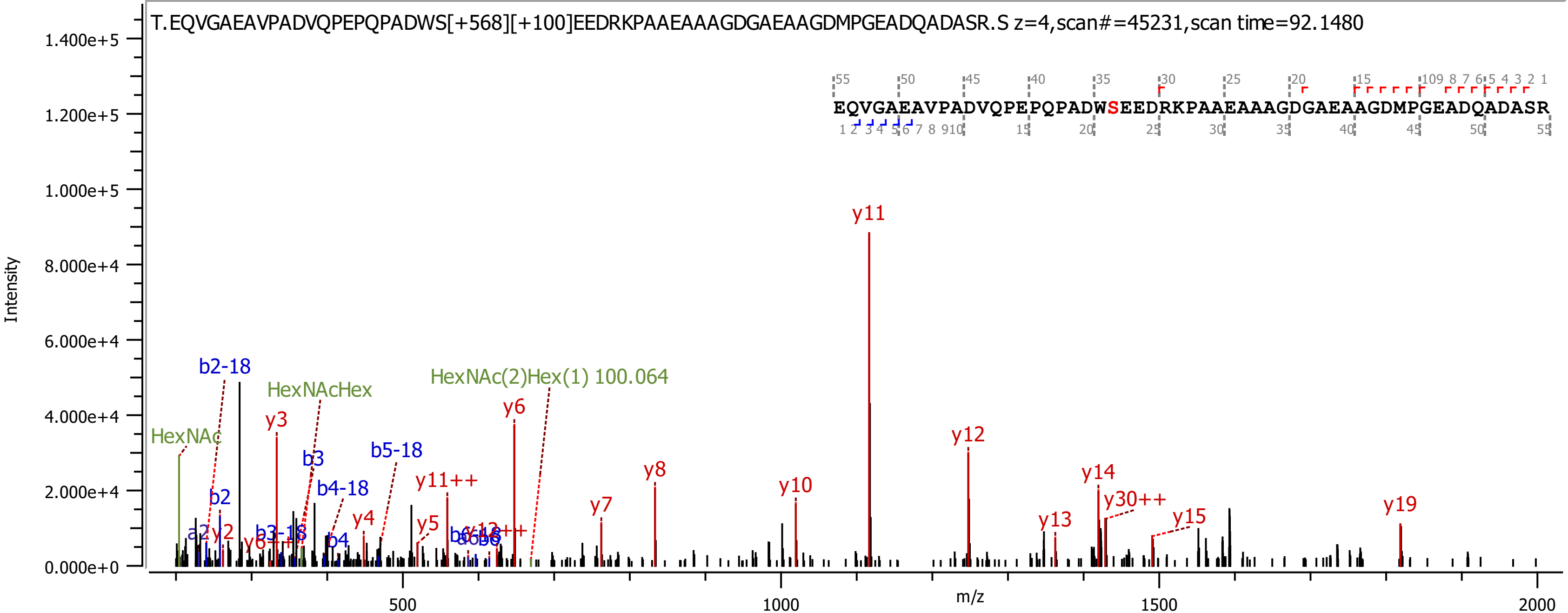

T.EQVGAEAVPADVQPEPQPADWS[+568][+100]EEDRKPAAEAAAGDGAEAAGDMPGEADQADASR.S z=4,scan#=45231,scan time=92.1480
EQVGAEAVPADVQPEPQPADWSEEDRKPAAEAAAGDGAEAAGDMPGEADQADASR
Intensity
m/z
HexNAc
b2-18
HexNAcHex
HexNAc(2)Hex(1) 100.064
y3
b3
b5-18
y6
y11
y12
b2
y2
b3-18
b4-18
b4
y4
y11++
y5
y7
y8
y10
y13
y14
y30++
y15
y19

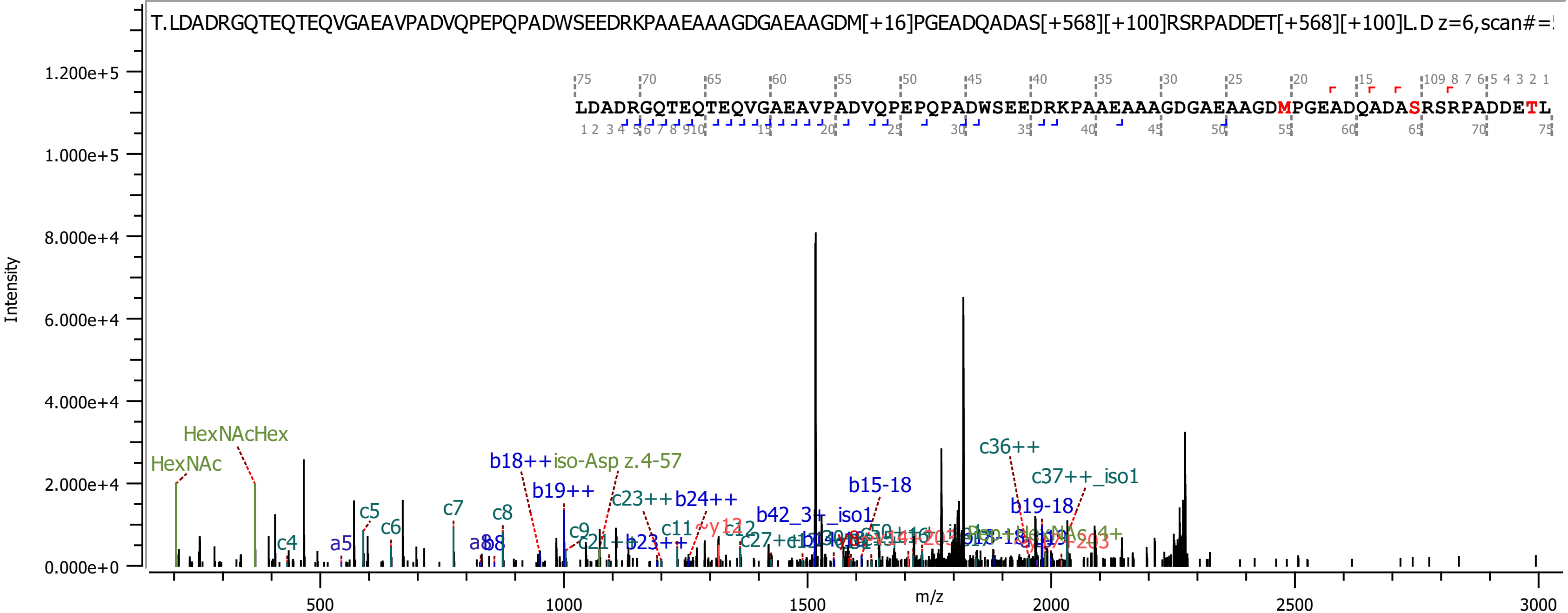
T.LDADRGQTEQTEQVGAEAVPADVQPEPQPADWSEEDRKPAAEAAAGDGAEAAGDM[+16]PGEADQADAS[+568][+100]RSRPADDET[+568][+100]L.D z=6,scan#=
LDADRGQTEQTEQVGAEAVPADVQPEPQPADWSEEDRKPAAEAAAGDGAEAAGDMPGEADQADASRSRPADDETL
Intensity
m/z
1.200e+5
1.000e+5
8.000e+4
6.000e+4
4.000e+4
2.000e+4
0.000e+0
500
1000
1500
2000
2500
3000
HexNAc
HexNAcHex
c4
a5
c5
c6
c7
a8
b8
c8
b18++iso-Asp z.4-57
b19++
c9
c21++
c23++
b23++
b24++
c11
c12
c27++
b42_3+_iso1
b15-18
c36++
c37++_iso1
b19-18

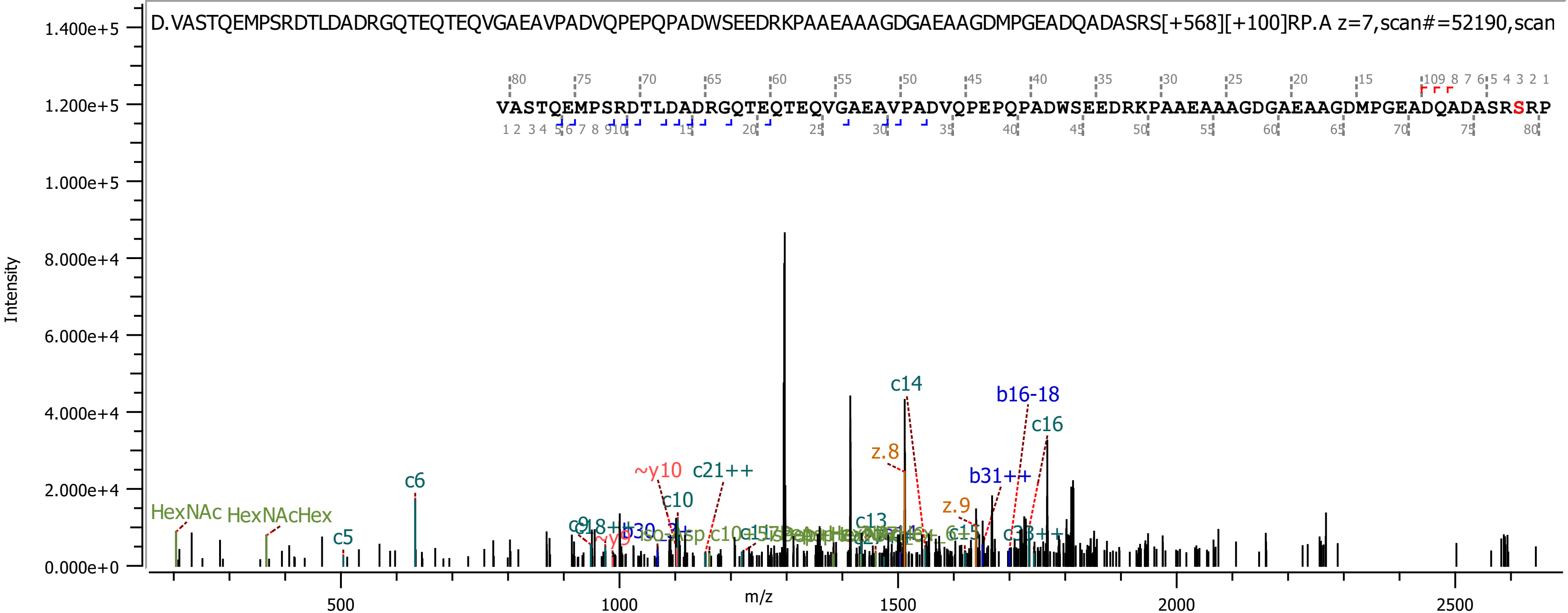

D.VASTQEMPSRDTLDADRGQTEQTEQVGAEAVPADVQPEPQPADWSEEDRKPAAEAAAGDGAEAAGDMPGEADQADASRS[+568][+100]RP.A z=7,scan#=52190,scan
VASTQEMPSRDTLDADRGQTEQTEQVGAEAVPADVQPEPQPADWSEEDRKPAAEAAAGDGAEAAGDMPGEADQADASRSRP
Intensity
m/z
1.400e+5
1.200e+5
1.000e+5
8.000e+4
6.000e+4
4.000e+4
2.000e+4
0.000e+0
500
1000
1500
2000
2500
HexNAc
HexNAcHex
c5
c6
~y10
c21++
c10
c14
z.8
b16-18
c16
b31++
z.9
c13

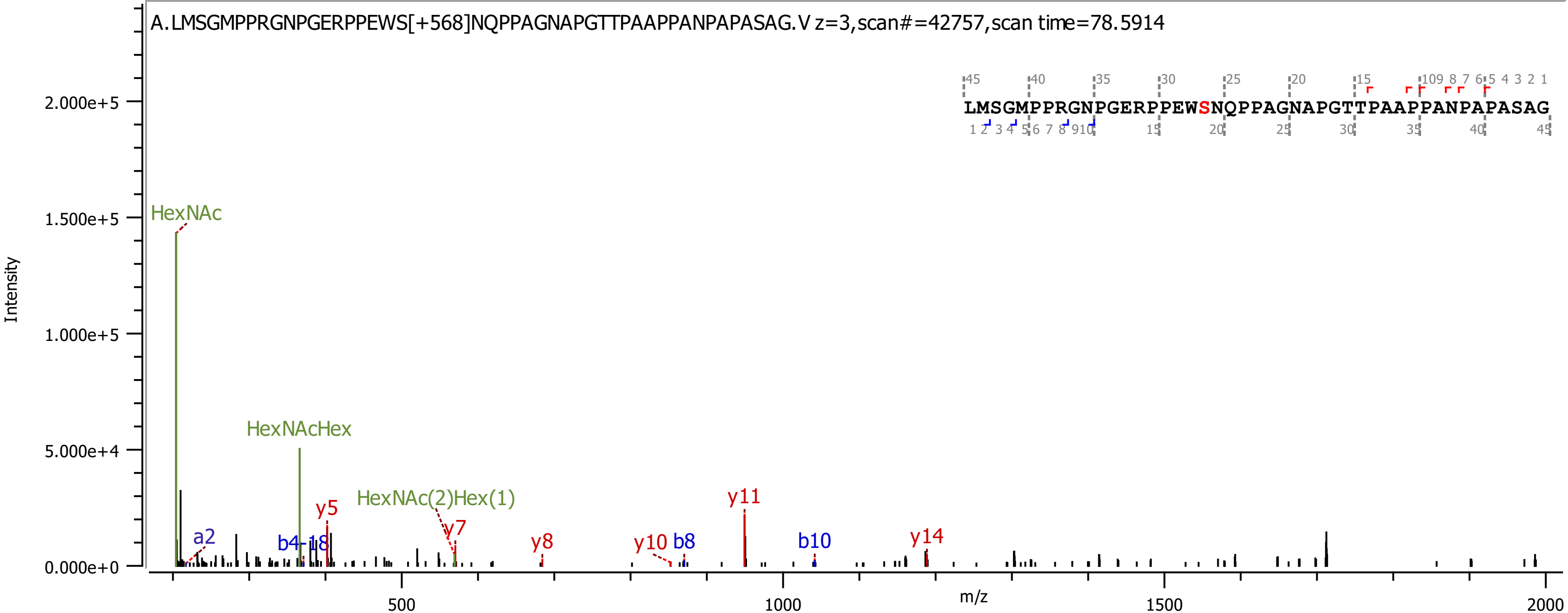

A.LMSGMPPRGNPGERPPEWS[+568]NQPPAGNAPGTTPAAPPANPAPASAG.V z=3,scan#=42757,scan time=78.5914
LMSGMPPRGNPGERPPEWSNQPPAGNAPGTTPAAPPANPAPASAG
Intensity
m/z
HexNAc
HexNAcHex
HexNAc(2)Hex(1)
a2
b4-18
y5
y7
y8
y10
b8
y11
b10
y14

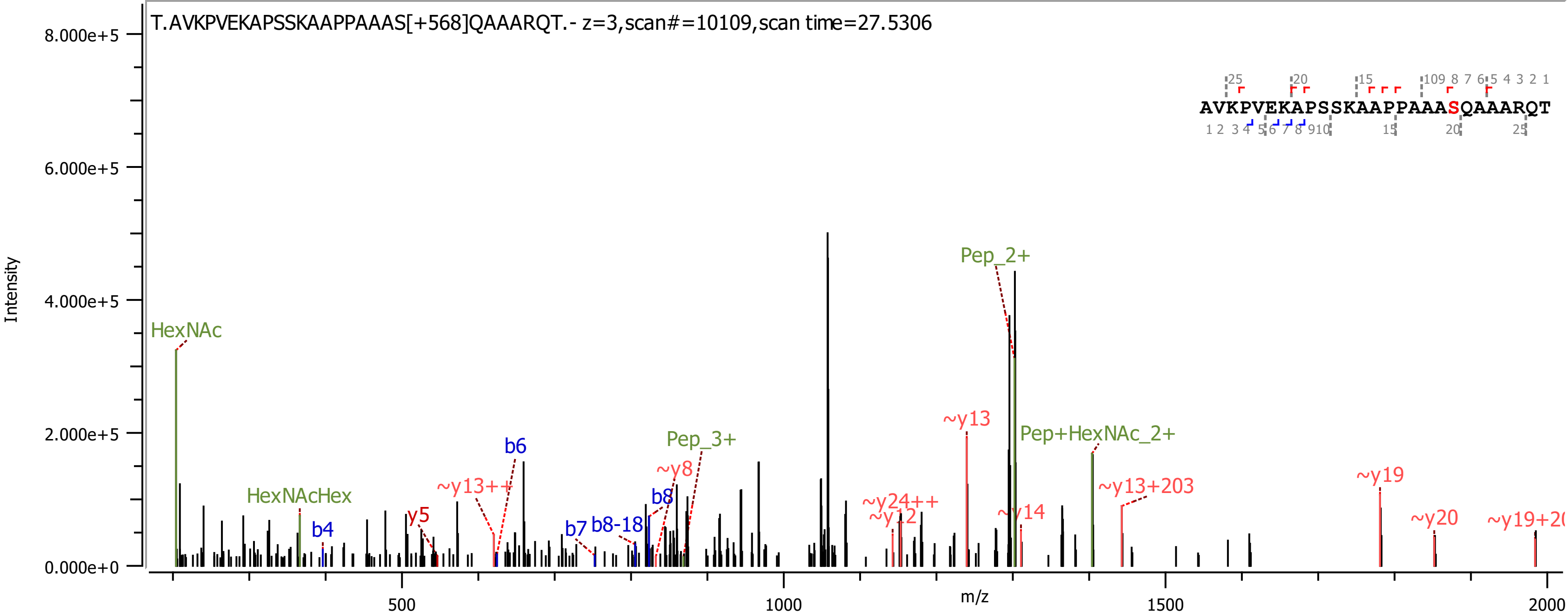

T.AVKPVEKAPSSKAAPPAAAS[+568]QAAARQT.- z=3,scan#=10109,scan time=27.5306
AVKPVEKAPSSKAAPPAAASQAAARQT
Intensity
m/z
HexNAc
HexNAcHex
b4
y5
~y13++
b6
b7
b8-18
b8
~y8
Pep_3+
~y24++
~y12
~y13
~y14
Pep_2+
Pep+HexNAc_2+
~y13+203
~y19
~y20
8.000e+5
6.000e+5
4.000e+5
2.000e+5
0.000e+0
500
1000
1500
2000

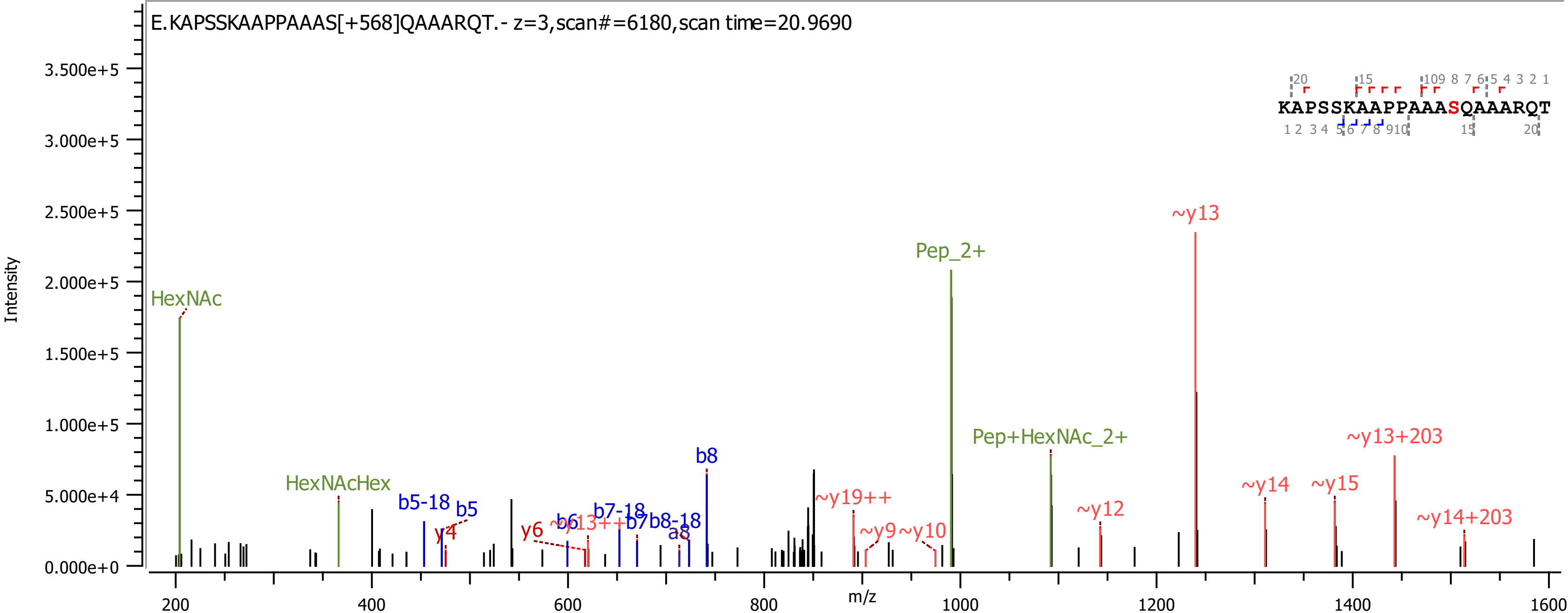
E.KAPSSKAAPPAAAS[+568]QAAARQT.- z=3,scan#=6180,scan time=20.9690
KAPSSKAAPPAAASQAAARQT
Intensity
m/z
HexNAc
HexNAcHex
b5-18
b5
y4
y6
b6
~y13++
b7-18
b7
b8-18
a8
b8
~y19++
~y9
~y10
Pep_2+
Pep+HexNAc_2+
~y12
~y13
~y14
~y15
~y13+203
~y14+203

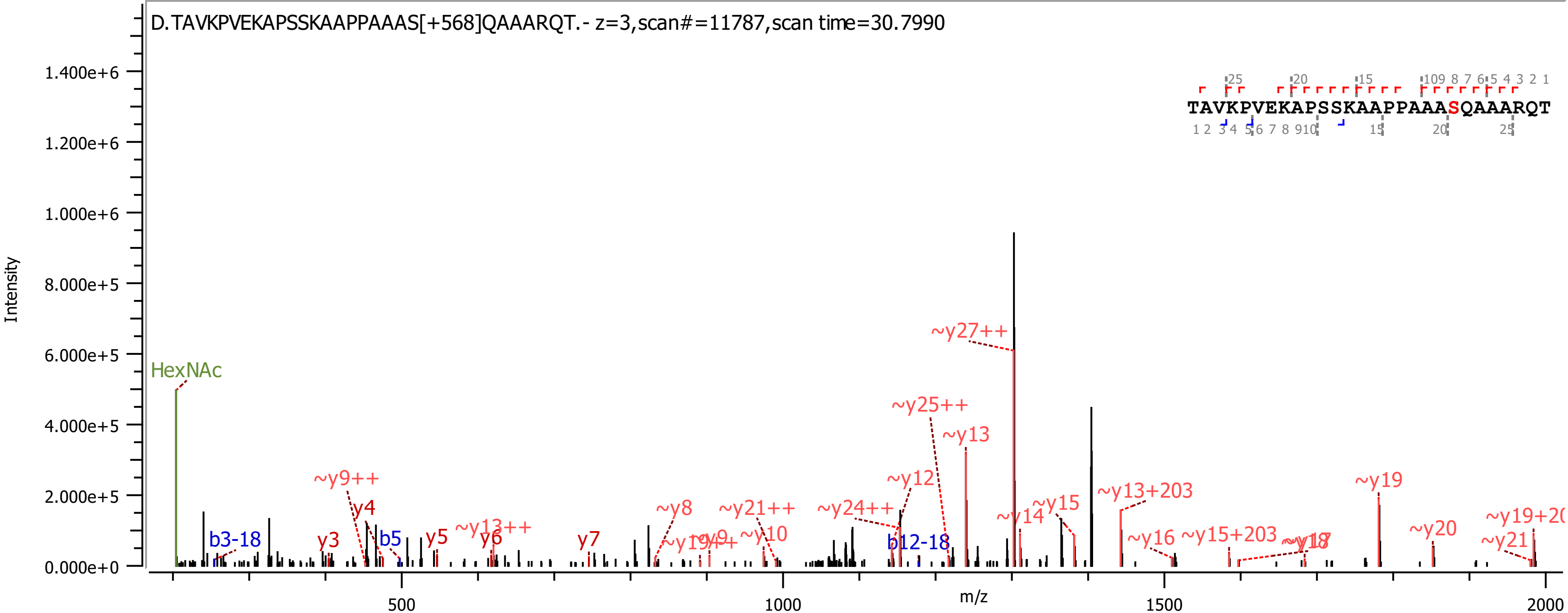
D.TAVKPVEKAPSSKAAPPAAAS[+568]QAAARQT.- z=3,scan#=11787,scan time=30.7990
TAVKPVEKAPSSKAAPPAAASQAAARQT
Intensity
m/z
HexNAc
b3-18
b5
b12-18
y3
y4
y5
y6
y7
~y8
~y9++
~y10
~y12
~y13
~y13++
~y13+203
~y14
~y15
~y15+203
~y16
~y19
~y20
~y21
~y21++
~y24++
~y25++
~y27++

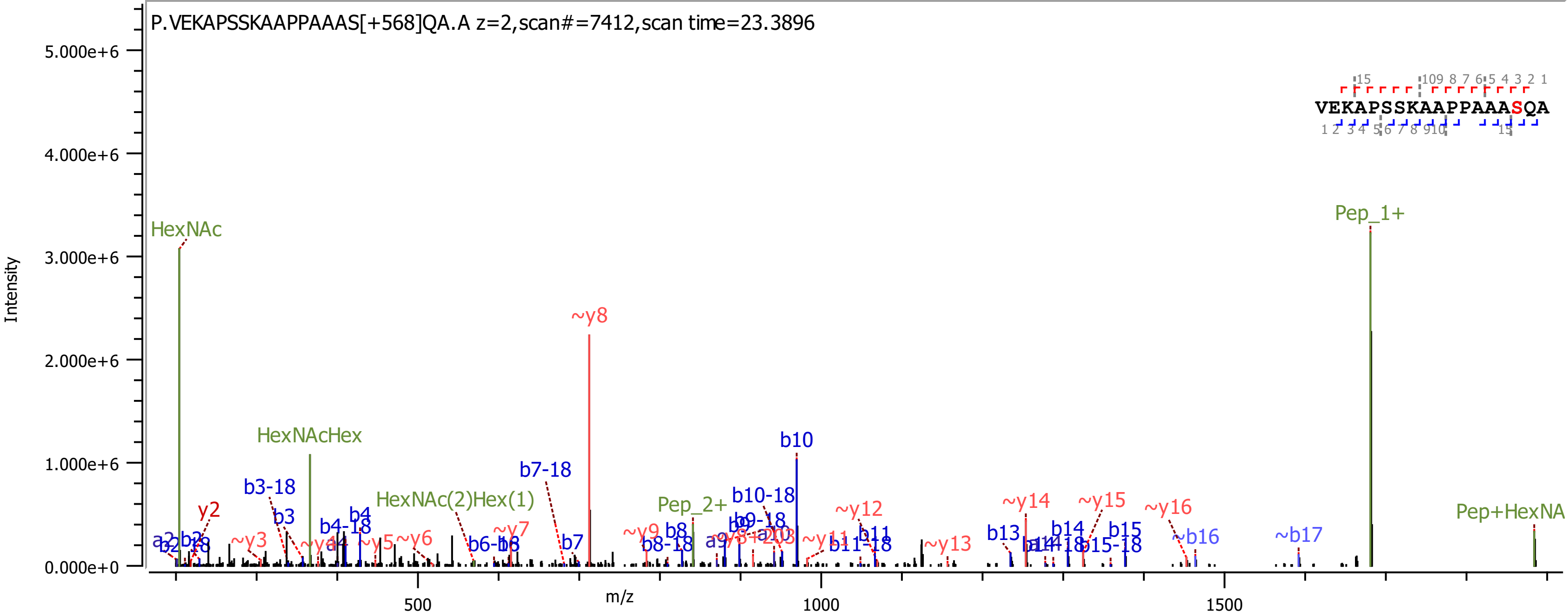

P.VEKAPSSKAAPPAAAS[+568]QA.A z=2,scan#=7412,scan time=23.3896
VEKAPSSKAAPPAAASQA
Intensity
m/z
HexNAc
HexNAcHex
HexNAc(2)Hex(1)
Pep_1+
Pep_2+
Pep+HexNA
~y8
b10
b7-18
b10-18
b3-18
~y14
~y15
~y16
~y12
~b16
~b17

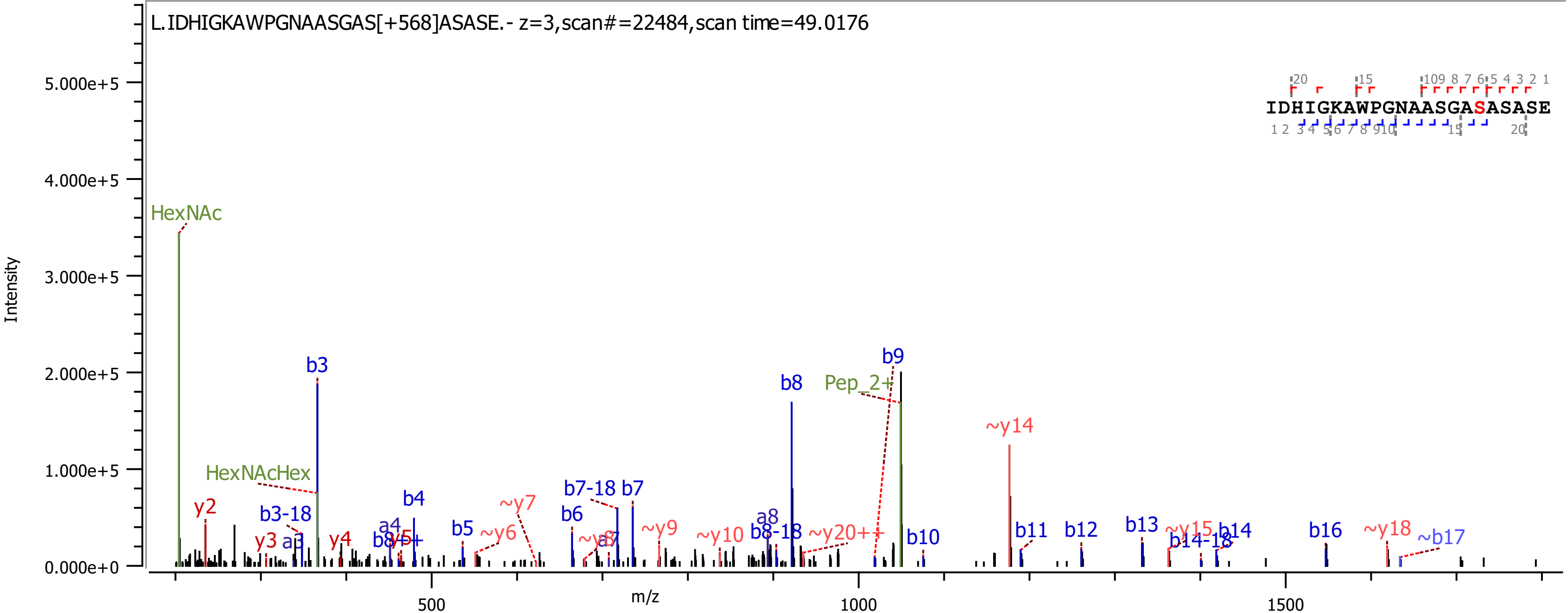

L.IDHIGKAWPGNAASGAS[+568]ASASE.- z=3,scan#=22484,scan time=49.0176
IDHIGKAWPGNAASGASASASE
Intensity
m/z
HexNAc
HexNAcHex
Pep_2+
b3
b4
b5
b6
b7
b8
b9
b10
b11
b12
b13
b14
b16
~b17
b3-18
b7-18
b8-18
b14-18
a3
a4
a8
y2
y3
y4
~y6
~y7
~y9
~y10
~y14
~y15
~y18
~y20++
5.000e+5
4.000e+5
3.000e+5
2.000e+5
1.000e+5
0.000e+0
500
1000
1500

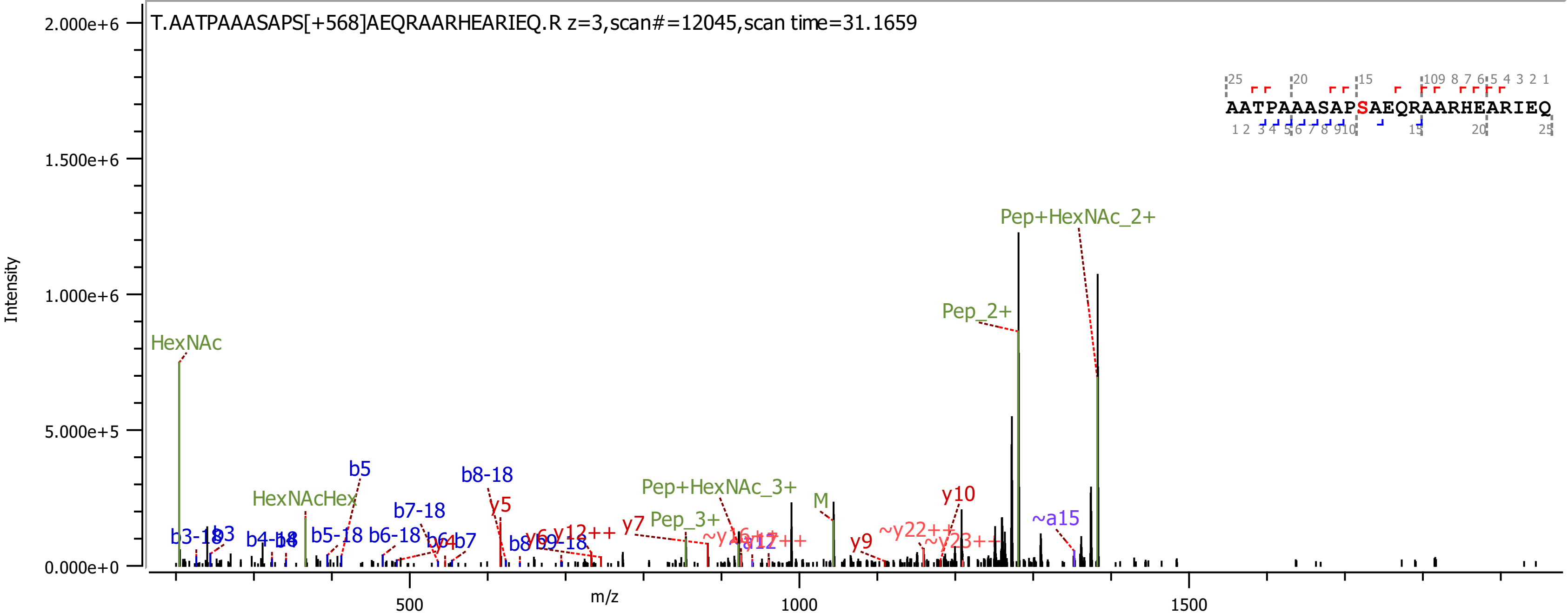
T.AATPAAASAPS[+568]AEQRAARHEARIEQ.R z=3,scan#=12045,scan time=31.1659
Intensity
m/z
AATPAAASAPSAEQRAARHEARIEQ
HexNAc
HexNAcHex
Pep+HexNAc_2+
Pep_2+
Pep+HexNAc_3+
Pep_3+
M
b5
b8-18
b7-18
y5
y10
~a15
y7
y9
~y22++
~y23++
y12++
b5-18
b6-18
b3-18
b3
b4-18
b6
b7
b8
b9-18
y6
~a12

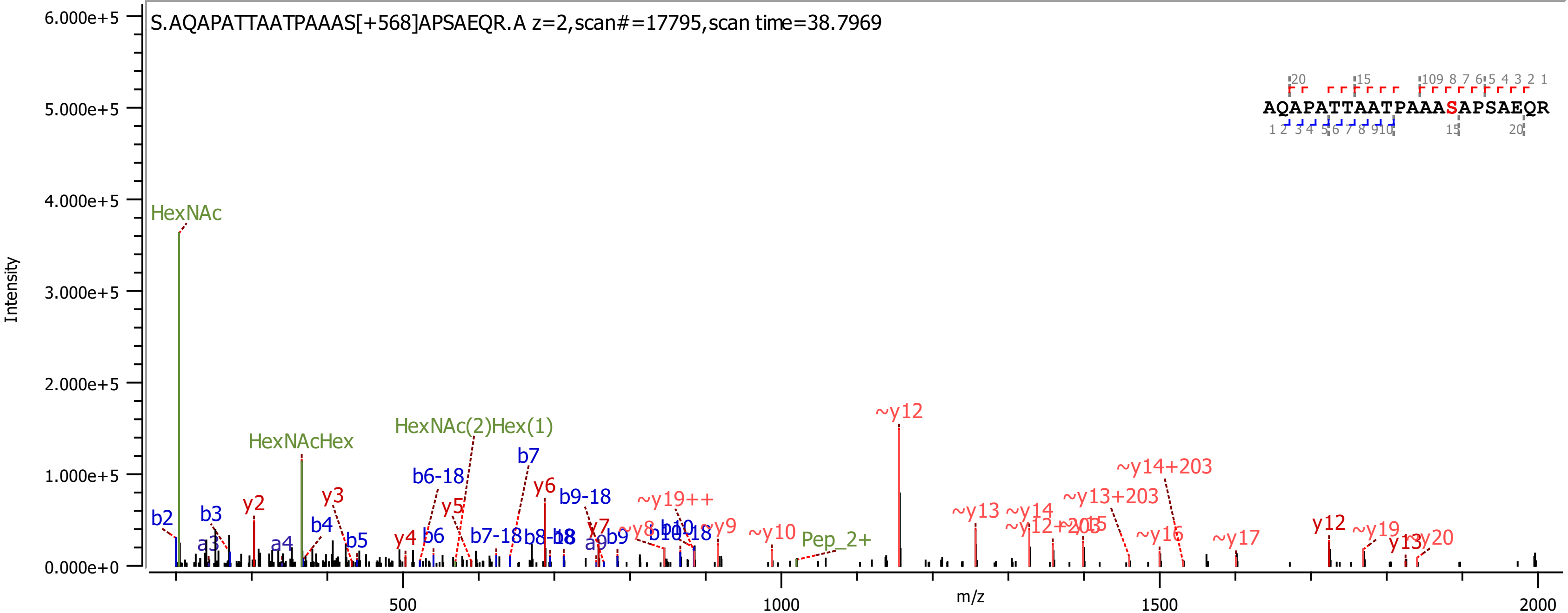
S.AQAPATTAATPAAAS[+568]APSAEQR.A z=2,scan#=17795,scan time=38.7969
AQAPATTAATPAAASAPSAEQR
Intensity
m/z
6.000e+5
5.000e+5
4.000e+5
3.000e+5
2.000e+5
1.000e+5
0.000e+0
500
1000
1500
2000
HexNAc
HexNAcHex
HexNAc(2)Hex(1)
Pep_2+
b2
b3
a3
a4
b4
b5
b6
b6-18
b7
b7-18
b9-18
b9
y2
y3
y4
y5
y6
y7
~y19++
~y9
~y10
~y12
~y13
~y14
~y13+203
~y14+203
~y16
~y17
y12
~y19
~y20

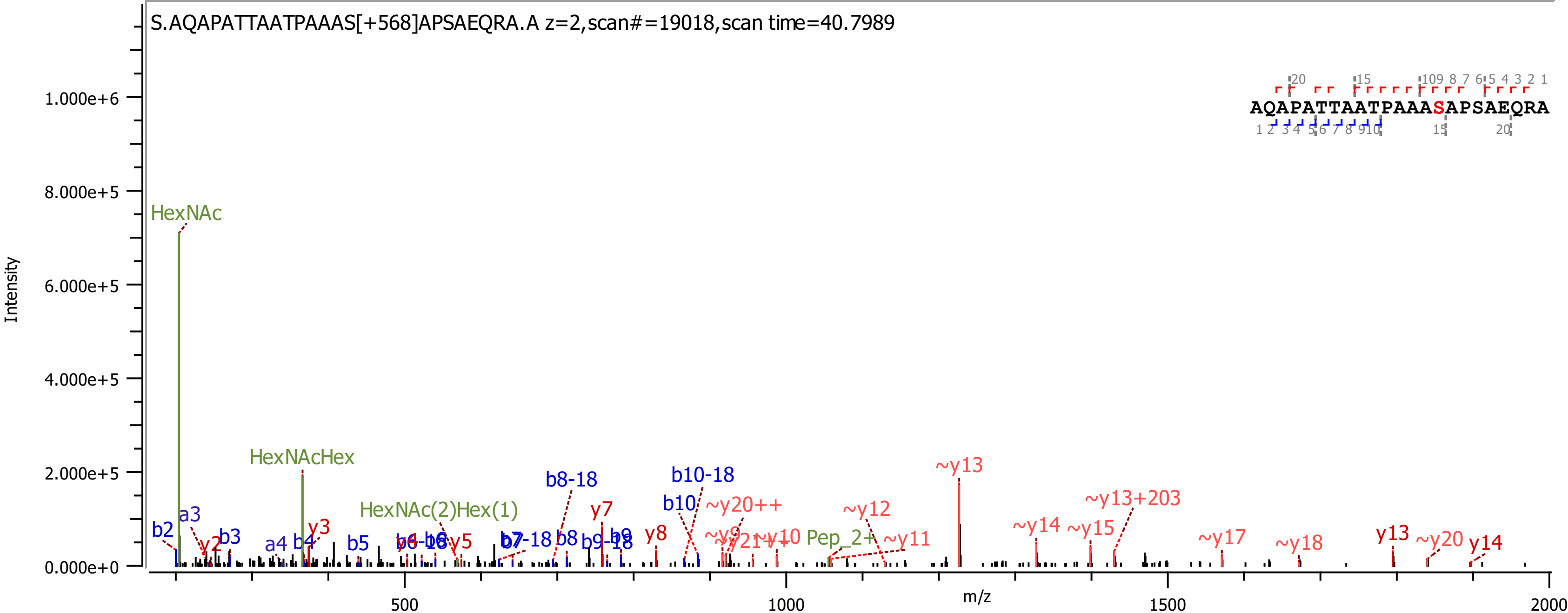
S.AQAPATTAATPAAAS[+568]APSAEQRA.A z=2,scan#=19018,scan time=40.7989
AQAPATTAATPAAASAPSAEQRA
Intensity
m/z
HexNAc
HexNAcHex
HexNAc(2)Hex(1)
b8-18
b10-18
~y13
~y13+203
~y20++
~y12
~y14
~y15
~y17
~y18
~y20
y13
y14
y7
y8
y3
y2
b10
Pep_2+
~y11
~y10
~y9
b2
a3
b3
a4
b4
b5
b7-18
b8
b9-18
b9
y5

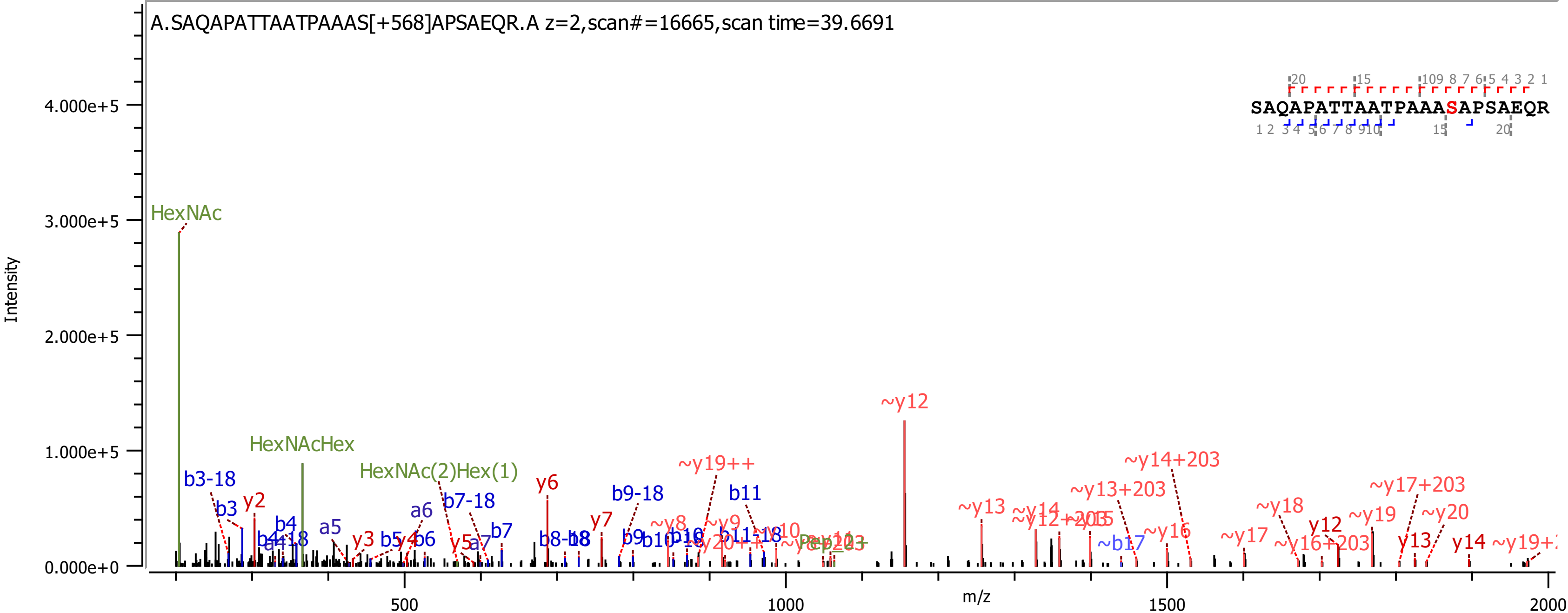

A.SAQAPATTAATPAAAS[+568]APSAEQR.A z=2,scan#=16665,scan time=39.6691
SAQAPATTAATPAAASAPSAEQR
Intensity
4.000e+5
3.000e+5
2.000e+5
1.000e+5
0.000e+0
m/z
500
1000
1500
2000
HexNAc
HexNAcHex
HexNAc(2)Hex(1)
b3-18
b3
y2
b4
a5
y3
b5
y4
b6
a6
b7-18
y5
a7
b7
y6
y7
b9-18
b9
b11
~y19++
~y8
~y9
~y12
~y13
~y14
~y13+203
~y14+203
~b17
~y16
~y17
~y18
~y16+203
y12
~y19
~y17+203
~y20
y13
y14

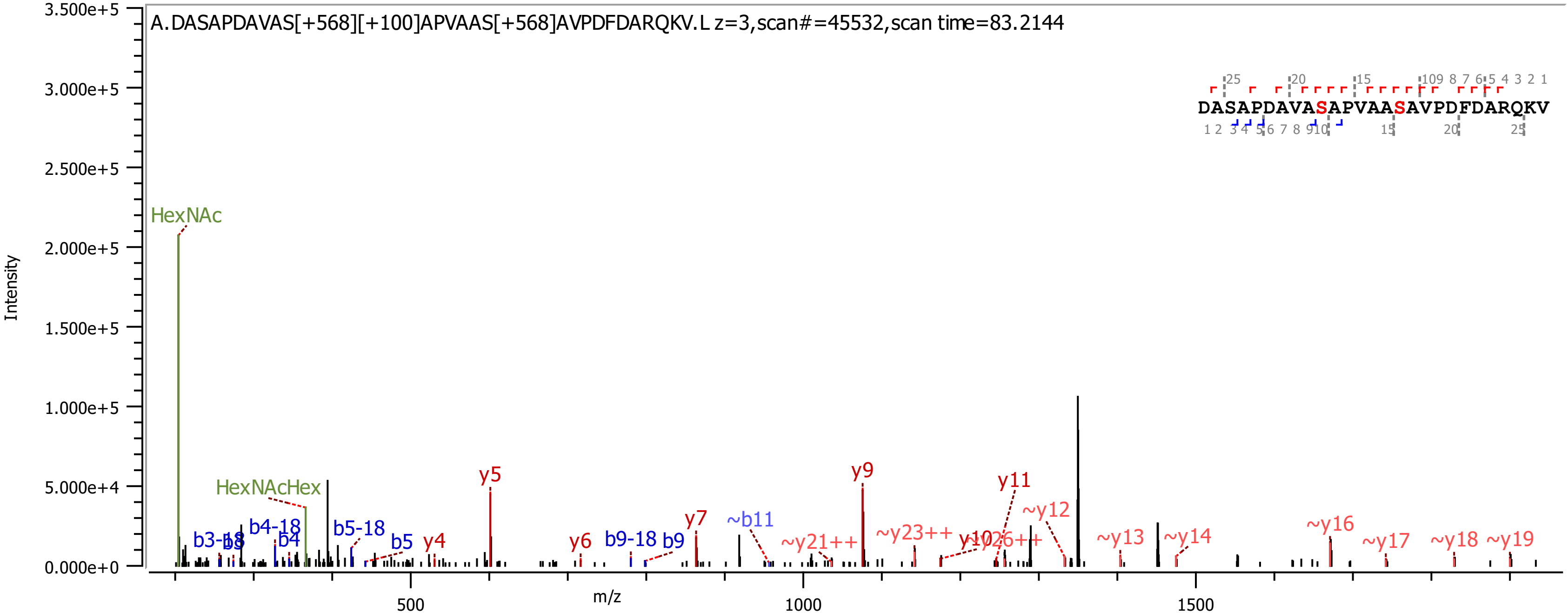
A.DASAPDAVAS[+568][+100]APVAAS[+568]AVPDFDARQKV.L z=3,scan#=45532,scan time=83.2144
DASAPDAVASAPVAASAVPDFDARQKV
Intensity
m/z
HexNAc
HexNAcHex
b3-18
b3
b4-18
b4
b5-18
b5
y4
y5
y6
b9-18
b9
y7
~b11
~y21++
y9
~y23++
y10
~y26++
y11
~y12
~y13
~y14
~y16
~y17
~y18
~y19
0.000e+0
5.000e+4
1.000e+5
1.500e+5
2.000e+5
2.500e+5
3.000e+5
3.500e+5
500
1000
1500

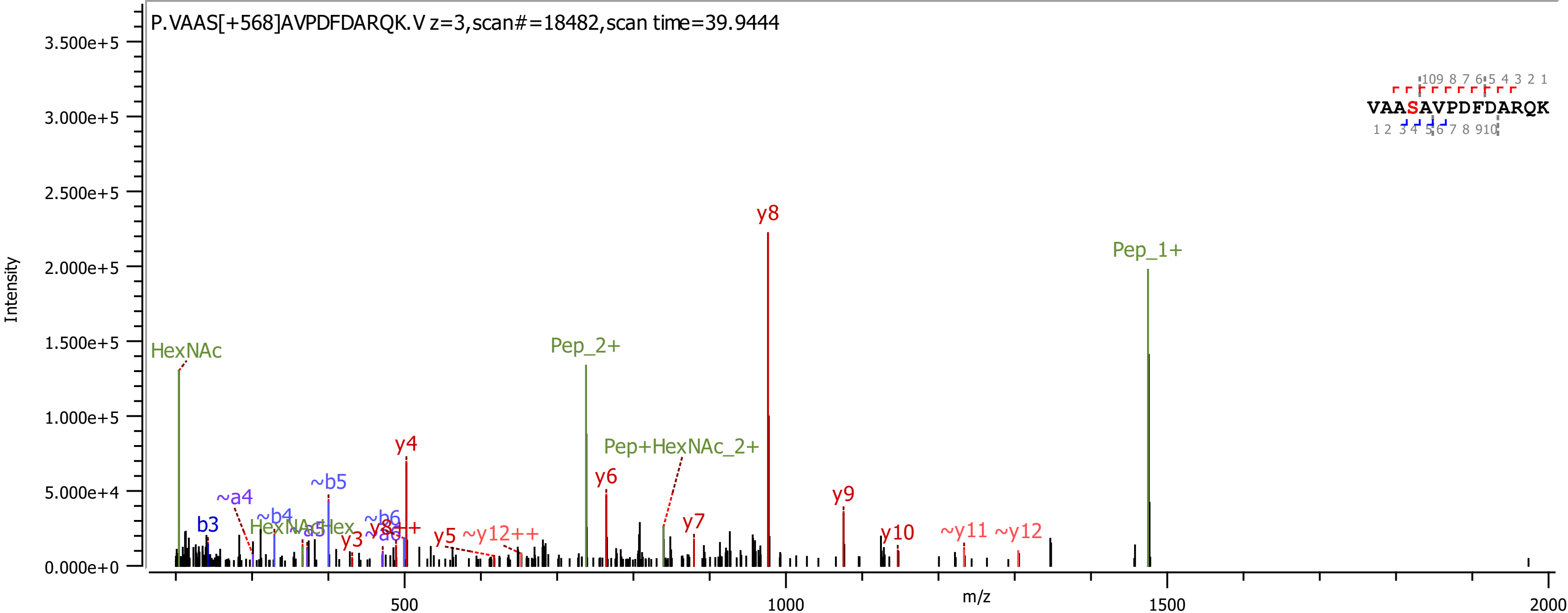

P.VAAS[+568]AVPDFDARQK.V z=3,scan#=18482,scan time=39.9444
Intensity
m/z
HexNAc
b3
~a4
~b4
HexNAc-Hex
~a5
~b5
~b6
~a6
y3
y8++
y4
y5
~y12++
Pep_2+
Pep+HexNAc_2+
y6
y7
y8
y9
y10
~y11
~y12
Pep_1+
VAASAVPDFDARQK

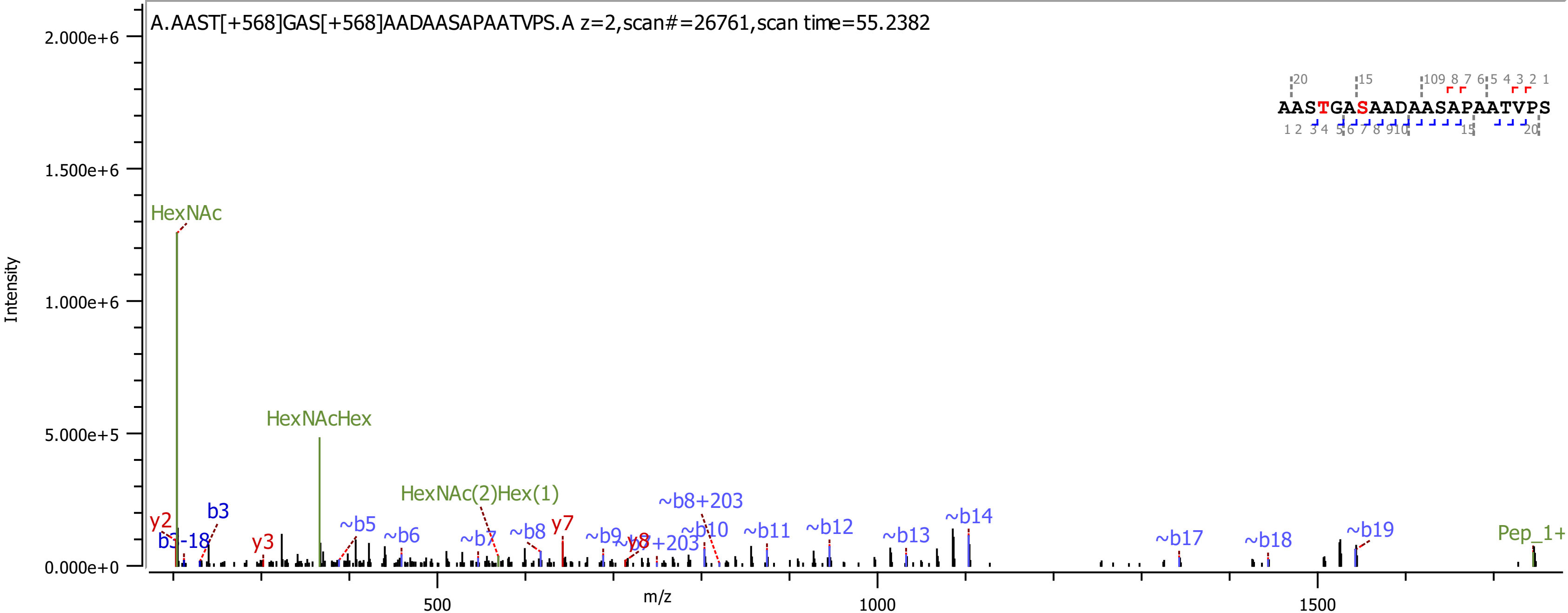

A.AAST[+568]GAS[+568]AADAASAPAATVPS.A z=2,scan#=26761,scan time=55.2382
AASTGASAADAASAPAATVPS
Intensity
m/z
HexNAc
HexNAcHex
HexNAc(2)Hex(1)
~b8+203
~b14
~b12
~b10
~b11
y7
~b5
~b6
~b7
~b8
~b9
~b13
~b17
~b18
~b19
Pep_1+
y2
b3
b3-18
y3
y8
~b7+203

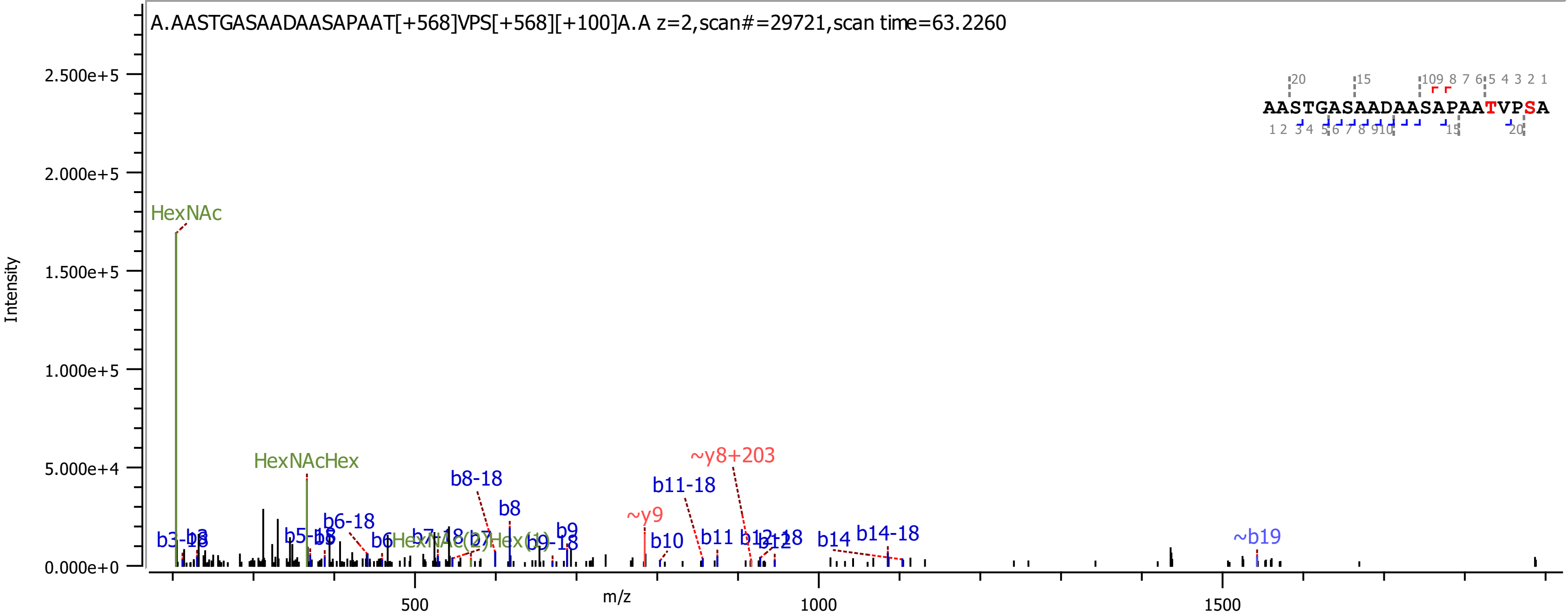

A.AASTGASAADAASAPAAT[+568]VPS[+568][+100]A.A z=2,scan#=29721,scan time=63.2260
AASTGASAADAASAPAATVPSA
Intensity
m/z
HexNAc
HexNAcHex
b8-18
b8
b6-18
b11-18
~y8+203
~y9
b10
b11
b14
b14-18
~b19
b9

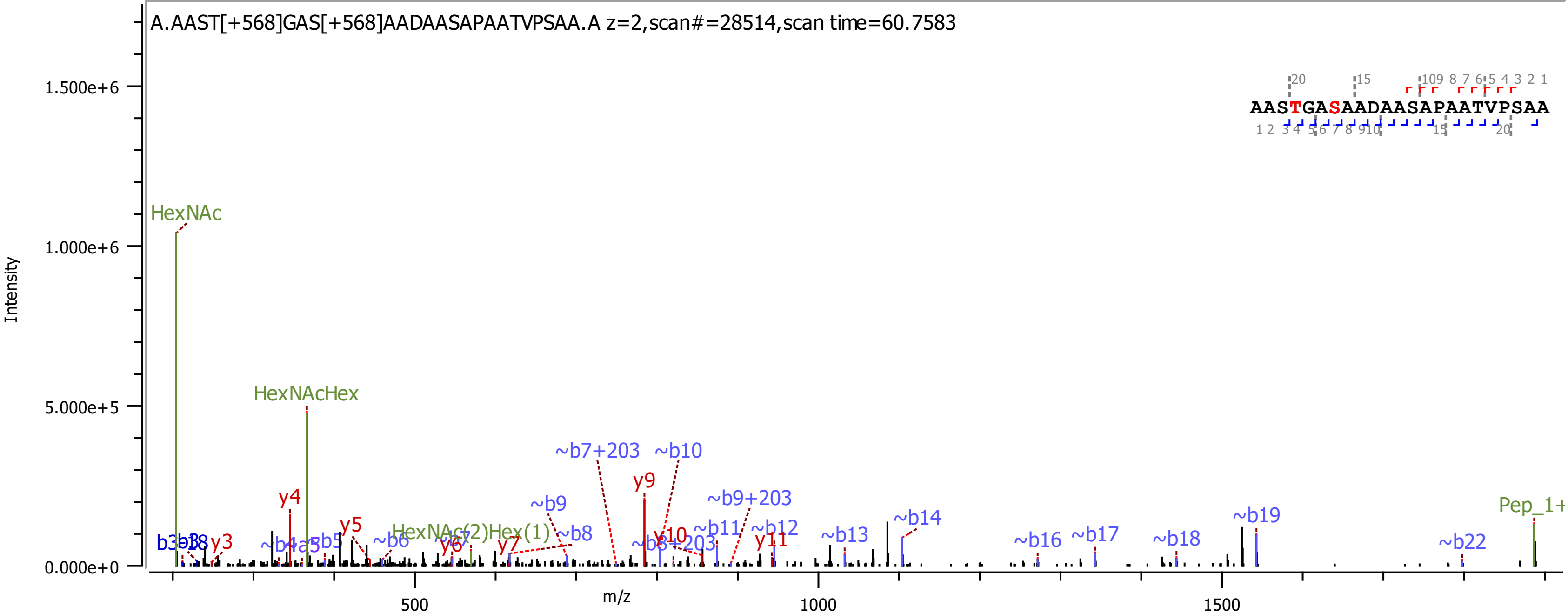
A.AAST[+568]GAS[+568]AADAASAPAATVPSAA.A z=2,scan#=28514,scan time=60.7583
AASTGASAADAASAPAATVPSAA
Intensity
m/z
1.500e+6
1.000e+6
5.000e+5
0.000e+0
500
1000
1500
HexNAc
HexNAcHex
HexNAc(2)Hex(1)
Pep_1+
y3
y4
y5
y6
y7
y9
y10
y11
~b5
~b6
~b8
~b9
~b10
~b7+203
~b9+203
~b11
~b12
~b13
~b14
~b16
~b17
~b18
~b19
~b22

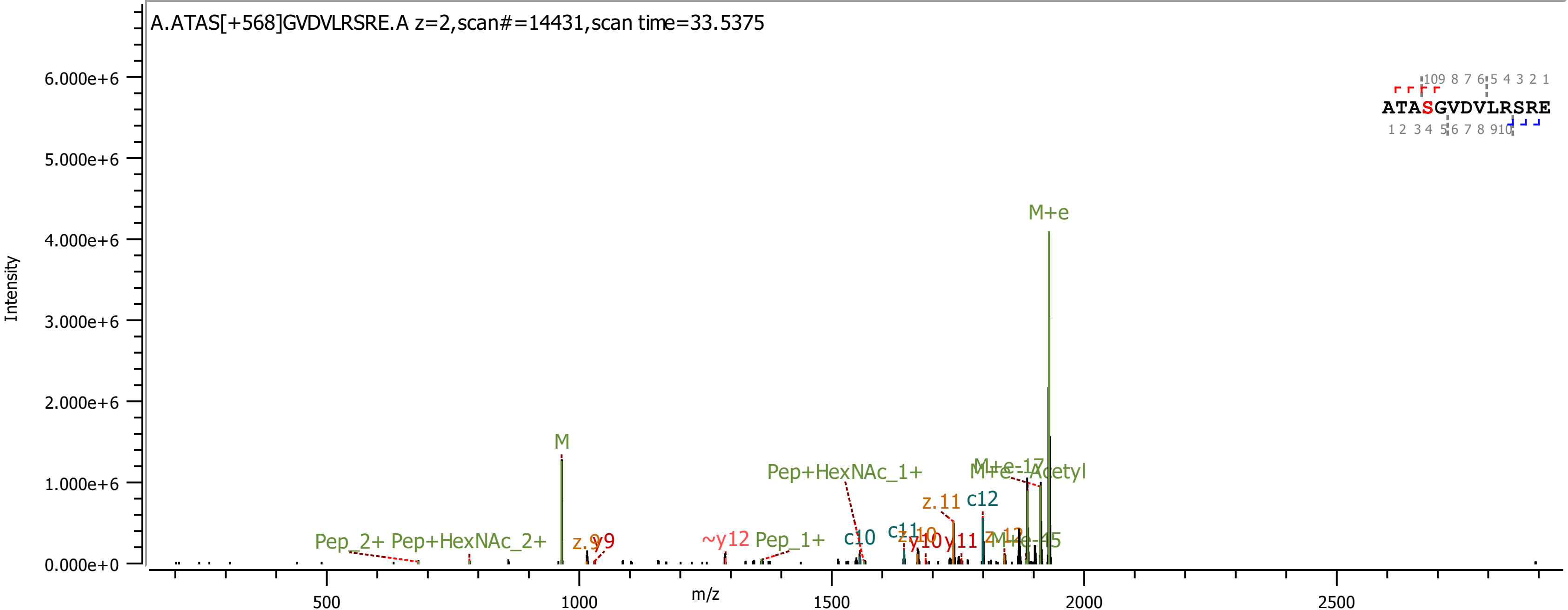

A.ATAS[+568]GVDVLRSRE.A z=2,scan#=14431,scan time=33.5375
ATASGVDVLRSRE
Intensity
6.000e+6
5.000e+6
4.000e+6
3.000e+6
2.000e+6
1.000e+6
0.000e+0
m/z
500
1000
1500
2000
2500
M+e
M
Pep+HexNAc_1+
M+e-Acetyl
z.11
c12
Pep_2+
Pep+HexNAc_2+
~y12
Pep_1+
c10
y10
y11

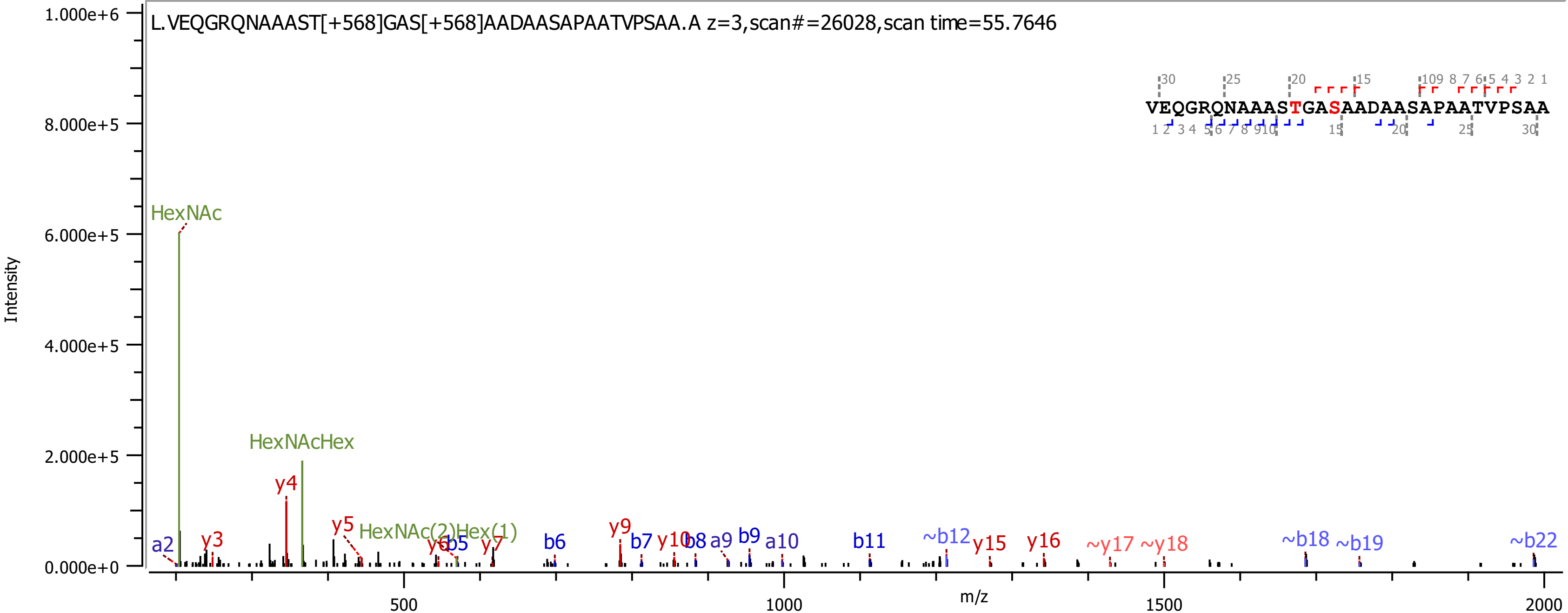
L.VEQGRQNAAAST[+568]GAS[+568]AADAASAPAATVPSAA.A z=3,scan#=26028,scan time=55.7646
VEQGRQNAAASTGASAADAASAPAATVPSAA
Intensity
m/z
HexNAc
HexNAcHex
HexNAc(2)Hex(1)
a2
y3
y4
y5
y6
b5
y7
b6
y9
b7
y10
b8
a9
b9
a10
b11
~b12
y15
y16
~y17
~y18
~b18
~b19
~b22
1.000e+6
8.000e+5
6.000e+5
4.000e+5
2.000e+5
0.000e+0
500
1000
1500
2000

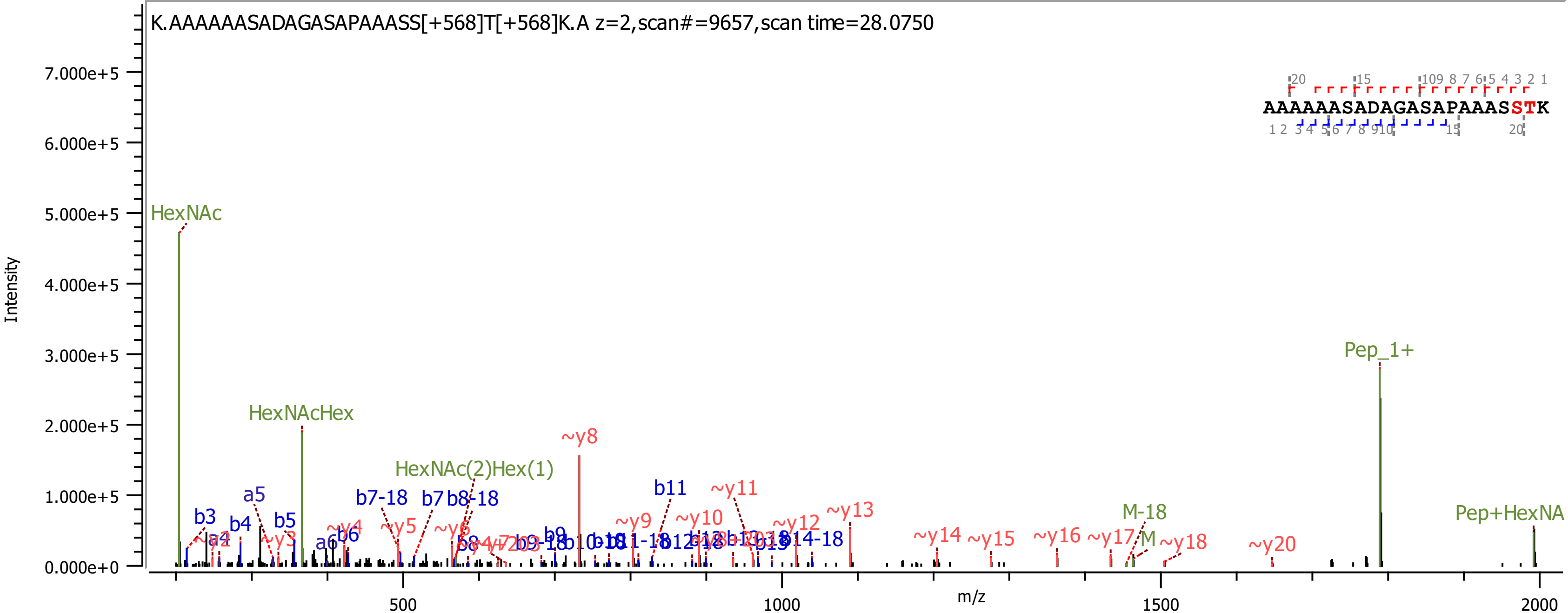
K.AAAAAASADAGASAPAAASS[+568]T[+568]K.A z=2,scan#=9657,scan time=28.0750
AAAAAASADAGASAPAAASSTK
Intensity
7.000e+5
6.000e+5
5.000e+5
4.000e+5
3.000e+5
2.000e+5
1.000e+5
0.000e+0
m/z
500
1000
1500
2000
HexNAc
HexNAcHex
HexNAc(2)Hex(1)
Pep_1+
Pep+HexNA
M-18
M
b3
b4
a5
b5
a6
b6
b7-18
b7
b8-18
b8
b9
b11
~y3
~y4
~y5
~y6
~y8
~y9
~y10
~y11
~y12
~y13
~y14
~y15
~y16
~y17
~y18
~y20

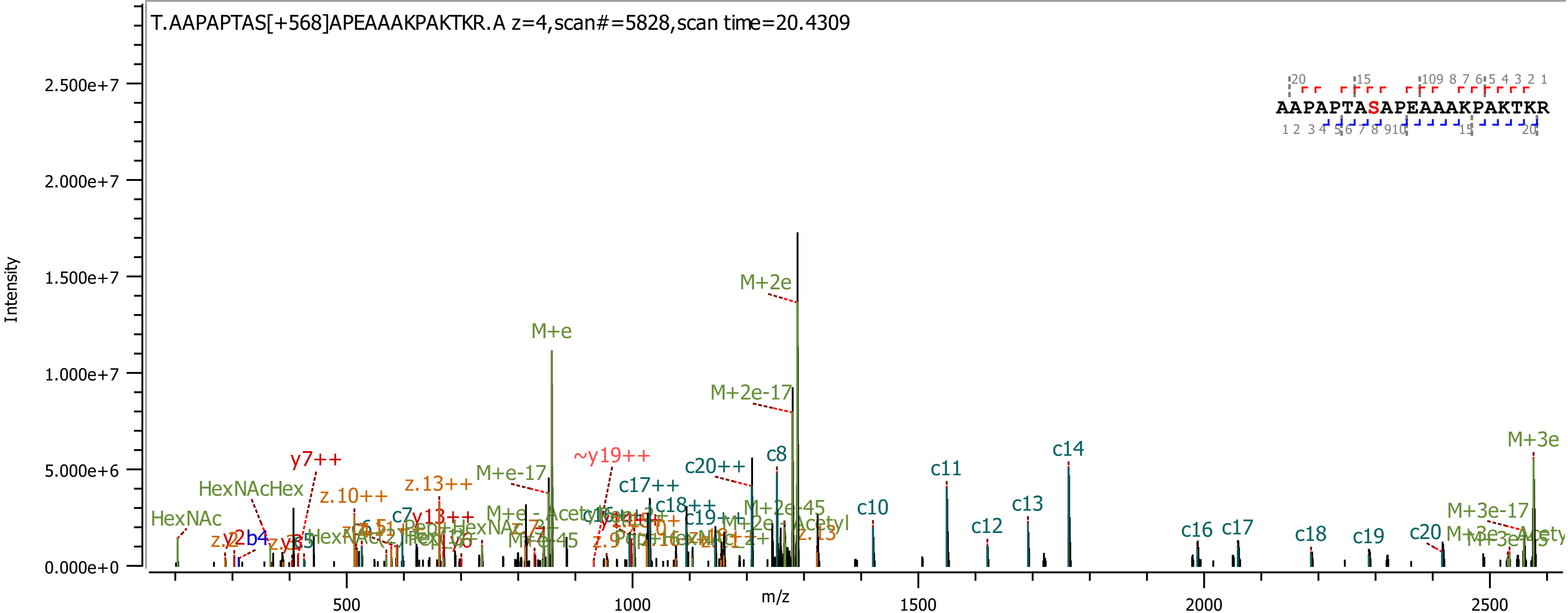
T.AAPAPTAS[+568]APEAAAKPAKTKR.A z=4,scan#=5828,scan time=20.4309
AAPAPTASAPEAAAKPAKTKR
Intensity
m/z
2.500e+7
2.000e+7
1.500e+7
1.000e+7
5.000e+6
0.000e+0
500
1000
1500
2000
2500
M+2e
M+e
M+2e-17
M+e-17
~y19++
c20++
c8
c17++
c18++
c10
c11
c12
c13
c14
c16
c17
c18
c19
c20
M+3e
M+3e-17
M+2e-45
y7++
z.10++
z.13++
y13++
HexNAcHex
HexNAc
b4

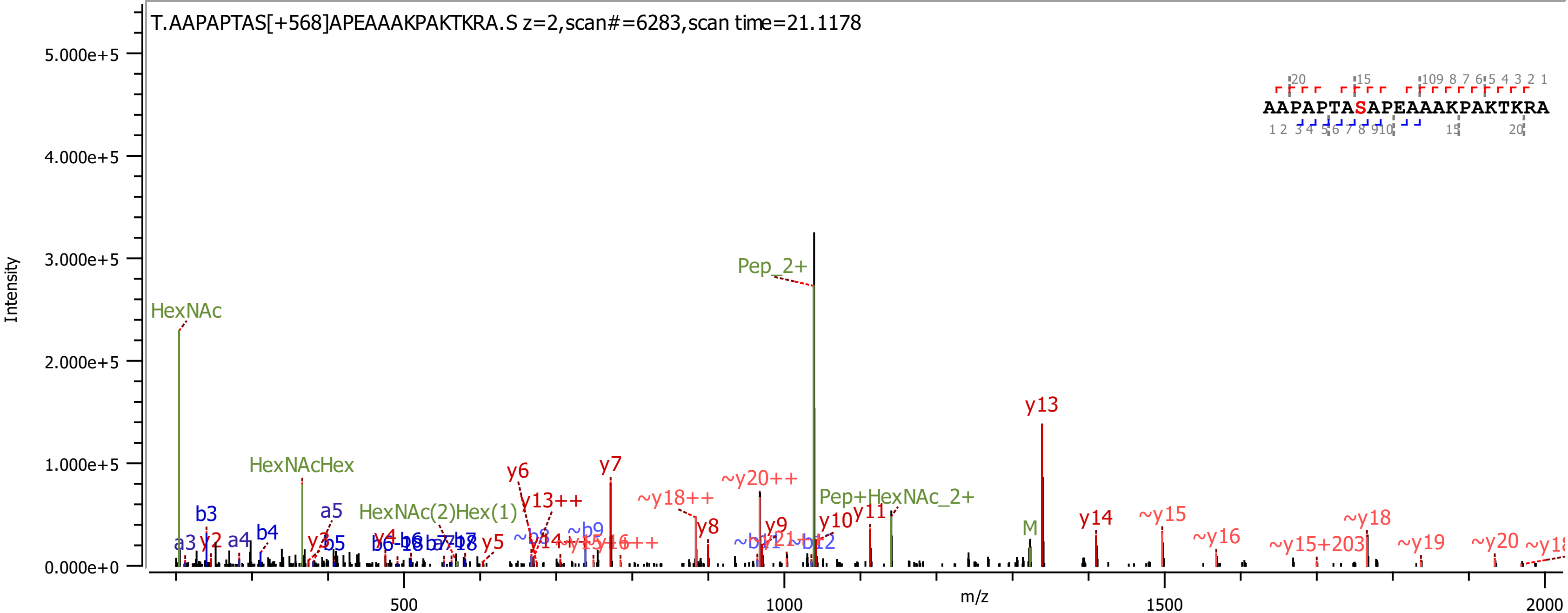

T.AAPAPTAS[+568]APEAAAKPAKTKRA.S z=2,scan#=6283,scan time=21.1178
AAPAPTASAPEAAAKPAKTKRA
Intensity
m/z
Pep_2+
HexNAc
HexNAcHex
HexNAc(2)Hex(1)
Pep+HexNAc_2+
y13
y7
y6
y13++
~y20++
~y18++
y8
y9
y10
y11
y14
~y15
~y18
~y16
~y15+203
~y19
~y20
b3
b4
a5
b5
a3
a4
y2
y3
y4
y5
b6
~b9
M

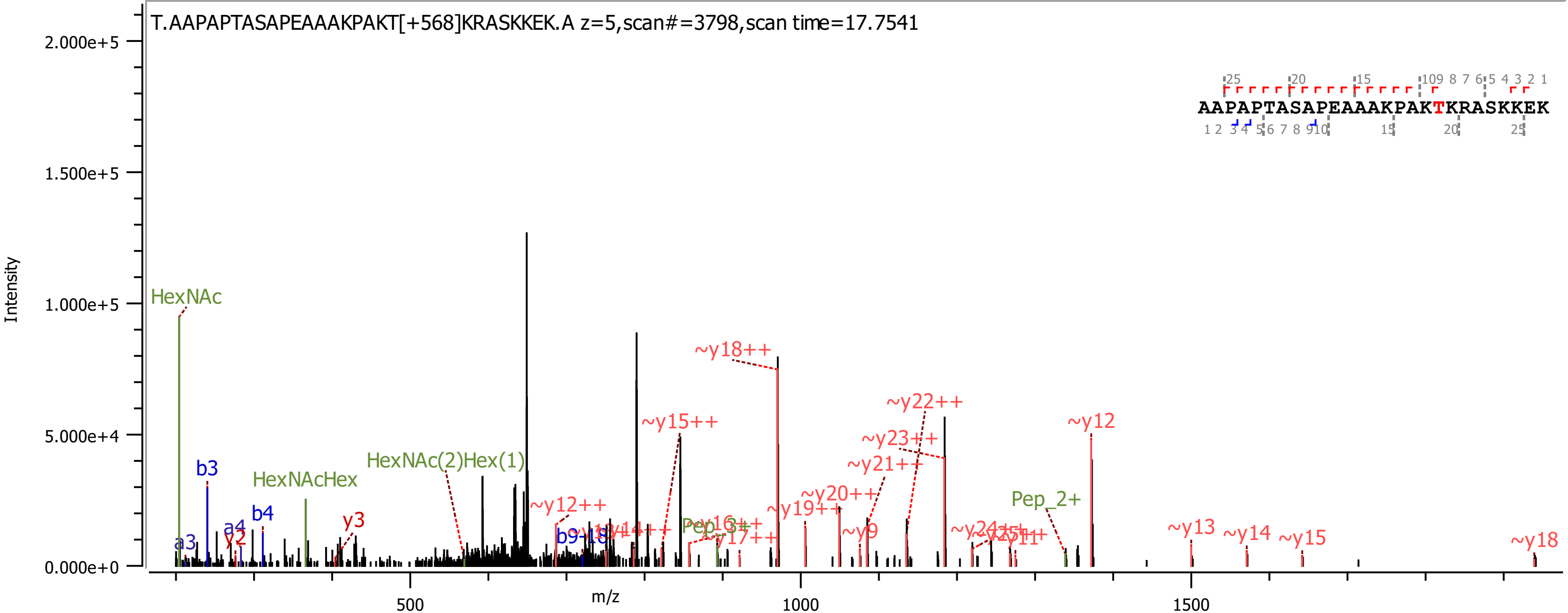

T.AAPAPTASAPEAAAKPAKT[+568]KRASKKEK.A z=5,scan#=3798,scan time=17.7541
AAPAPTASAPEAAAKPAKTKRASKKEK
Intensity
2.000e+5
1.500e+5
1.000e+5
5.000e+4
0.000e+0
m/z
500
1000
1500
HexNAc
b3
a3
a4
y2
b4
HexNAcHex
y3
HexNAc(2)Hex(1)
~y12++
~y15++
~y18++
~y19++
~y20++
~y21++
~y22++
~y23++
~y17++
~y9
Pep_2+
~y12
~y13
~y14
~y15
~y18

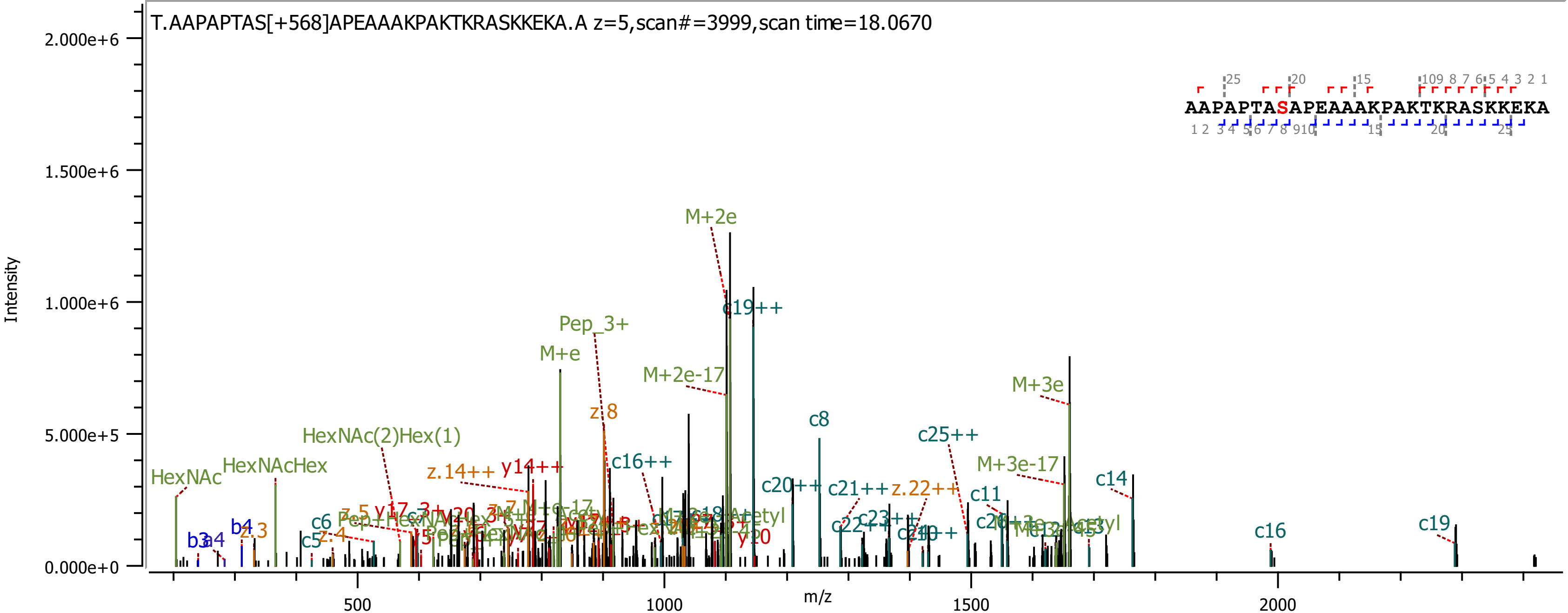
T.AAPAPTAS[+568]APEAAAKPAKTKRASKKEKA.A z=5,scan#=3999,scan time=18.0670
AAPAPTASAPEAAAKPAKTKRASKKEKA
Intensity
m/z
2.000e+6
1.500e+6
1.000e+6
5.000e+5
0.000e+0
500
1000
1500
2000
M+2e
c19++
Pep_3+
M+e
M+2e-17
z.8
c8
c25++
M+3e
M+3e-17
HexNAc(2)Hex(1)
HexNAcHex
HexNAc
z.14++
y14++
c16++
c20++
c21++
z.22++
c11
c14
c16
c19
b3
a4
b4
z.3
z.4
c5
c6
y20

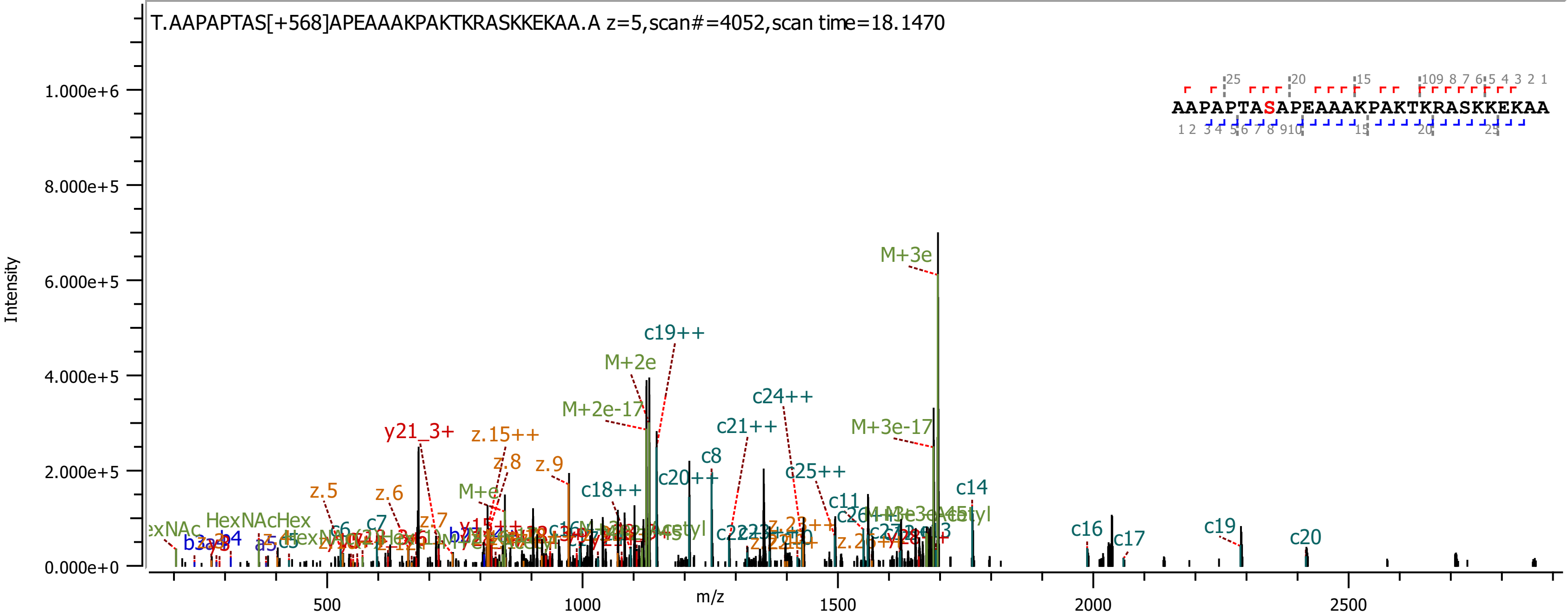

T.AAPAPTAS[+568]APEAAAKPAKTKRASKKEKAA.A z=5,scan#=4052,scan time=18.1470
AAPAPTASAPEAAAKPAKTKRASKKEKAA
Intensity
m/z
1.000e+6
8.000e+5
6.000e+5
4.000e+5
2.000e+5
0.000e+0
500
1000
1500
2000
2500
M+3e
c19++
M+2e
M+2e-17
c24++
c21++
M+3e-17
y21_3+
z.15++
z.8
z.9
c8
c20++
c25++
z.5
z.6
M+e
c18++
c14
c11
HexNAcHex
c16
c17
c19
c20
c6
c7
z.7
b4
a5

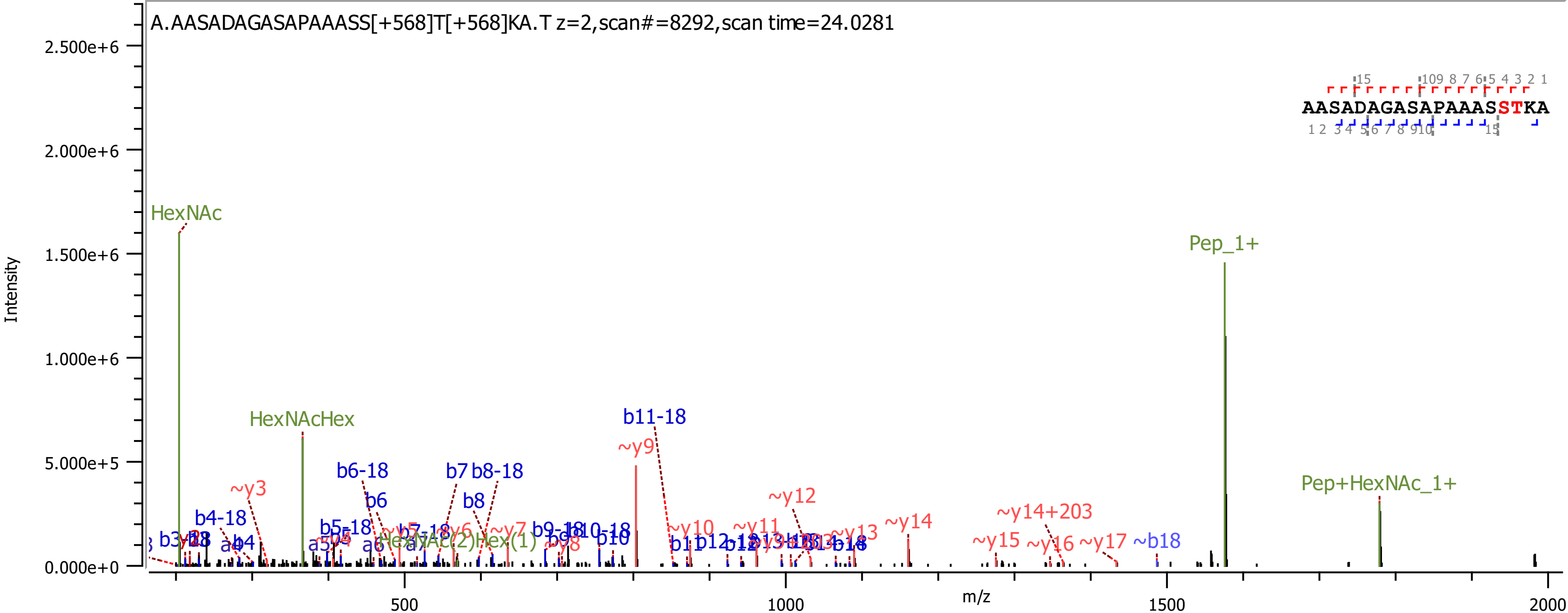
A.AASADAGASAPAAASS[+568]T[+568]KA.T z=2,scan#=8292,scan time=24.0281
AASADAGASAPAAASSTKA
Intensity
m/z
HexNAc
HexNAcHex
Pep_1+
Pep+HexNAc_1+
b11-18
~y9
b6-18
b7
b8-18
b6
b8
~y3
b4-18
~y12
~y14+203
~y14
~y10
~y11
~y13
~y15
~y16
~y17
~b18
b10
b11
b18

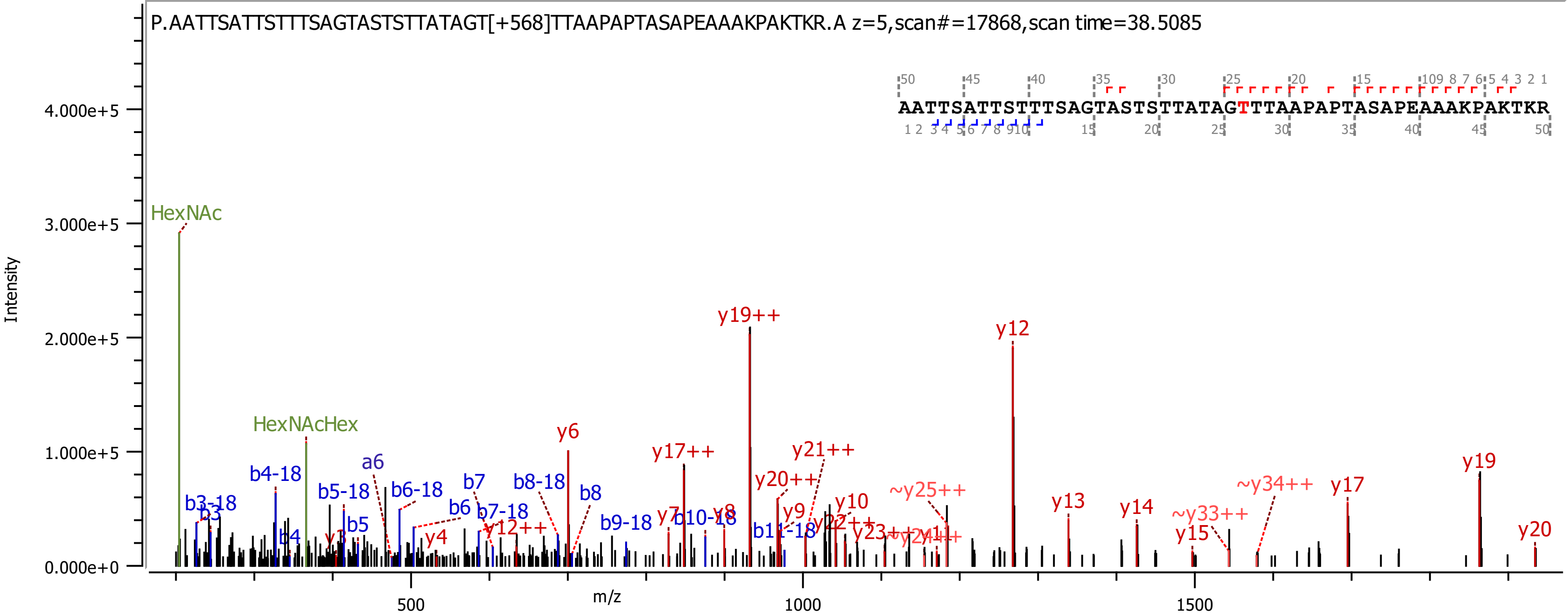

P.AATTSATTSTTTSAGTASTSTTATAGT[+568]TTAAPAPTASAPEAAAKPAKTKR.A z=5,scan#=17868,scan time=38.5085
AATTSATTSTTTSAGTASTSTTATAGTTTAAPAPTASAPEAAAKPAKTKR
Intensity
m/z
HexNAc
HexNAcHex
y19++
y12
y6
y17++
y21++
y20++
y19
y17
y20
y13
y14
y15
~y25++
~y34++
~y33++
y10
y9
b4-18
b5-18
b6-18
b8-18
b7-18
b9-18
b10-18
b3-18
a6
b7
b6
b8
b5
b4
b3
y4
y7
y8
y12++

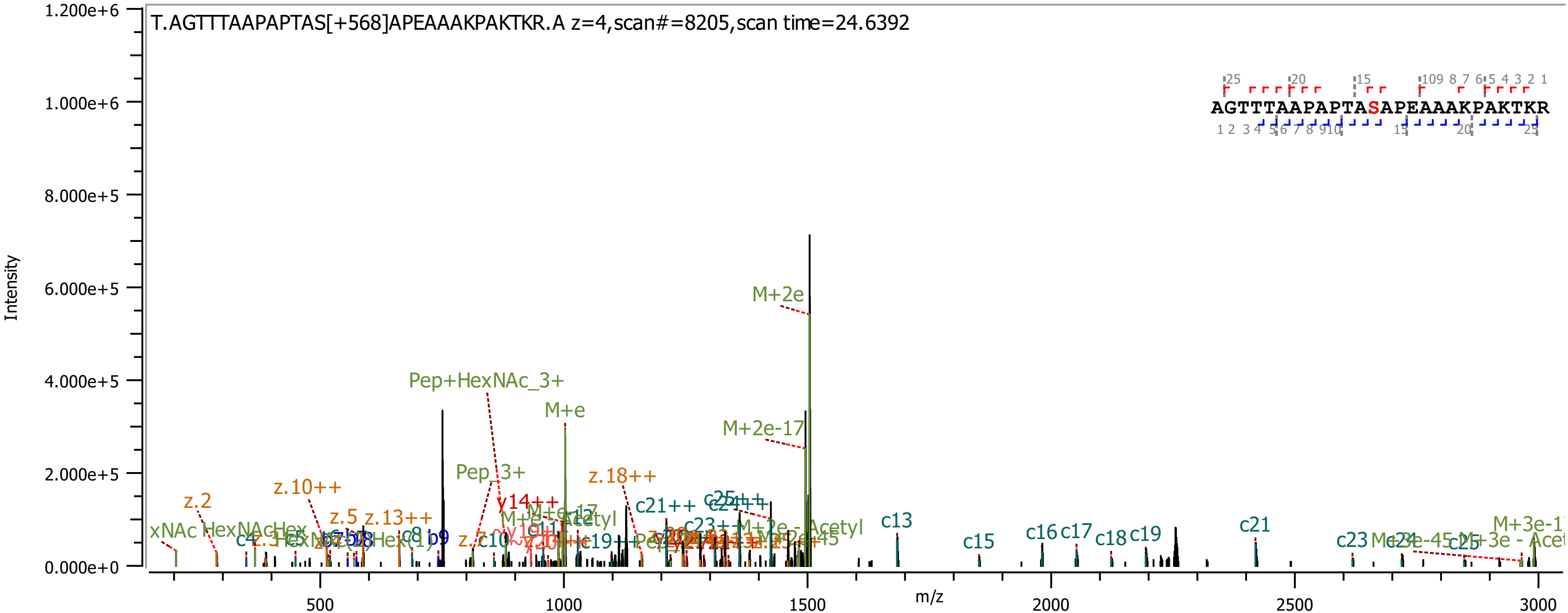

T.AGTTTAAPAPTAS[+568]APEAAAKPAKTKR.A z=4,scan#=8205,scan time=24.6392
AGTTTAAPAPTASAPEAAAKPAKTKR
Intensity
m/z
1.200e+6
1.000e+6
8.000e+5
6.000e+5
4.000e+5
2.000e+5
0.000e+0
500
1000
1500
2000
2500
3000
M+2e
M+e
M+2e-17
Pep+HexNAc_3+
Pep_3+
z.18++
z.10++
z.2
z.5
z.13++
y14++
c21++
c13
c15
c16
c17
c18
c19
c21
c23
c25
c8
b9
c4
c10

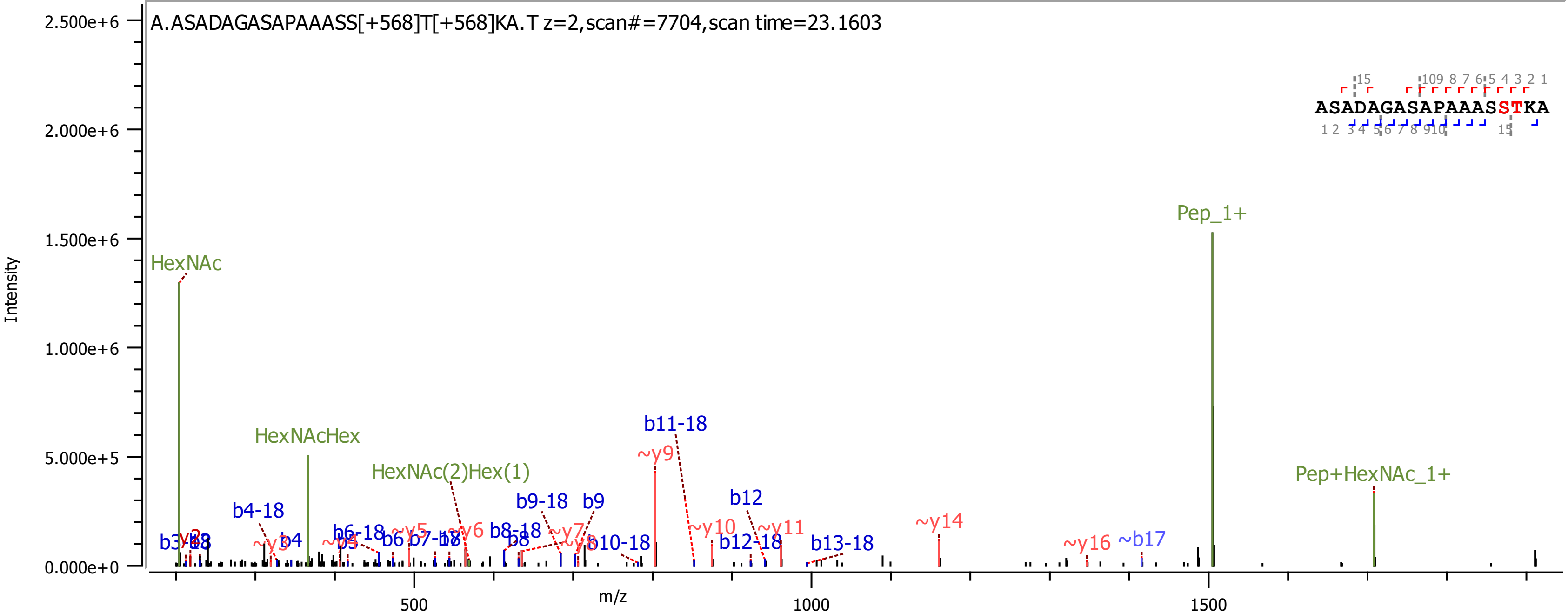
A.ASADAGASAPAAASS[+568]T[+568]KA.T z=2,scan#=7704,scan time=23.1603
ASADAGASAPAAASSTKA
Intensity
m/z
2.500e+6
2.000e+6
1.500e+6
1.000e+6
5.000e+5
0.000e+0
500
1000
1500
HexNAc
HexNAcHex
HexNAc(2)Hex(1)
Pep_1+
Pep+HexNAc_1+
b4-18
b4
b6-18
b6
b8-18
b8
b9-18
b9
b10-18
b11-18
b12
b12-18
b13-18
~b17
~y5
~y6
~y7
~y9
~y10
~y11
~y14
~y16

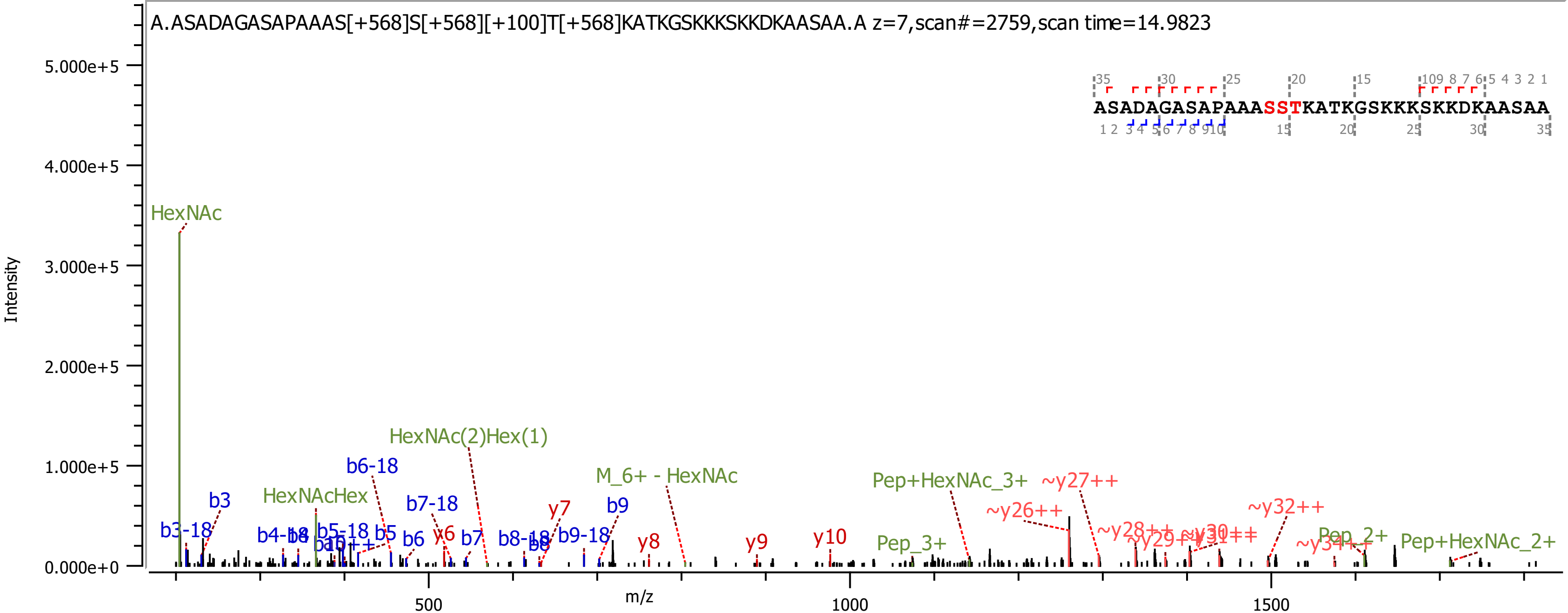
A.ASADAGASAPAAAS[+568]S[+568][+100]T[+568]KATKGSKKKSKKDKAASAA.A z=7,scan#=2759,scan time=14.9823
ASADAGASAPAAASSTKATKGSKKKSKKDKAASAA
Intensity
m/z
5.000e+5
4.000e+5
3.000e+5
2.000e+5
1.000e+5
0.000e+0
500
1000
1500
HexNAc
HexNAc(2)Hex(1)
b6-18
HexNAcHex
b3
b3-18
b7-18
y7
b9
M_6+ - HexNAc
Pep+HexNAc_3+
~y27++
~y26++
~y32++
b5-18
b5
b6
y6
b7
b8-18
b9-18
y8
y9
y10
Pep_3+
Pep_2+
Pep+HexNAc_2+

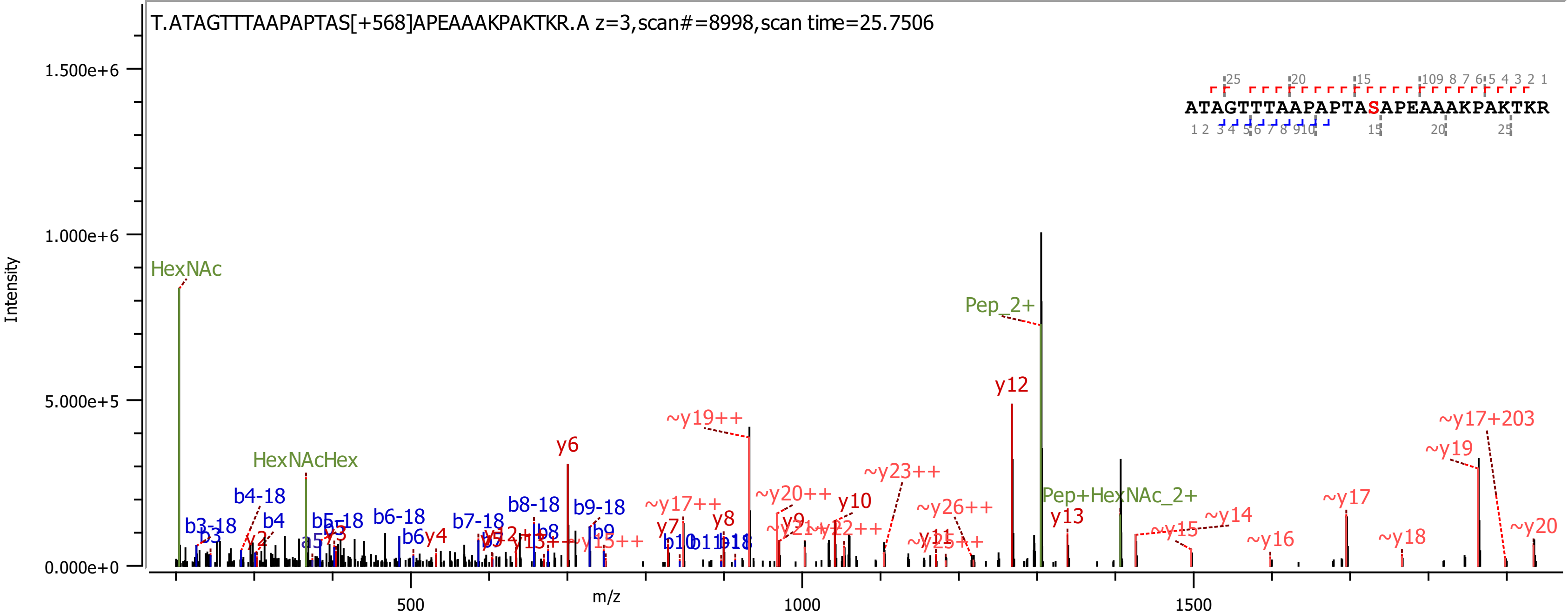

T.ATAGTTTAAPAPTAS[+568]APEAAAKPAKTKR.A z=3,scan#=8998,scan time=25.7506
ATAGTTTAAPAPTASAPEAAAKPAKTKR
Intensity
m/z
1.500e+6
1.000e+6
5.000e+5
0.000e+0
500
1000
1500
HexNAc
HexNAcHex
Pep_2+
Pep+HexNAc_2+
y6
y12
y13
~y19++
~y17+203
~y19
~y23++
~y20++
y10
~y26++
~y17
~y14
~y15
~y16
~y18
~y20
~y17++
y7
y8
y9
b3-18
b4-18
b4
b5-18
b6-18
b7-18
b8-18
b9-18
b6
y4
b8
b9
b10

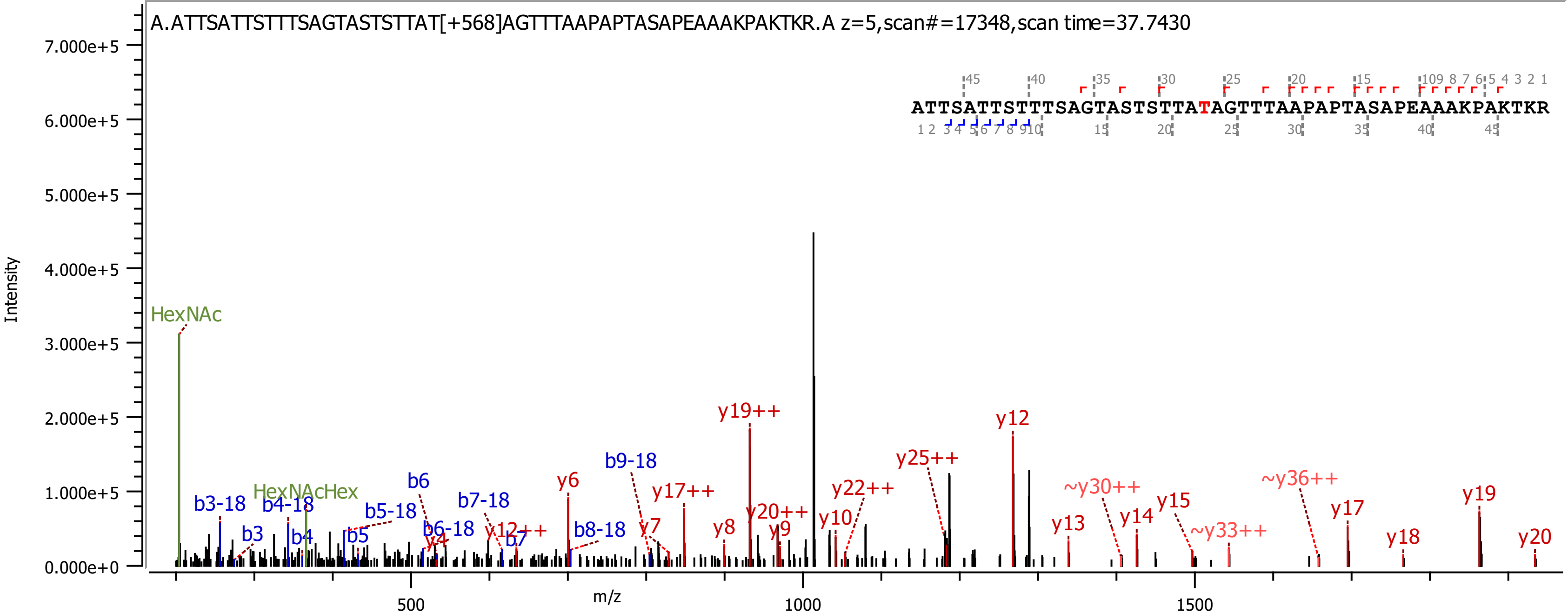
A.ATTSATTSTTTSAGTASTSTTAT[+568]AGTTTAAPAPTASAPEAAAKPAKTKR.A z=5,scan#=17348,scan time=37.7430
ATTSATTSTTTSAGTASTSTTATAGTTTAAPAPTASAPEAAAKPAKTKR
Intensity
m/z
HexNAc
HexNAcHex
b3-18
b3
b4-18
b4
b5-18
b5
b6
b6-18
b7-18
b7
b8-18
b9-18
y4
y6
y7
y8
y9
y10
y12
y13
y14
y15
y17
y18
y19
y20
y12++
y17++
y19++
y20++
y22++
y25++
~y30++
~y33++
~y36++

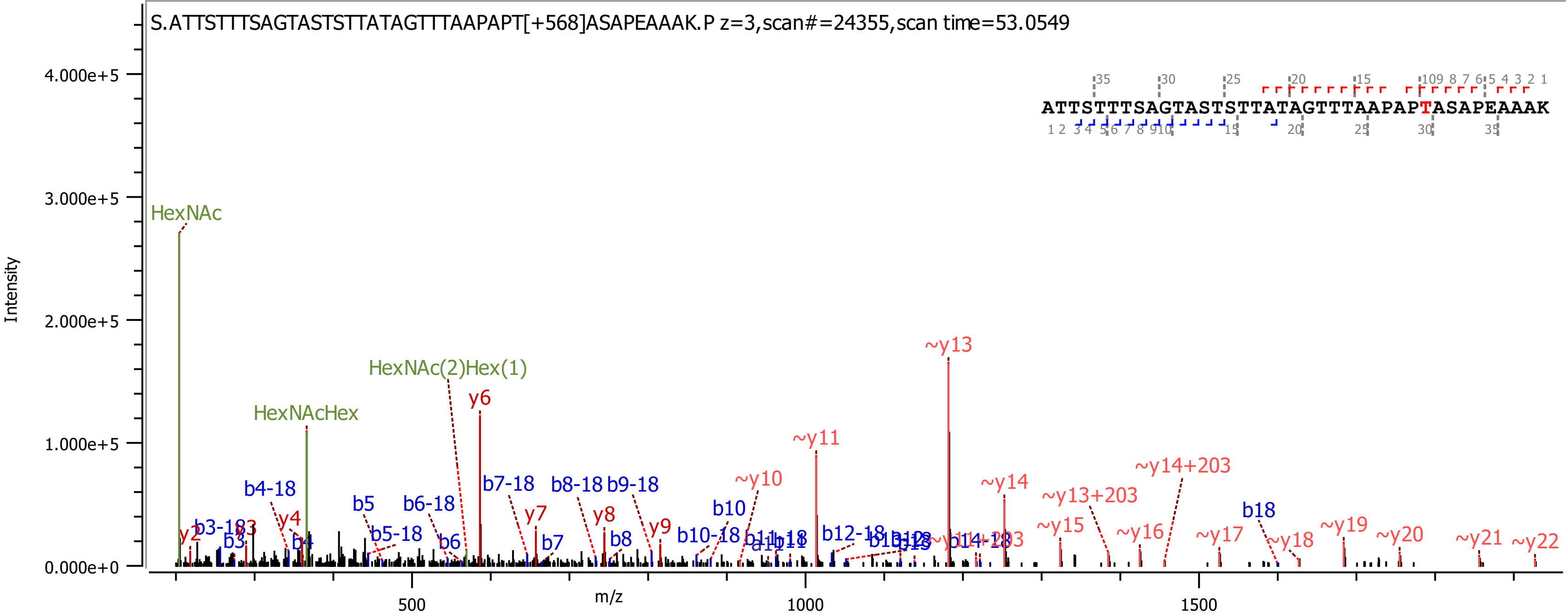
S.ATTSTTTSAGTASTSTTATAGTTTAAPAPT[+568]ASAPEAAAK.P z=3,scan#=24355,scan time=53.0549
ATTSTTTSAGTASTSTTATAGTTTAAPAPTASAPEAAAK
Intensity
m/z
HexNAc
HexNAcHex
HexNAc(2)Hex(1)
y6
~y13
~y11
~y10
~y14
~y14+203
~y13+203
b4-18
b5
b6-18
b7-18
b8-18
b9-18
b10
b18
y4
y7
y8
y9
~y15
~y16
~y17
~y18
~y19
~y20
~y21
~y22
b10-18
b12-18
b7
b8
b6
b5-18
b3
b4
y2

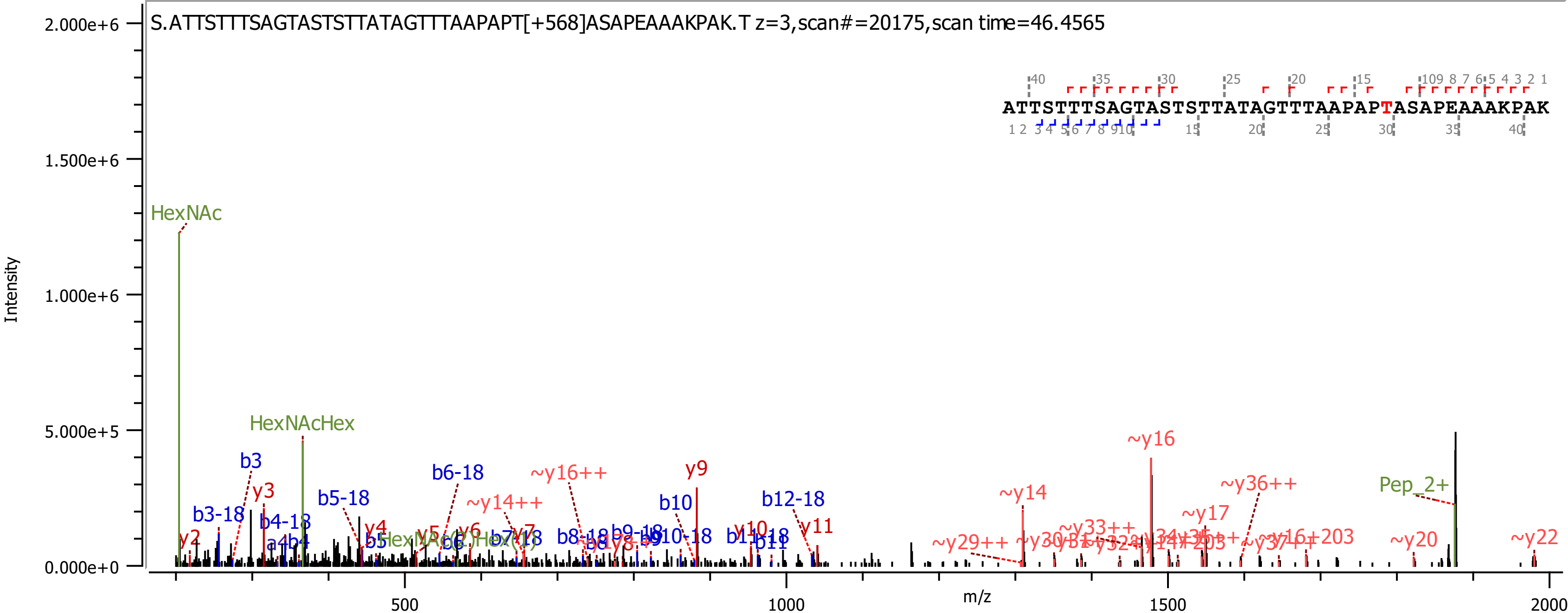

S.ATTSTTTSAGTASTSTTATAGTTTAAPAPT[+568]ASAPEAAAKPAK.T z=3,scan#=20175,scan time=46.4565
ATTSTTTSAGTASTSTTATAGTTTAAPAPTASAPEAAAKPAK
Intensity
m/z
HexNAc
HexNAcHex
Pep_2+
~y16
~y14
~y36++
~y17
~y20
~y22
~y16++
~y14++
y9
b10
b12-18
y11
y10
b3
b3-18
y3
b4-18
b5-18
b6-18
y2
y4
y6
y7

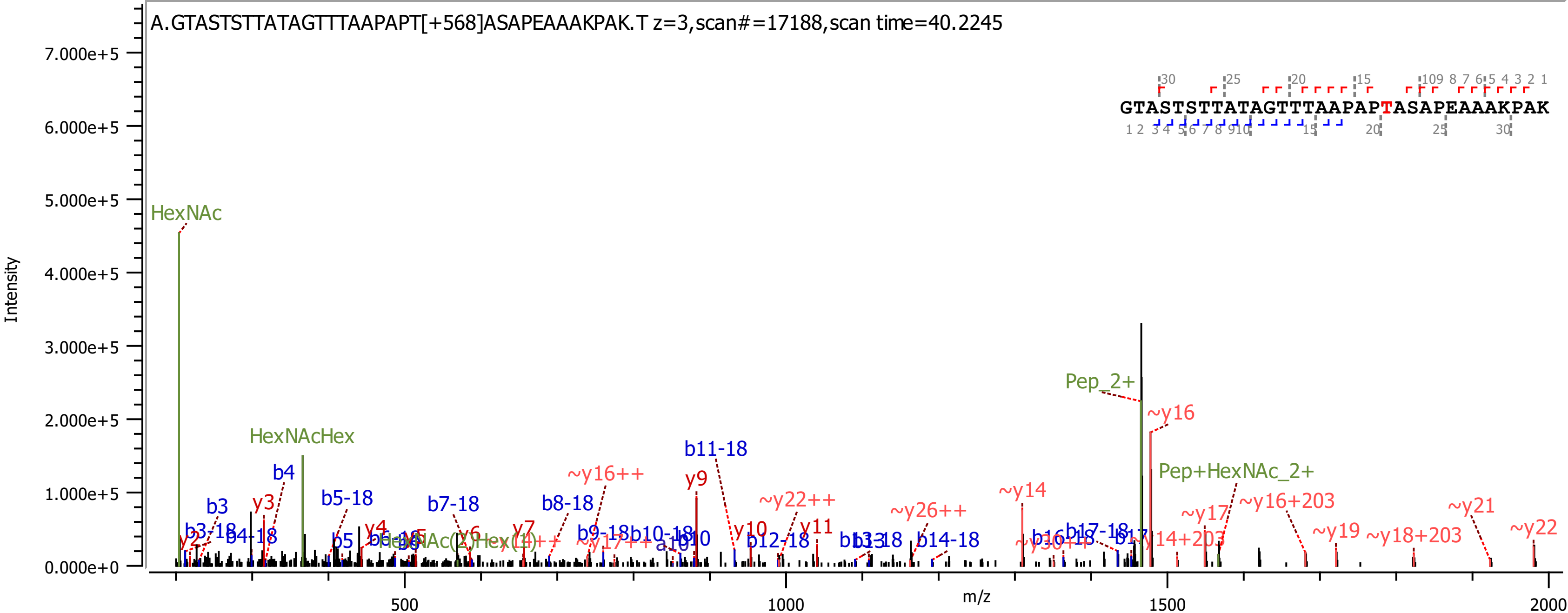
A.GTASTSTTATAGTTTAAPAPT[+568]ASAPEAAAKPAK.T z=3,scan#=17188,scan time=40.2245
GTASTSTTATAGTTTAAPAPTASAPEAAAKPAK
Intensity
m/z
7.000e+5
6.000e+5
5.000e+5
4.000e+5
3.000e+5
2.000e+5
1.000e+5
0.000e+0
500
1000
1500
2000
HexNAc
HexNAcHex
Pep_2+
Pep+HexNAc_2+
~y16
b11-18
y9
~y16++
~y22++
~y26++
~y14
~y17
~y16+203
~y21
~y22
~y19
~y18+203
b3
b4
y3
b5-18
b7-18
b8-18
b3-18
b4-18
b5
y4
y7
y10
y11
b12-18
b14-18
b9-18
b10-18
b17-18

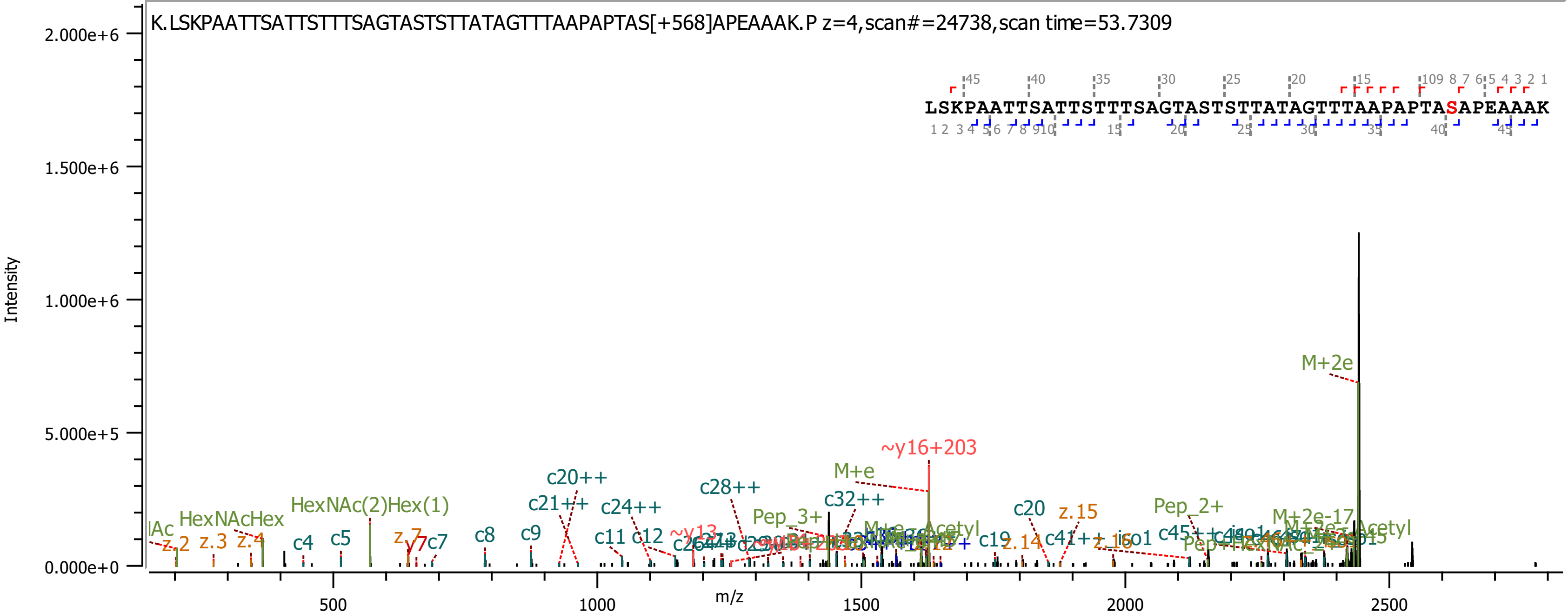
K.LSKPAATTSATTSTTTSAGTASTSTTATAGTTTAAPAPTAS[+568]APEAAAK.P z=4,scan#=24738,scan time=53.7309
LSKPAATTSATTSTTTSAGTASTSTTATAGTTTAAPAPTASAPEAAAK
Intensity
m/z
2.000e+6
1.500e+6
1.000e+6
5.000e+5
0.000e+0
500
1000
1500
2000
2500
M+2e
~y16+203
M+e
c20++
c28++
c21++
c24++
c32++
Pep_3+
HexNAc(2)Hex(1)
HexNAcHex
c20
z.15
Pep_2+
M+2e-17
c4
c5
c7
c8
c9
c11
c12
z.2
z.3
z.4
y7
c19
z.14
c41++
z.16
iso1
c45++

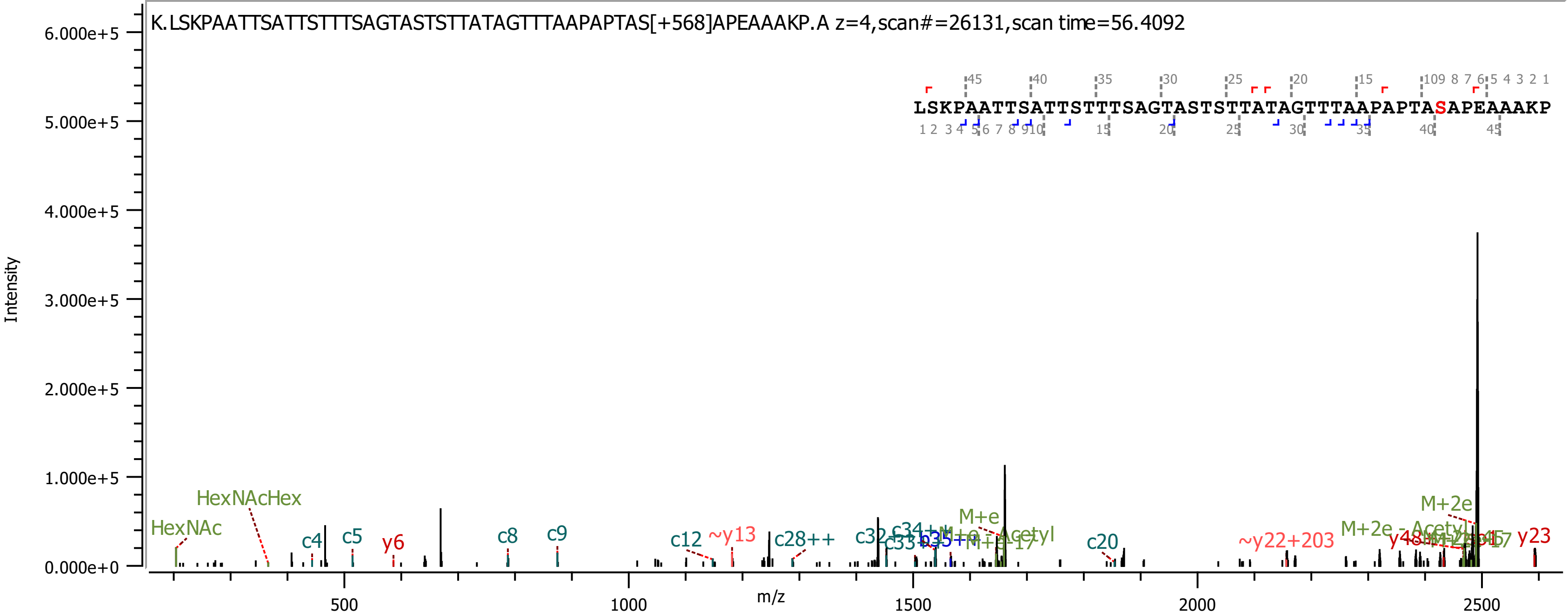
K.LSKPAATTSATTSTTTSAGTASTSTTATAGTTTAAPAPTAS[+568]APEAAAKP.A z=4,scan#=26131,scan time=56.4092
LSKPAATTSATTSTTTSAGTASTSTTATAGTTTAAPAPTASAPEAAAKP
Intensity
m/z
6.000e+5
5.000e+5
4.000e+5
3.000e+5
2.000e+5
1.000e+5
0.000e+0
500
1000
1500
2000
2500
HexNAc
HexNAcHex
c4
c5
y6
c8
c9
c12
~y13
c28++
c20
~y22+203
M+e
M+2e
M+2e - Acetyl
y23

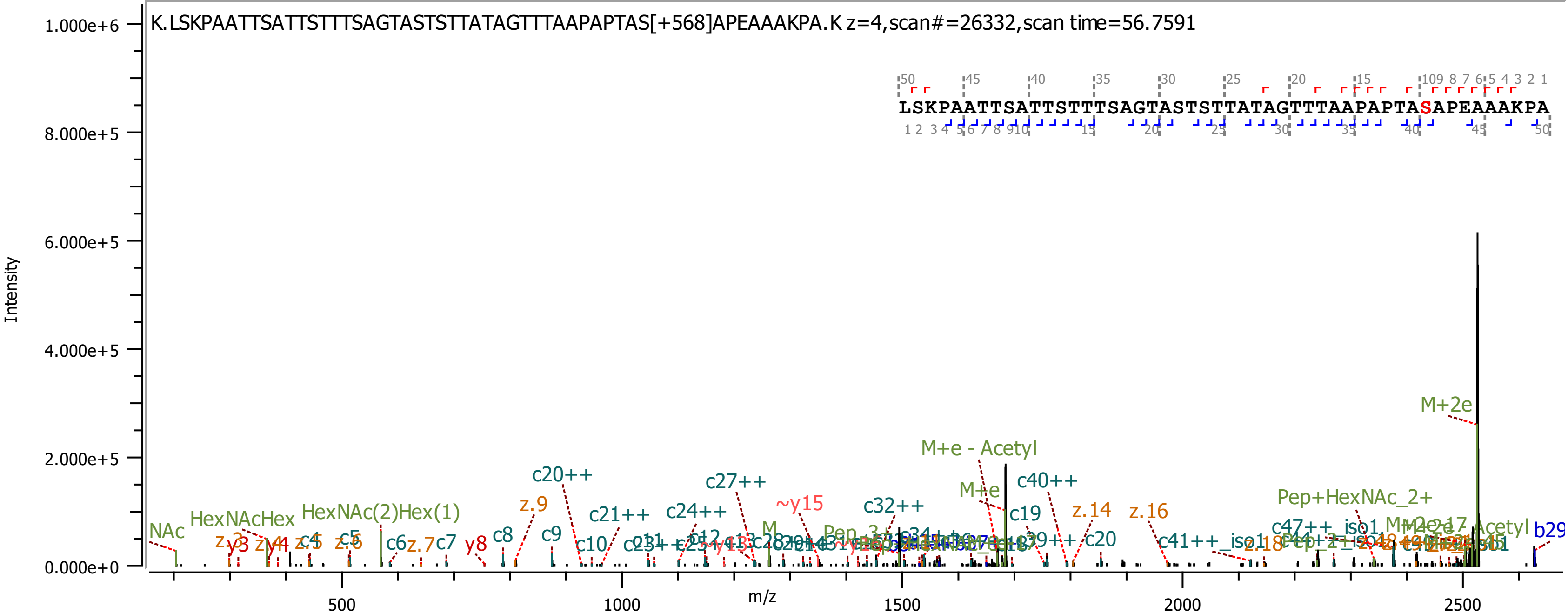
K.LSKPAATTSATTSTTTSAGTASTSTTATAGTTTAAPAPTAS[+568]APEAAAKPA.K z=4,scan#=26332,scan time=56.7591
LSKPAATTSATTSTTTSAGTASTSTTATAGTTTAAPAPTASAPEAAAKPA
Intensity
m/z
1.000e+6
8.000e+5
6.000e+5
4.000e+5
2.000e+5
0.000e+0
500
1000
1500
2000
2500
M+2e
M+e - Acetyl
M+e
c20++
c21++
c24++
c27++
~y15
c32++
c40++
c19
c20
z.9
z.14
z.16
z.18
c47++_iso1
c41++_iso1
Pep+HexNAc_2+
HexNAcHex
HexNAc(2)Hex(1)
NAc
c6
z.7
c7
y8
c8
c9
c10
b29

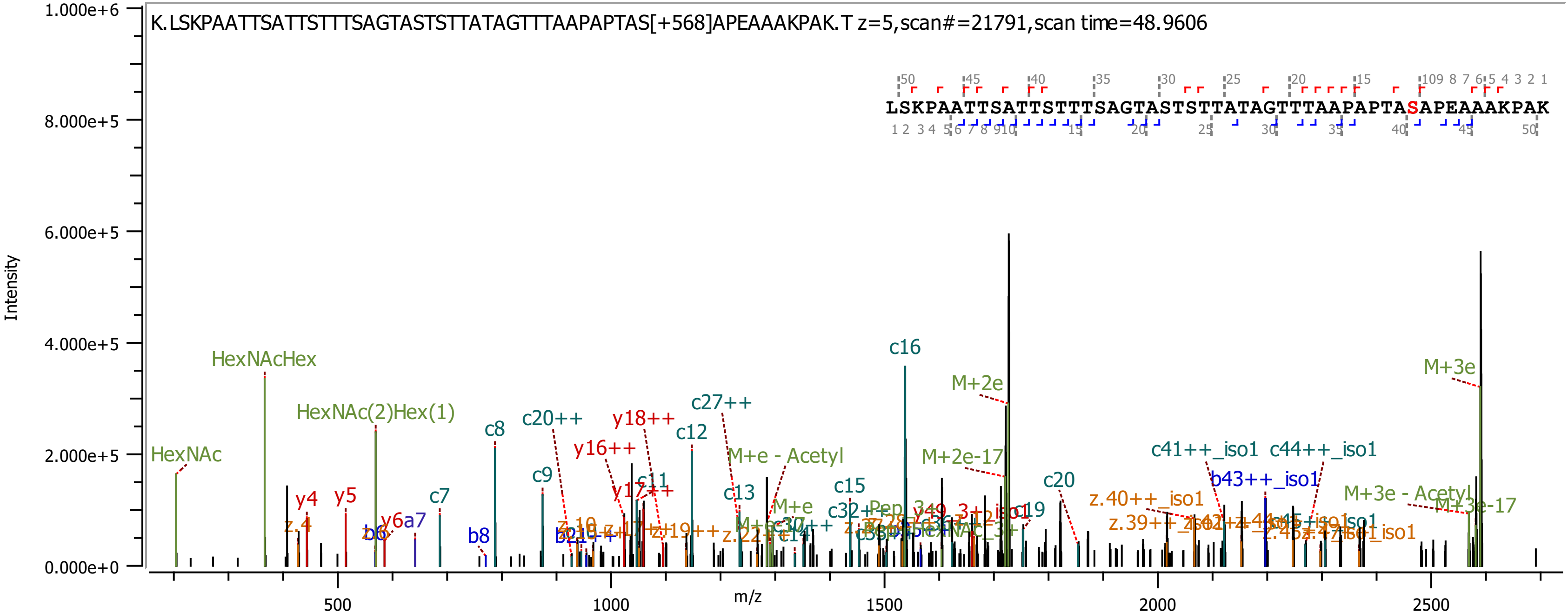
K.LSKPAATTSATTSTTTSAGTASTSTTATAGTTTAAPAPTAS[+568]APEAAAKPAK.T z=5,scan#=21791,scan time=48.9606
LSKPAATTSATTSTTTSAGTASTSTTATAGTTTAAPAPTASAPEAAAKPAK
Intensity
m/z
1.000e+6
8.000e+5
6.000e+5
4.000e+5
2.000e+5
0.000e+0
500
1000
1500
2000
2500
HexNAc
HexNAcHex
HexNAc(2)Hex(1)
y4
y5
c7
c8
c9
c20++
y16++
y18++
c12
c27++
c13
c15
c16
M+2e
M+2e-17
c20
M+e - Acetyl
c41++_iso1
c44++_iso1
b43++_iso1
z.40++_iso1
M+3e
M+3e - Acetyl
M+3e-17
b8

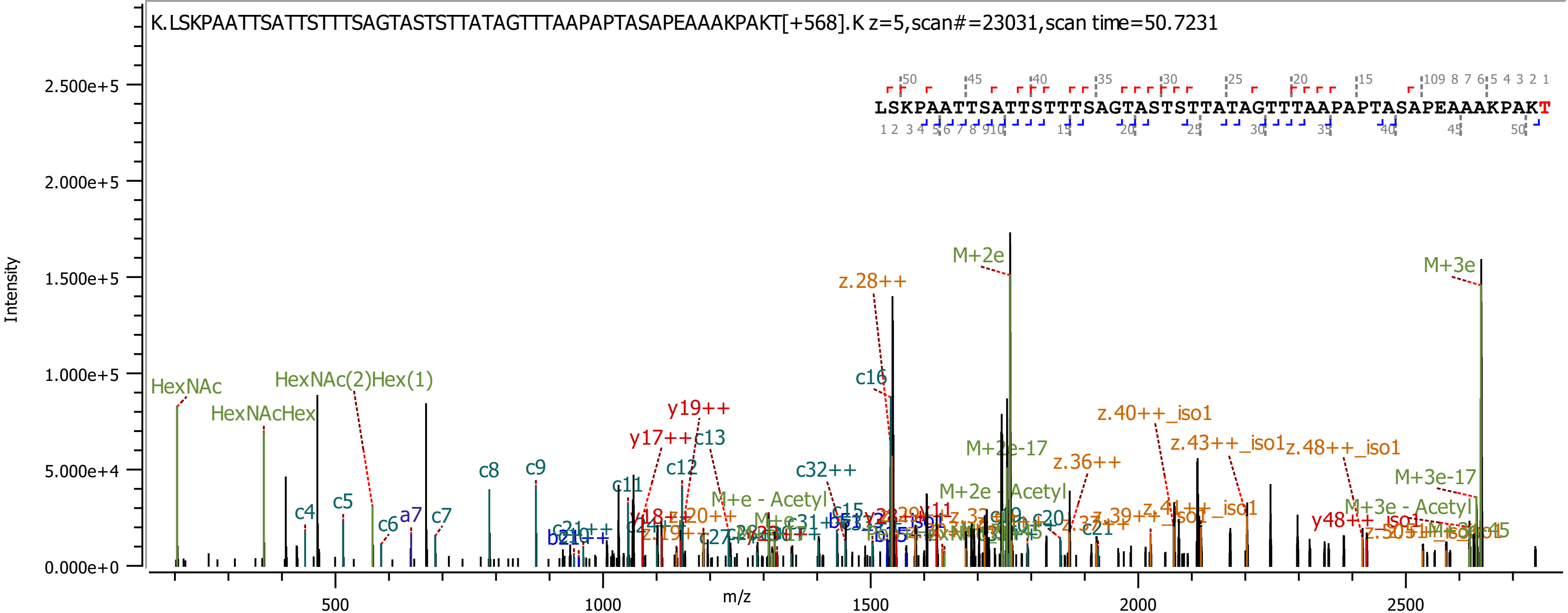
K.LSKPAATTSATTSTTTSAGTASTSTTATAGTTTAAPAPTASAPEAAAKPAKT[+568].K z=5,scan#=23031,scan time=50.7231
LSKPAATTSATTSTTTSAGTASTSTTATAGTTTAAPAPTASAPEAAAKPAKT
Intensity
m/z
HexNAc
HexNAcHex
HexNAc(2)Hex(1)
M+2e
M+3e
z.28++
c16
y19++
y17++
c13
c12
c11
c8
c9
c4
c5
c6
a7
c7
c32++
M+e - Acetyl
M+2e-17
M+2e - Acetyl
z.36++
z.40++_iso1
z.43++_iso1
z.48++_iso1
M+3e-17
M+3e - Acetyl
y48++
c15
c20
c21
z.19++
z.39++
z.41++_iso1
y18++
z.20++
c21++
b21++
c31++
y11
0.000e+0
5.000e+4
1.000e+5
1.500e+5
2.000e+5
2.500e+5
500
1000
1500
2000
2500

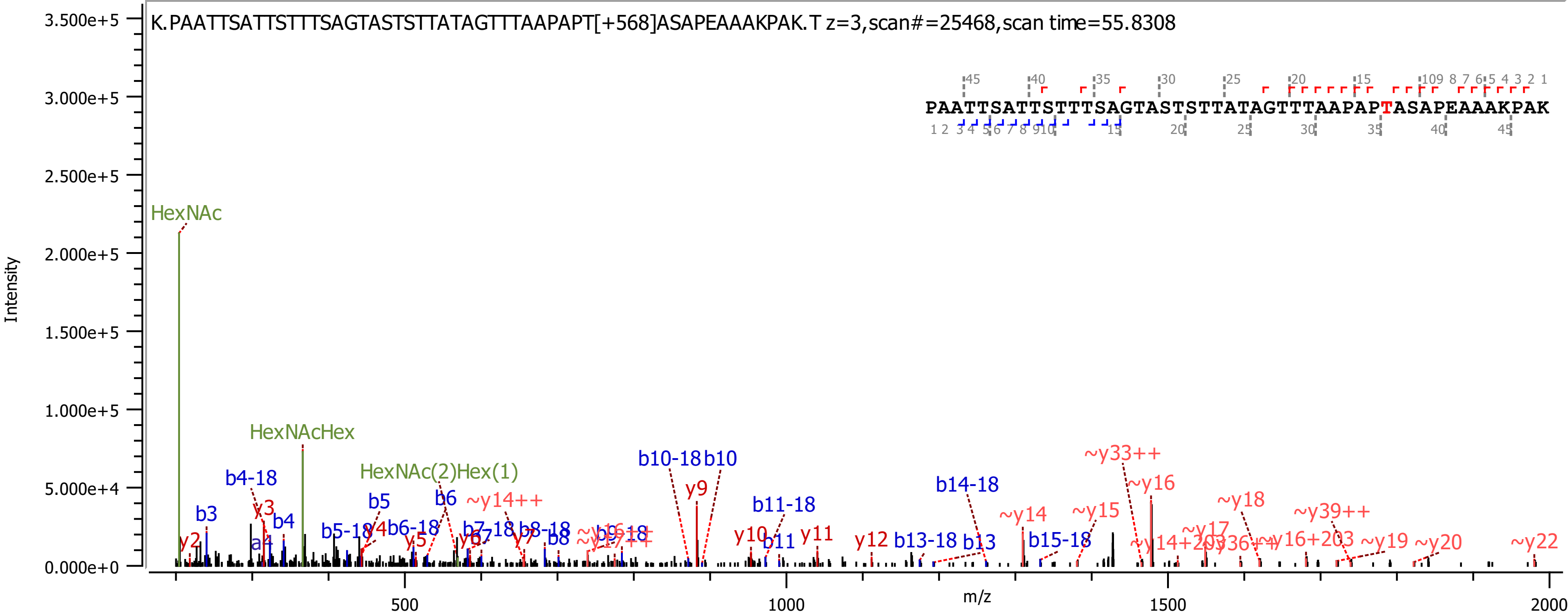
K.PAATTSATTSTTTSAGTASTSTTATAGTTTAAPAPT[+568]ASAPEAAAKPAK.T z=3,scan#=25468,scan time=55.8308
PAATTSATTSTTTSAGTASTSTTATAGTTTAAPAPTASAPEAAAKPAK
HexNAc
HexNAcHex
HexNAc(2)Hex(1)
b10-18 b10
y9
~y33++
~y16
b14-18
b11-18
~y18
~y14
~y15
~y39++
~y14++
b4-18
b3
y3
b4
b5
b6
y10
b11
y11
y12
b13-18
b13
b15-18
~y17
~y19
~y20
~y22
Intensity
m/z
3.500e+5
3.000e+5
2.500e+5
2.000e+5
1.500e+5
1.000e+5
5.000e+4
0.000e+0
500
1000
1500
2000

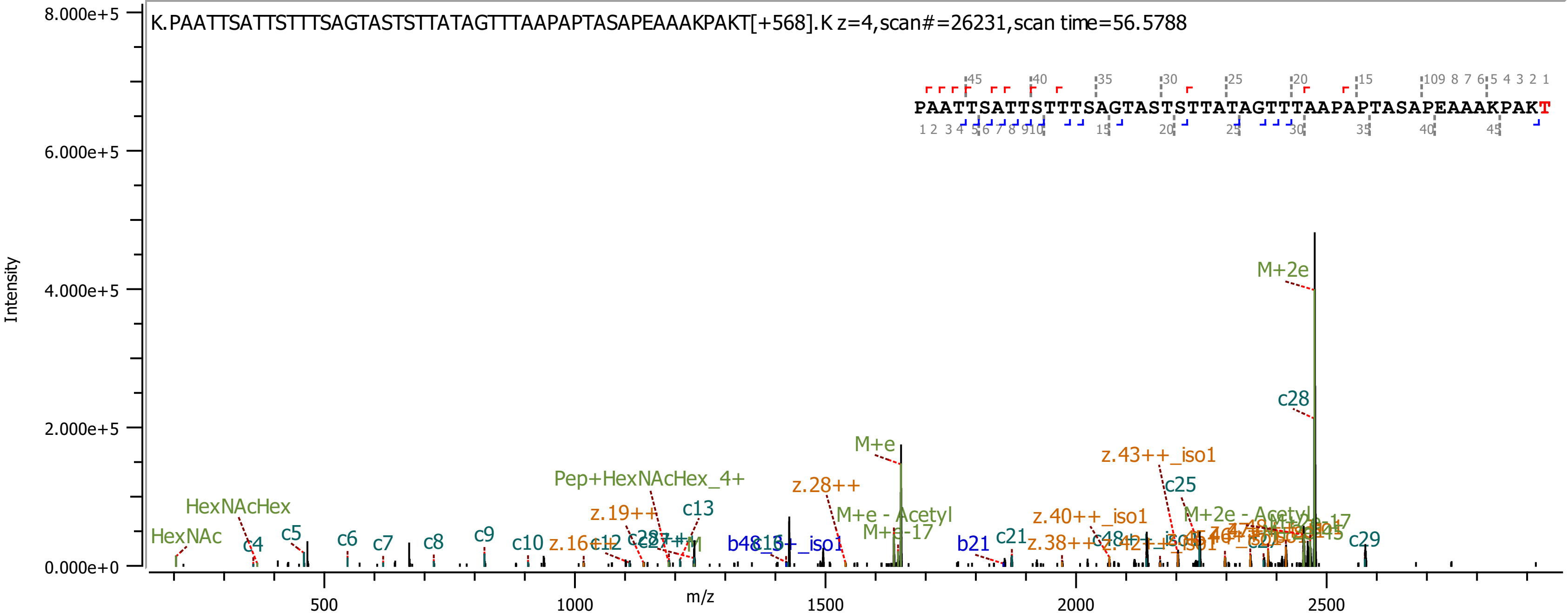
K.PAATTSATTSTTTSAGTASTSTTATAGTTTAAPAPTASAPEAAAKPAKT[+568].K z=4,scan#=26231,scan time=56.5788
PAATTSATTSTTTSAGTASTSTTATAGTTTAAPAPTASAPEAAAKPAKT
Intensity
m/z
8.000e+5
6.000e+5
4.000e+5
2.000e+5
0.000e+0
500
1000
1500
2000
2500
M+2e
c28
M+e
z.43++_iso1
c25
z.28++
M+e - Acetyl
M+e-17
Pep+HexNAcHex_4+
HexNAcHex
HexNAc
c4
c5
c6
c7
c8
c9
c10
z.16++
c12
z.19++
c13
z.40++_iso1
M+2e - Acetyl
b21
c21
z.38++
c29

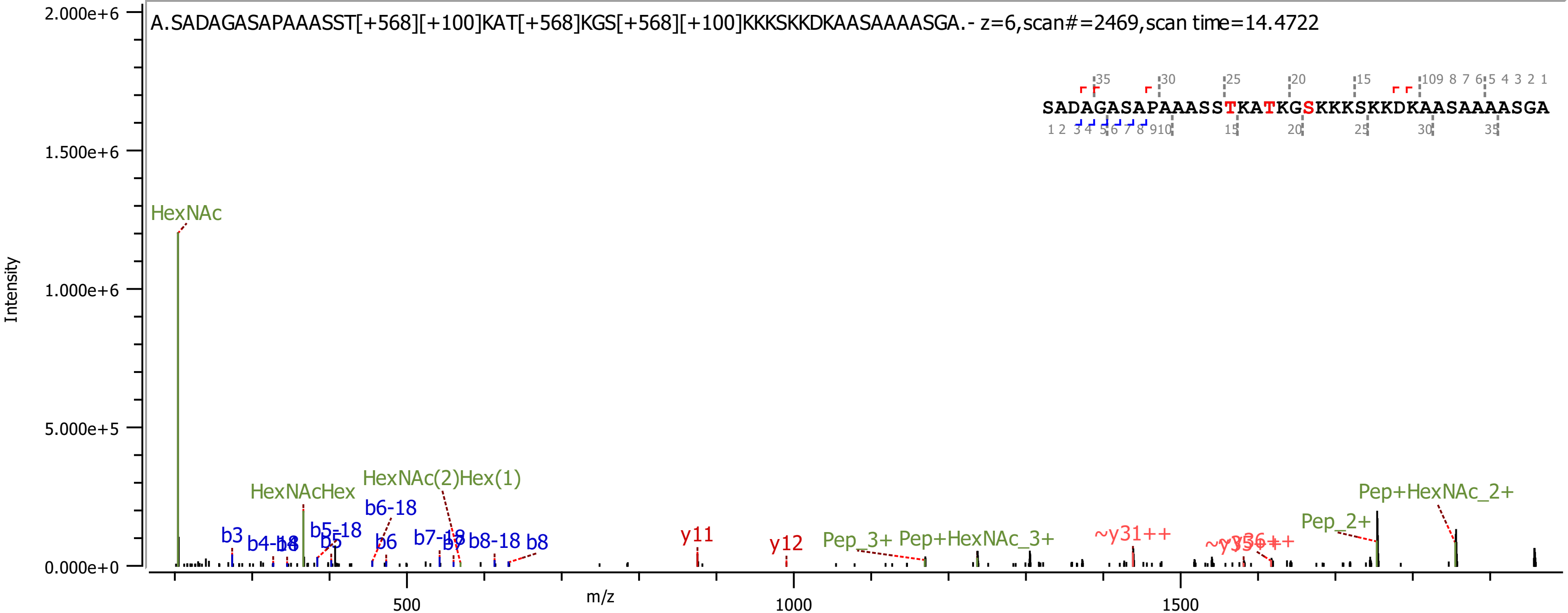

A.SADAGASAPAAASST[+568][+100]KAT[+568]KGS[+568][+100]KKKSKKDKAASAAAASGA.- z=6,scan#=2469,scan time=14.4722
SADAGASAPAAASSTKATKGSKKKSKKDKAASAAAASGA
HexNAc
HexNAcHex
HexNAc(2)Hex(1)
b3
b4-18
b4
b5-18
b5
b6-18
b6
b7-18
b7
b8-18
b8
y11
y12
Pep_3+
Pep+HexNAc_3+
~y31++
Pep_2+
Pep+HexNAc_2+
Intensity
m/z
2.000e+6
1.500e+6
1.000e+6
5.000e+5
0.000e+0
500
1000
1500

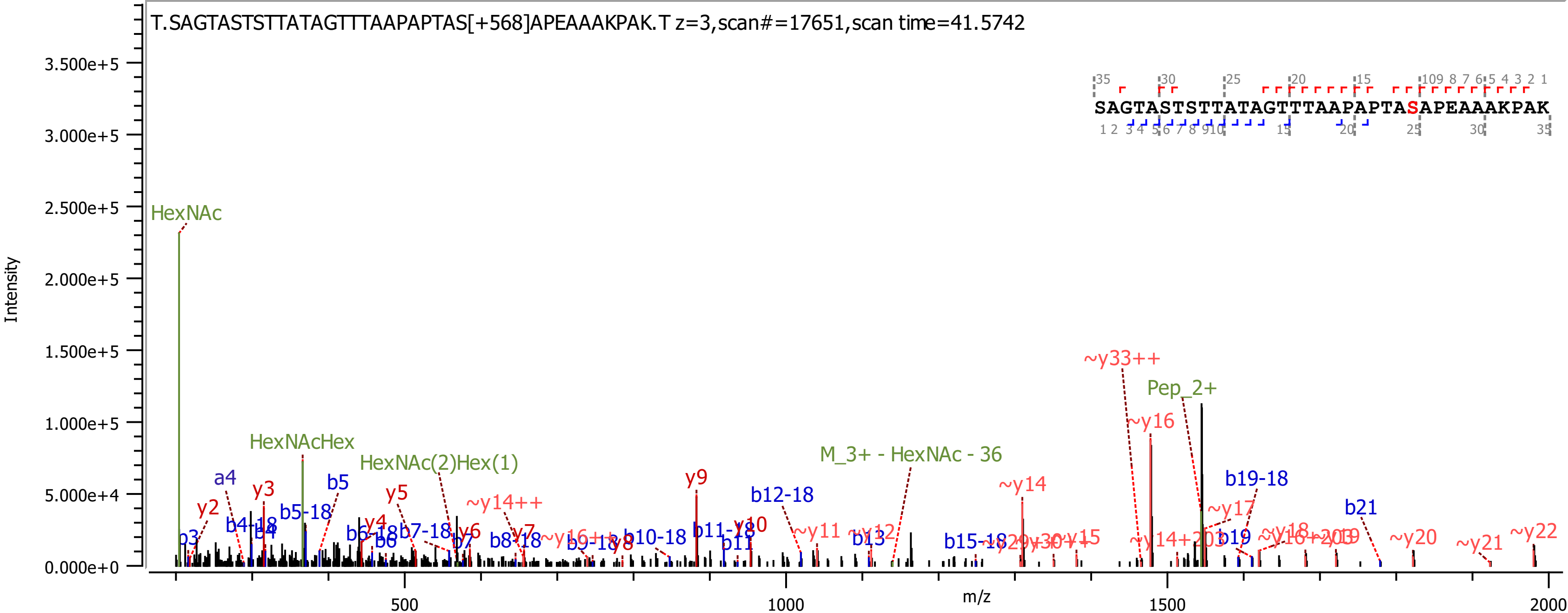

T.SAGTASTSTTATAGTTTAAPAPTAS[+568]APEAAAKPAK.T z=3,scan#=17651,scan time=41.5742
SAGTASTSTTATAGTTTAAPAPTASAPEAAAKPAK
Intensity
m/z
HexNAc
HexNAcHex
HexNAc(2)Hex(1)
M_3+ - HexNAc - 36
Pep_2+
~y33++
~y16
b19-18
b21
~y14
~y17
~y22
~y20
~y21
y9
b12-18
y10
b11-18
b11
b13
b15-18
~y15
b19
a4
y2
y3
b3
b4-18
b4
b5
b5-18
y4
y5
y6
y7
b6-18
b6
b7-18
b7
b8-18
~y14++
b9-18
y8
b10-18
~y11
~y12
500
1000
1500
2000
0.000e+0
5.000e+4
1.000e+5
1.500e+5
2.000e+5
2.500e+5
3.000e+5
3.500e+5

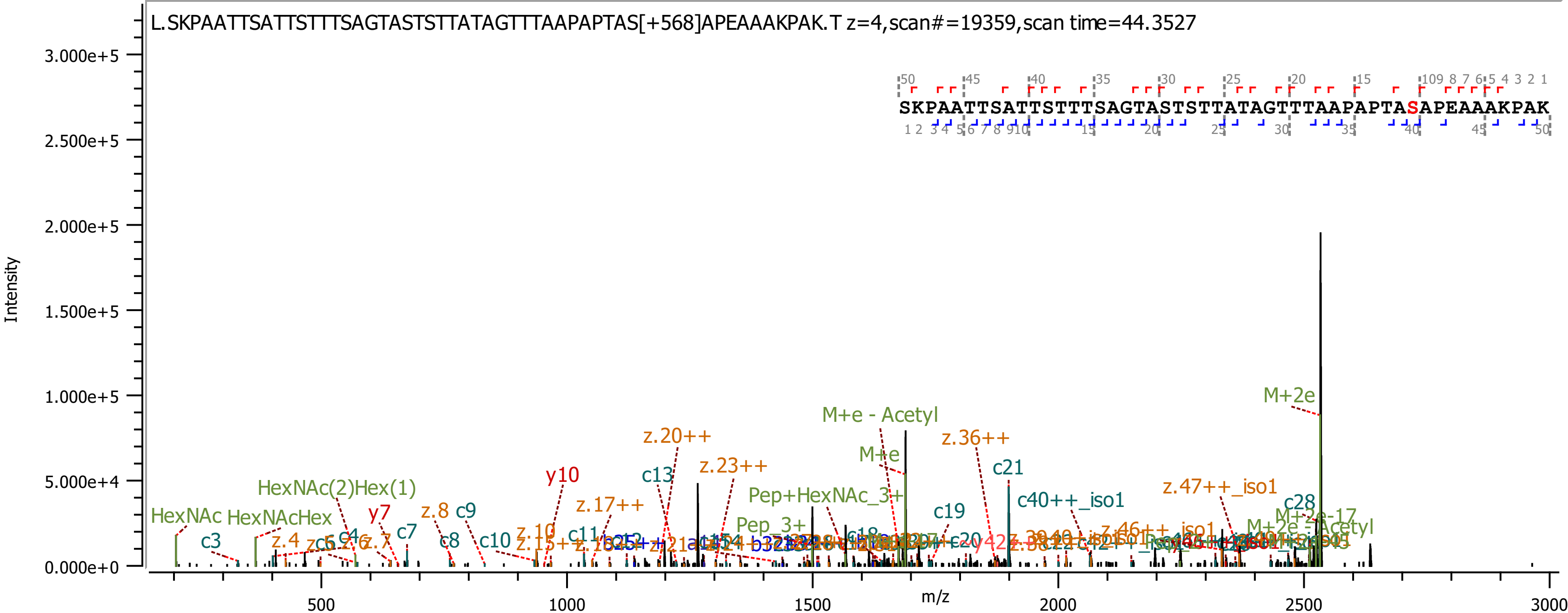

L.SKPAATTSATTSTTTSAGTASTSTTATAGTTTAAPAPTAS[+568]APEAAAKPAK.T z=4,scan#=19359,scan time=44.3527
SKPAATTSATTSTTTSAGTASTSTTATAGTTTAAPAPTASAPEAAAKPAK
Intensity
m/z
3.000e+5
2.500e+5
2.000e+5
1.500e+5
1.000e+5
5.000e+4
0.000e+0
500
1000
1500
2000
2500
3000
M+2e
M+e - Acetyl
M+e
z.36++
c21
z.20++
z.23++
c13
y10
HexNAc(2)Hex(1)
HexNAc
HexNAcHex
c3
z.4
c4
y7
c7
z.7
z.8
c9
c8
c10
z.10
z.17++
c11
Pep+HexNAc_3+
Pep_3+
c19
c40++_iso1
z.47++_iso1
c28
M+2e-17
M+2e - Acetyl

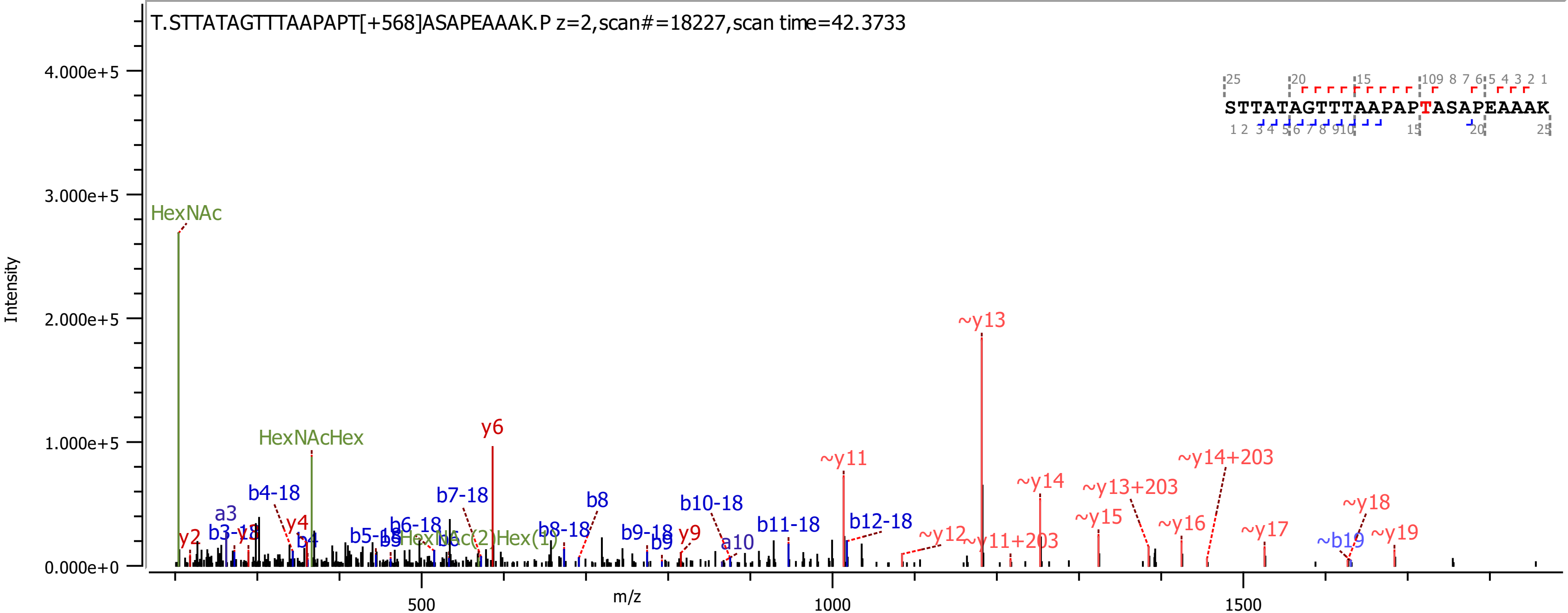
T.STTATAGTTTAAPAPT[+568]ASAPEAAAK.P z=2,scan#=18227,scan time=42.3733
STTATAGTTTAAPAPTASAPEAAAK
Intensity
m/z
HexNAc
HexNAcHex
y6
~y13
~y11
~y14
~y13+203
~y14+203
~y15
~y16
~y17
~y18
~y19
~y12
~y11+203
~b19
b4-18
b7-18
b8
b10-18
b11-18
b12-18
a3
b3-18
y2
y4
b4
b5-18
b6-18
b8-18
b9-18
b9
y9
a10
HexNAc(2)Hex(1)

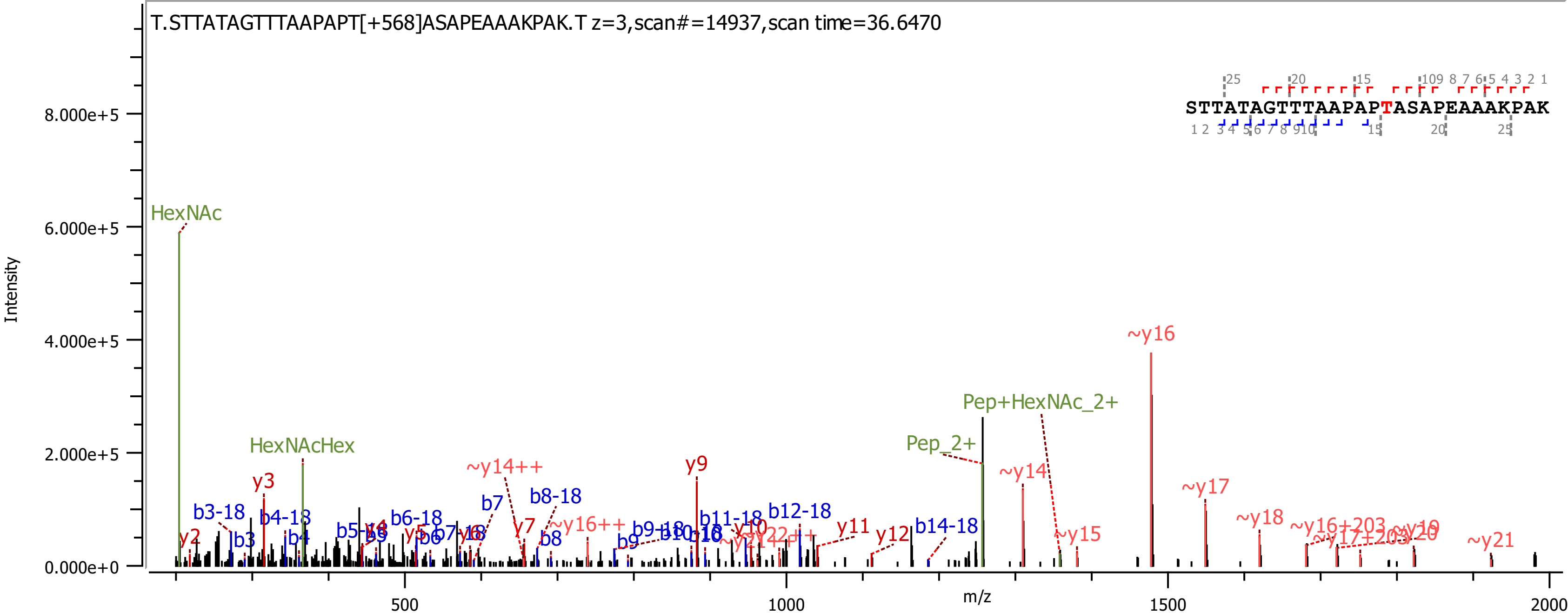

T.STTATAGTTTAAPAPT[+568]ASAPEAAAKPAK.T z=3,scan#=14937,scan time=36.6470
STTATAGTTTAAPAPTASAPEAAAKPAK
Intensity
m/z
HexNAc
HexNAcHex
Pep_2+
Pep+HexNAc_2+
~y16
~y14
~y17
~y18
~y15
~y21
~y14++
~y16++
y2
y3
y7
y9
y10
y11
y12
b3-18
b4-18
b6-18
b7
b8-18
b8
b9
b9-18
b11-18
b12-18
b14-18

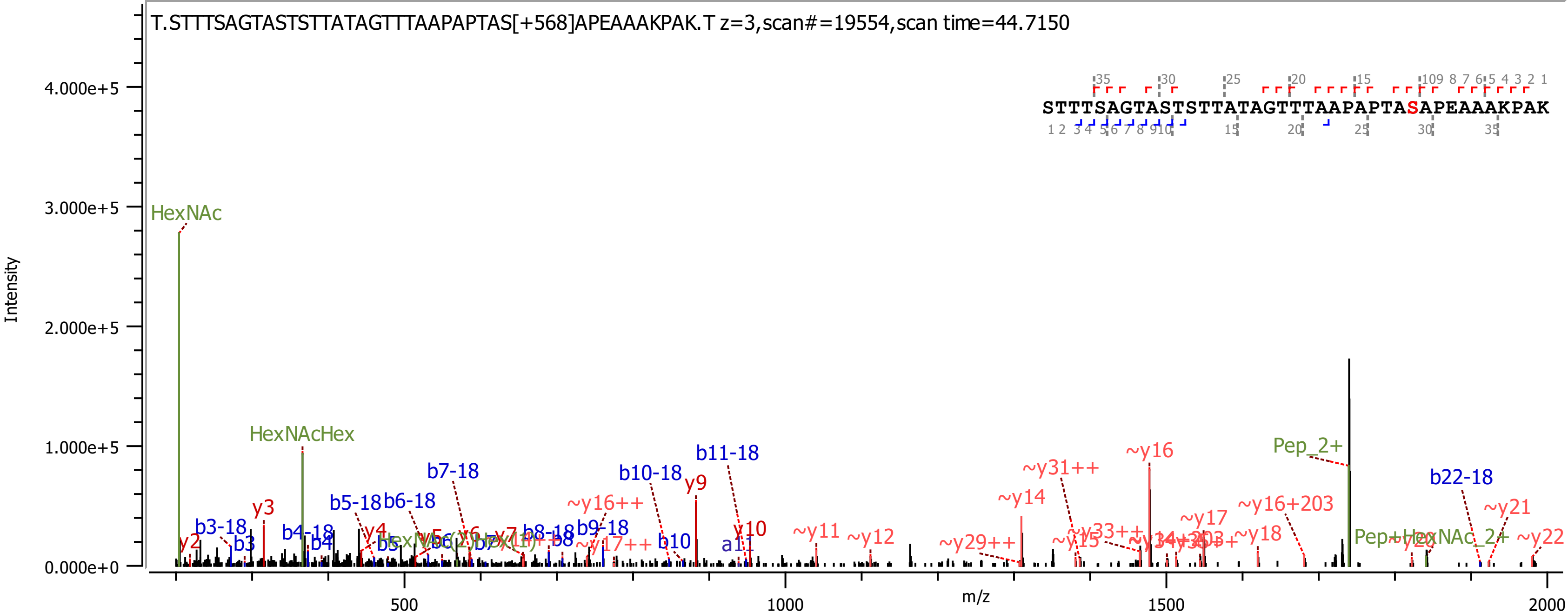

T.STTTSAGTASTSTTATAGTTTAAPAPTAS[+568]APEAAAKPAK.T z=3,scan#=19554,scan time=44.7150
STTTSAGTASTSTTATAGTTTAAPAPTASAPEAAAKPAK
Intensity
m/z
4.000e+5
3.000e+5
2.000e+5
1.000e+5
0.000e+0
500
1000
1500
2000
HexNAc
HexNAcHex
y2
b3-18
b3
y3
b4-18
b4
b5-18
b5
b6-18
y4
b7-18
y7
b8-18
b8
b9-18
~y16++
~y17++
b10-18
b10
y9
b11-18
a11
y10
~y11
~y12
~y29++
~y14
~y31++
~y15
~y33++
~y16
~y17
~y16+203
~y18
Pep_2+
Pep+HexNAc_2+
b22-18
~y21
~y22

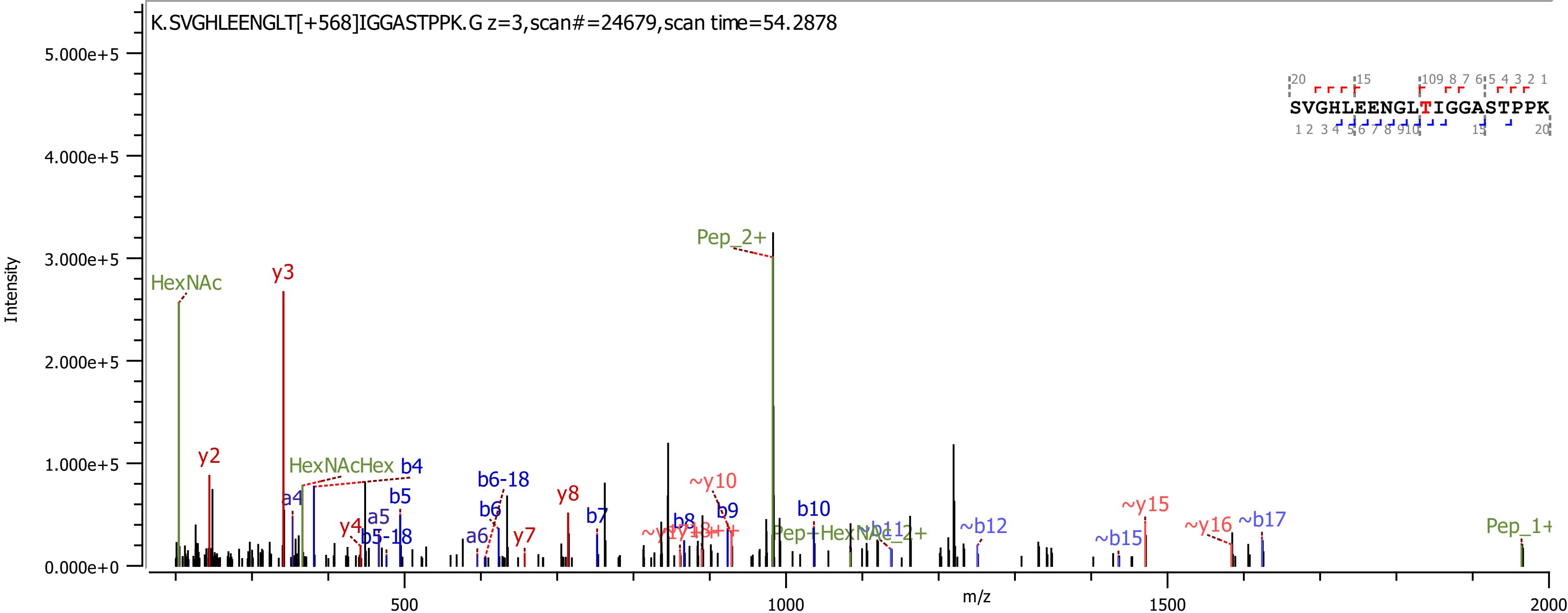
K.SVGHLEENGLT[+568]IGGASTPPK.G z=3,scan#=24679,scan time=54.2878
SVGHLEENGLTIGGASTPPK
Intensity
m/z
HexNAc
y3
y2
HexNAcHex
b4
a4
b5
a5
y4
b5-18
b6-18
b6
a6
y7
y8
b7
~y10
b8
b9
b10
Pep_2+
~b12
~y15
~b15
~y16
~b17
Pep_1+

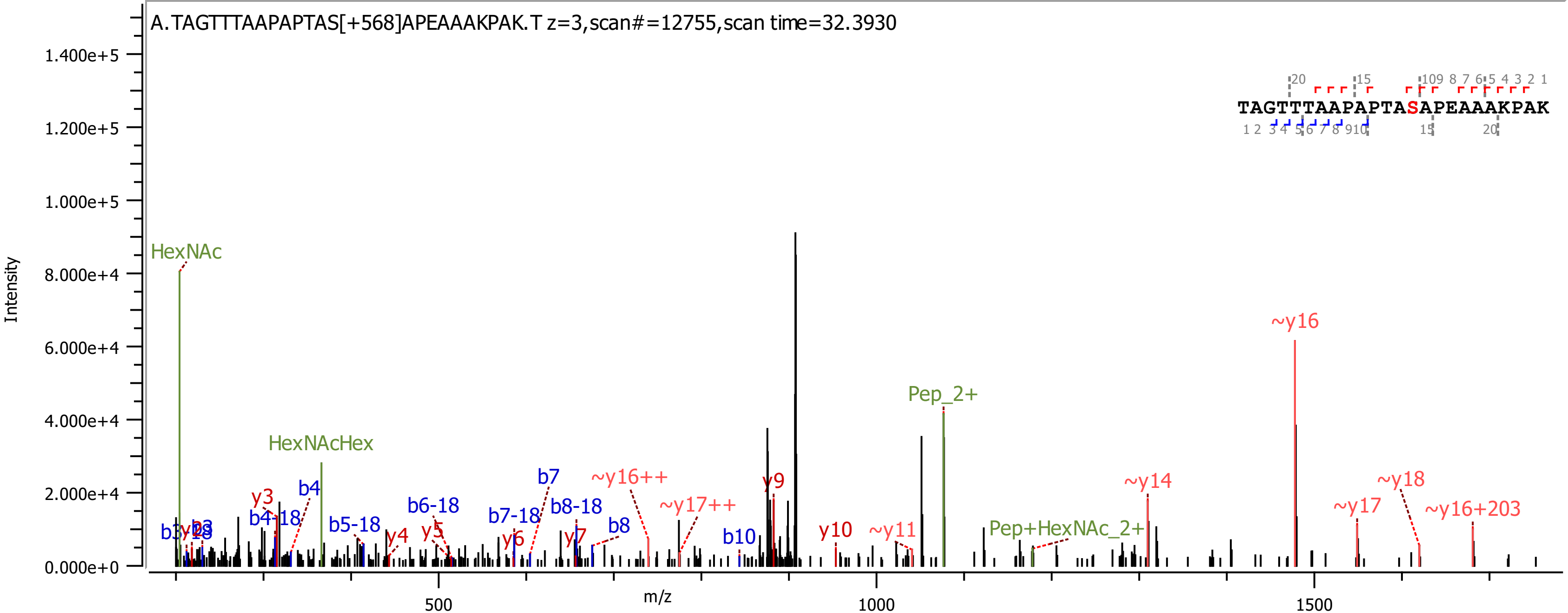
A.TAGTTTAAPAPTAS[+568]APEAAAKPAK.T z=3,scan#=12755,scan time=32.3930
TAGTTTAAPAPTASAPEAAAKPAK
Intensity
m/z
HexNAc
HexNAcHex
Pep_2+
Pep+HexNAc_2+
~y16
~y16++
~y17++
~y14
~y17
~y18
~y16+203
~y11
y3
y4
y5
y6
y7
y9
y10
b4
b4-18
b5-18
b6-18
b7
b7-18
b8-18
b8
b10

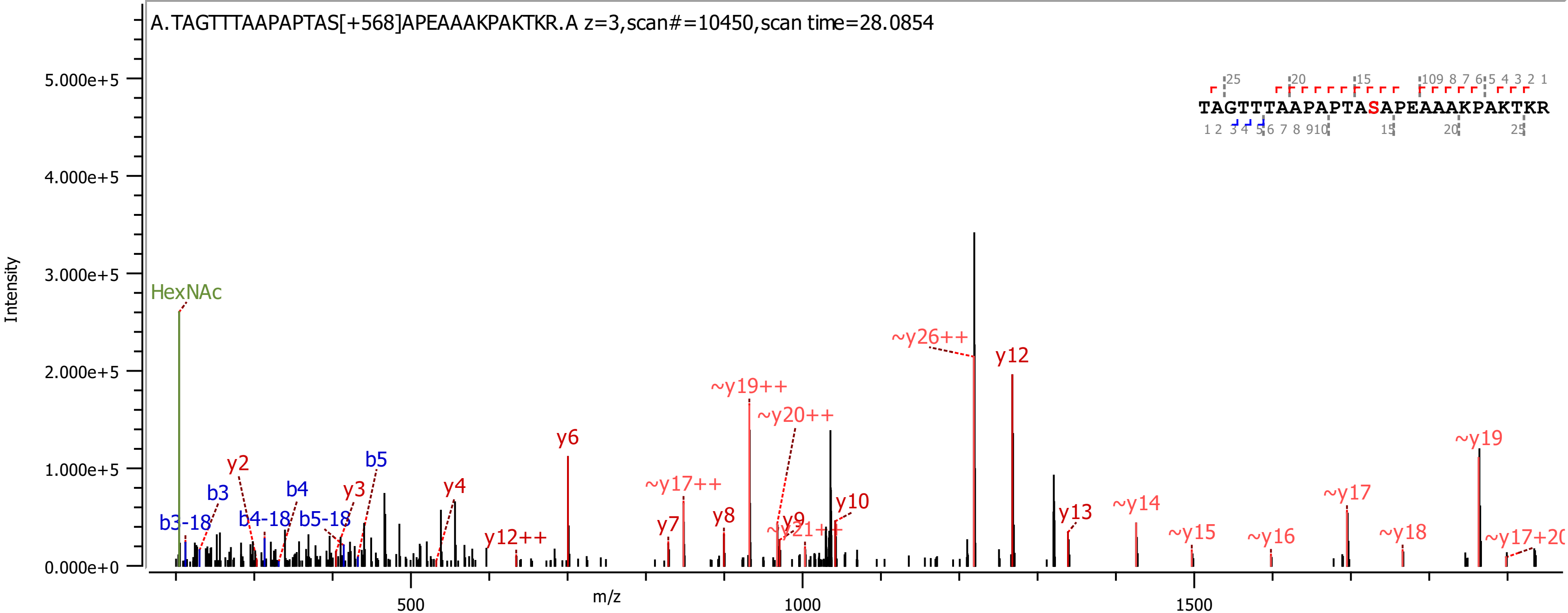

A.TAGTTTAAPAPTAS[+568]APEAAAKPAKTKR.A z=3,scan#=10450,scan time=28.0854
TAGTTTAAPAPTASAPEAAAKPAKTKR
Intensity
m/z
HexNAc
b3-18
b3
y2
b4-18
b4
b5-18
y3
b5
y4
y6
y12++
~y17++
y7
y8
~y19++
~y20++
y9
~y21++
y10
~y26++
y12
y13
~y14
~y15
~y16
~y17
~y18
~y19
~y17+20

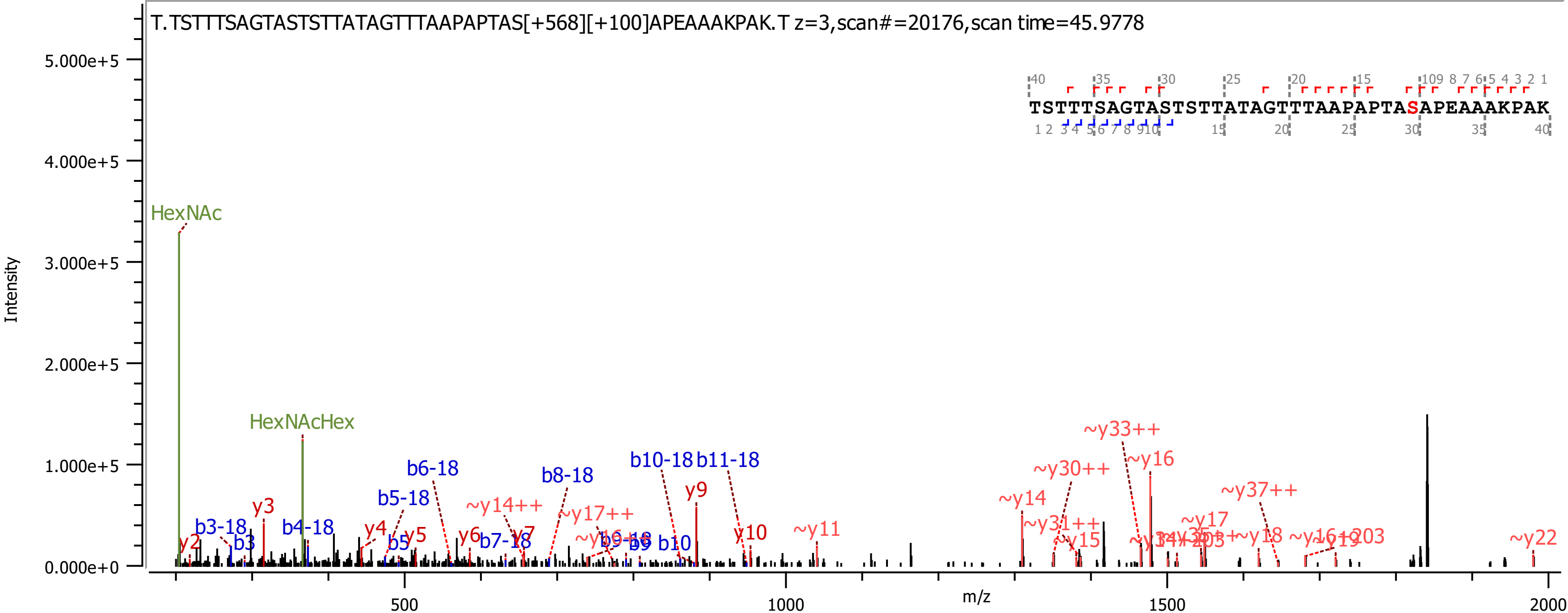

T.TSTTTSAGTASTSTTATAGTTTAAPAPTAS[+568][+100]APEAAAKPAK.T z=3,scan#=20176,scan time=45.9778
TSTTTSAGTASTSTTATAGTTTAAPAPTASAPEAAAKPAK
Intensity
m/z
5.000e+5
4.000e+5
3.000e+5
2.000e+5
1.000e+5
0.000e+0
500
1000
1500
2000
HexNAc
HexNAcHex
y2
b3-18
b3
y3
b4-18
y4
b5-18
b5
y5
b6-18
y6
b7-18
~y14++
y7
b8-18
~y17++
b9-18
b9
b10-18
b10
b11-18
y9
y10
~y11
~y14
~y30++
~y31++
~y15
~y33++
~y16
~y17
~y37++
~y18
~y22

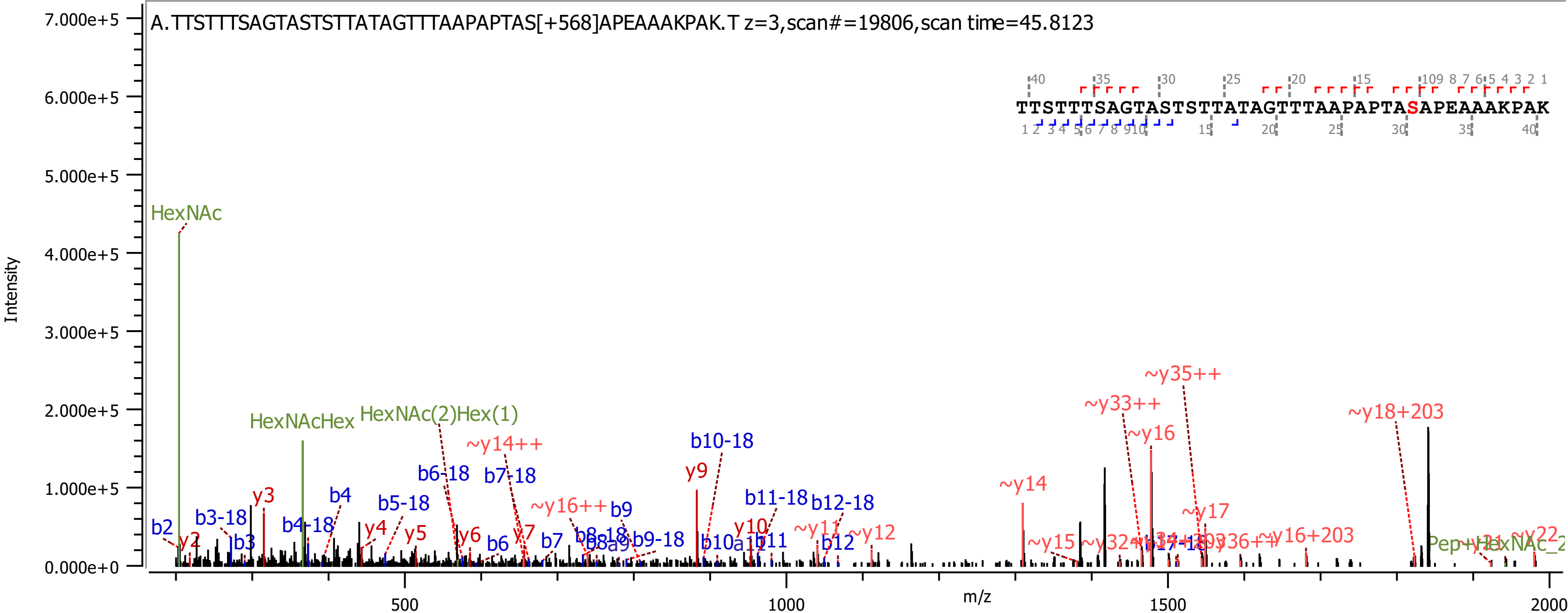
A.TTSTTTSAGTASTSTTATAGTTTAAPAPTAS[+568]APEAAAKPAK.T z=3,scan#=19806,scan time=45.8123
TTSTTTSAGTASTSTTATAGTTTAAPAPTASAPEAAAKPAK
Intensity
m/z
7.000e+5
6.000e+5
5.000e+5
4.000e+5
3.000e+5
2.000e+5
1.000e+5
0.000e+0
500
1000
1500
2000
HexNAc
HexNAcHex
HexNAc(2)Hex(1)
~y14++
~y16++
~y35++
~y33++
~y16
~y18+203
~y14
~y17
~y15
~y16+203
~y11
~y12
y2
y3
y4
y5
y6
y7
y9
y10
b2
b3-18
b3
b4-18
b4
b5-18
b6-18
b6
b7-18
b7
b9
b9-18
b10-18
b11-18
b11
b12-18
b12

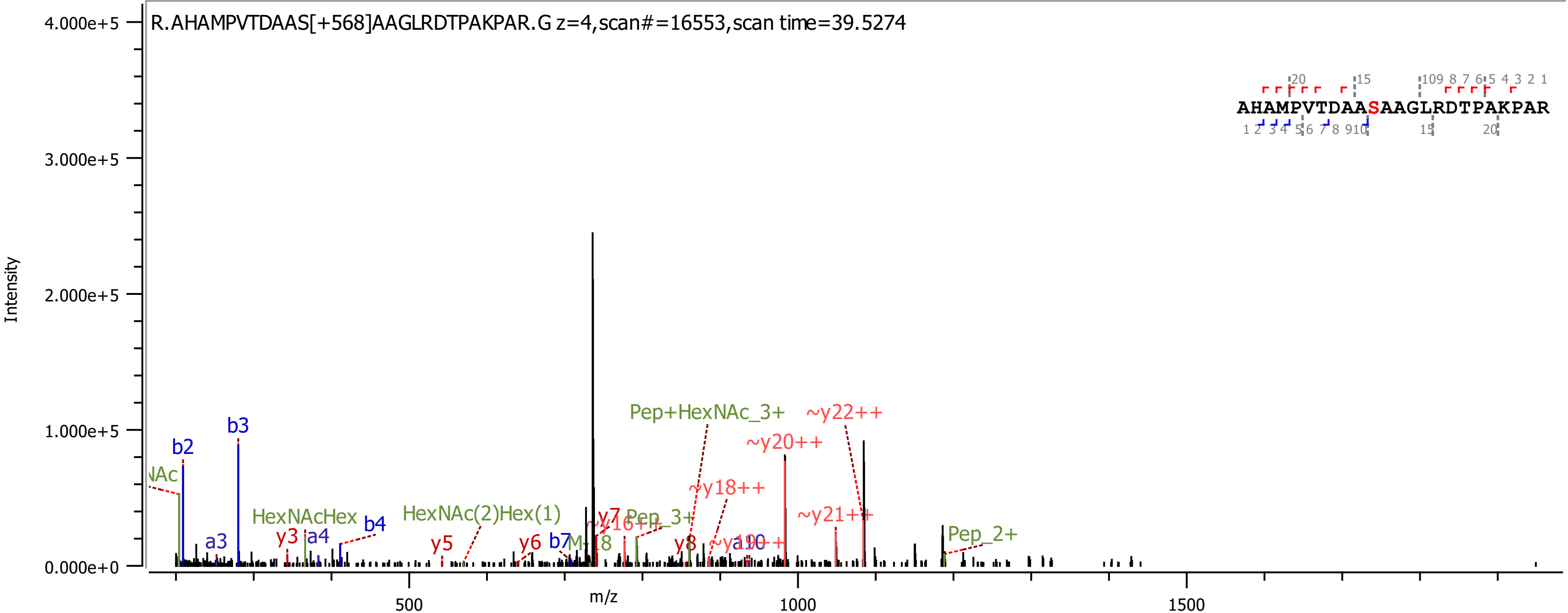
R.AHAMPVTDAAS[+568]AAGLRDTPAKPAR.G z=4,scan#=16553,scan time=39.5274
AHAMPVTDAASAAGLRDTPAKPAR
Intensity
m/z
Pep+HexNAc_3+
~y22++
~y20++
~y18++
~y21++
Pep_2+
Pep_3+
HexNAcHex
HexNAc(2)Hex(1)
b2
b3
b4
b7
a3
a4
y3
y5
y6
y7
y8
~y16++

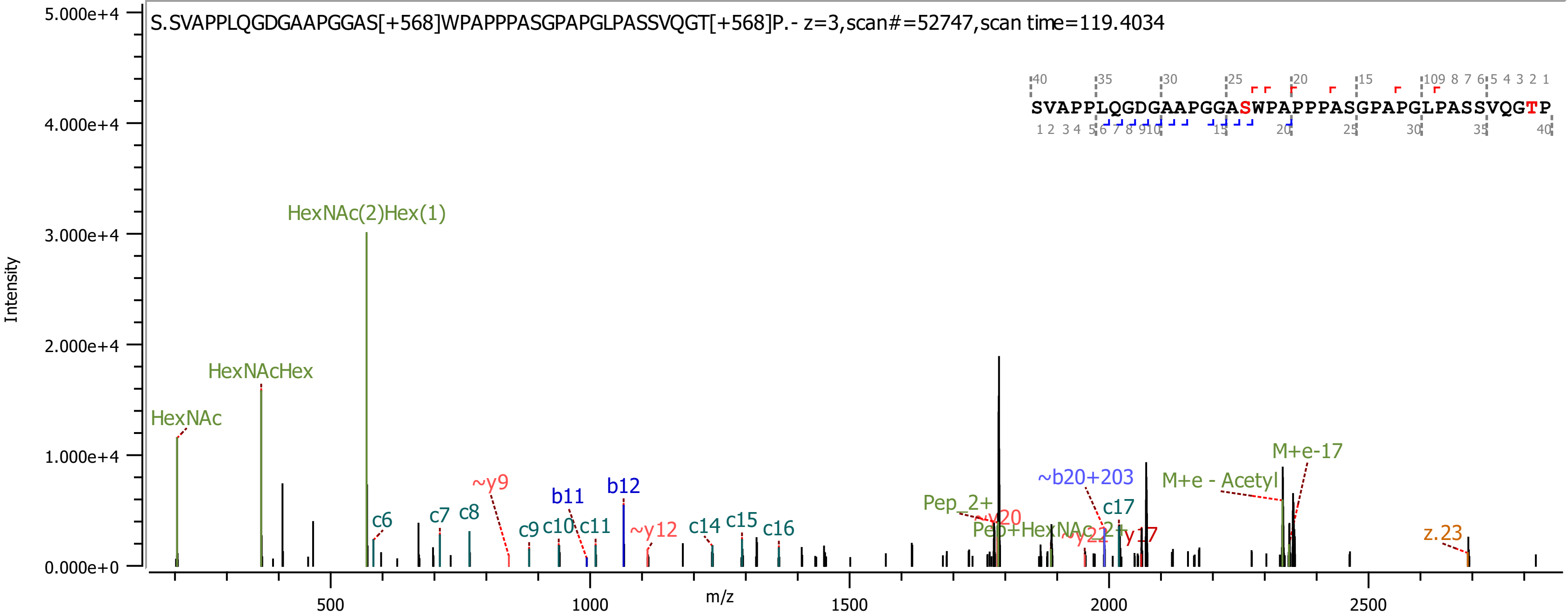
S.SVAPPLQGDGAAPGGAS[+568]WPAPPPASGPAPGLPASSVQGT[+568]P.- z=3,scan#=52747,scan time=119.4034
SVAPPLQGDGAAPGGASWPAPPPASGPAPGLPASSVQGTP
Intensity
m/z
HexNAc(2)Hex(1)
HexNAcHex
HexNAc
c6
c7
c8
~y9
b11
b12
c9
c10
c11
~y12
c14
c15
c16
Pep_2+
~y20
Pep+HexNAc_2+
~y22
y17
~b20+203
c17
M+e - Acetyl
M+e-17
z.23

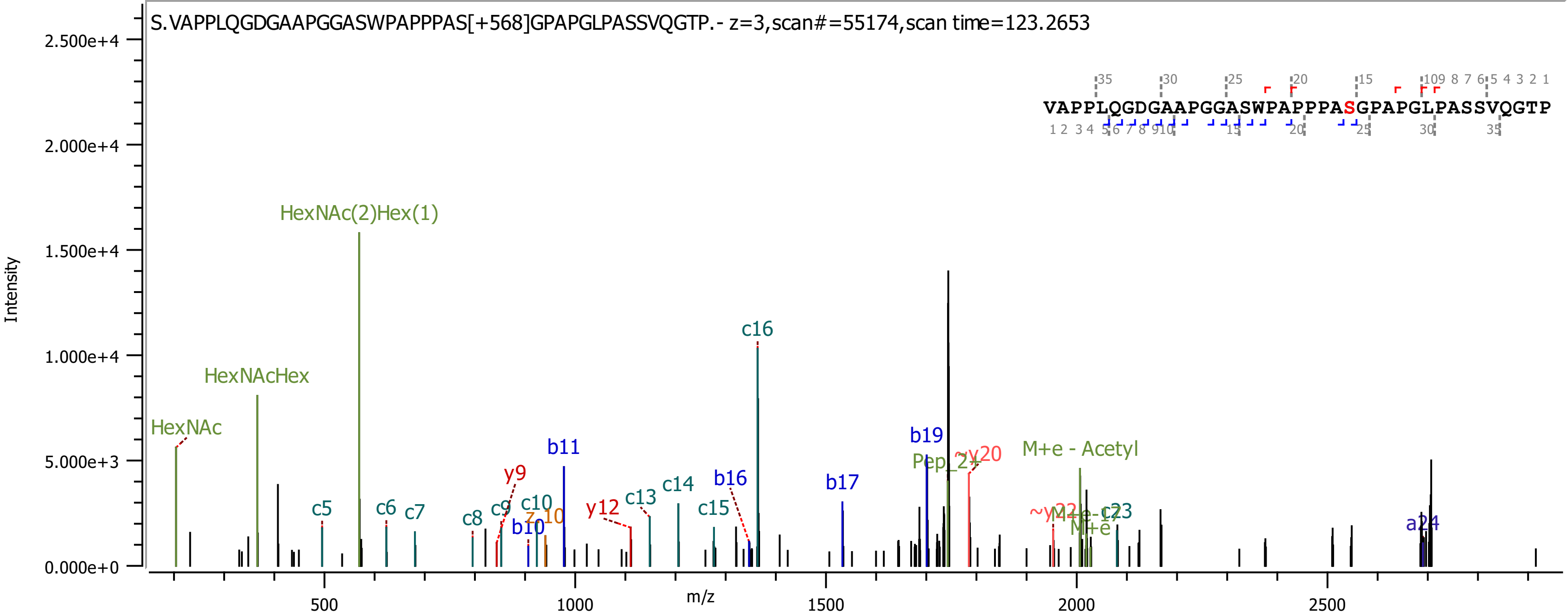
S.VAPPLQGDGAAPGGASWPAPPPAS[+568]GPAPGLPASSVQGTP.- z=3,scan#=55174,scan time=123.2653
VAPPLQGDGAAPGGASWPAPPPASGPAPGLPASSVQGTP
Intensity
m/z
HexNAc
HexNAcHex
HexNAc(2)Hex(1)
c5
c6
c7
c8
c9
c10
b10
z10
b11
y9
y12
c13
c14
c15
b16
c16
b17
b19
Pep_2+
~y20
M+e - Acetyl
~y22
M+e
c23
a24

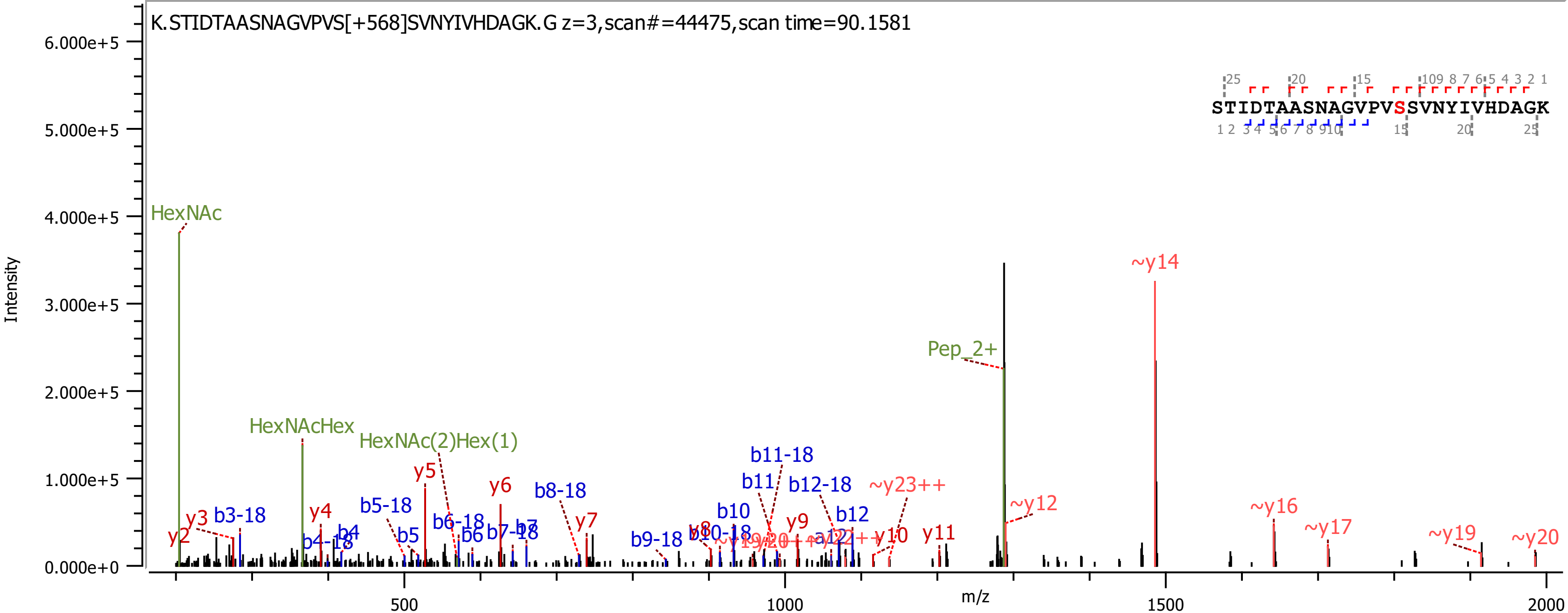

K.STIDTAASNAGVPVS[+568]SVNYIVHDAGK.G z=3,scan#=44475,scan time=90.1581
STIDTAASNAGVPVSSVNYIVHDAGK
Intensity
m/z
HexNAc
HexNAcHex
HexNAc(2)Hex(1)
Pep_2+
~y14
~y12
~y16
~y17
~y19
~y20
~y23++
y2
y3
y4
y5
y6
y7
y9
y10
y11
b3-18
b4-18
b4
b5-18
b5
b6-18
b6
b7-18
b8-18
b9-18
b10
b11
b11-18
b12-18
b12

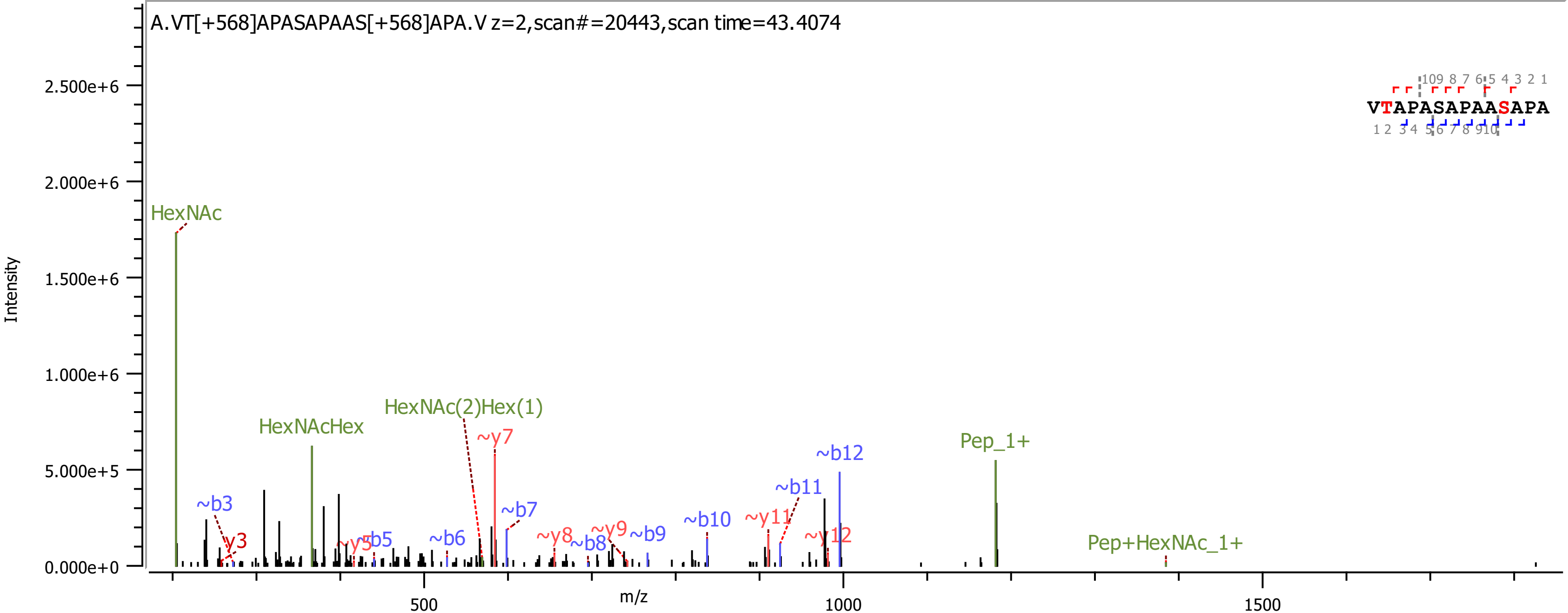
A.VT[+568]APASAPAAS[+568]APA.V z=2,scan#=20443,scan time=43.4074
Intensity
m/z
HexNAc
HexNAcHex
HexNAc(2)Hex(1)
Pep_1+
Pep+HexNAc_1+
~b3
~b5
~b6
~b7
~b8
~b9
~b10
~b11
~b12
y3
~y5
~y7
~y8
~y9
~y11
~y12
VTAPASAPAASAPA

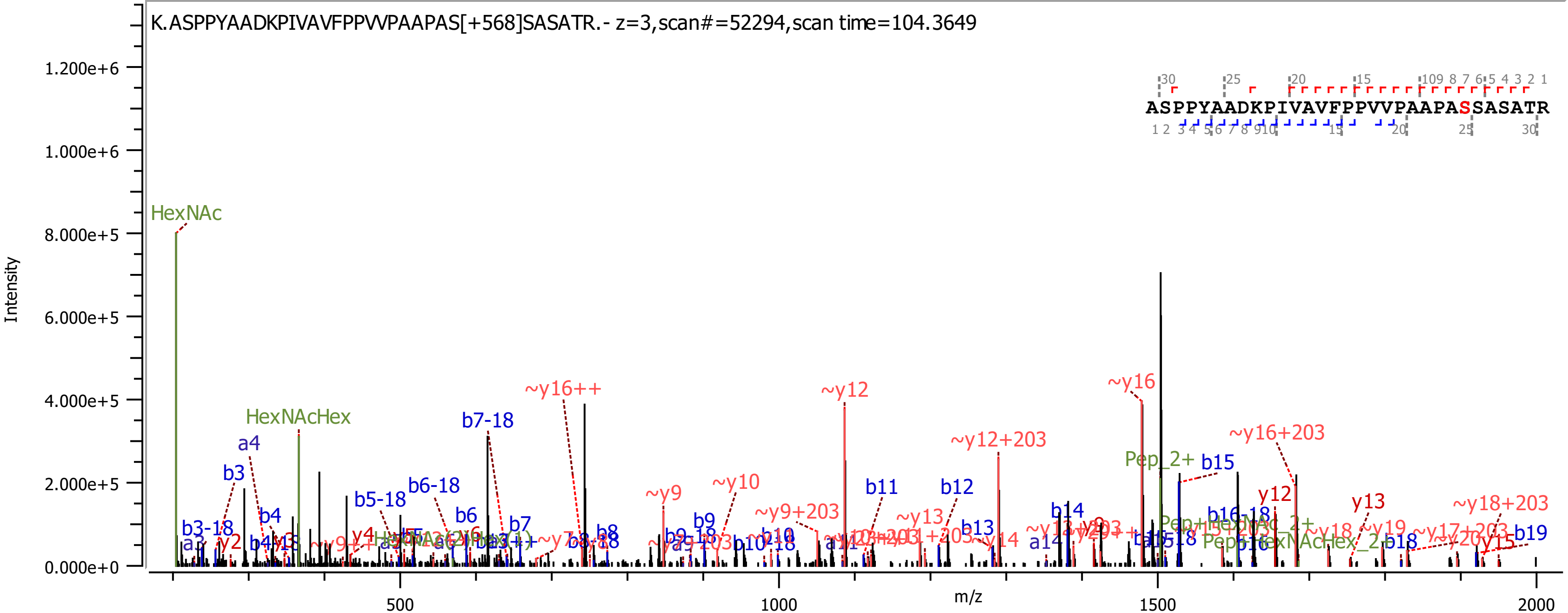

K.ASPPYAADKPIVAVFPPVVPAAPAS[+568]SASATR.- z=3,scan#=52294,scan time=104.3649
ASPPYAADKPIVAVFPPVVPAAPASSASATR
Intensity
m/z
1.200e+6
1.000e+6
8.000e+5
6.000e+5
4.000e+5
2.000e+5
0.000e+0
500
1000
1500
2000
HexNAc
HexNAcHex
a4
b3
b3-18
a3
y2
b4
b5-18
b6-18
b6
b7-18
b7
~y16++
b8
~y9
~y10
b9
~y9+203
b11
~y12
~y13
b12
~y12+203
b13
~y14
b14
a14
~y16
Pep_2+
b15
~y16+203
y12
b16-18
y13
~y18
~y19
b19
~y18+203
y15

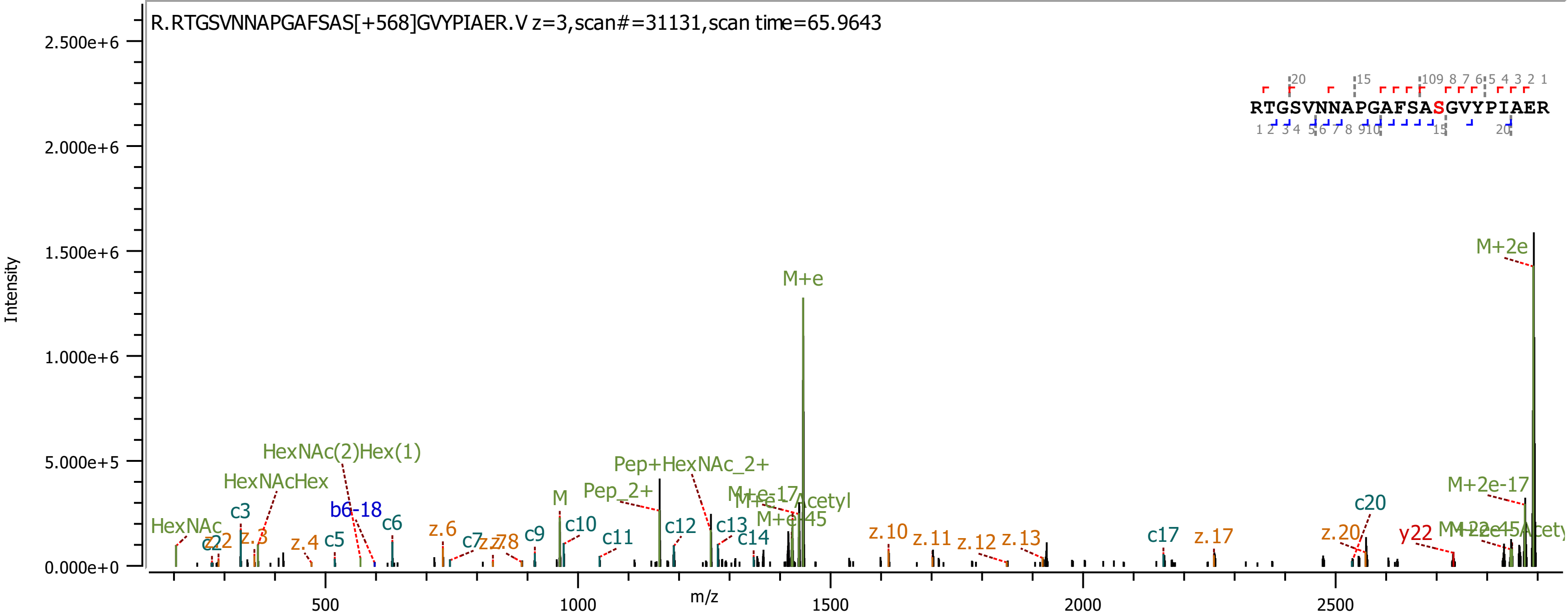

R.RTGSVNNAPGAFSAS[+568]GVYPIAER.V z=3,scan#=31131,scan time=65.9643
RTGSVNNAPGAFSASGVYPIAER
Intensity
m/z
M+e
M+2e
M+2e-17
HexNAc(2)Hex(1)
HexNAcHex
HexNAc
Pep+HexNAc_2+
Pep_2+
M
b6-18
y22
c20
c17
z.17
z.20
z.10
z.11
z.12
z.13

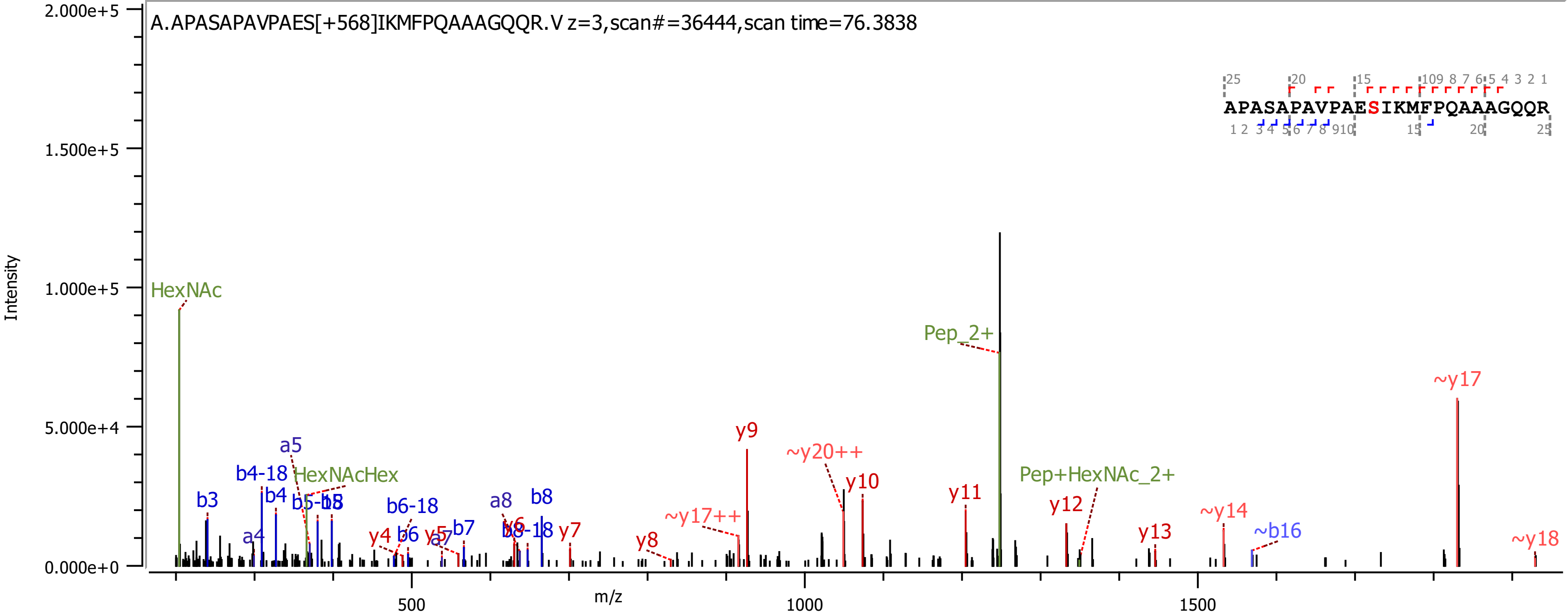
A.APASAPAVPAES[+568]IKMFPQAAAGQQR.V z=3,scan#=36444,scan time=76.3838
APASAPAVPAESIKMFPQAAAGQQR
Intensity
m/z
HexNAc
HexNAcHex
Pep_2+
Pep+HexNAc_2+
b3
b4-18
b4
a4
a5
b5-18
b6-18
b6
b7
a7
a8
b8
b8-18
y4
y5
y6
y7
y8
y9
y10
y11
y12
y13
~y17++
~y20++
~y14
~b16
~y17
~y18
2.000e+5
1.500e+5
1.000e+5
5.000e+4
0.000e+0
500
1000
1500

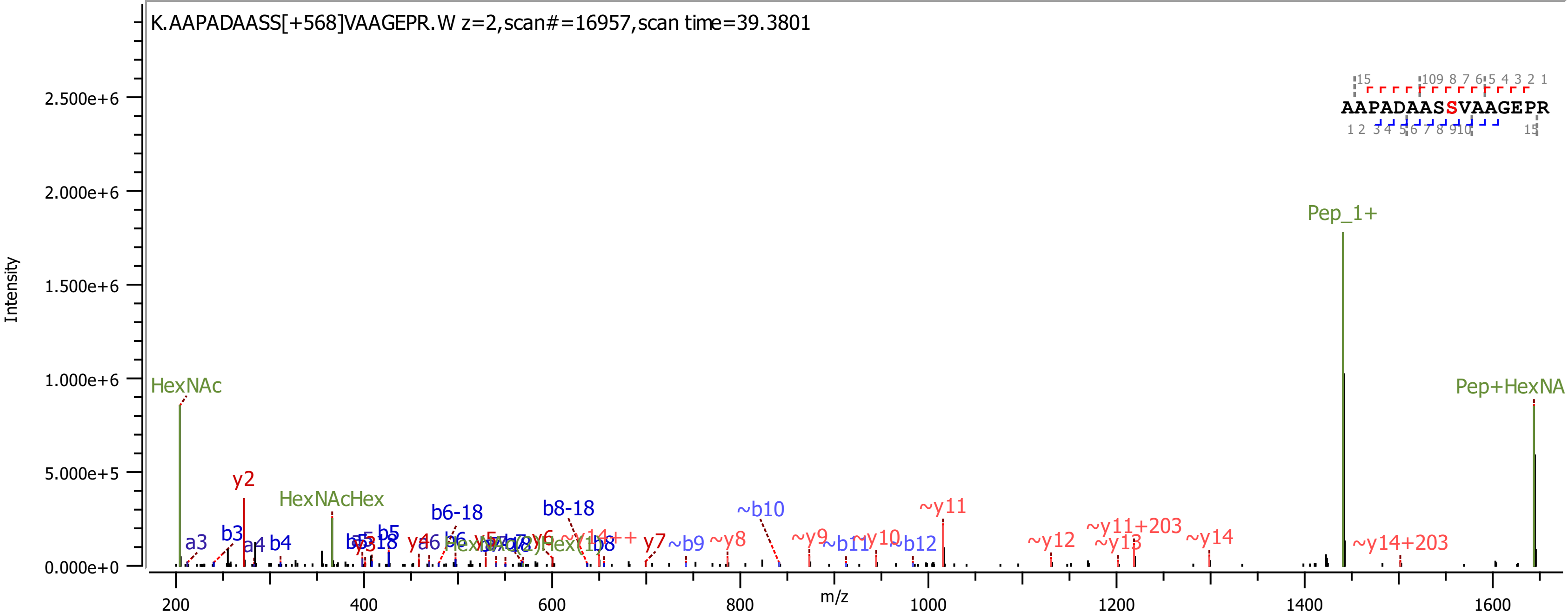

K.AAPADAASS[+568]VAAGEPR.W z=2,scan#=16957,scan time=39.3801
AAPADAASSVAAGEPR
Intensity
m/z
2.500e+6
2.000e+6
1.500e+6
1.000e+6
5.000e+5
0.000e+0
200
400
600
800
1000
1200
1400
1600
HexNAc
y2
a3
b3
a4
b4
HexNAcHex
b6-18
b8-18
b8
y7
~b9
~y8
~b10
~y9
~b11
~y10
~b12
~y11
~y12
~y11+203
~y13
~y14
~y14+203
Pep_1+
Pep+HexNA

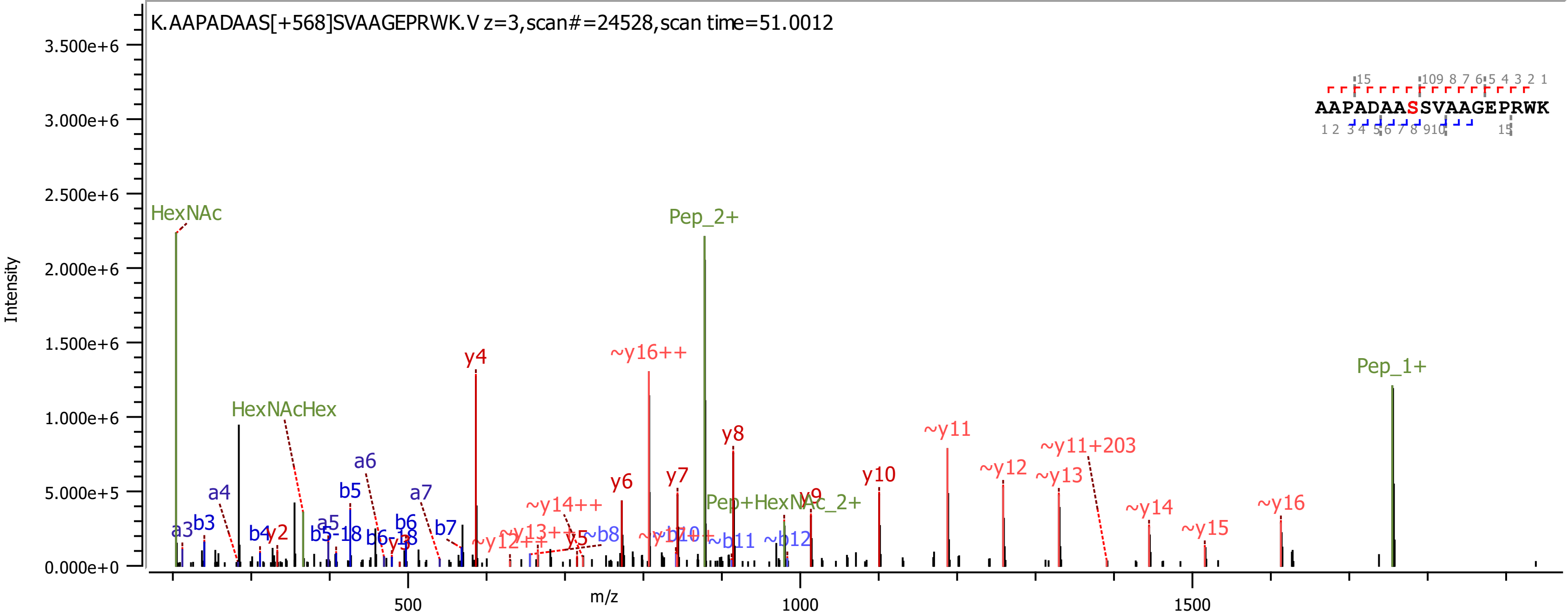

K.AAPADAAS[+568]SVAAGEPRWK.V z=3,scan#=24528,scan time=51.0012
AAPADAASSVAAGEPRWK
Intensity
m/z
HexNAc
HexNAcHex
Pep_2+
Pep_1+
Pep+HexNAc_2+
y4
~y16++
y8
~y11
~y11+203
~y12
~y13
y10
y6
y7
y9
~y14
~y15
~y16
~y14++
~y13++
~y12++
y5
y3
y2
a3
b3
a4
b4
a5
b5-18
b5
a6
b6-18
b6
a7
b7
~b8
~b10
~b11
~b12

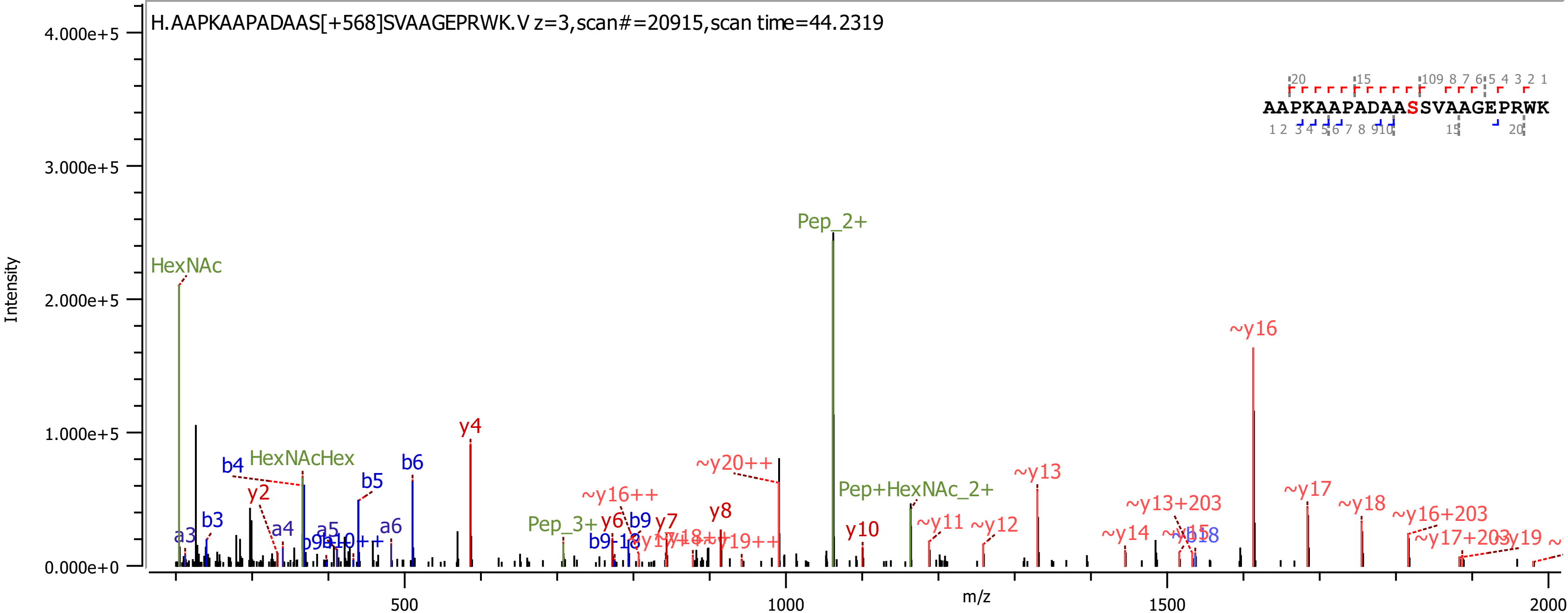
H.AAPKAAPADAAS[+568]SVAAGEPRWK.V z=3,scan#=20915,scan time=44.2319
AAPKAAPADAASSVAAGEPRWK
Intensity
m/z
4.000e+5
3.000e+5
2.000e+5
1.000e+5
0.000e+0
500
1000
1500
2000
HexNAc
HexNAcHex
Pep_2+
Pep_3+
Pep+HexNAc_2+
y4
~y16
~y13
~y17
~y18
~y20++
~y16++
~y11
~y12
~y14
~y13+203
~y16+203
~y17+203
~y19
y2
y6
y7
y8
y10
b3
b4
b5
b6
b9
a3
a4
a5
a6

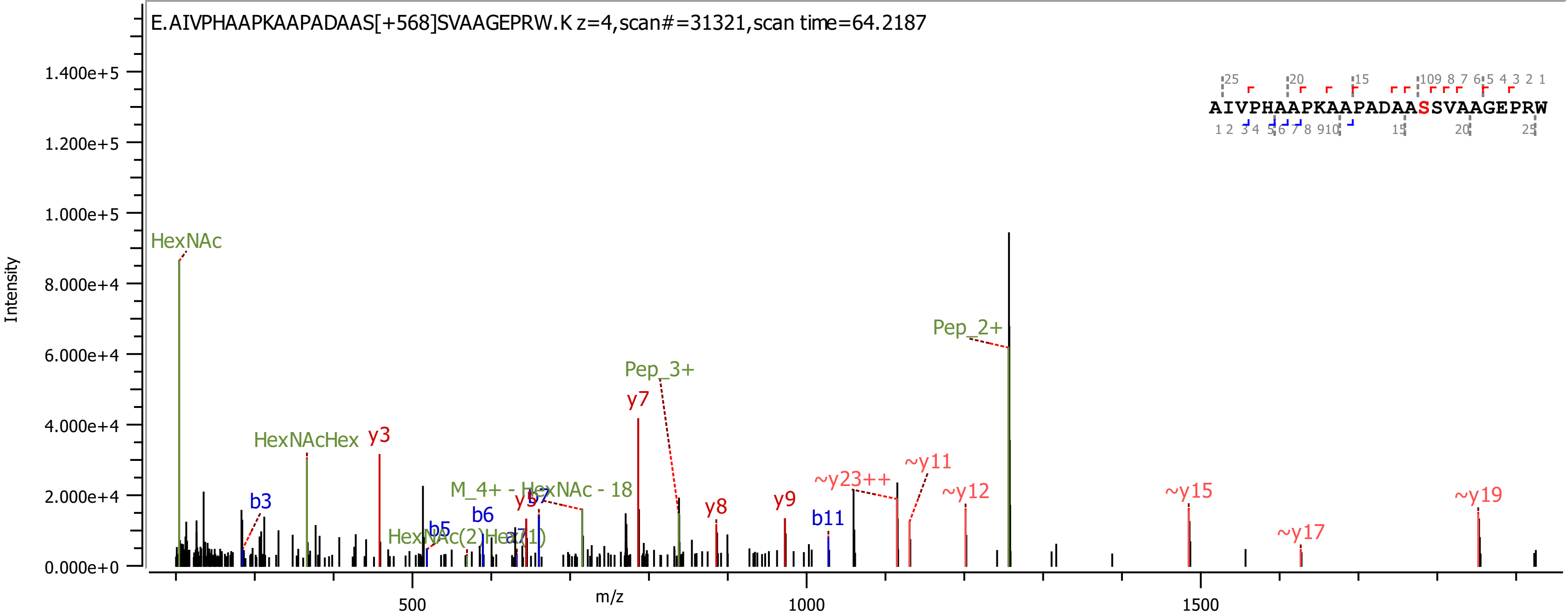

E.AIVPHAAPKAAPADAAS[+568]SVAAGEPRW.K z=4,scan#=31321,scan time=64.2187
AIVPHAAPKAAPADAASSVAAGEPRW
Intensity
m/z
HexNAc
HexNAcHex
y3
b3
b5
b6
y5
b7
HexNAc(2)Hex(1)
M_4+ - HexNAc - 18
y7
Pep_3+
y8
y9
b11
~y23++
~y11
~y12
Pep_2+
~y15
~y17
~y19

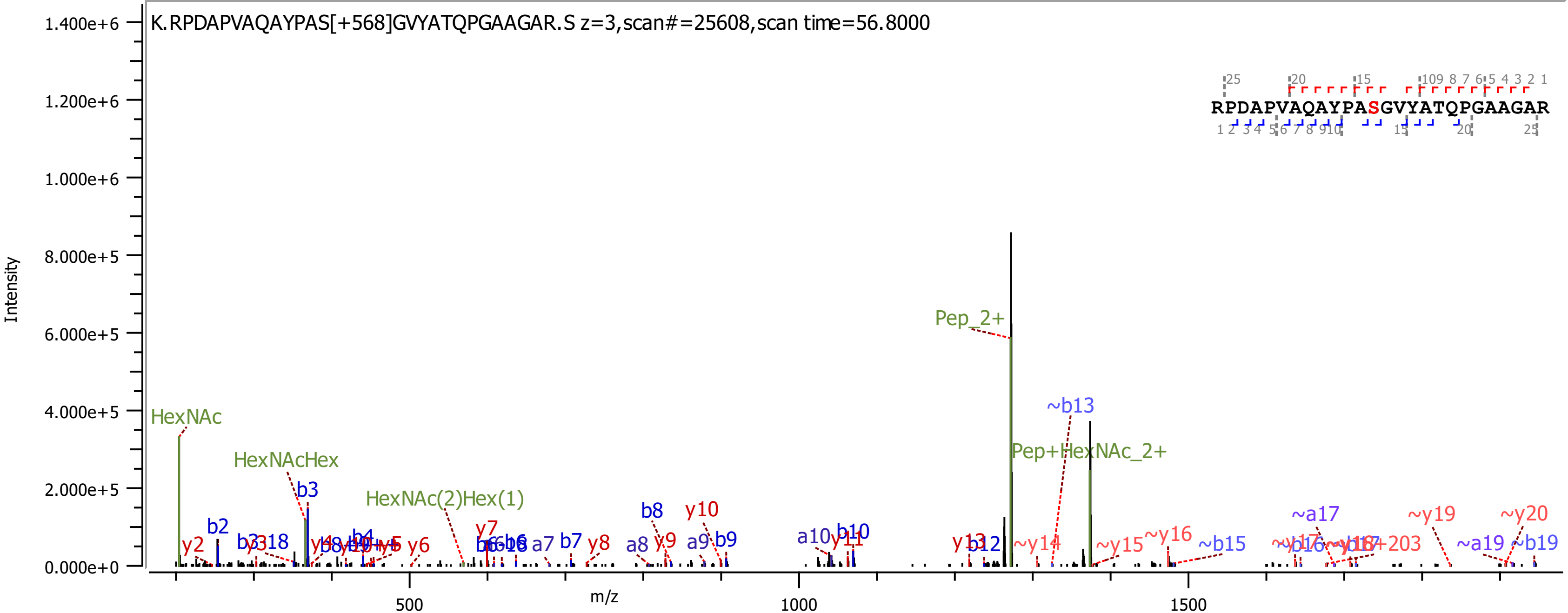
K.RPDAPVAQAYPAS[+568]GVYATQPGAAGAR.S z=3,scan#=25608,scan time=56.8000
RPDAPVAQAYPASGVYATQPGAAGAR
Intensity
m/z
0.000e+0
2.000e+5
4.000e+5
6.000e+5
8.000e+5
1.000e+6
1.200e+6
1.400e+6
500
1000
1500
HexNAc
HexNAcHex
HexNAc(2)Hex(1)
Pep_2+
Pep+HexNAc_2+
~b13
b2
b3
b4
b7
b8
b9
b10
b12
a7
a8
a9
a10
y2
y6
y7
y8
y9
y10
y11
y13
~y14
~y15
~y16
~b15
~a17
~y19
~y20
~a19
~b19

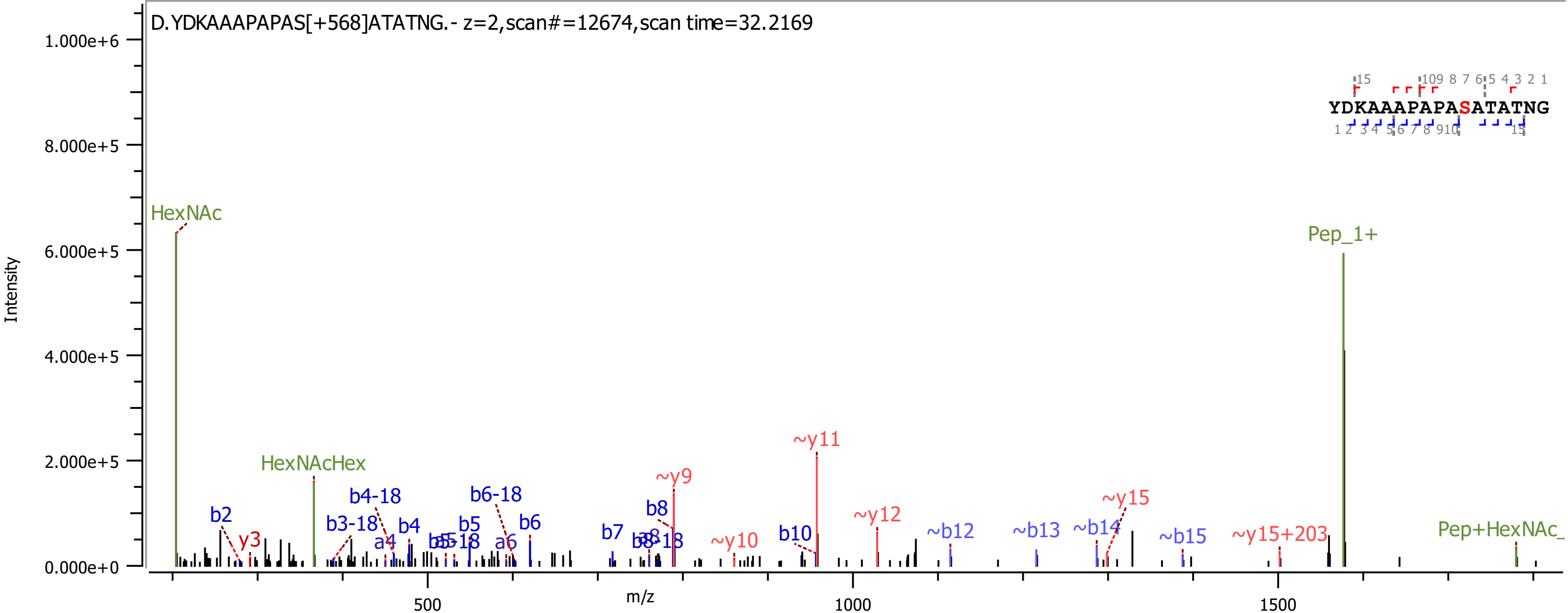

D.YDKAAAPAPAS[+568]ATATNG.- z=2,scan#=12674,scan time=32.2169
Intensity
m/z
1.000e+6
8.000e+5
6.000e+5
4.000e+5
2.000e+5
0.000e+0
500
1000
1500
HexNAc
HexNAcHex
Pep_1+
Pep+HexNAc_
b2
y3
b3-18
b4-18
a4
b4
b5
b6-18
a6
b6
b7
b8
b8-18
~y9
~y10
b10
~y11
~y12
~b12
~b13
~b14
~y15
~b15
~y15+203
YDKAAAPAPASATATNG

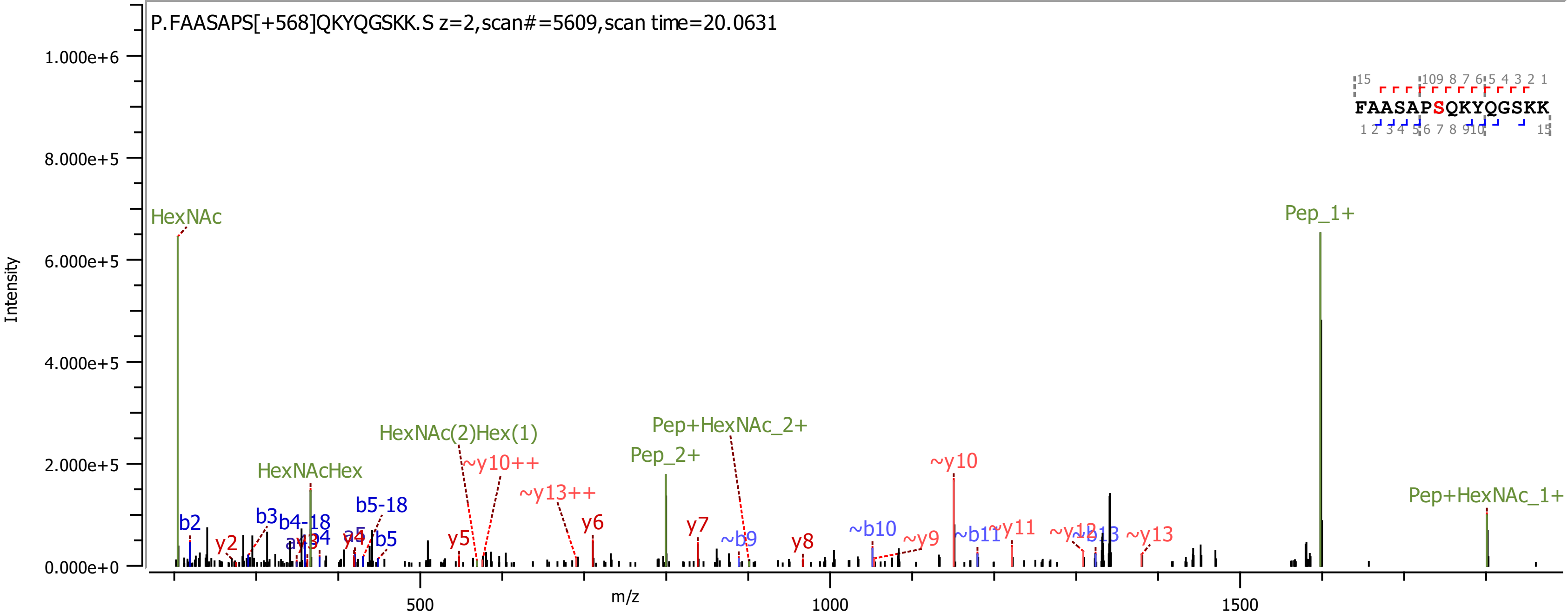
P.FAASAPS[+568]QKYQGSKK.S z=2,scan#=5609,scan time=20.0631
Intensity
m/z
HexNAc
Pep_1+
HexNAcHex
HexNAc(2)Hex(1)
Pep_2+
Pep+HexNAc_2+
Pep+HexNAc_1+
~y10++
~y13++
~y10
FAASAPSQKYQGSKK

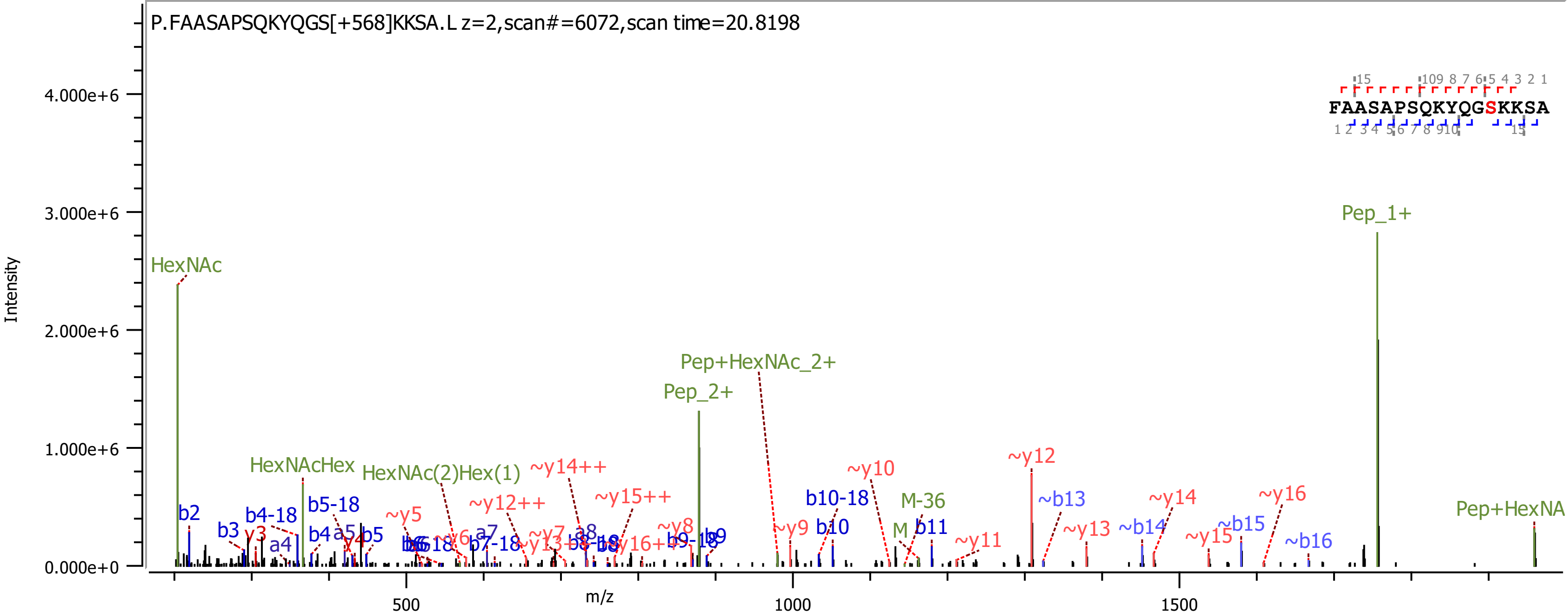
P.FAASAPSQKYQGS[+568]KKSA.L z=2,scan#=6072,scan time=20.8198
FAASAPSQKYQGSKKSA
Intensity
m/z
HexNAc
HexNAcHex
HexNAc(2)Hex(1)
Pep+HexNAc_2+
Pep_2+
Pep_1+
M-36
~y12
~y10
~y14++
~y12++
~y15++
~y5
~b13
~y14
~y16
~y13
~b14
~b15
~y15
~b16
~y11
~y9
~y8
b10-18
b10
b11
b2
b3
b4-18
b5-18
b4
b5
a4
a5
a7
a8
b7-18
b9-18
b9
4.000e+6
3.000e+6
2.000e+6
1.000e+6
0.000e+0
500
1000
1500

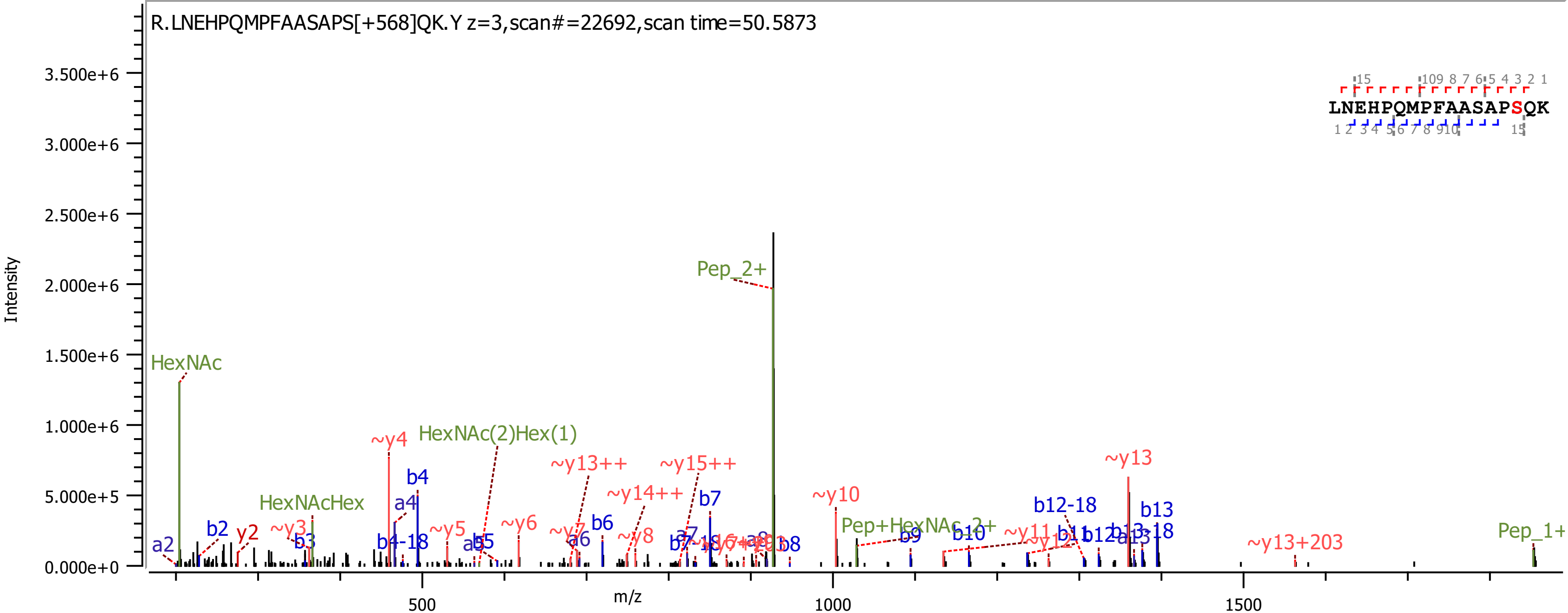
R.LNEHPQMPFAASAPS[+568]QK.Y z=3,scan#=22692,scan time=50.5873
LNEHPQMPFAASAPSQK
Intensity
m/z
HexNAc
HexNAcHex
HexNAc(2)Hex(1)
Pep_2+
Pep+HexNAc_2+
Pep_1+
~y4
~y13++
~y15++
~y14++
~y10
~y13
~y13+203
b4
a4
b7
b6
b12-18
b13
b2
y2
~y3
b3
b4-18
~y5
~y6
~y8
b9
b10
~y11
b13-18

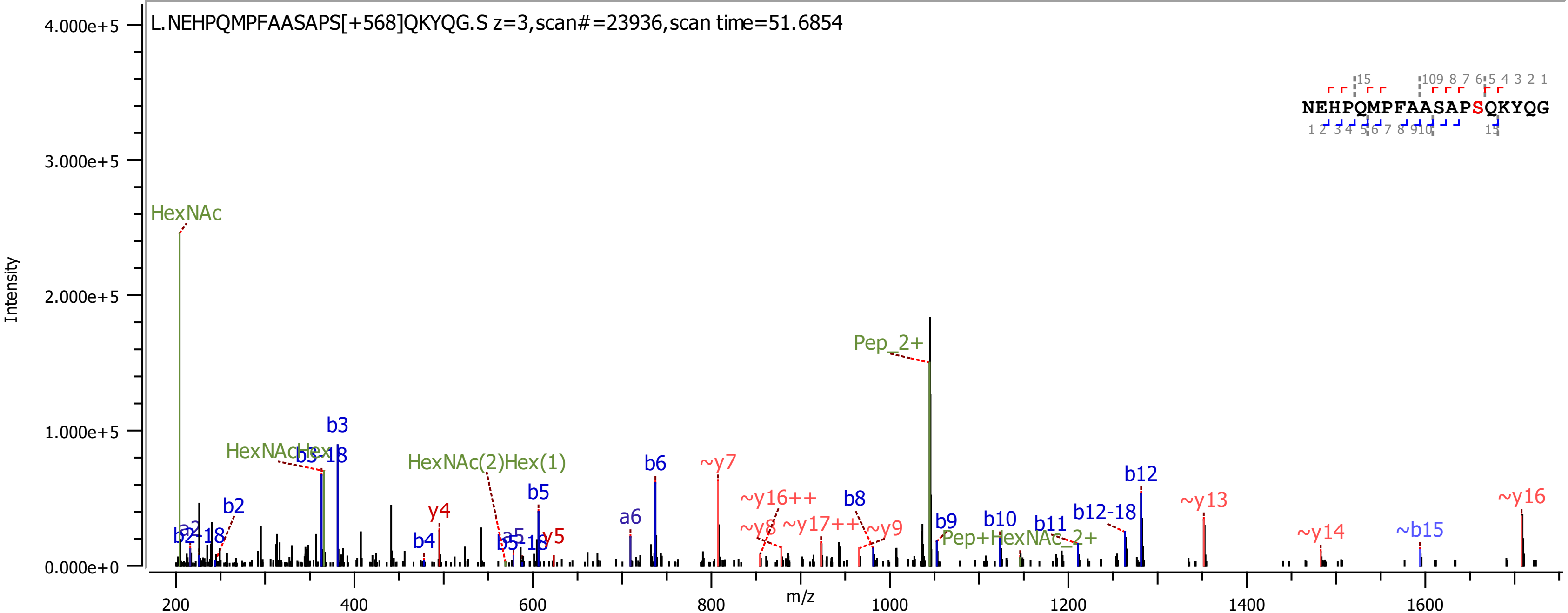

L.NEHPQMPFAASAPS[+568]QKYQG.S z=3,scan#=23936,scan time=51.6854
NEHPQMPFAASAPSQKYQG
Intensity
m/z
HexNAc
HexNAcHex
b3-18
b3
b2
a2
b2-18
HexNAc(2)Hex(1)
b4
y4
b5
a5
b5-18
y5
a6
b6
~y7
~y16++
~y8
~y17++
b8
~y9
Pep_2+
b9
b10
Pep+HexNAc_2+
b11
b12-18
b12
~y13
~y14
~b15
~y16

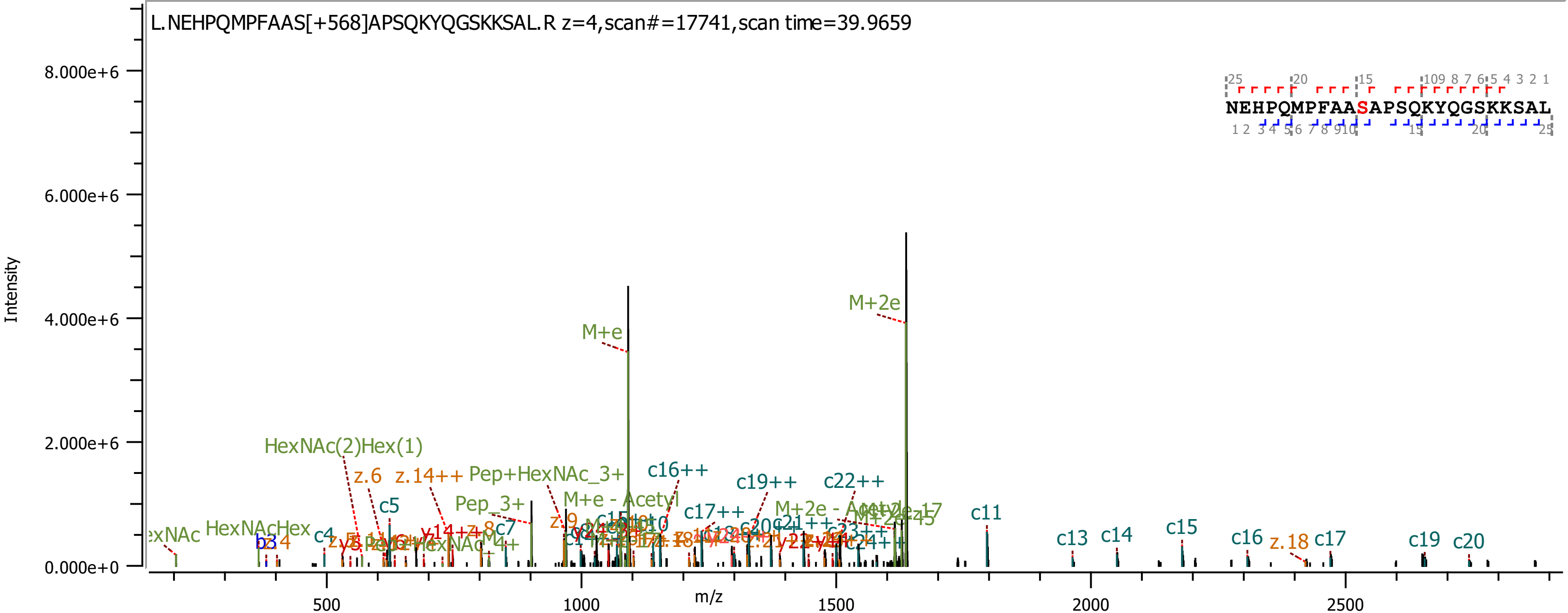

L.NEHPQMPFAAS[+568]APSQKYQGSKKSAL.R z=4,scan#=17741,scan time=39.9659
NEHPQMPFAASAPSQKYQGSKKSAL
Intensity
8.000e+6
6.000e+6
4.000e+6
2.000e+6
0.000e+0
m/z
500
1000
1500
2000
2500
M+e
M+2e
HexNAc(2)Hex(1)
z.6
z.14++
Pep+HexNAc_3+
c16++
c19++
c22++
c5
Pep_3+
M+e - Acetyl
c17++
HexNAcHex
c4
c7
c11
c13
c14
c15
c16
z.18
c17
c19
c20

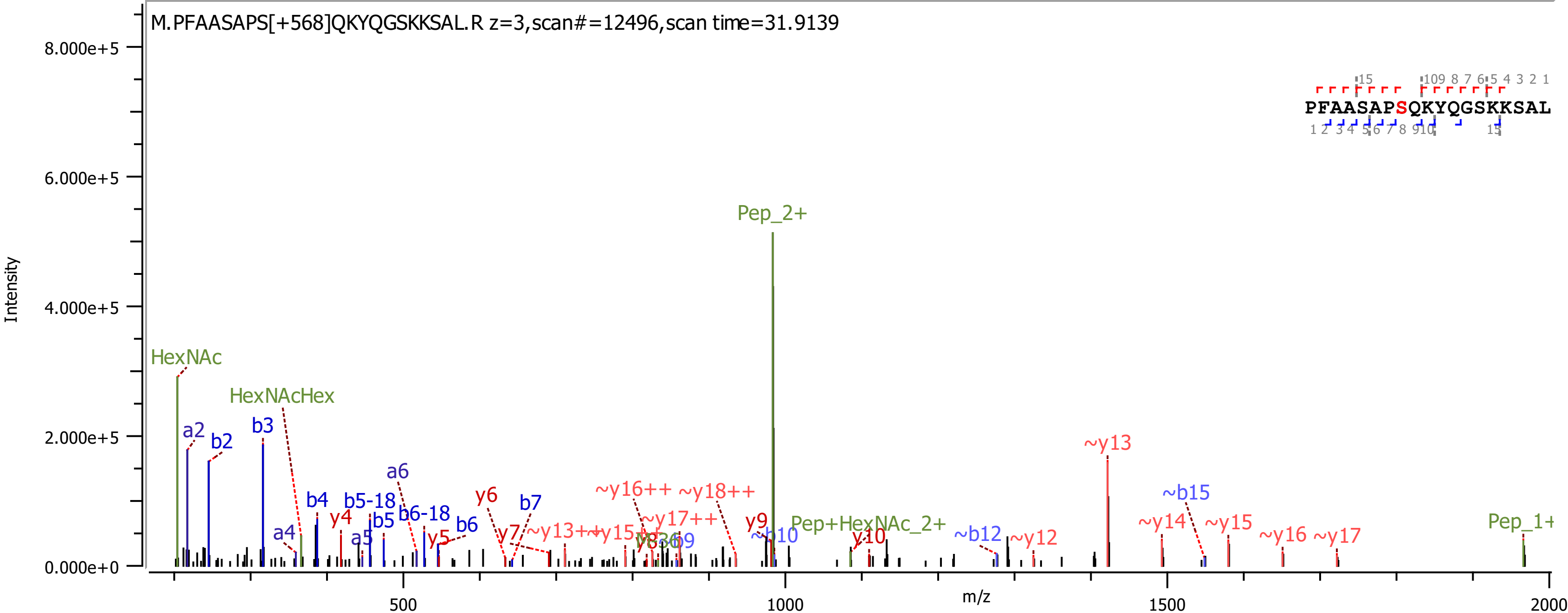

M.PFAASAPS[+568]QKYQGSKKSAL.R z=3,scan#=12496,scan time=31.9139
PFAASAPSQKYQGSKKSAL
Intensity
m/z
8.000e+5
6.000e+5
4.000e+5
2.000e+5
0.000e+0
500
1000
1500
2000
HexNAc
HexNAcHex
a2
b2
b3
a4
b4
b5-18
b5
a5
a6
b6-18
b6
y4
y5
y6
b7
y7
~y13++
~y15++
~y16++
~y18++
~y17++
y8
y9
~b9
~b10
y10
Pep_2+
Pep+HexNAc_2+
~b12
~y12
~y13
~y14
~b15
~y15
~y16
~y17
Pep_1+

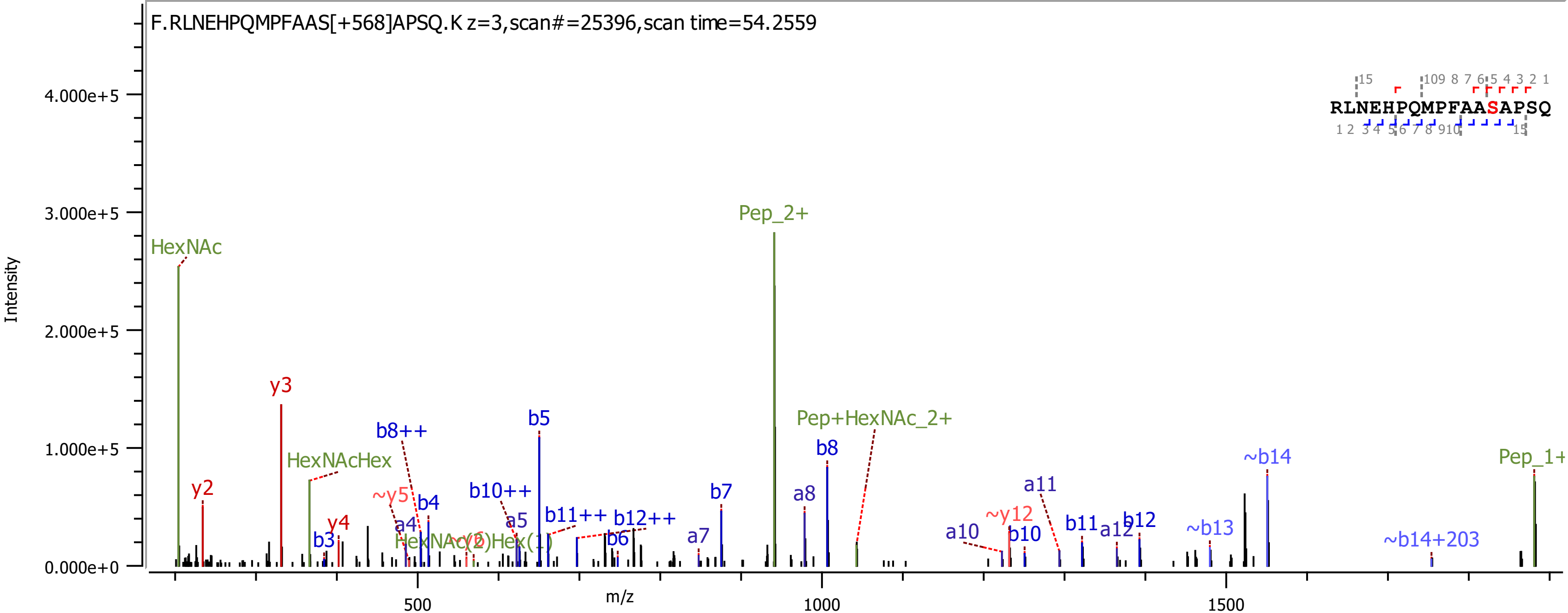

F.RLNEHPQMPFAAS[+568]APSQ.K z=3,scan#=25396,scan time=54.2559
RLNEHPQMPFAASAPSQ
Intensity
m/z
4.000e+5
3.000e+5
2.000e+5
1.000e+5
0.000e+0
500
1000
1500
HexNAc
y2
y3
HexNAcHex
y4
b3
~y5
a4
b8++
b4
HexNAc(2)Hex(1)
b10++
a5
b5
b11++
b12++
b6
a7
b7
Pep_2+
a8
b8
Pep+HexNAc_2+
a10
~y12
b10
a11
b11
a12
b12
~b13
~b14
~b14+203
Pep_1+

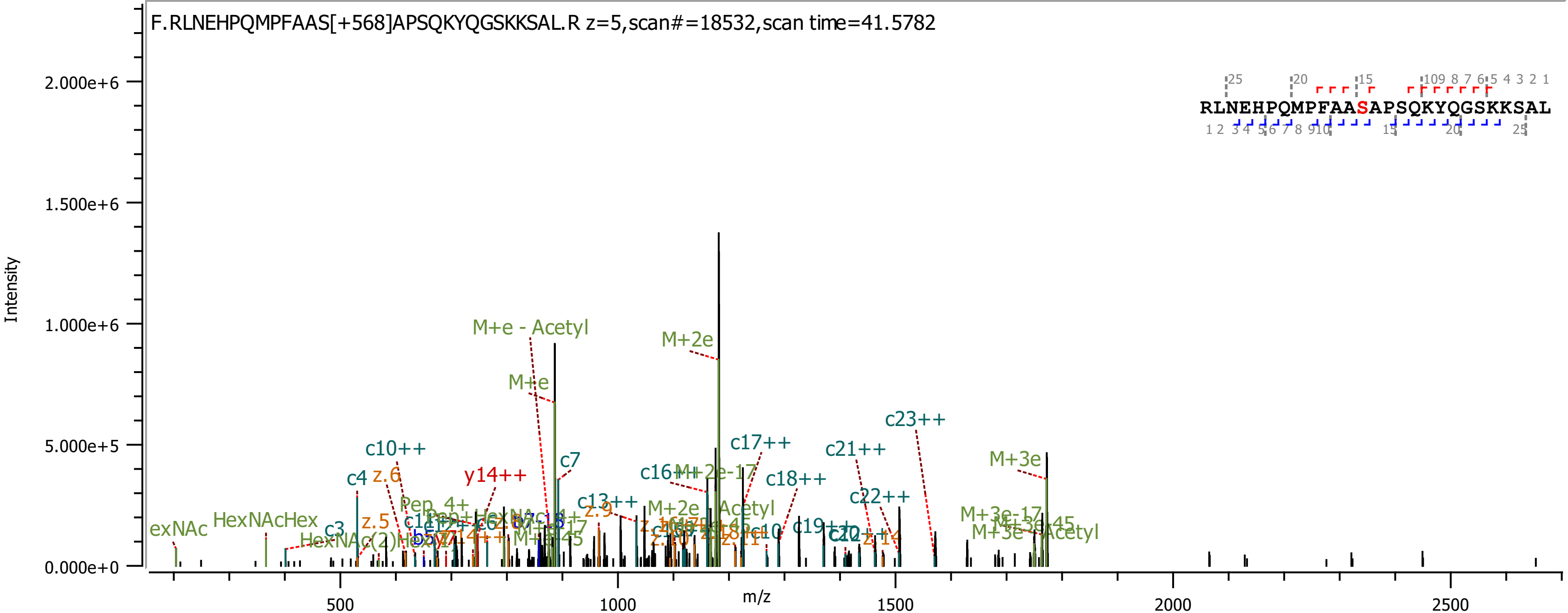
F.RLNEHPQMPFAAS[+568]APSQKYQGSKKSAL.R z=5,scan#=18532,scan time=41.5782
RLNEHPQMPFAASAPSQKYQGSKKSAL
Intensity
m/z
2.000e+6
1.500e+6
1.000e+6
5.000e+5
0.000e+0
500
1000
1500
2000
2500
M+e - Acetyl
M+2e
M+e
c7
c10++
c4
z.6
y14++
c16++
c17++
c18++
c21++
c23++
c22++
M+3e
c13++
z.9
M+2e - Acetyl
HexNAcHex
z.5
c3
c19++
M+3e-17
M+3e - Acetyl

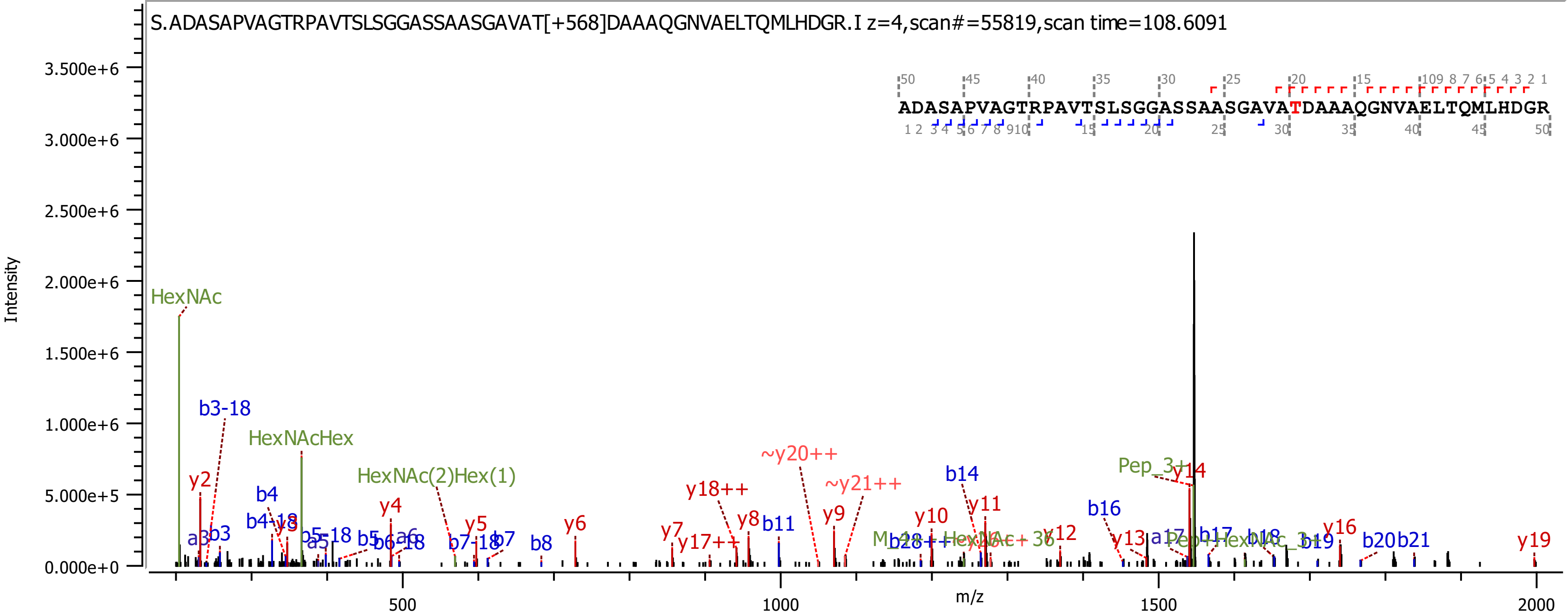
S.ADASAPVAGTRPAVTSLSGGASSAASGAVAT[+568]DAAAQGNVAELTQMLHDGR.I z=4,scan#=55819,scan time=108.6091
ADASAPVAGTRPAVTSLSGGASSAASGAVATDAAAQGNVAELTQMLHDGR
Intensity
m/z
HexNAc
HexNAcHex
HexNAc(2)Hex(1)
b3-18
y2
b4
y4
y5
y6
b7
b8
y7
y8
y9
y10
y11
y12
y13
y14
y16
y19
y18++
y17++
~y20++
~y21++
b11
b14
b16
b17
b18
b19
b20
b21
Pep_3+

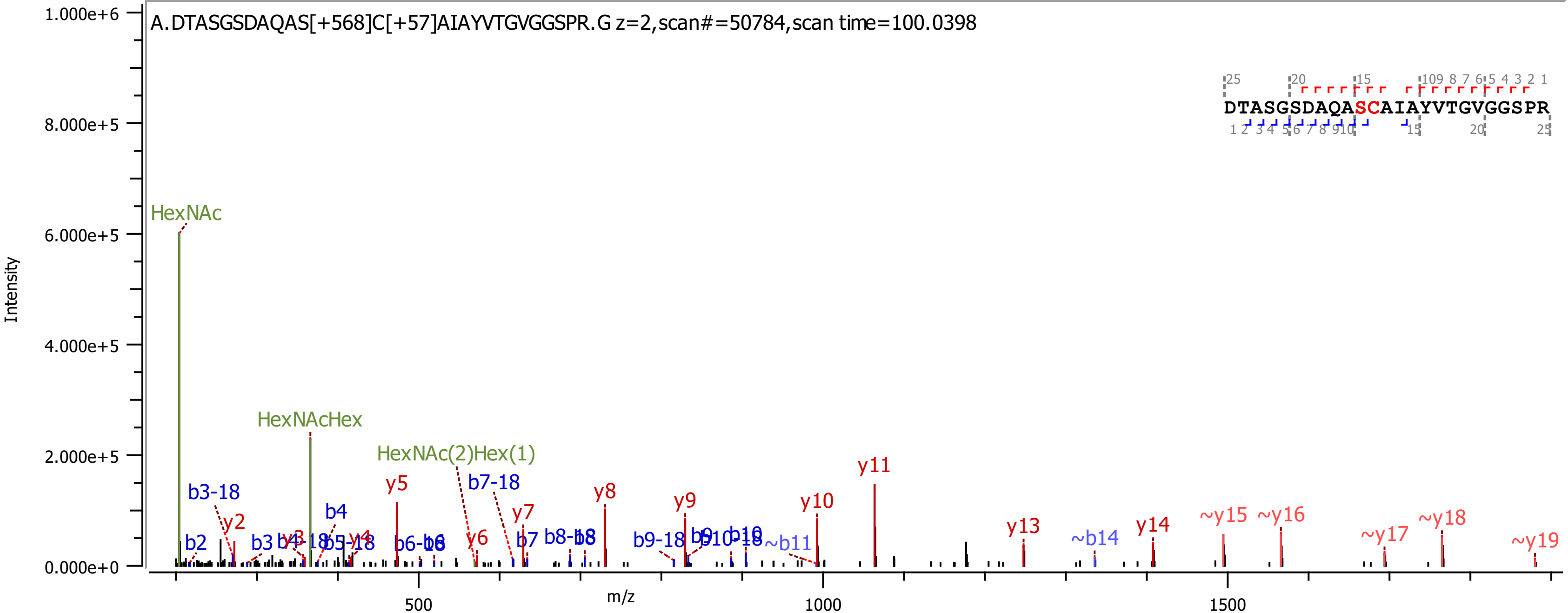
A.DTASGSDAQAS[+568]C[+57]AIAYVTGVGGSPR.G z=2,scan#=50784,scan time=100.0398
DTASGSDAQASCAIAYVTGVGGSPR
Intensity
m/z
HexNAc
HexNAcHex
HexNAc(2)Hex(1)
y2
y5
y6
y7
y8
y9
y10
y11
y13
y14
~y15
~y16
~y17
~y18
~y19
b2
b3
b4
b7
~b11
~b14
b3-18
b7-18
b9-18

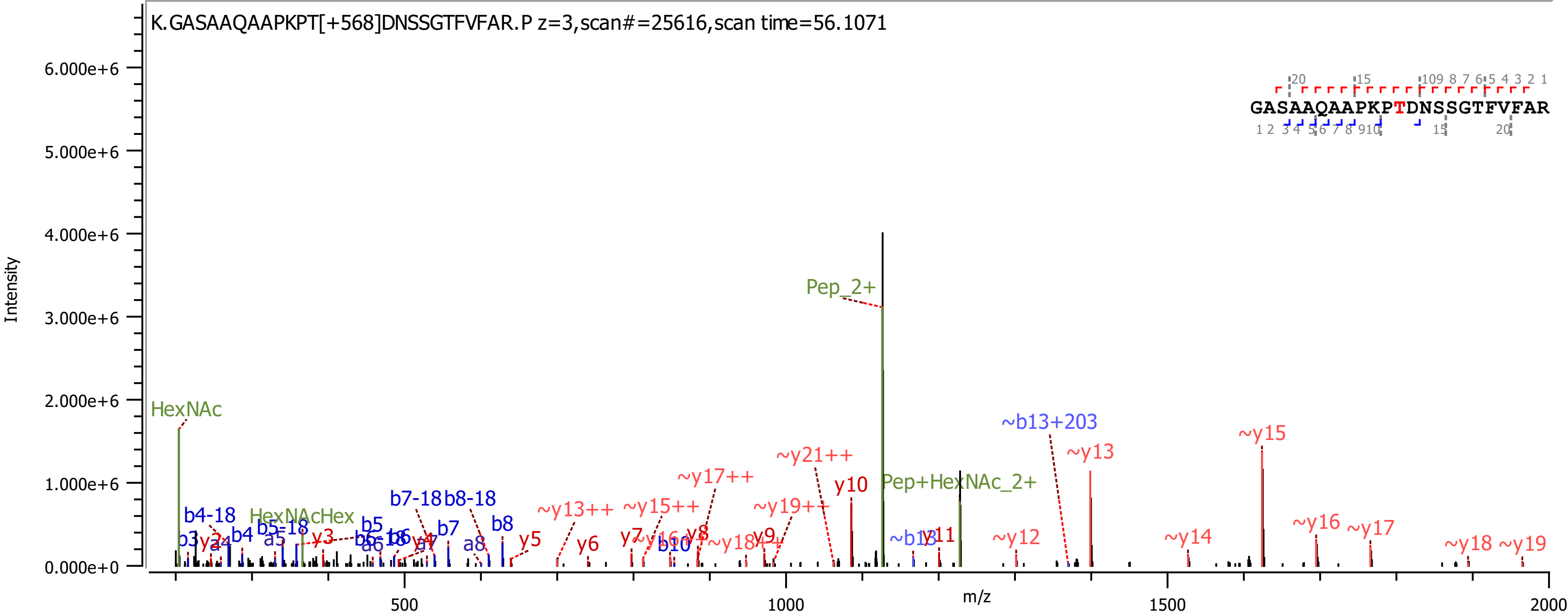
K.GASAAQAAPKPT[+568]DNSSGTFVFAR.P z=3,scan#=25616,scan time=56.1071
GASAAQAAPKPTDNSSGTFVFAR
Intensity
m/z
Pep_2+
HexNAc
HexNAcHex
Pep+HexNAc_2+
~b13+203
~y15
~y13
~y21++
~y17++
y10
~y19++
~y15++
~y13++
~y16
~y17
~y12
~y14
~y18
~y19
~b13
y11
b7-18
b8-18
b4-18
b5-18
b5
b6-18
b7
b8
y5
y6
y7
y8
y9
b10
~y18++
b3
b4
a5
a6
a8
y3
y4
0.000e+0
1.000e+6
2.000e+6
3.000e+6
4.000e+6
5.000e+6
6.000e+6
500
1000
1500
2000

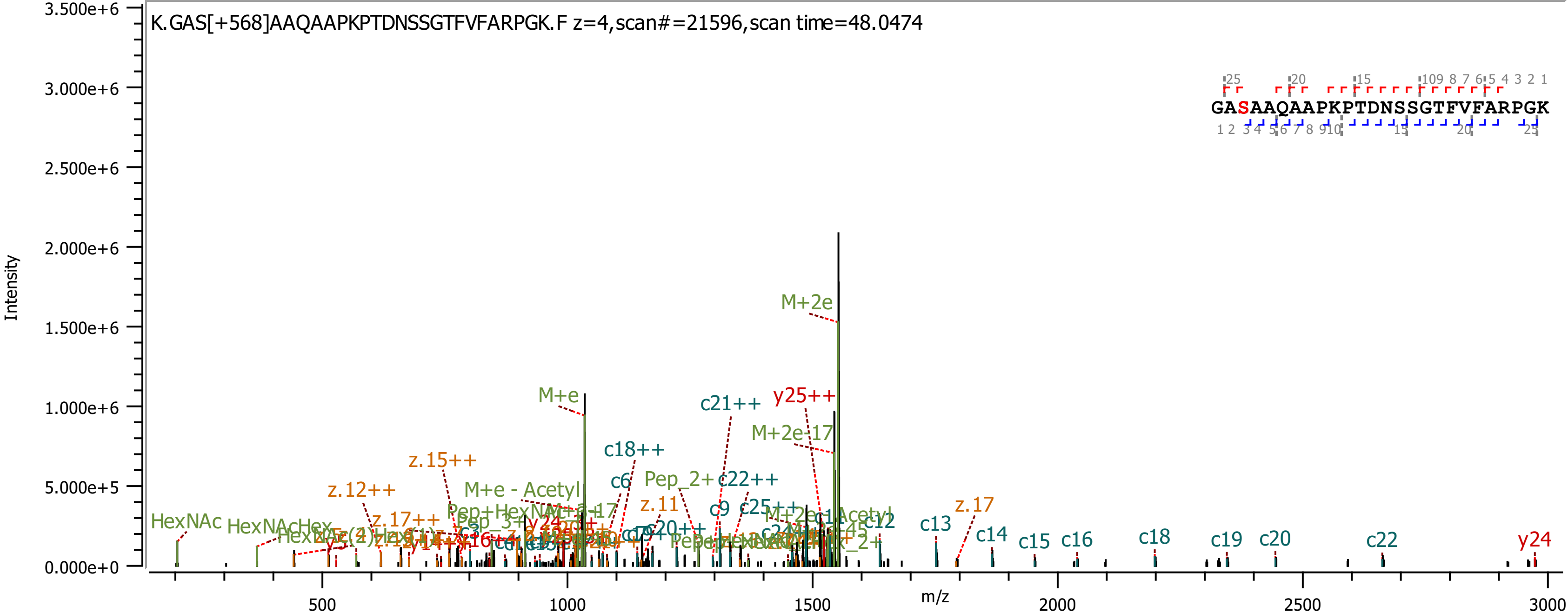

K.GAS[+568]AAQAAPKPTDNSSGTFVFARPGK.F z=4,scan#=21596,scan time=48.0474
GASAAQAAPKPTDNSSGTFVFARPGK
Intensity
m/z
3.500e+6
3.000e+6
2.500e+6
2.000e+6
1.500e+6
1.000e+6
5.000e+5
0.000e+0
500
1000
1500
2000
2500
3000
M+2e
M+e
c21++
y25++
M+2e-17
c18++
z.15++
M+e - Acetyl
c6
Pep_2+
c22++
z.12++
z.11
c9
c25++
HexNAc
HexNAcHex
z.17++
c12
c13
z.17
c14
c15
c16
c18
c19
c20
c22
y24

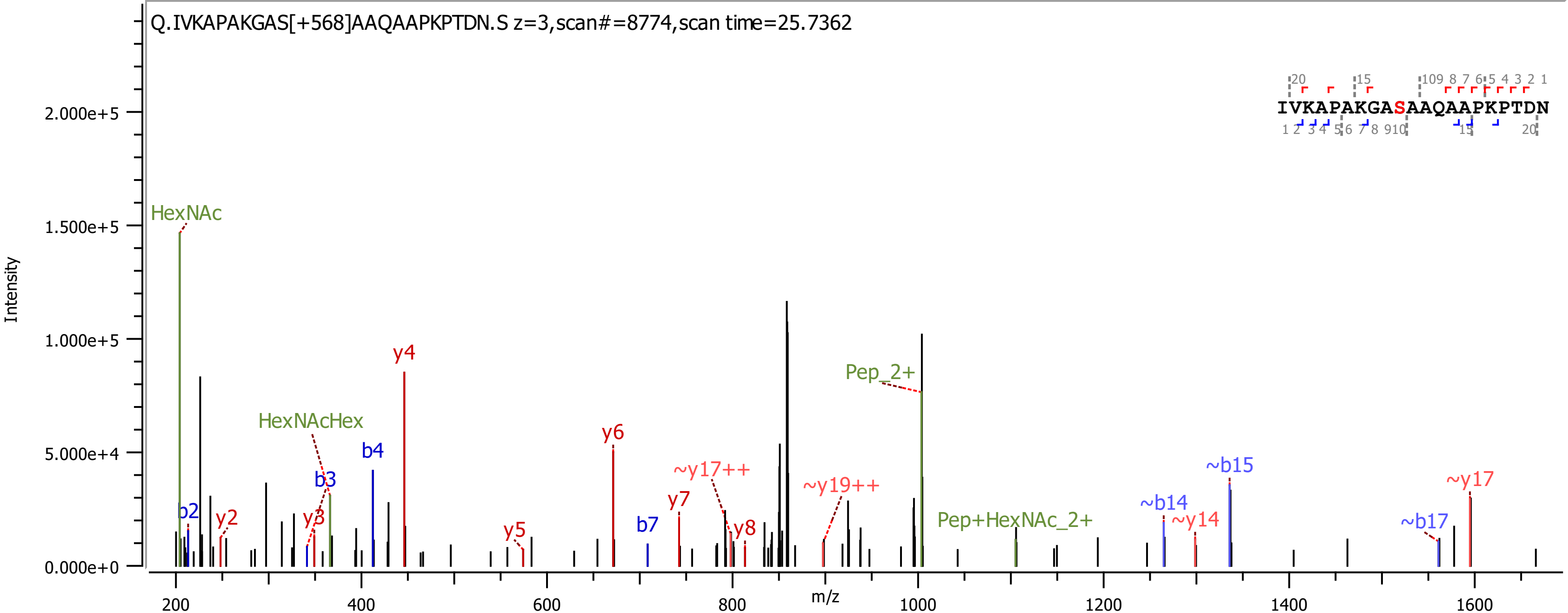

Q.IVKAPAKGAS[+568]AAQAAPKPTDN.S z=3,scan#=8774,scan time=25.7362
IVKAPAKGASAAQAAPKPTDN
Intensity
m/z
HexNAc
HexNAcHex
Pep_2+
Pep+HexNAc_2+
b2
y2
y3
b3
b4
y4
y5
y6
b7
y7
~y17++
y8
~y19++
~b14
~y14
~b15
~b17
~y17

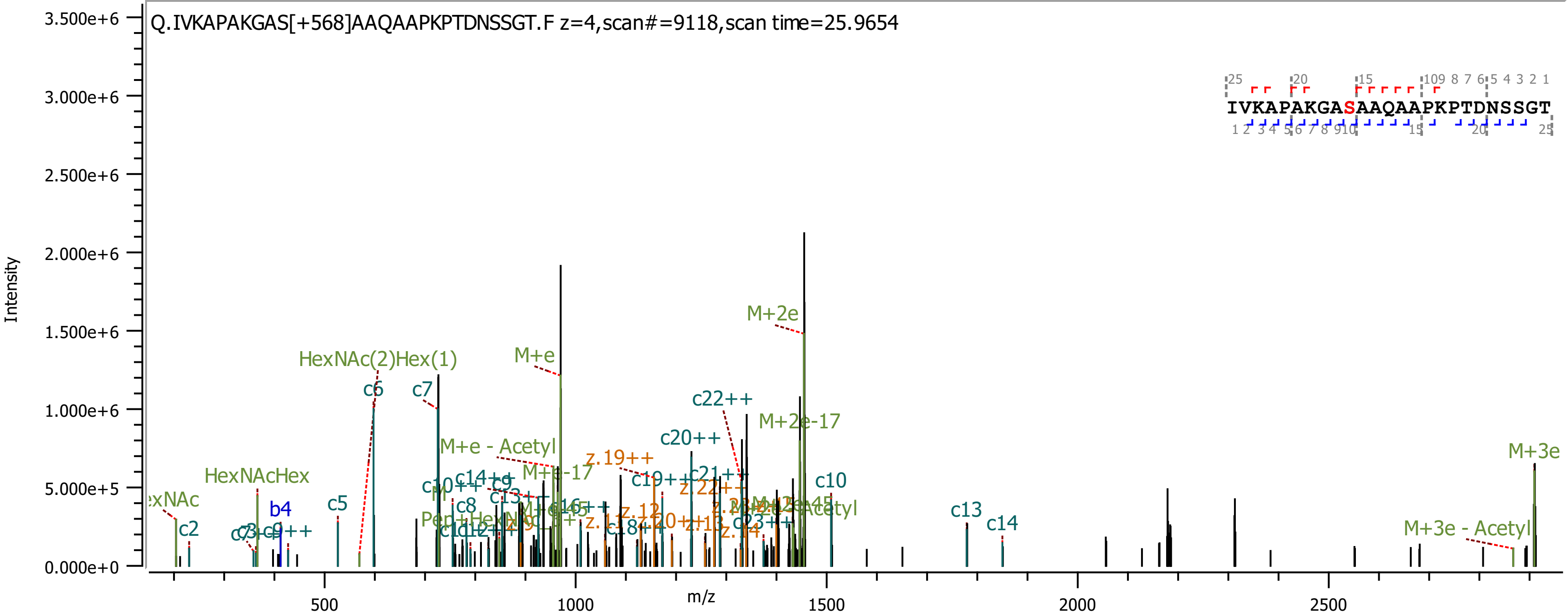
Q.IVKAPAKGAS[+568]AAQAAPKPTDNSSGT.F z=4,scan#=9118,scan time=25.9654
IVKAPAKGASAAQAAPKPTDNSSGT
Intensity
m/z
M+2e
HexNAc(2)Hex(1)
M+e
c6
c7
c22++
M+2e-17
c20++
M+e - Acetyl
z.19++
M+3e
HexNAcHex
c10
b4
c5
c2
c13
c14
M+3e - Acetyl

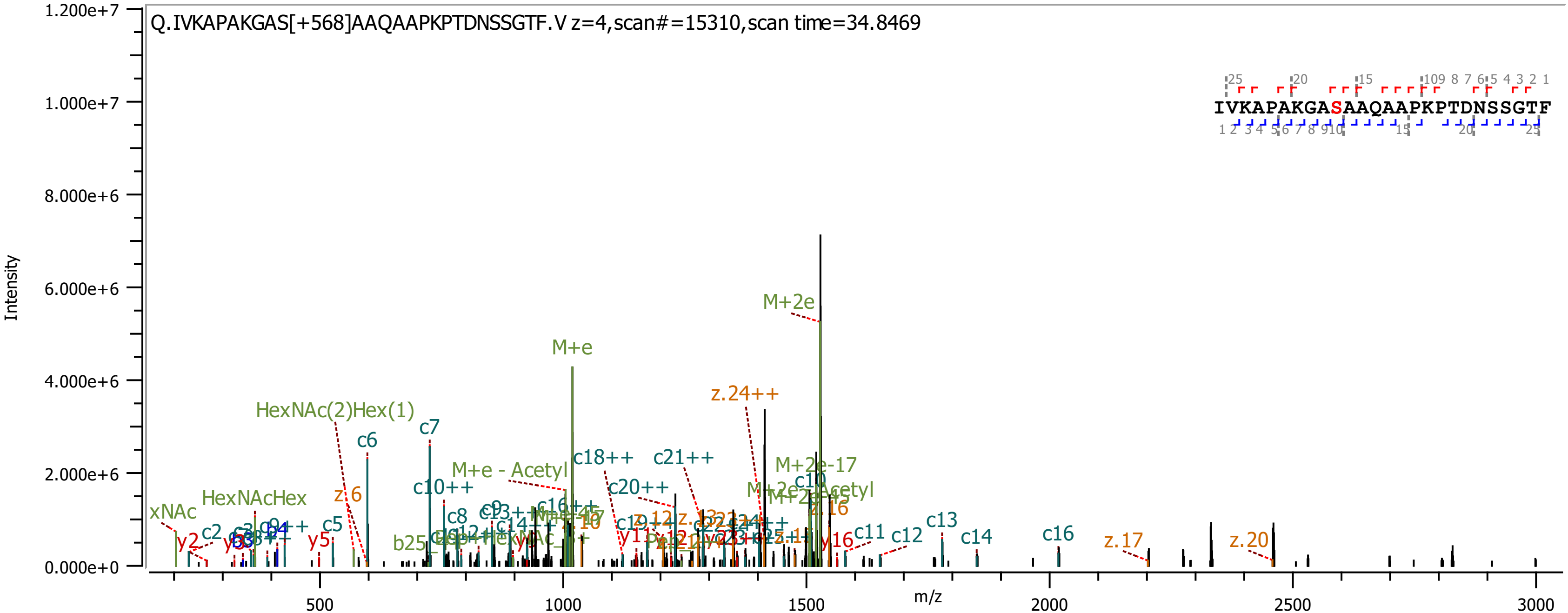

Q.IVKAPAKGAS[+568]AAQAAPKPTDNSSGTF.V z=4,scan#=15310,scan time=34.8469
IVKAPAKGASAAQAAPKPTDNSSGTF
Intensity
m/z
1.200e+7
1.000e+7
8.000e+6
6.000e+6
4.000e+6
2.000e+6
0.000e+0
500
1000
1500
2000
2500
3000
M+2e
M+e
z.24++
HexNAc(2)Hex(1)
c7
c6
c18++
c21++
M+e - Acetyl
M+2e-17
c10++
c20++
HexNAcHex
z.6
z.16
c5
c13
c11
c12
c14
c16
z.17
z.20
y2
c2
y5
y16

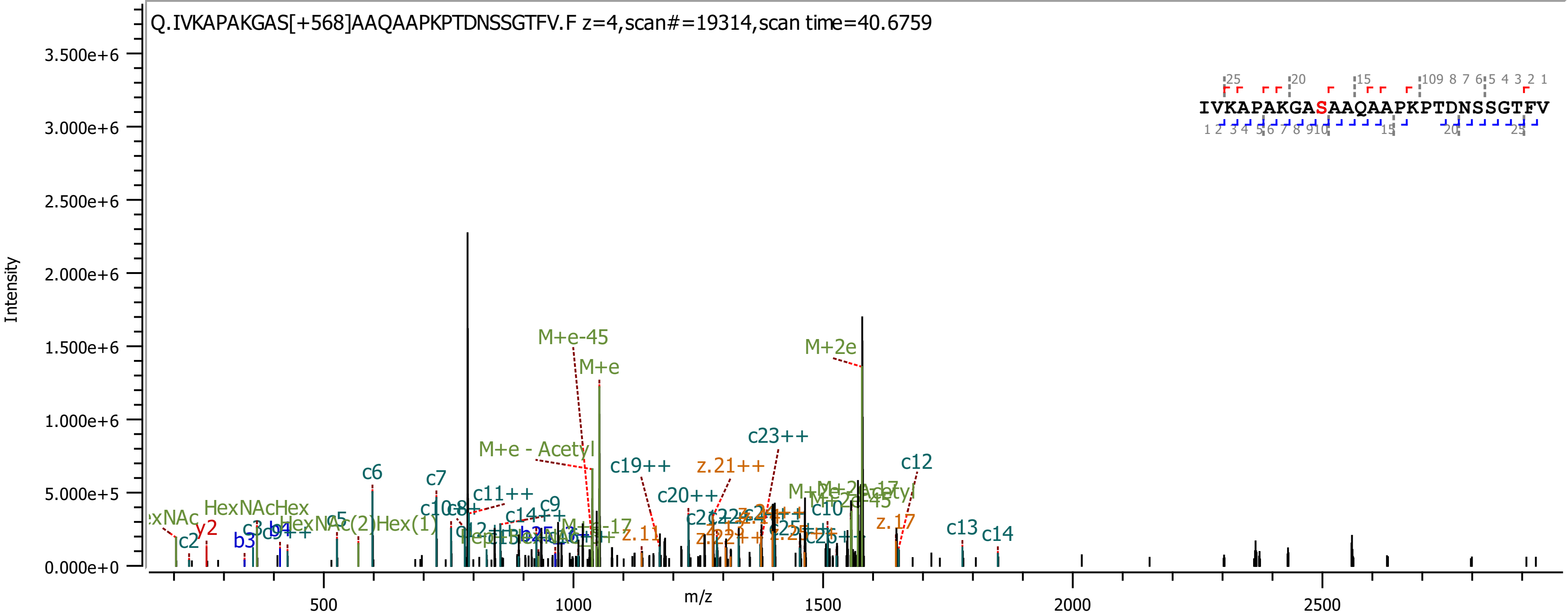
Q.IVKAPAKGAS[+568]AAQAAPKPTDNSSGTFV.F z=4,scan#=19314,scan time=40.6759
IVKAPAKGASAAQAAPKPTDNSSGTFV
Intensity
m/z
3.500e+6
3.000e+6
2.500e+6
2.000e+6
1.500e+6
1.000e+6
5.000e+5
0.000e+0
500
1000
1500
2000
2500
M+e-45
M+e
M+2e
M+e - Acetyl
c23++
c19++
z.21++
c12
c6
c7
c11++
c9
c14++
c20++
c10
z.17
c13
c14
HexNAcHex
HexNAc(2)Hex(1)
y2
c2
b3
c5
z.11

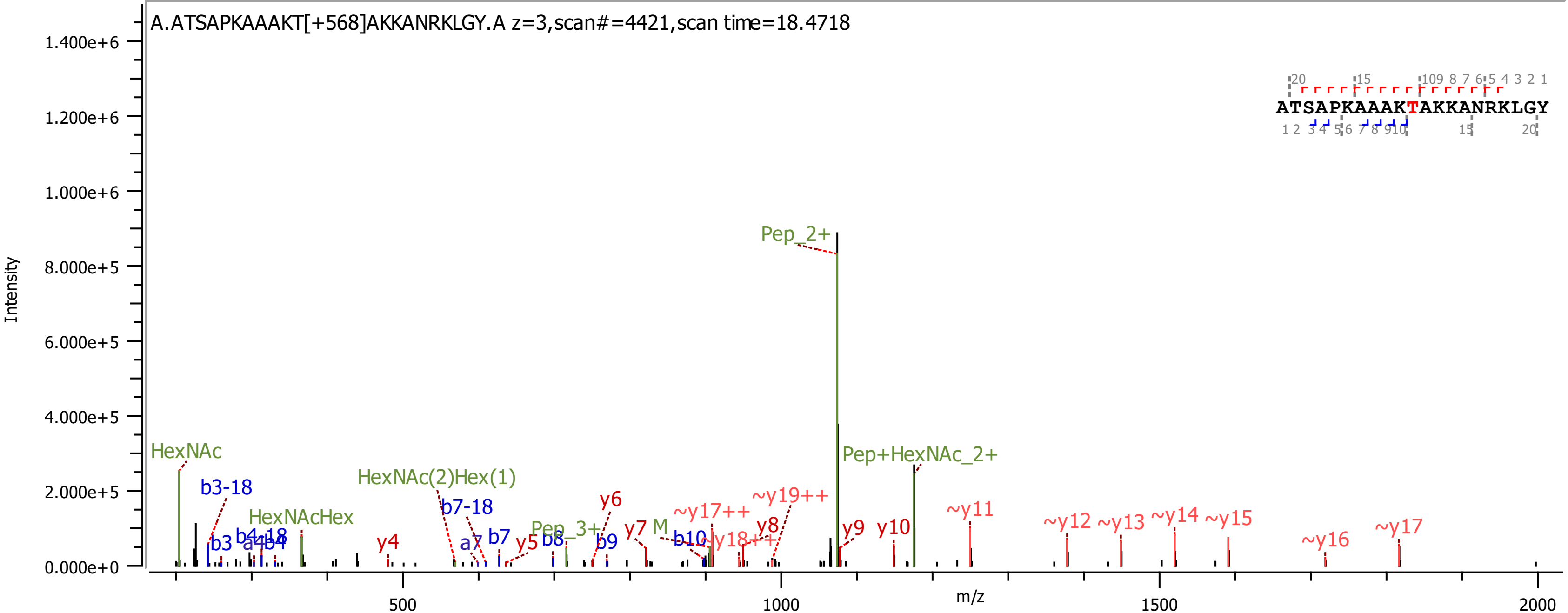
A.ATSAPKAAAKT[+568]AKKANRKLGY.A z=3,scan#=4421,scan time=18.4718
ATSAPKAAAKTAKKANRKLGY
Intensity
m/z
Pep_2+
Pep+HexNAc_2+
HexNAc
HexNAc(2)Hex(1)
HexNAcHex
Pep_3+
b3-18
b3
b4-18
a4
b4
b7-18
a7
b7
b8
b9
b10
y4
y5
y6
y7
M
y8
y9
y10
~y11
~y12
~y13
~y14
~y15
~y16
~y17
~y17++
~y18++
~y19++
0.000e+0
2.000e+5
4.000e+5
6.000e+5
8.000e+5
1.000e+6
1.200e+6
1.400e+6
500
1000
1500
2000

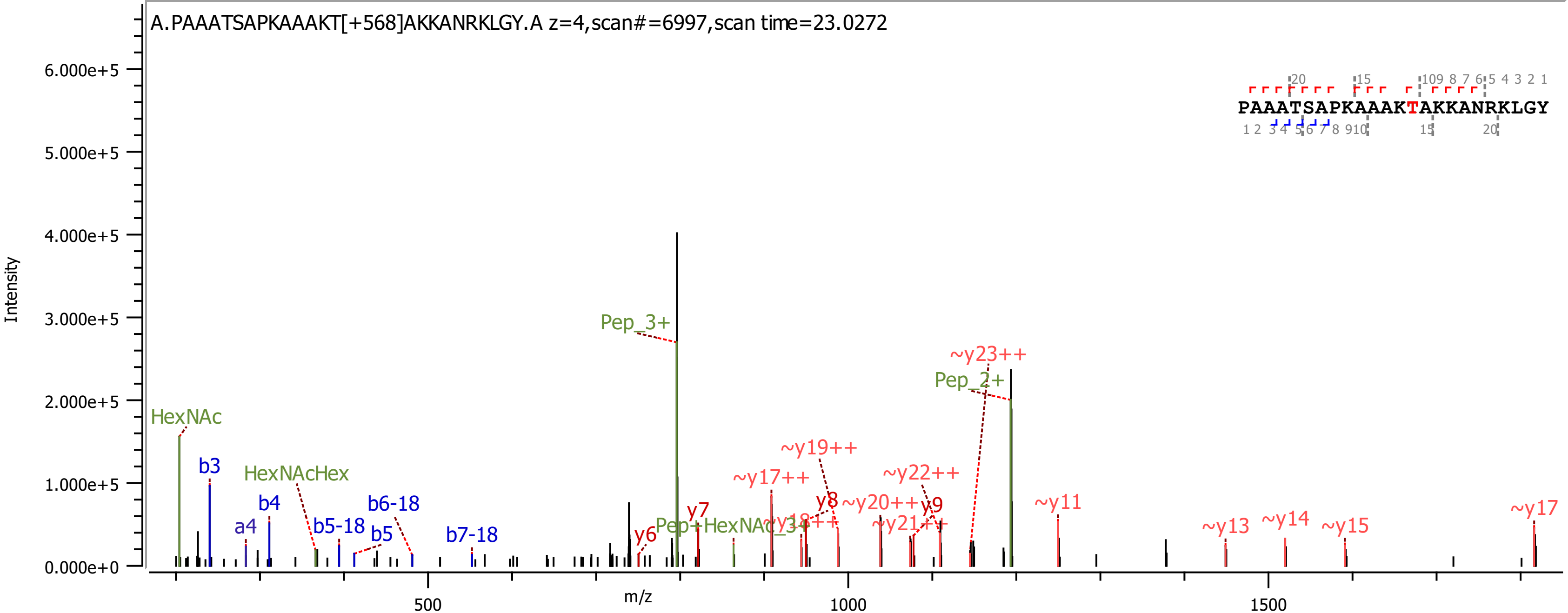
A.PAAATSAPKAAAKT[+568]AKKANRKLGY.A z=4,scan#=6997,scan time=23.0272
PAAATSAPKAAAKTAKKANRKLGY
Intensity
m/z
HexNAc
b3
a4
b4
HexNAcHex
b5-18
b5
b6-18
b7-18
y6
Pep_3+
y7
Pep+HexNAc_3+
~y17++
~y18++
~y19++
y8
~y20++
~y21++
~y22++
y9
~y23++
Pep_2+
~y11
~y13
~y14
~y15
~y17

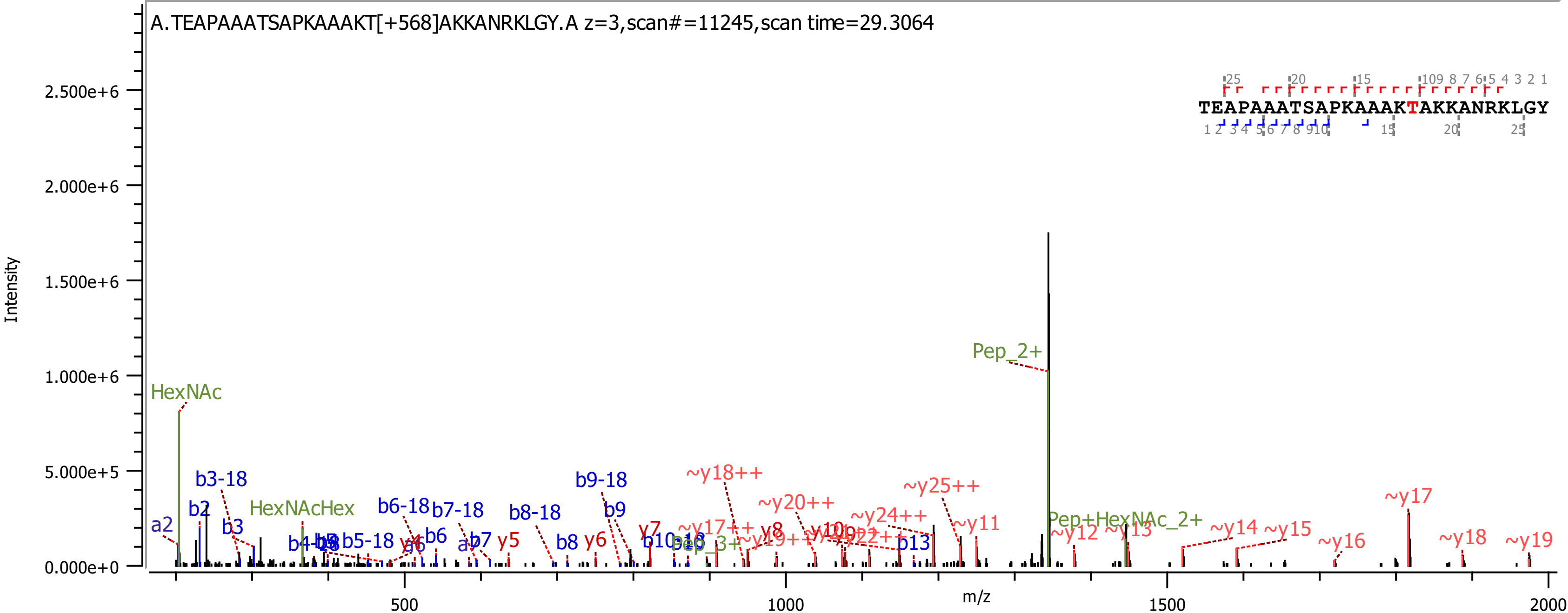

A.TEAPAAATSAPKAAAKT[+568]AKKANRKLGY.A z=3,scan#=11245,scan time=29.3064
TEAPAAATSAPKAAAKTAKKANRKLGY
Intensity
m/z
Pep_2+
HexNAc
HexNAcHex
Pep+HexNAc_2+
~y18++
~y20++
~y24++
~y25++
~y17
~y11
~y12
~y13
~y14
~y15
~y16
~y18
~y19
b3-18
b6-18
b7-18
b8-18
b9-18
b5-18
a2
b2
b3
b6
b7
b8
b9
b13
y5
y6
y7
y8

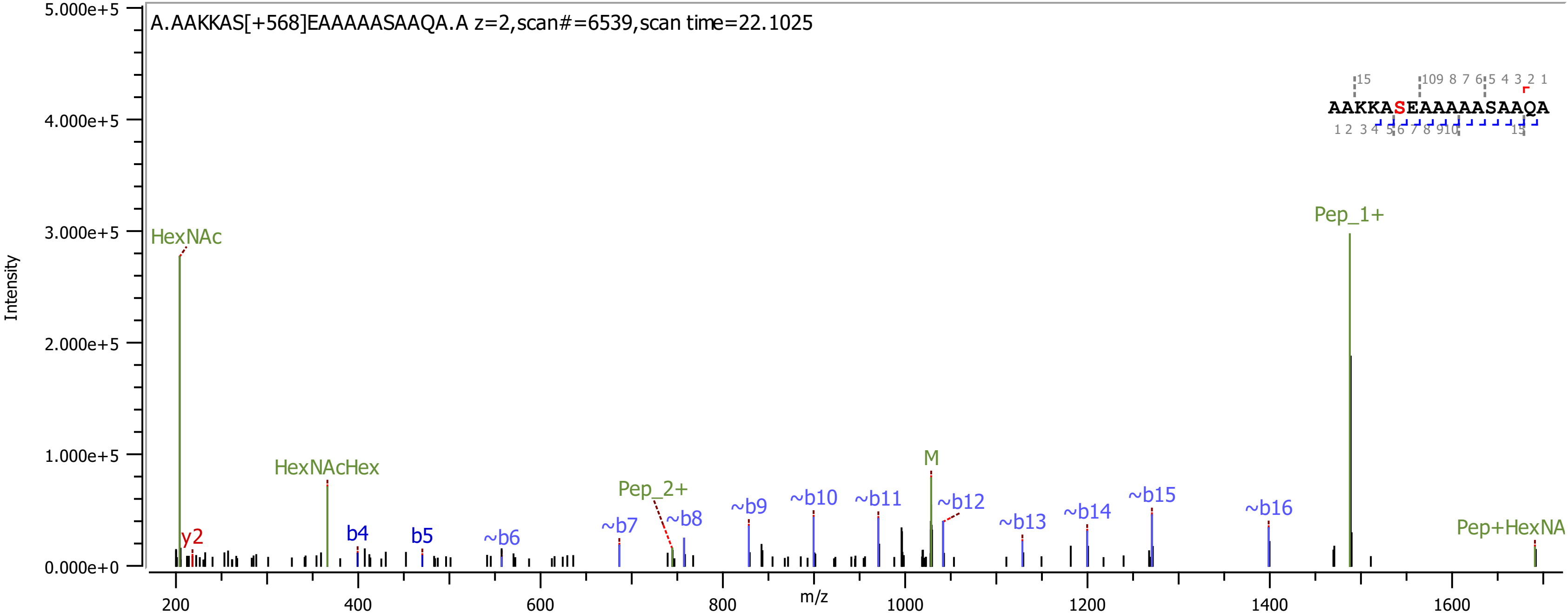
A.AAKKAS[+568]EAAAAASAAQA.A z=2,scan#=6539,scan time=22.1025
AAKKASEAAAAASAAQA
Intensity
m/z
HexNAc
HexNAcHex
y2
b4
b5
~b6
~b7
Pep_2+
~b8
~b9
~b10
~b11
M
~b12
~b13
~b14
~b15
~b16
Pep_1+
Pep+HexNA
5.000e+5
4.000e+5
3.000e+5
2.000e+5
1.000e+5
0.000e+0
200
400
600
800
1000
1200
1400
1600

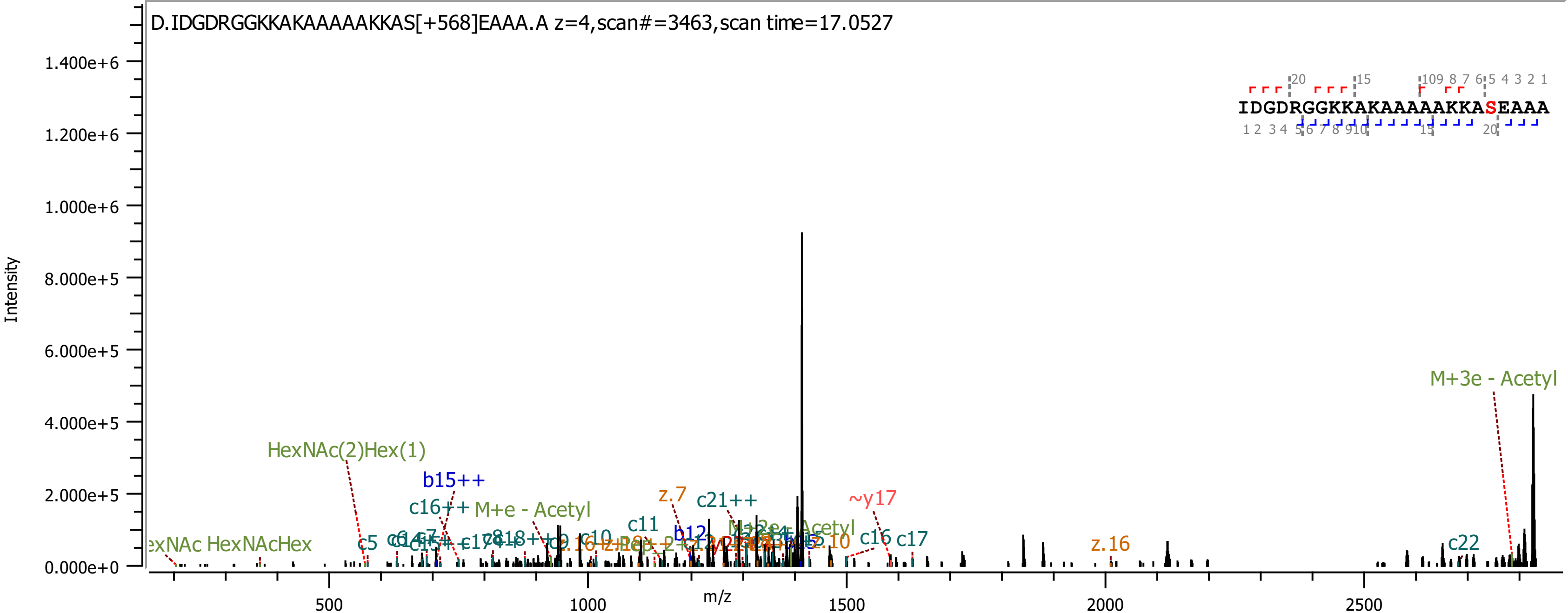
D.IDGDRGGKKAKAAAAAKKAS[+568]EAAA.A z=4,scan#=3463,scan time=17.0527
IDGDRGGKKAKAAAAAKKASEAAA
Intensity
m/z
M+3e - Acetyl
HexNAc(2)Hex(1)
b15++
c16++
M+e - Acetyl
z.7
c21++
~y17
c11
b12
c16
c17
z.16
c22
c5
HexNAcHex

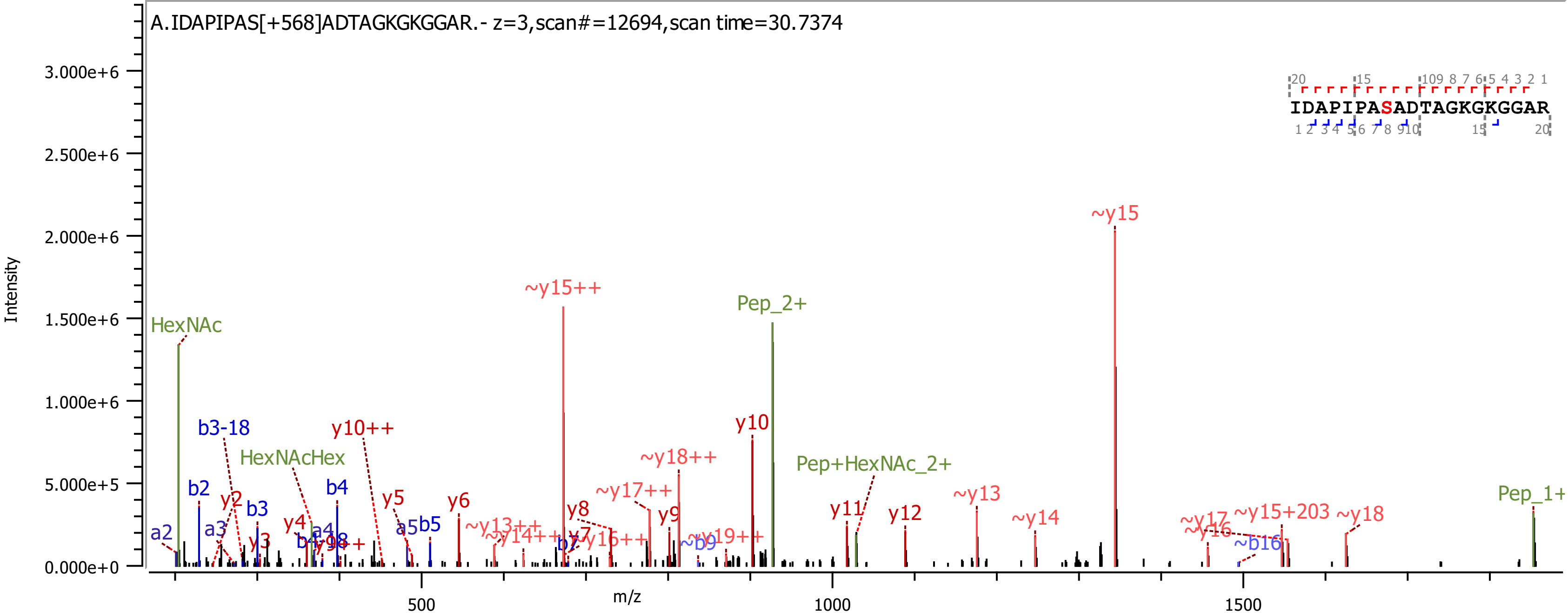
A.IDAPIPAS[+568]ADTAGKGKGGAR.- z=3,scan#=12694,scan time=30.7374
IDAPIPASADTAGKGKGGAR
Intensity
m/z
HexNAc
HexNAcHex
Pep_2+
Pep+HexNAc_2+
Pep_1+
~y15
~y15++
y10
y10++
~y18++
~y17++
b3-18
b2
b3
b4
y2
y3
y4
y5
y6
y7
y8
y9
y11
y12
~y13
~y14
~y17
~y16
~y15+203
~y18
~b16
~b9
~y19++
~y13++
~y14++
~y16++
a2
a3
a4
a5
b5
b7

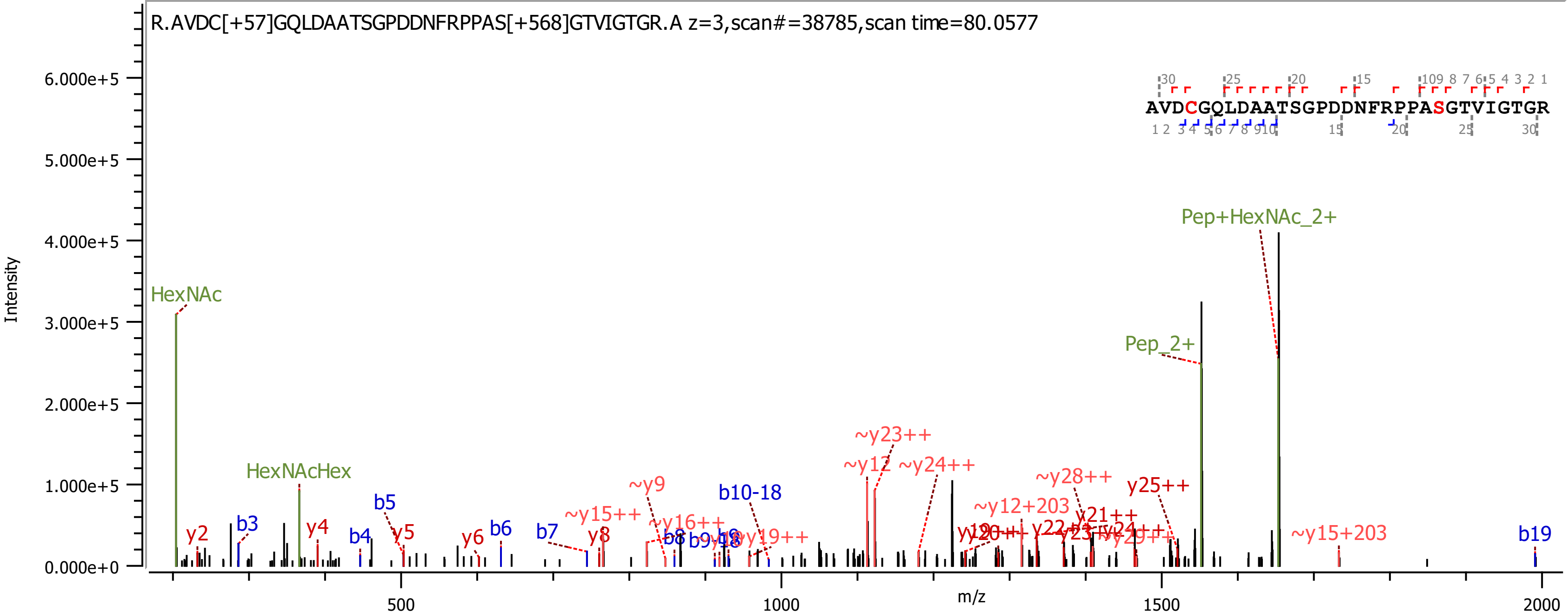
R.AVDC[+57]GQLDAATSGPDDNFRPPAS[+568]GTVIGTGR.A z=3,scan#=38785,scan time=80.0577
AVDCGQLDAATSGPDDNFRPPASGTVIGTGR
Intensity
m/z
6.000e+5
5.000e+5
4.000e+5
3.000e+5
2.000e+5
1.000e+5
0.000e+0
500
1000
1500
2000
HexNAc
HexNAcHex
Pep_2+
Pep+HexNAc_2+
y2
b3
y4
b4
b5
y5
y6
b6
b7
y8
~y15++
~y9
~y16++
b8
b10-18
~y19++
~y23++
~y12
~y24++
~y12+203
~y28++
y21++
y25++
~y15+203
b19

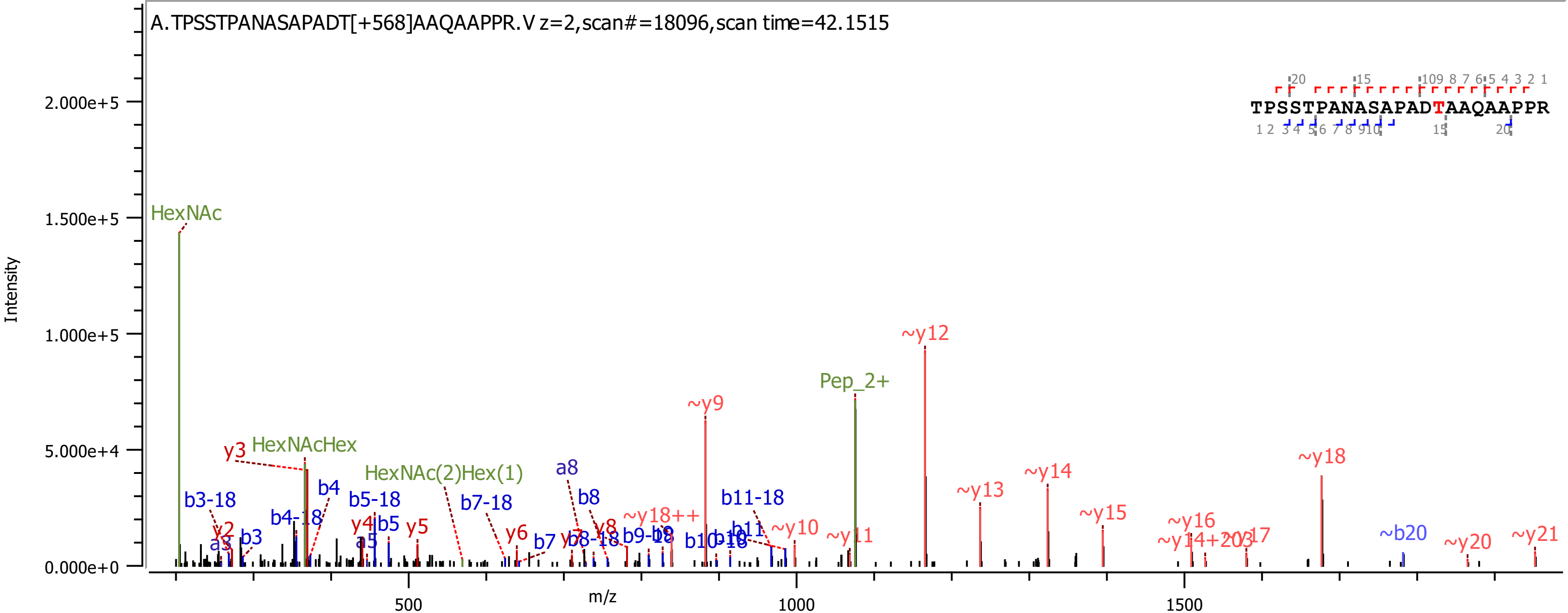
A.TPSSTPANASAPADT[+568]AAQAAPPR.V z=2,scan#=18096,scan time=42.1515
TPSSTPANASAPADTAAQAAPPR
Intensity
m/z
HexNAc
HexNAcHex
HexNAc(2)Hex(1)
Pep_2+
~y9
~y12
~y13
~y14
~y15
~y16
~y18
~b20
~y20
~y21
~y10
~y18++
b11-18
b11
b10-18
b9-18
b8
a8
b8-18
b7
b7-18
b5-18
b5
a5
y4
y5
y6
y3
b4
b4-18
b3-18
b3
a3
y2

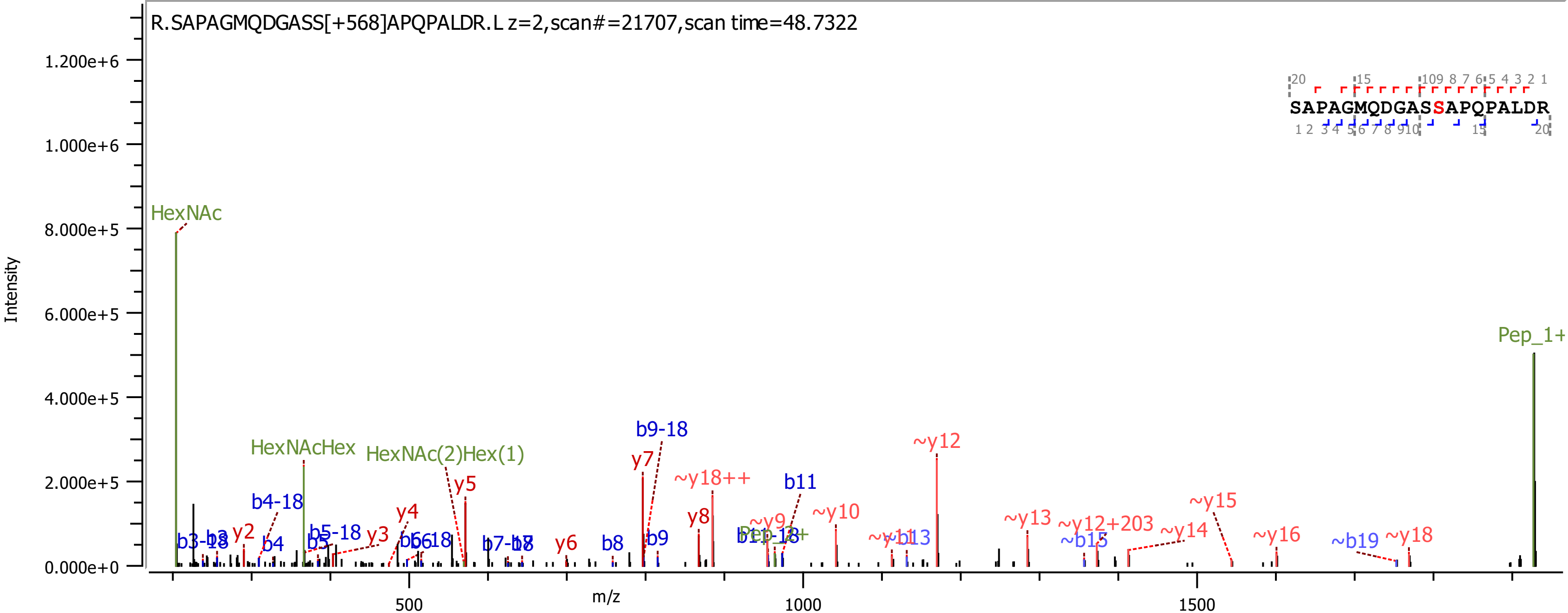
R.SAPAGMQDGASS[+568]APQPALDR.L z=2,scan#=21707,scan time=48.7322
SAPAGMQDGASSAPQPALDR
Intensity
m/z
HexNAc
HexNAcHex
HexNAc(2)Hex(1)
Pep_1+
b9-18
~y12
y7
~y18++
b11
y5
b4-18
y4
~y10
y8
~y15
~y9
~y13
~y12+203
~y14
~y16
~y18
~b19
~b15
~y11
~b13
y2
y3
b5-18
b4
b5
b6-18
b6
b7-18
b7
y6
b8
b9
b3-18
b3
1.200e+6
1.000e+6
8.000e+5
6.000e+5
4.000e+5
2.000e+5
0.000e+0
500
1000
1500

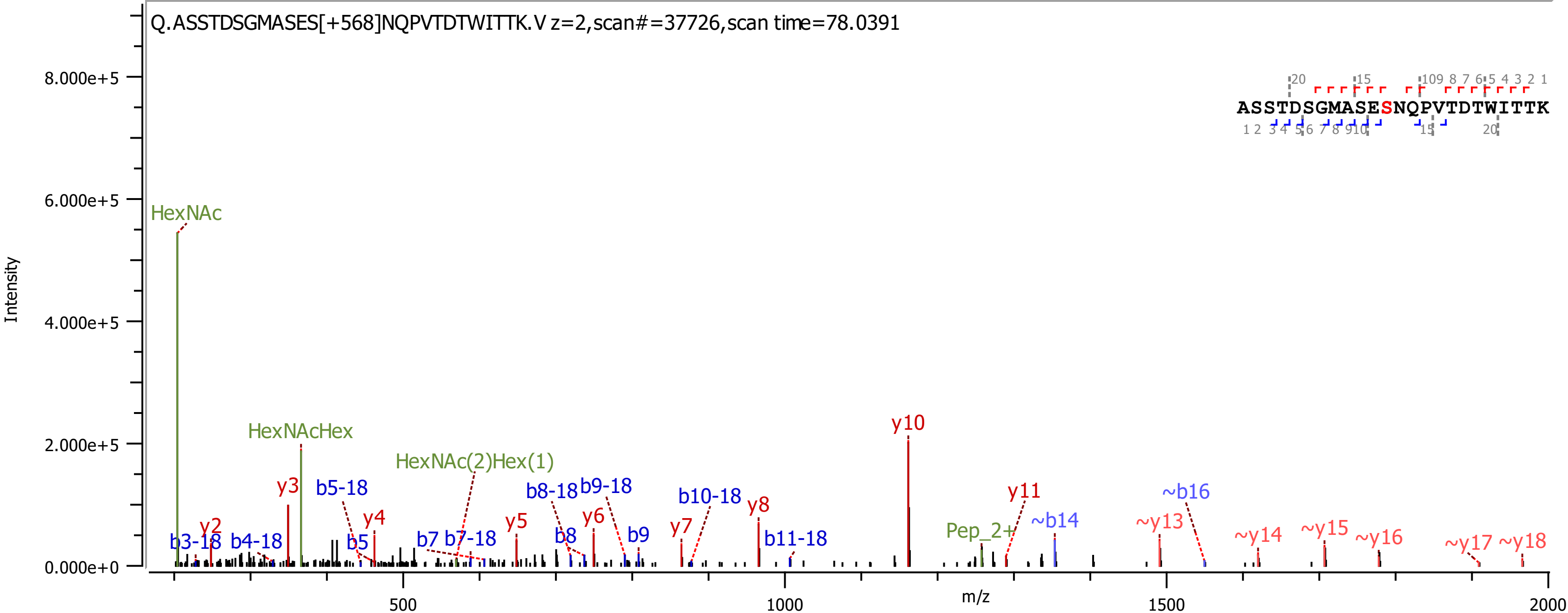
Q.ASSTDSGMASES[+568]NQPVTDTWITTK.V z=2,scan#=37726,scan time=78.0391
ASSTDSGMASESNQPVTDTWITTK
Intensity
m/z
HexNAc
HexNAcHex
HexNAc(2)Hex(1)
Pep_2+
y2
y3
y4
y5
y6
y7
y8
y10
y11
~y13
~y14
~y15
~y16
~y17
~y18
b3-18
b4-18
b5-18
b5
b7
b7-18
b8-18
b9-18
b8
b9
b10-18
b11-18
~b14
~b16
8.000e+5
6.000e+5
4.000e+5
2.000e+5
0.000e+0
500
1000
1500
2000

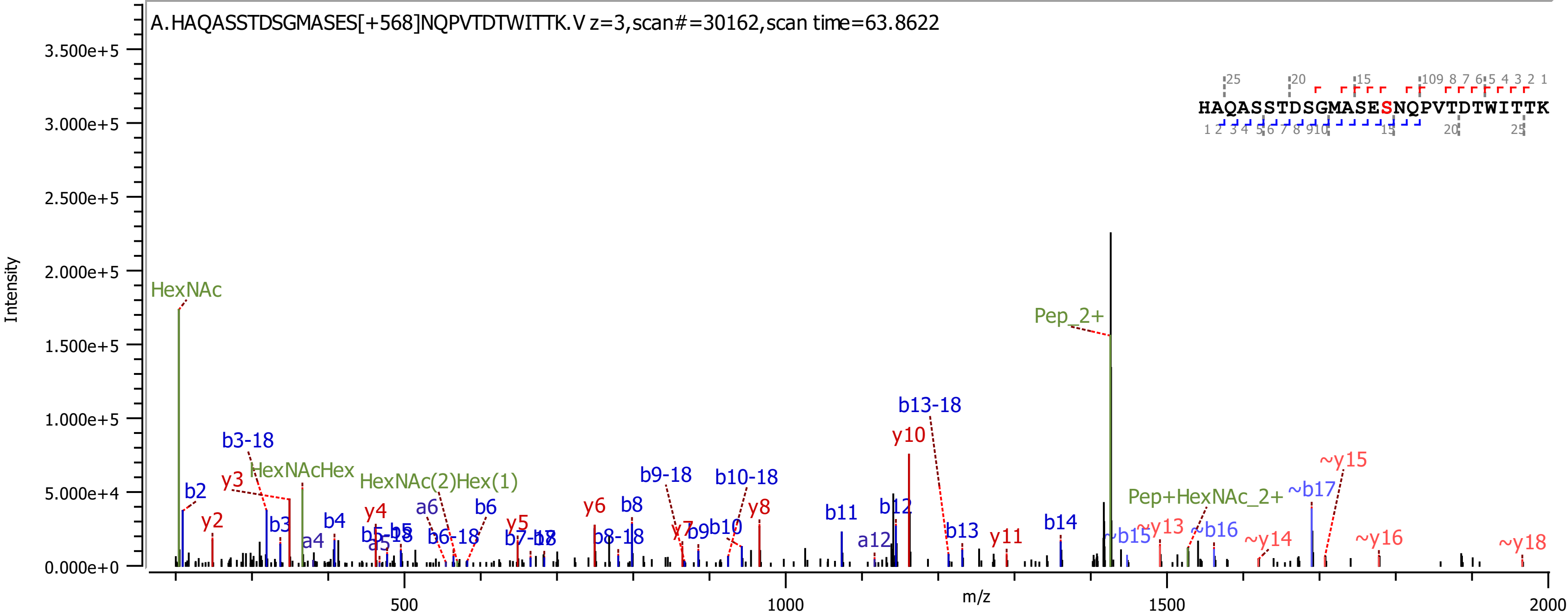
A.HAQASSTDSGMASES[+568]NQPVTDTWITTK.V z=3,scan#=30162,scan time=63.8622
HAQASSTDSGMASESNQPVTDTWITTK
Intensity
3.500e+5
3.000e+5
2.500e+5
2.000e+5
1.500e+5
1.000e+5
5.000e+4
0.000e+0
m/z
500
1000
1500
2000
HexNAc
Pep_2+
b13-18
y10
~y15
b3-18
HexNAcHex
HexNAc(2)Hex(1)
b9-18
b10-18
~b17
Pep+HexNAc_2+
y3
b2
y2
b3
a4
b4
y4
a6
b6
b5-18
a5
b5
b6-18
y5
b7-18
b7
y6
b8
b8-18
y7
b9
b10
y8
b11
b12
a12
b13
y11
b14
~b15
~y13
~b16
~y14
~y16
~y18

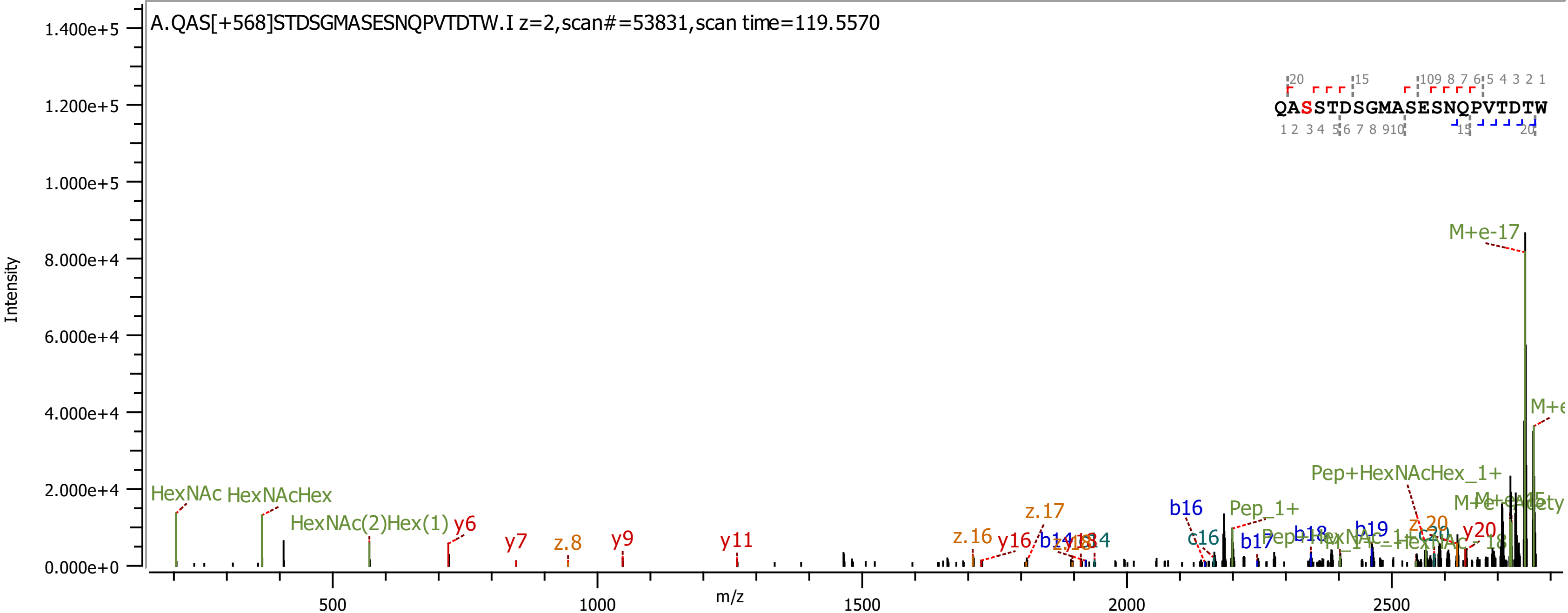
A.QAS[+568]STDSGMASESNQPVTDTW.I z=2,scan#=53831,scan time=119.5570
QASSTDSGMASESNQPVTDTW
Intensity
m/z
1.400e+5
1.200e+5
1.000e+5
8.000e+4
6.000e+4
4.000e+4
2.000e+4
0.000e+0
500
1000
1500
2000
2500
M+e-17
Pep+HexNAcHex_1+
Pep_1+
HexNAc
HexNAcHex
HexNAc(2)Hex(1)
y6
y7
z.8
y9
y11
z.16
y16
z.17
b16
c16
b17
b18
b19
z.20
y20

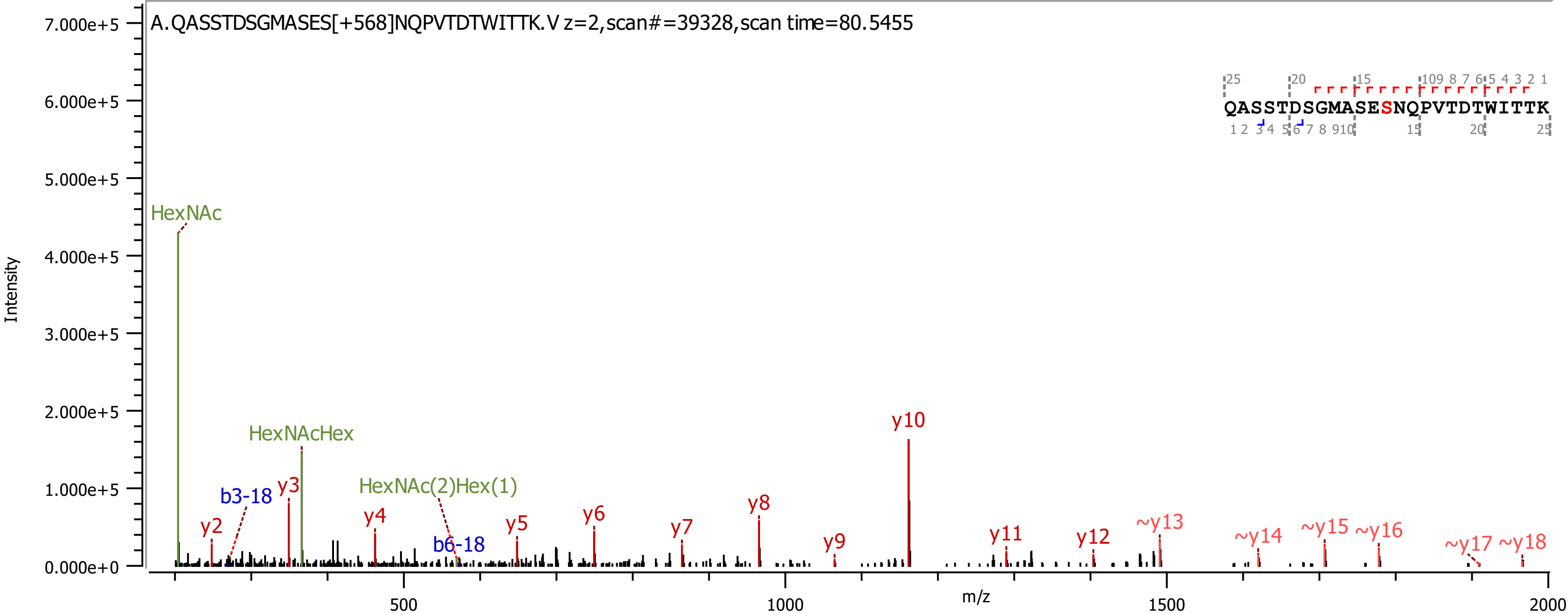

A.QASSTDSGMASES[+568]NQPVTDTWITTK.V z=2,scan#=39328,scan time=80.5455
QASSTDSGMASESNQPVTDTWITTK
Intensity
m/z
HexNAc
HexNAcHex
HexNAc(2)Hex(1)
b3-18
b6-18
y2
y3
y4
y5
y6
y7
y8
y9
y10
y11
y12
~y13
~y14
~y15
~y16
~y17
~y18

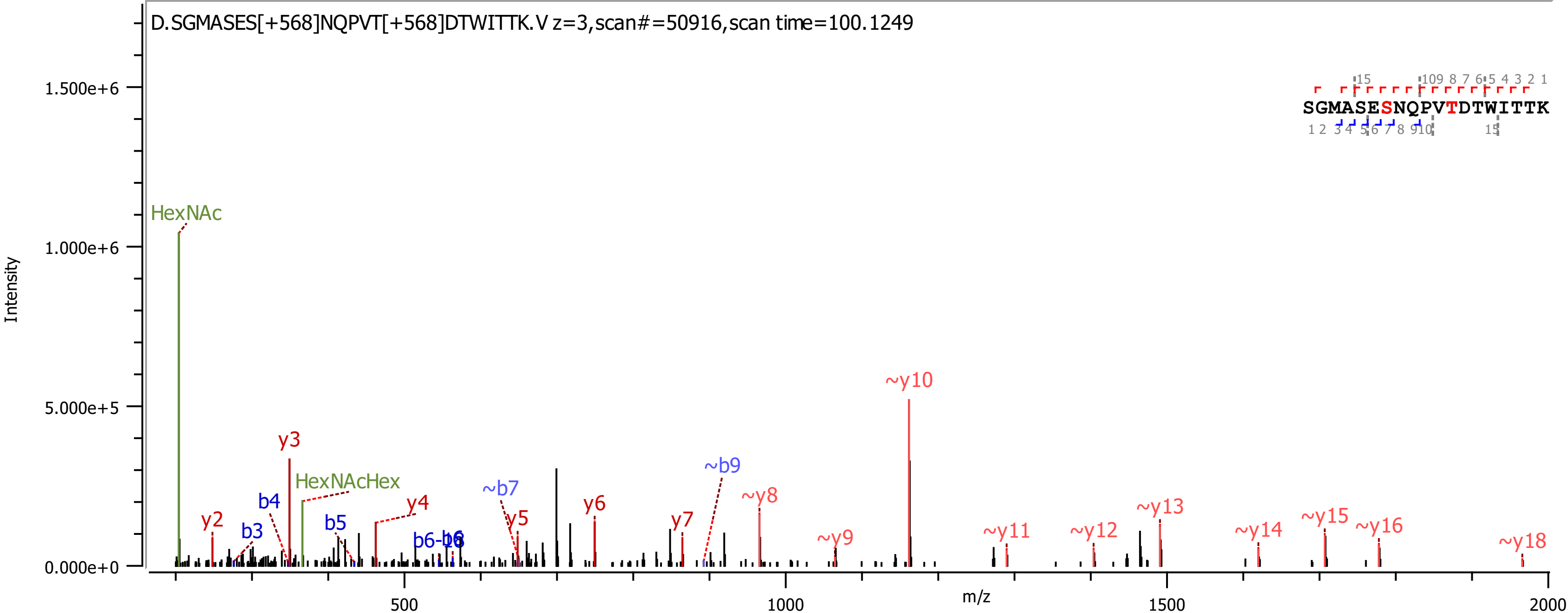
D.SGMASES[+568]NQPVT[+568]DTWITTK.V z=3,scan#=50916,scan time=100.1249
SGMASESNQPVTDTWITTK
Intensity
m/z
HexNAc
HexNAcHex
y2
y3
y4
y5
y6
y7
b3
b4
b5
b6-18
~b7
~b9
~y8
~y9
~y10
~y11
~y12
~y13
~y14
~y15
~y16
~y18
1.500e+6
1.000e+6
5.000e+5
0.000e+0
500
1000
1500
2000

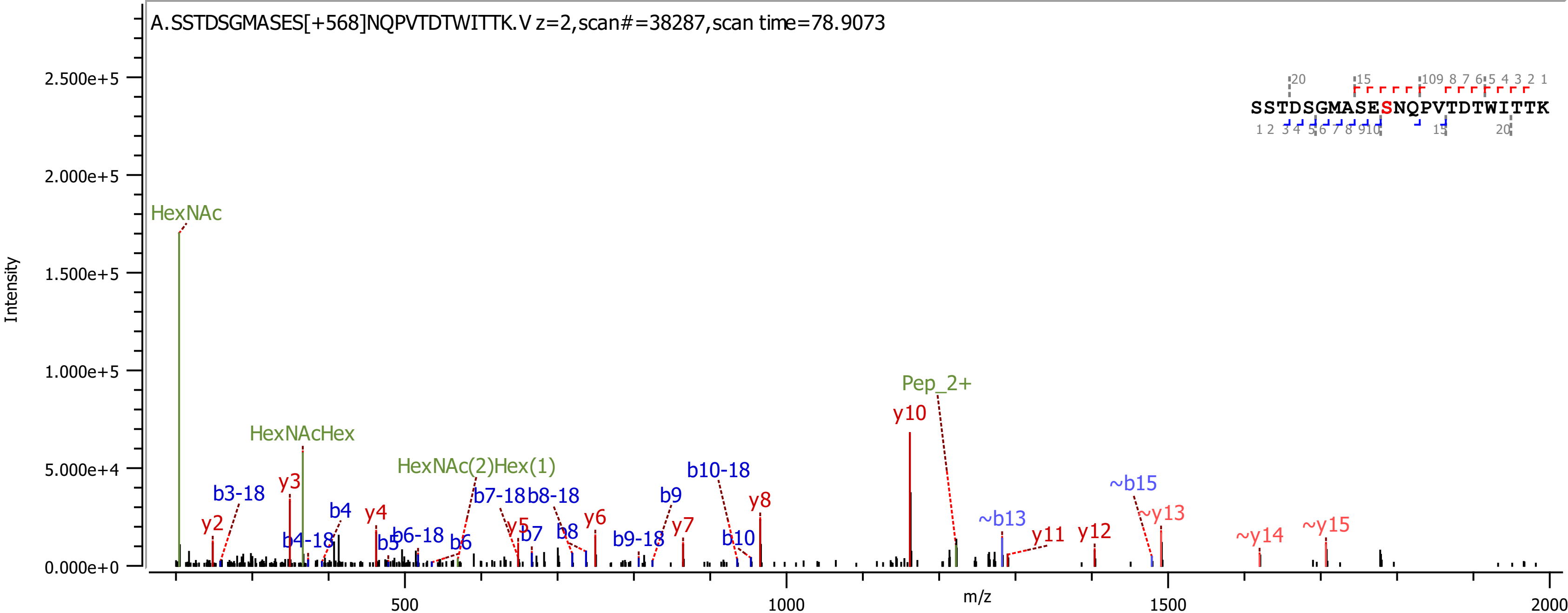

A.SSTDSGMASES[+568]NQPVTDTWITTK.V z=2,scan#=38287,scan time=78.9073
SSTDSGMASESNQPVTDTWITTK
Intensity
m/z
HexNAc
HexNAcHex
HexNAc(2)Hex(1)
Pep_2+
y2
y3
y4
y5
y6
y7
y8
y10
y11
y12
~y13
~y14
~y15
b3-18
b4-18
b4
b5
b6-18
b6
b7-18
b7
b8-18
b8
b9-18
b9
b10-18
b10
~b13
~b15

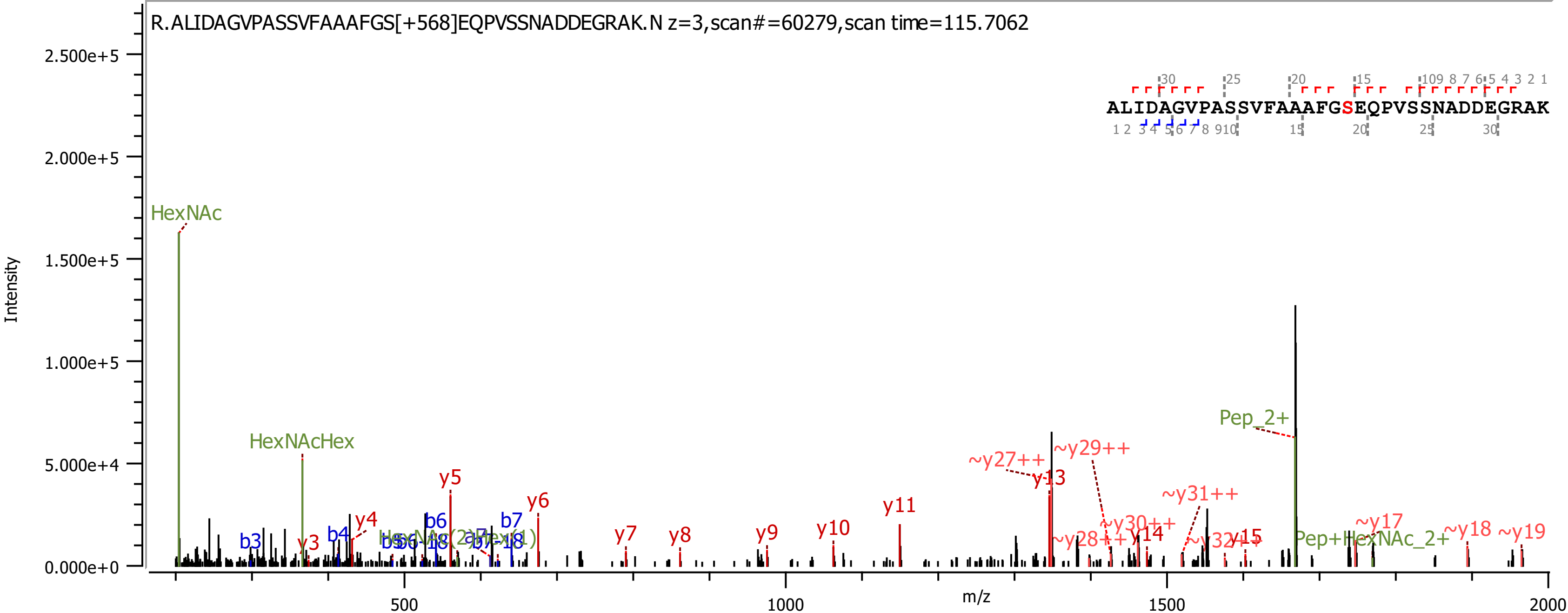

R.ALIDAGVPASSVFAAAFGS[+568]EQPVSSNADDEGRAK.N z=3,scan#=60279,scan time=115.7062
ALIDAGVPASSVFAAAFGSEQPVSSNADDEGRAK
Intensity
m/z
HexNAc
HexNAcHex
Pep_2+
Pep+HexNAc_2+
y3
y4
y5
y6
y7
y8
y9
y10
y11
y13
y14
y15
~y17
~y18
~y19
~y27++
~y28++
~y29++
~y30++
~y31++
~y32++
b3
b4
b6
b7

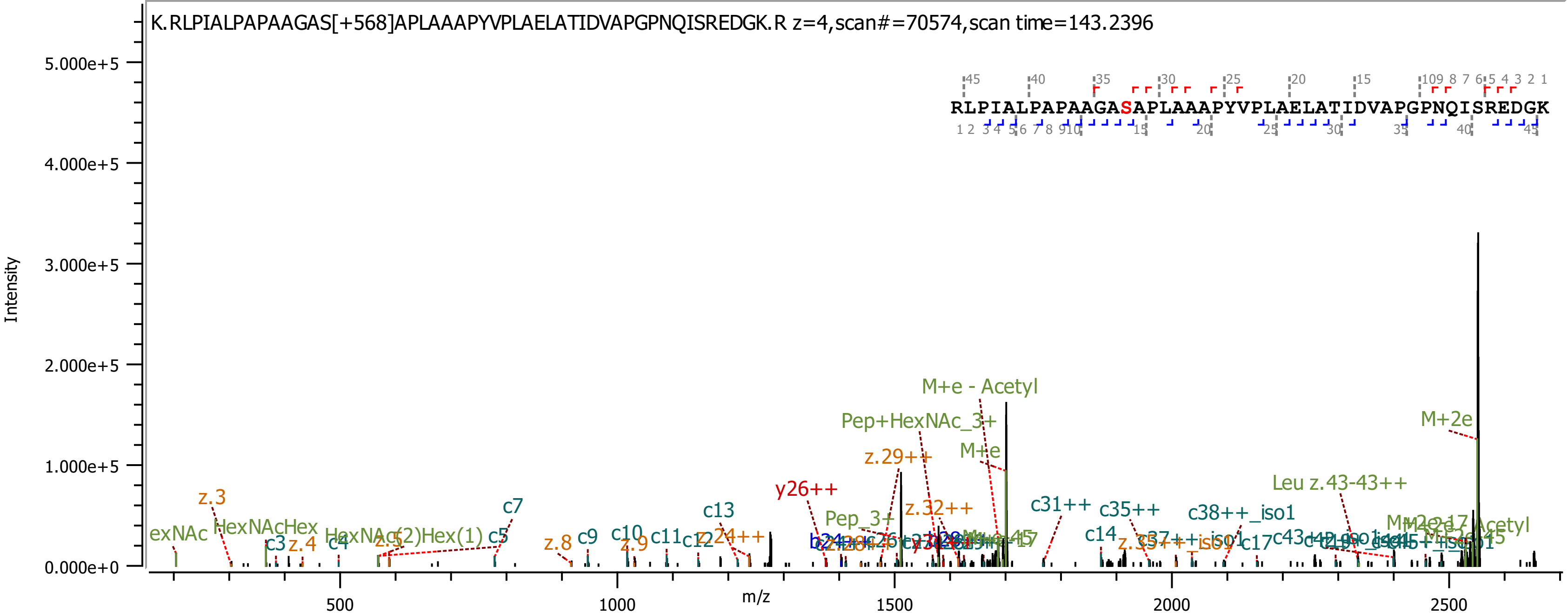

K.RLPIALPAPAAGAS[+568]APLAAAPYVPLAELATIDVAPGPNQISREDGK.R z=4,scan#=70574,scan time=143.2396
RLPIALPAPAAGASAPLAAAPYVPLAELATIDVAPGPNQISREDGK
Intensity
5.000e+5
4.000e+5
3.000e+5
2.000e+5
1.000e+5
0.000e+0
m/z
500
1000
1500
2000
2500
M+e - Acetyl
Pep+HexNAc_3+
M+e
M+2e
z.29++
y26++
z.32++
Pep_3+
Leu z.43-43++
c31++
c35++
c38++_iso1
c13
c14
c17
z.3
z.4
z.5
z.8
z.9
c3
c4
c5
c7
c9
c10
c11
c12
z.24++
HexNAcHex
HexNAc(2)Hex(1)
exNAc
z.35++
z.37++_iso1

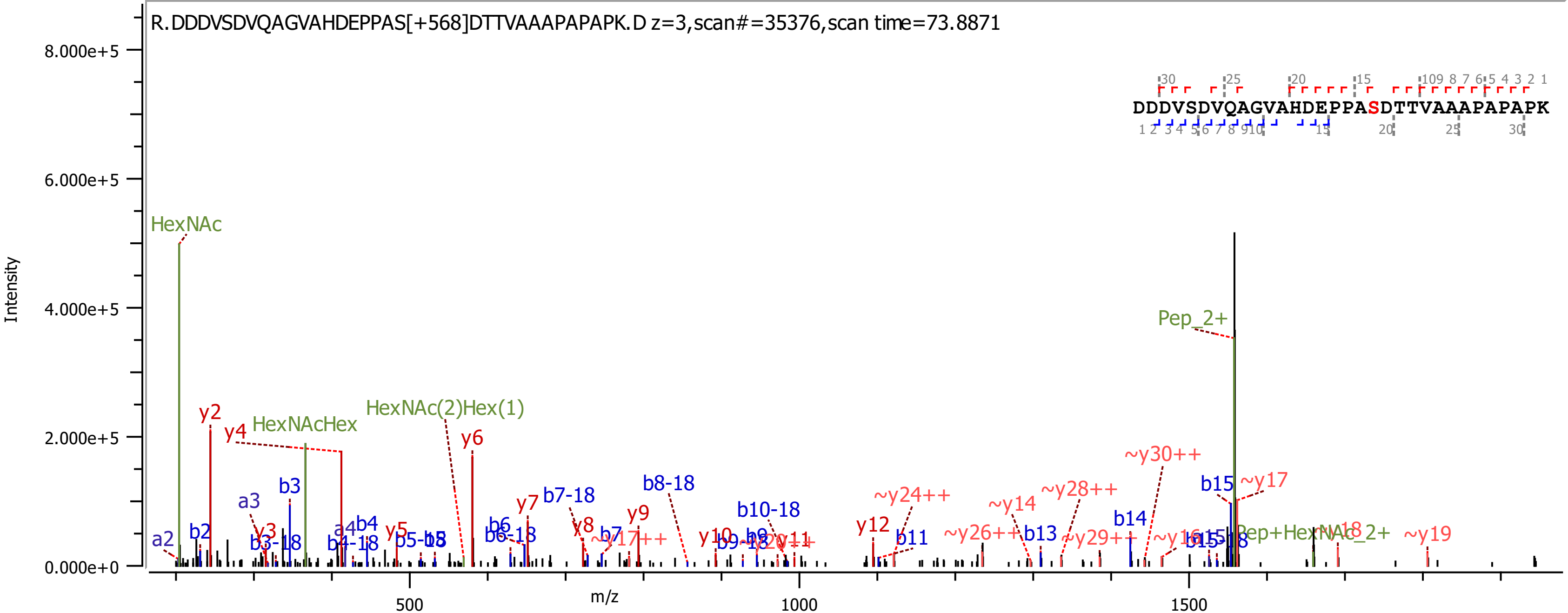

R.DDDVSDVQAGVAHDEPPAS[+568]DTTVAAAPAPAPK.D z=3,scan#=35376,scan time=73.8871
DDDVSDVQAGVAHDEPPASDTTVAAAPAPAPK
Intensity
m/z
HexNAc
HexNAcHex
HexNAc(2)Hex(1)
Pep_2+

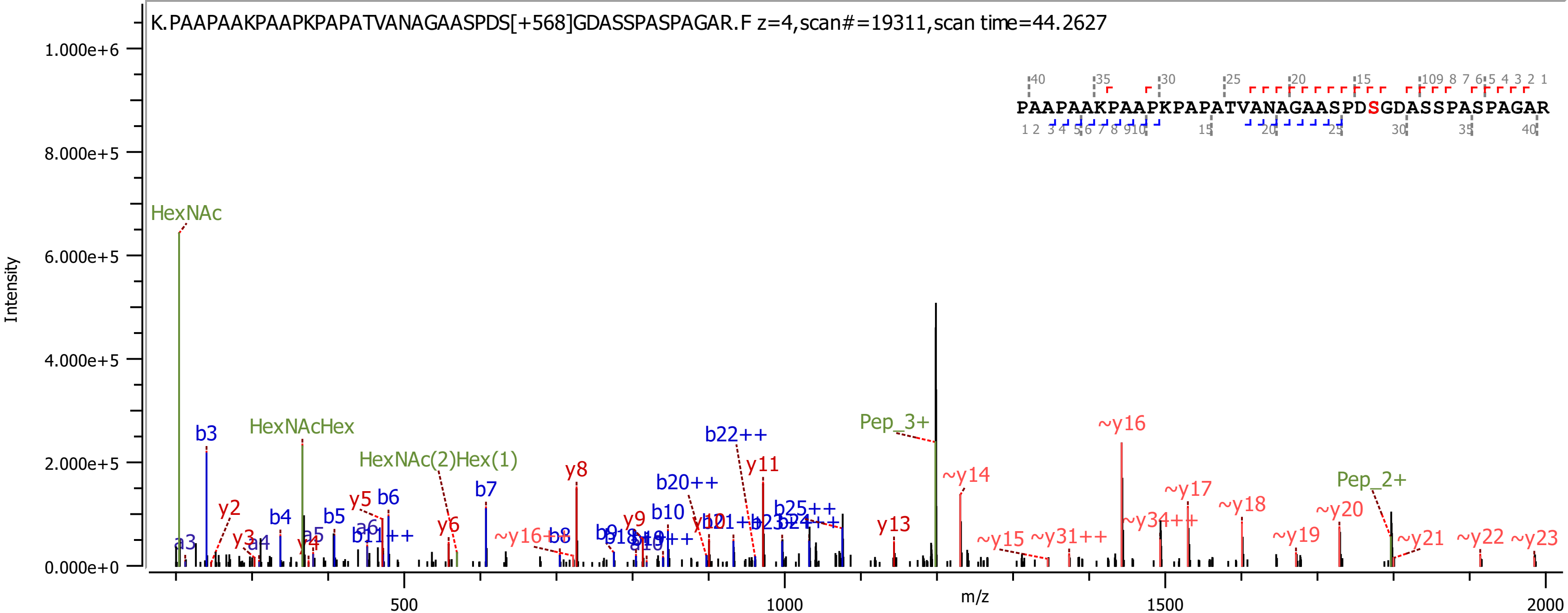
K.PAAPAAKPAAPKPAPATVANAGAASPDS[+568]GDASSPASPAGAR.F z=4,scan#=19311,scan time=44.2627
PAAPAAKPAAPKPAPATVANAGAASPDSGDASSPASPAGAR
Intensity
m/z
HexNAc
HexNAcHex
HexNAc(2)Hex(1)
Pep_3+
Pep_2+
~y16
~y14
~y17
~y18
~y20
~y19
~y21
~y22
~y23
~y15
~y31++
~y34++
~y16++
b22++
b20++
b25++
b3
b4
b5
b6
b7
b8
b10
b11++
y2
y3
y4
y5
y6
y8
y9
y11
y13
a3
a4
a5
a6

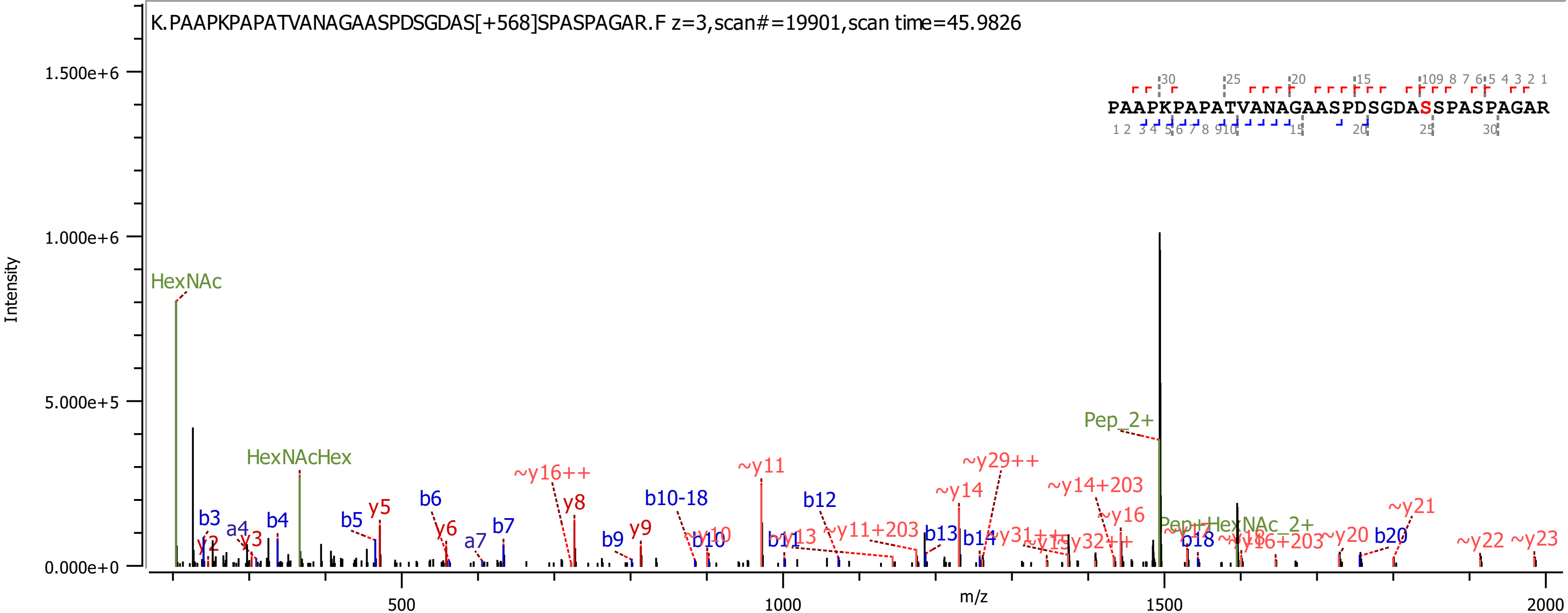
K.PAAPKPAPATVANAGAASPDSGDAS[+568]SPASPAGAR.F z=3,scan#=19901,scan time=45.9826
PAAPKPAPATVANAGAASPDSGDASSPASPAGAR
Intensity
m/z
1.500e+6
1.000e+6
5.000e+5
0.000e+0
500
1000
1500
2000
HexNAc
HexNAcHex
Pep_2+
Pep+HexNAc_2+
b3
a4
b4
b5
b6
a7
b7
b9
b10-18
b10
b11
b12
b13
b14
b18
b20
y2
y3
y5
y6
y8
y9
~y16++
~y10
~y11
~y13
~y11+203
~y14
~y29++
~y31++
~y15
~y32++
~y14+203
~y16
~y17
~y18
~y16+203
~y20
~y21
~y22
~y23

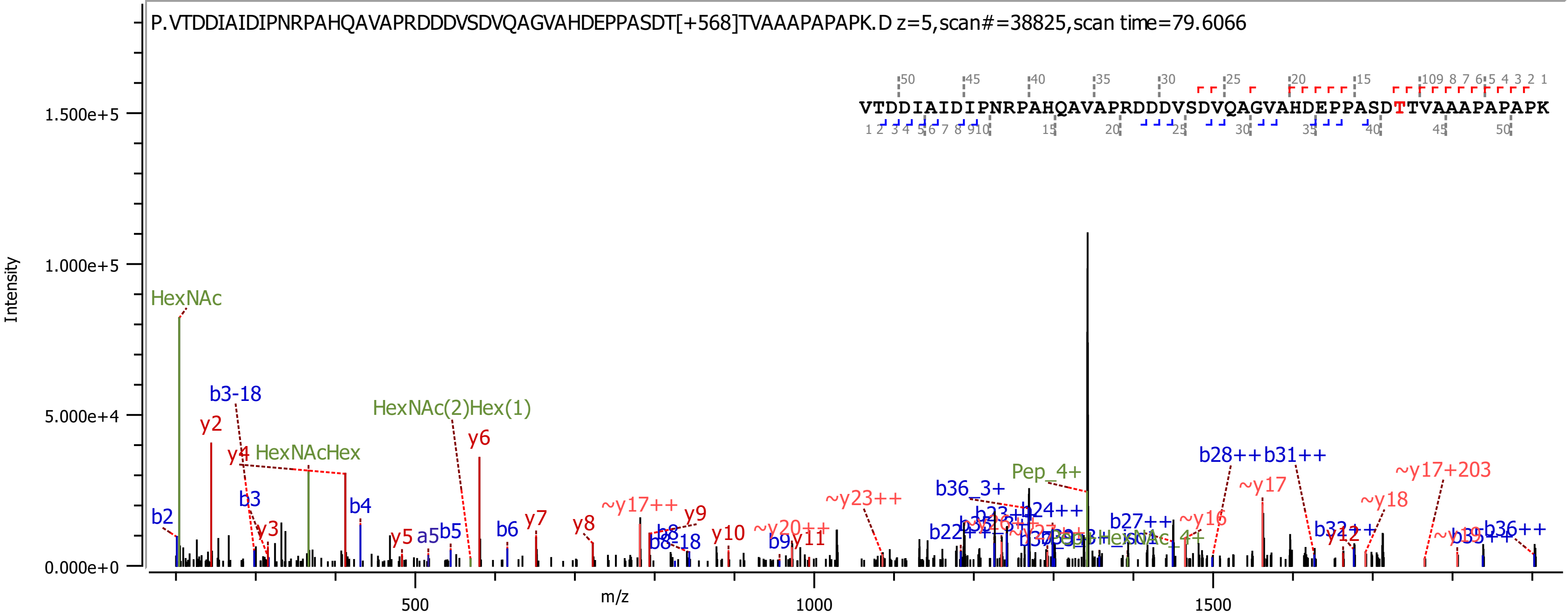
P.VTDDIAIDIPNRPAHQAVAPRDDDVSDVQAGVAHDEPPASDT[+568]TVAAAPAPAPK.D z=5,scan#=38825,scan time=79.6066
VTDDIAIDIPNRPAHQAVAPRDDDVSDVQAGVAHDEPPASDTTVAAAPAPAPK
Intensity
m/z
HexNAc
b3-18
y2
y4
HexNAcHex
HexNAc(2)Hex(1)
y6
b3
b4
y3
b2
y5
a5
b5
b6
y7
y8
~y17++
y9
b8
b8-18
y10
~y20++
b9
y11
~y23++
b36_3+
Pep_4+
b22++
b23++
b24++
b27++
~y16
b28++
b31++
~y17
~y18
~y17+203
b32++
y12
b35++
b36++

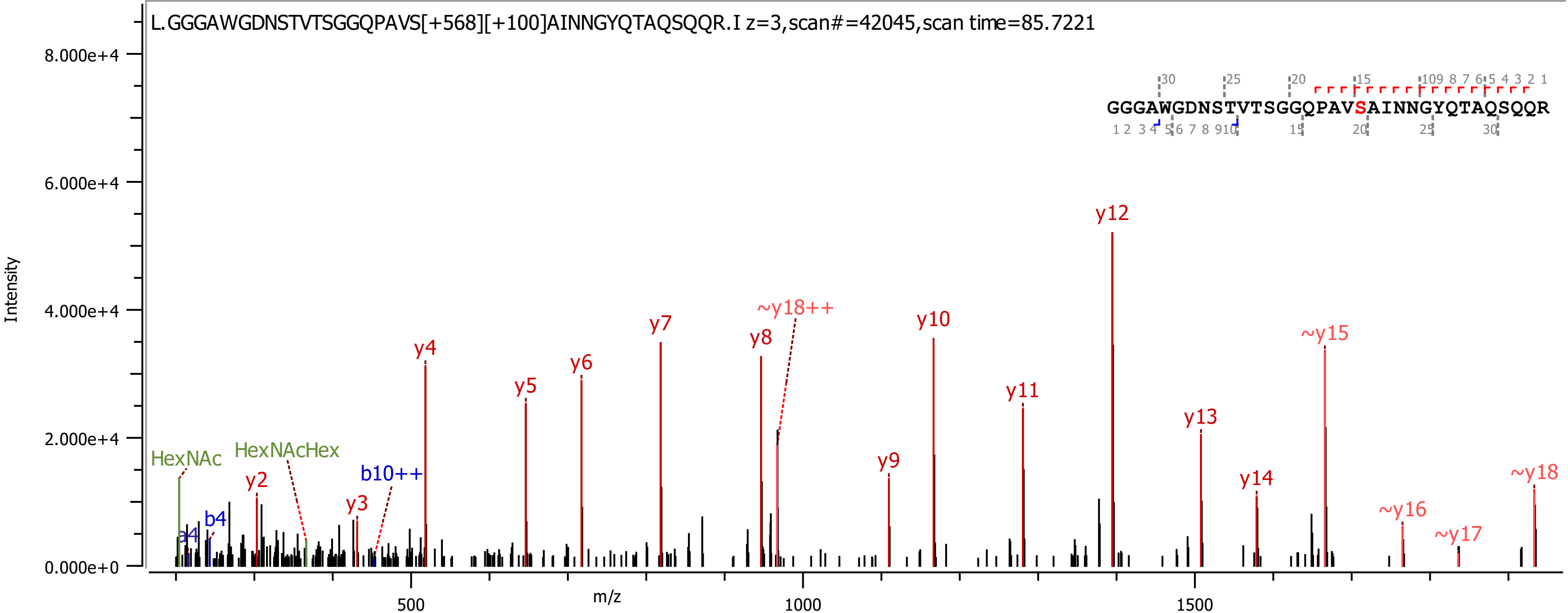
L.GGGAWGDNSTVTSGGQPAVS[+568][+100]AINNGYQTAQSQQR.I z=3,scan#=42045,scan time=85.7221
GGGAWGDNSTVTSGGQPAVSAINNGYQTAQSQQR
Intensity
m/z
8.000e+4
6.000e+4
4.000e+4
2.000e+4
0.000e+0
500
1000
1500
HexNAc
HexNAcHex
a4
b4
y2
y3
b10++
y4
y5
y6
y7
y8
~y18++
y9
y10
y11
y12
y13
y14
~y15
~y16
~y17
~y18

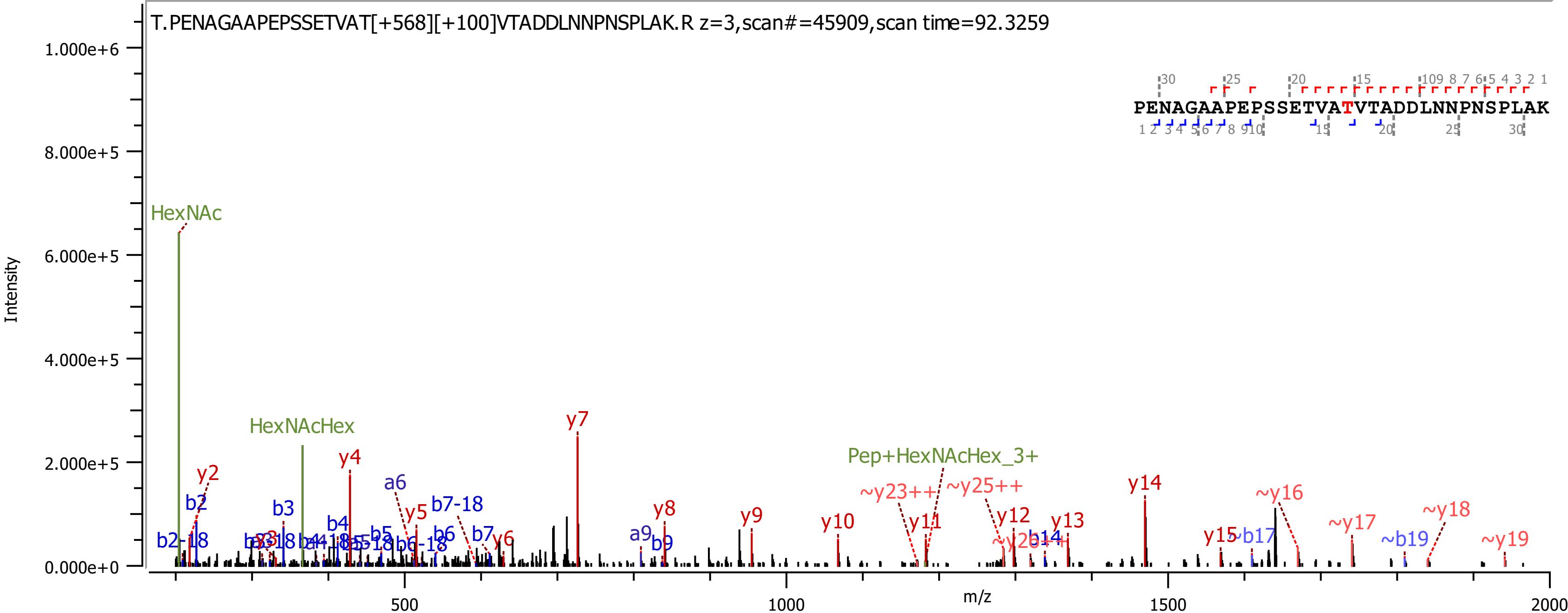
T.PENAGAAPEPSSETVAT[+568][+100]VTADDLNNPNSPLAK.R z=3,scan#=45909,scan time=92.3259
PENAGAAPEPSSETVATVTADDLNNPNSPLAK
Intensity
m/z
HexNAc
HexNAcHex
Pep+HexNAcHex_3+
y2
y4
y5
y6
y7
y8
y9
y10
y11
y12
y13
y14
y15
~y16
~y17
~y18
~y19
~y23++
~y25++
b2
b2-18
b3
b4
b5
b6
b6-18
b7
b7-18
b9
b14
~b17
~b19
a6
a9
0.000e+0
2.000e+5
4.000e+5
6.000e+5
8.000e+5
1.000e+6
500
1000
1500
2000

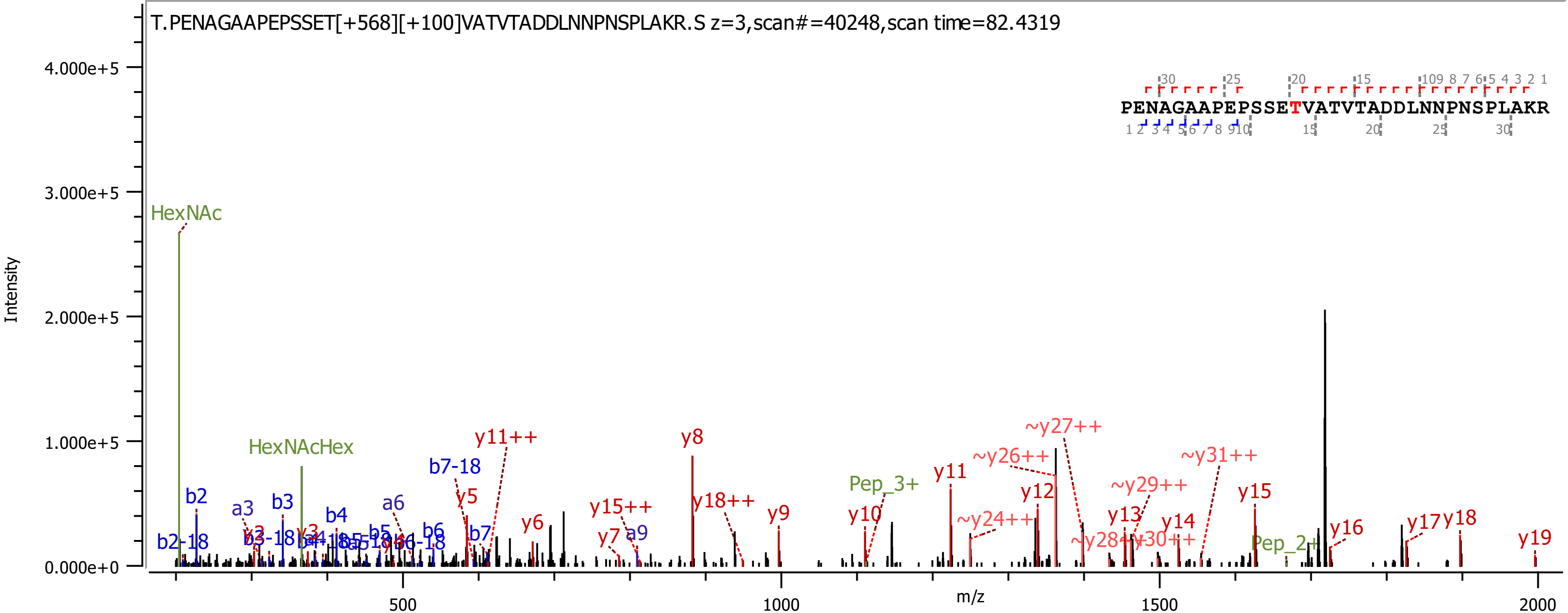
T.PENAGAAPEPSSET[+568][+100]VATVTADDLNNPNSPLAKR.S z=3,scan#=40248,scan time=82.4319
PENAGAAPEPSSETVATVTADDLNNPNSPLAKR
Intensity
m/z
HexNAc
HexNAcHex
Pep_3+
Pep_2+
b2
b2-18
a3
b3
b4
a6
b5
b6
b7
b7-18
a9
y5
y6
y7
y8
y9
y10
y11
y11++
y12
y13
y14
y15
y15++
y16
y17
y18
y18++
y19
~y24++
~y26++
~y27++
~y28++
~y29++
~y30++
~y31++

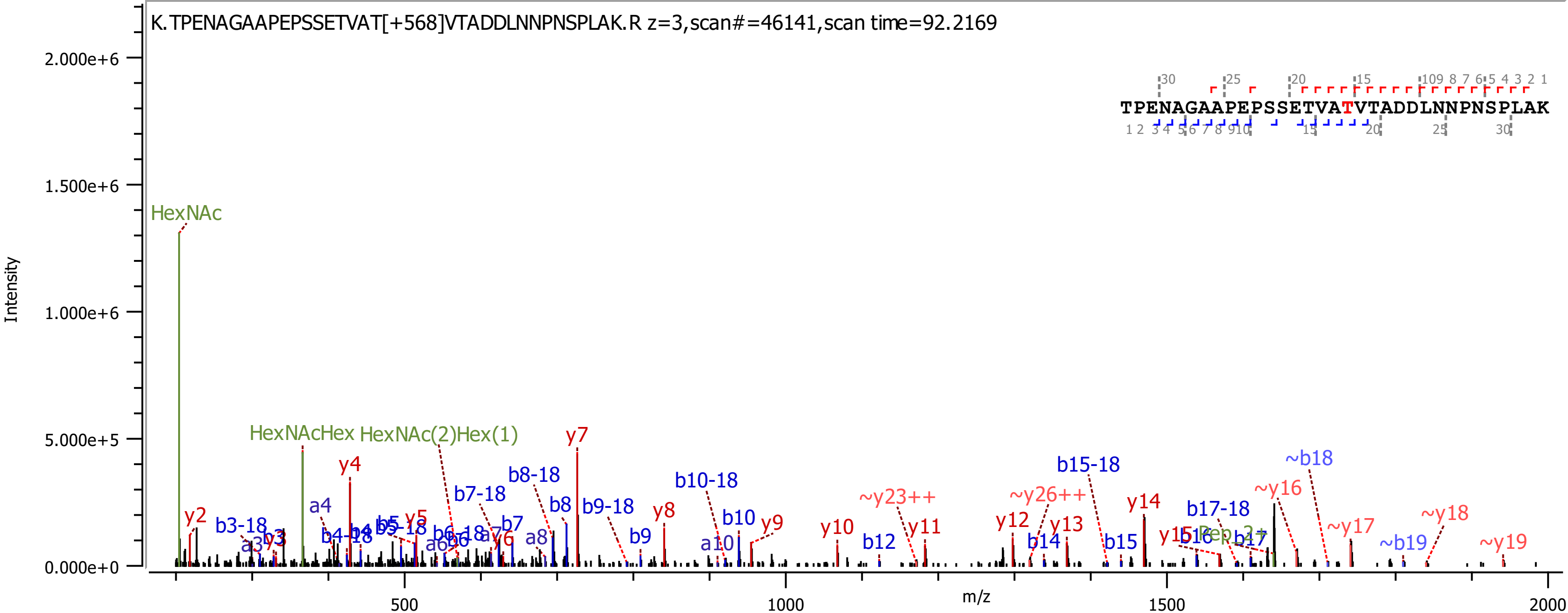

K.TPENAGAAPEPSSETVAT[+568]VTADDLNNPNSPLAK.R z=3,scan#=46141,scan time=92.2169
TPENAGAAPEPSSETVATVTADDLNNPNSPLAK
Intensity
m/z
HexNAc
HexNAcHex
HexNAc(2)Hex(1)
y2
y4
y5
y6
y7
y8
y9
y10
y11
y12
y13
y14
b3-18
b7-18
b8-18
b9-18
b10-18
b15-18
b17-18
b7
b8
b9
b10
b12
b14
b15
b16
b17
a4
a8
a10
~y23++
~y26++
~y16
~y17
~y18
~y19
~b18
~b19
2.000e+6
1.500e+6
1.000e+6
5.000e+5
0.000e+0
500
1000
1500
2000

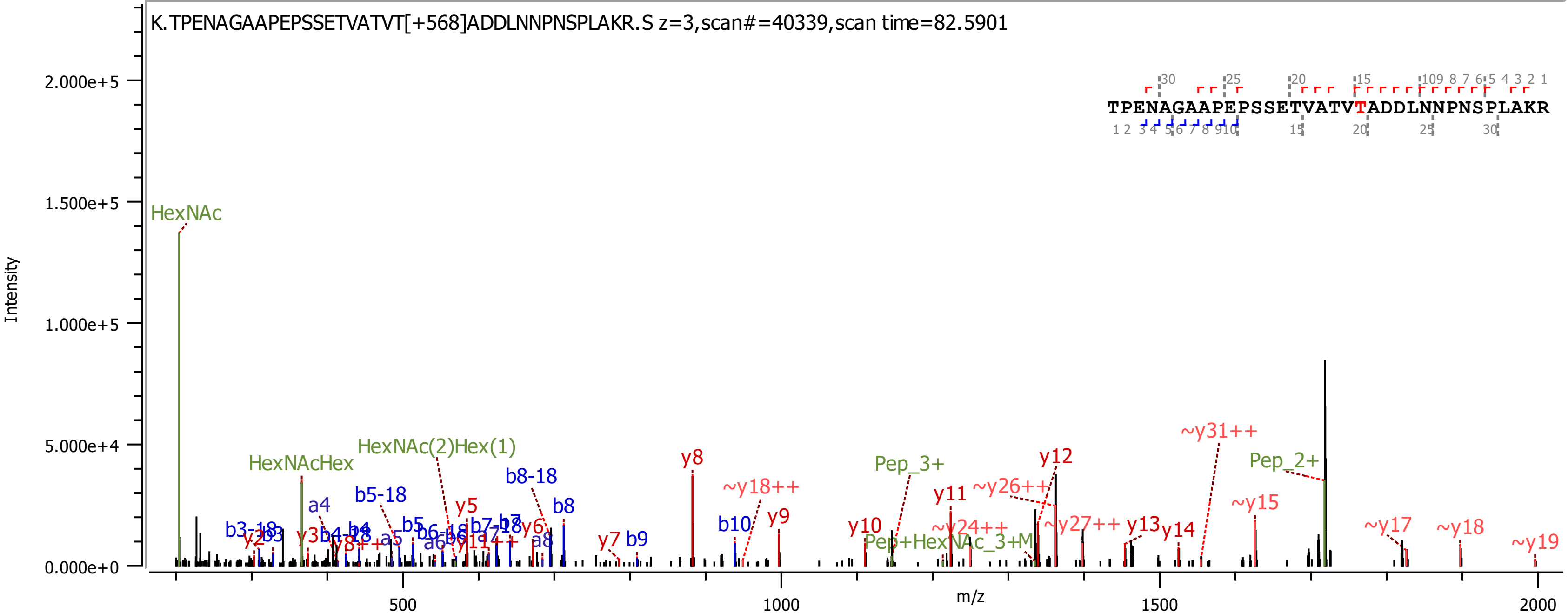
K.TPENAGAAPEPSSETVATVT[+568]ADDLNNPNSPLAKR.S z=3,scan#=40339,scan time=82.5901
TPENAGAAPEPSSETVATVTADDLNNPNSPLAKR
Intensity
m/z
2.000e+5
1.500e+5
1.000e+5
5.000e+4
0.000e+0
500
1000
1500
2000
HexNAc
HexNAcHex
HexNAc(2)Hex(1)
Pep_3+
Pep_2+
Pep+HexNAc_3+M
a4
a5
a6
a8
b3-18
b3
b4-18
b4
b5-18
b5
b6-18
b6
b7-18
b7
b8-18
b8
b9
b10
y2
y3
y5
y6
y7
y8
y8++
y9
y10
y11
y11++
y12
y13
y14
~y15
~y17
~y18
~y19
~y18++
~y24++
~y26++
~y27++
~y31++

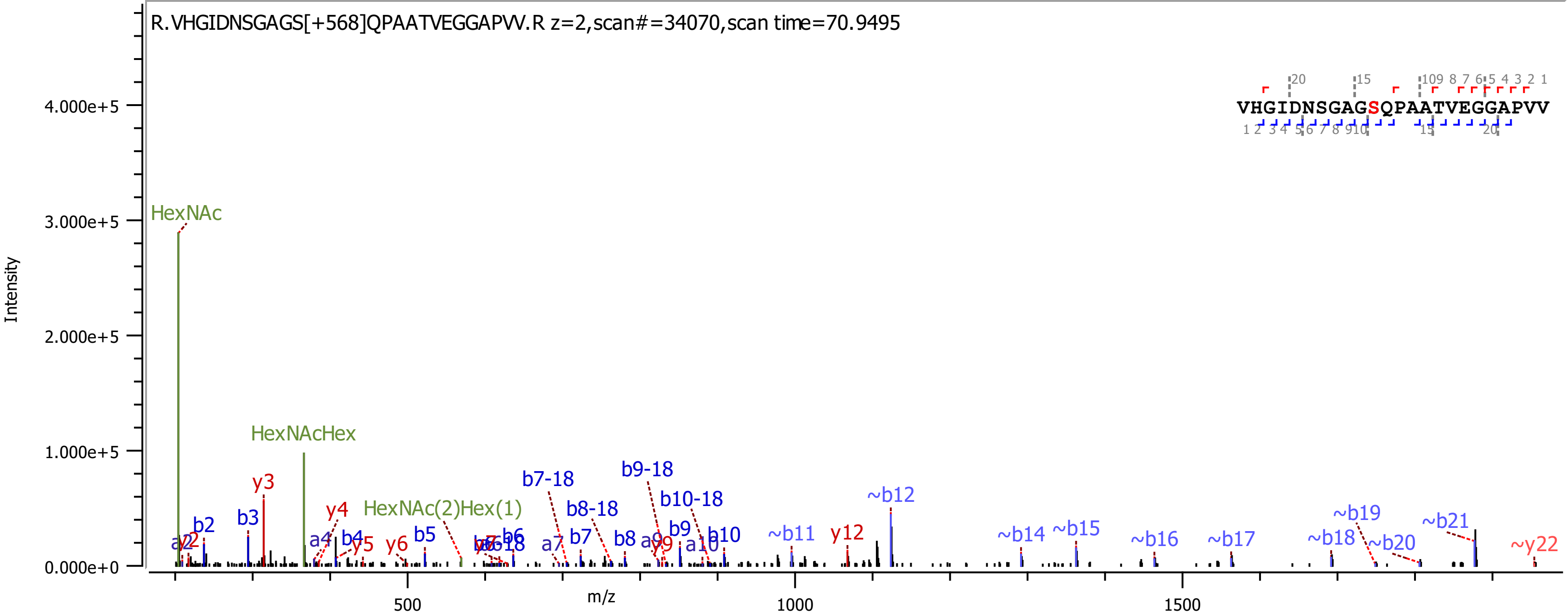
R.VHGIDNSGAGS[+568]QPAATVEGGAPVV.R z=2,scan#=34070,scan time=70.9495
VHGIDNSGAGSQPAATVEGGAPVV
Intensity
m/z
HexNAc
HexNAcHex
HexNAc(2)Hex(1)
y3
y4
y5
y6
y9
y12
~y22
b2
b3
b4
b5
b6
b7
b8
b9
b10
a4
a7
a9
a10
b7-18
b8-18
b9-18
b10-18
~b11
~b12
~b14
~b15
~b16
~b17
~b18
~b19
~b20
~b21

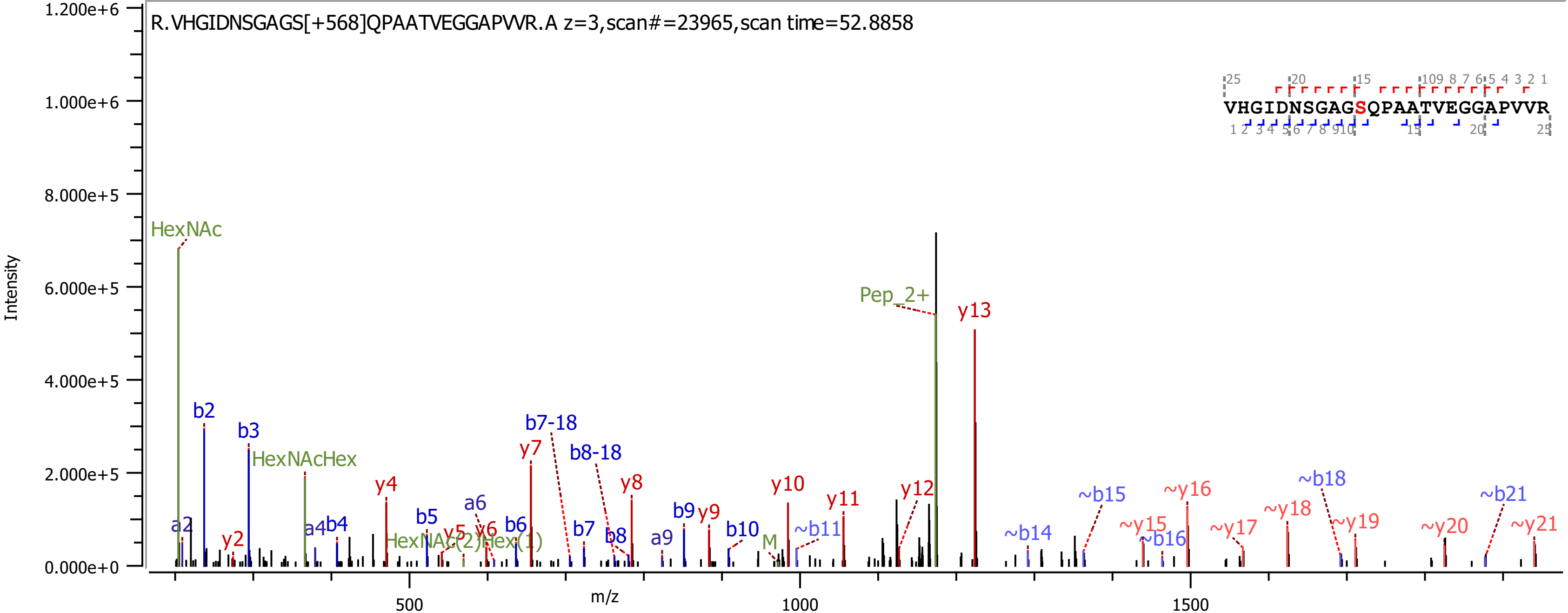

R.VHGIDNSGAGS[+568]QPAATVEGGAPVVR.A z=3,scan#=23965,scan time=52.8858
VHGIDNSGAGSQPAATVEGGAPVVR
Intensity
m/z
HexNAc
HexNAcHex
HexNAc(2)Hex(1)
Pep_2+
M
y13
y12
y11
y10
y9
y8
y7
y6
y5
y4
y2
b2
b3
b4
b5
b6
b7
b8
b9
b10
b7-18
b8-18
a2
a4
a6
a9
~b11
~b14
~b15
~b16
~b18
~b21
~y15
~y16
~y17
~y18
~y19
~y20
~y21

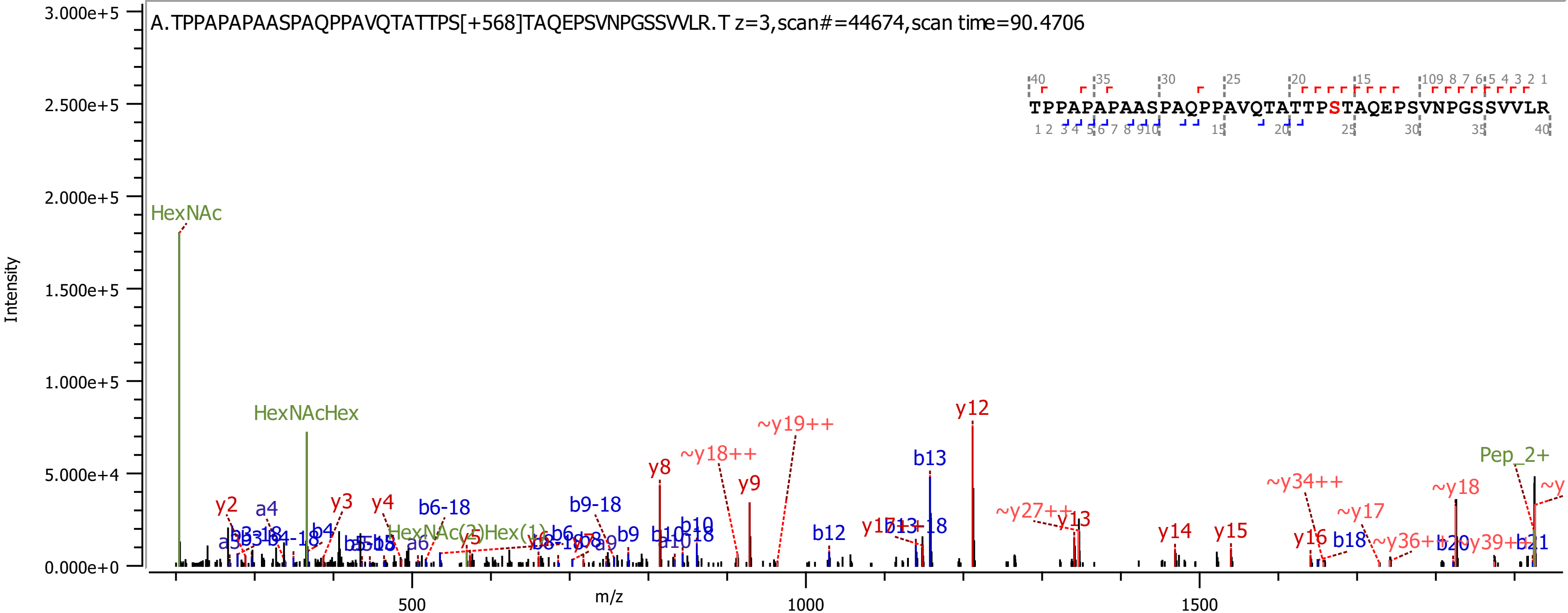
A.TPPAPAPAASPAQPPAVQTATTPS[+568]TAQEPSVNPGSSVVLR.T z=3,scan#=44674,scan time=90.4706
TPPAPAPAASPAQPPAVQTATTPSTAQEPSVNPGSSVVLR
Intensity
m/z
HexNAc
HexNAcHex
Pep_2+
y12
b13
~y19++
~y18++
y8
y9
b12
y14
y15
~y34++
~y18
~y17
b18
y16
y2
a4
y3
y4
b6-18
b9-18
b9
b10
y13
~y27++
3.000e+5
2.500e+5
2.000e+5
1.500e+5
1.000e+5
5.000e+4
0.000e+0
500
1000
1500

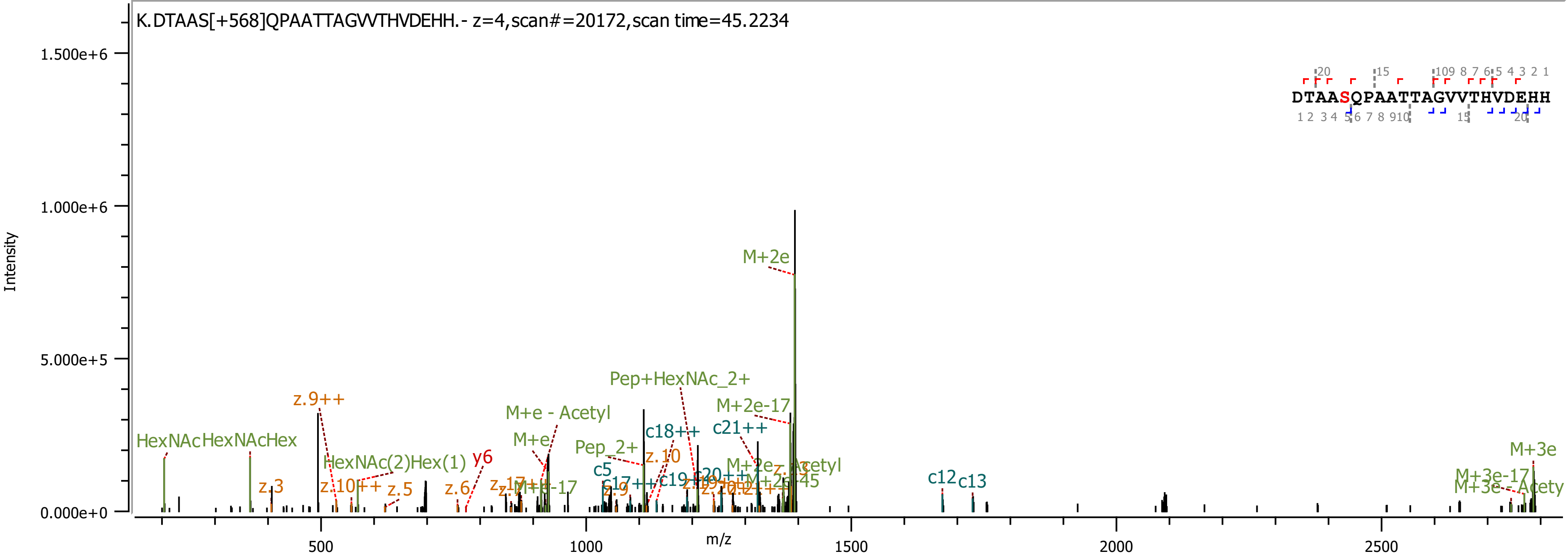

K.DTAAS[+568]QPAATTAGVVTHVDEHH.- z=4,scan#=20172,scan time=45.2234
Intensity
m/z
1.500e+6
1.000e+6
5.000e+5
0.000e+0
500
1000
1500
2000
2500
DTAASQPAATTAGVVTHVDEHH
M+2e
Pep+HexNAc_2+
M+2e-17
z.9++
M+e - Acetyl
c21++
c18++
M+e
Pep_2+
HexNAc
HexNAcHex
HexNAc(2)Hex(1)
y6
z.10
c5
z.3
z.10++
z.5
z.6
c12
c13
M+3e
M+3e-17

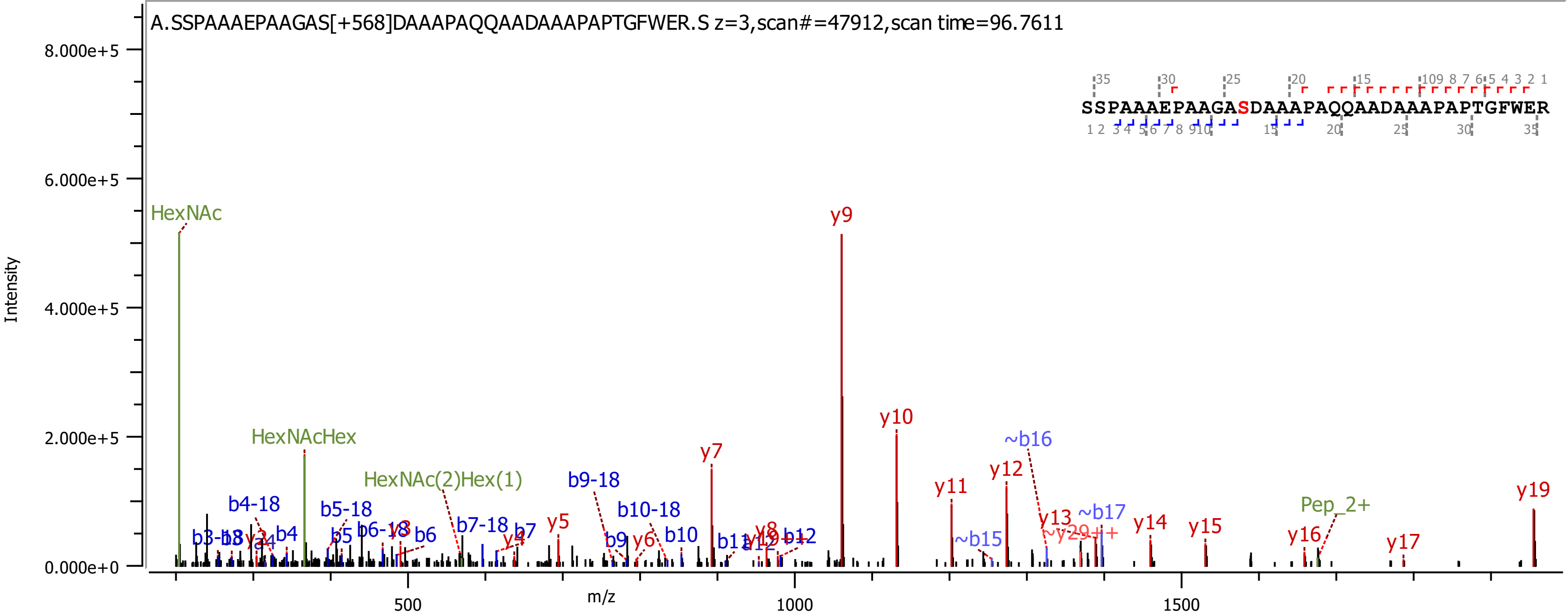
A.SSPAAAEPAAGAS[+568]DAAAPAQQAADAAAPAPTGFWER.S z=3,scan#=47912,scan time=96.7611
SSPAAAEPAAGASDAAAPAQQAADAAAPAPTGFWER
Intensity
m/z
8.000e+5
6.000e+5
4.000e+5
2.000e+5
0.000e+0
500
1000
1500
HexNAc
HexNAcHex
HexNAc(2)Hex(1)
Pep_2+
b4-18
b5-18
b6-18
b7-18
b9-18
b10-18
b4
b5
b6
b7
b9
b10
b12
~b15
~b16
~b17
y4
y5
y6
y7
y8
y9
y10
y11
y12
y13
y14
y15
y16
y17
y19

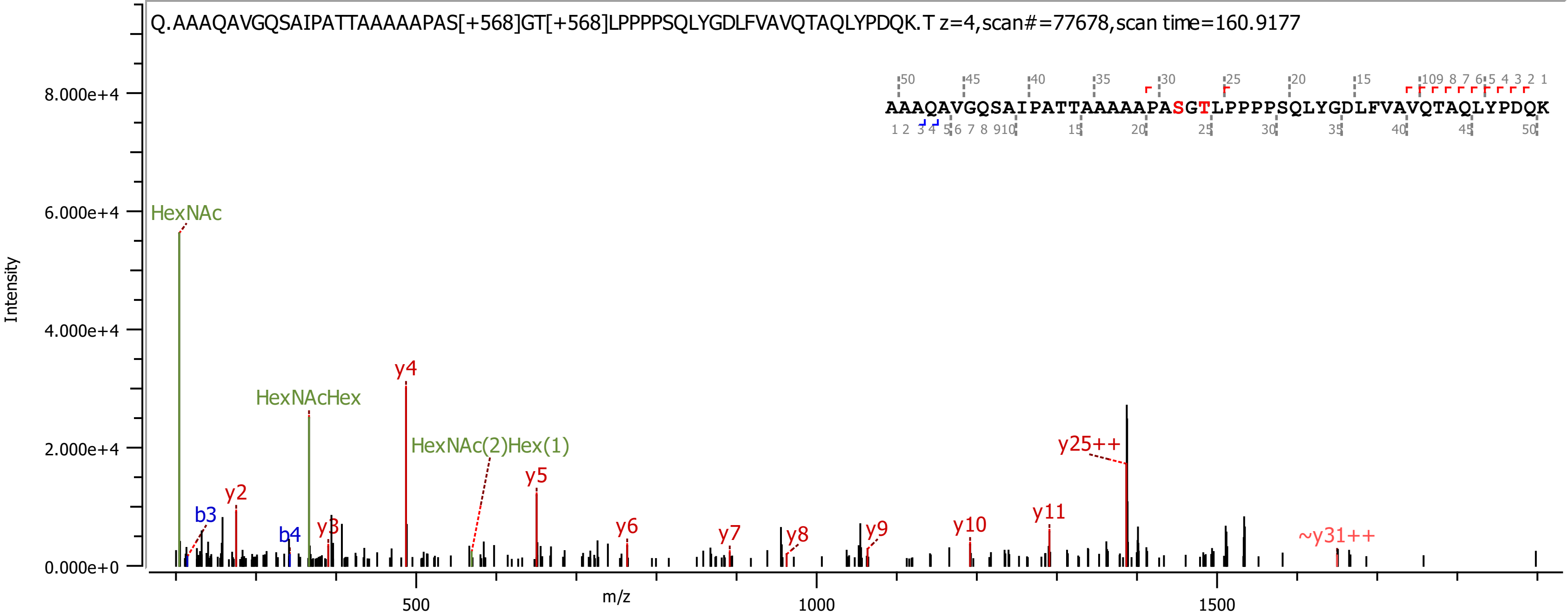

Q.AAAQAVGQSAIPATTAAAAAPAS[+568]GT[+568]LPPPPSQLYGDLFVAVQTAQLYPDQK.T z=4,scan#=77678,scan time=160.9177
AAAQAVGQSAIPATTAAAAAPASGTLPPPPSQLYGDLFVAVQTAQLYPDQK
Intensity
m/z
HexNAc
HexNAcHex
HexNAc(2)Hex(1)
b3
b4
y2
y3
y4
y5
y6
y7
y8
y9
y10
y11
y25++
~y31++

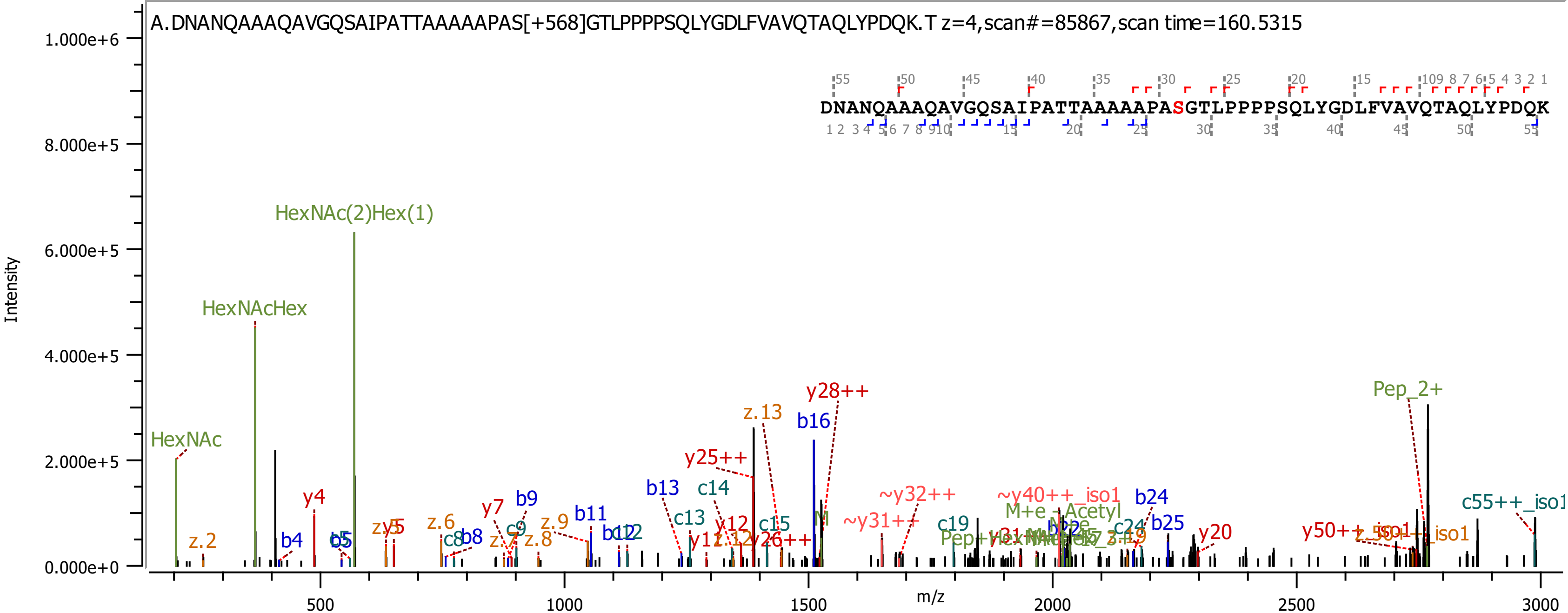

A.DNANQAAAQAVGQSAIPATTAAAAAPAS[+568]GTLPPPPSQLYGDLFVAVQTAQLYPDQK.T z=4,scan#=85867,scan time=160.5315
DNANQAAAQAVGQSAIPATTAAAAAPASGTLPPPPSQLYGDLFVAVQTAQLYPDQK
Intensity
m/z
1.000e+6
8.000e+5
6.000e+5
4.000e+5
2.000e+5
0.000e+0
500
1000
1500
2000
2500
3000
HexNAc(2)Hex(1)
HexNAcHex
HexNAc
Pep_2+
y28++
z.13
b16
y25++
b13
c14
~y32++
~y40++_iso1
b24
M+e -Acetyl
c55++_iso1
y4
b9
y7
c9
z.9
b11
b12
c13
y12
c15
~y31++
c19
c24
b25
y20
y50++
z.2
b4
b5
zy5
z.6
c8
b8
z.7
z.8
z.12
y26++

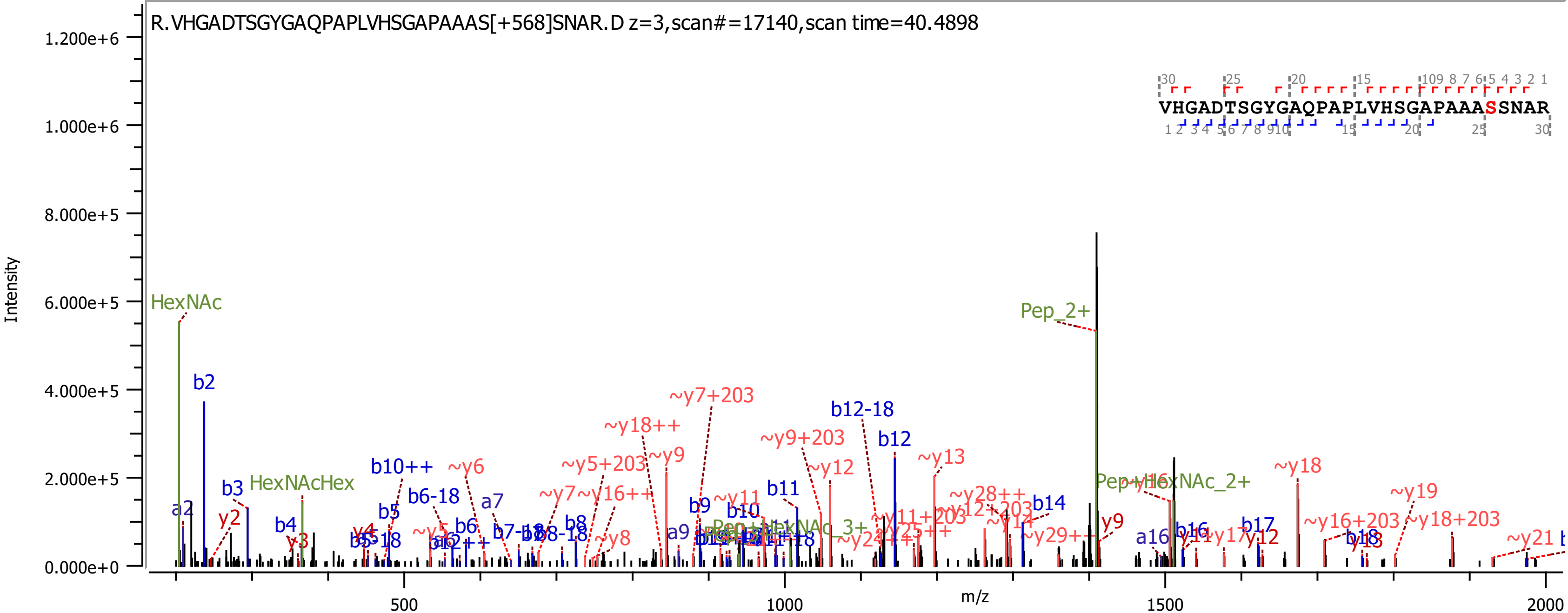
R.VHGADTSGYGAQPAPLVHSGAPAAAS[+568]SNAR.D z=3,scan#=17140,scan time=40.4898
VHGADTSGYGAQPAPLVHSGAPAAASSNAR
Intensity
m/z
1.200e+6
1.000e+6
8.000e+5
6.000e+5
4.000e+5
2.000e+5
0.000e+0
500
1000
1500
2000
HexNAc
b2
a2
b3
y2
HexNAcHex
b4
y3
b10++
b5
b6-18
~y6
a7
b6
~y5+203
~y9
~y18++
~y7+203
~y7
~y16++
b8
~y8
a9
b9
~y11
b10
b11
~y9+203
~y12
b12-18
b12
~y13
~y28++
~y12+203
b14
~y14
~y29++
y9
Pep_2+
Pep+HexNAc_2+
~y16
a16
b16
y11
~y17
b17
y12
~y18
~y16+203
b18
y18
~y19
~y18+203
~y21

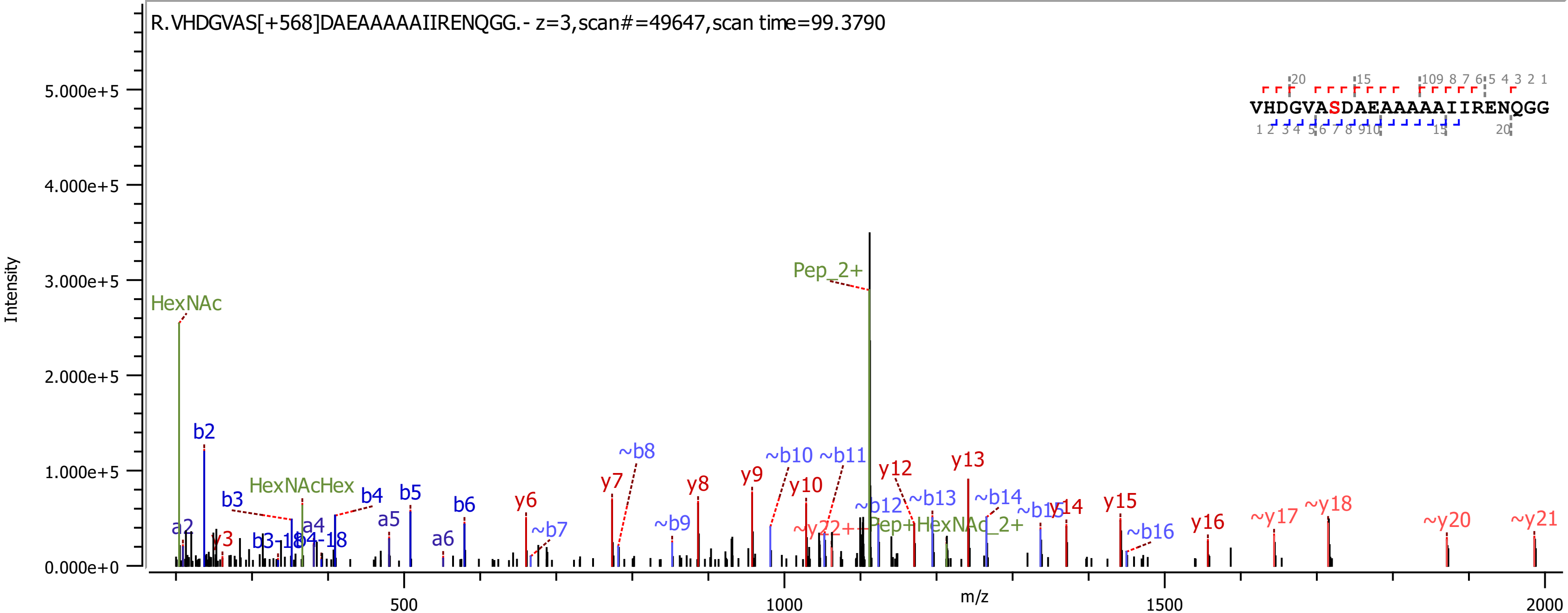
R.VHDGVAS[+568]DAEAAAAAIIRENQGG.- z=3,scan#=49647,scan time=99.3790
VHDGVASDAEAAAAAIIRENQGG
Intensity
5.000e+5
4.000e+5
3.000e+5
2.000e+5
1.000e+5
0.000e+0
m/z
500
1000
1500
2000
HexNAc
Pep_2+
HexNAcHex
Pep+HexNAc_2+
b2
b3
b4
b5
b6
a2
a4
a5
a6
y3
y6
y7
y8
y9
y10
y12
y13
y14
y15
y16
~b7
~b8
~b9
~b10
~b11
~b12
~b13
~b14
~b16
~y17
~y18
~y20
~y21
~y22+

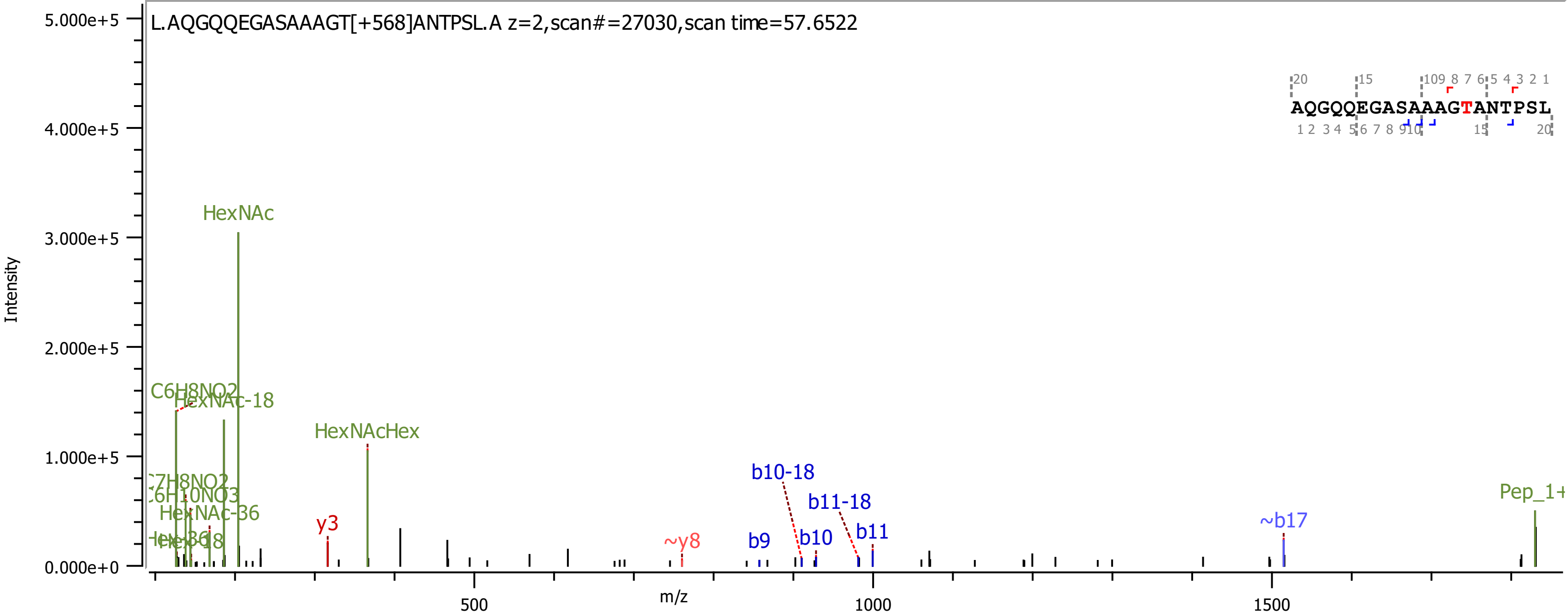
L.AQGQQEGASAAAGT[+568]ANTPSL.A z=2,scan#=27030,scan time=57.6522
AQGQQEGASAAAGTANTPSL
Intensity
m/z
HexNAc
HexNAc-18
C6H8NO2
HexNAcHex
C7H8NO2
C6H10NO3
HexNAc-36
y3
~y8
b9
b10
b10-18
b11-18
b11
~b17
Pep_1+
500
1000
1500
0.000e+0
1.000e+5
2.000e+5
3.000e+5
4.000e+5
5.000e+5

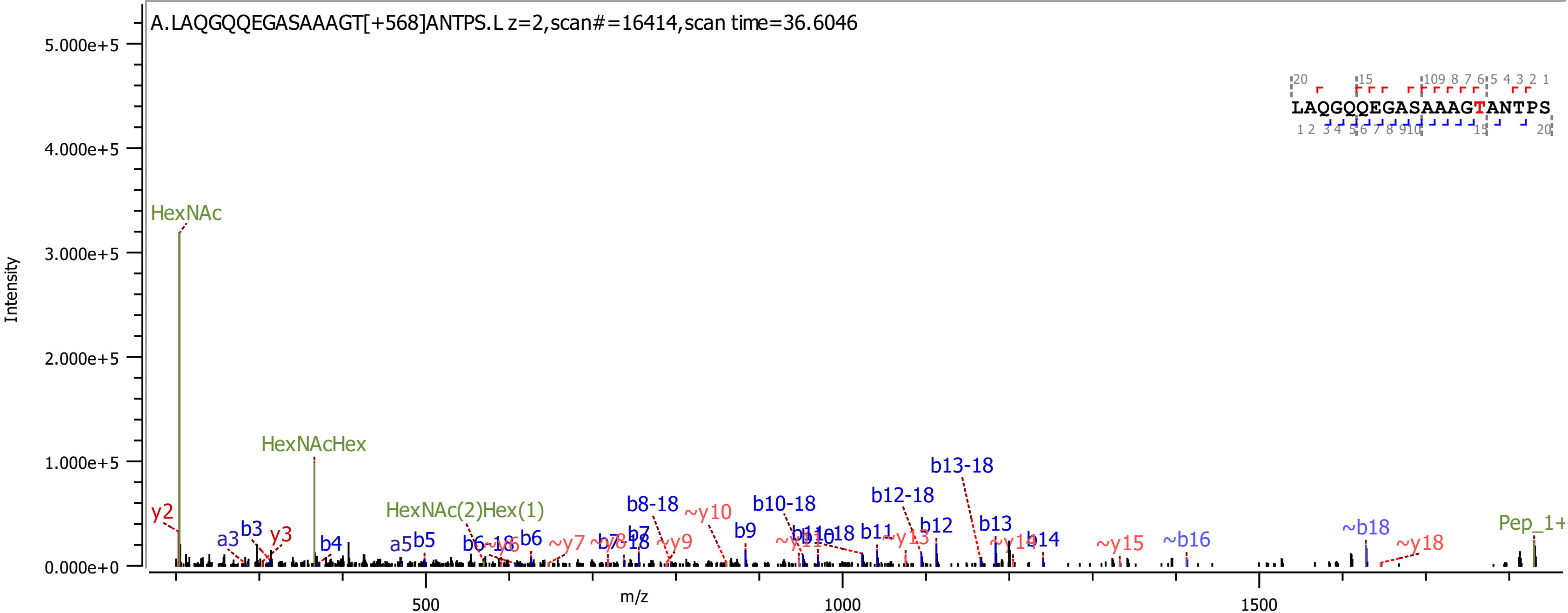
A.LAQGQQEGASAAAGT[+568]ANTPS.L z=2,scan#=16414,scan time=36.6046
Intensity
m/z
HexNAc
HexNAcHex
HexNAc(2)Hex(1)
Pep_1+
b8-18
b10-18
b12-18
b13-18
LAQGQQEGASAAAGTANTPS

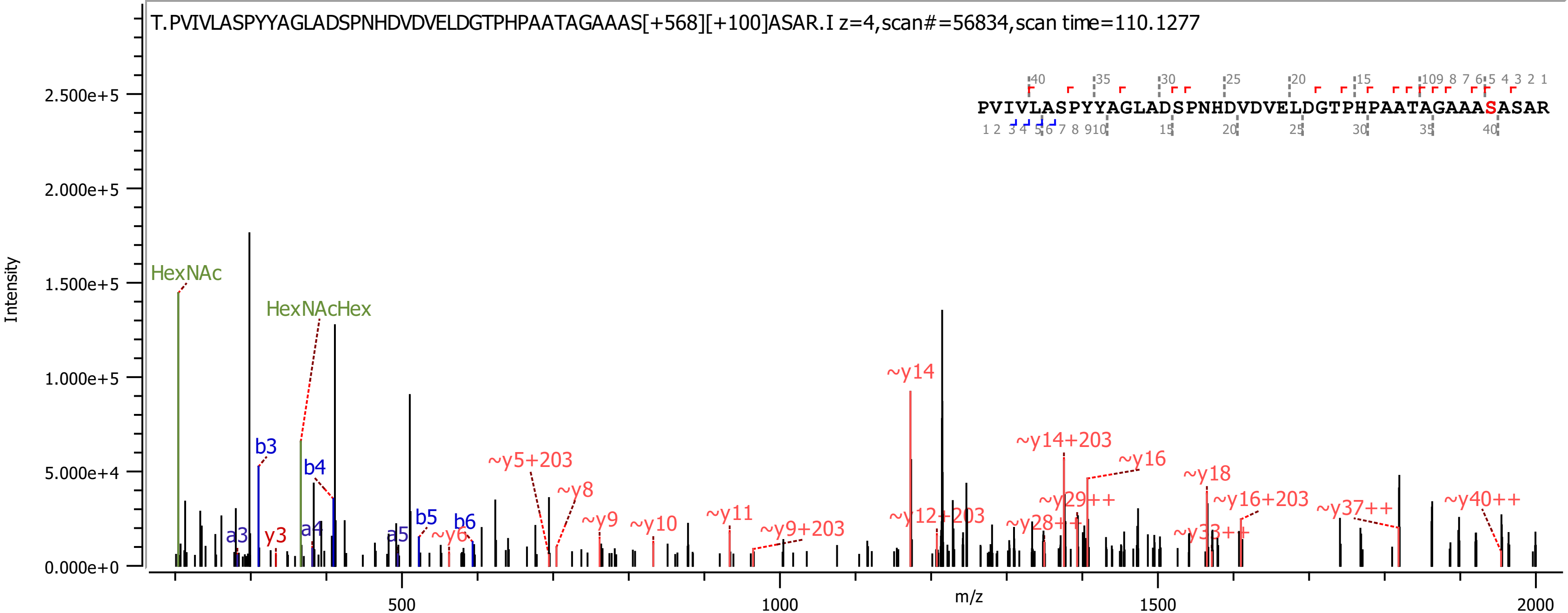

T.PVIVLASPYYAGLADSPNHDVDVELDGTPHPAATAGAAAS[+568][+100]ASAR.I z=4,scan#=56834,scan time=110.1277
PVIVLASPYYAGLADSPNHDVDVELDGTPHPAATAGAAASASAR
Intensity
m/z
HexNAc
HexNAcHex
b3
b4
a3
y3
a4
a5
b5
b6
~y6
~y5+203
~y8
~y9
~y10
~y11
~y9+203
~y14
~y12+203
~y14+203
~y16
~y29++
~y28++
~y18
~y16+203
~y33++
~y37++
~y40++

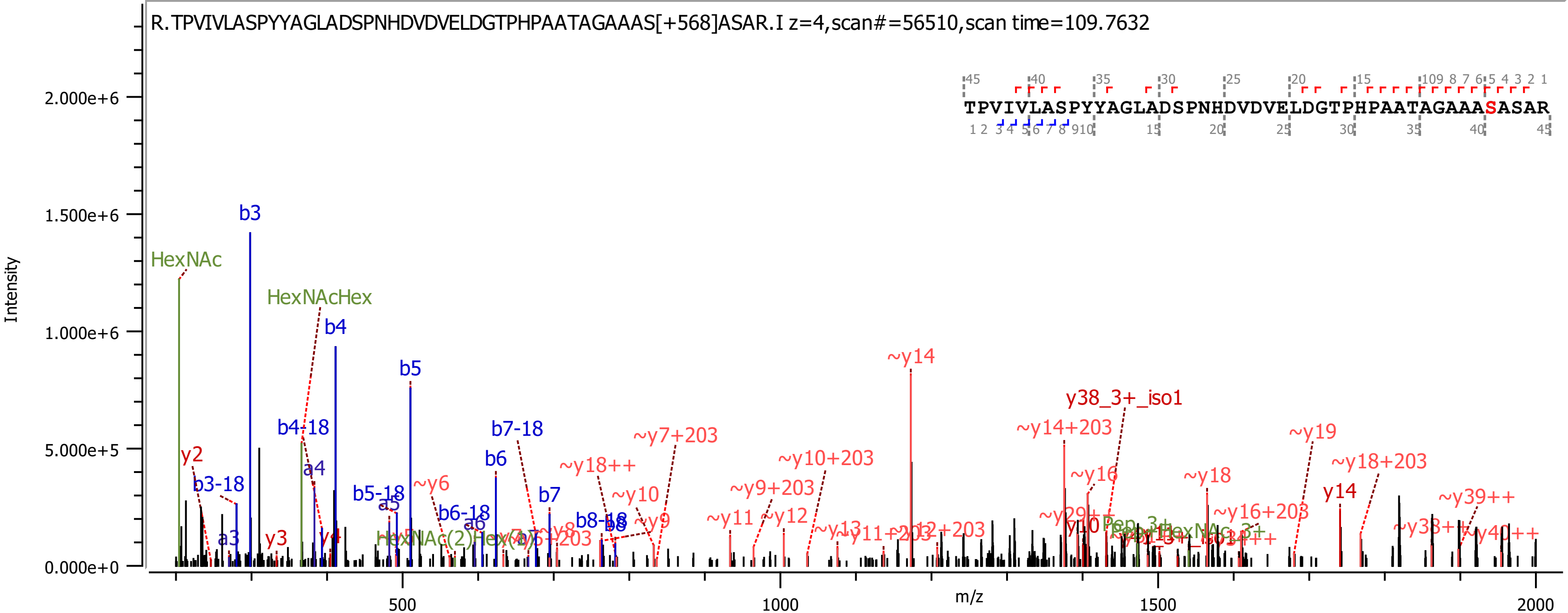

R.TPVIVLASPYYAGLADSPNHDVDVELDGTPHPAATAGAAAS[+568]ASAR.I z=4,scan#=56510,scan time=109.7632
TPVIVLASPYYAGLADSPNHDVDVELDGTPHPAATAGAAASASAR
Intensity
m/z
HexNAc
HexNAcHex
b3
b4
b5
b6
b7
b3-18
b4-18
b5-18
b6-18
b7-18
a3
a4
a5
a6
y2
y3
y4
~y6
~y9
~y10
~y11
~y12
~y13
~y14
~y16
~y18
~y19
y14
~y18++
~y7+203
~y9+203
~y10+203
~y14+203
~y16+203
~y18+203
~y39++
y38_3+_iso1

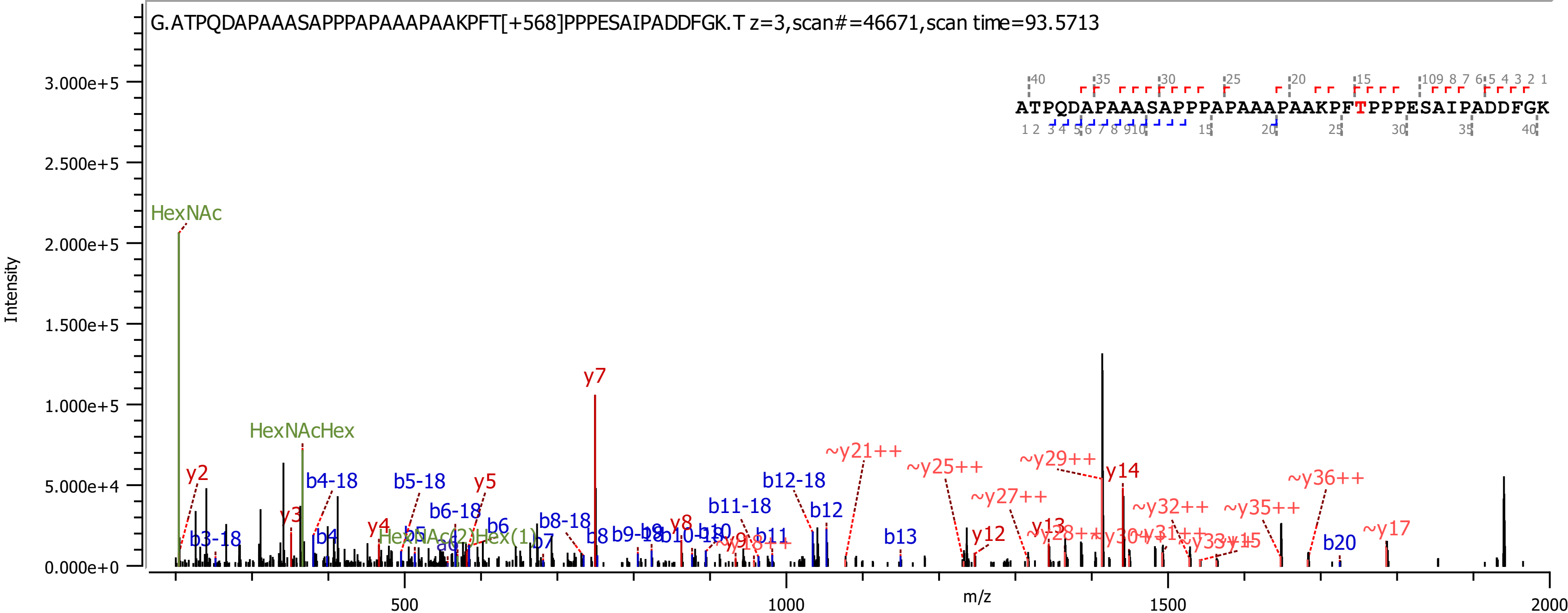

G.ATPQDAPAAASAPPPAPAAAPAAKPFT[+568]PPPESAIPADDFGK.T z=3,scan#=46671,scan time=93.5713
ATPQDAPAAASAPPPAPAAAPAAKPFTPPPESAIPADDFGK
Intensity
m/z
HexNAc
HexNAcHex
HexNAc(2)Hex(1)
y2
y3
y4
y5
y7
y8
y12
y13
y14
~y17
~y21++
~y25++
~y27++
~y29++
~y32++
~y35++
~y36++
b3-18
b4
b4-18
b5
b5-18
b6
b6-18
b7
b8
b8-18
b11
b11-18
b12
b12-18
b13
b20

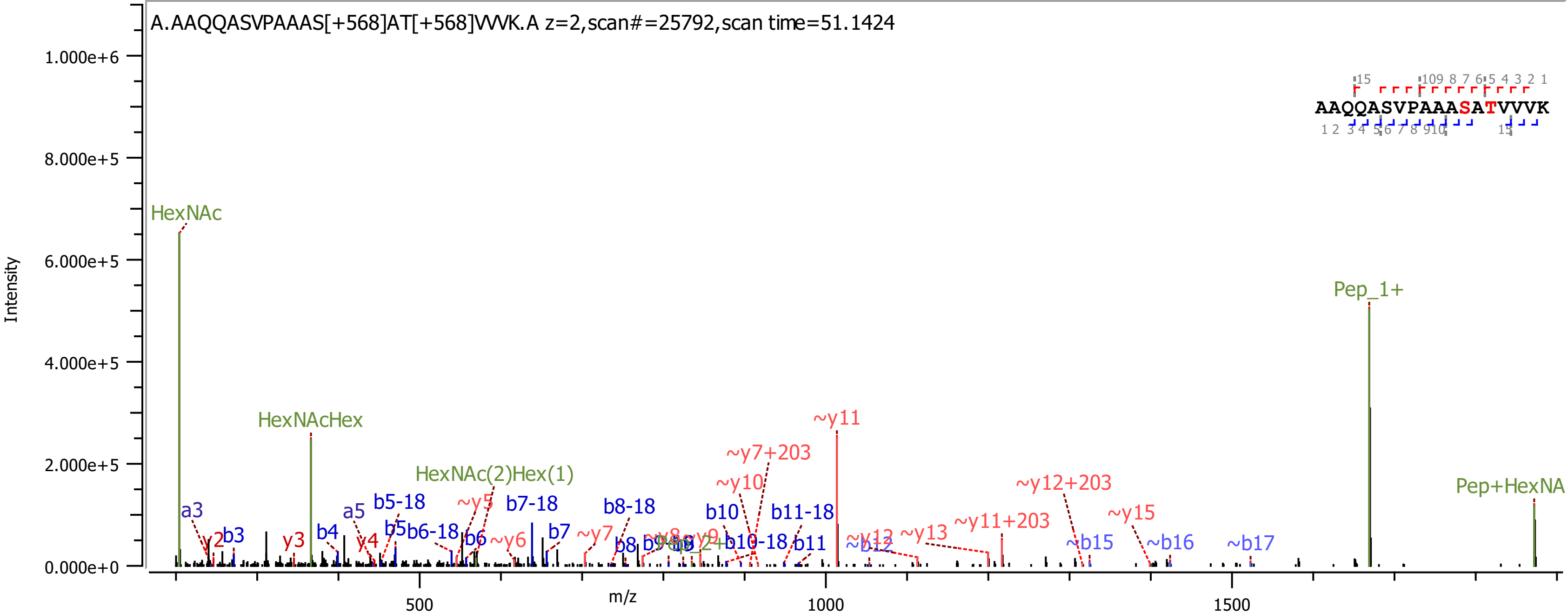
A.AAQQASVPAAAS[+568]AT[+568]VVVK.A z=2,scan#=25792,scan time=51.1424
AAQQASVPAAASATVVVK
Intensity
m/z
HexNAc
HexNAcHex
HexNAc(2)Hex(1)
Pep_1+
Pep+HexNA
~y11
~y7+203
~y10
~y12+203
~y11+203
~y15
~y13
~b15
~b16
~b17
b10
b11-18
b11
b10-18
b8-18
b7-18
b7
b5-18
b5
b6-18
b6
b4
b3
a3
a5
y2
y3
y4
~y5
~y6
~y7
b8
b9
~y12
~b12

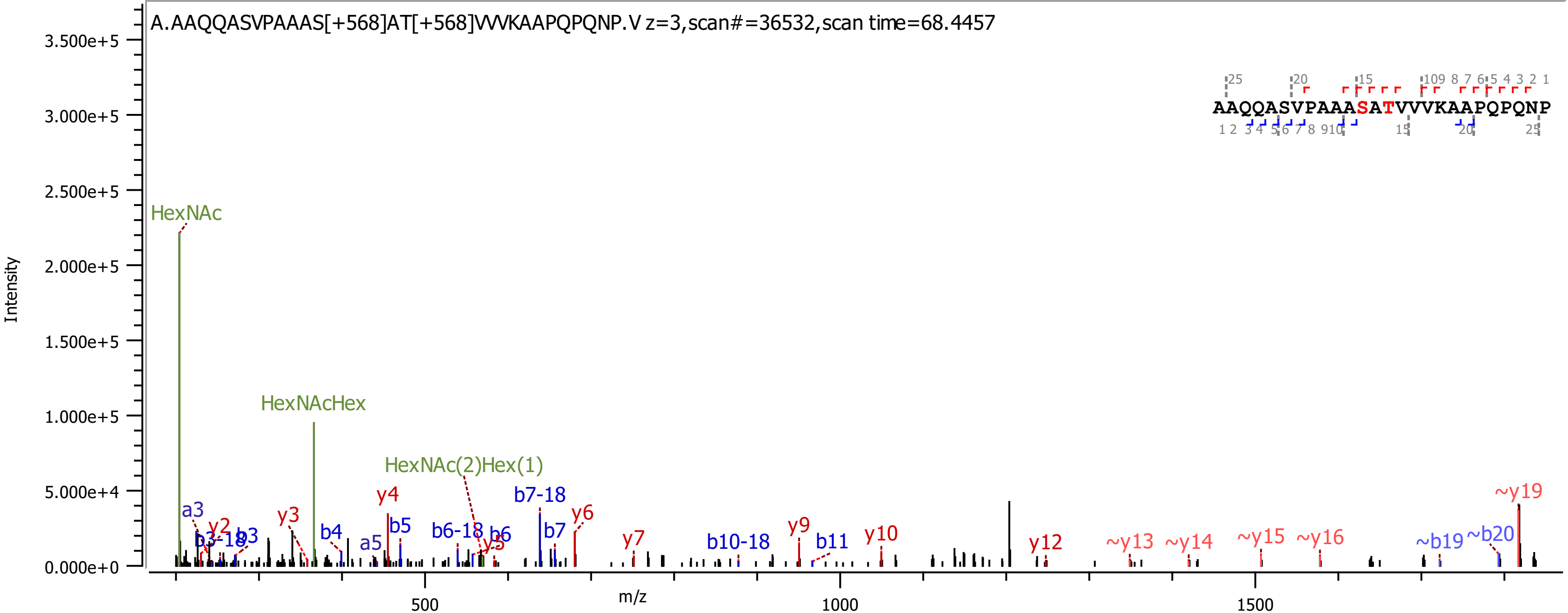
A.AAQQASVPAAAS[+568]AT[+568]VVVKAAPQPQNP.V z=3,scan#=36532,scan time=68.4457
AAQQASVPAAASATVVVKAAPQPQNP
Intensity
m/z
HexNAc
HexNAcHex
HexNAc(2)Hex(1)
a3
y2
b3-18
b3
y3
b4
a5
y4
b5
b6-18
b6
y5
b7-18
b7
y6
y7
b10-18
y9
b11
y10
y12
~y13
~y14
~y15
~y16
~b19
~b20
~y19
3.500e+5
3.000e+5
2.500e+5
2.000e+5
1.500e+5
1.000e+5
5.000e+4
0.000e+0
500
1000
1500

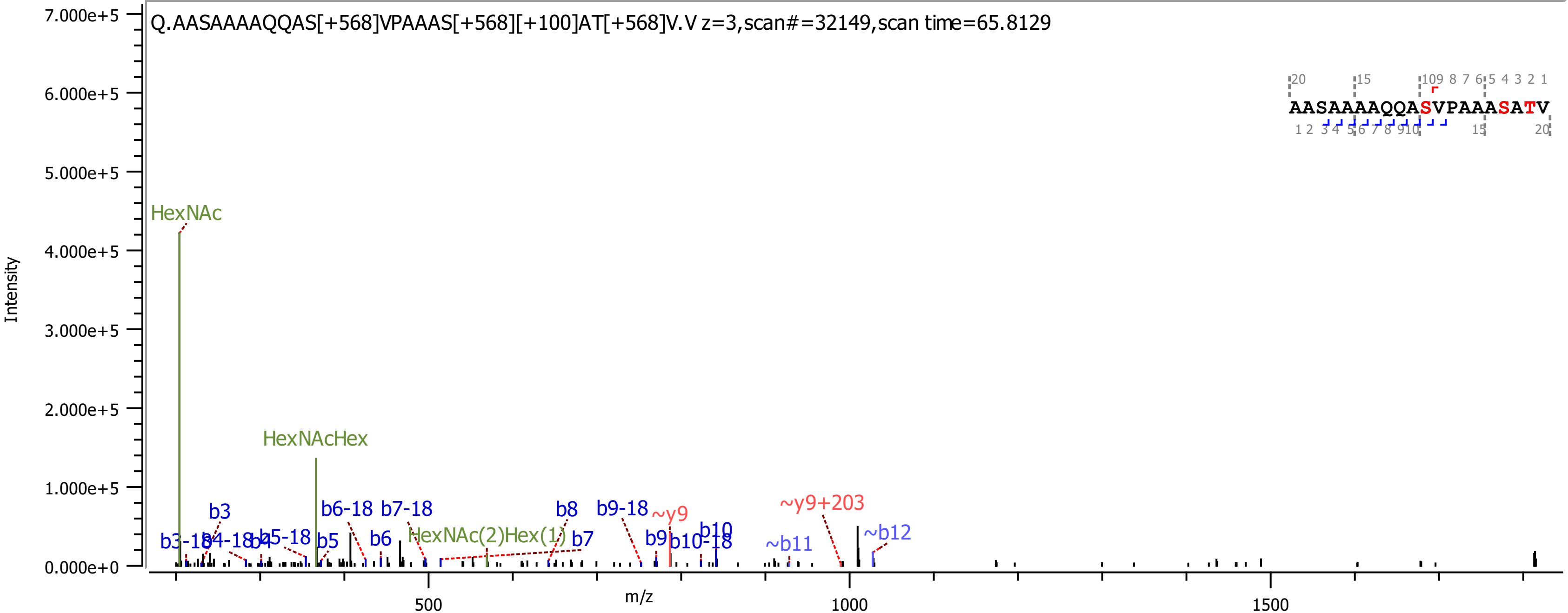

Q.AASAAAAQQAS[+568]VPAAAS[+568][+100]AT[+568]V.V z=3,scan#=32149,scan time=65.8129
AASAAAAQQASVPAAASATV
Intensity
m/z
HexNAc
HexNAcHex
HexNAc(2)Hex(1)
b3
b3-18
b4-18
b4
b5-18
b5
b6-18
b6
b7-18
b7
b8
b9-18
b9
b10
b10-18
~b11
~b12
~y9
~y9+203

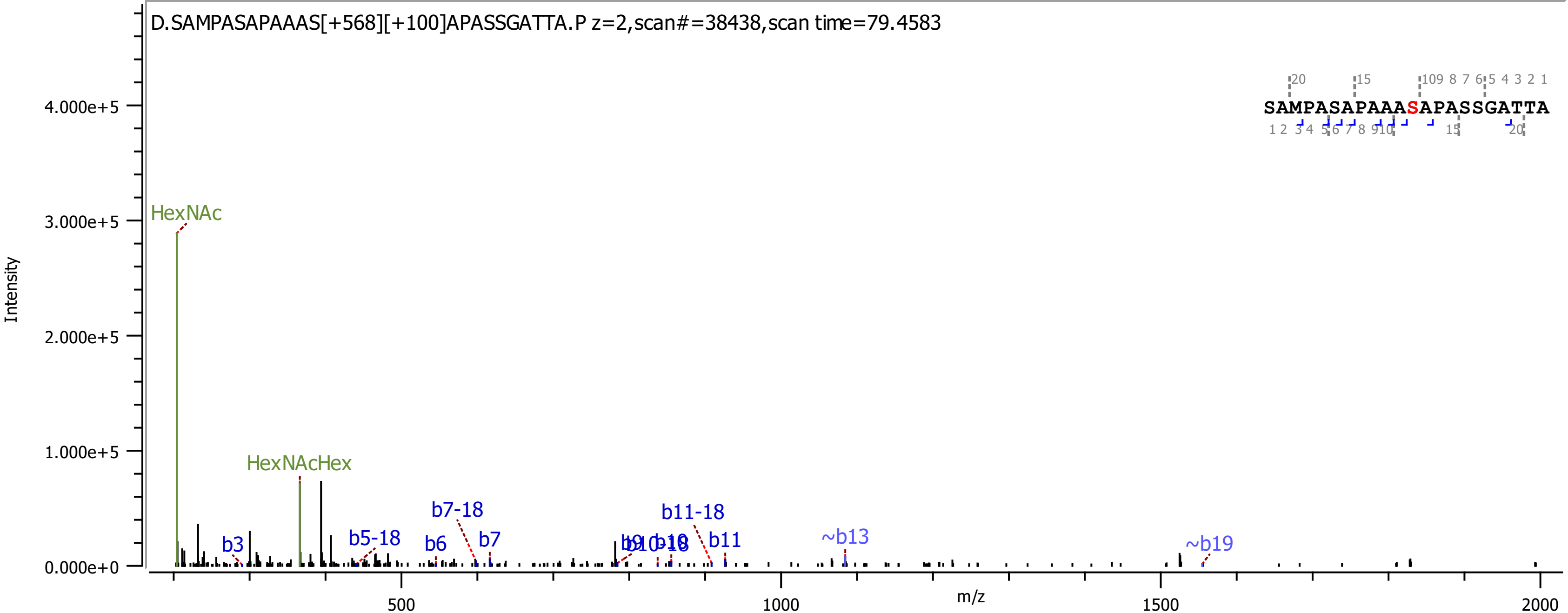
D.SAMPASAPAAAS[+568][+100]APASSGATTA.P z=2,scan#=38438,scan time=79.4583
SAMPASAPAAASAPASSGATTA
HexNAc
HexNAcHex
b3
b5-18
b6
b7-18
b7
b9
b10-18
b11-18
b11
~b13
~b19
Intensity
m/z
0.000e+0
1.000e+5
2.000e+5
3.000e+5
4.000e+5
500
1000
1500
2000

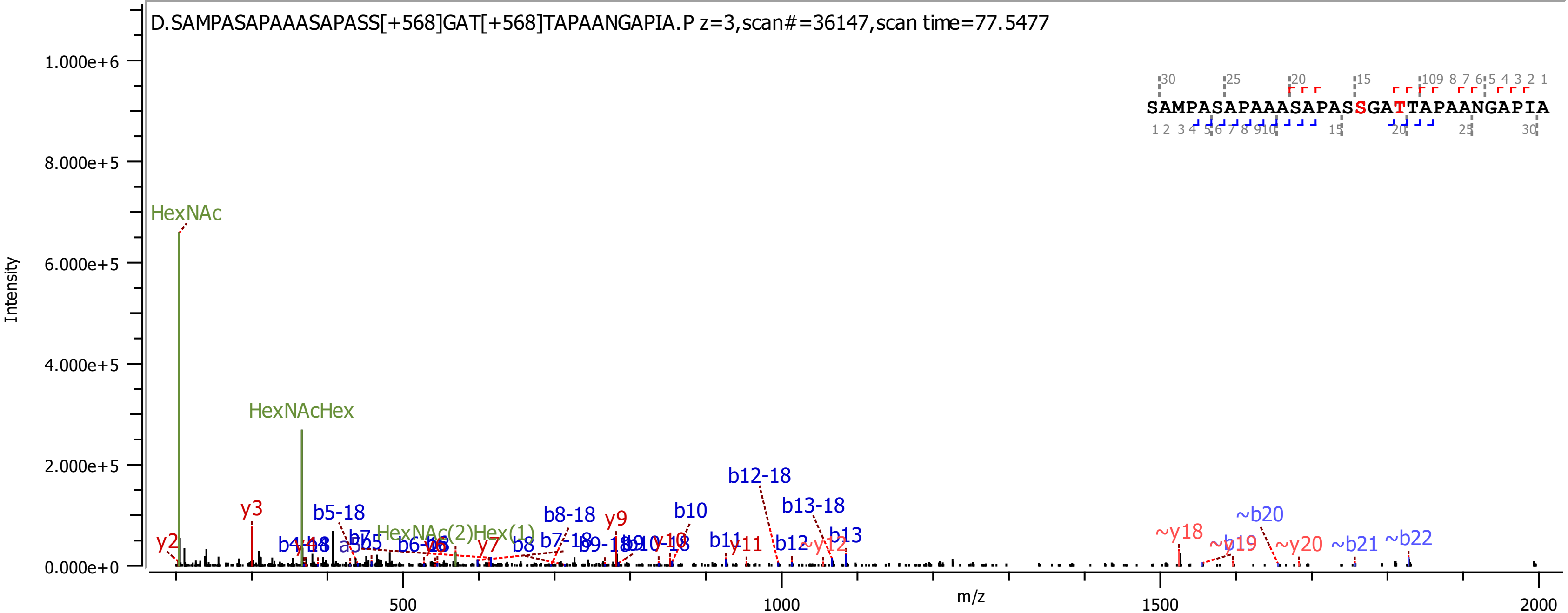

D.SAMPASAPAAASAPASS[+568]GAT[+568]TAPAANGAPIA.P z=3,scan#=36147,scan time=77.5477
SAMPASAPAAASAPASSGATTAPAANGAPIA
Intensity
m/z
1.000e+6
8.000e+5
6.000e+5
4.000e+5
2.000e+5
0.000e+0
500
1000
1500
2000
HexNAc
HexNAcHex
HexNAc(2)Hex(1)
y2
y3
y7
y9
y11
b5-18
b6
b8
b8-18
b7-18
b10
b11
b12
b12-18
b13-18
b13
~y12
~y18
~y19
~y20
~b20
~b21
~b22

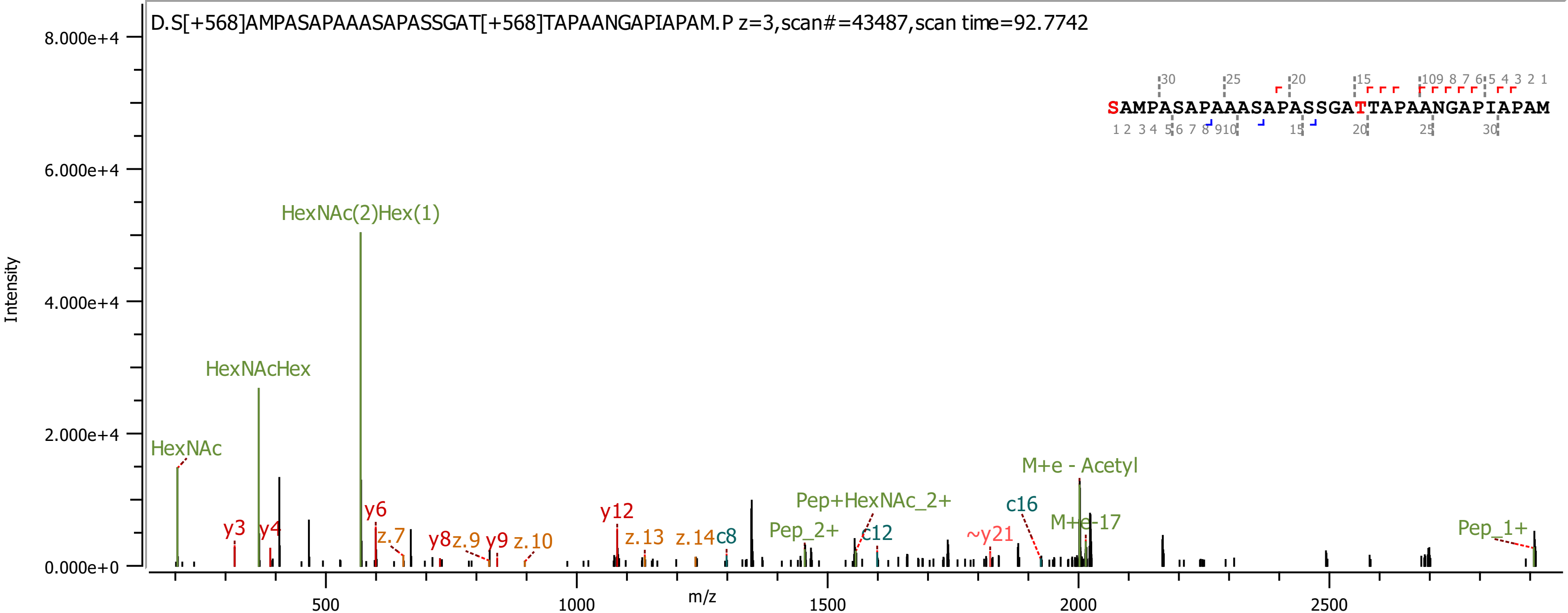
D.S[+568]AMPASAPAAASAPASSGAT[+568]TAPAANGAPIAPAM.P z=3,scan#=43487,scan time=92.7742
SAMPASAPAAASAPASSGATTAPAANGAPIAPAM
Intensity
m/z
8.000e+4
6.000e+4
4.000e+4
2.000e+4
0.000e+0
500
1000
1500
2000
2500
HexNAc(2)Hex(1)
HexNAcHex
HexNAc
y3
y4
y6
z.7
y8
z.9
y9
z.10
y12
z.13
z.14
c8
Pep_2+
Pep+HexNAc_2+
c12
~y21
c16
M+e - Acetyl
M+e-17
Pep_1+

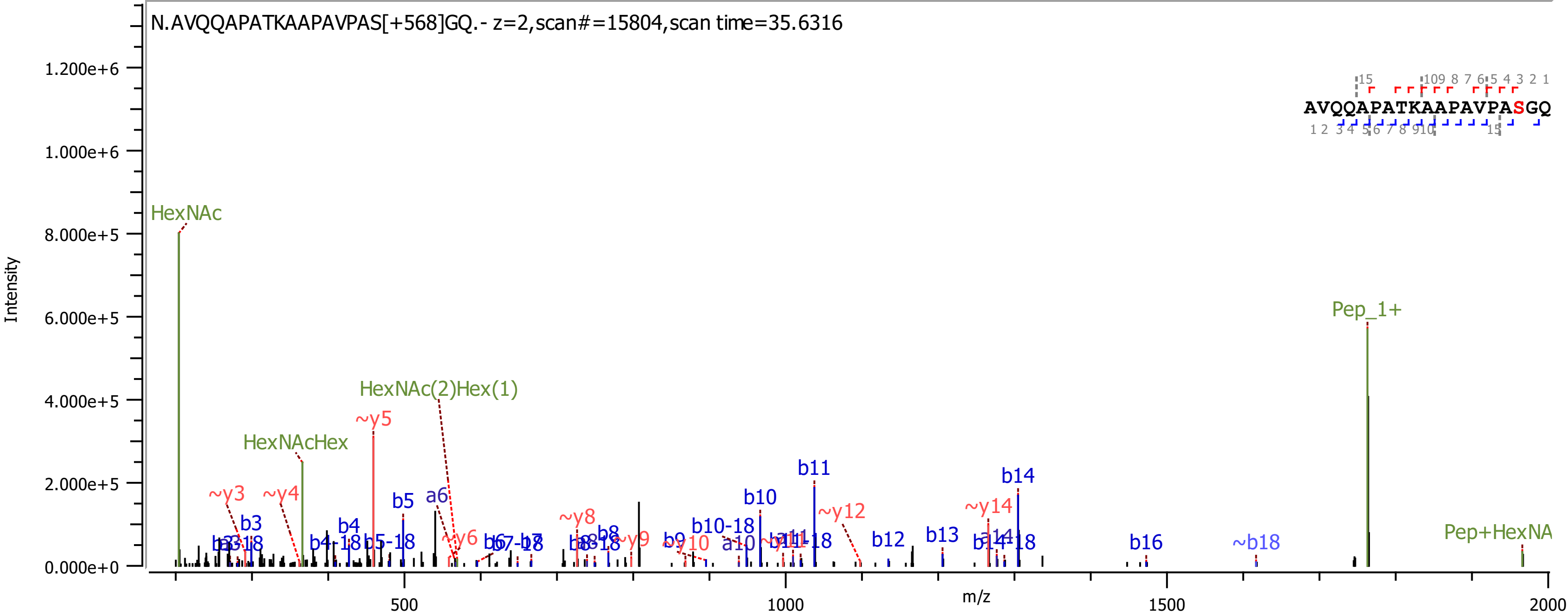
N.AVQQAPATKAAPAVPAS[+568]GQ.- z=2,scan#=15804,scan time=35.6316
AVQQAPATKAAPAVPASGQ
Intensity
m/z
0.000e+0
2.000e+5
4.000e+5
6.000e+5
8.000e+5
1.000e+6
1.200e+6
500
1000
1500
2000
HexNAc
HexNAcHex
HexNAc(2)Hex(1)
Pep_1+
Pep+HexNA
~y3
~y4
~y5
~y6
~y8
~y9
~y10
~y12
~y14
b3
b4
b4-18
b5
b5-18
a6
b10
b10-18
b11
b12
b13
b14
b14-18
b16
~b18
a10

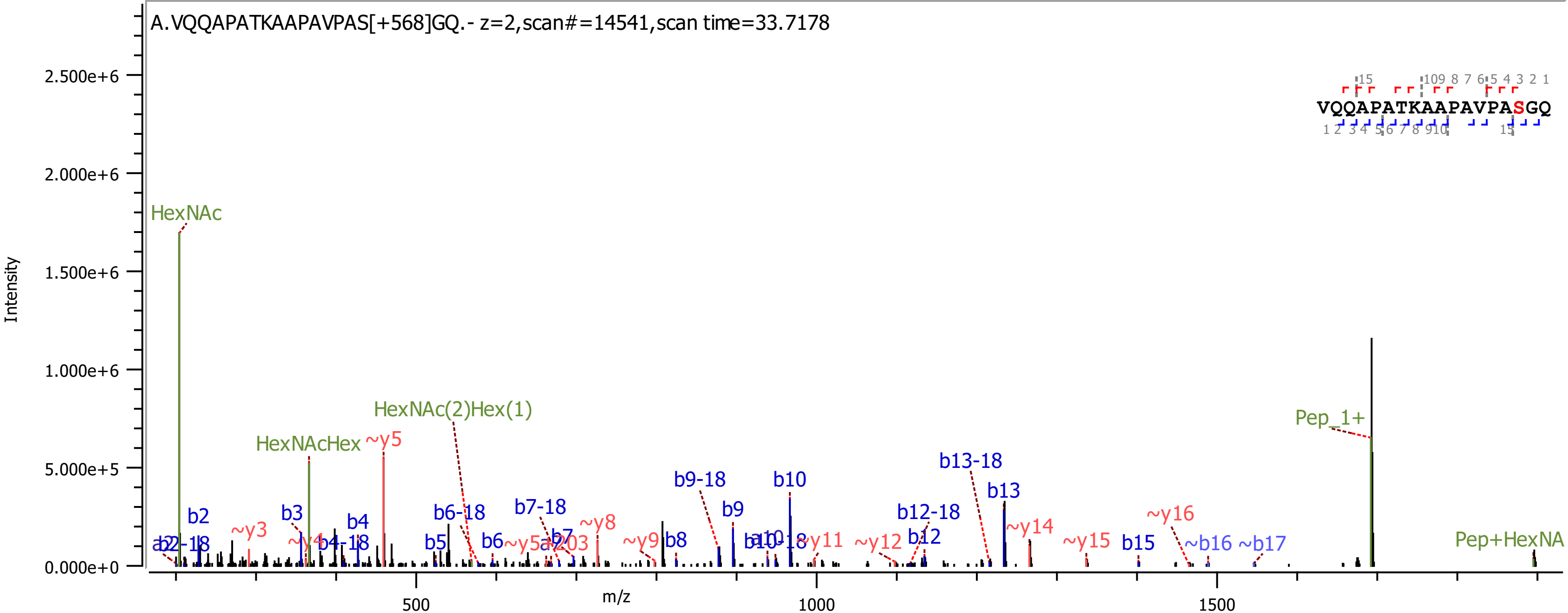
A.VQQAPATKAAPAVPAS[+568]GQ.- z=2,scan#=14541,scan time=33.7178
VQQAPATKAAPAVPASGQ
Intensity
m/z
HexNAc
HexNAcHex
HexNAc(2)Hex(1)
Pep_1+
Pep+HexNA
b2
b3
b4
b5
b6
b7
b8
b9
b10
b12
b13
b15
b6-18
b7-18
b9-18
b12-18
b13-18
~y3
~y5
~y8
~y9
~y11
~y12
~y14
~y15
~y16
~b16
~b17

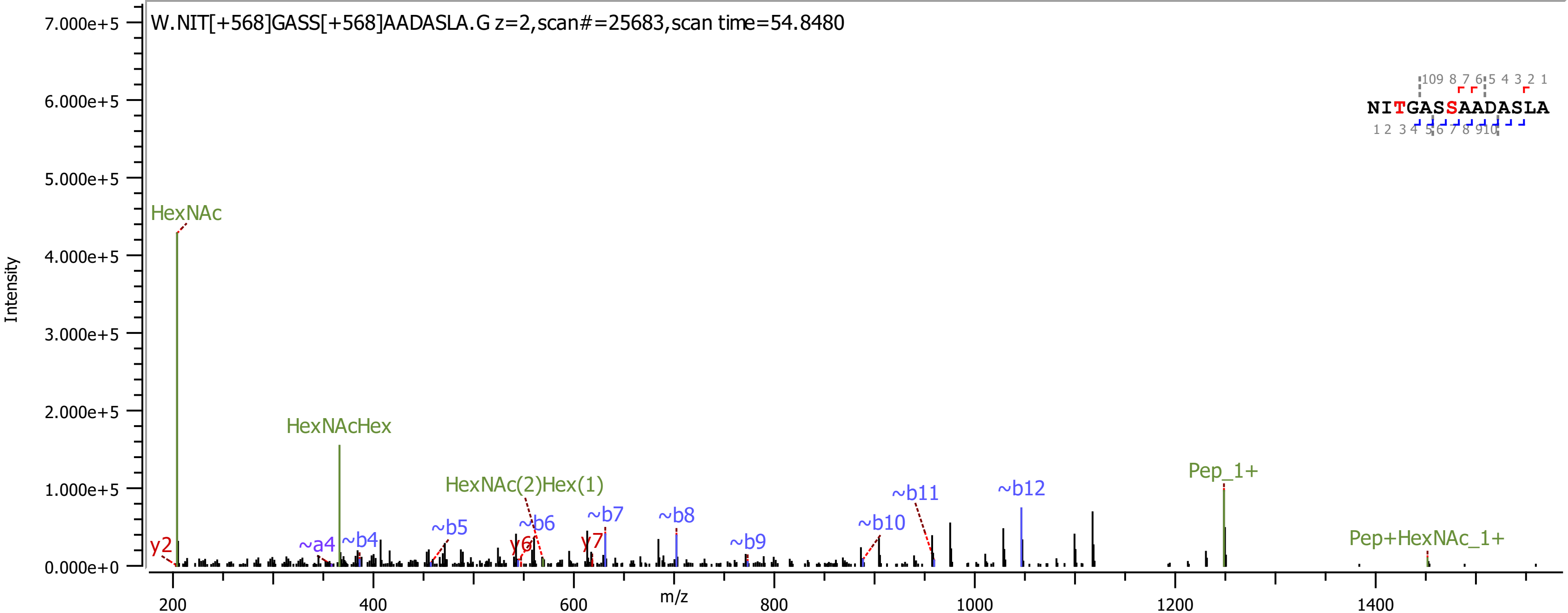

W.NIT[+568]GASS[+568]AADASLA.G z=2,scan#=25683,scan time=54.8480
NITGASSAADASLA
Intensity
m/z
HexNAc
HexNAcHex
HexNAc(2)Hex(1)
Pep_1+
Pep+HexNAc_1+
y2
~a4
~b4
~b5
~b6
y6
~b7
y7
~b8
~b9
~b10
~b11
~b12

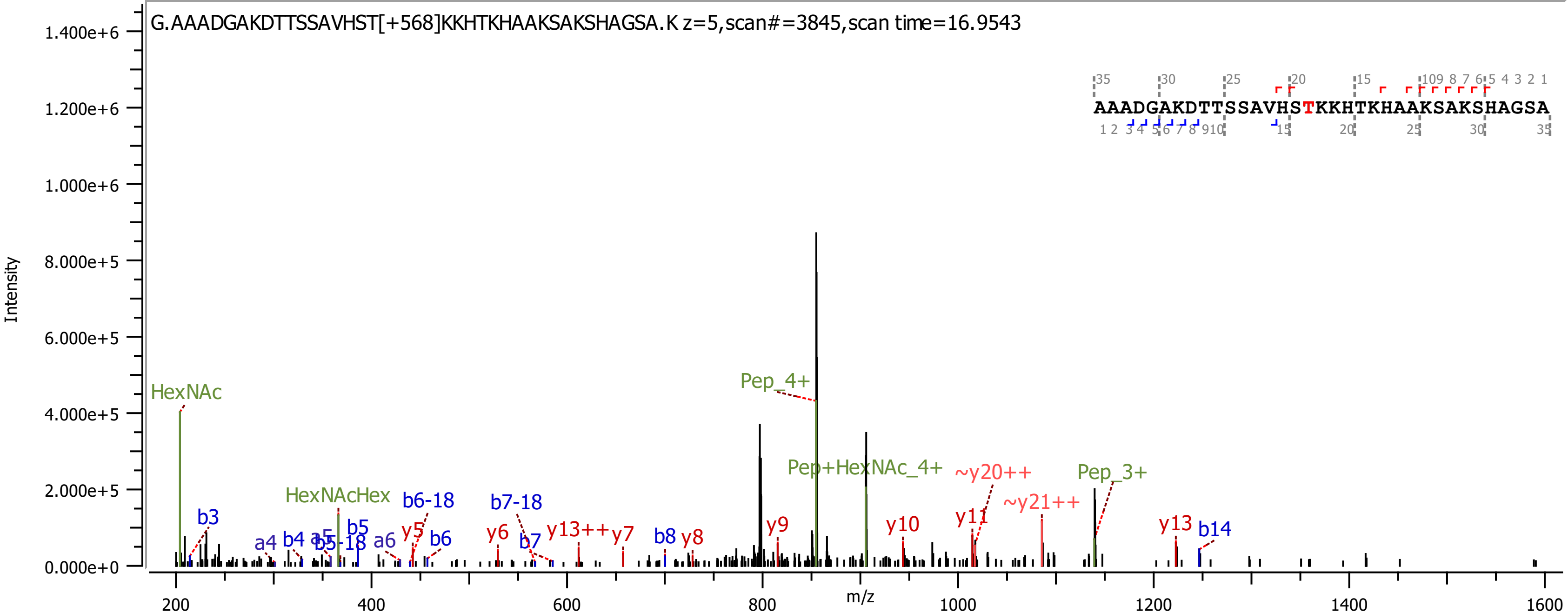

G.AAADGAKDTTSSAVHST[+568]KKHTKHAAKSAKSHAGSA.K z=5,scan#=3845,scan time=16.9543
AAADGAKDTTSSAVHSTKKHTKHAAKSAKSHAGSA
Intensity
m/z
HexNAc
HexNAcHex
Pep_4+
Pep+HexNAc_4+
Pep_3+
~y20++
~y21++
y13++
b3
a4
b4
a5
b5-18
b5
a6
b6-18
y5
b6
y6
b7-18
b7
y7
b8
y8
y9
y10
y11
y13
b14

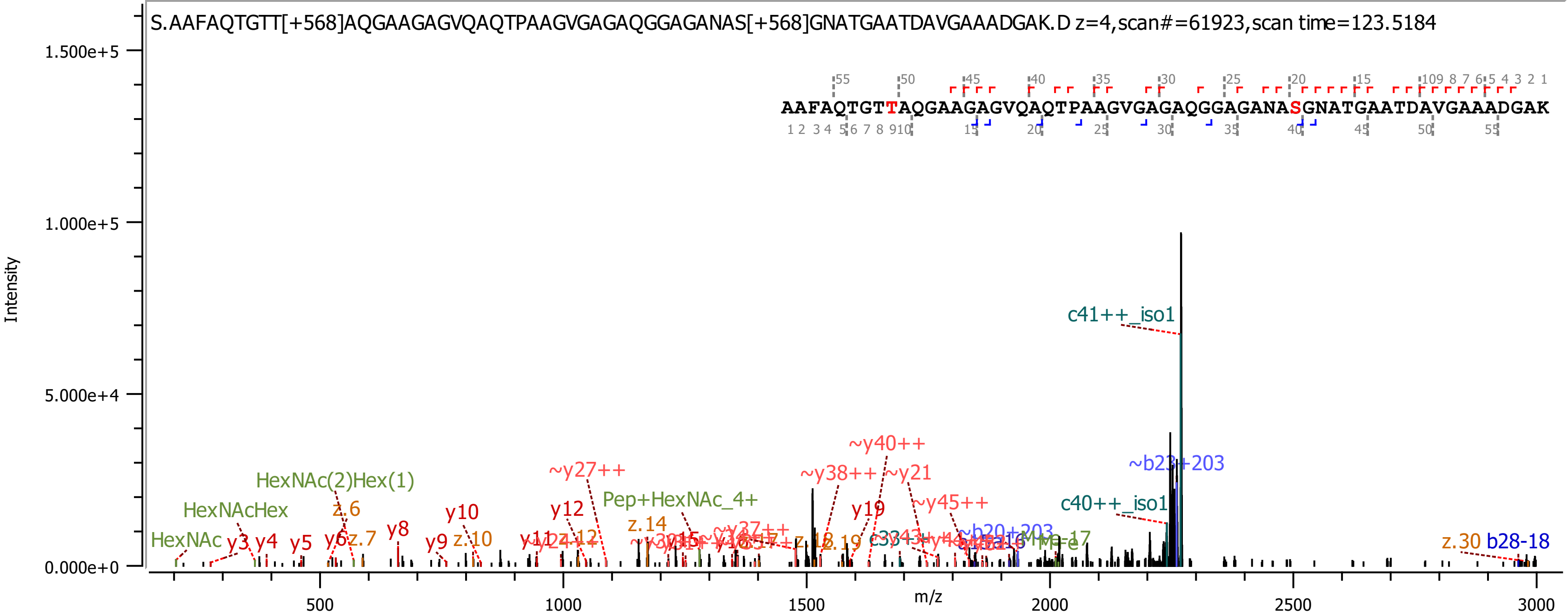
S.AAFAQTGTT[+568]AQGAAGAGVQAQTPAAGVGAGAQGGAGANAS[+568]GNATGAATDAVGAAADGAK.D z=4,scan#=61923,scan time=123.5184
AAFAQTGTTAQGAAGAGVQAQTPAAGVGAGAQGGAGANASGNATGAATDAVGAAADGAK
Intensity
m/z
1.500e+5
1.000e+5
5.000e+4
0.000e+0
500
1000
1500
2000
2500
3000
c41++_iso1
~y40++
~b23+203
~y27++
~y38++
~y21
HexNAc(2)Hex(1)
Pep+HexNAc_4+
~y45++
c40++_iso1
HexNAcHex
z.6
y10
y12
y19
y8
z.14
HexNAc
y3
y4
y5
y6
z.7
y9
z.10
z.12
z.19
z.30
b28-18

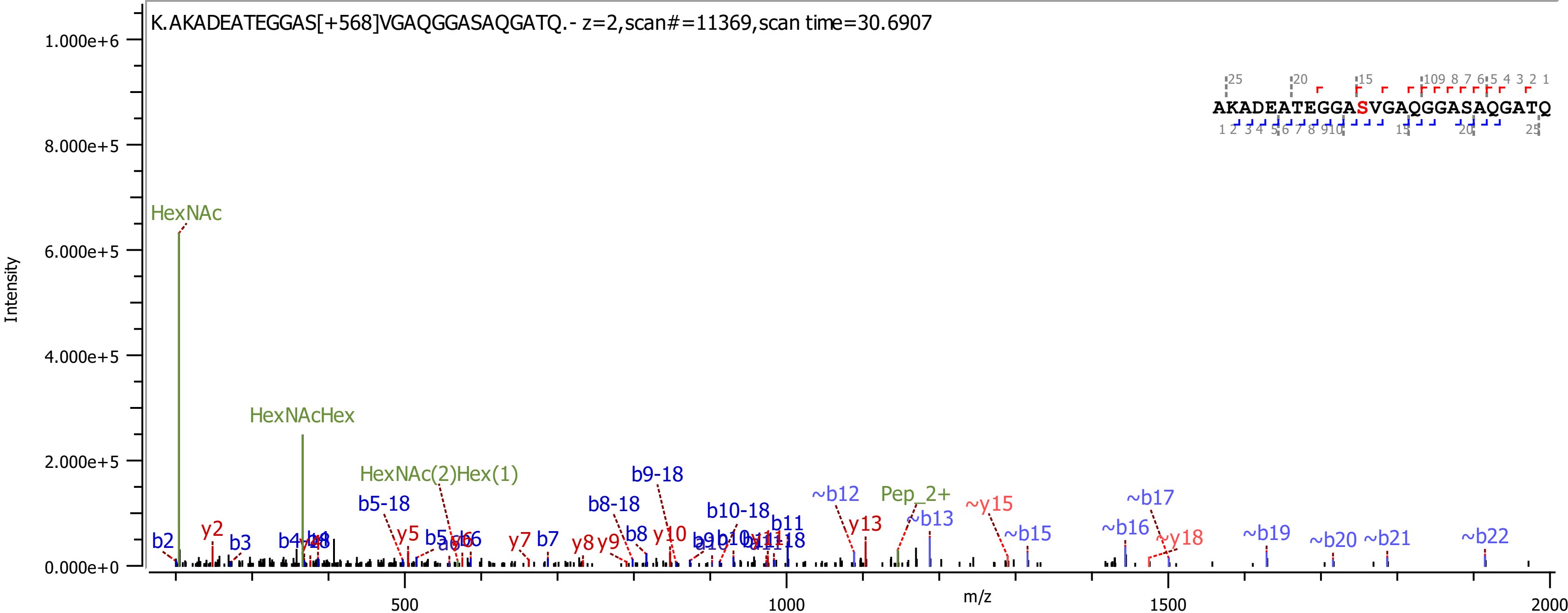
K.AKADEATEGGAS[+568]VGAQGGASAQGATQ.- z=2,scan#=11369,scan time=30.6907
Intensity
m/z
HexNAc
HexNAcHex
HexNAc(2)Hex(1)
Pep_2+
AKADEATEGGASVGAQGGASAQGATQ

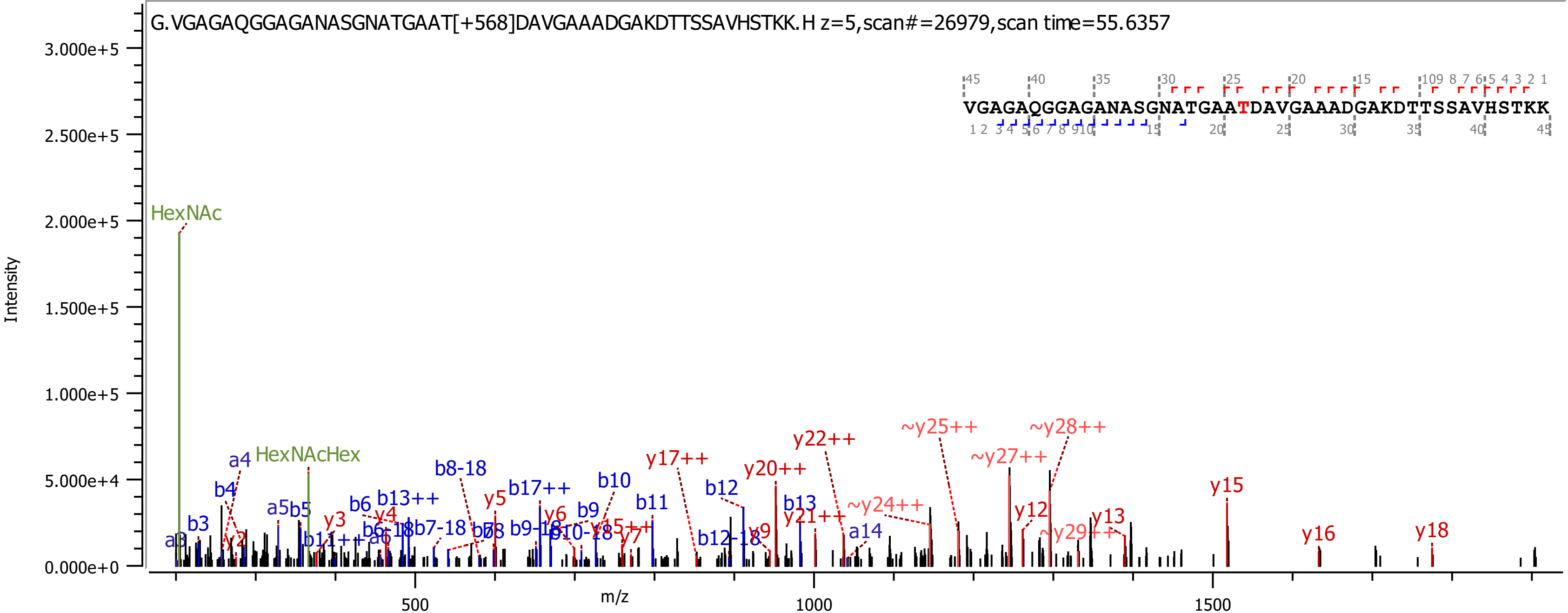
G.VGAGAQGGAGANASGNATGAAT[+568]DAVGAAADGAKDTTSSAVHSTKK.H z=5,scan#=26979,scan time=55.6357
VGAGAQGGAGANASGNATGAATDAVGAAADGAKDTTSSAVHSTKK
Intensity
m/z
HexNAc
HexNAcHex
y15
y16
y18
y22++
y20++
y17++
~y25++
~y28++
~y27++
~y24++
~y29++
y21++
y12
y13
b8-18
b17++
b10
b11
b12
b13
a14
b12-18
b13++
b6
b4
a4
b3
a3
a5
b5
y5
y6
y3
y4
y2
y7
y9
b9
b9-18
b10-18
b6-18
b7-18
a6
b11++
y15++

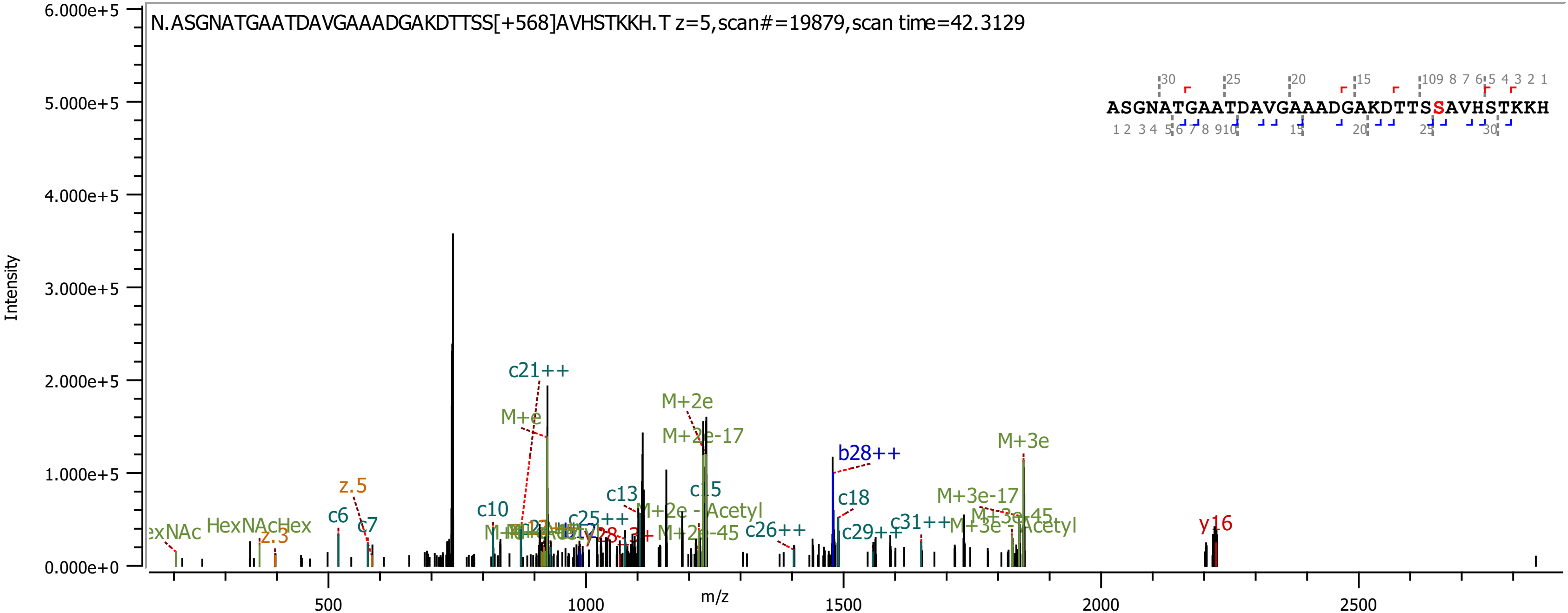

N.ASGNATGAATDAVGAAADGAKDTTSS[+568]AVHSTKKH.T z=5,scan#=19879,scan time=42.3129
ASGNATGAATDAVGAAADGAKDTTSSAVHSTKKH
Intensity
m/z
6.000e+5
5.000e+5
4.000e+5
3.000e+5
2.000e+5
1.000e+5
0.000e+0
500
1000
1500
2000
2500
c21++
M+e
M+2e
M+2e-17
b28++
M+3e
z.5
c6
c7
c10
c13
c15
c18
M+2e - Acetyl
M+2e-45
c25++
c26++
c29++
c31++
M+3e-17
M+3e-45
M+3e - Acetyl
y16
HexNAcHex
z.3

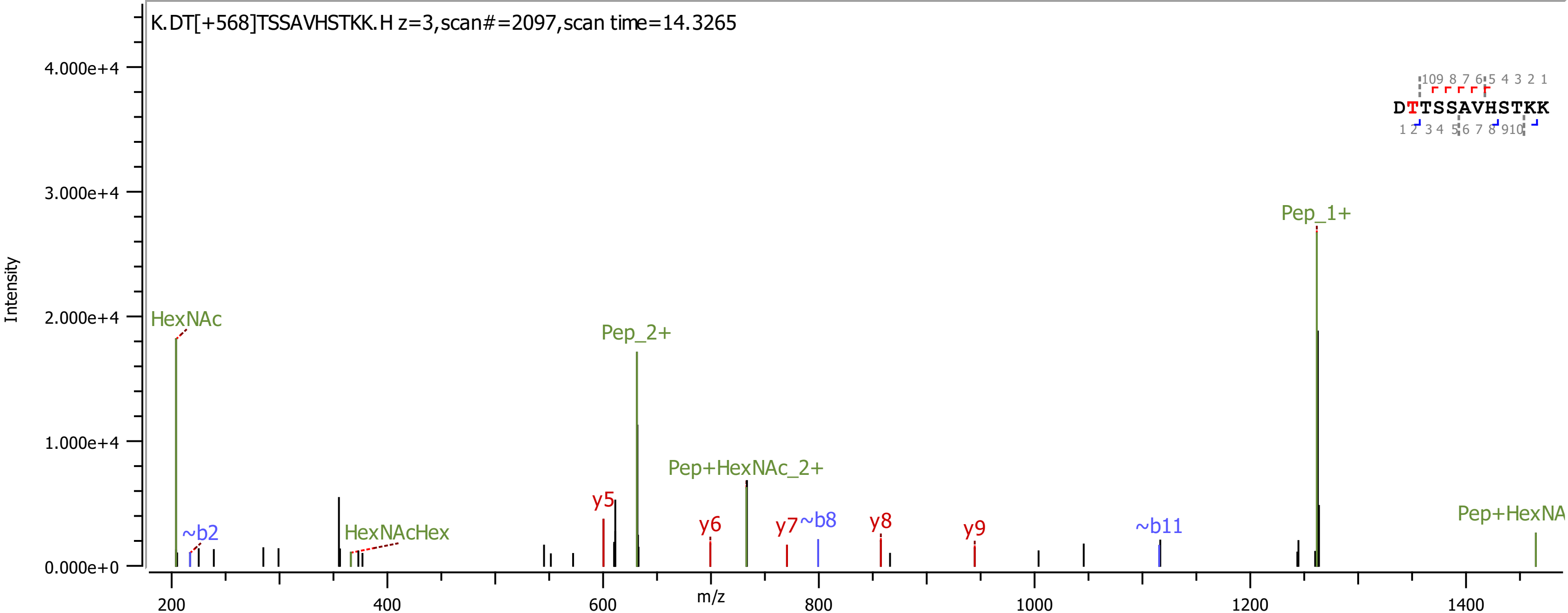
K.DT[+568]TSSAVHSTKK.H z=3,scan#=2097,scan time=14.3265
Intensity
4.000e+4
3.000e+4
2.000e+4
1.000e+4
0.000e+0
HexNAc
~b2
HexNAcHex
Pep_2+
y5
Pep+HexNAc_2+
y6
y7
~b8
y8
y9
~b11
Pep_1+
Pep+HexNA
DTTSSAVHSTKK
200
400
600
800
1000
1200
1400
m/z

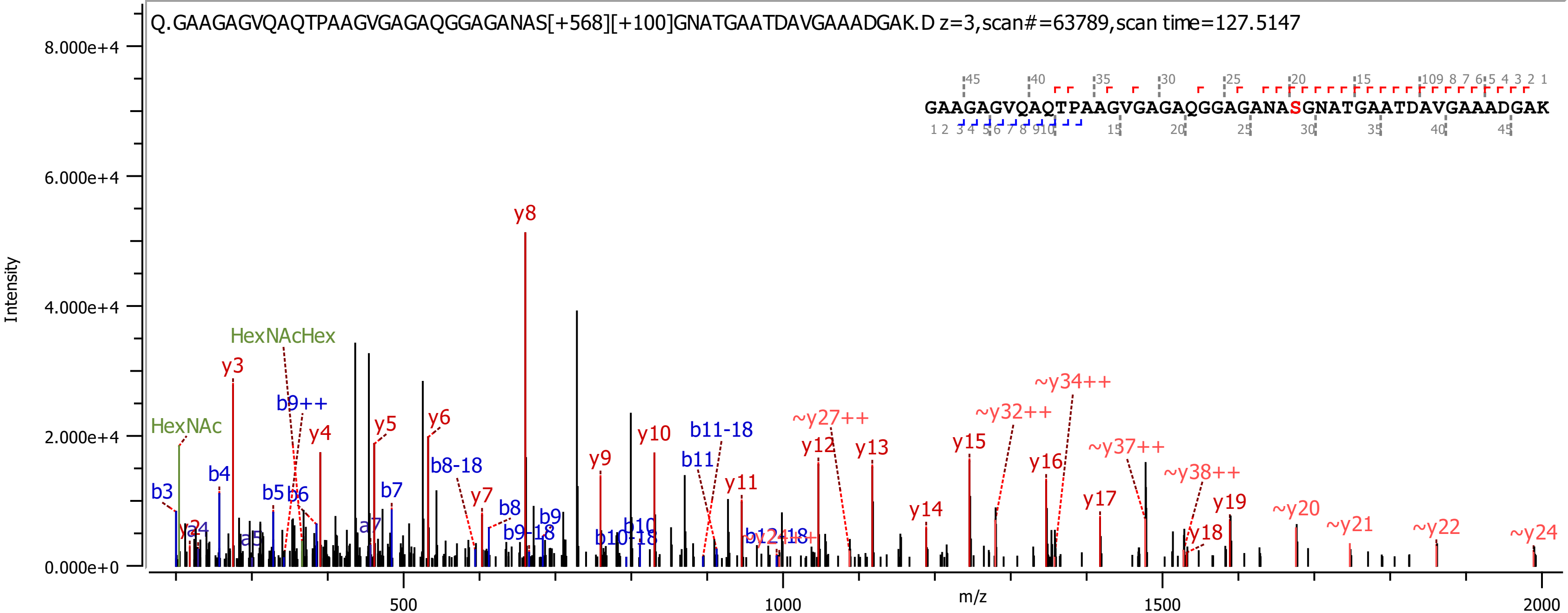

Q.GAAGAGVQAQTPAAGVGAGAQGGAGANAS[+568][+100]GNATGAATDAVGAAADGAK.D z=3,scan#=63789,scan time=127.5147
GAAGAGVQAQTPAAGVGAGAQGGAGANASGNATGAATDAVGAAADGAK
8.000e+4
6.000e+4
4.000e+4
2.000e+4
0.000e+0
Intensity
m/z
500
1000
1500
2000
HexNAcHex
HexNAc
y3
b9++
y4
y5
y6
y8
b8-18
y7
b8
b9
b9-18
b3
b4
b5
b6
b7
a7
a5
y9
y10
b11-18
b11
y11
b10
b10-18
~y27++
y12
y13
~y32++
~y34++
y14
y15
y16
~y37++
y17
~y38++
y19
y18
~y20
~y21
~y22
~y24

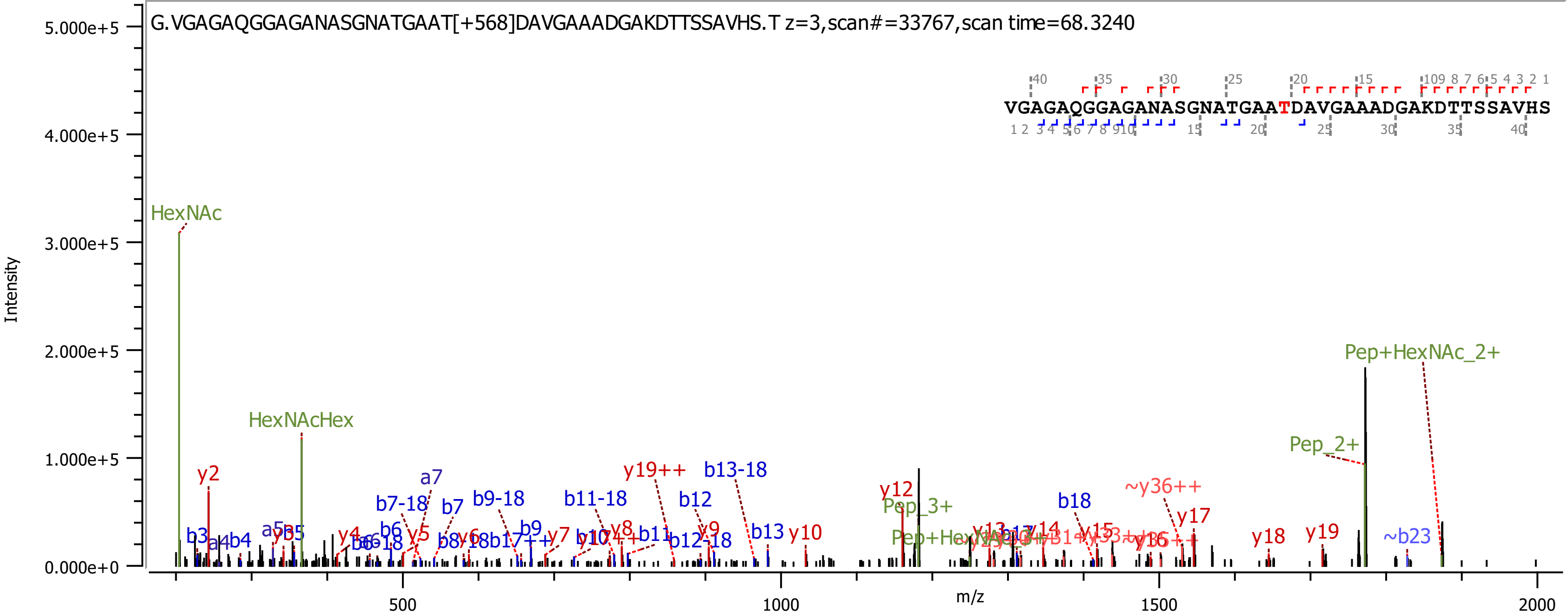
G.VGAGAQGGAGANASGNATGAAT[+568]DAVGAAADGAKDTTSSAVHS.T z=3,scan#=33767,scan time=68.3240
VGAGAQGGAGANASGNATGAATDAVGAAADGAKDTTSSAVHS
Intensity
5.000e+5
4.000e+5
3.000e+5
2.000e+5
1.000e+5
0.000e+0
m/z
500
1000
1500
2000
HexNAc
HexNAcHex
Pep+HexNAc_2+
Pep_2+
Pep_3+
y2
y10
y12
y17
y18
y19
b3
b7
b7-18
b9-18
b11-18
b12
b13
b13-18
b18
a7
y19++
~y36++
~b23

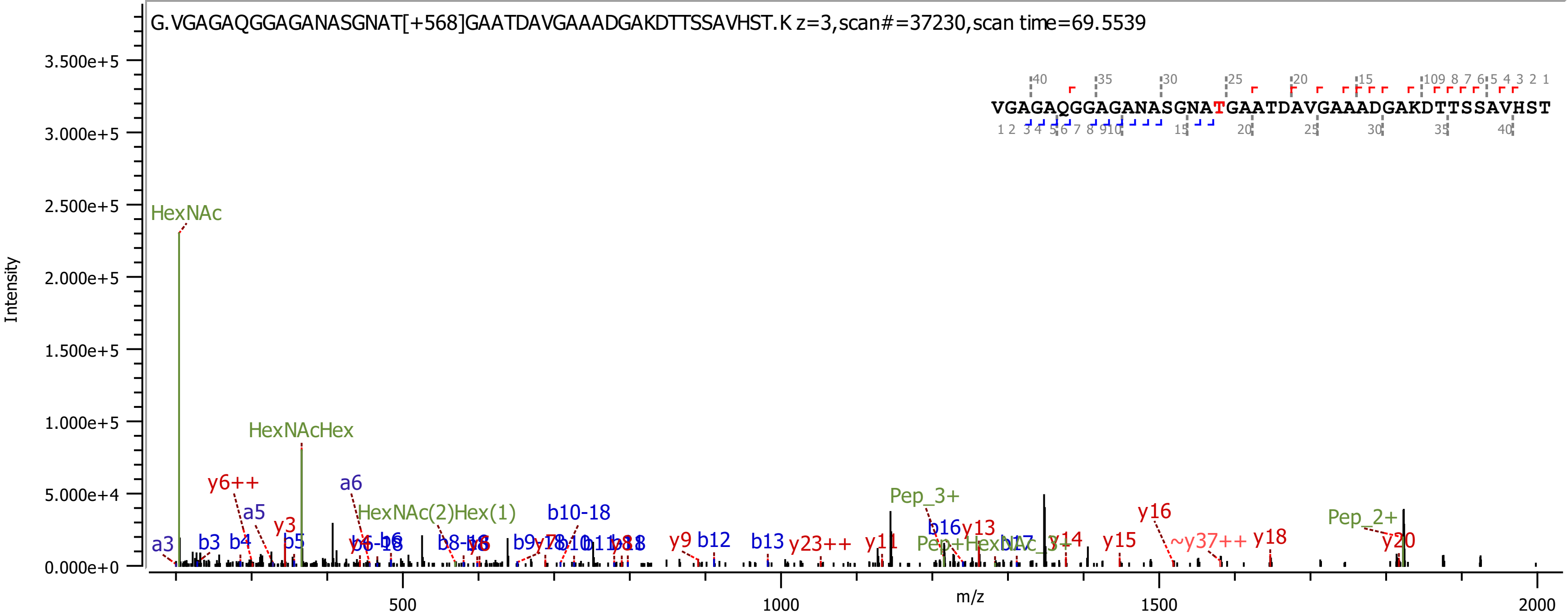
G.VGAGAQGGAGANASGNAT[+568]GAATDAVGAAADGAKDTTSSAVHST.K z=3,scan#=37230,scan time=69.5539
VGAGAQGGAGANASGNATGAATDAVGAAADGAKDTTSSAVHST
Intensity
m/z
HexNAc
HexNAcHex
HexNAc(2)Hex(1)
y6++
a3
b3
b4
a5
y3
b5
a6
b10-18
Pep_3+
b16
y13
y16
Pep_2+
b12
b13
y9
y23++
y11
y14
y15
~y37++
y18
y20
0.000e+0
5.000e+4
1.000e+5
1.500e+5
2.000e+5
2.500e+5
3.000e+5
3.500e+5
500
1000
1500
2000

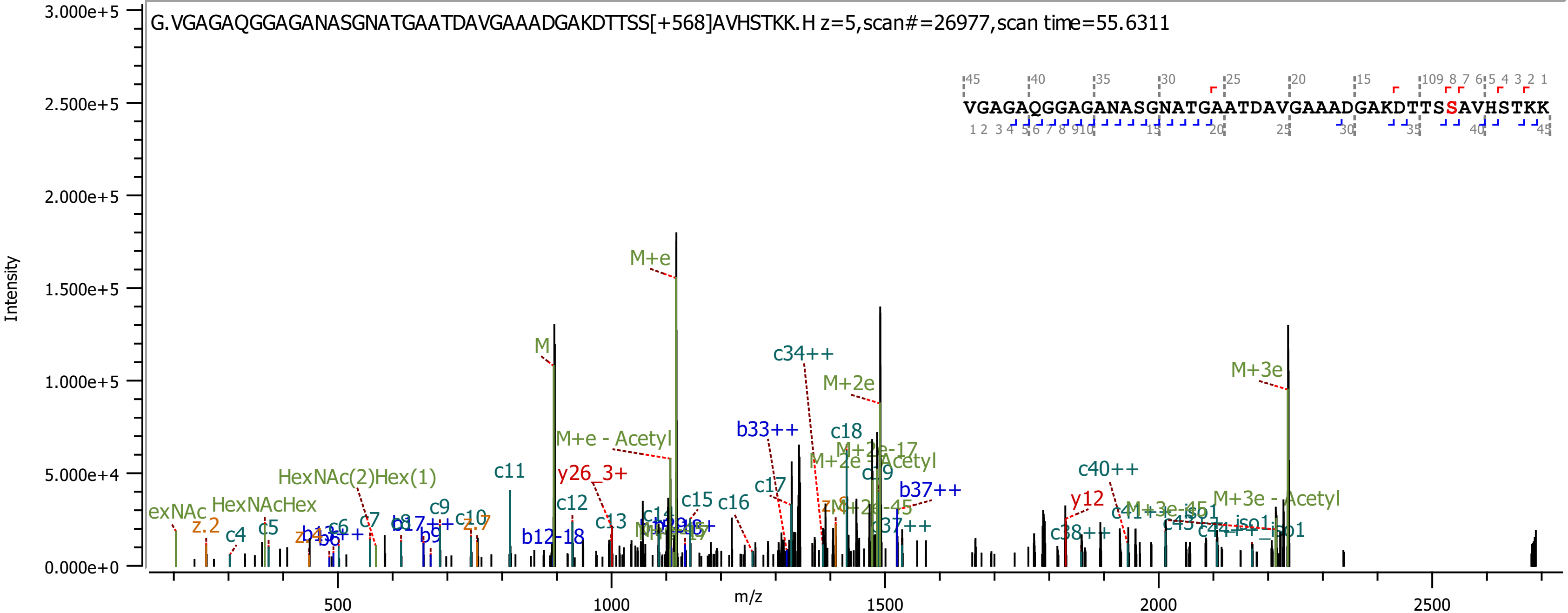
G.VGAGAQGGAGANASGNATGAATDAVGAAADGAKDTTSS[+568]AVHSTKK.H z=5,scan#=26977,scan time=55.6311
VGAGAQGGAGANASGNATGAATDAVGAAADGAKDTTSSAVHSTKK
Intensity
m/z
3.000e+5
2.500e+5
2.000e+5
1.500e+5
1.000e+5
5.000e+4
0.000e+0
500
1000
1500
2000
2500
M+e
M
M+3e
M+2e
c34++
M+e - Acetyl
b33++
c18
c11
y26_3+
HexNAc(2)Hex(1)
c40++
b37++
y12
M+3e - Acetyl
HexNAcHex
z.2
c4
c5
c7
c9
c12
c13
c15
c16
c17
b12-18
b9
c38++

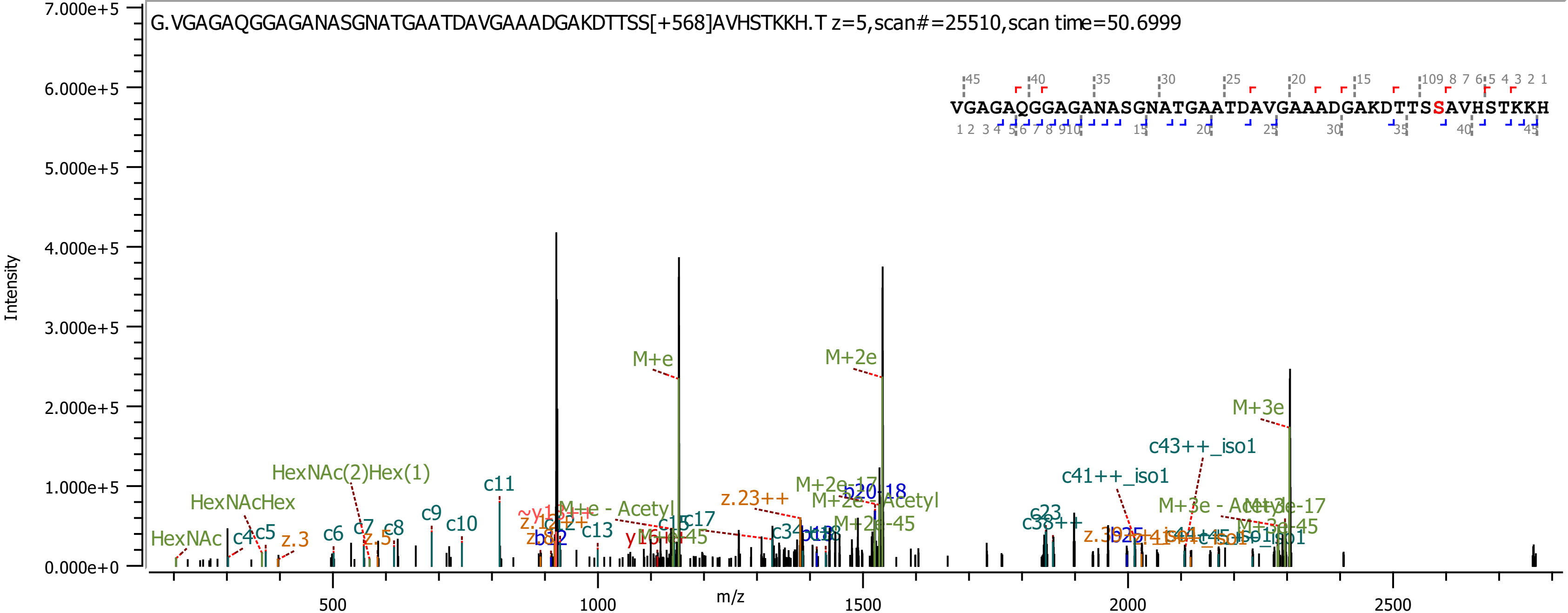
G.VGAGAQGGAGANASGNATGAATDAVGAAADGAKDTTSS[+568]AVHSTKKH.T z=5,scan#=25510,scan time=50.6999
VGAGAQGGAGANASGNATGAATDAVGAAADGAKDTTSSAVHSTKKH
Intensity
m/z
7.000e+5
6.000e+5
5.000e+5
4.000e+5
3.000e+5
2.000e+5
1.000e+5
0.000e+0
500
1000
1500
2000
2500
M+e
M+2e
M+3e
c43++_iso1
c41++_iso1
HexNAc(2)Hex(1)
HexNAcHex
HexNAc
c11
c9
c10
c4
c5
z.3
c6
c7
c8
z.5
c13
c17
z.23++
M+2e-17
b20-18
M+2e - Acetyl
M+2e-45
c23
c38++
M+3e - Acetyl
M+3e-17
M+3e-45

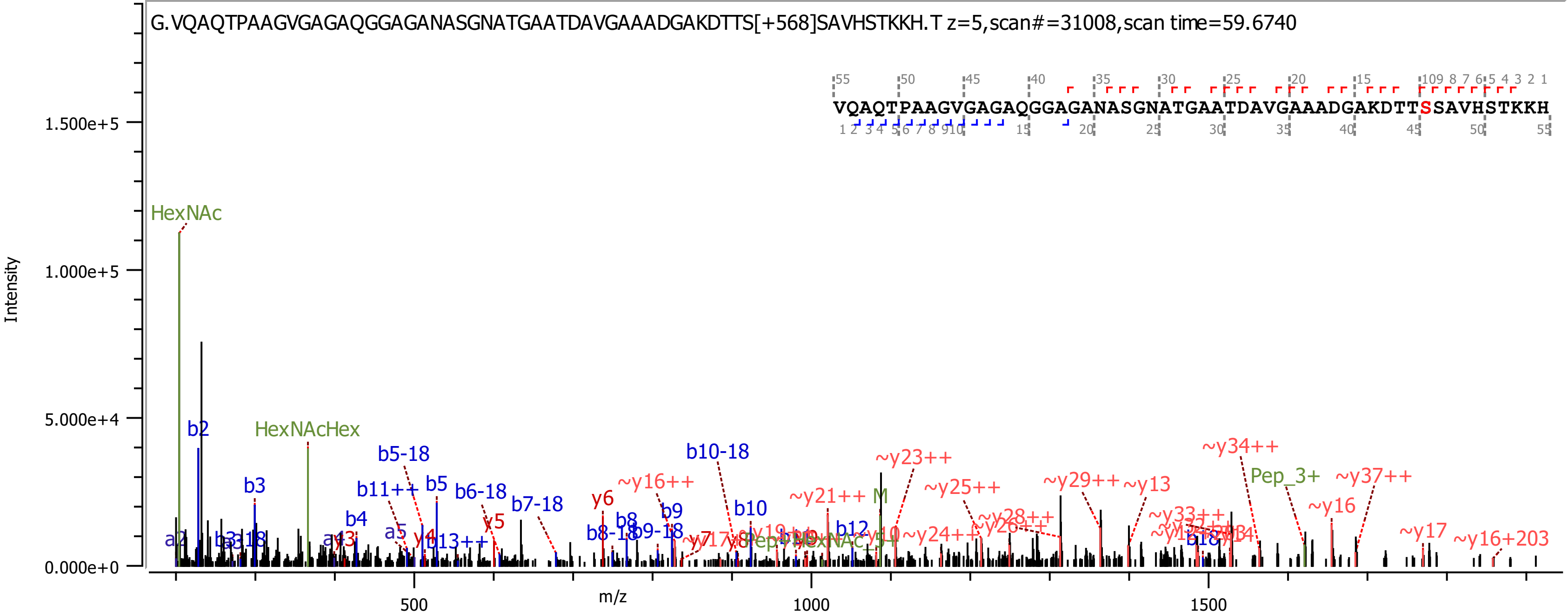
G.VQAQTPAAGVGAGAQGGAGANASGNATGAATDAVGAAADGAKDTTS[+568]SAVHSTKKH.T z=5,scan#=31008,scan time=59.6740
VQAQTPAAGVGAGAQGGAGANASGNATGAATDAVGAAADGAKDTTSSAVHSTKKH
Intensity
1.500e+5
1.000e+5
5.000e+4
0.000e+0
m/z
500
1000
1500
HexNAc
HexNAcHex
b2
b3
b4
b5
b5-18
b6-18
b7-18
b8
b9
b10
b10-18
b11++
b12
b13++
y4
y5
y6
~y16++
~y21++
~y23++
~y25++
~y29++
~y13
~y34++
Pep_3+
~y37++
~y16
~y17
~y16+203
~y24++
~y33++
M

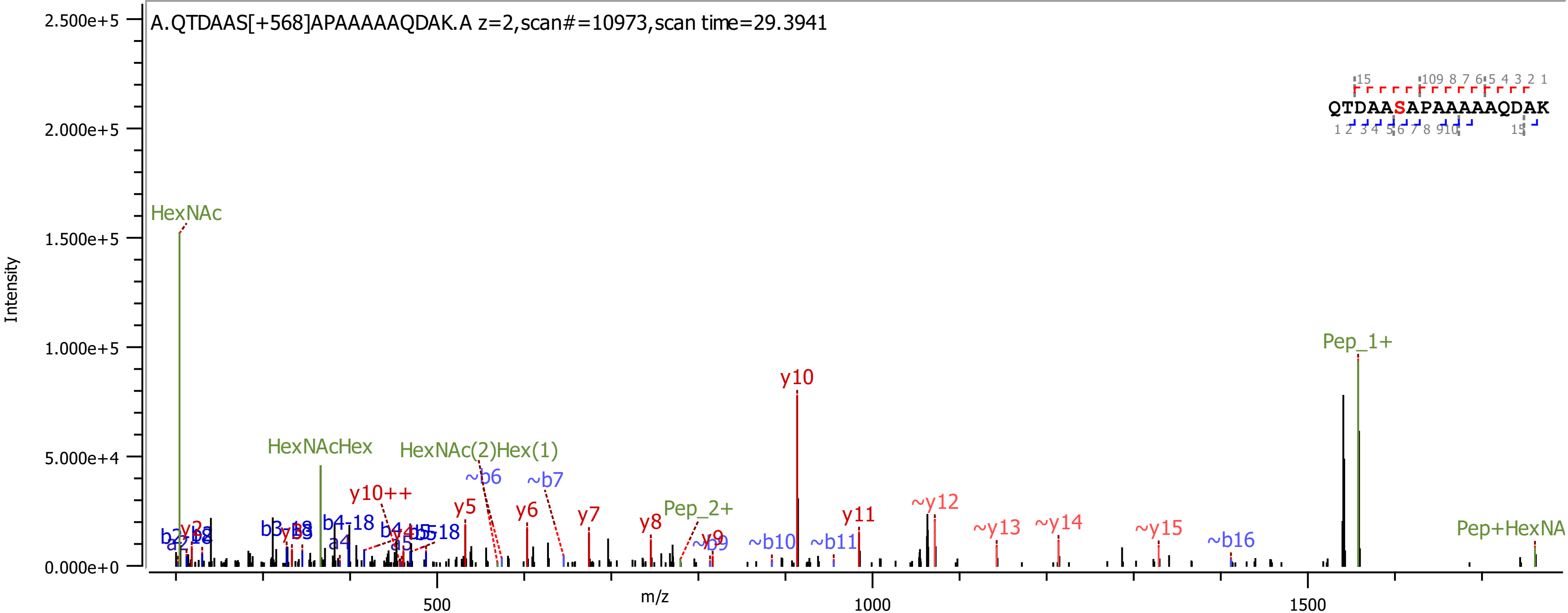
A.QTDAAS[+568]APAAAAAQDAK.A z=2,scan#=10973,scan time=29.3941
QTDAASAPAAAAAQDAK
Intensity
m/z
HexNAc
HexNAcHex
HexNAc(2)Hex(1)
Pep_1+
Pep_2+
Pep+HexNA
y10
y10++
y5
y6
y7
y8
y9
y11
~y12
~y13
~y14
~y15
~b6
~b7
~b9
~b10
~b11
~b16
b3-18
b4-18
b5-18
a4
a5
2.500e+5
2.000e+5
1.500e+5
1.000e+5
5.000e+4
0.000e+0
500
1000
1500
