## Supplementary Data 3 for "Burkholderia PglL enzymes are serine preferring oligosaccharidetransferases which target conserved proteins across the Burkholderia genus"

### *Burkholderia cenocepacia* H111 Best Localised Unique glycopeptides

<table>
<tr><th>J2315 numbers</th><th>Protein Name</th><th>Peptide < ProteinMetrics Confidential ></th><th>Glycans NHFAGNa</th><th>Localised site (Yes / No / Partial) ?</th><th>Observed m/z</th><th>z</th><th>Observed (M+H)</th><th>Calc. mass (M+H)</th><th>Mass error (ppm)</th><th>Score</th><th>Delta</th><th>Delta Mod</th><th>Comment</th><th>Scan Time</th><th>Enzyme</th><th>Replicate</th><th>Site</th><th>Page</th></tr>
<tr><td>BCAL0080</td><td>>tr|A0A1V2WZJ2|A0A1V2WZJ2_9BURK Class I cytochrome c OS=Burkholderia cenocepacia OX=95486 GN=A8E72_32710 PE=4 SV=1</td><td>Y.MANNDGANFPEPAAPAANAAQPAS[+568.212]GAPASGADASNAQA.A</td><td>HexNAc(2)Hex(1)</td><td>Yes</td><td>1345.2598</td><td>3</td><td>4033.7649</td><td>4033.7476</td><td>4.3</td><td>661.09</td><td>592.27</td><td>106.51</td><td>Nsco_20191108_BC_ZIC_HILIC_H111_thermolysin_B3.42890.42890.3</td><td>84.731</td><td>Thermolysin</td><td>B3</td><td>S176</td><td>16</td></tr>
<tr><td>BCAL0163</td><td>>tr|A0A144S4M8|A0A144S4M8_9BURK BON domain protein OS=Burkholderia cenocepacia OX=95486 GN=A8E72_32405 PE=4 SV=1</td><td>S.VQSTPLQPPAPISNSS[+568.212]SVHPGNPKAKAQ.-</td><td>HexNAc(2)Hex(1)</td><td>Yes</td><td>852.1834</td><td>4</td><td>3405.7117</td><td>3405.702</td><td>2.8</td><td>783.02</td><td>702.53</td><td>22.83</td><td>Nsco_20191108_BC_ZIC_HILIC_H111_pepsin_B2.21640.21640.4</td><td>45.203</td><td>Pepsin</td><td>B2</td><td>S262</td><td>17</td></tr>
<tr><td>BCAL0163</td><td>>tr|A0A144S4M8|A0A144S4M8_9BURK BON domain protein OS=Burkholderia cenocepacia OX=95486 GN=A8E72_32405 PE=4 SV=1</td><td>A.SVQSTPLQPPAPISNS[+568.212]SSVHPGNPKAKAQ.-</td><td>HexNAc(2)Hex(1)</td><td>Partial</td><td>873.9426</td><td>4</td><td>3492.7486</td><td>3492.734</td><td>4.2</td><td>310.05</td><td>310.05</td><td>1.6</td><td>Nsco_20191108_BC_ZIC_HILIC_H111_pepsin_B3.22644.22644.4</td><td>46.765</td><td>Pepsin</td><td>B3</td><td>NA</td><td>18</td></tr>
<tr><td>BCAL0163</td><td>>tr|A0A144S4M8|A0A144S4M8_9BURK BON domain protein OS=Burkholderia cenocepacia OX=95486 GN=A8E72_32405 PE=4 SV=1</td><td>A.LQDASPASGASGAQAAAAPADNATVGAVPDASVQSTPLQPPAPIS[+568.212]NSSSVHPGNPKAKAQ.-</td><td>HexNAc(2)Hex(1)</td><td>Partial</td><td>1549.7634</td><td>4</td><td>6196.0319</td><td>6196.0076</td><td>3.9</td><td>391.83</td><td>391.83</td><td>102.18</td><td>Nsco_20191108_BC_ZIC_HILIC_H111_thermolysin_B1.44265.44265.4</td><td>83.556</td><td>Thermolysin</td><td>B1</td><td>S224</td><td>19</td></tr>
<tr><td>BCAL0163</td><td>>tr|A0A144S4M8|A0A144S4M8_9BURK BON domain protein OS=Burkholderia cenocepacia OX=95486 GN=A8E72_32405 PE=4 SV=1</td><td>Y.VKPQDAQALQDASPAS[+568.212]GASGAQA.A</td><td>HexNAc(2)Hex(1)</td><td>Yes</td><td>1368.6424</td><td>2</td><td>2736.2775</td><td>2736.2694</td><td>3</td><td>280.25</td><td>83.45</td><td>62.99</td><td>Nsco_20191108_BC_ZIC_HILIC_H111_thermolysin_B1.26079.26079.2</td><td>51.977</td><td>Thermolysin</td><td>B1</td><td>S224</td><td>20</td></tr>
<tr><td>BCAL0163</td><td>>tr|A0A144S4M8|A0A144S4M8_9BURK BON domain protein OS=Burkholderia cenocepacia OX=95486 GN=A8E72_32405 PE=4 SV=1</td><td>Y.VKPQDAQALQDASPAS[+568.212]GASGAQ.A</td><td>HexNAc(2)Hex(1)</td><td>Yes</td><td>1333.1246</td><td>2</td><td>2665.242</td><td>2665.2323</td><td>3.6</td><td>191.34</td><td>46.96</td><td>34.32</td><td>Nsco_20191108_BC_ZIC_HILIC_H111_thermolysin_B3.23057.23057.2</td><td>49.251</td><td>Thermolysin</td><td>B3</td><td>S224</td><td>21</td></tr>
<tr><td>BCAL0163</td><td>>tr|A0A144S4M8|A0A144S4M8_9BURK BON domain protein OS=Burkholderia cenocepacia OX=95486 GN=A8E72_32405 PE=4 SV=1</td><td>Y.VKPQDAQALQDASPASGAS[+568.212]GAQAAAAPADNATVGAVPDASVQSTPLQPPAPISNSSSVHPGNPK.A</td><td>HexNAc(2)Hex(1)</td><td>Yes</td><td>1659.8156</td><td>4</td><td>6636.2404</td><td>6635.2143</td><td>3.4</td><td>901.28</td><td>550.13</td><td>85.05</td><td>Nsco_20191108_BC_ZIC_HILIC_H111_typsin_B2.47144.47144.4</td><td>91.282</td><td>Trypsin</td><td>B2</td><td>S224</td><td>22</td></tr>
<tr><td>BCAL0163</td><td>>tr|A0A144S4M8|A0A144S4M8_9BURK BON domain protein OS=Burkholderia cenocepacia OX=95486 GN=A8E72_32405 PE=4 SV=1</td><td>K.PQDAQALQDASPASGASGAQAAAAPADNATVGAVPDASVQST[+568.212]PLQPPAPISNSSSVHPGNPK.A</td><td>HexNAc(2)Hex(1)</td><td>Yes</td><td>1603.0234</td><td>4</td><td>6409.0719</td><td>6408.051</td><td>2.7</td><td>807.19</td><td>807.19</td><td>121.77</td><td>Nsco_20191108_BC_ZIC_HILIC_H111_typsin_B2.50883.50883.4</td><td>97.728</td><td>Trypsin</td><td>B2</td><td>S224</td><td>23</td></tr>
<tr><td>BCAL0163</td><td>>tr|A0A144S4M8|A0A144S4M8_9BURK BON domain protein OS=Burkholderia cenocepacia OX=95486 GN=A8E72_32405 PE=4 SV=1</td><td>K.VFQYVKPQDAQALQDASPASGASGAQAAAAPADNATVGAVPDASVQST[+568.212]PLQPPAPISNSSSVHPGNPK.A</td><td>HexNAc(2)Hex(1)</td><td>Yes</td><td>1436.1076</td><td>5</td><td>7176.5088</td><td>7172.4731</td><td>3.1</td><td>1109.8</td><td>690.41</td><td>100.78</td><td>Nsco_20191108_BC_ZIC_HILIC_H111_typsin_B2.52856.52856.5</td><td>101.38</td><td>Trypsin</td><td>B2</td><td>S224</td><td>24</td></tr>
<tr><td>BCAL0163</td><td>>tr|A0A144S4M8|A0A144S4M8_9BURK BON domain protein OS=Burkholderia cenocepacia OX=95486 GN=A8E72_32405 PE=4 SV=1</td><td>V.KVFQYVKPQDAQALQDASPASGAS[+568.212]GAQAAAAPADNATVGAVPDASVQSTPLQPPAPISNSSSVHPGNPK.A</td><td>HexNAc(2)Hex(1)</td><td>Yes</td><td>1460.9255</td><td>5</td><td>7300.5982</td><td>7300.568</td><td>4.1</td><td>1304.2</td><td>1304.2</td><td>54.21</td><td>Nsco_20191108_BC_ZIC_HILIC_H111_typsin_B1.47879.47879.5</td><td>93.267</td><td>Trypsin</td><td>B1</td><td>S224</td><td>25</td></tr>
</table>

| | | | | | | | | | | | | | | | | | | |
|---|---|---|---|---|---|---|---|---|---|---|---|---|---|---|---|---|---|---|
| BCAL0193 | >tr\|A0A071MDE2\|A0A071MDE2_9BURK Uncharacterized protein OS=Burkholderia cenocepacia OX=95486 GN=A8E72_00390 PE=4 SV=1 | Q.LKHHGSKKGQAKAAAAS[+568.212]AAGTNDAGTQN.- | HexNAc(2)Hex(1) | Yes | 652.5254 | 5 | 3258.5977 | 3258.5833 | 4.4 | 882.97 | 882.97 | 102.81 | Nsco_20191108_BC_ZIC_HILIC_H111_thermolysin_B1.2860.2860.5 | **16.086** | Thermolysin | B1 | S66 | 26 |
| BCAL0193 | >tr\|A0A071MDE2\|A0A071MDE2_9BURK Uncharacterized protein OS=Burkholderia cenocepacia OX=95486 GN=A8E72_00390 PE=4 SV=1 | Q.ASAPAADT[+568.212]SAAAPAPAKK.D | HexNAc(2)Hex(1) | Yes | 1082.5309 | 2 | 2164.0546 | 2164.0503 | 2 | 391.31 | 227.38 | 152.76 | Nsco_20191108_BC_ZIC_HILIC_H111_typsin_B1.6901.6901.2 | **22.631** | Trypsin | B1 | S25 | 27 |
| BCAL0272 | >tr\|A0A142PJB0\|A0A142PJB0_9BURK Lipoprotein OS=Burkholderia cenocepacia OX=95486 GN=A8E72_20920 PE=4 SV=1 | K.PIQPQNTPPSDVKPTDENASSDESPDTSGSPLTLSPELSSTSTMPAPAS[+568.212]GPAATK.- | HexNAc(2)Hex(1) | Yes | 1515.9615 | 4 | 6060.8244 | 6058.798 | 3.2 | 944.57 | 888.07 | 96.92 | Nsco_20191108_BC_ZIC_HILIC_H111_typsin_B2.51007.51007.4 | **97.936** | Trypsin | B2 | S85 | 28 |
| BCAL0303 | >tr\|A0A144S0T9\|A0A144S0T9_9BURK Outer membrane lipid asymmetry maintenance protein MlaD OS=Burkholderia cenocepacia OX=95486 GN=yrbD PE=4 SV=1 | S.KAADAGGAKPAAGAS[+568.212]AAPAAPAPVAV.P | HexNAc(2)Hex(1) | Yes | 909.4655 | 3 | 2726.3819 | 2726.3731 | 3.2 | 741.46 | 731.06 | 731.06 | Nsco_20191108_BC_ZIC_HILIC_H111_pepsin_B2.23564.23564.3 | 48.96 | Pepsin | B2 | S164 | 29 |
| BCAL0303 | >tr\|A0A144S0T9\|A0A144S0T9_9BURK Outer membrane lipid asymmetry maintenance protein MlaD OS=Burkholderia cenocepacia OX=95486 GN=yrbD PE=4 SV=1 | Y.SKAADAGGAKPAAGAS[+568.212]AAPAAPAPVAV.P | HexNAc(2)Hex(1) | Yes | 938.4753 | 3 | 2813.4114 | 2813.4051 | 2.2 | 386.22 | 312.9 | 280.27 | Nsco_20191108_BC_ZIC_HILIC_H111_pepsin_B2.23609.23609.3 | **49.036** | Pepsin | B2 | S164 | 30 |
| BCAL0303 | >tr\|A0A144S0T9\|A0A144S0T9_9BURK Outer membrane lipid asymmetry maintenance protein MlaD OS=Burkholderia cenocepacia OX=95486 GN=yrbD PE=4 SV=1 | L.YSKAADAGGAKPAAGAS[+568.212]AAPAAPAPVAV.P | HexNAc(2)Hex(1) | Yes | 744.8754 | 4 | 2976.48 | 2976.4684 | 3.9 | 843.54 | 806.59 | 509.24 | Nsco_20191108_BC_ZIC_HILIC_H111_pepsin_B3.25168.25168.4 | 51.945 | Pepsin | B3 | S164 | 31 |
| BCAL0303 | >tr\|A0A144S0T9\|A0A144S0T9_9BURK Outer membrane lipid asymmetry maintenance protein MlaD OS=Burkholderia cenocepacia OX=95486 GN=yrbD PE=4 SV=1 | K.PAAGASAAPAAPAPVAVPAS[+568.212][+100.064]AVS[+568.212]GSAGQ.- | HexNAc(2)Hex(1) 100.064,HexNAc(2)Hex(1) | Yes (1 of 2) | 1180.2254 | 3 | 3538.6616 | 3537.6705 | -3.4 | 247.45 | 206.29 | 1.76 | Nsco_20191108_BC_ZIC_HILIC_H111_typsin_B2.42457.42457.3 | 82.978 | Trypsin | B2 | S178 | 32 |
| BCAL0304 | >tr\|A0A2N9CYS8\|A0A2N9CYS8_9BURK Surface lipoprotein OS=Burkholderia cenocepacia OX=95486 GN=F01_50090 PE=4 SV=1 | A.VAAPASGTAESPNPAS[+568.212]ET[+568.212]NV.P | HexNAc(2)Hex(1),HexNAc(2)Hex(1) | Yes (1 of 2) | 1002.7773 | 3 | 3006.3172 | 3006.3056 | 3.9 | 162.23 | 111.11 | 25.82 | Nsco_20191108_BC_ZIC_HILIC_H111_pepsin_B3.27461.27461.3 | 56.738 | Pepsin | B3 | S287 | 33 |
| BCAL0304 | >tr\|A0A2N9CYS8\|A0A2N9CYS8_9BURK Surface lipoprotein OS=Burkholderia cenocepacia OX=95486 GN=F01_50090 PE=4 SV=1 | A.VAAPASGTAESPNPAS[+568.212]ET[+568.212]NVPAM.Q | HexNAc(2)Hex(1),HexNAc(2)Hex(1) | Yes (1 of 2) | 1102.4867 | 3 | 3305.4456 | 3305.436 | 2.9 | 194.56 | 117.32 | 9.93 | Nsco_20191108_BC_ZIC_HILIC_H111_pepsin_B2.36550.36550.3 | 76.601 | Pepsin | B2 | S297 | 34 |
| BCAL0304 | >tr\|A0A2N9CYS8\|A0A2N9CYS8_9BURK Surface lipoprotein OS=Burkholderia cenocepacia OX=95486 GN=F01_50090 PE=4 SV=1 | A.VAAPASGT[+568.212]AES[+568.212]PNPASETNVPAMQ.V | HexNAc(2)Hex(1),HexNAc(2)Hex(1) | Yes (1 of 2) | 1717.2566 | 2 | 3433.506 | 3433.4946 | 3.3 | 328.67 | 298.7 | 8.49 | Nsco_20191108_BC_ZIC_HILIC_H111_thermolysin_B3.36251.36251.2 | 72.229 | Thermolysin | B3 | S287 | 35 |
| BCAL0340 | >tr\|A0A1V2W5P4\|A0A1V2W5P4_9BURK Uncharacterized protein OS=Burkholderia cenocepacia OX=95486 GN=A8E72_11900 PE=4 SV=1 | K.AAAAPAAEAAASAAPT[+568.212]PPAAQK.G | HexNAc(2)Hex(1) | Yes | 1236.6064 | 2 | 2472.2055 | 2472.1988 | 2.7 | 548.17 | 380.52 | 57.59 | Nsco_20191108_BC_ZIC_HILIC_H111_typsin_B1.21380.21380.2 | 47.155 | Trypsin | B1 | S187 | 36 |
| BCAL0349 (type | >tr\|A0A2N9D2R1\|A0A2N9D2R1_9BURK Outer membrane protein OS=Burkholderia cenocepacia OX=95486 GN=F01_70026 PE=3 SV=1 | T.TALPQANPNAGGAS[+568.212]GTVVHGTAGTLTPPPANAAPGQ.V | HexNAc(2)Hex(1) | Yes | 958.7218 | 4 | 3831.8655 | 3831.8519 | 3.6 | 376.21 | 355.68 | 52.55 | Nsco_20191108_BC_ZIC_HILIC_H111_pepsin_B1.36442.36442.4 | **77.931** | Pepsin | B1 | S67 | 37 |
| BCAL0349 (type | >tr\|A0A2N9D2R1\|A0A2N9D2R1_9BURK Outer membrane protein OS=Burkholderia cenocepacia OX=95486 GN=F01_70026 PE=3 SV=1 | L.PQANPNAGGASGTVVHGT[+568.212]AGTLTPPPANAAPGQ.V | HexNAc(2)Hex(1) | Yes | 1182.9037 | 3 | 3546.6967 | 3546.6831 | 3.8 | 487.19 | 462.86 | 57.94 | Nsco_20191108_BC_ZIC_HILIC_H111_pepsin_B2.30908.30908.3 | 63.716 | Pepsin | B2 | S67 | 38 |

| | | | | | | | | | | | | | | | | | | |
|---|---|---|---|---|---|---|---|---|---|---|---|---|---|---|---|---|---|---|
| BCAL0349 (type | >tr\|A0A2N9D2R1\|A0A2N9D2R1_9BURK Outer membrane protein OS=Burkholderia cenocepacia OX=95486 GN=F01_70026 PE=3 SV=1 | A.TTALPQANPNAGGASGT[+568.212]VVHGTAGTLTPPPANAAPGQ.V | HexNAc(2)Hex(1) | Yes | 1311.6434 | 3 | 3932.9155 | 3932.8996 | 4.1 | 295.56 | 200.94 | 20.99 | Nsco_20191108_BC_ZIC_HILIC_H111_pepsin_B2.37365.37365.3 | 78.601 | Pepsin | B2 | S67 | 39 |
| BCAL0349 (type | >tr\|A0A2N9D2R1\|A0A2N9D2R1_9BURK Outer membrane protein OS=Burkholderia cenocepacia OX=95486 GN=F01_70026 PE=3 SV=1 | T.ALPQANPNAGGAS[+568.212]GTVVHGTAGTLTPPPANAAPGQVVVGGKVPDEATKA.A | HexNAc(2)Hex(1) | Yes | 1296.4121 | 4 | 5182.6266 | 5181.6189 | 0.8 | 199.81 | 199.81 | 91.51 | **Nsco_20191108_BC_ZIC_HILIC_H** | **78.307** | Thermolysin | B1 | S67 | 40 |
| BCAL0349 (type | >tr\|A0A2N9D2R1\|A0A2N9D2R1_9BURK Outer membrane protein OS=Burkholderia cenocepacia OX=95486 GN=F01_70026 PE=3 SV=1 | T.ALPQANPNAGGAS[+568.212][+100.064]GTVVHGTAGTLTPPPANAAPGQVVVGGKVPDEATKAAVLQK.L | HexNAc(2)Hex(1) 100.064 | Yes | 1145.1952 | 5 | 5721.9468 | 5720.9621 | -3.2 | 803.05 | 467.38 | 82.36 | Nsco_20191108_BC_ZIC_HILIC_H111_thermolysin_B1.44302.44302.5 | **83.627** | Thermolysin | B1 | S67 | 41 |
| BCAL0349 (type | >tr\|A0A2N9D2R1\|A0A2N9D2R1_9BURK Outer membrane protein OS=Burkholderia cenocepacia OX=95486 GN=F01_70026 PE=3 SV=1 | T.TALPQANPNAGGAS[+568.212]GTVVHGTAGTLTPPPANAAPGQVVVGGKVPDEATKA.A | HexNAc(2)Hex(1) | Yes | 1296.4056 | 4 | 5182.6007 | 5182.6026 | -0.4 | 747.56 | 107.6 | 99.01 | Nsco_20191108_BC_ZIC_HILIC_H111_thermolysin_B2.40555.40555.4 | 80.2 | Thermolysin | B2 | S67 | 42 |
| BCAL0349 (type | >tr\|A0A2N9D2R1\|A0A2N9D2R1_9BURK Outer membrane protein OS=Burkholderia cenocepacia OX=95486 GN=F01_70026 PE=3 SV=1 | T.VATTALPQANPNAGGAS[+568.212]GTVVHGTAGTLTPPPANAAPGQVVVGGKVPDEATKA.A | HexNAc(2)Hex(1) | Yes | 1364.2004 | 4 | 5453.7799 | 5453.7558 | 4.4 | 1043.8 | 1043.8 | 116.92 | Nsco_20191108_BC_ZIC_HILIC_H111_thermolysin_B3.42314.42314.4 | 83.58 | Thermolysin | B3 | S67 | 43 |
| BCAL0349 (type | >tr\|A0A2N9D2R1\|A0A2N9D2R1_9BURK Outer membrane protein OS=Burkholderia cenocepacia OX=95486 GN=F01_70026 PE=3 SV=1 | T.VATTALPQANPNAGGAS[+568.212]GTVVHGTAGTLTPPPAN.A | HexNAc(2)Hex(1) | Yes | 1227.2776 | 3 | 3679.8184 | 3678.7981 | 4.6 | 367.42 | 345.16 | 47.16 | Nsco_20191108_BC_ZIC_HILIC_H111_thermolysin_B2.39792.39792.3 | 78.721 | Thermolysin | B2 | S67 | 44 |
| BCAL0349 (type | >tr\|A0A2N9D2R1\|A0A2N9D2R1_9BURK Outer membrane protein OS=Burkholderia cenocepacia OX=95486 GN=F01_70026 PE=3 SV=1 | T.ALPQANPNAGGASGTVVHGTAGT[+568.212]LTPPPANAAPGQVVVGGK.V | HexNAc(2)Hex(1) | Yes | 1424.0583 | 3 | 4270.1603 | 4270.1474 | 3 | 208.24 | 167.3 | 47.75 | Nsco_20191108_BC_ZIC_HILIC_H111_typsin_B1.38426.38426.3 | **77.247** | Trypsin | B1 | S67 | 45 |
| BCAL0349 (type | >tr\|A0A2N9D2R1\|A0A2N9D2R1_9BURK Outer membrane protein OS=Burkholderia cenocepacia OX=95486 GN=F01_70026 PE=3 SV=1 | T.ALPQANPNAGGAS[+568.212]GTVVHGTAGTLTPPPANAAPGQVVVGGKVPDEATK.A | HexNAc(2)Hex(1) | Yes | 1253.3899 | 4 | 5010.5378 | 5010.5178 | 4 | 974.71 | 974.71 | 93.18 | Nsco_20191108_BC_ZIC_HILIC_H111_typsin_B2.39145.39145.4 | 77.158 | Trypsin | B2 | S67 | 46 |
| BCAL0426 | >tr\|A0A1V6KWZ1\|A0A1V6KWZ1_9BURK Membrane protein insertase YidC OS=Burkholderia cenocepacia OX=95486 GN=yidC PE=3 SV=1 | R.DHGRPSMFFPSATHTAPAAAGGAS[+568.212]GTGATTTAGDVPAAAAGAAPSTTAPAAQAQLVK.F | HexNAc(2)Hex(1) | Yes | 1446.9513 | 4 | 5784.7834 | 5784.757 | 4.6 | 1083.6 | 1083.6 | 48.54 | Nsco_20191108_BC_ZIC_HILIC_H111_typsin_B1.42478.42478.4 | 84.003 | Trypsin | B1 | S49 | 47 |
| BCAL0479 | >tr\|A0A2N9CSF4\|A0A2N9CSF4_9BURK Peptidoglycan D,D-transpeptidase MrdA OS=Burkholderia cenocepacia OX=95486 GN=mrdA PE=3 SV=1 | R.RPAS[+568.212]DAQPVVATPR.D | HexNAc(2)Hex(1) | Yes | 1017.0083 | 2 | 2033.0094 | 2033.0033 | 3 | 632.91 | 349.99 | 39.47 | Nsco_20191108_BC_ZIC_HILIC_H111_typsin_B1.9404.9404.2 | 27.408 | Trypsin | B1 | S681 | 48 |
| BCAL0525 | >tr\|A0A119PFA4\|A0A119PFA4_9BURK Flagellar M-ring protein OS=Burkholderia cenocepacia OX=95486 GN=fliF PE=3 SV=1 | L.SNTPPQPAS[+568.212]APIVAGNGQNAPQTTPVSD.R | HexNAc(2)Hex(1) | Yes | 1095.5181 | 3 | 3284.5397 | 3284.5288 | 3.3 | 328.95 | 301.17 | 24.3 | Nsco_20191108_BC_ZIC_HILIC_H111_pepsin_B1.36638.36638.3 | 78.434 | Pepsin | B1 | S358 | 49 |
| BCAL0525 | >tr\|A0A119PFA4\|A0A119PFA4_9BURK Flagellar M-ring protein OS=Burkholderia cenocepacia OX=95486 GN=fliF PE=3 SV=1 | R.SQQTSSATELAQGGASGVPGALS[568.212]NTPPQPAS[+568.212][+100.064]APIVAGNGQNAPQTTPVSDRK.D | HexNAc(2)Hex(1),HexNAc(2)Hex(1) 100.064 | Partial | 1560.2444 | 4 | 6237.9557 | 6234.9659 | -3.2 | 161.4 | 161.4 | 41.1 | Nsco_20191108_BC_ZIC_HILIC_H111_typsin_B3.41556.41556.4 | 86.147 | Trypsin | B3 | NA | 50 |
| BCAL0544 | >tr\|A0A2N9CS86\|A0A2N9CS86_9BURK Dipeptide transport system substrate-binding protein OS=Burkholderia cenocepacia OX=95486 GN=dppA PE=4 SV=1 | S.VYQGAGQAASAPMPPT[+568.212]QWSYDKN.L | HexNAc(2)Hex(1) | Partial | 1518.1818 | 2 | 3035.3563 | 3035.3463 | 3.3 | 336.38 | 270.08 | 45.08 | Nsco_20191108_BC_ZIC_HILIC_H111_thermolysin_B3.39157.39157.2 | 77.717 | Thermolysin | B3 | NA | 51 |

| | | | | | | | | | | | | | | | | | | |
|---|---|---|---|---|---|---|---|---|---|---|---|---|---|---|---|---|---|---|
| BCAL0544 | >tr\|A0A2N9CS86\|A0A2N9CS86_9BURK Dipeptide transport system substrate-binding protein OS=Burkholderia cenocepacia OX=95486 GN=dppA PE=4 SV=1 | K.KAILESVYQGAGQAAS[+568.212]APMPPTQWSYDK.N | HexNAc(2)Hex(1) | Yes | 1188.2358 | 3 | 3562.6928 | 3562.6782 | 4.1 | 568.83 | 554.97 | 117.74 | Nsco_20191108_BC_ZIC_HILIC_H111_typsin_B2.50148.50148.3 | 96.466 | Trypsin | B2 | S343 | 52 |
| BCAL0678 | >tr\|A0A142PL18\|A0A142PL18_9BURK Cell division protein OS=Burkholderia cenocepacia OX=95486 GN=A8E72_03155 PE=4 SV=1 | S.KVAPPPADNGAS[+568.212]QPQQFDPNRAL.Q | HexNAc(2)Hex(1) | Yes | 996.151 | 3 | 2986.4384 | 2986.4276 | 3.6 | 652.56 | 631.5 | 631.5 | Nsco_20191108_BC_ZIC_HILIC_H111_pepsin_B1.27669.27669.3 | 58.535 | Pepsin | B1 | S39 | 53 |
| BCAL0678 | >tr\|A0A142PL18\|A0A142PL18_9BURK Cell division protein OS=Burkholderia cenocepacia OX=95486 GN=A8E72_03155 PE=4 SV=1 | S.KVAPPPADNGAS[+568.212]QPQQFDPNRALQGKTPGQPVPQAAQPAPPNTAPGQA.A | HexNAc(2)Hex(1) | Yes | 1344.6699 | 4 | 5375.6579 | 5375.6415 | 3.1 | 753.23 | 753.23 | 297.94 | Nsco_20191108_BC_ZIC_HILIC_H1 | 73.703 | Pepsin | B1 | S39 | 54 |
| BCAL0678 | >tr\|A0A142PL18\|A0A142PL18_9BURK Cell division protein OS=Burkholderia cenocepacia OX=95486 GN=A8E72_03155 PE=4 SV=1 | S.KVAPPPADNGASQPQQFDPNRALQGKT[+568.212][+100.064]PGQPVPQAAQPAPPNTAPGQAANQ.T | HexNAc(2)Hex(1) 100.064 | Yes | 1422.9559 | 4 | 5688.8019 | 5688.7801 | 3.8 | 593.48 | 593.48 | 309.84 | Nsco_20191108_BC_ZIC_HILIC_H111_pepsin_B3.34118.34118.4 | 72.856 | Pepsin | B3 | S39 | 55 |
| BCAL0678 | >tr\|A0A142PL18\|A0A142PL18_9BURK Cell division protein OS=Burkholderia cenocepacia OX=95486 GN=A8E72_03155 PE=4 SV=1 | S.KVAPPPADNGAS[+568.212]QPQQFDPNRALQGKTPGQPVPQA.A | HexNAc(2)Hex(1) | Yes | 1044.523 | 4 | 4175.0701 | 4175.0527 | 4.2 | 776.89 | 776.89 | 543.75 | Nsco_20191108_BC_ZIC_HILIC_H111_pepsin_B2.31728.31728.4 | 65.589 | Pepsin | B2 | S39 | 56 |
| BCAL0678 | >tr\|A0A142PL18\|A0A142PL18_9BURK Cell division protein OS=Burkholderia cenocepacia OX=95486 GN=A8E72_03155 PE=4 SV=1 | S.KVAPPPADNGAS[+568.212]QPQQFDPNRALQGKTPGQPVPQAAQPAPPNTAPGQAAN.Q | HexNAc(2)Hex(1) | Yes | 1113.3546 | 5 | 5562.744 | 5560.7215 | 2.8 | 701.01 | 701.01 | 371.37 | Nsco_20191108_BC_ZIC_HILIC_H111_pepsin_B3.34070.34070.5 | 72.729 | Pepsin | B3 | S39 | 57 |
| BCAL0678 | >tr\|A0A142PL18\|A0A142PL18_9BURK Cell division protein OS=Burkholderia cenocepacia OX=95486 GN=A8E72_03155 PE=4 SV=1 | S.KVAPPPADNGAS[+568.212]QPQQFDPNRALQGKTPGQPVPQAAQPAPPNTAPGQ.A | HexNAc(2)Hex(1) | Yes | 1326.9117 | 4 | 5304.6252 | 5304.6044 | 3.9 | 691.37 | 691.37 | 332.32 | Nsco_20191108_BC_ZIC_HILIC_H111_pepsin_B1.34429.34429.4 | 73.067 | Pepsin | B1 | S39 | 58 |
| BCAL0678 | >tr\|A0A142PL18\|A0A142PL18_9BURK Cell division protein OS=Burkholderia cenocepacia OX=95486 GN=A8E72_03155 PE=4 SV=1 | S.KVAPPPADNGAS[+568.212]QPQQFDPNRALQGKTPGQPVPQAAQPAPPNTA.P | HexNAc(2)Hex(1) | Yes | 1256.3777 | 4 | 5022.4889 | 5022.4716 | 3.5 | 602.85 | 602.85 | 265.96 | Nsco_20191108_BC_ZIC_HILIC_H111_pepsin_B3.33636.33636.4 | 71.592 | Pepsin | B3 | S39 | 59 |
| BCAL0678 | >tr\|A0A142PL18\|A0A142PL18_9BURK Cell division protein OS=Burkholderia cenocepacia OX=95486 GN=A8E72_03155 PE=4 SV=1 | S.KVAPPPADNGAS[+568.212]QPQQFDPNRALQGKTPGQPVPQAAQPAPPNTAPGQAA.N | HexNAc(2)Hex(1) | Yes | 1362.4308 | 4 | 5446.7013 | 5446.6786 | 4.2 | 780.89 | 780.89 | 370.32 | Nsco_20191108_BC_ZIC_HILIC_H111_pepsin_B3.34360.34360.4 | 73.518 | Pepsin | B3 | S39 | 60 |
| BCAL0678 | >tr\|A0A142PL18\|A0A142PL18_9BURK Cell division protein OS=Burkholderia cenocepacia OX=95486 GN=A8E72_03155 PE=4 SV=1 | A.QKQQQQQQAANTPKPTSSAT[+568.212]AAA.A | HexNAc(2)Hex(1) | Partial | 984.4789 | 3 | 2951.4222 | 2951.4076 | 4.9 | 274.7 | 274.7 | 13.8 | Nsco_20191108_BC_ZIC_HILIC_H111_pepsin_B3.7849.7849.3 | 22.125 | Pepsin | B3 | NA | 61 |
| BCAL0678 | >tr\|A0A142PL18\|A0A142PL18_9BURK Cell division protein OS=Burkholderia cenocepacia OX=95486 GN=A8E72_03155 PE=4 SV=1 | K.QQQQQQAANTPKPTS[+568.212]SATAAAAAKPPTANDANTGYFLQVGAYK.T | HexNAc(2)Hex(1) | Yes | 1244.1075 | 4 | 4973.4084 | 4973.3923 | 3.2 | 1214.1 | 1143.8 | 28.53 | Nsco_20191108_BC_ZIC_HILIC_H111_typsin_B1.35749.35749.4 | 72.554 | Trypsin | B1 | S180 | 62 |
| BCAL0678 | >tr\|A0A142PL18\|A0A142PL18_9BURK Cell division protein OS=Burkholderia cenocepacia OX=95486 GN=A8E72_03155 PE=4 SV=1 | Q.KQQQQQQAANTPKPTS[+568.212]SATAAAAAKPPTANDANTGYFLQVGAYKTEGDAEQQR.A | HexNAc(2)Hex(1) | Yes | 1037.5025 | 6 | 6219.9785 | 6215.9879 | -3.7 | 1196.8 | 1084.2 | 19.24 | Nsco_20191108_BC_ZIC_HILIC_H111_typsin_B2.32751.32751.6 | 65.913 | Trypsin | B2 | S180 | 63 |

| | | | | | | | | | | | | | | | | | | |
|---|---|---|---|---|---|---|---|---|---|---|---|---|---|---|---|---|---|---|
| BCAL0678 | >tr\|A0A142PL18\|A0A142PL18_9BURK Cell division protein OS=Burkholderia cenocepacia OX=95486 GN=A8E72_03155 PE=4 SV=1 | Q.KQQQQQQAANTPKPTS[+568.212]SATAAAAAKPPTANDANTGYFLQVGAYK.T | HexNAc(2)Hex(1) | Yes | 1021.5052 | 5 | 5103.4971 | 5101.4873 | 0.6 | 1343.4 | 1328.3 | 42.28 | Nsco_20191108_BC_ZIC_HILIC_H111_typsin_B3.30947.30947.5 | 65.433 | Trypsin | B3 | S180 | 64 |
| BCAL0749 | >tr\|A0A2N9D2V0\|A0A2N9D2V0_9BURK Cytochrome c oxidase subunit 2 OS=Burkholderia cenocepacia OX=95486 GN=ctaC_1 PE=3 SV=1 | A.GAATAAAPAEAASAPAQAAS[+568.212]GAEQPAAAAS[+568.212][+100.064]AALSTIYFETGK.S | HexNAc(2)Hex(1),HexNAc(2)Hex(1) 100.064 | Partial | 1676.4614 | 3 | 5027.3696 | 5026.3437 | 4.5 | 318.21 | 294.02 | 30.22 | Nsco_20191108_BC_ZIC_HILIC_H111_typsin_B1.67136.67136.3 | 124.42 | Trypsin | B1 | NA | 65 |
| BCAL0749 | >tr\|A0A2N9D2V0\|A0A2N9D2V0_9BURK Cytochrome c oxidase subunit 2 OS=Burkholderia cenocepacia OX=95486 GN=ctaC_1 PE=3 SV=1 | G.KLPEDTAGAAT[+568.212][+100.064]AAAPAEAASAPAQAAS[+568.212]GAEQPAAAASAALSTIYFETGK.S | HexNAc(2)Hex(1) 100.064,HexNAc(2)Hex(1) | Yes (1 of 2) | 1447.1952 | 4 | 5785.7589 | 5780.7299 | 2.1 | 233.5 | 217.8 | 21.97 | Nsco_20191108_BC_ZIC_HILIC_H111_typsin_B2.63795.63795.4 | 123.3 | Trypsin | B2 | S410 | 66 |
| BCAL0749 | >tr\|A0A2N9D2V0\|A0A2N9D2V0_9BURK Cytochrome c oxidase subunit 2 OS=Burkholderia cenocepacia OX=95486 GN=ctaC_1 PE=3 SV=1 | L.PEDTAGAATAAAPAEAASAPAQAASGAEQPAAAASAALS[+568.212][+100.064]T[+568.212][+100.064]IYFETGK.S | HexNAc(2)Hex(1) 100.064,HexNAc(2)Hex(1) 100.064 | Yes (1 of 2) | 1411.1638 | 4 | 5641.6335 | 5639.6148 | 2.1 | 379.32 | 178.79 | 11.1 | Nsco_20191108_BC_ZIC_HILIC_H111_typsin_B3.58958.58958.4 | 120.69 | Trypsin | B3 | S410 | 67 |
| BCAL0786 | >tr\|A0A1V2XQK6\|A0A1V2XQK6_9BURK Membrane protein OS=Burkholderia cenocepacia OX=95486 GN=A8E72_26595 PE=4 SV=1 | K.LLSVPAPAST[+568.212]EGDHHDK.- | HexNAc(2)Hex(1) | Yes | 586.2802 | 4 | 2342.0991 | 2342.0882 | 4.7 | 445.92 | 357.63 | 52.5 | Nsco_20191108_BC_ZIC_HILIC_H111_typsin_B2.15426.15426.4 | **35.4** | Trypsin | B2 | S80 | 68 |
| BCAL1081 | >tr\|A0A1V2XQC3\|A0A1V2XQC3_9BURK Multidrug efflux system membrane fusion protein OS=Burkholderia cenocepacia OX=95486 GN=mdtA PE=3 SV=1 | K.RYQTLLSQDSIAS[+568.212]QTVDTQASLVK.Q | HexNAc(2)Hex(1) | Yes | 1074.5406 | 3 | 3221.6074 | 3220.5955 | 2.7 | 420.73 | 382.31 | 53.27 | Nsco_20191108_BC_ZIC_HILIC_H111_typsin_B3.39616.39616.3 | 82.427 | Trypsin | B3 | S183 | 69 |
| BCAL1086 | >tr\|A0A107L3U1\|A0A107L3U1_9BURK Putative lipoprotein OS=Burkholderia cenocepacia OX=95486 GN=A8E72_07105 PE=4 SV=1 | K.LQQWSQQSAAGAKPAS[+568.212]GE.- | HexNAc(2)Hex(1) | Yes | 1206.5597 | 2 | 2412.1122 | 2412.1049 | 3 | 260.84 | 151.11 | 64.75 | Nsco_20191108_BC_ZIC_HILIC_H111_typsin_B1.15214.15214.2 | 36.547 | Trypsin | B1 | S140 | 70 |
| BCAL1086 | >tr\|A0A107L3U1\|A0A107L3U1_9BURK Putative lipoprotein OS=Burkholderia cenocepacia OX=95486 GN=A8E72_07105 PE=4 SV=1 | K.KLQQWSQQS[+568.212]AAGAKPASGE.- | HexNAc(2)Hex(1) | Yes | 1270.6075 | 2 | 2540.2077 | 2540.1998 | 3.1 | 311.96 | 187.6 | 187.6 | Nsco_20191108_BC_ZIC_HILIC_H111_typsin_B3.11478.11478.2 | 29.282 | Trypsin | B3 | S140 | 71 |
| BCAL1086 | >tr\|A0A107L3U1\|A0A107L3U1_9BURK Putative lipoprotein OS=Burkholderia cenocepacia OX=95486 GN=A8E72_07105 PE=4 SV=1 | K.ALDQVAST[+568.212]VNQQINAAK.A | HexNAc(2)Hex(1) | Yes | 1170.0812 | 2 | 2339.1552 | 2339.146 | 3.9 | 468.08 | 292.44 | 38.51 | Nsco_20191108_BC_ZIC_HILIC_H111_typsin_B3.35313.35313.2 | **73.921** | Trypsin | B3 | S63 | 72 |
| BCAL1093 | >tr\|A0A2N9CGE7\|A0A2N9CGE7_9BURK Signal peptide transmembrane protein OS=Burkholderia cenocepacia OX=95486 GN=DFS07_119133 PE=4 SV=1 | K.HAYDEAHPAEAAS[+568.212]AASH.- | HexNAc(2)Hex(1) | Yes | 768.6602 | 3 | 2303.9659 | 2302.9582 | 1.9 | 353.8 | 272.78 | 95.02 | Nsco_20191108_BC_ZIC_HILIC_H111_typsin_B2.7957.7957.3 | 23.545 | Trypsin | B2 | S49 | 73 |
| BCAL1389 | >tr\|A0A2N9CG52\|A0A2N9CG52_9BURK Cyclic di-GMP-binding protein OS=Burkholderia cenocepacia OX=95486 GN=bcsB PE=3 SV=1 | P.AVPAAAS[+568.212]T[+568.212][+100.064]AAAGHAAAADVAPAAGPVAAPAAASAPAAGLPATTVHVPFASLGAFDPLR.L | HexNAc(2)Hex(1),HexNAc(2)Hex(1) 100.064 | Yes (1 of 2) | 1598.0516 | 4 | 6389.1847 | 6389.1838 | 0.1 | 770.16 | 495.83 | 33.82 | Nsco_20191108_BC_ZIC_HILIC_H111_typsin_B2.65504.65504.4 | 127.09 | Trypsin | B2 | S62 | 74 |
| BCAL1389 | >tr\|A0A2N9CG52\|A0A2N9CG52_9BURK Cyclic di-GMP-binding protein OS=Burkholderia cenocepacia OX=95486 GN=bcsB PE=3 SV=1 | A.APMPAVPAAAST[+568.212]AAAGHAAAADVAPAAGPVAAPAAAS[+568.212]APAAGLPATTVHVPFASLGAFDPLR.L | HexNAc(2)Hex(1),HexNAc(2)Hex(1) | Yes (1 of 2) | 1672.0869 | 4 | 6685.3257 | 6685.3029 | 3.4 | 1064.7 | 1061.8 | 159.46 | Nsco_20191108_BC_ZIC_HILIC_H111_typsin_B2.67298.67298.4 | 131.09 | Trypsin | B2 | S62 | 75 |

| | | | | | | | | | | | | | | | | | | |
|---|---|---|---|---|---|---|---|---|---|---|---|---|---|---|---|---|---|---|
| BCAL1496 | >tr\|A0A0M1IFG9\|A0A0M1IFG9_9BURK Signal peptide protein OS=Burkholderia cenocepacia OX=95486 GN=A8F55_16750 PE=4 SV=1 | K.KAGKKAKAADAAS[+568.212]Q.- | HexNAc(2)Hex(1) | Yes | 638.3309 | 3 | 1912.9781 | 1912.971 | 3.8 | 431.43 | 251.88 | 251.88 | Nsco_20191108_BC_ZIC_HILIC_H111_pepsin_B1.2102.2102.3 | **13.295** | Pepsin | B1 | S90 | 76 |
| BCAL1496 | >tr\|A0A0M1IFG9\|A0A0M1IFG9_9BURK Signal peptide protein OS=Burkholderia cenocepacia OX=95486 GN=A8F55_16750 PE=4 SV=1 | H.KAAAKKAGKKAKAADAAS[+568.212]Q.- | HexNAc(2)Hex(1) | Yes | 596.3257 | 4 | 2382.281 | 2382.2722 | 3.7 | 638.78 | 549.42 | 549.42 | Nsco_20191108_BC_ZIC_HILIC_H111_pepsin_B2.2867.2867.4 | 14.063 | Pepsin | B2 | S90 | 77 |
| BCAL1496 | >tr\|A0A0M1IFG9\|A0A0M1IFG9_9BURK Signal peptide protein OS=Burkholderia cenocepacia OX=95486 GN=A8F55_16750 PE=4 SV=1 | G.KKAKAADAAS[+568.212]Q.- | HexNAc(2)Hex(1) | Yes | 552.946 | 3 | 1656.8236 | 1656.8174 | 3.7 | 308.48 | 196.93 | 196.93 | Nsco_20191108_BC_ZIC_HILIC_H111_pepsin_B3.2523.2523.3 | 13.147 | Pepsin | B3 | S90 | 78 |
| BCAL1496 | >tr\|A0A0M1IFG9\|A0A0M1IFG9_9BURK Signal peptide protein OS=Burkholderia cenocepacia OX=95486 GN=A8F55_16750 PE=4 SV=1 | A.KKAGKKAKAADAAS[+568.212]Q.- | HexNAc(2)Hex(1) | Yes | 511.0242 | 4 | 2041.0751 | 2041.0659 | 4.5 | 308.02 | 306.43 | 306.43 | Nsco_20191108_BC_ZIC_HILIC_H111_pepsin_B3.2614.2614.4 | 13.491 | Pepsin | B3 | S90 | 79 |
| BCAL1496 | >tr\|A0A0M1IFG9\|A0A0M1IFG9_9BURK Signal peptide protein OS=Burkholderia cenocepacia OX=95486 GN=A8F55_16750 PE=4 SV=1 | A.AAKKAGKKAKAADAAS[+568.212]Q.- | HexNAc(2)Hex(1) | Yes | 546.5425 | 4 | 2183.1482 | 2183.1401 | 3.7 | 645.31 | 568.86 | 568.86 | Nsco_20191108_BC_ZIC_HILIC_H111_thermolysin_B1.2030.2030.4 | 13.248 | Thermolysin | B1 | S90 | 80 |
| BCAL1496 | >tr\|A0A0M1IFG9\|A0A0M1IFG9_9BURK Signal peptide protein OS=Burkholderia cenocepacia OX=95486 GN=A8F55_16750 PE=4 SV=1 | K.AAAKKAGKKAKAADAAS[+568.212]Q.- | HexNAc(2)Hex(1) | Yes | 564.3017 | 4 | 2254.1848 | 2254.1773 | 3.3 | 621.5 | 587.33 | 587.33 | Nsco_20191108_BC_ZIC_HILIC_H111_thermolysin_B2.2380.2380.4 | 13.86 | Thermolysin | B2 | S90 | 81 |
| BCAL1496 | >tr\|A0A0M1IFG9\|A0A0M1IFG9_9BURK Signal peptide protein OS=Burkholderia cenocepacia OX=95486 GN=A8F55_16750 PE=4 SV=1 | K.AKAADAAS[+568.212]Q.- | HexNAc(2)Hex(1) | Yes | 700.819 | 2 | 1400.6308 | 1400.6275 | 2.4 | 314.51 | 138.87 | 138.87 | Nsco_20191108_BC_ZIC_HILIC_H111_typsin_B1.2165.2165.2 | 14.283 | Trypsin | B1 | S90 | 82 |
| BCAL1674 | >tr\|A0A2N9CH15\|A0A2N9CH15_9BURK Multidrug efflux system OS=Burkholderia cenocepacia OX=95486 GN=acrA PE=3 SV=1 | Q.FAPDTAVKPVEKAPPS[+568.212]KAAPPAAASQA.A | HexNAc(2)Hex(1) | Yes | 1062.88 | 3 | 3186.6255 | 3185.61 | 3.8 | 693.94 | 656.34 | 21.6 | Nsco_20191108_BC_ZIC_HILIC_H111_thermolysin_B2.20980.20980.3 | 45.3 | Thermolysin | B2 | S404 | 83 |
| BCAL1746 | >tr\|A0A1V2Y428\|A0A1V2Y428_9BURK Uncharacterized protein OS=Burkholderia cenocepacia OX=95486 GN=A8F55_03700 PE=4 SV=1 | L.IDHIGKAWPGNAAS[+568.212]GASASE.- | HexNAc(2)Hex(1) | Yes | 836.3885 | 3 | 2507.1508 | 2507.142 | 3.5 | 313.93 | 234.39 | 59.12 | Nsco_20191108_BC_ZIC_HILIC_H111_pepsin_B3.22612.22612.3 | 46.702 | Pepsin | B3 | S140 | 84 |
| BCAL1917 | >tr\|A0A2N9CHU5\|A0A2N9CHU5_9BURK Uncharacterized protein OS=Burkholderia cenocepacia OX=95486 GN=F01_210059 PE=4 SV=1 | T.AAPAPTAS[+568.212]APEAAAKPAKTKRASKKEK.A | HexNAc(2)Hex(1) | Yes | 649.7541 | 5 | 3244.7412 | 3244.7271 | 4.4 | 796.81 | 784.32 | 121.44 | Nsco_20191108_BC_ZIC_HILIC_H111_thermolysin_B1.3500.3500.5 | 17.346 | Thermolysin | B1 | S151 | 85 |
| BCAL1917 | >tr\|A0A2N9CHU5\|A0A2N9CHU5_9BURK Uncharacterized protein OS=Burkholderia cenocepacia OX=95486 GN=F01_210059 PE=4 SV=1 | T.AAPAPTAS[+568.212]APEAAAKPAKTKRASKKEKA.A | HexNAc(2)Hex(1) | Yes | 663.9616 | 5 | 3315.7788 | 3315.7642 | 4.4 | 923.36 | 897.24 | 145.29 | Nsco_20191108_BC_ZIC_HILIC_H111_thermolysin_B1.3693.3693.5 | 17.662 | Thermolysin | B1 | S151 | 86 |
| BCAL1917 | >tr\|A0A2N9CHU5\|A0A2N9CHU5_9BURK Uncharacterized protein OS=Burkholderia cenocepacia OX=95486 GN=F01_210059 PE=4 SV=1 | T.AAPAPTAS[+568.212]APEAAAKPAKTKRASKKEKAAAA.A | HexNAc(2)Hex(1) | Yes | 882.9772 | 4 | 3528.887 | 3528.8755 | 3.3 | 580.06 | 580.06 | 133.19 | Nsco_20191108_BC_ZIC_HILIC_H111_thermolysin_B1.4372.4372.4 | 18.616 | Thermolysin | B1 | S151 | 87 |
| BCAL1917 | >tr\|A0A2N9CHU5\|A0A2N9CHU5_9BURK Uncharacterized protein OS=Burkholderia cenocepacia OX=95486 GN=F01_210059 PE=4 SV=1 | T.AAPAPTAS[+568.212]APEAAAKPAKTKRAS.K | HexNAc(2)Hex(1) | Yes | 683.6077 | 4 | 2731.4092 | 2731.3996 | 3.5 | 695.41 | 663.67 | 96.64 | Nsco_20191108_BC_ZIC_HILIC_H111_thermolysin_B2.5786.5786.4 | 20.941 | Thermolysin | B2 | S151 | 88 |

| | | | | | | | | | | | | | | | | | | |
|---|---|---|---|---|---|---|---|---|---|---|---|---|---|---|---|---|---|---|
| BCAL1917 | >tr\|A0A2N9CHU5\|A0A2N9CHU5_9BURK Uncharacterized protein OS=Burkholderia cenocepacia OX=95486 GN=F01_210059 PE=4 SV=1 | T.ATAGTTTAAPAPTAS[+568.212]APEAAAKPAKTKR.A | HexNAc(2)Hex(1) | Yes | 1059.8811 | 3 | 3177.6288 | 3176.6169 | 2.7 | 165.83 | 165.83 | 32.16 | Nsco_20191108_BC_ZIC_HILIC_H111_thermolysin_B2.9223.9223.3 | 25.928 | Thermolysin | B2 | S151 | 89 |
| BCAL1917 | >tr\|A0A2N9CHU5\|A0A2N9CHU5_9BURK Uncharacterized protein OS=Burkholderia cenocepacia OX=95486 GN=F01_210059 PE=4 SV=1 | T.AAPAPTAS[+568.212]APEAAAKPAKTKRASKKEKAAA.A | HexNAc(2)Hex(1) | Yes | 692.3761 | 5 | 3457.8512 | 3457.8384 | 3.7 | 749.03 | 741.46 | 142.57 | Nsco_20191108_BC_ZIC_HILIC_H111_thermolysin_B1.4098.4098.5 | 18.251 | Thermolysin | B1 | S151 | 90 |
| BCAL1917 | >tr\|A0A2N9CHU5\|A0A2N9CHU5_9BURK Uncharacterized protein OS=Burkholderia cenocepacia OX=95486 GN=F01_210059 PE=4 SV=1 | T.AAPAPTAS[+568.212]APEAAAKPAKTKRASKKEKAA.A | HexNAc(2)Hex(1) | Yes | 847.4583 | 4 | 3386.8115 | 3386.8013 | 3 | 800.88 | 800.88 | 134.79 | Nsco_20191108_BC_ZIC_HILIC_H111_thermolysin_B2.3946.3946.4 | 17.862 | Thermolysin | B2 | S151 | 91 |
| BCAL1917 | >tr\|A0A2N9CHU5\|A0A2N9CHU5_9BURK Uncharacterized protein OS=Burkholderia cenocepacia OX=95486 GN=F01_210059 PE=4 SV=1 | T.AAPAPTAS[+568.212]APEAAAKPAKTKR.A | HexNAc(2)Hex(1) | Yes | 644.0901 | 4 | 2573.3386 | 2573.3305 | 3.2 | 915.51 | 699.84 | 132.94 | Nsco_20191108_BC_ZIC_HILIC_H111_thermolysin_B2.5666.5666.4 | 20.778 | Thermolysin | B2 | S151 | 92 |
| BCAL1917 | >tr\|A0A2N9CHU5\|A0A2N9CHU5_9BURK Uncharacterized protein OS=Burkholderia cenocepacia OX=95486 GN=F01_210059 PE=4 SV=1 | T.STTATAGTTTAAPAPT[+568.212]ASAPEAAAKPAK.T | HexNAc(2)Hex(1) | Yes | 1027.5093 | 3 | 3080.5134 | 3080.5005 | 4.2 | 353.44 | 288.43 | 42.43 | Nsco_20191108_BC_ZIC_HILIC_H111_typsin_B1.15164.15164.3 | 36.476 | Trypsin | B1 | S151 | 93 |
| BCAL1917 | >tr\|A0A2N9CHU5\|A0A2N9CHU5_9BURK Uncharacterized protein OS=Burkholderia cenocepacia OX=95486 GN=F01_210059 PE=4 SV=1 | A.TAGTTTAAPAPTAS[+568.212]APEAAAKPAK.T | HexNAc(2)Hex(1) | Yes | 907.4542 | 3 | 2720.348 | 2720.336 | 4.4 | 344.84 | 260.24 | 49.3 | Nsco_20191108_BC_ZIC_HILIC_H111_typsin_B2.13385.13385.3 | 32.05 | Trypsin | B2 | S151 | 94 |
| BCAL1917 | >tr\|A0A2N9CHU5\|A0A2N9CHU5_9BURK Uncharacterized protein OS=Burkholderia cenocepacia OX=95486 GN=F01_210059 PE=4 SV=1 | T.SAGTASTSTTATAGTTTAAPAPTAS[+568.212]APEAAAKPAK.T | HexNAc(2)Hex(1) | Yes | 1219.262 | 3 | 3655.7715 | 3655.7556 | 4.3 | 335.99 | 302.12 | 43.51 | Nsco_20191108_BC_ZIC_HILIC_H111_typsin_B2.19049.19049.3 | 41.427 | Trypsin | B2 | S151 | 95 |
| BCAL1917 | >tr\|A0A2N9CHU5\|A0A2N9CHU5_9BURK Uncharacterized protein OS=Burkholderia cenocepacia OX=95486 GN=F01_210059 PE=4 SV=1 | S.KPAATTSATTSTTTSAGTASTSTTATAGTTTAAPAPTAS[+568.212]APEAAAKPAK.T | HexNAc(2)Hex(1) | Yes | 1244.8641 | 4 | 4976.4347 | 4975.4125 | 3.8 | 454.51 | 454.51 | 25.96 | Nsco_20191108_BC_ZIC_HILIC_H111_typsin_B2.20241.20241.4 | 43.54 | Trypsin | B2 | S151 | 96 |
| BCAL1917 | >tr\|A0A2N9CHU5\|A0A2N9CHU5_9BURK Uncharacterized protein OS=Burkholderia cenocepacia OX=95486 GN=F01_210059 PE=4 SV=1 | T.TSATTSTTTSAGTASTS[+568.212][+100.064]TTATAGTTTAAPAPT[+568.212][+100.064]ASAPEAAAKPAK.T | HexNAc(2)Hex(1) 100.064,HexNAc(2)Hex(1) 100.064 | Yes (1 of 2) | 1319.631 | 4 | 5275.5021 | 5275.4824 | 3.7 | 225.08 | 225.08 | 24.52 | Nsco_20191108_BC_ZIC_HILIC_H111_typsin_B2.43074.43074.4 | 84.038 | Trypsin | B2 | S151 | 97 |
| BCAL1917 | >tr\|A0A2N9CHU5\|A0A2N9CHU5_9BURK Uncharacterized protein OS=Burkholderia cenocepacia OX=95486 GN=F01_210059 PE=4 SV=1 | A.GTASTSTTATAGTTTAAPAPT[+568.212]ASAPEAAAKPAK.T | HexNAc(2)Hex(1) | Yes | 1166.5723 | 3 | 3497.7024 | 3497.6865 | 4.6 | 317.9 | 257.84 | 44.9 | Nsco_20191108_BC_ZIC_HILIC_H111_typsin_B3.17816.17816.3 | 40.101 | Trypsin | B3 | S151 | 98 |
| BCAL1917 | >tr\|A0A2N9CHU5\|A0A2N9CHU5_9BURK Uncharacterized protein OS=Burkholderia cenocepacia OX=95486 GN=F01_210059 PE=4 SV=1 | K.LSKPAATTSATTSTTTSAGTASTSTTATAGTTTAAPAPTAS[+568.212]APEAAAK.P | HexNAc(2)Hex(1) | Partial | 1220.5935 | 4 | 4879.3524 | 4879.3437 | 1.8 | 758.27 | 736.58 | 79.08 | Nsco_20191108_BC_ZIC_HILIC_H111_typsin_B2.25663.25663.4 | 53.316 | Trypsin | B2 | NA | 99 |
| BCAL1917 | >tr\|A0A2N9CHU5\|A0A2N9CHU5_9BURK Uncharacterized protein OS=Burkholderia cenocepacia OX=95486 GN=F01_210059 PE=4 SV=1 | K.PAATTSATTSTTTSAGTASTSTTATAGTTTAAPAPT[+568.212]ASAPEAAAKPAK.T | HexNAc(2)Hex(1) | Partial | 1616.4455 | 3 | 4847.3219 | 4847.3175 | 0.9 | 324.17 | 249.96 | 9.1 | Nsco_20191108_BC_ZIC_HILIC_H111_typsin_B2.26810.26810.3 | 55.497 | Trypsin | B2 | NA | 100 |

| | | | | | | | | | | | | | | | | | | |
|---|---|---|---|---|---|---|---|---|---|---|---|---|---|---|---|---|---|---|
| BCAL1917 | >tr\|A0A2N9CHU5\|A0A2N9CHU5_9BURK Uncharacterized protein OS=Burkholderia cenocepacia OX=95486 GN=F01_210059 PE=4 SV=1 | A.STSTTATAGTTTAAPAPT[+568.212]ASAPEAAAKPAK.T | HexNAc(2)Hex(1) | Yes | 1090.2005 | 3 | 3268.5869 | 3268.5802 | 2 | 292.48 | 253.17 | 40.26 | Nsco_20191108_BC_ZIC_HILIC_H111_typsin_B3.16579.16579.3 | 37.887 | Trypsin | B3 | S151 | 101 |
| BCAL1917 | >tr\|A0A2N9CHU5\|A0A2N9CHU5_9BURK Uncharacterized protein OS=Burkholderia cenocepacia OX=95486 GN=F01_210059 PE=4 SV=1 | T.STTTSAGTASTSTTATAGTTTAAPAT[+568.212]ASAPEAAAKPAK.T | HexNAc(2)Hex(1) | Yes | 1349.3188 | 3 | 4045.9418 | 4045.9307 | 2.7 | 382.44 | 294.2 | 50.7 | Nsco_20191108_BC_ZIC_HILIC_H111_typsin_B3.20202.20202.3 | 44.553 | Trypsin | B3 | S151 | 102 |
| BCAL1917 | >tr\|A0A2N9CHU5\|A0A2N9CHU5_9BURK Uncharacterized protein OS=Burkholderia cenocepacia OX=95486 GN=F01_210059 PE=4 SV=1 | A.TTSTTTSAGTASTSTTATAGTTTAAAPT[+568.212]ASAPEAAAKPAK.T | HexNAc(2)Hex(1) | Yes | 1417.6876 | 3 | 4251.0483 | 4248.026 | 2.9 | 231.98 | 192.01 | 32.92 | Nsco_20191108_BC_ZIC_HILIC_H111_typsin_B3.20807.20807.3 | 45.733 | Trypsin | B3 | S151 | 103 |
| BCAL1917 | >tr\|A0A2N9CHU5\|A0A2N9CHU5_9BURK Uncharacterized protein OS=Burkholderia cenocepacia OX=95486 GN=F01_210059 PE=4 SV=1 | S.ATTSTTTSAGTASTSTTATAGTTTAAPT[+568.212]ASAPEAAAKPAK.T | HexNAc(2)Hex(1) | Yes | 1440.3658 | 3 | 4319.0828 | 4319.0631 | 4.5 | 151.41 | 115.69 | 115.69 | Nsco_20191108_BC_ZIC_HILIC_H111_typsin_B3.21063.21063.3 | 46.286 | Trypsin | B3 | S151 | 104 |
| BCAL1917 | >tr\|A0A2N9CHU5\|A0A2N9CHU5_9BURK Uncharacterized protein OS=Burkholderia cenocepacia OX=95486 GN=F01_210059 PE=4 SV=1 | K.SVGHLEENGLT[+568.212]IGGASTPPK.G | HexNAc(2)Hex(1) | Yes | 844.7475 | 3 | 2532.2279 | 2532.2199 | 3.2 | 233.62 | 164.31 | 11.1 | Nsco_20191108_BC_ZIC_HILIC_H111_typsin_B3.25307.25307.3 | 54.409 | Trypsin | B3 | S104 | 105 |
| BCAL1917 | >tr\|A0A2N9CHU5\|A0A2N9CHU5_9BURK Uncharacterized protein OS=Burkholderia cenocepacia OX=95486 GN=F01_210059 PE=4 SV=1 | K.LSKPAATTSATTSTTTSAGTASTSTTTAGTTTAAPAPTAS[+568.212]APEAAAKPAK.T | HexNAc(2)Hex(1) | Partial | 1319.6501 | 4 | 5275.5788 | 5275.5926 | -2.6 | 591.05 | 591.05 | 205.17 | Nsco_20191108_BC_ZIC_HILIC_H111_typsin_B3.75172.75172.4 | 160.88 | Trypsin | B3 | NA | 106 |
| BCAL2107 | >tr\|A0A2N9CIG6\|A0A2N9CIG6_9BURK Fused putative transporter subunits of ABC superfamily: ATP-binding components OS=Burkholderia cenocepacia OX=95486 GN=yheS PE=4 SV=1 | A.DS[+568.212]AAAK.A | HexNAc(2)Hex(1) | Yes | 565.7523 | 2 | 1130.4974 | 1130.4947 | 2.4 | 241.82 | 23.12 | 23.12 | Nsco_20191108_BC_ZIC_HILIC_H111_typsin_B1.1961.1961.2 | 13.168 | Trypsin | B1 | NA | 107 |
| BCAL2161 | >tr\|A0A2N9CIP1\|A0A2N9CIP1_9BURK Putative exported protein OS=Burkholderia cenocepacia OX=95486 GN=F01_230051 PE=4 SV=1 | S.VAPPLQGDGAAPGGAS[+568.212]WPAPPPASGPAPGLPASSVQGT[+568.212]P.- | HexNAc(2)Hex(1),HexNAc(2)Hex(1) | Yes (1 of 2) | 1541.0685 | 3 | 4621.1911 | 4621.1727 | 4 | 465.76 | 361.02 | 34.77 | Nsco_20191108_BC_ZIC_HILIC_H111_thermolysin_B1.62166.62166.3 | 117.73 | Thermolysin | B1 | S391 | 108 |
| BCAL2276 | >tr\|A0A2N9CJ70\|A0A2N9CJ70_9BURK Putative membrane protein OS=Burkholderia cenocepacia OX=95486 GN=DFS07_11429 PE=4 SV=1 | K.STIDTAASNAGVPVS[+568.212]SVNYIVHDAGK.G | HexNAc(2)Hex(1) | Yes | 1048.1735 | 3 | 3142.5059 | 3141.4958 | 2.2 | 182.01 | 146.66 | 18.54 | Nsco_20191108_BC_ZIC_HILIC_H111_typsin_B1.45598.45598.3 | 89.504 | Trypsin | B1 | S353 | 109 |
| BCAL2398 | >tr\|A0A2N9CKD9\|A0A2N9CKD9_9BURK Periplasmic protein TonB, links inner and outer membranes OS=Burkholderia cenocepacia OX=95486 GN=F01_260105 PE=4 SV=1 | K.ASPPYAADKPIVAVFPPVVPAAPAS[+568.212]SASATR.- | HexNAc(2)Hex(1) | Yes | 1191.9487 | 3 | 3573.8314 | 3573.8211 | 2.9 | 382.56 | 364.3 | 20.38 | Nsco_20191108_BC_ZIC_HILIC_H111_typsin_B2.54241.54241.3 | 104.04 | Trypsin | B2 | S315 | 110 |
| BCAL2449 | >tr\|A0A144SYR9\|A0A144SYR9_9BURK Diguanylate cyclase (GGDEF) domain protein OS=Burkholderia cenocepacia OX=95486 GN=gmr_4 PE=4 SV=1 | R.RTGSVNNAPGAFS[+568.212]ASGVYPIAER.V | HexNAc(2)Hex(1) | Yes | 963.8006 | 3 | 2889.3871 | 2889.3749 | 4.3 | 382.29 | 269.92 | 29.81 | Nsco_20191108_BC_ZIC_HILIC_H111_typsin_B2.32822.32822.3 | 66.025 | Trypsin | B2 | S254 | 111 |
| BCAL2466 | >tr\|A0A144SXW2\|A0A144SXW2_9BURK Ecotin OS=Burkholderia cenocepacia OX=95486 GN=eco PE=4 SV=1 | A.APASAPAVPAES[+568.212]IKMFPQ.A | HexNAc(2)Hex(1) | Yes | 1190.5817 | 2 | 2380.1561 | 2380.1476 | 3.6 | 360.93 | 166 | 166 | Nsco_20191108_BC_ZIC_HILIC_H111_thermolysin_B1.47928.47928.2 | 89.914 | Thermolysin | B1 | S21 | 112 |

| | | | | | | | | | | | | | | | | | | |
|---|---|---|---|---|---|---|---|---|---|---|---|---|---|---|---|---|---|---|
| BCAL2466 | >tr\|A0A144SXW2\|A0A144SXW2_9BURK Ecotin OS=Burkholderia cenocepacia OX=95486 GN=eco PE=4 SV=1 | A.APASAPAVPAES[+568.212]IKMFPQA.A | HexNAc(2)Hex(1) | Yes | 1226.5984 | 2 | 2452.1896 | 2451.1847 | 0.6 | 272.18 | 109.18 | 109.18 | Nsco_20191108_BC_ZIC_HILIC_H111_thermolysin_B1.49472.49472.2 | 92.599 | Thermolysin | B1 | S21 | 113 |
| BCAL2466 | >tr\|A0A144SXW2\|A0A144SXW2_9BURK Ecotin OS=Burkholderia cenocepacia OX=95486 GN=eco PE=4 SV=1 | A.APASAPAVPAES[+568.212]IKMFPQAAAGQQR.V | HexNAc(2)Hex(1) | Yes | 1021.8416 | 3 | 3063.5102 | 3062.4987 | 2.7 | 298.81 | 255.28 | 113.68 | Nsco_20191108_BC_ZIC_HILIC_H111_typsin_B2.38632.38632.3 | 76.266 | Trypsin | B2 | S21 | 114 |
| BCAL2607 | >tr\|A0A1V2WZX0\|A0A1V2WZX0_9BURK DNA repair ATPase OS=Burkholderia cenocepacia OX=95486 GN=F01_260195 PE=4 SV=1 | M.KERWEQHRAAAKGAS[+568.212]APAQ.- | HexNAc(2)Hex(1) | Yes | 665.8308 | 4 | 2660.3013 | 2660.2911 | 3.8 | 659.69 | 477.63 | 477.63 | Nsco_20191108_BC_ZIC_HILIC_H111_pepsin_B2.4597.4597.4 | 17.427 | Pepsin | B2 | S180 | 115 |
| BCAL2640 | >tr\|A0A144SWF6\|A0A144SWF6_9BURK Putative exported protein OS=Burkholderia cenocepacia OX=95486 GN=A8E72_18590 PE=4 SV=1 | R.GAADRYAPPPAAVPVAAT[+568.212]SGAQGGAAAAAAPAGTKPANAPR.E | HexNAc(2)Hex(1) | Partial | 1052.5328 | 4 | 4207.1092 | 4207.0902 | 4.5 | 473.05 | 473.05 | 26.56 | Nsco_20191108_BC_ZIC_HILIC_H111_typsin_B1.29090.29090.4 | 60.942 | Trypsin | B1 | NA | 116 |
| BCAL2640 | >tr\|A0A144SWF6\|A0A144SWF6_9BURK Putative exported protein OS=Burkholderia cenocepacia OX=95486 GN=A8E72_18590 PE=4 SV=1 | R.YAPPPAAVPVAATSGAQGGAAAAAAPAGT[+568.212]KPANAPREPAVRR.V | HexNAc(2)Hex(1) | Yes | 890.0622 | 5 | 4446.282 | 4445.2695 | 2 | 378.48 | 378.48 | 33.62 | Nsco_20191108_BC_ZIC_HILIC_H111_typsin_B3.25543.25543.5 | 54.935 | Trypsin | B3 | S165 | 117 |
| BCAL2640 | >tr\|A0A144SWF6\|A0A144SWF6_9BURK Putative exported protein OS=Burkholderia cenocepacia OX=95486 GN=A8E72_18590 PE=4 SV=1 | R.GAADRYAPPPAAVPVAAT[+568.212]SGAQGGAAAAAAPAGTKPANAPREPAVR.R | HexNAc(2)Hex(1) | Yes | 952.6888 | 5 | 4759.4149 | 4759.3922 | 4.8 | 918.5 | 887.07 | 40.6 | Nsco_20191108_BC_ZIC_HILIC_H111_typsin_B3.28495.28495.5 | 60.698 | Trypsin | B3 | S165 | 118 |
| BCAL2640 | >tr\|A0A144SWF6\|A0A144SWF6_9BURK Putative exported protein OS=Burkholderia cenocepacia OX=95486 GN=A8E72_18590 PE=4 SV=1 | R.YAPPPAAVPVAATSGAQGGAAAAAAPAGT[+568.212]KPANAPREPAVR.R | HexNAc(2)Hex(1) | Yes | 1073.0513 | 4 | 4289.1832 | 4289.1684 | 3.5 | 611.4 | 611.4 | 13.02 | Nsco_20191108_BC_ZIC_HILIC_H111_typsin_B2.30519.30519.4 | 62.009 | Trypsin | B2 | S165 | 119 |
| BCAL2640 | >tr\|A0A144SWF6\|A0A144SWF6_9BURK Putative exported protein OS=Burkholderia cenocepacia OX=95486 GN=A8E72_18590 PE=4 SV=1 | R.GAADRYAPPPAAVPVAAT[+568.212]SGAQGGAAAAAAPAGTK.P | HexNAc(2)Hex(1) | Partial | 1200.9278 | 3 | 3600.7688 | 3600.7664 | 0.7 | 394.34 | 293.58 | 4.05 | Nsco_20191108_BC_ZIC_HILIC_H111_typsin_B2.32987.32987.5 | 66.318 | Trypsin | B2 | NA | 120 |
| BCAL2820 | >tr\|A0A1V2WC66\|A0A1V2WC66_9BURK Multidrug transporter OS=Burkholderia cenocepacia OX=95486 GN=oprM PE=3 SV=1 | D.YDKAAAPAPASAT[+568.212]ATNG.- | HexNAc(2)Hex(1) | Yes | 1072.9942 | 2 | 2144.9811 | 2144.9717 | 4.4 | 309.82 | 162.79 | 97.49 | Nsco_20191108_BC_ZIC_HILIC_H111_pepsin_B1.12911.12911.2 | 31.741 | Pepsin | B1 | S501 | 121 |
| BCAL2820 | >tr\|A0A1V2WC66\|A0A1V2WC66_9BURK Multidrug transporter OS=Burkholderia cenocepacia OX=95486 GN=oprM PE=3 SV=1 | A.LGGGWIQRAGETPRAPDAPVDYDKAAAPAPAS[+568.212]ATATNG.- | HexNAc(2)Hex(1) | Yes | 1073.021 | 4 | 4289.0623 | 4289.048 | 3.3 | 572.87 | 533.13 | 60.25 | Nsco_20191108_BC_ZIC_HILIC_H111_thermolysin_B1.40111.40111.4 | 76.557 | Thermolysin | B1 | S501 | 122 |
| BCAL2820 | >tr\|A0A1V2WC66\|A0A1V2WC66_9BURK Multidrug transporter OS=Burkholderia cenocepacia OX=95486 GN=oprM PE=3 SV=1 | P.VDYDKAAAPAPAS[+568.212]ATATNG.- | HexNAc(2)Hex(1) | Yes | 1180.5431 | 2 | 2360.0789 | 2359.0671 | 3.6 | 376.57 | 201.95 | 75.8 | Nsco_20191108_BC_ZIC_HILIC_H111_thermolysin_B2.20923.20923.2 | 45.214 | Thermolysin | B2 | S501 | 123 |
| BCAL2820 | >tr\|A0A1V2WC66\|A0A1V2WC66_9BURK Multidrug transporter OS=Burkholderia cenocepacia OX=95486 GN=oprM PE=3 SV=1 | W.IQRAGETPRAPDAPVDYDKAAAPAPAS[+568.212]ATATNG.- | HexNAc(2)Hex(1) | Yes | 955.4648 | 4 | 3818.8372 | 3818.8203 | 4.4 | 569.12 | 554.4 | 74.73 | Nsco_20191108_BC_ZIC_HILIC_H111_thermolysin_B3.25289.25289.4 | 53.253 | Thermolysin | B3 | S501 | 124 |
| BCAL2820 | >tr\|A0A1V2WC66\|A0A1V2WC66_9BURK Multidrug transporter OS=Burkholderia cenocepacia OX=95486 GN=oprM PE=3 SV=1 | K.AAAPAPASAT[+568.212]ATNG.- | HexNAc(2)Hex(1) | Yes | 869.9 | 2 | 1738.7927 | 1738.7865 | 3.5 | 211.91 | 115.51 | 82.44 | Nsco_20191108_BC_ZIC_HILIC_H111_typsin_B1.11030.11030.2 | 29.902 | Trypsin | B1 | S501 | 125 |
| BCAL2820 | >tr\|A0A1V2WC66\|A0A1V2WC66_9BURK Multidrug transporter OS=Burkholderia cenocepacia OX=95486 GN=oprM PE=3 SV=1 | K.RPDAPVAQAYPAS[+568.212]GVYATQPGAAGAR.S | HexNAc(2)Hex(1) | Yes | 1037.5055 | 3 | 3110.502 | 3110.4913 | 3.5 | 605.08 | 488.25 | 177.02 | Nsco_20191108_BC_ZIC_HILIC_H111_typsin_B1.26449.26449.3 | 56.523 | Trypsin | B1 | S39 | 126 |

| | | | | | | | | | | | | | | | | | | |
|---|---|---|---|---|---|---|---|---|---|---|---|---|---|---|---|---|---|---|
| BCAL2974 | >tr\|A0A144U3Q1\|A0A144U3Q1_9BURK Uncharacterized protein OS=Burkholderia cenocepacia OX=95486 GN=A8E72_12460 PE=4 SV=1 | M.PFAASAPS[+568.212]QKYQGSKKSAL.R | HexNAc(2)Hex(1) | Yes | 845.4247 | 3 | 2534.2597 | 2534.2508 | 3.5 | 281.45 | 203.96 | 50.36 | Nsco_20191108_BC_ZIC_HILIC_H111_pepsin_B1.12677.12677.3 | 31.412 | Pepsin | B1 | S91 | 127 |
| BCAL2974 | >tr\|A0A144U3Q1\|A0A144U3Q1_9BURK Uncharacterized protein OS=Burkholderia cenocepacia OX=95486 GN=A8E72_12460 PE=4 SV=1 | F.RLNEHPQMPFAAS[+568.212]APSQ.K | HexNAc(2)Hex(1) | Yes | 817.0482 | 3 | 2449.1302 | 2449.1188 | 4.7 | 170.39 | 100.54 | 34.72 | Nsco_20191108_BC_ZIC_HILIC_H111_pepsin_B1.26118.26118.3 | 55.491 | Pepsin | B1 | S91 | 128 |
| BCAL2974 | >tr\|A0A144U3Q1\|A0A144U3Q1_9BURK Uncharacterized protein OS=Burkholderia cenocepacia OX=95486 GN=A8E72_12460 PE=4 SV=1 | L.NEHPQMPFAASAPS[+568.212]QKYQG.S | HexNAc(2)Hex(1) | Yes | 886.0625 | 3 | 2656.1728 | 2656.1719 | 0.4 | 286.11 | 251.7 | 13.48 | Nsco_20191108_BC_ZIC_HILIC_H111_pepsin_B2.24696.24696.3 | 51.18 | Pepsin | B2 | S91 | 129 |
| BCAL2974 | >tr\|A0A144U3Q1\|A0A144U3Q1_9BURK Uncharacterized protein OS=Burkholderia cenocepacia OX=95486 GN=A8E72_12460 PE=4 SV=1 | F.RLNEHPQMPFAAS[+568.212]APSQKYQGSKKSAL.R | HexNAc(2)Hex(1) | Yes | 708.9552 | 5 | 3540.7467 | 3539.7323 | 3.1 | 1074.9 | 1068.2 | 124.4 | Nsco_20191108_BC_ZIC_HILIC_H111_pepsin_B1.18696.18696.5 | 41.048 | Pepsin | B1 | S91 | 130 |
| BCAL2974 | >tr\|A0A144U3Q1\|A0A144U3Q1_9BURK Uncharacterized protein OS=Burkholderia cenocepacia OX=95486 GN=A8E72_12460 PE=4 SV=1 | L.NEHPQMPFAAS[+568.212]APSQKYQGSKKSAL.R | HexNAc(2)Hex(1) | Yes | 818.3957 | 4 | 3270.5608 | 3270.5471 | 4.2 | 1164.9 | 1137.3 | 171.47 | Nsco_20191108_BC_ZIC_HILIC_H111_pepsin_B3.19176.19176.4 | 40.001 | Pepsin | B3 | S91 | 131 |
| BCAL2974 | >tr\|A0A144U3Q1\|A0A144U3Q1_9BURK Uncharacterized protein OS=Burkholderia cenocepacia OX=95486 GN=A8E72_12460 PE=4 SV=1 | P.FAASAPSQKYQGS[+568.212]KKSA.L | HexNAc(2)Hex(1) | Yes | 581.7866 | 4 | 2324.1247 | 2324.114 | 4.6 | 261.25 | 199.88 | 86.69 | Nsco_20191108_BC_ZIC_HILIC_H111_thermolysin_B1.1703.1703.4 | 11.115 | Thermolysin | B1 | S91 | 132 |
| BCAL2974 | >tr\|A0A144U3Q1\|A0A144U3Q1_9BURK Uncharacterized protein OS=Burkholderia cenocepacia OX=95486 GN=A8E72_12460 PE=4 SV=1 | R.LNEHPQMPFAASAPS[+568.212]QK.Y | HexNAc(2)Hex(1) | Yes | 1211.0634 | 2 | 2421.1194 | 2421.1126 | 2.8 | 435.34 | 227.13 | 75.7 | Nsco_20191108_BC_ZIC_HILIC_H111_typsin_B3.23325.23325.2 | 50.601 | Trypsin | B3 | S91 | 133 |
| BCAL2983A | >tr\|A0A2N9CL86\|A0A2N9CL86_9BURK Putative lipoprotein OS=Burkholderia cenocepacia OX=95486 GN=F01_310136 PE=4 SV=1 | S.ADASAPVAGTRPAVTSLSGGASSAASGAVAT[+568.212]DAAAQGNVAELTQMLHDGR.I | HexNAc(2)Hex(1) | Yes | 1301.8806 | 4 | 5204.5006 | 5204.4772 | 4.5 | 369.95 | 369.95 | 5.1 | Nsco_20191108_BC_ZIC_HILIC_H111_typsin_B3.53184.53184.5 | 108.58 | Trypsin | B3 | S55 | 134 |
| BCAL3033 | >tr\|A0A2N9CM16\|A0A2N9CM16_9BURK Outer-membrane lipoprotein carrier protein OS=Burkholderia cenocepacia OX=95486 GN=lolA PE=3 SV=1 | Q.IVKAPAKGAS[+568.212]AAQAAPKPTDN.S | HexNAc(2)Hex(1) | Yes | 644.5867 | 4 | 2575.3249 | 2574.3145 | 2.8 | 729.64 | 648.17 | 408.82 | Nsco_20191108_BC_ZIC_HILIC_H111_pepsin_B3.10005.10005.4 | 25.473 | Pepsin | B3 | S68 | 135 |
| BCAL3033 | >tr\|A0A2N9CM16\|A0A2N9CM16_9BURK Outer-membrane lipoprotein carrier protein OS=Burkholderia cenocepacia OX=95486 GN=lolA PE=3 SV=1 | Q.IVKAPAKGAS[+568.212]AAQAAPKPTDNSSGTF.V | HexNAc(2)Hex(1) | Yes | 764.1379 | 4 | 3053.5298 | 3053.5161 | 4.5 | 1082.9 | 1032.6 | 506.95 | Nsco_20191108_BC_ZIC_HILIC_H111_pepsin_B3.16451.16451.4 | 35.47 | Pepsin | B3 | S68 | 136 |
| BCAL3033 | >tr\|A0A2N9CM16\|A0A2N9CM16_9BURK Outer-membrane lipoprotein carrier protein OS=Burkholderia cenocepacia OX=95486 GN=lolA PE=3 SV=1 | Q.IVKAPAKGAS[+568.212]AAQAAPKPTDNSSGTFV.F | HexNAc(2)Hex(1) | Yes | 788.9051 | 4 | 3152.5986 | 3152.5845 | 4.5 | 805.9 | 805.9 | 378.72 | Nsco_20191108_BC_ZIC_HILIC_H111_pepsin_B3.19759.19759.4 | 41.093 | Pepsin | B3 | S68 | 137 |
| BCAL3033 | >tr\|A0A2N9CM16\|A0A2N9CM16_9BURK Outer-membrane lipoprotein carrier protein OS=Burkholderia cenocepacia OX=95486 GN=lolA PE=3 SV=1 | Q.IVKAPAKGAS[+568.212]AAQAAPKPTDNSSGT.F | HexNAc(2)Hex(1) | Yes | 727.371 | 4 | 2906.4621 | 2906.4477 | 5 | 727.77 | 543.13 | 403.44 | Nsco_20191108_BC_ZIC_HILIC_H111_thermolysin_B1.9738.9738.4 | 26.669 | Thermolysin | B1 | S68 | 138 |

| | | | | | | | | | | | | | | | | | | |
|---|---|---|---|---|---|---|---|---|---|---|---|---|---|---|---|---|---|---|
| BCAL3033 | >tr\|A0A2N9CM16\|A0A2N9CM16_9BURK Outer-membrane lipoprotein carrier protein OS=Burkholderia cenocepacia OX=95486 GN=lolA PE=3 SV=1 | Q.IVKAPAKGAS[+568.212]AAQAAPKPTDNSSGTFVF.A | HexNAc(2)Hex(1) | Yes | 825.6715 | 4 | 3299.6643 | 3299.6529 | 3.4 | 451.66 | 424.44 | 313.42 | Nsco_20191108_BC_ZIC_HILIC_H111_thermolysin_B3.26971.26971.4 | 55.985 | Thermolysin | B3 | S68 | 139 |
| BCAL3033 | >tr\|A0A2N9CM16\|A0A2N9CM16_9BURK Outer-membrane lipoprotein carrier protein OS=Burkholderia cenocepacia OX=95486 GN=lolA PE=3 SV=1 | K.GAS[+568.212]AAQAAPKPTDNSSGTFVFARPGK.F | HexNAc(2)Hex(1) | Yes | 776.3766 | 4 | 3102.4846 | 3101.4909 | -3.1 | 376.92 | 362.3 | 362.3 | Nsco_20191108_BC_ZIC_HILIC_H111_typsin_B3.24977.24977.4 | 53.711 | Trypsin | B3 | S68 | 140 |
| BCAL3033 | >tr\|A0A2N9CM16\|A0A2N9CM16_9BURK Outer-membrane lipoprotein carrier protein OS=Burkholderia cenocepacia OX=95486 GN=lolA PE=3 SV=1 | K.GASAAQAAPKPT[+568.212]DNSSGTFVFAR.P | HexNAc(2)Hex(1) | Yes | 940.448 | 3 | 2819.3293 | 2819.3218 | 2.7 | 537.54 | 363.25 | 188.19 | Nsco_20191108_BC_ZIC_HILIC_H111_typsin_B3.26102.26102.3 | 56.04 | Trypsin | B3 | S68 | 141 |
| BCAL3166 | >tr\|A0A1V2WZ65\|A0A1V2WZ65_9BURK BON domain-containing protein OS=Burkholderia cenocepacia OX=95486 GN=A8E72_33585 PE=4 SV=1 | A.TEAPAAATSAPKAAAKT[+568.212]AKKANRKLGY.A | HexNAc(2)Hex(1) | Yes | 1085.9139 | 3 | 3255.7273 | 3254.7114 | 3.8 | 197.65 | 97.64 | 15.49 | Nsco_20191108_BC_ZIC_HILIC_H111_pepsin_B3.12081.12081.3 | 28.676 | Pepsin | B3 | S40 | 142 |
| BCAL3377 | >tr\|A0A142PIH6\|A0A142PIH6_9BURK Outer membrane protein assembly factor BamE OS=Burkholderia cenocepacia OX=95486 GN=bamE PE=3 SV=1 | D.IDGDRGGKKAKAAAAAAKKAS[+568.212]EAAA.A | HexNAc(2)Hex(1) | Yes | 565.6979 | 5 | 2824.4605 | 2824.4534 | 2.5 | 649.03 | 597.74 | 597.74 | Nsco_20191108_BC_ZIC_HILIC_H111_pepsin_B2.4588.4588.5 | 17.418 | Pepsin | B2 | S145 | 143 |
| BCAL3377 | >tr\|A0A142PIH6\|A0A142PIH6_9BURK Outer membrane protein assembly factor BamE OS=Burkholderia cenocepacia OX=95486 GN=bamE PE=3 SV=1 | A.AAAASPATVPAS[+568.212]GAAVDQDANAQAARAANRATNQ.V | HexNAc(2)Hex(1) | Yes | 1249.5988 | 3 | 3746.7819 | 3746.77 | 3.2 | 149.53 | 95.59 | 48.93 | Nsco_20191108_BC_ZIC_HILIC_H111_thermolysin_B1.28240.28240.3 | 55.852 | Thermolysin | B1 | S168 | 144 |
| BCAL3377 | >tr\|A0A142PIH6\|A0A142PIH6_9BURK Outer membrane protein assembly factor BamE OS=Burkholderia cenocepacia OX=95486 GN=bamE PE=3 SV=1 | A.AASPATVPAS[+568.212]GAAVDQDANAQAARAANRATNQ.V | HexNAc(2)Hex(1) | Yes | 1202.2402 | 3 | 3604.706 | 3604.6957 | 2.8 | 151.2 | 73.1 | 18.05 | Nsco_20191108_BC_ZIC_HILIC_H111_thermolysin_B3.25905.25905.3 | 54.325 | Thermolysin | B3 | S168 | 145 |
| BCAL3377 | >tr\|A0A142PIH6\|A0A142PIH6_9BURK Outer membrane protein assembly factor BamE OS=Burkholderia cenocepacia OX=95486 GN=bamE PE=3 SV=1 | A.AAASPATVPAS[+568.212]GAAVDQDANAQAARAANR.A | HexNAc(2)Hex(1) | Yes | 1088.1919 | 3 | 3262.5612 | 3261.5466 | 3.5 | 183.94 | 140.81 | 140.81 | Nsco_20191108_BC_ZIC_HILIC_H111_thermolysin_B2.22542.22542.3 | 48.016 | Thermolysin | B2 | S168 | 146 |
| BCAL3377 | >tr\|A0A142PIH6\|A0A142PIH6_9BURK Outer membrane protein assembly factor BamE OS=Burkholderia cenocepacia OX=95486 GN=bamE PE=3 SV=1 | A.AAASPATVPAS[+568.212]GAAVDQDANAQAARAANRATNQ.V | HexNAc(2)Hex(1) | Yes | 1225.9203 | 3 | 3675.7465 | 3675.7329 | 3.7 | 169.36 | 108.65 | 25.51 | Nsco_20191108_BC_ZIC_HILIC_H111_thermolysin_B3.26466.26466.3 | 55.222 | Thermolysin | B3 | S168 | 147 |
| BCAL3377 | >tr\|A0A142PIH6\|A0A142PIH6_9BURK Outer membrane protein assembly factor BamE OS=Burkholderia cenocepacia OX=95486 GN=bamE PE=3 SV=1 | K.KASEAAAAAS[+568.212]AAQAAAAASPATVPASGAAVDQDANAQAAR.A | HexNAc(2)Hex(1) | Yes | 1172.5524 | 4 | 4687.1878 | 4687.1712 | 3.5 | 203.54 | 203.54 | 203.54 | Nsco_20191108_BC_ZIC_HILIC_H111_typsin_B1.36344.36344.4 | 73.494 | Trypsin | B1 | S152 | 148 |
| BCAL3377 | >tr\|A0A142PIH6\|A0A142PIH6_9BURK Outer membrane protein assembly factor BamE OS=Burkholderia cenocepacia OX=95486 GN=bamE PE=3 SV=1 | K.ASEAAAAASAAQAAAAASPAT[+568.212]VPAS[+568.212]GAAVDQDANAQAAR.A | HexNAc(2)Hex(1),HexNAc(2)Hex(1) | Yes (1 of 2) | 1520.3688 | 3 | 4559.0919 | 4559.0762 | 3.4 | 519.3 | 454.69 | 61 | Nsco_20191108_BC_ZIC_HILIC_H111_typsin_B2.43375.43375.3 | 84.583 | Trypsin | B2 | S145 | 149 |
| BCAL3377 | >tr\|A0A142PIH6\|A0A142PIH6_9BURK Outer membrane protein assembly factor BamE OS=Burkholderia cenocepacia OX=95486 GN=bamE PE=3 SV=1 | A.SEAAAAASAAQAAAAASPAT[+568.212][+100.064]VPAS[+568.212]GAAVDQDANAQAAR.A | HexNAc(2)Hex(1) 100.064,HexNAc(2)Hex(1) | Yes | 1340.6367 | 3 | 4019.8956 | 4019.8915 | 1 | 260.99 | 221.31 | 5.54 | Nsco_20191108_BC_ZIC_HILIC_H111_typsin_B3.45346.45346.3 | 93.292 | Trypsin | B3 | S145 | 150 |

| | | | | | | | | | | | | | | | | |
|---|---|---|---|---|---|---|---|---|---|---|---|---|---|---|---|---|
| BCAL3377 | >tr\|A0A142PIH6\|A0A142PIH6_9BURK Outer membrane protein assembly factor BamE OS=Burkholderia cenocepacia OX=95486 GN=bamE PE=3 SV=1 | A.KKASEAAAAAS[+568.212]AAQAAAAASPAT[+568.212]VPASGAAVDQDANAQAAR.A | HexNAc(2)Hex(1),HexNAc(2)Hex(1) | Yes (1 of 2) | 1204.5761 | 4 | 4815.2825 | 4815.2662 | 3.4 | 813.2 | 757.03 | 150.58 | Nsco_20191108_BC_ZIC_HILIC_H111_typsin_B3.30336.30336.4 | 64.258 | Trypsin | B3 | S145 | 151 |
| BCAL3377 | >tr\|A0A142PIH6\|A0A142PIH6_9BURK Outer membrane protein assembly factor BamE OS=Burkholderia cenocepacia OX=95486 GN=bamE PE=3 SV=1 | A.SAAQAAAAAS[+568.212]PAT[+568.212]VPAS[+568.212][+100.064]GAAVDQDANAQAAR.A | HexNAc(2)Hex(1),HexNAc(2)Hex(1),HexNAc(2)Hex(1) 100.064 | Partial | 1529.6973 | 3 | 4587.0774 | 4585.0545 | 3.5 | 201.77 | 105.38 | 0.38 | Nsco_20191108_BC_ZIC_HILIC_H111_typsin_B3.42021.42021.3 | 87.039 | Trypsin | B3 | NA | 152 |
| BCAL3469 | >tr\|A0A142PJ18\|A0A142PJ18_9BURK Cell division protein FtsL OS=Burkholderia cenocepacia OX=95486 GN=ftsL PE=3 SV=1 | A.IDAPIPAS[+568.212]ADTAGKGKGGAR.- | HexNAc(2)Hex(1) | Yes | 606.057 | 4 | 2421.206 | 2421.1991 | 2.9 | 540.75 | 477.57 | 169.63 | Nsco_20191108_BC_ZIC_HILIC_H111_thermolysin_B2.12795.12795.4 | 31.856 | Thermolysin | B2 | S101 | 153 |
| BCAM0505 | >tr\|A0A1V2WB29\|A0A1V2WB29_9BURK BON domain protein OS=Burkholderia cenocepacia OX=95486 GN=osmY_2 PE=4 SV=1 | A.QAS[+568.212]STDSGMASESNQPVTDTWITTKVKGELATTDGVKSTISVKT.V | HexNAc(2)Hex(1) | Partial | 1311.1262 | 4 | 5241.4831 | 5240.4897 | -1.9 | 319.81 | 319.81 | 1.4 | Nsco_20191108_BC_ZIC_HILIC_H111_thermolysin_B1.55291.55291.4 | 103.16 | Thermolysin | B1 | NA | 154 |
| BCAM0505 | >tr\|A0A1V2WB29\|A0A1V2WB29_9BURK BON domain protein OS=Burkholderia cenocepacia OX=95486 GN=osmY_2 PE=4 SV=1 | A.QASSTDSGMAS[+568.212]ESNQPVTDTWITTK.V | HexNAc(2)Hex(1) | Partial | 1606.2012 | 2 | 3211.3952 | 3210.4002 | -2.6 | 512.3 | 410.81 | 35.46 | Nsco_20191108_BC_ZIC_HILIC_H111_typsin_B2.41051.41051.2 | 80.551 | Trypsin | B2 | NA | 155 |
| BCAM0505 | >tr\|A0A1V2WB29\|A0A1V2WB29_9BURK BON domain protein OS=Burkholderia cenocepacia OX=95486 GN=osmY_2 PE=4 SV=1 | Q.ASSTDSGMASES[+568.212]NQPVTDTWITTK.V | HexNAc(2)Hex(1) | Yes | 1541.6788 | 2 | 3082.3502 | 3082.3416 | 2.8 | 589.8 | 443.36 | 24.29 | Nsco_20191108_BC_ZIC_HILIC_H111_typsin_B2.39568.39568.2 | 77.91 | Trypsin | B2 | S32 | 156 |
| BCAM0505 | >tr\|A0A1V2WB29\|A0A1V2WB29_9BURK BON domain protein OS=Burkholderia cenocepacia OX=95486 GN=osmY_2 PE=4 SV=1 | A.HAQASSTDSGMASES[+568.212]NQPVTDTWITTK.V | HexNAc(2)Hex(1) | Yes | 1140.1757 | 3 | 3418.5125 | 3418.4962 | 4.8 | 520.24 | 516.19 | 35.19 | Nsco_20191108_BC_ZIC_HILIC_H111_typsin_B3.29876.29876.3 | 63.391 | Trypsin | B3 | S32 | 157 |
| BCAM0690 | >tr\|A0A1V2Y2A4\|A0A1V2Y2A4_9BURK Flagellar motor protein OS=Burkholderia cenocepacia OX=95486 GN=A8F55_06330 PE=4 SV=1 | R.ALIDAGVPASSVFAAAFGS[+568.212]EQPVSSNADDEGRAK.N | HexNAc(2)Hex(1) | Yes | 1301.6209 | 3 | 3902.8482 | 3902.8302 | 4.6 | 167.6 | 138.68 | 22.81 | Nsco_20191108_BC_ZIC_HILIC_H111_typsin_B2.60235.60235.3 | 115.65 | Trypsin | B2 | S169 | 158 |
| BCAM0712 | >tr\|A0A2N9CPU2\|A0A2N9CPU2_9BURK Cobalt-zinc-cadmium resistance protein CzcB OS=Burkholderia cenocepacia OX=95486 GN=czcB PE=3 SV=1 | G.GGGASAPT[+568.212]AAEVAQPAAGAGAR.G | HexNAc(2)Hex(1) | Yes | 1204.0726 | 2 | 2407.138 | 2406.1267 | 3.3 | 387.01 | 160.83 | 160.83 | Nsco_20191108_BC_ZIC_HILIC_H111_typsin_B2.19918.19918.2 | 42.9 | Trypsin | B2 | S34 | 159 |
| BCAM0936 | >tr\|A0A1V2XJK6\|A0A1V2XJK6_9BURK Mechanosensitive ion channel protein OS=Burkholderia cenocepacia OX=95486 GN=A8F55_37840 PE=4 SV=1 | A.AAPAPAAASAAS[+568.212]DAAPALT[+568.212]PQQAR.Q | HexNAc(2)Hex(1),HexNAc(2)Hex(1) | Yes | 1656.7818 | 2 | 3312.5563 | 3311.5384 | 4.4 | 206.61 | 87.05 | 87.05 | Nsco_20191108_BC_ZIC_HILIC_H111_typsin_B2.28233.28233.2 | 57.819 | Trypsin | B2 | S28 | 160 |
| BCAM0988 | >tr\|A0A2N9CRH5\|A0A2N9CRH5_9BURK Putative exported protein OS=Burkholderia cenocepacia OX=95486 GN=F01_420505 PE=4 SV=1 | Q.PAAPTAGAS[+568.212]GPHVWSGA.I | HexNAc(2)Hex(1) | Yes | 1051.4849 | 2 | 2101.9625 | 2101.956 | 3.1 | 385.97 | 222.78 | 76.31 | Nsco_20191108_BC_ZIC_HILIC_H111_pepsin_B2.25602.25602.2 | 52.855 | Pepsin | B2 | S216 | 161 |
| BCAM0996 | >tr\|A0A2N9CRL1\|A0A2N9CRL1_9BURK Sporulation related OS=Burkholderia cenocepacia OX=95486 GN=F01_420513 PE=4 SV=1 | A.VAPRDDDVSDVQAGVAHDEPPAS[+568.212]DTT.V | HexNAc(2)Hex(1) | Yes | 1078.4798 | 3 | 3233.4248 | 3232.4135 | 2.5 | 215.74 | 117.92 | 27.68 | Nsco_20191108_BC_ZIC_HILIC_H111_thermolysin_B3.26931.26931.3 | 55.918 | Thermolysin | B3 | S104 | 162 |
| BCAM0996 | >tr\|A0A2N9CRL1\|A0A2N9CRL1_9BURK Sporulation related OS=Burkholderia cenocepacia OX=95486 GN=F01_420513 PE=4 SV=1 | K.DAAKPAAKPDTTTTASVTPPKPAPKPAAPAAKPAAPKPAPATVANAGPAS[+568.212]PDSGDASSPASPAGAR.F | HexNAc(2)Hex(1) | Partial | 1106.2348 | 6 | 6632.3723 | 6628.3541 | 0.7 | 431.46 | 429.91 | 31.05 | Nsco_20191108_BC_ZIC_HILIC_H111_typsin_B1.20631.20631.6 | 45.778 | Trypsin | B1 | NA | 163 |

| | | | | | | | | | | | | | | | | | | |
|---|---|---|---|---|---|---|---|---|---|---|---|---|---|---|---|---|---|---|
| BCAM0996 | >tr\|A0A2N9CRL1\|A0A2N9CRL1_9BURK Sporulation related OS=Burkholderia cenocepacia OX=95486 GN=F01_420513 PE=4 SV=1 | K.PAAPAAKPAAPKPAPAT[+568.212]VANAGPASPDSGDASSPASPAGAR.F | HexNAc(2)Hex(1) | Yes | 1048.2676 | 4 | 4190.0487 | 4186.0422 | -1.7 | 428.13 | 63.16 | 40.31 | Nsco_20191108_BC_ZIC_HILIC_H111_typsin_B1.19521.19521.4 | 43.77 | Trypsin | B1 | S178 | 164 |
| BCAM0996 | >tr\|A0A2N9CRL1\|A0A2N9CRL1_9BURK Sporulation related OS=Burkholderia cenocepacia OX=95486 GN=F01_420513 PE=4 SV=1 | R.DDDVSDVQAGVAHDEPPAS[+568.212]DTTVAAAPAPAPK.D | HexNAc(2)Hex(1) | Yes | 1228.2275 | 3 | 3682.6679 | 3682.6614 | 1.8 | 528.54 | 497.7 | 75.72 | Nsco_20191108_BC_ZIC_HILIC_H111_typsin_B2.37296.37296.3 | 73.805 | Trypsin | B2 | S104 | 165 |
| BCAM0996 | >tr\|A0A2N9CRL1\|A0A2N9CRL1_9BURK Sporulation related OS=Burkholderia cenocepacia OX=95486 GN=F01_420513 PE=4 SV=1 | P.AAPKPAPAT[+568.212][+100.064]VANAGPASPDSGDASSPASPAGAR.F | HexNAc(2)Hex(1) 100.064 | Yes | 1195.2382 | 3 | 3583.7001 | 3582.7045 | -2.2 | 221.21 | 171.96 | 42.4 | Nsco_20191108_BC_ZIC_HILIC_H111_typsin_B3.20792.20792.3 | 45.704 | Trypsin | B3 | T159 | 166 |
| BCAM0996 | >tr\|A0A2N9CRL1\|A0A2N9CRL1_9BURK Sporulation related OS=Burkholderia cenocepacia OX=95486 GN=F01_420513 PE=4 SV=1 | P.AAPAAKPAAPKPAPAT[+568.212][+100.064]VANAGPASPDSGDASSPASPAGAR.F | HexNAc(2)Hex(1) 100.064 | Partial | 1048.7652 | 4 | 4192.0389 | 4189.0535 | -5.9 | 255.61 | 226.89 | 37.83 | Nsco_20191108_BC_ZIC_HILIC_H111_typsin_B3.19863.19863.4 | 43.892 | Trypsin | B3 | NA | 167 |
| BCAM1550 | >tr\|A0A142PC85\|A0A142PC85_9BURK Peptidoglycan-associated protein OS=Burkholderia cenocepacia OX=95486 GN=pal PE=3 SV=1 | K.TPENAGAAPEPSSETVATVT[+568.212]ADDLNNPNSPLAKR.S | HexNAc(2)Hex(1) | Yes | 1335.3021 | 3 | 4003.8918 | 4002.8786 | 2.5 | 193.07 | 163.28 | 74.77 | Nsco_20191108_BC_ZIC_HILIC_H111_typsin_B1.41312.41312.3 | 82.212 | Trypsin | B1 | S37 | 168 |
| BCAM1550 | >tr\|A0A142PC85\|A0A142PC85_9BURK Peptidoglycan-associated protein OS=Burkholderia cenocepacia OX=95486 GN=pal PE=3 SV=1 | K.TPENAGAAPEPSSETVAT[+568.212]VTADDLNNPNSPLAK.R | HexNAc(2)Hex(1) | Yes | 1282.9354 | 3 | 3846.7915 | 3846.7775 | 3.7 | 737.69 | 370.44 | 26.1 | Nsco_20191108_BC_ZIC_HILIC_H111_typsin_B1.47218.47218.3 | 92.175 | Trypsin | B1 | S37 | 169 |
| BCAM1669 | >tr\|A0A1V2X7H1\|A0A1V2X7H1_9BURK Purine nucleoside phosphorylase OS=Burkholderia cenocepacia OX=95486 GN=A8E72_24650 PE=4 SV=1 | Q.ARVHGIDNS[+568.212]GAGSQPAAT.V | HexNAc(2)Hex(1) | Yes | 1139.0308 | 2 | 2277.0543 | 2277.0477 | 2.9 | 244.81 | 57.03 | 57.03 | Nsco_20191108_BC_ZIC_HILIC_H111_pepsin_B1.8943.8943.2 | 25.52 | Pepsin | B1 | S73 | 170 |
| BCAM1669 | >tr\|A0A1V2X7H1\|A0A1V2X7H1_9BURK Purine nucleoside phosphorylase OS=Burkholderia cenocepacia OX=95486 GN=A8E72_24650 PE=4 SV=1 | A.RVHGIDNSGAGS[+568.212]QPAA.T | HexNAc(2)Hex(1) | Yes | 1052.9897 | 2 | 2104.9721 | 2104.9629 | 4.3 | 181.24 | 18.99 | 18.99 | Nsco_20191108_BC_ZIC_HILIC_H111_pepsin_B3.8744.8744.2 | 23.545 | Pepsin | B3 | S73 | 171 |
| BCAM1669 | >tr\|A0A1V2X7H1\|A0A1V2X7H1_9BURK Purine nucleoside phosphorylase OS=Burkholderia cenocepacia OX=95486 GN=A8E72_24650 PE=4 SV=1 | R.VHGIDNSGAGSQPAAT[+568.212]VEGGAPVVRAQNPRDSVYF.G | HexNAc(2)Hex(1) | Yes | 1024.2471 | 4 | 4093.9665 | 4091.9429 | 4.1 | 402.09 | 402.09 | 43.31 | Nsco_20191108_BC_ZIC_HILIC_H111_thermolysin_B1.39133.39133.4 | 74.9 | Thermolysin | B1 | S73 | 172 |
| BCAM1669 | >tr\|A0A1V2X7H1\|A0A1V2X7H1_9BURK Purine nucleoside phosphorylase OS=Burkholderia cenocepacia OX=95486 GN=A8E72_24650 PE=4 SV=1 | R.VHGIDNSGAGSQPAAT[+568.212]VEGGAPVVRAQNPRDSVY.F | HexNAc(2)Hex(1) | Yes | 987.2284 | 4 | 3945.8917 | 3944.8744 | 3.5 | 888.69 | 843.46 | 140.13 | Nsco_20191108_BC_ZIC_HILIC_H111_thermolysin_B1.31465.31465.4 | 61.34 | Thermolysin | B1 | S73 | 173 |
| BCAM1669 | >tr\|A0A1V2X7H1\|A0A1V2X7H1_9BURK Purine nucleoside phosphorylase OS=Burkholderia cenocepacia OX=95486 GN=A8E72_24650 PE=4 SV=1 | R.VHGIDNSGAGS[+568.212]QPAATVEGGAP.V | HexNAc(2)Hex(1) | Partial | 1280.5865 | 2 | 2560.1657 | 2560.1533 | 4.8 | 240.26 | 124.47 | 124.47 | Nsco_20191108_BC_ZIC_HILIC_H111_thermolysin_B3.24497.24497.2 | 51.861 | Thermolysin | B3 | NA | 174 |
| BCAM1669 | >tr\|A0A1V2X7H1\|A0A1V2X7H1_9BURK Purine nucleoside phosphorylase OS=Burkholderia cenocepacia OX=95486 GN=A8E72_24650 PE=4 SV=1 | R.VHGIDNSGAGSQPAAT[+568.212]VEGGAPVVR.A | HexNAc(2)Hex(1) | Yes | 729.3552 | 4 | 2914.3992 | 2914.3912 | 2.7 | 958.01 | 894.26 | 149.49 | Nsco_20191108_BC_ZIC_HILIC_H111_typsin_B2.25408.25408.4 | 52.904 | Trypsin | B2 | S73 | 175 |

| | | | | | | | | | | | | | | | | | | |
|---|---|---|---|---|---|---|---|---|---|---|---|---|---|---|---|---|---|---|
| BCAM1737 | >tr\|A0A2N9CVE8\|A0A2N9CVE8_9BURK Putative exported glycoprotein OS=Burkholderia cenocepacia OX=95486 GN=F01_460192 PE=4 SV=1 | E.RAISWSQAGSAAPADT[+568.212]PASAAPSAS[+568.212]ATPATRA.A | HexNAc(2)Hex(1) | Yes (1 of 2) | 1378.3116 | 3 | 4132.9202 | 4132.9052 | 3.6 | 279.49 | 249.53 | 11.89 | Nsco_20191108_BC_ZIC_HILIC_H111_pepsin_B1.27720.27720.3 | 58.638 | Pepsin | B1 | S1820 | 176 |
| BCAM1737 | >tr\|A0A2N9CVE8\|A0A2N9CVE8_9BURK Putative exported glycoprotein OS=Burkholderia cenocepacia OX=95486 GN=F01_460192 PE=4 SV=1 | A.ISWSQAGSAAPADT[+568.212]PASAAPS[+568.212]ASATPATR.A | HexNAc(2)Hex(1),HexNAc(2)Hex(1) | Yes (1 of 2) | 1279.2545 | 3 | 3835.749 | 3834.7299 | 4.1 | 229.96 | 143.42 | 11.45 | Nsco_20191108_BC_ZIC_HILIC_H111_thermolysin_B2.34278.34278.3 | 68.064 | Thermolysin | B2 | S1820 | 177 |
| BCAM1737 | >tr\|A0A2N9CVE8\|A0A2N9CVE8_9BURK Putative exported glycoprotein OS=Burkholderia cenocepacia OX=95486 GN=F01_460192 PE=4 SV=1 | R.AISWSQAGSAAPADTPASAAPSAS[+568.212]ATPATR.A | HexNAc(2)Hex(1) | Yes | 1113.193 | 3 | 3337.5645 | 3337.5554 | 2.7 | 421.21 | 401.86 | 26.82 | Nsco_20191108_BC_ZIC_HILIC_H111_typsin_B1.36326.36326.3 | 73.457 | Trypsin | B1 | S1820 | 178 |
| BCAM2054 (typ | >tr\|A0A1V2XAF9\|A0A1V2XAF9_9BURK Peptide-binding protein OS=Burkholderia cenocepacia OX=95486 GN=bcscD PE=4 SV=1 | R.APEQAPAPPVAPLAS[+568.212]GAAAGVAQPVPGPTLLPREPAAGVSTK.E | HexNAc(2)Hex(1) | Yes | 1493.1215 | 3 | 4477.35 | 4477.3348 | 3.4 | 407.81 | 400.18 | 212.66 | Nsco_20191108_BC_ZIC_HILIC_H111_typsin_B1.55635.55635.3 | 105.77 | Trypsin | B1 | S416 | 179 |
| BCAM2055 | >tr\|A0A1V2VXC7\|A0A1V2VXC7_9BURK EscC/YscC/HrcC family type III secretion system outer membrane ring protein OS=Burkholderia cenocepacia OX=95486 GN=A8E72_28425 PE=3 SV=1 | D.VTVPLTPKPLPGTKFGPPALPPPKDTAAS[+568.212]QPAA.T | HexNAc(2)Hex(1) | Partial | 958.2687 | 4 | 3830.053 | 3830.0361 | 4.4 | 296.76 | 296.76 | 44.15 | Nsco_20191108_BC_ZIC_HILIC_H111_pepsin_B1.40237.40237.4 | 88.189 | Pepsin | B1 | NA | 180 |
| BCAM2063 | >tr\|A0A2N9CW94\|A0A2N9CW94_9BURK Porin OS=Burkholderia cenocepacia OX=95486 GN=F01_460501 PE=3 SV=1 | A.SSPAAAEPAAGASDAAAPAQQAADAAAPAPT[+568.212]GFWERSN.L | HexNAc(2)Hex(1) | Yes | 1374.963 | 3 | 4122.8745 | 4119.8538 | 2.6 | 470.77 | 441.26 | 113.52 | Nsco_20191108_BC_ZIC_HILIC_H111_thermolysin_B3.49036.49036.3 | 96.753 | Thermolysin | B3 | S46 | 181 |
| BCAM2064 | >tr\|A0A2N9CWA7\|A0A2N9CWA7_9BURK Periplasmic trehalase OS=Burkholderia cenocepacia OX=95486 GN=treA PE=3 SV=1 | A.DNANQAAAQAAGQSAIPATTAAAAAPASGT[+568.212]LPPPSQLYGDLFVAVQTAQLYPDQK.T | HexNAc(2)Hex(1) | Yes | 1494.9893 | 4 | 5976.9352 | 5976.9149 | 3.4 | 469.92 | 469.92 | 2.7 | Nsco_20191108_BC_ZIC_HILIC_H111_typsin_B2.79283.79283.4 | 160.56 | Trypsin | B2 | S80 | 182 |
| BCAM2289 | >tr\|A0A142PAR8\|A0A142PAR8_9BURK Purine nucleoside phosphorylase OS=Burkholderia cenocepacia OX=95486 GN=A8E72_27535 PE=4 SV=1 | D.TSGYGAQPAPLVHSGAPAAAS[+568.212]SNARD.S | HexNAc(2)Hex(1) | Yes | 1007.8045 | 3 | 3021.3989 | 3021.392 | 2.3 | 359.5 | 236.46 | 20.44 | Nsco_20191108_BC_ZIC_HILIC_H111_pepsin_B1.19533.19533.3 | 42.532 | Pepsin | B1 | S63 | 183 |
| BCAM2289 | >tr\|A0A142PAR8\|A0A142PAR8_9BURK Purine nucleoside phosphorylase OS=Burkholderia cenocepacia OX=95486 GN=A8E72_27535 PE=4 SV=1 | R.VHGADTSGYGAQPAPLVHSGAPAAAS[+568.212]SNAR.D | HexNAc(2)Hex(1) | Yes | 847.1534 | 4 | 3385.5918 | 3385.5779 | 4.1 | 1294.2 | 1230.2 | 71.23 | Nsco_20191108_BC_ZIC_HILIC_H111_typsin_B3.17844.17844.4 | 40.144 | Trypsin | B3 | S63 | 184 |
| BCAM2443 | >tr\|A0A2N9CXV2\|A0A2N9CXV2_9BURK Cytochrome c family protein OS=Burkholderia cenocepacia OX=95486 GN=F01_480355 PE=4 SV=1 | G.ATPQDAPAAAS[+568.212]APPPAPAAAAAPAAKPFTPPPES.A | HexNAc(2)Hex(1) | Yes | 1224.9368 | 3 | 3672.7959 | 3672.7803 | 4.3 | 356.59 | 69.23 | 69.23 | Nsco_20191108_BC_ZIC_HILIC_H111_pepsin_B2.38607.38607.3 | 81.685 | Pepsin | B2 | S38 | 185 |
| BCAM2681 | >tr\|A0A1V6L0G1\|A0A1V6L0G1_9BURK Putative lipoprotein OS=Burkholderia cenocepacia OX=95486 GN=F01_490209 PE=4 SV=1 | R.GASVAVHAGSAPSEAVGGGT[+568.212]PAEQVAALDPK.A | HexNAc(2)Hex(1) | Yes | 1124.2145 | 3 | 3370.6289 | 3369.618 | 2.2 | 344.3 | 294.43 | 32.01 | Nsco_20191108_BC_ZIC_HILIC_H111_typsin_B1.35984.35984.3 | 72.914 | Trypsin | B1 | S232 | 186 |
| BCAM2828 | >tr\|A0A2N9CZJ3\|A0A2N9CZJ3_9BURK Hopanoid biosynthesis associated RND transporter like protein HpnN OS=Burkholderia cenocepacia OX=95486 GN=hpnN PE=4 SV=1 | K.RAAIAAAAS[+568.212]ELLPALTQPAAPPATDAQR.V | HexNAc(2)Hex(1) | Yes | 1104.2439 | 3 | 3310.7172 | 3310.7013 | 4.8 | 435.85 | 424.66 | 122.25 | Nsco_20191108_BC_ZIC_HILIC_H111_typsin_B3.48545.48545.3 | 99.338 | Trypsin | B3 | S558 | 187 |
| BCAS0453 | >tr\|A0A142PSQ3\|A0A142PSQ3_9BURK Uncharacterized protein OS=Burkholderia cenocepacia OX=95486 GN=A8E72_07995 PE=4 SV=1 | R.DSLGNGVALDWPAS[+568.212]GVGGVADERQK.L | HexNAc(2)Hex(1) | Yes | 1023.1548 | 3 | 3067.4498 | 3066.4386 | 2.6 | 229.31 | 189.06 | 74.03 | Nsco_20191108_BC_ZIC_HILIC_H111_typsin_B1.47978.47978.3 | 93.418 | Trypsin | B1 | S98 | 188 |

| | | | | | | | | | | | | | | | | | | |
|---|---|---|---|---|---|---|---|---|---|---|---|---|---|---|---|---|---|---|
| I35_6143 | >tr\|A0A2N9CX34\|A0A2N9CX34_9BURK Uncharacterized protein OS=Burkholderia cenocepacia OX=95486 GN=F01_480128 PE=4 SV=1 | A.PAERSPFDTPDTRPLRVTS[+568.212]DAL.V | HexNAc(2)Hex(1) | Partial | 753.1206 | 4 | 3009.4607 | 3009.4535 | 2.4 | 684.21 | 615.43 | 14.88 | Nsco_20191108_BC_ZIC_HILIC_H111_pepsin_B1.34820.34820.4 | 73.999 | Pepsin | B1 | NA | 189 |
| I35_6143 | >tr\|A0A2N9CX34\|A0A2N9CX34_9BURK Uncharacterized protein OS=Burkholderia cenocepacia OX=95486 GN=F01_480128 PE=4 SV=1 | F.AAPAERSPFDTPDTRPLRVT[+568.212]SDAL.V | HexNAc(2)Hex(1) | Partial | 788.6396 | 4 | 3151.5366 | 3151.5277 | 2.8 | 221.14 | 221.14 | 8.73 | Nsco_20191108_BC_ZIC_HILIC_H111_pepsin_B1.35214.35214.4 | 74.891 | Pepsin | B1 | NA | 190 |

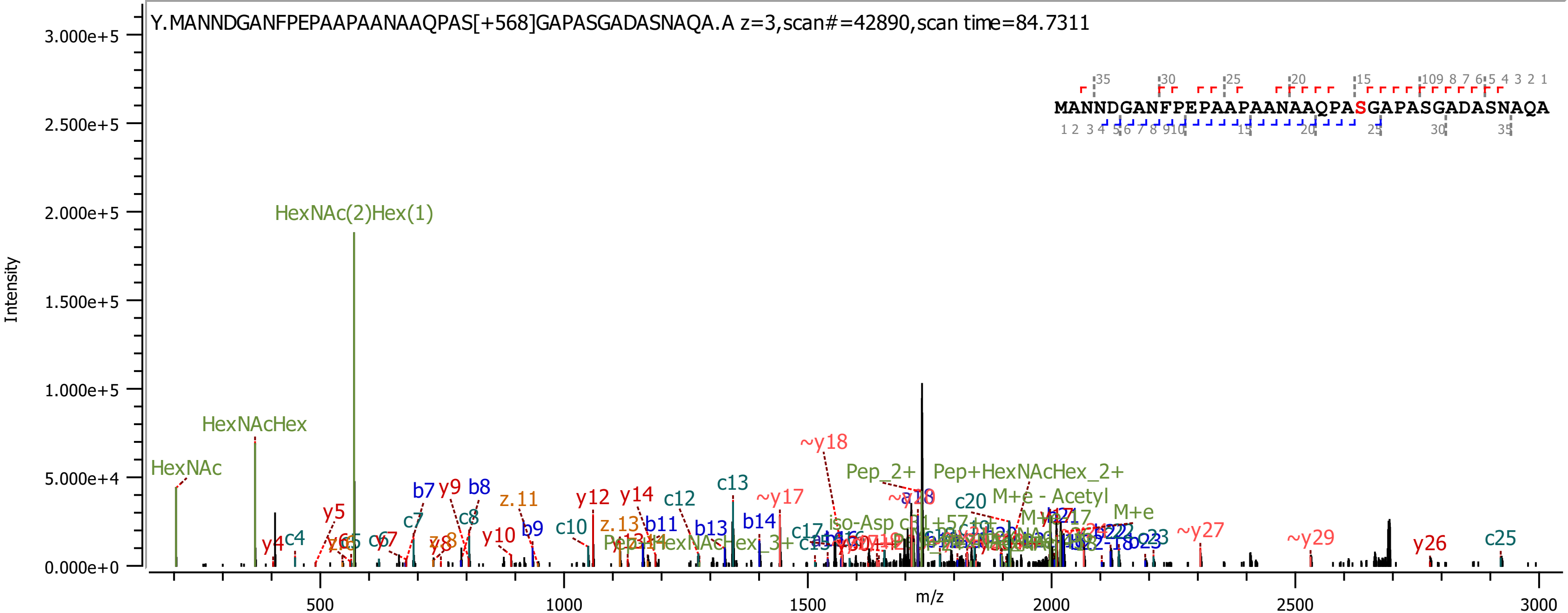

Y.MANNDGANFPEPAAPAANAAQPAS[+568]GAPASGADASNAQA.A z=3,scan#=42890,scan time=84.7311
MANNDGANFPEPAAPAANAAQPASGAPASGADASNAQA
Intensity
m/z
3.000e+5
2.500e+5
2.000e+5
1.500e+5
1.000e+5
5.000e+4
0.000e+0
500
1000
1500
2000
2500
3000
HexNAc(2)Hex(1)
HexNAcHex
HexNAc
y4
c4
y5
c7
b7
y9
b8
c8
z.11
y10
b9
c10
y12
z.13
y14
b11
c12
b13
c13
b14
~y17
~y18
Pep_2+
Pep+HexNAcHex_2+
c20
M+e - Acetyl
M+e
~y27
~y29
y26
c25

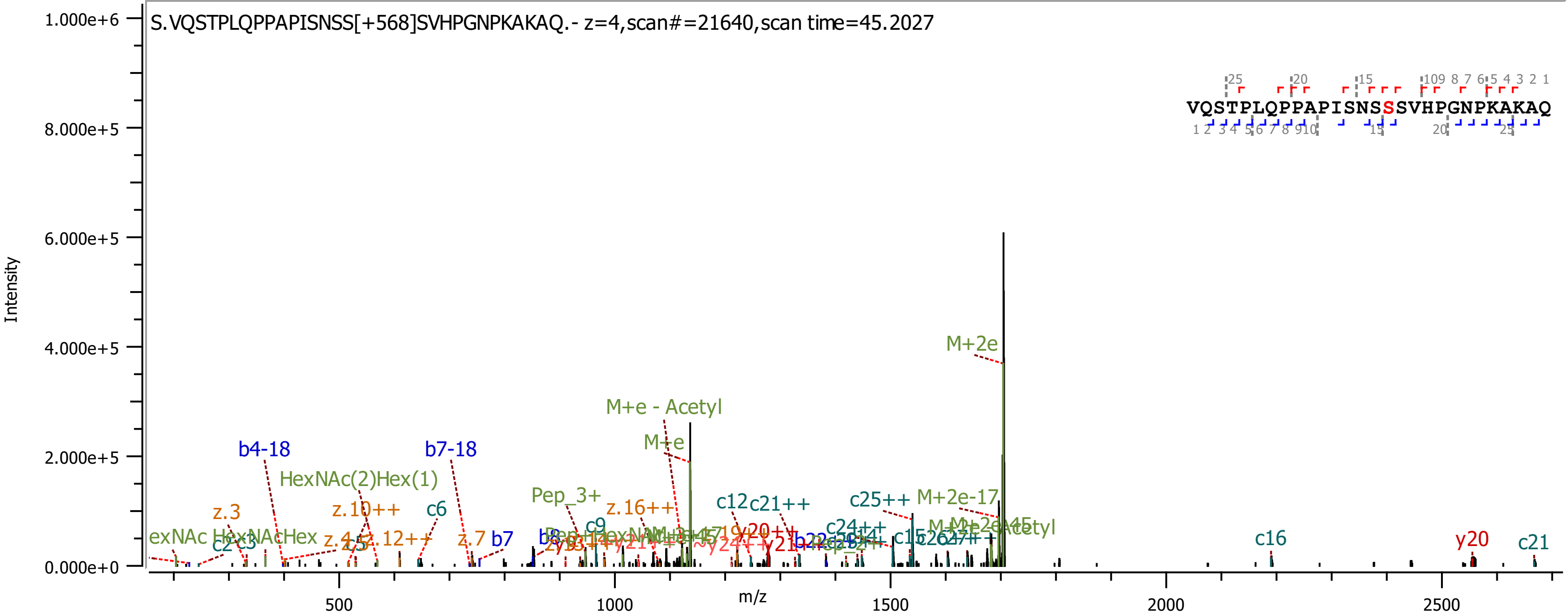

S.VQSTPLQPPAPISNSS[+568]SVHPGNPKAKAQ.- z=4,scan#=21640,scan time=45.2027
VQSTPLQPPAPISNSSSVHPGNPKAKAQ
Intensity
m/z
M+2e
M+e - Acetyl
M+e
b4-18
b7-18
HexNAc(2)Hex(1)
z.3
z.10++
c6
Pep_3+
z.16++
c12
c21++
c25++
M+2e-17
z.7
b7
c16
y20
c21

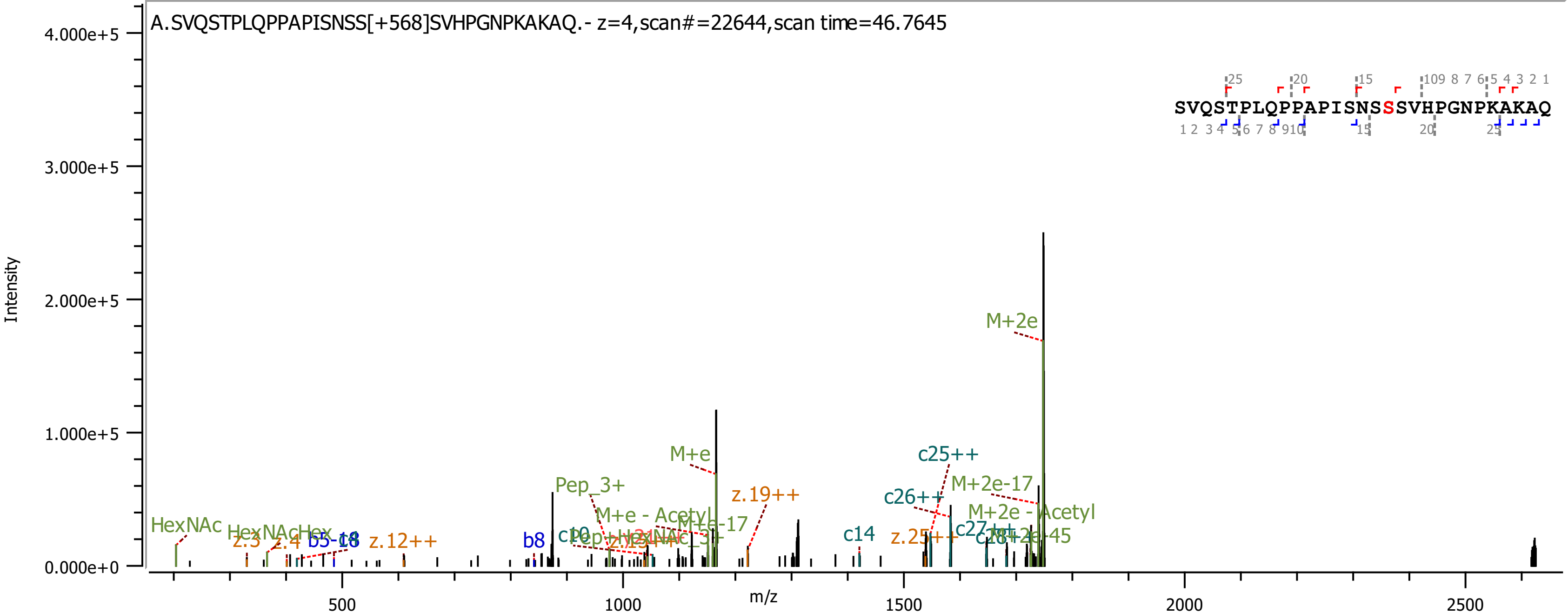

A.SVQSTPLQPPAPISNSS[+568]SVHPGNPKAKAQ.- z=4,scan#=22644,scan time=46.7645
SVQSTPLQPPAPISNSSSVHPGNPKAKAQ
Intensity
m/z
4.000e+5
3.000e+5
2.000e+5
1.000e+5
0.000e+0
500
1000
1500
2000
2500
M+2e
M+e
c25++
M+2e-17
Pep_3+
z.19++
c26++
M+e - Acetyl
M+e-17
M+2e - Acetyl
HexNAc
HexNAcHex
z.3
z.4
b5-18
z.12++
b8
c14
z.25++
c27++
M+2e-45

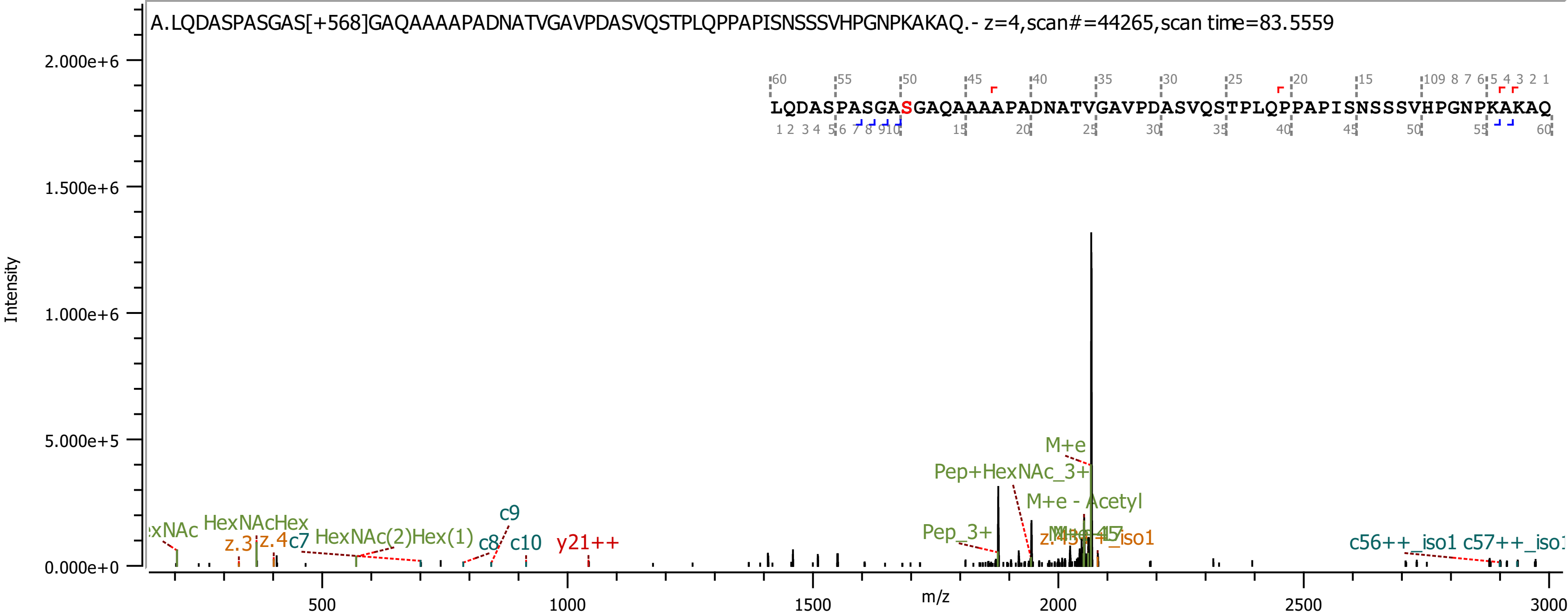
A.LQDASPASGAS[+568]GAQAAAAPADNATVGAVPDASVQSTPLQPPAPISNSSSVHPGNPKAKAQ.- z=4,scan#=44265,scan time=83.5559
LQDASPASGASGAQAAAAPADNATVGAVPDASVQSTPLQPPAPISNSSSVHPGNPKAKAQ
Intensity
2.000e+6
1.500e+6
1.000e+6
5.000e+5
0.000e+0
m/z
500
1000
1500
2000
2500
3000
HexNAcHex
z.3
z.4
c7
HexNAc(2)Hex(1)
c8
c9
c10
y21++
M+e
Pep+HexNAc_3+
M+e - Acetyl
Pep_3+
c56++_iso1

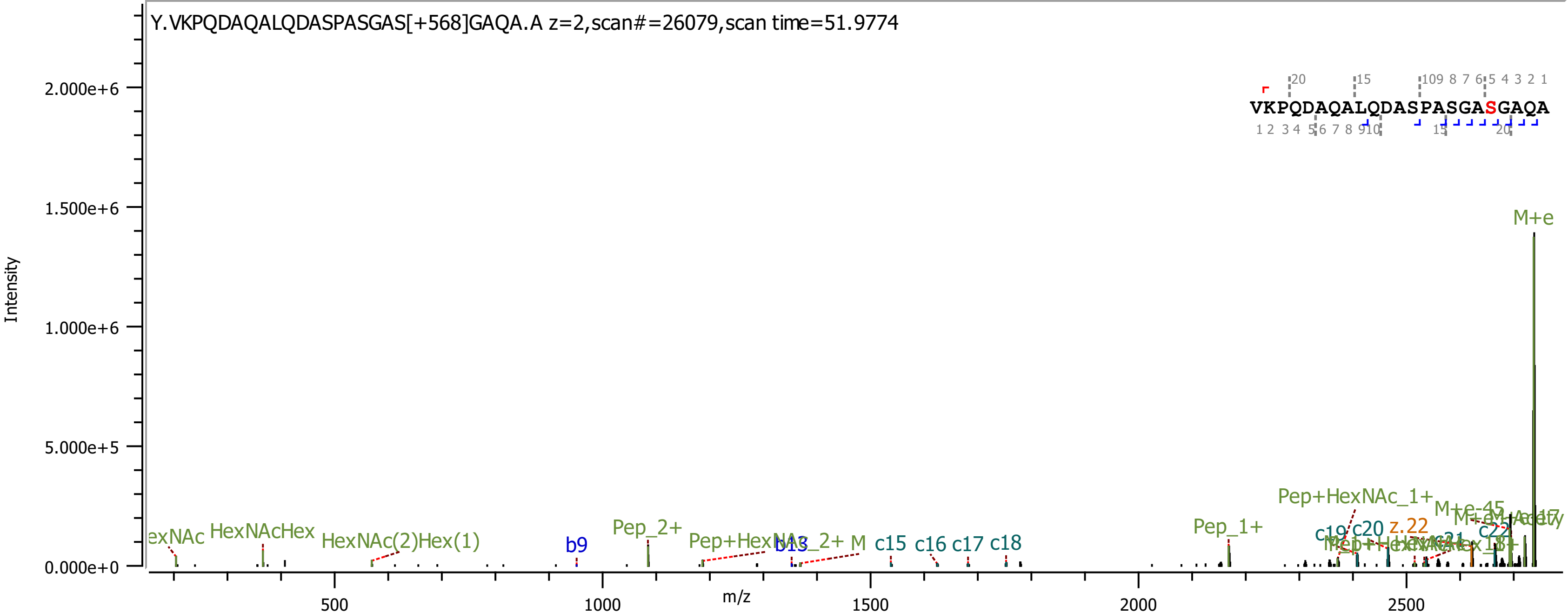
Y.VKPQDAQALQDASPASGAS[+568]GAQA.A z=2,scan#=26079,scan time=51.9774
VKPQDAQALQDASPASGASGAQA
Intensity
2.000e+6
1.500e+6
1.000e+6
5.000e+5
0.000e+0
m/z
500
1000
1500
2000
2500
M+e
HexNAcHex
HexNAc(2)Hex(1)
b9
Pep_2+
c15
c16
c17
c18
Pep_1+
Pep+HexNAc_1+
c20
z.22
M+e-45

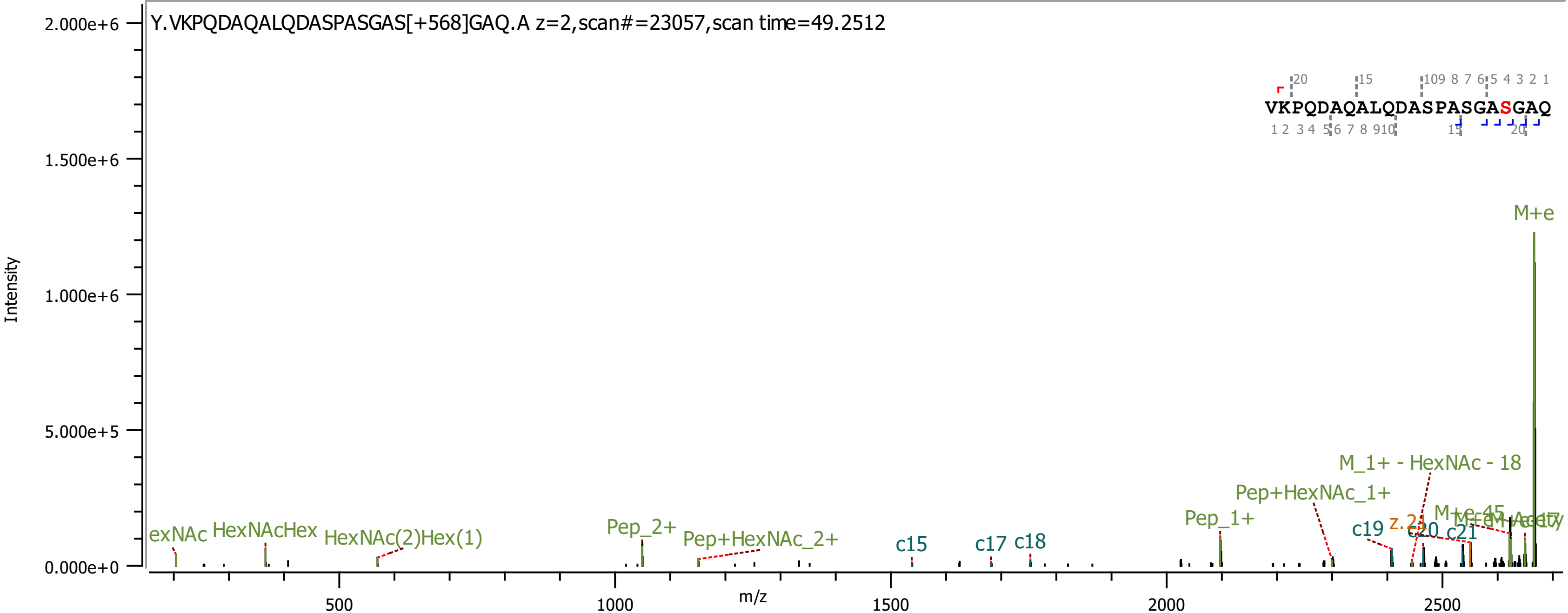
Y.VKPQDAQALQDASPASGAS[+568]GAQ.A z=2,scan#=23057,scan time=49.2512
Intensity
m/z
VKPQDAQALQDASPASGASGAQ
M+e
M_1+ - HexNAc - 18
Pep+HexNAc_1+
Pep_1+
Pep_2+
Pep+HexNAc_2+
HexNAcHex
HexNAc(2)Hex(1)
exNAc
c15
c17
c18
c19
c20
c21
z.21

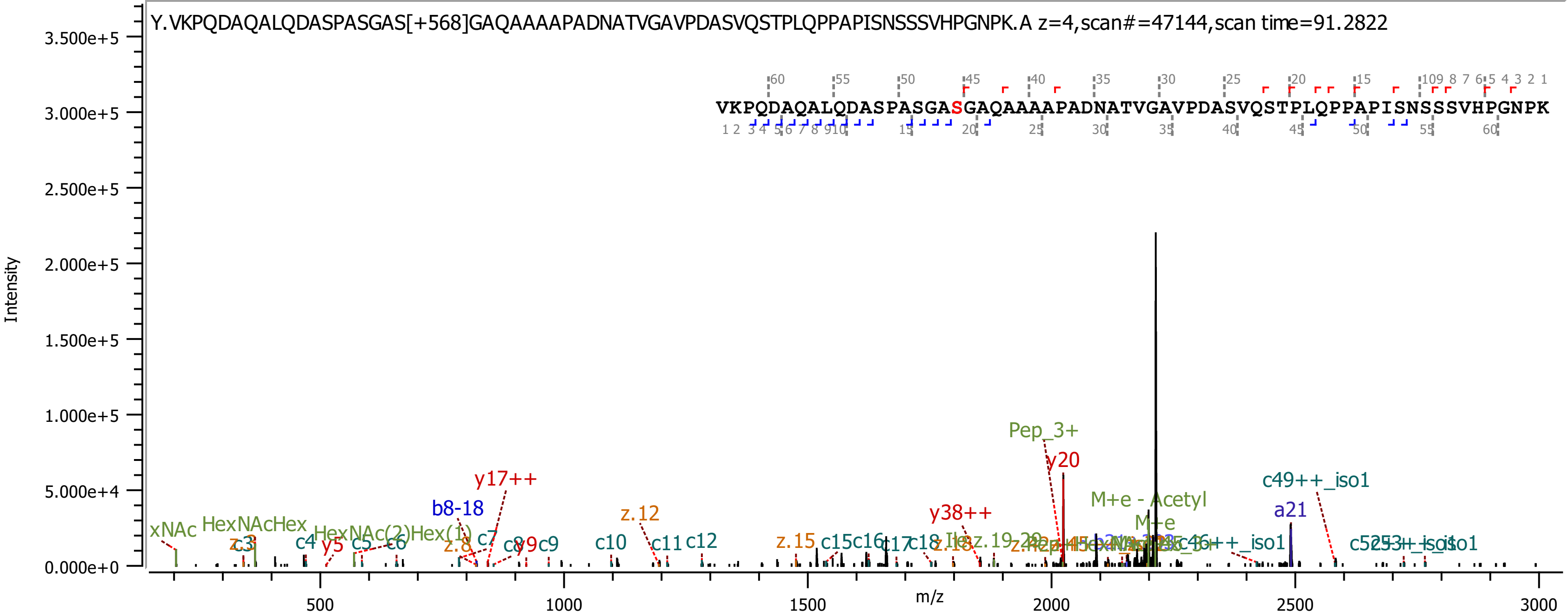

Y.VKPQDAQALQDASPASGAS[+568]GAQAAAAPADNATVGAVPDASVQSTPLQPPAPISNSSSVHPGNPK.A z=4,scan#=47144,scan time=91.2822
VKPQDAQALQDASPASGASGAQAAAAPADNATVGAVPDASVQSTPLQPPAPISNSSSVHPGNPK
Intensity
m/z
Pep_3+
y20
y17++
b8-18
z.12
y38++
c49++_iso1
a21
M+e - Acetyl
M+e
xNAc
HexNAcHex
HexNAc(2)Hex(1)
z.3
c3
c4
y5
c5
c6
c7
z.8
y9
c9
c10
c11
c12
z.15
c15
c16
c17
c18
z.18

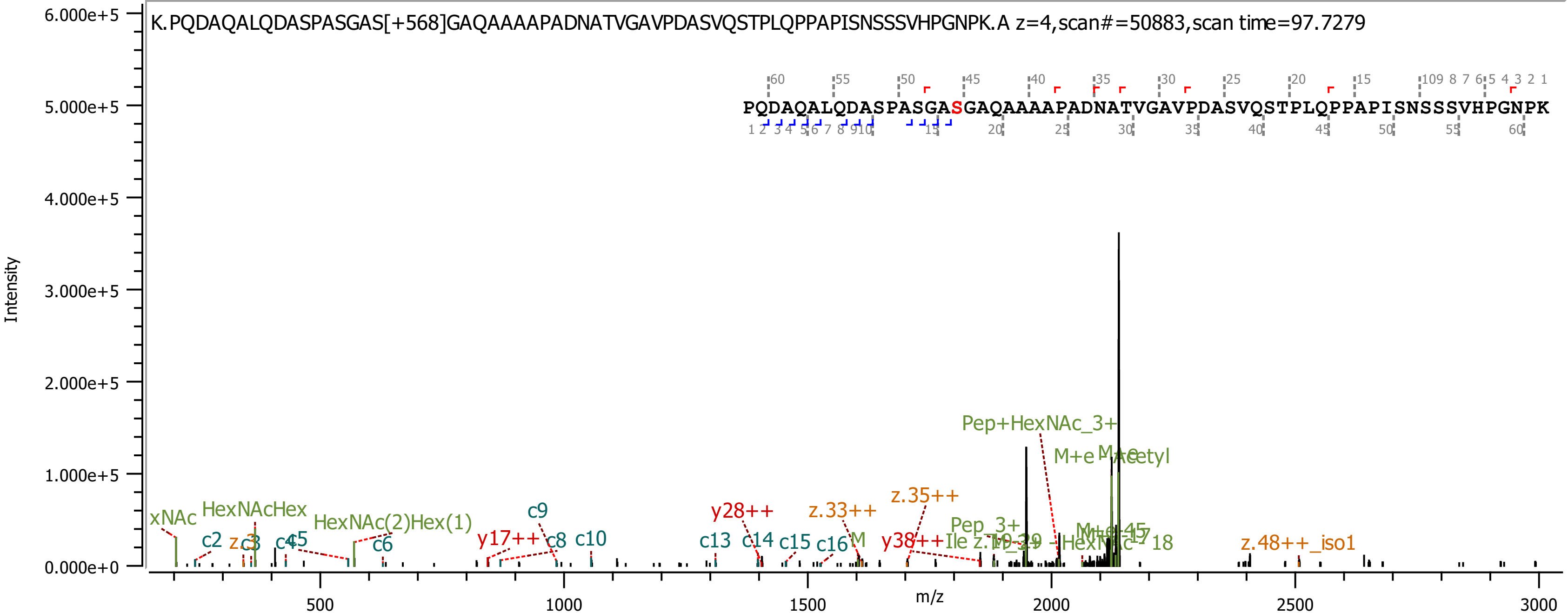

K.PQDAQALQDASPASGAS[+568]GAQAAAAPADNATVGAVPDASVQSTPLQPPAPISNSSSVHPGNPK.A z=4,scan#=50883,scan time=97.7279
PQDAQALQDASPASGASGAQAAAAPADNATVGAVPDASVQSTPLQPPAPISNSSSVHPGNPK
Intensity
m/z
Pep+HexNAc_3+
M+e
z.35++
z.33++
y28++
HexNAcHex
HexNAc(2)Hex(1)
c9
c10
c13
c14
c15
c16
y17++
c8
c6
c2
z.48++_iso1
Pep_3+

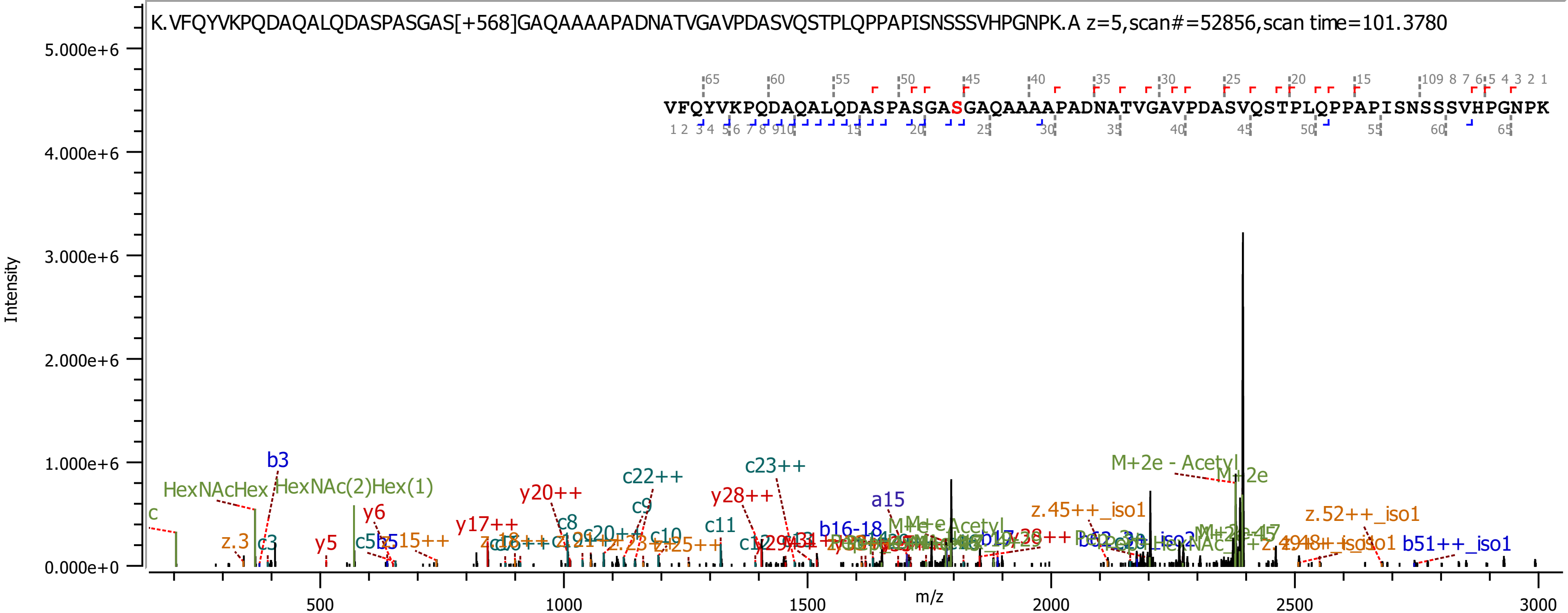
K.VFQYVKPQDAQALQDASPASGAS[+568]GAQAAAAPADNATVGAVPDASVQSTPLQPPAPISNSSSVHPGNPK.A z=5,scan#=52856,scan time=101.3780
VFQYVKPQDAQALQDASPASGASGAQAAAAPADNATVGAVPDASVQSTPLQPPAPISNSSSVHPGNPK
Intensity
m/z
5.000e+6
4.000e+6
3.000e+6
2.000e+6
1.000e+6
0.000e+0
500
1000
1500
2000
2500
3000
b3
HexNAcHex
HexNAc(2)Hex(1)
c
z.3
c3
y5
y6
c5
b5
z.15++
y17++
y20++
c8
c20++
c22++
c9
c10
c11
y28++
c23++
a15
b16-18
M+e - Acetyl
M+2e - Acetyl
M+2e
z.45++_iso1
z.52++_iso1
b51++_iso1

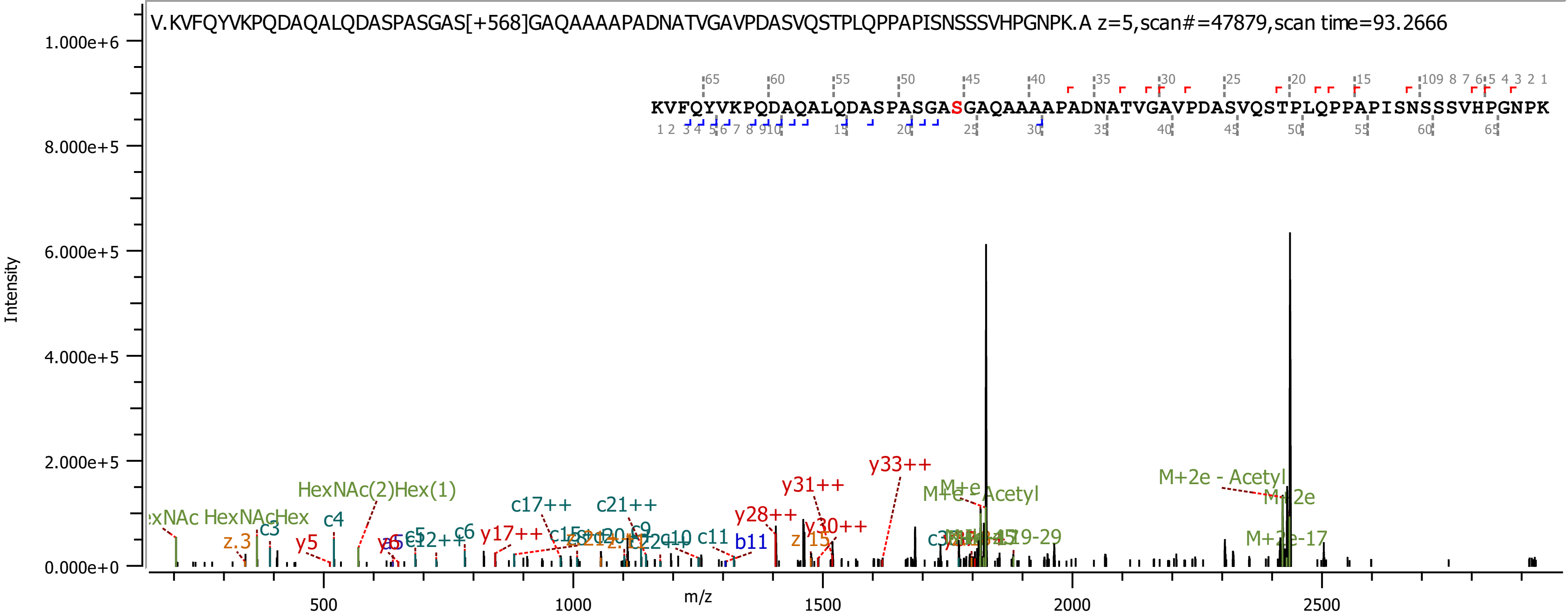

V.KVFQYVKPQDAQALQDASPASGAS[+568]GAQAAAAPADNATVGAVPDASVQSTPLQPPAPISNSSSVHPGNPK.A z=5,scan#=47879,scan time=93.2666
KVFQYVKPQDAQALQDASPASGASGAQAAAAPADNATVGAVPDASVQSTPLQPPAPISNSSSVHPGNPK
Intensity
m/z
1.000e+6
8.000e+5
6.000e+5
4.000e+5
2.000e+5
0.000e+0
500
1000
1500
2000
2500
HexNAc(2)Hex(1)
HexNAcHex
c3
c4
z.3
y5
c5
c6
y17++
c17++
c21++
c9
c10
c11
b11
y28++
y31++
y30++
z.15
y33++
M+e
M+e - Acetyl
M+2e - Acetyl
M+2e
M+2e-17

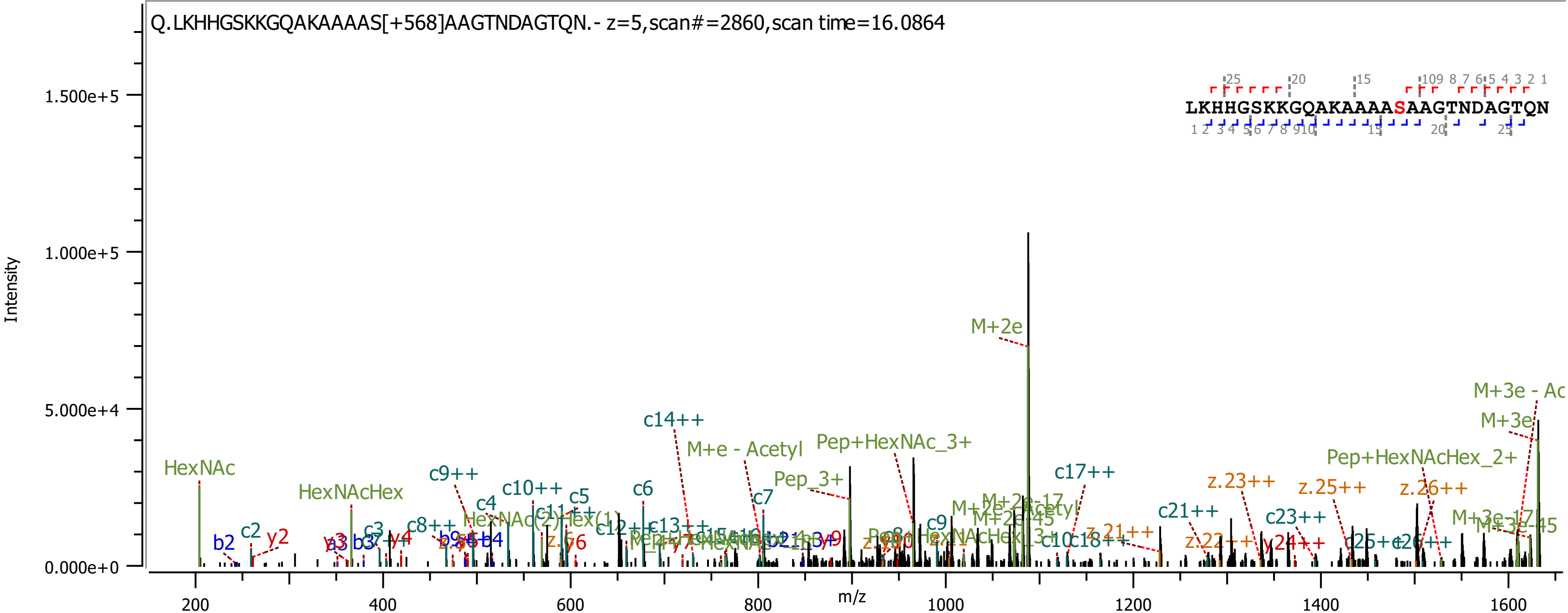
Q.LKHHGSKKGQAKAAAAS[+568]AAGTNDAGTQN.- z=5,scan#=2860,scan time=16.0864
LKHHGSKKGQAKAAAASAAGTNDAGTQN
Intensity
1.500e+5
1.000e+5
5.000e+4
0.000e+0
m/z
200
400
600
800
1000
1200
1400
1600
M+2e
HexNAc
HexNAcHex
c14++
M+e - Acetyl
Pep+HexNAc_3+
Pep_3+
c17++
Pep+HexNAcHex_2+
M+3e - Ac
M+3e
z.23++
z.25++
z.26++
c21++
c23++
c9++
c10++
c5
c6
c7
c4
c2
b2
y2
c9
y24++
M+3e-17
M+3e-45
M+2e-17
M+2e-45
z.21++
z.22++
c25++
c26++
c18++
c8++
y4
z.y6

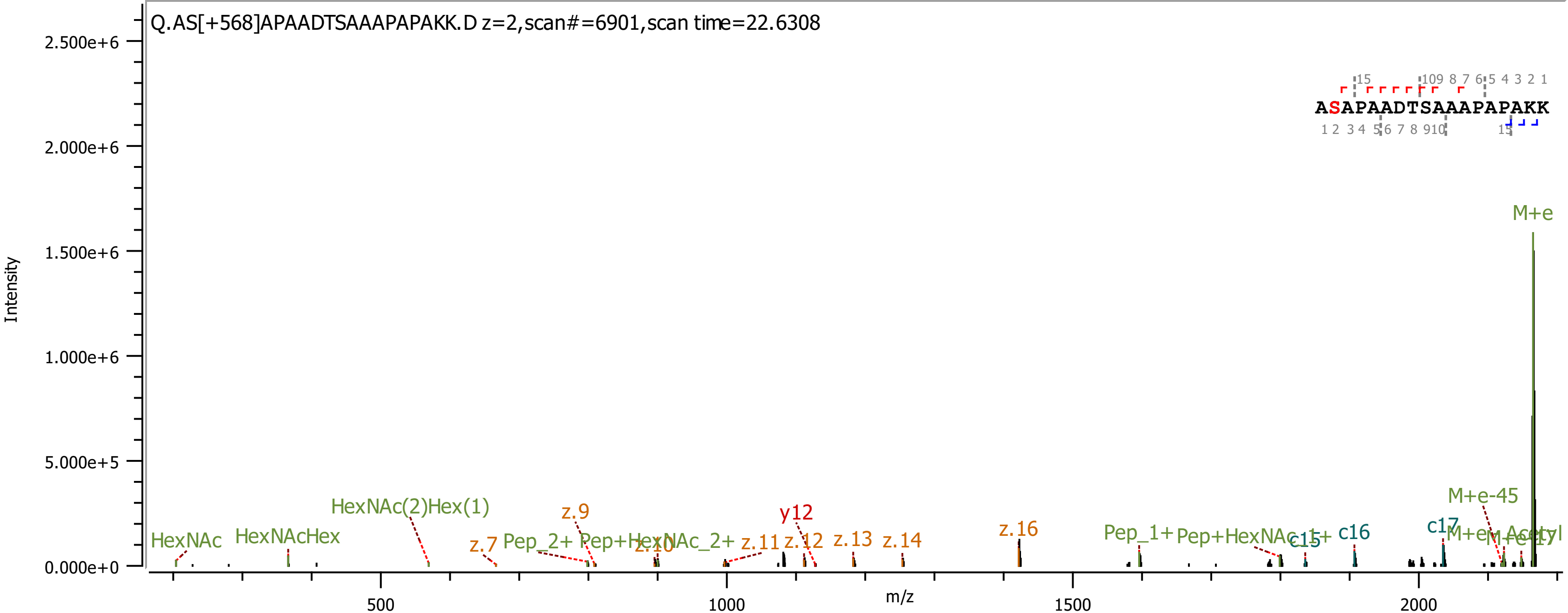

Q.AS[+568]APAADTSAAAPAPAKK.D z=2,scan#=6901,scan time=22.6308
ASAPAADTSAAAPAPAKK
Intensity
m/z
2.500e+6
2.000e+6
1.500e+6
1.000e+6
5.000e+5
0.000e+0
500
1000
1500
2000
M+e
M+e-45
HexNAc
HexNAcHex
HexNAc(2)Hex(1)
z.7
z.9
z.10
z.11
z.12
z.13
z.14
z.16
y12
Pep_2+
Pep+HexNAc_2+
Pep_1+
Pep+HexNAc_1+
c15
c16
c17

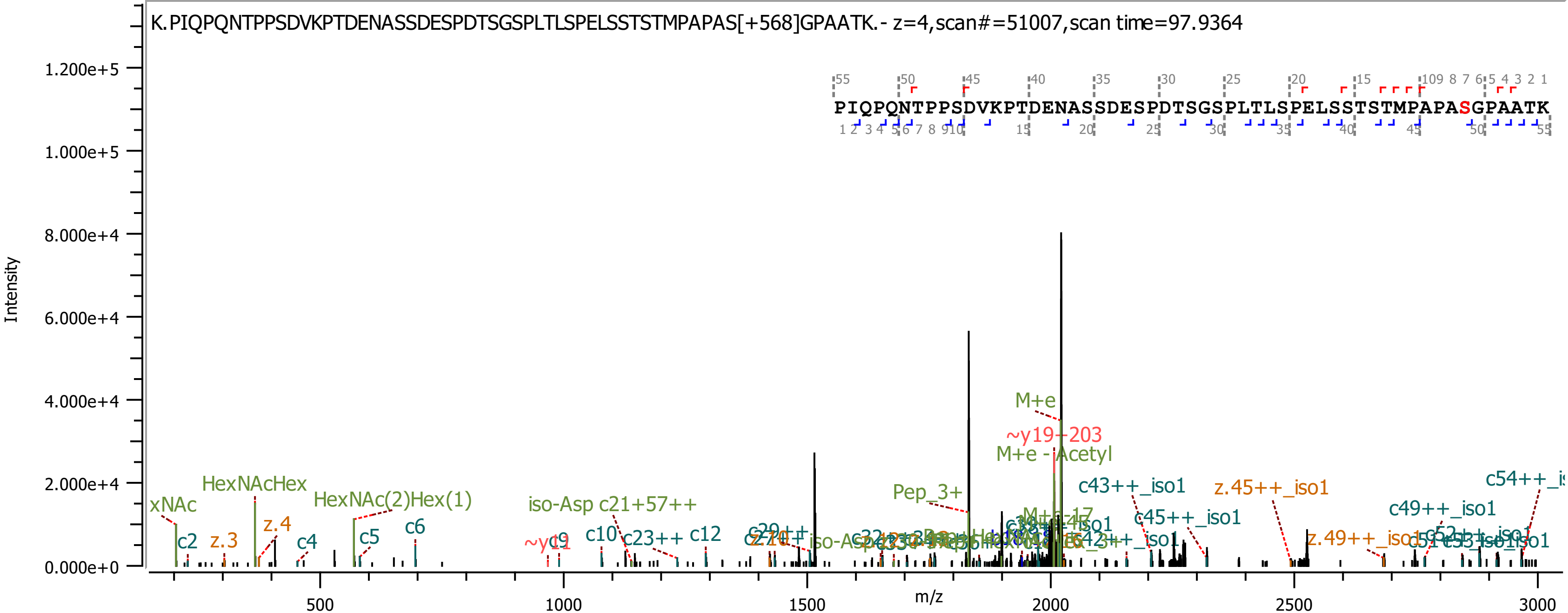
K.PIQPQNTPPSDVKPTDENASSDESPDTSGSPLTLSPELSSTSTMPAPAS[+568]GPAATK.- z=4,scan#=51007,scan time=97.9364
PIQPQNTPPSDVKPTDENASSDESPDTSGSPLTLSPELSSTSTMPAPASGPAATK
Intensity
m/z
1.200e+5
1.000e+5
8.000e+4
6.000e+4
4.000e+4
2.000e+4
0.000e+0
500
1000
1500
2000
2500
3000
xNAc
HexNAcHex
HexNAc(2)Hex(1)
iso-Asp c21+57++
Pep_3+
M+e
~y19-203
M+e - Acetyl
c43++_iso1
z.45++_iso1
c45++_iso1
c49++_iso1
c54++_i
c2
z.3
z.4
c4
c5
c6
c10
c23++
c12
z.49++_iso1

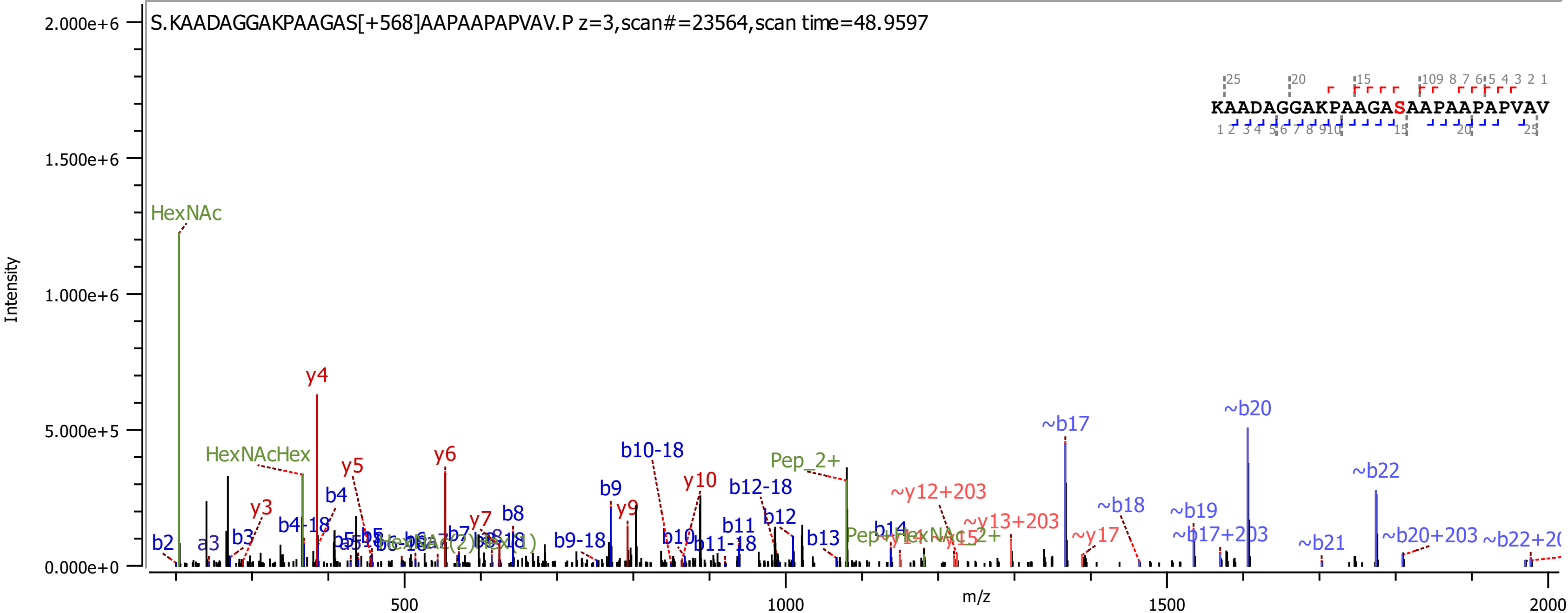

S.KAADAGGAKPAAGAS[+568]AAPAAPAPVAV.P z=3,scan#=23564,scan time=48.9597
KAADAGGAKPAAGASAAPAAPAPVAV
Intensity
m/z
HexNAc
HexNAcHex
y4
y5
y6
y3
b4
b4-18
y7
b8
b9
y9
b10-18
y10
b12-18
b12
b11
b11-18
b10
b13
b14
b9-18
Pep_2+
~y12+203
~y13+203
~b17
~b20
~b22
~b18
~b19
~b17+203
~b21
~b20+203
~y17
b2
a3
b3

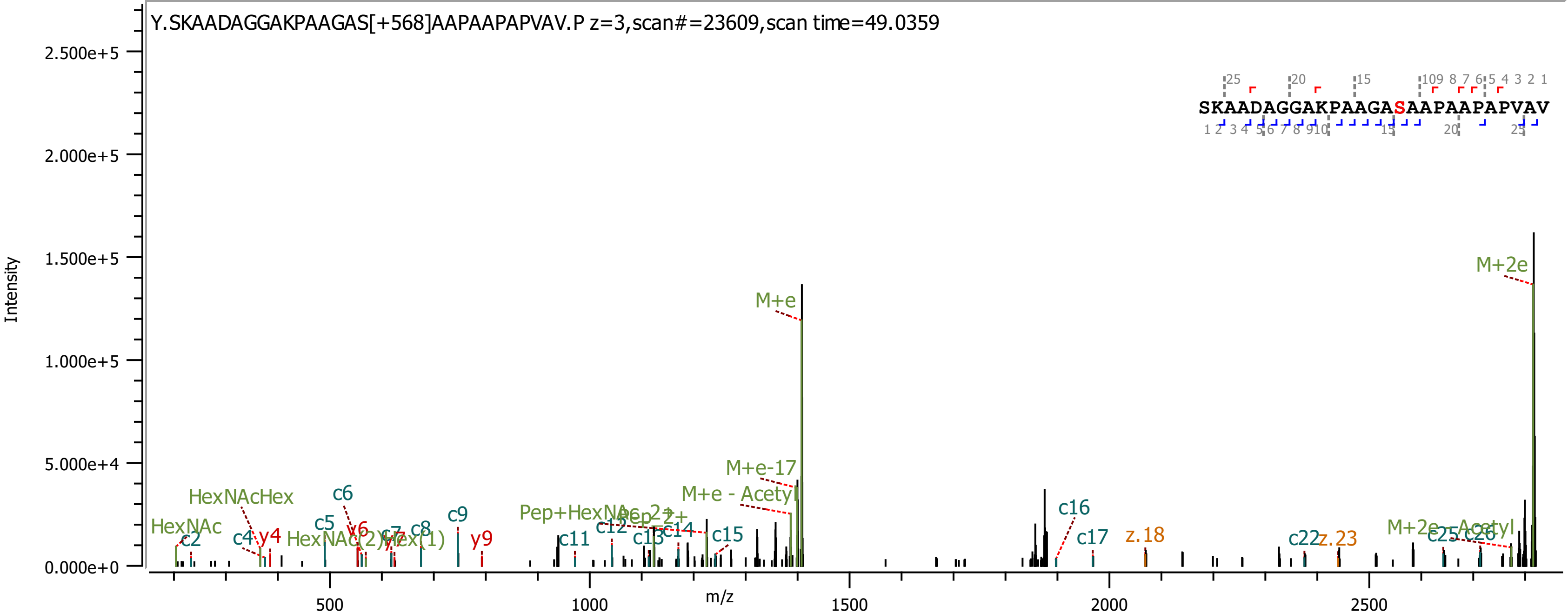

Y.SKAADAGGAKPAAGAS[+568]AAPAAPAPVAV.P z=3,scan#=23609,scan time=49.0359
SKAADAGGAKPAAGASAAPAAPAPVAV
Intensity
m/z
M+e
M+2e
M+e-17
M+e - Acetyl
M+2e - Acetyl
HexNAc
HexNAcHex
HexNAc(2)Hex(1)
Pep+HexNAc 2+
Pep 2+
c2
c4
c5
c6
c7
c8
c9
c11
c12
c13
c14
c15
c16
c17
c22
c25
c26
y4
y6
y7
y9
z.18
z.23

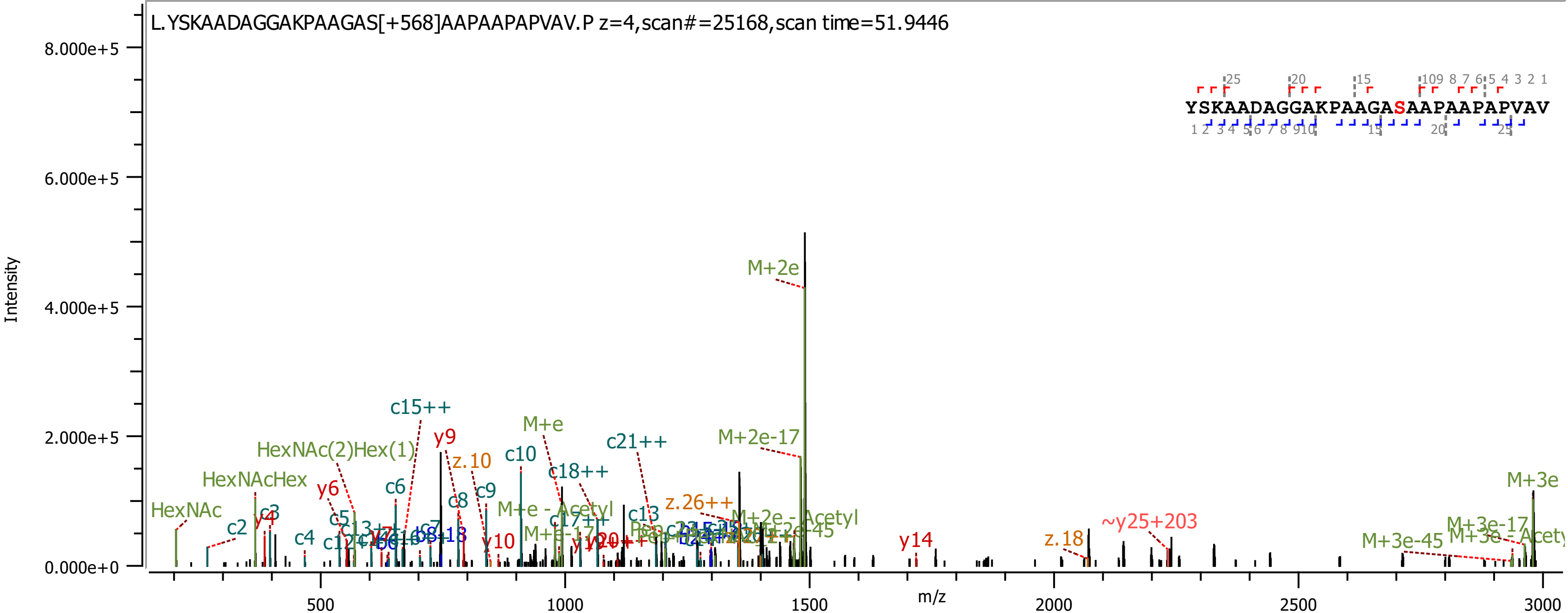
L.YSKAADAGGAKPAAGAS[+568]AAPAAPAPVAV.P z=4,scan#=25168,scan time=51.9446
Intensity
m/z
YSKAADAGGAKPAAGASAAPAAPAPVAV
M+2e
M+2e-17
M+3e
M+e
c15++
y9
z.10
c10
c21++
HexNAc(2)Hex(1)
HexNAcHex
HexNAc
c18++
c6
y6
c9
c8
M+e - Acetyl
z.26++
c13
c2
y4
c3
c4
c5
y10
y14
z.18
~y25+203
M+3e-45
M+3e-17
M+2e - Acetyl
0.000e+0
2.000e+5
4.000e+5
6.000e+5
8.000e+5
500
1000
1500
2000
2500
3000

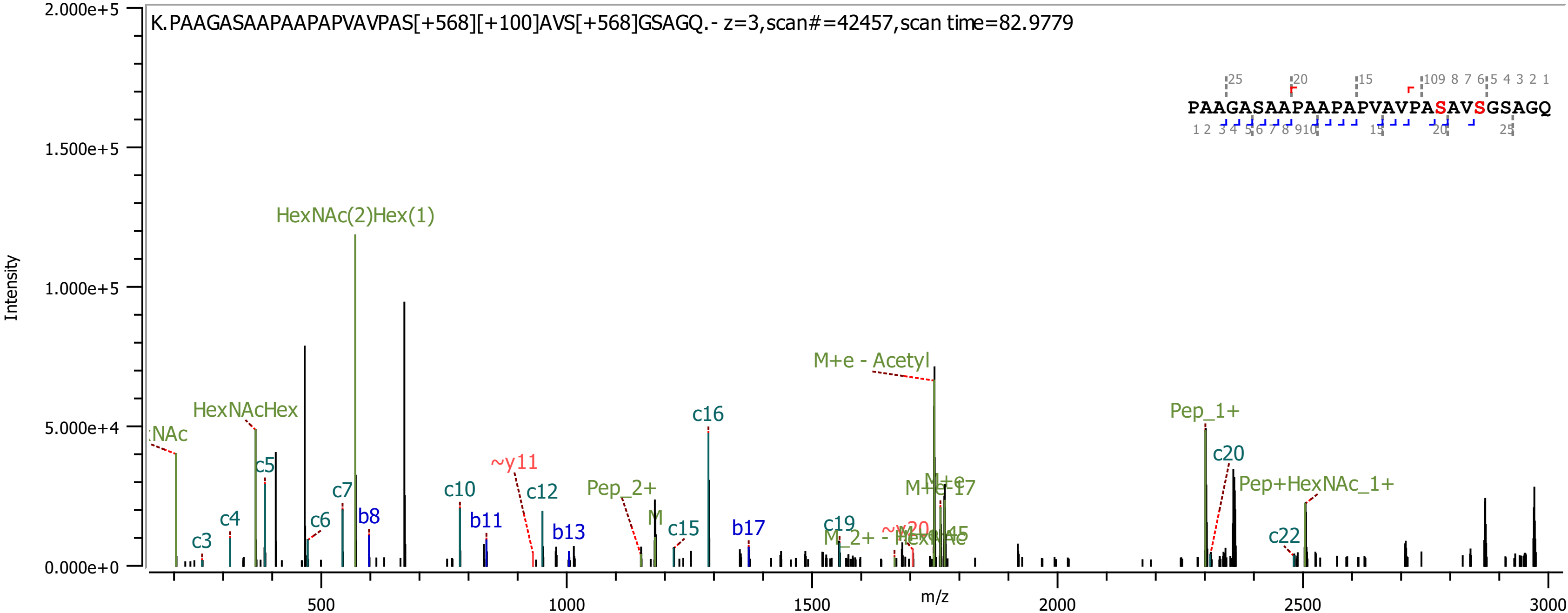

K.PAAGASAAPAAPAPVAVPAS[+568][+100]AVS[+568]GSAGQ.- z=3,scan#=42457,scan time=82.9779
PAAGASAAPAAPAPVAVPASAVSGSAGQ
Intensity
m/z
HexNAc(2)Hex(1)
HexNAcHex
M+e - Acetyl
Pep_1+
Pep+HexNAc_1+
Pep_2+
~y11
c3
c4
c5
c6
c7
b8
c10
b11
c12
b13
c15
c16
b17
c19
c20
c22

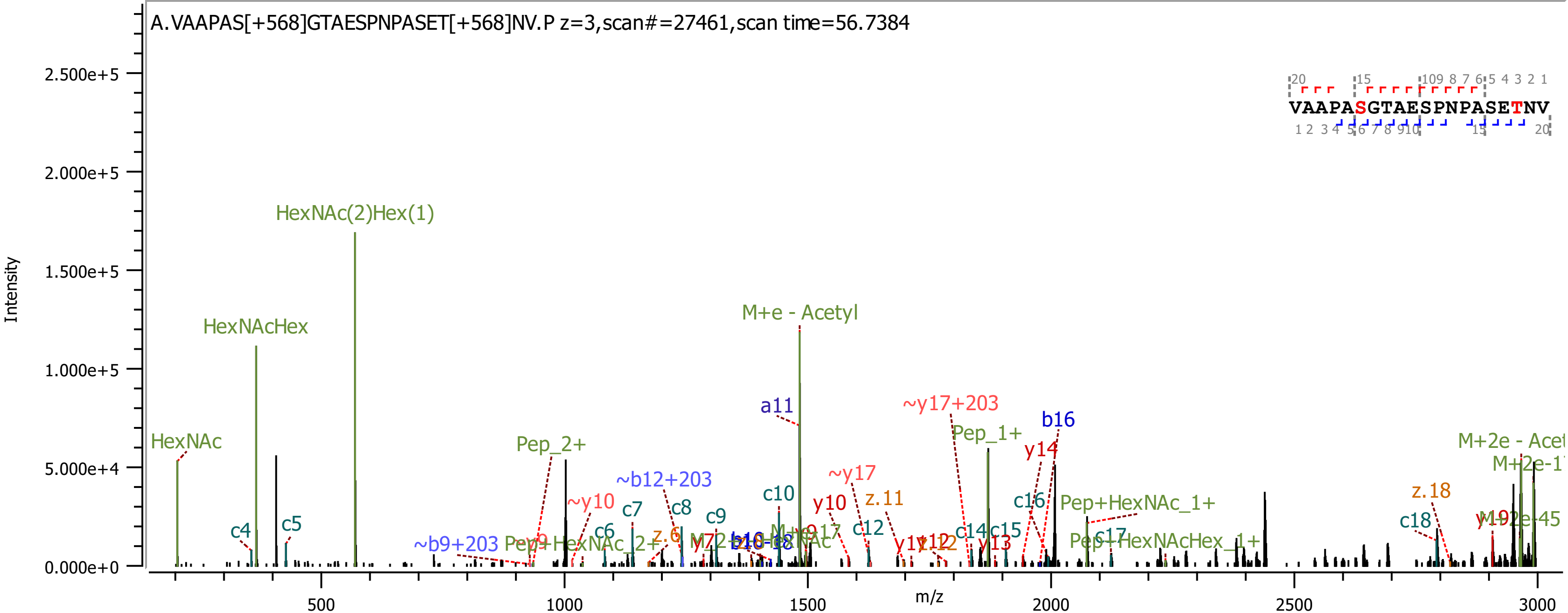
A.VAAPAS[+568]GTAESPNPASET[+568]NV.P z=3,scan#=27461,scan time=56.7384
VAAPASGTAESPNPASETNV
Intensity
2.500e+5
2.000e+5
1.500e+5
1.000e+5
5.000e+4
0.000e+0
m/z
500
1000
1500
2000
2500
3000
HexNAc(2)Hex(1)
HexNAcHex
HexNAc
M+e - Acetyl
a11
~y17+203
b16
Pep_1+
Pep_2+
y14
~b12+203
~y10
~y17
c4
c5
c6
c7
c8
c9
c10
c12
c14
c15
c16
c17
c18
y10
z.11
z.18
~b9+203
Pep+HexNAc_1+
Pep+HexNAcHex_1+
y13
M+2e - Acet
M+2e-1

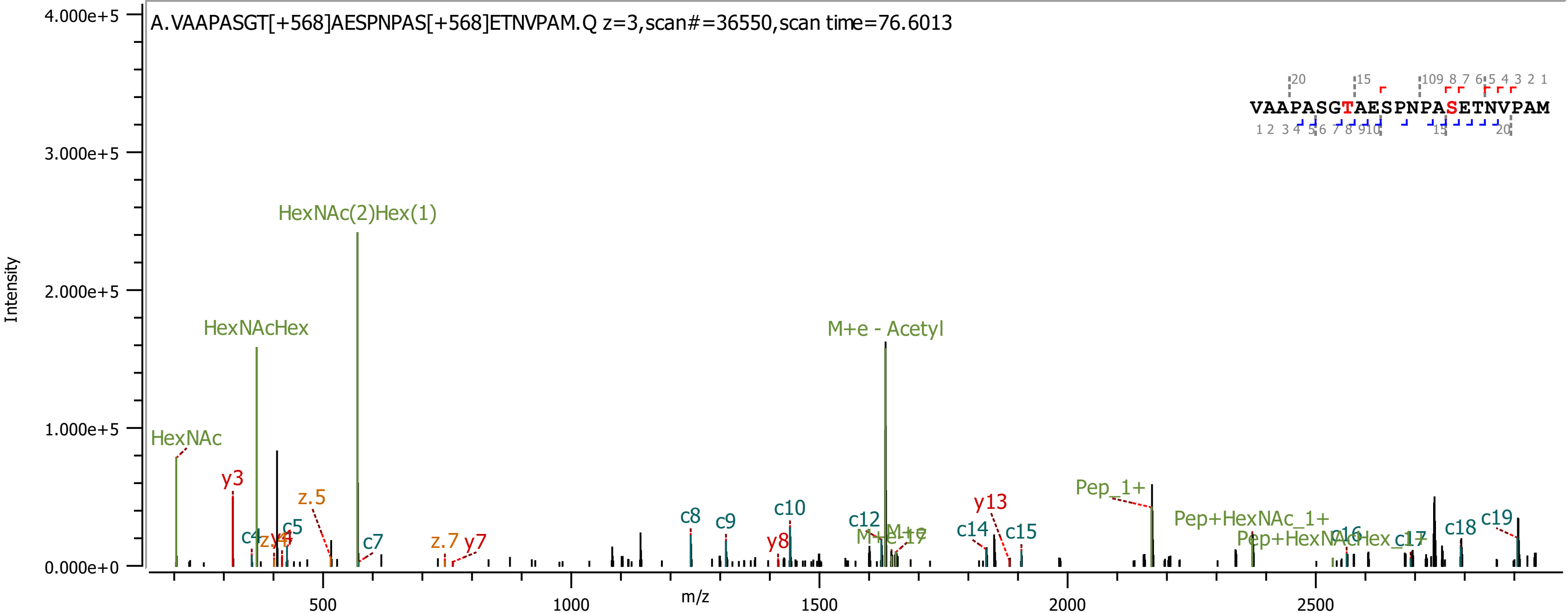
A.VAAPASGT[+568]AESPNPAS[+568]ETNVPAM.Q z=3,scan#=36550,scan time=76.6013
VAAPASGTAESPNPASETNVPAM
Intensity
m/z
HexNAc(2)Hex(1)
HexNAcHex
M+e - Acetyl
HexNAc
y3
z.5
c5
c7
z.7
y7
c8
c9
c10
y8
c12
c14
y13
c15
Pep_1+
Pep+HexNAc_1+
c18
c19
4.000e+5
3.000e+5
2.000e+5
1.000e+5
0.000e+0
500
1000
1500
2000
2500

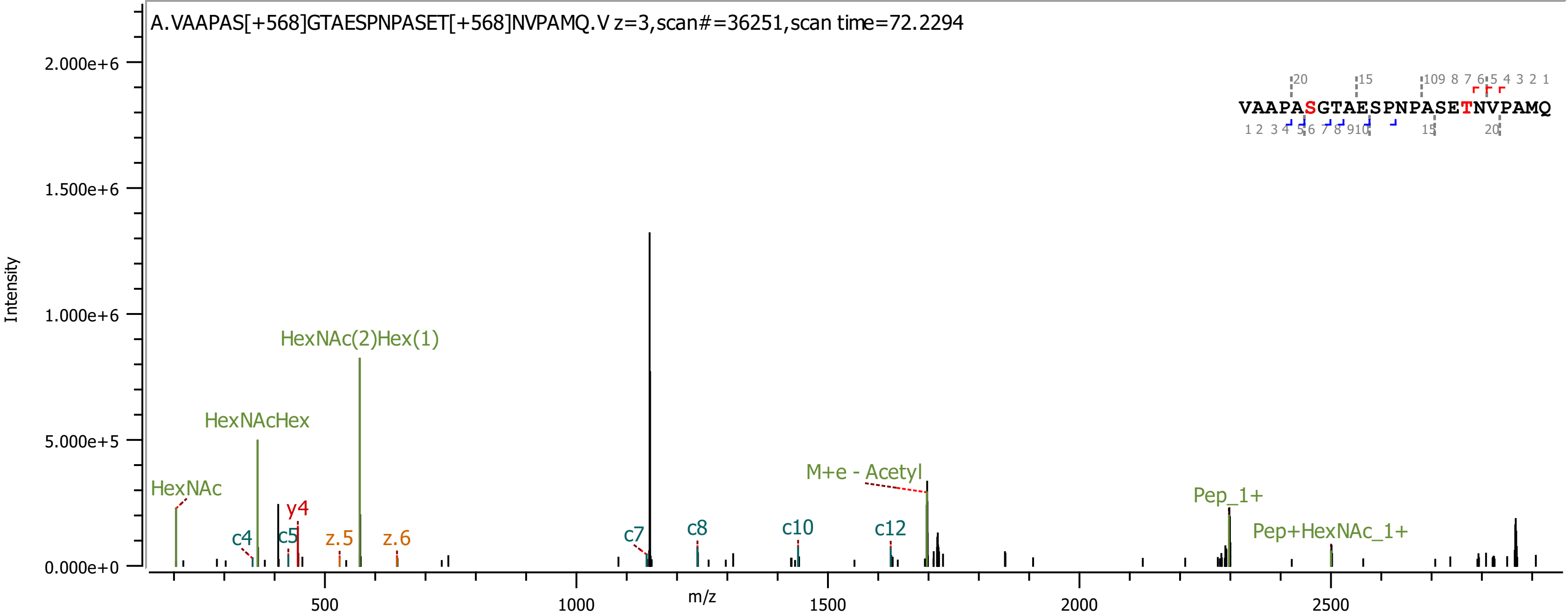

A.VAAPAS[+568]GTAESPNPASET[+568]NVPAMQ.V z=3,scan#=36251,scan time=72.2294
VAAPASGTAESPNPASETNVPAMQ
Intensity
m/z
HexNAc
HexNAcHex
HexNAc(2)Hex(1)
c4
c5
y4
z.5
z.6
c7
c8
c10
c12
M+e - Acetyl
Pep_1+
Pep+HexNAc_1+

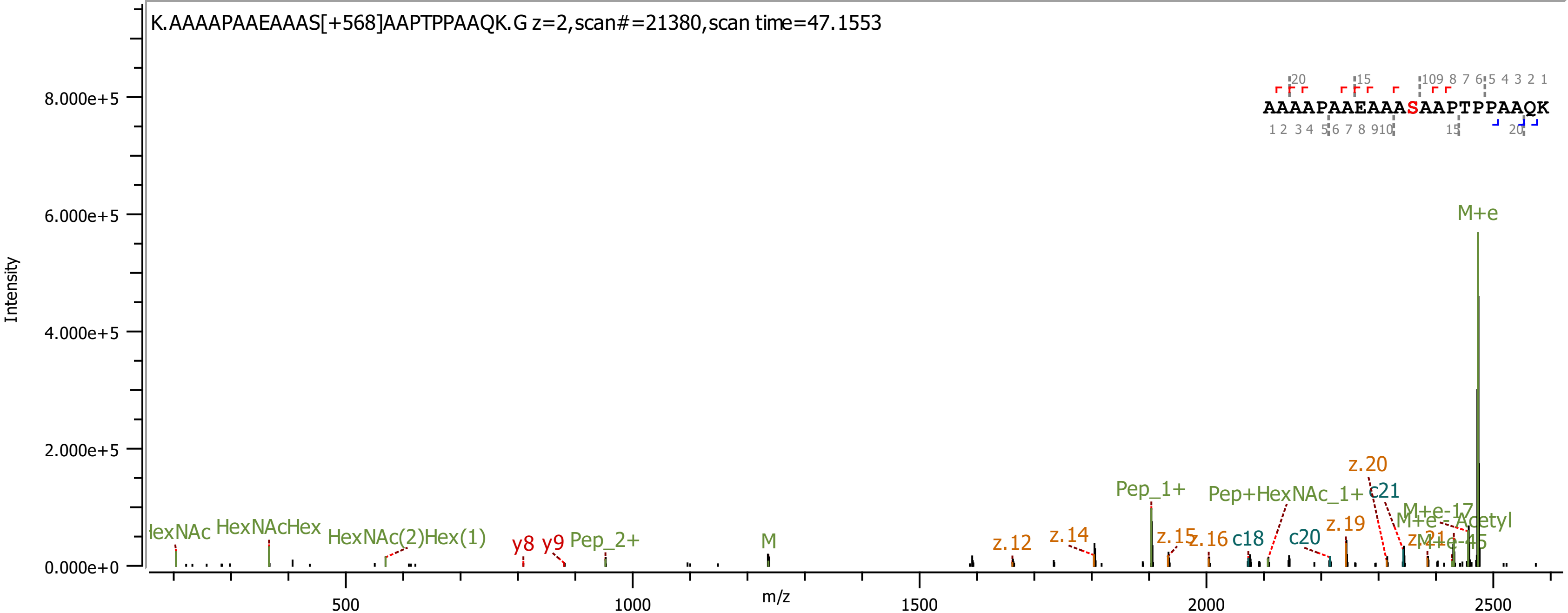
K.AAAAPAAEAAAS[+568]AAPTPPAAQK.G z=2,scan#=21380,scan time=47.1553
AAAAPAAEAAASAAPTPPAAQK
Intensity
8.000e+5
6.000e+5
4.000e+5
2.000e+5
0.000e+0
m/z
500
1000
1500
2000
2500
M+e
HexNAcHex
HexNAc(2)Hex(1)
y8
y9
Pep_2+
M
z.12
z.14
Pep_1+
z.15
z.16
c18
c20
Pep+HexNAc_1+
z.19
z.20
c21
M+e-17
M+e-Acetyl
M+e-45
z.21

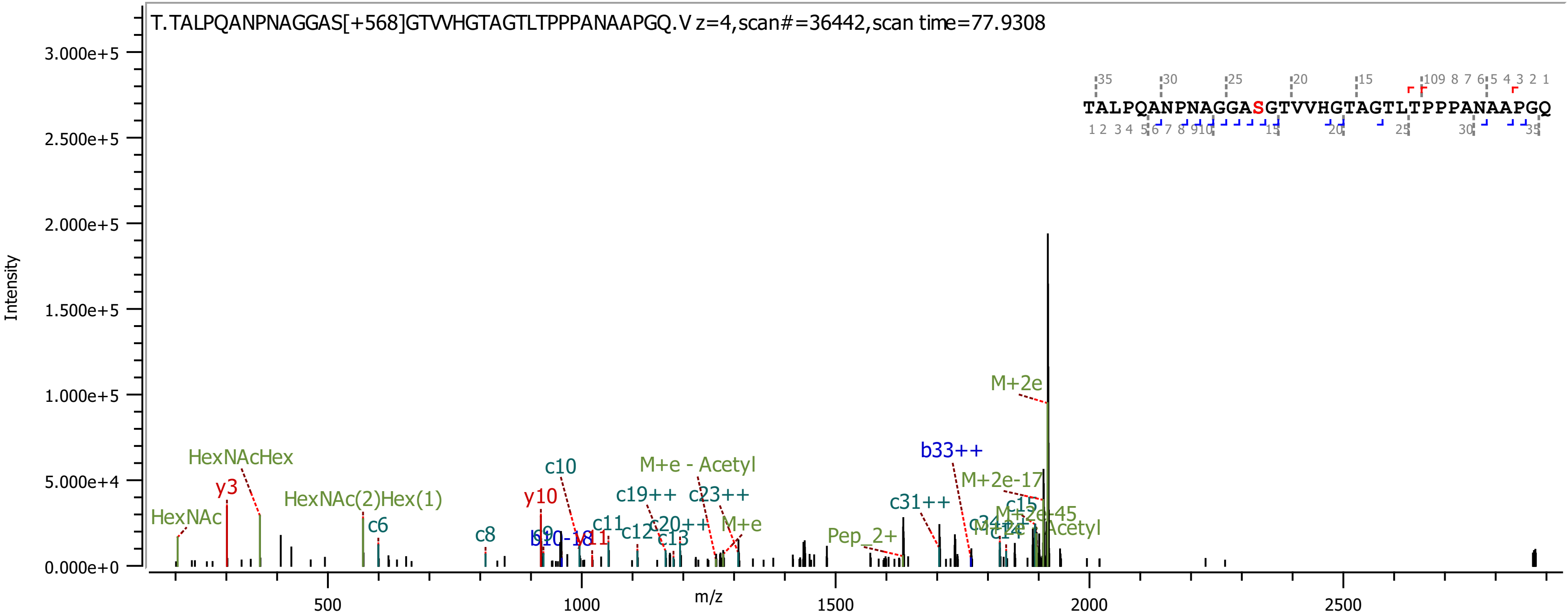

T.TALPQANPNAGGAS[+568]GTVVHGTAGTLTPPPANAAPGQ.V z=4,scan#=36442,scan time=77.9308
TALPQANPNAGGASGTVVHGTAGTLTPPPANAAPGQ
Intensity
m/z
3.000e+5
2.500e+5
2.000e+5
1.500e+5
1.000e+5
5.000e+4
0.000e+0
500
1000
1500
2000
2500
HexNAc
HexNAcHex
y3
HexNAc(2)Hex(1)
c6
c8
y10
c10
c11
c12
c13
c19++
c20++
c23++
M+e - Acetyl
M+e
Pep_2+
c31++
b33++
M+2e-17
M+2e
c15
c14
M+2e-45

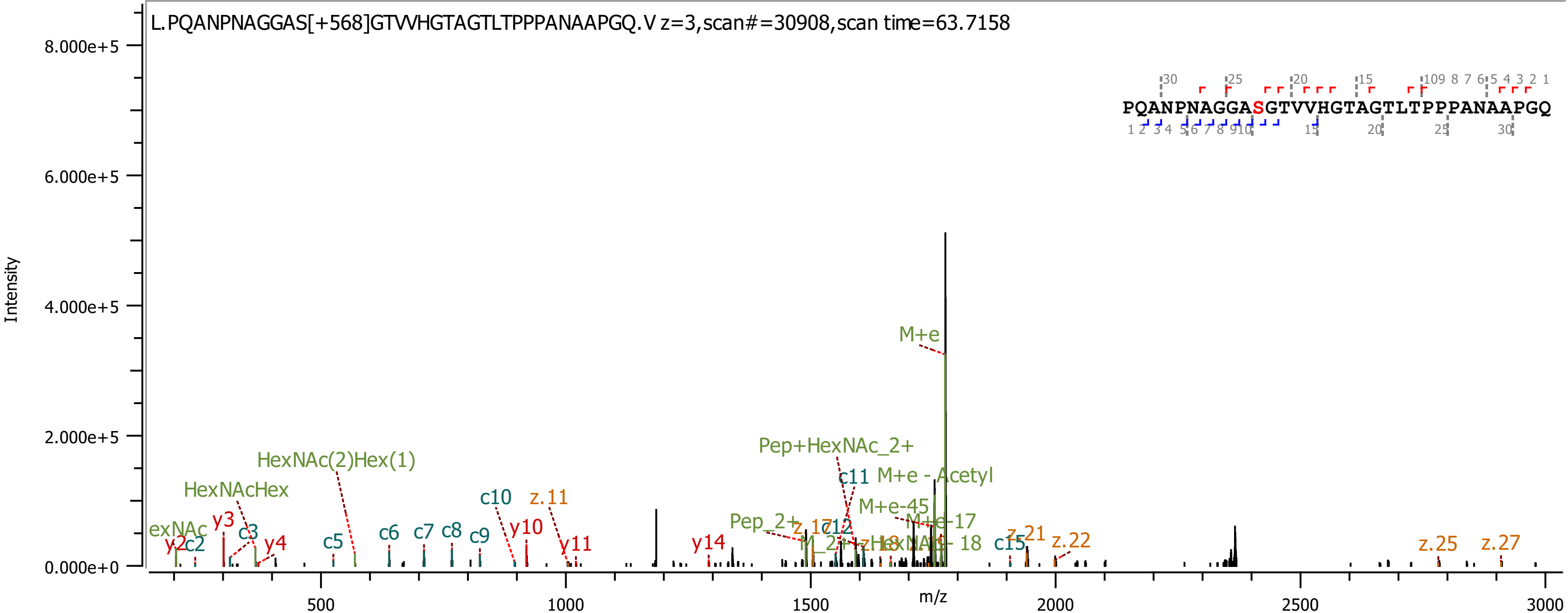

L.PQANPNAGGAS[+568]GTVVHGTAGTLTPPPANAAPGQ.V z=3,scan#=30908,scan time=63.7158
PQANPNAGGASGTVVHGTAGTLTPPPANAAPGQ
Intensity
m/z
8.000e+5
6.000e+5
4.000e+5
2.000e+5
0.000e+0
500
1000
1500
2000
2500
3000
M+e
Pep+HexNAc_2+
M+e - Acetyl
M+e-45
M+e-17
Pep_2+
HexNAc(2)Hex(1)
HexNAcHex
c10
z.11
y10
y11
y14
c11
c12
c15
z.21
z.22
z.25
z.27
y2
c2
y3
c3
y4
c5
c6
c7
c8
c9

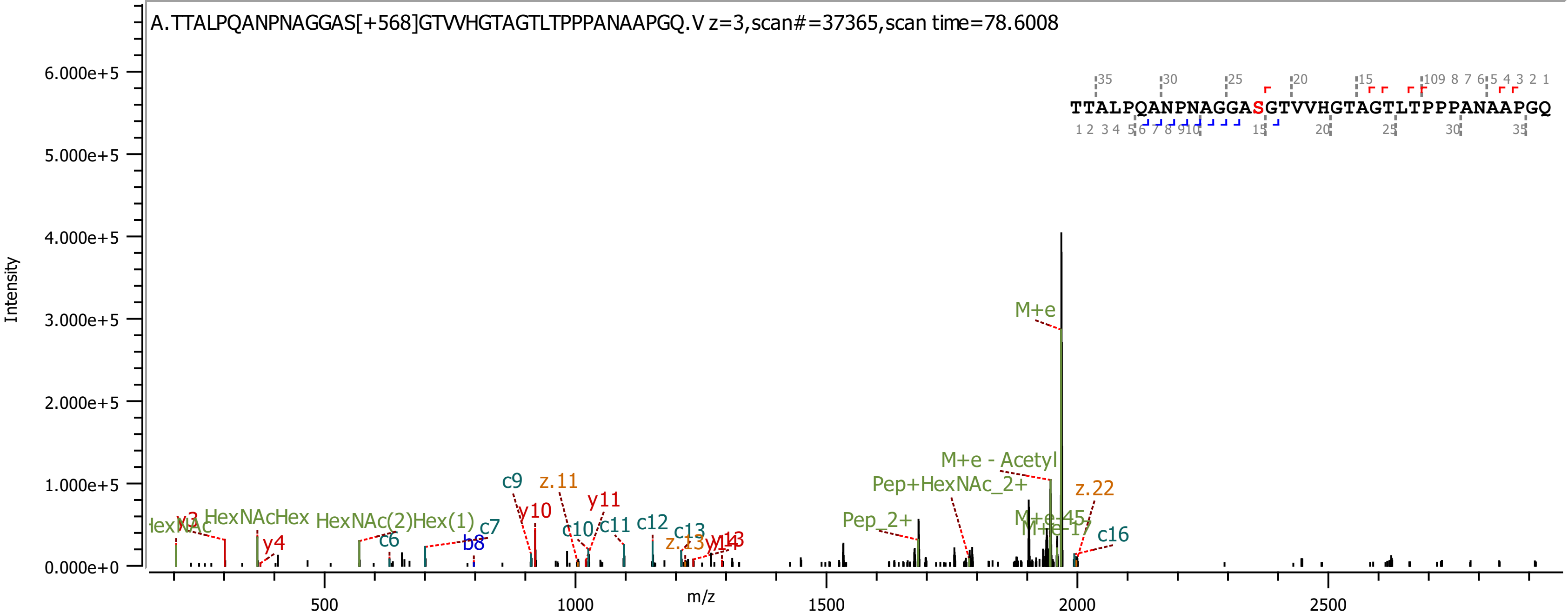
A.TTALPQANPNAGGAS[+568]GTVVHGTAGTLTPPPANAAPGQ.V z=3,scan#=37365,scan time=78.6008
TTALPQANPNAGGASGTVVHGTAGTLTPPPANAAPGQ
Intensity
m/z
6.000e+5
5.000e+5
4.000e+5
3.000e+5
2.000e+5
1.000e+5
0.000e+0
500
1000
1500
2000
2500
M+e
M+e - Acetyl
Pep+HexNAc_2+
Pep_2+
z.22
c16
HexNAcHex
HexNAc(2)Hex(1)
y4
c6
c7
b8
c9
z.11
y10
y11
c10
c11
c12
c13

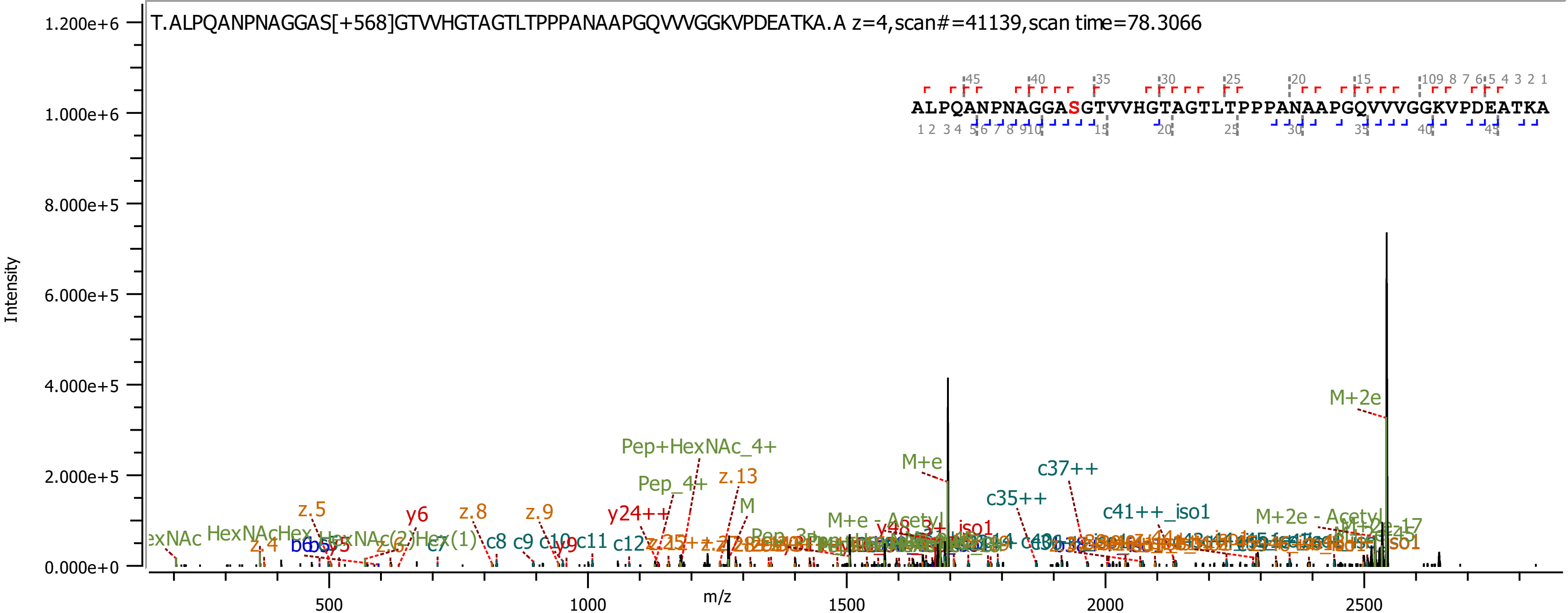

T.ALPQANPNAGGAS[+568]GTVVHGTAGTLTPPPANAAPGQVVVGGKVPDEATKA.A z=4,scan#=41139,scan time=78.3066
ALPQANPNAGGASGTVVHGTAGTLTPPPANAAPGQVVVGGKVPDEATKA
Intensity
m/z
1.200e+6
1.000e+6
8.000e+5
6.000e+5
4.000e+5
2.000e+5
0.000e+0
500
1000
1500
2000
2500
M+2e
Pep+HexNAc_4+
M+e
c37++
z.13
Pep_4+
c35++
z.5
z.8
z.9
y6
y24++
c41++_iso1
M+2e - Acetyl
M+e - Acetyl
HexNAcHex
c8
c9
c11
c12
z.4

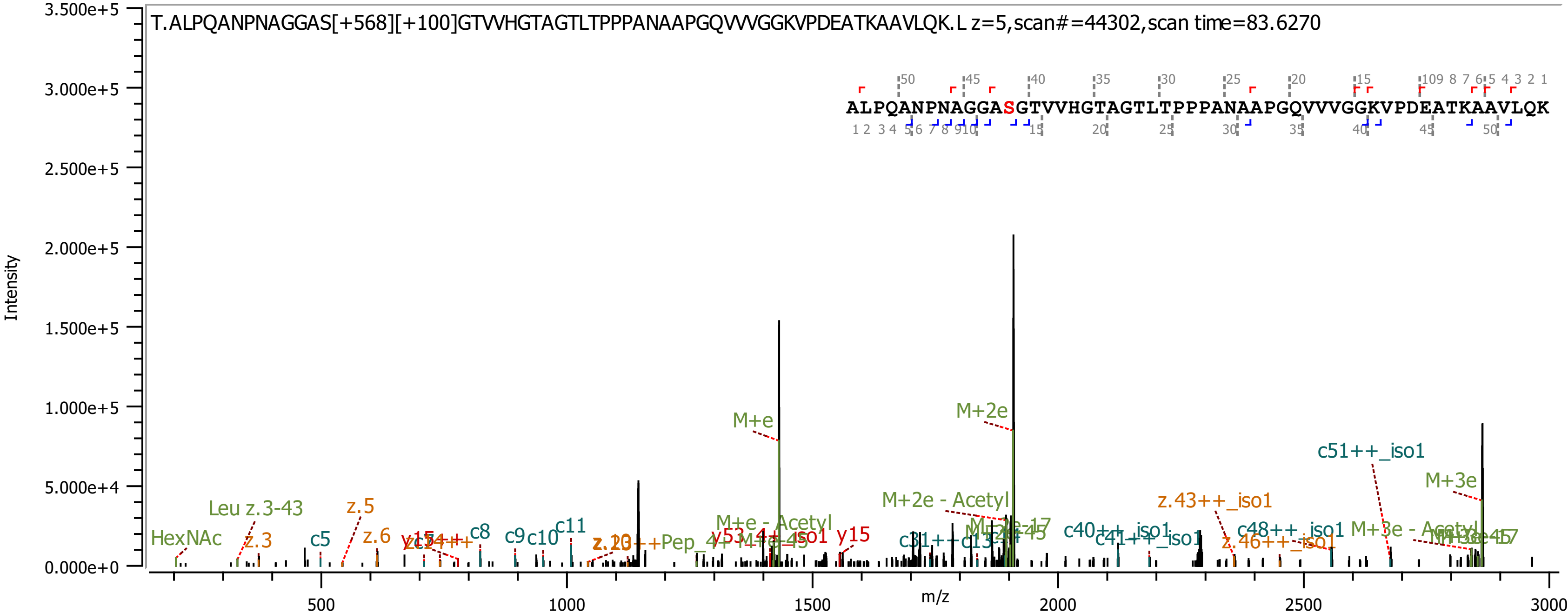
T.ALPQANPNAGGAS[+568][+100]GTVVHGTAGTLTPPPANAAPGQVVVGGKVPDEATKAAVLQK.L z=5,scan#=44302,scan time=83.6270
ALPQANPNAGGASGTVVHGTAGTLTPPPANAAPGQVVVGGKVPDEATKAAVLQK
Intensity
m/z
3.500e+5
3.000e+5
2.500e+5
2.000e+5
1.500e+5
1.000e+5
5.000e+4
0.000e+0
500
1000
1500
2000
2500
3000
HexNAc
Leu z.3-43
z.3
c5
z.5
z.6
c8
c9
c10
c11
M+e
M+2e
M+e - Acetyl
y15
M+2e - Acetyl
c51++_iso1
M+3e
z.43++_iso1
c40++_iso1
c41++_iso1
c48++_iso1
z.46++_iso1
M+3e - Acetyl

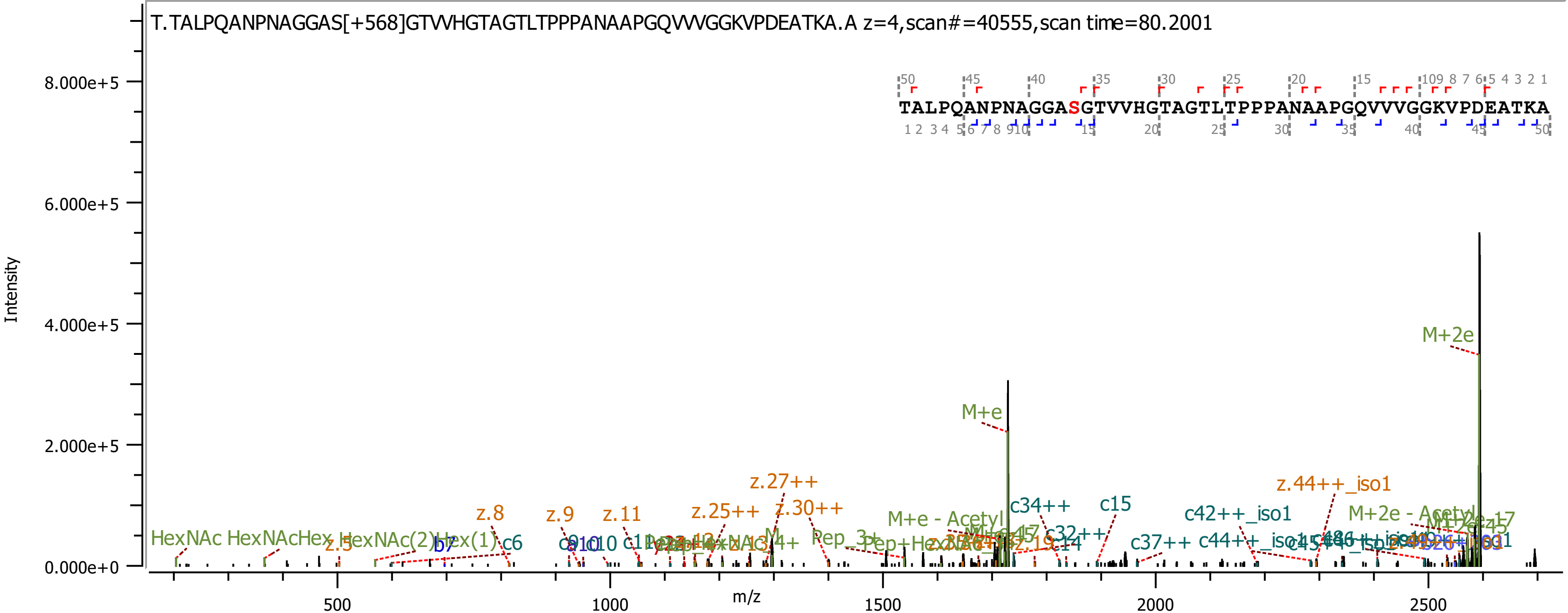

T.TALPQANPNAGGAS[+568]GTVVHGTAGTLTPPPANAAPGQVVVGGKVPDEATKA.A z=4,scan#=40555,scan time=80.2001
TALPQANPNAGGASGTVVHGTAGTLTPPPANAAPGQVVVGGKVPDEATKA
Intensity
m/z
8.000e+5
6.000e+5
4.000e+5
2.000e+5
0.000e+0
500
1000
1500
2000
2500
M+2e
M+e
z.27++
z.44++_iso1
z.25++
z.30++
z.8
z.9
z.11
c34++
c15
c42++_iso1
M+e - Acetyl
M+2e - Acetyl
HexNAc
HexNAcHex
HexNAc(2)Hex(1)
b7
c6
z.5
c37++
c32++
Pep_3+

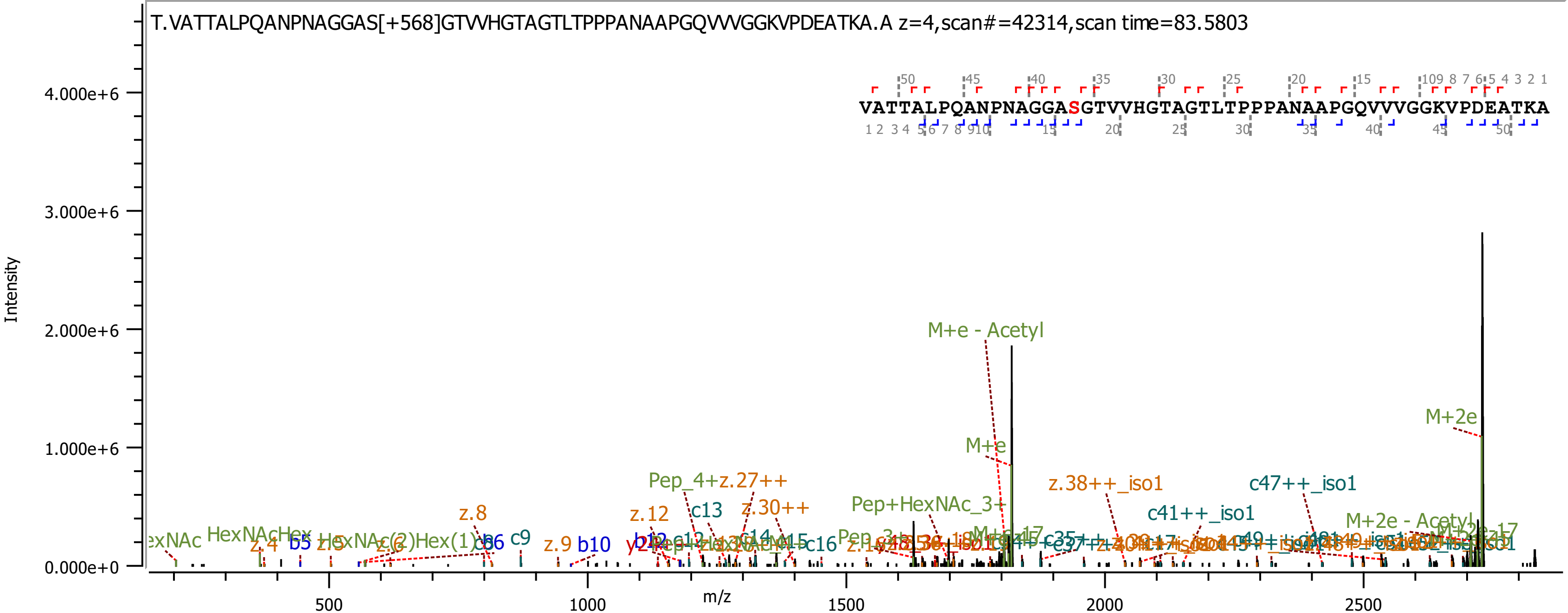
T.VATTALPQANPNAGGAS[+568]GTVVHGTAGTLTPPPANAAPGQVVVGGKVPDEATKA.A z=4,scan#=42314,scan time=83.5803
VATTALPQANPNAGGASGTVVHGTAGTLTPPPANAAPGQVVVGGKVPDEATKA
Intensity
4.000e+6
3.000e+6
2.000e+6
1.000e+6
0.000e+0
m/z
500
1000
1500
2000
2500
M+e - Acetyl
M+2e
M+e
Pep_4+z.27++
z.38++_iso1
c47++_iso1
Pep+HexNAc_3+
z.8
z.12
c13
z.30++
c41++_iso1
M+2e - Acetyl
HexNAcHex
HexNAc(2)Hex(1)
c9
z.9
b10
b5
z.4
z.5
z.6
c16

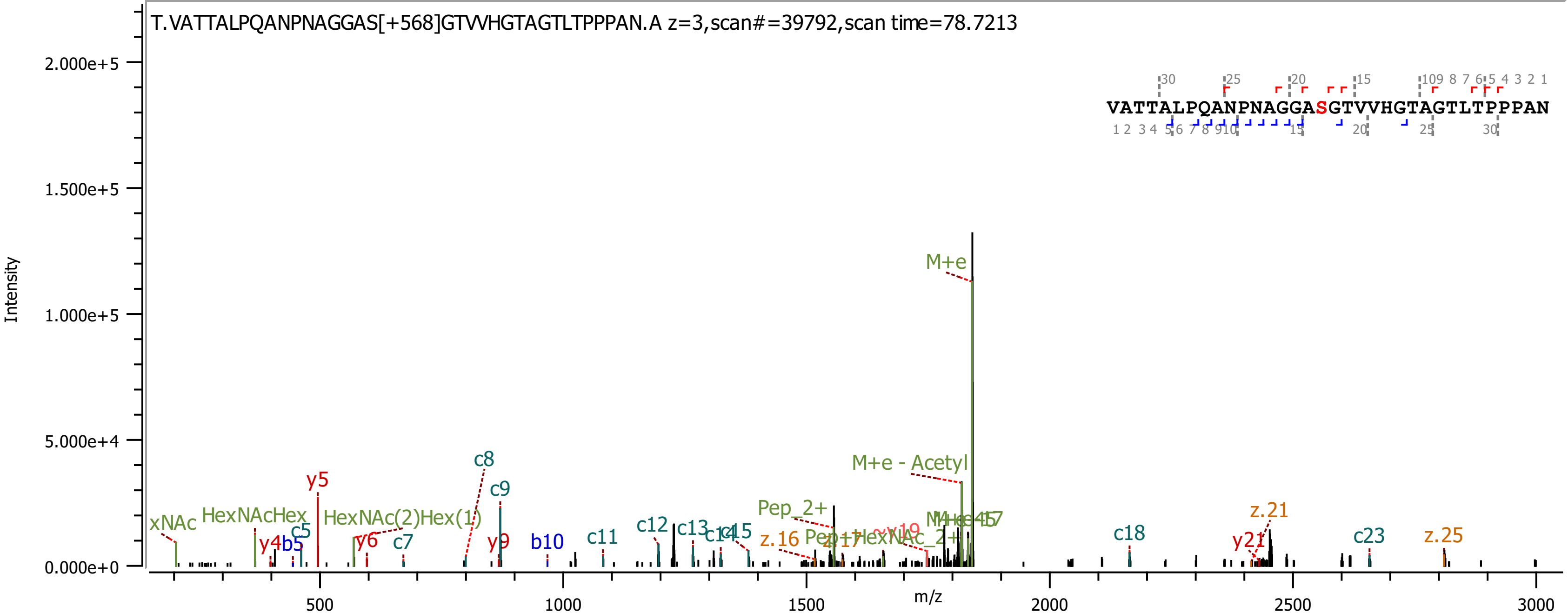

T.VATTALPQANPNAGGAS[+568]GTVVHGTAGTLTPPPAN.A z=3,scan#=39792,scan time=78.7213
VATTALPQANPNAGGASGTVVHGTAGTLTPPPAN
Intensity
m/z
2.000e+5
1.500e+5
1.000e+5
5.000e+4
0.000e+0
500
1000
1500
2000
2500
3000
M+e
M+e - Acetyl
Pep_2+
xNAc
HexNAcHex
HexNAc(2)Hex(1)
y4
b5
c5
y5
y6
c7
c8
c9
y9
b10
c11
c12
c13
c14
c15
z.16
z.17
c18
y21
z.21
c23
z.25

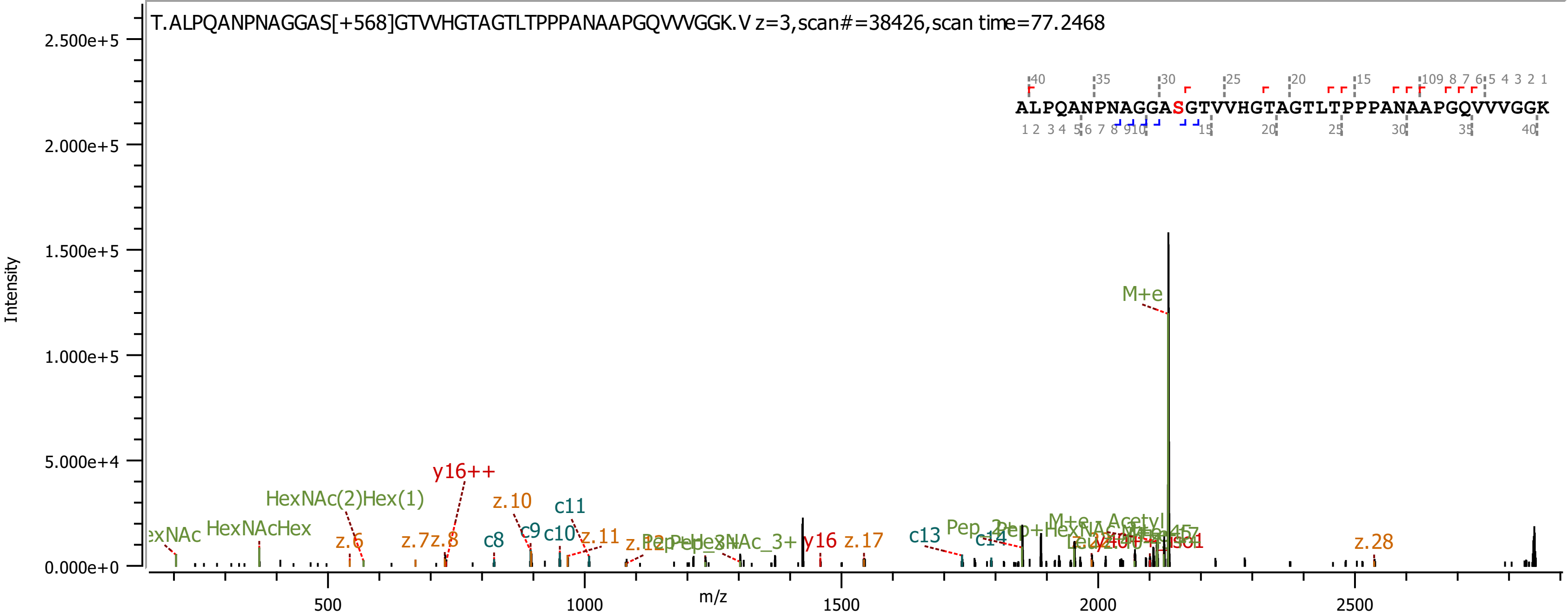
T.ALPQANPNAGGAS[+568]GTVVHGTAGTLTPPPANAAPGQVVVGGK.V z=3,scan#=38426,scan time=77.2468
ALPQANPNAGGASGTVVHGTAGTLTPPPANAAPGQVVVGGK
Intensity
m/z
2.500e+5
2.000e+5
1.500e+5
1.000e+5
5.000e+4
0.000e+0
500
1000
1500
2000
2500
M+e
y16++
HexNAc(2)Hex(1)
HexNAcHex
z.10
c11
z.6
z.7
z.8
c8
c9
c10
z.11
y16
z.17
c13
z.28

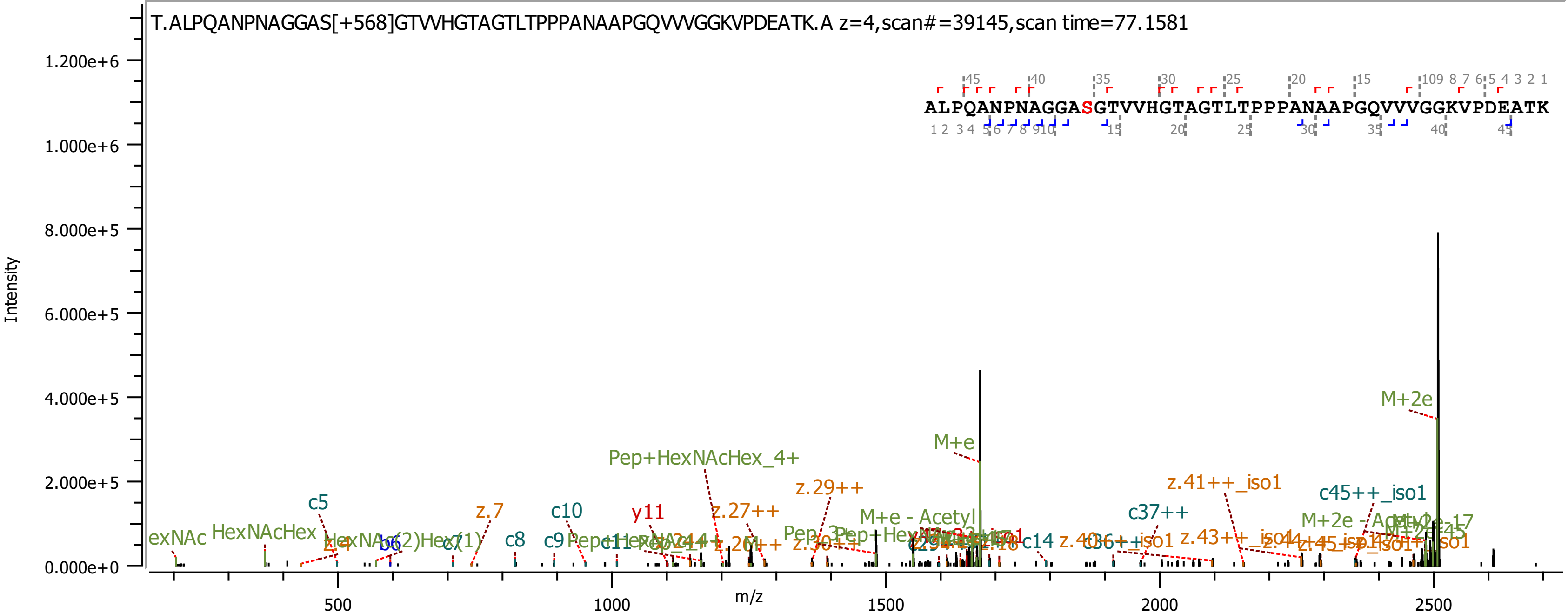
T.ALPQANPNAGGAS[+568]GTVVHGTAGTLTPPPANAAPGQVVVGGKVPDEATK.A z=4,scan#=39145,scan time=77.1581
ALPQANPNAGGASGTVVHGTAGTLTPPPANAAPGQVVVGGKVPDEATK
Intensity
m/z
M+2e
M+e
Pep+HexNAcHex_4+
z.29++
z.41++_iso1
c45++_iso1
c37++
M+e - Acetyl
z.27++
y11
c10
z.7
c5
HexNAcHex
c8
M+2e - Acetyl

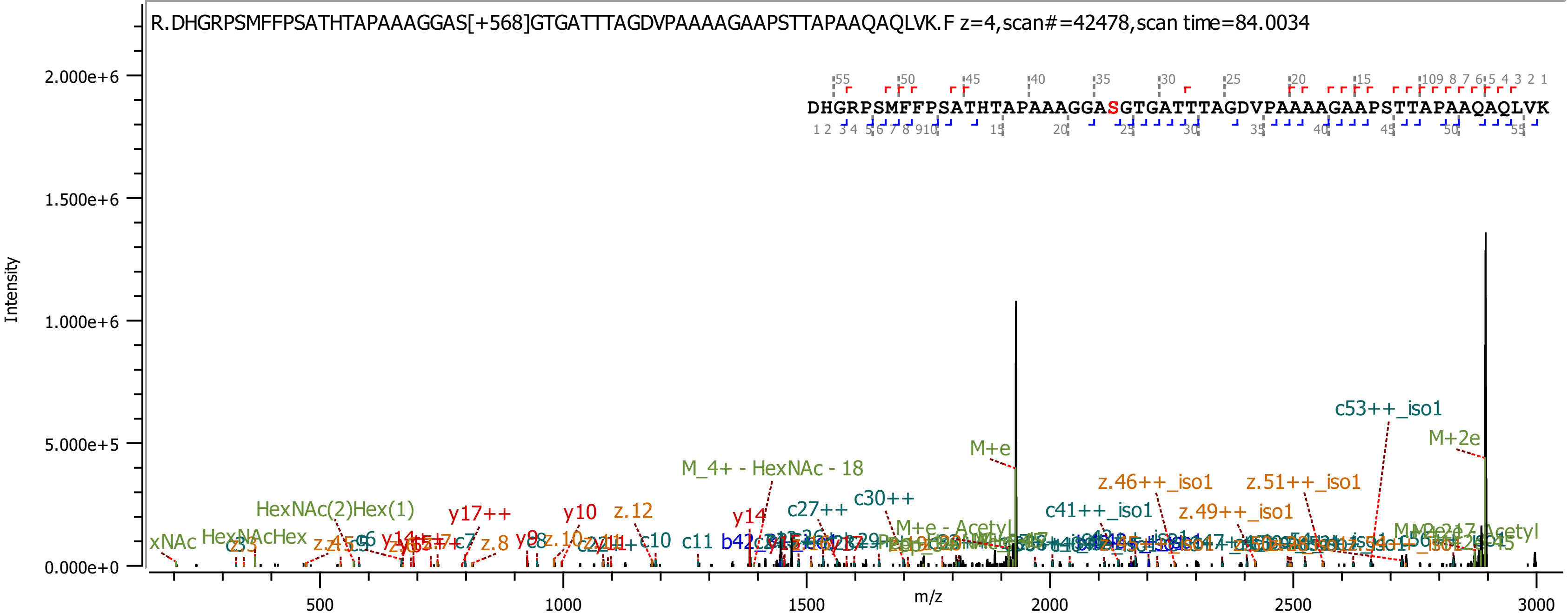
R.DHGRPSMFFPSATHTAPAAAGGAS[+568]GTGATTTAGDVPAAAAGAAPSTTAPAAQAQLVK.F z=4,scan#=42478,scan time=84.0034
DHGRPSMFFPSATHTAPAAAGGASGTGATTTAGDVPAAAAGAAPSTTAPAAQAQLVK
Intensity
2.000e+6
1.500e+6
1.000e+6
5.000e+5
0.000e+0
m/z
500
1000
1500
2000
2500
3000
c53++_iso1
M+2e
M+e
M_4+ - HexNAc - 18
z.46++_iso1
z.51++_iso1
c41++_iso1
z.49++_iso1
c30++
c27++
HexNAc(2)Hex(1)
y17++
y10
z.12
y14
HexNAcHex
xNAc
c6
c7
z.8
y9
c8
z.10
c10
c11
M+e - Acetyl

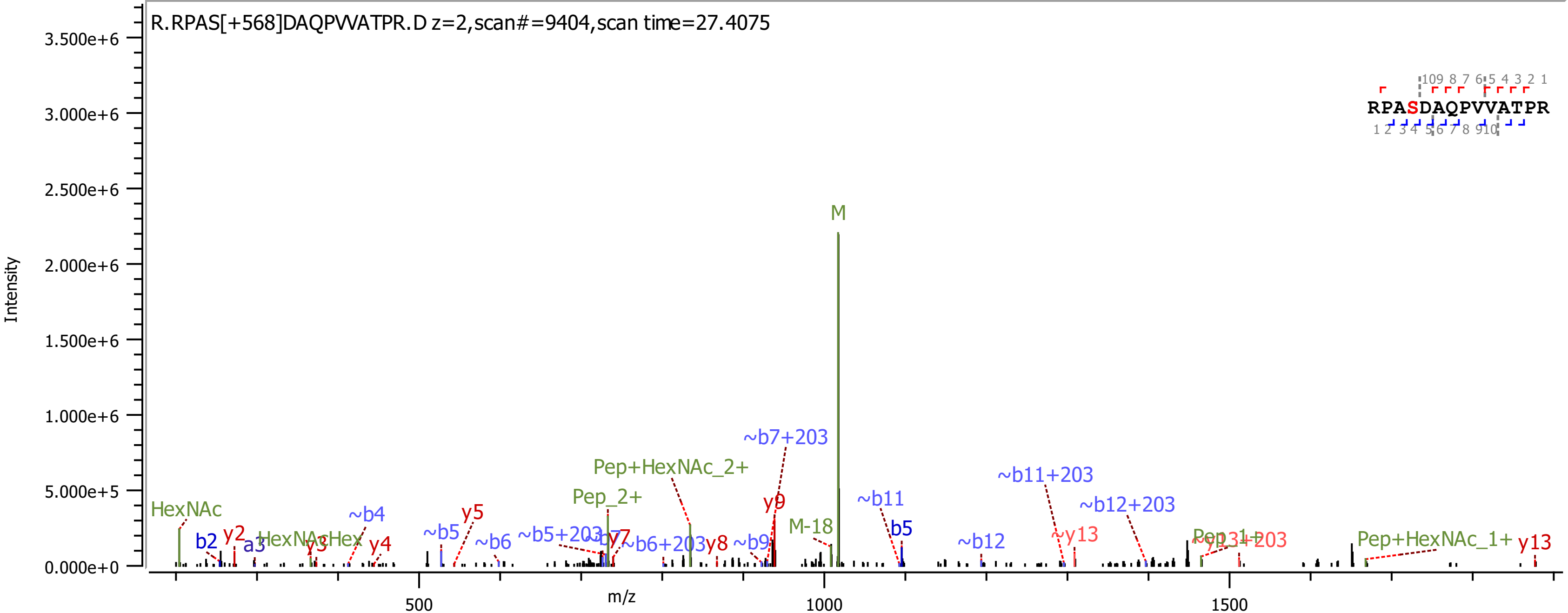
R.RPAS[+568]DAQPVVATPR.D z=2,scan#=9404,scan time=27.4075
RPASDAQPVVATPR
Intensity
m/z
M
~b7+203
Pep+HexNAc_2+
~b11+203
~b12+203
HexNAc
Pep_2+
~b4
y5
y9
~b11
M-18
b5
y2
b2
a3
HexNAcHex
y3
y4
~b5
~b6
~b5+203
y7
~b6+203
y8
~b9
~b12
~y13
Pep+HexNAc_1+
y13

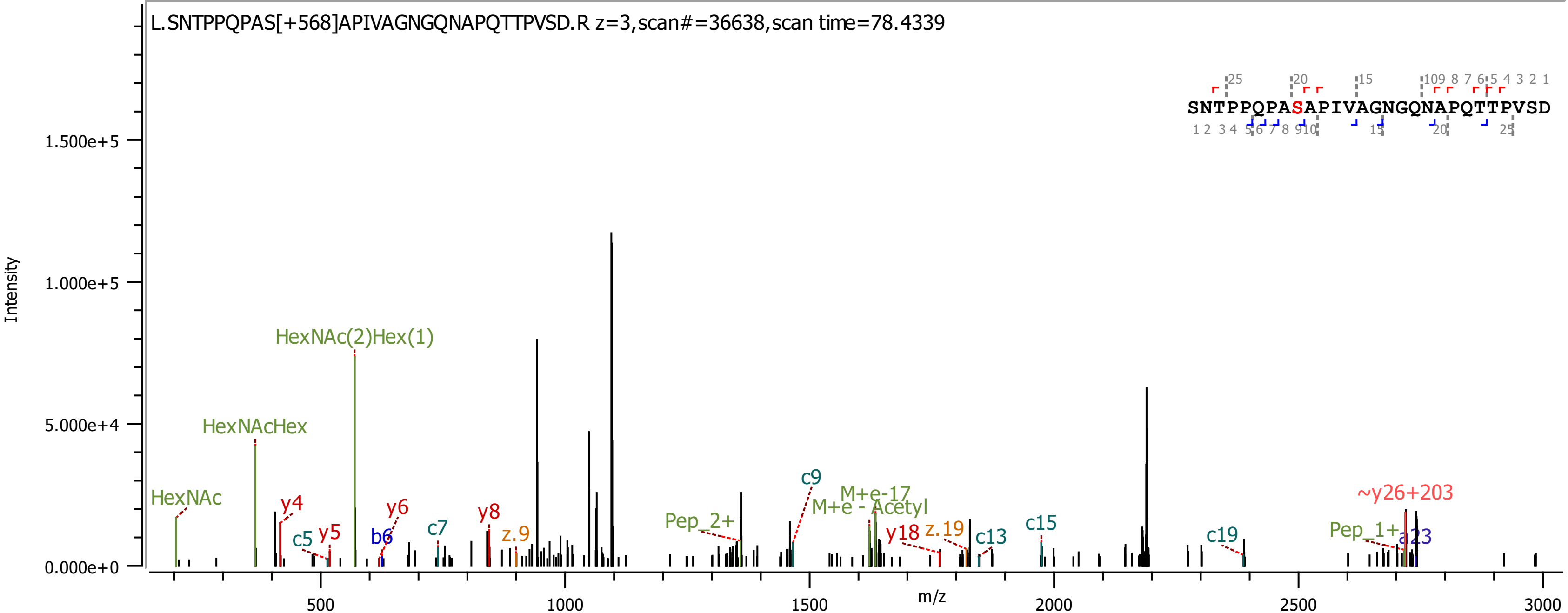

L.SNTPPQPAS[+568]APIVAGNGQNAPQTTPVSD.R z=3,scan#=36638,scan time=78.4339
SNTPPQPASAPIVAGNGQNAPQTTPVSD
Intensity
m/z
HexNAc(2)Hex(1)
HexNAcHex
HexNAc
y4
y5
c5
y6
b6
c7
y8
z.9
Pep_2+
c9
M+e-17
M+e - Acetyl
y18
z.19
c13
c15
c19
~y26+203
Pep_1+
a23

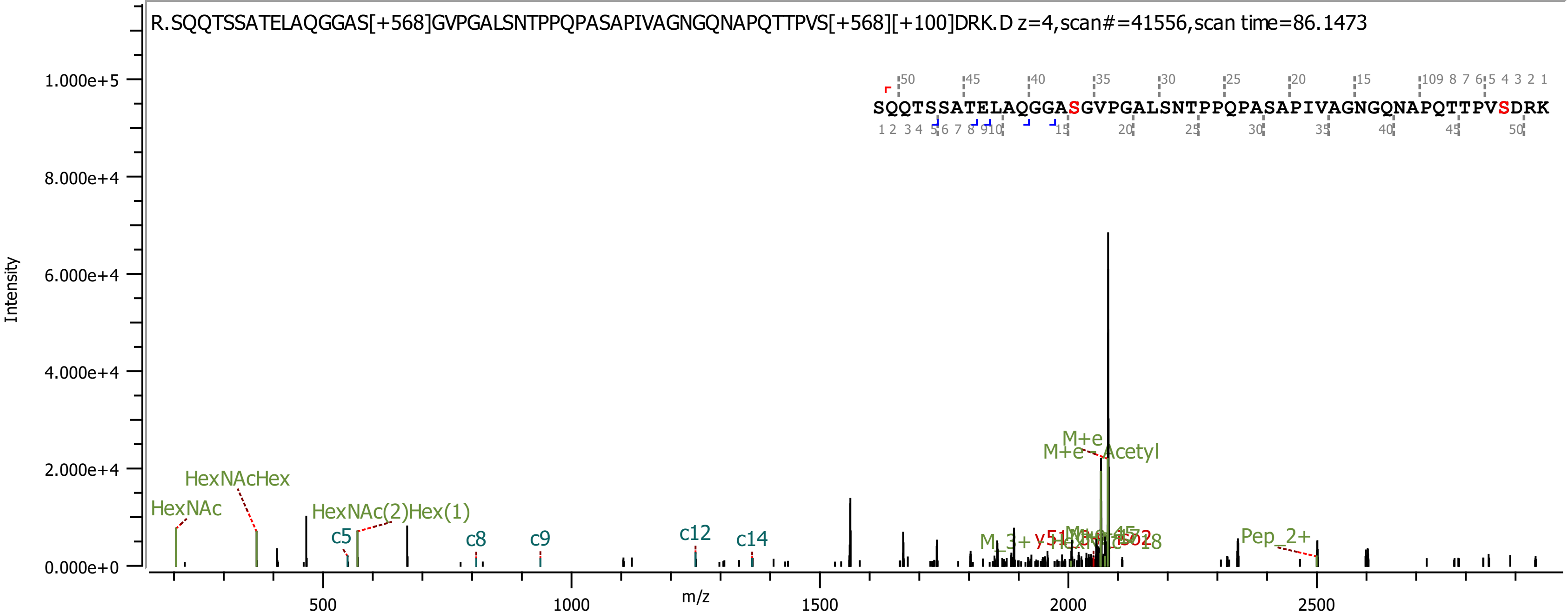

R.SQQTSSATELAQGGAS[+568]GVPGALSNTPPQPASAPIVAGNGQNAPQTTPVS[+568][+100]DRK.D z=4,scan#=41556,scan time=86.1473
SQQTSSATELAQGGASGVPGALSNTPPQPASAPIVAGNGQNAPQTTPVSDRK
Intensity
1.000e+5
8.000e+4
6.000e+4
4.000e+4
2.000e+4
0.000e+0
m/z
500
1000
1500
2000
2500
HexNAc
HexNAcHex
HexNAc(2)Hex(1)
c5
c8
c9
c12
c14
M+e
M+e - Acetyl
Pep_2+

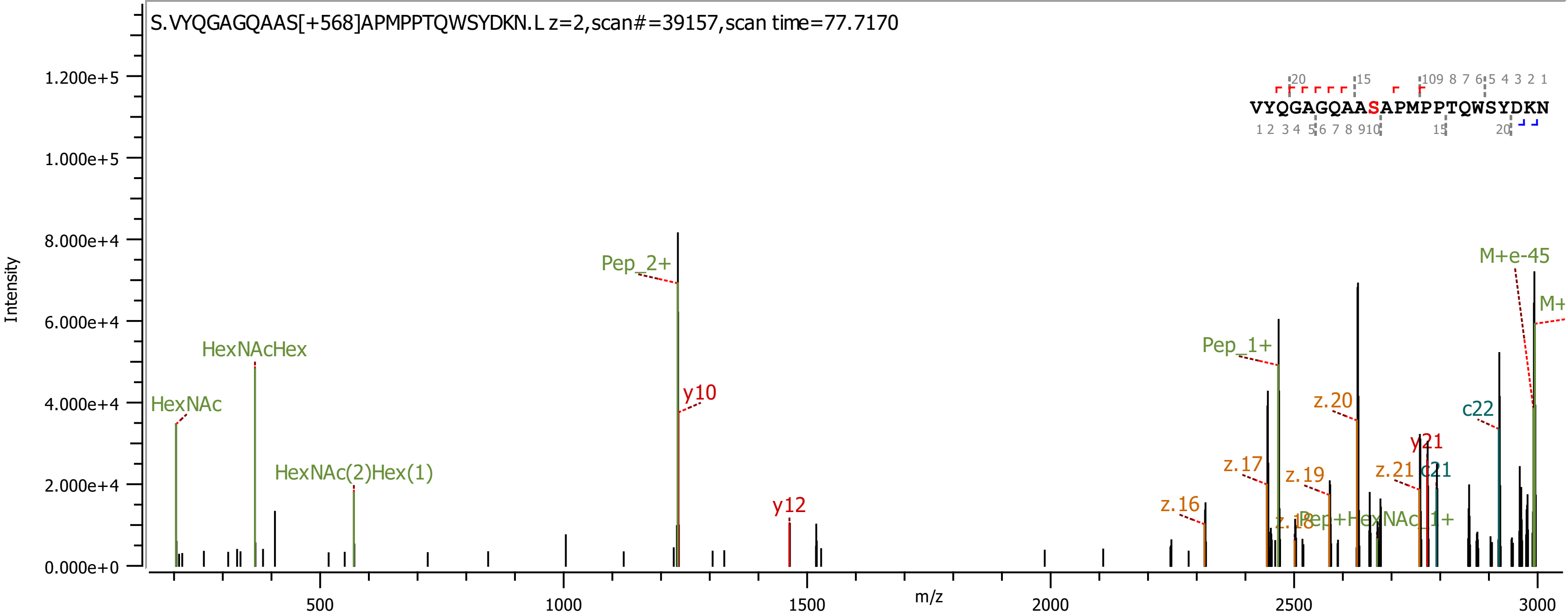

S.VYQGAGQAAS[+568]APMPPTQWSYDKN.L z=2,scan#=39157,scan time=77.7170
VYQGAGQAASAPMPPTQWSYDKN
Intensity
m/z
HexNAc
HexNAcHex
HexNAc(2)Hex(1)
Pep_2+
y10
y12
z.16
z.17
z.18
Pep_1+
z.19
Pep+HexNAc_1+
z.20
z.21
y21
c21
c22
M+e-45
M+

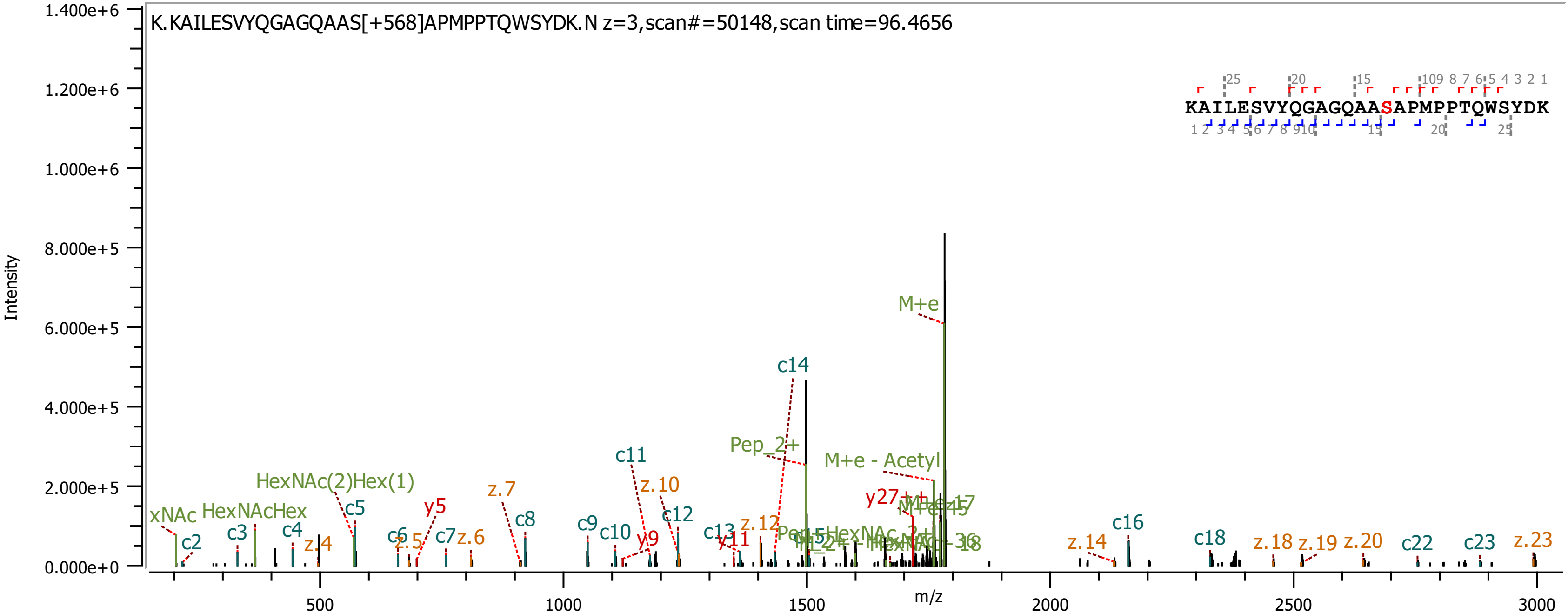

K.KAILESVYQGAGQAAS[+568]APMPPTQWSYDK.N z=3,scan#=50148,scan time=96.4656
KAILESVYQGAGQAASAPMPPTQWSYDK
Intensity
m/z
1.400e+6
1.200e+6
1.000e+6
8.000e+5
6.000e+5
4.000e+5
2.000e+5
0.000e+0
500
1000
1500
2000
2500
3000
M+e
c14
Pep_2+
M+e - Acetyl
c11
z.10
c12
HexNAc(2)Hex(1)
z.7
y5
c5
HexNAcHex
xNAc
c2
c3
c4
z.4
c6
z.5
c7
z.6
c8
c9
c10
y9
c13
y11
z.12
y27++
c16
c18
z.14
z.18
z.19
z.20
c22
c23
z.23

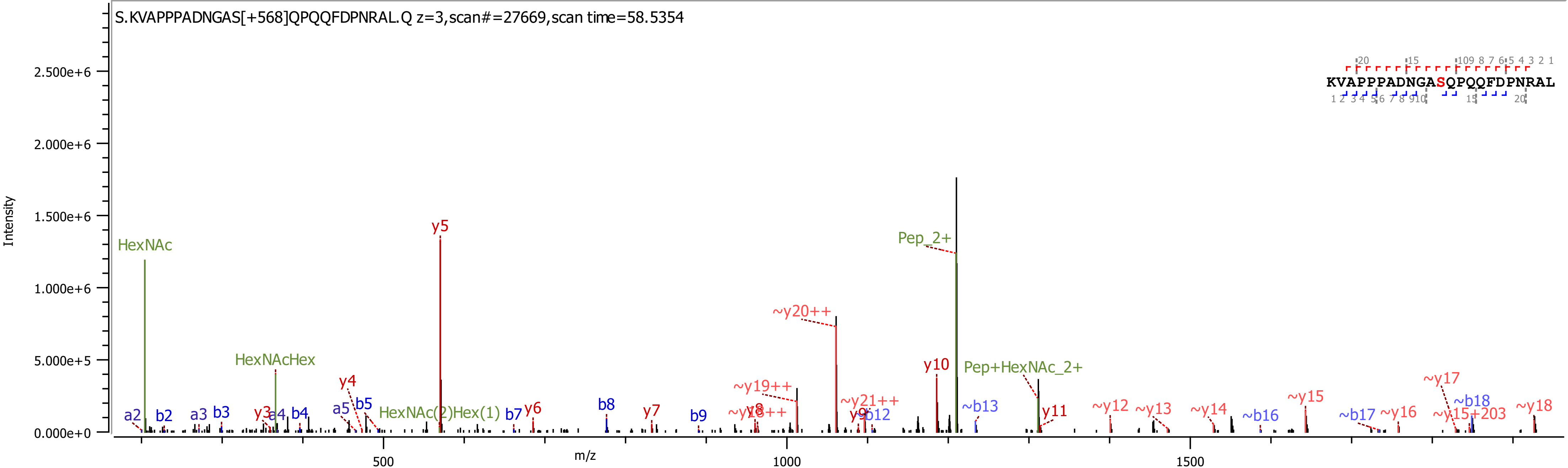

S.KVAPPPADNGAS[+568]QPQQFDPNRAL.Q z=3,scan#=27669,scan time=58.5354
KVAPPPADNGASQPQQFDPNRAL
Intensity
m/z
0.000e+0
5.000e+5
1.000e+6
1.500e+6
2.000e+6
2.500e+6
500
1000
1500
HexNAc
HexNAcHex
HexNAc(2)Hex(1)
Pep_2+
Pep+HexNAc_2+
y5
~y20++
y10
y4
y3
y6
y7
y8
y9
y11
~y19++
~y18++
~y21++
~y12
~y13
~y14
~y15
~y16
~y17
~y15+203
~y18
a2
a3
a4
a5
b2
b3
b4
b5
b7
b8
b9
b12
~b13
~b16
~b17
~b18

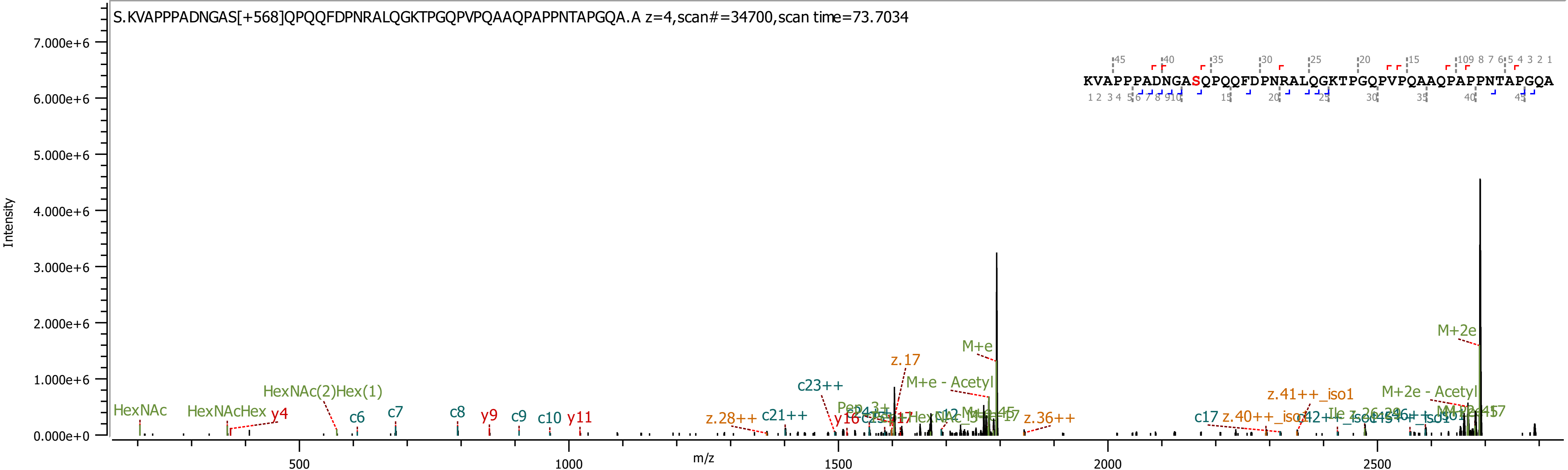
S.KVAPPPADNGAS[+568]QPQQFDPNRALQGKTPGQPVPQAAQPAPPNTAPGQA.A z=4,scan#=34700,scan time=73.7034
KVAPPPADNGASQPQQFDPNRALQGKTPGQPVPQAAQPAPPNTAPGQA
Intensity
m/z
7.000e+6
6.000e+6
5.000e+6
4.000e+6
3.000e+6
2.000e+6
1.000e+6
0.000e+0
500
1000
1500
2000
2500
HexNAc
HexNAcHex
y4
HexNAc(2)Hex(1)
c6
c7
c8
y9
c9
c10
y11
z.28++
c21++
c23++
z.17
M+e
M+e - Acetyl
z.36++
c17
z.40++_iso1
z.41++_iso1
M+2e - Acetyl
M+2e

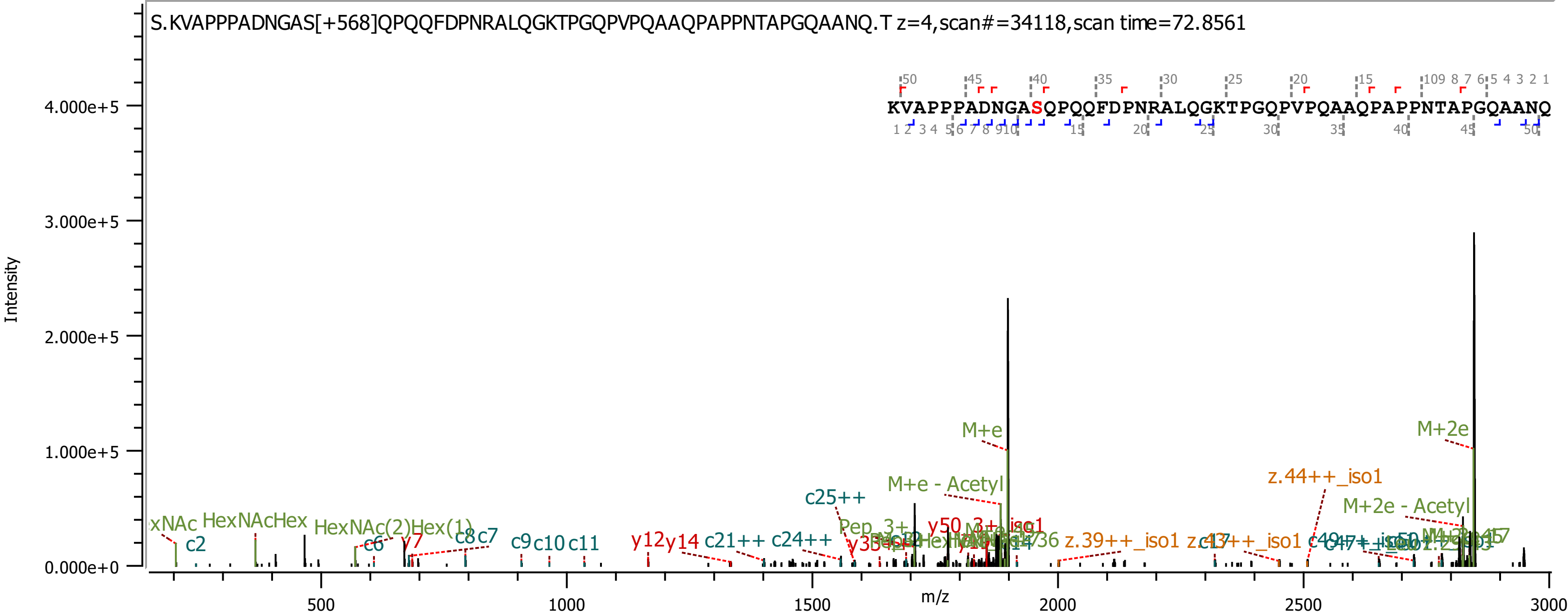

S.KVAPPPADNGAS[+568]QPQQFDPNRALQGKTPGQPVPQAAQPAPPNTAPGQAANQ.T z=4,scan#=34118,scan time=72.8561
KVAPPPADNGASQPQQFDPNRALQGKTPGQPVPQAAQPAPPNTAPGQAANQ
Intensity
m/z
0.000e+0
1.000e+5
2.000e+5
3.000e+5
4.000e+5
500
1000
1500
2000
2500
3000
M+e
M+2e
M+e - Acetyl
M+2e - Acetyl
z.44++_iso1
z.39++_iso1
c25++
c21++
c24++
y12
y14
y7
c2
c6
c8
c7
c9
c10
c11
HexNAcHex
HexNAc(2)Hex(1)

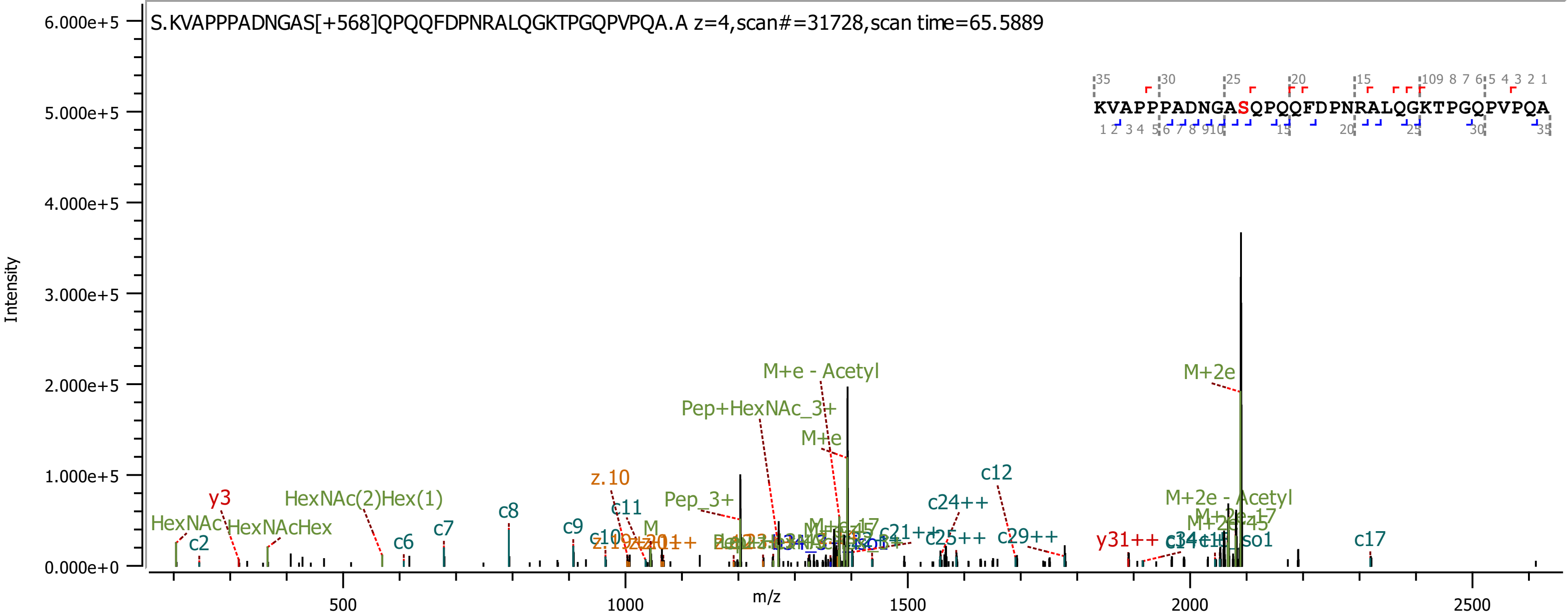

S.KVAPPPADNGAS[+568]QPQQFDPNRALQGKTPGQPVPQA.A z=4,scan#=31728,scan time=65.5889
KVAPPPADNGASQPQQFDPNRALQGKTPGQPVPQA
Intensity
m/z
6.000e+5
5.000e+5
4.000e+5
3.000e+5
2.000e+5
1.000e+5
0.000e+0
500
1000
1500
2000
2500
M+e - Acetyl
M+2e
Pep+HexNAc_3+
M+e
c12
z.10
y3
HexNAc(2)Hex(1)
c8
c11
Pep_3+
c24++
M+2e - Acetyl
HexNAc
HexNAcHex
c2
c6
c7
c9
c10
c21++
c25++
c29++
y31++
c17

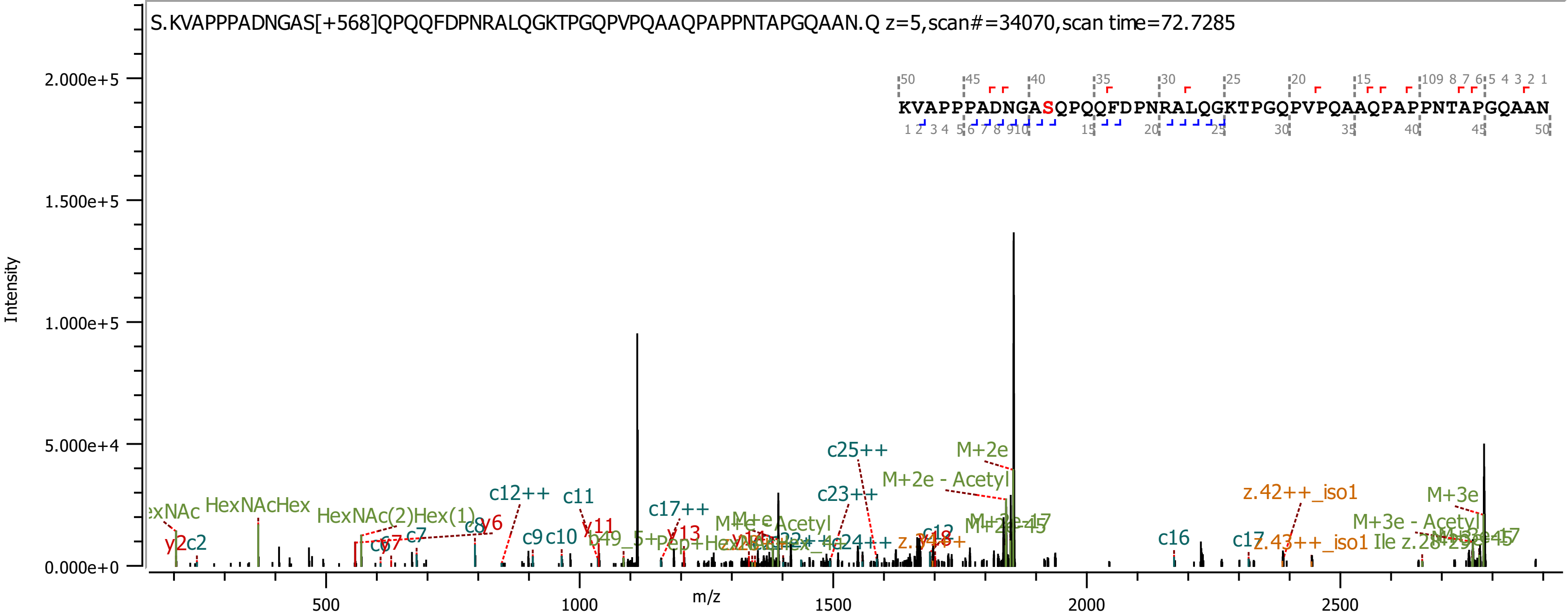
S.KVAPPPADNGAS[+568]QPQQFDPNRALQGKTPGQPVPQAAQPAPPNTAPGQAAN.Q z=5,scan#=34070,scan time=72.7285
KVAPPPADNGASQPQQFDPNRALQGKTPGQPVPQAAQPAPPNTAPGQAAN
Intensity
m/z
2.000e+5
1.500e+5
1.000e+5
5.000e+4
0.000e+0
500
1000
1500
2000
2500
HexNAcHex
HexNAc(2)Hex(1)
c12++
c11
c17++
c25++
c23++
M+2e
M+2e - Acetyl
z.42++_iso1
M+3e
M+3e - Acetyl
c16
c17
z.43++_iso1
y2
c2
y6
c7
c8
c9
c10
y11
y13
c12
c24++

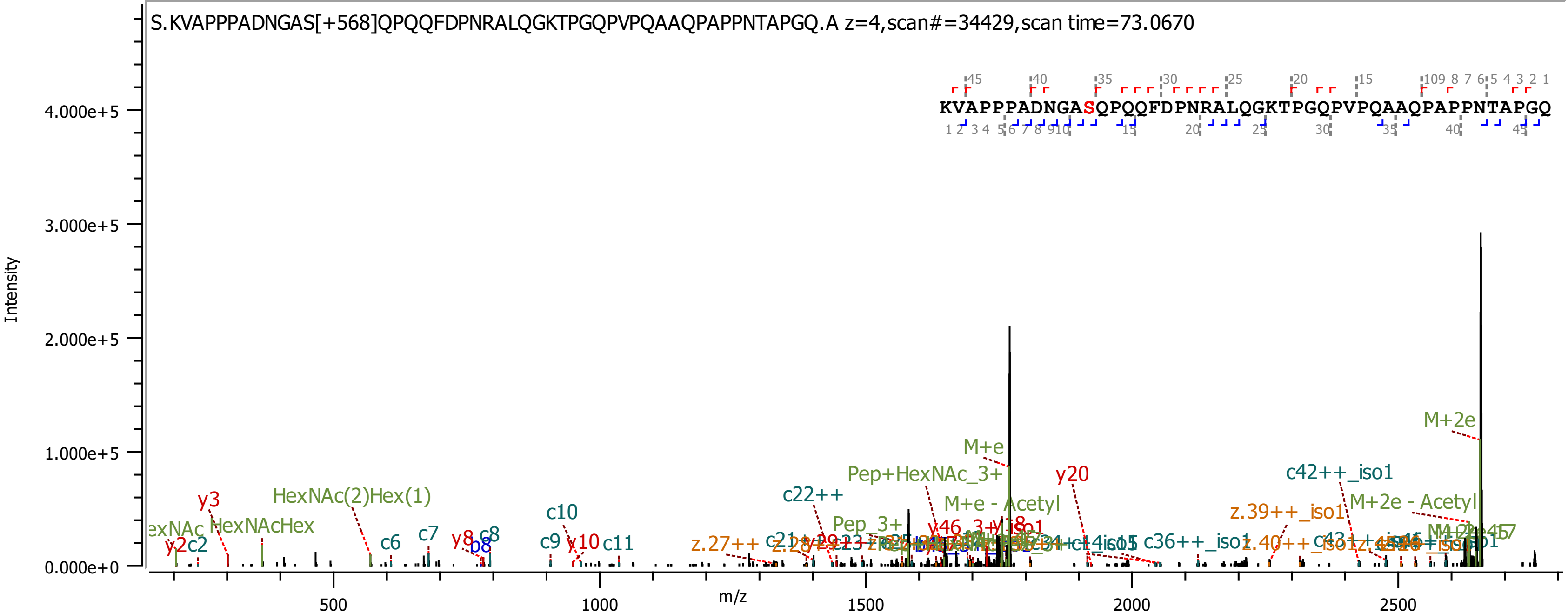
S.KVAPPPADNGAS[+568]QPQQFDPNRALQGKTPGQPVPQAAQPAPPNTAPGQ.A z=4,scan#=34429,scan time=73.0670
KVAPPPADNGASQPQQFDPNRALQGKTPGQPVPQAAQPAPPNTAPGQ
Intensity
m/z
4.000e+5
3.000e+5
2.000e+5
1.000e+5
0.000e+0
500
1000
1500
2000
2500
y3
HexNAc(2)Hex(1)
HexNAcHex
y2
c2
c6
c7
y8
b8
c8
c10
c9
y10
c11
z.27++
c22++
Pep+HexNAc_3+
M+e
M+e - Acetyl
Pep_3+
y46
y20
c36++_iso1
c42++_iso1
z.39++_iso1
M+2e
M+2e - Acetyl

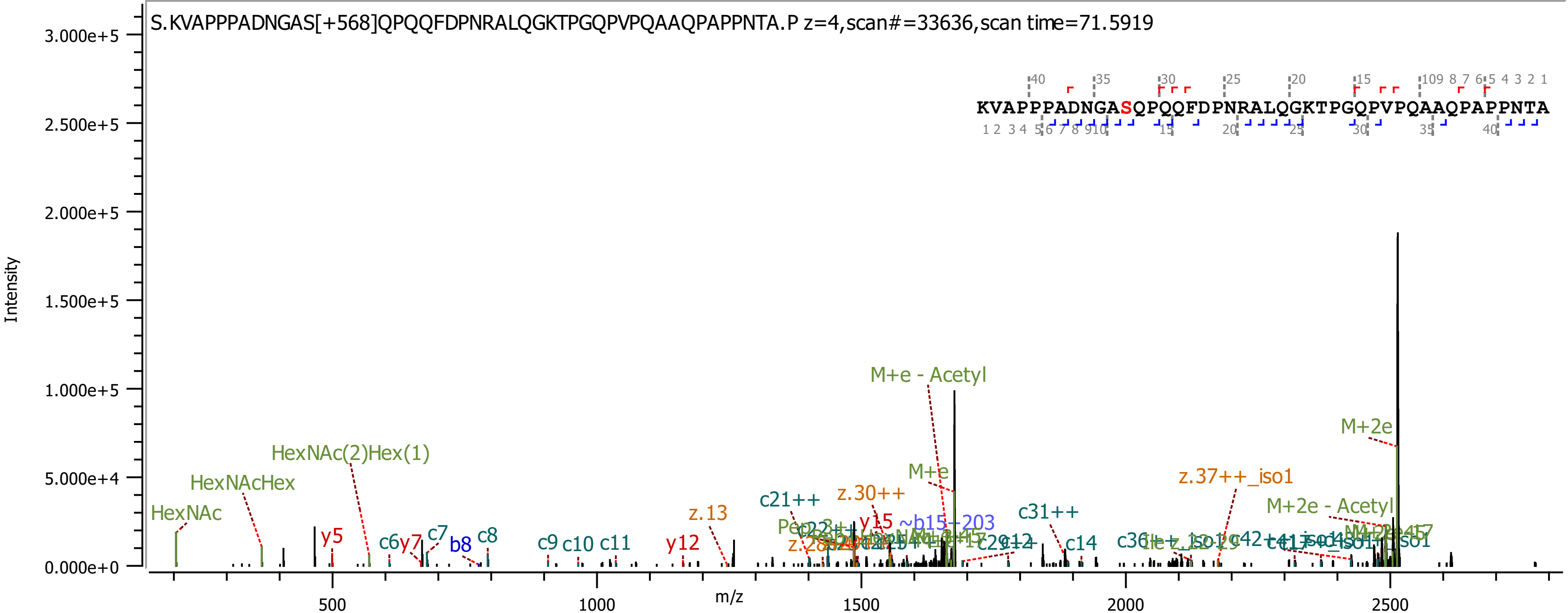
S.KVAPPPADNGAS[+568]QPQQFDPNRALQGKTPGQPVPQAAQPAPPNTA.P z=4,scan#=33636,scan time=71.5919
KVAPPPADNGASQPQQFDPNRALQGKTPGQPVPQAAQPAPPNTA
Intensity
m/z
M+e - Acetyl
M+2e
M+2e - Acetyl
M+e
HexNAc
HexNAcHex
HexNAc(2)Hex(1)
y5
c6
y7
c7
b8
c8
c9
c10
c11
y12
z.13
c21++
z.30++
y15
~b15--203
c31++
c14
z.37++_iso1

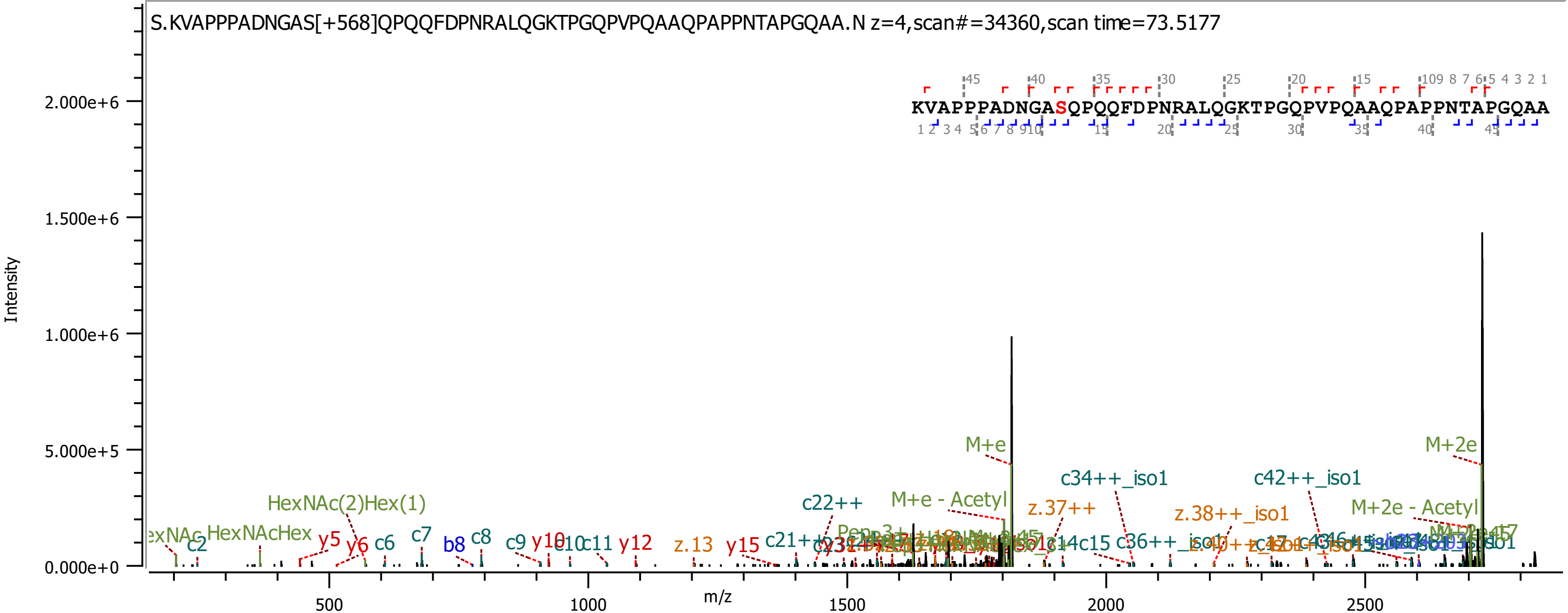

S.KVAPPPADNGAS[+568]QPQQFDPNRALQGKTPGQPVPQAAQPAPPNTAPGQAA.N z=4,scan#=34360,scan time=73.5177
KVAPPPADNGASQPQQFDPNRALQGKTPGQPVPQAAQPAPPNTAPGQAA
Intensity
m/z
0.000e+0
5.000e+5
1.000e+6
1.500e+6
2.000e+6
500
1000
1500
2000
2500
M+e
M+2e
M+e - Acetyl
M+2e - Acetyl
c34++_iso1
c42++_iso1
c22++
z.37++
z.38++_iso1
HexNAc(2)Hex(1)
HexNAcHex
c2
y5
y6
c6
c7
b8
c8
c9
y10
c11
y12
z.13
y15
c36++_iso1

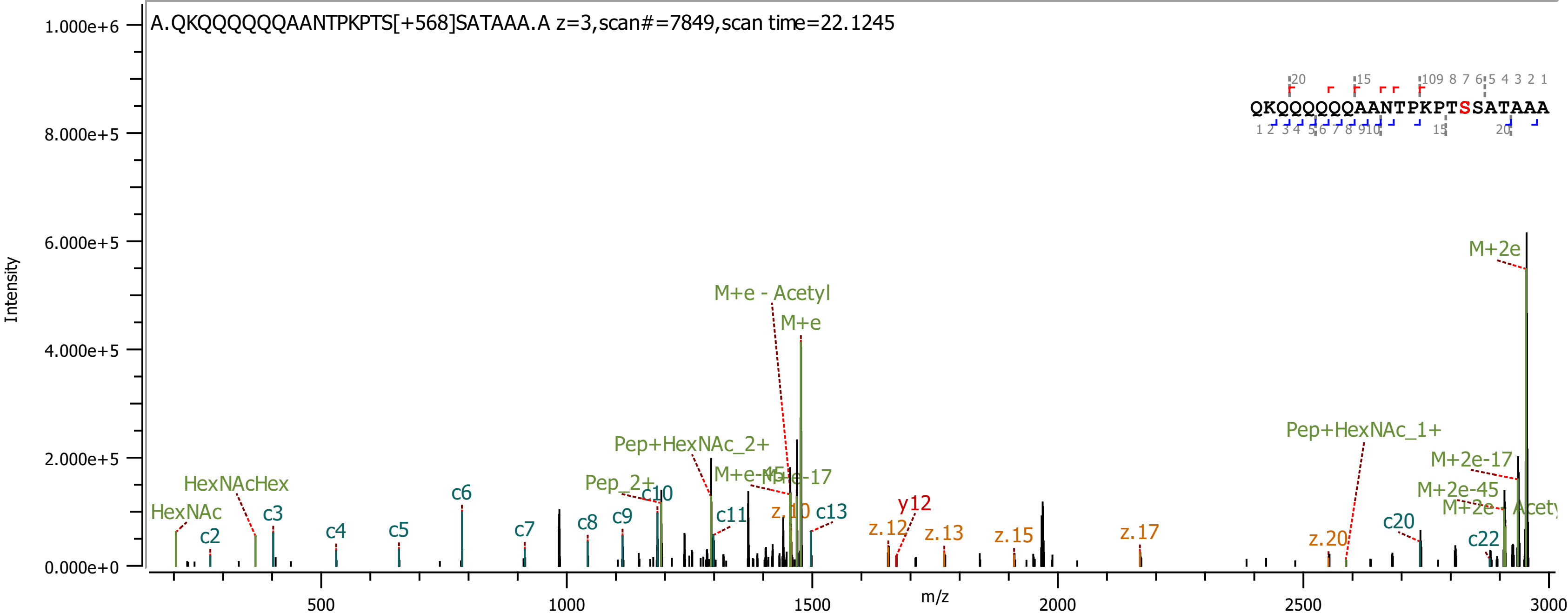
A.QKQQQQQQAANTPKPTS[+568]SATAAA.A z=3,scan#=7849,scan time=22.1245
QKQQQQQQAANTPKPTSSATAAA
Intensity
m/z
1.000e+6
8.000e+5
6.000e+5
4.000e+5
2.000e+5
0.000e+0
500
1000
1500
2000
2500
3000
M+2e
M+e - Acetyl
M+e
Pep+HexNAc_1+
M+2e-17
M+2e-45
Pep+HexNAc_2+
M+e-17
HexNAcHex
Pep_2+
HexNAc
c2
c3
c4
c5
c6
c7
c8
c9
c10
c11
c13
c20
c22
z.10
z.12
z.13
z.15
z.17
z.20
y12

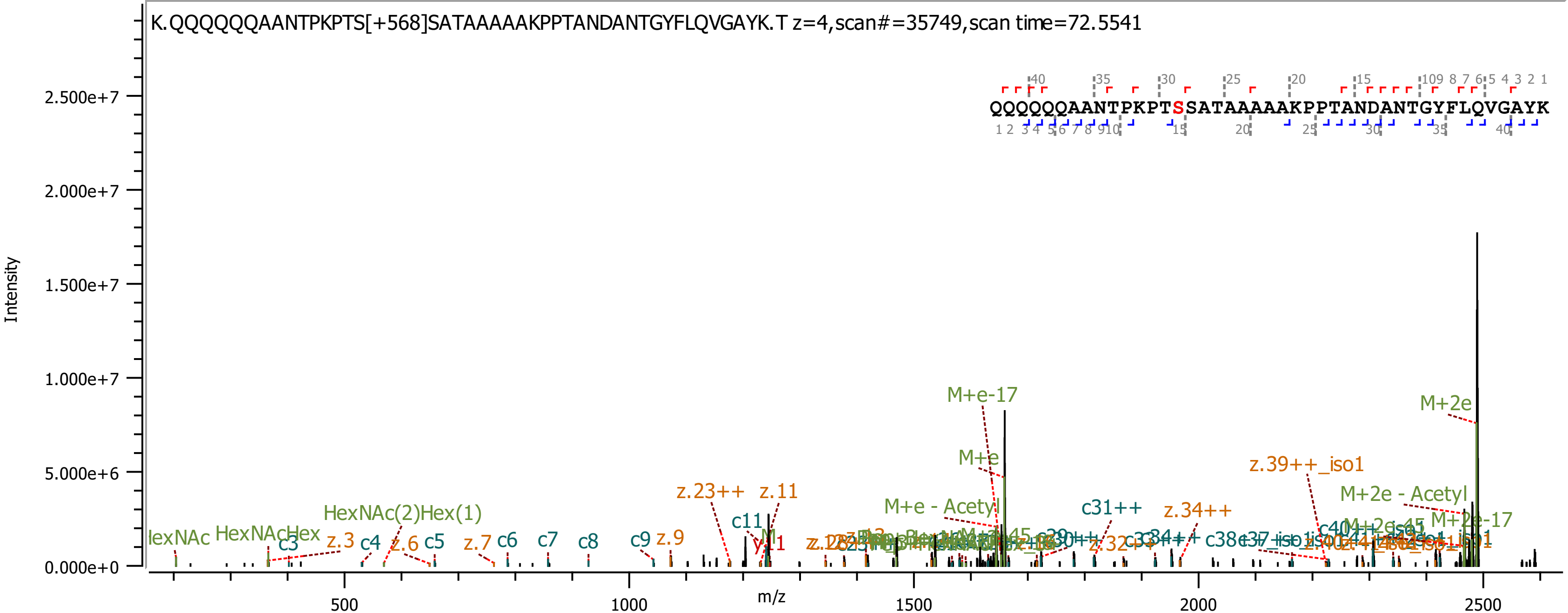

K.QQQQQQAANTPKPTS[+568]SATAAAAAKPPTANDANTGYFLQVGAYK.T z=4,scan#=35749,scan time=72.5541
QQQQQQAANTPKPTSSATAAAAAKPPTANDANTGYFLQVGAYK
Intensity
m/z
2.500e+7
2.000e+7
1.500e+7
1.000e+7
5.000e+6
0.000e+0
500
1000
1500
2000
2500
HexNAcHex
HexNAc(2)Hex(1)
c3
z.3
c4
z.6
c5
z.7
c6
c7
c8
c9
z.9
z.23++
z.11
c11
M+e-17
M+e
M+e - Acetyl
c31++
z.34++
z.32++
c38
z.39++_iso1
M+2e
M+2e - Acetyl
M+2e-17

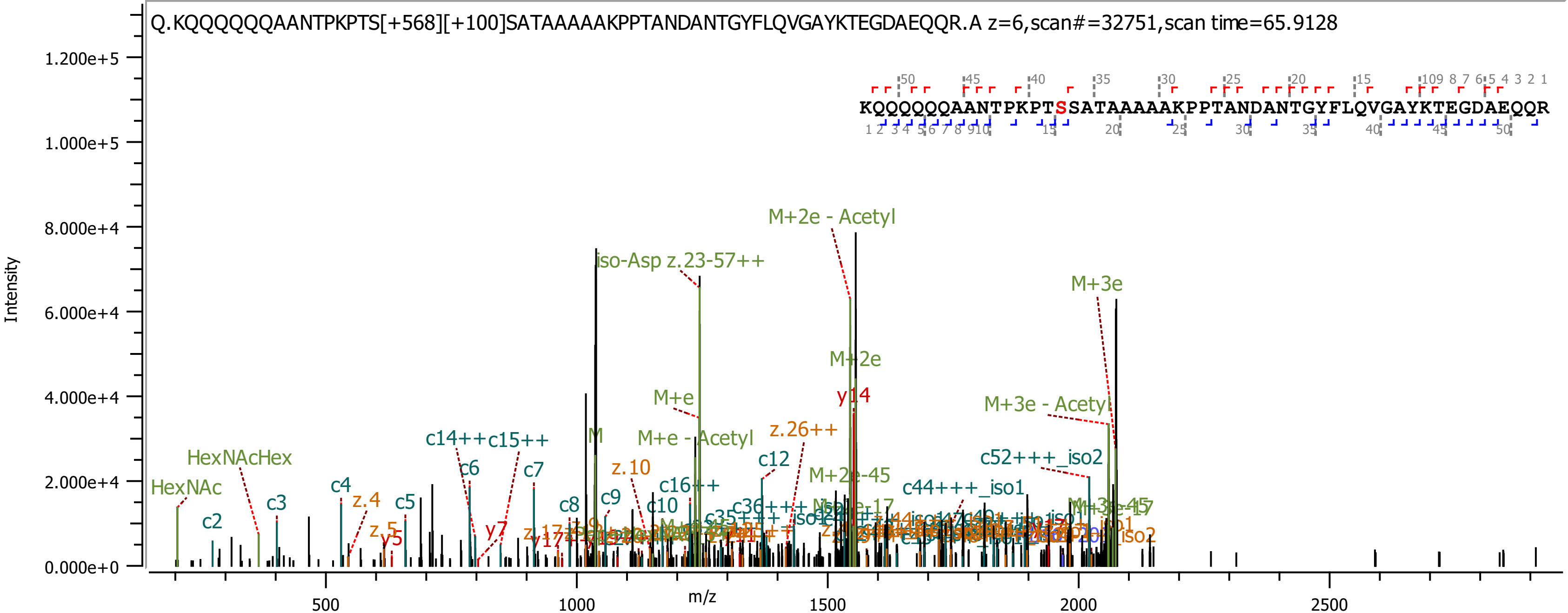

Q.KQQQQQQAANTPKPTS[+568][+100]SATAAAAAKPPTANDANTGYFLQVGAYKTEGDAEQQR.A z=6,scan#=32751,scan time=65.9128
KQQQQQQAANTPKPTSSATAAAAAKPPTANDANTGYFLQVGAYKTEGDAEQQR
Intensity
m/z
M+2e - Acetyl
iso-Asp z.23-57++
M+3e
M+2e
y14
M+e
M+3e - Acetyl
M+e - Acetyl
z.26++
c14++
c15++
HexNAcHex
HexNAc
c12
M+2e-45
c52+++_iso2
c44+++_iso1
z.10
c16++
c6
c7
c4
z.4
c5
c3
c2
c8
c9
c10
y7
y5

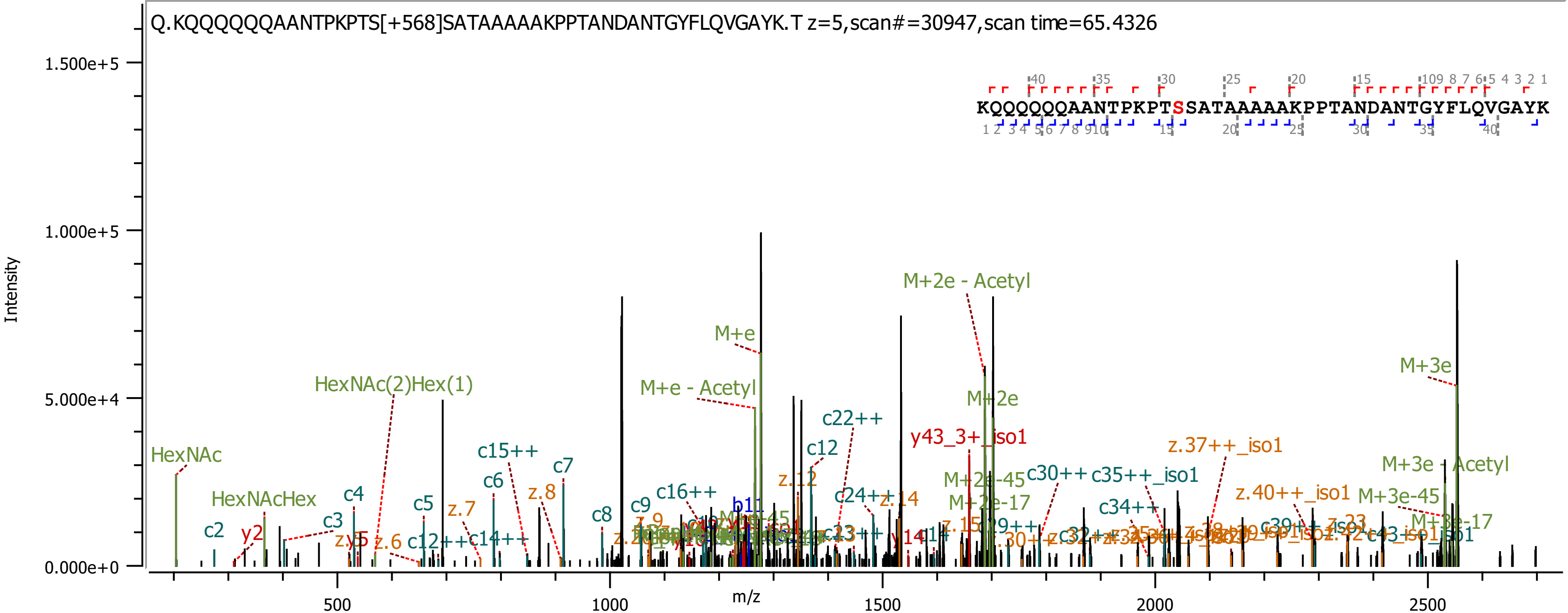

Q.KQQQQQQAANTPKPTS[+568]SATAAAAAKPPTANDANTGYFLQVGAYK.T z=5,scan#=30947,scan time=65.4326
KQQQQQQAANTPKPTSSATAAAAAKPPTANDANTGYFLQVGAYK
Intensity
m/z
1.500e+5
1.000e+5
5.000e+4
0.000e+0
500
1000
1500
2000
2500
HexNAc
HexNAcHex
HexNAc(2)Hex(1)
M+e
M+e - Acetyl
M+2e - Acetyl
M+2e
M+3e
M+3e - Acetyl
M+3e-45
M+3e-17
M+2e-45
M+2e-17
y43_3+_iso1
z.37++_iso1
z.40++_iso1
c2
c3
c4
c5
c6
c7
c8
c9
c12
c15++
c16++
c22++
c24++
c30++
c35++_iso1
c34++
y2
z.6
z.7
z.8
z.12
z.14

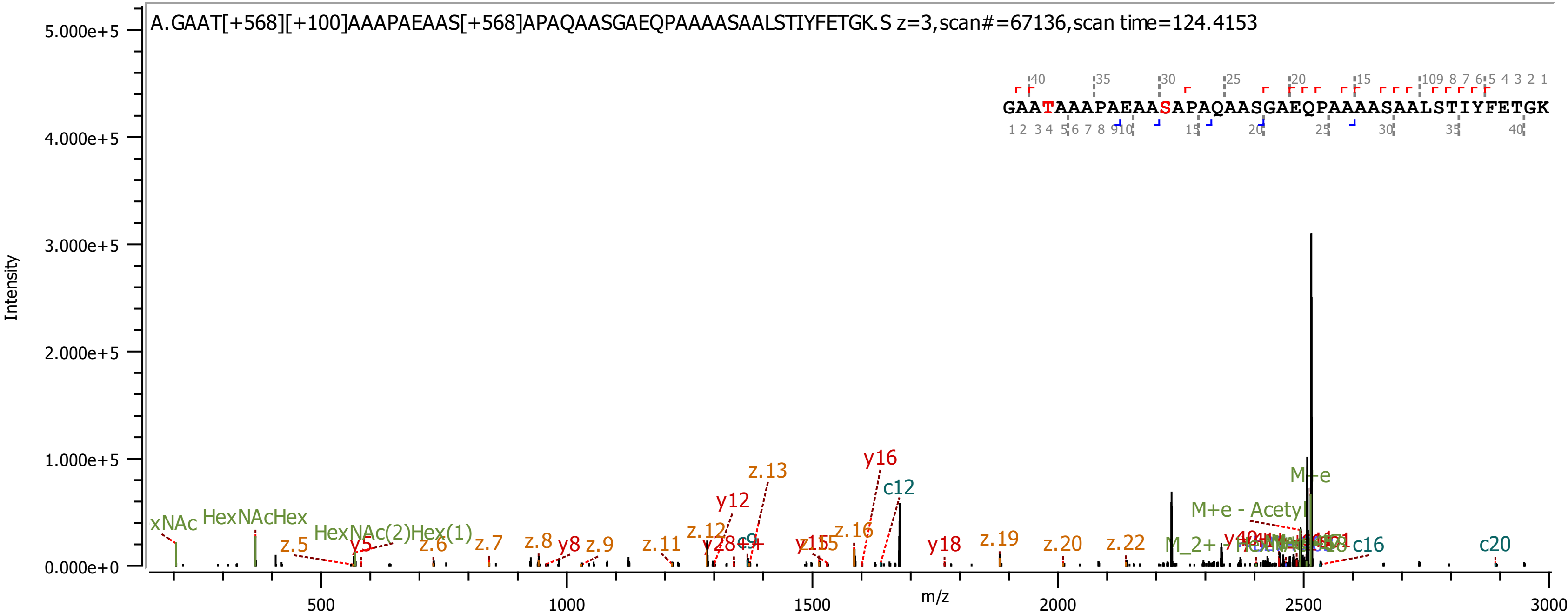

A.GAAT[+568][+100]AAAPAEAAS[+568]APAQAASGAEQPAAAASAALSTIYFETGK.S z=3,scan#=67136,scan time=124.4153
GAATAAAPAEAASAPAQAASGAEQPAAAASAALSTIYFETGK
Intensity
m/z
5.000e+5
4.000e+5
3.000e+5
2.000e+5
1.000e+5
0.000e+0
500
1000
1500
2000
2500
3000
HexNAcHex
HexNAc(2)Hex(1)
z.5
y5
z.6
z.7
z.8
y8
z.9
z.11
z.12
y12
z.13
y15
z.16
y16
c12
y18
z.19
z.20
z.22
M+e
M+e - Acetyl
c16
c20

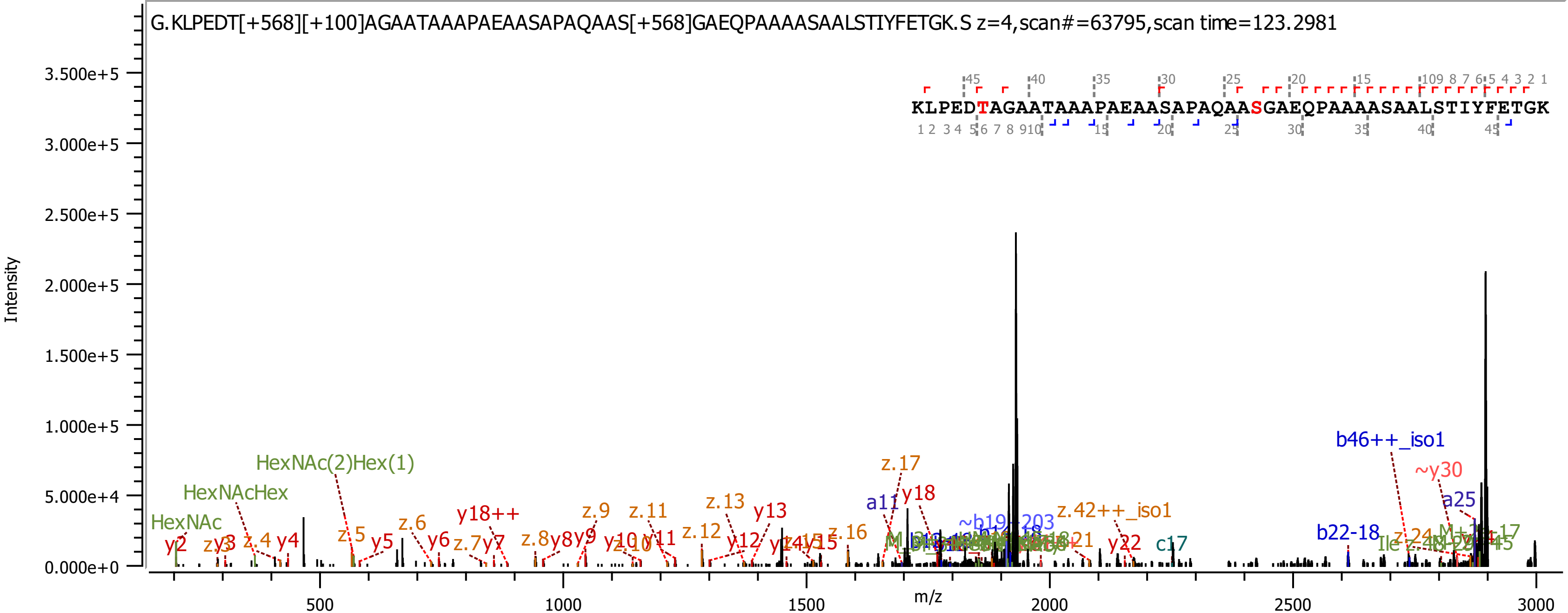
G.KLPEDT[+568][+100]AGAATAAAPAEAASAPAQAAS[+568]GAEQPAAAASAALSTIYFETGK.S z=4,scan#=63795,scan time=123.2981
KLPEDTAGAATAAAPAEAASAPAQAASGAEQPAAAASAALSTIYFETGK
Intensity
m/z
HexNAc(2)Hex(1)
HexNAcHex
HexNAc
b46++_iso1
~y30
a25
z.17
y18
a11
z.42++_iso1
y22
c17
b22-18
y13
z.13
z.11
z.9
y18++

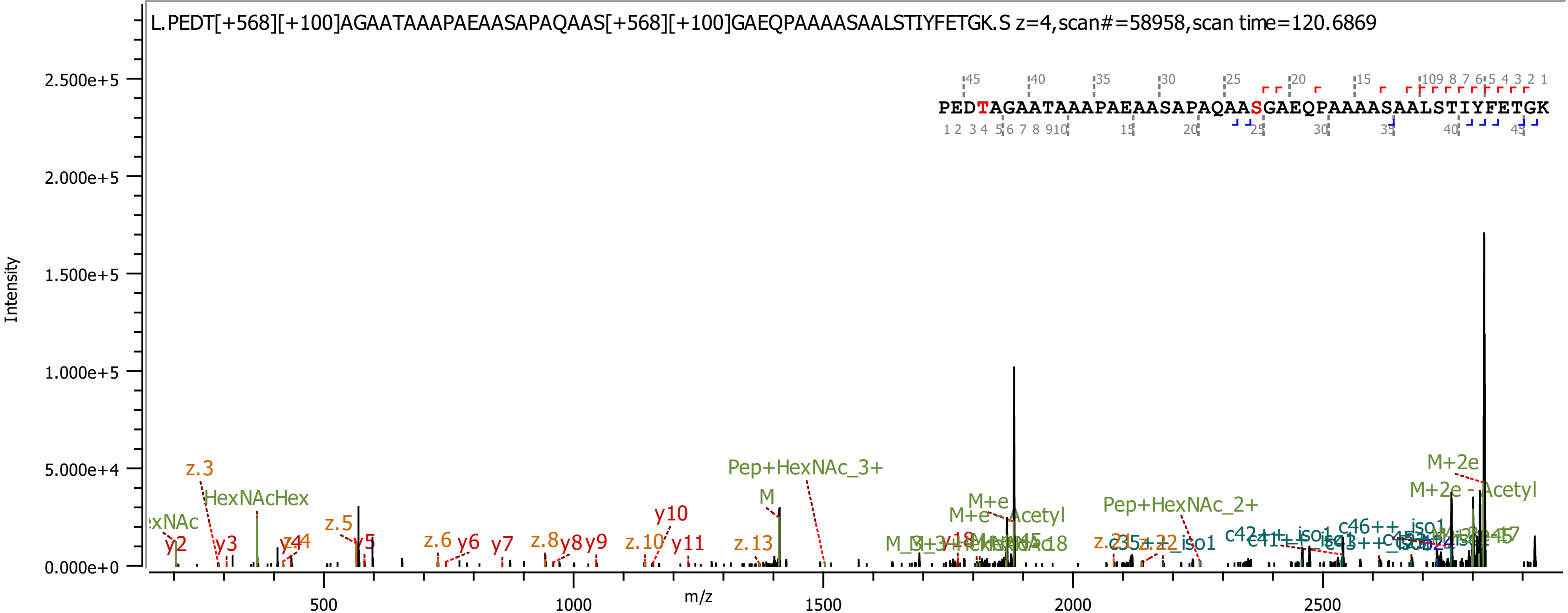
L.PEDT[+568][+100]AGAATAAAPAEAASAPAQAAS[+568][+100]GAEQPAAAASAALSTIYFETGK.S z=4,scan#=58958,scan time=120.6869
PEDTAGAATAAAPAEAASAPAQAASGAEQPAAAASAALSTIYFETGK
Intensity
m/z
2.500e+5
2.000e+5
1.500e+5
1.000e+5
5.000e+4
0.000e+0
500
1000
1500
2000
2500
z.3
HexNAcHex
y2
y3
y4
z.5
y5
z.6
y6
y7
z.8
y8
y9
z.10
y10
y11
z.13
M
Pep+HexNAc_3+
M+e
M+e - Acetyl
Pep+HexNAc_2+
M+2e
M+2e - Acetyl
c46++_iso1

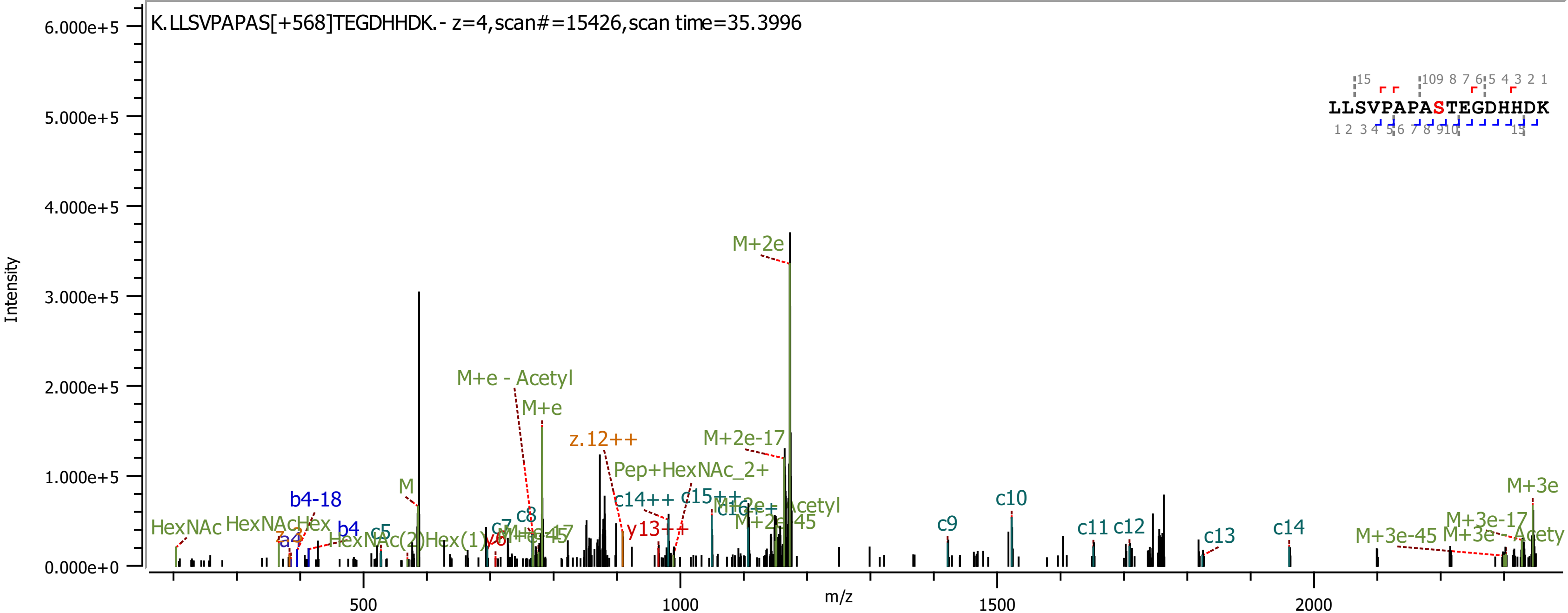
K.LLSVPAPAS[+568]TEGDHHDK.- z=4,scan#=15426,scan time=35.3996
Intensity
6.000e+5
5.000e+5
4.000e+5
3.000e+5
2.000e+5
1.000e+5
0.000e+0
m/z
500
1000
1500
2000
LLSVPAPASTEGDHHDK
M+2e
M+e - Acetyl
M+e
z.12++
M+2e-17
Pep+HexNAc_2+
c14++
c15++
y13++
M
b4-18
b4
c5
c7
c8
HexNAc
HexNAcHex
HexNAc(2)Hex(1)
c9
c10
c11
c12
c13
c14
M+3e
M+3e-17
M+3e-45

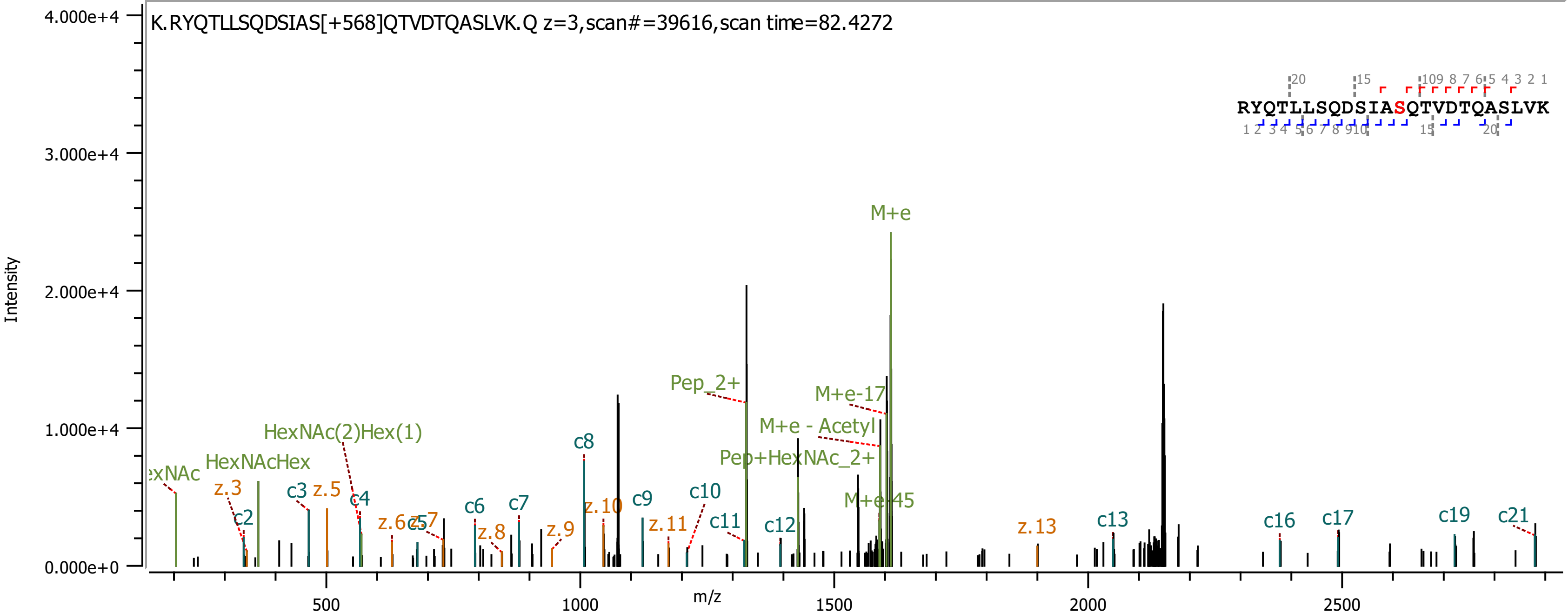
K.RYQTLLSQDSIAS[+568]QTVDTQASLVK.Q z=3,scan#=39616,scan time=82.4272
RYQTLLSQDSIASQTVDTQASLVK
Intensity
m/z
M+e
Pep_2+
M+e-17
M+e - Acetyl
Pep+HexNAc_2+
M+e-45
HexNAc(2)Hex(1)
HexNAcHex
exNAc
c2 c3 c4 c5 c6 c7 c8 c9 c10 c11 c12 c13 c16 c17 c19 c21
z.3 z.5 z.6 z.7 z.8 z.9 z.10 z.11 z.13

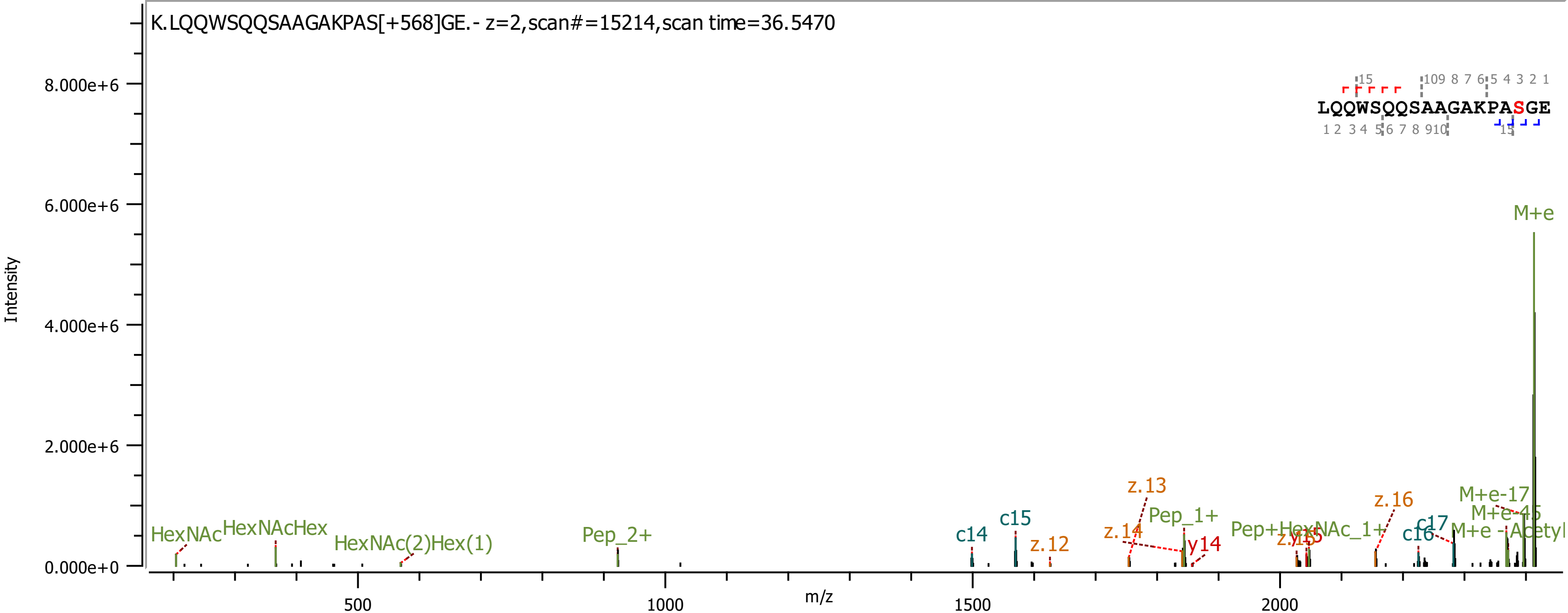
K.LQQWSQQSAAGAKPAS[+568]GE.- z=2,scan#=15214,scan time=36.5470
LQQWSQQSAAGAKPASGE
Intensity
m/z
M+e
M+e-17
M+e-45
M+e -Acetyl
HexNAc
HexNAcHex
HexNAc(2)Hex(1)
Pep_2+
Pep_1+
Pep+HexNAc_1+
c14
c15
c16
c17
z.12
z.13
z.14
z.16
y14

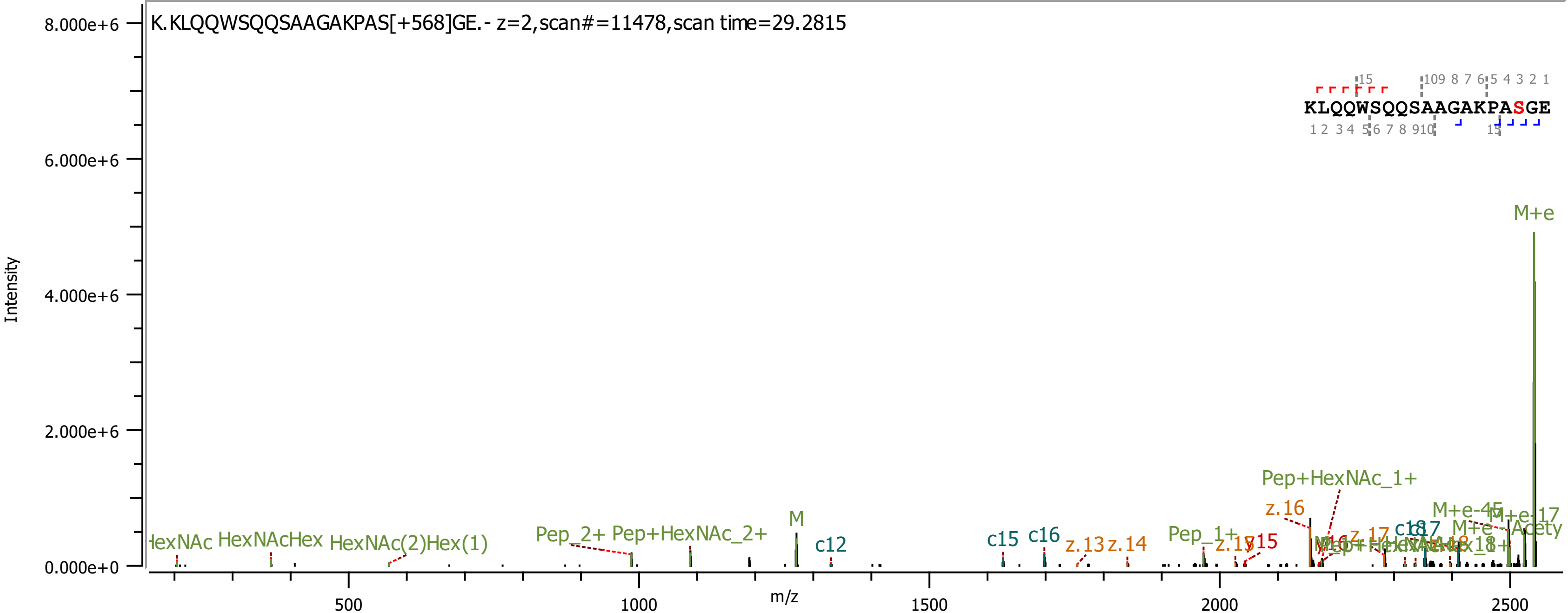
K.KLQQWSQQSAAGAKPAS[+568]GE.- z=2,scan#=11478,scan time=29.2815
KLQQWSQQSAAGAKPASGE
Intensity
m/z
8.000e+6
6.000e+6
4.000e+6
2.000e+6
0.000e+0
500
1000
1500
2000
2500
M+e
Pep+HexNAc_1+
z.16
M+e-45
M+e-17
c17
z.17
Pep_1+
z.15
y15
c15
c16
z.13
z.14
M
c12
Pep_2+
Pep+HexNAc_2+
HexNAcHex
HexNAc(2)Hex(1)

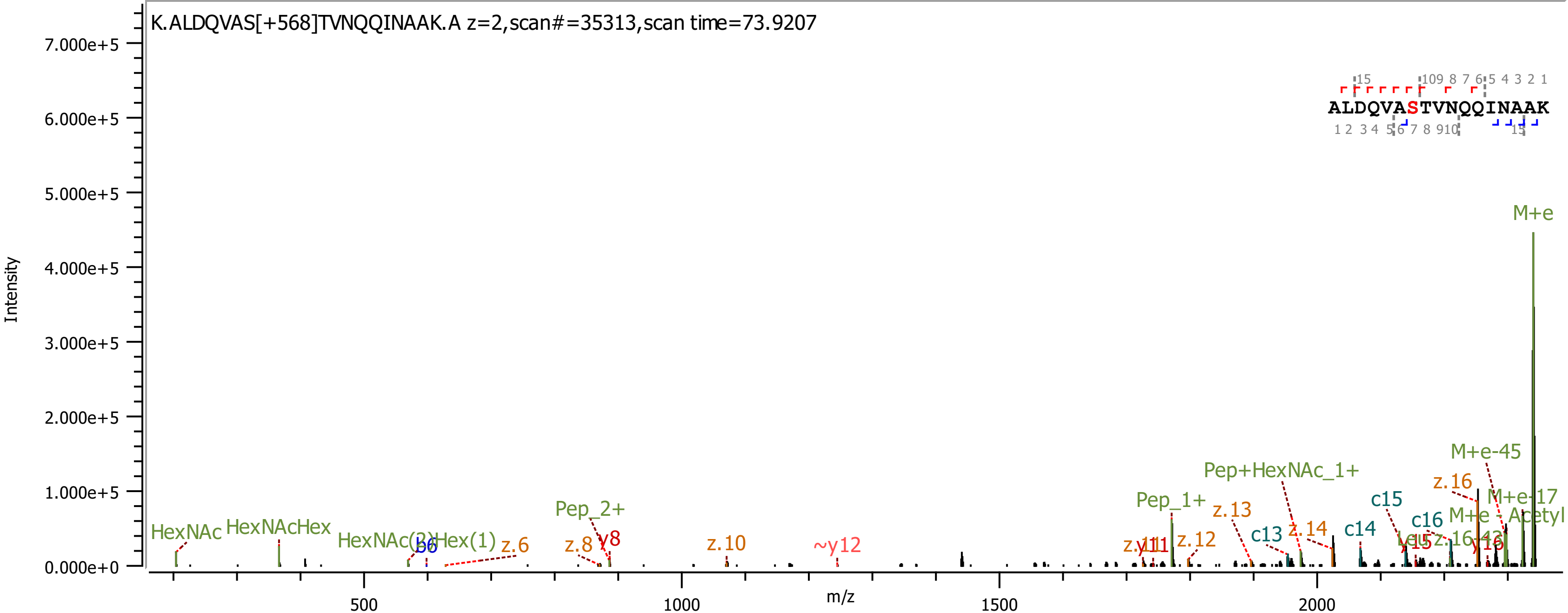
K.ALDQVAS[+568]TVNQQINAAK.A z=2,scan#=35313,scan time=73.9207
Intensity
7.000e+5
6.000e+5
5.000e+5
4.000e+5
3.000e+5
2.000e+5
1.000e+5
0.000e+0
m/z
500
1000
1500
2000
ALDQVASTVNQQINAAK
M+e
M+e-45
M+e-17
M+e - Acetyl
Pep+HexNAc_1+
Pep_1+
Pep_2+
HexNAc
HexNAcHex
HexNAc(2)Hex(1)
b6
z.6
z.8
y8
z.10
~y12
y11
z.12
z.13
c13
z.14
c14
c15
c16
y15
z.16
y16

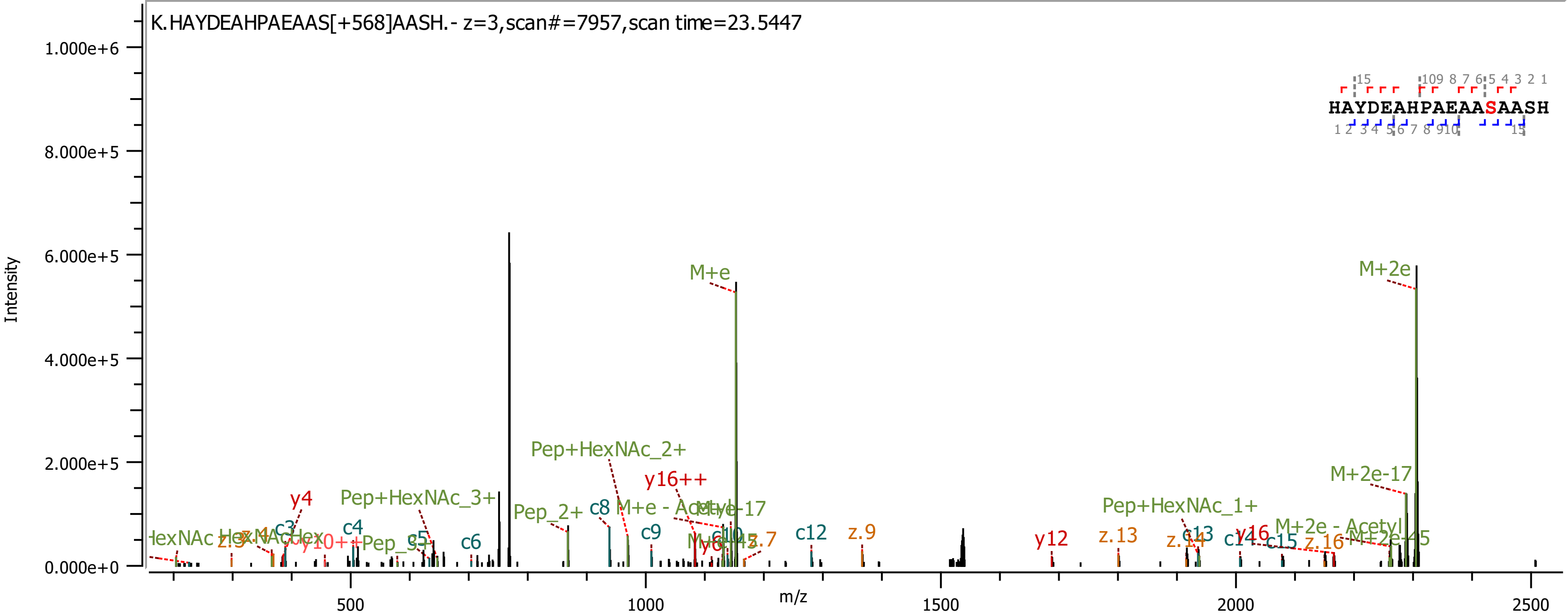

K.HAYDEAHPAEAAS[+568]AASH.- z=3,scan#=7957,scan time=23.5447
HAYDEAHPAEAASAASH
Intensity
m/z
M+e
M+2e
M+2e-17
Pep+HexNAc_2+
Pep+HexNAc_3+
Pep+HexNAc_1+
Pep_2+
y16++
y4
y12
y16
c3
c4
c5
c6
c8
c9
c12
c13
c14
c15
z.9
z.13
z.14
z.16

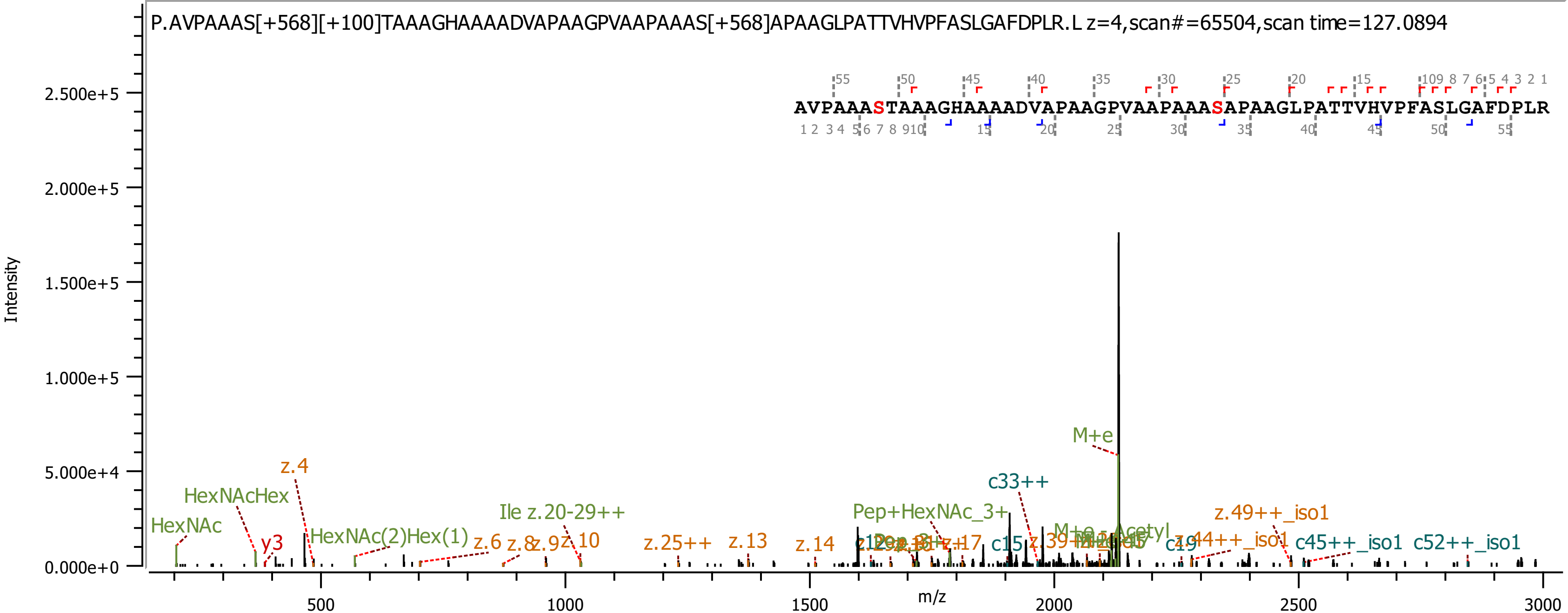

P.AVPAAAS[+568][+100]TAAAGHAAAADVAPAAGPVAAPAAAS[+568]APAAGLPATTVHVPFASLGAFDPLR.L z=4,scan#=65504,scan time=127.0894
AVPAAASTAAAGHAAAADVAPAAGPVAAPAAASAPAAGLPATTVHVPFASLGAFDPLR
Intensity
m/z
0.000e+0
5.000e+4
1.000e+5
1.500e+5
2.000e+5
2.500e+5
500
1000
1500
2000
2500
3000
HexNAc
HexNAcHex
y3
z.4
HexNAc(2)Hex(1)
z.6
z.8
z.9
z.10
Ile z.20-29++
z.25++
z.13
z.14
Pep+HexNAc_3+
z.17
c15
c33++
M+e
M+e - Acetyl
c19
z.44++_iso1
z.49++_iso1
c45++_iso1
c52++_iso1

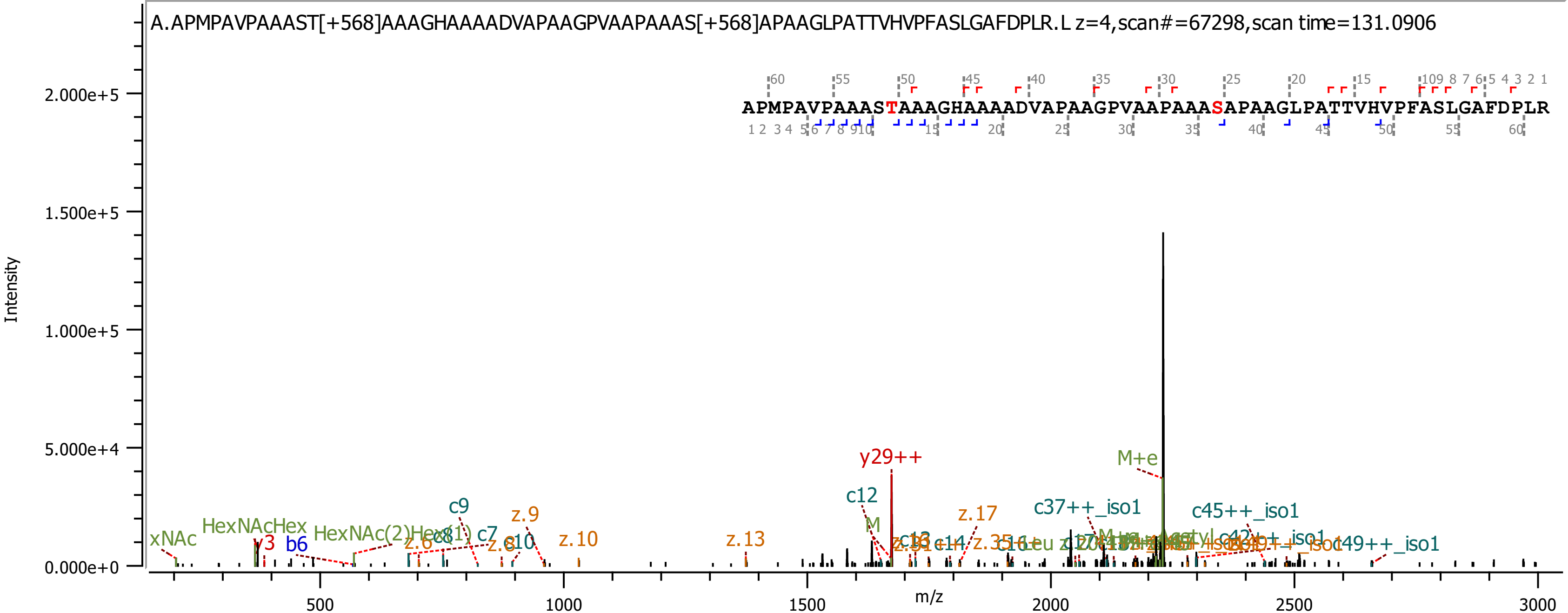
A.APMPAVPAAAST[+568]AAAGHAAAADVAPAAGPVAAPAAAS[+568]APAAGLPATTVHVPFASLGAFDPLR.L z=4,scan#=67298,scan time=131.0906
APMPAVPAAASTAAAGHAAAADVAPAAGPVAAPAAASAPAAGLPATTVHVPFASLGAFDPLR
Intensity
m/z
y29++
M+e
c12
c9
z.9
z.17
c37++_iso1
c45++_iso1
xNAc
HexNAcHex
y3
b6
c7
z.10
z.13
c49++_iso1

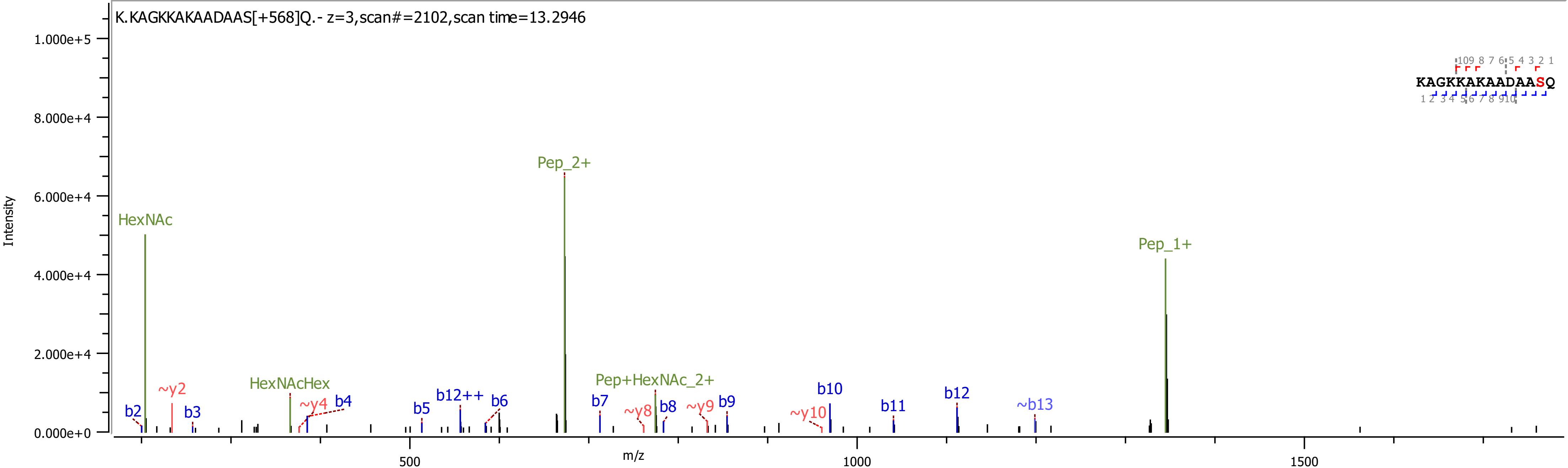

K.KAGKKAKAADAAS[+568]Q.- z=3,scan#=2102,scan time=13.2946
KAGKKAKAADAASQ
Intensity
m/z
HexNAc
Pep_2+
Pep_1+
HexNAcHex
Pep+HexNAc_2+
b2
~y2
b3
~y4
b4
b5
b12++
b6
b7
~y8
b8
~y9
b9
~y10
b10
b11
b12
~b13

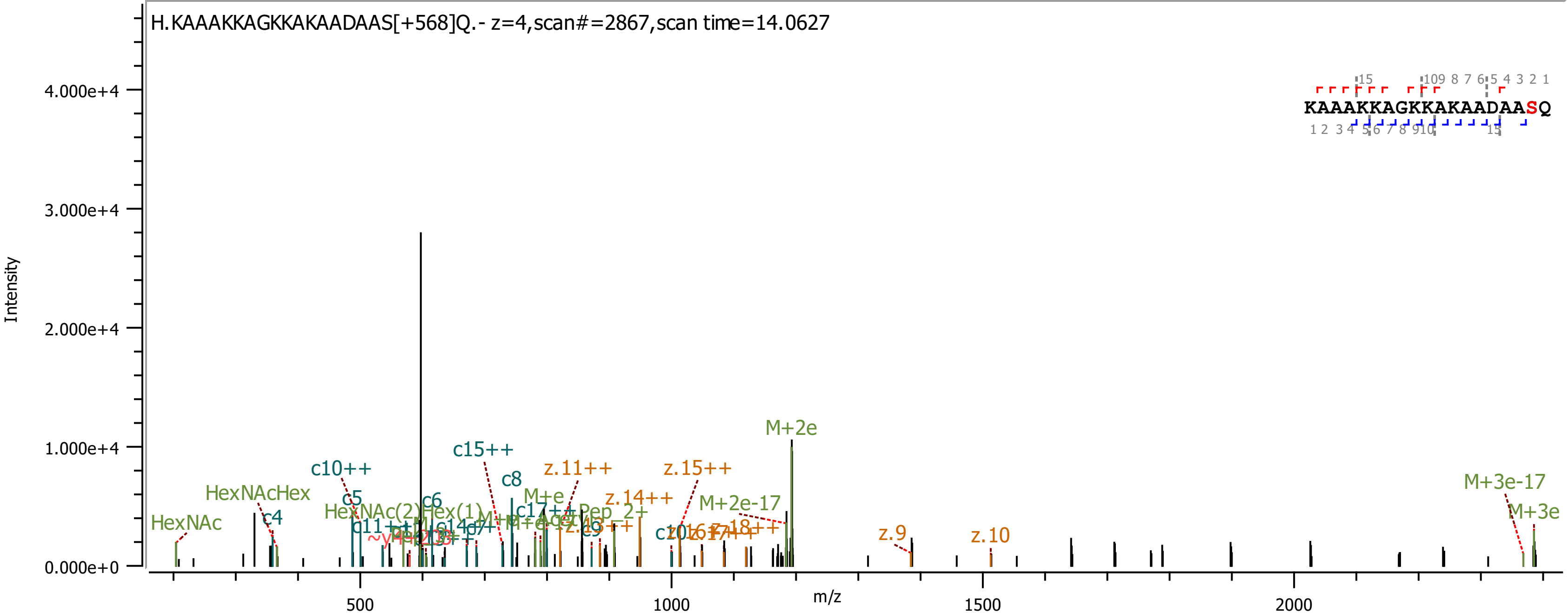
H.KAAAKKAGKKAKAADAAS[+568]Q.- z=4,scan#=2867,scan time=14.0627
KAAAKKAGKKAKAADAASQ
Intensity
m/z
4.000e+4
3.000e+4
2.000e+4
1.000e+4
0.000e+0
500
1000
1500
2000
HexNAc
HexNAcHex
c4
c5
c10++
c6
c8
c15++
M+e
z.11++
z.14++
z.15++
M+2e-17
M+2e
z.9
z.10
M+3e-17
M+3e

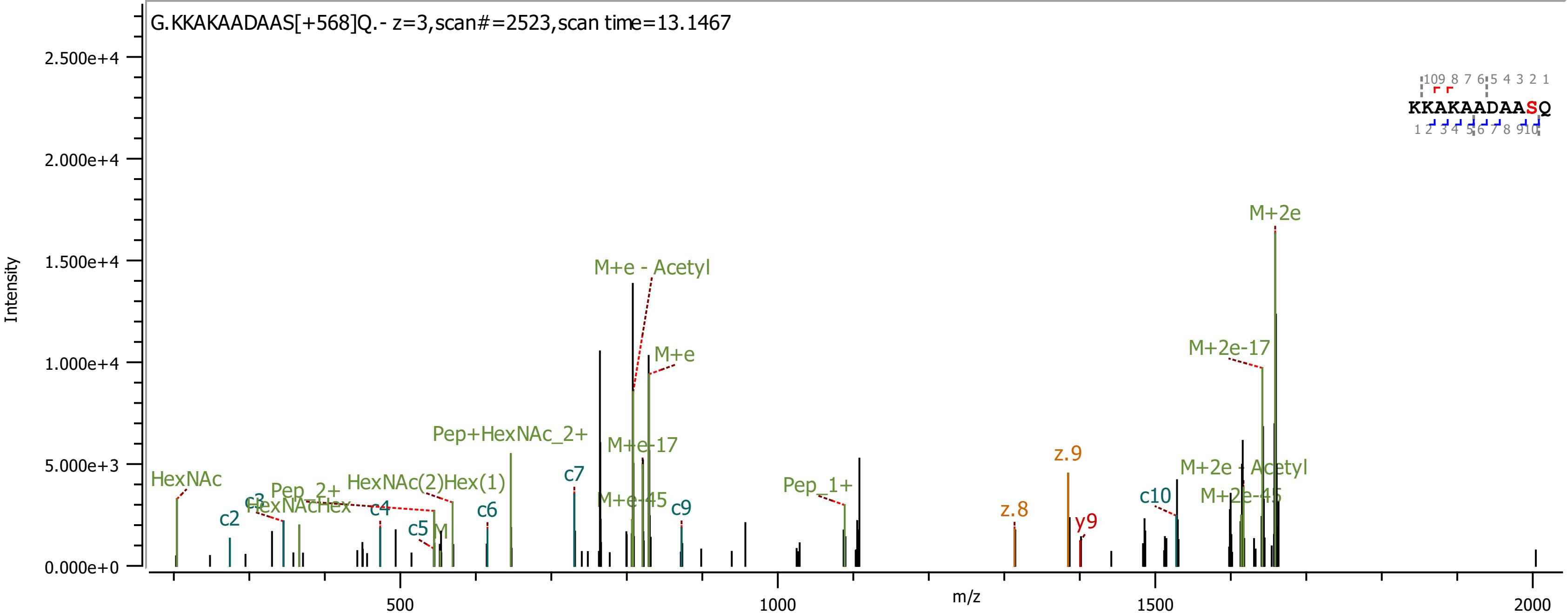

G.KKAKAADAAS[+568]Q.- z=3,scan#=2523,scan time=13.1467
Intensity
m/z
KKAKAADAASQ
HexNAc
c2
c3
HexNAcHex
Pep_2+
c4
HexNAc(2)Hex(1)
c5
M
c6
Pep+HexNAc_2+
c7
M+e - Acetyl
M+e
M+e-17
M+e-45
c9
Pep_1+
z.8
z.9
y9
c10
M+2e
M+2e-17
M+2e - Acetyl
M+2e-45

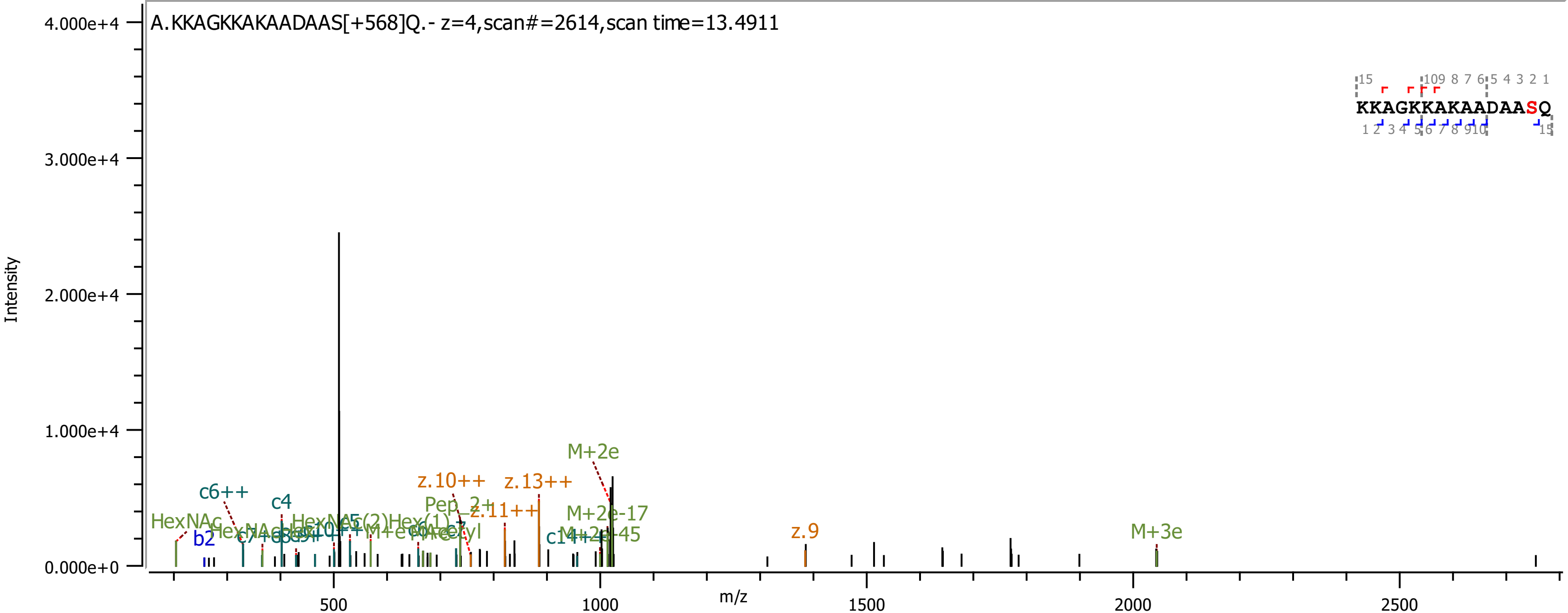
A.KKAGKKAKAADAAS[+568]Q.- z=4,scan#=2614,scan time=13.4911
Intensity
m/z
4.000e+4
3.000e+4
2.000e+4
1.000e+4
0.000e+0
500
1000
1500
2000
2500
KKAGKKAKAADAASQ
HexNAc
b2
c6++
c4
z.10++
z.13++
z.11++
Pep_2+
M+2e
M+2e-17
M+2e-45
z.9
M+3e

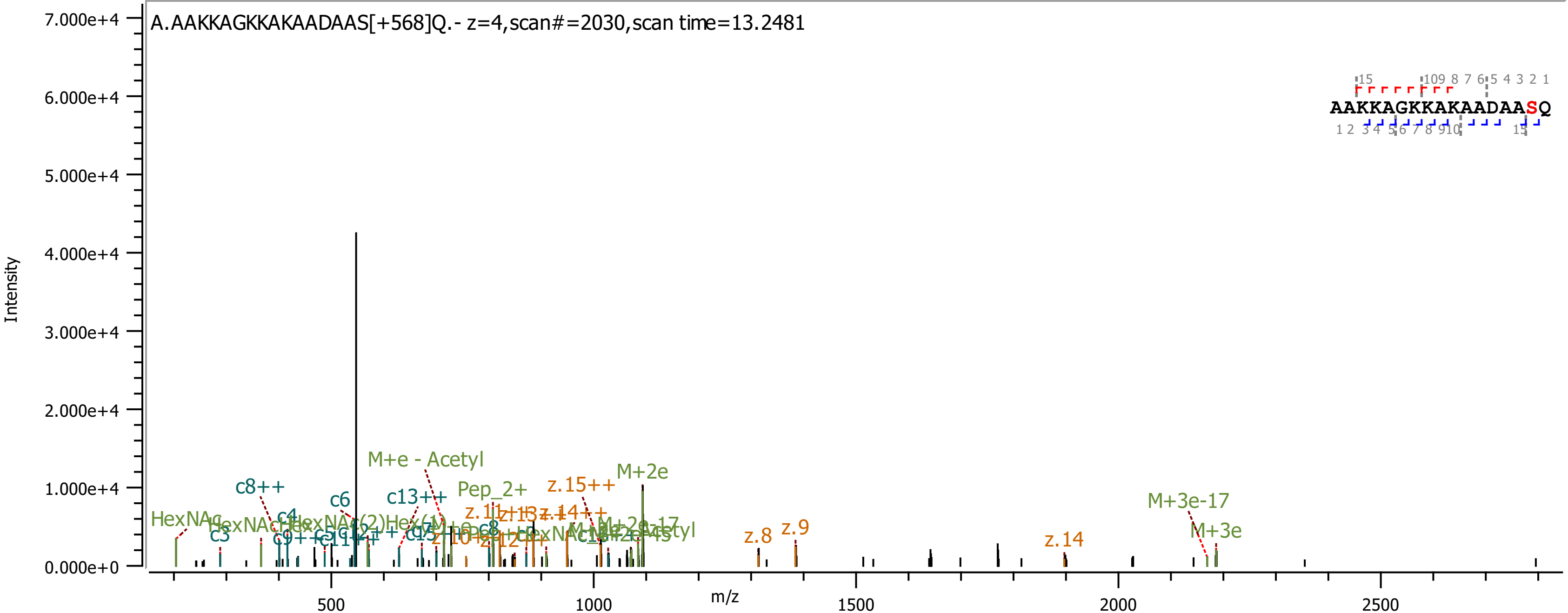
A.AAKKAGKKAKAADAAS[+568]Q.- z=4,scan#=2030,scan time=13.2481
AAKKAGKKAKAADAASQ
Intensity
7.000e+4
6.000e+4
5.000e+4
4.000e+4
3.000e+4
2.000e+4
1.000e+4
0.000e+0
m/z
500
1000
1500
2000
2500
HexNAc
c3
c8++
c4
c6
c13++
M+e - Acetyl
Pep_2+
z.15++
M+2e
z.8
z.9
z.14
M+3e-17
M+3e

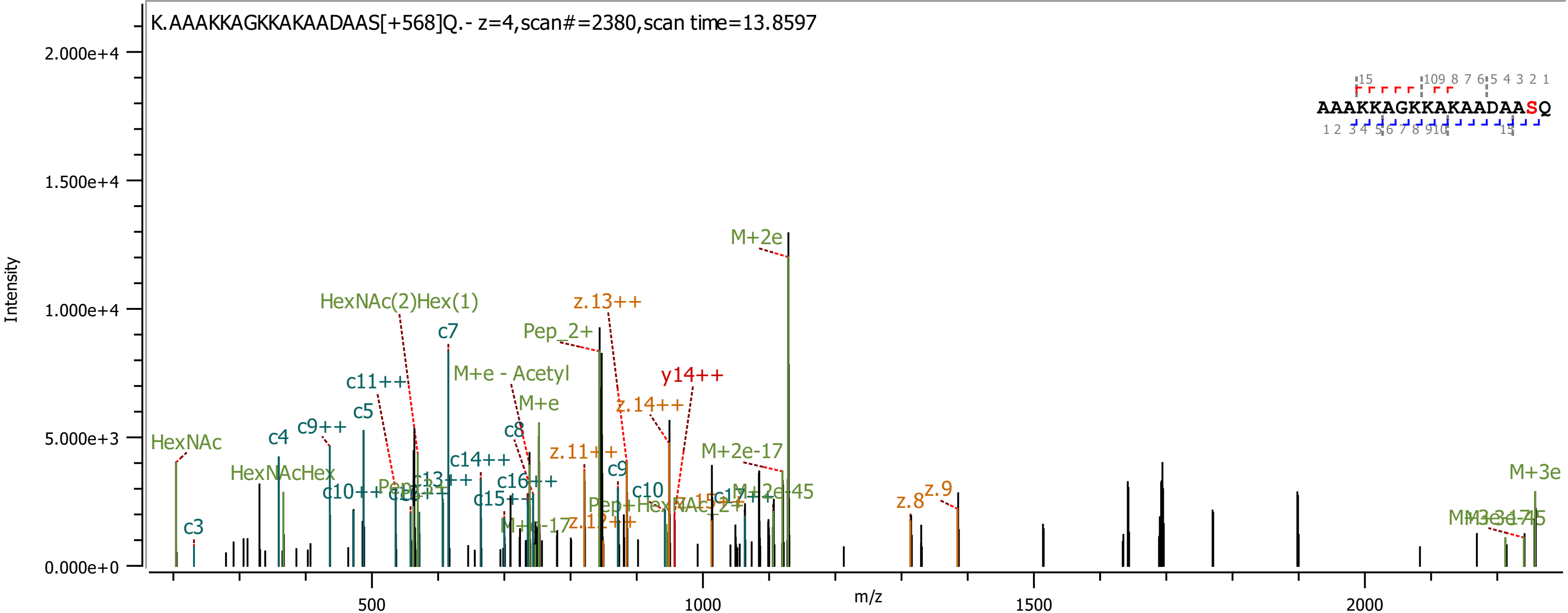
K.AAAKKAGKKAKAADAAS[+568]Q.- z=4,scan#=2380,scan time=13.8597
AAAKKAGKKAKAADAASQ
Intensity
m/z
2.000e+4
1.500e+4
1.000e+4
5.000e+3
0.000e+0
500
1000
1500
2000
M+2e
HexNAc(2)Hex(1)
z.13++
c7
Pep_2+
M+e - Acetyl
c11++
y14++
M+e
z.14++
c5
c9++
c8
c4
HexNAc
c14++
z.11++
M+2e-17
HexNAcHex
c9
c16++
c10
z.9
z.8
M+3e
c3

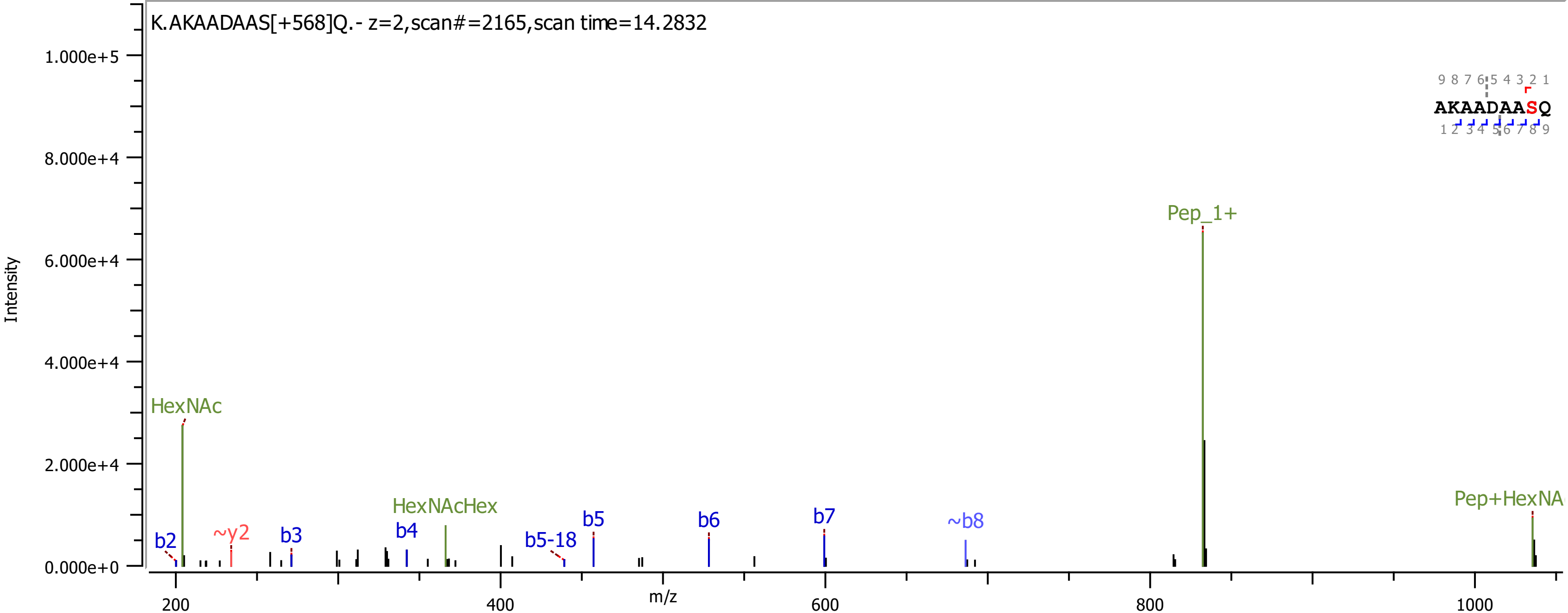

K.AKAADAAS[+568]Q.- z=2,scan#=2165,scan time=14.2832
Intensity
1.000e+5
8.000e+4
6.000e+4
4.000e+4
2.000e+4
0.000e+0
m/z
200
400
600
800
1000
AKAADAASQ
9 8 7 6 5 4 3 2 1
1 2 3 4 5 6 7 8 9
HexNAc
b2
~y2
b3
b4
HexNAcHex
b5-18
b5
b6
b7
~b8
Pep_1+
Pep+HexNA

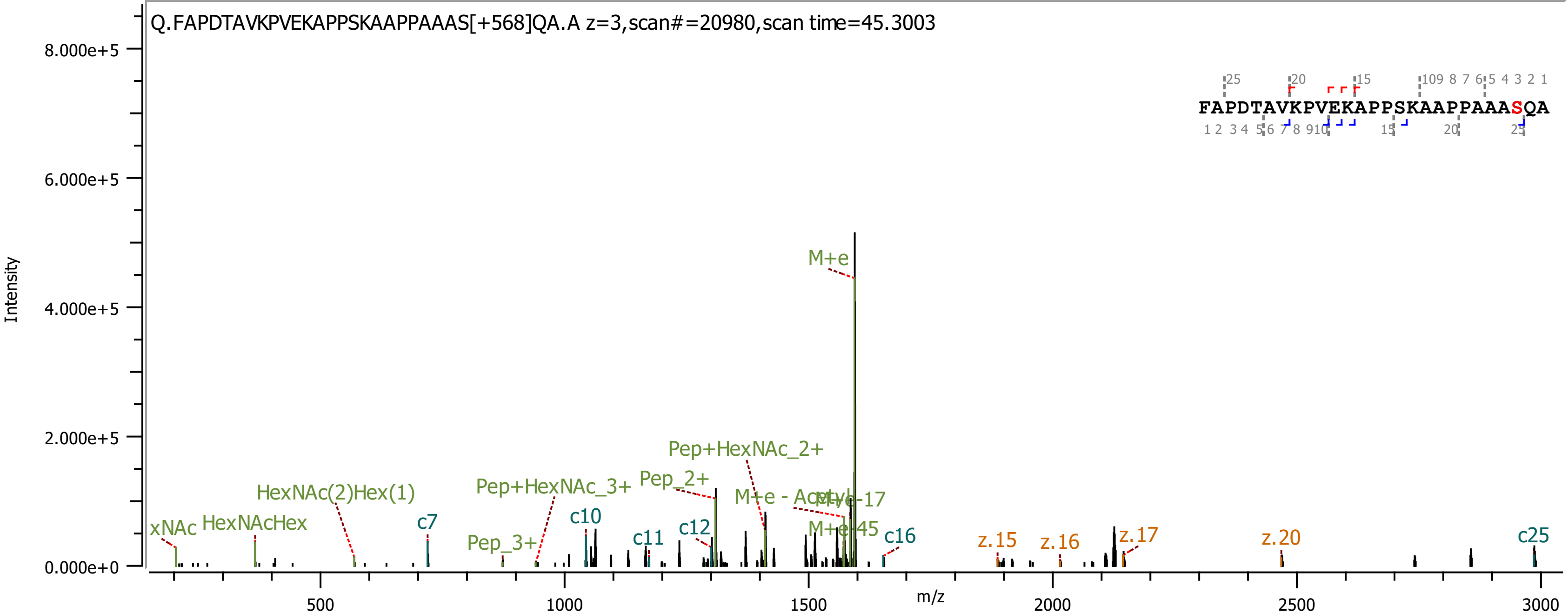
Q.FAPDTAVKPVEKAPPSKAAPPAAAS[+568]QA.A z=3,scan#=20980,scan time=45.3003
FAPDTAVKPVEKAPPSKAAPPAAASQA
Intensity
m/z
8.000e+5
6.000e+5
4.000e+5
2.000e+5
0.000e+0
500
1000
1500
2000
2500
3000
M+e
Pep+HexNAc_2+
Pep_2+
Pep+HexNAc_3+
HexNAc(2)Hex(1)
HexNAcHex
xNAc
Pep_3+
M+e - Acetyl
M+e-17
M+e-45
c7
c10
c11
c12
c16
c25
z.15
z.16
z.17
z.20

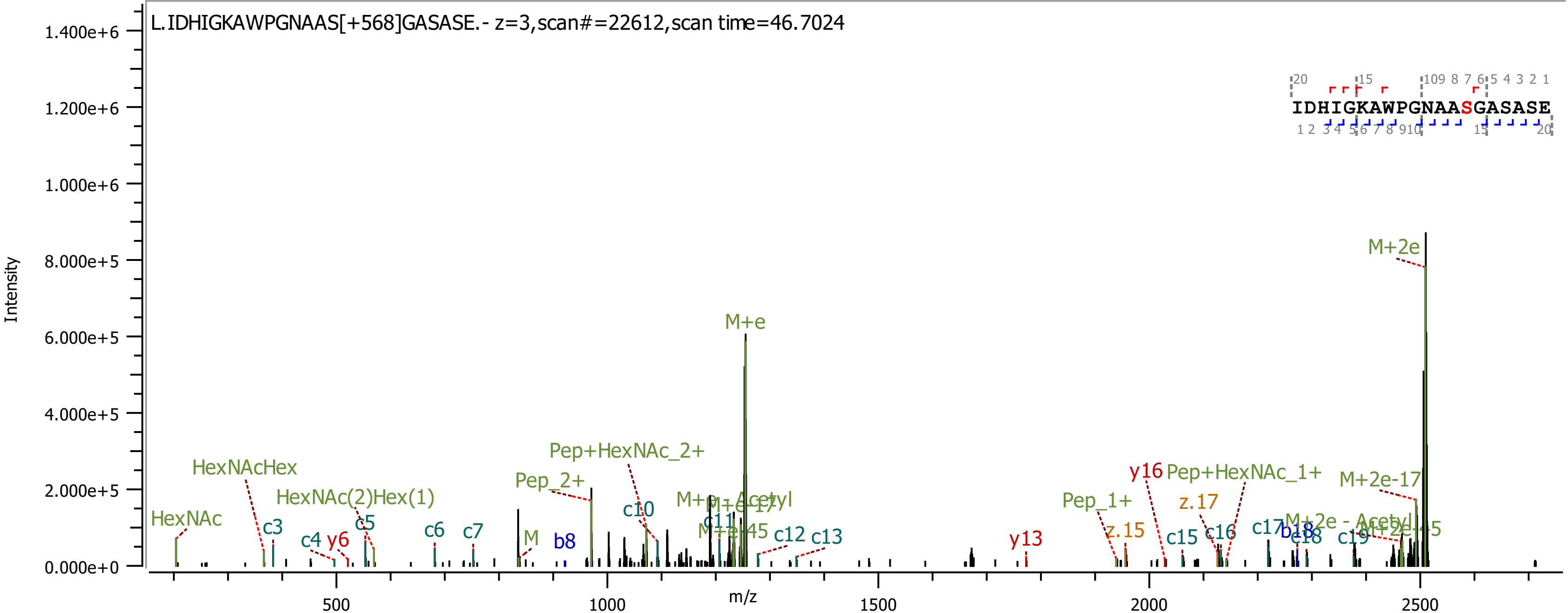

L.IDHIGKAWPGNAAS[+568]GASASE.- z=3,scan#=22612,scan time=46.7024
IDHIGKAWPGNAASGASASE
Intensity
m/z
0.000e+0
2.000e+5
4.000e+5
6.000e+5
8.000e+5
1.000e+6
1.200e+6
1.400e+6
500
1000
1500
2000
2500
HexNAc
HexNAcHex
HexNAc(2)Hex(1)
c3
c4
y6
c5
c6
c7
M
b8
Pep_2+
Pep+HexNAc_2+
c10
M+e
c11
c12
c13
y13
Pep_1+
z.15
y16
Pep+HexNAc_1+
z.17
c15
c16
c17
b18
c18
c19
M+2e-17
M+2e

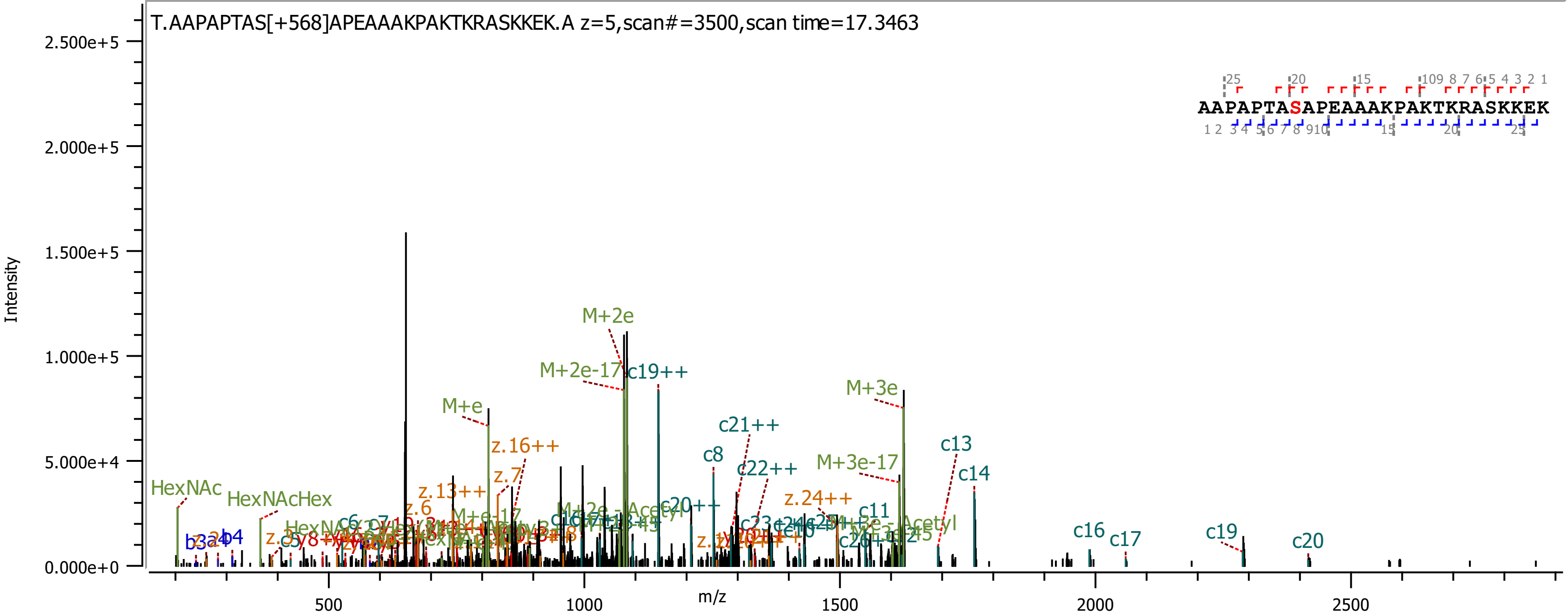
T.AAPAPTAS[+568]APEAAAKPAKTKRASKKEK.A z=5,scan#=3500,scan time=17.3463
Intensity
m/z
2.500e+5
2.000e+5
1.500e+5
1.000e+5
5.000e+4
0.000e+0
500
1000
1500
2000
2500
AAPAPTASAPEAAAKPAKTKRASKKEK
M+2e
M+2e-17
c19++
M+e
M+3e
c21++
z.16++
c8
c22++
M+3e-17
c13
c14
z.7
z.13++
z.24++
HexNAc
HexNAcHex
z.6
c20++
c11
c16
c17
c19
c20
b3
b4

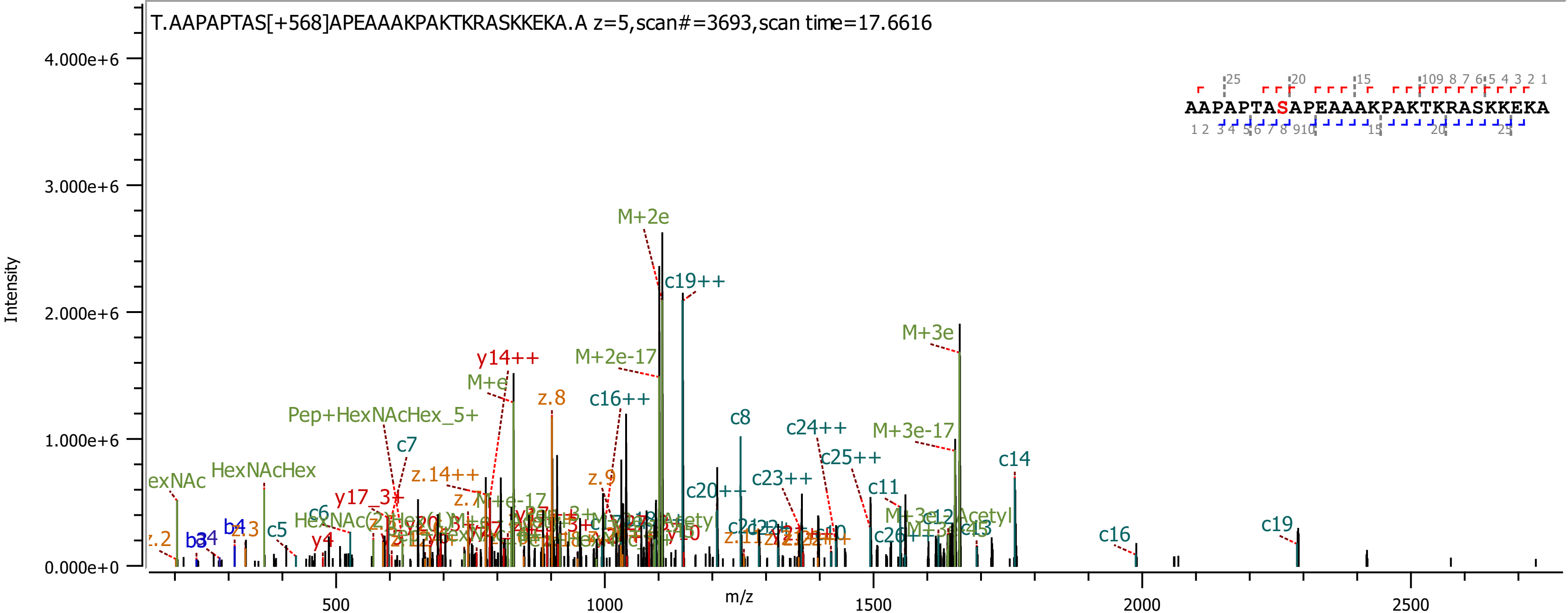

T.AAPAPTAS[+568]APEAAAKPAKTKRASKKEKA.A z=5,scan#=3693,scan time=17.6616
AAPAPTASAPEAAAKPAKTKRASKKEKA
Intensity
m/z
4.000e+6
3.000e+6
2.000e+6
1.000e+6
0.000e+0
500
1000
1500
2000
2500
M+2e
c19++
M+3e
M+2e-17
y14++
M+e
z.8
c16++
Pep+HexNAcHex_5+
c7
c8
c24++
M+3e-17
c25++
c14
HexNAcHex
z.14++
z.9
c20++
c23++
c11
y17_3+
c6
c5
b4
z.3
y4
c16
c19

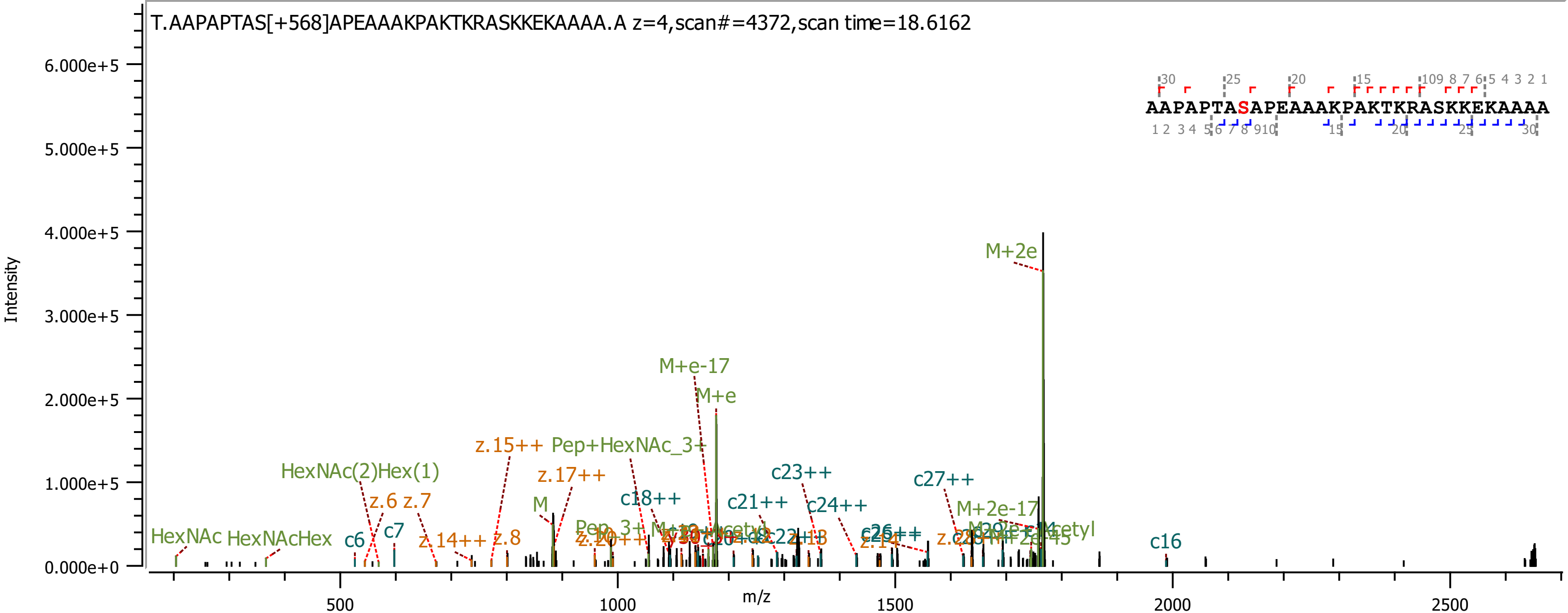
T.AAPAPTAS[+568]APEAAAKPAKTKRASKKEKAAAA.A z=4,scan#=4372,scan time=18.6162
AAPAPTASAPEAAAKPAKTKRASKKEKAAAA
Intensity
m/z
6.000e+5
5.000e+5
4.000e+5
3.000e+5
2.000e+5
1.000e+5
0.000e+0
500
1000
1500
2000
2500
M+2e
M+e-17
M+e
z.15++
Pep+HexNAc_3+
z.17++
HexNAc(2)Hex(1)
z.6
z.7
M
c18++
c21++
c23++
c24++
c27++
M+2e-17
c7
c6
HexNAc
HexNAcHex
z.14++
z.8
c16
z.14
c26++

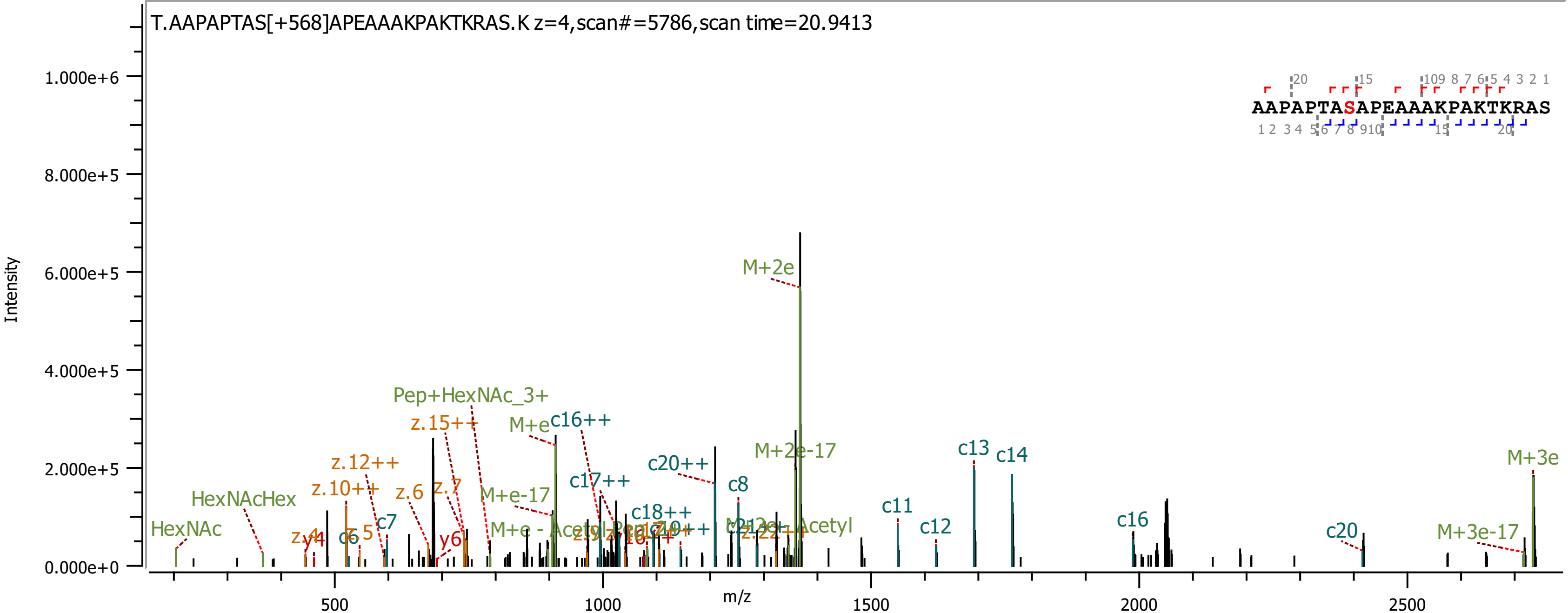

T.AAPAPTAS[+568]APEAAAKPAKTKRAS.K z=4,scan#=5786,scan time=20.9413
AAPAPTASAPEAAAKPAKTKRAS
Intensity
m/z
1.000e+6
8.000e+5
6.000e+5
4.000e+5
2.000e+5
0.000e+0
500
1000
1500
2000
2500
HexNAc
HexNAcHex
z.12++
z.10++
z.15++
Pep+HexNAc_3+
z.6
z.7
y6
c7
c6
M+e
c16++
M+e-17
c17++
c20++
c18++
c8
M+2e
M+2e-17
c11
c12
c13
c14
c16
c20
M+3e
M+3e-17

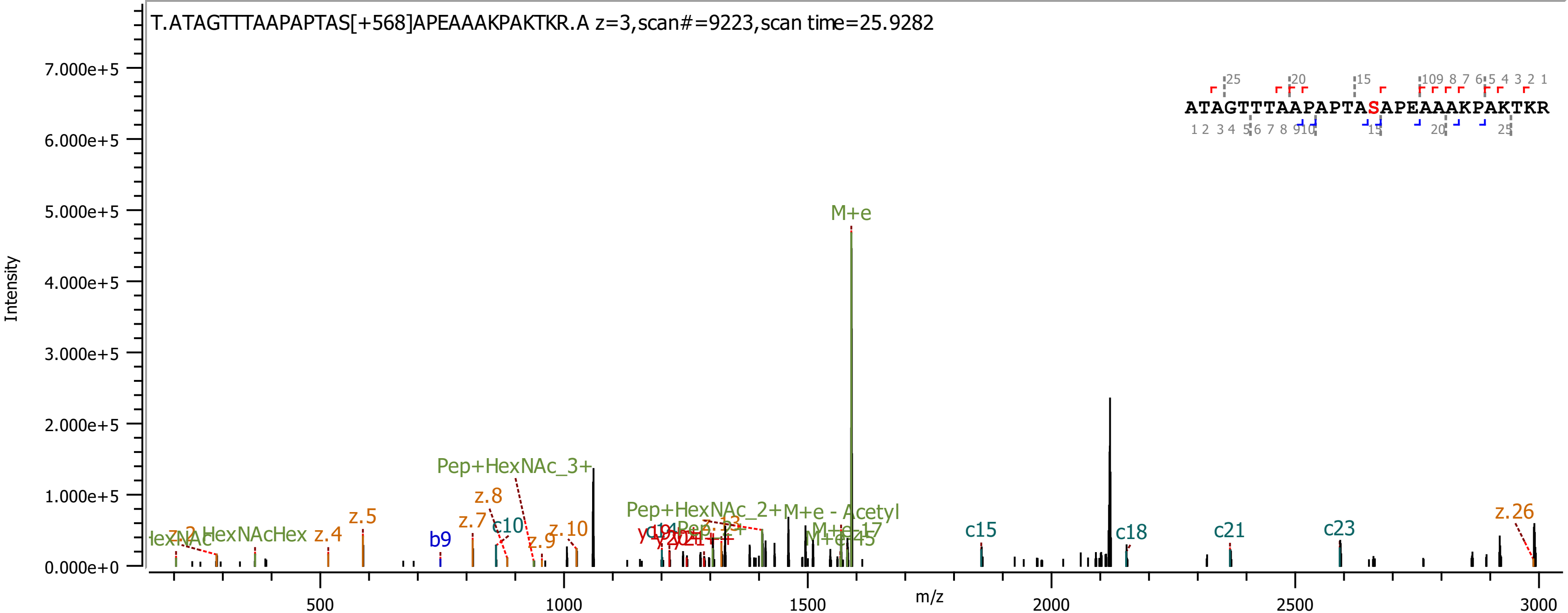

T.ATAGTTTAAPAPTAS[+568]APEAAAKPAKTKR.A z=3,scan#=9223,scan time=25.9282
ATAGTTTAAPAPTASAPEAAAKPAKTKR
Intensity
m/z
M+e
Pep+HexNAc_3+
Pep+HexNAc_2+
M+e - Acetyl
z.5
z.4
z.7
z.8
z.9
z.10
z.26
b9
c10
c15
c18
c21
c23

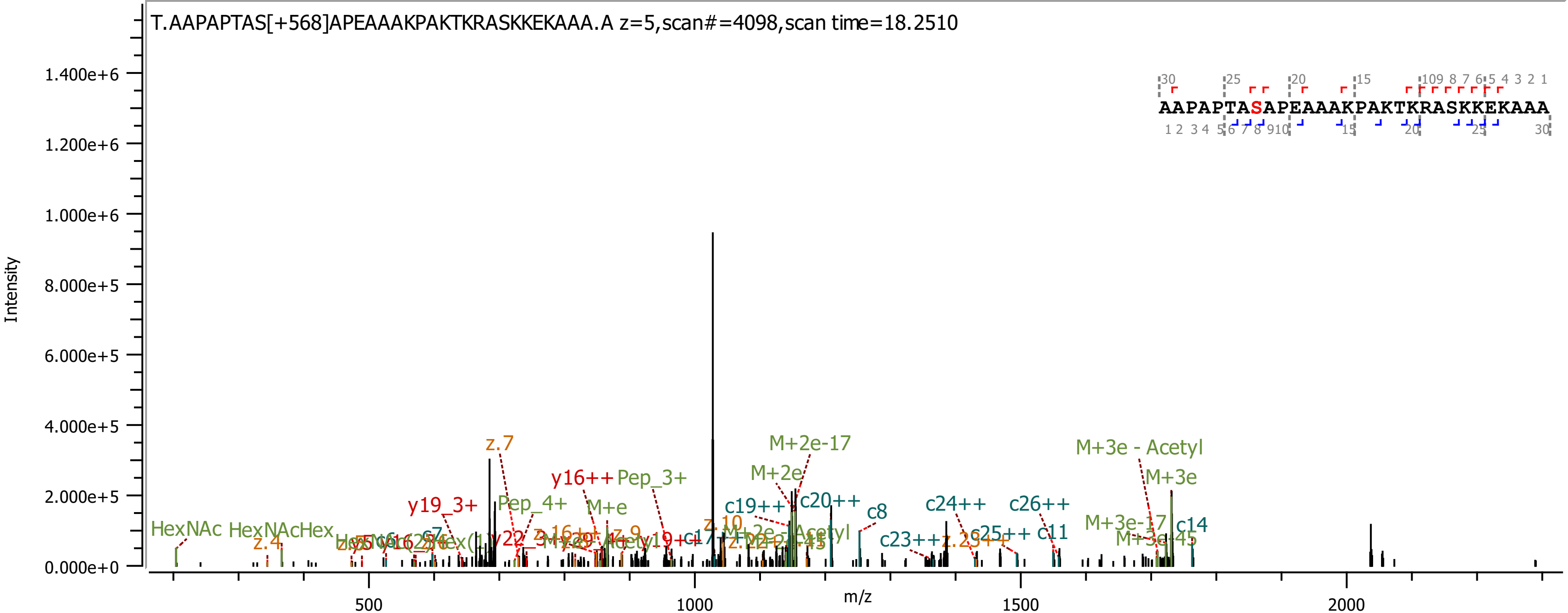
T.AAPAPTAS[+568]APEAAAKPAKTKRASKKEKAAA.A z=5,scan#=4098,scan time=18.2510
AAPAPTASAPEAAAKPAKTKRASKKEKAAA
Intensity
m/z
1.400e+6
1.200e+6
1.000e+6
8.000e+5
6.000e+5
4.000e+5
2.000e+5
0.000e+0
500
1000
1500
2000
z.7
y16++
Pep_3+
y19_3+
Pep_4+
M+e
M+2e-17
M+2e
c19++
c20++
c8
c24++
c26++
M+3e - Acetyl
M+3e
M+3e-17
c14
M+3e-45
c23++
c25++
c11
HexNAc
HexNAcHex
z.4
z.10

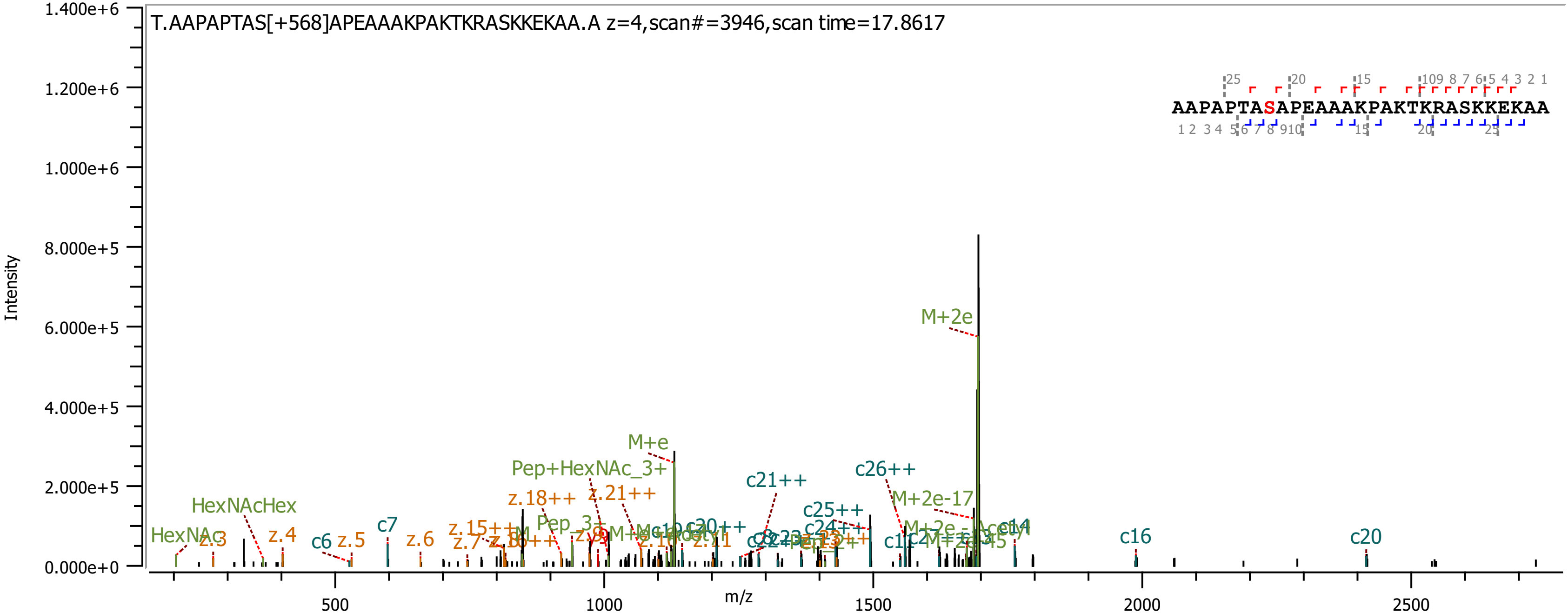

T.AAPAPTAS[+568]APEAAAKPAKTKRASKKEKAA.A z=4,scan#=3946,scan time=17.8617
AAPAPTASAPEAAAKPAKTKRASKKEKAA
Intensity
m/z
M+2e
M+e
Pep+HexNAc_3+
c21++
c26++
c25++
M+2e-17
z.18++
z.21++
HexNAcHex
HexNAc
z.3
z.4
c6
z.5
c7
z.6
c16
c20
c24++
1.400e+6
1.200e+6
1.000e+6
8.000e+5
6.000e+5
4.000e+5
2.000e+5
0.000e+0
500
1000
1500
2000
2500

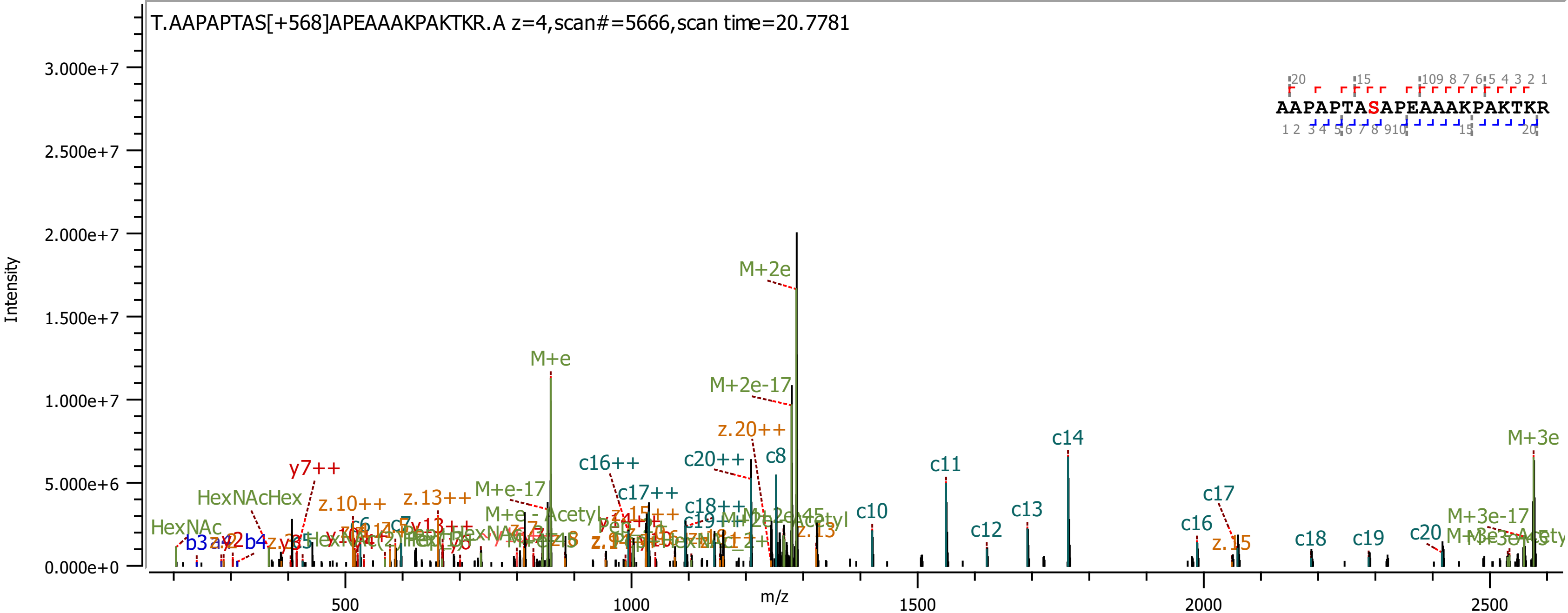
T.AAPAPTAS[+568]APEAAAKPAKTKR.A z=4,scan#=5666,scan time=20.7781
AAPAPTASAPEAAAKPAKTKR
Intensity
3.000e+7
2.500e+7
2.000e+7
1.500e+7
1.000e+7
5.000e+6
0.000e+0
m/z
500
1000
1500
2000
2500
M+2e
M+e
M+2e-17
z.20++
c8
c14
M+3e
c20++
c16++
c11
y7++
M+e-17
c17++
HexNAcHex
z.10++
z.13++
c18++
c10
c13
c17
c16
M+3e-17
HexNAc
c12
z.15
c18
c19
c20
b3
b4

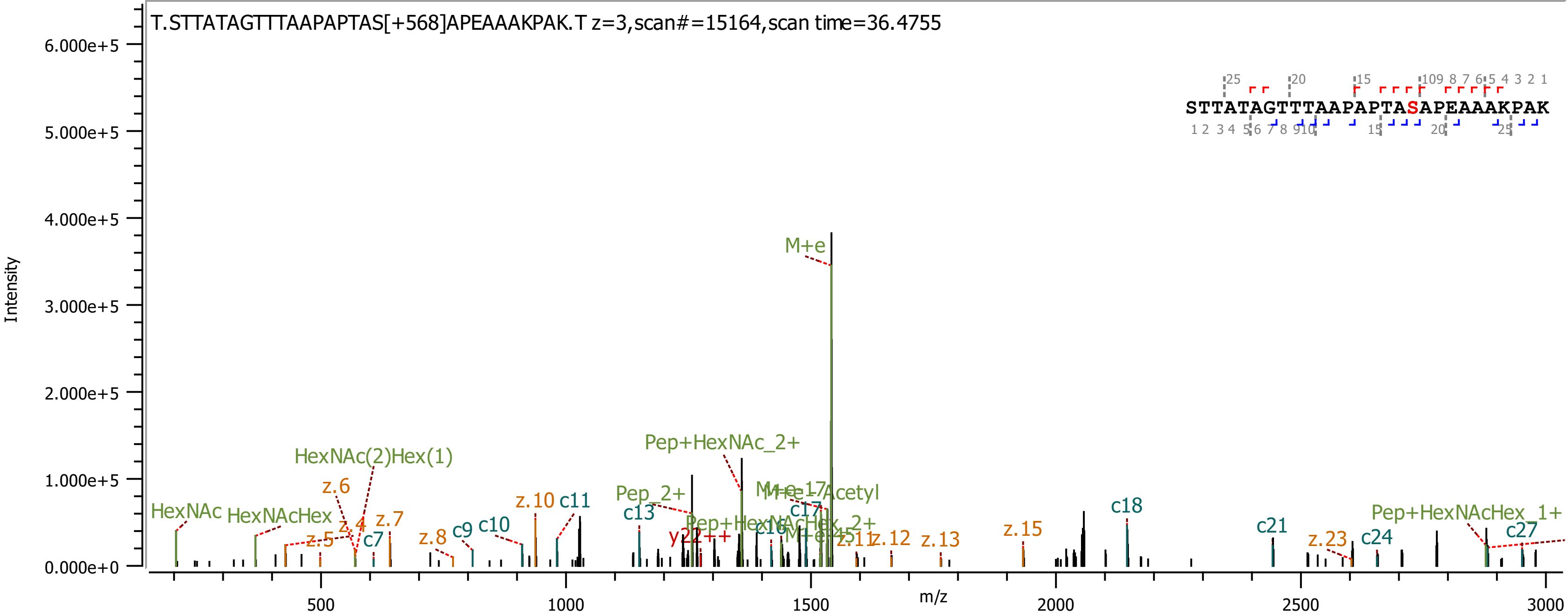

T.STTATAGTTTAAPAPTAS[+568]APEAAAKPAK.T z=3,scan#=15164,scan time=36.4755
STTATAGTTTAAPAPTASAPEAAAKPAK
Intensity
m/z
M+e
Pep+HexNAc_2+
HexNAc(2)Hex(1)
HexNAc
HexNAcHex
z.6
z.4
z.5
z.7
c7
z.8
c9
c10
z.10
c11
Pep_2+
c13
y22++
Pep+HexNAcHex_2+
c16
c17
M+e-Acetyl
z.11
z.12
z.13
z.15
c18
c21
z.23
c24
Pep+HexNAcHex_1+
c27

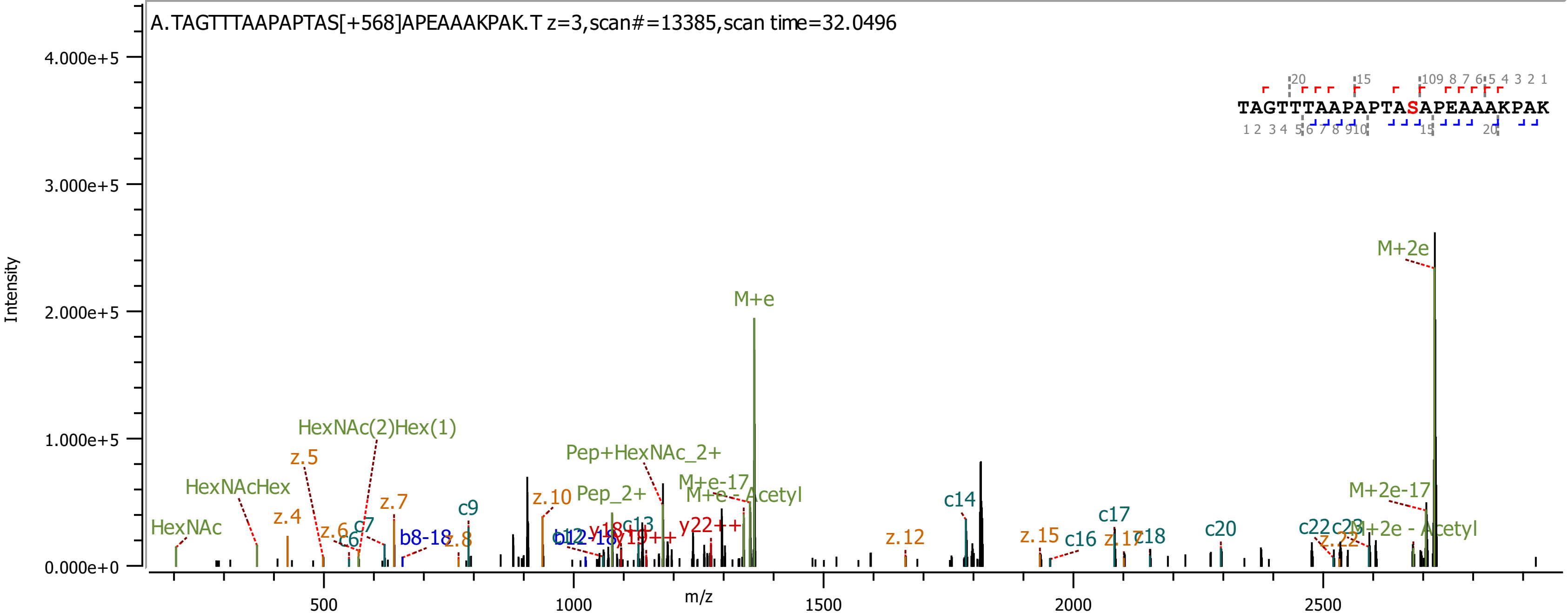
A.TAGTTTAAPAPTAS[+568]APEAAAKPAK.T z=3,scan#=13385,scan time=32.0496
TAGTTTAAPAPTASAPEAAAKPAK
Intensity
4.000e+5
3.000e+5
2.000e+5
1.000e+5
0.000e+0
m/z
500
1000
1500
2000
2500
M+2e
M+e
HexNAc(2)Hex(1)
HexNAcHex
HexNAc
Pep+HexNAc_2+
Pep_2+
M+e-17
M+e - Acetyl
M+2e-17
M+2e - Acetyl
z.4
z.5
z.6
z.7
z.8
z.10
z.12
z.15
z.17
z.22
c6
c7
c9
c13
c14
c16
c17
c18
c20
c22
b8-18
b12-18
y18
y19++
y22++

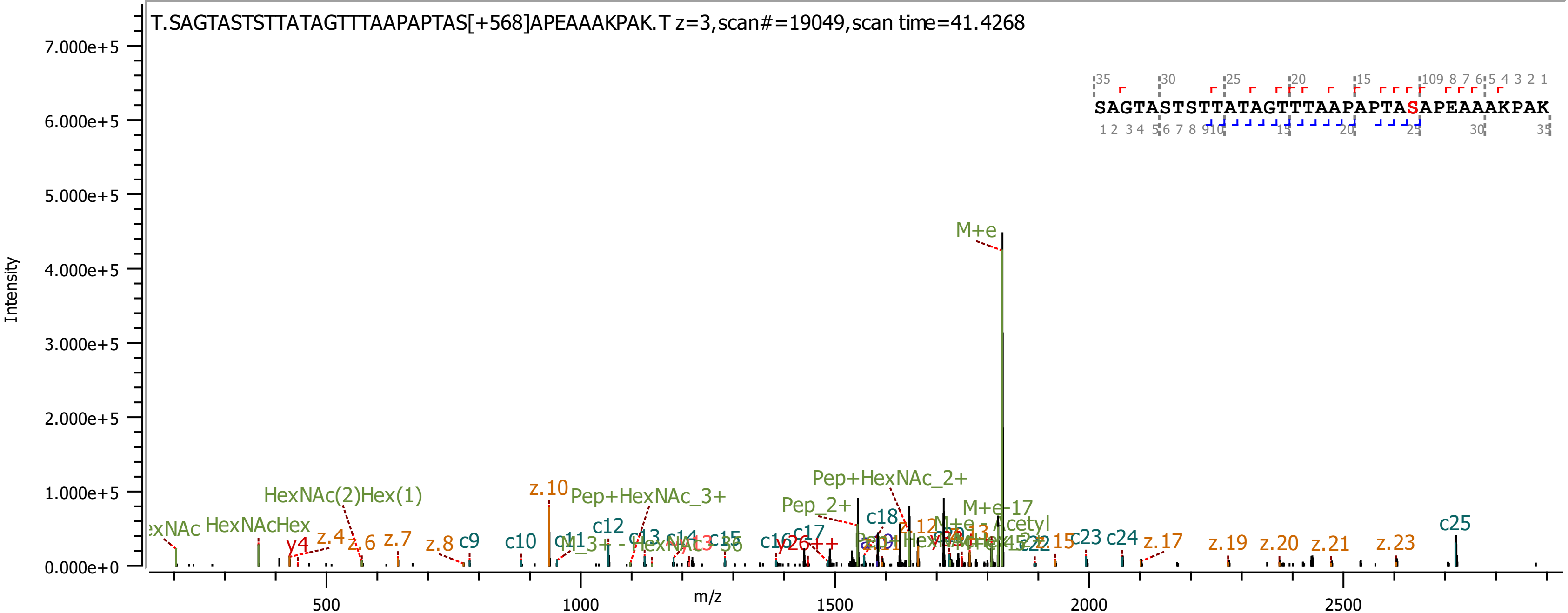
T.SAGTASTSTTATAGTTTAAPAPTAS[+568]APEAAAKPAK.T z=3,scan#=19049,scan time=41.4268
SAGTASTSTTATAGTTTAAPAPTASAPEAAAKPAK
Intensity
m/z
M+e
Pep+HexNAc_2+
Pep+HexNAc_3+
Pep_2+
HexNAc(2)Hex(1)
HexNAcHex
M+e-17
z.10
c25
c23
c24
z.17
z.19
z.20
z.21
z.23
c9
c10
c12
c15
c17
c18
y4
z.4
z.6
z.7
z.8

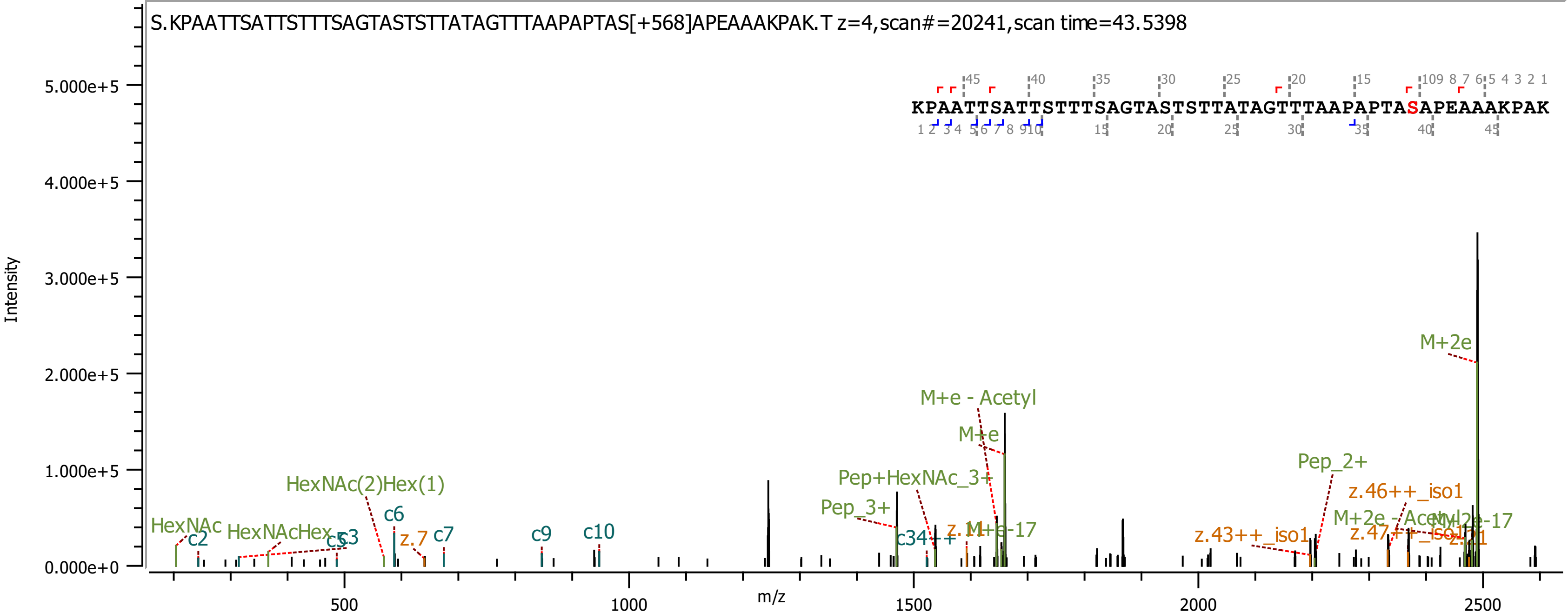
S.KPAATTSATTSTTTSAGTASTSTTATAGTTTAAPAPTAS[+568]APEAAAKPAK.T z=4,scan#=20241,scan time=43.5398
KPAATTSATTSTTTSAGTASTSTTATAGTTTAAPAPTASAPEAAAKPAK
Intensity
m/z
M+2e
M+e - Acetyl
M+e
Pep_2+
Pep+HexNAc_3+
Pep_3+
HexNAc(2)Hex(1)
HexNAc
HexNAcHex
c2
c3
c5
c6
c7
c9
c10
z.7
c34++
z.43++_iso1
z.46++_iso1
z.47++_iso1
M+2e - Acetyl
M+2e-17
M+e-17

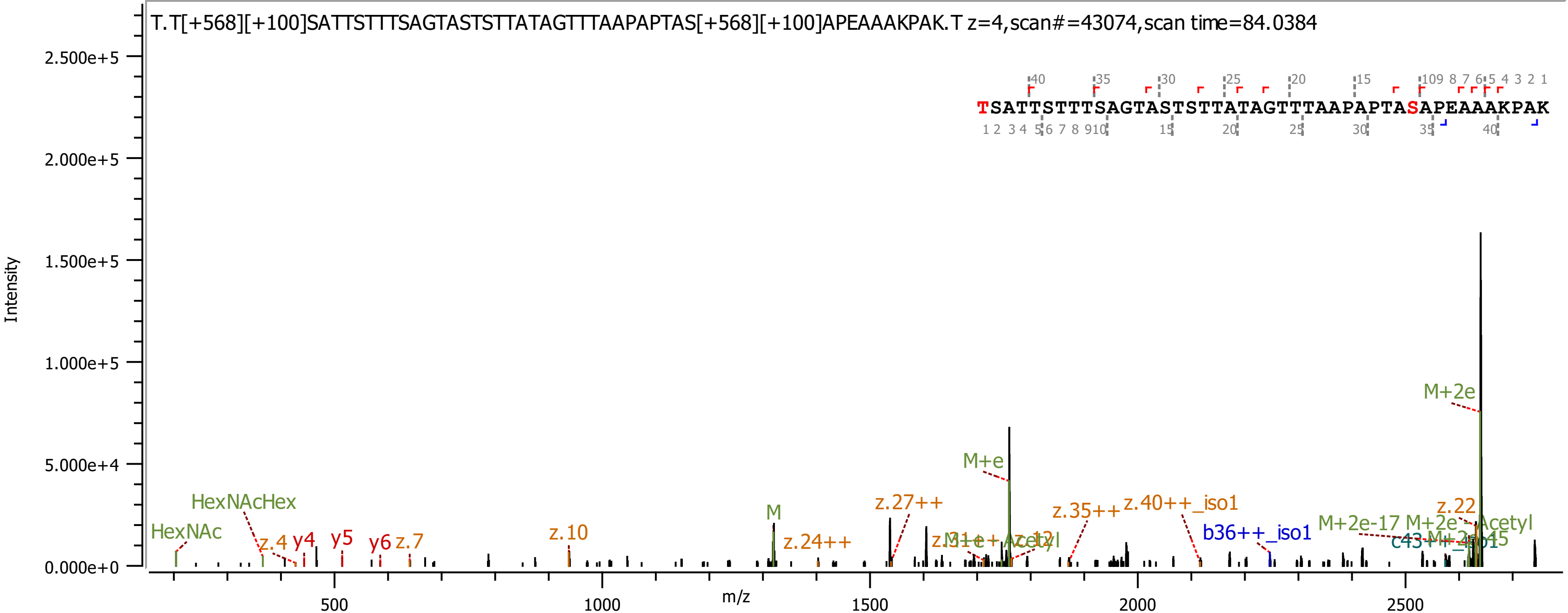
T.T[+568][+100]SATTSTTTSAGTASTSTTATAGTTTAAPAPTAS[+568][+100]APEAAAKPAK.T z=4,scan#=43074,scan time=84.0384
TSATTSTTTSAGTASTSTTATAGTTTAAPAPTASAPEAAAKPAK
Intensity
2.500e+5
2.000e+5
1.500e+5
1.000e+5
5.000e+4
0.000e+0
m/z
500
1000
1500
2000
2500
HexNAc
HexNAcHex
z.4
y4
y5
y6
z.7
z.10
M
z.24++
z.27++
M+e
z.35++
z.40++_iso1
b36++_iso1
M+2e-17
M+2e
z.22
M+2e

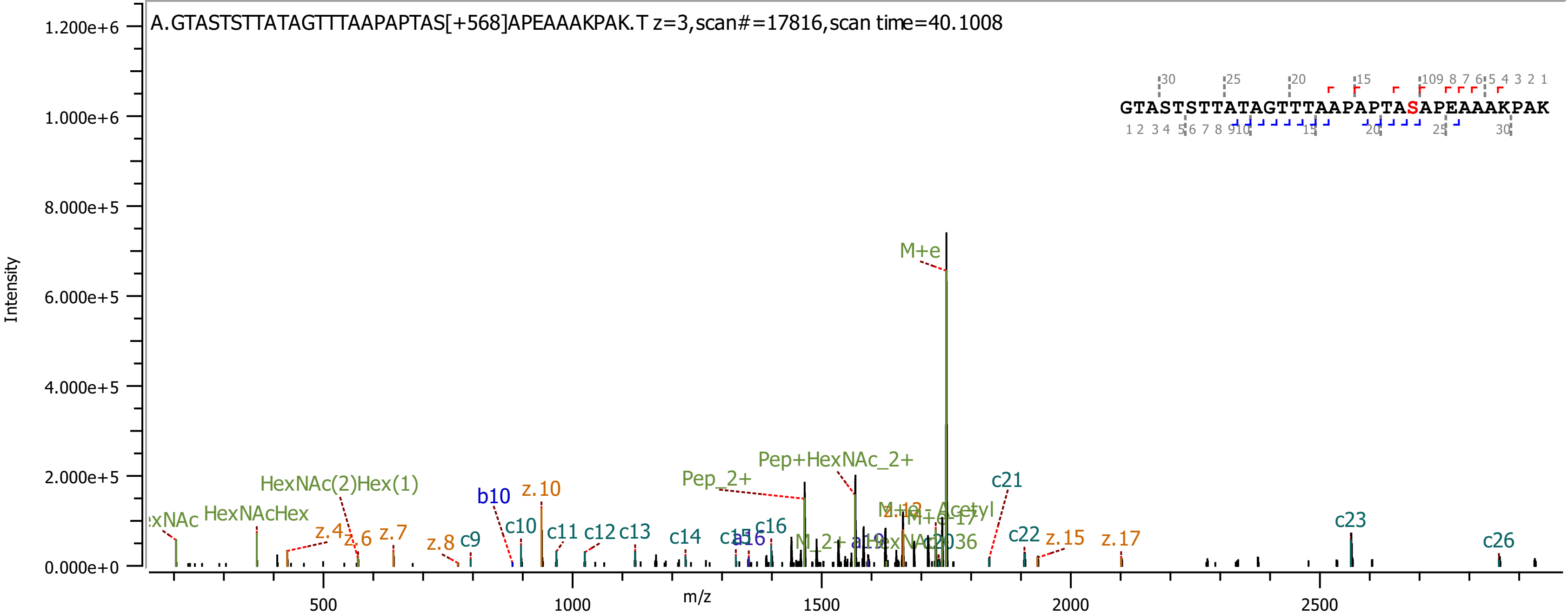

A.GTASTSTTATAGTTTAAPAPTAS[+568]APEAAAKPAK.T z=3,scan#=17816,scan time=40.1008
GTASTSTTATAGTTTAAPAPTASAPEAAAKPAK
Intensity
m/z
M+e
Pep+HexNAc_2+
Pep_2+
HexNAc(2)Hex(1)
HexNAcHex
b10
z.10
c21
c23
c26
c22
z.15
z.17
c10
c11
c12
c13
c14
c16
z.4
z.6
z.7
z.8
c9

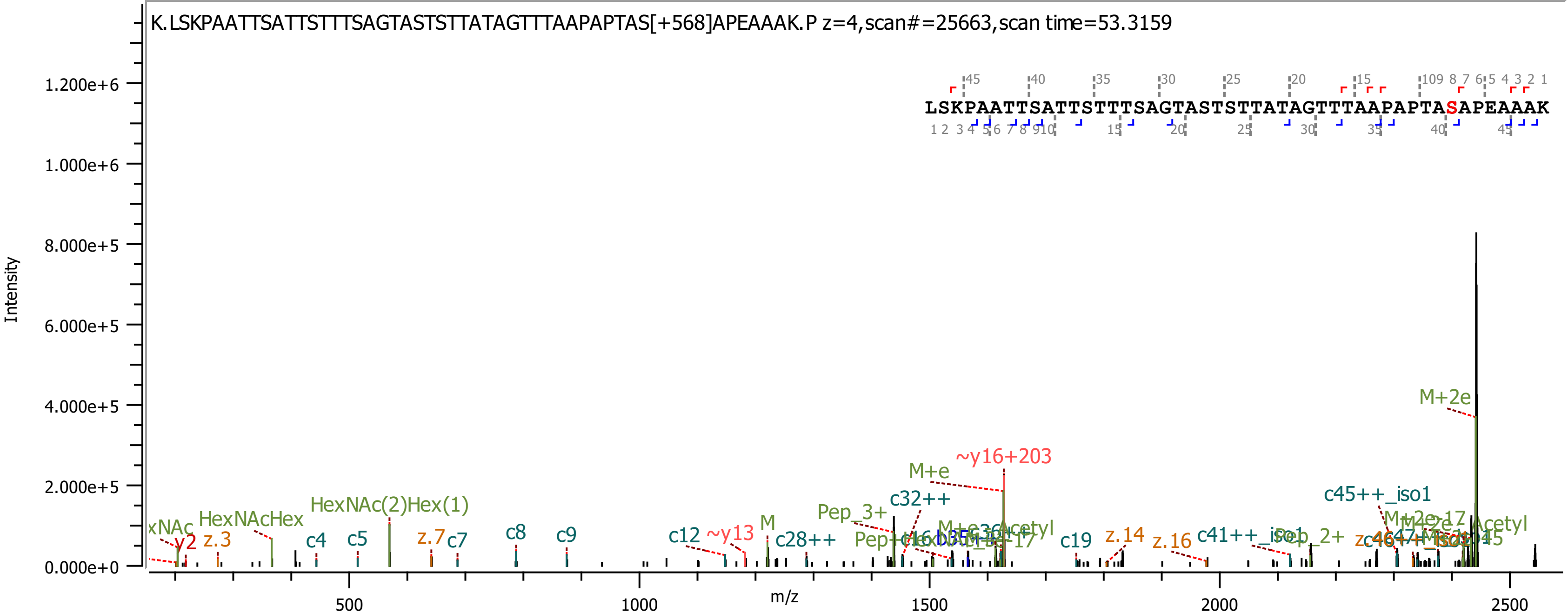
K.LSKPAATTSATTSTTTSAGTASTSTTATAGTTTAAPAPTAS[+568]APEAAAK.P z=4,scan#=25663,scan time=53.3159
LSKPAATTSATTSTTTSAGTASTSTTATAGTTTAAPAPTASAPEAAAK
Intensity
m/z
1.200e+6
1.000e+6
8.000e+5
6.000e+5
4.000e+5
2.000e+5
0.000e+0
500
1000
1500
2000
2500
M+2e
~y16+203
M+e
c32++
Pep_3+
c45++_iso1
HexNAc(2)Hex(1)
HexNAcHex
y2
z.3
c4
c5
z.7
c7
c8
c9
c12
~y13
M
c28++
c19
z.14
z.16
c41++_iso1

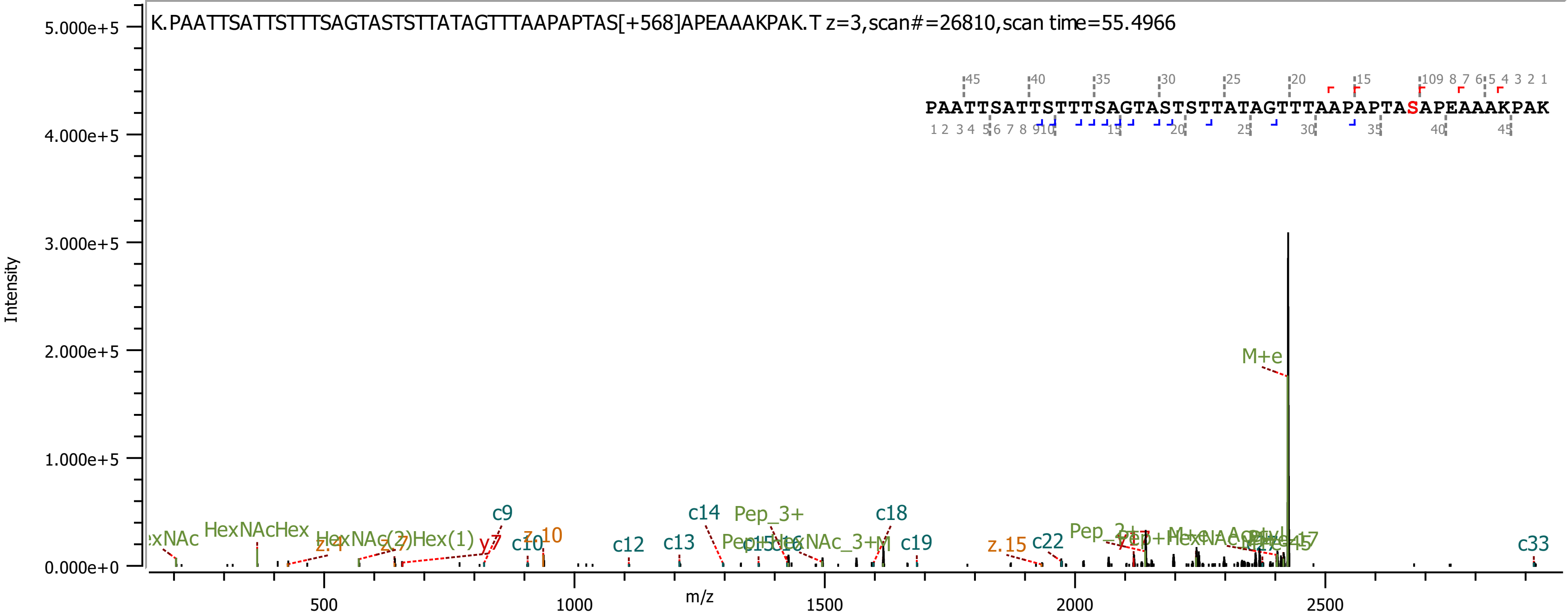
K.PAATTSATTSTTTSAGTASTSTTATAGTTTAAPAPTAS[+568]APEAAAKPAK.T z=3,scan#=26810,scan time=55.4966
PAATTSATTSTTTSAGTASTSTTATAGTTTAAPAPTASAPEAAAKPAK
Intensity
m/z
M+e
c9
c10
c12
c13
c14
c18
c19
c22
c33
z.10
z.15
y7
Pep_3+
HexNAcHex
HexNAc(2)Hex(1)

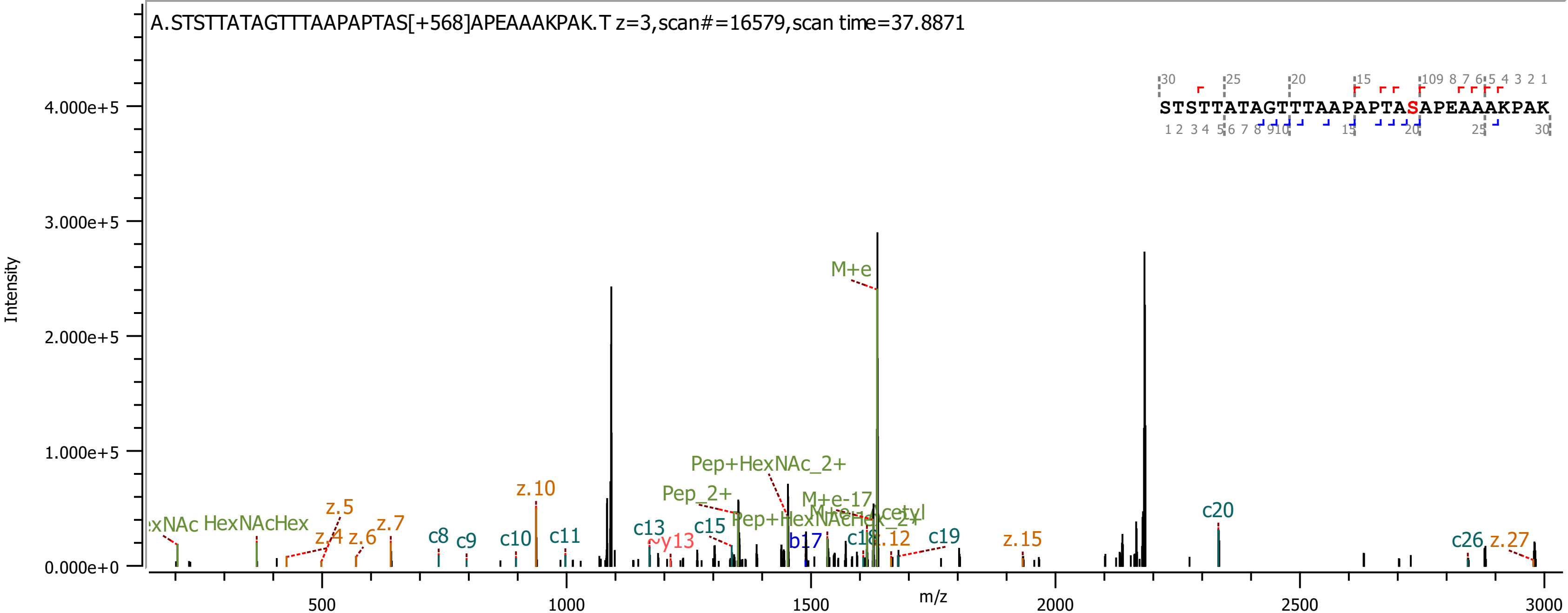

A.STSTTATAGTTTAAPAPTAS[+568]APEAAAKPAK.T z=3,scan#=16579,scan time=37.8871
STSTTATAGTTTAAPAPTASAPEAAAKPAK
Intensity
m/z
M+e
Pep+HexNAc_2+
Pep_2+
M+e-17
z.10
z.5
z.4
z.6
z.7
z.12
z.15
z.27
HexNAcHex
c8
c9
c10
c11
c13
c15
c19
c20
c26
y13
b17

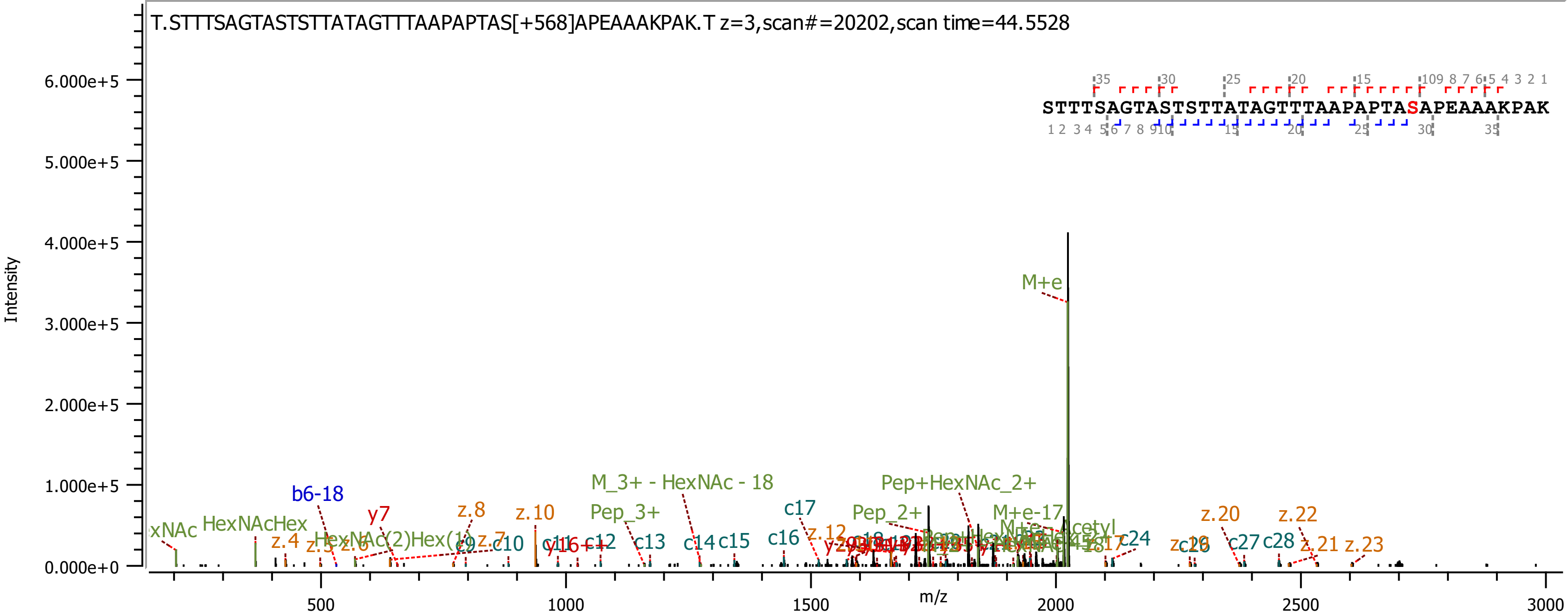

T.STTTSAGTASTSTTATAGTTTAAPAPTAS[+568]APEAAAKPAK.T z=3,scan#=20202,scan time=44.5528
STTTSAGTASTSTTATAGTTTAAPAPTASAPEAAAKPAK
Intensity
m/z
M+e
M_3+ - HexNAc - 18
Pep+HexNAc_2+
Pep_3+
Pep_2+
M+e-17
HexNAcHex
HexNAc(2)Hex(1)
b6-18
y7
y16++
z.4
z.5
z.6
z.7
z.8
z.10
z.12
z.20
z.21
z.22
z.23
c9
c10
c11
c12
c13
c14
c15
c16
c17
c24
c26
c27
c28

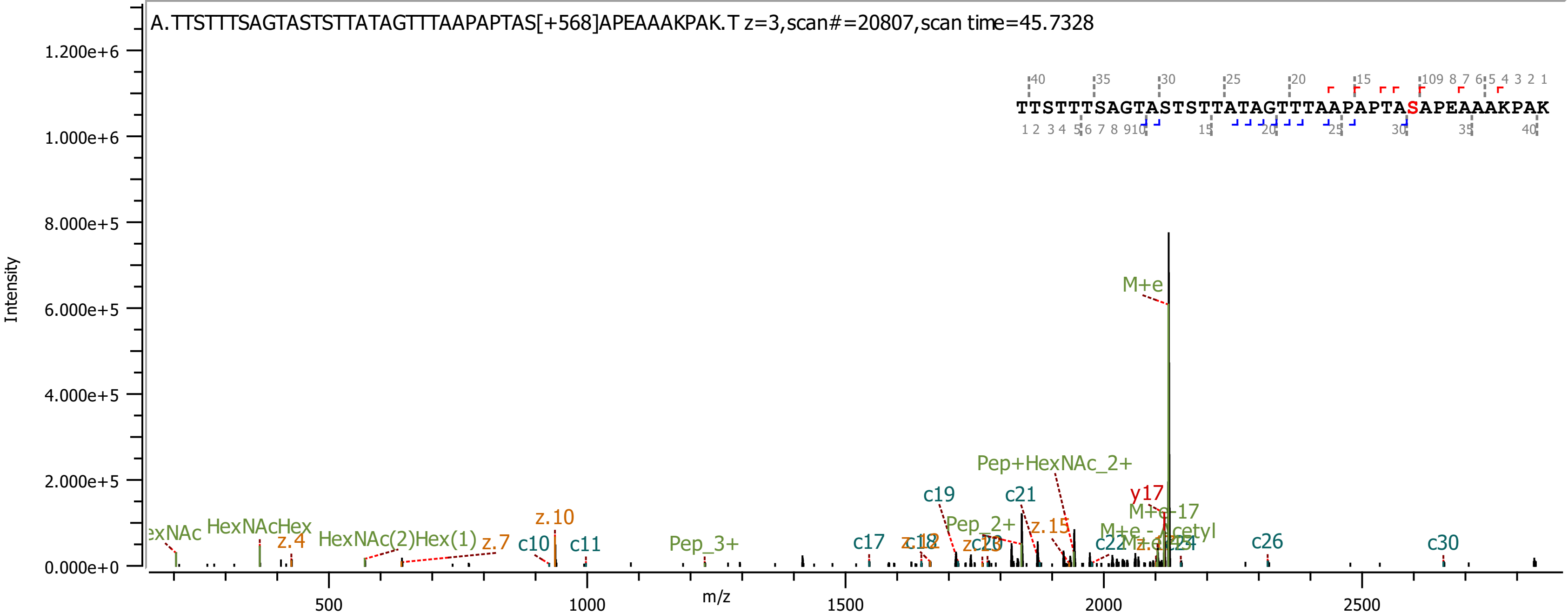

A.TTSTTTSAGTASTSTTATAGTTTAAPAPTAS[+568]APEAAAKPAK.T z=3,scan#=20807,scan time=45.7328
TTSTTTSAGTASTSTTATAGTTTAAPAPTASAPEAAAKPAK
Intensity
m/z
M+e
Pep+HexNAc_2+
c19
c21
y17
M+e-17
Pep_2+
z.15
M+e - Acetyl
c22
c26
c30
c17
c10
c11
z.10
z.4
z.7
HexNAcHex
HexNAc(2)Hex(1)
Pep_3+

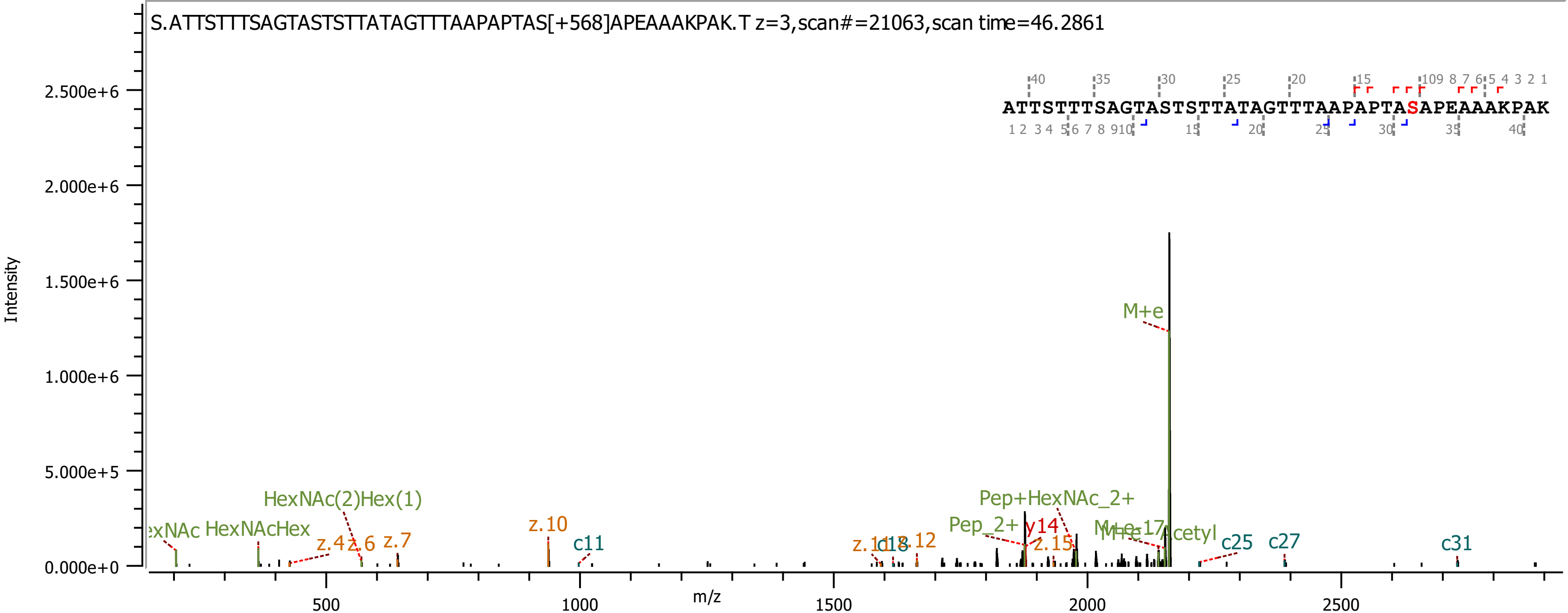

S.ATTSTTTSAGTASTSTTATAGTTTAAPAPTAS[+568]APEAAAKPAK.T z=3,scan#=21063,scan time=46.2861
ATTSTTTSAGTASTSTTATAGTTTAAPAPTASAPEAAAKPAK
Intensity
m/z
M+e
HexNAc(2)Hex(1)
HexNAcHex
Pep+HexNAc_2+
Pep_2+
y14
z.10
c11
z.4
z.6
z.7
z.15
c18
z.12
c25
c27
c31
cetyl

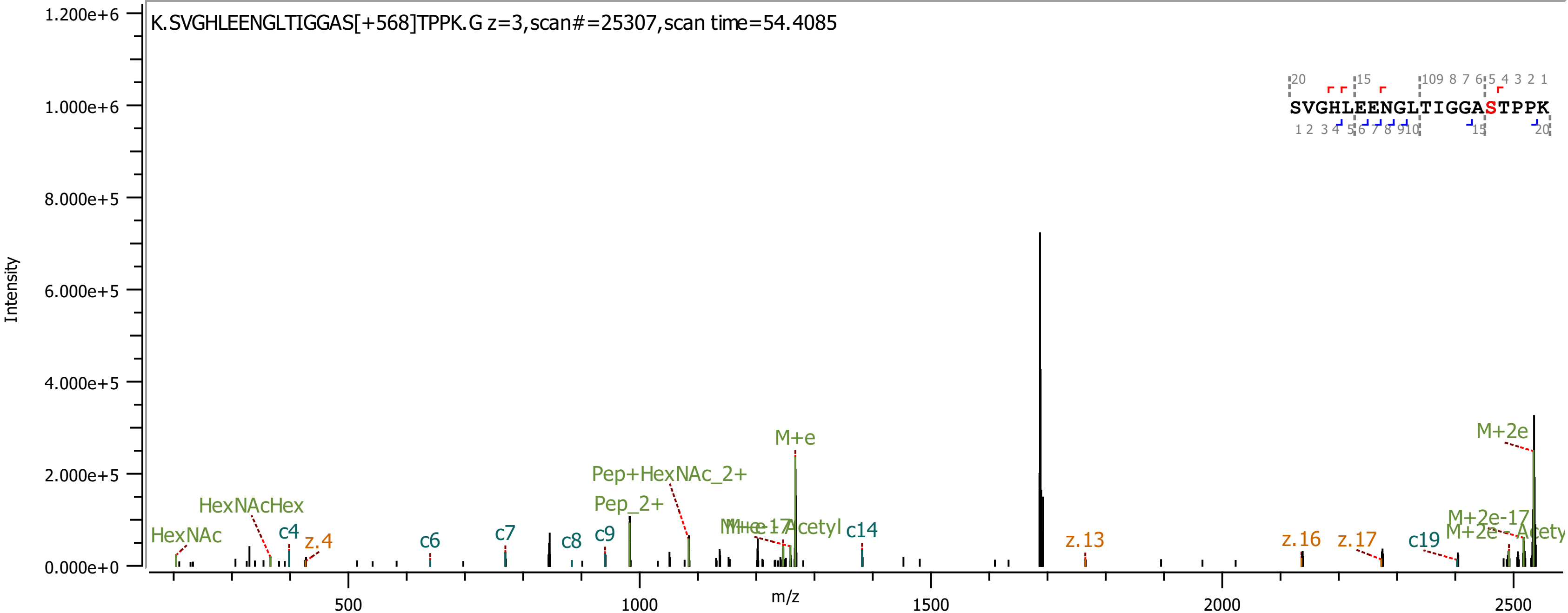

K.SVGHLEENGLTIGGAS[+568]TPPK.G z=3,scan#=25307,scan time=54.4085
SVGHLEENGLTIGGASTPPK
Intensity
m/z
HexNAc
HexNAcHex
c4
z.4
c6
c7
c8
c9
Pep_2+
Pep+HexNAc_2+
M+e
c14
z.13
z.16
z.17
c19
M+2e-17
M+2e

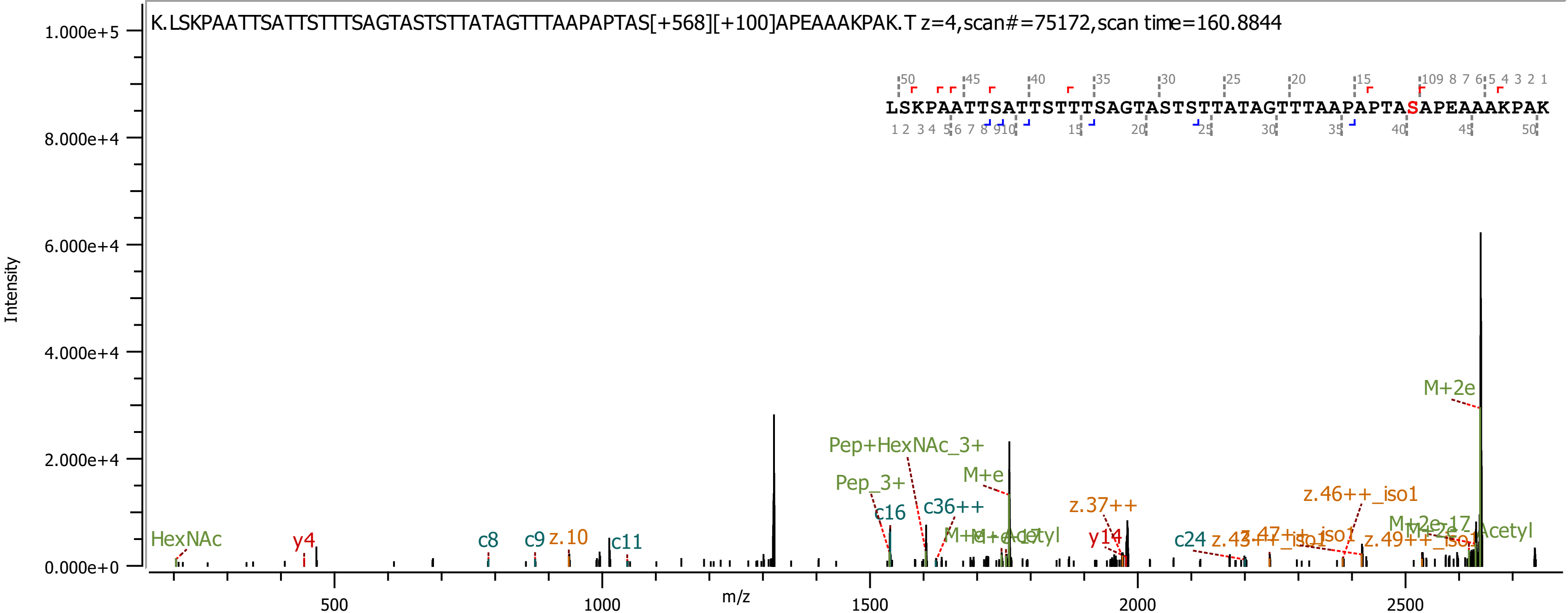

K.LSKPAATTSATTSTTTSAGTASTSTTATAGTTTAAPAPTAS[+568][+100]APEAAAKPAK.T z=4,scan#=75172,scan time=160.8844
LSKPAATTSATTSTTTSAGTASTSTTATAGTTTAAPAPTASAPEAAAKPAK
Intensity
m/z
HexNAc
y4
c8
c9
z.10
c11
c16
Pep_3+
Pep+HexNAc_3+
c36++
M+e
M+e-Acetyl
z.37++
y14
c24
z.46++_iso1
M+2e
M+2e-Acetyl

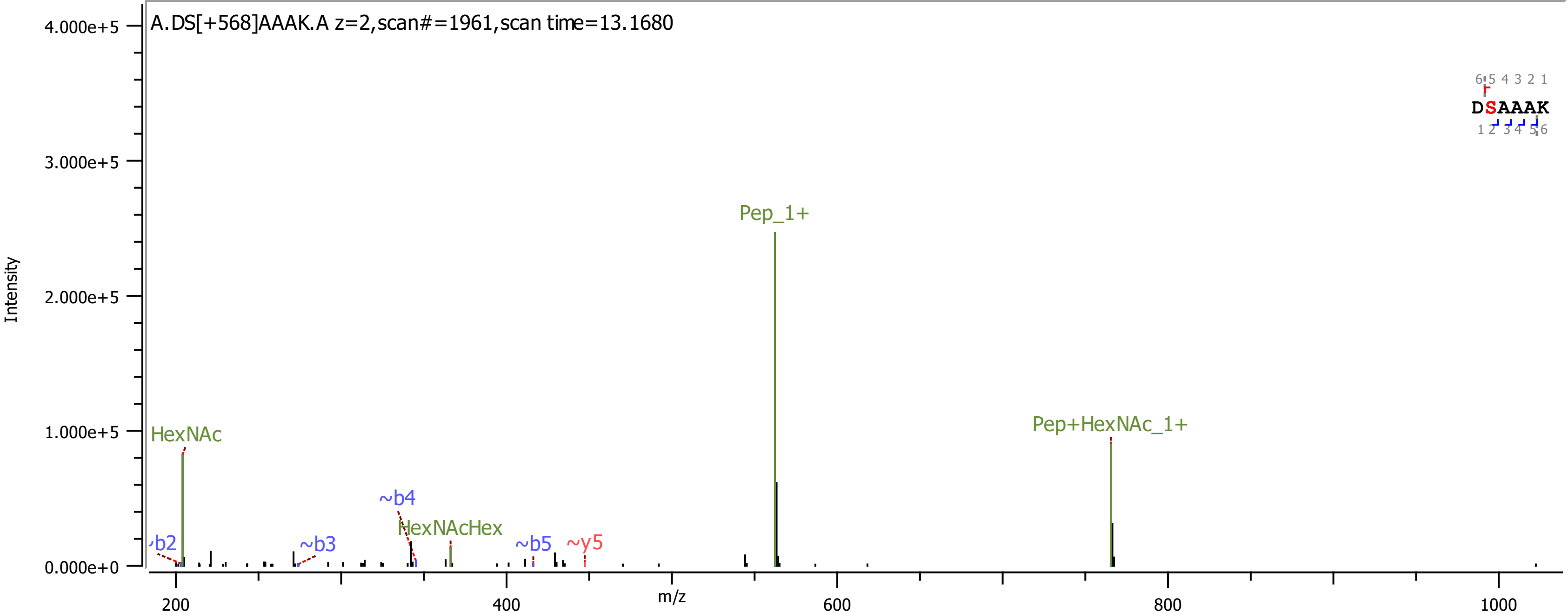
A.DS[+568]AAAK.A z=2,scan#=1961,scan time=13.1680
Intensity
4.000e+5
3.000e+5
2.000e+5
1.000e+5
0.000e+0
m/z
200
400
600
800
1000
HexNAc
~b2
~b3
~b4
HexNAcHex
~b5
~y5
Pep_1+
Pep+HexNAc_1+
DSAAAK
6 5 4 3 2 1
1 2 3 4 5 6

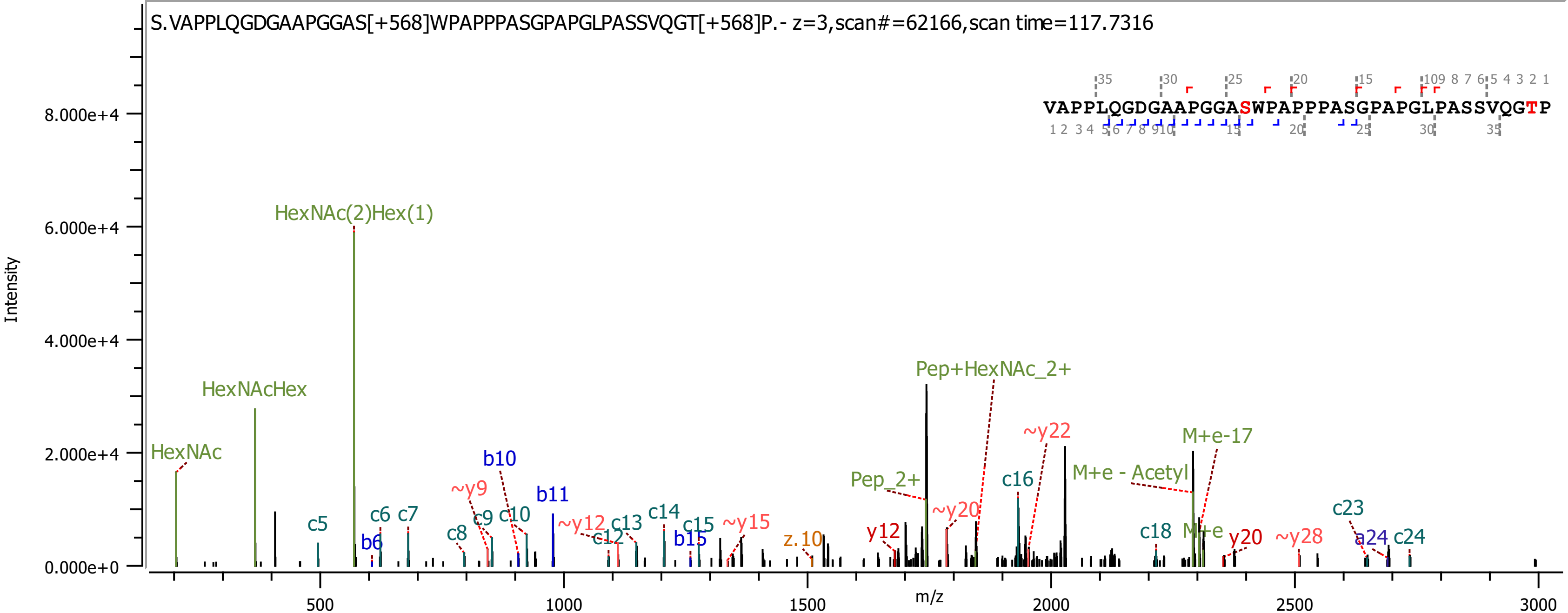

S.VAPPLQGDGAAPGGAS[+568]WPAPPPASGPAPGLPASSVQGT[+568]P.- z=3,scan#=62166,scan time=117.7316
VAPPLQGDGAAPGGASWPAPPPASGPAPGLPASSVQGTP
Intensity
m/z
HexNAc
HexNAcHex
HexNAc(2)Hex(1)
Pep+HexNAc_2+
Pep_2+
M+e-17
M+e - Acetyl
M+e
~y22
~y20
~y9
~y12
~y15
~y28
y12
y20
z.10
b6
b10
b11
b15
a24
c5
c6
c7
c8
c9
c10
c12
c13
c14
c15
c16
c18
c23
c24

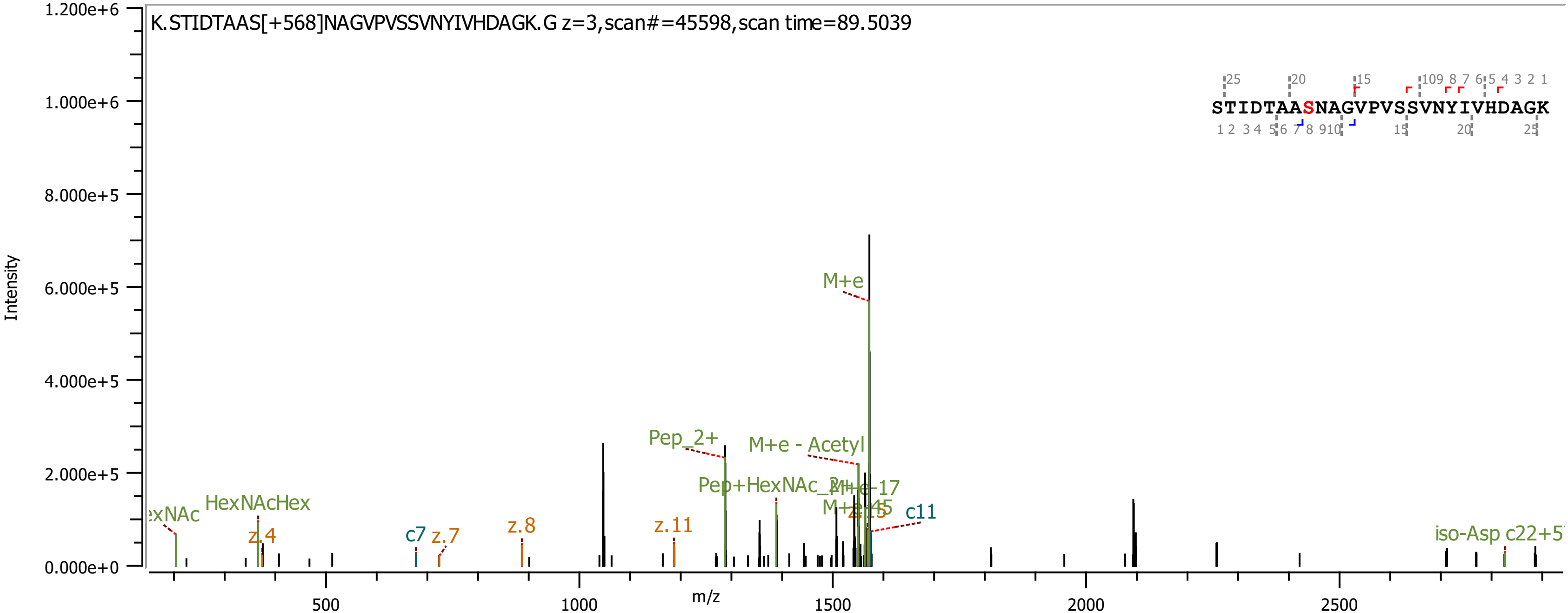
K.STIDTAAS[+568]NAGVPVSSVNYIVHDAGK.G z=3,scan#=45598,scan time=89.5039
STIDTAASNAGVPVSSVNYIVHDAGK
Intensity
m/z
M+e
Pep_2+
M+e - Acetyl
Pep+HexNAc_2+
M+e-17
M+e-45
z.15
c11
HexNAcHex
z.4
c7
z.7
z.8
z.11
iso-Asp c22+5
1.200e+6
1.000e+6
8.000e+5
6.000e+5
4.000e+5
2.000e+5
0.000e+0
500
1000
1500
2000
2500

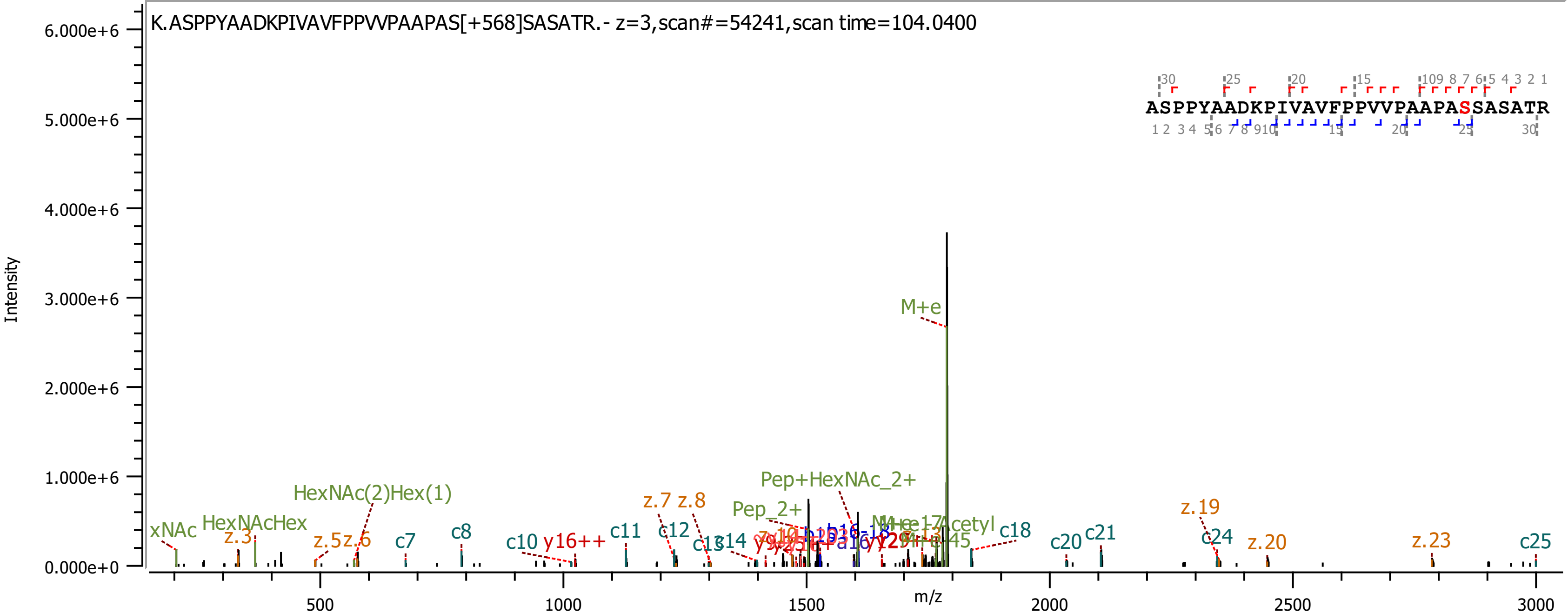
K.ASPPYAADKPIVAVFPPVVPAAPAS[+568]SASATR.- z=3,scan#=54241,scan time=104.0400
ASPPYAADKPIVAVFPPVVPAAPASSASATR
Intensity
m/z
6.000e+6
5.000e+6
4.000e+6
3.000e+6
2.000e+6
1.000e+6
0.000e+0
500
1000
1500
2000
2500
3000
M+e
Pep+HexNAc_2+
Pep_2+
HexNAc(2)Hex(1)
HexNAcHex
xNAc
z.3
z.5
z.6
c7
c8
c10
y16++
c11
z.7
z.8
c12
c13
c14
c18
c20
c21
z.19
c24
z.20
z.23
c25

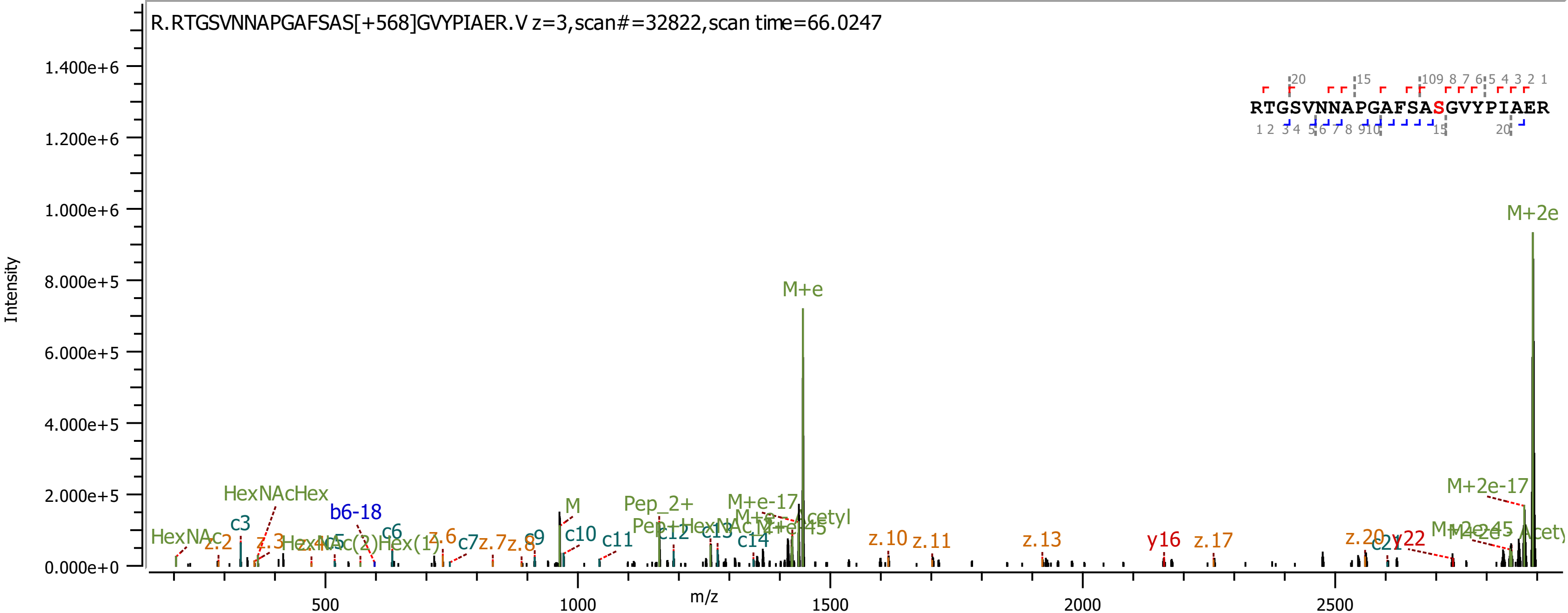

R.RTGSVNNAPGAFSAS[+568]GVYPIAER.V z=3,scan#=32822,scan time=66.0247
RTGSVNNAPGAFSASGVYPIAER
Intensity
m/z
M+2e
M+e
M+2e-17
M+e-17
HexNAcHex
HexNAc
b6-18
c3
c6
c7
c9
c10
c11
c12
c13
c14
c21
z.2
z.3
z.6
z.7
z.8
z.10
z.11
z.13
z.17
z.20
y16
y22
M
Pep_2+

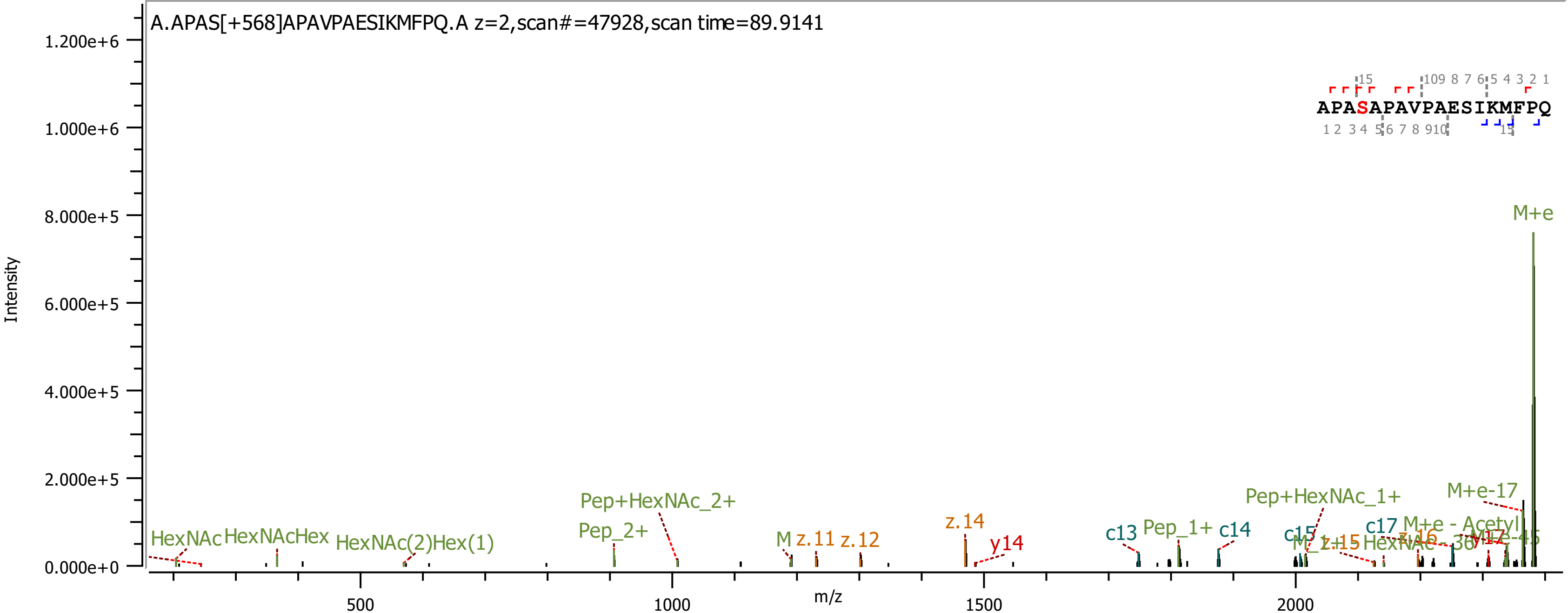
A.APAS[+568]APAVPAESIKMFPQ.A z=2,scan#=47928,scan time=89.9141
APASAPAVPAESIKMFPQ
Intensity
m/z
M+e
M+e-17
Pep+HexNAc_2+
Pep_2+
Pep+HexNAc_1+
Pep_1+
HexNAc
HexNAcHex
HexNAc(2)Hex(1)
z.11
z.12
z.14
y14
c13
c14
c15
c17
z.15
M+e - Acetyl

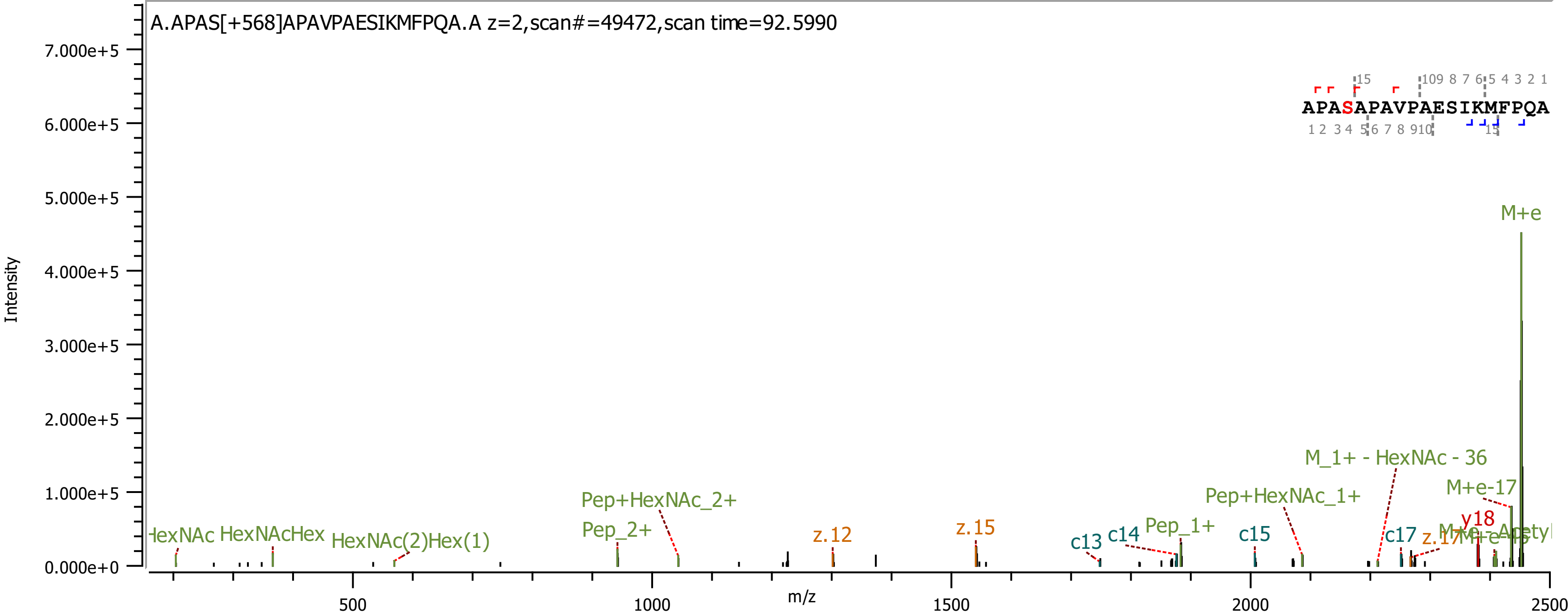

A.APAS[+568]APAVPAESIKMFPQA.A z=2,scan#=49472,scan time=92.5990
APASAPAVPAESIKMFPQA
Intensity
m/z
M+e
M_1+ - HexNAc - 36
M+e-17
Pep+HexNAc_1+
Pep+HexNAc_2+
Pep_2+
Pep_1+
HexNAcHex
HexNAc(2)Hex(1)
z.12
z.15
z.17
c13
c14
c15
c17
y18

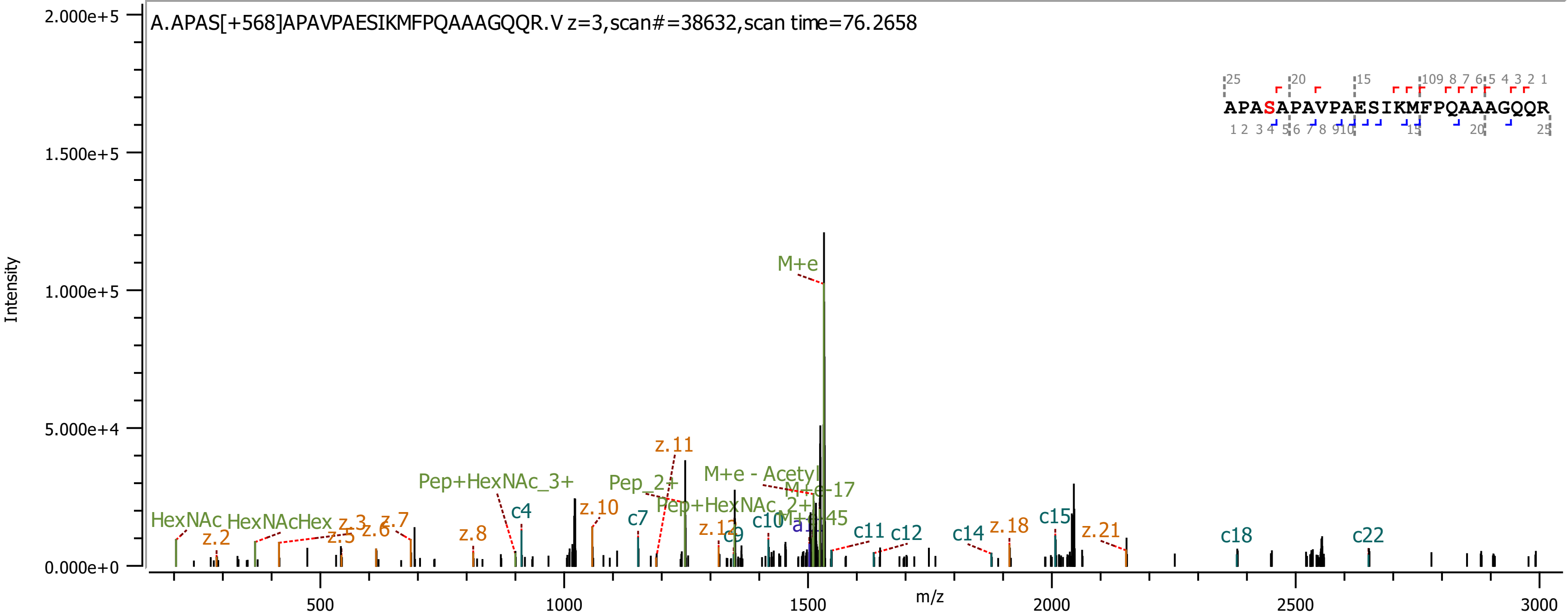

A.APAS[+568]APAVPAESIKMFPQAAAGQQR.V z=3,scan#=38632,scan time=76.2658
APASAPAVPAESIKMFPQAAAGQQR
Intensity
m/z
M+e
z.11
M+e - Acetyl
M+e-17
Pep+HexNAc_3+
Pep_2+
Pep+HexNAc_2+
HexNAc
HexNAcHex
c4
c7
c9
c10
c11
c12
c14
c15
c18
c22
z.2
z.3
z.5
z.6
z.7
z.8
z.10
z.12
z.18
z.21
2.000e+5
1.500e+5
1.000e+5
5.000e+4
0.000e+0
500
1000
1500
2000
2500
3000

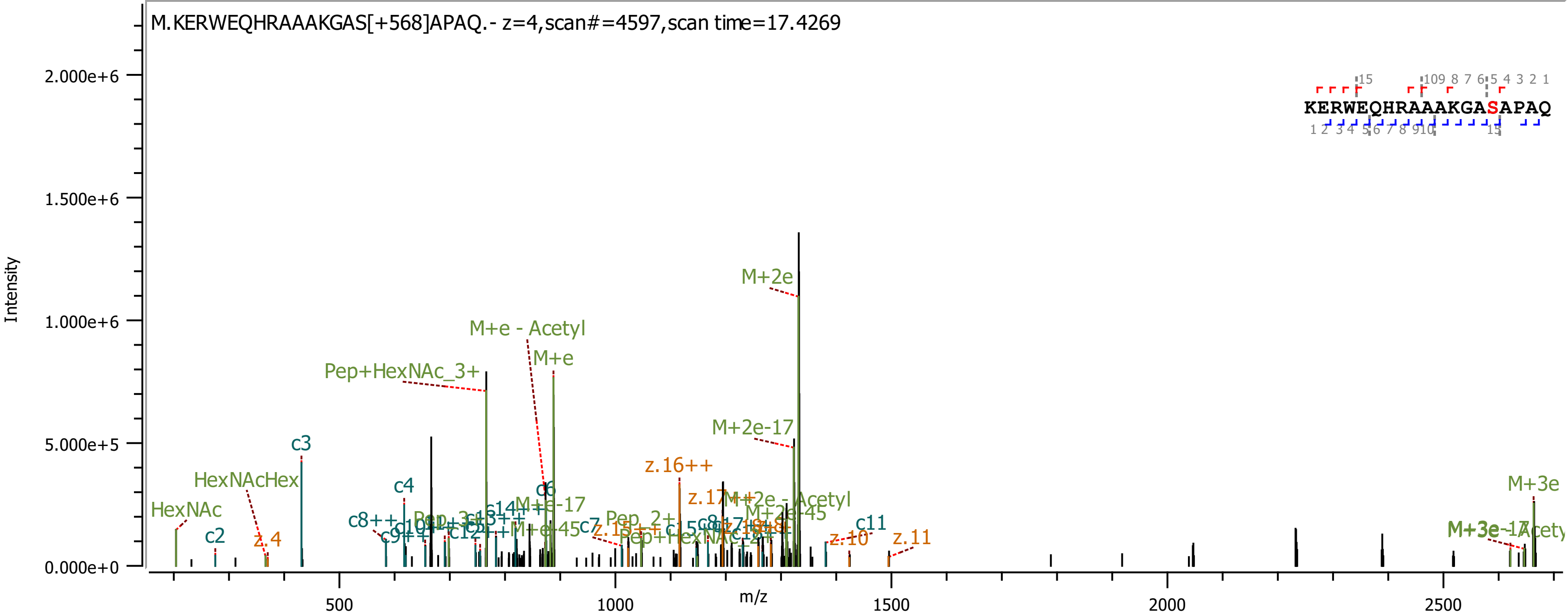

M.KERWEQHRAAAKGAS[+568]APAQ.- z=4,scan#=4597,scan time=17.4269
KERWEQHRAAAKGASAPAQ
Intensity
m/z
0.000e+0
5.000e+5
1.000e+6
1.500e+6
2.000e+6
500
1000
1500
2000
2500
M+2e
M+e - Acetyl
M+e
Pep+HexNAc_3+
M+2e-17
c3
HexNAcHex
HexNAc
c2
z.4
c4
c6
z.16++
c11
z.10
z.11
M+3e

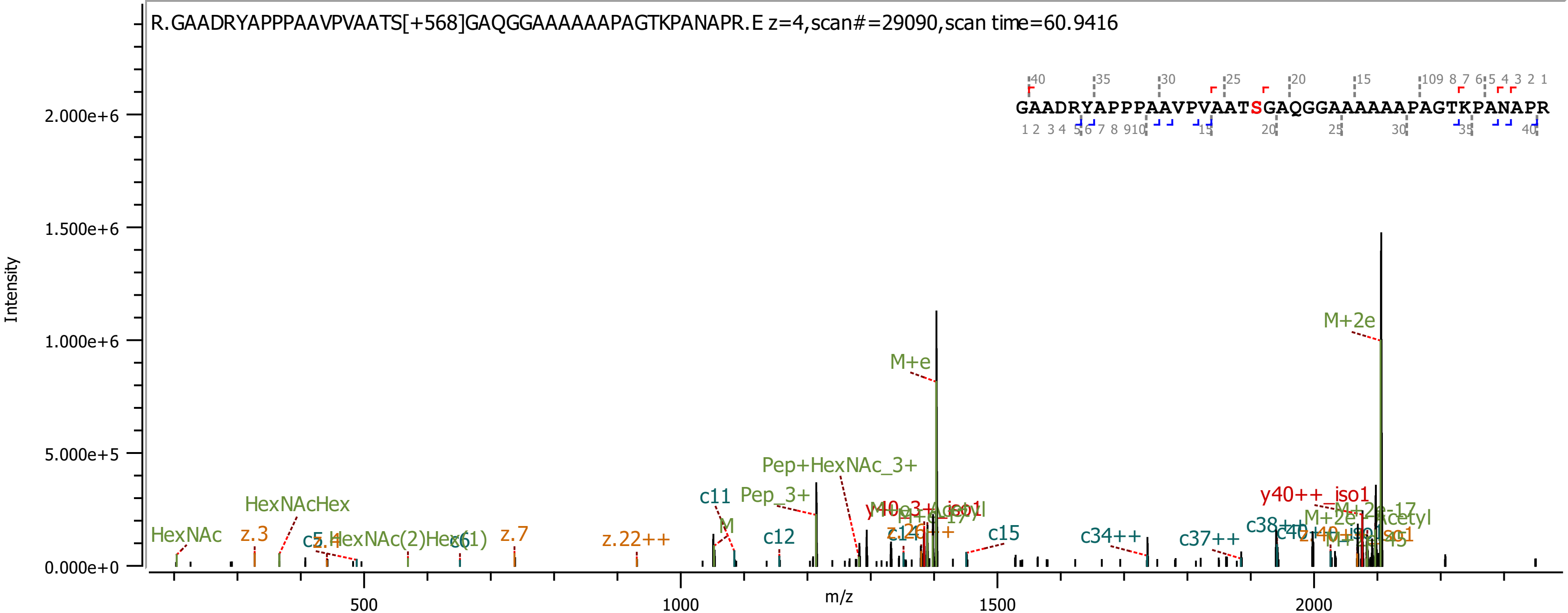

R.GAADRYAPPPAAVPVAATS[+568]GAQGGAAAAAAPAGTKPANAPR.E z=4,scan#=29090,scan time=60.9416
GAADRYAPPPAAVPVAATSGAQGGAAAAAAPAGTKPANAPR
Intensity
m/z
M+2e
M+e
Pep+HexNAc_3+
Pep_3+
c11
c12
c15
c34++
c37++
HexNAcHex
HexNAc
z.3
z.7
z.22++
y40++_iso1

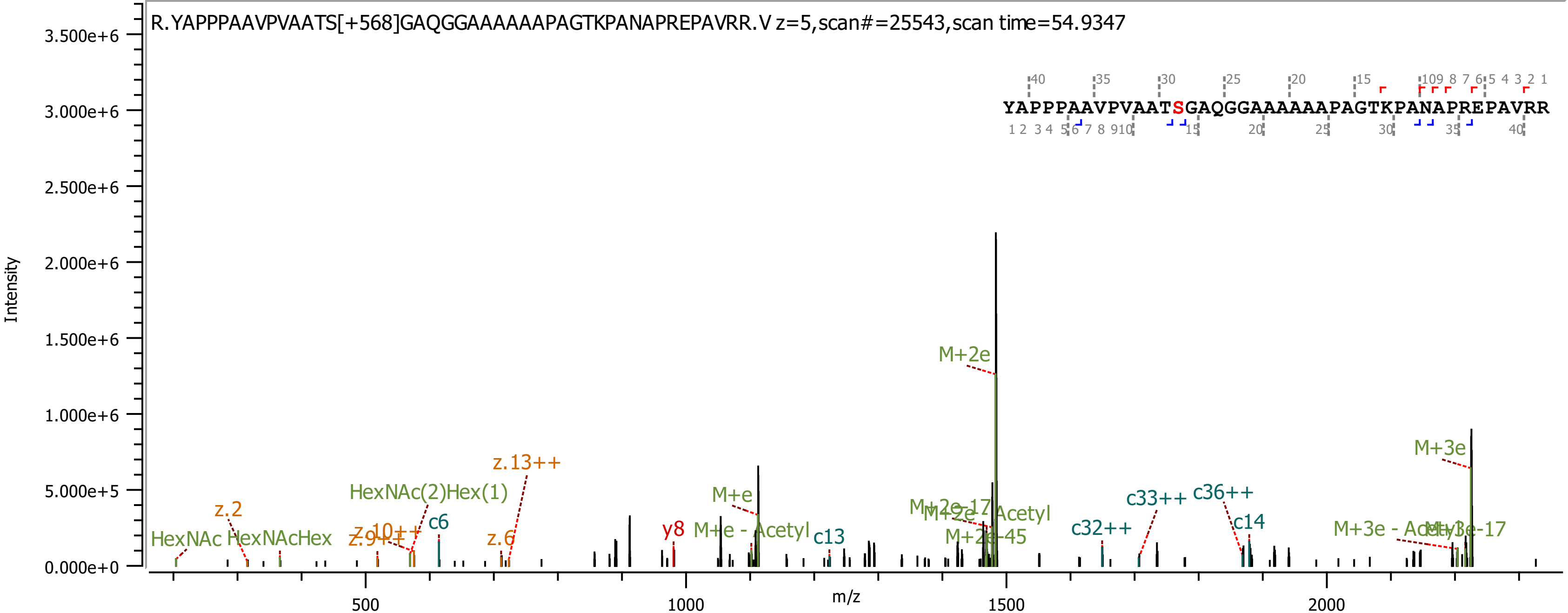

R.YAPPPAAVPVAATS[+568]GAQGGAAAAAAPAGTKPANAPREPAVRR.V z=5,scan#=25543,scan time=54.9347
YAPPPAAVPVAATSGAQGGAAAAAAPAGTKPANAPREPAVRR
Intensity
m/z
M+2e
M+3e
z.13++
HexNAc(2)Hex(1)
z.2
M+e
c6
HexNAc
HexNAcHex
z.9++
z.10++
z.6
y8
M+e - Acetyl
c13
M+2e-17
M+2e-Acetyl
M+2e-45
c32++
c33++
c36++
c14
M+3e - Acetyl
M+3e-17

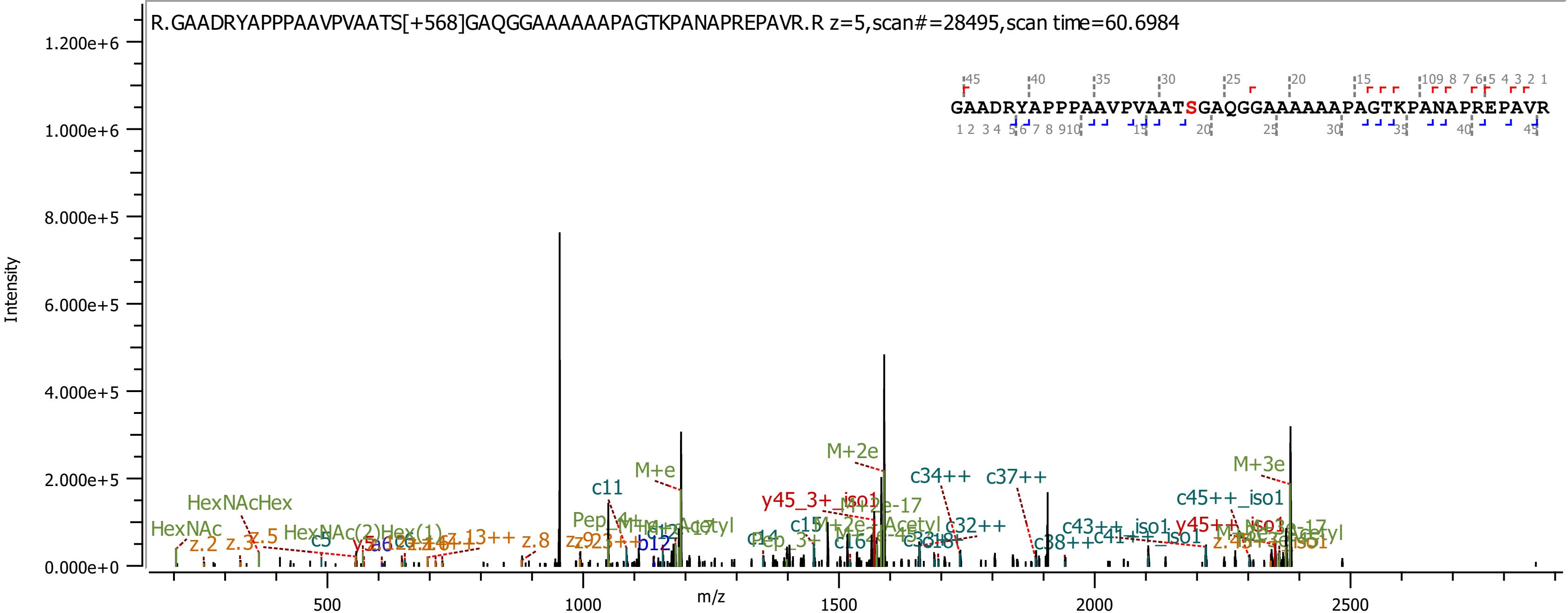
R.GAADRYAPPPAAVPVAATS[+568]GAQGGAAAAAAPAGTKPANAPREPAVR.R z=5,scan#=28495,scan time=60.6984
GAADRYAPPPAAVPVAATSGAQGGAAAAAAPAGTKPANAPREPAVR
Intensity
1.200e+6
1.000e+6
8.000e+5
6.000e+5
4.000e+5
2.000e+5
0.000e+0
m/z
500
1000
1500
2000
2500
HexNAc
HexNAcHex
HexNAc(2)Hex(1)
z.2
z.3
z.5
z.8
c5
c11
M+e
M+2e
M+3e
b12
c14
c15
c16
c32++
c34++
c37++
c38++
c43++_iso1
c45++_iso1
y45_3+_iso1
y45++_iso1

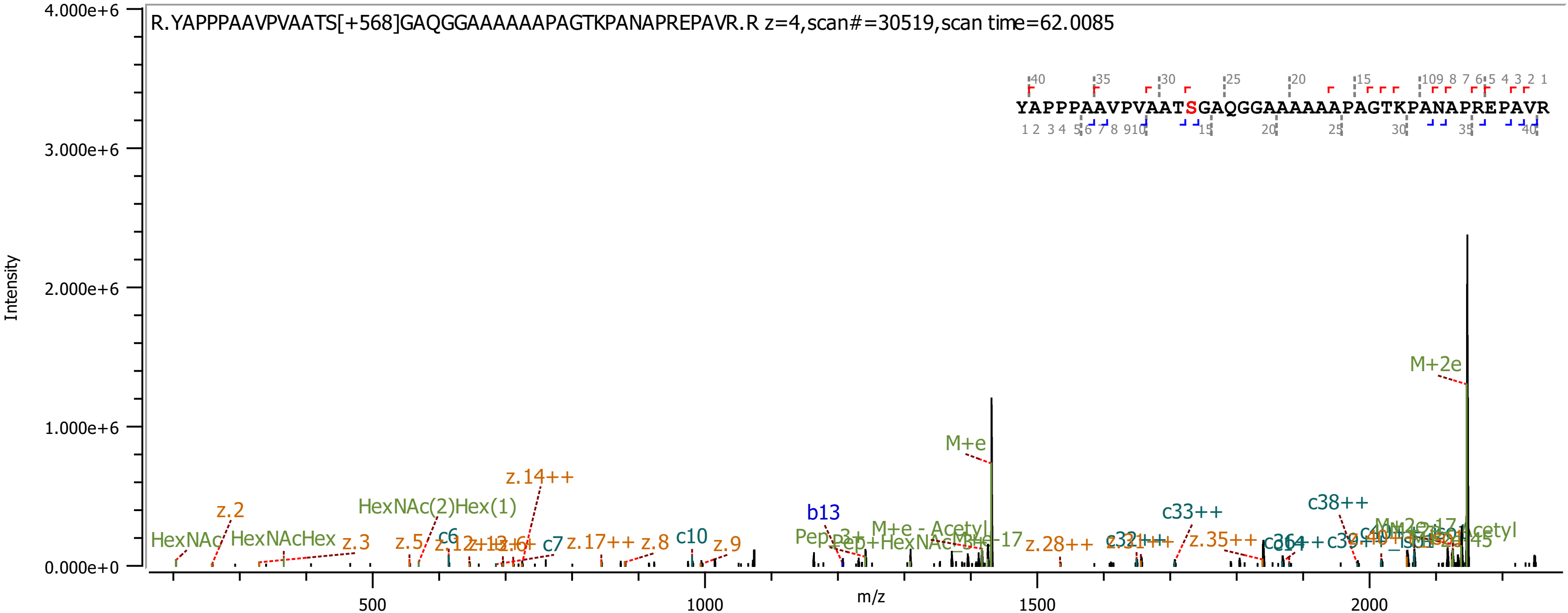

R.YAPPPAAVPVAATS[+568]GAQGGAAAAAAPAGTKPANAPREPAVR.R z=4,scan#=30519,scan time=62.0085
YAPPPAAVPVAATSGAQGGAAAAAAPAGTKPANAPREPAVR
Intensity
m/z
4.000e+6
3.000e+6
2.000e+6
1.000e+6
0.000e+0
500
1000
1500
2000
M+2e
M+e
z.14++
HexNAc(2)Hex(1)
z.2
HexNAc
HexNAcHex
z.3
z.5
c6
c7
z.17++
z.8
c10
z.9
b13
c33++
c38++
z.28++
z.35++

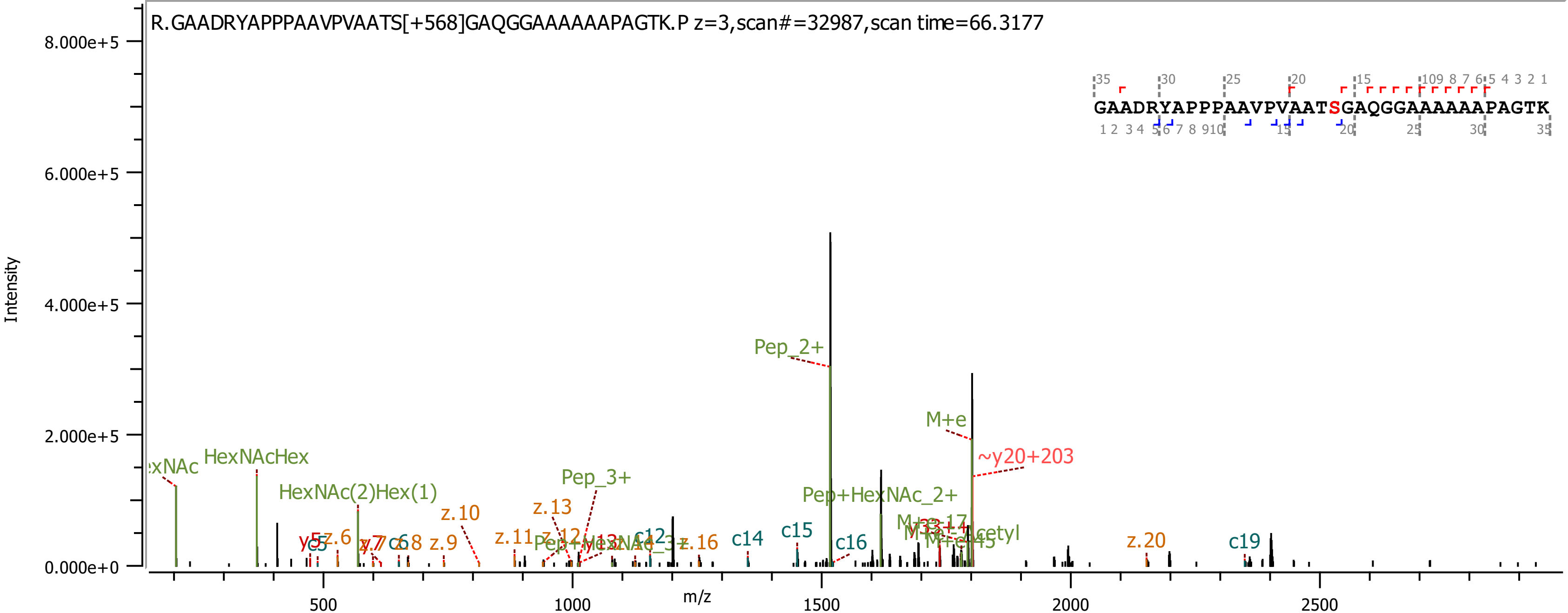

R.GAADRYAPPPAAVPVAATS[+568]GAQGGAAAAAAPAGTK.P z=3,scan#=32987,scan time=66.3177
GAADRYAPPPAAVPVAATSGAQGGAAAAAAPAGTK
Intensity
m/z
Pep_2+
M+e
~y20+203
Pep_3+
Pep+HexNAc_2+
HexNAcHex
HexNAc(2)Hex(1)
z.10
z.13
z.20
c14
c15
c16
c19

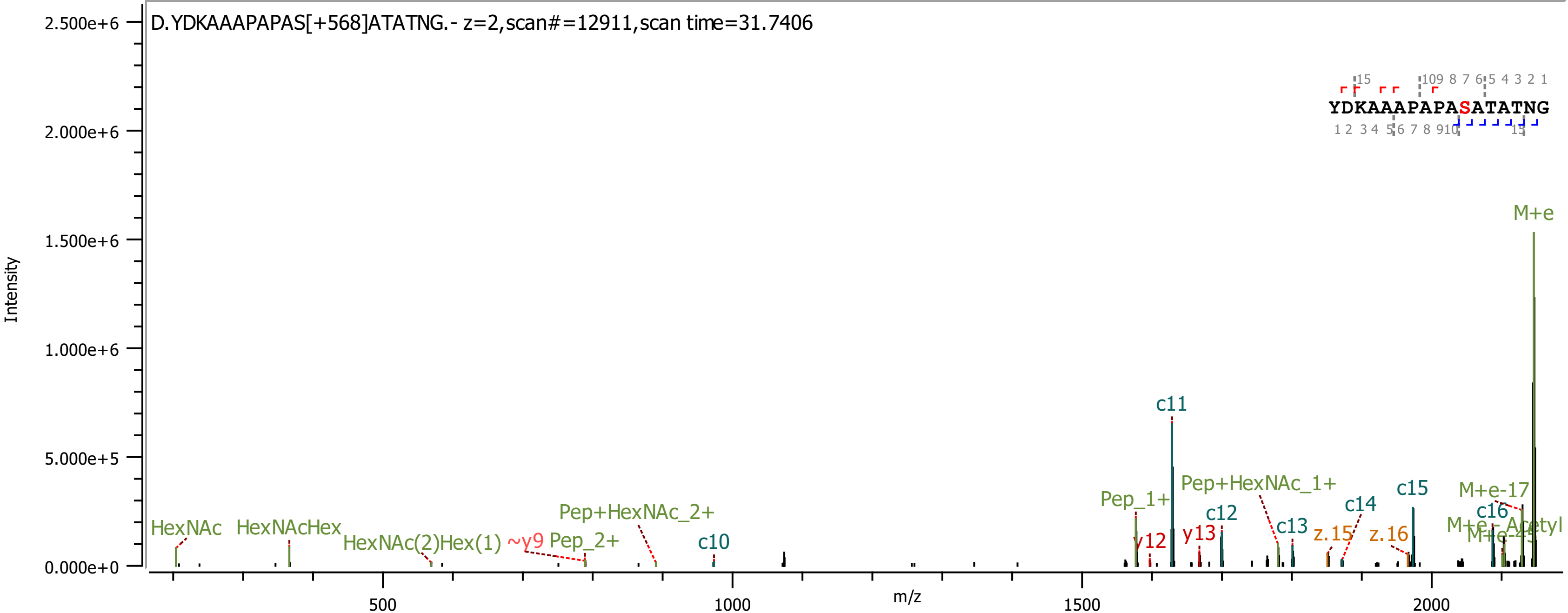

D.YDKAAAPAPAS[+568]ATATNG.- z=2,scan#=12911,scan time=31.7406
Intensity
m/z
M+e
c11
Pep+HexNAc_1+
Pep_1+
c15
M+e-17
c16
c12
c13
c14
y12
y13
z.15
z.16
HexNAc
HexNAcHex
HexNAc(2)Hex(1)
~y9
Pep_2+
Pep+HexNAc_2+
c10
YDKAAAPAPASATATNG

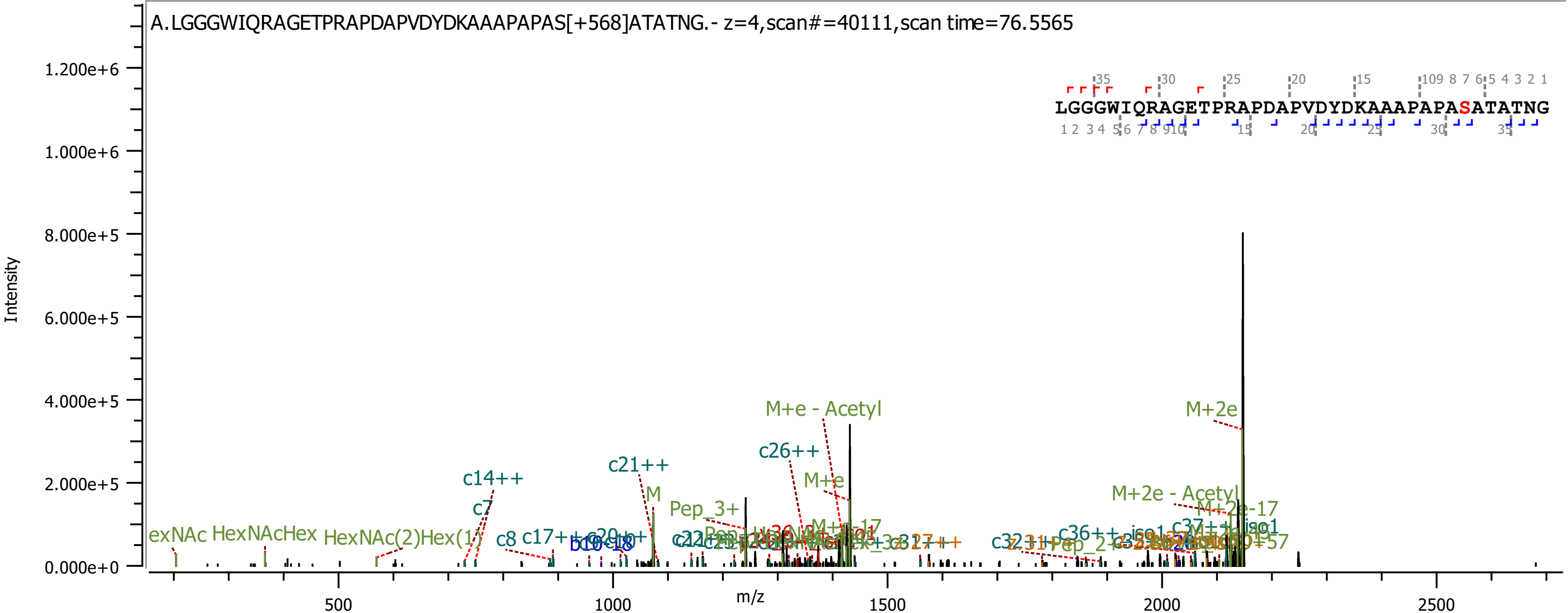
A.LGGGWIQRAGETPRAPDAPVDYDKAAAPAPAS[+568]ATATNG.- z=4,scan#=40111,scan time=76.5565
LGGGWIQRAGETPRAPDAPVDYDKAAAPAPASATATNG
Intensity
m/z
M+2e
M+e - Acetyl
c26++
M+e
c21++
c14++
c7
M
Pep_3+
M+2e - Acetyl
c8
c17++
c36++
c37++
HexNAcHex
HexNAc(2)Hex(1)

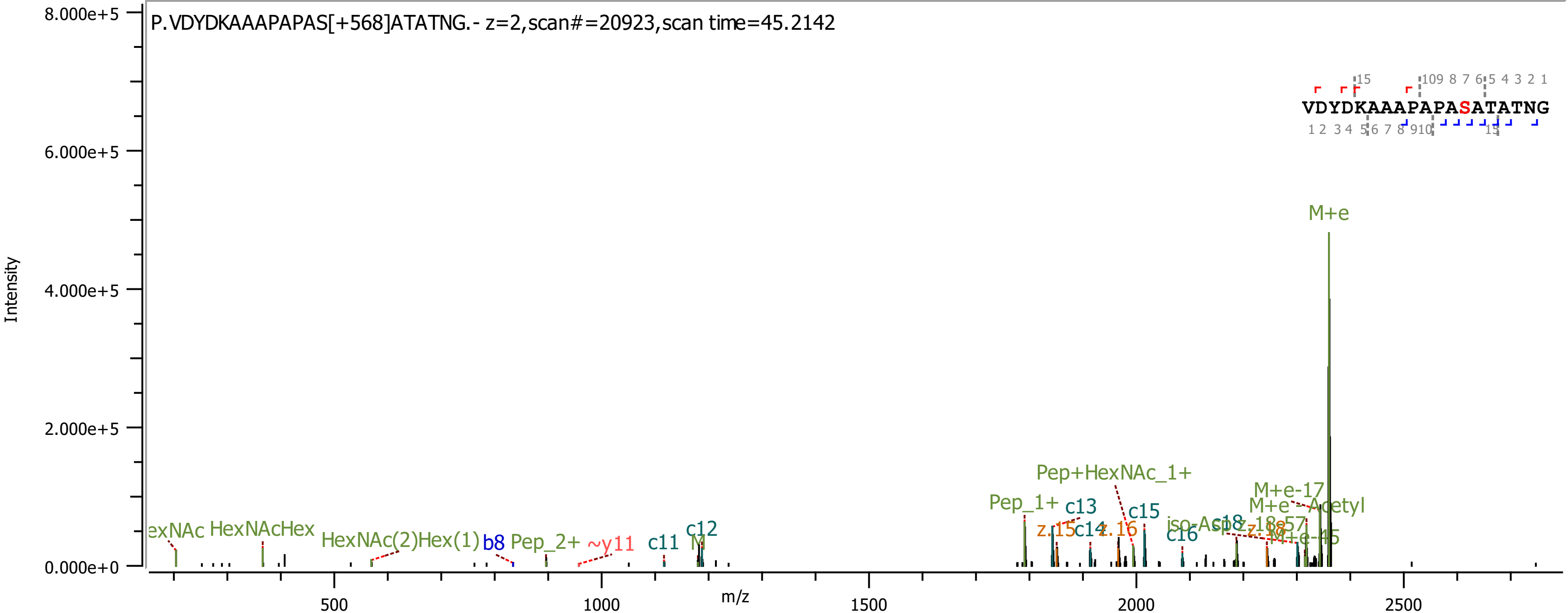

P.VDYDKAAAPAPAS[+568]ATATNG.- z=2,scan#=20923,scan time=45.2142
Intensity
m/z
VDYDKAAAPAPASATATNG
M+e
Pep+HexNAc_1+
M+e-17
M+e - Acetyl
Pep_1+
c13
c15
z.15
c14
z.16
iso-Asp
c18
c16
M+e-45
exNAc
HexNAcHex
HexNAc(2)Hex(1)
b8
Pep_2+
~y11
c11
c12
8.000e+5
6.000e+5
4.000e+5
2.000e+5
0.000e+0
500
1000
1500
2000
2500

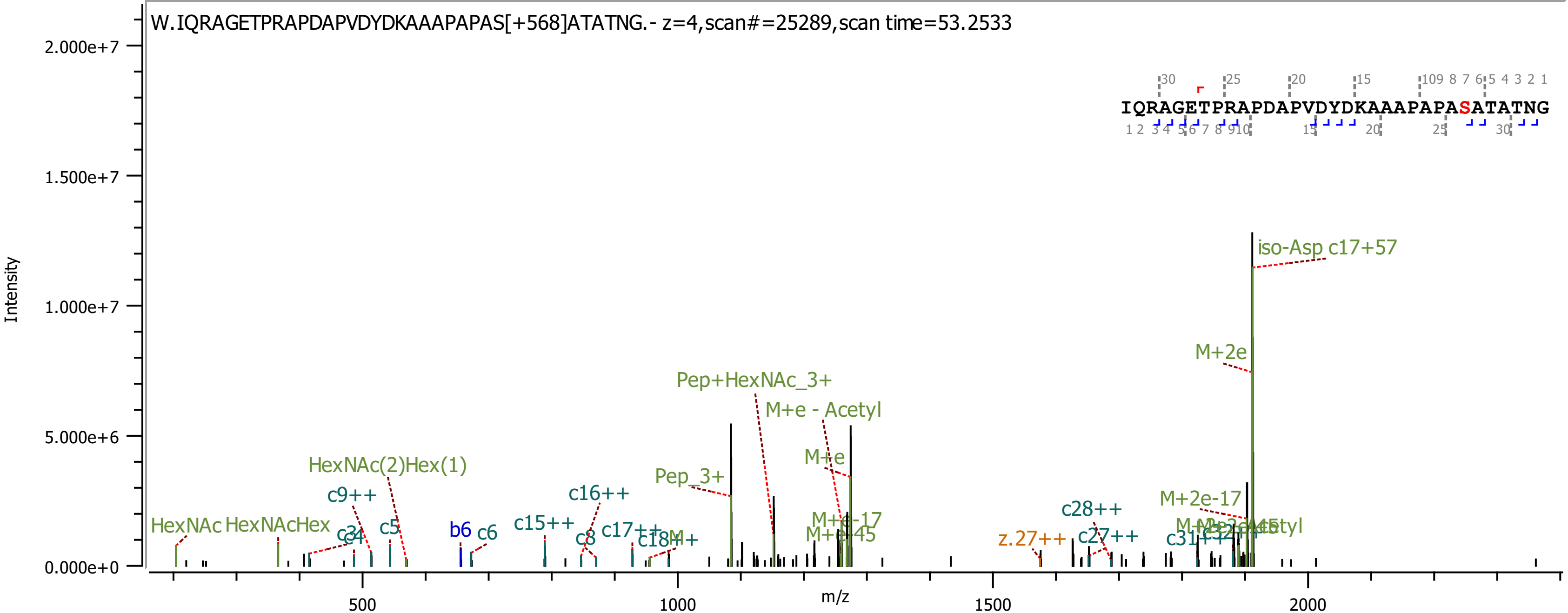

W.IQRAGETPRAPDAPVDYDKAAAPAPAS[+568]ATATNG.- z=4,scan#=25289,scan time=53.2533
IQRAGETPRAPDAPVDYDKAAAPAPASATATNG
Intensity
m/z
2.000e+7
1.500e+7
1.000e+7
5.000e+6
0.000e+0
500
1000
1500
2000
iso-Asp c17+57
M+2e
Pep+HexNAc_3+
M+e - Acetyl
M+e
Pep_3+
HexNAc(2)Hex(1)
c9++
c16++
c15++
c17++
c18++
M+e-17
M+e-45
c28++
M+2e-17
z.27++
c27++
c31++
HexNAc
HexNAcHex
c3
c4
c5
b6
c6
c8

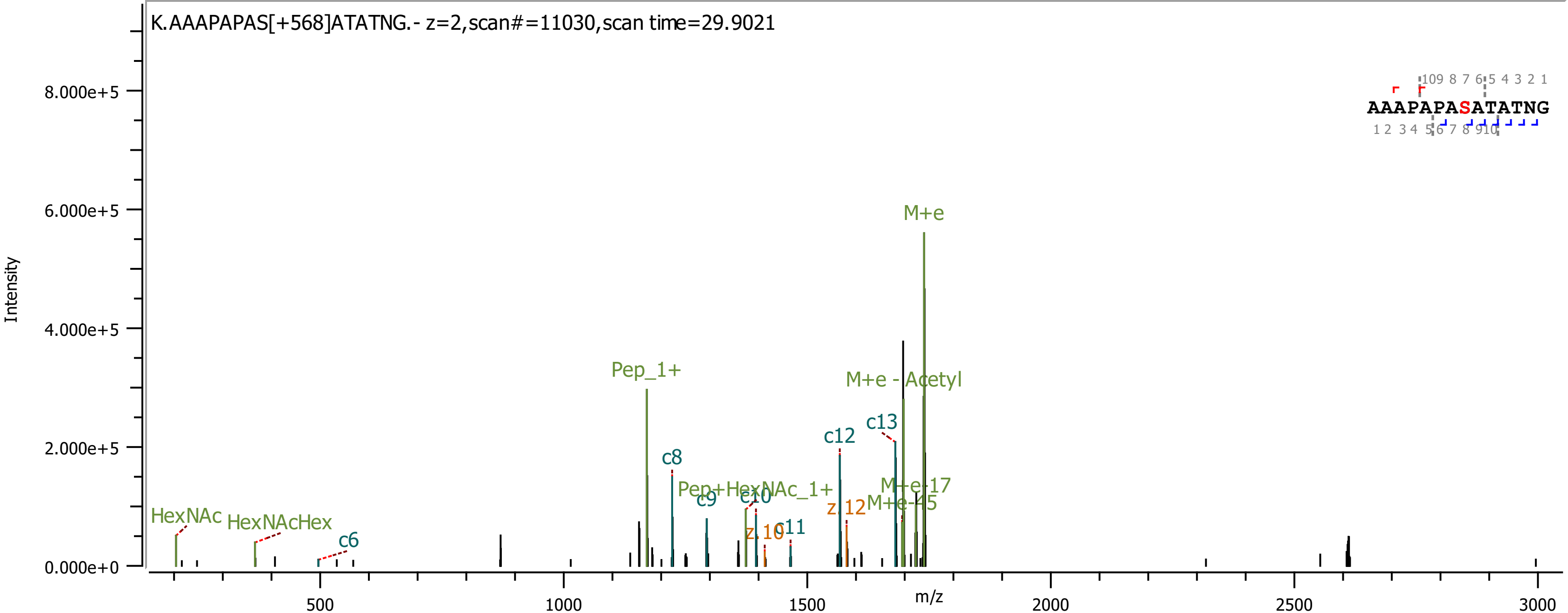
K.AAAPAPAS[+568]ATATNG.- z=2,scan#=11030,scan time=29.9021
Intensity
m/z
AAAPAPASATATNG
HexNAc
HexNAcHex
c6
Pep_1+
c8
c9
Pep+HexNAc_1+
c10
z 10
c11
c12
z 12
c13
M+e - Acetyl
M+e
M+e-17
M+e-45
8.000e+5
6.000e+5
4.000e+5
2.000e+5
0.000e+0
500
1000
1500
2000
2500
3000

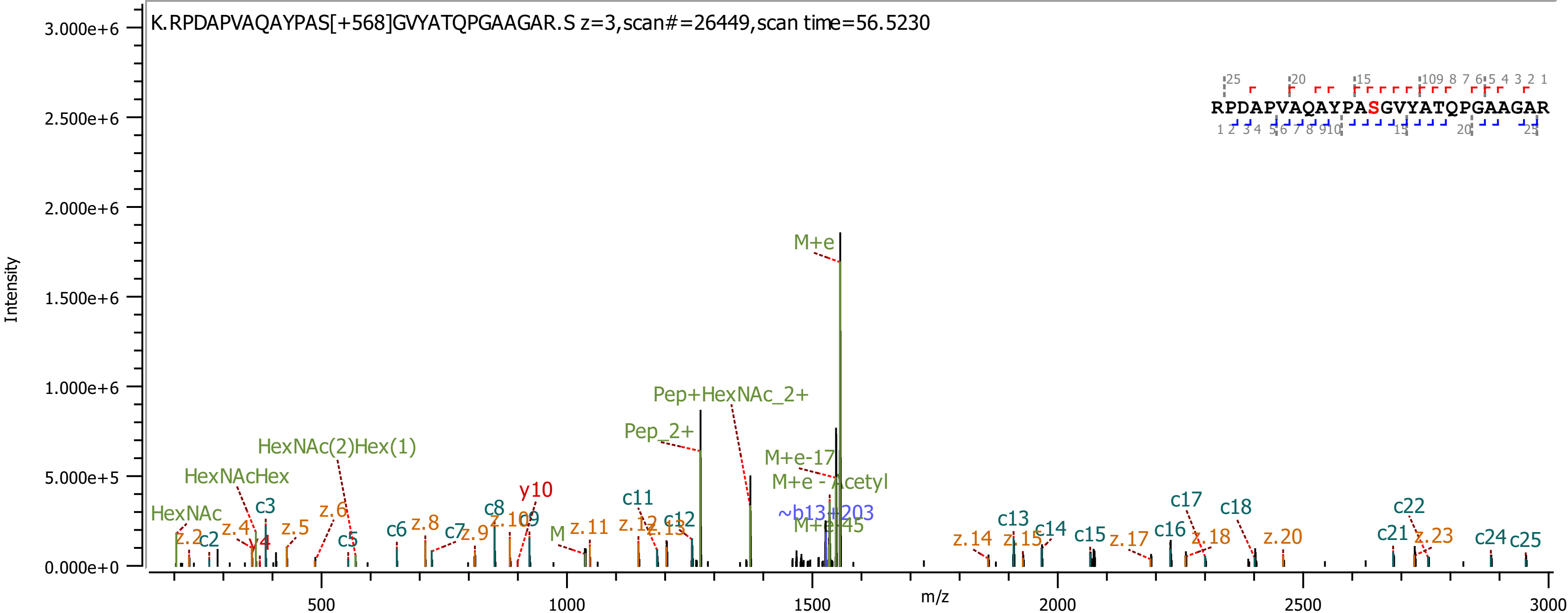
K.RPDAPVAQAYPAS[+568]GVYATQPGAAGAR.S z=3,scan#=26449,scan time=56.5230
RPDAPVAQAYPASGVYATQPGAAGAR
Intensity
m/z
M+e
Pep+HexNAc_2+
Pep_2+
HexNAc(2)Hex(1)
HexNAcHex
HexNAc
M+e-17
M+e - Acetyl
~b13+203
M+e-45
y10
y4
M
c2
c3
c5
c6
c7
c8
c9
c11
c12
c13
c14
c15
c16
c17
c18
c21
c22
c24
c25
z.2
z.4
z.5
z.6
z.8
z.9
z.10
z.11
z.12
z.13
z.14
z.15
z.17
z.18
z.20
z.23

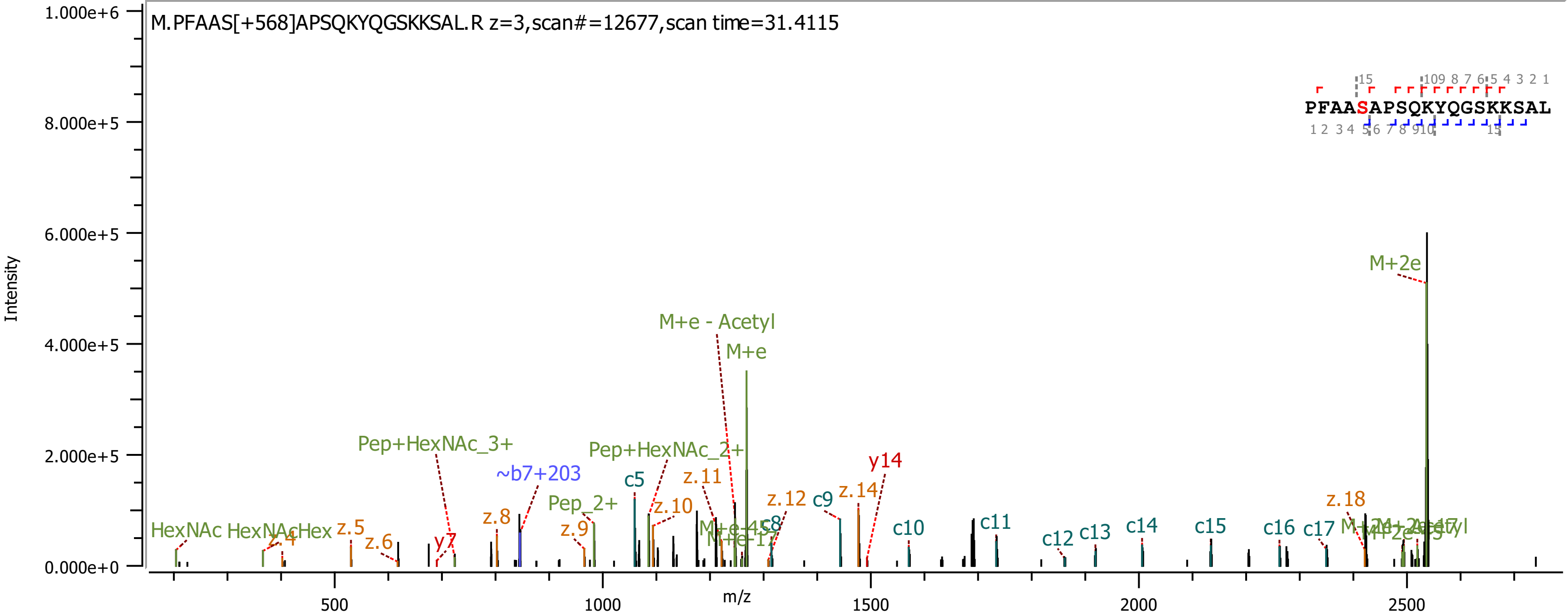
M.PFAAS[+568]APSQKYQGSKKSAL.R z=3,scan#=12677,scan time=31.4115
PFAASAPSQKYQGSKKSAL
Intensity
m/z
1.000e+6
8.000e+5
6.000e+5
4.000e+5
2.000e+5
0.000e+0
500
1000
1500
2000
2500
M+2e
M+e - Acetyl
M+e
Pep+HexNAc_3+
Pep+HexNAc_2+
~b7+203
Pep_2+
HexNAc
c5
z.11
z.10
y14
z.14
z.12
c9
z.8
z.9
z.5
z.6
z.18
c10
c11
c12
c13
c14
c15
c16
c17

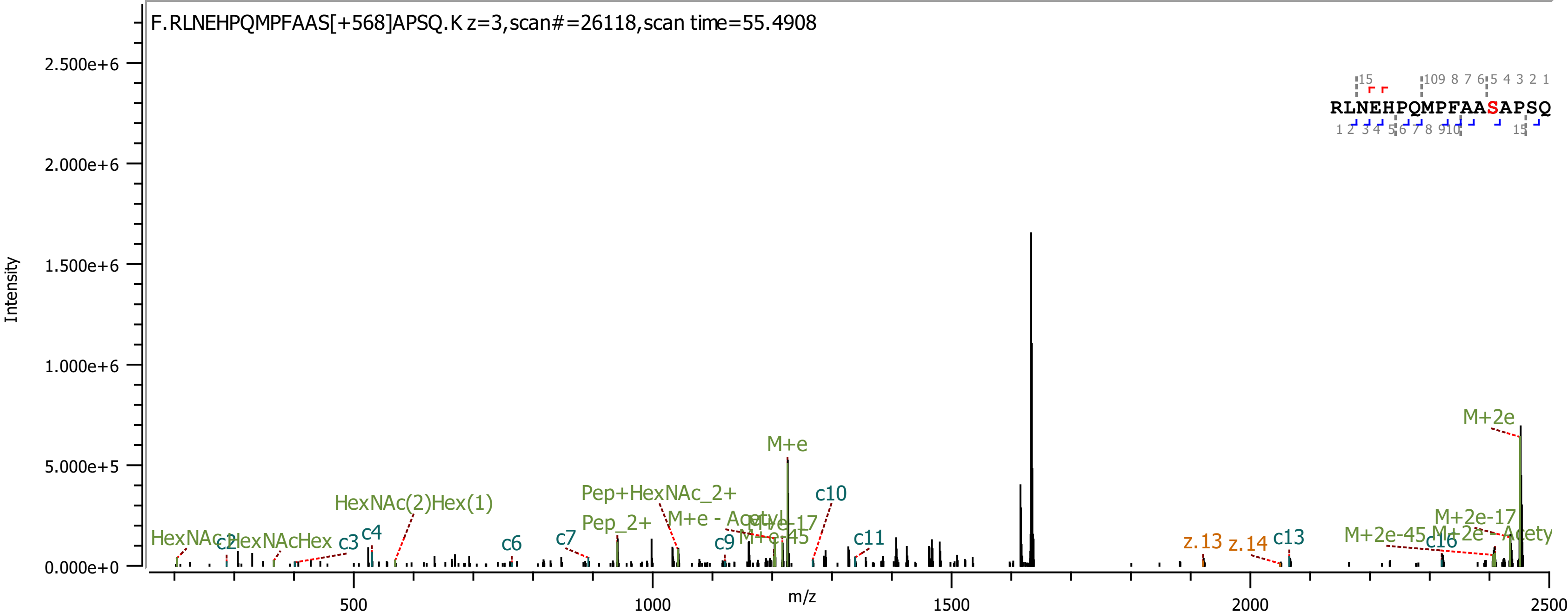

F.RLNEHPQMPFAAS[+568]APSQ.K z=3,scan#=26118,scan time=55.4908
RLNEHPQMPFAASAPSQ
Intensity
m/z
2.500e+6
2.000e+6
1.500e+6
1.000e+6
5.000e+5
0.000e+0
500
1000
1500
2000
2500
HexNAc
c2
HexNAcHex
c3
c4
HexNAc(2)Hex(1)
c6
c7
Pep+HexNAc_2+
Pep_2+
M+e - Acetyl
M+e-17
M+e-45
M+e
c9
c10
c11
z.13
z.14
c13
M+2e-45
c16
M+2e-17
M+2e - Acety
M+2e

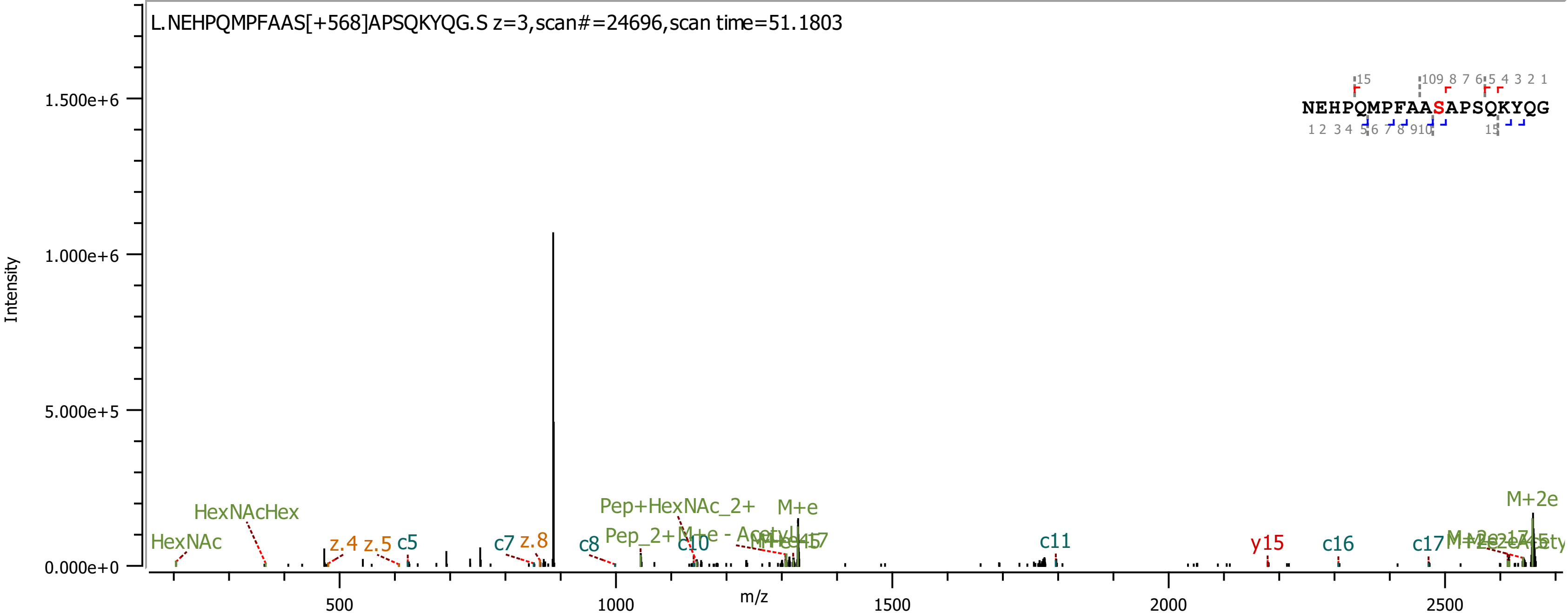
L.NEHPQMPFAAS[+568]APSQKYQG.S z=3,scan#=24696,scan time=51.1803
NEHPQMPFAASAPSQKYQG
Intensity
m/z
0.000e+0
5.000e+5
1.000e+6
1.500e+6
500
1000
1500
2000
2500
HexNAc
HexNAcHex
z.4
z.5
c5
c7
z.8
c8
Pep+HexNAc_2+
Pep_2+
M+e
c10
c11
y15
c16
c17
M+2e

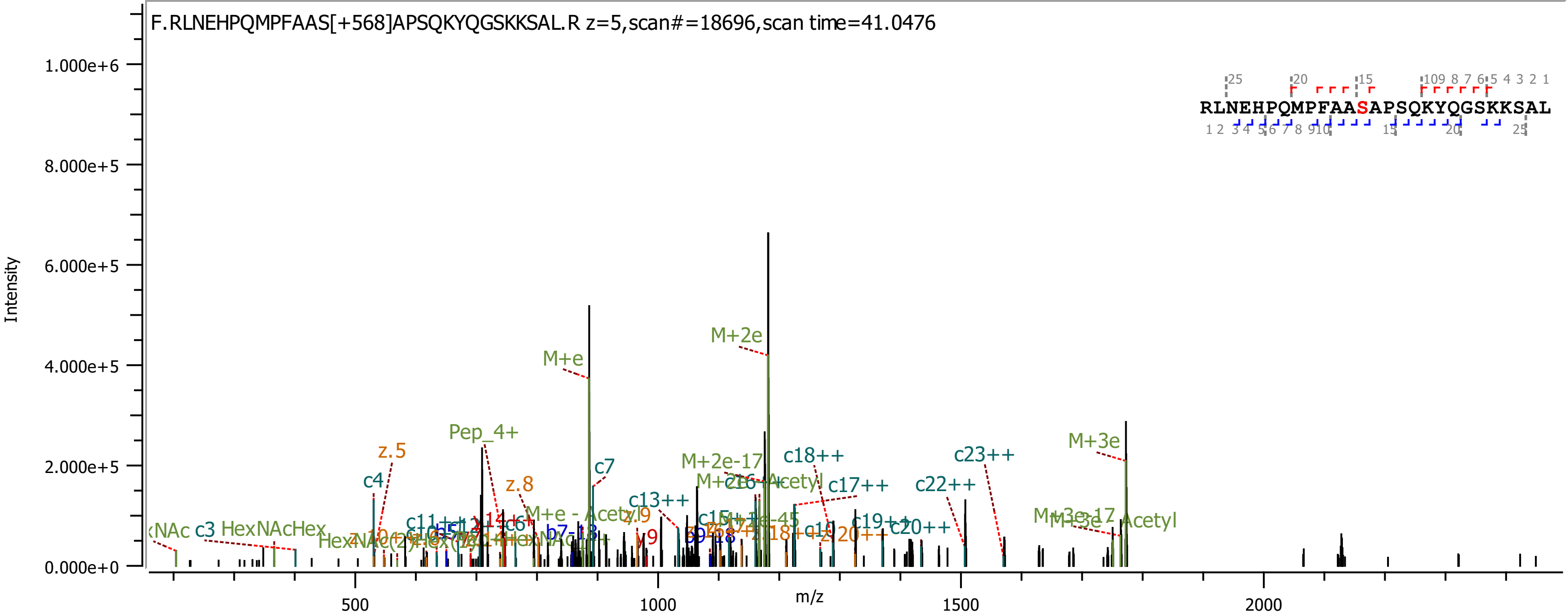
F.RLNEHPQMPFAAS[+568]APSQKYQGSKKSAL.R z=5,scan#=18696,scan time=41.0476
RLNEHPQMPFAASAPSQKYQGSKKSAL
Intensity
m/z
M+e
M+2e
M+3e
Pep_4+
c4
c7
z.5
z.8
c13++
c17++
c18++
c22++
c23++

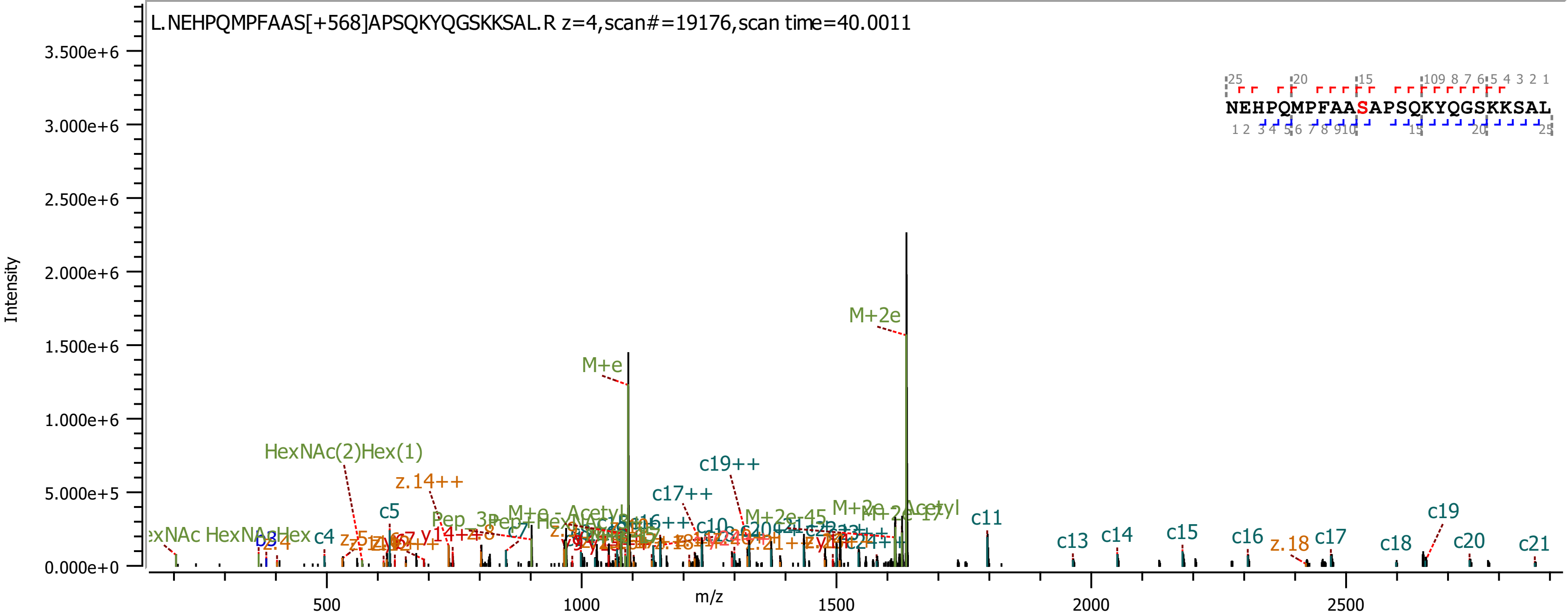
L.NEHPQMPFAAS[+568]APSQKYQGSKKSAL.R z=4,scan#=19176,scan time=40.0011
NEHPQMPFAASAPSQKYQGSKKSAL
Intensity
m/z
M+2e
M+e
HexNAc(2)Hex(1)
z.14++
c19++
c17++
c5
c4
c11
c13
c14
c15
c16
z.18
c17
c18
c19
c20
c21
M+2e-45
M+2e - Acetyl
M+e - Acetyl
HexNAc
HexNAcHex
z.4
c7
y14++
z.8
c10

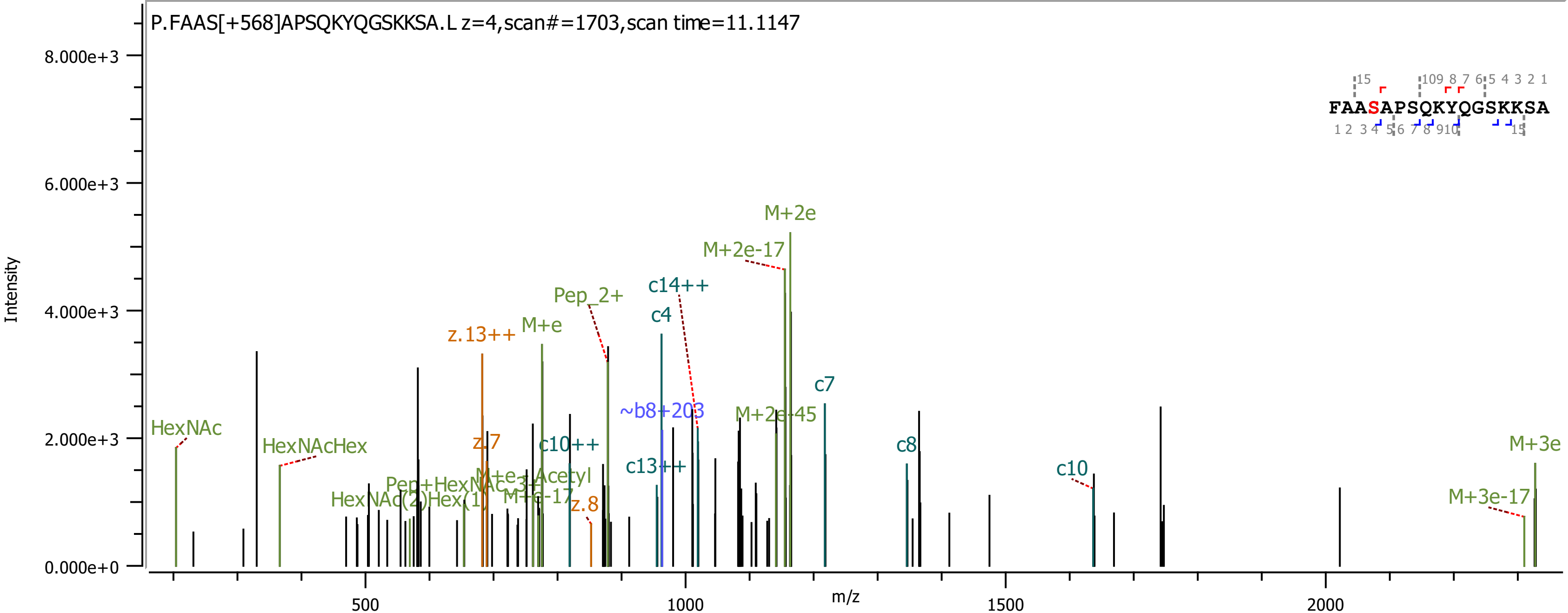
P.FAAS[+568]APSQKYQGSKKSA.L z=4,scan#=1703,scan time=11.1147
FAASAPSQKYQGSKKSA
Intensity
m/z
HexNAc
HexNAcHex
HexNAc(2)Hex(1)
Pep+HexNAc
z.13++
z.7
z.8
M+e
M+e-17
Pep_2+
c10++
c4
c14++
c13++
~b8+203
M+2e-17
M+2e
M+2e-45
c7
c8
c10
M+3e
M+3e-17

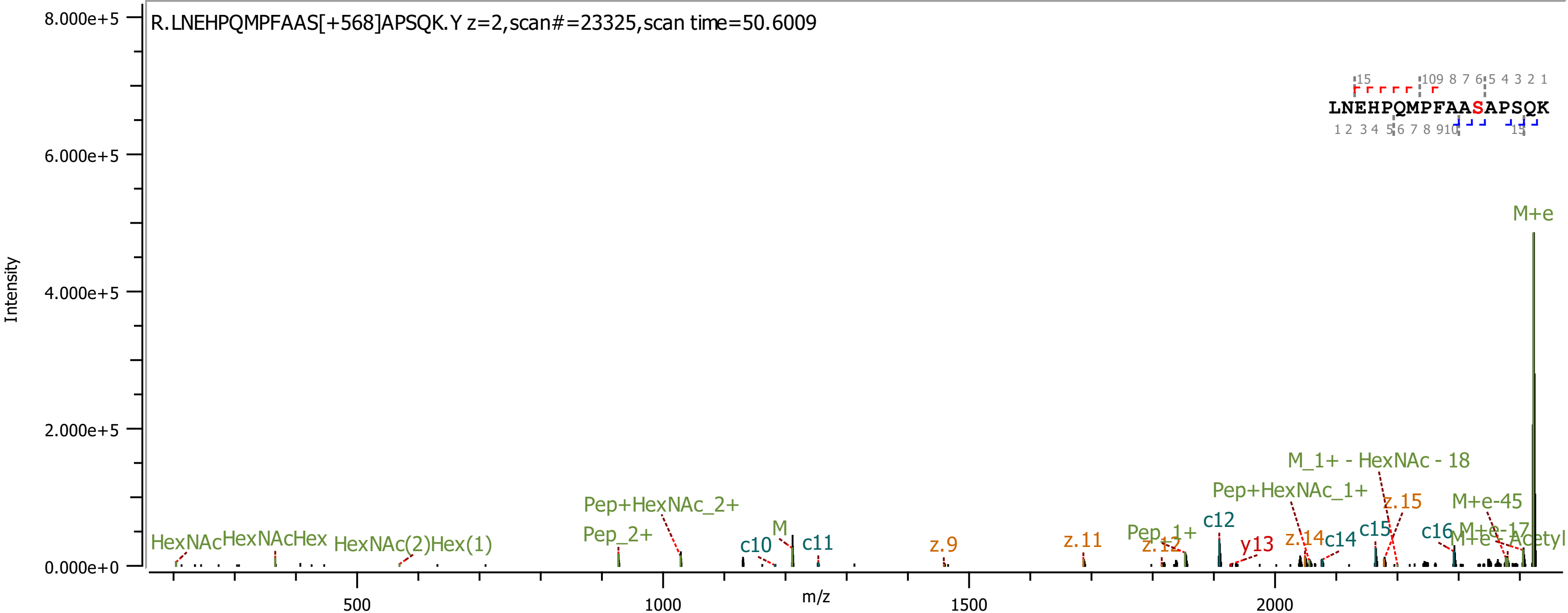
R.LNEHPQMPFAAS[+568]APSQK.Y z=2,scan#=23325,scan time=50.6009
LNEHPQMPFAASAPSQK
Intensity
m/z
M+e
M_1+ - HexNAc - 18
Pep+HexNAc_1+
z.15
M+e-45
Pep+HexNAc_2+
Pep_2+
c12
c15
c16
M+e-17
M+e-Acetyl
z.14
c14
y13
Pep_1+
z.12
z.11
z.9
c10
M
c11
HexNAc
HexNAcHex
HexNAc(2)Hex(1)

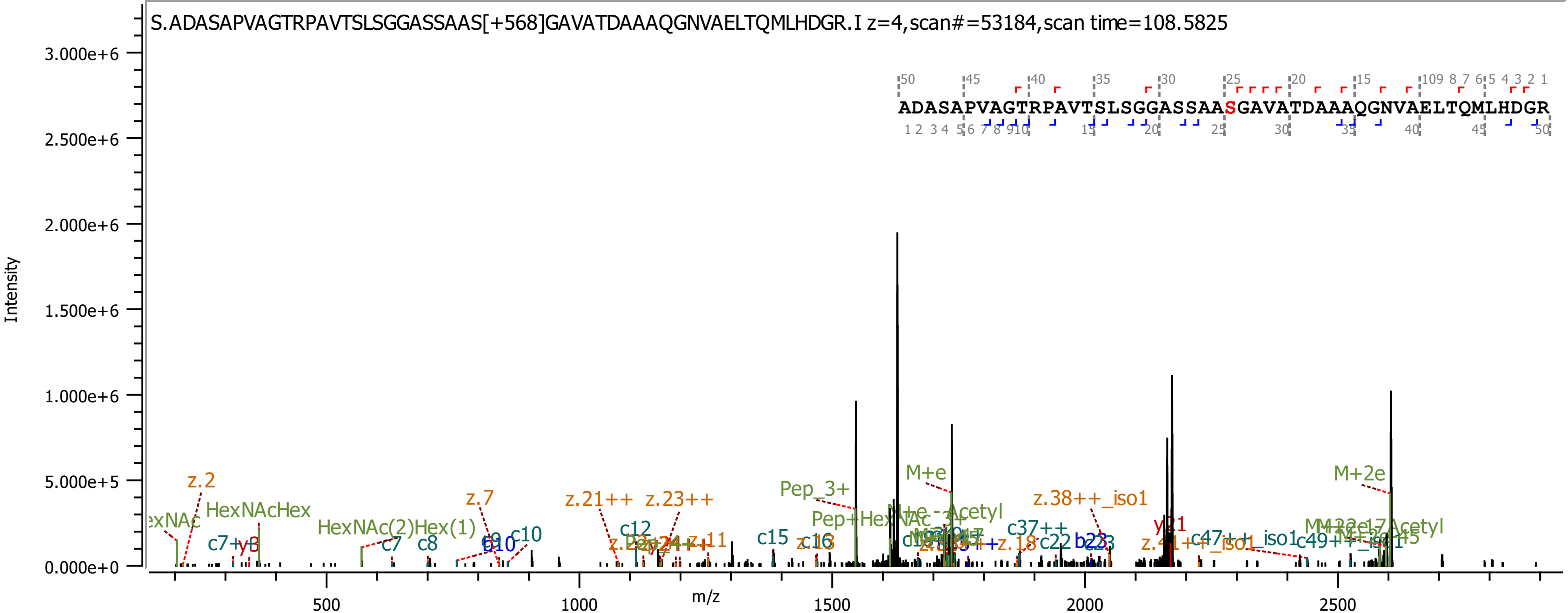
S.ADASAPVAGTRPAVTSLSGGASSAAS[+568]GAVATDAAAQGNVAELTQMLHDGR.I z=4,scan#=53184,scan time=108.5825
ADASAPVAGTRPAVTSLSGGASSAASGAVATDAAAQGNVAELTQMLHDGR
Intensity
m/z
z.2
HexNAcHex
HexNAc(2)Hex(1)
c7
c8
c10
z.7
z.21++
z.23++
c12
c15
Pep_3+
M+e
z.38++_iso1
y21
M+2e

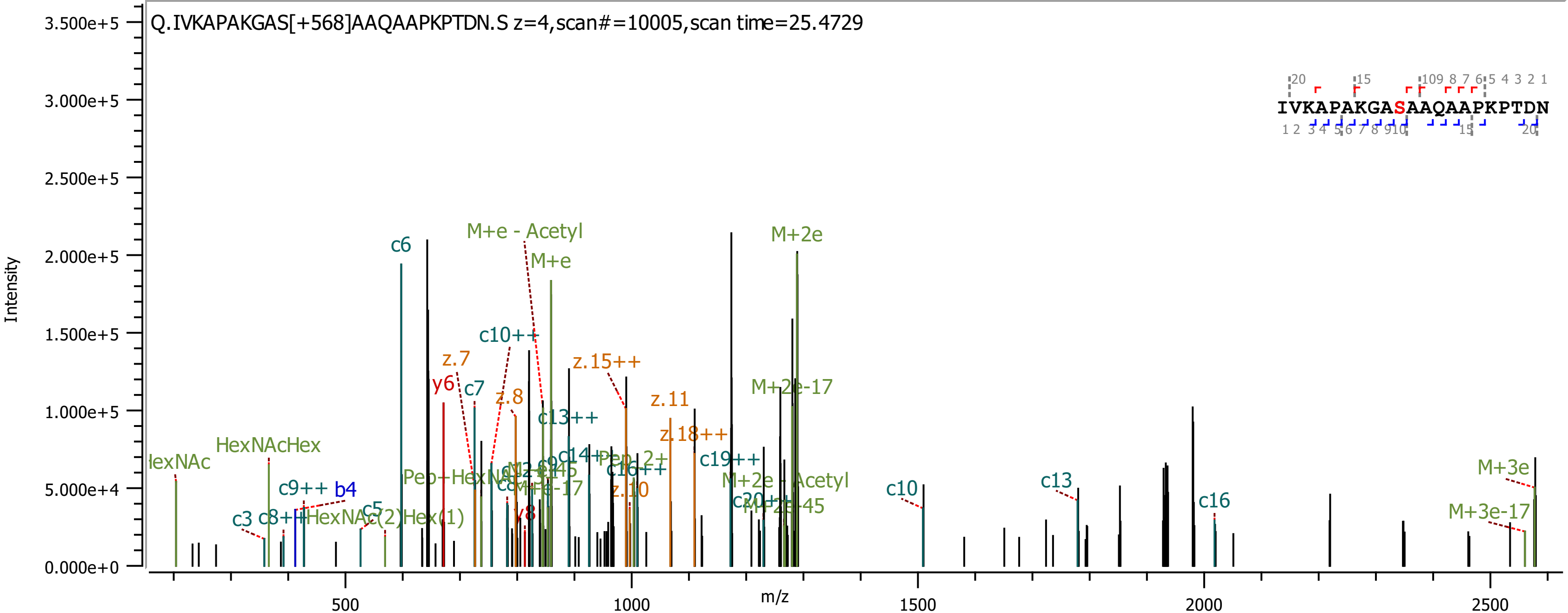
Q.IVKAPAKGAS[+568]AAQAAPKPTDN.S z=4,scan#=10005,scan time=25.4729
IVKAPAKGASAAQAAPKPTDN
Intensity
m/z
3.500e+5
3.000e+5
2.500e+5
2.000e+5
1.500e+5
1.000e+5
5.000e+4
0.000e+0
500
1000
1500
2000
2500
c6
M+e - Acetyl
M+e
M+2e
c10++
z.7
y6
c7
z.8
z.15++
M+2e-17
c13++
z.11
z.18++
HexNAcHex
c9++
b4
c3
c8++
HexNAc(2)Hex(1)
c5
c19++
M+2e - Acetyl
c10
c13
c16
M+3e
M+3e-17
z.10
c16++
c14++
y8

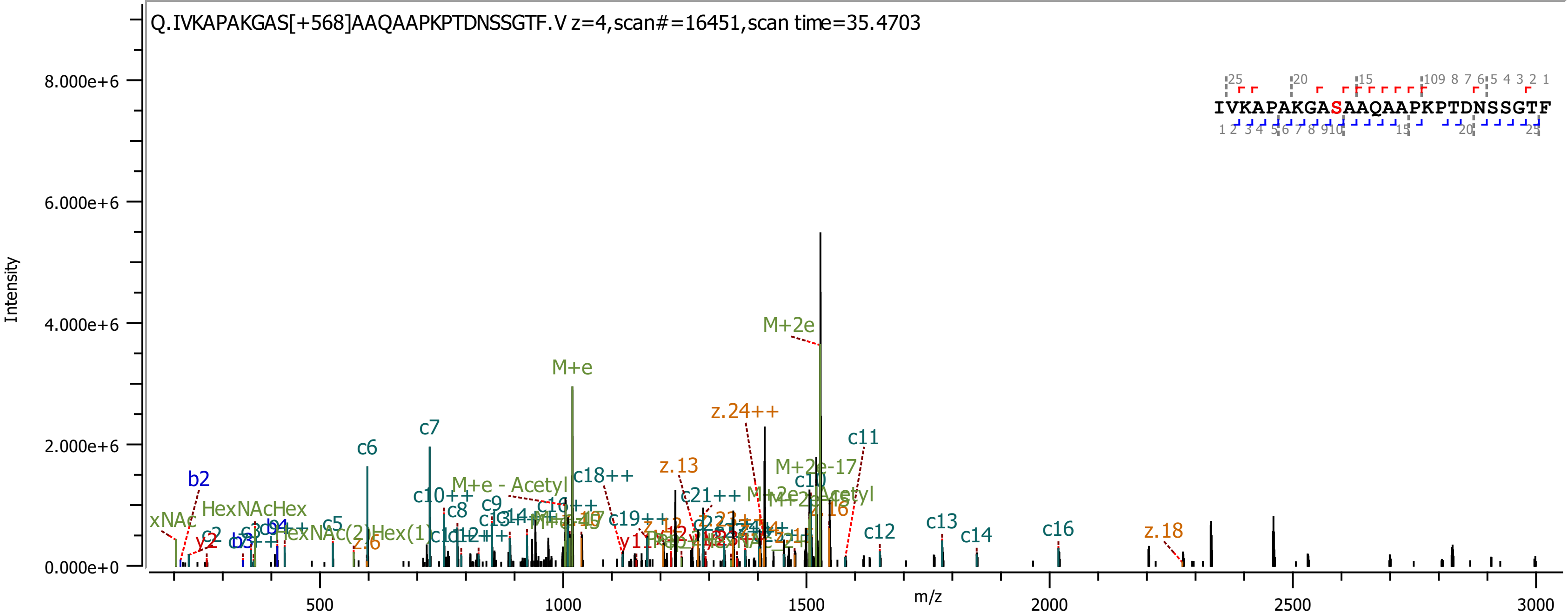

Q.IVKAPAKGAS[+568]AAQAAPKPTDNSSGTF.V z=4,scan#=16451,scan time=35.4703
IVKAPAKGASAAQAAPKPTDNSSGTF
Intensity
m/z
8.000e+6
6.000e+6
4.000e+6
2.000e+6
0.000e+0
500
1000
1500
2000
2500
3000
M+2e
M+e
z.24++
c11
c7
c6
b2
c18++
z.13
M+2e-17
M+e - Acetyl
HexNAcHex
c10
c10++
c8
c9
c5
z.16
c12
c13
c14
c16
z.18

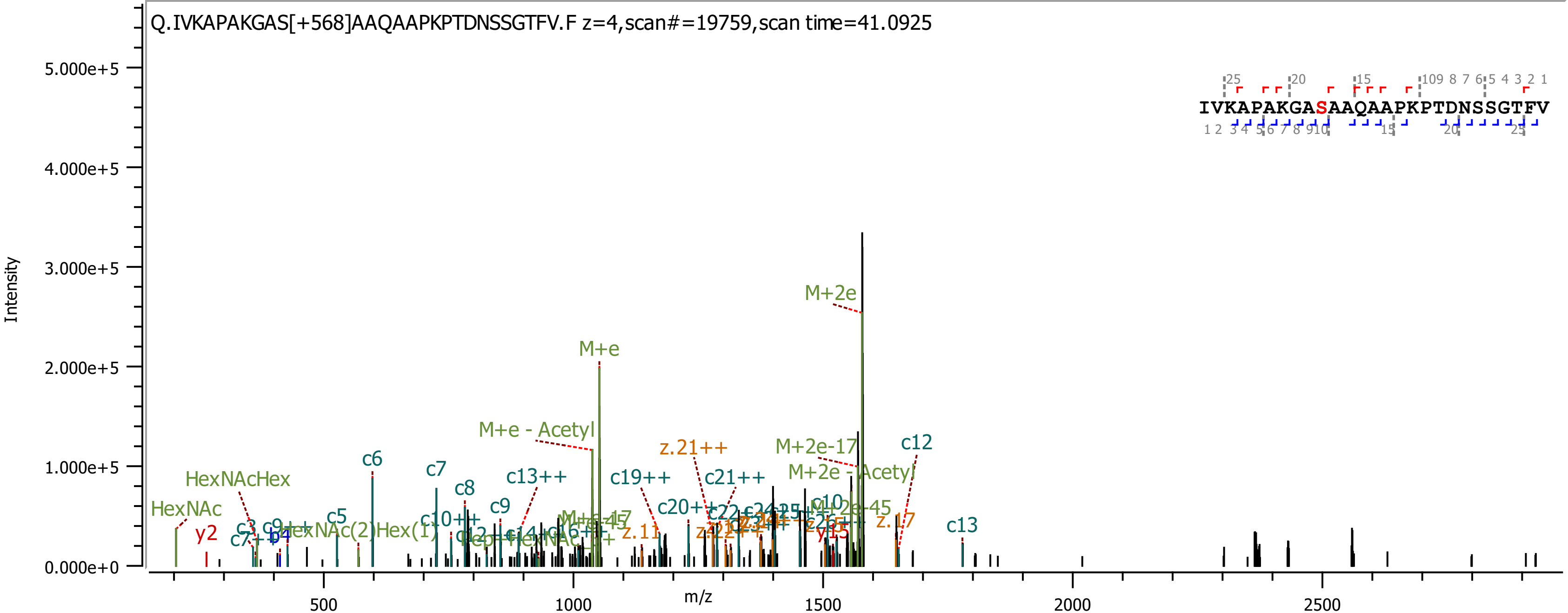
Q.IVKAPAKGAS[+568]AAQAAPKPTDNSSGTFV.F z=4,scan#=19759,scan time=41.0925
IVKAPAKGASAAQAAPKPTDNSSGTFV
Intensity
m/z
5.000e+5
4.000e+5
3.000e+5
2.000e+5
1.000e+5
0.000e+0
500
1000
1500
2000
2500
M+2e
M+e
M+e - Acetyl
z.21++
M+2e-17
c12
M+2e - Acetyl
c6
c7
HexNAcHex
c8
c13++
c19++
c21++
c9
HexNAc
c20++
c5
c10++
z.11
z.17
c13
y2
y15

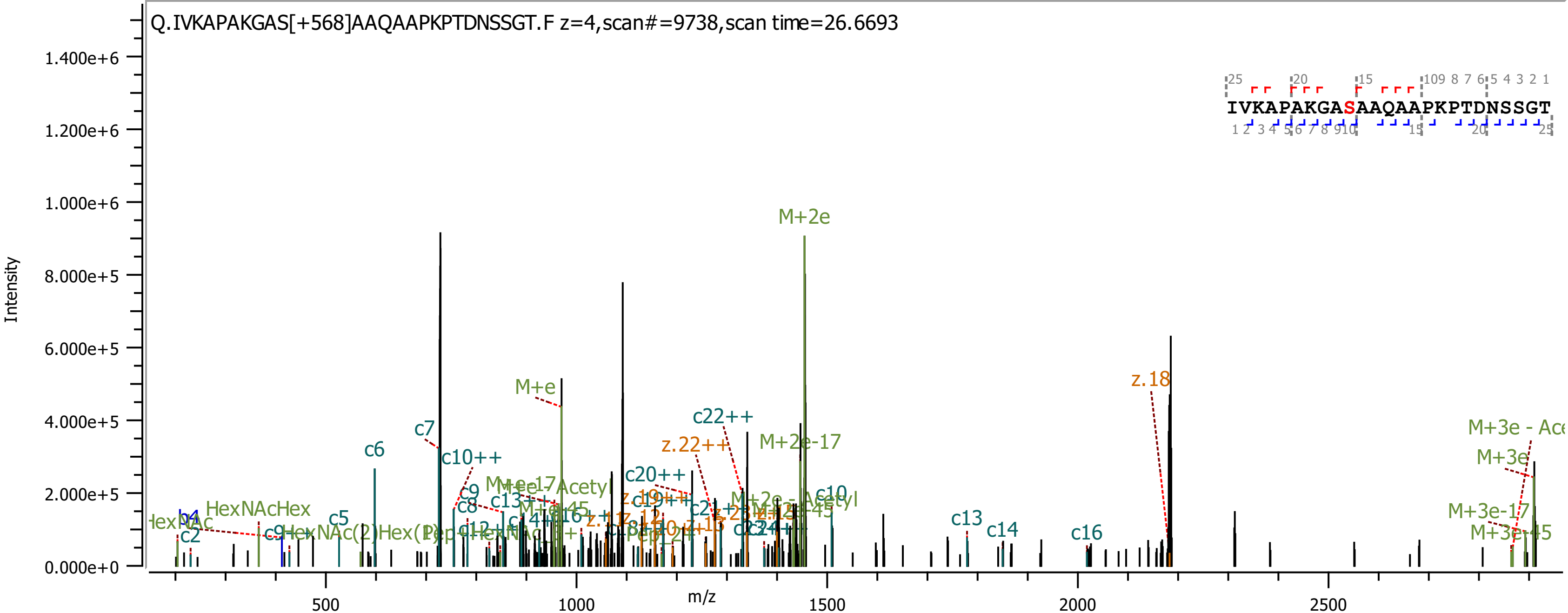

Q.IVKAPAKGAS[+568]AAQAAPKPTDNSSGT.F z=4,scan#=9738,scan time=26.6693
IVKAPAKGASAAQAAPKPTDNSSGT
Intensity
m/z
1.400e+6
1.200e+6
1.000e+6
8.000e+5
6.000e+5
4.000e+5
2.000e+5
0.000e+0
500
1000
1500
2000
2500
M+2e
M+e
z.18
c22++
z.22++
M+2e-17
c7
c6
c10++
c20++
M+3e
M+3e-17
M+3e-45
c5
c2
HexNAcHex
c13
c14
c16
c9
c8

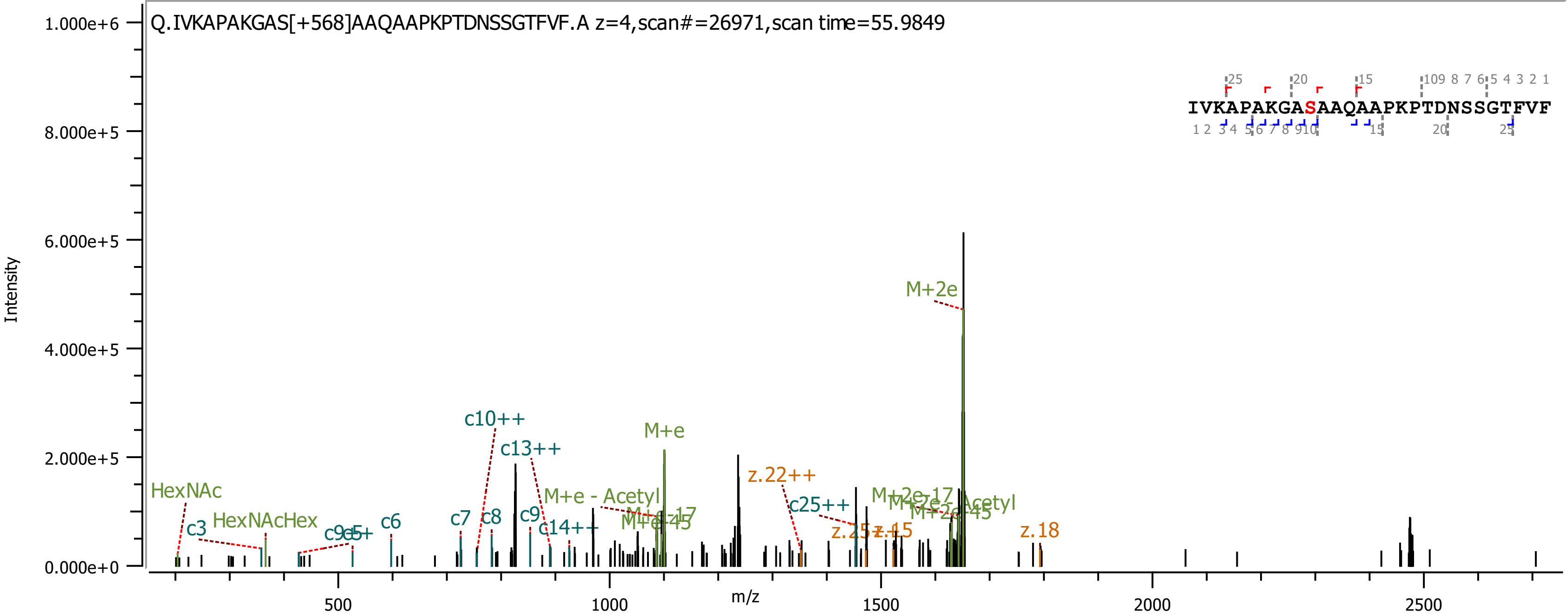

Q.IVKAPAKGAS[+568]AAQAAPKPTDNSSGTFVF.A z=4,scan#=26971,scan time=55.9849
IVKAPAKGASAAQAAPKPTDNSSGTFVF
Intensity
m/z
1.000e+6
8.000e+5
6.000e+5
4.000e+5
2.000e+5
0.000e+0
500
1000
1500
2000
2500
M+2e
M+e
c10++
c13++
z.22++
HexNAc
HexNAcHex
M+e - Acetyl
c25++
c3
c6
c7
c8
c9
c14++
z.18

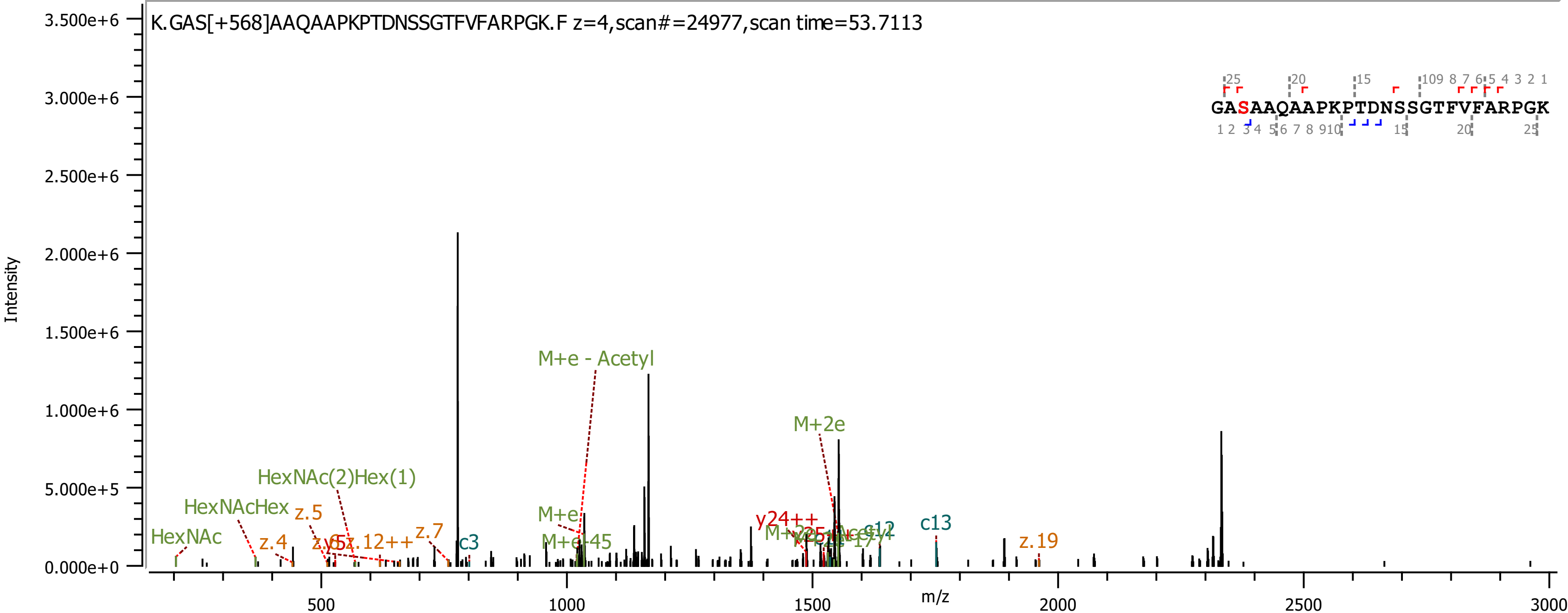
K.GAS[+568]AAQAAPKPTDNSSGTFVFARPGK.F z=4,scan#=24977,scan time=53.7113
GASAAQAAPKPTDNSSGTFVFARPGK
Intensity
m/z
3.500e+6
3.000e+6
2.500e+6
2.000e+6
1.500e+6
1.000e+6
5.000e+5
0.000e+0
500
1000
1500
2000
2500
3000
M+e - Acetyl
M+2e
HexNAc(2)Hex(1)
HexNAcHex
HexNAc
z.5
z.4
z.12++
z.7
c3
M+e
M+e-45
y24++
c12
c13
z.19

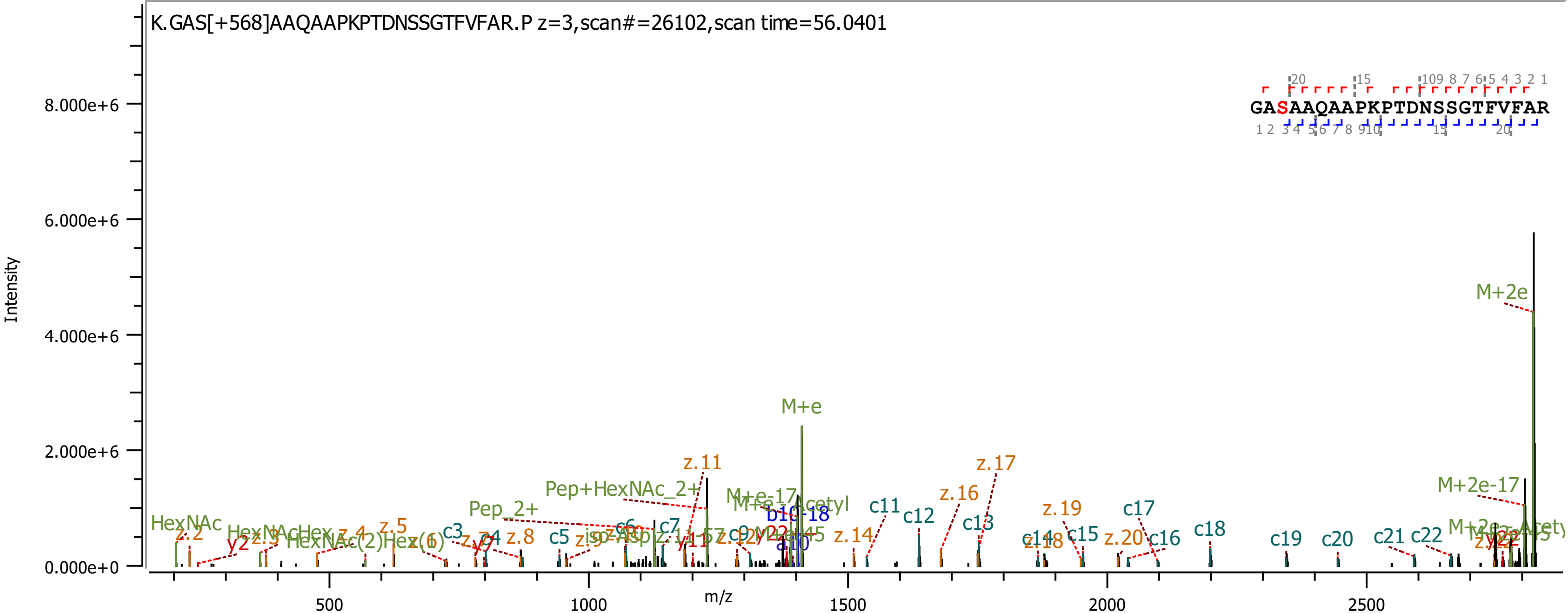
K.GAS[+568]AAQAAPKPTDNSSGTFVFAR.P z=3,scan#=26102,scan time=56.0401
GASAAQAAPKPTDNSSGTFVFAR
Intensity
m/z
8.000e+6
6.000e+6
4.000e+6
2.000e+6
0.000e+0
500
1000
1500
2000
2500
M+2e
M+e
M+2e-17
z.11
z.17
z.16
z.19
Pep+HexNAc_2+
Pep_2+
HexNAc
c11
c12
c13
c15
c16
c17
c18
c19
c20
c21
c22
z.14
z.20
z.8
z.5
c3
c5
c7
y2
b10-18
a10

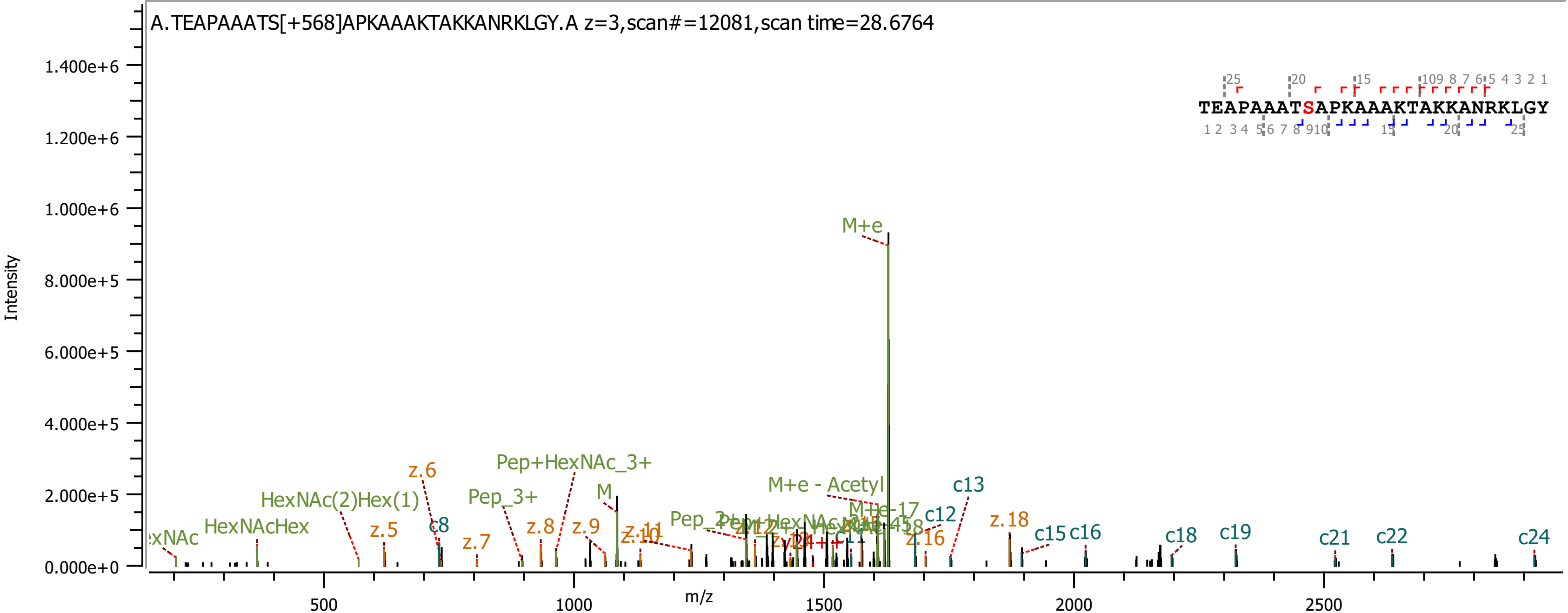

A.TEAPAAATS[+568]APKAAAKTAKKANRKLGY.A z=3,scan#=12081,scan time=28.6764
TEAPAAATSAPKAAAKTAKKANRKLGY
Intensity
m/z
M+e
Pep+HexNAc_3+
HexNAc(2)Hex(1)
HexNAcHex
Pep_3+
M
M+e - Acetyl
M+e-17
c8
c12
c13
c15
c16
c18
c19
c21
c22
c24
z.5
z.6
z.7
z.8
z.9
z.16
z.18

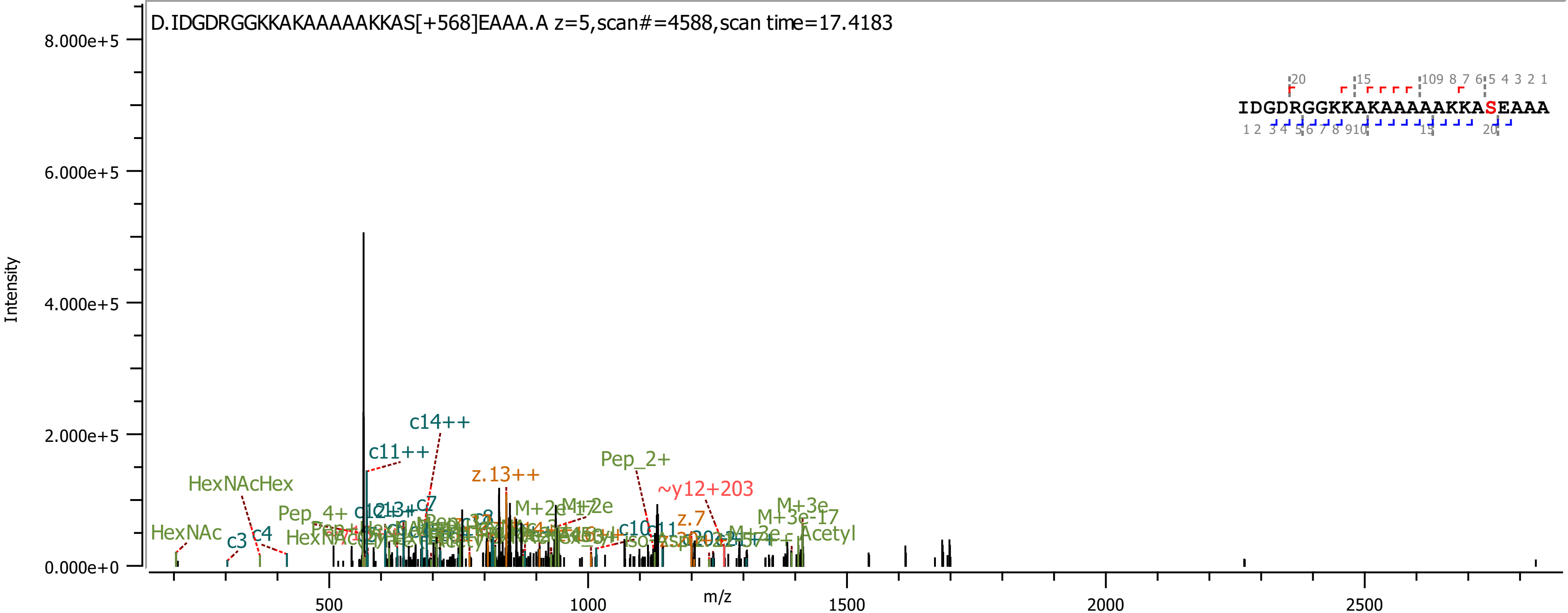

D.IDGDRGGKKAKAAAAAKKAS[+568]EAAA.A z=5,scan#=4588,scan time=17.4183
IDGDRGGKKAKAAAAAKKASEAAA
Intensity
m/z
8.000e+5
6.000e+5
4.000e+5
2.000e+5
0.000e+0
500
1000
1500
2000
2500
c14++
c11++
z.13++
Pep_2+
~y12+203
HexNAcHex
Pep_4+
c7
M+2e-17
M+2e
M+3e
M+3e-17
Acetyl
HexNAc
c3
c4
c10
c11
z.7

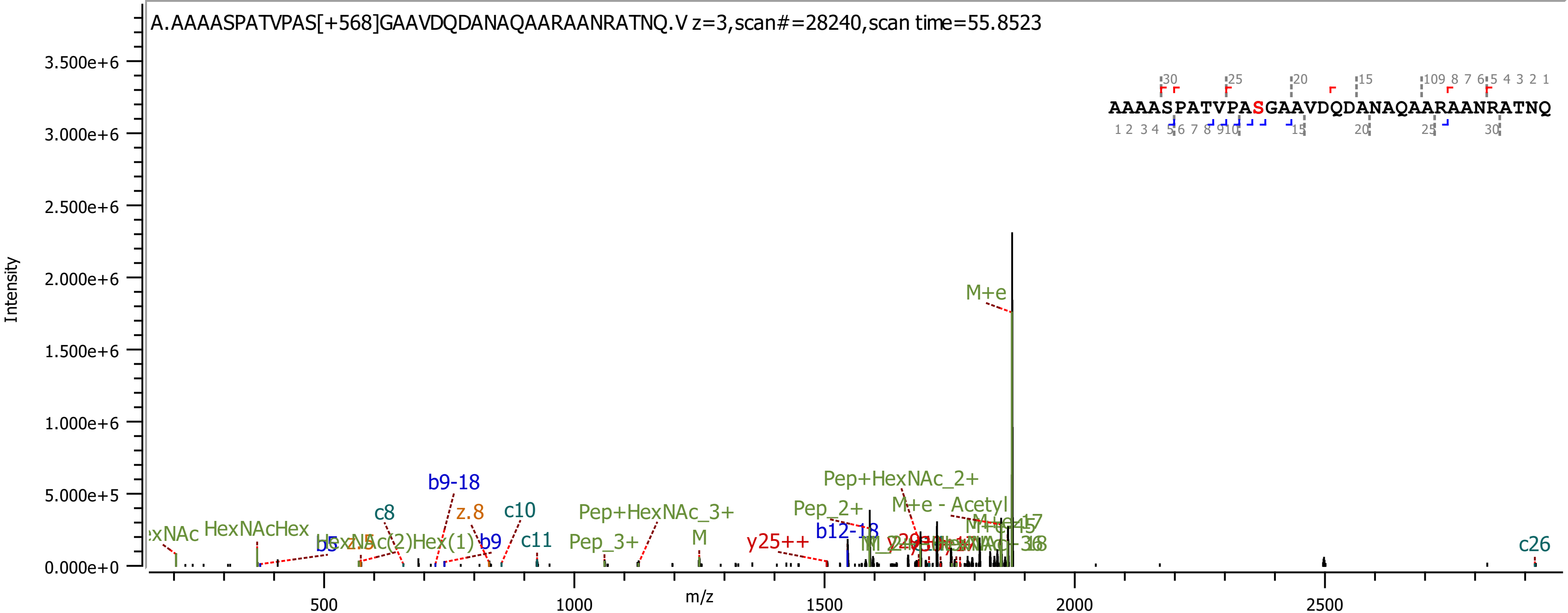
A.AAAASPATVPAS[+568]GAAVDQDANAQAARAANRATNQ.V z=3,scan#=28240,scan time=55.8523
AAAASPATVPASGAAVDQDANAQAARAANRATNQ
Intensity
m/z
M+e
Pep+HexNAc_2+
M+e - Acetyl
Pep_2+
Pep+HexNAc_3+
Pep_3+
b9-18
b12-18
z.8
c8
c10
c11
c26
b9
b5
y25++
M
HexNAcHex
HexNAc(2)Hex(1)

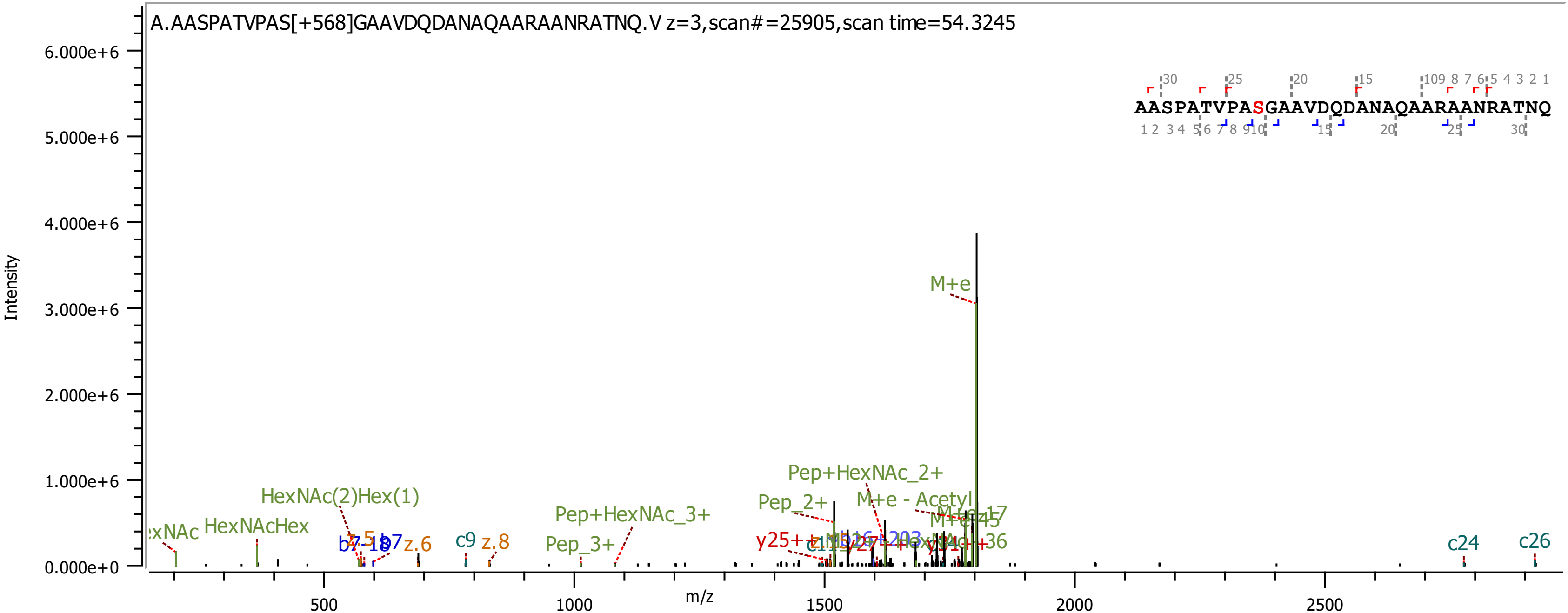

A.AASPATVPAS[+568]GAAVDQDANAQAARAANRATNQ.V z=3,scan#=25905,scan time=54.3245
AASPATVPASGAAVDQDANAQAARAANRATNQ
Intensity
m/z
M+e
Pep+HexNAc_2+
Pep_2+
M+e - Acetyl
HexNAc(2)Hex(1)
HexNAcHex
Pep+HexNAc_3+
Pep_3+
z.6
c9
z.8
c24
c26

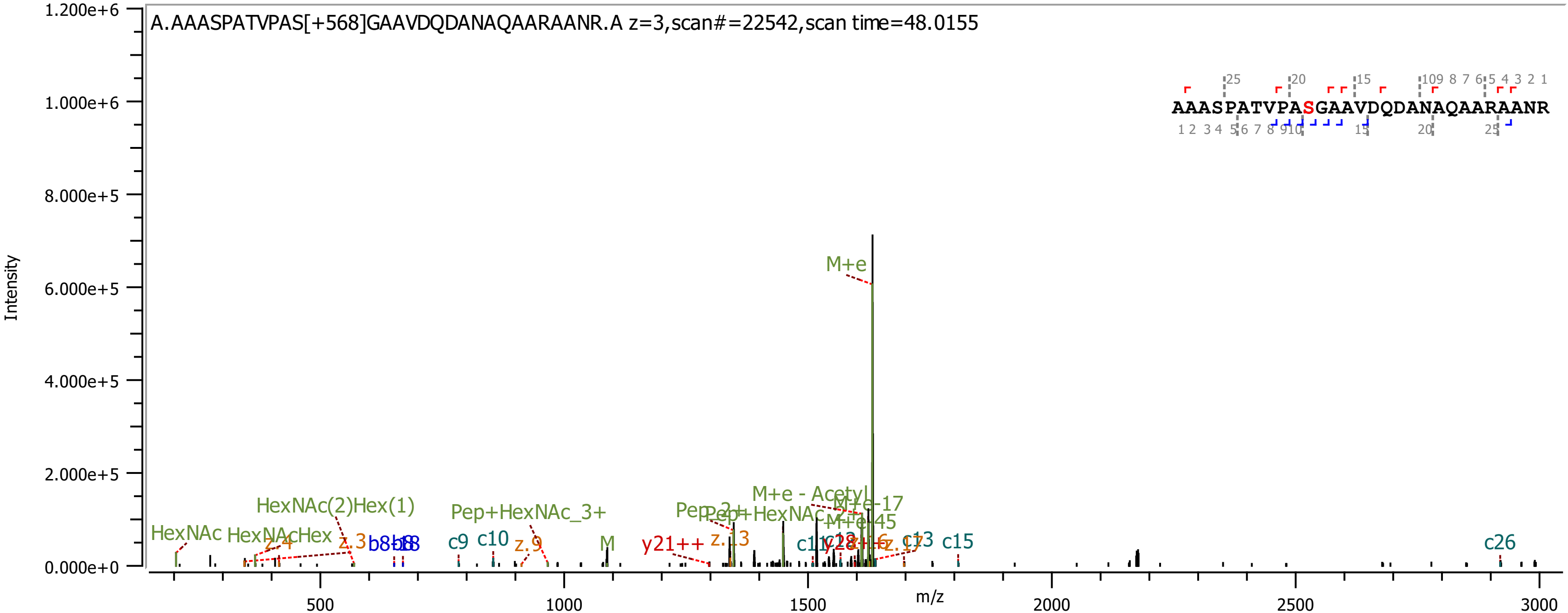

A.AAASPATVPAS[+568]GAAVDQDANAQAARAANR.A z=3,scan#=22542,scan time=48.0155
AAASPATVPASGAAVDQDANAQAARAANR
Intensity
m/z
M+e
M+e - Acetyl
M+e-17
M+e-45
HexNAc
HexNAcHex
HexNAc(2)Hex(1)
Pep+HexNAc_3+
Pep_2+
Pep+HexNAc
M
z.3
z.4
z.9
z.13
z.17
b8
c9
c10
c13
c15
c26
y21++
y28

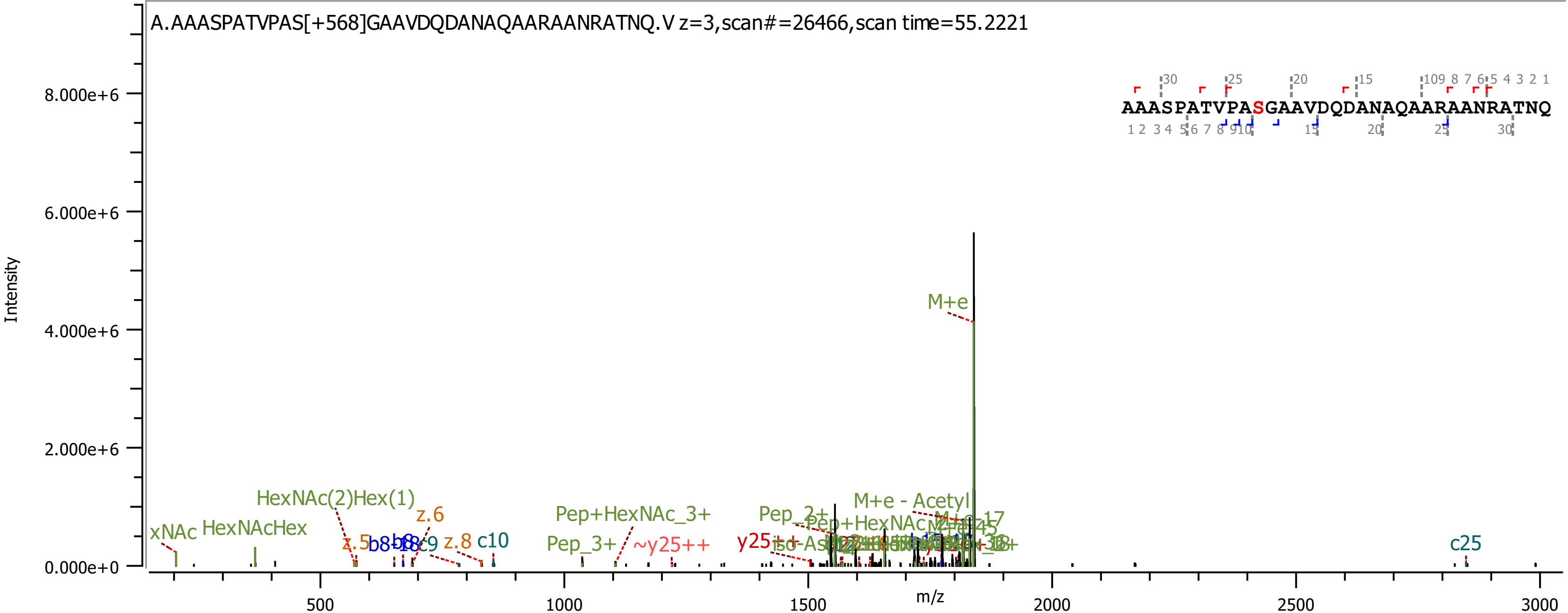
A.AAASPATVPAS[+568]GAAVDQDANAQAARAANRATNQ.V z=3,scan#=26466,scan time=55.2221
AAASPATVPASGAAVDQDANAQAARAANRATNQ
Intensity
m/z
8.000e+6
6.000e+6
4.000e+6
2.000e+6
0.000e+0
500
1000
1500
2000
2500
3000
M+e
HexNAc(2)Hex(1)
HexNAcHex
z.6
z.5
z.8
c10
c9
c25
Pep+HexNAc_3+
Pep_3+
~y25++
y25++
Pep_2+
M+e - Acetyl

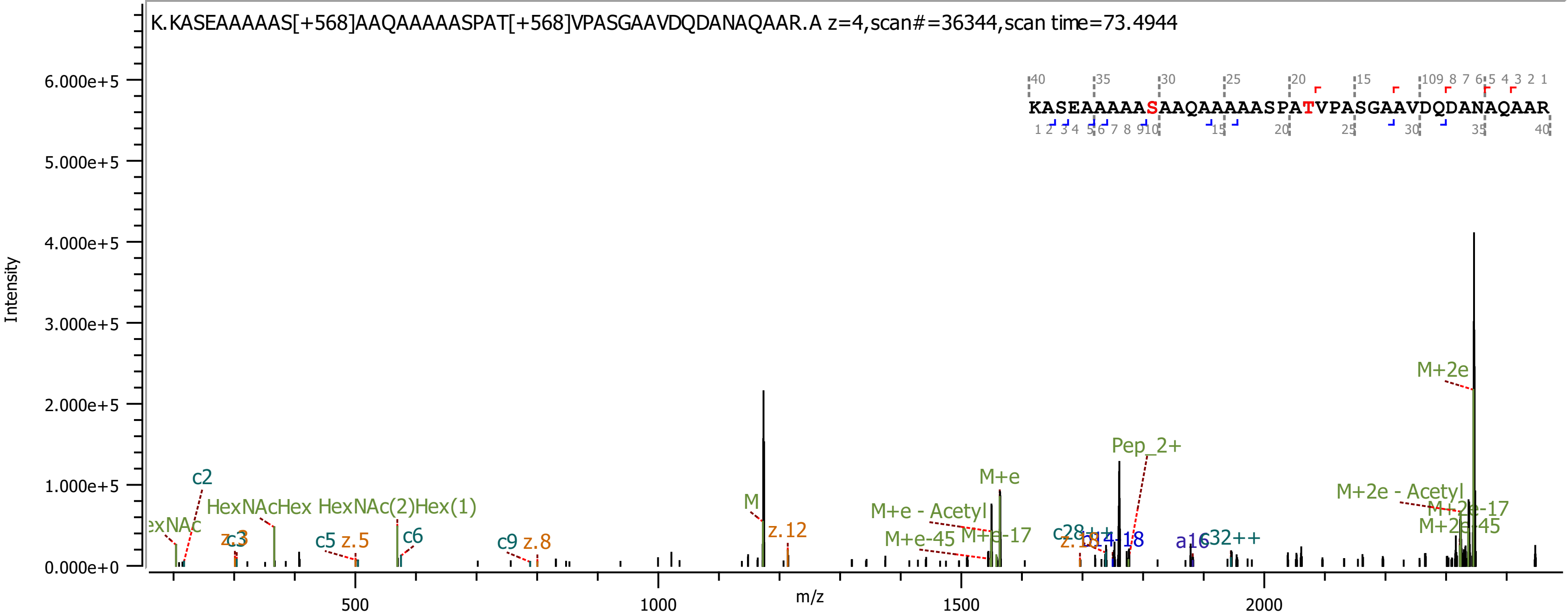

K.KASEAAAAAS[+568]AAQAAAAASPAT[+568]VPASGAAVDQDANAQAAR.A z=4,scan#=36344,scan time=73.4944
KASEAAAAASAAQAAAAASPATVPASGAAVDQDANAQAAR
Intensity
m/z
c2
HexNAcHex
HexNAc(2)Hex(1)
c5
z.5
c6
c9
z.8
M
z.12
M+e
M+e - Acetyl
M+e-45
M+e-17
Pep_2+
a16
c32++
M+2e
M+2e - Acetyl
M+2e-17
M+2e-45

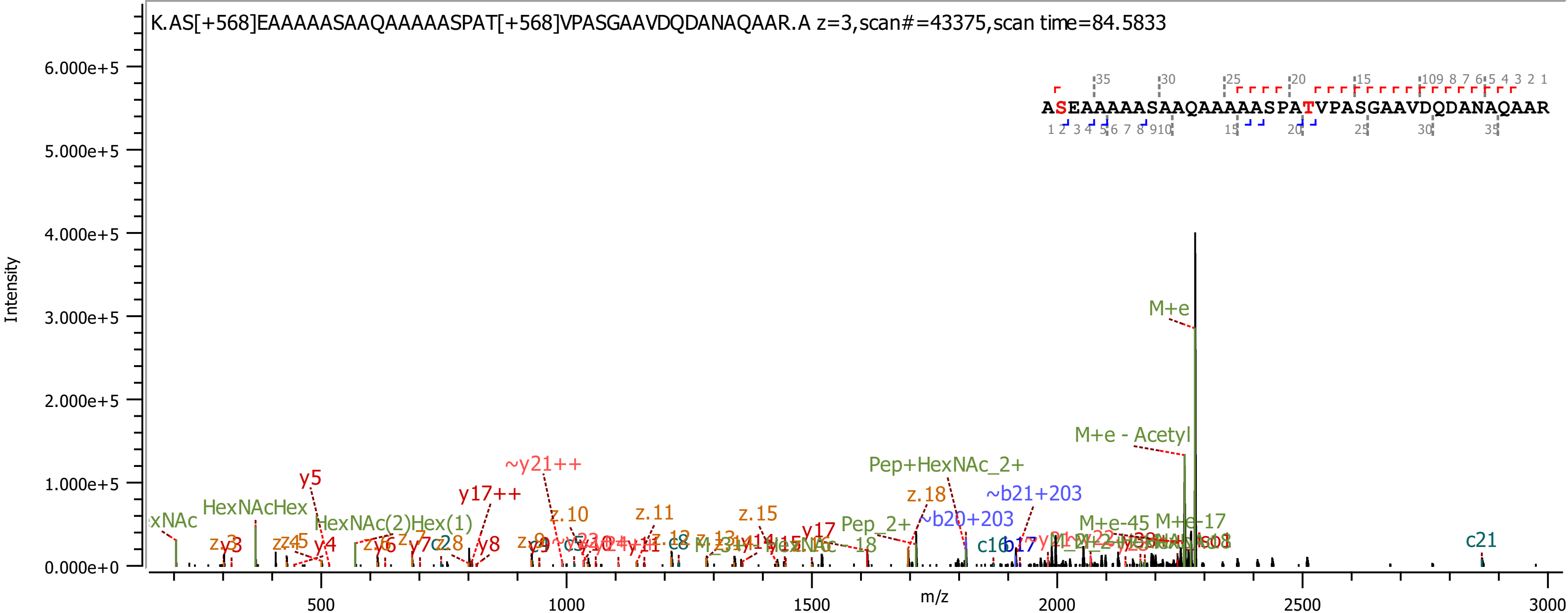
K.AS[+568]EAAAAASAAQAAAAASPAT[+568]VPASGAAVDQDANAQAAR.A z=3,scan#=43375,scan time=84.5833
ASEAAAAASAAQAAAAASPATVPASGAAVDQDANAQAAR
Intensity
m/z
6.000e+5
5.000e+5
4.000e+5
3.000e+5
2.000e+5
1.000e+5
0.000e+0
500
1000
1500
2000
2500
3000
M+e
M+e - Acetyl
Pep+HexNAc_2+
~y21++
y5
y17++
HexNAcHex
HexNAc(2)Hex(1)
z.10
z.11
z.15
z.18
~b21+203
~b20+203
Pep_2+
M+e-45
M+e-17
c21
c16
b17
y17
y4
y8
y9
c8
z.12
z.13
z.4
z.5
z.6
z.7
z.8
z.9
c28

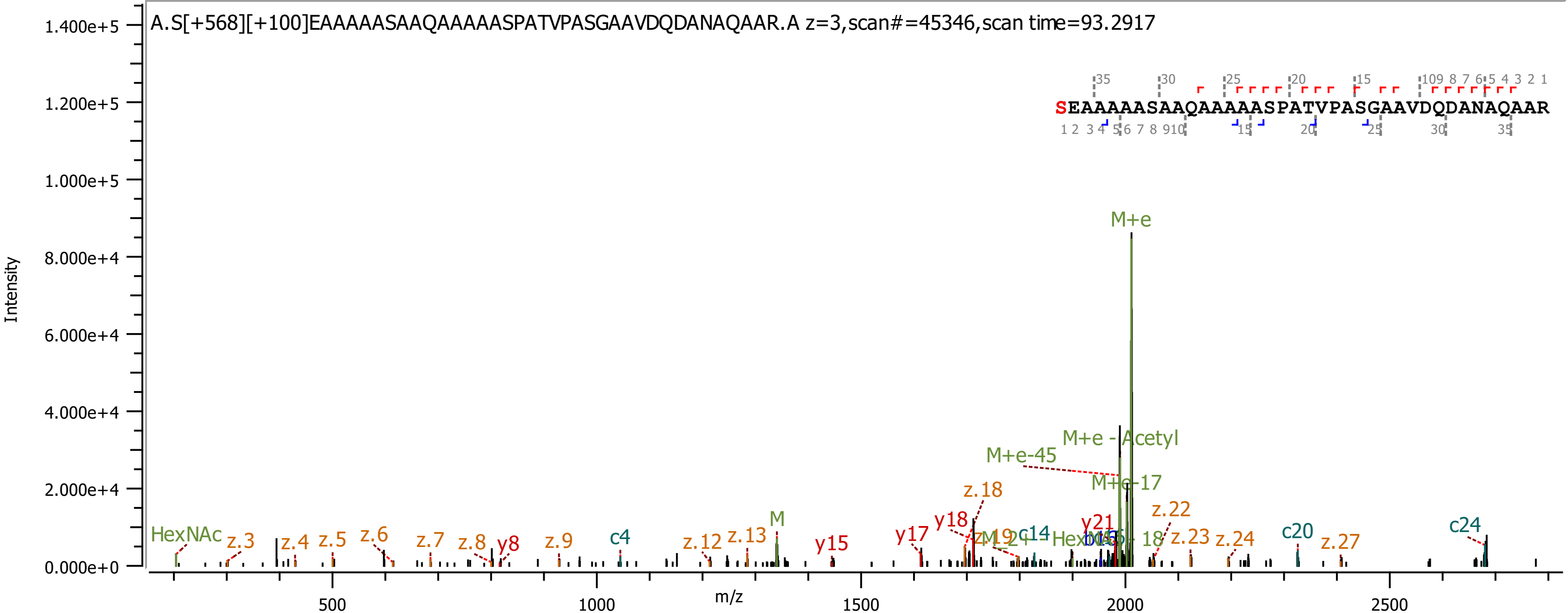
A.S[+568][+100]EAAAAASAAQAAAAASPATVPASGAAVDQDANAQAAR.A z=3,scan#=45346,scan time=93.2917
SEAAAAASAAQAAAAASPATVPASGAAVDQDANAQAAR
Intensity
m/z
M+e
M+e - Acetyl
M+e-45
M+e-17
HexNAc
M
z.3
z.4
z.5
z.6
z.7
z.8
y8
z.9
c4
z.12
z.13
y15
y17
y18
z.18
c14
y21
z.22
z.23
z.24
c20
z.27
c24
1.400e+5
1.200e+5
1.000e+5
8.000e+4
6.000e+4
4.000e+4
2.000e+4
0.000e+0
500
1000
1500
2000
2500

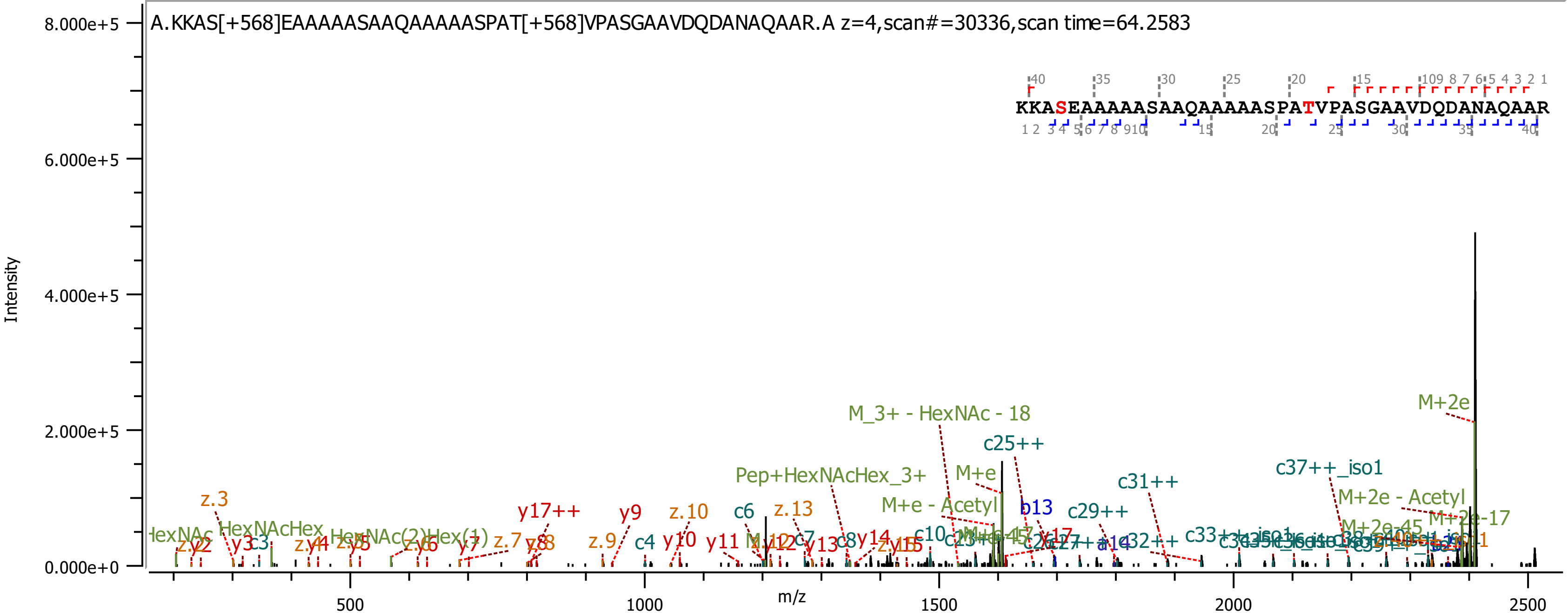

A.KKAS[+568]EAAAAASAAQAAAAASPAT[+568]VPASGAAVDQDANAQAAR.A z=4,scan#=30336,scan time=64.2583
KKASEAAAAASAAQAAAAASPATVPASGAAVDQDANAQAAR
Intensity
8.000e+5
6.000e+5
4.000e+5
2.000e+5
0.000e+0
m/z
500
1000
1500
2000
2500
M+2e
M_3+ - HexNAc - 18
c25++
c37++_iso1
M+e
Pep+HexNAcHex_3+
c31++
M+2e - Acetyl
M+e - Acetyl
b13
c29++
M+2e-17
M+2e-45
z.3
y17++
y9
z.10
c6
z.13
HexNAcHex
c3
z.7
y8
z.9
c4
y10
y11
c7
y13
c8
y14
c10
c32++
c33++_iso1

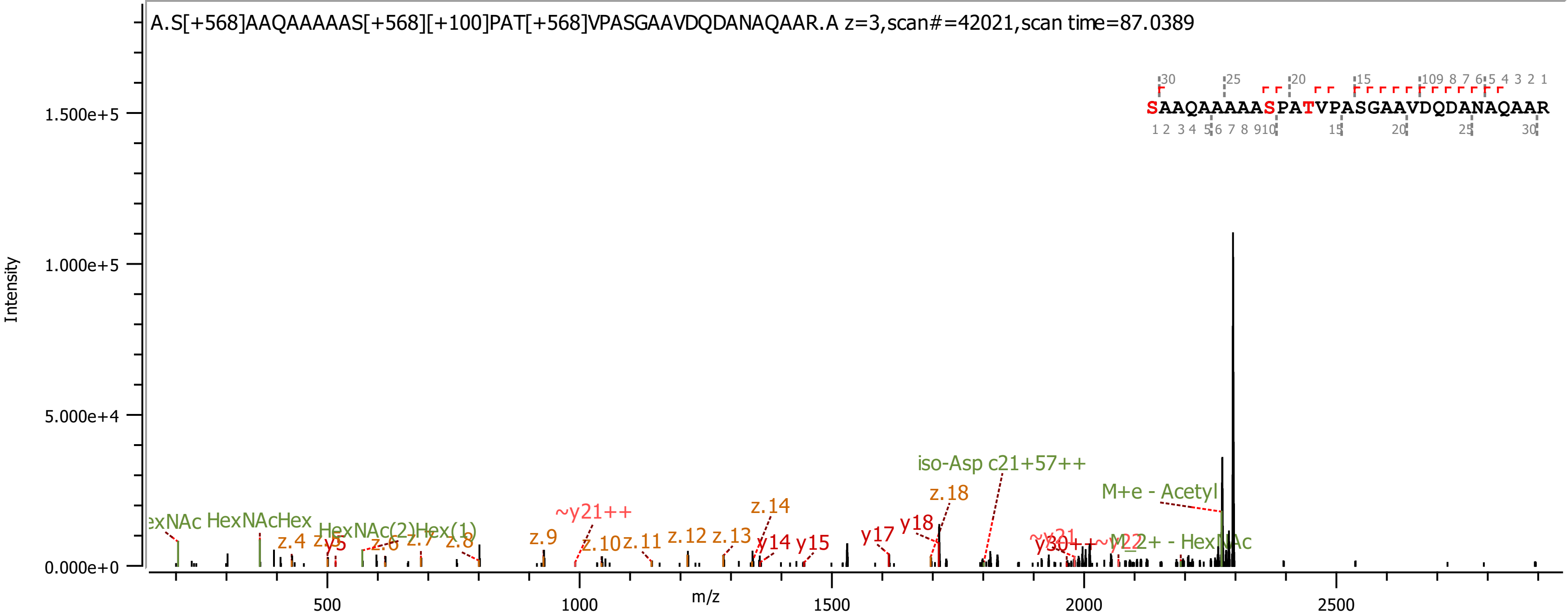
A.S[+568]AAQAAAAAS[+568][+100]PAT[+568]VPASGAAVDQDANAQAAR.A z=3,scan#=42021,scan time=87.0389
SAAQAAAAASPATVPASGAAVDQDANAQAAR
Intensity
m/z
1.500e+5
1.000e+5
5.000e+4
0.000e+0
500
1000
1500
2000
2500
iso-Asp c21+57++
M+e - Acetyl
z.18
y18
y17
z.14
y14
y15
~y21++
z.9
z.10
z.11
z.12
z.13
HexNAcHex
HexNAc(2)Hex(1)
z.4
y5
z.6
z.7
z.8
~y22++ - HexNAc

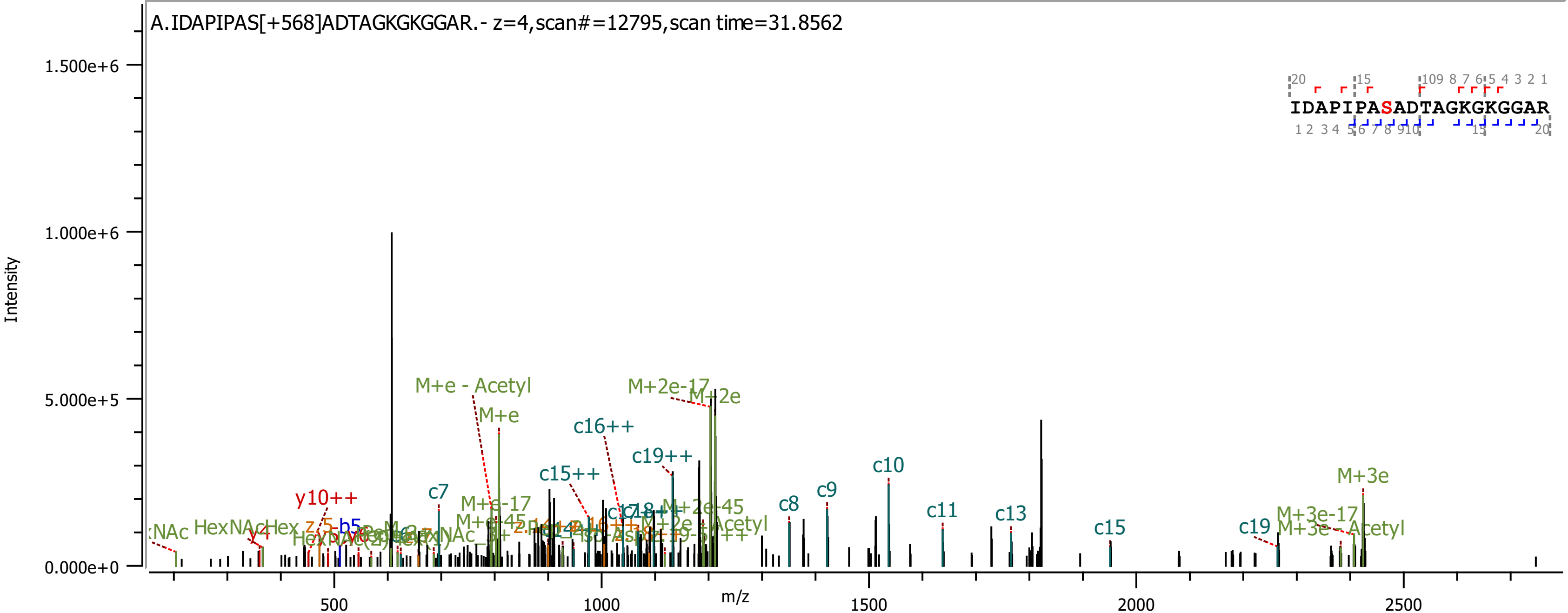
A.IDAPIPAS[+568]ADTAGKGKGGAR.- z=4,scan#=12795,scan time=31.8562
Intensity
1.500e+6
1.000e+6
5.000e+5
0.000e+0
m/z
500
1000
1500
2000
2500
IDAPIPASADTAGKGKGGAR
M+e - Acetyl
M+e
M+2e-17
M+2e
c16++
c19++
c15++
c7
y10++
M+e-17
c8
c9
c10
c11
c13
c15
c19
M+3e
M+3e-17
M+3e - Acetyl
HexNAcHex
b5
y4

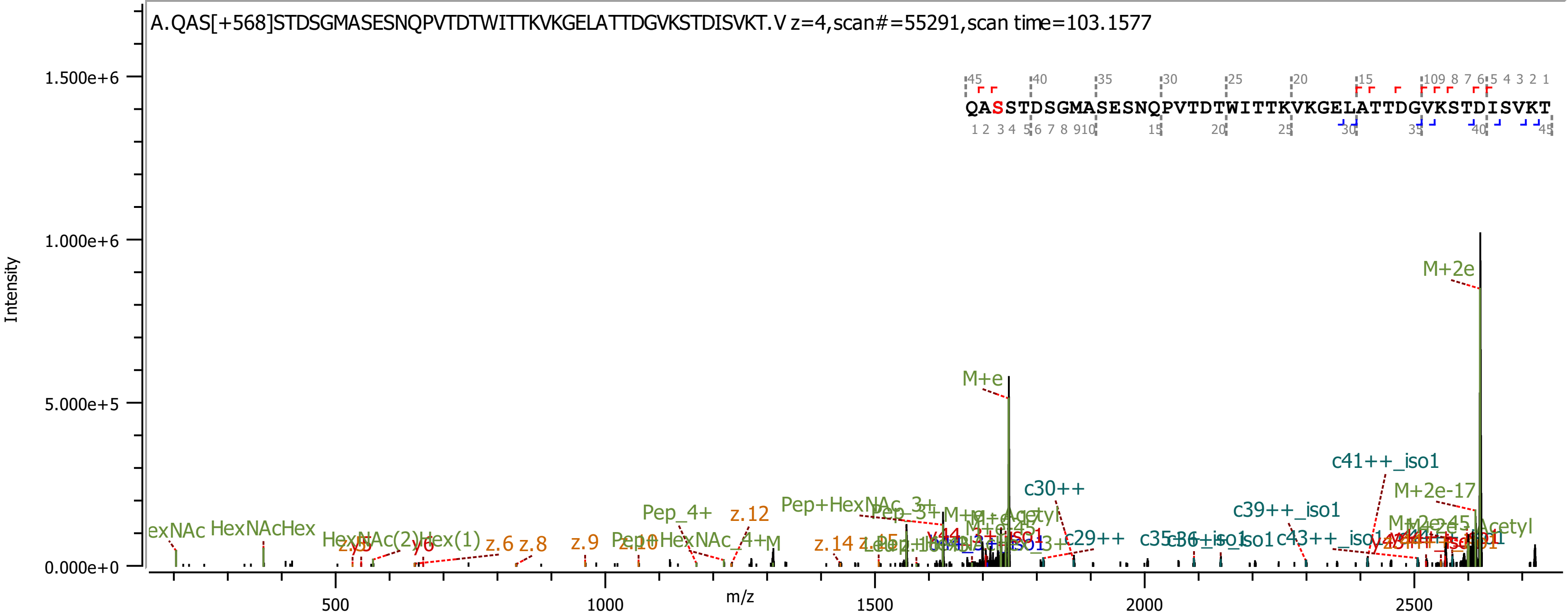

A.QAS[+568]STDSGMASESNQPVTDTWITTKVKGELATTDGVKSTDISVKT.V z=4,scan#=55291,scan time=103.1577
QASSTDSGMASESNQPVTDTWITTKVKGELATTDGVKSTDISVKT
Intensity
1.500e+6
1.000e+6
5.000e+5
0.000e+0
m/z
500
1000
1500
2000
2500
M+2e
M+e
c41++_iso1
M+2e-17
c30++
c39++_iso1
Pep_4+
z.12
Pep+HexNAc_3+
HexNAcHex
HexNAc(2)Hex(1)
y6
z.6
z.8
z.9
z.10
z.14
c29++
c43++_iso1

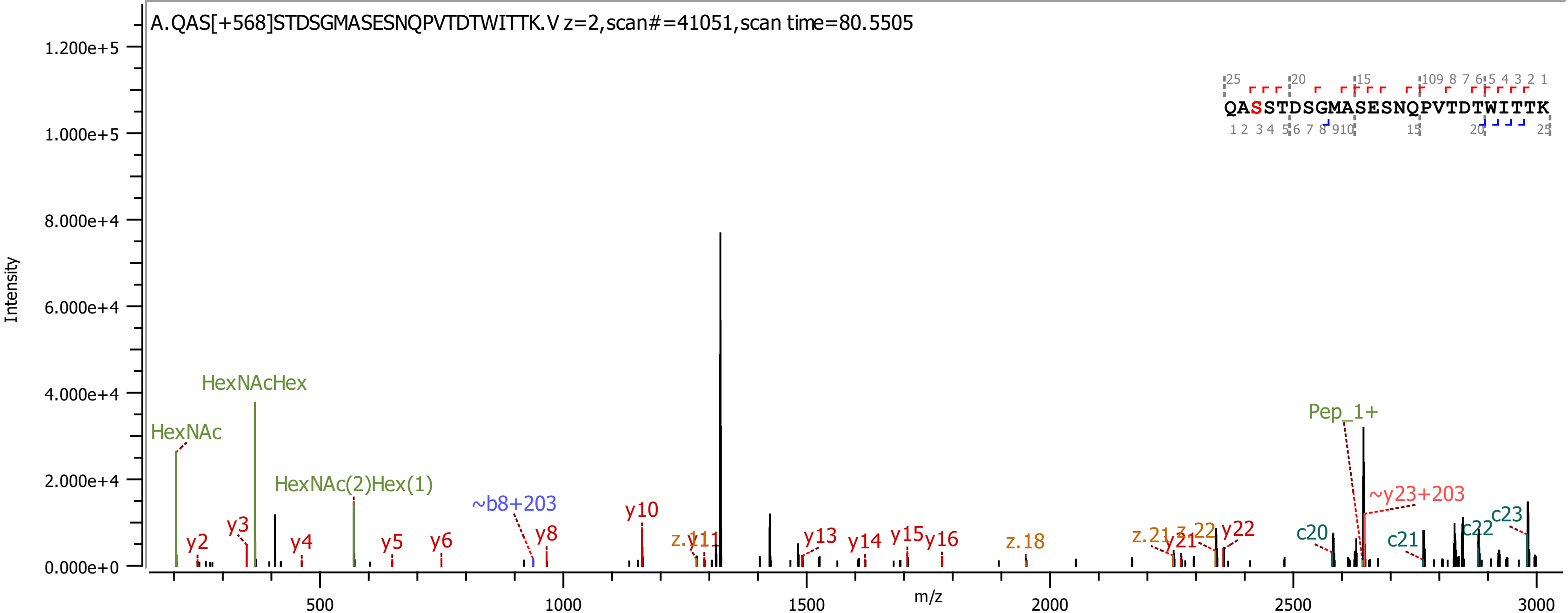

A.QAS[+568]STDSGMASESNQPVTDTWITTK.V z=2,scan#=41051,scan time=80.5505
QASSTDSGMASESNQPVTDTWITTK
Intensity
m/z
HexNAc
HexNAcHex
HexNAc(2)Hex(1)
~b8+203
Pep_1+
~y23+203
y2
y3
y4
y5
y6
y8
y10
y13
y14
y15
y16
y21
y22
z.y11
z.18
z.21
z.22
c20
c21
c22
c23

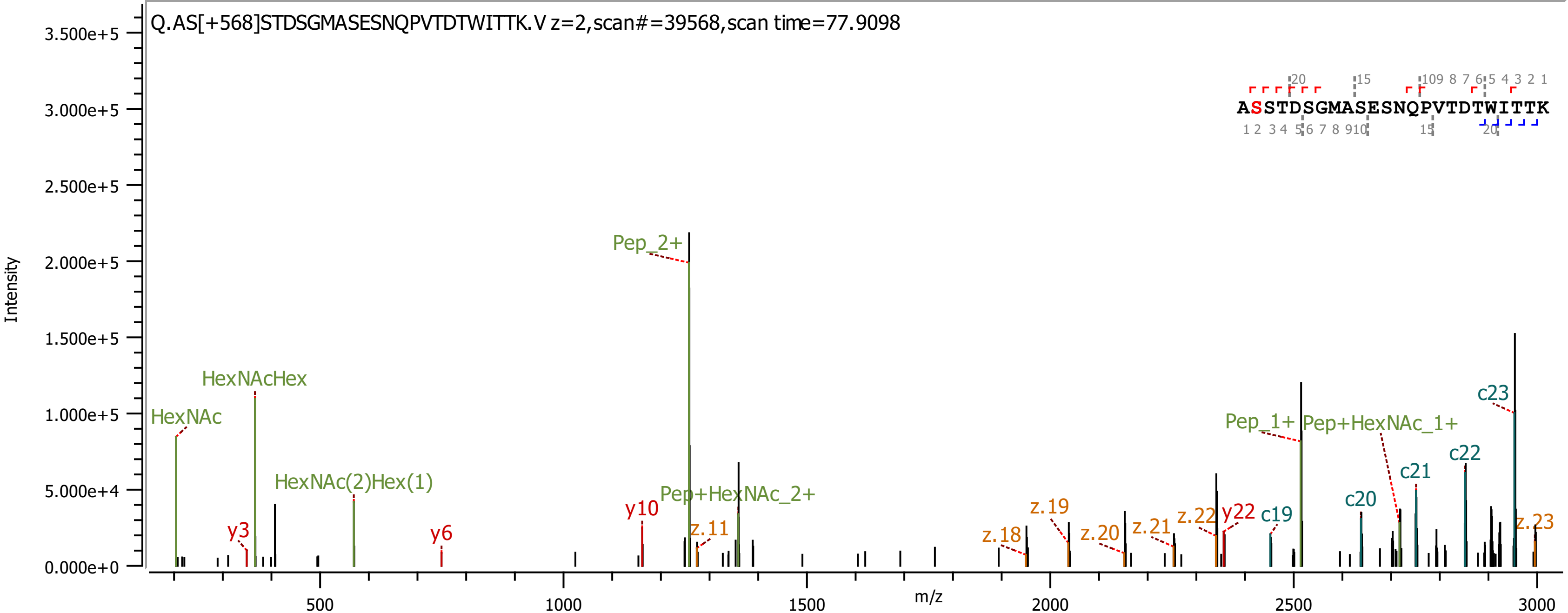

Q.AS[+568]STDSGMASESNQPVTDTWITTK.V z=2,scan#=39568,scan time=77.9098
ASSTDSGMASESNQPVTDTWITTK
Intensity
m/z
HexNAc
HexNAcHex
HexNAc(2)Hex(1)
y3
y6
y10
Pep_2+
Pep+HexNAc_2+
z.11
z.18
z.19
z.20
z.21
z.22
y22
c19
c20
c21
c22
c23
z.23
Pep_1+
Pep+HexNAc_1+

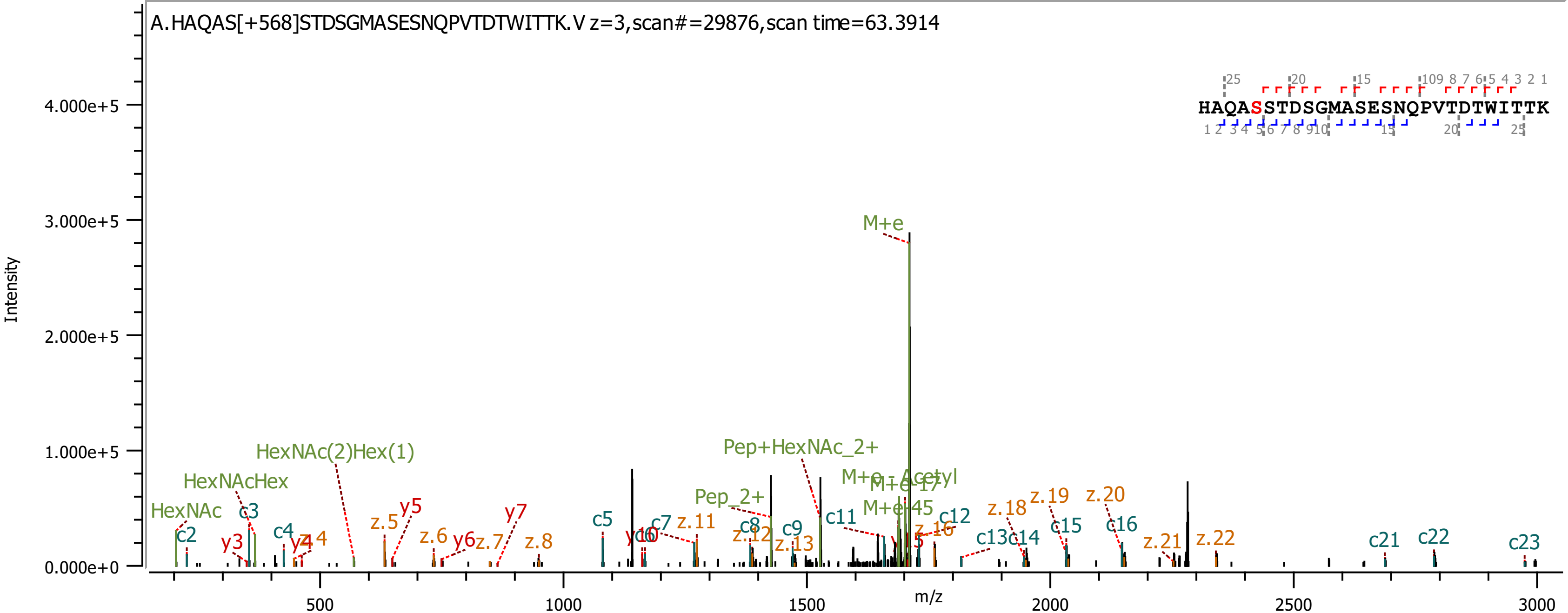
A.HAQAS[+568]STDSGMASESNQPVTDTWITTK.V z=3,scan#=29876,scan time=63.3914
HAQASSTDSGMASESNQPVTDTWITTK
Intensity
m/z
M+e
HexNAc(2)Hex(1)
HexNAcHex
HexNAc
Pep+HexNAc_2+
Pep_2+
M+e - Acetyl
M+e-17
M+e-45
c2
c3
c4
c5
c7
c8
c9
c11
c12
c13
c14
c15
c16
c21
c22
c23
y3
y4
y5
y6
y7
y10
z.4
z.5
z.6
z.7
z.8
z.11
z.12
z.13
z.16
z.18
z.19
z.20
z.21
z.22
0.000e+0
1.000e+5
2.000e+5
3.000e+5
4.000e+5
500
1000
1500
2000
2500
3000

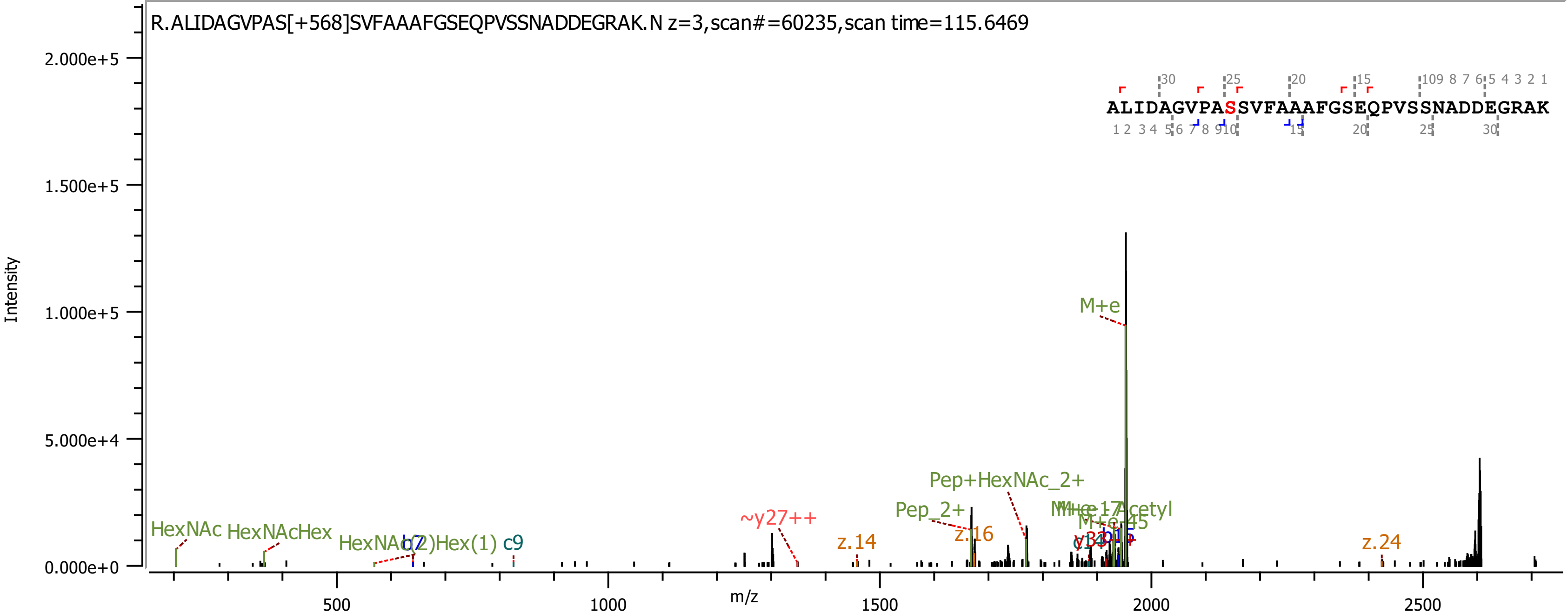
R.ALIDAGVPAS[+568]SVFAAAFGSEQPVSSNADDEGRAK.N z=3,scan#=60235,scan time=115.6469
ALIDAGVPASSVFAAAFGSEQPVSSNADDEGRAK
Intensity
m/z
2.000e+5
1.500e+5
1.000e+5
5.000e+4
0.000e+0
500
1000
1500
2000
2500
M+e
Pep+HexNAc_2+
Pep_2+
~y27++
HexNAc
HexNAcHex
HexNAc(2)Hex(1)
c9
z.14
z.16
z.24

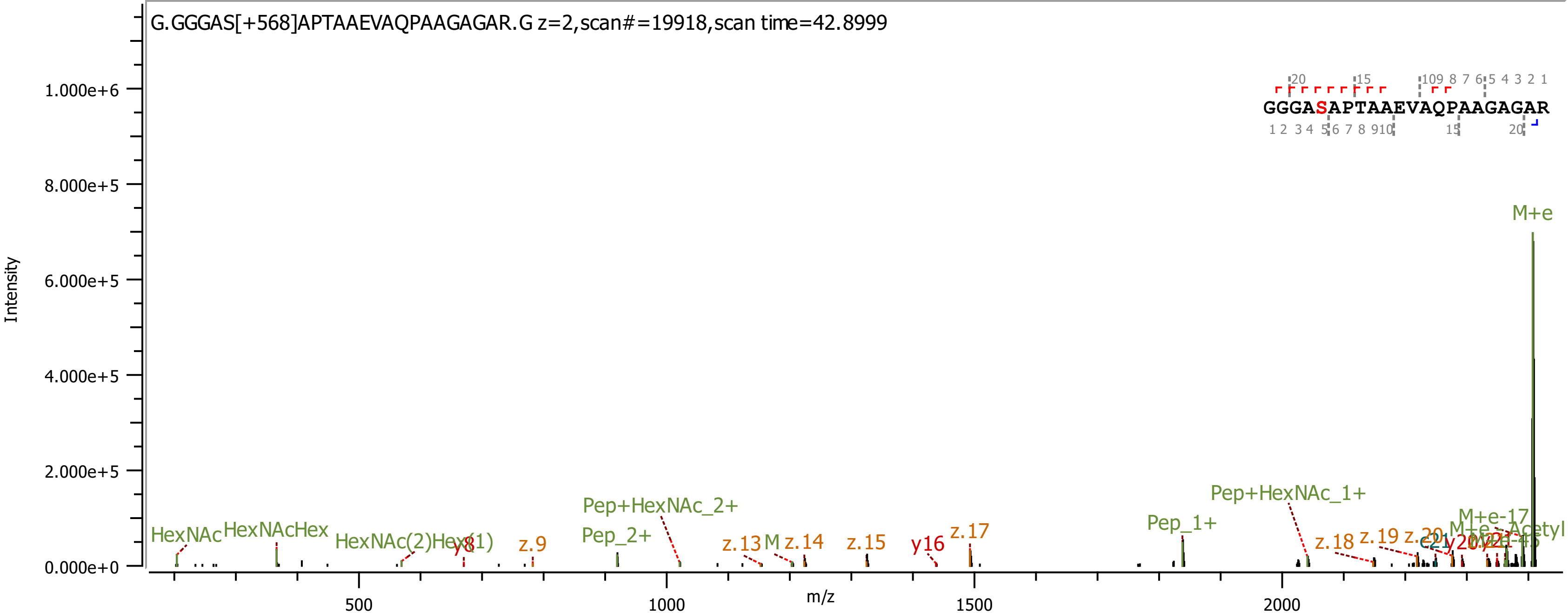
G.GGGAS[+568]APTAAEVAQPAAGAGAR.G z=2,scan#=19918,scan time=42.8999
GGGASAPTAAEVAQPAAGAGAR
Intensity
m/z
M+e
Pep+HexNAc_1+
Pep+HexNAc_2+
Pep_1+
Pep_2+
HexNAc
HexNAcHex
z.9
z.13
z.14
z.15
y16
z.17
z.18
z.19

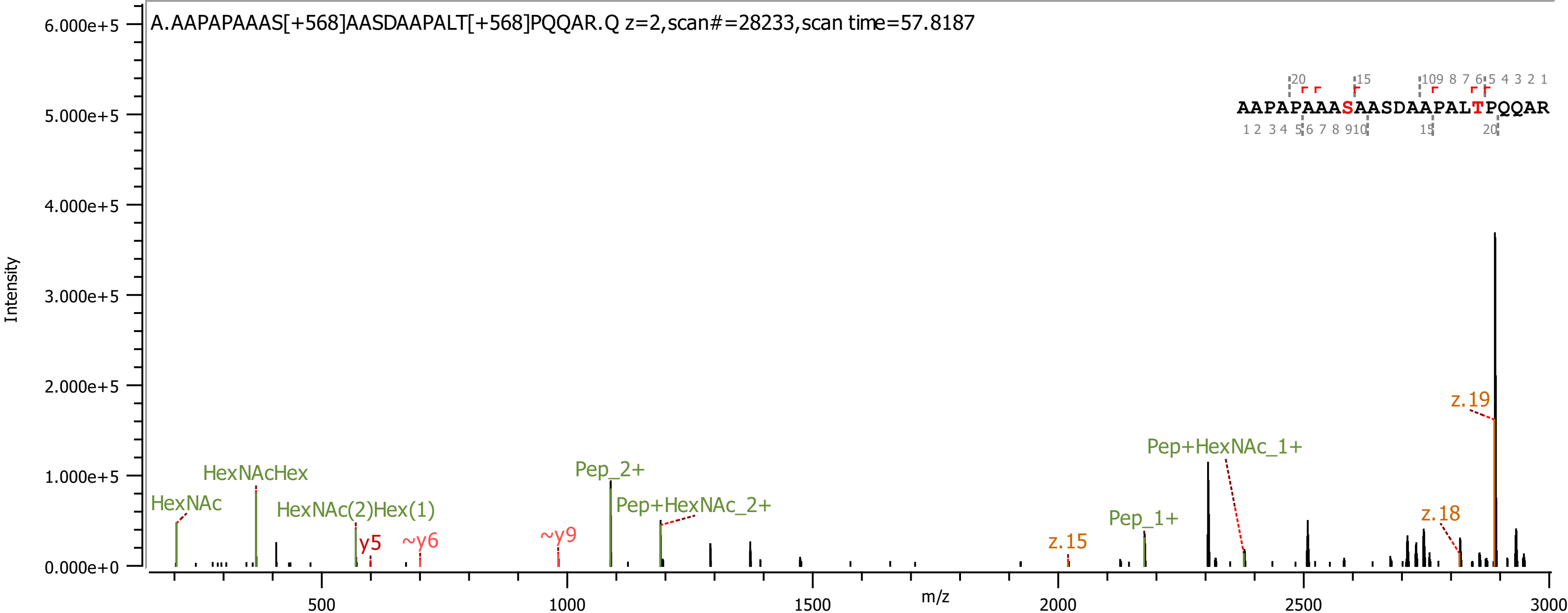

A.AAPAPAAAS[+568]AASDAAPALT[+568]PQQAR.Q z=2,scan#=28233,scan time=57.8187
AAPAPAAASAASDAAPALTPQQAR
Intensity
m/z
HexNAc
HexNAcHex
HexNAc(2)Hex(1)
y5
~y6
~y9
Pep_2+
Pep+HexNAc_2+
z.15
Pep_1+
Pep+HexNAc_1+
z.18
z.19

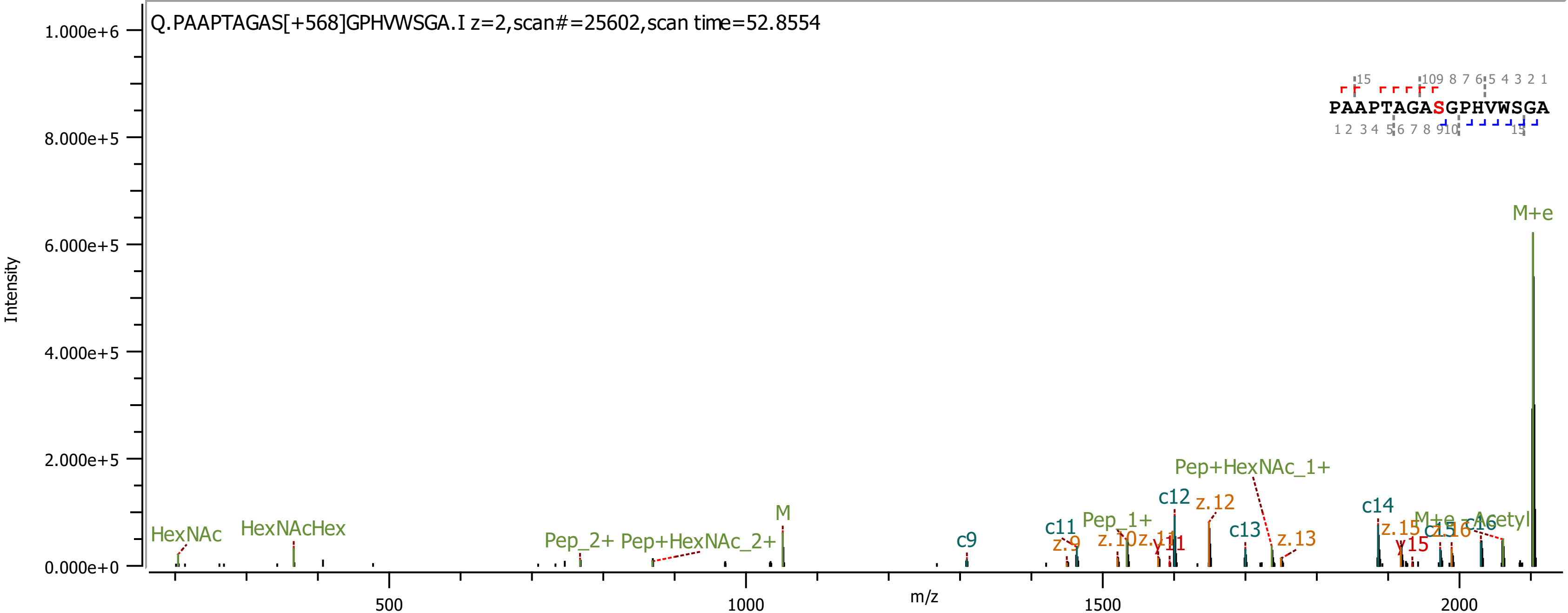
Q.PAAPTAGAS[+568]GPHVWSGA.I z=2,scan#=25602,scan time=52.8554
PAAPTAGASGPHVWSGA
Intensity
m/z
M+e
Pep+HexNAc_1+
c12
z.12
c14
M+e - Acetyl
c13
z.13
c11
Pep_1+
z.10
z.11
y11
z.9
c9
M
HexNAc
HexNAcHex
Pep_2+
Pep+HexNAc_2+
z.15
c15
z.16
c16
y15
1.000e+6
8.000e+5
6.000e+5
4.000e+5
2.000e+5
0.000e+0
500
1000
1500
2000

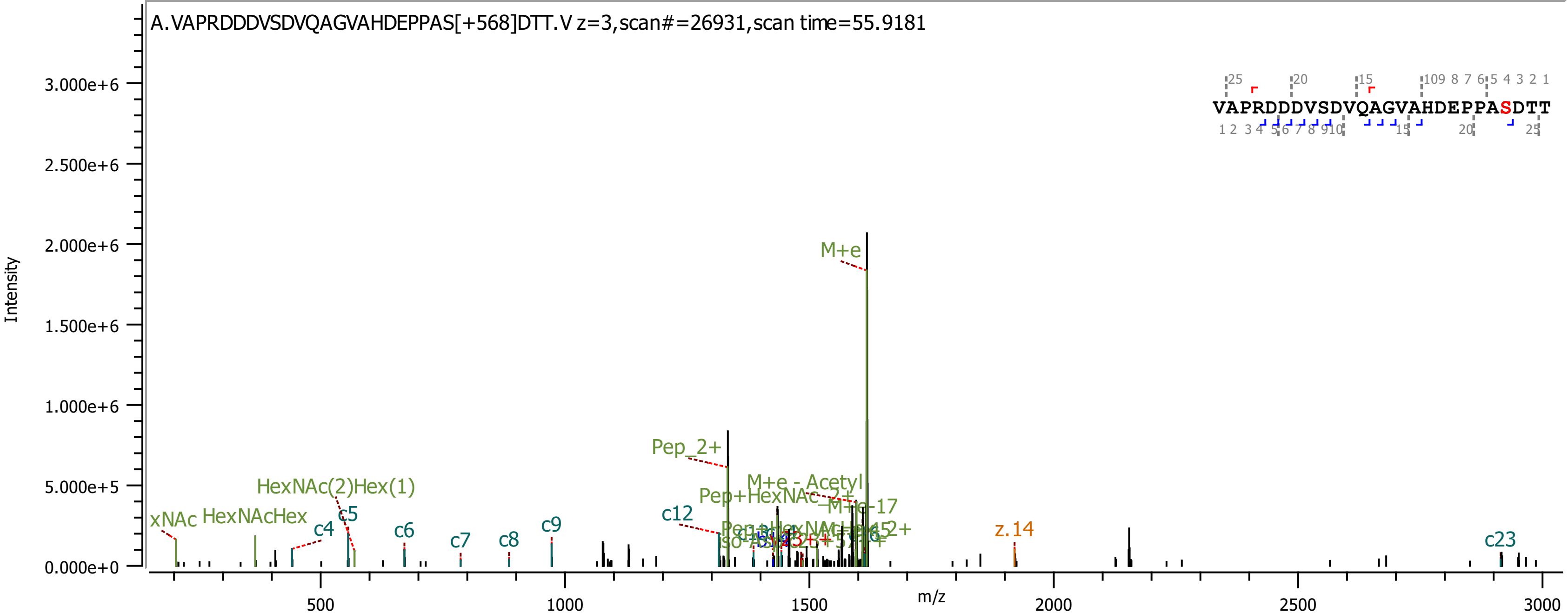
A.VAPRDDDVSDVQAGVAHDEPPAS[+568]DTT.V z=3,scan#=26931,scan time=55.9181
VAPRDDDVSDVQAGVAHDEPPASDTT
Intensity
m/z
M+e
Pep_2+
M+e - Acetyl
Pep+HexNAc_2+
M+e-17
HexNAc(2)Hex(1)
HexNAcHex
xNAc
c4
c5
c6
c7
c8
c9
c12
c23
z.14
0.000e+0
5.000e+5
1.000e+6
1.500e+6
2.000e+6
2.500e+6
3.000e+6
500
1000
1500
2000
2500
3000

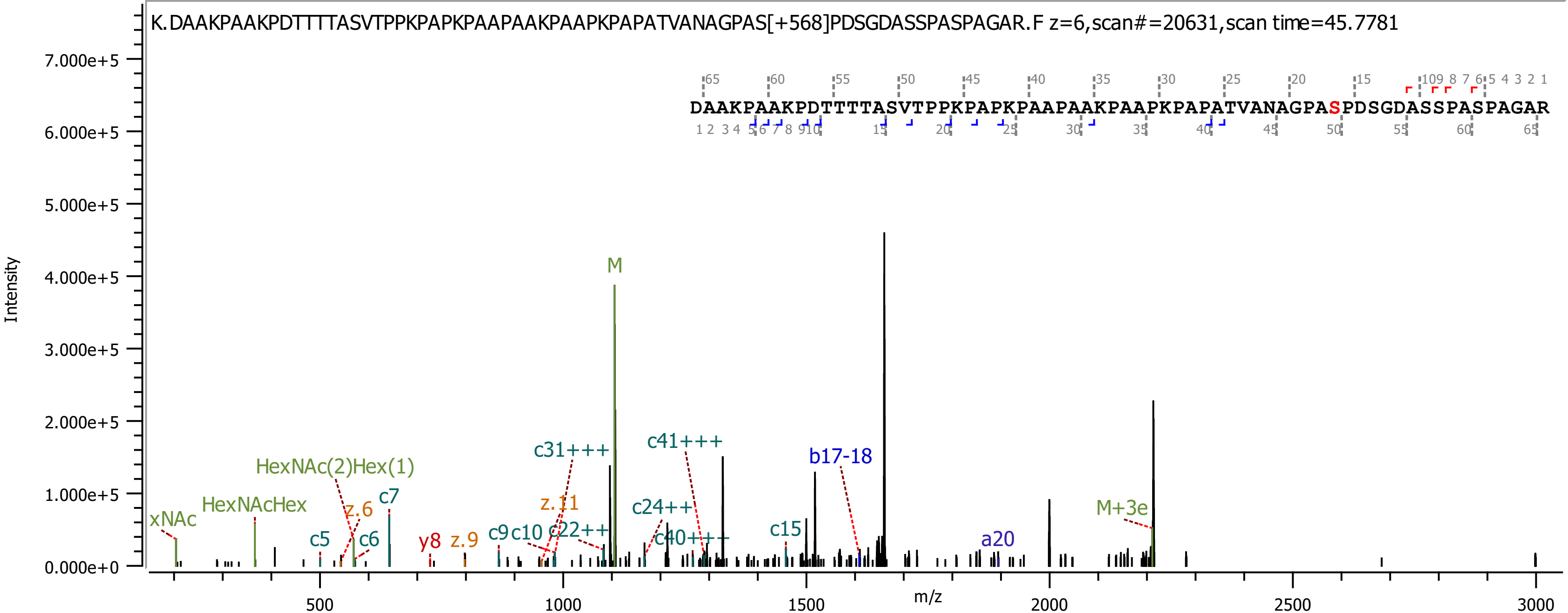
K.DAAKPAAKPDTTTTASVTPPKPAPKPAAPAAKPAAPKPAPATVANAGPAS[+568]PDSGDASSPASPAGAR.F z=6,scan#=20631,scan time=45.7781
DAAKPAAKPDTTTTASVTPPKPAPKPAAPAAKPAAPKPAPATVANAGPASPDSGDASSPASPAGAR
Intensity
m/z
M
M+3e
xNAc
HexNAcHex
HexNAc(2)Hex(1)
z.6
c5
c6
c7
y8
z.9
c9
c10
z.11
c22++
c31+++
c24++
c40+++
c41+++
c15
b17-18
a20

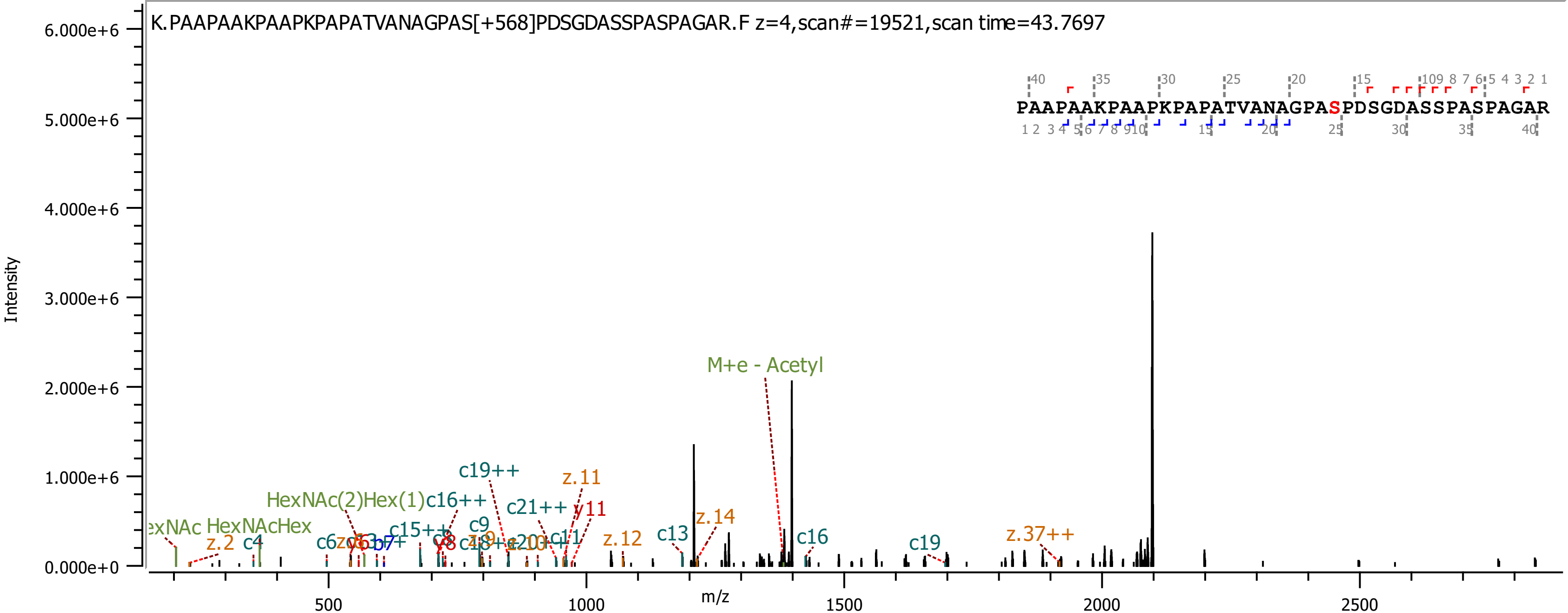

K.PAAPAAKPAAPKPAPATVANAGPAS[+568]PDSGDASSPASPAGAR.F z=4,scan#=19521,scan time=43.7697
PAAPAAKPAAPKPAPATVANAGPASPDSGDASSPASPAGAR
Intensity
m/z
M+e - Acetyl
HexNAc(2)Hex(1)c16++
HexNAcHex
c19++
z.11
c21++
y11
z.14
c13
c16
c19
z.37++
z.12
z.2
c4
c9
c15++

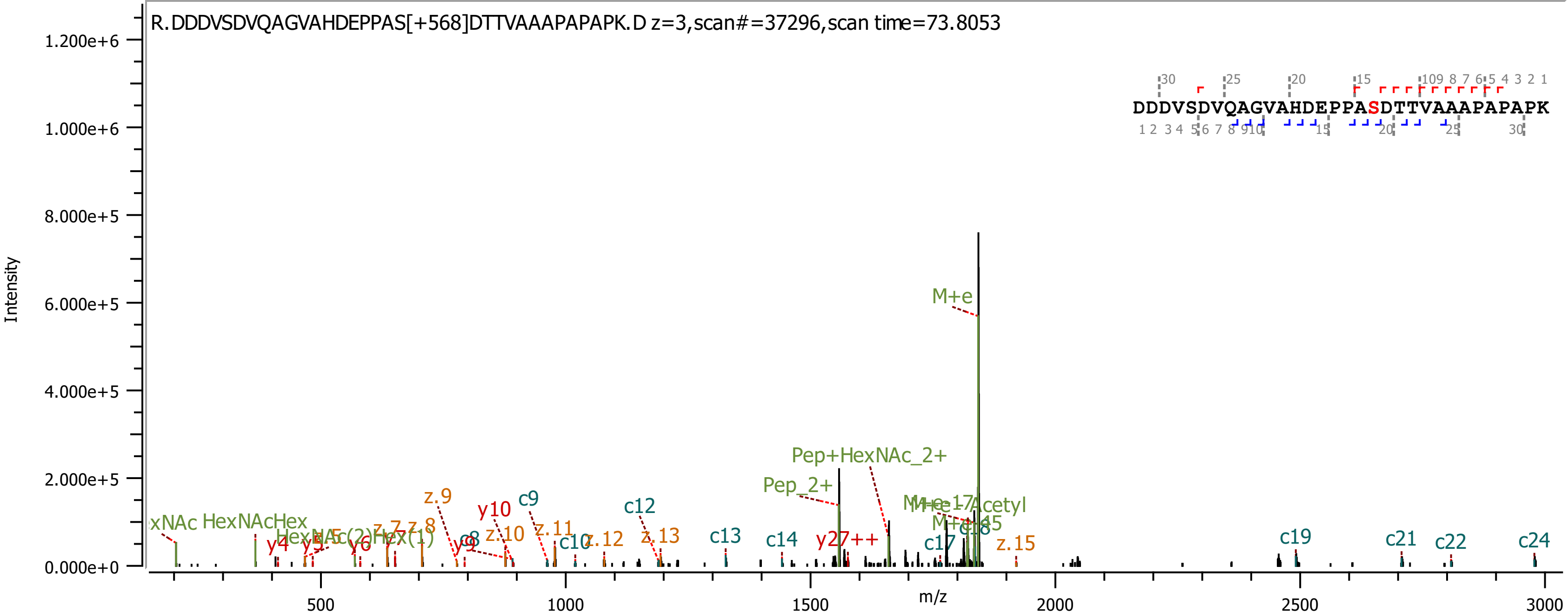
R.DDDVSDVQAGVAHDEPPAS[+568]DTTVAAAPAPAPK.D z=3,scan#=37296,scan time=73.8053
DDDVSDVQAGVAHDEPPASDTTVAAAPAPAPK
Intensity
m/z
M+e
Pep+HexNAc_2+
Pep_2+
HexNAcHex
y27++
c9
c12
c13
c14
c17
c19
c21
c22
c24
y4
y10
z.9
z.10
z.11
z.12
z.13
z.15

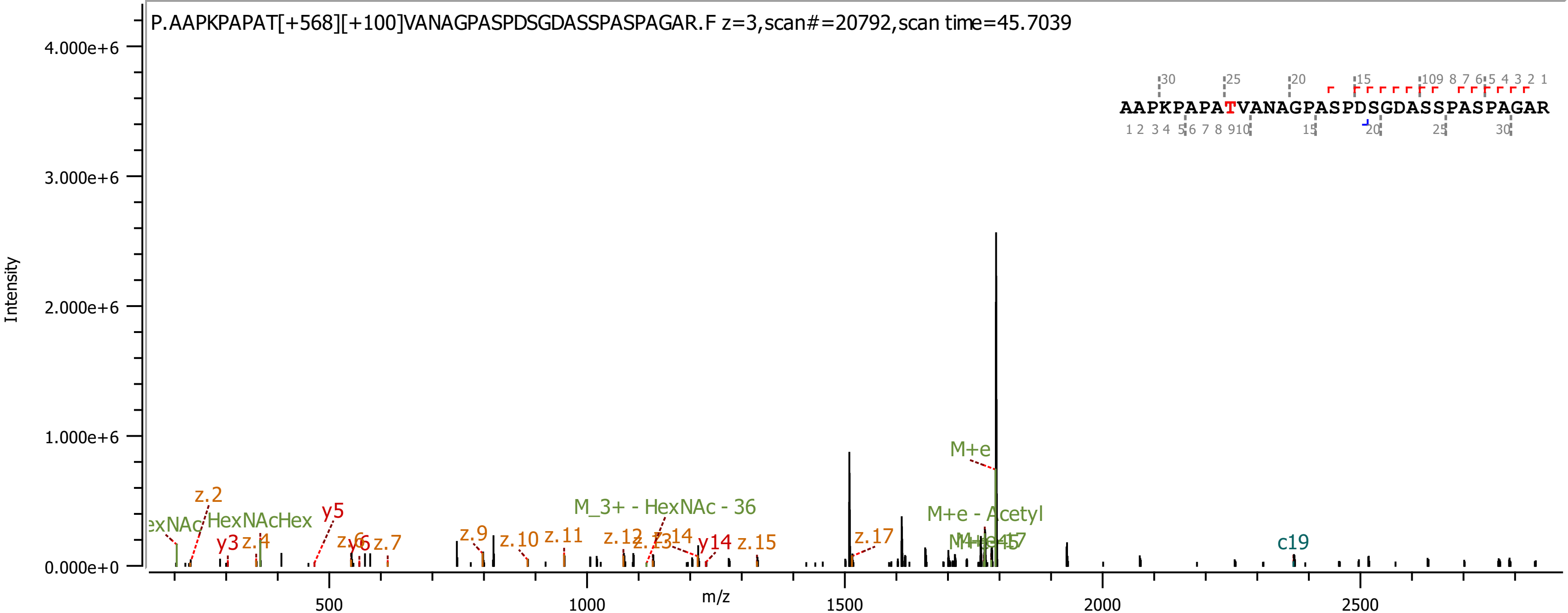
P.AAPKPAPAT[+568][+100]VANAGPASPDSGDASSPASPAGAR.F z=3,scan#=20792,scan time=45.7039
AAPKPAPATVANAGPASPDSGDASSPASPAGAR
Intensity
m/z
4.000e+6
3.000e+6
2.000e+6
1.000e+6
0.000e+0
500
1000
1500
2000
2500
M+e
M+e - Acetyl
M_3+ - HexNAc - 36
HexNAcHex
z.2
y5
y3
z.4
z.7
z.9
z.10
z.11
z.12
y14
z.15
z.17
c19

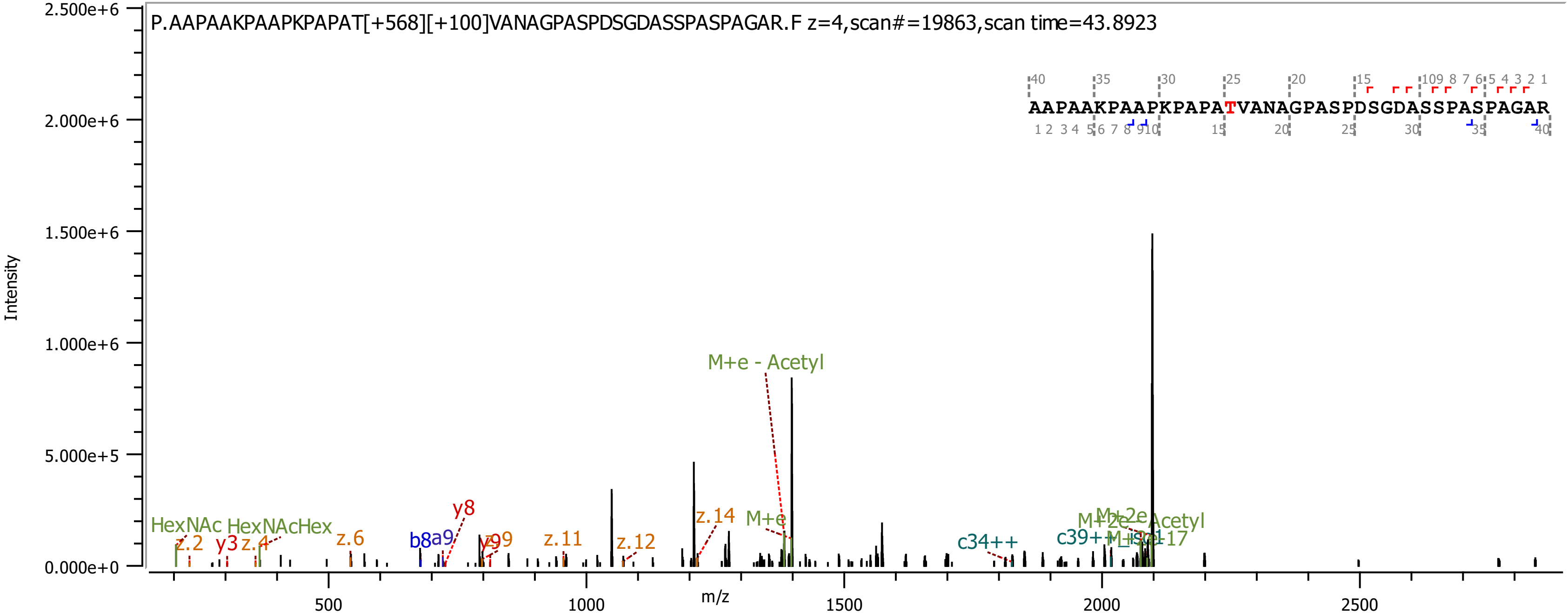

P.AAPAAKPAAPKPAPAT[+568][+100]VANAGPASPDSGDASSPASPAGAR.F z=4,scan#=19863,scan time=43.8923
AAPAAKPAAPKPAPATVANAGPASPDSGDASSPASPAGAR
Intensity
m/z
HexNAc
HexNAcHex
z.2
y3
z.4
z.6
b8
a9
y8
y9
z.11
z.12
z.14
M+e
M+e - Acetyl
c34++
c39++
M+2e
M+2e - Acetyl
M-H2O-17

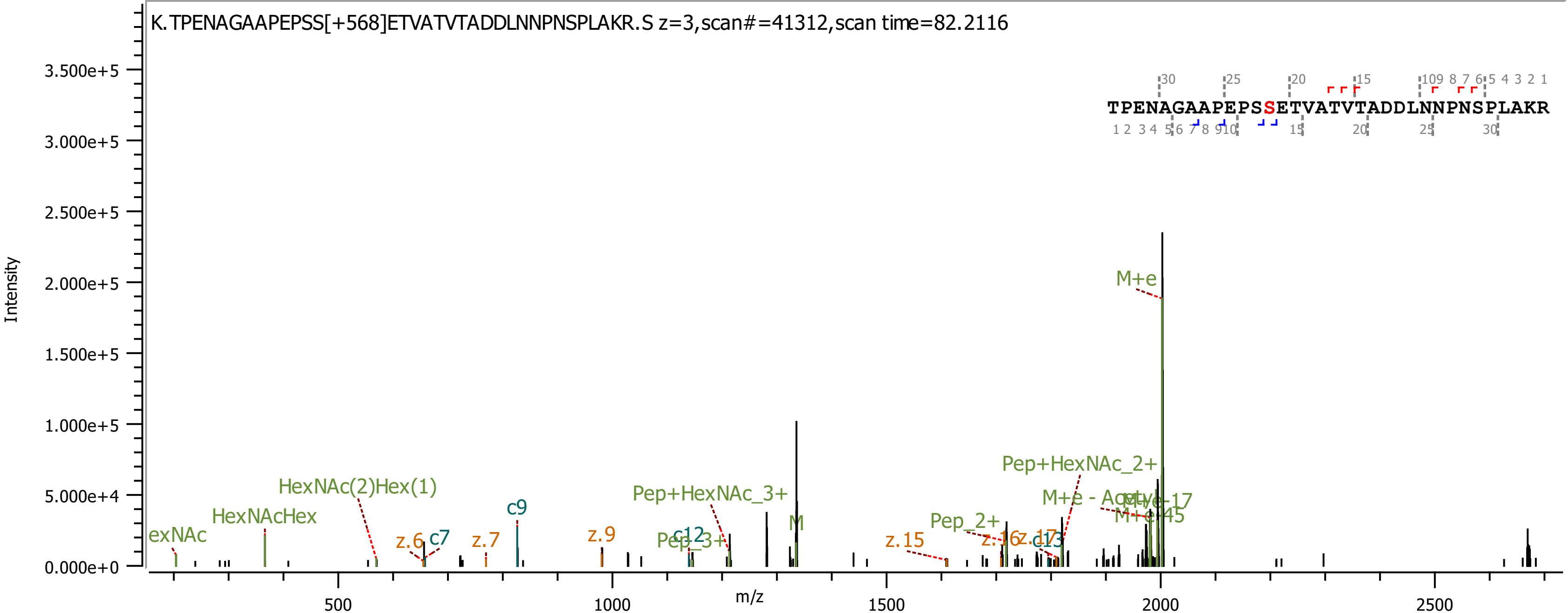
K.TPENAGAAPEPSS[+568]ETVATVTADDLNNPNSPLAKR.S z=3,scan#=41312,scan time=82.2116
TPENAGAAPEPSSETVATVTADDLNNPNSPLAKR
Intensity
m/z
M+e
Pep+HexNAc_2+
M+e - Acetyl
M+e-17
M+e-45
Pep_2+
z.15
z.16
z.17
c13
M
Pep+HexNAc_3+
Pep_3+
c12
z.9
c9
z.7
z.6
c7
HexNAc(2)Hex(1)
HexNAcHex
exNAc

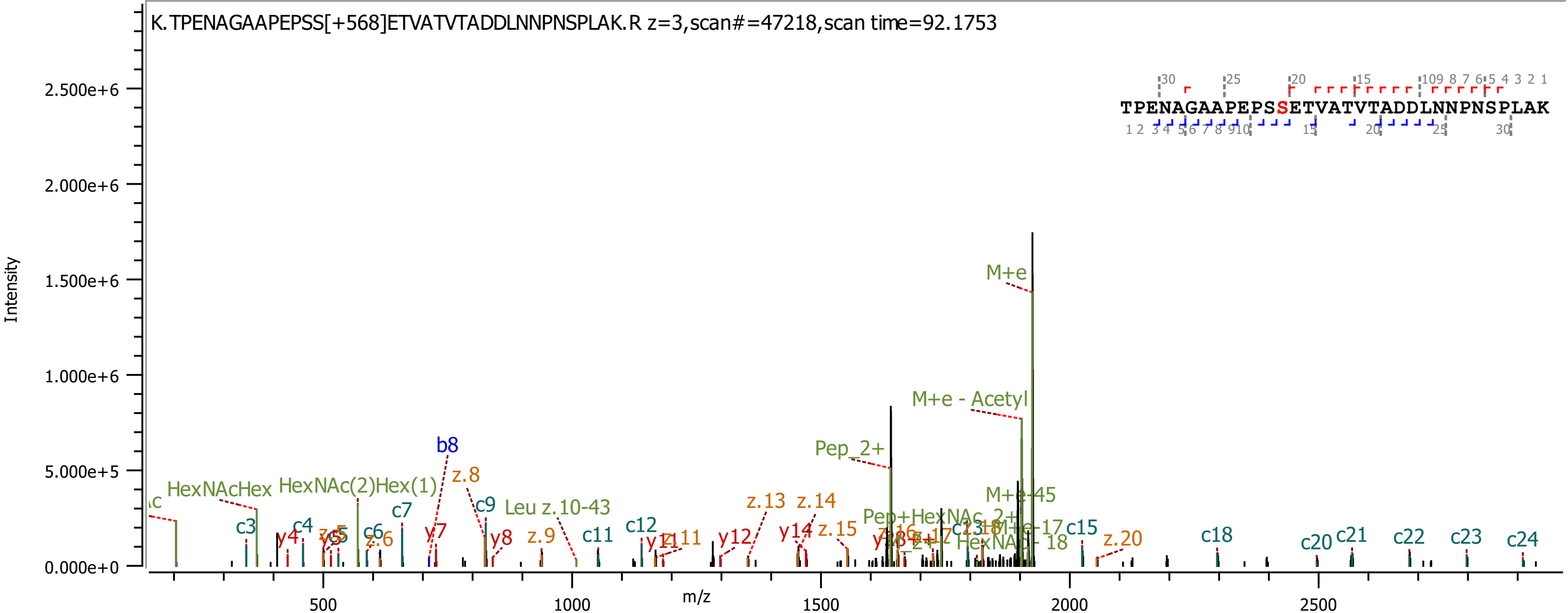

K.TPENAGAAPEPSS[+568]ETVATVTADDLNNPNSPLAK.R z=3,scan#=47218,scan time=92.1753
TPENAGAAPEPSSETVATVTADDLNNPNSPLAK
Intensity
m/z
2.500e+6
2.000e+6
1.500e+6
1.000e+6
5.000e+5
0.000e+0
500
1000
1500
2000
2500
M+e
M+e - Acetyl
Pep_2+
M+e-45
M+e-17
Pep+HexNAc_2+
HexNAc - 18
HexNAcHex
HexNAc(2)Hex(1)
Leu z.10-43
b8
z.8
z.9
z.13
z.14
z.15
z.20
c3
c4
c6
c7
c9
c11
c12
c15
c18
c20
c21
c22
c23
c24
y4
y7
y8
y12
y14

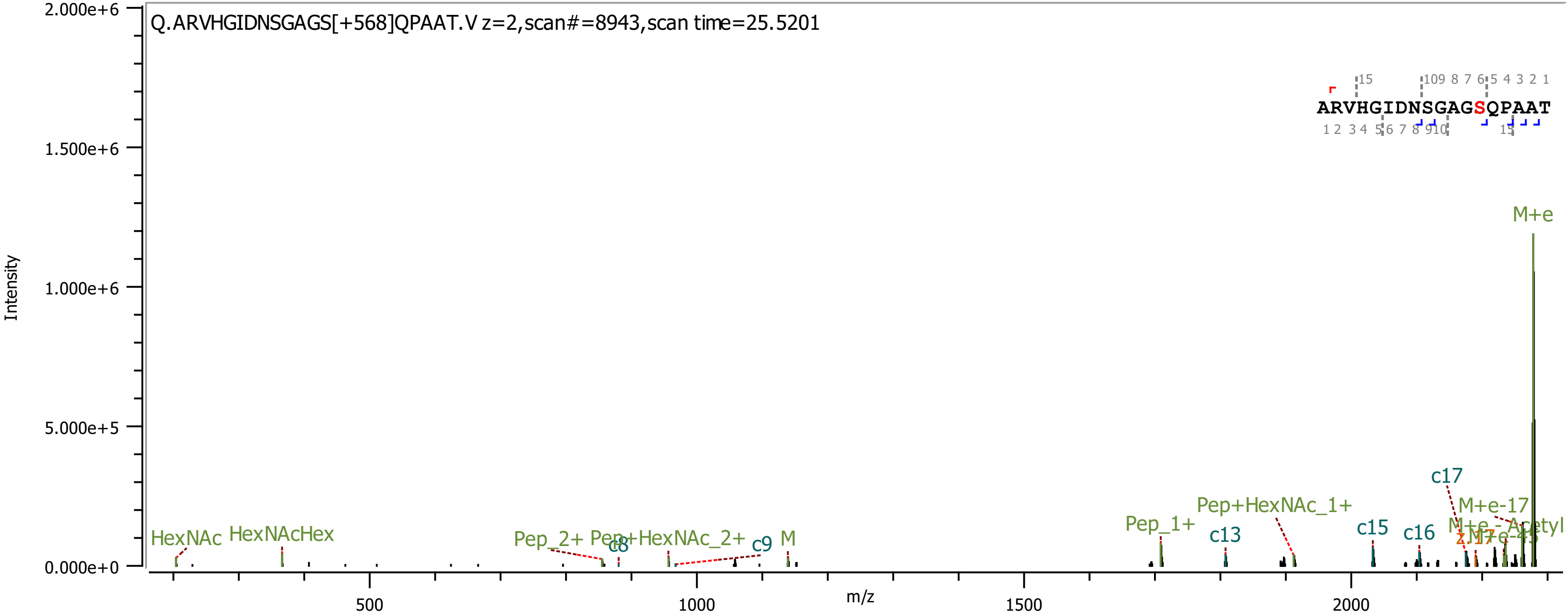

Q.ARVHGIDNSGAGS[+568]QPAAT.V z=2,scan#=8943,scan time=25.5201
ARVHGIDNSGAGSQPAAT
Intensity
m/z
2.000e+6
1.500e+6
1.000e+6
5.000e+5
0.000e+0
500
1000
1500
2000
HexNAc
HexNAcHex
Pep_2+
Pep+HexNAc_2+
c8
c9
M
Pep_1+
c13
Pep+HexNAc_1+
c15
c16
c17
M+e-17
M+e-Acetyl
M+e

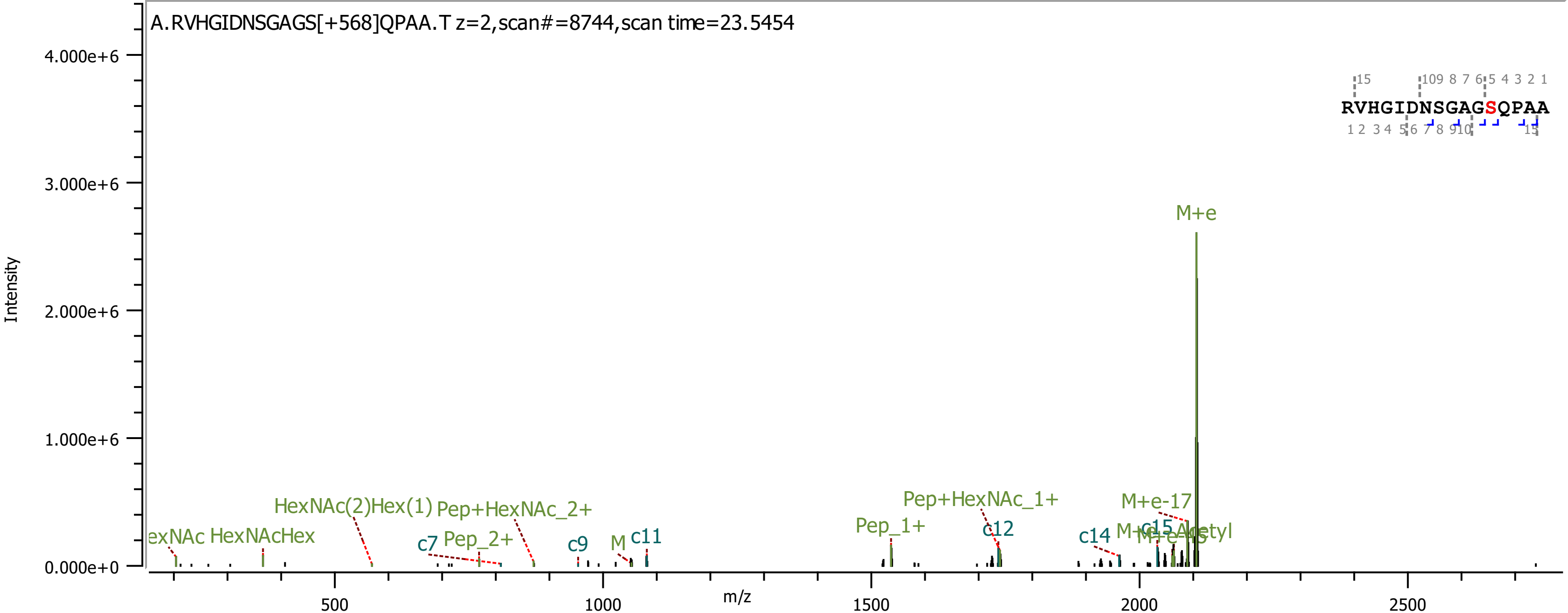

A.RVHGIDNSGAGS[+568]QPAA.T z=2,scan#=8744,scan time=23.5454
Intensity
4.000e+6
3.000e+6
2.000e+6
1.000e+6
0.000e+0
m/z
500
1000
1500
2000
2500
RVHGIDNSGAGSQPAA
M+e
M+e-17
Pep+HexNAc_1+
Pep_1+
c12
c14
c15
HexNAc(2)Hex(1)
Pep+HexNAc_2+
HexNAc
HexNAcHex
c7
Pep_2+
c9
M
c11

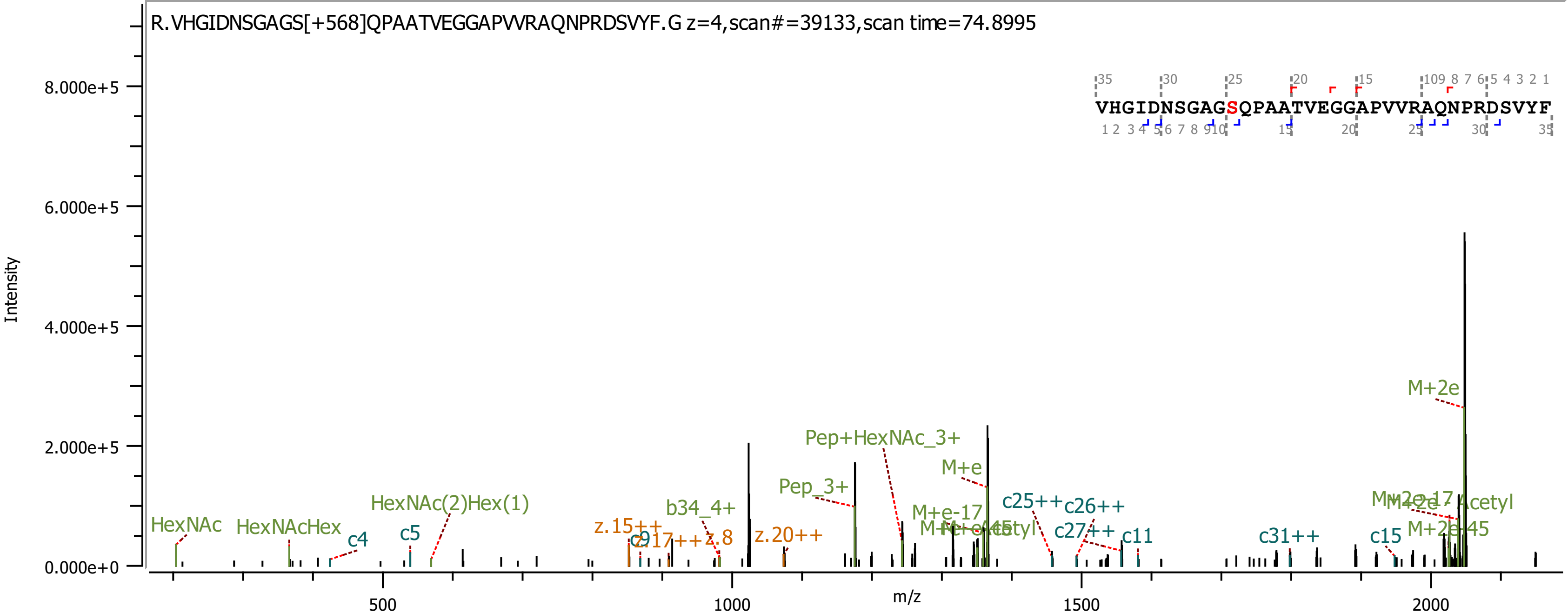

R.VHGIDNSGAGS[+568]QPAATVEGGAPVVRAQNPRDSVYF.G z=4,scan#=39133,scan time=74.8995
VHGIDNSGAGSQPAATVEGGAPVVRAQNPRDSVYF
Intensity
m/z
0.000e+0
2.000e+5
4.000e+5
6.000e+5
8.000e+5
500
1000
1500
2000
HexNAc
HexNAcHex
c4
c5
HexNAc(2)Hex(1)
z.15++
c9
z.8
b34_4+
z.20++
Pep_3+
Pep+HexNAc_3+
M+e
M+e-17
c25++
c26++
c27++
c11
c31++
c15
M+2e
M+2e-45

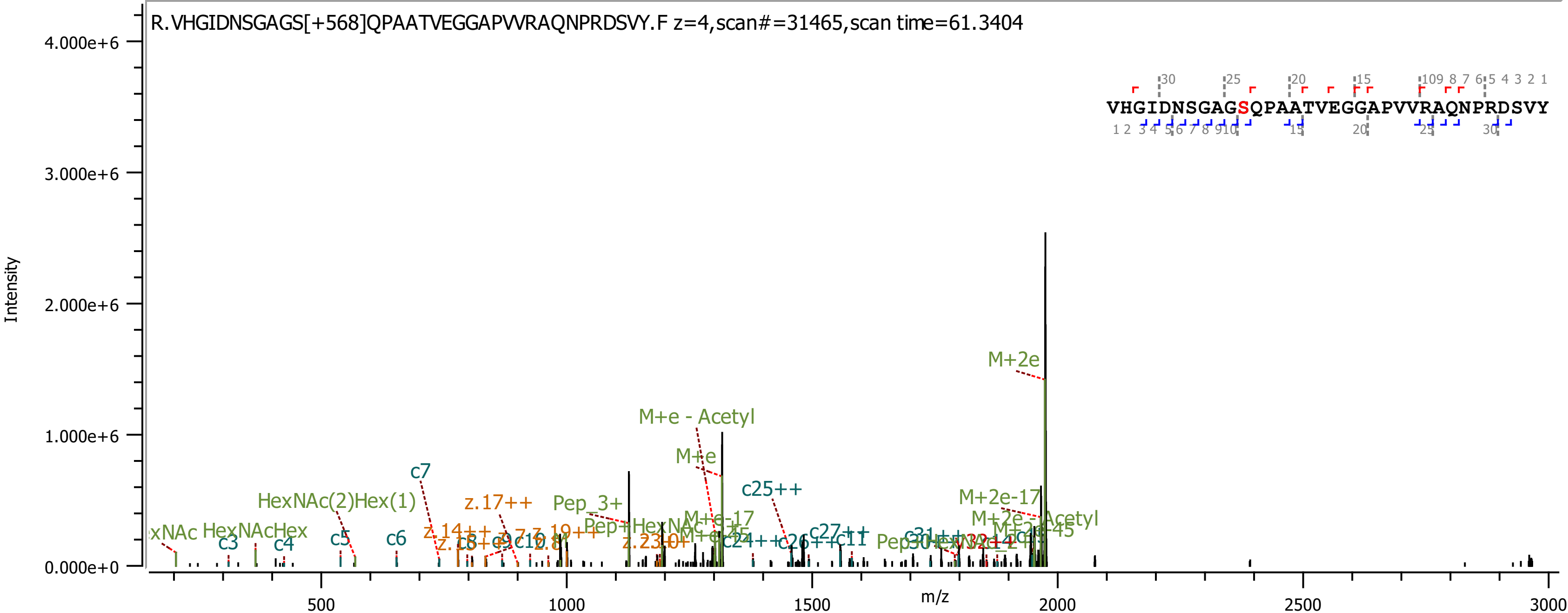
R.VHGIDNSGAGS[+568]QPAATVEGGAPVVRAQNPRDSVY.F z=4,scan#=31465,scan time=61.3404
VHGIDNSGAGSQPAATVEGGAPVVRAQNPRDSVY
Intensity
4.000e+6
3.000e+6
2.000e+6
1.000e+6
0.000e+0
m/z
500
1000
1500
2000
2500
3000
M+2e
M+e - Acetyl
M+e
c7
c25++
HexNAc(2)Hex(1)
z.17++
Pep_3+
M+2e-17
M+2e - Acetyl
HexNAcHex
c3
c4
c5
c6
c27++
c11

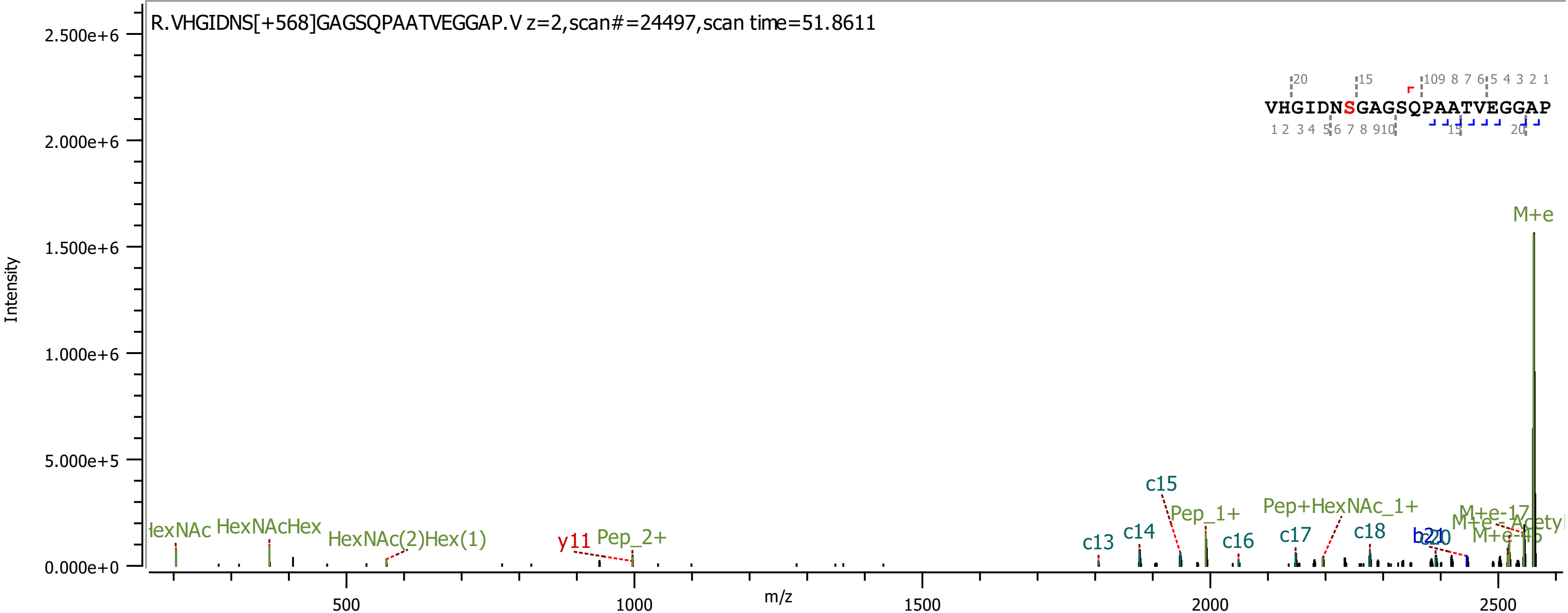
R.VHGIDNS[+568]GAGSQPAATVEGGAP.V z=2,scan#=24497,scan time=51.8611
VHGIDNSGAGSQPAATVEGGAP
Intensity
m/z
M+e
c13
c14
c15
c16
c17
c18
Pep_1+
Pep_2+
Pep+HexNAc_1+
HexNAcHex
HexNAc(2)Hex(1)
y11
M+e-17
M+e-45
2.500e+6
2.000e+6
1.500e+6
1.000e+6
5.000e+5
0.000e+0
500
1000
1500
2000
2500

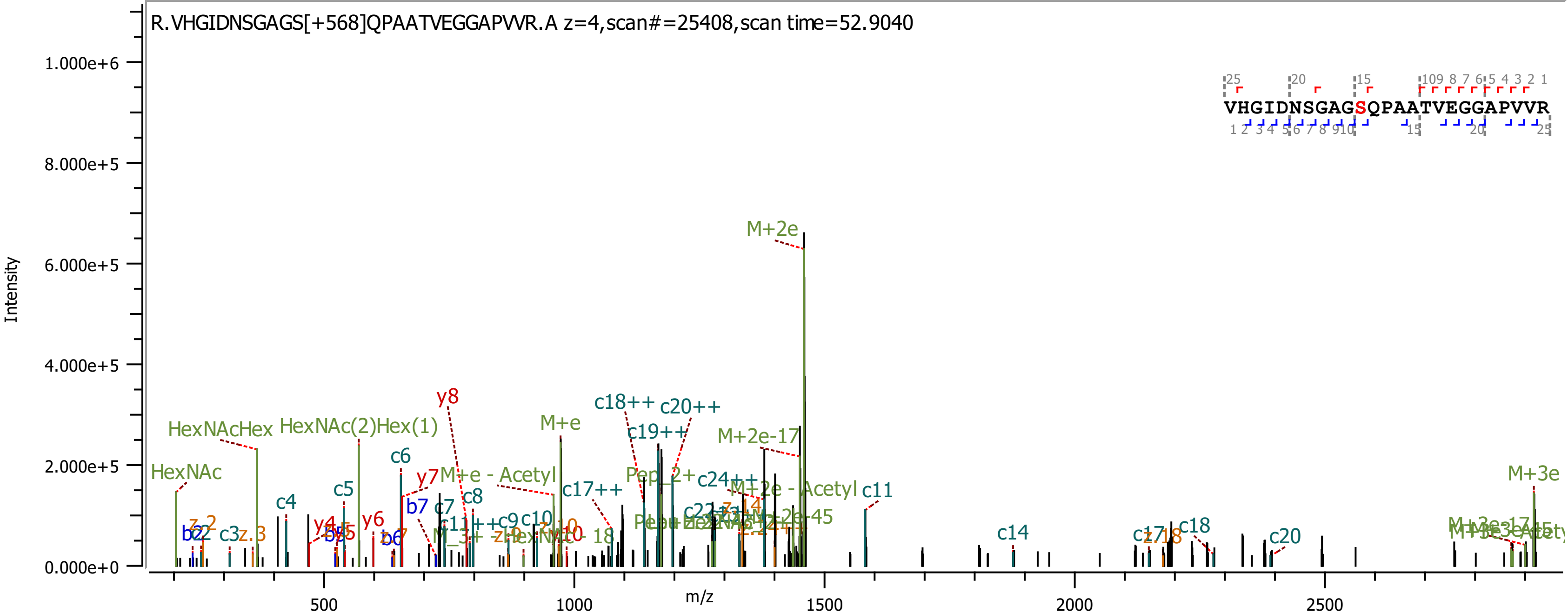
R.VHGIDNSGAGS[+568]QPAATVEGGAPVVR.A z=4,scan#=25408,scan time=52.9040
Intensity
m/z
1.000e+6
8.000e+5
6.000e+5
4.000e+5
2.000e+5
0.000e+0
500
1000
1500
2000
2500
VHGIDNSGAGSQPAATVEGGAPVVR
M+2e
HexNAcHex
HexNAc(2)Hex(1)
HexNAc
y8
M+e
c18++
c20++
c19++
M+2e-17
c6
y7
M+e - Acetyl
c5
c4
c8
b7
c7
c17++
c24++
M+2e - Acetyl
c11
y6
y4
b2
c3
c9
c10
c14
c17
c18
c20
M+3e

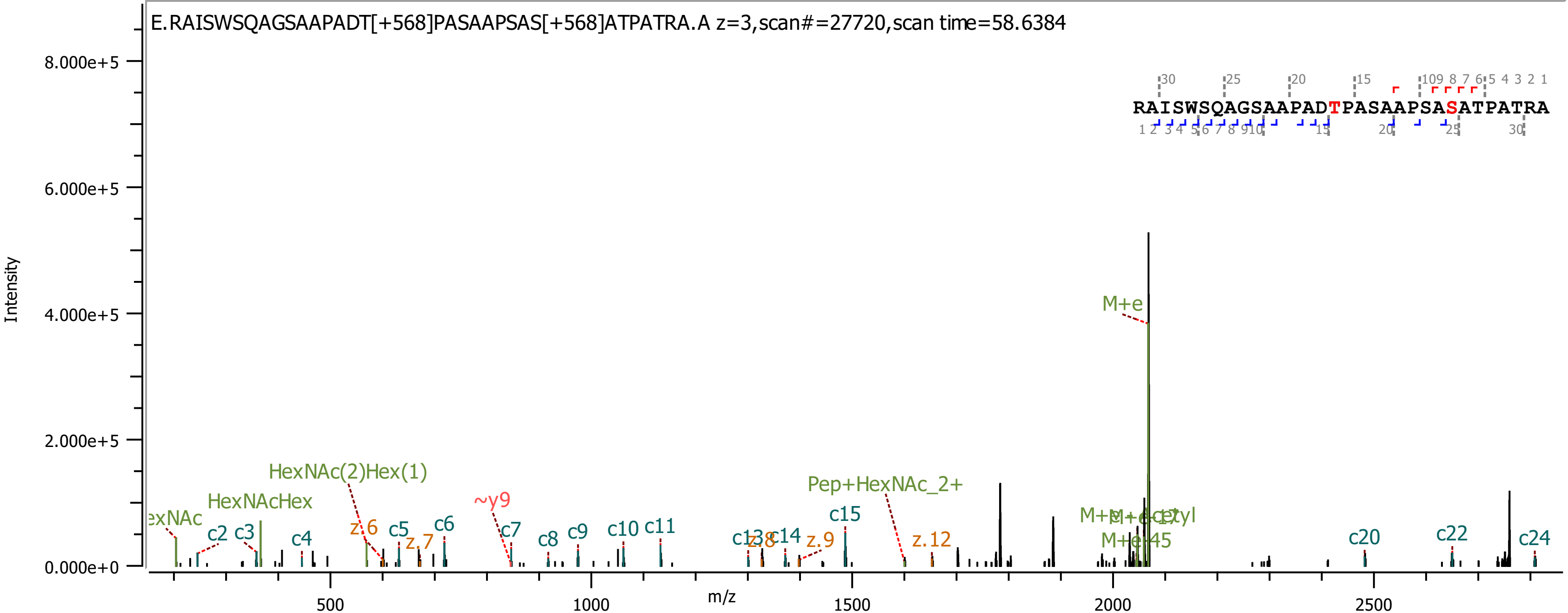
E.RAISWSQAGSAAPADT[+568]PASAAPSAS[+568]ATPATRA.A z=3,scan#=27720,scan time=58.6384
RAISWSQAGSAAPADTPASAAPSASATPATRA
Intensity
8.000e+5
6.000e+5
4.000e+5
2.000e+5
0.000e+0
m/z
500
1000
1500
2000
2500
M+e
HexNAc(2)Hex(1)
HexNAcHex
Pep+HexNAc_2+
~y9
c2
c3
c4
c5
c6
c7
c8
c9
c10
c11
c14
c15
c20
c22
c24
z.6
z.7
z.9
z.12
M+e-45

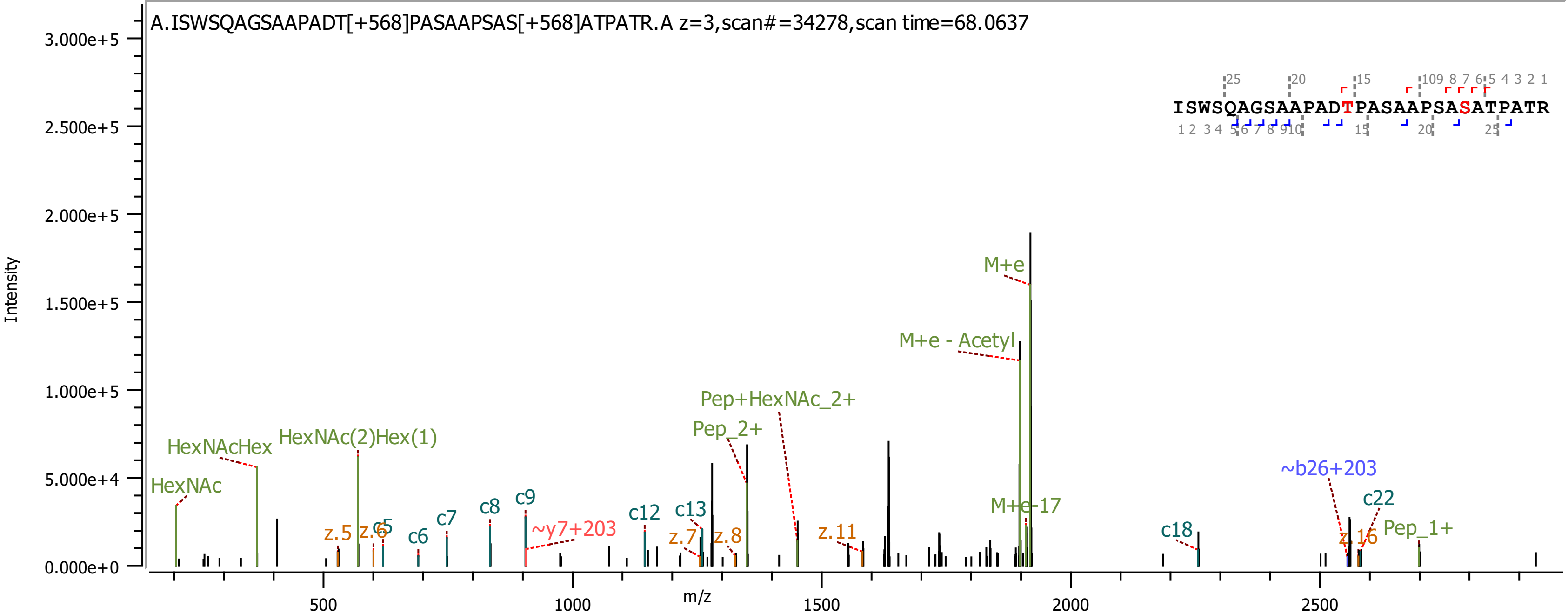
A.ISWSQAGSAAPADT[+568]PASAAPSAS[+568]ATPATR.A z=3,scan#=34278,scan time=68.0637
ISWSQAGSAAPADTPASAAPSASATPATR
Intensity
m/z
HexNAc
HexNAcHex
HexNAc(2)Hex(1)
z.5
z.6
c5
c6
c7
c8
c9
~y7+203
c12
c13
z.7
z.8
Pep_2+
Pep+HexNAc_2+
z.11
M+e - Acetyl
M+e
M+e-17
c18
~b26+203
c22
z.16
Pep_1+

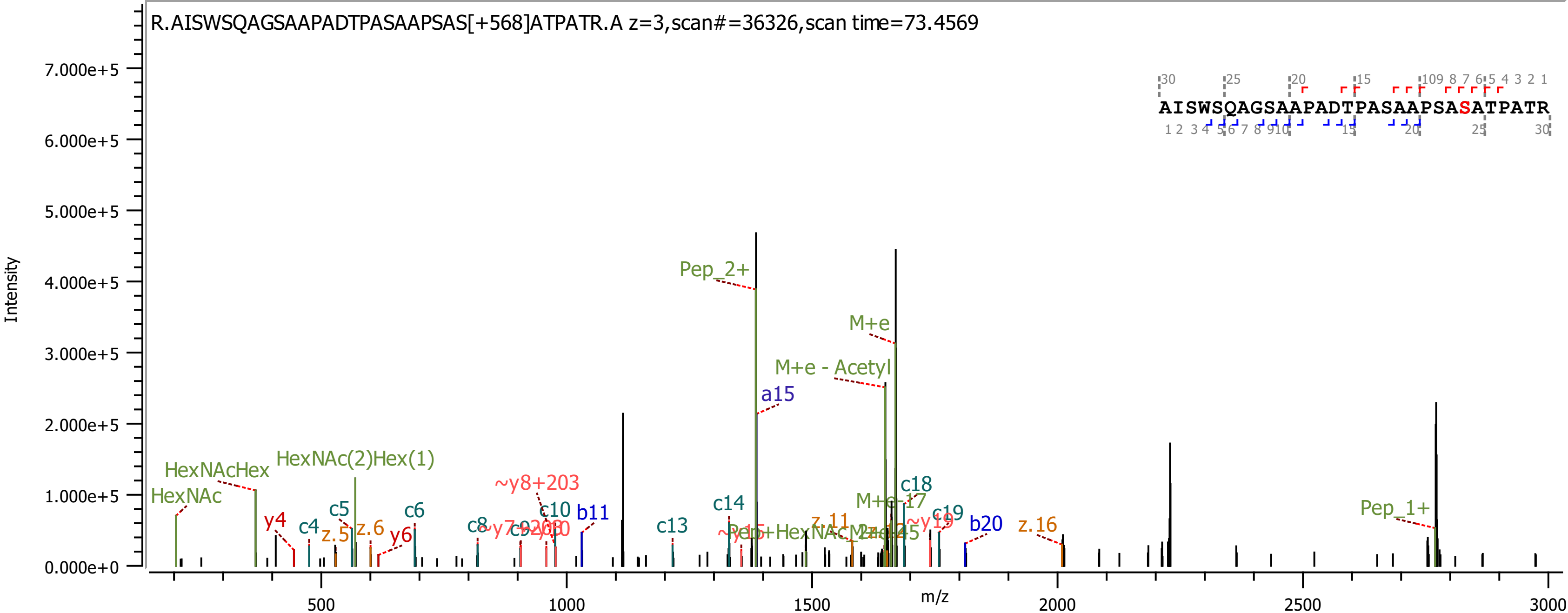

R.AISWSQAGSAAPADTPASAAPSAS[+568]ATPATR.A z=3,scan#=36326,scan time=73.4569
AISWSQAGSAAPADTPASAAPSASATPATR
Intensity
m/z
Pep_2+
M+e
M+e - Acetyl
a15
HexNAcHex
HexNAc(2)Hex(1)
HexNAc
y4
c4
c5
z.5
z.6
y6
c6
c8
~y8+203
c10
b11
c13
c14
c18
c19
b20
z.16
Pep_1+
z.11

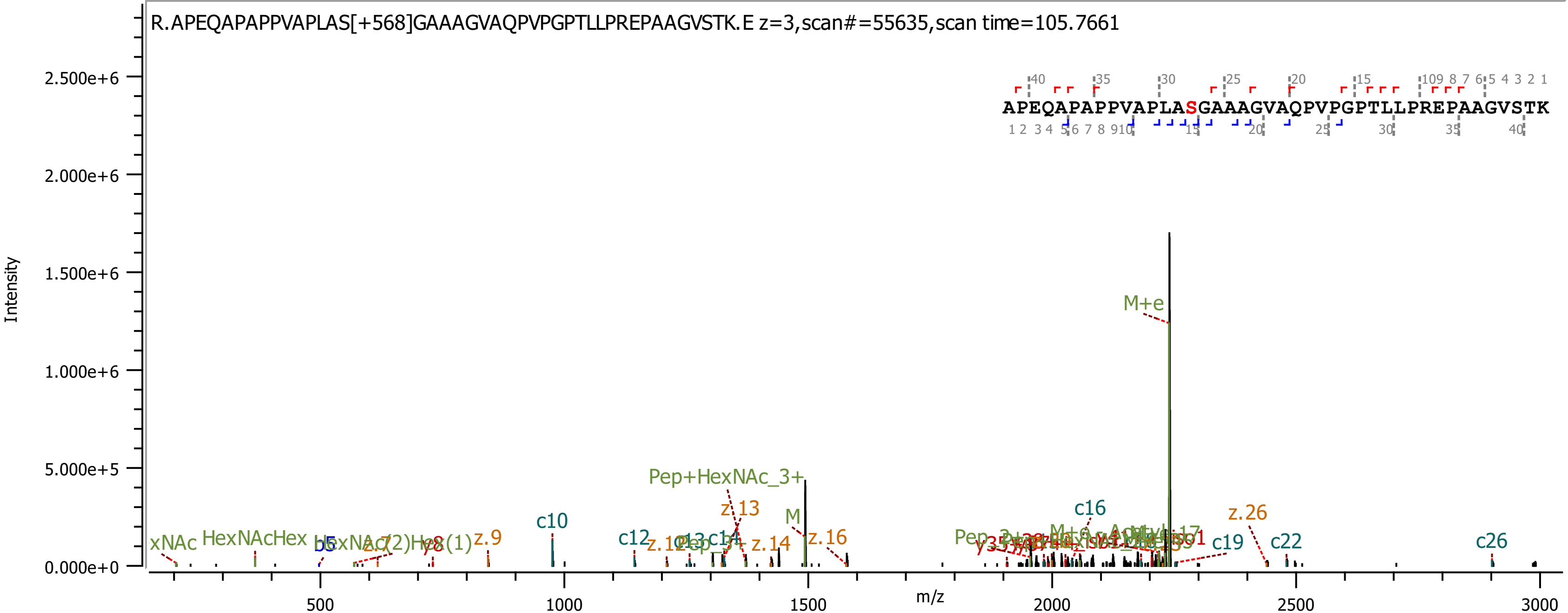

R.APEQAPAPPVAPLAS[+568]GAAAGVAQPVPGPTLLPREPAAGVSTK.E z=3,scan#=55635,scan time=105.7661
APEQAPAPPVAPLASGAAAGVAQPVPGPTLLPREPAAGVSTK
Intensity
m/z
M+e
Pep+HexNAc_3+
c10
c12
c16
c19
c22
c26
z.9
z.13
z.14
z.16
z.26
HexNAcHex
b5
M

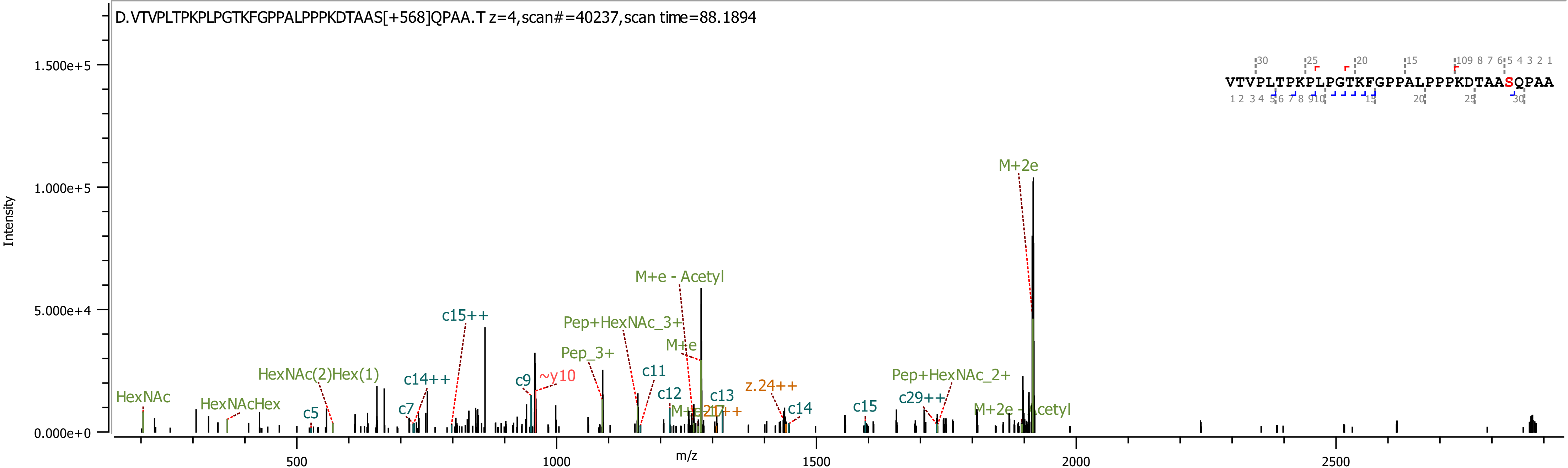
D.VTVPLTPKPLPGTKFGPPALPPPKDTAAS[+568]QPAA.T z=4,scan#=40237,scan time=88.1894
VTVPLTPKPLPGTKFGPPALPPPKDTAASQPAA
Intensity
1.500e+5
1.000e+5
5.000e+4
0.000e+0
m/z
500
1000
1500
2000
2500
HexNAc
HexNAcHex
c5
HexNAc(2)Hex(1)
c7
c14++
c15++
c9
~y10
Pep_3+
Pep+HexNAc_3+
M+e - Acetyl
M+e
c11
c12
c13
z.24++
c14
c15
Pep+HexNAc_2+
c29++
M+2e - Acetyl
M+2e

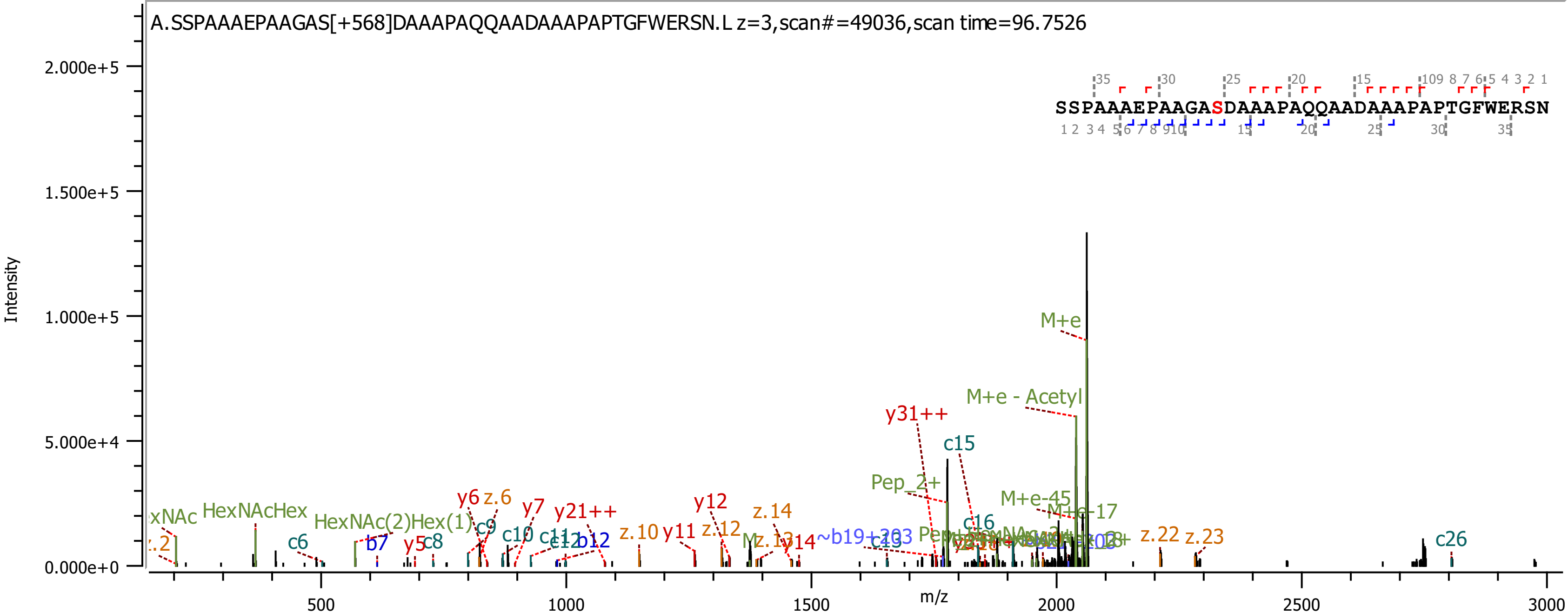
A.SSPAAAEPAAGAS[+568]DAAAPAQQAADAAAPAPTGFWERSN.L z=3,scan#=49036,scan time=96.7526
SSPAAAEPAAGASDAAAPAQQAADAAAPAPTGFWERSN
Intensity
2.000e+5
1.500e+5
1.000e+5
5.000e+4
0.000e+0
m/z
500
1000
1500
2000
2500
3000
M+e
M+e - Acetyl
y31++
c15
Pep_2+
M+e-45
M+e-17
c16
HexNAcHex
HexNAc(2)Hex(1)
c6
b7
y5
c8
y6
z.6
c9
y7
c10
y21++
c11
c12
b12
z.10
y11
y12
z.12
z.14
y14
~b19+203
c19
z.22
z.23
c26

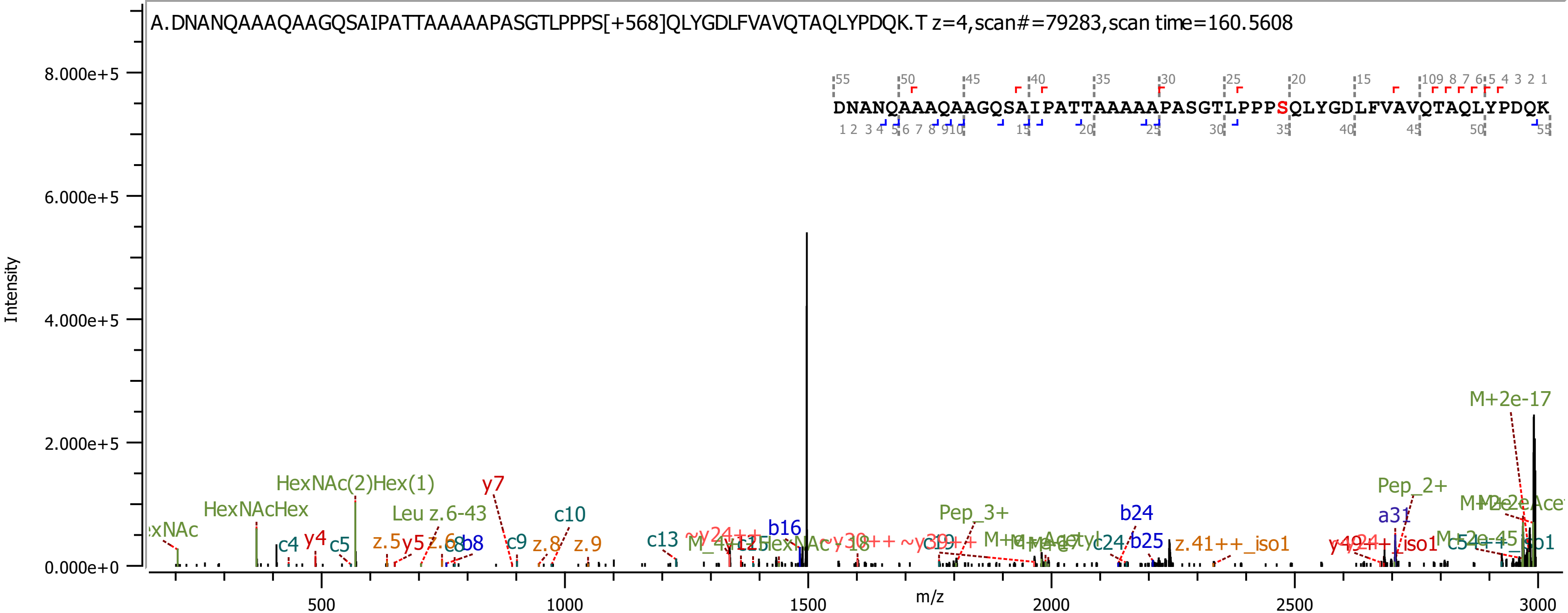
A.DNANQAAAQAAGQSAIPATTAAAAAPASGTLPPPS[+568]QLYGDLFVAVQTAQLYPDQK.T z=4,scan#=79283,scan time=160.5608
DNANQAAAQAAGQSAIPATTAAAAAPASGTLPPPSQLYGDLFVAVQTAQLYPDQK
Intensity
m/z
8.000e+5
6.000e+5
4.000e+5
2.000e+5
0.000e+0
500
1000
1500
2000
2500
3000
HexNAc(2)Hex(1)
HexNAcHex
Leu z.6-43
y7
c10
c4
y4
c5
z.5
y5
b8
c9
z.8
z.9
c13
b16
Pep_3+
b24
b25
z.41++_iso1
a31
Pep_2+
M+2e-17

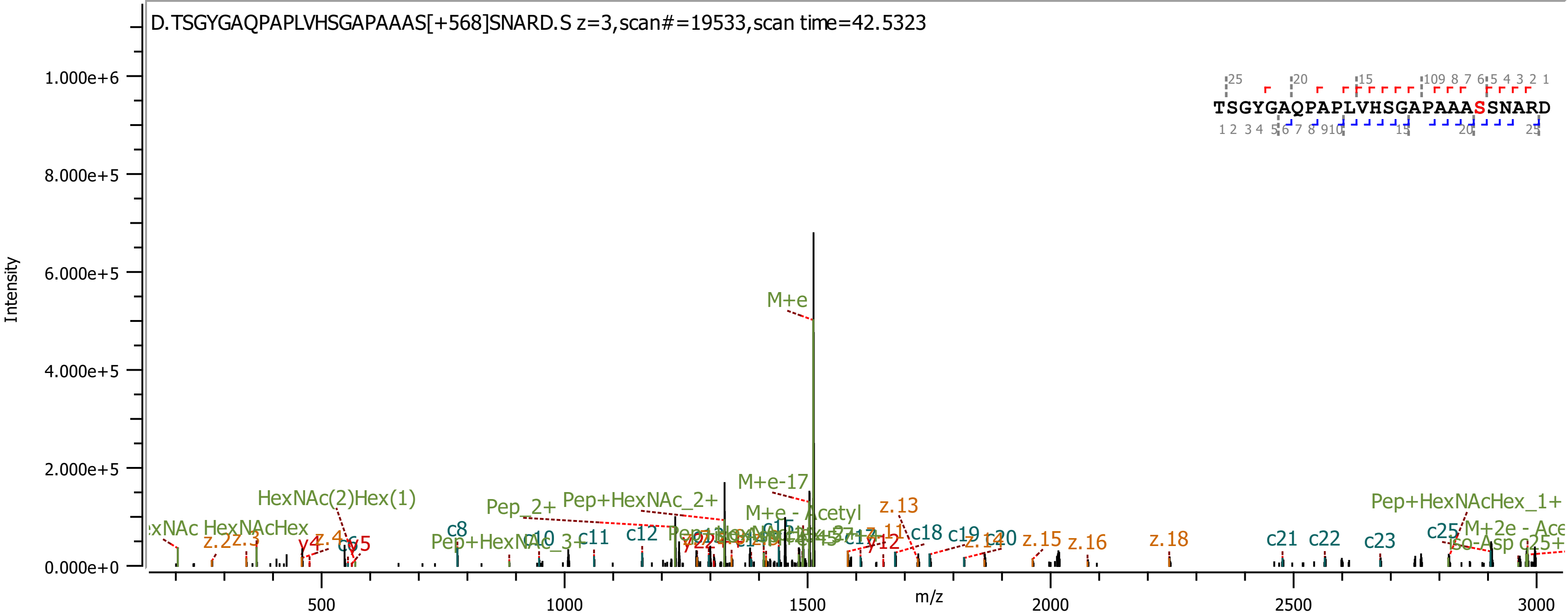
D.TSGYGAQPAPLVHSGAPAAAS[+568]SNARD.S z=3,scan#=19533,scan time=42.5323
TSGYGAQPAPLVHSGAPAAASSNARD
Intensity
m/z
1.000e+6
8.000e+5
6.000e+5
4.000e+5
2.000e+5
0.000e+0
500
1000
1500
2000
2500
3000
M+e
M+e-17
M+e - Acetyl
HexNAc(2)Hex(1)
HexNAcHex
Pep_2+
Pep+HexNAc_2+
Pep+HexNAc_3+
Pep+HexNAcHex_1+
c8
c10
c11
c12
c18
c19
c20
c21
c22
c23
c25
z.2
z.3
z.13
z.15
z.16
z.18
y12
Iso-Asp

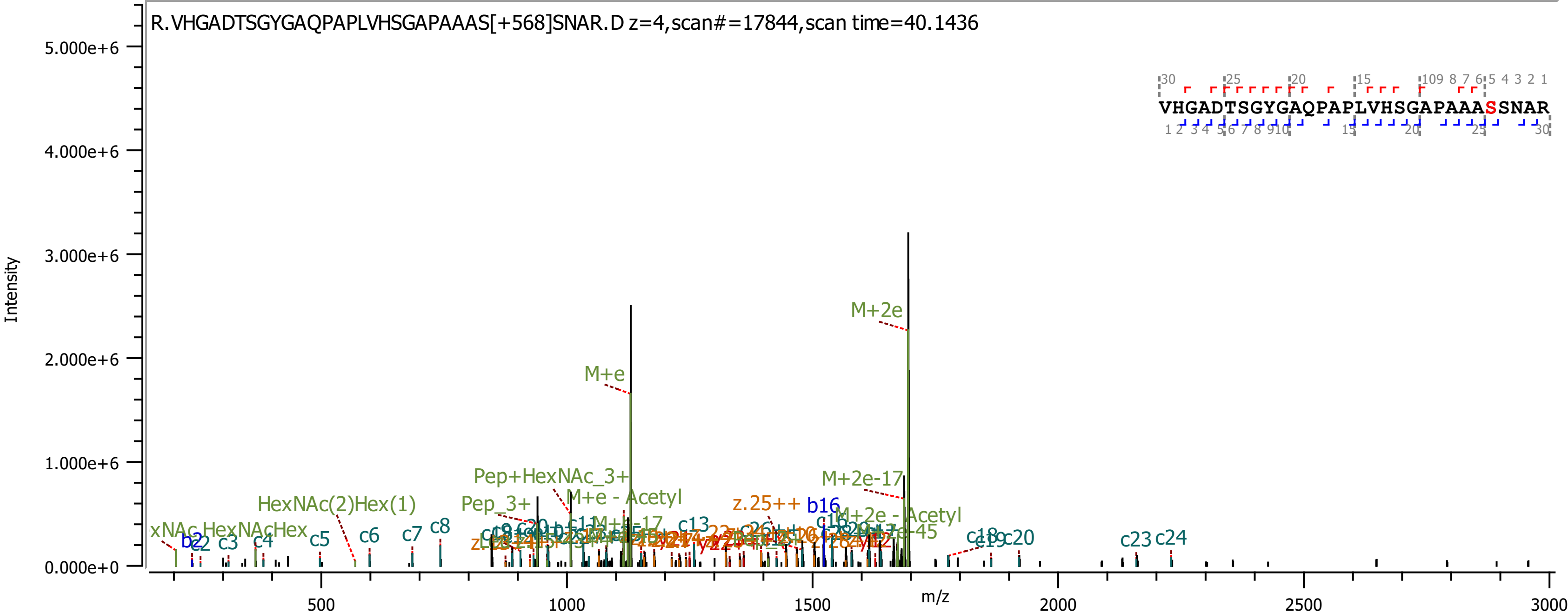
R.VHGADTSGYGAQPAPLVHSGAPAAAS[+568]SNAR.D z=4,scan#=17844,scan time=40.1436
VHGADTSGYGAQPAPLVHSGAPAAASSNAR
Intensity
m/z
5.000e+6
4.000e+6
3.000e+6
2.000e+6
1.000e+6
0.000e+0
500
1000
1500
2000
2500
3000
M+2e
M+e
Pep+HexNAc_3+
M+e - Acetyl
HexNAc(2)Hex(1)
Pep_3+
M+2e-17
z.25++
b16
M+2e - Acetyl
HexNAc
HexNAcHex
b2
c2
c3
c4
c5
c6
c7
c8
c13
c18
c19
c20
c23
c24

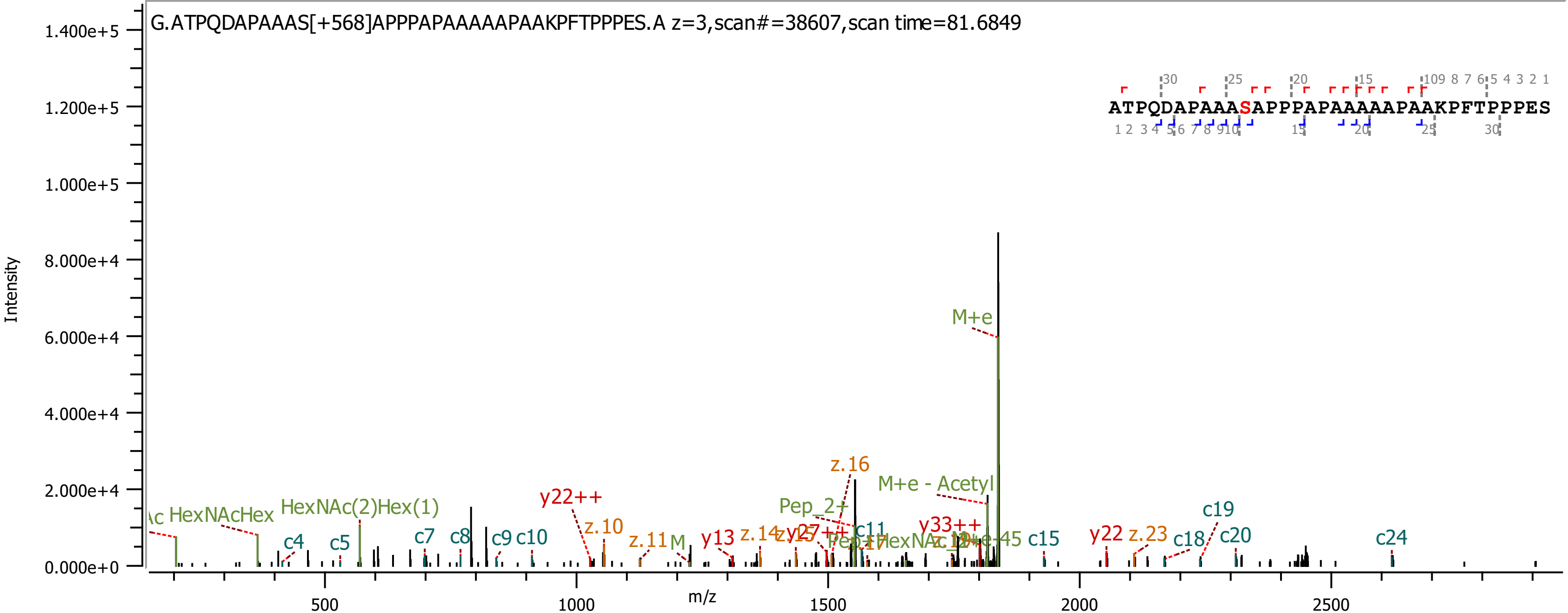

G.ATPQDAPAAAS[+568]APPPAPAAAAAPAAKPFTPPPES.A z=3,scan#=38607,scan time=81.6849
ATPQDAPAAASAPPPAPAAAAAPAAKPFTPPPES
Intensity
m/z
1.400e+5
1.200e+5
1.000e+5
8.000e+4
6.000e+4
4.000e+4
2.000e+4
0.000e+0
500
1000
1500
2000
2500
HexNAcHex
HexNAc(2)Hex(1)
c4
c5
c7
c8
c9
c10
y22++
z.10
z.11
y13
z.14
z.16
Pep_2+
M+e - Acetyl
M+e
y33++
c11
c15
y22
z.23
c18
c19
c20
c24

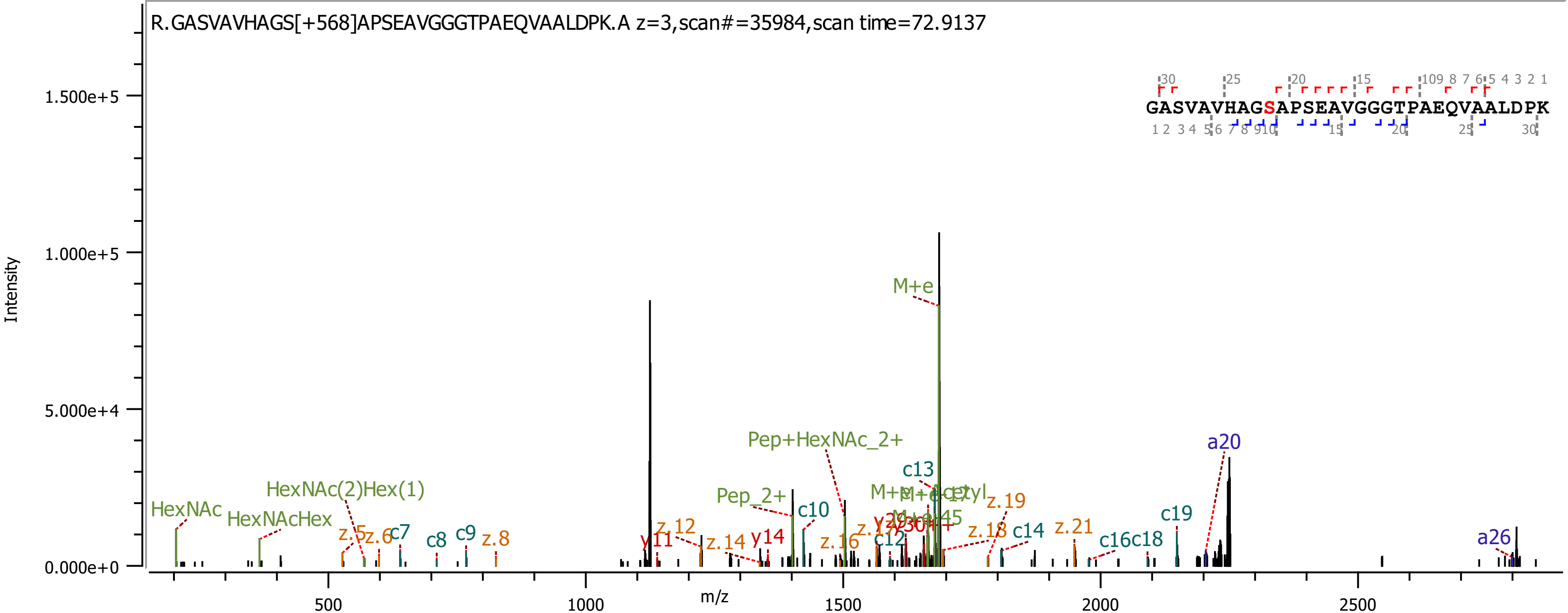
R.GASVAVHAGS[+568]APSEAVGGGTPAEQVAALDPK.A z=3,scan#=35984,scan time=72.9137
GASVAVHAGSAPSEAVGGGTPAEQVAALDPK
Intensity
m/z
0.000e+0
5.000e+4
1.000e+5
1.500e+5
500
1000
1500
2000
2500
HexNAc
HexNAcHex
HexNAc(2)Hex(1)
z.5
z.6
c7
c8
c9
z.8
y11
z.12
z.14
y14
Pep_2+
Pep+HexNAc_2+
c10
z.16
c12
c13
M+e
z.18
z.19
c14
z.21
c16
c18
c19
a20
a26

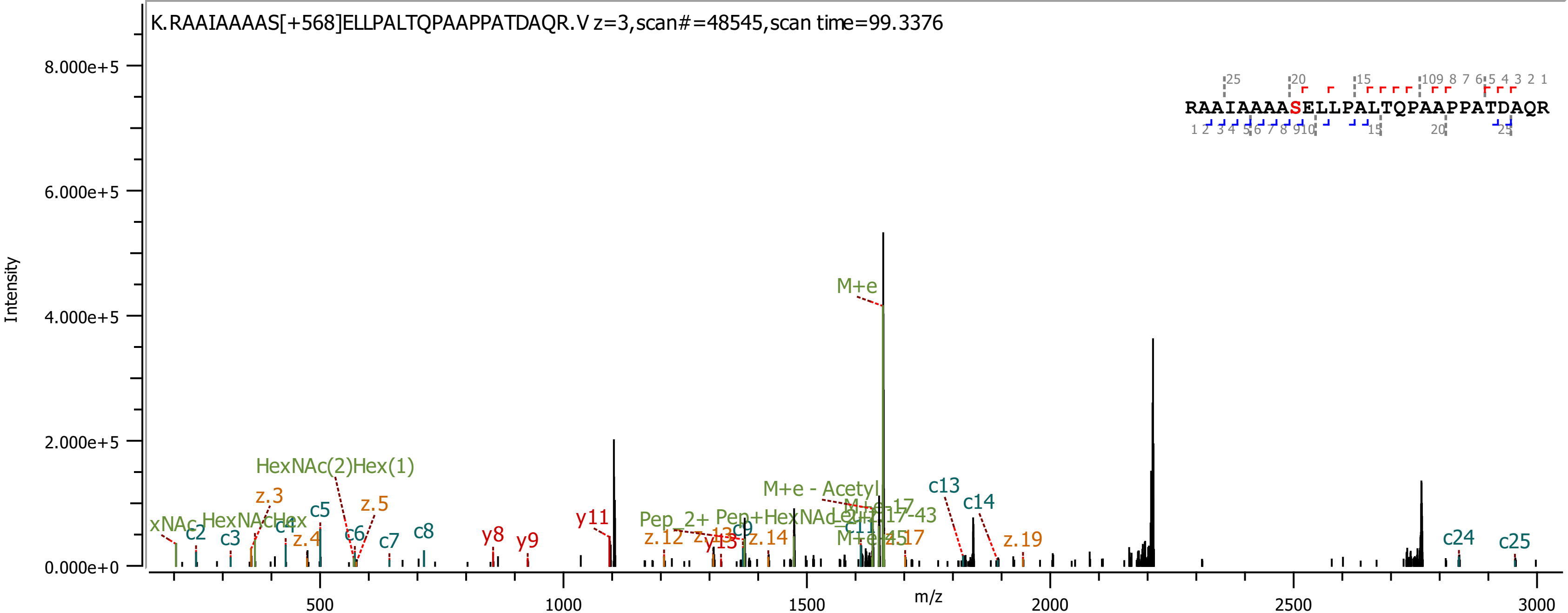

K.RAAIAAAAS[+568]ELLPALTQPAAPPATDAQR.V z=3,scan#=48545,scan time=99.3376
RAAIAAAASELLPALTQPAAPPATDAQR
Intensity
8.000e+5
6.000e+5
4.000e+5
2.000e+5
0.000e+0
m/z
500
1000
1500
2000
2500
3000
M+e
HexNAc(2)Hex(1)
M+e - Acetyl
c13
c14
z.3
z.5
c5
c2
c3
c4
z.4
c6
c7
c8
y8
y9
y11
z.12
z.14
c9
z.19
c24
c25

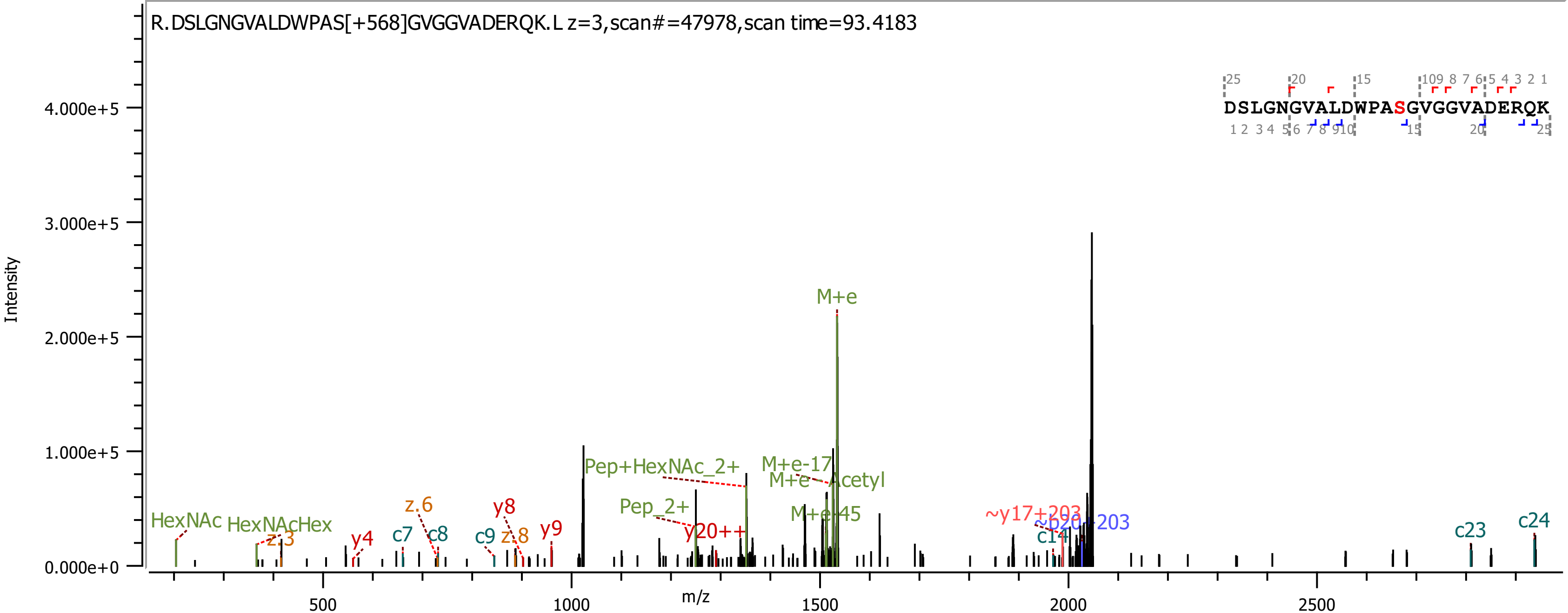

R.DSLGNGVALDWPAS[+568]GVGGVADERQK.L z=3,scan#=47978,scan time=93.4183
DSLGNGVALDWPASGVGGVADERQK
Intensity
m/z
0.000e+0
1.000e+5
2.000e+5
3.000e+5
4.000e+5
500
1000
1500
2000
2500
HexNAc
HexNAcHex
z.3
y4
c7
z.6
c8
c9
y8
z.8
y9
Pep+HexNAc_2+
Pep_2+
y20++
M+e-17
M+e-Acetyl
M+e
M+e-45
~y17+203
~b20+203
c14
c23
c24

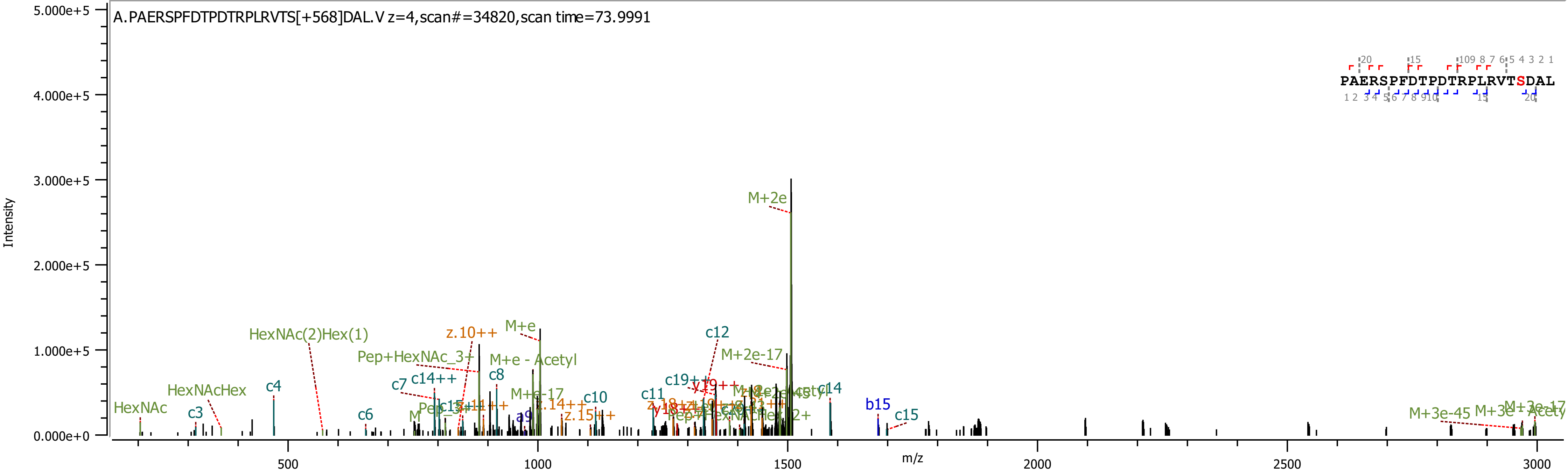
A.PAERSPFDTPDTRPLRVTS[+568]DAL.V z=4,scan#=34820,scan time=73.9991
PAERSPFDTPDTRPLRVTSDAL
Intensity
5.000e+5
4.000e+5
3.000e+5
2.000e+5
1.000e+5
0.000e+0
m/z
500
1000
1500
2000
2500
3000
M+2e
HexNAc(2)Hex(1)
z.10++
M+e
Pep+HexNAc_3+
M+e - Acetyl
c8
c12
M+2e-17
c7
c14++
c19++
c14
HexNAcHex
c4
M+e-17
c10
c11
HexNAc
c3
c6
z.14++
z.15++
b15
c15
a9
M+3e-45
M+3e-17

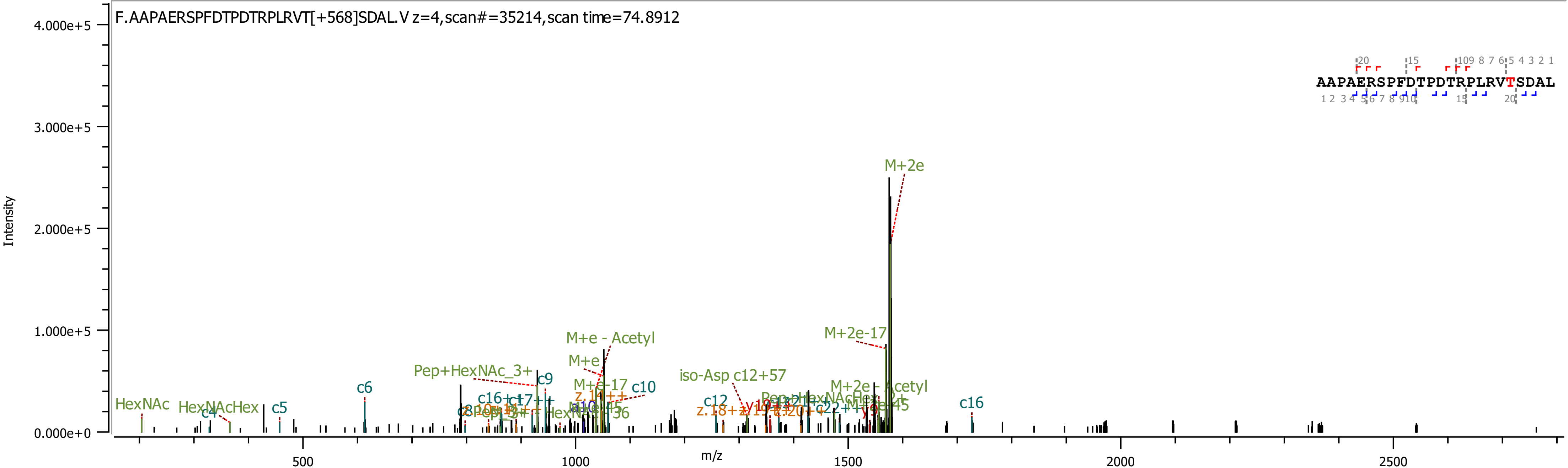
F.AAPAERSPFDTPDTRPLRVT[+568]SDAL.V z=4,scan#=35214,scan time=74.8912
AAPAERSPFDTPDTRPLRVTSDAL
Intensity
m/z
4.000e+5
3.000e+5
2.000e+5
1.000e+5
0.000e+0
500
1000
1500
2000
2500
M+2e
M+2e-17
M+e - Acetyl
M+e
M+e-17
Pep+HexNAc_3+
iso-Asp c12+57
M+2e- Acetyl
HexNAc
HexNAcHex
c4
c5
c6
c9
c10
c12
c16
