## Supplementary Data 4 for "Burkholderia PglL enzymes are serine preferring oligosaccharidetransferases which target conserved proteins across the Burkholderia genus"

### *Burkholderia cenocepacia* K56-2 Best Localised unique glycopeptides

| J2315 Gene name | Protein Name | Peptide < ProteinMetrics Confidential > | Glycans NHFAGNa | Localised site (Yes / No / Partial) ? | Observed m/z | z | Observed (M+H) | Calc. mass (M+H) | Mass error (ppm) | Cleavage | Score | Delta | Delta Mod | Comment | Scan Time | Enzyme | Replicate | Site | Page |
|---|---|---|---|---|---|---|---|---|---|---|---|---|---|---|---|---|---|---|---|
| BCAL0039 | >gi\|443605781\|gb\|ELT73605.1\| ABC transporter, substrate-binding protein, family 3 [Burkholderia cenocepacia K56-2Valvano] | T.AAPAS[+568.212]AATAGSRLDDVLARGA.L | HexNAc(2)Hex(1) | Yes | 837.0821 | 3 | 2509.2316 | 2509.226 | 2.1 | Non | 226.4 | 110.9 | 67.13 | Nsco_20191108_BC_ZIC_HILIC_K56_2_thermolysin_B1.32913.32913.3 | 66.64 | Thermolysin | B1 | S33 | 12 |
| BCAL0080 | >gi\|443593207\|gb\|ELT61961.1\| cytochrome C domain protein, partial [Burkholderia cenocepacia K56-2Valvano] | N.AAQPAS[+568.212]GAPASGADASNAQA.- | HexNAc(2)Hex(1) | Partial | 1141.0048 | 2 | 2281.0023 | 2280.995 | 3.2 | Non | 319.9 | 187.1 | 187.05 | Nsco_20191108_BC_ZIC_HILIC_K56_2_thermolysin_B1.12889.12889.2 | 31.01 | Thermolysin | B1 | NA | 13 |
| BCAL0080 | >gi\|443593207\|gb\|ELT61961.1\| cytochrome C domain protein, partial [Burkholderia cenocepacia K56-2Valvano] | Y.MANNDGANFPEPAAPAANAAQPAS[+568.212]GAPASGADASNAQ.A | HexNAc(2)Hex(1) | Yes | 1321.9149 | 3 | 3963.7302 | 3962.711 | 4.1 | Non | 498.8 | 403.1 | 58.83 | Nsco_20191108_BC_ZIC_HILIC_K56_2_thermolysin_B2_.45836.45836.3 | 83.76 | Thermolysin | B2 | S176 | 14 |
| BCAL0080 | >gi\|443593207\|gb\|ELT61961.1\| cytochrome C domain protein, partial [Burkholderia cenocepacia K56-2Valvano] | Y.MANNDGANFPEPAAPAANAAQPAS[+568.212]GAPASGADASNAQA.- | HexNAc(2)Hex(1) | Yes | 1345.9263 | 3 | 4035.7644 | 4033.748 | 2.5 | Non | 237.4 | 215.4 | 42.78 | Nsco_20191108_BC_ZIC_HILIC_K56_2_thermolysin_B1.42150.42150.3 | 86.91 | Thermolysin | B1 | S176 | 15 |
| BCAL0163 | >gi\|443595598\|gb\|ELT64173.1\| BON domain protein [Burkholderia cenocepacia K56-2Valvano] | V.KVFQYVKPQDAQALQDASPASGAS[+568.212]GAQAAAAPADNATVGAVPDASVQSTPLQPPAPISNSSSVHPGNPK.A | HexNAc(2)Hex(1) | Yes | 1460.9239 | 5 | 7300.5903 | 7300.568 | 3 | NRagged | 1310 | 1268 | 36.68 | Nsco_20191108_BC_ZIC_HILIC_K56_2_typsin_B2.46071.46071.5 | 93.66 | Trypsin | B2 | S224 | 16 |
| BCAL0163 | >gi\|443595598\|gb\|ELT64173.1\| BON domain protein [Burkholderia cenocepacia K56-2Valvano] | A.LQDASPASGAS[+568.212]GAQAAAAPADN.A | HexNAc(2)Hex(1) | Partial | 1255.0619 | 2 | 2509.1166 | 2509.106 | 4.2 | Non | 346.5 | 235.7 | 8.44 | Nsco_20191108_BC_ZIC_HILIC_K56_2_pepsin_B2.22492.22492.2 | 49.04 | Pepsin | B2 | Na | 17 |
| BCAL0163 | >gi\|443595598\|gb\|ELT64173.1\| BON domain protein [Burkholderia cenocepacia K56-2Valvano] | K.PQDAQALQDASPASGAS[+568.212]GAQAAAAPADNATVGAVPDASVQSTPLQPPAPISNSSSVHPGNPK.A | HexNAc(2)Hex(1) | Yes | 1602.7728 | 4 | 6408.0695 | 6408.051 | 2.9 | Specific | 830.5 | 830.5 | 81.53 | Nsco_20191108_BC_ZIC_HILIC_K56_2_typsin_B2.48571.48571.4 | 97.84 | Trypsin | B2 | S224 | 18 |
| BCAL0163 | >gi\|443595598\|gb\|ELT64173.1\| BON domain protein [Burkholderia cenocepacia K56-2Valvano] | K.VFQYVKPQDAQALQDASPASGAS[+568.212]GAQAAAAPADNATVGAVPDASVQSTPLQPPAPISNSSSVHPGNPK.A | HexNAc(2)Hex(1) | Yes | 1435.303 | 5 | 7172.486 | 7172.473 | 1.8 | Specific | 1194 | 1194 | 111.85 | Nsco_20191108_BC_ZIC_HILIC_K56_2_typsin_B1.51502.51502.5 | 101.1 | Trypsin | B1 | S224 | 19 |
| BCAL0163 | >gi\|443595598\|gb\|ELT64173.1\| BON domain protein [Burkholderia cenocepacia K56-2Valvano] | Y.VKPQDAQALQDASPASGAS[+568.212]GAQAAAAPADNATVGAVPDASVQSTPLQPPAPISNSSSVHPGNPK.A | HexNAc(2)Hex(1) | Yes | 1659.8165 | 4 | 6636.2443 | 6635.214 | 4 | NRagged | 963.7 | 963.7 | 93.72 | Nsco_20191108_BC_ZIC_HILIC_K56_2_typsin_B3.45475.45475.4 | 91.03 | Trypsin | B3 | S224 | 20 |
| BCAL0163 | >gi\|443595598\|gb\|ELT64173.1\| BON domain protein [Burkholderia cenocepacia K56-2Valvano] | S.VQSTPLQPPAPISNSSS[+568.212]VHPGNPKAKAQ.- | HexNAc(2)Hex(1) | Partial | 852.1829 | 4 | 3405.7097 | 3405.702 | 2.3 | Non | 314.8 | 314.8 | 170.78 | Nsco_20191108_BC_ZIC_HILIC_K56_2_pepsin_B1.20403.20403.4 | 44.79 | Pepsin | B1 | S262 | 21 |
| BCAL0193 | >gi\|443590944\|gb\|ELT59881.1\| hypothetical protein BURCENK562V_2433 [Burkholderia cenocepacia K56-2Valvano] | S.AAAPAPAKKDHSKPKHQLKHHGSKKGQAKAAAAS[+568.212][+100.064]AAGTNDAGTQN.- | HexNAc(2)Hex(1) 100.064 | Partial | 1025.3224 | 5 | 5122.5828 | 5121.606 | -5.3 | Non | 459.5 | 459.5 | 51.91 | Nsco_20191108_BC_ZIC_HILIC_K56_2_pepsin_B1.3536.3536.5 | 16.51 | Pepsin | B1 | Na | 22 |

| | | | | | | | | | | | | | | | | | | | |
|---|---|---|---|---|---|---|---|---|---|---|---|---|---|---|---|---|---|---|---|
| BCAL0193 | >gi\|443590944\|gb\|ELT59881.1\| hypothetical protein BURCENK562V_2433 [Burkholderia cenocepacia K56-2Valvano] | A.AAPAPAKKDHSKPKHQLKHHGSKKGQAKAAAAS[+568.212]AAGT[+568.212]ND.A | HexNAc(2)Hex(1),HexNAc(2)Hex(1) | Partial | 1010.7127 | 5 | 5049.5344 | 5047.509 | 3.7 | Non | 199.3 | 199.3 | 143.14 | Nsco_20191108_BC_ZIC_HILIC_K56_2_pepsin_B3_reinject.3350.3350.5 | 16.23 | Pepsin | B3 | Na | 23 |
| BCAL0193 | >gi\|443590944\|gb\|ELT59881.1\| hypothetical protein BURCENK562V_2433 [Burkholderia cenocepacia K56-2Valvano] | Q.AS[+568.212]APAADTSAAAPAPAKK.D | HexNAc(2)Hex(1) | Yes | 1082.5332 | 2 | 2164.0591 | 2164.05 | 4.1 | NRagged | 360 | 184.2 | 184.16 | Nsco_20191108_BC_ZIC_HILIC_K56_2_typsin_B1.6823.6823.2 | 22.75 | Trypsin | B1 | S25 | 24 |
| BCAL0193 | >gi\|443590944\|gb\|ELT59881.1\| hypothetical protein BURCENK562V_2433 [Burkholderia cenocepacia K56-2Valvano] | A.DTSAAAPAPAKKDHSKPKHQLKHHGSKKGQAKAAAASAAGTNDAGT[+568.212]QN.- | HexNAc(2)Hex(1) | Partial | 1066.1395 | 5 | 5326.6685 | 5324.649 | 2.4 | Non | 785 | 785 | 785.04 | Nsco_20191108_BC_ZIC_HILIC_K56_2_pepsin_B3_reinject.3443.3443.5 | 16.38 | Pepsin | B3 | NA | 25 |
| BCAL0193 | >gi\|443590944\|gb\|ELT59881.1\| hypothetical protein BURCENK562V_2433 [Burkholderia cenocepacia K56-2Valvano] | T.SAAAPAPAKKDHSKPKHQLKHHGSKKGQAKAAAAS[+568.212]AAGT[+568.212]ND.A | HexNAc(2)Hex(1),HexNAc(2)Hex(1) | Partial | 1042.5261 | 5 | 5208.6015 | 5205.578 | 2.5 | Non | 254.9 | 254.9 | 166.1 | Nsco_20191108_BC_ZIC_HILIC_K56_2_pepsin_B3_reinject.3735.3735.5 | 16.81 | Pepsin | B3 | NA | 26 |
| BCAL0193 | >gi\|443590944\|gb\|ELT59881.1\| hypothetical protein BURCENK562V_2433 [Burkholderia cenocepacia K56-2Valvano] | T.SAAAPAPAKKDHSKPKHQLKHHGSKKGQAKAAAASAAGTNDAGT[+568.212][+100.064]QN.- | HexNAc(2)Hex(1) | Partial | 1042.9348 | 5 | 5210.645 | 5208.638 | 0 | Non | 238.5 | 238.5 | 238.53 | Nsco_20191108_BC_ZIC_HILIC_K56_2_pepsin_B1.3351.3351.5 | 16.26 | Pepsin | B1 | NA | 27 |
| BCAL0303 | >gi\|443596149\|gb\|ELT64670.1\| hypothetical protein BURCENK562V_3632 [Burkholderia cenocepacia K56-2Valvano] | S.KAADAGGAKPAAGAS[+568.212]AAPAAPAPVAV.P | HexNAc(2)Hex(1) | Yes | 909.4652 | 3 | 2726.381 | 2726.373 | 2.9 | Non | 745.7 | 742.6 | 742.58 | Nsco_20191108_BC_ZIC_HILIC_K56_2_pepsin_B1.22517.22517.3 | 48.78 | Pepsin | B1 | S164 | 28 |
| BCAL0303 | >gi\|443596149\|gb\|ELT64670.1\| hypothetical protein BURCENK562V_3632 [Burkholderia cenocepacia K56-2Valvano] | L.YSKAADAGGAKPAAGAS[+568.212]AAPAAPAPVAV.P | HexNAc(2)Hex(1) | Yes | 992.8305 | 3 | 2976.4769 | 2976.468 | 2.8 | CRagged | 663.7 | 565.7 | 328.45 | Nsco_20191108_BC_ZIC_HILIC_K56_2_pepsin_B3_reinject.24722.24722.3 | 51.66 | Pepsin | B3 | S164 | 29 |
| BCAL0303 | >gi\|443596149\|gb\|ELT64670.1\| hypothetical protein BURCENK562V_3632 [Burkholderia cenocepacia K56-2Valvano] | L.YSKAADAGGAKPAAGAS[+568.212]AAPAAPAPVAVPASAVSG.S | HexNAc(2)Hex(1) | Partial | 1182.9243 | 3 | 3546.7582 | 3545.749 | 1.6 | CRagged | 275.2 | 256.1 | 19.2 | Nsco_20191108_BC_ZIC_HILIC_K56_2_pepsin_B3_reinject.31278.31278.3 | 64.13 | Pepsin | B3 | NA | 30 |
| BCAL0426 | >gi\|443602087\|gb\|ELT70191.1\| membrane protein insertase, YidC/Oxa1 family [Burkholderia cenocepacia K56-2Valvano] | R.DHGRPSMFFPSATHTAPAAAGGAS[+568.212]GTGATTAGDVPAAAAGAAPSTTAPAAQAQLVK.F | HexNAc(2)Hex(1) | Yes | 1422.4382 | 4 | 5686.7311 | 5683.709 | 2.1 | Specific | 1216 | 733.3 | 78.31 | Nsco_20191108_BC_ZIC_HILIC_K56_2_typsin_B2.40769.40769.4 | 84.14 | Trypsin | B2 | S49 | 31 |
| BCAL0426 | >gi\|443602087\|gb\|ELT70191.1\| membrane protein insertase, YidC/Oxa1 family [Burkholderia cenocepacia K56-2Valvano] | F.FPSATHTAPAAAGGAS[+568.212][+100.064]GTGATTAGDVPAAAAGAAPSTTAPAAQ.A | HexNAc(2)Hex(1) 100.064 | Partial | 1080.2585 | 4 | 4318.0124 | 4317.028 | -4.4 | CRagged | 416 | 416 | 20.65 | Nsco_20191108_BC_ZIC_HILIC_K56_2_pepsin_B2.36655.36655.4 | 78.73 | Pepsin | B2 | NA | 32 |
| BCAL0426 | >gi\|443602087\|gb\|ELT70191.1\| membrane protein insertase, YidC/Oxa1 family [Burkholderia cenocepacia K56-2Valvano] | T.HTAPAAAGGAS[+568.212]GTGATTAGDVPAAAAGAAPSTTAPAAQ.A | HexNAc(2)Hex(1) | Yes | 1238.5833 | 3 | 3713.7354 | 3713.726 | 2.5 | Non | 692.7 | 661.1 | 37.63 | Nsco_20191108_BC_ZIC_HILIC_K56_2_pepsin_B3_reinject.30627.30627.3 | 62.86 | Pepsin | B3 | S49 | 33 |
| BCAL0525 | >gi\|443597648\|gb\|ELT66063.1\| flagellar M-ring protein FliF, partial [Burkholderia cenocepacia K56-2Valvano] | L.SNTPPQPAS[+568.212]APIVAGNGQNAPQTTPVSD.R | HexNAc(2)Hex(1) | Yes | 1095.5191 | 3 | 3284.5427 | 3284.529 | 4.2 | CRagged | 535.3 | 449.8 | 35.51 | Nsco_20191108_BC_ZIC_HILIC_K56_2_pepsin_B3_reinject.37980.37980.3 | 78.38 | Pepsin | B3 | S358 | 34 |
| BCAL0525 | >gi\|443597648\|gb\|ELT66063.1\| flagellar M-ring protein FliF, partial [Burkholderia cenocepacia K56-2Valvano] | R.SQQTSSATELAQGGAS[+568.212]GVPGALSNTPPQPASAPIVAGNGQNAPQTT[+568.212]PVSDRK.D | HexNAc(2)Hex(1),HexNAc(2)Hex(1) | Partial | 1534.984 | 4 | 6136.9142 | 6134.902 | 0.9 | Specific | 328.6 | 328.6 | 81.93 | Nsco_20191108_BC_ZIC_HILIC_K56_2_typsin_B2.40291.40291.4 | 83.32 | Trypsin | B2 | NA | 35 |
| BCAL0544 | >gi\|443601095\|gb\|ELT69255.1\| ABC transporter, substrate-binding protein, family 5 [Burkholderia cenocepacia K56-2Valvano] | K.KAILESVYQGAGQAAS[+568.212]APMPPTQWSYDK.N | HexNAc(2)Hex(1) | Localised | 1188.2346 | 3 | 3562.6891 | 3562.678 | 3.1 | Non | 438.6 | 425.7 | 120.52 | Nsco_20191108_BC_ZIC_HILIC_K56_2_thermolysin_B3.48291.48291.3 | 96.95 | Thermolysin | B3 | S343 | 36 |
| BCAL0678 | >gi\|443605648\|gb\|ELT73487.1\| sporulation and cell division repeat protein [Burkholderia cenocepacia K56-2Valvano] | Q.AAAQKQQQQQQAANT[+568.212]PKPTSSAT.A | HexNAc(2)Hex(1) | Partial | 984.4776 | 3 | 2951.4183 | 2951.408 | 3.6 | Non | 418.1 | 342.9 | 11.7 | Nsco_20191108_BC_ZIC_HILIC_K56_2_thermolysin_B1.4855.4855.3 | 18.56 | Thermolysin | B1 | NA | 37 |

| | | | | | | | | | | | | | | | | | | | |
|---|---|---|---|---|---|---|---|---|---|---|---|---|---|---|---|---|---|---|---|
| BCAL0678 | >gi\|443605648\|gb\|ELT73487.1\| sporulation and cell division repeat protein [Burkholderia cenocepacia K56-2Valvano] | Q.AAAQKQQQQQQAANT[+568.212]PKPTSSATA.A | HexNAc(2)Hex(1) | Partial | 1008.1566 | 3 | 3022.4553 | 3022.445 | 3.5 | Non | 460.6 | 420 | 4.15 | Nsco_20191108_BC_ZIC_HILIC_K56_2_thermolysin_B2_.5227.5229.3 | 20.13 | Thermolysin | B2 | NA | 38 |
| BCAL0678 | >gi\|443605648\|gb\|ELT73487.1\| sporulation and cell division repeat protein [Burkholderia cenocepacia K56-2Valvano] | S.KVAPPPADNGAS[+568.212]QPQQFDPNRA.L | HexNAc(2)Hex(1) | Yes | 958.4554 | 3 | 2873.3517 | 2873.344 | 2.8 | Non | 630.6 | 630.6 | 630.58 | Nsco_20191108_BC_ZIC_HILIC_K56_2_pepsin_B3_reinject.18300.18300.3 | 40.82 | Pepsin | B3 | S39 | 39 |
| BCAL0678 | >gi\|443605648\|gb\|ELT73487.1\| sporulation and cell division repeat protein [Burkholderia cenocepacia K56-2Valvano] | S.KVAPPPADNGAS[+568.212]QPQQFDPNRALQG.K | HexNAc(2)Hex(1) | Yes | 1057.8449 | 3 | 3171.5201 | 3171.508 | 3.9 | Non | 621.4 | 611.9 | 611.92 | Nsco_20191108_BC_ZIC_HILIC_K56_2_pepsin_B1.27694.27694.3 | 59.06 | Pepsin | B1 | S39 | 40 |
| BCAL0678 | >gi\|443605648\|gb\|ELT73487.1\| sporulation and cell division repeat protein [Burkholderia cenocepacia K56-2Valvano] | S.KVAPPPADNGAS[+568.212]QPQQFDPNRALQGKTPGQPVPQA.A | HexNAc(2)Hex(1) | Yes | 1044.5223 | 4 | 4175.0672 | 4175.053 | 3.5 | Non | 743.2 | 743.2 | 488.41 | Nsco_20191108_BC_ZIC_HILIC_K56_2_pepsin_B3_reinject.31852.31852.4 | 65.21 | Pepsin | B3 | S39 | 41 |
| BCAL0678 | >gi\|443605648\|gb\|ELT73487.1\| sporulation and cell division repeat protein [Burkholderia cenocepacia K56-2Valvano] | S.KVAPPPADNGAS[+568.212]QPQQFDPNRALQGKTPGQPVPQAAQPAPPNTA.P | HexNAc(2)Hex(1) | Yes | 1256.3777 | 4 | 5022.4889 | 5022.472 | 3.5 | Non | 665.4 | 665.4 | 320.9 | Nsco_20191108_BC_ZIC_HILIC_K56_2_pepsin_B1.33145.33145.4 | 71.36 | Pepsin | B1 | S39 | 42 |
| BCAL0678 | >gi\|443605648\|gb\|ELT73487.1\| sporulation and cell division repeat protein [Burkholderia cenocepacia K56-2Valvano] | S.KVAPPPADNGAS[+568.212]QPQQFDPNRALQGKTPGQPVPQAAQPAPPNTAPGQ.A | HexNAc(2)Hex(1) | Yes | 1326.9113 | 4 | 5304.6232 | 5304.604 | 3.6 | Non | 836.8 | 836.8 | 393.47 | Nsco_20191108_BC_ZIC_HILIC_K56_2_pepsin_B3_reinject.35229.35229.4 | 72.36 | Pepsin | B3 | S39 | 43 |
| BCAL0678 | >gi\|443605648\|gb\|ELT73487.1\| sporulation and cell division repeat protein [Burkholderia cenocepacia K56-2Valvano] | S.KVAPPPADNGAS[+568.212]QPQQFDPNRALQGKTPGQPVPQAAQPAPPNTAPGQA.A | HexNAc(2)Hex(1) | Yes | 1075.9371 | 5 | 5375.6566 | 5375.642 | 2.8 | Non | 942 | 942 | 522.63 | Nsco_20191108_BC_ZIC_HILIC_K56_2_pepsin_B2.34589.34589.5 | 73.89 | Pepsin | B2 | S39 | 44 |
| BCAL0678 | >gi\|443605648\|gb\|ELT73487.1\| sporulation and cell division repeat protein [Burkholderia cenocepacia K56-2Valvano] | S.KVAPPPADNGAS[+568.212]QPQQFDPNRALQGKTPGQPVPQAAQPAPPNTAPGQAA.N | HexNAc(2)Hex(1) | Yes | 1362.4287 | 4 | 5446.693 | 5446.679 | 2.7 | Non | 807.9 | 807.9 | 435.58 | Nsco_20191108_BC_ZIC_HILIC_K56_2_pepsin_B1.33981.33981.4 | 73.55 | Pepsin | B1 | S39 | 45 |
| BCAL0678 | >gi\|443605648\|gb\|ELT73487.1\| sporulation and cell division repeat protein [Burkholderia cenocepacia K56-2Valvano] | S.KVAPPPADNGAS[+568.212]QPQQFDPNRALQGKTPGQPVPQAAQPAPPNTAPGQAAN.Q | HexNAc(2)Hex(1) | Yes | 1113.3529 | 5 | 5562.7354 | 5560.722 | 1.3 | Non | 1015 | 1001 | 483.05 | Nsco_20191108_BC_ZIC_HILIC_K56_2_pepsin_B1.33588.33588.5 | 72.44 | Pepsin | B1 | S39 | 46 |
| BCAL0678 | >gi\|443605648\|gb\|ELT73487.1\| sporulation and cell division repeat protein [Burkholderia cenocepacia K56-2Valvano] | S.KVAPPPADNGAS[+568.212]QPQQFDPNRALQGKTPGQPVPQAAQPAPPNTAPGQAANQ.T | HexNAc(2)Hex(1) | Yes | 1422.9518 | 4 | 5688.7853 | 5688.78 | 0.9 | Non | 680.9 | 672.1 | 302.36 | Nsco_20191108_BC_ZIC_HILIC_K56_2_pepsin_B3_reinject.35184.35184.4 | 72.26 | Pepsin | B3 | S39 | 47 |
| BCAL0678 | >gi\|443605648\|gb\|ELT73487.1\| sporulation and cell division repeat protein [Burkholderia cenocepacia K56-2Valvano] | K.PT[+568.212]SSATAAAAAKPPTANDANTGYFLQVGAYK.T | HexNAc(2)Hex(1) | Partial | 1208.9186 | 3 | 3624.7412 | 3622.728 | 1.7 | Specific | 447.8 | 406.4 | 1.8 | Nsco_20191108_BC_ZIC_HILIC_K56_2_typsin_B1.39530.39530.3 | 81.3 | Trypsin | B1 | NA | 48 |
| BCAL0678 | >gi\|443605648\|gb\|ELT73487.1\| sporulation and cell division repeat protein [Burkholderia cenocepacia K56-2Valvano] | Q.QAANT[+568.212][+100.064]PKPTSS[+568.212][+100.064]ATAAAAAKPPTANDANTGYFLQVGAYK.T | HexNAc(2)Hex(1)100.064,HexNAc(2)Hex(1) | Partial | 1276.3612 | 4 | 5102.423 | 5101.439 | -3.8 | NRagged | 178.7 | 178.7 | 178.68 | Nsco_20191108_BC_ZIC_HILIC_K56_2_typsin_B1.36658.36658.4 | 76.27 | Trypsin | B1 | NA | 49 |
| BCAL0678 | >gi\|443605648\|gb\|ELT73487.1\| sporulation and cell division repeat protein [Burkholderia cenocepacia K56-2Valvano] | Q.QAANT[+568.212][+100.064]PKPTSSATAAAAAKPPT[+568.212]ANDANTGYFLQVGAYKTEGDAEQQR.A | HexNAc(2)Hex(1)100.064,HexNAc(2)Hex(1)100.064 | Partial | 1204.7736 | 5 | 6019.8387 | 6015.812 | 2.3 | NRagged | 275.4 | 234.3 | 39.05 | Nsco_20191108_BC_ZIC_HILIC_K56_2_typsin_B1.34882.34882.5 | 73.1 | Trypsin | B1 | NA | 50 |
| BCAL0678 | >gi\|443605648\|gb\|ELT73487.1\| sporulation and cell division repeat protein [Burkholderia cenocepacia K56-2Valvano] | A.QKQQQQQQAANT[+568.212]PKPTSSATAAA.A | HexNAc(2)Hex(1) | Partial | 1476.2091 | 2 | 2951.4109 | 2951.408 | 1.1 | Non | 209.4 | 187.4 | 13.8 | Nsco_20191108_BC_ZIC_HILIC_K56_2_pepsin_B1.7370.7370.2 | 22.89 | Pepsin | B1 | NA | 51 |

| | | | | | | | | | | | | | | | | | | | |
|---|---|---|---|---|---|---|---|---|---|---|---|---|---|---|---|---|---|---|---|
| BCAL0678 | >gi\|443605648\|gb\|ELT73487.1\| sporulation and cell division repeat protein [Burkholderia cenocepacia K56-2Valvano] | Q.QQAANT[+568.212]PKPTS[+568.212]SATAAAAAKPPTANDANTGYFLQVGAYK.T | HexNAc(2)Hex(1),HexNAc(2)Hex(1) | Partial | 1258.1008 | 4 | 5029.3815 | 5029.37 | 2.4 | NRagged | 262.2 | 262.2 | 6.08 | Nsco_20191108_BC_ZIC_HILIC_K56_2_typsin_B2.35560.35560.4 | 74.65 | Trypsin | B2 | NA | 52 |
| BCAL0678 | >gi\|443605648\|gb\|ELT73487.1\| sporulation and cell division repeat protein [Burkholderia cenocepacia K56-2Valvano] | Q.QQAANTPKPTS[+568.212]SATAAAAAKPPT[+568.212]ANDANTGYFLQVGAYKTEGDAEQQR.A | HexNAc(2)Hex(1),HexNAc(2)Hex(1) | Partial | 1209.5704 | 5 | 6043.8231 | 6043.806 | 2.8 | NRagged | 538.3 | 520.2 | 22.54 | Nsco_20191108_BC_ZIC_HILIC_K56_2_typsin_B1.33924.33924.5 | 71.43 | Trypsin | B1 | NA | 53 |
| BCAL0678 | >gi\|443605648\|gb\|ELT73487.1\| sporulation and cell division repeat protein [Burkholderia cenocepacia K56-2Valvano] | K.QQQQQQAANTPKPTS[+568.212]SATAAAAAKPPTANDANTGYFLQV.G | HexNAc(2)Hex(1) | Yes | 1519.0698 | 3 | 4555.1947 | 4554.175 | 3.5 | Non | 206.8 | 132 | 1.06 | Nsco_20191108_BC_ZIC_HILIC_K56_2_thermolysin_B3.38670.38670.3 | 79.71 | Thermolysin | B3 | S180 | 54 |
| BCAL0678 | >gi\|443605648\|gb\|ELT73487.1\| sporulation and cell division repeat protein [Burkholderia cenocepacia K56-2Valvano] | K.QQQQQQAANTPKPTS[+568.212]SATAAAAAKPPTANDANTGYFLQVGAYK.T | HexNAc(2)Hex(1) | Yes | 1244.1025 | 4 | 4973.3883 | 4973.392 | -0.8 | Specific | 1231 | 607.5 | 26.24 | Nsco_20191108_BC_ZIC_HILIC_K56_2_typsin_B3.35416.35416.4 | 73.18 | Trypsin | B3 | S180 | 55 |
| BCAL0678 | >gi\|443605648\|gb\|ELT73487.1\| sporulation and cell division repeat protein [Burkholderia cenocepacia K56-2Valvano] | K.QQQQQQAANTPKPTSSATAAAAAKPPT[+568.212]ANDANTGYFLQVGAYKTEGDAEQQR.A | HexNAc(2)Hex(1) | Partial | 1198.5741 | 5 | 5988.8414 | 5987.829 | 1.5 | Specific | 798 | 508.6 | 34.71 | Nsco_20191108_BC_ZIC_HILIC_K56_2_typsin_B1.34007.34007.5 | 71.56 | Trypsin | B1 | NA | 56 |
| BCAL0678 | >gi\|443605648\|gb\|ELT73487.1\| sporulation and cell division repeat protein [Burkholderia cenocepacia K56-2Valvano] | K.VAPPPADNGAS[+568.212]QPQQFDPNR.A | HexNAc(2)Hex(1) | Yes | 1337.6141 | 2 | 2674.2208 | 2674.212 | 3.5 | Specific | 607.5 | 540 | 540.04 | Nsco_20191108_BC_ZIC_HILIC_K56_2_typsin_B2.21592.21592.2 | 48.6 | Trypsin | B2 | S39 | 57 |
| BCAL0678 | >gi\|443605648\|gb\|ELT73487.1\| sporulation and cell division repeat protein [Burkholderia cenocepacia K56-2Valvano] | K.VAPPPADNGAS[+568.212]QPQQFDPNRALQGK.T | HexNAc(2)Hex(1) | Yes | 1057.8439 | 3 | 3171.5171 | 3171.508 | 3 | Specific | 699.6 | 699.6 | 699.57 | Nsco_20191108_BC_ZIC_HILIC_K56_2_typsin_B3.27804.27804.3 | 60.07 | Trypsin | B3 | S39 | 58 |
| BCAL0678 | >gi\|443605648\|gb\|ELT73487.1\| sporulation and cell division repeat protein [Burkholderia cenocepacia K56-2Valvano] | F.VSKVAPPPADNGAS[+568.212]QPQQFDPNRA.L | HexNAc(2)Hex(1) | Yes | 1020.4881 | 3 | 3059.4496 | 3059.444 | 1.8 | CRagged | 233.1 | 216.8 | 126.3 | Nsco_20191108_BC_ZIC_HILIC_K56_2_pepsin_B1.20586.20586.3 | 45.12 | Pepsin | B1 | S39 | 59 |
| BCAL0738 | >gi\|443604531\|gb\|ELT72458.1\| peptidase, S41 family [Burkholderia cenocepacia K56-2Valvano] | K.ASGASAAKPAS[+568.212]APKPASAPK.- | HexNAc(2)Hex(1) | Yes | 778.3989 | 3 | 2333.1821 | 2333.172 | 4.4 | Specific | 265.2 | 187.9 | 46.49 | Nsco_20191108_BC_ZIC_HILIC_K56_2_typsin_B3.4910.4910.4 | 19.47 | Trypsin | B3 | S508 | 60 |
| BCAL0749 | >gi\|443604502\|gb\|ELT72429.1\| cytochrome c oxidase, subunit II [Burkholderia cenocepacia K56-2Valvano] | R.NGKLPEDTAGAAT[+568.212]AAAPAEAASAPAQAAS[+568.212]GAEQPAAAASAALSTIYFETGK.S | HexNAc(2)Hex(1),HexNAc(2)Hex(1) | Partial (1 of 2) | 1463.693 | 4 | 5851.7502 | 5851.73 | 3.4 | Specific | 1107 | 1018 | 111.99 | Nsco_20191108_BC_ZIC_HILIC_K56_2_typsin_B3.61271.61271.4 | 117.8 | Trypsin | B3 | S410 | 61 |
| BCAL0749 | >gi\|443604502\|gb\|ELT72429.1\| cytochrome c oxidase, subunit II [Burkholderia cenocepacia K56-2Valvano] | L.PEDTAGAAT[+568.212][+100.064]AAAPAEAASAPAQAAS[+568.212][+100.064]GAEQPAAAASAALSTIYFETGK.S | HexNAc(2)Hex(1)100.064,HexNAc(2)Hex(1)100.064 | Partial (1 of 2) | 1410.6604 | 4 | 5639.6198 | 5639.615 | 0.9 | CRagged | 273.1 | 268.3 | 3.02 | Nsco_20191108_BC_ZIC_HILIC_K56_2_thermolysin_B3.60729.60729.4 | 120.7 | Thermolysin | B3 | S410 | 62 |
| BCAL0749 | >gi\|443604502\|gb\|ELT72429.1\| cytochrome c oxidase, subunit II [Burkholderia cenocepacia K56-2Valvano] | K.QVADARNGKLPEDTAGAATAAAPAEAAS[+568.212][+100.064]APAQAASGAEQPAAAASAALSTIYFETGK.S | HexNAc(2)Hex(1) | Partial | 1506.9807 | 4 | 6024.901 | 6023.912 | -2.4 | Specific | 362.8 | 362.8 | 47.23 | Nsco_20191108_BC_ZIC_HILIC_K56_2_typsin_B2.61326.61326.4 | 122.3 | Trypsin | B2 | NA | 63 |
| BCAL0786 | >gi\|443598725\|gb\|ELT67055.1\| hypothetical protein BURCENK562V_1882 [Burkholderia cenocepacia K56-2Valvano] | K.LLSVPAPAS[+568.212]TEGDHHDK.- | HexNAc(2)Hex(1) | Yes | 586.2788 | 4 | 2342.0932 | 2342.088 | 2.2 | Specific | 499.2 | 439.7 | 35.82 | Nsco_20191108_BC_ZIC_HILIC_K56_2_typsin_B3.14440.14440.4 | 35.98 | Trypsin | B3 | S80 | 64 |
| BCAL1086 | >gi\|443597491\|gb\|ELT65913.1\| putative lipoprotein [Burkholderia cenocepacia K56-2Valvano] | K.ALDQVAS[+568.212]TVNQQINAAK.A | HexNAc(2)Hex(1) | Partial | 1170.0798 | 2 | 2339.1522 | 2339.146 | 2.7 | Specific | 339.4 | 163 | 163.02 | Nsco_20191108_BC_ZIC_HILIC_K56_2_typsin_B1.35454.35454.2 | 74 | Trypsin | B1 | NA | 65 |
| BCAL1086 | >gi\|443597491\|gb\|ELT65913.1\| putative lipoprotein [Burkholderia cenocepacia K56-2Valvano] | Q.IDAAAS[+568.212]AVVAH.A | HexNAc(2)Hex(1) | Yes | 796.8835 | 2 | 1592.7597 | 1592.754 | 3.7 | Non | 394.4 | 147.3 | 147.31 | Nsco_20191108_BC_ZIC_HILIC_K56_2_thermolysin_B2_.18308.18308.2 | 39.2 | Thermolysin | B2 | S104 | 66 |

| | | | | | | | | | | | | | | | | | | | |
|---|---|---|---|---|---|---|---|---|---|---|---|---|---|---|---|---|---|---|---|
| BCAL1086 | >gi\|443597491\|gb\|ELT65913.1\| putative lipoprotein [Burkholderia cenocepacia K56-2Valvano] | K.KLQQWSQQSAAGAKPAS[+568.212]GE.- | HexNAc(2)Hex(1) | Yes | 1270.6071 | 2 | 2540.2069 | 2540.2 | 2.8 | CRagged | 331.7 | 204.8 | 204.84 | Nsco_20191108_BC_ZIC_HILIC_K56_2_typsin_B2.11121.11121.2 | 29.91 | Trypsin | B2 | S140 | 67 |
| BCAL1086 | >gi\|443597491\|gb\|ELT65913.1\| putative lipoprotein [Burkholderia cenocepacia K56-2Valvano] | E.KSGTPVAQPDTAAS[+568.212]AAAD.A | HexNAc(2)Hex(1) | Yes | 1113.5132 | 2 | 2226.0191 | 2226.014 | 2.1 | Non | 253.6 | 143 | 143 | Nsco_20191108_BC_ZIC_HILIC_K56_2_pepsin_B1.13983.13983.2 | 33.95 | Pepsin | B1 | S45 | 68 |
| BCAL1086 | >gi\|443597491\|gb\|ELT65913.1\| putative lipoprotein [Burkholderia cenocepacia K56-2Valvano] | E.KSGTPVAQPDTAAS[+568.212]AAADAANNAAKAL.D | HexNAc(2)Hex(1) | Yes | 1017.4953 | 3 | 3050.4713 | 3050.465 | 2.1 | NRagged | 380.7 | 265.4 | 30.76 | Nsco_20191108_BC_ZIC_HILIC_K56_2_pepsin_B1.37649.37649.4 | 82.74 | Pepsin | B1 | S45 | 69 |
| BCAL1086 | >gi\|443597491\|gb\|ELT65913.1\| putative lipoprotein [Burkholderia cenocepacia K56-2Valvano] | K.LQQWSQQSAAGAKPAS[+568.212]GE.- | HexNAc(2)Hex(1) | Yes | 1206.559 | 2 | 2412.1107 | 2412.105 | 2.4 | CRagged | 283.9 | 121.9 | 67.95 | Nsco_20191108_BC_ZIC_HILIC_K56_2_typsin_B1.15007.15007.2 | 36.76 | Trypsin | B1 | S140 | 70 |
| BCAL1093 | >gi\|443599421\|gb\|ELT67702.1\| hypothetical protein BURCENK562V_2687 [Burkholderia cenocepacia K56-2Valvano] | K.HAYDEAHPAEAAS[+568.212]AASH.- | HexNAc(2)Hex(1) | Yes | 768.3271 | 3 | 2302.9666 | 2302.958 | 3.6 | CRagged | 304.4 | 251.1 | 92.55 | Nsco_20191108_BC_ZIC_HILIC_K56_2_typsin_B3.7340.7340.3 | 24.24 | Trypsin | B3 | S49 | 71 |
| BCAL1453 | >gi\|443604751\|gb\|ELT72660.1\| HlyD family secretion protein [Burkholderia cenocepacia K56-2Valvano] | R.ENAAHSAS[+568.212]SADAQYQQAIAALDAAK.L | HexNAc(2)Hex(1) | Yes | 1024.4741 | 3 | 3071.4078 | 3070.397 | 2.4 | Specific | 263.2 | 209.4 | 14.32 | Nsco_20191108_BC_ZIC_HILIC_K56_2_typsin_B2.39533.39533.3 | 82 | Trypsin | B2 | S130 | 72 |
| BCAL1453 | >gi\|443604751\|gb\|ELT72660.1\| HlyD family secretion protein [Burkholderia cenocepacia K56-2Valvano] | K.ENRENAAHSAS[+568.212]SADAQYQQAIAALDAAK.L | HexNAc(2)Hex(1) | Yes | 868.1537 | 4 | 3469.5931 | 3469.584 | 2.7 | Specific | 1112 | 987.1 | 56.26 | Nsco_20191108_BC_ZIC_HILIC_K56_2_typsin_B3.34661.34661.4 | 71.93 | Trypsin | B3 | S130 | 73 |
| BCAL1496 | >gi\|443597624\|gb\|ELT66041.1\| hypothetical protein BURCENK562V_3510 [Burkholderia cenocepacia K56-2Valvano] | K.AAAKKAGKKAKAADAAS[+568.212]Q.- | HexNAc(2)Hex(1) | Yes | 564.3015 | 4 | 2254.1841 | 2254.177 | 3 | Non | 480.3 | 448.6 | 448.6 | Nsco_20191108_BC_ZIC_HILIC_K56_2_thermolysin_B2_.2250.2250.4 | 13.92 | Thermolysin | B2 | S90 | 74 |
| BCAL1496 | >gi\|443597624\|gb\|ELT66041.1\| hypothetical protein BURCENK562V_3510 [Burkholderia cenocepacia K56-2Valvano] | A.AAKKAGKKAKAADAAS[+568.212]Q.- | HexNAc(2)Hex(1) | Yes | 546.5424 | 4 | 2183.1477 | 2183.14 | 3.5 | Non | 455.2 | 455.2 | 455.21 | Nsco_20191108_BC_ZIC_HILIC_K56_2_thermolysin_B2_.2204.2204.4 | 13.71 | Thermolysin | B2 | S90 | 75 |
| BCAL1496 | >gi\|443597624\|gb\|ELT66041.1\| hypothetical protein BURCENK562V_3510 [Burkholderia cenocepacia K56-2Valvano] | K.AGKKAKAADAAS[+568.212]Q.- | HexNAc(2)Hex(1) | Yes | 595.632 | 3 | 1784.8815 | 1784.876 | 3.1 | Non | 193.6 | 183.9 | 183.89 | Nsco_20191108_BC_ZIC_HILIC_K56_2_thermolysin_B1.2214.2214.3 | 13.4 | Thermolysin | B1 | S90 | 76 |
| BCAL1496 | >gi\|443597624\|gb\|ELT66041.1\| hypothetical protein BURCENK562V_3510 [Burkholderia cenocepacia K56-2Valvano] | K.AKAADAAS[+568.212]Q.- | HexNAc(2)Hex(1) | Yes | 700.8192 | 2 | 1400.631 | 1400.628 | 2.5 | CRagged | 403.4 | 208.7 | 208.72 | Nsco_20191108_BC_ZIC_HILIC_K56_2_typsin_B3.2167.2167.2 | 14.41 | Trypsin | B3 | S90 | 77 |
| BCAL1496 | >gi\|443597624\|gb\|ELT66041.1\| hypothetical protein BURCENK562V_3510 [Burkholderia cenocepacia K56-2Valvano] | H.KAAAKKAGKKAKAADAAS[+568.212]Q.- | HexNAc(2)Hex(1) | Yes | 596.3253 | 4 | 2382.2795 | 2382.272 | 3.1 | Non | 709.7 | 648.2 | 648.15 | Nsco_20191108_BC_ZIC_HILIC_K56_2_pepsin_B3_reinject.2004.2004.4 | 13.57 | Pepsin | B3 | S90 | 78 |
| BCAL1496 | >gi\|443597624\|gb\|ELT66041.1\| hypothetical protein BURCENK562V_3510 [Burkholderia cenocepacia K56-2Valvano] | K.KAGKKAKAADAAS[+568.212]Q.- | HexNAc(2)Hex(1) | Yes | 638.3299 | 3 | 1912.9752 | 1912.971 | 2.2 | Non | 346.5 | 250.2 | 250.15 | Nsco_20191108_BC_ZIC_HILIC_K56_2_pepsin_B3_reinject.1926.1926.3 | 13.28 | Pepsin | B3 | S90 | 79 |
| BCAL1496 | >gi\|443597624\|gb\|ELT66041.1\| hypothetical protein BURCENK562V_3510 [Burkholderia cenocepacia K56-2Valvano] | A.KKAGKKAKAADAAS[+568.212]Q.- | HexNAc(2)Hex(1) | Yes | 511.0234 | 4 | 2041.0719 | 2041.066 | 2.9 | Non | 349.8 | 268.4 | 268.41 | Nsco_20191108_BC_ZIC_HILIC_K56_2_pepsin_B2.1914.1914.4 | 13.27 | Pepsin | B2 | S90 | 80 |
| BCAL1496 | >gi\|443597624\|gb\|ELT66041.1\| hypothetical protein BURCENK562V_3510 [Burkholderia cenocepacia K56-2Valvano] | G.KKAKAADAAS[+568.212]Q.- | HexNAc(2)Hex(1) | Yes | 552.9457 | 3 | 1656.8225 | 1656.817 | 3.1 | Non | 367.4 | 216.1 | 216.13 | Nsco_20191108_BC_ZIC_HILIC_K56_2_pepsin_B1.1926.1926.3 | 13.11 | Pepsin | B1 | S90 | 81 |
| BCAL1674 | >gi\|443605593\|gb\|ELT73435.1\| periplasmic multidrug efflux lipoprotein [Burkholderia cenocepacia K56-2Valvano] | P.VEKAPSSKAAPPAAAS[+568.212]QA.A | HexNAc(2)Hex(1) | Yes | 1125.0594 | 2 | 2249.1115 | 2249.103 | 3.7 | Non | 388 | 231.4 | 187.03 | Nsco_20191108_BC_ZIC_HILIC_K56_2_thermolysin_B1.7802.7802.2 | 23.32 | Thermolysin | B1 | S404 | 82 |
| BCAL1746 | >gi\|443592790\|gb\|ELT61568.1\| putative lipoprotein [Burkholderia cenocepacia K56-2Valvano] | L.IDHIGKAWPGNAAS[+568.212]GASASASE.- | HexNAc(2)Hex(1) | Yes | 889.4119 | 3 | 2666.221 | 2665.211 | 2.5 | CRagged | 334.5 | 226.7 | 49.17 | Nsco_20191108_BC_ZIC_HILIC_K56_2_pepsin_B1.22279.22279.3 | 48.33 | Pepsin | B1 | S140 | 83 |

| | | | | | | | | | | | | | | | | | | | |
|---|---|---|---|---|---|---|---|---|---|---|---|---|---|---|---|---|---|---|---|
| BCAL1798 | >gi\|443598613\|gb\|ELT66949.1\| hypothetical protein BURCENK562V_2315 [Burkholderia cenocepacia K56-2Valvano] | T.AATPAAASAPS[+568.212]AEQRAARHEARIEQ.R | HexNAc(2)Hex(1) | Partial | 1043.5109 | 3 | 3128.5181 | 3128.509 | 2.9 | Non | 503.5 | 426 | 7.2 | Nsco_20191108_BC_ZIC_HILIC_K56_2_pepsin_B3_reinject.12565.12565.3 | 31.5 | Pepsin | B3 | NA | 84 |
| BCAL1869 | >gi\|443596292\|gb\|ELT64810.1\| hypothetical protein BURCENK562V_2581 [Burkholderia cenocepacia K56-2Valvano] | P.VAAS[+568.212]AVPDFDARQK.V | HexNAc(2)Hex(1) | Yes | 681.6658 | 3 | 2042.9828 | 2042.976 | 3.1 | Non | 394.4 | 175.1 | 175.06 | Nsco_20191108_BC_ZIC_HILIC_K56_2_thermolysin_B1.18482.18482.3 | 39.94 | Thermolysin | B1 | S29 | 85 |
| BCAL1877 | >gi\|443594658\|gb\|ELT63293.1\| HflK protein [Burkholderia cenocepacia K56-2Valvano] | A.ATAS[+568.212]GVDVLRSRE.A | HexNAc(2)Hex(1) | Yes | 964.9725 | 2 | 1928.9376 | 1928.93 | 4.2 | Non | 171.9 | 59.51 | 59.51 | Nsco_20191108_BC_ZIC_HILIC_K56_2_thermolysin_B1.14431.14431.2 | 33.54 | Thermolysin | B1 | S427 | 86 |
| BCAL1877 | >gi\|443594658\|gb\|ELT63293.1\| HflK protein [Burkholderia cenocepacia K56-2Valvano] | L.VEQGRQNAAAST[+568.212]GASAADAAS[+568.212]APAATVPSAA.A | HexNAc(2)Hex(1),HexNAc(2)Hex(1) | Partial | 1302.5969 | 3 | 3905.776 | 3905.763 | 3.3 | CRagged | 179.8 | 79.69 | 16.04 | Nsco_20191108_BC_ZIC_HILIC_K56_2_pepsin_B2.26026.26026.3 | 55.76 | Pepsin | B2 | NA | 87 |
| BCAL1917 | >gi\|443605668\|gb\|ELT73505.1\| hypothetical protein BURCENK562V_0636 [Burkholderia cenocepacia K56-2Valvano] | K.AAAAAASADAGAS[+568.212]APAAASST[+568.212]K.A | HexNAc(2)Hex(1),HexNAc(2)Hex(1) | Partial (1 of 2) | 975.7731 | 3 | 2925.3047 | 2925.295 | 3.2 | Specific | 399.5 | 259.7 | 93.53 | Nsco_20191108_BC_ZIC_HILIC_K56_2_typsin_B2.9570.9570.3 | 27.51 | Trypsin | B2 | S184 | 88 |
| BCAL1917 | >gi\|443605668\|gb\|ELT73505.1\| hypothetical protein BURCENK562V_0636 [Burkholderia cenocepacia K56-2Valvano] | T.AAPAPTAS[+568.212]APEAAAKPAKTKR.A | HexNAc(2)Hex(1) | Yes | 644.0907 | 4 | 2573.341 | 2573.331 | 4.1 | Non | 928.6 | 779.6 | 128.8 | Nsco_20191108_BC_ZIC_HILIC_K56_2_thermolysin_B1.5828.5828.4 | 20.43 | Thermolysin | B1 | S151 | 89 |
| BCAL1917 | >gi\|443605668\|gb\|ELT73505.1\| hypothetical protein BURCENK562V_0636 [Burkholderia cenocepacia K56-2Valvano] | T.AAPAPTAS[+568.212]APEAAAKPAKTKRA.S | HexNAc(2)Hex(1) | Yes | 882.1293 | 3 | 2644.3733 | 2644.368 | 2.2 | Non | 263.7 | 185 | 50.94 | Nsco_20191108_BC_ZIC_HILIC_K56_2_thermolysin_B2_.5922.5922.3 | 21.24 | Thermolysin | B2 | S151 | 90 |
| BCAL1917 | >gi\|443605668\|gb\|ELT73505.1\| hypothetical protein BURCENK562V_0636 [Burkholderia cenocepacia K56-2Valvano] | T.AAPAPTAS[+568.212]APEAAAKPAKTKRASKKEKA.A | HexNAc(2)Hex(1) | Yes | 663.9601 | 5 | 3315.7712 | 3315.764 | 2.1 | Non | 1029 | 993.3 | 140.99 | Nsco_20191108_BC_ZIC_HILIC_K56_2_thermolysin_B2_.3999.3999.5 | 18.07 | Thermolysin | B2 | S151 | 91 |
| BCAL1917 | >gi\|443605668\|gb\|ELT73505.1\| hypothetical protein BURCENK562V_0636 [Burkholderia cenocepacia K56-2Valvano] | T.AAPAPTAS[+568.212]APEAAAKPAKTKRASKKEKAA.A | HexNAc(2)Hex(1) | Yes | 678.167 | 5 | 3386.8058 | 3386.801 | 1.3 | Non | 940.1 | 902.6 | 127.47 | Nsco_20191108_BC_ZIC_HILIC_K56_2_thermolysin_B2_.4052.4052.4 | 18.15 | Thermolysin | B2 | S151 | 92 |
| BCAL1917 | >gi\|443605668\|gb\|ELT73505.1\| hypothetical protein BURCENK562V_0636 [Burkholderia cenocepacia K56-2Valvano] | P.AATTSATTSTTTSAGTASTSTATAGTTTAAPAPTAS[+568.212]APEAAAKPAKTKR.A | HexNAc(2)Hex(1) | Partial | 1027.9116 | 5 | 5135.529 | 5135.509 | 4 | Non | 652.6 | 652.6 | 1.99 | Nsco_20191108_BC_ZIC_HILIC_K56_2_thermolysin_B2_.17866.17866.5 | 38.5 | Thermolysin | B2 | NA | 93 |
| BCAL1917 | >gi\|443605668\|gb\|ELT73505.1\| hypothetical protein BURCENK562V_0636 [Burkholderia cenocepacia K56-2Valvano] | T.AGTTTAAPAPTAS[+568.212]APEAAAKPAKTKR.A | HexNAc(2)Hex(1) | Yes | 751.8917 | 4 | 3004.5449 | 3004.532 | 4.3 | Non | 907.5 | 830.6 | 99.82 | Nsco_20191108_BC_ZIC_HILIC_K56_2_thermolysin_B2_.8205.8205.4 | 24.64 | Thermolysin | B2 | S151 | 94 |
| BCAL1917 | >gi\|443605668\|gb\|ELT73505.1\| hypothetical protein BURCENK562V_0636 [Burkholderia cenocepacia K56-2Valvano] | T.ATAGTTTAAPAPTAS[+568.212]APEAAAKPAKTKR.A | HexNAc(2)Hex(1) | Yes | 1059.5459 | 3 | 3176.6232 | 3176.617 | 2 | Non | 242.4 | 240.3 | 40.27 | Nsco_20191108_BC_ZIC_HILIC_K56_2_thermolysin_B2_.8996.8996.3 | 25.75 | Thermolysin | B2 | S151 | 95 |
| BCAL1917 | >gi\|443605668\|gb\|ELT73505.1\| hypothetical protein BURCENK562V_0636 [Burkholderia cenocepacia K56-2Valvano] | A.ATTSATTSTTTSAGTASTSTTTAGTTTAAPAPTASAPEAAAKPAKT[+568.212]KR.A | HexNAc(2)Hex(1) | Partial | 1013.9047 | 5 | 5065.4942 | 5064.471 | 3.8 | Non | 375.1 | 375.1 | 38.72 | Nsco_20191108_BC_ZIC_HILIC_K56_2_thermolysin_B2_.17346.17346.5 | 37.74 | Thermolysin | B2 | NA | 96 |
| BCAL1917 | >gi\|443605668\|gb\|ELT73505.1\| hypothetical protein BURCENK562V_0636 [Burkholderia cenocepacia K56-2Valvano] | S.ATTSTTTSAGTASTSTTATACTTAAPAPTAS[+568.212]APEAAAK.P | HexNAc(2)Hex(1) | Yes | 1341.9662 | 3 | 4023.884 | 4022.878 | 0.6 | Non | 603.2 | 547.8 | 37.41 | Nsco_20191108_BC_ZIC_HILIC_K56_2_thermolysin_B3.24381.24381.3 | 53.09 | Thermolysin | B3 | S151 | 97 |
| BCAL1917 | >gi\|443605668\|gb\|ELT73505.1\| hypothetical protein BURCENK562V_0636 [Burkholderia cenocepacia K56-2Valvano] | S.ATTSTTTSAGTASTSTTATACTTAAPAPTAS[+568.212]APEAAAKPAK.T | HexNAc(2)Hex(1) | Yes | 1440.3646 | 3 | 4319.0791 | 4319.063 | 3.7 | NRagged | 349.6 | 270.5 | 51.3 | Nsco_20191108_BC_ZIC_HILIC_K56_2_typsin_B3.20286.20286.3 | 46.2 | Trypsin | B3 | S151 | 98 |
| BCAL1917 | >gi\|443605668\|gb\|ELT73505.1\| hypothetical protein BURCENK562V_0636 [Burkholderia cenocepacia K56-2Valvano] | A.GTASTSTTATAGTTTAAPAPTAS[+568.212]APEAAAKPAK.T | HexNAc(2)Hex(1) | Yes | 1166.5707 | 3 | 3497.6977 | 3497.687 | 3.2 | Non | 314.1 | 293.6 | 41.62 | Nsco_20191108_BC_ZIC_HILIC_K56_2_thermolysin_B3.17558.17558.3 | 40.46 | Thermolysin | B3 | S151 | 99 |
| BCAL1917 | >gi\|443605668\|gb\|ELT73505.1\| hypothetical protein BURCENK562V_0636 [Burkholderia cenocepacia K56-2Valvano] | K.LSKPAATTSATTSTTTSAGTASTSTTATAGTTTAAPAPTAS[+568.212]APEAAAK.P | HexNAc(2)Hex(1) | Yes | 1220.595 | 4 | 4879.3582 | 4879.344 | 3 | Non | 743.2 | 678.9 | 64.85 | Nsco_20191108_BC_ZIC_HILIC_K56_2_thermolysin_B3.24903.24903.4 | 54.02 | Thermolysin | B3 | S151 | 100 |

| | | | | | | | | | | | | | | | | | | | |
|---|---|---|---|---|---|---|---|---|---|---|---|---|---|---|---|---|---|---|---|
| BCAL1917 | >gi\|443605668\|gb\|ELT73505.1\| hypothetical protein BURCENK562V_0636 [Burkholderia cenocepacia K56-2Valvano] | K.LSKPAATTSATTSTTTSAGTA STSTTATAGTTTAAPAPTAS[+ 568.212]APEAAAKPA.K | HexNAc(2)Hex(1) | Yes | 1262.6184 | 4 | 5047.4518 | 5047.434 | 3.6 | Non | 1202 | 1202 | 99.55 | Nsco_20191108_BC_ZIC_HILIC_K56_2_thermolysin_B3.26332.26332.4 | 56.76 | Thermolysin | B3 | S151 | 101 |
| BCAL1917 | >gi\|443605668\|gb\|ELT73505.1\| hypothetical protein BURCENK562V_0636 [Burkholderia cenocepacia K56-2Valvano] | K.LSKPAATTSATTSTTTSAGTA STSTTATAGTTTAAPAPTAS[+ 568.212]APEAAAKPAK.T | HexNAc(2)Hex(1) | Yes | 1294.6409 | 4 | 5175.5417 | 5175.529 | 2.5 | Non | 1556 | 1556 | 95.66 | Nsco_20191108_BC_ZIC_HILIC_K56_2_thermolysin_B3.21919.21919.4 | 48.63 | Thermolysin | B3 | S151 | 102 |
| BCAL1917 | >gi\|443605668\|gb\|ELT73505.1\| hypothetical protein BURCENK562V_0636 [Burkholderia cenocepacia K56-2Valvano] | K.LSKPAATTSATTSTTTSAGTA STSTTATAGTTTAAPAPTASAP EAAAKPAKT[+568.212].K | HexNAc(2)Hex(1) | Partial | 1056.3196 | 5 | 5277.5688 | 5276.576 | -2.1 | Non | 1203 | 1203 | 118.24 | Nsco_20191108_BC_ZIC_HILIC_K56_2_thermolysin_B3.23031.23031.5 | 50.72 | Thermolysin | B3 | NA | 103 |
| BCAL1917 | >gi\|443605668\|gb\|ELT73505.1\| hypothetical protein BURCENK562V_0636 [Burkholderia cenocepacia K56-2Valvano] | K.PAATTSATTSTTTSAGTASTS TATAGTTTAAPAPTAS[+568. 212]APEAAAKPAK.T | HexNAc(2)Hex(1) | Yes | 1616.7836 | 3 | 4848.3362 | 4847.318 | 3.2 | Specific | 284.7 | 231.1 | 16.42 | Nsco_20191108_BC_ZIC_HILIC_K56_2_typsin_B1.23256.23256.3 | 52.28 | Trypsin | B1 | S151 | 104 |
| BCAL1917 | >gi\|443605668\|gb\|ELT73505.1\| hypothetical protein BURCENK562V_0636 [Burkholderia cenocepacia K56-2Valvano] | T.SAGTASTSTTATAGTTTAAP PTAS[+568.212]APEAAAK PAK.T | HexNAc(2)Hex(1) | Yes | 1219.5968 | 3 | 3656.7759 | 3655.756 | 4.6 | NRagged | 186.1 | 186.1 | 37.59 | Nsco_20191108_BC_ZIC_HILIC_K56_2_typsin_B1.17687.17687.3 | 41.65 | Trypsin | B1 | S151 | 105 |
| BCAL1917 | >gi\|443605668\|gb\|ELT73505.1\| hypothetical protein BURCENK562V_0636 [Burkholderia cenocepacia K56-2Valvano] | L.SKPAATTSATTSTTTSAGTAS STTATAGTTTAAPAPTAS[+56 8.212]APEAAAKPAK.T | HexNAc(2)Hex(1) | Yes | 1266.371 | 4 | 5062.4621 | 5062.445 | 3.5 | NRagged | 1230 | 1230 | 53.78 | Nsco_20191108_BC_ZIC_HILIC_K56_2_typsin_B2.19359.19359.4 | 44.35 | Trypsin | B2 | S151 | 106 |
| BCAL1917 | >gi\|443605668\|gb\|ELT73505.1\| hypothetical protein BURCENK562V_0636 [Burkholderia cenocepacia K56-2Valvano] | T.STTATAGTTTAAPAPTAS[+5 68.212]APEAAAKPAK.T | HexNAc(2)Hex(1) | Yes | 1027.8418 | 3 | 3081.5108 | 3080.501 | 2.3 | NRagged | 173.3 | 149.7 | 89.39 | Nsco_20191108_BC_ZIC_HILIC_K56_2_typsin_B3.14962.14962.3 | 36.8 | Trypsin | B3 | S151 | 107 |
| BCAL1917 | >gi\|443605668\|gb\|ELT73505.1\| hypothetical protein BURCENK562V_0636 [Burkholderia cenocepacia K56-2Valvano] | T.STTTSAGTASTSTTATAGTT APAPTAS[+568.212]APEA AAKPAK.T | HexNAc(2)Hex(1) | Yes | 1349.6518 | 3 | 4046.9408 | 4045.931 | 1.7 | NRagged | 342.1 | 258.8 | 47.02 | Nsco_20191108_BC_ZIC_HILIC_K56_2_typsin_B2.19517.19517.3 | 44.64 | Trypsin | B2 | S151 | 108 |
| BCAL1917 | >gi\|443605668\|gb\|ELT73505.1\| hypothetical protein BURCENK562V_0636 [Burkholderia cenocepacia K56-2Valvano] | K.SVGHLEENGLTIGGAS[+56 8.212]TPPK.G | HexNAc(2)Hex(1) | Yes | 844.7462 | 3 | 2532.224 | 2532.22 | 1.6 | Specific | 257.6 | 195.8 | 11.82 | Nsco_20191108_BC_ZIC_HILIC_K56_2_typsin_B2.24677.24677.3 | 54.28 | Trypsin | B2 | S104 | 109 |
| BCAL1917 | >gi\|443605668\|gb\|ELT73505.1\| hypothetical protein BURCENK562V_0636 [Burkholderia cenocepacia K56-2Valvano] | A.TAGTTTAAPAPT[+568.21 2]ASAPEAAAKPAK.T | HexNAc(2)Hex(1) | Partial | 907.4528 | 3 | 2720.3437 | 2720.336 | 2.8 | NRagged | 164 | 111.5 | 49.17 | Nsco_20191108_BC_ZIC_HILIC_K56_2_typsin_B2.12753.12753.3 | 32.39 | Trypsin | B2 | NA | 110 |
| BCAL1917 | >gi\|443605668\|gb\|ELT73505.1\| hypothetical protein BURCENK562V_0636 [Burkholderia cenocepacia K56-2Valvano] | A.TTSTTTSAGTASTSTTATAGT TAAPAPTAS[+568.212]AP EAAAKPAK.T | HexNAc(2)Hex(1) | Yes | 1416.6862 | 3 | 4248.0439 | 4248.026 | 4.2 | NRagged | 273.6 | 207.2 | 53 | Nsco_20191108_BC_ZIC_HILIC_K56_2_typsin_B2.20180.20180.3 | 45.99 | Trypsin | B2 | S151 | 111 |
| BCAL2161 | >gi\|443595154\|gb\|ELT63756.1\| hypothetical protein BURCENK562V_1535 [Burkholderia cenocepacia K56-2Valvano] | S.SVAPPLQGDGAAPGGAS[+ 568.212]WPAPPPASGPA PGLPASSVQGT[+568.212] P.- | HexNAc(2)Hex(1) ,HexNAc(2)Hex(1 ) | Yes | 1570.0799 | 3 | 4708.2251 | 4708.205 | 4.3 | Non | 281.7 | 234.9 | 31.41 | Nsco_20191108_BC_ZIC_HILIC_K56_2_pepsin_B3_reinject.52747.52747.3 | 119.4 | Pepsin | B3 | S391 | 112 |
| BCAL2161 | >gi\|443595154\|gb\|ELT63756.1\| hypothetical protein BURCENK562V_1535 [Burkholderia cenocepacia K56-2Valvano] | S.VAPPLQGDGAAPGGAS[+ 568.212]WPAPPPASGPA PGLPASSVQGT[+568.212] P.- | HexNAc(2)Hex(1) | Partial (1 of 2) | 1541.0661 | 3 | 4621.1837 | 4621.173 | 2.4 | Non | 295.7 | 239.6 | 17.02 | Nsco_20191108_BC_ZIC_HILIC_K56_2_pepsin_B3_reinject.52161.52161.3 | 117.7 | Pepsin | B3 | S391 | 113 |
| BCAL2276 | >gi\|443595650\|gb\|ELT64220.1\| hypothetical protein BURCENK562V_2512 [Burkholderia cenocepacia K56-2Valvano] | K.STIDTAAS[+568.212]NA GVPVSSVNYIVHDAGK.G | HexNAc(2)Hex(1) | Yes | 1047.8408 | 3 | 3141.508 | 3141.496 | 3.9 | Non | 346.6 | 279.4 | 18.21 | Nsco_20191108_BC_ZIC_HILIC_K56_2_thermolysin_B3.44492.44492.3 | 90.18 | Thermolysin | B3 | S353 | 114 |
| BCAL2398 | >gi\|443599855\|gb\|ELT68098.1\| PRC-barrel domain protein [Burkholderia cenocepacia K56-2Valvano] | K.ASPPYAADKPIVAVFPPV VPAAPAS[+568.212]SASA TR.- | HexNAc(2)Hex(1) | Yes | 1191.9507 | 3 | 3573.8377 | 3573.821 | 4.6 | Non | 288.5 | 248.9 | 19.9 | Nsco_20191108_BC_ZIC_HILIC_K56_2_thermolysin_B3.52470.52470.3 | 104.7 | Thermolysin | B3 | S315 | 115 |

| | | | | | | | | | | | | | | | | | | | |
|---|---|---|---|---|---|---|---|---|---|---|---|---|---|---|---|---|---|---|---|
| BCAL2449 | >gi\|443591735\|gb\|ELT60603.1\| diguanylate cyclase (GGDEF) domain protein [Burkholderia cenocepacia K56-2Valvano] | R.RTGSVNNAPGAFSAS[+568.212]GVYPIAER.V | HexNAc(2)Hex(1) | Yes | 964.1344 | 3 | 2890.3886 | 2889.375 | 3.6 | Specific | 405.8 | 312.8 | 33.91 | Nsco_20191108_BC_ZIC_HILIC_K56_2_typsin_B3.31102.31102.3 | 65.91 | Trypsin | B3 | S254 | 116 |
| BCAL2466 | >gi\|443591728\|gb\|ELT60596.1\| ecotin [Burkholderia cenocepacia K56-2Valvano] | A.APAS[+568.212]APAVPAESIKMFPQAAAGQQR.V | HexNAc(2)Hex(1) | Yes | 1021.842 | 3 | 3063.5114 | 3062.499 | 3.1 | NRagged | 168.4 | 159.9 | 159.94 | Nsco_20191108_BC_ZIC_HILIC_K56_2_typsin_B1.36665.36665.3 | 76.28 | Trypsin | B1 | S13 | 117 |
| BCAL2629 | >gi\|443603168\|gb\|ELT71192.1\| bifunctional uroporphyrinogen-III synthetase/uroporphyrin-III C-methyltransferase, partial [Burkholderia cenocepacia K56-2Valvano] | K.AAPADAAS[+568.212]SVAAGEPR.W | HexNAc(2)Hex(1) | Yes | 1004.9679 | 2 | 2008.9285 | 2008.919 | 4.6 | Non | 194.2 | 80.45 | 27.14 | Nsco_20191108_BC_ZIC_HILIC_K56_2_thermolysin_B3.16955.16955.2 | 39.37 | Thermolysin | B3 | S546 | 118 |
| BCAL2629 | >gi\|443603168\|gb\|ELT71192.1\| bifunctional uroporphyrinogen-III synthetase/uroporphyrin-III C-methyltransferase, partial [Burkholderia cenocepacia K56-2Valvano] | K.AAPADAAS[+568.212]SVAAGEPRWK.V | HexNAc(2)Hex(1) | Yes | 775.0395 | 3 | 2323.1038 | 2323.094 | 4.4 | Non | 255 | 123.2 | 34.56 | Nsco_20191108_BC_ZIC_HILIC_K56_2_thermolysin_B2_.25473.25473.3 | 50.65 | Thermolysin | B2 | S546 | 119 |
| BCAL2629 | >gi\|443603168\|gb\|ELT71192.1\| bifunctional uroporphyrinogen-III synthetase/uroporphyrin-III C-methyltransferase, partial [Burkholderia cenocepacia K56-2Valvano] | H.AAPKAAPADAAS[+568.212]SVAAGEPRWK.V | HexNAc(2)Hex(1) | Yes | 897.4465 | 3 | 2690.325 | 2690.316 | 3.5 | Non | 242.7 | 185.5 | 25.72 | Nsco_20191108_BC_ZIC_HILIC_K56_2_thermolysin_B1.20913.20913.3 | 44.23 | Thermolysin | B1 | S546 | 120 |
| BCAL2629 | >gi\|443603168\|gb\|ELT71192.1\| bifunctional uroporphyrinogen-III synthetase/uroporphyrin-III C-methyltransferase, partial [Burkholderia cenocepacia K56-2Valvano] | E.AIVPHAAPKAAPADAAS[+568.212]SVAAGEPRW.K | HexNAc(2)Hex(1) | Partial | 770.6379 | 4 | 3079.5298 | 3079.522 | 2.6 | Non | 198.5 | 198.5 | 17.54 | Nsco_20191108_BC_ZIC_HILIC_K56_2_pepsin_B3_reinject.31319.31319.4 | 64.21 | Pepsin | B3 | NA | 121 |
| BCAL2820 | >gi\|443594108\|gb\|ELT62784.1\| outer membrane multidrug efflux protein OprB [Burkholderia cenocepacia K56-2Valvano] | K.RPDAPVAQAYPAS[+568.212]GVYATQPGAAGAR.S | HexNAc(2)Hex(1) | Yes | 1037.84 | 3 | 3111.5053 | 3110.491 | 3.4 | Non | 590.7 | 510.1 | 170.93 | Nsco_20191108_BC_ZIC_HILIC_K56_2_thermolysin_B3.26573.26573.3 | 57.17 | Thermolysin | B3 | S39 | 122 |
| BCAL2820 | >gi\|443594108\|gb\|ELT62784.1\| outer membrane multidrug efflux protein OprB [Burkholderia cenocepacia K56-2Valvano] | D.YDKAAAPAPAS[+568.212]ATATNG.- | HexNAc(2)Hex(1) | Yes | 1072.9926 | 2 | 2144.9779 | 2144.972 | 2.9 | Non | 223.1 | 100.2 | 97.92 | Nsco_20191108_BC_ZIC_HILIC_K56_2_pepsin_B2.12672.12672.2 | 32.21 | Pepsin | B2 | S501 | 123 |
| BCAL2974 | >gi\|443594432\|gb\|ELT63086.1\| hypothetical protein BURCENK562V_0304 [Burkholderia cenocepacia K56-2Valvano] | P.FAAS[+568.212]APSQKYQGSKK.S | HexNAc(2)Hex(1) | Yes | 1083.5312 | 2 | 2166.0551 | 2166.045 | 4.7 | Non | 204.3 | 71.42 | 14.81 | Nsco_20191108_BC_ZIC_HILIC_K56_2_thermolysin_B1.5607.5607.2 | 20.06 | Thermolysin | B1 | S91 | 124 |
| BCAL2974 | >gi\|443594432\|gb\|ELT63086.1\| hypothetical protein BURCENK562V_0304 [Burkholderia cenocepacia K56-2Valvano] | P.FAAS[+568.212]APSQKYQGSKKSA.L | HexNAc(2)Hex(1) | Yes | 581.786 | 4 | 2324.1221 | 2324.114 | 3.5 | Non | 691.1 | 577.5 | 116.79 | Nsco_20191108_BC_ZIC_HILIC_K56_2_thermolysin_B2_.5731.5731.4 | 20.96 | Thermolysin | B2 | S91 | 125 |
| BCAL2974 | >gi\|443594432\|gb\|ELT63086.1\| hypothetical protein BURCENK562V_0304 [Burkholderia cenocepacia K56-2Valvano] | R.LNEHPQMPFAAS[+568.212]APSQK.Y | HexNAc(2)Hex(1) | Yes | 1211.0643 | 2 | 2421.1212 | 2421.113 | 3.6 | Specific | 515.3 | 320 | 93.9 | Nsco_20191108_BC_ZIC_HILIC_K56_2_typsin_B3.22620.22620.2 | 50.4 | Trypsin | B3 | S91 | 126 |
| BCAL2974 | >gi\|443594432\|gb\|ELT63086.1\| hypothetical protein BURCENK562V_0304 [Burkholderia cenocepacia K56-2Valvano] | L.NEHPQMPFAAS[+568.212]APSQKYQG.S | HexNAc(2)Hex(1) | Yes | 886.0642 | 3 | 2656.178 | 2656.172 | 2.3 | CRagged | 257.8 | 178.7 | 21.09 | Nsco_20191108_BC_ZIC_HILIC_K56_2_pepsin_B2.23934.23934.3 | 51.68 | Pepsin | B2 | S91 | 127 |
| BCAL2974 | >gi\|443594432\|gb\|ELT63086.1\| hypothetical protein BURCENK562V_0304 [Burkholderia cenocepacia K56-2Valvano] | L.NEHPQMPFAAS[+568.212]APSQKYQGSKKSAL.R | HexNAc(2)Hex(1) | Yes | 818.3949 | 4 | 3270.5576 | 3270.547 | 3.2 | Specific | 1143 | 1119 | 147.53 | Nsco_20191108_BC_ZIC_HILIC_K56_2_pepsin_B3_reinject.18154.18154.4 | 40.6 | Pepsin | B3 | S91 | 128 |
| BCAL2974 | >gi\|443594432\|gb\|ELT63086.1\| hypothetical protein BURCENK562V_0304 [Burkholderia cenocepacia K56-2Valvano] | M.PFAAS[+568.212]APSQKYQGSKKSAL.R | HexNAc(2)Hex(1) | Yes | 634.5717 | 4 | 2535.2651 | 2534.251 | 4.3 | NRagged | 432.1 | 412.2 | 96.69 | Nsco_20191108_BC_ZIC_HILIC_K56_2_pepsin_B1.12465.12465.4 | 31.43 | Pepsin | B1 | S91 | 129 |
| BCAL2974 | >gi\|443594432\|gb\|ELT63086.1\| hypothetical protein BURCENK562V_0304 [Burkholderia cenocepacia K56-2Valvano] | F.RLNEHPQMPFAAS[+568.212]APSQ.K | HexNAc(2)Hex(1) | Yes | 817.0473 | 3 | 2449.1274 | 2449.119 | 3.5 | CRagged | 263.8 | 162 | 63.15 | Nsco_20191108_BC_ZIC_HILIC_K56_2_pepsin_B1.25394.25394.3 | 54.25 | Pepsin | B1 | S91 | 130 |

| | | | | | | | | | | | | | | | | | | | |
|---|---|---|---|---|---|---|---|---|---|---|---|---|---|---|---|---|---|---|---|
| BCAL2974 | >gi\|443594432\|gb\|ELT63086.1\| hypothetical protein BURCENK562V_0304 [Burkholderia cenocepacia K56-2Valvano] | F.RLNEHPQMPFAAS[+568.212]APSQKYQGSKKSAL.R | HexNAc(2)Hex(1) | Yes | 708.7548 | 5 | 3539.7451 | 3539.732 | 3.6 | Specific | 1159 | 1138 | 125.6 | Nsco_20191108_BC_ZIC_HILIC_K56_2_pepsin_B3_reinject.18210.18210.5 | 40.69 | Pepsin | B3 | S91 | 131 |
| BCAL2983A | >gi\|443599451\|gb\|ELT67730.1\| PF11180 family protein [Burkholderia cenocepacia K56-2Valvano] | S.ADASAPVAGT[+568.212]RPAVTSLSGGASSAASGAVATDAAAQGNVAELTQMLHDGR.I | HexNAc(2)Hex(1) | Partial | 1301.8793 | 4 | 5204.4953 | 5204.477 | 3.5 | NRagged | 310 | 303.4 | 139.62 | Nsco_20191108_BC_ZIC_HILIC_K56_2_typsin_B3.55925.55925.4 | 108.8 | Trypsin | B3 | NA | 132 |
| BCAL3017 | >gi\|443591015\|gb\|ELT59947.1\| hypothetical protein BURCENK562V_1182 [Burkholderia cenocepacia K56-2Valvano] | A.DTASGS[+568.212]DAQASC[+57.021]AIAYVTGVGGSPR.G | HexNAc(2)Hex(1) | Partial | 1483.6553 | 2 | 2966.3033 | 2966.306 | -0.8 | NRagged | 294.6 | 196.1 | 51.03 | Nsco_20191108_BC_ZIC_HILIC_K56_2_typsin_B2.48932.48932.2 | 98.49 | Trypsin | B2 | NA | 133 |
| BCAL3033 | >gi\|443591005\|gb\|ELT59937.1\| outer-membrane lipoprotein carrier protein [Burkholderia cenocepacia K56-2Valvano] | K.GAS[+568.212]AAQAAPKPTDNSSGTFVFAR.P | HexNAc(2)Hex(1) | Yes | 1410.1691 | 2 | 2819.3309 | 2819.322 | 3.2 | Specific | 458.8 | 247.8 | 247.82 | Nsco_20191108_BC_ZIC_HILIC_K56_2_typsin_B1.25150.25150.2 | 55.93 | Trypsin | B1 | S68 | 134 |
| BCAL3033 | >gi\|443591005\|gb\|ELT59937.1\| outer-membrane lipoprotein carrier protein [Burkholderia cenocepacia K56-2Valvano] | K.GAS[+568.212]AAQAAPKPTDNSSGTFVFARPGK.F | HexNAc(2)Hex(1) | Yes | 776.1305 | 4 | 3101.5 | 3101.491 | 2.9 | Non | 655.3 | 655.3 | 655.26 | Nsco_20191108_BC_ZIC_HILIC_K56_2_thermolysin_B3.22747.22747.4 | 50.25 | Thermolysin | B3 | S68 | 135 |
| BCAL3033 | >gi\|443591005\|gb\|ELT59937.1\| outer-membrane lipoprotein carrier protein [Burkholderia cenocepacia K56-2Valvano] | Q.IVKAPAKGAS[+568.212]AAQAAPKPTDN.S | HexNAc(2)Hex(1) | Yes | 859.1134 | 3 | 2575.3257 | 2574.315 | 3.1 | Non | 184.6 | 144.6 | 144.61 | Nsco_20191108_BC_ZIC_HILIC_K56_2_pepsin_B2.8772.8772.3 | 25.73 | Pepsin | B2 | S68 | 136 |
| BCAL3033 | >gi\|443591005\|gb\|ELT59937.1\| outer-membrane lipoprotein carrier protein [Burkholderia cenocepacia K56-2Valvano] | Q.IVKAPAKGAS[+568.212]AAQAAPKPTDNSSGT.F | HexNAc(2)Hex(1) | Yes | 727.3692 | 4 | 2906.4548 | 2906.448 | 2.5 | Non | 907.1 | 606.5 | 449.61 | Nsco_20191108_BC_ZIC_HILIC_K56_2_thermolysin_B2_.9118.9118.4 | 25.97 | Thermolysin | B2 | S68 | 137 |
| BCAL3033 | >gi\|443591005\|gb\|ELT59937.1\| outer-membrane lipoprotein carrier protein [Burkholderia cenocepacia K56-2Valvano] | Q.IVKAPAKGAS[+568.212]AAQAAPKPTDNSSGTF.V | HexNAc(2)Hex(1) | Yes | 764.1369 | 4 | 3053.5259 | 3053.516 | 3.2 | NRagged | 1186 | 1133 | 528.59 | Nsco_20191108_BC_ZIC_HILIC_K56_2_pepsin_B2.15586.15586.4 | 36.98 | Pepsin | B2 | S68 | 138 |
| BCAL3033 | >gi\|443591005\|gb\|ELT59937.1\| outer-membrane lipoprotein carrier protein [Burkholderia cenocepacia K56-2Valvano] | Q.IVKAPAKGAS[+568.212]AAQAAPKPTDNSSGTFV.F | HexNAc(2)Hex(1) | Yes | 788.9039 | 4 | 3152.5937 | 3152.585 | 2.9 | Non | 855.2 | 855.2 | 397.28 | Nsco_20191108_BC_ZIC_HILIC_K56_2_thermolysin_B2_.19314.19314.4 | 40.68 | Thermolysin | B2 | S68 | 139 |
| BCAL3166 | >gi\|443602480\|gb\|ELT70558.1\| BON domain protein [Burkholderia cenocepacia K56-2Valvano] | A.ATS[+568.212]APKAAAKTAKKANRKLGY.A | HexNAc(2)Hex(1) | Yes | 905.826 | 3 | 2715.4635 | 2714.457 | 1.1 | Non | 186.7 | 115.5 | 15.17 | Nsco_20191108_BC_ZIC_HILIC_K56_2_pepsin_B2.4419.4419.3 | 18.47 | Pepsin | B2 | S40 | 140 |
| BCAL3166 | >gi\|443602480\|gb\|ELT70558.1\| BON domain protein [Burkholderia cenocepacia K56-2Valvano] | A.PAAATS[+568.212]APKAAAKTAKKANRKLGY.A | HexNAc(2)Hex(1) | Yes | 739.1546 | 4 | 2953.5967 | 2953.584 | 4.3 | Non | 392.3 | 392.3 | 68.75 | Nsco_20191108_BC_ZIC_HILIC_K56_2_pepsin_B2.6995.6995.4 | 23.02 | Pepsin | B2 | S40 | 141 |
| BCAL3166 | >gi\|443602480\|gb\|ELT70558.1\| BON domain protein [Burkholderia cenocepacia K56-2Valvano] | A.TEAPAAATS[+568.212]APKAAAKTAKKANRKLGY.A | HexNAc(2)Hex(1) | Yes | 1085.9101 | 3 | 3255.7156 | 3254.711 | 0.3 | Non | 285.4 | 157.3 | 27.28 | Nsco_20191108_BC_ZIC_HILIC_K56_2_pepsin_B3_reinject.11243.11243.3 | 29.3 | Pepsin | B3 | S40 | 142 |
| BCAL3377 | >gi\|443595379\|gb\|ELT63971.1\| SmpA/OmlA family protein, partial [Burkholderia cenocepacia K56-2Valvano] | A.AAKKASEAAAAAS[+568.212]AAQA.A | HexNAc(2)Hex(1) | Yes | 1028.5033 | 2 | 2055.9993 | 2055.993 | 3.2 | Non | 181.1 | 71.41 | 64.93 | Nsco_20191108_BC_ZIC_HILIC_K56_2_thermolysin_B2_.6537.6537.2 | 22.1 | Thermolysin | B2 | S152 | 143 |
| BCAL3377 | >gi\|443595379\|gb\|ELT63971.1\| SmpA/OmlA family protein, partial [Burkholderia cenocepacia K56-2Valvano] | D.IDGDRGGKKAKAAAAAKKAS[+568.212]EAAA.A | HexNAc(2)Hex(1) | Yes | 706.8718 | 4 | 2824.4656 | 2824.453 | 4.3 | Non | 240.3 | 240.3 | 240.3 | Nsco_20191108_BC_ZIC_HILIC_K56_2_pepsin_B2.3463.3463.3 | 17.05 | Pepsin | B2 | S145 | 144 |
| BCAL3469 | >gi\|443597454\|gb\|ELT65879.1\| cell division protein FtsL [Burkholderia cenocepacia K56-2Valvano] | A.IDAPIPAS[+568.212]ADTAGKGKGGAR.- | HexNAc(2)Hex(1) | Yes | 606.0576 | 4 | 2421.2087 | 2421.199 | 4 | Non | 780.2 | 649.5 | 197.28 | Nsco_20191108_BC_ZIC_HILIC_K56_2_thermolysin_B1.12654.12654.4 | 30.68 | Thermolysin | B1 | S101 | 145 |
| BCAM0505 | >gi\|443591075\|gb\|ELT60005.1\| BON domain protein [Burkholderia cenocepacia K56-2Valvano] | A.HAQAS[+568.212]STDSGMASESNQPVTDTWITTK.V | HexNAc(2)Hex(1) | Yes | 1140.1741 | 3 | 3418.5077 | 3418.496 | 3.4 | Non | 505.8 | 485.6 | 37.17 | Nsco_20191108_BC_ZIC_HILIC_K56_2_thermolysin_B3.30160.30160.3 | 63.86 | Thermolysin | B3 | S32 | 146 |
| BCAM0505 | >gi\|443591075\|gb\|ELT60005.1\| BON domain protein [Burkholderia cenocepacia K56-2Valvano] | A.QAS[+568.212]STDSGMASESNQPVTDTW.I | HexNAc(2)Hex(1) | Yes | 1384.0723 | 2 | 2767.1373 | 2767.126 | 4.2 | Non | 269.6 | 185.6 | 21.49 | Nsco_20191108_BC_ZIC_HILIC_K56_2_thermolysin_B1.53831.53831.2 | 119.6 | Thermolysin | B1 | S32 | 147 |

| | | | | | | | | | | | | | | | | | | | |
|---|---|---|---|---|---|---|---|---|---|---|---|---|---|---|---|---|---|---|---|
| BCAM0505 | >gi\|443591075\|gb\|ELT60005.1\| BON domain protein [Burkholderia cenocepacia K56-2Valvano] | A.QAS[+568.212]STDSGMASESNQPVTDTWITTK.V | HexNAc(2)Hex(1) | Yes | 1606.2004 | 2 | 3211.3934 | 3210.4 | -3.1 | Non | 327.9 | 214 | 41.1 | Nsco_20191108_BC_ZIC_HILIC_K56_2_thermolysin_B3.39356.39356.2 | 80.95 | Thermolysin | B3 | S32 | 148 |
| BCAM0690 | >gi\|443592239\|gb\|ELT61058.1\| OmpA family protein [Burkholderia cenocepacia K56-2Valvano] | R.ALIDAGVPAS[+568.212]SVFAAAFGSEQPVSSNADDE[illegible]RAK.N | HexNAc(2)Hex(1) | Partial | 1301.9476 | 3 | 3903.8281 | 3902.83 | -1.4 | Specific | 159.3 | 114.1 | 15.7 | Nsco_20191108_BC_ZIC_HILIC_K56_2_typsin_B2.58333.58333.3 | 115.8 | Trypsin | B2 | NA | 149 |
| BCAM0713 | >gi\|443605703\|gb\|ELT73537.1\| cation efflux system protein CzcA [Burkholderia cenocepacia K56-2Valvano] | K.RLPIALPAPAAGAS[+568.212]APLAAAPYVPLAELATIDVAPGPNQISREDGK.R | HexNAc(2)Hex(1) | Yes | 1275.4307 | 4 | 5098.7008 | 5098.683 | 3.4 | Specific | 1132 | 1132 | 379.2 | Nsco_20191108_BC_ZIC_HILIC_K56_2_typsin_B2.70574.70574.4 | 143.2 | Trypsin | B2 | S802 | 150 |
| BCAM0996 | >gi\|443592743\|gb\|ELT61524.1\| sporulation and cell division repeat protein [Burkholderia cenocepacia K56-2Valvano] | R.DDDVSDVQAGVAHDEPPAS[+568.212]DTTVAAAPAPAPK.D | HexNAc(2)Hex(1) | Yes | 1228.2304 | 3 | 3682.6767 | 3682.661 | 4.1 | Non | 472.5 | 446.1 | 66.25 | Nsco_20191108_BC_ZIC_HILIC_K56_2_thermolysin_B3.35728.35728.3 | 74.11 | Thermolysin | B3 | S104 | 151 |
| BCAM0996 | >gi\|443592743\|gb\|ELT61524.1\| sporulation and cell division repeat protein [Burkholderia cenocepacia K56-2Valvano] | K.PAAPAAKPAAPKPAPATVANAGAASPDSGDAS[+568.212]SPASPAGAR.F | HexNAc(2)Hex(1) | Yes | 1040.7653 | 4 | 4160.0394 | 4160.027 | 3.1 | Specific | 1003 | 1003 | 77.16 | Nsco_20191108_BC_ZIC_HILIC_K56_2_typsin_B2.19309.19309.4 | 44.26 | Trypsin | B2 | S174 | 152 |
| BCAM0996 | >gi\|443592743\|gb\|ELT61524.1\| sporulation and cell division repeat protein [Burkholderia cenocepacia K56-2Valvano] | K.PAAPKPAPATVANAGAASPDSGDAS[+568.212]SPASPAGAR.F | HexNAc(2)Hex(1) | Yes | 1185.2319 | 3 | 3553.6811 | 3553.678 | 1 | Specific | 456 | 436.8 | 46.57 | Nsco_20191108_BC_ZIC_HILIC_K56_2_typsin_B3.20127.20127.3 | 45.91 | Trypsin | B3 | S174 | 153 |
| BCAM0996 | >gi\|443592743\|gb\|ELT61524.1\| sporulation and cell division repeat protein [Burkholderia cenocepacia K56-2Valvano] | P.VTDDIAIDIPNRPAHQAVAPRDDDVSDVQAGVAHDEPPAS[+568.212]DTTVAAAPAPAPK.D | HexNAc(2)Hex(1) | Yes | 1187.7791 | 5 | 5934.8662 | 5932.848 | 1.9 | NRagged | 727.3 | 559.4 | 65.07 | Nsco_20191108_BC_ZIC_HILIC_K56_2_typsin_B3.38823.38823.5 | 79.6 | Trypsin | B3 | S104 | 154 |
| BCAM1550 | >gi\|443602195\|gb\|ELT70283.1\| peptidoglycan-associated lipoprotein [Burkholderia cenocepacia K56-2Valvano] | K.TPENAGAAPEPSS[+568.212]ETVATVTADDLNNPNSPLAK.R | HexNAc(2)Hex(1) | Yes | 1282.936 | 3 | 3846.7934 | 3846.778 | 4.1 | Specific | 824.7 | 381.5 | 28.07 | Nsco_20191108_BC_ZIC_HILIC_K56_2_typsin_B2.45430.45430.3 | 92.51 | Trypsin | B2 | S37 | 155 |
| BCAM1550 | >gi\|443602195\|gb\|ELT70283.1\| peptidoglycan-associated lipoprotein [Burkholderia cenocepacia K56-2Valvano] | K.TPENAGAAPEPSS[+568.212]ETVATVTADDLNNPNSPLAKR.S | HexNAc(2)Hex(1) | Yes | 1334.9677 | 3 | 4002.8884 | 4002.879 | 2.5 | Specific | 339.5 | 201.2 | 18.88 | Nsco_20191108_BC_ZIC_HILIC_K56_2_typsin_B1.40337.40337.3 | 82.58 | Trypsin | B1 | S37 | 156 |
| BCAM1669 | >gi\|443598083\|gb\|ELT66474.1\| hypothetical protein BURCENK562V_A2079 [Burkholderia cenocepacia K56-2Valvano] | R.VHGIDNSGAGS[+568.212]QPAATVEGGAPVV.R | HexNAc(2)Hex(1) | Yes | 1380.1554 | 2 | 2759.3035 | 2758.29 | 3.6 | Non | 426.9 | 259.6 | 9.99 | Nsco_20191108_BC_ZIC_HILIC_K56_2_thermolysin_B3.34068.34068.2 | 70.94 | Thermolysin | B3 | S73 | 157 |
| BCAM1669 | >gi\|443598083\|gb\|ELT66474.1\| hypothetical protein BURCENK562V_A2079 [Burkholderia cenocepacia K56-2Valvano] | R.VHGIDNSGAGS[+568.212]QPAATVEGGAPVVR.A | HexNAc(2)Hex(1) | Yes | 972.1378 | 3 | 2914.3988 | 2914.391 | 2.6 | Non | 621.3 | 532.4 | 132.54 | Nsco_20191108_BC_ZIC_HILIC_K56_2_thermolysin_B3.24364.24364.3 | 53.07 | Thermolysin | B3 | S73 | 158 |
| BCAM1789 | >gi\|443592202\|gb\|ELT61024.1\| putative esterase [Burkholderia cenocepacia K56-2Valvano] | A.TPPAPAPAAS[+568.212]PAQPPAVQTATTPSTAQEPSVNPGSSVVLR.T | HexNAc(2)Hex(1) | Yes | 1473.0701 | 3 | 4417.1957 | 4416.194 | -0.4 | Non | 570.4 | 217.4 | 66.06 | Nsco_20191108_BC_ZIC_HILIC_K56_2_thermolysin_B3.44672.44672.3 | 90.46 | Thermolysin | B3 | S32 | 159 |
| BCAM2055 | >gi\|443591388\|gb\|ELT60286.1\| type III secretion outer membrane pore, YscC/HrcC-like family protein [Burkholderia cenocepacia K56-2Valvano] | K.DTAAS[+568.212]QPAATTAGVVTHVDEHH.- | HexNAc(2)Hex(1) | Yes | 696.5697 | 4 | 2783.2568 | 2783.249 | 2.8 | Non | 373.3 | 327.1 | 26.78 | Nsco_20191108_BC_ZIC_HILIC_K56_2_thermolysin_B3.20172.20172.4 | 45.22 | Thermolysin | B3 | S664 | 160 |
| BCAM2063 | >gi\|443595086\|gb\|ELT63692.1\| carbohydrate-selective porin, OprB family [Burkholderia cenocepacia K56-2Valvano] | A.SSPAAAEPAAGAS[+568.212]DAAAPAQQAADAAAPAPTGFWER.S | HexNAc(2)Hex(1) | Yes | 1306.934 | 3 | 3918.7875 | 3918.779 | 2.2 | NRagged | 384.3 | 376.9 | 134.79 | Nsco_20191108_BC_ZIC_HILIC_K56_2_typsin_B3.48808.48808.3 | 96.67 | Trypsin | B3 | S46 | 161 |
| BCAM2064 | >gi\|443595102\|gb\|ELT63708.1\| alpha,alpha-trehalase [Burkholderia cenocepacia K56-2Valvano] | A.DNANQAAAQAVGQSAIPATTAAAAAPASGTLPPPPSQLYGDLFVAVQT[+568.212]AQLYPDQK.T | HexNAc(2)Hex(1) | Partial | 1526.2583 | 4 | 6102.0114 | 6101.999 | 2 | NRagged | 595.4 | 553.5 | 29.48 | Nsco_20191108_BC_ZIC_HILIC_K56_2_typsin_B2.77596.77596.4 | 160.7 | Trypsin | B2 | NA | 162 |
| BCAM2289 | >gi\|443599704\|gb\|ELT67963.1\| hypothetical protein BURCENK562V_A2967 [Burkholderia cenocepacia K56-2Valvano] | R.VHGADTSGYGAQPAPLVHSGAPAAAS[+568.212]SNAR.D | HexNAc(2)Hex(1) | Yes | 1129.2016 | 3 | 3385.5901 | 3385.578 | 3.6 | Specific | 471.9 | 429.5 | 40.53 | Nsco_20191108_BC_ZIC_HILIC_K56_2_typsin_B1.16993.16993.3 | 40.39 | Trypsin | B1 | S63 | 163 |

| | | | | | | | | | | | | | | | | | | | |
|---|---|---|---|---|---|---|---|---|---|---|---|---|---|---|---|---|---|---|---|
| BCAM2334 | >gi\|443606050\|gb\|ELT73855.1\| auxiliary transport protein, membrane fusion protein (MFP) domain protein, partial [Burkholderia cenocepacia K56-2Valvano] | R.VHDGVAS[+568.212]DAEAAAAAIIRENQGG.- | HexNAc(2)Hex(1) | Yes | 930.7708 | 3 | 2790.298 | 2790.291 | 2.4 | Non | 592.5 | 592.5 | 592.51 | Nsco_20191108_BC_ZIC_HILIC_K56_2_thermolysin_B3.49647.49647.3 | 99.38 | Thermolysin | B3 | S380 | 164 |
| BCAM2378 | >gi\|443594879\|gb\|ELT63498.1\| X-Pro dipeptidyl-peptidase (S15 family) [Burkholderia cenocepacia K56-2Valvano] | A.LAQGQQEGAS[+568.212]AAAGTANTPS.L | HexNAc(2)Hex(1) | Yes | 1199.5439 | 2 | 2398.0806 | 2398.074 | 2.8 | Non | 437.4 | 304.9 | 139.31 | Nsco_20191108_BC_ZIC_HILIC_K56_2_thermolysin_B1.16412.16412.2 | 36.59 | Thermolysin | B1 | S48 | 165 |
| BCAM2378 | >gi\|443594879\|gb\|ELT63498.1\| X-Pro dipeptidyl-peptidase (S15 family) [Burkholderia cenocepacia K56-2Valvano] | R.TPVIVLASPYYAGLADSPNHDVDVELDGTPHPAATAGAAAS[+568.212]ASAR.I | HexNAc(2)Hex(1) | Yes | 1246.8543 | 4 | 4984.3952 | 4984.386 | 1.9 | Specific | 1038 | 1038 | 76.39 | Nsco_20191108_BC_ZIC_HILIC_K56_2_typsin_B1.56596.56596.4 | 109.8 | Trypsin | B1 | S174 | 166 |
| BCAM2680 | >gi\|443603690\|gb\|ELT71680.1\| hypothetical protein BURCENK562V_A2523, partial [Burkholderia cenocepacia K56-2Valvano] | A.AAQQAS[+568.212]VPAAASAT[+568.212]VVVK.A | HexNAc(2)Hex(1),HexNAc(2)Hex(1) | Partial (1 of 2) | 1403.1835 | 2 | 2805.3598 | 2805.351 | 3.1 | Non | 351.2 | 187.3 | 187.32 | Nsco_20191108_BC_ZIC_HILIC_K56_2_thermolysin_B1.24775.24775.2 | 51.53 | Thermolysin | B1 | S858 | 167 |
| BCAM2680 | >gi\|443603690\|gb\|ELT71680.1\| hypothetical protein BURCENK562V_A2523, partial [Burkholderia cenocepacia K56-2Valvano] | A.AAQQASVPAAAS[+568.212]AT[+568.212]VVVKAAPQPQNP.V | HexNAc(2)Hex(1),HexNAc(2)Hex(1) | Partial (1 of 2) | 1203.9231 | 3 | 3609.7547 | 3608.744 | 2.1 | Non | 194.9 | 194.9 | 74.58 | Nsco_20191108_BC_ZIC_HILIC_K56_2_thermolysin_B2_.36530.36530.3 | 68.44 | Thermolysin | B2 | S858 | 168 |
| BCAM2829 | >gi\|443591146\|gb\|ELT60069.1\| hypothetical protein BURCENK562V_A2847 [Burkholderia cenocepacia K56-2Valvano] | D.SAMPAS[+568.212]APAAASAPASSGAT[+568.212]TAPAANGAPIA.P | HexNAc(2)Hex(1),HexNAc(2)Hex(1) | Partial (1 of 2) | 1248.9023 | 3 | 3744.6923 | 3744.69 | 0.5 | Non | 369.3 | 262.4 | 6.65 | Nsco_20191108_BC_ZIC_HILIC_K56_2_pepsin_B3_reinject.37501.37501.3 | 77.29 | Pepsin | B3 | S296 | 169 |
| BCAS0089 | >gi\|443598997\|gb\|ELT67309.1\| hypothetical protein BURCENK562V_B0207 [Burkholderia cenocepacia K56-2Valvano] | N.AVQQAPATKAAPAVPAS[+568.212]GQ.- | HexNAc(2)Hex(1) | Yes | 1166.0856 | 2 | 2331.164 | 2331.156 | 3.3 | Non | 298.1 | 168.3 | 78.72 | Nsco_20191108_BC_ZIC_HILIC_K56_2_thermolysin_B1.15802.15802.2 | 35.63 | Thermolysin | B1 | S99 | 170 |
| BCAS0089 | >gi\|443598997\|gb\|ELT67309.1\| hypothetical protein BURCENK562V_B0207 [Burkholderia cenocepacia K56-2Valvano] | A.VQQAPATKAAPAVPAS[+568.212]GQ.- | HexNAc(2)Hex(1) | Yes | 1130.5662 | 2 | 2260.1251 | 2260.119 | 2.7 | Non | 503.1 | 244.5 | 150.76 | Nsco_20191108_BC_ZIC_HILIC_K56_2_thermolysin_B1.14539.14539.2 | 33.71 | Thermolysin | B1 | S99 | 171 |
| BURCENK562V_3104 | >gi\|443598112\|gb\|ELT66499.1\| hypothetical protein BURCENK562V_3104 [Burkholderia cenocepacia K56-2Valvano] | G.AAADGAKDTTSS[+568.212]AVHSTKKHTKHAAKSAKSHAGSA.K | HexNAc(2)Hex(1) | Partial | 797.6005 | 5 | 3983.9733 | 3983.954 | 4.8 | Non | 613.4 | 594.4 | 35.12 | Nsco_20191108_BC_ZIC_HILIC_K56_2_pepsin_B3_reinject.3843.3843.5 | 16.95 | Pepsin | B3 | NA | 172 |
| BURCENK562V_3104 | >gi\|443598112\|gb\|ELT66499.1\| hypothetical protein BURCENK562V_3104 [Burkholderia cenocepacia K56-2Valvano] | N.ASGNATGAATDAVGAAADGAKDTTSS[+568.212]AVHS.T | HexNAc(2)Hex(1) | Yes | 1067.4822 | 3 | 3200.4321 | 3200.42 | 3.9 | Non | 302.2 | 245.4 | 8.44 | Nsco_20191108_BC_ZIC_HILIC_K56_2_thermolysin_B1.29677.29677.3 | 60.55 | Thermolysin | B1 | S83 | 173 |
| BURCENK562V_3104 | >gi\|443598112\|gb\|ELT66499.1\| hypothetical protein BURCENK562V_3104 [Burkholderia cenocepacia K56-2Valvano] | N.ASGNATGAATDAVGAAADGAKDTTSS[+568.212]AVHSTKKH.T | HexNAc(2)Hex(1) | Yes | 739.751 | 5 | 3694.7259 | 3694.716 | 2.6 | Non | 493.6 | 493.6 | 20.41 | Nsco_20191108_BC_ZIC_HILIC_K56_2_thermolysin_B1.19879.19879.5 | 42.31 | Thermolysin | B1 | S83 | 174 |
| BURCENK562V_3104 | >gi\|443598112\|gb\|ELT66499.1\| hypothetical protein BURCENK562V_3104 [Burkholderia cenocepacia K56-2Valvano] | G.VGAGAQGGAGANASGNATGAATDAVGAAADGAKDTTSS[+568.212]AVHSTKK.H | HexNAc(2)Hex(1) | Yes | 894.4257 | 5 | 4468.0996 | 4468.083 | 3.7 | Non | 1193 | 979.4 | 27.83 | Nsco_20191108_BC_ZIC_HILIC_K56_2_thermolysin_B1.26977.26977.5 | 55.63 | Thermolysin | B1 | S83 | 175 |
| BURCENK562V_3104 | >gi\|443598112\|gb\|ELT66499.1\| hypothetical protein BURCENK562V_3104 [Burkholderia cenocepacia K56-2Valvano] | G.VGAGAQGGAGANASGNATGAATDAVGAAADGAKDTTSS[+568.212]AVHSTKKH.T | HexNAc(2)Hex(1) | Partial | 768.5336 | 6 | 4606.165 | 4605.142 | 4.3 | Non | 957.8 | 881.9 | 18.94 | Nsco_20191108_BC_ZIC_HILIC_K56_2_thermolysin_B1.24815.24815.6 | 51.6 | Thermolysin | B1 | NA | 176 |
| BURCENK562V_3104 | >gi\|443598112\|gb\|ELT66499.1\| hypothetical protein BURCENK562V_3104 [Burkholderia cenocepacia K56-2Valvano] | G.VQAQTPAAGVGAGAQGGAGANASGNATGAATDAVGAAADGAKDTTSS[+568.212]AVHSTKKH.T | HexNAc(2)Hex(1) | Partial | 1086.9199 | 5 | 5430.5705 | 5428.561 | 0.6 | Non | 335.9 | 335.6 | 6.22 | Nsco_20191108_BC_ZIC_HILIC_K56_2_thermolysin_B2_.31006.31006.5 | 59.67 | Thermolysin | B2 | NA | 177 |
| BCAS0773 | >gi\|443593562\|gb\|ELT62293.1\| hypothetical protein BURCENK562V_B0326 [Burkholderia cenocepacia K56-2Valvano] | A.QTDAAS[+568.212]APAAAAAQDAK.A | HexNAc(2)Hex(1) | Yes | 1063.4875 | 2 | 2125.9678 | 2125.962 | 2.8 | Non | 295.6 | 96.23 | 71.49 | Nsco_20191108_BC_ZIC_HILIC_K56_2_thermolysin_B3.10971.10971.2 | 29.38 | Thermolysin | B3 | S33 | 178 |

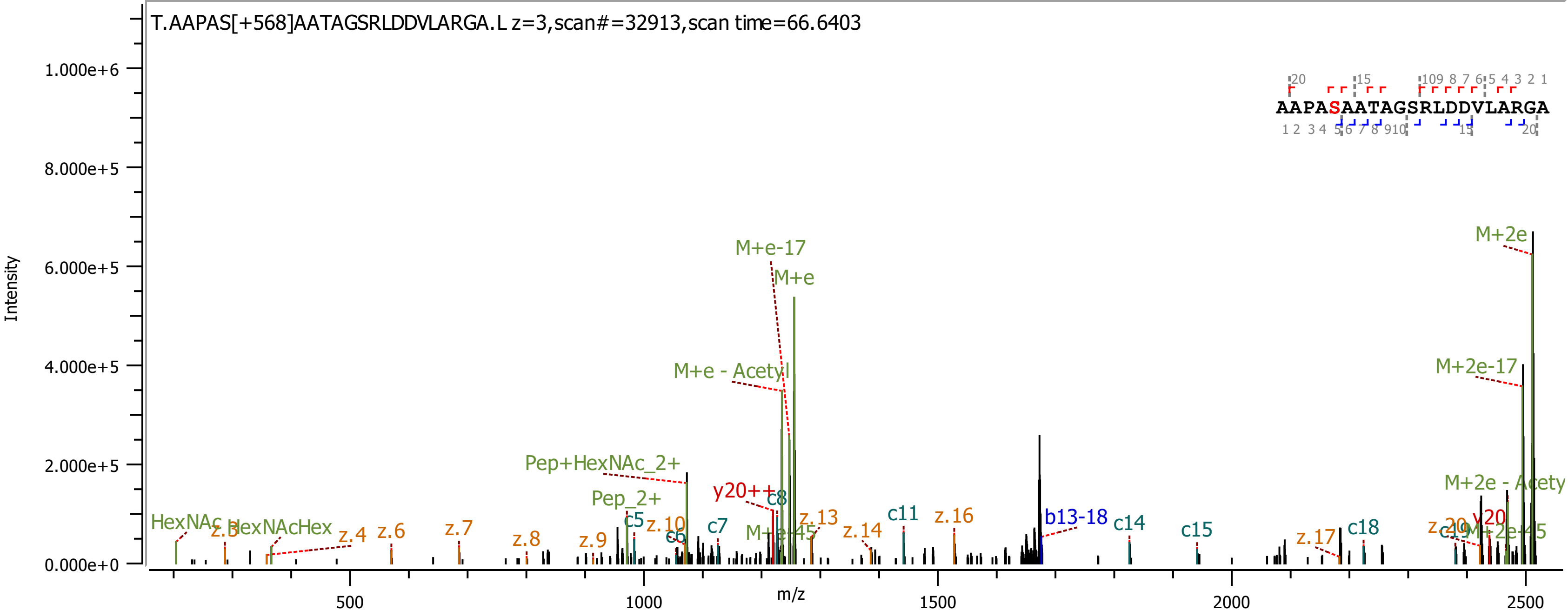
T.AAPAS[+568]AATAGSRLDDVLARGA.L z=3,scan#=32913,scan time=66.6403
AAPASAATAGSRLDDVLARGA
Intensity
m/z
M+2e
M+e-17
M+e
M+e - Acetyl
M+2e-17
Pep+HexNAc_2+
Pep_2+
y20++
c8
M+2e - Acety
HexNAc
z.3
HexNAcHex
z.4
z.6
z.7
z.8
z.9
c5
z.10
c6
c7
M+e-45
z.13
z.14
c11
z.16
b13-18
c14
c15
z.17
c18
y20
M+2e-45
0.000e+0
2.000e+5
4.000e+5
6.000e+5
8.000e+5
1.000e+6
500
1000
1500
2000
2500

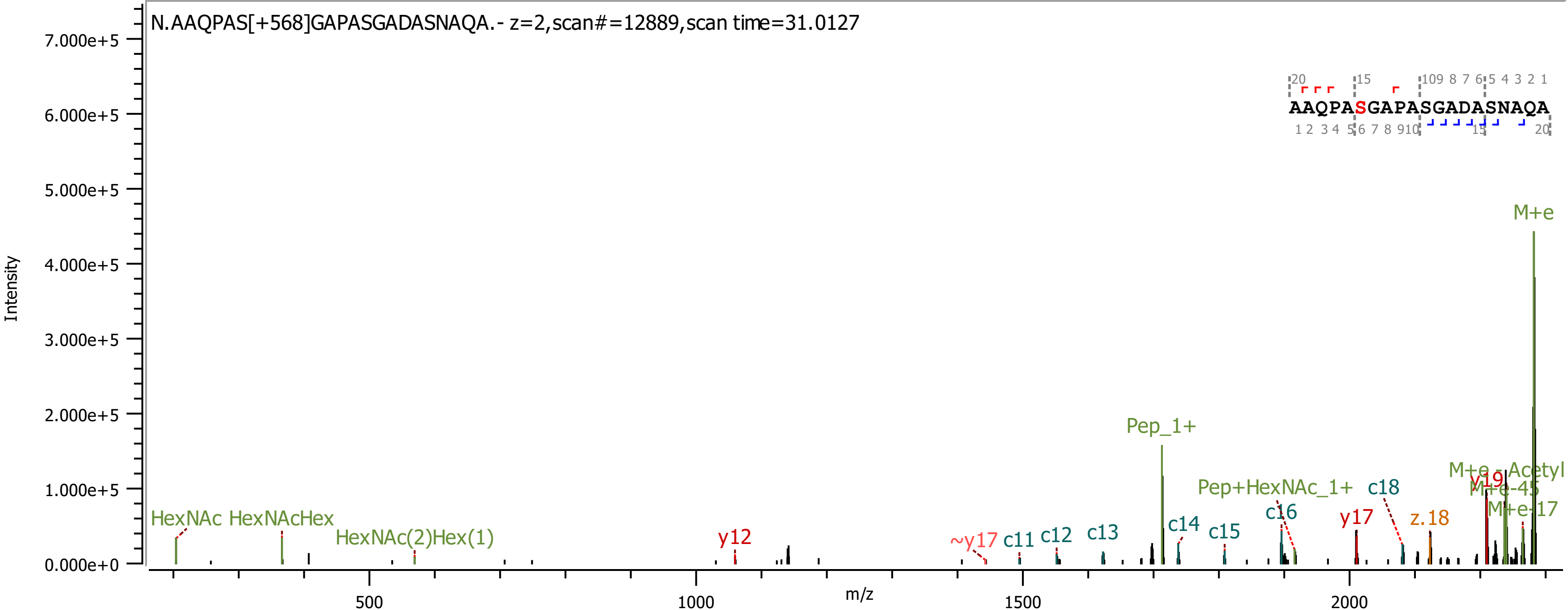
N.AAQPAS[+568]GAPASGADASNAQA.- z=2,scan#=12889,scan time=31.0127
AAQPASGAPASGADASNAQA
Intensity
m/z
HexNAc
HexNAcHex
HexNAc(2)Hex(1)
y12
~y17
c11
c12
c13
Pep_1+
c14
c15
c16
Pep+HexNAc_1+
c18
y17
z.18
y19
M+e-Acetyl
M+e-45
M+e-17
M+e

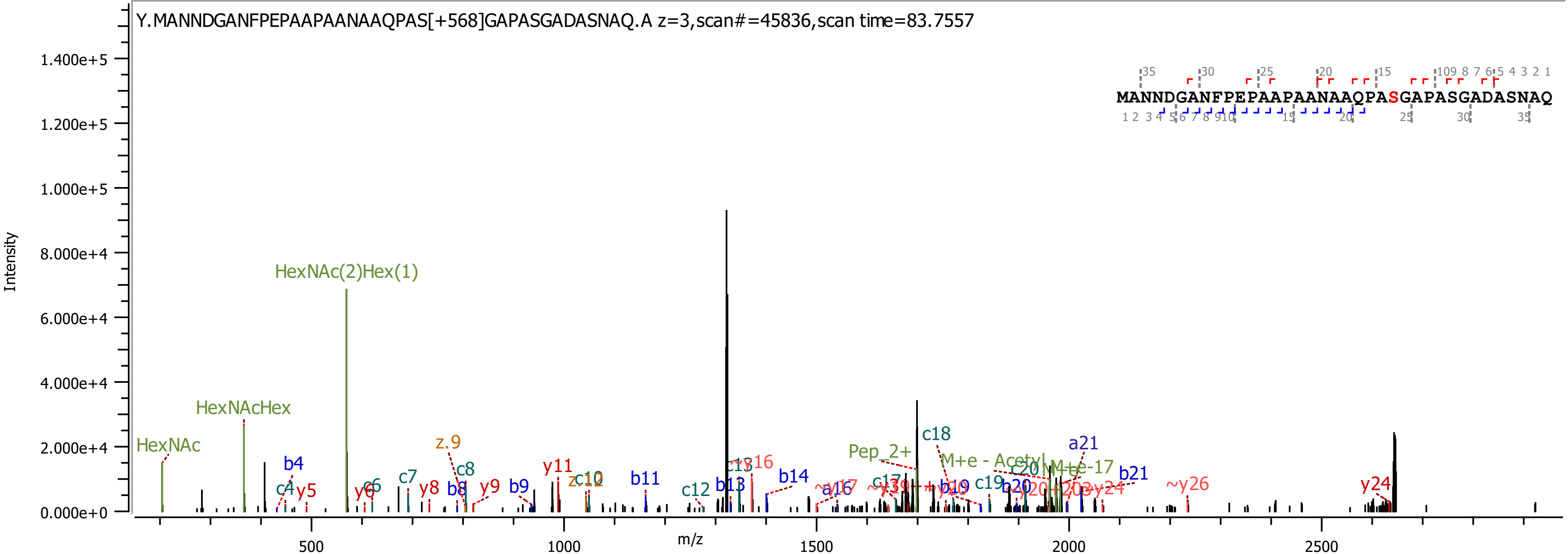

Y.MANNDGANFPEPAAPAANAAQPAS[+568]GAPASGADASNAQ.A z=3,scan#=45836,scan time=83.7557
MANNDGANFPEPAAPAANAAQPASGAPASGADASNAQ
Intensity
m/z
HexNAc(2)Hex(1)
HexNAcHex
HexNAc
Pep_2+
M+e - Acetyl
M+e-17

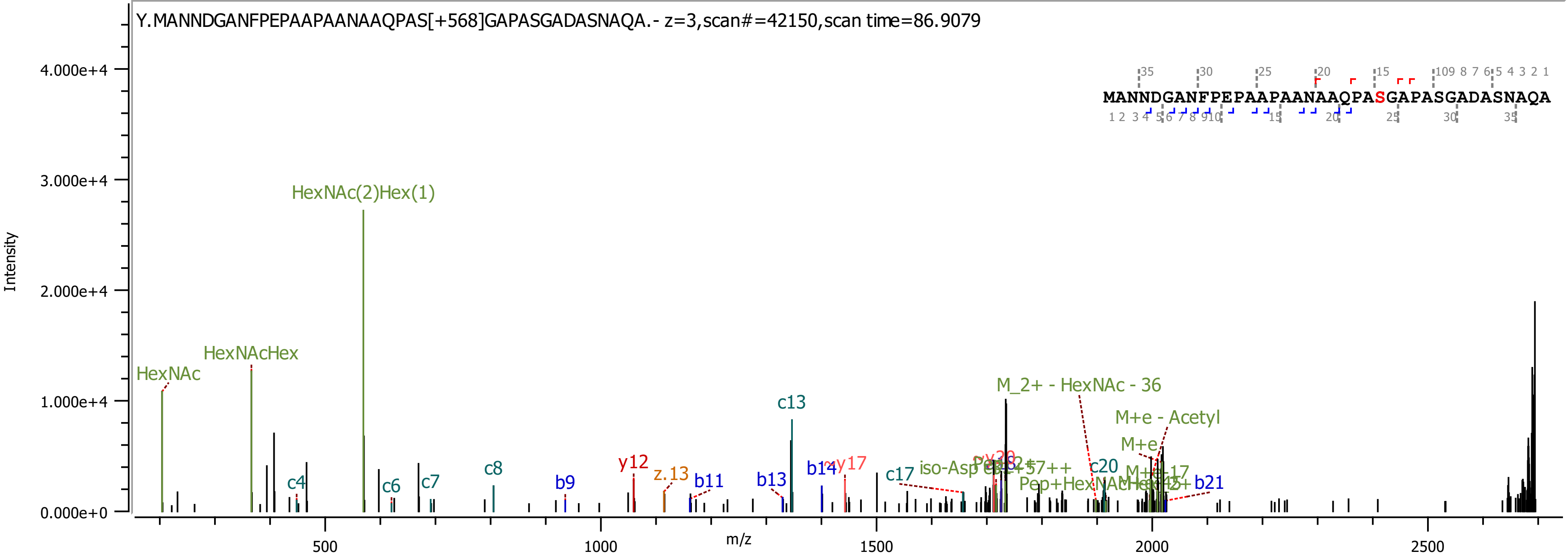

Y.MANNDGANFPEPAAPAANAAQPAS[+568]GAPASGADASNAQA.- z=3,scan#=42150,scan time=86.9079
MANNDGANFPEPAAPAANAAQPASGAPASGADASNAQA
Intensity
m/z
HexNAc(2)Hex(1)
HexNAcHex
HexNAc
c4
c6
c7
c8
b9
y12
z.13
b11
c13
b13
b14
y17
c17
M_2+ - HexNAc - 36
M+e - Acetyl
c20
b21

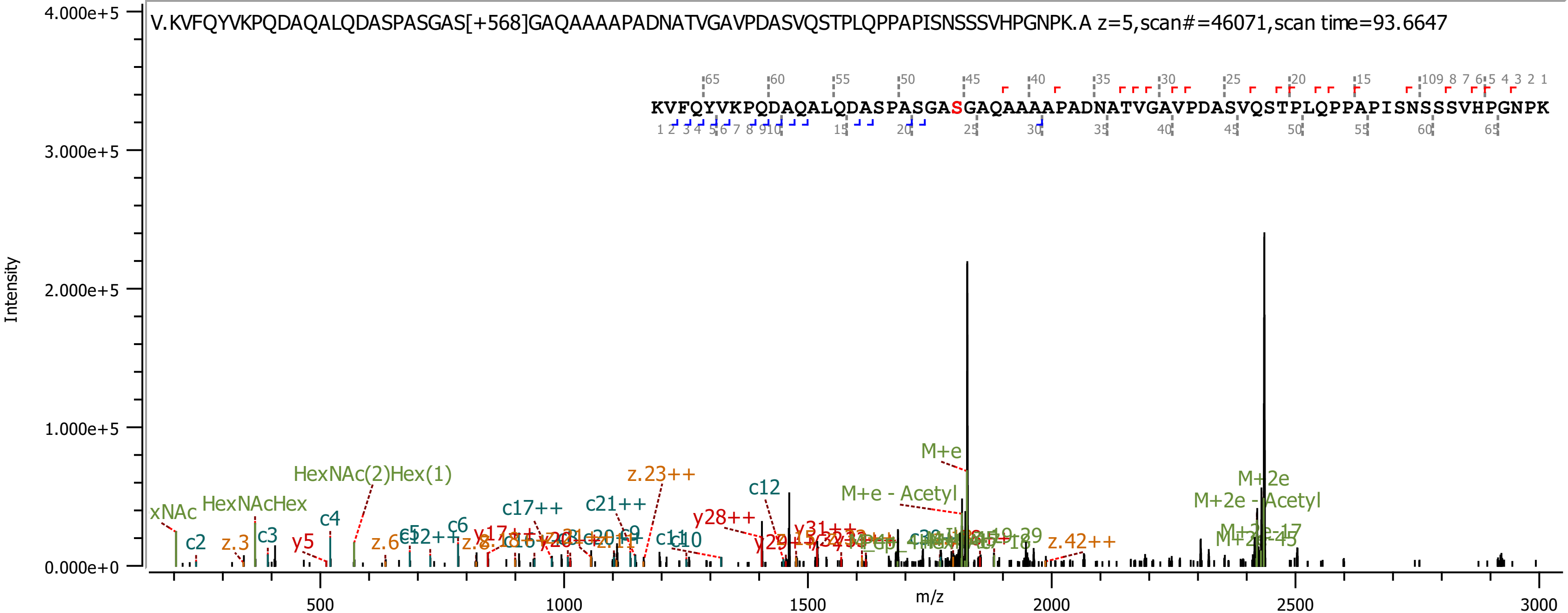

V.KVFQYVKPQDAQALQDASPASGAS[+568]GAQAAAAPADNATVGAVPDASVQSTPLQPPAPISNSSSVHPGNPK.A z=5,scan#=46071,scan time=93.6647
KVFQYVKPQDAQALQDASPASGASGAQAAAAPADNATVGAVPDASVQSTPLQPPAPISNSSSVHPGNPK
Intensity
m/z
4.000e+5
3.000e+5
2.000e+5
1.000e+5
0.000e+0
500
1000
1500
2000
2500
3000
M+e
M+e - Acetyl
M+2e
M+2e - Acetyl
HexNAc(2)Hex(1)
HexNAcHex
xNAc
z.23++
c12
c17++
c21++
y28++
z.42++
c2
c3
c4
c6
c9
c10
y5
z.3
z.6

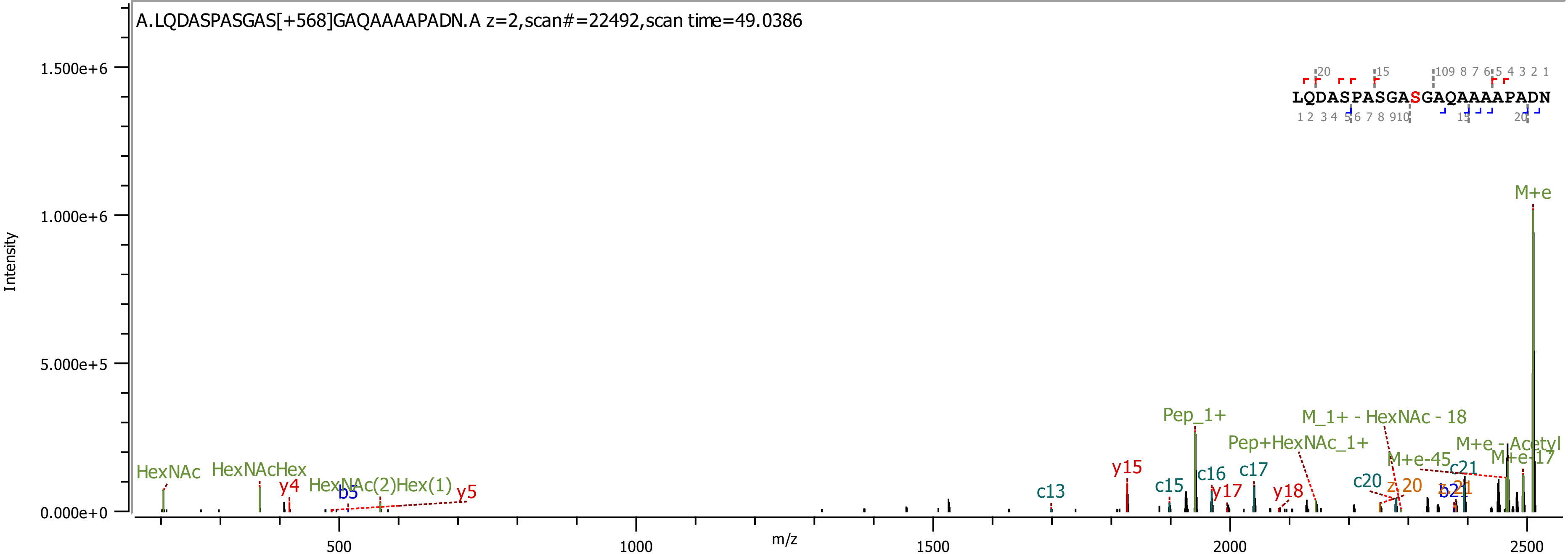

A.LQDASPASGAS[+568]GAQAAAAPADN.A z=2,scan#=22492,scan time=49.0386
LQDASPASGASGAQAAAAPADN
Intensity
m/z
M+e
Pep_1+
M_1+ - HexNAc - 18
Pep+HexNAc_1+
M+e - Acetyl
M+e-45
M+e-17
HexNAc
HexNAcHex
HexNAc(2)Hex(1)
y4
y5
y15
y17
y18
b5
b21
c13
c15
c16
c17
c20
c21
z 20
z 21

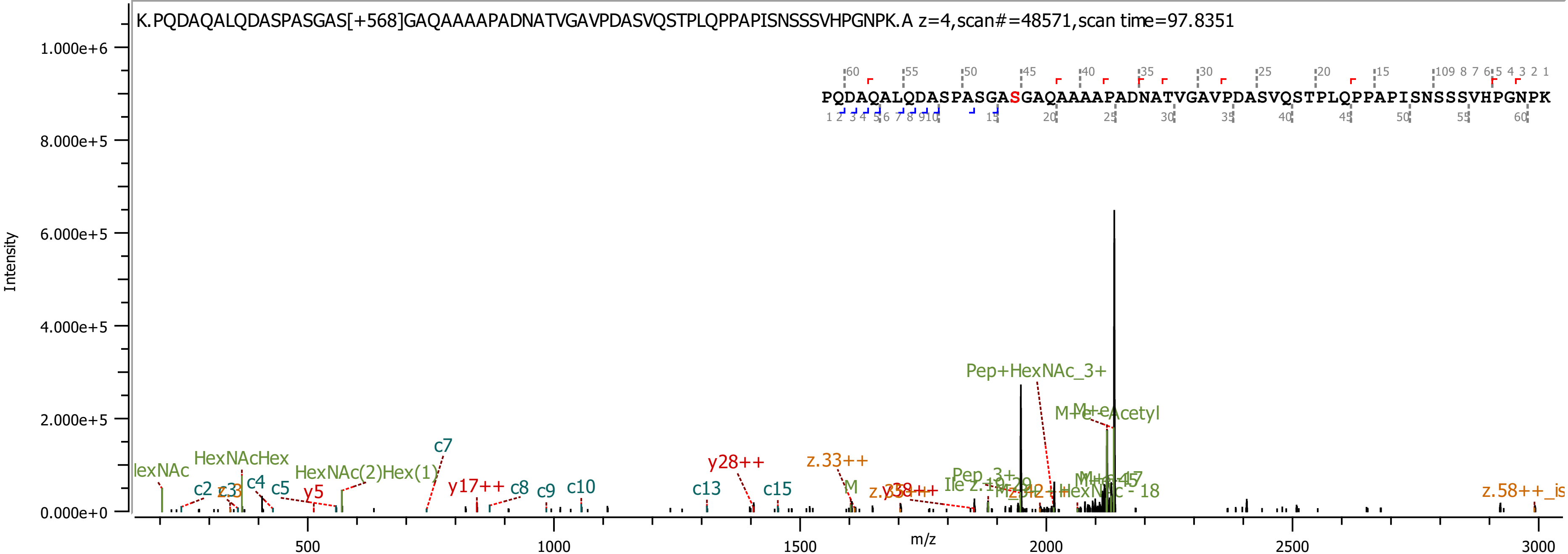
K.PQDAQALQDASPASGAS[+568]GAQAAAAPADNATVGAVPDASVQSTPLQPPAPISNSSSVHPGNPK.A z=4,scan#=48571,scan time=97.8351
PQDAQALQDASPASGASGAQAAAAPADNATVGAVPDASVQSTPLQPPAPISNSSSVHPGNPK
Intensity
1.000e+6
8.000e+5
6.000e+5
4.000e+5
2.000e+5
0.000e+0
m/z
500
1000
1500
2000
2500
3000
HexNAcHex
HexNAc(2)Hex(1)
c2
c4
c5
y5
c7
y17++
c8
c9
c10
y28++
c13
c15
z.33++
Pep+HexNAc_3+
M+e-Acetyl
z.58++

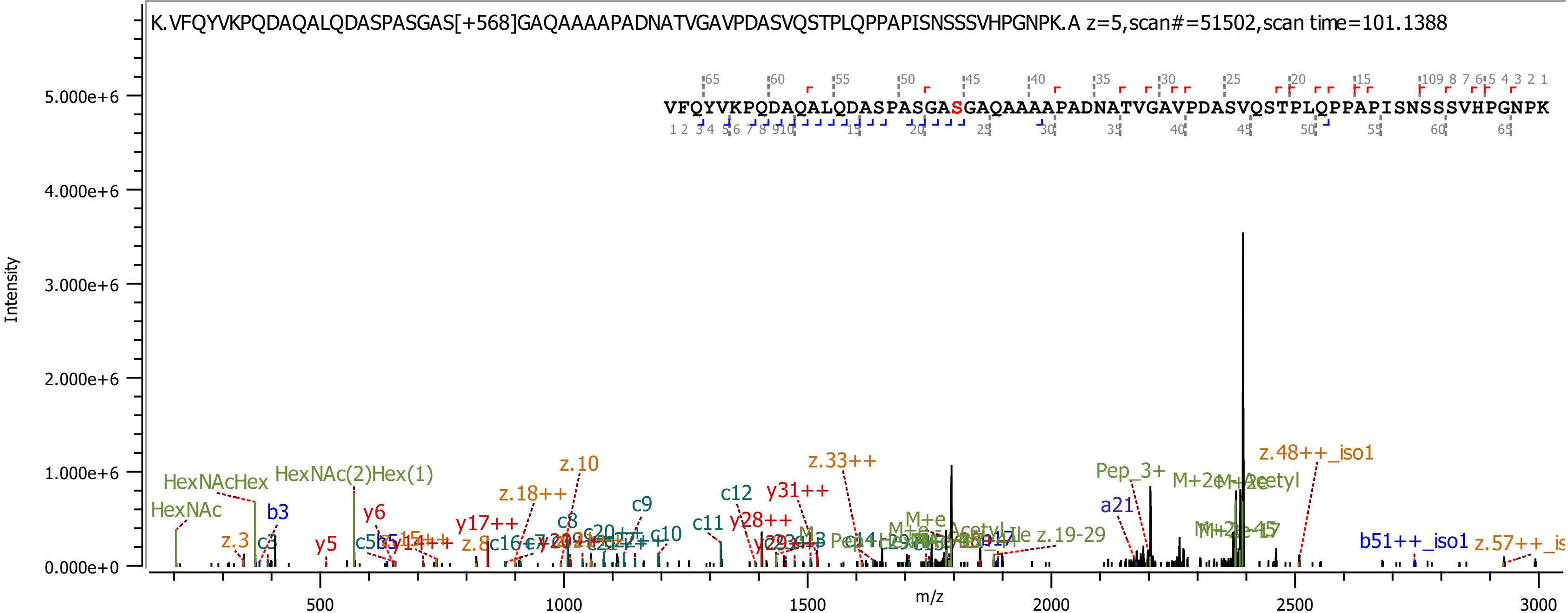

K.VFQYVKPQDAQALQDASPASGAS[+568]GAQAAAAPADNATVGAVPDASVQSTPLQPPAPISNSSSVHPGNPK.A z=5,scan#=51502,scan time=101.1388
VFQYVKPQDAQALQDASPASGASGAQAAAAPADNATVGAVPDASVQSTPLQPPAPISNSSSVHPGNPK
Intensity
5.000e+6
4.000e+6
3.000e+6
2.000e+6
1.000e+6
0.000e+0
m/z
500
1000
1500
2000
2500
3000
HexNAcHex
HexNAc(2)Hex(1)
HexNAc
b3
z.3
c3
y6
y5
z.10
z.18++
y17++
c8
c9
c12
y31++
z.33++
c11
y28++
c10
Pep_3+
a21
z.48++_iso1
z.19-29
b51++_iso1

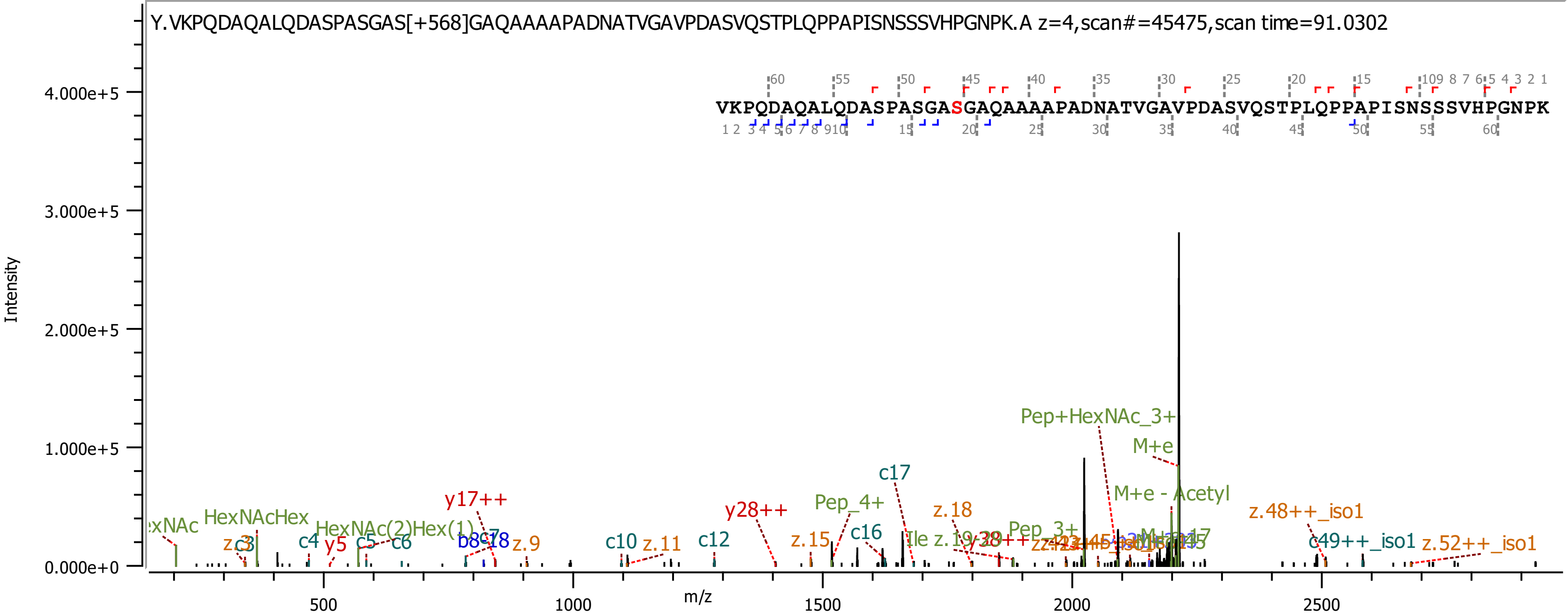
Y.VKPQDAQALQDASPASGAS[+568]GAQAAAAPADNATVGAVPDASVQSTPLQPPAPISNSSSVHPGNPK.A z=4,scan#=45475,scan time=91.0302
VKPQDAQALQDASPASGASGAQAAAAPADNATVGAVPDASVQSTPLQPPAPISNSSSVHPGNPK
Intensity
m/z
Pep+HexNAc_3+
M+e
M+e - Acetyl
c17
y17++
y28++
Pep_4+
z.18
z.48++_iso1
c49++_iso1
z.52++_iso1
HexNAcHex
HexNAc(2)Hex(1)
c16
c12
c10
z.11
z.15
z.9
c4
y5
c5
c6
Pep_3+

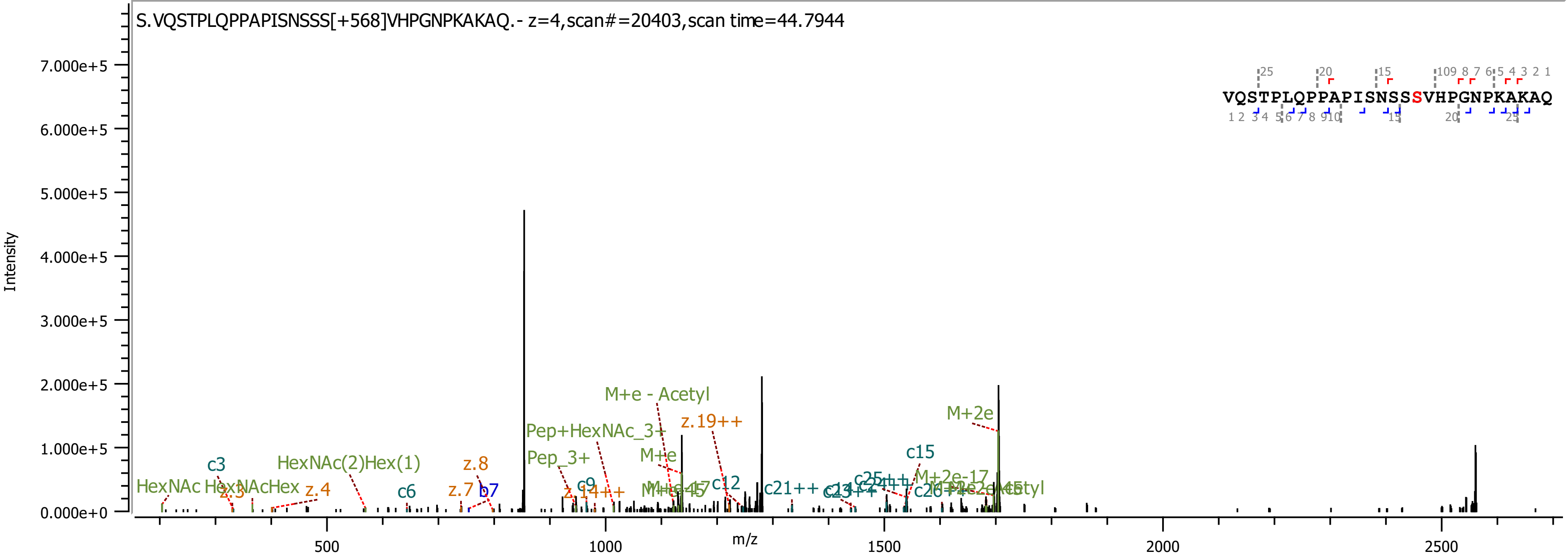
S.VQSTPLQPPAPISNSSS[+568]VHPGNPKAKAQ.- z=4,scan#=20403,scan time=44.7944
VQSTPLQPPAPISNSSSVHPGNPKAKAQ
Intensity
7.000e+5
6.000e+5
5.000e+5
4.000e+5
3.000e+5
2.000e+5
1.000e+5
0.000e+0
m/z
500
1000
1500
2000
2500
M+e - Acetyl
z.19++
M+2e
Pep+HexNAc_3+
M+e
c15
Pep_3+
HexNAc(2)Hex(1)
z.8
c3
HexNAc
HexNAcHex
z.4
z.3
c6
z.7
b7
c9
z.14++
c12
c21++
M+2e-17

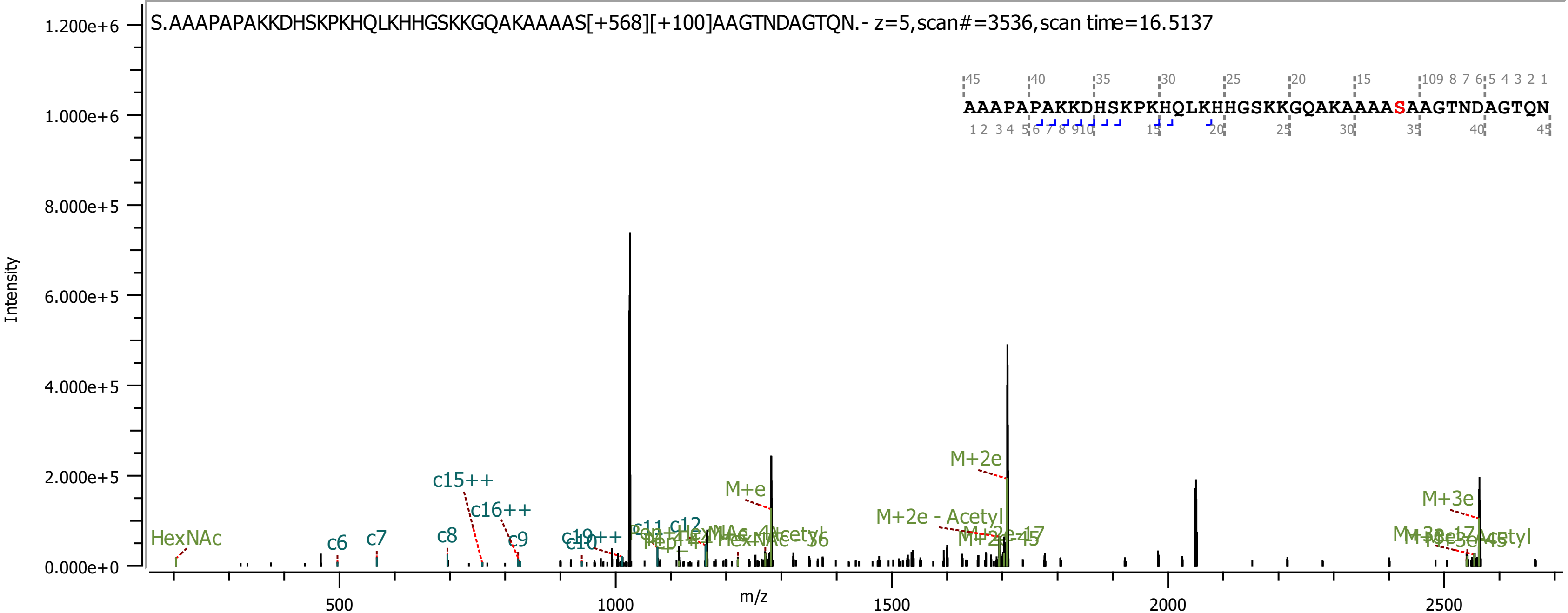

S.AAAPAPAKKDHSKPKHQLKHHGSKKGQAKAAAAS[+568][+100]AAGTNDAGTQN.- z=5,scan#=3536,scan time=16.5137
AAAPAPAKKDHSKPKHQLKHHGSKKGQAKAAAASAAGTNDAGTQN
Intensity
m/z
HexNAc
c6
c7
c8
c9
c10
c11
c12
c15++
c16++
c19++
M+e
M+2e
M+2e - Acetyl
M+3e
1.200e+6
1.000e+6
8.000e+5
6.000e+5
4.000e+5
2.000e+5
0.000e+0
500
1000
1500
2000
2500

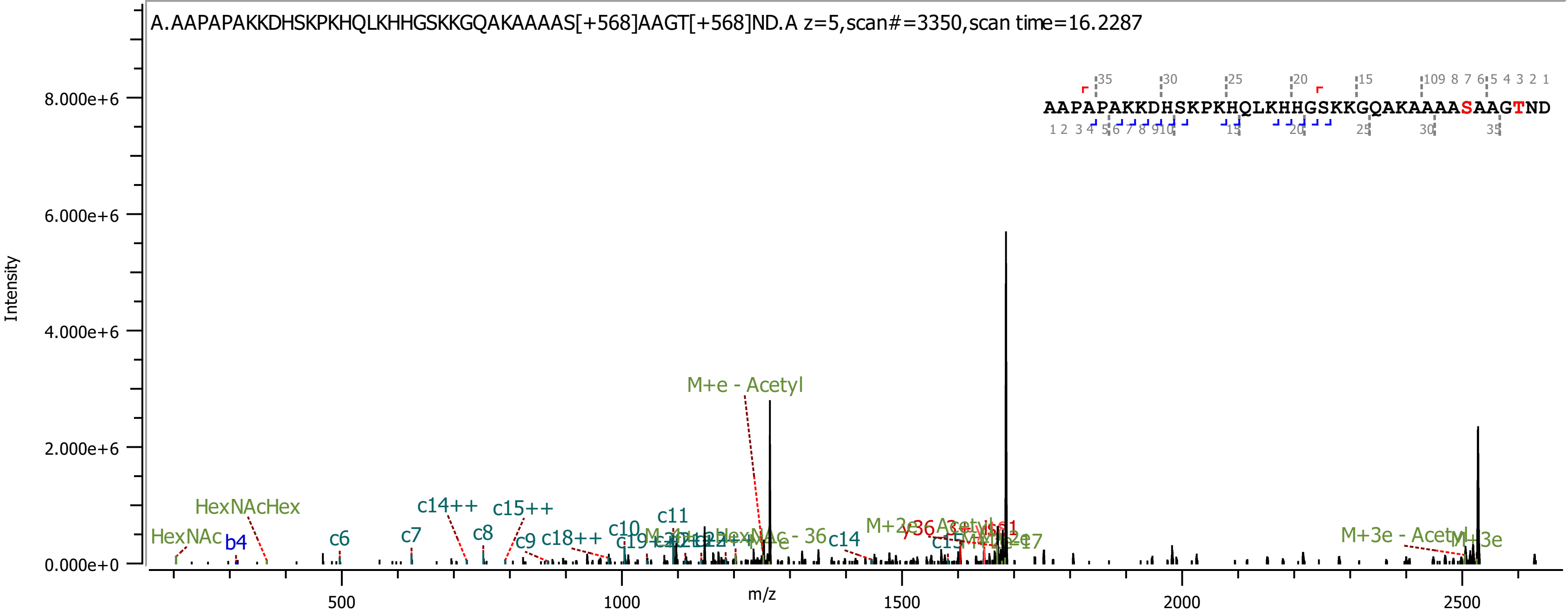
A.AAPAPAKKDHSKPKHQLKHHGSKKGQAKAAAAS[+568]AAGT[+568]ND.A z=5,scan#=3350,scan time=16.2287
AAPAPAKKDHSKPKHQLKHHGSKKGQAKAAAASAAGTND
Intensity
8.000e+6
6.000e+6
4.000e+6
2.000e+6
0.000e+0
m/z
500
1000
1500
2000
2500
M+e - Acetyl
HexNAcHex
HexNAc
b4
c6
c7
c8
c9
c14++
c15++
c18++
c10
c11
c14
M+3e - Acetyl
M+3e

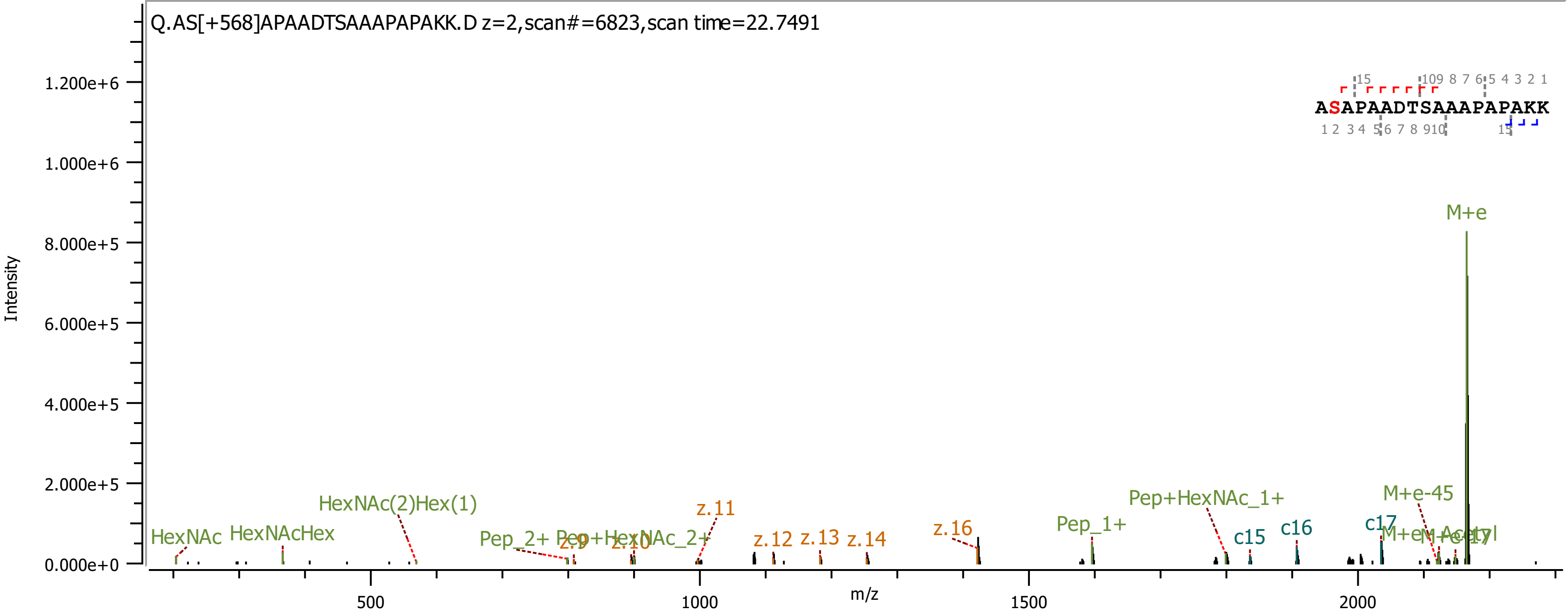

Q.AS[+568]APAADTSAAAPAPAKK.D z=2,scan#=6823,scan time=22.7491
Intensity
m/z
ASAPAADTSAAAPAPAKK
M+e
M+e-45
HexNAc
HexNAcHex
HexNAc(2)Hex(1)
Pep_2+
Pep+HexNAc_2+
Pep_1+
Pep+HexNAc_1+
z.9
z.10
z.11
z.12
z.13
z.14
z.16
c15
c16
c17

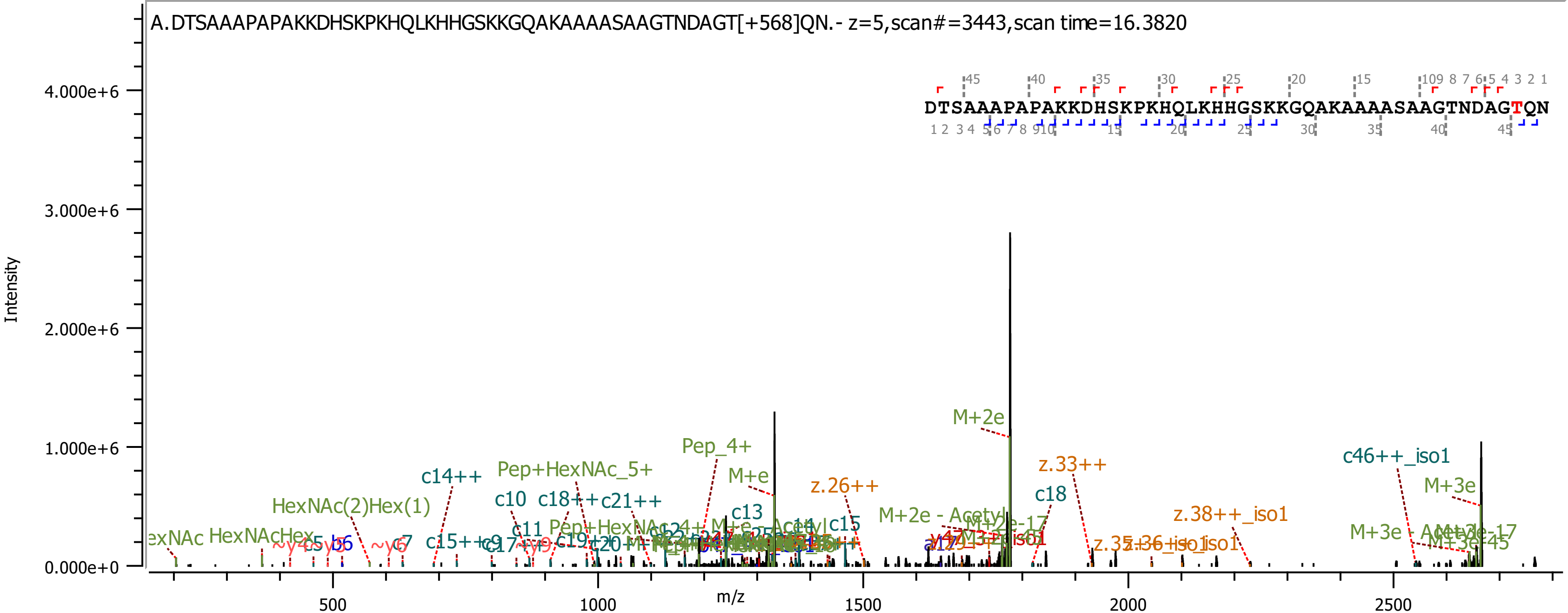
A.DTSAAAPAPAKKDHSKPKHQLKHHGSKKGQAKAAAASAAGTNDAGT[+568]QN.- z=5,scan#=3443,scan time=16.3820
DTSAAAPAPAKKDHSKPKHQLKHHGSKKGQAKAAAASAAGTNDAGTQN
Intensity
4.000e+6
3.000e+6
2.000e+6
1.000e+6
0.000e+0
m/z
500
1000
1500
2000
2500
M+2e
Pep_4+
M+e
c14++
Pep+HexNAc_5+
c46++_iso1
z.33++
z.26++
M+3e
c18
c13
c10
c18++
c21++
HexNAc(2)Hex(1)
z.38++_iso1
M+2e - Acetyl
M+3e - Acetyl
HexNAcHex
c11
c15
c7
b6

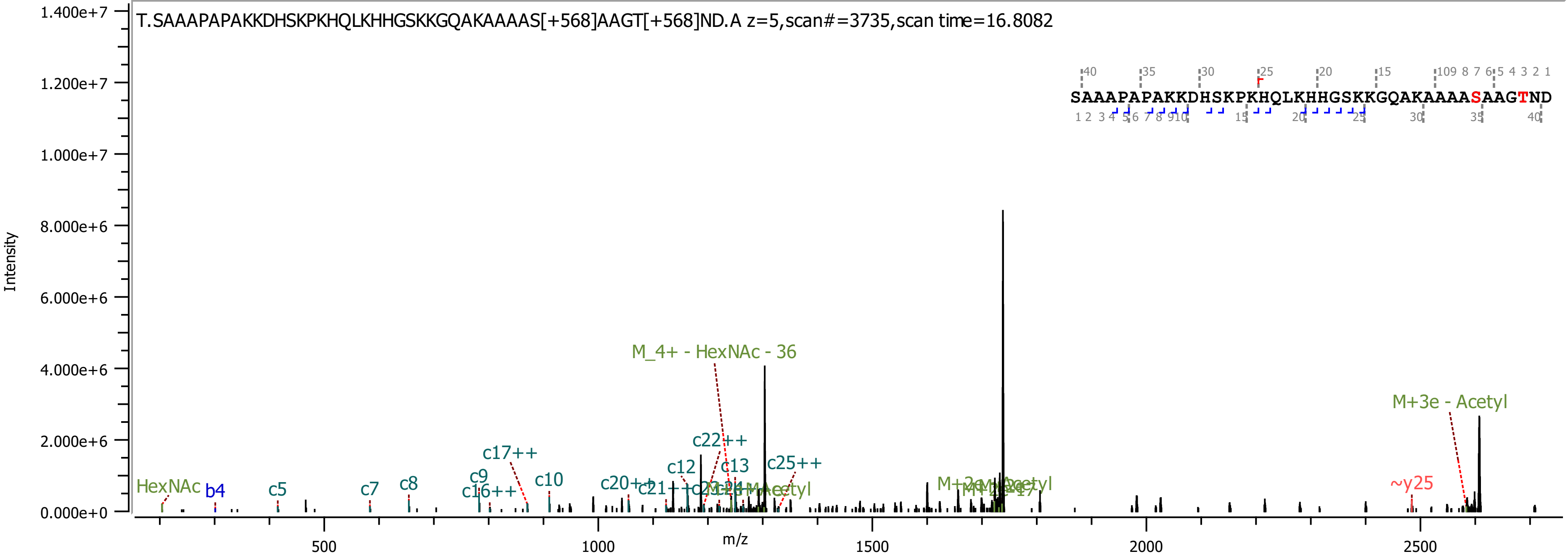

T.SAAAPAPAKKDHSKPKHQLKHHGSKKGQAKAAAAS[+568]AAGT[+568]ND.A z=5,scan#=3735,scan time=16.8082
SAAAPAPAKKDHSKPKHQLKHHGSKKGQAKAAAASAAGTND
Intensity
m/z
M_4+ - HexNAc - 36
M+3e - Acetyl
c22++
c17++
c25++
c12
c13
c9
c10
c16++
c20++
c8
c7
c5
b4
HexNAc
~y25

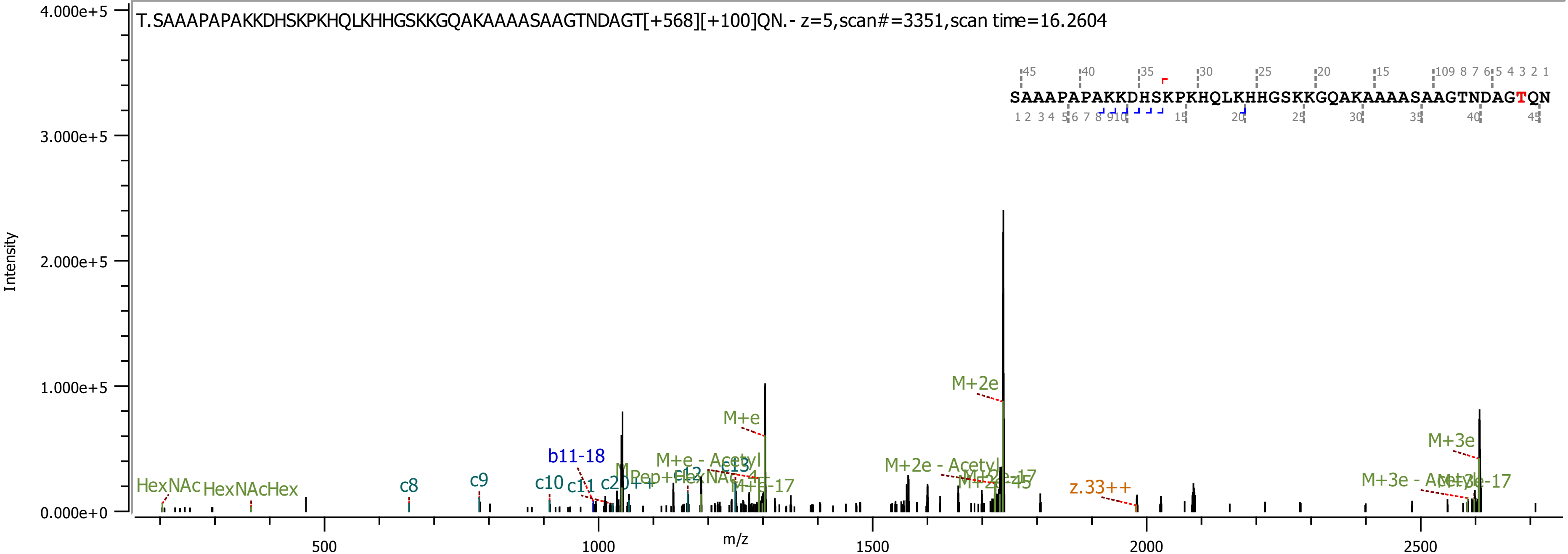

T.SAAAPAPAKKDHSKPKHQLKHHGSKKGQAKAAAASAAGTNDAGT[+568][+100]QN.- z=5,scan#=3351,scan time=16.2604
SAAAPAPAKKDHSKPKHQLKHHGSKKGQAKAAAASAAGTNDAGTQN
Intensity
m/z
4.000e+5
3.000e+5
2.000e+5
1.000e+5
0.000e+0
500
1000
1500
2000
2500
HexNAc
HexNAcHex
c8
c9
c10
c11
b11-18
c20++
c12
c13
M+e
M+e - Acetyl
M+e-17
M+2e
M+2e - Acetyl
M+2e-17
M+2e-45
z.33++
M+3e
M+3e - Acetyl
M+3e-17

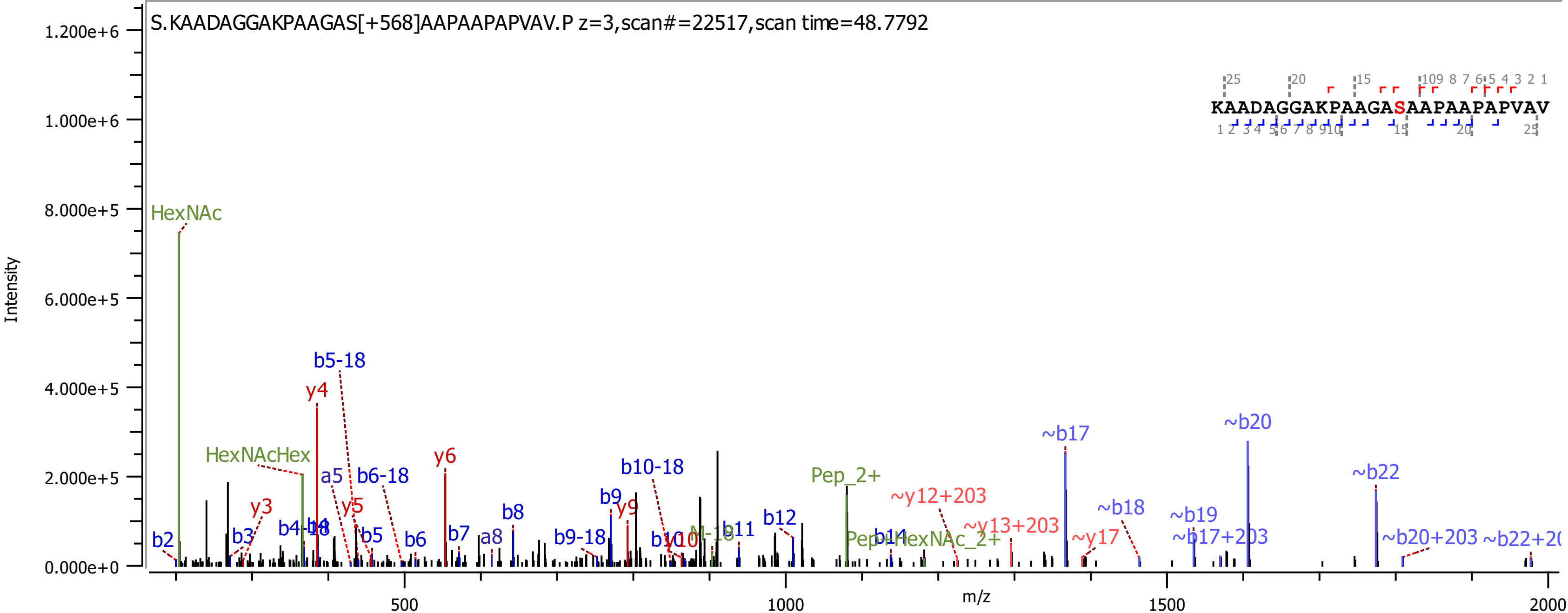
S.KAADAGGAKPAAGAS[+568]AAPAAPAPVAV.P z=3,scan#=22517,scan time=48.7792
KAADAGGAKPAAGASAAPAAPAPVAV
Intensity
m/z
HexNAc
HexNAcHex
b5-18
y4
y6
a5
b6-18
y3
y5
b2
b3
b4-18
b5
b6
b7
a8
b8
b9-18
b9
y9
b10-18
b11
b12
Pep_2+
Pep+HexNAc_2+
b14
~y12+203
~y13+203
~y17
~b17
~b18
~b19
~b17+203
~b20
~b22
~b20+203
0.000e+0
2.000e+5
4.000e+5
6.000e+5
8.000e+5
1.000e+6
1.200e+6
500
1000
1500
2000

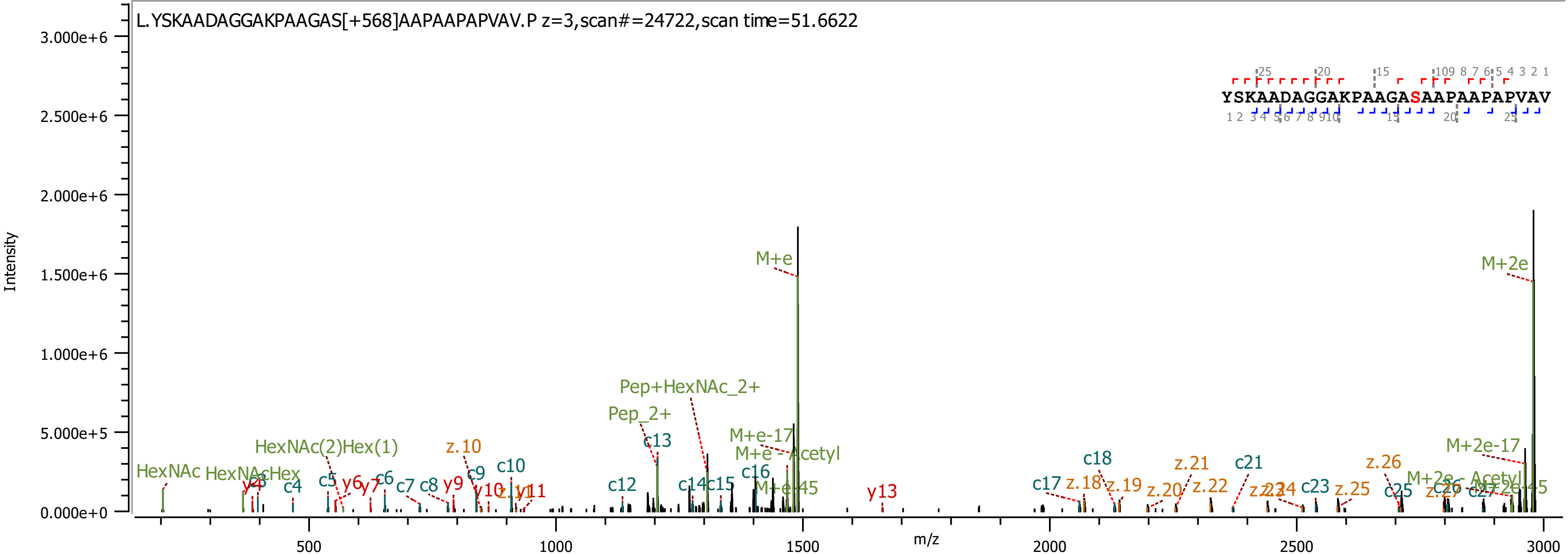
L.YSKAADAGGAKPAAGAS[+568]AAPAAPAPVAV.P z=3,scan#=24722,scan time=51.6622
YSKAADAGGAKPAAGASAAPAAPAPVAV
Intensity
m/z
M+e
M+2e
Pep+HexNAc_2+
Pep_2+
HexNAc(2)Hex(1)
HexNAc
HexNAcHex
M+e-17
M+e - Acetyl
M+e-45
M+2e-17
M+2e - Acetyl
M+2e-45
c4
c5
c6
c7
c8
c9
c10
c12
c13
c14
c15
c16
c17
c18
c21
c23
c25
c26
c27
y6
y7
y9
y10
y11
y13
z.10
z.18
z.19
z.20
z.21
z.22
z.25
z.26
z.27
0.000e+0
5.000e+5
1.000e+6
1.500e+6
2.000e+6
2.500e+6
3.000e+6
500
1000
1500
2000
2500
3000

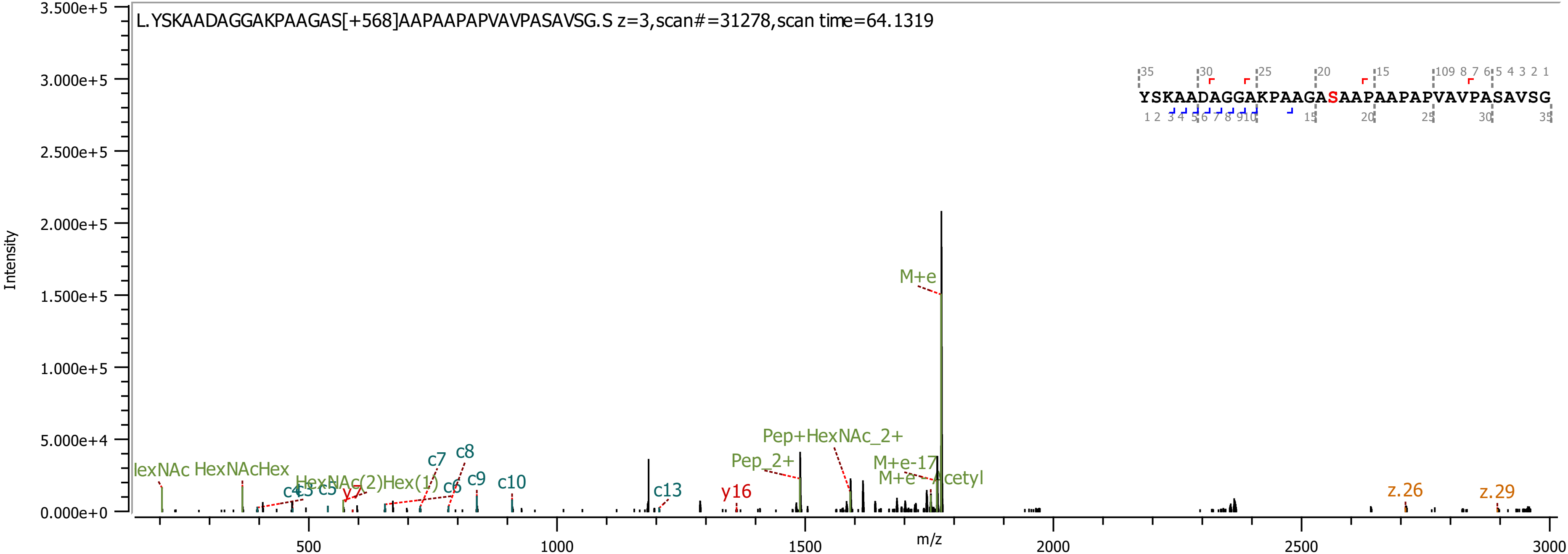

L.YSKAADAGGAKPAAGAS[+568]AAPAAPAPVAVPASAVSG.S z=3,scan#=31278,scan time=64.1319
YSKAADAGGAKPAAGASAAPAAPAPVAVPASAVSG
Intensity
m/z
M+e
Pep+HexNAc_2+
Pep_2+
M+e-17
M+e - Acetyl
HexNAc
HexNAcHex
HexNAc(2)Hex(1)
c4
c5
c5
c6
c7
c8
c9
c10
c13
y7
y16
z.26
z.29

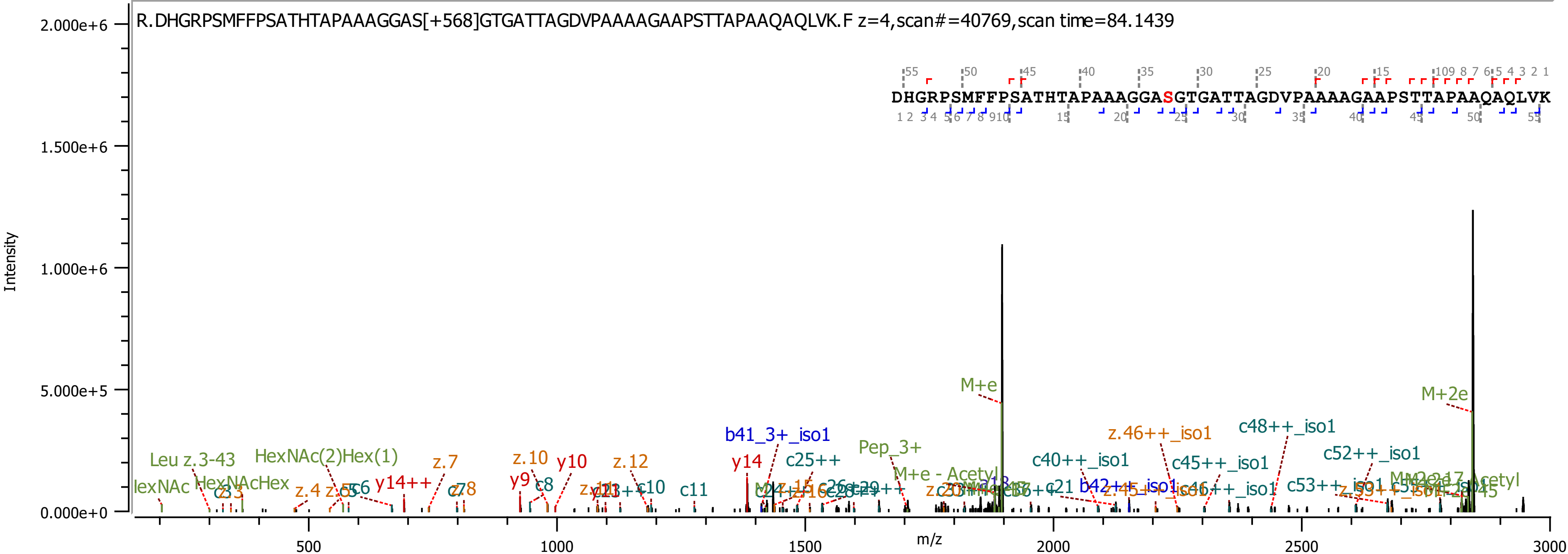
R.DHGRPSMFFPSATHTAPAAAGGAS[+568]GTGATTAGDVPAAAAGAAPSTTAPAAQAQLVK.F z=4,scan#=40769,scan time=84.1439
DHGRPSMFFPSATHTAPAAAGGASGTGATTAGDVPAAAAGAAPSTTAPAAQAQLVK
Intensity
m/z
2.000e+6
1.500e+6
1.000e+6
5.000e+5
0.000e+0
500
1000
1500
2000
2500
3000
M+e
M+2e
b41_3+_iso1
c25++
Pep_3+
z.46++_iso1
c48++_iso1
c40++_iso1
c45++_iso1
c52++_iso1
HexNAc(2)Hex(1)
Leu z.3-43
z.7
z.10
y10
z.12
y14
y14++
y9
c8
c11
HexNAcHex
z.4
c21
b42++_iso1
c53++_iso1
M+e - Acetyl

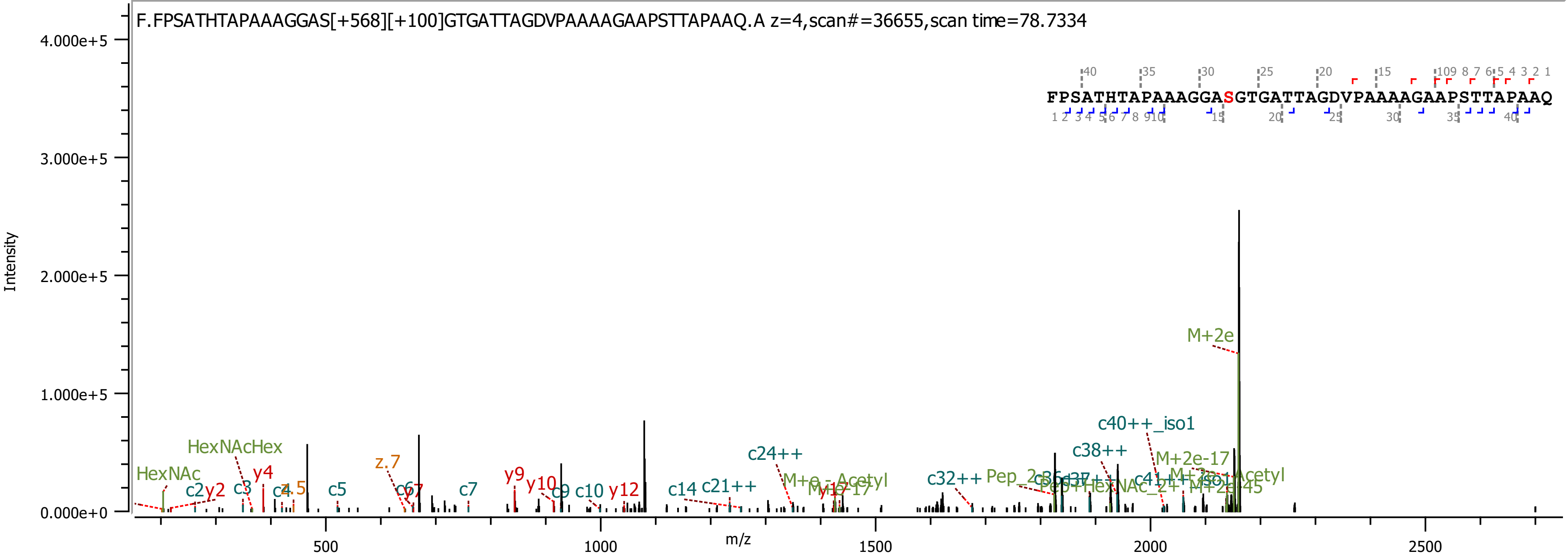

F.FPSATHTAPAAAGGAS[+568][+100]GTGATTAGDVPAAAAGAAPSTTAPAAQ.A z=4,scan#=36655,scan time=78.7334
FPSATHTAPAAAGGASGTGATTAGDVPAAAAGAAPSTTAPAAQ
Intensity
m/z
M+2e
c40++_iso1
c38++
M+2e-17
c24++
HexNAcHex
HexNAc
z.7
c32++
c21++
c14
y12
c10
y10
y9
c7
c5
y4
c3
c2
y2

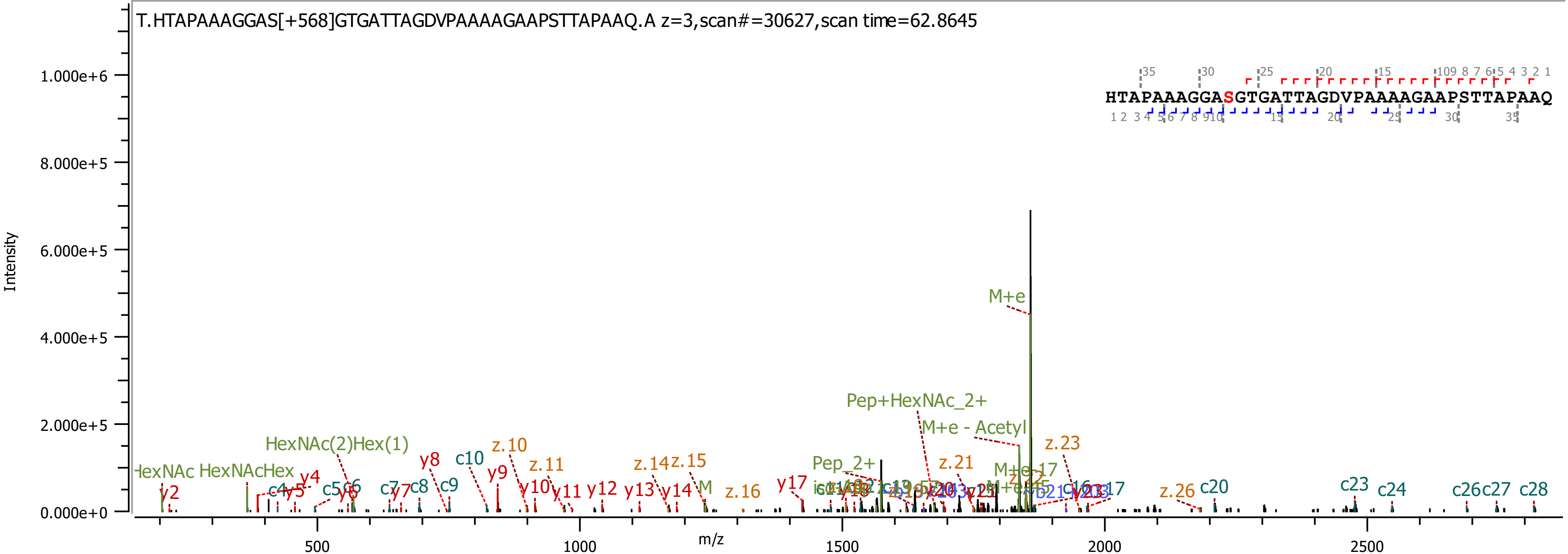
T.HTAPAAAGGAS[+568]GTGATTAGDVPAAAAGAAPSTTAPAAQ.A z=3,scan#=30627,scan time=62.8645
HTAPAAAGGASGTGATTAGDVPAAAAGAAPSTTAPAAQ
Intensity
m/z
1.000e+6
8.000e+5
6.000e+5
4.000e+5
2.000e+5
0.000e+0
500
1000
1500
2000
2500
M+e
Pep+HexNAc_2+
M+e - Acetyl
HexNAc(2)Hex(1)
HexNAcHex
Pep_2+
z.23
z.21
z.10
z.11
z.14
z.15
z.16
z.26
c20
c23
c24
c26
c27
c28
y2
y4
y5
y8
y9
y10
y11
y12
y13
y14
y17
c4
c5
c7
c8
c9
c10

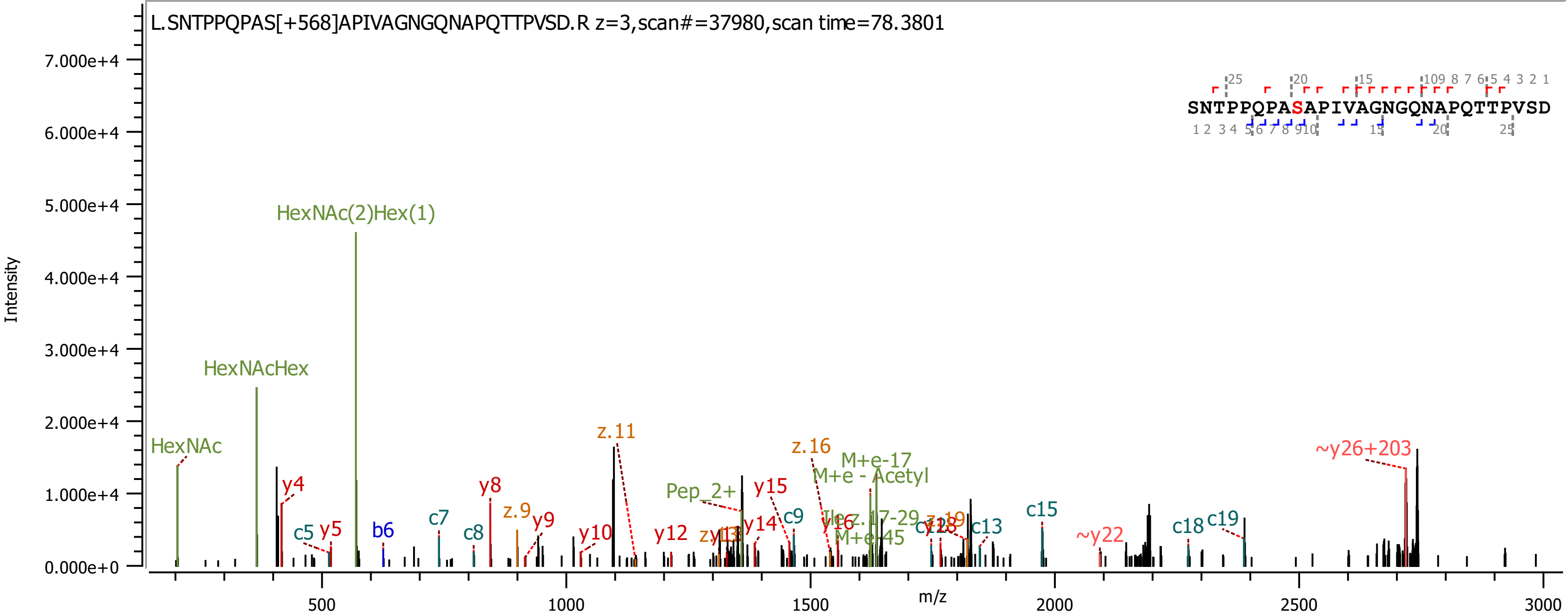

L.SNTPPQPAS[+568]APIVAGNGQNAPQTTPVSD.R z=3,scan#=37980,scan time=78.3801
SNTPPQPASAPIVAGNGQNAPQTTPVSD
Intensity
m/z
HexNAc(2)Hex(1)
HexNAcHex
HexNAc
y4
c5
y5
b6
c7
c8
y8
z.9
y9
y10
z.11
y12
y14
Pep_2+
y15
c9
z.16
M+e-17
M+e - Acetyl
y16
M+e-45
c13
c15
~y22
c18
c19
~y26+203

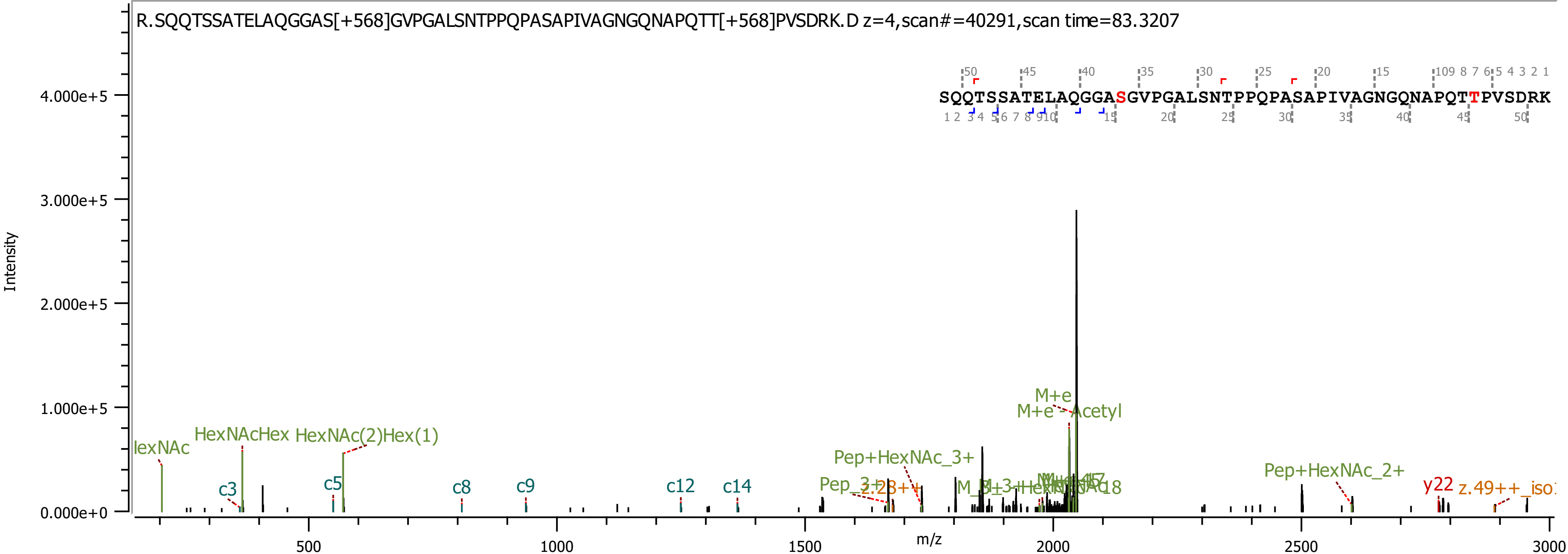

R.SQQTSSATELAQGGAS[+568]GVPGALSNTPPQPASAPIVAGNGQNAPQTT[+568]PVSDRK.D z=4,scan#=40291,scan time=83.3207
SQQTSSATELAQGGASGVPGALSNTPPQPASAPIVAGNGQNAPQTTPVSDRK
Intensity
m/z
M+e
M+e - Acetyl
HexNAcHex
HexNAc(2)Hex(1)
Pep+HexNAc_3+
Pep+HexNAc_2+
c3
c5
c8
c9
c12
c14
y22

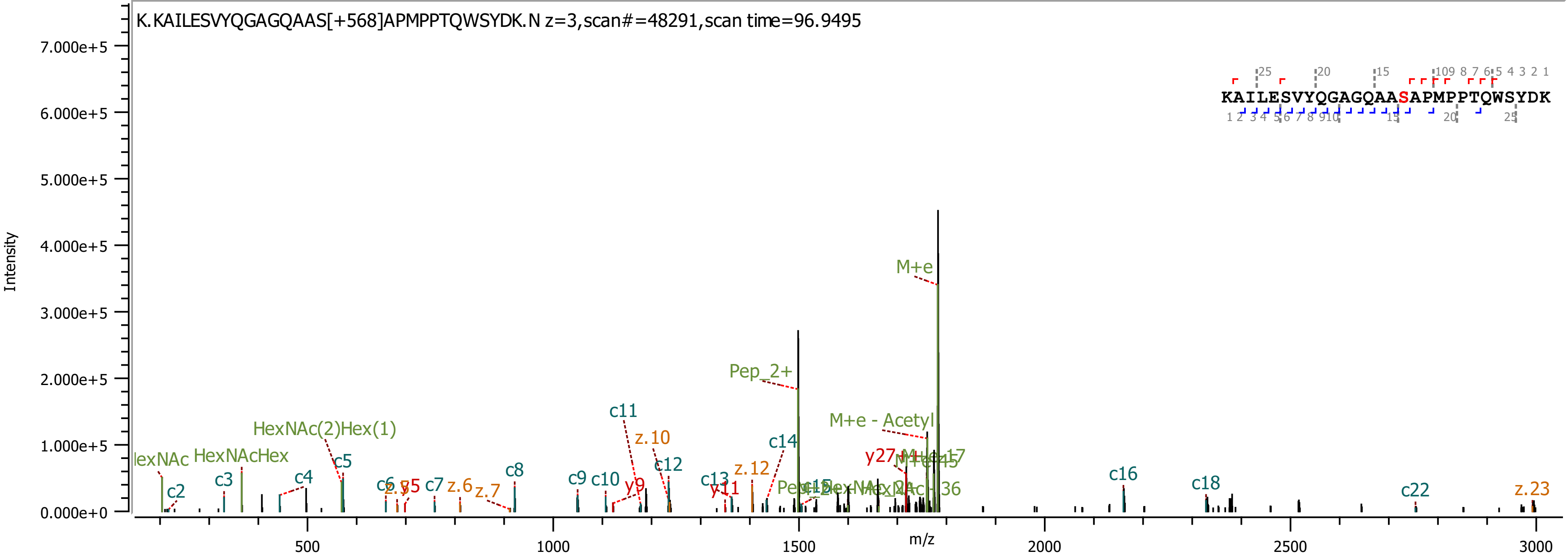
K.KAILESVYQGAGQAAS[+568]APMPPTQWSYDK.N z=3,scan#=48291,scan time=96.9495
KAILESVYQGAGQAASAPMPPTQWSYDK
Intensity
m/z
M+e
Pep_2+
M+e - Acetyl
HexNAc(2)Hex(1)
HexNAcHex
c2
c3
c4
c5
c7
c8
c9
c10
c11
c12
c13
c14
c16
c18
c22
z.6
z.7
z.10
z.12
z.23
y9
y11
y27

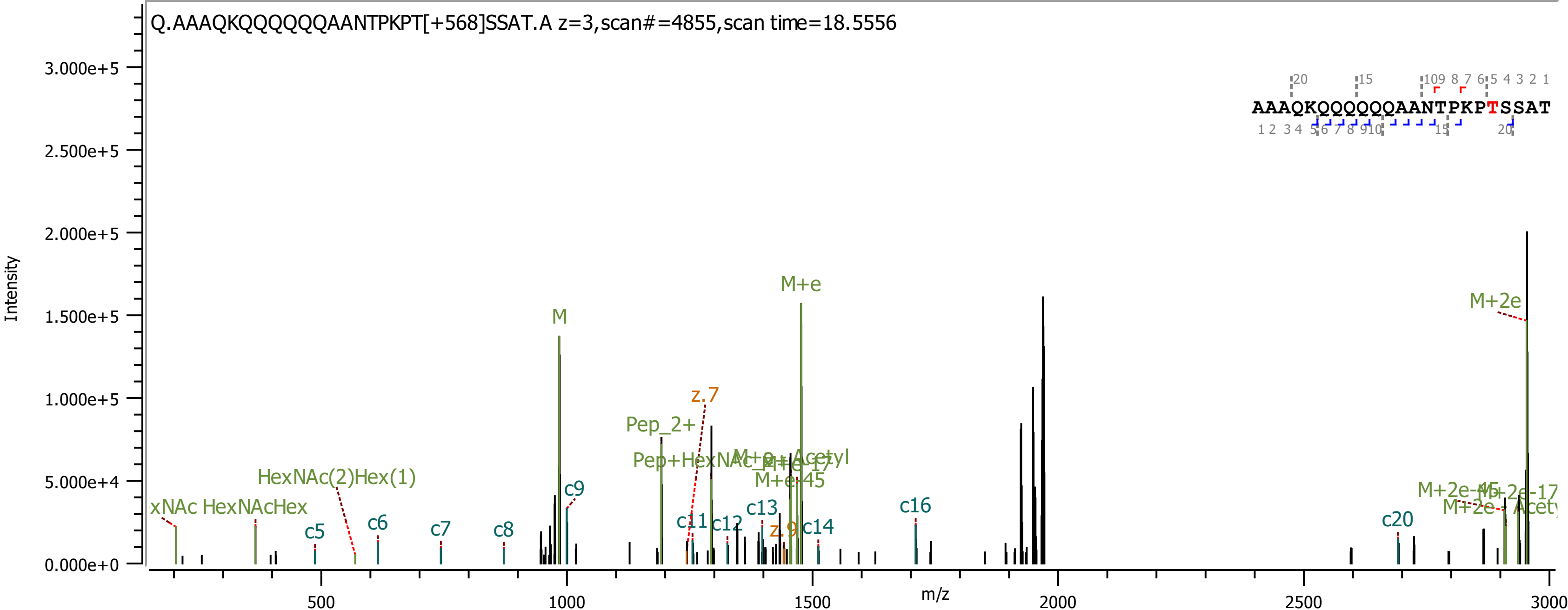

Q.AAAQKQQQQQQAANTPKPT[+568]SSAT.A z=3,scan#=4855,scan time=18.5556
Intensity
m/z
M
M+e
M+2e
Pep_2+
Pep+HexNAc
M+e-45
HexNAc(2)Hex(1)
HexNAcHex
c5
c6
c7
c8
c9
c11
c12
c13
c14
c16
c20
z.7
z.9
AAAQKQQQQQQAANTPKPTSSAT

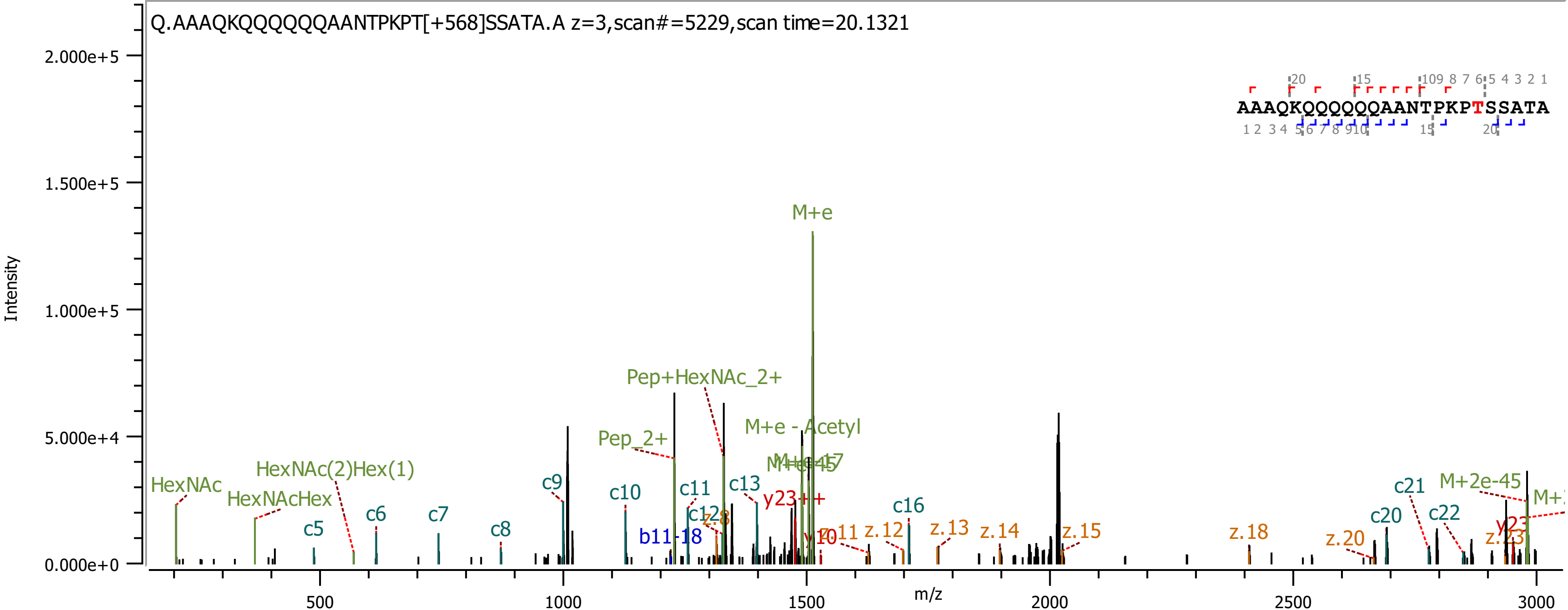

Q.AAAQKQQQQQQAANTPKPT[+568]SSATA.A z=3,scan#=5229,scan time=20.1321
AAAQKQQQQQQAANTPKPTSSATA
Intensity
m/z
M+e
Pep+HexNAc_2+
Pep_2+
M+e - Acetyl
HexNAc
HexNAcHex
HexNAc(2)Hex(1)
M+2e-45
c5
c6
c7
c8
c9
c10
c11
c13
c16
c20
c21
c22
b11-18
y23++
z.12
z.13
z.14
z.15
z.18
z.20

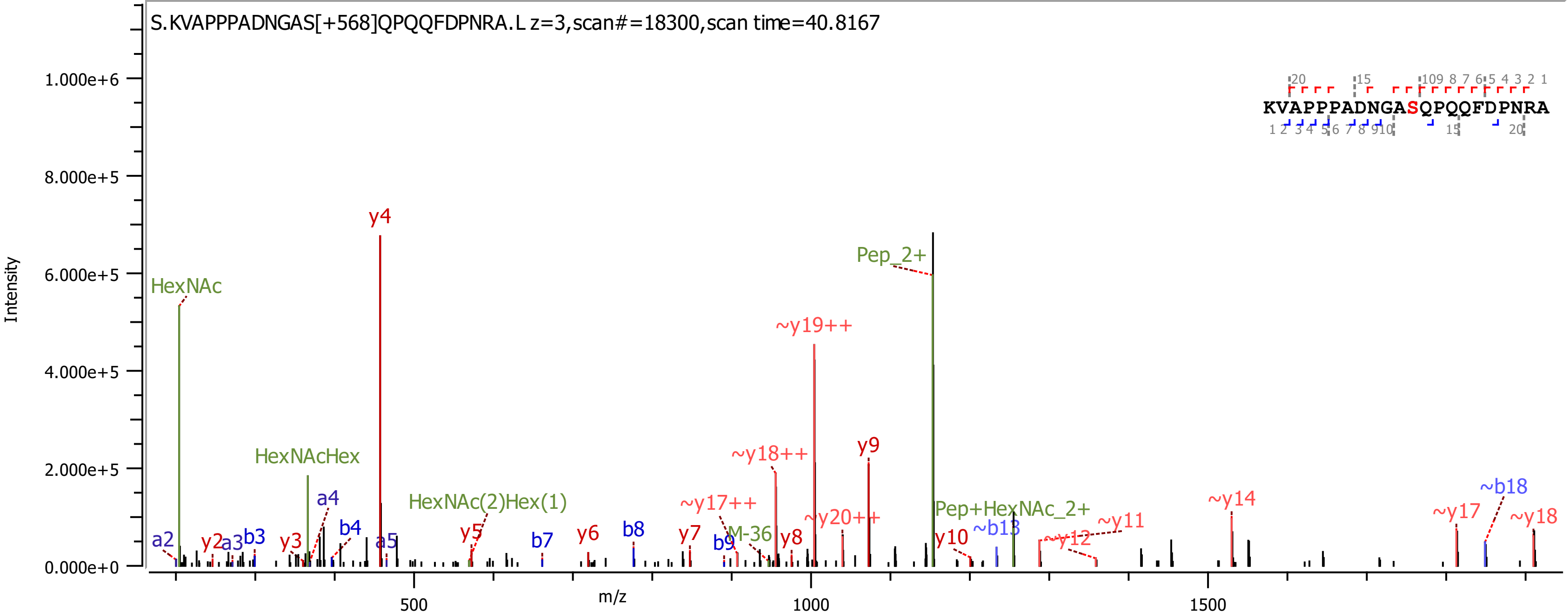
S.KVAPPPADNGAS[+568]QPQQFDPNRA.L z=3,scan#=18300,scan time=40.8167
KVAPPPADNGASQPQQFDPNRA
Intensity
m/z
HexNAc
HexNAcHex
HexNAc(2)Hex(1)
Pep_2+
Pep+HexNAc_2+
M-36
y4
~y19++
~y18++
y9
~y17++
~y20++
~y14
~b18
~y17
~y18
~y11
~y12
~b13
y10
y8
y7
y6
y5
y3
y2
a2
a3
b3
a4
b4
a5
b7
b8
b9

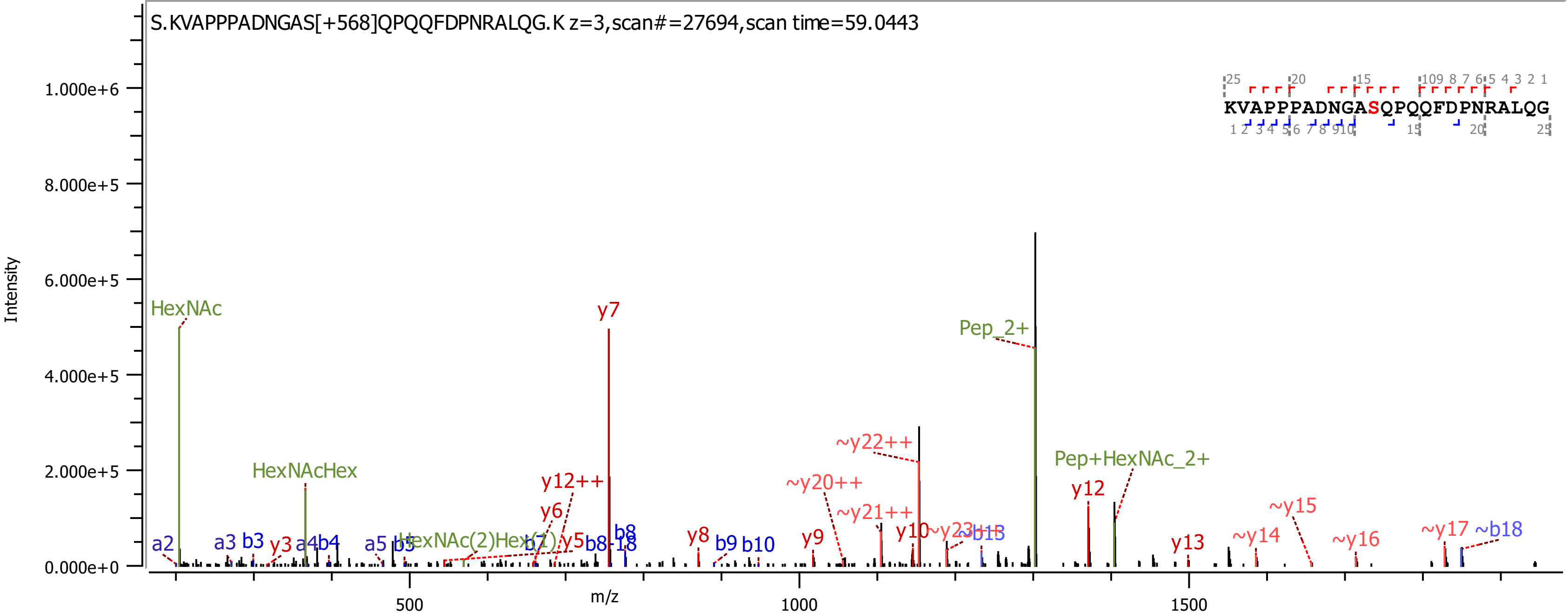

S.KVAPPPADNGAS[+568]QPQQFDPNRALQG.K z=3,scan#=27694,scan time=59.0443
KVAPPPADNGASQPQQFDPNRALQG
Intensity
m/z
1.000e+6
8.000e+5
6.000e+5
4.000e+5
2.000e+5
0.000e+0
500
1000
1500
HexNAc
HexNAcHex
HexNAc(2)Hex(1)
Pep_2+
Pep+HexNAc_2+
y7
y12++
y6
y5
y3
y8
y9
y10
y12
y13
~y14
~y15
~y16
~y17
~b18
~y20++
~y21++
~y22++
~y23++
~b13
a2
a3
b3
a4
b4
a5
b5
b7
b8
b8-18
b9
b10

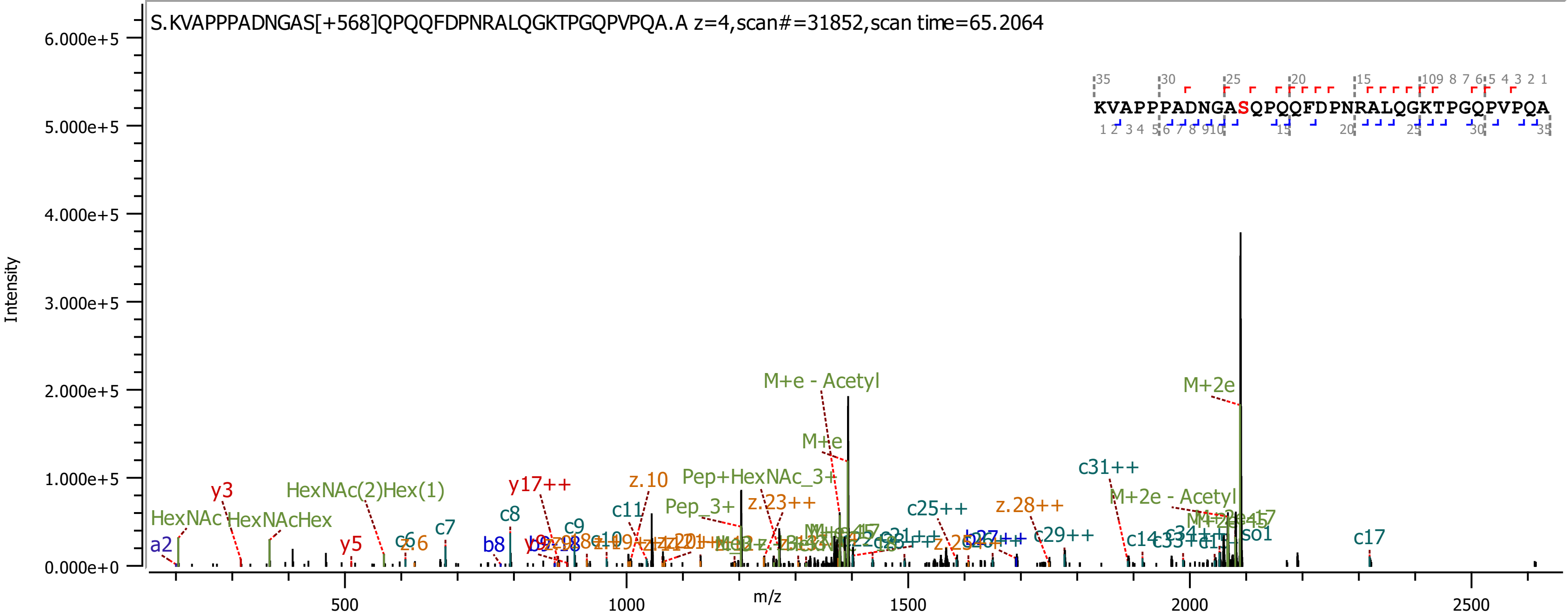
S.KVAPPPADNGAS[+568]QPQQFDPNRALQGKTPGQPVPQA.A z=4,scan#=31852,scan time=65.2064
KVAPPPADNGASQPQQFDPNRALQGKTPGQPVPQA
Intensity
m/z
6.000e+5
5.000e+5
4.000e+5
3.000e+5
2.000e+5
1.000e+5
0.000e+0
500
1000
1500
2000
2500
M+e - Acetyl
M+2e
M+e
Pep+HexNAc_3+
Pep_3+
c31++
M+2e - Acetyl
HexNAc(2)Hex(1)
y17++
z.10
z.23++
z.28++
c25++
c29++
HexNAc
HexNAcHex
y3
y5
c6
z.6
c7
c8
b8
c9
c11
c10
c17
c14
a2
iso1

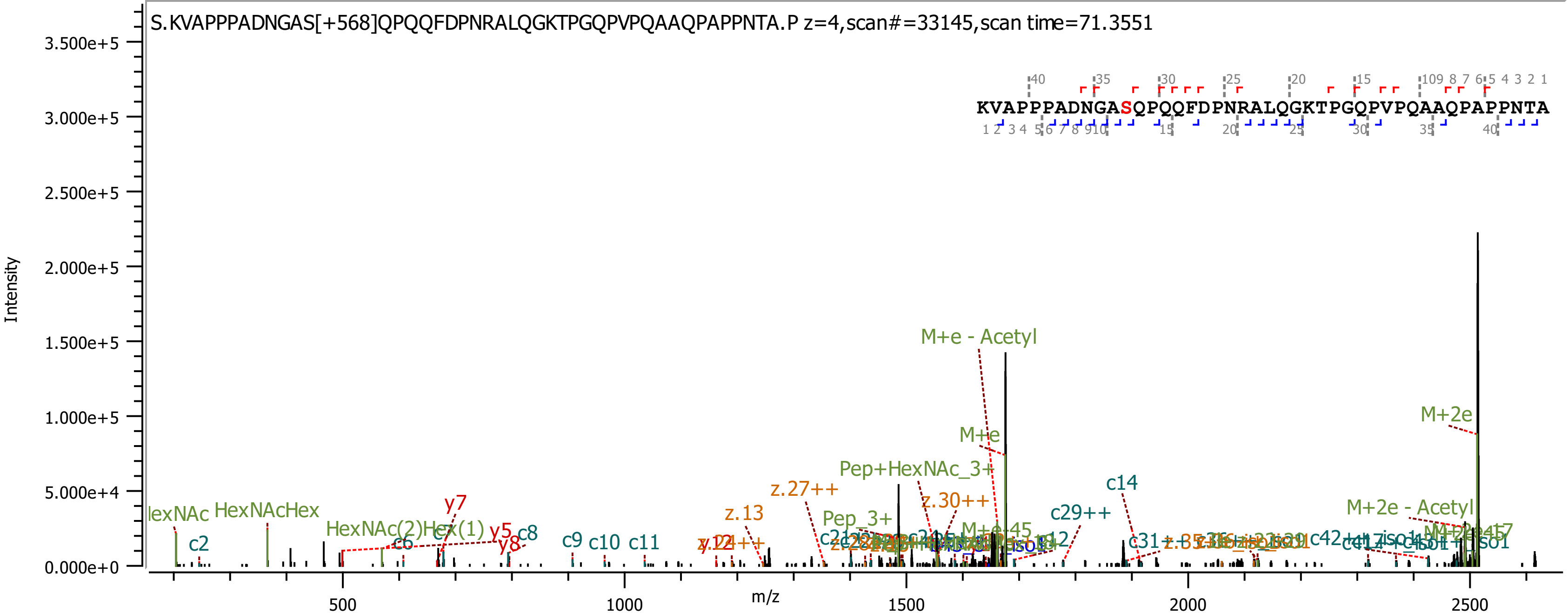

S.KVAPPPADNGAS[+568]QPQQFDPNRALQGKTPGQPVPQAAQPAPPNTA.P z=4,scan#=33145,scan time=71.3551
KVAPPPADNGASQPQQFDPNRALQGKTPGQPVPQAAQPAPPNTA
Intensity
m/z
M+e - Acetyl
M+2e
M+e
Pep+HexNAc_3+
M+2e - Acetyl
c14
c29++
z.30++
z.27++
z.13
Pep_3+
HexNAcHex
HexNAc(2)Hex(1)
y7
y5
y8
c8
c9
c10
c11
c2
c6

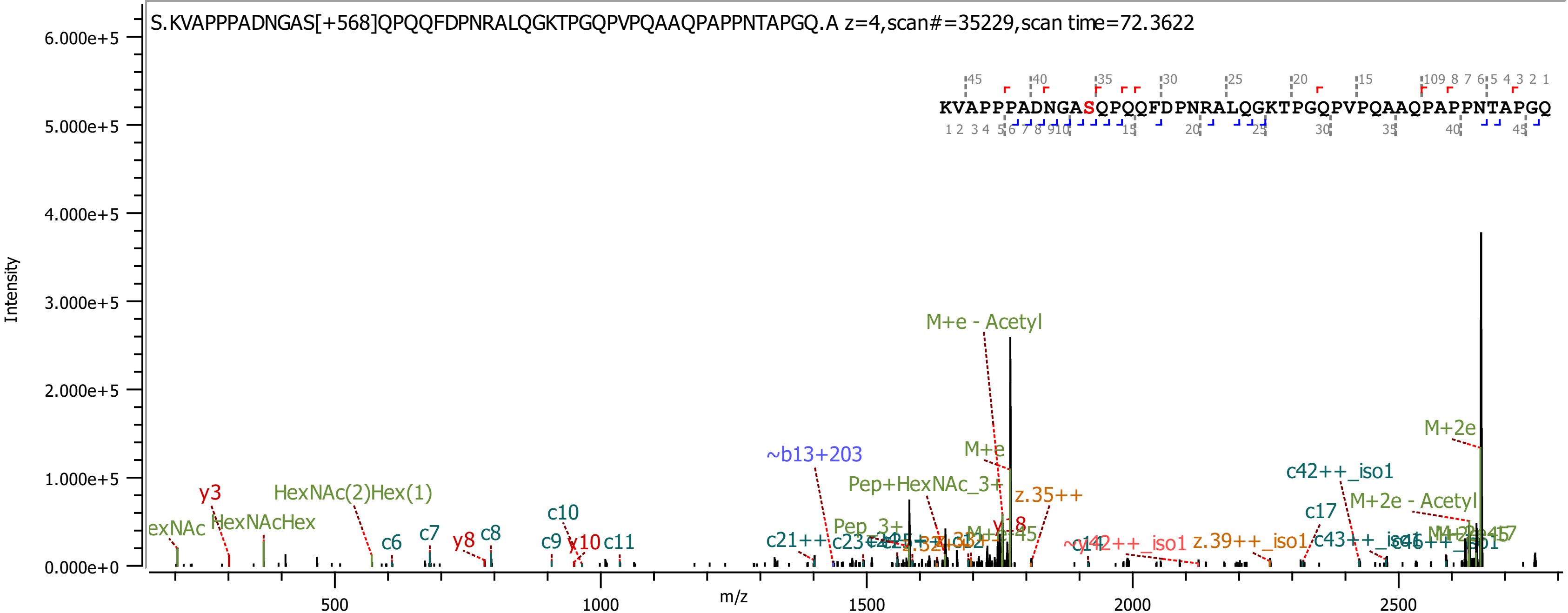

S.KVAPPPADNGAS[+568]QPQQFDPNRALQGKTPGQPVPQAAQPAPPNTAPGQ.A z=4,scan#=35229,scan time=72.3622
KVAPPPADNGASQPQQFDPNRALQGKTPGQPVPQAAQPAPPNTAPGQ
Intensity
m/z
M+e - Acetyl
M+2e
~b13+203
M+e
Pep+HexNAc_3+
z.35++
c42++_iso1
M+2e - Acetyl
c17
HexNAc(2)Hex(1)
y3
HexNAcHex
c10
c6
c7
y8
c8
c9
y10
c11
c21++
Pep_3+
y18
z.39++_iso1

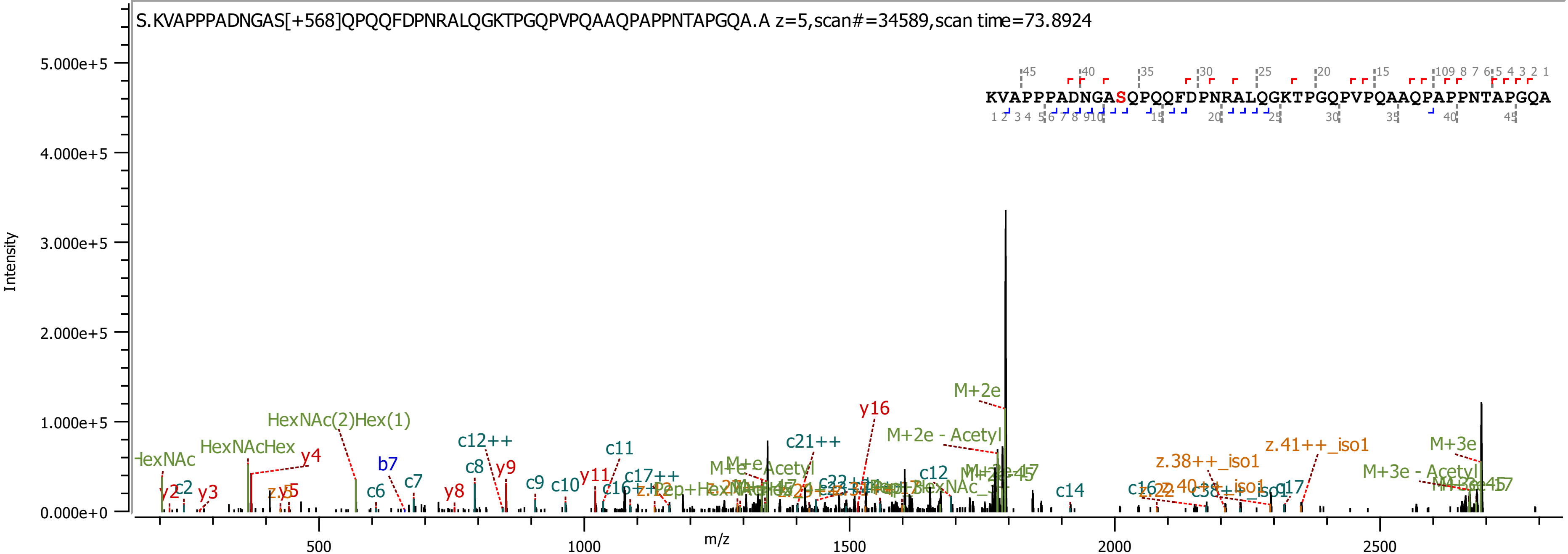

S.KVAPPPADNGAS[+568]QPQQFDPNRALQGKTPGQPVPQAAQPAPPNTAPGQA.A z=5,scan#=34589,scan time=73.8924
KVAPPPADNGASQPQQFDPNRALQGKTPGQPVPQAAQPAPPNTAPGQA
Intensity
m/z
5.000e+5
4.000e+5
3.000e+5
2.000e+5
1.000e+5
0.000e+0
500
1000
1500
2000
2500
HexNAcHex
HexNAc(2)Hex(1)
M+2e
M+2e - Acetyl
M+3e
M+3e - Acetyl
y16
c21++
c12++
c11
z.41++_iso1
z.38++_iso1

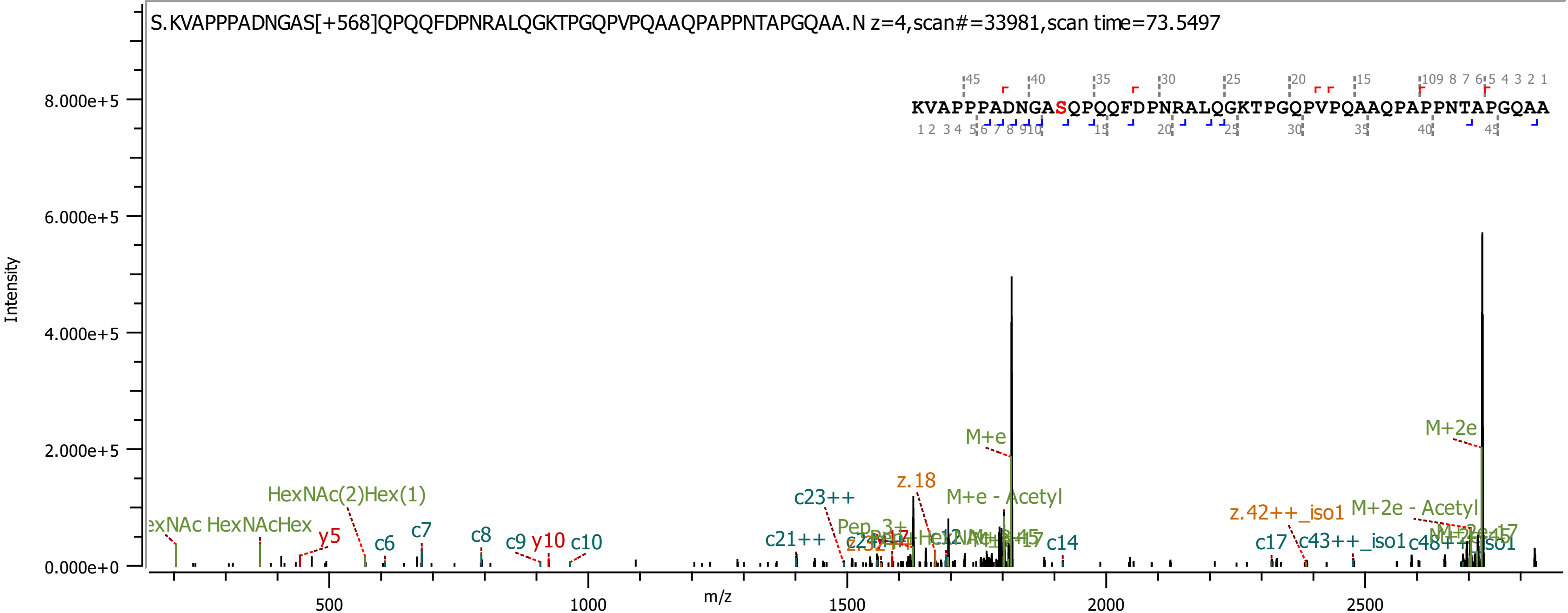

S.KVAPPPADNGAS[+568]QPQQFDPNRALQGKTPGQPVPQAAQPAPPNTAPGQAA.N z=4,scan#=33981,scan time=73.5497
KVAPPPADNGASQPQQFDPNRALQGKTPGQPVPQAAQPAPPNTAPGQAA
Intensity
8.000e+5
6.000e+5
4.000e+5
2.000e+5
0.000e+0
m/z
500
1000
1500
2000
2500
HexNAc(2)Hex(1)
HexNAcHex
y5
c6
c7
c8
c9
y10
c10
c21++
c23++
z.18
M+e
M+e - Acetyl
c14
c17
z.42++_iso1
c43++_iso1
M+2e - Acetyl
M+2e

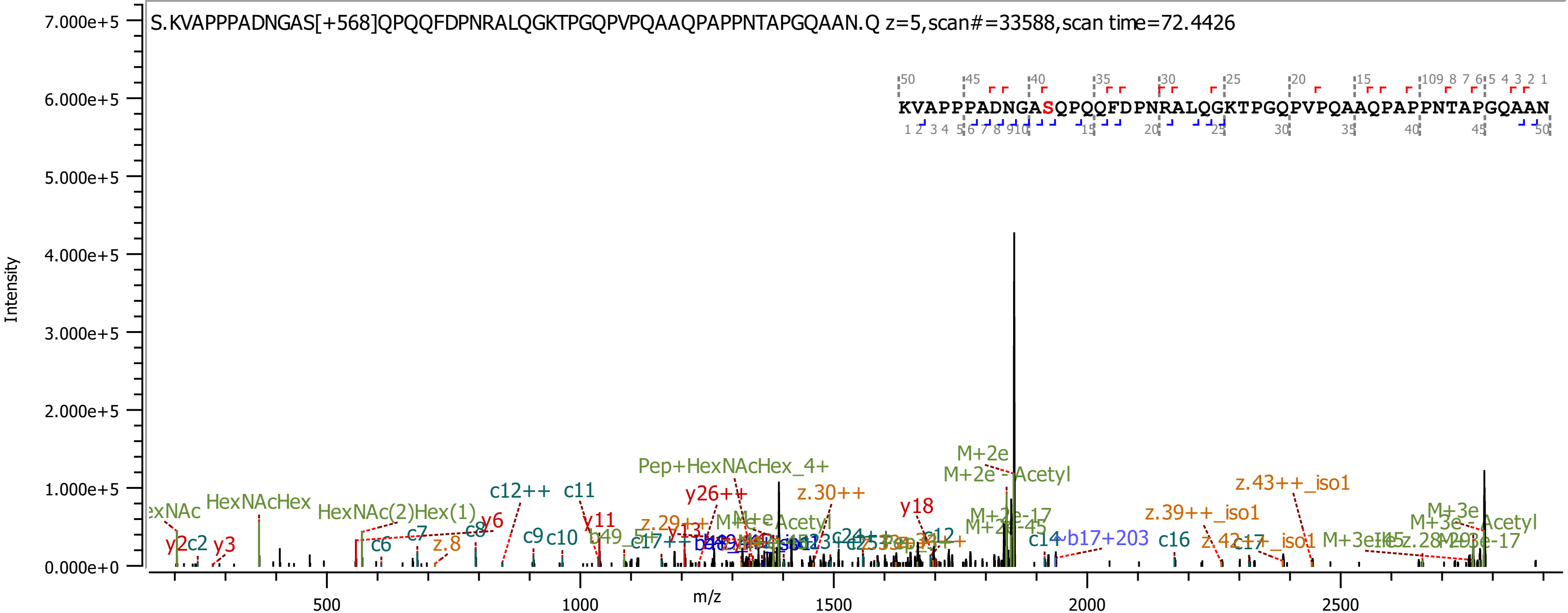
S.KVAPPPADNGAS[+568]QPQQFDPNRALQGKTPGQPVPQAAQPAPPNTAPGQAAN.Q z=5,scan#=33588,scan time=72.4426
KVAPPPADNGASQPQQFDPNRALQGKTPGQPVPQAAQPAPPNTAPGQAAN
Intensity
7.000e+5
6.000e+5
5.000e+5
4.000e+5
3.000e+5
2.000e+5
1.000e+5
0.000e+0
m/z
500
1000
1500
2000
2500
HexNAcHex
HexNAc(2)Hex(1)
Pep+HexNAcHex_4+
M+2e
M+2e-Acetyl
z.43++_iso1
z.39++_iso1
M+3e
M+3e-Acetyl
c12++
c11
y26++
z.30++
y18
y2
c2
y3
c6
c7
z.8
c8
y6
c9
c10
y11
z.29++
c12
M+2e-17
M+2e-45
c14
b17+203
c16
c17
M+3e-17

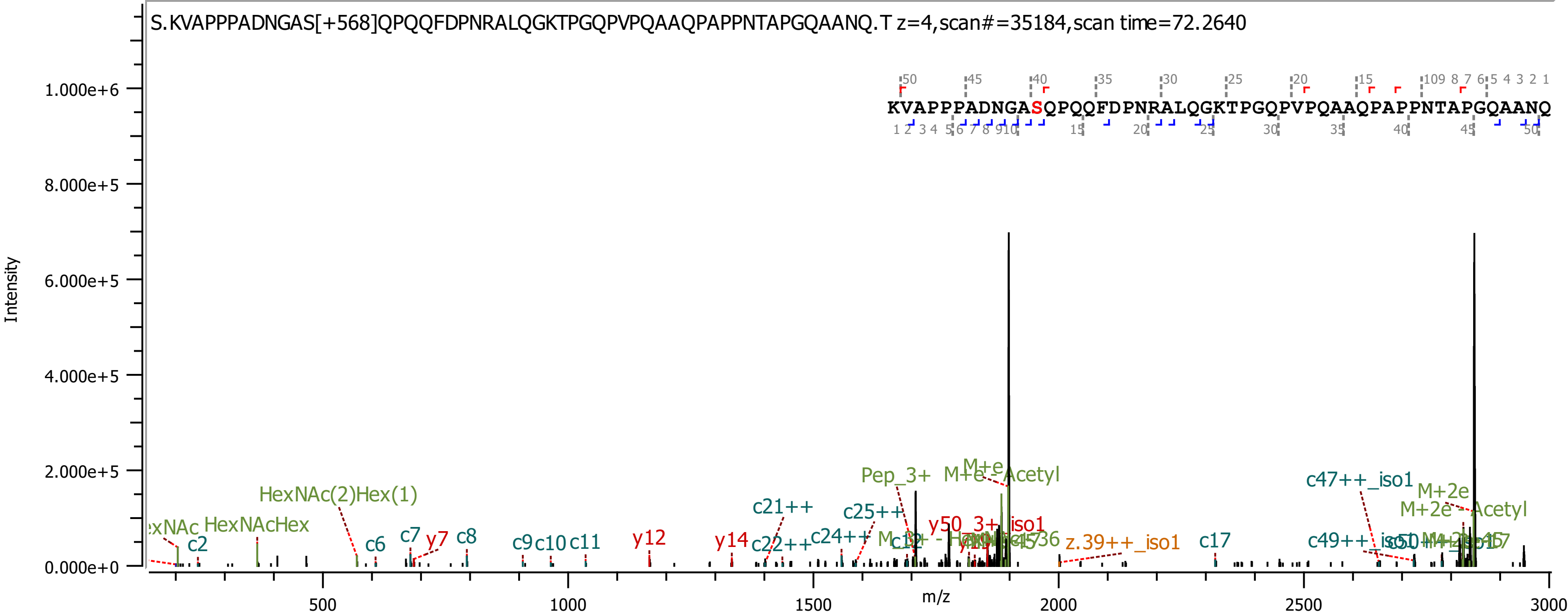

S.KVAPPPADNGAS[+568]QPQQFDPNRALQGKTPGQPVPQAAQPAPPNTAPGQAANQ.T z=4,scan#=35184,scan time=72.2640
KVAPPPADNGASQPQQFDPNRALQGKTPGQPVPQAAQPAPPNTAPGQAANQ
Intensity
m/z
HexNAcHex
HexNAc(2)Hex(1)
c2
c6
c7
y7
c8
c9
c10
c11
y12
y14
c21++
c22++
c24++
c25++
Pep_3+
M+e
M+e - Acetyl
y50_3+_iso1
z.39++_iso1
c17
c47++_iso1
c49++_iso1
M+2e
M+2e - Acetyl

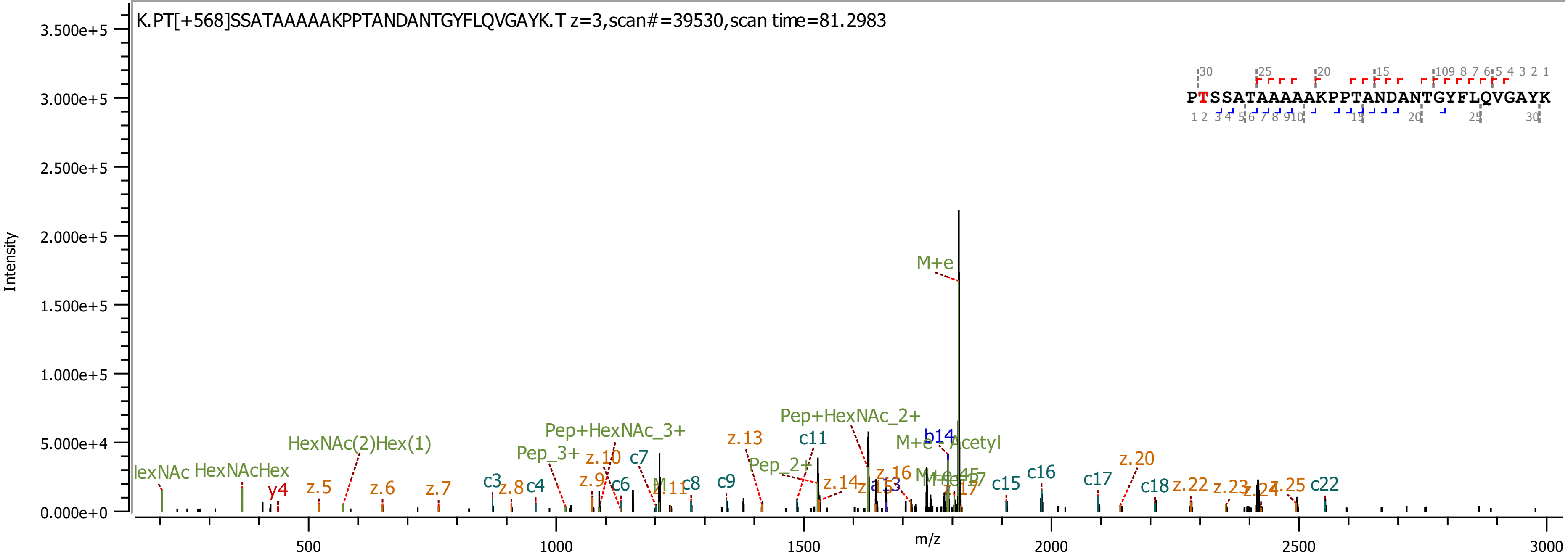
K.PT[+568]SSATAAAAAKPPTANDANTGYFLQVGAYK.T z=3,scan#=39530,scan time=81.2983
PTSSATAAAAAKPPTANDANTGYFLQVGAYK
Intensity
m/z
M+e
Pep+HexNAc_2+
Pep+HexNAc_3+
HexNAc(2)Hex(1)
HexNAcHex
Pep_3+
Pep_2+
M+e - Acetyl
b14
y4
c3
c4
c6
c7
c8
c9
c11
c15
c16
c17
c18
c22
z.5
z.6
z.7
z.8
z.9
z.10
z.13
z.14
z.16
z.20
z.22
z.23
z.25
0.000e+0
5.000e+4
1.000e+5
1.500e+5
2.000e+5
2.500e+5
3.000e+5
3.500e+5
500
1000
1500
2000
2500
3000

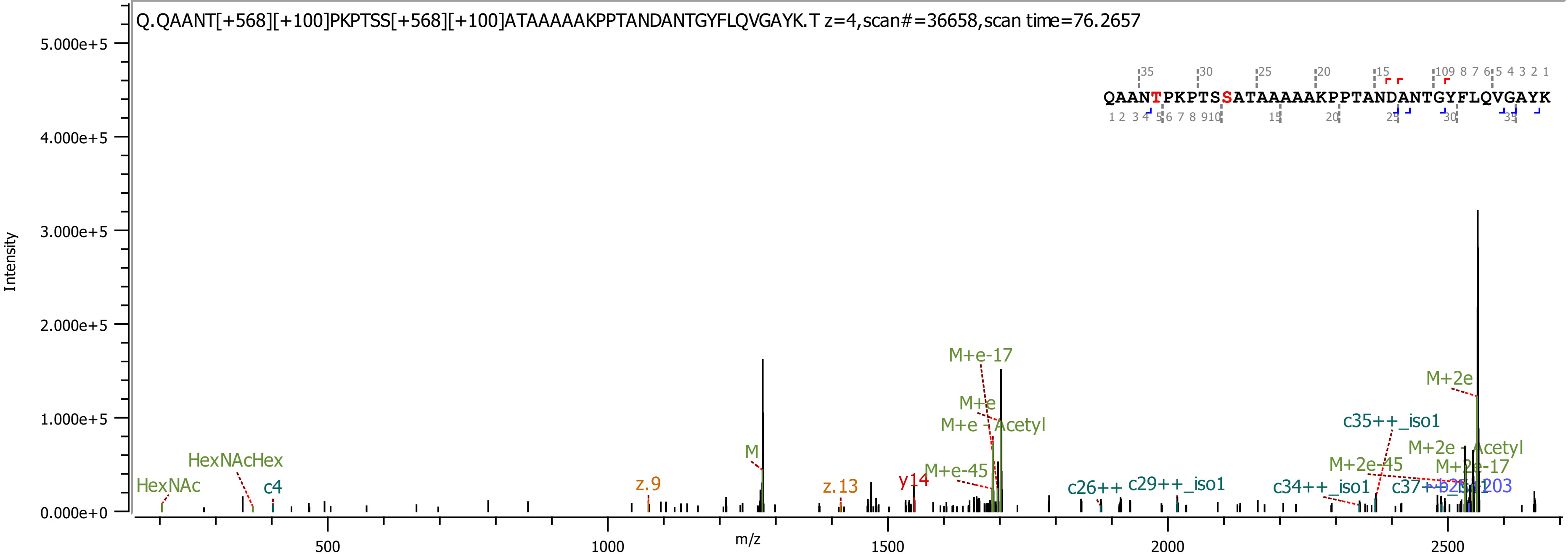

Q.QAANT[+568][+100]PKPTSS[+568][+100]ATAAAAAKPPTANDANTGYFLQVGAYK.T z=4,scan#=36658,scan time=76.2657
QAANTPKPTSSATAAAAAKPPTANDANTGYFLQVGAYK
Intensity
m/z
HexNAc
HexNAcHex
c4
z.9
M
z.13
y14
M+e-45
M+e-17
M+e
M+e - Acetyl
c26++
c29++_iso1
c34++_iso1
c35++_iso1
M+2e-45
M+2e
M+2e - Acetyl
M+2e-17

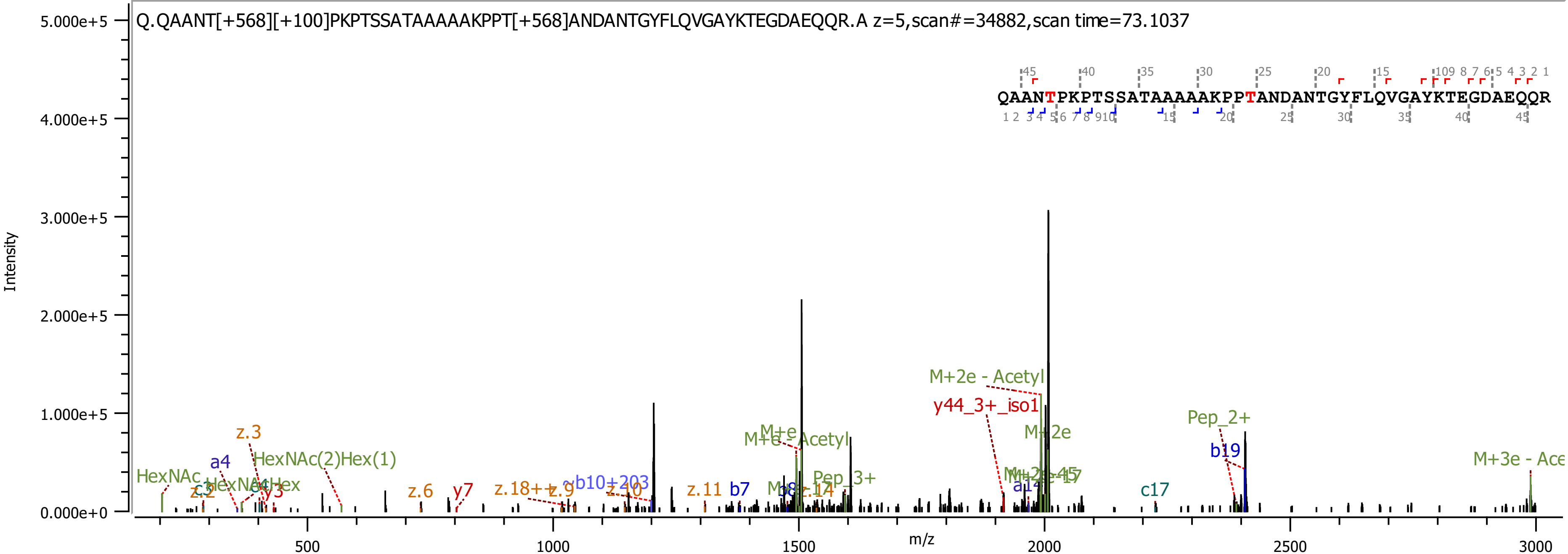
Q.QAANT[+568][+100]PKPTSSATAAAAAKPPT[+568]ANDANTGYFLQVGAYKTEGDAEQQR.A z=5,scan#=34882,scan time=73.1037
QAANTPKPTSSATAAAAAKPPTANDANTGYFLQVGAYKTEGDAEQQR
Intensity
m/z
M+2e - Acetyl
y44_3+_iso1
Pep_2+
b19
M+2e
M+3e - Ace
c17
M+e - Acetyl
HexNAc(2)Hex(1)
HexNAc
a4
z.3
z.6
y7
z.11
b7
Pep_3+

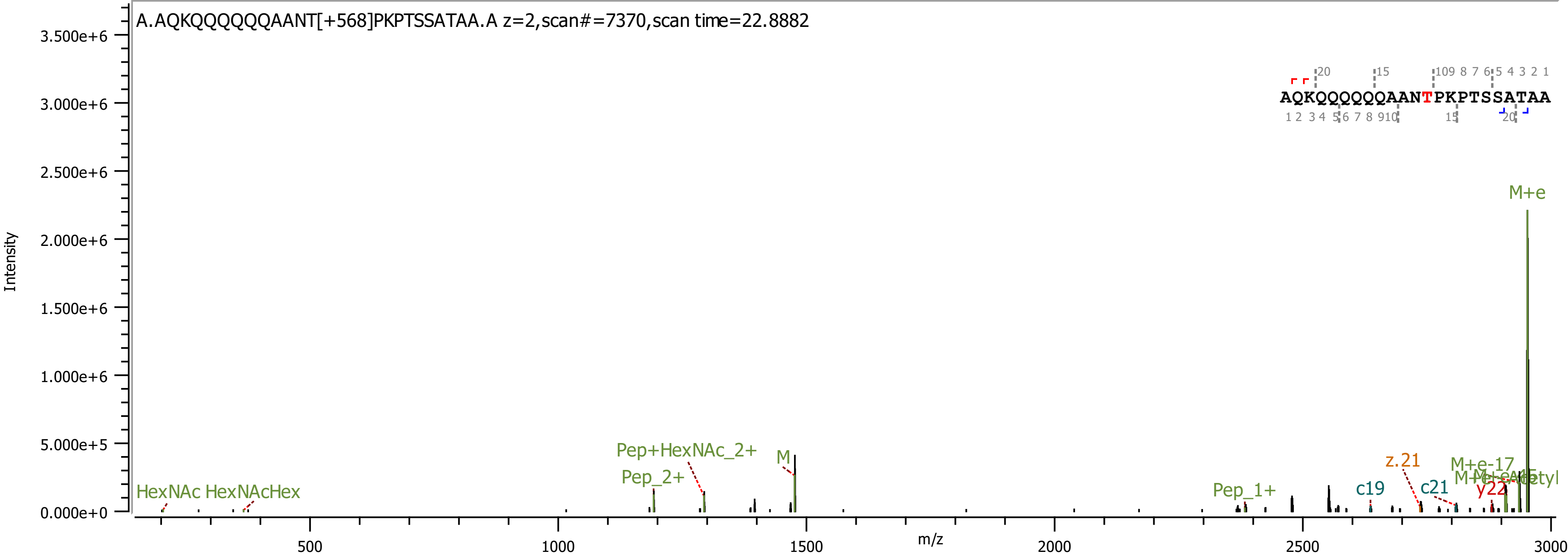
A.AQKQQQQQQAANT[+568]PKPTSSATAA.A z=2,scan#=7370,scan time=22.8882
Intensity
m/z
AQKQQQQQQAANTPKPTSSATAA
M+e
M+e-17
M
Pep+HexNAc_2+
Pep_2+
Pep_1+
HexNAc
HexNAcHex
z.21
c19
c21
y22

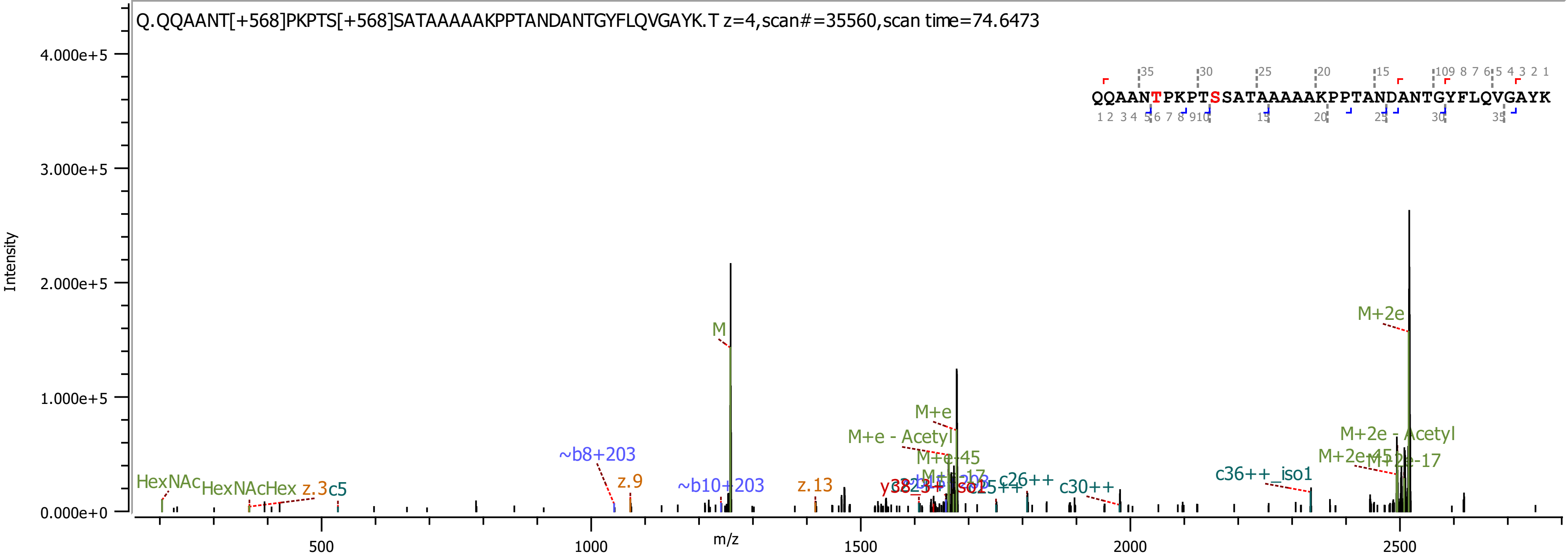
Q.QQAANT[+568]PKPTS[+568]SATAAAAAKPPTANDANTGYFLQVGAYK.T z=4,scan#=35560,scan time=74.6473
QQAANTPKPTSSATAAAAAKPPTANDANTGYFLQVGAYK
Intensity
m/z
4.000e+5
3.000e+5
2.000e+5
1.000e+5
0.000e+0
500
1000
1500
2000
2500
M
M+2e
M+e
M+e - Acetyl
M+e-45
M+2e - Acetyl
M+2e-45
M+2e-17
HexNAc
HexNAcHex
z.3
c5
~b8+203
z.9
~b10+203
z.13
c26++
c30++
c36++_iso1

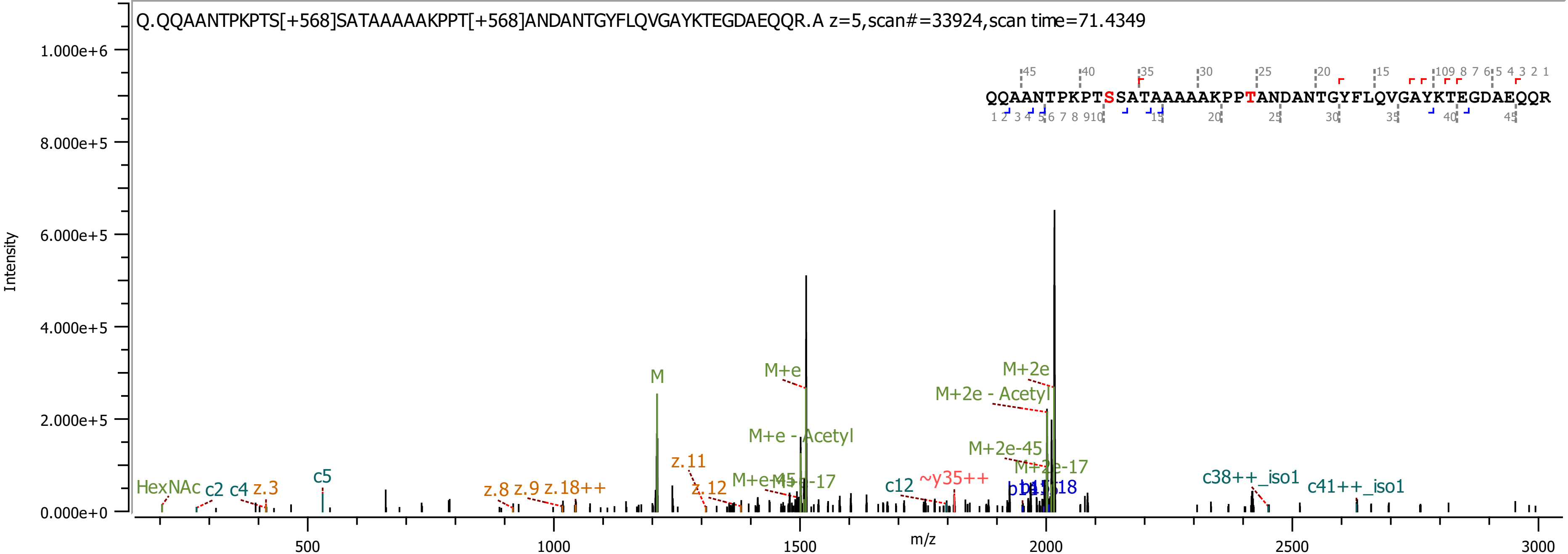
Q.QQAANTPKPTS[+568]SATAAAAAKPPT[+568]ANDANTGYFLQVGAYKTEGDAEQQR.A z=5,scan#=33924,scan time=71.4349
QQAANTPKPTSSATAAAAAKPPTANDANTGYFLQVGAYKTEGDAEQQR
Intensity
m/z
1.000e+6
8.000e+5
6.000e+5
4.000e+5
2.000e+5
0.000e+0
500
1000
1500
2000
2500
3000
HexNAc
c2
c4
z.3
c5
z.8
z.9
z.18++
M
z.11
z.12
M+e
M+e - Acetyl
M+e-45
M+e-17
c12
~y35++
M+2e
M+2e - Acetyl
M+2e-45
M+2e-17
c38++_iso1
c41++_iso1

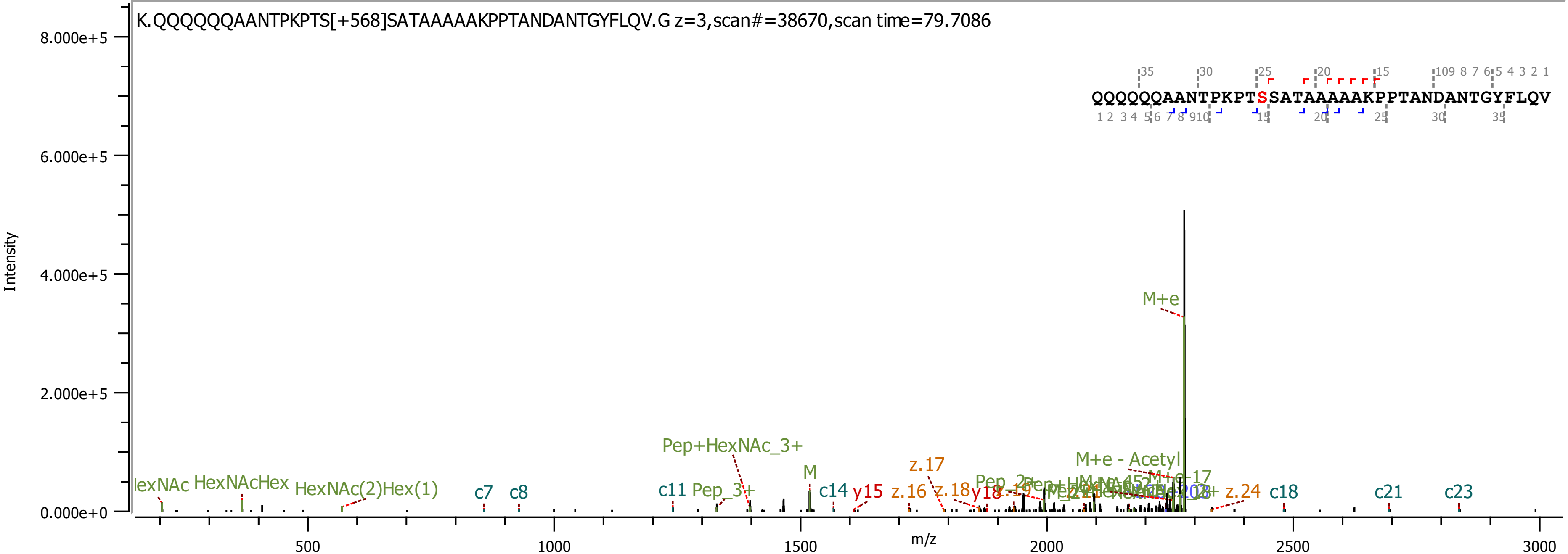

K.QQQQQQAANTPKPTS[+568]SATAAAAAKPPTANDANTGYFLQV.G z=3,scan#=38670,scan time=79.7086
QQQQQQAANTPKPTSSATAAAAAKPPTANDANTGYFLQV
Intensity
m/z
8.000e+5
6.000e+5
4.000e+5
2.000e+5
0.000e+0
500
1000
1500
2000
2500
3000
M+e
M+e - Acetyl
Pep+HexNAc_3+
HexNAcHex
HexNAc(2)Hex(1)
Pep_3+
M
c7
c8
c11
c14
c18
c21
c23
y15
z.16
z.17
z.18
z.24

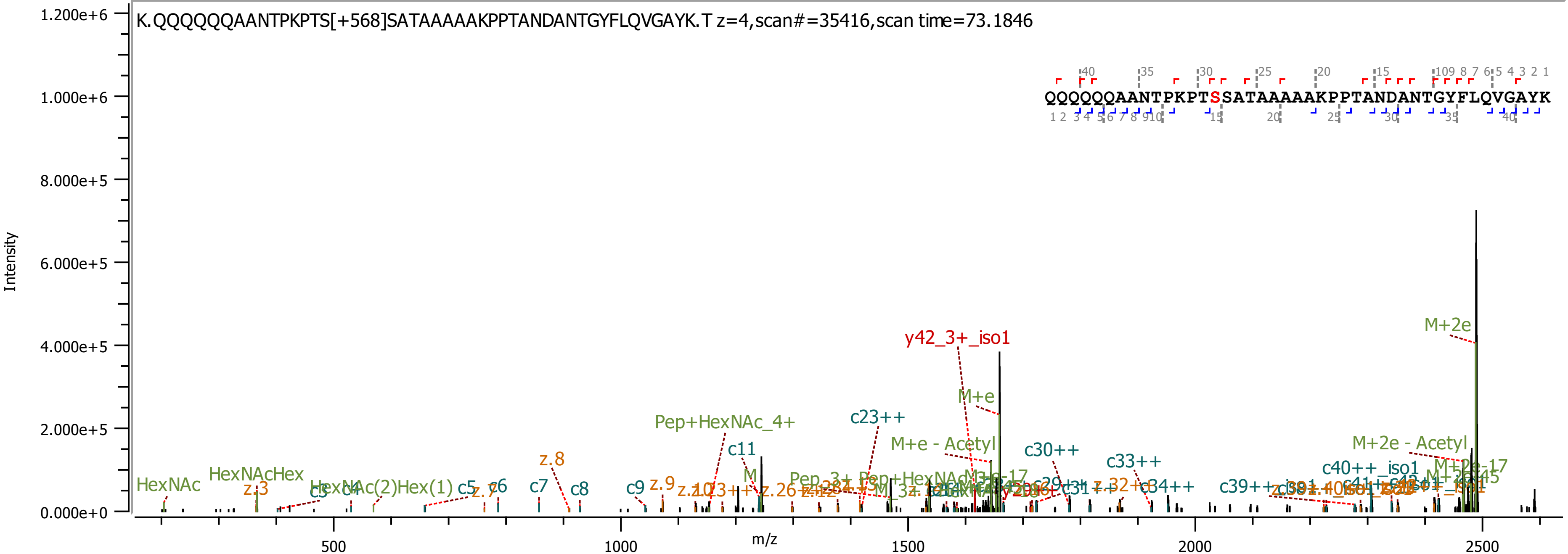

K.QQQQQQAANTPKPTS[+568]SATAAAAAKPPTANDANTGYFLQVGAYK.T z=4,scan#=35416,scan time=73.1846
QQQQQQAANTPKPTSSATAAAAAKPPTANDANTGYFLQVGAYK
Intensity
m/z
y42_3+_iso1
M+2e
M+e
c23++
Pep+HexNAc_4+
c11
M+e - Acetyl
c30++
c33++
M+2e - Acetyl
c40++_iso1
z.8
HexNAc
HexNAcHex
z.3
c5
c6
c7
c8
c9
z.9
c34++

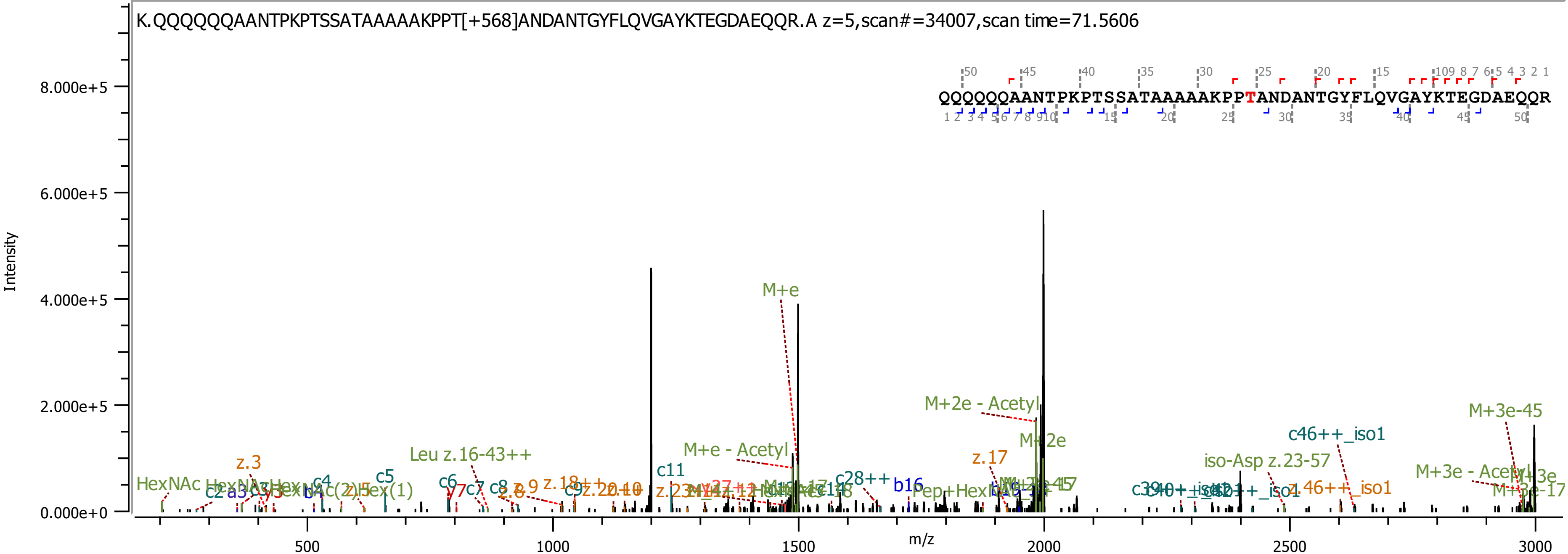

K.QQQQQQAANTPKPTSSATAAAAAKPPT[+568]ANDANTGYFLQVGAYKTEGDAEQQR.A z=5,scan#=34007,scan time=71.5606
QQQQQQAANTPKPTSSATAAAAAKPPTANDANTGYFLQVGAYKTEGDAEQQR
Intensity
m/z
M+e
M+2e - Acetyl
M+2e
M+e - Acetyl
M+3e-45
c46++_iso1
iso-Asp z.23-57
M+3e - Acetyl
z.17
Leu z.16-43++
c11
c28++
b16
HexNAc
z.3
c4
c5
c6
y7
c7
c8
z.46++_iso1

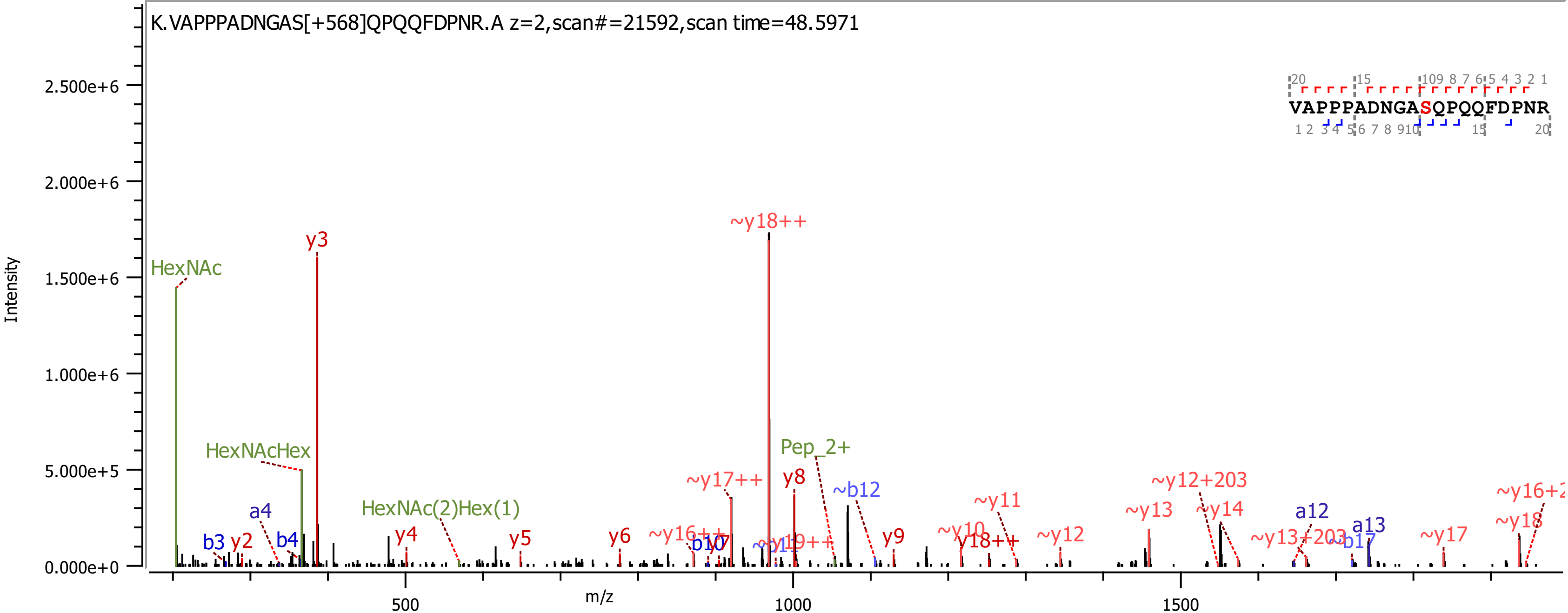
K.VAPPPADNGAS[+568]QPQQFDPNR.A z=2,scan#=21592,scan time=48.5971
Intensity
m/z
2.500e+6
2.000e+6
1.500e+6
1.000e+6
5.000e+5
0.000e+0
500
1000
1500
HexNAc
y3
~y18++
HexNAcHex
Pep_2+
~y17++
y8
~b12
HexNAc(2)Hex(1)
a4
b3
y2
b4
y4
y5
y6
~y11
~y12+203
~y13
~y14
a12
a13
~y17
~y18
~y12
y9
y18++
~y13+203
VAPPPADNGASQPQQFDPNR

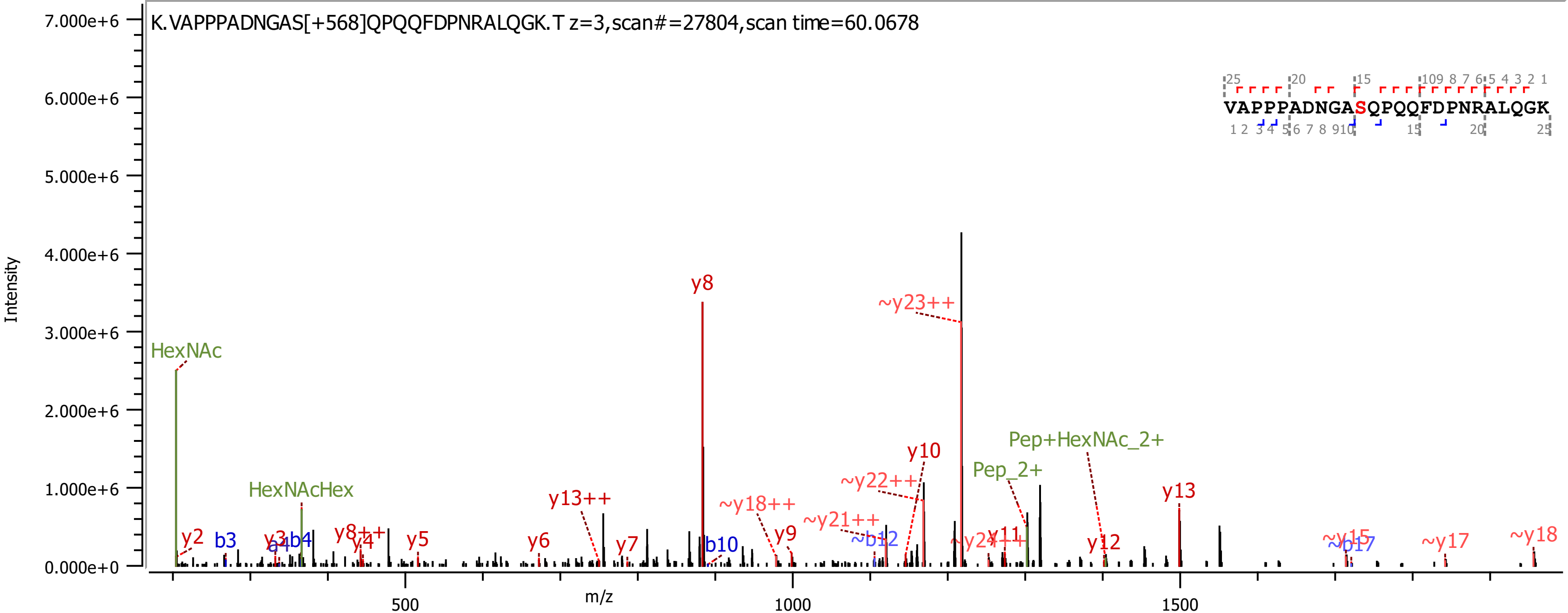
K.VAPPPADNGAS[+568]QPQQFDPNRALQGK.T z=3,scan#=27804,scan time=60.0678
Intensity
m/z
VAPPPADNGASQPQQFDPNRALQGK
HexNAc
HexNAcHex
y8
~y23++
y10
~y22++
Pep_2+
Pep+HexNAc_2+
y13
y13++
~y18++
~y21++
y2
b3
y3
b4
y8++
y4
y5
y6
y7
b10
y9
~b12
y11
y12
~y15
~b17
~y17
~y18

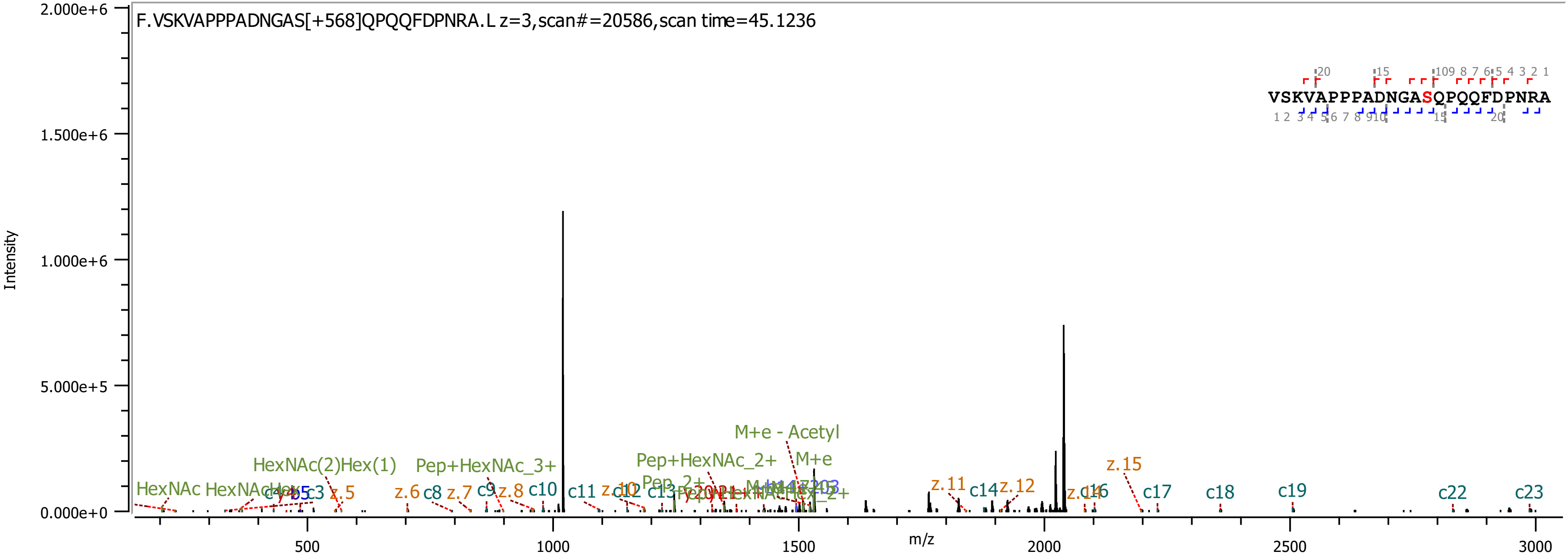
F.VSKVAPPPADNGAS[+568]QPQQFDPNRA.L z=3,scan#=20586,scan time=45.1236
VSKVAPPPADNGASQPQQFDPNRA
Intensity
m/z
M+e - Acetyl
M+e
Pep+HexNAc_2+
Pep+HexNAc_3+
HexNAc(2)Hex(1)
HexNAc
z.15
z.11
z.12
c14
c17
c18
c19
c22
c23
c10
c11
c8
z.6
z.7
z.8
z.5
c3
b5

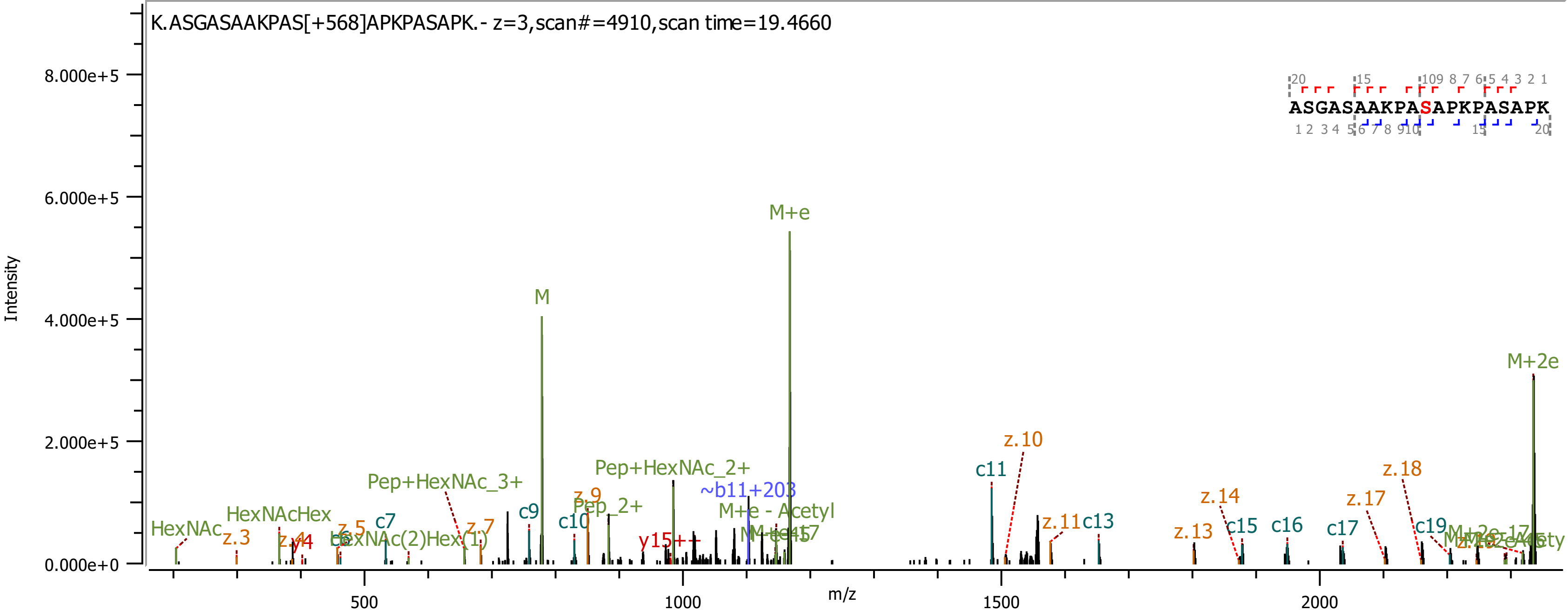

K.ASGASAAKPAS[+568]APKPASAPK.- z=3,scan#=4910,scan time=19.4660
Intensity
m/z
ASGASAAKPASAPKPASAPK
M+e
M
M+2e
c11
z.10
z.18
z.14
z.17
Pep+HexNAc_2+
Pep+HexNAc_3+
~b11+203
M+e - Acetyl
c9
c10
c13
c15
c16
c17
c19
c7
z.11
z.13
z.9
z.7
z.5
z.3
y15++
HexNAc
HexNAcHex
Pep_2+

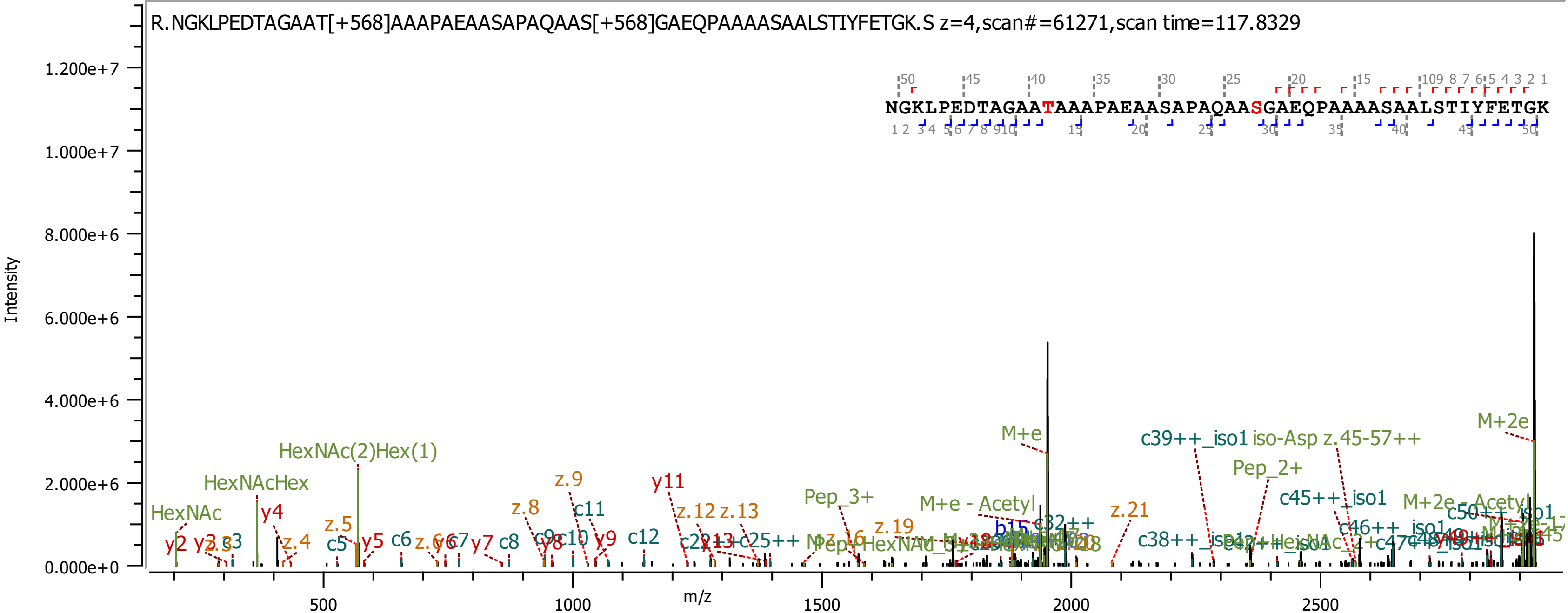

R.NGKLPEDTAGAAT[+568]AAAPAEAASAPAQAAS[+568]GAEQPAAAASAALSTIYFETGK.S z=4,scan#=61271,scan time=117.8329
NGKLPEDTAGAATAAAPAEAASAPAQAASGAEQPAAAASAALSTIYFETGK
Intensity
m/z
0.000e+0
2.000e+6
4.000e+6
6.000e+6
8.000e+6
1.000e+7
1.200e+7
500
1000
1500
2000
2500
HexNAc
HexNAcHex
HexNAc(2)Hex(1)
M+e
M+2e
M+e - Acetyl
M+2e - Acetyl
Pep_3+
Pep_2+
c39++_iso1
iso-Asp z.45-57++
c45++_iso1
c46++_iso1
z.21
z.19
z.16
z.13
z.12
z.9
z.8
z.5
z.4
y11
y9
y7
y5
y4
y2
c12
c11
c8
c6
c5
c38++_iso1

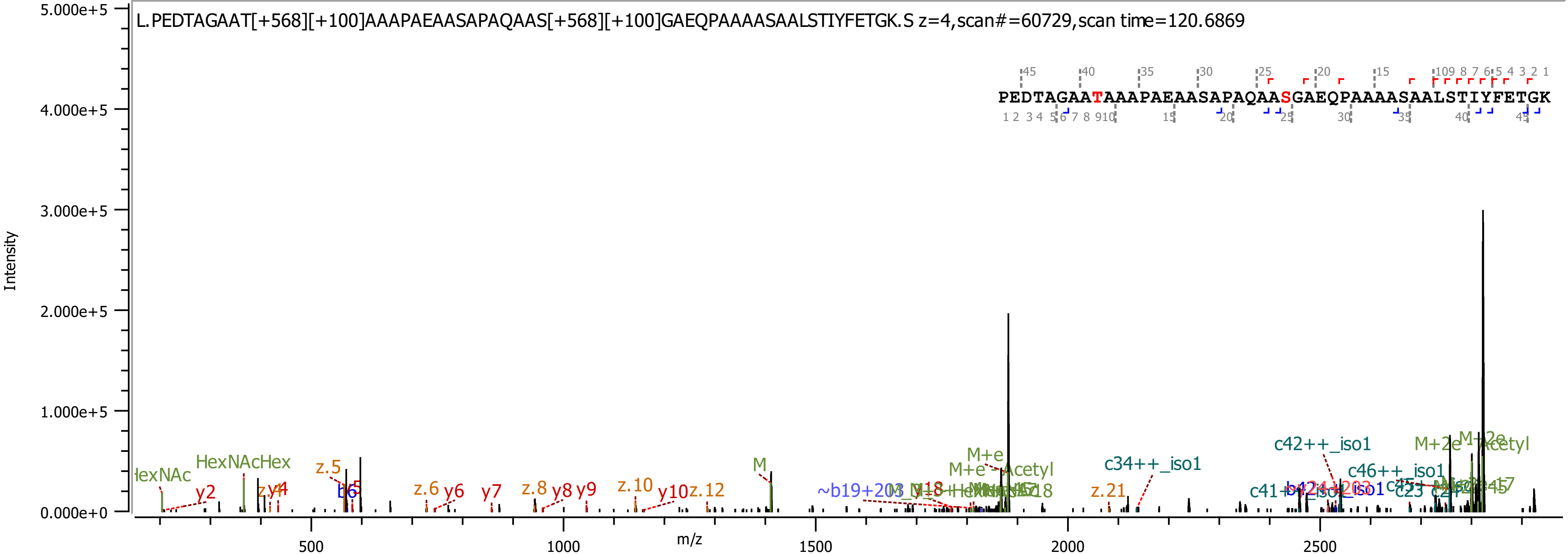
L.PEDTAGAAT[+568][+100]AAAPAEAASAPAQAAS[+568][+100]GAEQPAAAASAALSTIYFETGK.S z=4,scan#=60729,scan time=120.6869
PEDTAGAATAAAPAEAASAPAQAASGAEQPAAAASAALSTIYFETGK
Intensity
m/z
5.000e+5
4.000e+5
3.000e+5
2.000e+5
1.000e+5
0.000e+0
500
1000
1500
2000
2500
HexNAcHex
y2
z.5
b6
z.6
y6
y7
z.8
y8
y9
z.10
y10
z.12
M
M+e
M+e-Acetyl
c34++_iso1
z.21
c42++_iso1
c46++_iso1
M+2e
Acetyl

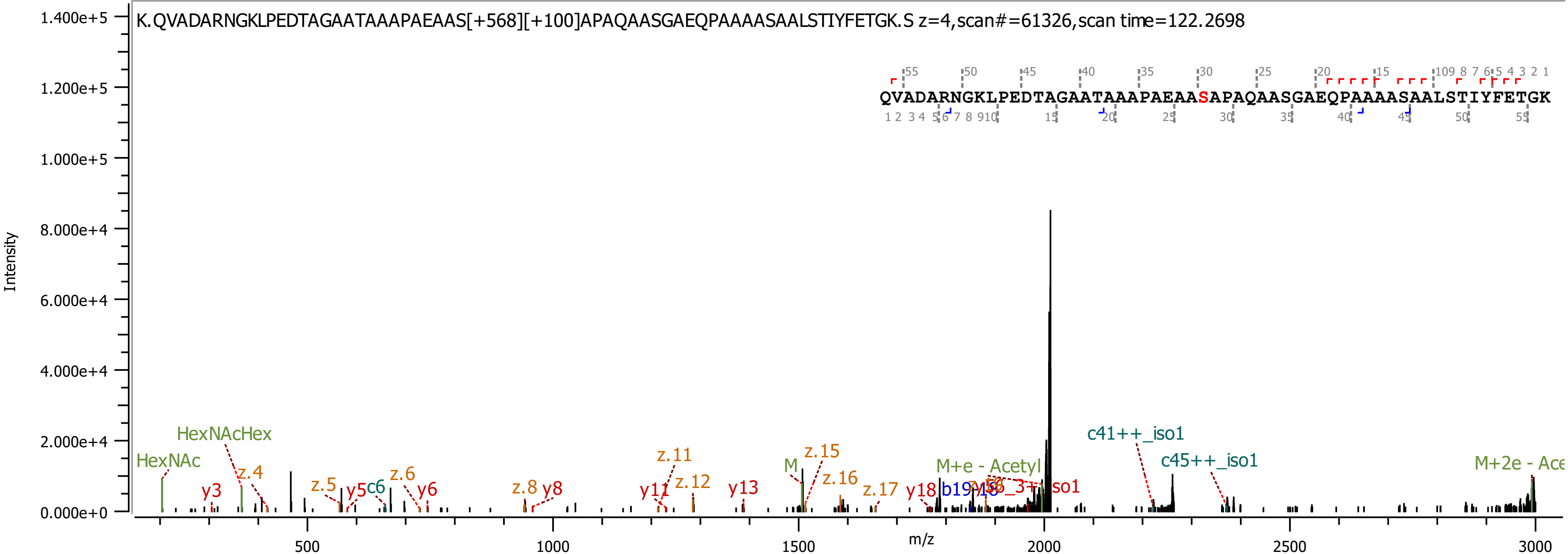
K.QVADARNGKLPEDTAGAATAAAPAEAAS[+568][+100]APAQAASGAEQPAAAASAALSTIYFETGK.S z=4,scan#=61326,scan time=122.2698
QVADARNGKLPEDTAGAATAAAPAEAASAPAQAASGAEQPAAAASAALSTIYFETGK
Intensity
m/z
1.400e+5
1.200e+5
1.000e+5
8.000e+4
6.000e+4
4.000e+4
2.000e+4
0.000e+0
500
1000
1500
2000
2500
3000
HexNAc
HexNAcHex
y3
z.4
z.5
y5
c6
z.6
y6
z.8
y8
y11
z.11
z.12
y13
M
z.15
z.16
z.17
y18
b19
M+e - Acetyl
c41++_iso1
c45++_iso1
M+2e - Ace

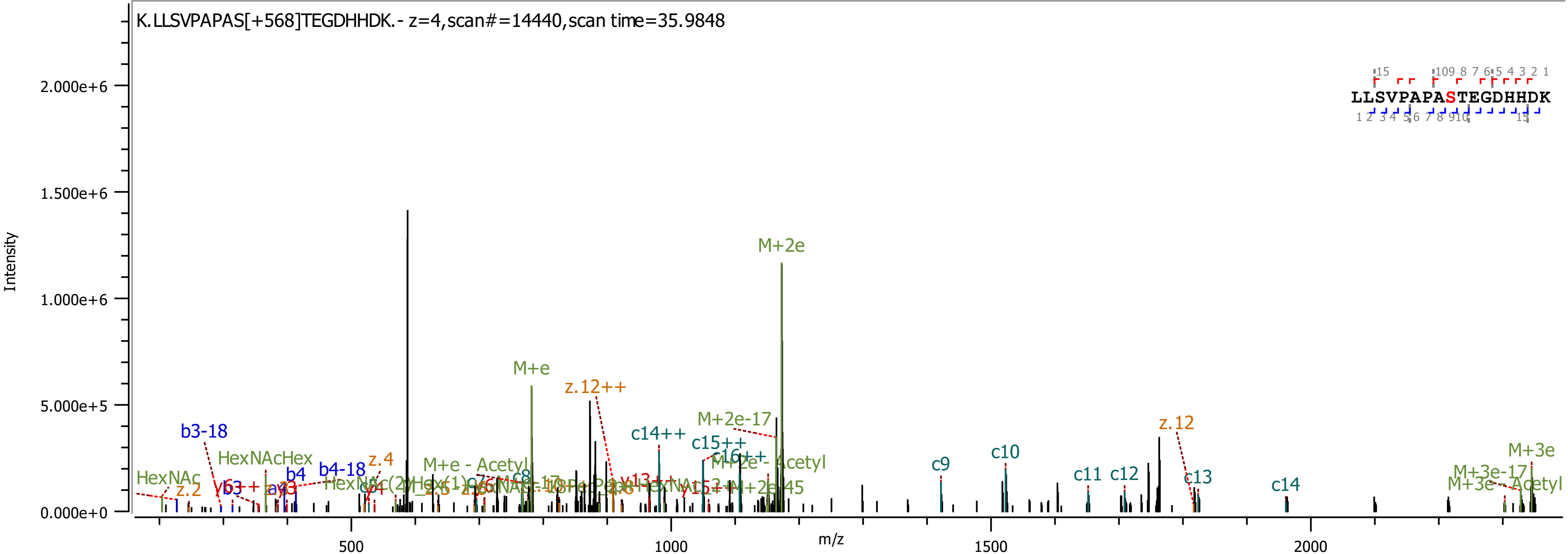

K.LLSVPAPAS[+568]TEGDHHDK.- z=4,scan#=14440,scan time=35.9848
Intensity
2.000e+6
1.500e+6
1.000e+6
5.000e+5
0.000e+0
m/z
500
1000
1500
2000
LLSVPAPASTEGDHHDK
M+2e
M+e
z.12++
M+2e-17
c14++
c15++
c16++
M+2e - Acetyl
b3-18
HexNAcHex
HexNAc
z.2
b3
b4
b4-18
z.4
M+e - Acetyl
c8
c9
c10
c11
c12
z.12
c13
c14
M+3e
M+3e-17
M+3e - Acetyl

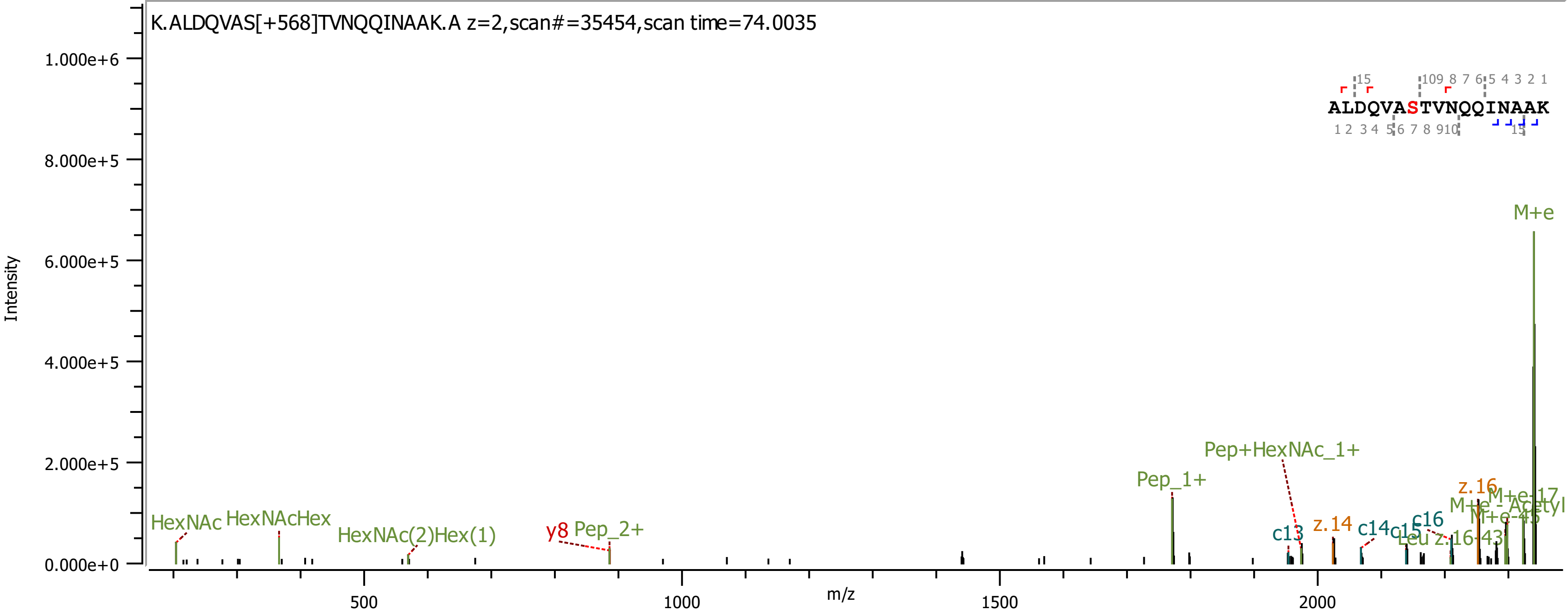

K.ALDQVAS[+568]TVNQQINAAK.A z=2,scan#=35454,scan time=74.0035
ALDQVASTVNQQINAAK
Intensity
m/z
M+e
Pep+HexNAc_1+
Pep_1+
HexNAc
HexNAcHex
HexNAc(2)Hex(1)
y8
Pep_2+
c13
z.14
c14
c15
c16
z.16

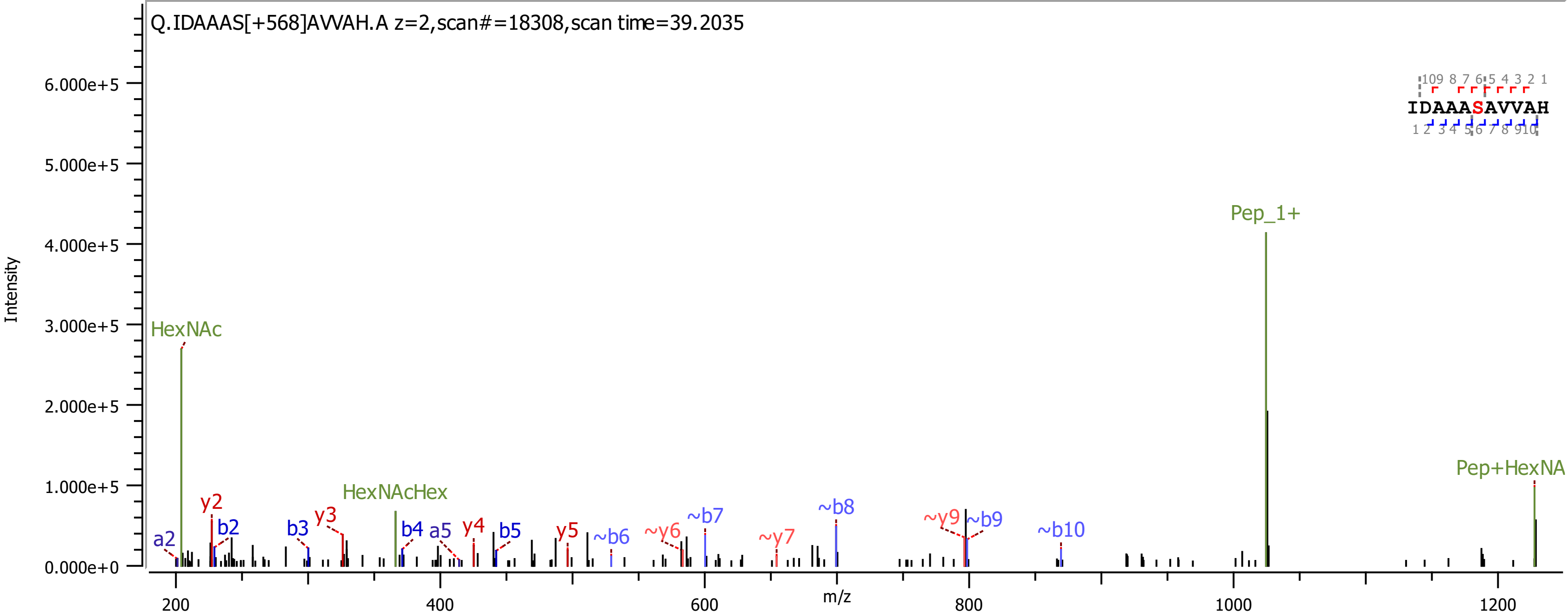

Q.IDAAAS[+568]AVVAH.A z=2,scan#=18308,scan time=39.2035
Intensity
m/z
6.000e+5
5.000e+5
4.000e+5
3.000e+5
2.000e+5
1.000e+5
0.000e+0
200
400
600
800
1000
1200
HexNAc
HexNAcHex
Pep_1+
Pep+HexNA
a2
y2
b2
b3
y3
b4
a5
y4
b5
y5
~b6
~y6
~b7
~y7
~b8
~y9
~b9
~b10
IDAAASAVVAH

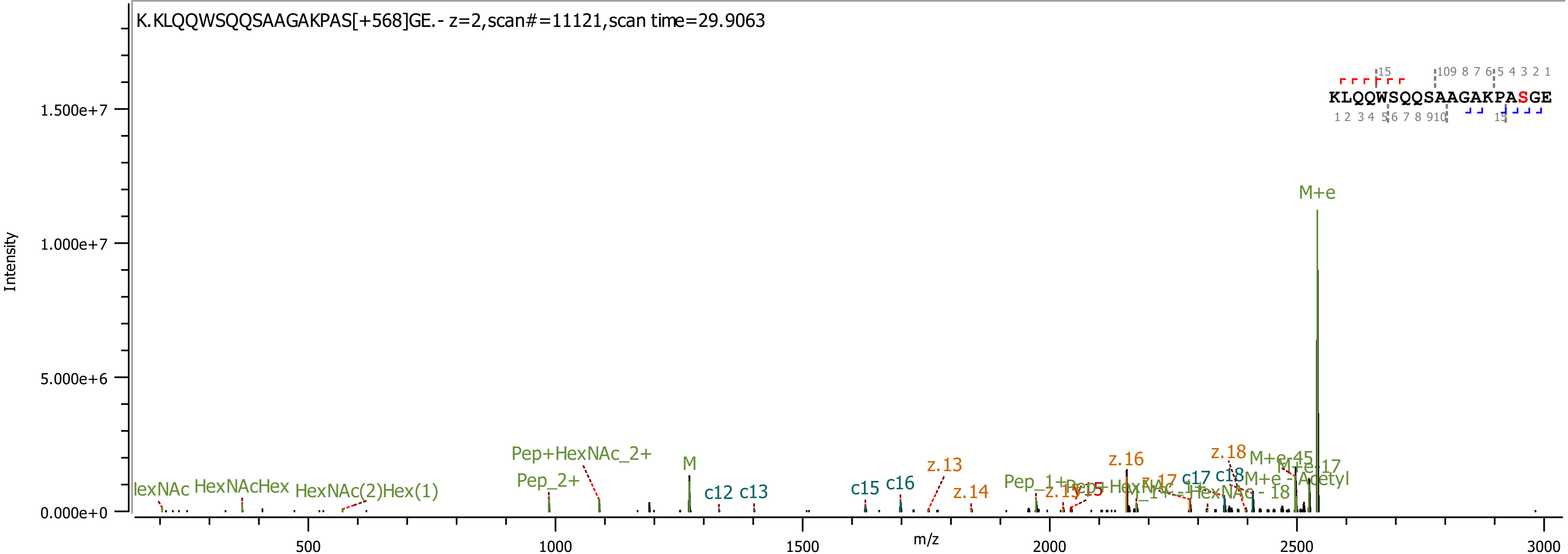
K.KLQQWSQQSAAGAKPAS[+568]GE.- z=2,scan#=11121,scan time=29.9063
KLQQWSQQSAAGAKPASGE
Intensity
m/z
0.000e+0
5.000e+6
1.000e+7
1.500e+7
500
1000
1500
2000
2500
3000
M+e
HexNAcHex
HexNAc(2)Hex(1)
Pep+HexNAc_2+
Pep_2+
M
c12
c13
c15
c16
z.13
z.14
z.16
z.18
c17
c18
M+e-45
M+e-17
M+e - Acetyl
M+e - 18
Pep_1+

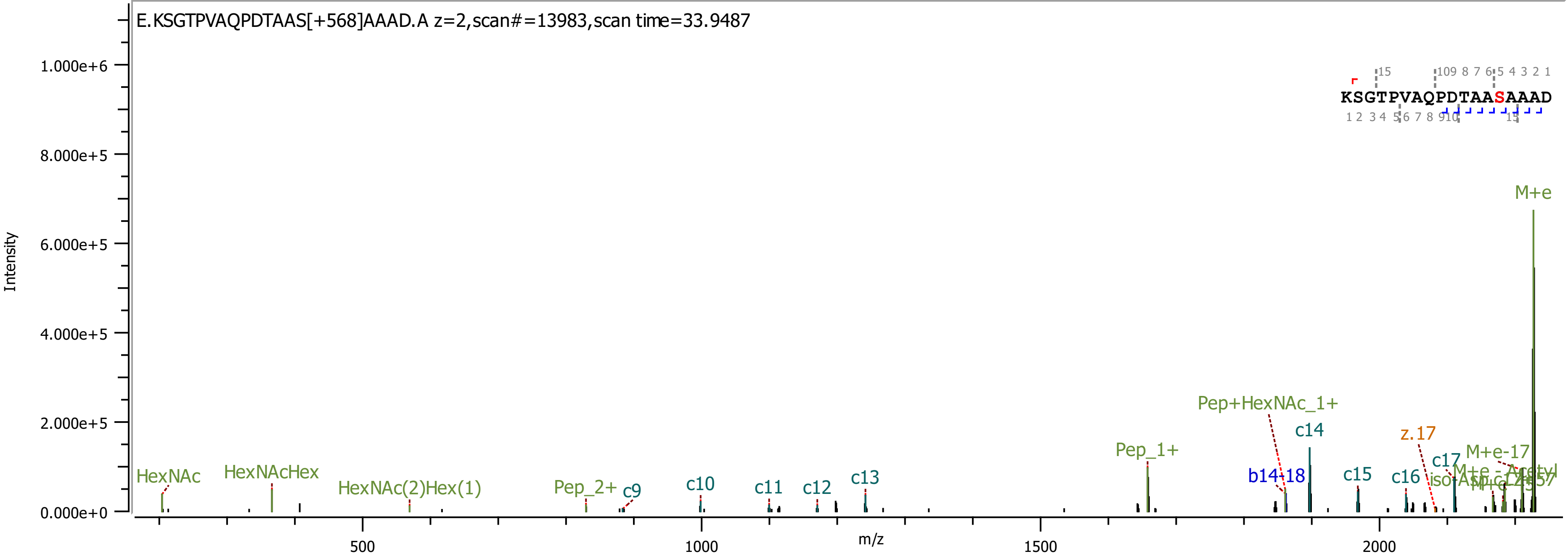

E.KSGTPVAQPDTAAS[+568]AAAD.A z=2,scan#=13983,scan time=33.9487
Intensity
m/z
KSGTPVAQPDTAASAAAD
HexNAc
HexNAcHex
HexNAc(2)Hex(1)
Pep_2+
c9
c10
c11
c12
c13
Pep_1+
Pep+HexNAc_1+
b14-18
c14
c15
c16
z.17
c17
M+e-17
M+e

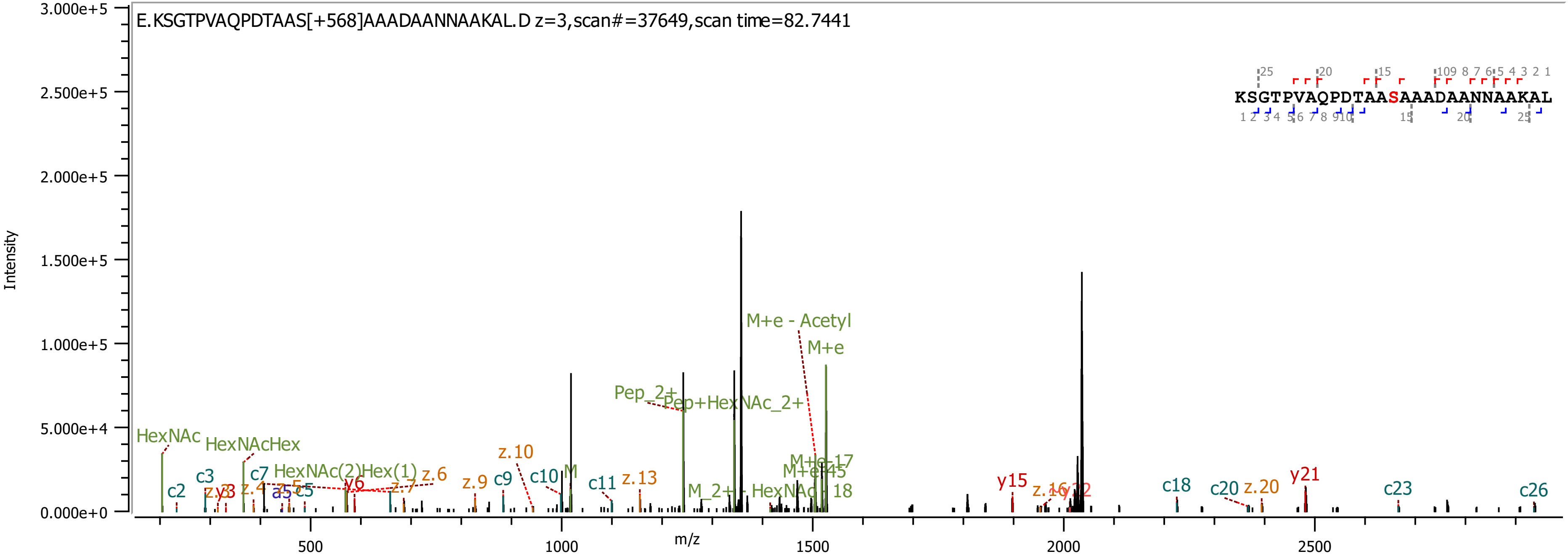
E.KSGTPVAQPDTAAS[+568]AAADAANNAAKAL.D z=3,scan#=37649,scan time=82.7441
KSGTPVAQPDTAASAAADAANNAAKAL
Intensity
m/z
3.000e+5
2.500e+5
2.000e+5
1.500e+5
1.000e+5
5.000e+4
0.000e+0
500
1000
1500
2000
2500
HexNAc
HexNAcHex
HexNAc(2)Hex(1)
c2
c3
c5
c7
c9
c10
c11
c18
c20
c23
c26
a5
y6
y15
y21
z.4
z.6
z.7
z.9
z.10
z.13
z.20
M
Pep_2+
Pep+HexNAc_2+
M+e - Acetyl
M+e
M+e-17
M+e-45
M_2+ - HexNAc - 18

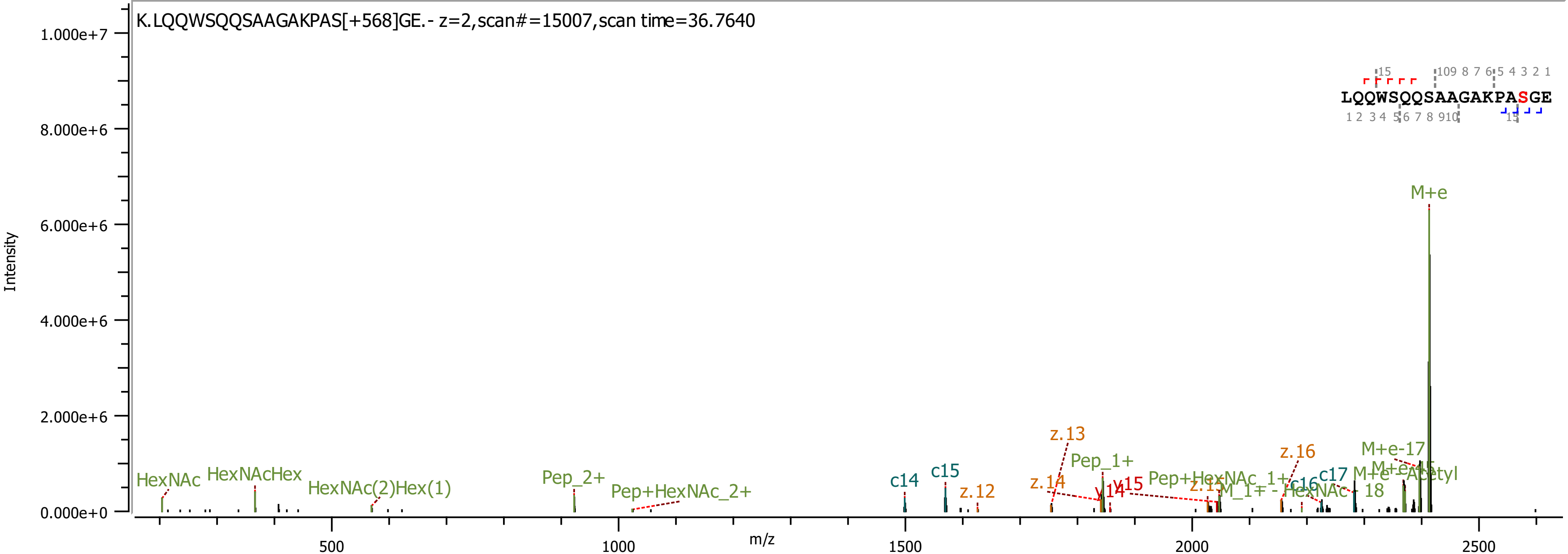
K.LQQWSQQSAAGAKPAS[+568]GE.- z=2,scan#=15007,scan time=36.7640
Intensity
m/z
LQQWSQQSAAGAKPASGE
M+e
HexNAc
HexNAcHex
HexNAc(2)Hex(1)
Pep_2+
Pep+HexNAc_2+
c14
c15
z.12
z.13
z.14
Pep_1+
y14
y15
Pep+HexNAc_1+
c16
c17
z.16
M+e-17
M+e-Acetyl
0.000e+0
2.000e+6
4.000e+6
6.000e+6
8.000e+6
1.000e+7
500
1000
1500
2000
2500

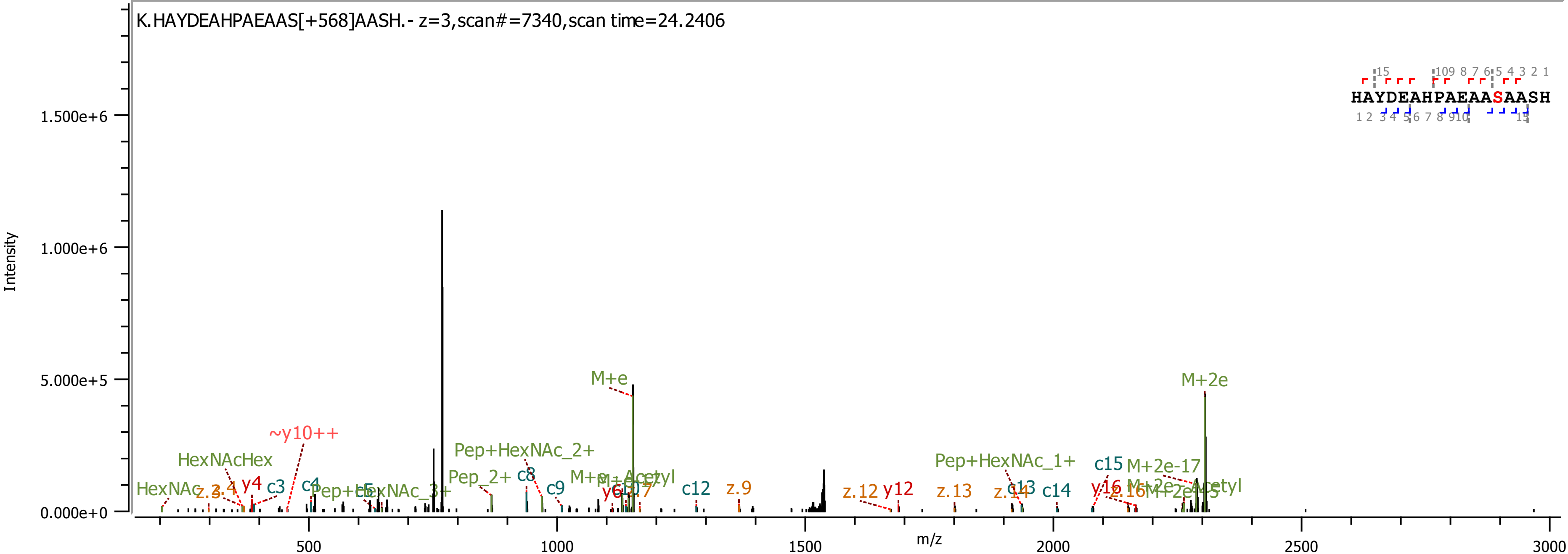
K.HAYDEAHPAEAAS[+568]AASH.- z=3,scan#=7340,scan time=24.2406
HAYDEAHPAEAASAASH
Intensity
m/z
1.500e+6
1.000e+6
5.000e+5
0.000e+0
500
1000
1500
2000
2500
3000
M+e
M+2e
~y10++
HexNAcHex
HexNAc
Pep+HexNAc_2+
Pep_2+
Pep+HexNAc_1+
c15
M+2e-17
c3
c4
c8
c9
c12
z.9
z.12
y12
z.13
c13
c14
y4
y16

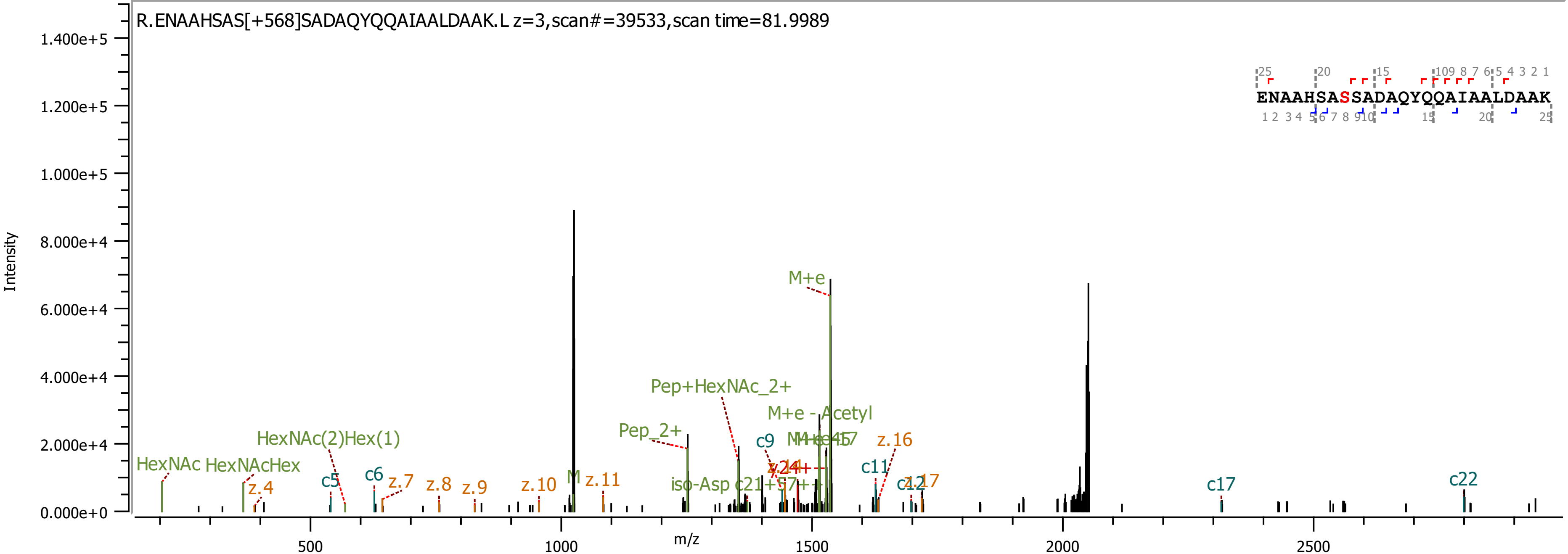

R.ENAAHSAS[+568]SADAQYQQAIAALDAAK.L z=3,scan#=39533,scan time=81.9989
ENAAHSASSADAQYQQAIAALDAAK
Intensity
m/z
M+e
Pep+HexNAc_2+
M+e - Acetyl
Pep_2+
HexNAc(2)Hex(1)
HexNAc
HexNAcHex
z.4
c5
c6
z.7
z.8
z.9
z.10
M
z.11
iso-Asp
c9
c11
z.16
c17
c22

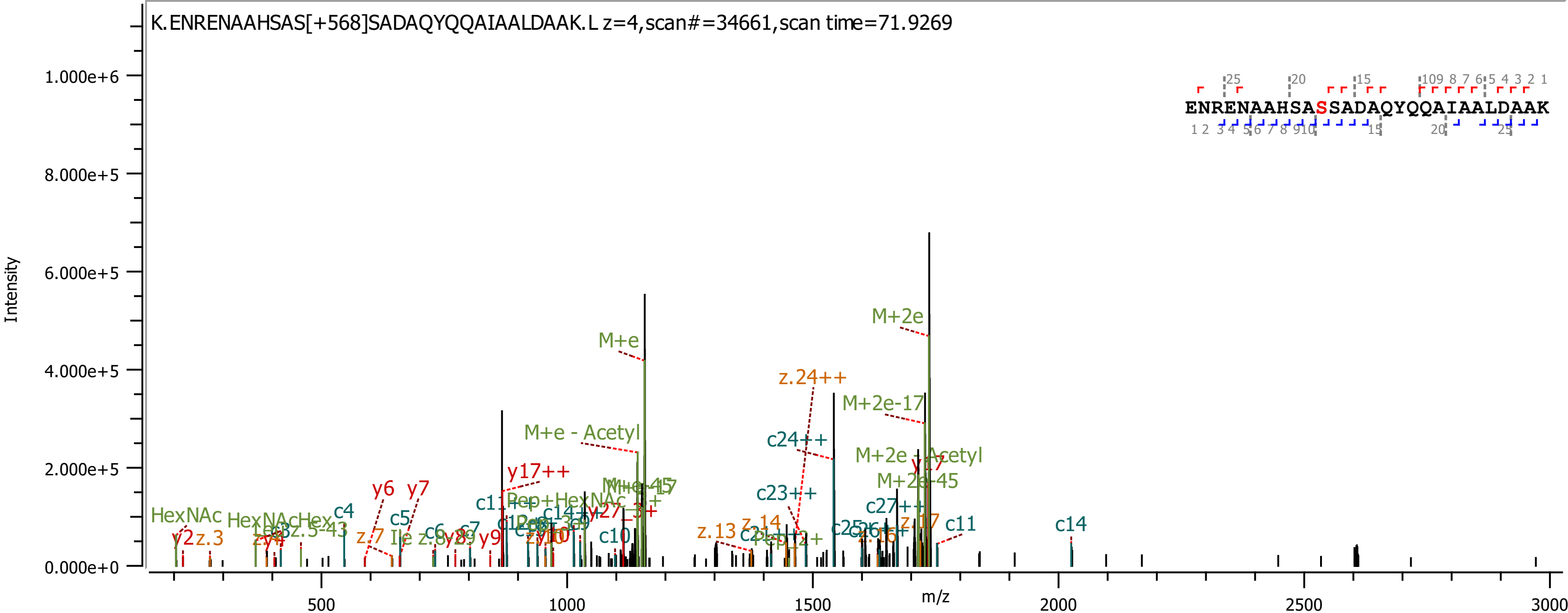
K.ENRENAAHSAS[+568]SADAQYQQAIAALDAAK.L z=4,scan#=34661,scan time=71.9269
ENRENAAHSASSADAQYQQAIAALDAAK
Intensity
m/z
1.000e+6
8.000e+5
6.000e+5
4.000e+5
2.000e+5
0.000e+0
500
1000
1500
2000
2500
3000
M+e
M+2e
M+2e-17
M+e - Acetyl
M+2e - Acetyl
M+2e-45
z.24++
c24++
c23++
c27++
y17++
y6
y7
y9
y2
z.3
HexNAc
c4
c5
c6
c7
c10
c11
c14
z.7
z.13
z.14
z.17

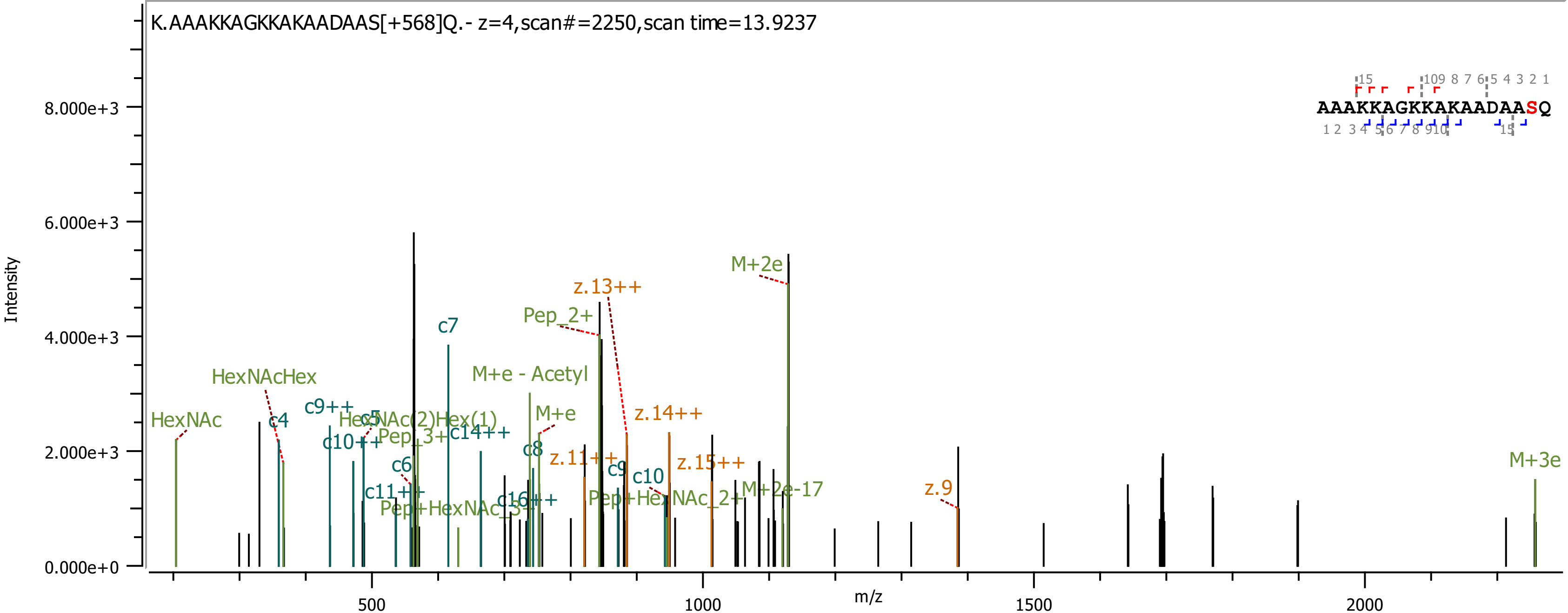
K.AAAKKAGKKAKAADAAS[+568]Q.- z=4,scan#=2250,scan time=13.9237
AAAKKAGKKAKAADAASQ
Intensity
m/z
HexNAc
HexNAcHex
c4
c9++
c5
HexNAc(2)Hex(1)
c10++
Pep_3+
c14++
c6
c11++
Pep+HexNAc_3+
c7
c16++
M+e - Acetyl
M+e
c8
Pep_2+
z.13++
z.11++
c9
c10
z.14++
Pep+HexNAc_2+
z.15++
M+2e-17
M+2e
z.9
M+3e

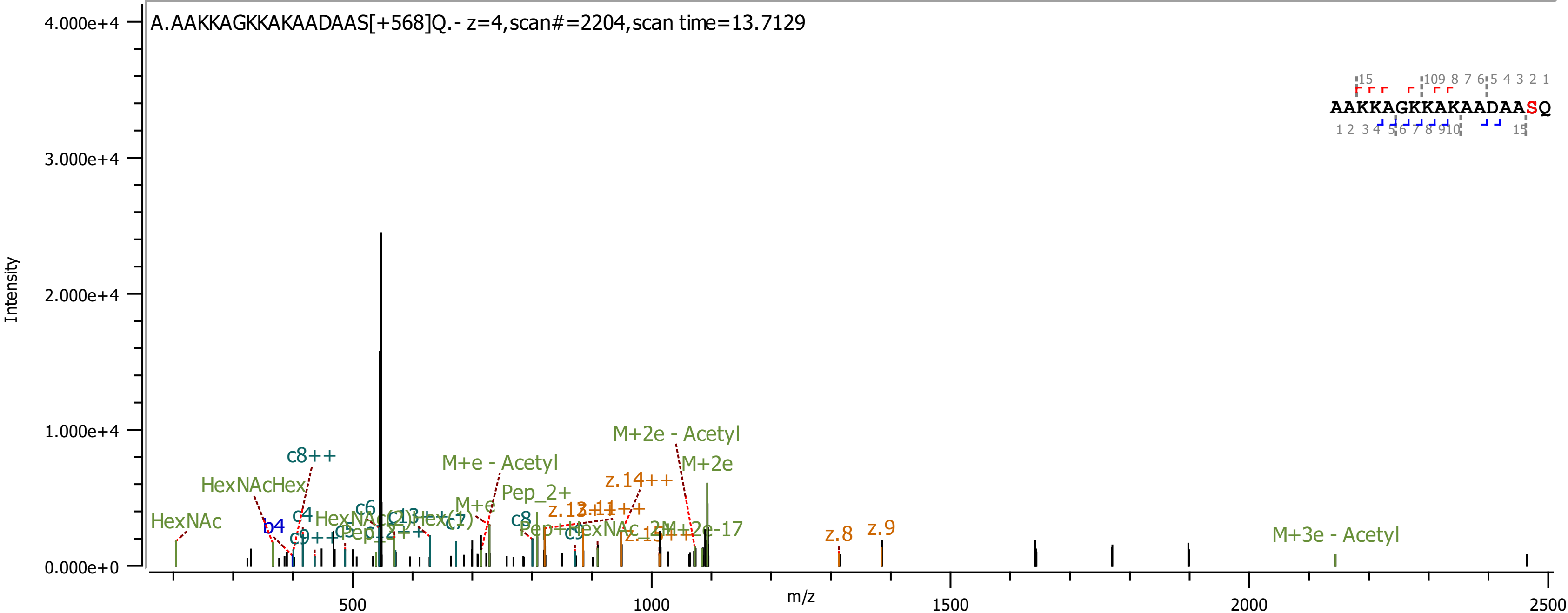
A.AAKKAGKKAKAADAAS[+568]Q.- z=4,scan#=2204,scan time=13.7129
Intensity
m/z
4.000e+4
3.000e+4
2.000e+4
1.000e+4
0.000e+0
500
1000
1500
2000
2500
AAKKAGKKAKAADAASQ
HexNAc
HexNAcHex
b4
c4
c8++
c6
c7
c8
c9
M+e - Acetyl
M+e
Pep_2+
M+2e - Acetyl
M+2e
M+2e-17
z.14++
z.13++
z.11++
z.15++
z.8
z.9
M+3e - Acetyl

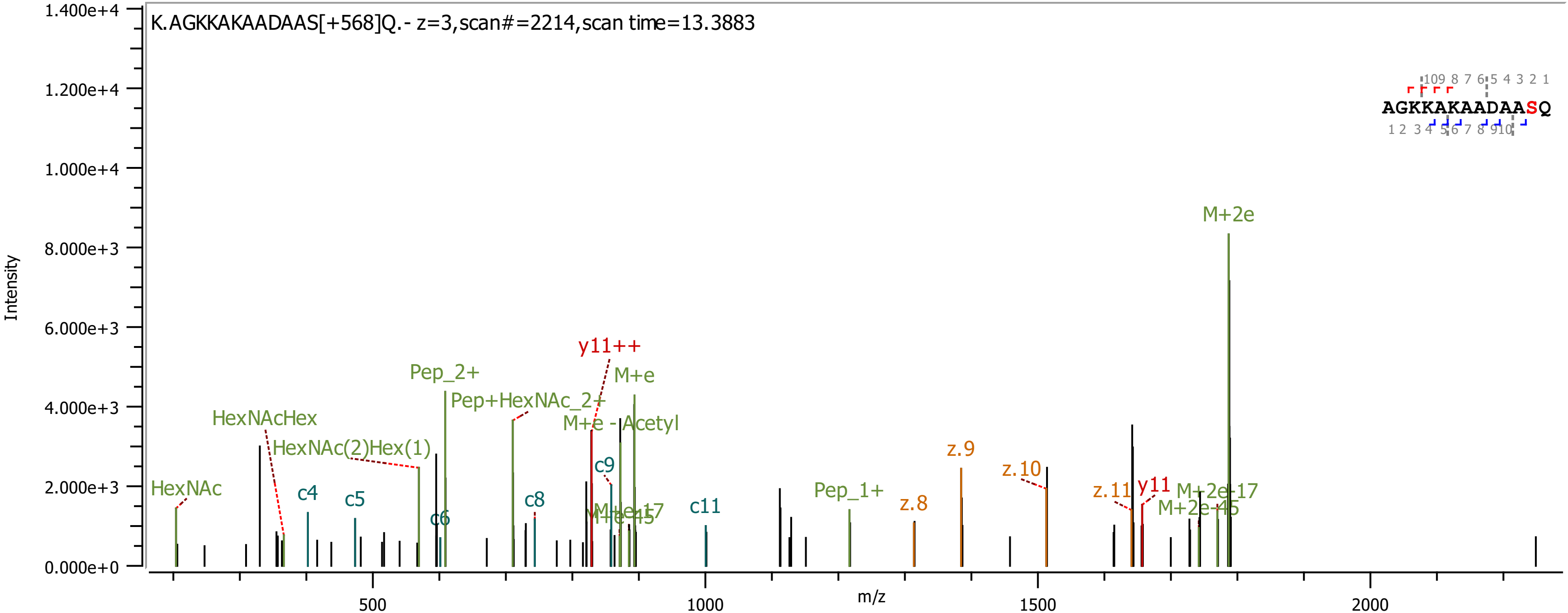

K.AGKKAKAADAAS[+568]Q.- z=3,scan#=2214,scan time=13.3883
Intensity
m/z
AGKKAKAADAASQ
HexNAc
HexNAcHex
HexNAc(2)Hex(1)
c4
c5
c6
Pep_2+
Pep+HexNAc_2+
c8
y11++
M+e
M+e - Acetyl
c9
M+e-17
M+e-45
c11
Pep_1+
z.8
z.9
z.10
z.11
y11
M+2e
M+2e-17
M+2e-45

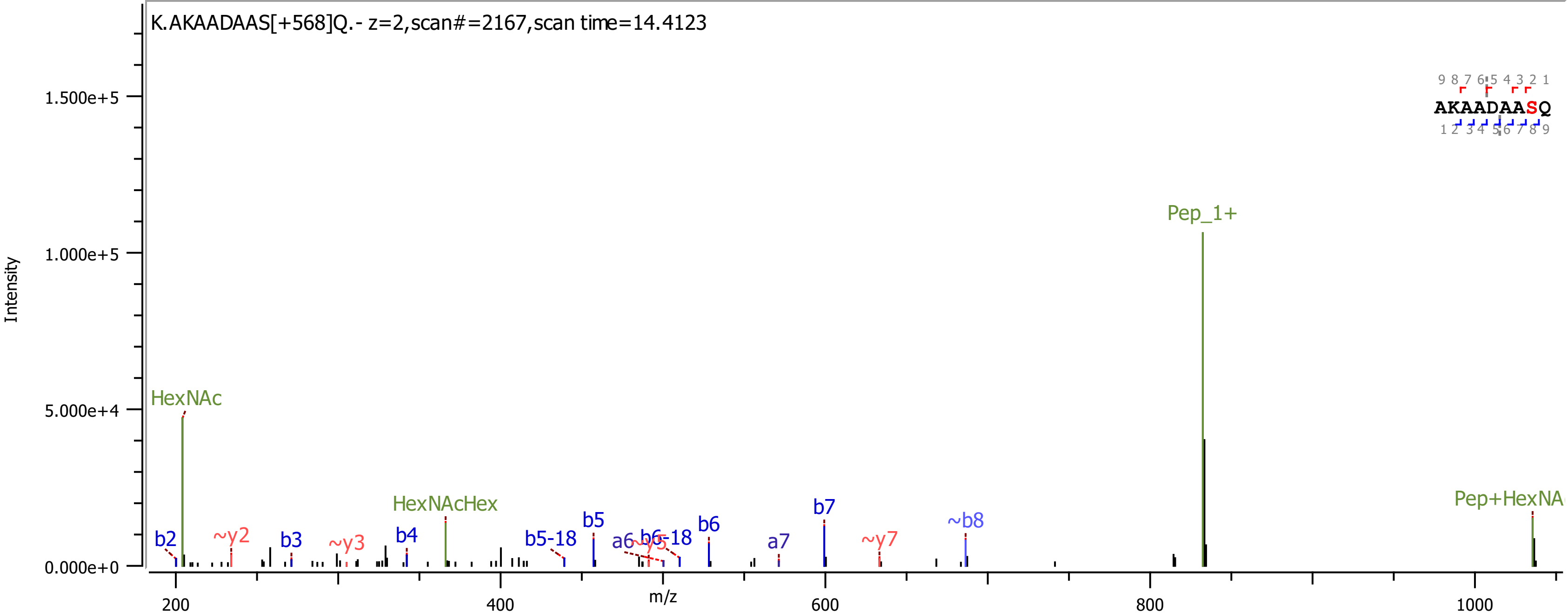
K.AKAADAAS[+568]Q.- z=2,scan#=2167,scan time=14.4123
Intensity
1.500e+5
1.000e+5
5.000e+4
0.000e+0
AKAADAASQ
HexNAc
~y2
b2
b3
~y3
b4
HexNAcHex
b5-18
b5
a6
~y5
b6-18
b6
a7
b7
~y7
~b8
Pep_1+
Pep+HexNA
200
400
600
800
1000
m/z

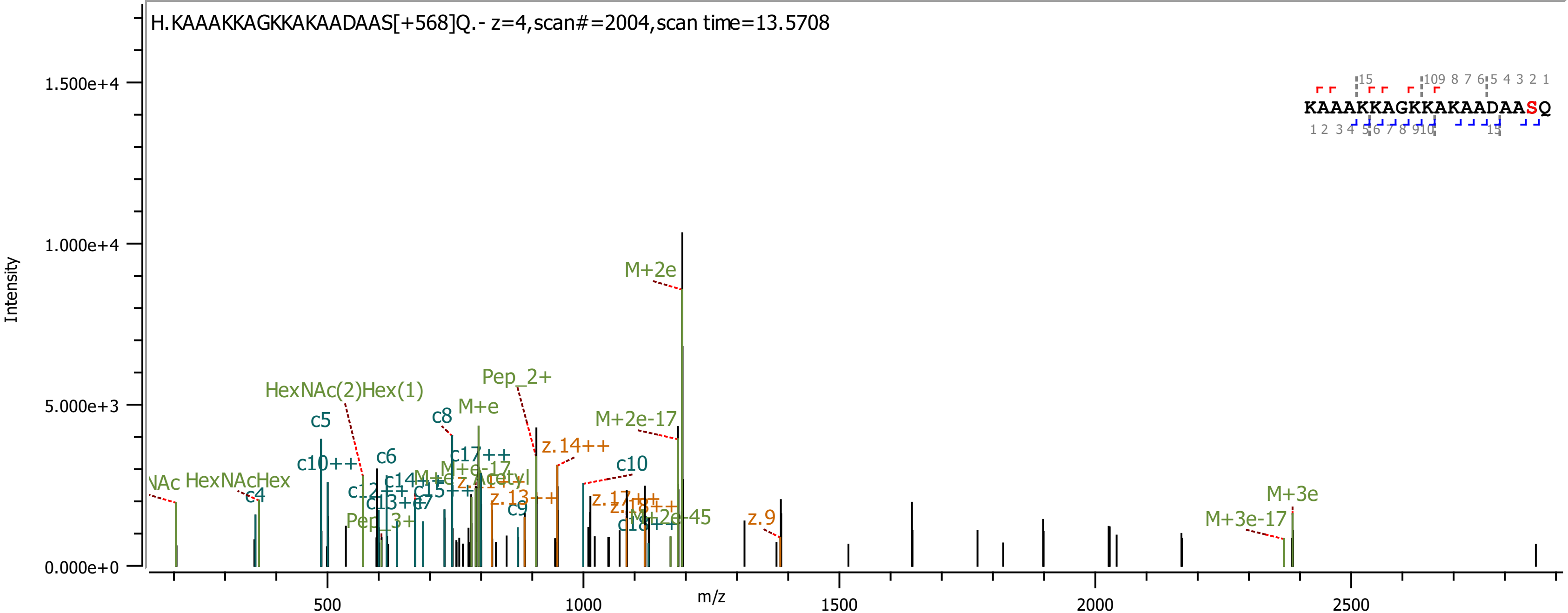

H.KAAAKKAGKKAKAADAAS[+568]Q.- z=4,scan#=2004,scan time=13.5708
Intensity
m/z
KAAAKKAGKKAKAADAASQ
M+2e
Pep_2+
HexNAc(2)Hex(1)
c5
c8
M+e
M+2e-17
z.14++
c10
c6
c10++
c17++
HexNAcHex
c4
z.9
M+3e
M+3e-17
c9
Pep_3+

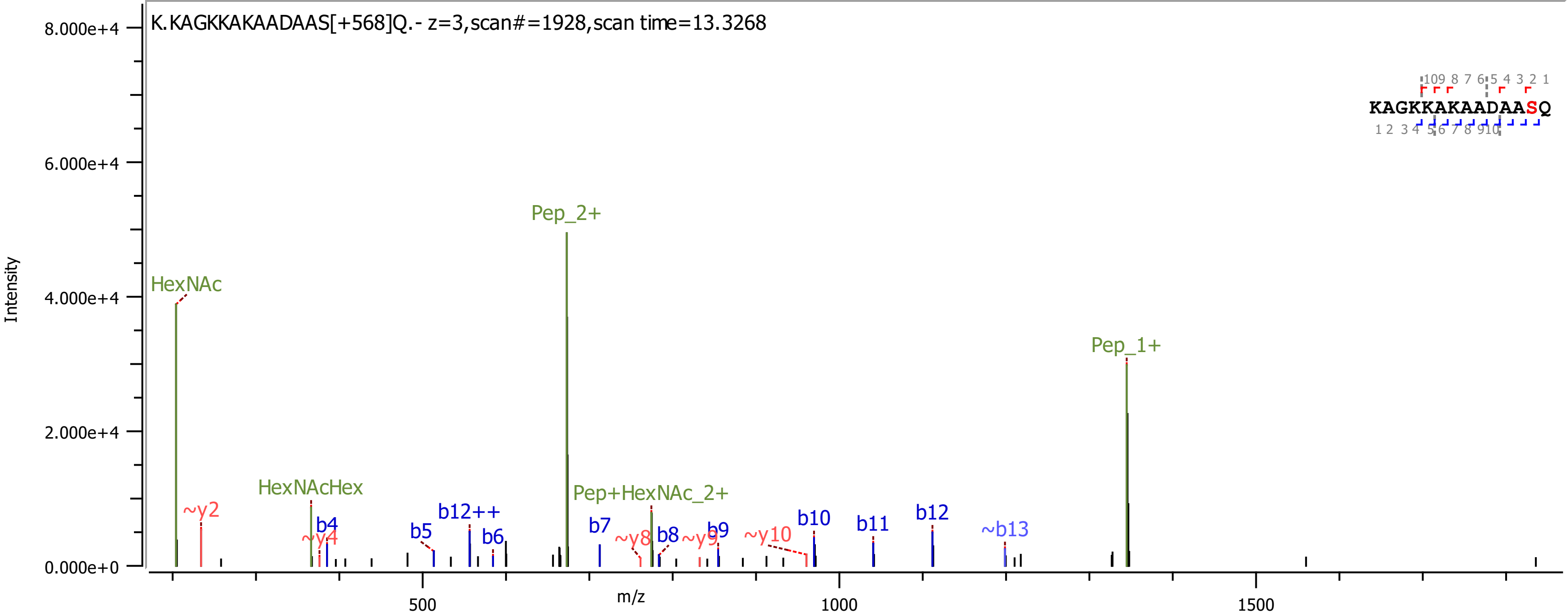
K.KAGKKAKAADAAS[+568]Q.- z=3,scan#=1928,scan time=13.3268
Intensity
8.000e+4
6.000e+4
4.000e+4
2.000e+4
0.000e+0
m/z
500
1000
1500
HexNAc
~y2
HexNAcHex
b4
~y4
b5
b12++
b6
Pep_2+
Pep+HexNAc_2+
b7
~y8
b8
~y9
b9
~y10
b10
b11
b12
~b13
Pep_1+
KAGKKAKAADAASQ

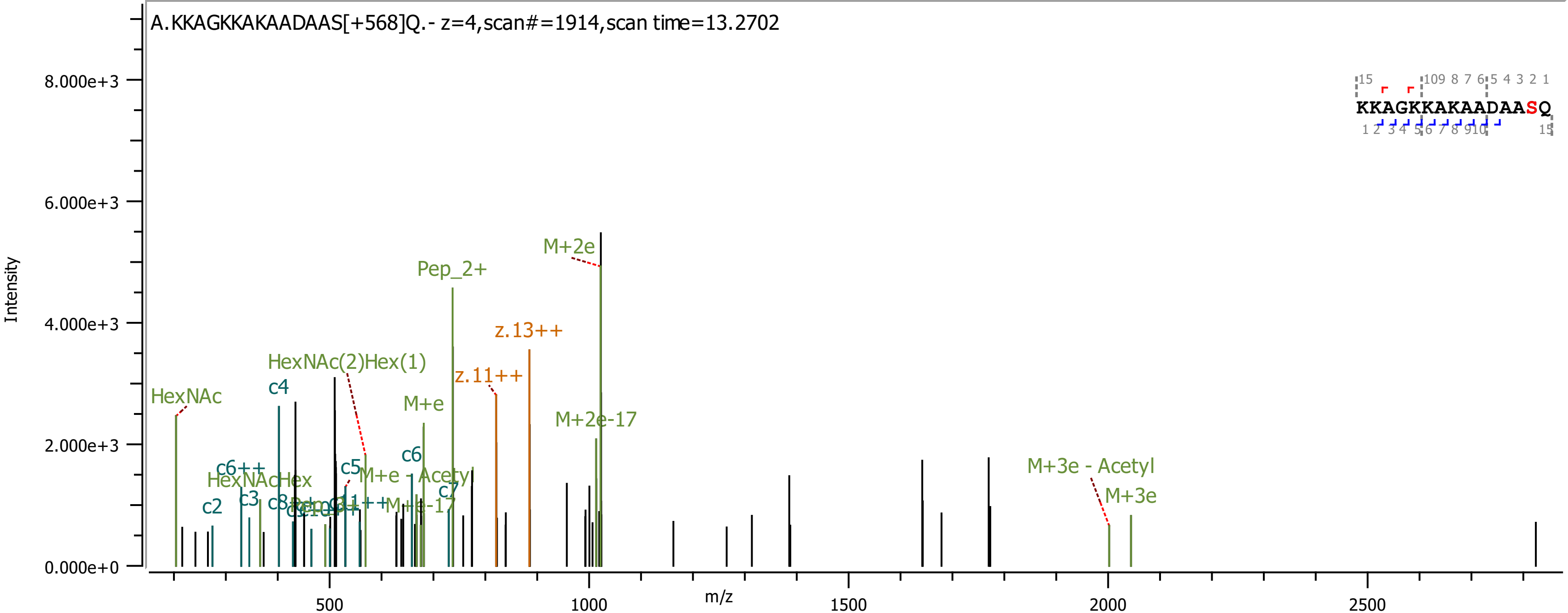
A.KKAGKKAKAADAAS[+568]Q.- z=4,scan#=1914,scan time=13.2702
Intensity
m/z
8.000e+3
6.000e+3
4.000e+3
2.000e+3
0.000e+0
500
1000
1500
2000
2500
KKAGKKAKAADAASQ
M+2e
Pep_2+
z.13++
z.11++
HexNAc(2)Hex(1)
HexNAc
c4
M+e
M+2e-17
c6
c5
c6++
HexNAcHex
M+e - Acetyl
c7
c3
c2
c8
M+e-17
M+3e - Acetyl
M+3e

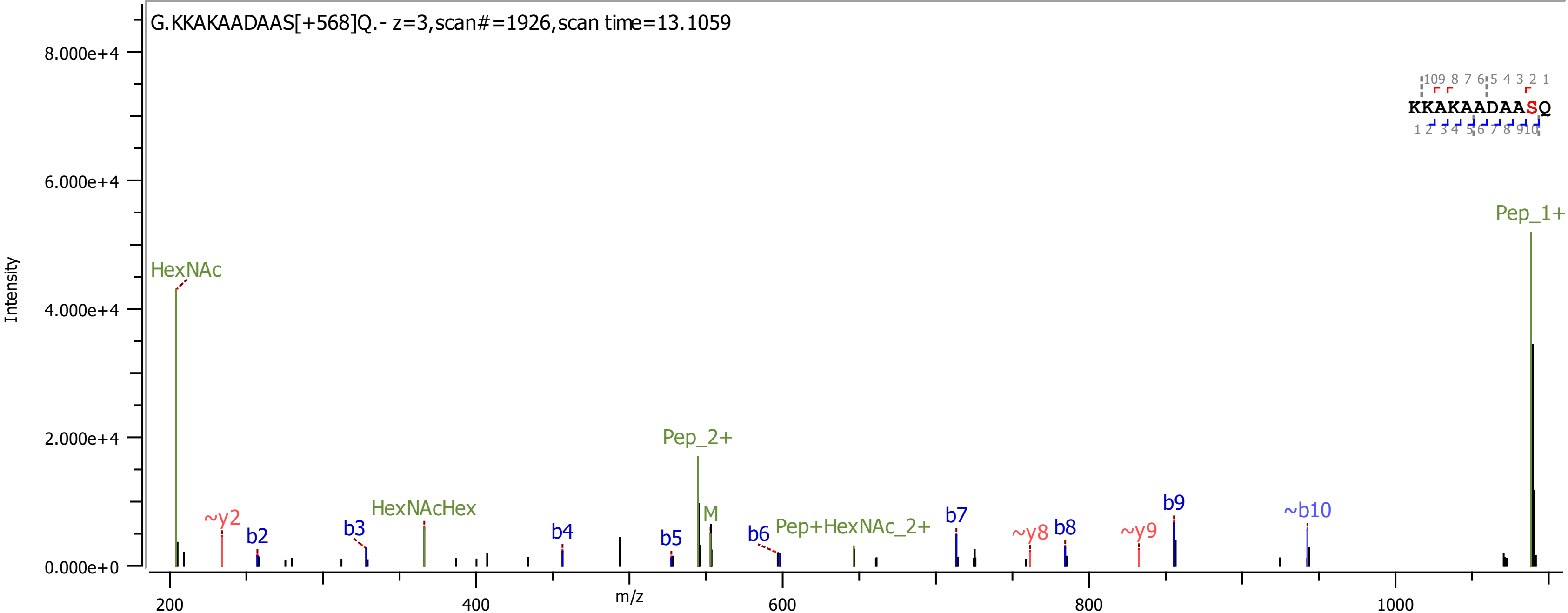

G.KKAKAADAAS[+568]Q.- z=3,scan#=1926,scan time=13.1059
Intensity
8.000e+4
6.000e+4
4.000e+4
2.000e+4
0.000e+0
m/z
200
400
600
800
1000
HexNAc
~y2
b2
b3
HexNAcHex
b4
b5
Pep_2+
M
b6
Pep+HexNAc_2+
b7
~y8
b8
~y9
b9
~b10
Pep_1+
KKAKAADAASQ

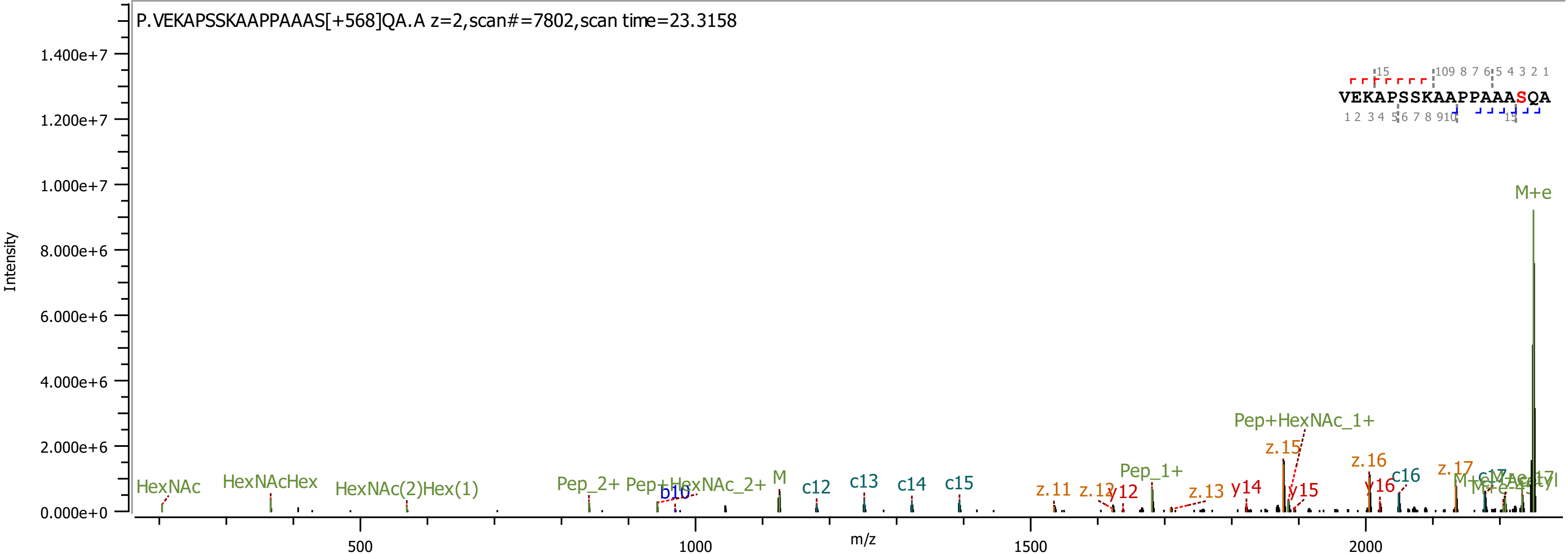

P.VEKAPSSKAAPPAAAS[+568]QA.A z=2,scan#=7802,scan time=23.3158
VEKAPSSKAAPPAAASQA
Intensity
m/z
M+e
Pep+HexNAc_1+
z.15
z.16
c16
z.17
Pep_1+
HexNAc
HexNAcHex
HexNAc(2)Hex(1)
Pep_2+
Pep+HexNAc_2+
b10
M
c12
c13
c14
c15
z.11
z.12
y12
z.13
y14
y15
y16

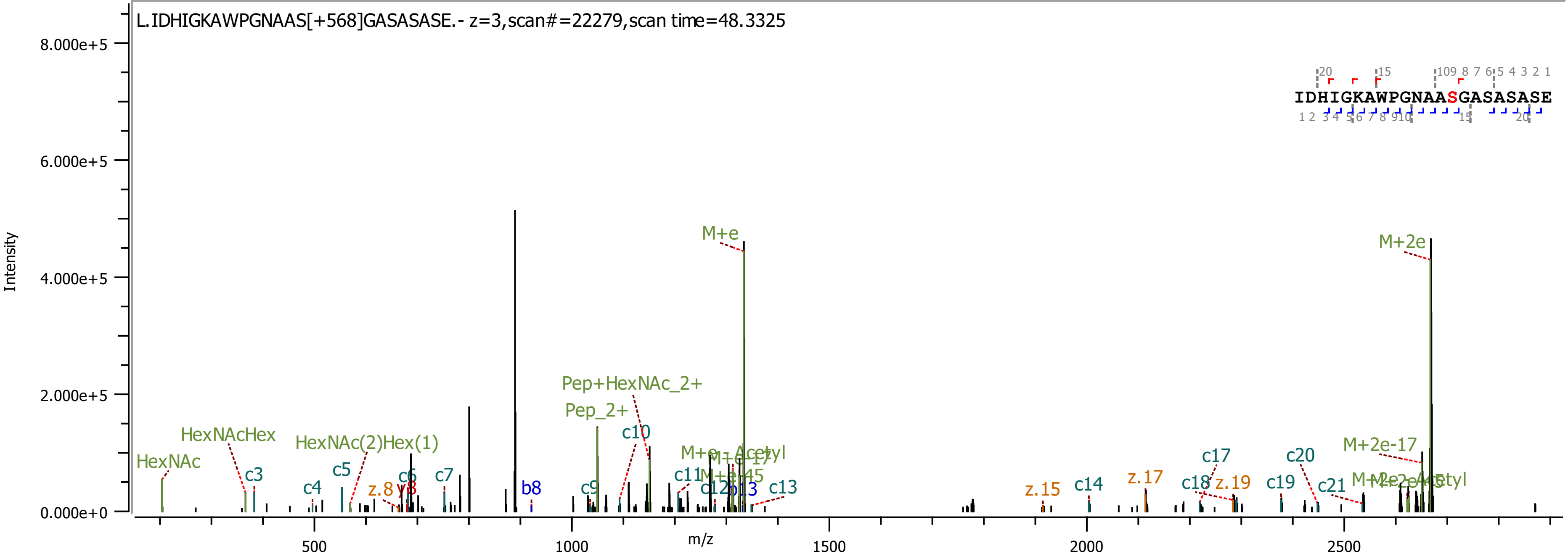

L.IDHIGKAWPGNAAS[+568]GASASASE.- z=3,scan#=22279,scan time=48.3325
Intensity
m/z
IDHIGKAWPGNAASGASASASE
M+e
M+2e
Pep+HexNAc_2+
Pep_2+
HexNAcHex
HexNAc(2)Hex(1)
HexNAc
M+2e-17
c3
c4
c5
c6
c7
c9
c10
c11
c12
c13
c14
c17
c18
c19
c20
c21
b8
b13
y8
z.8
z.15
z.17
z.19

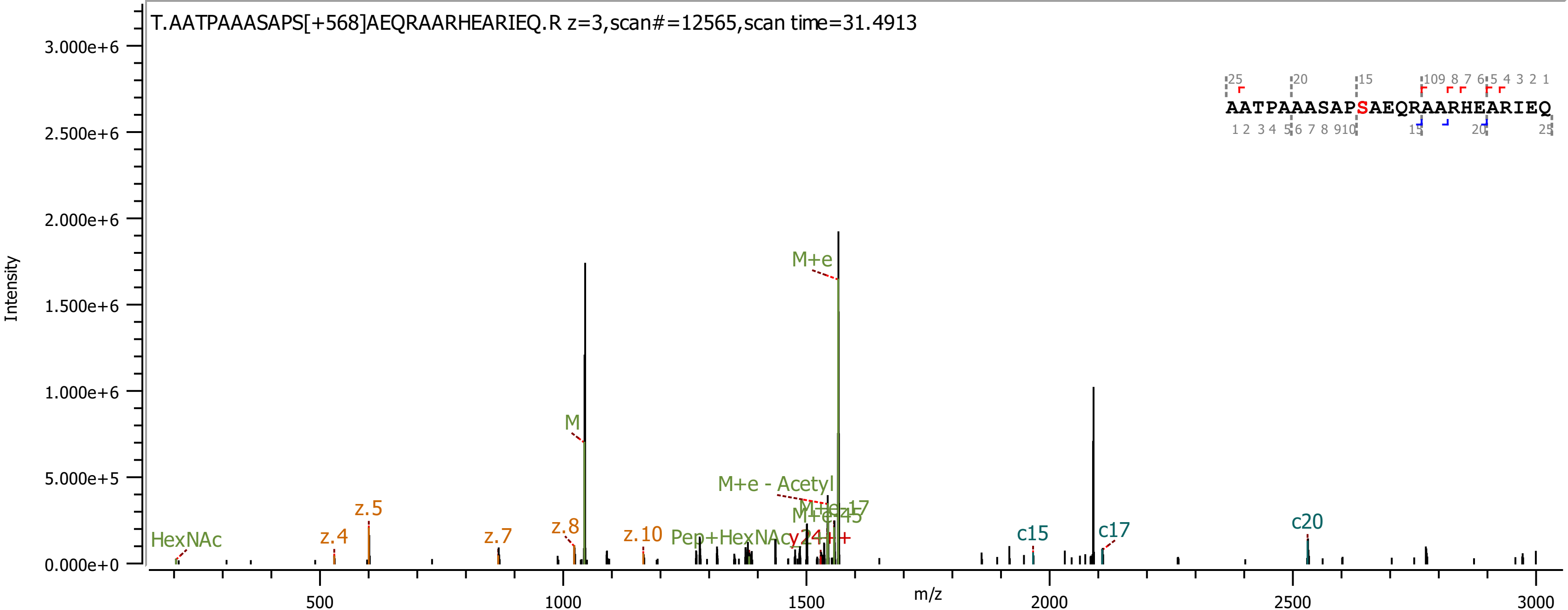
T.AATPAAASAPS[+568]AEQRAARHEARIEQ.R z=3,scan#=12565,scan time=31.4913
AATPAAASAPSAEQRAARHEARIEQ
Intensity
m/z
M+e
M
M+e - Acetyl
M+e-17
M+e-45
HexNAc
z.4
z.5
z.7
z.8
z.10
Pep+HexNAc
y24+++
c15
c17
c20

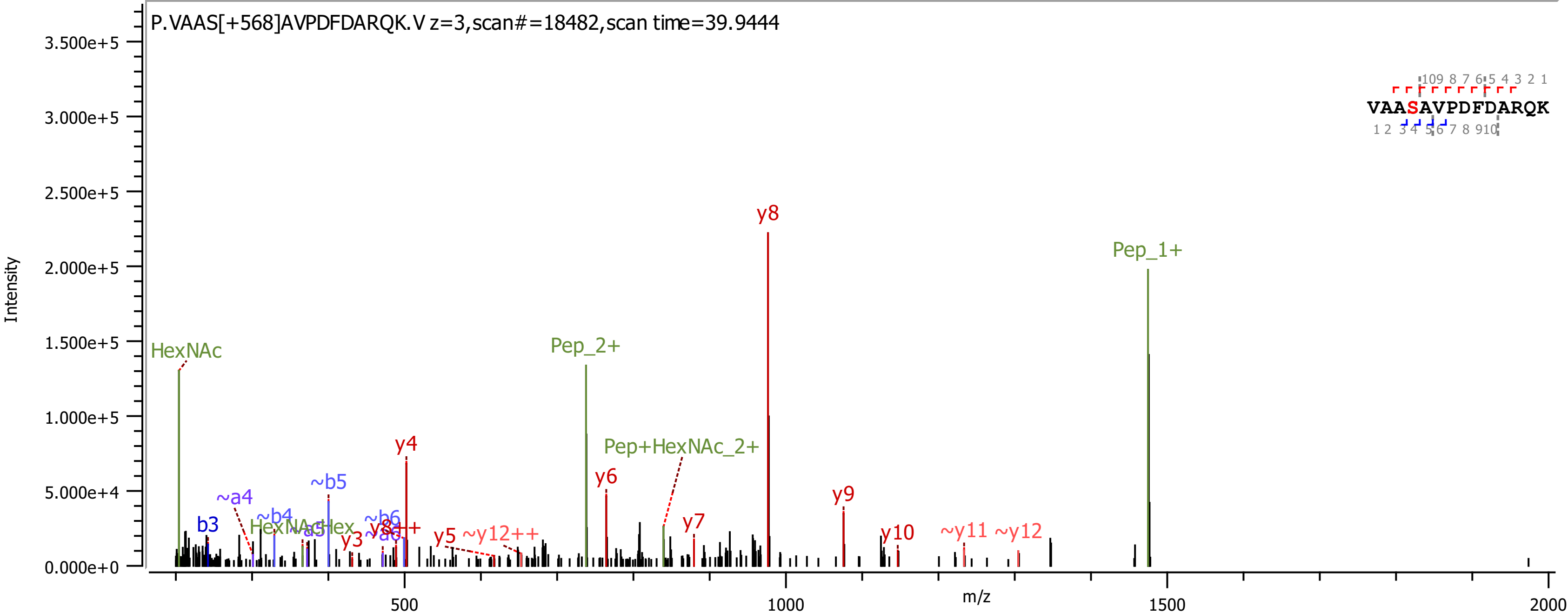
P.VAAS[+568]AVPDFDARQK.V z=3,scan#=18482,scan time=39.9444
VAASAVPDFDARQK
Intensity
m/z
HexNAc
Pep_2+
Pep_1+
Pep+HexNAc_2+
HexNAcHex
y8
y4
y6
y9
y7
y10
y3
y5
y8++
~y12++
~y11
~y12
b3
~a4
~b4
~a5
~b5
~b6
~a6

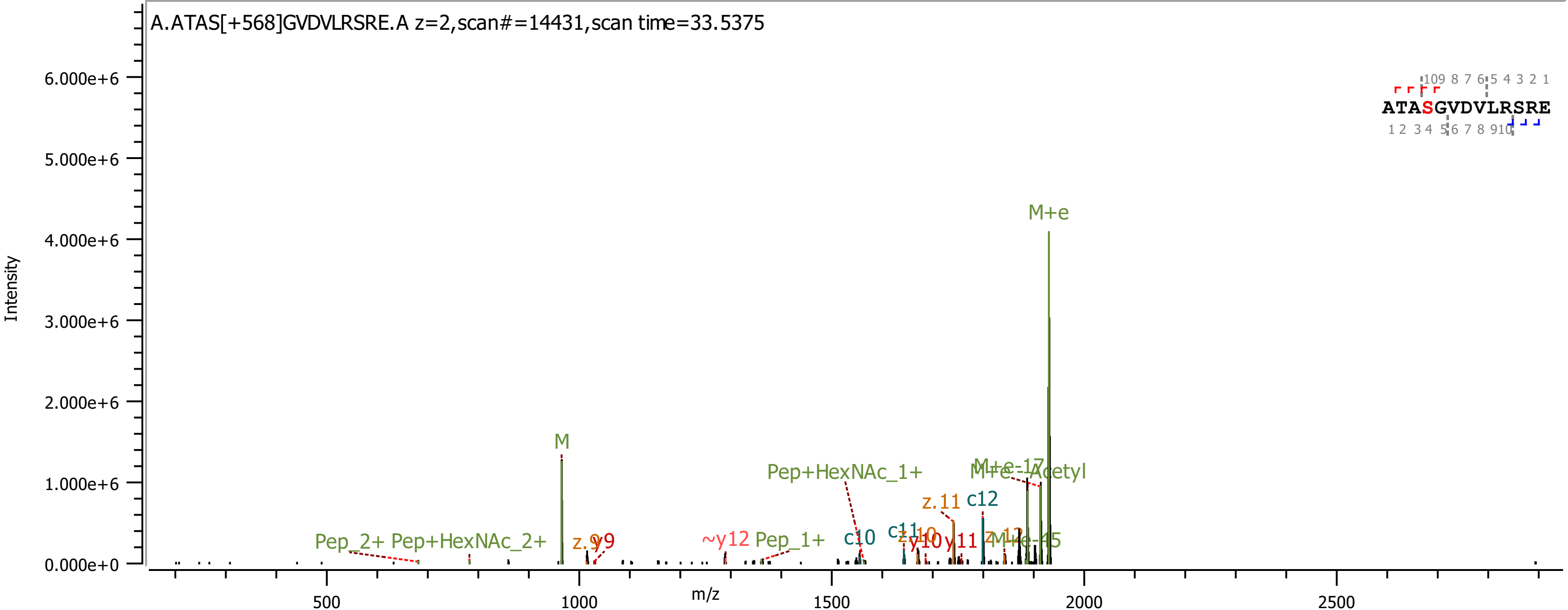

A.ATAS[+568]GVDVLRSRE.A z=2,scan#=14431,scan time=33.5375
Intensity
m/z
ATASGVDVLRSRE
M+e
M
Pep+HexNAc_1+
M+e-Acetyl
z.11
c12
c10
Pep_2+
Pep+HexNAc_2+
~y12
Pep_1+
y11

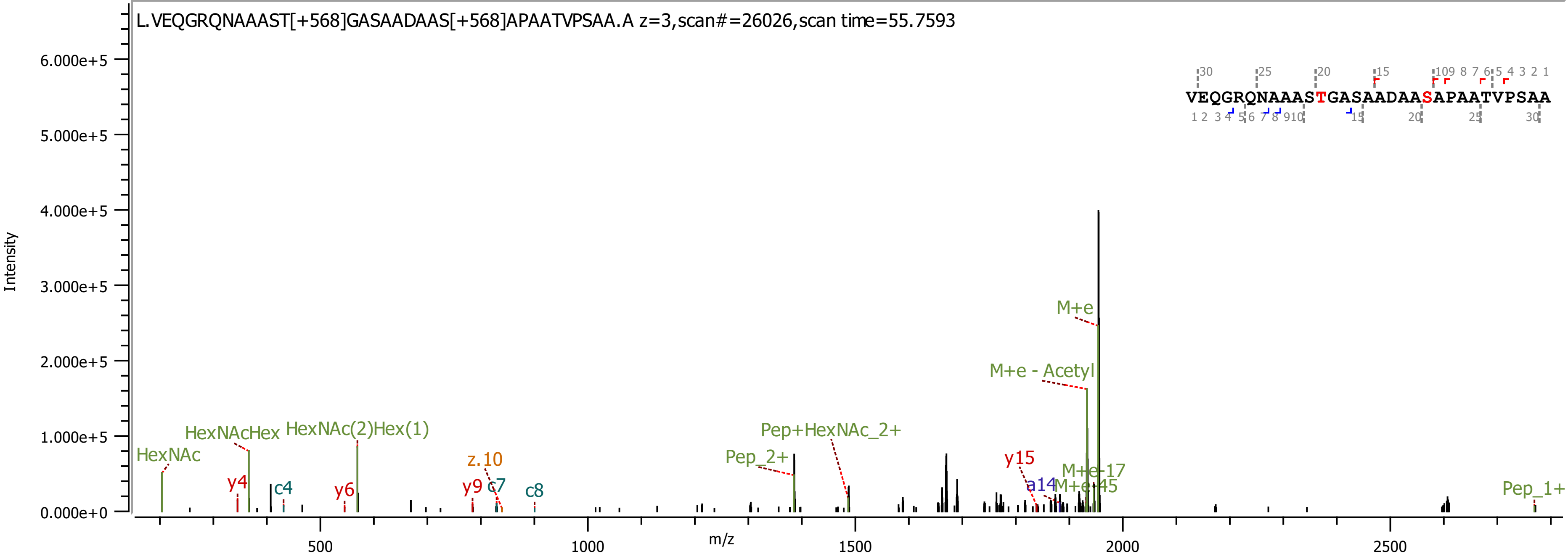

L.VEQGRQNAAAST[+568]GASAADAAS[+568]APAATVPSAA.A z=3,scan#=26026,scan time=55.7593
VEQGRQNAAASTGASAADAASAPAATVPSAA
Intensity
m/z
6.000e+5
5.000e+5
4.000e+5
3.000e+5
2.000e+5
1.000e+5
0.000e+0
500
1000
1500
2000
2500
HexNAc
HexNAcHex
HexNAc(2)Hex(1)
y4
c4
y6
z.10
y9
c7
c8
Pep_2+
Pep+HexNAc_2+
y15
a14
M+e-17
M+e-45
M+e - Acetyl
M+e
Pep_1+

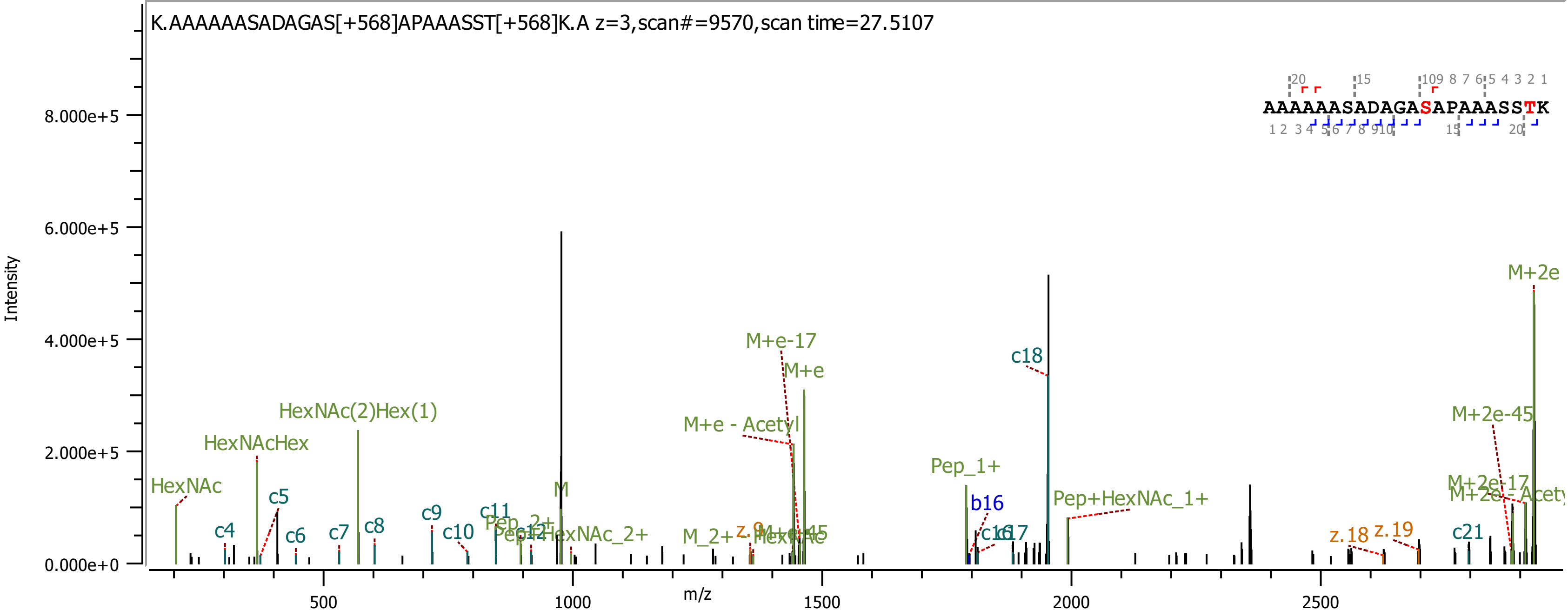
K.AAAAAASADAGAS[+568]APAAASST[+568]K.A z=3,scan#=9570,scan time=27.5107
AAAAAASADAGASAPAAASSTK
Intensity
m/z
8.000e+5
6.000e+5
4.000e+5
2.000e+5
0.000e+0
500
1000
1500
2000
2500
HexNAc
HexNAcHex
HexNAc(2)Hex(1)
c4
c5
c6
c7
c8
c9
c10
c11
c18
c21
M
M+e-17
M+e
M+e - Acetyl
Pep_1+
b16
Pep+HexNAc_1+
z.18
z.19
M+2e
M+2e-45
M+2e-17

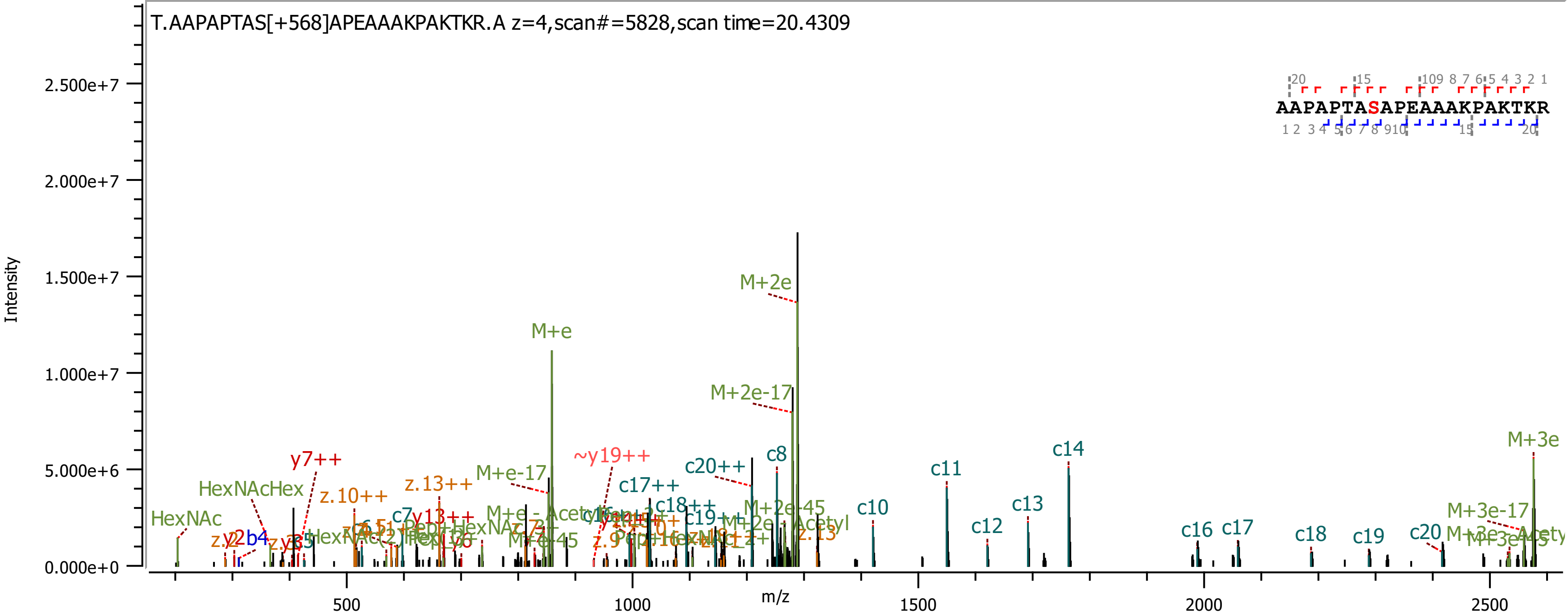
T.AAPAPTAS[+568]APEAAAKPAKTKR.A z=4,scan#=5828,scan time=20.4309
AAPAPTASAPEAAAKPAKTKR
Intensity
m/z
0.000e+0
5.000e+6
1.000e+7
1.500e+7
2.000e+7
2.500e+7
500
1000
1500
2000
2500
M+2e
M+2e-17
M+e
M+e-17
M+3e
M+3e-17
~y19++
y7++
c20++
c17++
c18++
c8
c10
c11
c12
c13
c14
c16
c17
c18
c19
c20
z.10++
z.13++
HexNAcHex
HexNAc
M+2e-45
b4
y13++

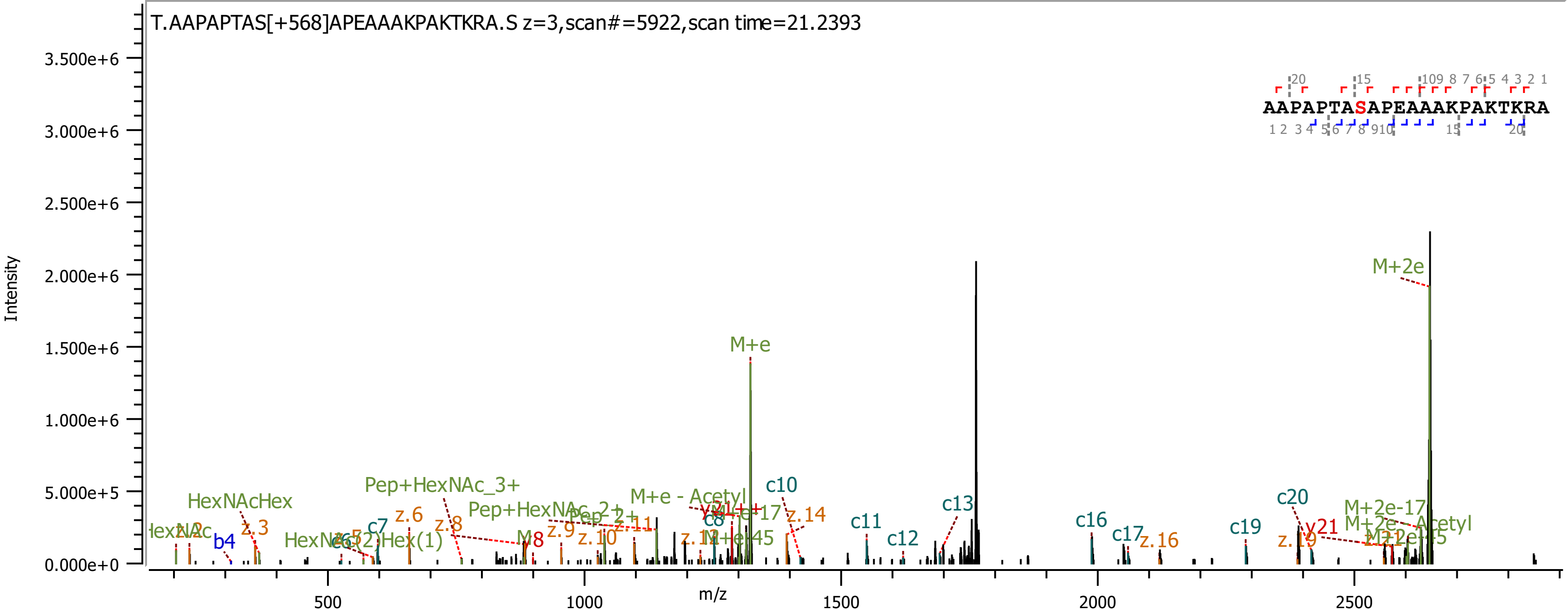

T.AAPAPTAS[+568]APEAAAKPAKTKRA.S z=3,scan#=5922,scan time=21.2393
AAPAPTASAPEAAAKPAKTKRA
Intensity
m/z
M+2e
M+e
c10
c11
c12
c13
c16
c17
c19
c20
y21
z.14
z.16
z.19
Pep+HexNAc_3+
Pep+HexNAc_2+
HexNAcHex
M+e - Acetyl
M+2e-17
M+2e - Acetyl
b4
c7
z.3
z.6
z.8
z.9
z.10
z.11

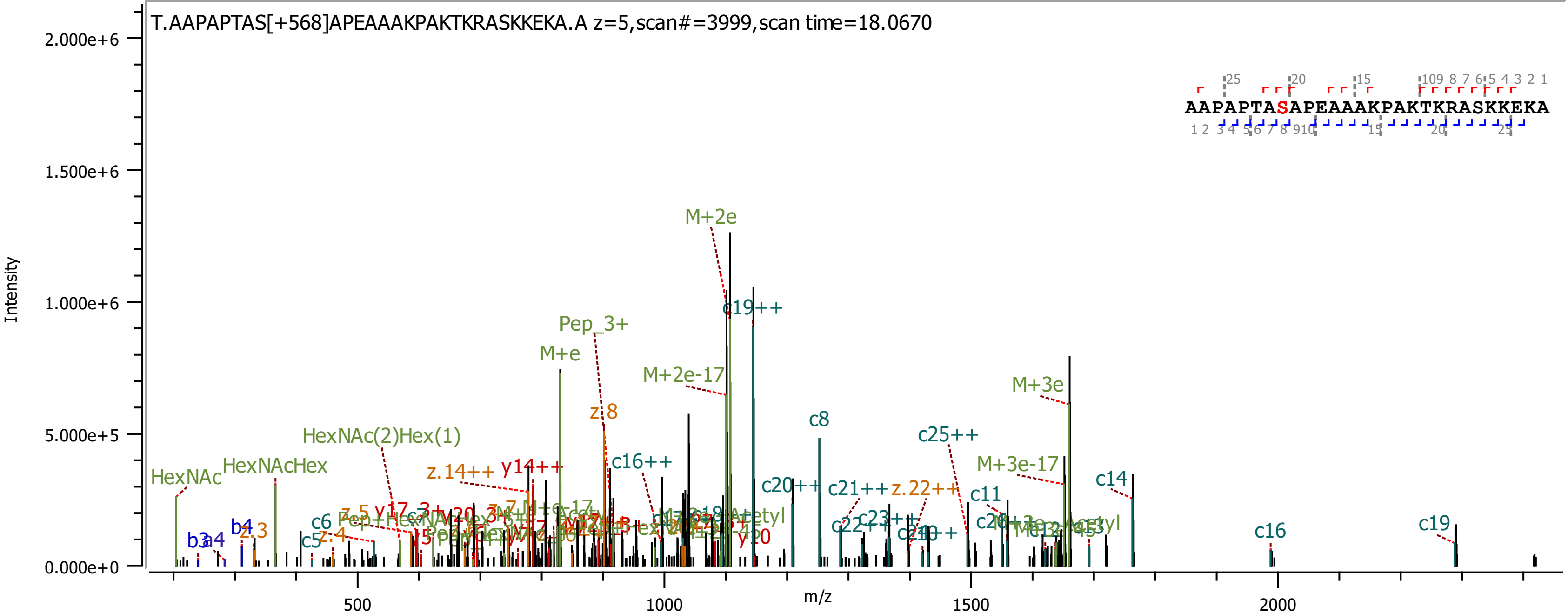

T.AAPAPTAS[+568]APEAAAKPAKTKRASKKEKA.A z=5,scan#=3999,scan time=18.0670
AAPAPTASAPEAAAKPAKTKRASKKEKA
Intensity
m/z
2.000e+6
1.500e+6
1.000e+6
5.000e+5
0.000e+0
500
1000
1500
2000
M+2e
c19++
Pep_3+
M+e
M+2e-17
z.8
c8
c25++
M+3e
M+3e-17
HexNAc(2)Hex(1)
HexNAcHex
HexNAc
z.14++
y14++
c16++
c20++
c21++
z.22++
c11
c14
c16
c19
b3
a4
b4
z.3
c5
z.4
c6
y20

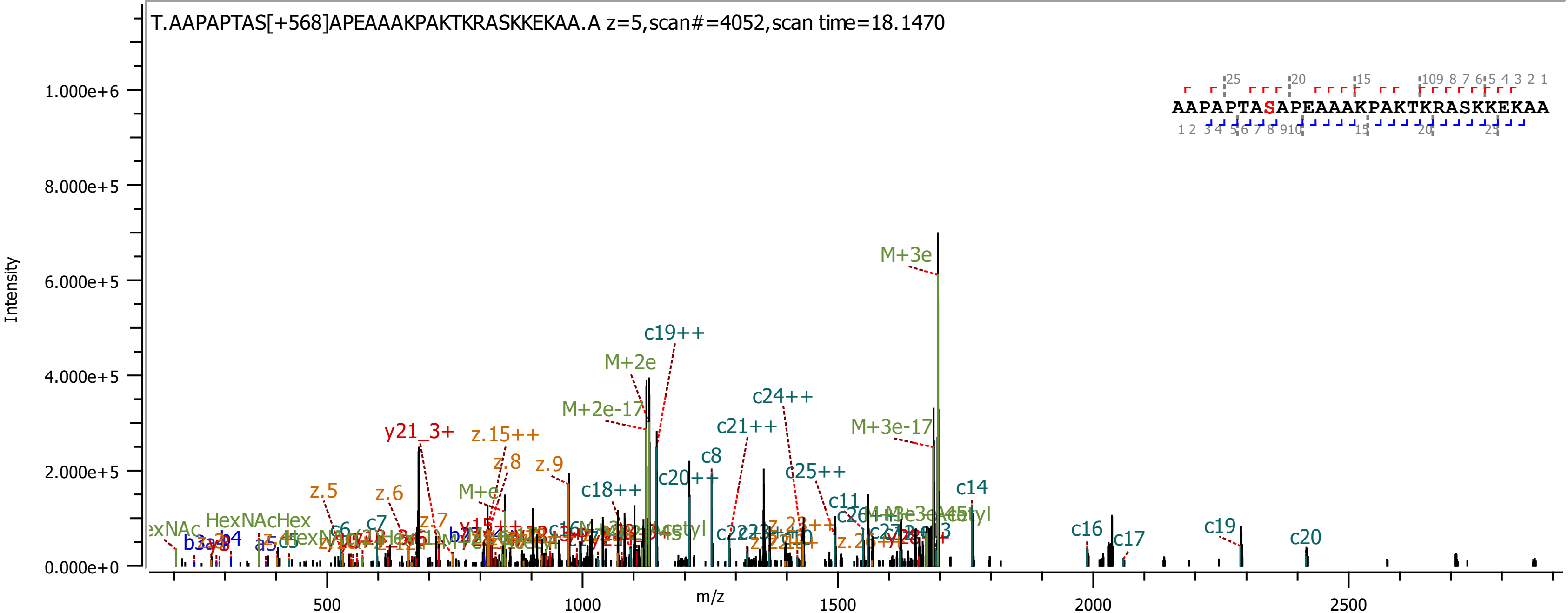
T.AAPAPTAS[+568]APEAAAKPAKTKRASKKEKAA.A z=5,scan#=4052,scan time=18.1470
AAPAPTASAPEAAAKPAKTKRASKKEKAA
Intensity
m/z
1.000e+6
8.000e+5
6.000e+5
4.000e+5
2.000e+5
0.000e+0
500
1000
1500
2000
2500
M+3e
c19++
M+2e
c24++
M+2e-17
M+3e-17
c21++
y21_3+
z.15++
z.8
z.9
c8
c20++
z.5
z.6
M+e
c18++
c25++
c14
c11
z.7
HexNAcHex
c6
c7
c16
c17
c19
c20
b4
a5
c5

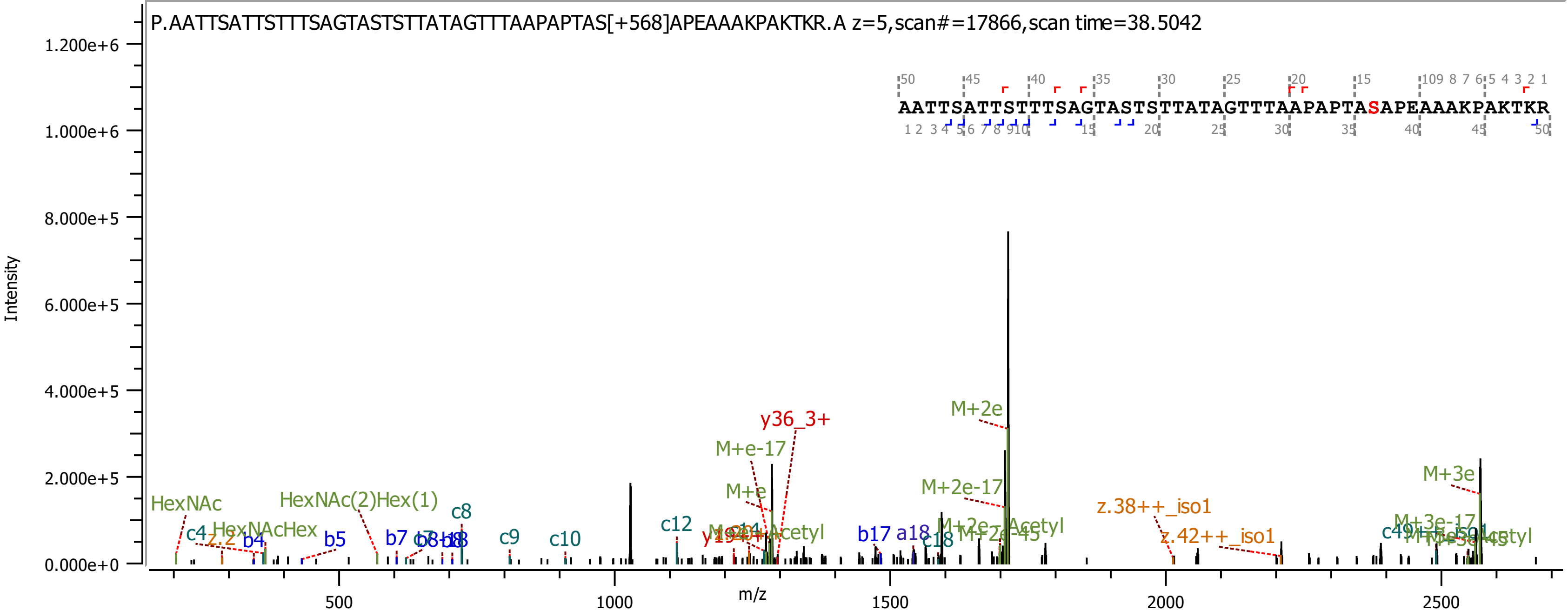
P.AATTSATTSTTTSAGTASTSTTATAGTTTAAPAPTAS[+568]APEAAAKPAKTKR.A z=5,scan#=17866,scan time=38.5042
AATTSATTSTTTSAGTASTSTTATAGTTTAAPAPTASAPEAAAKPAKTKR
Intensity
m/z
1.200e+6
1.000e+6
8.000e+5
6.000e+5
4.000e+5
2.000e+5
0.000e+0
500
1000
1500
2000
2500
HexNAc
c4
HexNAcHex
b4
b5
HexNAc(2)Hex(1)
b7
c8
c9
c10
c12
M+e-17
M+e
y36_3+
b17
a18
c18
M+2e
M+2e-17
z.38++_iso1
z.42++_iso1
M+3e
M+3e-17

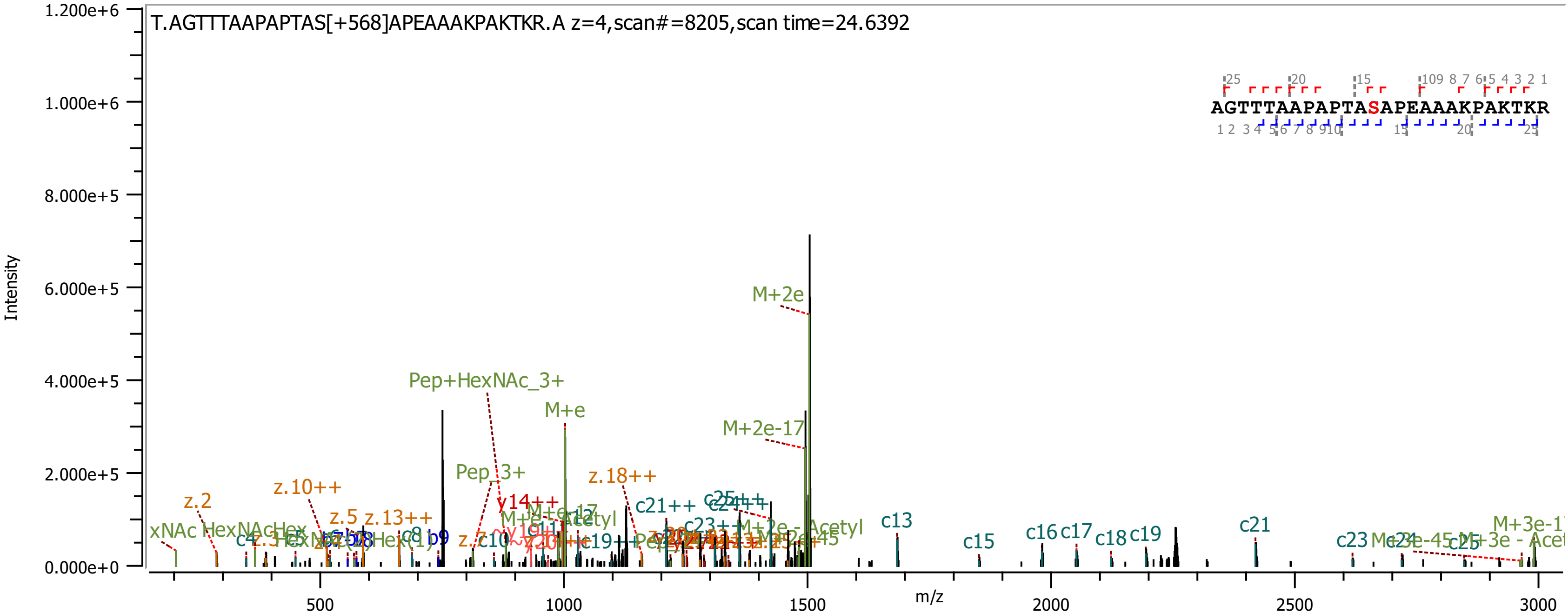

T.AGTTTAAPAPTAS[+568]APEAAAKPAKTKR.A z=4,scan#=8205,scan time=24.6392
AGTTTAAPAPTASAPEAAAKPAKTKR
Intensity
m/z
1.200e+6
1.000e+6
8.000e+5
6.000e+5
4.000e+5
2.000e+5
0.000e+0
500
1000
1500
2000
2500
3000
M+2e
Pep+HexNAc_3+
M+e
M+2e-17
Pep_3+
z.18++
z.10++
z.2
z.5
z.13++
y14++
c21++
c13
c15
c16
c17
c18
c19
c21
c23
c25
c8
b9
c4
c10
M+2e - Acetyl
M+3e-1

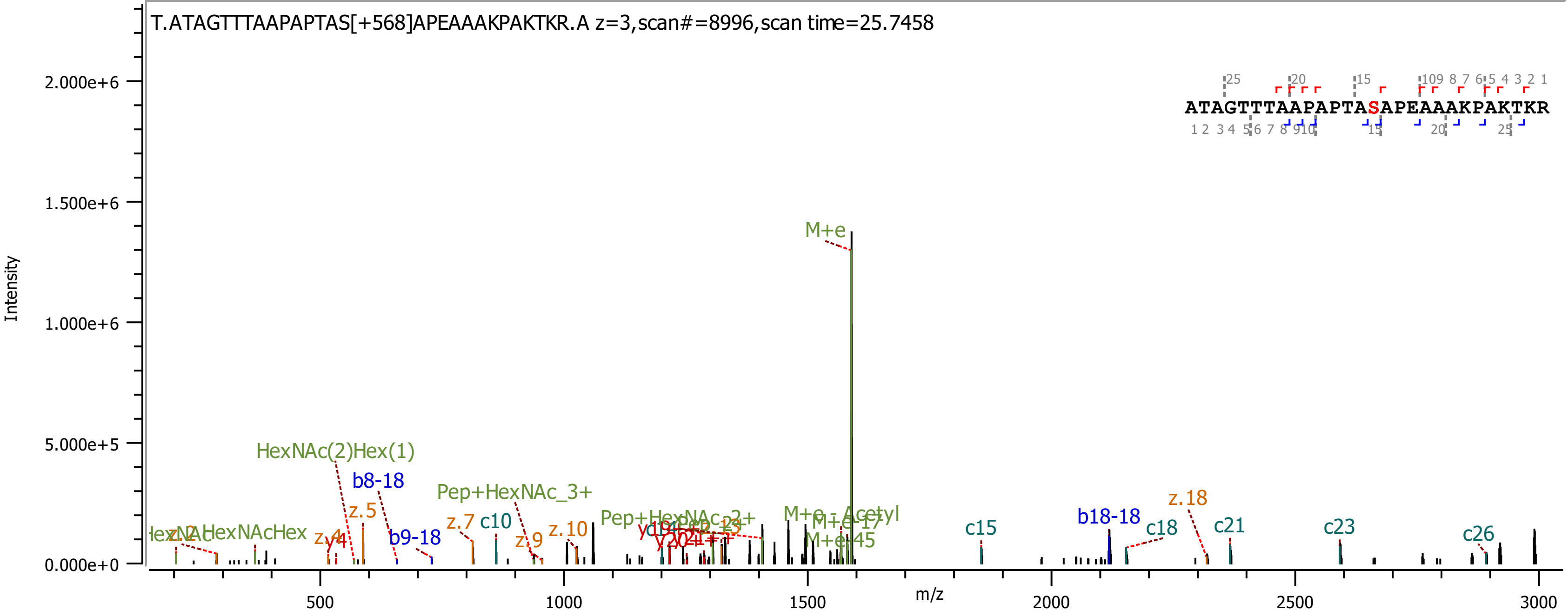

T.ATAGTTTAAPAPTAS[+568]APEAAAKPAKTKR.A z=3,scan#=8996,scan time=25.7458
ATAGTTTAAPAPTASAPEAAAKPAKTKR
Intensity
m/z
M+e
HexNAc(2)Hex(1)
b8-18
Pep+HexNAc_3+
z.5
z.7
c10
b9-18
z.9
z.10
z.4
M+e - Acetyl
M+e-17
M+e-45
c15
b18-18
c18
z.18
c21
c23
c26

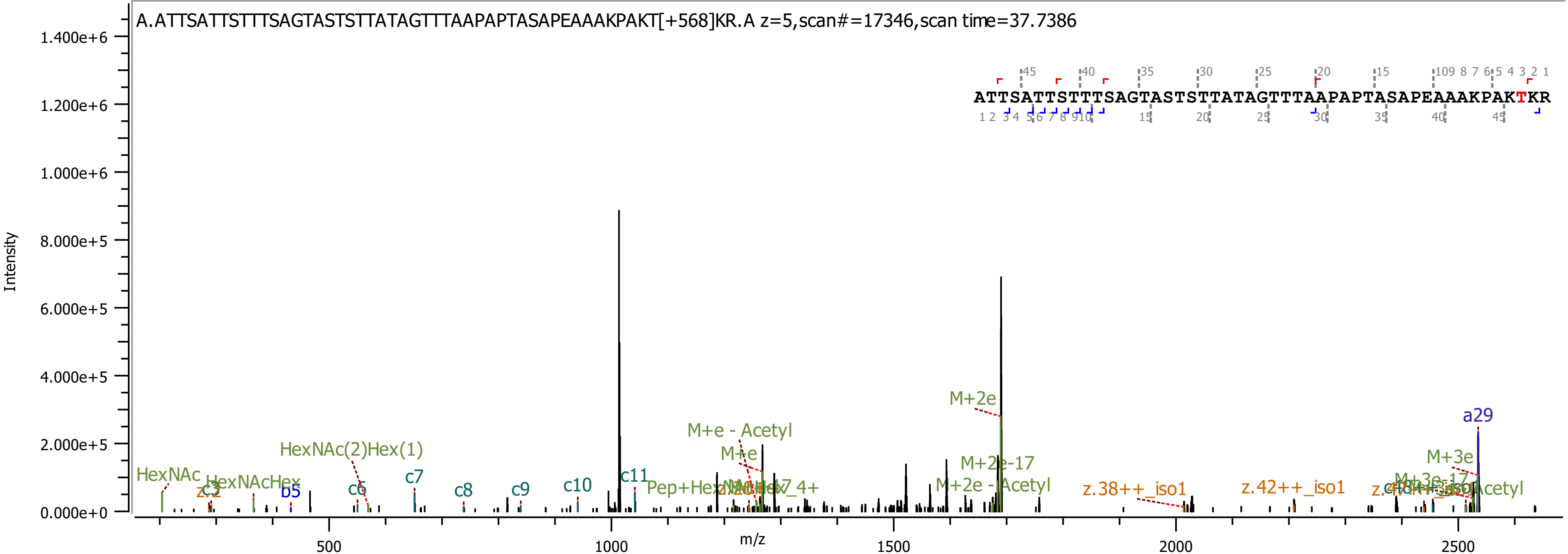

A.ATTSATTSTTTSAGTASTSTTATAGTTTAAPAPTASAPEAAAKPAKT[+568]KR.A z=5,scan#=17346,scan time=37.7386
ATTSATTSTTTSAGTASTSTTATAGTTTAAPAPTASAPEAAAKPAKTKR
Intensity
m/z
1.400e+6
1.200e+6
1.000e+6
8.000e+5
6.000e+5
4.000e+5
2.000e+5
0.000e+0
500
1000
1500
2000
2500
HexNAc
HexNAc(2)Hex(1)
b5
c6
c7
c8
c9
c10
c11
M+e - Acetyl
M+e
M+2e
M+2e-17
M+2e - Acetyl
z.38++_iso1
z.42++_iso1
M+3e
a29

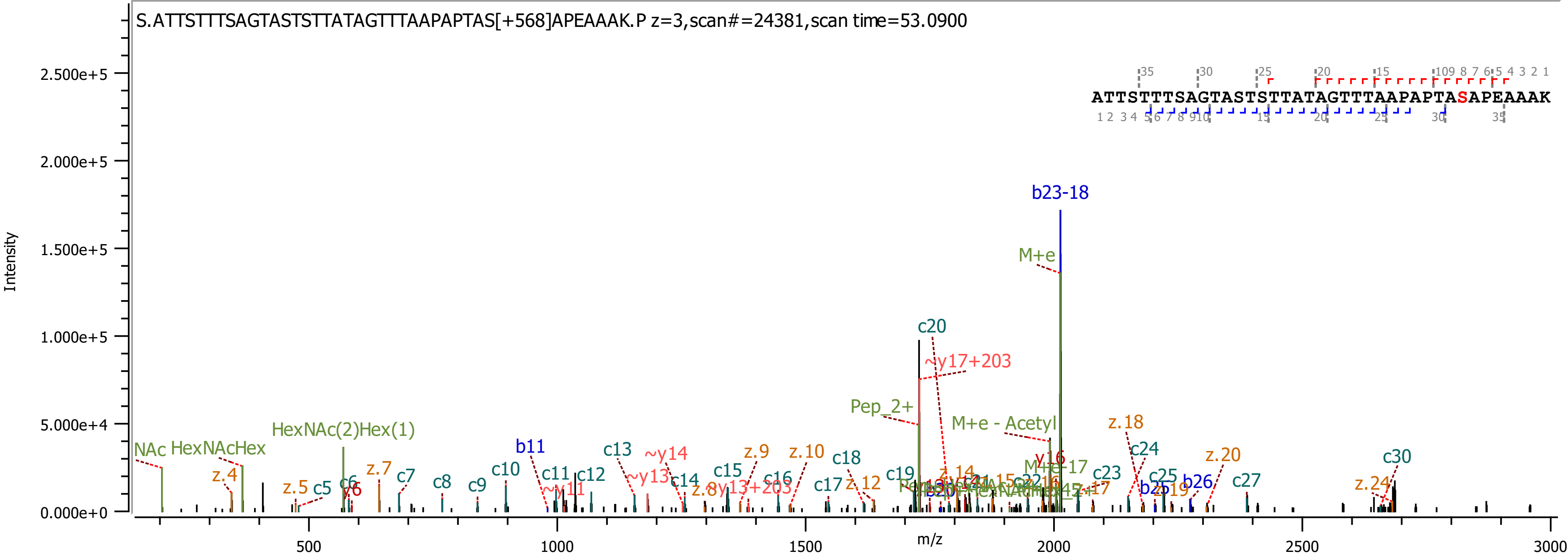

S.ATTSTTTSAGTASTSTTATAGTTTAAPAPTAS[+568]APEAAAK.P z=3,scan#=24381,scan time=53.0900
ATTSTTTSAGTASTSTTATAGTTTAAPAPTASAPEAAAK
Intensity
m/z
b23-18
M+e
c20
~y17+203
Pep_2+
M+e - Acetyl
z.18
HexNAc(2)Hex(1)
NAc
HexNAcHex
b11
c13
~y14
z.9
z.10
c18
c24
z.20
c30
z.4
z.7
c7
c8
c9
c10
c11
c12
c15
c16
c17
z.12
c19
c23
c25
b26
c27
z.24
z.5
c5
c6
y6
~y11
~y13
c14
y16

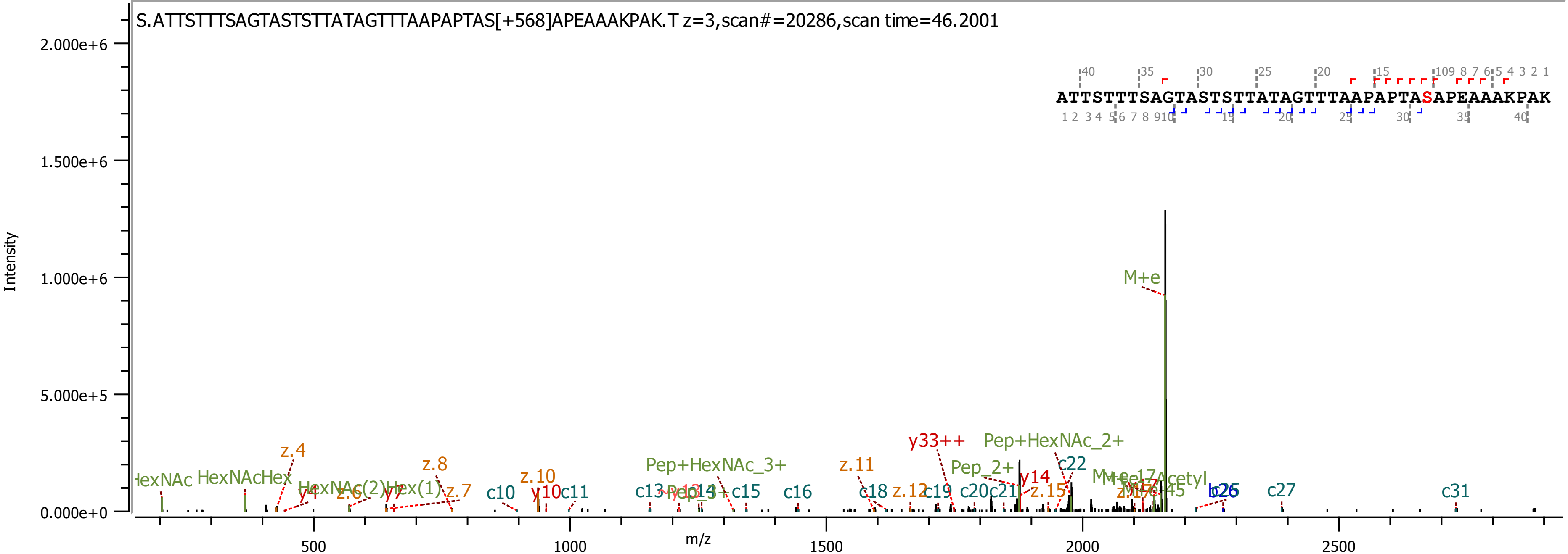
S.ATTSTTTSAGTASTSTTATAGTTTAAPAPTAS[+568]APEAAAKPAK.T z=3,scan#=20286,scan time=46.2001
ATTSTTTSAGTASTSTTATAGTTTAAPAPTASAPEAAAKPAK
Intensity
m/z
M+e
y33++
Pep+HexNAc_2+
Pep+HexNAc_3+
Pep_2+
c22
y14
z.15
z.11
z.12
c18
c19
c20
c21
c16
c15
c13
c10
y10
c11
z.10
z.8
z.7
z.4
y4
z.6
y7
HexNAcHex
HexNAc(2)Hex(1)
c27
c31
b26
M+e-17
Acetyl

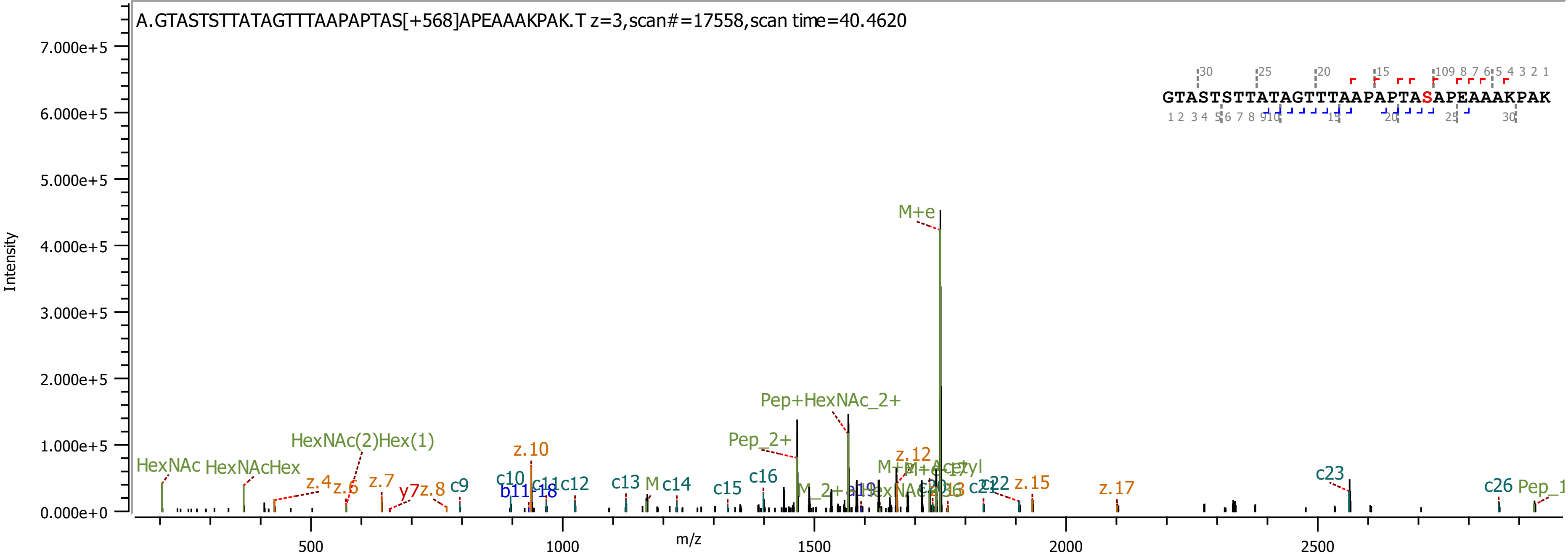

A.GTASTSTTATAGTTTAAPAPTAS[+568]APEAAAKPAK.T z=3,scan#=17558,scan time=40.4620
GTASTSTTATAGTTTAAPAPTASAPEAAAKPAK
Intensity
m/z
M+e
Pep+HexNAc_2+
Pep_2+
HexNAc(2)Hex(1)
HexNAc
HexNAcHex
z.10
z.12
z.4
z.6
z.7
y7
z.8
c9
c10
b11
c11
c12
c13
c14
c15
c16
c23
c26
z.15
z.17

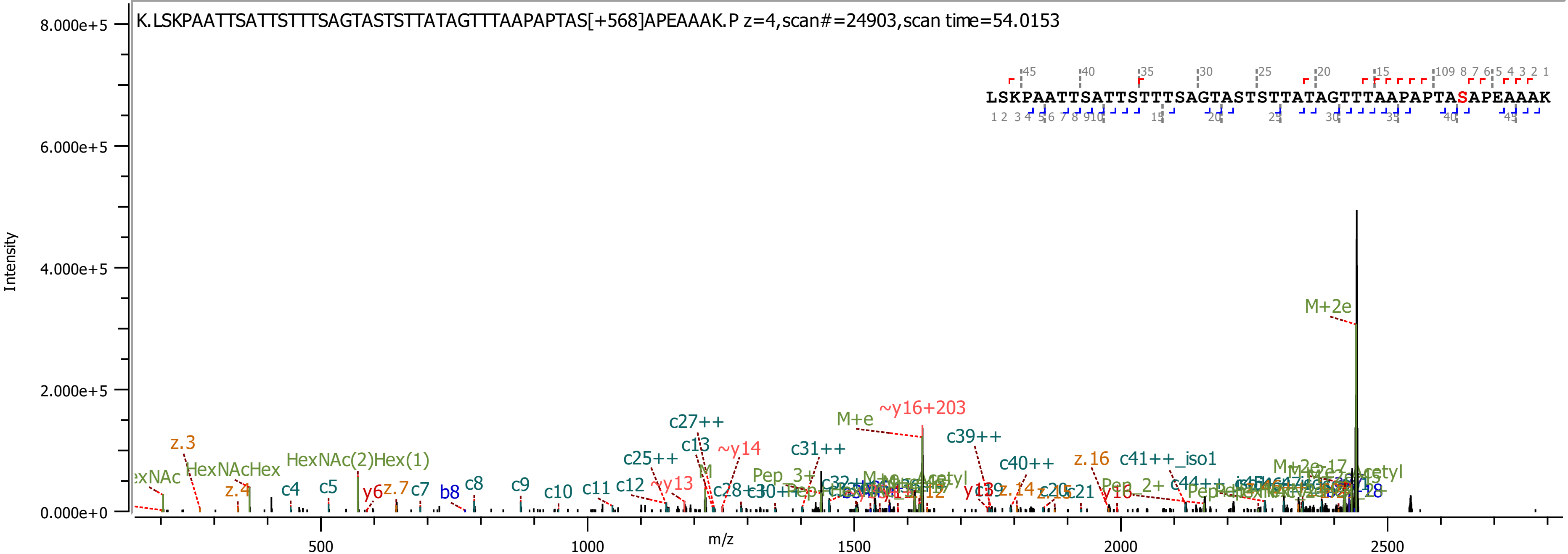
K.LSKPAATTSATTSTTTSAGTASTSTTATAGTTTAAPAPTAS[+568]APEAAAK.P z=4,scan#=24903,scan time=54.0153
LSKPAATTSATTSTTTSAGTASTSTTATAGTTTAAPAPTASAPEAAAK
Intensity
m/z
M+2e
~y16+203
M+e
c39++
c27++
c13
~y14
c31++
c25++
c40++
z.16
c41++_iso1
z.3
HexNAcHex
HexNAc(2)Hex(1)
z.4
c4
c5
y6
z.7
c7
b8
c8
c9
c10
c11
c12
~y13
c28++
y16
c21

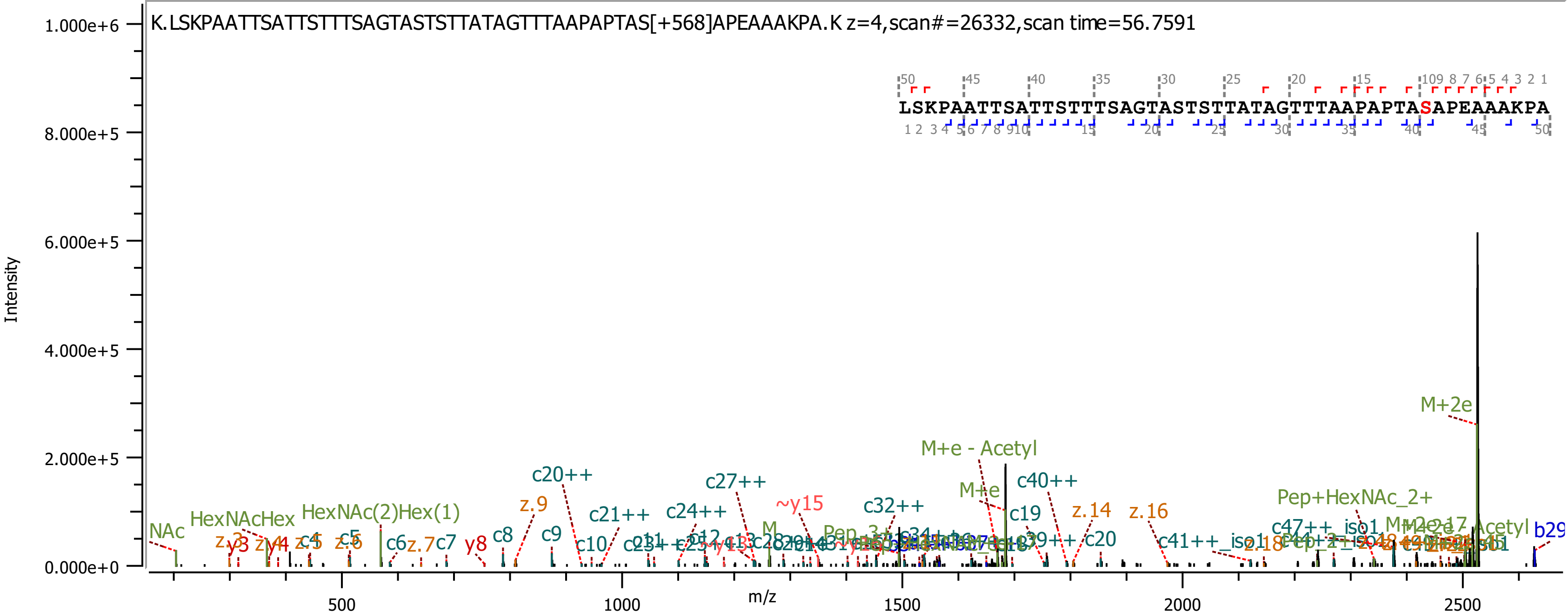
K.LSKPAATTSATTSTTTSAGTASTSTTATAGTTTAAPAPTAS[+568]APEAAAKPA.K z=4,scan#=26332,scan time=56.7591
LSKPAATTSATTSTTTSAGTASTSTTATAGTTTAAPAPTASAPEAAAKPA
Intensity
1.000e+6
8.000e+5
6.000e+5
4.000e+5
2.000e+5
0.000e+0
m/z
500
1000
1500
2000
2500
M+2e
M+e - Acetyl
M+e
c20++
c27++
c40++
c32++
~y15
c24++
c21++
z.9
z.14
z.16
c19
c20
c8
c9
c10
c6
c7
z.7
y8
NAc
HexNAcHex
HexNAc(2)Hex(1)
Pep+HexNAc_2+
c47++_iso1
c41++_iso1
z.18
b29

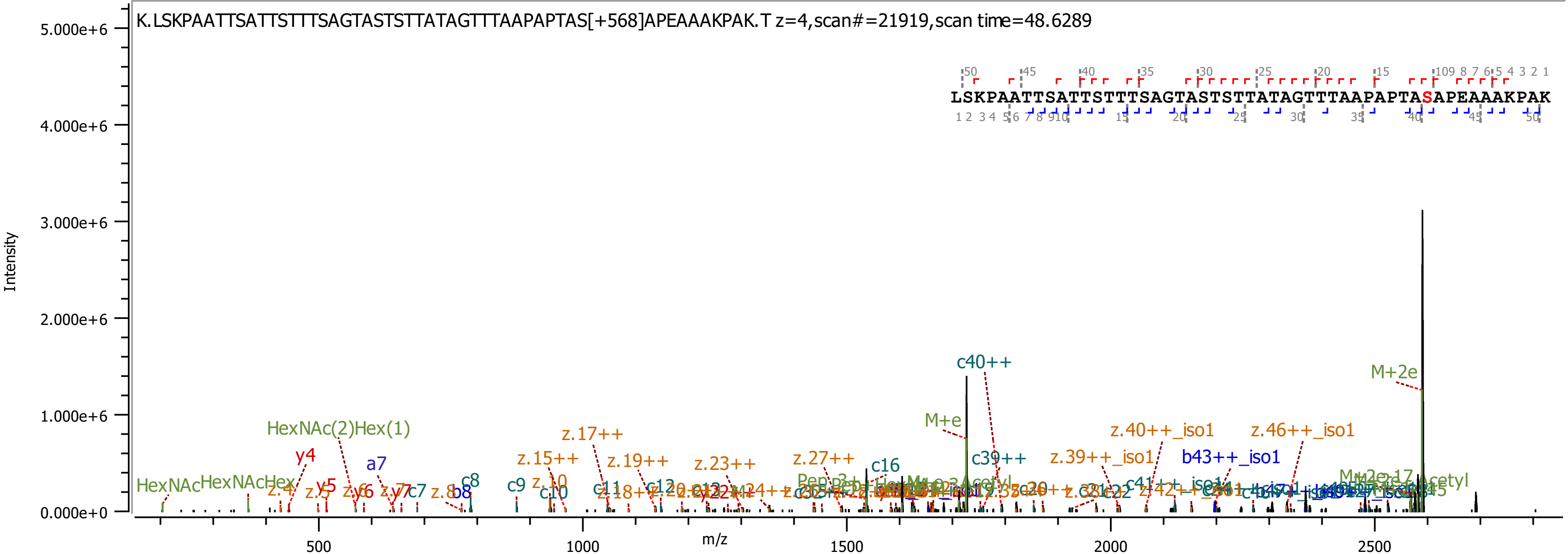

K.LSKPAATTSATTSTTTSAGTASTSTTATAGTTTAAPAPTAS[+568]APEAAAKPAK.T z=4,scan#=21919,scan time=48.6289
LSKPAATTSATTSTTTSAGTASTSTTATAGTTTAAPAPTASAPEAAAKPAK
Intensity
m/z
5.000e+6
4.000e+6
3.000e+6
2.000e+6
1.000e+6
0.000e+0
500
1000
1500
2000
2500
c40++
M+2e
M+e
HexNAc(2)Hex(1)
z.17++
z.15++
z.19++
z.23++
z.27++
c16
c39++
z.39++_iso1
z.40++_iso1
z.46++_iso1
b43++_iso1
y4
a7
y5
c7
c8
c9
b8
c10
c11
c12
HexNAc
HexNAcHex

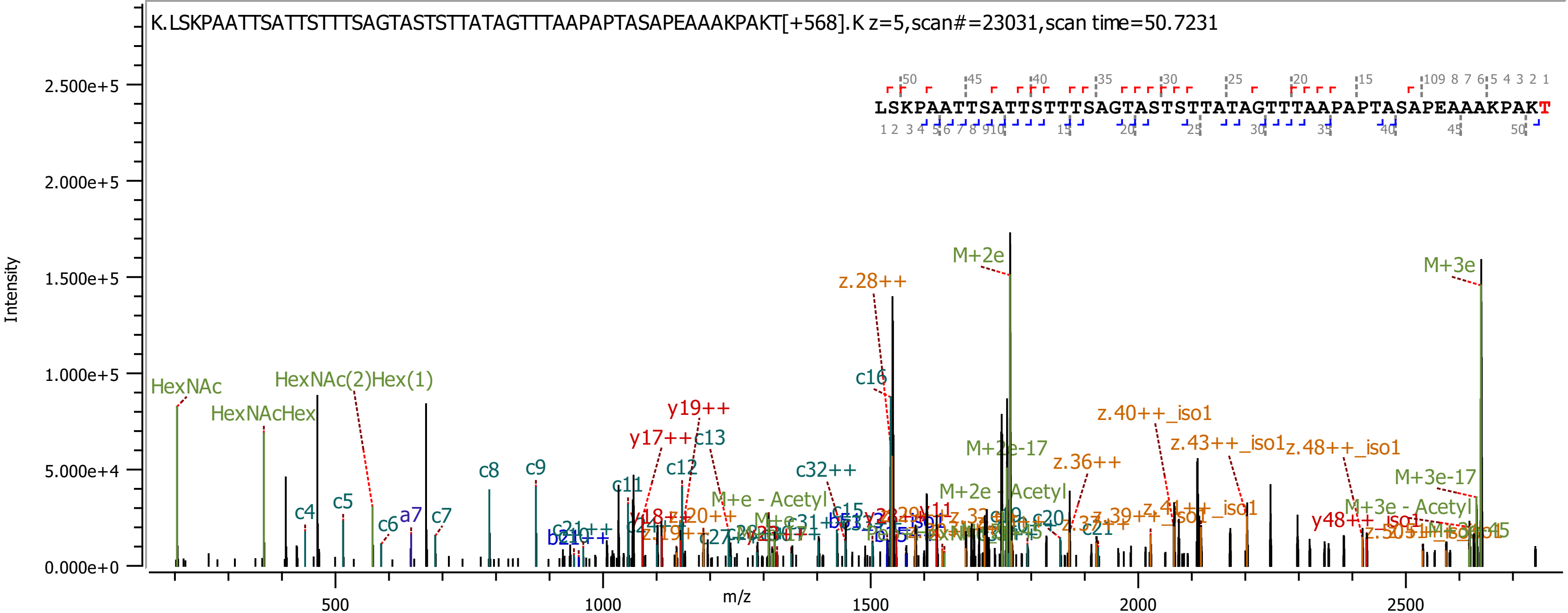
K.LSKPAATTSATTSTTTSAGTASTSTTATAGTTTAAPAPTASAPEAAAKPAKT[+568].K z=5,scan#=23031,scan time=50.7231
LSKPAATTSATTSTTTSAGTASTSTTATAGTTTAAPAPTASAPEAAAKPAKT
Intensity
m/z
2.500e+5
2.000e+5
1.500e+5
1.000e+5
5.000e+4
0.000e+0
500
1000
1500
2000
2500
HexNAc
HexNAcHex
HexNAc(2)Hex(1)
c4
c5
c6
a7
c7
c8
c9
c11
c12
y17++
y19++
c13
c32++
c15
c16
z.28++
M+2e
M+2e-17
M+2e - Acetyl
M+e - Acetyl
z.36++
z.40++_iso1
z.43++_iso1
z.48++_iso1
M+3e
M+3e-17
M+3e - Acetyl
y48++
c20
c21

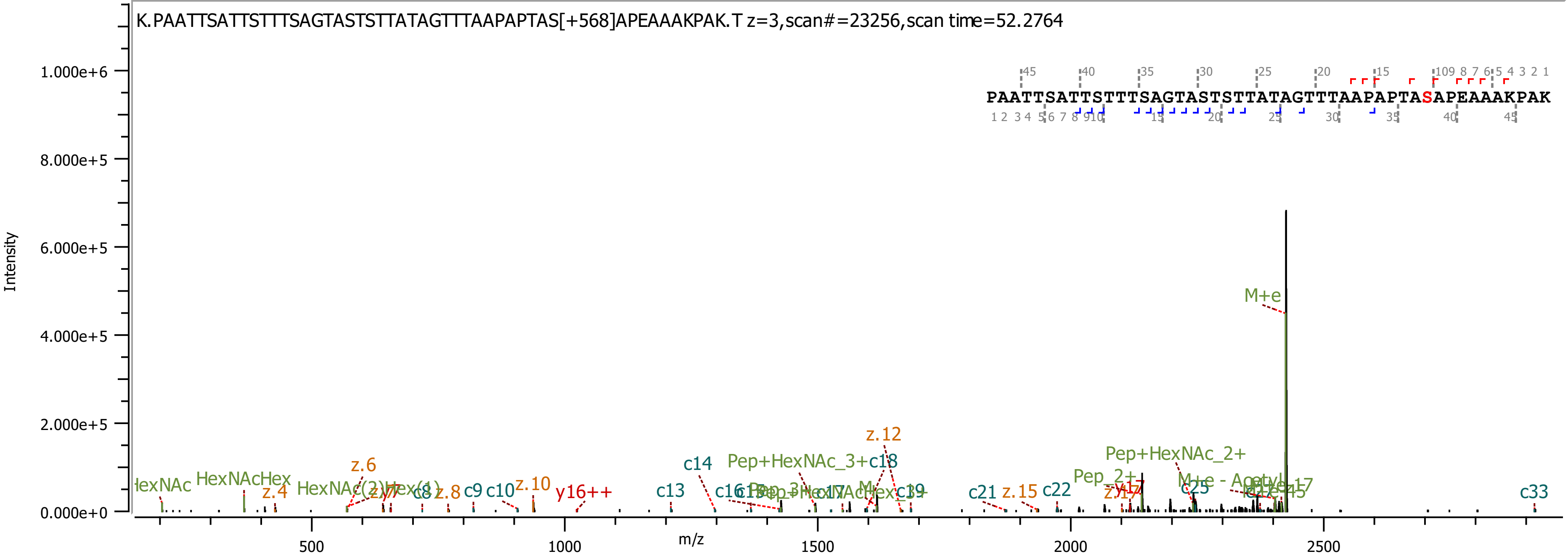
K.PAATTSATTSTTTSAGTASTSTTATAGTTTAAPAPTAS[+568]APEAAAKPAK.T z=3,scan#=23256,scan time=52.2764
PAATTSATTSTTTSAGTASTSTTATAGTTTAAPAPTASAPEAAAKPAK
Intensity
m/z
1.000e+6
8.000e+5
6.000e+5
4.000e+5
2.000e+5
0.000e+0
500
1000
1500
2000
2500
M+e
Pep+HexNAc_2+
Pep_2+
z.12
c14
c18
z.6
HexNAcHex
z.4
z.8
c9
c10
z.10
y16++
c13
c19
c21
z.15
c22
c25
c33

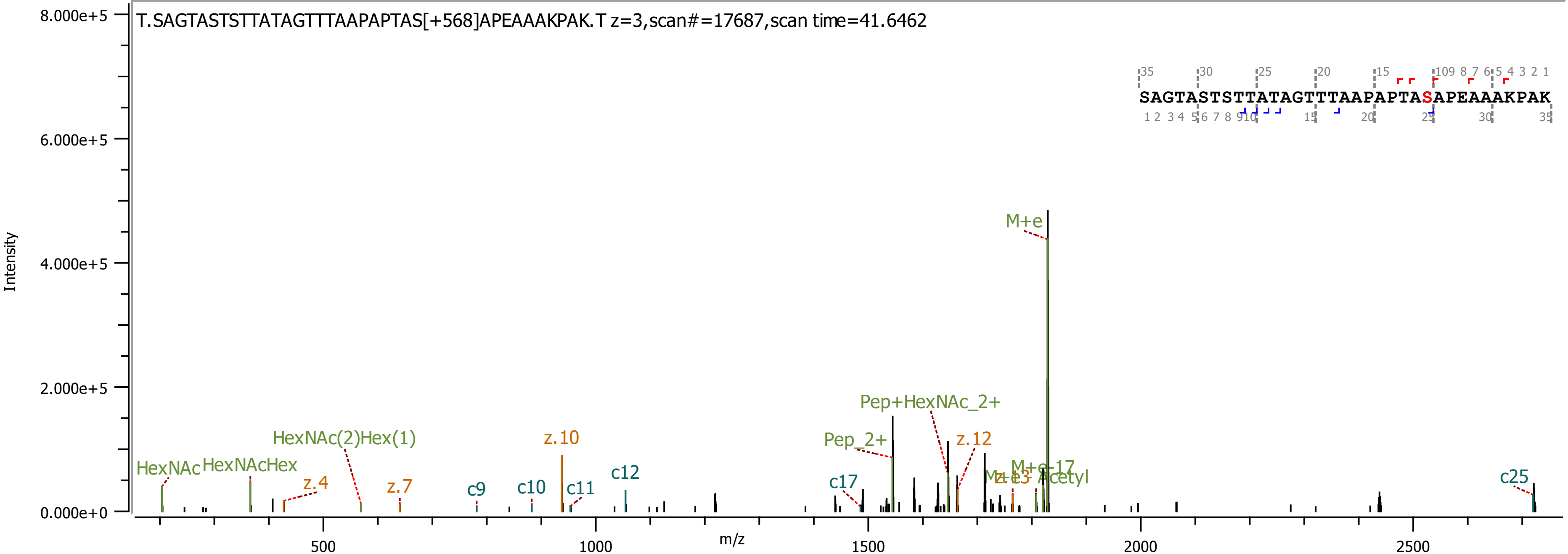

T.SAGTASTSTTATAGTTTAAPAPTAS[+568]APEAAAKPAK.T z=3,scan#=17687,scan time=41.6462
SAGTASTSTTATAGTTTAAPAPTASAPEAAAKPAK
Intensity
m/z
M+e
Pep+HexNAc_2+
Pep_2+
z.12
z.10
HexNAc(2)Hex(1)
HexNAc
HexNAcHex
z.4
z.7
c9
c10
c11
c12
c17
c25
M+e-17
Acetyl

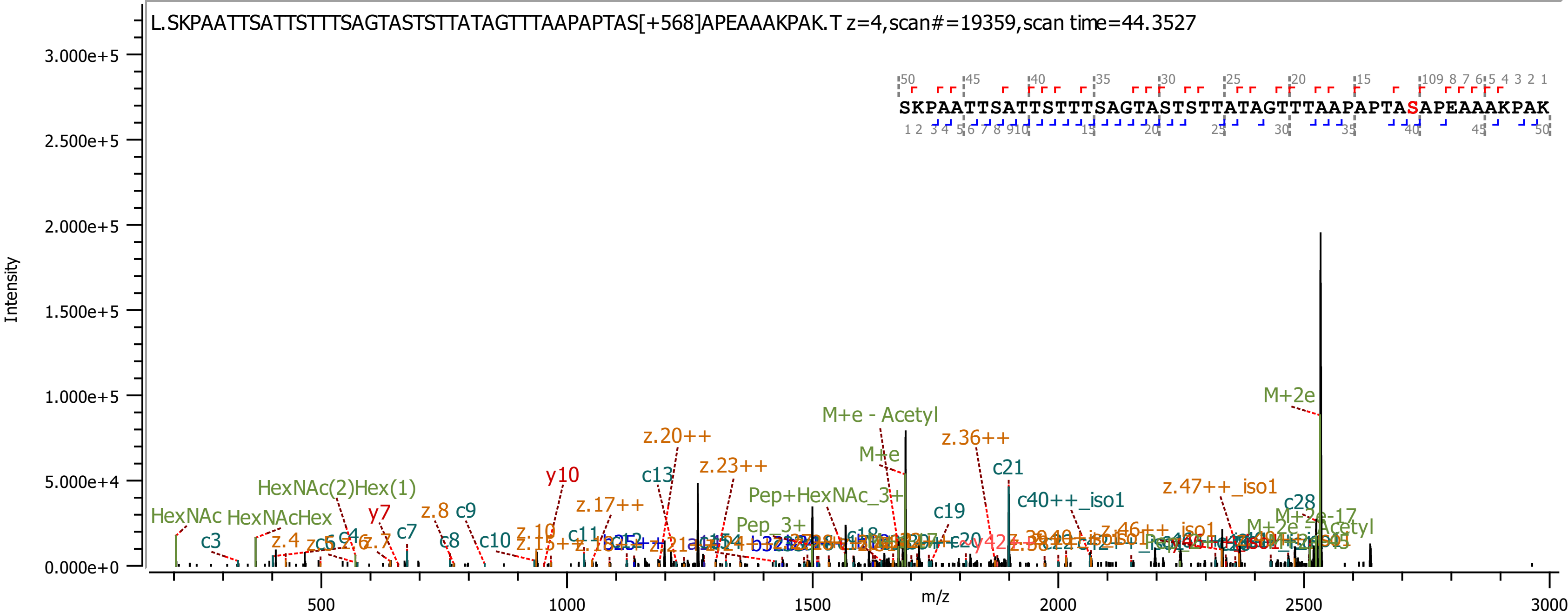
L.SKPAATTSATTSTTTSAGTASTSTTATAGTTTAAPAPTAS[+568]APEAAAKPAK.T z=4,scan#=19359,scan time=44.3527
SKPAATTSATTSTTTSAGTASTSTTATAGTTTAAPAPTASAPEAAAKPAK
Intensity
m/z
3.000e+5
2.500e+5
2.000e+5
1.500e+5
1.000e+5
5.000e+4
0.000e+0
500
1000
1500
2000
2500
3000
M+2e
M+e - Acetyl
M+e
z.20++
z.36++
z.23++
c21
c13
y10
c40++_iso1
z.47++_iso1
HexNAc(2)Hex(1)
Pep+HexNAc_3+
c19
z.17++
c28
HexNAc
HexNAcHex
y7
z.8
c9
c7
c3
z.4
c8
c10
z.10
c11
Pep_3+
M+2e-17
M+2e - Acetyl

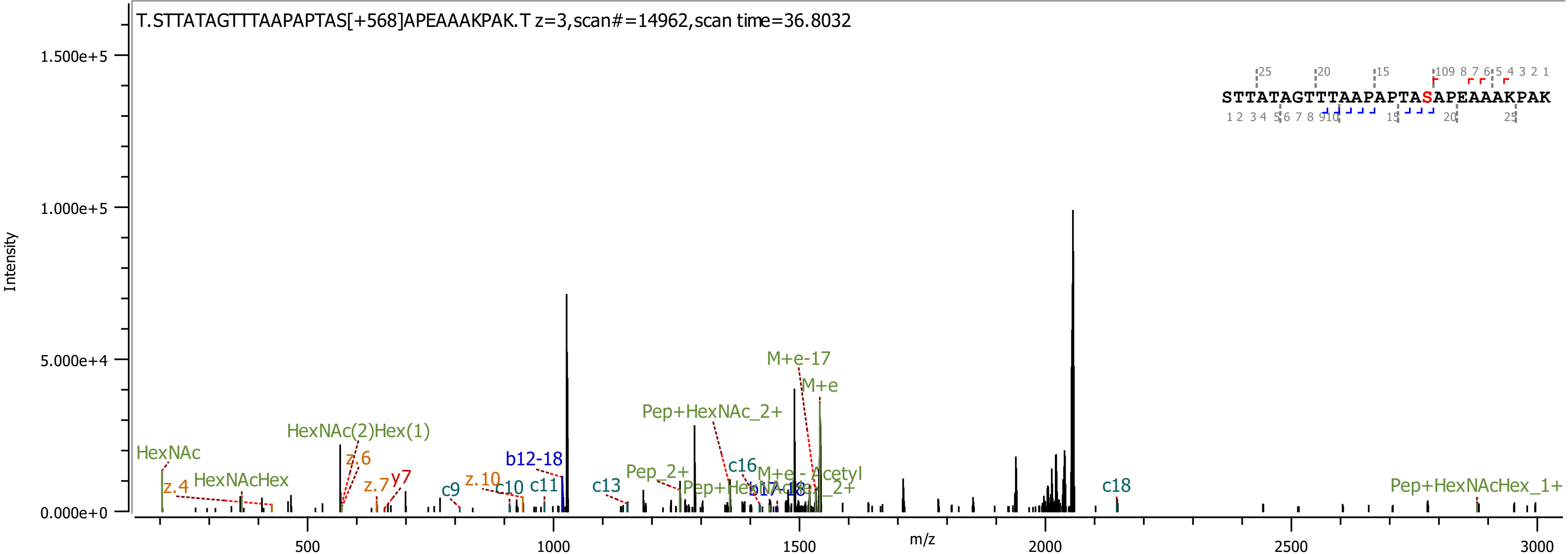
T.STTATAGTTTAAPAPTAS[+568]APEAAAKPAK.T z=3,scan#=14962,scan time=36.8032
STTATAGTTTAAPAPTASAPEAAAKPAK
Intensity
m/z
1.500e+5
1.000e+5
5.000e+4
0.000e+0
500
1000
1500
2000
2500
3000
HexNAc
z.4
HexNAcHex
HexNAc(2)Hex(1)
z.6
z.7
y7
c9
z.10
b12-18
c10
c11
c13
Pep_2+
Pep+HexNAc_2+
c16
M+e-17
M+e
M+e - Acetyl
c18
Pep+HexNAcHex_1+

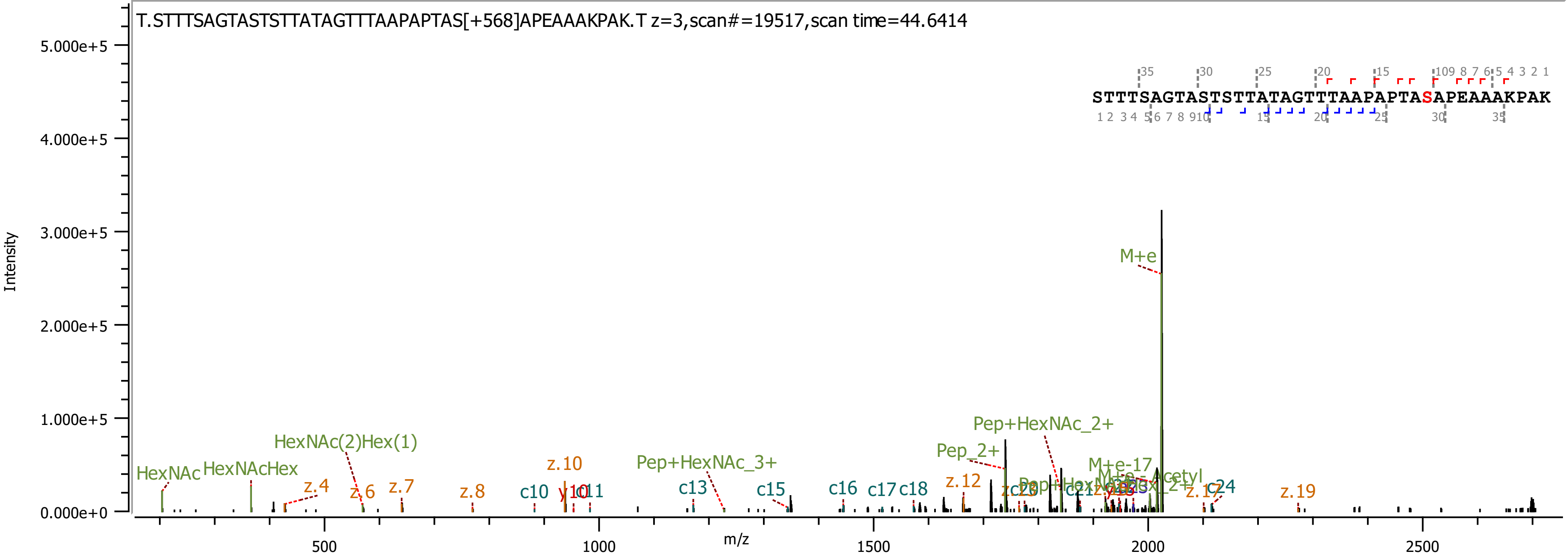

T.STTTSAGTASTSTTATAGTTTAAPAPTAS[+568]APEAAAKPAK.T z=3,scan#=19517,scan time=44.6414
STTTSAGTASTSTTATAGTTTAAPAPTASAPEAAAKPAK
Intensity
m/z
M+e
Pep+HexNAc_2+
Pep_2+
HexNAc(2)Hex(1)
HexNAc
HexNAcHex
Pep+HexNAc_3+
M+e-17
z.4
z.6
z.7
z.8
z.10
z.12
z.19
c10
y10
c11
c13
c15
c16
c17
c18
c24

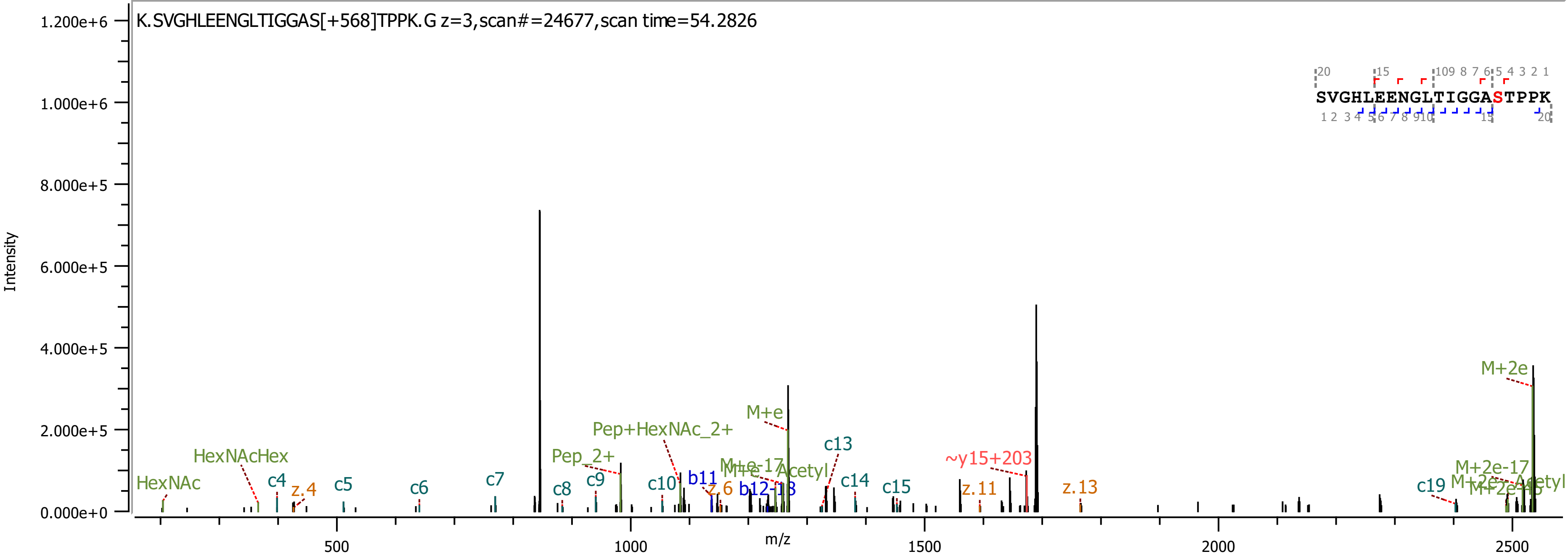
K.SVGHLEENGLTIGGAS[+568]TPPK.G z=3,scan#=24677,scan time=54.2826
SVGHLEENGLTIGGASTPPK
Intensity
m/z
1.200e+6
1.000e+6
8.000e+5
6.000e+5
4.000e+5
2.000e+5
0.000e+0
500
1000
1500
2000
2500
HexNAc
HexNAcHex
c4
z.4
c5
c6
c7
c8
c9
Pep_2+
Pep+HexNAc_2+
c10
b11
z.6
b12-18
M+e
M+e-17
M+e-Acetyl
c13
c14
c15
~y15+203
z.11
z.13
c19
M+2e
M+2e-17
M+2e-Acetyl

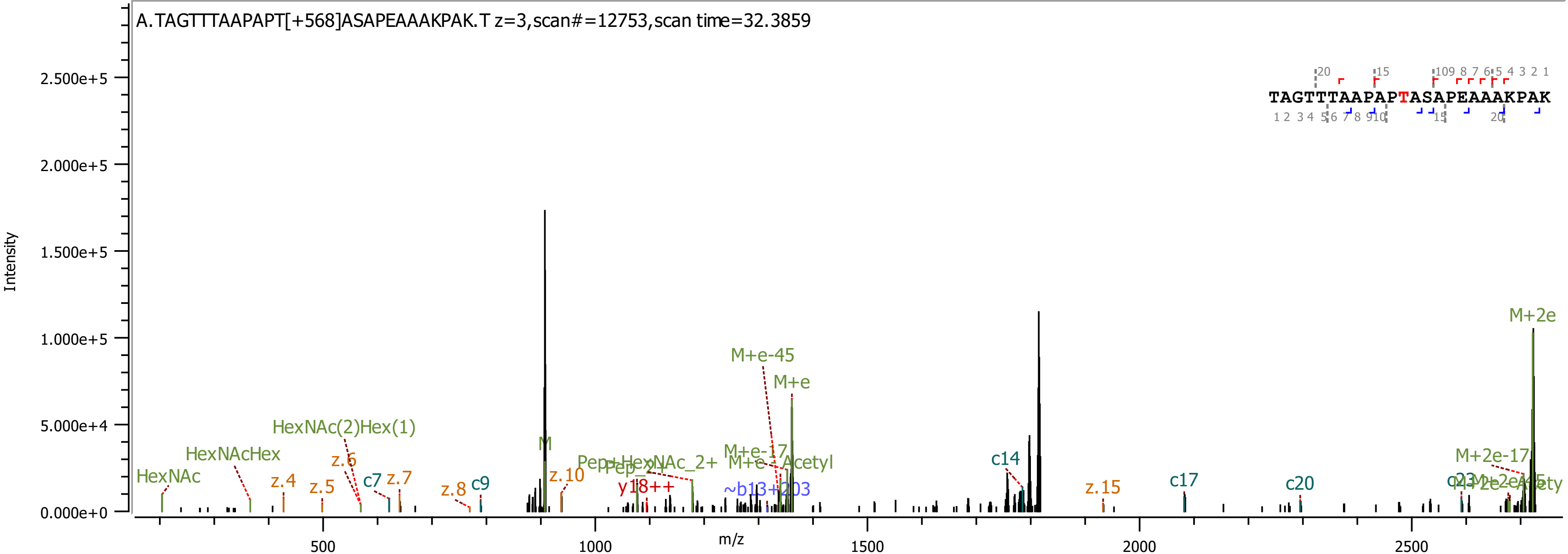

A.TAGTTTAAPAPT[+568]ASAPEAAAKPAK.T z=3,scan#=12753,scan time=32.3859
TAGTTTAAPAPTASAPEAAAKPAK
Intensity
m/z
HexNAc
HexNAcHex
HexNAc(2)Hex(1)
z.4
z.5
z.6
c7
z.7
z.8
c9
M
z.10
y18++
M+e-45
M+e
M+e-17
M+e-Acetyl
c14
z.15
c17
c20
M+2e-17
M+2e

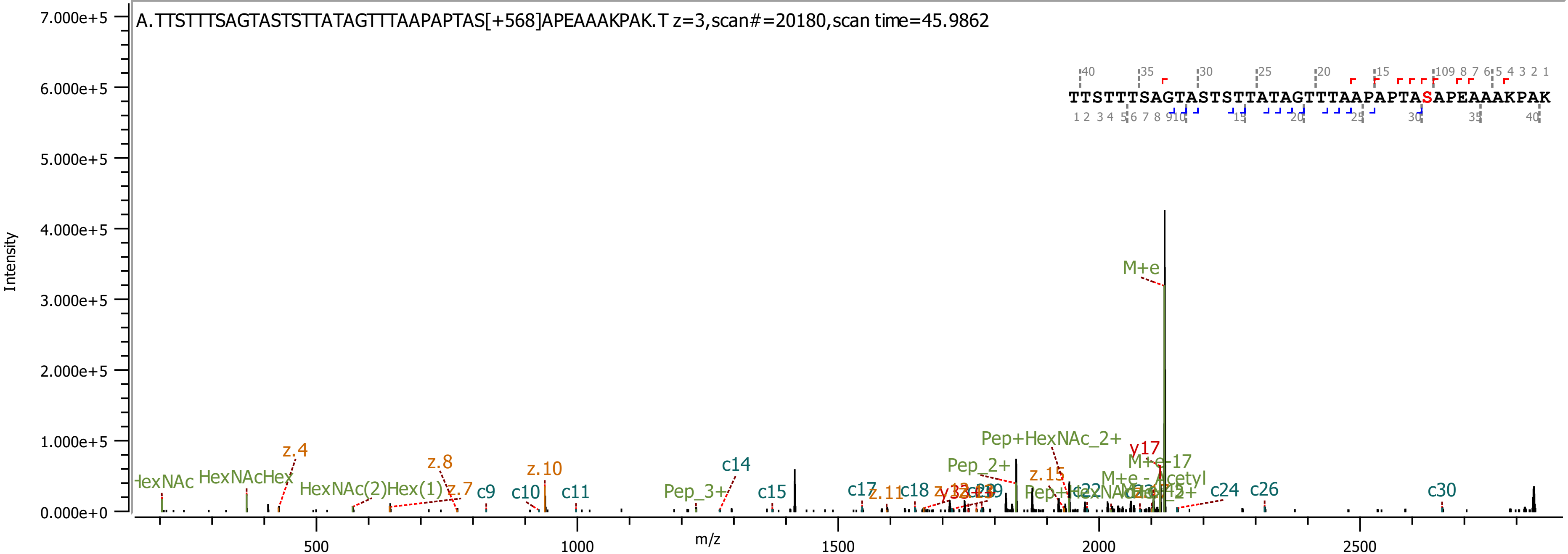

A.TTSTTTSAGTASTSTTATAGTTTAAPAPTAS[+568]APEAAAKPAK.T z=3,scan#=20180,scan time=45.9862
TTSTTTSAGTASTSTTATAGTTTAAPAPTASAPEAAAKPAK
Intensity
m/z
M+e
Pep+HexNAc_2+
Pep_2+
M+e-17
M+e - Acetyl
y17
z.15
Pep_3+
c14
c15
c17
c18
c24
c26
c30
z.4
z.8
z.7
z.10
z.11
c9
c10
c11
HexNAcHex
HexNAc(2)Hex(1)

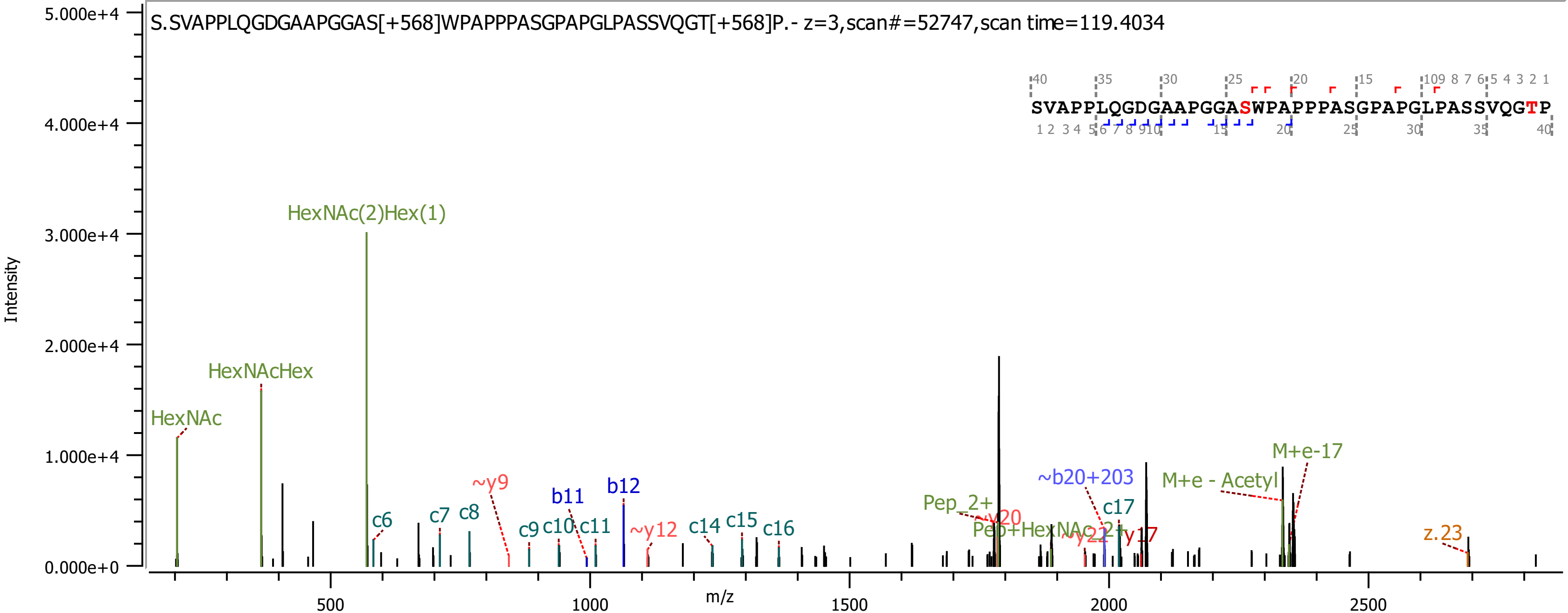

S.SVAPPLQGDGAAPGGAS[+568]WPAPPPASGPAPGLPASSVQGT[+568]P.- z=3,scan#=52747,scan time=119.4034
SVAPPLQGDGAAPGGASWPAPPPASGPAPGLPASSVQGTP
Intensity
m/z
HexNAc(2)Hex(1)
HexNAcHex
HexNAc
~y9
b11
b12
c6
c7
c8
c9
c10
c11
~y12
c14
c15
c16
Pep_2+
~y20
Pep+HexNAc_2+
~y22
y17
~b20+203
c17
M+e - Acetyl
M+e-17
z.23

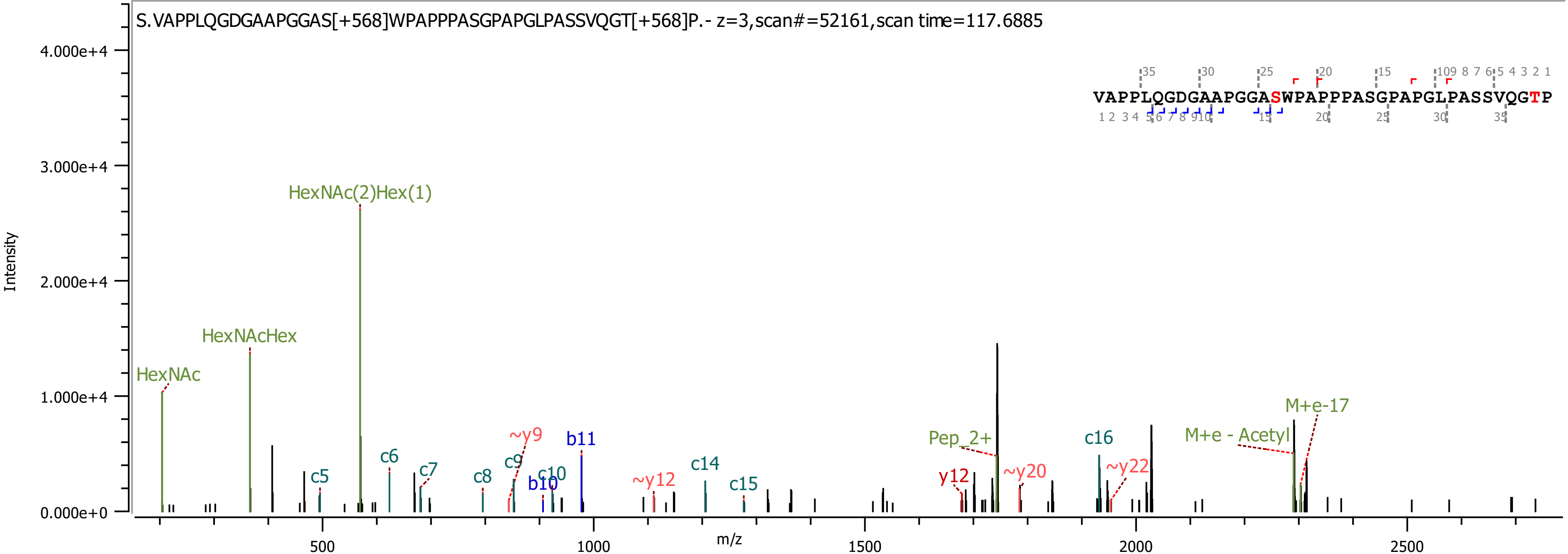

S.VAPPLQGDGAAPGGAS[+568]WPAPPPASGPAPGLPASSVQGT[+568]P.- z=3,scan#=52161,scan time=117.6885
VAPPLQGDGAAPGGASWPAPPPASGPAPGLPASSVQGTP
Intensity
m/z
HexNAc
HexNAcHex
HexNAc(2)Hex(1)
c5
c6
c7
c8
c9
~y9
b10
c10
b11
~y12
c14
c15
y12
Pep_2+
~y20
c16
~y22
M+e - Acetyl
M+e-17

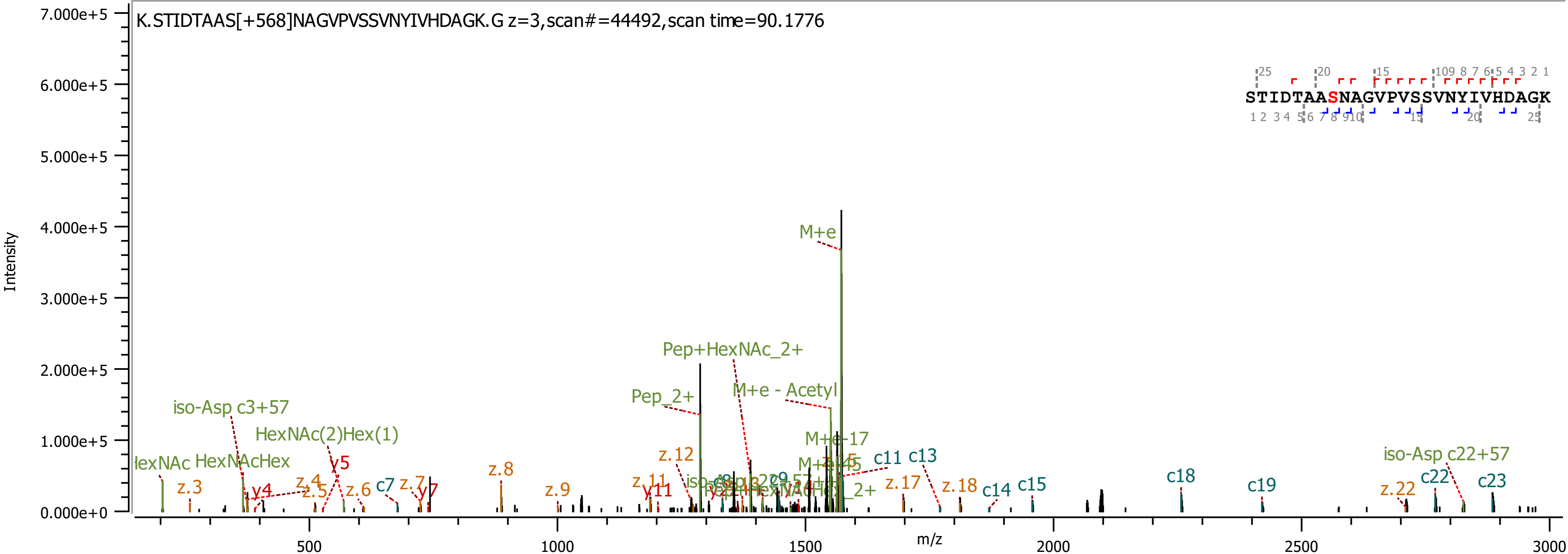

K.STIDTAAS[+568]NAGVPVSSVNYIVHDAGK.G z=3,scan#=44492,scan time=90.1776
Intensity
m/z
STIDTAASNAGVPVSSVNYIVHDAGK
M+e
Pep+HexNAc_2+
M+e - Acetyl
Pep_2+
iso-Asp c3+57
HexNAc(2)Hex(1)
HexNAcHex
y5
y4
y7
y11
z.3
z.4
z.5
z.6
z.7
z.8
z.9
z.11
z.12
z.17
z.18
z.22
c7
c11
c13
c14
c15
c18
c19
c22
c23
iso-Asp c22+57
M+e-17

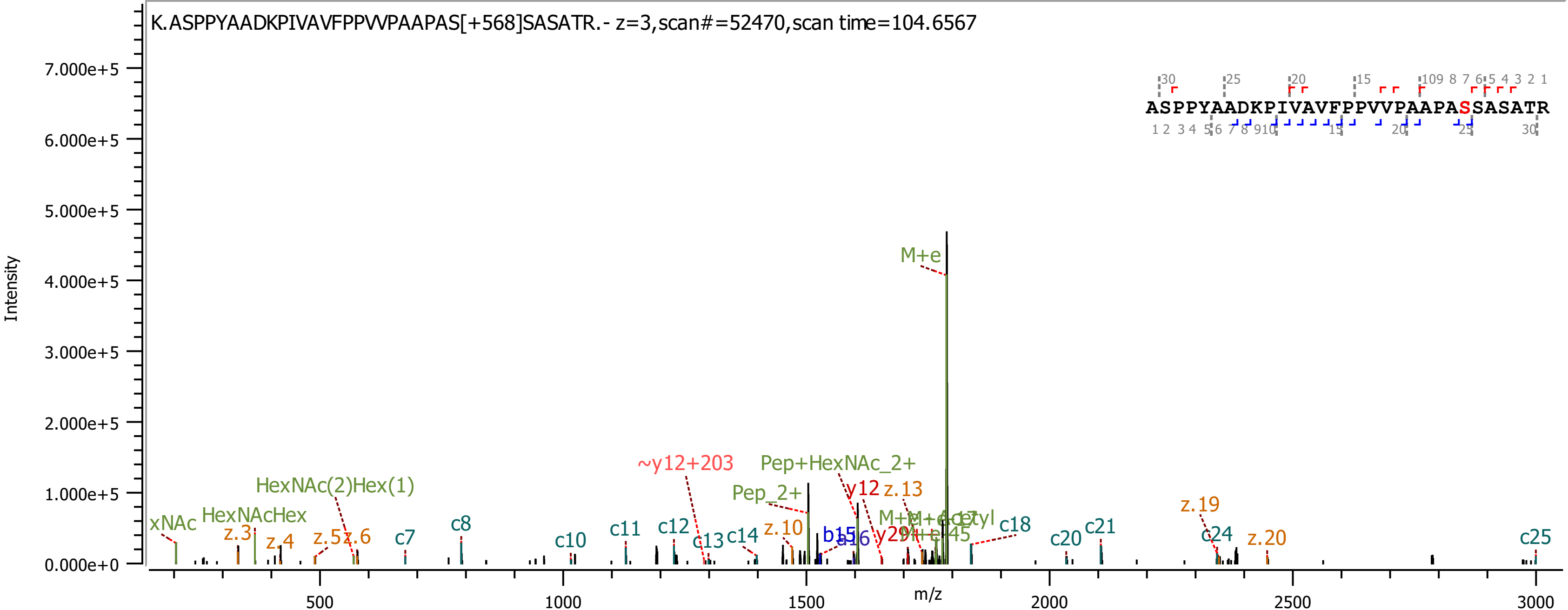

K.ASPPYAADKPIVAVFPPVVPAAPAS[+568]SASATR.- z=3,scan#=52470,scan time=104.6567
ASPPYAADKPIVAVFPPVVPAAPASSASATR
Intensity
m/z
M+e
Pep+HexNAc_2+
Pep_2+
~y12+203
y12
z.13
HexNAc(2)Hex(1)
HexNAcHex
xNAc
z.3
z.4
z.5
z.6
c7
c8
c10
c11
c12
c13
c14
z.10
b15
a16
c18
c20
c21
z.19
c24
z.20
c25

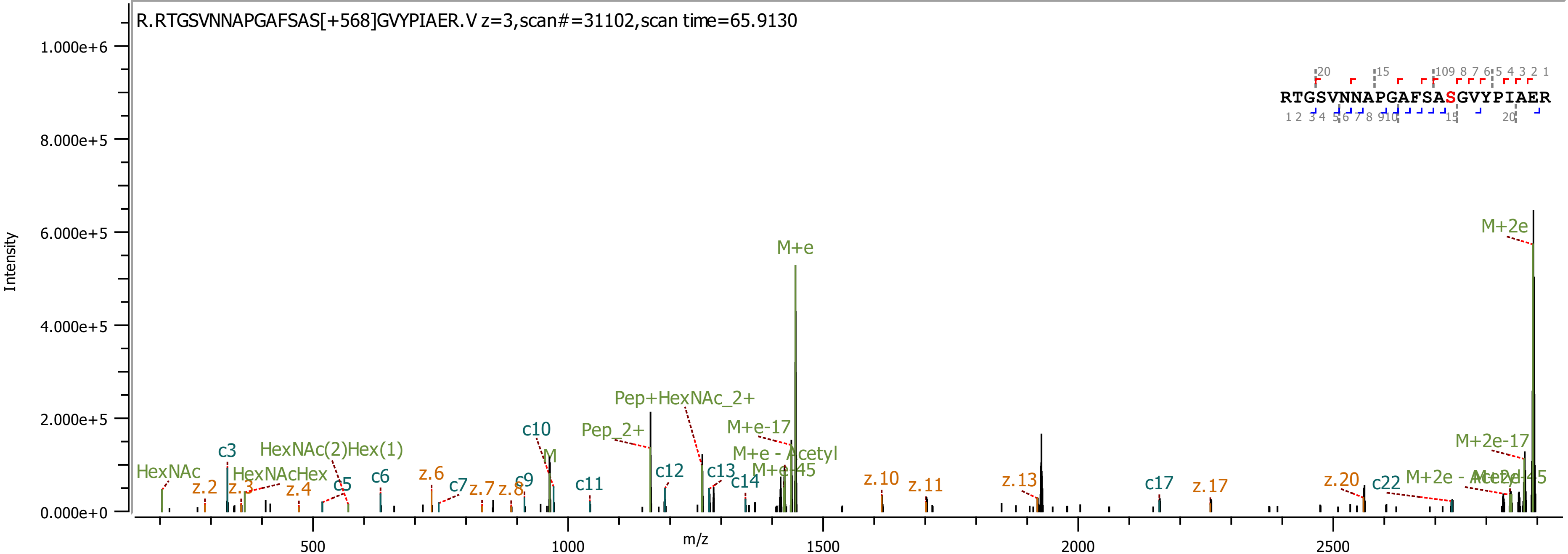

R.RTGSVNNAPGAFSAS[+568]GVYPIAER.V z=3,scan#=31102,scan time=65.9130
RTGSVNNAPGAFSASGVYPIAER
Intensity
m/z
M+2e
M+e
Pep+HexNAc_2+
Pep_2+
M+e-17
M+e - Acetyl
M+e-45
M+2e-17
M+2e - Acetyl
M+2e-45
HexNAc
HexNAcHex
HexNAc(2)Hex(1)
M
c3
c5
c6
c7
c9
c10
c11
c12
c13
c14
c17
c22
z.2
z.3
z.4
z.6
z.7
z.8
z.10
z.11
z.13
z.17
z.20

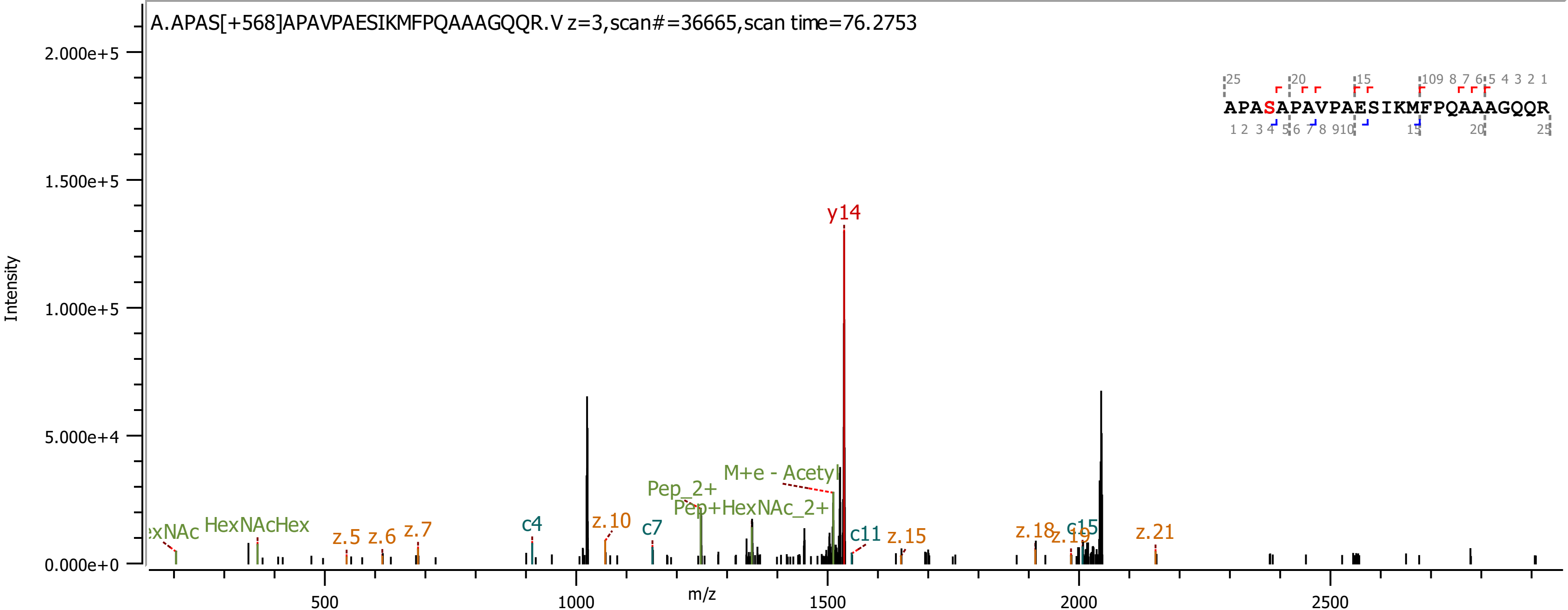
A.APAS[+568]APAVPAESIKMFPQAAAGQQR.V z=3,scan#=36665,scan time=76.2753
Intensity
m/z
2.000e+5
1.500e+5
1.000e+5
5.000e+4
0.000e+0
500
1000
1500
2000
2500
y14
M+e - Acetyl
Pep_2+
Pep+HexNAc_2+
HexNAcHex
z.5
z.6
z.7
c4
z.10
c7
c11
z.15
z.18
z.19
c15
z.21
APASAPAVPAESIKMFPQAAAGQQR

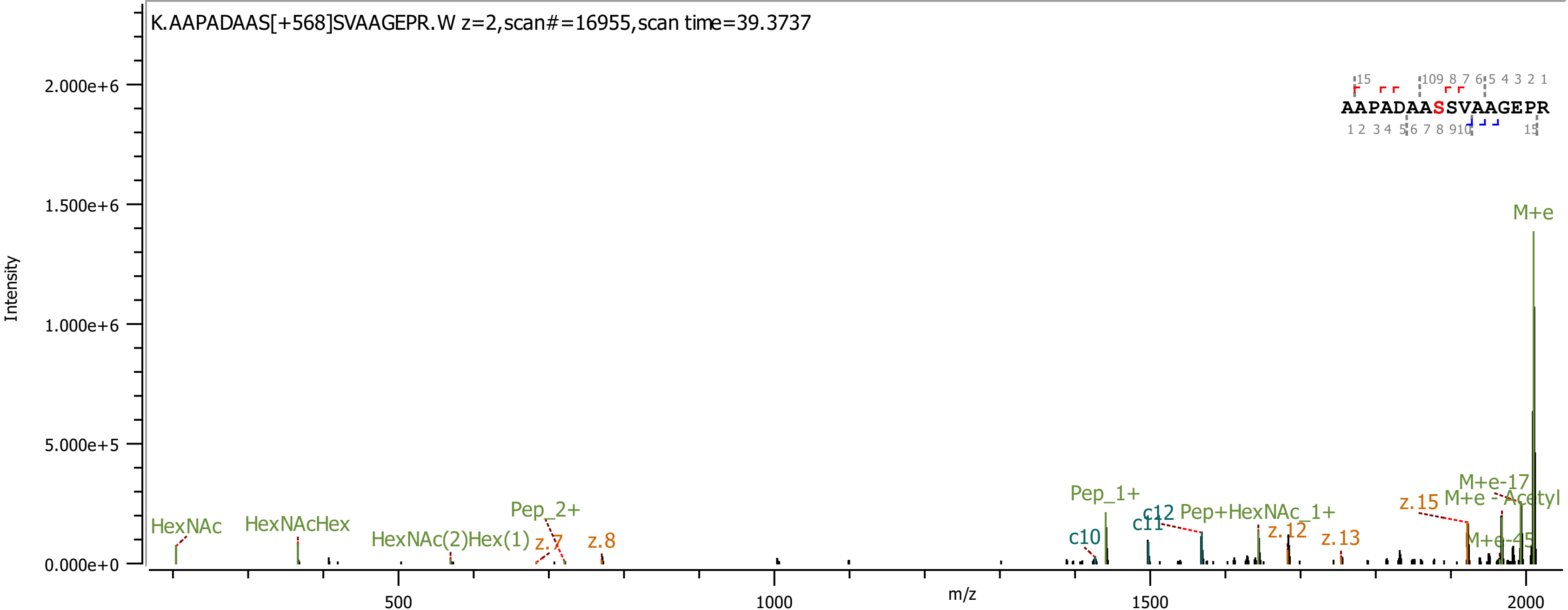
K.AAPADAAS[+568]SVAAGEPR.W z=2,scan#=16955,scan time=39.3737
AAPADAASSVAAGEPR
Intensity
m/z
M+e
M+e-17
M+e - Acetyl
M+e-45
z.15
Pep_1+
c12
c11
c10
Pep+HexNAc_1+
z.12
z.13
HexNAc
HexNAcHex
HexNAc(2)Hex(1)
Pep_2+
z.7
z.8

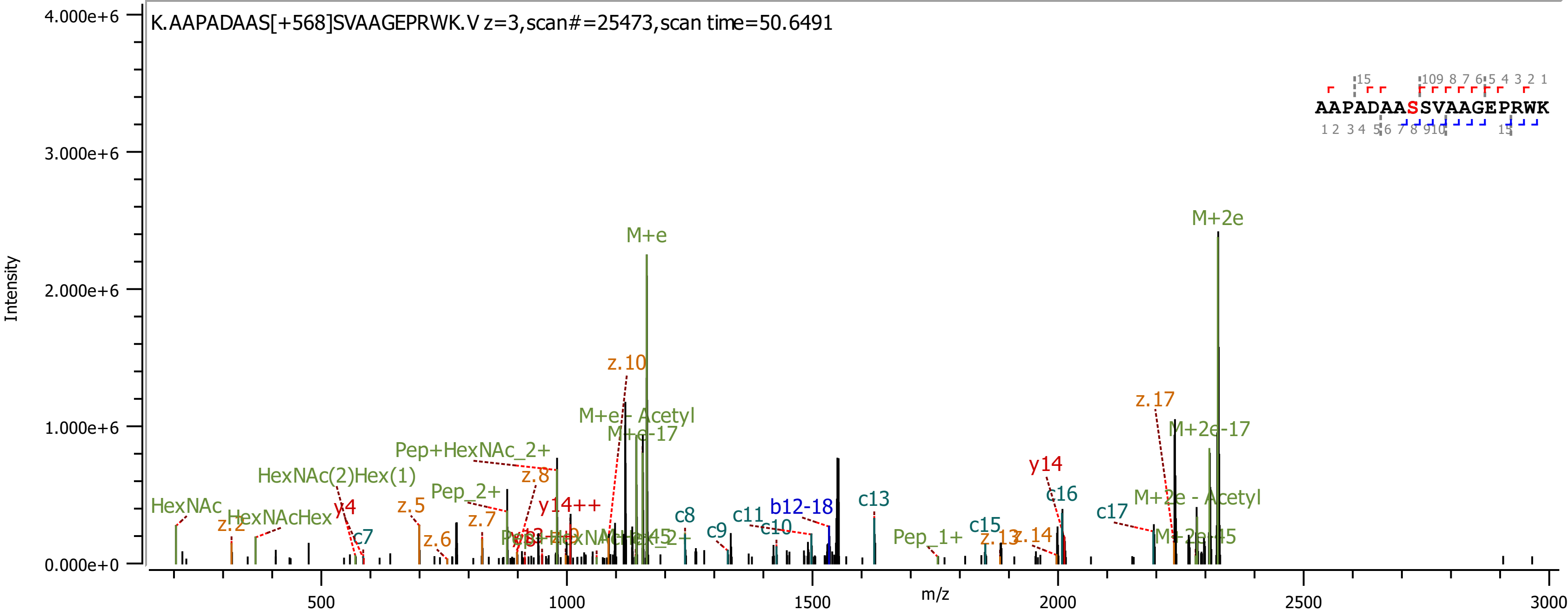
K.AAPADAAS[+568]SVAAGEPRWK.V z=3,scan#=25473,scan time=50.6491
AAPADAASSVAAGEPRWK
Intensity
m/z
4.000e+6
3.000e+6
2.000e+6
1.000e+6
0.000e+0
500
1000
1500
2000
2500
3000
M+e
M+2e
z.10
z.17
M+e - Acetyl
M+e-17
M+2e-17
Pep+HexNAc_2+
HexNAc(2)Hex(1)
Pep_2+
z.8
y14++
y14
c16
c17
M+2e - Acetyl
HexNAc
HexNAcHex
z.2
y4
z.5
z.7
z.6
c7
c8
c9
c11
c10
b12-18
c13
Pep_1+
c15
z.13
z.14
M+2e-45

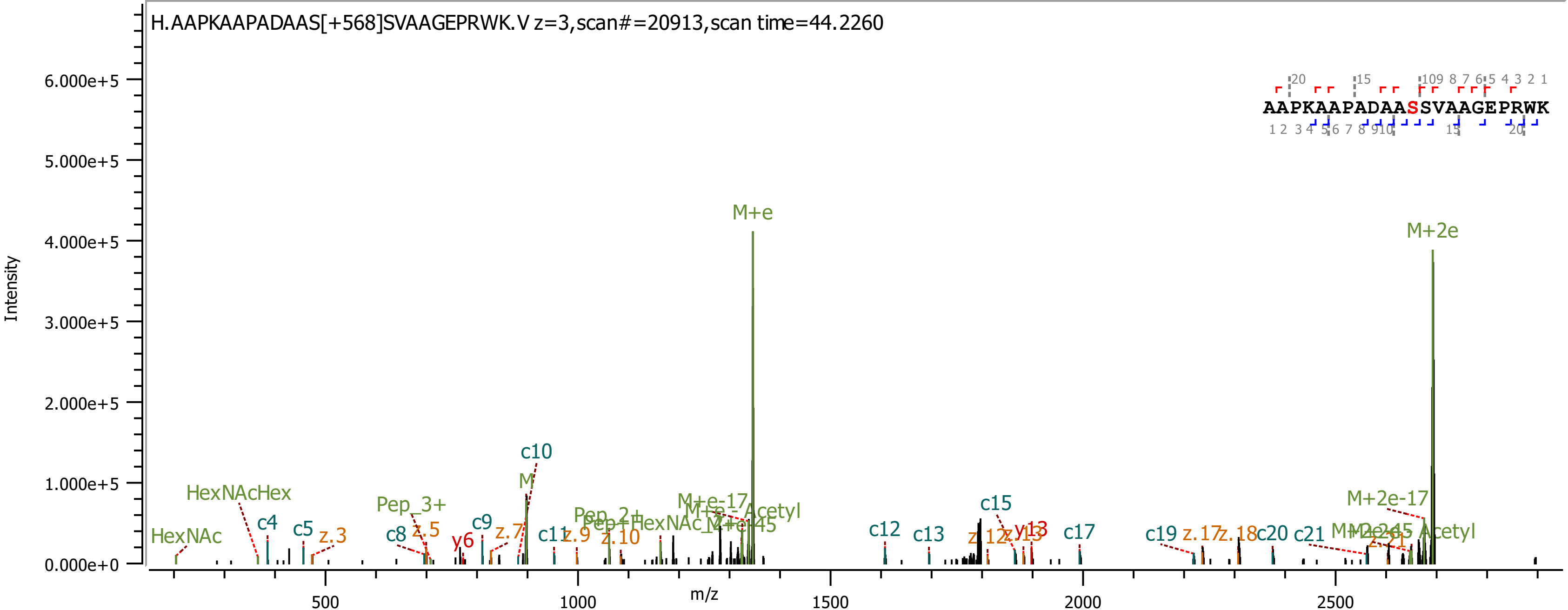
H.AAPKAAPADAAS[+568]SVAAGEPRWK.V z=3,scan#=20913,scan time=44.2260
AAPKAAPADAASSVAAGEPRWK
Intensity
m/z
M+e
M+2e
c10
M
HexNAcHex
HexNAc
c4
c5
z.3
Pep_3+
c8
z.5
y6
c9
z.7
c11
z.9
z.10
c12
c13
c15
y13
c17
c19
c20
c21
M+2e-17

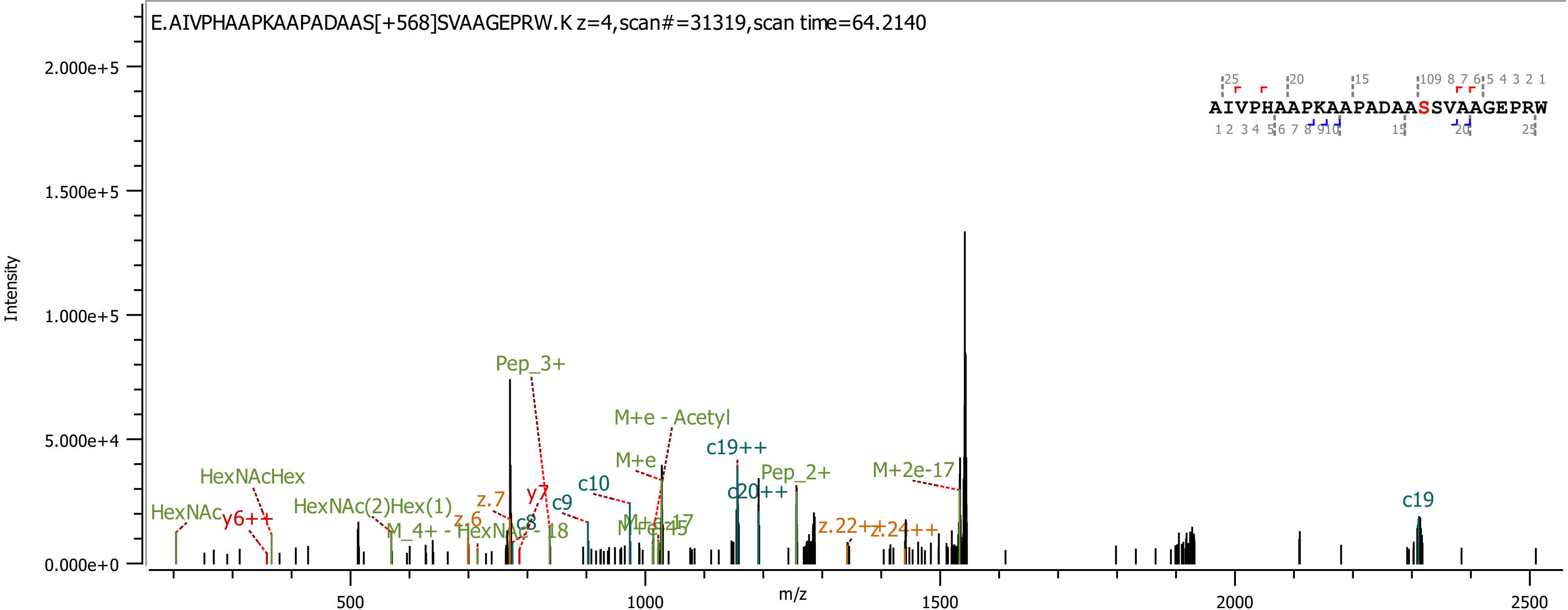

E.AIVPHAAPKAAPADAAS[+568]SVAAGEPRW.K z=4,scan#=31319,scan time=64.2140
AIVPHAAPKAAPADAASSVAAGEPRW
Intensity
m/z
Pep_3+
M+e - Acetyl
M+e
c19++
Pep_2+
M+2e-17
c20++
c10
c9
c8
y7
z.7
z.6
HexNAcHex
HexNAc
y6++
HexNAc(2)Hex(1)
M_4+ - HexNAc - 18
z.22++
z.24++
c19

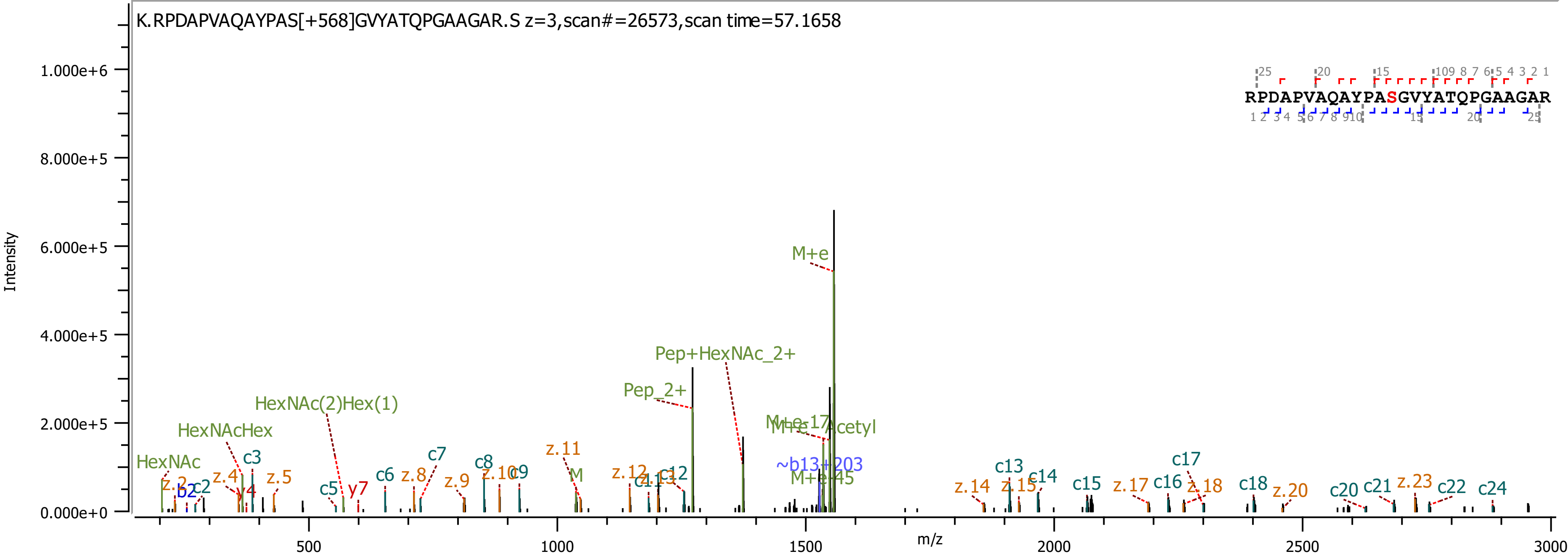
K.RPDAPVAQAYPAS[+568]GVYATQPGAAGAR.S z=3,scan#=26573,scan time=57.1658
RPDAPVAQAYPASGVYATQPGAAGAR
Intensity
m/z
M+e
Pep+HexNAc_2+
Pep_2+
HexNAc(2)Hex(1)
HexNAcHex
HexNAc
M+e-17
M+e-Acetyl
~b13+203
M+e-45
c2
c3
c5
c6
c7
c8
c9
c12
c13
c14
c15
c16
c17
c18
c20
c21
c22
c24
z.2
z.4
z.5
z.8
z.9
z.10
z.11
z.12
z.14
z.15
z.17
z.18
z.20
z.23
b2
y4
y7
M

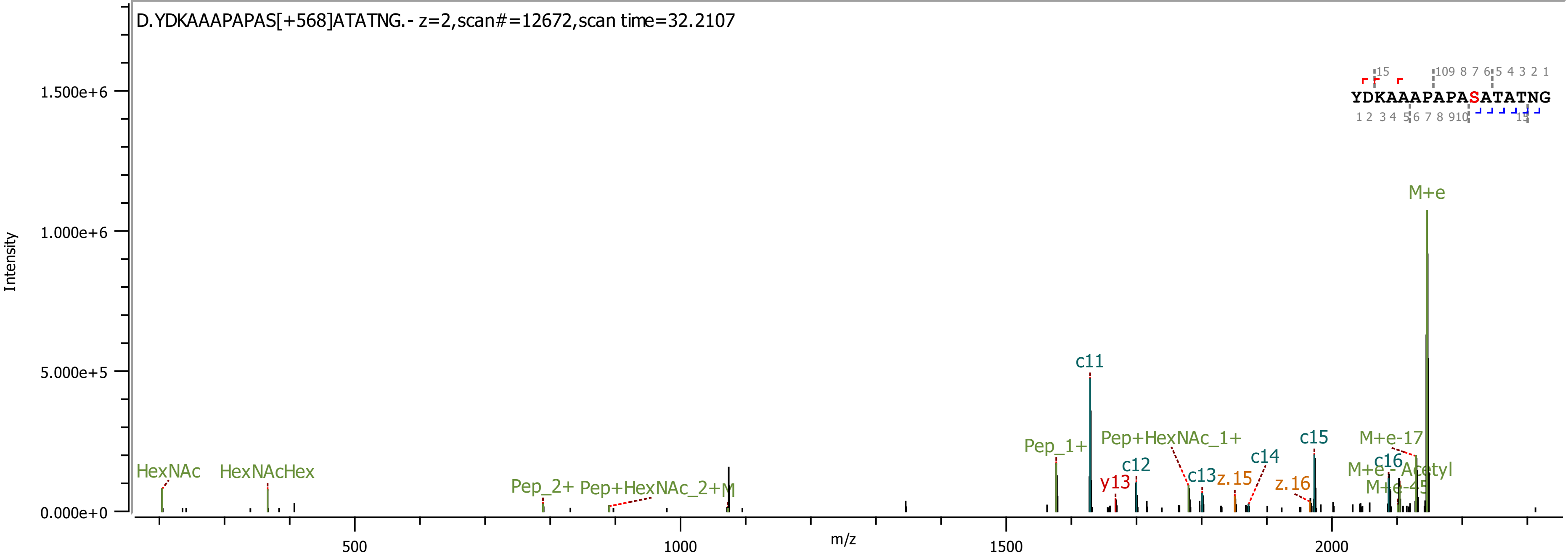

D.YDKAAAPAPAS[+568]ATATNG.- z=2,scan#=12672,scan time=32.2107
Intensity
m/z
YDKAAAPAPASATATNG
M+e
c11
Pep_1+
Pep+HexNAc_1+
c12
c13
c14
c15
c16
y13
z.15
z.16
M+e-17
M+e-Acetyl
M+e-45
HexNAc
HexNAcHex
Pep_2+
Pep+HexNAc_2+M

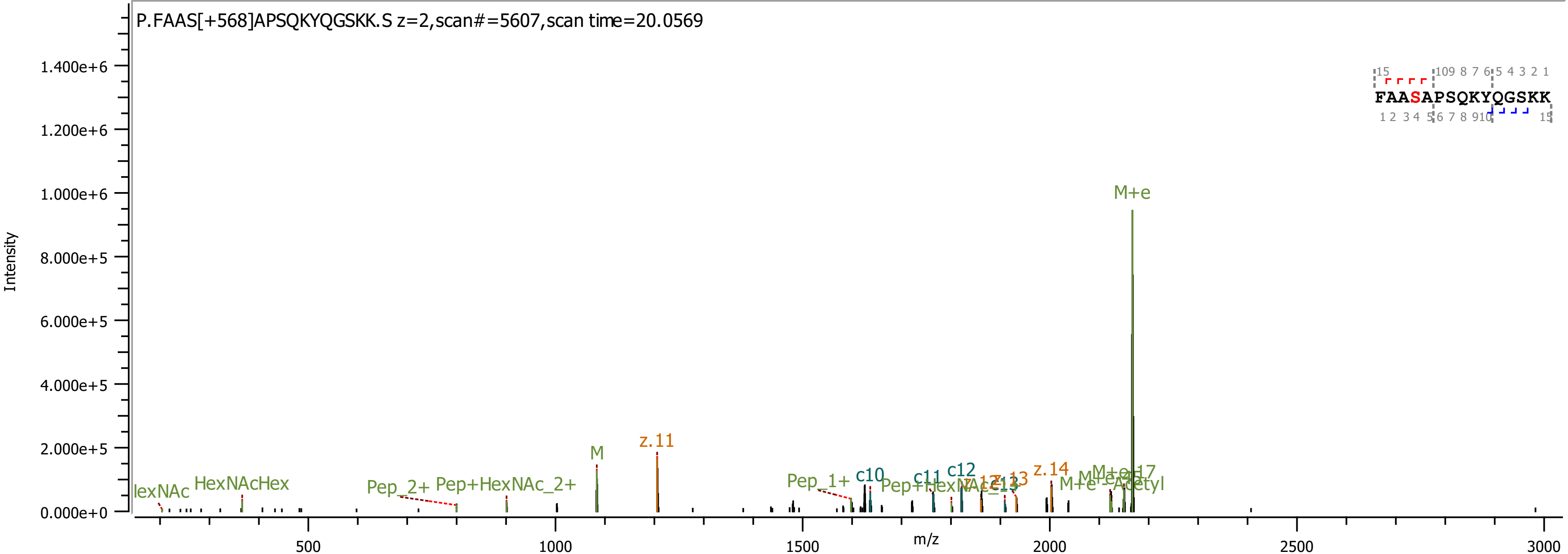

P.FAAS[+568]APSQKYQGSKK.S z=2,scan#=5607,scan time=20.0569
Intensity
m/z
1.400e+6
1.200e+6
1.000e+6
8.000e+5
6.000e+5
4.000e+5
2.000e+5
0.000e+0
500
1000
1500
2000
2500
3000
FAASAPSQKYQGSKK
M+e
z.11
M
HexNAcHex
Pep_2+
Pep+HexNAc_2+
Pep_1+
c10
c11
c12
z.14

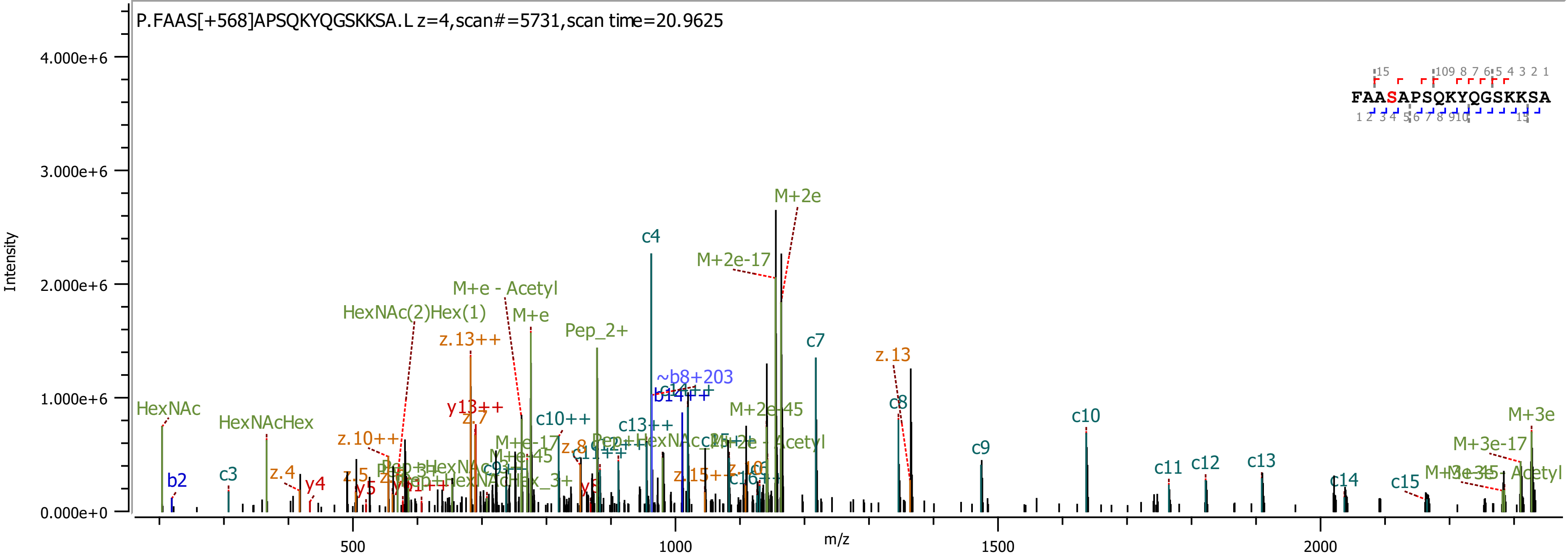
P.FAAS[+568]APSQKYQGSKKSA.L z=4,scan#=5731,scan time=20.9625
Intensity
m/z
4.000e+6
3.000e+6
2.000e+6
1.000e+6
0.000e+0
500
1000
1500
2000
FAASAPSQKYQGSKKSA
HexNAc
b2
c3
HexNAcHex
z.4
y4
z.10++
HexNAc(2)Hex(1)
z.13++
y13++
M+e - Acetyl
M+e
Pep_2+
c10++
c4
~b8+203
b14++
c13++
M+2e-17
M+2e
M+2e-45
c7
z.13
c8
c9
c10
c11
c12
c13
c14
c15
M+3e
M+3e-17

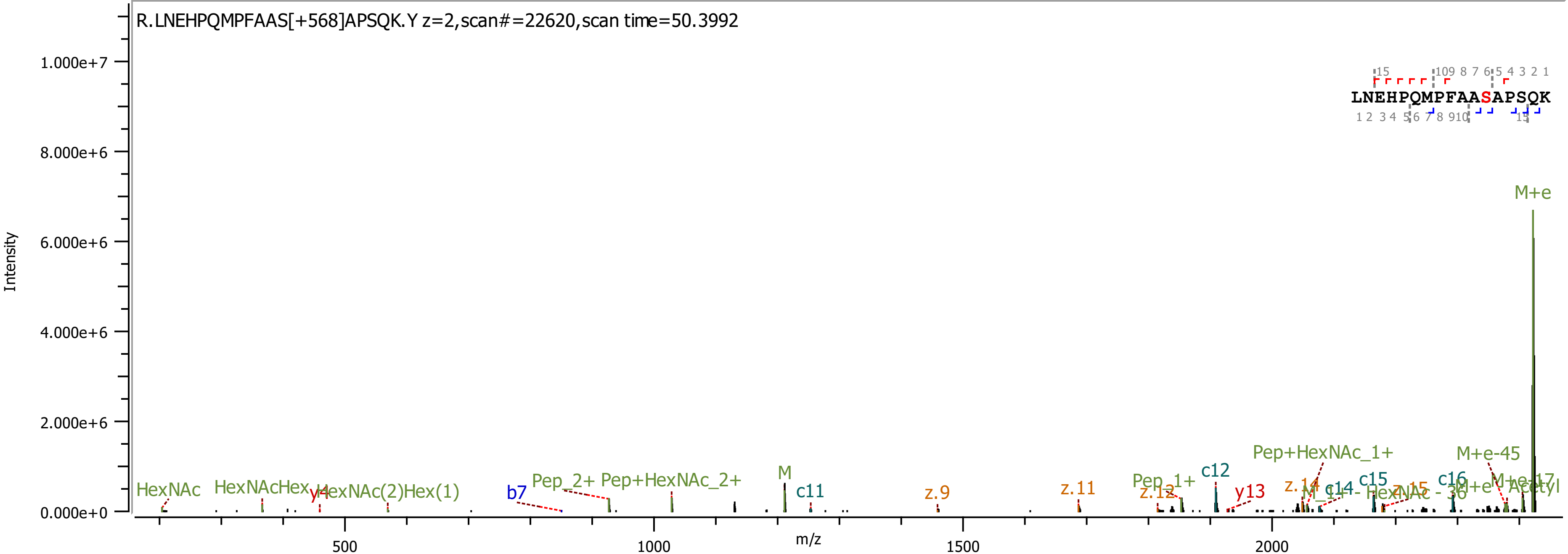

R.LNEHPQMPFAAS[+568]APSQK.Y z=2,scan#=22620,scan time=50.3992
LNEHPQMPFAASAPSQK
Intensity
m/z
1.000e+7
8.000e+6
6.000e+6
4.000e+6
2.000e+6
0.000e+0
500
1000
1500
2000
HexNAc
HexNAcHex
y4
HexNAc(2)Hex(1)
b7
Pep_2+
Pep+HexNAc_2+
M
c11
z.9
z.11
Pep_1+
z.12
c12
y13
Pep+HexNAc_1+
z.14
c14
c15
c16
M+e-45
M+e

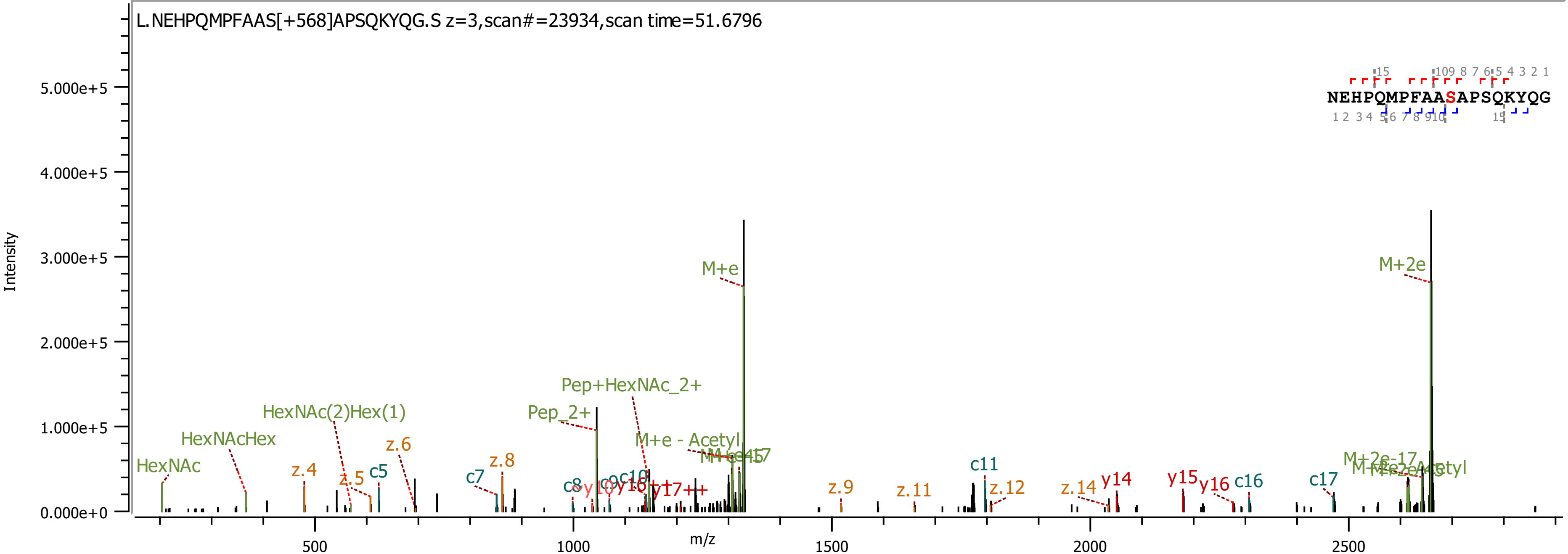

L.NEHPQMPFAAS[+568]APSQKYQG.S z=3,scan#=23934,scan time=51.6796
Intensity
m/z
NEHPQMPFAASAPSQKYQG
M+e
M+2e
Pep+HexNAc_2+
Pep_2+
HexNAc(2)Hex(1)
HexNAcHex
HexNAc
M+e - Acetyl
M+2e-17
c5
c7
c8
c11
c16
c17
z.4
z.5
z.6
z.8
z.9
z.11
z.12
z.14
y14
y15
y16
y17++

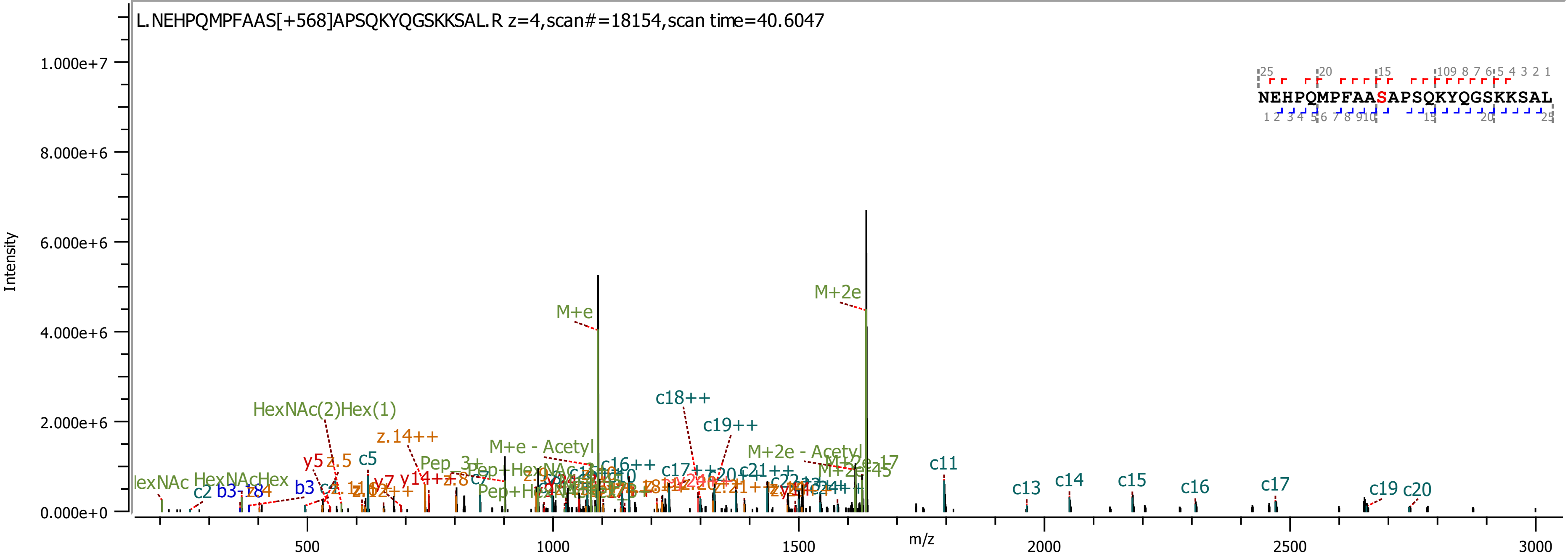

L.NEHPQMPFAAS[+568]APSQKYQGSKKSAL.R z=4,scan#=18154,scan time=40.6047
Intensity
m/z
NEHPQMPFAASAPSQKYQGSKKSAL
M+e
M+2e
c18++
c19++
HexNAc(2)Hex(1)
z.14++
M+e - Acetyl
M+2e - Acetyl
c11
c13
c14
c15
c16
c17
c19
c20
c5
y5
z.5
b3
c2

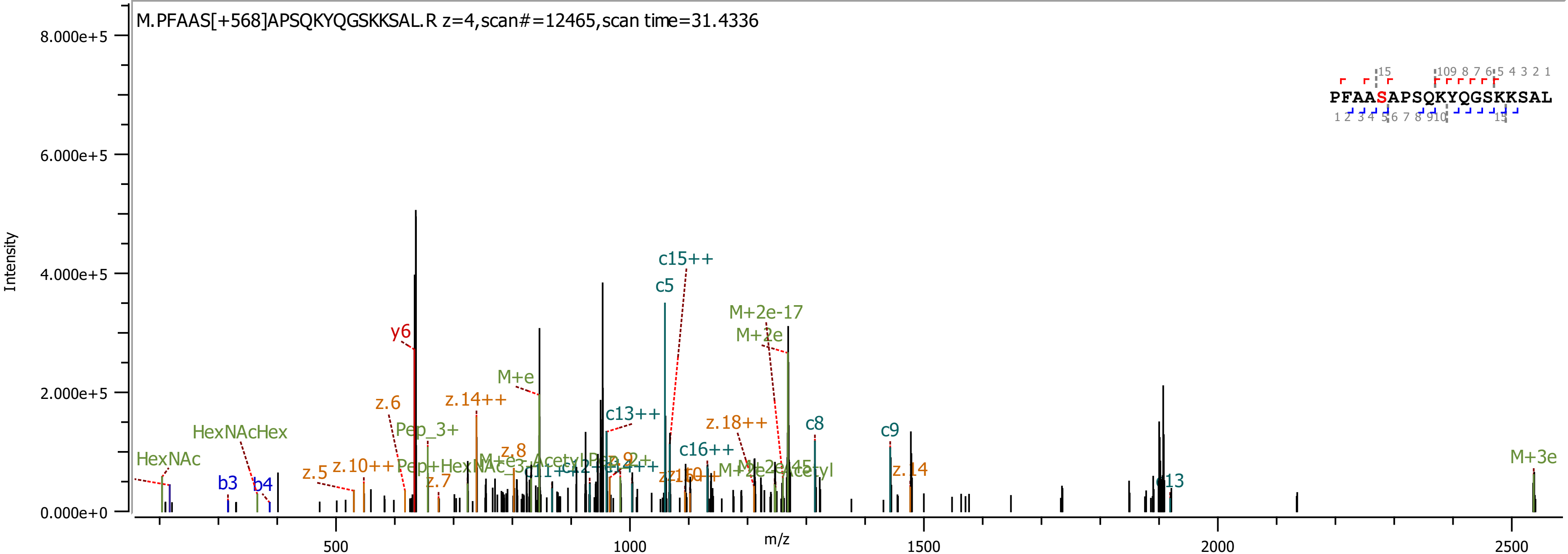

M.PFAAS[+568]APSQKYQGSKKSAL.R z=4,scan#=12465,scan time=31.4336
Intensity
m/z
PFAASAPSQKYQGSKKSAL
y6
c15++
c5
M+2e-17
M+2e
M+e
z.6
z.14++
c13++
z.18++
c8
c9
HexNAcHex
Pep_3+
HexNAc
b3
b4
z.5
z.10++
z.8
z.7
c16++
z.14
c13
M+3e

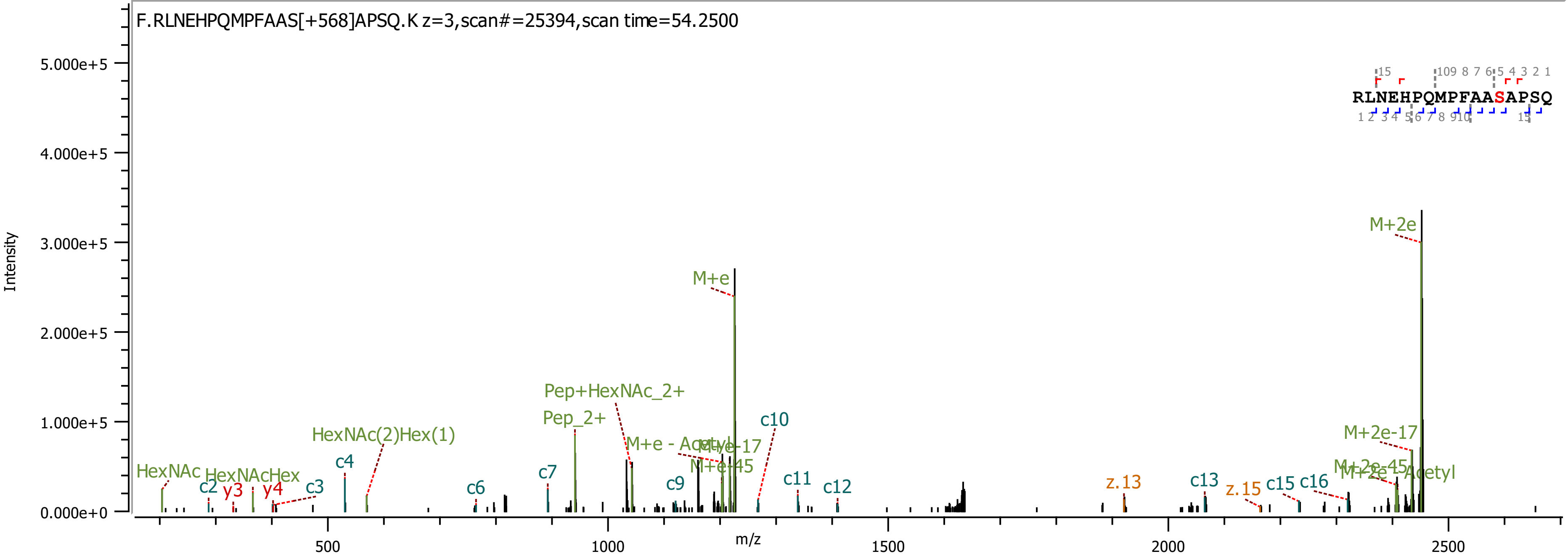

F.RLNEHPQMPFAAS[+568]APSQ.K z=3,scan#=25394,scan time=54.2500
RLNEHPQMPFAASAPSQ
Intensity
m/z
M+2e
M+e
Pep+HexNAc_2+
Pep_2+
HexNAc(2)Hex(1)
HexNAc
HexNAcHex
c2
y3
y4
c3
c4
c6
c7
c9
c10
c11
c12
c13
c15
c16
z.13
z.15
M+e - Acetyl
M+e-17
M+e-45
M+2e-17
M+2e-45

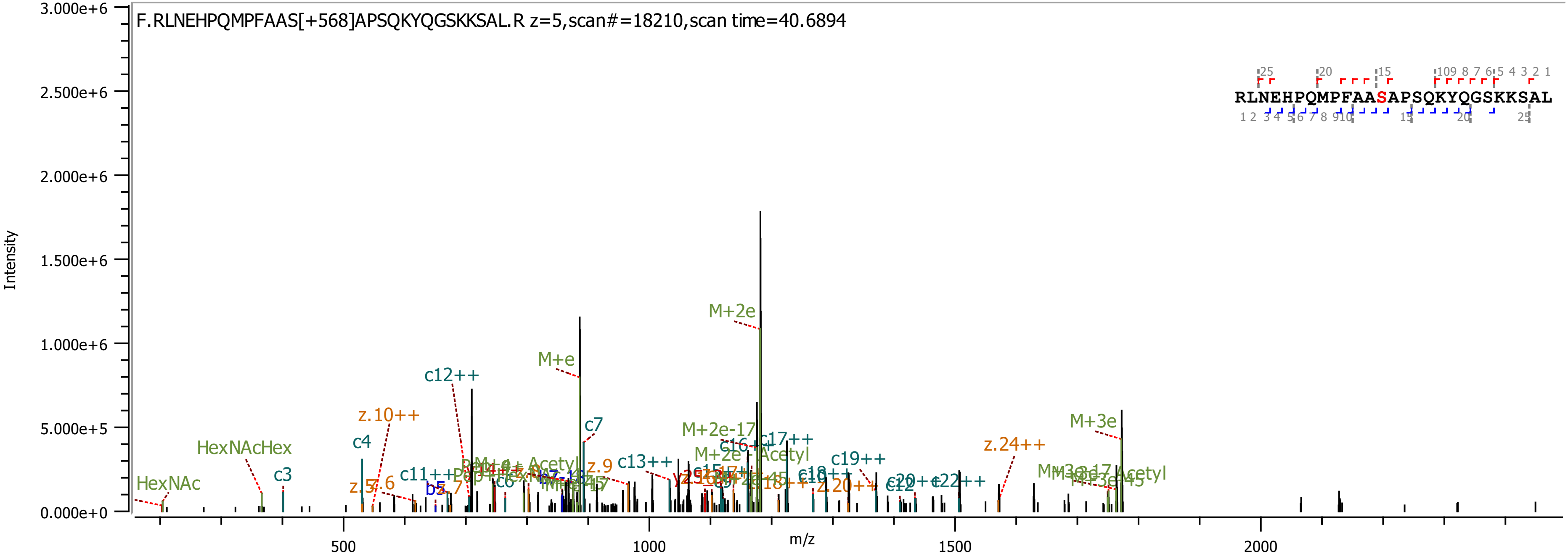

F.RLNEHPQMPFAAS[+568]APSQKYQGSKKSAL.R z=5,scan#=18210,scan time=40.6894
RLNEHPQMPFAASAPSQKYQGSKKSAL
Intensity
m/z
M+2e
M+e
M+3e
c12++
z.10++
HexNAcHex
HexNAc
c4
c3
c7
c13++
c17++
c19++
z.24++
M+2e-17
M+2e-Acetyl
c16
z.9
c11++
c6
z.7
z.6
b5
c18++
z.20++
c20++
c22++
c12

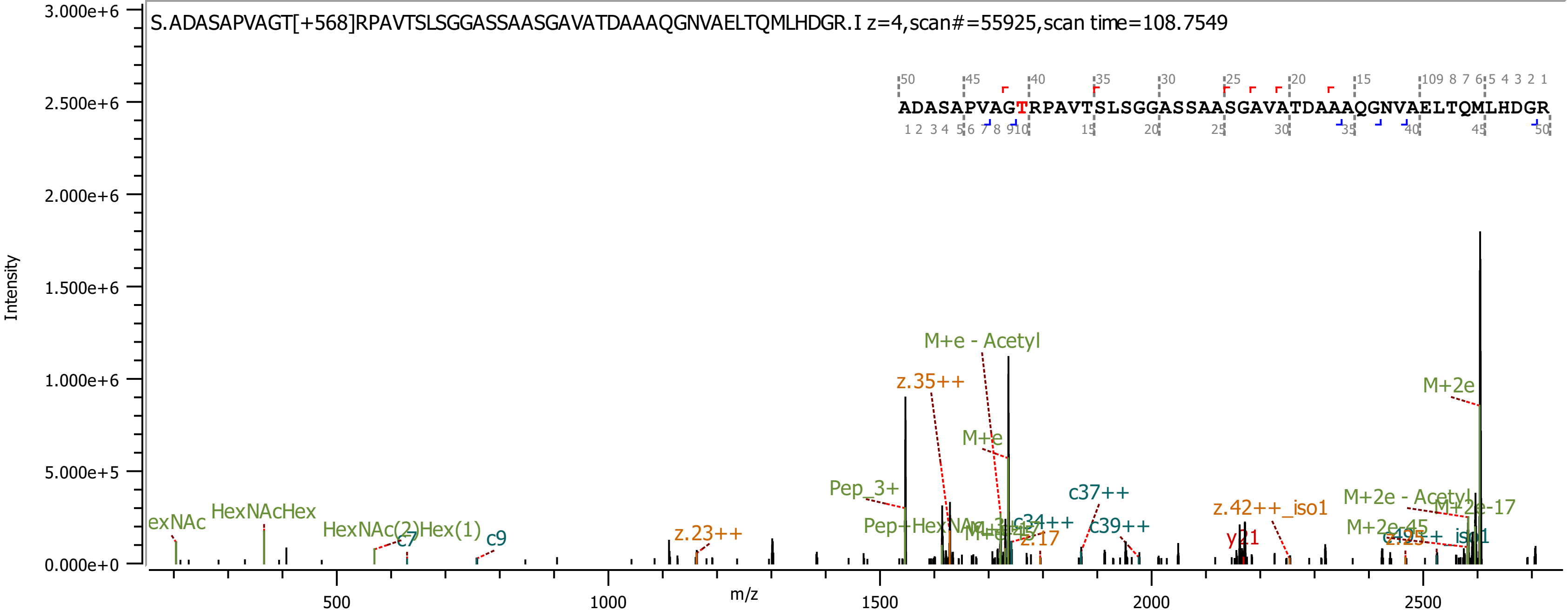
S.ADASAPVAGT[+568]RPAVTSLSGGASSAASGAVATDAAAQGNVAELTQMLHDGR.I z=4,scan#=55925,scan time=108.7549
ADASAPVAGTRPAVTSLSGGASSAASGAVATDAAAQGNVAELTQMLHDGR
Intensity
m/z
M+e - Acetyl
z.35++
M+2e
M+e
Pep_3+
c37++
z.42++_iso1
M+2e - Acetyl
M+2e-17
HexNAcHex
HexNAc(2)Hex(1)
c9
c7
z.23++
c39++
y21
z.17
M+2e-45
iso1
Pep+HexNAc
c34++
0.000e+0
5.000e+5
1.000e+6
1.500e+6
2.000e+6
2.500e+6
3.000e+6
500
1000
1500
2000
2500

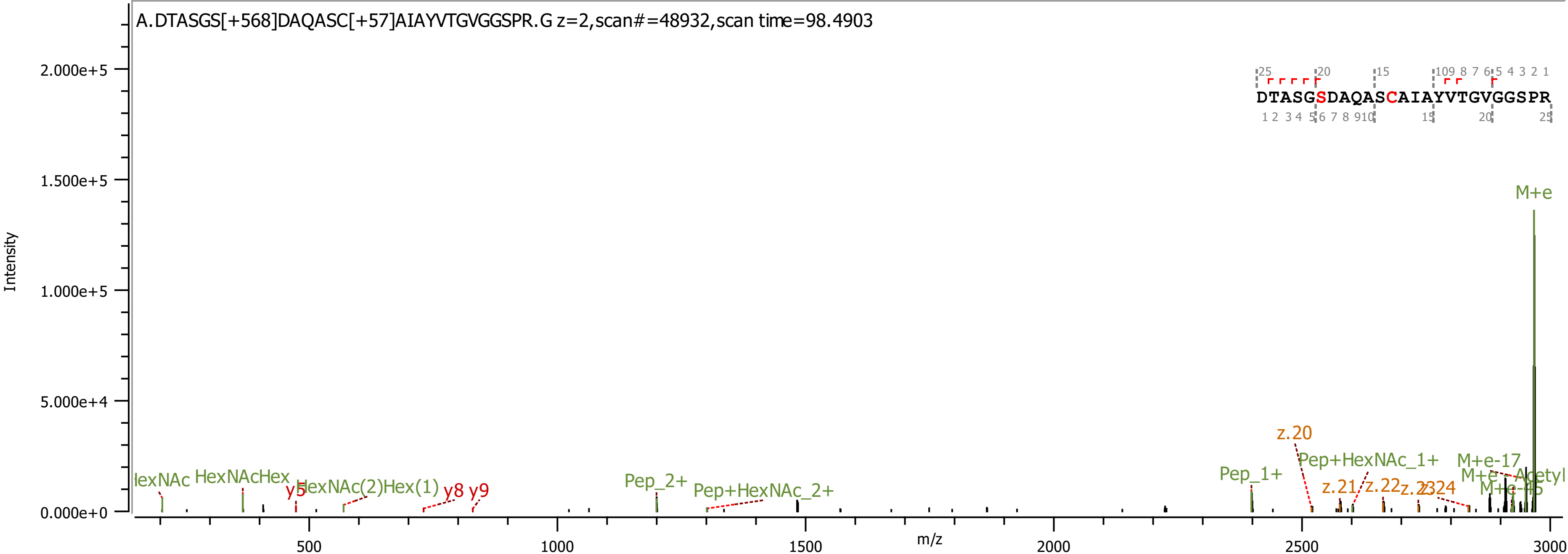

A.DTASGS[+568]DAQASC[+57]AIAYVTGVGGSPR.G z=2,scan#=48932,scan time=98.4903
DTASGSDAQASCAIAYVTGVGGSPR
Intensity
m/z
M+e
z.20
Pep+HexNAc_1+
M+e-17
Pep_1+
z.21
z.22
z.23
z.24
Pep_2+
Pep+HexNAc_2+
HexNAcHex
HexNAc(2)Hex(1)
y5
y8
y9

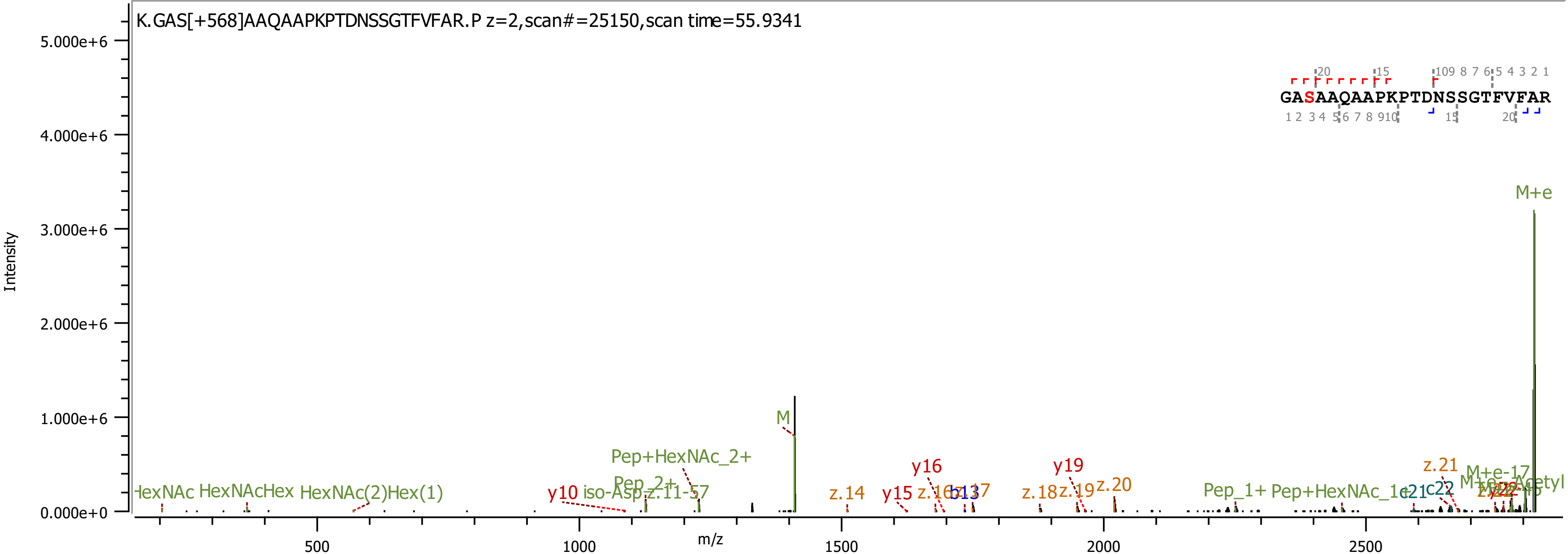

K.GAS[+568]AAQAAPKPTDNSSGTFVFAR.P z=2,scan#=25150,scan time=55.9341
GASAAQAAPKPTDNSSGTFVFAR
Intensity
m/z
5.000e+6
4.000e+6
3.000e+6
2.000e+6
1.000e+6
0.000e+0
500
1000
1500
2000
2500
M+e
M
Pep+HexNAc_2+
Pep_2+
HexNAcHex
HexNAc(2)Hex(1)
y10
z.14
y15
y16
y19
z.18
z.19
z.20
z.21
Pep_1+
Pep+HexNAc_1+
c21
c22
M+e-17

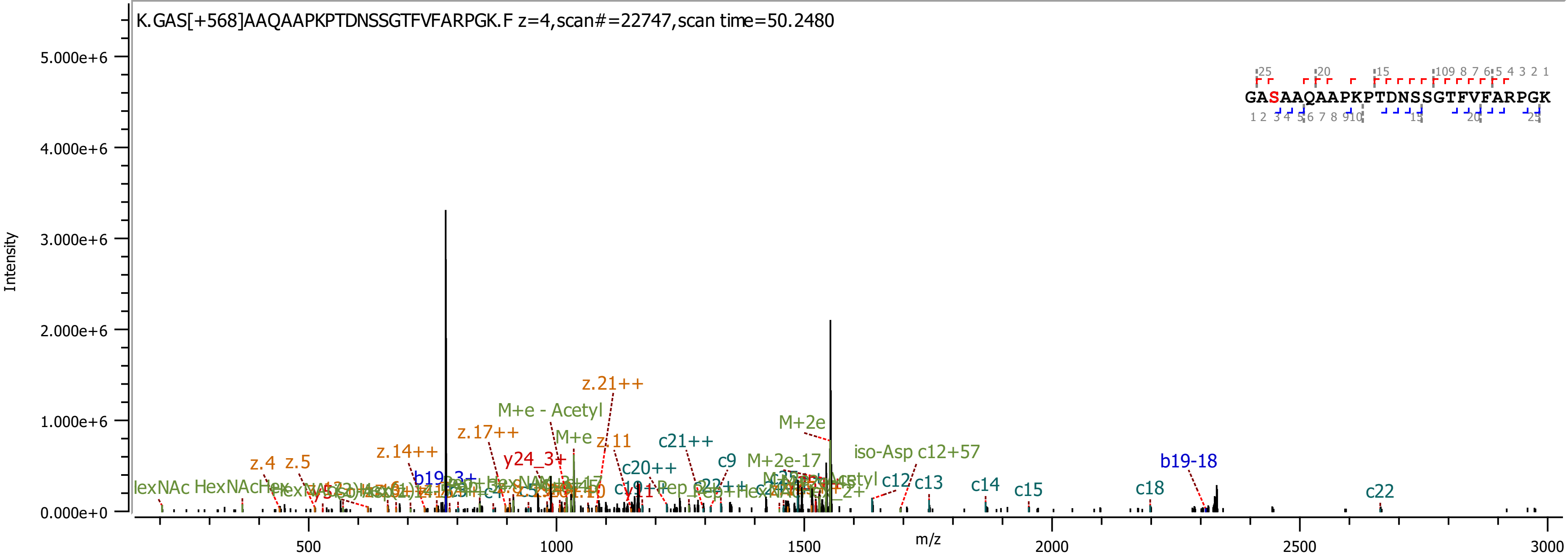
K.GAS[+568]AAQAAPKPTDNSSGTFVFARPGK.F z=4,scan#=22747,scan time=50.2480
GASAAQAAPKPTDNSSGTFVFARPGK
Intensity
m/z
z.21++
M+e - Acetyl
z.17++
M+e
z.11
c21++
M+2e
z.14++
y24_3+
c9
M+2e-17
iso-Asp c12+57
b19-18
z.4
z.5
c20++
c12
c13
c14
c15
c18
c22

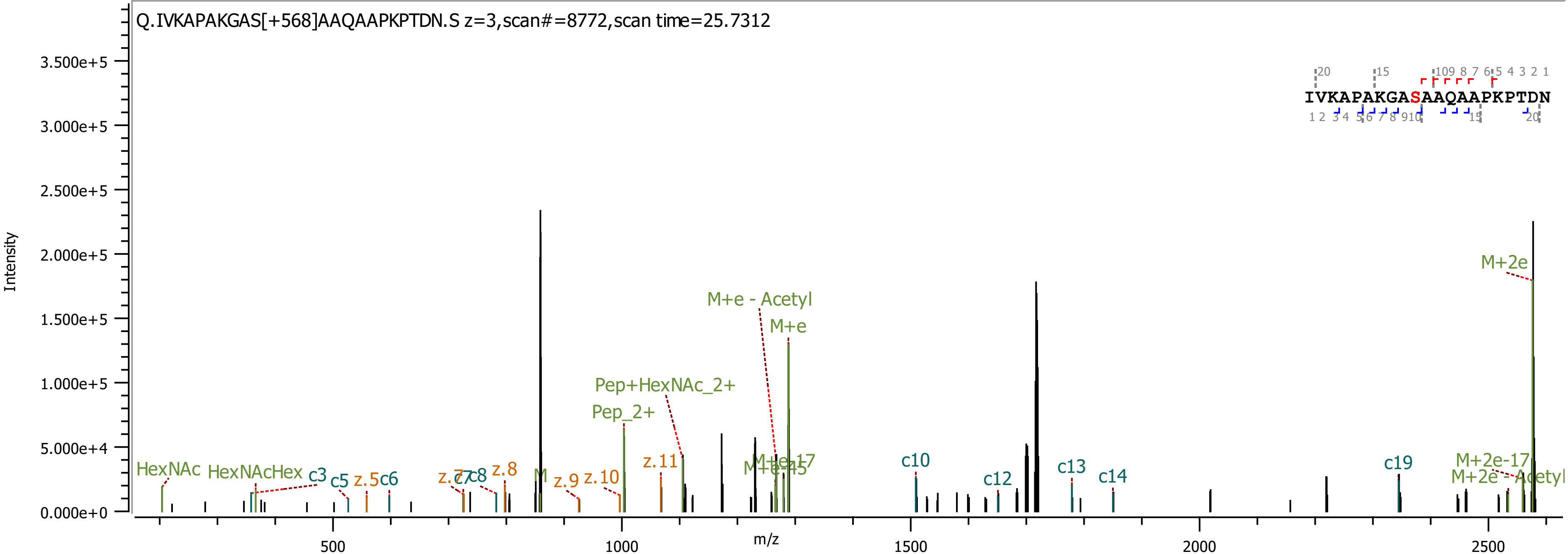

Q.IVKAPAKGAS[+568]AAQAAPKPTDN.S z=3,scan#=8772,scan time=25.7312
IVKAPAKGASAAQAAPKPTDN
Intensity
m/z
HexNAc
HexNAcHex
c3
c5
z.5
c6
z.7
c7
c8
z.8
M
z.9
z.10
Pep_2+
Pep+HexNAc_2+
z.11
M+e - Acetyl
M+e
M+e-17
M+e-45
c10
c12
c13
c14
c19
M+2e
M+2e-17
M+2e - Acetyl

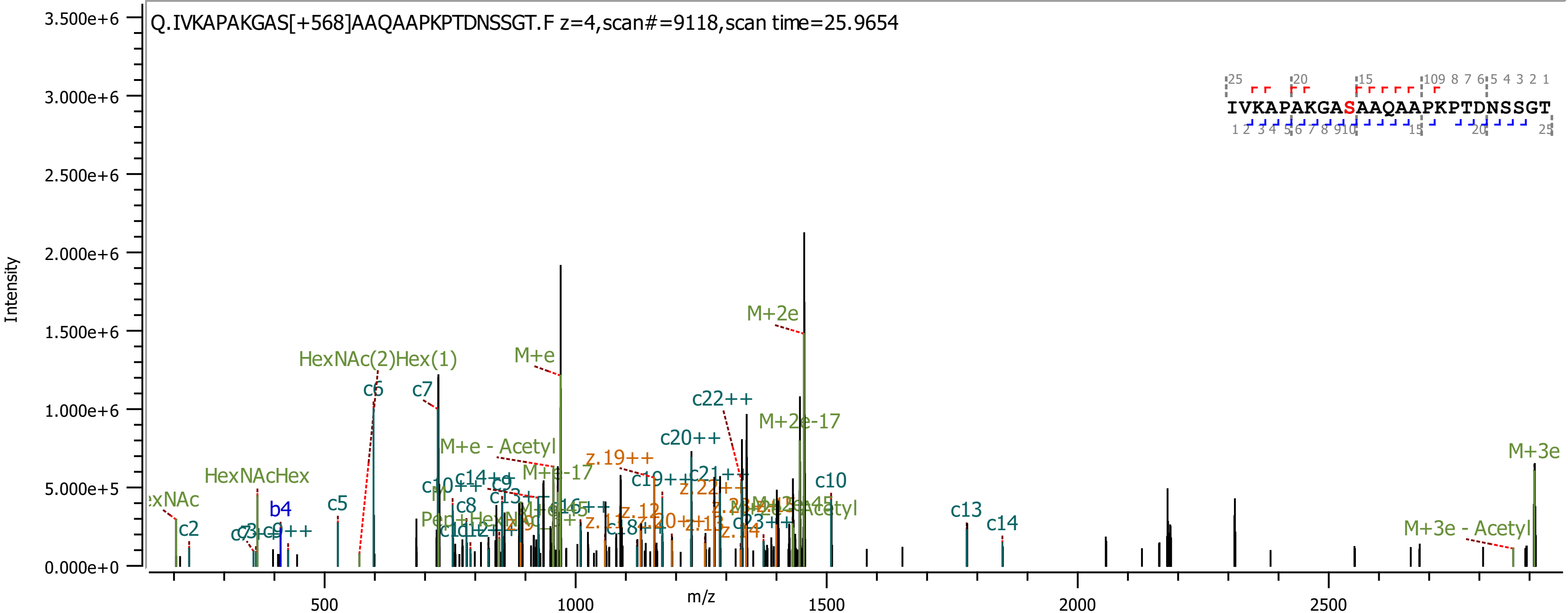
Q.IVKAPAKGAS[+568]AAQAAPKPTDNSSGT.F z=4,scan#=9118,scan time=25.9654
IVKAPAKGASAAQAAPKPTDNSSGT
Intensity
m/z
3.500e+6
3.000e+6
2.500e+6
2.000e+6
1.500e+6
1.000e+6
5.000e+5
0.000e+0
500
1000
1500
2000
2500
M+2e
M+e
HexNAc(2)Hex(1)
c6
c7
c22++
M+2e-17
c20++
M+e - Acetyl
z.19++
M+3e
HexNAcHex
c10
b4
c5
c2
c13
c14
M+3e - Acetyl

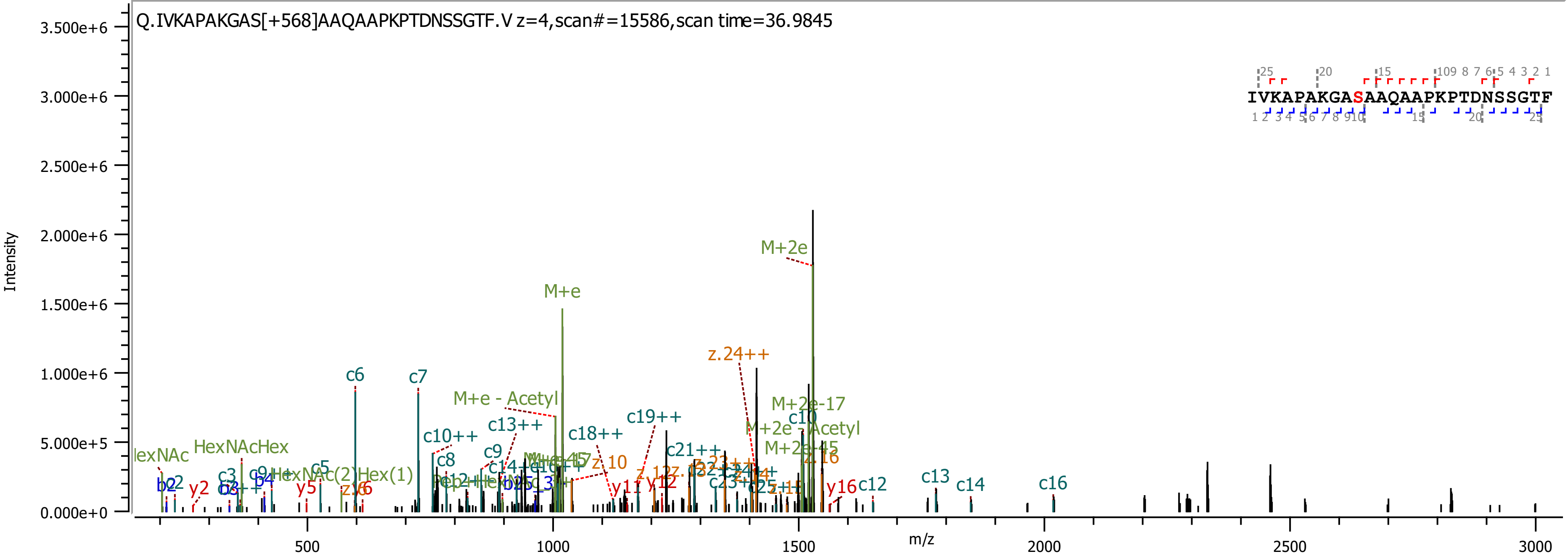
Q.IVKAPAKGAS[+568]AAQAAPKPTDNSSGTF.V z=4,scan#=15586,scan time=36.9845
IVKAPAKGASAAQAAPKPTDNSSGTF
Intensity
m/z
M+2e
M+e
c6
c7
z.24++
M+e - Acetyl
M+2e-17
c10++
c13++
c18++
c19++
c10
M+2e - Acetyl
M+2e-45
c21++
HexNAcHex
c8
c9
c5
HexNAc(2)Hex(1)
y5
y2
b2
c3
b4
y11
y12
y16
c12
c13
c14
c16
z.10
z.16
z.15

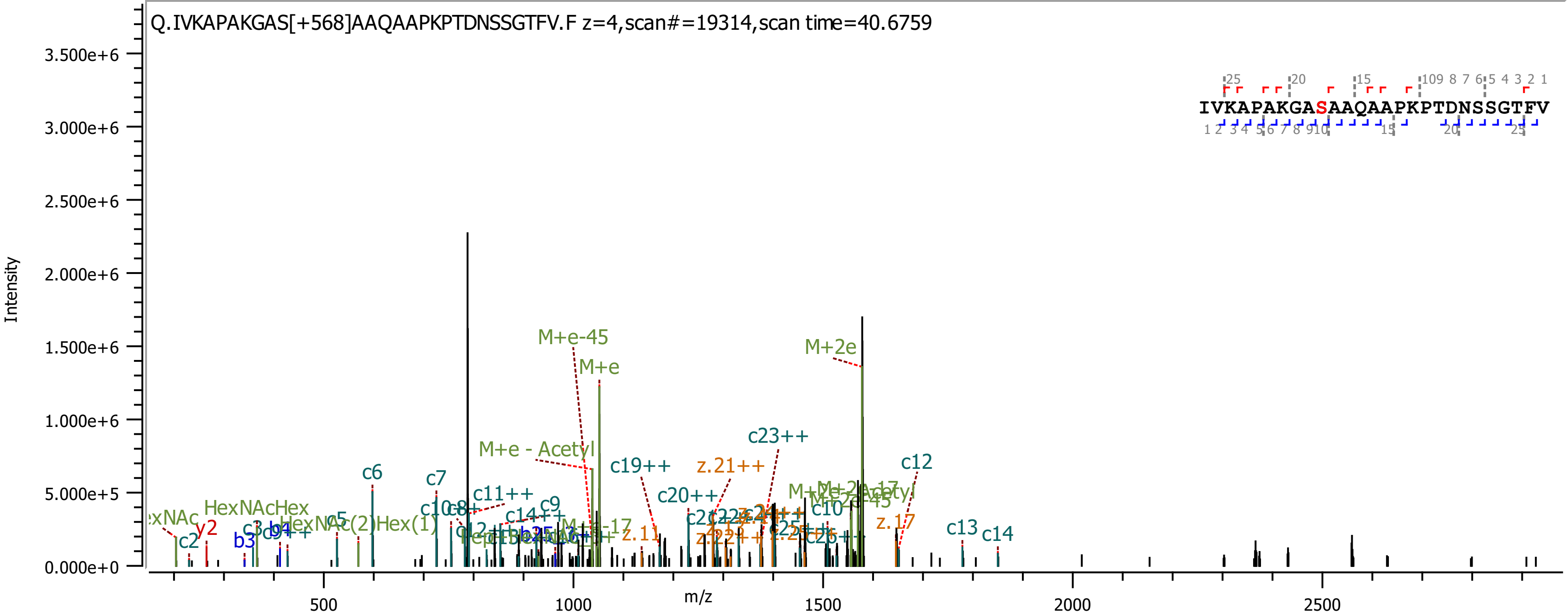
Q.IVKAPAKGAS[+568]AAQAAPKPTDNSSGTFV.F z=4,scan#=19314,scan time=40.6759
IVKAPAKGASAAQAAPKPTDNSSGTFV
Intensity
m/z
3.500e+6
3.000e+6
2.500e+6
2.000e+6
1.500e+6
1.000e+6
5.000e+5
0.000e+0
500
1000
1500
2000
2500
M+e-45
M+e
M+2e
M+e - Acetyl
c23++
c19++
z.21++
c12
c6
c7
c11++
c9
c20++
HexNAcHex
y2
c2
b3
c5
c13
c14
z.17
z.11
c10
c14++

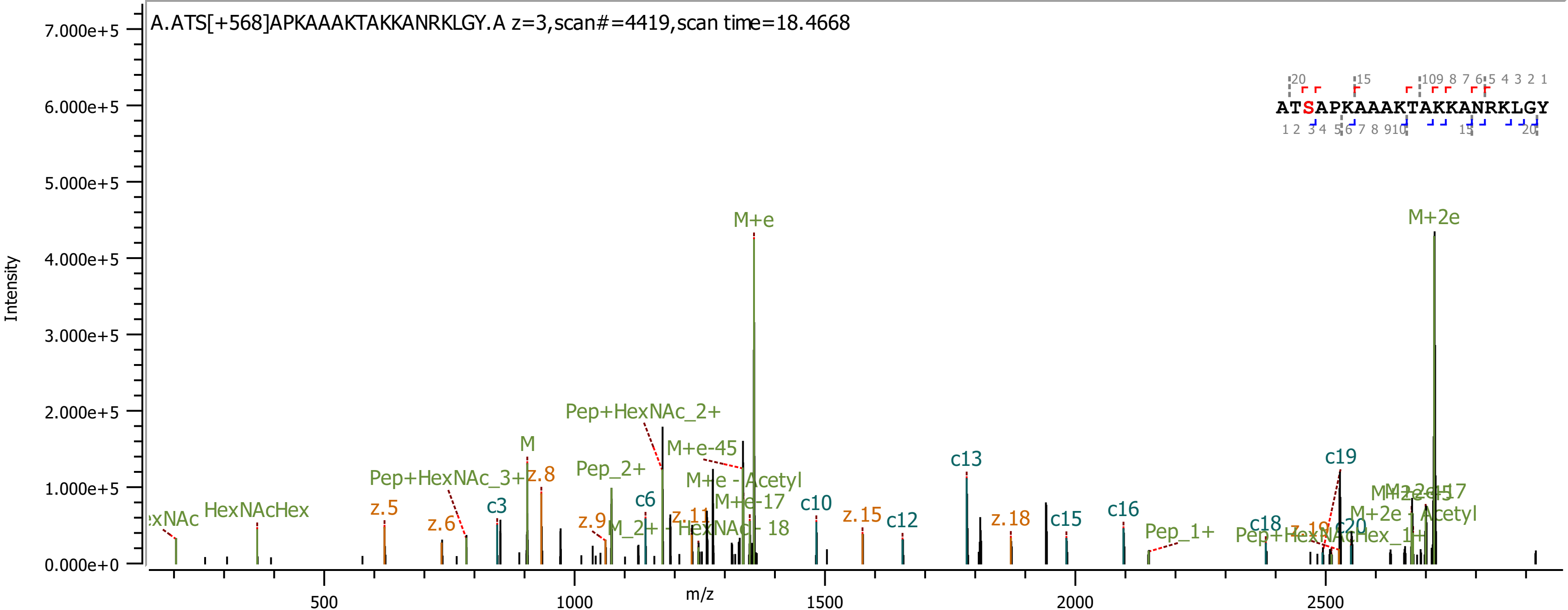

A.ATS[+568]APKAAAKTAKKANRKLGY.A z=3,scan#=4419,scan time=18.4668
Intensity
m/z
ATSAPKAAAKTAKKANRKLGY
M+e
M+2e
Pep+HexNAc_2+
M+e-45
M+e - Acetyl
M+e-17
M
z.8
Pep+HexNAc_3+
z.5
z.6
c3
Pep_2+
c6
z.9
z.11
M_2+ - HexNAc - 18
HexNAcHex
c10
z.15
c12
c13
z.18
c15
c16
Pep_1+
c18
c19
c20
M+2e - Acetyl
Pep+HexNAcHex_1+

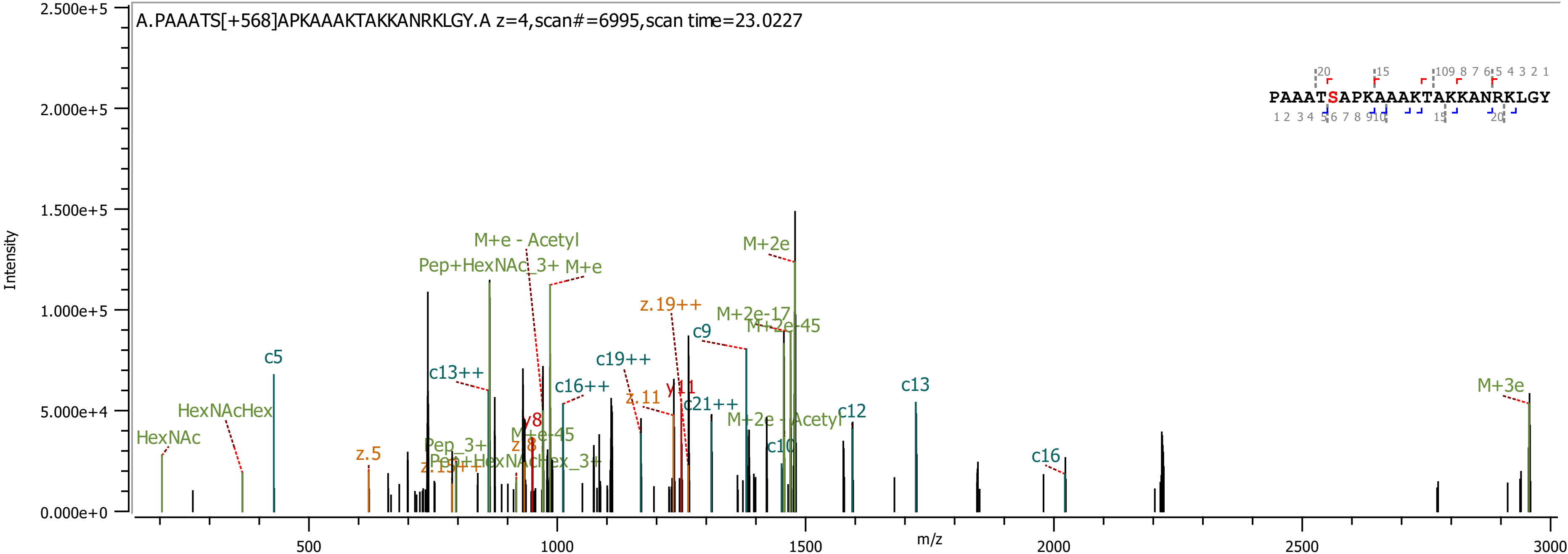

A.PAAATS[+568]APKAAAKTAKKANRKLGY.A z=4,scan#=6995,scan time=23.0227
PAAATSAPKAAAKTAKKANRKLGY
Intensity
m/z
HexNAc
HexNAcHex
c5
z.5
c13++
Pep+HexNAc_3+
M+e - Acetyl
M+e
M+e-45
y8
z.8
Pep_3+
Pep+HexNAcHex_3+
z.15++
c16++
c19++
z.11
z.19++
y11
c21++
c9
M+2e-17
M+2e-45
M+2e
M+2e - Acetyl
c10
c12
c13
c16
M+3e

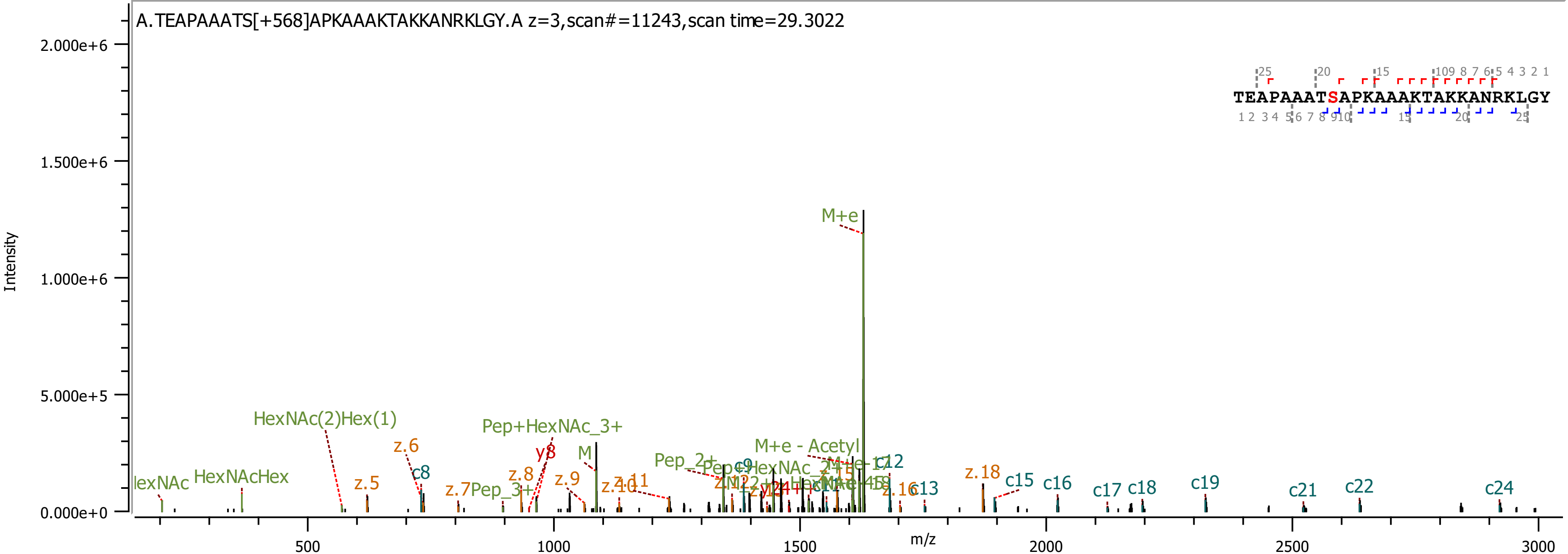

A.TEAPAAATS[+568]APKAAAKTAKKANRKLGY.A z=3,scan#=11243,scan time=29.3022
TEAPAAATSAPKAAAKTAKKANRKLGY
Intensity
m/z
2.000e+6
1.500e+6
1.000e+6
5.000e+5
0.000e+0
500
1000
1500
2000
2500
3000
M+e
HexNAc(2)Hex(1)
Pep+HexNAc_3+
M+e - Acetyl
HexNAcHex
z.5
z.6
c8
z.7
Pep_3+
z.8
y8
z.9
M
z.10
z.11
Pep_2+
c9
c12
c13
z.16
z.18
c15
c16
c17
c18
c19
c21
c22
c24

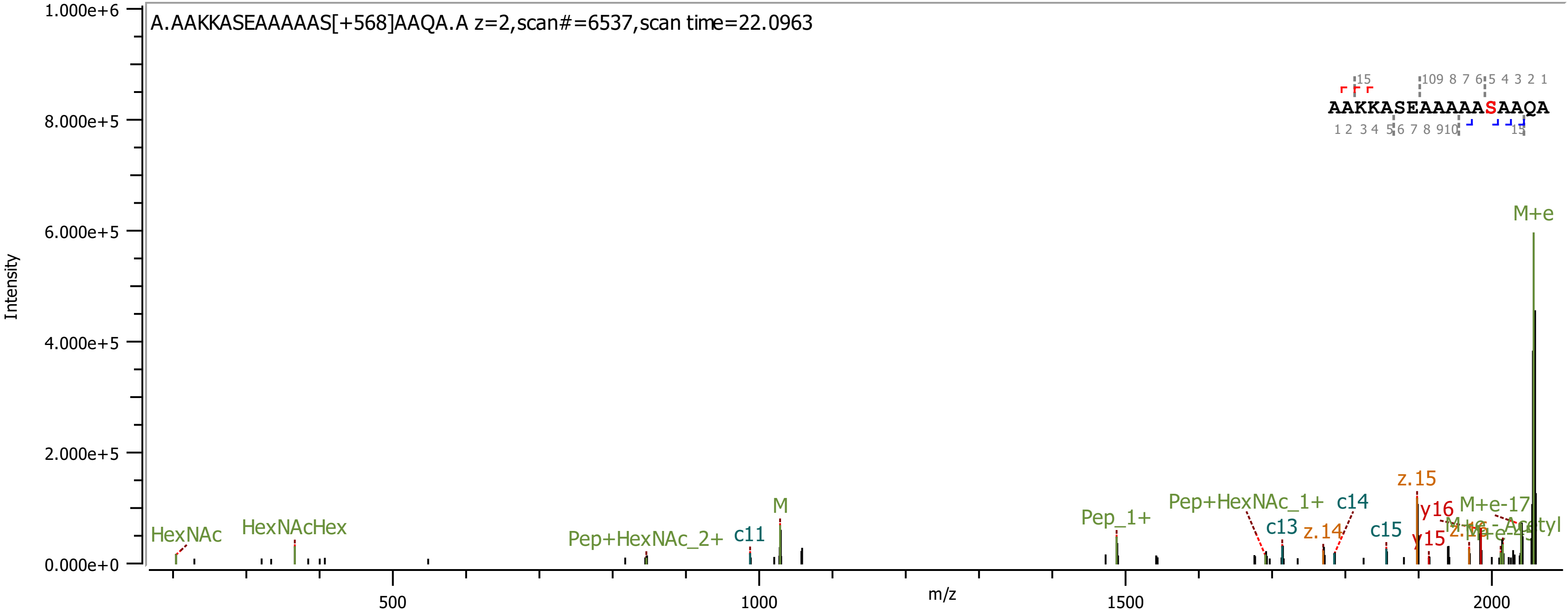
A.AAKKASEAAAAAS[+568]AAQA.A z=2,scan#=6537,scan time=22.0963
AAKKASEAAAAASAAQA
Intensity
m/z
M+e
z.15
M
HexNAc
HexNAcHex
Pep+HexNAc_2+
c11
Pep_1+
Pep+HexNAc_1+
c13
c14
z.14
c15
y16
y15
M+e-17
M+e-Acetyl

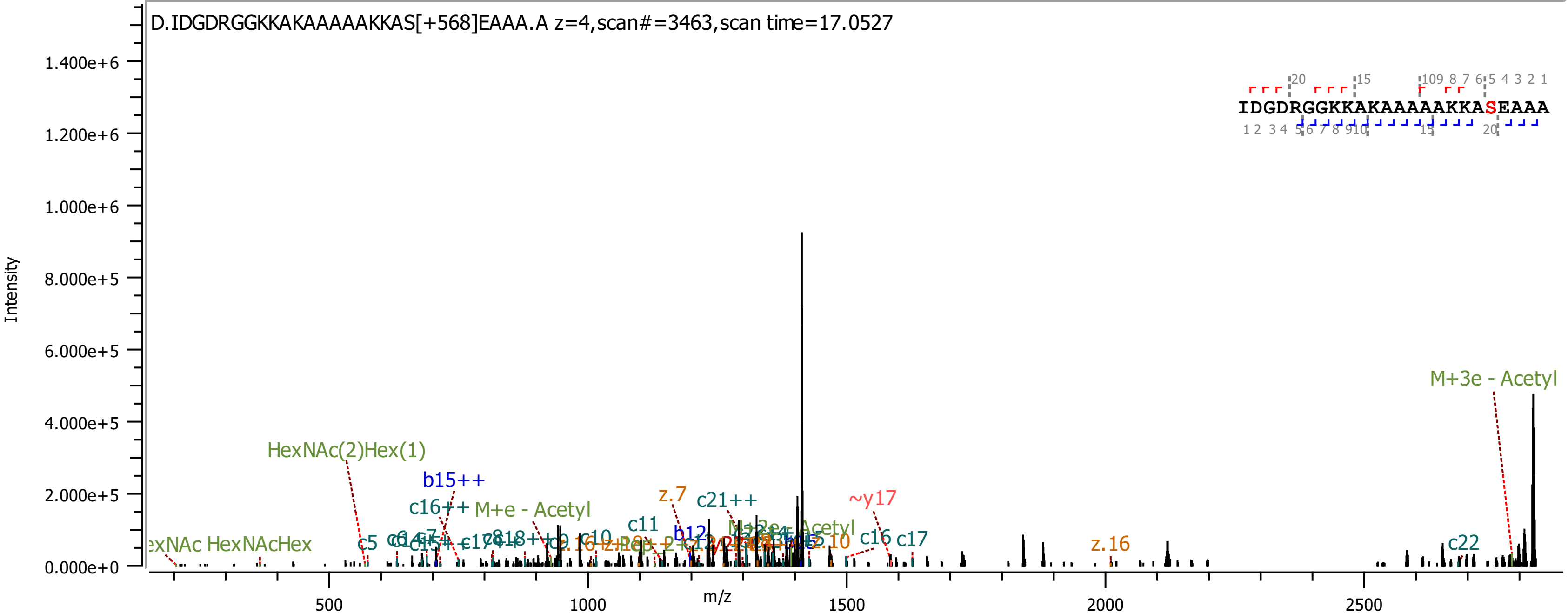

D.IDGDRGGKKAKAAAAAKKAS[+568]EAAA.A z=4,scan#=3463,scan time=17.0527
IDGDRGGKKAKAAAAAKKASEAAA
Intensity
m/z
1.400e+6
1.200e+6
1.000e+6
8.000e+5
6.000e+5
4.000e+5
2.000e+5
0.000e+0
500
1000
1500
2000
2500
M+3e - Acetyl
HexNAc(2)Hex(1)
b15++
c16++
M+e - Acetyl
z.7
c21++
~y17
c11
b12
c5
HexNAcHex
c16
c17
z.16
c22

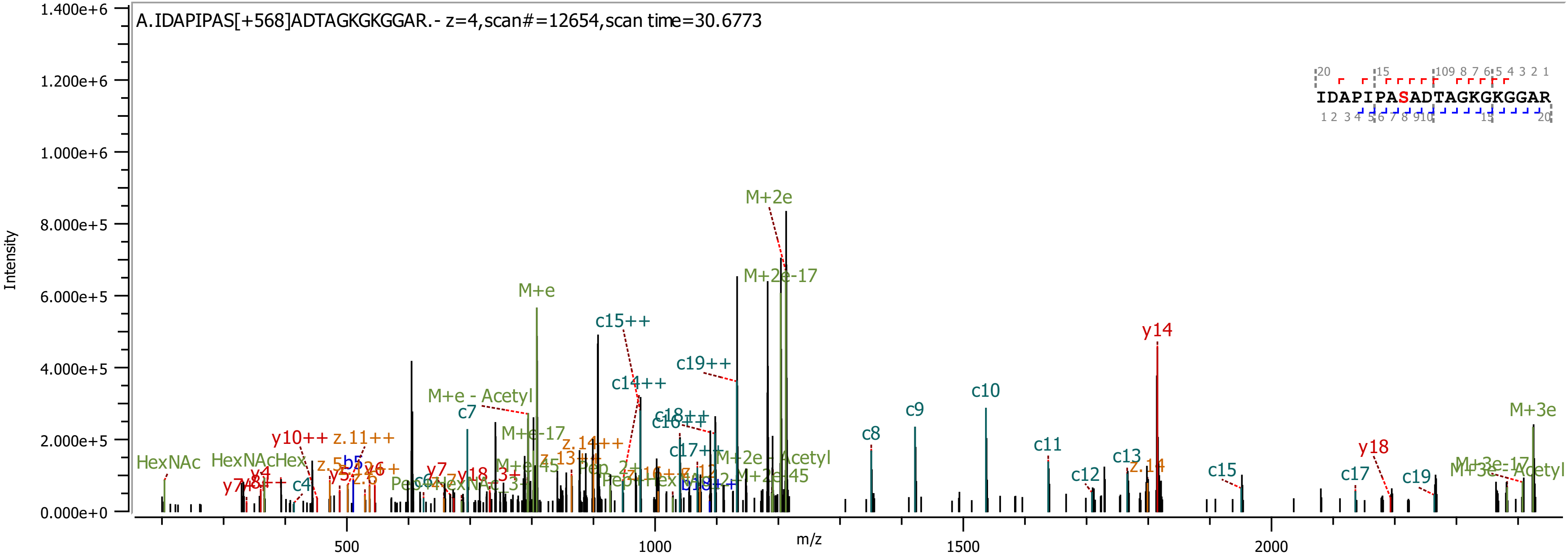
A.IDAPIPAS[+568]ADTAGKGKGGAR.- z=4,scan#=12654,scan time=30.6773
IDAPIPASADTAGKGKGGAR
Intensity
m/z
M+2e
M+2e-17
M+e
c15++
c19++
c14++
M+e - Acetyl
c7
y14
c10
c9
c8
M+3e
c11
y18
c13
c12
c15
c17
c19
HexNAc
HexNAcHex
y10++
z.11++
b5
c4
y4
M+e-17
z.14++
z.13++
c17++
c18++
c16++
M+2e - Acetyl
M+2e-45
M+e-45
M+3e-17
M+3e - Acetyl
z.14
y7
y18_3+
b18++
0.000e+0
2.000e+5
4.000e+5
6.000e+5
8.000e+5
1.000e+6
1.200e+6
1.400e+6
500
1000
1500
2000

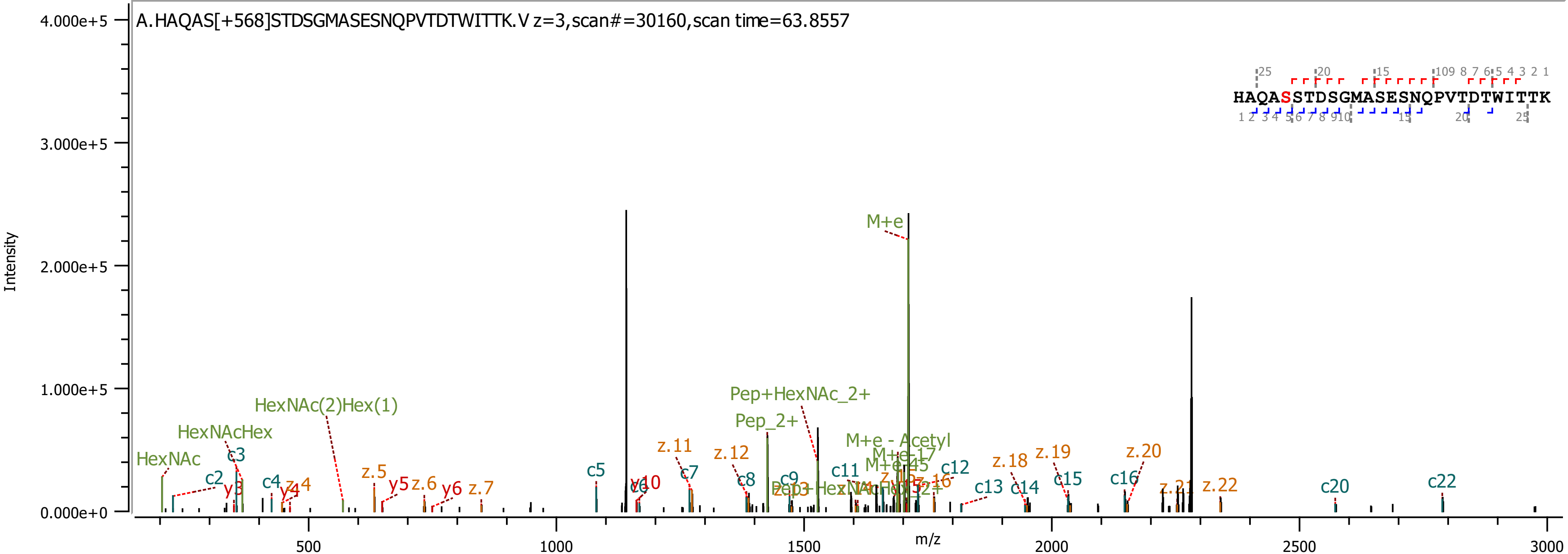
A.HAQAS[+568]STDSGMASESNQPVTDTWITTK.V z=3,scan#=30160,scan time=63.8557
Intensity
m/z
HAQASSTDSGMASESNQPVTDTWITTK
HexNAc
HexNAcHex
HexNAc(2)Hex(1)
Pep+HexNAc_2+
Pep_2+
M+e
M+e - Acetyl
c2
c3
c4
c5
c7
c8
c9
c11
c12
c13
c14
c15
c16
c20
c22
y3
y4
y5
y6
y10
z.4
z.5
z.6
z.7
z.11
z.12
z.18
z.19
z.20
z.22

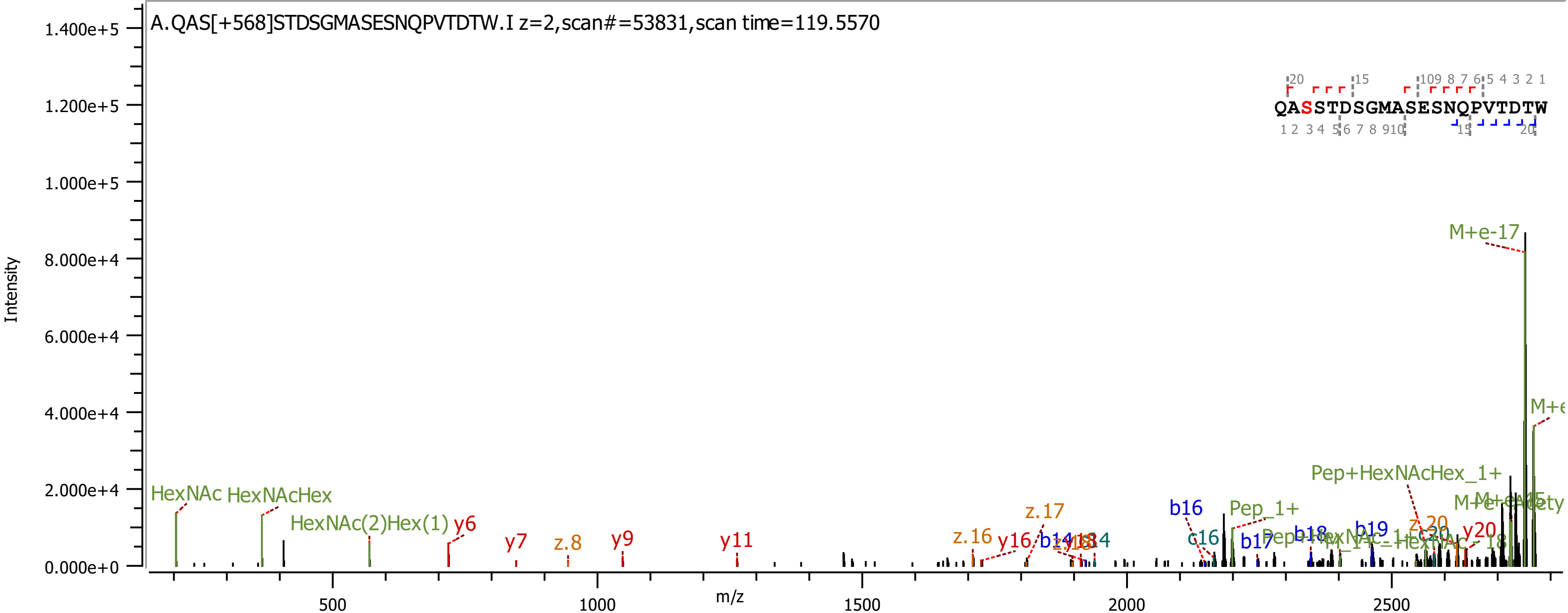
A.QAS[+568]STDSGMASESNQPVTDTW.I z=2,scan#=53831,scan time=119.5570
QASSTDSGMASESNQPVTDTW
Intensity
m/z
1.400e+5
1.200e+5
1.000e+5
8.000e+4
6.000e+4
4.000e+4
2.000e+4
0.000e+0
500
1000
1500
2000
2500
HexNAc
HexNAcHex
HexNAc(2)Hex(1)
y6
y7
z.8
y9
y11
z.16
y16
z.17
b16
c16
Pep_1+
b17
b18
b19
Pep+HexNAcHex_1+
y20
M+e-17

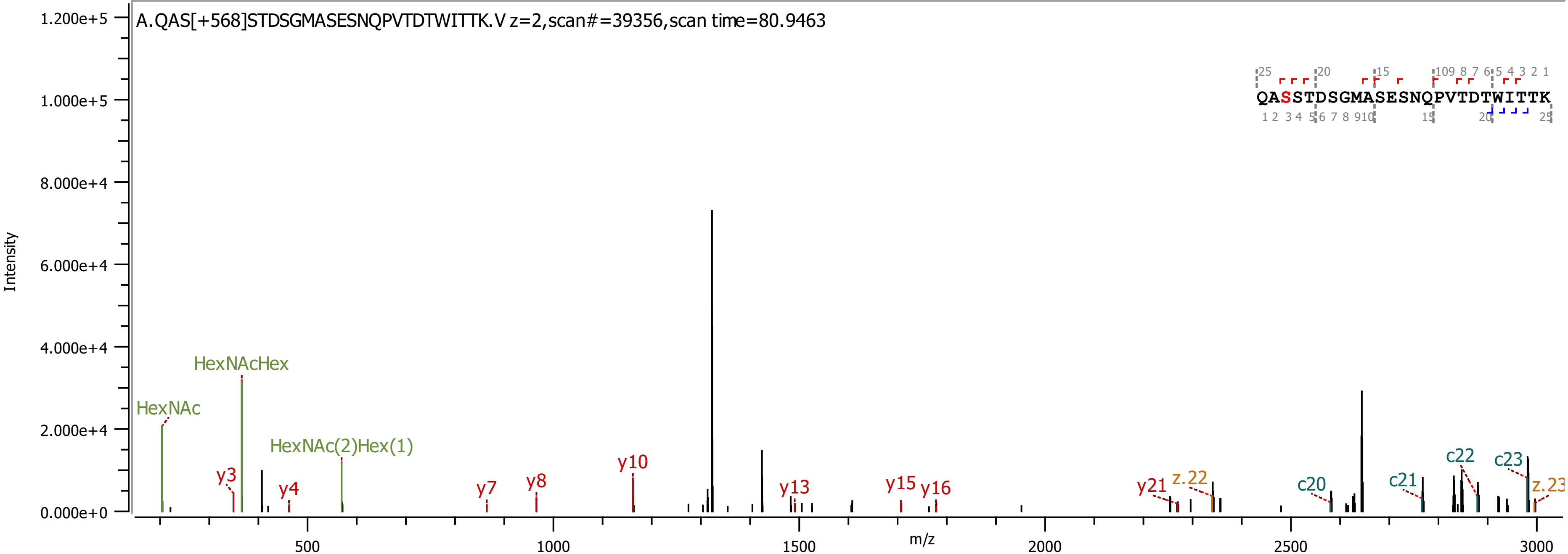

A.QAS[+568]STDSGMASESNQPVTDTWITTK.V z=2,scan#=39356,scan time=80.9463
QASSTDSGMASESNQPVTDTWITTK
Intensity
m/z
HexNAc
HexNAcHex
HexNAc(2)Hex(1)
y3
y4
y7
y8
y10
y13
y15
y16
y21
z.22
c20
c21
c22
c23
z.23

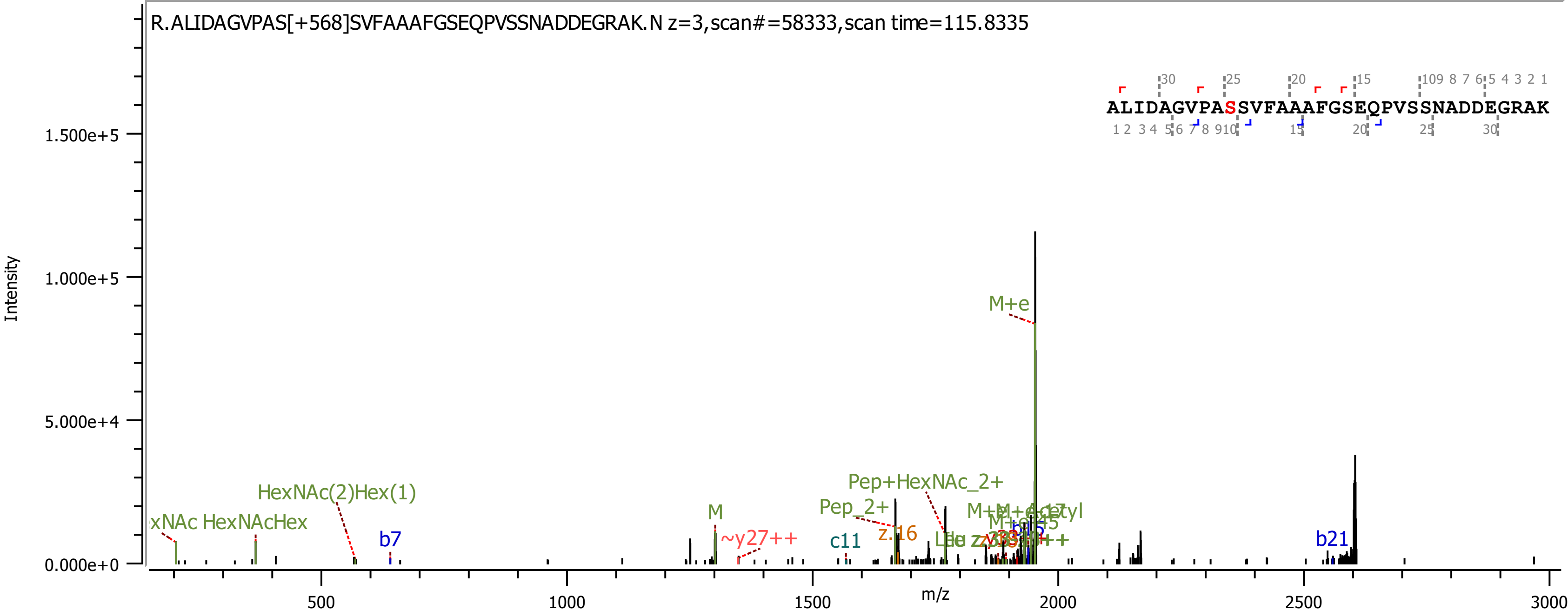
R.ALIDAGVPAS[+568]SVFAAAFGSEQPVSSNADDEGRAK.N z=3,scan#=58333,scan time=115.8335
ALIDAGVPASSVFAAAFGSEQPVSSNADDEGRAK
Intensity
m/z
1.500e+5
1.000e+5
5.000e+4
0.000e+0
500
1000
1500
2000
2500
3000
M+e
HexNAc(2)Hex(1)
HexNAcHex
M
b7
~y27++
c11
z.16
Pep_2+
Pep+HexNAc_2+
b21

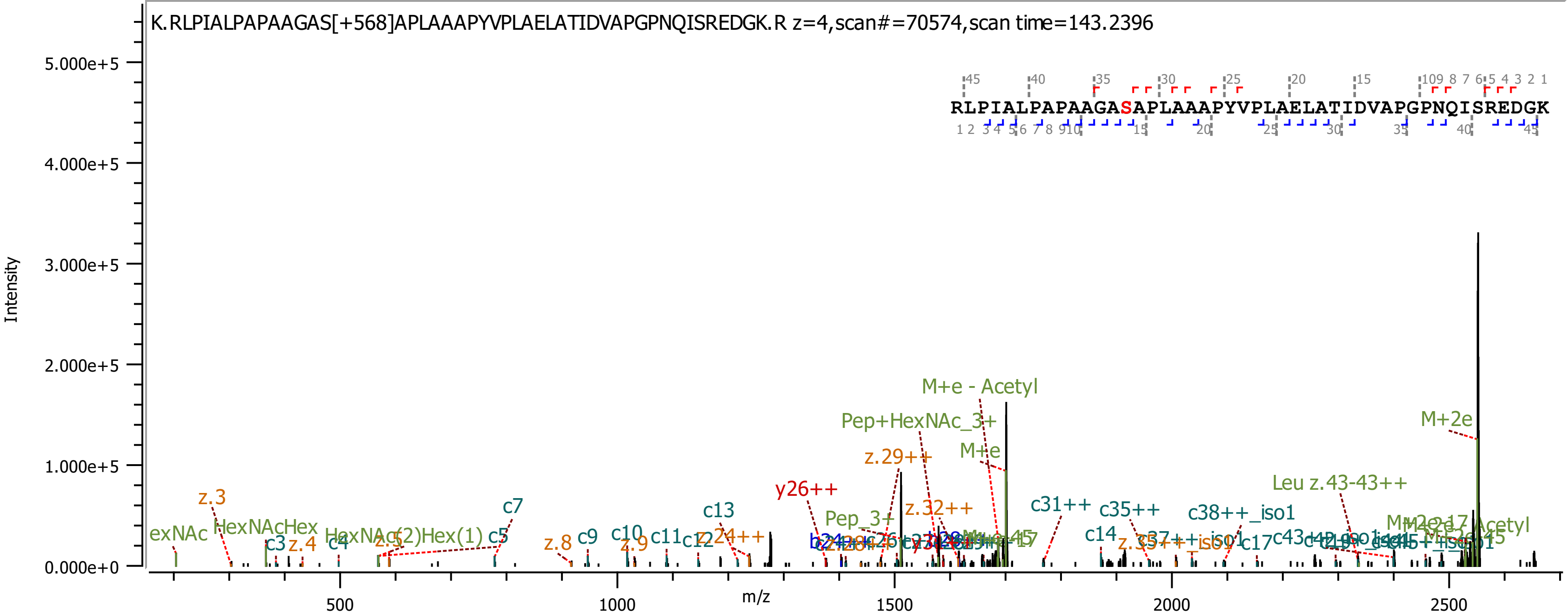
K.RLPIALPAPAAGAS[+568]APLAAAPYVPLAELATIDVAPGPNQISREDGK.R z=4,scan#=70574,scan time=143.2396
RLPIALPAPAAGASAPLAAAPYVPLAELATIDVAPGPNQISREDGK
Intensity
m/z
M+e - Acetyl
Pep+HexNAc_3+
M+e
z.29++
M+2e
y26++
Leu z.43-43++
c13
c31++
c35++
c38++_iso1
z.32++
Pep_3+
c7
z.3
HexNAcHex
c5
c9
c10
c11
c14
z.8
z.9
c3
z.4
c4
c17
exNAc

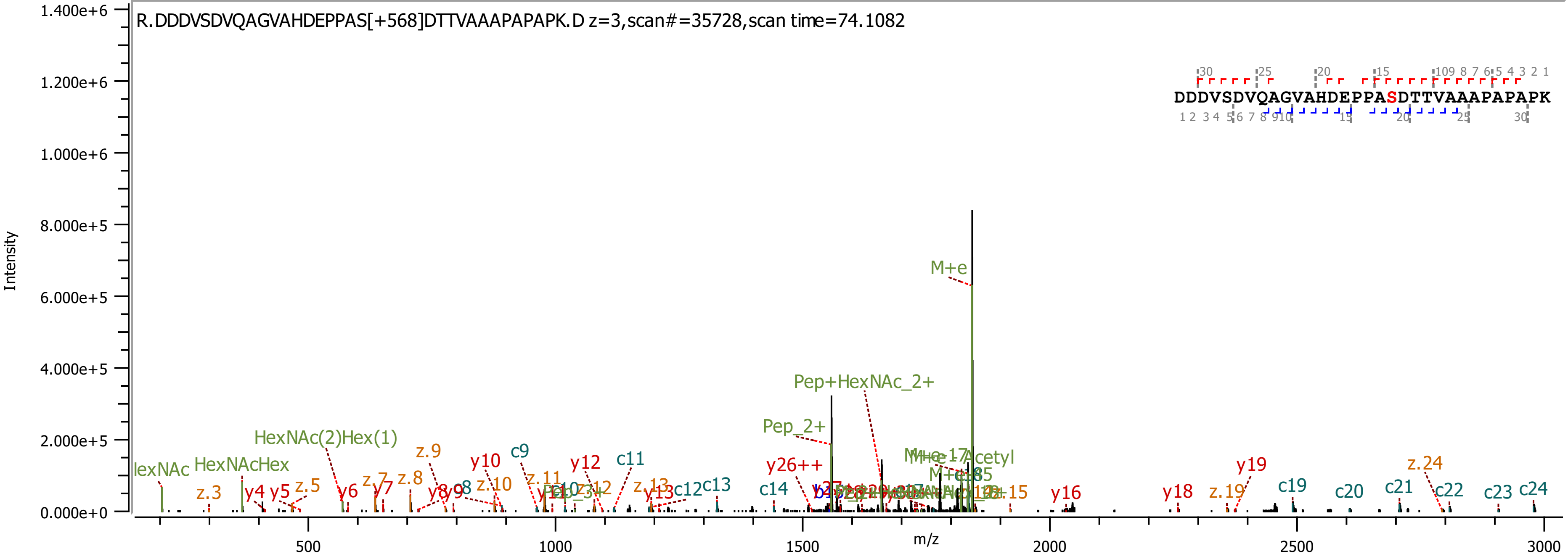

R.DDDVSDVQAGVAHDEPPAS[+568]DTTVAAAPAPAPK.D z=3,scan#=35728,scan time=74.1082
DDDVSDVQAGVAHDEPPASDTTVAAAPAPAPK
Intensity
m/z
M+e
Pep+HexNAc_2+
Pep_2+
HexNAc(2)Hex(1)
HexNAcHex
y26++
z.24
y19
c19
c20
c21
c22
c23
c24
y18
z.19
y16
c14
c13
c12
c11
c9
y10
y12
z.9
z.8
z.10
z.11
z.13
y13
y6
y5
y4
z.5
z.3

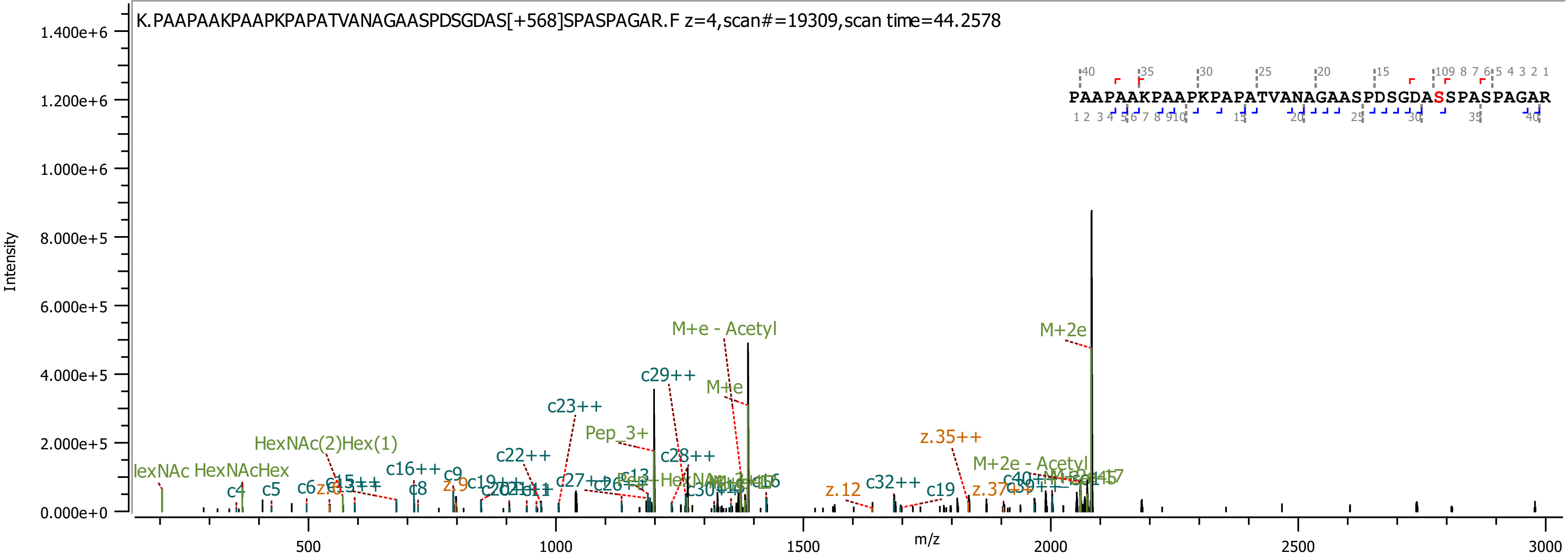
K.PAAPAAKPAAPKPAPATVANAGAASPDSGDAS[+568]SPASPAGAR.F z=4,scan#=19309,scan time=44.2578
Intensity
m/z
PAAPAAKPAAPKPAPATVANAGAASPDSGDASSPASPAGAR
M+2e
M+e - Acetyl
c29++
M+e
c23++
Pep_3+
c22++
c28++
z.35++
M+2e - Acetyl
HexNAc(2)Hex(1)
HexNAc HexNAcHex
c16++
c32++
c19
z.12
c4
c5
c8
c9

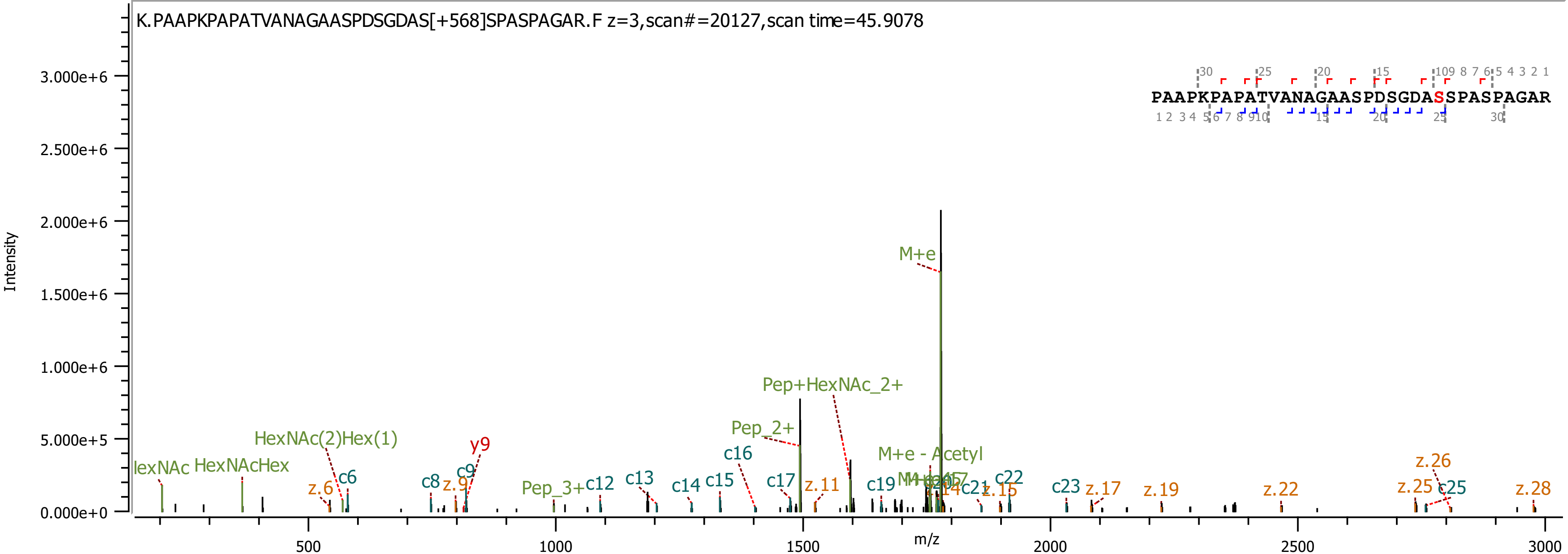
K.PAAPKPAPATVANAGAASPDSGDAS[+568]SPASPAGAR.F z=3,scan#=20127,scan time=45.9078
PAAPKPAPATVANAGAASPDSGDASSPASPAGAR
Intensity
m/z
3.000e+6
2.500e+6
2.000e+6
1.500e+6
1.000e+6
5.000e+5
0.000e+0
500
1000
1500
2000
2500
3000
M+e
Pep+HexNAc_2+
Pep_2+
M+e - Acetyl
HexNAc(2)Hex(1)
HexNAcHex
y9
c6
c8
c9
c12
c13
c14
c15
c16
c17
c19
c22
c23
c25
z.6
z.9
z.11
z.15
z.17
z.19
z.22
z.25
z.26
z.28
Pep_3+

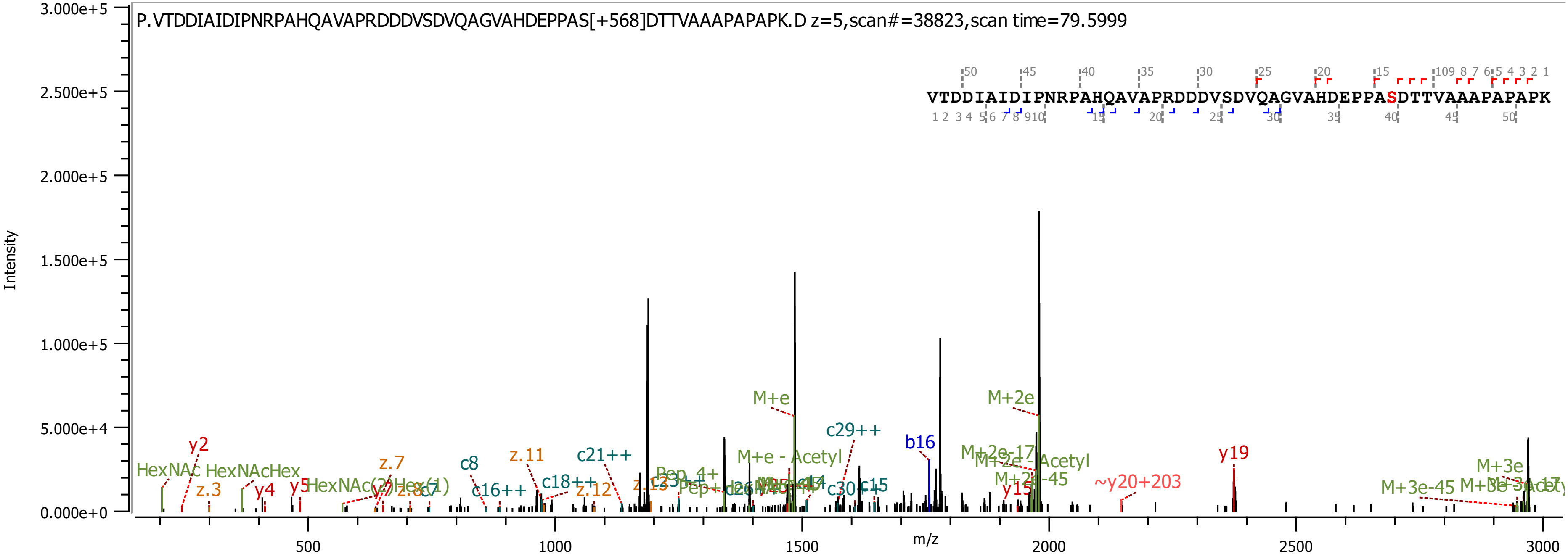

P.VTDDIAIDIPNRPAHQAVAPRDDDVSDVQAGVAHDEPPAS[+568]DTTVAAAPAPAPK.D z=5,scan#=38823,scan time=79.5999
VTDDIAIDIPNRPAHQAVAPRDDDVSDVQAGVAHDEPPASDTTVAAAPAPAPK
Intensity
m/z
M+e
M+2e
c29++
b16
y19
M+e - Acetyl
c21++
z.11
c8
y2
HexNAc
HexNAcHex
z.3
y4
y5
z.7
c16++
c18++
z.12
~y20+203
M+3e
M+3e-45
M+2e-45
y15
M+2e-17
c30++
c15

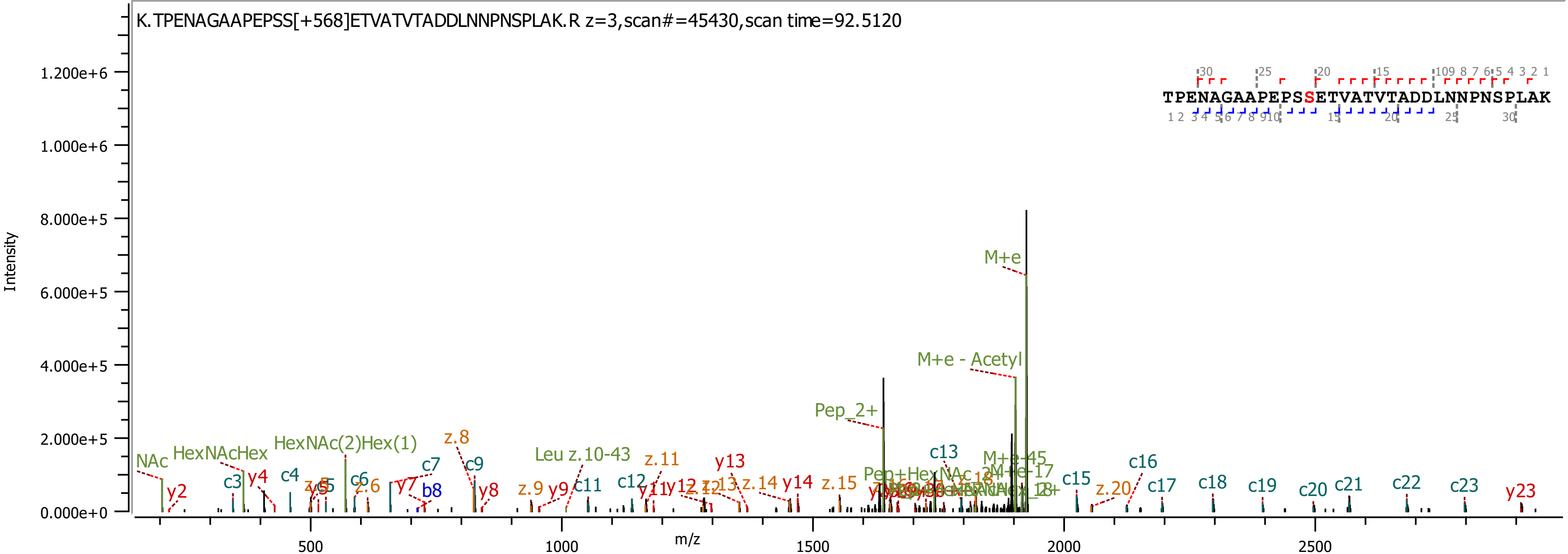
K.TPENAGAAPEPSS[+568]ETVATVTADDLNNPNSPLAK.R z=3,scan#=45430,scan time=92.5120
TPENAGAAPEPSSETVATVTADDLNNPNSPLAK
Intensity
m/z
M+e
M+e - Acetyl
Pep_2+
NAc
HexNAcHex
HexNAc(2)Hex(1)
Leu z.10-43
c3
c4
c6
c7
c9
c11
c12
c13
c15
c16
c17
c18
c19
c20
c21
c22
c23
y2
y4
y7
y8
y9
y11
y12
y13
y14
y23
b8
z.6
z.8
z.9
z.11
z.14
z.15
z.20

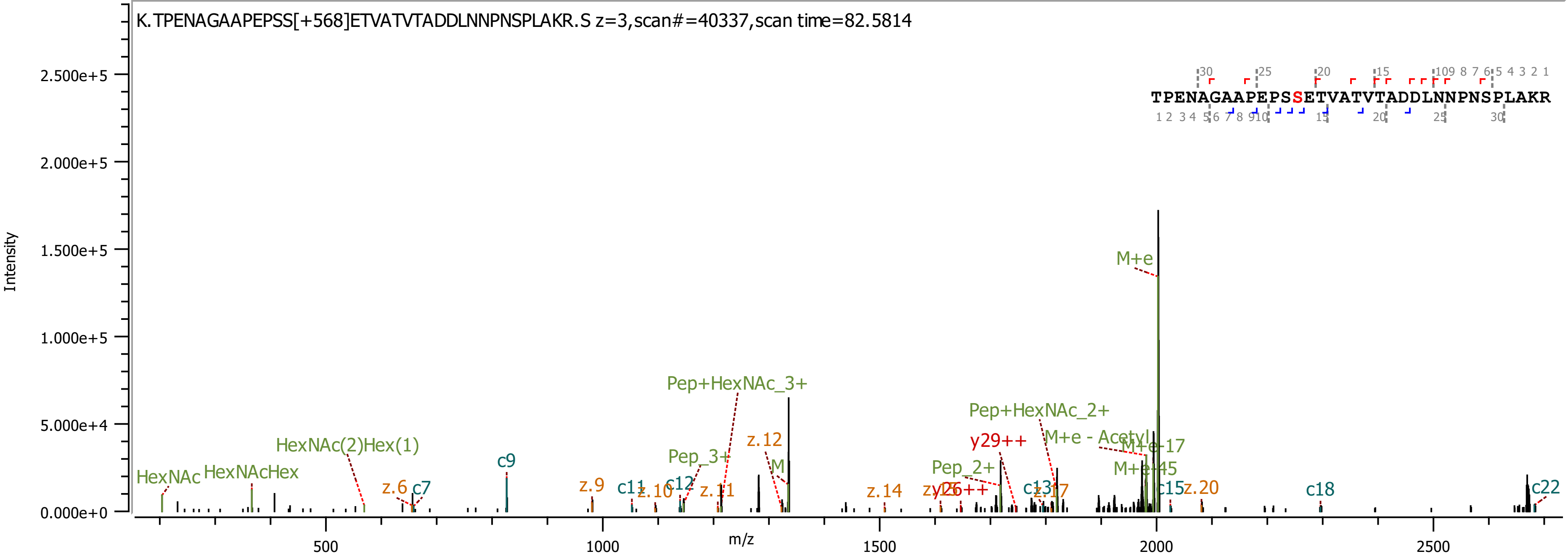
K.TPENAGAAPEPSS[+568]ETVATVTADDLNNPNSPLAKR.S z=3,scan#=40337,scan time=82.5814
TPENAGAAPEPSSETVATVTADDLNNPNSPLAKR
Intensity
m/z
M+e
Pep+HexNAc_3+
Pep+HexNAc_2+
HexNAc(2)Hex(1)
HexNAc
HexNAcHex
c9
z.6
c7
z.9
c11
z.10
c12
Pep_3+
z.11
z.12
y29++
M+e - Acetyl
M+e-17
M+e+45
Pep_2+
z.14
c13
z.17
c15
z.20
c18
c22

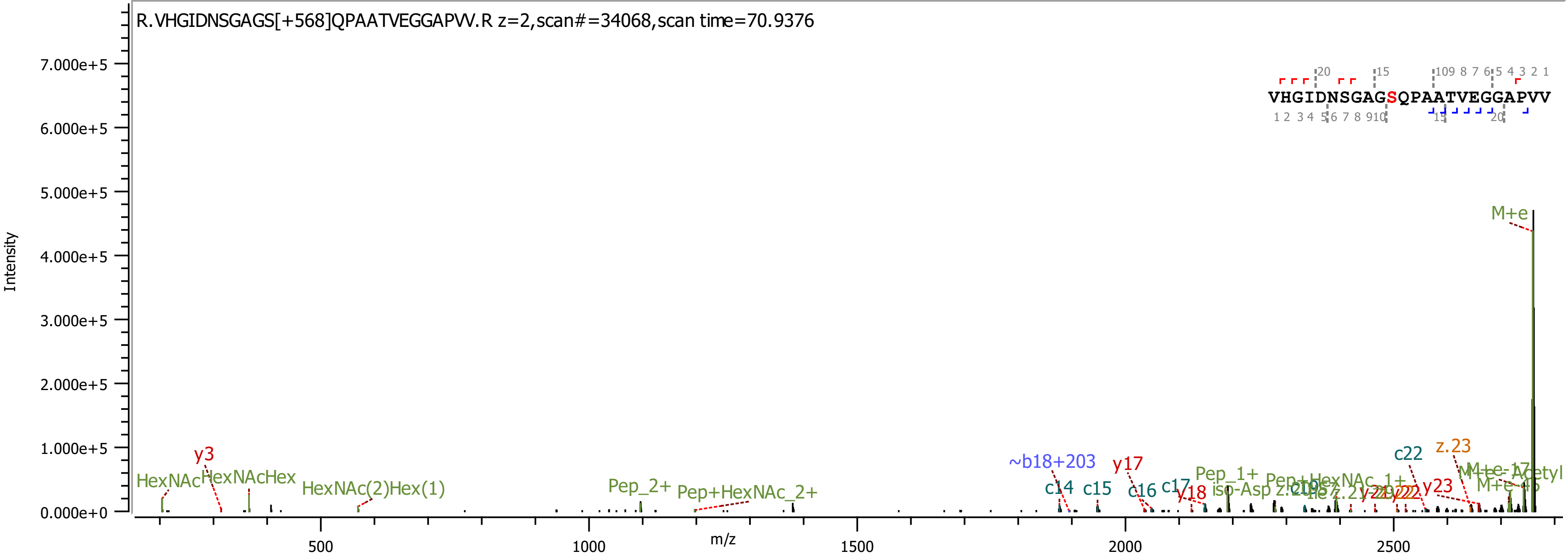
R.VHGIDNSGAGS[+568]QPAATVEGGAPVV.R z=2,scan#=34068,scan time=70.9376
VHGIDNSGAGSQPAATVEGGAPVV
Intensity
m/z
M+e
HexNAc
HexNAcHex
y3
HexNAc(2)Hex(1)
Pep_2+
Pep+HexNAc_2+
~b18+203
c14
c15
y17
c16
c17
y18
Pep_1+
c22
z.23
y23

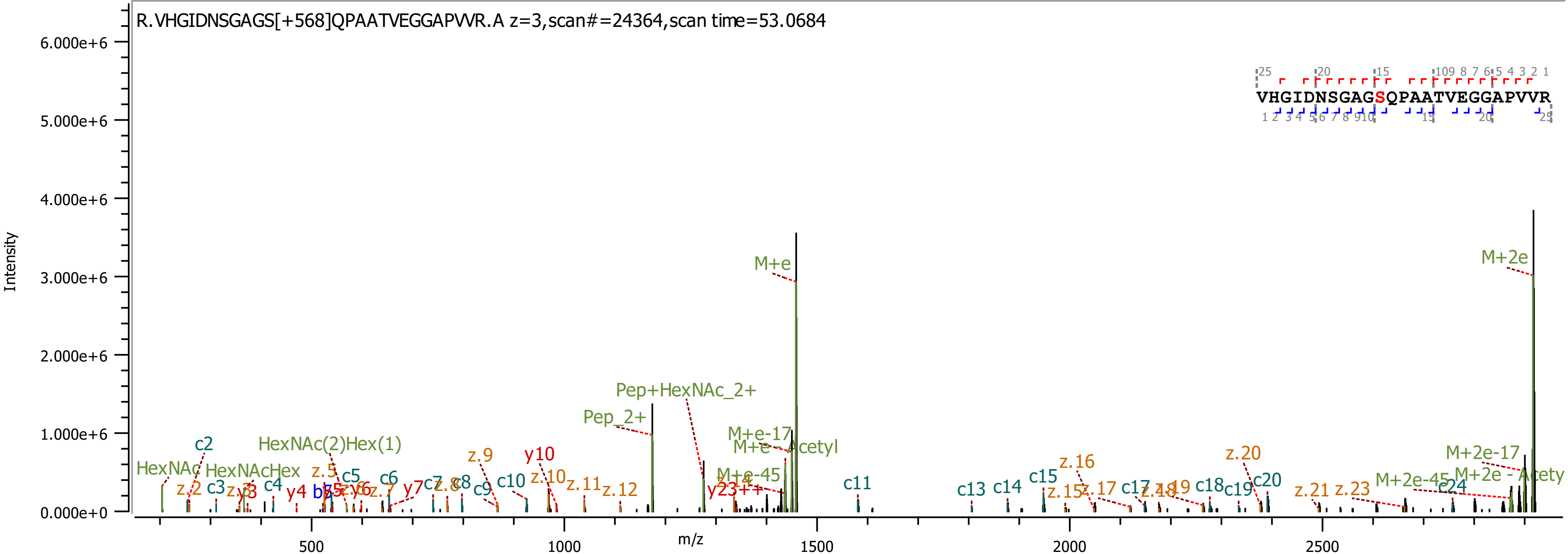
R.VHGIDNSGAGS[+568]QPAATVEGGAPVVR.A z=3,scan#=24364,scan time=53.0684
Intensity
m/z
VHGIDNSGAGSQPAATVEGGAPVVR
M+e
M+2e
Pep+HexNAc_2+
Pep_2+
HexNAc(2)Hex(1)
HexNAcHex
HexNAc
M+e-17
M+e - Acetyl
M+e-45
M+2e-17
M+2e - Acetyl
M+2e-45
y23++

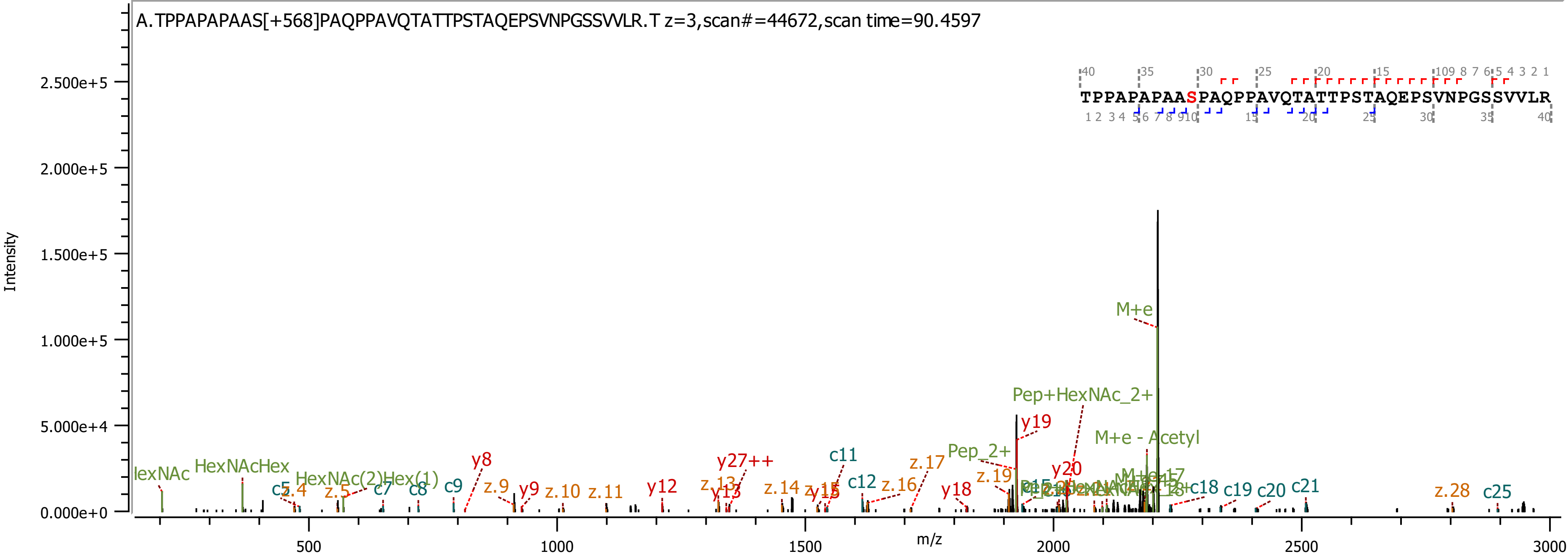

A.TPPAPAPAAS[+568]PAQPPAVQTATTPSTAQEPSVNPGSSVVLR.T z=3,scan#=44672,scan time=90.4597
TPPAPAPAASPAQPPAVQTATTPSTAQEPSVNPGSSVVLR
Intensity
m/z
M+e
Pep+HexNAc_2+
M+e - Acetyl
y19
Pep_2+
y20
c11
y27++
y8
z.17
HexNAcHex
HexNAc(2)Hex(1)
c12
z.16
y18
z.19
c18
c19
c20
c21
z.28
c25
z.9
y9
z.10
z.11
y12
z.13
y13
z.14
y15
c7
c8
c9
z.4
z.5

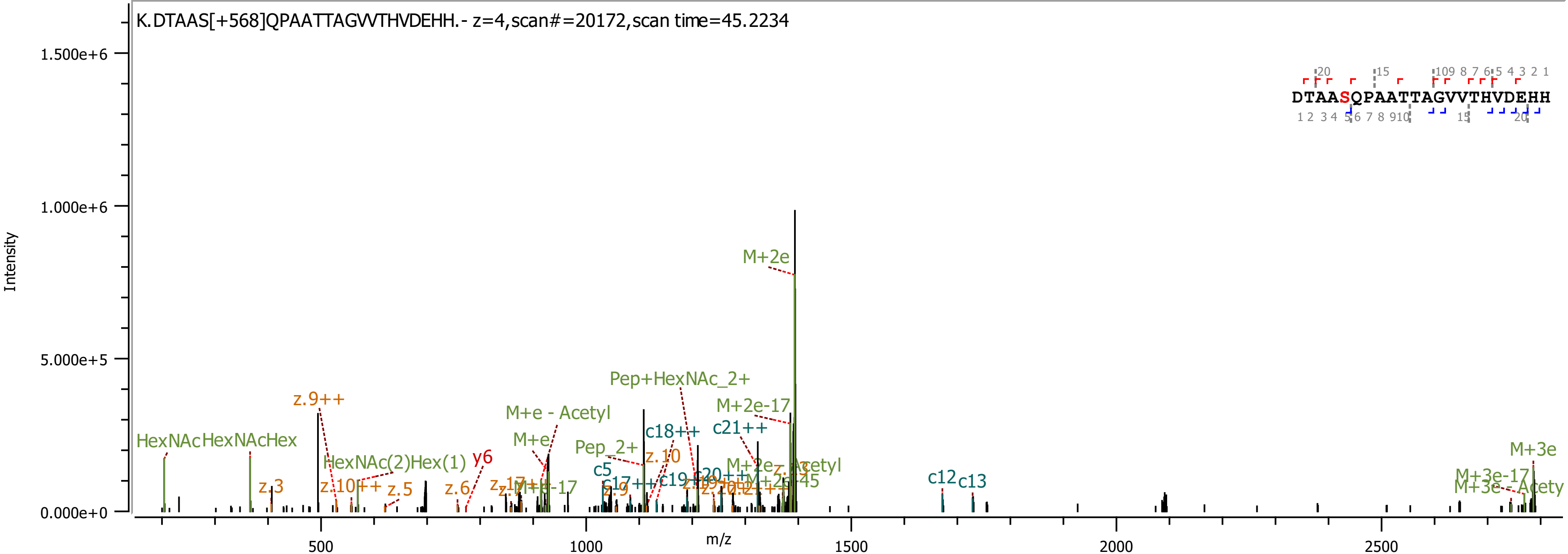

K.DTAAS[+568]QPAATTAGVVTHVDEHH.- z=4,scan#=20172,scan time=45.2234
DTAASQPAATTAGVVTHVDEHH
Intensity
m/z
1.500e+6
1.000e+6
5.000e+5
0.000e+0
500
1000
1500
2000
2500
M+2e
z.9++
Pep+HexNAc_2+
M+2e-17
M+e - Acetyl
c18++
c21++
HexNAc
HexNAcHex
M+e
Pep_2+
z.10
y6
HexNAc(2)Hex(1)
c5
z.3
z.10++
z.5
z.6
c12
c13
M+3e
M+3e-17

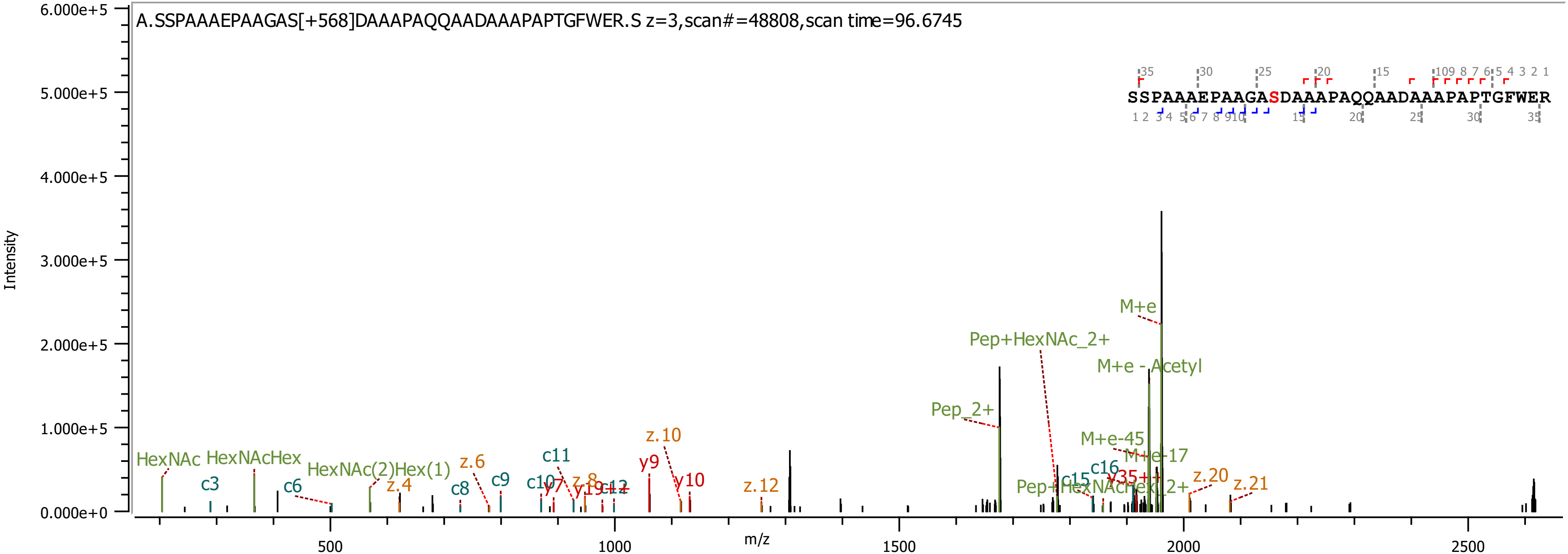

A.SSPAAAEPAAGAS[+568]DAAAPAQQAADAAAPAPTGFWER.S z=3,scan#=48808,scan time=96.6745
SSPAAAEPAAGASDAAAPAQQAADAAAPAPTGFWER
Intensity
m/z
M+e
Pep+HexNAc_2+
M+e - Acetyl
Pep_2+
M+e-45
M+e-17
HexNAc
HexNAcHex
HexNAc(2)Hex(1)
c3
c6
c8
c9
c11
c16
z.4
z.6
z.8
z.10
z.12
z.20
z.21
y9
y10

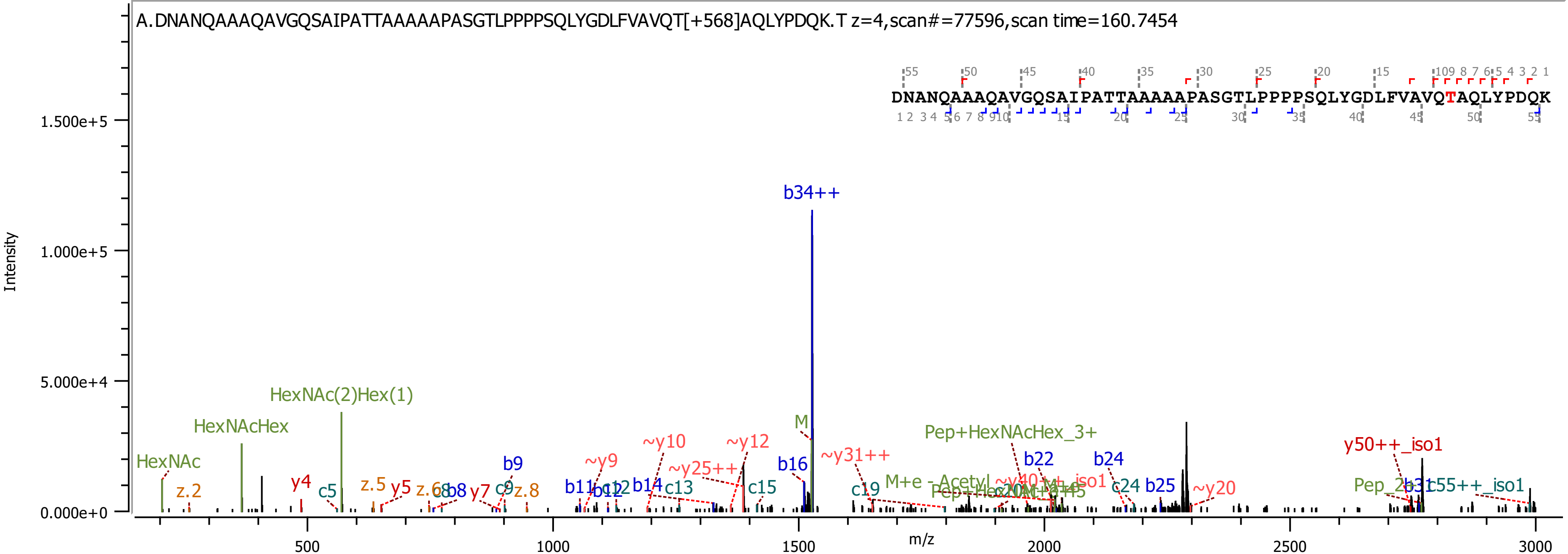
A.DNANQAAAQAVGQSAIPATTAAAAAPASGTLPPPPSQLYGDLFVAVQT[+568]AQLYPDQK.T z=4,scan#=77596,scan time=160.7454
DNANQAAAQAVGQSAIPATTAAAAAPASGTLPPPPSQLYGDLFVAVQTAQLYPDQK
Intensity
m/z
1.500e+5
1.000e+5
5.000e+4
0.000e+0
500
1000
1500
2000
2500
3000
b34++
HexNAc(2)Hex(1)
HexNAcHex
HexNAc
M
Pep+HexNAcHex_3+
y50++_iso1
b22
b24
~y10
~y12
~y9
~y25++
~y31++
b16
b9
y4
z.2
c5
z.5
y5
z.6
c8
b8
y7
c9
z.8
b11
c12
b12
b14
c13
c15
c19
c24
b25
~y20
Pep_2
b31
c55++_iso1

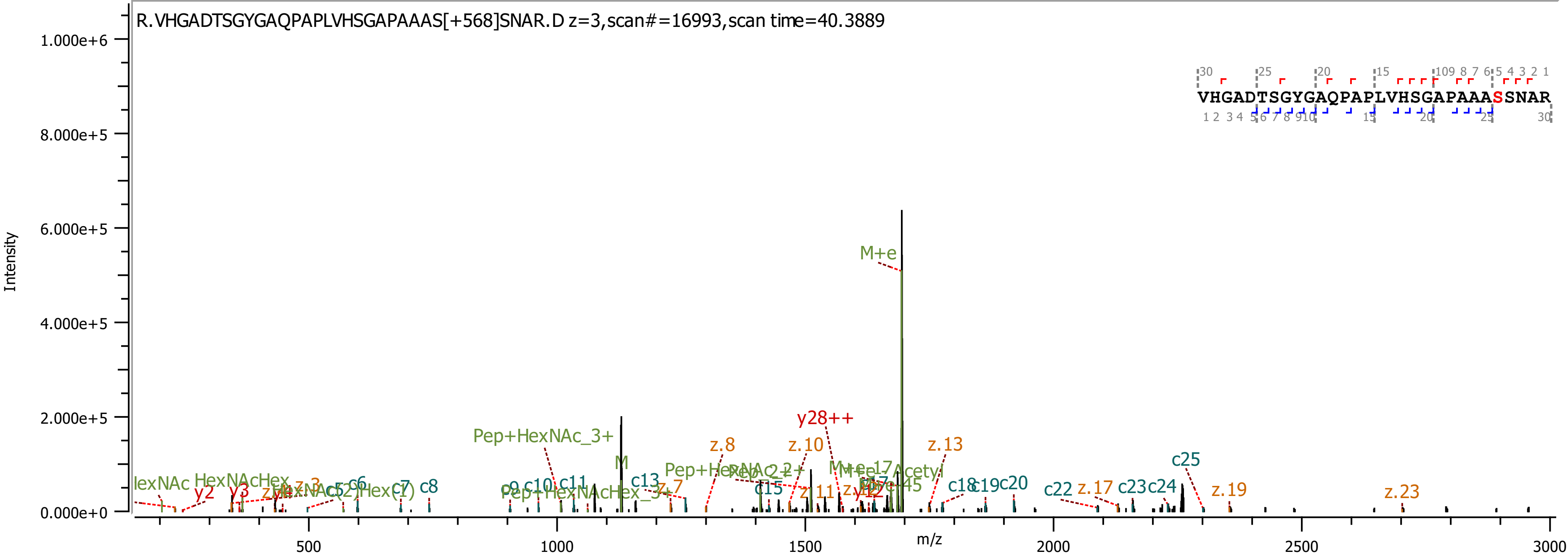
R.VHGADTSGYGAQPAPLVHSGAPAAAS[+568]SNAR.D z=3,scan#=16993,scan time=40.3889
VHGADTSGYGAQPAPLVHSGAPAAASSNAR
Intensity
m/z
1.000e+6
8.000e+5
6.000e+5
4.000e+5
2.000e+5
0.000e+0
500
1000
1500
2000
2500
3000
M+e
y28++
Pep+HexNAc_3+
z.8
z.10
z.13
c25
M
c13
z.7
HexNAcHex
y2
y3
c6
c7
c8
c10
c11
c15
z.11
c18
c19
c20
c22
z.17
c23
c24
z.19
z.23

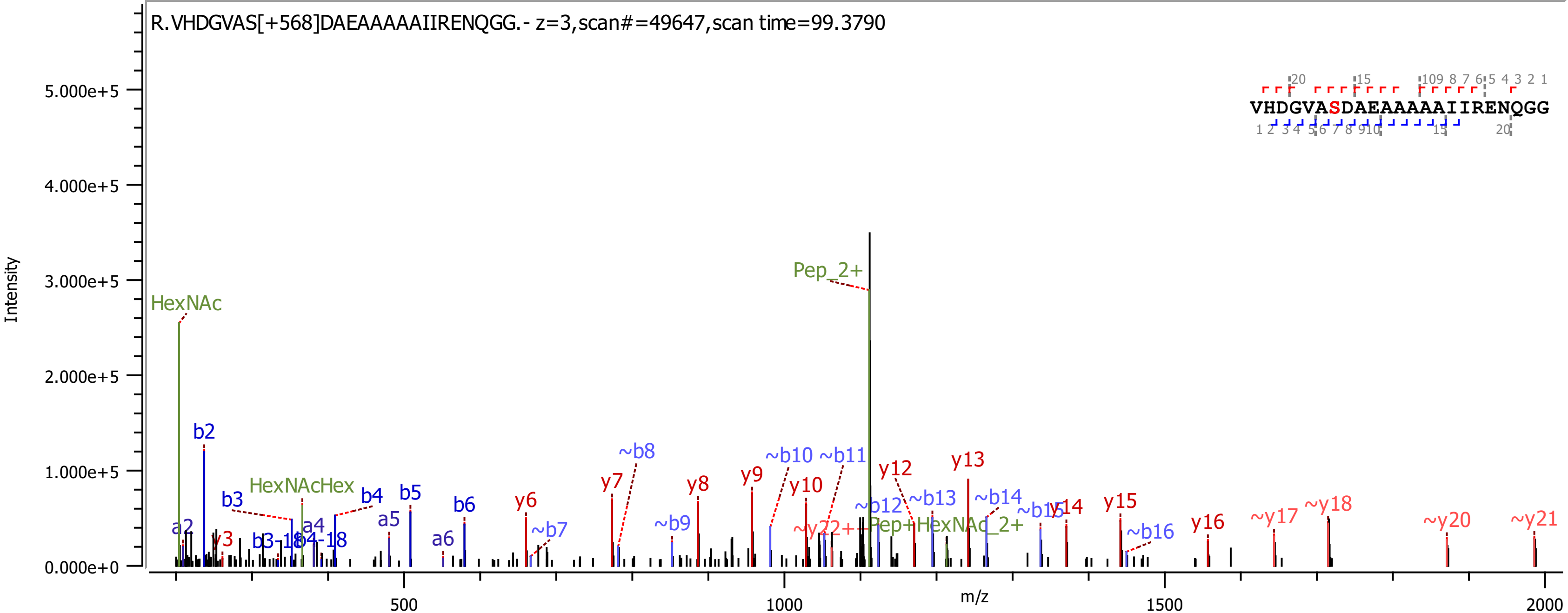

R.VHDGVAS[+568]DAEAAAAAIIRENQGG.- z=3,scan#=49647,scan time=99.3790
VHDGVASDAEAAAAAIIRENQGG
Intensity
m/z
5.000e+5
4.000e+5
3.000e+5
2.000e+5
1.000e+5
0.000e+0
500
1000
1500
2000
HexNAc
Pep_2+
HexNAcHex
Pep+HexNAc_2+
b2
b3
b4
b5
b6
a2
a4
a5
a6
y3
y6
y7
y8
y9
y10
y12
y13
y14
y15
y16
~b7
~b8
~b9
~b10
~b11
~b12
~b13
~b14
~b16
~y17
~y18
~y20
~y21

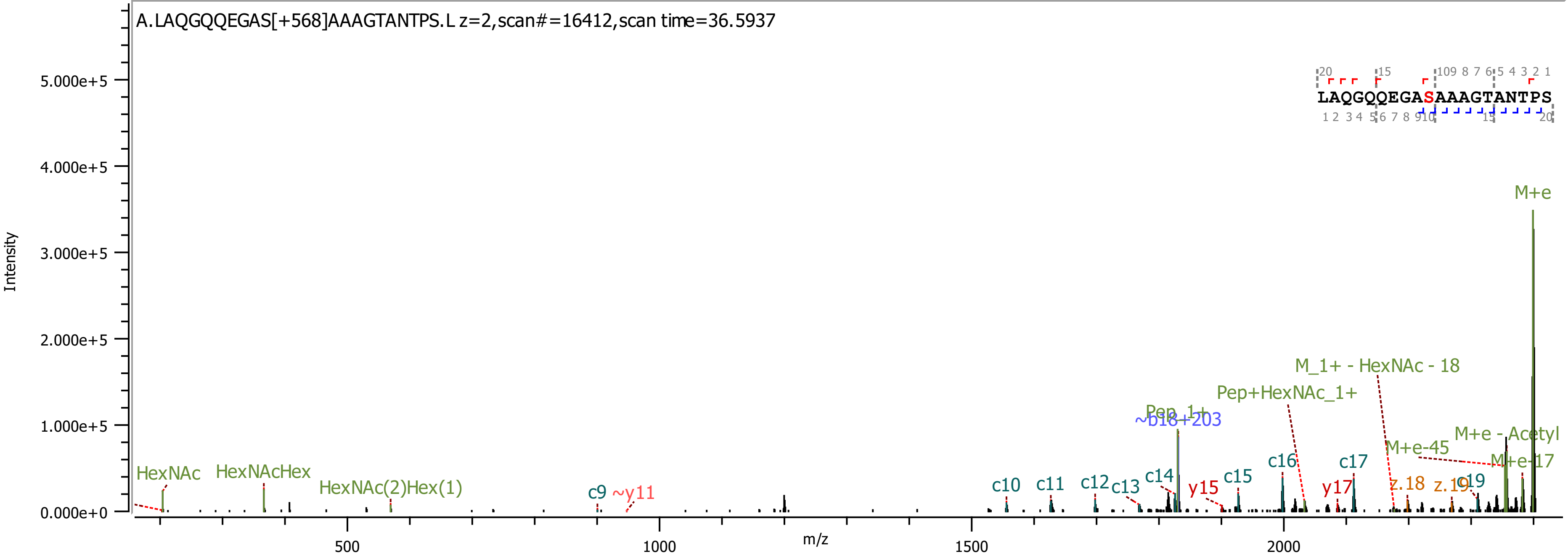
A.LAQGQQEGAS[+568]AAAGTANTPS.L z=2,scan#=16412,scan time=36.5937
Intensity
m/z
LAQGQQEGASAAAGTANTPS
M+e
M_1+ - HexNAc - 18
Pep+HexNAc_1+
Pep_1+
~b18+203
M+e - Acetyl
M+e-45
M+e-17
HexNAc
HexNAcHex
HexNAc(2)Hex(1)
c9
~y11
c10
c11
c12
c13
c14
y15
c15
c16
y17
c17
z.18
z.19
c19
0.000e+0
1.000e+5
2.000e+5
3.000e+5
4.000e+5
5.000e+5
500
1000
1500
2000

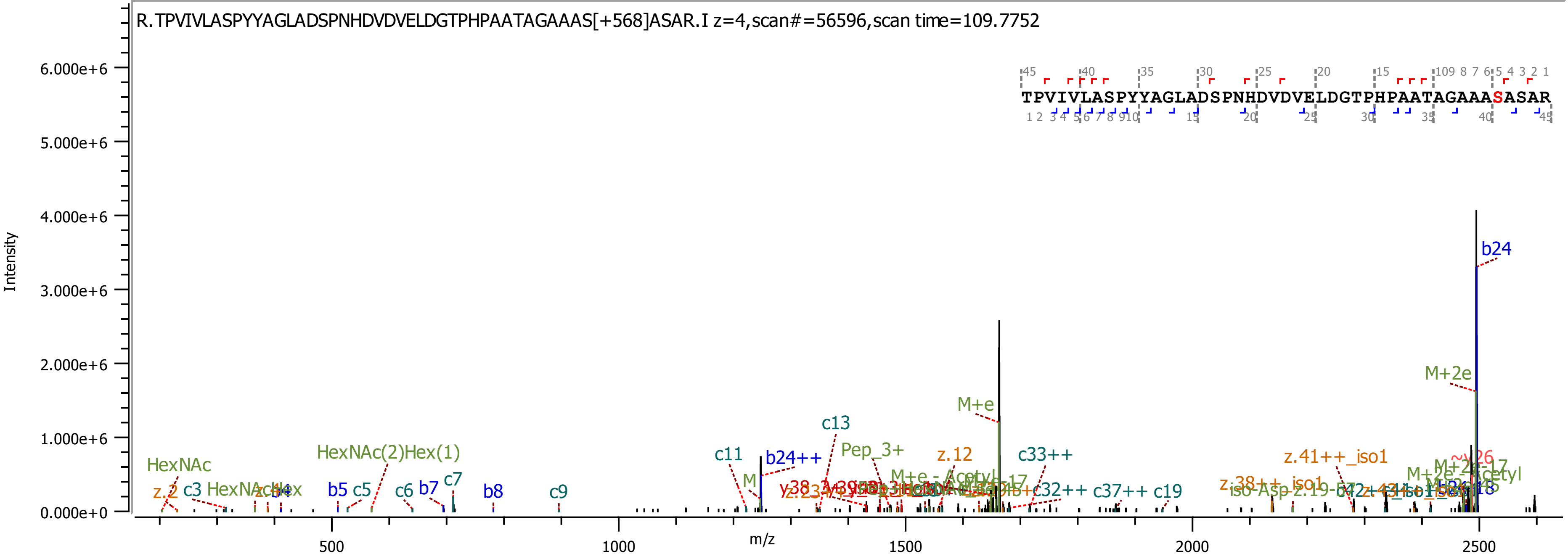
R.TPVIVLASPYYAGLADSPNHDVDVELDGTPHPAATAGAAAS[+568]ASAR.I z=4,scan#=56596,scan time=109.7752
TPVIVLASPYYAGLADSPNHDVDVELDGTPHPAATAGAAASASAR
Intensity
m/z
6.000e+6
5.000e+6
4.000e+6
3.000e+6
2.000e+6
1.000e+6
0.000e+0
500
1000
1500
2000
2500
b24
M+2e
M+e
c13
b24++
c11
Pep_3+
z.12
c33++
z.41++_iso1
HexNAc(2)Hex(1)
HexNAc
c3
b4
b5
c5
c6
b7
c7
b8
c9
c32++
c37++
c19
z.38++_iso1

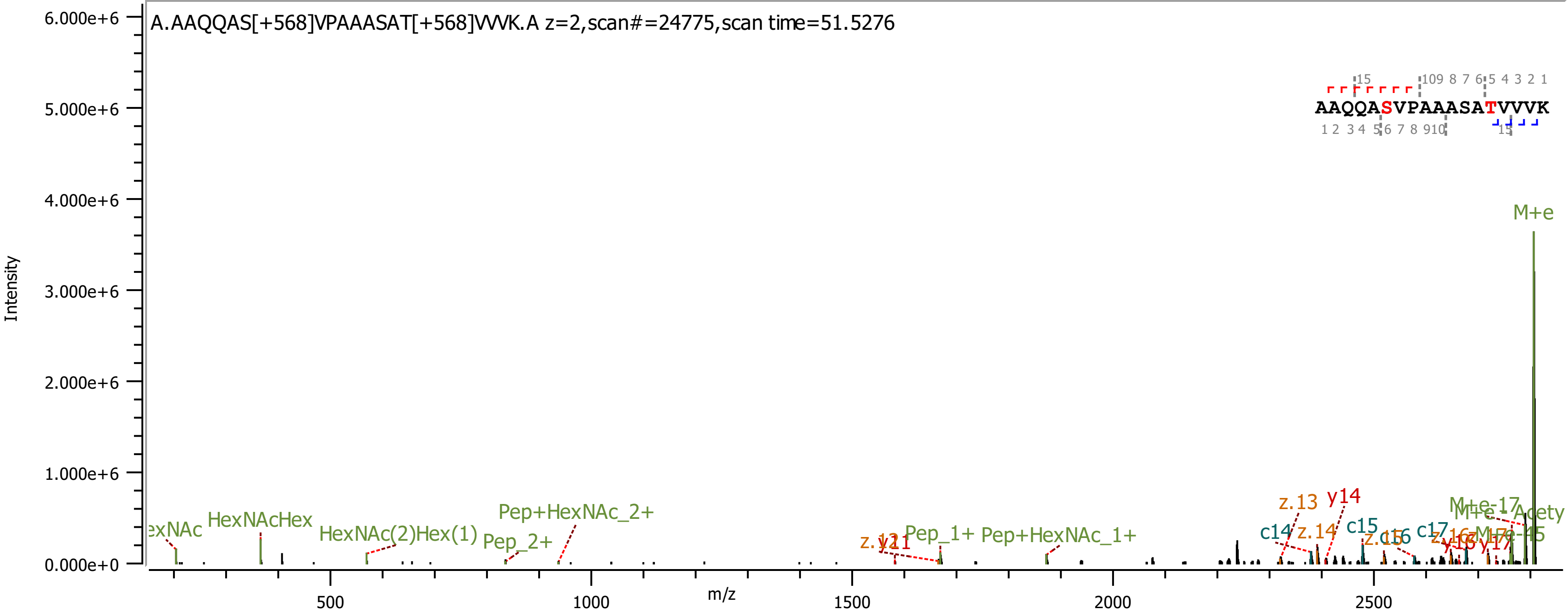

A.AAQQAS[+568]VPAAASAT[+568]VVVK.A z=2,scan#=24775,scan time=51.5276
AAQQASVPAAASATVVVK
Intensity
m/z
M+e
HexNAcHex
HexNAc(2)Hex(1)
Pep_2+
Pep+HexNAc_2+
Pep_1+
Pep+HexNAc_1+
z.13
y14
z.14
c14
c15
c16
c17
M+e-17
M+e-Acety

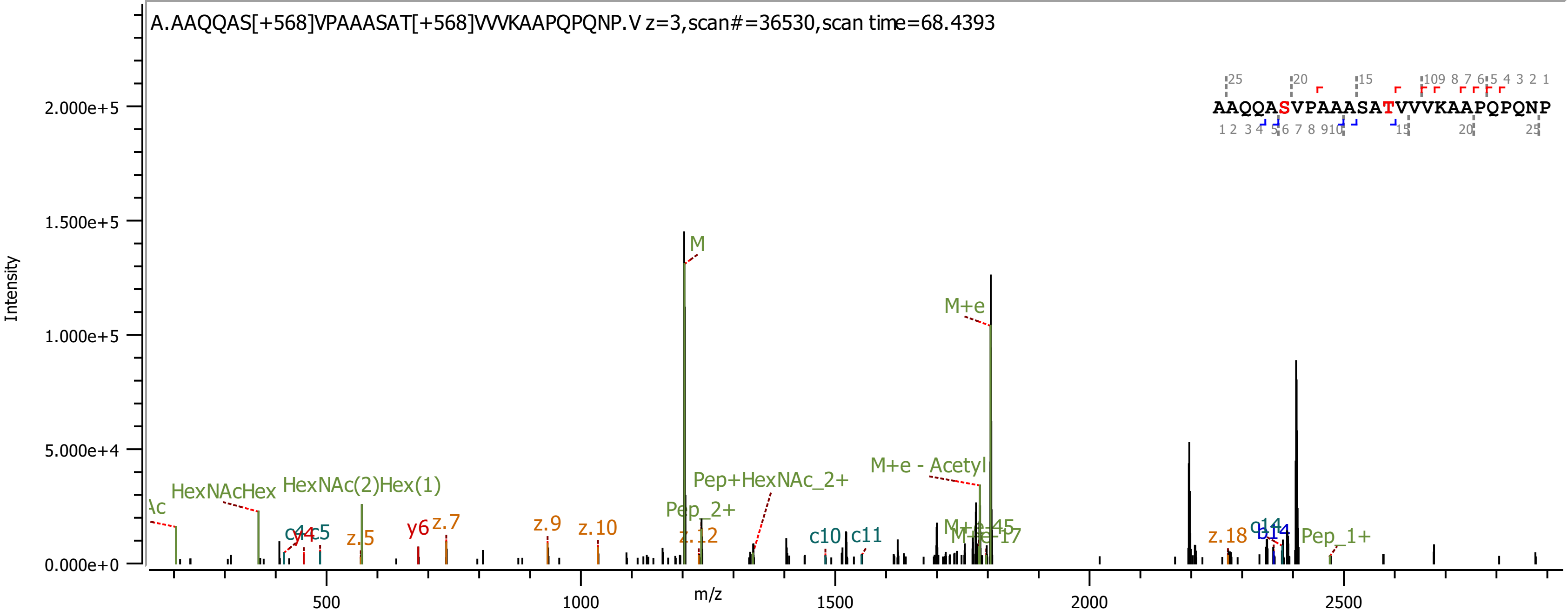

A.AAQQAS[+568]VPAAASAT[+568]VVVKAAPQPQNP.V z=3,scan#=36530,scan time=68.4393
AAQQASVPAAASATVVVKAAPQPQNP
Intensity
m/z
M
M+e
M+e - Acetyl
Pep+HexNAc_2+
Pep_2+
Pep_1+
HexNAcHex
HexNAc(2)Hex(1)
z.5
z.7
z.9
z.10
z.12
z.18
y6
c10
c11
c5

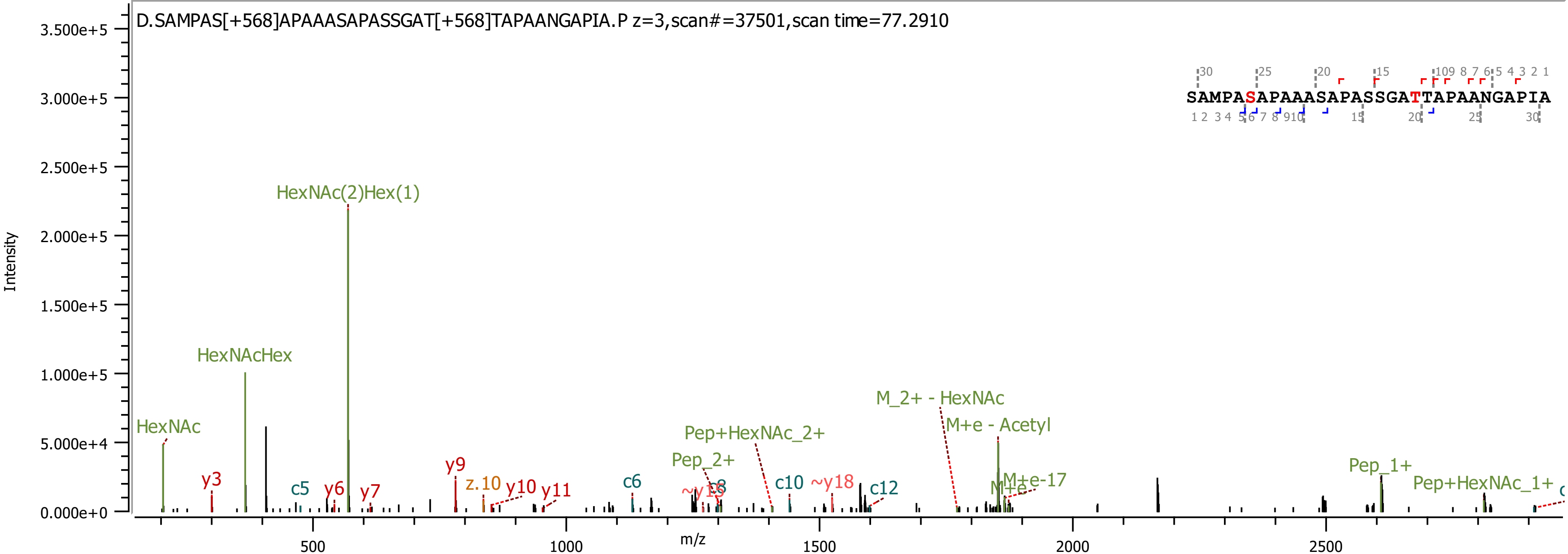

D.SAMPAS[+568]APAAASAPASSGAT[+568]TAPAANGAPIA.P z=3,scan#=37501,scan time=77.2910
SAMPASAPAAASAPASSGATTAPAANGAPIA
Intensity
m/z
HexNAc(2)Hex(1)
HexNAcHex
HexNAc
y3
c5
y6
y7
y9
z.10
y10
y11
c6
Pep_2+
Pep+HexNAc_2+
c10
~y18
c12
M_2+ - HexNAc
M+e - Acetyl
M+e-17
Pep_1+
Pep+HexNAc_1+

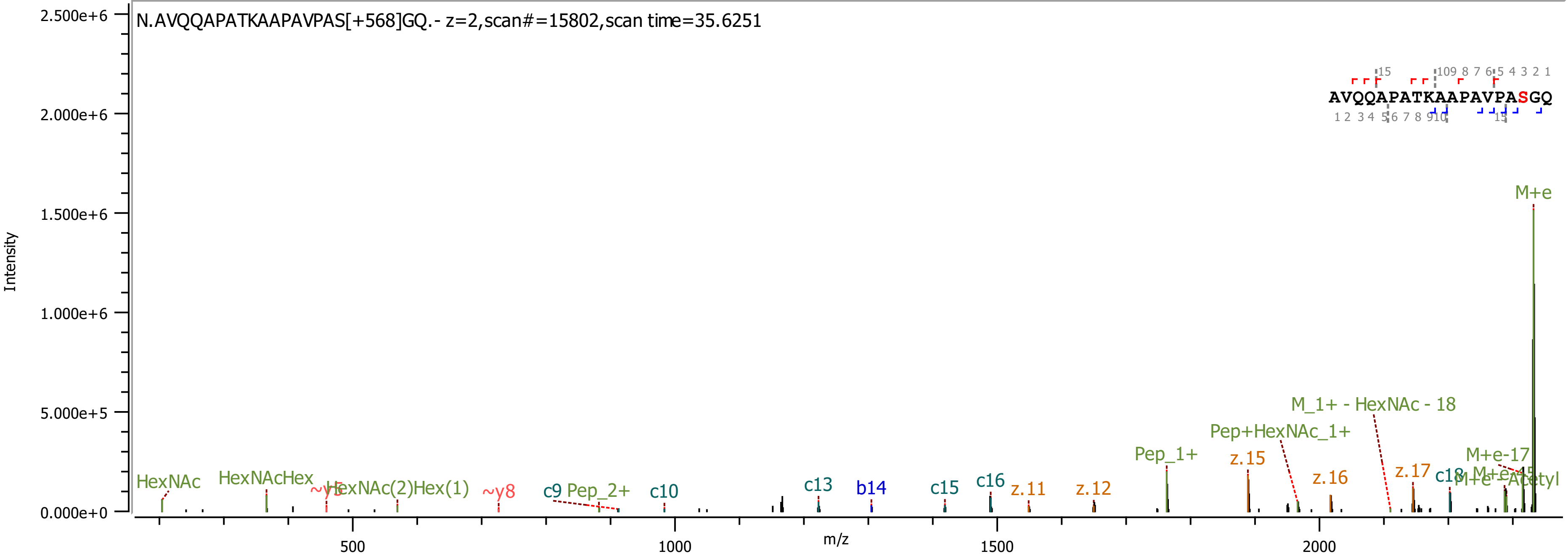
N.AVQQAPATKAAPAVPAS[+568]GQ.- z=2,scan#=15802,scan time=35.6251
AVQQAPATKAAPAVPASGQ
Intensity
m/z
2.500e+6
2.000e+6
1.500e+6
1.000e+6
5.000e+5
0.000e+0
500
1000
1500
2000
M+e
M_1+ - HexNAc - 18
Pep+HexNAc_1+
Pep_1+
z.15
M+e-17
z.16
z.17
c18
M+e-Acetyl
HexNAc
HexNAcHex
~y5
HexNAc(2)Hex(1)
~y8
c9
Pep_2+
c10
c13
b14
c15
c16
z.11
z.12

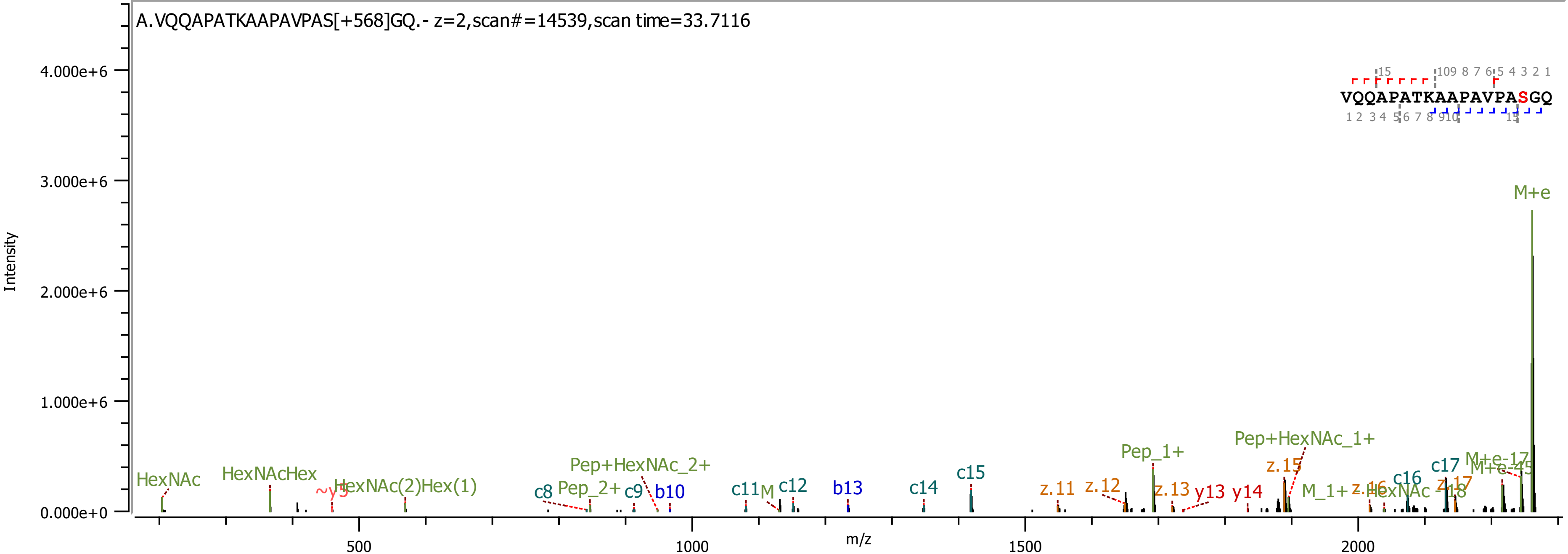
A.VQQAPATKAAPAVPAS[+568]GQ.- z=2,scan#=14539,scan time=33.7116
VQQAPATKAAPAVPASGQ
Intensity
m/z
4.000e+6
3.000e+6
2.000e+6
1.000e+6
0.000e+0
500
1000
1500
2000
M+e
HexNAc
HexNAcHex
HexNAc(2)Hex(1)
Pep+HexNAc_2+
Pep_2+
c8
c9
b10
c11
c12
b13
c14
c15
z.11
z.12
Pep_1+
z.13
y13
y14
Pep+HexNAc_1+
z.15
M_1+
c16
c17
M+e-17

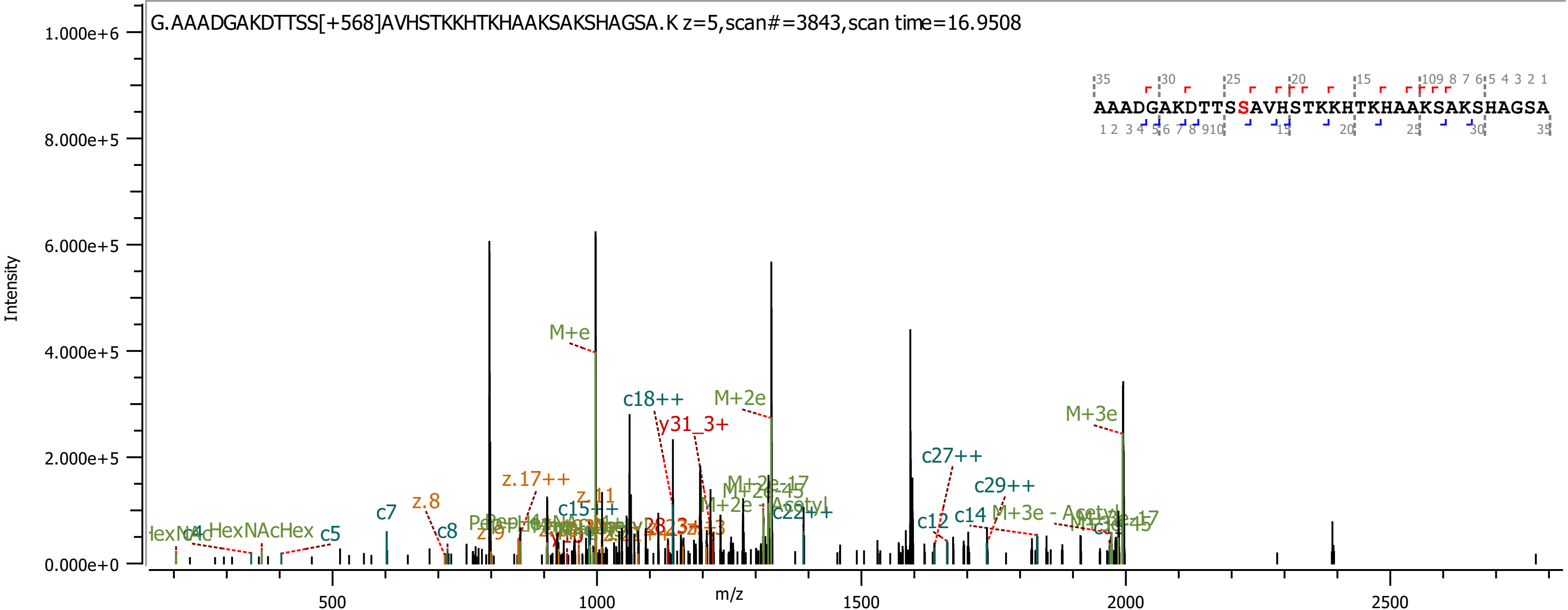
G.AAADGAKDTTSS[+568]AVHSTKKHTKHAAKSAKSHAGSA.K z=5,scan#=3843,scan time=16.9508
AAADGAKDTTSSAVHSTKKHTKHAAKSAKSHAGSA
Intensity
m/z
1.000e+6
8.000e+5
6.000e+5
4.000e+5
2.000e+5
0.000e+0
500
1000
1500
2000
2500
M+e
c18++
M+2e
y31_3+
M+3e
c27++
c29++
z.17++
z.8
c7
c8
c5
HexNAcHex
c15++
c22++
c12
c14
M+3e - Acetyl
M+2e - Acetyl

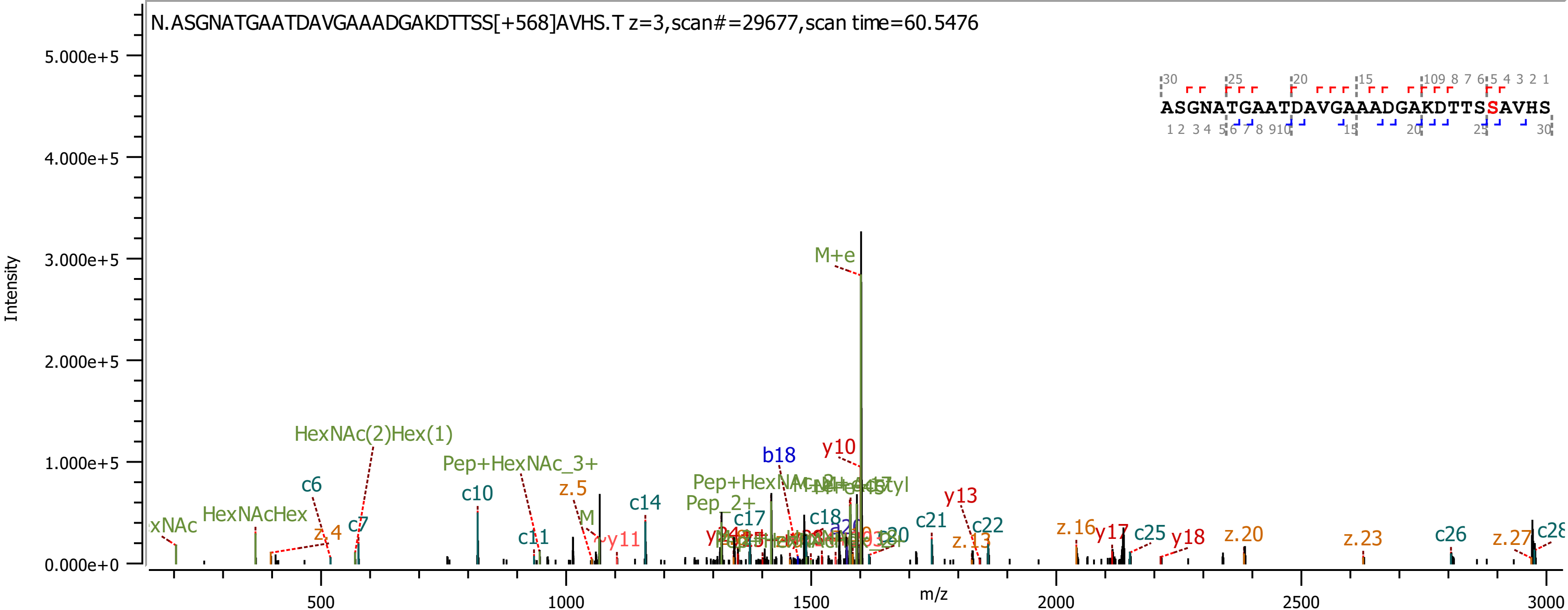
N.ASGNATGAATDAVGAAADGAKDTTSS[+568]AVHS.T z=3,scan#=29677,scan time=60.5476
ASGNATGAATDAVGAAADGAKDTTSSAVHS
Intensity
m/z
M+e
y10
b18
HexNAc(2)Hex(1)
Pep+HexNAc_3+
HexNAcHex
c6
c7
c10
c11
c14
c17
c18
c21
c22
c25
c26
z.4
z.5
z.13
z.16
z.20
z.23
z.27
y13
y17
y18
Pep_2+

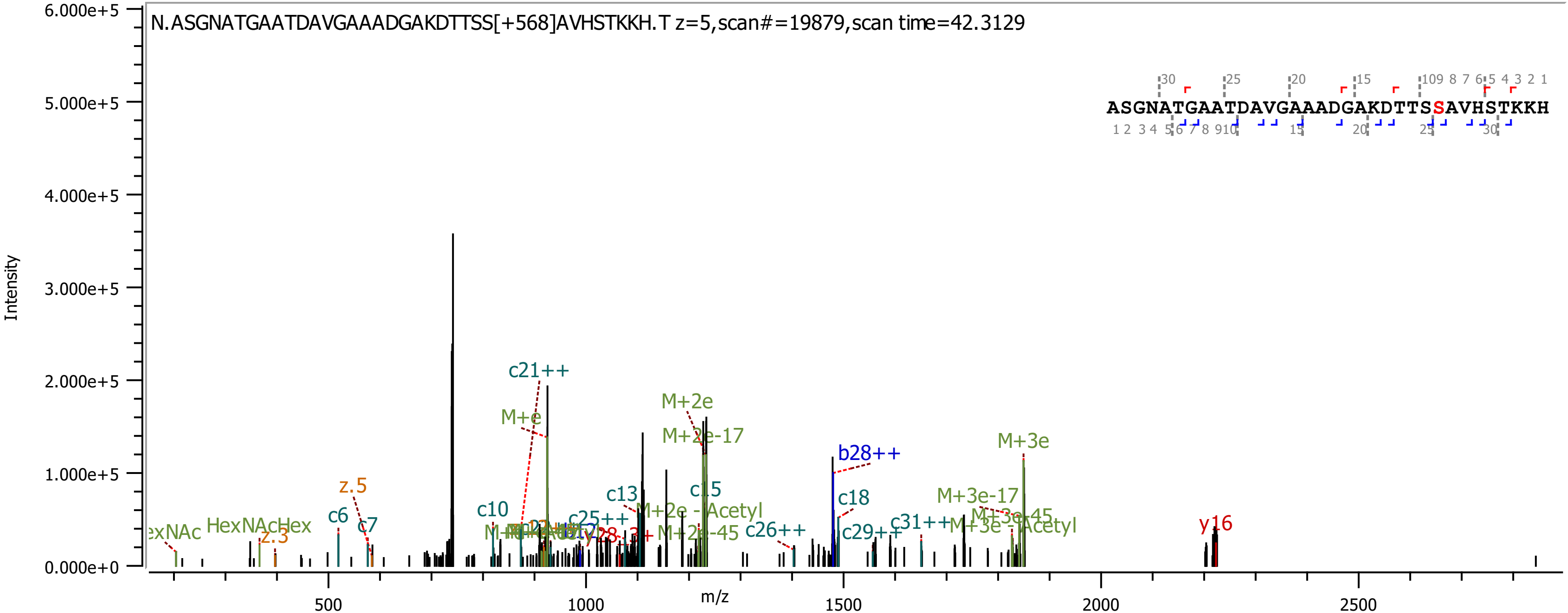
N.ASGNATGAATDAVGAAADGAKDTTSS[+568]AVHSTKKH.T z=5,scan#=19879,scan time=42.3129
ASGNATGAATDAVGAAADGAKDTTSSAVHSTKKH
Intensity
m/z
c21++
M+e
M+2e
M+2e-17
b28++
M+3e
c18
c15
c13
c10
c6
c7
z.5
z.3
HexNAcHex
M+3e-17
y16
c26++
c29++
c31++
c25++
M+2e - Acetyl
M+2e-45
M+3e - Acetyl
0.000e+0
1.000e+5
2.000e+5
3.000e+5
4.000e+5
5.000e+5
6.000e+5
500
1000
1500
2000
2500

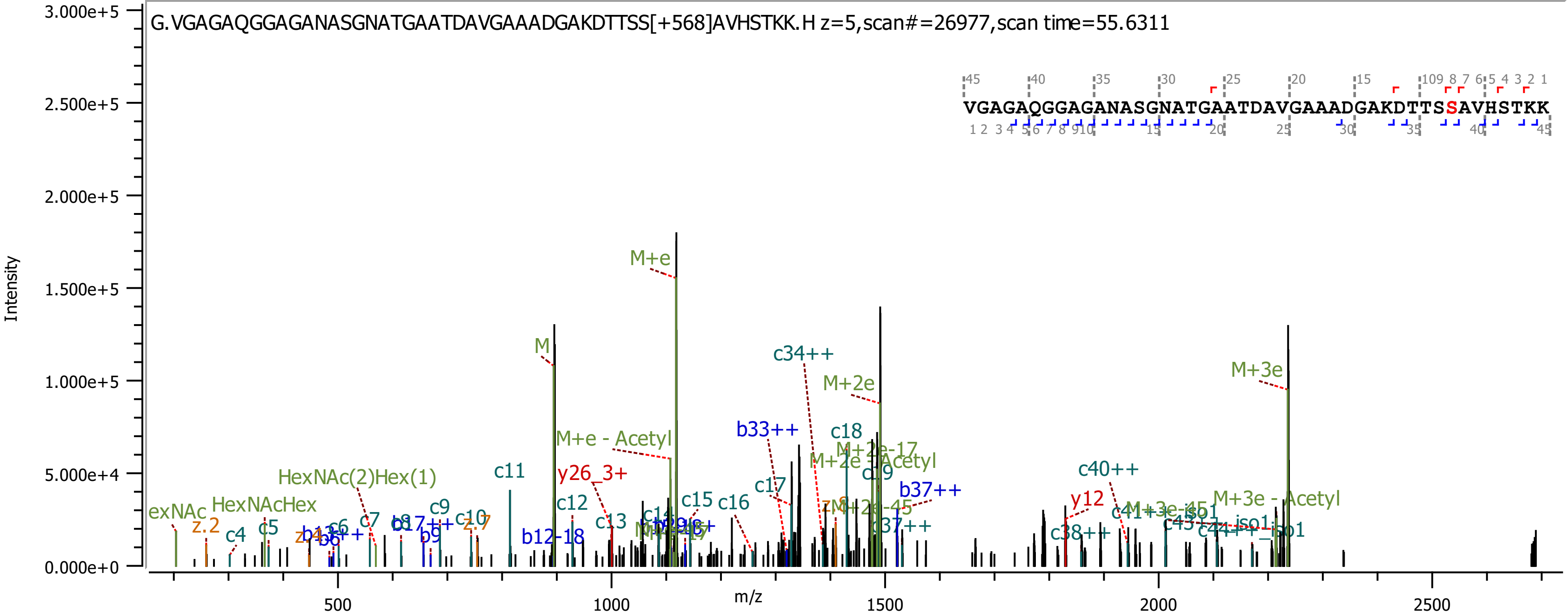
G.VGAGAQGGAGANASGNATGAATDAVGAAADGAKDTTSS[+568]AVHSTKK.H z=5,scan#=26977,scan time=55.6311
VGAGAQGGAGANASGNATGAATDAVGAAADGAKDTTSSAVHSTKK
Intensity
m/z
M+e
M
M+2e
M+3e
c34++
b33++
c18
c11
y26_3+
M+e - Acetyl
HexNAc(2)Hex(1)
HexNAcHex
c40++
y12
M+3e - Acetyl
b37++
c17
c16
c15
c13
c12
b12-18
c10
c9
c7
c5
c4
z.2
c38++
c37++

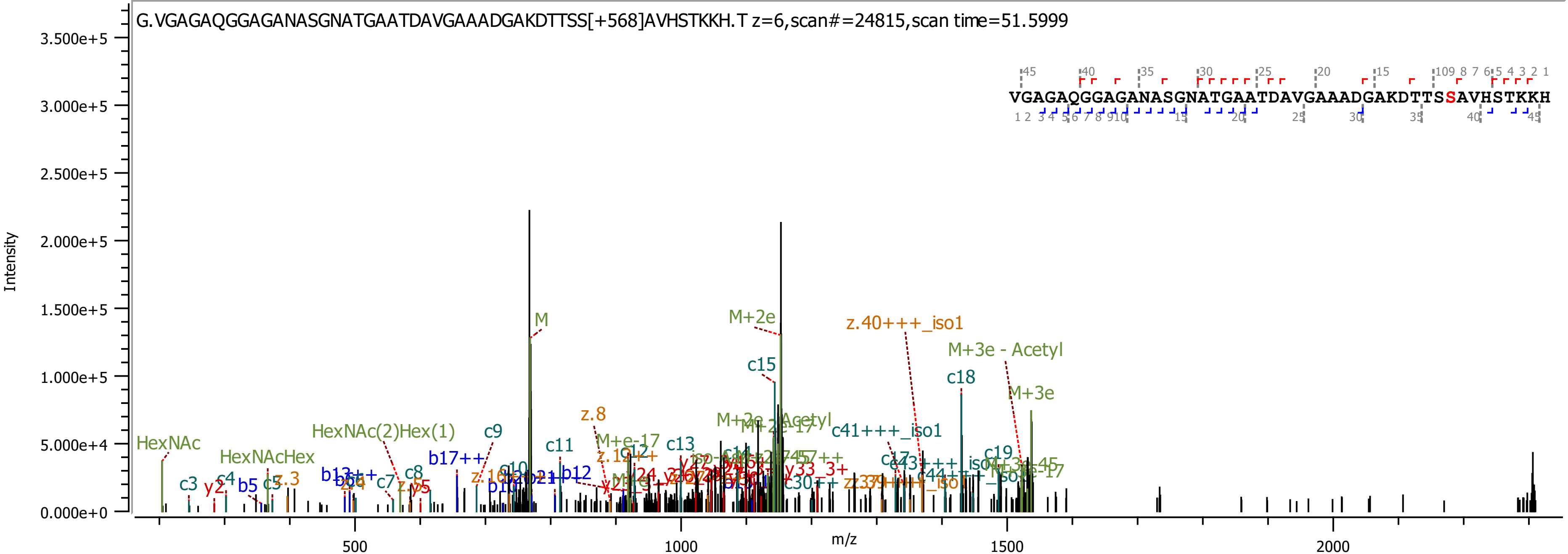
G.VGAGAQGGAGANASGNATGAATDAVGAAADGAKDTTSS[+568]AVHSTKKH.T z=6,scan#=24815,scan time=51.5999
VGAGAQGGAGANASGNATGAATDAVGAAADGAKDTTSSAVHSTKKH
Intensity
m/z
3.500e+5
3.000e+5
2.500e+5
2.000e+5
1.500e+5
1.000e+5
5.000e+4
0.000e+0
500
1000
1500
2000
M
M+2e
z.40+++_iso1
M+3e - Acetyl
c15
c18
M+3e
z.8
HexNAc(2)Hex(1)
c9
HexNAc
HexNAcHex
c11
c13
c41+++_iso1
c3
y2
c4
b5
c5
z.3
b17++
c8
c7
z.5
c19
y33_3+
b12
M+e-17

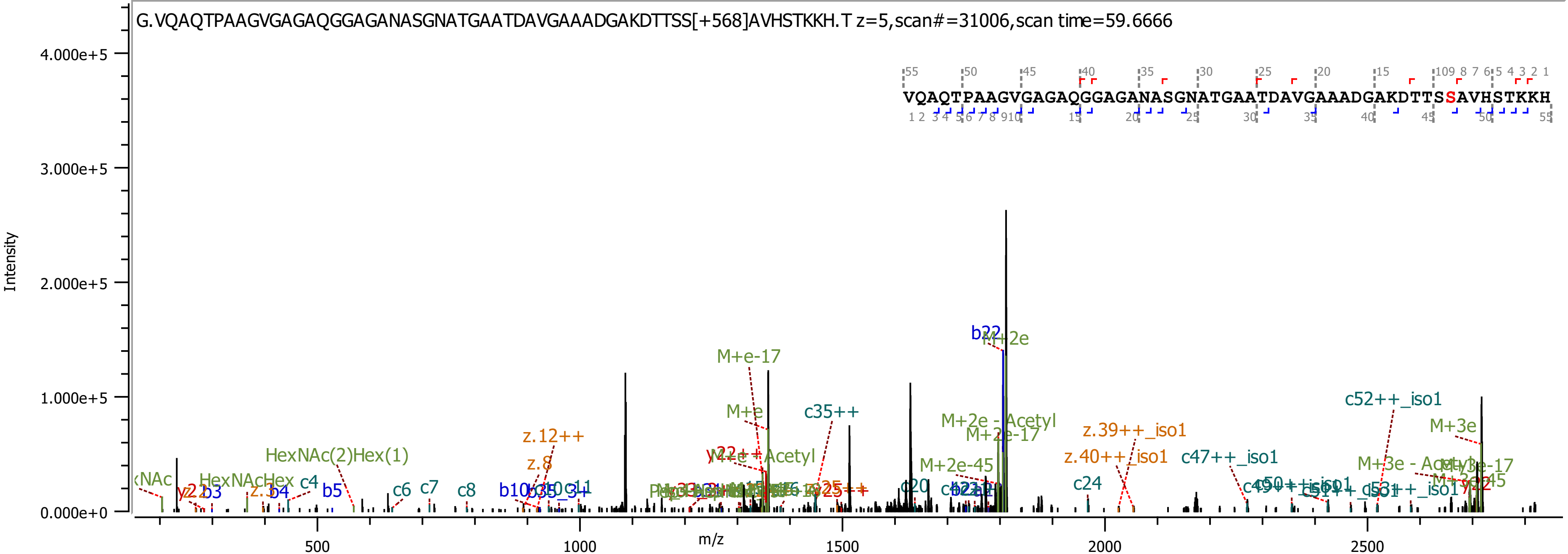

G.VQAQTPAAGVGAGAQGGAGANASGNATGAATDAVGAAADGAKDTTSS[+568]AVHSTKKH.T z=5,scan#=31006,scan time=59.6666
VQAQTPAAGVGAGAQGGAGANASGNATGAATDAVGAAADGAKDTTSSAVHSTKKH
Intensity
4.000e+5
3.000e+5
2.000e+5
1.000e+5
0.000e+0
m/z
500
1000
1500
2000
2500
b22
M+2e
M+e-17
M+e
c35++
M+2e - Acetyl
M+2e-17
M+2e-45
z.39++_iso1
z.40++_iso1
c47++_iso1
c52++_iso1
M+3e
M+3e - Acetyl
HexNAc(2)Hex(1)
HexNAcHex
z.12++
z.8
c4
b3
b4
b5
c6
c7
c8
c20
c24

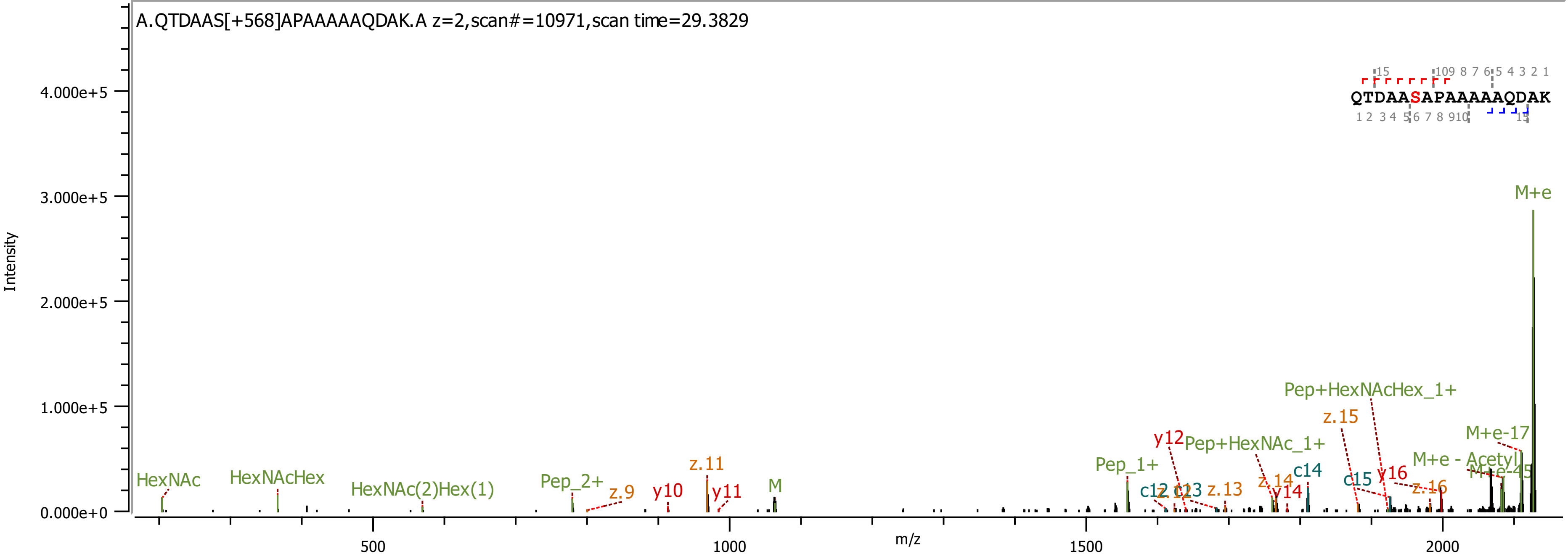
A.QTDAAS[+568]APAAAAAQDAK.A z=2,scan#=10971,scan time=29.3829
Intensity
m/z
QTDAASAPAAAAAQDAK
HexNAc
HexNAcHex
HexNAc(2)Hex(1)
Pep_2+
z.9
y10
z.11
y11
M
Pep_1+
y12
Pep+HexNAc_1+
c12
c13
z.13
z.14
y14
c14
z.15
c15
Pep+HexNAcHex_1+
y16
z.16
M+e - Acetyl
M+e-45
M+e-17
M+e
4.000e+5
3.000e+5
2.000e+5
1.000e+5
0.000e+0
500
1000
1500
2000
